# Sequence-specific trapping of EF-Tu/glycyl-tRNA complex on the ribosome by bottromycin

**DOI:** 10.1101/2025.08.17.670399

**Authors:** Dmitrii Y. Travin, Ritwika S. Basu, Madhura N. Paranjpe, Dorota Klepacki, Anna I. Zhurakovskaya, Nora Vázquez-Laslop, Alexander S. Mankin, Yury S. Polikanov, Matthieu G. Gagnon

## Abstract

The development of antibiotics with novel mechanisms of action is essential to address the growing threat of antimicrobial resistance. Protein synthesis-inhibiting antibiotic bottromycin (BOT), a ribosomally synthesized and posttranslationally modified peptide (RiPP), has long been known for its potent activity against Gram-positive bacteria but was largely neglected due in part to the lack of understanding of its mechanism of action. Here we uncover the unprecedented mode translation inhibition strategy employed by BOT. Using biochemical, microbiological, genetic, and structural approaches, we show that BOT acts by selectively trapping elongation factor-Tu (EF-Tu) in complex with glycyl-tRNA on the ribosome. BOT binds at the interface between EF-Tu and the CCA-end of Gly-tRNA, stabilizing the EF-Tu/Gly-tRNA complex in a pre-accommodated A/T-state on the ribosome, and specifically arresting translation at glycine codons. This mode of action is mechanistically distinct from that of other EF-Tu-targeting antibiotics, which act in a tRNA-agnostic fashion. Point mutations in EF-Tu confer high-level resistance to BOT, confirming EF-Tu as the direct and essential target of the drug. Our findings establish BOT as a founding member of a new class of antibiotics that stall the ribosome at defined mRNA sites by trapping a specific elongation factor-tRNA complex.

## INTRODUCTION

The relentless spread of antimicrobial resistance (AMR) among pathogenic bacteria continues to outpace the development of new drugs, posing a critical global health challenge^1^. Most antibiotics in clinical use today act through a limited number of well-characterized mechanisms, and resistance to these drugs is widely distributed in pathogenic microorganisms. Therefore, there is an urgent need for antibiotics with novel mechanisms of action. The prototypes of the majority of antimicrobial compounds currently used in clinic were discovered during the so-called “Golden Age of Antibiotics” (1940s-1960s), when the screening of soil actinobacteria yielded a wealth of bioactive natural products. Unfortunately, many of these compounds were shelved due to the lack of understanding of their mechanisms of action or unfavorable characteristics, such as poor solubility, low production yields, metabolic instability, etc. In the current era, however, these “forgotten” molecules represent a valuable reservoir of bioactivity, especially when the antibiotic discovery pipeline is draining, and because advances in synthetic biology and chemical derivatization of natural products could improve their properties. Yet, for many of them, the lack of mechanistic insight, particularly regarding their molecular targets and modes of action, hinders their further rational development into leads for preclinics.

Bottromycins (BOTs) are a striking example of such abandoned antibiotics. First isolated in the 1950s from *Streptomyces bottropensis*^2^, BOTs are active against a range of Gram-positive pathogens (e.g., *Streptococcus, Bacillus, Corynebacterium*), including multidrug-resistant strains such as methicillin-resistant *Staphylococcus aureus* and vancomycin-resistant enterococci, as well as certain *Mycobacterium* and *Mycoplasma* species^3–6^. BOTs also show activity against the devastating Gram-negative rice pathogen *Xanthomonas oryzae* pv. *oryzae*^7^, suggesting potential agricultural applications.

BOTs belong to ribosomally synthesized and posttranslationally modified peptides (RiPPs)^8,9^. BOTs produced by several *Streptomyces* strains differ in a single position in the sequence of the precursor peptide and a set of installed modifications. Among the naturally occurring variants, bottromycin A2 is the most studied and the only commercially available class representative (**Fig. 1a**, hereafter referred to as BOT). BOT is synthesized from an octapeptide GPVVVFDC and features multiple modifications, including a 12-membered macroamidine cycle at the N-terminus, several C-and O-methylations, epimerization of the Asp residue, and a thiazole ring at the C-terminus (**Fig. 1a**) (reviewed in ^10^), many of which are required for its antimicrobial activity **(Fig. 1b)**. Despite BOT’s high potency, the precise molecular mechanism by which it inhibits bacterial translation has remained elusive. Early reports showed that BOT inhibits protein synthesis both *in vitro* and *in vivo*^11,12^. Further studies suggested that BOT associates with the large (50S) ribosomal subunit^13^ and that it inhibits translocation or interferes with aminoacyl-tRNA (aa-tRNA) accommodation into the A site, but the data have often been contradictory and lacked a structural basis^14,15^.

**Figure 1.**
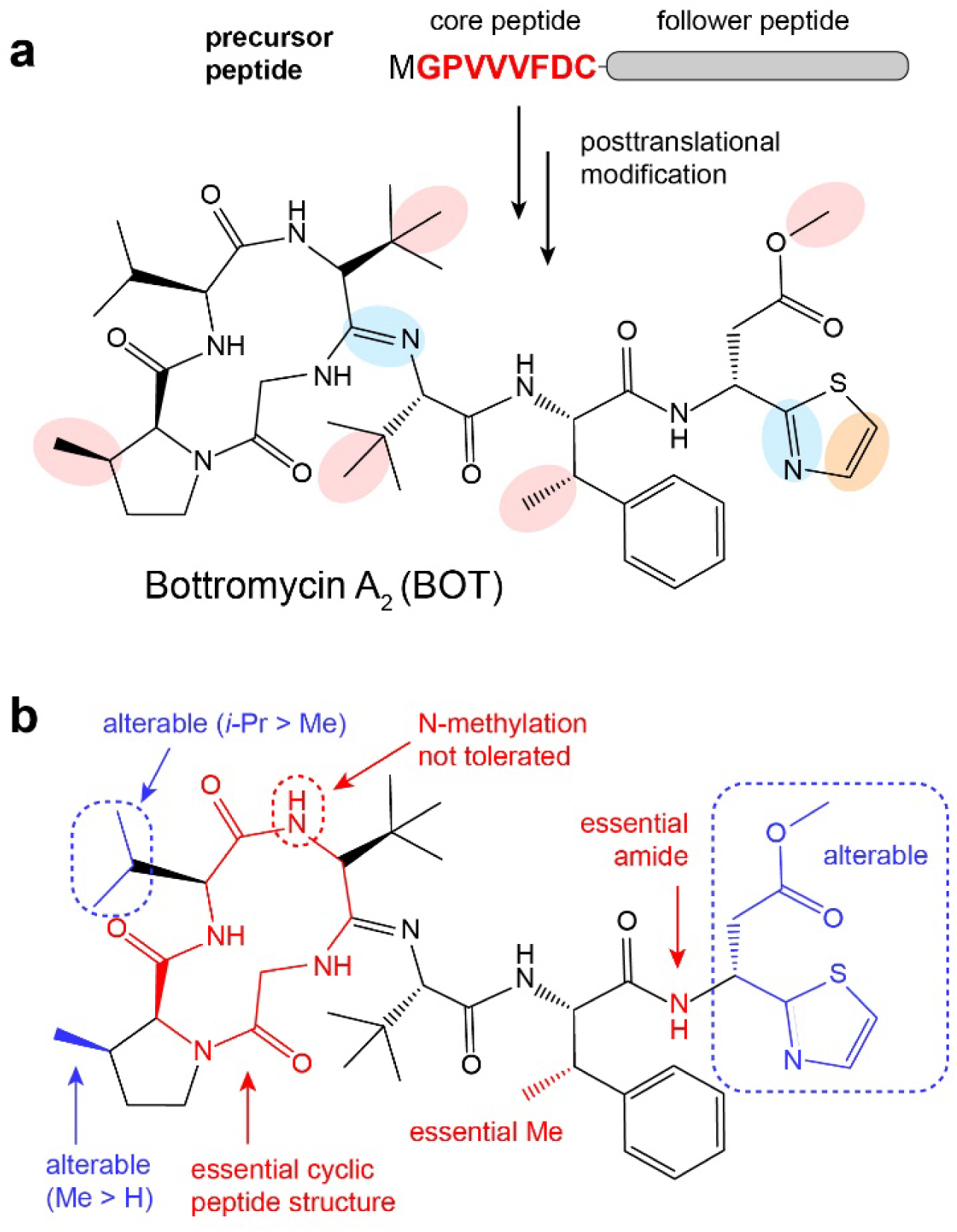
Bottromycins: chemical structure and structure-activity relationships. (**a**) Chemical structure of bottromycin A_2_. Posttranslational modifications installed on the core peptide are color-coded: side-chain methylations (red), YcaO enzyme-catalyzed cyclizations (blue), and oxidation of the C-terminal thiazoline into thiazole (orange). (**b**) Summary of structure-activity relationship (SAR) studies of naturally occurring and synthetic bottromycin variants.

Here, by employing a combination of biochemical, microbiological, genetic, and structural approaches, we uncovered the mechanism by which BOT inhibits translation. Contrary to earlier models implicating the A site of the peptidyl transferase center (PTC) on the 50S subunit as the drug’s binding site, we show that BOT does not target the ribosome *per se*. Instead, it binds to the EF-Tu/Gly-tRNA^Gly^ complex and traps it on the ribosome, preventing tRNA release and accommodation into the A site. As a result, the BOT stalls the translating ribosomes at the Gly codons of mRNA. This distinct mechanism establishes BOT as a first-in-class antibiotic that acts by targeting a key translation factor (EF-Tu) in a single-tRNA-specific manner. The redundancy of the EF-Tu genes in many pathogenic bacteria and the recessive nature of the resistance mutations account for the low frequency of appearance of resistant mutants. The exceptional and highly tRNA-dependent mode of BOT action not only expands our understanding of how translation can be selectively modulated but also opens new avenues for further rational development of BOTs into drugs with improved pharmacological profiles.

## RESULTS

### BOT specifically stalls ribosomes on Gly codons

We employed ribosome profiling (Ribo-seq)^16^ to obtain an unbiased, genome-wide view of the distribution of actively translating ribosomes throughout mRNAs in cells treated with BOT. As a model organism, we selected Gram-positive *Bacillus subtilis,* which is highly sensitive to BOT (minimal inhibitory concentration, MIC = 0.06 µg/mL)^5^. Exponentially growing cells were treated with BOT at 50×MIC for 5 minutes, a condition sufficient to shut down cellular protein synthesis to near completion (**Extended Data Fig. 1a**), polysomes were isolated, and ribosome-protected mRNA fragments were processed for Ribo-seq analysis.

Ribo-seq unexpectedly revealed a striking sequence specificity in BOT-induced ribosome stalling: ribosomes accumulated predominantly at sites where a Gly codon occupied the ribosomal A site (**Fig. 2a, b**). Such a specificity pattern is typically associated with depletion of charged Gly-tRNA^Gly^, as would be expected from a glycyl-tRNA-synthetase (GlyRS) inhibitor, whose activity decreases the amount of charged tRNA^Gly^ in the cell, ultimately leading to ribosome stalling at “hungry” Gly codons. To test this possibility, we monitored *in vitro* aminoacylation of tRNA^Gly^ with [^14^C]-glycine by *Escherichia coli* GlyRS in the presence and absence of 50 µM BOT. While the known GlyRS inhibitor Gly-AMS completely abolished aminoacylation, BOT had no detectable effect on tRNA charging (**Fig. 2c**), indicating that it does not inhibit GlyRS.

**Figure 2.**
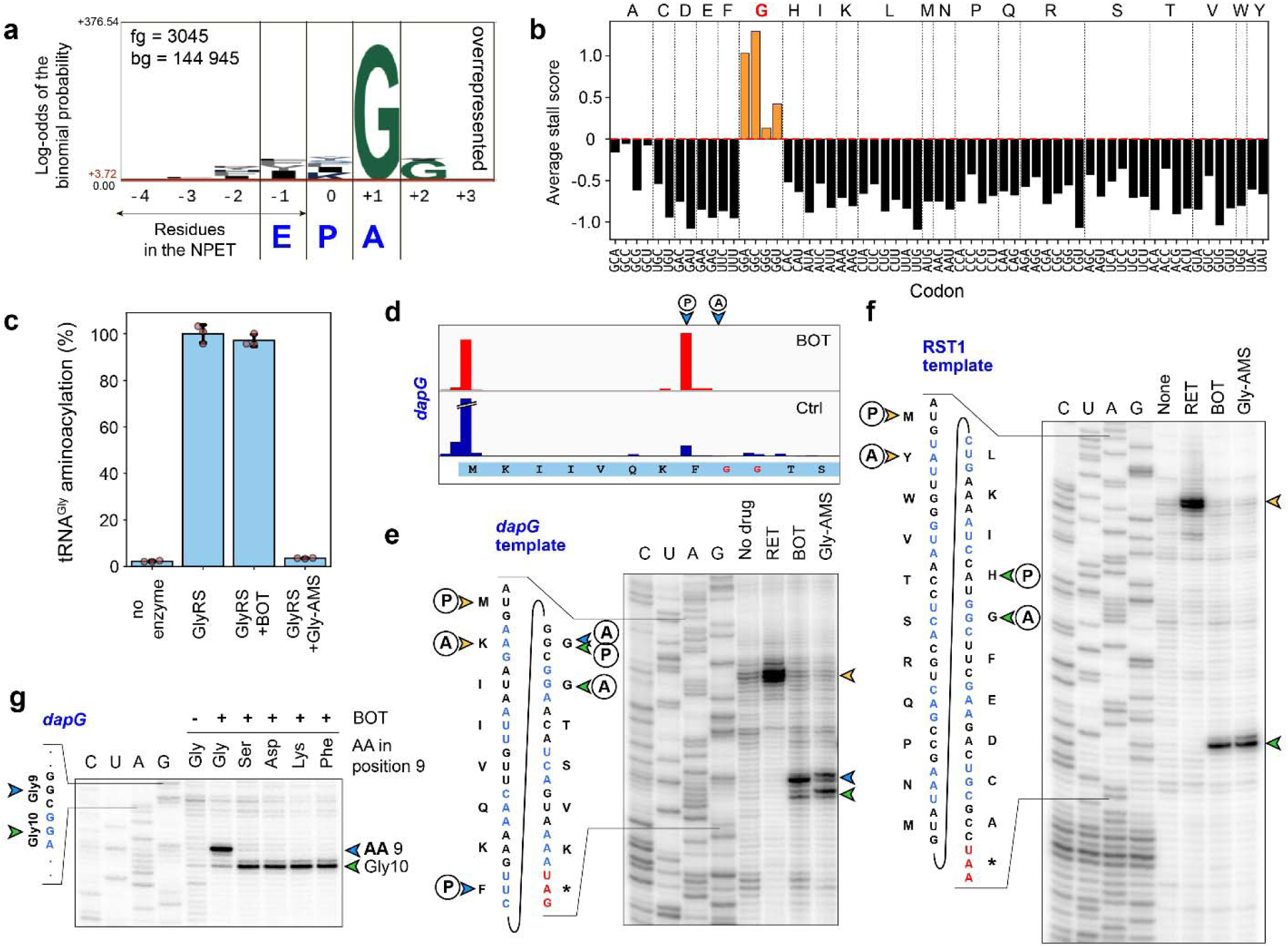
BOT inhibits bacterial translation in a sequence-specific manner. (**a**) pLOGO analysis of amino acid sequences corresponding to the mRNA fragments with increased ribosome occupancy (>5-fold change, n = 3045) in BOT-treated cells compared to untreated controls. (**b**) Average BOT stall scores calculated for each codon in the ribosomal A site. Note the high positive scores for Gly codons and negative scores for the codons of other amino acids. (**c**) High concentration of BOT (50 µM) does not inhibit aminoacylation of tRNA^Gly^ by glycyl-tRNA-synthetase *in vitro*. 5’-O-(glycylsulfamoyl)adenosine (Gly-AMS) serves as a positive control. (**d**) Ribosome stalling at the beginning of the *dapG* ORF as revealed by ribo-seq. (**e, f**) Toeprinting analysis of BOT-induced ribosome stalling on *dapG* (**e**) and RST1 (**f**) templates. Yellow arrowheads indicate start codon arrests; green and blue arrows mark drug-induced stalls within coding regions of the genes. RET, retapamulin (control). (**g**) Toeprinting analysis showing the effect of amino acid substitutions (Gly9→Ser, Gly9→Asp, Gly9→Lys and Gly9→Phe) on BOT-induced stalling on the *dapG* template.

To validate the sequence specificity of BOT action observed *in vivo*, we used toeprinting analysis, which reveals with high precision the sites of ribosome stalling during cell-free translation^17,18^. Guided by the Ribo-seq data, we selected the *rplQ* and *dapG* genes that showed strong BOT-induced ribosome stalling on Gly codons near the start of the open reading frame (ORF) (**Fig. 2d; Extended Data Fig. 1b, c**). In addition, we tested a model template RST1 encoding 20 proteinogenic amino acids^19^. Consistent with the Ribo-seq results, in all three mRNA templates, toeprinting revealed strong and specific ribosome stalling when Gly codons entered the A site (**Fig. 2e, f; Extended Data Fig. 1d**).

To reveal the determinants of the BOT-mediated specificity of ribosome stalling, we leveraged the presence of two sequential Gly codons, Gly9 and Gly10, in the *dapG* template: introducing stalling-disrupting mutations at the Gly9 codon allows trapping of the ribosomes that manage to proceed to the Gly10 codon. We first mutated the WT Gly9 codon (GGC) to the other three Gly codons (GGG, GGA, and GGU). Although our Ribo-seq data showed varying levels of BOT-induced stalling at the different Gly codons (**Fig. 2b**), BOT stalled the ribosome equally efficiently at all four Gly codons during cell-free translation (**Extended Data Fig. 1e**), possibly due to a higher excess of BOT over the components of the *in vitro* system. In stark contrast, mutating the Gly9 codon of *dapG* to Phe (UUC), Lys (AAA), Asp (GAU), or Ser (AGC) codons abolished stalling entirely (**Fig. 2g**), re-emphasizing the remarkable sequence specificity of BOT action. This result suggests that the size of the incoming amino acid’s side chain is the key factor modulating BOT activity: Gly, the smallest of all amino acids, allows efficient ribosome stalling, while larger incoming residues (such as Phe, Lys, Asp, or Ser) render the translation completely insensitive to BOT.

### BOT traps EF-Tu ternary complex containing Gly-tRNA^Gly^ on the ribosome

Earlier and, at times, conflicting reports suggested that BOT acts near or at the A site of the ribosomal PTC^14,15^. To verify those claims and directly visualize the BOT-stalled translation complex, we turned to single-particle cryo-electron microscopy (cryo-EM). The *dapG* mRNA template, where we had detected strong BOT-induced ribosome stalling at the early Gly codons *in vivo* and *in vitro* (**Fig. 2d, e**), was translated in the cell-free system in the presence of BOT, and the reaction mixture was applied directly to cryo-EM grids. 3D classification of the imaged dataset revealed ribosome particles in multiple states (**Extended Data Fig. 2a**). Two classes (32% and 9% of particles) showed density for tRNAs occupying the A and P sites or P and E sites, respectively, consistent with elongating ribosomes. Strikingly, however, these classes lacked any additional density within the ribosome, including the A site of the PTC, that could plausibly correspond to BOT.

Intriguingly, our analysis revealed an unexpected feature unique to this dataset: a well-defined predominant 3D subclass (51% of particles) in which EF-Tu was bound to the ribosome (**Extended Data Fig. 2a**). This is unusual, as EF-Tu typically interacts only transiently with the translating ribosomes during decoding and is rarely captured by cryo-EM in the steady-state translation reactions. The presence of a distinct EF-Tu-bound population suggested that BOT may stabilize an otherwise short-lived intermediate of the elongation cycle. Upon refining this EF-Tu-containing particle class to an overall resolution of 2.06 Å, we identified a prominent and well-resolved extra density within the ribosome-bound EF-Tu (**Fig. 3a-c; Extended Data Fig. 3a**). The shape and size of this density corresponded precisely to a single molecule of BOT. The overall high quality of the obtained map allowed unambiguous modeling of the compound and its interactions with both EF-Tu and the tRNA (**Fig. 3c; Extended Data Fig. 3a**).

**Figure 3.**
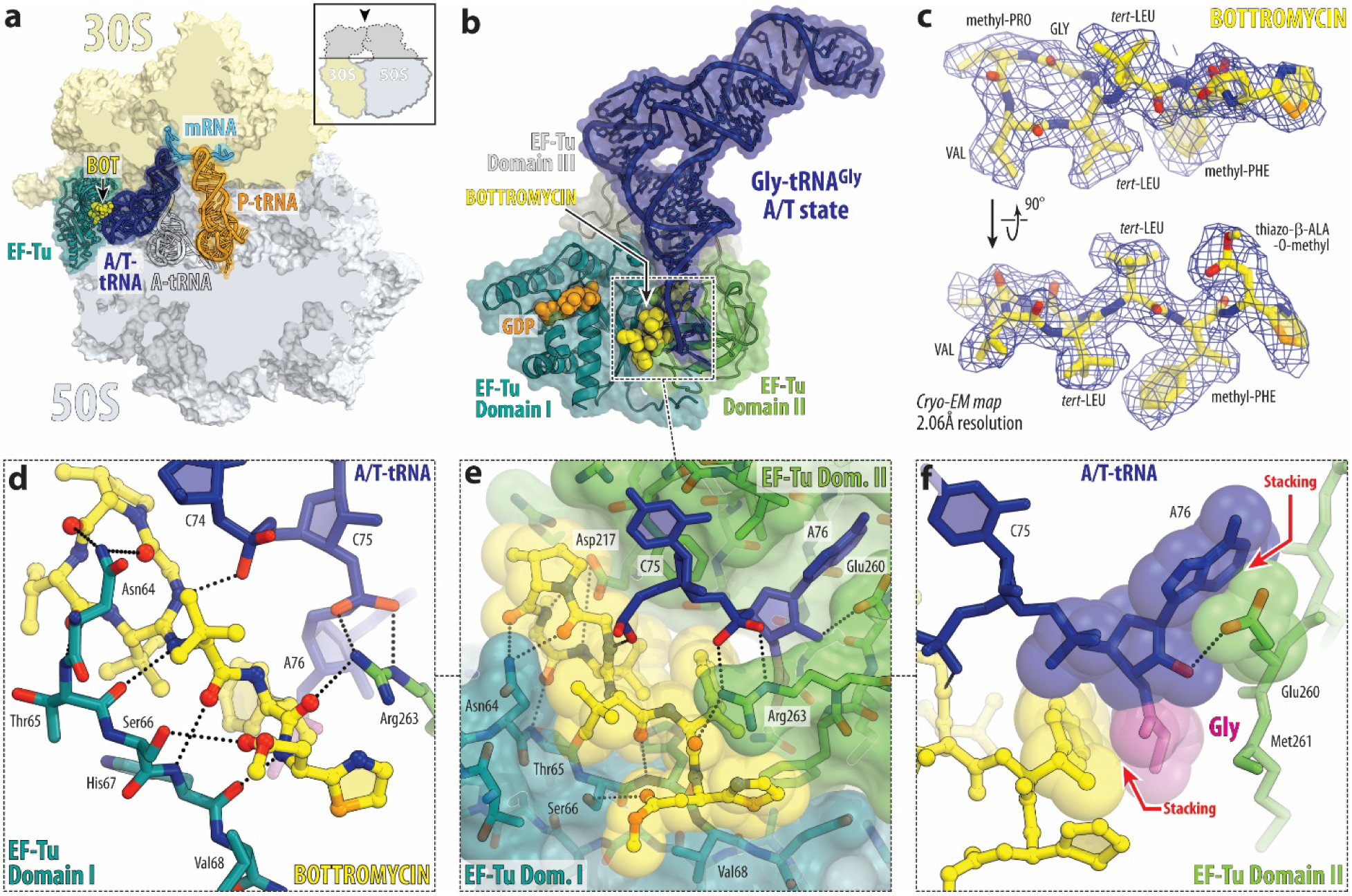
Cryo-EM structure of the EF-Tu•GDP•Gly-tRNA ternary complex stalled by BOT on the 70S ribosome during *in vitro* translation. (**a**) Overview of the structure of the *E. coli* 70S ribosome stalled by BOT (yellow) during *in vitro* translation of the *dapG* mRNA. The 30S and 50S subunits are shown in light yellow and light blue, respectively; EF-Tu is in teal; the mRNA is in blue, and the A-and P-site tRNAs are in dark blue and orange, respectively. BOT traps glycyl-tRNA in complex with EF-Tu on the ribosome in an A/T-state, preventing proper accommodation into the A site. The position of the fully accommodated A/A tRNA is indicated by a black outline. (**b**) Close-up view of the ribosome-bound EF-Tu•GDP•Gly-tRNA ternary complex with BOT bound at the interface between EF-Tu domains I and II (teal and green), adjacent to the CCA-end of the Gly-tRNA (dark blue). GDP (orange) in the GTPase center indicates a post-hydrolysis state of EF-Tu. (**c**) Cryo-EM density map (blue mesh) for BOT with the refined atomic model overlaid. Two orthogonal views reveal well-resolved density for all key chemical moieties of the drug, allowing confident placement and modeling. (**d, e**) Detailed views of the interaction network between BOT, EF-Tu, and the CCA-end of Gly-tRNA. BOT forms an extensive H-bonding interface with EF-Tu domain I (Asn64-Val68), domain II (Asp217 and Arg263) residues, and the phosphate of nucleotide C75. H-bonds are shown as black dotted lines. (**f**) CH-π stacking between the β-methyl-phenylalanine (mPhe) side chain of BOT and the glycine moiety of the aa-tRNA. This highly specific interaction explains the strict selectivity of BOT for glycyl-tRNA and its amino acid-dependent inhibitory activity.

The cryo-EM structure reveals that BOT binds snugly within a deep cleft formed between domains I and II of EF-Tu, directly adjacent to the CCA-end of the A-site tRNA (**Fig. 3a-c; Extended Data Fig. 3b**). As anticipated from our biochemical data, the captured tRNA in this complex is aminoacylated with glycine and adopts an A/T conformation, with its anticodon interacting with the A-site mRNA codon and the aminoacylated CCA-end bound to EF-Tu. Notably, in these particles, the ribosomal P site is occupied by a phenylalanine-specific tRNA carrying a clearly resolved QKF-tripeptide moiety (**Extended Data Fig. 4**), corresponding precisely to the nascent peptide sequence encoded by the *dapG* mRNA immediately upstream of the Gly9 codon, in full agreement with our toeprinting results (**Fig. 2e**). BOT is tightly coordinated at the EF-Tu/tRNA interface, engaging in an extensive network of hydrogen bonds (H-bonds) with both binding partners (**Fig. 3d-f**). Remarkably, the main-chain amino and carbonyl groups of BOT form multiple H-bonds with EF-Tu residues 65-68 in domain I, aligning in a β-strand-like fashion that extends the existing β-sheet of this domain (**Fig. 3d**). In addition to its interactions with EF-Tu, BOT forms an H-bond with the phosphate of the C75 residue of the tRNA (**Fig. 3d, e**). Most notably, the β-methyl-phenylalanine (mPhe) residue of BOT engages in extensive CH-π stacking interactions with the glycine moiety at the 3′-end of the tRNA (**Fig. 3f**). These contacts with the side chain-lacking Gly provide a compelling structural rationale for the strict Gly codon specificity of BOT-induced stalling.

Interestingly, the conformation of BOT in its binding site in the ribosome-bound EF-Tu contrasts with its structure in solution, determined by NMR, where BOT adopts a compact, curled-up conformation, folding nearly twice upon itself^20^. Bound to EF-Tu, BOT assumes an extended shape that fits precisely into the interdomain cleft of the Gly-tRNA-bound EF-Tu. The switch from a self-stabilized to target-stabilized conformation is likely driven by the abundance of H-bonds and stacking interactions that BOT molecule forms with EF-Tu and Gly-tRNA and is critical for maximizing its intermolecular contacts upon target engagement.

To better understand why BOT does not inhibit translation at other codons, we compared our BOT-bound EF-Tu/Gly-tRNA structure with the published structures of EF-Tu ternary complexes carrying tRNAs acylated with bulkier amino acids. Superposition with EF-Tu/Phe-tRNA or EF-Tu/Cys-tRNA complexes, both in solution and in the absence of the ribosome (**Extended Data Fig. 5a-d**), as well as with the EF-Tu/Trp-tRNA complex on the ribosome (**Extended Data Fig. 5e, f**), revealed severe steric clashes between the amino acid side chains and the mPhe moiety of BOT. In these structures, the side chains of the tRNA-bound amino acids occupy a pocket in EF-Tu that in our structure is occluded by BOT, rendering these ternary complexes incompatible with stable drug binding, and explaining why only glycine supports BOT-induced stalling. On the basis of this analysis, we reasoned that alanine, with its small side chain, could partially fit into the BOT-occupied pocket. Indeed, in the Ribo-seq dataset, positions with Ala codons in the A-site had higher average stall scores compared to all other non-Gly residues (**Fig. 2b**), and if Gly9 in the *dapG* mRNA is mutated to Ala, a residual BOT-induced stalling can be observed *in vitro* (**Extended Data Fig. 1f**). In contrast, larger side chain amino acids would clash with the BOT molecule on EF-Tu, likely resulting in BOT displacement from the complex and a failure to stall ribosomes at the respective codons (**Fig. 2g**).

Interestingly, in the BOT-stalled complex, EF-Tu is bound to GDP rather than GTP (**Fig. 3b; Extended Data Fig. 3c**), consistent with the state where GTP hydrolysis has already occurred following the decoding by Gly-tRNA and suggesting that BOT is unlikely to interfere with the GTPase activity of EF-Tu. During canonical translation elongation, GTP hydrolysis triggers the release of the amino-acid-carrying CCA-end of tRNA from EF-Tu, dissociation of the factor, and subsequent tRNA accommodation into the A site. However, in the presence of BOT, the CCA-end of Gly-tRNA remains “glued” to EF-Tu, trapping the translation complex in an A/T-state and providing the mechanistic basis for the highly selective inhibition of translation at Gly codons.

To rigorously test whether EF-Tu and Gly-tRNA are the only factors required for BOT-induced ribosome arrest, we reconstituted the complex using individually purified *E. coli* 70S ribosomes, recombinant EF-Tu, and a synthetic mini-mRNA encoding the Met-Gly sequence^21^. To stabilize the substrate and prevent any spontaneous hydrolysis of Gly-tRNA, we used non-hydrolyzable amide-linked Gly-NH-tRNA^Gly^ and fMet-tRNA_i_^Met^ as the A-and P-site substrates, respectively^21^. In addition, we included non-hydrolyzable GTP analog 5’-guanosyl-methylene-triphosphate (GDPCP). The assembled mixture was applied to the cryo-EM grids and subjected to single-particle cryo-EM analysis.

Cryo-EM data revealed that the assembled complex constituted the major particle population in the dataset (60%, **Extended Data Fig. 2b**). We refined this class to an overall resolution of 2.04 Å, enabling detailed structural interpretation. The architecture of this reconstituted complex was virtually indistinguishable from the cell-free translation-derived structure (**Extended Data Fig. 6**), except for the presence of GDPCP instead of GDP (**Extended Data Fig. 3e, g**) and the amide-linked (rather than ester-linked) aminoacyl-tRNA in the A site (**Extended Data Fig. 3e, f**). BOT was again observed sandwiched between domains I and II of EF-Tu, interacting simultaneously with both EF-Tu and the CCA-end of the Gly-tRNA (**Extended Data Fig. 3e, h**), as observed in our initial reconstruction of the translating ribosome (**Fig. 3**). This result demonstrated that EF-Tu and Gly-tRNA are not only necessary but also sufficient for BOT action on the ribosome. The new structure also revealed that BOT is compatible with the GTP-bound state of EF-Tu, suggesting that it may bind to the ternary complex prior to its interaction with the ribosome.

Notably, neither in the cell-free translation system-derived nor in the reconstituted structures, we observed interpretable cryo-EM density for the switch I region (residues 38-64) of EF-Tu domain I (**Extended Data Fig. 6c**) known to be critical for EF-Tu domains’ rearrangements required for tRNA release following GTP hydrolysis^22–24^. The absence of visible density for switch I in the G-domain of EF-Tu in the IVT-derived structure is consistent with a post-hydrolysis state and reflects the intrinsic flexibility of the switch I region, as observed in a previous ribosome/EF-Tu•Thr-tRNA^Thr^•GDP•kirromycin complex^25^. However, the absence of density for the same switch I in the structure of reconstituted complex harboring non-hydrolysable GTP analog, GDPCP, is intriguing because it is usually visible in the presence of GTP or its analogs^24,26–28^. An overlay of the current structure with a previous complex^27^ shows a potential steric clash between BOT and the switch I α-helical region at residue 49 (**Extended Data Fig. 6c**), which may explain the lack of visible density for this region in our structure. During tRNA selection, the engagement of switch I with the GTP binding pocket in the G-domain and the CCA-end of aminoacyl-tRNA keeps EF-Tu in a closed conformation. Upon GTP hydrolysis, switch I refolds to form a β-hairpin and eventually becomes disordered, transitioning EF-Tu to the open conformation and dissociating from aminoacyl-tRNA^24^. Our pre-GTP hydrolysis structure suggests that BOT acts independently of switch I engagement and that its inhibitory effect results from physically bridging EF-Tu and the glycyl-tRNA CCA-end, effectively freezing the complex in the closed state that precludes tRNA dissociation and accommodation.

### BOT is a bacteria-specific inhibitor of translation

While being a potent inhibitor of bacterial protein synthesis (**Extended Data Fig. 7a**), BOT showed only limited effect upon translation in the rabbit reticulocyte lysate even at a concentration 1000 times higher than that affording complete inhibition by cycloheximide, a classic inhibitor of eukaryotic protein synthesis (**Extended Data Fig. 7b**). To understand the structural basis of this selectivity, we aligned our cryo-EM structure of the *E. coli* ribosome associated with the BOT-bound EF-Tu•Gly-tRNA complex with the reported structure of the human 80S ribosome containing the eukaryotic EF-Tu homolog, elongation factor eEF1A (**Extended Data Fig. 7c, d)**^29^. Although eEF1A features a similar cleft between domains I and II, this region is structurally incompatible with BOT binding. Several residues unique to eEF1A, including Lys62, Asp74, Trp78, and Tyr254, protrude into the space occupied by BOT in its complex with EF-Tu, creating steric clashes with the drug. Among these, Trp78 intrudes the most into the possible BOT binding site on eEF1A (**Extended Data Fig. 7d**). While the precise contribution of individual residues to the eEF1A intrinsic resistance to BOT is unclear, our analysis provides a structural rationale for BOT selectivity as a specifically antibacterial antibiotic.

### Recessive point mutations in EF-Tu genes render bacteria resistant to BOT

Our biochemical and structural data showed that BOT inhibits translation by acting upon the EF-Tu/Gly-tRNA complex. However, they did not necessarily establish that BOT stops bacterial growth by acting upon EF-Tu. To identify the primary cellular target of BOT action in the bacterial cell, we set out to isolate BOT-resistant mutants. We carried out these experiments using *B. subtilis*, whose genome harbors a single EF-Tu-encoding gene *tuf*. After plating ∼10^8^ cells onto agar plates supplemented with 32x MIC of BOT, resistant colonies appeared with a frequency of 10^-7^. Sequencing of the *tuf* gene in six individual resistant colonies identified four mutants with the single nucleotide substitution resulting in the D218Y mutation in EF-Tu, and two mutants with the D218N change (**Fig. 4a**). Both D218Y and D218N mutants demonstrated a dramatic increase in BOT MIC (>512 fold) relative to the parental strain (**Fig. 4a**). In the cryo-EM structure of the *E. coli* BOT-containing EF-Tu complex, Asp217 (corresponding to Asp218 of the *B. subtilis* EF-Tu) (**Fig. 4b**) forms H-bonds with the macroamidine cycle of BOT (**Fig. 4c**). *In silico* modelling of a Tyr residue at this position reveals a direct steric clash between the bulky Tyr side chain and EF-Tu-bound BOT molecule, rationalizing the resistance mechanism in the D218Y *B. subtilis* mutant (**Fig. 4d**). While the basis for resistance of the D218N mutant is less apparent, the loss of the Asp negatively charged carboxyl group may impair the specific H-bonding network required for BOT binding, thereby reducing its affinity.

**Figure 4.**
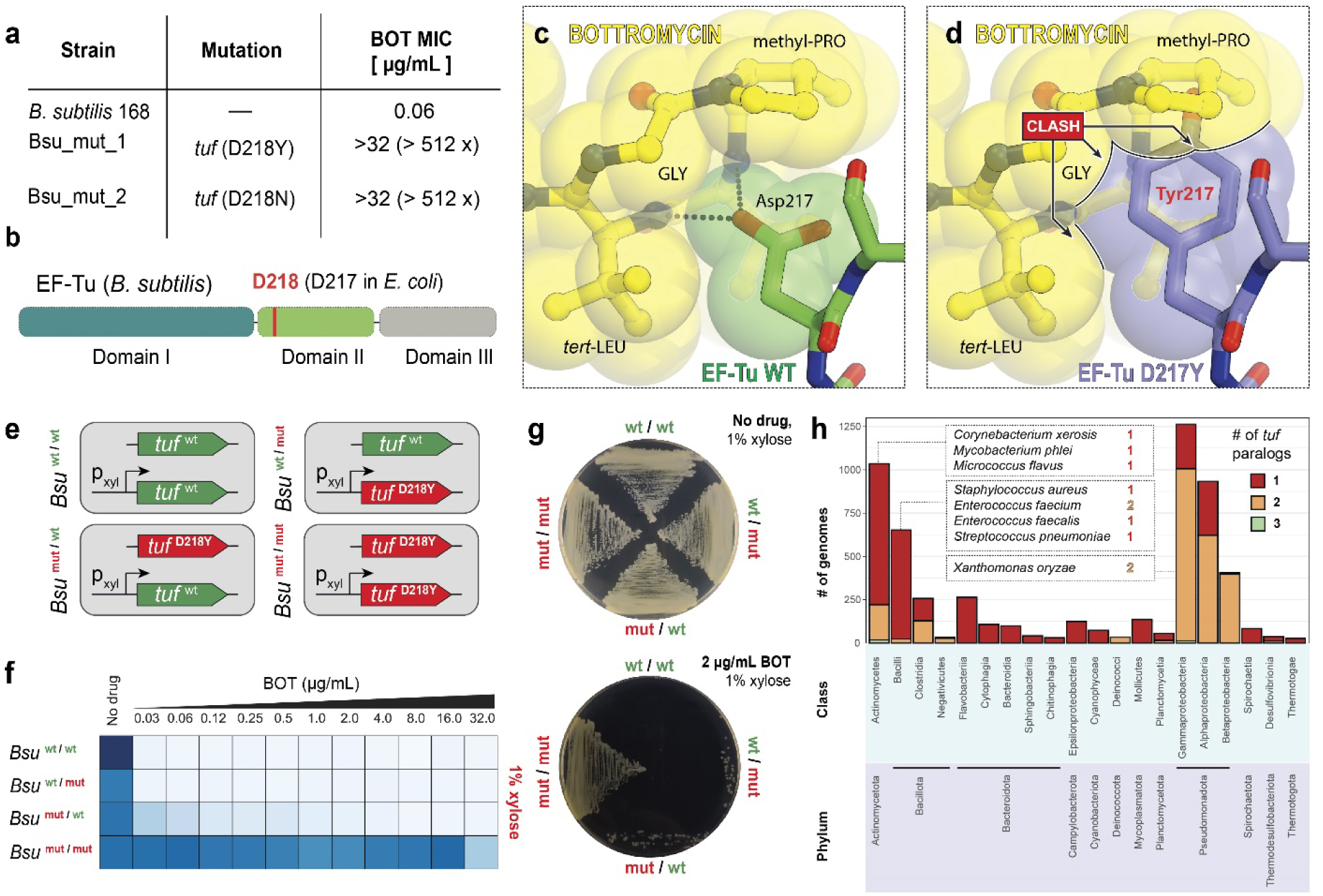
BOT resistance arises from point mutations in EF-Tu. (**a**) Characteristics of *B. subtilis* mutants selected on BOT-containing agar plates. All resistant clones carried point mutations at Asp218 residue in the *tuf* gene. (**b**) Location of residue Asp218 (*B. subtilis* numbering; equivalent to Asp217 in *E. coli*) within EF-Tu, mapped onto its domain organization. (**c, d**) *In silico* modeling of the D217Y (*E. coli* numbering) substitution in the structure of EF-Tu/Gly-tRNA/BOT complex reveals a steric clash between the bulky tyrosine side chain and the bound BOT molecule. (**e**) Schematic of engineered *B. subtilis* strains harboring a second copy of the *tuf* gene inserted at the chromosomal *lacA* locus under the control of an xylose-inducible promoter. (**f**) Heat map showing the growth (OD_600_) of the four strains shown in (**e**) in SM minimal medium supplemented with increasing concentrations of BOT and 1% xylose. (**g**) Growth of the same strains on SM minimal agar plates supplemented with 1% xylose, without the drug (top) or with 2 µg/mL BOT (bottom). (**h**) Distribution of *tuf* gene paralogs (encoding EF-Tu) across complete bacterial genomes from major classes/phyla. Selected species known to be highly sensitive to BOT are listed in the insets along with their respective *tuf* copy number.

Unlike with *B. subtilis*, similar experiments performed with *E. coli* strains yielded no BOT-resistant mutants. We reasoned that this is due to the presence of two EF-Tu encoding genes (*tufA* and *tufB*) in the *E. coli* genome: the probability of simultaneous spontaneous mutations in two gene copies would be too low to be detected by direct plating, whereas a single allele mutation would be manifested in the resistance phenotype only if it has a dominant nature. To test experimentally whether EF-Tu mutations are dominant or recessive in the two-allele setting, we constructed two pairs of *B. subtilis* strains, containing either WT or BOT-resistant (D218Y) genomic *tuf* variants and, in addition, harboring a second *tuf* allele, either WT or BOT-resistant (D218Y), integrated into a distinct genomic location under the control of a xylose-inducible promoter (**Fig. 4e**). In the absence of BOT, all four strains grew normally upon induction of the xylose-inducible *tuf* gene, indicating that expression of the second either WT or mutant EF-Tu allele had no detectable impact on cell viability (**Fig. 4g**, top panel). In the presence of BOT, only the strain carrying two copies of the genomic and inducible BOT-resistant (D218Y) *tuf* allele exhibited robust growth (**Fig. 4f, g**). In contrast, strains harboring either two copies of the WT *tuf* or a combination of WT and mutant *tuf* alleles remained sensitive to BOT. Notably, without xylose induction, the strain with the constitutively expressed genomic D218Y allele and inducible WT *tuf* grew in the presence of BOT (**Extended Data Fig. 8a, b**). These data firmly established EF-Tu as the primary target of BOT action in bacteria and confirmed that BOT resistance is recessive – only cells lacking any BOT-sensitive EF-Tu can survive in the presence of the compound. This also implies that resistance to BOT via point mutations in EF-Tu is unlikely to emerge easily in bacterial species with two or more genomic copies of EF-Tu-encoding genes.

Next, we surveyed the number of *tuf* gene copies in the 6096 completely assembled bacterial genomes from the RefSeq database^30^. This analysis revealed that while the majority of bacterial taxa have a single *tuf* gene, several groups, most notably Pseudomonadota (previously Proteobacteria), predominantly harbor two *tuf* paralogs (**Fig. 4h, Supplementary Data 1**). Importantly, some of the BOT-sensitive human (e.g., *Enterococcus faecium*) and plant (e.g., *Xanthomonas oryzae*) pathogens contain two *tuf* paralogs (**Fig. 4h**, inset), making them priority targets for further BOT development because the presence of multiple *tuf* alleles would reduce the probability of resistance development through target mutations.

## DISCUSSION

Protein synthesis is targeted by numerous antibiotics^31^. Historically, such inhibitors have been considered to indiscriminately halt the synthesis of all polypeptides. However, recent studies have revealed that the activity of many ribosome-targeting drugs is, in fact, context-dependent^32^. In the case of the ribosome-targeting drugs, such as macrolides^33^, phenicols^34^, and oxazolidinones^35^, context specificity emanates from interactions between the antibiotic and the emerging polypeptide chain. Other inhibitors, such as orthosomycins and paenilamycins, achieve context specificity through interactions with unique features of specific tRNAs^36,37^. Context-specificity of drug action can also arise from recognition of particular nucleotides within the mRNA by a ribosome-bound drug, as in the case of kasugamycin^38^. All these antibiotics directly interact with the ribosome. Our work identifies BOT as a context-specific inhibitor of translation that employs a fundamentally unique mechanism for achieving sequence selectivity, interacting with EF-Tu and trapping on the ribosome exclusively in complexes with Gly-tRNAs. Our structures show that BOT binds at the interface between EF-Tu and the CCA-end of the Gly-tRNA, preventing the release and subsequent accommodation of the tRNA into the A site. This results in ribosome arrest at Gly codons in a highly selective and codon-dependent manner. Remarkably, in stark contrast to other context-specific translation inhibitors, BOT does not make any direct contact with the ribosome itself.

Our findings attribute BOT to elfamycins – antibiotics targeting <u>el</u>ongation <u>fa</u>ctor Tu (hence “*<u>elfa</u>*”)^39^. This family includes kirromycin-like antibiotics (kirromycin, factumycin, aurodox, etc.), enacyloxin IIa, pulvomycin, and GE2270A. However, BOT stands apart from previously characterized elfamycins in both its binding site and mechanism of action. BOT engages a unique pocket at the interface between EF-Tu domains I and II and the CCA-end of the Gly-tRNA (**Extended Data Fig. 9a**), forming an extensive network of H-bonds and stacking interactions that are incompatible with tRNAs acylated with amino acids with larger side chains. The requirement for a specific EF-Tu conformation and a Gly-charged tRNA results in a remarkably high degree of substrate selectivity not seen in other translation inhibitors. While kirromycin-like antibiotics and enacyloxin IIa also stabilize EF-Tu•GDP•aa-tRNA ternary complexes on the ribosome, they do so by binding to a completely different site that is located between EF-Tu domains I and III (**Extended Data Fig. 9b**), and, importantly, without regard for the tRNA identity. Other elfamycins, such as pulvomycin and GE2270A, interact with EF-Tu domains I and II, locking the factor in an open conformation that sterically blocks any aa-tRNA engagement (**Extended Data Fig. 9c, d**). Thus, while BOT shares the general target with other elfamycins, it is the only known antibiotic that (i) binds a composite site at the EF-Tu/aa-tRNA interface and (ii) acts in a highly aminoacyl-tRNA-selective manner. These features distinguish BOT as a mechanistically unique EF-Tu inhibitor, making it the founding member of a new subclass of context-specific antibiotics.

An important question is whether BOT binds to the pre-assembled EF-Tu•GTP•Gly-tRNA complex or can engage even a tRNA-free EF-Tu? While we lack direct experimental evidence, our structural data suggest that the interaction of BOT with the vacant EF-Tu could be generally possible since BOT forms extensive contacts with the factor that could, in principle, sufficiently stabilize its association in the absence of Gly-tRNA. However, even if BOT can bind free EF-Tu, most aa-tRNAs would likely displace the drug during ternary complex formation due to the clash with the bulky side chains. Nevertheless, while generally conceivable, binding of BOT to the tRNA-free EF-Tu seems less favorable because, due to the rapid tRNA binding, only a small fraction of the cellular EF-Tu apparently exists in the tRNA-free form^40^. This is indirectly supported by the absence of BOT-induced ribosome stalling at non-glycine codons, which would be expected if BOT titrated out vacant EF-Tu. Therefore, we favor the model in which BOT scans the landscape of ternary complexes and neutralizes only the Gly-tRNA-loaded subset, thus explaining its amino acid-specific inhibition of translation *in vivo*.

Our findings have direct implications for antibiotic development. Previous medicinal chemistry efforts to improve BOT were undertaken in the absence of structural information^41,42^, and without knowing of BOT’s real molecular target, resulting in largely serendipitously empirical modifications (**Fig. 1b**). The high-resolution cryo-EM structures presented here provide a blueprint for rational, structure-guided optimization of BOT analogs. Chemical modifications aimed at enhancing interactions with domain I of EF-Tu or reinforcing glycine-specific contacts at the tRNA interface could improve potency and pharmacological properties. In particular, the mPhe moiety of BOT, which plays a pivotal role in enforcing selectivity by sterically excluding amino acids with bulkier side chains, emerges as a promising site for chemical tuning. Rational redesign of this residue to accommodate the side chains of other amino acids could, in principle, yield BOT derivatives with selectivity for other aa-tRNAs. This would establish a new platform for designing context-specific translation inhibitors tailored to distinct amino acid identities. This approach offers a new avenue for precision-targeted inhibition of protein synthesis, with the potential to minimize off-target effects and slow resistance evolution by exploiting molecular contexts unique to specific genes, pathways, or organisms.

## ACKNOWLEDGMENTS

We thank Drs. Alexander Voltz and Rolf Muller, as well as Dr. Ilya Osterman, for independently providing us samples of BOT used in our pilot experiments; Dr. Heather Feaga for providing the *B. subtilis* strain used in this work; Dr. Egor Syroegin and Elena Aleksandrova for preparing Gly-NH-tRNA^Gly^; Teresa Szal for purification of *E. coli* GlyRS; and Advita Sharma for performing the pilot mutant selection experiments in *E. coli*. We thank the Advanced Electron Microscopy Facility at the University of Chicago, and particularly Dr. Tera Lavoie, for training and support during initial cryo-EM data collection. We also thank Dr. Michael B. Sherman for advice and support with cryo-EM data collection, Drs. Ka-Yiu (Clem) Wong and John Perkyns for computational support, and the Sealy and Smith Foundation for supporting the Sealy Center for Structural Biology and Molecular Biophysics at the University of Texas Medical Branch.

This work was supported by the National Institute of General Medical Sciences of the National Institutes of Health [grants R01-GM132302 and R35-GM151957 to Y.S.P., R01-GM136936 and R35-GM158272 to M.G.G., and R35-GM127134 to A.S.M.], the National Institute of Allergy and Infectious Diseases of the National Institutes of Health [R01-AI162961 to A.S.M., N.V.-L., and Y.S.P.], Illinois State startup funds [to Y.S.P.], and the Welch Foundation [grant H-2032-20230405 to M.G.G.]. The funders had no role in study design, data collection and analysis, decision to publish, or manuscript preparation.

## AUTHOR CONTRIBUTIONS STATEMENT

D.Y.T., M.N.P., and D.K. conducted the *in vitro* biochemical experiments. R.S.B., M.N.P., M.G.G., and Y.S.P. designed and carried out the structural-biology studies. D.Y.T. and M.N.P. isolated BOT-resistant mutants. D.Y.T. performed genetic and microbiological experiments validating EF-Tu as the BOT target. A.I.Z. and D.Y.T. carried out the bioinformatic analysis of *tuf* gene distribution across bacterial taxa. A.S.M., N.V.-L., M.G.G., and Y.S.P. supervised the research. All authors contributed to data interpretation. D.Y.T., A.S.M., N.V.-L., M.G.G., and Y.S.P. wrote the manuscript.

## EXTENDED DATA FIGURES

**Extended Data Figure 1.**
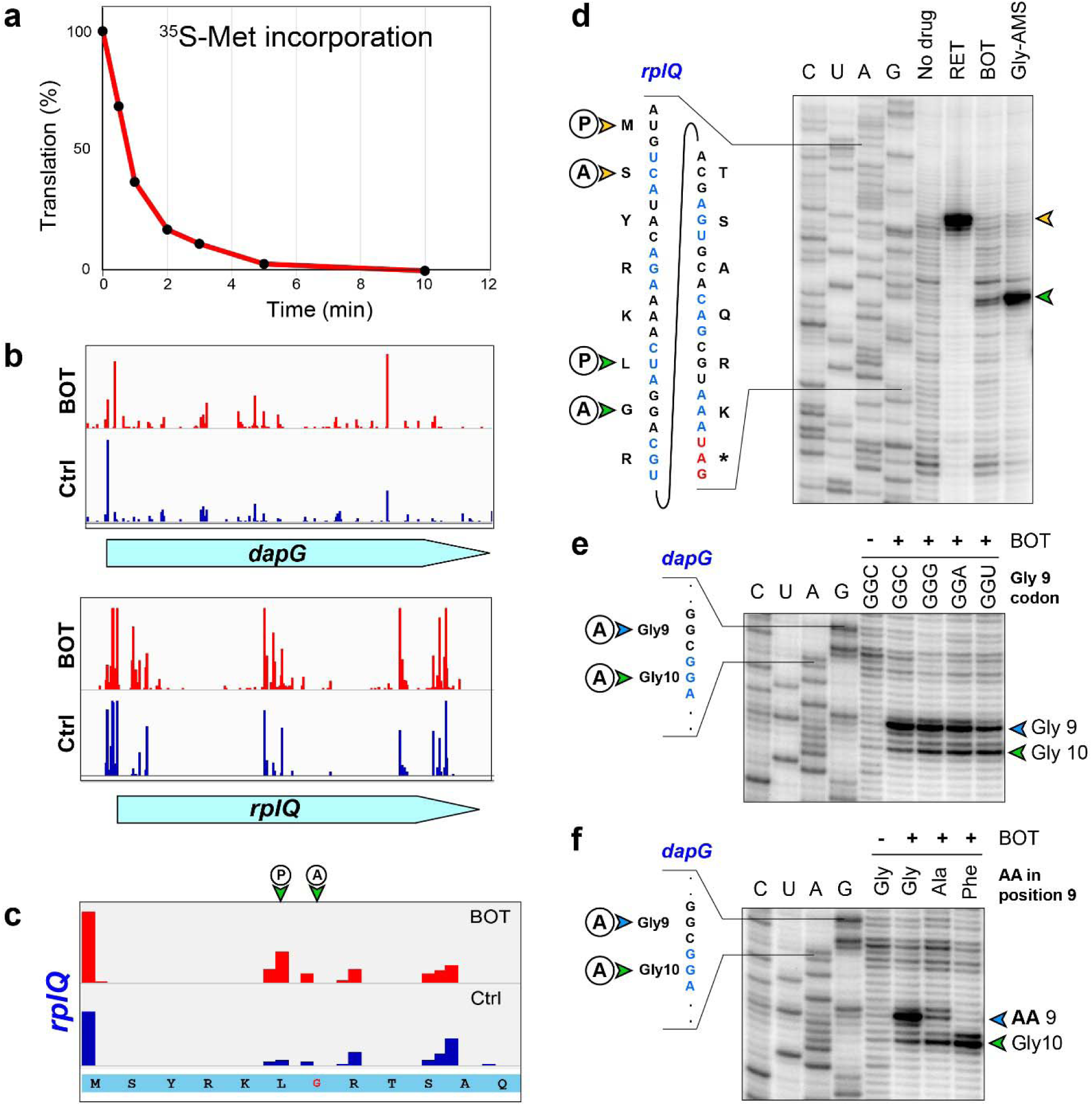
BOT acts as a sequence-specific inhibitor of translation. (**a**) Time course of ^35^S-Met incorporation in exponentially growing *B. subtilis* cultures following treatment with 3 µg/mL BOT (50× MIC). This experiment was done to determine the optimal 5-minute time point for the subsequent ribosome profiling. (**b**) Ribosome stalling on *dapG* and *rplQ* mRNAs in BOT-treated *versus* control ribo-seq datasets. (**c**) BOT-induced stalling at a Leu-Gly dipeptide near the start of the *rplQ* gene. (**d**) Toeprinting analysis of BOT-induced ribosome stalling on the *rplQ* gene. Yellow arrowheads indicate start codon arrest; green arrows mark drug-induced stalls within the coding region of the gene. RET, retapamulin (control). (**e**) Toeprinting analysis examining the impact of Gly codon identity on BOT-dependent ribosome stalling using the *dapG* template. (**f**) Toeprinting analysis showing the effect of amino acid substitutions (Gly9→Ala and Gly9→Phe) on BOT-induced stalling on the *dapG* template.

**Extended Data Figure 2.**
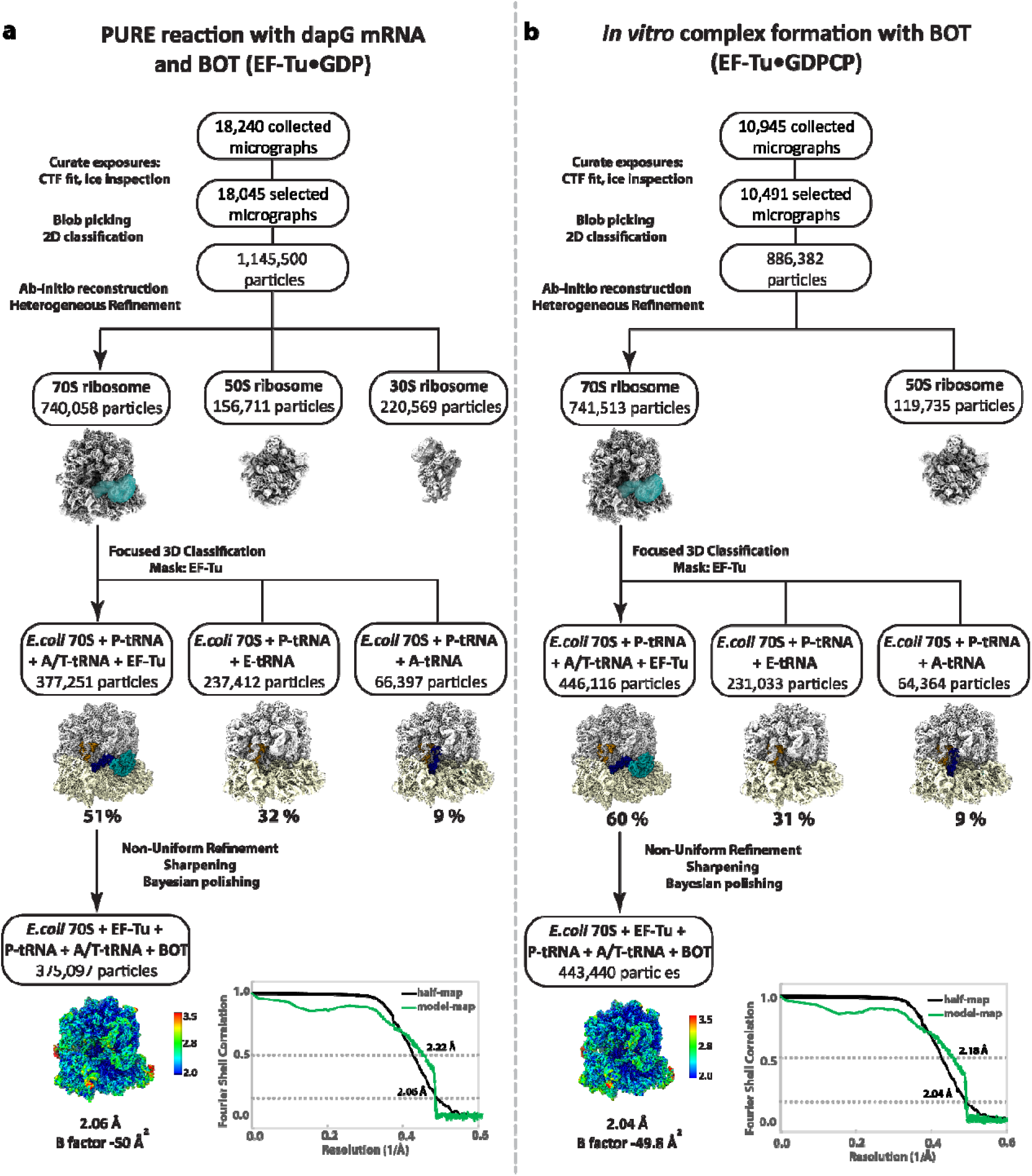
Cryo-EM data processing and particle classification workflow. Cryo-EM image processing and particle classification schemes for datasets obtained from (**a**) the PURE system reaction programmed with *dapG* mRNA, and (**b**) the reconstituted complex of 70S ribosomes, EF-Tu•GDPCP, glycyl-NH-tRNA, and BOT assembled *in vitro*. Final reconstructions were sharpened and polished, yielding high-resolution structures at 2.06 Å (a) and 2.04 Å (b).

**Extended Data Figure 3.**
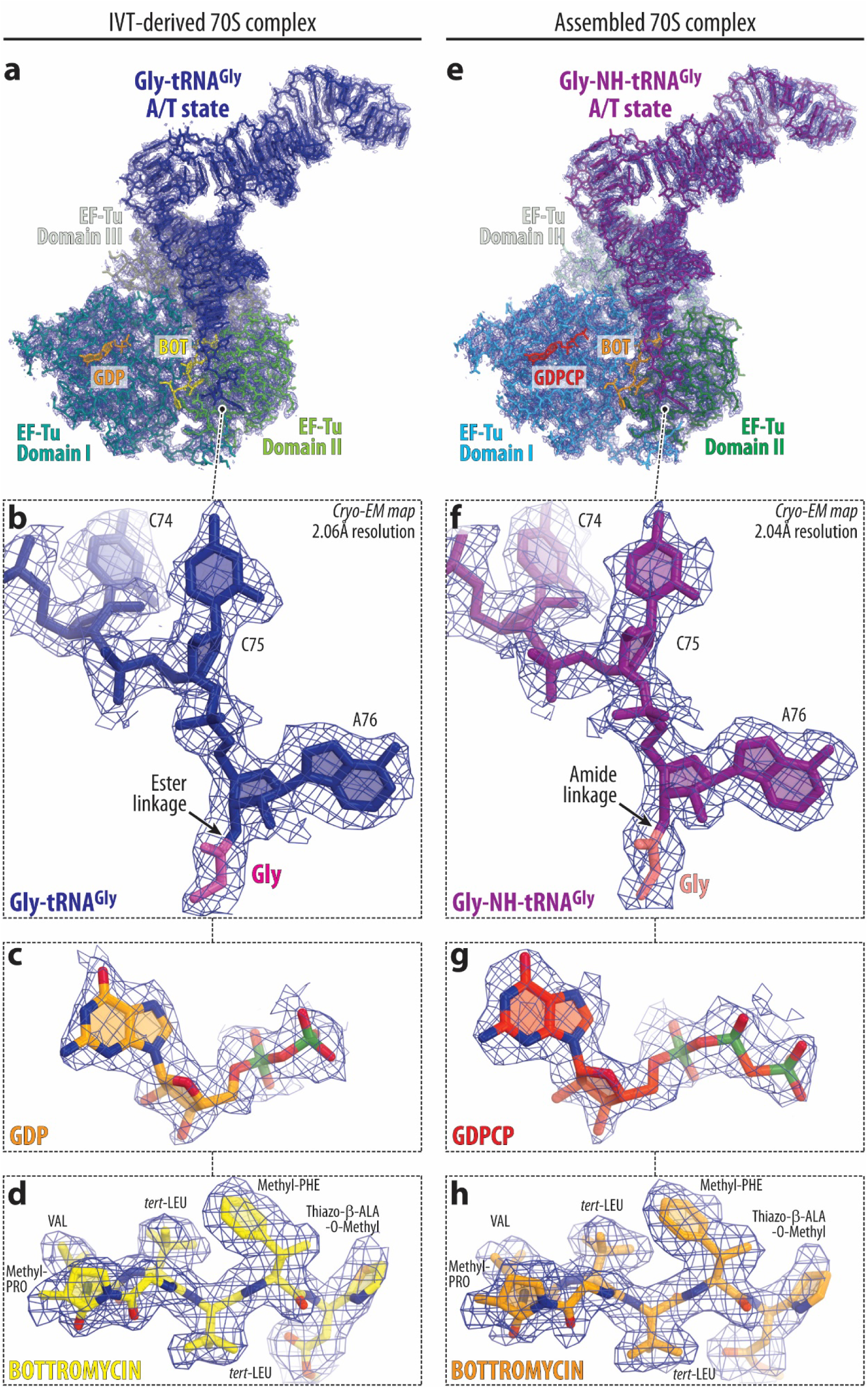
High-resolution cryo-EM maps of the ribosome-bound ternary complexes trapped by BOT. (**a-d**) Cryo-EM analysis of the *in vitro* translation (IVT)-derived 70S ribosome complex stalled by BOT. (**e-h**) Corresponding analysis of a reconstituted ribosome complex assembled from purified components. (**a, e**) Overall cryo-EM density maps (blue mesh) of the ribosome-bound EF-Tu•GDP•Gly-tRNA^Gly^ (a) or EF-Tu•GDPCP•Gly-NH-tRNA^Gly^ ternary complexes with tRNA in the A/T state stabilized by BOT (yellow or orange). EF-Tu domains I-III are colored teal, green, and artichoke, respectively, for the IVT-derived complex, and blue, dark green, and light teal for the assembled complex; Gly-tRNA^Gly^ is in dark blue, and GDP is in orange. (**b, f**) Close-up views of the CCA-ends of the A/T-state tRNAs, showing glycine residue (magenta) covalently linked to A76 via an ester (b) or an amide (f) bond. (**c, g**) Cryo-EM density for GDP (c) or GDPCP (g) in the GTPase center of EF-Tu domain I. (**d, h**) Cryo-EM density for BOT, revealing all major structural features and enabling confident model building.

**Extended Data Figure 4.**
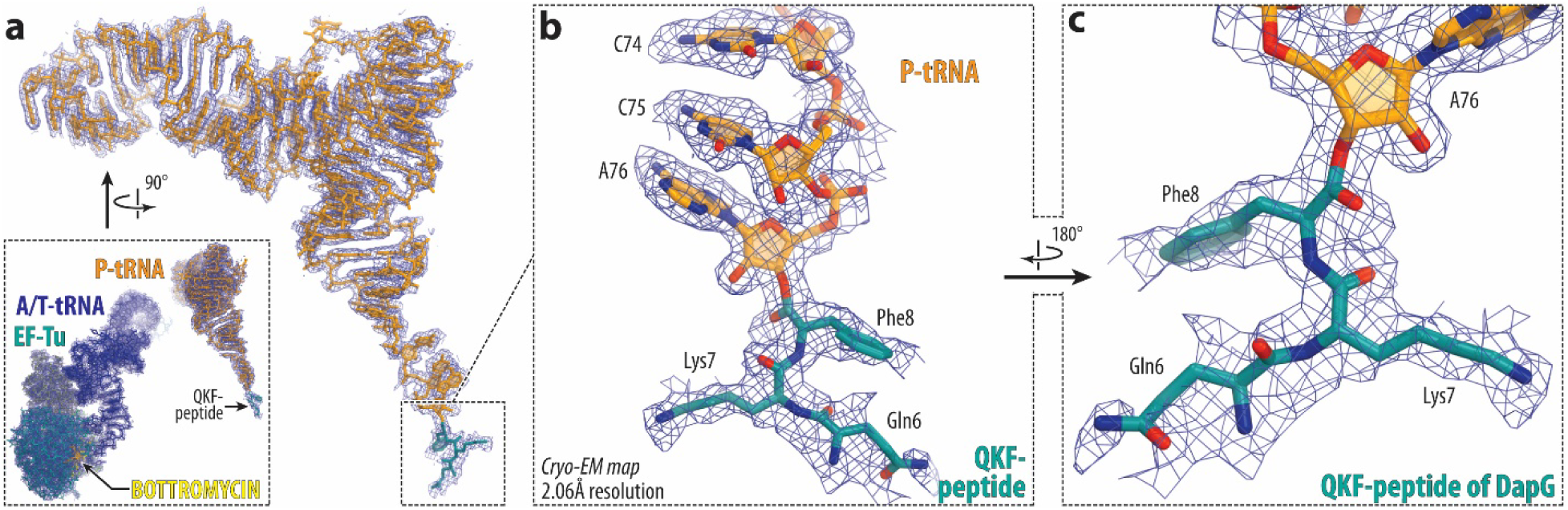
Visualization of the QKF-tripeptide nascent chain in the BOT-stalled ribosome complex. (**a**) Overview of the cryo-EM density map (blue mesh) of the ribosome-bound P-site peptidyl-tRNA (orange) carrying the QKF nascent peptide (teal) derived from the *dapG* mRNA template. The inset shows both A- and P-site substrates in the BOT-stalled ribosome complex, highlighting EF-Tu (teal), A/T-state tRNA (dark blue), and BOT (yellow). (**b, c**) Close-up views of the CCA-end of the P-site peptidyl-tRNA with the QKF-peptide modeled into the well-resolved continuous density. The observed density for the QKF-peptide confirms ribosome stalling at the Gly9 codon of the *dapG* ORF, in agreement with toeprinting and Ribo-seq data.

**Extended Data Figure 5.**
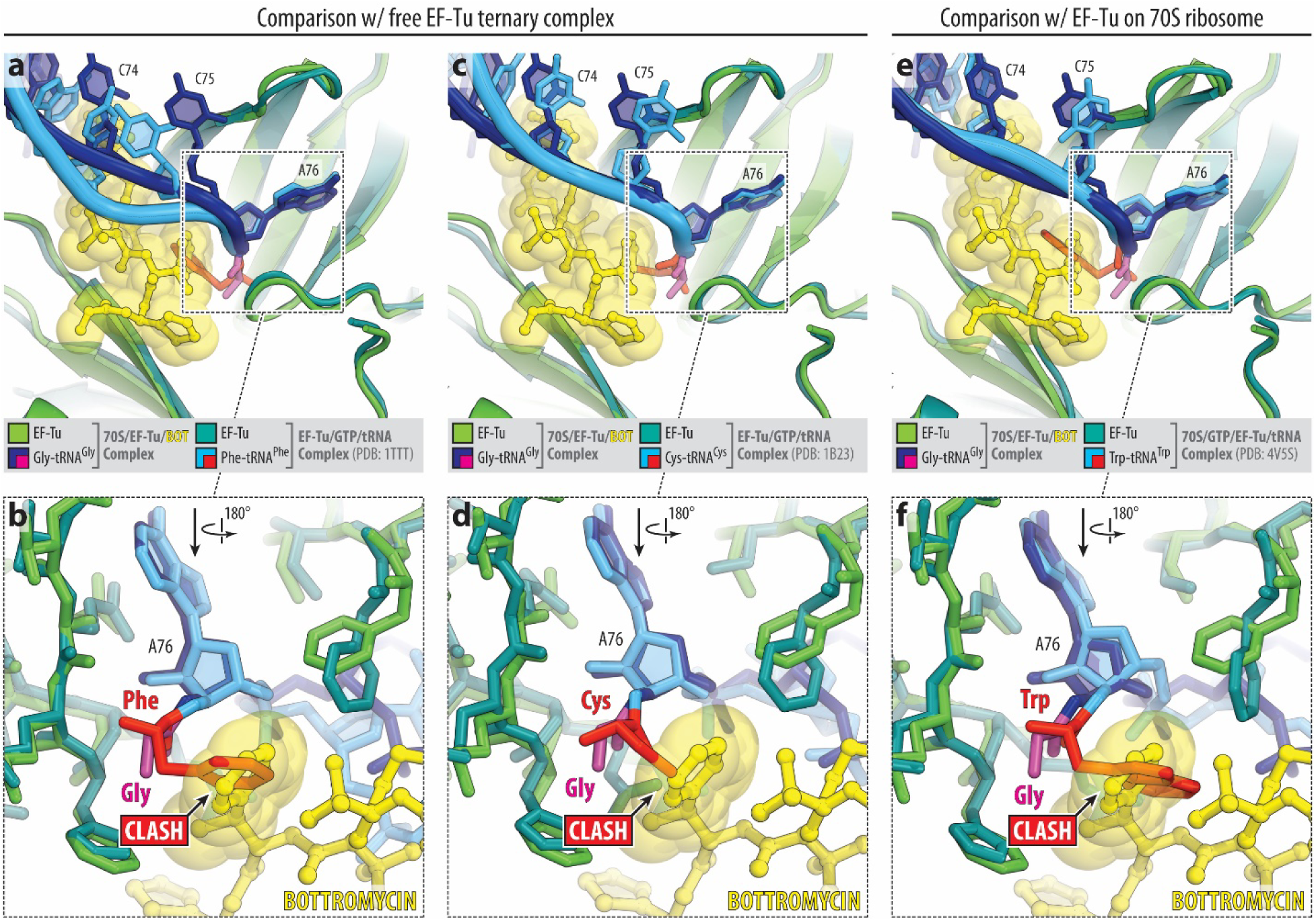
Structural basis for amino acid selectivity of BOT. Superposition of the structure of EF-Tu•GDP•Gly-tRNA^Gly^•BOT complex (this study) with previously determined structures of EF-Tu•GDPNP•aa-tRNA ternary complexes containing Phe-tRNA (**a, b**, PDB entry 1TTT^43^) or Cys-tRNA (**c, d**, PDB entry 1B23^44^), or EF-Tu•GDP•kirromycin•Trp-tRNA^Trp^ complex (**e, f**, PDB entry 4V5S^45^), analyzed either in isolation (a-d) or bound to the 70S ribosome (e, f). Overviews (a, c, e) and close-up views (rotated 180° from top panels) (b, d, f) show the CCA-ends of the aa-tRNAs within the EF-Tu. Bottromycin is shown in yellow; Gly-tRNA^Gly^ (this study) is shown in dark blue with glycine residue highlighted in magenta; comparison aa-tRNAs from previous studies are shown in light blue with Phe, Cys, or Trp residues highlighted in red. Note that the bulky side chains of these residues exhibit steric clashes with the mPhe moiety of BOT, providing a structural rationale for the strict selectivity of BOT for glycine-charged tRNA and its codon-specific inhibition of translation.

**Extended Data Figure 6.**
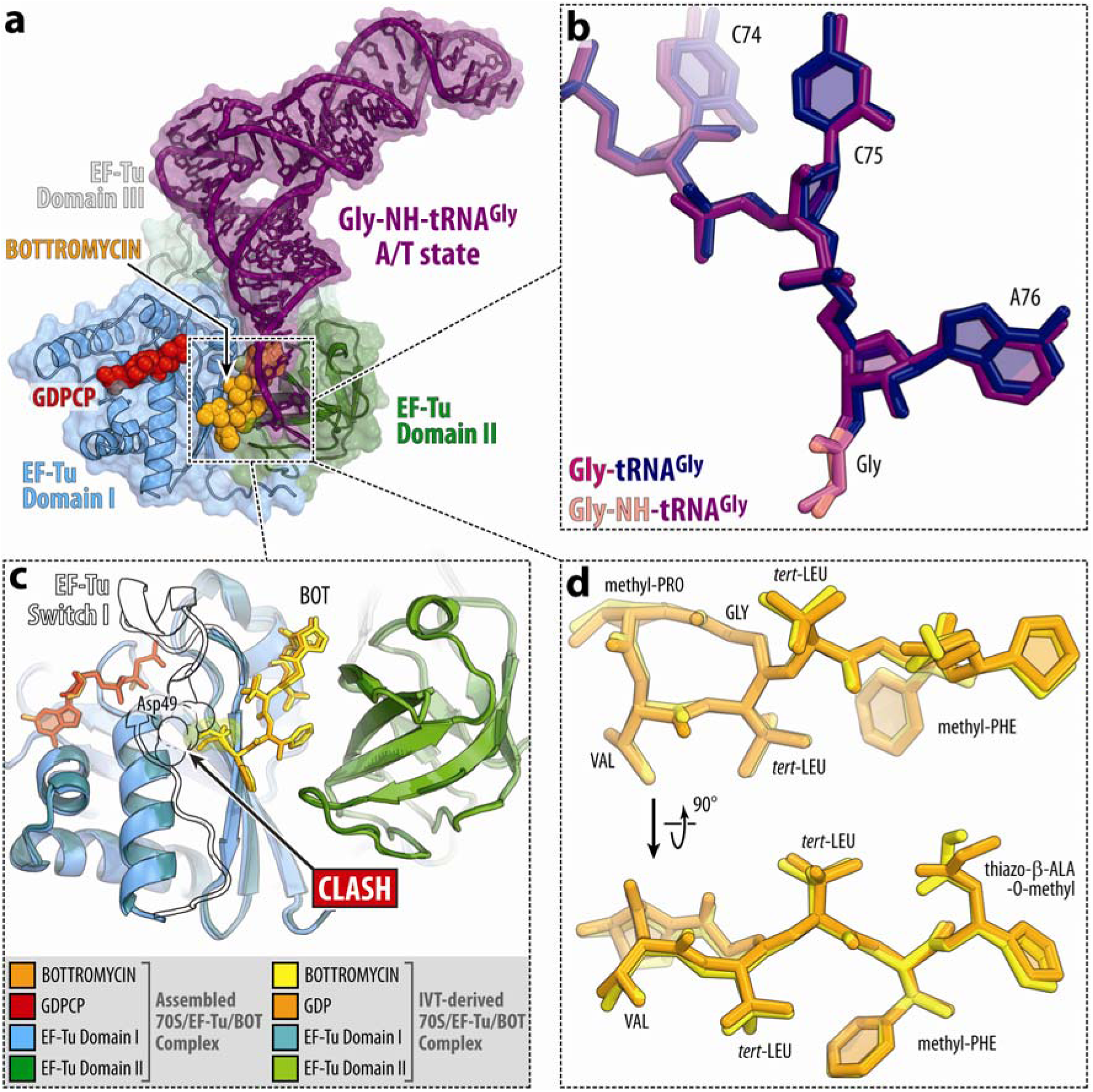
Cryo-EM structure of the EF-Tu•GDPCP•Gly-NH-tRNA ternary complex containing BOT on the 70S ribosome. (**a**) Cryo-EM structure of the *E. coli* 70S ribosome complex assembled from purified components containing BOT, EF-Tu (blue and dark green), non-hydrolyzable Gly-NH-tRNA^Gly^ (dark purple) in the A/T state, and non-hydrolyzable GTP analog GDPCP (red). (**b**) Superposition of the CCA-ends of A-site glycyl-tRNAs from the IVT-derived (ester-linked) and reconstituted (amide-linked) complexes reveals no significant structural differences. (**c**) Comparison of BOT-binding pockets between EF-Tu domains I and II in the IVT-derived and reconstituted complexes, demonstrating identical positioning of the drug. In both structures, switch I was not resolved in the cryo-EM density and was not modeled; its approximate position is shown as a black outline based on the EF-Tu structure from PDB entry 1B23. (**d**) Superposition of EF-Tu-bound BOT molecules in the two structures, highlighting their nearly indistinguishable conformations and interactions, confirming that BOT binds equivalently in both post- and pre-hydrolysis states of EF-Tu.

**Extended Data Figure 7.**
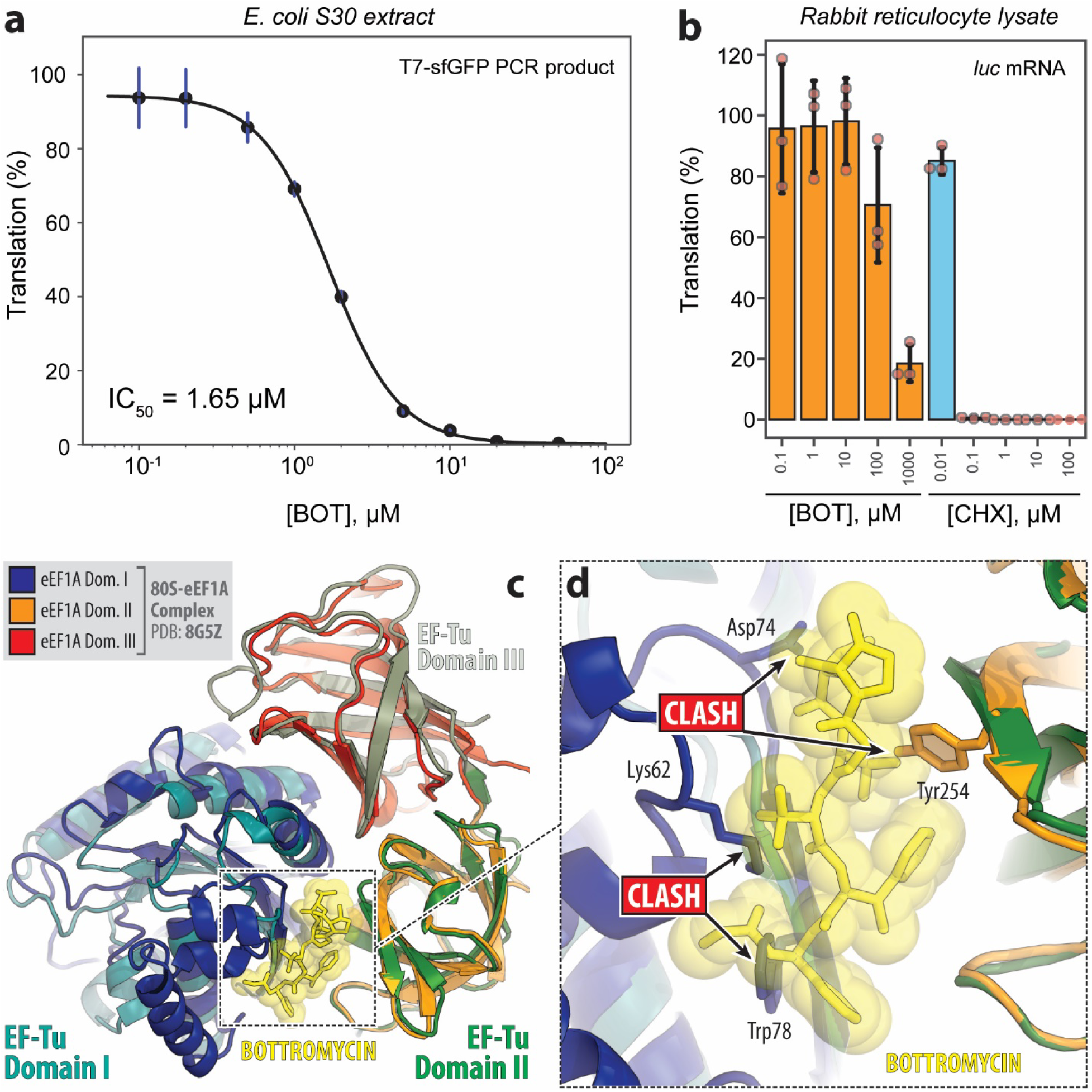
BOT is a selective inhibitor of bacterial translation. (**a**) Inhibition of protein synthesis by BOT in an *in vitro E. coli* cell-free transcription-translation coupled system programmed with a T7-*gfp* PCR template. Data represent three independent replicates. (**b**) Inhibition of luciferase mRNA translation by varying concentrations of BOT and cycloheximide (CHX) in rabbit reticulocyte lysate. Data represent three independent biological replicates. (**c, d**) Structural alignment of bacterial EF-Tu (from the BOT-arrested 70S ribosome complex in this study) and eukaryotic eEF1A (from a cryo-EM structure of the 80S ribosome taken from PDB entry 8G5Z^29^) based on domain II. While a similar interdomain cleft exists between domains I and II in eEF1A, multiple residues in the eukaryotic factor would sterically interfere with BOT binding, explaining the lack of inhibitory activity of BOT on eukaryotic translation.

**Extended Data Figure 8.**
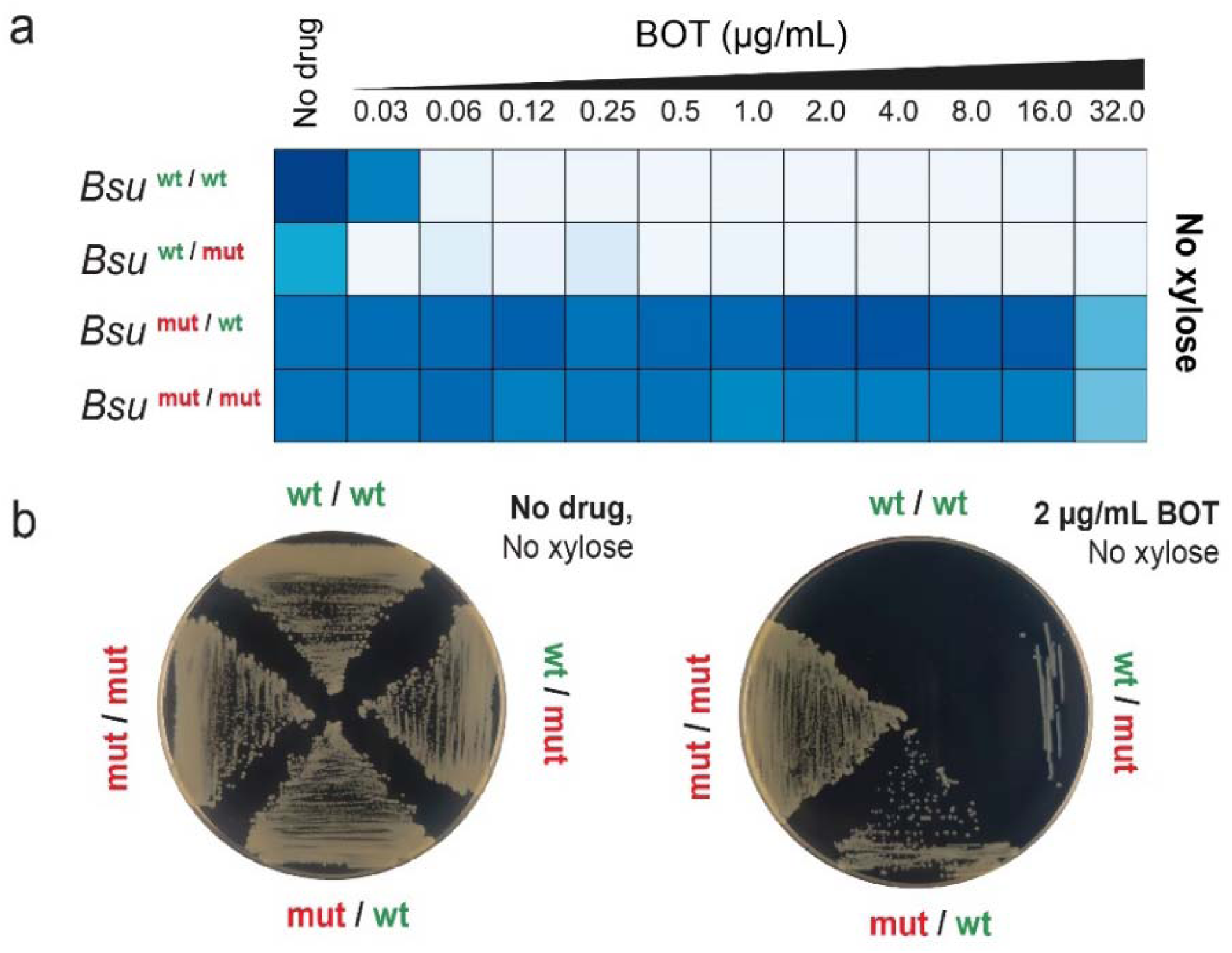
Point mutations in EF-Tu confer resistance to BOT. (**a**) Heat map showing the growth (OD_600_) of four *B. subtilus* strains (as in Fig. 4e) cultured in SM minimal medium with increasing concentrations of BOT, in the absence of xylose induction. (**b**) Growth of the same four strains on SM minimal agar plates without drug (left) or with 2 µg/mL BOT (right), also in the absence of xylose.

**Extended Data Figure 9.**
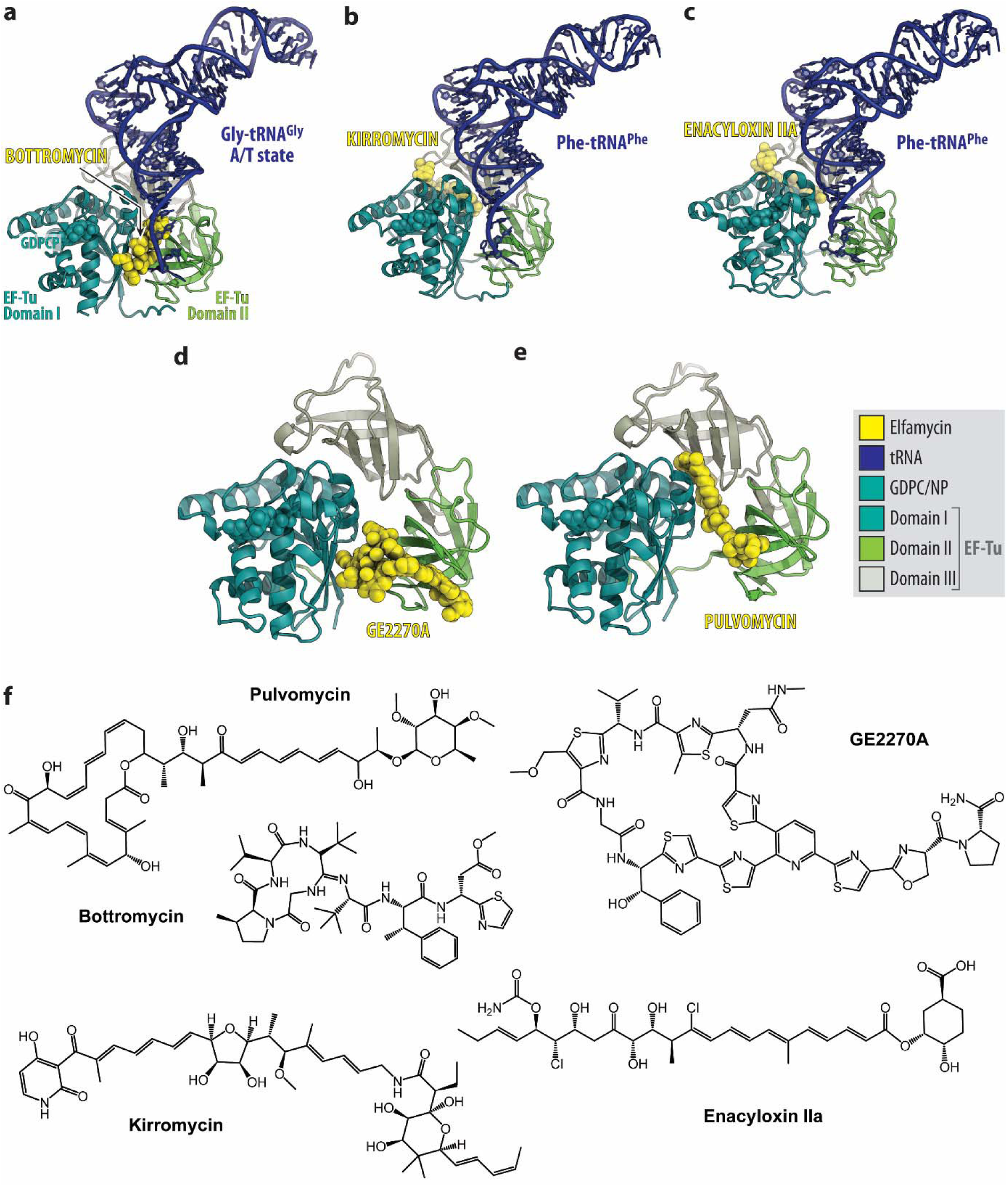
Comparison of BOT with other EF-Tu-targeting antibiotics (elfamycins). (**a-e**) Structural comparison of EF-Tu/tRNA complexes or EF-Tu bound to representative elfamycins: bottromycin (**a**, this study), kirromycin (**b**, PDB entry 1OB2), enacyloxin IIA (**c**, 1OB5^46^), GE2270A (**d**, PDB entry 2C77^47^), and pulvomycin (**e**, PDB entry 2C78^47^). Antibiotics are shown as yellow spheres. (**f**) Chemical structures of the antibiotics shown in panels **a-e**.

## SUPPLEMENTARY FIGURES

**Supplementary Fig. 1.**
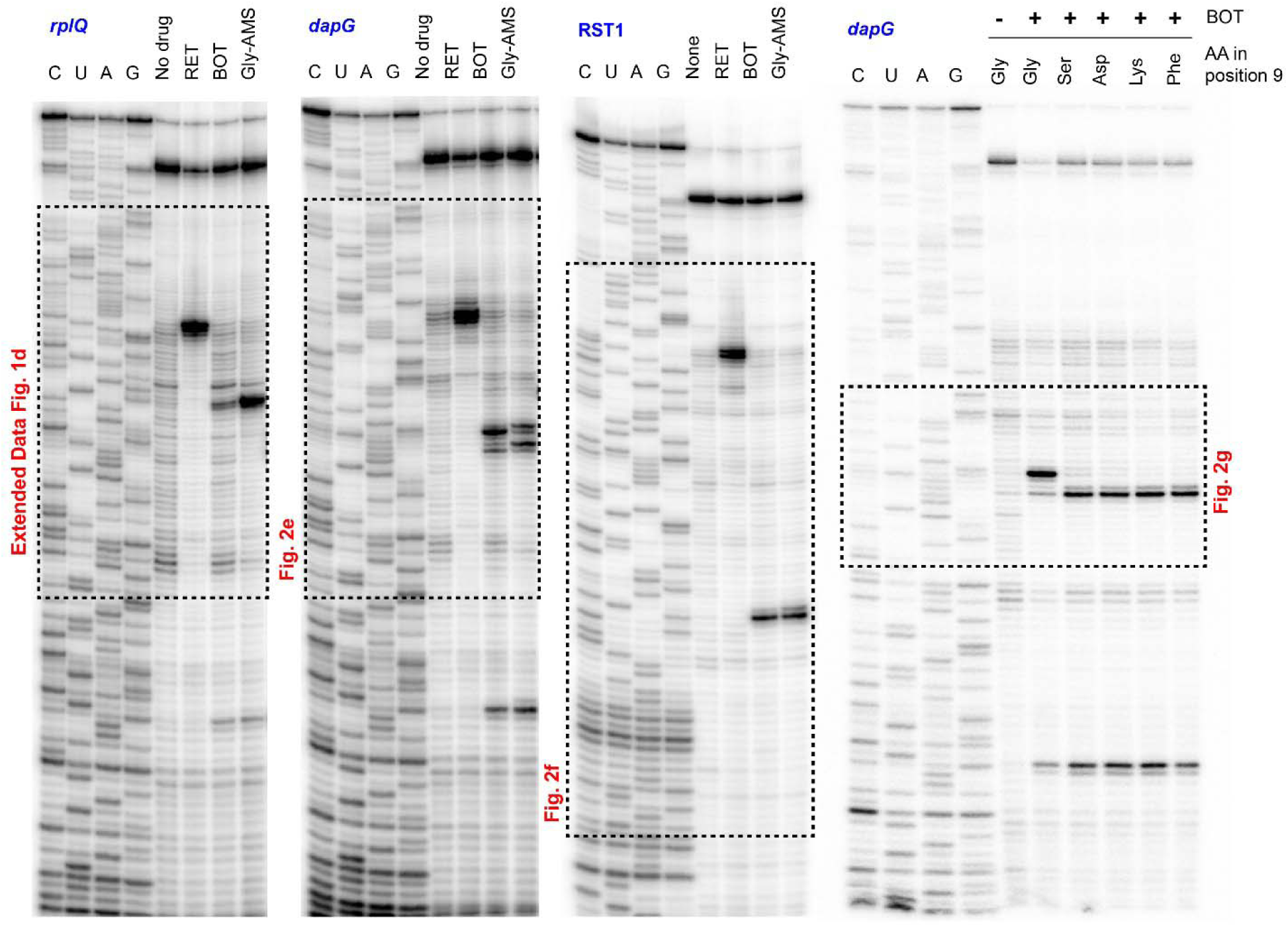
Full images of the toeprinting gels showing BOT-induced stalling on three different templates. Fragments of the gels used for making figures 2e, 2f, 2g, and Extended Data Figure 1d are framed.

**Supplementary Fig. 2.**
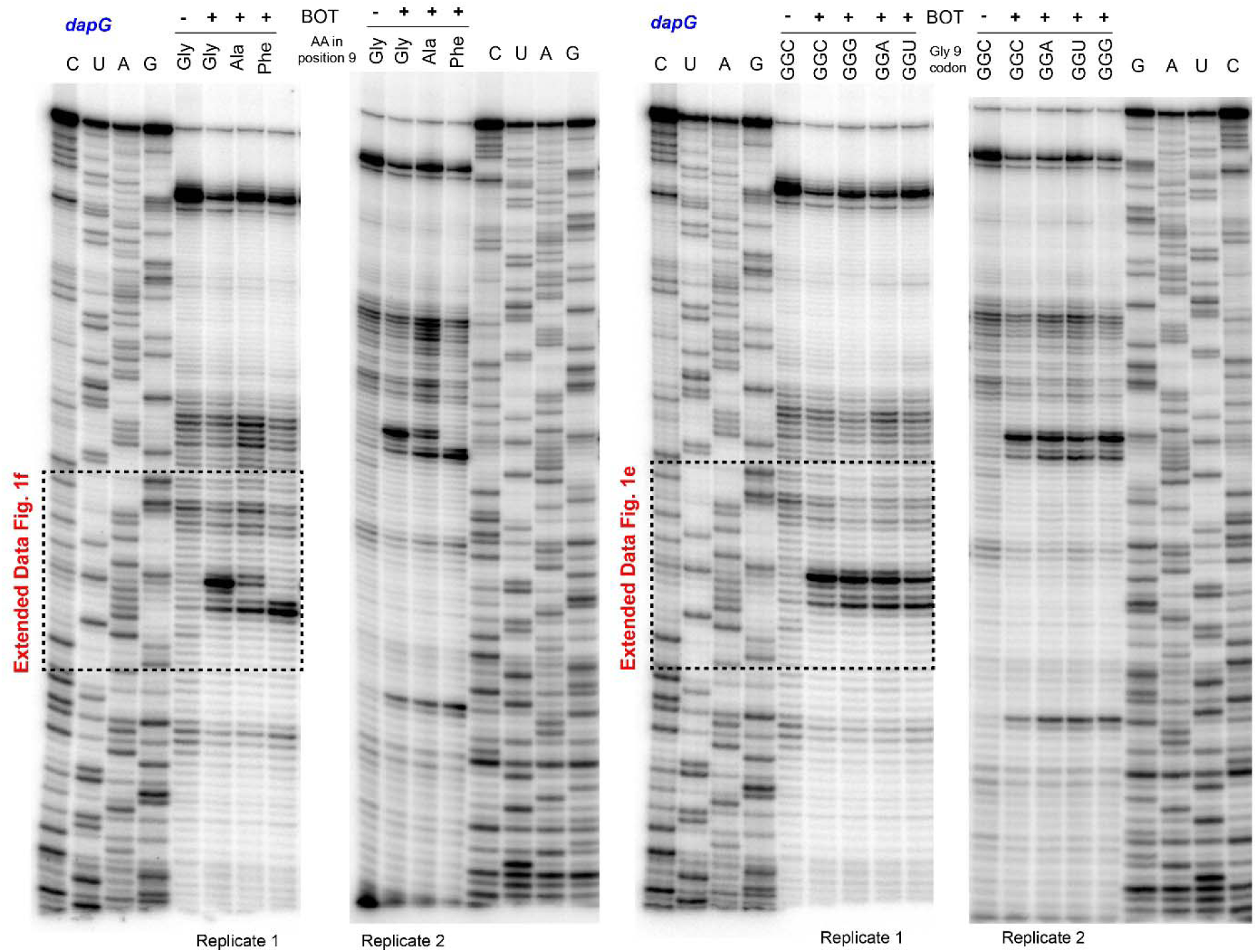
Full images of toeprinting gels showing BOT-induced stalling on the *dapG* template with varying codons in position 9. Two replicates for each experiment are shown. Fragments of the gels used for making Extended Data Figures 1e and 1f are framed.

## SUPPLEMENTARY TABLES

**Supplementary Table 1.**
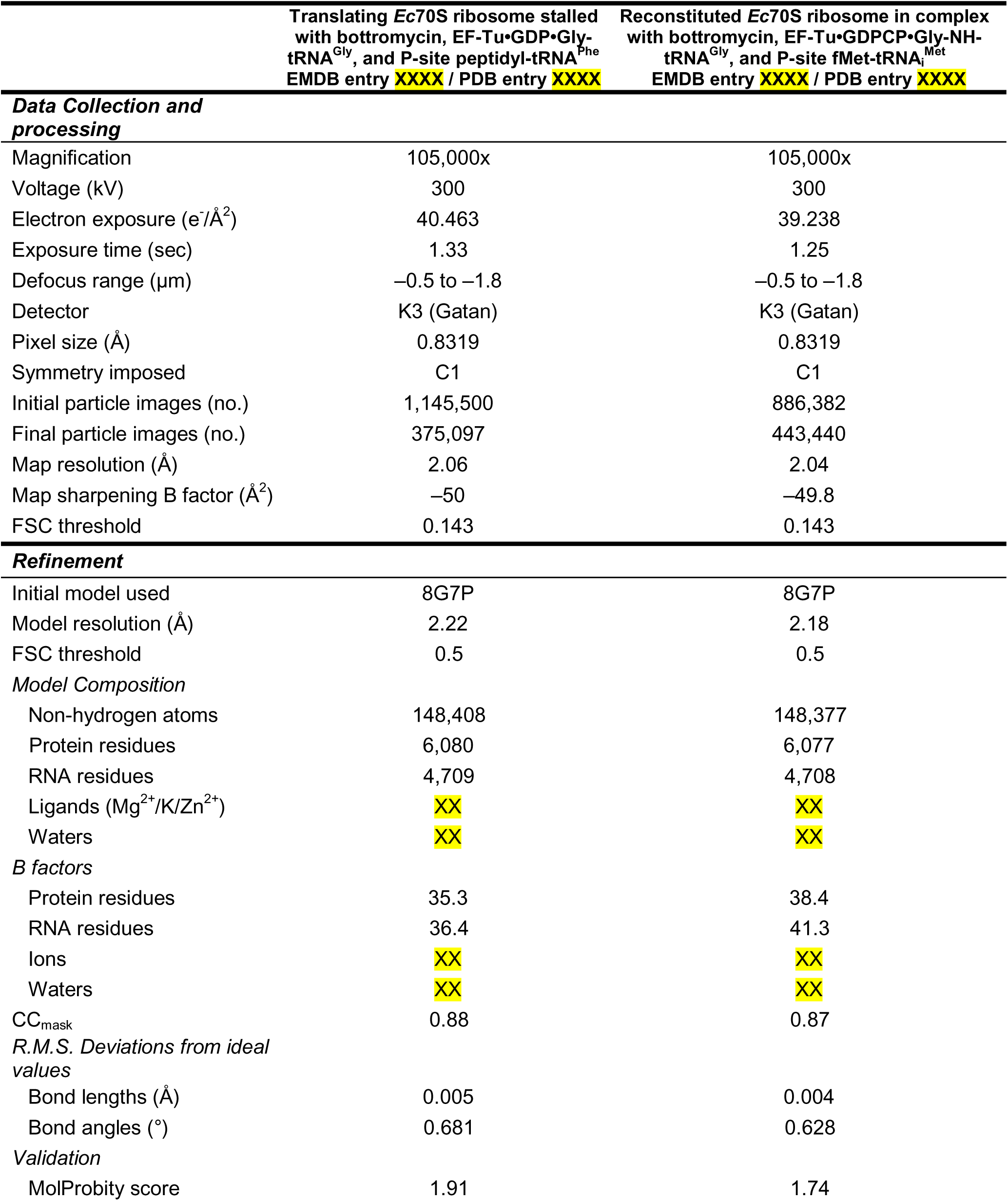
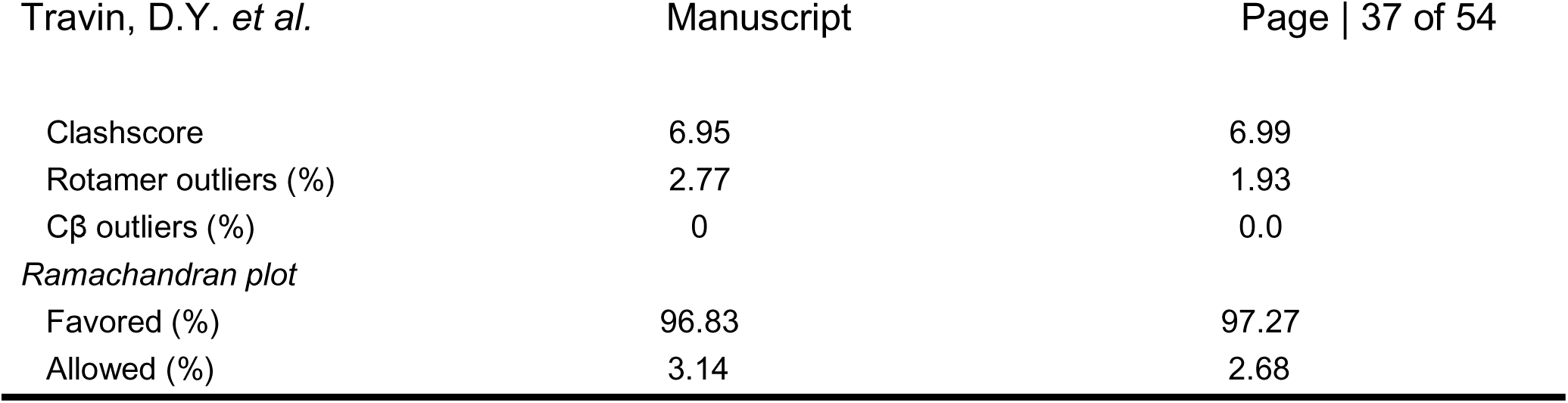
Cryo-EM data collection and refinement statistics.

|  | Translating *Ec*70S ribosome stalled with bottromycin, EF-Tu•GDP•Gly-tRNA<sup>Gly</sup>, and P-site peptidyl-tRNA<sup>Phe</sup><br>EMDB entry XXXX / PDB entry XXXX | Reconstituted *Ec*70S ribosome in complex with bottromycin, EF-Tu•GDPCP•Gly-NH-tRNA<sup>Gly</sup>, and P-site fMet-tRNA<sup>Met</sup><br>EMDB entry XXXX / PDB entry XXXX |
| --- | --- | --- |
| **Data Collection and processing** |  |  |
| Magnification | 105,000x | 105,000x |
| Voltage (kV) | 300 | 300 |
| Electron exposure (e<sup>-</sup>/Å<sup>2</sup>) | 40.463 | 39.238 |
| Exposure time (sec) | 1.33 | 1.25 |
| Defocus range (μm) | −0.5 to −1.8 | −0.5 to −1.8 |
| Detector | K3 (Gatan) | K3 (Gatan) |
| Pixel size (Å) | 0.8319 | 0.8319 |
| Symmetry imposed | C1 | C1 |
| Initial particle images (no.) | 1,145,500 | 886,382 |
| Final particle images (no.) | 375,097 | 443,440 |
| Map resolution (Å) | 2.06 | 2.04 |
| Map sharpening B factor (Å<sup>2</sup>) | −50 | −49.8 |
| FSC threshold | 0.143 | 0.143 |
| **Refinement** |  |  |
| Initial model used | 8G7P | 8G7P |
| Model resolution (Å) | 2.22 | 2.18 |
| FSC threshold | 0.5 | 0.5 |
| **Model Composition** |  |  |
| Non-hydrogen atoms | 148,408 | 148,377 |
| Protein residues | 6,080 | 6,077 |
| RNA residues | 4,709 | 4,708 |
| Ligands (Mg<sup>2+</sup>/K/Zn<sup>2+</sup>) | XX | XX |
| Waters | XX | XX |
| **B factors** |  |  |
| Protein residues | 35.3 | 38.4 |
| RNA residues | 36.4 | 41.3 |
| Ions | XX | XX |
| Waters | XX | XX |
| CC<sub>mask</sub> | 0.88 | 0.87 |
| **R.M.S. Deviations from ideal values** |  |  |
| Bond lengths (Å) | 0.005 | 0.004 |
| Bond angles (°) | 0.681 | 0.628 |
| **Validation** |  |  |
| MolProbity score | 1.91 | 1.74 |
| Clashscore | 6.95 | 6.99 |
| Rotamer outliers (%) | 2.77 | 1.93 |
| Cβ outliers (%) | 0 | 0.0 |
| *Ramachandran plot* |  |  |
| Favored (%) | 96.83 | 97.27 |
| Allowed (%) | 3.14 | 2.68 |

**Supplementary Table 2.**
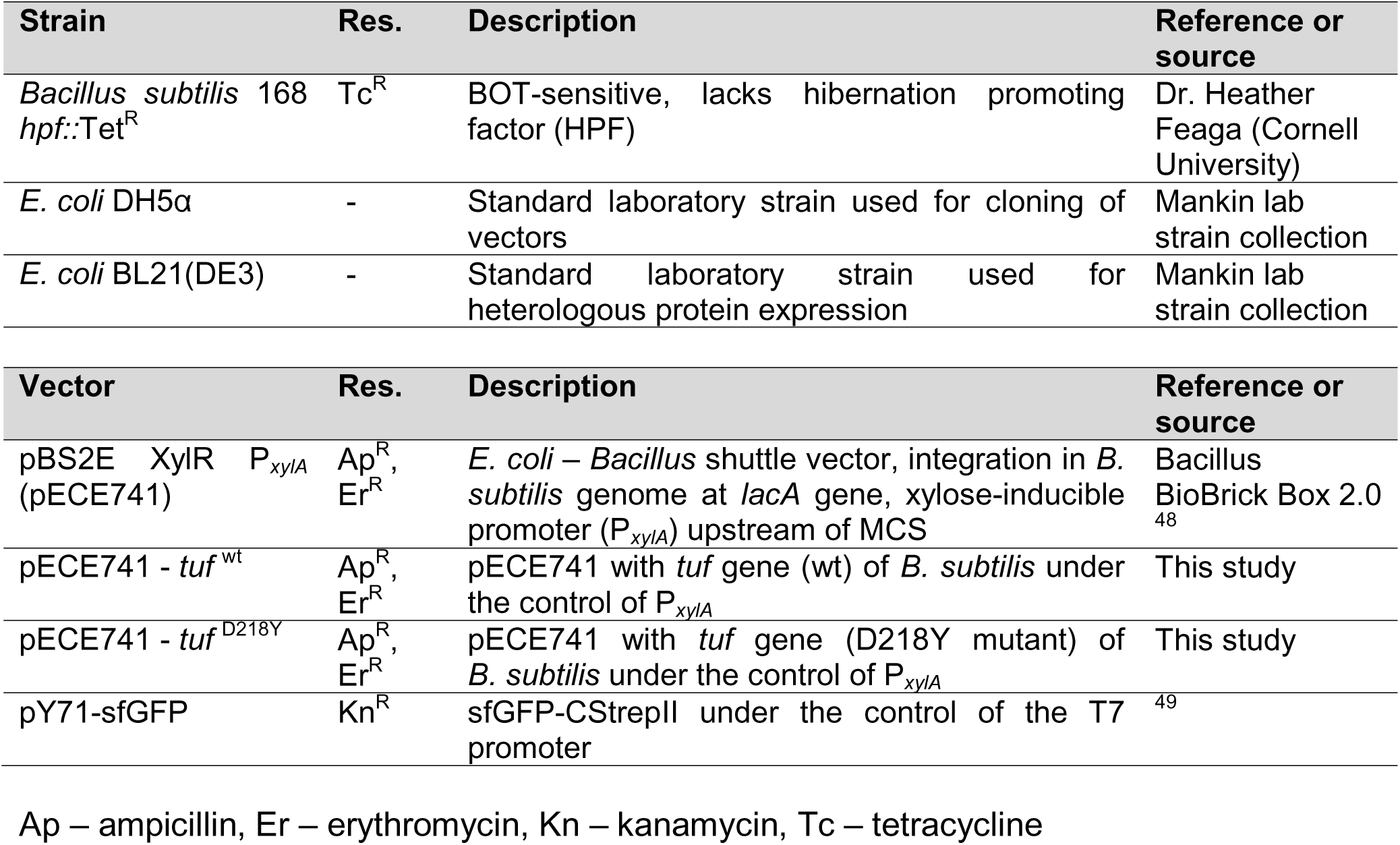
Bacterial strains and vectors used in the study.

**Supplementary Table 3.**
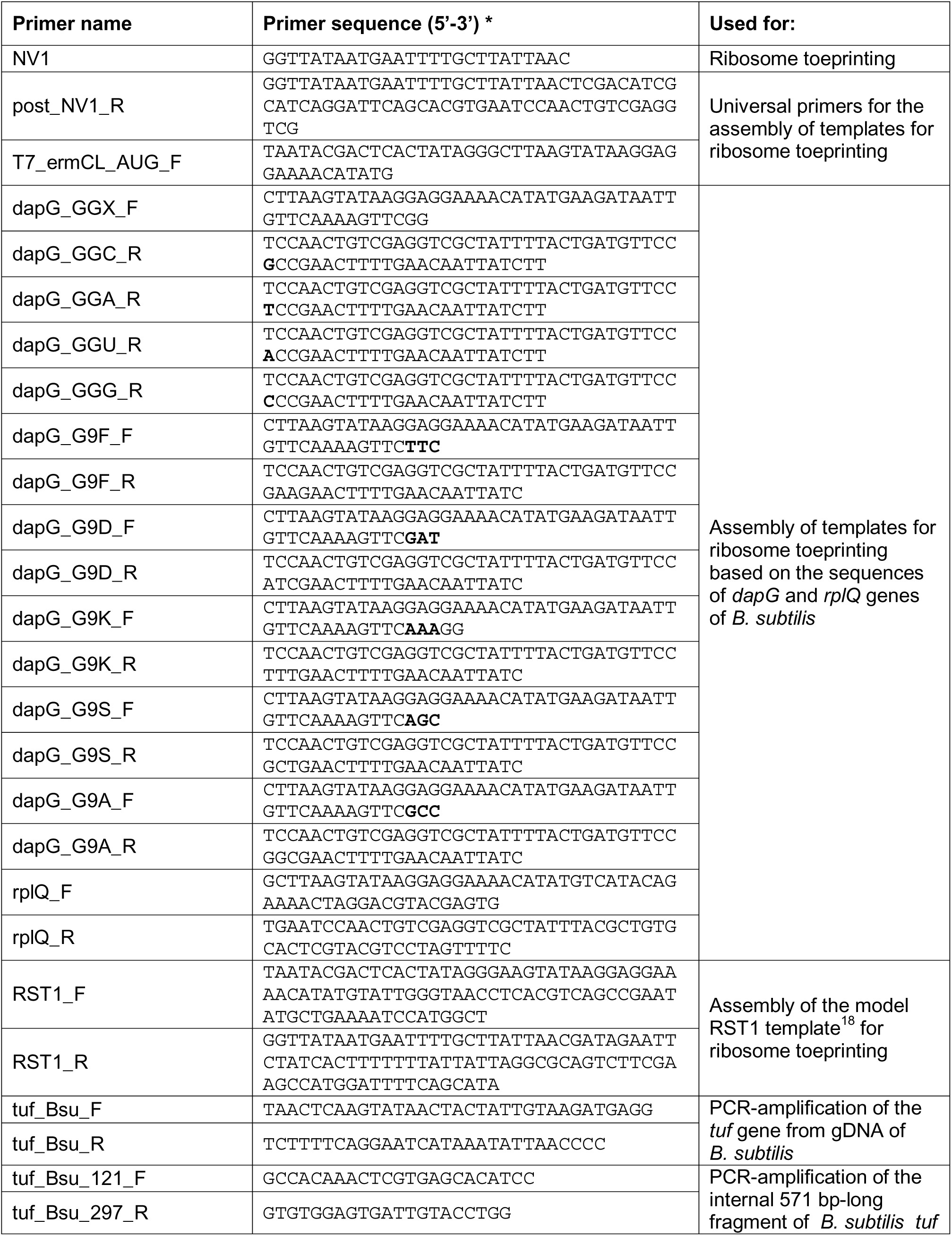
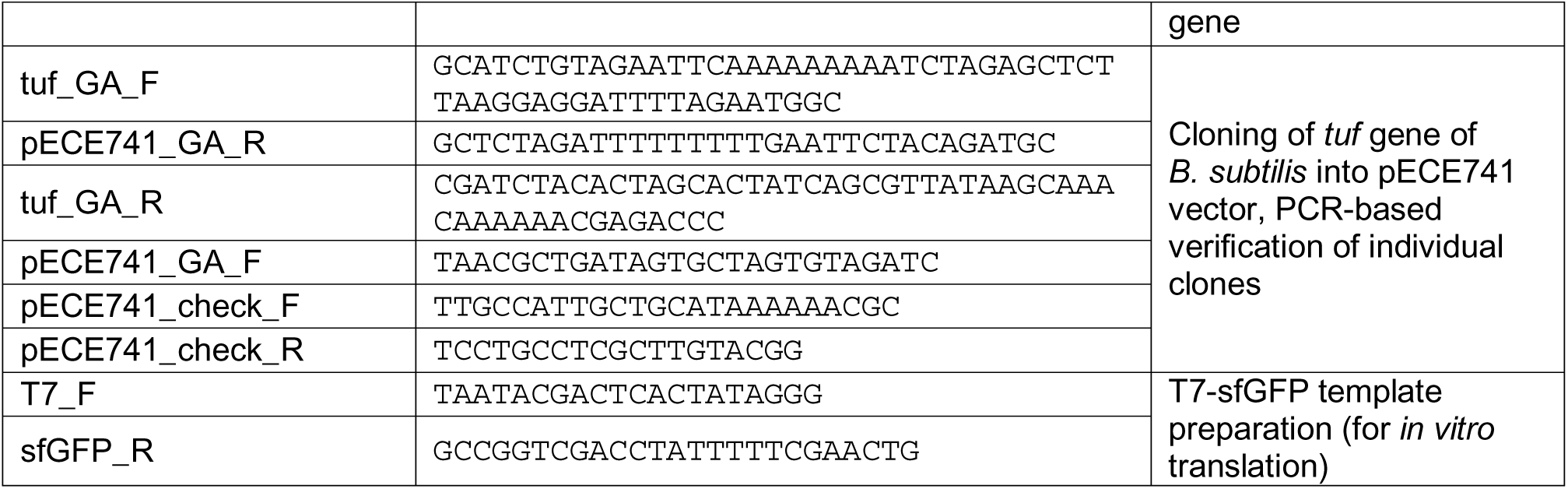
Nucleotide sequences of primers used in the study.

**Supplementary Table 4.**
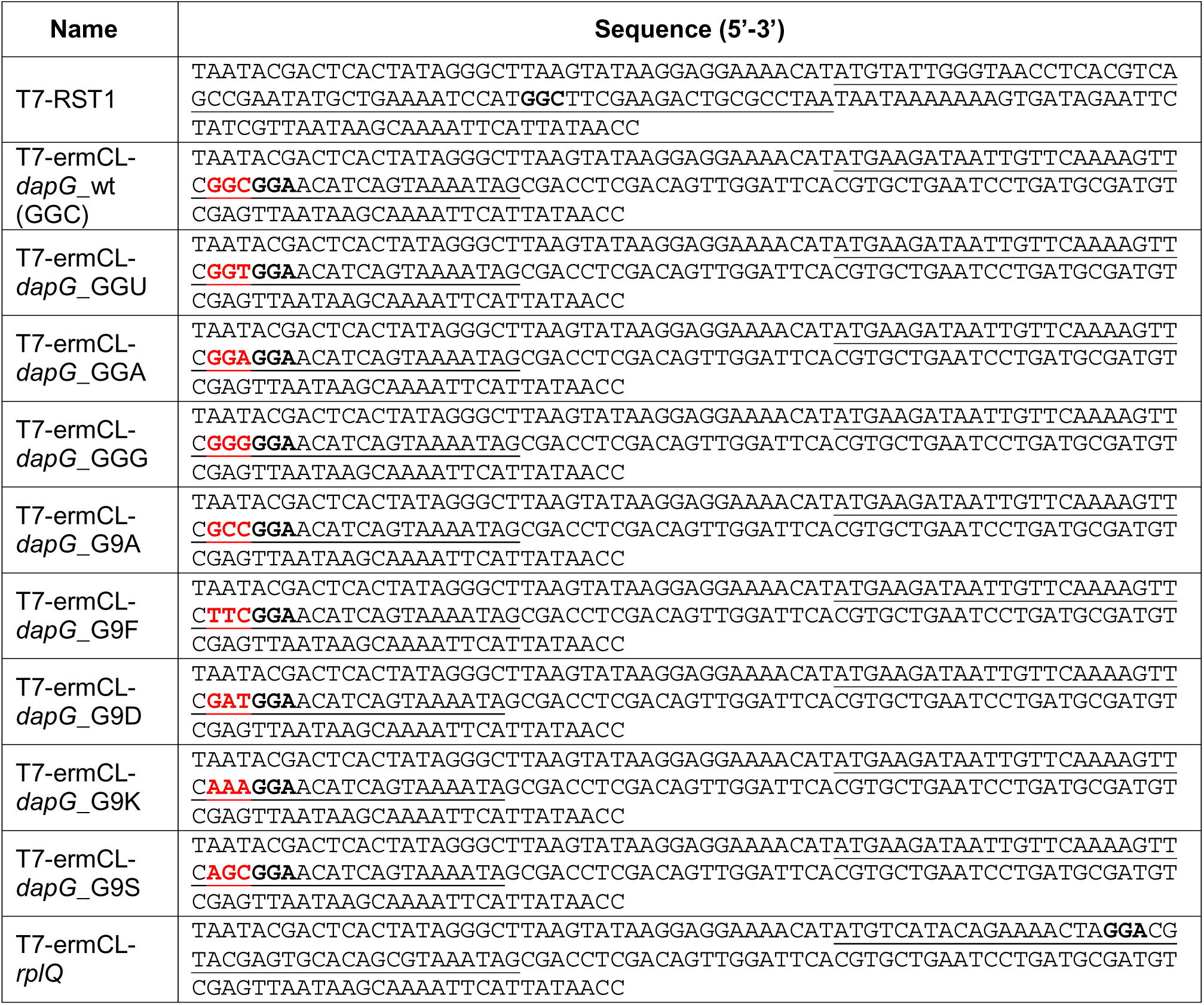
Nucleotide sequences of templates for *in vitro* translation.

## MATERIALS AND METHODS

### Reagents

Unless stated otherwise, all chemicals and reagents were obtained from MilliporeSigma (USA). All synthetic oligonucleotides, such as DNA primers and mRNA for structural studies, were obtained from Integrated DNA Technologies (USA).

### Antibiotics

Bottromycin A2 was purchased from Diagnoscine LLC (Cat. No. FNK-00592). GlyRS inhibitor Gly-AMS was purchased from MedChemExpress (Cat. No. HY-108940). Both compounds were dissolved in 100% dimethyl sulfoxide (DMSO) to a final stock concentration of 20 mM.

### Cultivation of bacteria

The list of bacterial strains and plasmids used in this study is provided in **Supplementary Table 2**. Lysogeny broth (LB, per 1L: 10 g tryptone, 5 g yeast extract, 10 g NaCl) was used for routine cultivation of *Escherichia coli* and *Bacillus subtilis*. Spizizen minimal (SM) medium (per 1L: 17.5 g K_2_HPO_4_, 7.5 g KH_2_PO_4_, 1.25 g of trisodium citrate × 2 H_2_O, 0.25 g MgSO_4_ × 7 H_2_O, 1% potassium glutamate, 0.5% glucose, 0.0055 g CaCl_2_, 0.001 g MnCl_2_ × 4 H_2_O, 0.0017 g ZnCl_2_, 0.00033 g CuCl_2_ × 2 H_2_O, 0.0006 g CoCl_2_ × 6 H_2_O, 0.0006 g Na_2_MoO_4_ × 2 H_2_O, 0.00135 g FeCl_3_ × 6 H_2_O) supplemented with 5 mg/mL of L-tryptophane was used for cultivation of *B. subtilis* where specified. When required antibiotics were added for selection (100 µg/mL ampicillin, 10 µg/mL erythromycin, 10 µg/mL tetracycline).

### Ribosome profiling

Ribosome profiling was performed essentially as described earlier^16^. Briefly, the overnight cultures of *Bacillus subtilis* 168 *hpf::*Tet^R^ (a strain, kindly provided by Dr. Heather Feaga, with increased proportion of actively translating ribosomes due to the lack of hibernation promoting factor, HPF^50^) were diluted 1:200 in 100 mL of fresh SM medium and grown with shaking at 37°C until reaching an OD_600_ of ∼0.4. For the BOT-treated sample, BOT was added to the culture to a final concentration of 3 µg/mL (50× MIC), and the culture was incubated with shaking for additional 5 min. The same procedure was performed for the control culture using the corresponding volume of 100% dimethyl sulfoxide (DMSO). Untreated (control) and BOT-treated cells were harvested by rapid filtration and flash frozen in liquid nitrogen. The cells were then resuspended in 650 µL lysis buffer (20 mM Tris pH 8.0, 10 mM MgCl_2_, 100 mM NH_4_Cl, 5 mM CaCl_2_, 0.4% Triton X-100, 0.1% NP-40) supplemented with 65 U RNase-free DNase I (Roche) and 208 U Superase•In RNase inhibitor (Invitrogen) and lysed using FastPrep-24™ bead beater (MP Biomedicals) (3 min, 6.5 beats/sec). The lysates were clarified by centrifugation for 10 min at 20,000×g (4°C). Fifteen OD_260_ units of obtained lysates were treated with 40U of S7 Micrococcal nuclease (Roche) per 1 OD_260_ of RNA for 1 h at 25°C with shaking. The reaction was quenched by the addition of ethylene glycol-bis(β-aminoethyl ether)-N,N,N′,N′-tetraacetic acid (EGTA) to the final concentration of 6 mM and lysates were layered over 2 mL of sucrose cushion (20% sucrose, 20 mM Tris/HCl pH 8.0, 10 mM MgCl_2_, 100 mM NH_4_Cl) in 4 mL tubes for S110AT rotor of Sorvall MX 120 Plus Micro-Ultracentrifuge (Thermo Fisher). Ribosomes were pelleted by centrifugation for 1 h at 422,000×g (100,000 RPM). The pellets were resuspended in 500 µL of resuspension buffer (20 mM Tris/HCl pH 8.0, 10 mM MgCl_2_, 100 mM NH_4_Cl, 1% SDS) and frozen in liquid nitrogen.

Total RNA was isolated from the obtained samples by the hot phenol-chloroform extraction procedure and precipitated for 30 min at -80°C following the addition of 1.1 volumes of ice-cold isopropanol. Subsequent steps, including size-selection of ribosome-protected fragments and preparation of sequencing libraries, followed the published protocol^51^.

### Ribosome profiling data analysis

A custom script (https://github.com/mmaiensc/RiboSeq) was used to demultiplex the samples, remove linker barcodes, and remove 2 nts from the 5’ end and 5 nts from the 3’ end, which were added as part of the library design. Bowtie2 (v2.2.9)^52^ within the Galaxy pipeline first aligned the trimmed reads to the non-coding RNA sequences, allowing filtering out the reads originating from rRNA and tRNA. The remaining unmapped reads were aligned to the reference genome of the *B. subtilis* strain 168 (GenBank ID: NC000964). The 24 to 46-nt-long reads that uniquely aligned to the genome were used in the subsequent analyses. The first position of the P-site codon was assigned 15 nt from the 3’ end of the read according to the 3’ assignment method used previously for Ribo-seq experiments in *Bacillus*^53^.

To analyze the sequence specificity of BOT-induced ribosome stalling, we first selected the codons in the bodies of the genes (the first 10 and the last 3 codons of the genes were excluded from analysis), for which the ribosome occupancy was at least 5 times higher in the BOT-treated sample compared to the untreated control. For each site, the corresponding encoded amino acid was determined, and the overrepresentation of amino acids for each position around the stall site was analyzed using the pLogo tool^54^ available online (https://plogo.uconn.edu/) with selected (n = 3045) and total (n = 144,945) samples of stalling sequences.

To assess the dependence of BOT-induced stalling efficiency on the identity of the A-site codon, we calculated BOT stall score on a codon-by-codon basis throughout the genome and determined average stall scores for each of 61 sense codons in the A-sites (**Fig. 2b**). For each codon BOT stall score was calculated as:

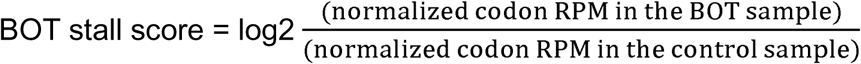

Each codon RPM value was normalized to the total gene RPM. Only the codons having more than 5 aligned reads in both the BOT and control samples were taken into the analysis. Genes having fewer than 100 aligned reads were excluded from the analysis.

### In vitro aminoacylation inhibition assay

To monitor aminoacylation of tRNA^Gly^ *in vitro,* the following components were combined in a final volume of 20 µL: 4 µM of deacylated tRNA^Gly^(CCC), 10 µM [^14^C]-glycine (100 mCi/mmol, American Radiolabeled Chemicals), 30 mM KCl, 60 mM HEPES pH 7.6, 10 mM MgCl_2_, 2.5 mM ATP, 0.5 mg/mL bovine serum albumin. The reaction contained either no drug, 50 µM BOT, or 50 µM 5’-O-(glycylsulfamoyl)adenosine (Gly-AMS). The reaction was then initiated by the addition of 0.5 µM *E. coli* Gly-tRNA synthetase (GlyRS, purified in-house) and incubated for 5 min at 37 °C. A reaction without enzyme addition was used as a background control. The experiment was performed in triplicate.

Each reaction (10 µL) was spotted on Whatman 3MM filter discs prerinsed with 25 µL of 10% trichloroacetic acid (TCA). The discs were immediately placed into a beaker with ice-cold 10% TCA, rinsed three times with 200 mL of cold 10% TCA, once with 100% acetone, and dried for 20 min under a fume hood. Dry disks were transferred to vials containing 5 mL of Ultima Gold liquid scintillation fluid (PerkinElmer), and the amount of radioactivity associated with precipitated tRNA was quantified in a Hidex 300 SL automatic liquid scintillation counter (**Fig. 2c**).

### MIC determination

Minimum inhibitory concentration (MIC) of BOT for *B. subtilis* 168 derivatives was determined in SM medium supplemented with 5 µg/mL L-tryptophane by the two-fold broth microdilution method in 96-well plates using a range of BOT concentrations of 0.03-32 µg/mL. Cells were added to the starting OD_600_ of 0.002, the plate was incubated for 18 hours at 37 °C, and the OD_600_ was determined in an Infinite M200 PRO microplate reader (TECAN).

### In vitro translation in bacterial cell lysate

The effect of BOT on bacterial translation was tested using NEBExpress® Cell-free *E. coli* Protein Synthesis System (NEB, Cat. No. E5360) programmed with a DNA fragment including the sfGFP gene under the control of T7 promoter. The fragment was PCR-amplified from the plasmid pY71-sfGFP^49^ using primers T7_F and sfGFP_R (**Supplementary Table 3**). Reactions were assembled according to the manufacturer’s protocol in a final volume of 5 μL and supplemented with 0.1–50 μM BOT. Reactions were placed into 384-well black/clear flat-bottom plate (BD, Cat. No. 353962). The plate was incubated at 37 °C in an Infinite M200 PRO microplate reader (TECAN), and fluorescence was read every 10 min for 2 hours with 488 nm excitation and 520 nm emission wavelengths. The signal at the 2-hour time point was used for data visualization (**Extended Data Fig. 7a**). The experiment was performed in triplicate.

### In vitro translation in the rabbit reticulocyte lysate

The effect of BOT on eukaryotic translation was tested using a Rabbit Reticulocyte Lysate System (Cat. No. L4960, Promega) programmed with firefly luciferase mRNA provided by the supplier. *In vitro* reactions were assembled according to the manufacturer’s protocol in a final volume of 10 μL and were supplemented with 0.1– 1000 μM of BOT, 0.01-100 μM of cycloheximide (CHX), or no antibiotic and incubated for 90 min at 30°C. Then, 2.5 μL from each reaction were mixed with 50 μL of prewarmed luciferase assay reagent (Cat. No. E1483, Promega), and the luminescence was measured in an Infinite M200 PRO plate reader (TECAN) for 10 min at 25°C (**Extended Data Fig. 7b**). The experiment was performed in triplicate.

### Toeprinting analysis

Toeprinting analysis was carried out using model mRNA templates listed in **Supplementary Table 4** and PURExpress *E. coli in vitro* transcription-translation coupled system (NEB) as described previously^19^. Reactions either contained no antibiotic or were supplemented with 50 μM retapamulin, 50 μM 5’-O-glycylsulfamoyl-adenosine (Gly-AMS), or 50 μM BOT. Model templates were generated by PCR using primers listed in **Supplementary Table 3**. Toeprinting experiments were performed in duplicates (**Supplementary Figs. 1, 2**).

### Cryo-EM sample preparation, data collection and processing

The *E. coli* 70S ribosomes were purified from MRE600 cells as described earlier^27^. The *in vitro* translation reaction was run using the PURExpress® i*n vitro* Protein Synthesis ΔRibosome kit (Cat. No. E3313S, NEB) supplemented with purified *E. coli* MRE600 70S ribosomes. The mRNA template was *in vitro* transcribed (HiScribe T7 RNA synthesis kit, Cat. No. E2040S, NEB) from a PCR fragment containing the *dapG* sequence, as described in the toeprinting assay. 5 µL PURE reaction containing 2 µM 70S ribosomes, 5 µM *dapG* mRNA template and 50 µM bottromycin A2 (Diagnoscine LLC) was incubated at 37°C for 30 minutes. The reaction was diluted to 300 nM final concentration of 70S ribosomes with the PURE system buffer (12 mM magnesium acetate, 5 mM potassium phosphate pH 7.3, 95 mM potassium glutamate, 5 mM NH_4_Cl, 0.5 mM CaCl_2_, 1 mM spermidine, 8 mM putrescine, 1 mM DTT) and applied directly onto EM grids.

Non-translating *E. coli* 70S ribosomes were reconstituted by mixing C-terminal 6×His-tagged *E. coli* EF-Tu, expressed and purified as described earlier^27^, with *E. coli* initiator fMet-tRNA_i_^Met^, purified according to established procedures^55^, and with 3’-amide-linked Gly-tRNA^Gly^, prepared as previously described^56^. The synthetic mRNA with sequence 5’-GGC AAG GAG GUA AAA <u>AUG GGG</u> UAA-3’ contained a strong Shine-Dalgarno sequence followed by the methionine (AUG), glycine (GGG) (underlined), and stop (UAA) codons. Briefly, the complex was assembled by incubating 600 nM ribosomes with 6 μM mRNA and 6 μM fMet-tRNA_i_^Met^ in 70S buffer containing 10 mM HEPES-KOH (pH 7.5), 60 mM KCl, 15 mM NH_4_Cl, 10 mM MgCl_2,_ and 6 mM β-mercaptoethanol, and incubated at 37°C for 10 minutes. The Gly-tRNA^Gly^-delivery ternary complex was assembled separately with 12 μM EF-Tu, 12 μM non-hydrolyzable 3’-amine-linked Gly-NH-tRNA^Gly^, 200 μM GDPCP, and 200 μM bottromycin A2 in 70S ribosome buffer at 37°C for 5 minutes and brought to room temperature. Equal volumes of 70S mixture and ternary complex were mixed and diluted with 70S ribosome buffer to a final concentration of 150 nM ribosomes and incubated at room temperature for 15 minutes before applying onto EM grids.

For both complexes, Quantifoil R2/1 Cu 200 mesh grids + 2nm C (Electron Microscopy Sciences, Hatfield, PA) were glow discharged at 15 mA for 15 seconds under 0.24 mBar vacuum in the PELCO easiGlow system (Ted Pella) and immediately used for freezing in the Leica EM GP2 cryoplunger. After the sample application, grids were blotted for 3 seconds in 22°C, 85% humidity and plunged into liquid ethane. Grids were then transferred into a Titan Krios G3i electron microscope (Thermo Fisher Scientific) operating at 300 keV and data acquired with a K3 direct electron detector (Gatan). The image stacks (movies) were acquired with pixel size of 0.8319 Å/pixel using the SerialEM software to record movies with 40 fractions and total accumulated dose of 40.4625 (PURE reaction) or 39.238 (reconstituted complex) e^-^/Å^2^/movie. A total of 18,240 (PURE reaction) or 10,945 (reconstituted complex) micrographs were collected with defocus values ranging from –0.5 to –1.8 µm.

Data processing was done in CryoSPARC 4.7.0^57^. The image stacks were motion-corrected with the patch motion correction job, followed by patch contrast transfer function (CTF) estimation. Based on relative ice thickness, CTF fit, and total frame motion, 18,045 micrographs were selected for the PURE reaction sample for further processing. 1,485,974 particles were filtered based on defocus adjusted power and pick scores, extracted with a box size of 512 x 512 pixels, and subjected to two rounds of reference-free two-dimensional (2D) classification. Selected 2D classes containing 1,145,500 particles were used in an ‘ab-initio reconstruction’ job to generate four 3D volumes that were further used for ‘heterogeneous refinement’. One main class average of 70S ribosomes containing 740,058 particles was obtained, in addition to 50S, 30S, and junk classes that were discarded **(Extended Data Fig. 2a)**. The 70S ribosome class average was bound to tRNA in the P site and EF-Tu with tRNA in the A/T conformation. These particles were classified using a soft mask on EF-Tu and A/T-tRNA using a focused 3D classification job resulting in 377,251 particles of 70S ribosomes bound to EF-Tu and A/T-tRNA corresponding to 51% of the total number of 70S particles, a class of 237,412 particles of 70S ribosomes bound to P- and E-tRNA, and a class of 66,397 particles of 70S ribosomes bound to A-, P- and E-tRNA. Non-uniform refinement with CTF refinement, per-particle defocus optimization was performed on the main class of 70S ribosomes in complex with EF-Tu and A/T-tRNA followed by reference-based motion correction of the particles, resulting in a final reconstruction with a resolution of 2.06 Å and sharpened with a B-factor of –50 Å^2^ from a final set of 375,097 particles.

Similarly, for the reconstituted complex sample, 10,491 micrographs were selected for further processing. 1,022,290 particles were selected based on defocus adjusted power and pick scores, extracted with a box size of 512 x 512 pixels, and subjected to two rounds of reference-free two-dimensional (2D) classification. Selected 2D classes containing 886,382 particles were used in an ‘ab-initio reconstruction’ job to generate four 3D volumes that were further used for ‘heterogeneous refinement’. One main class average of 70S ribosomes containing 741,513 particles was obtained, in addition to classes containing 50S subunits and junk particles, which were both discarded **(Extended Data Fig. 2b).** The 70S class average was bound to tRNA in the P site and EF-Tu with tRNA in the A/T conformation. These particles were further classified using a soft mask on EF-Tu and A/T-tRNA using a focused 3D classification job resulting in 446,116 particles of 70S ribosomes bound to EF-Tu and A/T-tRNA corresponding to 60% of the total number of 70S particles, a class of 231,033 particles of 70S bound to P- and E-tRNA, and a class of 64,364 particles of 70S bound to A-, P- and E-tRNA. Non-uniform refinement with CTF refinement and per-particle defocus optimization was performed on the main class of 70S in complex with EF-Tu and A/T-tRNA followed by reference-based motion correction of the particles, resulting in a final reconstruction with an overall resolution of 2.04 Å and sharpened with a B-factor of –49.8 Å^2^ from a final set of 443,440 particles.

### Model building and structure refinement

The initial model of *E. coli* 70S ribosome in complex with EF-Tu•GDPCP and Ile-tRNA_LAU_^Ile^ in the A/T conformation and fMet-tRNA_i_^Met^ in P site (PDB entry 8G7P^27^) was fitted into the refined volume of the reconstituted complex using UCSF Chimera (version 1.17.3)^58^ and manually adjusted in Coot (version 0.9.8.7)^59^. The model of Gly-tRNA^Gly^ was built by mutating the nucleotide sequence of Ile-tRNA_LAU_^Ile^ to that of Gly-tRNA^Gly^ CCC anticodon-isoform, real-space refined into the EM map, including the addition of modified nucleotides and the 3’-amide-linked non-hydrolysable glycine residue attached to nucleotide A76. The model and restraints for bottromycin were generated using phenix.elbow and fitted into the density clearly visible inside EF-Tu domain I near the CCA-end of A/T-tRNA. For the complex derived from the PURE translation reaction, the tRNA^Gly^ sequence was mutated to the GCC isoform, and three residues (QKF) of the nascent dapG peptide were added. In this complex, the glycine residue is 3’-ester linked to the Gly-tRNA^Gly^ and EF-Tu is bound to GDP. Magnesium ions and waters were added and the complete models were refined by phenix.real_space_refinement in PHENIX (version 1.19.2)^60^ using base pair restraints for rRNA, A-site tRNA in the A/T-conformation and bottromycin. The molecular models were validated using the comprehensive cryo-EM validation tool in PHENIX 1.19.2^61^. The statistics of data collection and refinement are compiled in **Supplementary Table 1**. All figures showing atomic models were rendered using the PyMol software (www.pymol.org).

### Selection of BOT-resistant mutants in B. subtilis

To identify mutations conferring resistance to BOT in the *tuf* gene, a culture of *B. subtilis* 168 *hpf*::Tet^R^ was grown overnight in LB supplemented with 10 µg/mL of tetracycline. Approximately 1.5 OD_600_ units of cells (∼1*10^8^ cells) were washed twice with 1 mL of SM medium and plated on a SM agar plate containing 10 µg/mL tetracycline and 1.6 μg/mL BOT (∼25×MIC). After 48 h of incubation at 37°C, nine colonies appeared on the plate. gDNA was extracted from ON cultures from six individual colonies; the *tuf* gene was PCR-amplified using the primers tuf_Bsu_F and tuf_Bsu_R (**Supplementary Table 3**) and sequenced by Oxford Nanopore Technology (ONT). BOT MIC in liquid SM medium was then determined for six isolated mutants along with the parent strain used for selection.

### Engineering of B. subtilis strains with two copies of the EF-Tu encoding genes

WT and mutant *tuf* genes were PCR amplified from gDNA of *B. subtilis* 168 *hpf*::TetR and its BOT-resistant derivative harbouring a D218Y mutation using primers tuf_GA_F and tuf_GA_R. PCR fragments were Gibson assembled with a PCR-amplified fragment of the pECE741 plasmid (primers pECE741_GA_R and pECE741_GA_F) and transformed into chemically competent *E. coli* DH5α. Correct assembly of plasmids was verified by whole-plasmid sequencing using ONT.

The obtained plasmids pECE741-*tuf* ^wt^ and pECE741-*tuf* ^D218Y^ harbouring EF-Tu encoding genes under the control of xylose-inducible promoter were transformed into *B. subtilis* 168 *hpf*::TetR or its BOT-resistant derivative following the procedure described as Method 3.8 in ^62^. Selection of transformants was performed on LB agar plates supplemented with 10 µg/mL erythromycin. Plasmid integrations into the *lacA* locus were confirmed by PCR. Additionally, using gDNA of the four engineered strains, the internal fragment of the *tuf* gene surrounding the site of single nucleotide substitution was amplified using primers tuf_Bsu_121_F and tuf_Bsu_297_R. The resulting fragments were sequenced by the Sanger method to assess the copy number of the two alleles in each strain.

### Bioinformatic analysis of tuf genes distribution across bacterial genomes

The list of reference genomes for analysis was retrieved from the Prokaryotic RefSeq Genomes dataset^30^ on May 22, 2025. Out of 21,014 genomes, 6,360 assembled to “Complete Genome” and “Chromosome” levels were used for subsequent analysis to avoid the bias potentially arising from incomplete assemblies. Complete genomes were downloaded using NCBI datasets software^63^, and the coordinates and descriptions of genes annotated as encoding EF-Tu (*tuf* genes) were extracted with a custom script. 115 *tuf* genes, which were either marked as “partial” or “pseudo” in the corresponding annotations or were shorter than 1000 bp (typical length of the *tuf* genes was 1160-1210 bp and was conserved across phyla) were filtered out. Additionally, from the analysis, we excluded 148 genomes of bacteria without assigned taxonomy, *i.e.,* no phylum or class was available in the NCBI Taxonomy database^64^. In addition, we manually checked the number of *tuf* paralogs encoded in the genomes of select BOT-sensitive bacteria if those were absent among the genomes in our global analysis.

## DATA AVAILABILITY STATEMENT

Sequencing data for ribosome profiling in *B. subtilis* were deposited in the NCBI Sequence Read Archive (SRA) under the accession number **PRJNA1256931**.

Coordinates were deposited in the RCSB Protein Data Bank with accession codes:

- **9XXX**for the IVT-derived *E. coli* 70S ribosome stalled with bottromycin, EF-Tu•GDP•Gly-tRNA^Gly^, and P-site peptidyl-tRNA^Phe^. The corresponding cryo-EM maps have been deposited in the Electron Microscopy Data Bank (EMDB) under accession code **EMD-XXXX**. The unaligned multi-frame cryo-EM micrographs have been deposited in the Electron Microscopy Public Image Archive (EMPIAR) with the accession code **EMPIAR-XXXX**.
- **9XXX**for the reconstituted *E. coli* 70S ribosome in complex with bottromycin, EF-Tu•GDPCP•Gly-NH-tRNA^Gly^, and P-site fMet-tRNA_i_^Met^. The corresponding cryo-EM maps have been deposited in the EMDB under accession code **EMD-XXXX**. The unaligned multi-frame cryo-EM micrographs have been deposited in the EMPIAR with the accession code **EMPIAR-XXXX**.

All previously published structures that were used in this work for structural comparisons were retrieved from the RCSB Protein Data Bank: PDB entries 1TTT, 1B23, 4V5S, 1OB2, 1OB5, 2C77, 2C78, 8G7P, and 8G5Z.

## REFERENCES

1. Miller, W. R. & Arias, C. A. ESKAPE pathogens: antimicrobial resistance, epidemiology, clinical impact and therapeutics. Nat. Rev. Microbiol. 22, 598–616 (2024).

2. M. Waisvisz, J., G. van der Hoeven, M., van Peppen, J. & C. M. Zwennis, W. Bottromycin. I. A New Sulfur-containing Antibiotic. J. Am. Chem. Soc. 79, 4520– 4521 (2002).

3. Kobayashi, Y. et al. Bottromycin derivatives: efficient chemical modifications of the ester moiety and evaluation of anti-MRSA and anti-VRE activities. Bioorg. Med. Chem. Lett. 20, 6116–20 (2010).

4. Tanaka, N., Nishimura, T., Nakamura, S., Umezawa, H. & Hayami, T. Activity of bottromycin against Mycoplasma gallisepticum. J. Antibiot. (Tokyo). 21, 75–76 (1968).

5. Nakamura, S., Yajima, T., Lin, Y. & Umezawa, H. Isolation and characterization of bottromycins A2, B2, C2. J. Antibiot. (Tokyo). 20, 1–5 (1967).

6. Lerchen, H.-G., et al. Cyclic iminopeptide derivatives. (2006).

7. Park, S. B., Lee, I. A., Suh, J.-W., Kim, J.-G. & Lee, C. H. Screening and identification of antimicrobial compounds from Streptomyces bottropensis suppressing rice bacterial blight. J. Microbiol. Biotechnol. 21, 1236–42 (2011).

8. Arnison, P. G. et al. Ribosomally synthesized and post-translationally modified peptide natural products: overview and recommendations for a universal nomenclature. Nat. Prod. Rep. 30, 108–60 (2013).

9. Gomez-Escribano, J. P., Song, L., Bibb, M. J. & Challis, G. L. Posttranslational [small beta]-methylation and macrolactamidination in the biosynthesis of the bottromycin complex of ribosomal peptide antibiotics. Chem. Sci. 3, 3522–3525 (2012).

10. Franz, L., Kazmaier, U., Truman, A. W. & Koehnke, J. Bottromycins - Biosynthesis, synthesis and activity. Nat. Prod. Rep. 38, 1659–1683 (2021).

11. Tanaka, N., Sashikata, K., Yamaguchi, H. & Umezawa, H. Inhibition of protein synthesis by bottromycin A2 and its hydrazide. J. Biochem. 60, 405–10 (1966).

12. Lin, Y. C. & Tanaka, N. Mechanism of action of bottromycin in polypeptide biosynthesis. J. Biochem. 63, 1–7 (1968).

13. Kinoshita, T. & Tanaka, N. On the site of action of bottromycin A2. J. Antibiot. (Tokyo). 23, 311–2 (1970).

14. Otaka, T. & Kaji, A. Mode of action of bottromycin A2. Release of aminoacyl- or peptidyl-tRNA from ribosomes. J. Biol. Chem. 251, 2299–306 (1976).

15. Otaka, T. & Kaji, A. Mode of action of bottromycin A2: effect on peptide bond formation. FEBS Lett. 123, 173–6 (1981).

16. Ingolia, N. T. Genome-wide translational profiling by ribosome footprinting. Methods Enzymol. 470, 119–42 (2010).

17. Vazquez-Laslop, N., Thum, C. & Mankin, A. S. Molecular mechanism of drug-dependent ribosome stalling. Mol. Cell 30, 190–202 (2008).

18. Orelle, C. et al. Tools for characterizing bacterial protein synthesis inhibitors. Antimicrob. Agents Chemother. 57, 5994–6004 (2013).

19. Orelle, C. et al. Identifying the targets of aminoacyl-tRNA synthetase inhibitors by primer extension inhibition. Nucleic Acids Res. 41, e144 (2013).

20. Gouda, H. et al. Three-dimensional solution structure of bottromycin A2: a potent antibiotic active against methicillin-resistant Staphylococcus aureus and vancomycin-resistant Enterococci. Chem. Pharm. Bull. (Tokyo). 60, 169–71 (2012).

21. Syroegin, E. A., Aleksandrova, E. V & Polikanov, Y. S. Structural basis for the inability of chloramphenicol to inhibit peptide bond formation in the presence of A-site glycine. Nucleic Acids Res. 50, 7669–7679 (2022).

22. Polekhina, G. et al. Helix unwinding in the effector region of elongation factor EF-Tu-GDP. Structure 4, 1141–51 (1996).

23. Abel, K., Yoder, M. D., Hilgenfeld, R. & Jurnak, F. An alpha to beta conformational switch in EF-Tu. Structure 4, 1153–9 (1996).

24. Loveland, A. B., Demo, G. & Korostelev, A. A. Cryo-EM of elongating ribosome with EF-Tu•GTP elucidates tRNA proofreading. Nature 584, 640–645 (2020).

25. Schmeing, T. M. et al. The crystal structure of the ribosome bound to EF-Tu and aminoacyl-tRNA. Science 326, 688–694 (2009).

26. Voorhees, R. M., Schmeing, T. M., Kelley, A. C. & Ramakrishnan, V. The mechanism for activation of GTP hydrolysis on the ribosome. Science 330, 835– 838 (2010).

27. Rybak, M. Y. & Gagnon, M. G. Structures of the ribosome bound to EF-Tu-isoleucine tRNA elucidate the mechanism of AUG avoidance. Nat. Struct. Mol. Biol. 31, 810–816 (2024).

28. Berchtold, H. et al. Crystal structure of active elongation factor Tu reveals major domain rearrangements. Nature 365, 126–32 (1993).

29. Holm, M. et al. mRNA decoding in human is kinetically and structurally distinct from bacteria. Nature 617, 200–207 (2023).

30. Goldfarb, T. et al. NCBI RefSeq: reference sequence standards through 25 years of curation and annotation. Nucleic Acids Res. 53, D243–D257 (2025).

31. Lin, J., Zhou, D., Steitz, T. A., Polikanov, Y. S. & Gagnon, M. G. Ribosome-Targeting Antibiotics: Modes of Action, Mechanisms of Resistance, and Implications for Drug Design. Annu. Rev. Biochem. 87, 451–478 (2018).

32. Vázquez-Laslop, N. & Mankin, A. S. Context-Specific Action of Ribosomal Antibiotics. Annu. Rev. Microbiol. 72, 185–207 (2018).

33. Beckert, B. et al. Structural and mechanistic basis for translation inhibition by macrolide and ketolide antibiotics. Nat. Commun. 12, 4466 (2021).

34. Syroegin, E. A. et al. Structural basis for the context-specific action of the classic peptidyl transferase inhibitor chloramphenicol. Nat. Struct. Mol. Biol. 29, 152–161 (2022).

35. Tsai, K. et al. Structural basis for context-specific inhibition of translation by oxazolidinone antibiotics. Nat. Struct. Mol. Biol. 29, 162–171 (2022).

36. Mangano, K. et al. Context-based sensing of orthosomycin antibiotics by the translating ribosome. Nat. Chem. Biol. 18, 1277–1286 (2022).

37. Koller, T. O. et al. Paenilamicins are context-specific translocation inhibitors of protein synthesis. Nat. Chem. Biol. 20, 1691–1700 (2024).

38. Zhang, Y. et al. The context of the ribosome binding site in mRNAs defines specificity of action of kasugamycin, an inhibitor of translation initiation. Proc. Natl. Acad. Sci. U. S. A. 119, (2022).

39. Prezioso, S. M., Brown, N. E. & Goldberg, J. B. Elfamycins: inhibitors of elongation factor-Tu. Mol. Microbiol. 106, 22–34 (2017).

40. Furano, A. V. Content of elongation factor Tu in Escherichia coli. Proc. Natl. Acad. Sci. U. S. A. 72, 4780–4 (1975).

41. Yamada, T. et al. Synthesis and Evaluation of Antibacterial Activity of Bottromycins. J. Org. Chem. 83, 7135–7149 (2018).

42. Bickel, E. & Kazmaier, U. Syntheses of bottromycin derivatives via Ugi-reactions and Matteson homologations. Org. Biomol. Chem. 22, 8811–8816 (2024).

43. Nissen, P. et al. Crystal structure of the ternary complex of Phe-tRNAPhe, EF-Tu, and a GTP analog. Science 270, 1464–72 (1995).

44. Nissen, P., Thirup, S., Kjeldgaard, M. & Nyborg, J. The crystal structure of Cys-tRNACys-EF-Tu-GDPNP reveals general and specific features in the ternary complex and in tRNA. Structure 7, 143–56 (1999).

45. Schmeing, T. M., Voorhees, R. M., Kelley, A. C. & Ramakrishnan, V. How mutations in tRNA distant from the anticodon affect the fidelity of decoding. Nat. Struct. Mol. Biol. 18, 432–6 (2011).

46. Parmeggiani, A. et al. Enacyloxin IIa pinpoints a binding pocket of elongation factor Tu for development of novel antibiotics. J. Biol. Chem. 281, 2893–900 (2006).

47. Parmeggiani, A. et al. Structural basis of the action of pulvomycin and GE2270 A on elongation factor Tu. Biochemistry 45, 6846–6857 (2006).

48. Popp, P. F., Dotzler, M., Radeck, J., Bartels, J. & Mascher, T. The Bacillus BioBrick Box 2.0: expanding the genetic toolbox for the standardized work with Bacillus subtilis. Sci. Rep. 7, 15058 (2017).

49. Bundy, B. C. & Swartz, J. R. Site-specific incorporation of p-propargyloxyphenylalanine in a cell-free environment for direct protein-protein click conjugation. Bioconjug. Chem. 21, 255–63 (2010).

50. Beckert, B. et al. Structure of the Bacillus subtilis hibernating 100S ribosome reveals the basis for 70S dimerization. EMBO J. 36, 2061–2072 (2017).

51. McGlincy, N. J. & Ingolia, N. T. Transcriptome-wide measurement of translation by ribosome profiling. Methods 126, 112–129 (2017).

52. Langmead, B. & Salzberg, S. L. Fast gapped-read alignment with Bowtie 2. Nat. Methods 9, 357–9 (2012).

53. Hummels, K. R. & Kearns, D. B. Suppressor mutations in ribosomal proteins and FliY restore Bacillus subtilis swarming motility in the absence of EF-P. PLoS Genet. 15, e1008179 (2019).

54. O’Shea, J. P. et al. pLogo: a probabilistic approach to visualizing sequence motifs. Nat. Methods 10, 1211–2 (2013).

55. Jünemann, R. et al. In vivo deuteration of transfer RNAs: overexpression and large-scale purification of deuterated specific tRNAs. Nucleic Acids Res. 24, 907– 13 (1996).

56. Gamper, H. & Hou, Y.-M. tRNA 3’-amino-tailing for stable amino acid attachment. RNA 24, 1878–1885 (2018).

57. Punjani, A., Rubinstein, J. L., Fleet, D. J. & Brubaker, M. A. cryoSPARC: algorithms for rapid unsupervised cryo-EM structure determination. Nat. Methods 14, 290–296 (2017).

58. Pettersen, E. F. et al. UCSF Chimera - a visualization system for exploratory research and analysis. J. Comput. Chem. 25, 1605–12 (2004).

59. Emsley, P., Lohkamp, B., Scott, W. G. & Cowtan, K. Features and development of Coot. Acta Crystallogr. D. Biol. Crystallogr. 66, 486–501 (2010).

60. Liebschner, D. et al. Macromolecular structure determination using X-rays, neutrons and electrons: recent developments in Phenix. Acta Crystallogr. Sect. D, Struct. Biol. 75, 861–877 (2019).

61. Afonine, P. V et al. New tools for the analysis and validation of cryo-EM maps and atomic models. Acta Crystallogr. Sect. D, Struct. Biol. 74, 814–840 (2018).

62. Harwood, C. R. & Cutting, S. M. Molecular biological methods for Bacillus. in (1990).

63. O’Leary, N. A. et al. Exploring and retrieving sequence and metadata for species across the tree of life with NCBI Datasets. Sci. data 11, 732 (2024).

64. Schoch, C. L. et al. NCBI Taxonomy: a comprehensive update on curation, resources and tools. Database (Oxford). 2020, (2020).

