## Supplementary Data 1 for "Sequence-specific trapping of EF-Tu/glycyl-tRNA complex on the ribosome by bottromycin"

| GeneBank_ID | RefSeq_ID | start | end | Length (nt) | strand | gene_name | protein_id |
|---|---|---|---|---|---|---|---|
| GCF_018304805.1 | NZ_CP072646.1 | 410452 | 411636 | 1184 | + | tuf | WP_015797865.1 |
| GCF_018304805.1 | NZ_CP072646.1 | 245731 | 246918 | 1187 | + | tuf | WP_153757942.1 |
| GCF_000226295.1 | NC_016024.1 | 263518 | 264705 | 1187 | + | tuf | WP_153757956.1 |
| GCF_000226295.1 | NC_016024.1 | 1675498 | 1676682 | 1184 | - | tuf | WP_421121315.1 |
| GCF_018304825.1 | NZ_CP072648.1 | 1714159 | 1715394 | 1235 | - | tuf | WP_421121364.1 |
| GCF_018304825.1 | NZ_CP072648.1 | 2358285 | 2359469 | 1184 | - | tuf | WP_227755646.1 |
| GCF_030295385.1 | NZ_AP027079.1 | 2377215 | 2378399 | 1184 | - | tuf | WP_227755646.1 |
| GCF_030295425.1 | NZ_AP027081.1 | 4107388 | 4108572 | 1184 | - | tuf | WP_272736202.1 |
| GCF_030295405.1 | NZ_AP027080.1 | 4126497 | 4127681 | 1184 | - | tuf | WP_272736202.1 |
| GCF_017498545.1 | NZ_CP071793.1 | 877386 | 878567 | 1181 | + | tuf | WP_015440347.1 |
| GCF_016592575.1 | NZ_AP017470.1 | 922388 | 923569 | 1181 | + | tuf | WP_015440347.1 |
| GCF_016592575.1 | NZ_AP017470.1 | 597609 | 598796 | 1187 | + | tuf | WP_004867925.1 |
| GCF_003330725.1 | NZ_CP030840.1 | 613620 | 614807 | 1187 | + | tuf | WP_004867925.1 |
| GCF_003330725.1 | NZ_CP030840.1 | 2582825 | 2584012 | 1187 | - | tuf | WP_423856148.1 |
| GCF_000022565.1 | NC_012483.1 | 2598847 | 2600034 | 1187 | - | tuf | WP_423856148.1 |
| GCF_000022565.1 | NC_012483.1 | 1060193 | 1061380 | 1187 | - | tuf | WP_113527398.1 |
| GCF_000014005.1 | NC_008009.1 | 1076208 | 1077395 | 1187 | - | tuf | WP_113527398.1 |
| GCF_000014005.1 | NC_008009.1 | 2213800 | 2214987 | 1187 | - | tuf | WP_064218407.1 |
| GCF_025264665.1 | NZ_CP073697.1 | 2229814 | 2231001 | 1187 | - | tuf | WP_064218407.1 |
| GCF_025264665.1 | NZ_CP073697.1 | 503388 | 504575 | 1187 | + | tuf | WP_014030034.1 |
| GCF_039993895.1 | NZ_CP121194.1 | 519402 | 520589 | 1187 | + | tuf | WP_014030034.1 |
| GCF_039993895.1 | NZ_CP121194.1 | 275372 | 276559 | 1187 | + | tuf | WP_009566505.1 |
| GCF_000178955.2 | NC_016631.1 | 291387 | 292574 | 1187 | + | tuf | WP_009566505.1 |
| GCF_000178955.2 | NC_016631.1 | 433726 | 434913 | 1187 | + | tuf | WP_009566505.1 |
| GCF_000178975.2 | NC_015064.1 | 449741 | 450928 | 1187 | + | tuf | WP_009566505.1 |
| GCF_000178975.2 | NC_015064.1 | 2552620 | 2553807 | 1187 | - | tuf | WP_215842731.1 |
| GCF_025264625.1 | NZ_CP093313.1 | 2568636 | 2569823 | 1187 | - | tuf | WP_215842720.1 |
| GCF_025264625.1 | NZ_CP093313.1 | 551722 | 552909 | 1187 | + | tuf | WP_123101140.1 |
| GCF_015277775.1 | NZ_CP063849.1 | 567778 | 568965 | 1187 | + | tuf | WP_123101140.1 |
| GCF_015277775.1 | NZ_CP063849.1 | 366596 | 367783 | 1187 | + | tuf | WP_010639711.1 |

| | | | | | | | |
|---|---|---|---|---|---|---|---|
| GCF_048629425.1 | NZ_CP184532.1 | 382615 | 383802 | 1187 | + | tuf | WP_010639711.1 |
| GCF_048629425.1 | NZ_CP184532.1 | 2248736 | 2249923 | 1187 | - | tuf | WP_418127834.1 |
| GCF_000265425.1 | NC_018014.1 | 2265122 | 2266309 | 1187 | - | tuf | WP_418127834.1 |
| GCF_000265425.1 | NC_018014.1 | 5516071 | 5517240 | 1169 | + | tuf | WP_080040436.1 |
| GCF_000179915.2 | NC_014963.1 | 9081979 | 9083169 | 1190 | - | tuf | WP_080043241.1 |
| GCF_000179915.2 | NC_014963.1 | 2965354 | 2966544 | 1190 | - | tuf | WP_014154597.1 |
| GCF_040358985.1 | NZ_CP132932.1 | 6324558 | 6325724 | 1166 | + | tuf | WP_329077392.1 |
| GCF_040358985.1 | NZ_CP132932.1 | 4053129 | 4054316 | 1187 | - | tuf | WP_308479191.1 |
| GCF_040358255.1 | NZ_CP132938.1 | 764292 | 765479 | 1187 | + | tuf | WP_308485647.1 |
| GCF_040358255.1 | NZ_CP132938.1 | 694619 | 695806 | 1187 | + | tuf | WP_308475587.1 |
| GCF_025264645.1 | NZ_CP073696.1 | 3958596 | 3959786 | 1190 | - | tuf | WP_078288431.1 |
| GCF_025264645.1 | NZ_CP073696.1 | 3838066 | 3839253 | 1187 | - | tuf | WP_316511540.1 |
| GCF_040359615.1 | NZ_CP132942.1 | 2069825 | 2071015 | 1190 | + | tuf | WP_015071256.1 |
| GCF_040359615.1 | NZ_CP132942.1 | 1794254 | 1795444 | 1190 | + | tuf | WP_028702624.1 |
| GCF_001618865.1 | NZ_CP015136.1 | 1044336 | 1045526 | 1190 | + | tuf | WP_114044179.1 |
| GCF_001618865.1 | NZ_CP015136.1 | 328542 | 329732 | 1190 | + | tuf | WP_011719141.1 |
| GCF_000023265.1 | NC_013124.1 | 4017836 | 4019026 | 1190 | - | tuf | WP_089226439.1 |
| GCF_009650215.1 | NZ_CP045851.1 | 7576850 | 7578019 | 1169 | + | tuf | WP_326578365.1 |
| GCF_009650215.1 | NZ_CP045851.1 | 5223392 | 5224582 | 1190 | + | tuf | WP_202235509.1 |
| GCF_049103755.1 | NZ_CP172265.1 | 644025 | 645215 | 1190 | + | tuf | WP_075738679.1 |
| GCF_049103755.1 | NZ_CP172265.1 | 621858 | 623048 | 1190 | + | tuf | WP_093939965.1 |
| GCF_020715685.1 | NZ_CP085840.1 | 621947 | 623137 | 1190 | + | tuf | WP_069846382.1 |
| GCF_020715685.1 | NZ_CP085840.1 | 2183031 | 2184221 | 1190 | + | tuf | WP_030446759.1 |
| GCF_028532485.1 | NZ_CP116942.1 | 5936812 | 5938002 | 1190 | - | tuf | WP_203964018.1 |
| GCF_028532485.1 | NZ_CP116942.1 | 7857155 | 7858345 | 1190 | - | tuf | WP_104477969.1 |
| GCF_000348785.1 | NC_020520.1 | 4573235 | 4574416 | 1181 | + | tuf | WP_231336329.1 |
| GCF_000348785.1 | NC_020520.1 | 5496370 | 5497560 | 1190 | + | tuf | WP_067454676.1 |
| GCF_002057455.1 | NZ_CP017717.1 | 7209494 | 7210684 | 1190 | - | tuf | WP_179837686.1 |
| GCF_002057455.1 | NZ_CP017717.1 | 8258884 | 8260062 | 1178 | + | tuf | WP_258940197.1 |
| GCF_036240075.1 | NZ_CP108452.1 | 6457513 | 6458691 | 1178 | - | tuf | WP_412515759.1 |
| GCF_036240075.1 | NZ_CP108452.1 | 7500602 | 7501792 | 1190 | + | tuf | WP_021597492.1 |

| | | | | | | | |
|---|---|---|---|---|---|---|---|
| GCF_963378095.1 | NZ_OY726397.1 | 2047816 | 2049003 | 1187 | + | tuf | WP_179900775.1 |
| GCF_963378105.1 | NZ_OY726398.1 | 2116656 | 2117843 | 1187 | - | tuf | WP_172120427.1 |
| GCF_963378075.1 | NZ_OY726394.1 | 1754792 | 1755979 | 1187 | - | tuf | WP_080461950.1 |
| GCF_002356335.1 | NZ_AP018165.1 | 1304415 | 1305602 | 1187 | - | tuf | WP_126382263.1 |
| GCF_963378085.1 | NZ_OY726395.1 | 2995379 | 2996566 | 1187 | - | tuf | WP_043558695.1 |
| GCF_001441165.1 | NZ_CP013126.1 | 735028 | 736215 | 1187 | + | tuf | WP_119837029.1 |
| GCF_005890135.1 | NZ_CP040635.1 | 556213 | 557400 | 1187 | + | tuf | WP_159524097.1 |
| GCF_003325455.1 | NZ_CP025198.1 | 3136369 | 3137556 | 1187 | - | tuf | WP_076067072.1 |
| GCF_000015025.1 | NC_008578.1 | 2062314 | 2063501 | 1187 | - | tuf | WP_070661478.1 |
| GCF_035918135.1 | NZ_CP109144.1 | 1716217 | 1717407 | 1190 | + | tuf | WP_072457385.1 |
| GCF_035918135.1 | NZ_CP109144.1 | 2774236 | 2775423 | 1187 | - | tuf | WP_136314048.1 |
| GCF_016756395.1 | NZ_AP018365.1 | 3120610 | 3121797 | 1187 | - | tuf | WP_128683484.1 |
| GCF_001941625.1 | NZ_CP016076.1 | 1706468 | 1707655 | 1187 | + | tuf | WP_067942040.1 |
| GCF_002234535.1 | NZ_CP022521.1 | 2298220 | 2299407 | 1187 | - | tuf | WP_166856090.1 |
| GCF_001747425.1 | NZ_CP014859.1 | 1269456 | 1270643 | 1187 | + | tuf | WP_026428010.1 |
| GCF_018324685.1 | NZ_AP023354.1 | 511913 | 513100 | 1187 | + | tuf | WP_194949786.1 |
| GCF_016865425.1 | NZ_AP023355.1 | 3402087 | 3403274 | 1187 | + | tuf | WP_126415184.1 |
| GCF_041897805.1 | NZ_CP154825.1 | 593226 | 594413 | 1187 | + | tuf | WP_200276696.1 |
| GCF_019175365.1 | NZ_CP059572.1 | 2223234 | 2224421 | 1187 | - | tuf | WP_127842560.1 |
| GCF_019175365.1 | NZ_CP059572.1 | 4567227 | 4568414 | 1187 | - | tuf | WP_092546112.1 |
| GCF_037081795.1 | NZ_CP146086.1 | 6889645 | 6890835 | 1190 | - | tuf | WP_092548818.1 |
| GCF_037081795.1 | NZ_CP146086.1 | 993745 | 994935 | 1190 | + | tuf | WP_023358166.1 |
| GCF_047130825.1 | NZ_CP083237.1 | 7048051 | 7049235 | 1184 | - | tuf | WP_023561080.1 |
| GCF_047130825.1 | NZ_CP083237.1 | 1622741 | 1623928 | 1187 | - | tuf | WP_189332741.1 |
| GCF_019974135.1 | NZ_AP025017.1 | 5413299 | 5414489 | 1190 | + | tuf | WP_189329191.1 |
| GCF_013184985.2 | NZ_CP063418.1 | 653091 | 654281 | 1190 | + | tuf | WP_014440721.1 |
| GCF_002076915.1 | NZ_CP020468.1 | 6194549 | 6195736 | 1187 | - | tuf | WP_014445897.1 |
| GCF_900637165.1 | NZ_LR134350.1 | 684642 | 685832 | 1190 | + | tuf | WP_284918482.1 |
| GCF_030253495.1 | NZ_CP124548.1 | 6864716 | 6865903 | 1187 | - | tuf | WP_284915390.1 |
| GCF_003606385.1 | NZ_CP032514.1 | 6796908 | 6798095 | 1187 | - | tuf | WP_317786866.1 |
| GCF_013177295.1 | NZ_CP053642.1 | 8851655 | 8852845 | 1190 | - | tuf | WP_093618762.1 |

| | | | | | | | |
|---|---|---|---|---|---|---|---|
| GCF_016127855.1 | NZ_CP066049.1 | 644265 | 645455 | 1190 | + | tuf | WP_122976224.1 |
| GCF_016127955.1 | NZ_CP066060.1 | 2963049 | 2964242 | 1193 | + | tuf | WP_092653801.1 |
| GCF_900120245.1 | NZ_LT635457.1 | 4560684 | 4561874 | 1190 | - | tuf | WP_043570386.1 |
| GCF_004798665.1 | NZ_CP039292.1 | 7747993 | 7749183 | 1190 | - | tuf | WP_015805292.1 |
| GCF_004102025.1 | NZ_CP025228.1 | 7591650 | 7592840 | 1190 | - | tuf | WP_096497128.1 |
| GCF_001553565.1 | NZ_CP014228.1 | 900312 | 901499 | 1187 | - | tuf | WP_026428288.1 |
| GCF_014595995.2 | NZ_CP063989.1 | 2085411 | 2086598 | 1187 | - | tuf | WP_087070535.1 |
| GCF_900637295.1 | NZ_LR134363.1 | 3074299 | 3075489 | 1190 | - | tuf | WP_108579730.1 |
| GCF_015711475.1 | NZ_CP065027.1 | 1014575 | 1015765 | 1190 | - | tuf | WP_078699133.1 |
| GCF_900637975.1 | NZ_LR134477.1 | 462650 | 463840 | 1190 | + | tuf | WP_187411723.1 |
| GCF_016598775.1 | NZ_CP066802.1 | 2878455 | 2879645 | 1190 | - | tuf | WP_067859832.1 |
| GCF_004010955.1 | NZ_CP025227.1 | 2053034 | 2054224 | 1190 | - | tuf | WP_007078639.1 |
| GCF_900104725.1 | NZ_LT629758.1 | 467852 | 469042 | 1190 | + | tuf | WP_187411723.1 |
| GCF_900104725.1 | NZ_LT629758.1 | 3060475 | 3061665 | 1190 | - | tuf | WP_153654355.1 |
| GCF_000494755.1 | NC_022657.1 | 558388 | 559578 | 1190 | + | tuf | WP_306233001.1 |
| GCF_000494755.1 | NC_022657.1 | 1934976 | 1936166 | 1190 | + | tuf | WP_092666756.1 |
| GCF_018324205.1 | NZ_AP023356.1 | 1066903 | 1068093 | 1190 | - | tuf | WP_092503052.1 |
| GCF_018324205.1 | NZ_AP023356.1 | 997061 | 998251 | 1190 | - | tuf | WP_210900442.1 |
| GCF_000284295.1 | NC_017093.1 | 3422683 | 3423873 | 1190 | - | tuf | WP_067878933.1 |
| GCF_000284295.1 | NC_017093.1 | 2434116 | 2435306 | 1190 | - | tuf | WP_108596010.1 |
| GCF_030252345.1 | NZ_CP126980.1 | 191201 | 192391 | 1190 | + | tuf | WP_092668510.1 |
| GCF_030252345.1 | NZ_CP126980.1 | 2755711 | 2756901 | 1190 | + | tuf | WP_149161161.1 |
| GCF_033097365.1 | NZ_AP028461.1 | 1138216 | 1139406 | 1190 | + | tuf | WP_139416362.1 |
| GCF_033097365.1 | NZ_AP028461.1 | 2159866 | 2161056 | 1190 | - | tuf | WP_243553565.1 |
| GCF_003711105.1 | NZ_CP023865.1 | 862164 | 863354 | 1190 | - | tuf | WP_286310356.1 |
| GCF_900104745.1 | NZ_LT629732.1 | 2527411 | 2528601 | 1190 | - | tuf | WP_234659723.1 |
| GCF_002263515.1 | NZ_CP022752.1 | 1525285 | 1526472 | 1187 | + | tuf | WP_129190136.1 |
| GCF_000023245.1 | NC_013093.1 | 1931898 | 1933088 | 1190 | + | tuf | WP_231430092.1 |
| GCF_002354875.1 | NZ_CP023445.1 | 3256098 | 3257279 | 1181 | - | tuf | WP_030429537.1 |
| GCF_000724605.1 | NZ_CP008802.1 | 5639177 | 5640367 | 1190 | - | tuf | WP_030427391.1 |
| GCF_002860685.3 | NZ_CP136960.1 | 4338258 | 4339448 | 1190 | - | tuf | WP_154078106.1 |

| | | | | | | | |
|---|---|---|---|---|---|---|---|
| GCF_003065605.1 | NZ_CP026952.1 | 754421 | 755608 | 1187 | + | tuf | WP_425307867.1 |
| GCF_900167475.1 | NZ_LT796768.1 | 629053 | 630243 | 1190 | + | tuf | WP_144643240.1 |
| GCF_024508295.1 | NZ_CP101990.1 | 6817793 | 6818983 | 1190 | - | tuf | WP_216210495.1 |
| GCF_001509405.1 | NZ_CP011502.1 | 7223282 | 7224472 | 1190 | + | tuf | WP_113695881.1 |
| GCF_000160775.2 | NZ_CM001024.1 | 643018 | 644208 | 1190 | + | tuf | WP_145938391.1 |
| GCF_014334735.1 | NZ_CP060587.1 | 862908 | 864098 | 1190 | - | tuf | WP_086675505.1 |
| GCF_009649075.1 | NZ_CP045737.1 | 6283025 | 6284215 | 1190 | - | tuf | WP_144586622.1 |
| GCF_030758955.1 | NZ_CP132360.1 | 7762133 | 7763323 | 1190 | - | tuf | WP_007031035.1 |
| GCF_900104705.1 | NZ_LT629734.1 | 678585 | 679775 | 1190 | + | tuf | WP_016331088.1 |
| GCF_900099705.1 | NZ_LT629695.1 | 743028 | 744218 | 1190 | + | tuf | WP_013222603.1 |
| GCF_018024495.1 | NZ_CP071696.1 | 4622263 | 4623447 | 1184 | + | tuf | WP_013226063.1 |
| GCF_001660485.1 | NZ_CP013979.1 | 661352 | 662542 | 1190 | + | tuf | WP_017986604.1 |
| GCF_003070885.1 | NZ_CP028913.1 | 1679932 | 1681122 | 1190 | - | tuf | WP_286000132.1 |
| GCF_900104685.1 | NZ_LT629755.1 | 7306176 | 7307360 | 1184 | + | tuf | WP_285995740.1 |
| GCF_008365295.1 | NZ_CP043505.1 | 1912532 | 1913722 | 1190 | - | tuf | WP_285456200.1 |
| GCF_006337065.1 | NZ_CP040872.1 | 8892997 | 8894181 | 1184 | - | tuf | WP_285452569.1 |
| GCF_022811705.1 | NZ_CP094528.1 | 8674083 | 8675273 | 1190 | - | tuf | WP_007031035.1 |
| GCF_030296695.1 | NZ_AP027737.1 | 8813728 | 8814918 | 1190 | + | tuf | WP_326568098.1 |
| GCF_021442325.1 | NZ_CP087879.1 | 4165871 | 4167061 | 1190 | - | tuf | WP_016331088.1 |
| GCF_004135405.1 | NZ_CP035491.1 | 1123837 | 1125027 | 1190 | - | tuf | WP_179495069.1 |
| GCF_022811605.1 | NZ_CP094533.1 | 137713 | 138900 | 1187 | + | tuf | WP_173493068.1 |
| GCF_900103775.1 | NZ_LT629701.1 | 2439500 | 2440696 | 1196 | - | tuf | WP_014847268.1 |
| GCF_900103775.1 | NZ_LT629701.1 | 1013236 | 1014432 | 1196 | + | tuf | WP_212325791.1 |
| GCF_009664835.1 | NZ_CP045929.1 | 1544219 | 1545406 | 1187 | - | tuf | WP_168918237.1 |
| GCF_039565995.1 | NZ_CP154795.1 | 470677 | 471864 | 1187 | + | tuf | WP_278013181.1 |
| GCF_021391495.1 | NZ_CP090063.1 | 692433 | 693620 | 1187 | + | tuf | WP_013170583.1 |
| GCF_018885265.1 | NZ_CP076538.1 | 975596 | 976783 | 1187 | + | tuf | WP_091280250.1 |
| GCF_003312875.1 | NZ_CP015163.1 | 1549473 | 1550660 | 1187 | - | tuf | WP_204423893.1 |
| GCF_046603725.1 | NZ_CP174150.1 | 1516432 | 1517619 | 1187 | - | tuf | WP_252672969.1 |
| GCF_030285625.1 | NZ_CP127294.1 | 559616 | 560803 | 1187 | + | tuf | WP_278059344.1 |
| GCF_026802195.1 | NZ_CP113836.1 | 172704 | 173894 | 1190 | - | tuf | WP_289032768.1 |

| | | | | | | | |
|---|---|---|---|---|---|---|---|
| GCF_000732925.1 | NZ_CP008953.1 | 2821113 | 2822303 | 1190 | - | tuf | WP_241452866.1 |
| GCF_000400635.2 | NC_021252.1 | 2699079 | 2700266 | 1187 | - | tuf | WP_274245070.1 |
| GCF_000220945.1 | NC_017186.1 | 3013493 | 3014680 | 1187 | - | tuf | WP_044576415.1 |
| GCF_000220945.1 | NC_017186.1 | 2962205 | 2963392 | 1187 | - | tuf | WP_043440601.1 |
| GCF_000739085.1 | NZ_CP009110.1 | 2787518 | 2788705 | 1187 | - | tuf | WP_104052739.1 |
| GCF_030285665.1 | NZ_CP127295.1 | 331378 | 332565 | 1187 | - | tuf | WP_074701282.1 |
| GCF_030285665.1 | NZ_CP127295.1 | 4278413 | 4279600 | 1187 | - | tuf | WP_113719168.1 |
| GCF_030264295.1 | NZ_CP127173.1 | 3003278 | 3004465 | 1187 | - | tuf | WP_227892319.1 |
| GCF_030264295.1 | NZ_CP127173.1 | 2878147 | 2879334 | 1187 | - | tuf | WP_210228057.1 |
| GCF_000943515.2 | NZ_CP016174.1 | 2860523 | 2861710 | 1187 | - | tuf | WP_227918747.1 |
| GCF_035918295.1 | NZ_CP142149.1 | 422246 | 423433 | 1187 | - | tuf | WP_009358310.1 |
| GCF_024628825.1 | NZ_CP102415.1 | 2978098 | 2979285 | 1187 | - | tuf | WP_233912370.1 |
| GCF_020535405.1 | NZ_CP085036.1 | 1075986 | 1077173 | 1187 | + | tuf | WP_127511058.1 |
| GCF_013283855.1 | NZ_CP054056.1 | 3185418 | 3186605 | 1187 | - | tuf | WP_104052739.1 |
| GCF_900637725.1 | NZ_LR134406.1 | 2749967 | 2751154 | 1187 | - | tuf | WP_227903865.1 |
| GCF_037055315.1 | NZ_CP146372.1 | 2727783 | 2728970 | 1187 | - | tuf | WP_230021153.1 |
| GCF_012563545.1 | NZ_CP050804.1 | 1264496 | 1265686 | 1190 | - | tuf | WP_096382562.1 |
| GCF_029625435.1 | NZ_CP121208.1 | 175267 | 176457 | 1190 | + | tuf | WP_110232718.1 |
| GCF_900475915.1 | NZ_LS483427.1 | 1722667 | 1723857 | 1190 | + | tuf | WP_090592200.1 |
| GCF_900105865.1 | NZ_LT629804.1 | 2535659 | 2536846 | 1187 | + | tuf | WP_110097930.1 |
| GCF_016904675.1 | NZ_CP070228.1 | 582329 | 583519 | 1190 | + | tuf | WP_142004343.1 |
| GCF_023973165.1 | NZ_CP099547.1 | 3496076 | 3497263 | 1187 | - | tuf | WP_015883629.1 |
| GCF_029632765.1 | NZ_CP121247.1 | 894909 | 896105 | 1196 | - | tuf | WP_033503735.1 |
| GCF_043162025.1 | NZ_CP171001.1 | 667935 | 669131 | 1196 | + | tuf | WP_003808607.1 |
| GCF_045782605.1 | NZ_CP124757.1 | 627007 | 628203 | 1196 | + | tuf | WP_003825777.1 |
| GCF_028736055.1 | NZ_CP117869.1 | 730186 | 731382 | 1196 | + | tuf | WP_004218878.1 |
| GCF_000755585.2 | NZ_CP013745.1 | 913676 | 914872 | 1196 | + | tuf | WP_369344810.1 |
| GCF_023100725.1 | NZ_CP095501.1 | 1245133 | 1246329 | 1196 | - | tuf | WP_015022138.1 |
| GCF_023921125.1 | NZ_CP099466.1 | 1494909 | 1496105 | 1196 | - | tuf | WP_003815489.1 |
| GCF_002849715.1 | NZ_CP018863.1 | 676880 | 678076 | 1196 | + | tuf | WP_003828654.1 |
| GCF_003268655.1 | NZ_CP029642.1 | 654136 | 655332 | 1196 | + | tuf | WP_003836467.1 |

| | | | | | | | |
|---|---|---|---|---|---|---|---|
| GCF_023022985.1 | NZ_CP095461.1 | 1397843 | 1399039 | 1196 | + | tuf | WP_024540765.1 |
| GCF_018622995.1 | NZ_CP076022.1 | 1204773 | 1205969 | 1196 | - | tuf | WP_033490410.1 |
| GCF_025244825.1 | NZ_CP104263.1 | 845776 | 846972 | 1196 | + | tuf | WP_003836844.1 |
| GCF_045833485.1 | NZ_CP174504.1 | 823353 | 824549 | 1196 | - | tuf | WP_072726493.1 |
| GCF_021398215.1 | NZ_CP071516.1 | 24239 | 25435 | 1196 | + | tuf | WP_369341627.1 |
| GCF_022637435.1 | NZ_CP093326.1 | 182904 | 184100 | 1196 | - | tuf | WP_033889609.1 |
| GCF_018866305.1 | NZ_CP076456.1 | 1944451 | 1945647 | 1196 | + | tuf | WP_072726493.1 |
| GCF_022869865.1 | NZ_CP094984.1 | 666574 | 667770 | 1196 | + | tuf | WP_007051202.1 |
| GCF_025244925.1 | NZ_CP104275.1 | 723922 | 725118 | 1196 | + | tuf | WP_003836467.1 |
| GCF_002355535.1 | NZ_AP017457.1 | 1500595 | 1501791 | 1196 | - | tuf | WP_022857769.1 |
| GCF_003194085.1 | NZ_CP023994.1 | 2126620 | 2127816 | 1196 | + | tuf | WP_033889609.1 |
| GCF_900101785.1 | NZ_LT629688.1 | 970386 | 971582 | 1196 | + | tuf | WP_033518543.1 |
| GCF_029853775.1 | NZ_CP122564.1 | 934051 | 935247 | 1196 | + | tuf | WP_024463698.1 |
| GCF_030296655.1 | NZ_AP027735.1 | 3750131 | 3751321 | 1190 | - | tuf | WP_324274421.1 |
| GCF_000023105.1 | NC_012669.1 | 3786468 | 3787658 | 1190 | - | tuf | WP_346620141.1 |
| GCF_001263395.1 | NZ_CP011786.1 | 1333636 | 1334832 | 1196 | - | tuf | WP_317642452.1 |
| GCF_000010425.1 | NC_008618.1 | 548217 | 549404 | 1187 | + | tuf | WP_071163690.1 |
| GCF_001025155.1 | NZ_AP012322.1 | 939579 | 940763 | 1184 | - | tuf | WP_089064427.1 |
| GCF_000022965.1 | NC_012815.1 | 1051466 | 1052650 | 1184 | + | tuf | WP_096798601.1 |
| GCF_041080795.1 | NZ_CP129674.1 | 3556618 | 3557802 | 1184 | + | tuf | WP_200501874.1 |
| GCF_002715865.1 | NZ_CP017696.1 | 144674 | 145858 | 1184 | - | tuf | WP_263594212.1 |
| GCF_017894325.1 | NZ_CP058603.1 | 5012 | 6196 | 1184 | + | tuf | WP_200501874.1 |
| GCF_001025175.1 | NZ_AP012324.1 | 2058123 | 2059307 | 1184 | - | tuf | WP_115413547.1 |
| GCF_001025195.1 | NZ_AP012325.1 | 1534806 | 1535990 | 1184 | + | tuf | WP_096802407.1 |
| GCF_002761235.1 | NZ_CP018044.1 | 1403446 | 1404636 | 1190 | + | tuf | WP_166970637.1 |
| GCF_000737865.1 | NZ_CP007287.1 | 1270176 | 1271366 | 1190 | + | tuf | WP_101639271.1 |
| GCF_001042595.1 | NZ_AP012326.1 | 66613 | 67803 | 1190 | + | tuf | WP_009379181.1 |
| GCF_014898155.1 | NZ_CP062938.1 | 1056491 | 1057681 | 1190 | + | tuf | WP_101621019.1 |
| GCF_041080835.1 | NZ_CP129683.1 | 2828321 | 2829511 | 1190 | - | tuf | WP_152348430.1 |
| GCF_017357045.1 | NZ_CP071591.1 | 1205758 | 1206948 | 1190 | + | tuf | WP_127364073.1 |
| GCF_014898175.1 | NZ_CP062948.1 | 1420975 | 1422165 | 1190 | + | tuf | WP_210604974.1 |

| | | | | | | | |
|---|---|---|---|---|---|---|---|
| GCF_000196555.1 | NC_015067.1 | 1170426 | 1171616 | 1190 | + | tuf | WP_167194667.1 |
| GCF_001025215.1 | NZ_AP012330.1 | 621786 | 622976 | 1190 | - | tuf | WP_092103017.1 |
| GCF_002706665.1 | NZ_CP017695.1 | 3345895 | 3347085 | 1190 | - | tuf | WP_092015663.1 |
| GCF_017378515.1 | NZ_CP071732.1 | 1349483 | 1350673 | 1190 | + | tuf | WP_265419747.1 |
| GCF_001042635.1 | NZ_AP012331.1 | 1482313 | 1483503 | 1190 | + | tuf | WP_101639271.1 |
| GCF_014898195.1 | NZ_CP062939.1 | 2390953 | 2392140 | 1187 | + | tuf | WP_286264887.1 |
| GCF_035320885.1 | NZ_CP141261.1 | 79537 | 80724 | 1187 | + | tuf | WP_371151599.1 |
| GCF_039679125.1 | NZ_CP155551.1 | 660370 | 661557 | 1187 | + | tuf | WP_219069221.1 |
| GCF_033095945.1 | NZ_AP026800.1 | 4627694 | 4628881 | 1187 | + | tuf | WP_285187245.1 |
| GCF_001856685.1 | NZ_CP017812.1 | 115109 | 116299 | 1190 | + | tuf | WP_095687710.1 |
| GCF_002216795.1 | NZ_CP022316.1 | 128587 | 129777 | 1190 | + | tuf | WP_095671983.1 |
| GCF_002407065.1 | NZ_CP023564.1 | 1097896 | 1099086 | 1190 | - | tuf | WP_095680228.1 |
| GCF_020792675.1 | NZ_CP086254.1 | 1104498 | 1105688 | 1190 | - | tuf | WP_095676573.1 |
| GCF_025725725.1 | NZ_CP107020.1 | 1135742 | 1136932 | 1190 | - | tuf | WP_095671538.1 |
| GCF_023373525.1 | NZ_CP097218.1 | 223777 | 224967 | 1190 | + | tuf | WP_095697587.1 |
| GCF_003355475.1 | NZ_CP031356.1 | 1099661 | 1100851 | 1190 | - | tuf | WP_095673928.1 |
| GCF_002407185.1 | NZ_CP023563.1 | 192542 | 193732 | 1190 | + | tuf | WP_095685399.1 |
| GCF_011617245.1 | NZ_CP050152.1 | 1083033 | 1084223 | 1190 | - | tuf | WP_095675224.1 |
| GCF_003932995.1 | NZ_CP025333.1 | 1205198 | 1206385 | 1187 | - | tuf | WP_070955008.1 |
| GCF_016026595.1 | NZ_CP065630.1 | 1076480 | 1077670 | 1190 | + | tuf | WP_012785151.1 |
| GCF_040256645.1 | NZ_CP158281.1 | 10277816 | 10279009 | 1193 | + | tuf | WP_015797402.1 |
| GCF_011617705.1 | NZ_CP050154.1 | 2941403 | 2942593 | 1190 | - | tuf | WP_227570304.1 |
| GCF_003999335.1 | NZ_CP026734.1 | 2934462 | 2935652 | 1190 | - | tuf | WP_208195836.1 |
| GCF_017948325.1 | NZ_CP072846.1 | 2915634 | 2916824 | 1190 | - | tuf | WP_056585863.1 |
| GCF_011617465.1 | NZ_CP050153.1 | 1104236 | 1105426 | 1190 | + | tuf | WP_013770163.1 |
| GCF_900105045.1 | NZ_LT629739.1 | 2958442 | 2959632 | 1190 | - | tuf | WP_013117880.1 |
| GCF_900105315.1 | NZ_LT629766.1 | 890190 | 891380 | 1190 | + | tuf | WP_013882844.1 |
| GCF_026168515.1 | NZ_CP093443.1 | 3619099 | 3620289 | 1190 | - | tuf | WP_236121553.1 |
| GCF_021117425.1 | NZ_CP088298.1 | 963131 | 964321 | 1190 | + | tuf | WP_135973047.1 |
| GCF_030297015.1 | NZ_AP028056.1 | 2809658 | 2810845 | 1187 | - | tuf | WP_169166353.1 |
| GCF_041380175.1 | NZ_CP159980.1 | 2918234 | 2919424 | 1190 | - | tuf | WP_207339057.1 |

| | | | | | | | |
|---|---|---|---|---|---|---|---|
| GCF_023515995.1 | NZ_CP097320.1 | 2989737 | 2990927 | 1190 | - | tuf | WP_227565746.1 |
| GCF_030252325.1 | NZ_CP126981.1 | 2936471 | 2937661 | 1190 | - | tuf | WP_227576136.1 |
| GCF_002288305.1 | NZ_CP016779.1 | 2411766 | 2412959 | 1193 | - | tuf | WP_021479872.1 |
| GCF_002288005.1 | NZ_CP016771.1 | 3871146 | 3872339 | 1193 | - | tuf | WP_168732021.1 |
| GCF_002287885.2 | NZ_CP016768.2 | 694018 | 695205 | 1187 | + | tuf | WP_182050250.1 |
| GCF_002287965.1 | NZ_CP016770.1 | 1081062 | 1082249 | 1187 | - | tuf | WP_345360709.1 |
| GCF_002288325.1 | NZ_CP016780.1 | 57558 | 58748 | 1190 | - | tuf | WP_012039309.1 |
| GCF_002288365.1 | NZ_CP016782.1 | 2688720 | 2689910 | 1190 | - | tuf | WP_053775260.1 |
| GCF_002288065.1 | NZ_CP016773.1 | 1935401 | 1936591 | 1190 | - | tuf | WP_012039309.1 |
| GCF_002288185.1 | NZ_CP016776.1 | 2719560 | 2720750 | 1190 | - | tuf | WP_015491193.1 |
| GCF_002288145.1 | NZ_CP016775.1 | 40777 | 41967 | 1190 | + | tuf | WP_012039309.1 |
| GCF_001854225.1 | NZ_CP015208.1 | 2877580 | 2878770 | 1190 | - | tuf | WP_043584326.1 |
| GCF_000024025.1 | NC_013131.1 | 322618 | 323808 | 1190 | + | tuf | WP_012297752.1 |
| GCF_000024025.1 | NC_013131.1 | 2763620 | 2764810 | 1190 | - | tuf | WP_191147464.1 |
| GCF_024508335.1 | NZ_CP101988.1 | 1000492 | 1001682 | 1190 | + | tuf | WP_085018728.1 |
| GCF_018623035.1 | NZ_CP076023.1 | 429014 | 430201 | 1187 | + | tuf | WP_005276884.1 |
| GCF_018388465.1 | NZ_CP074404.1 | 356269 | 357456 | 1187 | + | tuf | WP_194561138.1 |
| GCF_000212695.1 | NC_015514.1 | 484857 | 486044 | 1187 | + | tuf | WP_003848063.1 |
| GCF_000092865.1 | NC_014151.1 | 258528 | 259715 | 1187 | - | tuf | WP_038627641.1 |
| GCF_000218545.1 | NC_015671.1 | 1893783 | 1894970 | 1187 | - | tuf | WP_185770356.1 |
| GCF_021590045.1 | NZ_CP062221.1 | 284498 | 285685 | 1187 | + | tuf | WP_076598029.1 |
| GCF_004798685.1 | NZ_CP039291.1 | 2161415 | 2162602 | 1187 | - | tuf | WP_196825282.1 |
| GCF_012931845.1 | NZ_CP051884.1 | 388769 | 389959 | 1190 | + | tuf | WP_075724663.1 |
| GCF_018388445.1 | NZ_CP074405.1 | 252135 | 253325 | 1190 | + | tuf | WP_020975565.1 |
| GCF_024508275.1 | NZ_CP101989.1 | 385799 | 386989 | 1190 | + | tuf | WP_290219210.1 |
| GCF_024508315.1 | NZ_CP101987.1 | 411924 | 413111 | 1187 | + | tuf | WP_038604382.1 |
| GCF_018128265.1 | NZ_CP072387.1 | 420932 | 422119 | 1187 | + | tuf | WP_010189688.1 |
| GCF_009708005.2 | NZ_CP052757.1 | 382414 | 383601 | 1187 | + | tuf | WP_290342538.1 |
| GCF_014041995.1 | NZ_CP059492.1 | 2211155 | 2212342 | 1187 | - | tuf | WP_035113608.1 |
| GCF_047305825.1 | NZ_CP063070.1 | 439340 | 440527 | 1187 | + | tuf | WP_004566712.1 |
| GCF_021952865.1 | NZ_CP040792.1 | 274724 | 275911 | 1187 | + | tuf | WP_284825395.1 |

| | | | | | | | |
|---|---|---|---|---|---|---|---|
| GCF_011801245.1 | NZ_CP048047.1 | 525970 | 527157 | 1187 | + | tuf | WP_015650383.1 |
| GCF_021216655.1 | NZ_CP080437.1 | 352950 | 354137 | 1187 | + | tuf | WP_035106523.1 |
| GCF_000355695.1 | NC_020891.1 | 379979 | 381166 | 1187 | + | tuf | WP_146325690.1 |
| GCF_021922925.1 | NZ_CP040786.1 | 263839 | 265026 | 1187 | + | tuf | WP_018017468.1 |
| GCF_021919125.1 | NZ_CP083439.1 | 555570 | 556757 | 1187 | + | tuf | WP_006823212.1 |
| GCF_000069225.1 | NC_010407.1 | 1583610 | 1584797 | 1187 | - | tuf | WP_018340290.1 |
| GCF_014775655.1 | NZ_CP061274.1 | 2491043 | 2492233 | 1190 | - | tuf | WP_123929751.1 |
| GCF_002096055.1 | NZ_CP020715.1 | 318637 | 319824 | 1187 | + | tuf | WP_018018650.1 |
| GCF_023520795.1 | NZ_CP046605.1 | 446262 | 447449 | 1187 | + | tuf | WP_156226871.1 |
| GCF_030408375.1 | NZ_CP046530.1 | 396661 | 397848 | 1187 | + | tuf | WP_290224668.1 |
| GCF_001941425.1 | NZ_CP009244.1 | 1392251 | 1393438 | 1187 | + | tuf | WP_070422705.1 |
| GCF_016889425.1 | NZ_CP069513.1 | 549244 | 550431 | 1187 | + | tuf | WP_066564449.1 |
| GCF_014262665.1 | NZ_CP046883.1 | 2100080 | 2101267 | 1187 | - | tuf | WP_092256498.1 |
| GCF_030408415.1 | NZ_CP046976.1 | 548241 | 549428 | 1187 | + | tuf | WP_053544132.1 |
| GCF_030408395.1 | NZ_CP046980.1 | 381672 | 382859 | 1187 | + | tuf | WP_004566712.1 |
| GCF_001941445.1 | NZ_CP009245.1 | 530373 | 531560 | 1187 | + | tuf | WP_018021302.1 |
| GCF_000590555.1 | NC_022198.1 | 442983 | 444170 | 1187 | + | tuf | WP_006062917.1 |
| GCF_030408455.1 | NZ_CP046977.1 | 556746 | 557933 | 1187 | + | tuf | WP_006769810.1 |
| GCF_000732945.1 | NZ_CP008944.1 | 411117 | 412304 | 1187 | + | tuf | WP_136140510.1 |
| GCF_000022905.1 | NC_012590.1 | 388964 | 390151 | 1187 | + | tuf | WP_047239510.1 |
| GCF_030408575.1 | NZ_CP047047.1 | 458223 | 459410 | 1187 | + | tuf | WP_290278349.1 |
| GCF_030408435.1 | NZ_CP047046.1 | 68272 | 69459 | 1187 | - | tuf | WP_025403593.1 |
| GCF_041898185.1 | NZ_CP157823.1 | 367473 | 368660 | 1187 | + | tuf | WP_277105281.1 |
| GCF_030252165.1 | NZ_CP126969.1 | 454059 | 455246 | 1187 | + | tuf | WP_075729097.1 |
| GCF_000344785.1 | NC_020506.1 | 355047 | 356234 | 1187 | + | tuf | WP_085956816.1 |
| GCF_000980815.1 | NZ_CP011311.1 | 471427 | 472614 | 1187 | + | tuf | WP_075663214.1 |
| GCF_030408595.1 | NZ_CP047080.1 | 354216 | 355403 | 1187 | + | tuf | WP_027012651.1 |
| GCF_030440535.1 | NZ_CP047117.1 | 2298531 | 2299718 | 1187 | + | tuf | WP_070523233.1 |
| GCF_000550785.1 | NZ_CP004350.1 | 333003 | 334190 | 1187 | + | tuf | WP_005286783.1 |
| GCF_030440555.1 | NZ_CP047118.1 | 1968999 | 1970186 | 1187 | - | tuf | WP_123935309.1 |
| GCF_003813965.1 | NZ_CP033896.1 | 374643 | 375830 | 1187 | + | tuf | WP_290185955.1 |

| | | | | | | | |
|---|---|---|---|---|---|---|---|
| GCF_030440575.1 | NZ_CP047189.1 | 477683 | 478870 | 1187 | + | tuf | WP_005390564.1 |
| GCF_009734405.1 | NZ_CP046453.1 | 526376 | 527563 | 1187 | + | tuf | WP_011013696.1 |
| GCF_030408715.1 | NZ_CP047202.1 | 755551 | 756738 | 1187 | + | tuf | WP_038546237.1 |
| GCF_019048165.1 | NZ_CP077302.1 | 374064 | 375251 | 1187 | + | tuf | WP_095277977.1 |
| GCF_001643015.1 | NZ_CP015622.1 | 505073 | 506260 | 1187 | + | tuf | WP_290197336.1 |
| GCF_900187295.1 | NZ_LT906473.1 | 528008 | 529195 | 1187 | + | tuf | WP_015399921.1 |
| GCF_001277995.1 | NZ_CP009220.1 | 406090 | 407277 | 1187 | + | tuf | WP_048740066.1 |
| GCF_001457455.1 | NZ_LN831026.1 | 2012549 | 2013736 | 1187 | - | tuf | WP_112768812.1 |
| GCF_000767055.1 | NZ_CP006764.1 | 393256 | 394443 | 1187 | + | tuf | WP_070521778.1 |
| GCF_030408675.1 | NZ_CP047200.1 | 1129712 | 1130899 | 1187 | - | tuf | WP_182386981.1 |
| GCF_000011305.1 | NC_004369.1 | 362325 | 363512 | 1187 | + | tuf | WP_034997213.1 |
| GCF_004795735.1 | NZ_CP039247.1 | 464046 | 465233 | 1187 | + | tuf | WP_038588218.1 |
| GCF_001021025.1 | NZ_CP011541.1 | 1600278 | 1601465 | 1187 | + | tuf | WP_185175286.1 |
| GCF_030408735.1 | NZ_CP047204.1 | 411884 | 413071 | 1187 | + | tuf | WP_042406241.1 |
| GCF_020099275.1 | NZ_CP083646.1 | 2208660 | 2209847 | 1187 | - | tuf | WP_005291995.1 |
| GCF_030408755.1 | NZ_CP047209.1 | 2120412 | 2121599 | 1187 | - | tuf | WP_156193241.1 |
| GCF_001941465.1 | NZ_CP009246.1 | 2283098 | 2284285 | 1187 | - | tuf | WP_023020538.1 |
| GCF_030408775.1 | NZ_CP047210.1 | 2152512 | 2153699 | 1187 | - | tuf | WP_012732455.1 |
| GCF_001941485.1 | NZ_CP009247.1 | 327883 | 329070 | 1187 | + | tuf | WP_046438628.1 |
| GCF_030408815.1 | NZ_CP047355.1 | 465834 | 467021 | 1187 | + | tuf | WP_053411458.1 |
| GCF_019047805.1 | NZ_CP077258.1 | 357401 | 358588 | 1187 | + | tuf | WP_108403352.1 |
| GCF_024453835.1 | NZ_CP072935.1 | 463804 | 464991 | 1187 | + | tuf | WP_165008659.1 |
| GCF_003813985.1 | NZ_CP033897.1 | 346403 | 347590 | 1187 | + | tuf | WP_171194361.1 |
| GCF_030408855.1 | NZ_CP047358.1 | 432109 | 433296 | 1187 | + | tuf | WP_121910652.1 |
| GCF_030440595.1 | NZ_CP047452.1 | 378628 | 379815 | 1187 | + | tuf | WP_042620703.1 |
| GCF_000011325.1 | NC_003450.3 | 541138 | 542325 | 1187 | + | tuf | WP_020933913.1 |
| GCF_000626675.1 | NZ_CP006842.1 | 338312 | 339499 | 1187 | + | tuf | WP_022863448.1 |
| GCF_030408875.1 | NZ_CP047454.1 | 2212454 | 2213641 | 1187 | + | tuf | WP_342318995.1 |
| GCF_030408915.1 | NZ_CP047654.1 | 2308593 | 2309780 | 1187 | - | tuf | WP_039673679.1 |
| GCF_000341345.1 | NC_020302.1 | 447904 | 449094 | 1190 | + | tuf | WP_047261124.1 |
| GCF_030408795.1 | NZ_CP047211.1 | 1938350 | 1939537 | 1187 | + | tuf | WP_092151531.1 |

| | | | | | | | |
|---|---|---|---|---|---|---|---|
| GCF_028609845.1 | NZ_CP063191.1 | 1544365 | 1545552 | 1187 | - | tuf | WP_010120799.1 |
| GCF_037850265.1 | NZ_CP137211.1 | 485688 | 486875 | 1187 | + | tuf | WP_156229970.1 |
| GCF_014117265.1 | NZ_CP059833.1 | 2065444 | 2066631 | 1187 | - | tuf | WP_128890505.1 |
| GCF_028609825.1 | NZ_CP063190.1 | 2316937 | 2318124 | 1187 | - | tuf | WP_075735645.1 |
| GCF_900187215.1 | NZ_LT906467.1 | 474689 | 475876 | 1187 | + | tuf | WP_187975034.1 |
| GCF_014217255.1 | NZ_CP059404.1 | 788036 | 789223 | 1187 | - | tuf | WP_018121316.1 |
| GCF_028609865.1 | NZ_CP063194.1 | 413758 | 414945 | 1187 | + | tuf | WP_070363113.1 |
| GCF_028609885.1 | NZ_CP063195.1 | 2209009 | 2210196 | 1187 | - | tuf | WP_012732455.1 |
| GCF_009734385.1 | NZ_CP046452.1 | 2048726 | 2049913 | 1187 | - | tuf | WP_123960927.1 |
| GCF_016599755.1 | NZ_CP067012.1 | 360153 | 361340 | 1187 | + | tuf | WP_013241194.1 |
| GCF_000023145.1 | NC_012704.1 | 302291 | 303478 | 1187 | + | tuf | WP_165003463.1 |
| GCF_000980835.1 | NZ_CP011312.1 | 411401 | 412588 | 1187 | + | tuf | WP_013241194.1 |
| GCF_001274895.1 | NZ_CP006841.1 | 1284732 | 1285919 | 1187 | + | tuf | WP_048379947.1 |
| GCF_003070865.1 | NZ_CP026948.1 | 2271085 | 2272272 | 1187 | + | tuf | WP_013889249.1 |
| GCF_011038655.2 | NZ_CP064954.1 | 1675748 | 1676935 | 1187 | - | tuf | WP_052205408.1 |
| GCF_014490555.1 | NZ_CP061032.1 | 466756 | 467943 | 1187 | + | tuf | WP_004566712.1 |
| GCF_016889465.1 | NZ_CP069516.1 | 498145 | 499332 | 1187 | - | tuf | WP_136653350.1 |
| GCF_000835165.1 | NZ_CP007790.1 | 441586 | 442773 | 1187 | + | tuf | WP_013241194.1 |
| GCF_000442645.1 | NC_021915.1 | 1137677 | 1138864 | 1187 | - | tuf | WP_061920476.1 |
| GCF_028609805.1 | NZ_CP063189.1 | 460758 | 461945 | 1187 | + | tuf | WP_042529275.1 |
| GCF_038452275.1 | NZ_CP141643.1 | 321864 | 323051 | 1187 | + | tuf | WP_075691167.1 |
| GCF_900478045.1 | NZ_LS483460.1 | 482393 | 483580 | 1187 | + | tuf | WP_066792420.1 |
| GCF_001020985.1 | NZ_CP011542.1 | 1034407 | 1035594 | 1187 | - | tuf | WP_005527719.1 |
| GCF_900103625.1 | NZ_LT629700.1 | 1172837 | 1174024 | 1187 | + | tuf | WP_284875869.1 |
| GCF_007970465.1 | NZ_CP042429.1 | 692644 | 693831 | 1187 | + | tuf | WP_003854225.1 |
| GCF_009734425.1 | NZ_CP046455.1 | 2260537 | 2261724 | 1187 | - | tuf | WP_020441970.1 |
| GCF_004114895.1 | NZ_CP035299.1 | 404704 | 405894 | 1190 | + | tuf | WP_047252302.1 |
| GCF_001941565.1 | NZ_CP009249.1 | 1492375 | 1493562 | 1187 | + | tuf | WP_179387370.1 |
| GCF_014522205.1 | NZ_CP046884.1 | 387757 | 388944 | 1187 | + | tuf | WP_227280887.1 |
| GCF_016728665.1 | NZ_CP068161.1 | 432282 | 433469 | 1187 | + | tuf | WP_013241194.1 |
| GCF_024453815.1 | NZ_CP072934.1 | 2073257 | 2074444 | 1187 | + | tuf | WP_012359571.1 |

| | | | | | | | |
|---|---|---|---|---|---|---|---|
| GCF_030486685.2 | NZ_CP137757.1 | 293424 | 294611 | 1187 + | tuf | | WP_038609604.1 |
| GCF_003814005.1 | NZ_CP033898.1 | 2001325 | 2002512 | 1187 - | tuf | | WP_151903405.1 |
| GCF_002155265.1 | NZ_CP021251.1 | 330662 | 331849 | 1187 + | tuf | | WP_047258944.1 |
| GCF_011038645.2 | NZ_CP064955.1 | 452340 | 453527 | 1187 + | tuf | | WP_025251885.1 |
| GCF_000767685.1 | NZ_CP009622.1 | 364739 | 365926 | 1187 + | tuf | | WP_108430829.1 |
| GCF_900478035.1 | NZ_LS483464.1 | 1824229 | 1825416 | 1187 - | tuf | | WP_151842186.1 |
| GCF_000177535.2 | NC_015673.1 | 3444953 | 3446143 | 1190 + | tuf | | WP_066597682.1 |
| GCF_001263755.1 | NZ_CP012342.1 | 3119423 | 3120607 | 1184 + | tuf | | WP_216259451.1 |
| GCF_902702935.1 | NZ_LR738855.1 | 3533521 | 3534705 | 1184 - | tuf | | WP_027466734.1 |
| GCF_007641235.1 | NZ_CP038157.1 | 3352231 | 3353415 | 1184 - | tuf | | WP_017885537.1 |
| GCF_002162115.2 | NZ_CP021417.2 | 1976996 | 1978186 | 1190 - | tuf | | WP_002514878.1 |
| GCF_001586235.1 | NZ_CP014635.1 | 355634 | 356824 | 1190 + | tuf | | WP_004809063.1 |
| GCF_000833575.1 | NZ_CP010827.1 | 1870195 | 1871385 | 1190 - | tuf | | WP_271417744.1 |
| GCF_001941505.1 | NZ_CP009248.1 | 1669414 | 1670604 | 1190 - | tuf | | WP_065860439.1 |
| GCF_001941345.1 | NZ_CP009251.1 | 9702828 | 9704012 | 1184 - | tuf | | WP_033362545.1 |
| GCF_016728105.1 | NZ_CP068158.1 | 10790482 | 10791672 | 1190 - | tuf | | WP_033357668.1 |
| GCF_030252185.1 | NZ_CP126970.1 | 8558118 | 8559302 | 1184 - | tuf | | WP_259858886.1 |
| GCF_002355155.1 | NZ_AP017369.1 | 9630066 | 9631256 | 1190 - | tuf | | WP_259859814.1 |
| GCF_000418365.1 | NC_021663.1 | 6389094 | 6390278 | 1184 - | tuf | | WP_260724631.1 |
| GCF_001021045.1 | NZ_CP011545.1 | 7332033 | 7333223 | 1190 - | tuf | | WP_260725397.1 |
| GCF_016728365.1 | NZ_CP068156.1 | 9939158 | 9940342 | 1184 - | tuf | | WP_223097092.1 |
| GCF_020616335.1 | NZ_CP085051.1 | 11085251 | 11086441 | 1190 - | tuf | | WP_223099115.1 |
| GCF_000215645.1 | NC_017317.1 | 2479013 | 2480200 | 1187 - | tuf | | WP_313497459.1 |
| GCF_900187235.1 | NZ_LT906481.1 | 1261489 | 1262673 | 1184 + | tuf | | WP_016664017.1 |
| GCF_000747315.1 | NZ_CP009215.1 | 621125 | 622309 | 1184 + | tuf | | WP_065247383.1 |
| GCF_009026825.1 | NZ_CP045032.1 | 801239 | 802432 | 1193 + | tuf | | WP_006946303.1 |
| GCF_001021065.1 | NZ_CP011546.1 | 2307686 | 2308876 | 1190 - | tuf | | WP_028326551.1 |
| GCF_000550805.1 | NZ_CP004353.1 | 2482865 | 2484049 | 1184 + | tuf | | WP_058235316.1 |
| GCF_003065405.1 | NZ_CP026947.1 | 2544200 | 2545387 | 1187 - | tuf | | WP_007628137.1 |
| GCF_014490575.1 | NZ_CP061033.1 | 2381188 | 2382375 | 1187 - | tuf | | WP_120284055.1 |
| GCF_003611035.1 | NZ_CP030033.1 | 2537469 | 2538656 | 1187 - | tuf | | WP_108847901.1 |

| | | | | | | | |
|---|---|---|---|---|---|---|---|
| GCF_018885305.1 | NZ_CP076544.1 | 2731345 | 2732532 | 1187 | - | tuf | WP_107746266.1 |
| GCF_024925425.1 | NZ_CP103853.1 | 2602148 | 2603335 | 1187 | - | tuf | WP_067475790.1 |
| GCF_047226435.1 | NZ_CP178434.1 | 2945169 | 2946362 | 1193 | + | tuf | WP_159546452.1 |
| GCF_006739385.1 | NZ_AP019723.1 | 630875 | 632065 | 1190 | + | tuf | WP_182173453.1 |
| GCF_000367205.1 | NC_021064.1 | 2550056 | 2551246 | 1190 | + | tuf | WP_126042105.1 |
| GCF_028021195.1 | NZ_CP115668.1 | 1158163 | 1159353 | 1190 | + | tuf | WP_009740529.1 |
| GCF_030295285.1 | NZ_AP026710.1 | 675819 | 677009 | 1190 | + | tuf | WP_011435038.1 |
| GCF_025264705.1 | NZ_CP073767.1 | 3746198 | 3747388 | 1190 | - | tuf | WP_091414456.1 |
| GCF_025264705.1 | NZ_CP073767.1 | 1212044 | 1213234 | 1190 | + | tuf | WP_286345939.1 |
| GCF_025137375.1 | NZ_CP073720.1 | 980128 | 981324 | 1196 | - | tuf | WP_004118785.1 |
| GCF_025137375.1 | NZ_CP073720.1 | 445805 | 447001 | 1196 | + | tuf | WP_004112314.1 |
| GCF_025264685.1 | NZ_CP073721.1 | 4726239 | 4727429 | 1190 | - | tuf | WP_012950484.1 |
| GCF_025264685.1 | NZ_CP073721.1 | 2860324 | 2861514 | 1190 | - | tuf | WP_122824928.1 |
| GCF_019904175.1 | NZ_CP061913.1 | 3021552 | 3022742 | 1190 | - | tuf | WP_139071847.1 |
| GCF_019904175.1 | NZ_CP061913.1 | 3494086 | 3495276 | 1190 | - | tuf | WP_139930142.1 |
| GCF_032102965.1 | NZ_CP134880.1 | 1247623 | 1248813 | 1190 | - | tuf | WP_168582116.1 |
| GCF_902381725.1 | NZ_LR698963.1 | 1674971 | 1676158 | 1187 | - | tuf | WP_016444141.1 |
| GCF_001678905.1 | NZ_CP012117.1 | 2653668 | 2654855 | 1187 | - | tuf | WP_013349691.1 |
| GCF_000725405.1 | NZ_CP008889.1 | 2343532 | 2344719 | 1187 | - | tuf | WP_138177181.1 |
| GCF_900187045.1 | NZ_LT906453.1 | 2480854 | 2482041 | 1187 | - | tuf | WP_345470202.1 |
| GCF_900070355.1 | NZ_LN849456.1 | 2475672 | 2476859 | 1187 | - | tuf | WP_309406197.1 |
| GCF_036287335.1 | NZ_CP143053.1 | 2631086 | 2632273 | 1187 | - | tuf | WP_060702225.1 |
| GCF_045689975.1 | NZ_CP143297.1 | 639048 | 640235 | 1187 | - | tuf | WP_022874140.1 |
| GCF_003096075.1 | NZ_CP015449.1 | 2582178 | 2583365 | 1187 | - | tuf | WP_204675222.1 |
| GCF_003096095.1 | NZ_CP015453.1 | 1946902 | 1948089 | 1187 | + | tuf | WP_005199938.1 |
| GCF_001659785.1 | NZ_CP015961.1 | 4171568 | 4172755 | 1187 | - | tuf | WP_040516240.1 |
| GCF_009834025.1 | NZ_CP047156.1 | 4838687 | 4839874 | 1187 | - | tuf | WP_006434719.1 |
| GCF_014069875.1 | NZ_CP059676.1 | 4088230 | 4089417 | 1187 | - | tuf | WP_012835485.1 |
| GCF_003952265.1 | NZ_CP034438.1 | 4061992 | 4063179 | 1187 | - | tuf | WP_004020593.1 |
| GCF_000058485.1 | NC_008278.1 | 2600025 | 2601212 | 1187 | + | tuf | WP_066164529.1 |
| GCF_000013345.1 | NC_007777.1 | 5285356 | 5286543 | 1187 | + | tuf | WP_124710518.1 |

| | | | | | | | |
|---|---|---|---|---|---|---|---|
| GCF_900105065.1 | NZ_LT629749.1 | 3616533 | 3617720 | 1187 | - | tuf | WP_105943287.1 |
| GCF_030296615.1 | NZ_AP027732.1 | 4051851 | 4053038 | 1187 | - | tuf | WP_188328762.1 |
| GCF_003293675.1 | NZ_CP029984.1 | 772155 | 773342 | 1187 | - | tuf | WP_160900584.1 |
| GCF_001042655.1 | NZ_AP012332.1 | 3421744 | 3422931 | 1187 | + | tuf | WP_007236639.1 |
| GCF_000025345.1 | NC_013757.1 | 3449294 | 3450481 | 1187 | - | tuf | WP_062393821.1 |
| GCF_003710105.1 | NZ_CP033325.1 | 4183664 | 4184851 | 1187 | - | tuf | WP_006367846.1 |
| GCF_006349365.1 | NZ_CP040899.1 | 4021901 | 4023088 | 1187 | - | tuf | WP_213244645.1 |
| GCF_006352065.1 | NZ_CP040915.1 | 74924 | 76111 | 1187 | - | tuf | WP_005199938.1 |
| GCF_012277275.1 | NZ_CP051155.1 | 473757 | 474944 | 1187 | + | tuf | WP_006894480.1 |
| GCF_002871995.2 | NZ_CP126966.1 | 4431852 | 4433039 | 1187 | - | tuf | WP_004020593.1 |
| GCF_000197735.1 | NC_014550.1 | 2608814 | 2610001 | 1187 | - | tuf | WP_132992108.1 |
| GCF_006094275.1 | NZ_CP034412.1 | 1040767 | 1041957 | 1190 | - | tuf | WP_120788524.1 |
| GCF_039602135.1 | NZ_CP125942.1 | 3172988 | 3174175 | 1187 | + | tuf | WP_026937631.1 |
| GCF_048457885.1 | NZ_CP174200.1 | 1628051 | 1629235 | 1184 | - | tuf | WP_317699932.1 |
| GCF_001302565.1 | NZ_CP012750.1 | 462967 | 464157 | 1190 | + | tuf | WP_013805260.1 |
| GCF_012221945.1 | NZ_CP032549.1 | 2106268 | 2107458 | 1190 | - | tuf | WP_146320429.1 |
| GCF_016894325.1 | NZ_CP069806.1 | 3073014 | 3074207 | 1193 | - | tuf | WP_013493572.1 |
| GCF_004011905.1 | NZ_CP027114.1 | 1731771 | 1732964 | 1193 | - | tuf | WP_068202533.1 |
| GCF_044358485.1 | NZ_CP045810.1 | 712777 | 713970 | 1193 | + | tuf | WP_013837836.1 |
| GCF_022059885.1 | NZ_CP091855.1 | 564237 | 565427 | 1190 | + | tuf | WP_338750495.1 |
| GCF_000024785.1 | NC_013441.1 | 1763542 | 1764732 | 1190 | - | tuf | WP_123092236.1 |
| GCF_029639525.1 | NZ_CP121270.1 | 1138499 | 1139689 | 1190 | + | tuf | WP_068326060.1 |
| GCF_038024885.1 | NZ_CP136137.1 | 1842411 | 1843601 | 1190 | - | tuf | WP_269445413.1 |
| GCF_003855095.1 | NZ_CP033972.1 | 3713176 | 3714366 | 1190 | - | tuf | WP_249771090.1 |
| GCF_002993285.1 | NZ_CP027433.1 | 7694328 | 7695521 | 1193 | - | tuf | WP_046772615.1 |
| GCF_014041935.1 | NZ_CP059491.1 | 637864 | 639051 | 1187 | + | tuf | WP_015770878.1 |
| GCF_024734075.1 | NZ_CP102850.1 | 2693369 | 2694547 | 1178 | + | tuf | WP_054289559.1 |
| GCF_020735545.1 | NZ_CP085974.1 | 11108794 | 11109984 | 1190 | - | tuf | WP_054295770.1 |
| GCF_001305675.1 | NZ_CP011853.1 | 1589261 | 1590451 | 1190 | - | tuf | WP_012084984.1 |
| GCF_000247715.1 | NC_016906.1 | 3311194 | 3312384 | 1190 | - | tuf | WP_354641044.1 |
| GCF_025273675.1 | NZ_CP045809.1 | 5099029 | 5100219 | 1190 | + | tuf | WP_030280743.1 |

| | | | | | | | |
|---|---|---|---|---|---|---|---|
| GCF_014070455.1 | NZ_CP059694.1 | 3206738 | 3207928 | 1190 | - | tuf | WP_282205979.1 |
| GCF_029632925.1 | NZ_CP121248.1 | 4979560 | 4980750 | 1190 | + | tuf | WP_030457430.1 |
| GCF_001698225.1 | NZ_CP016594.1 | 3880764 | 3881954 | 1190 | - | tuf | WP_190210874.1 |
| GCF_007559005.1 | NZ_CP041763.1 | 4251220 | 4252410 | 1190 | - | tuf | WP_266269924.1 |
| GCF_003627055.1 | NZ_CP032624.1 | 8310673 | 8311851 | 1178 | - | tuf | WP_266273035.1 |
| GCF_003010915.2 | NZ_CP028426.1 | 3575660 | 3576850 | 1190 | - | tuf | WP_014136300.1 |
| GCF_033106145.1 | NZ_CP136959.1 | 3138445 | 3139632 | 1187 | - | tuf | WP_058859366.1 |
| GCF_000214175.1 | NC_015564.1 | 2035251 | 2036438 | 1187 | + | tuf | WP_019309651.1 |
| GCF_007859675.1 | NZ_CP042305.1 | 1243472 | 1244659 | 1187 | - | tuf | WP_062734957.1 |
| GCF_000184685.1 | NC_014830.1 | 6395308 | 6396492 | 1184 | + | tuf | WP_012923507.1 |
| GCF_001636295.1 | NZ_CP014209.1 | 6567970 | 6569160 | 1190 | - | tuf | WP_012923689.1 |
| GCF_000215105.1 | NC_015588.1 | 9219868 | 9221058 | 1190 | - | tuf | WP_025361384.1 |
| GCF_037198315.1 | NZ_CP144913.1 | 6235933 | 6237123 | 1190 | + | tuf | WP_116172401.1 |
| GCF_003721155.3 | NZ_CP044548.2 | 6722888 | 6724078 | 1190 | - | tuf | WP_273936043.1 |
| GCF_039636765.1 | NZ_CP144167.1 | 1721544 | 1722734 | 1190 | - | tuf | WP_015780021.1 |
| GCF_027247405.1 | NZ_CP097463.1 | 637225 | 638412 | 1187 | - | tuf | WP_053962667.1 |
| GCF_023516435.1 | NZ_CP097332.1 | 384523 | 385713 | 1190 | + | tuf | WP_021754069.1 |
| GCF_900105925.1 | NZ_LT629791.1 | 3492170 | 3493360 | 1190 | + | tuf | WP_065915969.1 |
| GCF_900475835.1 | NZ_LS483423.1 | 4057704 | 4058882 | 1178 | - | tuf | WP_065916409.1 |
| GCF_001302585.1 | NZ_CP012752.1 | 820103 | 821293 | 1190 | + | tuf | WP_244693238.1 |
| GCF_001302585.1 | NZ_CP012752.1 | 1012190 | 1013380 | 1190 | + | tuf | WP_166329437.1 |
| GCF_000017305.1 | NC_009664.2 | 2271842 | 2273032 | 1190 | + | tuf | WP_187555198.1 |
| GCF_040571325.1 | NZ_CP159872.1 | 1324534 | 1325724 | 1190 | + | tuf | WP_166322728.1 |
| GCF_027627435.1 | NZ_CP115450.1 | 861723 | 862913 | 1190 | + | tuf | WP_017885332.1 |
| GCF_948472415.1 | NZ_OX419519.1 | 1150967 | 1152157 | 1190 | - | tuf | WP_244687812.1 |
| GCF_044999035.1 | NZ_CP173360.1 | 1045850 | 1047040 | 1190 | - | tuf | WP_223044600.1 |
| GCF_036243155.1 | NZ_CP108460.1 | 890172 | 891362 | 1190 | - | tuf | WP_130109263.1 |
| GCF_036010405.1 | NZ_CP108561.1 | 3250514 | 3251704 | 1190 | + | tuf | WP_166292549.1 |
| GCF_036010405.1 | NZ_CP108561.1 | 952243 | 953436 | 1193 | + | tuf | WP_036953300.1 |
| GCF_000269985.1 | NC_016109.1 | 4484078 | 4485268 | 1190 | - | tuf | WP_052594854.1 |
| GCF_001482365.1 | NZ_CP013254.1 | 975731 | 976921 | 1190 | + | tuf | WP_161885392.1 |

| | | | | | | | |
|---|---|---|---|---|---|---|---|
| GCF_035990065.1 | NZ_CP108528.1 | 731360 | 732547 | 1187 | + | tuf | WP_194421673.1 |
| GCF_001580365.1 | NZ_CP014480.1 | 2362720 | 2363910 | 1190 | + | tuf | WP_124291715.1 |
| GCF_000024345.1 | NC_013729.1 | 1995282 | 1996472 | 1190 | + | tuf | WP_165136180.1 |
| GCF_000024345.1 | NC_013729.1 | 2973997 | 2975187 | 1190 | - | tuf | WP_023954029.1 |
| GCF_000525635.1 | NZ_CP007155.1 | 605991 | 607181 | 1190 | + | tuf | WP_127818193.1 |
| GCF_043575855.1 | NZ_CP144375.1 | 723926 | 725113 | 1187 | + | tuf | WP_047524110.1 |
| GCF_028622115.1 | NZ_CP097263.1 | 2950664 | 2951854 | 1190 | - | tuf | WP_076691663.1 |
| GCF_016127495.1 | NZ_CP066015.1 | 2923362 | 2924552 | 1190 | - | tuf | WP_045249929.1 |
| GCF_001293125.1 | NZ_CP009312.1 | 2867255 | 2868445 | 1190 | - | tuf | WP_317138878.1 |
| GCF_000470775.1 | NC_022438.1 | 453921 | 455111 | 1190 | + | tuf | WP_150970983.1 |
| GCF_001701025.1 | NZ_CP016793.1 | 1208539 | 1209729 | 1190 | + | tuf | WP_053546381.1 |
| GCF_001701025.1 | NZ_CP016793.1 | 3049083 | 3050273 | 1190 | - | tuf | WP_127818193.1 |
| GCF_022919155.1 | NZ_CP095045.1 | 984709 | 985899 | 1190 | - | tuf | WP_165139820.1 |
| GCF_011382985.1 | NZ_CP049933.1 | 2797153 | 2798340 | 1187 | - | tuf | WP_045253854.1 |
| GCF_014396385.1 | NZ_CP060716.1 | 3280101 | 3281288 | 1187 | + | tuf | WP_042538710.1 |
| GCF_011382965.1 | NZ_CP049934.1 | 567609 | 568796 | 1187 | - | tuf | WP_042538710.1 |
| GCF_004028235.1 | NZ_CP035037.1 | 691231 | 692421 | 1190 | - | tuf | WP_060960249.1 |
| GCF_022919175.1 | NZ_CP095043.1 | 531001 | 532188 | 1187 | + | tuf | WP_275278763.1 |
| GCF_019884725.1 | NZ_CP082322.1 | 468850 | 470040 | 1190 | + | tuf | WP_127094611.1 |
| GCF_004208635.1 | NZ_CP035806.1 | 2377222 | 2378412 | 1190 | - | tuf | WP_330170441.1 |
| GCF_011299575.1 | NZ_CP049863.1 | 535342 | 536532 | 1190 | + | tuf | WP_135063311.1 |
| GCF_009429125.1 | NZ_CP045529.1 | 2886651 | 2887841 | 1190 | - | tuf | WP_071044915.1 |
| GCF_001190945.1 | NZ_CP011112.1 | 2980758 | 2981945 | 1187 | - | tuf | WP_017829210.1 |
| GCF_009930795.1 | NZ_CP017146.1 | 2142323 | 2143513 | 1190 | - | tuf | WP_259606612.1 |
| GCF_015277895.1 | NZ_CP063815.1 | 2552764 | 2553951 | 1187 | - | tuf | WP_036309808.1 |
| GCF_030347395.1 | NZ_CP128170.1 | 1689273 | 1690463 | 1190 | - | tuf | WP_156242094.1 |
| GCF_011046975.1 | NZ_CP049253.1 | 1325553 | 1326743 | 1190 | - | tuf | WP_025102410.1 |
| GCF_030369635.1 | NZ_CP128474.1 | 2896801 | 2897991 | 1190 | - | tuf | WP_023954029.1 |
| GCF_015277815.1 | NZ_CP063813.1 | 2112903 | 2114090 | 1187 | + | tuf | WP_129389245.1 |
| GCF_023101205.1 | NZ_CP078078.1 | 53246 | 54436 | 1190 | - | tuf | WP_091484849.1 |
| GCF_001974985.1 | NZ_CP018762.1 | 3692919 | 3694109 | 1190 | + | tuf | WP_067247212.1 |

| | | | | | | | |
|---|---|---|---|---|---|---|---|
| GCF_025583665.1 | NZ_CP099706.1 | 1013004 | 1014194 | 1190 | + | tuf | WP_320943250.1 |
| GCF_032878875.1 | NZ_CP118157.1 | 2422940 | 2424130 | 1190 | - | tuf | WP_067024829.1 |
| GCF_008727755.1 | NZ_CP044231.1 | 233106 | 234293 | 1187 | - | tuf | WP_136028873.1 |
| GCF_001652465.1 | NZ_CP015810.1 | 2660692 | 2661882 | 1190 | - | tuf | WP_263796465.1 |
| GCF_015277855.1 | NZ_CP063812.1 | 579027 | 580217 | 1190 | + | tuf | WP_308492915.1 |
| GCF_011047135.1 | NZ_CP049255.1 | 560501 | 561691 | 1190 | + | tuf | WP_135063311.1 |
| GCF_003367705.1 | NZ_CP031425.1 | 284137 | 285327 | 1190 | - | tuf | WP_346225640.1 |
| GCF_031885415.1 | NZ_CP116871.1 | 1356671 | 1357861 | 1190 | + | tuf | WP_267738837.1 |
| GCF_023101185.1 | NZ_CP078077.1 | 345596 | 346786 | 1190 | + | tuf | WP_168914897.1 |
| GCF_014725695.1 | NZ_CP061344.1 | 1376408 | 1377598 | 1190 | + | tuf | WP_255158176.1 |
| GCF_029094505.1 | NZ_CP119108.1 | 215472 | 216662 | 1190 | + | tuf | WP_179872980.1 |
| GCF_003991875.1 | NZ_CP031423.1 | 1125514 | 1126701 | 1187 | - | tuf | WP_248116035.1 |
| GCF_036324365.1 | NZ_CP137080.1 | 412515 | 413702 | 1187 | - | tuf | WP_010080381.1 |
| GCF_008727775.1 | NZ_CP044232.1 | 1138460 | 1139647 | 1187 | + | tuf | WP_224356573.1 |
| GCF_015277875.1 | NZ_CP063814.1 | 1336364 | 1337554 | 1190 | + | tuf | WP_143985476.1 |
| GCF_024362265.1 | NZ_CP101471.1 | 1202543 | 1203733 | 1190 | + | tuf | WP_013861987.1 |
| GCF_024971095.1 | NZ_CP069717.1 | 3106084 | 3107274 | 1190 | - | tuf | WP_091075641.1 |
| GCF_001975955.2 | NZ_CP031421.1 | 4018514 | 4019704 | 1190 | + | tuf | WP_091526922.1 |
| GCF_009735645.1 | NZ_CP032550.1 | 6183901 | 6185091 | 1190 | - | tuf | WP_013288635.1 |
| GCF_900105335.1 | NZ_LT629770.1 | 4277789 | 4278979 | 1190 | - | tuf | WP_091666211.1 |
| GCF_035231185.1 | NZ_CP141117.1 | 6597153 | 6598343 | 1190 | - | tuf | WP_043965499.1 |
| GCF_004135285.1 | NZ_CP035494.1 | 551394 | 552584 | 1190 | - | tuf | WP_275032063.1 |
| GCF_900100885.1 | NZ_LT629692.1 | 4054604 | 4055794 | 1190 | - | tuf | WP_013288635.1 |
| GCF_021172025.1 | NZ_CP082781.1 | 2006122 | 2007312 | 1190 | + | tuf | WP_088987573.1 |
| GCF_034120055.1 | NZ_CP139368.1 | 3426752 | 3427942 | 1190 | + | tuf | WP_089018942.1 |
| GCF_004564075.1 | NZ_CP038256.1 | 2787131 | 2788321 | 1190 | - | tuf | WP_088976126.1 |
| GCF_023101165.1 | NZ_CP078076.1 | 4208373 | 4209563 | 1190 | + | tuf | WP_117226257.1 |
| GCF_025758395.1 | NZ_CP107222.1 | 7178474 | 7179664 | 1190 | + | tuf | WP_088997372.1 |
| GCF_030866805.1 | NZ_CP133019.1 | 303238 | 304428 | 1190 | + | tuf | WP_088998349.1 |
| GCF_004564355.1 | NZ_CP038266.1 | 6284881 | 6286071 | 1190 | + | tuf | WP_120330660.1 |
| GCF_039649665.1 | NZ_JBDJAW0100000( | 3196812 | 3198002 | 1190 | + | tuf | WP_262283035.1 |

| | | | | | | | |
|---|---|---|---|---|---|---|---|
| GCF_026625045.1 | NZ_CP113089.1 | 505118 | 506308 | 1190 | + | tuf | WP_043965499.1 |
| GCF_012530535.1 | NZ_CP051299.1 | 2649530 | 2650720 | 1190 | + | tuf | WP_089012497.1 |
| GCF_024362325.1 | NZ_CP101497.1 | 5545478 | 5546668 | 1190 | + | tuf | WP_091594933.1 |
| GCF_013414345.1 | NZ_CP058670.1 | 5932895 | 5934085 | 1190 | - | tuf | WP_013735955.1 |
| GCF_044872765.1 | NZ_CP085253.1 | 4666113 | 4667303 | 1190 | - | tuf | WP_091198304.1 |
| GCF_900475555.1 | NZ_LS483396.1 | 1384293 | 1385483 | 1190 | + | tuf | WP_088960251.1 |
| GCF_020097155.1 | NZ_CP083691.1 | 5260639 | 5261829 | 1190 | + | tuf | WP_067301408.1 |
| GCF_007362775.1 | NZ_CP041692.1 | 6217088 | 6218278 | 1190 | + | tuf | WP_181570341.1 |
| GCF_000270245.1 | NC_015635.1 | 2111095 | 2112285 | 1190 | - | tuf | WP_145815457.1 |
| GCF_900105785.1 | NZ_LT629799.1 | 3913028 | 3914212 | 1184 | + | tuf | WP_088971625.1 |
| GCF_900105385.1 | NZ_LT629772.1 | 6111079 | 6112269 | 1190 | - | tuf | WP_088973245.1 |
| GCF_000145235.1 | NC_014391.1 | 4836629 | 4837819 | 1190 | - | tuf | WP_091262807.1 |
| GCF_900089595.1 | NZ_LT594323.1 | 1075975 | 1077165 | 1190 | + | tuf | WP_089005210.1 |
| GCF_013389765.1 | NZ_CP058322.1 | 5535780 | 5536970 | 1190 | - | tuf | WP_088950239.1 |
| GCF_028993575.1 | NZ_CP118615.1 | 428406 | 429596 | 1190 | + | tuf | WP_105184678.1 |
| GCF_028768405.1 | NZ_CP118233.1 | 1070389 | 1071579 | 1190 | + | tuf | WP_083363048.1 |
| GCF_900091505.1 | NZ_LT607409.1 | 403316 | 404509 | 1193 | - | tuf | WP_004008360.1 |
| GCF_900091455.1 | NZ_LT607412.1 | 4908623 | 4909813 | 1190 | - | tuf | WP_014742860.1 |
| GCF_900090295.1 | NZ_LT607753.1 | 3864397 | 3865587 | 1190 | - | tuf | WP_370616232.1 |
| GCF_014764405.1 | NZ_CP061725.1 | 2504988 | 2506178 | 1190 | + | tuf | WP_226081060.1 |
| GCF_900090235.1 | NZ_LT607750.1 | 2414689 | 2415879 | 1190 | - | tuf | WP_108391780.1 |
| GCF_900091445.1 | NZ_LT607733.1 | 5331196 | 5332383 | 1187 | - | tuf | WP_068000509.1 |
| GCF_036184225.1 | NZ_CP108084.1 | 4390335 | 4391522 | 1187 | - | tuf | WP_003873538.1 |
| GCF_032108445.1 | NZ_CP134876.1 | 974210 | 975397 | 1187 | + | tuf | WP_158015502.1 |
| GCF_045348645.1 | NZ_CP160047.1 | 589226 | 590413 | 1187 | - | tuf | WP_045382598.1 |
| GCF_900090285.1 | NZ_LT607754.1 | 796635 | 797822 | 1187 | + | tuf | WP_003403463.1 |
| GCF_900091425.1 | NZ_LT598496.1 | 608496 | 609683 | 1187 | + | tuf | WP_085233424.1 |
| GCF_000204155.1 | NC_015434.1 | 4462287 | 4463474 | 1187 | + | tuf | WP_163779725.1 |
| GCF_900089605.1 | NZ_LT594324.1 | 5915245 | 5916432 | 1187 | - | tuf | WP_087079323.1 |
| GCF_900091515.1 | NZ_LT607410.1 | 3877766 | 3878953 | 1187 | - | tuf | WP_067921884.1 |
| GCF_900090265.1 | NZ_LT607752.1 | 2723139 | 2724326 | 1187 | - | tuf | WP_085222925.1 |

| | | | | | | | |
|---|---|---|---|---|---|---|---|
| GCF_013694245.2 | NZ_CP059322.2 | 5504513 | 5505700 | 1187 | + | tuf | WP_163735367.1 |
| GCF_014680085.1 | NZ_AP023438.1 | 748192 | 749379 | 1187 | + | tuf | WP_047313536.1 |
| GCF_900090305.1 | NZ_LT607751.1 | 4016230 | 4017417 | 1187 | - | tuf | WP_003923354.1 |
| GCF_900090305.1 | NZ_LT607751.1 | 1229726 | 1230913 | 1187 | - | tuf | WP_083073965.1 |
| GCF_009671205.1 | NZ_CP045309.1 | 1169542 | 1170729 | 1187 | + | tuf | WP_008260847.1 |
| GCF_900091545.1 | NZ_LT607411.1 | 4285034 | 4286221 | 1187 | + | tuf | WP_023370882.1 |
| GCF_900090275.1 | NZ_LT607755.1 | 1026879 | 1028066 | 1187 | + | tuf | WP_068033909.1 |
| GCF_900289045.1 | NZ_LT985188.1 | 3722724 | 3723911 | 1187 | - | tuf | WP_003403463.1 |
| GCF_900104895.1 | NZ_LT629742.1 | 952675 | 953862 | 1187 | + | tuf | WP_036466681.1 |
| GCF_000196535.1 | NC_014246.1 | 2200761 | 2201948 | 1187 | - | tuf | WP_010908588.1 |
| GCF_000306785.1 | NC_017955.1 | 2297920 | 2299107 | 1187 | - | tuf | WP_045843458.1 |
| GCF_041320275.1 | NZ_CP162023.1 | 903554 | 904741 | 1187 | + | tuf | WP_071509425.1 |
| GCF_020424085.1 | NZ_CP080203.1 | 3508215 | 3509402 | 1187 | - | tuf | WP_064883866.1 |
| GCF_003065425.1 | NZ_CP026949.1 | 1184739 | 1185926 | 1187 | + | tuf | WP_011739036.1 |
| GCF_001644575.1 | NZ_CP015596.1 | 4459126 | 4460313 | 1187 | - | tuf | WP_067166249.1 |
| GCF_022175585.2 | NZ_CP018019.2 | 2696594 | 2697781 | 1187 | + | tuf | WP_083089163.1 |
| GCF_900292015.1 | NZ_LR130759.1 | 1230777 | 1231964 | 1187 | - | tuf | WP_013827639.1 |
| GCF_010728725.1 | NZ_AP022606.1 | 4532692 | 4533879 | 1187 | - | tuf | WP_023370882.1 |
| GCF_000253375.1 | NC_015848.1 | 1192209 | 1193396 | 1187 | + | tuf | WP_055578873.1 |
| GCF_010730195.1 | NZ_AP022613.1 | 4713287 | 4714474 | 1187 | - | tuf | WP_008260847.1 |
| GCF_010727945.1 | NZ_AP022569.1 | 4964387 | 4965574 | 1187 | + | tuf | WP_083171590.1 |
| GCF_002157835.1 | NZ_CP020809.1 | 3017561 | 3018748 | 1187 | + | tuf | WP_067334850.1 |
| GCF_034555895.2 | NZ_CP157315.1 | 3350446 | 3351633 | 1187 | - | tuf | WP_085271926.1 |
| GCF_010730355.1 | NZ_AP022576.1 | 1263732 | 1264919 | 1187 | + | tuf | WP_011739036.1 |
| GCF_010726765.1 | NZ_AP022601.1 | 1062516 | 1063703 | 1187 | + | tuf | WP_085249133.1 |
| GCF_000340435.2 | NZ_CP011883.2 | 4683334 | 4684521 | 1187 | + | tuf | WP_085258094.1 |
| GCF_016861545.1 | NZ_AP024310.1 | 727965 | 729152 | 1187 | - | tuf | WP_044483827.1 |
| GCF_010730745.1 | NZ_AP022615.1 | 5164705 | 5165892 | 1187 | + | tuf | WP_163683851.1 |
| GCF_020736005.1 | NZ_CP085945.1 | 899606 | 900793 | 1187 | + | tuf | WP_096437398.1 |
| GCF_000157895.3 | NC_022663.1 | 3014668 | 3015855 | 1187 | - | tuf | WP_083045950.1 |
| GCF_014263315.1 | NZ_CP045081.1 | 2699114 | 2700301 | 1187 | + | tuf | WP_011739036.1 |

| | | | | | | | |
|---|---|---|---|---|---|---|---|
| GCF_010731535.1 | NZ_AP022581.1 | 1734283 | 1735470 | 1187 | - | tuf | WP_044509807.1 |
| GCF_022374895.2 | NZ_CP092423.2 | 1067650 | 1068837 | 1187 | + | tuf | WP_211697885.1 |
| GCF_003253775.1 | NZ_CP029543.1 | 3005043 | 3006230 | 1187 | - | tuf | WP_163790664.1 |
| GCF_000975265.2 | NZ_CP083405.1 | 784821 | 786008 | 1187 | + | tuf | NP_215199.1 |
| GCF_019645855.1 | NZ_CP080999.1 | 4687182 | 4688369 | 1187 | - | tuf | WP_011739036.1 |
| GCF_010731775.1 | NZ_AP022590.1 | 1160717 | 1161904 | 1187 | + | tuf | WP_180916919.1 |
| GCF_000723425.2 | NZ_HG917972.2 | 3895077 | 3896267 | 1190 | - | tuf | WP_005055596.1 |
| GCF_002285715.1 | NZ_CP023147.1 | 3944302 | 3945492 | 1190 | - | tuf | WP_046254679.1 |
| GCF_010731635.1 | NZ_AP022583.1 | 4404299 | 4405489 | 1190 | - | tuf | WP_043076432.1 |
| GCF_010726505.1 | NZ_AP022562.1 | 3664204 | 3665394 | 1190 | - | tuf | WP_078288431.1 |
| GCF_021183725.1 | NZ_CP089224.1 | 3599591 | 3600781 | 1190 | - | tuf | WP_070909368.1 |
| GCF_003614435.1 | NZ_CP025546.1 | 2299300 | 2300487 | 1187 | - | tuf | WP_069392792.1 |
| GCF_000276825.1 | NC_016948.1 | 3159212 | 3160399 | 1187 | - | tuf | WP_240170443.1 |
| GCF_010731655.1 | NZ_AP022619.1 | 817405 | 818592 | 1187 | + | tuf | WP_047317600.1 |
| GCF_022430545.2 | NZ_CP092488.2 | 211638 | 212825 | 1187 | + | tuf | WP_085134912.1 |
| GCF_010730575.1 | NZ_AP022614.1 | 2526314 | 2527501 | 1187 | + | tuf | WP_083027766.1 |
| GCF_003584745.1 | NZ_AP018410.1 | 741963 | 743150 | 1187 | + | tuf | WP_013827639.1 |
| GCF_963853645.1 | NZ_OY970456.1 | 3530682 | 3531869 | 1187 | - | tuf | WP_085258910.1 |
| GCF_010729105.1 | NZ_AP022573.1 | 794223 | 795410 | 1187 | + | tuf | WP_046282637.1 |
| GCF_019668465.1 | NZ_AP024828.1 | 4721780 | 4722967 | 1187 | - | tuf | WP_115321986.1 |
| GCF_010731595.1 | NZ_AP022582.1 | 2494732 | 2495919 | 1187 | - | tuf | WP_163664246.1 |
| GCF_002356315.1 | NZ_AP018164.1 | 5200715 | 5201902 | 1187 | + | tuf | WP_163806937.1 |
| GCF_010730055.1 | NZ_AP022575.1 | 2451891 | 2453078 | 1187 | - | tuf | WP_163918991.1 |
| GCF_010728525.1 | NZ_AP022572.1 | 2233417 | 2234604 | 1187 | - | tuf | WP_138232222.1 |
| GCF_010727605.1 | NZ_AP022568.1 | 1099551 | 1100738 | 1187 | + | tuf | WP_048630018.1 |
| GCF_018278905.1 | NZ_CP046600.1 | 1080667 | 1081854 | 1187 | + | tuf | WP_011778548.1 |
| GCF_010731715.1 | NZ_AP022587.1 | 4808432 | 4809619 | 1187 | - | tuf | WP_193046244.1 |
| GCF_000195955.2 | NC_000962.3 | 6478690 | 6479877 | 1187 | - | tuf | WP_003881870.1 |
| GCF_002355775.1 | NZ_AP017624.1 | 3350094 | 3351281 | 1187 | - | tuf | WP_090588362.1 |
| GCF_013466425.1 | NZ_CP059165.1 | 2360456 | 2361643 | 1187 | - | tuf | WP_067217723.1 |
| GCF_000069185.1 | NC_010397.1 | 4010717 | 4011904 | 1187 | - | tuf | WP_126335204.1 |

| | | | | | | | |
|---|---|---|---|---|---|---|---|
| GCF_001632805.1 | NZ_CP007220.1 | 4041809 | 4042996 | 1187 | - | tuf | WP_085149707.1 |
| GCF_001605725.1 | NZ_CP011530.1 | 1199584 | 1200771 | 1187 | + | tuf | WP_115321986.1 |
| GCF_004924335.1 | NZ_CP024633.1 | 1825186 | 1826457 | 1271 | + | tuf | WP_262871761.1 |
| GCF_001456355.1 | NZ_CP010271.1 | 1233457 | 1234644 | 1187 | + | tuf | WP_073856275.1 |
| GCF_010731835.1 | NZ_AP022594.1 | 615922 | 617109 | 1187 | + | tuf | WP_011558404.1 |
| GCF_022370835.2 | NZ_CP092365.1 | 3566966 | 3568153 | 1187 | - | tuf | WP_098002444.1 |
| GCF_019645815.1 | NZ_CP080997.1 | 2236546 | 2237733 | 1187 | - | tuf | WP_085096354.1 |
| GCF_010729485.1 | NZ_AP022609.1 | 5350839 | 5352026 | 1187 | - | tuf | WP_003881870.1 |
| GCF_010731755.1 | NZ_AP022589.1 | 1907632 | 1908819 | 1187 | + | tuf | WP_090359967.1 |
| GCF_000214155.1 | NC_015576.1 | 1511570 | 1512757 | 1187 | + | tuf | WP_003881870.1 |
| GCF_010727125.1 | NZ_AP022564.1 | 1351617 | 1352804 | 1187 | - | tuf | WP_168141740.1 |
| GCF_022374935.2 | NZ_CP092430.2 | 2871813 | 2873000 | 1187 | - | tuf | WP_163687026.1 |
| GCF_010726245.1 | NZ_AP022561.1 | 4647165 | 4648352 | 1187 | - | tuf | WP_011895899.1 |
| GCF_010727325.1 | NZ_AP022565.1 | 4475182 | 4476369 | 1187 | - | tuf | WP_214397222.1 |
| GCF_010731855.1 | NZ_AP022620.1 | 2375118 | 2376305 | 1187 | + | tuf | WP_070352555.1 |
| GCF_010731815.2 | NZ_AP022593.1 | 239525 | 240712 | 1187 | + | tuf | WP_005625383.1 |
| GCF_010730955.1 | NZ_AP022577.1 | 1304563 | 1305750 | 1187 | - | tuf | WP_163746689.1 |
| GCF_900637195.1 | NZ_LR134356.1 | 4421654 | 4422841 | 1187 | - | tuf | WP_069407796.1 |
| GCF_016919645.1 | NZ_CP070380.1 | 1358540 | 1359727 | 1187 | + | tuf | WP_083033768.1 |
| GCF_016313185.1 | NZ_CP066218.1 | 3301463 | 3302650 | 1187 | - | tuf | WP_134054324.1 |
| GCF_010731295.1 | NZ_AP022579.1 | 1171356 | 1172543 | 1187 | + | tuf | WP_163733616.1 |
| GCF_025215495.1 | NZ_CP104302.1 | 4366851 | 4368038 | 1187 | + | tuf | WP_036429167.1 |
| GCF_010731795.1 | NZ_AP022591.1 | 4288872 | 4290059 | 1187 | - | tuf | WP_197382466.1 |
| GCF_900637205.1 | NZ_LR134355.1 | 3776537 | 3777724 | 1187 | - | tuf | WP_011558404.1 |
| GCF_010729895.1 | NZ_AP022612.1 | 5510924 | 5512111 | 1187 | - | tuf | WP_083156089.1 |
| GCF_022370635.2 | NZ_CP092362.2 | 4737868 | 4739055 | 1187 | - | tuf | WP_020101374.1 |
| GCF_022370635.2 | NZ_CP092362.2 | 1357860 | 1359047 | 1187 | + | tuf | WP_019514020.1 |
| GCF_019456655.1 | NZ_CP080332.1 | 5141819 | 5143006 | 1187 | - | tuf | WP_090424593.1 |
| GCF_010728155.1 | NZ_AP022605.1 | 1059546 | 1060733 | 1187 | + | tuf | WP_071947504.1 |
| GCF_010726645.1 | NZ_AP022563.1 | 306556 | 307743 | 1187 | - | tuf | WP_011895899.1 |
| GCF_010726955.1 | NZ_AP022603.1 | 5005212 | 5006399 | 1187 | + | tuf | WP_020101374.1 |

| | | | | | | | |
|---|---|---|---|---|---|---|---|
| GCF_019739075.1 | NZ_CP081673.1 | 1638202 | 1639389 | 1187 | + | tuf | WP_152515676.1 |
| GCF_014295435.1 | NZ_CP059894.1 | 5559877 | 5561064 | 1187 | - | tuf | WP_057151156.1 |
| GCF_022179545.1 | NZ_AP025518.1 | 1150271 | 1151458 | 1187 | - | tuf | WP_163898030.1 |
| GCF_012223425.1 | NZ_CP038799.1 | 1895067 | 1896254 | 1187 | - | tuf | WP_043405705.1 |
| GCF_010728925.1 | NZ_AP022608.1 | 27886 | 29073 | 1187 | - | tuf | WP_083405429.1 |
| GCF_000184435.1 | NC_014814.1 | 2478972 | 2480159 | 1187 | + | tuf | WP_163697373.1 |
| GCF_022370755.2 | NZ_CP092364.2 | 2439318 | 2440505 | 1187 | + | tuf | WP_163797072.1 |
| GCF_008329645.1 | NZ_CP043474.1 | 1351365 | 1352552 | 1187 | + | tuf | WP_003881870.1 |
| GCF_900603025.1 | NZ_LR026975.1 | 2492068 | 2493255 | 1187 | + | tuf | WP_029106828.1 |
| GCF_010731895.1 | NZ_AP022596.1 | 1444586 | 1445773 | 1187 | + | tuf | WP_003892789.1 |
| GCF_019645835.1 | NZ_CP080998.1 | 976375 | 977562 | 1187 | + | tuf | WP_003926032.1 |
| GCF_010731615.1 | NZ_AP022618.1 | 3250918 | 3252105 | 1187 | + | tuf | WP_068915586.1 |
| GCF_010731695.1 | NZ_AP022586.1 | 1132672 | 1133859 | 1187 | + | tuf | WP_003929599.1 |
| GCF_010729665.1 | NZ_AP022610.1 | 1374740 | 1375927 | 1187 | + | tuf | WP_011778548.1 |
| GCF_010727475.1 | NZ_AP022567.1 | 861749 | 862939 | 1190 | - | tuf | WP_286278460.1 |
| GCF_015710575.1 | NZ_CP065373.1 | 2608486 | 2609676 | 1190 | - | tuf | WP_124799605.1 |
| GCF_010731575.1 | NZ_AP022617.1 | 1247817 | 1249004 | 1187 | + | tuf | WP_015746440.1 |
| GCF_010726085.1 | NZ_AP022560.1 | 2845497 | 2846684 | 1187 | + | tuf | WP_090476333.1 |
| GCF_005670685.2 | NZ_CP062008.1 | 2848610 | 2849800 | 1190 | - | tuf | WP_213170541.1 |
| GCF_000317305.3 | NC_023036.2 | 872215 | 873405 | 1190 | + | tuf | WP_239677695.1 |
| GCF_003855255.1 | NZ_CP034072.1 | 6965148 | 6966335 | 1187 | - | tuf | WP_019049356.1 |
| GCF_019456675.1 | NZ_CP080333.1 | 7393236 | 7394423 | 1187 | + | tuf | WP_029901625.1 |
| GCF_010725485.1 | NZ_AP022598.1 | 8042366 | 8043541 | 1175 | + | tuf | WP_042264133.1 |
| GCF_010731115.1 | NZ_AP022616.1 | 2454828 | 2456015 | 1187 | + | tuf | WP_014353061.1 |
| GCF_010728325.1 | NZ_AP022570.1 | 5373831 | 5375018 | 1187 | - | tuf | WP_011211609.1 |
| GCF_010729305.1 | NZ_AP022574.1 | 1817194 | 1818381 | 1187 | - | tuf | WP_043698828.1 |
| GCF_010725725.1 | NZ_AP022599.1 | 1976718 | 1977905 | 1187 | + | tuf | WP_181583544.1 |
| GCF_022374875.2 | NZ_CP092427.2 | 3531362 | 3532555 | 1193 | + | tuf | WP_181584644.1 |
| GCF_900108565.1 | NZ_LT629971.1 | 4215063 | 4216235 | 1172 | + | tuf | WP_181578776.1 |
| GCF_010731875.1 | NZ_AP022595.1 | 7461663 | 7462838 | 1175 | - | tuf | WP_218471308.1 |
| GCF_010731735.1 | NZ_AP022588.1 | 8123417 | 8124604 | 1187 | - | tuf | WP_218471834.1 |

| | | | | | | | |
|---|---|---|---|---|---|---|---|
| GCF_019645875.1 | NZ_CP081000.1 | 6550713 | 6551900 | 1187 | - | tuf | WP_071930486.1 |
| GCF_016919345.1 | NZ_CP070349.1 | 115115 | 116302 | 1187 | + | tuf | WP_029933056.1 |
| GCF_022370415.1 | NZ_CP080274.1 | 4640389 | 4641564 | 1175 | + | tuf | WP_143981991.1 |
| GCF_900187065.1 | NZ_LT906483.1 | 1640867 | 1642042 | 1175 | + | tuf | WP_067509814.1 |
| GCF_010725885.1 | NZ_AP022600.1 | 2625411 | 2626598 | 1187 | + | tuf | WP_067506332.1 |
| GCF_001655245.1 | NZ_CP011491.1 | 5344123 | 5345310 | 1187 | - | tuf | WP_056814209.1 |
| GCF_000015305.1 | NC_008726.1 | 6228003 | 6229190 | 1187 | - | tuf | WP_063024777.1 |
| GCF_030296595.1 | NZ_AP027731.1 | 2313918 | 2315093 | 1175 | + | tuf | WP_098698508.1 |
| GCF_003860405.1 | NZ_CP034170.1 | 8171004 | 8172191 | 1187 | - | tuf | WP_067594060.1 |
| GCF_000024365.1 | NC_013235.1 | 770925 | 772103 | 1178 | + | tuf | WP_187686570.1 |
| GCF_900104535.1 | NZ_LT629710.1 | 7099876 | 7101063 | 1187 | - | tuf | WP_187685391.1 |
| GCF_016925535.1 | NZ_CP070496.1 | 3197941 | 3199113 | 1172 | - | tuf | WP_233040665.1 |
| GCF_016925615.1 | NZ_CP070499.1 | 3605615 | 3606808 | 1193 | - | tuf | WP_067723546.1 |
| GCF_900637185.1 | NZ_LR134352.1 | 5120381 | 5121568 | 1187 | - | tuf | WP_067716365.1 |
| GCF_002209125.2 | NZ_CP022088.2 | 1458331 | 1459518 | 1187 | + | tuf | WP_120735569.1 |
| GCF_002209125.2 | NZ_CP022088.2 | 4527063 | 4528232 | 1169 | + | tuf | WP_120738981.1 |
| GCF_002949635.1 | NZ_CP026746.1 | 7070324 | 7071517 | 1193 | + | tuf | WP_120742727.1 |
| GCF_000009805.1 | NC_006361.1 | 315549 | 316739 | 1190 | + | tuf | WP_165228177.1 |
| GCF_021183645.1 | NZ_CP089225.1 | 1047466 | 1048659 | 1193 | + | tuf | WP_214058205.1 |
| GCF_013744875.1 | NZ_CP059399.1 | 2210989 | 2212179 | 1190 | + | tuf | WP_125569281.1 |
| GCF_013744875.1 | NZ_CP059399.1 | 3681968 | 3683158 | 1190 | - | tuf | WP_322456217.1 |
| GCF_013744875.1 | NZ_CP059399.1 | 4200234 | 4201418 | 1184 | - | tuf | WP_322937299.1 |
| GCF_019222765.1 | NZ_CP078145.1 | 3263938 | 3265128 | 1190 | - | tuf | WP_181312626.1 |
| GCF_019222765.1 | NZ_CP078145.1 | 2936048 | 2937238 | 1190 | - | tuf | WP_222895241.1 |
| GCF_001886715.1 | NZ_CP018082.1 | 3636910 | 3638100 | 1190 | + | tuf | WP_151084436.1 |
| GCF_007362295.1 | NZ_CP041695.1 | 487751 | 488941 | 1190 | - | tuf | WP_068105950.1 |
| GCF_007362295.1 | NZ_CP041695.1 | 3569384 | 3570574 | 1190 | - | tuf | WP_182524566.1 |
| GCF_036051655.1 | NZ_CP107943.1 | 3219152 | 3220342 | 1190 | + | tuf | WP_135079034.1 |
| GCF_036051655.1 | NZ_CP107943.1 | 3196274 | 3197464 | 1190 | - | tuf | WP_205291087.1 |
| GCF_036142455.1 | NZ_CP108014.1 | 1677919 | 1679109 | 1190 | - | tuf | WP_141004032.1 |
| GCF_027924405.1 | NZ_AP026978.1 | 4547365 | 4548549 | 1184 | - | tuf | WP_141006311.1 |

| | | | | | | | |
|---|---|---|---|---|---|---|---|
| GCF_002568625.1 | NZ_CP023778.1 | 3696616 | 3697806 | 1190 | - | tuf | WP_171896990.1 |
| GCF_002568625.1 | NZ_CP023778.1 | 2558008 | 2559198 | 1190 | + | tuf | WP_187577230.1 |
| GCF_014466955.1 | NZ_AP023396.1 | 639938 | 641128 | 1190 | + | tuf | WP_153322086.1 |
| GCF_014466955.1 | NZ_AP023396.1 | 3535488 | 3536678 | 1190 | + | tuf | WP_211733383.1 |
| GCF_021183565.1 | NZ_CP089226.1 | 3252062 | 3253252 | 1190 | - | tuf | WP_216938423.1 |
| GCF_021183565.1 | NZ_CP089226.1 | 3394835 | 3396025 | 1190 | - | tuf | WP_122818660.1 |
| GCF_021183565.1 | NZ_CP089226.1 | 823180 | 824370 | 1190 | + | tuf | WP_172269841.1 |
| GCF_003626895.1 | NZ_CP032568.1 | 1049274 | 1050467 | 1193 | + | tuf | WP_223397717.1 |
| GCF_003626895.1 | NZ_CP032568.1 | 42229 | 43419 | 1190 | + | tuf | WP_141012505.1 |
| GCF_003626895.1 | NZ_CP032568.1 | 2259600 | 2260790 | 1190 | - | tuf | WP_091729479.1 |
| GCF_011046535.1 | NZ_CP049257.1 | 857383 | 858573 | 1190 | - | tuf | WP_135266723.1 |
| GCF_018459925.1 | NZ_CP075371.1 | 2690130 | 2691320 | 1190 | - | tuf | WP_128221547.1 |
| GCF_003945325.1 | NZ_AP019307.1 | 6111615 | 6112805 | 1190 | - | tuf | WP_212641596.1 |
| GCF_034661235.1 | NZ_CP141059.1 | 6330071 | 6331261 | 1190 | - | tuf | WP_220563401.1 |
| GCF_034661235.1 | NZ_CP141059.1 | 4856358 | 4857548 | 1190 | + | tuf | WP_013156109.1 |
| GCF_013624435.2 | NZ_CP061768.1 | 1168982 | 1170172 | 1190 | + | tuf | WP_017578644.1 |
| GCF_019880385.1 | NZ_CP082273.1 | 5780128 | 5781318 | 1190 | - | tuf | WP_017621535.1 |
| GCF_008761635.1 | NZ_CP044344.1 | 822986 | 824176 | 1190 | + | tuf | WP_020542312.1 |
| GCF_001653335.1 | NZ_CP015079.1 | 3934043 | 3935200 | 1157 | - | tuf | WP_020539947.1 |
| GCF_014127485.1 | NZ_CP059903.1 | 3808384 | 3809574 | 1190 | + | tuf | WP_364068663.1 |
| GCF_004564375.1 | NZ_CP038267.1 | 6662433 | 6663599 | 1166 | - | tuf | WP_378788218.1 |
| GCF_018388425.1 | NZ_CP074406.1 | 5357981 | 5359186 | 1205 | + | tuf | WP_106237253.1 |
| GCF_006494775.1 | NZ_CP041146.1 | 9285867 | 9287057 | 1190 | - | tuf | WP_106253412.1 |
| GCF_006494775.1 | NZ_CP041146.1 | 906611 | 907801 | 1190 | + | tuf | WP_043623429.1 |
| GCF_013177455.1 | NZ_CP053660.1 | 4663739 | 4664911 | 1172 | - | tuf | WP_225274749.1 |
| GCF_014395785.1 | NZ_CP060713.1 | 3969894 | 3971084 | 1190 | + | tuf | WP_327108822.1 |
| GCF_011420305.2 | NZ_CP061769.1 | 7201054 | 7202217 | 1163 | - | tuf | WP_327103854.1 |
| GCF_020991065.1 | NZ_CP087710.1 | 941362 | 942552 | 1190 | + | tuf | WP_139628812.1 |
| GCF_019039255.1 | NZ_CP077062.1 | 1230670 | 1231857 | 1187 | + | tuf | WP_154792915.1 |
| GCF_003710085.1 | NZ_CP033324.1 | 3666862 | 3668052 | 1190 | + | tuf | WP_114930051.1 |
| GCF_034661175.1 | NZ_CP141058.1 | 1115298 | 1116491 | 1193 | + | tuf | WP_143782469.1 |

| | | | | | | | |
|---|---|---|---|---|---|---|---|
| GCF_019931675.1 | NZ_CP082922.1 | 767005 | 768198 | 1193 | + | tuf | WP_252621713.1 |
| GCF_006494815.1 | NZ_CP041091.1 | 971998 | 973191 | 1193 | + | tuf | WP_252594291.1 |
| GCF_900104965.1 | NZ_LT629757.1 | 1791072 | 1792253 | 1181 | - | tuf | WP_252595158.1 |
| GCF_004683965.1 | NZ_CP038436.1 | 2816628 | 2817818 | 1190 | - | tuf | WP_134773984.1 |
| GCF_004014875.1 | NZ_CP034929.1 | 2597431 | 2598621 | 1190 | - | tuf | WP_158061699.1 |
| GCF_018316795.1 | NZ_CP074132.1 | 3156001 | 3157191 | 1190 | - | tuf | WP_131105564.1 |
| GCF_018316655.1 | NZ_CP074133.1 | 3301883 | 3303076 | 1193 | - | tuf | WP_392543049.1 |
| GCF_900638215.1 | NZ_LR134501.1 | 2987824 | 2989017 | 1193 | + | tuf | WP_377640630.1 |
| GCF_024134545.1 | NZ_CP099837.1 | 3004182 | 3005369 | 1187 | - | tuf | WP_373091518.1 |
| GCF_002263495.1 | NZ_CP022753.1 | 3055099 | 3056286 | 1187 | - | tuf | WP_018776689.1 |
| GCF_019397265.1 | NZ_CP068985.1 | 1969242 | 1970429 | 1187 | + | tuf | WP_021472401.1 |
| GCF_019397265.1 | NZ_CP068985.1 | 691067 | 692257 | 1190 | + | tuf | WP_166988242.1 |
| GCF_042751225.1 | NZ_CP170570.1 | 812186 | 813376 | 1190 | + | tuf | WP_166984476.1 |
| GCF_042751225.1 | NZ_CP170570.1 | 2676204 | 2677397 | 1193 | - | tuf | WP_029252168.1 |
| GCF_035917475.1 | NZ_CP109074.1 | 485144 | 486337 | 1193 | - | tuf | WP_286218418.1 |
| GCF_035917475.1 | NZ_CP109074.1 | 1656120 | 1657316 | 1196 | - | tuf | WP_006288641.1 |
| GCF_020215645.1 | NZ_CP084058.1 | 1086652 | 1087836 | 1184 | - | tuf | WP_009743583.1 |
| GCF_020215645.1 | NZ_CP084058.1 | 3923131 | 3924324 | 1193 | + | tuf | WP_091788921.1 |
| GCF_035920485.1 | NZ_CP109166.1 | 2672937 | 2674127 | 1190 | + | tuf | WP_407341403.1 |
| GCF_035920485.1 | NZ_CP109166.1 | 3850267 | 3851457 | 1190 | - | tuf | WP_227423348.1 |
| GCF_009497075.1 | NZ_CP045572.1 | 2723089 | 2724279 | 1190 | - | tuf | WP_111489775.1 |
| GCF_009708215.1 | NZ_CP046175.1 | 3688340 | 3689533 | 1193 | - | tuf | WP_030526824.1 |
| GCF_003351765.1 | NZ_CP031229.1 | 1383645 | 1384829 | 1184 | - | tuf | WP_173154833.1 |
| GCF_007197575.1 | NZ_CP041616.1 | 4370798 | 4371988 | 1190 | + | tuf | WP_173158440.1 |
| GCF_023923205.1 | NZ_CP099490.1 | 588697 | 589881 | 1184 | + | tuf | WP_038676485.1 |
| GCF_023923225.1 | NZ_CP099489.1 | 1254648 | 1255838 | 1190 | + | tuf | WP_038677270.1 |
| GCF_023923225.1 | NZ_CP099489.1 | 479116 | 480303 | 1187 | - | tuf | WP_212821055.1 |
| GCF_004526345.1 | NZ_CP038213.1 | 1710927 | 1712117 | 1190 | - | tuf | WP_212823058.1 |
| GCF_008843165.1 | NZ_CP044427.1 | 288194 | 289384 | 1190 | + | tuf | WP_104912937.1 |
| GCF_004322775.1 | NZ_CP036403.1 | 638266 | 639456 | 1190 | + | tuf | WP_091802124.1 |
| GCF_043099285.1 | NZ_CP170723.1 | 4084763 | 4085950 | 1187 | - | tuf | WP_005518275.1 |

| | | | | | | | |
|---|---|---|---|---|---|---|---|
| GCF_042756445.1 | NZ_CP170578.1 | 4233510 | 4234697 | 1187 | + | tuf | WP_348604181.1 |
| GCF_041549525.1 | NZ_CP157456.1 | 2295599 | 2296786 | 1187 | + | tuf | WP_021797163.1 |
| GCF_021919345.1 | NZ_CP089293.1 | 2169135 | 2170322 | 1187 | - | tuf | WP_119161001.1 |
| GCF_017357445.1 | NZ_CP014574.1 | 1876293 | 1877480 | 1187 | - | tuf | WP_055345899.1 |
| GCF_011751765.2 | NZ_CP061169.1 | 614518 | 615708 | 1190 | - | tuf | WP_166231877.1 |
| GCF_011751745.2 | NZ_CP061170.1 | 1011284 | 1012474 | 1190 | + | tuf | WP_232549018.1 |
| GCF_900105445.1 | NZ_LT629776.1 | 892404 | 893594 | 1190 | - | tuf | WP_120761920.1 |
| GCF_030296575.1 | NZ_AP027729.1 | 854853 | 856043 | 1190 | + | tuf | WP_149324665.1 |
| GCF_001042675.1 | NZ_AP012333.1 | 3012686 | 3013873 | 1187 | - | tuf | WP_015937856.1 |
| GCF_001746855.1 | NZ_CP017298.1 | 3415490 | 3416677 | 1187 | - | tuf | WP_056075385.1 |
| GCF_900104525.1 | NZ_LT629711.1 | 1095775 | 1096962 | 1187 | + | tuf | WP_056334246.1 |
| GCF_045689905.1 | NZ_CP144210.1 | 349665 | 350840 | 1175 | - | tuf | WP_013421487.1 |
| GCF_017569225.1 | NZ_CP071868.1 | 7666085 | 7667275 | 1190 | - | tuf | WP_013427328.1 |
| GCF_003258605.2 | NZ_CP031264.1 | 5720756 | 5721946 | 1190 | + | tuf | WP_185721380.1 |
| GCF_044797865.1 | NZ_CP085209.1 | 6233734 | 6234924 | 1190 | - | tuf | WP_085912591.1 |
| GCF_011764565.1 | NZ_AP022871.1 | 2447854 | 2449044 | 1190 | + | tuf | WP_172157753.1 |
| GCF_011764565.1 | NZ_AP022871.1 | 5787064 | 5788254 | 1190 | - | tuf | WP_013677466.1 |
| GCF_000785495.1 | NZ_CP009896.1 | 2405599 | 2406789 | 1190 | + | tuf | WP_185721380.1 |
| GCF_000785495.1 | NZ_CP009896.1 | 2081415 | 2082602 | 1187 | - | tuf | WP_045075284.1 |
| GCF_018324255.1 | NZ_AP023359.1 | 158927 | 160117 | 1190 | + | tuf | WP_123445037.1 |
| GCF_018324255.1 | NZ_AP023359.1 | 2026666 | 2027856 | 1190 | - | tuf | WP_104265235.1 |
| GCF_002950575.1 | NZ_CP026923.1 | 313867 | 315057 | 1190 | + | tuf | WP_068251449.1 |
| GCF_002240355.1 | NZ_CP016353.1 | 864038 | 865228 | 1190 | + | tuf | WP_308467483.1 |
| GCF_003013675.1 | NZ_CP027793.1 | 2558499 | 2559689 | 1190 | - | tuf | WP_068209460.1 |
| GCF_040024445.1 | NZ_CP157276.1 | 1959521 | 1960711 | 1190 | - | tuf | WP_027692588.1 |
| GCF_003798325.1 | NZ_CP033719.1 | 639593 | 640783 | 1190 | + | tuf | WP_068251449.1 |
| GCF_900637915.1 | NZ_LR134442.1 | 1913345 | 1914532 | 1187 | - | tuf | WP_012245569.1 |
| GCF_900087655.1 | NZ_LT593929.1 | 2153077 | 2154264 | 1187 | + | tuf | WP_006936215.1 |
| GCF_011393335.1 | NZ_CP049865.1 | 444651 | 445841 | 1190 | + | tuf | WP_265383427.1 |
| GCF_038502305.1 | NZ_CP115965.1 | 3932938 | 3934125 | 1187 | + | tuf | WP_072704529.1 |
| GCF_003627075.1 | NZ_CP032630.1 | 5080518 | 5081705 | 1187 | - | tuf | WP_019744283.1 |

| | | | | | | | |
|---|---|---|---|---|---|---|---|
| GCF_008365275.1 | NZ_CP043504.1 | 2152157 | 2153344 | 1187 | - | tuf | WP_033236419.1 |
| GCF_000022025.1 | NC_011886.1 | 3904676 | 3905863 | 1187 | - | tuf | WP_003937665.1 |
| GCF_900105535.1 | NZ_LT629779.1 | 2119280 | 2120467 | 1187 | - | tuf | WP_072940147.1 |
| GCF_045291495.1 | NZ_CP148033.1 | 6319138 | 6320325 | 1187 | - | tuf | WP_005252203.1 |
| GCF_000166135.1 | NC_014666.1 | 1767388 | 1768575 | 1187 | + | tuf | WP_109328041.1 |
| GCF_000166135.1 | NC_014666.1 | 6746140 | 6747327 | 1187 | + | tuf | WP_087557087.1 |
| GCF_019263705.2 | NZ_JADQDK0100000( | 1934731 | 1935918 | 1187 | - | tuf | WP_016690893.1 |
| GCF_003945385.1 | NZ_AP018920.1 | 1821829 | 1823016 | 1187 | + | tuf | WP_003941926.1 |
| GCF_013155125.1 | NZ_CP053564.1 | 3790761 | 3791948 | 1187 | - | tuf | WP_016690893.1 |
| GCF_000196675.2 | NC_015312.1 | 4280104 | 4281291 | 1187 | - | tuf | WP_003937665.1 |
| GCF_014235185.1 | NZ_CP060131.1 | 3421519 | 3422706 | 1187 | - | tuf | WP_264151749.1 |
| GCF_000950575.1 | NZ_CP011005.1 | 3560082 | 3561269 | 1187 | - | tuf | WP_338886298.1 |
| GCF_004011135.1 | NZ_CP028137.1 | 3079012 | 3080199 | 1187 | - | tuf | WP_072738341.1 |
| GCF_004011115.1 | NZ_CP028130.1 | 2938273 | 2939460 | 1187 | + | tuf | WP_005252203.1 |
| GCF_004011095.1 | NZ_CP028129.1 | 1241010 | 1242197 | 1187 | - | tuf | WP_038503185.1 |
| GCF_030866825.1 | NZ_CP133020.1 | 174612 | 175799 | 1187 | + | tuf | WP_138315159.1 |
| GCF_009834165.1 | NZ_CP047186.1 | 2084272 | 2085459 | 1187 | + | tuf | WP_006887686.1 |
| GCF_014770185.1 | NZ_CP061742.1 | 664482 | 665669 | 1187 | + | tuf | WP_068169718.1 |
| GCF_001641005.1 | NZ_CP015515.1 | 316757 | 317944 | 1187 | - | tuf | WP_013397618.1 |
| GCF_000018885.1 | NC_010168.1 | 304301 | 305488 | 1187 | - | tuf | WP_055684589.1 |
| GCF_025722975.1 | NZ_CP106982.1 | 688547 | 689734 | 1187 | + | tuf | WP_168614565.1 |
| GCF_026153295.1 | NZ_CP110615.1 | 3433194 | 3434381 | 1187 | - | tuf | WP_193496682.1 |
| GCF_900478115.1 | NZ_LS483468.1 | 3372014 | 3373201 | 1187 | - | tuf | WP_231064658.1 |
| GCF_000696675.2 | NZ_CP007255.1 | 2862840 | 2864027 | 1187 | - | tuf | WP_231053372.1 |
| GCF_019334125.1 | NZ_CP079698.1 | 4225765 | 4226952 | 1187 | - | tuf | WP_147918565.1 |
| GCF_047782815.1 | NZ_CP180630.1 | 3385507 | 3386697 | 1190 | + | tuf | WP_005443241.1 |
| GCF_017068375.1 | NZ_CP070609.1 | 473691 | 474881 | 1190 | + | tuf | WP_005453235.1 |
| GCF_020542785.1 | NZ_CP080954.1 | 412987 | 414177 | 1190 | + | tuf | WP_005461350.1 |
| GCF_003130705.1 | NZ_CP021354.1 | 714188 | 715378 | 1190 | + | tuf | WP_009152375.1 |
| GCF_017068395.1 | NZ_CP070619.1 | 396822 | 398012 | 1190 | + | tuf | WP_012795906.1 |
| GCF_000511305.1 | NC_023150.1 | 7632855 | 7634045 | 1190 | - | tuf | WP_009948624.1 |

| | | | | | | | |
|---|---|---|---|---|---|---|---|
| GCF_015099595.1 | NZ_AP023172.1 | 5881215 | 5882405 | 1190 | - | tuf | WP_224956669.1 |
| GCF_029457455.1 | NZ_CP120356.1 | 7119563 | 7120753 | 1190 | + | tuf | WP_190820737.1 |
| GCF_016804345.1 | NZ_CP044211.1 | 6074188 | 6075378 | 1190 | - | tuf | WP_263249843.1 |
| GCF_025837095.1 | NZ_CP107551.1 | 8756485 | 8757675 | 1190 | - | tuf | WP_015105265.1 |
| GCF_037327425.1 | NZ_CP147846.1 | 10407121 | 10408311 | 1190 | - | tuf | WP_033428574.1 |
| GCF_014217765.1 | NZ_CP048813.1 | 4474735 | 4475925 | 1190 | - | tuf | WP_012015126.1 |
| GCF_041228765.1 | NZ_CP150488.1 | 3316157 | 3317344 | 1187 | - | tuf | WP_056128674.1 |
| GCF_000699505.1 | NZ_CP007490.1 | 3303130 | 3304317 | 1187 | - | tuf | WP_012867948.1 |
| GCF_005845365.1 | NZ_CP040509.1 | 2591824 | 2593014 | 1190 | - | tuf | WP_349637686.1 |
| GCF_016726365.1 | NZ_CP068102.1 | 233253 | 234449 | 1196 | + | tuf | WP_006292611.1 |
| GCF_014705945.2 | NZ_CP061538.1 | 367869 | 369053 | 1184 | + | tuf | WP_005962253.1 |
| GCF_000164695.2 | NC_014643.1 | 1365831 | 1367015 | 1184 | - | tuf | WP_050694285.1 |
| GCF_016028855.1 | NZ_CP065738.1 | 1617544 | 1618728 | 1184 | + | tuf | WP_111823850.1 |
| GCF_014705925.2 | NZ_CP061539.1 | 1812049 | 1813236 | 1187 | - | tuf | WP_058237169.1 |
| GCF_014960965.1 | NZ_CP063169.1 | 1851064 | 1852251 | 1187 | - | tuf | WP_350257872.1 |
| GCF_021049285.1 | NZ_CP088017.1 | 524517 | 525707 | 1190 | + | tuf | WP_013137487.1 |
| GCF_021049265.1 | NZ_CP088018.1 | 2480878 | 2482068 | 1190 | - | tuf | WP_130014039.1 |
| GCF_008000995.1 | NZ_CP042828.1 | 3168968 | 3170158 | 1190 | + | tuf | WP_066641554.1 |
| GCF_000231055.2 | NZ_CM001466.1 | 3250488 | 3251675 | 1187 | - | tuf | WP_066499550.1 |
| GCF_000244975.1 | NZ_CM001440.1 | 3129920 | 3131107 | 1187 | - | tuf | WP_229229897.1 |
| GCF_000243395.2 | NZ_CM001484.1 | 3028737 | 3029924 | 1187 | - | tuf | WP_307955630.1 |
| GCF_000244955.1 | NZ_CM001439.1 | 777660 | 778847 | 1187 | + | tuf | WP_066466674.1 |
| GCF_000023865.1 | NC_013159.1 | 2958584 | 2959774 | 1190 | + | tuf | WP_271632383.1 |
| GCF_000062885.1 | NC_009142.1 | 161882 | 163051 | 1169 | - | tuf | WP_424527953.1 |
| GCF_024734405.1 | NZ_CP059556.1 | 1148424 | 1149614 | 1190 | + | tuf | WP_405144533.1 |
| GCF_014697215.1 | NZ_CP031142.1 | 4222403 | 4223593 | 1190 | + | tuf | WP_236539360.1 |
| GCF_025643595.1 | NZ_CP102826.1 | 1272630 | 1273820 | 1190 | + | tuf | WP_013016464.1 |
| GCF_000328705.1 | NC_019673.1 | 3354709 | 3355899 | 1190 | - | tuf | WP_014143915.1 |
| GCF_009498035.1 | NZ_CP034550.1 | 3790740 | 3791930 | 1190 | + | tuf | WP_014144292.1 |
| GCF_000016425.1 | NC_009380.1 | 4868641 | 4869831 | 1190 | - | tuf | WP_131100478.1 |
| GCF_033798665.1 | NZ_CP138359.1 | 4985872 | 4987062 | 1190 | + | tuf | WP_030601270.1 |

| | | | | | | | |
|---|---|---|---|---|---|---|---|
| GCF_000024925.1 | NC_013521.1 | 4070629 | 4071819 | 1190 | - | tuf | WP_033528329.1 |
| GCF_025787225.1 | NZ_CP090958.1 | 8712274 | 8713440 | 1166 | + | tuf | WP_220650152.1 |
| GCF_001042695.1 | NZ_AP012334.1 | 1571827 | 1572996 | 1169 | - | tuf | WP_073895152.1 |
| GCF_005886435.1 | NZ_CP040505.1 | 3697707 | 3698897 | 1190 | - | tuf | WP_073901448.1 |
| GCF_016127995.1 | NZ_CP066065.1 | 4487531 | 4488721 | 1190 | + | tuf | WP_003948649.1 |
| GCF_031191545.1 | NZ_CP133472.1 | 6501599 | 6502768 | 1169 | + | tuf | WP_008412796.1 |
| GCF_900106055.1 | NZ_LT629792.1 | 1106842 | 1108008 | 1166 | - | tuf | WP_055532280.1 |
| GCF_040256745.1 | NZ_CP138335.1 | 4676691 | 4677881 | 1190 | + | tuf | WP_055527121.1 |
| GCF_000092825.1 | NC_014168.1 | 3017131 | 3018321 | 1190 | - | tuf | WP_016473456.1 |
| GCF_006337125.1 | NZ_CP040887.1 | 1489279 | 1490445 | 1166 | - | tuf | WP_198502080.1 |
| GCF_001685415.1 | NZ_CP014989.1 | 5044764 | 5045954 | 1190 | + | tuf | WP_062775123.1 |
| GCF_001577305.1 | NZ_CP014518.1 | 3091192 | 3092382 | 1190 | - | tuf | WP_033272475.1 |
| GCF_020886775.1 | NZ_AP024525.1 | 6372919 | 6374085 | 1166 | + | tuf | WP_037638438.1 |
| GCF_041022165.1 | NZ_CP163302.1 | 1418861 | 1420048 | 1187 | - | tuf | WP_053127763.1 |
| GCF_019285775.1 | NZ_CP079105.1 | 4745393 | 4746583 | 1190 | + | tuf | WP_053135585.1 |
| GCF_025790165.1 | NZ_CP094970.1 | 668023 | 669192 | 1169 | - | tuf | WP_086717753.1 |
| GCF_049674955.1 | NZ_CP187232.1 | 4544315 | 4545505 | 1190 | + | tuf | WP_086717603.1 |
| GCF_049674955.1 | NZ_CP187232.1 | 3361478 | 3362668 | 1190 | - | tuf | WP_011029795.1 |
| GCF_018437505.1 | NZ_CP072467.1 | 7005394 | 7006569 | 1175 | + | tuf | WP_003977508.1 |
| GCF_000024545.1 | NC_013947.1 | 6183514 | 6184704 | 1190 | + | tuf | WP_037952879.1 |
| GCF_000237305.1 | NC_016111.1 | 8869329 | 8870507 | 1178 | + | tuf | WP_086705547.1 |
| GCF_000237305.1 | NC_016111.1 | 1023653 | 1024825 | 1172 | - | tuf | WP_057661547.1 |
| GCF_004323735.1 | NZ_CP036455.1 | 5222055 | 5223245 | 1190 | + | tuf | WP_007445914.1 |
| GCF_045160875.1 | NZ_CP108164.1 | 1973649 | 1974815 | 1166 | - | tuf | WP_126270601.1 |
| GCF_019599145.1 | NZ_CP080647.1 | 6286875 | 6288065 | 1190 | + | tuf | WP_079308664.1 |
| GCF_019599145.1 | NZ_CP080647.1 | 9007077 | 9008267 | 1190 | + | tuf | WP_126275774.1 |
| GCF_025642175.1 | NZ_CP106795.1 | 3198583 | 3199773 | 1190 | - | tuf | WP_030778468.1 |
| GCF_025642175.1 | NZ_CP106795.1 | 6591065 | 6592231 | 1166 | + | tuf | WP_171158408.1 |
| GCF_045039635.1 | NZ_CP108647.1 | 1170375 | 1171565 | 1190 | - | tuf | WP_208876130.1 |
| GCF_045039635.1 | NZ_CP108647.1 | 3459251 | 3460417 | 1166 | - | tuf | WP_208878808.1 |
| GCF_008704395.1 | NZ_CP023695.1 | 1591417 | 1592583 | 1166 | - | tuf | WP_114243352.1 |

| | | | | | | | |
|---|---|---|---|---|---|---|---|
| GCF_008704395.1 | NZ_CP023695.1 | 5367677 | 5368867 | 1190 | + | tuf | WP_093897236.1 |
| GCF_024752535.1 | NZ_CP103060.1 | 5654295 | 5655485 | 1190 | + | tuf | WP_055517363.1 |
| GCF_018141485.1 | NZ_CP060742.1 | 8036833 | 8038008 | 1175 | + | tuf | WP_190853889.1 |
| GCF_018141485.1 | NZ_CP060742.1 | 3484831 | 3486021 | 1190 | - | tuf | WP_006604872.1 |
| GCF_045276965.1 | NZ_CP109234.1 | 7130935 | 7132104 | 1169 | + | tuf | WP_006601669.1 |
| GCF_045276965.1 | NZ_CP109234.1 | 5981306 | 5982496 | 1190 | + | tuf | WP_010986337.1 |
| GCF_001267885.1 | NZ_CP012382.1 | 8388106 | 8389275 | 1169 | + | tuf | WP_010988423.1 |
| GCF_001267885.1 | NZ_CP012382.1 | 3869592 | 3870782 | 1190 | - | tuf | WP_189970634.1 |
| GCF_019933235.1 | NZ_CP082945.1 | 7121357 | 7122535 | 1178 | + | tuf | WP_405441891.1 |
| GCF_019933235.1 | NZ_CP082945.1 | 2821762 | 2822952 | 1190 | - | tuf | WP_028433855.1 |
| GCF_035917655.1 | NZ_CP109102.1 | 7106721 | 7107893 | 1172 | + | tuf | WP_197353853.1 |
| GCF_035917655.1 | NZ_CP109102.1 | 3849316 | 3850506 | 1190 | - | tuf | WP_086767450.1 |
| GCF_045161845.1 | NZ_CP108856.1 | 7694225 | 7695391 | 1166 | + | tuf | WP_086770480.1 |
| GCF_045161845.1 | NZ_CP108856.1 | 780093 | 781265 | 1172 | - | tuf | WP_419993009.1 |
| GCF_001434355.1 | NZ_CM003601.1 | 5812355 | 5813545 | 1190 | + | tuf | WP_419997205.1 |
| GCF_001434355.1 | NZ_CM003601.1 | 324078 | 325268 | 1190 | - | tuf | WP_158916993.1 |
| GCF_003955715.1 | NZ_CP034463.1 | 1991108 | 1992283 | 1175 | - | tuf | WP_158919144.1 |
| GCF_003955715.1 | NZ_CP034463.1 | 5615280 | 5616470 | 1190 | + | tuf | WP_158923752.1 |
| GCF_003955715.1 | NZ_CP034463.1 | 8593884 | 8595074 | 1190 | + | tuf | WP_158927947.1 |
| GCF_013046785.1 | NZ_CP053189.1 | 3506551 | 3507741 | 1190 | - | tuf | WP_176163497.1 |
| GCF_013046785.1 | NZ_CP053189.1 | 3971875 | 3973065 | 1190 | - | tuf | WP_053759521.1 |
| GCF_003355155.1 | NZ_CP031320.1 | 760381 | 761553 | 1172 | - | tuf | WP_030118935.1 |
| GCF_003355155.1 | NZ_CP031320.1 | 4865756 | 4866946 | 1190 | + | tuf | WP_030112819.1 |
| GCF_003330865.1 | NZ_CP027306.1 | 4325507 | 4326697 | 1190 | + | tuf | WP_142193697.1 |
| GCF_003330865.1 | NZ_CP027306.1 | 6740771 | 6741937 | 1166 | + | tuf | WP_142233568.1 |
| GCF_014701115.1 | NZ_AP023440.1 | 3787971 | 3789161 | 1190 | - | tuf | WP_225100422.1 |
| GCF_014701115.1 | NZ_AP023440.1 | 7674714 | 7675889 | 1175 | + | tuf | WP_270084991.1 |
| GCF_000280865.2 | NZ_CP072931.1 | 3914204 | 3915394 | 1190 | - | tuf | WP_159481495.1 |
| GCF_000280865.2 | NZ_CP072931.1 | 8213783 | 8214952 | 1169 | + | tuf | WP_328691864.1 |
| GCF_000009765.2 | NC_003155.5 | 4003735 | 4004925 | 1190 | - | tuf | WP_045559444.1 |
| GCF_000009765.2 | NC_003155.5 | 8547756 | 8548922 | 1166 | + | tuf | WP_086802747.1 |

| | | | | | | | |
|---|---|---|---|---|---|---|---|
| GCF_045161815.1 | NZ_CP108863.1 | 4251229 | 4252419 | 1190 | - | tuf | WP_328780959.1 |
| GCF_045161815.1 | NZ_CP108863.1 | 8660016 | 8661182 | 1166 | + | tuf | WP_328784167.1 |
| GCF_015910445.1 | NZ_CP048882.1 | 3119243 | 3120421 | 1178 | - | tuf | WP_189785065.1 |
| GCF_015910445.1 | NZ_CP048882.1 | 7486587 | 7487777 | 1190 | - | tuf | WP_115906452.1 |
| GCF_036172715.1 | NZ_CP108038.1 | 619102 | 620277 | 1175 | - | tuf | WP_347227097.1 |
| GCF_036172715.1 | NZ_CP108038.1 | 2613272 | 2614462 | 1190 | - | tuf | WP_019433401.1 |
| GCF_048572725.1 | NZ_CP183837.1 | 791889 | 793055 | 1166 | - | tuf | WP_306051560.1 |
| GCF_048572725.1 | NZ_CP183837.1 | 4756697 | 4757887 | 1190 | + | tuf | WP_018550640.1 |
| GCF_009796285.1 | NZ_CP047020.1 | 3310328 | 3311518 | 1190 | - | tuf | WP_279334448.1 |
| GCF_009796285.1 | NZ_CP047020.1 | 826843 | 828009 | 1166 | - | tuf | WP_058941252.1 |
| GCF_009796285.1 | NZ_CP047020.1 | 4007721 | 4008911 | 1190 | + | tuf | WP_058942712.1 |
| GCF_009796285.1 | NZ_CP047020.1 | 4109436 | 4110626 | 1190 | - | tuf | WP_030046358.1 |
| GCF_014295035.1 | NZ_CP060404.1 | 8306831 | 8307997 | 1166 | + | tuf | WP_125525669.1 |
| GCF_003261055.1 | NZ_CP030073.1 | 1055370 | 1056536 | 1166 | - | tuf | WP_290348093.1 |
| GCF_016906185.1 | NZ_CP070242.1 | 4650370 | 4651560 | 1190 | + | tuf | WP_017949669.1 |
| GCF_016906185.1 | NZ_CP070242.1 | 1269278 | 1270447 | 1169 | - | tuf | WP_189697147.1 |
| GCF_006716135.1 | NZ_CP022310.1 | 4742034 | 4743224 | 1190 | + | tuf | WP_189700577.1 |
| GCF_006716135.1 | NZ_CP022310.1 | 4548960 | 4550150 | 1190 | + | tuf | WP_030208973.1 |
| GCF_027625935.1 | NZ_CP115300.1 | 1440113 | 1441279 | 1166 | - | tuf | WP_269657823.1 |
| GCF_027625935.1 | NZ_CP115300.1 | 5324525 | 5325715 | 1190 | + | tuf | WP_269661065.1 |
| GCF_036178885.1 | NZ_CP108073.1 | 8137315 | 8138505 | 1190 | + | tuf | WP_269663310.1 |
| GCF_036178885.1 | NZ_CP108073.1 | 1201664 | 1202842 | 1178 | - | tuf | WP_338694159.1 |
| GCF_014852565.2 | NZ_CP119182.1 | 3530125 | 3531315 | 1190 | - | tuf | WP_338697311.1 |
| GCF_014852565.2 | NZ_CP119182.1 | 1032547 | 1033713 | 1166 | - | tuf | WP_056795811.1 |
| GCF_036177905.1 | NZ_CP108065.1 | 4493290 | 4494480 | 1190 | + | tuf | WP_056796624.1 |
| GCF_036177905.1 | NZ_CP108065.1 | 2517989 | 2519179 | 1190 | - | tuf | WP_003956470.1 |
| GCF_002214185.1 | NZ_CP022161.1 | 3399861 | 3401051 | 1190 | - | tuf | WP_007443930.1 |
| GCF_002214185.1 | NZ_CP022161.1 | 7018106 | 7019293 | 1187 | + | tuf | WP_153180862.1 |
| GCF_025133285.1 | NZ_CP104005.1 | 1315668 | 1316834 | 1166 | - | tuf | WP_184843278.1 |
| GCF_025133285.1 | NZ_CP104005.1 | 5325778 | 5326968 | 1190 | + | tuf | WP_030839706.1 |
| GCF_030719295.1 | NZ_CP120997.1 | 830454 | 831644 | 1190 | - | tuf | WP_126388458.1 |

| | | | | | | | |
|---|---|---|---|---|---|---|---|
| GCF_030719295.1 | NZ_CP120997.1 | 1680519 | 1681685 | 1166 | - | tuf | WP_126389802.1 |
| GCF_029760955.1 | NZ_CP121682.1 | 6044044 | 6045234 | 1190 | + | tuf | WP_030046358.1 |
| GCF_024600715.1 | NZ_CP102332.1 | 3156364 | 3157554 | 1190 | - | tuf | WP_018490814.1 |
| GCF_024600715.1 | NZ_CP102332.1 | 7252039 | 7253211 | 1172 | + | tuf | WP_317822988.1 |
| GCF_008704715.1 | NZ_CP023689.1 | 1653348 | 1654514 | 1166 | - | tuf | WP_208031027.1 |
| GCF_008704715.1 | NZ_CP023689.1 | 5185980 | 5187170 | 1190 | + | tuf | WP_014674366.1 |
| GCF_030406905.1 | NZ_CP129137.1 | 1527776 | 1528942 | 1166 | - | tuf | WP_263228778.1 |
| GCF_030406905.1 | NZ_CP129137.1 | 5198072 | 5199262 | 1190 | + | tuf | WP_103841183.1 |
| GCF_015160875.1 | NZ_CP063374.1 | 3263511 | 3264701 | 1190 | - | tuf | WP_121787379.1 |
| GCF_015160875.1 | NZ_CP063374.1 | 1188585 | 1189775 | 1190 | - | tuf | WP_015655872.1 |
| GCF_009299385.1 | NZ_CP023693.1 | 4195187 | 4196377 | 1190 | - | tuf | WP_015658472.1 |
| GCF_027270315.1 | NZ_CP114413.1 | 8147026 | 8148192 | 1166 | + | tuf | WP_015661980.1 |
| GCF_027270315.1 | NZ_CP114413.1 | 1721463 | 1722629 | 1166 | - | tuf | WP_239752283.1 |
| GCF_027270315.1 | NZ_CP114413.1 | 6274890 | 6276080 | 1190 | + | tuf | WP_115906452.1 |
| GCF_036967205.1 | NZ_CP146022.1 | 941318 | 942487 | 1169 | - | tuf | WP_250057664.1 |
| GCF_036967205.1 | NZ_CP146022.1 | 5119167 | 5120357 | 1190 | + | tuf | WP_006604872.1 |
| GCF_036345975.1 | NZ_CP107793.1 | 710403 | 711593 | 1190 | - | tuf | WP_099501388.1 |
| GCF_036345975.1 | NZ_CP107793.1 | 3576890 | 3578080 | 1190 | - | tuf | WP_099500619.1 |
| GCF_005519465.1 | NZ_CP027858.1 | 7504493 | 7505659 | 1166 | + | tuf | WP_099505544.1 |
| GCF_045041415.1 | NZ_CP108218.1 | 4117341 | 4118531 | 1190 | - | tuf | WP_160506655.1 |
| GCF_045041415.1 | NZ_CP108218.1 | 7948906 | 7950072 | 1166 | + | tuf | WP_249591290.1 |
| GCF_031348265.1 | NZ_CP133771.1 | 3533686 | 3534876 | 1190 | - | tuf | WP_030653602.1 |
| GCF_031348265.1 | NZ_CP133771.1 | 6837209 | 6838387 | 1178 | + | tuf | WP_031149938.1 |
| GCF_003963535.1 | NZ_CP034539.1 | 1723928 | 1725094 | 1166 | - | tuf | WP_086749758.1 |
| GCF_003963535.1 | NZ_CP034539.1 | 6386984 | 6388174 | 1190 | + | tuf | WP_037694088.1 |
| GCF_003963535.1 | NZ_CP034539.1 | 4038328 | 4039518 | 1190 | - | tuf | WP_153289127.1 |
| GCF_033109485.1 | NZ_CP137002.1 | 7734628 | 7735806 | 1178 | + | tuf | WP_153291824.1 |
| GCF_033109485.1 | NZ_CP137002.1 | 3539435 | 3540601 | 1166 | + | tuf | WP_194048385.1 |
| GCF_017526105.1 | NZ_CP071839.1 | 9197753 | 9198943 | 1190 | - | tuf | WP_194042007.1 |
| GCF_017526105.1 | NZ_CP071839.1 | 2981164 | 2982354 | 1190 | - | tuf | WP_156692845.1 |
| GCF_025642135.1 | NZ_CP106793.1 | 6311930 | 6313096 | 1166 | + | tuf | WP_156695504.1 |

| | | | | | | | |
|---|---|---|---|---|---|---|---|
| GCF_025642135.1 | NZ_CP106793.1 | 4219949 | 4221139 | 1190 | - | tuf | WP_069779222.1 |
| GCF_003675325.1 | NZ_CP033073.1 | 8087816 | 8088994 | 1178 | + | tuf | WP_069782123.1 |
| GCF_000349325.1 | NC_020504.1 | 806022 | 807212 | 1190 | + | tuf | WP_242329288.1 |
| GCF_000349325.1 | NC_020504.1 | 4965115 | 4966281 | 1166 | - | tuf | WP_242332653.1 |
| GCF_000349325.1 | NC_020504.1 | 740649 | 741821 | 1172 | - | tuf | WP_031132976.1 |
| GCF_022385335.1 | NZ_CP092431.1 | 4037797 | 4038987 | 1190 | + | tuf | WP_031134103.1 |
| GCF_022385335.1 | NZ_CP092431.1 | 1166727 | 1167896 | 1169 | - | tuf | WP_176116527.1 |
| GCF_045208505.1 | NZ_CP109339.1 | 4588665 | 4589855 | 1190 | + | tuf | WP_004927329.1 |
| GCF_045208505.1 | NZ_CP109339.1 | 5301884 | 5303074 | 1190 | + | tuf | WP_026058274.1 |
| GCF_002946835.1 | NZ_CP026652.1 | 918080 | 919252 | 1172 | - | tuf | WP_187739379.1 |
| GCF_002946835.1 | NZ_CP026652.1 | 4292403 | 4293593 | 1190 | + | tuf | WP_187741868.1 |
| GCF_002946835.1 | NZ_CP026652.1 | 4214477 | 4215667 | 1190 | - | tuf | WP_046927441.1 |
| GCF_023498005.1 | NZ_CP097289.1 | 8246674 | 8247843 | 1169 | + | tuf | WP_083109133.1 |
| GCF_023498005.1 | NZ_CP097289.1 | 1031514 | 1032686 | 1172 | - | tuf | WP_010062273.1 |
| GCF_036170895.1 | NZ_CP108036.1 | 4842159 | 4843349 | 1190 | + | tuf | WP_010058805.1 |
| GCF_036170895.1 | NZ_CP108036.1 | 3367757 | 3368947 | 1190 | - | tuf | WP_014154597.1 |
| GCF_036346855.1 | NZ_CP107841.1 | 7257829 | 7258995 | 1166 | + | tuf | WP_147962508.1 |
| GCF_036346855.1 | NZ_CP107841.1 | 3173569 | 3174756 | 1187 | + | tuf | WP_232269631.1 |
| GCF_009498275.1 | NZ_CP045643.1 | 3466766 | 3467956 | 1190 | + | tuf | WP_232269877.1 |
| GCF_009498275.1 | NZ_CP045643.1 | 5564741 | 5565913 | 1172 | + | tuf | WP_232271590.1 |
| GCF_015160855.1 | NZ_CP063373.1 | 4400466 | 4401656 | 1190 | + | tuf | WP_037798043.1 |
| GCF_015160855.1 | NZ_CP063373.1 | 6086019 | 6087197 | 1178 | + | tuf | WP_037798572.1 |
| GCF_009739905.1 | NZ_CP034279.1 | 4115284 | 4116474 | 1190 | + | tuf | WP_100452283.1 |
| GCF_009739905.1 | NZ_CP034279.1 | 5954347 | 5955516 | 1169 | + | tuf | WP_100455547.1 |
| GCF_001735805.1 | NZ_CP017248.1 | 8011761 | 8012951 | 1190 | + | tuf | WP_286256221.1 |
| GCF_001735805.1 | NZ_CP017248.1 | 10531557 | 10532723 | 1166 | + | tuf | WP_286258639.1 |
| GCF_022647665.1 | NZ_CP071872.1 | 11239791 | 11240981 | 1190 | + | tuf | WP_286259382.1 |
| GCF_022647665.1 | NZ_CP071872.1 | 3066592 | 3067782 | 1190 | - | tuf | WP_006140924.1 |
| GCF_008704425.1 | NZ_CP023696.1 | 6367636 | 6368847 | 1211 | + | tuf | WP_367513424.1 |
| GCF_008704425.1 | NZ_CP023696.1 | 878029 | 879219 | 1190 | + | tuf | WP_067299840.1 |
| GCF_020783575.1 | NZ_CP086173.1 | 7656111 | 7657280 | 1169 | - | tuf | WP_067311995.1 |

| | | | | | | | |
|---|---|---|---|---|---|---|---|
| GCF_020783575.1 | NZ_CP086173.1 | 3674378 | 3675568 | 1190 | - | tuf | WP_037657541.1 |
| GCF_015710995.1 | NZ_CP059991.1 | 7064214 | 7065389 | 1175 | + | tuf | WP_037658066.1 |
| GCF_014489615.1 | NZ_CP060825.1 | 3974453 | 3975643 | 1190 | - | tuf | WP_007384061.1 |
| GCF_014489615.1 | NZ_CP060825.1 | 8032315 | 8033481 | 1166 | + | tuf | WP_062026168.1 |
| GCF_002082195.1 | NZ_CP020569.1 | 714014 | 715183 | 1169 | - | tuf | WP_248861818.1 |
| GCF_002082195.1 | NZ_CP020569.1 | 5494309 | 5495499 | 1190 | + | tuf | WP_248865552.1 |
| GCF_000261345.2 | NZ_CP013738.1 | 3289911 | 3291101 | 1190 | - | tuf | WP_028439262.1 |
| GCF_000261345.2 | NZ_CP013738.1 | 6854717 | 6855883 | 1166 | + | tuf | WP_037875452.1 |
| GCF_030719215.1 | NZ_CP120983.1 | 515912 | 517087 | 1175 | - | tuf | WP_019434668.1 |
| GCF_030719215.1 | NZ_CP120983.1 | 2091031 | 2092221 | 1190 | - | tuf | WP_019433401.1 |
| GCF_021216675.1 | NZ_CP086120.1 | 1473488 | 1474654 | 1166 | - | tuf | WP_175431301.1 |
| GCF_021216675.1 | NZ_CP086120.1 | 5244856 | 5246046 | 1190 | + | tuf | WP_030839706.1 |
| GCF_021216675.1 | NZ_CP086120.1 | 1226090 | 1227256 | 1166 | - | tuf | WP_326751547.1 |
| GCF_027942015.1 | NZ_CP115936.1 | 5056542 | 5057732 | 1190 | + | tuf | WP_326754257.1 |
| GCF_027942015.1 | NZ_CP115936.1 | 1039661 | 1040827 | 1166 | - | tuf | WP_230899593.1 |
| GCF_045039555.1 | NZ_CP108653.1 | 4812226 | 4813416 | 1190 | + | tuf | WP_127910221.1 |
| GCF_045039555.1 | NZ_CP108653.1 | 5453697 | 5454887 | 1190 | + | tuf | WP_030835078.1 |
| GCF_030294945.1 | NZ_AP018448.1 | 8023170 | 8024348 | 1178 | + | tuf | WP_060945410.1 |
| GCF_030294945.1 | NZ_AP018448.1 | 1408066 | 1409232 | 1166 | - | tuf | WP_033307318.1 |
| GCF_030294945.1 | NZ_AP018448.1 | 5420101 | 5421291 | 1190 | + | tuf | WP_033308978.1 |
| GCF_037099765.1 | NZ_CP146837.1 | 3581028 | 3582218 | 1190 | - | tuf | WP_208899320.1 |
| GCF_037099765.1 | NZ_CP146837.1 | 7209402 | 7210571 | 1169 | + | tuf | WP_208902093.1 |
| GCF_001542625.1 | NZ_CP016279.1 | 4483364 | 4484554 | 1190 | - | tuf | WP_238573626.1 |
| GCF_001542625.1 | NZ_CP016279.1 | 8782840 | 8784009 | 1169 | + | tuf | WP_238576793.1 |
| GCF_014673495.1 | NZ_CP051006.1 | 2228254 | 2229426 | 1172 | - | tuf | WP_044568208.1 |
| GCF_014673495.1 | NZ_CP051006.1 | 5535106 | 5536296 | 1190 | - | tuf | WP_037952879.1 |
| GCF_003595235.1 | NZ_CP032427.1 | 3616421 | 3617611 | 1190 | - | tuf | WP_193482362.1 |
| GCF_003595235.1 | NZ_CP032427.1 | 7855865 | 7857031 | 1166 | + | tuf | WP_193482110.1 |
| GCF_023277745.1 | NZ_CP086322.1 | 1060544 | 1061740 | 1196 | - | tuf | WP_328921649.1 |
| GCF_023277745.1 | NZ_CP086322.1 | 4364458 | 4365648 | 1190 | + | tuf | WP_006140924.1 |
| GCF_045207185.1 | NZ_CP109044.1 | 4173265 | 4174455 | 1190 | - | tuf | WP_055553487.1 |

| | | | | | | | |
|---|---|---|---|---|---|---|---|
| GCF_045207185.1 | NZ_CP109044.1 | 8605488 | 8606669 | 1181 | - | tuf | WP_150494475.1 |
| GCF_013364095.1 | NZ_CP054938.1 | 3231836 | 3233026 | 1190 | - | tuf | WP_193482362.1 |
| GCF_013364095.1 | NZ_CP054938.1 | 7270745 | 7271911 | 1166 | + | tuf | WP_215122779.1 |
| GCF_004803895.1 | NZ_CP021978.1 | 4555073 | 4556263 | 1190 | + | tuf | WP_047138873.1 |
| GCF_004803895.1 | NZ_CP021978.1 | 3648640 | 3649830 | 1190 | - | tuf | WP_073732686.1 |
| GCF_035918035.1 | NZ_CP109134.1 | 7641432 | 7642598 | 1166 | + | tuf | WP_306091754.1 |
| GCF_035918035.1 | NZ_CP109134.1 | 3689197 | 3690387 | 1190 | - | tuf | WP_030230628.1 |
| GCF_021028635.1 | NZ_CP086119.1 | 6910666 | 6911841 | 1175 | + | tuf | WP_030236668.1 |
| GCF_021028635.1 | NZ_CP086119.1 | 3565542 | 3566732 | 1190 | - | tuf | WP_029386959.1 |
| GCF_037126205.1 | NZ_CP146988.1 | 6761149 | 6762315 | 1166 | + | tuf | WP_029383553.1 |
| GCF_037126205.1 | NZ_CP146988.1 | 3149004 | 3150194 | 1190 | - | tuf | WP_161308202.1 |
| GCF_036287475.1 | NZ_CP143088.1 | 6817585 | 6818760 | 1175 | + | tuf | WP_167034477.1 |
| GCF_036287475.1 | NZ_CP143088.1 | 894736 | 895905 | 1169 | - | tuf | WP_109895044.1 |
| GCF_001027185.1 | NZ_CP011497.1 | 4873593 | 4874783 | 1190 | + | tuf | WP_018090904.1 |
| GCF_001027185.1 | NZ_CP011497.1 | 705050 | 706240 | 1190 | + | tuf | WP_159329772.1 |
| GCF_022221585.1 | NZ_CP084681.1 | 5164929 | 5166104 | 1175 | - | tuf | WP_271414785.1 |
| GCF_022221585.1 | NZ_CP084681.1 | 6451157 | 6452347 | 1190 | + | tuf | WP_055610510.1 |
| GCF_042466515.1 | NZ_CP136563.1 | 9082914 | 9084092 | 1178 | + | tuf | WP_200399672.1 |
| GCF_042466515.1 | NZ_CP136563.1 | 1825896 | 1827062 | 1166 | - | tuf | WP_067428449.1 |
| GCF_031593255.1 | NZ_CP134213.1 | 6016831 | 6018021 | 1190 | + | tuf | WP_067437927.1 |
| GCF_031593255.1 | NZ_CP134213.1 | 9206271 | 9207461 | 1190 | + | tuf | WP_067443430.1 |
| GCF_036189845.1 | NZ_CP108326.1 | 2061264 | 2062439 | 1175 | - | tuf | WP_227293504.1 |
| GCF_036189845.1 | NZ_CP108326.1 | 5699590 | 5700780 | 1190 | + | tuf | WP_116511070.1 |
| GCF_008704495.1 | NZ_CP023699.1 | 1358972 | 1360147 | 1175 | - | tuf | WP_108147482.1 |
| GCF_008704495.1 | NZ_CP023699.1 | 5159882 | 5161072 | 1190 | + | tuf | WP_108151498.1 |
| GCF_018604545.1 | NZ_CP075896.1 | 1411680 | 1412867 | 1187 | - | tuf | WP_311034310.1 |
| GCF_018604545.1 | NZ_CP075896.1 | 5741068 | 5742258 | 1190 | + | tuf | WP_030835078.1 |
| GCF_003428925.1 | NZ_CP031742.1 | 3974059 | 3975249 | 1190 | - | tuf | WP_372408336.1 |
| GCF_030719275.1 | NZ_CP120992.1 | 3876082 | 3877272 | 1190 | - | tuf | WP_033269709.1 |
| GCF_030719275.1 | NZ_CP120992.1 | 7749085 | 7750254 | 1169 | + | tuf | WP_077194279.1 |
| GCF_002803845.1 | NZ_CP024985.1 | 5574010 | 5575200 | 1190 | + | tuf | WP_069865218.1 |

| | | | | | | | |
|---|---|---|---|---|---|---|---|
| GCF_002803845.1 | NZ_CP024985.1 | 8373025 | 8374194 | 1169 | + | tuf | WP_079259781.1 |
| GCF_001013905.1 | NZ_LN831790.1 | 1239713 | 1240891 | 1178 | - | tuf | WP_365494949.1 |
| GCF_001013905.1 | NZ_LN831790.1 | 4378805 | 4379995 | 1190 | + | tuf | WP_030653602.1 |
| GCF_011694815.1 | NZ_CP050177.1 | 4317216 | 4318406 | 1190 | + | tuf | WP_086158608.1 |
| GCF_011694815.1 | NZ_CP050177.1 | 916179 | 917351 | 1172 | - | tuf | WP_031121751.1 |
| GCF_027497475.1 | NZ_CP114202.1 | 4780075 | 4781265 | 1190 | + | tuf | WP_015610719.1 |
| GCF_027497475.1 | NZ_CP114202.1 | 2334108 | 2335286 | 1178 | - | tuf | WP_014181375.1 |
| GCF_027947595.1 | NZ_CP116257.1 | 5618500 | 5619690 | 1190 | - | tuf | WP_014178750.1 |
| GCF_027947595.1 | NZ_CP116257.1 | 1467389 | 1468579 | 1190 | - | tuf | WP_328508851.1 |
| GCF_016598615.1 | NZ_CP066831.1 | 2647986 | 2649182 | 1196 | - | tuf | WP_060895448.1 |
| GCF_016598615.1 | NZ_CP066831.1 | 6953748 | 6954938 | 1190 | + | tuf | WP_060900339.1 |
| GCF_001685355.1 | NZ_CP016438.1 | 4235439 | 4236629 | 1190 | + | tuf | WP_185943959.1 |
| GCF_001685355.1 | NZ_CP016438.1 | 693587 | 694777 | 1190 | + | tuf | WP_037657541.1 |
| GCF_001685355.1 | NZ_CP016438.1 | 5653306 | 5654481 | 1175 | - | tuf | WP_128789076.1 |
| GCF_020616555.1 | NZ_CP085173.1 | 1238561 | 1239727 | 1166 | - | tuf | WP_107478422.1 |
| GCF_020616555.1 | NZ_CP085173.1 | 4640751 | 4641941 | 1190 | + | tuf | WP_043381065.1 |
| GCF_003054555.1 | NZ_CP026304.1 | 3472273 | 3473463 | 1190 | - | tuf | WP_130878980.1 |
| GCF_003054555.1 | NZ_CP026304.1 | 1014209 | 1015375 | 1166 | - | tuf | WP_108705852.1 |
| GCF_031679605.1 | NZ_CP117522.1 | 4457415 | 4458605 | 1190 | + | tuf | WP_108708255.1 |
| GCF_031679605.1 | NZ_CP117522.1 | 935656 | 936825 | 1169 | - | tuf | WP_109895044.1 |
| GCF_041519305.1 | NZ_CP167927.1 | 5330948 | 5332138 | 1190 | + | tuf | WP_018090904.1 |
| GCF_001984445.1 | NZ_CP019457.1 | 3073158 | 3074348 | 1190 | - | tuf | WP_150488209.1 |
| GCF_001984445.1 | NZ_CP019457.1 | 6776558 | 6777724 | 1166 | + | tuf | WP_150490975.1 |
| GCF_002591335.1 | NZ_CP023992.1 | 860692 | 861867 | 1175 | - | tuf | WP_329081163.1 |
| GCF_002591335.1 | NZ_CP023992.1 | 5018649 | 5019839 | 1190 | + | tuf | WP_069627353.1 |
| GCF_049570255.1 | NZ_CP186950.1 | 1386089 | 1387255 | 1166 | - | tuf | WP_043438288.1 |
| GCF_049570255.1 | NZ_CP186950.1 | 4420278 | 4421468 | 1190 | + | tuf | WP_043443001.1 |
| GCF_020410765.1 | NZ_CP084541.1 | 2075154 | 2076332 | 1178 | - | tuf | WP_189746167.1 |
| GCF_013364315.1 | NZ_CP054926.1 | 5412463 | 5413653 | 1190 | + | tuf | WP_030008455.1 |
| GCF_013364315.1 | NZ_CP054926.1 | 4616062 | 4617252 | 1190 | - | tuf | WP_205365570.1 |
| GCF_025908395.1 | NZ_CP109672.1 | 8769252 | 8770424 | 1172 | - | tuf | WP_205368584.1 |

| | | | | | | | |
|---|---|---|---|---|---|---|---|
| GCF_025908395.1 | NZ_CP109672.1 | 3623509 | 3624699 | 1190 | - | tuf | WP_031045706.1 |
| GCF_036073155.1 | NZ_CP107957.1 | 7253896 | 7255083 | 1187 | + | tuf | WP_194278527.1 |
| GCF_036073155.1 | NZ_CP107957.1 | 549749 | 550915 | 1166 | - | tuf | WP_114658086.1 |
| GCF_036073155.1 | NZ_CP107957.1 | 2775997 | 2777187 | 1190 | - | tuf | WP_114659679.1 |
| GCF_020099395.1 | NZ_CP083590.1 | 1053961 | 1055139 | 1178 | - | tuf | WP_064726775.1 |
| GCF_025231465.1 | NZ_CP046623.1 | 4322435 | 4323625 | 1190 | + | tuf | WP_064729336.1 |
| GCF_025231465.1 | NZ_CP046623.1 | 2134984 | 2136153 | 1169 | - | tuf | WP_055617114.1 |
| GCF_046628145.1 | NZ_CP176874.1 | 4764581 | 4765771 | 1190 | - | tuf | WP_055610510.1 |
| GCF_046628145.1 | NZ_CP176874.1 | 1271846 | 1273024 | 1178 | - | tuf | WP_152167343.1 |
| GCF_037414505.1 | NZ_CP147867.1 | 6576367 | 6577557 | 1190 | + | tuf | WP_037694088.1 |
| GCF_003074055.1 | NZ_CP029043.1 | 807582 | 808772 | 1190 | + | tuf | WP_073491913.1 |
| GCF_003074055.1 | NZ_CP029043.1 | 6094883 | 6096049 | 1166 | - | tuf | WP_252552894.1 |
| GCF_027626975.1 | NZ_CP114203.1 | 2440237 | 2441427 | 1190 | - | tuf | WP_121518463.1 |
| GCF_027626975.1 | NZ_CP114203.1 | 3968518 | 3969708 | 1190 | - | tuf | WP_006604872.1 |
| GCF_008704555.1 | NZ_CP023702.1 | 7911972 | 7913141 | 1169 | + | tuf | WP_085922196.1 |
| GCF_008704555.1 | NZ_CP023702.1 | 724616 | 725785 | 1169 | - | tuf | WP_039654124.1 |
| GCF_036227585.1 | NZ_CP109483.1 | 2792998 | 2794188 | 1190 | - | tuf | WP_039656254.1 |
| GCF_036227585.1 | NZ_CP109483.1 | 1253965 | 1255131 | 1166 | - | tuf | WP_219567911.1 |
| GCF_008704995.1 | NZ_CP023747.1 | 5463120 | 5464310 | 1190 | + | tuf | WP_219573674.1 |
| GCF_008704995.1 | NZ_CP023747.1 | 1229623 | 1230789 | 1166 | - | tuf | WP_055609984.1 |
| GCF_017639205.1 | NZ_CP071139.1 | 4680889 | 4682079 | 1190 | + | tuf | WP_055605682.1 |
| GCF_017639205.1 | NZ_CP071139.1 | 788633 | 789823 | 1190 | + | tuf | WP_033297591.1 |
| GCF_016917755.1 | NZ_CP070326.1 | 5095364 | 5096530 | 1166 | - | tuf | WP_203184889.1 |
| GCF_016917755.1 | NZ_CP070326.1 | 1307698 | 1308864 | 1166 | - | tuf | WP_005320257.1 |
| GCF_015244315.1 | NZ_CP043317.1 | 4788783 | 4789973 | 1190 | + | tuf | WP_005313609.1 |
| GCF_015244315.1 | NZ_CP043317.1 | 108690 | 109862 | 1172 | - | tuf | WP_228077585.1 |
| GCF_003344965.1 | NZ_CP031194.1 | 2513787 | 2514977 | 1190 | - | tuf | WP_062207500.1 |
| GCF_003344965.1 | NZ_CP031194.1 | 1681830 | 1682996 | 1166 | - | tuf | WP_405946601.1 |
| GCF_001660045.1 | NZ_CP015866.1 | 6490923 | 6492113 | 1190 | + | tuf | WP_019055493.1 |
| GCF_001660045.1 | NZ_CP015866.1 | 10058098 | 10059288 | 1190 | + | tuf | WP_366093629.1 |
| GCF_035957715.1 | NZ_CP108382.1 | 4217187 | 4218377 | 1190 | - | tuf | WP_007384061.1 |

| | | | | | | | |
|---|---|---|---|---|---|---|---|
| GCF_035957715.1 | NZ_CP108382.1 | 8130446 | 8131612 | 1166 | + | tuf | WP_031059312.1 |
| GCF_009184865.1 | NZ_CP045096.1 | 1501935 | 1503101 | 1166 | - | tuf | WP_405505022.1 |
| GCF_009184865.1 | NZ_CP045096.1 | 5648865 | 5650055 | 1190 | + | tuf | WP_189728731.1 |
| GCF_023923245.1 | NZ_CP099468.1 | 1510075 | 1511241 | 1166 | - | tuf | WP_062925618.1 |
| GCF_023923245.1 | NZ_CP099468.1 | 5434881 | 5436071 | 1190 | + | tuf | WP_030839706.1 |
| GCF_040746085.1 | NZ_CP160458.1 | 892149 | 893318 | 1169 | - | tuf | WP_146479038.1 |
| GCF_008704855.1 | NZ_CP023691.1 | 4736517 | 4737707 | 1190 | + | tuf | WP_006347239.1 |
| GCF_008704855.1 | NZ_CP023691.1 | 2008324 | 2009496 | 1172 | - | tuf | WP_020866656.1 |
| GCF_000802245.2 | NZ_CP021080.1 | 5240153 | 5241343 | 1190 | - | tuf | WP_020869272.1 |
| GCF_000802245.2 | NZ_CP021080.1 | 3282340 | 3283530 | 1190 | - | tuf | WP_058046830.1 |
| GCF_030719255.1 | NZ_CP120990.1 | 1066296 | 1067468 | 1172 | - | tuf | WP_003987131.1 |
| GCF_030719255.1 | NZ_CP120990.1 | 4167527 | 4168717 | 1190 | - | tuf | WP_004571815.1 |
| GCF_008704445.1 | NZ_CP023697.1 | 1499611 | 1500786 | 1175 | - | tuf | WP_030976330.1 |
| GCF_008704445.1 | NZ_CP023697.1 | 4853879 | 4855069 | 1190 | + | tuf | WP_306692931.1 |
| GCF_016804005.1 | NZ_CP051486.1 | 3911256 | 3912446 | 1190 | - | tuf | WP_187747791.1 |
| GCF_016804005.1 | NZ_CP051486.1 | 7868402 | 7869571 | 1169 | + | tuf | WP_187750904.1 |
| GCF_014070435.1 | NZ_CP059696.1 | 8478102 | 8479292 | 1190 | + | tuf | WP_187751392.1 |
| GCF_014070435.1 | NZ_CP059696.1 | 3768425 | 3769615 | 1190 | - | tuf | WP_023547407.1 |
| GCF_020740535.1 | NZ_CP082362.1 | 8180785 | 8181957 | 1172 | + | tuf | WP_023552392.1 |
| GCF_020740535.1 | NZ_CP082362.1 | 2877893 | 2879068 | 1175 | + | tuf | WP_109030204.1 |
| GCF_045162825.1 | NZ_CP108676.1 | 7657671 | 7658861 | 1190 | - | tuf | WP_011029795.1 |
| GCF_045162825.1 | NZ_CP108676.1 | 763359 | 764528 | 1169 | - | tuf | WP_185392813.1 |
| GCF_045162825.1 | NZ_CP108676.1 | 2592940 | 2594130 | 1190 | - | tuf | WP_100452283.1 |
| GCF_036345035.1 | NZ_CP107755.1 | 4175900 | 4177090 | 1190 | - | tuf | WP_013001430.1 |
| GCF_036345035.1 | NZ_CP107755.1 | 8586665 | 8587831 | 1166 | + | tuf | WP_013005184.1 |
| GCF_045040895.1 | NZ_CP108341.1 | 575079 | 576326 | 1247 | - | tuf | WP_051871267.1 |
| GCF_045040895.1 | NZ_CP108341.1 | 3843770 | 3844960 | 1190 | - | tuf | WP_030619950.1 |
| GCF_001611795.1 | NZ_CP015098.1 | 3479077 | 3480267 | 1190 | - | tuf | WP_030351429.1 |
| GCF_001611795.1 | NZ_CP015098.1 | 7251531 | 7252697 | 1166 | + | tuf | WP_326613065.1 |
| GCF_007856155.1 | NZ_CP042266.1 | 751046 | 752212 | 1166 | - | tuf | WP_031181948.1 |
| GCF_007856155.1 | NZ_CP042266.1 | 3677897 | 3679087 | 1190 | + | tuf | WP_031180867.1 |

| | | | | | | | |
|---|---|---|---|---|---|---|---|
| GCF_024298965.1 | NZ_CP085193.1 | 4175762 | 4176952 | 1190 | - | tuf | WP_052859607.1 |
| GCF_024298965.1 | NZ_CP085193.1 | 8220334 | 8221503 | 1169 | + | tuf | WP_407288918.1 |
| GCF_013394065.1 | NZ_CP040752.1 | 875250 | 876416 | 1166 | - | tuf | WP_406859557.1 |
| GCF_000331185.2 | NZ_CP048261.1 | 4213083 | 4214273 | 1190 | + | tuf | WP_406862541.1 |
| GCF_000331185.2 | NZ_CP048261.1 | 3369301 | 3370491 | 1190 | - | tuf | WP_318103828.1 |
| GCF_030863865.1 | NZ_CP121271.1 | 6784877 | 6786043 | 1166 | + | tuf | WP_318108338.1 |
| GCF_030863865.1 | NZ_CP121271.1 | 6461333 | 6462523 | 1190 | + | tuf | WP_069865218.1 |
| GCF_014489635.1 | NZ_CP060828.1 | 9180836 | 9182005 | 1169 | + | tuf | WP_206333821.1 |
| GCF_014489635.1 | NZ_CP060828.1 | 1768693 | 1769868 | 1175 | - | tuf | WP_150509700.1 |
| GCF_014489635.1 | NZ_CP060828.1 | 5723478 | 5724668 | 1190 | + | tuf | WP_030685484.1 |
| GCF_000497445.1 | NZ_CM002285.1 | 1326736 | 1327902 | 1166 | - | tuf | WP_108009991.1 |
| GCF_000497445.1 | NZ_CM002285.1 | 3782144 | 3783334 | 1190 | - | tuf | WP_030839706.1 |
| GCF_027947575.1 | NZ_CP116256.1 | 7808757 | 7809947 | 1190 | + | tuf | WP_108015654.1 |
| GCF_027947575.1 | NZ_CP116256.1 | 2484491 | 2485657 | 1166 | - | tuf | WP_109294293.1 |
| GCF_014216335.1 | NZ_CP045704.1 | 4274918 | 4276108 | 1190 | + | tuf | WP_109295541.1 |
| GCF_014216335.1 | NZ_CP045704.1 | 1168308 | 1169489 | 1181 | + | tuf | WP_150516823.1 |
| GCF_000091305.1 | NC_013929.1 | 3226301 | 3227491 | 1190 | - | tuf | WP_150518320.1 |
| GCF_000091305.1 | NC_013929.1 | 1130793 | 1131983 | 1190 | - | tuf | WP_010474158.1 |
| GCF_040907265.1 | NZ_JBFOHP0100000( | 1609304 | 1610470 | 1166 | - | tuf | WP_007380508.1 |
| GCF_040907265.1 | NZ_JBFOHP0100000( | 5596820 | 5598010 | 1190 | + | tuf | WP_007384061.1 |
| GCF_035917595.1 | NZ_CP109091.1 | 1251627 | 1252796 | 1169 | + | tuf | WP_353941189.1 |
| GCF_035917595.1 | NZ_CP109091.1 | 5016245 | 5017435 | 1190 | + | tuf | WP_353944180.1 |
| GCF_004328625.1 | NZ_CP032229.1 | 3748407 | 3749597 | 1190 | - | tuf | WP_041662773.1 |
| GCF_004328625.1 | NZ_CP032229.1 | 1356263 | 1357453 | 1190 | - | tuf | WP_248776023.1 |
| GCF_045690365.1 | NZ_CP147982.1 | 5951480 | 5952670 | 1190 | + | tuf | WP_107020991.1 |
| GCF_045690365.1 | NZ_CP147982.1 | 8340052 | 8341218 | 1166 | + | tuf | WP_189779621.1 |
| GCF_045348655.1 | NZ_CP160404.1 | 903705 | 904871 | 1166 | - | tuf | WP_028959843.1 |
| GCF_045348655.1 | NZ_CP160404.1 | 4233693 | 4234883 | 1190 | + | tuf | WP_019525214.1 |
| GCF_033343995.1 | NZ_CP137573.1 | 3371632 | 3372822 | 1190 | + | tuf | WP_182313923.1 |
| GCF_033343995.1 | NZ_CP137573.1 | 2434929 | 2436119 | 1190 | - | tuf | WP_125496524.1 |
| GCF_015689475.1 | NZ_CP065050.1 | 2428839 | 2430005 | 1166 | - | tuf | WP_108906278.1 |

| | | | | | | | |
|---|---|---|---|---|---|---|---|
| GCF_015689475.1 | NZ_CP065050.1 | 4045155 | 4046345 | 1190 | + | tuf | WP_017949669.1 |
| GCF_008704795.1 | NZ_CP023690.1 | 751500 | 752666 | 1166 | - | tuf | WP_418955834.1 |
| GCF_008704795.1 | NZ_CP023690.1 | 4035836 | 4037026 | 1190 | + | tuf | WP_418958557.1 |
| GCF_021441825.1 | NZ_CP090447.1 | 1074148 | 1075317 | 1169 | - | tuf | WP_006350493.1 |
| GCF_021441825.1 | NZ_CP090447.1 | 4871533 | 4872723 | 1190 | + | tuf | WP_006347239.1 |
| GCF_021441825.1 | NZ_CP090447.1 | 4569840 | 4571030 | 1190 | + | tuf | WP_242753366.1 |
| GCF_003122365.1 | NZ_CP029254.1 | 494887 | 496056 | 1169 | - | tuf | WP_159741969.1 |
| GCF_003122365.1 | NZ_CP029254.1 | 4573183 | 4574373 | 1190 | + | tuf | WP_159744864.1 |
| GCF_008704535.1 | NZ_CP023701.1 | 917414 | 918580 | 1166 | - | tuf | WP_190897146.1 |
| GCF_008704535.1 | NZ_CP023701.1 | 4540622 | 4541812 | 1190 | + | tuf | WP_016434619.1 |
| GCF_023614315.1 | NZ_CP095474.1 | 5662969 | 5664159 | 1190 | + | tuf | WP_073792895.1 |
| GCF_000154965.1 | NZ_CM000951.1 | 7947862 | 7949028 | 1166 | + | tuf | WP_073794845.1 |
| GCF_000154965.1 | NZ_CM000951.1 | 4727771 | 4728961 | 1190 | + | tuf | WP_041662773.1 |
| GCF_040529335.1 | NZ_CP159534.1 | 5062909 | 5064099 | 1190 | + | tuf | WP_041130770.1 |
| GCF_040529335.1 | NZ_CP159534.1 | 2903421 | 2904587 | 1166 | + | tuf | WP_137990675.1 |
| GCF_026427415.1 | NZ_CP084204.1 | 6846290 | 6847480 | 1190 | - | tuf | WP_137993946.1 |
| GCF_036204065.1 | NZ_CP108133.1 | 3143754 | 3144944 | 1190 | - | tuf | WP_132759734.1 |
| GCF_036204065.1 | NZ_CP108133.1 | 6221867 | 6223045 | 1178 | + | tuf | WP_150530379.1 |
| GCF_036204065.1 | NZ_CP108133.1 | 3451636 | 3452826 | 1190 | - | tuf | WP_011029795.1 |
| GCF_024760485.1 | NZ_CP058273.1 | 7103025 | 7104200 | 1175 | + | tuf | WP_003977508.1 |
| GCF_024760485.1 | NZ_CP058273.1 | 3577753 | 3578943 | 1190 | - | tuf | WP_193482362.1 |
| GCF_044987285.1 | NZ_CP173276.1 | 7629426 | 7630592 | 1166 | + | tuf | WP_328415954.1 |
| GCF_045209695.1 | NZ_CP109280.1 | 1607824 | 1609002 | 1178 | - | tuf | WP_004990992.1 |
| GCF_003097515.1 | NZ_CP029188.1 | 4947038 | 4948228 | 1190 | + | tuf | WP_004984544.1 |
| GCF_003097515.1 | NZ_CP029188.1 | 3188676 | 3189866 | 1190 | - | tuf | WP_188337386.1 |
| GCF_048500665.1 | NZ_CP183183.1 | 6749025 | 6750206 | 1181 | + | tuf | WP_188340168.1 |
| GCF_048500665.1 | NZ_CP183183.1 | 1615524 | 1616705 | 1181 | - | tuf | WP_031142225.1 |
| GCF_003932715.1 | NZ_CP020700.1 | 4876304 | 4877494 | 1190 | + | tuf | WP_136229671.1 |
| GCF_003932715.1 | NZ_CP020700.1 | 2262970 | 2264160 | 1190 | - | tuf | WP_030736332.1 |
| GCF_022699385.1 | NZ_CP093846.1 | 5744811 | 5745986 | 1175 | + | tuf | WP_030737604.1 |
| GCF_027497495.1 | NZ_CP114205.1 | 214787 | 215956 | 1169 | - | tuf | WP_095756425.1 |

| | | | | | | | |
|---|---|---|---|---|---|---|---|
| GCF_027497495.1 | NZ_CP114205.1 | 2946834 | 2948024 | 1190 | - | tuf | WP_019707818.1 |
| GCF_014701095.1 | NZ_AP023439.1 | 1216820 | 1217998 | 1178 | - | tuf | WP_183068926.1 |
| GCF_014701095.1 | NZ_AP023439.1 | 4387502 | 4388692 | 1190 | + | tuf | WP_030653602.1 |
| GCF_036250755.1 | NZ_CP109583.1 | 1747750 | 1748928 | 1178 | - | tuf | WP_211122314.1 |
| GCF_036250755.1 | NZ_CP109583.1 | 4592456 | 4593646 | 1190 | - | tuf | WP_037952879.1 |
| GCF_008639165.1 | NZ_CP029197.1 | 1422688 | 1423860 | 1172 | - | tuf | WP_275306627.1 |
| GCF_000830005.1 | NZ_CP010407.1 | 5424045 | 5425235 | 1190 | + | tuf | WP_039635340.1 |
| GCF_005517195.1 | NZ_CP040244.1 | 5382744 | 5383934 | 1190 | + | tuf | WP_026058274.1 |
| GCF_005517195.1 | NZ_CP040244.1 | 1111301 | 1112491 | 1190 | + | tuf | WP_012887829.1 |
| GCF_008704935.1 | NZ_CP023692.1 | 4706723 | 4707913 | 1190 | + | tuf | WP_336413038.1 |
| GCF_008704935.1 | NZ_CP023692.1 | 94452 | 95639 | 1187 | + | tuf | WP_188675408.1 |
| GCF_033406955.1 | NZ_CP137734.1 | 1933003 | 1934199 | 1196 | - | tuf | WP_077685959.1 |
| GCF_033406955.1 | NZ_CP137734.1 | 661083 | 662279 | 1196 | - | tuf | WP_077347858.1 |
| GCF_036014075.1 | NZ_CP107911.1 | 675949 | 677139 | 1190 | + | tuf | WP_077340474.1 |
| GCF_036014075.1 | NZ_CP107911.1 | 820633 | 821829 | 1196 | + | tuf | WP_281145573.1 |
| GCF_008704515.1 | NZ_CP023700.1 | 2291689 | 2292885 | 1196 | - | tuf | WP_219080780.1 |
| GCF_008704515.1 | NZ_CP023700.1 | 288558 | 289754 | 1196 | - | tuf | WP_108870159.1 |
| GCF_014621695.1 | NZ_CP061281.1 | 3093446 | 3094636 | 1190 | - | tuf | WP_248590341.1 |
| GCF_014621695.1 | NZ_CP061281.1 | 3109135 | 3110325 | 1190 | - | tuf | WP_011293071.1 |
| GCF_028421465.1 | NZ_CP116355.1 | 3640948 | 3642138 | 1190 | - | tuf | WP_068688277.1 |
| GCF_028421465.1 | NZ_CP116355.1 | 624694 | 625884 | 1190 | + | tuf | WP_013130820.1 |
| GCF_000993785.3 | NZ_CP009922.3 | 2789504 | 2790694 | 1190 | + | tuf | WP_119728429.1 |
| GCF_000993785.3 | NZ_CP009922.3 | 4912713 | 4913903 | 1190 | - | tuf | WP_012854633.1 |
| GCF_000220705.2 | NZ_CP023202.1 | 799239 | 800426 | 1187 | + | tuf | WP_143906427.1 |
| GCF_000220705.2 | NZ_CP023202.1 | 943323 | 944510 | 1187 | + | tuf | WP_182347946.1 |
| GCF_024666385.1 | NZ_CP102514.1 | 136972 | 138162 | 1190 | + | tuf | WP_011096092.1 |
| GCF_024666385.1 | NZ_CP102514.1 | 423398 | 424585 | 1187 | + | tuf | WP_068538836.1 |
| GCF_018069625.1 | NZ_CP072941.1 | 587334 | 588521 | 1187 | - | tuf | WP_126416032.1 |
| GCF_018069625.1 | NZ_CP072941.1 | 453231 | 454418 | 1187 | + | tuf | WP_193327100.1 |
| GCF_029011865.1 | NZ_CP095749.1 | 1281016 | 1282203 | 1187 | + | tuf | WP_024963335.1 |
| GCF_029011865.1 | NZ_CP095749.1 | 3546601 | 3547788 | 1187 | - | tuf | WP_013128044.1 |

| | | | | | | | |
|---|---|---|---|---|---|---|---|
| GCF_036035005.1 | NZ_CP107933.1 | 1095921 | 1097108 | 1187 | - | tuf | WP_076389811.1 |
| GCF_000024865.1 | NC_013595.1 | 1853894 | 1855081 | 1187 | - | tuf | WP_215523694.1 |
| GCF_047456005.1 | NZ_CP179755.1 | 1705900 | 1707087 | 1187 | - | tuf | WP_045823745.1 |
| GCF_021044565.1 | NZ_CP087997.1 | 3499641 | 3500843 | 1202 | + | tuf | WP_129205577.1 |
| GCF_001997345.1 | NZ_CP019606.1 | 690821 | 692023 | 1202 | + | tuf | WP_012877408.1 |
| GCF_001998865.1 | NZ_CP019607.1 | 1860516 | 1861718 | 1202 | + | tuf | WP_129187592.1 |
| GCF_001997295.1 | NZ_CP019605.1 | 1359289 | 1360476 | 1187 | + | tuf | WP_275248347.1 |
| GCF_029917005.1 | NZ_CP123967.1 | 1336578 | 1337750 | 1172 | + | tuf | WP_274129365.1 |
| GCF_019316905.1 | NZ_CP079216.1 | 1376750 | 1377922 | 1172 | + | tuf | WP_274129365.1 |
| GCF_900343145.1 | NZ_LT996886.1 | 313551 | 314738 | 1187 | - | tuf | WP_019090046.1 |
| GCF_023208015.1 | NZ_CP051627.1 | 1010420 | 1011607 | 1187 | + | tuf | WP_006116052.1 |
| GCF_000012405.1 | NC_007333.1 | 880404 | 881591 | 1187 | + | tuf | WP_042786750.1 |
| GCF_003574835.2 | NZ_CP063196.1 | 313723 | 314910 | 1187 | - | tuf | WP_003622701.1 |
| GCF_000092645.1 | NC_014165.1 | 166524 | 167711 | 1187 | - | tuf | WP_039903569.1 |
| GCF_003589885.1 | NZ_CP032402.1 | 135179 | 136366 | 1187 | - | tuf | WP_149277881.1 |
| GCF_000024385.1 | NC_013510.1 | 1146936 | 1148123 | 1187 | + | tuf | WP_114911462.1 |
| GCF_007559025.1 | NZ_CP041765.1 | 2413525 | 2414709 | 1184 | - | tuf | WP_007424171.1 |
| GCF_014126825.1 | NZ_CP059900.1 | 4222672 | 4223859 | 1187 | + | tuf | WP_284945261.1 |
| GCF_000196075.1 | NC_004551.1 | 2359775 | 2360959 | 1184 | - | tuf | WP_169655151.1 |
| GCF_030246965.1 | NZ_CP127099.1 | 2379348 | 2380532 | 1184 | - | tuf | WP_169655151.1 |
| GCF_900637955.1 | NZ_LR134476.1 | 3444711 | 3445898 | 1187 | - | tuf | WP_259780403.1 |
| GCF_014926385.1 | NZ_CP053291.1 | 3462694 | 3463881 | 1187 | - | tuf | WP_259780403.1 |
| GCF_001281085.1 | NZ_CP012649.1 | 2503138 | 2504325 | 1187 | - | tuf | WP_012562832.1 |
| GCF_000092225.1 | NC_014158.1 | 1701559 | 1702731 | 1172 | - | tuf | WP_080824790.1 |
| GCF_900155645.1 | NZ_LT706985.1 | 1721586 | 1722758 | 1172 | - | tuf | WP_080824790.1 |
| GCF_943169825.2 | NZ_OW968402.1 | 1916032 | 1917204 | 1172 | - | tuf | WP_010971964.1 |
| GCF_036153515.1 | NZ_CP108021.1 | 1936385 | 1937557 | 1172 | - | tuf | WP_010971964.1 |
| GCF_004135345.1 | NZ_CP035495.1 | 1350203 | 1351375 | 1172 | + | tuf | WP_136954323.1 |
| GCF_000024965.1 | NC_013530.1 | 1377075 | 1378247 | 1172 | + | tuf | WP_136954323.1 |
| GCF_004135385.1 | NZ_CP035493.1 | 1621077 | 1622249 | 1172 | - | tuf | WP_003525183.1 |
| GCF_000478885.1 | NC_022567.1 | 1641299 | 1642471 | 1172 | - | tuf | WP_003525183.1 |

| | | | | | | | |
|---|---|---|---|---|---|---|---|
| GCF_011405655.1 | NZ_AP022829.1 | 1540328 | 1541500 | 1172 | - | tuf | WP_004442617.1 |
| GCF_902386925.1 | NZ_LR698996.1 | 1560673 | 1561845 | 1172 | - | tuf | WP_173614730.1 |
| GCF_000814825.1 | NZ_CP009302.1 | 1553358 | 1554530 | 1172 | - | tuf | WP_013636434.1 |
| GCF_013201155.1 | NZ_CP053716.1 | 1573743 | 1574915 | 1172 | - | tuf | WP_013636434.1 |
| GCF_010509075.1 | NZ_CP048433.1 | 1046201 | 1047373 | 1172 | + | tuf | WP_111841877.1 |
| GCF_025149625.1 | NZ_CP102276.1 | 1066210 | 1067382 | 1172 | + | tuf | WP_111841877.1 |
| GCF_000195315.1 | NC_015389.1 | 130346 | 131518 | 1172 | + | tuf | WP_161860369.1 |
| GCF_000023845.1 | NC_013170.1 | 2340923 | 2342095 | 1172 | - | tuf | WP_161860369.1 |
| GCF_001643775.1 | NZ_CP011402.1 | 250128 | 251300 | 1172 | + | tuf | WP_173485861.1 |
| GCF_009834925.2 | NZ_CP063310.1 | 269032 | 270204 | 1172 | + | tuf | WP_173485861.1 |
| GCF_021378605.1 | NZ_CP089331.1 | 753241 | 754413 | 1172 | + | tuf | WP_406646644.1 |
| GCF_016026575.1 | NZ_CP065631.1 | 4367603 | 4368775 | 1172 | - | tuf | WP_406646644.1 |
| GCF_000024225.1 | NC_013203.1 | 1509896 | 1511068 | 1172 | + | tuf | WP_015915721.1 |
| GCF_023168405.1 | NZ_AP025285.1 | 1530874 | 1532046 | 1172 | + | tuf | WP_015915721.1 |
| GCF_900119915.1 | NZ_LT635455.1 | 3266499 | 3267671 | 1172 | - | tuf | WP_165220168.1 |
| GCF_000143845.1 | NC_014363.1 | 3284945 | 3286117 | 1172 | - | tuf | WP_165220190.1 |
| GCF_900105025.1 | NZ_LT629759.1 | 1477842 | 1479029 | 1187 | + | tuf | WP_085218205.1 |
| GCF_003966955.1 | NZ_AP019367.1 | 190791 | 191981 | 1190 | - | tuf | WP_342262178.1 |
| GCF_900143685.1 | NZ_LT671675.1 | 2350450 | 2351637 | 1187 | + | tuf | WP_278015412.1 |
| GCF_900169485.1 | NZ_LT821227.1 | 1497494 | 1498681 | 1187 | - | tuf | WP_061924931.1 |
| GCF_022845715.1 | NZ_AP025564.1 | 1375854 | 1377026 | 1172 | + | tuf | WP_067598853.1 |
| GCF_900637575.1 | NZ_LR134379.1 | 1099344 | 1100531 | 1187 | + | tuf | WP_285976861.1 |
| GCF_015277515.1 | NZ_CP063767.1 | 1893478 | 1894650 | 1172 | + | tuf | WP_290134765.1 |
| GCF_017945865.1 | NZ_CP072829.1 | 2336443 | 2337630 | 1187 | - | tuf | WP_317080615.1 |
| GCF_004322855.1 | NZ_CP036402.1 | 3393560 | 3394732 | 1172 | - | tuf | WP_108608191.1 |
| GCF_004322855.1 | NZ_CP036402.1 | 3415198 | 3416370 | 1172 | - | tuf | WP_108608191.1 |
| GCF_004300825.1 | NZ_CP036250.1 | 403680 | 404858 | 1178 | + | tuf | WP_012880528.1 |
| GCF_004300825.1 | NZ_CP036250.1 | 1037393 | 1038571 | 1178 | - | tuf | WP_012880528.1 |
| GCF_003344865.1 | NZ_CP031165.1 | 210328 | 211506 | 1178 | + | tuf | WP_011114180.1 |
| GCF_003344865.1 | NZ_CP031165.1 | 834676 | 835854 | 1178 | - | tuf | WP_011114180.1 |
| GCF_003568865.1 | NZ_CP031115.1 | 172052 | 173230 | 1178 | + | tuf | WP_011114180.1 |

| | | | | | | | |
|---|---|---|---|---|---|---|---|
| GCF_003568865.1 | NZ_CP031115.1 | 810762 | 811940 | 1178 | - | tuf | WP_011114180.1 |
| GCF_000014185.1 | NC_008148.1 | 965868 | 967046 | 1178 | - | tuf | WP_169193403.1 |
| GCF_000014185.1 | NC_008148.1 | 1493649 | 1494836 | 1187 | + | tuf | WP_013166243.1 |
| GCF_028416635.1 | NZ_CP114013.1 | 1515451 | 1516638 | 1187 | + | tuf | WP_013166243.1 |
| GCF_007970665.1 | NZ_CP042430.1 | 113417 | 114604 | 1187 | - | tuf | WP_163073317.1 |
| GCF_021172045.1 | NZ_CP087164.1 | 134135 | 135322 | 1187 | - | tuf | WP_163073317.1 |
| GCF_000025265.1 | NC_013739.1 | 1011990 | 1013162 | 1172 | + | tuf | WP_187971225.1 |
| GCF_018141525.1 | NZ_CP064655.1 | 1977033 | 1978205 | 1172 | - | tuf | WP_187971225.1 |
| GCF_018141545.1 | NZ_CP064356.1 | 2303384 | 2304556 | 1172 | + | tuf | WP_072604130.1 |
| GCF_024220175.1 | NZ_CP098502.1 | 3312335 | 3313507 | 1172 | - | tuf | WP_072604130.1 |
| GCF_024666565.1 | NZ_CP088295.1 | 984630 | 985817 | 1187 | + | tuf | WP_219894728.1 |
| GCF_000008625.1 | NC_000918.1 | 633668 | 634855 | 1187 | - | tuf | WP_160618982.1 |
| GCF_000008625.1 | NC_000918.1 | 811083 | 812270 | 1187 | + | tuf | WP_023979005.1 |
| GCF_000191045.1 | NC_015185.1 | 996513 | 997700 | 1187 | + | tuf | WP_013478455.1 |
| GCF_000191045.1 | NC_015185.1 | 1456681 | 1457853 | 1172 | + | tuf | WP_048885356.1 |
| GCF_000164905.1 | NC_017161.1 | 1531960 | 1533132 | 1172 | + | tuf | WP_047005405.1 |
| GCF_000164905.1 | NC_017161.1 | 1661399 | 1662571 | 1172 | + | tuf | WP_185982939.1 |
| GCF_000021565.1 | NC_012440.1 | 2288787 | 2289959 | 1172 | - | tuf | WP_185982939.1 |
| GCF_000021565.1 | NC_012440.1 | 3454485 | 3455657 | 1172 | - | tuf | WP_090957917.1 |
| GCF_000021545.1 | NC_012438.1 | 3473199 | 3474371 | 1172 | - | tuf | WP_090957917.1 |
| GCF_000025605.1 | NC_013894.1 | 2612173 | 2613345 | 1172 | + | tuf | WP_209736196.1 |
| GCF_000025605.1 | NC_013894.1 | 2630888 | 2632060 | 1172 | + | tuf | WP_209736196.1 |
| GCF_900142435.1 | NZ_LT670846.1 | 982046 | 983236 | 1190 | + | tuf | WP_012169411.1 |
| GCF_900142435.1 | NZ_LT670846.1 | 2881685 | 2882875 | 1190 | - | tuf | WP_012169411.1 |
| GCF_000512735.1 | NZ_CP007028.1 | 750667 | 751854 | 1187 | + | tuf | WP_014241041.1 |
| GCF_000512735.1 | NZ_CP007028.1 | 768236 | 769423 | 1187 | + | tuf | WP_035675071.1 |
| GCF_000185805.1 | NC_014926.1 | 1219727 | 1220914 | 1187 | + | tuf | WP_014241041.1 |
| GCF_000185805.1 | NC_014926.1 | 1237295 | 1238482 | 1187 | + | tuf | WP_014241041.1 |
| GCF_003574315.2 | NZ_AP025739.1 | 394460 | 395647 | 1187 | + | tuf | WP_063633859.1 |
| GCF_003574315.2 | NZ_AP025739.1 | 412539 | 413675 | 1136 | + | tuf | WP_063633866.1 |
| GCF_000427095.1 | NC_021487.1 | 2481484 | 2482671 | 1187 | - | tuf | WP_136705163.1 |

| | | | | | | | |
|---|---|---|---|---|---|---|---|
| GCF_000427095.1 | NC_021487.1 | 2498368 | 2499555 | 1187 | - | tuf | WP_136705163.1 |
| GCF_000724625.1 | NZ_CP007139.1 | 335711 | 336898 | 1187 | + | tuf | WP_111065615.1 |
| GCF_015775515.1 | NZ_CP065383.1 | 352591 | 353778 | 1187 | + | tuf | WP_111065615.1 |
| GCF_015775515.1 | NZ_CP065383.1 | 901339 | 902526 | 1187 | + | tuf | WP_109326847.1 |
| GCF_045864955.1 | NZ_AP028803.1 | 918160 | 919347 | 1187 | + | tuf | WP_109326847.1 |
| GCF_045864955.1 | NZ_AP028803.1 | 318267 | 319454 | 1187 | - | tuf | WP_045581796.1 |
| GCF_038396615.1 | NZ_CP121689.1 | 335126 | 336313 | 1187 | - | tuf | WP_045581796.1 |
| GCF_038396615.1 | NZ_CP121689.1 | 233463 | 234635 | 1172 | - | tuf | WP_005865050.1 |
| GCF_050267035.1 | NZ_CP187957.1 | 1572008 | 1573180 | 1172 | + | tuf | WP_005865050.1 |
| GCF_050267035.1 | NZ_CP187957.1 | 917853 | 919025 | 1172 | + | tuf | WP_053944181.1 |
| GCF_000093085.1 | NC_014219.1 | 1197355 | 1198527 | 1172 | - | tuf | WP_053944181.1 |
| GCF_013267415.1 | NZ_CP053988.1 | 654982 | 656154 | 1172 | + | tuf | WP_015397909.1 |
| GCF_001989575.1 | NZ_CP019573.1 | 979988 | 981160 | 1172 | - | tuf | WP_015397909.1 |
| GCF_004359375.1 | NZ_CP034726.1 | 572688 | 573860 | 1172 | + | tuf | WP_005766708.1 |
| GCF_030762805.1 | NZ_CP129113.1 | 683896 | 685068 | 1172 | + | tuf | WP_005766892.1 |
| GCF_031760945.1 | NZ_CP134501.1 | 710052 | 711224 | 1172 | + | tuf | WP_005766892.1 |
| GCF_001543105.1 | NZ_CP014159.1 | 1001347 | 1002519 | 1172 | - | tuf | WP_013545217.1 |
| GCF_001543145.1 | NZ_CP014160.1 | 1030579 | 1031751 | 1172 | - | tuf | WP_013545217.1 |
| GCF_008726475.3 | NZ_CP145134.1 | 962849 | 964021 | 1172 | - | tuf | WP_005773266.1 |
| GCF_001543175.1 | NZ_CP014161.1 | 1086840 | 1088012 | 1172 | - | tuf | WP_005773266.1 |
| GCF_001543205.1 | NZ_CP014162.1 | 151428 | 152600 | 1172 | + | tuf | WP_078691533.1 |
| GCF_001543245.1 | NZ_CP014163.1 | 580413 | 581585 | 1172 | - | tuf | WP_078691533.1 |
| GCF_001543285.1 | NZ_CP014164.1 | 828796 | 829968 | 1172 | + | tuf | WP_012755022.1 |
| GCF_047943045.1 | NZ_CP181347.1 | 1484814 | 1485986 | 1172 | - | tuf | WP_012755022.1 |
| GCF_000024285.1 | NC_013205.1 | 438393 | 439565 | 1172 | + | tuf | WP_144756285.1 |
| GCF_022674245.1 | NZ_CP080467.1 | 621359 | 622531 | 1172 | + | tuf | WP_144756285.1 |
| GCF_028751525.1 | NZ_CP067097.1 | 559328 | 560500 | 1172 | + | tuf | WP_078691533.1 |
| GCF_026651605.1 | NZ_CP104064.1 | 1074254 | 1075426 | 1172 | - | tuf | WP_078691533.1 |
| GCF_026651875.1 | NZ_CP104067.1 | 503922 | 505094 | 1172 | + | tuf | WP_011180527.1 |
| GCF_017298635.1 | NZ_CP071182.1 | 956113 | 957285 | 1172 | - | tuf | WP_011180527.1 |
| GCF_040959045.1 | NZ_CP162615.1 | 826763 | 827935 | 1172 | + | tuf | WP_005773266.1 |

| | | | | | | | |
|---|---|---|---|---|---|---|---|
| GCF_008014775.2 | NZ_CP144914.1 | 1376724 | 1377896 | 1172 | + | tuf | WP_005773266.1 |
| GCF_029094545.1 | NZ_CP117835.1 | 606626 | 607798 | 1172 | + | tuf | WP_120122039.1 |
| GCF_028743495.1 | NZ_CP117968.1 | 1200051 | 1201223 | 1172 | - | tuf | WP_120122039.1 |
| GCF_000307165.1 | NC_018704.1 | 455996 | 457168 | 1172 | + | tuf | WP_144756285.1 |
| GCF_001936335.1 | NZ_CP018888.1 | 1027626 | 1028798 | 1172 | + | tuf | WP_144756285.1 |
| GCF_001866075.4 | NZ_CP063356.2 | 564889 | 566061 | 1172 | + | tuf | WP_011179553.1 |
| GCF_002355375.1 | NZ_AP017312.1 | 678707 | 679879 | 1172 | + | tuf | WP_011179553.1 |
| GCF_019603355.1 | NZ_CP080764.1 | 315089 | 316261 | 1172 | - | tuf | WP_007477058.1 |
| GCF_028471905.1 | NZ_CP116902.1 | 634055 | 635227 | 1172 | - | tuf | WP_007477058.1 |
| GCF_000019045.1 | NC_011567.1 | 147567 | 148739 | 1172 | + | tuf | WP_078691533.1 |
| GCF_001187595.1 | NZ_CP012152.1 | 520665 | 521837 | 1172 | - | tuf | WP_078691533.1 |
| GCF_001634285.1 | NZ_CP015438.1 | 949421 | 950593 | 1172 | + | tuf | WP_012231465.1 |
| GCF_946888465.1 | NZ_OX335302.1 | 1607759 | 1608931 | 1172 | - | tuf | WP_012231465.1 |
| GCF_023380225.1 | NZ_CP093362.1 | 1286303 | 1287475 | 1172 | - | tuf | WP_004861849.1 |
| GCF_003627035.1 | NZ_CP032626.1 | 1885589 | 1886761 | 1172 | + | tuf | WP_004861849.1 |
| GCF_019575995.1 | NZ_CP080568.1 | 1567399 | 1568586 | 1187 | + | tuf | WP_012384349.1 |
| GCF_030292135.1 | NZ_CP128152.1 | 2220506 | 2221693 | 1187 | - | tuf | WP_012384349.1 |
| GCF_001191605.1 | NZ_CP009108.1 | 1917126 | 1918313 | 1187 | - | tuf | WP_126399461.1 |
| GCF_019396925.1 | NZ_CP072120.1 | 1506096 | 1507283 | 1187 | + | tuf | WP_055036992.1 |
| GCF_000008445.1 | NC_007530.2 | 95067 | 96254 | 1187 | - | tuf | WP_054134239.1 |
| GCF_019915265.1 | NZ_CP082780.1 | 374286 | 375458 | 1172 | + | tuf | WP_188641343.1 |
| GCF_030285945.1 | NZ_CP127376.1 | 5037564 | 5038751 | 1187 | - | tuf | WP_069692156.1 |
| GCF_039519175.1 | NZ_CP154443.1 | 5059136 | 5060323 | 1187 | - | tuf | WP_069692171.1 |
| GCF_004124315.2 | NZ_CP096889.1 | 4427546 | 4428733 | 1187 | - | tuf | WP_008136415.1 |
| GCF_000017425.1 | NC_009674.1 | 6286498 | 6287685 | 1187 | - | tuf | WP_027564408.1 |
| GCF_012955605.1 | NZ_CP052135.1 | 3411776 | 3412963 | 1187 | + | tuf | WP_063982577.1 |
| GCF_004103615.1 | NZ_CP035232.1 | 6120844 | 6122031 | 1187 | + | tuf | WP_008136415.1 |
| GCF_001278705.1 | NZ_CP012600.1 | 4043010 | 4044197 | 1187 | + | tuf | WP_018272140.1 |
| GCF_004006435.1 | NZ_CP029364.1 | 3202152 | 3203339 | 1187 | + | tuf | WP_128926343.1 |
| GCF_014042035.1 | NZ_CP059494.1 | 3069876 | 3071063 | 1187 | + | tuf | WP_195803894.1 |
| GCF_023238325.1 | NZ_CP096592.1 | 5977446 | 5978648 | 1202 | + | tuf | WP_195799912.1 |

| | | | | | | | |
|---|---|---|---|---|---|---|---|
| GCF_040703065.1 | NZ_CP160000.1 | 2987193 | 2988380 | 1187 | + | tuf | WP_008136415.1 |
| GCF_037201805.1 | NZ_CP147404.1 | 5991797 | 5992984 | 1187 | - | tuf | WP_028177965.1 |
| GCF_034478925.1 | NZ_CP140161.1 | 5256653 | 5257840 | 1187 | - | tuf | WP_011088152.1 |
| GCF_009739945.1 | NZ_CP040336.1 | 4165723 | 4166910 | 1187 | + | tuf | WP_018272140.1 |
| GCF_040500475.1 | NZ_CP159358.1 | 5752329 | 5753516 | 1187 | - | tuf | WP_008136415.1 |
| GCF_012648005.1 | NZ_CP051464.1 | 4722183 | 4723370 | 1187 | - | tuf | WP_128966831.1 |
| GCF_018739485.1 | NZ_CP035997.1 | 3971924 | 3973111 | 1187 | - | tuf | WP_128926343.1 |
| GCF_020861345.1 | NZ_CP086328.1 | 3021249 | 3022436 | 1187 | + | tuf | WP_057745503.1 |
| GCF_002993925.1 | NZ_CP023665.1 | 3843600 | 3844787 | 1187 | + | tuf | WP_080137239.1 |
| GCF_024296885.1 | NZ_CP101135.1 | 3589523 | 3590710 | 1187 | + | tuf | WP_065728816.1 |
| GCF_000161455.1 | NZ_CM000745.1 | 5595102 | 5596289 | 1187 | - | tuf | WP_014494497.1 |
| GCF_000017885.4 | NC_009848.4 | 2682223 | 2683410 | 1187 | + | tuf | WP_018456453.1 |
| GCF_023238245.1 | NZ_CP096590.1 | 5139434 | 5140621 | 1187 | - | tuf | WP_006611837.1 |
| GCF_008244765.1 | NZ_CP043404.1 | 2025176 | 2026363 | 1187 | - | tuf | WP_231325640.1 |
| GCF_020519665.1 | NZ_CP084703.1 | 3562398 | 3563585 | 1187 | + | tuf | WP_008136415.1 |
| GCF_001050115.1 | NZ_CP012024.1 | 5529655 | 5530842 | 1187 | - | tuf | WP_018272140.1 |
| GCF_034055115.1 | NZ_CP139190.1 | 3004222 | 3005409 | 1187 | + | tuf | WP_065748163.1 |
| GCF_000227465.1 | NC_016047.1 | 5919156 | 5920343 | 1187 | + | tuf | WP_315831967.1 |
| GCF_029772985.1 | NZ_CP121861.1 | 5950754 | 5951941 | 1187 | + | tuf | WP_315831967.1 |
| GCF_000009045.1 | NC_000964.3 | 603860 | 605047 | 1187 | - | tuf | WP_176530645.1 |
| GCF_030908325.1 | NZ_CP133085.1 | 2967954 | 2969141 | 1187 | + | tuf | WP_300947269.1 |
| GCF_012225885.1 | NZ_CP048852.1 | 2489911 | 2491098 | 1187 | + | tuf | WP_215612320.1 |
| GCF_000161615.1 | NZ_CM000753.1 | 1184600 | 1185802 | 1202 | + | tuf | WP_166214273.1 |
| GCF_016605985.1 | NZ_CP064875.1 | 8714686 | 8715873 | 1187 | - | tuf | WP_166205926.1 |
| GCF_031357045.1 | NZ_CP119875.1 | 4340946 | 4342133 | 1187 | - | tuf | WP_008136415.1 |
| GCF_003667885.1 | NZ_CP033052.1 | 3421076 | 3422263 | 1187 | + | tuf | WP_027574053.1 |
| GCF_000015785.2 | NC_009725.2 | 4879506 | 4880693 | 1187 | - | tuf | WP_270160728.1 |
| GCF_001889165.1 | NZ_CP016020.1 | 4441522 | 4442709 | 1187 | - | tuf | WP_025036994.1 |
| GCF_008807735.1 | NZ_CP032365.1 | 4192152 | 4193339 | 1187 | + | tuf | WP_027574053.1 |
| GCF_001857925.1 | NZ_CP017786.1 | 1643628 | 1644815 | 1187 | - | tuf | WP_201216432.1 |
| GCF_964340835.1 | NZ_OZ204251.1 | 1882031 | 1883218 | 1187 | - | tuf | WP_201216432.1 |

| | | | | | | | |
|---|---|---|---|---|---|---|---|
| GCF_023380265.1 | NZ_CP093366.1 | 913334 | 914521 | 1187 | + | tuf | WP_114818832.1 |
| GCF_023380245.1 | NZ_CP093365.1 | 954927 | 956114 | 1187 | + | tuf | WP_114818832.1 |
| GCF_004117055.1 | NZ_CP026363.1 | 1764012 | 1765199 | 1187 | + | tuf | WP_017506879.1 |
| GCF_961514395.1 | NZ_OY569118.1 | 2449778 | 2450965 | 1187 | - | tuf | WP_017506879.1 |
| GCF_036700025.1 | NZ_CP144922.1 | 377696 | 378883 | 1187 | + | tuf | WP_207870893.1 |
| GCF_900637055.1 | NZ_LR134338.1 | 1198708 | 1199895 | 1187 | + | tuf | WP_207870893.1 |
| GCF_046622605.1 | NZ_CP176856.1 | 1157206 | 1158393 | 1187 | + | tuf | WP_008260274.1 |
| GCF_016406105.1 | NZ_CP066308.1 | 2177498 | 2178685 | 1187 | - | tuf | WP_008260274.1 |
| GCF_020923435.1 | NZ_CP087263.1 | 531104 | 532291 | 1187 | + | tuf | WP_201100531.1 |
| GCF_002706795.1 | NZ_CP017705.1 | 570923 | 572110 | 1187 | + | tuf | WP_201100531.1 |
| GCF_003963515.1 | NZ_CP034541.1 | 1424176 | 1425363 | 1187 | + | tuf | WP_338578415.1 |
| GCF_022701015.1 | NZ_CP064090.1 | 2573895 | 2575082 | 1187 | - | tuf | WP_338576128.1 |
| GCF_023746555.1 | NZ_CP098755.1 | 1151860 | 1153047 | 1187 | + | tuf | WP_064108282.1 |
| GCF_046058945.1 | NZ_CP175511.1 | 1191582 | 1192769 | 1187 | + | tuf | WP_064108282.1 |
| GCF_001703515.1 | NZ_CP016839.1 | 2437871 | 2439058 | 1187 | - | tuf | WP_374036239.1 |
| GCF_019856495.1 | NZ_CP082237.1 | 3220905 | 3222092 | 1187 | - | tuf | WP_374036239.1 |
| GCF_039654145.1 | NZ_CP155467.1 | 1559028 | 1560215 | 1187 | + | tuf | WP_013269004.1 |
| GCF_002140975.2 | NZ_CP147247.1 | 1781208 | 1782395 | 1187 | + | tuf | WP_013269004.1 |
| GCF_002140975.2 | NZ_CP147247.1 | 260293 | 261480 | 1187 | - | tuf | WP_055753383.1 |
| GCF_002174455.2 | NZ_CP147246.1 | 2971478 | 2972665 | 1187 | + | tuf | WP_055753383.1 |
| GCF_002174455.2 | NZ_CP147246.1 | 89230 | 90417 | 1187 | + | tuf | WP_201102799.1 |
| GCF_002148425.2 | NZ_CP147248.1 | 2826812 | 2827999 | 1187 | + | tuf | WP_201102799.1 |
| GCF_002148425.2 | NZ_CP147248.1 | 1435627 | 1436799 | 1172 | - | tuf | WP_012091906.1 |
| GCF_017426705.2 | NZ_CP147251.1 | 1453650 | 1454822 | 1172 | - | tuf | WP_012091906.1 |
| GCF_017426705.2 | NZ_CP147251.1 | 1199691 | 1200863 | 1172 | - | tuf | WP_002970090.1 |
| GCF_017316305.2 | NZ_CP147250.1 | 1217652 | 1218824 | 1172 | - | tuf | WP_002970090.1 |
| GCF_017316305.2 | NZ_CP147250.1 | 157402 | 158574 | 1172 | + | tuf | WP_002970090.1 |
| GCF_002141365.2 | NZ_CP147244.1 | 175363 | 176535 | 1172 | + | tuf | WP_002970090.1 |
| GCF_002141365.2 | NZ_CP147244.1 | 1486850 | 1488022 | 1172 | + | tuf | WP_002970090.1 |
| GCF_000493735.1 | NC_022606.1 | 1504808 | 1505980 | 1172 | + | tuf | WP_002970090.1 |
| GCF_949790605.1 | NZ_OX460976.1 | 768090 | 769262 | 1172 | + | tuf | WP_002970090.1 |

| | | | | | | | |
|---|---|---|---|---|---|---|---|
| GCF_949790605.1 | NZ_OX460976.1 | 786042 | 787214 | 1172 | + | tuf | WP_002970090.1 |
| GCF_949790605.1 | NZ_OX460976.1 | 1207116 | 1208288 | 1172 | - | tuf | WP_002970090.1 |
| GCF_043182735.1 | NZ_CP171117.1 | 1225095 | 1226267 | 1172 | - | tuf | WP_002970090.1 |
| GCF_004295585.1 | NZ_AP019400.1 | 1206589 | 1207761 | 1172 | - | tuf | WP_002970090.1 |
| GCF_003713065.1 | NZ_CP033433.1 | 1224532 | 1225704 | 1172 | - | tuf | WP_002970090.1 |
| GCF_014107495.1 | NZ_CP041969.1 | 786816 | 787988 | 1172 | - | tuf | WP_151682093.1 |
| GCF_012849095.1 | NZ_CP051680.1 | 805385 | 806557 | 1172 | - | tuf | WP_151682093.1 |
| GCF_002849895.1 | NZ_CP018867.1 | 892242 | 893414 | 1172 | + | tuf | WP_007875417.1 |
| GCF_001971585.1 | NZ_CP019323.1 | 910199 | 911371 | 1172 | + | tuf | WP_007875417.1 |
| GCF_001951175.1 | NZ_CP017996.1 | 1200801 | 1201973 | 1172 | - | tuf | WP_002970090.1 |
| GCF_002706745.1 | NZ_CP017702.1 | 1218761 | 1219933 | 1172 | - | tuf | WP_002970090.1 |
| GCF_006007945.1 | NZ_CP040736.1 | 215149 | 216321 | 1172 | - | tuf | WP_021689147.1 |
| GCF_001050475.1 | NZ_CP012034.1 | 296706 | 297884 | 1178 | + | tuf | WP_282472236.1 |
| GCF_000831645.3 | NZ_CP012559.1 | 429753 | 430931 | 1178 | + | tuf | WP_282472236.1 |
| GCF_040688005.1 | NZ_CP159983.1 | 658094 | 659269 | 1175 | - | tuf | WP_323732500.1 |
| GCF_014058425.1 | NZ_CP049366.1 | 863273 | 864448 | 1175 | + | tuf | WP_323732500.1 |
| GCF_963227125.1 | NZ_OY724131.1 | 1565258 | 1566445 | 1187 | + | tuf | WP_331256166.1 |
| GCF_003606365.2 | NZ_CP031933.2 | 1761083 | 1762270 | 1187 | - | tuf | WP_331256166.1 |
| GCF_007995155.1 | NZ_CP042593.1 | 72490 | 73677 | 1187 | + | tuf | WP_350332078.1 |
| GCF_018582665.1 | NZ_CP071709.1 | 181807 | 183003 | 1196 | + | tuf | WP_148980766.1 |
| GCF_044790165.1 | NZ_CP085254.1 | 594281 | 595480 | 1199 | - | tuf | WP_148971539.1 |
| GCF_038398625.1 | NZ_CP151651.1 | 796780 | 797958 | 1178 | + | tuf | WP_146820828.1 |
| GCF_006704205.1 | NZ_CP041305.1 | 237217 | 238407 | 1190 | - | tuf | WP_015087949.1 |
| GCF_021390075.1 | NZ_CP089997.1 | 215259 | 216431 | 1172 | + | tuf | WP_341760426.1 |
| GCF_039830095.1 | NZ_CP142435.1 | 264042 | 265220 | 1178 | + | tuf | WP_323722394.1 |
| GCF_015097055.1 | NZ_AP023366.1 | 170395 | 171573 | 1178 | + | tuf | WP_108673014.1 |
| GCF_000157355.2 | NC_020995.1 | 396923 | 398107 | 1184 | + | tuf | WP_099340218.1 |
| GCF_000157355.2 | NC_020995.1 | 521707 | 522933 | 1226 | + | tuf | WP_231158845.1 |
| GCF_900474605.1 | NZ_LS483306.1 | 594 | 1763 | 1169 | + | tuf | WP_146656917.1 |
| GCF_002277935.1 | NZ_CP022930.1 | 21409 | 22578 | 1169 | - | tuf | WP_146656963.1 |
| GCF_002277935.1 | NZ_CP022930.1 | 814984 | 816174 | 1190 | + | tuf | WP_236865047.1 |

| | | | | | | | |
|---|---|---|---|---|---|---|---|
| GCF_009734005.1 | NZ_CP038996.1 | 180750 | 181937 | 1187 | + | tuf | WP_319807058.1 |
| GCF_009734005.1 | NZ_CP038996.1 | 407097 | 408269 | 1172 | + | tuf | WP_339052400.1 |
| GCF_021496385.1 | NZ_CP050816.1 | 490093 | 491265 | 1172 | + | tuf | WP_339042023.1 |
| GCF_021496385.1 | NZ_CP050816.1 | 198277 | 199455 | 1178 | - | tuf | WP_047263908.1 |
| GCF_003343305.1 | NZ_CP030932.1 | 908225 | 909403 | 1178 | + | tuf | WP_047263908.1 |
| GCF_003343305.1 | NZ_CP030932.1 | 517123 | 518301 | 1178 | + | tuf | WP_007557022.1 |
| GCF_002278015.2 | NZ_CP023011.2 | 1072806 | 1073984 | 1178 | - | tuf | WP_007557022.1 |
| GCF_002278015.2 | NZ_CP023011.2 | 25545 | 26720 | 1175 | - | tuf | WP_012778370.1 |
| GCF_022846515.1 | NZ_AP025635.1 | 137259 | 138434 | 1175 | - | tuf | WP_012778370.1 |
| GCF_022846515.1 | NZ_AP025635.1 | 1074294 | 1075472 | 1178 | - | tuf | WP_013462409.1 |
| GCF_019343125.1 | NZ_CP079880.1 | 1145409 | 1146587 | 1178 | - | tuf | WP_013462409.1 |
| GCF_019343125.1 | NZ_CP079880.1 | 590224 | 591399 | 1175 | + | tuf | WP_282486697.1 |
| GCF_029983095.1 | NZ_CP120467.1 | 1298997 | 1300178 | 1181 | - | tuf | WP_339045596.1 |
| GCF_029983095.1 | NZ_CP120467.1 | 1391902 | 1393083 | 1181 | - | tuf | WP_339045687.1 |
| GCF_002813755.1 | NZ_CP018061.1 | 472127 | 473308 | 1181 | + | tuf | WP_339047451.1 |
| GCF_002813755.1 | NZ_CP018061.1 | 1024787 | 1025968 | 1181 | - | tuf | WP_339047451.1 |
| GCF_019175485.1 | NZ_CP072888.1 | 522732 | 523907 | 1175 | + | tuf | WP_013951074.1 |
| GCF_019175485.1 | NZ_CP072888.1 | 126894 | 128102 | 1208 | + | tuf | WP_148972308.1 |
| GCF_001465345.1 | NZ_CP013655.1 | 1666939 | 1668126 | 1187 | + | tuf | WP_085784866.1 |
| GCF_001465345.1 | NZ_CP013655.1 | 1683107 | 1684294 | 1187 | + | tuf | WP_085784866.1 |
| GCF_029023825.1 | NZ_CP118957.1 | 1776990 | 1778177 | 1187 | - | tuf | WP_038465458.1 |
| GCF_011397115.1 | NZ_AP022822.1 | 1793141 | 1794328 | 1187 | - | tuf | WP_038465458.1 |
| GCF_011397115.1 | NZ_AP022822.1 | 63329 | 64516 | 1187 | + | tuf | WP_168606156.1 |
| GCF_002290025.1 | NZ_CP023074.1 | 1085519 | 1086706 | 1187 | - | tuf | WP_006996798.1 |
| GCF_002290025.1 | NZ_CP023074.1 | 892516 | 893697 | 1181 | - | tuf | WP_106875037.1 |
| GCF_002197645.1 | NZ_CP021874.1 | 2590924 | 2592111 | 1187 | + | tuf | WP_013047245.1 |
| GCF_002197645.1 | NZ_CP021874.1 | 2610377 | 2611564 | 1187 | + | tuf | WP_013047245.1 |
| GCF_000177235.2 | NC_014829.1 | 493795 | 494973 | 1178 | + | tuf | WP_148952126.1 |
| GCF_018604565.1 | NZ_CP075897.1 | 1413917 | 1415098 | 1181 | + | tuf | WP_341763930.1 |
| GCF_018138125.1 | NZ_CP073101.1 | 534886 | 536067 | 1181 | - | tuf | WP_341749203.1 |
| GCF_000019905.1 | NC_010556.1 | 484821 | 486002 | 1181 | + | tuf | WP_341748170.1 |

| | | | | | | | |
|---|---|---|---|---|---|---|---|
| GCF_026547225.1 | NZ_CP106878.1 | 599871 | 601052 | 1181 | - | tuf | WP_341750556.1 |
| GCF_026547205.1 | NZ_CP106877.1 | 478594 | 479775 | 1181 | - | tuf | WP_341750556.1 |
| GCF_022818495.1 | NZ_CP089291.1 | 1642026 | 1643207 | 1181 | - | tuf | WP_341757726.1 |
| GCF_024029675.1 | NZ_CP097121.1 | 674041 | 675222 | 1181 | - | tuf | WP_341748170.1 |
| GCF_024029595.1 | NZ_CP097119.1 | 1285576 | 1286757 | 1181 | - | tuf | WP_341748170.1 |
| GCF_024029435.1 | NZ_CP097118.1 | 204625 | 205806 | 1181 | + | tuf | WP_341748170.1 |
| GCF_024029915.1 | NZ_CP097478.1 | 589691 | 590872 | 1181 | + | tuf | WP_417905409.1 |
| GCF_001702115.1 | NZ_CP014872.1 | 856535 | 857716 | 1181 | + | tuf | WP_341748170.1 |
| GCF_024029335.1 | NZ_CP097116.1 | 821099 | 822280 | 1181 | + | tuf | WP_341749203.1 |
| GCF_029023685.1 | NZ_CP118925.1 | 272410 | 273591 | 1181 | - | tuf | WP_367364419.1 |
| GCF_047324295.1 | NZ_CP137138.1 | 939120 | 940301 | 1181 | + | tuf | WP_341756194.1 |
| GCF_024029775.1 | NZ_CP097122.1 | 1309290 | 1310471 | 1181 | - | tuf | WP_375330011.1 |
| GCF_024661895.1 | NZ_CP102453.1 | 269568 | 270749 | 1181 | + | tuf | WP_375359097.1 |
| GCF_013394005.1 | NZ_CP040586.1 | 773126 | 774307 | 1181 | - | tuf | WP_375331465.1 |
| GCF_900476045.1 | NZ_LS483440.1 | 1774511 | 1775692 | 1181 | + | tuf | WP_342271298.1 |
| GCF_009730295.1 | NZ_CP046313.1 | 1068695 | 1069876 | 1181 | - | tuf | WP_341751363.1 |
| GCF_000009785.1 | NC_006510.1 | 1533151 | 1534332 | 1181 | + | tuf | WP_341748170.1 |
| GCF_001618685.1 | NZ_CP014342.1 | 1242673 | 1243854 | 1181 | - | tuf | WP_341748170.1 |
| GCF_002072065.1 | NZ_CP017694.1 | 410433 | 411614 | 1181 | - | tuf | WP_341749203.1 |
| GCF_001610955.1 | NZ_CP014335.1 | 1186390 | 1187571 | 1181 | - | tuf | WP_425360985.1 |
| GCF_030414215.1 | NZ_CP129217.1 | 191856 | 193037 | 1181 | + | tuf | WP_341748170.1 |
| GCF_002847845.2 | NZ_CP126962.1 | 82672 | 83853 | 1181 | - | tuf | WP_341748170.1 |
| GCF_022919115.1 | NZ_CP095072.1 | 812119 | 813300 | 1181 | + | tuf | WP_375333702.1 |
| GCF_022919575.1 | NZ_CP095071.1 | 1551234 | 1552415 | 1181 | + | tuf | WP_316354982.1 |
| GCF_009650095.1 | NZ_CP045915.1 | 3670732 | 3671919 | 1187 | + | tuf | WP_066774275.1 |
| GCF_019931005.1 | NZ_CP082858.1 | 2838930 | 2840117 | 1187 | + | tuf | WP_058346798.1 |
| GCF_005671335.1 | NZ_CP040441.1 | 5297196 | 5298383 | 1187 | - | tuf | WP_058346798.1 |
| GCF_022921115.1 | NZ_CP095075.1 | 1320324 | 1321511 | 1187 | + | tuf | WP_062145518.1 |
| GCF_000284515.1 | NC_017668.1 | 3153533 | 3154720 | 1187 | - | tuf | WP_062145518.1 |
| GCF_002097535.1 | NZ_CP020772.1 | 640527 | 641714 | 1187 | + | tuf | WP_099620739.1 |
| GCF_029714185.1 | NZ_CP121671.1 | 1867012 | 1868199 | 1187 | + | tuf | WP_099620739.1 |

| | | | | | | | |
|---|---|---|---|---|---|---|---|
| GCF_022919095.1 | NZ_CP095073.1 | 3566026 | 3567213 | 1187 | - | tuf | WP_163231737.1 |
| GCF_022919835.1 | NZ_CP095074.1 | 4826192 | 4827379 | 1187 | - | tuf | WP_035072114.1 |
| GCF_000832905.1 | NZ_CP009709.1 | 794842 | 796029 | 1187 | + | tuf | WP_013077929.1 |
| GCF_040267745.1 | NZ_CP158453.1 | 1587710 | 1588897 | 1187 | + | tuf | WP_013078638.1 |
| GCF_016772275.1 | NZ_CP065425.1 | 854157 | 855344 | 1187 | + | tuf | WP_165184995.1 |
| GCF_023195815.2 | NZ_CP096206.2 | 1721162 | 1722349 | 1187 | + | tuf | WP_165184995.1 |
| GCF_011100465.1 | NZ_CP049740.1 | 3043509 | 3044681 | 1172 | + | tuf | WP_066707018.1 |
| GCF_003955755.1 | NZ_CP034465.1 | 3065517 | 3066689 | 1172 | + | tuf | WP_066707018.1 |
| GCF_002005405.1 | NZ_CP019728.1 | 638877 | 640049 | 1172 | + | tuf | WP_043868431.1 |
| GCF_011299095.1 | NZ_CP049889.1 | 656400 | 657572 | 1172 | + | tuf | WP_043868431.1 |
| GCF_000756715.1 | NZ_CCSE01000001.1 | 1454589 | 1455761 | 1172 | - | tuf | WP_062217321.1 |
| GCF_013282215.1 | NZ_CP048103.1 | 1476210 | 1477382 | 1172 | - | tuf | WP_062217321.1 |
| GCF_013265585.1 | NZ_CP048104.1 | 2315605 | 2316777 | 1172 | - | tuf | WP_011909488.1 |
| GCF_002804065.1 | NZ_CP024955.1 | 2333551 | 2334723 | 1172 | - | tuf | WP_011909488.1 |
| GCF_000092905.1 | NC_014098.1 | 287813 | 288985 | 1172 | + | tuf | WP_002722461.1 |
| GCF_024927925.1 | NZ_CP103866.1 | 306490 | 307662 | 1172 | + | tuf | WP_002722461.1 |
| GCF_000829055.1 | NZ_AP012544.1 | 1935809 | 1936981 | 1172 | - | tuf | WP_182304381.1 |
| GCF_029726355.1 | NZ_CP120687.1 | 1955583 | 1956755 | 1172 | - | tuf | WP_182304381.1 |
| GCF_009184845.1 | NZ_CP045068.1 | 486559 | 487746 | 1187 | + | tuf | WP_319568407.1 |
| GCF_028736235.1 | NZ_CP117884.1 | 505256 | 506443 | 1187 | + | tuf | WP_319568407.1 |
| GCF_000155515.2 | NC_022112.1 | 1542324 | 1543511 | 1187 | - | tuf | WP_110438408.1 |
| GCF_030770265.1 | NZ_CP132482.1 | 291067 | 292254 | 1187 | - | tuf | WP_350334709.1 |
| GCF_006151905.1 | NZ_CP040780.1 | 309857 | 311044 | 1187 | - | tuf | WP_350334709.1 |
| GCF_030770295.1 | NZ_CP132483.1 | 1873031 | 1874218 | 1187 | + | tuf | WP_305095623.1 |
| GCF_030770335.1 | NZ_CP132484.1 | 66054 | 67226 | 1172 | + | tuf | WP_046902245.1 |
| GCF_037290055.1 | NZ_CP147647.1 | 2405304 | 2406491 | 1187 | - | tuf | WP_066843138.1 |
| GCF_045862825.1 | NZ_AP027463.1 | 1126330 | 1127502 | 1172 | + | tuf | WP_047820270.1 |
| GCF_947539765.1 | NZ_OX443570.1 | 1091068 | 1092255 | 1187 | - | tuf | WP_190262331.1 |
| GCF_029025825.1 | NZ_CP118745.1 | 1107937 | 1109124 | 1187 | - | tuf | WP_190262331.1 |
| GCF_003641185.1 | NZ_CP032757.1 | 3517173 | 3518360 | 1187 | + | tuf | WP_193335629.1 |
| GCF_009913655.1 | NZ_CP028221.1 | 1379839 | 1381026 | 1187 | - | tuf | WP_146289260.1 |

| | | | | | | | |
|---|---|---|---|---|---|---|---|
| GCF_963243495.1 | NZ_OY725316.1 | 2305484 | 2306671 | 1187 | - | tuf | WP_240229340.1 |
| GCF_008831485.1 | NZ_CP044496.1 | 3166481 | 3167668 | 1187 | + | tuf | WP_035102615.1 |
| GCF_034298135.1 | NZ_CP139575.1 | 2156097 | 2157284 | 1187 | + | tuf | WP_201653348.1 |
| GCF_003999355.1 | NZ_CP031835.1 | 1093769 | 1094956 | 1187 | + | tuf | WP_201635787.1 |
| GCF_000194115.1 | NC_017470.1 | 1747344 | 1748531 | 1187 | + | tuf | WP_284580812.1 |
| GCF_003150935.1 | NZ_CP029476.1 | 76155 | 77342 | 1187 | - | tuf | WP_282211488.1 |
| GCF_009769205.1 | NZ_CP039266.1 | 1978152 | 1979339 | 1187 | - | tuf | WP_220307217.1 |
| GCF_001888925.1 | NZ_CP018218.1 | 220923 | 222095 | 1172 | + | tuf | WP_012176905.1 |
| GCF_000014425.1 | NC_008530.1 | 277686 | 278858 | 1172 | + | tuf | WP_012176905.1 |
| GCF_003173695.1 | NZ_CP029544.1 | 244136 | 245320 | 1184 | + | tuf | WP_011304291.1 |
| GCF_046109915.1 | NZ_CP151471.1 | 863868 | 865052 | 1184 | - | tuf | WP_011304291.1 |
| GCF_011058775.1 | NZ_CP049230.1 | 396752 | 397936 | 1184 | + | tuf | WP_006011505.1 |
| GCF_024397795.1 | NZ_CP072983.1 | 983266 | 984450 | 1184 | - | tuf | WP_006011505.1 |
| GCF_947539375.1 | NZ_OX443569.1 | 387293 | 388477 | 1184 | + | tuf | WP_044194392.1 |
| GCF_001936235.1 | NZ_CP018809.1 | 965297 | 966481 | 1184 | - | tuf | WP_044194392.1 |
| GCF_014058685.1 | NZ_CP059055.1 | 191510 | 192694 | 1184 | + | tuf | WP_024071794.1 |
| GCF_014656585.1 | NZ_CP061341.1 | 789696 | 790880 | 1184 | - | tuf | WP_024071794.1 |
| GCF_003151025.1 | NZ_CP029477.1 | 283090 | 284274 | 1184 | + | tuf | WP_011154850.1 |
| GCF_042997415.1 | NZ_CP160088.1 | 1022635 | 1023819 | 1184 | - | tuf | WP_011154850.1 |
| GCF_019469265.1 | NZ_CP048268.1 | 2529648 | 2530835 | 1187 | + | tuf | WP_218284964.1 |
| GCF_003584685.1 | NZ_AP018549.1 | 2545812 | 2546999 | 1187 | + | tuf | WP_218284964.1 |
| GCF_017894345.1 | NZ_CP059276.1 | 736436 | 737608 | 1172 | - | tuf | WP_038539026.1 |
| GCF_002762335.1 | NZ_CP024610.1 | 1227700 | 1228872 | 1172 | - | tuf | WP_038539026.1 |
| GCF_016647595.1 | NZ_CP059829.1 | 1142679 | 1143851 | 1172 | + | tuf | WP_034795467.1 |
| GCF_033096005.1 | NZ_AP026803.1 | 1161217 | 1162389 | 1172 | + | tuf | WP_034795467.1 |
| GCF_003627095.1 | NZ_CP032627.1 | 1105479 | 1106651 | 1172 | + | tuf | WP_025426651.1 |
| GCF_006770265.1 | NZ_CP017194.1 | 1124027 | 1125199 | 1172 | + | tuf | WP_025426651.1 |
| GCF_000468955.1 | NC_022369.1 | 62053 | 63240 | 1187 | - | tuf | WP_203413821.1 |
| GCF_037892375.1 | NZ_CP141721.1 | 2553152 | 2554339 | 1187 | - | tuf | WP_252258123.1 |
| GCF_016026695.1 | NZ_CP065637.1 | 2428092 | 2429279 | 1187 | - | tuf | WP_115417106.1 |
| GCF_003176835.1 | NZ_CP028160.1 | 1260704 | 1261891 | 1187 | + | tuf | WP_034905026.1 |

| | | | | | | | |
|---|---|---|---|---|---|---|---|
| GCF_006770285.1 | NZ_CP017195.1 | 2276020 | 2277207 | 1187 | - | tuf | WP_173215006.1 |
| GCF_023499275.1 | NZ_CP094882.1 | 2979903 | 2981090 | 1187 | - | tuf | WP_068352939.1 |
| GCF_006965445.1 | NZ_CP041356.1 | 856101 | 857276 | 1175 | - | tuf | WP_062628133.1 |
| GCF_010669245.1 | NZ_CP048686.1 | 878072 | 879247 | 1175 | - | tuf | WP_062628136.1 |
| GCF_017068355.1 | NZ_CP070872.1 | 1984054 | 1985226 | 1172 | + | tuf | WP_211783765.1 |
| GCF_008831385.1 | NZ_CP044499.1 | 2001743 | 2002915 | 1172 | + | tuf | WP_211783765.1 |
| GCF_004101845.1 | NZ_CP026116.1 | 3169163 | 3170350 | 1187 | + | tuf | WP_144069569.1 |
| GCF_900258455.1 | NZ_LT984417.1 | 3567585 | 3568772 | 1187 | - | tuf | WP_144069569.1 |
| GCF_009025875.1 | NZ_CP045007.1 | 1396488 | 1397660 | 1172 | + | tuf | WP_145214922.1 |
| GCF_009676365.1 | NZ_CP046037.1 | 1421599 | 1422771 | 1172 | + | tuf | WP_145214922.1 |
| GCF_900478165.1 | NZ_LS483476.1 | 432197 | 433384 | 1187 | - | tuf | WP_141442821.1 |
| GCF_001307805.1 | NZ_CP013862.1 | 654009 | 655181 | 1172 | + | tuf | WP_371397445.1 |
| GCF_027186265.1 | NZ_CP114176.1 | 898055 | 899227 | 1172 | + | tuf | WP_371397659.1 |
| GCF_018314255.1 | NZ_CP073066.1 | 401669 | 402841 | 1172 | + | tuf | WP_066809175.1 |
| GCF_000785105.2 | NZ_CP018906.1 | 421639 | 422811 | 1172 | + | tuf | WP_066809175.1 |
| GCF_009832765.1 | NZ_CP047121.1 | 3229078 | 3230250 | 1172 | + | tuf | WP_281465230.1 |
| GCF_009739535.1 | NZ_CP029971.1 | 3249643 | 3250815 | 1172 | + | tuf | WP_281465230.1 |
| GCF_020616595.1 | NZ_CP085278.1 | 2481152 | 2482324 | 1172 | + | tuf | WP_215504664.1 |
| GCF_014879295.1 | NZ_CP062473.1 | 2503212 | 2504384 | 1172 | + | tuf | WP_215504664.1 |
| GCF_026913205.2 | NZ_CP168151.1 | 2526916 | 2528088 | 1172 | + | tuf | WP_126010234.1 |
| GCF_007954525.1 | NZ_CP042374.1 | 2544728 | 2545900 | 1172 | + | tuf | WP_126010234.1 |
| GCF_007954785.1 | NZ_CP042410.1 | 1637946 | 1639133 | 1187 | + | tuf | WP_289505465.1 |
| GCF_045801155.1 | NZ_CP174386.1 | 1655071 | 1656258 | 1187 | + | tuf | WP_289505465.1 |
| GCF_000196855.1 | NC_014319.1 | 3250838 | 3252025 | 1187 | + | tuf | WP_012227963.1 |
| GCF_000298875.1 | NC_018631.1 | 1960001 | 1961188 | 1187 | - | tuf | WP_062029287.1 |
| GCF_000092505.1 | NC_014136.1 | 420393 | 421580 | 1187 | - | tuf | WP_041110853.1 |
| GCF_007954605.1 | NZ_CP042387.1 | 468801 | 469988 | 1187 | - | tuf | WP_141351583.1 |
| GCF_000014445.1 | NC_008531.1 | 221900 | 223072 | 1172 | + | tuf | WP_371393812.1 |
| GCF_016127255.1 | NZ_CP065993.1 | 244843 | 246015 | 1172 | + | tuf | WP_371393812.1 |
| GCF_001891125.1 | NZ_CP015247.1 | 877307 | 878479 | 1172 | + | tuf | WP_127565786.1 |
| GCF_900475625.1 | NZ_LS483405.1 | 897449 | 898621 | 1172 | + | tuf | WP_127565797.1 |

| | | | | | | | |
|---|---|---|---|---|---|---|---|
| GCF_964062745.1 | NZ_OZ061257.1 | 644270 | 645457 | 1187 | + | tuf | WP_011631260.1 |
| GCF_001050435.1 | NZ_CP012033.1 | 1861165 | 1862337 | 1172 | - | tuf | WP_035715313.1 |
| GCF_044051715.1 | NZ_CP168688.1 | 1882887 | 1884059 | 1172 | - | tuf | WP_035715313.1 |
| GCF_014069315.1 | NZ_CP059603.1 | 187429 | 188616 | 1187 | - | tuf | WP_066132245.1 |
| GCF_963243745.1 | NZ_OY725319.1 | 204637 | 205824 | 1187 | - | tuf | WP_066132245.1 |
| GCF_900116935.1 | NZ_LT630287.1 | 2229901 | 2231088 | 1187 | + | tuf | WP_222872877.1 |
| GCF_025311475.1 | NZ_CP104397.1 | 1623404 | 1624591 | 1187 | + | tuf | WP_099555611.1 |
| GCF_009933595.1 | NZ_CP047141.1 | 2697671 | 2698858 | 1187 | - | tuf | WP_099555611.1 |
| GCF_030389495.1 | NZ_CP127832.1 | 2222506 | 2223696 | 1190 | - | tuf | WP_015827918.1 |
| GCF_029889745.1 | NZ_CP123639.1 | 1996881 | 1998053 | 1172 | + | tuf | WP_007197090.1 |
| GCF_003288115.1 | NZ_CP023565.1 | 2016701 | 2017873 | 1172 | + | tuf | WP_007197090.1 |
| GCF_000224985.1 | NC_015975.1 | 2070038 | 2071225 | 1187 | + | tuf | WP_151116732.1 |
| GCF_036689625.1 | NZ_CP144759.1 | 2087170 | 2088357 | 1187 | + | tuf | WP_151116732.1 |
| GCF_035231985.1 | NZ_CP117983.1 | 3199425 | 3200612 | 1187 | - | tuf | WP_151177881.1 |
| GCF_029961225.1 | NZ_CP124737.1 | 3216546 | 3217733 | 1187 | - | tuf | WP_151177881.1 |
| GCF_008876665.1 | NZ_CP044534.1 | 2854240 | 2855412 | 1172 | + | tuf | WP_274492808.1 |
| GCF_013423865.1 | NZ_CP058954.1 | 779716 | 780888 | 1172 | + | tuf | WP_411818051.1 |
| GCF_025311495.1 | NZ_CP104398.1 | 1074696 | 1075901 | 1205 | + | tuf | WP_013215044.1 |
| GCF_009428965.1 | NZ_CP045530.1 | 2924696 | 2925901 | 1205 | - | tuf | WP_023788139.1 |
| GCF_028607025.1 | NZ_CP117296.1 | 2729184 | 2730371 | 1187 | - | tuf | WP_272983375.1 |
| GCF_025311515.1 | NZ_CP104399.1 | 3444654 | 3445841 | 1187 | - | tuf | WP_272983375.1 |
| GCF_019443985.1 | NZ_CP049303.1 | 1776506 | 1777693 | 1187 | + | tuf | WP_011646739.1 |
| GCF_009184705.1 | NZ_CP045035.1 | 3004401 | 3005588 | 1187 | - | tuf | WP_011647828.1 |
| GCF_019444005.1 | NZ_CP049304.1 | 376624 | 377808 | 1184 | + | tuf | WP_119677612.1 |
| GCF_900638015.1 | NZ_LR134483.1 | 394582 | 395766 | 1184 | + | tuf | WP_119677612.1 |
| GCF_028596125.1 | NZ_CP117229.1 | 505749 | 506921 | 1172 | - | tuf | WP_279965948.1 |
| GCF_000763515.1 | NZ_CP009577.1 | 528344 | 529516 | 1172 | - | tuf | WP_279965948.1 |
| GCF_000196035.1 | NC_003210.1 | 62489 | 63661 | 1172 | + | tuf | WP_213162543.1 |
| GCF_026871175.1 | NZ_CP113980.1 | 79709 | 80881 | 1172 | + | tuf | WP_213162555.1 |
| GCF_000027145.1 | NC_013891.1 | 2193099 | 2194271 | 1172 | - | tuf | WP_085786935.1 |
| GCF_031843265.2 | NZ_CP156021.1 | 2212929 | 2214101 | 1172 | - | tuf | WP_085786935.1 |

| | | | | | | | |
|---|---|---|---|---|---|---|---|
| GCF_003534205.1 | NZ_CP011102.1 | 2603093 | 2604265 | 1172 | - | tuf | WP_013385324.1 |
| GCF_900187315.1 | NZ_LT906444.1 | 2624851 | 2626023 | 1172 | - | tuf | WP_013385324.1 |
| GCF_001663675.1 | NZ_CP014873.1 | 1705286 | 1706473 | 1187 | - | tuf | WP_010506739.1 |
| GCF_009650015.1 | NZ_CP045872.1 | 1639452 | 1640639 | 1187 | - | tuf | WP_010506739.1 |
| GCF_002706425.1 | NZ_CP017713.1 | 1219670 | 1220857 | 1187 | - | tuf | WP_014105031.1 |
| GCF_016638705.1 | NZ_CP067341.1 | 1031588 | 1032775 | 1187 | - | tuf | WP_078524560.1 |
| GCF_020217405.1 | NZ_CP084108.1 | 2449536 | 2450723 | 1187 | - | tuf | WP_007399618.1 |
| GCF_000724775.3 | NZ_CP010820.1 | 1366430 | 1367617 | 1187 | - | tuf | WP_102323163.1 |
| GCF_028877475.1 | NZ_CP113527.1 | 701628 | 702815 | 1187 | + | tuf | WP_010506739.1 |
| GCF_033880585.1 | NZ_CP137624.1 | 1365350 | 1366537 | 1187 | - | tuf | WP_061271396.1 |
| GCF_016026755.1 | NZ_CP065643.1 | 1993043 | 1994230 | 1187 | + | tuf | WP_194215668.1 |
| GCF_030123245.1 | NZ_CP126101.1 | 2010084 | 2011271 | 1187 | + | tuf | WP_194215668.1 |
| GCF_003049605.2 | NZ_CP073713.1 | 1759675 | 1760862 | 1187 | - | tuf | WP_029604134.1 |
| GCF_002982115.1 | NZ_CP019980.1 | 3016060 | 3017247 | 1187 | + | tuf | WP_284317121.1 |
| GCF_039654155.1 | NZ_CP155468.1 | 3037369 | 3038556 | 1187 | + | tuf | WP_284317121.1 |
| GCF_900291985.1 | NZ_LT985980.1 | 2932973 | 2934145 | 1172 | - | tuf | WP_008555140.1 |
| GCF_000600105.1 | NZ_CP006837.1 | 2957645 | 2958817 | 1172 | - | tuf | WP_008555140.1 |
| GCF_016865215.1 | NZ_CP054482.1 | 209272 | 210444 | 1172 | + | tuf | WP_027235550.1 |
| GCF_002119805.1 | NZ_CP021059.1 | 3281712 | 3282884 | 1172 | - | tuf | WP_027235550.1 |
| GCF_016028795.1 | NZ_CP065729.1 | 169445 | 170617 | 1172 | + | tuf | WP_024088515.1 |
| GCF_020097135.1 | NZ_CP083608.1 | 192704 | 193876 | 1172 | + | tuf | WP_024088515.1 |
| GCF_030376725.1 | NZ_CP128470.1 | 224710 | 225882 | 1172 | + | tuf | WP_264212159.1 |
| GCF_022343725.1 | NZ_CP092179.1 | 249150 | 250322 | 1172 | + | tuf | WP_264212159.1 |
| GCF_024205965.1 | NZ_CP073819.1 | 517771 | 518943 | 1172 | + | tuf | WP_015272827.1 |
| GCF_024205945.1 | NZ_CP073809.1 | 987564 | 988736 | 1172 | - | tuf | WP_015272827.1 |
| GCF_038095555.1 | NZ_CP124591.1 | 1146458 | 1147645 | 1187 | + | tuf | WP_171834513.1 |
| GCF_947036595.1 | NZ_OX344834.1 | 2463987 | 2465159 | 1172 | + | tuf | WP_183469893.1 |
| GCF_900474615.1 | NZ_LS483305.1 | 2486255 | 2487427 | 1172 | + | tuf | WP_183469893.1 |
| GCF_900187075.1 | NZ_LT906462.1 | 157169 | 158341 | 1172 | + | tuf | WP_254248016.1 |
| GCF_016725245.1 | NZ_CP068061.1 | 176919 | 178091 | 1172 | + | tuf | WP_254248021.1 |
| GCF_015482585.1 | NZ_CP049742.1 | 957157 | 958344 | 1187 | + | tuf | WP_138379486.1 |

| | | | | | | | |
|---|---|---|---|---|---|---|---|
| GCF_003966875.1 | NZ_AP018492.1 | 974186 | 975373 | 1187 | + | tuf | WP_138379486.1 |
| GCF_002874535.1 | NZ_CP025025.1 | 1036850 | 1038037 | 1187 | + | tuf | WP_011712511.1 |
| GCF_044803185.1 | NZ_CP085225.1 | 1054110 | 1055297 | 1187 | + | tuf | WP_011712511.1 |
| GCF_019933155.2 | NZ_CP082944.1 | 22870 | 24057 | 1187 | + | tuf | WP_024078283.1 |
| GCF_014217835.1 | NZ_CP055263.1 | 42752 | 43939 | 1187 | + | tuf | WP_024078283.1 |
| GCF_030127205.1 | NZ_CP126116.1 | 984746 | 985933 | 1187 | - | tuf | WP_303618850.1 |
| GCF_046603695.1 | NZ_CP176757.1 | 1005509 | 1006696 | 1187 | - | tuf | WP_303618850.1 |
| GCF_037201795.1 | NZ_CP147403.1 | 1319621 | 1320793 | 1172 | - | tuf | WP_206340921.1 |
| GCF_009720625.1 | NZ_CP046266.1 | 1344427 | 1345599 | 1172 | - | tuf | WP_206340921.1 |
| GCF_037201815.1 | NZ_CP147407.1 | 1718450 | 1719637 | 1187 | + | tuf | WP_117395607.1 |
| GCF_048572935.1 | NZ_CP183883.1 | 1798606 | 1799793 | 1187 | + | tuf | WP_117395607.1 |
| GCF_013267435.1 | NZ_CP053989.1 | 4069425 | 4070597 | 1172 | - | tuf | WP_045683516.1 |
| GCF_036670025.1 | NZ_CP137640.1 | 4292275 | 4293447 | 1172 | - | tuf | WP_045683812.1 |
| GCF_023380205.1 | NZ_CP093361.1 | 2546334 | 2547506 | 1172 | + | tuf | WP_018063709.1 |
| GCF_019357495.1 | NZ_CP079110.1 | 4259560 | 4260732 | 1172 | - | tuf | WP_018063709.1 |
| GCF_002005165.1 | NZ_CP019699.1 | 4215840 | 4217012 | 1172 | - | tuf | WP_006202147.1 |
| GCF_000011245.1 | NC_004193.1 | 4239952 | 4241124 | 1172 | - | tuf | WP_006202147.1 |
| GCF_044790185.1 | NZ_CP085256.1 | 4310690 | 4311862 | 1172 | - | tuf | WP_015317986.1 |
| GCF_000340475.1 | NZ_CM001792.1 | 4332760 | 4333932 | 1172 | - | tuf | WP_015317986.1 |
| GCF_003352005.1 | NZ_CP024848.1 | 4148028 | 4149200 | 1172 | - | tuf | WP_010909256.1 |
| GCF_000241055.1 | NZ_CM001398.1 | 4171182 | 4172354 | 1172 | - | tuf | WP_010909256.1 |
| GCF_020510245.1 | NZ_CP084701.1 | 3252970 | 3254142 | 1172 | + | tuf | WP_010909256.1 |
| GCF_004102045.2 | NZ_CP029684.2 | 3277676 | 3278848 | 1172 | + | tuf | WP_010909256.1 |
| GCF_006965545.2 | NZ_CP041372.2 | 4028172 | 4029344 | 1172 | - | tuf | WP_010909256.1 |
| GCF_018604145.1 | NZ_CP075889.1 | 4050639 | 4051811 | 1172 | - | tuf | WP_010909256.1 |
| GCF_012676905.1 | NZ_CP051428.1 | 4786523 | 4787695 | 1172 | - | tuf | WP_010909256.1 |
| GCF_003952225.1 | NZ_CP034437.1 | 4808612 | 4809784 | 1172 | - | tuf | WP_010909256.1 |
| GCF_005577435.1 | NZ_CP040396.1 | 3803013 | 3804185 | 1172 | - | tuf | WP_311028980.1 |
| GCF_036894225.1 | NZ_CP145892.1 | 3820638 | 3821810 | 1172 | - | tuf | WP_311028980.1 |
| GCF_025369935.1 | NZ_CP104467.1 | 4596439 | 4597611 | 1172 | - | tuf | WP_126106702.1 |
| GCF_008369725.1 | NZ_CP043611.1 | 4618129 | 4619301 | 1172 | - | tuf | WP_126106702.1 |

| | | | | | | | |
|---|---|---|---|---|---|---|---|
| GCF_032268605.1 | NZ_CP130318.1 | 4557956 | 4559128 | 1172 | - | tuf | WP_258117360.1 |
| GCF_021654775.1 | NZ_AP025343.1 | 4582105 | 4583277 | 1172 | - | tuf | WP_258117360.1 |
| GCF_003945345.1 | NZ_AP019308.1 | 4745205 | 4746377 | 1172 | - | tuf | WP_013895701.1 |
| GCF_013347305.1 | NZ_CP054614.1 | 4769653 | 4770825 | 1172 | - | tuf | WP_013895701.1 |
| GCF_000961095.1 | NZ_CP011058.1 | 1432144 | 1433331 | 1187 | + | tuf | WP_018044231.1 |
| GCF_000758665.1 | NZ_CP009285.1 | 1454760 | 1455947 | 1187 | + | tuf | WP_018044231.1 |
| GCF_001421015.2 | NZ_CP013023.1 | 2966366 | 2967553 | 1187 | + | tuf | WP_099953741.1 |
| GCF_009363115.1 | NZ_CP045298.1 | 2986026 | 2987213 | 1187 | + | tuf | WP_099953741.1 |
| GCF_013347265.1 | NZ_CP054613.1 | 6596930 | 6598117 | 1187 | - | tuf | WP_109895584.1 |
| GCF_009363095.1 | NZ_CP045295.1 | 6618919 | 6620106 | 1187 | - | tuf | WP_109895584.1 |
| GCF_004117095.1 | NZ_CP026520.1 | 5628810 | 5629997 | 1187 | - | tuf | WP_056480538.1 |
| GCF_001857945.1 | NZ_CP017770.1 | 5650596 | 5651783 | 1187 | - | tuf | WP_056480538.1 |
| GCF_021654795.1 | NZ_AP025344.1 | 1356909 | 1358096 | 1187 | - | tuf | WP_058617276.1 |
| GCF_002192415.1 | NZ_CP021780.1 | 3378860 | 3380047 | 1187 | - | tuf | WP_058617276.1 |
| GCF_000756615.1 | NZ_CP009288.1 | 1993175 | 1994362 | 1187 | + | tuf | WP_015928572.1 |
| GCF_029625475.1 | NZ_CP121215.1 | 2012769 | 2013956 | 1187 | + | tuf | WP_015928572.1 |
| GCF_037333485.1 | NZ_CP147845.1 | 3454239 | 3455426 | 1187 | + | tuf | WP_238315046.1 |
| GCF_000758705.1 | NZ_CP009287.1 | 4947469 | 4948656 | 1187 | + | tuf | WP_238315046.1 |
| GCF_009363075.1 | NZ_CP045293.1 | 2675594 | 2676781 | 1187 | + | tuf | WP_043348276.1 |
| GCF_021513275.1 | NZ_CP090978.1 | 4411003 | 4412190 | 1187 | - | tuf | WP_043348276.1 |
| GCF_034055775.1 | NZ_CP139194.1 | 1962999 | 1964186 | 1187 | + | tuf | WP_043348276.1 |
| GCF_002741055.1 | NZ_CP016809.1 | 3547731 | 3548918 | 1187 | - | tuf | WP_043348276.1 |
| GCF_010450915.1 | NZ_CP048429.1 | 4004379 | 4005566 | 1187 | - | tuf | WP_066923801.1 |
| GCF_002240415.1 | NZ_CP020028.1 | 4760311 | 4761498 | 1187 | - | tuf | WP_066923801.1 |
| GCF_028606985.1 | NZ_CP117416.1 | 2354156 | 2355343 | 1187 | + | tuf | WP_012319185.1 |
| GCF_002951935.1 | NZ_CP019659.1 | 4097901 | 4099088 | 1187 | - | tuf | WP_012319185.1 |
| GCF_003931855.1 | NZ_CP034248.1 | 4102326 | 4103513 | 1187 | - | tuf | WP_109960477.1 |
| GCF_003991425.1 | NZ_CP034346.1 | 4122209 | 4123396 | 1187 | - | tuf | WP_109960477.1 |
| GCF_010119935.1 | NZ_CP048209.1 | 3574602 | 3575789 | 1187 | + | tuf | WP_407338808.1 |
| GCF_028771845.1 | NZ_CP118270.1 | 2383672 | 2384859 | 1187 | - | tuf | WP_045367563.1 |
| GCF_000258535.2 | NC_017672.3 | 630504 | 631691 | 1187 | - | tuf | WP_012589624.1 |

| | | | | | | | |
|---|---|---|---|---|---|---|---|
| GCF_001465255.1 | NZ_CP013652.1 | 1336249 | 1337436 | 1187 | - | tuf | WP_134488375.1 |
| GCF_963311185.1 | NZ_OY725677.1 | 1442658 | 1443845 | 1187 | + | tuf | WP_085772126.1 |
| GCF_000758725.1 | NZ_CP009428.1 | 1756859 | 1758046 | 1187 | + | tuf | WP_085772126.1 |
| GCF_031599675.1 | NZ_CP134223.1 | 1824696 | 1825883 | 1187 | + | tuf | WP_363348256.1 |
| GCF_002240225.1 | NZ_CP022584.1 | 3387702 | 3388889 | 1187 | + | tuf | WP_363348256.1 |
| GCF_030263935.1 | NZ_CP127162.1 | 1453869 | 1455056 | 1187 | - | tuf | WP_136495663.1 |
| GCF_022811565.1 | NZ_CP024795.1 | 3927214 | 3928401 | 1187 | + | tuf | WP_136495663.1 |
| GCF_004135365.1 | NZ_CP035492.1 | 2143715 | 2144902 | 1187 | - | tuf | WP_202071278.1 |
| GCF_009728935.1 | NZ_CP034235.1 | 3518556 | 3519743 | 1187 | - | tuf | WP_202071278.1 |
| GCF_010365285.1 | NZ_CP048286.1 | 846870 | 848057 | 1187 | - | tuf | WP_016917710.1 |
| GCF_000981585.1 | NZ_LN831776.1 | 2078144 | 2079331 | 1187 | - | tuf | WP_016917710.1 |
| GCF_032271945.1 | NZ_CP130319.1 | 921936 | 923123 | 1187 | + | tuf | WP_407050625.1 |
| GCF_000612505.1 | NZ_CP004078.1 | 941309 | 942496 | 1187 | + | tuf | WP_407050625.1 |
| GCF_030866765.1 | NZ_CP133017.1 | 2866004 | 2867191 | 1187 | + | tuf | WP_012253644.1 |
| GCF_018966525.1 | NZ_CP076607.1 | 5370323 | 5371510 | 1187 | - | tuf | WP_012255488.1 |
| GCF_024734895.1 | NZ_CP091430.1 | 2282068 | 2283255 | 1187 | + | tuf | WP_012454025.1 |
| GCF_000758685.1 | NZ_CP009286.1 | 5238990 | 5240177 | 1187 | - | tuf | WP_012456596.1 |
| GCF_001644605.1 | NZ_CP011388.1 | 3220754 | 3221941 | 1187 | - | tuf | WP_115836066.1 |
| GCF_000235585.1 | NC_016641.1 | 129753 | 130940 | 1187 | - | tuf | WP_009764297.1 |
| GCF_007066085.1 | NZ_CP041405.1 | 154141 | 155328 | 1187 | - | tuf | WP_009764297.1 |
| GCF_017086365.1 | NZ_CP070969.1 | 4412588 | 4413775 | 1187 | + | tuf | WP_099511374.1 |
| GCF_018207515.1 | NZ_CP073365.1 | 4433279 | 4434466 | 1187 | + | tuf | WP_099511374.1 |
| GCF_036884255.1 | NZ_CP145605.1 | 1937700 | 1938887 | 1187 | + | tuf | WP_259060525.1 |
| GCF_900177425.1 | NZ_LT840184.1 | 1956906 | 1958093 | 1187 | + | tuf | WP_259060525.1 |
| GCF_028747985.1 | NZ_CP118108.1 | 1547004 | 1548188 | 1184 | - | tuf | WP_426958299.1 |
| GCF_030122845.1 | NZ_CP126084.1 | 1566230 | 1567414 | 1184 | - | tuf | WP_426958299.1 |
| GCF_001908275.1 | NZ_CP018620.1 | 1328167 | 1329354 | 1187 | - | tuf | WP_077806599.1 |
| GCF_001685395.1 | NZ_CP014167.1 | 122416 | 123597 | 1181 | - | tuf | WP_218193899.1 |
| GCF_004367585.1 | NZ_CP038015.1 | 724026 | 725207 | 1181 | + | tuf | WP_218193899.1 |
| GCF_003595195.1 | NZ_CP032418.1 | 363779 | 364966 | 1187 | - | tuf | WP_141492040.1 |
| GCF_001700985.1 | NZ_CP016622.1 | 1246433 | 1247605 | 1172 | + | tuf | WP_038585819.1 |

| | | | | | | | |
|---|---|---|---|---|---|---|---|
| GCF_016939435.1 | NZ_CP070511.1 | 1264926 | 1266098 | 1172 | + | tuf | WP_038585819.1 |
| GCF_003268595.1 | NZ_CP029797.1 | 1402045 | 1403217 | 1172 | + | tuf | WP_037154426.1 |
| GCF_000829395.1 | NZ_AP014680.1 | 1423996 | 1425168 | 1172 | + | tuf | WP_037154426.1 |
| GCF_008632615.1 | NZ_CP043939.1 | 606601 | 607890 | 1289 | - | tuf | WP_067979822.1 |
| GCF_900015005.1 | NZ_LN898144.1 | 613858 | 615141 | 1283 | - | tuf | WP_038559816.1 |
| GCF_040409965.1 | NZ_CP158670.1 | 623098 | 624387 | 1289 | - | tuf | WP_015816524.1 |
| GCF_013127755.1 | NZ_CP053421.1 | 605265 | 606554 | 1289 | - | tuf | WP_011452069.1 |
| GCF_001702215.1 | NZ_CP014933.1 | 124191 | 125363 | 1172 | - | tuf | WP_220664386.1 |
| GCF_001611155.1 | NZ_CP012294.1 | 2753450 | 2754622 | 1172 | - | tuf | WP_220660990.1 |
| GCF_034049535.1 | NZ_CP139080.1 | 1837957 | 1839129 | 1172 | + | tuf | WP_048600220.1 |
| GCF_964063155.1 | NZ_OZ061292.1 | 1859095 | 1860267 | 1172 | + | tuf | WP_048600220.1 |
| GCF_000014505.1 | NC_008525.1 | 4347814 | 4349001 | 1187 | + | tuf | WP_176638759.1 |
| GCF_047220215.1 | NZ_CP178375.1 | 3160662 | 3161849 | 1187 | + | tuf | WP_257767150.1 |
| GCF_022394675.1 | NZ_CP091882.1 | 3177422 | 3178609 | 1187 | + | tuf | WP_257767150.1 |
| GCF_045159795.1 | NZ_CP048037.1 | 646765 | 647937 | 1172 | + | tuf | WP_206547510.1 |
| GCF_016724865.1 | NZ_CP068053.1 | 665861 | 667033 | 1172 | + | tuf | WP_206547510.1 |
| GCF_002243645.1 | NZ_CP017704.1 | 3770907 | 3772079 | 1172 | - | tuf | WP_223020762.1 |
| GCF_004358325.1 | NZ_CP037940.1 | 3788971 | 3790143 | 1172 | - | tuf | WP_223020762.1 |
| GCF_001687565.2 | NZ_CP016534.2 | 937126 | 938298 | 1172 | + | tuf | WP_146298239.1 |
| GCF_001687665.2 | NZ_CP016543.2 | 956120 | 957292 | 1172 | + | tuf | WP_146298239.1 |
| GCF_002009235.1 | NZ_CP019401.1 | 3969719 | 3970891 | 1172 | + | tuf | WP_338531531.1 |
| GCF_013368775.1 | NZ_CP051177.1 | 3998910 | 4000082 | 1172 | + | tuf | WP_338531531.1 |
| GCF_001687585.2 | NZ_CP016537.2 | 1757400 | 1758587 | 1187 | + | tuf | WP_011510053.1 |
| GCF_001465835.2 | NZ_CP013661.2 | 1498402 | 1499589 | 1187 | + | tuf | WP_011314640.1 |
| GCF_001999905.1 | NZ_CP019640.1 | 858595 | 859782 | 1187 | + | tuf | WP_088871000.1 |
| GCF_030413595.1 | NZ_CP129238.1 | 2005684 | 2006871 | 1187 | - | tuf | WP_088871000.1 |
| GCF_001687625.2 | NZ_CP016538.2 | 3235812 | 3236996 | 1184 | + | tuf | WP_102113007.1 |
| GCF_001687605.2 | NZ_CP016539.2 | 3256918 | 3258102 | 1184 | + | tuf | WP_102113007.1 |
| GCF_001465795.2 | NZ_CP013659.2 | 1189499 | 1190686 | 1187 | + | tuf | WP_048858902.1 |
| GCF_001186155.3 | NZ_CP016540.2 | 1296706 | 1297893 | 1187 | + | tuf | WP_011444905.1 |
| GCF_018326425.1 | NZ_AP024601.1 | 4171265 | 4172452 | 1187 | + | tuf | WP_039388092.1 |

| | | | | | | |
|---|---|---|---|---|---|---|
| GCF_002688605.1 | NZ_CP024035.1 | 3084313 | 3085500 | 1187 - | tuf | WP_147091457.1 |
| GCF_021441905.1 | NZ_CP090431.1 | 2454218 | 2455405 | 1187 - | tuf | WP_168603832.1 |
| GCF_006094495.1 | NZ_CP035094.1 | 2037841 | 2039028 | 1187 - | tuf | WP_232495571.1 |
| GCF_008973485.1 | NZ_CP031223.1 | 888059 | 889246 | 1187 + | tuf | WP_169492433.1 |
| GCF_007301515.1 | NZ_CP041666.1 | 3190009 | 3191196 | 1187 + | tuf | WP_007011876.1 |
| GCF_021049245.1 | NZ_CP088020.1 | 1991088 | 1992275 | 1187 - | tuf | WP_008828240.1 |
| GCF_022170785.1 | NZ_CP081870.1 | 114778 | 115965 | 1187 + | tuf | WP_187335650.1 |
| GCF_006542275.1 | NZ_CP041217.1 | 3541690 | 3542862 | 1172 + | tuf | WP_306051289.1 |
| GCF_019272935.1 | NZ_CP040553.1 | 3564113 | 3565285 | 1172 + | tuf | WP_306051289.1 |
| GCF_016495725.1 | NZ_CP054705.1 | 1300791 | 1301963 | 1172 - | tuf | WP_095446817.1 |
| GCF_016495865.1 | NZ_CP054706.1 | 1318841 | 1320013 | 1172 - | tuf | WP_095446817.1 |
| GCF_006740705.1 | NZ_CP035485.1 | 4356452 | 4357624 | 1172 - | tuf | WP_015501705.1 |
| GCF_003336365.1 | NZ_CP031092.1 | 246839 | 248011 | 1172 + | tuf | WP_015493755.1 |
| GCF_001685435.2 | NZ_CP011361.2 | 269890 | 271062 | 1172 + | tuf | WP_015493755.1 |
| GCF_038561955.2 | NZ_CP138333.2 | 218006 | 219178 | 1172 + | tuf | WP_049833262.1 |
| GCF_001721685.1 | NZ_CP012502.1 | 243156 | 244328 | 1172 + | tuf | WP_049833262.1 |
| GCF_008694105.1 | NZ_CP041364.1 | 411569 | 412756 | 1187 - | tuf | WP_141450933.1 |
| GCF_008033155.1 | NZ_CP042371.1 | 1459255 | 1460427 | 1172 - | tuf | WP_175276136.1 |
| GCF_001702155.1 | NZ_CP014912.1 | 1477010 | 1478182 | 1172 - | tuf | WP_175276136.1 |
| GCF_017792025.1 | NZ_CP046956.1 | 754163 | 755344 | 1181 - | tuf | WP_109227340.1 |
| GCF_034479305.1 | NZ_CP140150.1 | 1900402 | 1901583 | 1181 + | tuf | WP_012462217.1 |
| GCF_028735875.1 | NZ_CP117834.1 | 2979663 | 2980835 | 1172 - | tuf | WP_088665145.1 |
| GCF_007725405.1 | NZ_CP042163.1 | 3000819 | 3001991 | 1172 - | tuf | WP_088665145.1 |
| GCF_036700015.1 | NZ_CP144921.1 | 4561514 | 4562686 | 1172 + | tuf | WP_306753150.1 |
| GCF_028747805.1 | NZ_CP114887.1 | 1085417 | 1086589 | 1172 + | tuf | WP_271112670.1 |
| GCF_019930545.1 | NZ_CP082837.1 | 2222778 | 2223950 | 1172 - | tuf | WP_271112670.1 |
| GCF_033304595.1 | NZ_CP116341.1 | 434839 | 436011 | 1172 + | tuf | WP_271072128.1 |
| GCF_033304615.1 | NZ_CP129118.1 | 1143022 | 1144194 | 1172 + | tuf | WP_271072128.1 |
| GCF_041295575.1 | NZ_CP160452.1 | 12545 | 13717 | 1172 - | tuf | WP_090616862.1 |
| GCF_001590685.1 | NZ_CP014616.1 | 3157167 | 3158339 | 1172 - | tuf | WP_090616862.1 |
| GCF_030406775.1 | NZ_CP129119.1 | 3320618 | 3321790 | 1172 - | tuf | WP_020951885.1 |

| | | | | | | | |
|---|---|---|---|---|---|---|---|
| GCF_002081995.1 | NZ_CP015108.1 | 3343930 | 3345102 | 1172 | - | tuf | WP_020951885.1 |
| GCF_001753205.1 | NZ_CP017560.1 | 579466 | 580638 | 1172 | + | tuf | WP_074966919.1 |
| GCF_011466855.1 | NZ_CP045927.1 | 604493 | 605665 | 1172 | + | tuf | WP_074966919.1 |
| GCF_000236925.1 | NC_016941.1 | 391378 | 392550 | 1172 | - | tuf | WP_085376690.1 |
| GCF_025560845.1 | NZ_CP094803.1 | 1590628 | 1591800 | 1172 | + | tuf | WP_085376690.1 |
| GCF_016028295.1 | NZ_CP065712.1 | 2311335 | 2312507 | 1172 | + | tuf | WP_011747079.1 |
| GCF_040739365.1 | NZ_CP145258.1 | 2336771 | 2337943 | 1172 | + | tuf | WP_011747079.1 |
| GCF_003966625.1 | NZ_AP018587.1 | 2488929 | 2490101 | 1172 | - | tuf | WP_234853844.1 |
| GCF_016028275.1 | NZ_CP065711.1 | 2515831 | 2517003 | 1172 | - | tuf | WP_234853844.1 |
| GCF_030294405.1 | NZ_CP125718.1 | 3089919 | 3091091 | 1172 | + | tuf | WP_272833720.1 |
| GCF_007814525.1 | NZ_CP031274.1 | 3118111 | 3119283 | 1172 | + | tuf | WP_272833720.1 |
| GCF_022557135.1 | NZ_CP092966.1 | 412163 | 413335 | 1172 | + | tuf | WP_101498624.1 |
| GCF_900637235.1 | NZ_LR134360.1 | 1669569 | 1670741 | 1172 | - | tuf | WP_101498624.1 |
| GCF_003718735.1 | NZ_CP033460.1 | 1803192 | 1804364 | 1172 | - | tuf | WP_010401000.1 |
| GCF_025558825.1 | NZ_CP094740.1 | 1827185 | 1828357 | 1172 | - | tuf | WP_010401000.1 |
| GCF_006094375.1 | NZ_CP035288.1 | 599118 | 600290 | 1172 | + | tuf | WP_135312074.1 |
| GCF_016127455.1 | NZ_CP066013.1 | 2717139 | 2718311 | 1172 | + | tuf | WP_135313800.1 |
| GCF_003012915.1 | NZ_CP027770.1 | 531267 | 532439 | 1172 | + | tuf | WP_045983513.1 |
| GCF_029774255.1 | NZ_CP069082.1 | 2445505 | 2446677 | 1172 | - | tuf | WP_045983513.1 |
| GCF_006094395.1 | NZ_CP035291.1 | 547265 | 548437 | 1172 | + | tuf | WP_205294585.1 |
| GCF_003812505.1 | NZ_CP033732.1 | 571626 | 572798 | 1172 | + | tuf | WP_205294585.1 |
| GCF_038789205.1 | NZ_CP128355.1 | 1529306 | 1530478 | 1172 | - | tuf | WP_024843763.1 |
| GCF_000816085.1 | NZ_CP008747.1 | 1554207 | 1555379 | 1172 | - | tuf | WP_024843763.1 |
| GCF_003019255.1 | NZ_CP027846.1 | 3235910 | 3237082 | 1172 | - | tuf | WP_076526689.1 |
| GCF_015775975.1 | NZ_CP064056.1 | 3259355 | 3260527 | 1172 | - | tuf | WP_076526689.1 |
| GCF_001558775.1 | NZ_CP014022.1 | 1946376 | 1947548 | 1172 | + | tuf | WP_036703025.1 |
| GCF_002101335.1 | NZ_CP020773.1 | 1975499 | 1976671 | 1172 | + | tuf | WP_036703025.1 |
| GCF_043186315.1 | NZ_CP171119.1 | 227568 | 228740 | 1172 | + | tuf | WP_271116819.1 |
| GCF_900187005.1 | NZ_LT906464.1 | 798493 | 799665 | 1172 | + | tuf | WP_271116819.1 |
| GCF_002442935.1 | NZ_CP017466.1 | 635070 | 636242 | 1172 | + | tuf | WP_089345724.1 |
| GCF_016599795.1 | NZ_CP067014.1 | 656861 | 658033 | 1172 | + | tuf | WP_089345724.1 |

| | | | | | | | |
|---|---|---|---|---|---|---|---|
| GCF_002208805.2 | NZ_CP022096.2 | 3082857 | 3084029 | 1172 | - | tuf | WP_272858444.1 |
| GCF_900186985.1 | NZ_LT906447.1 | 3107367 | 3108539 | 1172 | - | tuf | WP_272858444.1 |
| GCF_016026635.1 | NZ_CP065635.1 | 1288265 | 1289437 | 1172 | - | tuf | WP_114075655.1 |
| GCF_044794325.1 | NZ_CP085024.1 | 1313522 | 1314694 | 1172 | + | tuf | WP_114075655.1 |
| GCF_020883535.1 | NZ_CP086654.1 | 999623 | 1000795 | 1172 | - | tuf | WP_101459508.1 |
| GCF_031297465.1 | NZ_CP113107.1 | 2076218 | 2077390 | 1172 | + | tuf | WP_101459508.1 |
| GCF_016127715.1 | NZ_CP066042.1 | 1117436 | 1118608 | 1172 | - | tuf | WP_036754146.1 |
| GCF_000010125.1 | NC_007350.1 | 1142629 | 1143801 | 1172 | - | tuf | WP_036754146.1 |
| GCF_025558325.1 | NZ_CP094712.1 | 353684 | 354856 | 1172 | - | tuf | WP_080621080.1 |
| GCF_900636685.1 | NZ_LR134304.1 | 649843 | 651015 | 1172 | - | tuf | WP_080621080.1 |
| GCF_900187055.1 | NZ_LT906460.1 | 1837026 | 1838198 | 1172 | - | tuf | WP_101752314.1 |
| GCF_025561465.1 | NZ_CP095100.1 | 2074218 | 2075390 | 1172 | - | tuf | WP_101752314.1 |
| GCF_029024945.1 | NZ_CP118976.1 | 3527383 | 3528570 | 1187 | + | tuf | WP_035038342.1 |
| GCF_025558845.1 | NZ_CP094825.1 | 2748011 | 2749198 | 1187 | - | tuf | WP_049705631.1 |
| GCF_003571725.1 | NZ_CP032159.1 | 385326 | 386498 | 1172 | + | tuf | WP_108434239.1 |
| GCF_000709415.1 | NZ_CP008724.1 | 411878 | 413050 | 1172 | + | tuf | WP_108434260.1 |
| GCF_003856495.1 | NZ_CP034118.1 | 3377436 | 3378623 | 1187 | - | tuf | WP_011385499.1 |
| GCF_001552035.1 | NZ_CP012480.1 | 3397443 | 3398630 | 1187 | - | tuf | WP_011385499.1 |
| GCF_900636475.1 | NZ_LR134283.1 | 2022858 | 2024030 | 1172 | + | tuf | WP_275790763.1 |
| GCF_900476055.1 | NZ_LS483444.1 | 2043628 | 2044800 | 1172 | + | tuf | WP_275790763.1 |
| GCF_900636575.1 | NZ_LR134293.1 | 3037709 | 3038881 | 1172 | - | tuf | WP_276118530.1 |
| GCF_003086355.1 | NZ_CP031733.1 | 3057053 | 3058225 | 1172 | - | tuf | WP_276118533.1 |
| GCF_016725005.1 | NZ_CP068213.1 | 245178 | 246350 | 1172 | + | tuf | WP_127747093.1 |
| GCF_900475445.1 | NZ_LS483383.1 | 3669863 | 3671035 | 1172 | - | tuf | WP_127747093.1 |
| GCF_034137685.1 | NZ_CP139418.1 | 1232453 | 1233625 | 1172 | - | tuf | WP_286018247.1 |
| GCF_016128095.1 | NZ_CP066069.1 | 2115201 | 2116373 | 1172 | + | tuf | WP_286018247.1 |
| GCF_015689395.1 | NZ_CP065054.1 | 3052012 | 3053199 | 1187 | + | tuf | WP_176870551.1 |
| GCF_900475025.1 | NZ_LS483343.1 | 3295304 | 3296491 | 1187 | + | tuf | WP_207987477.1 |
| GCF_013267695.1 | NZ_CP054015.1 | 1650774 | 1651961 | 1187 | - | tuf | WP_168819134.1 |
| GCF_019046945.1 | NZ_CP077224.1 | 2488057 | 2489244 | 1187 | - | tuf | WP_229953893.1 |
| GCF_003627155.1 | NZ_CP032621.1 | 1317653 | 1318825 | 1172 | + | tuf | WP_067786961.1 |

| | | | | | | | |
|---|---|---|---|---|---|---|---|
| GCF_019774635.1 | NZ_CP081502.1 | 3283418 | 3284605 | 1187 | - | tuf | WP_099470873.1 |
| GCF_001598035.1 | NZ_CP014835.1 | 3300356 | 3301543 | 1187 | - | tuf | WP_099470873.1 |
| GCF_041081565.1 | NZ_CP163513.1 | 386233 | 387405 | 1172 | - | tuf | WP_143972233.1 |
| GCF_001708305.1 | NZ_CP016953.1 | 404096 | 405268 | 1172 | - | tuf | WP_143972233.1 |
| GCF_000479335.1 | NC_022584.1 | 2928299 | 2929486 | 1187 | + | tuf | WP_012111645.1 |
| GCF_032595075.1 | NZ_CP118735.1 | 2945395 | 2946582 | 1187 | + | tuf | WP_012111645.1 |
| GCF_016127275.1 | NZ_CP065994.1 | 1892790 | 1893962 | 1172 | + | tuf | WP_013300913.1 |
| GCF_000831485.1 | NZ_CP010783.1 | 526297 | 527469 | 1172 | + | tuf | WP_047805732.1 |
| GCF_900475975.1 | NZ_LS483436.1 | 1023742 | 1024914 | 1172 | + | tuf | WP_160732480.1 |
| GCF_003627135.1 | NZ_CP032620.1 | 1852927 | 1854114 | 1187 | + | tuf | WP_264227481.1 |
| GCF_018127725.1 | NZ_CP072329.1 | 1871336 | 1872523 | 1187 | + | tuf | WP_264227481.1 |
| GCF_034134805.1 | NZ_CP139419.1 | 1728995 | 1730182 | 1187 | + | tuf | WP_014130838.1 |
| GCF_900475675.1 | NZ_LS483403.1 | 1747407 | 1748594 | 1187 | + | tuf | WP_014130838.1 |
| GCF_029866925.1 | NZ_CP119172.1 | 1670377 | 1671564 | 1187 | + | tuf | WP_338610390.1 |
| GCF_001623565.1 | NZ_CP015196.1 | 1688667 | 1689854 | 1187 | + | tuf | WP_338610390.1 |
| GCF_900187085.1 | NZ_LT906439.1 | 3253891 | 3255063 | 1172 | - | tuf | WP_140196245.1 |
| GCF_001281025.1 | NZ_CP012646.1 | 3277105 | 3278277 | 1172 | - | tuf | WP_140196245.1 |
| GCF_019048645.1 | NZ_CP077404.1 | 577201 | 578373 | 1172 | + | tuf | WP_245137060.1 |
| GCF_900637025.1 | NZ_LR134336.1 | 596129 | 597301 | 1172 | + | tuf | WP_245137060.1 |
| GCF_018137985.1 | NZ_CP073084.1 | 2455094 | 2456266 | 1172 | + | tuf | WP_179028205.1 |
| GCF_001642085.1 | NZ_CP014699.1 | 2473519 | 2474691 | 1172 | + | tuf | WP_179028205.1 |
| GCF_037076355.1 | NZ_AP026968.1 | 268850 | 270022 | 1172 | + | tuf | WP_024095804.1 |
| GCF_000164675.2 | NC_015678.1 | 295361 | 296533 | 1172 | + | tuf | WP_024095804.1 |
| GCF_021654455.1 | NZ_AP024276.1 | 3148871 | 3150043 | 1172 | - | tuf | WP_014876013.1 |
| GCF_000187935.1 | NZ_AEUT02000001.1 | 3177759 | 3178931 | 1172 | - | tuf | WP_014876013.1 |
| GCF_004843545.1 | NZ_CP039457.1 | 3135132 | 3136304 | 1172 | - | tuf | WP_024095804.1 |
| GCF_003963555.1 | NZ_CP034543.1 | 3160239 | 3161411 | 1172 | - | tuf | WP_024095804.1 |
| GCF_002953735.1 | NZ_CP025536.1 | 363934 | 365106 | 1172 | + | tuf | WP_072503619.1 |
| GCF_001457635.1 | NZ_LN831051.1 | 390602 | 391774 | 1172 | + | tuf | WP_072503619.1 |
| GCF_900475415.1 | NZ_LS483388.1 | 5130725 | 5131912 | 1187 | + | tuf | WP_211938006.1 |
| GCF_000221985.1 | NC_015875.1 | 5151846 | 5153033 | 1187 | + | tuf | WP_211938023.1 |

| | | | | | | | |
|---|---|---|---|---|---|---|---|
| GCF_900637075.1 | NZ_LR134341.1 | 1105449 | 1106636 | 1187 | + | tuf | WP_110449801.1 |
| GCF_900475035.1 | NZ_LS483338.1 | 1126460 | 1127647 | 1187 | + | tuf | WP_110449801.1 |
| GCF_008803015.1 | NZ_CP043405.1 | 1394298 | 1395485 | 1187 | + | tuf | WP_012521769.1 |
| GCF_003595525.1 | NZ_CP022680.1 | 1416764 | 1417951 | 1187 | + | tuf | WP_012521769.1 |
| GCF_003609975.1 | NZ_AP018400.1 | 3350852 | 3352039 | 1187 | + | tuf | WP_137100437.1 |
| GCF_011387195.1 | NZ_CP046919.1 | 3369703 | 3370890 | 1187 | + | tuf | WP_137100451.1 |
| GCF_000253315.1 | NC_017595.1 | 1250073 | 1251260 | 1187 | + | tuf | WP_106747996.1 |
| GCF_900475505.1 | NZ_LS483385.1 | 1268260 | 1269447 | 1187 | + | tuf | WP_106747996.1 |
| GCF_003172995.1 | NZ_CP029490.1 | 7040935 | 7042122 | 1187 | - | tuf | WP_136964105.1 |
| GCF_000026745.1 | NC_012926.1 | 7060051 | 7061238 | 1187 | - | tuf | WP_136964105.1 |
| GCF_032594915.1 | NZ_CP118733.1 | 1929838 | 1931010 | 1172 | - | tuf | WP_099999166.1 |
| GCF_903886475.1 | NZ_LR822015.1 | 1948832 | 1950004 | 1172 | - | tuf | WP_099999166.1 |
| GCF_024346585.1 | NZ_AP024523.1 | 3515971 | 3517143 | 1172 | + | tuf | WP_165100936.1 |
| GCF_002355215.1 | NZ_AP014612.1 | 4250120 | 4251292 | 1172 | + | tuf | WP_165102735.1 |
| GCF_900475595.1 | NZ_LS483397.1 | 1837049 | 1838236 | 1187 | + | tuf | WP_216708527.1 |
| GCF_900636885.1 | NZ_LR134323.1 | 1822045 | 1823232 | 1187 | + | tuf | WP_013652468.1 |
| GCF_900636445.1 | NZ_LR134275.1 | 1839389 | 1840576 | 1187 | + | tuf | WP_013652468.1 |
| GCF_017639855.1 | NZ_CP072115.1 | 489478 | 490650 | 1172 | + | tuf | WP_039615500.1 |
| GCF_003546865.1 | NZ_CP023434.1 | 3556542 | 3557714 | 1172 | + | tuf | WP_039615500.1 |
| GCF_002250055.1 | NZ_CP018866.1 | 963695 | 964867 | 1172 | + | tuf | WP_166189173.1 |
| GCF_002157855.1 | NZ_CP020880.1 | 983646 | 984818 | 1172 | + | tuf | WP_166189173.1 |
| GCF_037949505.1 | NZ_CP149939.1 | 2378281 | 2379453 | 1172 | + | tuf | WP_196101674.1 |
| GCF_000725365.1 | NZ_CP008876.1 | 2398382 | 2399554 | 1172 | + | tuf | WP_196101674.1 |
| GCF_001712815.1 | NZ_CP012047.1 | 125083 | 126255 | 1172 | + | tuf | WP_118941211.1 |
| GCF_003795145.1 | NZ_CP027786.1 | 3343139 | 3344311 | 1172 | + | tuf | WP_118941211.1 |
| GCF_003795125.1 | NZ_CP027783.1 | 636841 | 638028 | 1187 | + | tuf | WP_249037192.1 |
| GCF_015326725.1 | NZ_CP064060.1 | 653584 | 654771 | 1187 | + | tuf | WP_249037192.1 |
| GCF_005886755.1 | NZ_CP039710.1 | 1959431 | 1960603 | 1172 | - | tuf | WP_022710937.1 |
| GCF_000227705.2 | NC_019897.1 | 1977406 | 1978578 | 1172 | - | tuf | WP_022710937.1 |
| GCF_963662535.1 | NZ_OY759718.1 | 2637133 | 2638305 | 1172 | + | tuf | WP_108644067.1 |
| GCF_963666195.1 | NZ_OY762653.1 | 2657065 | 2658237 | 1172 | + | tuf | WP_108644067.1 |

| | | | | | | | |
|---|---|---|---|---|---|---|---|
| GCF_002243515.1 | NZ_CP022657.1 | 647526 | 648713 | 1187 | + | tuf | WP_115688386.1 |
| GCF_002162355.1 | NZ_CP021434.1 | 2179459 | 2180631 | 1172 | - | tuf | WP_196941842.1 |
| GCF_044795995.1 | NZ_CP085009.1 | 2200239 | 2201411 | 1172 | - | tuf | WP_196941842.1 |
| GCF_004331915.1 | NZ_CP036528.1 | 2897711 | 2898883 | 1172 | + | tuf | WP_043752481.1 |
| GCF_002370195.2 | NZ_AP018335.1 | 2933021 | 2934193 | 1172 | + | tuf | WP_043752481.1 |
| GCF_014397115.1 | NZ_CP060720.1 | 987088 | 988260 | 1172 | + | tuf | WP_276718893.1 |
| GCF_014397115.1 | NZ_CP060720.1 | 4148051 | 4149223 | 1172 | + | tuf | WP_193081177.1 |
| GCF_011303955.1 | NZ_CP049886.1 | 4169292 | 4170464 | 1172 | + | tuf | WP_193081201.1 |
| GCF_011304195.1 | NZ_CP049887.1 | 1511036 | 1512208 | 1172 | + | tuf | WP_052638114.1 |
| GCF_029144185.1 | NZ_CP110232.1 | 1530022 | 1531194 | 1172 | + | tuf | WP_052638114.1 |
| GCF_035918315.1 | NZ_CP142146.1 | 1161861 | 1163033 | 1172 | - | tuf | WP_137192969.1 |
| GCF_024637875.1 | NZ_CP102451.1 | 580751 | 581923 | 1172 | - | tuf | WP_137192969.1 |
| GCF_023299185.1 | NZ_CP097017.1 | 4388294 | 4389481 | 1187 | + | tuf | WP_086089731.1 |
| GCF_001870205.1 | NZ_CP017267.1 | 738039 | 739211 | 1172 | + | tuf | WP_073194192.1 |
| GCF_004792515.1 | NZ_CP038865.1 | 764776 | 765948 | 1172 | + | tuf | WP_073194193.1 |
| GCF_005145005.1 | NZ_CP039712.1 | 3024751 | 3025923 | 1172 | + | tuf | WP_075698795.1 |
| GCF_002849835.1 | NZ_CP018622.1 | 3042264 | 3043436 | 1172 | + | tuf | WP_075698795.1 |
| GCF_001878675.1 | NZ_CP017962.1 | 2090115 | 2091287 | 1172 | + | tuf | WP_106472375.1 |
| GCF_026013645.1 | NZ_CP110224.1 | 2528659 | 2529831 | 1172 | + | tuf | WP_106472375.1 |
| GCF_002224365.1 | NZ_CP022437.1 | 1075977 | 1077164 | 1187 | + | tuf | WP_170080183.1 |
| GCF_018075365.1 | NZ_CP073011.1 | 1094348 | 1095535 | 1187 | + | tuf | WP_170080183.1 |
| GCF_002216775.1 | NZ_CP022315.1 | 137144 | 138316 | 1172 | + | tuf | WP_146362762.1 |
| GCF_018394055.1 | NZ_CP074441.1 | 155505 | 156677 | 1172 | + | tuf | WP_146362762.1 |
| GCF_001951095.1 | NZ_CP013934.1 | 2292474 | 2293646 | 1172 | + | tuf | WP_040963560.1 |
| GCF_011304355.1 | NZ_CP049888.1 | 2668766 | 2669953 | 1187 | - | tuf | WP_067464776.1 |
| GCF_004771075.1 | NZ_CP027563.1 | 546857 | 548029 | 1172 | - | tuf | WP_221431384.1 |
| GCF_014397255.1 | NZ_CP060724.1 | 1628919 | 1630106 | 1187 | + | tuf | WP_253515080.1 |
| GCF_001932615.1 | NZ_CP014332.1 | 1512451 | 1513638 | 1187 | - | tuf | WP_221421664.1 |
| GCF_000219805.1 | NC_015759.1 | 1244680 | 1245867 | 1187 | - | tuf | WP_133365668.1 |
| GCF_036327715.1 | NZ_CP130559.1 | 236881 | 238053 | 1172 | - | tuf | WP_100259031.1 |
| GCF_001761545.1 | NZ_CP017326.1 | 1568449 | 1569636 | 1187 | - | tuf | WP_200980826.1 |

| | | | | | | | |
|---|---|---|---|---|---|---|---|
| GCF_018502505.1 | NZ_CP075544.1 | 1569216 | 1570388 | 1172 | + | tuf | WP_265557387.1 |
| GCF_014789145.1 | NZ_CP061835.1 | 1327040 | 1328212 | 1172 | + | tuf | WP_003573803.1 |
| GCF_033095985.1 | NZ_AP026802.1 | 1346274 | 1347446 | 1172 | + | tuf | WP_003573803.1 |
| GCF_033095965.1 | NZ_AP026801.1 | 2026049 | 2027221 | 1172 | + | tuf | WP_028757605.1 |
| GCF_025149125.1 | NZ_CP102272.1 | 2045314 | 2046486 | 1172 | + | tuf | WP_028757605.1 |
| GCF_940677205.1 | NZ_OW712331.1 | 1716015 | 1717187 | 1172 | + | tuf | WP_064706966.1 |
| GCF_022845755.1 | NZ_AP025566.1 | 1735192 | 1736364 | 1172 | + | tuf | WP_064706966.1 |
| GCF_000144625.1 | NC_014376.1 | 1761459 | 1762631 | 1172 | + | tuf | WP_207600958.1 |
| GCF_004295125.1 | NZ_CP036170.1 | 1780697 | 1781869 | 1172 | + | tuf | WP_207600958.1 |
| GCF_025152405.1 | NZ_CP102292.1 | 1425708 | 1426880 | 1172 | + | tuf | WP_020049533.1 |
| GCF_044591705.1 | NZ_CP172594.1 | 1444923 | 1446095 | 1172 | + | tuf | WP_020049533.1 |
| GCF_017357865.1 | NZ_CP060096.1 | 1304886 | 1306058 | 1172 | + | tuf | WP_113438755.1 |
| GCF_017357865.1 | NZ_CP060096.1 | 1324582 | 1325754 | 1172 | + | tuf | WP_113438755.1 |
| GCF_000237085.1 | NC_016627.1 | 3806606 | 3807778 | 1172 | + | tuf | WP_163904961.1 |
| GCF_002844395.1 | NZ_CP025197.1 | 3826565 | 3827737 | 1172 | + | tuf | WP_163904961.1 |
| GCF_000015865.1 | NC_009012.1 | 1760534 | 1761706 | 1172 | + | tuf | WP_011424970.1 |
| GCF_000196455.1 | NC_014614.1 | 1779706 | 1780878 | 1172 | + | tuf | WP_011424970.1 |
| GCF_000196455.1 | NC_014614.1 | 1747562 | 1748734 | 1172 | + | tuf | WP_011424970.1 |
| GCF_047456275.1 | NZ_CP173357.1 | 1766733 | 1767908 | 1175 | + | tuf | WP_011424980.1 |
| GCF_047456275.1 | NZ_CP173357.1 | 1536172 | 1537344 | 1172 | + | tuf | WP_037069361.1 |
| GCF_047456285.1 | NZ_CP174122.1 | 1555811 | 1556983 | 1172 | + | tuf | WP_037069361.1 |
| GCF_047456285.1 | NZ_CP174122.1 | 1560686 | 1561858 | 1172 | + | tuf | WP_039844630.1 |
| GCF_023700225.1 | NZ_CP097897.1 | 1580349 | 1581521 | 1172 | + | tuf | WP_039844638.1 |
| GCF_023700225.1 | NZ_CP097897.1 | 2634326 | 2635498 | 1172 | - | tuf | WP_003573803.1 |
| GCF_000247605.1 | NC_016894.1 | 2653544 | 2654716 | 1172 | - | tuf | WP_003573803.1 |
| GCF_000247605.1 | NC_016894.1 | 787157 | 788329 | 1172 | - | tuf | WP_020049533.1 |
| GCF_000144695.1 | NC_014378.1 | 806363 | 807535 | 1172 | - | tuf | WP_020049533.1 |
| GCF_000144695.1 | NC_014378.1 | 1426737 | 1427909 | 1172 | + | tuf | WP_018241634.1 |
| GCF_902729405.1 | NZ_LR746496.1 | 1445950 | 1447122 | 1172 | + | tuf | WP_018241634.1 |
| GCF_002201475.1 | NZ_CP021422.1 | 1756253 | 1757425 | 1172 | + | tuf | WP_120703906.1 |
| GCF_022453685.1 | NZ_CP092643.1 | 1778183 | 1779355 | 1172 | + | tuf | WP_120703906.1 |

| | | | | | | | |
|---|---|---|---|---|---|---|---|
| GCF_017352215.1 | NZ_CP071444.1 | 1856153 | 1857325 | 1172 | + | tuf | WP_003558535.1 |
| GCF_017352215.1 | NZ_CP071444.1 | 1875369 | 1876541 | 1172 | + | tuf | WP_003558535.1 |
| GCF_014467015.1 | NZ_CP058559.1 | 3592516 | 3593688 | 1172 | + | tuf | WP_207241574.1 |
| GCF_014467015.1 | NZ_CP058559.1 | 3611714 | 3612886 | 1172 | + | tuf | WP_207241574.1 |
| GCF_000016985.1 | NC_009633.1 | 45772 | 46944 | 1172 | + | tuf | WP_003495154.1 |
| GCF_000016985.1 | NC_009633.1 | 66121 | 67293 | 1172 | + | tuf | WP_003495154.1 |
| GCF_000018325.1 | NC_009922.1 | 1756442 | 1757614 | 1172 | + | tuf | WP_004669088.1 |
| GCF_000018325.1 | NC_009922.1 | 1775695 | 1776867 | 1172 | + | tuf | WP_004669088.1 |
| GCF_010669305.1 | NZ_CP048649.1 | 1080662 | 1081834 | 1172 | + | tuf | WP_012651262.1 |
| GCF_010669305.1 | NZ_CP048649.1 | 1102109 | 1103281 | 1172 | + | tuf | WP_034518454.1 |
| GCF_004103735.1 | NZ_CP035281.1 | 2873330 | 2874502 | 1172 | - | tuf | WP_137136476.1 |
| GCF_004103735.1 | NZ_CP035281.1 | 2892119 | 2893291 | 1172 | - | tuf | WP_137136476.1 |
| GCF_010120715.1 | NZ_CP047591.1 | 1056458 | 1057630 | 1172 | + | tuf | WP_142822264.1 |
| GCF_010120715.1 | NZ_CP047591.1 | 1110986 | 1112158 | 1172 | + | tuf | WP_142822264.1 |
| GCF_000024605.1 | NC_013385.1 | 1993569 | 1994741 | 1172 | + | tuf | WP_017963897.1 |
| GCF_000024605.1 | NC_013385.1 | 2012784 | 2013956 | 1172 | + | tuf | WP_027685040.1 |
| GCF_029101545.1 | NZ_CP118866.1 | 1354885 | 1356057 | 1172 | + | tuf | WP_009996414.1 |
| GCF_018140635.1 | NZ_CP073280.1 | 1374117 | 1375289 | 1172 | + | tuf | WP_009996414.1 |
| GCF_014218335.1 | NZ_AP023367.1 | 1978284 | 1979456 | 1172 | - | tuf | WP_027508614.1 |
| GCF_014218355.1 | NZ_AP023368.1 | 1998243 | 1999415 | 1172 | - | tuf | WP_027508614.1 |
| GCF_009931695.1 | NZ_CP048000.1 | 1497604 | 1498776 | 1172 | + | tuf | WP_015339584.1 |
| GCF_016591975.1 | NZ_AP024169.1 | 1519548 | 1520720 | 1172 | + | tuf | WP_015339584.1 |
| GCF_940670685.1 | NZ_OW711693.1 | 1102392 | 1103564 | 1172 | + | tuf | WP_111220656.1 |
| GCF_020181435.1 | NZ_CP084016.1 | 1129236 | 1130408 | 1172 | + | tuf | WP_111220656.1 |
| GCF_030864025.1 | NZ_CP132968.1 | 1667421 | 1668608 | 1187 | + | tuf | WP_066966372.1 |
| GCF_005280655.1 | NZ_CP040058.1 | 3929942 | 3931129 | 1187 | + | tuf | WP_165324458.1 |
| GCF_001561955.1 | NZ_CP014223.1 | 5552277 | 5553464 | 1187 | + | tuf | WP_011952198.1 |
| GCF_023008305.1 | NZ_CP094682.1 | 158427 | 159599 | 1172 | + | tuf | WP_013065894.1 |
| GCF_003991135.1 | NZ_CP016379.1 | 339356 | 340528 | 1172 | + | tuf | WP_013065894.1 |
| GCF_003991135.1 | NZ_CP016379.1 | 2835672 | 2836844 | 1172 | - | tuf | WP_071166712.1 |
| GCF_018128945.1 | NZ_CP068393.1 | 2856019 | 2857191 | 1172 | - | tuf | WP_071166712.1 |

| | | | | | | | |
|---|---|---|---|---|---|---|---|
| GCF_018128945.1 | NZ_CP068393.1 | 3247407 | 3248594 | 1187 | + | tuf | WP_013420551.1 |
| GCF_018118585.1 | NZ_CP069421.1 | 3679123 | 3680310 | 1187 | - | tuf | WP_233022165.1 |
| GCF_018118585.1 | NZ_CP069421.1 | 3715557 | 3716744 | 1187 | - | tuf | WP_233022165.1 |
| GCF_039719005.1 | NZ_CP133584.1 | 3648716 | 3649903 | 1187 | - | tuf | WP_011158796.1 |
| GCF_003691585.1 | NZ_CP033169.1 | 3690851 | 3692038 | 1187 | - | tuf | WP_011158796.1 |
| GCF_003691585.1 | NZ_CP033169.1 | 739538 | 740722 | 1184 | + | tuf | WP_012565917.1 |
| GCF_003287895.1 | NZ_CP030280.1 | 760621 | 761805 | 1184 | + | tuf | WP_012565917.1 |
| GCF_034355335.1 | NZ_CP136422.1 | 3108376 | 3109563 | 1187 | - | tuf | WP_011390500.1 |
| GCF_002222595.2 | NZ_CP022413.2 | 3125607 | 3126794 | 1187 | - | tuf | WP_011390512.1 |
| GCF_034356035.1 | NZ_CP136423.1 | 3166761 | 3167945 | 1184 | + | tuf | WP_150040696.1 |
| GCF_015159595.1 | NZ_CP063304.1 | 238488 | 239660 | 1172 | - | tuf | WP_042456853.1 |
| GCF_033096465.1 | NZ_AP028156.1 | 258011 | 259183 | 1172 | - | tuf | WP_042456853.1 |
| GCF_025147765.1 | NZ_CP102265.1 | 914016 | 915200 | 1184 | - | tuf | WP_012149907.1 |
| GCF_020215665.1 | NZ_CP084061.1 | 1002556 | 1003737 | 1181 | - | tuf | WP_147143209.1 |
| GCF_010669205.1 | NZ_CP048626.1 | 501246 | 502427 | 1181 | + | tuf | WP_014412445.1 |
| GCF_001689125.2 | NZ_CP015405.2 | 1153900 | 1155084 | 1184 | - | tuf | WP_011477723.1 |
| GCF_025148125.1 | NZ_CP102267.1 | 869023 | 870204 | 1181 | - | tuf | WP_012148996.1 |
| GCF_037113525.1 | NZ_CP146963.1 | 934454 | 935635 | 1181 | - | tuf | WP_010977589.1 |
| GCF_001858005.1 | NZ_CP017831.1 | 1597955 | 1599136 | 1181 | + | tuf | WP_253308023.1 |
| GCF_000145035.1 | NC_014387.1 | 394562 | 395743 | 1181 | + | tuf | WP_341787577.1 |
| GCF_000007085.1 | NC_003869.1 | 322442 | 323623 | 1181 | + | tuf | WP_253308023.1 |
| GCF_000007085.1 | NC_003869.1 | 1335308 | 1336492 | 1184 | - | tuf | WP_341791125.1 |
| GCF_025999835.1 | NZ_AP025628.1 | 1152692 | 1153876 | 1184 | - | tuf | WP_341788298.1 |
| GCF_003932015.1 | NZ_CP034260.1 | 277511 | 278695 | 1184 | + | tuf | WP_395476545.1 |
| GCF_003932015.1 | NZ_CP034260.1 | 1920094 | 1921275 | 1181 | - | tuf | WP_250310946.1 |
| GCF_030708075.1 | NZ_CP117982.1 | 1268923 | 1270107 | 1184 | - | tuf | WP_395476545.1 |
| GCF_002777255.1 | NZ_CP017634.1 | 1396813 | 1397994 | 1181 | - | tuf | WP_341789528.1 |
| GCF_002777255.1 | NZ_CP017634.1 | 1293956 | 1295140 | 1184 | - | tuf | WP_341794459.1 |
| GCF_015708635.1 | NZ_CP051754.1 | 1126311 | 1127495 | 1184 | - | tuf | WP_342169915.1 |
| GCF_015708635.1 | NZ_CP051754.1 | 79587 | 80768 | 1181 | - | tuf | WP_010421650.1 |
| GCF_009734445.1 | NZ_CP046457.1 | 954043 | 955224 | 1181 | - | tuf | WP_004997779.1 |

| | | | | | | | |
|---|---|---|---|---|---|---|---|
| GCF_009734445.1 | NZ_CP046457.1 | 1297434 | 1298615 | 1181 | - | tuf | WP_023507966.1 |
| GCF_014303955.1 | NZ_CP060286.1 | 818497 | 819678 | 1181 | - | tuf | WP_004596195.1 |
| GCF_014467055.1 | NZ_CP060696.1 | 1035827 | 1037008 | 1181 | - | tuf | WP_014408924.1 |
| GCF_029211325.2 | NZ_CP135996.1 | 931652 | 932833 | 1181 | - | tuf | WP_004997779.1 |
| GCF_013315815.1 | NZ_CP046051.1 | 993297 | 994478 | 1181 | + | tuf | WP_004997779.1 |
| GCF_000012865.1 | NC_007503.1 | 944291 | 945472 | 1181 | - | tuf | WP_004997779.1 |
| GCF_000012865.1 | NC_007503.1 | 1078746 | 1079927 | 1181 | - | tuf | WP_202068916.1 |
| GCF_000178835.2 | NC_015275.1 | 828639 | 829820 | 1181 | - | tuf | WP_011191093.1 |
| GCF_030067835.1 | NZ_CP092441.1 | 1044169 | 1045356 | 1187 | + | tuf | WP_125216271.1 |
| GCF_019355395.1 | NZ_CP048829.1 | 1113954 | 1115141 | 1187 | + | tuf | WP_125216271.1 |
| GCF_040409835.1 | NZ_CP117826.1 | 506663 | 507850 | 1187 | + | tuf | WP_008188955.1 |
| GCF_040409835.1 | NZ_CP117826.1 | 526219 | 527406 | 1187 | + | tuf | WP_008188955.1 |
| GCF_003628755.1 | NZ_CP029256.1 | 5274877 | 5276064 | 1187 | + | tuf | WP_006936456.1 |
| GCF_003628755.1 | NZ_CP029256.1 | 5292510 | 5293697 | 1187 | + | tuf | WP_006936456.1 |
| GCF_030296775.1 | NZ_AP027742.1 | 2877639 | 2878826 | 1187 | + | tuf | WP_152501357.1 |
| GCF_018885085.1 | NZ_CP076401.1 | 2895296 | 2896483 | 1187 | + | tuf | WP_152501357.1 |
| GCF_018885085.1 | NZ_CP076401.1 | 1503477 | 1504649 | 1172 | + | tuf | WP_025311745.1 |
| GCF_001042715.1 | NZ_CP009687.1 | 1529093 | 1530265 | 1172 | + | tuf | WP_025311745.1 |
| GCF_001042715.1 | NZ_CP009687.1 | 1680516 | 1681688 | 1172 | + | tuf | WP_131616265.1 |
| GCF_000218855.1 | NC_015687.1 | 1704979 | 1706151 | 1172 | + | tuf | WP_131616265.1 |
| GCF_000218855.1 | NC_015687.1 | 1889286 | 1890458 | 1172 | - | tuf | WP_011567656.1 |
| GCF_002074155.1 | NZ_CP014176.1 | 3495917 | 3497089 | 1172 | + | tuf | WP_011567656.1 |
| GCF_002074155.1 | NZ_CP014176.1 | 3265355 | 3266527 | 1172 | - | tuf | WP_187429016.1 |
| GCF_001484725.1 | NZ_CP012395.1 | 3503139 | 3504311 | 1172 | - | tuf | WP_187429016.1 |
| GCF_001484725.1 | NZ_CP012395.1 | 457263 | 458435 | 1172 | + | tuf | WP_013960398.1 |
| GCF_001991075.2 | NZ_CP014204.2 | 3455419 | 3456591 | 1172 | - | tuf | WP_013960398.1 |
| GCF_001991075.2 | NZ_CP014204.2 | 568203 | 569375 | 1172 | + | tuf | WP_169639388.1 |
| GCF_000577895.1 | NZ_HG917868.1 | 589026 | 590198 | 1172 | + | tuf | WP_169639388.1 |
| GCF_000577895.1 | NZ_HG917868.1 | 3698454 | 3699638 | 1184 | + | tuf | WP_191083921.1 |
| GCF_000063585.1 | NC_009495.1 | 3714093 | 3715277 | 1184 | + | tuf | WP_191083928.1 |
| GCF_000063585.1 | NC_009495.1 | 1371228 | 1372412 | 1184 | + | tuf | WP_230449014.1 |

| | | | | | | | |
|---|---|---|---|---|---|---|---|
| GCF_014131795.1 | NZ_CP040626.1 | 3227474 | 3228658 | 1184 | - | tuf | WP_230449014.1 |
| GCF_014131795.1 | NZ_CP040626.1 | 4276255 | 4277439 | 1184 | - | tuf | WP_244408372.1 |
| GCF_020911725.1 | NZ_CP076620.1 | 4295475 | 4296659 | 1184 | - | tuf | WP_244408372.1 |
| GCF_020911725.1 | NZ_CP076620.1 | 3019173 | 3020357 | 1184 | - | tuf | WP_075798816.1 |
| GCF_001038625.1 | NZ_CP011803.1 | 3039505 | 3040689 | 1184 | - | tuf | WP_075798816.1 |
| GCF_001038625.1 | NZ_CP011803.1 | 224436 | 225620 | 1184 | - | tuf | WP_207417764.1 |
| GCF_000145275.1 | NC_014393.1 | 239746 | 240930 | 1184 | - | tuf | WP_207417764.1 |
| GCF_000145275.1 | NC_014393.1 | 997116 | 998300 | 1184 | + | tuf | WP_207448639.1 |
| GCF_002327205.1 | NZ_CP018630.1 | 1013240 | 1014424 | 1184 | + | tuf | WP_207448477.1 |
| GCF_002327205.1 | NZ_CP018630.1 | 3752598 | 3753782 | 1184 | - | tuf | WP_019463282.1 |
| GCF_900187165.1 | NZ_LT906477.1 | 3772927 | 3774111 | 1184 | - | tuf | WP_019463282.1 |
| GCF_900187165.1 | NZ_LT906477.1 | 916144 | 917316 | 1172 | + | tuf | WP_157706269.1 |
| GCF_008705175.1 | NZ_CP043998.1 | 213193 | 214365 | 1172 | + | tuf | WP_151175212.1 |
| GCF_008705175.1 | NZ_CP043998.1 | 3991591 | 3992763 | 1172 | - | tuf | WP_151175212.1 |
| GCF_003096175.1 | NZ_CP020953.1 | 1268406 | 1269578 | 1172 | - | tuf | WP_165196847.1 |
| GCF_003096175.1 | NZ_CP020953.1 | 1292396 | 1293568 | 1172 | - | tuf | WP_165196847.1 |
| GCF_030585465.1 | NZ_CP077615.1 | 3080164 | 3081336 | 1172 | - | tuf | WP_317057050.1 |
| GCF_030585465.1 | NZ_CP077615.1 | 3138458 | 3139630 | 1172 | - | tuf | WP_317057050.1 |
| GCF_002006355.2 | NZ_CP096980.1 | 2165846 | 2167018 | 1172 | + | tuf | WP_317057008.1 |
| GCF_002006355.2 | NZ_CP096980.1 | 2213152 | 2214324 | 1172 | + | tuf | WP_317057050.1 |
| GCF_003600355.1 | NZ_CP032416.1 | 935800 | 936972 | 1172 | + | tuf | WP_274789771.1 |
| GCF_003600355.1 | NZ_CP032416.1 | 2678486 | 2679658 | 1172 | - | tuf | WP_274789771.1 |
| GCF_002080475.1 | NZ_CP020559.1 | 756638 | 757810 | 1172 | + | tuf | WP_011046478.1 |
| GCF_002080475.1 | NZ_CP020559.1 | 3701729 | 3702901 | 1172 | - | tuf | WP_011046478.1 |
| GCF_017348895.1 | NZ_CP071376.1 | 4419646 | 4420818 | 1172 | - | tuf | WP_005862789.1 |
| GCF_017348895.1 | NZ_CP071376.1 | 4446459 | 4447631 | 1172 | - | tuf | WP_005862789.1 |
| GCF_019977655.1 | NZ_AP024849.1 | 1248944 | 1250116 | 1172 | - | tuf | WP_075996142.1 |
| GCF_019977655.1 | NZ_AP024849.1 | 1318914 | 1320086 | 1172 | + | tuf | WP_075996184.1 |
| GCF_035905475.1 | NZ_CP121166.1 | 866761 | 867933 | 1172 | - | tuf | WP_076696759.1 |
| GCF_035905475.1 | NZ_CP121166.1 | 891367 | 892539 | 1172 | - | tuf | WP_076696798.1 |
| GCF_002285495.1 | NZ_CP016786.1 | 2024808 | 2025980 | 1172 | + | tuf | WP_017467314.1 |

| | | | | | | | |
|---|---|---|---|---|---|---|---|
| GCF_002285495.1 | NZ_CP016786.1 | 2047112 | 2048284 | 1172 | + | tuf | WP_017467314.1 |
| GCF_000010265.1 | NC_011837.1 | 271186 | 272373 | 1187 | + | tuf | WP_182296794.1 |
| GCF_000010265.1 | NC_011837.1 | 1638795 | 1639979 | 1184 | + | tuf | WP_318650744.1 |
| GCF_000143685.1 | NC_014328.1 | 1660110 | 1661294 | 1184 | + | tuf | WP_318650744.1 |
| GCF_000143685.1 | NC_014328.1 | 1180019 | 1181191 | 1172 | + | tuf | WP_134711005.1 |
| GCF_004015185.1 | NZ_CP025746.1 | 1198786 | 1199958 | 1172 | + | tuf | WP_134711005.1 |
| GCF_004015185.1 | NZ_CP025746.1 | 1576817 | 1577989 | 1172 | - | tuf | WP_134711005.1 |
| GCF_003614235.1 | NZ_CP029458.1 | 1595681 | 1596853 | 1172 | - | tuf | WP_134711005.1 |
| GCF_003614235.1 | NZ_CP029458.1 | 1323851 | 1325023 | 1172 | + | tuf | WP_160784683.1 |
| GCF_001856645.1 | NZ_CP013018.1 | 1341804 | 1342976 | 1172 | + | tuf | WP_160784683.1 |
| GCF_001856645.1 | NZ_CP013018.1 | 95129 | 96301 | 1172 | + | tuf | WP_126918448.1 |
| GCF_016027375.1 | NZ_CP065681.1 | 4089076 | 4090248 | 1172 | - | tuf | WP_126918621.1 |
| GCF_016027375.1 | NZ_CP065681.1 | 1079626 | 1080798 | 1172 | + | tuf | WP_069456381.1 |
| GCF_000473995.1 | NC_022571.1 | 1099923 | 1101095 | 1172 | + | tuf | WP_069456381.1 |
| GCF_000473995.1 | NC_022571.1 | 1478997 | 1480169 | 1172 | + | tuf | WP_037382919.1 |
| GCF_000340885.1 | NC_020291.1 | 1498116 | 1499288 | 1172 | + | tuf | WP_037382919.1 |
| GCF_000340885.1 | NC_020291.1 | 1227712 | 1228884 | 1172 | + | tuf | WP_153442062.1 |
| GCF_000968375.1 | NZ_CP009933.1 | 1247618 | 1248790 | 1172 | + | tuf | WP_153442062.1 |
| GCF_000968375.1 | NZ_CP009933.1 | 1265792 | 1266964 | 1172 | + | tuf | WP_012707739.1 |
| GCF_003606265.1 | NZ_CP023671.1 | 1284702 | 1285874 | 1172 | + | tuf | WP_012707739.1 |
| GCF_003606265.1 | NZ_CP023671.1 | 3971412 | 3972584 | 1172 | - | tuf | WP_280659423.1 |
| GCF_000973705.1 | NZ_CP009225.1 | 3991290 | 3992462 | 1172 | - | tuf | WP_280659423.1 |
| GCF_000973705.1 | NZ_CP009225.1 | 87375 | 88547 | 1172 | + | tuf | WP_010969189.1 |
| GCF_001735765.2 | NZ_CP017253.2 | 107894 | 109066 | 1172 | + | tuf | WP_010969189.1 |
| GCF_001735765.2 | NZ_CP017253.2 | 1848422 | 1849594 | 1172 | + | tuf | WP_010969189.1 |
| GCF_026650945.1 | NZ_CP086155.1 | 1868942 | 1870114 | 1172 | + | tuf | WP_010969189.1 |
| GCF_026650945.1 | NZ_CP086155.1 | 1510971 | 1512143 | 1172 | + | tuf | WP_180940833.1 |
| GCF_000007625.1 | NC_004557.1 | 1530869 | 1532041 | 1172 | + | tuf | WP_180940833.1 |
| GCF_000007625.1 | NC_004557.1 | 1426079 | 1427251 | 1172 | + | tuf | WP_280732814.1 |
| GCF_006351925.1 | NZ_CP040924.1 | 1445885 | 1447057 | 1172 | + | tuf | WP_280732814.1 |
| GCF_006351925.1 | NZ_CP040924.1 | 1047740 | 1048912 | 1172 | + | tuf | WP_034859927.1 |

| | | | | | | | |
|---|---|---|---|---|---|---|---|
| GCF_002896855.1 | NZ_CP021850.1 | 1144496 | 1145668 | 1172 | + | tuf | WP_034859927.1 |
| GCF_001642655.1 | NZ_CP014170.1 | 1030330 | 1031502 | 1172 | + | tuf | WP_153442062.1 |
| GCF_001642655.1 | NZ_CP014170.1 | 1050252 | 1051424 | 1172 | + | tuf | WP_153442062.1 |
| GCF_025149785.1 | NZ_CP102277.1 | 1620196 | 1621383 | 1187 | + | tuf | WP_201073040.1 |
| GCF_025149915.1 | NZ_CP102278.1 | 4310325 | 4311512 | 1187 | - | tuf | WP_201073040.1 |
| GCF_019203185.1 | NZ_CP078093.1 | 1320167 | 1321354 | 1187 | + | tuf | WP_201073040.1 |
| GCF_019203185.1 | NZ_CP078093.1 | 4305719 | 4306906 | 1187 | - | tuf | WP_201073040.1 |
| GCF_019884785.1 | NZ_CP082326.1 | 1239765 | 1240952 | 1187 | + | tuf | WP_158044268.1 |
| GCF_019884785.1 | NZ_CP082326.1 | 4035317 | 4036504 | 1187 | - | tuf | WP_192499022.1 |
| GCF_008152325.1 | NZ_CP042243.1 | 1274487 | 1275674 | 1187 | + | tuf | WP_201073040.1 |
| GCF_008152325.1 | NZ_CP042243.1 | 4020710 | 4021897 | 1187 | - | tuf | WP_201073040.1 |
| GCF_038396635.1 | NZ_CP121687.1 | 2813214 | 2814401 | 1187 | - | tuf | WP_267177726.1 |
| GCF_009796305.1 | NZ_CP046996.1 | 2829936 | 2831123 | 1187 | - | tuf | WP_267177726.1 |
| GCF_002224645.1 | NZ_CP022121.1 | 2880361 | 2881548 | 1187 | - | tuf | WP_251933092.1 |
| GCF_002224645.1 | NZ_CP022121.1 | 2897090 | 2898277 | 1187 | - | tuf | WP_251933092.1 |
| GCF_000243155.2 | NC_018017.1 | 2919984 | 2921171 | 1187 | - | tuf | WP_066700956.1 |
| GCF_000243135.2 | NC_019903.1 | 2137705 | 2138892 | 1187 | - | tuf | WP_066520453.1 |
| GCF_000021925.1 | NC_011830.1 | 2540921 | 2542108 | 1187 | - | tuf | WP_025547259.1 |
| GCF_000231405.2 | NZ_CP007032.1 | 641575 | 642762 | 1187 | + | tuf | WP_007686588.1 |
| GCF_000024205.1 | NC_013216.1 | 1961674 | 1962861 | 1187 | - | tuf | WP_394437375.1 |
| GCF_000024205.1 | NC_013216.1 | 306035 | 307222 | 1187 | + | tuf | WP_070156847.1 |
| GCF_000214705.1 | NC_015573.1 | 1756060 | 1757247 | 1187 | + | tuf | WP_017184028.1 |
| GCF_000214705.1 | NC_015573.1 | 4084520 | 4085707 | 1187 | - | tuf | WP_004208725.1 |
| GCF_002005145.1 | NZ_CP019698.1 | 891820 | 893007 | 1187 | - | tuf | WP_245110852.1 |
| GCF_002005145.1 | NZ_CP019698.1 | 1455971 | 1457158 | 1187 | - | tuf | WP_260482428.1 |
| GCF_000016165.1 | NC_009253.1 | 877043 | 878230 | 1187 | + | tuf | WP_270077966.1 |
| GCF_000016165.1 | NC_009253.1 | 1411088 | 1412278 | 1190 | + | tuf | WP_031393767.1 |
| GCF_000215085.1 | NC_015589.1 | 2766558 | 2767748 | 1190 | + | tuf | WP_031393767.1 |
| GCF_000215085.1 | NC_015589.1 | 1295719 | 1296909 | 1190 | + | tuf | WP_202095162.1 |
| GCF_000233715.2 | NC_021184.1 | 1074363 | 1075550 | 1187 | + | tuf | WP_066606490.1 |
| GCF_000233715.2 | NC_021184.1 | 3862265 | 3863452 | 1187 | - | tuf | WP_233783925.1 |

| | | | | | | | |
|---|---|---|---|---|---|---|---|
| GCF_000255115.2 | NC_018068.1 | 3208526 | 3209719 | 1193 | + | tuf | WP_112381155.1 |
| GCF_000231385.2 | NC_018515.1 | 45958 | 47145 | 1187 | - | tuf | WP_160590894.1 |
| GCF_000235605.1 | NC_016584.1 | 522806 | 523999 | 1193 | + | tuf | WP_250753004.1 |
| GCF_045689965.1 | NZ_CP144211.1 | 3700965 | 3702155 | 1190 | + | tuf | WP_260598034.1 |
| GCF_000244895.1 | NZ_CM001441.1 | 1115593 | 1116780 | 1187 | - | tuf | WP_308517693.1 |
| GCF_000214435.1 | NC_015565.1 | 1188878 | 1190065 | 1187 | - | tuf | WP_114226649.1 |
| GCF_000214435.1 | NC_015565.1 | 2731790 | 2732977 | 1187 | + | tuf | WP_249503702.1 |
| GCF_025150245.1 | NZ_CP102279.1 | 3457749 | 3458939 | 1190 | - | tuf | WP_044333399.1 |
| GCF_025150085.1 | NZ_CP102280.1 | 3304377 | 3305567 | 1190 | - | tuf | WP_037534177.1 |
| GCF_012273195.1 | NZ_CP050964.1 | 3742270 | 3743460 | 1190 | - | tuf | WP_066574475.1 |
| GCF_025148445.1 | NZ_CP102270.1 | 1116405 | 1117592 | 1187 | + | tuf | WP_169944545.1 |
| GCF_000178115.2 | NC_014828.1 | 2122309 | 2123496 | 1187 | - | tuf | WP_187537832.1 |
| GCF_000152245.2 | NC_014624.2 | 2229457 | 2230644 | 1187 | + | tuf | WP_252165655.1 |
| GCF_000152245.2 | NC_014624.2 | 3142827 | 3144014 | 1187 | - | tuf | WP_273686935.1 |
| GCF_000807675.2 | NZ_CP019962.1 | 4087322 | 4088509 | 1187 | - | tuf | WP_223797639.1 |
| GCF_000807675.2 | NZ_CP019962.1 | 746464 | 747651 | 1187 | - | tuf | WP_121152713.1 |
| GCF_002441855.2 | NZ_CP029487.1 | 4891477 | 4892667 | 1190 | - | tuf | WP_069206812.1 |
| GCF_002441855.2 | NZ_CP029487.1 | 1128457 | 1129650 | 1193 | + | tuf | WP_146570117.1 |
| GCF_045002895.1 | NZ_CP173382.1 | 904051 | 905241 | 1190 | - | tuf | WP_007406059.1 |
| GCF_010509575.1 | NZ_CP048437.1 | 2105167 | 2106354 | 1187 | + | tuf | WP_166411679.1 |
| GCF_036632915.2 | NZ_CP155552.1 | 312834 | 314021 | 1187 | + | tuf | WP_156677899.1 |
| GCF_042846195.1 | NZ_AP031411.1 | 4566634 | 4567824 | 1190 | + | tuf | WP_100283858.1 |
| GCF_002998295.1 | NZ_CP027226.1 | 257078 | 258265 | 1187 | + | tuf | WP_256506758.1 |
| GCF_023347705.1 | NZ_CP096649.1 | 655117 | 656307 | 1190 | - | tuf | WP_228947341.1 |
| GCF_943169635.2 | NZ_OW968186.1 | 1662050 | 1663237 | 1187 | - | tuf | WP_187541384.1 |
| GCF_000163895.2 | NC_016630.1 | 1025014 | 1026201 | 1187 | - | tuf | WP_187480675.1 |
| GCF_000163895.2 | NC_016630.1 | 4316678 | 4317865 | 1187 | - | tuf | WP_025293893.1 |
| GCF_010508875.1 | NZ_CP048436.1 | 5970904 | 5972088 | 1184 | + | tuf | WP_025295363.1 |
| GCF_001750685.1 | NZ_CP017269.1 | 1825917 | 1827104 | 1187 | - | tuf | WP_187707971.1 |
| GCF_001750685.1 | NZ_CP017269.1 | 811897 | 813084 | 1187 | + | tuf | WP_166093276.1 |
| GCF_000166415.1 | NC_014654.1 | 241079 | 242269 | 1190 | + | tuf | WP_145844357.1 |

| | | | | | | | |
|---|---|---|---|---|---|---|---|
| GCF_000166415.1 | NC_014654.1 | 2598306 | 2599496 | 1190 | - | tuf | WP_038663408.1 |
| GCF_000328625.1 | NC_019978.1 | 677832 | 679019 | 1187 | - | tuf | WP_147493551.1 |
| GCF_000328625.1 | NC_019978.1 | 2973664 | 2974854 | 1190 | - | tuf | WP_011543090.1 |
| GCF_000020485.1 | NC_011899.1 | 3617566 | 3618756 | 1190 | - | tuf | WP_179497634.1 |
| GCF_000020485.1 | NC_011899.1 | 4409526 | 4410716 | 1190 | + | tuf | WP_039578268.1 |
| GCF_901482605.1 | NZ_LR590481.1 | 317270 | 318457 | 1187 | + | tuf | WP_067180121.1 |
| GCF_901482605.1 | NZ_LR590481.1 | 288638 | 289825 | 1187 | - | tuf | WP_089216047.1 |
| GCF_033097505.1 | NZ_AP028654.1 | 1569274 | 1570464 | 1190 | - | tuf | WP_338427021.1 |
| GCF_033097505.1 | NZ_AP028654.1 | 2293812 | 2294999 | 1187 | + | tuf | WP_105999103.1 |
| GCF_000019165.1 | NC_010337.2 | 4881996 | 4883186 | 1190 | + | tuf | WP_054732125.1 |
| GCF_009649955.1 | NZ_CP045875.1 | 3058094 | 3059281 | 1187 | + | tuf | WP_158902165.1 |
| GCF_900070325.1 | NZ_LN879430.1 | 2447454 | 2448641 | 1187 | - | tuf | WP_072559987.1 |
| GCF_025149285.1 | NZ_CP102274.1 | 1344428 | 1345615 | 1187 | - | tuf | WP_200970646.1 |
| GCF_025148965.1 | NZ_CP102273.1 | 3387012 | 3388202 | 1190 | - | tuf | WP_165328198.1 |
| GCF_025148965.1 | NZ_CP102273.1 | 1361945 | 1363132 | 1187 | + | tuf | WP_313917814.1 |
| GCF_004154955.1 | NZ_CP011524.1 | 1020586 | 1021758 | 1172 | - | tuf | WP_158192886.1 |
| GCF_017751145.1 | NZ_CP046640.1 | 1037996 | 1039168 | 1172 | - | tuf | WP_158192886.1 |
| GCF_017751145.1 | NZ_CP046640.1 | 890972 | 892159 | 1187 | + | tuf | WP_142235648.1 |
| GCF_039134705.1 | NZ_CP146256.1 | 907804 | 908991 | 1187 | + | tuf | WP_142235648.1 |
| GCF_017301615.1 | NZ_CP059066.1 | 2482456 | 2483628 | 1172 | - | tuf | WP_212703905.1 |
| GCF_017301615.1 | NZ_CP059066.1 | 2510576 | 2511748 | 1172 | - | tuf | WP_212703905.1 |
| GCF_003589745.1 | NZ_CP032364.1 | 368782 | 369954 | 1172 | + | tuf | WP_071969790.1 |
| GCF_000018685.1 | NC_010001.1 | 392591 | 393763 | 1172 | + | tuf | WP_071969790.1 |
| GCF_000146185.1 | NC_012778.1 | 533983 | 535155 | 1172 | + | tuf | WP_067260569.1 |
| GCF_026723765.1 | NZ_CP113524.1 | 2952571 | 2953743 | 1172 | - | tuf | WP_322328393.1 |
| GCF_000025225.2 | NC_013895.2 | 522104 | 523276 | 1172 | + | tuf | WP_007118020.1 |
| GCF_000213255.1 | NC_015520.1 | 2972299 | 2973471 | 1172 | - | tuf | WP_007118020.1 |
| GCF_000213255.1 | NC_015520.1 | 2165348 | 2166520 | 1172 | - | tuf | WP_037911852.1 |
| GCF_025148285.1 | NZ_CP102268.1 | 2194361 | 2195533 | 1172 | - | tuf | WP_037911852.1 |
| GCF_042852765.1 | NZ_AP031414.1 | 689934 | 691106 | 1172 | - | tuf | WP_005849852.1 |
| GCF_009831375.1 | NZ_CP027002.1 | 709410 | 710582 | 1172 | - | tuf | WP_197145831.1 |

| | | | | | | | |
|---|---|---|---|---|---|---|---|
| GCF_035573695.1 | NZ_CP121694.1 | 416809 | 417996 | 1187 | - | tuf | WP_141459484.1 |
| GCF_035573695.1 | NZ_CP121694.1 | 1677099 | 1678271 | 1172 | - | tuf | WP_125325003.1 |
| GCF_002998925.1 | NZ_CP027228.1 | 1698028 | 1699200 | 1172 | - | tuf | WP_125325003.1 |
| GCF_002998925.1 | NZ_CP027228.1 | 3257370 | 3258557 | 1187 | + | tuf | WP_072598406.1 |
| GCF_030644205.1 | NZ_CP128647.1 | 3747064 | 3748251 | 1187 | - | tuf | WP_211909031.1 |
| GCF_030644205.1 | NZ_CP128647.1 | 3799556 | 3800743 | 1187 | - | tuf | WP_211909031.1 |
| GCF_002874775.1 | NZ_CP020991.1 | 35380 | 36567 | 1187 | - | tuf | WP_310798633.1 |
| GCF_000020005.1 | NC_010718.1 | 52451 | 53638 | 1187 | - | tuf | WP_310798633.1 |
| GCF_000020005.1 | NC_010718.1 | 5027507 | 5028679 | 1172 | - | tuf | WP_065998099.1 |
| GCF_022539405.1 | NZ_CP054394.1 | 5050776 | 5051948 | 1172 | - | tuf | WP_065998099.1 |
| GCF_022539405.1 | NZ_CP054394.1 | 1798993 | 1800180 | 1187 | + | tuf | WP_158765896.1 |
| GCF_045278715.1 | NZ_CP136420.1 | 1877884 | 1879071 | 1187 | - | tuf | WP_222874876.1 |
| GCF_045278715.1 | NZ_CP136420.1 | 1897181 | 1898368 | 1187 | - | tuf | WP_222874876.1 |
| GCF_009735625.1 | NZ_CP046244.1 | 1172851 | 1174038 | 1187 | + | tuf | WP_342361919.1 |
| GCF_009735625.1 | NZ_CP046244.1 | 1880742 | 1881929 | 1187 | + | tuf | WP_129542700.1 |
| GCF_037477945.1 | NZ_CP136421.1 | 1897592 | 1898779 | 1187 | + | tuf | WP_120225142.1 |
| GCF_037477945.1 | NZ_CP136421.1 | 1285062 | 1286249 | 1187 | + | tuf | WP_101284269.1 |
| GCF_001267435.1 | NZ_CP012370.1 | 1301876 | 1303063 | 1187 | + | tuf | WP_101284269.1 |
| GCF_018866245.1 | NZ_CP076458.1 | 1285020 | 1286207 | 1187 | + | tuf | WP_040059651.1 |
| GCF_014334055.1 | NZ_CP060490.1 | 1301853 | 1303040 | 1187 | + | tuf | WP_040059651.1 |
| GCF_045865535.1 | NZ_AP031406.1 | 1613558 | 1614730 | 1172 | - | tuf | WP_074646985.1 |
| GCF_019916025.1 | NZ_CP079737.1 | 2781678 | 2782850 | 1172 | + | tuf | WP_058262003.1 |
| GCF_019916025.1 | NZ_CP079737.1 | 3839455 | 3840627 | 1172 | - | tuf | WP_078571262.1 |
| GCF_048568265.1 | NZ_CP183813.1 | 3879030 | 3880202 | 1172 | - | tuf | WP_078571262.1 |
| GCF_048568265.1 | NZ_CP183813.1 | 694803 | 695975 | 1172 | - | tuf | WP_339108092.1 |
| GCF_023653455.2 | NZ_CP161000.1 | 715191 | 716363 | 1172 | - | tuf | WP_339108092.1 |
| GCF_023653455.2 | NZ_CP161000.1 | 3709583 | 3710755 | 1172 | - | tuf | WP_077701325.1 |
| GCF_000953675.1 | NZ_LN679998.1 | 3749185 | 3750357 | 1172 | - | tuf | WP_075773810.1 |
| GCF_000953675.1 | NZ_LN679998.1 | 603015 | 604202 | 1187 | + | tuf | WP_188577922.1 |
| GCF_008151785.1 | NZ_CP036523.1 | 620966 | 622153 | 1187 | + | tuf | WP_188577943.1 |
| GCF_008151785.1 | NZ_CP036523.1 | 4024845 | 4026032 | 1187 | - | tuf | WP_372050751.1 |

| | | | | | | | |
|---|---|---|---|---|---|---|---|
| GCF_000597865.1 | NZ_CP007452.1 | 4042760 | 4043947 | 1187 | - | tuf | WP_372050752.1 |
| GCF_000597865.1 | NZ_CP007452.1 | 179076 | 180248 | 1172 | + | tuf | WP_009176153.1 |
| GCF_027125355.1 | NZ_CP114052.1 | 202788 | 203960 | 1172 | + | tuf | WP_065317462.1 |
| GCF_027125355.1 | NZ_CP114052.1 | 2856127 | 2857299 | 1172 | - | tuf | WP_317384588.1 |
| GCF_900538275.1 | NZ_LR130778.1 | 3848643 | 3849815 | 1172 | + | tuf | WP_317385267.1 |
| GCF_040515995.1 | NZ_CP159485.1 | 1366111 | 1367283 | 1172 | + | tuf | WP_282301826.1 |
| GCF_040267685.1 | NZ_CP158367.1 | 1391049 | 1392221 | 1172 | + | tuf | WP_282301826.1 |
| GCF_018408705.1 | NZ_AP023420.1 | 441468 | 442640 | 1172 | - | tuf | WP_120715975.1 |
| GCF_014337195.1 | NZ_CP060634.1 | 1312782 | 1313954 | 1172 | + | tuf | WP_067677476.1 |
| GCF_900015215.1 | NZ_LN555523.1 | 1128264 | 1129451 | 1187 | - | tuf | WP_095011835.1 |
| GCF_900015215.1 | NZ_LN555523.1 | 3664973 | 3666160 | 1187 | - | tuf | WP_096357782.1 |
| GCF_000225345.1 | NC_015977.1 | 3697892 | 3699079 | 1187 | - | tuf | WP_096357823.1 |
| GCF_900537995.1 | NZ_LR027880.1 | 2703979 | 2705166 | 1187 | + | tuf | WP_276157034.1 |
| GCF_014287515.2 | NZ_CP092473.1 | 2720825 | 2722012 | 1187 | + | tuf | WP_276157034.1 |
| GCF_000022065.1 | NC_011898.1 | 899709 | 900878 | 1169 | - | tuf | WP_253303165.1 |
| GCF_005473905.2 | NZ_CP061336.1 | 1086984 | 1088153 | 1169 | - | tuf | WP_253303378.1 |
| GCF_029318685.1 | NZ_CP119677.1 | 543082 | 544251 | 1169 | + | tuf | WP_265025679.1 |
| GCF_000179635.2 | NC_014833.1 | 790325 | 791494 | 1169 | - | tuf | WP_265025679.1 |
| GCF_005601135.1 | NZ_CP039381.1 | 421587 | 422756 | 1169 | + | tuf | WP_010962759.1 |
| GCF_025151995.1 | NZ_CP102290.1 | 1128657 | 1129826 | 1169 | - | tuf | WP_010962309.1 |
| GCF_045038405.1 | NZ_CP173660.1 | 312188 | 313357 | 1169 | + | tuf | WP_253303378.1 |
| GCF_045061075.1 | NZ_CP173694.1 | 1019393 | 1020562 | 1169 | - | tuf | WP_353271140.1 |
| GCF_018141405.1 | NZ_CP058648.1 | 321145 | 322314 | 1169 | + | tuf | WP_010962759.1 |
| GCF_018141405.1 | NZ_CP058648.1 | 952216 | 953385 | 1169 | - | tuf | WP_264705807.1 |
| GCF_014337175.1 | NZ_CP060633.1 | 377268 | 378437 | 1169 | + | tuf | WP_010962309.1 |
| GCF_015140235.1 | NZ_AP023321.1 | 1076308 | 1077477 | 1169 | - | tuf | WP_010962759.1 |
| GCF_025152575.1 | NZ_CP102293.1 | 62379 | 63548 | 1169 | + | tuf | WP_010962759.1 |
| GCF_000009905.1 | NC_006177.1 | 852040 | 853209 | 1169 | - | tuf | WP_010962309.1 |
| GCF_000190635.1 | NC_015172.1 | 355018 | 356187 | 1169 | + | tuf | WP_010962309.1 |
| GCF_000014725.1 | NC_008346.1 | 1064293 | 1065462 | 1169 | - | tuf | WP_010962759.1 |
| GCF_000014725.1 | NC_008346.1 | 491042 | 492211 | 1169 | + | tuf | WP_264722989.1 |

| | | | | | | | |
|---|---|---|---|---|---|---|---|
| GCF_000092405.1 | NC_014220.1 | 1148798 | 1149967 | 1169 | + | tuf | WP_341817940.1 |
| GCF_934924865.1 | NZ_OW235315.1 | 495736 | 496905 | 1169 | - | tuf | WP_010962759.1 |
| GCF_934924865.1 | NZ_OW235315.1 | 1081709 | 1082878 | 1169 | - | tuf | WP_010962759.1 |
| GCF_029542625.1 | NZ_CP120733.1 | 402175 | 403344 | 1169 | + | tuf | WP_353285501.1 |
| GCF_029542625.1 | NZ_CP120733.1 | 1326775 | 1327944 | 1169 | - | tuf | WP_010962309.1 |
| GCF_036812735.1 | NZ_CP117523.1 | 313145 | 314314 | 1169 | + | tuf | WP_264329094.1 |
| GCF_036812735.1 | NZ_CP117523.1 | 1204183 | 1205352 | 1169 | + | tuf | WP_264329094.1 |
| GCF_020809405.1 | NZ_CP081135.1 | 491222 | 492391 | 1169 | + | tuf | WP_265037154.1 |
| GCF_020809405.1 | NZ_CP081135.1 | 597016 | 598185 | 1169 | + | tuf | WP_265037313.1 |
| GCF_031202285.1 | NZ_CP101637.1 | 280307 | 281476 | 1169 | - | tuf | WP_265016035.1 |
| GCF_031202285.1 | NZ_CP101637.1 | 464161 | 465330 | 1169 | + | tuf | WP_264329094.1 |
| GCF_032598785.1 | NZ_CP102984.1 | 279686 | 280855 | 1169 | - | tuf | WP_264329094.1 |
| GCF_032598785.1 | NZ_CP102984.1 | 814530 | 815699 | 1169 | + | tuf | WP_353286675.1 |
| GCF_000305935.1 | NC_018870.1 | 615837 | 617006 | 1169 | + | tuf | WP_265025679.1 |
| GCF_900176005.1 | NZ_LT838272.1 | 863080 | 864249 | 1169 | - | tuf | WP_265025679.1 |
| GCF_000746025.2 | NZ_CP045798.1 | 290412 | 291581 | 1169 | + | tuf | WP_012673185.1 |
| GCF_000746025.2 | NZ_CP045798.1 | 806386 | 807555 | 1169 | + | tuf | WP_012673037.1 |
| GCF_000175295.2 | NC_014964.1 | 316524 | 317693 | 1169 | + | tuf | WP_341810069.1 |
| GCF_000175295.2 | NC_014964.1 | 1025371 | 1026540 | 1169 | - | tuf | WP_265041119.1 |
| GCF_003722315.1 | NZ_CP033580.1 | 490205 | 491374 | 1169 | + | tuf | WP_425385771.1 |
| GCF_003722315.1 | NZ_CP033580.1 | 803996 | 805165 | 1169 | + | tuf | WP_425385771.1 |
| GCF_000025645.1 | NC_013921.1 | 579190 | 580359 | 1169 | + | tuf | WP_265025679.1 |
| GCF_000025645.1 | NC_013921.1 | 824875 | 826044 | 1169 | - | tuf | WP_265025679.1 |
| GCF_000763575.1 | NZ_CP009170.1 | 698249 | 699418 | 1169 | + | tuf | WP_265025460.1 |
| GCF_000763575.1 | NZ_CP009170.1 | 939694 | 940863 | 1169 | - | tuf | WP_265025679.1 |
| GCF_000092965.1 | NC_014209.1 | 443556 | 444725 | 1169 | + | tuf | WP_265027314.1 |
| GCF_000019085.1 | NC_010321.1 | 551038 | 552207 | 1169 | + | tuf | WP_010962759.1 |
| GCF_000019085.1 | NC_010321.1 | 163299 | 164468 | 1169 | + | tuf | WP_010962309.1 |
| GCF_000147695.2 | NC_015958.1 | 973122 | 974291 | 1169 | - | tuf | WP_010962759.1 |
| GCF_000147695.2 | NC_015958.1 | 296719 | 297888 | 1169 | + | tuf | WP_265027922.1 |
| GCF_000307585.2 | NC_017992.1 | 1192208 | 1193377 | 1169 | - | tuf | WP_010962759.1 |

| | | | | | | | |
|---|---|---|---|---|---|---|---|
| GCF_000307585.2 | NC_017992.1 | 401012 | 402181 | 1169 | + | tuf | WP_236514841.1 |
| GCF_000145615.1 | NC_014410.1 | 1074302 | 1075471 | 1169 | - | tuf | WP_341810597.1 |
| GCF_000145615.1 | NC_014410.1 | 296530 | 297699 | 1169 | + | tuf | WP_010962759.1 |
| GCF_000189775.2 | NC_015555.1 | 1047355 | 1048524 | 1169 | - | tuf | WP_010962759.1 |
| GCF_000189775.2 | NC_015555.1 | 404594 | 405763 | 1169 | + | tuf | WP_182158996.1 |
| GCF_024498295.1 | NZ_CP101827.1 | 563329 | 564498 | 1169 | + | tuf | WP_182158996.1 |
| GCF_000331995.1 | NC_020134.1 | 303688 | 304857 | 1169 | + | tuf | WP_010962309.1 |
| GCF_000144645.1 | NC_014377.1 | 562226 | 563395 | 1169 | + | tuf | WP_010962309.1 |
| GCF_000144645.1 | NC_014377.1 | 296860 | 298029 | 1169 | + | tuf | WP_141456826.1 |
| GCF_018141425.1 | NZ_CP058561.1 | 1041043 | 1042212 | 1169 | - | tuf | WP_264378096.1 |
| GCF_018141445.1 | NZ_CP058649.1 | 286243 | 287412 | 1169 | + | tuf | WP_264329094.1 |
| GCF_018408575.1 | NZ_AP023418.1 | 1043423 | 1044592 | 1169 | + | tuf | WP_341822803.1 |
| GCF_018326305.1 | NZ_AP023416.1 | 301015 | 302184 | 1169 | - | tuf | WP_264329094.1 |
| GCF_014337215.1 | NZ_CP060635.1 | 996667 | 997836 | 1169 | + | tuf | WP_264329094.1 |
| GCF_014337155.1 | NZ_CP060632.1 | 498009 | 499178 | 1169 | - | tuf | WP_264329094.1 |
| GCF_004353525.1 | NZ_CP085524.1 | 874404 | 875573 | 1169 | - | tuf | WP_264329094.1 |
| GCF_004353525.1 | NZ_CP085524.1 | 297747 | 298916 | 1169 | + | tuf | WP_265018017.1 |
| GCF_022558445.1 | NZ_CP054699.1 | 550002 | 551171 | 1169 | + | tuf | WP_010962759.1 |
| GCF_012317185.1 | NZ_CP048838.1 | 734485 | 735654 | 1169 | - | tuf | WP_010962759.1 |
| GCF_022701195.1 | NZ_CP078089.1 | 913594 | 914763 | 1169 | - | tuf | WP_010962309.1 |
| GCF_024399475.1 | NZ_CP101620.1 | 293893 | 295062 | 1169 | + | tuf | WP_010962309.1 |
| GCF_010537335.1 | NZ_AP019711.1 | 938050 | 939219 | 1169 | - | tuf | WP_010962759.1 |
| GCF_025148785.1 | NZ_CP102271.1 | 498357 | 499526 | 1169 | - | tuf | WP_010962759.1 |
| GCF_940143175.1 | NZ_OW659496.1 | 592209 | 593378 | 1169 | - | tuf | WP_010962309.1 |
| GCF_035066605.1 | NZ_CP125807.1 | 571291 | 572460 | 1169 | - | tuf | WP_341814532.1 |
| GCF_014396165.1 | NZ_CP060715.1 | 1007440 | 1008609 | 1169 | - | tuf | WP_010962309.1 |
| GCF_001545095.1 | NZ_CP013213.1 | 696715 | 697884 | 1169 | - | tuf | WP_236514841.1 |
| GCF_003931795.1 | NZ_CP034234.1 | 900283 | 901452 | 1169 | - | tuf | WP_236514501.1 |
| GCF_000270085.1 | NC_015601.1 | 1035583 | 1036752 | 1169 | - | tuf | WP_264953849.1 |
| GCF_015097455.1 | NZ_AP024085.1 | 1142049 | 1143218 | 1169 | - | tuf | WP_010962759.1 |
| GCF_001564455.1 | NZ_CP011391.1 | 638308 | 639477 | 1169 | - | tuf | WP_264683056.1 |

| | | | | | | | |
|---|---|---|---|---|---|---|---|
| GCF_003925875.1 | NZ_AP019309.1 | 1240408 | 1241577 | 1169 | + | tuf | WP_264683056.1 |
| GCF_018406465.1 | NZ_AP024510.1 | 518776 | 519945 | 1169 | + | tuf | WP_265024216.1 |
| GCF_042850985.1 | NZ_AP031415.1 | 1378359 | 1379528 | 1169 | - | tuf | WP_265023969.1 |
| GCF_036323475.1 | NZ_AP028934.1 | 295319 | 296488 | 1169 | + | tuf | WP_141456826.1 |
| GCF_021054785.1 | NZ_CP088239.1 | 871681 | 872850 | 1169 | - | tuf | WP_141456826.1 |
| GCF_016728785.1 | NZ_CP068170.1 | 637776 | 638945 | 1169 | - | tuf | WP_265016035.1 |
| GCF_025149465.1 | NZ_CP102275.1 | 1286489 | 1287658 | 1169 | + | tuf | WP_253303165.1 |
| GCF_024499545.1 | NZ_CP071251.1 | 537155 | 538324 | 1169 | + | tuf | WP_010962309.1 |
| GCF_037076425.1 | NZ_AP028127.1 | 944877 | 946046 | 1169 | - | tuf | WP_341812695.1 |
| GCF_013046825.1 | NZ_CP053187.1 | 292831 | 294000 | 1169 | + | tuf | WP_010962309.1 |
| GCF_035593305.1 | NZ_CP141615.1 | 1146799 | 1147968 | 1169 | - | tuf | WP_010962759.1 |
| GCF_035593265.1 | NZ_CP141614.1 | 475373 | 476542 | 1169 | + | tuf | WP_253303378.1 |
| GCF_001544015.1 | NZ_AP014924.1 | 1038013 | 1039182 | 1169 | - | tuf | WP_236514841.1 |
| GCF_001544015.1 | NZ_AP014924.1 | 406914 | 408083 | 1169 | + | tuf | WP_253303378.1 |
| GCF_000025305.1 | NC_013740.1 | 855658 | 856827 | 1169 | + | tuf | WP_264704984.1 |
| GCF_000025305.1 | NC_013740.1 | 518991 | 520160 | 1169 | + | tuf | WP_353270736.1 |
| GCF_025144545.1 | NZ_CP102249.1 | 1341805 | 1342974 | 1169 | - | tuf | WP_353270736.1 |
| GCF_025144545.1 | NZ_CP102249.1 | 360588 | 361757 | 1169 | + | tuf | WP_264722989.1 |
| GCF_007164725.1 | NZ_AP019697.1 | 625615 | 626784 | 1169 | + | tuf | WP_236514841.1 |
| GCF_007164725.1 | NZ_AP019697.1 | 468986 | 470155 | 1169 | + | tuf | WP_265021207.1 |
| GCF_900343095.1 | NZ_LT996885.1 | 724295 | 725464 | 1169 | - | tuf | WP_265021207.1 |
| GCF_900343095.1 | NZ_LT996885.1 | 338632 | 339801 | 1169 | + | tuf | WP_010962309.1 |
| GCF_928721695.1 | NZ_OV788641.1 | 1014214 | 1015383 | 1169 | - | tuf | WP_010962759.1 |
| GCF_928721695.1 | NZ_OV788641.1 | 384970 | 386139 | 1169 | + | tuf | WP_353284683.1 |
| GCF_010669225.1 | NZ_CP048627.1 | 890506 | 891675 | 1169 | - | tuf | WP_353284171.1 |
| GCF_900187035.1 | NZ_LT906446.1 | 587890 | 589059 | 1169 | + | tuf | WP_010962759.1 |
| GCF_003010495.1 | NZ_CP027570.1 | 893404 | 894573 | 1169 | - | tuf | WP_010962309.1 |
| GCF_003010495.1 | NZ_CP027570.1 | 445374 | 446543 | 1169 | + | tuf | WP_010962309.1 |
| GCF_043520715.1 | NZ_CP171361.1 | 956735 | 957904 | 1169 | - | tuf | WP_010962759.1 |
| GCF_043520715.1 | NZ_CP171361.1 | 263642 | 264811 | 1169 | - | tuf | WP_010962759.1 |
| GCF_003367905.1 | NZ_CP029462.1 | 433178 | 434347 | 1169 | - | tuf | WP_010962309.1 |

| | | | | | | | |
|---|---|---|---|---|---|---|---|
| GCF_003367905.1 | NZ_CP029462.1 | 770525 | 771694 | 1169 | - | tuf | WP_374699506.1 |
| GCF_050219315.1 | NZ_CP116939.1 | 1497834 | 1499003 | 1169 | + | tuf | WP_374699506.1 |
| GCF_050219315.1 | NZ_CP116939.1 | 217521 | 218690 | 1169 | - | tuf | WP_010962309.1 |
| GCF_003966895.1 | NZ_AP018449.1 | 451272 | 452441 | 1169 | - | tuf | WP_010962309.1 |
| GCF_003966895.1 | NZ_AP018449.1 | 610937 | 612106 | 1169 | + | tuf | WP_353283866.1 |
| GCF_014076555.1 | NZ_CP045424.1 | 1498909 | 1500078 | 1169 | - | tuf | WP_010962759.1 |
| GCF_003945365.1 | NZ_AP019004.1 | 464286 | 465455 | 1169 | + | tuf | WP_182158996.1 |
| GCF_003945365.1 | NZ_AP019004.1 | 708656 | 709825 | 1169 | - | tuf | WP_182158996.1 |
| GCF_023497905.1 | NZ_CP097284.1 | 524151 | 525320 | 1169 | + | tuf | WP_182158996.1 |
| GCF_023497905.1 | NZ_CP097284.1 | 682233 | 683402 | 1169 | + | tuf | WP_182158996.1 |
| GCF_030273445.1 | NZ_CP120678.1 | 212980 | 214149 | 1169 | + | tuf | WP_010962309.1 |
| GCF_030644225.1 | NZ_CP128650.1 | 491664 | 492833 | 1169 | - | tuf | WP_425383564.1 |
| GCF_000208405.1 | NC_015437.1 | 316594 | 317763 | 1169 | + | tuf | WP_010962309.1 |
| GCF_014250475.1 | NZ_CP060204.1 | 948810 | 949979 | 1169 | - | tuf | WP_010962309.1 |
| GCF_002257695.2 | NZ_CP155571.1 | 788963 | 790132 | 1169 | - | tuf | WP_236514841.1 |
| GCF_002257695.2 | NZ_CP155571.1 | 1168837 | 1170006 | 1169 | - | tuf | WP_236514501.1 |
| GCF_041428895.1 | NZ_CP156926.1 | 167231 | 168400 | 1169 | - | tuf | WP_341820594.1 |
| GCF_041428895.1 | NZ_CP156926.1 | 651077 | 652246 | 1169 | - | tuf | WP_341820594.1 |
| GCF_041428575.1 | NZ_CP155570.1 | 309077 | 310246 | 1169 | + | tuf | WP_010962309.1 |
| GCF_041428575.1 | NZ_CP155570.1 | 1029777 | 1030946 | 1169 | - | tuf | WP_010962759.1 |
| GCF_041428715.1 | NZ_CP155572.1 | 128954 | 130123 | 1169 | + | tuf | WP_010962759.1 |
| GCF_041428715.1 | NZ_CP155572.1 | 793272 | 794441 | 1169 | - | tuf | WP_010962309.1 |
| GCF_000445445.2 | NZ_CP146301.1 | 840108 | 841277 | 1169 | - | tuf | WP_341814063.1 |
| GCF_000445445.2 | NZ_CP146301.1 | 1091633 | 1092802 | 1169 | + | tuf | WP_341814072.1 |
| GCF_041428605.1 | NZ_CP155574.1 | 299191 | 300360 | 1169 | + | tuf | WP_010962309.1 |
| GCF_041428605.1 | NZ_CP155574.1 | 884734 | 885903 | 1169 | - | tuf | WP_010962759.1 |
| GCF_041428845.1 | NZ_CP156925.1 | 770364 | 771533 | 1169 | - | tuf | WP_182158996.1 |
| GCF_041428845.1 | NZ_CP156925.1 | 929076 | 930245 | 1169 | - | tuf | WP_182158996.1 |
| GCF_002257705.2 | NZ_CP155573.1 | 656941 | 658110 | 1169 | - | tuf | WP_010962759.1 |
| GCF_002257705.2 | NZ_CP155573.1 | 1290226 | 1291395 | 1169 | + | tuf | WP_010962309.1 |
| GCF_001941975.2 | NZ_CP146991.1 | 304870 | 306039 | 1169 | + | tuf | WP_010962309.1 |

| | | | | | | | |
|---|---|---|---|---|---|---|---|
| GCF_001941975.2 | NZ_CP146991.1 | 972656 | 973825 | 1169 | - | tuf | WP_010962759.1 |
| GCF_007641255.1 | NZ_CP036259.1 | 102573 | 103742 | 1169 | + | tuf | WP_264329094.1 |
| GCF_007641255.1 | NZ_CP036259.1 | 622972 | 624141 | 1169 | - | tuf | WP_264329094.1 |
| GCF_900637515.1 | NZ_LR134375.1 | 420614 | 421783 | 1169 | + | tuf | WP_253303378.1 |
| GCF_900637515.1 | NZ_LR134375.1 | 1073029 | 1074198 | 1169 | - | tuf | WP_253303165.1 |
| GCF_013393365.1 | NZ_AP022321.1 | 369775 | 370944 | 1169 | + | tuf | WP_253303378.1 |
| GCF_013393365.1 | NZ_AP022321.1 | 1051900 | 1053069 | 1169 | - | tuf | WP_236514841.1 |
| GCF_018406505.1 | NZ_AP022322.1 | 113967 | 115136 | 1169 | + | tuf | WP_010962759.1 |
| GCF_018406505.1 | NZ_AP022322.1 | 843371 | 844540 | 1169 | - | tuf | WP_010962309.1 |
| GCF_900186885.1 | NZ_LT906445.1 | 891088 | 892257 | 1169 | - | tuf | WP_341811184.1 |
| GCF_900186885.1 | NZ_LT906445.1 | 1057728 | 1058897 | 1169 | - | tuf | WP_341811243.1 |
| GCF_900187285.1 | NZ_LT906470.1 | 736129 | 737298 | 1169 | - | tuf | WP_010962759.1 |
| GCF_900187285.1 | NZ_LT906470.1 | 1111352 | 1112521 | 1169 | - | tuf | WP_010962309.1 |
| GCF_004103715.1 | NZ_CP035282.1 | 262454 | 263623 | 1169 | + | tuf | WP_010962759.1 |
| GCF_004103715.1 | NZ_CP035282.1 | 903179 | 904348 | 1169 | - | tuf | WP_264723768.1 |
| GCF_900128415.1 | NZ_LT635772.1 | 550627 | 551796 | 1169 | + | tuf | WP_353280111.1 |
| GCF_900128415.1 | NZ_LT635772.1 | 1129507 | 1130676 | 1169 | - | tuf | WP_353280111.1 |
| GCF_040364225.1 | NZ_AP031484.1 | 299263 | 300432 | 1169 | + | tuf | WP_265018017.1 |
| GCF_040364225.1 | NZ_AP031484.1 | 1022037 | 1023206 | 1169 | - | tuf | WP_010962759.1 |
| GCF_016599815.1 | NZ_CP067016.1 | 1097228 | 1098397 | 1169 | - | tuf | WP_341817467.1 |
| GCF_016599815.1 | NZ_CP067016.1 | 1178894 | 1180063 | 1169 | + | tuf | WP_010962309.1 |
| GCF_000024105.1 | NC_013171.1 | 248152 | 249321 | 1169 | - | tuf | WP_182158996.1 |
| GCF_000024105.1 | NC_013171.1 | 921701 | 922870 | 1169 | - | tuf | WP_182158996.1 |
| GCF_016127475.1 | NZ_CP066014.1 | 348494 | 349663 | 1169 | + | tuf | WP_236514841.1 |
| GCF_016127475.1 | NZ_CP066014.1 | 1480970 | 1482139 | 1169 | - | tuf | WP_253303378.1 |
| GCF_900120165.1 | NZ_LT635475.1 | 379658 | 380827 | 1169 | - | tuf | WP_264329094.1 |
| GCF_900120165.1 | NZ_LT635475.1 | 676498 | 677667 | 1169 | + | tuf | WP_264329094.1 |
| GCF_000299355.1 | NC_018664.1 | 30589 | 31758 | 1169 | + | tuf | WP_264329094.1 |
| GCF_000299355.1 | NC_018664.1 | 807812 | 808981 | 1169 | + | tuf | WP_264329094.1 |
| GCF_004101785.1 | NZ_CP035130.1 | 274640 | 275809 | 1169 | - | tuf | WP_264329094.1 |
| GCF_004101785.1 | NZ_CP035130.1 | 941949 | 943118 | 1169 | + | tuf | WP_353273072.1 |

| | | | | | | | |
|---|---|---|---|---|---|---|---|
| GCF_004524775.2 | NZ_CP119081.1 | 298245 | 299414 | 1169 | + | tuf | WP_265018017.1 |
| GCF_016767175.1 | NZ_CP068564.1 | 1023503 | 1024672 | 1169 | - | tuf | WP_010962759.1 |
| GCF_016767175.1 | NZ_CP068564.1 | 77275 | 78444 | 1169 | + | tuf | WP_264329094.1 |
| GCF_036321625.1 | NZ_CP143259.1 | 846924 | 848093 | 1169 | - | tuf | WP_264329094.1 |
| GCF_036321625.1 | NZ_CP143259.1 | 354082 | 355251 | 1169 | - | tuf | WP_353271467.1 |
| GCF_016458825.1 | NZ_CP066744.1 | 983611 | 984780 | 1169 | + | tuf | WP_010962309.1 |
| GCF_016458825.1 | NZ_CP066744.1 | 327163 | 328332 | 1169 | + | tuf | WP_265043073.1 |
| GCF_900119705.1 | NZ_LT632322.1 | 863413 | 864582 | 1169 | - | tuf | WP_265043073.1 |
| GCF_900637905.1 | NZ_LR134472.1 | 273651 | 274820 | 1169 | + | tuf | WP_007301992.1 |
| GCF_045866535.1 | NZ_AP038371.1 | 522669 | 523838 | 1169 | + | tuf | WP_007301992.1 |
| GCF_027921445.1 | NZ_CP115667.1 | 464105 | 465274 | 1169 | + | tuf | WP_010962309.1 |
| GCF_027921445.1 | NZ_CP115667.1 | 1101636 | 1102805 | 1169 | - | tuf | WP_265041119.1 |
| GCF_023614525.1 | NZ_CP097885.1 | 506639 | 507808 | 1169 | + | tuf | WP_264329094.1 |
| GCF_023614525.1 | NZ_CP097885.1 | 996845 | 998014 | 1169 | - | tuf | WP_264329094.1 |
| GCF_016726485.1 | NZ_CP068103.1 | 412131 | 413300 | 1169 | + | tuf | WP_264329094.1 |
| GCF_016726485.1 | NZ_CP068103.1 | 1065831 | 1067000 | 1169 | + | tuf | WP_264329094.1 |
| GCF_000166695.1 | NC_014721.1 | 298777 | 299946 | 1169 | + | tuf | WP_010962309.1 |
| GCF_000022325.1 | NC_012034.1 | 820977 | 822146 | 1169 | - | tuf | WP_010962759.1 |
| GCF_003999255.1 | NZ_CP034791.1 | 307388 | 308557 | 1169 | + | tuf | WP_010962309.1 |
| GCF_017347585.1 | NZ_AP024480.1 | 952430 | 953599 | 1169 | - | tuf | WP_010962759.1 |
| GCF_000166355.1 | NC_014652.1 | 75659 | 76828 | 1169 | - | tuf | WP_264329094.1 |
| GCF_000166775.1 | NC_014720.1 | 1058940 | 1060109 | 1169 | + | tuf | WP_264329094.1 |
| GCF_026810225.1 | NZ_CP113865.1 | 244322 | 245491 | 1169 | + | tuf | WP_010962309.1 |
| GCF_026914285.1 | NZ_CP113864.1 | 735873 | 737042 | 1169 | + | tuf | WP_010962759.1 |
| GCF_000145215.1 | NC_014392.1 | 308788 | 309957 | 1169 | - | tuf | WP_010962759.1 |
| GCF_000166335.1 | NC_014657.1 | 874484 | 875653 | 1169 | - | tuf | WP_264329094.1 |
| GCF_000016545.1 | NC_009437.1 | 813665 | 814834 | 1169 | - | tuf | WP_010962759.1 |
| GCF_001584725.1 | NZ_CP014606.1 | 984901 | 986070 | 1169 | - | tuf | WP_010962309.1 |
| GCF_001584725.1 | NZ_CP014606.1 | 561520 | 562689 | 1169 | - | tuf | WP_265039989.1 |
| GCF_900120375.1 | NZ_LT635480.1 | 1133036 | 1134205 | 1169 | - | tuf | WP_265040235.1 |
| GCF_900120375.1 | NZ_LT635480.1 | 448821 | 449990 | 1169 | + | tuf | WP_141456826.1 |

| | | | | | | | |
|---|---|---|---|---|---|---|---|
| GCF_006542665.1 | NZ_AP019735.1 | 1273867 | 1275036 | 1169 | - | tuf | WP_264378096.1 |
| GCF_006542685.1 | NZ_AP019736.1 | 1110816 | 1111985 | 1169 | + | tuf | WP_353272616.1 |
| GCF_025144665.1 | NZ_CP102294.1 | 1204702 | 1205871 | 1169 | - | tuf | WP_353272616.1 |
| GCF_025144995.1 | NZ_CP102250.1 | 431939 | 433108 | 1169 | + | tuf | WP_236514501.1 |
| GCF_900604385.1 | NZ_LR027382.1 | 507030 | 508199 | 1169 | + | tuf | WP_236514841.1 |
| GCF_006542645.1 | NZ_AP019734.1 | 756100 | 757269 | 1169 | + | tuf | WP_010962759.1 |
| GCF_025145845.1 | NZ_CP102253.1 | 1167820 | 1168989 | 1169 | + | tuf | WP_010962759.1 |
| GCF_002201795.1 | NZ_CP021904.1 | 239249 | 240418 | 1169 | + | tuf | WP_395460931.1 |
| GCF_009689585.1 | NZ_AP018694.1 | 932707 | 933876 | 1169 | - | tuf | WP_395460931.1 |
| GCF_018292205.1 | NZ_CP072258.1 | 145948 | 147117 | 1169 | - | tuf | WP_007301992.1 |
| GCF_001688725.2 | NZ_CP015401.2 | 768494 | 769663 | 1169 | + | tuf | WP_007301992.1 |
| GCF_018292125.1 | NZ_CP072251.1 | 355233 | 356402 | 1169 | + | tuf | WP_264374619.1 |
| GCF_048319815.1 | NZ_CP182814.1 | 1546502 | 1547671 | 1169 | - | tuf | WP_264374619.1 |
| GCF_025146565.1 | NZ_CP102258.1 | 356798 | 357967 | 1169 | + | tuf | WP_007301992.1 |
| GCF_020091505.1 | NZ_CP081916.1 | 1179941 | 1181110 | 1169 | - | tuf | WP_007301992.1 |
| GCF_012113595.1 | NZ_CP050831.1 | 311200 | 312369 | 1169 | + | tuf | WP_007301992.1 |
| GCF_000025985.1 | NC_003228.3 | 951678 | 952847 | 1169 | - | tuf | WP_007301992.1 |
| GCF_963666085.1 | NZ_OY762556.1 | 273651 | 274820 | 1169 | + | tuf | WP_007301992.1 |
| GCF_000186225.1 | NC_014933.1 | 983179 | 984348 | 1169 | - | tuf | WP_007301992.1 |
| GCF_002998535.1 | NZ_CP027234.1 | 317740 | 318909 | 1169 | + | tuf | WP_007301992.1 |
| GCF_033042785.2 | NZ_CP178829.1 | 1024266 | 1025435 | 1169 | - | tuf | WP_007301992.1 |
| GCF_017309675.2 | NZ_CP084680.1 | 446212 | 447381 | 1169 | + | tuf | WP_007301992.1 |
| GCF_009193295.2 | NZ_CP059973.1 | 1237509 | 1238678 | 1169 | - | tuf | WP_007301992.1 |
| GCF_036416795.1 | NZ_CP143867.1 | 162122 | 163291 | 1169 | - | tuf | WP_007301992.1 |
| GCF_019930665.1 | NZ_CP082886.1 | 763424 | 764593 | 1169 | + | tuf | WP_007301992.1 |
| GCF_001314995.1 | NZ_CP012938.1 | 38519 | 39688 | 1169 | + | tuf | WP_264686636.1 |
| GCF_048595595.1 | NZ_CP184347.1 | 814868 | 816037 | 1169 | - | tuf | WP_264686636.1 |
| GCF_031172035.1 | NZ_CP133452.1 | 290507 | 291676 | 1169 | + | tuf | WP_264374619.1 |
| GCF_040365225.1 | NZ_AP028055.1 | 1214601 | 1215770 | 1169 | - | tuf | WP_264374619.1 |
| GCF_014131755.1 | NZ_CP040530.1 | 385747 | 386916 | 1169 | + | tuf | WP_264706756.1 |
| GCF_044361425.1 | NZ_CP129143.1 | 1559532 | 1560701 | 1169 | - | tuf | WP_264706756.1 |

| | | | | | | | |
|---|---|---|---|---|---|---|---|
| GCF_018289135.1 | NZ_CP072216.1 | 373767 | 374936 | 1169 | + | tuf | WP_341815302.1 |
| GCF_009193325.2 | NZ_CP059856.1 | 1346343 | 1347512 | 1169 | - | tuf | WP_007301992.1 |
| GCF_002998435.1 | NZ_CP027231.1 | 324641 | 325810 | 1169 | + | tuf | WP_265030537.1 |
| GCF_000512915.1 | NZ_CP007034.1 | 1069971 | 1071140 | 1169 | - | tuf | WP_265030537.1 |
| GCF_003991565.1 | NZ_CP032819.1 | 352215 | 353384 | 1169 | + | tuf | WP_007301992.1 |
| GCF_032878895.1 | NZ_CP043836.1 | 1190955 | 1192124 | 1169 | - | tuf | WP_007301992.1 |
| GCF_032878955.1 | NZ_CP043839.1 | 1209526 | 1210695 | 1169 | - | tuf | WP_265014698.1 |
| GCF_025148635.1 | NZ_CP102269.1 | 1357789 | 1358958 | 1169 | - | tuf | WP_265014793.1 |
| GCF_000010645.1 | NC_011565.1 | 647713 | 648882 | 1169 | + | tuf | WP_264688705.1 |
| GCF_030296935.1 | NZ_AP028032.1 | 1347097 | 1348266 | 1169 | - | tuf | WP_007301992.1 |
| GCF_015097275.1 | NZ_AP023322.1 | 300006 | 301175 | 1169 | + | tuf | WP_264683056.1 |
| GCF_010448835.1 | NZ_CP048409.1 | 1160078 | 1161247 | 1169 | - | tuf | WP_264682335.1 |
| GCF_004803915.1 | NZ_CP039396.1 | 417031 | 418200 | 1169 | - | tuf | WP_264336206.1 |
| GCF_036511535.1 | NZ_AP028867.1 | 600237 | 601406 | 1169 | + | tuf | WP_264336932.1 |
| GCF_025151385.1 | NZ_CP102287.1 | 488298 | 489467 | 1169 | - | tuf | WP_007301992.1 |
| GCF_001444445.1 | NZ_CP013195.1 | 823846 | 825015 | 1169 | - | tuf | WP_007301992.1 |
| GCF_002356295.1 | NZ_AP018042.1 | 376534 | 377703 | 1169 | + | tuf | WP_211907845.1 |
| GCF_041902845.1 | NZ_CP159486.1 | 1359149 | 1360318 | 1169 | - | tuf | WP_211907845.1 |
| GCF_009762775.1 | NZ_CP046401.1 | 351660 | 352829 | 1169 | + | tuf | WP_007301992.1 |
| GCF_963677365.1 | NZ_OY781962.1 | 1295262 | 1296431 | 1169 | - | tuf | WP_265024966.1 |
| GCF_004803695.1 | NZ_CP039393.1 | 714743 | 715912 | 1169 | - | tuf | WP_264374619.1 |
| GCF_001688845.2 | NZ_CP015402.2 | 1388741 | 1389910 | 1169 | + | tuf | WP_264374619.1 |
| GCF_900187175.1 | NZ_LT906459.1 | 276924 | 278093 | 1169 | + | tuf | WP_007301992.1 |
| GCF_000183135.1 | NC_014734.1 | 1069899 | 1071068 | 1169 | - | tuf | WP_007301992.1 |
| GCF_023744375.2 | NZ_CP146284.1 | 288925 | 290094 | 1169 | + | tuf | WP_264374619.1 |
| GCF_029581355.1 | NZ_CP120849.1 | 1307119 | 1308288 | 1169 | - | tuf | WP_264374619.1 |
| GCF_018279895.1 | NZ_CP042285.1 | 664307 | 665476 | 1169 | + | tuf | WP_341818208.1 |
| GCF_020091425.1 | NZ_CP081906.1 | 914437 | 915606 | 1169 | - | tuf | WP_341818208.1 |
| GCF_025151045.1 | NZ_CP102285.1 | 233313 | 234482 | 1169 | + | tuf | WP_264729962.1 |
| GCF_030759015.1 | NZ_CP098825.1 | 906095 | 907264 | 1169 | - | tuf | WP_264729962.1 |
| GCF_900683745.1 | NZ_LR215980.1 | 320892 | 322061 | 1169 | - | tuf | WP_007301992.1 |

| | | | | | | | |
|---|---|---|---|---|---|---|---|
| GCF_900095795.1 | NZ_LT608328.1 | 920291 | 921460 | 1169 | + | tuf | WP_007301992.1 |
| GCF_016888945.1 | NZ_CP069440.1 | 323029 | 324198 | 1169 | + | tuf | WP_264719577.1 |
| GCF_013009555.1 | NZ_CP046176.1 | 997947 | 999116 | 1169 | + | tuf | WP_264719577.1 |
| GCF_000190575.1 | NC_015164.1 | 353401 | 354570 | 1169 | + | tuf | WP_007301992.1 |
| GCF_000012825.1 | NC_009614.1 | 959293 | 960462 | 1169 | - | tuf | WP_007301992.1 |
| GCF_000212375.1 | NC_015501.1 | 380962 | 382131 | 1169 | + | tuf | WP_264683056.1 |
| GCF_900638305.1 | NZ_LR134506.1 | 1022375 | 1023544 | 1169 | + | tuf | WP_264682335.1 |
| GCF_900476255.1 | NZ_LS483447.1 | 1405934 | 1407103 | 1169 | - | tuf | WP_425384121.1 |
| GCF_020097375.1 | NZ_CP083629.1 | 1509182 | 1510351 | 1169 | + | tuf | WP_425384121.1 |
| GCF_000010505.1 | NC_010729.1 | 273651 | 274820 | 1169 | + | tuf | WP_007301992.1 |
| GCF_036320655.1 | NZ_CP143258.1 | 1012955 | 1014124 | 1169 | - | tuf | WP_007301992.1 |
| GCF_022845615.1 | NZ_AP025559.1 | 277640 | 278809 | 1169 | + | tuf | WP_007301992.1 |
| GCF_022024115.1 | NZ_CP091792.1 | 354626 | 355795 | 1169 | + | tuf | WP_007301992.1 |
| GCF_000242335.1 | NC_019960.1 | 711636 | 712805 | 1169 | + | tuf | WP_353287553.1 |
| GCF_018128205.1 | NZ_CP072374.1 | 1450704 | 1451873 | 1169 | + | tuf | WP_353287553.1 |
| GCF_001262015.1 | NZ_CP012074.1 | 557863 | 559032 | 1169 | - | tuf | WP_015587867.1 |
| GCF_017347605.1 | NZ_AP024484.1 | 1170457 | 1171626 | 1169 | + | tuf | WP_015587867.1 |
| GCF_018128125.1 | NZ_CP072367.1 | 333572 | 334741 | 1169 | + | tuf | WP_353281258.1 |
| GCF_002849795.1 | NZ_CP023863.1 | 1160040 | 1161209 | 1169 | - | tuf | WP_353281258.1 |
| GCF_000144405.1 | NC_014370.1 | 311514 | 312683 | 1169 | + | tuf | WP_007301992.1 |
| GCF_018127985.1 | NZ_CP072357.1 | 995069 | 996238 | 1169 | - | tuf | WP_007301992.1 |
| GCF_018127825.1 | NZ_CP072339.1 | 412745 | 413914 | 1169 | + | tuf | WP_264374619.1 |
| GCF_018127785.1 | NZ_CP072390.1 | 1466796 | 1467965 | 1169 | - | tuf | WP_264374619.1 |
| GCF_018127765.1 | NZ_CP072330.1 | 754832 | 756001 | 1169 | + | tuf | WP_265023047.1 |
| GCF_022267555.1 | NZ_CP073586.1 | 1198156 | 1199325 | 1169 | + | tuf | WP_265022257.1 |
| GCF_900095135.1 | NZ_LT605205.1 | 468429 | 469598 | 1169 | + | tuf | WP_374698952.1 |
| GCF_003265305.2 | NZ_CP033459.1 | 811052 | 812221 | 1169 | - | tuf | WP_374698332.1 |
| GCF_001443605.1 | NZ_CP013118.1 | 1271464 | 1272633 | 1169 | + | tuf | WP_213863921.1 |
| GCF_022024215.1 | NZ_CP091798.1 | 1413221 | 1414390 | 1169 | - | tuf | WP_213863921.1 |
| GCF_020735445.1 | NZ_CP085932.1 | 284964 | 286133 | 1169 | + | tuf | WP_007301992.1 |
| GCF_900637655.1 | NZ_LR134384.1 | 999100 | 1000269 | 1169 | - | tuf | WP_007301992.1 |

| | | | | | | | |
|---|---|---|---|---|---|---|---|
| GCF_009676955.1 | NZ_CP045696.1 | 680104 | 681273 | 1169 | - | tuf | WP_007301992.1 |
| GCF_000238215.1 | NC_016610.1 | 1338117 | 1339286 | 1169 | + | tuf | WP_007301992.1 |
| GCF_001717525.2 | NZ_CP017038.2 | 550481 | 551650 | 1169 | - | tuf | WP_265041690.1 |
| GCF_013265555.1 | NZ_CP041345.1 | 1258092 | 1259261 | 1169 | + | tuf | WP_265041690.1 |
| GCF_023674465.1 | NZ_CP098400.1 | 118244 | 119413 | 1169 | + | tuf | WP_007301992.1 |
| GCF_000025925.1 | NC_014033.1 | 944851 | 946020 | 1169 | - | tuf | WP_007301992.1 |
| GCF_003600625.1 | NZ_CP032489.1 | 281663 | 282832 | 1169 | + | tuf | WP_007301992.1 |
| GCF_020042345.1 | NZ_CP083387.1 | 925920 | 927089 | 1169 | - | tuf | WP_007301992.1 |
| GCF_010093065.1 | NZ_CP048113.1 | 247671 | 248840 | 1169 | - | tuf | WP_007301992.1 |
| GCF_002557795.1 | NZ_CP023777.1 | 848068 | 849237 | 1169 | + | tuf | WP_007301992.1 |
| GCF_038396765.1 | NZ_CP150096.1 | 289365 | 290534 | 1169 | + | tuf | WP_265025099.1 |
| GCF_022867795.2 | NZ_CP107006.1 | 844386 | 845555 | 1169 | - | tuf | WP_265025099.1 |
| GCF_012516355.1 | NZ_CP051205.1 | 303977 | 305146 | 1169 | + | tuf | WP_007301992.1 |
| GCF_020386615.1 | NZ_CP077769.1 | 1182653 | 1183822 | 1169 | - | tuf | WP_007301992.1 |
| GCF_000024005.1 | NC_013132.1 | 366445 | 367614 | 1169 | + | tuf | WP_264374619.1 |
| GCF_038396755.1 | NZ_CP149822.1 | 1227948 | 1229117 | 1169 | - | tuf | WP_264374619.1 |
| GCF_034424315.1 | NZ_CP139972.1 | 384775 | 385944 | 1169 | + | tuf | WP_007301992.1 |
| GCF_020042285.1 | NZ_CP083388.1 | 1021345 | 1022514 | 1169 | - | tuf | WP_007301992.1 |
| GCF_020783315.1 | NZ_CP086063.1 | 188443 | 189612 | 1169 | - | tuf | WP_007301992.1 |
| GCF_002355595.1 | NZ_AP017422.1 | 776784 | 777953 | 1169 | + | tuf | WP_007301992.1 |
| GCF_018595675.2 | NZ_CP092333.1 | 462977 | 464146 | 1169 | + | tuf | WP_007301992.1 |
| GCF_018595685.2 | NZ_CP092334.1 | 1060214 | 1061383 | 1169 | - | tuf | WP_007301992.1 |
| GCF_007970805.1 | NZ_CP042433.1 | 309114 | 310283 | 1169 | + | tuf | WP_007301992.1 |
| GCF_001644645.1 | NZ_CP011390.1 | 954974 | 956143 | 1169 | - | tuf | WP_007301992.1 |
| GCF_014168535.1 | NZ_CP060007.1 | 169999 | 171168 | 1169 | + | tuf | WP_007301992.1 |
| GCF_028485475.1 | NZ_CP144143.1 | 1119507 | 1120676 | 1169 | - | tuf | WP_007301992.1 |
| GCF_001654455.1 | NZ_CP015772.1 | 525732 | 526901 | 1169 | - | tuf | WP_007301992.1 |
| GCF_039652435.2 | NZ_CP190246.1 | 829565 | 830734 | 1169 | - | tuf | WP_007301992.1 |
| GCF_000243115.2 | NZ_CP007035.1 | 366634 | 367803 | 1169 | + | tuf | WP_007301992.1 |
| GCF_034424215.1 | NZ_CP139960.1 | 510176 | 511345 | 1169 | + | tuf | WP_265024966.1 |
| GCF_000246855.1 | NC_016609.1 | 320512 | 321681 | 1169 | + | tuf | WP_007301992.1 |

| | | | | | | | |
|---|---|---|---|---|---|---|---|
| GCF_007971225.1 | NZ_CP042435.1 | 977990 | 979159 | 1169 | - | tuf | WP_007301992.1 |
| GCF_003555545.1 | NZ_CP032157.1 | 1372639 | 1373808 | 1169 | - | tuf | WP_264686636.1 |
| GCF_032395985.1 | NZ_CP135920.1 | 1429298 | 1430467 | 1169 | - | tuf | WP_264686636.1 |
| GCF_004347775.1 | NZ_CP037755.1 | 351067 | 352236 | 1169 | + | tuf | WP_007301992.1 |
| GCF_004337365.2 | NZ_CP044016.1 | 1099452 | 1100621 | 1169 | - | tuf | WP_007301992.1 |
| GCF_014075995.1 | NZ_CP055153.1 | 653633 | 654802 | 1169 | - | tuf | WP_007301992.1 |
| GCF_014217805.1 | NZ_CP055156.1 | 1250262 | 1251431 | 1169 | + | tuf | WP_007301992.1 |
| GCF_026013705.1 | NZ_CP110226.1 | 292753 | 293922 | 1169 | - | tuf | WP_007301992.1 |
| GCF_000166275.1 | NZ_CM001023.1 | 913606 | 914775 | 1169 | + | tuf | WP_007301992.1 |
| GCF_001593605.1 | NZ_CP012836.1 | 435174 | 436343 | 1169 | - | tuf | WP_007301992.1 |
| GCF_900176595.1 | NZ_LT838813.1 | 1026746 | 1027915 | 1169 | + | tuf | WP_007301992.1 |
| GCF_023921165.1 | NZ_CP049835.1 | 385120 | 386289 | 1169 | + | tuf | WP_007301992.1 |
| GCF_019916085.1 | NZ_CP049834.1 | 1021669 | 1022838 | 1169 | - | tuf | WP_007301992.1 |
| GCF_003143535.1 | NZ_CP029346.1 | 253400 | 254569 | 1169 | + | tuf | WP_353276541.1 |
| GCF_003258705.1 | NZ_CP029480.1 | 1105298 | 1106467 | 1169 | - | tuf | WP_353276541.1 |
| GCF_036492635.1 | NZ_AP025305.1 | 460368 | 461537 | 1169 | + | tuf | WP_007301992.1 |
| GCF_000265405.1 | NC_018010.1 | 1339774 | 1340943 | 1169 | - | tuf | WP_007301992.1 |
| GCF_000265505.1 | NC_018018.1 | 271529 | 272698 | 1169 | + | tuf | WP_353279424.1 |
| GCF_000020565.1 | NC_010830.1 | 984605 | 985774 | 1169 | - | tuf | WP_353279424.1 |
| GCF_003176915.1 | NZ_CP029619.1 | 351216 | 352385 | 1169 | + | tuf | WP_007301992.1 |
| GCF_000304455.1 | NC_018605.1 | 1055528 | 1056697 | 1169 | - | tuf | WP_007301992.1 |
| GCF_965196625.1 | NZ_OZ239769.1 | 310153 | 311322 | 1169 | + | tuf | WP_007301992.1 |
| GCF_936981045.1 | NZ_OW441264.1 | 701024 | 702193 | 1169 | + | tuf | WP_007301992.1 |
| GCF_964030745.1 | NZ_OZ034918.1 | 249399 | 250568 | 1169 | - | tuf | WP_007301992.1 |
| GCF_003351905.1 | NZ_CP022339.1 | 848110 | 849279 | 1169 | + | tuf | WP_007301992.1 |
| GCF_964020025.1 | NZ_OZ026476.1 | 411946 | 413115 | 1169 | + | tuf | WP_341808635.1 |
| GCF_030506285.1 | NZ_CP119422.1 | 462755 | 463924 | 1169 | + | tuf | WP_341808635.1 |
| GCF_026250525.1 | NZ_CP111043.1 | 16118 | 17287 | 1169 | + | tuf | WP_010962309.1 |
| GCF_003589925.1 | NZ_CP032382.1 | 657839 | 659008 | 1169 | - | tuf | WP_010962759.1 |
| GCF_001050135.1 | NZ_CP012040.1 | 38326 | 39495 | 1169 | + | tuf | WP_096617024.1 |
| GCF_000222485.1 | NC_015914.1 | 980641 | 981810 | 1169 | + | tuf | WP_096617024.1 |

| | | | | | | | |
|---|---|---|---|---|---|---|---|
| GCF_000014145.1 | NC_008255.1 | 275762 | 276931 | 1169 | + | tuf | WP_250296088.1 |
| GCF_023821985.2 | NZ_CP098805.1 | 1136155 | 1137324 | 1169 | + | tuf | WP_250296088.1 |
| GCF_022869785.1 | NZ_CP094997.1 | 268931 | 270100 | 1169 | + | tuf | WP_349967923.1 |
| GCF_023822005.2 | NZ_CP098806.1 | 568775 | 569944 | 1169 | + | tuf | WP_349967923.1 |
| GCF_000023125.1 | NC_013037.1 | 308328 | 309497 | 1169 | - | tuf | WP_127463721.1 |
| GCF_026625565.1 | NZ_CP112998.1 | 917410 | 918579 | 1169 | + | tuf | WP_127463721.1 |
| GCF_016894445.1 | NZ_CP056775.1 | 430303 | 431499 | 1196 | + | tuf | WP_167895909.1 |
| GCF_020463795.1 | NZ_CP080025.1 | 768763 | 769932 | 1169 | - | tuf | WP_167896116.1 |
| GCF_005281475.1 | NZ_CP040106.1 | 285193 | 286362 | 1169 | - | tuf | WP_395463318.1 |
| GCF_006575665.1 | NZ_CP041253.1 | 1038492 | 1039682 | 1190 | - | tuf | WP_395462874.1 |
| GCF_003260975.1 | NZ_CP030041.1 | 562738 | 563907 | 1169 | - | tuf | WP_141456826.1 |
| GCF_000325705.1 | NC_019904.1 | 750134 | 751303 | 1169 | + | tuf | WP_010962759.1 |
| GCF_000263195.1 | NC_018748.1 | 468059 | 469228 | 1169 | + | tuf | WP_010962309.1 |
| GCF_000331105.1 | NC_020054.1 | 1008791 | 1009960 | 1169 | - | tuf | WP_010962309.1 |
| GCF_018736065.1 | NZ_CP076128.1 | 48224 | 49393 | 1169 | - | tuf | WP_007301992.1 |
| GCF_003970675.1 | NZ_CP034562.1 | 708119 | 709288 | 1169 | + | tuf | WP_007301992.1 |
| GCF_018736045.1 | NZ_CP076132.1 | 215955 | 217127 | 1172 | + | tuf | WP_041045201.1 |
| GCF_036492835.1 | NZ_AP025314.1 | 900772 | 901944 | 1172 | + | tuf | WP_041045201.1 |
| GCF_021389935.1 | NZ_CP089979.1 | 352746 | 353915 | 1169 | + | tuf | WP_175814040.1 |
| GCF_017068455.1 | NZ_CP070608.1 | 767776 | 768945 | 1169 | - | tuf | WP_175814040.1 |
| GCF_021389955.1 | NZ_CP089980.1 | 548956 | 550128 | 1172 | + | tuf | WP_168455695.1 |
| GCF_021389975.1 | NZ_CP089981.1 | 1131009 | 1132202 | 1193 | - | tuf | WP_168456167.1 |
| GCF_022921095.1 | NZ_CP095053.1 | 128111 | 129280 | 1169 | + | tuf | WP_168463869.1 |
| GCF_008728635.1 | NZ_CP044285.1 | 730060 | 731244 | 1184 | - | tuf | WP_168464297.1 |
| GCF_027359925.1 | NZ_CP114767.1 | 605969 | 607138 | 1169 | - | tuf | WP_007301992.1 |
| GCF_022919135.1 | NZ_CP095049.1 | 1252115 | 1253284 | 1169 | + | tuf | WP_007301992.1 |
| GCF_006337165.1 | NZ_CP040896.1 | 296920 | 298089 | 1169 | - | tuf | WP_236514841.1 |
| GCF_022811645.1 | NZ_CP094534.1 | 1383042 | 1384211 | 1169 | - | tuf | WP_236514501.1 |
| GCF_003149515.1 | NZ_CP029145.1 | 332357 | 333526 | 1169 | - | tuf | WP_017532483.1 |
| GCF_003574965.1 | NZ_CP032317.1 | 953756 | 954925 | 1169 | - | tuf | WP_017532483.1 |
| GCF_004355185.1 | NZ_CP037922.1 | 230614 | 231783 | 1169 | - | tuf | WP_175809394.1 |

| | | | | | | | |
|---|---|---|---|---|---|---|---|
| GCF_001507645.1 | NZ_CP013909.1 | 633659 | 634828 | 1169 | - | tuf | WP_175809394.1 |
| GCF_014250515.1 | NZ_CP060202.1 | 265042 | 266211 | 1169 | + | tuf | WP_175818227.1 |
| GCF_023101345.1 | NZ_CP095848.1 | 695758 | 696927 | 1169 | + | tuf | WP_175818532.1 |
| GCF_000576555.1 | NZ_CP007145.1 | 139468 | 140637 | 1169 | - | tuf | WP_010962309.1 |
| GCF_013137895.1 | NZ_CP053538.1 | 942587 | 943756 | 1169 | + | tuf | WP_317051348.1 |
| GCF_022827545.1 | NZ_CP094669.1 | 16395 | 17564 | 1169 | + | tuf | WP_012673037.1 |
| GCF_022921055.1 | NZ_CP095061.1 | 549661 | 550830 | 1169 | - | tuf | WP_012673185.1 |
| GCF_027625995.1 | NZ_CP115396.1 | 16274 | 17443 | 1169 | + | tuf | WP_010962309.1 |
| GCF_032999765.1 | NZ_CP136051.1 | 658037 | 659206 | 1169 | - | tuf | WP_070356882.1 |
| GCF_026248825.1 | NZ_CP110973.1 | 307673 | 308842 | 1169 | + | tuf | WP_015587867.1 |
| GCF_000166395.1 | NC_014655.1 | 1028043 | 1029212 | 1169 | - | tuf | WP_015587867.1 |
| GCF_005166025.1 | NZ_CP028923.1 | 16118 | 17287 | 1169 | + | tuf | WP_010962309.1 |
| GCF_020149935.1 | NZ_CP083757.1 | 657839 | 659008 | 1169 | - | tuf | WP_010962759.1 |
| GCF_030503875.2 | NZ_CP129968.2 | 16159 | 17328 | 1169 | + | tuf | WP_253303378.1 |
| GCF_030534335.1 | NZ_CP130565.1 | 549800 | 550969 | 1169 | + | tuf | WP_253303165.1 |
| GCF_030503855.1 | NZ_CP129971.1 | 16232 | 17401 | 1169 | + | tuf | WP_010962309.1 |
| GCF_000183425.1 | NC_014759.1 | 498884 | 500053 | 1169 | - | tuf | WP_010962759.1 |
| GCF_021596945.1 | NZ_CP063779.1 | 16350 | 17519 | 1169 | + | tuf | WP_010962309.1 |
| GCF_009913235.1 | NZ_CP047897.1 | 691830 | 692999 | 1169 | - | tuf | WP_010962759.1 |
| GCF_036492425.1 | NZ_AP025292.1 | 393546 | 394715 | 1169 | + | tuf | WP_095742604.1 |
| GCF_003585765.1 | NZ_CP021235.1 | 1032467 | 1033636 | 1169 | - | tuf | WP_095742716.1 |
| GCF_001611675.1 | NZ_CP014766.1 | 16232 | 17401 | 1169 | + | tuf | WP_010962309.1 |
| GCF_026278245.1 | NZ_CP111079.1 | 498892 | 500061 | 1169 | - | tuf | WP_010962759.1 |
| GCF_000973725.1 | NZ_CP009621.1 | 188574 | 189743 | 1169 | + | tuf | WP_264336932.1 |
| GCF_010092985.1 | NZ_CP048106.1 | 806189 | 807358 | 1169 | + | tuf | WP_264336932.1 |
| GCF_009931655.1 | NZ_CP047984.1 | 62435 | 63604 | 1169 | + | tuf | WP_110409550.1 |
| GCF_030506305.1 | NZ_CP119421.1 | 164456 | 165625 | 1169 | + | tuf | WP_110409475.1 |
| GCF_025502585.1 | NZ_CP106679.1 | 566833 | 568002 | 1169 | + | tuf | WP_280143440.1 |
| GCF_025639805.1 | NZ_CP106735.1 | 1173613 | 1174782 | 1169 | - | tuf | WP_280143440.1 |
| GCF_010119975.1 | NZ_CP048222.1 | 321744 | 322913 | 1169 | + | tuf | WP_259535456.1 |
| GCF_034376045.1 | NZ_CP139724.1 | 14901 | 16070 | 1169 | + | tuf | WP_010962759.1 |

| | | | | | | | |
|---|---|---|---|---|---|---|---|
| GCF_001078055.1 | NZ_CP010777.1 | 1003534 | 1004703 | 1169 | + | tuf | WP_010962309.1 |
| GCF_001310085.1 | NZ_CP012643.1 | 663943 | 665112 | 1169 | - | tuf | WP_175838037.1 |
| GCF_003325355.1 | NZ_CP030850.1 | 262699 | 263868 | 1169 | + | tuf | WP_250296088.1 |
| GCF_000218895.1 | NC_015703.1 | 1168915 | 1170084 | 1169 | - | tuf | WP_250296088.1 |
| GCF_002056795.1 | NZ_CP020104.1 | 138673 | 139842 | 1169 | + | tuf | WP_014868768.1 |
| GCF_011604685.1 | NZ_CP050063.1 | 941244 | 942413 | 1169 | - | tuf | WP_014869309.1 |
| GCF_010233585.1 | NZ_CP045997.1 | 579213 | 580382 | 1169 | - | tuf | WP_025264205.1 |
| GCF_014117325.1 | NZ_CP059732.1 | 734797 | 735966 | 1169 | + | tuf | WP_025264278.1 |
| GCF_000024525.1 | NC_013730.1 | 932935 | 934104 | 1169 | - | tuf | WP_007301992.1 |
| GCF_001988955.1 | NZ_CP014263.1 | 1724343 | 1725512 | 1169 | + | tuf | WP_007301992.1 |
| GCF_021233055.1 | NZ_CP089538.1 | 344106 | 345275 | 1169 | + | tuf | WP_250120085.1 |
| GCF_002831565.1 | NZ_CP025096.1 | 1078720 | 1079889 | 1169 | - | tuf | WP_250120085.1 |
| GCF_000974425.1 | NZ_CP010429.1 | 344221 | 345390 | 1169 | + | tuf | WP_250120085.1 |
| GCF_012849055.1 | NZ_CP051677.1 | 1074625 | 1075794 | 1169 | - | tuf | WP_250120085.1 |
| GCF_002067135.1 | NZ_CP020105.1 | 44537 | 45706 | 1169 | + | tuf | WP_253303378.1 |
| GCF_013127955.1 | NZ_CP053435.1 | 1312876 | 1314045 | 1169 | + | tuf | WP_253303165.1 |
| GCF_021484805.1 | NZ_CP090557.1 | 181720 | 182889 | 1169 | - | tuf | WP_253302495.1 |
| GCF_026278345.1 | NZ_CP111085.1 | 438756 | 439925 | 1169 | - | tuf | WP_253302495.1 |
| GCF_030506325.1 | NZ_CP120683.1 | 16174 | 17343 | 1169 | + | tuf | WP_010962309.1 |
| GCF_022807975.1 | NZ_CP094358.1 | 303479 | 304648 | 1169 | + | tuf | WP_010962759.1 |
| GCF_015476115.1 | NZ_CP064295.1 | 531344 | 532513 | 1169 | - | tuf | WP_211907845.1 |
| GCF_004022485.1 | NZ_CP034951.1 | 1127956 | 1129125 | 1169 | + | tuf | WP_211907845.1 |
| GCF_016757735.1 | NZ_CP068439.1 | 28078 | 29247 | 1169 | + | tuf | WP_007301992.1 |
| GCF_029814825.1 | NZ_CP122379.1 | 742813 | 743982 | 1169 | - | tuf | WP_007301992.1 |
| GCF_046352995.1 | NZ_CP130281.1 | 483260 | 484429 | 1169 | - | tuf | WP_068651682.1 |
| GCF_000265385.1 | NC_018013.1 | 1063390 | 1064559 | 1169 | + | tuf | WP_068651682.1 |
| GCF_032926325.1 | NZ_CP136709.1 | 358582 | 359751 | 1169 | - | tuf | WP_010962759.1 |
| GCF_005943945.1 | NZ_CP040710.1 | 417540 | 418709 | 1169 | + | tuf | WP_010962309.1 |
| GCF_030344935.1 | NZ_CP128313.1 | 666684 | 667853 | 1169 | + | tuf | WP_010962309.1 |
| GCF_000224085.1 | NC_015945.1 | 1141990 | 1143159 | 1169 | - | tuf | WP_010962759.1 |
| GCF_045788535.1 | NZ_CP074074.1 | 430785 | 431981 | 1196 | - | tuf | WP_011256542.1 |

| | | | | | | | |
|---|---|---|---|---|---|---|---|
| GCF_007993795.1 | NZ_CP042476.1 | 853811 | 854980 | 1169 | - | tuf | WP_011256851.1 |
| GCF_006159205.1 | NZ_CP040812.1 | 16270 | 17439 | 1169 | + | tuf | WP_010962309.1 |
| GCF_004014855.1 | NZ_CP034930.1 | 657991 | 659160 | 1169 | - | tuf | WP_010962759.1 |
| GCF_002234495.1 | NZ_CP022515.1 | 3481028 | 3482215 | 1187 | + | tuf | WP_203191581.1 |
| GCF_041320605.1 | NZ_CP166679.1 | 3837579 | 3838766 | 1187 | + | tuf | WP_203191581.1 |
| GCF_006065315.1 | NZ_CP040749.1 | 3114630 | 3115802 | 1172 | + | tuf | WP_342069681.1 |
| GCF_009914665.1 | NZ_CP029149.1 | 3456468 | 3457640 | 1172 | + | tuf | WP_342069681.1 |
| GCF_003268615.1 | NZ_CP029844.1 | 592180 | 593352 | 1172 | - | tuf | WP_341368310.1 |
| GCF_014251515.1 | NZ_CP059188.1 | 625105 | 626277 | 1172 | - | tuf | WP_341368310.1 |
| GCF_003226695.1 | NZ_CP029808.1 | 3094216 | 3095388 | 1172 | + | tuf | WP_342077052.1 |
| GCF_910589745.1 | NZ_OU342399.1 | 3117774 | 3118946 | 1172 | + | tuf | WP_342077052.1 |
| GCF_020084865.1 | NZ_CP071953.1 | 852763 | 853953 | 1190 | - | tuf | WP_011240421.1 |
| GCF_001447915.1 | NZ_CP013212.1 | 1115644 | 1116843 | 1199 | + | tuf | WP_013559427.1 |
| GCF_951229915.1 | NZ_OX579588.1 | 2136742 | 2137941 | 1199 | - | tuf | WP_013559427.1 |
| GCF_033372575.1 | NZ_CP104094.1 | 2248031 | 2249230 | 1199 | + | tuf | WP_087862848.1 |
| GCF_964030875.1 | NZ_OZ034981.1 | 11708 | 12907 | 1199 | + | tuf | WP_116226398.1 |
| GCF_000709555.1 | NZ_CP006873.1 | 2902930 | 2904129 | 1199 | - | tuf | WP_116226398.1 |
| GCF_002302565.1 | NZ_CP022382.1 | 2117 | 3331 | 1214 | + | tuf | WP_010879925.1 |
| GCF_043187445.1 | NZ_CP171121.1 | 1356932 | 1358146 | 1214 | + | tuf | WP_010881259.1 |
| GCF_002302475.1 | NZ_CP022378.1 | 153243 | 154436 | 1193 | - | tuf | WP_013637778.1 |
| GCF_002209445.1 | NZ_CP022022.1 | 185152 | 186345 | 1193 | + | tuf | WP_013637778.1 |
| GCF_002302415.1 | NZ_CP022386.1 | 790054 | 791268 | 1214 | - | tuf | WP_012963494.1 |
| GCF_900187255.1 | NZ_LT906449.1 | 806837 | 808051 | 1214 | - | tuf | WP_012963494.1 |
| GCF_000023285.1 | NC_013162.1 | 1122719 | 1123906 | 1187 | + | tuf | WP_012675691.1 |
| GCF_002302495.1 | NZ_CP022379.1 | 1140047 | 1141234 | 1187 | + | tuf | WP_012675691.1 |
| GCF_002302635.1 | NZ_CP022387.1 | 1623156 | 1624343 | 1187 | - | tuf | WP_012674290.1 |
| GCF_000186265.1 | NC_014934.1 | 587438 | 588652 | 1214 | - | tuf | WP_012991661.1 |
| GCF_000468615.2 | NZ_CP009976.1 | 604236 | 605450 | 1214 | - | tuf | WP_012991674.1 |
| GCF_000190595.1 | NC_015167.1 | 589835 | 591049 | 1214 | - | tuf | WP_079653812.1 |
| GCF_027915275.1 | NZ_CP115819.1 | 606603 | 607817 | 1214 | - | tuf | WP_079653825.1 |
| GCF_006974145.1 | NZ_CP039456.1 | 1307296 | 1308510 | 1214 | + | tuf | WP_025306573.1 |

| | | | | | | | |
|---|---|---|---|---|---|---|---|
| GCF_001951155.1 | NZ_CP016359.1 | 1324012 | 1325226 | 1214 | + | tuf | WP_025306573.1 |
| GCF_000060345.1 | NC_008571.1 | 241771 | 242964 | 1193 | + | tuf | WP_013537014.1 |
| GCF_003024155.1 | NZ_CP028136.1 | 1552968 | 1554161 | 1193 | - | tuf | WP_013537014.1 |
| GCF_001889005.1 | NZ_CP018153.1 | 3755570 | 3756769 | 1199 | + | tuf | WP_095044541.1 |
| GCF_020872015.1 | NZ_CP073074.1 | 827627 | 828826 | 1199 | + | tuf | WP_095044541.1 |
| GCF_003971235.1 | NZ_CP034661.1 | 3994289 | 3995479 | 1190 | + | tuf | WP_119322401.1 |
| GCF_003815995.1 | NZ_CP033934.1 | 4503349 | 4504539 | 1190 | + | tuf | WP_119322401.1 |
| GCF_003815975.1 | NZ_CP033932.1 | 325049 | 326248 | 1199 | - | tuf | WP_218112352.1 |
| GCF_002770595.1 | NZ_CP022986.1 | 2899104 | 2900303 | 1199 | - | tuf | WP_218111870.1 |
| GCF_003815695.1 | NZ_CP033920.1 | 302229 | 303428 | 1199 | + | tuf | WP_412709935.1 |
| GCF_026314945.1 | NZ_CP111121.1 | 1532747 | 1533946 | 1199 | - | tuf | WP_369017822.1 |
| GCF_001021975.1 | NZ_CP009928.1 | 604176 | 605375 | 1199 | + | tuf | WP_369017822.1 |
| GCF_027920505.1 | NZ_CP115857.1 | 1781146 | 1782345 | 1199 | - | tuf | WP_369017822.1 |
| GCF_001648155.1 | NZ_CP015199.1 | 136696 | 137883 | 1187 | + | tuf | WP_013171091.1 |
| GCF_016766875.1 | NZ_CP068486.1 | 994061 | 995245 | 1184 | + | tuf | WP_023392493.1 |
| GCF_030012525.1 | NZ_CP124855.1 | 1545590 | 1546774 | 1184 | - | tuf | WP_077140921.1 |
| GCF_003815915.1 | NZ_CP033929.1 | 932801 | 933988 | 1187 | - | tuf | WP_133441945.1 |
| GCF_003815775.1 | NZ_CP033926.1 | 126785 | 127972 | 1187 | + | tuf | WP_348028206.1 |
| GCF_003815875.1 | NZ_CP033924.1 | 145063 | 146247 | 1184 | + | tuf | WP_063387156.1 |
| GCF_014623485.1 | NZ_CP060203.1 | 955329 | 956513 | 1184 | + | tuf | WP_060776838.1 |
| GCF_900637665.1 | NZ_LR134386.1 | 702892 | 704076 | 1184 | - | tuf | WP_067973078.1 |
| GCF_023195755.1 | NZ_CP096203.1 | 1549258 | 1550439 | 1181 | - | tuf | WP_060778677.1 |
| GCF_022811665.1 | NZ_CP094529.1 | 1529086 | 1530267 | 1181 | - | tuf | WP_060778677.1 |
| GCF_025403485.1 | NZ_CP099966.1 | 1292809 | 1293993 | 1184 | + | tuf | WP_026465803.1 |
| GCF_002285635.2 | NZ_CP023049.2 | 824499 | 825683 | 1184 | - | tuf | WP_067978347.1 |
| GCF_004359195.1 | NZ_CP037954.1 | 408245 | 409429 | 1184 | + | tuf | WP_003143114.1 |
| GCF_003815615.1 | NZ_CP033914.1 | 1502546 | 1503733 | 1187 | - | tuf | WP_416352996.1 |
| GCF_022811685.1 | NZ_CP094532.1 | 2777440 | 2778624 | 1184 | - | tuf | WP_012811933.1 |
| GCF_900683625.1 | NZ_LR215974.1 | 364458 | 365642 | 1184 | + | tuf | WP_021296614.1 |
| GCF_900187185.1 | NZ_LT906465.1 | 1682876 | 1684060 | 1184 | + | tuf | WP_274457443.1 |
| GCF_907163105.1 | NZ_OU015321.1 | 459320 | 460504 | 1184 | + | tuf | WP_268044875.1 |

| | | | | | | | |
|---|---|---|---|---|---|---|---|
| GCF_003860565.1 | NZ_CP034157.1 | 389625 | 390809 | 1184 | + | tuf | WP_268006155.1 |
| GCF_014295415.1 | NZ_CP052909.1 | 648900 | 650084 | 1184 | + | tuf | WP_206657326.1 |
| GCF_013402795.1 | NZ_CP058595.1 | 197124 | 198305 | 1181 | + | tuf | WP_368645621.1 |
| GCF_000196315.1 | NC_014230.1 | 141051 | 142238 | 1187 | + | tuf | WP_147804599.1 |
| GCF_014524565.1 | NZ_CP060139.1 | 140398 | 141585 | 1187 | + | tuf | WP_012957135.1 |
| GCF_014961145.1 | NZ_CP063145.1 | 2693664 | 2694851 | 1187 | + | tuf | WP_144163614.1 |
| GCF_001653755.1 | NZ_CP015125.1 | 137080 | 138267 | 1187 | + | tuf | WP_015008860.1 |
| GCF_002023665.2 | NZ_CP023401.1 | 1469611 | 1470798 | 1187 | + | tuf | WP_054745160.1 |
| GCF_002024805.1 | NZ_CP014337.1 | 179228 | 180415 | 1187 | + | tuf | WP_071315269.1 |
| GCF_900475375.1 | NZ_LS483376.1 | 182890 | 184077 | 1187 | - | tuf | WP_096463195.1 |
| GCF_002022125.1 | NZ_CP016377.1 | 188078 | 189265 | 1187 | + | tuf | WP_057897068.1 |
| GCF_013488205.1 | NZ_CP040908.1 | 3693676 | 3694863 | 1187 | - | tuf | WP_272561102.1 |
| GCF_014621655.1 | NZ_CP053698.1 | 119051 | 120235 | 1184 | + | tuf | WP_003397655.1 |
| GCF_003860525.1 | NZ_CP034161.1 | 118148 | 119332 | 1184 | + | tuf | WP_035066573.1 |
| GCF_003687485.1 | NZ_CP032050.1 | 61675 | 62859 | 1184 | - | tuf | WP_066322072.1 |
| GCF_017948305.1 | NZ_CP072842.1 | 954986 | 956170 | 1184 | - | tuf | WP_053950214.1 |
| GCF_000963865.1 | NZ_CP011071.1 | 531646 | 532830 | 1184 | - | tuf | WP_249511479.1 |
| GCF_003269425.1 | NZ_CP030104.1 | 561471 | 562655 | 1184 | + | tuf | WP_120784237.1 |
| GCF_011068285.1 | NZ_CP049616.1 | 979079 | 980263 | 1184 | - | tuf | WP_054450182.1 |
| GCF_030540775.2 | NZ_CP141266.1 | 124706 | 125890 | 1184 | + | tuf | WP_001029614.1 |
| GCF_002886045.1 | NZ_CP025791.1 | 1101697 | 1102884 | 1187 | + | tuf | WP_007496302.1 |
| GCF_020911885.1 | NZ_CP087132.1 | 1946345 | 1947532 | 1187 | + | tuf | WP_007410400.1 |
| GCF_026870175.2 | NZ_CP113977.2 | 119376 | 120560 | 1184 | + | tuf | WP_001029614.1 |
| GCF_025919725.1 | NZ_CP081495.1 | 3591070 | 3592254 | 1184 | + | tuf | WP_163241616.1 |
| GCF_003096035.1 | NZ_CP029186.1 | 126107 | 127294 | 1187 | + | tuf | WP_026593164.1 |
| GCF_008000935.1 | NZ_CP042831.1 | 1391183 | 1392370 | 1187 | + | tuf | WP_003328291.1 |
| GCF_020886115.1 | NZ_AP025185.1 | 133781 | 134968 | 1187 | + | tuf | WP_019257280.1 |
| GCF_020886055.1 | NZ_AP025184.1 | 125691 | 126875 | 1184 | + | tuf | WP_011983244.1 |
| GCF_001705175.1 | NZ_CP016907.1 | 74828 | 76015 | 1187 | + | tuf | WP_170005905.1 |
| GCF_027111335.1 | NZ_CP114288.1 | 138740 | 139924 | 1184 | + | tuf | WP_046128850.1 |
| GCF_003344925.1 | NZ_CP031188.1 | 3708339 | 3709526 | 1187 | + | tuf | WP_053605170.1 |

| | | | | | | | |
|---|---|---|---|---|---|---|---|
| GCF_033547075.1 | NZ_CP137891.1 | 1875637 | 1876824 | 1187 | - | tuf | WP_010332768.1 |
| GCF_023195775.1 | NZ_CP096205.1 | 2903522 | 2904709 | 1187 | - | tuf | WP_003178321.1 |
| GCF_024271895.1 | NZ_CP091861.1 | 132343 | 133530 | 1187 | + | tuf | WP_003242008.1 |
| GCF_000253275.1 | NC_016001.1 | 4780790 | 4781974 | 1184 | + | tuf | WP_009791331.1 |
| GCF_032098525.1 | NZ_CP134890.1 | 125247 | 126431 | 1184 | + | tuf | WP_338752609.1 |
| GCF_021172165.1 | NZ_CP089096.1 | 483746 | 484933 | 1187 | - | tuf | WP_003178321.1 |
| GCF_946903285.1 | NZ_OX336425.1 | 4993616 | 4994803 | 1187 | - | tuf | WP_001029613.1 |
| GCF_014844295.1 | NZ_CP054494.1 | 2060145 | 2061332 | 1187 | - | tuf | WP_353854008.1 |
| GCF_001857965.1 | NZ_CP017774.1 | 132593 | 133780 | 1187 | + | tuf | WP_010332768.1 |
| GCF_001534645.1 | NZ_CP013992.1 | 2654477 | 2655664 | 1187 | - | tuf | WP_002009778.1 |
| GCF_001831475.1 | NZ_CP017688.1 | 4968997 | 4970181 | 1184 | - | tuf | WP_001029615.1 |
| GCF_003122385.1 | NZ_CP029255.1 | 164475 | 165662 | 1187 | + | tuf | WP_023855913.1 |
| GCF_020911925.1 | NZ_CP087134.1 | 124732 | 125916 | 1184 | + | tuf | WP_001029615.1 |
| GCF_021172145.1 | NZ_CP089095.1 | 98743 | 99930 | 1187 | - | tuf | WP_003205504.1 |
| GCF_004151275.2 | NZ_CP130821.1 | 131135 | 132322 | 1187 | + | tuf | WP_012008701.1 |
| GCF_042182265.1 | NZ_CP097869.1 | 132492 | 133679 | 1187 | + | tuf | WP_024714047.1 |
| GCF_027111355.2 | NZ_CP114291.2 | 1411547 | 1412734 | 1187 | - | tuf | WP_024425595.1 |
| GCF_017352115.1 | NZ_CP071448.1 | 154653 | 155840 | 1187 | + | tuf | WP_226516366.1 |
| GCF_003076455.1 | NZ_CP020918.1 | 136176 | 137360 | 1184 | + | tuf | WP_048622373.1 |
| GCF_027111255.2 | NZ_CP114285.2 | 137466 | 138650 | 1184 | + | tuf | WP_006640316.1 |
| GCF_020911905.1 | NZ_CP087133.1 | 132960 | 134147 | 1187 | + | tuf | WP_003225787.1 |
| GCF_003312915.1 | NZ_CP030261.1 | 4234377 | 4235564 | 1187 | - | tuf | WP_003242008.1 |
| GCF_020911945.1 | NZ_CP087135.1 | 132882 | 134069 | 1187 | + | tufA | NP_387994.1 |
| GCF_027111295.1 | NZ_CP114287.1 | 137219 | 138406 | 1187 | + | tuf | WP_307893103.1 |
| GCF_001761465.1 | NZ_CP017479.1 | 132820 | 134007 | 1187 | + | tuf | WP_024714047.1 |
| GCF_037478105.1 | NZ_CP147988.1 | 87826 | 89010 | 1184 | - | tuf | WP_001029617.1 |
| GCF_029625295.1 | NZ_CP121112.1 | 124820 | 126004 | 1184 | + | tuf | WP_001029617.1 |
| GCF_036840915.1 | NZ_CP085725.1 | 5138302 | 5139486 | 1184 | - | tuf | WP_001029614.1 |
| GCF_023272795.1 | NZ_CP096829.1 | 268869 | 270056 | 1187 | + | tuf | WP_087991688.1 |
| GCF_000455605.1 | NC_017025.1 | 133588 | 134775 | 1187 | + | tuf | WP_007410400.1 |
| GCF_013389455.1 | NZ_CP058278.1 | 4049029 | 4050216 | 1187 | - | tuf | WP_072581462.1 |

| | | | | | | | |
|---|---|---|---|---|---|---|---|
| GCF_000016645.1 | NC_009441.1 | 124738 | 125922 | 1184 | + | tuf | WP_001029617.1 |
| GCF_021650975.2 | NZ_CP091285.1 | 1494478 | 1495665 | 1187 | - | tuf | WP_008356525.1 |
| GCF_029866485.1 | NZ_CP092332.1 | 861809 | 862996 | 1187 | + | tuf | WP_118901988.1 |
| GCF_003076475.1 | NZ_CP020919.1 | 707524 | 708711 | 1187 | + | tuf | WP_249515026.1 |
| GCF_027474525.2 | NZ_CP114882.2 | 122121 | 123308 | 1187 | - | tuf | WP_249512934.1 |
| GCF_027111315.2 | NZ_CP114289.2 | 292104 | 293291 | 1187 | - | tuf | WP_005828864.1 |
| GCF_019613795.1 | NZ_CP080429.1 | 3915841 | 3917028 | 1187 | - | tuf | WP_171567158.1 |
| GCF_027111275.1 | NZ_CP114286.1 | 5545722 | 5546909 | 1187 | - | tuf | WP_003391992.1 |
| GCF_003055625.1 | NZ_CP028811.1 | 263690 | 264877 | 1187 | + | tuf | WP_106656573.1 |
| GCF_026870155.1 | NZ_CP113975.1 | 236066 | 237253 | 1187 | + | tuf | WP_092276973.1 |
| GCF_004355225.1 | NZ_CP037933.1 | 247757 | 248944 | 1187 | + | tuf | WP_198828248.1 |
| GCF_031326205.1 | NZ_CP133721.1 | 4524815 | 4526002 | 1187 | + | tuf | WP_230044101.1 |
| GCF_029625335.1 | NZ_CP121111.1 | 2318695 | 2319882 | 1187 | + | tuf | WP_018673901.1 |
| GCF_008369745.1 | NZ_CP043612.1 | 220822 | 222009 | 1187 | + | tuf | WP_126424787.1 |
| GCF_019702945.1 | NZ_AP024749.1 | 641623 | 642810 | 1187 | - | tuf | WP_122966804.1 |
| GCF_040703125.1 | NZ_CP160035.1 | 287302 | 288489 | 1187 | + | tuf | WP_251873129.1 |
| GCF_003097535.1 | NZ_CP029187.1 | 106888 | 108069 | 1181 | + | tuf | WP_035313873.1 |
| GCF_030378165.1 | NZ_CP119766.1 | 1329691 | 1330872 | 1181 | - | tuf | WP_029091545.1 |
| GCF_024499525.1 | NZ_CP100437.1 | 649300 | 650487 | 1187 | + | tuf | WP_007505929.1 |
| GCF_024638035.1 | NZ_CP100442.1 | 142451 | 143635 | 1184 | + | tuf | WP_034767019.1 |
| GCF_026810365.1 | NZ_CP113948.1 | 516475 | 517659 | 1184 | - | tuf | WP_086347683.1 |
| GCF_013343195.2 | NZ_CP039120.2 | 2321750 | 2322934 | 1184 | - | tuf | WP_086349654.1 |
| GCF_003403075.1 | NZ_CP031557.1 | 348654 | 349838 | 1184 | - | tuf | WP_087640137.1 |
| GCF_030643905.1 | NZ_CP121446.1 | 2451247 | 2452431 | 1184 | + | tuf | WP_087642001.1 |
| GCF_004797125.1 | NZ_CP038810.1 | 610718 | 611902 | 1184 | - | tuf | WP_086444423.1 |
| GCF_023008245.1 | NZ_CP090145.1 | 1900499 | 1901683 | 1184 | - | tuf | WP_086278226.1 |
| GCF_003148385.1 | NZ_CP029463.1 | 700566 | 701750 | 1184 | + | tuf | WP_207942221.1 |
| GCF_045348165.1 | NZ_CP150845.1 | 2759693 | 2760877 | 1184 | - | tuf | WP_207941794.1 |
| GCF_000194605.1 | NC_015321.1 | 1327137 | 1328321 | 1184 | + | tuf | WP_010734472.1 |
| GCF_000723205.1 | NZ_HG315671.1 | 2383897 | 2385081 | 1184 | + | tuf | WP_206858827.1 |
| GCF_007197735.1 | NZ_CP041637.1 | 641057 | 642241 | 1184 | + | tuf | WP_086313014.1 |

| | | | | | | | |
|---|---|---|---|---|---|---|---|
| GCF_032814565.1 | NZ_CP136521.1 | 2967191 | 2968375 | 1184 | + | tuf | WP_010761693.1 |
| GCF_002369955.1 | NZ_AP014564.1 | 2266673 | 2267860 | 1187 | - | tuf | WP_023179572.1 |
| GCF_028743515.2 | NZ_CP117969.2 | 244897 | 246081 | 1184 | + | tuf | WP_010052044.1 |
| GCF_900637895.1 | NZ_LR134441.1 | 916255 | 917439 | 1184 | + | tuf | WP_010051635.1 |
| GCF_003860585.1 | NZ_CP034159.1 | 1620723 | 1621907 | 1184 | + | tuf | WP_087064524.1 |
| GCF_020985365.1 | NZ_CP087583.1 | 264283 | 265467 | 1184 | + | tuf | WP_204591485.1 |
| GCF_900638245.1 | NZ_LR134503.1 | 6138460 | 6139647 | 1187 | - | tuf | WP_130615208.1 |
| GCF_020410745.1 | NZ_CP084528.1 | 3636569 | 3637756 | 1187 | - | tuf | WP_123042132.1 |
| GCF_009901525.1 | NZ_CP019288.1 | 2210475 | 2211665 | 1190 | - | tuf | WP_182302921.1 |
| GCF_026625145.1 | NZ_CP113088.1 | 4499793 | 4500983 | 1190 | - | tuf | WP_169281355.1 |
| GCF_001971745.1 | NZ_CP019352.1 | 362899 | 364086 | 1187 | + | tuf | WP_057739334.1 |
| GCF_020310025.1 | NZ_CP084318.1 | 2188624 | 2189811 | 1187 | - | tuf | WP_076617490.1 |
| GCF_001543325.1 | NZ_CP013355.1 | 1264583 | 1265770 | 1187 | - | tuf | WP_057867224.1 |
| GCF_030503675.1 | NZ_CP129964.1 | 1599205 | 1600392 | 1187 | - | tuf | WP_010020061.1 |
| GCF_037126215.1 | NZ_CP136924.1 | 1489395 | 1490582 | 1187 | - | tuf | WP_057811984.1 |
| GCF_027474845.2 | NZ_CP107031.2 | 368506 | 369693 | 1187 | + | tuf | WP_048702836.1 |
| GCF_002269385.1 | NZ_CP022957.1 | 1065425 | 1066612 | 1187 | + | tuf | WP_041499424.1 |
| GCF_900105105.1 | NZ_LT629754.1 | 1555962 | 1557149 | 1187 | + | tuf | WP_119326889.1 |
| GCF_027474825.1 | NZ_CP107030.1 | 851111 | 852298 | 1187 | + | tuf | WP_059074196.1 |
| GCF_001913155.1 | NZ_CP018760.1 | 1565540 | 1566727 | 1187 | - | tuf | WP_054643611.1 |
| GCF_025244665.1 | NZ_CP104205.1 | 1634712 | 1635899 | 1187 | - | tuf | WP_120142302.1 |
| GCF_031128465.2 | NZ_CP155618.1 | 135608 | 136792 | 1184 | + | tuf | WP_057776536.1 |
| GCF_014764305.1 | NZ_CP061703.1 | 139301 | 140485 | 1184 | + | tuf | WP_066441586.1 |
| GCF_004139715.1 | NZ_CP035544.1 | 129424 | 130608 | 1184 | + | tuf | WP_095371226.1 |
| GCF_024722315.1 | NZ_CP102754.1 | 141261 | 142445 | 1184 | + | tuf | WP_009336569.1 |
| GCF_009792295.1 | NZ_CP039934.1 | 1757444 | 1758628 | 1184 | - | tuf | WP_053478266.1 |
| GCF_004337635.1 | NZ_CP037427.1 | 139986 | 141170 | 1184 | + | tuf | WP_233809073.1 |
| GCF_016726985.1 | NZ_CP068108.1 | 1035324 | 1036505 | 1181 | + | tuf | WP_347297691.1 |
| GCF_000833025.1 | NZ_CP010817.1 | 2212756 | 2213943 | 1187 | - | tuf | WP_200757224.1 |
| GCF_000332115.1 | NC_020156.1 | 247904 | 249088 | 1184 | + | tuf | WP_005226902.1 |
| GCF_000831385.1 | NZ_AP014548.1 | 2375612 | 2376796 | 1184 | - | tuf | WP_005229289.1 |

| | | | | | | | |
|---|---|---|---|---|---|---|---|
| GCF_003966335.1 | NZ_CP034549.1 | 230643 | 231827 | 1184 | + | tuf | WP_016250285.1 |
| GCF_002117125.1 | NZ_CP019344.1 | 1104825 | 1106009 | 1184 | + | tuf | WP_005880819.1 |
| GCF_002117085.1 | NZ_CP019342.1 | 2066924 | 2068108 | 1184 | + | tuf | WP_005878630.1 |
| GCF_032922765.1 | NZ_CP136694.1 | 72500 | 73684 | 1184 | + | tuf | WP_002290442.1 |
| GCF_000265465.1 | NC_018016.1 | 1013673 | 1014857 | 1184 | + | tuf | WP_002289463.1 |
| GCF_000236705.1 | NC_016599.1 | 879698 | 880882 | 1184 | + | tuf | WP_003127959.1 |
| GCF_024628805.1 | NZ_CP102382.1 | 1569938 | 1571122 | 1184 | + | tuf | WP_003126794.1 |
| GCF_019740355.2 | NZ_CP091819.1 | 1320685 | 1321869 | 1184 | - | tuf | WP_086303686.1 |
| GCF_907165195.1 | NZ_OU015584.1 | 2749632 | 2750816 | 1184 | + | tuf | WP_114525277.1 |
| GCF_038023995.1 | NZ_CP150660.1 | 599876 | 601060 | 1184 | - | tuf | WP_010720459.1 |
| GCF_017498485.1 | NZ_CP071795.1 | 2463101 | 2464285 | 1184 | - | tuf | WP_010719593.1 |
| GCF_038024105.1 | NZ_CP150661.1 | 2245711 | 2246895 | 1184 | + | tuf | WP_005226902.1 |
| GCF_017569185.1 | NZ_CP071869.1 | 3017250 | 3018434 | 1184 | + | tuf | WP_005237121.1 |
| GCF_024362345.1 | NZ_CP101505.1 | 61811 | 62995 | 1184 | + | tuf | WP_002290442.1 |
| GCF_038024125.1 | NZ_CP150662.1 | 1657040 | 1658224 | 1184 | - | tuf | WP_002289463.1 |
| GCF_014784055.1 | NZ_CP061813.1 | 285183 | 286367 | 1184 | + | tuf | WP_206903369.1 |
| GCF_030444335.1 | NZ_CP128595.1 | 1061343 | 1062527 | 1184 | + | tuf | WP_206905417.1 |
| GCF_019968605.1 | NZ_CP082966.1 | 974591 | 975775 | 1184 | + | tuf | WP_019723509.1 |
| GCF_038019025.1 | NZ_CP150496.1 | 3154860 | 3156044 | 1184 | + | tuf | WP_010734472.1 |
| GCF_038024035.1 | NZ_CP150664.1 | 246533 | 247717 | 1184 | + | tuf | WP_010747219.1 |
| GCF_014352875.1 | NZ_CP060695.1 | 1767103 | 1768287 | 1184 | - | tuf | WP_010745022.1 |
| GCF_001975665.1 | NZ_CP019419.1 | 685124 | 686308 | 1184 | - | tuf | WP_010761693.1 |
| GCF_024918935.1 | NZ_CP103460.1 | 3058093 | 3059277 | 1184 | - | tuf | WP_208927711.1 |
| GCF_038024045.1 | NZ_CP150665.1 | 258587 | 259771 | 1184 | + | tuf | WP_016176306.1 |
| GCF_038024095.1 | NZ_CP150668.1 | 1705697 | 1706881 | 1184 | - | tuf | WP_173103257.1 |
| GCF_001761365.1 | NZ_CP017477.1 | 2443208 | 2444392 | 1184 | - | tuf | WP_173103879.1 |
| GCF_001310225.1 | NZ_CP012898.1 | 64923 | 66107 | 1184 | + | tuf | WP_067483550.1 |
| GCF_002893765.1 | NZ_CP025938.1 | 866638 | 867822 | 1184 | + | tuf | WP_067484820.1 |
| GCF_000153485.2 | NC_018721.1 | 692093 | 693277 | 1184 | - | tuf | WP_088268797.1 |
| GCF_023556315.2 | NZ_CP116221.1 | 2065639 | 2066823 | 1184 | - | tuf | WP_087642001.1 |
| GCF_002966125.1 | NZ_CP027062.1 | 147784 | 148971 | 1187 | + | tuf | WP_013486748.1 |

| | | | | | | | |
|---|---|---|---|---|---|---|---|
| GCF_011044175.1 | NZ_CP049057.1 | 122528 | 123712 | 1184 | + | tuf | WP_023466627.1 |
| GCF_000183155.1 | NC_014738.1 | 447755 | 448939 | 1184 | + | tuf | WP_034817130.1 |
| GCF_000024125.1 | NC_013222.1 | 123206 | 124390 | 1184 | + | tuf | WP_012369007.1 |
| GCF_020310005.1 | NZ_CP084316.1 | 2667745 | 2668929 | 1184 | - | tuf | WP_275417215.1 |
| GCF_900142975.1 | NZ_LT670848.1 | 2712901 | 2714085 | 1184 | + | tuf | WP_275420530.1 |
| GCF_026109735.1 | NZ_CP069620.1 | 1391904 | 1393091 | 1187 | + | tuf | WP_347438581.1 |
| GCF_030845195.1 | NZ_CP132543.1 | 299019 | 300203 | 1184 | + | tuf | WP_252795429.1 |
| GCF_000941055.1 | NZ_CP007202.1 | 277772 | 278956 | 1184 | + | tuf | WP_252767049.1 |
| GCF_021183825.1 | NZ_CP089211.1 | 808177 | 809361 | 1184 | + | tuf | WP_252796597.1 |
| GCF_964507685.1 | NZ_OZ208414.1 | 886261 | 887445 | 1184 | - | tuf | WP_252779625.1 |
| GCF_964036635.1 | NZ_OZ038524.1 | 912950 | 914134 | 1184 | + | tuf | WP_054646868.1 |
| GCF_964512435.1 | NZ_OZ208512.1 | 416285 | 417469 | 1184 | - | tuf | WP_252750379.1 |
| GCF_900198195.1 | NZ_LT899436.1 | 988706 | 989890 | 1184 | - | tuf | WP_014082224.1 |
| GCF_900119795.1 | NZ_LT634361.1 | 944269 | 945453 | 1184 | + | tuf | WP_413627113.1 |
| GCF_009362255.1 | NZ_CP045192.1 | 1321346 | 1322530 | 1184 | + | tuf | WP_059393490.1 |
| GCF_027941775.1 | NZ_CP115917.1 | 1776563 | 1777747 | 1184 | - | tuf | WP_313792375.1 |
| GCF_964036595.1 | NZ_OZ038523.1 | 1577500 | 1578684 | 1184 | - | tuf | WP_017262024.1 |
| GCF_964048965.1 | NZ_OZ040189.1 | 1164997 | 1166181 | 1184 | - | tuf | WP_003144664.1 |
| GCF_027941755.1 | NZ_CP115916.1 | 330167 | 331351 | 1184 | - | tuf | WP_006364510.1 |
| GCF_003867015.1 | NZ_CP032548.1 | 126202 | 127386 | 1184 | + | tuf | WP_011229619.1 |
| GCF_029853675.1 | NZ_CP122539.1 | 3101537 | 3102721 | 1184 | - | tuf | WP_063166994.1 |
| GCF_001889045.1 | NZ_CP018155.1 | 1715025 | 1716209 | 1184 | - | tuf | WP_008881946.1 |
| GCF_964036645.1 | NZ_OZ038525.1 | 107604 | 108788 | 1184 | + | tuf | WP_011229619.1 |
| GCF_031826515.1 | NZ_CP134536.1 | 123937 | 125121 | 1184 | + | tuf | WP_011229619.1 |
| GCF_001761325.1 | NZ_CP017478.1 | 1610671 | 1611855 | 1184 | + | tuf | WP_066124406.1 |
| GCF_900637795.1 | NZ_LR134404.1 | 1005548 | 1006735 | 1187 | - | tuf | WP_244721600.1 |
| GCF_001697185.1 | NZ_CP014224.1 | 3085864 | 3087051 | 1187 | + | tuf | WP_244741179.1 |
| GCF_013394595.1 | NZ_CP053348.1 | 184410 | 185597 | 1187 | + | tuf | WP_100362626.1 |
| GCF_013394655.1 | NZ_CP053351.1 | 661003 | 662187 | 1184 | - | tuf | WP_005607487.1 |
| GCF_900105425.1 | NZ_LT629774.1 | 170844 | 172031 | 1187 | + | tuf | WP_010896315.1 |
| GCF_022760175.1 | NZ_CP094326.1 | 2543770 | 2544957 | 1187 | - | tuf | WP_245029798.1 |

| | | | | | | | |
|---|---|---|---|---|---|---|---|
| GCF_029323795.1 | NZ_CP119758.1 | 142194 | 143381 | 1187 | + | tuf | WP_014641419.1 |
| GCF_000973105.1 | NC_015844.1 | 58308 | 59495 | 1187 | - | tuf | WP_085026897.1 |
| GCF_044597075.1 | NZ_CP139000.1 | 154186 | 155373 | 1187 | + | tuf | WP_283076986.1 |
| GCF_000023465.1 | NC_014041.1 | 196602 | 197789 | 1187 | - | tuf | WP_244710721.1 |
| GCF_026000115.1 | NZ_AP026867.1 | 2536040 | 2537227 | 1187 | + | tuf | WP_244751035.1 |
| GCF_000212735.1 | NC_015510.1 | 2337996 | 2339180 | 1184 | + | tuf | WP_029141997.1 |
| GCF_000250635.1 | NC_016940.1 | 3485105 | 3486289 | 1184 | - | tuf | WP_014096502.1 |
| GCF_029625215.1 | NZ_CP121197.1 | 129083 | 130267 | 1184 | + | tuf | WP_202778551.1 |
| GCF_003576455.2 | NZ_CP032869.1 | 198570 | 199751 | 1181 | + | tuf | WP_248416808.1 |
| GCF_020783335.1 | NZ_CP086068.1 | 583357 | 584541 | 1184 | - | tuf | WP_166161208.1 |
| GCF_009754905.2 | NZ_CP066775.1 | 764804 | 765988 | 1184 | - | tuf | WP_126108911.1 |
| GCF_007971025.1 | NZ_CP042436.1 | 1479394 | 1480578 | 1184 | - | tuf | WP_062470809.1 |
| GCF_007971525.1 | NZ_CP042437.1 | 1658946 | 1660130 | 1184 | + | tuf | WP_166063117.1 |
| GCF_017569325.2 | NZ_CP071878.2 | 1787882 | 1789063 | 1181 | + | tuf | WP_035810527.1 |
| GCF_002355435.1 | NZ_AP017313.1 | 234595 | 235782 | 1187 | + | tuf | WP_009710894.1 |
| GCF_028596025.1 | NZ_CP117167.1 | 3082558 | 3083742 | 1184 | - | tuf | WP_173224399.1 |
| GCF_013283875.1 | NZ_CP054139.1 | 145256 | 146443 | 1187 | + | tuf | WP_013074222.1 |
| GCF_900105165.1 | NZ_LT629740.1 | 151848 | 153035 | 1187 | + | tuf | WP_013074222.1 |
| GCF_000166195.2 | NZ_CM001403.1 | 247720 | 248904 | 1184 | + | tuf | WP_132223232.1 |
| GCF_012849215.1 | NZ_CP051682.1 | 1233515 | 1234702 | 1187 | + | tuf | WP_010488554.1 |
| GCF_034262375.1 | NZ_CP139558.1 | 2594172 | 2595359 | 1187 | + | tuf | WP_010488554.1 |
| GCF_002257585.1 | NZ_CP022743.1 | 1408379 | 1409566 | 1187 | + | tuf | WP_056962329.1 |
| GCF_008195825.1 | NZ_CP043329.1 | 1427170 | 1428357 | 1187 | - | tuf | WP_274258522.1 |
| GCF_015679185.1 | NZ_CP064939.1 | 1319837 | 1321024 | 1187 | + | tuf | WP_003565275.1 |
| GCF_044501175.1 | NZ_CP162529.1 | 1313578 | 1314765 | 1187 | + | tuf | WP_010488554.1 |
| GCF_000023825.1 | NC_013061.1 | 1791418 | 1792605 | 1187 | + | tuf | WP_005685646.1 |
| GCF_040026395.1 | NZ_CP157278.1 | 1362443 | 1363630 | 1187 | + | tuf | WP_010488554.1 |
| GCF_022200785.1 | NZ_CP087585.1 | 1832336 | 1833523 | 1187 | - | tuf | WP_010488554.1 |
| GCF_014692875.1 | NZ_CP061171.1 | 1742614 | 1743798 | 1184 | - | tuf | WP_003640798.1 |
| GCF_014395225.1 | NZ_CP060723.1 | 1712181 | 1713365 | 1184 | - | tuf | WP_308703309.1 |
| GCF_900564155.1 | NZ_LS999839.1 | 2888888 | 2890072 | 1184 | + | tuf | WP_021732063.1 |

| | | | | | | | |
|---|---|---|---|---|---|---|---|
| GCF_007833675.1 | NZ_CP031708.1 | 1890856 | 1892040 | 1184 | - | tuf | WP_003640798.1 |
| GCF_000190735.1 | NC_015177.1 | 2153701 | 2154885 | 1184 | - | tuf | WP_050338445.1 |
| GCF_000242635.2 | NC_017770.1 | 51992 | 53176 | 1184 | - | tuf | WP_003640798.1 |
| GCF_022014595.1 | NZ_CP091740.1 | 1830962 | 1832146 | 1184 | - | tuf | WP_057707446.1 |
| GCF_033955345.1 | NZ_CP138332.1 | 1009251 | 1010441 | 1190 | - | tuf | WP_054681243.1 |
| GCF_043187465.1 | NZ_CP171136.1 | 987101 | 988288 | 1187 | - | tuf | WP_003546862.1 |
| GCF_008274825.1 | NZ_CP030848.1 | 775092 | 776279 | 1187 | - | tuf | WP_080881452.1 |
| GCF_011046555.1 | NZ_CP049246.1 | 829335 | 830522 | 1187 | + | tuf | WP_014565740.1 |
| GCF_900187125.1 | NZ_LT906468.1 | 733497 | 734684 | 1187 | + | tuf | WP_046306787.1 |
| GCF_016894225.1 | NZ_CP069793.1 | 855367 | 856554 | 1187 | + | tuf | WP_013086177.1 |
| GCF_028736175.1 | NZ_CP117880.1 | 1030120 | 1031307 | 1187 | + | tuf | WP_002879079.1 |
| GCF_014109745.1 | NZ_CP058555.1 | 1168693 | 1169880 | 1187 | - | tuf | WP_003648636.1 |
| GCF_004421025.1 | NZ_CP038029.1 | 861384 | 862571 | 1187 | + | tuf | WP_038522433.1 |
| GCF_023277505.1 | NZ_CP080574.1 | 847645 | 848832 | 1187 | - | tuf | WP_014919004.1 |
| GCF_016725645.1 | NZ_CP068084.1 | 576838 | 578025 | 1187 | + | tuf | WP_006730011.1 |
| GCF_901482695.1 | NZ_LR590484.1 | 1023089 | 1024276 | 1187 | - | tuf | WP_004042801.1 |
| GCF_009557615.1 | NZ_CP045652.1 | 956459 | 957646 | 1187 | - | tuf | WP_282806338.1 |
| GCF_020149915.1 | NZ_CP083760.1 | 1165391 | 1166578 | 1187 | - | tuf | WP_006584557.1 |
| GCF_003353065.1 | NZ_CP027806.1 | 1214518 | 1215705 | 1187 | - | tuf | WP_004894320.1 |
| GCF_003353065.1 | NZ_CP027806.1 | 1288088 | 1289275 | 1187 | - | tuf | WP_013854292.1 |
| GCF_002872415.1 | NZ_CP025704.1 | 929086 | 930273 | 1187 | + | tuf | WP_045928030.1 |
| GCF_002872415.1 | NZ_CP025704.1 | 958803 | 959990 | 1187 | + | tuf | WP_006584557.1 |
| GCF_000210915.2 | NC_016620.1 | 996902 | 998089 | 1187 | + | tuf | WP_103751563.1 |
| GCF_000210915.2 | NC_016620.1 | 1209913 | 1211100 | 1187 | - | tuf | WP_113576145.1 |
| GCF_034259205.1 | NZ_CP139487.1 | 1170422 | 1171609 | 1187 | - | tuf | WP_057719388.1 |
| GCF_034259205.1 | NZ_CP139487.1 | 919876 | 921063 | 1187 | + | tuf | WP_099974685.1 |
| GCF_000196175.1 | NC_005363.1 | 1385643 | 1386830 | 1187 | - | tuf | WP_007125426.1 |
| GCF_022814725.1 | NZ_CP093442.1 | 723226 | 724413 | 1187 | + | tuf | WP_317638138.1 |
| GCF_000348725.1 | NC_020813.1 | 1573101 | 1574285 | 1184 | - | tuf | WP_120772293.1 |
| GCF_004295685.1 | NZ_AP019368.1 | 1577664 | 1578857 | 1193 | - | tuf | WP_047915022.1 |
| GCF_001907975.1 | NZ_CP017834.1 | 2001654 | 2002838 | 1184 | - | tuf | WP_003132374.1 |

| | | | | | | | |
|---|---|---|---|---|---|---|---|
| GCF_037081835.1 | NZ_CP146516.1 | 1085179 | 1086363 | 1184 | + | tuf | WP_017370373.1 |
| GCF_000284335.1 | NC_017096.1 | 754158 | 755342 | 1184 | + | tuf | WP_003133474.1 |
| GCF_000284335.1 | NC_017096.1 | 2021429 | 2022613 | 1184 | - | tuf | WP_003132374.1 |
| GCF_001886815.1 | NZ_CP018099.1 | 1695536 | 1696729 | 1193 | - | tuf | WP_047915022.1 |
| GCF_001886815.1 | NZ_CP018099.1 | 1734859 | 1736043 | 1184 | - | tuf | WP_003133474.1 |
| GCF_000194135.1 | NC_015318.1 | 2642860 | 2644044 | 1184 | - | tuf | WP_142767354.1 |
| GCF_000194135.1 | NC_015318.1 | 1187110 | 1188303 | 1193 | - | tuf | WP_061774772.1 |
| GCF_000014025.1 | NC_009850.1 | 1781266 | 1782450 | 1184 | - | tuf | WP_075525307.1 |
| GCF_013372265.1 | NZ_CP054051.1 | 1467156 | 1468343 | 1187 | - | tuf | WP_057757390.1 |
| GCF_003660105.1 | NZ_CP032823.1 | 348323 | 349510 | 1187 | + | tuf | WP_039099648.1 |
| GCF_013201705.1 | NZ_CP053837.1 | 1290460 | 1291647 | 1187 | - | tuf | WP_025082375.1 |
| GCF_013201625.1 | NZ_CP053839.1 | 1111095 | 1112282 | 1187 | - | tuf | WP_057907881.1 |
| GCF_003544835.1 | NZ_CP032099.1 | 1865699 | 1866886 | 1187 | - | tuf | WP_011374763.1 |
| GCF_004214815.1 | NZ_CP035926.1 | 126248 | 127432 | 1184 | + | tuf | WP_066145207.1 |
| GCF_003355515.1 | NZ_CP031367.1 | 2544985 | 2546172 | 1187 | - | tuf | WP_068446123.1 |
| GCF_048320235.1 | NZ_CP182882.1 | 125224 | 126411 | 1187 | + | tuf | WP_269410877.1 |
| GCF_013177675.1 | NZ_CP042652.1 | 2056973 | 2058157 | 1184 | + | tuf | WP_013727577.1 |
| GCF_013177635.1 | NZ_CP030944.1 | 88024 | 89208 | 1184 | - | tuf | WP_035168468.1 |
| GCF_013201935.1 | NZ_CP053833.1 | 1284380 | 1285564 | 1184 | + | tuf | WP_003555271.1 |
| GCF_013201725.1 | NZ_CP053835.1 | 263207 | 264391 | 1184 | - | tuf | WP_054769507.1 |
| GCF_003544915.1 | NZ_CP032097.1 | 383060 | 384244 | 1184 | + | tuf | WP_227361373.1 |
| GCF_000092245.1 | NC_014166.1 | 146162 | 147346 | 1184 | - | tuf | WP_057909557.1 |
| GCF_003711085.1 | NZ_CP032363.1 | 1467819 | 1469003 | 1184 | - | tuf | WP_268913837.1 |
| GCF_004299785.2 | NZ_CP036246.2 | 1541558 | 1542742 | 1184 | + | tuf | WP_014974550.1 |
| GCF_024267655.1 | NZ_CP100595.1 | 1119382 | 1120566 | 1184 | - | tuf | WP_004900335.1 |
| GCF_003544815.1 | NZ_CP032100.1 | 661336 | 662520 | 1184 | + | tuf | WP_080519256.1 |
| GCF_013201665.1 | NZ_CP053840.1 | 1275336 | 1276520 | 1184 | - | tuf | WP_010383678.1 |
| GCF_005843985.1 | NZ_CP040463.1 | 1274920 | 1276104 | 1184 | - | tuf | WP_010017300.1 |
| GCF_005083985.2 | NZ_CP027432.2 | 388528 | 389712 | 1184 | - | tuf | WP_013102558.1 |
| GCF_013372105.1 | NZ_CP053825.1 | 134528 | 135712 | 1184 | - | tuf | WP_010001907.1 |
| GCF_002238335.1 | NZ_CP022347.1 | 594404 | 595588 | 1184 | + | tuf | WP_011679439.1 |

| | | | | | | | |
|---|---|---|---|---|---|---|---|
| GCF_041735015.1 | NZ_CP168723.1 | 715042 | 716226 | 1184 | + | tuf | WP_080519256.1 |
| GCF_013201895.1 | NZ_CP053841.1 | 586424 | 587608 | 1184 | + | tuf | WP_072613296.1 |
| GCF_004803835.1 | NZ_CP012545.1 | 1417149 | 1418336 | 1187 | - | tuf | WP_024526778.1 |
| GCF_013177655.1 | NZ_CP035946.1 | 1153381 | 1154568 | 1187 | + | tuf | WP_125579748.1 |
| GCF_009730395.1 | NZ_CP046317.1 | 2582419 | 2583606 | 1187 | + | tuf | WP_048735869.1 |
| GCF_001298465.1 | NZ_CP012541.1 | 1259138 | 1260325 | 1187 | + | tuf | WP_020090257.1 |
| GCF_013201645.1 | NZ_CP053842.1 | 1807013 | 1808200 | 1187 | + | tuf | WP_182150618.1 |
| GCF_002104335.1 | NZ_CP020867.1 | 1192747 | 1193934 | 1187 | + | tuf | WP_057730629.1 |
| GCF_013372125.1 | NZ_CP053826.1 | 1455655 | 1456845 | 1190 | - | tuf | WP_010494526.1 |
| GCF_011600945.2 | NZ_CP059443.1 | 1363847 | 1365031 | 1184 | - | tuf | WP_089144350.1 |
| GCF_013201685.1 | NZ_CP053844.1 | 761443 | 762627 | 1184 | + | tuf | WP_010688796.1 |
| GCF_001190745.1 | NZ_CP012196.1 | 744335 | 745519 | 1184 | - | tuf | WP_290034168.1 |
| GCF_002080395.1 | NZ_CP020478.1 | 2241752 | 2242936 | 1184 | - | tuf | WP_280606114.1 |
| GCF_001687475.2 | NZ_CP031611.1 | 934432 | 935616 | 1184 | - | tuf | WP_112194775.1 |
| GCF_000017585.1 | NC_009714.1 | 1299213 | 1300397 | 1184 | - | tuf | WP_014073681.1 |
| GCF_013372165.1 | NZ_CP053828.1 | 886214 | 887398 | 1184 | - | tuf | WP_027826984.1 |
| GCF_000736415.1 | NZ_CP009043.1 | 729748 | 730932 | 1184 | + | tuf | WP_003706071.1 |
| GCF_000816185.1 | NZ_CP007770.1 | 837757 | 838944 | 1187 | - | tuf | WP_014562272.1 |
| GCF_000009085.1 | NC_002163.1 | 706899 | 708086 | 1187 | + | tuf | WP_057750281.1 |
| GCF_002139935.1 | NZ_CP015578.1 | 635908 | 637095 | 1187 | + | tuf | WP_006499744.1 |
| GCF_008245025.1 | NZ_CP043426.1 | 91695 | 92882 | 1187 | + | tuf | WP_003711740.1 |
| GCF_013372205.1 | NZ_CP053831.1 | 691432 | 692619 | 1187 | + | tuf | WP_057808178.1 |
| GCF_018968685.1 | NZ_CP076657.1 | 1081213 | 1082400 | 1187 | - | tuf | WP_102169071.1 |
| GCF_013201605.1 | NZ_CP053848.1 | 744627 | 745814 | 1187 | + | tuf | WP_003716883.1 |
| GCF_000816785.1 | NZ_CP007766.1 | 1677634 | 1678818 | 1184 | - | tuf | WP_057868978.1 |
| GCF_002021985.1 | NZ_CP017018.1 | 1491456 | 1492640 | 1184 | - | tuf | WP_003689860.1 |
| GCF_002139855.1 | NZ_CP018789.1 | 486186 | 487370 | 1184 | + | tuf | WP_057885763.1 |
| GCF_004803795.1 | NZ_CP012543.1 | 2648018 | 2649202 | 1184 | - | tuf | WP_036105441.1 |
| GCF_004803815.1 | NZ_CP012544.1 | 2587988 | 2589172 | 1184 | - | tuf | WP_003739865.1 |
| GCF_008245005.1 | NZ_CP043427.1 | 2751188 | 2752372 | 1184 | - | tuf | WP_003753951.1 |
| GCF_000816305.1 | NZ_CP007773.1 | 2726011 | 2727195 | 1184 | - | tuf | NP_466175.1 |

| | | | | | | | |
|---|---|---|---|---|---|---|---|
| GCF_013372245.1 | NZ_CP053849.1 | 1787087 | 1788271 | 1184 | + | tuf | WP_036094372.1 |
| GCF_013372225.1 | NZ_CP053832.1 | 2620445 | 2621629 | 1184 | - | tuf | WP_012986510.1 |
| GCF_002139825.2 | NZ_CP144916.1 | 2616197 | 2617381 | 1184 | - | tuf | WP_185639342.1 |
| GCF_008245045.1 | NZ_CP043428.1 | 368621 | 369805 | 1184 | + | tuf | WP_036060245.1 |
| GCF_014217995.1 | NZ_CP041617.1 | 2630773 | 2631957 | 1184 | - | tuf | WP_003723640.1 |
| GCF_013416015.1 | NZ_CP049075.1 | 2239103 | 2240287 | 1184 | - | tuf | WP_068224739.1 |
| GCF_015265455.1 | NZ_CP054492.1 | 297730 | 298914 | 1184 | + | tuf | WP_057905110.1 |
| GCF_015265475.1 | NZ_CP054493.1 | 1325542 | 1326726 | 1184 | + | tuf | WP_004562580.1 |
| GCF_006459125.1 | NZ_CP041070.1 | 5110238 | 5111422 | 1184 | - | tuf | WP_053596053.1 |
| GCF_003346815.1 | NZ_CP031217.1 | 4569956 | 4571140 | 1184 | - | tuf | WP_004233614.1 |
| GCF_013201825.1 | NZ_CP053836.1 | 191627 | 192811 | 1184 | + | tuf | WP_008181778.1 |
| GCF_000009305.1 | NC_008229.1 | 4285316 | 4286500 | 1184 | - | tuf | WP_274794957.1 |
| GCF_003097575.1 | NZ_CP021886.1 | 1183706 | 1184890 | 1184 | + | tuf | WP_293924102.1 |
| GCF_000162575.1 | NZ_CM000776.2 | 4034913 | 4036097 | 1184 | + | tuf | WP_053997228.1 |
| GCF_000259275.1 | NC_017735.1 | 5070585 | 5071769 | 1184 | - | tuf | WP_283870073.1 |
| GCF_900638485.1 | NZ_LR134518.1 | 4680612 | 4681796 | 1184 | - | tuf | WP_107925082.1 |
| GCF_000284635.1 | NC_017761.1 | 235475 | 236659 | 1184 | + | tuf | WP_024364376.1 |
| GCF_001693335.1 | NZ_CP016503.1 | 4412975 | 4414159 | 1184 | - | tuf | WP_126292326.1 |
| GCF_000200595.1 | NC_014810.2 | 2194329 | 2195513 | 1184 | - | tuf | WP_106781840.1 |
| GCF_030296895.1 | NZ_AP028022.1 | 4508071 | 4509255 | 1184 | - | tuf | WP_024364376.1 |
| GCF_019703305.1 | NZ_AP024814.1 | 2236210 | 2237394 | 1184 | - | tuf | WP_203545863.1 |
| GCF_000007905.1 | NC_004917.1 | 2230072 | 2231256 | 1184 | - | tuf | WP_086043509.1 |
| GCF_001602095.1 | NZ_CP014991.1 | 1619478 | 1620662 | 1184 | - | tuf | WP_086039297.1 |
| GCF_039555295.1 | NZ_CP145316.1 | 2074748 | 2075932 | 1184 | - | tuf | WP_224183607.1 |
| GCF_900476215.1 | NZ_LS483446.1 | 1952739 | 1953923 | 1184 | - | tuf | WP_289650164.1 |
| GCF_900638355.1 | NZ_LR134509.1 | 1986031 | 1987215 | 1184 | - | tuf | WP_133431848.1 |
| GCF_025998455.1 | NZ_AP026446.1 | 2341282 | 2342466 | 1184 | - | tuf | WP_111717039.1 |
| GCF_013374215.1 | NZ_AP023046.1 | 2007835 | 2009019 | 1184 | - | tuf | WP_149459284.1 |
| GCF_001460635.1 | NZ_LN907858.1 | 1908417 | 1909601 | 1184 | - | tuf | WP_144781510.1 |
| GCF_028751035.1 | NZ_CP063533.1 | 2652163 | 2653350 | 1187 | - | tuf | WP_016999427.1 |
| GCF_030296055.1 | NZ_AP027370.1 | 2637235 | 2638422 | 1187 | - | tuf | WP_048541361.1 |

| | | | | | | | |
|---|---|---|---|---|---|---|---|
| GCF_040436365.1 | NZ_AP026369.1 | 2261468 | 2262655 | 1187 | - | tuf | WP_095089740.1 |
| GCF_032850805.1 | NZ_CP136593.1 | 2415969 | 2417156 | 1187 | + | tuf | WP_107536373.1 |
| GCF_016593255.1 | NZ_AP023212.1 | 2847006 | 2848193 | 1187 | + | tuf | WP_239672714.1 |
| GCF_008000835.1 | NZ_CP042812.1 | 254616 | 255800 | 1184 | + | tuf | WP_013773176.1 |
| GCF_003346755.1 | NZ_CP031218.1 | 4606809 | 4607993 | 1184 | + | tuf | WP_102264854.1 |
| GCF_003544855.1 | NZ_CP032101.1 | 135743 | 136927 | 1184 | + | tuf | WP_404330703.1 |
| GCF_003544935.1 | NZ_CP032098.1 | 140625 | 141812 | 1187 | + | tuf | WP_223438939.1 |
| GCF_003346775.1 | NZ_CP031219.1 | 2098853 | 2100040 | 1187 | + | tuf | WP_066330429.1 |
| GCF_004214795.1 | NZ_CP035928.1 | 140606 | 141793 | 1187 | + | tuf | WP_223438939.1 |
| GCF_000021725.1 | NC_012115.1 | 142436 | 143623 | 1187 | + | tuf | WP_029282835.1 |
| GCF_000186245.1 | NC_014935.1 | 542674 | 543861 | 1187 | - | tuf | WP_066330429.1 |
| GCF_027946175.1 | NZ_AP026671.1 | 142426 | 143610 | 1184 | + | tuf | WP_136357926.1 |
| GCF_015100395.1 | NZ_AP022847.1 | 145075 | 146262 | 1187 | + | tuf | WP_035406786.1 |
| GCF_027943725.1 | NZ_AP025696.1 | 723455 | 724639 | 1184 | - | tuf | WP_034676069.1 |
| GCF_014466985.1 | NZ_AP022826.1 | 248205 | 249389 | 1184 | + | tuf | WP_047943714.1 |
| GCF_013283835.1 | NZ_CP054052.1 | 128702 | 129886 | 1184 | + | tuf | WP_338450520.1 |
| GCF_001956695.1 | NZ_CP019070.1 | 1121277 | 1122461 | 1184 | + | tuf | WP_260116278.1 |
| GCF_000183725.1 | NC_014762.1 | 1387737 | 1388921 | 1184 | - | tuf | WP_183673694.1 |
| GCF_017357825.1 | NZ_CP046072.1 | 126042 | 127229 | 1187 | + | tuf | WP_077718346.1 |
| GCF_000147355.1 | NC_014506.1 | 139729 | 140913 | 1184 | + | tuf | WP_011064520.1 |
| GCF_000012965.1 | NC_007575.1 | 140786 | 141973 | 1187 | + | tuf | WP_379560383.1 |
| GCF_039645995.1 | NZ_CP147920.1 | 30967 | 32154 | 1187 | + | tuf | WP_017795118.1 |
| GCF_000242915.1 | NZ_AFRZ01000001.1 | 128457 | 129641 | 1184 | + | tuf | WP_114914887.1 |
| GCF_009068765.1 | NZ_CP035534.1 | 1607383 | 1608570 | 1187 | - | tuf | WP_007746813.1 |
| GCF_009258225.1 | NZ_CP043617.1 | 1213060 | 1214247 | 1187 | - | tuf | WP_002818673.1 |
| GCF_014905095.1 | NZ_CP041165.1 | 1247205 | 1248392 | 1187 | + | tuf | WP_174813062.1 |
| GCF_014905135.1 | NZ_CP041406.1 | 136899 | 138086 | 1187 | + | tuf | WP_176007715.1 |
| GCF_014905115.1 | NZ_CP041235.1 | 7558920 | 7560107 | 1187 | + | tuf | WP_087570305.1 |
| GCF_009258045.1 | NZ_CP041166.1 | 706207 | 707394 | 1187 | + | tuf | WP_168906331.1 |
| GCF_000265295.1 | NC_018002.1 | 231010 | 232197 | 1187 | - | tuf | WP_126011539.1 |
| GCF_000024885.1 | NC_013512.1 | 3938872 | 3940059 | 1187 | - | tuf | WP_138227148.1 |

| | | | | | | | |
|---|---|---|---|---|---|---|---|
| GCF_002309535.1 | NZ_CP023275.1 | 6057167 | 6058354 | 1187 | - | tuf | WP_017692073.1 |
| GCF_001723605.1 | NZ_CP017111.1 | 7198827 | 7200014 | 1187 | - | tuf | WP_145669490.1 |
| GCF_000568815.1 | NZ_CP007201.1 | 1745367 | 1746554 | 1187 | + | tuf | WP_068650942.1 |
| GCF_014931715.1 | NZ_CP063164.1 | 279912 | 281099 | 1187 | + | tuf | WP_315605515.1 |
| GCF_000987835.1 | NZ_CP011308.1 | 6656154 | 6657341 | 1187 | - | tuf | WP_194234534.1 |
| GCF_020911965.1 | NZ_CP087124.1 | 6049025 | 6050212 | 1187 | + | tuf | WP_125663898.1 |
| GCF_900637325.1 | NZ_LR134366.1 | 2954265 | 2955452 | 1187 | - | tuf | WP_110899278.1 |
| GCF_021057185.1 | NZ_CP054059.1 | 2784713 | 2785900 | 1187 | + | tuf | WP_045670702.1 |
| GCF_000146065.2 | NC_020449.1 | 7406335 | 7407522 | 1187 | - | tuf | WP_039298612.1 |
| GCF_030520105.1 | NZ_CP130454.1 | 5137912 | 5139099 | 1187 | - | tuf | WP_046213348.1 |
| GCF_004102945.1 | NZ_CP019384.1 | 1537712 | 1538899 | 1187 | - | tuf | WP_007432584.1 |
| GCF_047159735.1 | NZ_AP038758.1 | 1617712 | 1618899 | 1187 | - | tuf | WP_110046859.1 |
| GCF_005697565.1 | NZ_CP040004.1 | 4384871 | 4386058 | 1187 | + | tuf | WP_145030007.1 |
| GCF_000803625.1 | NZ_CP007496.1 | 6394473 | 6395660 | 1187 | - | tuf | WP_042235513.1 |
| GCF_000392435.1 | NC_021219.1 | 3251933 | 3253120 | 1187 | + | tuf | WP_068657358.1 |
| GCF_030253535.1 | NZ_CP124550.1 | 5598554 | 5599744 | 1190 | - | tuf | WP_168182396.1 |
| GCF_025273655.1 | NZ_CP045921.1 | 6286772 | 6287959 | 1187 | - | tuf | WP_087918352.1 |
| GCF_001653975.1 | NZ_CP014639.1 | 5301508 | 5302695 | 1187 | - | tuf | WP_042208399.1 |
| GCF_000011565.2 | NC_005861.2 | 295411 | 296598 | 1187 | + | tuf | WP_054975995.1 |
| GCF_001499655.1 | NZ_LN879502.1 | 735448 | 736635 | 1187 | + | tuf | WP_006212927.1 |
| GCF_019453995.1 | NZ_CP075587.1 | 6440473 | 6441660 | 1187 | - | tuf | WP_020427072.1 |
| GCF_015356815.2 | NZ_CP075585.1 | 4847288 | 4848475 | 1187 | - | tuf | WP_152395417.1 |
| GCF_040235295.1 | NZ_CP158101.1 | 6552073 | 6553260 | 1187 | + | tuf | WP_235120084.1 |
| GCF_019056495.1 | NZ_CP050318.1 | 4990190 | 4991377 | 1187 | - | tuf | WP_188777913.1 |
| GCF_000007605.1 | NC_003361.3 | 3980453 | 3981640 | 1187 | + | tuf | WP_077568289.1 |
| GCF_018343815.1 | NZ_CP060791.1 | 1134923 | 1136110 | 1187 | - | tuf | WP_020427072.1 |
| GCF_000009945.1 | NC_007899.1 | 658617 | 659804 | 1187 | + | tuf | WP_007432584.1 |
| GCF_000471025.2 | NZ_CP015840.1 | 4421263 | 4422450 | 1187 | - | tuf | WP_204827267.1 |
| GCF_000006685.1 | NC_002620.2 | 3994662 | 3995849 | 1187 | - | tuf | WP_077996267.1 |
| GCF_020549965.1 | NZ_CP080403.1 | 1632489 | 1633679 | 1190 | + | tuf | WP_125082107.1 |
| GCF_000007205.1 | NC_005043.1 | 5705801 | 5706988 | 1187 | - | tuf | WP_127003008.1 |

| | | | | | | | |
|---|---|---|---|---|---|---|---|
| GCF_900239975.1 | NZ_LS992154.1 | 6318592 | 6319779 | 1187 | - | tuf | WP_162359832.1 |
| GCF_000204255.1 | NC_017287.1 | 320275 | 321462 | 1187 | - | tuf | WP_024633588.1 |
| GCF_900239945.1 | NZ_LT993738.1 | 8378642 | 8379829 | 1187 | - | tuf | WP_013921164.1 |
| GCF_900149385.2 | NZ_LT999999.1 | 4575813 | 4577000 | 1187 | + | tuf | WP_054818710.1 |
| GCF_000008725.1 | NC_000117.1 | 5042954 | 5044141 | 1187 | + | tuf | WP_069328352.1 |
| GCF_902807005.1 | NZ_LR777658.1 | 6161393 | 6162580 | 1187 | - | tuf | WP_036680523.1 |
| GCF_902806995.1 | NZ_LR777654.1 | 5801038 | 5802225 | 1187 | - | tuf | WP_236337404.1 |
| GCF_000253035.1 | NC_015702.1 | 2534250 | 2535437 | 1187 | - | tuf | WP_094094351.1 |
| GCF_000237205.1 | NC_015713.1 | 4390200 | 4391387 | 1187 | - | tuf | WP_160036605.1 |
| GCF_000092785.1 | NC_014225.1 | 2913870 | 2915057 | 1187 | - | tuf | WP_013373267.1 |
| GCF_001747405.1 | NZ_CP017305.1 | 3718612 | 3719799 | 1187 | + | tuf | WP_129443063.1 |
| GCF_000020505.1 | NC_011027.1 | 242755 | 243942 | 1187 | + | tuf | WP_155698562.1 |
| GCF_000006985.1 | NC_002932.3 | 4216018 | 4217205 | 1187 | + | tuf | WP_162642518.1 |
| GCF_000020465.1 | NC_010803.1 | 7142092 | 7143279 | 1187 | - | tuf | WP_020427072.1 |
| GCF_000015125.1 | NC_008639.1 | 5494928 | 5496115 | 1187 | - | tuf | WP_314799778.1 |
| GCF_000020525.1 | NC_011026.1 | 4694497 | 4695684 | 1187 | - | tuf | WP_025336677.1 |
| GCF_000012485.1 | NC_007512.1 | 6247327 | 6248514 | 1187 | - | tuf | WP_116191465.1 |
| GCF_000020645.1 | NC_011060.1 | 22549 | 23736 | 1187 | - | tuf | WP_036589455.1 |
| GCF_000020625.1 | NC_011059.1 | 6961889 | 6963076 | 1187 | - | tuf | WP_258385896.1 |
| GCF_000199675.1 | NC_014960.1 | 5024066 | 5025253 | 1187 | - | tuf | WP_038699020.1 |
| GCF_000199675.1 | NC_014960.1 | 2604034 | 2605221 | 1187 | + | tuf | WP_068606585.1 |
| GCF_900184705.1 | NZ_LT859958.1 | 786565 | 787752 | 1187 | - | tuf | WP_010348813.1 |
| GCF_003966975.1 | NZ_AP018437.1 | 3087368 | 3088558 | 1190 | - | tuf | WP_087444231.1 |
| GCF_003966975.1 | NZ_AP018437.1 | 5621546 | 5622733 | 1187 | - | tuf | WP_019908221.1 |
| GCF_900066015.1 | NZ_LN890655.1 | 6907235 | 6908422 | 1187 | - | tuf | WP_173140462.1 |
| GCF_900066015.1 | NZ_LN890656.1 | 5681768 | 5682955 | 1187 | - | tuf | WP_145053340.1 |
| GCF_000281175.1 | NC_017079.1 | 540217 | 541404 | 1187 | + | tuf | WP_208917602.1 |
| GCF_000281175.1 | NC_017079.1 | 4767168 | 4768355 | 1187 | - | tuf | WP_047914091.1 |
| GCF_015500095.1 | NZ_CP062983.1 | 4638155 | 4639345 | 1190 | - | tuf | WP_283925951.1 |
| GCF_030389965.1 | NZ_CP128399.1 | 6933541 | 6934728 | 1187 | - | tuf | WP_017692073.1 |
| GCF_030389965.1 | NZ_CP128399.1 | 3627208 | 3628395 | 1187 | - | tuf | WP_068698244.1 |

| | | | | | | | |
|---|---|---|---|---|---|---|---|
| GCF_000021945.1 | NC_011831.1 | 157205 | 158389 | 1184 | + | tuf | WP_017381021.1 |
| GCF_000021945.1 | NC_011831.1 | 2403248 | 2404432 | 1184 | - | tuf | WP_119884329.1 |
| GCF_000018865.1 | NC_010175.1 | 484316 | 485500 | 1184 | - | tuf | WP_003247557.1 |
| GCF_000018865.1 | NC_010175.1 | 350994 | 352178 | 1184 | - | tuf | WP_062756063.1 |
| GCF_000017805.1 | NC_009767.1 | 175780 | 176967 | 1187 | + | tuf | WP_112179271.1 |
| GCF_000017805.1 | NC_009767.1 | 1318975 | 1320162 | 1187 | - | tuf | WP_041093791.1 |
| GCF_037094495.1 | NZ_CP146612.1 | 1453357 | 1454544 | 1187 | - | tuf | WP_150204210.1 |
| GCF_029593895.1 | NZ_CP046146.1 | 1154455 | 1155642 | 1187 | - | tuf | WP_057890380.1 |
| GCF_000011905.1 | NC_002936.3 | 1687573 | 1688760 | 1187 | - | tuf | WP_010622374.1 |
| GCF_014672715.2 | NZ_CP058566.2 | 1246498 | 1247682 | 1184 | - | tuf | WP_002830466.1 |
| GCF_001953175.1 | NZ_CP018258.1 | 1699502 | 1700686 | 1184 | + | tuf | WP_014215321.1 |
| GCF_004208415.1 | NZ_CP035758.1 | 96935 | 98119 | 1184 | - | tuf | WP_062913065.1 |
| GCF_004208415.1 | NZ_CP035758.1 | 881723 | 882907 | 1184 | + | tuf | WP_320532066.1 |
| GCF_008838325.1 | NZ_CP042829.1 | 1002176 | 1003360 | 1184 | + | tuf | WP_057784497.1 |
| GCF_008838325.1 | NZ_CP042829.1 | 1142648 | 1143832 | 1184 | - | tuf | WP_002833446.1 |
| GCF_000025005.1 | NC_013525.1 | 823931 | 825115 | 1184 | + | tuf | WP_412990136.1 |
| GCF_000025005.1 | NC_013525.1 | 133393 | 134577 | 1184 | + | tuf | WP_048677883.1 |
| GCF_000177635.2 | NC_014836.1 | 112564 | 113748 | 1184 | + | tuf | WP_241582232.1 |
| GCF_000020945.1 | NC_011295.1 | 2700048 | 2701232 | 1184 | - | tuf | WP_040372538.1 |
| GCF_000020945.1 | NC_011295.1 | 2441297 | 2442481 | 1184 | - | tuf | WP_063235794.1 |
| GCF_000018105.1 | NC_009925.1 | 2530933 | 2532117 | 1184 | + | tuf | WP_133364068.1 |
| GCF_000317515.1 | NC_019738.1 | 133737 | 134921 | 1184 | + | tuf | WP_006828917.1 |
| GCF_000317695.1 | NC_019771.1 | 149280 | 150464 | 1184 | + | tuf | WP_065527574.1 |
| GCF_015245355.1 | NZ_CP063311.1 | 129285 | 130469 | 1184 | + | tuf | WP_058386734.1 |
| GCF_018389385.1 | NZ_CP062698.1 | 1602314 | 1603498 | 1184 | + | tuf | WP_036803876.1 |
| GCF_006968745.1 | NZ_CP030118.1 | 129144 | 130328 | 1184 | + | tuf | WP_065528393.1 |
| GCF_000025125.1 | NC_013771.1 | 1632409 | 1633593 | 1184 | - | tuf | WP_058386734.1 |
| GCF_047276495.1 | NZ_CP178543.1 | 113001 | 114185 | 1184 | + | tuf | WP_077587635.1 |
| GCF_000317145.1 | NC_019697.1 | 127660 | 128844 | 1184 | + | tuf | WP_074511495.1 |
| GCF_000317125.1 | NC_019695.1 | 126678 | 127862 | 1184 | + | tuf | WP_068459676.1 |
| GCF_000317495.1 | NC_019753.1 | 126064 | 127248 | 1184 | + | tuf | WP_068871334.1 |

| | | | | | | | |
|---|---|---|---|---|---|---|---|
| GCF_000017845.1 | NC_010546.1 | 3314844 | 3316028 | 1184 | + | tuf | WP_058382174.1 |
| GCF_000317675.1 | NC_019776.1 | 117135 | 118319 | 1184 | + | tuf | WP_049694537.1 |
| GCF_000316515.1 | NC_019675.1 | 184994 | 186181 | 1187 | + | tuf | WP_212773787.1 |
| GCF_963920485.1 | NZ_OY986431.1 | 126551 | 127738 | 1187 | + | tuf | WP_013081371.1 |
| GCF_014489415.1 | NZ_CP060822.1 | 3873481 | 3874668 | 1187 | + | tuf | WP_025909723.1 |
| GCF_003367075.2 | NZ_CP065936.1 | 126805 | 127992 | 1187 | + | tuf | WP_013081371.1 |
| GCF_000317535.1 | NC_019757.1 | 4137789 | 4138973 | 1184 | - | tuf | WP_151701737.1 |
| GCF_000317615.1 | NC_019780.1 | 149017 | 150204 | 1187 | + | tuf | WP_143891636.1 |
| GCF_002368115.1 | NZ_AP018316.1 | 85972 | 87159 | 1187 | + | tuf | WP_231095133.1 |
| GCF_012516395.1 | NZ_CP051206.1 | 144682 | 145869 | 1187 | + | tuf | WP_053429952.1 |
| GCF_024584745.1 | NZ_CP099464.1 | 1813120 | 1814307 | 1187 | - | tuf | WP_141447291.1 |
| GCF_007904085.1 | NZ_CP042326.1 | 1589420 | 1590604 | 1184 | - | tuf | WP_017434018.1 |
| GCF_000332235.1 | NZ_CM001775.1 | 207294 | 208481 | 1187 | + | tuf | WP_200126445.1 |
| GCF_000484535.1 | NC_022600.1 | 164532 | 165719 | 1187 | + | tuf | WP_200086907.1 |
| GCF_021018745.1 | NZ_CP063845.1 | 2081979 | 2083166 | 1187 | - | tuf | WP_142089673.1 |
| GCF_000011385.1 | NC_005125.1 | 2352291 | 2353478 | 1187 | + | tuf | WP_114373631.1 |
| GCF_032163395.1 | NZ_CP119968.1 | 2579483 | 2580670 | 1187 | - | tuf | WP_008592520.1 |
| GCF_001870225.1 | NZ_CP017675.1 | 2479303 | 2480487 | 1184 | - | tuf | WP_342388020.1 |
| GCF_000021825.1 | NC_011729.1 | 3446917 | 3448104 | 1187 | - | tuf | WP_069366434.1 |
| GCF_000147335.1 | NC_014501.1 | 706503 | 707690 | 1187 | + | tuf | WP_027827358.1 |
| GCF_002075285.3 | NZ_CP021983.2 | 138947 | 140131 | 1184 | - | tuf | WP_010620049.1 |
| GCF_029953635.1 | NZ_CP124543.1 | 2514455 | 2515639 | 1184 | + | tuf | WP_054707779.1 |
| GCF_021379005.1 | NZ_CP075171.1 | 21132 | 22319 | 1187 | + | tuf | WP_026569042.1 |
| GCF_002142475.1 | NZ_AP014638.1 | 128508 | 129695 | 1187 | + | tuf | WP_062751231.1 |
| GCF_019891175.1 | NZ_CP072600.1 | 1660889 | 1662076 | 1187 | + | tuf | WP_038475783.1 |
| GCF_000973065.1 | NZ_FO818640.1 | 173528 | 174715 | 1187 | + | tuf | WP_038475783.1 |
| GCF_042465725.1 | NZ_CP120853.1 | 2017060 | 2018247 | 1187 | + | tuf | WP_095327382.1 |
| GCF_022376295.1 | NZ_CP060140.1 | 3992034 | 3993218 | 1184 | - | tuf | WP_241370734.1 |
| GCF_003054475.1 | NZ_CP020114.1 | 2919856 | 2921040 | 1184 | + | tuf | WP_028977338.1 |
| GCF_000196515.1 | NC_014248.1 | 2916016 | 2917200 | 1184 | - | tuf | WP_039043316.1 |
| GCF_014023275.1 | NZ_CP054698.1 | 3186896 | 3188080 | 1184 | - | tuf | WP_317967275.1 |

| | | | | | | | |
|---|---|---|---|---|---|---|---|
| GCF_002813575.1 | NZ_CP024785.1 | 3118994 | 3120178 | 1184 | - | tuf | WP_115363647.1 |
| GCF_001298445.1 | NZ_CP012036.1 | 327421 | 328605 | 1184 | + | tuf | WP_067204738.1 |
| GCF_000020025.1 | NC_010628.1 | 2647672 | 2648856 | 1184 | + | tuf | WP_040285480.1 |
| GCF_003443655.1 | NZ_CP031941.1 | 2111846 | 2113030 | 1184 | + | tuf | WP_029053279.1 |
| GCF_038050375.1 | NZ_CP150886.1 | 426537 | 427721 | 1184 | + | tuf | WP_075526622.1 |
| GCF_000317105.1 | NC_019693.1 | 262339 | 263523 | 1184 | + | tuf | WP_037566322.1 |
| GCF_000317475.1 | NC_019729.1 | 569660 | 570841 | 1181 | + | tuf | WP_001040566.1 |
| GCF_000317475.1 | NC_019729.1 | 2370576 | 2371760 | 1184 | - | tuf | WP_002509426.1 |
| GCF_012295525.1 | NZ_CP051167.1 | 388517 | 389701 | 1184 | + | tuf | WP_059107364.1 |
| GCF_000195975.1 | NC_005070.1 | 2250136 | 2251317 | 1181 | - | tuf | WP_002432994.1 |
| GCF_002754935.1 | NZ_CP018092.1 | 2387456 | 2388637 | 1181 | - | tuf | WP_002432994.1 |
| GCF_023983615.1 | NZ_CP098611.1 | 1964396 | 1965580 | 1184 | + | tuf | WP_012664235.1 |
| GCF_040564635.1 | NZ_CP159837.1 | 2389884 | 2391068 | 1184 | - | tuf | WP_046837709.1 |
| GCF_904830935.1 | NZ_LR882952.1 | 2121141 | 2122325 | 1184 | - | tuf | WP_105985711.1 |
| GCF_904830955.1 | NZ_LR882970.1 | 2305181 | 2306365 | 1184 | - | tuf | WP_016424306.1 |
| GCF_000015665.1 | NC_008817.1 | 2406993 | 2408177 | 1184 | - | tuf | WP_047131407.1 |
| GCF_029910235.1 | NZ_CP112874.1 | 2658964 | 2660148 | 1184 | - | tuf | WP_123145595.1 |
| GCF_030323025.1 | NZ_CP097329.1 | 2371005 | 2372189 | 1184 | - | tuf | WP_019166641.1 |
| GCF_019056575.1 | NZ_CP021056.1 | 2257321 | 2258502 | 1181 | - | tuf | WP_001832289.1 |
| GCF_000021805.1 | NC_011726.1 | 2003221 | 2004405 | 1184 | - | tuf | WP_002508563.1 |
| GCF_019598945.1 | NZ_CP080598.1 | 381728 | 382912 | 1184 | - | tuf | WP_103207207.1 |
| GCF_000317575.1 | NC_019748.1 | 2492789 | 2493973 | 1184 | - | tuf | WP_042737971.1 |
| GCF_022984195.1 | NZ_CP085785.1 | 2270373 | 2271554 | 1181 | - | tuf | WP_011276713.1 |
| GCF_026802175.1 | NZ_CP113797.1 | 671657 | 672838 | 1181 | - | tuf | WP_002449023.1 |
| GCF_032705245.1 | NZ_CP053540.1 | 880753 | 881937 | 1184 | + | tuf | WP_251518605.1 |
| GCF_013177315.1 | NZ_CP053661.1 | 2235122 | 2236306 | 1184 | - | tuf | WP_039647031.1 |
| GCF_003555505.1 | NZ_CP032152.1 | 2583704 | 2584888 | 1184 | - | tuf | WP_061855460.1 |
| GCF_000011345.1 | NC_004113.1 | 2250200 | 2251384 | 1184 | - | tuf | WP_048792370.1 |
| GCF_030409055.1 | NZ_CP116619.1 | 1200592 | 1201773 | 1181 | - | tuf | WP_002459931.1 |
| GCF_000014265.1 | NC_008312.1 | 1561068 | 1562252 | 1184 | + | tuf | WP_085237570.1 |
| GCF_009856605.1 | NZ_CP047242.1 | 240954 | 242138 | 1184 | - | tuf | WP_009383851.1 |

| | | | | | | | |
|---|---|---|---|---|---|---|---|
| GCF_026240635.1 | NZ_CP110848.1 | 232309 | 233493 | 1184 | + | tuf | WP_095115234.1 |
| GCF_029919255.1 | NZ_CP123991.1 | 2598902 | 2600086 | 1184 | - | tuf | WP_096810913.1 |
| GCF_000829235.1 | NZ_AP012549.1 | 2239603 | 2240784 | 1181 | - | tuf | WP_017637541.1 |
| GCF_003574135.1 | NZ_AP018341.1 | 2040731 | 2041915 | 1184 | + | tuf | WP_002471493.1 |
| GCF_000183405.1 | NC_014758.1 | 2403693 | 2404877 | 1184 | - | tuf | WP_095106857.1 |
| GCF_000183405.1 | NC_014758.1 | 1683973 | 1685157 | 1184 | + | tuf | WP_014614744.1 |
| GCF_000010985.1 | NC_013939.1 | 2509416 | 2510600 | 1184 | - | tuf | WP_107530605.1 |
| GCF_000010985.1 | NC_013939.1 | 2086339 | 2087523 | 1184 | - | tuf | WP_229292422.1 |
| GCF_000025725.1 | NC_013943.1 | 1582321 | 1583505 | 1184 | + | tuf | WP_103358484.1 |
| GCF_000025725.1 | NC_013943.1 | 1129659 | 1130840 | 1181 | + | tuf | WP_115313946.1 |
| GCF_000218625.1 | NC_015672.1 | 2265099 | 2266283 | 1184 | - | tuf | WP_002484153.1 |
| GCF_000218625.1 | NC_015672.1 | 2294471 | 2295655 | 1184 | - | tuf | WP_016424306.1 |
| GCF_004087915.1 | NZ_CP035108.1 | 553420 | 554601 | 1181 | + | tuf | WP_047428308.1 |
| GCF_004087915.1 | NZ_CP035108.1 | 557832 | 559013 | 1181 | + | tuf | WP_002465098.1 |
| GCF_000487995.2 | NZ_CP097562.1 | 2413809 | 2414993 | 1184 | - | tuf | WP_002481337.1 |
| GCF_000092125.1 | NC_014212.1 | 2324945 | 2326129 | 1184 | - | tuf | WP_046837709.1 |
| GCF_000092125.1 | NC_014212.1 | 273371 | 274555 | 1184 | + | tuf | WP_046207677.1 |
| GCF_025997855.1 | NZ_AP026560.1 | 2256949 | 2258130 | 1181 | - | tuf | WP_002451225.1 |
| GCF_025997855.1 | NZ_AP026560.1 | 2377229 | 2378413 | 1184 | - | tuf | WP_042363297.1 |
| GCF_045784255.1 | NZ_CP129415.1 | 175950 | 177134 | 1184 | + | tuf | WP_124726396.1 |
| GCF_045784255.1 | NZ_CP129415.1 | 785075 | 786268 | 1193 | + | tuf | WP_001040730.1 |
| GCF_028622095.1 | NZ_CP115165.1 | 266480 | 267673 | 1193 | + | tuf | WP_003023932.1 |
| GCF_028622095.1 | NZ_CP115165.1 | 1430232 | 1431425 | 1193 | - | tuf | WP_006595927.1 |
| GCF_000020685.1 | NC_012526.1 | 1988750 | 1989943 | 1193 | - | tuf | WP_129545031.1 |
| GCF_000020685.1 | NC_012526.1 | 1695631 | 1696824 | 1193 | - | tuf | WP_116878219.1 |
| GCF_003444775.1 | NZ_CP021081.1 | 800130 | 801323 | 1193 | - | tuf | WP_003023932.1 |
| GCF_003444775.1 | NZ_CP021081.1 | 683671 | 684864 | 1193 | + | tuf | WP_005590656.1 |
| GCF_000196275.1 | NC_008025.1 | 1071279 | 1072472 | 1193 | + | tuf | WP_031574016.1 |
| GCF_000196275.1 | NC_008025.1 | 1873562 | 1874755 | 1193 | - | tuf | WP_003054233.1 |
| GCF_000252445.1 | NC_017790.1 | 1462522 | 1463715 | 1193 | - | tuf | WP_012515923.1 |
| GCF_000252445.1 | NC_017790.1 | 1297319 | 1298512 | 1193 | - | tuf | WP_018030176.1 |

| | | | | | | | |
|---|---|---|---|---|---|---|---|
| GCF_003173015.1 | NZ_CP029494.1 | 120876 | 122069 | 1193 | - | tuf | WP_009854603.1 |
| GCF_003173015.1 | NZ_CP029494.1 | 348816 | 350009 | 1193 | + | tuf | WP_060554003.1 |
| GCF_000186385.1 | NC_014958.1 | 972618 | 973811 | 1193 | - | tuf | WP_120770296.1 |
| GCF_000186385.1 | NC_014958.1 | 1138640 | 1139833 | 1193 | + | tuf | WP_159545289.1 |
| GCF_004758605.1 | NZ_CP038512.1 | 25893 | 27086 | 1193 | + | tuf | WP_062707346.1 |
| GCF_004758605.1 | NZ_CP038512.1 | 1721500 | 1722693 | 1193 | - | tuf | WP_121836359.1 |
| GCF_000317835.1 | NC_019793.1 | 1204719 | 1205912 | 1193 | + | tuf | WP_068991479.1 |
| GCF_000317835.1 | NC_019793.1 | 853248 | 854441 | 1193 | - | tuf | WP_023023854.1 |
| GCF_000190555.1 | NC_015161.1 | 1753966 | 1755159 | 1193 | - | tuf | WP_024531546.1 |
| GCF_000190555.1 | NC_015161.1 | 1716783 | 1717976 | 1193 | + | tuf | WP_043878441.1 |
| GCF_003860465.1 | NZ_CP034183.1 | 1295574 | 1296767 | 1193 | - | tuf | WP_003101188.1 |
| GCF_003860465.1 | NZ_CP034183.1 | 704860 | 706053 | 1193 | + | tuf | WP_003073108.1 |
| GCF_001644565.1 | NZ_CP011387.1 | 1267209 | 1268402 | 1193 | + | tuf | WP_023023854.1 |
| GCF_001644565.1 | NZ_CP011387.1 | 1481897 | 1483090 | 1193 | - | tuf | WP_187540490.1 |
| GCF_045784305.1 | NZ_CP129906.1 | 848960 | 850153 | 1193 | + | tuf | WP_003008055.1 |
| GCF_045784305.1 | NZ_CP129906.1 | 1238768 | 1239961 | 1193 | - | tuf | WP_020917184.1 |
| GCF_000800395.1 | NZ_CP010028.1 | 1563791 | 1564984 | 1193 | - | tuf | WP_014294966.1 |
| GCF_000800395.1 | NZ_CP010028.1 | 1602502 | 1603695 | 1193 | + | tuf | WP_067089207.1 |
| GCF_025244745.1 | NZ_CP104213.1 | 425756 | 426949 | 1193 | + | tuf | WP_018373820.1 |
| GCF_025244745.1 | NZ_CP104213.1 | 1490629 | 1491822 | 1193 | - | tuf | WP_001040726.1 |
| GCF_040256395.1 | NZ_CP158299.1 | 1145693 | 1146886 | 1193 | - | tuf | WP_002263314.1 |
| GCF_040256395.1 | NZ_CP158299.1 | 788341 | 789534 | 1193 | + | tuf | WP_001040720.1 |
| GCF_020229735.1 | NZ_CP083455.1 | 2151200 | 2152393 | 1193 | + | tuf | WP_212570723.1 |
| GCF_020229735.1 | NZ_CP083455.1 | 565399 | 566592 | 1193 | + | tuf | WP_067060855.1 |
| GCF_011067105.1 | NZ_CP049357.1 | 786721 | 787914 | 1193 | + | tuf | WP_001040724.1 |
| GCF_011067105.1 | NZ_CP049357.1 | 889712 | 890905 | 1193 | - | tuf | WP_003002543.1 |
| GCF_046774955.1 | NZ_CP177252.1 | 614354 | 615547 | 1193 | + | tuf | WP_024391225.1 |
| GCF_046774955.1 | NZ_CP177252.1 | 138567 | 139760 | 1193 | + | tuf | WP_003102427.1 |
| GCF_000195335.1 | NC_015387.1 | 1460424 | 1461617 | 1193 | - | tuf | WP_003065688.1 |
| GCF_000195335.1 | NC_015387.1 | 1214746 | 1215939 | 1193 | - | tuf | WP_003023932.1 |
| GCF_000024425.1 | NC_013946.1 | 1370270 | 1371463 | 1193 | - | tuf | WP_104968187.1 |

| | | | | | | | |
|---|---|---|---|---|---|---|---|
| GCF_000024425.1 | NC_013946.1 | 1238181 | 1239374 | 1193 | + | tuf | WP_001040724.1 |
| GCF_003568585.1 | NZ_CP021130.1 | 1328722 | 1329915 | 1193 | - | tuf | WP_003083585.1 |
| GCF_003568585.1 | NZ_CP021130.1 | 1395788 | 1396981 | 1193 | - | tuf | WP_001040724.1 |
| GCF_018771645.1 | NZ_CP076325.1 | 1935866 | 1937059 | 1193 | + | tuf | WP_003083585.1 |
| GCF_018771645.1 | NZ_CP076325.1 | 1257397 | 1258590 | 1193 | - | tuf | WP_003054233.1 |
| GCF_001399775.1 | NZ_CP010822.1 | 1367223 | 1368416 | 1193 | + | tuf | WP_003087525.1 |
| GCF_001399775.1 | NZ_CP010822.1 | 1740170 | 1741363 | 1193 | - | tuf | WP_067089207.1 |
| GCF_022846375.1 | NZ_AP025593.1 | 1522571 | 1523764 | 1193 | - | tuf | WP_024531546.1 |
| GCF_022846375.1 | NZ_AP025593.1 | 285625 | 286818 | 1193 | - | tuf | WP_021142772.1 |
| GCF_004684245.1 | NZ_CP038452.1 | 462960 | 464153 | 1193 | + | tuf | WP_002886182.1 |
| GCF_004684245.1 | NZ_CP038452.1 | 1490521 | 1491714 | 1193 | - | tuf | WP_002895003.1 |
| GCF_000309885.1 | NC_019386.1 | 1593346 | 1594539 | 1193 | - | tuf | WP_019769452.1 |
| GCF_000309885.1 | NC_019386.1 | 593671 | 594864 | 1193 | + | tuf | WP_012775011.1 |
| GCF_001535545.1 | NZ_CP014141.1 | 587332 | 588525 | 1193 | + | tuf | WP_024531546.1 |
| GCF_001535545.1 | NZ_CP014141.1 | 466926 | 468119 | 1193 | + | tuf | WP_002949971.1 |
| GCF_000091545.1 | NC_006461.1 | 865773 | 866966 | 1193 | + | tuf | WP_001040721.1 |
| GCF_000091545.1 | NC_006461.1 | 1427915 | 1429108 | 1193 | - | tuf | WP_002263314.1 |
| GCF_000092425.1 | NC_014221.1 | 622622 | 623815 | 1193 | + | tuf | WP_037592336.1 |
| GCF_000092425.1 | NC_014221.1 | 1360920 | 1362113 | 1193 | - | tuf | WP_006739206.1 |
| GCF_000020965.1 | NC_011297.1 | 496430 | 497623 | 1193 | + | tuf | WP_003095670.1 |
| GCF_000020965.1 | NC_011297.1 | 1362394 | 1363587 | 1193 | - | tuf | WP_067089207.1 |
| GCF_000021645.1 | NC_011661.1 | 2389679 | 2390863 | 1184 | - | tuf | WP_118991381.1 |
| GCF_000021645.1 | NC_011661.1 | 4733742 | 4734929 | 1187 | + | tuf | WP_066414023.1 |
| GCF_000020145.1 | NC_010644.1 | 131748 | 132935 | 1187 | + | tuf | WP_010191482.1 |
| GCF_000020145.1 | NC_010644.1 | 2263482 | 2264669 | 1187 | - | tuf | WP_339060639.1 |
| GCF_002355835.1 | NZ_AP017459.1 | 3447430 | 3448617 | 1187 | + | tuf | WP_038565169.1 |
| GCF_002355835.1 | NZ_AP017459.1 | 2162316 | 2163500 | 1184 | - | tuf | WP_061840770.1 |
| GCF_001027545.1 | NZ_CP009498.1 | 1668947 | 1670134 | 1187 | - | tuf | WP_124006386.1 |
| GCF_001027545.1 | NZ_CP009498.1 | 1149009 | 1150196 | 1187 | + | tuf | WP_123935341.1 |
| GCF_000146505.1 | NC_017448.1 | 3262617 | 3263801 | 1184 | + | tuf | WP_181557436.1 |
| GCF_000146505.1 | NC_017448.1 | 55003 | 56187 | 1184 | - | tuf | WP_037997041.1 |

| | | | | | | | |
|---|---|---|---|---|---|---|---|
| GCF_022430525.1 | NZ_CP092519.1 | 306783 | 307973 | 1190 | + | tuf | WP_015253285.1 |
| GCF_022430525.1 | NZ_CP092519.1 | 2676927 | 2678111 | 1184 | - | tuf | WP_321433426.1 |
| GCF_001296145.1 | NZ_CP012715.1 | 953830 | 955014 | 1184 | - | tuf | WP_319996104.1 |
| GCF_016724785.1 | NZ_CP068114.1 | 4371425 | 4372609 | 1184 | - | tuf | WP_094238155.1 |
| GCF_003019695.1 | NZ_CP028106.1 | 1785411 | 1786595 | 1184 | + | tuf | WP_087456447.1 |
| GCF_003019695.1 | NZ_CP028106.1 | 3787441 | 3788625 | 1184 | - | tuf | WP_107932289.1 |
| GCF_014337255.1 | NZ_CP060637.1 | 629540 | 630724 | 1184 | + | tuf | WP_208651257.1 |
| GCF_014337255.1 | NZ_CP060637.1 | 739489 | 740673 | 1184 | + | tuf | WP_096550061.1 |
| GCF_001455085.1 | NZ_CP013336.1 | 197759 | 198943 | 1184 | + | tuf | WP_126796029.1 |
| GCF_003019315.1 | NZ_CP028102.1 | 2824398 | 2825582 | 1184 | - | tuf | WP_126792996.1 |
| GCF_003019315.1 | NZ_CP028102.1 | 401593 | 402777 | 1184 | - | tuf | WP_166006969.1 |
| GCF_003732525.1 | NZ_CP019306.1 | 1130262 | 1131446 | 1184 | + | tuf | WP_166034181.1 |
| GCF_003732525.1 | NZ_CP019306.1 | 43517 | 44701 | 1184 | + | tuf | WP_275469174.1 |
| GCF_003019295.1 | NZ_CP028101.1 | 40335 | 41519 | 1184 | + | tuf | WP_326717159.1 |
| GCF_001457555.1 | NZ_LN831027.1 | 543232 | 544416 | 1184 | + | tuf | WP_257701974.1 |
| GCF_003019675.1 | NZ_CP028105.1 | 1409827 | 1411011 | 1184 | + | tuf | WP_023606957.1 |
| GCF_003019675.1 | NZ_CP028105.1 | 2126324 | 2127508 | 1184 | - | tuf | WP_071457768.1 |
| GCF_003019655.1 | NZ_CP028103.1 | 1943838 | 1945022 | 1184 | - | tuf | WP_135253812.1 |
| GCF_003019655.1 | NZ_CP028103.1 | 66544 | 67728 | 1184 | + | tuf | WP_136952357.1 |
| GCF_019552005.1 | NZ_CP071093.1 | 1375788 | 1376975 | 1187 | - | tuf | WP_073012567.1 |
| GCF_965119615.1 | NZ_OZ221964.1 | 145023 | 146210 | 1187 | + | tuf | WP_019375536.1 |
| GCF_030295325.1 | NZ_AP027059.1 | 233161 | 234348 | 1187 | + | tuf | WP_209463952.1 |
| GCF_030295325.1 | NZ_AP027059.1 | 173868 | 175055 | 1187 | - | tuf | WP_089530257.1 |
| GCF_000165505.1 | NC_014632.1 | 189962 | 191149 | 1187 | + | tuf | WP_050349640.1 |
| GCF_000165505.1 | NC_014632.1 | 1065008 | 1066195 | 1187 | - | tuf | WP_089060920.1 |
| GCF_000023905.1 | NC_013192.1 | 464307 | 465491 | 1184 | + | tuf | WP_213409394.1 |
| GCF_000023905.1 | NC_013192.1 | 980605 | 981789 | 1184 | + | tuf | WP_010369738.1 |
| GCF_007990525.1 | NZ_AP019823.1 | 1234905 | 1236095 | 1190 | - | tuf | WP_206212036.1 |
| GCF_007990525.1 | NZ_AP019823.1 | 890038 | 891222 | 1184 | + | tuf | WP_003609958.1 |
| GCF_007990635.1 | NZ_AP019846.1 | 215454 | 216638 | 1184 | - | tuf | WP_187529353.1 |
| GCF_007990635.1 | NZ_AP019846.1 | 1307387 | 1308571 | 1184 | - | tuf | WP_075269867.1 |

| | | | | | | | |
|---|---|---|---|---|---|---|---|
| GCF_041199995.1 | NZ_CP165646.1 | 1193959 | 1195143 | 1184 | + | tuf | WP_006845167.1 |
| GCF_041199995.1 | NZ_CP165646.1 | 1158637 | 1159821 | 1184 | - | tuf | WP_002828990.1 |
| GCF_007990365.1 | NZ_AP019831.1 | 833224 | 834408 | 1184 | - | tuf | WP_070230055.1 |
| GCF_007990365.1 | NZ_AP019831.1 | 478608 | 479792 | 1184 | + | tuf | WP_009765485.1 |
| GCF_007990545.2 | NZ_AP019829.2 | 687263 | 688447 | 1184 | + | tuf | WP_057747191.1 |
| GCF_007990545.2 | NZ_AP019829.2 | 1779189 | 1780376 | 1187 | - | tuf | WP_317635057.1 |
| GCF_000973085.1 | NZ_CP011280.1 | 1857560 | 1858747 | 1187 | - | tuf | WP_317696056.1 |
| GCF_000973085.1 | NZ_CP011280.1 | 3657106 | 3658293 | 1187 | - | tuf | WP_102245281.1 |
| GCF_000024565.1 | NC_013515.1 | 3673505 | 3674692 | 1187 | - | tuf | WP_102245281.1 |
| GCF_000024565.1 | NC_013515.1 | 3290363 | 3291550 | 1187 | - | tuf | WP_014245821.1 |
| GCF_000010305.1 | NC_012489.1 | 3306942 | 3308129 | 1187 | - | tuf | WP_014245821.1 |
| GCF_013004105.1 | NZ_CP053085.1 | 116326 | 117513 | 1187 | - | tuf | WP_321395441.1 |
| GCF_000695095.2 | NZ_CP011454.1 | 132524 | 133711 | 1187 | - | tuf | WP_321395441.1 |
| GCF_000522985.1 | NZ_CP007128.1 | 866328 | 867512 | 1184 | - | tuf | WP_019131491.1 |
| GCF_030623445.1 | NZ_CP130613.1 | 924356 | 925540 | 1184 | - | tuf | WP_141428012.1 |
| GCF_038095605.1 | NZ_CP144380.1 | 1488540 | 1489724 | 1184 | - | tuf | WP_026089672.1 |
| GCF_038095605.1 | NZ_CP144380.1 | 1544136 | 1545320 | 1184 | + | tuf | WP_009133745.1 |
| GCF_000258405.1 | NC_017464.1 | 1726002 | 1727186 | 1184 | + | tuf | WP_087309943.1 |
| GCF_000279145.1 | NC_018178.1 | 658575 | 659759 | 1184 | - | tuf | WP_018696712.1 |
| GCF_001017655.1 | NZ_CP010904.1 | 1010320 | 1011504 | 1184 | - | tuf | WP_015548066.1 |
| GCF_009905215.1 | NZ_CP047593.1 | 3072824 | 3074008 | 1184 | + | tuf | WP_079556527.1 |
| GCF_028728065.1 | NZ_CP117812.1 | 3863732 | 3864916 | 1184 | + | tuf | WP_318347545.1 |
| GCF_014023125.1 | NZ_CP048914.1 | 527187 | 528368 | 1181 | - | tuf | WP_005675411.1 |
| GCF_013745115.1 | NZ_CP051151.1 | 3864673 | 3865854 | 1181 | - | tuf | WP_065539685.1 |
| GCF_900660755.1 | NZ_LR215050.1 | 446717 | 447898 | 1181 | - | tuf | WP_007212053.1 |
| GCF_000018785.1 | NC_010163.1 | 669947 | 671128 | 1181 | - | tuf | WP_055279243.1 |
| GCF_000968055.1 | NC_022538.1 | 283316 | 284497 | 1181 | - | tuf | WP_004291289.1 |
| GCF_000582535.1 | NZ_CP006932.1 | 507149 | 508330 | 1181 | - | tuf | WP_008762026.1 |
| GCF_030550815.1 | NZ_CP114583.1 | 561626 | 562807 | 1181 | - | tuf | WP_167959840.1 |
| GCF_038505995.1 | NZ_CP128414.1 | 4725505 | 4726686 | 1181 | - | tuf | WP_005782155.1 |
| GCF_038024965.1 | NZ_CP146843.1 | 1331877 | 1333058 | 1181 | + | tuf | WP_319504455.1 |

| | | | | | | | |
|---|---|---|---|---|---|---|---|
| GCF_018024475.1 | NZ_CP054393.1 | 1339556 | 1340737 | 1181 | + | tuf | WP_013546664.1 |
| GCF_000026205.1 | NC_011047.1 | 1552628 | 1553809 | 1181 | - | tuf | WP_106069097.1 |
| GCF_026821955.1 | NZ_CP114006.1 | 671420 | 672601 | 1181 | - | tuf | WP_005782155.1 |
| GCF_040126175.1 | NZ_CP155828.1 | 1511461 | 1512642 | 1181 | + | tuf | WP_117601000.1 |
| GCF_003640545.1 | NZ_CP025121.1 | 684265 | 685446 | 1181 | - | tuf | WP_004307553.1 |
| GCF_002804205.1 | NZ_CP024962.1 | 2330619 | 2331800 | 1181 | + | tuf | WP_073314903.1 |
| GCF_900660745.1 | NZ_LR215048.1 | 4062110 | 4063291 | 1181 | - | tuf | WP_007487391.1 |
| GCF_016865445.1 | NZ_AP024412.1 | 3427453 | 3428634 | 1181 | - | tuf | WP_004296364.1 |
| GCF_002290085.1 | NZ_CP023173.1 | 450925 | 452106 | 1181 | - | tuf | WP_021645843.1 |
| GCF_002804245.1 | NZ_CP024968.1 | 536574 | 537755 | 1181 | - | tuf | WP_005928979.1 |
| GCF_002749675.1 | NZ_CP024411.1 | 2664663 | 2665847 | 1184 | + | tuf | WP_353330558.1 |
| GCF_000008305.1 | NC_006055.1 | 696670 | 697851 | 1181 | - | tuf | WP_008762026.1 |
| GCF_002441935.1 | NZ_CP023668.1 | 1726493 | 1727674 | 1181 | + | tuf | WP_005825727.1 |
| GCF_002804105.1 | NZ_CP024964.1 | 3843003 | 3844184 | 1181 | - | tuf | WP_004307553.1 |
| GCF_002843565.1 | NZ_CP025257.1 | 4443064 | 4444245 | 1181 | + | tuf | WP_055279243.1 |
| GCF_002804025.1 | NZ_CP024969.1 | 1404366 | 1405547 | 1181 | - | tuf | WP_106040872.1 |
| GCF_000723365.1 | NZ_HG937516.1 | 2588576 | 2589757 | 1181 | + | tuf | WP_025279249.1 |
| GCF_007859615.1 | NZ_CP042295.1 | 2211102 | 2212286 | 1184 | - | tuf | WP_106480531.1 |
| GCF_048074525.1 | NZ_CP182208.1 | 4843441 | 4844625 | 1184 | + | tuf | WP_151411504.1 |
| GCF_024338845.1 | NZ_CP101414.1 | 4074064 | 4075248 | 1184 | + | tuf | WP_087420800.1 |
| GCF_000012765.1 | NC_007633.1 | 702258 | 703442 | 1184 | - | tuf | WP_027200804.1 |
| GCF_045276875.1 | NZ_CP174163.1 | 18524 | 19708 | 1184 | - | tuf | WP_012573037.1 |
| GCF_024918975.1 | NZ_CP103424.1 | 1911327 | 1912511 | 1184 | - | tuf | WP_009318002.1 |
| GCF_000025845.1 | NC_014014.1 | 3242902 | 3244086 | 1184 | + | tuf | WP_200754963.1 |
| GCF_029677705.1 | NZ_CP104009.1 | 4306170 | 4307351 | 1181 | - | tuf | WP_163348369.1 |
| GCF_049942695.1 | NZ_CP187940.1 | 1520848 | 1522029 | 1181 | + | tuf | WP_123613574.1 |
| GCF_024722375.1 | NZ_CP102734.1 | 3953284 | 3954468 | 1184 | + | tuf | WP_026626453.1 |
| GCF_000183365.1 | NC_014751.1 | 2899131 | 2900324 | 1193 | + | tuf | WP_004350087.1 |
| GCF_013008635.1 | NZ_CP053096.1 | 184049 | 185239 | 1190 | - | tuf | WP_025065911.1 |
| GCF_013008815.1 | NZ_CP053097.1 | 3346113 | 3347297 | 1184 | + | tuf | WP_096429747.1 |
| GCF_018389705.1 | NZ_CP065581.1 | 959674 | 960858 | 1184 | + | tuf | WP_418894499.1 |

| | | | | | | | |
|---|---|---|---|---|---|---|---|
| GCF_006228185.1 | NZ_CP040825.1 | 1284829 | 1286013 | 1184 | - | tuf | WP_158863882.1 |
| GCF_000508245.1 | NC_023062.1 | 2884790 | 2885974 | 1184 | - | tuf | WP_321279232.1 |
| GCF_000477415.1 | NC_022575.1 | 334533 | 335714 | 1181 | + | tuf | WP_123395280.1 |
| GCF_012934855.1 | NZ_CP051481.1 | 2506465 | 2507646 | 1181 | + | tuf | WP_068961376.1 |
| GCF_012934885.1 | NZ_CP051480.1 | 2358952 | 2360136 | 1184 | - | tuf | WP_013612174.1 |
| GCF_900476175.1 | NZ_LS991954.1 | 2057151 | 2058335 | 1184 | + | tuf | WP_013445226.1 |
| GCF_030718845.1 | NZ_CP132191.1 | 1899940 | 1901124 | 1184 | - | tuf | WP_251967775.1 |
| GCF_003855455.1 | NZ_CP034044.1 | 287396 | 288580 | 1184 | - | tuf | WP_122362432.1 |
| GCF_000179035.2 | NC_015155.1 | 899884 | 901068 | 1184 | - | tuf | WP_048927526.1 |
| GCF_016925555.1 | NZ_CP070479.1 | 6893987 | 6895171 | 1184 | + | tuf | WP_128133172.1 |
| GCF_014068355.1 | NZ_CP059674.1 | 4709193 | 4710377 | 1184 | + | tuf | WP_008152453.1 |
| GCF_000277795.1 | NC_018149.1 | 2438171 | 2439352 | 1181 | - | tuf | WP_107035015.1 |
| GCF_000277795.1 | NC_018149.1 | 2216082 | 2217260 | 1178 | - | tuf | WP_008628203.1 |
| GCF_000875755.1 | NZ_CP007520.1 | 2933457 | 2934641 | 1184 | + | tuf | WP_071137551.1 |
| GCF_000967915.1 | NC_022549.1 | 3298887 | 3300068 | 1181 | - | tuf | WP_008140167.1 |
| GCF_003363775.1 | NZ_CP031376.1 | 4930471 | 4931652 | 1181 | - | tuf | WP_007838973.1 |
| GCF_000500935.1 | NC_022998.1 | 2919862 | 2921043 | 1181 | + | tuf | WP_013618472.1 |
| GCF_001281045.1 | NZ_CP012622.1 | 1118824 | 1120005 | 1181 | - | tuf | WP_005844822.1 |
| GCF_008086545.1 | NZ_CP043026.1 | 930353 | 931537 | 1184 | - | tuf | WP_004331700.1 |
| GCF_000400935.1 | NC_021280.1 | 2060575 | 2061759 | 1184 | + | tuf | WP_025836724.1 |
| GCF_001886855.1 | NZ_CP013197.1 | 254130 | 255311 | 1181 | - | tuf | WP_023936385.1 |
| GCF_002795265.1 | NZ_CP024870.1 | 802518 | 803705 | 1187 | + | tuf | WP_004335339.1 |
| GCF_002237575.1 | NZ_CP022535.1 | 1768510 | 1769694 | 1184 | - | tuf | WP_012458382.1 |
| GCF_000565175.1 | NZ_CP006681.1 | 2726651 | 2727835 | 1184 | - | tuf | WP_329904094.1 |
| GCF_000439455.1 | NC_021833.1 | 2597916 | 2599100 | 1184 | + | tuf | WP_021902511.1 |
| GCF_964031275.1 | NZ_OZ035019.1 | 435876 | 437057 | 1181 | + | tuf | WP_091814739.1 |
| GCF_964019485.1 | NZ_OZ026448.1 | 274641 | 275843 | 1202 | + | tuf | WP_005844206.1 |
| GCF_964031135.1 | NZ_OZ035008.1 | 1336237 | 1337430 | 1193 | - | tuf | WP_013671024.1 |
| GCF_964020965.1 | NZ_OZ026611.1 | 1171242 | 1172435 | 1193 | - | tuf | WP_025078268.1 |
| GCF_964030965.1 | NZ_OZ034999.1 | 1166760 | 1167947 | 1187 | - | tuf | WP_018462903.1 |
| GCF_964019575.1 | NZ_OZ026455.1 | 1657838 | 1659031 | 1193 | - | tuf | WP_008821962.1 |

| | | | | | | | |
|---|---|---|---|---|---|---|---|
| GCF_042465095.1 | NZ_CP102059.1 | 390813 | 392006 | 1193 | + | tuf | WP_009012371.1 |
| GCF_964020205.1 | NZ_OZ026543.1 | 412212 | 413405 | 1193 | + | tuf | WP_013264904.1 |
| GCF_964031085.1 | NZ_OZ035001.1 | 937578 | 938771 | 1193 | - | tuf | WP_211813686.1 |
| GCF_964031045.1 | NZ_OZ034994.1 | 2218066 | 2219253 | 1187 | - | tuf | WP_004362495.1 |
| GCF_964020005.1 | NZ_OZ026474.1 | 352767 | 353960 | 1193 | + | tuf | WP_025838806.1 |
| GCF_964019925.1 | NZ_OZ026470.1 | 1630604 | 1631797 | 1193 | - | tuf | WP_004383925.1 |
| GCF_964019665.1 | NZ_OZ026459.1 | 477735 | 478919 | 1184 | + | tuf | WP_238867992.1 |
| GCF_964031115.1 | NZ_OZ035006.1 | 3679882 | 3681066 | 1184 | + | tuf | WP_076931508.1 |
| GCF_964020095.1 | NZ_OZ026536.1 | 2326640 | 2327818 | 1178 | + | tuf | WP_111898107.1 |
| GCF_964020035.1 | NZ_OZ026475.1 | 1352896 | 1354080 | 1184 | - | tuf | WP_057952290.1 |
| GCF_964019495.1 | NZ_OZ026447.1 | 2120466 | 2121680 | 1214 | - | tuf | WP_074803322.1 |
| GCF_964030685.1 | NZ_OZ034912.1 | 134946 | 136139 | 1193 | - | tuf | WP_006848849.1 |
| GCF_964030865.1 | NZ_OZ034984.1 | 2876118 | 2877311 | 1193 | + | tuf | WP_004373959.1 |
| GCF_964019625.1 | NZ_OZ026458.1 | 1091180 | 1092370 | 1190 | - | tuf | WP_154327241.1 |
| GCF_964031155.1 | NZ_OZ035010.1 | 2062015 | 2063199 | 1184 | + | tuf | WP_014225269.1 |
| GCF_964019525.1 | NZ_OZ026453.1 | 182793 | 183986 | 1193 | + | tuf | WP_037984105.1 |
| GCF_049669535.1 | NZ_CP187162.1 | 1065772 | 1066962 | 1190 | + | tuf | WP_173073353.1 |
| GCF_964019845.1 | NZ_OZ026465.1 | 2105609 | 2106793 | 1184 | - | tuf | WP_250722467.1 |
| GCF_964020065.1 | NZ_OZ026534.1 | 2569414 | 2570595 | 1181 | - | tuf | WP_013063031.1 |
| GCF_964020195.1 | NZ_OZ026539.1 | 1030886 | 1032070 | 1184 | + | tuf | WP_114983388.1 |
| GCF_964019715.1 | NZ_OZ026463.1 | 1256363 | 1257547 | 1184 | + | tuf | WP_114983388.1 |
| GCF_964031175.1 | NZ_OZ035012.1 | 2890463 | 2891650 | 1187 | - | tuf | WP_011165367.1 |
| GCF_964019975.1 | NZ_OZ026473.1 | 3431726 | 3432913 | 1187 | + | tuf | WP_243537559.1 |
| GCF_964030895.1 | NZ_OZ034986.1 | 1944779 | 1945966 | 1187 | - | tuf | WP_015470630.1 |
| GCF_018831625.1 | NZ_CP065193.1 | 289235 | 290422 | 1187 | + | tuf | WP_012655361.1 |
| GCF_964030835.1 | NZ_OZ034982.1 | 3471728 | 3472915 | 1187 | - | tuf | WP_012655361.1 |
| GCF_964030795.1 | NZ_OZ034962.1 | 908737 | 909924 | 1187 | + | tuf | WP_062680839.1 |
| GCF_964277355.1 | NZ_OZ195517.1 | 932980 | 934167 | 1187 | + | tuf | WP_062680839.1 |
| GCF_964030985.1 | NZ_OZ034993.1 | 6393202 | 6394389 | 1187 | - | tuf | WP_042795889.1 |
| GCF_964019505.1 | NZ_OZ026449.1 | 6417424 | 6418611 | 1187 | - | tuf | WP_042795889.1 |
| GCF_964019765.1 | NZ_OZ026461.1 | 5599937 | 5601124 | 1187 | - | tuf | WP_063953628.1 |

| | | | | | | | |
|---|---|---|---|---|---|---|---|
| GCF_964019705.1 | NZ_OZ026460.1 | 5624170 | 5625357 | 1187 | - | tuf | WP_062839263.1 |
| GCF_918697755.1 | NZ_OU953854.1 | 6289836 | 6291023 | 1187 | - | tuf | WP_020925806.1 |
| GCF_964030655.1 | NZ_OZ034915.1 | 6314210 | 6315397 | 1187 | - | tuf | WP_020925806.1 |
| GCF_964019755.1 | NZ_OZ026469.1 | 4881224 | 4882411 | 1187 | + | tuf | WP_306940817.1 |
| GCF_964263245.1 | NZ_OZ181656.1 | 4905428 | 4906615 | 1187 | + | tuf | WP_306940817.1 |
| GCF_964030825.1 | NZ_OZ034964.1 | 1384704 | 1385891 | 1187 | + | tuf | WP_006221899.1 |
| GCF_964019595.1 | NZ_OZ026456.1 | 1410461 | 1411648 | 1187 | + | tuf | WP_006221899.1 |
| GCF_964020235.1 | NZ_OZ026545.1 | 935477 | 936664 | 1187 | - | tuf | WP_287753809.1 |
| GCF_964031205.1 | NZ_OZ035018.1 | 959746 | 960933 | 1187 | - | tuf | WP_287753809.1 |
| GCF_964019735.1 | NZ_OZ026464.1 | 836872 | 838059 | 1187 | - | tuf | WP_020925806.1 |
| GCF_964020225.1 | NZ_OZ026544.1 | 861224 | 862411 | 1187 | - | tuf | WP_020925806.1 |
| GCF_964264785.1 | NZ_OZ183484.1 | 279828 | 281015 | 1187 | + | tuf | WP_086911058.1 |
| GCF_964019535.1 | NZ_OZ026450.1 | 321265 | 322452 | 1187 | + | tuf | WP_086911058.1 |
| GCF_964030925.1 | NZ_OZ034992.1 | 467090 | 468277 | 1187 | + | tuf | WP_011793521.1 |
| GCF_964264745.1 | NZ_OZ183478.1 | 5250246 | 5251433 | 1187 | - | tuf | WP_011793521.1 |
| GCF_964030805.1 | NZ_OZ034965.1 | 4048908 | 4050095 | 1187 | - | tuf | WP_207373633.1 |
| GCF_001029265.1 | NZ_CP011856.1 | 4085829 | 4087016 | 1187 | - | tuf | WP_207373633.1 |
| GCF_002813555.1 | NZ_CP025057.1 | 4611642 | 4612829 | 1187 | - | tuf | WP_025374750.1 |
| GCF_004379335.1 | NZ_CP038013.1 | 4683692 | 4684879 | 1187 | - | tuf | WP_025374750.1 |
| GCF_001715535.1 | NZ_CP017015.1 | 3714132 | 3715319 | 1187 | - | tuf | WP_059318443.1 |
| GCF_030316605.1 | NZ_CP127039.1 | 3737184 | 3738371 | 1187 | - | tuf | WP_059318443.1 |
| GCF_001274875.1 | NZ_CP010899.1 | 3767393 | 3768580 | 1187 | - | tuf | WP_094198179.1 |
| GCF_001267155.1 | NZ_CP012357.1 | 3790448 | 3791635 | 1187 | - | tuf | WP_094198179.1 |
| GCF_001029245.1 | NZ_CP011855.1 | 381731 | 382918 | 1187 | + | tuf | WP_013517320.1 |
| GCF_002865545.1 | NZ_CP025543.1 | 420914 | 422101 | 1187 | + | tuf | WP_013517320.1 |
| GCF_003339775.1 | NZ_CP031088.1 | 1444984 | 1446165 | 1181 | + | tuf | WP_115723606.1 |
| GCF_021496725.1 | NZ_CP051215.1 | 1462196 | 1463377 | 1181 | - | tuf | WP_115723606.1 |
| GCF_009866525.1 | NZ_CM020866.1 | 1189342 | 1190523 | 1181 | - | tuf | WP_115723606.1 |
| GCF_000565215.1 | NZ_CP006934.1 | 1210321 | 1211502 | 1181 | - | tuf | WP_115723606.1 |
| GCF_000400955.1 | NC_021284.1 | 3298532 | 3299719 | 1187 | - | tuf | WP_424192846.1 |
| GCF_009730595.1 | NZ_CP046276.1 | 3315027 | 3316214 | 1187 | - | tuf | WP_424192846.1 |

| | | | | | | | |
|---|---|---|---|---|---|---|---|
| GCF_000439435.1 | NC_021846.1 | 191538 | 192725 | 1187 | + | tuf | WP_109033866.1 |
| GCF_001262715.1 | NZ_CP012328.1 | 215456 | 216643 | 1187 | + | tuf | WP_109033866.1 |
| GCF_000397185.1 | NC_021236.1 | 436090 | 437277 | 1187 | + | tuf | WP_102950428.1 |
| GCF_002803985.1 | NZ_CP024963.1 | 2439155 | 2440342 | 1187 | - | tuf | WP_102950428.1 |
| GCF_002804005.1 | NZ_CP024965.1 | 3477705 | 3478892 | 1187 | - | tuf | WP_250198098.1 |
| GCF_003285045.1 | NZ_CP030140.1 | 3495345 | 3496532 | 1187 | - | tuf | WP_250198098.1 |
| GCF_042984175.1 | NZ_CP155569.1 | 1553477 | 1554664 | 1187 | + | tuf | WP_137009171.1 |
| GCF_037482225.1 | NZ_CP148066.1 | 1570069 | 1571256 | 1187 | + | tuf | WP_137009173.1 |
| GCF_003332325.1 | NZ_CP029295.1 | 3963842 | 3965029 | 1187 | - | tuf | WP_159879912.1 |
| GCF_900660615.1 | NZ_LR215023.1 | 3980439 | 3981626 | 1187 | - | tuf | WP_159879912.1 |
| GCF_046774425.1 | NZ_CP177248.1 | 2243032 | 2244219 | 1187 | + | tuf | WP_011237959.1 |
| GCF_000011225.1 | NC_004432.1 | 2259484 | 2260671 | 1187 | + | tuf | WP_011237959.1 |
| GCF_000524555.1 | NZ_CP007154.1 | 2430004 | 2431191 | 1187 | + | tuf | WP_169201689.1 |
| GCF_000026765.1 | NC_012806.1 | 2446466 | 2447653 | 1187 | + | tuf | WP_169201689.1 |
| GCF_000941075.1 | NZ_CP007229.1 | 4767863 | 4769050 | 1187 | - | tuf | WP_169127209.1 |
| GCF_000815065.1 | NZ_CP007585.1 | 4784322 | 4785509 | 1187 | - | tuf | WP_169127224.1 |
| GCF_000008205.1 | NC_007295.1 | 2089164 | 2090351 | 1187 | - | tuf | WP_169127224.1 |
| GCF_900476065.1 | NZ_LS991950.1 | 2105625 | 2106812 | 1187 | - | tuf | WP_169127224.1 |
| GCF_029854295.1 | NZ_CP122979.1 | 3829631 | 3830818 | 1187 | - | tuf | WP_011767142.1 |
| GCF_045276915.1 | NZ_CP174166.1 | 3846064 | 3847251 | 1187 | - | tuf | WP_011767142.1 |
| GCF_024918955.1 | NZ_CP103423.1 | 1163562 | 1164749 | 1187 | - | tuf | WP_216131374.1 |
| GCF_900660485.1 | NZ_LR214951.1 | 1180261 | 1181448 | 1187 | - | tuf | WP_216131374.1 |
| GCF_028885435.1 | NZ_CP118522.1 | 355840 | 357027 | 1187 | - | tuf | WP_014235481.1 |
| GCF_900476125.1 | NZ_LS991949.1 | 372610 | 373797 | 1187 | - | tuf | WP_014235481.1 |
| GCF_900660715.1 | NZ_LR215047.1 | 1233182 | 1234369 | 1187 | + | tuf | WP_203388413.1 |
| GCF_046774455.1 | NZ_CP177249.1 | 1249925 | 1251112 | 1187 | + | tuf | WP_203388413.1 |
| GCF_000828855.1 | NZ_AP014631.1 | 80292 | 81479 | 1187 | + | tuf | WP_038497853.1 |
| GCF_003269445.1 | NZ_CP030103.1 | 1948578 | 1949765 | 1187 | - | tuf | WP_038497853.1 |
| GCF_033546855.1 | NZ_CP137845.1 | 45495 | 46676 | 1181 | + | tuf | WP_066081383.1 |
| GCF_045160055.1 | NZ_CP088155.1 | 92582 | 93763 | 1181 | + | tuf | WP_066081383.1 |
| GCF_036352135.1 | NZ_CP143578.1 | 3669721 | 3670908 | 1187 | - | tuf | WP_012415717.1 |

| | | | | | | | |
|---|---|---|---|---|---|---|---|
| GCF_000767725.1 | NZ_CP009652.1 | 3693788 | 3694975 | 1187 | - | tuf | WP_012415717.1 |
| GCF_024297005.1 | NZ_CP101127.1 | 5803936 | 5805123 | 1187 | - | tuf | WP_066357411.1 |
| GCF_900660435.1 | NZ_LR214940.1 | 5832878 | 5834065 | 1187 | - | tuf | WP_066357411.1 |
| GCF_003383595.3 | NZ_CP033058.2 | 5138593 | 5139780 | 1187 | - | tuf | WP_003806883.1 |
| GCF_900660445.2 | NZ_LR214938.2 | 5164197 | 5165384 | 1187 | - | tuf | WP_003806883.1 |
| GCF_041026625.1 | NZ_CP163393.1 | 5756433 | 5757620 | 1187 | - | tuf | WP_066664207.1 |
| GCF_045276925.1 | NZ_CP174167.1 | 5788277 | 5789464 | 1187 | - | tuf | WP_066664267.1 |
| GCF_033546815.1 | NZ_CP137846.1 | 3500656 | 3501843 | 1187 | + | tuf | WP_029579994.1 |
| GCF_000008365.1 | NC_006908.1 | 3524707 | 3525894 | 1187 | + | tuf | WP_029579994.1 |
| GCF_024498275.1 | NZ_CP101809.1 | 3420123 | 3421310 | 1187 | - | tuf | WP_005017271.1 |
| GCF_900476085.1 | NZ_LS991952.1 | 3444195 | 3445382 | 1187 | - | tuf | WP_005017230.1 |
| GCF_040556925.1 | NZ_CP159789.1 | 4317952 | 4319139 | 1187 | + | tuf | WP_003806883.1 |
| GCF_900660465.1 | NZ_LR214945.1 | 4343497 | 4344684 | 1187 | + | tuf | WP_003806883.1 |
| GCF_045276205.1 | NZ_CP174162.1 | 6722 | 7909 | 1187 | + | tuf | WP_003806883.1 |
| GCF_000063605.1 | NC_009497.1 | 243624 | 244811 | 1187 | - | tuf | WP_003806883.1 |
| GCF_003285065.1 | NZ_CP030141.1 | 5212533 | 5213720 | 1187 | - | tuf | WP_012251771.1 |
| GCF_001547975.1 | NZ_AP014657.1 | 5236788 | 5237975 | 1187 | - | tuf | WP_012251771.1 |
| GCF_900660525.1 | NZ_LR214970.1 | 4430577 | 4431764 | 1187 | - | tuf | WP_043207328.1 |
| GCF_900660515.1 | NZ_LR214972.1 | 4454627 | 4455814 | 1187 | - | tuf | WP_043207328.1 |
| GCF_000183385.1 | NC_014760.1 | 314477 | 315664 | 1187 | + | tuf | WP_011350666.1 |
| GCF_000695835.1 | NZ_CP007521.1 | 333724 | 334911 | 1187 | + | tuf | WP_011350666.1 |
| GCF_001553195.1 | NZ_CP014281.1 | 1365855 | 1367042 | 1187 | - | tuf | WP_011350666.1 |
| GCF_024498215.1 | NZ_CP101806.1 | 1385033 | 1386220 | 1187 | - | tuf | WP_011350666.1 |
| GCF_900660645.1 | NZ_LR215036.1 | 319528 | 320715 | 1187 | + | tuf | WP_011350666.1 |
| GCF_900660685.1 | NZ_LR215041.1 | 338734 | 339921 | 1187 | + | tuf | WP_011350666.1 |
| GCF_900660705.1 | NZ_LR215043.1 | 307820 | 309007 | 1187 | + | tuf | WP_006482910.1 |
| GCF_900660675.1 | NZ_LR215039.1 | 327041 | 328228 | 1187 | + | tuf | WP_006482910.1 |
| GCF_000328725.1 | NC_019949.1 | 303814 | 305001 | 1187 | + | tuf | WP_006482910.1 |
| GCF_900476105.1 | NZ_LS991951.1 | 323069 | 324256 | 1187 | + | tuf | WP_057056448.1 |
| GCF_024498255.1 | NZ_CP101808.1 | 2666403 | 2667590 | 1187 | - | tuf | WP_011350666.1 |
| GCF_034647275.2 | NZ_CP141030.2 | 2685536 | 2686723 | 1187 | - | tuf | WP_011350666.1 |

| | | | | | | | |
|---|---|---|---|---|---|---|---|
| GCF_000186005.1 | NC_014921.1 | 976602 | 977789 | 1187 | + | tuf | WP_045552492.1 |
| GCF_900660495.1 | NZ_LR214950.1 | 996726 | 997913 | 1187 | + | tuf | WP_045552492.1 |
| GCF_900660635.1 | NZ_LR215031.1 | 319279 | 320466 | 1187 | + | tuf | WP_013696349.1 |
| GCF_900660635.1 | NZ_LR215031.1 | 349737 | 350924 | 1187 | + | tuf | WP_013696349.1 |
| GCF_900660605.1 | NZ_LR215024.1 | 1296080 | 1297267 | 1187 | + | tuf | WP_012734343.1 |
| GCF_045276885.1 | NZ_CP174164.1 | 1316054 | 1317241 | 1187 | + | tuf | WP_012734343.1 |
| GCF_045276895.1 | NZ_CP174165.1 | 317227 | 318414 | 1187 | + | tuf | WP_015601108.1 |
| GCF_900660665.1 | NZ_LR215037.1 | 336320 | 337507 | 1187 | + | tuf | WP_015601108.1 |
| GCF_900660695.1 | NZ_LR215042.1 | 301228 | 302415 | 1187 | + | tuf | WP_011350666.1 |
| GCF_004216495.1 | NZ_CP034841.1 | 320417 | 321604 | 1187 | + | tuf | WP_011350666.1 |
| GCF_900660575.1 | NZ_LR215008.1 | 2730355 | 2731542 | 1187 | - | tuf | WP_004198356.1 |
| GCF_000969765.1 | NZ_CP011096.1 | 2750765 | 2751952 | 1187 | - | tuf | WP_004198356.1 |
| GCF_033546915.1 | NZ_CP137850.1 | 330195 | 331382 | 1187 | + | tuf | WP_038802783.1 |
| GCF_000941095.1 | NZ_CP009770.1 | 349285 | 350472 | 1187 | + | tuf | WP_038802782.1 |
| GCF_000019345.1 | NC_010503.1 | 295526 | 296713 | 1187 | + | tuf | WP_011350666.1 |
| GCF_000022145.1 | NC_011891.1 | 314691 | 315878 | 1187 | + | tuf | WP_011350666.1 |
| GCF_000022145.1 | NC_011891.1 | 267304 | 268491 | 1187 | + | tuf | WP_006400660.1 |
| GCF_023169945.1 | NZ_AP025591.1 | 286467 | 287654 | 1187 | + | tuf | WP_006400660.1 |
| GCF_023169945.1 | NZ_AP025591.1 | 1979354 | 1980541 | 1187 | + | tuf | WP_038802783.1 |
| GCF_023169965.1 | NZ_AP025592.1 | 1998444 | 1999631 | 1187 | + | tuf | WP_038802782.1 |
| GCF_023169965.1 | NZ_AP025592.1 | 288092 | 289279 | 1187 | + | tuf | WP_246796495.1 |
| GCF_001027285.1 | NZ_CP011509.1 | 308053 | 309240 | 1187 | + | tuf | WP_246796495.1 |
| GCF_001027285.1 | NZ_CP011509.1 | 293925 | 295112 | 1187 | + | tuf | WP_012734343.1 |
| GCF_001027285.1 | NZ_CP011509.1 | 313887 | 315074 | 1187 | + | tuf | WP_012734343.1 |
| GCF_033547035.1 | NZ_CP137851.1 | 3692377 | 3693564 | 1187 | - | tuf | WP_004198356.1 |
| GCF_033547035.1 | NZ_CP137851.1 | 3711474 | 3712661 | 1187 | - | tuf | WP_004198356.1 |
| GCF_033547035.1 | NZ_CP137851.1 | 1597287 | 1598474 | 1187 | + | tuf | WP_017333531.1 |
| GCF_000255295.1 | NC_017030.1 | 1616516 | 1617703 | 1187 | + | tuf | WP_017333531.1 |
| GCF_000255295.1 | NC_017030.1 | 96441 | 97628 | 1187 | + | tuf | WP_059577396.1 |
| GCF_002305895.1 | NZ_CP022203.1 | 115547 | 116734 | 1187 | + | tuf | WP_059577396.1 |
| GCF_002305895.1 | NZ_CP022203.1 | 310342 | 311529 | 1187 | + | tuf | WP_006482910.1 |

| | | | | | | | |
|---|---|---|---|---|---|---|---|
| GCF_002305855.1 | NZ_CP022163.1 | 329551 | 330738 | 1187 | + | tuf | WP_006482910.1 |
| GCF_002305855.1 | NZ_CP022163.1 | 562521 | 563708 | 1187 | + | tuf | WP_069233656.1 |
| GCF_017301635.1 | NZ_CP071091.1 | 582961 | 584148 | 1187 | + | tuf | WP_069233656.1 |
| GCF_017301635.1 | NZ_CP071091.1 | 2288225 | 2289412 | 1187 | - | tuf | WP_011350666.1 |
| GCF_000331735.1 | NC_020126.1 | 2307471 | 2308658 | 1187 | - | tuf | WP_011350666.1 |
| GCF_000331735.1 | NC_020126.1 | 175457 | 176647 | 1190 | - | tuf | WP_347468078.1 |
| GCF_000012685.1 | NC_008095.1 | 194580 | 195767 | 1187 | - | tuf | WP_011882996.1 |
| GCF_000012685.1 | NC_008095.1 | 302873 | 304060 | 1187 | + | tuf | WP_034209732.1 |
| GCF_037157815.1 | NZ_CP089984.1 | 322063 | 323250 | 1187 | + | tuf | WP_034209732.1 |
| GCF_037157815.1 | NZ_CP089984.1 | 3502159 | 3503346 | 1187 | - | tuf | WP_004198356.1 |
| GCF_037157775.1 | NZ_CP089982.1 | 3521254 | 3522441 | 1187 | - | tuf | WP_004198356.1 |
| GCF_037157775.1 | NZ_CP089982.1 | 346080 | 347267 | 1187 | + | tuf | WP_129513512.1 |
| GCF_037157775.1 | NZ_CP089982.1 | 366577 | 367764 | 1187 | + | tuf | WP_129513512.1 |
| GCF_037157805.1 | NZ_CP089983.1 | 324660 | 325847 | 1187 | + | tuf | WP_006482910.1 |
| GCF_037157805.1 | NZ_CP089983.1 | 343816 | 345003 | 1187 | + | tuf | WP_017333531.1 |
| GCF_000280925.3 | NZ_CP012109.1 | 516119 | 517306 | 1187 | - | tuf | WP_011882996.1 |
| GCF_000280925.3 | NZ_CP012109.1 | 535335 | 536522 | 1187 | - | tuf | WP_011882996.1 |
| GCF_017301735.1 | NZ_CP071090.1 | 260948 | 262135 | 1187 | + | tuf | WP_061138394.1 |
| GCF_017301735.1 | NZ_CP071090.1 | 280099 | 281286 | 1187 | + | tuf | WP_061138394.1 |
| GCF_001263175.1 | NZ_CP012332.1 | 2680131 | 2681318 | 1187 | - | tuf | WP_016346480.1 |
| GCF_001263175.1 | NZ_CP012332.1 | 2699307 | 2700494 | 1187 | - | tuf | WP_016346480.1 |
| GCF_001189295.1 | NZ_CP012159.1 | 2684120 | 2685307 | 1187 | - | tuf | WP_008343820.1 |
| GCF_001189295.1 | NZ_CP012159.1 | 2703265 | 2704452 | 1187 | - | tuf | WP_008343820.1 |
| GCF_000024805.1 | NC_013440.1 | 2942792 | 2943979 | 1187 | - | tuf | WP_264892270.1 |
| GCF_000024805.1 | NC_013440.1 | 2976304 | 2977491 | 1187 | - | tuf | WP_264892270.1 |
| GCF_001263205.1 | NZ_CP012333.1 | 5741970 | 5743157 | 1187 | - | tuf | WP_047196718.1 |
| GCF_001263205.1 | NZ_CP012333.1 | 5760394 | 5761581 | 1187 | - | tuf | WP_047196718.1 |
| GCF_026965555.1 | NZ_CP114040.1 | 3360344 | 3361531 | 1187 | - | tuf | WP_104357075.1 |
| GCF_026965555.1 | NZ_CP114040.1 | 3378560 | 3379747 | 1187 | - | tuf | WP_104357075.1 |
| GCF_001931505.1 | NZ_CP016908.1 | 4003736 | 4004923 | 1187 | - | tuf | WP_015767927.1 |
| GCF_005144635.2 | NZ_CP079217.1 | 4020535 | 4021722 | 1187 | - | tuf | WP_015767927.1 |

| | | | | | | | |
|---|---|---|---|---|---|---|---|
| GCF_005144635.2 | NZ_CP079217.1 | 492590 | 493771 | 1181 | + | tuf | WP_092343884.1 |
| GCF_000737325.1 | NZ_CP011125.1 | 508718 | 509899 | 1181 | + | tuf | WP_092343884.1 |
| GCF_000737325.1 | NZ_CP011125.1 | 145743 | 146930 | 1187 | - | tuf | WP_015237709.1 |
| GCF_000067165.1 | NC_010162.1 | 162272 | 163459 | 1187 | - | tuf | WP_015237709.1 |
| GCF_000067165.1 | NC_010162.1 | 796866 | 798053 | 1187 | - | tuf | WP_015238841.1 |
| GCF_946900835.1 | NZ_OX336137.1 | 813342 | 814529 | 1187 | - | tuf | WP_015238841.1 |
| GCF_032050975.1 | NZ_CP116967.1 | 807686 | 808873 | 1187 | - | tuf | WP_015396695.1 |
| GCF_032050975.1 | NZ_CP116967.1 | 824174 | 825361 | 1187 | - | tuf | WP_015396707.1 |
| GCF_001458695.1 | NZ_LN885086.1 | 717598 | 718785 | 1187 | - | tuf | WP_108674188.1 |
| GCF_001458695.1 | NZ_LN885086.1 | 784954 | 786141 | 1187 | - | tuf | WP_015397386.1 |
| GCF_032051675.1 | NZ_CP116968.1 | 801510 | 802697 | 1187 | - | tuf | WP_015397386.1 |
| GCF_032051675.1 | NZ_CP116968.1 | 166699 | 167886 | 1187 | + | tuf | WP_046486930.1 |
| GCF_000284315.1 | NC_017094.1 | 182971 | 184158 | 1187 | + | tuf | WP_046486930.1 |
| GCF_000284315.1 | NC_017094.1 | 1116382 | 1117569 | 1187 | - | tuf | WP_139873840.1 |
| GCF_900169565.1 | NZ_LT828648.1 | 1132658 | 1133845 | 1187 | - | tuf | WP_139873840.1 |
| GCF_900169565.1 | NZ_LT828648.1 | 155114 | 156301 | 1187 | + | tuf | WP_139866848.1 |
| GCF_001273775.1 | NZ_CP011801.1 | 171391 | 172578 | 1187 | + | tuf | WP_139866869.1 |
| GCF_001273775.1 | NZ_CP011801.1 | 92101 | 93291 | 1190 | - | tuf | WP_259286621.1 |
| GCF_014701235.1 | NZ_AP022873.1 | 3023594 | 3024781 | 1187 | - | tuf | WP_239289265.1 |
| GCF_014701235.1 | NZ_AP022873.1 | 3040094 | 3041281 | 1187 | - | tuf | WP_239289265.1 |
| GCF_040450955.1 | NZ_CP144373.1 | 14955 | 16142 | 1187 | + | tuf | WP_020915288.1 |
| GCF_040450955.1 | NZ_CP144373.1 | 426710 | 427897 | 1187 | + | tuf | WP_022771327.1 |
| GCF_040451425.1 | NZ_CP144374.1 | 1131074 | 1132261 | 1187 | - | tuf | WP_022771327.1 |
| GCF_040451425.1 | NZ_CP144374.1 | 64165 | 65352 | 1187 | + | tuf | WP_226809876.1 |
| GCF_000020985.1 | NC_011296.1 | 83564 | 84751 | 1187 | + | tuf | WP_226809876.1 |
| GCF_000020985.1 | NC_011296.1 | 67065 | 68252 | 1187 | + | tuf | WP_226806517.1 |
| GCF_011066545.1 | NZ_CP049055.1 | 86871 | 88058 | 1187 | + | tuf | WP_226806517.1 |
| GCF_011066545.1 | NZ_CP049055.1 | 1060747 | 1061934 | 1187 | - | tuf | WP_216797272.1 |
| GCF_009002475.1 | NZ_AP019860.1 | 1080857 | 1082044 | 1187 | - | tuf | WP_216797272.1 |
| GCF_041508025.1 | NZ_CP165719.1 | 106453 | 107640 | 1187 | - | tuf | WP_339045084.1 |
| GCF_002007645.1 | NZ_CP019791.1 | 1000401 | 1001588 | 1187 | + | tuf | WP_156861654.1 |

| | | | | | | | |
|---|---|---|---|---|---|---|---|
| GCF_015169395.1 | NZ_CP063458.1 | 1016947 | 1018134 | 1187 | + | tuf | WP_156861654.1 |
| GCF_015169395.1 | NZ_CP063458.1 | 133413 | 134600 | 1187 | - | tuf | WP_043679011.1 |
| GCF_001999965.1 | NZ_CP019646.1 | 171186 | 172373 | 1187 | - | tuf | WP_043679088.1 |
| GCF_007748075.1 | NZ_CP036280.1 | 1146320 | 1147507 | 1187 | + | tuf | WP_368639903.1 |
| GCF_000284115.1 | NC_017080.1 | 1172070 | 1173257 | 1187 | + | tuf | WP_368639903.1 |
| GCF_007747445.1 | NZ_CP036425.1 | 2309255 | 2310442 | 1187 | - | tuf | WP_179355217.1 |
| GCF_001997385.1 | NZ_CP019633.1 | 2325908 | 2327095 | 1187 | - | tuf | WP_179355217.1 |
| GCF_002117005.1 | NZ_CP021023.1 | 2190449 | 2191636 | 1187 | + | tuf | WP_180306008.1 |
| GCF_007741515.1 | NZ_CP036263.1 | 2207124 | 2208311 | 1187 | + | tuf | WP_180306008.1 |
| GCF_007748035.1 | NZ_CP036278.1 | 1863566 | 1864753 | 1187 | - | tuf | WP_144277782.1 |
| GCF_007743815.1 | NZ_CP036265.1 | 1880415 | 1881602 | 1187 | - | tuf | WP_144277782.1 |
| GCF_007747655.1 | NZ_CP036274.1 | 4341214 | 4342401 | 1187 | - | tuf | WP_136773613.1 |
| GCF_008087625.1 | NZ_CP042997.1 | 4357723 | 4358910 | 1187 | - | tuf | WP_136773613.1 |
| GCF_007752135.1 | NZ_CP036298.1 | 1318125 | 1319312 | 1187 | + | tuf | WP_263125923.1 |
| GCF_007752135.1 | NZ_CP036298.1 | 1334660 | 1335847 | 1187 | + | tuf | WP_263125923.1 |
| GCF_007753265.1 | NZ_CP036349.1 | 1060490 | 1061677 | 1187 | + | tuf | WP_148714527.1 |
| GCF_007748115.1 | NZ_CP036289.1 | 1077017 | 1078204 | 1187 | + | tuf | WP_148714527.1 |
| GCF_008065115.1 | NZ_CP042913.1 | 408819 | 410006 | 1187 | + | tuf | WP_081555062.1 |
| GCF_007745435.1 | NZ_CP036316.1 | 425474 | 426661 | 1187 | + | tuf | WP_081574305.1 |
| GCF_007745175.1 | NZ_CP036271.1 | 4058085 | 4059272 | 1187 | + | tuf | WP_149299336.1 |
| GCF_007752935.1 | NZ_CP036319.1 | 4074682 | 4075869 | 1187 | + | tuf | WP_149299336.1 |
| GCF_007752935.1 | NZ_CP036319.1 | 2190712 | 2191899 | 1187 | + | tuf | WP_114061405.1 |
| GCF_047300955.1 | NZ_CP178615.1 | 2207313 | 2208500 | 1187 | + | tuf | WP_114061413.1 |
| GCF_047300955.1 | NZ_CP178615.1 | 555071 | 556258 | 1187 | + | tuf | WP_227106574.1 |
| GCF_013128195.2 | NZ_CP053452.2 | 571722 | 572909 | 1187 | + | tuf | WP_227106574.1 |
| GCF_001983935.1 | NZ_CP017641.1 | 4528003 | 4529190 | 1187 | - | tuf | WP_101527487.1 |
| GCF_901538265.1 | NZ_LR593886.1 | 4544622 | 4545809 | 1187 | - | tuf | WP_101527486.1 |
| GCF_901538265.1 | NZ_LR593886.1 | 3547402 | 3548589 | 1187 | + | tuf | WP_046166914.1 |
| GCF_007744675.1 | NZ_CP036269.1 | 3563998 | 3565185 | 1187 | + | tuf | WP_046166914.1 |
| GCF_007744675.1 | NZ_CP036269.1 | 1975167 | 1976354 | 1187 | + | tuf | WP_011137735.1 |
| GCF_007746795.1 | NZ_CP036343.1 | 1991760 | 1992947 | 1187 | + | tuf | WP_011137735.1 |

| | | | | | | | |
|---|---|---|---|---|---|---|---|
| GCF_007748195.1 | NZ_CP037920.1 | 303898 | 305088 | 1190 | + | tuf | WP_061532002.1 |
| GCF_007748195.1 | NZ_CP037920.1 | 323459 | 324652 | 1193 | + | tuf | WP_061532011.1 |
| GCF_009720525.1 | NZ_CP043930.1 | 689666 | 690859 | 1193 | + | tuf | WP_014004379.1 |
| GCF_009720525.1 | NZ_CP043930.1 | 709954 | 711147 | 1193 | + | tuf | WP_014004379.1 |
| GCF_007754425.1 | NZ_CP037452.1 | 415593 | 416786 | 1193 | + | tuf | WP_061936342.1 |
| GCF_007754425.1 | NZ_CP037452.1 | 435839 | 437032 | 1193 | + | tuf | WP_061936342.1 |
| GCF_008298035.1 | NZ_CP042910.1 | 3841935 | 3843122 | 1187 | - | tuf | WP_175505395.1 |
| GCF_007748155.1 | NZ_CP037421.1 | 3874029 | 3875216 | 1187 | - | tuf | WP_175505395.1 |
| GCF_007748155.1 | NZ_CP037421.1 | 3758761 | 3759948 | 1187 | + | tuf | WP_045265651.1 |
| GCF_000186345.1 | NC_014962.1 | 3794292 | 3795479 | 1187 | + | tuf | WP_045265651.1 |
| GCF_007746075.1 | NZ_CP036339.1 | 1595893 | 1597080 | 1187 | - | tuf | WP_231043118.1 |
| GCF_009177095.1 | NZ_AP021861.1 | 1628717 | 1629904 | 1187 | - | tuf | WP_231043086.1 |
| GCF_007751035.1 | NZ_CP036433.1 | 1111968 | 1113155 | 1187 | - | tuf | WP_146912112.1 |
| GCF_008254045.1 | NZ_CP042425.1 | 1305837 | 1307024 | 1187 | - | tuf | WP_146912112.1 |
| GCF_008254045.1 | NZ_CP042425.1 | 266510 | 267697 | 1187 | + | tuf | WP_218242018.1 |
| GCF_007747795.1 | NZ_CP036275.1 | 316311 | 317498 | 1187 | + | tuf | WP_218242018.1 |
| GCF_008087665.1 | NZ_CP042912.1 | 1006589 | 1007776 | 1187 | - | tuf | WP_054065871.1 |
| GCF_001956985.1 | NZ_CP019082.1 | 1041680 | 1042867 | 1187 | - | tuf | WP_054065871.1 |
| GCF_007750855.1 | NZ_CP036291.1 | 453234 | 454421 | 1187 | + | tuf | WP_224151095.1 |
| GCF_030622045.1 | NZ_CP130886.1 | 511270 | 512457 | 1187 | + | tuf | WP_224151095.1 |
| GCF_007752345.1 | NZ_CP036299.1 | 99106 | 100293 | 1187 | - | tuf | WP_182325873.1 |
| GCF_000092105.1 | NC_014148.1 | 141953 | 143140 | 1187 | - | tuf | WP_182325873.1 |
| GCF_007750395.1 | NZ_CP036281.1 | 322061 | 323248 | 1187 | + | tuf | WP_283486951.1 |
| GCF_000196115.1 | NC_005027.1 | 361429 | 362616 | 1187 | + | tuf | WP_283486951.1 |
| GCF_007741495.1 | NZ_CP036262.1 | 235385 | 236572 | 1187 | + | tuf | WP_087276052.1 |
| GCF_008065135.1 | NZ_CP042914.1 | 4492694 | 4493881 | 1187 | - | tuf | WP_087276052.1 |
| GCF_007753095.1 | NZ_CP036348.1 | 3150688 | 3151875 | 1187 | - | tuf | WP_116926819.1 |
| GCF_007751715.1 | NZ_CP036292.1 | 3193742 | 3194929 | 1187 | - | tuf | WP_116926819.1 |
| GCF_007741475.1 | NZ_CP036261.1 | 273564 | 274751 | 1187 | + | tuf | WP_098066016.1 |
| GCF_000165715.2 | NC_015174.1 | 343652 | 344839 | 1187 | + | tuf | WP_098066324.1 |
| GCF_007741535.1 | NZ_CP036525.1 | 347572 | 348759 | 1187 | + | tuf | WP_012836789.1 |

| | | | | | | | |
|---|---|---|---|---|---|---|---|
| GCF_000242455.2 | NC_019892.1 | 386552 | 387739 | 1187 | + | tuf | WP_012836789.1 |
| GCF_008035925.1 | NZ_CP036264.1 | 295510 | 296691 | 1181 | + | tuf | WP_182078444.1 |
| GCF_008035925.1 | NZ_CP036264.1 | 323809 | 324990 | 1181 | - | tuf | WP_124794465.1 |
| GCF_007754155.1 | NZ_CP037423.1 | 714182 | 715369 | 1187 | - | tuf | WP_115432508.1 |
| GCF_007754155.1 | NZ_CP037423.1 | 730810 | 731997 | 1187 | - | tuf | WP_115432508.1 |
| GCF_038443385.1 | NZ_CP151726.1 | 4145090 | 4146277 | 1187 | - | tuf | WP_006164373.1 |
| GCF_038443385.1 | NZ_CP151726.1 | 152944 | 154131 | 1187 | + | tuf | WP_053822680.1 |
| GCF_007744515.1 | NZ_CP036268.1 | 173413 | 174600 | 1187 | + | tuf | WP_053822680.1 |
| GCF_007752535.1 | NZ_CP036426.1 | 4342517 | 4343704 | 1187 | - | tuf | WP_071010149.1 |
| GCF_018398935.1 | NZ_CP074694.1 | 4362322 | 4363509 | 1187 | - | tuf | WP_071010149.1 |
| GCF_007744255.1 | NZ_CP036267.1 | 3573659 | 3574846 | 1187 | - | tuf | WP_010810470.1 |
| GCF_002277955.1 | NZ_CP018477.1 | 3593381 | 3594568 | 1187 | - | tuf | WP_010810470.1 |
| GCF_901538355.1 | NZ_LR593887.1 | 3318761 | 3319948 | 1187 | - | tuf | WP_018004885.1 |
| GCF_007747215.1 | NZ_CP036273.1 | 3338463 | 3339650 | 1187 | - | tuf | WP_018004885.1 |
| GCF_007747215.1 | NZ_CP036273.1 | 3577615 | 3578802 | 1187 | - | tuf | WP_063242169.1 |
| GCF_007748095.1 | NZ_CP036287.1 | 3597346 | 3598533 | 1187 | - | tuf | WP_063242169.1 |
| GCF_007748055.1 | NZ_CP036279.1 | 1432063 | 1433250 | 1187 | - | tuf | WP_061956567.1 |
| GCF_007750655.1 | NZ_CP036290.1 | 1451770 | 1452957 | 1187 | - | tuf | WP_223505860.1 |
| GCF_007751475.1 | NZ_CP036434.1 | 3575770 | 3576957 | 1187 | - | tuf | WP_011299314.1 |
| GCF_007745635.1 | NZ_CP036272.1 | 3596105 | 3597292 | 1187 | - | tuf | WP_011299314.1 |
| GCF_007753675.1 | NZ_CP036432.1 | 3112211 | 3113398 | 1187 | - | tuf | WP_012354145.1 |
| GCF_007753675.1 | NZ_CP036432.1 | 3131918 | 3133105 | 1187 | - | tuf | WP_012354145.1 |
| GCF_007741555.1 | NZ_CP036526.1 | 3144955 | 3146142 | 1187 | - | tuf | WP_371323575.1 |
| GCF_000221025.1 | NC_015850.1 | 3161511 | 3162698 | 1187 | - | tuf | WP_371323575.1 |
| GCF_000221025.1 | NC_015850.1 | 816356 | 817543 | 1187 | - | tuf | WP_173532451.1 |
| GCF_010378095.2 | NZ_CP127523.1 | 833034 | 834221 | 1187 | - | tuf | WP_173532462.1 |
| GCF_010378095.2 | NZ_CP127523.1 | 2666843 | 2668030 | 1187 | - | tuf | WP_221005482.1 |
| GCF_003966655.1 | NZ_AP018795.1 | 2683548 | 2684735 | 1187 | - | tuf | WP_221005482.1 |
| GCF_003966655.1 | NZ_AP018795.1 | 432152 | 433339 | 1187 | + | tuf | WP_012202328.1 |
| GCF_028215975.1 | NZ_CP116458.1 | 534475 | 535662 | 1187 | + | tuf | WP_012202435.1 |
| GCF_028215975.1 | NZ_CP116458.1 | 550332 | 551519 | 1187 | + | tuf | WP_016448514.1 |

| | | | | | | | |
|---|---|---|---|---|---|---|---|
| GCF_016250455.1 | NZ_CP059488.1 | 587297 | 588484 | 1187 | + | tuf | WP_016448514.1 |
| GCF_016250455.1 | NZ_CP059488.1 | 6325057 | 6326244 | 1187 | - | tuf | WP_016448514.1 |
| GCF_000021485.1 | NC_011761.1 | 6362013 | 6363200 | 1187 | - | tuf | WP_016448514.1 |
| GCF_000021485.1 | NC_011761.1 | 3259859 | 3261046 | 1187 | - | tuf | WP_183148266.1 |
| GCF_018854495.2 | NZ_CP130946.1 | 3276133 | 3277320 | 1187 | - | tuf | WP_183148266.1 |
| GCF_018854495.2 | NZ_CP130946.1 | 2496846 | 2498033 | 1187 | - | tuf | WP_187722815.1 |
| GCF_018853465.2 | NZ_CP127526.1 | 3311058 | 3312245 | 1187 | + | tuf | WP_187722815.1 |
| GCF_018853465.2 | NZ_CP127526.1 | 2304768 | 2305955 | 1187 | - | tuf | WP_317700467.1 |
| GCF_003721225.2 | NZ_CP127527.1 | 2337272 | 2338459 | 1187 | - | tuf | WP_317700467.1 |
| GCF_003721225.2 | NZ_CP127527.1 | 2952705 | 2953892 | 1187 | - | tuf | WP_011806991.1 |
| GCF_009662475.1 | NZ_CP045571.1 | 3561111 | 3562298 | 1187 | + | tuf | WP_011806991.1 |
| GCF_009662475.1 | NZ_CP045571.1 | 170701 | 171888 | 1187 | + | tuf | WP_187597420.1 |
| GCF_048300925.1 | NZ_CP133759.1 | 4944126 | 4945313 | 1187 | + | tuf | WP_187597420.1 |
| GCF_048300925.1 | NZ_CP133759.1 | 2781729 | 2782916 | 1187 | + | tuf | WP_110846314.1 |
| GCF_029030725.1 | NZ_CP113162.1 | 2879230 | 2880417 | 1187 | + | tuf | WP_110846314.1 |
| GCF_020447305.1 | NZ_CP095732.1 | 5543161 | 5544348 | 1187 | - | tuf | WP_019923500.1 |
| GCF_020447305.1 | NZ_CP095732.1 | 5626465 | 5627652 | 1187 | - | tuf | WP_019923500.1 |
| GCF_002173775.1 | NZ_CP021524.1 | 1344374 | 1345561 | 1187 | - | tuf | WP_074008611.1 |
| GCF_002220195.1 | NZ_CP022374.1 | 1364397 | 1365584 | 1187 | - | tuf | WP_074008611.1 |
| GCF_004014775.2 | NZ_CP042808.1 | 1127787 | 1128968 | 1181 | + | tuf | WP_003825050.1 |
| GCF_009914215.2 | NZ_CP039846.2 | 1130842 | 1132023 | 1181 | + | tuf | WP_003825050.1 |
| GCF_001499615.1 | NZ_LN606600.1 | 17630 | 18811 | 1181 | + | tuf | WP_067442335.1 |
| GCF_008365315.1 | NZ_CP043506.1 | 20701 | 21882 | 1181 | + | tuf | WP_067442335.1 |
| GCF_003352165.1 | NZ_CP029176.1 | 3923770 | 3924957 | 1187 | + | tuf | WP_301256521.1 |
| GCF_000202835.1 | NC_015186.1 | 3943653 | 3944840 | 1187 | + | tuf | WP_301256521.1 |
| GCF_030252425.1 | NZ_CP126994.1 | 129052 | 130239 | 1187 | - | tuf | WP_114561733.1 |
| GCF_012913305.1 | NZ_CP051775.1 | 262997 | 264184 | 1187 | - | tuf | WP_114561733.1 |
| GCF_012913305.1 | NZ_CP051775.1 | 342226 | 343413 | 1187 | + | tuf | WP_117607634.1 |
| GCF_025136295.1 | NZ_CP062788.1 | 358801 | 359988 | 1187 | + | tuf | WP_117607634.1 |
| GCF_025136295.1 | NZ_CP062788.1 | 1981129 | 1982316 | 1187 | - | tuf | WP_212785422.1 |
| GCF_000218565.1 | NC_015684.1 | 1997459 | 1998646 | 1187 | - | tuf | WP_212785434.1 |

| | | | | | | | |
|---|---|---|---|---|---|---|---|
| GCF_030036535.1 | NZ_CP080387.1 | 491090 | 492277 | 1187 | + | tuf | WP_062625514.1 |
| GCF_030036535.1 | NZ_CP080387.1 | 507421 | 508608 | 1187 | + | tuf | WP_062625514.1 |
| GCF_000092025.1 | NC_003062.2 | 1839000 | 1840187 | 1187 | + | tuf | WP_062187353.1 |
| GCF_000092025.1 | NC_003062.2 | 1855212 | 1856399 | 1187 | + | tuf | WP_062187353.1 |
| GCF_005144425.2 | NZ_CP124733.1 | 1916016 | 1917203 | 1187 | - | tuf | WP_162048921.1 |
| GCF_005144425.2 | NZ_CP124733.1 | 1932873 | 1934060 | 1187 | - | tuf | WP_162048921.1 |
| GCF_005144435.2 | NZ_CP120211.1 | 591325 | 592512 | 1187 | + | tuf | WP_013292530.1 |
| GCF_005144435.2 | NZ_CP120211.1 | 607810 | 608997 | 1187 | + | tuf | WP_013292530.1 |
| GCF_013285525.1 | NZ_CP053856.1 | 55281 | 56468 | 1187 | - | tuf | WP_034312043.1 |
| GCF_013285525.1 | NZ_CP053856.1 | 76538 | 77725 | 1187 | - | tuf | WP_034312043.1 |
| GCF_013318015.2 | NZ_CP115841.1 | 4785500 | 4786693 | 1193 | - | tuf | WP_016836040.1 |
| GCF_013318015.2 | NZ_CP115841.1 | 4806880 | 4808073 | 1193 | - | tuf | WP_016836040.1 |
| GCF_021310995.1 | NZ_CP054150.1 | 149616 | 150803 | 1187 | + | tuf | WP_008334097.1 |
| GCF_021310995.1 | NZ_CP054150.1 | 171037 | 172224 | 1187 | + | tuf | WP_008334097.1 |
| GCF_009931615.1 | NZ_CP046620.1 | 3417518 | 3418705 | 1187 | - | tuf | WP_079217281.1 |
| GCF_009931615.1 | NZ_CP046620.1 | 3438937 | 3440124 | 1187 | - | tuf | WP_079217292.1 |
| GCF_025137635.1 | NZ_CP080772.1 | 129130 | 130317 | 1187 | + | tuf | WP_034312043.1 |
| GCF_025137635.1 | NZ_CP080772.1 | 149193 | 150380 | 1187 | + | tuf | WP_034312043.1 |
| GCF_045277565.1 | NZ_CP123584.1 | 110427 | 111614 | 1187 | + | tuf | WP_008334097.1 |
| GCF_045277565.1 | NZ_CP123584.1 | 131765 | 132952 | 1187 | + | tuf | WP_008334097.1 |
| GCF_000016285.1 | NC_011989.1 | 2757337 | 2758524 | 1187 | + | tuf | WP_076593020.1 |
| GCF_000016285.1 | NC_011989.1 | 2778600 | 2779787 | 1187 | + | tuf | WP_076593020.1 |
| GCF_011046245.1 | NZ_CP049241.1 | 398782 | 399969 | 1187 | + | tuf | WP_070263901.1 |
| GCF_011046245.1 | NZ_CP049241.1 | 4639932 | 4641119 | 1187 | - | tuf | WP_070263901.1 |
| GCF_900177405.1 | NZ_LT840185.1 | 3409627 | 3410814 | 1187 | - | tuf | WP_166228711.1 |
| GCF_964031065.1 | NZ_OZ034995.1 | 3951306 | 3952493 | 1187 | + | tuf | WP_166228711.1 |
| GCF_029623755.1 | NZ_CP121106.1 | 404368 | 405555 | 1187 | + | tuf | WP_079364208.1 |
| GCF_001281485.1 | NZ_CP012669.1 | 4511586 | 4512773 | 1187 | - | tuf | WP_079364208.1 |
| GCF_001663155.1 | NZ_CP015963.1 | 3325942 | 3327129 | 1187 | - | tuf | WP_151123650.1 |
| GCF_030284385.1 | NZ_CP127221.1 | 3343807 | 3344994 | 1187 | - | tuf | WP_151123650.1 |
| GCF_030406725.1 | NZ_CP129107.1 | 635454 | 636641 | 1187 | + | tuf | WP_138855352.1 |

| | | | | | | | |
|---|---|---|---|---|---|---|---|
| GCF_032852305.1 | NZ_CP136594.1 | 652950 | 654137 | 1187 | + | tuf | WP_138855352.1 |
| GCF_946995915.1 | NZ_OX341517.1 | 2622370 | 2623557 | 1187 | - | tuf | WP_125974279.1 |
| GCF_946995915.1 | NZ_OX341517.1 | 2639280 | 2640467 | 1187 | - | tuf | WP_125974293.1 |
| GCF_000024505.1 | NC_013532.1 | 707618 | 708805 | 1187 | + | tuf | WP_034753125.1 |
| GCF_000024505.1 | NC_013532.1 | 773277 | 774464 | 1187 | + | tuf | WP_034753125.1 |
| GCF_000020305.1 | NC_012026.1 | 4 | 1203 | 1199 | + | tuf | WP_038498161.1 |
| GCF_000020305.1 | NC_012026.1 | 649053 | 650240 | 1187 | + | tuf | WP_034753125.1 |
| GCF_002214625.1 | NZ_CP015994.1 | 712735 | 713922 | 1187 | + | tuf | WP_034753125.1 |
| GCF_002214625.1 | NZ_CP015994.1 | 656965 | 658152 | 1187 | + | tuf | WP_034753125.1 |
| GCF_012790675.1 | NZ_CP046391.1 | 722323 | 723510 | 1187 | + | tuf | WP_034753125.1 |
| GCF_000092925.1 | NC_014217.1 | 688711 | 689898 | 1187 | + | tuf | WP_096233366.1 |
| GCF_000092925.1 | NC_014217.1 | 751313 | 752500 | 1187 | + | tuf | WP_096233366.1 |
| GCF_010669125.1 | NZ_CP048630.1 | 4547621 | 4548808 | 1187 | + | tuf | WP_070345915.1 |
| GCF_010669125.1 | NZ_CP048630.1 | 4609239 | 4610426 | 1187 | + | tuf | WP_070345915.1 |
| GCF_014495845.1 | NZ_CP061080.1 | 3675509 | 3676696 | 1187 | - | tuf | WP_068375540.1 |
| GCF_014495845.1 | NZ_CP061080.1 | 3828886 | 3830073 | 1187 | - | tuf | WP_068375540.1 |
| GCF_001889605.1 | NZ_CP018171.1 | 166374 | 167555 | 1181 | - | tuf | WP_003784924.1 |
| GCF_001889605.1 | NZ_CP018171.1 | 234422 | 235603 | 1181 | + | tuf | WP_003784924.1 |
| GCF_019443285.1 | NZ_CP080327.1 | 365416 | 366597 | 1181 | - | tuf | WP_038302692.1 |
| GCF_022459015.1 | NZ_CP092734.1 | 512053 | 513234 | 1181 | + | tuf | WP_038302692.1 |
| GCF_001547995.1 | NZ_AP014690.1 | 1430893 | 1432074 | 1181 | + | tuf | WP_085815394.1 |
| GCF_000175215.2 | NC_014816.1 | 1552601 | 1553782 | 1181 | + | tuf | WP_085815394.1 |
| GCF_001077815.2 | NZ_CP011310.1 | 894043 | 895230 | 1187 | + | tuf | WP_126348071.1 |
| GCF_001886695.1 | NZ_CP018097.1 | 912697 | 913884 | 1187 | + | tuf | WP_126348071.1 |
| GCF_014058705.1 | NZ_CP059692.1 | 4332224 | 4333411 | 1187 | + | tuf | WP_012348865.1 |
| GCF_014058705.1 | NZ_CP059692.1 | 4346757 | 4347944 | 1187 | - | tuf | WP_012348865.1 |
| GCF_047943035.1 | NZ_CP181348.1 | 378575 | 379765 | 1190 | + | tuf | WP_171097401.1 |
| GCF_047943035.1 | NZ_CP181348.1 | 395041 | 396231 | 1190 | + | tuf | WP_171097401.1 |
| GCF_017815515.1 | NZ_CP072611.1 | 3136028 | 3137218 | 1190 | - | tuf | WP_130557569.1 |
| GCF_017815515.1 | NZ_CP072611.1 | 3152470 | 3153660 | 1190 | - | tuf | WP_130557569.1 |
| GCF_000010525.1 | NC_009937.1 | 2155604 | 2156806 | 1202 | + | tuf | WP_054265806.1 |

| | | | | | | | |
|---|---|---|---|---|---|---|---|
| GCF_000010525.1 | NC_009937.1 | 2256688 | 2257890 | 1202 | + | tuf | WP_054265806.1 |
| GCF_000632475.1 | NZ_CP007793.1 | 784798 | 786000 | 1202 | + | tuf | WP_404932179.1 |
| GCF_000632475.1 | NZ_CP007793.1 | 897836 | 899038 | 1202 | + | tuf | WP_404932179.1 |
| GCF_003119195.2 | NZ_CP022253.1 | 6467114 | 6468301 | 1187 | - | tuf | WP_160406738.1 |
| GCF_003119195.2 | NZ_CP022253.1 | 6573572 | 6574759 | 1187 | - | tuf | WP_160406738.1 |
| GCF_001639105.2 | NZ_CP015285.1 | 4423365 | 4424552 | 1187 | - | tuf | WP_231766970.1 |
| GCF_001639105.2 | NZ_CP015285.1 | 5155064 | 5156251 | 1187 | + | tuf | WP_231767596.1 |
| GCF_013347285.1 | NZ_CP054619.1 | 1475709 | 1476896 | 1187 | - | tuf | WP_169434656.1 |
| GCF_013347285.1 | NZ_CP054619.1 | 1566239 | 1567426 | 1187 | - | tuf | WP_169434704.1 |
| GCF_003233655.1 | NZ_CP029829.1 | 888642 | 889829 | 1187 | - | tuf | WP_193687473.1 |
| GCF_003233655.1 | NZ_CP029829.1 | 1645536 | 1646723 | 1187 | + | tuf | WP_193687473.1 |
| GCF_003130795.1 | NZ_CP029353.1 | 2582758 | 2583945 | 1187 | - | tuf | WP_109345418.1 |
| GCF_003130795.1 | NZ_CP029353.1 | 2684102 | 2685289 | 1187 | - | tuf | WP_109345418.1 |
| GCF_001305595.1 | NZ_CP012401.1 | 4333345 | 4334532 | 1187 | + | tuf | WP_075794180.1 |
| GCF_001305595.1 | NZ_CP012401.1 | 4418880 | 4420067 | 1187 | + | tuf | WP_075794180.1 |
| GCF_013388295.1 | NZ_CP058235.1 | 794481 | 795683 | 1202 | + | tuf | WP_099873741.1 |
| GCF_013388295.1 | NZ_CP058235.1 | 923602 | 924804 | 1202 | + | tuf | WP_099873741.1 |
| GCF_001281405.1 | NZ_CP010401.1 | 254495 | 255682 | 1187 | + | tuf | WP_209423174.1 |
| GCF_001281405.1 | NZ_CP010401.1 | 3333584 | 3334771 | 1187 | - | tuf | WP_209423174.1 |
| GCF_000341355.1 | NC_020300.1 | 1596388 | 1597575 | 1187 | - | tuf | WP_106683576.1 |
| GCF_000341355.1 | NC_020300.1 | 2226598 | 2227785 | 1187 | + | tuf | WP_106683576.1 |
| GCF_000015445.1 | NC_008783.1 | 3649643 | 3650830 | 1187 | - | tuf | WP_011831019.1 |
| GCF_000015445.1 | NC_008783.1 | 3667375 | 3668562 | 1187 | - | tuf | WP_011831019.1 |
| GCF_000015445.1 | NC_008783.1 | 270103 | 271290 | 1187 | + | tuf | WP_011478633.1 |
| GCF_000253015.1 | NC_014932.1 | 286472 | 287659 | 1187 | + | tuf | WP_011478645.1 |
| GCF_000253015.1 | NC_014932.1 | 338470 | 339657 | 1187 | + | tuf | WP_140002317.1 |
| GCF_900638615.1 | NZ_LR134527.1 | 355126 | 356313 | 1187 | + | tuf | WP_140002317.1 |
| GCF_900638615.1 | NZ_LR134527.1 | 268873 | 270060 | 1187 | + | tuf | WP_221764568.1 |
| GCF_045277155.1 | NZ_CP123962.1 | 285511 | 286698 | 1187 | + | tuf | WP_221764568.1 |
| GCF_045277155.1 | NZ_CP123962.1 | 357255 | 358442 | 1187 | + | tuf | WP_015831283.1 |
| GCF_000022725.1 | NC_012846.1 | 373999 | 375186 | 1187 | + | tuf | WP_015831283.1 |

| | | | | | | | |
|---|---|---|---|---|---|---|---|
| GCF_000022725.1 | NC_012846.1 | 357829 | 359016 | 1187 | + | tuf | WP_013147026.1 |
| GCF_024297065.1 | NZ_CP101114.1 | 374516 | 375703 | 1187 | + | tuf | WP_013147026.1 |
| GCF_024297065.1 | NZ_CP101114.1 | 303664 | 304851 | 1187 | + | tuf | WP_013441116.1 |
| GCF_046245125.1 | NZ_CP176404.1 | 320052 | 321239 | 1187 | + | tuf | WP_013441116.1 |
| GCF_046245125.1 | NZ_CP176404.1 | 1065398 | 1066585 | 1187 | - | tuf | WP_107888850.1 |
| GCF_019930925.1 | NZ_CP082885.1 | 1082049 | 1083236 | 1187 | - | tuf | WP_107888851.1 |
| GCF_019930925.1 | NZ_CP082885.1 | 76626 | 77807 | 1181 | + | tuf | WP_003742643.1 |
| GCF_003606325.2 | NZ_CP031843.2 | 93551 | 94732 | 1181 | + | tuf | WP_003742643.1 |
| GCF_003606325.2 | NZ_CP031843.2 | 267094 | 268281 | 1187 | + | tuf | WP_104077832.1 |
| GCF_003606345.3 | NZ_CP031844.2 | 289176 | 290363 | 1187 | + | tuf | WP_104077832.1 |
| GCF_003606345.3 | NZ_CP031844.2 | 283738 | 284925 | 1187 | + | tuf | WP_013434762.1 |
| GCF_022559585.1 | NZ_CP087114.1 | 305716 | 306903 | 1187 | + | tuf | WP_013434762.1 |
| GCF_022559585.1 | NZ_CP087114.1 | 268222 | 269412 | 1190 | + | tuf | WP_026922084.1 |
| GCF_009936175.1 | NZ_AP019773.1 | 1043990 | 1045171 | 1181 | + | tuf | WP_126325827.1 |
| GCF_009936175.1 | NZ_AP019773.1 | 1095184 | 1096365 | 1181 | + | tuf | WP_126325827.1 |
| GCF_039555305.1 | NZ_CP154603.1 | 1672883 | 1674064 | 1181 | + | tuf | WP_085390775.1 |
| GCF_039555305.1 | NZ_CP154603.1 | 1737442 | 1738623 | 1181 | + | tuf | WP_085390775.1 |
| GCF_023920085.1 | NZ_CP083444.1 | 177699 | 178880 | 1181 | + | tuf | WP_047761939.1 |
| GCF_023920085.1 | NZ_CP083444.1 | 2293727 | 2294908 | 1181 | - | tuf | WP_047761939.1 |
| GCF_000196435.1 | NC_010161.1 | 456971 | 458158 | 1187 | + | tuf | WP_182109863.1 |
| GCF_000196435.1 | NC_010161.1 | 523723 | 524910 | 1187 | + | tuf | WP_182109863.1 |
| GCF_900638635.1 | NZ_LR134529.1 | 1241345 | 1242526 | 1181 | + | tuf | WP_095502524.1 |
| GCF_900638635.1 | NZ_LR134529.1 | 1260823 | 1262004 | 1181 | + | tuf | WP_095502524.1 |
| GCF_000019845.1 | NC_010581.1 | 179382 | 180563 | 1181 | - | tuf | WP_126326477.1 |
| GCF_000019845.1 | NC_010581.1 | 1143604 | 1144785 | 1181 | + | tuf | WP_126326477.1 |
| GCF_003966715.1 | NZ_AP018907.1 | 923903 | 925084 | 1181 | - | tuf | WP_089035815.1 |
| GCF_001402875.1 | NZ_CP012946.1 | 1045831 | 1047012 | 1181 | - | tuf | WP_089035815.1 |
| GCF_003431825.1 | NZ_CP020083.1 | 1672146 | 1673327 | 1181 | + | tuf | WP_002215366.1 |
| GCF_034374265.1 | NZ_CP139718.1 | 1688855 | 1690036 | 1181 | + | tuf | WP_002215366.1 |
| GCF_001741865.1 | NZ_CP017147.1 | 2308011 | 2309192 | 1181 | - | tuf | WP_115336235.1 |
| GCF_001741865.1 | NZ_CP017147.1 | 2679377 | 2680558 | 1181 | + | tuf | WP_115336235.1 |

| | | | | | | | |
|---|---|---|---|---|---|---|---|
| GCF_002266435.3 | NZ_CP029426.2 | 244436 | 245617 | 1181 | + | tuf | WP_085360440.1 |
| GCF_015291705.1 | NZ_CP030050.1 | 2220642 | 2221823 | 1181 | - | tuf | WP_085360440.1 |
| GCF_011068405.5 | NZ_CP088280.1 | 1930234 | 1931415 | 1181 | + | tuf | WP_002215366.1 |
| GCF_008932115.1 | NZ_CP044543.1 | 1950509 | 1951690 | 1181 | + | tuf | WP_002215366.1 |
| GCF_029714345.1 | NZ_CP121669.1 | 1946269 | 1947450 | 1181 | + | tuf | WP_003690212.1 |
| GCF_045571495.1 | NZ_CP120703.1 | 1965787 | 1966968 | 1181 | + | tuf | WP_003690212.1 |
| GCF_015624505.1 | NZ_CP061379.1 | 1917345 | 1918526 | 1181 | - | tuf | WP_013448204.1 |
| GCF_015624505.1 | NZ_CP061379.1 | 1934372 | 1935553 | 1181 | - | tuf | WP_013448204.1 |
| GCF_000284275.1 | NC_017082.1 | 1388855 | 1390036 | 1181 | - | tuf | WP_338577843.1 |
| GCF_021044685.1 | NZ_CP088014.1 | 1454205 | 1455386 | 1181 | - | tuf | WP_338577843.1 |
| GCF_001642675.1 | NZ_CP011360.1 | 1865915 | 1867096 | 1181 | + | tuf | WP_237090465.1 |
| GCF_023278185.1 | NZ_CP066356.1 | 1883470 | 1884651 | 1181 | + | tuf | WP_237090465.1 |
| GCF_049949285.1 | NZ_CP181318.1 | 346893 | 348074 | 1181 | + | tuf | WP_003742643.1 |
| GCF_004114975.1 | NZ_CP030051.1 | 363869 | 365050 | 1181 | + | tuf | WP_003742643.1 |
| GCF_004114915.1 | NZ_CP022219.1 | 1342032 | 1343213 | 1181 | + | tuf | WP_002215366.1 |
| GCF_004114955.1 | NZ_CP030053.1 | 1359358 | 1360539 | 1181 | + | tuf | WP_002215366.1 |
| GCF_025200885.1 | NZ_CP104172.1 | 1955396 | 1956577 | 1181 | - | tuf | WP_187000175.1 |
| GCF_001693385.1 | NZ_CP016428.1 | 2118741 | 2119922 | 1181 | - | tuf | WP_187000175.1 |
| GCF_013752735.1 | NZ_CP058354.1 | 2111506 | 2112687 | 1181 | - | tuf | WP_182121945.1 |
| GCF_042691665.1 | NZ_CP170504.1 | 2276530 | 2277711 | 1181 | - | tuf | WP_182121945.1 |
| GCF_000344805.1 | NC_020453.1 | 2318493 | 2319674 | 1181 | + | tuf | WP_003742643.1 |
| GCF_021088345.1 | NZ_CP088156.1 | 2335316 | 2336497 | 1181 | + | tuf | WP_003742643.1 |
| GCF_002278135.3 | NZ_CP029425.2 | 1795451 | 1796632 | 1181 | - | tuf | WP_049350946.1 |
| GCF_029714545.1 | NZ_CP121667.1 | 1812250 | 1813431 | 1181 | - | tuf | WP_049350946.1 |
| GCF_001693515.2 | NZ_CP042968.1 | 292289 | 293470 | 1181 | - | tuf | WP_107689181.1 |
| GCF_032397745.1 | NZ_CP135921.1 | 1193847 | 1195028 | 1181 | + | tuf | WP_107689181.1 |
| GCF_032397745.1 | NZ_CP135921.1 | 1342326 | 1343507 | 1181 | + | tuf | WP_004284587.1 |
| GCF_017493175.3 | NZ_CP088282.1 | 1359286 | 1360467 | 1181 | + | tuf | WP_064130938.1 |
| GCF_030413175.1 | NZ_CP129212.1 | 99601 | 100782 | 1181 | + | tuf | WP_165090209.1 |
| GCF_018736085.1 | NZ_CP076134.1 | 119084 | 120265 | 1181 | + | tuf | WP_165090209.1 |
| GCF_011516645.4 | NZ_CP088285.1 | 582058 | 583239 | 1181 | + | tuf | WP_151049960.1 |

| | | | | | | | |
|---|---|---|---|---|---|---|---|
| GCF_011516645.4 | NZ_CP088285.1 | 756054 | 757235 | 1181 | + | tuf | WP_151049960.1 |
| GCF_011604665.2 | NZ_CP050066.2 | 38190 | 39371 | 1181 | - | tuf | WP_085360686.1 |
| GCF_900324035.1 | NZ_LS398110.1 | 790225 | 791406 | 1181 | + | tuf | WP_085360686.1 |
| GCF_027594865.1 | NZ_CP089391.1 | 1517852 | 1519039 | 1187 | - | tuf | WP_289175498.1 |
| GCF_025200905.1 | NZ_CP104173.1 | 1546906 | 1548093 | 1187 | - | tuf | WP_289175536.1 |
| GCF_004114935.1 | NZ_CP022221.1 | 3941022 | 3942209 | 1187 | - | tuf | WP_227816688.1 |
| GCF_023573565.1 | NZ_CP097649.1 | 3957588 | 3958775 | 1187 | - | tuf | WP_227816688.1 |
| GCF_023573565.1 | NZ_CP097649.1 | 3536830 | 3538017 | 1187 | - | tuf | WP_173767314.1 |
| GCF_034424665.1 | NZ_CP139996.1 | 3553383 | 3554570 | 1187 | - | tuf | WP_173767314.1 |
| GCF_034424665.1 | NZ_CP139996.1 | 303076 | 304263 | 1187 | - | tuf | WP_046848862.1 |
| GCF_017086445.1 | NZ_CP070968.1 | 3695804 | 3696991 | 1187 | - | tuf | WP_046851109.1 |
| GCF_017086445.1 | NZ_CP070968.1 | 451898 | 453085 | 1187 | + | tuf | WP_011111034.1 |
| GCF_017487405.1 | NZ_CP062222.1 | 2228140 | 2229327 | 1187 | - | tuf | WP_011111034.1 |
| GCF_017487405.1 | NZ_CP062222.1 | 578796 | 579983 | 1187 | + | tuf | WP_011633671.1 |
| GCF_041892135.1 | NZ_CP169067.1 | 1911854 | 1913041 | 1187 | - | tuf | WP_011634842.1 |
| GCF_041892135.1 | NZ_CP169067.1 | 1888411 | 1889598 | 1187 | + | tuf | WP_013647103.1 |
| GCF_019395145.1 | NZ_CP080034.1 | 2194827 | 2196014 | 1187 | + | tuf | WP_013647103.1 |
| GCF_019395145.1 | NZ_CP080034.1 | 941181 | 942368 | 1187 | - | tuf | WP_025040225.1 |
| GCF_037055245.1 | NZ_CP146369.1 | 959918 | 961105 | 1187 | - | tuf | WP_025040225.1 |
| GCF_037055245.1 | NZ_CP146369.1 | 1363218 | 1364405 | 1187 | + | tuf | WP_040853253.1 |
| GCF_029636725.1 | NZ_CP116770.1 | 1381983 | 1383170 | 1187 | + | tuf | WP_040853253.1 |
| GCF_029636725.1 | NZ_CP116770.1 | 875181 | 876368 | 1187 | + | tuf | WP_011380109.1 |
| GCF_041893865.1 | NZ_CP169082.1 | 893921 | 895108 | 1187 | + | tuf | WP_011380109.1 |
| GCF_041893865.1 | NZ_CP169082.1 | 711641 | 712828 | 1187 | + | tuf | WP_206255063.1 |
| GCF_000144605.1 | NC_014375.1 | 728264 | 729451 | 1187 | + | tuf | WP_206255063.1 |
| GCF_000144605.1 | NC_014375.1 | 1379495 | 1380682 | 1187 | - | tuf | WP_018574320.1 |
| GCF_002208825.2 | NZ_CP022048.2 | 1430984 | 1432171 | 1187 | - | tuf | WP_018574320.1 |
| GCF_002208825.2 | NZ_CP022048.2 | 167891 | 169078 | 1187 | - | tuf | WP_243478918.1 |
| GCF_016656965.1 | NZ_CP067977.1 | 184658 | 185845 | 1187 | - | tuf | WP_243478918.1 |
| GCF_016656965.1 | NZ_CP067977.1 | 86171 | 87358 | 1187 | - | tuf | WP_067758359.1 |
| GCF_015326295.1 | NZ_CP064063.1 | 138433 | 139620 | 1187 | - | tuf | WP_067758359.1 |

| | | | | | | | |
|---|---|---|---|---|---|---|---|
| GCF_015326295.1 | NZ_CP064063.1 | 3095267 | 3096454 | 1187 | - | tuf | WP_108622091.1 |
| GCF_000018525.1 | NC_010103.1 | 3112019 | 3113206 | 1187 | - | tuf | WP_108622091.1 |
| GCF_000018525.1 | NC_010103.1 | 3089992 | 3091179 | 1187 | - | tuf | WP_149426143.1 |
| GCF_047324105.1 | NZ_CP178745.1 | 3106868 | 3108055 | 1187 | - | tuf | WP_149426143.1 |
| GCF_047324105.1 | NZ_CP178745.1 | 356781 | 357968 | 1187 | - | tuf | WP_106701453.1 |
| GCF_900095155.1 | NZ_LT605585.1 | 534403 | 535590 | 1187 | - | tuf | WP_106701453.1 |
| GCF_900095155.1 | NZ_LT605585.1 | 63556 | 64743 | 1187 | - | tuf | WP_086131724.1 |
| GCF_000007125.1 | NC_003317.1 | 80509 | 81696 | 1187 | - | tuf | WP_086131724.1 |
| GCF_000007125.1 | NC_003317.1 | 1569281 | 1570468 | 1187 | + | tuf | WP_269308170.1 |
| GCF_000022745.1 | NC_013119.1 | 1586333 | 1587520 | 1187 | + | tuf | WP_269308170.1 |
| GCF_000022745.1 | NC_013119.1 | 2670484 | 2671671 | 1187 | + | tuf | WP_269900973.1 |
| GCF_000016845.1 | NC_009505.1 | 2691207 | 2692394 | 1187 | + | tuf | WP_269900973.1 |
| GCF_000016845.1 | NC_009505.1 | 3842900 | 3844087 | 1187 | - | tuf | WP_215796421.1 |
| GCF_025118245.1 | NZ_CP099967.1 | 3862033 | 3863220 | 1187 | - | tuf | WP_215796421.1 |
| GCF_025118245.1 | NZ_CP099967.1 | 4164987 | 4166174 | 1187 | - | tuf | WP_028536828.1 |
| GCF_022024335.1 | NZ_CP091783.1 | 4181556 | 4182743 | 1187 | - | tuf | WP_215796677.1 |
| GCF_022024335.1 | NZ_CP091783.1 | 314745 | 315932 | 1187 | + | tuf | WP_048628233.1 |
| GCF_000007505.1 | NC_004310.3 | 354422 | 355627 | 1205 | + | tuf | WP_042113424.1 |
| GCF_000007505.1 | NC_004310.3 | 5567287 | 5568492 | 1205 | - | tuf | WP_257958814.1 |
| GCF_003860345.1 | NZ_CP034179.1 | 5615279 | 5616484 | 1205 | - | tuf | WP_257958831.1 |
| GCF_029981845.1 | NZ_CP104164.1 | 4858268 | 4859473 | 1205 | - | tuf | WP_047906206.1 |
| GCF_029981845.1 | NZ_CP104164.1 | 4900728 | 4901915 | 1187 | - | tuf | WP_047906231.1 |
| GCF_034359465.1 | NZ_CP110820.1 | 1104692 | 1105879 | 1187 | + | tuf | WP_039374987.1 |
| GCF_034359465.1 | NZ_CP110820.1 | 1146997 | 1148202 | 1205 | + | tuf | WP_039374948.1 |
| GCF_036670005.1 | NZ_CP133270.1 | 5805097 | 5806284 | 1187 | - | tuf | WP_058376384.1 |
| GCF_036670005.1 | NZ_CP133270.1 | 5844951 | 5846138 | 1187 | - | tuf | WP_058376384.1 |
| GCF_014896945.1 | NZ_CP054719.1 | 5222701 | 5223906 | 1205 | - | tuf | WP_046290034.1 |
| GCF_008189285.1 | NZ_CP043315.1 | 5270978 | 5272183 | 1205 | - | tuf | WP_046290065.1 |
| GCF_951805245.1 | NZ_OX638200.1 | 5020280 | 5021485 | 1205 | - | tuf | WP_023593823.1 |
| GCF_007993655.1 | NZ_CP029077.1 | 5066305 | 5067492 | 1187 | - | tuf | WP_023593795.1 |
| GCF_000319385.1 | NC_019566.1 | 5494391 | 5495596 | 1205 | - | tuf | WP_039406364.1 |

| | | | | | | | |
|---|---|---|---|---|---|---|---|
| GCF_964026585.1 | NZ_OZ032160.1 | 5535172 | 5536359 | 1187 | - | tuf | WP_039406415.1 |
| GCF_034359305.1 | NZ_CP110343.1 | 5376705 | 5377910 | 1205 | - | tuf | WP_039393730.1 |
| GCF_003072485.1 | NZ_CP025989.1 | 5423726 | 5424931 | 1205 | - | tuf | WP_039393730.1 |
| GCF_002688585.1 | NZ_CP024034.1 | 227654 | 228841 | 1187 | + | tuf | WP_047215714.1 |
| GCF_021065005.1 | NZ_CP069247.1 | 244612 | 245799 | 1187 | + | tuf | WP_047215714.1 |
| GCF_002774925.1 | NZ_NXGO01000052.1 | 5255270 | 5256475 | 1205 | - | tuf | WP_044457329.1 |
| GCF_002774925.1 | NZ_NXGO01000070.1 | 5300535 | 5301722 | 1187 | - | tuf | WP_044457352.1 |
| GCF_021654655.1 | NZ_AP025225.1 | 1469121 | 1470308 | 1187 | + | tuf | WP_007180138.1 |
| GCF_025736875.1 | NZ_CP107052.1 | 1488318 | 1489505 | 1187 | + | tuf | WP_007180138.1 |
| GCF_964019805.1 | NZ_OZ026472.1 | 3199533 | 3200720 | 1187 | - | tuf | WP_013090760.1 |
| GCF_964020185.1 | NZ_OZ026538.1 | 3218723 | 3219910 | 1187 | - | tuf | WP_013090760.1 |
| GCF_001021085.1 | NZ_CP004021.1 | 4304629 | 4305816 | 1187 | - | tuf | WP_111932934.1 |
| GCF_001021085.1 | NZ_CP004021.1 | 4323889 | 4325076 | 1187 | - | tuf | WP_111932934.1 |
| GCF_000496595.1 | NC_022793.1 | 1815053 | 1816240 | 1187 | - | tuf | WP_115781224.1 |
| GCF_000496595.1 | NC_022793.1 | 1834228 | 1835415 | 1187 | - | tuf | WP_115781224.1 |
| GCF_000590865.3 | NZ_CP010804.2 | 1074845 | 1076032 | 1187 | + | tuf | WP_028371383.1 |
| GCF_000590865.3 | NZ_CP010804.2 | 1094026 | 1095213 | 1187 | + | tuf | WP_028371383.1 |
| GCF_000183665.1 | NC_014774.1 | 4159784 | 4160971 | 1187 | - | tuf | WP_007180138.1 |
| GCF_000183665.1 | NC_014774.1 | 4178971 | 4180158 | 1187 | - | tuf | WP_007180138.1 |
| GCF_029982035.1 | NZ_CP104166.1 | 2804407 | 2805594 | 1187 | - | tuf | WP_129563002.1 |
| GCF_964019585.1 | NZ_OZ026454.1 | 2841089 | 2842276 | 1187 | - | tuf | WP_129563002.1 |
| GCF_964019585.1 | NZ_OZ026454.1 | 666272 | 667459 | 1187 | + | tuf | WP_219798700.1 |
| GCF_964020175.1 | NZ_OZ026541.1 | 688037 | 689224 | 1187 | + | tuf | WP_219798700.1 |
| GCF_964020175.1 | NZ_OZ026541.1 | 1992255 | 1993442 | 1187 | + | tuf | WP_025496714.1 |
| GCF_000219355.1 | NC_015722.1 | 2011480 | 2012667 | 1187 | + | tuf | WP_025496714.1 |
| GCF_960610765.1 | NZ_OY490516.1 | 2970316 | 2971503 | 1187 | + | tuf | WP_198488102.1 |
| GCF_002117145.1 | NZ_CP008743.1 | 2989477 | 2990664 | 1187 | + | tuf | WP_198488102.1 |
| GCF_002117145.1 | NZ_CP008743.1 | 3693348 | 3694535 | 1187 | - | tuf | WP_012434572.1 |
| GCF_000742835.1 | NZ_CP008941.1 | 3712509 | 3713696 | 1187 | - | tuf | WP_012434572.1 |
| GCF_000742835.1 | NZ_CP008941.1 | 3698584 | 3699771 | 1187 | - | tuf | WP_063497539.1 |
| GCF_012276695.1 | NZ_CP038852.1 | 3717764 | 3718951 | 1187 | - | tuf | WP_063497539.1 |

| | | | | | | | |
|---|---|---|---|---|---|---|---|
| GCF_000012345.1 | NC_007205.1 | 2481030 | 2482217 | 1187 | - | tuf | WP_012434572.1 |
| GCF_003015145.1 | NZ_CP027845.1 | 2500208 | 2501395 | 1187 | - | tuf | WP_012434572.1 |
| GCF_000024465.1 | NC_014010.1 | 3736038 | 3737225 | 1187 | - | tuf | WP_017777140.1 |
| GCF_000024465.1 | NC_014010.1 | 3755270 | 3756457 | 1187 | - | tuf | WP_017777140.1 |
| GCF_008189685.1 | NZ_CP043312.1 | 3402403 | 3403590 | 1187 | - | tuf | WP_028371383.1 |
| GCF_964026555.1 | NZ_OZ032159.1 | 3421578 | 3422765 | 1187 | - | tuf | WP_028371383.1 |
| GCF_964030735.1 | NZ_OZ034917.1 | 926231 | 927418 | 1187 | + | tuf | WP_028371383.1 |
| GCF_964026405.1 | NZ_OZ032146.1 | 963048 | 964235 | 1187 | + | tuf | WP_028371383.1 |
| GCF_964026545.1 | NZ_OZ032156.1 | 3181530 | 3182717 | 1187 | - | tuf | WP_012402197.1 |
| GCF_964026595.1 | NZ_OZ032163.1 | 3200708 | 3201895 | 1187 | - | tuf | WP_012402197.1 |
| GCF_964026425.1 | NZ_OZ032143.1 | 4075481 | 4076668 | 1187 | - | tuf | WP_012434572.1 |
| GCF_964031035.1 | NZ_OZ034997.1 | 4094667 | 4095854 | 1187 | - | tuf | WP_012434572.1 |
| GCF_964026575.1 | NZ_OZ032161.1 | 6271337 | 6272524 | 1187 | - | tuf | WP_028371383.1 |
| GCF_964030715.1 | NZ_OZ034919.1 | 6290521 | 6291708 | 1187 | - | tuf | WP_028371383.1 |
| GCF_965178575.1 | NZ_OZ238268.1 | 3335004 | 3336191 | 1187 | - | tuf | WP_027199087.1 |
| GCF_964031105.1 | NZ_OZ035005.1 | 3354195 | 3355382 | 1187 | - | tuf | WP_027199087.1 |
| GCF_964031095.1 | NZ_OZ035004.1 | 3298179 | 3299366 | 1187 | - | tuf | WP_028371383.1 |
| GCF_964059235.1 | NZ_OZ060452.1 | 3317360 | 3318547 | 1187 | - | tuf | WP_028371383.1 |
| GCF_964026475.1 | NZ_OZ032151.1 | 3711091 | 3712278 | 1187 | - | tuf | WP_012434572.1 |
| GCF_964264755.1 | NZ_OZ183481.1 | 3730256 | 3731443 | 1187 | - | tuf | WP_012434572.1 |
| GCF_964264975.1 | NZ_OZ183649.1 | 2961510 | 2962697 | 1187 | - | tuf | WP_011350666.1 |
| GCF_964264865.1 | NZ_OZ183486.1 | 2981876 | 2983063 | 1187 | - | tuf | WP_011350666.1 |
| GCF_964030955.1 | NZ_OZ035003.1 | 4514874 | 4516061 | 1187 | - | tuf | WP_007180138.1 |
| GCF_964026415.1 | NZ_OZ032144.1 | 4534073 | 4535260 | 1187 | - | tuf | WP_007180138.1 |
| GCF_964264915.1 | NZ_OZ183487.1 | 446929 | 448116 | 1187 | + | tuf | WP_011793521.1 |
| GCF_964026605.1 | NZ_OZ032164.1 | 5096420 | 5097607 | 1187 | - | tuf | WP_011793521.1 |
| GCF_964026535.1 | NZ_OZ032153.1 | 765547 | 766734 | 1187 | - | tuf | WP_011793521.1 |
| GCF_964030695.1 | NZ_OZ034911.1 | 1446639 | 1447826 | 1187 | + | tuf | WP_011793521.1 |
| GCF_964264675.1 | NZ_OZ183472.1 | 775987 | 777174 | 1187 | + | tuf | WP_230339842.1 |
| GCF_964026395.1 | NZ_OZ032145.1 | 801280 | 802467 | 1187 | + | tuf | WP_230339842.1 |
| GCF_964264705.1 | NZ_OZ183473.1 | 2223528 | 2224715 | 1187 | + | tuf | WP_285235081.1 |

| | | | | | | | |
|---|---|---|---|---|---|---|---|
| GCF_963422225.1 | NZ_OY728607.1 | 2241975 | 2243162 | 1187 | + | tuf | WP_285235081.1 |
| GCF_001679665.1 | NZ_CP013244.1 | 1585442 | 1586629 | 1187 | - | tuf | WP_159991453.1 |
| GCF_003722335.1 | NZ_CP026100.1 | 1617989 | 1619176 | 1187 | - | tuf | WP_159991453.1 |
| GCF_003722335.1 | NZ_CP026100.1 | 5850326 | 5851513 | 1187 | - | tuf | WP_148818257.1 |
| GCF_001414055.1 | NZ_CP013002.1 | 5856204 | 5857391 | 1187 | + | tuf | WP_148818257.1 |
| GCF_001414055.1 | NZ_CP013002.1 | 587160 | 588347 | 1187 | + | tuf | WP_085749122.1 |
| GCF_002749615.1 | NZ_CP024201.1 | 605047 | 606234 | 1187 | + | tuf | WP_085749122.1 |
| GCF_002749615.1 | NZ_CP024201.1 | 2258516 | 2259703 | 1187 | - | tuf | WP_418317390.1 |
| GCF_010977555.1 | NZ_CP048815.1 | 2276594 | 2277781 | 1187 | - | tuf | WP_418317390.1 |
| GCF_010977555.1 | NZ_CP048815.1 | 378781 | 379968 | 1187 | + | tuf | WP_377304296.1 |
| GCF_003015125.1 | NZ_CP027850.1 | 3579225 | 3580412 | 1187 | - | tuf | WP_377304296.1 |
| GCF_003015125.1 | NZ_CP027850.1 | 210311 | 211498 | 1187 | + | tuf | WP_011799636.1 |
| GCF_011045195.1 | NZ_CP049199.1 | 583152 | 584339 | 1187 | + | tuf | WP_011799636.1 |
| GCF_011045195.1 | NZ_CP049199.1 | 213103 | 214290 | 1187 | + | tuf | WP_011799636.1 |
| GCF_002407265.1 | NZ_CP022196.1 | 3881986 | 3883173 | 1187 | - | tuf | WP_011799636.1 |
| GCF_002407265.1 | NZ_CP022196.1 | 166456 | 167643 | 1187 | + | tuf | WP_168920827.1 |
| GCF_000819565.1 | NZ_CP004393.1 | 3628837 | 3630024 | 1187 | - | tuf | WP_168920827.1 |
| GCF_000819565.1 | NZ_CP004393.1 | 2972992 | 2974179 | 1187 | + | tuf | WP_128355916.1 |
| GCF_001308265.1 | NZ_CP012023.1 | 2993657 | 2994844 | 1187 | + | tuf | WP_128355930.1 |
| GCF_001308265.1 | NZ_CP012023.1 | 33459 | 34646 | 1187 | + | tuf | WP_108507569.1 |
| GCF_021391315.1 | NZ_CP090021.1 | 50089 | 51276 | 1187 | + | tuf | WP_108507569.1 |
| GCF_021391315.1 | NZ_CP090021.1 | 33296 | 34483 | 1187 | + | tuf | WP_173941800.1 |
| GCF_000012905.2 | NC_007493.2 | 50111 | 51298 | 1187 | + | tuf | WP_173941800.1 |
| GCF_000012905.2 | NC_007493.2 | 33127 | 34314 | 1187 | + | tuf | WP_173959374.1 |
| GCF_014126615.1 | NZ_CP059896.1 | 49914 | 51101 | 1187 | + | tuf | WP_173959374.1 |
| GCF_014126615.1 | NZ_CP059896.1 | 42390 | 43577 | 1187 | + | tuf | WP_011901887.1 |
| GCF_963665895.1 | NZ_OY762534.1 | 59194 | 60381 | 1187 | + | tuf | WP_011901887.1 |
| GCF_963665895.1 | NZ_OY762534.1 | 37806 | 38993 | 1187 | + | tuf | WP_011901887.1 |
| GCF_003691365.1 | NZ_CP033087.1 | 54621 | 55808 | 1187 | + | tuf | WP_011901887.1 |
| GCF_030845175.1 | NZ_CP132542.1 | 33704 | 34891 | 1187 | + | tuf | WP_108468228.1 |
| GCF_030845175.1 | NZ_CP132542.1 | 50494 | 51681 | 1187 | + | tuf | WP_108468228.1 |

| | | | | | | | |
|---|---|---|---|---|---|---|---|
| GCF_030657335.1 | NZ_CP131039.1 | 37806 | 38993 | 1187 | + | tuf | WP_046329368.1 |
| GCF_001008165.2 | NZ_CP011452.2 | 54581 | 55768 | 1187 | + | tuf | WP_046329368.1 |
| GCF_014211955.1 | NZ_CP060052.1 | 39196 | 40383 | 1187 | + | tuf | WP_071464430.1 |
| GCF_001028705.1 | NZ_CP011770.1 | 56003 | 57190 | 1187 | + | tuf | WP_071464430.1 |
| GCF_014672695.1 | NZ_CP053923.1 | 1556726 | 1557913 | 1187 | + | tuf | WP_114639423.1 |
| GCF_014672695.1 | NZ_CP053923.1 | 1573474 | 1574661 | 1187 | + | tuf | WP_114639423.1 |
| GCF_014926585.1 | NZ_CP045422.1 | 38339 | 39526 | 1187 | + | tuf | WP_215336684.1 |
| GCF_007859655.1 | NZ_CP042304.1 | 55142 | 56329 | 1187 | + | tuf | WP_215336684.1 |
| GCF_022429625.1 | NZ_CP092483.1 | 38255 | 39442 | 1187 | + | tuf | WP_112202573.1 |
| GCF_025452235.1 | NZ_CP104965.1 | 55057 | 56244 | 1187 | + | tuf | WP_112202573.1 |
| GCF_016698645.1 | NZ_CP068047.1 | 39521 | 40708 | 1187 | + | tuf | WP_173954841.1 |
| GCF_016698665.1 | NZ_CP068046.1 | 56327 | 57514 | 1187 | + | tuf | WP_173954841.1 |
| GCF_030177755.1 | NZ_CP126539.1 | 39391 | 40578 | 1187 | + | tuf | WP_068947683.1 |
| GCF_028816475.1 | NZ_CP118247.1 | 56208 | 57395 | 1187 | + | tuf | WP_068947683.1 |
| GCF_019504385.1 | NZ_CP080590.1 | 39534 | 40721 | 1187 | + | tuf | WP_215348176.1 |
| GCF_000018145.1 | NC_009952.1 | 56337 | 57524 | 1187 | + | tuf | WP_215348176.1 |
| GCF_000018145.1 | NC_009952.1 | 616671 | 617858 | 1187 | + | tuf | WP_102246059.1 |
| GCF_000012565.1 | NC_007354.1 | 633195 | 634382 | 1187 | + | tuf | WP_102246059.1 |
| GCF_000012565.1 | NC_007354.1 | 6474352 | 6475539 | 1187 | - | tuf | WP_130186564.1 |
| GCF_000013145.1 | NC_007799.1 | 6538334 | 6539521 | 1187 | - | tuf | WP_130186564.1 |
| GCF_000013145.1 | NC_007799.1 | 820791 | 821978 | 1187 | + | tuf | WP_161071440.1 |
| GCF_000632845.1 | NZ_CP007474.1 | 912467 | 913654 | 1187 | + | tuf | WP_161071440.1 |
| GCF_000632845.1 | NZ_CP007474.1 | 6050277 | 6051464 | 1187 | - | tuf | WP_107144103.1 |
| GCF_000508225.1 | NC_023063.1 | 6144082 | 6145269 | 1187 | - | tuf | WP_107144103.1 |
| GCF_000508225.1 | NC_023063.1 | 5351179 | 5352366 | 1187 | + | tuf | WP_277415161.1 |
| GCF_000026005.1 | NC_005295.2 | 5446017 | 5447204 | 1187 | + | tuf | WP_277415161.1 |
| GCF_000026005.1 | NC_005295.2 | 4168098 | 4169285 | 1187 | + | tuf | WP_158206780.1 |
| GCF_019203965.1 | NZ_CP076448.1 | 4252352 | 4253539 | 1187 | + | tuf | WP_158206780.1 |
| GCF_019203965.1 | NZ_CP076448.1 | 2638168 | 2639355 | 1187 | - | tuf | WP_130186564.1 |
| GCF_000730245.1 | NZ_CP004403.1 | 2701119 | 2702306 | 1187 | - | tuf | WP_130186564.1 |
| GCF_000730245.1 | NZ_CP004403.1 | 4138846 | 4140033 | 1187 | + | tuf | WP_134386401.1 |

| | | | | | | | |
|---|---|---|---|---|---|---|---|
| GCF_000697965.2 | NZ_CP015880.1 | 4204898 | 4206085 | 1187 | + | tuf | WP_134386401.1 |
| GCF_000697965.2 | NZ_CP015880.1 | 5334070 | 5335257 | 1187 | + | tuf | WP_130186564.1 |
| GCF_017488845.2 | NZ_CP083370.1 | 5398533 | 5399720 | 1187 | + | tuf | WP_130186564.1 |
| GCF_017488845.2 | NZ_CP083370.1 | 464728 | 465915 | 1187 | + | tuf | WP_102040831.1 |
| GCF_014672835.1 | NZ_CP060244.1 | 481258 | 482445 | 1187 | + | tuf | WP_102040831.1 |
| GCF_023823125.1 | NZ_CP090949.1 | 594429 | 595616 | 1187 | + | tuf | WP_106845249.1 |
| GCF_003355455.1 | NZ_CP031357.1 | 705503 | 706690 | 1187 | + | tuf | WP_106845249.1 |
| GCF_001719165.1 | NZ_CP017057.1 | 332610 | 333797 | 1187 | + | tuf | WP_226442088.1 |
| GCF_013260645.1 | NZ_CP053921.1 | 349177 | 350364 | 1187 | + | tuf | WP_226442088.1 |
| GCF_001663175.1 | NZ_CP016033.1 | 3381295 | 3382500 | 1205 | - | tuf | WP_021197867.1 |
| GCF_000812665.2 | NZ_CP014327.1 | 3414873 | 3416078 | 1205 | - | tuf | WP_021197867.1 |
| GCF_000812665.2 | NZ_CP014327.1 | 136257 | 137444 | 1187 | - | tuf | WP_045785052.1 |
| GCF_018279165.1 | NZ_CP047289.1 | 169741 | 170928 | 1187 | - | tuf | WP_045785052.1 |
| GCF_018279165.1 | NZ_CP047289.1 | 382718 | 383905 | 1187 | + | tuf | WP_011002918.1 |
| GCF_007197755.1 | NZ_CP041636.1 | 414012 | 415199 | 1187 | + | tuf | WP_011002918.1 |
| GCF_007197755.1 | NZ_CP041636.1 | 1579409 | 1580596 | 1187 | - | tuf | WP_013204939.1 |
| GCF_030407165.1 | NZ_CP129111.1 | 1610694 | 1611881 | 1187 | - | tuf | WP_013204939.1 |
| GCF_030407165.1 | NZ_CP129111.1 | 363928 | 365115 | 1187 | + | tuf | WP_013211201.1 |
| GCF_006542395.2 | NZ_CP038231.1 | 395187 | 396374 | 1187 | + | tuf | WP_013211201.1 |
| GCF_041429785.1 | NZ_CP163423.1 | 2957841 | 2959046 | 1205 | - | tuf | WP_102067425.1 |
| GCF_041429785.1 | NZ_CP163423.1 | 2991367 | 2992554 | 1187 | - | tuf | WP_015855767.1 |
| GCF_001620265.1 | NZ_CP012661.1 | 332450 | 333637 | 1187 | + | tuf | WP_013899627.1 |
| GCF_001620265.1 | NZ_CP012661.1 | 3817343 | 3818530 | 1187 | - | tuf | WP_013899627.1 |
| GCF_029953595.1 | NZ_CP124535.1 | 645007 | 646194 | 1187 | - | tuf | WP_075586132.1 |
| GCF_029953595.1 | NZ_CP124535.1 | 3859807 | 3860994 | 1187 | - | tuf | WP_075585500.1 |
| GCF_018798885.1 | NZ_CP076361.1 | 4341492 | 4342679 | 1187 | - | tuf | WP_142813687.1 |
| GCF_018798885.1 | NZ_CP076361.1 | 4457277 | 4458464 | 1187 | - | tuf | WP_142813687.1 |
| GCF_003952725.1 | NZ_CP032509.1 | 4009647 | 4010834 | 1187 | - | tuf | WP_011465866.1 |
| GCF_003952725.1 | NZ_CP032509.1 | 4236862 | 4238049 | 1187 | - | tuf | WP_011466060.1 |
| GCF_028463845.1 | NZ_CP116805.1 | 409044 | 410231 | 1187 | + | tuf | WP_076196153.1 |
| GCF_028463845.1 | NZ_CP116805.1 | 783754 | 784941 | 1187 | - | tuf | WP_076196153.1 |

| | | | | | | | |
|---|---|---|---|---|---|---|---|
| GCF_000021325.1 | NC_011365.1 | 151574 | 152761 | 1187 | - | tuf | WP_313867780.1 |
| GCF_002005485.1 | NZ_CP014689.1 | 325263 | 326450 | 1187 | - | tuf | WP_313867780.1 |
| GCF_000583855.1 | NZ_CP004373.1 | 308424 | 309611 | 1187 | + | tuf | WP_076069245.1 |
| GCF_016757795.1 | NZ_CP068419.1 | 489748 | 490935 | 1187 | + | tuf | WP_076069245.1 |
| GCF_041429775.1 | NZ_CP163421.1 | 292322 | 293509 | 1187 | + | tuf | WP_142817246.1 |
| GCF_041429775.1 | NZ_CP163421.1 | 4191468 | 4192655 | 1187 | - | tuf | WP_142817246.1 |
| GCF_004000605.1 | NZ_CP018911.1 | 3431116 | 3432303 | 1187 | - | tuf | WP_024902012.1 |
| GCF_004000605.1 | NZ_CP018911.1 | 3453267 | 3454454 | 1187 | - | tuf | WP_024902012.1 |
| GCF_001889445.1 | NZ_CP018193.1 | 689416 | 690603 | 1187 | + | tuf | WP_261758742.1 |
| GCF_004328555.1 | NZ_CP035510.1 | 707490 | 708677 | 1187 | + | tuf | WP_261758742.1 |
| GCF_004328555.1 | NZ_CP035510.1 | 626939 | 628126 | 1187 | + | tuf | WP_058933581.1 |
| GCF_001611975.1 | NZ_CP014525.1 | 645484 | 646671 | 1187 | + | tuf | WP_058933581.1 |
| GCF_001611975.1 | NZ_CP014525.1 | 5019388 | 5020575 | 1187 | - | tuf | WP_200222282.1 |
| GCF_005144965.1 | NZ_CP039704.1 | 5037102 | 5038289 | 1187 | - | tuf | WP_200222282.1 |
| GCF_900231165.1 | NZ_LT960614.1 | 2568383 | 2569570 | 1187 | - | tuf | WP_202635128.1 |
| GCF_900231165.1 | NZ_LT960614.1 | 2584721 | 2585908 | 1187 | - | tuf | WP_202635128.1 |
| GCF_000023785.1 | NC_012982.1 | 2219549 | 2220736 | 1187 | - | tuf | WP_045536781.1 |
| GCF_000154705.2 | NZ_CM002917.1 | 2312483 | 2313670 | 1187 | - | tuf | WP_045538050.1 |
| GCF_000154705.2 | NZ_CM002917.1 | 2183467 | 2184654 | 1187 | - | tuf | WP_045532507.1 |
| GCF_008728835.1 | NZ_CP042582.1 | 2278154 | 2279341 | 1187 | - | tuf | WP_045532620.1 |
| GCF_008728835.1 | NZ_CP042582.1 | 705771 | 706958 | 1187 | + | tuf | WP_237248073.1 |
| GCF_008728855.1 | NZ_CP042906.1 | 722046 | 723233 | 1187 | + | tuf | WP_237248073.1 |
| GCF_008728855.1 | NZ_CP042906.1 | 753089 | 754276 | 1187 | + | tuf | WP_013028891.1 |
| GCF_028748065.1 | NZ_CP118166.1 | 769424 | 770611 | 1187 | + | tuf | WP_013028891.1 |
| GCF_046736345.1 | NZ_CP157210.1 | 3104206 | 3105393 | 1187 | + | tuf | WP_283148192.1 |
| GCF_000143145.1 | NC_014313.1 | 3120899 | 3122086 | 1187 | + | tuf | WP_283148192.1 |
| GCF_000503895.1 | NC_022997.1 | 2303167 | 2304348 | 1181 | + | tuf | WP_002642354.1 |
| GCF_017795365.1 | NZ_CP051254.1 | 2340856 | 2342037 | 1181 | + | tuf | WP_002642323.1 |
| GCF_017795365.1 | NZ_CP051254.1 | 1252549 | 1253736 | 1187 | - | tuf | WP_106445868.1 |
| GCF_000013025.1 | NC_008358.1 | 1281639 | 1282826 | 1187 | - | tuf | WP_106445868.1 |
| GCF_000013025.1 | NC_008358.1 | 295515 | 296702 | 1187 | + | tuf | WP_025329891.1 |

| | | | | | | | |
|---|---|---|---|---|---|---|---|
| GCF_003568845.1 | NZ_CP030126.1 | 312243 | 313430 | 1187 | + | tuf | WP_025329891.1 |
| GCF_003568845.1 | NZ_CP030126.1 | 1197220 | 1198407 | 1187 | + | tuf | WP_251972229.1 |
| GCF_029853695.1 | NZ_CP122537.1 | 1213593 | 1214780 | 1187 | - | tuf | WP_251972239.1 |
| GCF_029853695.1 | NZ_CP122537.1 | 133722 | 134909 | 1187 | + | tuf | WP_149502220.1 |
| GCF_015482775.1 | NZ_CP058214.1 | 3390745 | 3391932 | 1187 | + | tuf | WP_149502220.1 |
| GCF_015482775.1 | NZ_CP058214.1 | 620332 | 621519 | 1187 | + | tuf | WP_009206587.1 |
| GCF_002117445.1 | NZ_CP019937.1 | 636855 | 638042 | 1187 | + | tuf | WP_009206587.1 |
| GCF_002117445.1 | NZ_CP019937.1 | 524711 | 525898 | 1187 | + | tuf | WP_126444092.1 |
| GCF_001399515.1 | NZ_CP012908.1 | 541047 | 542234 | 1187 | + | tuf | WP_126444092.1 |
| GCF_001399515.1 | NZ_CP012908.1 | 547370 | 548557 | 1187 | + | tuf | WP_131110469.1 |
| GCF_043995075.1 | NZ_CP171836.1 | 563708 | 564895 | 1187 | + | tuf | WP_131110469.1 |
| GCF_002173515.1 | NZ_CP021467.1 | 546448 | 547635 | 1187 | + | tuf | WP_162083851.1 |
| GCF_000182745.2 | NC_016027.1 | 562842 | 564029 | 1187 | + | tuf | WP_162083851.1 |
| GCF_002009295.1 | NZ_CP019875.1 | 672147 | 673334 | 1187 | + | tuf | WP_147075184.1 |
| GCF_011611525.1 | NZ_CP050139.1 | 688685 | 689872 | 1187 | + | tuf | WP_147075184.1 |
| GCF_004348195.1 | NZ_CP036404.1 | 545021 | 546208 | 1187 | + | tuf | WP_173059960.1 |
| GCF_040581405.1 | NZ_CP137157.1 | 561576 | 562763 | 1187 | + | tuf | WP_173059960.1 |
| GCF_004006375.1 | NZ_CP025269.1 | 663742 | 664929 | 1187 | + | tuf | WP_041097028.1 |
| GCF_015240255.1 | NZ_CP061205.1 | 680138 | 681325 | 1187 | + | tuf | WP_041097028.1 |
| GCF_015240255.1 | NZ_CP061205.1 | 404116 | 405303 | 1187 | + | tuf | WP_126458950.1 |
| GCF_001787355.1 | NZ_CP014681.1 | 420535 | 421722 | 1187 | + | tuf | WP_126458950.1 |
| GCF_047345005.1 | NZ_CP178864.1 | 1997953 | 1999140 | 1187 | + | tuf | WP_139688388.1 |
| GCF_047345005.1 | NZ_CP178864.1 | 2015509 | 2016696 | 1187 | + | tuf | WP_139688388.1 |
| GCF_006740765.1 | NZ_CP041159.1 | 305075 | 306262 | 1187 | + | tuf | WP_120176064.1 |
| GCF_006740765.1 | NZ_CP041159.1 | 322516 | 323703 | 1187 | + | tuf | WP_120176064.1 |
| GCF_025139665.1 | NZ_CP081070.1 | 2513321 | 2514508 | 1187 | - | tuf | WP_016475357.1 |
| GCF_025139665.1 | NZ_CP081070.1 | 2530770 | 2531957 | 1187 | - | tuf | WP_016475357.1 |
| GCF_000511355.1 | NC_023135.1 | 1596227 | 1597414 | 1187 | - | tuf | WP_013521790.1 |
| GCF_000511355.1 | NC_023135.1 | 1628011 | 1629198 | 1187 | - | tuf | WP_013521790.1 |
| GCF_025857195.1 | NZ_CP109787.1 | 1661906 | 1663093 | 1187 | - | tuf | WP_013521790.1 |
| GCF_025857195.1 | NZ_CP109787.1 | 1693806 | 1694993 | 1187 | - | tuf | WP_013521790.1 |

| | | | | | | | |
|---|---|---|---|---|---|---|---|
| GCF_001543305.1 | NZ_CP010522.1 | 6008547 | 6009935 | 1388 | + | tuf | WP_306394437.1 |
| GCF_001543305.1 | NZ_CP010522.1 | 6012831 | 6014018 | 1187 | + | tuf | WP_109345418.1 |
| GCF_013201075.1 | NZ_CP053708.1 | 1613816 | 1615003 | 1187 | - | tuf | WP_259448231.1 |
| GCF_034371605.1 | NZ_CP139691.1 | 1714789 | 1715976 | 1187 | - | tuf | WP_259448231.1 |
| GCF_034371605.1 | NZ_CP139691.1 | 2809910 | 2811097 | 1187 | + | tuf | WP_143898650.1 |
| GCF_024206795.1 | NZ_CP072994.1 | 2864044 | 2865231 | 1187 | + | tuf | WP_143898650.1 |
| GCF_024206795.1 | NZ_CP072994.1 | 3644102 | 3645289 | 1187 | - | tuf | WP_004298398.1 |
| GCF_005849285.1 | NZ_CP040517.1 | 3660748 | 3661935 | 1187 | - | tuf | WP_004298398.1 |
| GCF_005849285.1 | NZ_CP040517.1 | 2912777 | 2913964 | 1187 | - | tuf | WP_107221693.1 |
| GCF_000014865.1 | NC_008576.1 | 2929356 | 2930543 | 1187 | - | tuf | WP_107221693.1 |
| GCF_000014865.1 | NC_008576.1 | 3828711 | 3829898 | 1187 | - | tuf | WP_418647214.1 |
| GCF_000513295.1 | NC_023065.1 | 3845322 | 3846509 | 1187 | - | tuf | WP_418647214.1 |
| GCF_000513295.1 | NC_023065.1 | 1873708 | 1874895 | 1187 | + | tuf | WP_075148051.1 |
| GCF_021611515.1 | NZ_CP090960.1 | 1890273 | 1891460 | 1187 | + | tuf | WP_075148051.1 |
| GCF_021611515.1 | NZ_CP090960.1 | 2463185 | 2464372 | 1187 | - | tuf | WP_004258139.1 |
| GCF_017161425.1 | NZ_CP071057.1 | 2479771 | 2480958 | 1187 | - | tuf | WP_004258139.1 |
| GCF_017161425.1 | NZ_CP071057.1 | 475147 | 476334 | 1187 | + | tuf | WP_324780245.1 |
| GCF_003433515.1 | NZ_CP021330.1 | 491549 | 492736 | 1187 | + | tuf | WP_324780245.1 |
| GCF_003433515.1 | NZ_CP021330.1 | 3343103 | 3344290 | 1187 | - | tuf | WP_013107281.1 |
| GCF_000960975.1 | NZ_CP010803.1 | 3628470 | 3629657 | 1187 | - | tuf | WP_013107281.1 |
| GCF_000960975.1 | NZ_CP010803.1 | 1477398 | 1478585 | 1187 | - | tuf | WP_079417354.1 |
| GCF_002043005.1 | NZ_CP020330.1 | 1659789 | 1660976 | 1187 | - | tuf | WP_079417354.1 |
| GCF_002043005.1 | NZ_CP020330.1 | 1756715 | 1757902 | 1187 | - | tuf | WP_085228323.1 |
| GCF_001686985.1 | NZ_CP015318.1 | 1789427 | 1790614 | 1187 | - | tuf | WP_085228323.1 |
| GCF_001686985.1 | NZ_CP015318.1 | 4401896 | 4403083 | 1187 | - | tuf | WP_102645092.1 |
| GCF_000230995.2 | NC_019973.1 | 4421072 | 4422259 | 1187 | - | tuf | WP_102645092.1 |
| GCF_000230995.2 | NC_019973.1 | 1576279 | 1577466 | 1187 | + | tuf | WP_066594855.1 |
| GCF_013170725.1 | NZ_CP033361.1 | 1592986 | 1594173 | 1187 | + | tuf | WP_066594855.1 |
| GCF_013170725.1 | NZ_CP033361.1 | 665978 | 667165 | 1187 | + | tuf | WP_309482688.1 |
| GCF_034421895.1 | NZ_CP139858.1 | 855387 | 856574 | 1187 | + | tuf | WP_309482688.1 |
| GCF_034421895.1 | NZ_CP139858.1 | 1396262 | 1397452 | 1190 | - | tuf | WP_126126972.1 |

| | | | | | | | |
|---|---|---|---|---|---|---|---|
| GCF_012913625.1 | NZ_CP051772.1 | 2620262 | 2621452 | 1190 | + | tuf | WP_126126972.1 |
| GCF_012913625.1 | NZ_CP051772.1 | 2475136 | 2476323 | 1187 | - | tuf | WP_244802573.1 |
| GCF_013170785.1 | NZ_CP033507.1 | 2491948 | 2493135 | 1187 | - | tuf | WP_244802573.1 |
| GCF_013170785.1 | NZ_CP033507.1 | 2807727 | 2808914 | 1187 | - | tuf | WP_244785051.1 |
| GCF_031656215.1 | NZ_CP134228.1 | 2824633 | 2825820 | 1187 | - | tuf | WP_244785051.1 |
| GCF_031656215.1 | NZ_CP134228.1 | 3266991 | 3268178 | 1187 | - | tuf | WP_171164371.1 |
| GCF_030272285.1 | NZ_CP088151.1 | 3283206 | 3284393 | 1187 | - | tuf | WP_171164393.1 |
| GCF_030272285.1 | NZ_CP088151.1 | 3886765 | 3887952 | 1187 | - | tuf | WP_171094875.1 |
| GCF_024707545.1 | NZ_CP062229.1 | 3903110 | 3904297 | 1187 | - | tuf | WP_171094875.1 |
| GCF_024707545.1 | NZ_CP062229.1 | 638225 | 639415 | 1190 | + | tuf | WP_012745744.1 |
| GCF_000176035.2 | NC_015675.1 | 5121792 | 5122982 | 1190 | - | tuf | WP_012745744.1 |
| GCF_000176035.2 | NC_015675.1 | 1404246 | 1405433 | 1187 | + | tuf | WP_011809041.1 |
| GCF_044501185.1 | NZ_CP162530.1 | 2557928 | 2559115 | 1187 | + | tuf | WP_011809041.1 |
| GCF_044501185.1 | NZ_CP162530.1 | 842213 | 843400 | 1187 | + | tuf | WP_089415880.1 |
| GCF_003058325.1 | NZ_CP028843.1 | 1319533 | 1320720 | 1187 | + | tuf | WP_089415880.1 |
| GCF_003058325.1 | NZ_CP028843.1 | 2371176 | 2372363 | 1187 | - | tuf | WP_058305387.1 |
| GCF_003173715.1 | NZ_CP029550.1 | 2387912 | 2389099 | 1187 | - | tuf | WP_058305387.1 |
| GCF_003173715.1 | NZ_CP029550.1 | 105776 | 106957 | 1181 | - | tuf | WP_249442883.1 |
| GCF_025813715.1 | NZ_CP107532.1 | 285967 | 287148 | 1181 | + | tuf | WP_249442883.1 |
| GCF_025813715.1 | NZ_CP107532.1 | 2566136 | 2567323 | 1187 | - | tuf | WP_160552150.1 |
| GCF_017347565.1 | NZ_AP024145.1 | 3138615 | 3139802 | 1187 | + | tuf | WP_160552150.1 |
| GCF_017347565.1 | NZ_AP024145.1 | 272172 | 273356 | 1184 | + | tuf | WP_211425379.1 |
| GCF_000022085.1 | NC_011894.1 | 1772025 | 1773209 | 1184 | - | tuf | WP_211425379.1 |
| GCF_000022085.1 | NC_011894.1 | 282683 | 283867 | 1184 | + | tuf | WP_014099979.1 |
| GCF_022533465.1 | NZ_CP092848.1 | 1794280 | 1795464 | 1184 | - | tuf | WP_014099979.1 |
| GCF_022533465.1 | NZ_CP092848.1 | 431182 | 432366 | 1184 | + | tuf | WP_211429092.1 |
| GCF_000757795.1 | NZ_CP003811.1 | 776786 | 777970 | 1184 | + | tuf | WP_211429092.1 |
| GCF_000757795.1 | NZ_CP003811.1 | 514060 | 515256 | 1196 | - | tuf | WP_014432758.1 |
| GCF_001936175.1 | NZ_CP015367.1 | 1860548 | 1861744 | 1196 | + | tuf | WP_014432758.1 |
| GCF_001936175.1 | NZ_CP015367.1 | 1071183 | 1072391 | 1208 | - | tuf | WP_014453623.1 |
| GCF_003173735.1 | NZ_CP029551.1 | 1097137 | 1098345 | 1208 | - | tuf | WP_014453623.1 |

| | | | | | | | |
|---|---|---|---|---|---|---|---|
| GCF_003173735.1 | NZ_CP029551.1 | 1059010 | 1060197 | 1187 | + | tuf | WP_006928897.1 |
| GCF_000019725.1 | NC_010505.1 | 1074649 | 1075836 | 1187 | + | tuf | WP_006928897.1 |
| GCF_000019725.1 | NC_010505.1 | 451528 | 452715 | 1187 | - | tuf | WP_023791762.1 |
| GCF_003173755.1 | NZ_CP029553.1 | 349970 | 351157 | 1187 | - | tuf | WP_115585485.1 |
| GCF_003173755.1 | NZ_CP029553.1 | 1220773 | 1221972 | 1199 | + | tuf | WP_099325874.1 |
| GCF_045689225.1 | NZ_CP136862.1 | 1236629 | 1237828 | 1199 | + | tuf | WP_099325874.1 |
| GCF_000828475.1 | NZ_AP014648.1 | 525367 | 526548 | 1181 | - | tuf | WP_044278825.1 |
| GCF_000021745.1 | NC_011666.1 | 461119 | 462315 | 1196 | - | tuf | WP_066060581.1 |
| GCF_038024855.1 | NZ_CP139089.1 | 476965 | 478161 | 1196 | - | tuf | WP_066060581.1 |
| GCF_027925445.1 | NZ_AP027149.1 | 1306760 | 1307947 | 1187 | + | tuf | WP_258877158.1 |
| GCF_027925445.1 | NZ_AP027149.1 | 1104732 | 1105913 | 1181 | - | tuf | WP_312031116.1 |
| GCF_040687965.1 | NZ_CP156084.1 | 470525 | 471718 | 1193 | + | tuf | WP_412732970.1 |
| GCF_040687965.1 | NZ_CP156084.1 | 381456 | 382637 | 1181 | - | tuf | WP_138078991.1 |
| GCF_004802635.2 | NZ_CP046052.1 | 364933 | 366114 | 1181 | - | tuf | WP_039327427.1 |
| GCF_004802635.2 | NZ_CP046052.1 | 1004056 | 1005237 | 1181 | - | tuf | WP_015642185.1 |
| GCF_027925385.1 | NZ_AP027142.1 | 444134 | 445315 | 1181 | - | tuf | WP_376753648.1 |
| GCF_027925385.1 | NZ_AP027142.1 | 2479779 | 2480975 | 1196 | + | tuf | WP_195172862.1 |
| GCF_027571405.1 | NZ_CP092968.1 | 5363289 | 5364479 | 1190 | + | tuf | WP_151969663.1 |
| GCF_027571405.1 | NZ_CP092968.1 | 648580 | 649770 | 1190 | - | tuf | WP_372368593.1 |
| GCF_040269895.1 | NZ_CP158568.1 | 741208 | 742392 | 1184 | - | tuf | WP_119985113.1 |
| GCF_040269895.1 | NZ_CP158568.1 | 1249075 | 1250259 | 1184 | + | tuf | WP_224070960.1 |
| GCF_000083545.1 | NC_012988.1 | 5942450 | 5943634 | 1184 | + | tuf | WP_162334157.1 |
| GCF_000083545.1 | NC_012988.1 | 2386267 | 2387451 | 1184 | + | tuf | WP_098193879.1 |
| GCF_000019945.1 | NC_010725.1 | 6266771 | 6267955 | 1184 | - | tuf | WP_126246845.1 |
| GCF_000019945.1 | NC_010725.1 | 309681 | 310865 | 1184 | + | tuf | WP_264281769.1 |
| GCF_004135935.1 | NZ_CP025086.1 | 5338132 | 5339316 | 1184 | - | tuf | WP_078667074.1 |
| GCF_034627025.1 | NZ_CP141048.1 | 5020207 | 5021391 | 1184 | - | tuf | WP_095838008.1 |
| GCF_034627025.1 | NZ_CP141048.1 | 2044162 | 2045346 | 1184 | + | tuf | WP_012789372.1 |
| GCF_002741015.1 | NZ_CP016616.1 | 4449984 | 4451168 | 1184 | + | tuf | WP_341834980.1 |
| GCF_002741015.1 | NZ_CP016616.1 | 6603657 | 6604841 | 1184 | + | tuf | WP_072360731.1 |
| GCF_013307435.2 | NZ_CP102845.1 | 1676649 | 1677833 | 1184 | + | tuf | WP_224019020.1 |

| | | | | | | | |
|---|---|---|---|---|---|---|---|
| GCF_013307435.2 | NZ_CP102845.1 | 2394229 | 2395413 | 1184 | + | tuf | WP_229759990.1 |
| GCF_050154705.1 | NZ_CP189561.1 | 809726 | 810910 | 1184 | + | tuf | WP_076381965.1 |
| GCF_050154705.1 | NZ_CP189561.1 | 2453598 | 2454782 | 1184 | - | tuf | WP_214458176.1 |
| GCF_002005465.1 | NZ_CP014691.1 | 1847899 | 1849083 | 1184 | - | tuf | WP_214447726.1 |
| GCF_024299005.1 | NZ_CP089285.1 | 4969893 | 4971077 | 1184 | + | tuf | WP_146791566.1 |
| GCF_024299005.1 | NZ_CP089285.1 | 5209757 | 5210941 | 1184 | + | tuf | WP_066407891.1 |
| GCF_006542335.1 | NZ_CP032485.1 | 1479182 | 1480369 | 1187 | + | tuf | WP_182805217.1 |
| GCF_000731315.1 | NZ_HG938353.1 | 1493595 | 1494779 | 1184 | + | tuf | WP_409966752.1 |
| GCF_000731315.1 | NZ_HG938353.1 | 3820691 | 3821875 | 1184 | - | tuf | WP_067758148.1 |
| GCF_029854435.1 | NZ_CP123000.1 | 2956202 | 2957386 | 1184 | - | tuf | WP_346237100.1 |
| GCF_029854435.1 | NZ_CP123000.1 | 2434426 | 2435610 | 1184 | - | tuf | WP_008585570.1 |
| GCF_009856525.1 | NZ_CP047224.1 | 1879202 | 1880386 | 1184 | - | tuf | WP_114789363.1 |
| GCF_000632985.1 | NZ_CP007481.1 | 6103484 | 6104668 | 1184 | - | tuf | WP_014221094.1 |
| GCF_000022525.1 | NC_013009.1 | 871732 | 872916 | 1184 | - | tuf | WP_147188228.1 |
| GCF_000013165.1 | NC_007798.1 | 4363537 | 4364721 | 1184 | - | tuf | WP_119051310.1 |
| GCF_019623905.1 | NZ_CP069370.1 | 6596872 | 6598056 | 1184 | - | tuf | WP_315816827.1 |
| GCF_019623905.1 | NZ_CP069370.1 | 4097569 | 4098753 | 1184 | - | tuf | WP_114235055.1 |
| GCF_047114695.1 | NZ_CP112885.1 | 3733765 | 3734949 | 1184 | - | tuf | WP_131331042.1 |
| GCF_047114695.1 | NZ_CP112885.1 | 767965 | 769146 | 1181 | + | tuf | WP_066482532.1 |
| GCF_038719015.1 | NZ_CP152276.1 | 754070 | 755251 | 1181 | + | tuf | WP_011175145.1 |
| GCF_024662015.1 | NZ_CP102480.1 | 2028028 | 2029209 | 1181 | - | tuf | WP_032124106.1 |
| GCF_024662015.1 | NZ_CP102480.1 | 815581 | 816855 | 1274 | + | tuf | WP_215216594.1 |
| GCF_046773475.1 | NZ_CP177239.1 | 506727 | 507905 | 1178 | + | tuf | WP_194844923.1 |
| GCF_046773475.1 | NZ_CP177239.1 | 272285 | 273466 | 1181 | - | tuf | WP_006343364.1 |
| GCF_019891395.1 | NZ_CP078143.1 | 462480 | 463661 | 1181 | + | tuf | WP_131743633.1 |
| GCF_019891395.1 | NZ_CP078143.1 | 806454 | 807635 | 1181 | - | tuf | WP_011006655.1 |
| GCF_007922615.2 | NZ_CP042301.2 | 161133 | 162314 | 1181 | - | tuf | WP_213241254.1 |
| GCF_007922615.2 | NZ_CP042301.2 | 359800 | 360981 | 1181 | + | tuf | WP_011457865.1 |
| GCF_036621415.1 | NZ_CP030941.1 | 85701 | 86882 | 1181 | - | tuf | WP_021828487.1 |
| GCF_036621415.1 | NZ_CP030941.1 | 712354 | 713535 | 1181 | - | tuf | WP_010904371.1 |
| GCF_000013885.1 | NC_007964.1 | 1068756 | 1069937 | 1181 | - | tuf | WP_013713068.1 |

| | | | | | | |
|---|---|---|---|---|---|---|
| GCF_000012725.1 | NC_007406.1 | 87268 | 88449 | 1181 + | tuf | WP_010882724.1 |
| GCF_000730165.1 | NZ_CP022110.1 | 790013 | 791194 | 1181 - | tuf | WP_117274461.1 |
| GCF_000730165.1 | NZ_CP022110.1 | 804559 | 805740 | 1181 - | tuf | WP_006343364.1 |
| GCF_002868735.1 | NZ_CP025611.1 | 66349 | 67530 | 1181 + | tuf | WP_108896231.1 |
| GCF_002868735.1 | NZ_CP025611.1 | 388432 | 389613 | 1181 - | tuf | WP_035406303.1 |
| GCF_014843995.1 | NZ_CP062147.1 | 361983 | 363164 | 1181 - | tufA | NP_219827.1 |
| GCF_000013325.1 | NC_007794.1 | 146948 | 148129 | 1181 + | tuf | WP_213318203.1 |
| GCF_018417475.1 | NZ_CP054856.1 | 246591 | 247772 | 1181 + | tuf | WP_213318203.1 |
| GCF_007954425.1 | NZ_CP042345.1 | 2198959 | 2200143 | 1184 - | tuf | WP_006340068.1 |
| GCF_028607105.1 | NZ_CP117417.1 | 1585140 | 1586321 | 1181 - | tuf | WP_013943862.1 |
| GCF_021227995.1 | NZ_CP089301.1 | 631037 | 632218 | 1181 + | tuf | WP_013181678.1 |
| GCF_037076535.1 | NZ_AP029021.1 | 2679081 | 2680259 | 1178 - | tuf | WP_069811009.1 |
| GCF_000767465.1 | NZ_CP009291.1 | 188146 | 189324 | 1178 + | tuf | WP_012501437.1 |
| GCF_001742225.1 | NZ_CP017075.1 | 2064827 | 2066005 | 1178 - | tuf | WP_010933844.1 |
| GCF_014334095.1 | NZ_CP060495.1 | 2457517 | 2458695 | 1178 - | tuf | WP_012467122.1 |
| GCF_965223445.1 | NZ_OZ244994.1 | 2798987 | 2800165 | 1178 - | tuf | WP_011746195.1 |
| GCF_965223445.1 | NZ_OZ244994.1 | 1292738 | 1293916 | 1178 + | tuf | WP_012499634.1 |
| GCF_002278035.1 | NZ_CP022604.1 | 186385 | 187563 | 1178 + | tuf | WP_011356944.1 |
| GCF_002278035.1 | NZ_CP022604.1 | 286197 | 287375 | 1178 + | tuf | WP_012507116.1 |
| GCF_000155675.2 | NC_020911.1 | 2249067 | 2250245 | 1178 - | tuf | WP_012506608.1 |
| GCF_000155735.2 | NC_020908.1 | 1636902 | 1638104 | 1202 + | tuf | WP_341467502.1 |
| GCF_000155735.2 | NC_020908.1 | 1648122 | 1649324 | 1202 + | tuf | WP_341467502.1 |
| GCF_001187845.1 | NZ_CP012160.1 | 1812590 | 1813792 | 1202 - | tuf | WP_015940243.1 |
| GCF_001187845.1 | NZ_CP012160.1 | 3681588 | 3682790 | 1202 - | tuf | WP_015940243.1 |
| GCF_006542375.1 | NZ_CP038143.1 | 2978443 | 2979645 | 1202 + | tuf | WP_012258047.1 |
| GCF_013358405.1 | NZ_CP054836.1 | 3210940 | 3212142 | 1202 + | tuf | WP_012258047.1 |
| GCF_013358405.1 | NZ_CP054836.1 | 5226055 | 5227257 | 1202 - | tuf | WP_012122483.1 |
| GCF_900327255.1 | NZ_LS398552.1 | 5571914 | 5573116 | 1202 - | tuf | WP_012122768.1 |
| GCF_900327255.1 | NZ_LS398552.1 | 2140570 | 2141754 | 1184 - | tuf | WP_013506558.1 |
| GCF_002504165.1 | NZ_CP021404.1 | 668149 | 669345 | 1196 - | tuf | WP_016481989.1 |
| GCF_002504165.1 | NZ_CP021404.1 | 3372433 | 3373683 | 1250 + | tuf | WP_016484149.1 |

| | | | | | | | |
|---|---|---|---|---|---|---|---|
| GCF_030863885.1 | NZ_CP132956.1 | 976102 | 977292 | 1190 | + | tuf | WP_007717083.1 |
| GCF_027913215.1 | NZ_CP115784.1 | 508781 | 509965 | 1184 | + | tuf | WP_250278558.1 |
| GCF_027913215.1 | NZ_CP115784.1 | 3486872 | 3488062 | 1190 | - | tuf | WP_006443176.1 |
| GCF_027913035.1 | NZ_CP115775.1 | 4081337 | 4082527 | 1190 | - | tuf | WP_013274353.1 |
| GCF_027913035.1 | NZ_CP115775.1 | 743732 | 744922 | 1190 | - | tuf | WP_004606376.1 |
| GCF_028553725.1 | NZ_CP067124.1 | 2304780 | 2305970 | 1190 | - | tuf | WP_005608983.1 |
| GCF_028553725.1 | NZ_CP067124.1 | 2556139 | 2557329 | 1190 | - | tuf | WP_009243125.1 |
| GCF_000444995.1 | NC_022041.1 | 2030689 | 2031888 | 1199 | - | tuf | WP_284679850.1 |
| GCF_000444995.1 | NC_022041.1 | 2046378 | 2047577 | 1199 | - | tuf | WP_284679850.1 |
| GCF_900005615.1 | NZ_LN832559.1 | 775296 | 776495 | 1199 | + | tuf | WP_014253999.1 |
| GCF_900005615.1 | NZ_LN832559.1 | 2899395 | 2900594 | 1199 | - | tuf | WP_101303003.1 |
| GCF_002105555.1 | NZ_CP020612.1 | 3222531 | 3223730 | 1199 | + | tuf | WP_020457902.1 |
| GCF_002105555.1 | NZ_CP020612.1 | 2314667 | 2315857 | 1190 | - | tuf | WP_013362418.1 |
| GCF_004063735.1 | NZ_CP035090.1 | 2330334 | 2331524 | 1190 | - | tuf | WP_013362418.1 |
| GCF_004063735.1 | NZ_CP035090.1 | 714124 | 715314 | 1190 | + | tuf | WP_111889120.1 |
| GCF_021491915.1 | NZ_CP090836.1 | 3264321 | 3265511 | 1190 | - | tuf | WP_111889120.1 |
| GCF_021491915.1 | NZ_CP090836.1 | 731973 | 733163 | 1190 | + | tuf | WP_414149961.1 |
| GCF_028553785.1 | NZ_CP067136.1 | 3424776 | 3425966 | 1190 | - | tuf | WP_414149961.1 |
| GCF_028553785.1 | NZ_CP067136.1 | 666278 | 667468 | 1190 | + | tuf | WP_070369929.1 |
| GCF_002865605.1 | NZ_CP025583.1 | 2915722 | 2916912 | 1190 | - | tuf | WP_070372670.1 |
| GCF_002865605.1 | NZ_CP025583.1 | 1066213 | 1067403 | 1190 | + | tuf | WP_014355310.1 |
| GCF_009301415.1 | NZ_CP045072.1 | 3321968 | 3323158 | 1190 | - | tuf | WP_014355310.1 |
| GCF_009301415.1 | NZ_CP045072.1 | 163627 | 164820 | 1193 | + | tuf | WP_013277145.1 |
| GCF_004683865.2 | NZ_CP038439.1 | 178653 | 179846 | 1193 | + | tuf | WP_013277145.1 |
| GCF_004683865.2 | NZ_CP038439.1 | 874499 | 875698 | 1199 | + | tuf | WP_240983856.1 |
| GCF_019141545.1 | NZ_CP065892.1 | 357529 | 358728 | 1199 | + | tuf | WP_066536463.1 |
| GCF_019141545.1 | NZ_CP065892.1 | 351331 | 352515 | 1184 | + | tuf | WP_012741295.1 |
| GCF_016919705.1 | NZ_CP070368.1 | 1293423 | 1294613 | 1190 | - | tuf | WP_207298835.1 |
| GCF_016919705.1 | NZ_CP070368.1 | 1309340 | 1310530 | 1190 | - | tuf | WP_207298835.1 |
| GCF_008824185.1 | NZ_CP044426.1 | 1696917 | 1698116 | 1199 | - | tuf | WP_213168783.1 |
| GCF_008824185.1 | NZ_CP044426.1 | 1714162 | 1715361 | 1199 | - | tuf | WP_213168810.1 |

| | | | | | | | |
|---|---|---|---|---|---|---|---|
| GCF_028553805.1 | NZ_CP067140.1 | 4578584 | 4579774 | 1190 | - | tuf | WP_012065443.1 |
| GCF_028553805.1 | NZ_CP067140.1 | 4595000 | 4596190 | 1190 | - | tuf | WP_041721235.1 |
| GCF_012689545.1 | NZ_CP051542.1 | 541105 | 542295 | 1190 | + | tuf | WP_012158353.1 |
| GCF_012689545.1 | NZ_CP051542.1 | 557737 | 558927 | 1190 | + | tuf | WP_012158353.1 |
| GCF_027912835.1 | NZ_CP115768.1 | 2827454 | 2828644 | 1190 | - | tuf | WP_163067470.1 |
| GCF_027912835.1 | NZ_CP115768.1 | 2848546 | 2849736 | 1190 | - | tuf | WP_163067470.1 |
| GCF_028553745.1 | NZ_CP067129.1 | 2136718 | 2137908 | 1190 | - | tuf | WP_128746293.1 |
| GCF_028553745.1 | NZ_CP067129.1 | 2157701 | 2158891 | 1190 | - | tuf | WP_128746293.1 |
| GCF_028553765.1 | NZ_CP067134.1 | 2273586 | 2274776 | 1190 | + | tuf | WP_162362584.1 |
| GCF_028553765.1 | NZ_CP067134.1 | 2296416 | 2297606 | 1190 | + | tuf | WP_162362600.1 |
| GCF_003324675.1 | NZ_CP030918.1 | 417898 | 419097 | 1199 | + | tuf | WP_015738440.1 |
| GCF_003324675.1 | NZ_CP030918.1 | 1516980 | 1518179 | 1199 | - | tuf | WP_015739498.1 |
| GCF_002847305.1 | NZ_CP025408.1 | 540310 | 541509 | 1199 | + | tuf | WP_315570449.1 |
| GCF_002847305.1 | NZ_CP025408.1 | 553306 | 554496 | 1190 | + | tuf | WP_005344951.1 |
| GCF_029909795.1 | NZ_CP038197.1 | 5079173 | 5080363 | 1190 | - | tuf | WP_184091980.1 |
| GCF_029909795.1 | NZ_CP038197.1 | 403187 | 404377 | 1190 | + | tuf | WP_185257752.1 |
| GCF_002749495.1 | NZ_CP024422.1 | 874751 | 875941 | 1190 | - | tuf | WP_161836859.1 |
| GCF_002749495.1 | NZ_CP024422.1 | 4790427 | 4791617 | 1190 | - | tuf | WP_271713791.1 |
| GCF_002847445.1 | NZ_CP025430.1 | 3159049 | 3160239 | 1190 | - | tuf | WP_250228285.1 |
| GCF_002847445.1 | NZ_CP025430.1 | 1316369 | 1317559 | 1190 | - | tuf | WP_006568224.1 |
| GCF_016801025.1 | NZ_CP068983.1 | 375333 | 376523 | 1190 | + | tuf | WP_008393909.1 |
| GCF_008000755.1 | NZ_CP041690.1 | 3454786 | 3455976 | 1190 | - | tuf | WP_137330058.1 |
| GCF_003060865.1 | NZ_CP028918.1 | 2678842 | 2680032 | 1190 | - | tuf | WP_066052497.1 |
| GCF_003060865.1 | NZ_CP028918.1 | 2343809 | 2345008 | 1199 | + | tuf | WP_024729920.1 |
| GCF_000009985.1 | NC_007626.1 | 3517066 | 3518256 | 1190 | + | tuf | WP_127018070.1 |
| GCF_000009985.1 | NC_007626.1 | 3533905 | 3535095 | 1190 | + | tuf | WP_127018070.1 |
| GCF_029223885.1 | NZ_CP119409.1 | 1361295 | 1362506 | 1211 | - | tuf | WP_304583099.1 |
| GCF_029223885.1 | NZ_CP119409.1 | 2948036 | 2949247 | 1211 | - | tuf | WP_304583099.1 |
| GCF_029277445.1 | NZ_CP119566.1 | 1244091 | 1245302 | 1211 | - | tuf | WP_304588086.1 |
| GCF_029277445.1 | NZ_CP119566.1 | 2849662 | 2850873 | 1211 | - | tuf | WP_304588086.1 |
| GCF_004006175.1 | NZ_CP033219.1 | 3525296 | 3526492 | 1196 | + | tuf | WP_409968151.1 |

| | | | | | | | |
|---|---|---|---|---|---|---|---|
| GCF_004006175.1 | NZ_CP033219.1 | 287259 | 288461 | 1202 | + | tuf | WP_122013879.1 |
| GCF_030285785.1 | NZ_CP127247.1 | 3123767 | 3124969 | 1202 | - | tuf | WP_122013879.1 |
| GCF_030285785.1 | NZ_CP127247.1 | 309969 | 311159 | 1190 | + | tuf | WP_111917899.1 |
| GCF_013378075.1 | NZ_CP051131.1 | 376213 | 377403 | 1190 | + | tuf | WP_115624166.1 |
| GCF_017498565.1 | NZ_CP071794.1 | 2855866 | 2857056 | 1190 | - | tuf | WP_003022245.1 |
| GCF_012516475.1 | NZ_CP051217.1 | 196146 | 197336 | 1190 | + | tuf | WP_005947864.1 |
| GCF_020906275.1 | NZ_CP086727.1 | 261561 | 262751 | 1190 | + | tuf | WP_193735956.1 |
| GCF_001687545.1 | NZ_CP016545.1 | 3333691 | 3334881 | 1190 | - | tuf | WP_118509158.1 |
| GCF_006385135.1 | NZ_CP041025.1 | 3220982 | 3222172 | 1190 | - | tuf | WP_005423313.1 |
| GCF_006385135.1 | NZ_CP041025.1 | 3657258 | 3658448 | 1190 | - | tuf | WP_033139946.1 |
| GCF_005887635.2 | NZ_CP040818.1 | 3504905 | 3506095 | 1190 | + | tuf | WP_018596934.1 |
| GCF_005887635.2 | NZ_CP040818.1 | 3838142 | 3839332 | 1190 | - | tuf | WP_065543566.1 |
| GCF_000017565.1 | NC_009719.1 | 4092563 | 4093753 | 1190 | - | tuf | WP_019163091.1 |
| GCF_000017565.1 | NC_009719.1 | 3997541 | 3998728 | 1187 | - | tuf | WP_073388378.1 |
| GCF_000152825.2 | NC_014414.1 | 690119 | 691306 | 1187 | + | tuf | WP_071175402.1 |
| GCF_001028625.1 | NZ_CP011805.1 | 2886140 | 2887327 | 1187 | - | tuf | WP_013281750.1 |
| GCF_034375205.1 | NZ_CP139719.1 | 2176300 | 2177499 | 1199 | - | tuf | WP_011026340.1 |
| GCF_025854335.1 | NZ_CP107716.1 | 2191779 | 2192978 | 1199 | - | tuf | WP_011026354.1 |
| GCF_025854335.1 | NZ_CP107716.1 | 3385630 | 3386814 | 1184 | + | tuf | WP_264842812.1 |
| GCF_013004005.1 | NZ_CP053086.1 | 236652 | 237848 | 1196 | + | tuf | WP_012301361.1 |
| GCF_013004005.1 | NZ_CP053086.1 | 255915 | 257114 | 1199 | + | tuf | WP_012301376.1 |
| GCF_037044555.1 | NZ_CP146275.1 | 1891034 | 1892224 | 1190 | - | tuf | WP_010166217.1 |
| GCF_037044555.1 | NZ_CP146275.1 | 4889037 | 4890239 | 1202 | + | tuf | WP_148136444.1 |
| GCF_016134045.1 | NZ_CP065915.1 | 4904832 | 4906034 | 1202 | + | tuf | WP_148136458.1 |
| GCF_016134045.1 | NZ_CP065915.1 | 2091431 | 2092621 | 1190 | - | tuf | WP_058991536.1 |
| GCF_030369675.1 | NZ_CP128477.1 | 2107238 | 2108428 | 1190 | - | tuf | WP_058991536.1 |
| GCF_030369675.1 | NZ_CP128477.1 | 2229773 | 2230972 | 1199 | + | tuf | WP_156204511.1 |
| GCF_005860795.2 | NZ_CP058350.1 | 2245997 | 2247196 | 1199 | + | tuf | WP_156204511.1 |
| GCF_005860795.2 | NZ_CP058350.1 | 606308 | 607507 | 1199 | - | tuf | WP_187036618.1 |
| GCF_000511385.1 | NC_023137.1 | 1687075 | 1688274 | 1199 | - | tuf | WP_212505990.1 |
| GCF_000511385.1 | NC_023137.1 | 1653950 | 1655149 | 1199 | - | tuf | WP_275844009.1 |

| | | | | | | | |
|---|---|---|---|---|---|---|---|
| GCF_002892185.1 | NZ_CP010650.1 | 1641405 | 1642604 | 1199 | - | tuf | WP_174193475.1 |
| GCF_002892185.1 | NZ_CP010650.1 | 2073887 | 2075086 | 1199 | - | tuf | WP_011345194.1 |
| GCF_000826835.2 | NZ_CP080275.1 | 2089332 | 2090531 | 1199 | - | tuf | WP_011345209.1 |
| GCF_000826835.2 | NZ_CP080275.1 | 3862194 | 3863384 | 1190 | - | tuf | WP_013658508.1 |
| GCF_001888185.1 | NZ_CP016364.1 | 2235664 | 2236854 | 1190 | - | tuf | WP_317854119.1 |
| GCF_001888185.1 | NZ_CP016364.1 | 2239611 | 2240804 | 1193 | + | tuf | WP_274939746.1 |
| GCF_018135625.1 | NZ_CP073078.1 | 953833 | 955044 | 1211 | - | tuf | WP_079546096.1 |
| GCF_018135625.1 | NZ_CP073078.1 | 2013259 | 2014470 | 1211 | + | tuf | WP_079546096.1 |
| GCF_003150835.1 | NZ_CP029479.1 | 1419685 | 1420893 | 1208 | - | tuf | WP_066523778.1 |
| GCF_003150835.1 | NZ_CP029479.1 | 2145659 | 2146867 | 1208 | - | tuf | WP_066523778.1 |
| GCF_000017265.1 | NC_011144.1 | 549842 | 551032 | 1190 | + | tuf | WP_316266153.1 |
| GCF_000017265.1 | NC_011144.1 | 88967 | 90157 | 1190 | + | tuf | WP_009887863.1 |
| GCF_005160265.1 | NZ_CP039865.1 | 106159 | 107349 | 1190 | + | tuf | WP_009887863.1 |
| GCF_005160265.1 | NZ_CP039865.1 | 3824338 | 3825528 | 1190 | - | tuf | WP_044826578.1 |
| GCF_003008515.1 | NZ_CP027668.1 | 3840936 | 3842126 | 1190 | - | tuf | WP_044826578.1 |
| GCF_003008515.1 | NZ_CP027668.1 | 3271930 | 3273120 | 1190 | - | tuf | WP_010966415.1 |
| GCF_005144885.1 | NZ_CP039690.1 | 3288139 | 3289329 | 1190 | - | tuf | WP_010966415.1 |
| GCF_005144885.1 | NZ_CP039690.1 | 630342 | 631532 | 1190 | + | tuf | WP_039629580.1 |
| GCF_002764115.1 | NZ_CP017940.1 | 646362 | 647552 | 1190 | + | tuf | WP_039629580.1 |
| GCF_002764115.1 | NZ_CP017940.1 | 2079197 | 2080387 | 1190 | - | tuf | WP_013240693.1 |
| GCF_011044155.1 | NZ_CP049056.1 | 2095132 | 2096322 | 1190 | - | tuf | WP_013240693.1 |
| GCF_011044155.1 | NZ_CP049056.1 | 261022 | 262212 | 1190 | + | tuf | WP_039312760.1 |
| GCF_018982885.2 | NZ_CP081848.1 | 276957 | 278147 | 1190 | + | tuf | WP_039312760.1 |
| GCF_000192745.1 | NC_015259.1 | 2663007 | 2664197 | 1190 | - | tuf | WP_044039390.1 |
| GCF_000192745.1 | NC_015259.1 | 2679849 | 2681039 | 1190 | - | tuf | WP_044039404.1 |
| GCF_016806125.1 | NZ_CP047166.1 | 3707977 | 3709167 | 1190 | - | tuf | WP_003357263.1 |
| GCF_016806125.1 | NZ_CP047166.1 | 3723874 | 3725064 | 1190 | - | tuf | WP_003357263.1 |
| GCF_011290545.1 | NZ_CP049811.1 | 3711547 | 3712737 | 1190 | - | tuf | WP_002582604.1 |
| GCF_011290545.1 | NZ_CP049811.1 | 3727510 | 3728700 | 1190 | - | tuf | WP_002582604.1 |
| GCF_015679265.1 | NZ_CP064942.1 | 3223079 | 3224269 | 1190 | - | tuf | WP_074845260.1 |
| GCF_015679265.1 | NZ_CP064942.1 | 3239112 | 3240302 | 1190 | - | tuf | WP_074845260.1 |

| | | | | | | | |
|---|---|---|---|---|---|---|---|
| GCF_003544895.1 | NZ_CP032125.1 | 4439829 | 4441019 | 1190 | - | tuf | WP_048030358.1 |
| GCF_003544895.1 | NZ_CP032125.1 | 4455749 | 4456939 | 1190 | - | tuf | WP_048030358.1 |
| GCF_028869115.1 | NZ_CP079800.1 | 4503034 | 4504224 | 1190 | - | tuf | WP_013291937.1 |
| GCF_028869115.1 | NZ_CP079800.1 | 4519096 | 4520286 | 1190 | - | tuf | WP_013291937.1 |
| GCF_018436245.1 | NZ_CP075361.1 | 226916 | 228106 | 1190 | + | tuf | WP_079481046.1 |
| GCF_018436245.1 | NZ_CP075361.1 | 242875 | 244065 | 1190 | + | tuf | WP_079481046.1 |
| GCF_003071405.1 | NZ_CP020470.1 | 2267773 | 2268963 | 1190 | - | tuf | WP_089866438.1 |
| GCF_003071405.1 | NZ_CP020470.1 | 2283743 | 2284933 | 1190 | - | tuf | WP_089866438.1 |
| GCF_003367395.1 | NZ_CP031417.1 | 173074 | 174264 | 1190 | + | tuf | WP_031276475.1 |
| GCF_003590145.2 | NZ_CP060436.1 | 188995 | 190185 | 1190 | + | tuf | WP_031276475.1 |
| GCF_003590145.2 | NZ_CP060436.1 | 5290208 | 5291398 | 1190 | - | tuf | WP_032079783.1 |
| GCF_017795125.1 | NZ_CP051248.1 | 5306129 | 5307319 | 1190 | - | tuf | WP_032079783.1 |
| GCF_017795125.1 | NZ_CP051248.1 | 362485 | 363675 | 1190 | - | tuf | WP_071614634.1 |
| GCF_017795185.1 | NZ_CP051251.1 | 378560 | 379750 | 1190 | - | tuf | WP_071614634.1 |
| GCF_014899185.1 | NZ_CP045201.1 | 820790 | 821980 | 1190 | + | tuf | WP_077836261.1 |
| GCF_014899185.1 | NZ_CP045201.1 | 836981 | 838171 | 1190 | + | tuf | WP_077836261.1 |
| GCF_902502825.2 | NZ_LR723670.1 | 248685 | 249875 | 1190 | + | tuf | WP_119969948.1 |
| GCF_902502825.2 | NZ_LR723670.1 | 264691 | 265881 | 1190 | + | tuf | WP_119969948.1 |
| GCF_005234135.1 | NZ_CP039964.1 | 4184896 | 4186086 | 1190 | - | tuf | WP_070964306.1 |
| GCF_005234135.1 | NZ_CP039965.1 | 4201418 | 4202608 | 1190 | - | tuf | WP_070964306.1 |
| GCF_002119765.1 | NZ_CP021112.1 | 981034 | 982224 | 1190 | - | tuf | WP_089969057.1 |
| GCF_013347125.1 | NZ_CP054599.1 | 997170 | 998360 | 1190 | - | tuf | WP_089969034.1 |
| GCF_013347125.1 | NZ_CP054599.1 | 190254 | 191444 | 1190 | + | tuf | WP_224035801.1 |
| GCF_018282095.1 | NZ_CP074126.1 | 206131 | 207321 | 1190 | + | tuf | WP_224035801.1 |
| GCF_018282095.1 | NZ_CP074126.1 | 3094249 | 3095439 | 1190 | - | tuf | WP_021800139.1 |
| GCF_003008555.2 | NZ_CP027665.1 | 3110162 | 3111352 | 1190 | - | tuf | WP_021800139.1 |
| GCF_003008555.2 | NZ_CP027665.1 | 233640 | 234830 | 1190 | + | tuf | WP_119864272.1 |
| GCF_012848855.1 | NZ_CP051630.1 | 249558 | 250748 | 1190 | + | tuf | WP_119864272.1 |
| GCF_012848855.1 | NZ_CP051630.1 | 195777 | 196967 | 1190 | + | tuf | WP_011988788.1 |
| GCF_007855645.1 | NZ_CP042261.1 | 211876 | 213066 | 1190 | + | tuf | WP_011988788.1 |
| GCF_007855645.1 | NZ_CP042261.1 | 4446427 | 4447617 | 1190 | - | tuf | WP_013240693.1 |

| | | | | | | |
|---|---|---|---|---|---|---|
| GCF_019711375.1 | NZ_CP081295.1 | 4462362 | 4463552 | 1190 - | tuf | WP_013240693.1 |
| GCF_008932225.1 | NZ_CP032228.1 | 808167 | 809360 | 1193 + | tuf | WP_128211256.1 |
| GCF_019711315.1 | NZ_CP081294.1 | 827852 | 829045 | 1193 + | tuf | WP_128211256.1 |
| GCF_033956045.1 | NZ_CP138853.1 | 624661 | 625839 | 1178 - | tuf | WP_003375253.1 |
| GCF_019711355.1 | NZ_CP081297.1 | 640701 | 641879 | 1178 - | tuf | WP_003375253.1 |
| GCF_004358425.1 | NZ_CP037948.1 | 3938935 | 3940125 | 1190 - | tuf | WP_003447962.1 |
| GCF_002795865.1 | NZ_CP024920.1 | 3954912 | 3956102 | 1190 - | tuf | WP_003447962.1 |
| GCF_015529805.1 | NZ_CP064654.1 | 585277 | 586467 | 1190 + | tuf | WP_003452162.1 |
| GCF_026168555.1 | NZ_CP092471.1 | 601288 | 602478 | 1190 + | tuf | WP_003452162.1 |
| GCF_002531755.2 | NZ_CP034998.1 | 201155 | 202345 | 1190 + | tuf | WP_022743536.1 |
| GCF_002531755.2 | NZ_CP034998.1 | 217072 | 218262 | 1190 + | tuf | WP_022743536.1 |
| GCF_024346605.1 | NZ_CP098803.1 | 176098 | 177288 | 1190 + | tuf | WP_015390326.1 |
| GCF_024346605.1 | NZ_CP098803.1 | 192007 | 193197 | 1190 + | tuf | WP_015390326.1 |
| GCF_017357245.1 | NZ_CP071612.1 | 4616887 | 4618077 | 1190 + | tuf | WP_046066033.1 |
| GCF_017357245.1 | NZ_CP071612.1 | 4632804 | 4633994 | 1190 + | tuf | WP_046066033.1 |
| GCF_017357225.1 | NZ_CP071604.1 | 1990355 | 1991545 | 1190 + | tuf | WP_120140791.1 |
| GCF_017357225.1 | NZ_CP071604.1 | 2006309 | 2007499 | 1190 + | tuf | WP_120140791.1 |
| GCF_000769405.2 | NZ_CP053439.1 | 3959138 | 3960328 | 1190 - | tuf | WP_003490638.1 |
| GCF_000769405.2 | NZ_CP053439.1 | 3975062 | 3976252 | 1190 - | tuf | WP_003490638.1 |
| GCF_047553425.1 | NZ_CP180217.1 | 215553 | 216743 | 1190 + | tuf | WP_069678553.1 |
| GCF_047553425.1 | NZ_CP180217.1 | 231412 | 232602 | 1190 + | tuf | WP_069678553.1 |
| GCF_010450895.1 | NZ_CP048427.1 | 4546631 | 4547821 | 1190 - | tuf | WP_216276048.1 |
| GCF_010450895.1 | NZ_CP048427.1 | 4562648 | 4563838 | 1190 - | tuf | WP_216276048.1 |
| GCF_001664265.1 | NZ_CP013500.1 | 2748400 | 2749590 | 1190 - | tuf | WP_011100718.1 |
| GCF_001664265.1 | NZ_CP013500.1 | 2764341 | 2765531 | 1190 - | tuf | WP_011100718.1 |
| GCF_000092045.1 | NC_007761.1 | 3352769 | 3353959 | 1190 - | tuf | WP_139905874.1 |
| GCF_000092045.1 | NC_007761.1 | 3368653 | 3369843 | 1190 - | tuf | WP_139905874.1 |
| GCF_000577275.2 | NZ_HG916852.1 | 3537278 | 3538477 | 1199 - | tuf | WP_103080995.1 |
| GCF_000577275.2 | NZ_HG916852.1 | 2882219 | 2883409 | 1190 - | tuf | WP_017752389.1 |
| GCF_000816845.1 | NZ_CP006877.1 | 2898279 | 2899469 | 1190 - | tuf | WP_017752389.1 |
| GCF_000816845.1 | NZ_CP006877.1 | 2830942 | 2832132 | 1190 - | tuf | WP_044999124.1 |

| | | | | | | | |
|---|---|---|---|---|---|---|---|
| GCF_005862185.2 | NZ_CP054027.1 | 2779583 | 2780767 | 1184 | - | tuf | WP_004850603.1 |
| GCF_005862185.2 | NZ_CP054027.1 | 1341468 | 1342658 | 1190 | + | tuf | WP_218283981.1 |
| GCF_005862305.2 | NZ_CP054021.1 | 1357901 | 1359091 | 1190 | + | tuf | WP_218283981.1 |
| GCF_005862305.2 | NZ_CP054021.1 | 3566971 | 3568161 | 1190 | - | tuf | WP_223037135.1 |
| GCF_034554815.1 | NZ_CP140635.1 | 3583440 | 3584630 | 1190 | - | tuf | WP_223037135.1 |
| GCF_034554815.1 | NZ_CP140635.1 | 343183 | 344373 | 1190 | + | tuf | WP_148808226.1 |
| GCF_003627755.1 | NZ_CP032694.1 | 359678 | 360868 | 1190 | + | tuf | WP_148808226.1 |
| GCF_003627755.1 | NZ_CP032694.1 | 1572608 | 1573798 | 1190 | - | tuf | WP_341875722.1 |
| GCF_000009265.1 | NC_008380.1 | 554936 | 556135 | 1199 | + | tuf | WP_015043944.1 |
| GCF_000009265.1 | NC_008380.1 | 1152435 | 1153637 | 1202 | + | tuf | WP_089609453.1 |
| GCF_017352135.1 | NZ_CP071454.1 | 1168300 | 1169502 | 1202 | + | tuf | WP_089609453.1 |
| GCF_017352135.1 | NZ_CP071454.1 | 436002 | 437201 | 1199 | + | tuf | WP_014792382.1 |
| GCF_010669145.1 | NZ_CP048632.1 | 459936 | 461084 | 1148 | + | tuf | WP_242825458.1 |
| GCF_010669145.1 | NZ_CP048632.1 | 467678 | 468877 | 1199 | + | tuf | WP_005810165.1 |
| GCF_001664385.1 | NZ_CP013532.1 | 300587 | 301786 | 1199 | + | tuf | WP_006716334.1 |
| GCF_001664385.1 | NZ_CP013532.1 | 295055 | 296254 | 1199 | + | tuf | WP_015755944.1 |
| GCF_016864595.1 | NZ_CP044122.1 | 311554 | 312753 | 1199 | + | tuf | WP_015755958.1 |
| GCF_016864595.1 | NZ_CP044122.1 | 3062329 | 3063528 | 1199 | - | tuf | WP_013824065.1 |
| GCF_011046895.1 | NZ_CP049250.1 | 3078067 | 3079266 | 1199 | - | tuf | WP_013824079.1 |
| GCF_011046895.1 | NZ_CP049250.1 | 368538 | 369737 | 1199 | + | tuf | WP_077712910.1 |
| GCF_007000325.2 | NZ_CP117267.1 | 384526 | 385725 | 1199 | + | tuf | WP_077712923.1 |
| GCF_007000325.2 | NZ_CP117267.1 | 214459 | 215658 | 1199 | + | tuf | WP_011876589.1 |
| GCF_017357285.1 | NZ_CP071678.1 | 230469 | 231668 | 1199 | + | tuf | WP_011876589.1 |
| GCF_017357285.1 | NZ_CP071678.1 | 286310 | 287509 | 1199 | + | tuf | WP_013840356.1 |
| GCF_025200795.1 | NZ_CP104152.1 | 302500 | 303699 | 1199 | + | tuf | WP_013840356.1 |
| GCF_025200795.1 | NZ_CP104152.1 | 326282 | 327481 | 1199 | + | tuf | WP_006522873.1 |
| GCF_025200715.1 | NZ_CP104143.1 | 342049 | 343248 | 1199 | + | tuf | WP_006522873.1 |
| GCF_025200715.1 | NZ_CP104143.1 | 498567 | 499766 | 1199 | + | tuf | WP_014825567.1 |
| GCF_000330885.1 | NC_020059.1 | 236907 | 238109 | 1202 | + | tuf | WP_014901233.1 |
| GCF_000330885.1 | NC_020059.1 | 235662 | 236861 | 1199 | + | tuf | WP_014182790.1 |
| GCF_003240565.2 | NZ_CP117255.1 | 2397237 | 2398436 | 1199 | - | tuf | WP_407313880.1 |

| | | | | | | | |
|---|---|---|---|---|---|---|---|
| GCF_003240565.2 | NZ_CP117255.1 | 262173 | 263372 | 1199 | + | tuf | WP_007778406.1 |
| GCF_002355275.1 | NZ_CP023449.1 | 215469 | 216668 | 1199 | + | tuf | WP_013809477.1 |
| GCF_011044255.1 | NZ_CP049107.1 | 231553 | 232752 | 1199 | + | tuf | WP_013809477.1 |
| GCF_017607425.1 | NZ_CP059319.1 | 157712 | 158902 | 1190 | + | tuf | WP_005331928.1 |
| GCF_000021865.1 | NC_014034.1 | 2557795 | 2558985 | 1190 | - | tuf | WP_028087889.1 |
| GCF_000021865.1 | NC_014034.1 | 3973137 | 3974327 | 1190 | + | tuf | WP_002585860.1 |
| GCF_001856665.1 | NZ_CP017781.1 | 2210367 | 2211551 | 1184 | - | tuf | WP_005601267.1 |
| GCF_001856665.1 | NZ_CP017781.1 | 1189004 | 1190203 | 1199 | + | tuf | WP_013484988.1 |
| GCF_000166055.1 | NC_014664.1 | 3762481 | 3763671 | 1190 | - | tuf | WP_013382292.1 |
| GCF_021184025.1 | NZ_CP036537.1 | 3889184 | 3890374 | 1190 | - | tuf | WP_013382292.1 |
| GCF_021184025.1 | NZ_CP036537.1 | 1243558 | 1244748 | 1190 | + | tuf | WP_081571111.1 |
| GCF_016584445.1 | NZ_CP066699.1 | 1372774 | 1373964 | 1190 | + | tuf | WP_081571111.1 |
| GCF_016584445.1 | NZ_CP066699.1 | 2170398 | 2171588 | 1190 | - | tuf | WP_058693315.1 |
| GCF_000016185.1 | NC_011420.2 | 2299830 | 2301020 | 1190 | - | tuf | WP_058693315.1 |
| GCF_000016185.1 | NC_011420.2 | 2296893 | 2298077 | 1184 | - | tuf | WP_044964739.1 |
| GCF_000013085.1 | NC_007643.1 | 2071598 | 2072800 | 1202 | - | tuf | WP_005930548.1 |
| GCF_000013085.1 | NC_007643.1 | 1411422 | 1412624 | 1202 | + | tuf | WP_005940209.1 |
| GCF_937425535.1 | NZ_OW485601.1 | 546262 | 547452 | 1190 | + | tuf | WP_009263102.1 |
| GCF_001633165.1 | NZ_CP015418.1 | 1034459 | 1035658 | 1199 | - | tuf | WP_106012479.1 |
| GCF_001633165.1 | NZ_CP015418.1 | 1329566 | 1330756 | 1190 | - | tuf | WP_249242433.1 |
| GCF_000018205.1 | NC_009881.1 | 243078 | 244268 | 1190 | - | tuf | WP_215491777.1 |
| GCF_007989425.1 | NZ_AP019563.1 | 693702 | 694892 | 1190 | - | tuf | WP_014262273.1 |
| GCF_000284155.1 | NC_017058.1 | 1398925 | 1400115 | 1190 | - | tuf | WP_014262273.1 |
| GCF_000012385.1 | NC_007940.1 | 1904335 | 1905531 | 1196 | + | tuf | WP_007495837.1 |
| GCF_000283915.1 | NC_016929.1 | 3697263 | 3698453 | 1190 | + | tuf | WP_069978673.1 |
| GCF_000007025.1 | NC_003103.1 | 3717131 | 3718321 | 1190 | + | tuf | WP_069978673.1 |
| GCF_964030775.1 | NZ_OZ034920.1 | 2113265 | 2114455 | 1190 | - | tuf | WP_013406412.1 |
| GCF_964026445.1 | NZ_OZ032150.1 | 2128508 | 2129698 | 1190 | - | tuf | WP_013406412.1 |
| GCF_964026565.1 | NZ_OZ032162.1 | 180960 | 182141 | 1181 | + | tuf | WP_015325914.1 |
| GCF_964026435.1 | NZ_OZ032147.1 | 196476 | 197657 | 1181 | + | tuf | WP_015325914.1 |
| GCF_964026455.1 | NZ_OZ032152.1 | 119180 | 120370 | 1190 | + | tuf | WP_012635061.1 |

| | | | | | | | |
|---|---|---|---|---|---|---|---|
| GCF_965196655.1 | NZ_OZ239771.1 | 134987 | 136177 | 1190 | + | tuf | WP_012635061.1 |
| GCF_936269705.1 | NZ_OW370493.1 | 2428755 | 2429945 | 1190 | - | tuf | WP_138210969.1 |
| GCF_964291875.1 | NZ_OZ198175.1 | 2444810 | 2446000 | 1190 | - | tuf | WP_138210969.1 |
| GCF_964026385.1 | NZ_OZ032142.1 | 2747574 | 2748761 | 1187 | - | tuf | WP_338535729.1 |
| GCF_964026465.1 | NZ_OZ032155.1 | 2764617 | 2765804 | 1187 | - | tuf | WP_338535743.1 |
| GCF_964030815.1 | NZ_OZ034963.1 | 1400714 | 1401913 | 1199 | + | tuf | WP_012282469.1 |
| GCF_963970025.1 | NZ_OZ018776.1 | 2963970 | 2965169 | 1199 | - | tuf | WP_153726220.1 |
| GCF_000283595.1 | NC_016050.1 | 2365402 | 2366592 | 1190 | - | tuf | WP_058258946.1 |
| GCF_000499665.2 | NZ_LN794217.1 | 5924185 | 5925375 | 1190 | - | tuf | WP_055650430.1 |
| GCF_000277165.1 | NC_017049.1 | 271490 | 272680 | 1190 | + | tuf | WP_337979326.1 |
| GCF_000284075.1 | NC_017042.1 | 287371 | 288561 | 1190 | + | tuf | WP_007285284.1 |
| GCF_000018225.1 | NC_009882.1 | 1048666 | 1049862 | 1196 | + | tuf | WP_075704221.1 |
| GCF_000166935.1 | NZ_AABW01000001.1 | 3275897 | 3277087 | 1190 | - | tuf | WP_230867677.1 |
| GCF_000237845.1 | NC_016639.1 | 3291264 | 3292454 | 1190 | - | tuf | WP_230867677.1 |
| GCF_016743795.2 | NZ_CP060138.2 | 878362 | 879552 | 1190 | + | tuf | WP_342758530.1 |
| GCF_000008045.1 | NC_006142.1 | 1384361 | 1385560 | 1199 | + | tuf | WP_206709159.1 |
| GCF_003932735.1 | NZ_CP033611.1 | 1400363 | 1401562 | 1199 | + | tuf | WP_206709159.1 |
| GCF_003932735.1 | NZ_CP033611.1 | 95966 | 97162 | 1196 | + | tuf | WP_111526075.1 |
| GCF_000158095.2 | NZ_CM011002.1 | 295200 | 296390 | 1190 | + | tuf | WP_012198268.1 |
| GCF_000158095.2 | NZ_CM011002.1 | 322473 | 323663 | 1190 | + | tuf | WP_012738561.1 |
| GCF_001999245.1 | NZ_CP019630.1 | 3922524 | 3923714 | 1190 | - | tuf | WP_024834856.1 |
| GCF_001999245.1 | NZ_CP019630.1 | 505875 | 507071 | 1196 | + | tuf | WP_012993097.1 |
| GCF_026191725.2 | NZ_CP120863.1 | 2287744 | 2288940 | 1196 | - | tuf | WP_013781758.1 |
| GCF_026191725.2 | NZ_CP120863.1 | 2303485 | 2304681 | 1196 | - | tuf | WP_013781758.1 |
| GCF_000590925.1 | NZ_CP004372.1 | 2832785 | 2833975 | 1190 | + | tuf | WP_006863836.1 |
| GCF_000590925.1 | NZ_CP004372.1 | 2813552 | 2814736 | 1184 | - | tuf | WP_009266662.1 |
| GCF_004331955.1 | NZ_CP036532.1 | 1733095 | 1734285 | 1190 | - | tuf | WP_004842437.1 |
| GCF_004331955.1 | NZ_CP036532.1 | 114985 | 116184 | 1199 | - | tuf | WP_366923265.1 |
| GCF_002983865.1 | NZ_CP027407.1 | 131109 | 132308 | 1199 | - | tuf | WP_366923265.1 |
| GCF_002983865.1 | NZ_CP027407.1 | 84340 | 85530 | 1190 | + | tuf | WP_106056140.1 |
| GCF_014337925.2 | NZ_CP143423.1 | 1330158 | 1331348 | 1190 | - | tuf | WP_106056140.1 |

| | | | | | | | |
|---|---|---|---|---|---|---|---|
| GCF_014337925.2 | NZ_CP143423.1 | 371339 | 372529 | 1190 | + | tuf | WP_304989574.1 |
| GCF_000154785.2 | NC_015730.1 | 1572312 | 1573502 | 1190 | - | tuf | WP_304989574.1 |
| GCF_000154785.2 | NC_015730.1 | 1447579 | 1448778 | 1199 | + | tuf | WP_102365640.1 |
| GCF_012932215.1 | NZ_CP048788.1 | 200993 | 202192 | 1199 | + | tuf | WP_012446667.1 |
| GCF_012932215.1 | NZ_CP048788.1 | 218420 | 219619 | 1199 | + | tuf | WP_012446667.1 |
| GCF_014764365.1 | NZ_CP061718.1 | 3167257 | 3168456 | 1199 | + | tuf | WP_241079349.1 |
| GCF_014764365.1 | NZ_CP061718.1 | 3183379 | 3184578 | 1199 | + | tuf | WP_241079349.1 |
| GCF_020991085.1 | NZ_CP087719.1 | 2712884 | 2714083 | 1199 | - | tuf | WP_406677325.1 |
| GCF_020991085.1 | NZ_CP087719.1 | 2728556 | 2729755 | 1199 | - | tuf | WP_406677336.1 |
| GCF_022846615.1 | NZ_AP025637.1 | 1186531 | 1187730 | 1199 | - | tuf | WP_156272499.1 |
| GCF_022846615.1 | NZ_AP025637.1 | 1202023 | 1203222 | 1199 | - | tuf | WP_156272499.1 |
| GCF_001941945.1 | NZ_CP015583.1 | 2357125 | 2358324 | 1199 | - | tuf | WP_106006347.1 |
| GCF_001941945.1 | NZ_CP015583.1 | 2372381 | 2373580 | 1199 | - | tuf | WP_106006347.1 |
| GCF_017355405.1 | NZ_CP061177.1 | 2561170 | 2562369 | 1199 | - | tuf | WP_011393939.1 |
| GCF_017355405.1 | NZ_CP061177.1 | 373382 | 374572 | 1190 | + | tuf | WP_230398946.1 |
| GCF_017654485.1 | NZ_CP061091.1 | 318194 | 319393 | 1199 | + | tuf | WP_187333365.1 |
| GCF_017654485.1 | NZ_CP061091.1 | 3211072 | 3212271 | 1199 | + | tuf | WP_349216421.1 |
| GCF_006384915.1 | NZ_CP025060.1 | 64498 | 65688 | 1190 | + | tuf | WP_021428171.1 |
| GCF_006384915.1 | NZ_CP025060.1 | 80299 | 81489 | 1190 | + | tuf | WP_021428171.1 |
| GCF_009762325.1 | NZ_CP034348.1 | 3153867 | 3155057 | 1190 | + | tuf | WP_021428171.1 |
| GCF_008728195.1 | NZ_CP031598.1 | 3169668 | 3170858 | 1190 | + | tuf | WP_021428171.1 |
| GCF_008728195.1 | NZ_CP031598.1 | 67505 | 68695 | 1190 | + | tuf | WP_250672937.1 |
| GCF_025639885.1 | NZ_CP106738.1 | 83304 | 84494 | 1190 | + | tuf | WP_250672929.1 |
| GCF_025639885.1 | NZ_CP106738.1 | 29435 | 30625 | 1190 | + | tuf | WP_054632066.1 |
| GCF_037023735.1 | NZ_CP146069.1 | 45226 | 46416 | 1190 | + | tuf | WP_054632066.1 |
| GCF_037023735.1 | NZ_CP146069.1 | 37949 | 39139 | 1190 | - | tuf | WP_118549961.1 |
| GCF_032842945.2 | NZ_CP146606.1 | 54027 | 55217 | 1190 | - | tuf | WP_118549961.1 |
| GCF_032842945.2 | NZ_CP146606.1 | 124191 | 125381 | 1190 | + | tuf | WP_025434519.1 |
| GCF_028885395.1 | NZ_CP118494.1 | 140150 | 141340 | 1190 | + | tuf | WP_025434519.1 |
| GCF_028885395.1 | NZ_CP118494.1 | 1909411 | 1910601 | 1190 | - | tuf | WP_269311528.1 |
| GCF_000011965.2 | NC_003911.12 | 1946085 | 1947275 | 1190 | - | tuf | WP_269311528.1 |

| | | | | | | | |
|---|---|---|---|---|---|---|---|
| GCF_000011965.2 | NC_003911.12 | 2726972 | 2728162 | 1190 | + | tuf | WP_125137687.1 |
| GCF_039724765.1 | NZ_CP155729.1 | 170862 | 172058 | 1196 | + | tuf | WP_353893430.1 |
| GCF_039724765.1 | NZ_CP155729.1 | 162611 | 163807 | 1196 | + | tuf | WP_350343807.1 |
| GCF_001953055.1 | NZ_CP019044.1 | 620208 | 621410 | 1202 | - | tuf | WP_187032085.1 |
| GCF_001953055.1 | NZ_CP019044.1 | 1308018 | 1309202 | 1184 | - | tuf | WP_249304370.1 |
| GCF_001975705.1 | NZ_CP015093.1 | 2531818 | 2533008 | 1190 | + | tuf | WP_180702534.1 |
| GCF_001975705.1 | NZ_CP015093.1 | 2547609 | 2548799 | 1190 | + | tuf | WP_180702534.1 |
| GCF_001969385.1 | NZ_CP014796.1 | 244746 | 245930 | 1184 | + | tuf | WP_014078427.1 |
| GCF_001969385.1 | NZ_CP014796.1 | 3926305 | 3927489 | 1184 | - | tuf | WP_006859046.1 |
| GCF_014117445.1 | NZ_CP059851.1 | 555264 | 556448 | 1184 | + | tuf | WP_006859046.1 |
| GCF_033547095.1 | NZ_CP137852.1 | 368924 | 370123 | 1199 | + | tuf | WP_012634770.1 |
| GCF_033547095.1 | NZ_CP137852.1 | 4226558 | 4227757 | 1199 | - | tuf | WP_137697343.1 |
| GCF_023038235.1 | NZ_CP081987.1 | 425090 | 426289 | 1199 | + | tuf | WP_004619849.1 |
| GCF_023038235.1 | NZ_CP081987.1 | 3270859 | 3272055 | 1196 | - | tuf | WP_013499486.1 |
| GCF_030733865.1 | NZ_CP132302.1 | 1398339 | 1399538 | 1199 | + | tuf | WP_138157103.1 |
| GCF_030733865.1 | NZ_CP132302.1 | 1753439 | 1754629 | 1190 | - | tuf | WP_028529583.1 |
| GCF_020883495.1 | NZ_CP086610.1 | 1592030 | 1593220 | 1190 | + | tuf | WP_087199795.1 |
| GCF_020883495.1 | NZ_CP086610.1 | 2670698 | 2671888 | 1190 | - | tuf | WP_024733721.1 |
| GCF_003970735.1 | NZ_CP034588.1 | 1267118 | 1268308 | 1190 | + | tuf | WP_213050339.1 |
| GCF_003970735.1 | NZ_CP034588.1 | 1283330 | 1284520 | 1190 | + | tuf | WP_213050339.1 |
| GCF_008932245.1 | NZ_CP034909.1 | 2761168 | 2762352 | 1184 | - | tuf | WP_118549993.1 |
| GCF_008932245.1 | NZ_CP034909.1 | 1875322 | 1876521 | 1199 | - | tuf | WP_090264528.1 |
| GCF_001889105.1 | NZ_CP013107.1 | 2978543 | 2979745 | 1202 | + | tuf | WP_007046520.1 |
| GCF_001889105.1 | NZ_CP013107.1 | 3275811 | 3276995 | 1184 | - | tuf | WP_011197192.1 |
| GCF_036488675.1 | NZ_CP133148.1 | 404335 | 405534 | 1199 | + | tuf | WP_013623627.1 |
| GCF_036488675.1 | NZ_CP133148.1 | 2665505 | 2666704 | 1199 | - | tuf | WP_011641710.1 |
| GCF_003177055.1 | NZ_CP029451.1 | 2681824 | 2683023 | 1199 | - | tuf | WP_011641725.1 |
| GCF_003177055.1 | NZ_CP029451.1 | 2261299 | 2262498 | 1199 | - | tuf | WP_013176382.1 |
| GCF_029892065.1 | NZ_CP120373.1 | 3398531 | 3399721 | 1190 | - | tuf | WP_248481116.1 |
| GCF_029892065.1 | NZ_CP120373.1 | 3414570 | 3415760 | 1190 | - | tuf | WP_248481116.1 |
| GCF_030064585.1 | NZ_CP120365.1 | 3997774 | 3998964 | 1190 | + | tuf | WP_277732348.1 |

| | | | | | | | |
|---|---|---|---|---|---|---|---|
| GCF_030064585.1 | NZ_CP120365.1 | 4013815 | 4015005 | 1190 | + | tuf | WP_277732357.1 |
| GCF_037023865.1 | NZ_CP146208.1 | 71918 | 73108 | 1190 | + | tuf | WP_103980790.1 |
| GCF_037023865.1 | NZ_CP146208.1 | 87812 | 89002 | 1190 | + | tuf | WP_103980790.1 |
| GCF_013488225.1 | NZ_CP041238.1 | 1805075 | 1806265 | 1190 | - | tuf | WP_228417295.1 |
| GCF_013488225.1 | NZ_CP041238.1 | 1820986 | 1822176 | 1190 | - | tuf | WP_228417295.1 |
| GCF_029892045.1 | NZ_CP120368.1 | 2938157 | 2939347 | 1190 | + | tuf | WP_228106565.1 |
| GCF_029892045.1 | NZ_CP120368.1 | 2954066 | 2955256 | 1190 | + | tuf | WP_228106565.1 |
| GCF_002288525.1 | NZ_CP023067.1 | 72152 | 73342 | 1190 | + | tuf | WP_228108692.1 |
| GCF_002288525.1 | NZ_CP023067.1 | 88063 | 89253 | 1190 | + | tuf | WP_228108685.1 |
| GCF_029714365.1 | NZ_CP121659.1 | 2599316 | 2600515 | 1199 | - | tuf | WP_015051622.1 |
| GCF_029714365.1 | NZ_CP121659.1 | 59935 | 61134 | 1199 | + | tuf | WP_084662972.1 |
| GCF_016653635.2 | NZ_CP067420.1 | 2917399 | 2918586 | 1187 | + | tuf | WP_034424880.1 |
| GCF_016653635.2 | NZ_CP067420.1 | 2933328 | 2934515 | 1187 | + | tuf | WP_034424880.1 |
| GCF_016765655.2 | NZ_CP086106.1 | 397263 | 398462 | 1199 | + | tuf | WP_003870265.1 |
| GCF_016765655.2 | NZ_CP086106.1 | 412926 | 414125 | 1199 | + | tuf | WP_012268992.1 |
| GCF_008843145.1 | NZ_CP030265.1 | 2765371 | 2766570 | 1199 | - | tuf | WP_003870265.1 |
| GCF_008843145.1 | NZ_CP030265.1 | 2781038 | 2782237 | 1199 | - | tuf | WP_003870265.1 |
| GCF_016806835.2 | NZ_CP086111.1 | 2015383 | 2016582 | 1199 | - | tuf | WP_012995915.1 |
| GCF_016806835.2 | NZ_CP086111.1 | 2031045 | 2032244 | 1199 | - | tuf | WP_012995915.1 |
| GCF_026409565.1 | NZ_CP112881.1 | 1964764 | 1965963 | 1199 | - | tuf | WP_049685822.1 |
| GCF_026409565.1 | NZ_CP112881.1 | 1980417 | 1981616 | 1199 | - | tuf | WP_049685822.1 |
| GCF_023746535.1 | NZ_CP098747.1 | 1944539 | 1945738 | 1199 | - | tuf | WP_012995915.1 |
| GCF_023746535.1 | NZ_CP098747.1 | 401847 | 403046 | 1199 | + | tuf | WP_003870265.1 |
| GCF_003967075.1 | NZ_AP018664.1 | 417510 | 418709 | 1199 | + | tuf | WP_012268992.1 |
| GCF_002355855.1 | NZ_AP017655.1 | 2295063 | 2296262 | 1199 | - | tuf | WP_003870265.1 |
| GCF_015074805.1 | NZ_CP060035.1 | 2310725 | 2311924 | 1199 | - | tuf | WP_003870265.1 |
| GCF_000091125.1 | NC_014006.1 | 1042829 | 1044028 | 1199 | + | tuf | WP_014757913.1 |
| GCF_043589935.1 | NZ_CP171454.1 | 1058351 | 1059550 | 1199 | + | tuf | WP_014757922.1 |
| GCF_018286635.1 | NZ_CP073910.1 | 412274 | 413473 | 1199 | + | tuf | WP_013296961.1 |
| GCF_002288285.1 | NZ_CP022745.1 | 427815 | 429014 | 1199 | + | tuf | WP_013296975.1 |
| GCF_002504085.1 | NZ_CP023741.1 | 336104 | 337303 | 1199 | + | tuf | WP_013787134.1 |

| | | | | | | | |
|---|---|---|---|---|---|---|---|
| GCF_024721585.1 | NZ_CP102629.1 | 351635 | 352834 | 1199 | + | tuf | WP_013787148.1 |
| GCF_024721605.1 | NZ_CP102630.1 | 1060082 | 1061281 | 1199 | - | tuf | WP_283109536.1 |
| GCF_027625475.1 | NZ_CP115174.1 | 2843766 | 2844971 | 1205 | - | tuf | WP_015360289.1 |
| GCF_023935825.1 | NZ_CP098762.1 | 126428 | 127627 | 1199 | + | tuf | WP_013274960.1 |
| GCF_044501755.1 | NZ_CP162536.1 | 2112245 | 2113444 | 1199 | - | tuf | WP_013274960.1 |
| GCF_016743815.1 | NZ_CP061035.1 | 1896648 | 1897835 | 1187 | - | tuf | WP_212693111.1 |
| GCF_024349785.1 | NZ_AP018817.1 | 938687 | 939877 | 1190 | + | tuf | WP_212696946.1 |
| GCF_021391395.1 | NZ_CP090059.1 | 1665232 | 1666434 | 1202 | - | tuf | WP_213540515.1 |
| GCF_047223455.1 | NZ_CP178397.1 | 697302 | 698501 | 1199 | + | tuf | WP_212821738.1 |
| GCF_009913435.1 | NZ_CP047895.1 | 3537482 | 3538672 | 1190 | + | tuf | WP_118644900.1 |
| GCF_023674425.1 | NZ_CP098401.1 | 2228451 | 2229635 | 1184 | - | tuf | WP_118544680.1 |
| GCF_025231525.1 | NZ_CP103057.1 | 2953917 | 2955116 | 1199 | + | tuf | WP_227764085.1 |
| GCF_030866745.1 | NZ_CP133016.1 | 2970108 | 2971307 | 1199 | + | tuf | WP_227764108.1 |
| GCF_009363895.1 | NZ_CP045434.1 | 856517 | 857731 | 1214 | - | tuf | WP_012544048.1 |
| GCF_023380025.1 | NZ_CP097253.1 | 892637 | 893851 | 1214 | - | tuf | WP_012544210.1 |
| GCF_000935025.1 | NZ_CP010836.1 | 1876808 | 1878007 | 1199 | - | tuf | WP_022740306.1 |
| GCF_010450875.1 | NZ_CP048422.1 | 1119199 | 1120398 | 1199 | + | tuf | WP_114539934.1 |
| GCF_001922385.1 | NZ_CP018820.1 | 3300534 | 3301733 | 1199 | + | tuf | WP_080803363.1 |
| GCF_012979535.1 | NZ_CP053015.1 | 1426620 | 1427819 | 1199 | - | tuf | WP_039690697.1 |
| GCF_021497585.1 | NZ_CP071573.1 | 1754934 | 1756133 | 1199 | + | tuf | WP_172165954.1 |
| GCF_023822065.1 | NZ_CP084930.1 | 1220480 | 1221667 | 1187 | - | tuf | WP_035137538.1 |
| GCF_028607085.1 | NZ_CP117411.1 | 245667 | 246854 | 1187 | - | tuf | WP_006720938.1 |
| GCF_019930585.1 | NZ_CP082839.1 | 1598828 | 1600015 | 1187 | - | tuf | WP_013709218.1 |
| GCF_003660165.1 | NZ_CP032829.1 | 1112368 | 1113567 | 1199 | - | tuf | WP_012803351.1 |
| GCF_001717955.1 | NZ_CP014168.1 | 1683051 | 1684250 | 1199 | - | tuf | WP_066663213.1 |
| GCF_007859635.1 | NZ_CP042306.1 | 2852934 | 2854133 | 1199 | - | tuf | WP_160942775.1 |
| GCF_016027095.1 | NZ_CP065670.1 | 2404430 | 2405629 | 1199 | - | tuf | WP_009306296.1 |
| GCF_011300455.1 | NZ_CP049869.1 | 1295544 | 1296746 | 1202 | + | tuf | WP_006302344.1 |
| GCF_009739515.1 | NZ_CP046535.1 | 1317773 | 1318975 | 1202 | - | tuf | WP_012809262.1 |
| GCF_002796605.1 | NZ_CP024923.1 | 527548 | 528735 | 1187 | + | tuf | WP_136012364.1 |
| GCF_024496245.1 | NZ_CP101740.1 | 69335 | 70537 | 1202 | + | tuf | WP_075575820.1 |

| | | | | | | | |
|---|---|---|---|---|---|---|---|
| GCF_020866845.1 | NZ_CP086594.1 | 1861033 | 1862235 | 1202 | - | tuf | WP_013252478.1 |
| GCF_014396585.1 | NZ_CP060717.1 | 1928867 | 1930069 | 1202 | - | tuf | WP_090845484.1 |
| GCF_014352855.1 | NZ_CP060697.1 | 205700 | 206887 | 1187 | + | tuf | WP_126420797.1 |
| GCF_000512205.2 | NZ_CP006644.1 | 1715524 | 1716711 | 1187 | - | tuf | WP_073296341.1 |
| GCF_000512205.2 | NZ_CP006644.1 | 509910 | 511103 | 1193 | + | tuf | WP_079535498.1 |
| GCF_014489515.1 | NZ_CP060782.1 | 2541363 | 2542562 | 1199 | - | tuf | WP_102377746.1 |
| GCF_011302055.1 | NZ_CP049871.1 | 894146 | 895345 | 1199 | + | tuf | WP_012798017.1 |
| GCF_007833215.1 | NZ_CP042239.1 | 1962445 | 1963647 | 1202 | - | tuf | WP_194371035.1 |
| GCF_000764535.1 | NZ_CP009571.1 | 1516799 | 1517998 | 1199 | - | tuf | WP_165058867.1 |
| GCF_007998985.1 | NZ_CP041659.1 | 1783829 | 1785055 | 1226 | - | tuf | WP_012162324.1 |
| GCF_000013985.1 | NC_008048.1 | 7267042 | 7268268 | 1226 | - | tuf | WP_015185939.1 |
| GCF_035930445.1 | NZ_CP142394.1 | 2822784 | 2824010 | 1226 | - | tuf | WP_015214570.1 |
| GCF_000803645.1 | NZ_CP009122.1 | 1915474 | 1916700 | 1226 | + | tuf | WP_200989780.1 |
| GCF_001559015.1 | NZ_CP012199.1 | 1242869 | 1244086 | 1217 | + | tuf | WP_218082158.1 |
| GCF_032879645.1 | NZ_CP076395.1 | 1569336 | 1570562 | 1226 | - | tuf | WP_169263501.1 |
| GCF_036687315.1 | NZ_CP144683.1 | 944372 | 945598 | 1226 | + | tuf | WP_012954204.1 |
| GCF_002993885.1 | NZ_CM009578.1 | 7388599 | 7389825 | 1226 | + | tuf | WP_413163748.1 |
| GCF_001314325.1 | NZ_CP009429.1 | 4510122 | 4511348 | 1226 | - | tuf | WP_015161332.1 |
| GCF_009768975.1 | NZ_CP035733.1 | 5379877 | 5381103 | 1226 | - | tuf | WP_015156707.1 |
| GCF_001889025.1 | NZ_CP018154.1 | 3399422 | 3400648 | 1226 | + | tuf | WP_015203969.1 |
| GCF_016025255.1 | NZ_CP065592.1 | 4206500 | 4207726 | 1226 | - | tuf | WP_009543830.1 |
| GCF_011057975.1 | NZ_CP049109.1 | 518416 | 519642 | 1226 | - | tuf | WP_015218284.1 |
| GCF_032229225.1 | NZ_CP135076.1 | 1811878 | 1813104 | 1226 | + | tuf | WP_015109451.1 |
| GCF_009789575.1 | NZ_CP046908.1 | 2024888 | 2026114 | 1226 | + | tuf | WP_322764240.1 |
| GCF_009789575.1 | NZ_CP046908.1 | 3407584 | 3408810 | 1226 | + | tuf | WP_141302906.1 |
| GCF_006738645.1 | NZ_AP019700.1 | 1766961 | 1768187 | 1226 | + | tuf | WP_115538209.1 |
| GCF_006738645.1 | NZ_AP019700.1 | 5668061 | 5669287 | 1226 | - | tuf | WP_015210286.1 |
| GCF_018200035.1 | NZ_CP073581.1 | 589923 | 591149 | 1226 | - | tuf | WP_015228368.1 |
| GCF_018200035.1 | NZ_CP073581.1 | 4829394 | 4830620 | 1226 | + | tuf | WP_096670707.1 |
| GCF_001886735.1 | NZ_CP018076.1 | 726081 | 727307 | 1226 | - | tuf | WP_168695010.1 |
| GCF_001886735.1 | NZ_CP018076.1 | 4518396 | 4519622 | 1226 | + | tuf | WP_027402056.1 |

| | | | | | | | |
|---|---|---|---|---|---|---|---|
| GCF_034376865.1 | NZ_CP139725.1 | 2381923 | 2383149 | 1226 | + | tuf | WP_146296399.1 |
| GCF_034376865.1 | NZ_CP139725.1 | 1033184 | 1034410 | 1226 | + | tuf | WP_017293371.1 |
| GCF_022788655.1 | NZ_CP084951.1 | 2787230 | 2788456 | 1226 | + | tuf | WP_023174090.1 |
| GCF_022788655.1 | NZ_CP084951.1 | 3625357 | 3626583 | 1226 | + | tuf | WP_230840580.1 |
| GCF_016801755.1 | NZ_CP068998.1 | 4122116 | 4123342 | 1226 | + | tuf | WP_011143916.1 |
| GCF_016801755.1 | NZ_CP068998.1 | 1766571 | 1767797 | 1226 | - | tuf | WP_105219137.1 |
| GCF_040790665.1 | NZ_CP160849.1 | 1256506 | 1257732 | 1226 | - | tuf | WP_071454104.1 |
| GCF_040790665.1 | NZ_CP160849.1 | 58051 | 59280 | 1229 | - | tuf | WP_012597473.1 |
| GCF_006542355.1 | NZ_CP038141.1 | 2374884 | 2376113 | 1229 | + | tuf | WP_013322224.1 |
| GCF_003940805.1 | NZ_CP034328.1 | 935885 | 937111 | 1226 | - | tuf | WP_080809384.1 |
| GCF_003940805.1 | NZ_CP034328.1 | 589412 | 590638 | 1226 | - | tuf | WP_281483677.1 |
| GCF_001890385.1 | NZ_CP018221.1 | 2393964 | 2395190 | 1226 | + | tuf | WP_233748621.1 |
| GCF_018279705.1 | NZ_CP036498.1 | 3506617 | 3507843 | 1226 | - | tuf | WP_017288041.1 |
| GCF_018279705.1 | NZ_CP036498.1 | 3696457 | 3697683 | 1226 | + | tuf | WP_223046198.1 |
| GCF_031598375.1 | NZ_CP123872.1 | 4730481 | 4731707 | 1226 | - | tuf | WP_006623906.1 |
| GCF_031598375.1 | NZ_CP123872.1 | 5806418 | 5807644 | 1226 | + | tuf | WP_377481705.1 |
| GCF_008727715.1 | NZ_CP044218.1 | 2641183 | 2642409 | 1226 | + | tuf | WP_239731860.1 |
| GCF_008727715.1 | NZ_CP044218.1 | 2319321 | 2320547 | 1226 | - | tuf | WP_006196403.1 |
| GCF_009792355.1 | NZ_CP047045.1 | 699067 | 700293 | 1226 | + | tuf | WP_013190149.1 |
| GCF_014191545.1 | NZ_AP023361.1 | 2947624 | 2948850 | 1226 | - | tuf | WP_181931916.1 |
| GCF_014191545.1 | NZ_AP023361.1 | 4183101 | 4184327 | 1226 | - | tuf | WP_100899834.1 |
| GCF_022059845.1 | NZ_CP091851.1 | 6332771 | 6333997 | 1226 | + | tuf | WP_062297078.1 |
| GCF_003403095.1 | NZ_CP031555.1 | 4901872 | 4903098 | 1226 | + | tuf | WP_012410239.1 |
| GCF_003403095.1 | NZ_CP031555.1 | 5519932 | 5521158 | 1226 | - | tuf | WP_118168986.1 |
| GCF_002844375.1 | NZ_CP024199.1 | 400733 | 401962 | 1229 | + | tuf | WP_353931348.1 |
| GCF_002844375.1 | NZ_CP024199.1 | 5332325 | 5333572 | 1247 | + | tuf | WP_015150336.1 |
| GCF_000300235.2 | NZ_CP004388.1 | 74794 | 76020 | 1226 | - | tuf | WP_015174047.1 |
| GCF_000300235.2 | NZ_CP004388.1 | 462604 | 463830 | 1226 | + | tuf | WP_015174343.1 |
| GCF_012295595.1 | NZ_CP051181.1 | 4966562 | 4967788 | 1226 | - | tuf | WP_168570730.1 |
| GCF_012295595.1 | NZ_CP051181.1 | 2028879 | 2030075 | 1196 | + | tuf | WP_011128994.1 |
| GCF_002085925.2 | NZ_CP053562.1 | 801090 | 802316 | 1226 | - | tuf | WP_099798387.1 |

| | | | | | | | |
|---|---|---|---|---|---|---|---|
| GCF_002085925.2 | NZ_CP053562.1 | 571307 | 572533 | 1226 | + | tuf | WP_252663613.1 |
| GCF_045278885.1 | NZ_CP135443.1 | 5974885 | 5976111 | 1226 | - | tuf | WP_190877107.1 |
| GCF_045278885.1 | NZ_CP135443.1 | 1142690 | 1143916 | 1226 | - | tuf | WP_042152457.1 |
| GCF_001940525.2 | NZ_CP019437.1 | 4477170 | 4478396 | 1226 | + | tuf | WP_072721121.1 |
| GCF_001940525.2 | NZ_CP019437.1 | 1483883 | 1485079 | 1196 | - | tuf | WP_011820988.1 |
| GCF_041504375.1 | NZ_CP121009.1 | 3014117 | 3015346 | 1229 | - | tuf | WP_281005783.1 |
| GCF_041504375.1 | NZ_CP121009.1 | 2376531 | 2377760 | 1229 | + | tuf | WP_286390236.1 |
| GCF_041475515.1 | NZ_CP121045.1 | 5542029 | 5543255 | 1226 | + | tuf | WP_190601984.1 |
| GCF_041475515.1 | NZ_CP121045.1 | 1609575 | 1610801 | 1226 | - | tuf | WP_012594862.1 |
| GCF_000376545.2 | NZ_CP015230.1 | 1795713 | 1796939 | 1226 | + | tuf | WP_220611073.1 |
| GCF_000376545.2 | NZ_CP015230.1 | 2587617 | 2588843 | 1226 | - | tuf | WP_015193496.1 |
| GCF_032924845.1 | NZ_CP136704.1 | 889185 | 890411 | 1226 | - | tuf | WP_011242968.1 |
| GCF_032924845.1 | NZ_CP136704.1 | 1406451 | 1407677 | 1226 | - | tuf | WP_268611552.1 |
| GCF_029958925.1 | NZ_CP124616.1 | 2915205 | 2916431 | 1226 | + | tuf | WP_316786620.1 |
| GCF_029958925.1 | NZ_CP124616.1 | 4800050 | 4801276 | 1226 | - | tuf | WP_172358194.1 |
| GCF_003626915.1 | NZ_CP032570.1 | 2048562 | 2049788 | 1226 | + | tuf | WP_181494896.1 |
| GCF_001698205.1 | NZ_CP016591.1 | 1831732 | 1832958 | 1226 | + | tuf | WP_011057587.1 |
| GCF_002269345.1 | NZ_CP022889.1 | 791199 | 792425 | 1226 | + | tuf | WP_290226551.1 |
| GCF_002355335.1 | NZ_AP014946.1 | 755363 | 756589 | 1226 | + | tuf | WP_011610327.1 |
| GCF_002355335.1 | NZ_AP014946.1 | 2867276 | 2868502 | 1226 | - | tuf | WP_011318125.1 |
| GCF_029318635.1 | NZ_CP119676.1 | 2677327 | 2678556 | 1229 | - | tuf | WP_265900621.1 |
| GCF_029318635.1 | NZ_CP119676.1 | 5162554 | 5163738 | 1184 | - | tuf | WP_182412737.1 |
| GCF_947250015.1 | NZ_OX366272.1 | 540547 | 541731 | 1184 | - | tuf | WP_185272598.1 |
| GCF_947250015.1 | NZ_OX366272.1 | 3829948 | 3831132 | 1184 | + | tuf | WP_264808571.1 |
| GCF_964026805.1 | NZ_OZ034699.1 | 4423344 | 4424528 | 1184 | - | tuf | WP_008203052.1 |
| GCF_964026805.1 | NZ_OZ034699.1 | 1231417 | 1232601 | 1184 | + | tuf | WP_067544555.1 |
| GCF_964028465.1 | NZ_OZ034782.1 | 205152 | 206336 | 1184 | - | tuf | WP_084118536.1 |
| GCF_964028465.1 | NZ_OZ034782.1 | 226613 | 227797 | 1184 | - | tuf | WP_130894758.1 |
| GCF_964028505.1 | NZ_OZ034793.1 | 229478 | 230662 | 1184 | - | tuf | WP_223130981.1 |
| GCF_964028505.1 | NZ_OZ034793.1 | 374208 | 375392 | 1184 | - | tuf | WP_109321965.1 |
| GCF_947250785.1 | NZ_OX366348.1 | 3541920 | 3543104 | 1184 | + | tuf | WP_111372620.1 |

| | | | | | | | |
|---|---|---|---|---|---|---|---|
| GCF_947250785.1 | NZ_OX366348.1 | 573811 | 574995 | 1184 | + | tuf | WP_309937416.1 |
| GCF_964026715.1 | NZ_OZ034690.1 | 1802977 | 1804161 | 1184 | + | tuf | WP_014772281.1 |
| GCF_964026715.1 | NZ_OZ034690.1 | 3187392 | 3188573 | 1181 | + | tuf | WP_014798455.1 |
| GCF_964027085.1 | NZ_OZ034726.1 | 1786977 | 1788161 | 1184 | - | tuf | WP_012473441.1 |
| GCF_964027085.1 | NZ_OZ034726.1 | 25654 | 26838 | 1184 | + | tuf | WP_109996828.1 |
| GCF_947251765.1 | NZ_OX366389.1 | 337360 | 338547 | 1187 | + | tuf | WP_014934481.1 |
| GCF_947251765.1 | NZ_OX366389.1 | 607810 | 608994 | 1184 | + | tuf | WP_419241012.1 |
| GCF_964026815.1 | NZ_OZ034701.1 | 790738 | 791922 | 1184 | - | tuf | WP_250237859.1 |
| GCF_964026815.1 | NZ_OZ034701.1 | 279474 | 280658 | 1184 | - | tuf | WP_342265531.1 |
| GCF_947250565.1 | NZ_OX366326.1 | 1039036 | 1040220 | 1184 | + | tuf | WP_114910403.1 |
| GCF_947250565.1 | NZ_OX366326.1 | 907288 | 908472 | 1184 | - | tuf | WP_339043904.1 |
| GCF_964028415.1 | NZ_OZ034779.1 | 2238933 | 2240117 | 1184 | - | tuf | WP_302255548.1 |
| GCF_964028415.1 | NZ_OZ034779.1 | 1470575 | 1471759 | 1184 | - | tuf | WP_284685191.1 |
| GCF_964028445.1 | NZ_OZ034776.1 | 2784084 | 2785268 | 1184 | + | tuf | WP_073133806.1 |
| GCF_964028445.1 | NZ_OZ034776.1 | 3027351 | 3028535 | 1184 | + | tuf | WP_048642211.1 |
| GCF_947251545.1 | NZ_OX366362.1 | 3016515 | 3017699 | 1184 | + | tuf | WP_014020607.1 |
| GCF_947251545.1 | NZ_OX366362.1 | 3614244 | 3615428 | 1184 | - | tuf | WP_011586525.1 |
| GCF_947251795.1 | NZ_OX366385.1 | 4800142 | 4801326 | 1184 | + | tuf | WP_026631774.1 |
| GCF_947251795.1 | NZ_OX366385.1 | 5867843 | 5869027 | 1184 | + | tuf | WP_234605594.1 |
| GCF_964028115.1 | NZ_OZ034756.1 | 5059347 | 5060531 | 1184 | + | tuf | WP_026631774.1 |
| GCF_964028115.1 | NZ_OZ034756.1 | 4232363 | 4233547 | 1184 | - | tuf | WP_015812910.1 |
| GCF_947251615.1 | NZ_OX366367.1 | 5674219 | 5675403 | 1184 | - | tuf | WP_244822403.1 |
| GCF_947251615.1 | NZ_OX366367.1 | 5020462 | 5021646 | 1184 | + | tuf | WP_204658821.1 |
| GCF_964277365.1 | NZ_OZ195518.1 | 249037 | 250221 | 1184 | - | tuf | WP_226334355.1 |
| GCF_964277365.1 | NZ_OZ195518.1 | 272255 | 273439 | 1184 | - | tuf | WP_137400930.1 |
| GCF_964027025.1 | NZ_OZ034721.1 | 273216 | 274400 | 1184 | - | tuf | WP_141613022.1 |
| GCF_964027025.1 | NZ_OZ034721.1 | 2052149 | 2053333 | 1184 | - | tuf | WP_112783441.1 |
| GCF_964030885.1 | NZ_OZ034985.1 | 274686 | 275870 | 1184 | - | tuf | WP_015264131.1 |
| GCF_964030885.1 | NZ_OZ034985.1 | 1432723 | 1433907 | 1184 | - | tuf | WP_015028368.1 |
| GCF_964028185.1 | NZ_OZ034760.1 | 1355957 | 1357141 | 1184 | + | tuf | WP_015330198.1 |
| GCF_964028185.1 | NZ_OZ034760.1 | 3728158 | 3729333 | 1175 | + | tuf | WP_144074027.1 |

| | | | | | | | |
|---|---|---|---|---|---|---|---|
| GCF_947251645.1 | NZ_OX366364.1 | 3572034 | 3573209 | 1175 | - | tuf | WP_126615805.1 |
| GCF_947251645.1 | NZ_OX366364.1 | 4579593 | 4580768 | 1175 | - | tuf | WP_169667294.1 |
| GCF_947251895.1 | NZ_OX366395.1 | 4335271 | 4336455 | 1184 | - | tuf | WP_338392488.1 |
| GCF_947251895.1 | NZ_OX366395.1 | 209140 | 210324 | 1184 | - | tuf | WP_233771713.1 |
| GCF_964026995.1 | NZ_OZ034720.1 | 5294 | 6478 | 1184 | - | tuf | WP_205722027.1 |
| GCF_964026995.1 | NZ_OZ034720.1 | 1202252 | 1203436 | 1184 | - | tuf | WP_233779193.1 |
| GCF_947251935.1 | NZ_OX366397.1 | 1405030 | 1406214 | 1184 | - | tuf | WP_155170770.1 |
| GCF_947251935.1 | NZ_OX366397.1 | 4976526 | 4977710 | 1184 | + | tuf | WP_149069446.1 |
| GCF_964026755.1 | NZ_OZ034695.1 | 408134 | 409321 | 1187 | - | tuf | WP_151086072.1 |
| GCF_964026755.1 | NZ_OZ034695.1 | 2843556 | 2844740 | 1184 | - | tuf | WP_269558676.1 |
| GCF_964026895.1 | NZ_OZ034713.1 | 229093 | 230277 | 1184 | - | tuf | WP_100337265.1 |
| GCF_964026895.1 | NZ_OZ034713.1 | 2825393 | 2826580 | 1187 | + | tuf | WP_139515888.1 |
| GCF_964031185.1 | NZ_OZ035014.1 | 516556 | 517743 | 1187 | - | tuf | WP_196286000.1 |
| GCF_964031185.1 | NZ_OZ035014.1 | 2891980 | 2893167 | 1187 | - | tuf | WP_109656661.1 |
| GCF_964026765.1 | NZ_OZ034697.1 | 3741870 | 3743054 | 1184 | + | tuf | WP_119445973.1 |
| GCF_964026765.1 | NZ_OZ034697.1 | 651024 | 652211 | 1187 | + | tuf | WP_133271644.1 |
| GCF_947250625.1 | NZ_OX366332.1 | 1906980 | 1908167 | 1187 | + | tuf | WP_068191696.1 |
| GCF_947250625.1 | NZ_OX366332.1 | 799638 | 800822 | 1184 | + | tuf | WP_185888775.1 |
| GCF_964026865.1 | NZ_OZ034707.1 | 3584691 | 3585875 | 1184 | - | tuf | WP_185888775.1 |
| GCF_964026865.1 | NZ_OZ034707.1 | 4068956 | 4070140 | 1184 | - | tuf | WP_044003269.1 |
| GCF_964264715.1 | NZ_OZ183471.1 | 949865 | 951049 | 1184 | - | tuf | WP_135431220.1 |
| GCF_964264715.1 | NZ_OZ183471.1 | 4857923 | 4859110 | 1187 | - | tuf | WP_243797598.1 |
| GCF_964027015.1 | NZ_OZ034722.1 | 4448455 | 4449642 | 1187 | - | tuf | WP_245119664.1 |
| GCF_964027015.1 | NZ_OZ034722.1 | 3290837 | 3292021 | 1184 | - | tuf | WP_044015999.1 |
| GCF_947251955.1 | NZ_OX366399.1 | 4534799 | 4535983 | 1184 | - | tuf | WP_317487974.1 |
| GCF_947251955.1 | NZ_OX366399.1 | 2416324 | 2417508 | 1184 | + | tuf | WP_265993608.1 |
| GCF_964028475.1 | NZ_OZ034785.1 | 2621522 | 2622706 | 1184 | - | tuf | WP_013409093.1 |
| GCF_964028475.1 | NZ_OZ034785.1 | 930145 | 931329 | 1184 | + | tuf | WP_137089610.1 |
| GCF_964028495.1 | NZ_OZ034792.1 | 2299380 | 2300564 | 1184 | - | tuf | WP_261452293.1 |
| GCF_964028495.1 | NZ_OZ034792.1 | 1163971 | 1165155 | 1184 | + | tuf | WP_308357494.1 |
| GCF_964026745.1 | NZ_OZ034694.1 | 2113100 | 2114284 | 1184 | + | tuf | WP_303271996.1 |

| | | | | | | | |
|---|---|---|---|---|---|---|---|
| GCF_964026745.1 | NZ_OZ034694.1 | 4421288 | 4422472 | 1184 | - | tuf | WP_308349502.1 |
| GCF_964026835.1 | NZ_OZ034702.1 | 3575227 | 3576411 | 1184 | + | tuf | WP_013455137.1 |
| GCF_964026835.1 | NZ_OZ034702.1 | 2124407 | 2125591 | 1184 | + | tuf | WP_236139535.1 |
| GCF_947251695.1 | NZ_OX366378.1 | 3625577 | 3626761 | 1184 | + | tuf | WP_160693389.1 |
| GCF_947251695.1 | NZ_OX366378.1 | 2806257 | 2807441 | 1184 | - | tuf | WP_332921331.1 |
| GCF_947250805.1 | NZ_OX366352.1 | 4471856 | 4473040 | 1184 | + | tuf | WP_025609038.1 |
| GCF_947250805.1 | NZ_OX366352.1 | 2494520 | 2495704 | 1184 | + | tuf | WP_068838118.1 |
| GCF_964028535.1 | NZ_OZ034790.1 | 4694558 | 4695742 | 1184 | + | tuf | WP_266203992.1 |
| GCF_964028535.1 | NZ_OZ034790.1 | 1955137 | 1956321 | 1184 | + | tuf | WP_046310159.1 |
| GCF_947251775.1 | NZ_OX366387.1 | 2826116 | 2827300 | 1184 | - | tuf | WP_162427727.1 |
| GCF_947251775.1 | NZ_OX366387.1 | 3262969 | 3264153 | 1184 | + | tuf | WP_161889978.1 |
| GCF_947251475.1 | NZ_OX366359.1 | 335173 | 336357 | 1184 | - | tuf | WP_302240190.1 |
| GCF_947251475.1 | NZ_OX366359.1 | 630657 | 631841 | 1184 | - | tuf | WP_262310265.1 |
| GCF_947251435.1 | NZ_OX366363.1 | 2157081 | 2158265 | 1184 | + | tuf | WP_263052870.1 |
| GCF_947251435.1 | NZ_OX366363.1 | 5240801 | 5241985 | 1184 | + | tuf | WP_162445055.1 |
| GCF_964026915.1 | NZ_OZ034712.1 | 2298649 | 2299833 | 1184 | + | tuf | WP_068221192.1 |
| GCF_964026915.1 | NZ_OZ034712.1 | 255359 | 256543 | 1184 | + | tuf | WP_048919285.1 |
| GCF_947251595.1 | NZ_OX366369.1 | 2855891 | 2857075 | 1184 | + | tuf | WP_062543967.1 |
| GCF_947251595.1 | NZ_OX366369.1 | 4279526 | 4280710 | 1184 | + | tuf | WP_114068263.1 |
| GCF_964028265.1 | NZ_OZ034763.1 | 2053759 | 2054943 | 1184 | + | tuf | WP_013927548.1 |
| GCF_964028265.1 | NZ_OZ034763.1 | 6621671 | 6622855 | 1184 | + | tuf | WP_020598178.1 |
| GCF_947250765.1 | NZ_OX366347.1 | 204135 | 205319 | 1184 | - | tuf | WP_167204329.1 |
| GCF_947250765.1 | NZ_OX366347.1 | 6209592 | 6210776 | 1184 | - | tuf | WP_162388508.1 |
| GCF_964028325.1 | NZ_OZ034755.1 | 5973881 | 5975065 | 1184 | + | tuf | WP_182458540.1 |
| GCF_964028325.1 | NZ_OZ034755.1 | 5433180 | 5434364 | 1184 | - | tuf | WP_012928968.1 |
| GCF_947251915.1 | NZ_OX366396.1 | 5003998 | 5005182 | 1184 | + | tuf | WP_077133091.1 |
| GCF_947251915.1 | NZ_OX366396.1 | 5531305 | 5532489 | 1184 | - | tuf | WP_232560330.1 |
| GCF_947251635.1 | NZ_OX366357.1 | 1108332 | 1109516 | 1184 | + | tuf | WP_100986724.1 |
| GCF_947251635.1 | NZ_OX366357.1 | 5968223 | 5969407 | 1184 | - | tuf | WP_046577464.1 |
| GCF_964026985.1 | NZ_OZ034715.1 | 1849662 | 1850846 | 1184 | + | tuf | WP_169550262.1 |
| GCF_964026985.1 | NZ_OZ034715.1 | 671120 | 672304 | 1184 | - | tuf | WP_080237288.1 |

| | | | | | | | |
|---|---|---|---|---|---|---|---|
| GCF_964028605.1 | NZ_OZ034798.1 | 52607 | 53791 | 1184 | + | tuf | WP_171737752.1 |
| GCF_964028605.1 | NZ_OZ034798.1 | 1643378 | 1644562 | 1184 | + | tuf | WP_234736397.1 |
| GCF_947251945.1 | NZ_OX366398.1 | 3030426 | 3031610 | 1184 | + | tuf | WP_266362298.1 |
| GCF_947251945.1 | NZ_OX366398.1 | 5776365 | 5777549 | 1184 | + | tuf | WP_302205964.1 |
| GCF_947250585.1 | NZ_OX366327.1 | 1557611 | 1558798 | 1187 | - | tuf | WP_013451628.1 |
| GCF_947250585.1 | NZ_OX366327.1 | 1585144 | 1586331 | 1187 | - | tuf | WP_013451655.1 |
| GCF_964026955.1 | NZ_OZ034716.1 | 1785598 | 1786785 | 1187 | - | tuf | WP_013008475.1 |
| GCF_964026955.1 | NZ_OZ034716.1 | 1801122 | 1802309 | 1187 | - | tuf | WP_013008475.1 |
| GCF_964263225.1 | NZ_OZ181654.1 | 374237 | 375418 | 1181 | + | tuf | WP_013009702.1 |
| GCF_964263225.1 | NZ_OZ181654.1 | 390168 | 391349 | 1181 | + | tuf | WP_013009702.1 |
| GCF_963931955.1 | NZ_OZ008337.1 | 477125 | 478312 | 1187 | + | tuf | WP_013885646.1 |
| GCF_963931955.1 | NZ_OZ008337.1 | 492787 | 493974 | 1187 | + | tuf | WP_013885646.1 |
| GCF_964028575.1 | NZ_OZ034796.1 | 1062451 | 1063638 | 1187 | + | tuf | WP_128466050.1 |
| GCF_964028575.1 | NZ_OZ034796.1 | 1079085 | 1080272 | 1187 | + | tuf | WP_128466050.1 |
| GCF_964027095.1 | NZ_OZ034727.1 | 1015858 | 1017045 | 1187 | - | tuf | WP_023275261.1 |
| GCF_964027095.1 | NZ_OZ034727.1 | 1211731 | 1212930 | 1199 | + | tuf | WP_338736988.1 |
| GCF_947251555.1 | NZ_OX366372.1 | 627208 | 628404 | 1196 | + | tuf | WP_342825267.1 |
| GCF_947251555.1 | NZ_OX366372.1 | 912113 | 913312 | 1199 | - | tuf | WP_010936699.1 |
| GCF_964031125.1 | NZ_OZ035007.1 | 1156354 | 1157553 | 1199 | - | tuf | WP_102330188.1 |
| GCF_964031125.1 | NZ_OZ035007.1 | 1167110 | 1168309 | 1199 | - | tuf | WP_076004225.1 |
| GCF_947251675.1 | NZ_OX366375.1 | 2729563 | 2730777 | 1214 | - | tuf | WP_013159074.1 |
| GCF_947251675.1 | NZ_OX366375.1 | 2955607 | 2956821 | 1214 | - | tuf | WP_013159074.1 |
| GCF_964031025.1 | NZ_OZ034998.1 | 383497 | 384711 | 1214 | + | tuf | WP_264774903.1 |
| GCF_964031025.1 | NZ_OZ034998.1 | 2151162 | 2152376 | 1214 | + | tuf | WP_264774903.1 |
| GCF_964027115.1 | NZ_OZ034729.1 | 68988 | 70202 | 1214 | + | tuf | WP_407569228.1 |
| GCF_964027115.1 | NZ_OZ034729.1 | 2169270 | 2170484 | 1214 | - | tuf | WP_407569228.1 |
| GCF_947250795.1 | NZ_OX366351.1 | 373835 | 375049 | 1214 | - | tuf | WP_273987408.1 |
| GCF_947250795.1 | NZ_OX366351.1 | 1814310 | 1815524 | 1214 | - | tuf | WP_273987408.1 |
| GCF_964027105.1 | NZ_OZ034728.1 | 718688 | 719902 | 1214 | - | tuf | WP_012692578.1 |
| GCF_964027105.1 | NZ_OZ034728.1 | 2286439 | 2287653 | 1214 | - | tuf | WP_012692578.1 |
| GCF_964027055.1 | NZ_OZ034725.1 | 440028 | 441242 | 1214 | + | tuf | WP_022800164.1 |

| | | | | | | | |
|---|---|---|---|---|---|---|---|
| GCF_964027055.1 | NZ_OZ034725.1 | 592044 | 593258 | 1214 | - | tuf | WP_022800164.1 |
| GCF_964028175.1 | NZ_OZ034762.1 | 698608 | 699822 | 1214 | - | tuf | WP_011529789.1 |
| GCF_964028175.1 | NZ_OZ034762.1 | 1965805 | 1967019 | 1214 | - | tuf | WP_011529789.1 |
| GCF_964026705.1 | NZ_OZ034693.1 | 1250143 | 1251357 | 1214 | - | tuf | WP_014685999.1 |
| GCF_964026705.1 | NZ_OZ034693.1 | 2742312 | 2743526 | 1214 | - | tuf | WP_014685999.1 |
| GCF_963966195.1 | NZ_OZ014602.1 | 299835 | 301049 | 1214 | + | tuf | WP_109824827.1 |
| GCF_963966195.1 | NZ_OZ014602.1 | 1068732 | 1069946 | 1214 | - | tuf | WP_109824827.1 |
| GCF_964277465.1 | NZ_OZ195525.1 | 921765 | 922979 | 1214 | - | tuf | WP_013555989.1 |
| GCF_964277465.1 | NZ_OZ195525.1 | 3100905 | 3102119 | 1214 | - | tuf | WP_013555989.1 |
| GCF_964273535.1 | NZ_OZ187006.1 | 2007080 | 2008294 | 1214 | + | tuf | WP_146719929.1 |
| GCF_964273535.1 | NZ_OZ187006.1 | 2447342 | 2448556 | 1214 | - | tuf | WP_146719929.1 |
| GCF_963969245.1 | NZ_OZ017714.1 | 997480 | 998694 | 1214 | + | tuf | WP_015234848.1 |
| GCF_963969245.1 | NZ_OZ017714.1 | 3055511 | 3056725 | 1214 | + | tuf | WP_015234848.1 |
| GCF_964019365.1 | NZ_OZ025779.1 | 416838 | 418052 | 1214 | + | tuf | WP_013614171.1 |
| GCF_964019365.1 | NZ_OZ025779.1 | 1526691 | 1527905 | 1214 | + | tuf | WP_013614171.1 |
| GCF_964264765.1 | NZ_OZ183482.1 | 430222 | 431436 | 1214 | - | tuf | WP_124867456.1 |
| GCF_964264765.1 | NZ_OZ183482.1 | 1764754 | 1765968 | 1214 | - | tuf | WP_124870145.1 |
| GCF_964027005.1 | NZ_OZ034719.1 | 2205912 | 2207126 | 1214 | + | tuf | WP_064015142.1 |
| GCF_964027005.1 | NZ_OZ034719.1 | 2474169 | 2475383 | 1214 | - | tuf | WP_064015142.1 |
| GCF_964028485.1 | NZ_OZ034784.1 | 614672 | 615886 | 1214 | - | tuf | WP_407540428.1 |
| GCF_964028485.1 | NZ_OZ034784.1 | 1674492 | 1675706 | 1214 | + | tuf | WP_407540428.1 |
| GCF_947250505.1 | NZ_OX366319.1 | 219071 | 220285 | 1214 | - | tuf | WP_039681653.1 |
| GCF_947250505.1 | NZ_OX366319.1 | 1150971 | 1152185 | 1214 | + | tuf | WP_039682845.1 |
| GCF_964028125.1 | NZ_OZ034764.1 | 594905 | 596119 | 1214 | + | tuf | WP_260560932.1 |
| GCF_964028125.1 | NZ_OZ034764.1 | 875417 | 876631 | 1214 | - | tuf | WP_260561212.1 |
| GCF_947251565.1 | NZ_OX366379.1 | 816797 | 818011 | 1214 | - | tuf | WP_350243532.1 |
| GCF_947251565.1 | NZ_OX366379.1 | 2808946 | 2810160 | 1214 | + | tuf | WP_350243532.1 |
| GCF_964026875.1 | NZ_OZ034706.1 | 495747 | 496961 | 1214 | + | tuf | WP_225475552.1 |
| GCF_964026875.1 | NZ_OZ034706.1 | 2008671 | 2009885 | 1214 | + | tuf | WP_225475552.1 |
| GCF_964028305.1 | NZ_OZ034757.1 | 190689 | 191903 | 1214 | + | tuf | WP_164993950.1 |
| GCF_964028305.1 | NZ_OZ034757.1 | 2078669 | 2079883 | 1214 | - | tuf | WP_017872078.1 |

| | | | | | | | |
|---|---|---|---|---|---|---|---|
| GCF_947251755.1 | NZ_OX366388.1 | 1860593 | 1861807 | 1214 | + | tuf | WP_322472781.1 |
| GCF_947251755.1 | NZ_OX366388.1 | 2262222 | 2263436 | 1214 | - | tuf | WP_322473048.1 |
| GCF_964028555.1 | NZ_OZ034787.1 | 419407 | 420621 | 1214 | + | tuf | WP_013703243.1 |
| GCF_964028555.1 | NZ_OZ034787.1 | 1725621 | 1726835 | 1214 | - | tuf | WP_013703243.1 |
| GCF_964028245.1 | NZ_OZ034770.1 | 1193351 | 1194565 | 1214 | - | tuf | WP_013013466.1 |
| GCF_964028245.1 | NZ_OZ034770.1 | 2656553 | 2657767 | 1214 | - | tuf | WP_013014869.1 |
| GCF_964028525.1 | NZ_OZ034786.1 | 526088 | 527302 | 1214 | + | tuf | WP_063843310.1 |
| GCF_964028525.1 | NZ_OZ034786.1 | 1785986 | 1787200 | 1214 | + | tuf | WP_187388530.1 |
| GCF_947251425.1 | NZ_OX366368.1 | 509319 | 510536 | 1217 | - | tuf | WP_273106246.1 |
| GCF_947251425.1 | NZ_OX366368.1 | 913967 | 915184 | 1217 | + | tuf | WP_273107077.1 |
| GCF_947250775.1 | NZ_OX366349.1 | 61898 | 63115 | 1217 | + | tuf | WP_003043841.1 |
| GCF_947250775.1 | NZ_OX366349.1 | 318218 | 319435 | 1217 | + | tuf | WP_003043841.1 |
| GCF_964028425.1 | NZ_OZ034769.1 | 1781309 | 1782526 | 1217 | - | tuf | WP_071677189.1 |
| GCF_964028425.1 | NZ_OZ034769.1 | 1847227 | 1848444 | 1217 | - | tuf | WP_071677189.1 |
| GCF_947251485.1 | NZ_OX366356.1 | 1566516 | 1567733 | 1217 | - | tuf | WP_135257046.1 |
| GCF_947251485.1 | NZ_OX366356.1 | 1979608 | 1980825 | 1217 | + | tuf | WP_135257392.1 |
| GCF_964031055.1 | NZ_OZ034991.1 | 256749 | 257966 | 1217 | + | tuf | WP_016328551.1 |
| GCF_964031055.1 | NZ_OZ034991.1 | 1853986 | 1855203 | 1217 | - | tuf | WP_016328551.1 |
| GCF_964030675.1 | NZ_OZ034913.1 | 24893 | 26110 | 1217 | - | tuf | WP_060383976.1 |
| GCF_964030675.1 | NZ_OZ034913.1 | 407945 | 409162 | 1217 | + | tuf | WP_060383976.1 |
| GCF_947251515.1 | NZ_OX366370.1 | 238950 | 240167 | 1217 | - | tuf | WP_011227805.1 |
| GCF_947251515.1 | NZ_OX366370.1 | 1591099 | 1592316 | 1217 | - | tuf | WP_011228847.1 |
| GCF_947250735.1 | NZ_OX366344.1 | 1131510 | 1132724 | 1214 | + | tuf | WP_013177534.1 |
| GCF_947250735.1 | NZ_OX366344.1 | 2711658 | 2712872 | 1214 | - | tuf | WP_013177534.1 |
| GCF_947251965.1 | NZ_OX366400.1 | 285310 | 286503 | 1193 | + | tuf | WP_111331316.1 |
| GCF_947251965.1 | NZ_OX366400.1 | 4763538 | 4764731 | 1193 | + | tuf | WP_111331316.1 |
| GCF_964030905.1 | NZ_OZ034988.1 | 751622 | 752809 | 1187 | + | tuf | WP_237889265.1 |
| GCF_964030905.1 | NZ_OZ034988.1 | 878835 | 880022 | 1187 | + | tuf | WP_237889265.1 |
| GCF_947250715.1 | NZ_OX366342.1 | 4539248 | 4540441 | 1193 | + | tuf | WP_146962450.1 |
| GCF_947250715.1 | NZ_OX366342.1 | 4805519 | 4806709 | 1190 | + | tuf | WP_141198970.1 |
| GCF_964277345.1 | NZ_OZ195515.1 | 4992245 | 4993435 | 1190 | + | tuf | WP_141199100.1 |

| | | | | | | | |
|---|---|---|---|---|---|---|---|
| GCF_964277345.1 | NZ_OZ195515.1 | 3236738 | 3237934 | 1196 | - | tuf | WP_013259731.1 |
| GCF_947251725.1 | NZ_OX366380.1 | 3253899 | 3255095 | 1196 | - | tuf | WP_013259731.1 |
| GCF_947251725.1 | NZ_OX366380.1 | 3866753 | 3867949 | 1196 | - | tuf | WP_338602517.1 |
| GCF_964028565.1 | NZ_OZ034791.1 | 3883687 | 3884883 | 1196 | - | tuf | WP_338602517.1 |
| GCF_964028565.1 | NZ_OZ034791.1 | 1659430 | 1660620 | 1190 | - | tuf | WP_013706424.1 |
| GCF_947251665.1 | NZ_OX366374.1 | 1676266 | 1677456 | 1190 | - | tuf | WP_013706424.1 |
| GCF_947251665.1 | NZ_OX366374.1 | 2492666 | 2493856 | 1190 | - | tuf | WP_012611045.1 |
| GCF_964028365.1 | NZ_OZ034773.1 | 2507530 | 2508720 | 1190 | - | tuf | WP_012611045.1 |
| GCF_964028365.1 | NZ_OZ034773.1 | 2090080 | 2091270 | 1190 | + | tuf | WP_111956861.1 |
| GCF_964028435.1 | NZ_OZ034774.1 | 2653757 | 2654947 | 1190 | + | tuf | WP_004073802.1 |
| GCF_964028435.1 | NZ_OZ034774.1 | 4067223 | 4068413 | 1190 | - | tuf | WP_014958831.1 |
| GCF_964026965.1 | NZ_OZ034717.1 | 4081694 | 4082884 | 1190 | - | tuf | WP_014958831.1 |
| GCF_964026965.1 | NZ_OZ034717.1 | 3347846 | 3349036 | 1190 | - | tuf | WP_070962341.1 |
| GCF_947251575.1 | NZ_OX366353.1 | 3362608 | 3363798 | 1190 | - | tuf | WP_070962341.1 |
| GCF_947251575.1 | NZ_OX366353.1 | 5184506 | 5185699 | 1193 | + | tuf | WP_236889854.1 |
| GCF_964277395.1 | NZ_OZ195519.1 | 5199256 | 5200449 | 1193 | + | tuf | WP_236889864.1 |
| GCF_964277395.1 | NZ_OZ195519.1 | 1189965 | 1191155 | 1190 | + | tuf | WP_207690645.1 |
| GCF_964028545.1 | NZ_OZ034789.1 | 1204623 | 1205813 | 1190 | + | tuf | WP_207690655.1 |
| GCF_964028545.1 | NZ_OZ034789.1 | 1030403 | 1031593 | 1190 | + | tuf | WP_207681444.1 |
| GCF_947250755.1 | NZ_OX366345.1 | 4076478 | 4077668 | 1190 | - | tuf | WP_041273359.1 |
| GCF_947250755.1 | NZ_OX366345.1 | 4091089 | 4092279 | 1190 | - | tuf | WP_015905455.1 |
| GCF_964026845.1 | NZ_OZ034705.1 | 2379550 | 2380740 | 1190 | - | tuf | WP_155320377.1 |
| GCF_964026845.1 | NZ_OZ034705.1 | 2394397 | 2395587 | 1190 | - | tuf | WP_155320377.1 |
| GCF_964028455.1 | NZ_OZ034765.1 | 6962102 | 6963292 | 1190 | + | tuf | WP_155313290.1 |
| GCF_964028455.1 | NZ_OZ034765.1 | 6976948 | 6978138 | 1190 | + | tuf | WP_155313290.1 |
| GCF_964028275.1 | NZ_OZ034751.1 | 2191274 | 2192464 | 1190 | + | tuf | WP_419661508.1 |
| GCF_964028275.1 | NZ_OZ034751.1 | 2206167 | 2207357 | 1190 | + | tuf | WP_419661508.1 |
| GCF_964030705.1 | NZ_OZ034914.1 | 2416997 | 2418187 | 1190 | - | tuf | WP_155303870.1 |
| GCF_964030705.1 | NZ_OZ034914.1 | 2431858 | 2433048 | 1190 | - | tuf | WP_155303870.1 |
| GCF_964028375.1 | NZ_OZ034783.1 | 824525 | 825715 | 1190 | + | tuf | WP_012174124.1 |
| GCF_964028375.1 | NZ_OZ034783.1 | 839167 | 840357 | 1190 | + | tuf | WP_012174124.1 |

| | | | | | | | |
|---|---|---|---|---|---|---|---|
| GCF_947250485.1 | NZ_OX366322.1 | 1667781 | 1668968 | 1187 | + | tuf | WP_199261106.1 |
| GCF_947250485.1 | NZ_OX366322.1 | 1684253 | 1685440 | 1187 | + | tuf | WP_199261106.1 |
| GCF_947250475.1 | NZ_OX366320.1 | 2424820 | 2426007 | 1187 | - | tuf | WP_104937110.1 |
| GCF_947250475.1 | NZ_OX366320.1 | 2441319 | 2442506 | 1187 | - | tuf | WP_104937110.1 |
| GCF_947251835.1 | NZ_OX366384.1 | 862847 | 864034 | 1187 | + | tuf | WP_015723495.1 |
| GCF_947251835.1 | NZ_OX366384.1 | 879338 | 880525 | 1187 | + | tuf | WP_015723495.1 |
| GCF_947250705.1 | NZ_OX366341.1 | 2296532 | 2297719 | 1187 | + | tuf | WP_015404268.1 |
| GCF_947250705.1 | NZ_OX366341.1 | 2313577 | 2314764 | 1187 | + | tuf | WP_015404268.1 |
| GCF_964026975.1 | NZ_OZ034718.1 | 520698 | 521885 | 1187 | + | tuf | WP_284153214.1 |
| GCF_964026975.1 | NZ_OZ034718.1 | 537385 | 538572 | 1187 | + | tuf | WP_284153227.1 |
| GCF_947251845.1 | NZ_OX366383.1 | 2596558 | 2597745 | 1187 | - | tuf | WP_267926716.1 |
| GCF_947251845.1 | NZ_OX366383.1 | 2613370 | 2614557 | 1187 | - | tuf | WP_267926716.1 |
| GCF_947250595.1 | NZ_OX366330.1 | 517823 | 519010 | 1187 | + | tuf | WP_228855950.1 |
| GCF_947250595.1 | NZ_OX366330.1 | 534524 | 535708 | 1184 | + | tuf | WP_228855963.1 |
| GCF_947251825.1 | NZ_OX366391.1 | 508094 | 509281 | 1187 | + | tuf | WP_136810561.1 |
| GCF_947251825.1 | NZ_OX366391.1 | 524889 | 526076 | 1187 | + | tuf | WP_136810573.1 |
| GCF_947250515.1 | NZ_OX366321.1 | 5068625 | 5069812 | 1187 | - | tuf | WP_136798084.1 |
| GCF_947250515.1 | NZ_OX366321.1 | 5085435 | 5086622 | 1187 | - | tuf | WP_136798096.1 |
| GCF_964026945.1 | NZ_OZ034711.1 | 1254233 | 1255420 | 1187 | + | tuf | WP_011188352.1 |
| GCF_964026945.1 | NZ_OZ034711.1 | 1271102 | 1272289 | 1187 | + | tuf | WP_011188365.1 |
| GCF_947251815.1 | NZ_OX366390.1 | 1698265 | 1699452 | 1187 | - | tuf | WP_013163677.1 |
| GCF_947251815.1 | NZ_OX366390.1 | 1715531 | 1716718 | 1187 | - | tuf | WP_013163677.1 |
| GCF_947250525.1 | NZ_OX366323.1 | 6054108 | 6055298 | 1190 | + | tuf | WP_014812936.1 |
| GCF_947250525.1 | NZ_OX366323.1 | 6072157 | 6073347 | 1190 | + | tuf | WP_014812936.1 |
| GCF_964028235.1 | NZ_OZ034749.1 | 382272 | 383462 | 1190 | - | tuf | WP_096399336.1 |
| GCF_964028235.1 | NZ_OZ034749.1 | 250365 | 251555 | 1190 | - | tuf | WP_014258449.1 |
| GCF_964026695.1 | NZ_OZ034689.1 | 2446933 | 2448123 | 1190 | + | tuf | WP_014258449.1 |
| GCF_964026695.1 | NZ_OZ034689.1 | 2353221 | 2354414 | 1193 | - | tuf | WP_015752493.1 |
| GCF_964026825.1 | NZ_OZ034703.1 | 2369360 | 2370553 | 1193 | - | tuf | WP_015752493.1 |
| GCF_964026825.1 | NZ_OZ034703.1 | 198144 | 199334 | 1190 | + | tuf | WP_176629177.1 |
| GCF_947251605.1 | NZ_OX366361.1 | 1480006 | 1481196 | 1190 | - | tuf | WP_176629177.1 |

| | | | | | | | |
|---|---|---|---|---|---|---|---|
| GCF_947251605.1 | NZ_OX366361.1 | 1257011 | 1258201 | 1190 | + | tuf | WP_176630472.1 |
| GCF_947251735.1 | NZ_OX366382.1 | 1717502 | 1718692 | 1190 | + | tuf | WP_176630472.1 |
| GCF_947251735.1 | NZ_OX366382.1 | 3152179 | 3153369 | 1190 | - | tuf | WP_015774941.1 |
| GCF_964026905.1 | NZ_OZ034708.1 | 3168262 | 3169452 | 1190 | - | tuf | WP_015774953.1 |
| GCF_964026905.1 | NZ_OZ034708.1 | 225899 | 227089 | 1190 | - | tuf | WP_066602121.1 |
| GCF_947250575.1 | NZ_OX366325.1 | 242027 | 243298 | 1271 | - | tuf | WP_083517780.1 |
| GCF_947250575.1 | NZ_OX366325.1 | 3437773 | 3438963 | 1190 | + | tuf | WP_027175311.1 |
| GCF_947250545.1 | NZ_OX366331.1 | 2332258 | 2333448 | 1190 | + | tuf | WP_022658438.1 |
| GCF_947250545.1 | NZ_OX366331.1 | 866064 | 867254 | 1190 | - | tuf | WP_008684736.1 |
| GCF_947250645.1 | NZ_OX366334.1 | 414609 | 415799 | 1190 | + | tuf | WP_126376003.1 |
| GCF_947250645.1 | NZ_OX366334.1 | 438945 | 440135 | 1190 | + | tuf | WP_126376003.1 |
| GCF_947250695.1 | NZ_OX366337.1 | 3619174 | 3620364 | 1190 | - | tuf | WP_265826566.1 |
| GCF_947250695.1 | NZ_OX366337.1 | 3429618 | 3430808 | 1190 | - | tuf | WP_174404409.1 |
| GCF_947250535.1 | NZ_OX366324.1 | 1163815 | 1165005 | 1190 | - | tuf | WP_011527018.1 |
| GCF_947250535.1 | NZ_OX366324.1 | 3427016 | 3428206 | 1190 | - | tuf | WP_015337874.1 |
| GCF_964030915.1 | NZ_OZ034987.1 | 1303108 | 1304298 | 1190 | + | tuf | WP_015851053.1 |
| GCF_964030915.1 | NZ_OZ034987.1 | 1883660 | 1884850 | 1190 | + | tuf | WP_021760441.1 |
| GCF_964028595.1 | NZ_OZ034799.1 | 571590 | 572780 | 1190 | - | tuf | WP_010940180.1 |
| GCF_964028595.1 | NZ_OZ034799.1 | 308428 | 309618 | 1190 | + | tuf | WP_171266355.1 |
| GCF_964026725.1 | NZ_OZ034691.1 | 3855214 | 3856404 | 1190 | + | tuf | WP_171266355.1 |
| GCF_964026725.1 | NZ_OZ034691.1 | 2969589 | 2970779 | 1190 | + | tuf | WP_011368756.1 |
| GCF_964028355.1 | NZ_OZ034761.1 | 2474791 | 2475981 | 1190 | - | tuf | WP_013515186.1 |
| GCF_964028355.1 | NZ_OZ034761.1 | 2586341 | 2587531 | 1190 | + | tuf | WP_158948336.1 |
| GCF_964028515.1 | NZ_OZ034788.1 | 3781898 | 3783088 | 1190 | + | tuf | WP_014324013.1 |
| GCF_964028515.1 | NZ_OZ034788.1 | 1911446 | 1912636 | 1190 | + | tuf | WP_338670015.1 |
| GCF_964028625.1 | NZ_OZ034797.1 | 3692531 | 3693721 | 1190 | + | tuf | WP_281761508.1 |
| GCF_964028625.1 | NZ_OZ034797.1 | 3577214 | 3578404 | 1190 | + | tuf | WP_015416558.1 |
| GCF_947251975.1 | NZ_OX366401.1 | 3009417 | 3010607 | 1190 | + | tuf | WP_264982166.1 |
| GCF_947251975.1 | NZ_OX366401.1 | 4100701 | 4101891 | 1190 | + | tuf | WP_097013534.1 |
| GCF_964026785.1 | NZ_OZ034700.1 | 3696822 | 3698012 | 1190 | + | tuf | WP_229592175.1 |
| GCF_964026785.1 | NZ_OZ034700.1 | 1246951 | 1248141 | 1190 | + | tuf | WP_269941392.1 |

| | | | | | | | |
|---|---|---|---|---|---|---|---|
| GCF_947251585.1 | NZ_OX366371.1 | 131497 | 132687 | 1190 | + | tuf | WP_243545550.1 |
| GCF_947251585.1 | NZ_OX366371.1 | 3259769 | 3260959 | 1190 | - | tuf | WP_147822077.1 |
| GCF_964263235.1 | NZ_OZ181655.1 | 1493091 | 1494281 | 1190 | + | tuf | WP_015859931.1 |
| GCF_964263235.1 | NZ_OZ181655.1 | 3396580 | 3397770 | 1190 | - | tuf | WP_015859931.1 |
| GCF_964028135.1 | NZ_OZ034758.1 | 329478 | 330668 | 1190 | + | tuf | WP_009179754.1 |
| GCF_964028135.1 | NZ_OZ034758.1 | 3641767 | 3642957 | 1190 | + | tuf | WP_009182549.1 |
| GCF_947251875.1 | NZ_OX366392.1 | 1339276 | 1340466 | 1190 | + | tuf | WP_015859931.1 |
| GCF_947251875.1 | NZ_OX366392.1 | 3537718 | 3538908 | 1190 | - | tuf | WP_015859931.1 |
| GCF_964030625.1 | NZ_OZ034910.1 | 2124905 | 2126095 | 1190 | + | tuf | WP_286438244.1 |
| GCF_964030625.1 | NZ_OZ034910.1 | 1351864 | 1353045 | 1181 | - | tuf | WP_013682367.1 |
| GCF_947251465.1 | NZ_OX366366.1 | 1367975 | 1369156 | 1181 | - | tuf | WP_013682367.1 |
| GCF_947251465.1 | NZ_OX366366.1 | 1080005 | 1081192 | 1187 | + | tuf | WP_012529351.1 |
| GCF_964277435.1 | NZ_OZ195524.1 | 1096444 | 1097631 | 1187 | + | tuf | WP_012529364.1 |
| GCF_964277435.1 | NZ_OZ195524.1 | 3688538 | 3689725 | 1187 | - | tuf | WP_085815242.1 |
| GCF_947250465.1 | NZ_OX366317.1 | 3705011 | 3706198 | 1187 | - | tuf | WP_085815242.1 |
| GCF_947250465.1 | NZ_OX366317.1 | 3112958 | 3114154 | 1196 | - | tuf | WP_053551585.1 |
| GCF_965178645.1 | NZ_OZ238265.1 | 3129232 | 3130428 | 1196 | - | tuf | WP_053551598.1 |
| GCF_965178645.1 | NZ_OZ238265.1 | 1518444 | 1519631 | 1187 | + | tuf | WP_092076061.1 |
| GCF_947251495.1 | NZ_OX366360.1 | 1534626 | 1535813 | 1187 | + | tuf | WP_092076061.1 |
| GCF_947251495.1 | NZ_OX366360.1 | 1261531 | 1262727 | 1196 | + | tuf | WP_221251454.1 |
| GCF_947250555.1 | NZ_OX366328.1 | 1277913 | 1279109 | 1196 | + | tuf | WP_221251454.1 |
| GCF_947250555.1 | NZ_OX366328.1 | 3258719 | 3259906 | 1187 | + | tuf | WP_260747309.1 |
| GCF_947250635.1 | NZ_OX366335.1 | 3274984 | 3276171 | 1187 | + | tuf | WP_260747320.1 |
| GCF_947250635.1 | NZ_OX366335.1 | 2702015 | 2703202 | 1187 | - | tuf | WP_040201096.1 |
| GCF_947250725.1 | NZ_OX366343.1 | 2718320 | 2719507 | 1187 | - | tuf | WP_040201096.1 |
| GCF_947250725.1 | NZ_OX366343.1 | 1853545 | 1854732 | 1187 | + | tuf | WP_207161844.1 |
| GCF_947251625.1 | NZ_OX366358.1 | 1869850 | 1871037 | 1187 | + | tuf | WP_207161844.1 |
| GCF_947251625.1 | NZ_OX366358.1 | 682481 | 683668 | 1187 | + | tuf | WP_011365695.1 |
| GCF_964027035.1 | NZ_OZ034723.1 | 698775 | 699962 | 1187 | + | tuf | WP_011365695.1 |
| GCF_964027035.1 | NZ_OZ034723.1 | 725861 | 727048 | 1187 | + | tuf | WP_039740461.1 |
| GCF_947251655.1 | NZ_OX366373.1 | 742184 | 743371 | 1187 | + | tuf | WP_039740461.1 |

| | | | | | | | |
|---|---|---|---|---|---|---|---|
| GCF_947251655.1 | NZ_OX366373.1 | 3129543 | 3130730 | 1187 | - | tuf | WP_010943488.1 |
| GCF_964028155.1 | NZ_OZ034754.1 | 3145859 | 3147046 | 1187 | - | tuf | WP_010943488.1 |
| GCF_964028155.1 | NZ_OZ034754.1 | 3547703 | 3548890 | 1187 | - | tuf | WP_216500028.1 |
| GCF_947250745.1 | NZ_OX366346.1 | 3564158 | 3565345 | 1187 | - | tuf | WP_216500028.1 |
| GCF_947250745.1 | NZ_OX366346.1 | 3560785 | 3561972 | 1187 | - | tuf | WP_216799378.1 |
| GCF_947251865.1 | NZ_OX366394.1 | 3577195 | 3578382 | 1187 | - | tuf | WP_216799378.1 |
| GCF_947251865.1 | NZ_OX366394.1 | 1017714 | 1018901 | 1187 | + | tuf | WP_248646986.1 |
| GCF_947250605.1 | NZ_OX366329.1 | 1034177 | 1035364 | 1187 | + | tuf | WP_183350961.1 |
| GCF_947250605.1 | NZ_OX366329.1 | 1993485 | 1994672 | 1187 | - | tuf | WP_217287374.1 |
| GCF_947250615.1 | NZ_OX366333.1 | 2009892 | 2011079 | 1187 | - | tuf | WP_217287374.1 |
| GCF_947250615.1 | NZ_OX366333.1 | 4016709 | 4017896 | 1187 | - | tuf | WP_012648695.1 |
| GCF_964188425.1 | NZ_OZ077190.1 | 4032909 | 4034096 | 1187 | - | tuf | WP_012648695.1 |
| GCF_964188425.1 | NZ_OZ077190.1 | 1255557 | 1256744 | 1187 | + | tuf | WP_011937983.1 |
| GCF_964028615.1 | NZ_OZ034795.1 | 1271835 | 1273022 | 1187 | + | tuf | WP_011937983.1 |
| GCF_964028615.1 | NZ_OZ034795.1 | 704085 | 705272 | 1187 | + | tuf | WP_011734609.1 |
| GCF_964028405.1 | NZ_OZ034777.1 | 721135 | 722322 | 1187 | + | tuf | WP_011734609.1 |
| GCF_964028405.1 | NZ_OZ034777.1 | 718318 | 719514 | 1196 | - | tuf | WP_072286011.1 |
| GCF_947251505.1 | NZ_OX366365.1 | 734569 | 735765 | 1196 | - | tuf | WP_072286011.1 |
| GCF_947251505.1 | NZ_OX366365.1 | 1657214 | 1658410 | 1196 | - | tuf | WP_072283666.1 |
| GCF_964028225.1 | NZ_OZ034752.1 | 1673484 | 1674680 | 1196 | - | tuf | WP_072283679.1 |
| GCF_964028225.1 | NZ_OZ034752.1 | 835370 | 836566 | 1196 | + | tuf | WP_011340388.1 |
| GCF_964028285.1 | NZ_OZ034766.1 | 851687 | 852883 | 1196 | + | tuf | WP_011340401.1 |
| GCF_964028285.1 | NZ_OZ034766.1 | 816992 | 818179 | 1187 | + | tuf | WP_305731492.1 |
| GCF_947250655.1 | NZ_OX366336.1 | 833890 | 835077 | 1187 | + | tuf | WP_305731492.1 |
| GCF_947250655.1 | NZ_OX366336.1 | 1376905 | 1378092 | 1187 | + | tuf | WP_012469398.1 |
| GCF_964028585.1 | NZ_OZ034794.1 | 1393820 | 1395007 | 1187 | + | tuf | WP_012469398.1 |
| GCF_964028585.1 | NZ_OZ034794.1 | 399352 | 400551 | 1199 | + | tuf | WP_241422196.1 |
| GCF_947250675.1 | NZ_OX366340.1 | 845661 | 846875 | 1214 | + | tuf | WP_012547866.1 |
| GCF_947250675.1 | NZ_OX366340.1 | 1208974 | 1210188 | 1214 | - | tuf | WP_012547745.1 |
| GCF_964028205.1 | NZ_OZ034771.1 | 1013046 | 1014260 | 1214 | + | tuf | WP_012583343.1 |
| GCF_964028205.1 | NZ_OZ034771.1 | 1389279 | 1390493 | 1214 | - | tuf | WP_012583343.1 |

| | | | | | | | |
|---|---|---|---|---|---|---|---|
| GCF_964026795.1 | NZ_OZ034698.1 | 1511550 | 1512734 | 1184 | - | tuf | WP_012415587.1 |
| GCF_964026795.1 | NZ_OZ034698.1 | 1591337 | 1592521 | 1184 | - | tuf | WP_012415587.1 |
| GCF_964026735.1 | NZ_OZ034692.1 | 146020 | 147216 | 1196 | + | tuf | WP_015423092.1 |
| GCF_964026735.1 | NZ_OZ034692.1 | 338850 | 340046 | 1196 | + | tuf | WP_015423092.1 |
| GCF_017896245.1 | NZ_CP072669.1 | 963236 | 964432 | 1196 | - | tuf | WP_052570717.1 |
| GCF_017896245.1 | NZ_CP072669.1 | 1547373 | 1548569 | 1196 | + | tuf | WP_052570717.1 |
| GCF_024804185.1 | NZ_CP101657.1 | 1896573 | 1897778 | 1205 | - | tuf | WP_004511262.1 |
| GCF_024804185.1 | NZ_CP101657.1 | 583272 | 584477 | 1205 | + | tuf | WP_024775592.1 |
| GCF_042182995.1 | NZ_CP102058.1 | 446598 | 447803 | 1205 | + | tuf | WP_105918105.1 |
| GCF_042182995.1 | NZ_CP102058.1 | 317190 | 318395 | 1205 | + | tuf | WP_026806437.1 |
| GCF_040207875.1 | NZ_CP157942.1 | 1796175 | 1797380 | 1205 | - | tuf | WP_026803894.1 |
| GCF_040207875.1 | NZ_CP157942.1 | 459194 | 460399 | 1205 | + | tuf | WP_066350498.1 |
| GCF_003999585.1 | NZ_CP016430.1 | 1490492 | 1491697 | 1205 | - | tuf | WP_066171420.1 |
| GCF_003999585.1 | NZ_CP016430.1 | 1503993 | 1505198 | 1205 | - | tuf | WP_115428875.1 |
| GCF_012030695.1 | NZ_CP050521.1 | 917494 | 918699 | 1205 | - | tuf | WP_113909562.1 |
| GCF_012030695.1 | NZ_CP050521.1 | 2517585 | 2518790 | 1205 | - | tuf | WP_172127527.1 |
| GCF_964277405.1 | NZ_OZ195521.1 | 2137235 | 2138440 | 1205 | - | tuf | WP_129095361.1 |
| GCF_964277405.1 | NZ_OZ195521.1 | 2262484 | 2263689 | 1205 | - | tuf | WP_129013497.1 |
| GCF_006542295.1 | NZ_CP041215.1 | 2606088 | 2607293 | 1205 | - | tuf | WP_129012191.1 |
| GCF_006542295.1 | NZ_CP041215.1 | 2415711 | 2416916 | 1205 | - | tuf | WP_118918229.1 |
| GCF_017869155.1 | NZ_CP054598.1 | 2637827 | 2639032 | 1205 | - | tuf | WP_013136418.1 |
| GCF_017869155.1 | NZ_CP054598.1 | 2514020 | 2515225 | 1205 | + | tuf | WP_076088264.1 |
| GCF_008245065.1 | NZ_CP021120.1 | 1579273 | 1580478 | 1205 | - | tuf | WP_066171420.1 |
| GCF_008245065.1 | NZ_CP021120.1 | 2611240 | 2612445 | 1205 | - | tuf | WP_254576230.1 |
| GCF_000829315.1 | NZ_AP013028.1 | 2242880 | 2244085 | 1205 | - | tuf | WP_118887240.1 |
| GCF_000829315.1 | NZ_AP013028.1 | 2679654 | 2680859 | 1205 | - | tuf | WP_128359744.1 |
| GCF_013365475.1 | NZ_CP046579.1 | 21497 | 22693 | 1196 | - | tuf | WP_138322676.1 |
| GCF_013365475.1 | NZ_CP046579.1 | 1569576 | 1570772 | 1196 | - | tuf | WP_123352908.1 |
| GCF_012277315.1 | NZ_CP051157.1 | 408892 | 410088 | 1196 | + | tuf | WP_139427114.1 |
| GCF_012277315.1 | NZ_CP051157.1 | 1107702 | 1108898 | 1196 | - | tuf | WP_094325565.1 |
| GCF_012277295.1 | NZ_CP051156.1 | 340822 | 342018 | 1196 | + | tuf | WP_158334875.1 |

| | | | | | | | |
|---|---|---|---|---|---|---|---|
| GCF_012277295.1 | NZ_CP051156.1 | 405932 | 407128 | 1196 | + | tuf | WP_106871538.1 |
| GCF_000073005.1 | NC_010981.1 | 1272800 | 1273996 | 1196 | - | tuf | WP_169937767.1 |
| GCF_000073005.1 | NC_010981.1 | 48919 | 50115 | 1196 | + | tuf | WP_172229688.1 |
| GCF_021609905.1 | NZ_CP091195.1 | 1016677 | 1017873 | 1196 | - | tuf | WP_002779448.1 |
| GCF_021609905.1 | NZ_CP091195.1 | 674693 | 675889 | 1196 | + | tuf | WP_021090663.1 |
| GCF_013458815.1 | NZ_CP051608.1 | 354379 | 355575 | 1196 | + | tuf | WP_025803105.1 |
| GCF_013458815.1 | NZ_CP051608.1 | 619752 | 620948 | 1196 | + | tuf | WP_027306289.1 |
| GCF_013365495.1 | NZ_CP046580.1 | 1409588 | 1410784 | 1196 | - | tuf | WP_009650564.1 |
| GCF_013365495.1 | NZ_CP046580.1 | 1318238 | 1319434 | 1196 | - | tuf | WP_002850129.1 |
| GCF_013365455.1 | NZ_CP046578.1 | 505444 | 506640 | 1196 | + | tuf | WP_075494174.1 |
| GCF_013365455.1 | NZ_CP046578.1 | 1668863 | 1670059 | 1196 | - | tuf | WP_040304280.1 |
| GCF_032848195.1 | NZ_CP136586.1 | 314983 | 316179 | 1196 | + | tuf | WP_082199256.1 |
| GCF_032848195.1 | NZ_CP136586.1 | 53031 | 54227 | 1196 | + | tuf | WP_066777275.1 |
| GCF_008033215.1 | NZ_CP042904.1 | 1594186 | 1595382 | 1196 | - | tuf | WP_012109494.1 |
| GCF_008033215.1 | NZ_CP042904.1 | 446835 | 448031 | 1196 | + | tuf | WP_059431330.1 |
| GCF_001758565.1 | NZ_CP011148.1 | 1337099 | 1338295 | 1196 | - | tuf | WP_038454838.1 |
| GCF_001758565.1 | NZ_CP011148.1 | 389910 | 391106 | 1196 | + | tuf | WP_039649445.1 |
| GCF_004795975.1 | NZ_CP034335.1 | 434408 | 435604 | 1196 | + | tuf | YP_002343904.1 |
| GCF_004795975.1 | NZ_CP034335.1 | 491027 | 492223 | 1196 | + | tuf | WP_096013476.1 |
| GCF_016584425.1 | NZ_CP046925.1 | 411164 | 412360 | 1196 | + | tuf | WP_039668122.1 |
| GCF_016584425.1 | NZ_CP046925.1 | 445721 | 446917 | 1196 | + | tuf | WP_034967874.1 |
| GCF_026015925.1 | NZ_CP110359.1 | 506179 | 507375 | 1196 | + | tuf | WP_131163605.1 |
| GCF_026015925.1 | NZ_CP110359.1 | 421277 | 422473 | 1196 | + | tuf | WP_043019496.1 |
| GCF_018467135.1 | NZ_CP069060.1 | 451788 | 452984 | 1196 | + | tuf | WP_044598446.1 |
| GCF_018467135.1 | NZ_CP069060.1 | 1334941 | 1336137 | 1196 | - | tuf | WP_078423731.1 |
| GCF_000376605.1 | NC_021089.1 | 571448 | 572644 | 1196 | + | tuf | WP_086237856.1 |
| GCF_000376605.1 | NC_021089.1 | 1916079 | 1917275 | 1196 | - | tuf | WP_039887818.1 |
| GCF_040084705.1 | NZ_CP157591.1 | 509416 | 510612 | 1196 | + | tuf | WP_002947618.1 |
| GCF_040084705.1 | NZ_CP157591.1 | 454286 | 455482 | 1196 | + | tuf | WP_033916502.1 |
| GCF_018467115.1 | NZ_CP069053.1 | 426362 | 427558 | 1196 | + | tuf | WP_043019496.1 |
| GCF_018467115.1 | NZ_CP069053.1 | 1309326 | 1310522 | 1196 | - | tuf | WP_004277763.1 |

| | | | | | | | |
|---|---|---|---|---|---|---|---|
| GCF_039540065.1 | NZ_CP154278.1 | 1435699 | 1436895 | 1196 | - | tuf | WP_018712753.1 |
| GCF_039540065.1 | NZ_CP154278.1 | 411724 | 412920 | 1196 | + | tuf | WP_086246781.1 |
| GCF_001931755.2 | NZ_CP015510.2 | 399901 | 401097 | 1196 | + | tuf | WP_039665247.1 |
| GCF_001931755.2 | NZ_CP015510.2 | 317193 | 318389 | 1196 | + | tuf | WP_099461586.1 |
| GCF_029856955.1 | NZ_CP097148.1 | 729841 | 731037 | 1196 | + | tuf | WP_179975809.1 |
| GCF_029856955.1 | NZ_CP097148.1 | 2263603 | 2264799 | 1196 | - | tuf | WP_194369124.1 |
| GCF_025021925.1 | NZ_CP092368.1 | 272939 | 274135 | 1196 | + | tuf | WP_194366985.1 |
| GCF_042928915.1 | NZ_CP170632.1 | 2490389 | 2491594 | 1205 | - | tuf | WP_129082174.1 |
| GCF_042928915.1 | NZ_CP170632.1 | 363694 | 364899 | 1205 | + | tuf | WP_114838271.1 |
| GCF_013365435.1 | NZ_CP046577.1 | 2554629 | 2555834 | 1205 | - | tuf | WP_129088074.1 |
| GCF_936270145.1 | NZ_OW370537.1 | 1378421 | 1379617 | 1196 | - | tuf | WP_011578381.1 |
| GCF_936270145.1 | NZ_OW370537.1 | 1651259 | 1652455 | 1196 | + | tuf | WP_034553869.1 |
| GCF_000306885.1 | NC_018267.1 | 210224 | 211420 | 1196 | + | tuf | WP_006655954.1 |
| GCF_000306885.1 | NC_018267.1 | 621942 | 623138 | 1196 | - | tuf | WP_014659111.1 |
| GCF_000530755.1 | NZ_HG810405.1 | 804854 | 806050 | 1196 | - | tuf | WP_104724713.1 |
| GCF_000530755.1 | NZ_HG810405.1 | 1507845 | 1509041 | 1196 | - | tuf | WP_002955418.1 |
| GCF_025617515.1 | NZ_CP092526.1 | 932784 | 933980 | 1196 | - | tuf | WP_066340225.1 |
| GCF_025617515.1 | NZ_CP092526.1 | 503165 | 504361 | 1196 | + | tuf | WP_013468981.1 |
| GCF_023559125.1 | NZ_CP096925.1 | 683294 | 684490 | 1196 | + | tuf | WP_286349256.1 |
| GCF_023559125.1 | NZ_CP096925.1 | 1141986 | 1143182 | 1196 | + | tuf | WP_221279194.1 |
| GCF_023559145.1 | NZ_CP096926.1 | 359584 | 360780 | 1196 | - | tuf | WP_011115211.1 |
| GCF_023559145.1 | NZ_CP096926.1 | 754958 | 756154 | 1196 | + | tuf | WP_066388108.1 |
| GCF_918697765.1 | NZ_OU953855.1 | 230389 | 231585 | 1196 | - | tuf | WP_295699162.1 |
| GCF_918697765.1 | NZ_OU953855.1 | 369590 | 370786 | 1196 | + | tuf | WP_013022666.1 |
| GCF_918342435.1 | NZ_OU909470.1 | 1532398 | 1533594 | 1196 | - | tuf | WP_005021383.1 |
| GCF_918342435.1 | NZ_OU909470.1 | 1449261 | 1450457 | 1196 | + | tuf | WP_001040572.1 |
| GCF_048537655.1 | NZ_CP175927.1 | 1430709 | 1431905 | 1196 | + | tuf | WP_006564788.1 |
| GCF_048537655.1 | NZ_CP175927.1 | 1195358 | 1196554 | 1196 | - | tuf | WP_034326281.1 |
| GCF_018141665.1 | NZ_CP067976.1 | 182673 | 183872 | 1199 | + | tuf | WP_006802146.1 |
| GCF_018141665.1 | NZ_CP067976.1 | 1526078 | 1527274 | 1196 | + | tuf | WP_286336210.1 |
| GCF_046529255.1 | NZ_CP114181.1 | 1471544 | 1472740 | 1196 | - | tuf | WP_353661427.1 |

| | | | | | | | |
|---|---|---|---|---|---|---|---|
| GCF_046529255.1 | NZ_CP114181.1 | 170385 | 171581 | 1196 | + | tuf | WP_092910249.1 |
| GCF_001439985.1 | NZ_CM003641.1 | 1317510 | 1318706 | 1196 | - | tuf | WP_201351947.1 |
| GCF_001439985.1 | NZ_CM003641.1 | 2334509 | 2335714 | 1205 | - | tuf | WP_099334243.1 |
| GCF_017869285.1 | NZ_CP054557.1 | 2274634 | 2275839 | 1205 | - | tuf | WP_101184936.1 |
| GCF_017869285.1 | NZ_CP054557.1 | 2387506 | 2388711 | 1205 | - | tuf | WP_099310468.1 |
| GCF_032849005.1 | NZ_CP136587.1 | 2309632 | 2310837 | 1205 | - | tuf | WP_099343445.1 |
| GCF_032849005.1 | NZ_CP136587.1 | 2332358 | 2333563 | 1205 | - | tuf | WP_114842665.1 |
| GCF_000008385.1 | NC_006833.1 | 2133200 | 2134405 | 1205 | - | tuf | WP_130234113.1 |
| GCF_000008385.1 | NC_006833.1 | 147102 | 148298 | 1196 | + | tuf | WP_015902389.1 |
| GCF_007971685.1 | NZ_CP042446.1 | 1943436 | 1944641 | 1205 | - | tuf | WP_013554900.1 |
| GCF_007971685.1 | NZ_CP042446.1 | 1375179 | 1376375 | 1196 | - | tuf | WP_084275194.1 |
| GCF_016807805.1 | NZ_CP063362.1 | 177136 | 178332 | 1196 | + | tuf | WP_200763219.1 |
| GCF_016807805.1 | NZ_CP063362.1 | 819455 | 820651 | 1196 | + | tuf | WP_281951956.1 |
| GCF_038443675.1 | NZ_CP151762.1 | 160626 | 161822 | 1196 | + | tuf | WP_187647783.1 |
| GCF_038443675.1 | NZ_CP151762.1 | 3026092 | 3027297 | 1205 | - | tuf | WP_071626842.1 |
| GCF_038069655.2 | NZ_CP150951.2 | 2365833 | 2367038 | 1205 | - | tuf | WP_076088264.1 |
| GCF_038069655.2 | NZ_CP150951.2 | 2304929 | 2306125 | 1196 | - | tuf | WP_013461135.1 |
| GCF_038443705.2 | NZ_CP151767.2 | 334677 | 335873 | 1196 | + | tuf | WP_207562143.1 |
| GCF_038443705.2 | NZ_CP151767.2 | 285960 | 287156 | 1196 | + | tuf | WP_013326116.1 |
| GCF_000498655.1 | NC_022900.1 | 343130 | 344326 | 1196 | + | tuf | WP_011371980.1 |
| GCF_000022305.1 | NC_011992.1 | 368835 | 370031 | 1196 | + | tuf | WP_345970563.1 |
| GCF_000022305.1 | NC_011992.1 | 943684 | 944880 | 1196 | - | tuf | WP_008339112.1 |
| GCF_013267395.1 | NZ_CP053987.1 | 2021853 | 2023049 | 1196 | - | tuf | WP_151901157.1 |
| GCF_013267395.1 | NZ_CP053987.1 | 2017825 | 2019021 | 1196 | - | tuf | WP_152308005.1 |
| GCF_900637265.1 | NZ_LR134361.1 | 246439 | 247635 | 1196 | + | tuf | WP_193113881.1 |
| GCF_900637265.1 | NZ_LR134361.1 | 223128 | 224324 | 1196 | + | tuf | WP_193111245.1 |
| GCF_029866465.1 | NZ_CP123364.1 | 324076 | 325272 | 1196 | + | tuf | WP_193151096.1 |
| GCF_029866465.1 | NZ_CP123364.1 | 239260 | 240456 | 1196 | + | tuf | WP_152298704.1 |
| GCF_002082135.1 | NZ_CP017433.1 | 349513 | 350709 | 1196 | + | tuf | WP_014768580.1 |
| GCF_002082135.1 | NZ_CP017433.1 | 325495 | 326691 | 1196 | + | tuf | WP_012856148.1 |
| GCF_030864005.1 | NZ_CP132976.1 | 401592 | 402788 | 1196 | + | tuf | WP_096045766.1 |

| | | | | | | | |
|---|---|---|---|---|---|---|---|
| GCF_030864005.1 | NZ_CP132976.1 | 476983 | 478179 | 1196 | + | tuf | WP_069477217.1 |
| GCF_002812705.1 | NZ_CP025030.1 | 452362 | 453558 | 1196 | + | tuf | WP_025343665.1 |
| GCF_002812705.1 | NZ_CP025030.1 | 2062072 | 2063277 | 1205 | - | tuf | WP_197548492.1 |
| GCF_037619525.1 | NZ_CP148753.1 | 234144 | 235349 | 1205 | + | tuf | WP_046550269.1 |
| GCF_037619525.1 | NZ_CP148753.1 | 138670 | 139878 | 1208 | + | tuf | WP_229937294.1 |
| GCF_016728825.1 | NZ_CP068211.1 | 1484360 | 1485556 | 1196 | + | tuf | WP_041571700.1 |
| GCF_016728825.1 | NZ_CP068211.1 | 758927 | 760108 | 1181 | + | tuf | WP_002607408.1 |
| GCF_002157165.1 | NZ_CP021359.1 | 1950255 | 1951436 | 1181 | + | tuf | WP_230226609.1 |
| GCF_002157165.1 | NZ_CP021359.1 | 1614731 | 1615912 | 1181 | + | tuf | WP_290137900.1 |
| GCF_042847045.1 | NZ_AP035783.1 | 2322205 | 2323386 | 1181 | + | tuf | WP_115715796.1 |
| GCF_042847045.1 | NZ_AP035783.1 | 655617 | 656798 | 1181 | + | tuf | WP_006505718.1 |
| GCF_028596105.1 | NZ_CP117193.1 | 1334457 | 1335638 | 1181 | - | tuf | WP_003775179.1 |
| GCF_028596105.1 | NZ_CP117193.1 | 124143 | 125324 | 1181 | - | tuf | WP_323611372.1 |
| GCF_000521505.1 | NZ_CP003915.1 | 1232089 | 1233270 | 1181 | + | tuf | WP_187533056.1 |
| GCF_000521505.1 | NZ_CP003915.1 | 738766 | 739947 | 1181 | + | tuf | WP_067631459.1 |
| GCF_019343455.1 | NZ_CP049362.1 | 848987 | 850168 | 1181 | + | tuf | WP_018579501.1 |
| GCF_019343455.1 | NZ_CP049362.1 | 1244753 | 1245934 | 1181 | - | tuf | WP_013853154.1 |
| GCF_023373785.1 | NZ_CP094619.1 | 465342 | 466523 | 1181 | + | tuf | WP_022002420.1 |
| GCF_023373785.1 | NZ_CP094619.1 | 404156 | 405337 | 1181 | + | tuf | WP_067554867.1 |
| GCF_000204645.1 | NC_015422.1 | 749550 | 750731 | 1181 | + | tuf | WP_125118706.1 |
| GCF_000204645.1 | NC_015422.1 | 332401 | 333582 | 1181 | + | tuf | WP_008690123.1 |
| GCF_022871045.2 | NZ_CP091519.2 | 2228327 | 2229508 | 1181 | - | tuf | WP_087357336.1 |
| GCF_022871045.2 | NZ_CP091519.2 | 2401377 | 2402567 | 1190 | - | tuf | WP_028078654.1 |
| GCF_020162295.1 | NZ_CP083931.1 | 2245630 | 2246811 | 1181 | + | tuf | WP_249029468.1 |
| GCF_020162295.1 | NZ_CP083931.1 | 3165879 | 3167060 | 1181 | + | tuf | WP_003539660.1 |
| GCF_049675895.1 | NZ_CP183834.1 | 2400540 | 2401721 | 1181 | - | tuf | WP_004610979.1 |
| GCF_049675895.1 | NZ_CP183834.1 | 958475 | 959656 | 1181 | + | tuf | WP_055275531.1 |
| GCF_003100395.1 | NZ_CP029210.1 | 93945 | 95126 | 1181 | + | tuf | WP_161831759.1 |
| GCF_003100395.1 | NZ_CP029210.1 | 1139944 | 1141125 | 1181 | + | tuf | WP_132942678.1 |
| GCF_002892535.1 | NZ_CP024847.1 | 2026520 | 2027701 | 1181 | - | tuf | WP_014545994.1 |
| GCF_002892535.1 | NZ_CP024847.1 | 2183085 | 2184266 | 1181 | - | tuf | WP_014545994.1 |

| | | | | | | | |
|---|---|---|---|---|---|---|---|
| GCF_023573145.1 | NZ_CP097636.1 | 4793851 | 4795059 | 1208 | - | tuf | WP_025228350.1 |
| GCF_023573145.1 | NZ_CP097636.1 | 3364051 | 3365235 | 1184 | + | tuf | WP_255842115.1 |
| GCF_005155025.1 | NZ_CP039731.1 | 299452 | 300636 | 1184 | - | tuf | WP_228852242.1 |
| GCF_005155025.1 | NZ_CP039731.1 | 5496 | 6680 | 1184 | + | tuf | WP_128248619.1 |
| GCF_009856625.1 | NZ_CP047241.1 | 2025463 | 2026647 | 1184 | + | tuf | WP_202335376.1 |
| GCF_009856625.1 | NZ_CP047241.1 | 1930029 | 1931213 | 1184 | + | tuf | WP_279447269.1 |
| GCF_000025965.1 | NC_006513.1 | 245661 | 246845 | 1184 | - | tuf | WP_410005931.1 |
| GCF_000025965.1 | NC_006513.1 | 2151193 | 2152377 | 1184 | + | tuf | WP_014782724.1 |
| GCF_017894365.1 | NZ_CP059467.1 | 424394 | 425578 | 1184 | - | tuf | WP_240571641.1 |
| GCF_017894365.1 | NZ_CP059467.1 | 829213 | 830397 | 1184 | + | tuf | WP_138851582.1 |
| GCF_045689615.1 | NZ_CP141259.1 | 2994634 | 2995818 | 1184 | - | tuf | WP_019388698.1 |
| GCF_045689615.1 | NZ_CP141259.1 | 2362802 | 2363986 | 1184 | + | tuf | WP_014033477.1 |
| GCF_017894385.1 | NZ_CP059560.1 | 46906 | 48090 | 1184 | - | tuf | WP_139697922.1 |
| GCF_017894385.1 | NZ_CP059560.1 | 1765888 | 1767072 | 1184 | - | tuf | WP_146833245.1 |
| GCF_001682385.1 | NZ_CP016210.1 | 3522690 | 3523874 | 1184 | + | tuf | WP_139067232.1 |
| GCF_001682385.1 | NZ_CP016210.1 | 70352 | 71536 | 1184 | + | tuf | WP_128148704.1 |
| GCF_018905915.1 | NZ_CP064782.1 | 20733 | 21917 | 1184 | - | tuf | WP_031444833.1 |
| GCF_018905915.1 | NZ_CP064782.1 | 2733957 | 2735141 | 1184 | + | tuf | WP_251809299.1 |
| GCF_000236665.1 | NC_016616.1 | 3568528 | 3569709 | 1181 | - | tuf | WP_138950528.1 |
| GCF_000236665.1 | NC_016616.1 | 840219 | 841403 | 1184 | - | tuf | WP_120488296.1 |
| GCF_016858125.1 | NZ_CP064781.1 | 171718 | 172902 | 1184 | + | tuf | WP_113738178.1 |
| GCF_016858125.1 | NZ_CP064781.1 | 484012 | 485196 | 1184 | - | tuf | WP_185849665.1 |
| GCF_000743945.1 | NZ_CP009238.1 | 168257 | 169441 | 1184 | + | tuf | WP_110548600.1 |
| GCF_000743945.1 | NZ_CP009238.1 | 1645816 | 1647000 | 1184 | - | tuf | WP_252158710.1 |
| GCF_047839125.1 | NZ_CP181246.1 | 1488362 | 1489546 | 1184 | - | tuf | WP_224134260.1 |
| GCF_047839125.1 | NZ_CP181246.1 | 183682 | 184866 | 1184 | - | tuf | WP_020931606.1 |
| GCF_034424645.1 | NZ_CP139969.1 | 404435 | 405619 | 1184 | + | tuf | WP_119058163.1 |
| GCF_034424645.1 | NZ_CP139969.1 | 5723 | 6895 | 1172 | - | tuf | WP_318165776.1 |
| GCF_001676705.1 | NZ_CP016170.1 | 68446 | 69630 | 1184 | - | tuf | WP_342226899.1 |
| GCF_001676705.1 | NZ_CP016170.1 | 167952 | 169136 | 1184 | - | tuf | WP_038436163.1 |
| GCF_900636925.1 | NZ_LR134326.1 | 977226 | 978410 | 1184 | - | tuf | WP_041913636.1 |

| | | | | | | | |
|---|---|---|---|---|---|---|---|
| GCF_900636925.1 | NZ_LR134326.1 | 2226391 | 2227575 | 1184 | + | tuf | WP_042344887.1 |
| GCF_001676725.1 | NZ_CP016172.1 | 1999127 | 2000311 | 1184 | + | tuf | WP_041989985.1 |
| GCF_001676725.1 | NZ_CP016172.1 | 42149 | 43333 | 1184 | - | tuf | WP_034575764.1 |
| GCF_900637615.1 | NZ_LR134382.1 | 1467862 | 1469046 | 1184 | - | tuf | WP_002666853.1 |
| GCF_900637615.1 | NZ_LR134382.1 | 1351951 | 1353135 | 1184 | - | tuf | WP_066428030.1 |
| GCF_020735985.1 | NZ_CP085973.1 | 1419368 | 1420552 | 1184 | + | tuf | WP_015782377.1 |
| GCF_020735985.1 | NZ_CP085973.1 | 1936795 | 1937979 | 1184 | + | tuf | WP_002677857.1 |
| GCF_020735925.1 | NZ_CP085971.1 | 730478 | 731662 | 1184 | - | tuf | WP_095895093.1 |
| GCF_020735925.1 | NZ_CP085971.1 | 2691154 | 2692338 | 1184 | + | tuf | WP_013551135.1 |
| GCF_004008975.1 | NZ_CP025371.1 | 2132513 | 2133697 | 1184 | + | tuf | WP_029446957.1 |
| GCF_004008975.1 | NZ_CP025371.1 | 588000 | 589184 | 1184 | - | tuf | WP_013620114.1 |
| GCF_000067205.1 | NC_010170.1 | 612781 | 613965 | 1184 | - | tuf | WP_077400567.1 |
| GCF_000067205.1 | NC_010170.1 | 624394 | 625578 | 1184 | - | tuf | WP_142783707.1 |
| GCF_001698185.1 | NZ_CP016440.1 | 2755506 | 2756690 | 1184 | + | tuf | WP_083644856.1 |
| GCF_001698185.1 | NZ_CP016440.1 | 2526752 | 2527936 | 1184 | - | tuf | WP_011710244.1 |
| GCF_048627025.1 | NZ_CP184468.1 | 2484847 | 2486031 | 1184 | + | tuf | WP_107012519.1 |
| GCF_048627025.1 | NZ_CP184468.1 | 1404134 | 1405318 | 1184 | - | tuf | WP_072552713.1 |
| GCF_000959545.1 | NZ_CP009798.1 | 2885288 | 2886472 | 1184 | - | tuf | WP_062697363.1 |
| GCF_000959545.1 | NZ_CP009798.1 | 2599075 | 2600283 | 1208 | - | tuf | WP_126652417.1 |
| GCF_038723655.1 | NZ_CP109821.1 | 10811 | 12022 | 1211 | + | tuf | WP_076452082.1 |
| GCF_038723655.1 | NZ_CP109821.1 | 529433 | 530635 | 1202 | - | tuf | WP_123868707.1 |
| GCF_001718895.1 | NZ_CP013450.1 | 2155521 | 2156705 | 1184 | - | tuf | WP_027380911.1 |
| GCF_001718895.1 | NZ_CP013450.1 | 2079301 | 2080509 | 1208 | + | tuf | WP_034749673.1 |
| GCF_009586235.1 | NZ_CP045235.1 | 2503544 | 2504752 | 1208 | - | tuf | WP_202088548.1 |
| GCF_009586235.1 | NZ_CP045235.1 | 1093082 | 1094266 | 1184 | - | tuf | WP_002976407.1 |
| GCF_040215475.1 | NZ_CP157953.1 | 2188687 | 2189889 | 1202 | - | tuf | WP_076390704.1 |
| GCF_040215475.1 | NZ_CP157953.1 | 2717570 | 2718778 | 1208 | + | tuf | WP_066754958.1 |
| GCF_019930785.1 | NZ_CP082882.1 | 1535544 | 1536728 | 1184 | + | tuf | WP_002976407.1 |
| GCF_019930785.1 | NZ_CP082882.1 | 2876002 | 2877186 | 1184 | + | tuf | WP_282904706.1 |
| GCF_016698705.1 | NZ_CP068049.1 | 4018133 | 4019341 | 1208 | - | tuf | WP_115596457.1 |
| GCF_016698705.1 | NZ_CP068049.1 | 2593046 | 2594254 | 1208 | - | tuf | WP_034687396.1 |

| | | | | | | | |
|---|---|---|---|---|---|---|---|
| GCF_000960995.1 | NZ_CP009435.1 | 519960 | 521168 | 1208 | - | tuf | WP_103291895.1 |
| GCF_000960995.1 | NZ_CP009435.1 | 2730194 | 2731378 | 1184 | + | tuf | WP_123248723.1 |
| GCF_001513745.1 | NZ_CP013380.1 | 5099325 | 5100527 | 1202 | - | tuf | WP_123857806.1 |
| GCF_001513745.1 | NZ_CP013380.1 | 2027642 | 2028844 | 1202 | - | tuf | WP_042721320.1 |
| GCF_000012945.1 | NC_007510.1 | 2659976 | 2661178 | 1202 | - | tuf | WP_209390828.1 |
| GCF_000012945.1 | NC_007510.1 | 2850694 | 2851902 | 1208 | - | tuf | WP_214827758.1 |
| GCF_033956065.1 | NZ_CP124608.1 | 3883894 | 3885102 | 1208 | + | tuf | WP_034687396.1 |
| GCF_033956065.1 | NZ_CP124608.1 | 1554099 | 1555283 | 1184 | + | tuf | WP_133439733.1 |
| GCF_001523745.2 | NZ_CP013386.1 | 555962 | 557146 | 1184 | + | tuf | WP_029295578.1 |
| GCF_001523745.2 | NZ_CP013386.1 | 611073 | 612257 | 1184 | + | tuf | WP_243550409.1 |
| GCF_001718555.1 | NZ_CP013401.1 | 3039345 | 3040553 | 1208 | + | tuf | WP_089745515.1 |
| GCF_001718555.1 | NZ_CP013401.1 | 1276748 | 1277932 | 1184 | - | tuf | WP_095071474.1 |
| GCF_036352035.1 | NZ_CP143552.1 | 1797125 | 1798309 | 1184 | - | tuf | WP_213196721.1 |
| GCF_036352035.1 | NZ_CP143552.1 | 204271 | 205455 | 1184 | + | tuf | WP_069799579.1 |
| GCF_000959365.1 | NZ_CP009555.1 | 1877494 | 1878678 | 1184 | + | tuf | WP_186987866.1 |
| GCF_000959365.1 | NZ_CP009555.1 | 522920 | 524104 | 1184 | - | tuf | WP_179240578.1 |
| GCF_022870505.1 | NZ_CP045094.1 | 39835 | 41019 | 1184 | - | tuf | WP_013185783.1 |
| GCF_022870505.1 | NZ_CP045094.1 | 670215 | 671399 | 1184 | - | tuf | WP_210759362.1 |
| GCF_001411805.1 | NZ_CP007212.1 | 976688 | 977872 | 1184 | - | tuf | WP_193440689.1 |
| GCF_001411805.1 | NZ_CP007212.1 | 3052208 | 3053392 | 1184 | - | tuf | WP_035325197.1 |
| GCF_030297255.1 | NZ_AP028077.1 | 2910541 | 2911725 | 1184 | + | tuf | WP_009085650.1 |
| GCF_030297255.1 | NZ_AP028077.1 | 3308533 | 3309717 | 1184 | - | tuf | WP_034869614.1 |
| GCF_001028665.1 | NZ_CP011503.1 | 2299765 | 2300949 | 1184 | + | tuf | WP_016198713.1 |
| GCF_001028665.1 | NZ_CP011503.1 | 2447347 | 2448531 | 1184 | + | tuf | WP_034869614.1 |
| GCF_001524445.2 | NZ_CP013417.1 | 328452 | 329636 | 1184 | + | tuf | WP_038331067.1 |
| GCF_001524445.2 | NZ_CP013417.1 | 14374 | 15558 | 1184 | + | tuf | WP_171622822.1 |
| GCF_001718535.1 | NZ_CP013398.1 | 1222616 | 1223824 | 1208 | + | tuf | WP_089769837.1 |
| GCF_001718535.1 | NZ_CP013398.1 | 1559512 | 1560696 | 1184 | + | tuf | WP_121848083.1 |
| GCF_001718875.1 | NZ_CP013448.1 | 1411771 | 1412955 | 1184 | + | tuf | WP_230477822.1 |
| GCF_001718875.1 | NZ_CP013448.1 | 2380395 | 2381579 | 1184 | + | tuf | WP_045802446.1 |
| GCF_900240005.1 | NZ_LR025742.1 | 2741656 | 2742840 | 1184 | + | tuf | WP_112378851.1 |

| | | | | | | | |
|---|---|---|---|---|---|---|---|
| GCF_900240005.1 | NZ_LR025742.1 | 352412 | 353596 | 1184 | + | tuf | WP_108247122.1 |
| GCF_039848545.1 | NZ_CP156685.1 | 3014920 | 3016104 | 1184 | + | tuf | WP_303316168.1 |
| GCF_039848545.1 | NZ_CP156685.1 | 1418970 | 1420154 | 1184 | - | tuf | WP_102754920.1 |
| GCF_001718335.1 | NZ_CP013366.1 | 1673985 | 1675169 | 1184 | + | tuf | WP_230064136.1 |
| GCF_001718335.1 | NZ_CP013366.1 | 1790520 | 1791704 | 1184 | + | tuf | WP_268849072.1 |
| GCF_000012365.1 | NC_007651.1 | 1663727 | 1664911 | 1184 | - | tuf | WP_264432389.1 |
| GCF_000012365.1 | NC_007651.1 | 1983296 | 1984480 | 1184 | + | tuf | WP_108777911.1 |
| GCF_048565265.1 | NZ_CP183461.1 | 122847 | 124031 | 1184 | - | tuf | WP_129751645.1 |
| GCF_048565265.1 | NZ_CP183461.1 | 39855 | 41036 | 1181 | - | tuf | WP_229317261.1 |
| GCF_002097715.1 | NZ_CP020737.1 | 38806 | 39987 | 1181 | - | tuf | WP_229317261.1 |
| GCF_002097715.1 | NZ_CP020737.1 | 791278 | 792462 | 1184 | - | tuf | WP_008466928.1 |
| GCF_000959445.1 | NZ_CP009631.1 | 4539771 | 4540955 | 1184 | - | tuf | WP_281226532.1 |
| GCF_000959445.1 | NZ_CP009631.1 | 174909 | 176093 | 1184 | - | tuf | WP_114676604.1 |
| GCF_022879615.1 | NZ_CP084271.1 | 74882 | 76072 | 1190 | - | tuf | WP_318641048.1 |
| GCF_022879615.1 | NZ_CP084271.1 | 515426 | 516610 | 1184 | - | tuf | WP_248435026.1 |
| GCF_000402035.1 | NC_021287.1 | 2308600 | 2309784 | 1184 | + | tuf | WP_017496723.1 |
| GCF_000402035.1 | NC_021287.1 | 912980 | 914161 | 1181 | + | tuf | WP_014083367.1 |
| GCF_022879815.1 | NZ_CP084283.1 | 1527446 | 1528630 | 1184 | - | tuf | WP_313323585.1 |
| GCF_022879815.1 | NZ_CP084283.1 | 2125392 | 2126576 | 1184 | - | tuf | WP_231835104.1 |
| GCF_026013905.1 | NZ_CP110257.1 | 2282089 | 2283273 | 1184 | + | tuf | WP_007807448.1 |
| GCF_026013905.1 | NZ_CP110257.1 | 545407 | 546591 | 1184 | - | tuf | WP_014166167.1 |
| GCF_001017435.1 | NZ_CP011371.1 | 2611852 | 2613036 | 1184 | + | tuf | WP_071185144.1 |
| GCF_001017435.1 | NZ_CP011371.1 | 2076319 | 2077503 | 1184 | - | tuf | WP_060382900.1 |
| GCF_015476235.1 | NZ_CP064338.1 | 39357 | 40538 | 1181 | - | tuf | WP_066332625.1 |
| GCF_015476235.1 | NZ_CP064338.1 | 2278600 | 2279784 | 1184 | + | tuf | WP_109192121.1 |
| GCF_000024165.1 | NC_013194.1 | 2240500 | 2241684 | 1184 | + | tuf | WP_047777914.1 |
| GCF_000024165.1 | NC_013194.1 | 759933 | 761117 | 1184 | + | tuf | WP_231835104.1 |
| GCF_001460935.1 | NZ_LN906597.1 | 2516938 | 2518122 | 1184 | - | tuf | WP_129758209.1 |
| GCF_001460935.1 | NZ_LN906597.1 | 61352 | 62536 | 1184 | - | tuf | WP_060382900.1 |
| GCF_000319245.1 | NC_019814.1 | 3247451 | 3248635 | 1184 | - | tuf | WP_281336406.1 |
| GCF_000319245.1 | NC_019814.1 | 1385290 | 1386474 | 1184 | - | tuf | WP_012024011.1 |

| | | | | | | | |
|---|---|---|---|---|---|---|---|
| GCF_000340825.1 | NC_020283.1 | 47412 | 48596 | 1184 | - | tuf | WP_108739020.1 |
| GCF_000340825.1 | NC_020283.1 | 4170400 | 4171584 | 1184 | + | tuf | WP_026730432.1 |
| GCF_000340795.1 | NC_020294.1 | 2178730 | 2179914 | 1184 | + | tuf | WP_059116012.1 |
| GCF_000340795.1 | NC_020294.1 | 2073425 | 2074609 | 1184 | + | tuf | WP_057117058.1 |
| GCF_003072465.1 | NZ_CP025628.1 | 162903 | 164087 | 1184 | - | tuf | WP_210644253.1 |
| GCF_000340865.1 | NC_020299.1 | 2325137 | 2326321 | 1184 | - | tuf | WP_012024011.1 |
| GCF_000340865.1 | NC_020299.1 | 1915080 | 1916264 | 1184 | + | tuf | WP_035639372.1 |
| GCF_006364455.1 | NZ_CP040979.1 | 1078198 | 1079382 | 1184 | - | tuf | WP_057123935.1 |
| GCF_006364455.1 | NZ_CP040979.1 | 3209252 | 3210436 | 1184 | + | tuf | WP_026730432.1 |
| GCF_006364615.1 | NZ_CP040986.1 | 1877694 | 1878878 | 1184 | - | tuf | WP_264543454.1 |
| GCF_006364615.1 | NZ_CP040986.1 | 2242459 | 2243643 | 1184 | + | tuf | WP_026730432.1 |
| GCF_006363895.1 | NZ_CP040953.1 | 2292990 | 2294174 | 1184 | - | tuf | WP_014389199.1 |
| GCF_006363895.1 | NZ_CP040953.1 | 64157 | 65341 | 1184 | - | tuf | WP_177729869.1 |
| GCF_024820135.1 | NZ_CP094197.1 | 2238888 | 2240072 | 1184 | + | tuf | WP_012024011.1 |
| GCF_918378365.1 | NZ_OU912926.1 | 2565532 | 2566716 | 1184 | + | tuf | WP_236453026.1 |
| GCF_918378365.1 | NZ_OU912926.1 | 42833 | 44014 | 1181 | - | tuf | WP_264532587.1 |
| GCF_000441555.1 | NC_021885.1 | 36995 | 38179 | 1184 | - | tuf | WP_108735424.1 |
| GCF_000477435.1 | NC_022576.1 | 244904 | 246085 | 1181 | - | tuf | WP_269685524.1 |
| GCF_000477435.1 | NC_022576.1 | 36183 | 37367 | 1184 | + | tuf | WP_269241669.1 |
| GCF_020541325.1 | NZ_CP047399.1 | 159289 | 160473 | 1184 | - | tuf | WP_220640787.1 |
| GCF_020541325.1 | NZ_CP047399.1 | 3200609 | 3201793 | 1184 | + | tuf | WP_109192121.1 |
| GCF_020541305.1 | NZ_CP047396.1 | 1555826 | 1557016 | 1190 | - | tuf | WP_108370320.1 |
| GCF_020541305.1 | NZ_CP047396.1 | 4152804 | 4153988 | 1184 | - | tuf | WP_194640762.1 |
| GCF_910593725.1 | NZ_OU343031.1 | 3436228 | 3437409 | 1181 | + | tuf | WP_133277512.1 |
| GCF_910593725.1 | NZ_OU343031.1 | 573683 | 574867 | 1184 | + | tuf | WP_298655819.1 |
| GCF_964019745.1 | NZ_OZ026462.1 | 2745328 | 2746512 | 1184 | + | tuf | WP_278035058.1 |
| GCF_009746125.1 | NZ_CP041352.1 | 597433 | 598617 | 1184 | - | tuf | WP_109192121.1 |
| GCF_009746125.1 | NZ_CP041352.1 | 1461187 | 1462371 | 1184 | + | tuf | WP_221257561.1 |
| GCF_000612685.1 | NZ_HG916765.1 | 44846 | 46030 | 1184 | - | tuf | WP_366184660.1 |
| GCF_000612685.1 | NZ_HG916765.1 | 1674507 | 1675697 | 1190 | - | tuf | WP_108903399.1 |
| GCF_040955415.1 | NZ_CP158257.1 | 2334124 | 2335308 | 1184 | + | tuf | WP_115889623.1 |

| | | | | | | | |
|---|---|---|---|---|---|---|---|
| GCF_040955415.1 | NZ_CP158257.1 | 2394293 | 2395477 | 1184 | + | tuf | WP_053472306.1 |
| GCF_013403565.1 | NZ_CP058627.1 | 2257748 | 2258932 | 1184 | + | tuf | WP_089059268.1 |
| GCF_013403565.1 | NZ_CP058627.1 | 2187215 | 2188399 | 1184 | - | tuf | WP_269221969.1 |
| GCF_013423785.1 | NZ_CP058952.1 | 1846746 | 1847930 | 1184 | - | tuf | WP_011963319.1 |
| GCF_013423785.1 | NZ_CP058952.1 | 3096413 | 3097597 | 1184 | - | tuf | WP_116789263.1 |
| GCF_007431345.1 | NZ_CP041730.1 | 3108850 | 3110034 | 1184 | - | tuf | WP_171222207.1 |
| GCF_007431345.1 | NZ_CP041730.1 | 106375 | 107559 | 1184 | - | tuf | WP_136150607.1 |
| GCF_042140075.1 | NZ_CP156074.1 | 2060595 | 2061779 | 1184 | + | tuf | WP_045967694.1 |
| GCF_042140075.1 | NZ_CP156074.1 | 2352695 | 2353879 | 1184 | + | tuf | WP_109569654.1 |
| GCF_025642115.1 | NZ_CP106753.1 | 2295155 | 2296339 | 1184 | + | tuf | WP_109192121.1 |
| GCF_025642115.1 | NZ_CP106753.1 | 2479835 | 2481022 | 1187 | + | tuf | WP_013686909.1 |
| GCF_008033135.1 | NZ_CP041335.1 | 3001692 | 3002876 | 1184 | - | tuf | WP_038531091.1 |
| GCF_008033135.1 | NZ_CP041335.1 | 106316 | 107500 | 1184 | - | tuf | WP_143379434.1 |
| GCF_009936155.1 | NZ_AP019312.1 | 2546384 | 2547568 | 1184 | + | tuf | WP_316982276.1 |
| GCF_009936155.1 | NZ_AP019312.1 | 157117 | 158301 | 1184 | - | tuf | WP_096684748.1 |
| GCF_008275125.1 | NZ_CP043473.1 | 154436 | 155620 | 1184 | + | tuf | WP_008238257.1 |
| GCF_008275125.1 | NZ_CP043473.1 | 1843159 | 1844343 | 1184 | - | tuf | WP_034719150.1 |
| GCF_003325475.1 | NZ_CP029495.1 | 2824594 | 2825778 | 1184 | + | tuf | WP_125025684.1 |
| GCF_003325475.1 | NZ_CP029495.1 | 1816091 | 1817275 | 1184 | + | tuf | WP_218248548.1 |
| GCF_020544465.1 | NZ_CP066126.1 | 2690008 | 2691192 | 1184 | + | tuf | WP_039353714.1 |
| GCF_020544465.1 | NZ_CP066126.1 | 2286652 | 2287836 | 1184 | + | tuf | WP_226063900.1 |
| GCF_036320555.1 | NZ_CP143257.1 | 4806538 | 4807722 | 1184 | + | tuf | WP_160131285.1 |
| GCF_036320555.1 | NZ_CP143257.1 | 1292675 | 1293859 | 1184 | - | tuf | WP_267677902.1 |
| GCF_001855275.1 | NZ_CP017707.1 | 2529341 | 2530525 | 1184 | - | tuf | WP_076733804.1 |
| GCF_001855275.1 | NZ_CP017707.1 | 807963 | 809147 | 1184 | - | tuf | WP_072983799.1 |
| GCF_016890085.1 | NZ_CP069587.1 | 1008653 | 1009837 | 1184 | - | tuf | WP_068206653.1 |
| GCF_016890085.1 | NZ_CP069587.1 | 2664927 | 2666111 | 1184 | - | tuf | WP_302043563.1 |
| GCF_001584205.1 | NZ_CP013235.1 | 2821774 | 2822958 | 1184 | + | tuf | WP_338731143.1 |
| GCF_001584205.1 | NZ_CP013235.1 | 579385 | 580569 | 1184 | - | tuf | WP_281540829.1 |
| GCF_001584145.1 | NZ_CP013232.1 | 3231774 | 3232958 | 1184 | + | tuf | WP_094997908.1 |
| GCF_001584145.1 | NZ_CP013232.1 | 566515 | 567699 | 1184 | + | tuf | WP_091602376.1 |

| | | | | | | | |
|---|---|---|---|---|---|---|---|
| GCF_001584225.1 | NZ_CP013236.1 | 663278 | 664462 | 1184 | - | tuf | WP_273567716.1 |
| GCF_001584225.1 | NZ_CP013236.1 | 773895 | 775079 | 1184 | - | tuf | WP_068484278.1 |
| GCF_013363755.1 | NZ_CP054840.1 | 2522162 | 2523346 | 1184 | - | tuf | WP_260571691.1 |
| GCF_013363755.1 | NZ_CP054840.1 | 3031988 | 3033172 | 1184 | + | tuf | WP_308990828.1 |
| GCF_018064205.1 | NZ_CP072916.1 | 303099 | 304283 | 1184 | - | tuf | WP_106688304.1 |
| GCF_018064205.1 | NZ_CP072916.1 | 596824 | 598008 | 1184 | + | tuf | WP_129602652.1 |
| GCF_023634805.2 | NZ_CP106881.1 | 1108785 | 1109969 | 1184 | - | tuf | WP_155091303.1 |
| GCF_023634805.2 | NZ_CP106881.1 | 1991260 | 1992444 | 1184 | + | tuf | WP_158962437.1 |
| GCF_007954405.1 | NZ_CP042344.1 | 606545 | 607729 | 1184 | - | tuf | WP_131540232.1 |
| GCF_007954405.1 | NZ_CP042344.1 | 3353094 | 3354278 | 1184 | - | tuf | WP_002987571.1 |
| GCF_019186805.1 | NZ_CP066783.1 | 3582028 | 3583212 | 1184 | - | tuf | WP_006259596.1 |
| GCF_019186805.1 | NZ_CP066783.1 | 2247290 | 2248474 | 1184 | + | tuf | WP_015362696.1 |
| GCF_002056725.1 | NZ_CP020121.1 | 1556285 | 1557469 | 1184 | + | tuf | WP_041496079.1 |
| GCF_002056725.1 | NZ_CP020121.1 | 1146288 | 1147472 | 1184 | + | tuf | WP_126446310.1 |
| GCF_020080045.1 | NZ_CP083451.1 | 3269790 | 3270974 | 1184 | + | tuf | WP_085767996.1 |
| GCF_020080045.1 | NZ_CP083451.1 | 1999597 | 2000781 | 1184 | - | tuf | WP_042279745.1 |
| GCF_014109725.1 | NZ_CP058554.1 | 3168468 | 3169652 | 1184 | + | tuf | WP_036582240.1 |
| GCF_014109725.1 | NZ_CP058554.1 | 1564283 | 1565467 | 1184 | + | tuf | WP_014791257.1 |
| GCF_030064165.1 | NZ_CP125947.1 | 1605836 | 1607020 | 1184 | - | tuf | WP_014201829.1 |
| GCF_030064165.1 | NZ_CP125947.1 | 1520767 | 1521951 | 1184 | + | tuf | WP_257498067.1 |
| GCF_002158865.1 | NZ_CP021455.1 | 3328583 | 3329767 | 1184 | + | tuf | WP_221832972.1 |
| GCF_002158865.1 | NZ_CP021455.1 | 2999097 | 3000275 | 1178 | - | tuf | WP_258542897.1 |
| GCF_025309455.1 | NZ_CP104377.1 | 1265565 | 1266749 | 1184 | - | tuf | WP_068450828.1 |
| GCF_025309455.1 | NZ_CP104377.1 | 3187475 | 3188659 | 1184 | + | tuf | WP_207971408.1 |
| GCF_006740045.1 | NZ_AP019749.1 | 1597303 | 1598487 | 1184 | - | tuf | WP_105050517.1 |
| GCF_006740045.1 | NZ_AP019749.1 | 3694329 | 3695513 | 1184 | + | tuf | WP_208078162.1 |
| GCF_020886175.1 | NZ_AP025193.1 | 1208594 | 1209778 | 1184 | - | tuf | WP_053972786.1 |
| GCF_020886175.1 | NZ_AP025193.1 | 1078388 | 1079572 | 1184 | - | tuf | WP_105045973.1 |
| GCF_014054725.1 | NZ_CP059563.1 | 3477798 | 3478982 | 1184 | + | tuf | WP_088354342.1 |
| GCF_014054725.1 | NZ_CP059563.1 | 1700845 | 1702026 | 1181 | + | tuf | WP_301399907.1 |
| GCF_003355495.1 | NZ_CP031337.1 | 730035 | 731219 | 1184 | + | tuf | WP_223445311.1 |

| | | | | | | | |
|---|---|---|---|---|---|---|---|
| GCF_003355495.1 | NZ_CP031337.1 | 1006900 | 1008084 | 1184 | - | tuf | WP_340934429.1 |
| GCF_008801925.2 | NZ_CP062803.1 | 1036284 | 1037465 | 1181 | - | tuf | WP_188597338.1 |
| GCF_013347325.1 | NZ_CP054624.1 | 454849 | 456033 | 1184 | + | tuf | WP_187482840.1 |
| GCF_013347325.1 | NZ_CP054624.1 | 2422016 | 2423200 | 1184 | + | tuf | WP_068357682.1 |
| GCF_001854325.1 | NZ_CP017754.1 | 1736128 | 1737312 | 1184 | - | tuf | WP_166385072.1 |
| GCF_001854325.1 | NZ_CP017754.1 | 1349025 | 1350209 | 1184 | - | tuf | WP_105050517.1 |
| GCF_000219215.1 | NC_015726.1 | 641392 | 642576 | 1184 | - | tuf | WP_088323156.1 |
| GCF_000219215.1 | NC_015726.1 | 1170572 | 1171756 | 1184 | - | tuf | WP_065318947.1 |
| GCF_900250125.1 | NZ_LT984806.1 | 1348685 | 1349869 | 1184 | - | tuf | WP_054725766.1 |
| GCF_900250125.1 | NZ_LT984806.1 | 2407103 | 2408287 | 1184 | + | tuf | WP_102995890.1 |
| GCF_016894385.1 | NZ_CP069812.1 | 2763434 | 2764615 | 1181 | + | tuf | WP_015024891.1 |
| GCF_016894385.1 | NZ_CP069812.1 | 2715940 | 2717124 | 1184 | - | tuf | WP_249995792.1 |
| GCF_019931045.1 | NZ_CP082862.1 | 329261 | 330445 | 1184 | - | tuf | WP_105214275.1 |
| GCF_019931045.1 | NZ_CP082862.1 | 1249396 | 1250580 | 1184 | - | tuf | WP_164679079.1 |
| GCF_019602855.1 | NZ_CP080767.1 | 1640448 | 1641632 | 1184 | + | tuf | WP_004916760.1 |
| GCF_019602855.1 | NZ_CP080767.1 | 3284087 | 3285271 | 1184 | - | tuf | WP_015755618.1 |
| GCF_000069785.1 | NC_010528.1 | 344080 | 345264 | 1184 | - | tuf | WP_224470590.1 |
| GCF_000069785.1 | NC_010528.1 | 3465806 | 3466990 | 1184 | + | tuf | WP_079736068.1 |
| GCF_041378895.1 | NZ_CP144212.1 | 870131 | 871315 | 1184 | - | tuf | WP_265164415.1 |
| GCF_041378895.1 | NZ_CP144212.1 | 578386 | 579570 | 1184 | + | tuf | WP_306642486.1 |
| GCF_013284055.1 | NZ_CP054143.1 | 138443 | 139627 | 1184 | - | tuf | WP_044637043.1 |
| GCF_013284055.1 | NZ_CP054143.1 | 1062739 | 1063923 | 1184 | - | tuf | WP_233030496.1 |
| GCF_019665765.1 | NZ_CP081150.1 | 984980 | 986164 | 1184 | - | tuf | WP_418650910.1 |
| GCF_019665765.1 | NZ_CP081150.1 | 2096338 | 2097522 | 1184 | - | tuf | WP_101902917.1 |
| GCF_000018665.1 | NC_010002.1 | 526875 | 528059 | 1184 | - | tuf | WP_303555686.1 |
| GCF_000018665.1 | NC_010002.1 | 1331483 | 1332667 | 1184 | + | tuf | WP_095070388.1 |
| GCF_040267515.1 | NZ_CP141274.1 | 1001965 | 1003149 | 1184 | + | tuf | WP_100210908.1 |
| GCF_040267515.1 | NZ_CP141274.1 | 873907 | 875091 | 1184 | - | tuf | WP_047790103.1 |
| GCF_029215875.1 | NZ_CP118775.1 | 666722 | 667909 | 1187 | + | tuf | WP_271232118.1 |
| GCF_029215875.1 | NZ_CP118775.1 | 1519861 | 1521045 | 1184 | - | tuf | WP_100923982.1 |
| GCF_902813185.1 | NZ_LR778301.1 | 3115463 | 3116647 | 1184 | - | tuf | WP_348714112.1 |

| | | | | | | | |
|---|---|---|---|---|---|---|---|
| GCF_902813185.1 | NZ_LR778301.1 | 2184007 | 2185191 | 1184 | - | tuf | WP_275687476.1 |
| GCF_014489535.1 | NZ_CP060783.1 | 1383003 | 1384187 | 1184 | + | tuf | WP_125066964.1 |
| GCF_014489535.1 | NZ_CP060783.1 | 1556564 | 1557748 | 1184 | - | tuf | WP_279652693.1 |
| GCF_033100095.1 | NZ_CP136921.1 | 1367626 | 1368810 | 1184 | - | tuf | WP_072555291.1 |
| GCF_033100095.1 | NZ_CP136921.1 | 2635847 | 2637031 | 1184 | - | tuf | WP_348702416.1 |
| GCF_002214645.1 | NZ_CP016278.1 | 2758968 | 2760152 | 1184 | + | tuf | WP_415861541.1 |
| GCF_002214645.1 | NZ_CP016278.1 | 1245282 | 1246463 | 1181 | - | tuf | WP_070236335.1 |
| GCF_014395975.1 | NZ_CP060714.1 | 81464 | 82648 | 1184 | + | tuf | WP_013597833.1 |
| GCF_014395975.1 | NZ_CP060714.1 | 507899 | 509080 | 1181 | + | tuf | WP_068824280.1 |
| GCF_012849375.1 | NZ_CP051684.1 | 3051799 | 3052983 | 1184 | - | tuf | WP_179008537.1 |
| GCF_012849375.1 | NZ_CP051684.1 | 2980613 | 2981797 | 1184 | - | tuf | WP_178989340.1 |
| GCF_034479515.1 | NZ_CP140152.1 | 2738642 | 2739826 | 1184 | + | tuf | WP_092446933.1 |
| GCF_034479515.1 | NZ_CP140152.1 | 218251 | 219435 | 1184 | - | tuf | WP_242937346.1 |
| GCF_928721995.1 | NZ_OV788642.1 | 3286910 | 3288094 | 1184 | + | tuf | WP_276165801.1 |
| GCF_928721995.1 | NZ_OV788642.1 | 809602 | 810786 | 1184 | - | tuf | WP_013992033.1 |
| GCF_900187105.1 | NZ_LT906482.1 | 2068915 | 2070099 | 1184 | - | tuf | WP_215441894.1 |
| GCF_900187105.1 | NZ_LT906482.1 | 1266529 | 1267713 | 1184 | + | tuf | WP_013070653.1 |
| GCF_008805035.1 | NZ_CP038018.1 | 1672305 | 1673486 | 1181 | - | tuf | WP_040406695.1 |
| GCF_008805035.1 | NZ_CP038018.1 | 1778391 | 1779572 | 1181 | - | tuf | WP_040406695.1 |
| GCF_030419005.1 | NZ_CP106951.1 | 127772 | 128953 | 1181 | + | tuf | WP_008693350.1 |
| GCF_030419005.1 | NZ_CP106951.1 | 2254801 | 2255982 | 1181 | - | tuf | WP_198480673.1 |
| GCF_003339525.1 | NZ_CP031124.1 | 48935 | 50116 | 1181 | - | tuf | WP_048911094.1 |
| GCF_003339525.1 | NZ_CP031124.1 | 205049 | 206230 | 1181 | - | tuf | WP_048911094.1 |
| GCF_020510565.1 | NZ_CP075189.1 | 1715561 | 1716742 | 1181 | + | tuf | WP_101474504.1 |
| GCF_020510565.1 | NZ_CP075189.1 | 1820123 | 1821304 | 1181 | + | tuf | WP_101474504.1 |
| GCF_018324385.1 | NZ_AP019536.1 | 848702 | 849883 | 1181 | + | tuf | WP_029490858.1 |
| GCF_018324385.1 | NZ_AP019536.1 | 413064 | 414245 | 1181 | - | tuf | WP_005885863.1 |
| GCF_000974685.2 | NZ_AP018738.1 | 716663 | 717844 | 1181 | + | tuf | WP_005885863.1 |
| GCF_000974685.2 | NZ_AP018738.1 | 86362 | 87543 | 1181 | + | tuf | WP_005959409.1 |
| GCF_013181415.1 | NZ_CP053676.1 | 260700 | 261881 | 1181 | + | tuf | WP_005959409.1 |
| GCF_013181415.1 | NZ_CP053676.1 | 76345 | 77526 | 1181 | - | tuf | WP_005903497.1 |

| | | | | | | | |
|---|---|---|---|---|---|---|---|
| GCF_016110345.1 | NZ_AP019011.1 | 2192616 | 2193797 | 1181 | - | tuf | WP_005894517.1 |
| GCF_016110345.1 | NZ_AP019011.1 | 1114207 | 1115388 | 1181 | - | tuf | WP_005979589.1 |
| GCF_000145255.1 | NC_014394.1 | 1448909 | 1450090 | 1181 | + | tuf | WP_005979589.1 |
| GCF_000145255.1 | NC_014394.1 | 816139 | 817320 | 1181 | - | tuf | WP_005948793.1 |
| GCF_047325095.1 | NZ_CP168558.1 | 1124248 | 1125429 | 1181 | + | tuf | WP_005948793.1 |
| GCF_047325095.1 | NZ_CP168558.1 | 661064 | 662245 | 1181 | - | tuf | WP_220278817.1 |
| GCF_001267925.1 | NZ_CP011409.1 | 721487 | 722668 | 1181 | + | tuf | WP_150152296.1 |
| GCF_001267925.1 | NZ_CP011409.1 | 116853 | 118034 | 1181 | + | tuf | WP_307904473.1 |
| GCF_004367745.1 | NZ_CP037993.1 | 712240 | 713421 | 1181 | + | tuf | WP_307904473.1 |
| GCF_004367745.1 | NZ_CP037993.1 | 1602430 | 1603611 | 1181 | - | tuf | WP_013387933.1 |
| GCF_002025725.1 | NZ_CP018845.1 | 1930638 | 1931819 | 1181 | - | tuf | WP_013387933.1 |
| GCF_002025725.1 | NZ_CP018845.1 | 241703 | 242884 | 1181 | + | tuf | WP_012806348.1 |
| GCF_003719195.1 | NZ_CP024996.1 | 2035149 | 2036330 | 1181 | + | tuf | WP_015770120.1 |
| GCF_003719195.1 | NZ_CP024996.1 | 331082 | 332263 | 1181 | + | tuf | WP_026745237.1 |
| GCF_001040945.1 | NZ_CP011930.1 | 2402706 | 2403887 | 1181 | - | tuf | WP_026745851.1 |
| GCF_001040945.1 | NZ_CP011930.1 | 123477 | 124658 | 1181 | + | tuf | WP_026745237.1 |
| GCF_900130075.1 | NZ_LT671418.1 | 1837943 | 1839124 | 1181 | - | tuf | WP_026745237.1 |
| GCF_900130075.1 | NZ_LT671418.1 | 142287 | 143468 | 1181 | + | tuf | WP_369713009.1 |
| GCF_001761385.1 | NZ_CP017476.1 | 295303 | 296484 | 1181 | + | tuf | WP_369713111.1 |
| GCF_001761385.1 | NZ_CP017476.1 | 2492237 | 2493418 | 1181 | - | tuf | WP_026748143.1 |
| GCF_011388215.1 | NZ_CP049989.1 | 2667460 | 2668641 | 1181 | - | tuf | WP_026748143.1 |
| GCF_011388215.1 | NZ_CP049989.1 | 1959572 | 1960753 | 1181 | - | tuf | WP_060917639.1 |
| GCF_004353865.1 | NZ_CP037867.1 | 2135560 | 2136741 | 1181 | - | tuf | WP_060917639.1 |
| GCF_004353865.1 | NZ_CP037867.1 | 109073 | 110254 | 1181 | + | tuf | WP_046328181.1 |
| GCF_021049305.1 | NZ_CP088081.1 | 840020 | 841201 | 1181 | + | tuf | WP_046328181.1 |
| GCF_021049305.1 | NZ_CP088081.1 | 235764 | 236945 | 1181 | + | tuf | WP_012858235.1 |
| GCF_005952805.1 | NZ_CP040709.1 | 1144214 | 1145395 | 1181 | - | tuf | WP_012858235.1 |
| GCF_005952805.1 | NZ_CP040709.1 | 421507 | 422688 | 1181 | + | tuf | WP_013364443.1 |
| GCF_003952345.1 | NZ_CP034433.1 | 4516943 | 4518124 | 1181 | - | tuf | WP_013368129.1 |
| GCF_003952345.1 | NZ_CP034433.1 | 555317 | 556498 | 1181 | - | tuf | WP_064082225.1 |
| GCF_036885605.1 | NZ_CP142523.1 | 737752 | 738933 | 1181 | + | tuf | WP_064082225.1 |

| | | | | | | | |
|---|---|---|---|---|---|---|---|
| GCF_036885605.1 | NZ_CP142523.1 | 141198 | 142379 | 1181 | + | tuf | WP_011199348.1 |
| GCF_000723165.1 | NZ_HG322949.1 | 2124538 | 2125719 | 1181 | - | tuf | WP_011199348.1 |
| GCF_013372045.1 | NZ_CP048832.1 | 404961 | 406142 | 1181 | + | tuf | WP_242080467.1 |
| GCF_013372045.1 | NZ_CP048832.1 | 3813463 | 3814644 | 1181 | - | tuf | WP_154325460.1 |
| GCF_029690045.1 | NZ_CP121464.1 | 1003018 | 1004205 | 1187 | - | tuf | WP_201340880.1 |
| GCF_029690045.1 | NZ_CP121464.1 | 1227106 | 1228293 | 1187 | - | tuf | WP_201340880.1 |
| GCF_002327145.1 | NZ_CP023422.1 | 531510 | 532697 | 1187 | + | tuf | WP_065971275.1 |
| GCF_002327145.1 | NZ_CP023422.1 | 547724 | 548911 | 1187 | + | tuf | WP_065971275.1 |
| GCF_006517255.1 | NZ_CP041185.1 | 2484448 | 2485635 | 1187 | - | tuf | WP_070073148.1 |
| GCF_006517255.1 | NZ_CP041185.1 | 2501001 | 2502188 | 1187 | - | tuf | WP_070073148.1 |
| GCF_048595715.1 | NZ_CP169556.1 | 2978344 | 2979531 | 1187 | - | tuf | WP_076837710.1 |
| GCF_048595715.1 | NZ_CP169556.1 | 2995014 | 2996201 | 1187 | - | tuf | WP_076837722.1 |
| GCF_016127355.1 | NZ_CP065999.1 | 2631373 | 2632560 | 1187 | - | tuf | WP_070079064.1 |
| GCF_016127355.1 | NZ_CP065999.1 | 2647939 | 2649126 | 1187 | - | tuf | WP_070079064.1 |
| GCF_900475905.1 | NZ_LS483426.1 | 354792 | 355979 | 1187 | + | tuf | WP_001029610.1 |
| GCF_900475905.1 | NZ_LS483426.1 | 907942 | 909129 | 1187 | + | tuf | WP_001029610.1 |
| GCF_030177895.1 | NZ_CP123448.1 | 297344 | 298531 | 1187 | + | tuf | WP_004922118.1 |
| GCF_030177895.1 | NZ_CP123448.1 | 868301 | 869488 | 1187 | + | tuf | WP_004922118.1 |
| GCF_900637555.1 | NZ_LR134378.1 | 3815549 | 3816736 | 1187 | - | tuf | WP_004718491.1 |
| GCF_900637555.1 | NZ_LR134378.1 | 4396464 | 4397651 | 1187 | - | tuf | WP_004718491.1 |
| GCF_000019785.1 | NC_010524.1 | 777317 | 778504 | 1187 | + | tuf | WP_005006977.1 |
| GCF_000019785.1 | NC_010524.1 | 3142807 | 3143994 | 1187 | - | tuf | WP_005006977.1 |
| GCF_013004065.1 | NZ_CP053084.1 | 298586 | 299773 | 1187 | + | tuf | WP_118998476.1 |
| GCF_013004065.1 | NZ_CP053084.1 | 781368 | 782555 | 1187 | + | tuf | WP_118998476.1 |
| GCF_036323495.1 | NZ_AP028947.1 | 1992835 | 1994022 | 1187 | - | tuf | WP_283266036.1 |
| GCF_036323495.1 | NZ_AP028947.1 | 3235146 | 3236333 | 1187 | + | tuf | WP_283266921.1 |
| GCF_015689335.1 | NZ_CP065053.1 | 2724328 | 2725515 | 1187 | - | tuf | WP_137287916.1 |
| GCF_015689335.1 | NZ_CP065053.1 | 3176977 | 3178164 | 1187 | - | tuf | WP_137287908.1 |
| GCF_045016065.1 | NZ_CP051166.1 | 426174 | 427361 | 1187 | + | tuf | WP_065995613.1 |
| GCF_045016065.1 | NZ_CP051166.1 | 2678435 | 2679622 | 1187 | - | tuf | WP_065995613.1 |
| GCF_047325105.1 | NZ_CP168562.1 | 2149560 | 2150747 | 1187 | - | tuf | WP_004802222.1 |

| | | | | | | | |
|---|---|---|---|---|---|---|---|
| GCF_047325105.1 | NZ_CP168562.1 | 2768548 | 2769735 | 1187 | - | tuf | WP_004802222.1 |
| GCF_021165955.1 | NZ_CP088952.1 | 594849 | 596036 | 1187 | - | tuf | WP_054580458.1 |
| GCF_021165955.1 | NZ_CP088952.1 | 1479127 | 1480314 | 1187 | + | tuf | WP_054580458.1 |
| GCF_012849555.1 | NZ_CP051685.1 | 265152 | 266339 | 1187 | + | tuf | WP_004723653.1 |
| GCF_012849555.1 | NZ_CP051685.1 | 3744502 | 3745689 | 1187 | - | tuf | WP_004723653.1 |
| GCF_015101885.1 | NZ_CP062941.1 | 2580836 | 2582023 | 1187 | - | tuf | WP_005081540.1 |
| GCF_015101885.1 | NZ_CP062941.1 | 3162771 | 3163958 | 1187 | - | tuf | WP_005081540.1 |
| GCF_003143515.1 | NZ_CP029343.1 | 1796296 | 1797483 | 1187 | - | tuf | WP_160241040.1 |
| GCF_003143515.1 | NZ_CP029343.1 | 2773174 | 2774361 | 1187 | + | tuf | WP_104513317.1 |
| GCF_001941825.1 | NZ_CP019038.1 | 855392 | 856579 | 1187 | + | tuf | WP_004913952.1 |
| GCF_001941825.1 | NZ_CP019038.1 | 3080473 | 3081660 | 1187 | - | tuf | WP_004913952.1 |
| GCF_002752675.1 | NZ_CP024608.1 | 3069696 | 3070883 | 1187 | - | tuf | WP_005065785.1 |
| GCF_002752675.1 | NZ_CP024608.1 | 3662794 | 3663981 | 1187 | - | tuf | WP_005065785.1 |
| GCF_021249385.1 | NZ_CP089742.1 | 2641545 | 2642732 | 1187 | - | tuf | WP_166326224.1 |
| GCF_021249385.1 | NZ_CP089742.1 | 3113396 | 3114583 | 1187 | - | tuf | WP_166326224.1 |
| GCF_003008575.1 | NZ_CP027667.1 | 605746 | 606933 | 1187 | + | tuf | WP_067552282.1 |
| GCF_003008575.1 | NZ_CP027667.1 | 3403174 | 3404361 | 1187 | - | tuf | WP_067558061.1 |
| GCF_000015725.1 | NC_008825.1 | 1535653 | 1536840 | 1187 | - | tuf | WP_004646112.1 |
| GCF_000015725.1 | NC_008825.1 | 2461205 | 2462392 | 1187 | + | tuf | WP_004646112.1 |
| GCF_000013705.1 | NC_007947.1 | 329666 | 330853 | 1187 | + | tuf | WP_002157016.1 |
| GCF_000013705.1 | NC_007947.1 | 921463 | 922650 | 1187 | + | tuf | WP_002157016.1 |
| GCF_006363915.1 | NZ_CP040946.1 | 3270524 | 3271711 | 1187 | - | tuf | WP_004794160.1 |
| GCF_006363915.1 | NZ_CP040946.1 | 3844648 | 3845835 | 1187 | - | tuf | WP_004794160.1 |
| GCF_019703875.1 | NZ_AP024110.1 | 139409 | 140596 | 1187 | + | tuf1 | YP_004994400.1 |
| GCF_019703875.1 | NZ_AP024110.1 | 3419505 | 3420692 | 1187 | + | tufA | YP_004997453.1 |
| GCF_000023705.1 | NC_012968.1 | 211194 | 212381 | 1187 | + | tuf | WP_005017071.1 |
| GCF_000023705.1 | NC_012968.1 | 2398849 | 2400036 | 1187 | - | tuf | WP_005017071.1 |
| GCF_000093025.1 | NC_014207.1 | 2384221 | 2385408 | 1187 | - | tuf | WP_004811166.1 |
| GCF_000093025.1 | NC_014207.1 | 2839877 | 2841064 | 1187 | - | tuf | WP_004811166.1 |
| GCF_000023745.1 | NC_012969.1 | 3200299 | 3201486 | 1187 | - | tuf | WP_002157016.1 |
| GCF_000023745.1 | NC_012969.1 | 3758680 | 3759867 | 1187 | - | tuf | WP_002157016.1 |

| | | | | | | | |
|---|---|---|---|---|---|---|---|
| GCF_003051865.1 | NZ_CP028519.1 | 1820076 | 1821263 | 1187 | - | tuf | WP_004991043.1 |
| GCF_003051865.1 | NZ_CP028519.1 | 2893106 | 2894293 | 1187 | + | tuf | WP_004991043.1 |
| GCF_022749515.1 | NZ_CP094242.1 | 103953 | 105140 | 1187 | + | tuf | WP_004932358.1 |
| GCF_022749515.1 | NZ_CP094242.1 | 2263513 | 2264700 | 1187 | - | tuf | WP_004932358.1 |
| GCF_037478035.1 | NZ_CP062168.1 | 2476837 | 2478024 | 1187 | - | tuf | WP_150026484.1 |
| GCF_037478035.1 | NZ_CP062168.1 | 3065800 | 3066987 | 1187 | - | tuf | WP_150026484.1 |
| GCF_037478055.1 | NZ_CP062174.1 | 731932 | 733119 | 1187 | + | tuf | WP_347453254.1 |
| GCF_037478055.1 | NZ_CP062174.1 | 2805343 | 2806530 | 1187 | - | tuf | WP_347453254.1 |
| GCF_003966915.1 | NZ_AP018150.1 | 687898 | 689085 | 1187 | + | tuf | WP_116763782.1 |
| GCF_008806995.1 | NZ_CP031699.1 | 3093643 | 3094830 | 1187 | - | tuf | WP_116763782.1 |
| GCF_008806995.1 | NZ_CP031699.1 | 788525 | 789712 | 1187 | + | tuf | WP_004975768.1 |
| GCF_900637855.1 | NZ_LR134440.1 | 2501319 | 2502506 | 1187 | - | tuf | WP_004975768.1 |
| GCF_900637855.1 | NZ_LR134440.1 | 2425372 | 2426559 | 1187 | - | tuf | WP_004781663.1 |
| GCF_022870905.1 | NZ_CP091510.1 | 2908529 | 2909716 | 1187 | - | tuf | WP_179991967.1 |
| GCF_022870905.1 | NZ_CP091510.1 | 226402 | 227589 | 1187 | + | tuf | WP_068972734.1 |
| GCF_014055025.1 | NZ_CP059571.1 | 894397 | 895584 | 1187 | + | tuf | WP_068972734.1 |
| GCF_014055025.1 | NZ_CP059571.1 | 850265 | 851446 | 1181 | + | tuf | WP_126599201.1 |
| GCF_009671065.1 | NZ_CP046027.1 | 1415216 | 1416397 | 1181 | + | tuf | WP_126599201.1 |
| GCF_009671065.1 | NZ_CP046027.1 | 1350278 | 1351459 | 1181 | + | tuf | WP_015674055.1 |
| GCF_900636765.1 | NZ_LR134313.1 | 1434640 | 1435821 | 1181 | + | tuf | WP_015674055.1 |
| GCF_900636765.1 | NZ_LR134313.1 | 1268701 | 1269885 | 1184 | + | tuf | WP_162856707.1 |
| GCF_002216145.1 | NZ_CP022278.1 | 1309261 | 1310445 | 1184 | - | tuf | WP_162856707.1 |
| GCF_002216145.1 | NZ_CP022278.1 | 1740771 | 1741952 | 1181 | + | tuf | WP_009875214.1 |
| GCF_900475315.1 | NZ_LS483369.1 | 1772412 | 1773593 | 1181 | - | tuf | WP_009875214.1 |
| GCF_900475315.1 | NZ_LS483369.1 | 366006 | 367187 | 1181 | - | tuf | WP_108922804.1 |
| GCF_014055005.1 | NZ_CP059570.1 | 751420 | 752601 | 1181 | + | tuf | WP_108922804.1 |
| GCF_014055005.1 | NZ_CP059570.1 | 110605 | 111786 | 1181 | - | tuf | WP_011978750.1 |
| GCF_022870885.1 | NZ_CP091509.1 | 313709 | 314890 | 1181 | - | tuf | WP_011978750.1 |
| GCF_022870885.1 | NZ_CP091509.1 | 1940975 | 1942156 | 1181 | - | tuf | WP_015674055.1 |
| GCF_005221285.1 | NZ_CP039886.1 | 2293500 | 2294681 | 1181 | + | tuf | WP_015674055.1 |
| GCF_005221285.1 | NZ_CP039886.1 | 410172 | 411392 | 1220 | + | tuf | WP_160195571.1 |

| | | | | | | | |
|---|---|---|---|---|---|---|---|
| GCF_013030075.1 | NZ_AP023069.1 | 426884 | 428104 | 1220 | + | tuf | WP_160195571.1 |
| GCF_013030075.1 | NZ_AP023069.1 | 3794734 | 3795915 | 1181 | + | tuf | WP_042657281.1 |
| GCF_003351565.1 | NZ_CP031253.1 | 3813934 | 3815115 | 1181 | + | tuf | WP_042657281.1 |
| GCF_003351565.1 | NZ_CP031253.1 | 4403417 | 4404598 | 1181 | - | tuf | WP_011898272.1 |
| GCF_028776105.2 | NZ_CP145606.1 | 4422618 | 4423799 | 1181 | - | tuf | WP_011898272.1 |
| GCF_028776105.2 | NZ_CP145606.1 | 285607 | 286788 | 1181 | + | tuf | WP_005306325.1 |
| GCF_028463985.1 | NZ_CP116766.1 | 304812 | 305993 | 1181 | + | tuf | WP_005306325.1 |
| GCF_028463985.1 | NZ_CP116766.1 | 442824 | 444005 | 1181 | - | tuf | WP_005306325.1 |
| GCF_022749495.1 | NZ_CP094241.1 | 462020 | 463201 | 1181 | - | tuf | WP_005306325.1 |
| GCF_022749495.1 | NZ_CP094241.1 | 252596 | 253777 | 1181 | + | tuf | WP_126623228.1 |
| GCF_022869645.1 | NZ_CP064367.1 | 271795 | 272976 | 1181 | + | tuf | WP_126623228.1 |
| GCF_022869645.1 | NZ_CP064367.1 | 1661993 | 1663174 | 1181 | + | tuf | WP_061476754.1 |
| GCF_014297595.2 | NZ_CP060414.2 | 1681197 | 1682378 | 1181 | + | tuf | WP_061476754.1 |
| GCF_014297595.2 | NZ_CP060414.2 | 4455504 | 4456685 | 1181 | - | tuf | WP_011707694.1 |
| GCF_014054965.1 | NZ_CP059567.1 | 4474711 | 4475892 | 1181 | - | tuf | WP_011707694.1 |
| GCF_014054965.1 | NZ_CP059567.1 | 1999433 | 2000614 | 1181 | - | tuf | WP_041209577.1 |
| GCF_017753665.1 | NZ_CP072524.1 | 2018629 | 2019810 | 1181 | - | tuf | WP_074042664.1 |
| GCF_017753665.1 | NZ_CP072524.1 | 4177103 | 4178284 | 1181 | - | tuf | WP_041206539.1 |
| GCF_005221305.1 | NZ_CP039887.1 | 4196303 | 4197484 | 1181 | - | tuf | WP_025328607.1 |
| GCF_005221305.1 | NZ_CP039887.1 | 455723 | 456904 | 1181 | + | tuf | WP_033135315.1 |
| GCF_014054885.1 | NZ_CP059565.1 | 474934 | 476115 | 1181 | + | tuf | WP_042036023.1 |
| GCF_014054885.1 | NZ_CP059565.1 | 231496 | 232677 | 1181 | + | tuf | WP_224431767.1 |
| GCF_900638685.1 | NZ_LR134533.1 | 250682 | 251863 | 1181 | + | tuf | WP_042644447.1 |
| GCF_900638685.1 | NZ_LR134533.1 | 4481021 | 4482202 | 1181 | - | tuf | WP_011898272.1 |
| GCF_014898075.1 | NZ_CP062976.1 | 4500221 | 4501402 | 1181 | - | tuf | WP_011898272.1 |
| GCF_014898075.1 | NZ_CP062976.1 | 3891415 | 3892596 | 1181 | + | tuf | WP_042048519.1 |
| GCF_008807015.1 | NZ_CP031700.1 | 3910568 | 3911749 | 1181 | + | tuf | WP_193002862.1 |
| GCF_008807015.1 | NZ_CP031700.1 | 3547828 | 3549009 | 1181 | + | tuf | WP_005341825.1 |
| GCF_900187305.1 | NZ_LT906434.1 | 3567031 | 3568212 | 1181 | + | tuf | WP_005339879.1 |
| GCF_900187305.1 | NZ_LT906434.1 | 2147930 | 2149117 | 1187 | - | tuf | WP_126538823.1 |
| GCF_030344915.1 | NZ_CP128233.1 | 2165192 | 2166379 | 1187 | - | tuf | WP_126538823.1 |

| | | | | | | | |
|---|---|---|---|---|---|---|---|
| GCF_030344915.1 | NZ_CP128233.1 | 5199412 | 5200632 | 1220 | - | tuf | WP_086933207.1 |
| GCF_017309585.1 | NZ_CP071321.1 | 5219319 | 5220539 | 1220 | - | tuf | WP_086933207.1 |
| GCF_017309585.1 | NZ_CP071321.1 | 5236926 | 5238146 | 1220 | - | tuf | WP_086933207.1 |
| GCF_010983895.1 | NZ_CP048836.1 | 4037235 | 4038416 | 1181 | - | tuf | WP_221074962.1 |
| GCF_010983895.1 | NZ_CP048836.1 | 4055498 | 4056679 | 1181 | - | tuf | WP_221074962.1 |
| GCF_001007935.1 | NZ_CP011451.1 | 4132345 | 4133526 | 1181 | - | tuf | WP_221077417.1 |
| GCF_001007935.1 | NZ_CP011451.1 | 4150608 | 4151789 | 1181 | - | tuf | WP_221077417.1 |
| GCF_000009145.1 | NC_004757.1 | 4168871 | 4170052 | 1181 | - | tuf | WP_221077417.1 |
| GCF_000009145.1 | NC_004757.1 | 372193 | 373374 | 1181 | - | tuf | WP_014167500.1 |
| GCF_000014765.1 | NC_008344.1 | 819018 | 820199 | 1181 | - | tuf | WP_014167500.1 |
| GCF_000014765.1 | NC_008344.1 | 101439 | 102620 | 1181 | + | tuf | WP_005702465.1 |
| GCF_001455205.1 | NZ_CP013341.1 | 1311004 | 1312185 | 1181 | - | tuf | WP_005702465.1 |
| GCF_001455205.1 | NZ_CP013341.1 | 133411 | 134592 | 1181 | + | tuf | WP_109092123.1 |
| GCF_000619905.2 | NZ_CP012371.1 | 392351 | 393532 | 1181 | - | tuf | WP_109092123.1 |
| GCF_000619905.2 | NZ_CP012371.1 | 2245514 | 2246701 | 1187 | + | tuf | WP_187711347.1 |
| GCF_000355765.4 | NZ_CP021106.3 | 2263322 | 2264509 | 1187 | + | tuf | WP_187711347.1 |
| GCF_000355765.4 | NZ_CP021106.3 | 4179485 | 4180672 | 1187 | - | tuf | WP_106892548.1 |
| GCF_000196355.1 | NC_007614.1 | 4196286 | 4197473 | 1187 | - | tuf | WP_106892548.1 |
| GCF_000196355.1 | NC_007614.1 | 414014 | 415201 | 1187 | + | tuf | WP_011587666.1 |
| GCF_017161445.1 | NZ_CP071060.1 | 430828 | 432015 | 1187 | + | tuf | WP_011587666.1 |
| GCF_017161445.1 | NZ_CP071060.1 | 4777290 | 4778510 | 1220 | - | tuf | WP_261842608.1 |
| GCF_016028735.1 | NZ_CP065725.1 | 4793833 | 4795053 | 1220 | - | tuf | WP_261842608.1 |
| GCF_016028735.1 | NZ_CP065725.1 | 4766308 | 4767528 | 1220 | - | tuf | WP_271271947.1 |
| GCF_022811525.1 | NZ_CP063982.1 | 4782839 | 4784059 | 1220 | - | tuf | WP_271271947.1 |
| GCF_022811525.1 | NZ_CP063982.1 | 251632 | 252813 | 1181 | + | tuf | WP_005417220.1 |
| GCF_900089455.2 | NZ_LT907988.1 | 2707877 | 2709058 | 1181 | - | tuf | WP_005417220.1 |
| GCF_900089455.2 | NZ_LT907988.1 | 357073 | 358254 | 1181 | + | tuf | WP_012549148.1 |
| GCF_003058465.1 | NZ_CP028901.1 | 3112534 | 3113715 | 1181 | - | tuf | WP_012549148.1 |
| GCF_003058465.1 | NZ_CP028901.1 | 499981 | 501168 | 1187 | + | tuf | WP_011628194.1 |
| GCF_008274805.1 | NZ_CP022579.1 | 516615 | 517802 | 1187 | + | tuf | WP_011628194.1 |
| GCF_008274805.1 | NZ_CP022579.1 | 3883754 | 3884974 | 1220 | - | tuf | WP_265047316.1 |

| | | | | | | | |
|---|---|---|---|---|---|---|---|
| GCF_003008535.1 | NZ_CP027666.1 | 4551976 | 4553196 | 1220 | + | tuf | WP_250656534.1 |
| GCF_003008535.1 | NZ_CP027666.1 | 441636 | 442856 | 1220 | + | tuf | WP_251812750.1 |
| GCF_002127965.1 | NZ_CP018787.1 | 4203116 | 4204306 | 1190 | - | tuf | WP_041717128.1 |
| GCF_002127965.1 | NZ_CP018787.1 | 4219838 | 4221028 | 1190 | - | tuf | WP_041717128.1 |
| GCF_027118995.1 | NZ_CP098242.1 | 3417933 | 3419123 | 1190 | - | tuf | WP_142949704.1 |
| GCF_027118995.1 | NZ_CP098242.1 | 3434669 | 3435859 | 1190 | - | tuf | WP_142949704.1 |
| GCF_027557445.1 | NZ_CP106841.1 | 3795788 | 3796978 | 1190 | - | tuf | WP_026949537.1 |
| GCF_027557445.1 | NZ_CP106841.1 | 3812524 | 3813714 | 1190 | - | tuf | WP_026949537.1 |
| GCF_018802605.1 | NZ_CP069161.1 | 1389903 | 1391090 | 1187 | + | tuf | WP_213381006.1 |
| GCF_018802605.1 | NZ_CP069161.1 | 1407264 | 1408451 | 1187 | + | tuf | WP_213381006.1 |
| GCF_018802645.1 | NZ_CP069162.1 | 2680046 | 2681233 | 1187 | - | tuf | WP_012971553.1 |
| GCF_018802645.1 | NZ_CP069162.1 | 2697431 | 2698618 | 1187 | - | tuf | WP_012971553.1 |
| GCF_001465595.2 | NZ_CP013481.2 | 1384629 | 1385810 | 1181 | - | tuf | WP_039123801.1 |
| GCF_001465595.2 | NZ_CP013481.2 | 545960 | 547141 | 1181 | + | tuf | WP_039123801.1 |
| GCF_024704525.1 | NZ_CP102780.1 | 281922 | 283103 | 1181 | + | tuf | WP_133942155.1 |
| GCF_024704525.1 | NZ_CP102780.1 | 3958622 | 3959803 | 1181 | - | tuf | WP_014977498.1 |
| GCF_001029105.3 | NZ_CP011807.3 | 4164096 | 4165277 | 1181 | - | tuf | WP_014977498.1 |
| GCF_001029105.3 | NZ_CP011807.3 | 54739 | 55917 | 1178 | + | tuf | WP_013782545.1 |
| GCF_000807775.2 | NZ_CP047385.1 | 4411881 | 4413059 | 1178 | - | tuf | WP_013782545.1 |
| GCF_000807775.2 | NZ_CP047385.1 | 2704816 | 2705997 | 1181 | + | tuf | WP_075610371.1 |
| GCF_001465545.3 | NZ_CP013480.3 | 2850394 | 2851575 | 1181 | + | tuf | WP_075610371.1 |
| GCF_001465545.3 | NZ_CP013480.3 | 4209910 | 4211145 | 1235 | - | tuf | WP_231701243.1 |
| GCF_000972785.3 | NZ_CP011253.3 | 4504076 | 4505296 | 1220 | - | tuf | WP_261832956.1 |
| GCF_000972785.3 | NZ_CP011253.3 | 4520753 | 4521973 | 1220 | - | tuf | WP_261832956.1 |
| GCF_000504585.2 | NC_023018.2 | 3582778 | 3583998 | 1220 | - | tuf | WP_019622814.1 |
| GCF_000504585.2 | NC_023018.2 | 3599337 | 3600557 | 1220 | - | tuf | WP_019622814.1 |
| GCF_000815105.2 | NZ_CP010310.2 | 1718296 | 1719483 | 1187 | - | tuf | WP_114833746.1 |
| GCF_000815105.2 | NZ_CP010310.2 | 1735416 | 1736603 | 1187 | - | tuf | WP_114833746.1 |
| GCF_900187205.1 | NZ_LT906435.1 | 457625 | 458812 | 1187 | + | tuf | WP_148338180.1 |
| GCF_900187205.1 | NZ_LT906435.1 | 474319 | 475506 | 1187 | + | tuf | WP_148338201.1 |
| GCF_001017775.3 | NZ_CP011568.3 | 4282142 | 4283332 | 1190 | - | tuf | WP_021702710.1 |

| | | | | | | | |
|---|---|---|---|---|---|---|---|
| GCF_001017775.3 | NZ_CP011568.3 | 4298688 | 4299878 | 1190 | - | tuf | WP_021702710.1 |
| GCF_000934605.2 | NZ_CP010897.2 | 561094 | 562284 | 1190 | + | tuf | WP_173203010.1 |
| GCF_000934605.2 | NZ_CP010897.2 | 577592 | 578782 | 1190 | + | tuf | WP_173203010.1 |
| GCF_002278075.1 | NZ_CP022989.1 | 1837735 | 1838922 | 1187 | - | tuf | WP_114898932.1 |
| GCF_002278075.1 | NZ_CP022989.1 | 2502262 | 2503449 | 1187 | - | tuf | WP_114898932.1 |
| GCF_000092885.1 | NC_014117.1 | 3068340 | 3069521 | 1181 | - | tuf | WP_225506352.1 |
| GCF_000092885.1 | NC_014117.1 | 3107094 | 3108275 | 1181 | - | tuf | WP_225506447.1 |
| GCF_028473685.1 | NZ_CP099394.1 | 577684 | 578865 | 1181 | + | tuf | WP_121463410.1 |
| GCF_028473685.1 | NZ_CP099394.1 | 389741 | 390922 | 1181 | + | tuf | WP_180816921.1 |
| GCF_003368325.1 | NZ_CP031467.1 | 480441 | 481622 | 1181 | - | tuf | WP_180816921.1 |
| GCF_003368325.1 | NZ_CP031467.1 | 1084237 | 1085418 | 1181 | + | tuf | WP_032116381.1 |
| GCF_000522545.2 | NZ_CP012746.1 | 3769251 | 3770432 | 1181 | - | tuf | WP_032116381.1 |
| GCF_000522545.2 | NZ_CP012746.1 | 752097 | 753278 | 1181 | - | tuf | WP_066283224.1 |
| GCF_902459535.1 | NZ_LR699553.1 | 3792530 | 3793711 | 1181 | - | tuf | WP_135678127.1 |
| GCF_902459535.1 | NZ_LR699553.1 | 3833259 | 3834440 | 1181 | - | tuf | WP_135678135.1 |
| GCF_003286395.1 | NZ_CP029640.1 | 205317 | 206498 | 1181 | + | tuf | WP_348666479.1 |
| GCF_003286395.1 | NZ_CP029640.1 | 126460 | 127641 | 1181 | + | tuf | WP_148051980.1 |
| GCF_019428685.1 | NZ_CP080095.1 | 3951795 | 3952976 | 1181 | - | tuf | WP_125317950.1 |
| GCF_019428685.1 | NZ_CP080095.1 | 389014 | 390195 | 1181 | + | tuf | WP_125317950.1 |
| GCF_000961515.1 | NZ_CP010026.1 | 4421219 | 4422400 | 1181 | - | tuf | WP_125317950.1 |
| GCF_000961515.1 | NZ_CP010026.1 | 2042664 | 2043845 | 1181 | - | tuf | WP_035689535.1 |
| GCF_016128195.1 | NZ_CP066075.1 | 2369553 | 2370734 | 1181 | + | tuf | WP_035689535.1 |
| GCF_016128195.1 | NZ_CP066075.1 | 242706 | 243887 | 1181 | - | tuf | WP_115248696.1 |
| GCF_003330785.1 | NZ_CP024934.1 | 447941 | 449122 | 1181 | + | tuf | WP_046097610.1 |
| GCF_003330785.1 | NZ_CP024934.1 | 3870154 | 3871344 | 1190 | - | tuf | WP_377968817.1 |
| GCF_002902965.1 | NZ_CP026105.1 | 3886625 | 3887815 | 1190 | - | tuf | WP_377968817.1 |
| GCF_002902965.1 | NZ_CP026105.1 | 3961614 | 3962804 | 1190 | - | tuf | WP_089169348.1 |
| GCF_032806925.1 | NZ_CP136513.1 | 3978095 | 3979285 | 1190 | - | tuf | WP_089169348.1 |
| GCF_032806925.1 | NZ_CP136513.1 | 3397289 | 3398479 | 1190 | + | tuf | WP_152388454.1 |
| GCF_965118185.1 | NZ_OZ221290.1 | 3413772 | 3414962 | 1190 | + | tuf | WP_152388462.1 |
| GCF_965118185.1 | NZ_OZ221290.1 | 588990 | 590180 | 1190 | + | tuf | WP_012699289.1 |

| | | | | | | | |
|---|---|---|---|---|---|---|---|
| GCF_013426895.1 | NZ_AP023174.1 | 605463 | 606653 | 1190 | + | tuf | WP_012699289.1 |
| GCF_013426895.1 | NZ_AP023174.1 | 1716481 | 1717662 | 1181 | - | tuf | WP_011199348.1 |
| GCF_004524855.1 | NZ_CP038148.1 | 2014649 | 2015830 | 1181 | + | tuf | WP_011199348.1 |
| GCF_004524855.1 | NZ_CP038148.1 | 634847 | 636034 | 1187 | + | tuf | WP_122951122.1 |
| GCF_000020045.1 | NC_010622.1 | 2521925 | 2523112 | 1187 | - | tuf | WP_122951122.1 |
| GCF_000020045.1 | NC_010622.1 | 2328668 | 2329855 | 1187 | - | tuf | WP_062152299.1 |
| GCF_000020125.1 | NC_010681.1 | 3303828 | 3305015 | 1187 | - | tuf | WP_062153804.1 |
| GCF_000020125.1 | NC_010681.1 | 3116023 | 3117243 | 1220 | - | tuf | WP_007018319.1 |
| GCF_030412785.1 | NZ_CP125295.1 | 3134017 | 3135237 | 1220 | - | tuf | WP_007018319.1 |
| GCF_030412785.1 | NZ_CP125295.1 | 2008746 | 2009930 | 1184 | + | tuf | WP_015431789.1 |
| GCF_001865575.2 | NZ_CP017561.2 | 2112489 | 2113673 | 1184 | - | tuf | WP_025267414.1 |
| GCF_001865575.2 | NZ_CP017561.2 | 3962902 | 3964092 | 1190 | - | tuf | WP_197448635.1 |
| GCF_002902925.1 | NZ_CP026111.1 | 233288 | 234469 | 1181 | + | tuf | WP_132023998.1 |
| GCF_002902925.1 | NZ_CP026111.1 | 1923817 | 1924998 | 1181 | - | tuf | WP_132023998.1 |
| GCF_003330825.1 | NZ_CP024941.1 | 684635 | 685816 | 1181 | - | tuf | WP_046305253.1 |
| GCF_003330825.1 | NZ_CP024941.1 | 673661 | 674842 | 1181 | + | tuf | WP_250241513.1 |
| GCF_014171495.1 | NZ_CP049134.1 | 498923 | 500104 | 1181 | + | tuf | WP_159715096.1 |
| GCF_014171495.1 | NZ_CP049134.1 | 278810 | 279991 | 1181 | + | tuf | WP_250231781.1 |
| GCF_000013645.1 | NC_007951.1 | 752687 | 753868 | 1181 | + | tuf | WP_250226908.1 |
| GCF_000013645.1 | NC_007951.1 | 580445 | 581626 | 1181 | + | tuf | WP_192380149.1 |
| GCF_000176855.2 | NC_015138.1 | 666052 | 667233 | 1181 | - | tuf | WP_046289005.1 |
| GCF_000176855.2 | NC_015138.1 | 3941468 | 3942649 | 1181 | - | tuf | WP_048636590.1 |
| GCF_022493915.1 | NZ_CP042323.1 | 4794063 | 4795244 | 1181 | + | tuf | WP_048637241.1 |
| GCF_022493915.1 | NZ_CP042323.1 | 484024 | 485205 | 1181 | + | tuf | WP_208229545.1 |
| GCF_016894345.1 | NZ_CP069798.1 | 5006260 | 5007441 | 1181 | - | tuf | WP_208228844.1 |
| GCF_016894345.1 | NZ_CP069798.1 | 482249 | 483430 | 1181 | + | tuf | WP_009111200.1 |
| GCF_030254645.1 | NZ_CP116346.1 | 4525452 | 4526633 | 1181 | - | tuf | WP_009114822.1 |
| GCF_030254645.1 | NZ_CP116346.1 | 407103 | 408284 | 1181 | + | tuf | WP_021013352.1 |
| GCF_009833965.1 | NZ_CP047165.1 | 4709307 | 4710488 | 1181 | - | tuf | WP_021013352.1 |
| GCF_009833965.1 | NZ_CP047165.1 | 745612 | 746793 | 1181 | + | tuf | WP_014107696.1 |
| GCF_008119665.1 | NZ_CP043046.1 | 3373991 | 3375172 | 1181 | - | tuf | WP_014107696.1 |

| | | | | | | | |
|---|---|---|---|---|---|---|---|
| GCF_008119665.1 | NZ_CP043046.1 | 561330 | 562511 | 1181 | - | tuf | WP_011054016.1 |
| GCF_002762215.1 | NZ_CP024645.1 | 410056 | 411237 | 1181 | + | tuf | WP_121814555.1 |
| GCF_002762215.1 | NZ_CP024645.1 | 5044605 | 5045786 | 1181 | - | tuf | WP_121814555.1 |
| GCF_048320135.1 | NZ_CP182947.1 | 405933 | 407114 | 1181 | + | tuf | WP_309876750.1 |
| GCF_048320135.1 | NZ_CP182947.1 | 4931250 | 4932431 | 1181 | - | tuf | WP_309876750.1 |
| GCF_047549485.1 | NZ_CP180191.1 | 689429 | 690616 | 1187 | + | tuf | WP_166269962.1 |
| GCF_047549485.1 | NZ_CP180191.1 | 706635 | 707822 | 1187 | + | tuf | WP_166269962.1 |
| GCF_040105105.1 | NZ_CP157675.1 | 170263 | 171444 | 1181 | + | tuf | WP_126071556.1 |
| GCF_040105105.1 | NZ_CP157675.1 | 26823 | 28004 | 1181 | + | tuf | WP_226799142.1 |
| GCF_000015505.1 | NC_008781.1 | 344495 | 345676 | 1181 | - | tuf | WP_066283224.1 |
| GCF_000015505.1 | NC_008781.1 | 2760278 | 2761459 | 1181 | - | tuf | WP_032116381.1 |
| GCF_012584515.1 | NZ_CP051461.1 | 2794933 | 2796114 | 1181 | - | tuf | WP_032116381.1 |
| GCF_012584515.1 | NZ_CP051461.1 | 264975 | 266162 | 1187 | + | tuf | WP_176605036.1 |
| GCF_004022565.1 | NZ_CP022987.1 | 611522 | 612703 | 1181 | - | tuf | WP_250247217.1 |
| GCF_004022565.1 | NZ_CP022987.1 | 700395 | 701576 | 1181 | - | tuf | WP_015344803.1 |
| GCF_003065385.1 | NZ_CP023277.1 | 700813 | 701994 | 1181 | - | tuf | WP_011283101.1 |
| GCF_003065385.1 | NZ_CP023277.1 | 645968 | 647149 | 1181 | - | tuf | WP_013516869.1 |
| GCF_013307245.1 | NZ_CP028941.1 | 141481 | 142668 | 1187 | - | tuf | WP_014887324.1 |
| GCF_013307245.1 | NZ_CP028941.1 | 740180 | 741358 | 1178 | - | tuf | WP_258568084.1 |
| GCF_013307205.1 | NZ_CP028940.1 | 761530 | 762702 | 1172 | - | tuf | WP_258568100.1 |
| GCF_013307205.1 | NZ_CP028940.1 | 1461762 | 1462952 | 1190 | - | tuf | WP_048875579.1 |
| GCF_000016345.1 | NC_009379.1 | 1478332 | 1479522 | 1190 | - | tuf | WP_048875579.1 |
| GCF_000016345.1 | NC_009379.1 | 671669 | 672850 | 1181 | + | tuf | WP_395479650.1 |
| GCF_018688255.1 | NZ_CP061314.1 | 454592 | 455773 | 1181 | - | tuf | WP_067565843.1 |
| GCF_018688255.1 | NZ_CP061314.1 | 362340 | 363527 | 1187 | + | tuf | WP_251859242.1 |
| GCF_003065365.1 | NZ_CP023276.1 | 378891 | 380078 | 1187 | + | tuf | WP_251859242.1 |
| GCF_003065365.1 | NZ_CP023276.1 | 1364453 | 1365634 | 1181 | + | tuf | WP_096619358.1 |
| GCF_000973625.1 | NZ_CP007501.1 | 435503 | 436684 | 1181 | + | tuf | WP_157992549.1 |
| GCF_000973625.1 | NZ_CP007501.1 | 1032235 | 1033416 | 1181 | - | tuf | WP_040009155.1 |
| GCF_018687955.1 | NZ_CP061309.1 | 343107 | 344288 | 1181 | + | tuf | WP_072550019.1 |
| GCF_018687955.1 | NZ_CP061309.1 | 59447 | 60628 | 1181 | + | tuf | WP_067497389.1 |

| | | | | | | | |
|---|---|---|---|---|---|---|---|
| GCF_900096765.1 | NZ_LT615228.1 | 587683 | 588864 | 1181 | + | tuf | WP_067497389.1 |
| GCF_900096765.1 | NZ_LT615228.1 | 711985 | 713166 | 1181 | - | tuf | WP_015873272.1 |
| GCF_018687595.1 | NZ_CP061298.1 | 339779 | 340960 | 1181 | - | tuf | WP_067567847.1 |
| GCF_018687595.1 | NZ_CP061298.1 | 300367 | 301548 | 1181 | + | tuf | WP_041069078.1 |
| GCF_018685455.1 | NZ_CP049682.1 | 604859 | 606040 | 1181 | - | tuf | WP_041069693.1 |
| GCF_018685455.1 | NZ_CP049682.1 | 154227 | 155414 | 1187 | - | tuf | WP_100115026.1 |
| GCF_013307225.1 | NZ_CP028942.1 | 2277193 | 2278380 | 1187 | + | tuf | WP_348757530.1 |
| GCF_013307225.1 | NZ_CP028942.1 | 2530811 | 2531998 | 1187 | + | tuf | WP_348757530.1 |
| GCF_001659725.1 | NZ_CP015922.1 | 3172695 | 3173882 | 1187 | - | tuf | WP_153249646.1 |
| GCF_001659725.1 | NZ_CP015922.1 | 3280811 | 3281998 | 1187 | - | tuf | WP_153249731.1 |
| GCF_027924165.1 | NZ_AP026973.1 | 105408 | 106592 | 1184 | - | tuf | WP_067569158.1 |
| GCF_027924165.1 | NZ_AP026973.1 | 307745 | 308926 | 1181 | - | tuf | WP_013975810.1 |
| GCF_002872475.1 | NZ_CP025682.1 | 338003 | 339190 | 1187 | + | tuf | WP_197744704.1 |
| GCF_002872475.1 | NZ_CP025682.1 | 354265 | 355452 | 1187 | + | tuf | WP_197744704.1 |
| GCF_004322755.1 | NZ_CP036401.1 | 685190 | 686377 | 1187 | + | tuf | WP_096526604.1 |
| GCF_004322755.1 | NZ_CP036401.1 | 701741 | 702928 | 1187 | + | tuf | WP_096526604.1 |
| GCF_020865585.1 | NZ_CP086370.1 | 557559 | 558740 | 1181 | + | tuf | WP_038498984.1 |
| GCF_020865585.1 | NZ_CP086370.1 | 980021 | 981202 | 1181 | + | tuf | WP_022564217.1 |
| GCF_003028855.1 | NZ_CP028324.1 | 208412 | 209593 | 1181 | + | tuf | WP_145886281.1 |
| GCF_003028855.1 | NZ_CP028324.1 | 3403842 | 3405023 | 1181 | - | tuf | WP_145886281.1 |
| GCF_029028125.1 | NZ_CP119083.1 | 1005400 | 1006620 | 1220 | + | tuf | WP_279242885.1 |
| GCF_029028125.1 | NZ_CP119083.1 | 97217 | 98413 | 1196 | + | tuf | WP_014894988.1 |
| GCF_009789595.1 | NZ_CP046904.1 | 780739 | 781920 | 1181 | + | tuf | WP_226813170.1 |
| GCF_009789595.1 | NZ_CP046904.1 | 769195 | 770376 | 1181 | + | tuf | WP_216782210.1 |
| GCF_004209755.1 | NZ_CP035913.1 | 245405 | 246586 | 1181 | - | tuf | WP_341765196.1 |
| GCF_004209755.1 | NZ_CP035913.1 | 491840 | 493030 | 1190 | + | tuf | WP_129210747.1 |
| GCF_004421005.1 | NZ_CP038026.1 | 274114 | 275301 | 1187 | + | tuf | WP_020026356.1 |
| GCF_004421005.1 | NZ_CP038026.1 | 434531 | 435718 | 1187 | + | tuf | WP_020026356.1 |
| GCF_005280315.1 | NZ_CP040017.1 | 219698 | 220879 | 1181 | - | tuf | WP_115956018.1 |
| GCF_005280315.1 | NZ_CP040017.1 | 1754796 | 1755980 | 1184 | + | tuf | WP_339056396.1 |
| GCF_003345255.1 | NZ_CP029331.1 | 1246320 | 1247507 | 1187 | + | tuf | WP_174672959.1 |

| | | | | | | | |
|---|---|---|---|---|---|---|---|
| GCF_003345255.1 | NZ_CP029331.1 | 1262900 | 1264087 | 1187 | + | tuf | WP_174672960.1 |
| GCF_003013695.1 | NZ_CP027792.1 | 1159479 | 1160678 | 1199 | - | tuf | WP_126323073.1 |
| GCF_003013695.1 | NZ_CP027792.1 | 409818 | 410999 | 1181 | - | tuf | WP_013087723.1 |
| GCF_020510525.1 | NZ_CP075188.1 | 156687 | 157868 | 1181 | + | tuf | WP_039719495.1 |
| GCF_020510525.1 | NZ_CP075188.1 | 566 | 1753 | 1187 | + | tuf | WP_174605066.1 |
| GCF_001663855.1 | NZ_CP016022.1 | 714995 | 716182 | 1187 | - | tuf | WP_174605066.1 |
| GCF_001663855.1 | NZ_CP016022.1 | 190320 | 191507 | 1187 | + | tuf | WP_011737581.1 |
| GCF_000954135.1 | NZ_CP010799.1 | 874131 | 875318 | 1187 | - | tuf | WP_011738164.1 |
| GCF_000954135.1 | NZ_CP010799.1 | 15168 | 16349 | 1181 | - | tuf | WP_223138644.1 |
| GCF_018243235.1 | NZ_CP046674.1 | 213950 | 215131 | 1181 | - | tuf | WP_025244013.1 |
| GCF_018243235.1 | NZ_CP046674.1 | 3901893 | 3903074 | 1181 | - | tuf | WP_025246566.1 |
| GCF_001587155.1 | NZ_CP012943.1 | 657618 | 658799 | 1181 | + | tuf | WP_226811221.1 |
| GCF_001587155.1 | NZ_CP012943.1 | 360712 | 361893 | 1181 | + | tuf | WP_041062952.1 |
| GCF_037101365.1 | NZ_AP029014.1 | 406839 | 408020 | 1181 | - | tuf | WP_041062952.1 |
| GCF_037101365.1 | NZ_AP029014.1 | 3681283 | 3682503 | 1220 | - | tuf | WP_369855604.1 |
| GCF_021173085.1 | NZ_CP076413.1 | 3725183 | 3726403 | 1220 | - | tuf | WP_369855604.1 |
| GCF_021173085.1 | NZ_CP076413.1 | 1309723 | 1310910 | 1187 | + | tuf | WP_100918275.1 |
| GCF_000215705.1 | NC_015677.1 | 5624359 | 5625546 | 1187 | - | tuf | WP_100918275.1 |
| GCF_000215705.1 | NC_015677.1 | 1403108 | 1404295 | 1187 | + | tuf | WP_369150815.1 |
| GCF_001955735.1 | NZ_CP019240.1 | 1499354 | 1500541 | 1187 | + | tuf | WP_369177340.1 |
| GCF_001955735.1 | NZ_CP019240.1 | 977418 | 978605 | 1187 | + | tuf | WP_053951621.1 |
| GCF_006974105.1 | NZ_CP036282.1 | 1103880 | 1105067 | 1187 | + | tuf | WP_053951621.1 |
| GCF_006974105.1 | NZ_CP036282.1 | 1439946 | 1441133 | 1187 | + | tuf | WP_210229715.1 |
| GCF_000013605.1 | NC_007908.1 | 1456263 | 1457450 | 1187 | + | tuf | WP_210229715.1 |
| GCF_000013605.1 | NC_007908.1 | 179972 | 181159 | 1187 | + | tuf | WP_011929567.1 |
| GCF_001955695.1 | NZ_CP019236.1 | 786415 | 787602 | 1187 | - | tuf | WP_011929567.1 |
| GCF_001955695.1 | NZ_CP019236.1 | 120450 | 121631 | 1181 | + | tuf | WP_281263974.1 |
| GCF_032191775.1 | NZ_CP132507.1 | 1699412 | 1700593 | 1181 | + | tuf | WP_015873272.1 |
| GCF_032191775.1 | NZ_CP132507.1 | 814294 | 815481 | 1187 | + | tuf | WP_079541704.1 |
| GCF_001955715.1 | NZ_CP019239.1 | 830538 | 831725 | 1187 | + | tuf | WP_126316002.1 |
| GCF_001955715.1 | NZ_CP019239.1 | 494217 | 495398 | 1181 | + | tuf | WP_268074995.1 |

| | | | | | | | |
|---|---|---|---|---|---|---|---|
| GCF_006970865.1 | NZ_CP035503.1 | 510735 | 511916 | 1181 | + | tuf | WP_268074995.1 |
| GCF_006970865.1 | NZ_CP035503.1 | 2148907 | 2150088 | 1181 | + | tuf | WP_143871240.1 |
| GCF_034047095.1 | NZ_CP139170.1 | 2165401 | 2166582 | 1181 | + | tuf | WP_143871251.1 |
| GCF_034047095.1 | NZ_CP139170.1 | 246338 | 247519 | 1181 | + | tuf | WP_115586309.1 |
| GCF_025398155.2 | NZ_CP104562.2 | 4566734 | 4567915 | 1181 | - | tuf | WP_115586309.1 |
| GCF_025398155.2 | NZ_CP104562.2 | 233201 | 234382 | 1181 | + | tuf | WP_038478205.1 |
| GCF_001483865.1 | NZ_CP013729.1 | 4384385 | 4385566 | 1181 | - | tuf | WP_038478205.1 |
| GCF_001483865.1 | NZ_CP013729.1 | 228694 | 229875 | 1181 | + | tuf | WP_038478205.1 |
| GCF_047946925.1 | NZ_CP181386.1 | 4809134 | 4810315 | 1181 | - | tuf | WP_038476797.1 |
| GCF_047946925.1 | NZ_CP181386.1 | 3742849 | 3744030 | 1181 | - | tuf | WP_131912229.1 |
| GCF_000934545.1 | NZ_CP010554.1 | 3762481 | 3763662 | 1181 | - | tuf | WP_131912213.1 |
| GCF_000934545.1 | NZ_CP010554.1 | 816845 | 818065 | 1220 | + | tuf | WP_012486350.1 |
| GCF_000828915.1 | NZ_AP014569.1 | 833725 | 834945 | 1220 | + | tuf | WP_012486350.1 |
| GCF_000828915.1 | NZ_AP014569.1 | 597723 | 598904 | 1181 | - | tuf | WP_025297190.1 |
| GCF_000828895.1 | NZ_AP014568.1 | 1379229 | 1380410 | 1181 | + | tuf | WP_025297183.1 |
| GCF_000828895.1 | NZ_AP014568.1 | 1954550 | 1955740 | 1190 | + | tuf | WP_097024033.1 |
| GCF_021654335.1 | NZ_AP023423.1 | 478004 | 479194 | 1190 | + | tuf | WP_011505727.1 |
| GCF_021654335.1 | NZ_AP023423.1 | 3616739 | 3617920 | 1181 | - | tuf | WP_042998968.1 |
| GCF_000025705.1 | NC_013959.1 | 4391674 | 4392855 | 1181 | + | tuf | WP_003031109.1 |
| GCF_000025705.1 | NC_013959.1 | 278866 | 280047 | 1181 | + | tuf | WP_016157942.1 |
| GCF_030035605.1 | NZ_CP106736.1 | 4793819 | 4795000 | 1181 | - | tuf | WP_003031109.1 |
| GCF_030035605.1 | NZ_CP106736.1 | 706505 | 707686 | 1181 | - | tuf | WP_003031109.1 |
| GCF_002951835.1 | NZ_CP019448.1 | 1384732 | 1385913 | 1181 | + | tuf | WP_003031109.1 |
| GCF_002951835.1 | NZ_CP019448.1 | 882293 | 883474 | 1181 | - | tuf | WP_003031109.1 |
| GCF_003008595.1 | NZ_CP027669.1 | 1540028 | 1541209 | 1181 | + | tuf | WP_016157942.1 |
| GCF_003008595.1 | NZ_CP027669.1 | 409692 | 410873 | 1181 | + | tuf | WP_003031109.1 |
| GCF_000600005.1 | NZ_CP007446.1 | 4837013 | 4838194 | 1181 | - | tuf | WP_047407352.1 |
| GCF_000600005.1 | NZ_CP007446.1 | 76640 | 77821 | 1181 | + | tuf | WP_042998968.1 |
| GCF_023734135.1 | NZ_AP025730.1 | 4384981 | 4386162 | 1181 | - | tuf | WP_003031109.1 |
| GCF_023734135.1 | NZ_AP025730.1 | 397813 | 398994 | 1181 | + | tuf | WP_003031109.1 |
| GCF_008329925.1 | NZ_CP035708.1 | 4633571 | 4634752 | 1181 | - | tuf | WP_003031109.1 |

| | | | | | | | |
|---|---|---|---|---|---|---|---|
| GCF_008329925.1 | NZ_CP035708.1 | 2790389 | 2791570 | 1181 | - | tuf | WP_012133818.1 |
| GCF_000297055.2 | NC_022357.1 | 4353425 | 4354606 | 1181 | - | tuf | WP_024131004.1 |
| GCF_000297055.2 | NC_022357.1 | 449946 | 451127 | 1181 | + | tuf | WP_003031109.1 |
| GCF_003966565.1 | NZ_AP018718.1 | 5006814 | 5007995 | 1181 | - | tuf | WP_003031109.1 |
| GCF_003966565.1 | NZ_AP018718.1 | 26047 | 27228 | 1181 | - | tuf | WP_003031109.1 |
| GCF_004323595.1 | NZ_AP019373.1 | 676252 | 677433 | 1181 | + | tuf | WP_003031109.1 |
| GCF_004323595.1 | NZ_AP019373.1 | 422516 | 423697 | 1181 | + | tuf | WP_003031109.1 |
| GCF_009937995.1 | NZ_AP021881.1 | 4830776 | 4831957 | 1181 | - | tuf | WP_003031109.1 |
| GCF_009937995.1 | NZ_AP021881.1 | 865840 | 867021 | 1181 | - | tuf | WP_012907761.1 |
| GCF_009938015.1 | NZ_AP021884.1 | 1631637 | 1632818 | 1181 | + | tuf | WP_012907761.1 |
| GCF_009938015.1 | NZ_AP021884.1 | 465053 | 466234 | 1181 | + | tuf | WP_181822206.1 |
| GCF_011764585.1 | NZ_AP022853.1 | 4477252 | 4478433 | 1181 | - | tuf | WP_181822206.1 |
| GCF_011764585.1 | NZ_AP022853.1 | 3158652 | 3159833 | 1181 | - | tuf | WP_135323597.1 |
| GCF_000828635.1 | NZ_AP012547.1 | 3823014 | 3824195 | 1181 | + | tuf | WP_135324094.1 |
| GCF_000828635.1 | NZ_AP012547.1 | 448603 | 449784 | 1181 | + | tuf | WP_016157942.1 |
| GCF_003967275.1 | NZ_AP018721.1 | 4684624 | 4685805 | 1181 | - | tuf | WP_003031109.1 |
| GCF_003967275.1 | NZ_AP018721.1 | 152817 | 154007 | 1190 | - | tuf | WP_043334873.1 |
| GCF_006337085.1 | NZ_CP040882.1 | 142478 | 143668 | 1190 | + | tuf | WP_065392570.1 |
| GCF_006337085.1 | NZ_CP040882.1 | 2175499 | 2176680 | 1181 | + | tuf | WP_081151013.1 |
| GCF_003609995.1 | NZ_AP018786.1 | 2195998 | 2197179 | 1181 | + | tuf | WP_081151013.1 |
| GCF_003609995.1 | NZ_AP018786.1 | 5042156 | 5043337 | 1181 | - | tuf | WP_011045489.1 |
| GCF_016724805.1 | NZ_CP068055.1 | 5064629 | 5065810 | 1181 | - | tuf | WP_011045489.1 |
| GCF_016724805.1 | NZ_CP068055.1 | 3593417 | 3594637 | 1220 | - | tuf | WP_407327396.1 |
| GCF_000226625.1 | NC_016043.1 | 3611700 | 3612920 | 1220 | - | tuf | WP_407327396.1 |
| GCF_000226625.1 | NC_016043.1 | 985739 | 986959 | 1220 | + | tuf | WP_008292621.1 |
| GCF_028868935.1 | NZ_CP076516.1 | 1004001 | 1005221 | 1220 | + | tuf | WP_008292621.1 |
| GCF_028868935.1 | NZ_CP076516.1 | 3519292 | 3520512 | 1220 | + | tuf | WP_407347854.1 |
| GCF_030770395.1 | NZ_CP132480.1 | 3537495 | 3538715 | 1220 | + | tuf | WP_407347854.1 |
| GCF_030770395.1 | NZ_CP132480.1 | 223400 | 224590 | 1190 | + | tuf-2 | NP_819280.1 |
| GCF_029223865.1 | NZ_CP119520.1 | 329771 | 330961 | 1190 | + | tuf | WP_039669934.1 |
| GCF_029223865.1 | NZ_CP119520.1 | 346616 | 347806 | 1190 | + | tuf | WP_039669934.1 |

| | | | | | | | |
|---|---|---|---|---|---|---|---|
| GCF_020162115.1 | NZ_CP083911.1 | 333096 | 334280 | 1184 | + | tuf | WP_211923999.1 |
| GCF_020162115.1 | NZ_CP083911.1 | 349755 | 350939 | 1184 | + | tuf | WP_211923999.1 |
| GCF_000021765.1 | NC_011662.2 | 329628 | 330818 | 1190 | + | tuf | WP_159747926.1 |
| GCF_000021765.1 | NC_011662.2 | 329775 | 330965 | 1190 | + | tuf | WP_039669934.1 |
| GCF_003030465.1 | NZ_CP028339.1 | 346620 | 347810 | 1190 | + | tuf | WP_039669934.1 |
| GCF_003030465.1 | NZ_CP028339.1 | 349605 | 350786 | 1181 | + | tuf | WP_007872666.1 |
| GCF_048398245.1 | NZ_CP183038.1 | 3970742 | 3971923 | 1181 | - | tuf | WP_053532071.1 |
| GCF_048398245.1 | NZ_CP183038.1 | 343485 | 344666 | 1181 | + | tuf | WP_038869252.1 |
| GCF_001922305.1 | NZ_CP018839.1 | 4060629 | 4061810 | 1181 | - | tuf | WP_038869252.1 |
| GCF_001922305.1 | NZ_CP018839.1 | 219365 | 220546 | 1181 | + | tuf | WP_007872666.1 |
| GCF_001051995.2 | NZ_CP014646.1 | 3816118 | 3817299 | 1181 | - | tuf | WP_007872666.1 |
| GCF_001051995.2 | NZ_CP014646.1 | 73349 | 74530 | 1181 | + | tuf | WP_038869252.1 |
| GCF_035621355.1 | NZ_CP141769.1 | 3693302 | 3694483 | 1181 | - | tuf | WP_038869252.1 |
| GCF_035621355.1 | NZ_CP141769.1 | 3095657 | 3096838 | 1181 | - | tuf | WP_007872666.1 |
| GCF_000253115.1 | NC_014145.1 | 3792109 | 3793290 | 1181 | + | tuf | WP_007872666.1 |
| GCF_000253115.1 | NC_014145.1 | 367566 | 368747 | 1181 | + | tuf | WP_015742710.1 |
| GCF_002028405.1 | NZ_CP020046.1 | 4046175 | 4047356 | 1181 | - | tuf | WP_175573005.1 |
| GCF_002028405.1 | NZ_CP020046.1 | 334486 | 335667 | 1181 | + | tuf | WP_007872666.1 |
| GCF_034424545.1 | NZ_CP139970.1 | 3893389 | 3894570 | 1181 | - | tuf | WP_007872666.1 |
| GCF_034424545.1 | NZ_CP139970.1 | 288922 | 290103 | 1181 | + | tuf | WP_180706188.1 |
| GCF_005280735.1 | NZ_CP040077.1 | 4049280 | 4050461 | 1181 | - | tuf | WP_180706188.1 |
| GCF_005280735.1 | NZ_CP040077.1 | 373480 | 374661 | 1181 | + | tuf | WP_012768098.1 |
| GCF_016696765.1 | NZ_CP065313.1 | 4398219 | 4399400 | 1181 | - | tuf | WP_012768098.1 |
| GCF_016696765.1 | NZ_CP065313.1 | 252949 | 254130 | 1181 | + | tuf | WP_013315919.1 |
| GCF_031326225.1 | NZ_CP133720.1 | 4439807 | 4440988 | 1181 | - | tuf | WP_013315919.1 |
| GCF_031326225.1 | NZ_CP133720.1 | 250031 | 251212 | 1181 | + | tuf | WP_013315919.1 |
| GCF_003955735.1 | NZ_CP034464.1 | 4429902 | 4431083 | 1181 | - | tuf | WP_013315919.1 |
| GCF_003955735.1 | NZ_CP034464.1 | 247869 | 249050 | 1181 | + | tuf | WP_012882957.1 |
| GCF_022870845.1 | NZ_CP091507.1 | 4516253 | 4517434 | 1181 | - | tuf | WP_012882957.1 |
| GCF_022870845.1 | NZ_CP091507.1 | 419200 | 420381 | 1181 | + | tuf | WP_012882957.1 |
| GCF_022870865.1 | NZ_CP091508.1 | 4459202 | 4460383 | 1181 | - | tuf | WP_012882957.1 |

| | | | | | | | |
|---|---|---|---|---|---|---|---|
| GCF_022870865.1 | NZ_CP091508.1 | 240662 | 241843 | 1181 | + | tuf | WP_012882957.1 |
| GCF_013003985.1 | NZ_CP053073.1 | 4269798 | 4270979 | 1181 | - | tuf | WP_012882957.1 |
| GCF_013003985.1 | NZ_CP053073.1 | 252016 | 253197 | 1181 | + | tuf | WP_019845203.1 |
| GCF_013003965.1 | NZ_CP053069.1 | 470207 | 471388 | 1181 | + | tuf | WP_019845203.1 |
| GCF_013003965.1 | NZ_CP053069.1 | 1649735 | 1650916 | 1181 | - | tuf | WP_013315919.1 |
| GCF_048305125.1 | NZ_CP157626.1 | 2411720 | 2412901 | 1181 | + | tuf | WP_022631747.1 |
| GCF_048305125.1 | NZ_CP157626.1 | 252954 | 254135 | 1181 | + | tuf | WP_019845203.1 |
| GCF_000015565.1 | NC_008786.1 | 4193120 | 4194301 | 1181 | - | tuf | WP_012882957.1 |
| GCF_000015565.1 | NC_008786.1 | 3009676 | 3010863 | 1187 | - | tuf | WP_067648358.1 |
| GCF_002222655.1 | NZ_CP022423.1 | 3026485 | 3027672 | 1187 | - | tuf | WP_067648392.1 |
| GCF_002222655.1 | NZ_CP022423.1 | 4490987 | 4492168 | 1181 | + | tuf | WP_108652423.1 |
| GCF_022870945.1 | NZ_CP091512.1 | 214121 | 215302 | 1181 | + | tuf | WP_187488813.1 |
| GCF_022870945.1 | NZ_CP091512.1 | 3314414 | 3315595 | 1181 | - | tuf | WP_067427337.1 |
| GCF_022354465.1 | NZ_CP092361.1 | 1013574 | 1014761 | 1187 | + | tuf | WP_329742737.1 |
| GCF_022354465.1 | NZ_CP092361.1 | 1030702 | 1031889 | 1187 | + | tuf | WP_329742737.1 |
| GCF_009906855.1 | NZ_CP047650.1 | 3304449 | 3305636 | 1187 | - | tuf | WP_188799894.1 |
| GCF_009906855.1 | NZ_CP047650.1 | 3321479 | 3322666 | 1187 | - | tuf | WP_188799894.1 |
| GCF_000164865.1 | NC_014618.1 | 1046105 | 1047292 | 1187 | - | tuf | WP_187057920.1 |
| GCF_000164865.1 | NC_014618.1 | 1063180 | 1064367 | 1187 | - | tuf | WP_187057920.1 |
| GCF_001647695.1 | NZ_CP015425.1 | 2051801 | 2052988 | 1187 | - | tuf | WP_017463590.1 |
| GCF_001647695.1 | NZ_CP015425.1 | 2068818 | 2070005 | 1187 | - | tuf | WP_017463590.1 |
| GCF_000007745.1 | NC_006300.1 | 4566731 | 4567921 | 1190 | - | tuf | WP_252576691.1 |
| GCF_000007745.1 | NC_006300.1 | 4583278 | 4584468 | 1190 | - | tuf | WP_004373450.1 |
| GCF_022647505.1 | NZ_CP071409.1 | 4578699 | 4579889 | 1190 | - | tuf | WP_134675806.1 |
| GCF_022647505.1 | NZ_CP071409.1 | 4595245 | 4596435 | 1190 | - | tuf | WP_134675806.1 |
| GCF_016592595.1 | NZ_AP012977.1 | 719784 | 720974 | 1190 | + | tuf | WP_013717348.1 |
| GCF_016592595.1 | NZ_AP012977.1 | 736314 | 737504 | 1190 | + | tuf | WP_036994748.1 |
| GCF_001705075.2 | NZ_CP080624.1 | 1474120 | 1475310 | 1190 | - | tuf | WP_017675622.1 |
| GCF_001705075.2 | NZ_CP080624.1 | 1490663 | 1491853 | 1190 | - | tuf | WP_017675622.1 |
| GCF_001753165.1 | NZ_CP017448.1 | 1421861 | 1423042 | 1181 | - | tuf | WP_038631315.1 |
| GCF_001753165.1 | NZ_CP017448.1 | 1930990 | 1932171 | 1181 | + | tuf | WP_038631315.1 |

| | | | | | | | |
|---|---|---|---|---|---|---|---|
| GCF_001975725.1 | NZ_CP019434.1 | 376636 | 377817 | 1181 | - | tuf | WP_024522593.1 |
| GCF_001975725.1 | NZ_CP019434.1 | 893810 | 894991 | 1181 | + | tuf | WP_024522593.1 |
| GCF_001753245.1 | NZ_CP017415.1 | 159930 | 161111 | 1181 | + | tuf | WP_012847048.1 |
| GCF_001753245.1 | NZ_CP017415.1 | 3501151 | 3502332 | 1181 | - | tuf | WP_012847048.1 |
| GCF_009035845.1 | NZ_CP045110.1 | 180103 | 181284 | 1181 | + | tuf | WP_012847048.1 |
| GCF_009035845.1 | NZ_CP045110.1 | 3405047 | 3406228 | 1181 | - | tuf | WP_012847048.1 |
| GCF_000046845.1 | NC_005966.1 | 38365 | 39546 | 1181 | - | tuf | WP_024522593.1 |
| GCF_000046845.1 | NC_005966.1 | 589320 | 590501 | 1181 | + | tuf | WP_024522593.1 |
| GCF_016576965.1 | NZ_CP066119.1 | 266739 | 267926 | 1187 | + | tuf | WP_066042991.1 |
| GCF_016576965.1 | NZ_CP066119.1 | 409040 | 410227 | 1187 | + | tuf | WP_066042991.1 |
| GCF_018324105.1 | NZ_AP024595.1 | 168266 | 169447 | 1181 | - | tuf | WP_130624143.1 |
| GCF_018324105.1 | NZ_AP024595.1 | 88102 | 89283 | 1181 | + | tuf | WP_130579433.1 |
| GCF_002165375.2 | NZ_CP032134.1 | 126618 | 127799 | 1181 | + | tuf | WP_148708428.1 |
| GCF_002165375.2 | NZ_CP032134.1 | 113986 | 115167 | 1181 | - | tuf | WP_130616901.1 |
| GCF_030053775.1 | NZ_CP125669.1 | 184867 | 186054 | 1187 | + | tuf | WP_043106870.1 |
| GCF_030053775.1 | NZ_CP125669.1 | 201614 | 202801 | 1187 | + | tuf | WP_043106870.1 |
| GCF_003024525.3 | NZ_CP035934.2 | 3324203 | 3325423 | 1220 | + | tuf | WP_262595310.1 |
| GCF_003024525.3 | NZ_CP035934.2 | 5030864 | 5032084 | 1220 | - | tuf | WP_034877901.1 |
| GCF_001704615.3 | NZ_CP029397.2 | 207010 | 208191 | 1181 | + | tuf | WP_014830236.1 |
| GCF_001704615.3 | NZ_CP029397.2 | 4234356 | 4235537 | 1181 | - | tuf | WP_023326212.1 |
| GCF_009884975.1 | NZ_CP041970.1 | 209792 | 210973 | 1181 | + | tuf | WP_023309456.1 |
| GCF_009884975.1 | NZ_CP041970.1 | 4230795 | 4231976 | 1181 | - | tuf | WP_014068448.1 |
| GCF_001307195.1 | NZ_CP012808.1 | 215242 | 216423 | 1181 | + | tuf | WP_014068448.1 |
| GCF_001307195.1 | NZ_CP012808.1 | 4289217 | 4290398 | 1181 | - | tuf | WP_014068448.1 |
| GCF_002370525.2 | NZ_AP014630.1 | 29169 | 30350 | 1181 | - | tuf | WP_220710802.1 |
| GCF_002370525.2 | NZ_AP014630.1 | 4538832 | 4540013 | 1181 | - | tuf | WP_039274549.1 |
| GCF_003323815.1 | NZ_CP030880.1 | 227863 | 229044 | 1181 | + | tuf | WP_014068448.1 |
| GCF_003323815.1 | NZ_CP030880.1 | 4634800 | 4635981 | 1181 | - | tuf | WP_014068448.1 |
| GCF_009914475.1 | NZ_CP045198.1 | 222835 | 224016 | 1181 | + | tuf | WP_023309456.1 |
| GCF_009914475.1 | NZ_CP045198.1 | 4224268 | 4225449 | 1181 | - | tuf | WP_023309456.1 |
| GCF_018336855.1 | NZ_CP059558.1 | 237768 | 238949 | 1181 | + | tuf | WP_013094997.1 |

| | | | | | | | |
|---|---|---|---|---|---|---|---|
| GCF_018336855.1 | NZ_CP059558.1 | 4411860 | 4413041 | 1181 | - | tuf | WP_023620081.1 |
| GCF_013122135.1 | NZ_CP053391.1 | 197334 | 198515 | 1181 | + | tuf | WP_023309456.1 |
| GCF_013122135.1 | NZ_CP053391.1 | 4128425 | 4129606 | 1181 | - | tuf | WP_014068448.1 |
| GCF_011578285.1 | NZ_CP049916.1 | 150536 | 151717 | 1181 | + | tuf | WP_039274549.1 |
| GCF_011578285.1 | NZ_CP049916.1 | 4156264 | 4157445 | 1181 | - | tuf | WP_023323999.1 |
| GCF_001704115.1 | NZ_CP016895.1 | 419622 | 420803 | 1181 | + | tuf | WP_243544814.1 |
| GCF_001704115.1 | NZ_CP016895.1 | 4611415 | 4612596 | 1181 | - | tuf | WP_243544814.1 |
| GCF_019343495.1 | NZ_CP072549.1 | 237361 | 238542 | 1181 | + | tuf | WP_017693880.1 |
| GCF_019343495.1 | NZ_CP072549.1 | 4247859 | 4249040 | 1181 | - | tuf | WP_017693880.1 |
| GCF_041021905.1 | NZ_CP157432.1 | 192623 | 193804 | 1181 | + | tuf | WP_014168151.1 |
| GCF_041021905.1 | NZ_CP157432.1 | 4229752 | 4230933 | 1181 | - | tuf | WP_014171903.1 |
| GCF_000196795.1 | NC_014259.1 | 2861243 | 2862424 | 1181 | - | tuf | WP_014171903.1 |
| GCF_000196795.1 | NC_014259.1 | 3440026 | 3441207 | 1181 | + | tuf | WP_014830236.1 |
| GCF_000191145.1 | NC_016603.1 | 210096 | 211277 | 1181 | + | tuf | WP_014068448.1 |
| GCF_000191145.1 | NC_016603.1 | 4271076 | 4272257 | 1181 | - | tuf | WP_014068448.1 |
| GCF_003258335.1 | NZ_CP030031.1 | 2031600 | 2032781 | 1181 | - | tuf | WP_152082495.1 |
| GCF_003258335.1 | NZ_CP030031.1 | 2712909 | 2714090 | 1181 | + | tuf | WP_013094997.1 |
| GCF_010918895.1 | NZ_CP044483.1 | 208345 | 209526 | 1181 | + | tuf | WP_023326212.1 |
| GCF_010918895.1 | NZ_CP044483.1 | 4365795 | 4366976 | 1181 | - | tuf | WP_023326212.1 |
| GCF_016064815.1 | NZ_CP065820.1 | 166030 | 167211 | 1181 | - | tuf | WP_014068448.1 |
| GCF_016064815.1 | NZ_CP065820.1 | 805430 | 806611 | 1181 | + | tuf | WP_014830236.1 |
| GCF_019355975.1 | NZ_CP079898.1 | 213029 | 214210 | 1181 | + | tuf | WP_014830236.1 |
| GCF_019355975.1 | NZ_CP079898.1 | 4298683 | 4299864 | 1181 | - | tuf | WP_014830236.1 |
| GCF_018139125.1 | NZ_CP073241.1 | 215121 | 216302 | 1181 | + | tuf | WP_014830236.1 |
| GCF_018139125.1 | NZ_CP073241.1 | 4294584 | 4295765 | 1181 | - | tuf | WP_014830236.1 |
| GCF_008630915.1 | NZ_CP043909.1 | 226872 | 228053 | 1181 | + | tuf | WP_014068448.1 |
| GCF_008630915.1 | NZ_CP043909.1 | 4313147 | 4314328 | 1181 | - | tuf | WP_014068448.1 |
| GCF_039865415.1 | NZ_CP156787.1 | 204609 | 205790 | 1181 | + | tuf | WP_014068448.1 |
| GCF_039865415.1 | NZ_CP156787.1 | 4051188 | 4052369 | 1181 | - | tuf | WP_014068448.1 |
| GCF_023824315.1 | NZ_CP098732.1 | 276853 | 278034 | 1181 | + | tuf | WP_326469424.1 |
| GCF_023824315.1 | NZ_CP098732.1 | 4368773 | 4369954 | 1181 | - | tuf | WP_326468683.1 |

| | | | | | | | |
|---|---|---|---|---|---|---|---|
| GCF_017498585.1 | NZ_CP071766.1 | 329542 | 330723 | 1181 | - | tuf | WP_168918948.1 |
| GCF_017498585.1 | NZ_CP071766.1 | 327110 | 328291 | 1181 | - | tuf | WP_168893972.1 |
| GCF_030518655.1 | NZ_CP130328.1 | 333431 | 334612 | 1181 | - | tuf | WP_168894397.1 |
| GCF_030518655.1 | NZ_CP130328.1 | 327370 | 328551 | 1181 | - | tuf | WP_168919752.1 |
| GCF_001696605.3 | NZ_CP031716.1 | 330473 | 331654 | 1181 | - | tuf | WP_169786034.1 |
| GCF_001696605.3 | NZ_CP031716.1 | 328751 | 329932 | 1181 | - | tuf | WP_168920138.1 |
| GCF_900638385.1 | NZ_LR134510.1 | 331418 | 332599 | 1181 | - | tuf | WP_208753800.1 |
| GCF_900638385.1 | NZ_LR134510.1 | 331490 | 332671 | 1181 | - | tuf | WP_168893158.1 |
| GCF_900636745.1 | NZ_LR134310.1 | 331625 | 332806 | 1181 | - | tuf | WP_169780480.1 |
| GCF_900636745.1 | NZ_LR134310.1 | 325870 | 327051 | 1181 | - | tuf | WP_168821868.1 |
| GCF_004519515.1 | NZ_CP038145.1 | 330533 | 331714 | 1181 | - | tuf | WP_169767749.1 |
| GCF_004519515.1 | NZ_CP038145.1 | 328943 | 330124 | 1181 | - | tuf | WP_168868505.1 |
| GCF_003290385.1 | NZ_CP030753.1 | 331610 | 332791 | 1181 | - | tuf | WP_168893571.1 |
| GCF_003290385.1 | NZ_CP030753.1 | 335898 | 337079 | 1181 | - | tuf | WP_168821209.1 |
| GCF_003101015.1 | NZ_CP029206.1 | 331221 | 332402 | 1181 | - | tuf | WP_168919343.1 |
| GCF_003101015.1 | NZ_CP029206.1 | 350662 | 351843 | 1181 | - | tuf | WP_168895208.1 |
| GCF_000017245.1 | NC_009655.1 | 328801 | 329982 | 1181 | - | tuf | WP_168894798.1 |
| GCF_000017245.1 | NC_009655.1 | 330754 | 331935 | 1181 | - | tuf | WP_168868104.1 |
| GCF_000739435.1 | NZ_CP009159.1 | 338664 | 339845 | 1181 | - | tuf | WP_168920548.1 |
| GCF_000739435.1 | NZ_CP009159.1 | 339488 | 340669 | 1181 | - | tuf | WP_168895680.1 |
| GCF_009910365.1 | NZ_CP036423.1 | 343916 | 345097 | 1181 | - | tuf | WP_208754488.1 |
| GCF_009910365.1 | NZ_CP036423.1 | 364482 | 365663 | 1181 | - | tuf | WP_168867667.1 |
| GCF_016026615.1 | NZ_CP065745.1 | 365522 | 366703 | 1181 | - | tuf | WP_169785614.1 |
| GCF_016026615.1 | NZ_CP065745.1 | 366879 | 368060 | 1181 | - | tuf | WP_168892729.1 |
| GCF_028751155.1 | NZ_CP064746.1 | 354030 | 355211 | 1181 | - | tuf | WP_168819697.1 |
| GCF_028751155.1 | NZ_CP064746.1 | 368483 | 369664 | 1181 | - | tuf | WP_168867220.1 |
| GCF_900476005.1 | NZ_LS483441.1 | 2868726 | 2869916 | 1190 | - | tuf | WP_201091844.1 |
| GCF_900476005.1 | NZ_LS483441.1 | 3024412 | 3025602 | 1190 | + | tuf | WP_127164546.1 |
| GCF_002285935.1 | NZ_CP023141.1 | 345362 | 346543 | 1181 | + | tuf | WP_004160596.1 |
| GCF_002285935.1 | NZ_CP023141.1 | 3555401 | 3556582 | 1181 | - | tuf | WP_004154983.1 |
| GCF_900637545.1 | NZ_LR134376.1 | 293345 | 294526 | 1181 | + | tuf | WP_013200290.1 |

| | | | | | | | |
|---|---|---|---|---|---|---|---|
| GCF_900637545.1 | NZ_LR134376.1 | 4592615 | 4593796 | 1181 | - | tuf | WP_013204139.1 |
| GCF_020097295.1 | NZ_CP083625.1 | 386128 | 387309 | 1181 | + | tuf | WP_118665375.1 |
| GCF_020097295.1 | NZ_CP083625.1 | 4364314 | 4365495 | 1181 | - | tuf | WP_222681782.1 |
| GCF_000014805.1 | NC_008570.1 | 432110 | 433291 | 1181 | + | tuf | WP_306209981.1 |
| GCF_000014805.1 | NC_008570.1 | 4220613 | 4221794 | 1181 | - | tuf | WP_306208812.1 |
| GCF_016127195.1 | NZ_CP066092.1 | 310887 | 312068 | 1181 | + | tuf | WP_015899096.1 |
| GCF_016127195.1 | NZ_CP066092.1 | 3712006 | 3713187 | 1181 | - | tuf | WP_012666622.1 |
| GCF_020423125.1 | NZ_CP075564.1 | 467873 | 469054 | 1181 | + | tuf | WP_118665375.1 |
| GCF_020423125.1 | NZ_CP075564.1 | 4730714 | 4731895 | 1181 | - | tuf | WP_118664988.1 |
| GCF_037818705.1 | NZ_CP149129.1 | 408719 | 409900 | 1181 | + | tuf | WP_156286830.1 |
| GCF_037818705.1 | NZ_CP149129.1 | 4374328 | 4375509 | 1181 | - | tuf | WP_154754597.1 |
| GCF_020149575.1 | NZ_CP079742.1 | 173301 | 174482 | 1181 | + | tuf | WP_012439918.1 |
| GCF_020149575.1 | NZ_CP079742.1 | 3533409 | 3534590 | 1181 | - | tuf | WP_012442846.1 |
| GCF_028355655.1 | NZ_CP110645.1 | 279764 | 280945 | 1181 | + | tuf | WP_016191423.1 |
| GCF_028355655.1 | NZ_CP110645.1 | 564404 | 565585 | 1181 | + | tuf | WP_020322785.1 |
| GCF_014892695.1 | NZ_CP040449.1 | 2474717 | 2475898 | 1181 | - | tuf | WP_000031783.1 |
| GCF_014892695.1 | NZ_CP040449.1 | 3252184 | 3253365 | 1181 | + | tuf | WP_025237921.1 |
| GCF_008693705.1 | NZ_CP044060.1 | 4190182 | 4191363 | 1181 | - | tufB | NP_312217.1 |
| GCF_008693705.1 | NZ_CP044060.1 | 4985088 | 4986269 | 1181 | + | tufB | NP_312930.1 |
| GCF_003967035.1 | NZ_AP018560.1 | 3470148 | 3471329 | 1181 | - | tufA | NP_417798.1 |
| GCF_003967035.1 | NZ_AP018560.1 | 4175944 | 4177125 | 1181 | + | tufB | NP_418407.1 |
| GCF_002157225.2 | NZ_CP020038.1 | 3340787 | 3341968 | 1181 | - | tuf | WP_000031784.1 |
| GCF_002157225.2 | NZ_CP020038.1 | 4104685 | 4105866 | 1181 | + | tuf | WP_000031783.1 |
| GCF_002157225.2 | NZ_CP020038.1 | 217210 | 218391 | 1181 | + | tuf | WP_000031784.1 |
| GCF_019670125.1 | NZ_AP023033.1 | 4066857 | 4068038 | 1181 | - | tuf | WP_000031784.1 |
| GCF_019670125.1 | NZ_AP023033.1 | 149293 | 150480 | 1187 | + | tuf | WP_410472501.1 |
| GCF_019670105.1 | NZ_AP023032.1 | 637973 | 639160 | 1187 | - | tuf | WP_410472809.1 |
| GCF_019670105.1 | NZ_AP023032.1 | 273822 | 275003 | 1181 | + | tuf | WP_013343764.1 |
| GCF_019670105.1 | NZ_AP023032.1 | 290492 | 291673 | 1181 | + | tuf | WP_013343776.1 |
| GCF_023518055.1 | NZ_CP085096.1 | 3728266 | 3729447 | 1181 | + | tuf | WP_168662391.1 |
| GCF_023518055.1 | NZ_CP085096.1 | 3745012 | 3746193 | 1181 | + | tuf | WP_168662391.1 |

| | | | | | | | |
|---|---|---|---|---|---|---|---|
| GCF_900636915.1 | NZ_LR134327.1 | 711700 | 712893 | 1193 | - | tuf | WP_228347456.1 |
| GCF_900636915.1 | NZ_LR134327.1 | 728400 | 729593 | 1193 | - | tuf | WP_228347456.1 |
| GCF_900476035.1 | NZ_LS483443.1 | 512744 | 513925 | 1181 | + | tuf | WP_199610662.1 |
| GCF_900476035.1 | NZ_LS483443.1 | 3473065 | 3474246 | 1181 | - | tuf | WP_199609801.1 |
| GCF_014489475.1 | NZ_CP060820.1 | 386320 | 387501 | 1181 | + | tuf | WP_112869428.1 |
| GCF_014489475.1 | NZ_CP060820.1 | 653757 | 654938 | 1181 | + | tuf | WP_071663629.1 |
| GCF_003015185.1 | NZ_CP027860.1 | 230734 | 231915 | 1181 | + | tuf | WP_088771753.1 |
| GCF_003015185.1 | NZ_CP027860.1 | 1702340 | 1703521 | 1181 | - | tuf | WP_014548920.1 |
| GCF_000009365.1 | NC_008260.1 | 2055119 | 2056300 | 1181 | + | tuf | WP_013922406.1 |
| GCF_000009365.1 | NC_008260.1 | 1600934 | 1602115 | 1181 | - | tuf | WP_071629749.1 |
| GCF_024347215.1 | NZ_AP025282.1 | 1690991 | 1692172 | 1181 | - | tuf | WP_014714676.1 |
| GCF_024347215.1 | NZ_AP025282.1 | 44095 | 45276 | 1181 | + | tuf | WP_064460677.1 |
| GCF_024347195.1 | NZ_AP025281.1 | 1641109 | 1642290 | 1181 | - | tuf | WP_004287053.1 |
| GCF_024347195.1 | NZ_AP025281.1 | 1691432 | 1692613 | 1181 | - | tuf | WP_013922406.1 |
| GCF_000011805.1 | NC_006840.2 | 639297 | 640478 | 1181 | + | tuf | WP_013922406.1 |
| GCF_000011805.1 | NC_006840.2 | 1119390 | 1120571 | 1181 | + | tuf | WP_014548920.1 |
| GCF_000196495.1 | NC_011312.1 | 1952242 | 1953423 | 1181 | - | tuf | WP_072713270.1 |
| GCF_000196495.1 | NC_011312.1 | 709387 | 710574 | 1187 | + | tuf | WP_014402118.1 |
| GCF_000014785.1 | NC_008340.1 | 726286 | 727473 | 1187 | + | tuf | WP_014402130.1 |
| GCF_000014785.1 | NC_008340.1 | 844109 | 845296 | 1187 | + | tuf | WP_231324151.1 |
| GCF_026016465.1 | NZ_CP100390.1 | 861548 | 862735 | 1187 | + | tuf | WP_231324151.1 |
| GCF_023650515.1 | NZ_CP096016.1 | 702854 | 704041 | 1187 | + | tuf | WP_231324151.1 |
| GCF_026427595.1 | NZ_CP101527.1 | 720317 | 721504 | 1187 | + | tuf | WP_231324151.1 |
| GCF_000300005.1 | NC_018691.1 | 778497 | 779678 | 1181 | + | tuf | WP_123957622.1 |
| GCF_000300005.1 | NC_018691.1 | 781096 | 782277 | 1181 | - | tuf | WP_123957622.1 |
| GCF_047495515.1 | NZ_CP179934.1 | 2265231 | 2266412 | 1181 | + | tuf | WP_039105709.1 |
| GCF_047495515.1 | NZ_CP179934.1 | 2273486 | 2274667 | 1181 | - | tuf | WP_039105709.1 |
| GCF_032810065.1 | NZ_CP136538.1 | 335374 | 336555 | 1181 | + | tuf | WP_301497104.1 |
| GCF_032810065.1 | NZ_CP136538.1 | 352065 | 353246 | 1181 | + | tuf | WP_301497127.1 |
| GCF_018409545.1 | NZ_AP024563.1 | 280749 | 281930 | 1181 | + | tuf | WP_115717125.1 |
| GCF_018409545.1 | NZ_AP024563.1 | 297449 | 298630 | 1181 | + | tuf | WP_115717125.1 |

| | | | | | | | |
|---|---|---|---|---|---|---|---|
| GCF_000025485.1 | NC_013851.1 | 2254127 | 2255308 | 1181 | - | tuf | WP_302501071.1 |
| GCF_000025485.1 | NC_013851.1 | 2737212 | 2738393 | 1181 | + | tuf | WP_302501071.1 |
| GCF_012222825.1 | NZ_CP038017.1 | 1349933 | 1351129 | 1196 | + | tuf | WP_090213021.1 |
| GCF_000815225.1 | NZ_CP010427.1 | 2235988 | 2237169 | 1181 | - | tuf | WP_095846273.1 |
| GCF_012222965.1 | NZ_CP038241.1 | 2896139 | 2897320 | 1181 | + | tuf | WP_095846273.1 |
| GCF_002849875.1 | NZ_CP018321.1 | 409739 | 410920 | 1181 | + | tuf | WP_025314614.1 |
| GCF_002849875.1 | NZ_CP018321.1 | 2896964 | 2898145 | 1181 | + | tuf | WP_025314614.1 |
| GCF_000213655.1 | NC_015554.1 | 387725 | 388906 | 1181 | - | tuf | WP_034900851.1 |
| GCF_000213655.1 | NC_015554.1 | 621449 | 622630 | 1181 | + | tuf | WP_034900851.1 |
| GCF_002499975.2 | NZ_CP052766.1 | 651479 | 652702 | 1223 | + | tuf | WP_382413938.1 |
| GCF_002499975.2 | NZ_CP052766.1 | 668220 | 669443 | 1223 | + | tuf | WP_382413938.1 |
| GCF_001562195.1 | NZ_CP014322.1 | 250281 | 251465 | 1184 | - | tuf | WP_015939681.1 |
| GCF_024347255.1 | NZ_AP025284.1 | 554895 | 556079 | 1184 | - | tuf | WP_015939681.1 |
| GCF_024347255.1 | NZ_AP025284.1 | 1908439 | 1909626 | 1187 | + | tuf | WP_088917154.1 |
| GCF_024347275.1 | NZ_AP025761.1 | 1968143 | 1969330 | 1187 | + | tuf | WP_088917154.1 |
| GCF_024347275.1 | NZ_AP025761.1 | 978041 | 979222 | 1181 | - | tuf | WP_005501288.1 |
| GCF_902459475.1 | NZ_LR699114.1 | 1444062 | 1445243 | 1181 | + | tuf | WP_005501217.1 |
| GCF_902459475.1 | NZ_LR699114.1 | 251961 | 253142 | 1181 | + | tuf | WP_251877228.1 |
| GCF_902459485.1 | NZ_LR699119.1 | 3578455 | 3579636 | 1181 | - | tuf | WP_251876817.1 |
| GCF_902459485.1 | NZ_LR699119.1 | 2320860 | 2322047 | 1187 | - | tuf | WP_322521218.1 |
| GCF_001597285.1 | NZ_CP014784.1 | 2337415 | 2338602 | 1187 | - | tuf | WP_322521227.1 |
| GCF_001597285.1 | NZ_CP014784.1 | 426356 | 427537 | 1181 | - | tuf | WP_005654658.1 |
| GCF_013200955.2 | NZ_CP053697.2 | 518943 | 520124 | 1181 | + | tuf | WP_005654658.1 |
| GCF_013200955.2 | NZ_CP053697.2 | 398177 | 399358 | 1181 | - | tuf | WP_005646706.1 |
| GCF_003351745.1 | NZ_CP031222.1 | 583357 | 584538 | 1181 | + | tuf | WP_005646706.1 |
| GCF_003351745.1 | NZ_CP031222.1 | 634021 | 635202 | 1181 | - | tuf | WP_005667564.1 |
| GCF_020268605.1 | NZ_CP084222.1 | 675327 | 676508 | 1181 | + | tuf | WP_005667564.1 |
| GCF_020268605.1 | NZ_CP084222.1 | 1430941 | 1432122 | 1181 | - | tuf | WP_078208097.1 |
| GCF_900343015.1 | NZ_LR025108.1 | 1740012 | 1741193 | 1181 | + | tuf | WP_078177606.1 |
| GCF_013460135.1 | NZ_CP038155.1 | 1807790 | 1808971 | 1181 | - | tuf | WP_005635637.1 |
| GCF_013460135.1 | NZ_CP038155.1 | 2015860 | 2017041 | 1181 | + | tuf | WP_005635637.1 |

| | | | | | | | |
|---|---|---|---|---|---|---|---|
| GCF_964020125.1 | NZ_OZ026540.1 | 325419 | 326600 | 1181 | - | tuf | WP_005635637.1 |
| GCF_964020125.1 | NZ_OZ026540.1 | 612436 | 613617 | 1181 | + | tuf | WP_005635637.1 |
| GCF_964277425.1 | NZ_OZ195522.1 | 280836 | 282017 | 1181 | + | tuf | WP_038501901.1 |
| GCF_004768525.1 | NZ_CP038613.1 | 4261967 | 4263148 | 1181 | - | tuf | WP_038502512.1 |
| GCF_004768525.1 | NZ_CP038613.1 | 273790 | 274971 | 1181 | + | tuf | WP_008815921.1 |
| GCF_964026495.1 | NZ_OZ032148.1 | 4329564 | 4330745 | 1181 | - | tuf | WP_008815465.1 |
| GCF_016027855.1 | NZ_CP065700.1 | 6345649 | 6346839 | 1190 | - | tuf | WP_011399922.1 |
| GCF_016027855.1 | NZ_CP065700.1 | 6362041 | 6363231 | 1190 | - | tuf | WP_011399922.1 |
| GCF_024224615.1 | NZ_CP100494.1 | 703103 | 704323 | 1220 | + | tuf | WP_152660830.1 |
| GCF_024224615.1 | NZ_CP100494.1 | 726281 | 727501 | 1220 | + | tuf | WP_152660830.1 |
| GCF_011765605.1 | NZ_CP050316.1 | 889188 | 890408 | 1220 | + | tuf | WP_390517762.1 |
| GCF_011765605.1 | NZ_CP050316.1 | 909344 | 910564 | 1220 | + | tuf | WP_390517762.1 |
| GCF_900635775.1 | NZ_LR134167.1 | 1961483 | 1962673 | 1190 | + | tuf | WP_071943789.1 |
| GCF_900635775.1 | NZ_LR134167.1 | 167241 | 168431 | 1190 | + | tuf | WP_280105344.1 |
| GCF_043587585.1 | NZ_CP171449.1 | 3583931 | 3585121 | 1190 | - | tuf | WP_169959409.1 |
| GCF_043587585.1 | NZ_CP171449.1 | 3644495 | 3645685 | 1190 | - | tuf | WP_097790623.1 |
| GCF_002220155.1 | NZ_CP011835.1 | 443752 | 444942 | 1190 | + | tuf | WP_149283006.1 |
| GCF_002220155.1 | NZ_CP011835.1 | 126438 | 127628 | 1190 | + | tuf | WP_066443631.1 |
| GCF_009363155.1 | NZ_CP045302.1 | 282912 | 284102 | 1190 | + | tuf | WP_013330991.1 |
| GCF_009363155.1 | NZ_CP045302.1 | 3279637 | 3280827 | 1190 | - | tuf | WP_035596342.1 |
| GCF_000021045.1 | NC_012560.1 | 366917 | 368107 | 1190 | + | tuf | WP_021819601.1 |
| GCF_000021045.1 | NC_012560.1 | 203687 | 204877 | 1190 | + | tuf | WP_172419592.1 |
| GCF_011455875.1 | NZ_CP015031.1 | 3139838 | 3141028 | 1190 | - | tuf | WP_346796667.1 |
| GCF_011455875.1 | NZ_CP015031.1 | 479934 | 481124 | 1190 | + | tuf | WP_311884220.1 |
| GCF_003711265.1 | NZ_CP024634.1 | 199789 | 200979 | 1190 | + | tuf | WP_009723905.1 |
| GCF_003711265.1 | NZ_CP024634.1 | 1431592 | 1432782 | 1190 | - | tuf | WP_009723905.1 |
| GCF_001305575.3 | NZ_CP012373.2 | 4014798 | 4015988 | 1190 | - | tuf | WP_209474204.1 |
| GCF_001305575.3 | NZ_CP012373.2 | 1476065 | 1477255 | 1190 | + | tuf | WP_228247484.1 |
| GCF_012295615.1 | NZ_CP051183.1 | 1492540 | 1493730 | 1190 | + | tuf | WP_228247484.1 |
| GCF_012295615.1 | NZ_CP051183.1 | 3800282 | 3801478 | 1196 | + | tuf | WP_090275798.1 |
| GCF_000521725.1 | NZ_CP006954.1 | 3816761 | 3817957 | 1196 | + | tuf | WP_090275798.1 |

| | | | | | | | |
|---|---|---|---|---|---|---|---|
| GCF_000521725.1 | NZ_CP006954.1 | 1542357 | 1543547 | 1190 | - | tuf | WP_238873023.1 |
| GCF_017868775.1 | NZ_CP053381.1 | 1558827 | 1560017 | 1190 | - | tuf | WP_238873023.1 |
| GCF_013377195.1 | NZ_CP016605.1 | 3353852 | 3355042 | 1190 | - | tuf | WP_093395601.1 |
| GCF_013377195.1 | NZ_CP016605.1 | 3370311 | 3371501 | 1190 | - | tuf | WP_093395601.1 |
| GCF_000973545.1 | NZ_CP010049.1 | 915761 | 916951 | 1190 | + | tuf | WP_321350145.1 |
| GCF_023585785.1 | NZ_CP097765.1 | 932229 | 933419 | 1190 | + | tuf | WP_321350145.1 |
| GCF_009827135.1 | NZ_CP046534.1 | 738343 | 739533 | 1190 | - | tuf | WP_080048723.1 |
| GCF_023585765.1 | NZ_CP097760.1 | 754829 | 756019 | 1190 | - | tuf | WP_080048723.1 |
| GCF_023585865.1 | NZ_CP097752.1 | 3024716 | 3025906 | 1190 | + | tuf | WP_092287457.1 |
| GCF_014857065.1 | NZ_CP046533.1 | 3041171 | 3042361 | 1190 | + | tuf | WP_092287457.1 |
| GCF_000973505.1 | NZ_CP010048.1 | 2994968 | 2996158 | 1190 | - | tuf | WP_092387918.1 |
| GCF_002291445.1 | NZ_CP014137.1 | 3011413 | 3012603 | 1190 | - | tuf | WP_092387918.1 |
| GCF_002291445.1 | NZ_CP014137.1 | 2820278 | 2821468 | 1190 | - | tuf | WP_093395601.1 |
| GCF_017569925.1 | NZ_CP050854.1 | 2836729 | 2837919 | 1190 | - | tuf | WP_093395623.1 |
| GCF_017569925.1 | NZ_CP050854.1 | 2434411 | 2435598 | 1187 | - | tuf | WP_096410219.1 |
| GCF_005484965.1 | NZ_CP034036.1 | 2451255 | 2452442 | 1187 | - | tuf | WP_096410231.1 |
| GCF_005484965.1 | NZ_CP034036.1 | 927839 | 929026 | 1187 | - | tuf | WP_011813659.1 |
| GCF_032395385.1 | NZ_CP135917.1 | 944629 | 945816 | 1187 | - | tuf | WP_011813671.1 |
| GCF_032395385.1 | NZ_CP135917.1 | 2371250 | 2372440 | 1190 | + | tuf | WP_066101216.1 |
| GCF_000226565.1 | NC_016041.1 | 2387763 | 2388953 | 1190 | + | tuf | WP_066101216.1 |
| GCF_000226565.1 | NC_016041.1 | 319633 | 320820 | 1187 | + | tuf | WP_012823208.1 |
| GCF_000007365.1 | NC_004061.1 | 336147 | 337334 | 1187 | + | tuf | WP_012823220.1 |
| GCF_022637515.1 | NZ_CP093332.1 | 55441 | 56622 | 1181 | - | tuf | WP_011608348.1 |
| GCF_022637515.1 | NZ_CP093332.1 | 2057102 | 2058283 | 1181 | - | tuf | WP_011608348.1 |
| GCF_031432975.1 | NZ_CP133838.1 | 598194 | 599384 | 1190 | + | tuf | WP_136546657.1 |
| GCF_031432975.1 | NZ_CP133838.1 | 614740 | 615930 | 1190 | + | tuf | WP_136546657.1 |
| GCF_011290485.1 | NZ_CP048029.1 | 329135 | 330322 | 1187 | + | tuf | WP_135795007.1 |
| GCF_011290485.1 | NZ_CP048029.1 | 345459 | 346646 | 1187 | + | tuf | WP_135795007.1 |
| GCF_003956045.1 | NZ_CP026513.1 | 361012 | 362193 | 1181 | + | tuf | WP_029910557.1 |
| GCF_020541245.1 | NZ_CP045876.1 | 377291 | 378472 | 1181 | + | tuf | WP_029910557.1 |
| GCF_001534665.1 | NZ_CP013920.1 | 427539 | 428726 | 1187 | + | tuf | WP_029939562.1 |

| | | | | | | | |
|---|---|---|---|---|---|---|---|
| GCF_045346035.1 | NZ_CP158507.1 | 443843 | 445030 | 1187 | + | tuf | WP_029939562.1 |
| GCF_045346035.1 | NZ_CP158507.1 | 303532 | 304713 | 1181 | + | tuf | WP_199495133.1 |
| GCF_902860225.1 | NZ_LR794158.1 | 320284 | 321465 | 1181 | + | tuf | WP_199495133.1 |
| GCF_023585725.1 | NZ_CP097750.1 | 348574 | 349755 | 1181 | + | tuf | WP_011233600.1 |
| GCF_000331065.1 | NC_020075.1 | 365324 | 366505 | 1181 | + | tuf | WP_011233600.1 |
| GCF_000011745.1 | NC_007292.1 | 246128 | 247315 | 1187 | + | tuf | WP_109188571.1 |
| GCF_000185985.2 | NC_014909.2 | 1979480 | 1980667 | 1187 | - | tuf | WP_109188571.1 |
| GCF_000287275.1 | NC_018416.1 | 1971435 | 1972622 | 1187 | - | tuf | WP_242147423.1 |
| GCF_024172705.1 | NZ_CP092900.1 | 2991974 | 2993161 | 1187 | + | tuf | WP_242147423.1 |
| GCF_024172705.1 | NZ_CP092900.1 | 2435643 | 2436830 | 1187 | + | tuf | WP_068805581.1 |
| GCF_001077715.1 | NZ_CP011126.1 | 2452080 | 2453267 | 1187 | + | tuf | WP_068805615.1 |
| GCF_001077715.1 | NZ_CP011126.1 | 597073 | 598263 | 1190 | + | tuf | WP_041025923.1 |
| GCF_964275155.1 | NZ_OZ193413.1 | 614021 | 615211 | 1190 | + | tuf | WP_041025923.1 |
| GCF_900039485.1 | NZ_LN999833.1 | 115988 | 117175 | 1187 | + | tuf | WP_201339494.1 |
| GCF_023733635.1 | NZ_CP090569.1 | 1353701 | 1354888 | 1187 | - | tuf | WP_201339494.1 |
| GCF_023733635.1 | NZ_CP090569.1 | 717823 | 719004 | 1181 | - | tuf | WP_138094439.1 |
| GCF_002381345.1 | NZ_CP020660.1 | 1506688 | 1507869 | 1181 | + | tuf | WP_138095288.1 |
| GCF_900698925.1 | NZ_LR217713.1 | 2133515 | 2134702 | 1187 | - | tuf | WP_046561889.1 |
| GCF_000764555.1 | NZ_CP009574.1 | 2150629 | 2151816 | 1187 | - | tuf | WP_046561889.1 |
| GCF_964030845.1 | NZ_OZ034983.1 | 445477 | 446664 | 1187 | + | tuf | WP_012800358.1 |
| GCF_900048035.1 | NZ_LN999832.1 | 462274 | 463461 | 1187 | + | tuf | WP_012800358.1 |
| GCF_900048035.1 | NZ_LN999832.1 | 2132815 | 2134002 | 1187 | - | tuf | WP_068993330.1 |
| GCF_000021705.1 | NC_012751.1 | 2149929 | 2151116 | 1187 | - | tuf | WP_068993330.1 |
| GCF_900044015.1 | NZ_LN999835.1 | 1478659 | 1479846 | 1187 | - | tuf | WP_101893403.1 |
| GCF_000828515.1 | NZ_AP010872.1 | 1495270 | 1496457 | 1187 | - | tuf | WP_101893403.1 |
| GCF_000828515.1 | NZ_AP010872.1 | 52627 | 53808 | 1181 | + | tuf | WP_015368828.1 |
| GCF_002776555.1 | NZ_CP021497.1 | 4655357 | 4656538 | 1181 | - | tuf | WP_015369418.1 |
| GCF_964023245.1 | NZ_OZ026884.1 | 433722 | 434903 | 1181 | + | tuf | WP_004174069.1 |
| GCF_964023245.1 | NZ_OZ026884.1 | 4977617 | 4978798 | 1181 | - | tuf | WP_004174069.1 |
| GCF_009498235.1 | NZ_CP044205.1 | 417972 | 419153 | 1181 | + | tuf | WP_047663733.1 |
| GCF_009498235.1 | NZ_CP044205.1 | 5027290 | 5028471 | 1181 | - | tuf | WP_047663733.1 |

| | | | | | | | |
|---|---|---|---|---|---|---|---|
| GCF_900048045.1 | NZ_LN999831.1 | 450386 | 451567 | 1181 | + | tuf | WP_004097636.1 |
| GCF_000364725.1 | NC_021057.1 | 5419734 | 5420915 | 1181 | - | tuf | WP_004097636.1 |
| GCF_902810445.1 | NZ_LR778175.1 | 539233 | 540414 | 1181 | + | tuf | WP_004097636.1 |
| GCF_902810445.1 | NZ_LR778175.1 | 5837231 | 5838412 | 1181 | - | tuf | WP_130624411.1 |
| GCF_002356115.1 | NZ_AP014836.1 | 468949 | 470130 | 1181 | + | tuf | WP_004097636.1 |
| GCF_002356115.1 | NZ_AP014836.1 | 5623687 | 5624868 | 1181 | - | tuf | WP_004097636.1 |
| GCF_000754265.1 | NZ_CP008985.1 | 39036 | 40217 | 1181 | - | tuf | WP_004106368.1 |
| GCF_000971765.1 | NZ_CP010907.1 | 771437 | 772618 | 1181 | + | tuf | WP_004115955.1 |
| GCF_007833795.1 | NZ_CP032702.1 | 3437650 | 3438831 | 1181 | - | tuf | WP_004097636.1 |
| GCF_007833795.1 | NZ_CP032702.1 | 4223393 | 4224574 | 1181 | + | tuf | WP_004097636.1 |
| GCF_026230195.1 | NZ_CP036501.1 | 2201045 | 2202226 | 1181 | - | tuf | WP_004174069.1 |
| GCF_000292685.1 | NC_018507.1 | 2901229 | 2902410 | 1181 | + | tuf | WP_004174069.1 |
| GCF_020541265.1 | NZ_CP045925.1 | 437685 | 438866 | 1181 | + | tuf | WP_004174069.1 |
| GCF_904848675.1 | NZ_LR890047.1 | 5221994 | 5223175 | 1181 | - | tuf | WP_004174069.1 |
| GCF_964026685.1 | NZ_OZ034688.1 | 428574 | 429755 | 1181 | + | tuf | WP_012540432.1 |
| GCF_004124255.1 | NZ_CP026512.1 | 5263211 | 5264392 | 1181 | - | tuf | WP_012540432.1 |
| GCF_001281385.1 | NZ_CP006911.1 | 433158 | 434339 | 1181 | + | tuf | WP_046491013.1 |
| GCF_001281385.1 | NZ_CP006911.1 | 4780551 | 4781732 | 1181 | - | tuf | WP_046491013.1 |
| GCF_003391335.1 | NZ_CP028374.1 | 454066 | 455247 | 1181 | + | tuf | WP_126440962.1 |
| GCF_964020115.1 | NZ_OZ026542.1 | 4430353 | 4431534 | 1181 | - | tuf | WP_126440962.1 |
| GCF_013343005.1 | NZ_CP054491.1 | 3271327 | 3272508 | 1181 | - | tuf | WP_046491013.1 |
| GCF_013343005.1 | NZ_CP054491.1 | 3995606 | 3996787 | 1181 | + | tuf | WP_115074753.1 |
| GCF_003966755.1 | NZ_AP018005.1 | 2907856 | 2909037 | 1181 | - | tuf | WP_090127228.1 |
| GCF_000093065.1 | NC_014109.1 | 3554072 | 3555253 | 1181 | + | tuf | WP_090127228.1 |
| GCF_002073895.1 | NZ_CP012839.1 | 482534 | 483715 | 1181 | + | tuf | WP_342322705.1 |
| GCF_013342985.1 | NZ_CP054490.1 | 4494282 | 4495463 | 1181 | - | tuf | WP_342322705.1 |
| GCF_013342985.1 | NZ_CP054490.1 | 418072 | 419253 | 1181 | + | tuf | WP_094419090.1 |
| GCF_000015105.1 | NC_008610.1 | 4431633 | 4432814 | 1181 | - | tuf | WP_094419090.1 |
| GCF_000015105.1 | NC_008610.1 | 446426 | 447607 | 1181 | + | tuf | WP_064563291.1 |
| GCF_019923565.1 | NZ_CP074435.1 | 5148525 | 5149706 | 1181 | - | tuf | WP_064563291.1 |
| GCF_000517405.1 | NZ_CP006568.1 | 494785 | 495966 | 1181 | + | tuf | WP_061492922.1 |

| | | | | | | |
|---|---|---|---|---|---|---|
| GCF_000517405.1 | NZ_CP006568.1 | 4855748 | 4856929 | 1181 - | tuf | WP_263098500.1 |
| GCF_020541225.1 | NZ_CP045889.1 | 228223 | 229404 | 1181 + | tuf | WP_017460031.1 |
| GCF_000828815.1 | NZ_AP014521.1 | 4449670 | 4450851 | 1181 - | tuf | WP_015742710.1 |
| GCF_000828815.1 | NZ_AP014521.1 | 438328 | 439509 | 1181 + | tuf | WP_017460031.1 |
| GCF_041222825.1 | NZ_CP139482.1 | 5384022 | 5385203 | 1181 - | tuf | WP_017460031.1 |
| GCF_041222825.1 | NZ_CP139482.1 | 421679 | 422860 | 1181 + | tuf | WP_015742710.1 |
| GCF_002813775.1 | NZ_CP020370.1 | 4638586 | 4639767 | 1181 - | tuf | WP_017460031.1 |
| GCF_002813775.1 | NZ_CP020370.1 | 766779 | 767969 | 1190 - | tuf | WP_086620795.1 |
| GCF_964212635.1 | NZ_OZ156463.1 | 2531531 | 2532721 | 1190 + | tuf | WP_086620795.1 |
| GCF_964212635.1 | NZ_OZ156463.1 | 3033236 | 3034426 | 1190 - | tuf | WP_070977996.1 |
| GCF_001293165.1 | NZ_CP010552.1 | 442200 | 443381 | 1181 + | tuf | WP_059307073.1 |
| GCF_001293165.1 | NZ_CP010552.1 | 4589590 | 4590771 | 1181 - | tuf | WP_059307073.1 |
| GCF_017901155.1 | NZ_CP072800.1 | 192021 | 193202 | 1181 + | tuf | WP_040078350.1 |
| GCF_017901155.1 | NZ_CP072800.1 | 3909006 | 3910187 | 1181 - | tuf | WP_040078350.1 |
| GCF_000010405.1 | NC_009465.1 | 3960028 | 3961215 | 1187 - | tuf | WP_019233360.1 |
| GCF_000010405.1 | NC_009465.1 | 3976350 | 3977537 | 1187 - | tuf | WP_019233360.1 |
| GCF_001242845.1 | NZ_LN774881.1 | 357669 | 358856 | 1187 + | tuf | WP_173235947.1 |
| GCF_964019775.1 | NZ_OZ026471.1 | 373941 | 375128 | 1187 + | tuf | WP_173235947.1 |
| GCF_900637305.1 | NZ_LR134365.1 | 2761300 | 2762487 | 1187 - | tuf | WP_275088455.1 |
| GCF_900637305.1 | NZ_LR134365.1 | 2777531 | 2778718 | 1187 - | tuf | WP_275088455.1 |
| GCF_026725475.1 | NZ_CP109965.1 | 2970909 | 2972096 | 1187 - | tuf | WP_094091987.1 |
| GCF_026725475.1 | NZ_CP109965.1 | 2987151 | 2988338 | 1187 - | tuf | WP_094091987.1 |
| GCF_007289895.1 | NZ_CP041660.1 | 3088653 | 3089840 | 1187 - | tuf | WP_045096555.1 |
| GCF_007289895.1 | NZ_CP041660.1 | 3104963 | 3106150 | 1187 - | tuf | WP_045096555.1 |
| GCF_044095835.1 | NZ_CP172054.1 | 1691525 | 1692712 | 1187 + | tuf | WP_115152860.1 |
| GCF_044095835.1 | NZ_CP172054.1 | 1707893 | 1709080 | 1187 + | tuf | WP_028386052.1 |
| GCF_900635955.1 | NZ_LR134201.1 | 319009 | 320196 | 1187 + | tuf | WP_045104964.1 |
| GCF_900635955.1 | NZ_LR134201.1 | 335248 | 336435 | 1187 + | tuf | WP_045104964.1 |
| GCF_002393445.1 | NZ_CP023525.1 | 242121 | 243308 | 1187 - | tuf | WP_058502161.1 |
| GCF_002393445.1 | NZ_CP023525.1 | 258324 | 259511 | 1187 - | tuf | WP_058502161.1 |
| GCF_914590485.1 | NZ_OU594967.1 | 270201 | 271388 | 1187 + | tuf | WP_064108440.1 |

| | | | | | | | |
|---|---|---|---|---|---|---|---|
| GCF_914590485.1 | NZ_OU594967.1 | 286431 | 287618 | 1187 | + | tuf | WP_064108440.1 |
| GCF_000019225.1 | NC_010995.1 | 237388 | 238575 | 1187 | + | tuf | WP_028373276.1 |
| GCF_000019225.1 | NC_010995.1 | 253631 | 254818 | 1187 | + | tuf | WP_028373276.1 |
| GCF_000520015.2 | NZ_CP007044.2 | 545756 | 546943 | 1187 | + | tuf | WP_004450463.1 |
| GCF_000520015.2 | NZ_CP007044.2 | 562041 | 563228 | 1187 | + | tuf | WP_004450463.1 |
| GCF_034479555.1 | NZ_CP140151.1 | 325835 | 327022 | 1187 | + | tuf | WP_252580409.1 |
| GCF_000055785.1 | NC_007963.1 | 342155 | 343342 | 1187 | + | tuf | WP_252580409.1 |
| GCF_020099335.1 | NZ_CP083651.1 | 427859 | 429046 | 1187 | + | tuf | WP_045098297.1 |
| GCF_020099335.1 | NZ_CP083651.1 | 444138 | 445325 | 1187 | + | tuf | WP_045098297.1 |
| GCF_004353845.1 | NZ_CP037864.1 | 365434 | 366621 | 1187 | + | tuf | WP_010946066.1 |
| GCF_004353845.1 | NZ_CP037864.1 | 381716 | 382903 | 1187 | + | tuf | WP_010946066.1 |
| GCF_009648935.1 | NZ_CP045771.1 | 4315848 | 4317035 | 1187 | + | tuf | WP_058390692.1 |
| GCF_009648935.1 | NZ_CP045771.1 | 4332131 | 4333318 | 1187 | + | tuf | WP_058390692.1 |
| GCF_016893685.1 | NZ_CP069763.1 | 407141 | 408328 | 1187 | + | tuf | WP_058483732.1 |
| GCF_016893685.1 | NZ_CP069763.1 | 423464 | 424651 | 1187 | + | tuf | WP_058483732.1 |
| GCF_037482245.1 | NZ_CP148084.1 | 633968 | 635155 | 1187 | + | tuf | WP_058481016.1 |
| GCF_037482245.1 | NZ_CP148084.1 | 650231 | 651418 | 1187 | + | tuf | WP_058481016.1 |
| GCF_019048065.1 | NZ_CP077291.1 | 1353958 | 1355139 | 1181 | - | tuf | WP_096759551.1 |
| GCF_019048065.1 | NZ_CP077291.1 | 2066313 | 2067494 | 1181 | + | tuf | WP_096759551.1 |
| GCF_904859905.1 | NZ_LR890181.1 | 203927 | 205108 | 1181 | + | tuf | WP_100778110.1 |
| GCF_904859905.1 | NZ_LR890181.1 | 4228178 | 4229359 | 1181 | - | tuf | WP_100778647.1 |
| GCF_000018045.1 | NC_009792.1 | 422961 | 424142 | 1181 | + | tuf | WP_036038139.1 |
| GCF_000018045.1 | NC_009792.1 | 3809395 | 3810576 | 1181 | - | tuf | WP_036038195.1 |
| GCF_041320245.1 | NZ_CP101036.1 | 367211 | 368392 | 1181 | + | tuf | WP_111739027.1 |
| GCF_041320245.1 | NZ_CP101036.1 | 3498824 | 3500005 | 1181 | - | tuf | WP_111741586.1 |
| GCF_019047765.1 | NZ_CP077262.1 | 363229 | 364410 | 1181 | + | tuf | WP_108900585.1 |
| GCF_019047765.1 | NZ_CP077262.1 | 3335897 | 3337078 | 1181 | - | tuf | WP_108900585.1 |
| GCF_023374935.1 | NZ_CP092466.1 | 4026303 | 4027484 | 1181 | - | tuf | WP_130593059.1 |
| GCF_023374935.1 | NZ_CP092466.1 | 4150565 | 4151746 | 1181 | - | tuf | WP_130593148.1 |
| GCF_021278985.1 | NZ_CP082833.1 | 2200419 | 2201600 | 1181 | - | tuf | WP_130601696.1 |
| GCF_021278985.1 | NZ_CP082833.1 | 2217898 | 2219079 | 1181 | - | tuf | WP_130601718.1 |

| | | | | | | | |
|---|---|---|---|---|---|---|---|
| GCF_018128425.1 | NZ_CP071070.1 | 1934029 | 1935216 | 1187 | - | tuf | WP_153714367.1 |
| GCF_018128425.1 | NZ_CP071070.1 | 1950503 | 1951690 | 1187 | - | tuf | WP_153714367.1 |
| GCF_004684345.1 | NZ_CP038469.1 | 135058 | 136239 | 1181 | + | tuf | WP_132299727.1 |
| GCF_004684345.1 | NZ_CP038469.1 | 2297498 | 2298679 | 1181 | + | tuf | WP_132299727.1 |
| GCF_008693645.1 | NZ_CP044101.1 | 752883 | 754064 | 1181 | - | tuf | WP_027063854.1 |
| GCF_008693645.1 | NZ_CP044101.1 | 1321787 | 1322968 | 1181 | + | tuf | WP_118894785.1 |
| GCF_032923285.1 | NZ_CP136695.1 | 312316 | 313497 | 1181 | + | tuf | WP_197669175.1 |
| GCF_001720485.1 | NZ_CP017114.1 | 3619869 | 3621050 | 1181 | - | tuf | WP_197669175.1 |
| GCF_002076895.1 | NZ_CP020465.1 | 1255312 | 1256499 | 1187 | + | tuf | WP_247333630.1 |
| GCF_002076895.1 | NZ_CP020465.1 | 1272147 | 1273334 | 1187 | + | tuf | WP_247333702.1 |
| GCF_000012325.1 | NC_003910.7 | 3731948 | 3733135 | 1187 | - | tuf | WP_247369575.1 |
| GCF_000012325.1 | NC_003910.7 | 3748797 | 3749984 | 1187 | - | tuf | WP_247369619.1 |
| GCF_045689245.1 | NZ_CP136865.1 | 1136992 | 1138179 | 1187 | + | tuf | WP_247433799.1 |
| GCF_045689245.1 | NZ_CP136865.1 | 1153845 | 1155032 | 1187 | + | tuf | WP_247433817.1 |
| GCF_000153125.2 | NZ_CM002299.1 | 452079 | 453266 | 1187 | - | tuf | WP_166951192.1 |
| GCF_000153125.2 | NZ_CM002299.1 | 468921 | 470108 | 1187 | - | tuf | WP_166951174.1 |
| GCF_045689235.1 | NZ_CP136864.1 | 872312 | 873499 | 1187 | + | tuf | WP_250339853.1 |
| GCF_045689235.1 | NZ_CP136864.1 | 889177 | 890364 | 1187 | + | tuf | WP_250339853.1 |
| GCF_000007765.2 | NC_002971.4 | 1102088 | 1103275 | 1187 | + | tuf | WP_139980268.1 |
| GCF_000815025.1 | NZ_CP007541.1 | 1118950 | 1120137 | 1187 | + | tuf | WP_139980268.1 |
| GCF_000815025.1 | NZ_CP007541.1 | 2664496 | 2665683 | 1187 | + | tuf | WP_071924949.1 |
| GCF_018107685.1 | NZ_CP064834.1 | 2681283 | 2682470 | 1187 | + | tuf | WP_071924950.1 |
| GCF_018107685.1 | NZ_CP064834.1 | 1340449 | 1341636 | 1187 | + | tuf | WP_096297397.1 |
| GCF_009883795.1 | NZ_CP033868.1 | 1357938 | 1359125 | 1187 | + | tuf | WP_096297397.1 |
| GCF_002850495.1 | NZ_CP021379.1 | 857315 | 858502 | 1187 | - | tuf | WP_232580488.1 |
| GCF_002850495.1 | NZ_CP021379.1 | 874807 | 875994 | 1187 | - | tuf | WP_232134451.1 |
| GCF_001277255.1 | NZ_CP012264.1 | 2307143 | 2308330 | 1187 | + | tuf | WP_144892832.1 |
| GCF_001277255.1 | NZ_CP012264.1 | 2324454 | 2325641 | 1187 | + | tuf | WP_144892832.1 |
| GCF_001277235.1 | NZ_CP012266.1 | 2347226 | 2348413 | 1187 | - | tuf | WP_134673967.1 |
| GCF_001277235.1 | NZ_CP012266.1 | 2364696 | 2365883 | 1187 | - | tuf | WP_134673967.1 |
| GCF_001277215.2 | NZ_CP013940.1 | 4499928 | 4501115 | 1187 | - | tuf | WP_057918900.1 |

| | | | | | | | |
|---|---|---|---|---|---|---|---|
| GCF_001277215.2 | NZ_CP013940.1 | 4517719 | 4518906 | 1187 | - | tuf | WP_057918900.1 |
| GCF_001277195.1 | NZ_CP012268.1 | 1882467 | 1883654 | 1187 | - | tuf | WP_200606508.1 |
| GCF_001277195.1 | NZ_CP012268.1 | 1900062 | 1901249 | 1187 | - | tuf | WP_200606508.1 |
| GCF_003516125.1 | NZ_CP027107.1 | 1046656 | 1047843 | 1187 | + | tuf | WP_115843879.1 |
| GCF_003516125.1 | NZ_CP027107.1 | 1064355 | 1065542 | 1187 | + | tuf | WP_115843879.1 |
| GCF_041222865.1 | NZ_CP165997.1 | 4436131 | 4437318 | 1187 | - | tuf | WP_247024592.1 |
| GCF_041222865.1 | NZ_CP165997.1 | 4454230 | 4455417 | 1187 | - | tuf | WP_046657641.1 |
| GCF_001277175.1 | NZ_CP012257.1 | 1537774 | 1538961 | 1187 | + | tuf | WP_096377136.1 |
| GCF_001277175.1 | NZ_CP012257.1 | 1555917 | 1557104 | 1187 | + | tuf | WP_096377136.1 |
| GCF_900095885.1 | NZ_LT615367.1 | 2814753 | 2815940 | 1187 | + | tuf | WP_064747322.1 |
| GCF_900095885.1 | NZ_LT615367.1 | 2832801 | 2833988 | 1187 | + | tuf | WP_064747322.1 |
| GCF_000023565.1 | NC_012912.1 | 4218113 | 4219300 | 1187 | - | tuf | WP_057943879.1 |
| GCF_000023565.1 | NC_012912.1 | 4236249 | 4237436 | 1187 | - | tuf | WP_057943879.1 |
| GCF_003049785.1 | NZ_CP023467.1 | 505170 | 506357 | 1187 | - | tuf | WP_213435476.1 |
| GCF_003049785.1 | NZ_CP023467.1 | 522131 | 523318 | 1187 | - | tuf | WP_213435476.1 |
| GCF_003403135.1 | NZ_CP031560.1 | 1060066 | 1061253 | 1187 | + | tuf | WP_115843879.1 |
| GCF_003403135.1 | NZ_CP031560.1 | 1077596 | 1078783 | 1187 | + | tuf | WP_115843879.1 |
| GCF_002812485.1 | NZ_CP025003.1 | 705296 | 706483 | 1187 | + | tuf | WP_220380357.1 |
| GCF_002812485.1 | NZ_CP025003.1 | 722506 | 723693 | 1187 | + | tuf | WP_220380357.1 |
| GCF_020406815.2 | NZ_CP162670.1 | 3281641 | 3282828 | 1187 | + | tuf | WP_057948556.1 |
| GCF_020406815.2 | NZ_CP162670.1 | 3299650 | 3300837 | 1187 | + | tuf | WP_057948556.1 |
| GCF_000025065.1 | NC_013592.1 | 3916867 | 3918048 | 1181 | - | tuf | WP_353613075.1 |
| GCF_000025065.1 | NC_013592.1 | 5433168 | 5434349 | 1181 | - | tuf | WP_353613075.1 |
| GCF_007858975.2 | NZ_CP042220.2 | 172897 | 174078 | 1181 | + | tuf | WP_027073920.1 |
| GCF_007858975.2 | NZ_CP042220.2 | 599515 | 600696 | 1181 | + | tuf | WP_027073920.1 |
| GCF_001644705.1 | NZ_CP015137.1 | 139096 | 140277 | 1181 | - | tuf | WP_006249832.1 |
| GCF_001644705.1 | NZ_CP015137.1 | 652118 | 653299 | 1181 | + | tuf | WP_006249832.1 |
| GCF_002887555.1 | NZ_CP025799.1 | 427892 | 429073 | 1181 | + | tuf | WP_006249832.1 |
| GCF_002887555.1 | NZ_CP025799.1 | 2627574 | 2628755 | 1181 | - | tuf | WP_006249832.1 |
| GCF_001632775.1 | NZ_CP015249.1 | 314181 | 315362 | 1181 | - | tuf | WP_176807589.1 |
| GCF_001632775.1 | NZ_CP015249.1 | 768109 | 769290 | 1181 | + | tuf | WP_176807589.1 |

| | | | | | | | |
|---|---|---|---|---|---|---|---|
| GCF_003072645.1 | NZ_CP028897.1 | 1060991 | 1062172 | 1181 | - | tuf | WP_025218017.1 |
| GCF_020342335.1 | NZ_CP073262.1 | 1253075 | 1254256 | 1181 | - | tuf | WP_025218017.1 |
| GCF_020342335.1 | NZ_CP073262.1 | 795406 | 796593 | 1187 | + | tuf | WP_005224655.1 |
| GCF_036320485.1 | NZ_CP132911.1 | 812626 | 813813 | 1187 | + | tuf | WP_005224655.1 |
| GCF_036320485.1 | NZ_CP132911.1 | 477432 | 478652 | 1220 | + | tuf | WP_390617142.1 |
| GCF_016865235.1 | NZ_CP064030.1 | 494637 | 495857 | 1220 | + | tuf | WP_390617142.1 |
| GCF_016865235.1 | NZ_CP064030.1 | 2691155 | 2692342 | 1187 | - | tuf | WP_141624027.1 |
| GCF_014297575.1 | NZ_CP060412.1 | 2708544 | 2709731 | 1187 | - | tuf | WP_141624039.1 |
| GCF_014297575.1 | NZ_CP060412.1 | 2874149 | 2875336 | 1187 | - | tuf | WP_111267080.1 |
| GCF_001641285.1 | NZ_CP014841.1 | 2891393 | 2892580 | 1187 | - | tuf | WP_111267080.1 |
| GCF_001641285.1 | NZ_CP014841.1 | 463090 | 464310 | 1220 | + | tuf | WP_236985640.1 |
| GCF_021495345.1 | NZ_CP099397.1 | 1009992 | 1011212 | 1220 | - | tuf | WP_422380500.1 |
| GCF_021495345.1 | NZ_CP099397.1 | 4141383 | 4142603 | 1220 | - | tuf | WP_422380500.1 |
| GCF_017915135.1 | NZ_CP068551.1 | 1298874 | 1300067 | 1193 | + | tuf | WP_041644923.1 |
| GCF_017915135.1 | NZ_CP068551.1 | 1315454 | 1316647 | 1193 | + | tuf | WP_008174823.1 |
| GCF_000733715.2 | NZ_CP013124.1 | 603674 | 604867 | 1193 | + | tuf | WP_068351550.1 |
| GCF_000733715.2 | NZ_CP013124.1 | 620240 | 621433 | 1193 | + | tuf | WP_068351550.1 |
| GCF_017167985.1 | NZ_CP070505.1 | 641536 | 642729 | 1193 | + | tuf | WP_286221773.1 |
| GCF_017167985.1 | NZ_CP070505.1 | 658112 | 659305 | 1193 | + | tuf | WP_286221773.1 |
| GCF_000264765.2 | NZ_CP006664.1 | 4392233 | 4393426 | 1193 | + | tuf | WP_008174851.1 |
| GCF_000264765.2 | NZ_CP006664.1 | 4408812 | 4410005 | 1193 | + | tuf | WP_008174851.1 |
| GCF_016026395.1 | NZ_CP065626.1 | 786276 | 787469 | 1193 | + | tuf | WP_148861585.1 |
| GCF_016026395.1 | NZ_CP065626.1 | 802774 | 803967 | 1193 | + | tuf | WP_148861585.1 |
| GCF_003074995.2 | NZ_CP084521.1 | 3464699 | 3465892 | 1193 | + | tuf | WP_117619468.1 |
| GCF_003074995.2 | NZ_CP084521.1 | 3481279 | 3482472 | 1193 | + | tuf | WP_117619468.1 |
| GCF_021733145.1 | NZ_CP090968.1 | 600250 | 601443 | 1193 | + | tuf | WP_011784245.1 |
| GCF_021733145.1 | NZ_CP090968.1 | 616824 | 618017 | 1193 | + | tuf | WP_011784245.1 |
| GCF_019933175.1 | NZ_CP082939.1 | 703921 | 705114 | 1193 | + | tuf | WP_014869911.1 |
| GCF_019933175.1 | NZ_CP082939.1 | 720526 | 721719 | 1193 | + | tuf | WP_014869911.1 |
| GCF_001547755.1 | NZ_AP013042.1 | 3794452 | 3795645 | 1193 | - | tuf | WP_085681648.1 |
| GCF_001547755.1 | NZ_AP013042.1 | 3811048 | 3812241 | 1193 | - | tuf | WP_007154018.1 |

| | | | | | | | |
|---|---|---|---|---|---|---|---|
| GCF_004296535.1 | NZ_AP018159.1 | 1374933 | 1376126 | 1193 | + | tuf | WP_206645217.1 |
| GCF_004296515.1 | NZ_AP018160.1 | 1391495 | 1392688 | 1193 | + | tuf | WP_206645217.1 |
| GCF_004296495.1 | NZ_AP018161.1 | 2678958 | 2680151 | 1193 | - | tuf | WP_070970532.1 |
| GCF_004296475.1 | NZ_AP018162.1 | 2695563 | 2696756 | 1193 | - | tuf | WP_070970562.1 |
| GCF_000801295.1 | NZ_AP012978.1 | 761269 | 762462 | 1193 | + | tuf | WP_068351550.1 |
| GCF_000801295.1 | NZ_AP012978.1 | 777835 | 779028 | 1193 | + | tuf | WP_068351550.1 |
| GCF_025562755.1 | NZ_CP103300.1 | 3973653 | 3974837 | 1184 | + | tuf | WP_041343366.1 |
| GCF_001583435.1 | NZ_CP013251.1 | 4883056 | 4884276 | 1220 | - | tuf | WP_067386410.1 |
| GCF_041937165.1 | NZ_CP154353.1 | 4899698 | 4900918 | 1220 | - | tuf | WP_067386410.1 |
| GCF_041937165.1 | NZ_CP154353.1 | 242940 | 244130 | 1190 | + | tuf | WP_206838263.1 |
| GCF_007035805.1 | NZ_AP019630.1 | 259467 | 260657 | 1190 | + | tuf | WP_091826387.1 |
| GCF_007035805.1 | NZ_AP019630.1 | 227607 | 228827 | 1220 | + | tuf | WP_191601890.1 |
| GCF_900324475.1 | NZ_LT992502.1 | 4312891 | 4314111 | 1220 | - | tuf | WP_111608627.1 |
| GCF_900324475.1 | NZ_LT992502.1 | 1779553 | 1780773 | 1220 | - | tuf | WP_114412704.1 |
| GCF_019665745.1 | NZ_CP081105.1 | 249841 | 251061 | 1220 | + | tuf | WP_013659447.1 |
| GCF_019665745.1 | NZ_CP081105.1 | 3348696 | 3349916 | 1220 | - | tuf | WP_072841835.1 |
| GCF_001984825.2 | NZ_CP043318.1 | 3306640 | 3307860 | 1220 | - | tuf | WP_265727223.1 |
| GCF_001984825.2 | NZ_CP043318.1 | 3683857 | 3685077 | 1220 | - | tuf | WP_013797843.1 |
| GCF_039718975.1 | NZ_CP097173.1 | 3848619 | 3849839 | 1220 | - | tuf | WP_176336546.1 |
| GCF_039718975.1 | NZ_CP097173.1 | 1496926 | 1498146 | 1220 | - | tuf | WP_168826626.1 |
| GCF_905331265.2 | NZ_OW968328.1 | 5644 | 6825 | 1181 | - | tuf | WP_261919790.1 |
| GCF_905331265.2 | NZ_OW968328.1 | 245332 | 246513 | 1181 | + | tuf | WP_261919790.1 |
| GCF_008364625.2 | NZ_CP126604.1 | 2368870 | 2370060 | 1190 | + | tuf | WP_215377555.1 |
| GCF_008364625.2 | NZ_CP126604.1 | 2385406 | 2386596 | 1190 | + | tuf | WP_215377555.1 |
| GCF_019048245.1 | NZ_CP077308.1 | 604328 | 605518 | 1190 | + | tuf | WP_096368001.1 |
| GCF_019048245.1 | NZ_CP077308.1 | 620868 | 622058 | 1190 | + | tuf | WP_096368001.1 |
| GCF_003594935.2 | NZ_CP043342.1 | 606816 | 608006 | 1190 | + | tuf | WP_103102850.1 |
| GCF_003594935.2 | NZ_CP043342.1 | 623370 | 624560 | 1190 | + | tuf | WP_103102850.1 |
| GCF_048568405.1 | NZ_CP183772.1 | 693085 | 694275 | 1190 | + | tuf | WP_044410941.1 |
| GCF_048568405.1 | NZ_CP183772.1 | 709649 | 710839 | 1190 | + | tuf | WP_044410941.1 |
| GCF_023023125.1 | NZ_CP083828.1 | 2984561 | 2985748 | 1187 | - | tuf | WP_119630037.1 |

| | | | | | | | |
|---|---|---|---|---|---|---|---|
| GCF_023023125.1 | NZ_CP083828.1 | 3634216 | 3635403 | 1187 | - | tuf | WP_119630481.1 |
| GCF_001750725.1 | NZ_CP017279.1 | 2700752 | 2701939 | 1187 | - | tuf | WP_026609161.1 |
| GCF_001750725.1 | NZ_CP017279.1 | 3469917 | 3471104 | 1187 | + | tuf | WP_026609161.1 |
| GCF_022014715.1 | NZ_CP091779.1 | 1109771 | 1110958 | 1187 | + | tuf | WP_010960359.1 |
| GCF_022014715.1 | NZ_CP091779.1 | 2553521 | 2554708 | 1187 | - | tuf | WP_010960359.1 |
| GCF_009176645.1 | NZ_AP019007.1 | 2745969 | 2747156 | 1187 | - | tuf | WP_169604020.1 |
| GCF_009176645.1 | NZ_AP019007.1 | 3142084 | 3143271 | 1187 | - | tuf | WP_169604020.1 |
| GCF_030406165.1 | NZ_CP129026.1 | 3546568 | 3547755 | 1187 | + | tuf | WP_266021805.1 |
| GCF_030406165.1 | NZ_CP129026.1 | 3836847 | 3838034 | 1187 | + | tuf | WP_266021805.1 |
| GCF_043972695.1 | NZ_CP143745.1 | 375893 | 377080 | 1187 | + | tuf | WP_221048041.1 |
| GCF_043972695.1 | NZ_CP143745.1 | 392377 | 393564 | 1187 | + | tuf | WP_221048041.1 |
| GCF_001729805.1 | NZ_CP017184.1 | 1022643 | 1023830 | 1187 | + | tuf | WP_221065815.1 |
| GCF_001729805.1 | NZ_CP017184.1 | 4122059 | 4123300 | 1241 | - | tuf | WP_343223273.1 |
| GCF_009036245.1 | NZ_CP027986.1 | 92982 | 94172 | 1190 | - | tuf | WP_317705514.1 |
| GCF_009036245.1 | NZ_CP027986.1 | 109404 | 110594 | 1190 | - | tuf | WP_317705514.1 |
| GCF_000224675.1 | NC_015968.1 | 1202452 | 1203642 | 1190 | + | tuf | WP_286290982.1 |
| GCF_000224675.1 | NC_015968.1 | 1218873 | 1220063 | 1190 | + | tuf | WP_286290982.1 |
| GCF_040834095.1 | NZ_CP162152.1 | 266285 | 267472 | 1187 | - | tuf | WP_305906359.1 |
| GCF_040834095.1 | NZ_CP162152.1 | 282579 | 283766 | 1187 | - | tuf | WP_305906359.1 |
| GCF_035835995.1 | NZ_CP142124.1 | 3713325 | 3714512 | 1187 | - | tuf | WP_005373968.1 |
| GCF_035835995.1 | NZ_CP142124.1 | 3729628 | 3730815 | 1187 | - | tuf | WP_005373968.1 |
| GCF_012567955.1 | NZ_CP046186.1 | 3175242 | 3176429 | 1187 | + | tuf | WP_020484790.1 |
| GCF_012568845.1 | NZ_CP046194.1 | 3191443 | 3192630 | 1187 | + | tuf | WP_020484790.1 |
| GCF_012569925.1 | NZ_CP046204.1 | 4058666 | 4059853 | 1187 | - | tuf | WP_054763846.1 |
| GCF_012570365.1 | NZ_CP046208.1 | 4074844 | 4076031 | 1187 | - | tuf | WP_054763846.1 |
| GCF_012569785.1 | NZ_CP046202.1 | 3697908 | 3699095 | 1187 | - | tuf | WP_301671687.1 |
| GCF_012570745.1 | NZ_CP046212.1 | 3714113 | 3715300 | 1187 | - | tuf | WP_301671687.1 |
| GCF_012567545.1 | NZ_CP046182.1 | 2915130 | 2916317 | 1187 | - | tuf | WP_215581294.1 |
| GCF_012567685.1 | NZ_CP046184.1 | 2931329 | 2932516 | 1187 | - | tuf | WP_215581314.1 |
| GCF_012569525.1 | NZ_CP046200.1 | 3900113 | 3901300 | 1187 | - | tuf | WP_255189293.1 |
| GCF_012570145.1 | NZ_CP046206.1 | 3916314 | 3917501 | 1187 | - | tuf | WP_255189293.1 |

| | | | | | | | |
|---|---|---|---|---|---|---|---|
| GCF_012569285.1 | NZ_CP046198.1 | 528361 | 529548 | 1187 | + | tuf | WP_150046429.1 |
| GCF_012570535.1 | NZ_CP046210.1 | 544522 | 545709 | 1187 | + | tuf | WP_150046429.1 |
| GCF_012568645.1 | NZ_CP046192.1 | 2382633 | 2383820 | 1187 | + | tuf | WP_014704932.1 |
| GCF_012569045.1 | NZ_CP046196.1 | 2398963 | 2400150 | 1187 | + | tuf | WP_014704932.1 |
| GCF_012568245.1 | NZ_CP046188.1 | 991868 | 993055 | 1187 | - | tuf | WP_286305549.1 |
| GCF_012571135.1 | NZ_CP046216.1 | 1008250 | 1009437 | 1187 | - | tuf | WP_286305555.1 |
| GCF_012570965.1 | NZ_CP046214.1 | 2917948 | 2919135 | 1187 | - | tuf | WP_014707495.1 |
| GCF_012568445.1 | NZ_CP046190.1 | 2934306 | 2935493 | 1187 | - | tuf | WP_014707495.1 |
| GCF_012571605.1 | NZ_CP046220.1 | 2843290 | 2844477 | 1187 | - | tuf | WP_247363583.1 |
| GCF_012571345.1 | NZ_CP046218.1 | 2859645 | 2860832 | 1187 | - | tuf | WP_247363583.1 |
| GCF_012571795.1 | NZ_CP046222.1 | 3863878 | 3865065 | 1187 | - | tuf | WP_014149775.1 |
| GCF_012563325.1 | NZ_CP046232.1 | 3880126 | 3881313 | 1187 | - | tuf | WP_014149775.1 |
| GCF_012563145.1 | NZ_CP046230.1 | 810768 | 811955 | 1187 | + | tuf | WP_017840760.1 |
| GCF_012562765.1 | NZ_CP046226.1 | 827027 | 828214 | 1187 | + | tuf | WP_017840760.1 |
| GCF_012562965.1 | NZ_CP046228.1 | 1165525 | 1166712 | 1187 | - | tuf | WP_088618399.1 |
| GCF_012562605.1 | NZ_CP046224.1 | 1181895 | 1183082 | 1187 | - | tuf | WP_088618399.1 |
| GCF_016653615.1 | NZ_CP067393.1 | 3548106 | 3549296 | 1190 | - | tuf | WP_077406707.1 |
| GCF_003991975.1 | NZ_CP029822.1 | 3565302 | 3566492 | 1190 | - | tuf | WP_077406707.1 |
| GCF_043228865.1 | NZ_CP171267.1 | 1553501 | 1554691 | 1190 | + | tuf | WP_069946910.1 |
| GCF_043228865.1 | NZ_CP171267.1 | 1570628 | 1571818 | 1190 | + | tuf | WP_069946920.1 |
| GCF_000196615.1 | NC_014306.1 | 1916408 | 1917598 | 1190 | - | tuf | WP_280322258.1 |
| GCF_000196615.1 | NC_014306.1 | 1933607 | 1934797 | 1190 | - | tuf | WP_280322258.1 |
| GCF_019844095.1 | NZ_CP082141.1 | 2840111 | 2841301 | 1190 | - | tuf | WP_230435682.1 |
| GCF_019844095.1 | NZ_CP082141.1 | 2857404 | 2858594 | 1190 | - | tuf | WP_230435702.1 |
| GCF_030758455.1 | NZ_CP132353.1 | 1494665 | 1495885 | 1220 | + | tuf | WP_231759496.1 |
| GCF_030758455.1 | NZ_CP132353.1 | 1511919 | 1513139 | 1220 | + | tuf | WP_231759505.1 |
| GCF_002952315.1 | NZ_CP023567.1 | 3448297 | 3449517 | 1220 | - | tuf | WP_161859424.1 |
| GCF_002952315.1 | NZ_CP023567.1 | 3465554 | 3466744 | 1190 | - | tuf | WP_161859433.1 |
| GCF_020683125.1 | NZ_CP085627.1 | 1423205 | 1424395 | 1190 | - | tuf | WP_318955203.1 |
| GCF_020683125.1 | NZ_CP085627.1 | 1440355 | 1441545 | 1190 | - | tuf | WP_318955203.1 |
| GCF_009738185.1 | NZ_CP046509.1 | 3894461 | 3895651 | 1190 | - | tuf | WP_301415178.1 |

| | | | | | | | |
|---|---|---|---|---|---|---|---|
| GCF_009738185.1 | NZ_CP046509.1 | 3911547 | 3912737 | 1190 | - | tuf | WP_301415178.1 |
| GCF_000026185.1 | NC_010694.1 | 575161 | 576351 | 1190 | + | tuf | WP_067150963.1 |
| GCF_000026185.1 | NC_010694.1 | 592255 | 593445 | 1190 | + | tuf | WP_067150963.1 |
| GCF_021365465.1 | NZ_CP089932.1 | 472314 | 473495 | 1181 | + | tuf | WP_038629750.1 |
| GCF_021365465.1 | NZ_CP089932.1 | 4274108 | 4275289 | 1181 | - | tuf | WP_104958836.1 |
| GCF_028622335.1 | NZ_CP117568.1 | 434283 | 435464 | 1181 | + | tuf | WP_249893123.1 |
| GCF_028622335.1 | NZ_CP117568.1 | 3982654 | 3983835 | 1181 | - | tuf | WP_249892531.1 |
| GCF_000008865.2 | NC_002695.2 | 3407968 | 3409149 | 1181 | - | tuf | WP_160251210.1 |
| GCF_000008865.2 | NC_002695.2 | 4221290 | 4222471 | 1181 | + | tuf | WP_160623185.1 |
| GCF_000005845.2 | NC_000913.3 | 288374 | 289555 | 1181 | + | tuf | WP_241501149.1 |
| GCF_000005845.2 | NC_000913.3 | 3449800 | 3450981 | 1181 | - | tuf | WP_241501149.1 |
| GCF_020097475.1 | NZ_CP083638.1 | 332952 | 334139 | 1187 | + | tuf | WP_029102688.1 |
| GCF_020097475.1 | NZ_CP083638.1 | 2492634 | 2493821 | 1187 | - | tuf | WP_029102688.1 |
| GCF_029962465.1 | NZ_CP072689.1 | 1753256 | 1754443 | 1187 | - | tuf | WP_036363988.1 |
| GCF_029962465.1 | NZ_CP072689.1 | 1791930 | 1793117 | 1187 | - | tuf | WP_036363988.1 |
| GCF_046523945.1 | NZ_CP171363.1 | 1360723 | 1361910 | 1187 | + | tuf | WP_078256771.1 |
| GCF_046523945.1 | NZ_CP171363.1 | 1380352 | 1381539 | 1187 | + | tuf | WP_049237885.1 |
| GCF_000148645.1 | NC_014541.1 | 1474957 | 1476144 | 1187 | - | tuf | WP_003656211.1 |
| GCF_000148645.1 | NC_014541.1 | 1492625 | 1493812 | 1187 | - | tuf | WP_003656211.1 |
| GCF_012295575.1 | NZ_CP051180.1 | 822306 | 823493 | 1187 | + | tuf | WP_076554779.1 |
| GCF_012295575.1 | NZ_CP051180.1 | 842347 | 843534 | 1187 | + | tuf | WP_076554779.1 |
| GCF_016918765.2 | NZ_CP092104.1 | 288973 | 290160 | 1187 | + | tuf | WP_284673437.1 |
| GCF_016918765.2 | NZ_CP092104.1 | 1805145 | 1806332 | 1187 | - | tuf | WP_284673437.1 |
| GCF_016469335.1 | NZ_CP059888.1 | 537660 | 538847 | 1187 | + | tuf | WP_338755227.1 |
| GCF_016469335.1 | NZ_CP059888.1 | 1620107 | 1621294 | 1187 | - | tuf | WP_338755227.1 |
| GCF_003290445.1 | NZ_CP021781.1 | 270557 | 271744 | 1187 | + | tuf | WP_277808322.1 |
| GCF_001880225.1 | NZ_CP009654.1 | 1952770 | 1953957 | 1187 | - | tuf | WP_277808322.1 |
| GCF_002211785.1 | NZ_CP022132.1 | 778669 | 779856 | 1187 | - | tuf | WP_263075701.1 |
| GCF_000195555.1 | NC_017449.1 | 1336303 | 1337490 | 1187 | + | tuf | WP_263075701.1 |
| GCF_008369785.1 | NZ_CP043552.1 | 753439 | 754626 | 1187 | - | tuf | WP_242746475.1 |
| GCF_003347135.1 | NZ_CP022377.1 | 1288413 | 1289600 | 1187 | + | tuf | WP_242746475.1 |

| | | | | | | | |
|---|---|---|---|---|---|---|---|
| GCF_000505725.1 | NC_023029.1 | 484069 | 485256 | 1187 | + | tuf | WP_066886983.1 |
| GCF_001275365.1 | NZ_CP012505.1 | 2183186 | 2184373 | 1187 | - | tuf | WP_066886983.1 |
| GCF_018135955.1 | NZ_CP063138.1 | 852871 | 854058 | 1187 | + | tuf | WP_062331575.1 |
| GCF_018972105.1 | NZ_CP076680.1 | 2058053 | 2059240 | 1187 | + | tuf | WP_062331575.1 |
| GCF_000219045.1 | NC_015696.1 | 1860909 | 1862096 | 1187 | - | tuf | WP_063514600.1 |
| GCF_000833355.1 | NZ_CP009607.1 | 1897133 | 1898320 | 1187 | - | tuf | WP_063514600.1 |
| GCF_001895265.1 | NZ_CP016796.1 | 169167 | 170354 | 1187 | + | tuf | WP_064609853.1 |
| GCF_000242255.2 | NC_017033.1 | 187373 | 188560 | 1187 | + | tuf | WP_064609853.1 |
| GCF_000242255.2 | NC_017033.1 | 215700 | 216881 | 1181 | + | tuf | WP_036413223.1 |
| GCF_021117405.1 | NZ_CP088251.1 | 3689374 | 3690555 | 1181 | - | tuf | WP_015422337.1 |
| GCF_021117405.1 | NZ_CP088251.1 | 3630101 | 3631282 | 1181 | - | tuf | WP_019443188.1 |
| GCF_021117385.1 | NZ_CP088252.1 | 2718730 | 2719911 | 1181 | - | tuf | WP_112715686.1 |
| GCF_021117385.1 | NZ_CP088252.1 | 448155 | 449336 | 1181 | + | tuf | WP_012764037.1 |
| GCF_011455495.1 | NZ_CP015029.1 | 4332024 | 4333205 | 1181 | - | tuf | WP_012764037.1 |
| GCF_011455495.1 | NZ_CP015029.1 | 1440101 | 1441321 | 1220 | - | tuf | WP_286240384.1 |
| GCF_000807275.1 | NZ_CP009056.1 | 1456651 | 1457871 | 1220 | - | tuf | WP_286240384.1 |
| GCF_000807275.1 | NZ_CP009056.1 | 1798 | 3018 | 1220 | - | tuf | WP_067862880.1 |
| GCF_030450055.1 | NZ_CP118480.1 | 18355 | 19575 | 1220 | - | tuf | WP_067862880.1 |
| GCF_030450055.1 | NZ_CP118480.1 | 3309616 | 3310836 | 1220 | - | tuf | WP_138988116.1 |
| GCF_003367375.1 | NZ_CP031416.1 | 3326131 | 3327351 | 1220 | - | tuf | WP_138988116.1 |
| GCF_003367375.1 | NZ_CP031416.1 | 66869 | 68089 | 1220 | + | tuf | WP_178967740.1 |
| GCF_030517105.1 | NZ_CP103875.1 | 83387 | 84607 | 1220 | + | tuf | WP_178967740.1 |
| GCF_030517105.1 | NZ_CP103875.1 | 232821 | 234011 | 1190 | + | tuf | WP_151053272.1 |
| GCF_900105885.1 | NZ_LT629780.1 | 249347 | 250537 | 1190 | + | tuf | WP_151053272.1 |
| GCF_002291425.1 | NZ_CP014136.1 | 2444499 | 2445686 | 1187 | + | tuf | WP_013033320.1 |
| GCF_002291425.1 | NZ_CP014136.1 | 2461151 | 2462338 | 1187 | + | tuf | WP_013033320.1 |
| GCF_000599985.1 | NZ_CP007445.1 | 2665198 | 2666385 | 1187 | - | tuf | WP_011330929.1 |
| GCF_000599985.1 | NZ_CP007445.1 | 2681846 | 2683033 | 1187 | - | tuf | WP_011330929.1 |
| GCF_030758635.1 | NZ_CP132382.1 | 1873393 | 1874580 | 1187 | + | tuf | WP_134357843.1 |
| GCF_030758635.1 | NZ_CP132382.1 | 1890047 | 1891234 | 1187 | + | tuf | WP_134357843.1 |
| GCF_042847215.1 | NZ_AP031500.1 | 2346930 | 2348117 | 1187 | - | tuf | WP_013221080.1 |

| | | | | | | | |
|---|---|---|---|---|---|---|---|
| GCF_042847215.1 | NZ_AP031500.1 | 2363601 | 2364788 | 1187 | - | tuf | WP_013221080.1 |
| GCF_017352235.1 | NZ_CP071487.1 | 2380272 | 2381459 | 1187 | - | tuf | WP_013221080.1 |
| GCF_017352235.1 | NZ_CP071487.1 | 505172 | 506359 | 1187 | - | tuf | WP_213435476.1 |
| GCF_002215215.1 | NZ_CP018632.1 | 522133 | 523320 | 1187 | - | tuf | WP_213435476.1 |
| GCF_002215215.1 | NZ_CP018632.1 | 556629 | 557816 | 1187 | + | tuf | WP_043959529.1 |
| GCF_001558255.2 | NZ_CP014056.2 | 573808 | 574995 | 1187 | + | tuf | WP_043959529.1 |
| GCF_001558255.2 | NZ_CP014056.2 | 533967 | 535154 | 1187 | + | tuf | WP_193985918.1 |
| GCF_023746615.1 | NZ_CP082275.1 | 551163 | 552350 | 1187 | + | tuf | WP_193985918.1 |
| GCF_023746615.1 | NZ_CP082275.1 | 2193803 | 2194990 | 1187 | - | tuf | WP_215218602.1 |
| GCF_034479635.1 | NZ_CP140153.1 | 2210987 | 2212174 | 1187 | - | tuf | WP_215218602.1 |
| GCF_034479635.1 | NZ_CP140153.1 | 1051702 | 1052922 | 1220 | - | tuf | WP_085757595.1 |
| GCF_900475885.1 | NZ_LS483429.1 | 2232134 | 2233354 | 1220 | + | tuf | WP_085757595.1 |
| GCF_900475885.1 | NZ_LS483429.1 | 190178 | 191359 | 1181 | + | tuf | WP_306762213.1 |
| GCF_900477945.1 | NZ_LS483458.1 | 3006870 | 3008051 | 1181 | - | tuf | WP_306761451.1 |
| GCF_900477945.1 | NZ_LS483458.1 | 1867569 | 1868750 | 1181 | - | tuf | WP_086964071.1 |
| GCF_020736045.1 | NZ_CP085952.1 | 1267478 | 1268659 | 1181 | - | tuf | WP_087035258.1 |
| GCF_020736045.1 | NZ_CP085952.1 | 744783 | 745973 | 1190 | + | tuf | WP_087459942.1 |
| GCF_016889385.1 | NZ_CP069510.1 | 2345431 | 2346612 | 1181 | + | tuf | WP_392558953.1 |
| GCF_016889385.1 | NZ_CP069510.1 | 2385081 | 2386262 | 1181 | + | tuf | WP_392558953.1 |
| GCF_016127215.1 | NZ_CP065991.1 | 2203942 | 2205123 | 1181 | + | tuf | WP_392561417.1 |
| GCF_016127215.1 | NZ_CP065991.1 | 2214257 | 2215438 | 1181 | - | tuf | WP_392561417.1 |
| GCF_900186995.1 | NZ_LT906463.1 | 2792098 | 2793279 | 1181 | + | tuf | WP_392552214.1 |
| GCF_900186995.1 | NZ_LT906463.1 | 2799629 | 2800810 | 1181 | - | tuf | WP_392552214.1 |
| GCF_011617105.1 | NZ_CP050150.1 | 249789 | 250970 | 1181 | + | tuf | WP_121120754.1 |
| GCF_011617105.1 | NZ_CP050150.1 | 1827880 | 1829061 | 1181 | - | tuf | WP_121120754.1 |
| GCF_020150375.1 | NZ_CP083737.1 | 370091 | 371272 | 1181 | - | tuf | WP_010261012.1 |
| GCF_020150375.1 | NZ_CP083737.1 | 998514 | 999695 | 1181 | + | tuf | WP_010253018.1 |
| GCF_000012985.1 | NC_007645.1 | 452672 | 453853 | 1181 | + | tuf | WP_105099977.1 |
| GCF_000012985.1 | NC_007645.1 | 3783414 | 3784595 | 1181 | - | tuf | WP_222924737.1 |
| GCF_009388985.1 | NZ_CP036422.1 | 1192556 | 1193737 | 1181 | - | tuf | WP_085068484.1 |
| GCF_009388985.1 | NZ_CP036422.1 | 1885121 | 1886302 | 1181 | + | tuf | WP_085069541.1 |

| | | | | | | | |
|---|---|---|---|---|---|---|---|
| GCF_042850945.1 | NZ_AP031496.1 | 464525 | 465706 | 1181 | + | tuf | WP_013027471.1 |
| GCF_042850945.1 | NZ_AP031496.1 | 4420578 | 4421759 | 1181 | - | tuf | WP_013027471.1 |
| GCF_001886615.1 | NZ_CP018139.1 | 451810 | 452991 | 1181 | + | tuf | WP_013027471.1 |
| GCF_029854215.1 | NZ_CP122961.1 | 4439323 | 4440504 | 1181 | - | tuf | WP_013027471.1 |
| GCF_030505695.1 | NZ_CP130143.1 | 448996 | 450177 | 1181 | + | tuf | WP_009087599.1 |
| GCF_002549795.1 | NZ_CP021435.1 | 3721645 | 3722826 | 1181 | - | tuf | WP_167429651.1 |
| GCF_008329985.2 | NZ_CP038437.2 | 440296 | 441477 | 1181 | + | tuf | WP_242281354.1 |
| GCF_001545155.1 | NZ_CP014226.1 | 3814404 | 3815585 | 1181 | - | tuf | WP_167429651.1 |
| GCF_000196875.2 | NC_014532.2 | 316915 | 318096 | 1181 | + | tuf | WP_222951547.1 |
| GCF_030406665.1 | NZ_CP129121.1 | 3715506 | 3716687 | 1181 | - | tuf | WP_150059788.1 |
| GCF_001431725.1 | NZ_CP013106.1 | 451309 | 452490 | 1181 | + | tuf | WP_105099977.1 |
| GCF_011399095.1 | NZ_AP022843.1 | 3868461 | 3869642 | 1181 | - | tuf | WP_013359730.1 |
| GCF_039691755.1 | NZ_CP137552.1 | 159458 | 160639 | 1181 | + | tuf | WP_039380832.1 |
| GCF_031886125.1 | NZ_CP119391.1 | 3068987 | 3070168 | 1181 | - | tuf | WP_039380832.1 |
| GCF_025917315.1 | NZ_CP080627.1 | 2686659 | 2687840 | 1181 | - | tuf | WP_215845418.1 |
| GCF_025820815.1 | NZ_CP096973.1 | 3324813 | 3325994 | 1181 | + | tuf | WP_409075112.1 |
| GCF_017868935.1 | NZ_CP053383.1 | 2452994 | 2454175 | 1181 | - | tuf | WP_006120578.1 |
| GCF_030295685.1 | NZ_AP027273.1 | 3049271 | 3050452 | 1181 | + | tuf | WP_006120578.1 |
| GCF_030295685.1 | NZ_AP027273.1 | 461448 | 462629 | 1181 | + | tuf | WP_135907598.1 |
| GCF_900105005.1 | NZ_LT629748.1 | 3878629 | 3879810 | 1181 | - | tuf | WP_013359730.1 |
| GCF_900105005.1 | NZ_LT629748.1 | 406891 | 408072 | 1181 | + | tuf | WP_173632468.1 |
| GCF_021545785.1 | NZ_CP079801.1 | 4301314 | 4302495 | 1181 | - | tuf | WP_173635814.1 |
| GCF_021545785.1 | NZ_CP079801.1 | 462883 | 464064 | 1181 | - | tuf | WP_089072914.1 |
| GCF_020025155.1 | NZ_CP073751.1 | 720269 | 721450 | 1181 | + | tuf | WP_089072914.1 |
| GCF_020025155.1 | NZ_CP073751.1 | 146951 | 148132 | 1181 | - | tuf | WP_133309412.1 |
| GCF_963677335.1 | NZ_OY781932.1 | 5379435 | 5380616 | 1181 | - | tuf | WP_133309412.1 |
| GCF_963677335.1 | NZ_OY781932.1 | 98188 | 99369 | 1181 | + | tuf | WP_133405295.1 |
| GCF_002056295.1 | NZ_CP020100.1 | 124709 | 125890 | 1181 | + | tuf | WP_133405295.1 |
| GCF_002056295.1 | NZ_CP020100.1 | 332044 | 333225 | 1181 | - | tuf | WP_306347534.1 |
| GCF_900105255.1 | NZ_LT629763.1 | 724675 | 725856 | 1181 | + | tuf | WP_090921093.1 |
| GCF_900105255.1 | NZ_LT629763.1 | 1660290 | 1661471 | 1181 | + | tuf | WP_049214773.1 |

| | | | | | | | |
|---|---|---|---|---|---|---|---|
| GCF_900105655.1 | NZ_LT629787.1 | 2062895 | 2064076 | 1181 | + | tuf | WP_049214773.1 |
| GCF_900105655.1 | NZ_LT629787.1 | 1635615 | 1636796 | 1181 | - | tuf | WP_095075298.1 |
| GCF_900104945.1 | NZ_LT629736.1 | 1780034 | 1781215 | 1181 | + | tuf | WP_095075298.1 |
| GCF_900104945.1 | NZ_LT629736.1 | 660662 | 661843 | 1181 | - | tuf | WP_005751979.1 |
| GCF_002356555.2 | NZ_AP017372.2 | 1078591 | 1079772 | 1181 | - | tuf | WP_005751979.1 |
| GCF_002356555.2 | NZ_AP017372.2 | 210767 | 211948 | 1181 | + | tuf | WP_102116828.1 |
| GCF_000015585.1 | NC_008789.1 | 3954093 | 3955274 | 1181 | - | tuf | WP_102116828.1 |
| GCF_000015585.1 | NC_008789.1 | 399164 | 400345 | 1181 | + | tuf | WP_261846679.1 |
| GCF_001663815.1 | NZ_CP016027.1 | 4236886 | 4238067 | 1181 | - | tuf | WP_261846679.1 |
| GCF_001663815.1 | NZ_CP016027.1 | 515269 | 516450 | 1181 | + | tuf | WP_012772931.1 |
| GCF_000024765.1 | NC_013422.1 | 4682170 | 4683351 | 1181 | - | tuf | WP_012772931.1 |
| GCF_000024765.1 | NC_013422.1 | 253250 | 254431 | 1181 | + | tuf | WP_011091854.1 |
| GCF_000019405.1 | NC_010519.1 | 4408236 | 4409417 | 1181 | - | tuf | WP_011091854.1 |
| GCF_000019405.1 | NC_010519.1 | 1856996 | 1858177 | 1181 | + | tuf | WP_010277443.1 |
| GCF_004851605.1 | NZ_CP031093.1 | 2132443 | 2133624 | 1181 | + | tuf | WP_010277443.1 |
| GCF_004851605.1 | NZ_CP031093.1 | 328145 | 329326 | 1181 | + | tuf | WP_264496896.1 |
| GCF_004786015.1 | NZ_CP032096.1 | 3686326 | 3687507 | 1181 | - | tuf | WP_264496896.1 |
| GCF_004786015.1 | NZ_CP032096.1 | 474222 | 475403 | 1181 | + | tuf | WP_014913795.1 |
| GCF_013340845.1 | NZ_AP020335.1 | 4589136 | 4590317 | 1181 | - | tuf | WP_014913795.1 |
| GCF_013340845.1 | NZ_AP020335.1 | 222789 | 223970 | 1181 | + | tuf | WP_225086522.1 |
| GCF_004028275.1 | NZ_CP035033.1 | 4364307 | 4365488 | 1181 | - | tuf | WP_012772931.1 |
| GCF_004028275.1 | NZ_CP035033.1 | 239637 | 240818 | 1181 | + | tuf | WP_010277443.1 |
| GCF_019797805.1 | NZ_CP081832.1 | 4398773 | 4399954 | 1181 | - | tuf | WP_010277443.1 |
| GCF_019797805.1 | NZ_CP081832.1 | 3829818 | 3830999 | 1181 | - | tuf | WP_005969581.1 |
| GCF_000008465.1 | NC_006512.1 | 4730488 | 4731669 | 1181 | + | tuf | WP_005969581.1 |
| GCF_000008465.1 | NC_006512.1 | 436469 | 437650 | 1181 | + | tuf | WP_039512662.1 |
| GCF_038500265.1 | NZ_CP150637.1 | 726601 | 727782 | 1181 | + | tuf | WP_039512662.1 |
| GCF_038500265.1 | NZ_CP150637.1 | 504343 | 505524 | 1181 | + | tuf | WP_011091854.1 |
| GCF_022655595.1 | NZ_CP093379.1 | 4638262 | 4639443 | 1181 | - | tuf | WP_368350107.1 |
| GCF_022655595.1 | NZ_CP093379.1 | 3211578 | 3212759 | 1181 | - | tuf | WP_095700750.1 |
| GCF_001697225.1 | NZ_CP014671.1 | 4016681 | 4017862 | 1181 | + | tuf | WP_095700750.1 |

| | | | | | | | |
|---|---|---|---|---|---|---|---|
| GCF_001697225.1 | NZ_CP014671.1 | 262983 | 264164 | 1181 | + | tuf | WP_014913795.1 |
| GCF_000299335.2 | NZ_CP004387.1 | 4238336 | 4239517 | 1181 | - | tuf | WP_014913795.1 |
| GCF_000299335.2 | NZ_CP004387.1 | 221364 | 222545 | 1181 | + | tuf | WP_102116828.1 |
| GCF_016592615.1 | NZ_AP013043.1 | 3951110 | 3952291 | 1181 | - | tuf | WP_102116828.1 |
| GCF_016592615.1 | NZ_AP013043.1 | 269026 | 270207 | 1181 | + | tuf | WP_095700750.1 |
| GCF_005671395.1 | NZ_CP040428.1 | 4432999 | 4434180 | 1181 | - | tuf | WP_095700750.1 |
| GCF_005671395.1 | NZ_CP040428.1 | 1139962 | 1141143 | 1181 | - | tuf | WP_005969581.1 |
| GCF_000981765.1 | NZ_CP010975.1 | 1435752 | 1436933 | 1181 | - | tuf | WP_005969581.1 |
| GCF_000981765.1 | NZ_CP010975.1 | 94870 | 96051 | 1181 | + | tuf | WP_090921093.1 |
| GCF_000024085.1 | NC_013166.1 | 1962522 | 1963703 | 1181 | - | tuf | WP_090920738.1 |
| GCF_000024085.1 | NC_013166.1 | 263699 | 264880 | 1181 | + | tuf | WP_255389246.1 |
| GCF_001708405.1 | NZ_CP012418.1 | 3257031 | 3258212 | 1181 | - | tuf | WP_255388828.1 |
| GCF_001708405.1 | NZ_CP012418.1 | 243626 | 244807 | 1181 | + | tuf | WP_069107610.1 |
| GCF_002863865.1 | NZ_CP022684.1 | 3012315 | 3013496 | 1181 | - | tuf | WP_237033314.1 |
| GCF_002863865.1 | NZ_CP022684.1 | 262878 | 264059 | 1181 | + | tuf | WP_107193913.1 |
| GCF_019048125.1 | NZ_CP077293.1 | 3314533 | 3315714 | 1181 | - | tuf | WP_107193928.1 |
| GCF_019048125.1 | NZ_CP077293.1 | 1164888 | 1166069 | 1181 | - | tuf | WP_107184525.1 |
| GCF_020526085.1 | NZ_CP084874.1 | 1625834 | 1627015 | 1181 | + | tuf | WP_107184525.1 |
| GCF_020526085.1 | NZ_CP084874.1 | 141735 | 142916 | 1181 | - | tuf | WP_107277200.1 |
| GCF_006711645.1 | NZ_CP041247.1 | 615391 | 616572 | 1181 | + | tuf | WP_107276051.1 |
| GCF_006711645.1 | NZ_CP041247.1 | 268615 | 269796 | 1181 | + | tuf | WP_261857707.1 |
| GCF_042137965.1 | NZ_CP169721.1 | 3509175 | 3510356 | 1181 | - | tuf | WP_261857554.1 |
| GCF_042137965.1 | NZ_CP169721.1 | 287390 | 288571 | 1181 | + | tuf | WP_107302837.1 |
| GCF_003261575.2 | NZ_CP036175.1 | 3598896 | 3600077 | 1181 | - | tuf | WP_107303053.1 |
| GCF_003261575.2 | NZ_CP036175.1 | 247942 | 249123 | 1181 | + | tuf | WP_107276051.1 |
| GCF_015139575.1 | NZ_AP022547.1 | 2796018 | 2797199 | 1181 | - | tuf | WP_256383349.1 |
| GCF_015139575.1 | NZ_AP022547.1 | 453334 | 454515 | 1181 | + | tuf | WP_049584863.1 |
| GCF_900636985.1 | NZ_LR134333.1 | 5507129 | 5508310 | 1181 | - | tuf | WP_217470518.1 |
| GCF_900636985.1 | NZ_LR134333.1 | 370622 | 371803 | 1181 | + | tuf | WP_012776534.1 |
| GCF_018139045.1 | NZ_CP073236.1 | 4880029 | 4881210 | 1181 | - | tuf | WP_015836288.1 |
| GCF_018139045.1 | NZ_CP073236.1 | 459157 | 460338 | 1181 | + | tuf | WP_011144818.1 |

| | | | | | | | |
|---|---|---|---|---|---|---|---|
| GCF_020099175.1 | NZ_CP083633.1 | 5499056 | 5500237 | 1181 | - | tuf | WP_011148798.1 |
| GCF_020099175.1 | NZ_CP083633.1 | 1494011 | 1495192 | 1181 | - | tuf | WP_046974281.1 |
| GCF_020525665.1 | NZ_CP084768.1 | 1688430 | 1689611 | 1181 | - | tuf | WP_046974414.1 |
| GCF_020525665.1 | NZ_CP084768.1 | 4310250 | 4311431 | 1181 | - | tuf | WP_047372260.1 |
| GCF_020525545.1 | NZ_CP084765.1 | 5134701 | 5135882 | 1181 | + | tuf | WP_047372876.1 |
| GCF_020525545.1 | NZ_CP084765.1 | 2844636 | 2845823 | 1187 | - | tuf | WP_016209759.1 |
| GCF_023195735.1 | NZ_CP096201.1 | 524925 | 526115 | 1190 | - | tuf | WP_147183123.1 |
| GCF_023195735.1 | NZ_CP096201.1 | 2052034 | 2053215 | 1181 | - | tuf | WP_309204340.1 |
| GCF_009649915.1 | NZ_CP045845.1 | 2068806 | 2069987 | 1181 | - | tuf | WP_309204340.1 |
| GCF_009649915.1 | NZ_CP045845.1 | 944101 | 945282 | 1181 | - | tuf | WP_064978262.1 |
| GCF_040930405.1 | NZ_CP162271.1 | 1556820 | 1558001 | 1181 | + | tuf | WP_230433021.1 |
| GCF_040930405.1 | NZ_CP162271.1 | 515407 | 516588 | 1181 | - | tuf | WP_043081015.1 |
| GCF_009363135.1 | NZ_CP045300.1 | 1174844 | 1176025 | 1181 | + | tuf | WP_043081015.1 |
| GCF_009363135.1 | NZ_CP045300.1 | 5406049 | 5407239 | 1190 | - | tuf | WP_110185682.1 |
| GCF_038449125.1 | NZ_CP151800.1 | 300965 | 302146 | 1181 | + | tuf | WP_047779688.1 |
| GCF_038449125.1 | NZ_CP151800.1 | 3533814 | 3534995 | 1181 | - | tuf | WP_047782102.1 |
| GCF_025758125.1 | NZ_CP107077.1 | 521528 | 522709 | 1181 | - | tuf | WP_021013352.1 |
| GCF_025758125.1 | NZ_CP107077.1 | 1012838 | 1014019 | 1181 | + | tuf | WP_021013352.1 |
| GCF_001658025.2 | NZ_CP014007.2 | 166962 | 168143 | 1181 | + | tuf | WP_151436284.1 |
| GCF_001658025.2 | NZ_CP014007.2 | 3752270 | 3753451 | 1181 | - | tuf | WP_151436284.1 |
| GCF_027886705.1 | NZ_CP115659.1 | 1214522 | 1215703 | 1181 | - | tuf | WP_129038369.1 |
| GCF_027886705.1 | NZ_CP115659.1 | 1701240 | 1702421 | 1181 | + | tuf | WP_129038369.1 |
| GCF_015167415.1 | NZ_CP063425.1 | 3065935 | 3067116 | 1181 | - | tuf | WP_012368508.1 |
| GCF_015167415.1 | NZ_CP063425.1 | 3579887 | 3581068 | 1181 | + | tuf | WP_012368688.1 |
| GCF_008330085.1 | NZ_CP040392.1 | 3170440 | 3171621 | 1181 | - | tuf | WP_006535489.1 |
| GCF_008330085.1 | NZ_CP040392.1 | 3676414 | 3677595 | 1181 | + | tuf | WP_006535489.1 |
| GCF_000300455.3 | NZ_CP007215.2 | 131820 | 133001 | 1181 | + | tuf | WP_036933947.1 |
| GCF_000300455.3 | NZ_CP007215.2 | 3599540 | 3600721 | 1181 | - | tuf | WP_036933947.1 |
| GCF_002155145.1 | NZ_CP021323.1 | 331467 | 332648 | 1181 | + | tuf | WP_036933947.1 |
| GCF_002157205.1 | NZ_CP021358.1 | 3796460 | 3797641 | 1181 | - | tuf | WP_036933947.1 |
| GCF_008247605.1 | NZ_CP043420.1 | 358938 | 360119 | 1181 | + | tuf | WP_004264373.1 |

| | | | | | | | |
|---|---|---|---|---|---|---|---|
| GCF_001518835.1 | NZ_CP013990.1 | 4334831 | 4336012 | 1181 | - | tuf | WP_004264373.1 |
| GCF_001518835.1 | NZ_CP013990.1 | 432379 | 433560 | 1181 | + | tuf | WP_068438404.1 |
| GCF_017348915.1 | NZ_CP071383.1 | 4108530 | 4109711 | 1181 | - | tuf | WP_068438404.1 |
| GCF_017348915.1 | NZ_CP071383.1 | 422829 | 424010 | 1181 | + | tuf | WP_116068795.1 |
| GCF_003176875.1 | NZ_CP029563.1 | 4191805 | 4192986 | 1181 | - | tuf | WP_116068795.1 |
| GCF_003176875.1 | NZ_CP029563.1 | 469205 | 470386 | 1181 | + | tuf | WP_337979750.1 |
| GCF_011764505.1 | NZ_AP022839.1 | 4169185 | 4170366 | 1181 | - | tuf | WP_337979645.1 |
| GCF_011764505.1 | NZ_AP022839.1 | 252089 | 253270 | 1181 | - | tuf | WP_004264373.1 |
| GCF_029026145.1 | NZ_CP119078.1 | 801770 | 802951 | 1181 | + | tuf | WP_004264373.1 |
| GCF_029026145.1 | NZ_CP119078.1 | 422276 | 423457 | 1181 | + | tuf | WP_112836004.1 |
| GCF_002240035.1 | NZ_CP016397.1 | 3720002 | 3721183 | 1181 | - | tuf | WP_112836004.1 |
| GCF_002240035.1 | NZ_CP016397.1 | 765593 | 766774 | 1181 | + | tuf | WP_008914507.1 |
| GCF_000953135.1 | NZ_LN614827.1 | 412574 | 413755 | 1181 | + | tuf | WP_036942149.1 |
| GCF_000953135.1 | NZ_LN614827.1 | 4158484 | 4159665 | 1181 | - | tuf | WP_036942149.1 |
| GCF_004571195.1 | NZ_CP038271.1 | 381139 | 382320 | 1181 | + | tuf | WP_318825957.1 |
| GCF_004571195.1 | NZ_CP038271.1 | 4285090 | 4286271 | 1181 | - | tuf | WP_318825957.1 |
| GCF_000953655.1 | NZ_LN681225.1 | 394050 | 395231 | 1181 | + | tuf | WP_154624778.1 |
| GCF_000953655.1 | NZ_LN681225.1 | 4070926 | 4072107 | 1181 | - | tuf | WP_154624778.1 |
| GCF_007361795.1 | NZ_CP041668.1 | 331599 | 332780 | 1181 | + | tuf | WP_096759607.1 |
| GCF_007361795.1 | NZ_CP041668.1 | 3629299 | 3630480 | 1181 | - | tuf | WP_096759607.1 |
| GCF_900637635.1 | NZ_LR134383.1 | 215286 | 216467 | 1181 | - | tuf | WP_115084474.1 |
| GCF_900637635.1 | NZ_LR134383.1 | 1687629 | 1688810 | 1181 | - | tuf | WP_024589451.1 |
| GCF_900187355.1 | NZ_LT906451.1 | 2168949 | 2170130 | 1181 | + | tuf | WP_024589451.1 |
| GCF_900187355.1 | NZ_LT906451.1 | 1502047 | 1503228 | 1181 | - | tuf | WP_054564107.1 |
| GCF_000091785.1 | NC_013861.1 | 2072853 | 2074034 | 1181 | + | tuf | WP_054564107.1 |
| GCF_000091785.1 | NC_013861.1 | 130320 | 131501 | 1181 | + | tuf | WP_024589451.1 |
| GCF_023921225.1 | NZ_CP071527.1 | 291403 | 292584 | 1181 | + | tuf | WP_024589451.1 |
| GCF_023921225.1 | NZ_CP071527.1 | 127193 | 128374 | 1181 | + | tuf | WP_058548756.1 |
| GCF_000953635.1 | NZ_LN614830.1 | 3304805 | 3305986 | 1181 | - | tuf | WP_058548756.1 |
| GCF_000953635.1 | NZ_LN614830.1 | 12114 | 13295 | 1181 | + | tuf | WP_002958888.1 |
| GCF_001941585.1 | NZ_CP013742.1 | 3583619 | 3584800 | 1181 | + | tuf | WP_039488657.1 |

| | | | | | | | |
|---|---|---|---|---|---|---|---|
| GCF_001941585.1 | NZ_CP013742.1 | 294622 | 295803 | 1181 | + | tuf | WP_004589660.1 |
| GCF_002848365.2 | NZ_CP025491.2 | 3547966 | 3549147 | 1181 | - | tuf | WP_004589660.1 |
| GCF_002848365.2 | NZ_CP025491.2 | 261812 | 262993 | 1181 | + | tuf | WP_013463851.1 |
| GCF_900186965.1 | NZ_LT906457.1 | 3251495 | 3252676 | 1181 | - | tuf | WP_013463851.1 |
| GCF_900186965.1 | NZ_LT906457.1 | 426161 | 427342 | 1181 | + | tuf | WP_036971551.1 |
| GCF_900187095.1 | NZ_LT906442.1 | 3506225 | 3507406 | 1181 | - | tuf | WP_036971551.1 |
| GCF_900187095.1 | NZ_LT906442.1 | 3528710 | 3529891 | 1181 | - | tuf | WP_063359275.1 |
| GCF_019047465.1 | NZ_CP077236.1 | 3678659 | 3679840 | 1181 | - | tuf | WP_063359275.1 |
| GCF_019047465.1 | NZ_CP077236.1 | 289384 | 290565 | 1181 | + | tuf | WP_119230648.1 |
| GCF_041464075.1 | NZ_CP167172.1 | 412036 | 413217 | 1181 | + | tuf | WP_119230648.1 |
| GCF_041464075.1 | NZ_CP167172.1 | 257258 | 258439 | 1181 | + | tuf | WP_011326944.1 |
| GCF_039789245.1 | NZ_CP146357.1 | 3385146 | 3386327 | 1181 | - | tuf | WP_011329538.1 |
| GCF_039789245.1 | NZ_CP146357.1 | 310249 | 311430 | 1181 | + | tuf | WP_210798819.1 |
| GCF_900478135.1 | NZ_LS483470.1 | 3654318 | 3655499 | 1181 | - | tuf | WP_210798819.1 |
| GCF_900478135.1 | NZ_LS483470.1 | 165433 | 166614 | 1181 | + | tuf | WP_058028464.1 |
| GCF_003096015.2 | NZ_CP029185.2 | 225500 | 226681 | 1181 | + | tuf | WP_058028464.1 |
| GCF_003096015.2 | NZ_CP029185.2 | 844517 | 845698 | 1181 | + | tuf | WP_038638961.1 |
| GCF_004295645.1 | NZ_CP034752.1 | 863812 | 864993 | 1181 | + | tuf | WP_038638961.1 |
| GCF_004295645.1 | NZ_CP034752.1 | 254276 | 255457 | 1181 | + | tuf | WP_039494551.1 |
| GCF_004295665.1 | NZ_CP034759.1 | 376611 | 377792 | 1181 | + | tuf | WP_039494551.1 |
| GCF_004295665.1 | NZ_CP034759.1 | 61 | 1242 | 1181 | - | tuf | WP_064668786.1 |
| GCF_009650135.1 | NZ_CP045871.1 | 311560 | 312741 | 1181 | - | tuf | WP_064668786.1 |
| GCF_009650135.1 | NZ_CP045871.1 | 29469 | 30650 | 1181 | + | tuf | WP_064668786.1 |
| GCF_042926695.1 | NZ_CP170642.1 | 171222 | 172403 | 1181 | - | tuf | WP_182758002.1 |
| GCF_042926695.1 | NZ_CP170642.1 | 631874 | 633055 | 1181 | + | tuf | WP_257325261.1 |
| GCF_003515985.1 | NZ_CP023009.1 | 126599 | 127780 | 1181 | + | tuf | WP_010562291.1 |
| GCF_003515985.1 | NZ_CP023009.1 | 145910 | 147091 | 1181 | + | tuf | WP_010562291.1 |
| GCF_015999465.1 | NZ_CP065534.1 | 239981 | 241162 | 1181 | + | tuf | WP_011326944.1 |
| GCF_015999465.1 | NZ_CP065534.1 | 3079386 | 3080567 | 1181 | - | tuf | WP_011329538.1 |
| GCF_023078475.1 | NZ_CP095741.1 | 2944374 | 2945555 | 1181 | - | tuf | WP_009840797.1 |
| GCF_023078475.1 | NZ_CP095741.1 | 2963723 | 2964904 | 1181 | - | tuf | WP_009840797.1 |

| | | | | | | | |
|---|---|---|---|---|---|---|---|
| GCF_023078455.1 | NZ_CP095740.1 | 126515 | 127696 | 1181 | + | tuf | WP_109875886.1 |
| GCF_023078455.1 | NZ_CP095740.1 | 2343441 | 2344622 | 1181 | - | tuf | WP_109875886.1 |
| GCF_023078495.1 | NZ_CP095742.1 | 1067564 | 1068745 | 1181 | - | tuf | WP_054017727.1 |
| GCF_023078495.1 | NZ_CP095742.1 | 1661695 | 1662876 | 1181 | + | tuf | WP_054017727.1 |
| GCF_023699965.1 | NZ_CP077072.1 | 3610753 | 3611934 | 1181 | - | tuf | WP_003031109.1 |
| GCF_023699965.1 | NZ_CP077072.1 | 4298072 | 4299253 | 1181 | + | tuf | WP_003031109.1 |
| GCF_023612255.1 | NZ_CP063231.1 | 2662211 | 2663404 | 1193 | - | tuf | WP_058022517.1 |
| GCF_023612255.1 | NZ_CP063231.1 | 2678802 | 2679995 | 1193 | - | tuf | WP_058022517.1 |
| GCF_006385595.1 | NZ_CP041046.1 | 3507514 | 3508701 | 1187 | - | tuf | WP_129834707.1 |
| GCF_006385595.1 | NZ_CP041046.1 | 3524481 | 3525668 | 1187 | - | tuf | WP_129834707.1 |
| GCF_001887595.1 | NZ_CP017480.1 | 3237496 | 3238686 | 1190 | + | tuf | WP_249679516.1 |
| GCF_001887595.1 | NZ_CP017480.1 | 4767814 | 4769004 | 1190 | - | tufA | NP_252955.1 |
| GCF_002327105.1 | NZ_CP023406.1 | 4784319 | 4785509 | 1190 | - | tufB | NP_252967.1 |
| GCF_002327105.1 | NZ_CP023406.1 | 446461 | 447651 | 1190 | + | tuf | WP_133860552.1 |
| GCF_021233355.1 | NZ_CP089507.1 | 462948 | 464138 | 1190 | + | tuf | WP_133860552.1 |
| GCF_021233355.1 | NZ_CP089507.1 | 669281 | 670471 | 1190 | + | tuf | WP_010220303.1 |
| GCF_007795095.1 | NZ_CP042218.1 | 685890 | 687080 | 1190 | + | tuf | WP_010220303.1 |
| GCF_007795095.1 | NZ_CP042218.1 | 5862001 | 5863191 | 1190 | - | tuf | WP_024014521.1 |
| GCF_004803715.2 | NZ_CP039383.2 | 5878556 | 5879746 | 1190 | - | tuf | WP_024014521.1 |
| GCF_004803715.2 | NZ_CP039383.2 | 511896 | 513086 | 1190 | + | tuf | WP_003255489.1 |
| GCF_001442745.1 | NZ_CP011129.1 | 528486 | 529676 | 1190 | + | tuf | WP_010951775.1 |
| GCF_001442745.1 | NZ_CP011129.1 | 5836532 | 5837722 | 1190 | - | tuf | WP_003186103.1 |
| GCF_016613475.2 | NZ_CP071517.1 | 5853085 | 5854275 | 1190 | - | tuf | WP_003186103.1 |
| GCF_016613475.2 | NZ_CP071517.1 | 5403079 | 5404269 | 1190 | - | tuf | WP_003369027.1 |
| GCF_027924565.1 | NZ_AP027041.1 | 1506751 | 1507941 | 1190 | + | tuf | WP_003176426.1 |
| GCF_027924565.1 | NZ_AP027041.1 | 485620 | 486810 | 1190 | + | tuf | WP_170027986.1 |
| GCF_014779555.2 | NZ_CP094357.1 | 502215 | 503405 | 1190 | + | tuf | WP_170027986.1 |
| GCF_014779555.2 | NZ_CP094357.1 | 1764090 | 1765280 | 1190 | + | tuf | WP_027924378.1 |
| GCF_002355975.1 | NZ_AP014940.1 | 5667482 | 5668672 | 1190 | - | tuf | WP_003176426.1 |
| GCF_002355975.1 | NZ_AP014940.1 | 545457 | 546647 | 1190 | + | tuf | WP_013970626.1 |
| GCF_040687735.1 | NZ_CP159925.1 | 562054 | 563244 | 1190 | + | tuf | WP_013970626.1 |

| | | | | | | | |
|---|---|---|---|---|---|---|---|
| GCF_040687735.1 | NZ_CP159925.1 | 2324189 | 2325379 | 1190 | + | tuf | WP_090204646.1 |
| GCF_001442805.1 | NZ_CP011131.1 | 781470 | 782660 | 1190 | + | tuf | WP_024014521.1 |
| GCF_001442805.1 | NZ_CP011131.1 | 798028 | 799218 | 1190 | + | tuf | WP_024014521.1 |
| GCF_018406645.1 | NZ_AP024546.1 | 5769334 | 5770524 | 1190 | - | tuf | WP_007970421.1 |
| GCF_018406645.1 | NZ_AP024546.1 | 4043483 | 4044673 | 1190 | - | tuf | WP_003176426.1 |
| GCF_009755585.1 | NZ_CP046603.1 | 4500715 | 4501905 | 1190 | + | tuf | WP_338522482.1 |
| GCF_009755585.1 | NZ_CP046603.1 | 4517287 | 4518477 | 1190 | + | tuf | WP_338522484.1 |
| GCF_019551435.1 | NZ_CP080544.1 | 5808930 | 5810120 | 1190 | - | tuf | WP_003186071.1 |
| GCF_019551435.1 | NZ_CP080544.1 | 5825482 | 5826672 | 1190 | - | tuf | WP_003186103.1 |
| GCF_031199375.1 | NZ_CP133568.1 | 3855617 | 3856807 | 1190 | - | tuf | WP_318643776.1 |
| GCF_031199375.1 | NZ_CP133568.1 | 3872146 | 3873336 | 1190 | - | tuf | WP_318643776.1 |
| GCF_040429585.1 | NZ_CP134782.1 | 4591607 | 4592797 | 1190 | - | tuf | WP_160921986.1 |
| GCF_040429585.1 | NZ_CP134782.1 | 4608094 | 4609284 | 1190 | - | tuf | WP_338545365.1 |
| GCF_014541205.1 | NZ_CP061280.1 | 4767607 | 4768797 | 1190 | - | tuf | WP_182368693.1 |
| GCF_014541205.1 | NZ_CP061280.1 | 4784156 | 4785346 | 1190 | - | tuf | WP_182368693.1 |
| GCF_046267875.1 | NZ_CP176502.1 | 6196212 | 6197402 | 1190 | - | tuf | WP_003186103.1 |
| GCF_046267875.1 | NZ_CP176502.1 | 6212764 | 6213954 | 1190 | - | tuf | WP_003186103.1 |
| GCF_002285575.1 | NZ_CP023044.1 | 6064488 | 6065678 | 1190 | - | tuf | WP_003186071.1 |
| GCF_002285575.1 | NZ_CP023044.1 | 6081040 | 6082230 | 1190 | - | tuf | WP_003186103.1 |
| GCF_013377995.1 | NZ_CP055305.1 | 1228704 | 1229894 | 1190 | + | tuf | WP_016779174.1 |
| GCF_013377995.1 | NZ_CP055305.1 | 5631092 | 5632282 | 1190 | - | tuf | WP_003176426.1 |
| GCF_013377235.1 | NZ_CP016226.1 | 507542 | 508732 | 1190 | + | tuf | WP_007253311.1 |
| GCF_013377235.1 | NZ_CP016226.1 | 596739 | 597929 | 1190 | + | tuf | WP_119891999.1 |
| GCF_000224005.2 | NZ_CP007031.1 | 613203 | 614393 | 1190 | + | tuf | WP_119892010.1 |
| GCF_000224005.2 | NZ_CP007031.1 | 5023155 | 5024345 | 1190 | - | tuf | WP_003176426.1 |
| GCF_042854845.1 | NZ_AP031498.1 | 6340482 | 6341672 | 1190 | - | tuf | WP_007924207.1 |
| GCF_042854845.1 | NZ_AP031498.1 | 6357052 | 6358242 | 1190 | - | tuf | WP_007924207.1 |
| GCF_006547045.1 | NZ_CP041242.1 | 5315608 | 5316798 | 1190 | - | tuf | WP_025262054.1 |
| GCF_006547045.1 | NZ_CP041242.1 | 801624 | 802814 | 1190 | + | tuf | WP_009617114.1 |
| GCF_003233695.1 | NZ_CP029843.1 | 818142 | 819332 | 1190 | + | tuf | WP_009617114.1 |
| GCF_003233695.1 | NZ_CP029843.1 | 4857276 | 4858466 | 1190 | - | tuf | WP_005887082.1 |

| | | | | | | | |
|---|---|---|---|---|---|---|---|
| GCF_021655555.1 | NZ_AP023086.1 | 2875020 | 2876210 | 1190 | - | tuf | WP_053191465.1 |
| GCF_045862395.1 | NZ_AP023087.1 | 2891575 | 2892765 | 1190 | - | tuf | WP_053191465.1 |
| GCF_045862395.1 | NZ_AP023087.1 | 5524738 | 5525928 | 1190 | - | tuf | WP_003186071.1 |
| GCF_018972165.1 | NZ_CP076686.1 | 5541309 | 5542499 | 1190 | - | tuf | WP_007896619.1 |
| GCF_018972165.1 | NZ_CP076686.1 | 2269171 | 2270361 | 1190 | + | tuf | WP_060841339.1 |
| GCF_038784545.1 | NZ_CP152380.1 | 2285769 | 2286959 | 1190 | + | tuf | WP_060841339.1 |
| GCF_038784545.1 | NZ_CP152380.1 | 5199744 | 5200934 | 1190 | - | tuf | WP_010220303.1 |
| GCF_030297175.1 | NZ_AP028070.1 | 5216353 | 5217543 | 1190 | - | tuf | WP_010220292.1 |
| GCF_030297175.1 | NZ_AP028070.1 | 498083 | 499273 | 1190 | + | tuf | WP_010951775.1 |
| GCF_045057155.1 | NZ_CP171359.1 | 514677 | 515867 | 1190 | + | tuf | WP_010951775.1 |
| GCF_045057155.1 | NZ_CP171359.1 | 5770367 | 5771557 | 1190 | - | tuf | WP_010220303.1 |
| GCF_008122265.1 | NZ_CP043042.1 | 5571523 | 5572713 | 1190 | + | tuf | WP_010566854.1 |
| GCF_008122265.1 | NZ_CP043042.1 | 5712972 | 5714162 | 1190 | - | tuf | WP_010206920.1 |
| GCF_038098645.1 | NZ_CP101118.1 | 581129 | 582319 | 1190 | + | tuf | WP_010951775.1 |
| GCF_038098645.1 | NZ_CP101118.1 | 597723 | 598913 | 1190 | + | tuf | WP_010951775.1 |
| GCF_000284615.1 | NC_017067.1 | 723559 | 724749 | 1190 | + | tuf | WP_003369027.1 |
| GCF_000284615.1 | NC_017067.1 | 5997748 | 5998938 | 1190 | - | tuf | WP_010220303.1 |
| GCF_001043175.1 | NZ_CP011494.1 | 6014292 | 6015482 | 1190 | - | tuf | WP_064382246.1 |
| GCF_001043175.1 | NZ_CP011494.1 | 1469597 | 1470787 | 1190 | - | tuf | WP_003176426.1 |
| GCF_002116735.1 | NZ_CP020931.1 | 542273 | 543463 | 1190 | + | tuf | WP_016484647.1 |
| GCF_002116735.1 | NZ_CP020931.1 | 558860 | 560050 | 1190 | + | tuf | WP_016484647.1 |
| GCF_017301335.1 | NZ_CP071247.1 | 4736287 | 4737477 | 1190 | - | tuf | WP_016779174.1 |
| GCF_017301335.1 | NZ_CP071247.1 | 3556954 | 3558144 | 1190 | + | tuf | WP_019693885.1 |
| GCF_001854125.1 | NZ_CP017715.1 | 3573521 | 3574711 | 1190 | + | tuf | WP_071553149.1 |
| GCF_001854125.1 | NZ_CP017715.1 | 4361755 | 4362945 | 1190 | - | tuf | WP_038411791.1 |
| GCF_030297115.1 | NZ_AP028062.1 | 4378346 | 4379536 | 1190 | - | tuf | WP_038411791.1 |
| GCF_030297115.1 | NZ_AP028062.1 | 6232520 | 6233710 | 1190 | - | tuf | WP_064361246.1 |
| GCF_000830985.1 | NZ_CP007151.1 | 6249078 | 6250268 | 1190 | - | tuf | WP_064361246.1 |
| GCF_001651805.1 | NZ_CP015839.1 | 6010952 | 6012142 | 1190 | - | tuf | WP_010220303.1 |
| GCF_001651805.1 | NZ_CP015839.1 | 6252104 | 6253294 | 1190 | - | tuf | WP_010566854.1 |
| GCF_017310015.1 | NZ_CP022297.1 | 5852501 | 5853691 | 1190 | - | tuf | WP_010220303.1 |

| | | | | | | | |
|---|---|---|---|---|---|---|---|
| GCF_017310015.1 | NZ_CP022297.1 | 5869062 | 5870252 | 1190 | - | tuf | WP_010220303.1 |
| GCF_014805825.1 | NZ_CP061941.1 | 4912086 | 4913276 | 1190 | - | tuf | WP_024014521.1 |
| GCF_014623465.1 | NZ_CP061081.1 | 4928645 | 4929835 | 1190 | - | tuf | WP_024014521.1 |
| GCF_016906065.1 | NZ_CP070273.1 | 6120281 | 6121471 | 1190 | - | tuf | WP_347909399.1 |
| GCF_000192865.1 | NC_015276.1 | 4774949 | 4776139 | 1190 | - | tuf | WP_099593972.1 |
| GCF_041227765.1 | NZ_CP165966.1 | 4791551 | 4792741 | 1190 | - | tuf | WP_099593972.1 |
| GCF_036322735.1 | NZ_AP027271.1 | 5926506 | 5927696 | 1190 | - | tuf | WP_003176426.1 |
| GCF_000214215.1 | NC_015559.1 | 5943064 | 5944254 | 1190 | - | tuf | WP_003176426.1 |
| GCF_013372285.1 | NZ_CP054301.1 | 483777 | 484967 | 1190 | + | tuf | WP_048373848.1 |
| GCF_020694005.1 | NZ_CP073013.1 | 5687523 | 5688713 | 1190 | - | tuf | WP_054594277.1 |
| GCF_003265225.1 | NZ_CP022013.1 | 5704088 | 5705278 | 1190 | - | tuf | WP_347928143.1 |
| GCF_003265225.1 | NZ_CP022013.1 | 5049439 | 5050629 | 1190 | - | tuf | WP_007896660.1 |
| GCF_042232855.1 | NZ_CP113889.1 | 5171651 | 5172841 | 1190 | - | tuf | WP_268798242.1 |
| GCF_042232855.1 | NZ_CP113889.1 | 6111176 | 6112366 | 1190 | - | tuf | WP_060841339.1 |
| GCF_002355475.1 | NZ_AP014862.1 | 6127774 | 6128964 | 1190 | - | tuf | WP_060841339.1 |
| GCF_002355475.1 | NZ_AP014862.1 | 3152458 | 3153648 | 1190 | + | tuf | WP_013970626.1 |
| GCF_008807375.1 | NZ_CP043311.1 | 3169048 | 3170238 | 1190 | + | tuf | WP_013970626.1 |
| GCF_008807375.1 | NZ_CP043311.1 | 5607077 | 5608267 | 1190 | - | tuf | WP_024014521.1 |
| GCF_011397855.1 | NZ_AP022642.1 | 5623635 | 5624825 | 1190 | - | tuf | WP_024014521.1 |
| GCF_011397855.1 | NZ_AP022642.1 | 6104571 | 6105761 | 1190 | - | tuf | WP_008018195.1 |
| GCF_003584645.1 | NZ_AP017928.1 | 480594 | 481784 | 1190 | + | tuf | WP_054881963.1 |
| GCF_003584645.1 | NZ_AP017928.1 | 497184 | 498374 | 1190 | + | tuf | WP_054881963.1 |
| GCF_949769195.1 | NZ_OX458333.1 | 531696 | 532886 | 1190 | + | tuf | WP_016484647.1 |
| GCF_949769195.1 | NZ_OX458333.1 | 548285 | 549475 | 1190 | + | tuf | WP_016484647.1 |
| GCF_000008325.1 | NC_002977.6 | 5878186 | 5879376 | 1190 | - | tuf | WP_003176426.1 |
| GCF_000008325.1 | NC_002977.6 | 2826225 | 2827415 | 1190 | + | tuf | WP_010467251.1 |
| GCF_012769535.1 | NZ_CP046565.1 | 2842794 | 2843984 | 1190 | + | tuf | WP_150582439.1 |
| GCF_012769535.1 | NZ_CP046565.1 | 674876 | 676066 | 1190 | + | tuf | WP_043248914.1 |
| GCF_026247885.1 | NZ_CP110921.1 | 691384 | 692574 | 1190 | + | tuf | WP_043248914.1 |
| GCF_026247885.1 | NZ_CP110921.1 | 5282679 | 5283869 | 1190 | - | tuf | WP_024014521.1 |
| GCF_019669985.1 | NZ_AP019782.1 | 5299235 | 5300425 | 1190 | - | tuf | WP_024014521.1 |

| | | | | | | | |
|---|---|---|---|---|---|---|---|
| GCF_019669985.1 | NZ_AP019782.1 | 5645852 | 5647042 | 1190 | - | tuf | WP_010220303.1 |
| GCF_019670005.1 | NZ_AP019783.1 | 5662414 | 5663604 | 1190 | - | tuf | WP_010220303.1 |
| GCF_019670005.1 | NZ_AP019783.1 | 581786 | 582976 | 1190 | + | tuf | WP_003444379.1 |
| GCF_033126985.1 | NZ_AP024714.1 | 579949 | 581139 | 1190 | + | tuf | WP_010951775.1 |
| GCF_033126985.1 | NZ_AP024714.1 | 596539 | 597729 | 1190 | + | tuf | WP_013970626.1 |
| GCF_030294985.1 | NZ_AP024718.1 | 3090842 | 3092032 | 1190 | - | tuf | WP_163933278.1 |
| GCF_030294985.1 | NZ_AP024718.1 | 3107429 | 3108619 | 1190 | - | tuf | WP_102051728.1 |
| GCF_030717995.2 | NZ_CP157743.1 | 4628441 | 4629631 | 1190 | + | tuf | WP_136665592.1 |
| GCF_030717995.2 | NZ_CP157743.1 | 4644935 | 4646125 | 1190 | + | tuf | WP_415754459.1 |
| GCF_000214275.2 | NZ_CM001475.1 | 5159800 | 5160990 | 1190 | - | tuf | WP_025262054.1 |
| GCF_000214275.2 | NZ_CM001475.1 | 1832049 | 1833239 | 1190 | + | tuf | WP_007896619.1 |
| GCF_000785705.2 | NZ_CP014476.1 | 1848614 | 1849804 | 1190 | + | tuf | WP_007896660.1 |
| GCF_000785705.2 | NZ_CP014476.1 | 621089 | 622279 | 1190 | + | tuf | WP_005791980.1 |
| GCF_019669905.1 | NZ_AP019777.1 | 3011136 | 3012326 | 1190 | + | tuf | WP_010467251.1 |
| GCF_019669905.1 | NZ_AP019777.1 | 3027707 | 3028897 | 1190 | + | tuf | WP_007896660.1 |
| GCF_030490285.1 | NZ_CP129884.1 | 6124390 | 6125580 | 1190 | - | tuf | WP_003176426.1 |
| GCF_030490285.1 | NZ_CP129884.1 | 749003 | 750193 | 1190 | + | tuf | WP_069901943.1 |
| GCF_018734325.1 | NZ_CP073754.1 | 765515 | 766705 | 1190 | + | tuf | WP_069901953.1 |
| GCF_018734325.1 | NZ_CP073754.1 | 510976 | 512166 | 1190 | + | tuf | WP_217868085.1 |
| GCF_024360925.2 | NZ_CP113517.1 | 527565 | 528755 | 1190 | + | tuf | WP_217868085.1 |
| GCF_024360925.2 | NZ_CP113517.1 | 1306888 | 1308078 | 1190 | + | tuf | WP_047702780.1 |
| GCF_008632455.1 | NZ_CP043929.1 | 1323438 | 1324628 | 1190 | + | tuf | WP_047702780.1 |
| GCF_008632455.1 | NZ_CP043929.1 | 5540109 | 5541299 | 1190 | - | tuf | WP_008053875.1 |
| GCF_000260965.1 | NC_017856.1 | 5914238 | 5915428 | 1190 | - | tuf | WP_024164425.1 |
| GCF_000260965.1 | NC_017856.1 | 5930796 | 5931986 | 1190 | - | tuf | WP_186746080.1 |
| GCF_030296755.1 | NZ_AP027741.1 | 590055 | 591245 | 1190 | + | tuf | WP_013970626.1 |
| GCF_030296755.1 | NZ_AP027741.1 | 606645 | 607835 | 1190 | + | tuf | WP_013970626.1 |
| GCF_000260985.4 | NC_017857.3 | 659840 | 661030 | 1190 | - | tuf | WP_090325641.1 |
| GCF_000260985.4 | NC_017857.3 | 5184892 | 5186082 | 1190 | - | tuf | WP_062575362.1 |
| GCF_020447175.2 | NZ_CP095733.1 | 5201485 | 5202675 | 1190 | - | tuf | WP_062575362.1 |
| GCF_020447175.2 | NZ_CP095733.1 | 5180281 | 5181471 | 1190 | - | tuf | WP_029299269.1 |

| | | | | | | | |
|---|---|---|---|---|---|---|---|
| GCF_000968535.2 | NC_016112.1 | 6422510 | 6423700 | 1190 | - | tuf | WP_054910987.1 |
| GCF_000968535.2 | NC_016112.1 | 6439036 | 6440226 | 1190 | - | tuf | WP_054910987.1 |
| GCF_005931095.1 | NZ_CP035467.1 | 109929 | 111119 | 1190 | - | tuf | WP_182323863.1 |
| GCF_005931095.1 | NZ_CP035467.1 | 126519 | 127709 | 1190 | - | tuf | WP_104960689.1 |
| GCF_002209385.1 | NZ_CP022129.1 | 6941379 | 6942569 | 1190 | + | tuf | WP_007896619.1 |
| GCF_002209385.1 | NZ_CP022129.1 | 6957950 | 6959140 | 1190 | + | tuf | WP_008145495.1 |
| GCF_001999945.1 | NZ_CP019650.1 | 6073069 | 6074259 | 1190 | - | tuf | WP_003186071.1 |
| GCF_001999945.1 | NZ_CP019650.1 | 6089621 | 6090811 | 1190 | - | tuf | WP_003186103.1 |
| GCF_001750105.1 | NZ_CP014143.1 | 5255304 | 5256494 | 1190 | - | tuf | WP_003176426.1 |
| GCF_001750105.1 | NZ_CP014143.1 | 3502230 | 3503420 | 1190 | - | tuf | WP_090350222.1 |
| GCF_029869925.1 | NZ_CP118605.1 | 3518705 | 3519895 | 1190 | - | tuf | WP_090350222.1 |
| GCF_029869925.1 | NZ_CP118605.1 | 512463 | 513653 | 1190 | + | tuf | WP_016497741.1 |
| GCF_020991125.1 | NZ_CP087715.1 | 529052 | 530242 | 1190 | + | tuf | WP_016497741.1 |
| GCF_020991125.1 | NZ_CP087715.1 | 552345 | 553535 | 1190 | + | tuf | WP_079204508.1 |
| GCF_021165935.1 | NZ_CP088953.1 | 568870 | 570060 | 1190 | + | tuf | WP_079204519.1 |
| GCF_021165935.1 | NZ_CP088953.1 | 1191469 | 1192659 | 1190 | + | tuf | WP_060755286.1 |
| GCF_009931115.1 | NZ_CP047491.1 | 615717 | 616907 | 1190 | + | tuf | WP_005791980.1 |
| GCF_009931115.1 | NZ_CP047491.1 | 680592 | 681782 | 1190 | + | tuf | WP_003115146.1 |
| GCF_033723955.1 | NZ_CP137555.1 | 697097 | 698287 | 1190 | + | tuf | WP_003115146.1 |
| GCF_033723955.1 | NZ_CP137555.1 | 5718028 | 5719218 | 1190 | - | tuf | WP_095109688.1 |
| GCF_030440425.1 | NZ_CP098023.1 | 5934617 | 5935807 | 1190 | - | tuf | WP_010206920.1 |
| GCF_030440425.1 | NZ_CP098023.1 | 6908605 | 6909795 | 1190 | - | tuf | WP_003176426.1 |
| GCF_001617625.1 | NZ_CP014864.1 | 1642651 | 1643841 | 1190 | + | tuf | WP_138300984.1 |
| GCF_001617625.1 | NZ_CP014864.1 | 1659102 | 1660292 | 1190 | + | tuf | WP_138300984.1 |
| GCF_002953195.1 | NZ_CP026377.1 | 2912913 | 2914103 | 1190 | - | tuf | WP_010951775.1 |
| GCF_002953195.1 | NZ_CP026377.1 | 2929504 | 2930694 | 1190 | - | tuf | WP_010951775.1 |
| GCF_023517775.1 | NZ_CP082904.1 | 4113623 | 4114813 | 1190 | - | tuf | WP_003232413.1 |
| GCF_023517775.1 | NZ_CP082904.1 | 183530 | 184720 | 1190 | + | tuf | WP_090192642.1 |
| GCF_009914055.1 | NZ_CP028271.1 | 199969 | 201159 | 1190 | + | tuf | WP_090192642.1 |
| GCF_009914055.1 | NZ_CP028271.1 | 488183 | 489373 | 1190 | + | tuf | WP_010951775.1 |
| GCF_022592395.1 | NZ_CP093242.1 | 504776 | 505966 | 1190 | + | tuf | WP_010951775.1 |

| | | | | | | | |
|---|---|---|---|---|---|---|---|
| GCF_022592395.1 | NZ_CP093242.1 | 3537145 | 3538335 | 1190 | + | tuf | WP_092277391.1 |
| GCF_025985205.1 | NZ_CP087840.1 | 3553716 | 3554906 | 1190 | + | tuf | WP_008145495.1 |
| GCF_025985205.1 | NZ_CP087840.1 | 6156032 | 6157222 | 1190 | - | tuf | WP_015637231.1 |
| GCF_000988745.2 | NZ_CP011381.2 | 6172629 | 6173819 | 1190 | - | tuf | WP_011063776.1 |
| GCF_000988745.2 | NZ_CP011381.2 | 1235949 | 1237139 | 1190 | + | tuf | WP_003444379.1 |
| GCF_034424565.1 | NZ_CP139961.1 | 5181390 | 5182580 | 1190 | - | tuf | WP_008145495.1 |
| GCF_034424565.1 | NZ_CP139961.1 | 5197961 | 5199151 | 1190 | - | tuf | WP_007896619.1 |
| GCF_002080125.1 | NZ_CP018059.1 | 558139 | 559329 | 1190 | + | tuf | WP_016497741.1 |
| GCF_002080125.1 | NZ_CP018059.1 | 574726 | 575916 | 1190 | + | tuf | WP_016497741.1 |
| GCF_900637105.1 | NZ_LR134343.1 | 5297066 | 5298256 | 1190 | - | tuf | WP_043861932.1 |
| GCF_900637105.1 | NZ_LR134343.1 | 5313668 | 5314858 | 1190 | - | tuf | WP_213606403.1 |
| GCF_030177935.1 | NZ_CP089974.1 | 372254 | 373444 | 1190 | + | tuf | WP_008374172.1 |
| GCF_030177935.1 | NZ_CP089974.1 | 3419767 | 3420957 | 1190 | + | tuf | WP_008065716.1 |
| GCF_037201985.1 | NZ_CP147511.1 | 5200961 | 5202151 | 1190 | - | tuf | WP_107322989.1 |
| GCF_037201985.1 | NZ_CP147511.1 | 4168838 | 4170028 | 1190 | + | tuf | WP_043192520.1 |
| GCF_029581575.1 | NZ_CP089975.1 | 5324444 | 5325634 | 1190 | - | tuf | WP_034139514.1 |
| GCF_029581575.1 | NZ_CP089975.1 | 2855876 | 2857066 | 1190 | - | tuf | WP_003176426.1 |
| GCF_025643275.1 | NZ_CP089977.1 | 5294358 | 5295548 | 1190 | - | tuf | WP_003369027.1 |
| GCF_025643275.1 | NZ_CP089977.1 | 1019803 | 1020993 | 1190 | + | tuf | WP_084331690.1 |
| GCF_022701215.1 | NZ_CP089976.1 | 1036346 | 1037536 | 1190 | + | tuf | WP_084331690.1 |
| GCF_022701215.1 | NZ_CP089976.1 | 2920803 | 2921993 | 1190 | - | tuf | WP_064361246.1 |
| GCF_016028775.1 | NZ_CP065728.1 | 2937359 | 2938549 | 1190 | - | tuf | WP_064361246.1 |
| GCF_016028775.1 | NZ_CP065728.1 | 4345383 | 4346573 | 1190 | + | tuf | WP_011063776.1 |
| GCF_001553955.1 | NZ_CP014234.1 | 4361978 | 4363168 | 1190 | + | tuf | WP_011063776.1 |
| GCF_001553955.1 | NZ_CP014234.1 | 5597528 | 5598718 | 1190 | - | tuf | WP_003176426.1 |
| GCF_001636015.1 | NZ_CP011158.1 | 526424 | 527614 | 1190 | + | tuf | WP_101322346.1 |
| GCF_001636015.1 | NZ_CP011158.1 | 543018 | 544208 | 1190 | + | tuf | WP_101322346.1 |
| GCF_044740715.1 | NZ_CP170124.1 | 4758985 | 4760175 | 1190 | - | tuf | WP_013970626.1 |
| GCF_044740715.1 | NZ_CP170124.1 | 4775582 | 4776772 | 1190 | - | tuf | WP_013970626.1 |
| GCF_006094455.1 | NZ_CP034944.1 | 536978 | 538168 | 1190 | + | tuf | WP_258644205.1 |
| GCF_006094455.1 | NZ_CP034944.1 | 553567 | 554757 | 1190 | + | tuf | WP_016497741.1 |

| | | | | | | | |
|---|---|---|---|---|---|---|---|
| GCF_008931805.1 | NZ_CP044399.1 | 3293546 | 3294736 | 1190 | - | tuf | WP_092377958.1 |
| GCF_900465055.1 | NZ_LS483250.1 | 3310088 | 3311278 | 1190 | - | tuf | WP_017675622.1 |
| GCF_000023545.1 | NC_012880.1 | 6142415 | 6143605 | 1190 | - | tuf | WP_007896660.1 |
| GCF_000023545.1 | NC_012880.1 | 5276086 | 5277276 | 1190 | - | tuf | WP_064361246.1 |
| GCF_030295765.1 | NZ_AP027292.1 | 5292644 | 5293834 | 1190 | - | tuf | WP_064361246.1 |
| GCF_030295765.1 | NZ_AP027292.1 | 5483833 | 5485023 | 1190 | - | tuf | WP_010206920.1 |
| GCF_965223595.1 | NZ_OZ245004.1 | 5637867 | 5639057 | 1190 | - | tuf | WP_003176426.1 |
| GCF_965223595.1 | NZ_OZ245004.1 | 775781 | 776971 | 1190 | + | tuf | WP_184492125.1 |
| GCF_008630635.1 | NZ_CP043869.1 | 792342 | 793532 | 1190 | + | tuf | WP_184492125.1 |
| GCF_008630635.1 | NZ_CP043869.1 | 501464 | 502654 | 1190 | + | tuf | WP_038707241.1 |
| GCF_013394205.1 | NZ_CP041336.1 | 518053 | 519243 | 1190 | + | tuf | WP_038707241.1 |
| GCF_013394205.1 | NZ_CP041336.1 | 5638337 | 5639527 | 1190 | - | tuf | WP_134423355.1 |
| GCF_008727795.1 | NZ_CP044222.1 | 5654908 | 5656098 | 1190 | - | tuf | WP_019693885.1 |
| GCF_008727795.1 | NZ_CP044222.1 | 5858327 | 5859517 | 1190 | - | tuf | WP_003176426.1 |
| GCF_000024725.1 | NC_013960.1 | 4983697 | 4984887 | 1190 | - | tuf | WP_003344210.1 |
| GCF_000024725.1 | NC_013960.1 | 5190747 | 5191937 | 1190 | - | tuf | WP_003369027.1 |
| GCF_000012805.1 | NC_007484.1 | 690156 | 691346 | 1190 | + | tuf | WP_007245455.1 |
| GCF_000012805.1 | NC_007484.1 | 530537 | 531727 | 1190 | + | tuf | WP_048378426.1 |
| GCF_004421105.1 | NZ_CP038033.1 | 6151853 | 6153043 | 1190 | - | tuf | WP_095122212.1 |
| GCF_004421105.1 | NZ_CP038033.1 | 6168409 | 6169599 | 1190 | - | tuf | WP_095122212.1 |
| GCF_000143085.1 | NC_014315.1 | 868206 | 869396 | 1190 | + | tuf | WP_175384238.1 |
| GCF_000143085.1 | NC_014315.1 | 884749 | 885939 | 1190 | + | tuf | WP_175384238.1 |
| GCF_000143085.1 | NC_014315.1 | 729282 | 730472 | 1190 | + | tuf | WP_024670793.1 |
| GCF_018406605.1 | NZ_AP024545.1 | 692064 | 693254 | 1190 | + | tuf | WP_065327628.1 |
| GCF_018406605.1 | NZ_AP024545.1 | 708571 | 709761 | 1190 | + | tuf | WP_065327628.1 |
| GCF_015209745.1 | NZ_CP063657.1 | 5789197 | 5790387 | 1190 | - | tuf | WP_003176426.1 |
| GCF_015209745.1 | NZ_CP063657.1 | 4832548 | 4833738 | 1190 | - | tuf | WP_057010371.1 |
| GCF_015209725.1 | NZ_CP063656.1 | 5404639 | 5405829 | 1190 | - | tuf | WP_130266518.1 |
| GCF_015209725.1 | NZ_CP063656.1 | 5421245 | 5422435 | 1190 | - | tuf | WP_130266525.1 |
| GCF_907164845.1 | NZ_OU015430.1 | 6213504 | 6214694 | 1190 | - | tuf | WP_008065716.1 |
| GCF_907164845.1 | NZ_OU015430.1 | 6124977 | 6126167 | 1190 | + | tuf | WP_093229673.1 |

| | | | | | | | |
|---|---|---|---|---|---|---|---|
| GCF_002117105.1 | NZ_CP019343.1 | 6527662 | 6528852 | 1190 | - | tuf | WP_026144793.1 |
| GCF_002117105.1 | NZ_CP019343.1 | 6189718 | 6190908 | 1190 | - | tuf | WP_017848801.1 |
| GCF_030849025.1 | NZ_CP118224.1 | 5050645 | 5051835 | 1190 | + | tuf | WP_003444379.1 |
| GCF_030849025.1 | NZ_CP118224.1 | 6016881 | 6018071 | 1190 | - | tuf | WP_003186071.1 |
| GCF_002157875.1 | NZ_CP021376.1 | 6033434 | 6034624 | 1190 | - | tuf | WP_003186103.1 |
| GCF_002157895.1 | NZ_CP021377.1 | 690944 | 692134 | 1190 | + | tuf | WP_025995600.1 |
| GCF_002162375.1 | NZ_CP021425.1 | 5492710 | 5493900 | 1190 | + | tuf | WP_100634181.1 |
| GCF_036251205.1 | NZ_CP133958.1 | 5509318 | 5510508 | 1190 | + | tuf | WP_105642145.1 |
| GCF_036251205.1 | NZ_CP133958.1 | 3940612 | 3941802 | 1190 | - | tuf | WP_147810603.1 |
| GCF_036251875.1 | NZ_CP133960.1 | 3957142 | 3958332 | 1190 | - | tuf | WP_230925669.1 |
| GCF_036251875.1 | NZ_CP133960.1 | 693863 | 695053 | 1190 | + | tuf | WP_007896619.1 |
| GCF_036251935.1 | NZ_CP133967.1 | 710434 | 711624 | 1190 | + | tuf | WP_305424907.1 |
| GCF_036251935.1 | NZ_CP133967.1 | 530209 | 531399 | 1190 | + | tuf | WP_182323863.1 |
| GCF_009684715.1 | NZ_CP016604.1 | 546799 | 547989 | 1190 | + | tuf | WP_182323863.1 |
| GCF_009684715.1 | NZ_CP016604.1 | 1295405 | 1296595 | 1190 | + | tuf | WP_010951775.1 |
| GCF_019048385.1 | NZ_CP077366.1 | 1311999 | 1313189 | 1190 | + | tuf | WP_010951775.1 |
| GCF_019048385.1 | NZ_CP077366.1 | 4133383 | 4134573 | 1190 | + | tuf | WP_003210085.1 |
| GCF_019880205.1 | NZ_CP082292.1 | 6145082 | 6146272 | 1190 | - | tuf | WP_064361246.1 |
| GCF_019880205.1 | NZ_CP082292.1 | 6161640 | 6162830 | 1190 | - | tuf | WP_064361246.1 |
| GCF_002101395.1 | NZ_CP019706.1 | 547564 | 548751 | 1187 | - | tuf | WP_162410744.1 |
| GCF_002101395.1 | NZ_CP019706.1 | 564801 | 565988 | 1187 | - | tuf | WP_162410744.1 |
| GCF_046529425.1 | NZ_CP125958.1 | 3059923 | 3061110 | 1187 | - | tuf | WP_162338029.1 |
| GCF_046529425.1 | NZ_CP125958.1 | 3077372 | 3078559 | 1187 | - | tuf | WP_213600565.1 |
| GCF_000233595.1 | NC_017554.1 | 1835460 | 1836647 | 1187 | + | tuf | WP_162109794.1 |
| GCF_000233595.1 | NC_017554.1 | 1852765 | 1853952 | 1187 | + | tuf | WP_162109794.1 |
| GCF_026104315.1 | NZ_CP110473.1 | 2585283 | 2586470 | 1187 | - | tuf | WP_014161384.1 |
| GCF_026104315.1 | NZ_CP110473.1 | 2602881 | 2604068 | 1187 | - | tuf | WP_014161396.1 |
| GCF_022647325.1 | NZ_CP071405.1 | 1203152 | 1204339 | 1187 | + | tuf | WP_130515940.1 |
| GCF_022647325.1 | NZ_CP071405.1 | 1220541 | 1221728 | 1187 | + | tuf | WP_130515940.1 |
| GCF_019890955.1 | NZ_CP082346.1 | 588117 | 589304 | 1187 | + | tuf | WP_062843867.1 |
| GCF_019890955.1 | NZ_CP082346.1 | 2955372 | 2956559 | 1187 | - | tuf | WP_062843867.1 |

| | | | | | | | |
|---|---|---|---|---|---|---|---|
| GCF_009646115.1 | NZ_CP045720.1 | 479294 | 480481 | 1187 | + | tuf | WP_011279689.1 |
| GCF_009646115.1 | NZ_CP045720.1 | 2338445 | 2339632 | 1187 | - | tuf | WP_011279689.1 |
| GCF_020079945.1 | NZ_CP083448.1 | 1364480 | 1365667 | 1187 | - | tuf | WP_201497832.1 |
| GCF_020079945.1 | NZ_CP083448.1 | 2525360 | 2526547 | 1187 | + | tuf | WP_201497832.1 |
| GCF_046118715.1 | NZ_CP173698.1 | 519410 | 520597 | 1187 | + | tuf | WP_011512793.1 |
| GCF_046118715.1 | NZ_CP173698.1 | 2683943 | 2685130 | 1187 | - | tuf | WP_011512793.1 |
| GCF_011044475.1 | NZ_CP049115.1 | 409863 | 411050 | 1187 | + | tuf | WP_186473561.1 |
| GCF_011044475.1 | NZ_CP049115.1 | 2457935 | 2459122 | 1187 | - | tuf | WP_071002260.1 |
| GCF_004792415.1 | NZ_CP038853.1 | 2414237 | 2415424 | 1187 | - | tuf | WP_011961077.1 |
| GCF_004792415.1 | NZ_CP038853.1 | 2663475 | 2664662 | 1187 | - | tuf | WP_011961077.1 |
| GCF_013285385.1 | NZ_CP054212.1 | 4249457 | 4250638 | 1181 | - | tuf | WP_011771673.1 |
| GCF_013285385.1 | NZ_CP054212.1 | 70566 | 71747 | 1181 | - | tuf | WP_210560552.1 |
| GCF_002216855.1 | NZ_CP022355.1 | 92025 | 93206 | 1181 | - | tuf | WP_210560552.1 |
| GCF_002216855.1 | NZ_CP022355.1 | 1895931 | 1897112 | 1181 | - | tuf | WP_208830321.1 |
| GCF_004358345.1 | NZ_CP037952.1 | 1917511 | 1918692 | 1181 | - | tuf | WP_208830321.1 |
| GCF_004358345.1 | NZ_CP037952.1 | 3253209 | 3254390 | 1181 | - | tuf | WP_013573670.1 |
| GCF_004358445.1 | NZ_CP037951.1 | 3956385 | 3957566 | 1181 | + | tuf | WP_013573670.1 |
| GCF_004358445.1 | NZ_CP037951.1 | 368834 | 370015 | 1181 | + | tuf | WP_013573670.1 |
| GCF_963693435.1 | NZ_OY856306.1 | 4524735 | 4525916 | 1181 | - | tuf | WP_013573670.1 |
| GCF_963693435.1 | NZ_OY856306.1 | 83680 | 84861 | 1181 | - | tuf | WP_213051554.1 |
| GCF_020810675.1 | NZ_CP085871.1 | 770525 | 771706 | 1181 | + | tuf | WP_213051554.1 |
| GCF_020810675.1 | NZ_CP085871.1 | 3250546 | 3251727 | 1181 | - | tuf | WP_104923588.1 |
| GCF_900186835.1 | NZ_LT906448.1 | 3968082 | 3969263 | 1181 | + | tuf | WP_104923588.1 |
| GCF_900186835.1 | NZ_LT906448.1 | 1336553 | 1337734 | 1181 | - | tuf | WP_095924283.1 |
| GCF_002073255.2 | NZ_CP020405.2 | 2034294 | 2035475 | 1181 | + | tuf | WP_095923767.1 |
| GCF_002073255.2 | NZ_CP020405.2 | 455994 | 457175 | 1181 | + | tuf | WP_004118409.1 |
| GCF_003382565.3 | NZ_CP086253.1 | 5292264 | 5293445 | 1181 | - | tuf | WP_004118409.1 |
| GCF_003382565.3 | NZ_CP086253.1 | 1259270 | 1260451 | 1181 | - | tuf | WP_004118409.1 |
| GCF_037076465.1 | NZ_AP028908.1 | 1984846 | 1986027 | 1181 | + | tuf | WP_047663733.1 |
| GCF_037076465.1 | NZ_AP028908.1 | 419534 | 420715 | 1181 | + | tuf | WP_045854065.1 |
| GCF_015689195.1 | NZ_CP065044.1 | 5347316 | 5348497 | 1181 | - | tuf | WP_041142972.1 |

| | | | | | | | |
|---|---|---|---|---|---|---|---|
| GCF_015689195.1 | NZ_CP065044.1 | 3413884 | 3415074 | 1190 | - | tuf | WP_100258443.1 |
| GCF_000740965.1 | NZ_CP009125.1 | 298537 | 299718 | 1181 | + | tuf | WP_127019486.1 |
| GCF_000740965.1 | NZ_CP009125.1 | 315243 | 316424 | 1181 | + | tuf | WP_127019503.1 |
| GCF_009873295.1 | NZ_CP047495.1 | 1074440 | 1075627 | 1187 | + | tuf | WP_007514312.1 |
| GCF_009873295.1 | NZ_CP047495.1 | 1091740 | 1092927 | 1187 | + | tuf | WP_007514312.1 |
| GCF_036885195.1 | NZ_CP133656.1 | 1578784 | 1579983 | 1199 | + | tuf | WP_342147784.1 |
| GCF_036885195.1 | NZ_CP133656.1 | 432575 | 433774 | 1199 | + | tuf | WP_218814451.1 |
| GCF_013488025.1 | NZ_CP051652.1 | 1080678 | 1081877 | 1199 | - | tuf | WP_071662576.1 |
| GCF_013488025.1 | NZ_CP051652.1 | 1194346 | 1195545 | 1199 | - | tuf | WP_342219684.1 |
| GCF_020181655.1 | NZ_CP084032.1 | 1208570 | 1209769 | 1199 | - | tuf | WP_342227282.1 |
| GCF_020181655.1 | NZ_CP084032.1 | 1255050 | 1256249 | 1199 | - | tuf | WP_339049903.1 |
| GCF_009931295.1 | NZ_CP034938.1 | 1496683 | 1497864 | 1181 | - | tuf | WP_059368590.1 |
| GCF_009931295.1 | NZ_CP034938.1 | 1742292 | 1743473 | 1181 | + | tuf | WP_018356115.1 |
| GCF_001742145.1 | NZ_CP015749.1 | 1234922 | 1236103 | 1181 | - | tuf | WP_018356115.1 |
| GCF_001742145.1 | NZ_CP015749.1 | 1249000 | 1250181 | 1181 | - | tuf | WP_018356115.1 |
| GCF_020971565.1 | NZ_CP087392.1 | 3774366 | 3775547 | 1181 | - | tuf | WP_192504208.1 |
| GCF_020971565.1 | NZ_CP087392.1 | 4385242 | 4386423 | 1181 | + | tuf | WP_192504208.1 |
| GCF_003312355.2 | NZ_CP161826.1 | 2791739 | 2792920 | 1181 | + | tuf | WP_108602772.1 |
| GCF_003312355.2 | NZ_CP161826.1 | 2808700 | 2809881 | 1181 | + | tuf | WP_108602772.1 |
| GCF_002288545.1 | NZ_CP017482.1 | 1199589 | 1200809 | 1220 | + | tuf | WP_011467412.1 |
| GCF_002288545.1 | NZ_CP017482.1 | 1203474 | 1204694 | 1220 | + | tuf | WP_011467412.1 |
| GCF_012427845.1 | NZ_CP038498.1 | 524426 | 525616 | 1190 | + | tuf | WP_108127778.1 |
| GCF_012427845.1 | NZ_CP038498.1 | 541283 | 542473 | 1190 | + | tuf | WP_108127778.1 |
| GCF_014946775.2 | NZ_CP065177.1 | 450935 | 452116 | 1181 | + | tuf | WP_139755069.1 |
| GCF_014946775.2 | NZ_CP065177.1 | 585638 | 586819 | 1181 | + | tuf | WP_139755069.1 |
| GCF_003932035.1 | NZ_CP034276.1 | 533260 | 534441 | 1181 | + | tuf | WP_137165714.1 |
| GCF_003932035.1 | NZ_CP034276.1 | 3545088 | 3546269 | 1181 | - | tuf | WP_137168108.1 |
| GCF_001742185.1 | NZ_CP015750.1 | 311897 | 313078 | 1181 | + | tuf | WP_195811017.1 |
| GCF_001742185.1 | NZ_CP015750.1 | 443929 | 445110 | 1181 | + | tuf | WP_195811017.1 |
| GCF_013377295.1 | NZ_CP016180.1 | 3769136 | 3770317 | 1181 | - | tuf | WP_117318059.1 |
| GCF_013377295.1 | NZ_CP016180.1 | 3952352 | 3953533 | 1181 | - | tuf | WP_117318200.1 |

| | | | | | | | |
|---|---|---|---|---|---|---|---|
| GCF_024380015.1 | NZ_CP101508.1 | 406412 | 407593 | 1181 | + | tuf | WP_109338602.1 |
| GCF_024380015.1 | NZ_CP101508.1 | 424175 | 425356 | 1181 | + | tuf | WP_109338602.1 |
| GCF_038086725.1 | NZ_CP146970.1 | 289815 | 290996 | 1181 | + | tuf | WP_269587314.1 |
| GCF_038086725.1 | NZ_CP146970.1 | 2675734 | 2676915 | 1181 | - | tuf | WP_269587150.1 |
| GCF_017329545.1 | NZ_CP071325.1 | 291694 | 292875 | 1181 | + | tuf | WP_096630667.1 |
| GCF_017329545.1 | NZ_CP071325.1 | 2828373 | 2829554 | 1181 | - | tuf | WP_096630185.1 |
| GCF_048537465.1 | NZ_CP183250.1 | 3329653 | 3330834 | 1181 | - | tuf | WP_000031748.1 |
| GCF_048537465.1 | NZ_CP183250.1 | 4007751 | 4008932 | 1181 | + | tuf | WP_000031748.1 |
| GCF_046058925.1 | NZ_CP175534.1 | 3598312 | 3599493 | 1181 | - | tufA | NP_462349.1 |
| GCF_046058925.1 | NZ_CP175534.1 | 4360616 | 4361797 | 1181 | + | tufB | NP_463015.1 |
| GCF_024346675.1 | NZ_AP024850.1 | 369505 | 370686 | 1181 | + | tuf | WP_174081212.1 |
| GCF_024346675.1 | NZ_AP024850.1 | 4326212 | 4327393 | 1181 | - | tuf | WP_174081212.1 |
| GCF_024346715.1 | NZ_AP024852.1 | 589770 | 590951 | 1181 | - | tuf | WP_014888098.1 |
| GCF_024346715.1 | NZ_AP024852.1 | 798728 | 799909 | 1181 | + | tuf | WP_014889039.1 |
| GCF_024529955.1 | NZ_AP024854.1 | 280803 | 281984 | 1181 | - | tuf | WP_083172596.1 |
| GCF_024529955.1 | NZ_AP024854.1 | 3135163 | 3136350 | 1187 | + | tuf | WP_046860283.1 |
| GCF_019090985.1 | NZ_CP020335.1 | 3151966 | 3153153 | 1187 | + | tuf | WP_046860295.1 |
| GCF_019090985.1 | NZ_CP020335.1 | 372973 | 374154 | 1181 | + | tuf | WP_380676595.1 |
| GCF_000196475.1 | NC_012962.1 | 4979545 | 4980726 | 1181 | - | tuf | WP_380676818.1 |
| GCF_000196475.1 | NC_012962.1 | 228598 | 229779 | 1181 | + | tuf | WP_234587542.1 |
| GCF_003343245.1 | NZ_CP024901.1 | 4581497 | 4582678 | 1181 | - | tuf | WP_234586705.1 |
| GCF_003343245.1 | NZ_CP024901.1 | 275633 | 276814 | 1181 | + | tuf | WP_095095107.1 |
| GCF_001010285.1 | NZ_CP011104.1 | 4812665 | 4813846 | 1181 | - | tuf | WP_095099556.1 |
| GCF_001010285.1 | NZ_CP011104.1 | 1321190 | 1322371 | 1181 | - | tuf | WP_046808028.1 |
| GCF_001022135.1 | NZ_CP011602.1 | 2222512 | 2223693 | 1181 | + | tuf | WP_046808053.1 |
| GCF_001022135.1 | NZ_CP011602.1 | 4825749 | 4826930 | 1181 | - | tuf | WP_020828747.1 |
| GCF_009708955.1 | NZ_CP039032.1 | 33676 | 34857 | 1181 | - | tuf | WP_020828747.1 |
| GCF_007954585.1 | NZ_CP042382.1 | 261607 | 262788 | 1181 | + | tuf | WP_015376315.1 |
| GCF_031198435.1 | NZ_CP133548.1 | 4627104 | 4628285 | 1181 | - | tuf | WP_015376315.1 |
| GCF_031198435.1 | NZ_CP133548.1 | 245120 | 246301 | 1181 | + | tuf | WP_015379267.1 |
| GCF_020991025.1 | NZ_CP087711.1 | 4848367 | 4849548 | 1181 | - | tuf | WP_015379267.1 |

| | | | | | | | |
|---|---|---|---|---|---|---|---|
| GCF_020991025.1 | NZ_CP087711.1 | 245088 | 246269 | 1181 | + | tuf | WP_015376315.1 |
| GCF_003019925.1 | NZ_CP020388.1 | 4939903 | 4941084 | 1181 | - | tuf | WP_015376315.1 |
| GCF_003019925.1 | NZ_CP020388.1 | 263115 | 264296 | 1181 | + | tuf | WP_013811124.1 |
| GCF_046844695.1 | NZ_CP177354.1 | 5015146 | 5016327 | 1181 | - | tuf | WP_013811124.1 |
| GCF_900638655.1 | NZ_LR134531.1 | 283400 | 284581 | 1181 | + | tuf | WP_208718403.1 |
| GCF_900638655.1 | NZ_LR134531.1 | 4983467 | 4984648 | 1181 | - | tuf | WP_208718396.1 |
| GCF_000463345.2 | NZ_CP025084.1 | 279791 | 280972 | 1181 | + | tuf | WP_126501225.1 |
| GCF_000463345.2 | NZ_CP025084.1 | 4943102 | 4944283 | 1181 | - | tuf | WP_112363683.1 |
| GCF_030271835.1 | NZ_CP127389.1 | 1814880 | 1816061 | 1181 | - | tuf | WP_105229426.1 |
| GCF_030271835.1 | NZ_CP127389.1 | 2486126 | 2487307 | 1181 | + | tuf | WP_105229426.1 |
| GCF_004116975.1 | NZ_CP026364.1 | 1121377 | 1122558 | 1181 | - | tuf | WP_015344220.1 |
| GCF_004116975.1 | NZ_CP026364.1 | 1817156 | 1818337 | 1181 | + | tuf | WP_015344220.1 |
| GCF_000069965.1 | NC_010554.1 | 253693 | 254874 | 1181 | + | tuf | WP_015376315.1 |
| GCF_000069965.1 | NC_010554.1 | 4646581 | 4647762 | 1181 | - | tuf | WP_015376315.1 |
| GCF_022369495.1 | NZ_CP059690.1 | 4076466 | 4077647 | 1181 | - | tuf | WP_055314187.1 |
| GCF_022369495.1 | NZ_CP059690.1 | 4730660 | 4731841 | 1181 | + | tuf | WP_015376315.1 |
| GCF_011045835.1 | NZ_CP047349.1 | 2938244 | 2939425 | 1181 | - | tuf | WP_040263589.1 |
| GCF_011045835.1 | NZ_CP047349.1 | 3282226 | 3283407 | 1181 | + | tuf | WP_040264637.1 |
| GCF_025200655.1 | NZ_CP104121.1 | 333237 | 334418 | 1181 | + | tuf | WP_015376315.1 |
| GCF_025200655.1 | NZ_CP104121.1 | 4786958 | 4788139 | 1181 | - | tuf | WP_015376315.1 |
| GCF_029193595.2 | NZ_CP135052.1 | 153255 | 154436 | 1181 | + | tuf | WP_220050889.1 |
| GCF_029193595.2 | NZ_CP135052.1 | 170038 | 171219 | 1181 | + | tuf | WP_220050889.1 |
| GCF_900475855.1 | NZ_LS483422.1 | 160206 | 161387 | 1181 | + | tuf | WP_011863965.1 |
| GCF_900475855.1 | NZ_LS483422.1 | 176979 | 178160 | 1181 | + | tuf | WP_011863965.1 |
| GCF_036898185.1 | NZ_CP145931.1 | 244981 | 246162 | 1181 | + | tuf | WP_071239068.1 |
| GCF_036898185.1 | NZ_CP145931.1 | 261724 | 262905 | 1181 | + | tuf | WP_071239068.1 |
| GCF_016618195.1 | NZ_CP067099.1 | 161048 | 162229 | 1181 | + | tuf | WP_011863965.1 |
| GCF_016618195.1 | NZ_CP067099.1 | 177821 | 179002 | 1181 | + | tuf | WP_011863965.1 |
| GCF_019048105.1 | NZ_CP077317.1 | 233618 | 234799 | 1181 | + | tuf | WP_011758321.1 |
| GCF_019048105.1 | NZ_CP077317.1 | 250344 | 251525 | 1181 | + | tuf | WP_011758321.1 |
| GCF_900637755.1 | NZ_LR134396.1 | 3939138 | 3940319 | 1181 | - | tuf | WP_207354695.1 |

| | | | | | | |
|---|---|---|---|---|---|---|
| GCF_900637755.1 | NZ_LR134396.1 | 3955868 | 3957049 | 1181 - | tuf | WP_207354695.1 |
| GCF_000314895.2 | NZ_CM001773.1 | 229794 | 230975 | 1181 + | tuf | WP_006087508.1 |
| GCF_035747985.1 | NZ_CP142095.1 | 246558 | 247739 | 1181 + | tuf | WP_006083602.1 |
| GCF_035747985.1 | NZ_CP142095.1 | 205672 | 206853 | 1181 + | tuf | WP_221050213.1 |
| GCF_041953405.1 | NZ_CP169565.1 | 222413 | 223594 | 1181 + | tuf | WP_221050213.1 |
| GCF_041953405.1 | NZ_CP169565.1 | 208318 | 209499 | 1181 + | tuf | WP_165564289.1 |
| GCF_041953975.1 | NZ_CP169566.1 | 4378314 | 4379495 | 1181 - | tuf | WP_165564289.1 |
| GCF_041953975.1 | NZ_CP169566.1 | 222369 | 223550 | 1181 + | tuf | WP_207321607.1 |
| GCF_030315915.2 | NZ_CP135990.1 | 239113 | 240294 | 1181 + | tuf | WP_207321619.1 |
| GCF_030315915.2 | NZ_CP135990.1 | 4416038 | 4417219 | 1181 - | tuf | WP_133182414.1 |
| GCF_902164725.1 | NZ_LR607338.1 | 194785 | 195966 | 1181 + | tuf | WP_011494634.1 |
| GCF_001999225.1 | NZ_CP019628.1 | 3397057 | 3398238 | 1181 + | tuf | WP_144211908.1 |
| GCF_001999225.1 | NZ_CP019628.1 | 3413804 | 3414985 | 1181 + | tuf | WP_144211908.1 |
| GCF_030297135.1 | NZ_AP028065.1 | 5591200 | 5592381 | 1181 - | tuf | WP_142873441.1 |
| GCF_030297135.1 | NZ_AP028065.1 | 5607969 | 5609150 | 1181 - | tuf | WP_142873441.1 |
| GCF_000238395.3 | NZ_CP011025.1 | 152246 | 153427 | 1181 + | tuf | WP_011635626.1 |
| GCF_000238395.3 | NZ_CP011025.1 | 169019 | 170200 | 1181 + | tuf | WP_011635638.1 |
| GCF_900239935.1 | NZ_LT965928.1 | 4283078 | 4284259 | 1181 - | tuf | WP_226411993.1 |
| GCF_900239935.1 | NZ_LT965928.1 | 4299857 | 4301038 | 1181 - | tuf | WP_226411993.1 |
| GCF_005877035.1 | NZ_CP040558.1 | 195225 | 196406 | 1181 + | tuf | WP_220755189.1 |
| GCF_005877035.1 | NZ_CP040558.1 | 211956 | 213137 | 1181 + | tuf | WP_220755197.1 |
| GCF_002221525.1 | NZ_CP011028.1 | 5025443 | 5026624 | 1181 - | tuf | WP_012279183.1 |
| GCF_002221525.1 | NZ_CP011028.1 | 5042231 | 5043412 | 1181 - | tuf | WP_012279183.1 |
| GCF_002310795.1 | NZ_CP011030.1 | 159954 | 161135 | 1181 + | tuf | WP_011863953.1 |
| GCF_002310795.1 | NZ_CP011030.1 | 176727 | 177908 | 1181 + | tuf | WP_011863953.1 |
| GCF_003515105.1 | NZ_CP032090.1 | 618942 | 620123 | 1181 + | tuf | WP_188740705.1 |
| GCF_003515105.1 | NZ_CP032090.1 | 635724 | 636905 | 1181 + | tuf | WP_188740705.1 |
| GCF_006704145.1 | NZ_VIGJ01000001.1 | 4747315 | 4748496 | 1181 - | tuf | WP_080917157.1 |
| GCF_006704145.1 | NZ_VIGJ01000001.1 | 4764038 | 4765219 | 1181 - | tuf | WP_080917157.1 |
| GCF_024746855.1 | NZ_CP024625.1 | 4392757 | 4393938 | 1181 + | tuf | WP_126169070.1 |
| GCF_024746855.1 | NZ_CP024625.1 | 4409483 | 4410664 | 1181 + | tuf | WP_126169070.1 |

| | | | | | | | |
|---|---|---|---|---|---|---|---|
| GCF_002221505.1 | NZ_CP011036.1 | 203777 | 204958 | 1181 | + | tuf | WP_011758321.1 |
| GCF_002221505.1 | NZ_CP011036.1 | 220502 | 221683 | 1181 | + | tuf | WP_011758321.1 |
| GCF_029023665.1 | NZ_CP118916.1 | 404450 | 405631 | 1181 | + | tuf | WP_124729261.1 |
| GCF_029023665.1 | NZ_CP118916.1 | 421218 | 422399 | 1181 | + | tuf | WP_124729272.1 |
| GCF_001444405.1 | NZ_CP013187.1 | 160907 | 162088 | 1181 | + | tuf | WP_011863953.1 |
| GCF_001444405.1 | NZ_CP013187.1 | 177680 | 178861 | 1181 | + | tuf | WP_011863965.1 |
| GCF_000788395.1 | NZ_CP009888.1 | 277646 | 278827 | 1181 | + | tuf | WP_011715454.1 |
| GCF_000788395.1 | NZ_CP009888.1 | 294432 | 295613 | 1181 | + | tuf | WP_011715454.1 |
| GCF_019797925.1 | NZ_CP081860.1 | 149617 | 150798 | 1181 | + | tuf | WP_088903540.1 |
| GCF_019797925.1 | NZ_CP081860.1 | 166385 | 167566 | 1181 | + | tuf | WP_088903540.1 |
| GCF_014925355.1 | NZ_AQHH01000004.1 | 2595936 | 2597117 | 1181 | + | tuf | WP_130600035.1 |
| GCF_014925355.1 | NZ_AQHH01000009.1 | 2612677 | 2613858 | 1181 | + | tuf | WP_130600035.1 |
| GCF_014925355.1 | NZ_AQHH01000014.1 | 139630 | 140811 | 1181 | + | tuf | WP_220062677.1 |
| GCF_028885455.1 | NZ_CP118496.1 | 156417 | 157598 | 1181 | + | tuf | WP_220062677.1 |
| GCF_028885455.1 | NZ_CP118496.1 | 230308 | 231489 | 1181 | + | tuf | WP_105251028.1 |
| GCF_000238255.3 | NZ_CP011039.1 | 247068 | 248249 | 1181 | + | tuf | WP_105251028.1 |
| GCF_000238255.3 | NZ_CP011039.1 | 227280 | 228461 | 1181 | + | tuf | WP_011070604.1 |
| GCF_000026085.1 | NC_007481.1 | 244057 | 245238 | 1181 | + | tuf | WP_011070615.1 |
| GCF_000026085.1 | NC_007481.1 | 192087 | 193268 | 1181 | + | tuf | WP_012153445.1 |
| GCF_003568825.1 | NZ_CP031961.1 | 208877 | 210058 | 1181 | + | tuf | WP_012153445.1 |
| GCF_003568825.1 | NZ_CP031961.1 | 2072970 | 2074151 | 1181 | + | tuf | WP_020912125.1 |
| GCF_000238275.3 | NZ_AHCF03000002.1 | 2089774 | 2090955 | 1181 | + | tuf | WP_020912136.1 |
| GCF_000238275.3 | NZ_AHCF03000003.1 | 4439863 | 4441044 | 1181 | - | tuf | WP_137224523.1 |
| GCF_017742995.1 | NZ_CP072425.1 | 4456633 | 4457814 | 1181 | - | tuf | WP_137224523.1 |
| GCF_017742995.1 | NZ_CP072425.1 | 1324698 | 1325879 | 1181 | + | tuf | WP_101033683.1 |
| GCF_021172065.1 | NZ_CP087880.1 | 1341465 | 1342646 | 1181 | + | tuf | WP_101033683.1 |
| GCF_021172065.1 | NZ_CP087880.1 | 6132153 | 6133334 | 1181 | - | tuf | WP_144048512.1 |
| GCF_001444425.1 | NZ_CP013189.1 | 6148913 | 6150094 | 1181 | - | tuf | WP_144048512.1 |
| GCF_001444425.1 | NZ_CP013189.1 | 147036 | 148217 | 1181 | + | tuf | WP_220069694.1 |
| GCF_004168365.1 | NZ_CP035704.1 | 163825 | 165006 | 1181 | + | tuf | WP_220069694.1 |
| GCF_004168365.1 | NZ_CP035704.1 | 175182 | 176363 | 1181 | + | tuf | WP_011787562.1 |

| | | | | | | | |
|---|---|---|---|---|---|---|---|
| GCF_023509015.1 | NZ_CP094975.1 | 191952 | 193133 | 1181 | + | tuf | WP_011787562.1 |
| GCF_000006765.1 | NC_002516.2 | 160941 | 162122 | 1181 | + | tuf | WP_011863965.1 |
| GCF_000006765.1 | NC_002516.2 | 177714 | 178895 | 1181 | + | tuf | WP_011863965.1 |
| GCF_039908605.1 | NZ_CP150916.1 | 229673 | 230854 | 1181 | + | tuf | WP_207380691.1 |
| GCF_039908605.1 | NZ_CP150916.1 | 246420 | 247601 | 1181 | + | tuf | WP_207380691.1 |
| GCF_009755645.1 | NZ_CP046621.1 | 160804 | 161985 | 1181 | + | tuf | WP_011863953.1 |
| GCF_009755645.1 | NZ_CP046621.1 | 177577 | 178758 | 1181 | + | tuf | WP_011863953.1 |
| GCF_014863605.1 | NZ_CP062252.1 | 4755387 | 4756568 | 1181 | - | tuf | WP_013053420.1 |
| GCF_014863605.1 | NZ_CP062252.1 | 4772150 | 4773331 | 1181 | - | tuf | WP_013053420.1 |
| GCF_021283545.2 | NZ_CP128540.1 | 5656905 | 5658086 | 1181 | - | tuf | WP_012327261.1 |
| GCF_021283545.2 | NZ_CP128540.1 | 5673656 | 5674837 | 1181 | - | tuf | WP_012327272.1 |
| GCF_019141525.1 | NZ_CP077080.1 | 4807435 | 4808616 | 1181 | - | tuf | WP_055647561.1 |
| GCF_019141525.1 | NZ_CP077080.1 | 4824207 | 4825388 | 1181 | - | tuf | WP_055647561.1 |
| GCF_000145945.2 | NZ_CP042804.1 | 4207570 | 4208751 | 1181 | - | tuf | WP_310472776.1 |
| GCF_900103795.1 | NZ_LT629704.1 | 4224365 | 4225546 | 1181 | - | tuf | WP_310472776.1 |
| GCF_014269225.2 | NZ_CP077097.1 | 183144 | 184325 | 1181 | + | tuf | WP_212595092.1 |
| GCF_014269225.2 | NZ_CP077097.1 | 199852 | 201033 | 1181 | + | tuf | WP_212595092.1 |
| GCF_900103875.1 | NZ_LT629705.1 | 224873 | 226054 | 1181 | + | tuf | WP_011758321.1 |
| GCF_019139815.1 | NZ_CP077079.1 | 241599 | 242780 | 1181 | + | tuf | WP_011758321.1 |
| GCF_034358925.1 | NZ_CP139776.1 | 104945 | 106126 | 1181 | + | tuf | WP_000031784.1 |
| GCF_034358925.1 | NZ_CP139776.1 | 4354174 | 4355355 | 1181 | - | tuf | WP_000031783.1 |
| GCF_900105475.1 | NZ_LT629777.1 | 3439425 | 3440606 | 1181 | - | tuf | NP_709113.2 |
| GCF_017167965.1 | NZ_CP070503.1 | 4195696 | 4196877 | 1181 | + | tuf | NP_709775.1 |
| GCF_017167965.1 | NZ_CP070503.1 | 3482553 | 3483734 | 1181 | - | tuf | WP_000031784.1 |
| GCF_019139795.1 | NZ_CP077078.1 | 4261981 | 4263162 | 1181 | + | tuf | WP_000031783.1 |
| GCF_900103345.1 | NZ_LT629702.1 | 306563 | 307744 | 1181 | + | tuf | WP_014716237.1 |
| GCF_036585405.1 | NZ_CP144470.1 | 3862876 | 3864057 | 1181 | - | tuf | WP_014716237.1 |
| GCF_036585405.1 | NZ_CP144470.1 | 346094 | 347275 | 1181 | + | tuf | WP_264384974.1 |
| GCF_030687875.1 | NZ_CP117451.1 | 3785409 | 3786590 | 1181 | - | tuf | WP_264384974.1 |
| GCF_030687875.1 | NZ_CP117451.1 | 543296 | 544489 | 1193 | + | tuf | WP_390593289.1 |
| GCF_033547155.1 | NZ_CP137892.1 | 560107 | 561300 | 1193 | + | tuf | WP_390593289.1 |

| | | | | | | | |
|---|---|---|---|---|---|---|---|
| GCF_033547155.1 | NZ_CP137892.1 | 133013 | 134194 | 1181 | - | tuf | WP_097032089.1 |
| GCF_036884265.1 | NZ_CP145723.1 | 212377 | 213558 | 1181 | + | tuf | WP_011409992.1 |
| GCF_036884265.1 | NZ_CP145723.1 | 3935041 | 3936222 | 1181 | - | tuf | WP_011409992.1 |
| GCF_014109765.1 | NZ_CP059139.1 | 108196 | 109377 | 1181 | + | tuf | WP_367670683.1 |
| GCF_014109765.1 | NZ_CP059139.1 | 312797 | 313984 | 1187 | + | tuf | WP_112925754.1 |
| GCF_023330585.1 | NZ_CP097108.1 | 330176 | 331363 | 1187 | + | tuf | WP_112925754.1 |
| GCF_023330585.1 | NZ_CP097108.1 | 5644099 | 5645289 | 1190 | - | tuf | WP_163835727.1 |
| GCF_008370715.1 | NZ_CP034725.1 | 5660687 | 5661877 | 1190 | - | tuf | WP_163836601.1 |
| GCF_008370715.1 | NZ_CP034725.1 | 3151653 | 3152840 | 1187 | - | tuf | WP_109679292.1 |
| GCF_037023725.1 | NZ_CP146077.1 | 3168700 | 3169887 | 1187 | - | tuf | WP_109679292.1 |
| GCF_030980095.1 | NZ_CP133298.1 | 1522215 | 1523402 | 1187 | - | tuf | WP_144347953.1 |
| GCF_016599635.1 | NZ_CP067022.1 | 1538590 | 1539777 | 1187 | - | tuf | WP_148122436.1 |
| GCF_003595175.1 | NZ_CP032419.1 | 1522946 | 1524133 | 1187 | - | tuf | WP_016353898.1 |
| GCF_003595175.1 | NZ_CP032419.1 | 3133263 | 3134483 | 1220 | - | tuf | WP_198569037.1 |
| GCF_900104915.1 | NZ_LT629753.1 | 3149919 | 3151139 | 1220 | - | tuf | WP_198569037.1 |
| GCF_036689615.1 | NZ_CP144767.1 | 1635304 | 1636524 | 1220 | - | tuf | WP_251089157.1 |
| GCF_036689615.1 | NZ_CP144767.1 | 1651978 | 1653198 | 1220 | - | tuf | WP_251089157.1 |
| GCF_018343775.1 | NZ_CP074349.1 | 1640757 | 1641944 | 1187 | + | tuf | WP_182336326.1 |
| GCF_001586155.1 | NZ_CP014158.1 | 1658085 | 1659272 | 1187 | + | tuf | WP_054663274.1 |
| GCF_001586155.1 | NZ_CP014158.1 | 861042 | 862229 | 1187 | + | tuf | WP_311183892.1 |
| GCF_000156995.2 | NZ_CP046035.1 | 878278 | 879465 | 1187 | + | tuf | WP_311183892.1 |
| GCF_001708425.1 | NZ_CP014262.1 | 840128 | 841315 | 1187 | + | tuf | WP_004154360.1 |
| GCF_001708425.1 | NZ_CP014262.1 | 857554 | 858741 | 1187 | + | tuf | WP_004154360.1 |
| GCF_030687935.1 | NZ_CP117454.1 | 1059131 | 1060318 | 1187 | - | tuf | WP_014036096.1 |
| GCF_030687935.1 | NZ_CP117454.1 | 1076553 | 1077740 | 1187 | - | tuf | WP_014036096.1 |
| GCF_045013515.1 | NZ_CP120725.1 | 900916 | 902103 | 1187 | + | tuf | WP_032977440.1 |
| GCF_045013515.1 | NZ_CP120725.1 | 918391 | 919578 | 1187 | + | tuf | WP_032977440.1 |
| GCF_000696345.2 | NZ_CP071706.1 | 3596868 | 3598055 | 1187 | - | tuf | WP_089238985.1 |
| GCF_000696345.2 | NZ_CP071706.1 | 3614324 | 3615511 | 1187 | - | tuf | WP_089238985.1 |
| GCF_000026105.1 | NC_008027.1 | 897212 | 898399 | 1187 | + | tuf | WP_004154360.1 |
| GCF_000026105.1 | NC_008027.1 | 914734 | 915921 | 1187 | + | tuf | WP_004154360.1 |

| | | | | | | | |
|---|---|---|---|---|---|---|---|
| GCF_013374995.1 | NZ_CP056030.1 | 2600344 | 2601531 | 1187 | + | tuf | WP_311190285.1 |
| GCF_900102035.1 | NZ_LT629689.1 | 2617691 | 2618878 | 1187 | + | tuf | WP_311190285.1 |
| GCF_024584605.1 | NZ_CP089519.1 | 1170555 | 1171742 | 1187 | + | tuf | WP_303749021.1 |
| GCF_019139895.1 | NZ_CP077076.1 | 1188024 | 1189211 | 1187 | + | tuf | WP_303749021.1 |
| GCF_019139895.1 | NZ_CP077076.1 | 3619013 | 3620200 | 1187 | - | tuf | WP_038690301.1 |
| GCF_024584585.1 | NZ_CP085684.1 | 3636502 | 3637689 | 1187 | - | tuf | WP_038690301.1 |
| GCF_021018765.1 | NZ_CP075567.1 | 817132 | 818319 | 1187 | + | tuf | WP_004154360.1 |
| GCF_021018765.1 | NZ_CP075567.1 | 834593 | 835780 | 1187 | + | tuf | WP_049463838.1 |
| GCF_900215245.1 | NZ_LT907842.1 | 820942 | 822129 | 1187 | + | tuf | WP_065178958.1 |
| GCF_026898135.2 | NZ_CP114035.2 | 838440 | 839627 | 1187 | + | tuf | WP_065178958.1 |
| GCF_026898135.2 | NZ_CP114035.2 | 1179038 | 1180225 | 1187 | + | tuf | WP_066919279.1 |
| GCF_015476275.1 | NZ_CP064354.1 | 1196202 | 1197389 | 1187 | + | tuf | WP_066919279.1 |
| GCF_001874645.1 | NZ_CP017886.1 | 3731610 | 3732800 | 1190 | - | tuf | WP_043222062.1 |
| GCF_001874645.1 | NZ_CP017886.1 | 3748047 | 3749237 | 1190 | - | tuf | WP_043222066.1 |
| GCF_039634495.1 | NZ_CP132347.1 | 3682011 | 3683201 | 1190 | - | tuf | WP_014821633.1 |
| GCF_039634495.1 | NZ_CP132347.1 | 3698469 | 3699659 | 1190 | - | tuf | WP_014821633.1 |
| GCF_019614655.1 | NZ_CP071586.1 | 3846877 | 3848067 | 1190 | - | tuf | WP_038663376.1 |
| GCF_019614655.1 | NZ_CP071586.1 | 3863334 | 3864524 | 1190 | - | tuf | WP_038663376.1 |
| GCF_001594225.2 | NZ_CP014205.2 | 2516608 | 2517798 | 1190 | - | tuf | WP_165594100.1 |
| GCF_000508205.1 | NC_023064.1 | 2533056 | 2534246 | 1190 | - | tuf | WP_165594100.1 |
| GCF_014863585.1 | NZ_CP062253.1 | 1867221 | 1868411 | 1190 | + | tuf | WP_063544037.1 |
| GCF_014863585.1 | NZ_CP062253.1 | 1883701 | 1884891 | 1190 | + | tuf | WP_063544037.1 |
| GCF_900105485.1 | NZ_LT629778.1 | 740006 | 741196 | 1190 | + | tuf | WP_011911986.1 |
| GCF_900105485.1 | NZ_LT629778.1 | 756488 | 757678 | 1190 | + | tuf | WP_011911986.1 |
| GCF_039909015.1 | NZ_CP150919.1 | 955727 | 956917 | 1190 | + | tuf | WP_073301046.1 |
| GCF_040822035.1 | NZ_CP162012.1 | 972226 | 973416 | 1190 | + | tuf | WP_073301046.1 |
| GCF_040822035.1 | NZ_CP162012.1 | 103430 | 104611 | 1181 | - | tuf | WP_078929370.1 |
| GCF_014268595.2 | NZ_CP077091.1 | 122404 | 123585 | 1181 | - | tuf | WP_078929370.1 |
| GCF_014268595.2 | NZ_CP077091.1 | 2340738 | 2341925 | 1187 | - | tuf | WP_096361314.1 |
| GCF_045830615.1 | NZ_CP174501.1 | 2357318 | 2358505 | 1187 | - | tuf | WP_096361314.1 |
| GCF_039908645.1 | NZ_CP150917.1 | 2402797 | 2403984 | 1187 | - | tuf | WP_126456454.1 |

| | | | | | | | |
|---|---|---|---|---|---|---|---|
| GCF_039908645.1 | NZ_CP150917.1 | 2419347 | 2420534 | 1187 | - | tuf | WP_126456454.1 |
| GCF_018502625.1 | NZ_CP075566.1 | 3398789 | 3399976 | 1187 | - | tuf | WP_096462258.1 |
| GCF_026810445.1 | NZ_CP113949.1 | 3415521 | 3416708 | 1187 | - | tuf | WP_096462258.1 |
| GCF_037903935.1 | NZ_CP149586.1 | 618799 | 619986 | 1187 | + | tuf | WP_127475816.1 |
| GCF_037903935.1 | NZ_CP149586.1 | 635235 | 636422 | 1187 | + | tuf | WP_127475816.1 |
| GCF_032164165.1 | NZ_CP134401.1 | 4575466 | 4576647 | 1181 | - | tuf | WP_222158739.1 |
| GCF_032164165.1 | NZ_CP134401.1 | 4856105 | 4857286 | 1181 | - | tuf | WP_222158937.1 |
| GCF_014268585.2 | NZ_CP077092.1 | 5603209 | 5604396 | 1187 | - | tuf | WP_261694362.1 |
| GCF_014268585.2 | NZ_CP077092.1 | 5619953 | 5621140 | 1187 | - | tuf | WP_261694362.1 |
| GCF_002356535.2 | NZ_AP017423.2 | 100281 | 101462 | 1181 | + | tuf | WP_087486823.1 |
| GCF_021560075.1 | NZ_CP091088.1 | 3923543 | 3924724 | 1181 | - | tuf | WP_087486823.1 |
| GCF_021560075.1 | NZ_CP091088.1 | 206609 | 207829 | 1220 | + | tuf | WP_075186033.1 |
| GCF_029215585.1 | NZ_CP119382.1 | 4441060 | 4442280 | 1220 | - | tuf | WP_045860746.1 |
| GCF_029215585.1 | NZ_CP119382.1 | 4296552 | 4297772 | 1220 | - | tuf | WP_018274087.1 |
| GCF_036704165.1 | NZ_CP129946.1 | 3585258 | 3586478 | 1220 | - | tuf | WP_260997559.1 |
| GCF_030505375.1 | NZ_CP130043.1 | 3589133 | 3590353 | 1220 | - | tuf | WP_260997559.1 |
| GCF_030505375.1 | NZ_CP130043.1 | 3605575 | 3606795 | 1220 | - | tuf | WP_260997559.1 |
| GCF_009911755.1 | NZ_CP047698.1 | 3613996 | 3615216 | 1220 | - | tuf | WP_015488545.1 |
| GCF_009911755.1 | NZ_CP047698.1 | 3630638 | 3631858 | 1220 | - | tuf | WP_015488557.1 |
| GCF_001654515.1 | NZ_CP015852.1 | 3634398 | 3635618 | 1220 | - | tuf | WP_015488559.1 |
| GCF_001654515.1 | NZ_CP015852.1 | 6140435 | 6141616 | 1181 | - | tuf | WP_274055300.1 |
| GCF_003352185.1 | NZ_CP029608.1 | 6159358 | 6160539 | 1181 | - | tuf | WP_274055300.1 |
| GCF_003352185.1 | NZ_CP029608.1 | 5960135 | 5961316 | 1181 | - | tuf | WP_274051678.1 |
| GCF_963681935.1 | NZ_OY817088.1 | 5978309 | 5979490 | 1181 | - | tuf | WP_274051678.1 |
| GCF_030291635.1 | NZ_CP127871.1 | 382759 | 383940 | 1181 | + | tuf | WP_274038487.1 |
| GCF_030291635.1 | NZ_CP127871.1 | 400243 | 401424 | 1181 | + | tuf | WP_274038487.1 |
| GCF_963678005.1 | NZ_OY782661.1 | 350643 | 351824 | 1181 | + | tuf | WP_286219620.1 |
| GCF_963678005.1 | NZ_OY782661.1 | 451916 | 453097 | 1181 | + | tuf | WP_286219620.1 |
| GCF_047758745.1 | NZ_CP180477.1 | 2895261 | 2896442 | 1181 | - | tuf | WP_371374513.1 |
| GCF_047758745.1 | NZ_CP180477.1 | 2900468 | 2901649 | 1181 | - | tuf | WP_371374513.1 |
| GCF_018968705.1 | NZ_CP076668.1 | 2918362 | 2919543 | 1181 | - | tuf | WP_371374513.1 |

| | | | | | | | |
|---|---|---|---|---|---|---|---|
| GCF_900104735.1 | NZ_LT629746.1 | 508094 | 509275 | 1181 | + | tuf | WP_286264720.1 |
| GCF_900104735.1 | NZ_LT629746.1 | 526803 | 527984 | 1181 | + | tuf | WP_286264720.1 |
| GCF_015377145.2 | NZ_CP062158.2 | 273795 | 274976 | 1181 | + | tuf | WP_070795866.1 |
| GCF_900106065.1 | NZ_LT629796.1 | 467808 | 468989 | 1181 | + | tuf | WP_070795875.1 |
| GCF_900106065.1 | NZ_LT629796.1 | 3978706 | 3979887 | 1181 | + | tuf | WP_348395720.1 |
| GCF_034011435.1 | NZ_CP125381.1 | 3998024 | 3999205 | 1181 | + | tuf | WP_348395720.1 |
| GCF_900682675.2 | NZ_LR215729.2 | 4023597 | 4024778 | 1181 | + | tuf | WP_348395720.1 |
| GCF_900682675.2 | NZ_LR215729.2 | 418994 | 420175 | 1181 | + | tuf | WP_286271148.1 |
| GCF_019139675.1 | NZ_CP077077.1 | 437871 | 439052 | 1181 | + | tuf | WP_286271148.1 |
| GCF_019139675.1 | NZ_CP077077.1 | 3032319 | 3033500 | 1181 | - | tuf | WP_371187036.1 |
| GCF_900106005.1 | NZ_LT629790.1 | 3041663 | 3042844 | 1181 | - | tuf | WP_371187036.1 |
| GCF_900106005.1 | NZ_LT629790.1 | 3062337 | 3063518 | 1181 | - | tuf | WP_371187036.1 |
| GCF_047345015.1 | NZ_CP178857.1 | 480009 | 481190 | 1181 | + | tuf | WP_348388146.1 |
| GCF_014268495.2 | NZ_CP077087.1 | 498411 | 499592 | 1181 | + | tuf | WP_348388146.1 |
| GCF_014268495.2 | NZ_CP077087.1 | 523479 | 524660 | 1181 | + | tuf | WP_348388146.1 |
| GCF_003671975.1 | NZ_CP022562.1 | 3558635 | 3559816 | 1181 | - | tuf | WP_286289064.1 |
| GCF_003671975.1 | NZ_CP022562.1 | 3576749 | 3577930 | 1181 | - | tuf | WP_286289064.1 |
| GCF_042846945.1 | NZ_AP035776.1 | 2993856 | 2995037 | 1181 | - | tuf | WP_371189930.1 |
| GCF_002736065.1 | NZ_CP024159.1 | 3008327 | 3009508 | 1181 | - | tuf | WP_371189930.1 |
| GCF_002736065.1 | NZ_CP024159.1 | 3028267 | 3029448 | 1181 | - | tuf | WP_371189930.1 |
| GCF_900475945.1 | NZ_LS483433.1 | 4161876 | 4163057 | 1181 | - | tuf | WP_348390994.1 |
| GCF_012971725.1 | NZ_CP048833.1 | 4186196 | 4187377 | 1181 | - | tuf | WP_348391007.1 |
| GCF_012971725.1 | NZ_CP048833.1 | 4205191 | 4206372 | 1181 | - | tuf | WP_348390994.1 |
| GCF_019139535.1 | NZ_CP077073.1 | 3419316 | 3420497 | 1181 | - | tuf | WP_286233792.1 |
| GCF_019139535.1 | NZ_CP077073.1 | 3440183 | 3441364 | 1181 | - | tuf | WP_286233792.1 |
| GCF_024296925.1 | NZ_CP101125.1 | 255168 | 256349 | 1181 | + | tuf | WP_087017546.1 |
| GCF_024296925.1 | NZ_CP101125.1 | 3428447 | 3429628 | 1181 | - | tuf | WP_087017546.1 |
| GCF_014268695.2 | NZ_CP077086.1 | 1591044 | 1592231 | 1187 | - | tuf | WP_153974998.1 |
| GCF_014268695.2 | NZ_CP077086.1 | 1608400 | 1609587 | 1187 | - | tuf | WP_153974998.1 |
| GCF_003852045.1 | NZ_CP027723.1 | 1000413 | 1001600 | 1187 | - | tuf | WP_139715750.1 |
| GCF_900104805.1 | NZ_LT629751.1 | 1017734 | 1018921 | 1187 | - | tuf | WP_139715750.1 |

| | | | | | | | |
|---|---|---|---|---|---|---|---|
| GCF_900104805.1 | NZ_LT629751.1 | 2599452 | 2600639 | 1187 | + | tuf | WP_187569674.1 |
| GCF_003940825.1 | NZ_CP034338.1 | 2616749 | 2617936 | 1187 | + | tuf | WP_187569674.1 |
| GCF_003940825.1 | NZ_CP034338.1 | 536123 | 537310 | 1187 | + | tuf | WP_256646399.1 |
| GCF_035621475.1 | NZ_CP141799.1 | 553457 | 554644 | 1187 | + | tuf | WP_256646399.1 |
| GCF_035621475.1 | NZ_CP141799.1 | 2658600 | 2659787 | 1187 | - | tuf | WP_015259329.1 |
| GCF_022836895.1 | NZ_CP092411.1 | 2674912 | 2676099 | 1187 | - | tuf | WP_015259329.1 |
| GCF_046272155.1 | NZ_CP176709.1 | 3093508 | 3094695 | 1187 | - | tuf | WP_006747220.1 |
| GCF_003025345.2 | NZ_CP020560.1 | 3109804 | 3110991 | 1187 | - | tuf | WP_006747208.1 |
| GCF_003025345.2 | NZ_CP020560.1 | 2444246 | 2445433 | 1187 | - | tuf | WP_012638882.1 |
| GCF_963681955.1 | NZ_OY817130.1 | 2460736 | 2461923 | 1187 | - | tuf | WP_012638882.1 |
| GCF_024112395.2 | NZ_CP078013.2 | 483800 | 484987 | 1187 | + | tuf | WP_018168213.1 |
| GCF_045037875.1 | NZ_CP173614.1 | 500137 | 501324 | 1187 | + | tuf | WP_018168213.1 |
| GCF_021432765.1 | NZ_CP058908.1 | 4590203 | 4591390 | 1187 | + | tuf | WP_232407326.1 |
| GCF_021432765.1 | NZ_CP058908.1 | 4607340 | 4608527 | 1187 | + | tuf | WP_232407326.1 |
| GCF_003391255.1 | NZ_CP031146.1 | 4553296 | 4554483 | 1187 | + | tuf | WP_014780383.1 |
| GCF_003391255.1 | NZ_CP031146.1 | 4570513 | 4571700 | 1187 | + | tuf | WP_014780383.1 |
| GCF_000336465.1 | NC_020209.1 | 3142511 | 3143698 | 1187 | - | tuf | WP_015281698.1 |
| GCF_900105995.1 | NZ_LT629785.1 | 3159537 | 3160724 | 1187 | - | tuf | WP_015281698.1 |
| GCF_900105995.1 | NZ_LT629785.1 | 341386 | 342573 | 1187 | + | tuf | WP_127470077.1 |
| GCF_014269025.2 | NZ_CP077094.1 | 357724 | 358911 | 1187 | + | tuf | WP_127470077.1 |
| GCF_014269025.2 | NZ_CP077094.1 | 411994 | 413181 | 1187 | + | tuf | WP_237262248.1 |
| GCF_900105155.1 | NZ_LT629762.1 | 428330 | 429517 | 1187 | + | tuf | WP_237262248.1 |
| GCF_900105155.1 | NZ_LT629762.1 | 435792 | 436973 | 1181 | + | tuf | WP_130536562.1 |
| GCF_900560965.1 | NZ_LS999205.1 | 452216 | 453397 | 1181 | + | tuf | WP_130536562.1 |
| GCF_900560965.1 | NZ_LS999205.1 | 406123 | 407310 | 1187 | + | tuf | WP_275595249.1 |
| GCF_011040435.1 | NZ_CP049044.1 | 422478 | 423665 | 1187 | + | tuf | WP_275595261.1 |
| GCF_039908635.1 | NZ_CP150918.1 | 344152 | 345339 | 1187 | + | tuf | WP_138563715.1 |
| GCF_039908635.1 | NZ_CP150918.1 | 360500 | 361687 | 1187 | + | tuf | WP_138563715.1 |
| GCF_000412675.1 | NC_021505.1 | 417604 | 418791 | 1187 | + | tuf | WP_173284012.1 |
| GCF_000412675.1 | NC_021505.1 | 433970 | 435157 | 1187 | + | tuf | WP_173284012.1 |
| GCF_018398475.1 | NZ_CP074676.1 | 149751 | 150938 | 1187 | + | tuf | WP_025299247.1 |

| | | | | | | | |
|---|---|---|---|---|---|---|---|
| GCF_018398475.1 | NZ_CP074676.1 | 165977 | 167164 | 1187 | + | tuf | WP_006459613.1 |
| GCF_026410085.1 | NZ_CP112866.1 | 114249 | 115436 | 1187 | + | tuf | WP_013834672.1 |
| GCF_900104125.1 | NZ_LT629709.1 | 130478 | 131665 | 1187 | + | tuf | WP_013834672.1 |
| GCF_003033885.1 | NZ_CP024081.1 | 2052120 | 2053316 | 1196 | + | tuf | WP_053103335.1 |
| GCF_000761155.1 | NZ_CP009533.1 | 2068499 | 2069695 | 1196 | + | tuf | WP_053103335.1 |
| GCF_013285305.1 | NZ_CP054205.1 | 1887683 | 1888870 | 1187 | + | tuf | WP_009148299.1 |
| GCF_014268375.2 | NZ_CP077083.1 | 1905414 | 1906601 | 1187 | + | tuf | WP_009148299.1 |
| GCF_020917325.1 | NZ_CP076652.1 | 3734525 | 3735712 | 1187 | - | tuf | WP_201064603.1 |
| GCF_013409125.2 | NZ_CP060009.1 | 3752039 | 3753226 | 1187 | - | tuf | WP_201064603.1 |
| GCF_013409125.2 | NZ_CP060009.1 | 1852895 | 1854082 | 1187 | + | tuf | WP_328987257.1 |
| GCF_028830055.1 | NZ_CP101655.1 | 1872406 | 1873593 | 1187 | + | tuf | WP_328987257.1 |
| GCF_028830055.1 | NZ_CP101655.1 | 4337321 | 4338508 | 1187 | - | tuf | WP_078121434.1 |
| GCF_003850565.1 | NZ_CP027706.1 | 4354197 | 4355384 | 1187 | - | tuf | WP_078121434.1 |
| GCF_003850565.1 | NZ_CP027706.1 | 2402410 | 2403591 | 1181 | - | tuf | WP_173273900.1 |
| GCF_014268455.2 | NZ_CP077085.1 | 2418779 | 2419960 | 1181 | - | tuf | WP_173273900.1 |
| GCF_046619455.1 | NZ_CP177041.1 | 558789 | 559976 | 1187 | + | tuf | WP_173290349.1 |
| GCF_046619455.1 | NZ_CP177041.1 | 575093 | 576280 | 1187 | + | tuf | WP_173290349.1 |
| GCF_015168335.1 | NZ_CP063456.1 | 4073133 | 4074320 | 1187 | + | tuf | WP_210222532.1 |
| GCF_015168335.1 | NZ_CP063456.1 | 4089478 | 4090665 | 1187 | + | tuf | WP_210222539.1 |
| GCF_024749065.1 | NZ_CP087185.1 | 1364783 | 1365970 | 1187 | - | tuf | WP_202718508.1 |
| GCF_024749065.1 | NZ_CP087185.1 | 1445292 | 1446479 | 1187 | + | tuf | WP_236501160.1 |
| GCF_900106015.1 | NZ_LT629797.1 | 1461642 | 1462829 | 1187 | + | tuf | WP_236501172.1 |
| GCF_900106015.1 | NZ_LT629797.1 | 3047572 | 3048753 | 1181 | - | tuf | WP_015879816.1 |
| GCF_001661075.1 | NZ_CP014870.1 | 3065830 | 3067011 | 1181 | - | tuf | WP_015879832.1 |
| GCF_024072275.1 | NZ_CP099598.1 | 348255 | 349442 | 1187 | + | tuf | WP_283282465.1 |
| GCF_024072275.1 | NZ_CP099598.1 | 4374831 | 4376012 | 1181 | - | tuf | WP_283281603.1 |
| GCF_000934565.1 | NZ_CP010896.1 | 602037 | 603218 | 1181 | + | tuf | WP_392566526.1 |
| GCF_021391435.1 | NZ_CP090029.1 | 611880 | 613061 | 1181 | - | tuf | WP_392566526.1 |
| GCF_040281505.1 | NZ_CP158373.1 | 210326 | 211546 | 1220 | + | tuf | WP_228345651.1 |
| GCF_040281505.1 | NZ_CP158373.1 | 226978 | 228198 | 1220 | + | tuf | WP_228345651.1 |
| GCF_021283165.2 | NZ_CP128543.1 | 230864 | 232084 | 1220 | + | tuf | WP_228345651.1 |

| | | | | | | |
|---|---|---|---|---|---|---|
| GCF_021283165.2 | NZ_CP128543.1 | 562158 | 563339 | 1181 - | tuf | WP_176672521.1 |
| GCF_030053115.1 | NZ_CP125619.1 | 1123585 | 1124766 | 1181 + | tuf | WP_176672521.1 |
| GCF_030053115.1 | NZ_CP125619.1 | 3549245 | 3550426 | 1181 - | tuf | WP_073603663.1 |
| GCF_003851555.1 | NZ_CP027756.1 | 3597479 | 3598660 | 1181 - | tuf | WP_073603663.1 |
| GCF_018394375.1 | NZ_CP074578.1 | 2839386 | 2840567 | 1181 - | tuf | WP_221068231.1 |
| GCF_022557255.1 | NZ_CP092923.1 | 3003283 | 3004464 | 1181 - | tuf | WP_221068231.1 |
| GCF_000007805.1 | NC_004578.1 | 103290 | 104471 | 1181 + | tuf | WP_153445645.1 |
| GCF_900475285.1 | NZ_LS483370.1 | 2305539 | 2306720 | 1181 - | tuf | WP_153447924.1 |
| GCF_014268445.2 | NZ_CP077089.1 | 146769 | 147950 | 1181 + | tuf | WP_005381361.1 |
| GCF_014268445.2 | NZ_CP077089.1 | 305818 | 306999 | 1181 + | tuf | WP_005381361.1 |
| GCF_012767755.2 | NZ_AP023189.1 | 589166 | 590347 | 1181 + | tuf | WP_012841309.1 |
| GCF_012767755.2 | NZ_AP023189.1 | 752206 | 753387 | 1181 + | tuf | WP_012841343.1 |
| GCF_023278125.1 | NZ_CP066270.1 | 276562 | 277743 | 1181 + | tuf | WP_086981798.1 |
| GCF_026686735.1 | NZ_CP113432.1 | 2434994 | 2436175 | 1181 - | tuf | WP_086981798.1 |
| GCF_026686735.1 | NZ_CP113432.1 | 3437610 | 3438791 | 1181 - | tuf | WP_012604914.1 |
| GCF_014268275.3 | NZ_CP077084.1 | 3597575 | 3598756 | 1181 - | tuf | WP_012604914.1 |
| GCF_036864995.1 | NZ_CP142151.1 | 269579 | 270760 | 1181 + | tuf | WP_152429398.1 |
| GCF_004214895.1 | NZ_CP035952.1 | 316031 | 317212 | 1181 + | tuf | WP_152429398.1 |
| GCF_004214895.1 | NZ_CP035952.1 | 2720388 | 2721569 | 1181 - | tuf | WP_261838232.1 |
| GCF_012647205.1 | NZ_CP051487.1 | 2862671 | 2863852 | 1181 - | tuf | WP_261838232.1 |
| GCF_900105825.1 | NZ_LT629803.1 | 2786759 | 2787940 | 1181 - | tuf | WP_164649258.1 |
| GCF_014268755.2 | NZ_CP077093.1 | 2832832 | 2834013 | 1181 - | tuf | WP_164649258.1 |
| GCF_002028325.1 | NZ_CP018420.1 | 3100945 | 3102126 | 1181 - | tuf | WP_012604914.1 |
| GCF_001294575.1 | NZ_CP012676.1 | 3251958 | 3253139 | 1181 - | tuf | WP_012604914.1 |
| GCF_004786035.1 | NZ_CP035088.1 | 2786872 | 2788053 | 1181 - | tuf | WP_101903900.1 |
| GCF_004786035.1 | NZ_CP035088.1 | 2929169 | 2930350 | 1181 - | tuf | WP_101903902.1 |
| GCF_900184295.1 | NZ_LT855380.1 | 3286833 | 3288014 | 1181 - | tuf | WP_012604914.1 |
| GCF_040438935.1 | NZ_CP159259.1 | 3434733 | 3435914 | 1181 - | tuf | WP_012604914.1 |
| GCF_040438935.1 | NZ_CP159259.1 | 114278 | 115459 | 1181 + | tuf | WP_065210307.1 |
| GCF_021029445.1 | NZ_CP072610.1 | 2603968 | 2605149 | 1181 - | tuf | WP_261825416.1 |
| GCF_021029445.1 | NZ_CP072610.1 | 156192 | 157373 | 1181 + | tuf | WP_005438393.1 |

| | | | | | | | |
|---|---|---|---|---|---|---|---|
| GCF_030687395.1 | NZ_CP117430.1 | 317535 | 318716 | 1181 | + | tuf | WP_005438393.1 |
| GCF_030687395.1 | NZ_CP117430.1 | 314339 | 315520 | 1181 | + | tuf | WP_353497269.1 |
| GCF_014268885.2 | NZ_CP077095.1 | 3360083 | 3361264 | 1181 | - | tuf | WP_353497269.1 |
| GCF_014268885.2 | NZ_CP077095.1 | 3072890 | 3074071 | 1181 | - | tuf | WP_105057636.1 |
| GCF_019139835.1 | NZ_CP077075.1 | 3220598 | 3221779 | 1181 | - | tuf | WP_105057636.1 |
| GCF_019139835.1 | NZ_CP077075.1 | 2615656 | 2616837 | 1181 | - | tuf | WP_000031644.1 |
| GCF_900105735.1 | NZ_LT629793.1 | 2662943 | 2664124 | 1181 | - | tuf | WP_000031644.1 |
| GCF_014268485.2 | NZ_CP077090.1 | 1876076 | 1877257 | 1181 | + | tuf | WP_061900646.1 |
| GCF_014268485.2 | NZ_CP077090.1 | 2023634 | 2024815 | 1181 | + | tuf | WP_065819404.1 |
| GCF_022637495.1 | NZ_CP093331.1 | 114945 | 116126 | 1181 | + | tuf | WP_337972447.1 |
| GCF_022637495.1 | NZ_CP093331.1 | 2755833 | 2757014 | 1181 | - | tuf | WP_337972447.1 |
| GCF_043632995.1 | NZ_CP171632.1 | 873884 | 875065 | 1181 | + | tuf | WP_038509321.1 |
| GCF_043632995.1 | NZ_CP171632.1 | 922747 | 923928 | 1181 | + | tuf | WP_038509321.1 |
| GCF_014397415.1 | NZ_CP060731.1 | 310537 | 311718 | 1181 | + | tuf | WP_261888623.1 |
| GCF_014397415.1 | NZ_CP060731.1 | 2929418 | 2930599 | 1181 | - | tuf | WP_261888623.1 |
| GCF_000233915.3 | NC_016147.2 | 3140979 | 3142160 | 1181 | - | tuf | WP_010435158.1 |
| GCF_000233915.3 | NC_016147.2 | 3287618 | 3288799 | 1181 | - | tuf | WP_010435158.1 |
| GCF_030322905.1 | NZ_CP095475.1 | 159152 | 160333 | 1181 | + | tuf | WP_010435158.1 |
| GCF_030322905.1 | NZ_CP095475.1 | 306505 | 307686 | 1181 | + | tuf | WP_010435158.1 |
| GCF_001606025.1 | NZ_CP014945.1 | 176058 | 177239 | 1181 | + | tuf | WP_012841309.1 |
| GCF_001606025.1 | NZ_CP014945.1 | 335549 | 336730 | 1181 | + | tuf | WP_012841343.1 |
| GCF_000012305.1 | NC_007204.1 | 2974581 | 2975762 | 1181 | - | tuf | WP_337164282.1 |
| GCF_000012305.1 | NC_007204.1 | 3024414 | 3025595 | 1181 | - | tuf | WP_342291494.1 |
| GCF_904848165.1 | NZ_LR884459.1 | 164887 | 166068 | 1181 | + | tuf | WP_135457668.1 |
| GCF_904848165.1 | NZ_LR884459.1 | 312199 | 313380 | 1181 | + | tuf | WP_135457668.1 |
| GCF_000013905.1 | NC_007969.1 | 2460706 | 2461887 | 1181 | - | tuf | WP_061056950.1 |
| GCF_000013905.1 | NC_007969.1 | 2509279 | 2510460 | 1181 | - | tuf | WP_020332747.1 |
| GCF_018336615.1 | NZ_CP048752.1 | 2931222 | 2932403 | 1181 | - | tuf | WP_261903221.1 |
| GCF_018336615.1 | NZ_CP048752.1 | 3070301 | 3071482 | 1181 | - | tuf | WP_261903221.1 |
| GCF_022631235.3 | NZ_CP093310.2 | 737121 | 738302 | 1181 | + | tuf | WP_004728252.1 |
| GCF_022631235.3 | NZ_CP093310.2 | 785771 | 786952 | 1181 | + | tuf | WP_004728252.1 |

| | | | | | | | |
|---|---|---|---|---|---|---|---|
| GCF_000015285.1 | NC_008709.1 | 2954183 | 2955364 | 1181 | - | tuf | WP_137375015.1 |
| GCF_017948405.1 | NZ_CP072844.1 | 3099071 | 3100252 | 1181 | - | tuf | WP_137375015.1 |
| GCF_017948405.1 | NZ_CP072844.1 | 204563 | 205744 | 1181 | + | tuf | WP_261816238.1 |
| GCF_017638545.1 | NZ_CP072110.1 | 2386002 | 2387183 | 1181 | - | tuf | WP_261816238.1 |
| GCF_017638545.1 | NZ_CP072110.1 | 73406 | 74587 | 1181 | + | tuf | WP_264923430.1 |
| GCF_022637475.1 | NZ_CP093328.1 | 2341963 | 2343144 | 1181 | - | tuf | WP_264923430.1 |
| GCF_022637475.1 | NZ_CP093328.1 | 3142476 | 3143657 | 1181 | - | tuf | WP_252518335.1 |
| GCF_000241955.1 | NC_016818.1 | 3187202 | 3188383 | 1181 | - | tuf | WP_252518335.1 |
| GCF_000241955.1 | NC_016818.1 | 3110206 | 3111387 | 1181 | - | tuf | WP_010435158.1 |
| GCF_018288855.1 | NZ_CP065024.1 | 3263278 | 3264459 | 1181 | - | tuf | WP_010435158.1 |
| GCF_018288855.1 | NZ_CP065024.1 | 3253848 | 3255029 | 1181 | + | tuf | WP_005451119.1 |
| GCF_002951615.1 | NZ_CP019062.1 | 3417197 | 3418378 | 1181 | + | tuf | WP_005438393.1 |
| GCF_002951615.1 | NZ_CP019062.1 | 2358530 | 2359711 | 1181 | + | tuf | WP_103415287.1 |
| GCF_021276285.1 | NZ_CP089919.1 | 2526233 | 2527414 | 1181 | + | tuf | WP_103415287.1 |
| GCF_021276285.1 | NZ_CP089919.1 | 114564 | 115745 | 1181 | + | tuf | WP_337983893.1 |
| GCF_030505655.1 | NZ_CP130153.1 | 2613454 | 2614635 | 1181 | - | tuf | WP_337983893.1 |
| GCF_030505655.1 | NZ_CP130153.1 | 229738 | 230919 | 1181 | + | tuf | WP_261834263.1 |
| GCF_000783935.2 | NZ_CP026047.1 | 2823766 | 2824947 | 1181 | - | tuf | WP_261834263.1 |
| GCF_000783935.2 | NZ_CP026047.1 | 248892 | 250073 | 1181 | + | tuf | WP_257084556.1 |
| GCF_012029655.1 | NZ_CP050508.1 | 298448 | 299629 | 1181 | + | tuf | WP_257084556.1 |
| GCF_012029655.1 | NZ_CP050508.1 | 2410989 | 2412170 | 1181 | + | tuf | WP_042605518.1 |
| GCF_002795845.1 | NZ_CP011797.1 | 2577569 | 2578750 | 1181 | + | tuf | WP_042605518.1 |
| GCF_003990335.1 | NZ_CP034683.1 | 153542 | 154723 | 1181 | + | tuf | WP_077680933.1 |
| GCF_003990335.1 | NZ_CP034683.1 | 323089 | 324270 | 1181 | + | tuf | WP_077680933.1 |
| GCF_021545825.1 | NZ_CP088980.1 | 358963 | 360144 | 1181 | + | tuf | WP_087481459.1 |
| GCF_021545825.1 | NZ_CP088980.1 | 403683 | 404864 | 1181 | + | tuf | WP_087481459.1 |
| GCF_964030935.1 | NZ_OZ034990.1 | 2634631 | 2635812 | 1181 | - | tuf | WP_000031640.1 |
| GCF_019285595.1 | NZ_CP079094.1 | 2682607 | 2683788 | 1181 | - | tuf | WP_000031640.1 |
| GCF_964277335.1 | NZ_OZ195516.1 | 217816 | 218997 | 1181 | + | tuf | WP_278182950.1 |
| GCF_964031245.1 | NZ_OZ035017.1 | 264509 | 265690 | 1181 | + | tuf | WP_278182950.1 |
| GCF_964031165.1 | NZ_OZ035011.1 | 11619 | 12800 | 1181 | + | tuf | WP_014233170.1 |

| | | | | | | | |
|---|---|---|---|---|---|---|---|
| GCF_964019545.1 | NZ_OZ026451.1 | 3197386 | 3198567 | 1181 | + | tuf | WP_061778638.1 |
| GCF_010587025.1 | NZ_CP040863.1 | 222843 | 224024 | 1181 | + | tuf | WP_023404590.1 |
| GCF_010587025.1 | NZ_CP040863.1 | 2601609 | 2602790 | 1181 | - | tuf | WP_023404590.1 |
| GCF_015356115.1 | NZ_CP063056.1 | 185290 | 186471 | 1181 | + | tuf | WP_022551812.1 |
| GCF_015356115.1 | NZ_CP063056.1 | 336670 | 337851 | 1181 | + | tuf | WP_281870126.1 |
| GCF_026967515.2 | NZ_CP114062.1 | 2429837 | 2431018 | 1181 | - | tuf | WP_218562431.1 |
| GCF_026967515.2 | NZ_CP114062.1 | 2477717 | 2478898 | 1181 | - | tuf | WP_218562431.1 |
| GCF_003063625.1 | NZ_CP026604.1 | 1993095 | 1994276 | 1181 | + | tuf | WP_005438393.1 |
| GCF_003063625.1 | NZ_CP026604.1 | 2156216 | 2157397 | 1181 | + | tuf | WP_005438393.1 |
| GCF_000013665.1 | NC_007912.1 | 236944 | 238125 | 1181 | + | tuf | WP_077315602.1 |
| GCF_000013665.1 | NC_007912.1 | 281296 | 282477 | 1181 | + | tuf | WP_077315602.1 |
| GCF_003367315.1 | NZ_CP031415.1 | 215538 | 216719 | 1181 | - | tuf | WP_000031640.1 |
| GCF_003367315.1 | NZ_CP031415.1 | 262338 | 263519 | 1181 | - | tuf | WP_000031640.1 |
| GCF_006228385.1 | NZ_CP039852.1 | 2938517 | 2939698 | 1181 | - | tuf | WP_005480487.1 |
| GCF_006228385.1 | NZ_CP039852.1 | 3120895 | 3122076 | 1181 | - | tuf | WP_005480487.1 |
| GCF_005222225.1 | NZ_CP036536.1 | 294082 | 295263 | 1181 | + | tuf | WP_125319293.1 |
| GCF_005222225.1 | NZ_CP036536.1 | 340949 | 342130 | 1181 | + | tuf | WP_125319293.1 |
| GCF_015644725.1 | NZ_CP064795.1 | 2855493 | 2856674 | 1181 | - | tuf | WP_261882166.1 |
| GCF_015644725.1 | NZ_CP064795.1 | 2990780 | 2991961 | 1181 | - | tuf | WP_261882166.1 |
| GCF_003429285.1 | NZ_CP031769.1 | 234065 | 235246 | 1181 | + | tuf | WP_221768496.1 |
| GCF_003429285.1 | NZ_CP031769.1 | 406375 | 407556 | 1181 | + | tuf | WP_101111190.1 |
| GCF_003143555.1 | NZ_CP029347.1 | 3293450 | 3294631 | 1181 | - | tuf | WP_010435158.1 |
| GCF_003143555.1 | NZ_CP029347.1 | 3440918 | 3442099 | 1181 | - | tuf | WP_010435158.1 |
| GCF_027286345.1 | NZ_CP114590.1 | 2578512 | 2579693 | 1181 | - | tuf | WP_075652508.1 |
| GCF_027286345.1 | NZ_CP114590.1 | 2627110 | 2628291 | 1181 | - | tuf | WP_161987050.1 |
| GCF_027286365.1 | NZ_CP114584.1 | 302105 | 303286 | 1181 | + | tuf | WP_261894522.1 |
| GCF_027286365.1 | NZ_CP114584.1 | 347664 | 348845 | 1181 | + | tuf | WP_261894522.1 |
| GCF_000252995.1 | NC_015761.1 | 200126 | 201307 | 1181 | + | tuf | WP_013857920.1 |
| GCF_000252995.1 | NC_015761.1 | 247669 | 248850 | 1181 | + | tuf | WP_013857920.1 |
| GCF_000006945.2 | NC_003197.2 | 352611 | 353792 | 1181 | + | tuf | WP_073579957.1 |
| GCF_000006945.2 | NC_003197.2 | 403421 | 404602 | 1181 | + | tuf | WP_073579957.1 |

| | | | | | | | |
|---|---|---|---|---|---|---|---|
| GCF_003935895.2 | NZ_CP054058.1 | 221493 | 222674 | 1181 | + | tuf | WP_261873467.1 |
| GCF_003935895.2 | NZ_CP054058.1 | 2686360 | 2687541 | 1181 | - | tuf | WP_261873467.1 |
| GCF_000287335.1 | NC_018419.1 | 3153110 | 3154291 | 1181 | - | tuf | WP_038181313.1 |
| GCF_000287355.1 | NC_018420.1 | 3197913 | 3199094 | 1181 | - | tuf | WP_038181313.1 |
| GCF_900090215.1 | NZ_LT594522.1 | 123250 | 124431 | 1181 | + | tuf | WP_088880292.1 |
| GCF_001007875.1 | NZ_CP011412.1 | 3289303 | 3290484 | 1181 | + | tuf | WP_088880292.1 |
| GCF_001007875.1 | NZ_CP011412.1 | 3099486 | 3100667 | 1181 | - | tuf | WP_038181313.1 |
| GCF_044865145.1 | NZ_CP173186.1 | 3144305 | 3145486 | 1181 | - | tuf | WP_038181313.1 |
| GCF_044865145.1 | NZ_CP173186.1 | 311468 | 312649 | 1181 | + | tuf | WP_089137635.1 |
| GCF_021462285.1 | NZ_CP082787.1 | 2706533 | 2707714 | 1181 | - | tuf | WP_089137635.1 |
| GCF_021462285.1 | NZ_CP082787.1 | 1090479 | 1091660 | 1181 | - | tuf | WP_065431012.1 |
| GCF_900187015.1 | NZ_LT906479.1 | 1140028 | 1141209 | 1181 | - | tuf | WP_065431012.1 |
| GCF_900187015.1 | NZ_LT906479.1 | 3566420 | 3567601 | 1181 | - | tuf | WP_182287811.1 |
| GCF_001006005.1 | NZ_CP011254.1 | 3611369 | 3612550 | 1181 | - | tuf | WP_182287820.1 |
| GCF_001006005.1 | NZ_CP011254.1 | 2097618 | 2098799 | 1181 | + | tuf | WP_012604914.1 |
| GCF_000422085.1 | NC_021741.1 | 2251054 | 2252235 | 1181 | + | tuf | WP_012604914.1 |
| GCF_000422085.1 | NC_021742.1 | 161737 | 162918 | 1181 | + | tuf | WP_012604914.1 |
| GCF_030291735.1 | NZ_CP127881.1 | 312654 | 313835 | 1181 | + | tuf | WP_012604914.1 |
| GCF_030291735.1 | NZ_CP127881.1 | 170327 | 171508 | 1181 | + | tuf | WP_012604914.1 |
| GCF_004768745.1 | NZ_CP038662.1 | 326793 | 327974 | 1181 | + | tuf | WP_012604914.1 |
| GCF_004768745.1 | NZ_CP038662.1 | 2832685 | 2833866 | 1181 | - | tuf | WP_060468717.1 |
| GCF_037948395.1 | NZ_CP149940.1 | 2979312 | 2980493 | 1181 | - | tuf | WP_060468717.1 |
| GCF_037948395.1 | NZ_CP149940.1 | 291442 | 292623 | 1181 | + | tuf | WP_068711691.1 |
| GCF_000214235.1 | NC_015567.1 | 337143 | 338324 | 1181 | + | tuf | WP_068711691.1 |
| GCF_000214235.1 | NC_015567.1 | 2953492 | 2954673 | 1181 | - | tuf | WP_038550586.1 |
| GCF_949794035.1 | NZ_OX461105.1 | 3003694 | 3004875 | 1181 | - | tuf | WP_038550586.1 |
| GCF_949794035.1 | NZ_OX461105.1 | 1668280 | 1669461 | 1181 | + | tuf | WP_011079194.1 |
| GCF_900638135.1 | NZ_LR134494.1 | 1810780 | 1811961 | 1181 | + | tuf | WP_011079318.1 |
| GCF_900638135.1 | NZ_LR134494.1 | 240700 | 241881 | 1181 | + | tuf | WP_123015191.1 |
| GCF_009817885.1 | NZ_CP041764.1 | 286071 | 287252 | 1181 | + | tuf | WP_123015191.1 |
| GCF_009817885.1 | NZ_CP041764.1 | 2869741 | 2870922 | 1181 | - | tuf | WP_165312425.1 |

| | | | | | | | |
|---|---|---|---|---|---|---|---|
| GCF_016026735.1 | NZ_CP065640.1 | 2927495 | 2928676 | 1181 | - | tuf | WP_165312469.1 |
| GCF_016026735.1 | NZ_CP065640.1 | 122621 | 123811 | 1190 | + | tuf | WP_009288143.1 |
| GCF_035749905.1 | NZ_CP142089.1 | 209259 | 210449 | 1190 | + | tuf | WP_139527870.1 |
| GCF_035749905.1 | NZ_CP142089.1 | 101775 | 102965 | 1190 | + | tuf | WP_235040994.1 |
| GCF_001642805.2 | NZ_CP016948.1 | 910085 | 911275 | 1190 | - | tuf | WP_016915773.1 |
| GCF_001642805.2 | NZ_CP016948.1 | 1695169 | 1696359 | 1190 | - | tuf | WP_079552768.1 |
| GCF_000821185.2 | NZ_CP050855.1 | 707065 | 708255 | 1190 | - | tuf | WP_009288143.1 |
| GCF_000821185.2 | NZ_CP050855.1 | 203148 | 204338 | 1190 | + | tuf | WP_206048526.1 |
| GCF_017309605.1 | NZ_CP071320.1 | 2709188 | 2710375 | 1187 | - | tuf | WP_049726200.1 |
| GCF_017309605.1 | NZ_CP071320.1 | 2733334 | 2734521 | 1187 | - | tuf | WP_049726200.1 |
| GCF_019457475.1 | NZ_CP080413.1 | 236377 | 237558 | 1181 | - | tuf | WP_014354021.1 |
| GCF_019457475.1 | NZ_CP080413.1 | 417946 | 419127 | 1181 | + | tuf | WP_313854325.1 |
| GCF_019457495.1 | NZ_CP080416.1 | 4629069 | 4630250 | 1181 | - | tuf | WP_313854325.1 |
| GCF_019457495.1 | NZ_CP080416.1 | 1136928 | 1138115 | 1187 | + | tuf | WP_068613575.1 |
| GCF_009183365.2 | NZ_CP068230.1 | 1154111 | 1155298 | 1187 | + | tuf | WP_068613575.1 |
| GCF_009183365.2 | NZ_CP068230.1 | 497420 | 498607 | 1187 | + | tuf | WP_012915046.1 |
| GCF_019457575.1 | NZ_CP080422.1 | 514882 | 516069 | 1187 | + | tuf | WP_012915046.1 |
| GCF_019457575.1 | NZ_CP080422.1 | 1187596 | 1188783 | 1187 | + | tuf | WP_024938571.1 |
| GCF_000015245.1 | NC_008700.1 | 1204978 | 1206165 | 1187 | + | tuf | WP_024938571.1 |
| GCF_000015245.1 | NC_008700.1 | 1051219 | 1052406 | 1187 | + | tuf | WP_003486726.1 |
| GCF_017354945.1 | NZ_CP071503.1 | 1068573 | 1069760 | 1187 | + | tuf | WP_003486726.1 |
| GCF_017354945.1 | NZ_CP071503.1 | 4177384 | 4178571 | 1187 | - | tuf | WP_369976718.1 |
| GCF_000178875.2 | NC_016901.1 | 4194761 | 4195948 | 1187 | - | tuf | WP_369976728.1 |
| GCF_000178875.2 | NC_016901.1 | 3779638 | 3780825 | 1187 | - | tuf | WP_014506811.1 |
| GCF_019670705.1 | NZ_AP024618.1 | 3796974 | 3798161 | 1187 | - | tuf | WP_014506811.1 |
| GCF_019670705.1 | NZ_AP024618.1 | 1169518 | 1170705 | 1187 | + | tuf | WP_047695376.1 |
| GCF_037953065.1 | NZ_CP149966.1 | 1186896 | 1188083 | 1187 | + | tuf | WP_166752379.1 |
| GCF_037953065.1 | NZ_CP149966.1 | 105856 | 107043 | 1187 | + | tuf | WP_029218063.1 |
| GCF_017354985.1 | NZ_CP071501.1 | 3452214 | 3453401 | 1187 | - | tuf | WP_142742534.1 |
| GCF_017354985.1 | NZ_CP071501.1 | 3469647 | 3470834 | 1187 | - | tuf | WP_142742534.1 |
| GCF_004354305.1 | NZ_CP037898.1 | 1116065 | 1117252 | 1187 | + | tuf | WP_003486726.1 |

| | | | | | | | |
|---|---|---|---|---|---|---|---|
| GCF_000013765.1 | NC_007954.1 | 1133463 | 1134650 | 1187 | + | tuf | WP_003486726.1 |
| GCF_007567505.1 | NZ_CP041783.1 | 874507 | 875694 | 1187 | + | tuf | WP_159406665.1 |
| GCF_007567505.1 | NZ_CP041783.1 | 891962 | 893149 | 1187 | + | tuf | WP_159406666.1 |
| GCF_007004545.3 | NZ_CP045503.2 | 1267508 | 1268695 | 1187 | + | tuf | WP_104617719.1 |
| GCF_007004545.3 | NZ_CP045503.2 | 1284891 | 1286078 | 1187 | + | tuf | WP_104617719.1 |
| GCF_000014705.1 | NC_008345.1 | 1112704 | 1113891 | 1187 | + | tuf | WP_104582262.1 |
| GCF_000014705.1 | NC_008345.1 | 1130082 | 1131269 | 1187 | + | tuf | WP_104582262.1 |
| GCF_020511155.1 | NZ_CP041216.1 | 1158132 | 1159319 | 1187 | + | tuf | WP_003486726.1 |
| GCF_020511155.1 | NZ_CP041216.1 | 1175493 | 1176680 | 1187 | + | tuf | WP_003486726.1 |
| GCF_030252245.1 | NZ_CP126972.1 | 3139211 | 3140398 | 1187 | - | tuf | WP_002811703.1 |
| GCF_030252245.1 | NZ_CP126972.1 | 3156615 | 3157802 | 1187 | - | tuf | WP_002811703.1 |
| GCF_000019185.1 | NC_010334.1 | 930760 | 931947 | 1187 | + | tuf | WP_068832715.1 |
| GCF_000019185.1 | NC_010334.1 | 948201 | 949388 | 1187 | + | tuf | WP_068832715.1 |
| GCF_019457535.1 | NZ_CP080417.1 | 958153 | 959340 | 1187 | + | tuf | WP_048490041.1 |
| GCF_019457535.1 | NZ_CP080417.1 | 975615 | 976802 | 1187 | + | tuf | WP_048490041.1 |
| GCF_019931735.1 | NZ_CP082926.1 | 2416225 | 2417412 | 1187 | - | tuf | WP_074056769.1 |
| GCF_019931735.1 | NZ_CP082926.1 | 2433610 | 2434797 | 1187 | - | tuf | WP_046934093.1 |
| GCF_002075795.1 | NZ_CP020472.1 | 825926 | 827113 | 1187 | - | tuf | WP_104557641.1 |
| GCF_002075795.1 | NZ_CP020472.1 | 843305 | 844492 | 1187 | - | tuf | WP_104557641.1 |
| GCF_003957745.1 | NZ_CP020373.1 | 3289957 | 3291144 | 1187 | + | tuf | WP_316693967.1 |
| GCF_003957745.1 | NZ_CP020373.1 | 3307347 | 3308534 | 1187 | + | tuf | WP_316693967.1 |
| GCF_016834455.1 | NZ_CP069213.1 | 1704644 | 1705831 | 1187 | + | tuf | WP_242081676.1 |
| GCF_016834455.1 | NZ_CP069213.1 | 1722102 | 1723289 | 1187 | + | tuf | WP_242081676.1 |
| GCF_003855395.1 | NZ_CP034015.1 | 4238438 | 4239625 | 1187 | - | tuf | WP_369940045.1 |
| GCF_003855395.1 | NZ_CP034015.1 | 4255901 | 4257088 | 1187 | - | tuf | WP_369940045.1 |
| GCF_000016065.1 | NC_009092.1 | 1098938 | 1100125 | 1187 | + | tuf | WP_064634192.1 |
| GCF_000016065.1 | NC_009092.1 | 1116363 | 1117550 | 1187 | + | tuf | WP_064634192.1 |
| GCF_019457635.1 | NZ_CP080412.1 | 3808366 | 3809553 | 1187 | - | tuf | WP_014502146.1 |
| GCF_019457635.1 | NZ_CP080412.1 | 3825787 | 3826974 | 1187 | - | tuf | WP_014502146.1 |
| GCF_002215585.1 | NZ_CP022272.1 | 958165 | 959352 | 1187 | + | tuf | WP_150145591.1 |
| GCF_002215585.1 | NZ_CP022272.1 | 975587 | 976774 | 1187 | + | tuf | WP_003486726.1 |

| | | | | | | |
|---|---|---|---|---|---|---|
| GCF_004295345.1 | NZ_CP036200.1 | 1450825 | 1452012 | 1187 - | tuf | WP_053051265.1 |
| GCF_004295345.1 | NZ_CP036200.1 | 1468251 | 1469438 | 1187 - | tuf | WP_053051265.1 |
| GCF_019457515.1 | NZ_CP080421.1 | 3964657 | 3965844 | 1187 - | tuf | WP_160966994.1 |
| GCF_019457515.1 | NZ_CP080421.1 | 3982116 | 3983303 | 1187 - | tuf | WP_160966994.1 |
| GCF_030848765.1 | NZ_CP132914.1 | 4163014 | 4164201 | 1187 - | tuf | WP_043093035.1 |
| GCF_030848765.1 | NZ_CP132914.1 | 4180473 | 4181660 | 1187 - | tuf | WP_043093035.1 |
| GCF_000146165.2 | NC_004347.2 | 4740892 | 4742079 | 1187 - | tuf | WP_148828348.1 |
| GCF_000146165.2 | NC_004347.2 | 4758351 | 4759538 | 1187 - | tuf | WP_043093035.1 |
| GCF_000018285.1 | NC_009901.1 | 4046270 | 4047457 | 1187 - | tuf | WP_184411725.1 |
| GCF_000018285.1 | NC_009901.1 | 4063648 | 4064835 | 1187 - | tuf | WP_184411725.1 |
| GCF_000014885.1 | NC_011566.1 | 3173713 | 3174900 | 1187 - | tuf | WP_128421991.1 |
| GCF_000014885.1 | NC_011566.1 | 3191157 | 3192344 | 1187 - | tuf | WP_128421991.1 |
| GCF_006385555.1 | NZ_CP041036.1 | 1514551 | 1515738 | 1187 - | tuf | WP_047324579.1 |
| GCF_006385555.1 | NZ_CP041036.1 | 1532008 | 1533195 | 1187 - | tuf | WP_047324579.1 |
| GCF_003855155.1 | NZ_CP034073.1 | 1092938 | 1094125 | 1187 + | tuf | WP_010376339.1 |
| GCF_003855155.1 | NZ_CP034073.1 | 1110365 | 1111552 | 1187 + | tuf | WP_010376339.1 |
| GCF_007197555.1 | NZ_CP041614.1 | 4086461 | 4087648 | 1187 + | tuf | WP_074059437.1 |
| GCF_007197555.1 | NZ_CP041614.1 | 4103825 | 4105012 | 1187 + | tuf | WP_005993360.1 |
| GCF_019457595.1 | NZ_CP080419.1 | 2817016 | 2818197 | 1181 - | tuf | WP_047770768.1 |
| GCF_019457595.1 | NZ_CP080419.1 | 3261875 | 3263056 | 1181 + | tuf | WP_047770768.1 |
| GCF_016406325.1 | NZ_CP066370.1 | 355554 | 356735 | 1181 + | tuf | WP_209027467.1 |
| GCF_016406325.1 | NZ_CP066370.1 | 4198426 | 4199607 | 1181 - | tuf | WP_209027467.1 |
| GCF_019457655.1 | NZ_CP080420.1 | 268715 | 269896 | 1181 + | tuf | WP_045968037.1 |
| GCF_019457655.1 | NZ_CP080420.1 | 3866707 | 3867888 | 1181 - | tuf | WP_045968037.1 |
| GCF_017354965.1 | NZ_CP071502.1 | 361605 | 362786 | 1181 + | tuf | WP_282885534.1 |
| GCF_017354965.1 | NZ_CP071502.1 | 4371004 | 4372185 | 1181 - | tuf | WP_282885569.1 |
| GCF_019457555.1 | NZ_CP080418.1 | 3167363 | 3168544 | 1181 - | tuf | WP_069317339.1 |
| GCF_019457555.1 | NZ_CP080418.1 | 3648125 | 3649306 | 1181 + | tuf | WP_069317694.1 |
| GCF_000091325.1 | NC_014012.1 | 258325 | 259506 | 1181 + | tuf | WP_013183218.1 |
| GCF_000091325.1 | NC_014012.1 | 3100842 | 3102023 | 1181 - | tuf | WP_045959653.1 |
| GCF_000019525.1 | NC_010506.1 | 3492505 | 3493686 | 1181 - | tuf | WP_045959653.1 |

| | | | | | | | |
|---|---|---|---|---|---|---|---|
| GCF_000019525.1 | NC_010506.1 | 308370 | 309551 | 1181 | + | tuf | WP_374052009.1 |
| GCF_024971755.1 | NZ_CP079718.1 | 3840389 | 3841570 | 1181 | - | tuf | WP_374052009.1 |
| GCF_024971755.1 | NZ_CP079718.1 | 2389785 | 2390972 | 1187 | - | tuf | WP_004089480.1 |
| GCF_033180165.1 | NZ_CP136522.1 | 2407172 | 2408359 | 1187 | - | tuf | WP_004089480.1 |
| GCF_033180165.1 | NZ_CP136522.1 | 2449164 | 2450351 | 1187 | - | tuf | WP_038273292.1 |
| GCF_018223345.1 | NZ_CP073587.1 | 2466558 | 2467745 | 1187 | - | tuf | WP_038273292.1 |
| GCF_018223345.1 | NZ_CP073587.1 | 2361983 | 2363164 | 1181 | - | tuf | WP_004704670.1 |
| GCF_019457615.1 | NZ_CP080414.1 | 3000459 | 3001640 | 1181 | + | tuf | WP_004702310.1 |
| GCF_019457615.1 | NZ_CP080414.1 | 299336 | 300517 | 1181 | + | tuf | WP_050110014.1 |
| GCF_022354085.1 | NZ_CP055055.1 | 4533793 | 4534974 | 1181 | - | tuf | WP_050111023.1 |
| GCF_022354085.1 | NZ_CP055055.1 | 4011380 | 4012561 | 1181 | - | tuf | WP_032898571.1 |
| GCF_000006925.2 | NC_004337.2 | 4050536 | 4051717 | 1181 | - | tuf | WP_005271793.1 |
| GCF_000006925.2 | NC_004337.2 | 4333452 | 4334633 | 1181 | - | tuf | WP_145553661.1 |
| GCF_002950395.1 | NZ_CP026802.1 | 4370546 | 4371727 | 1181 | - | tuf | WP_004391430.1 |
| GCF_002950395.1 | NZ_CP026802.1 | 3405816 | 3406997 | 1181 | + | tuf | WP_005159831.1 |
| GCF_000262305.1 | NC_017910.1 | 3614139 | 3615320 | 1181 | + | tuf | WP_005165809.1 |
| GCF_000262305.1 | NC_017910.1 | 1529630 | 1530811 | 1181 | - | tuf | WP_064514258.1 |
| GCF_025914095.1 | NZ_CP074352.1 | 2472058 | 2473239 | 1181 | - | tuf | WP_064515581.1 |
| GCF_025914095.1 | NZ_CP074352.1 | 319110 | 320291 | 1181 | + | tuf | WP_100273993.1 |
| GCF_042853465.1 | NZ_AP031497.1 | 4384647 | 4385828 | 1181 | - | tuf | WP_050076773.1 |
| GCF_042853465.1 | NZ_AP031497.1 | 133833 | 135014 | 1181 | + | tuf | WP_005186230.1 |
| GCF_900161835.1 | NZ_LT897836.1 | 4361283 | 4362464 | 1181 | - | tuf | WP_005192344.1 |
| GCF_000010085.1 | NC_007712.1 | 859542 | 860723 | 1181 | - | tuf | WP_004876753.1 |
| GCF_000010085.1 | NC_007712.1 | 1561031 | 1562212 | 1181 | + | tuf | WP_004875106.1 |
| GCF_964059955.1 | NZ_OZ060924.1 | 885460 | 886641 | 1181 | - | tuf | WP_002212326.1 |
| GCF_003293695.1 | NZ_CP029556.1 | 1550879 | 1552060 | 1181 | + | tuf | WP_002210669.1 |
| GCF_003293695.1 | NZ_CP029556.1 | 304740 | 305921 | 1181 | + | tuf | WP_002212326.1 |
| GCF_011009015.1 | NZ_CP048878.1 | 4243661 | 4244842 | 1181 | - | tuf | WP_002210669.1 |
| GCF_011009015.1 | NZ_CP048878.1 | 4040571 | 4041752 | 1181 | - | tuf | WP_038638412.1 |
| GCF_020883455.1 | NZ_CP086615.1 | 4077933 | 4079114 | 1181 | - | tuf | WP_038637113.1 |
| GCF_020883455.1 | NZ_CP086615.1 | 2596309 | 2597490 | 1181 | - | tuf | WP_002212326.1 |

| | | | | | | | |
|---|---|---|---|---|---|---|---|
| GCF_002813635.1 | NZ_CP016382.1 | 3205441 | 3206622 | 1181 | + | tuf | WP_004716473.1 |
| GCF_002813635.1 | NZ_CP016382.1 | 269083 | 270264 | 1181 | + | tuf | WP_004718571.1 |
| GCF_000319575.2 | NC_021291.1 | 3432732 | 3433913 | 1181 | - | tuf | WP_038244641.1 |
| GCF_016132545.1 | NZ_CP066167.1 | 1270302 | 1271483 | 1181 | - | tuf | WP_002212326.1 |
| GCF_016132545.1 | NZ_CP066167.1 | 1956311 | 1957492 | 1181 | + | tuf | WP_025382234.1 |
| GCF_023702635.1 | NZ_CP098455.1 | 287664 | 288845 | 1181 | + | tuf | WP_050076773.1 |
| GCF_023702635.1 | NZ_CP098455.1 | 4392495 | 4393676 | 1181 | - | tuf | WP_004712401.1 |
| GCF_014109845.1 | NZ_CP043567.1 | 482488 | 483708 | 1220 | + | tuf | WP_008248373.1 |
| GCF_014109845.1 | NZ_CP043567.1 | 499153 | 500373 | 1220 | + | tuf | WP_008248373.1 |
| GCF_031834615.1 | NZ_CP115543.1 | 262971 | 264152 | 1181 | + | tuf | WP_096778336.1 |
| GCF_031834615.1 | NZ_CP115543.1 | 359579 | 360760 | 1181 | + | tuf | WP_096778336.1 |
| GCF_028752115.1 | NZ_CP082270.1 | 757867 | 759048 | 1181 | + | tuf | WP_166914980.1 |
| GCF_028752115.1 | NZ_CP082270.1 | 766853 | 768034 | 1181 | - | tuf | WP_166914980.1 |
| GCF_034508895.1 | NZ_CP140571.1 | 324834 | 326024 | 1190 | + | tuf | WP_027704510.1 |
| GCF_034508895.1 | NZ_CP140571.1 | 1046448 | 1047647 | 1199 | + | tuf | WP_012682360.1 |
| GCF_004551575.1 | NZ_CP037883.1 | 2482760 | 2483956 | 1196 | - | tuf | WP_171225343.1 |
| GCF_004551575.1 | NZ_CP037883.1 | 1244691 | 1245890 | 1199 | + | tuf | WP_026849972.1 |
| GCF_041956365.1 | NZ_CP167120.1 | 3234191 | 3235390 | 1199 | + | tuf | WP_025411781.1 |
| GCF_041956365.1 | NZ_CP167120.1 | 806351 | 807550 | 1199 | + | tuf | WP_367887158.1 |
| GCF_900186865.1 | NZ_LT906480.1 | 2923389 | 2924579 | 1190 | - | tuf | WP_286354204.1 |
| GCF_900186865.1 | NZ_LT906480.1 | 275760 | 276950 | 1190 | + | tuf | WP_243330169.1 |
| GCF_031834595.1 | NZ_CP115541.1 | 280512 | 281702 | 1190 | + | tuf | WP_316413953.1 |
| GCF_031834595.1 | NZ_CP115541.1 | 3708420 | 3709601 | 1181 | - | tuf | WP_237383031.1 |
| GCF_030549615.1 | NZ_CP130832.1 | 1485172 | 1486371 | 1199 | - | tuf | WP_201327234.1 |
| GCF_030549615.1 | NZ_CP130832.1 | 1501351 | 1502550 | 1199 | - | tuf | WP_201327234.1 |
| GCF_000661955.1 | NZ_CP007597.1 | 412484 | 413671 | 1187 | + | tuf | WP_119334464.1 |
| GCF_000661955.1 | NZ_CP007597.1 | 428701 | 429888 | 1187 | + | tuf | WP_119334476.1 |
| GCF_040256885.1 | NZ_CP155568.1 | 1018551 | 1019753 | 1202 | + | tuf | WP_014559777.1 |
| GCF_040256885.1 | NZ_CP155568.1 | 245013 | 246215 | 1202 | - | tuf | WP_014854899.1 |
| GCF_041531565.1 | NZ_CP168059.1 | 2009697 | 2010899 | 1202 | - | tuf | WP_052882247.1 |
| GCF_041531565.1 | NZ_CP168059.1 | 7045466 | 7046668 | 1202 | + | tuf | WP_129891152.1 |

| | | | | | | |
|---|---|---|---|---|---|---|
| GCF_001579945.1 | NZ_CP011971.1 | 7310825 | 7312027 | 1202 + | tuf | WP_129891152.1 |
| GCF_001579945.1 | NZ_CP011971.1 | 325689 | 326882 | 1193 - | tuf | WP_274153596.1 |
| GCF_000818015.1 | NZ_CP007511.1 | 337239 | 338438 | 1199 + | tuf | WP_324716984.1 |
| GCF_000818015.1 | NZ_CP007511.1 | 2529453 | 2530652 | 1199 - | tuf | WP_324668460.1 |
| GCF_020783375.1 | NZ_CP086067.1 | 3438648 | 3439847 | 1199 - | tuf | WP_068140403.1 |
| GCF_020783375.1 | NZ_CP086067.1 | 3454839 | 3456038 | 1199 - | tuf | WP_068140434.1 |
| GCF_000661915.1 | NZ_CP007509.1 | 1006734 | 1007915 | 1181 + | tuf | WP_009059040.1 |
| GCF_000661915.1 | NZ_CP007509.1 | 504509 | 505690 | 1181 - | tuf | WP_134438989.1 |
| GCF_016028635.1 | NZ_CP065721.1 | 755527 | 756708 | 1181 + | tuf | WP_012463134.1 |
| GCF_016028635.1 | NZ_CP065721.1 | 849370 | 850551 | 1181 + | tuf | WP_039721804.1 |
| GCF_016028515.1 | NZ_CP065720.1 | 951890 | 953071 | 1181 - | tuf | WP_035368549.1 |
| GCF_016028515.1 | NZ_CP065720.1 | 181291 | 182475 | 1184 + | tuf | WP_012242145.1 |
| GCF_019704535.1 | NZ_AP024722.1 | 1377396 | 1378577 | 1181 - | tuf | WP_030003721.1 |
| GCF_019704535.1 | NZ_AP024722.1 | 544324 | 545508 | 1184 - | tuf | WP_025208900.1 |
| GCF_900108535.1 | NZ_LT629970.1 | 237730 | 238914 | 1184 + | tuf | WP_303717360.1 |
| GCF_900108535.1 | NZ_LT629970.1 | 372164 | 373345 | 1181 + | tuf | WP_011160639.1 |
| GCF_016747875.1 | NZ_CP068345.1 | 412389 | 413573 | 1184 - | tuf | WP_341266427.1 |
| GCF_016747875.1 | NZ_CP068345.1 | 283640 | 284827 | 1187 - | tuf | WP_210954883.1 |
| GCF_002355735.1 | NZ_AP014879.1 | 474914 | 476089 | 1175 + | tuf | WP_012504541.1 |
| GCF_002355735.1 | NZ_AP014879.1 | 573354 | 574514 | 1160 - | tuf | WP_268850211.1 |
| GCF_003967195.1 | NZ_AP018725.1 | 268326 | 269507 | 1181 + | tuf | WP_349402149.1 |
| GCF_003967195.1 | NZ_AP018725.1 | 478755 | 479915 | 1160 + | tuf | WP_121464061.1 |
| GCF_002355415.1 | NZ_AP014936.1 | 129071 | 130255 | 1184 + | tuf | WP_100609095.1 |
| GCF_002355415.1 | NZ_AP014936.1 | 572050 | 573231 | 1181 + | tuf | WP_026390661.1 |
| GCF_004001245.1 | NZ_AP018724.1 | 319524 | 320705 | 1181 + | tuf | WP_176239666.1 |
| GCF_004001245.1 | NZ_AP018724.1 | 783827 | 785008 | 1181 - | tuf | WP_027875536.1 |
| GCF_019797845.1 | NZ_CP081864.1 | 740466 | 741647 | 1181 - | tuf | WP_100671436.1 |
| GCF_019797845.1 | NZ_CP081864.1 | 788053 | 789234 | 1181 - | tuf | WP_099651467.1 |
| GCF_025398735.1 | NZ_CP104694.1 | 727925 | 729106 | 1181 - | tuf | WP_011183518.1 |
| GCF_025398735.1 | NZ_CP104694.1 | 792200 | 793384 | 1184 + | tuf | WP_096863078.1 |
| GCF_002163585.1 | NZ_CP015579.1 | 777080 | 778261 | 1181 - | tuf | WP_028124553.1 |

| | | | | | | | |
|---|---|---|---|---|---|---|---|
| GCF_002163585.1 | NZ_CP015579.1 | 130880 | 132064 | 1184 | + | tuf | WP_027048400.1 |
| GCF_014211975.1 | NZ_CP060084.1 | 772660 | 773841 | 1181 | - | tuf | WP_100679781.1 |
| GCF_014217335.1 | NZ_CP060092.1 | 31888 | 33069 | 1181 | - | tuf | WP_343251503.1 |
| GCF_037935975.1 | NZ_CP149817.1 | 898133 | 899317 | 1184 | - | tuf | WP_146367676.1 |
| GCF_025345565.1 | NZ_CP054475.1 | 102103 | 103287 | 1184 | + | tuf | WP_416754641.1 |
| GCF_025345565.1 | NZ_CP054475.1 | 263660 | 264841 | 1181 | + | tuf | WP_255034860.1 |
| GCF_025345565.1 | NZ_CP054475.1 | 183880 | 185064 | 1184 | + | tuf | WP_011166362.1 |
| GCF_000355675.1 | NC_020888.1 | 841531 | 842715 | 1184 | - | tuf | WP_406614085.1 |
| GCF_000355675.1 | NC_020888.1 | 114704 | 115888 | 1184 | + | tuf | WP_259430275.1 |
| GCF_000355675.1 | NC_020888.1 | 42789 | 43979 | 1190 | + | tuf | WP_013054124.1 |
| GCF_000948975.2 | NZ_CP059735.1 | 167788 | 168972 | 1184 | + | tuf | WP_008362398.1 |
| GCF_000948975.2 | NZ_CP059735.1 | 888708 | 889892 | 1184 | + | tuf | WP_426461223.1 |
| GCF_028657945.1 | NZ_CP059693.1 | 562825 | 564012 | 1187 | - | tuf | WP_258210765.1 |
| GCF_028657945.1 | NZ_CP059693.1 | 232076 | 233260 | 1184 | + | tuf | WP_011166362.1 |
| GCF_000948985.2 | NZ_CP059733.1 | 731275 | 732462 | 1187 | - | tuf | WP_171111904.1 |
| GCF_000948985.2 | NZ_CP059733.1 | 605013 | 606197 | 1184 | + | tuf | WP_171113237.1 |
| GCF_030295955.1 | NZ_AP027363.1 | 872816 | 874000 | 1184 | - | tuf | WP_011166362.1 |
| GCF_030295955.1 | NZ_AP027363.1 | 712926 | 714110 | 1184 | + | tuf | WP_139592409.1 |
| GCF_041410465.1 | NZ_CP166961.1 | 151272 | 152453 | 1181 | + | tuf | WP_024070985.1 |
| GCF_041410465.1 | NZ_CP166961.1 | 477907 | 479088 | 1181 | - | tuf | WP_022770413.1 |
| GCF_041410465.1 | NZ_CP166961.1 | 178427 | 179611 | 1184 | + | tuf | WP_169604890.1 |
| GCF_030295975.1 | NZ_AP027364.1 | 98722 | 99912 | 1190 | - | tuf | WP_169580029.1 |
| GCF_030295975.1 | NZ_AP027364.1 | 768659 | 769843 | 1184 | - | tuf | WP_014035359.1 |
| GCF_001831495.1 | NZ_CP017689.1 | 963827 | 965011 | 1184 | - | tuf | WP_305937913.1 |
| GCF_001831495.1 | NZ_CP017689.1 | 457797 | 458987 | 1190 | + | tuf | WP_124724406.1 |
| GCF_032911225.1 | NZ_CP136600.1 | 232079 | 233260 | 1181 | + | tuf | WP_013608939.1 |
| GCF_032911225.1 | NZ_CP136600.1 | 361011 | 362198 | 1187 | + | tuf | WP_205517621.1 |
| GCF_032911225.1 | NZ_CP136600.1 | 256384 | 257565 | 1181 | + | tuf | WP_182078992.1 |
| GCF_030295995.1 | NZ_AP027365.1 | 302268 | 303449 | 1181 | + | tuf | WP_014849796.1 |
| GCF_030295995.1 | NZ_AP027365.1 | 488474 | 489655 | 1181 | - | tuf | WP_014849796.1 |
| GCF_041379765.1 | NZ_CP166919.1 | 117237 | 118421 | 1184 | + | tuf | WP_004428385.1 |

| | | | | | | | |
|---|---|---|---|---|---|---|---|
| GCF_041379765.1 | NZ_CP166919.1 | 341171 | 342352 | 1181 | + | tuf | WP_030004194.1 |
| GCF_041379765.1 | NZ_CP166919.1 | 311260 | 312444 | 1184 | + | tuf | WP_115557861.1 |
| GCF_031583575.1 | NZ_CP134146.1 | 1105392 | 1106576 | 1184 | - | tuf | WP_023790227.1 |
| GCF_031583575.1 | NZ_CP134146.1 | 1114088 | 1115272 | 1184 | - | tuf | WP_053946606.1 |
| GCF_031583575.1 | NZ_CP134146.1 | 1297910 | 1299094 | 1184 | - | tuf | WP_166508713.1 |
| GCF_030295935.1 | NZ_AP027362.1 | 74743 | 75927 | 1184 | + | tuf | WP_016338472.1 |
| GCF_030295935.1 | NZ_AP027362.1 | 96868 | 98055 | 1187 | + | tuf | WP_071890513.1 |
| GCF_041379045.1 | NZ_CP166871.1 | 203626 | 204810 | 1184 | + | tuf | WP_100254065.1 |
| GCF_041379045.1 | NZ_CP166871.1 | 1107721 | 1108905 | 1184 | - | tuf | WP_094049719.1 |
| GCF_041379045.1 | NZ_CP166871.1 | 1106525 | 1107709 | 1184 | - | tuf | WP_025363567.1 |
| GCF_031583595.1 | NZ_CP134145.1 | 879941 | 881125 | 1184 | - | tuf | WP_020836730.1 |
| GCF_031583595.1 | NZ_CP134145.1 | 61441 | 62694 | 1253 | + | tuf | WP_338961656.1 |
| GCF_031583595.1 | NZ_CP134145.1 | 1095901 | 1097085 | 1184 | - | tuf | WP_339024329.1 |
| GCF_030295915.1 | NZ_AP027361.1 | 177387 | 178640 | 1253 | - | tuf | WP_425377202.1 |
| GCF_030295915.1 | NZ_AP027361.1 | 179051 | 180238 | 1187 | + | tuf | WP_348736040.1 |
| GCF_019703835.1 | NZ_AP023054.1 | 412901 | 414154 | 1253 | + | tuf | WP_342218735.1 |
| GCF_019703835.1 | NZ_AP023054.1 | 630081 | 631334 | 1253 | + | tuf | WP_286641788.1 |
| GCF_009664085.1 | NZ_CP039268.1 | 440235 | 441419 | 1184 | + | tuf | WP_381414809.1 |
| GCF_009664085.1 | NZ_CP039268.1 | 68427 | 69680 | 1253 | + | tuf | WP_339042031.1 |
| GCF_006337105.1 | NZ_CP040871.1 | 1227663 | 1228847 | 1184 | - | tuf | WP_342223976.1 |
| GCF_006337105.1 | NZ_CP040871.1 | 1090589 | 1091773 | 1184 | - | tuf | WP_342268807.1 |
| GCF_014395425.1 | NZ_CP060711.1 | 1001951 | 1003135 | 1184 | - | tuf | WP_339020784.1 |
| GCF_014395425.1 | NZ_CP060711.1 | 1124780 | 1125964 | 1184 | + | tuf | WP_339030065.1 |
| GCF_024498955.1 | NZ_CP093311.1 | 1178181 | 1179365 | 1184 | - | tuf | WP_339034047.1 |
| GCF_024498955.1 | NZ_CP093311.1 | 1945348 | 1946532 | 1184 | - | tuf | WP_342264066.1 |
| GCF_000321415.2 | NC_019902.2 | 372297 | 373550 | 1253 | + | tuf | WP_286641788.1 |
| GCF_000321415.2 | NC_019902.2 | 648337 | 649521 | 1184 | - | tuf | WP_339022686.1 |
| GCF_000227685.2 | NZ_CP007029.1 | 381155 | 382408 | 1253 | + | tuf | WP_338988011.1 |
| GCF_000227685.2 | NZ_CP007029.1 | 838569 | 839753 | 1184 | - | tuf | WP_338969016.1 |
| GCF_000021985.1 | NC_011901.1 | 1120457 | 1121641 | 1184 | + | tuf | WP_342258788.1 |
| GCF_000021985.1 | NC_011901.1 | 918302 | 919486 | 1184 | - | tuf | WP_338984976.1 |

| | | | | | | | |
|---|---|---|---|---|---|---|---|
| GCF_001020955.1 | NZ_CP011367.1 | 1271055 | 1272239 | 1184 | - | tuf | WP_342262727.1 |
| GCF_001020955.1 | NZ_CP011367.1 | 996424 | 997608 | 1184 | - | tuf | WP_338982691.1 |
| GCF_021228795.1 | NZ_CP089309.1 | 96624 | 97811 | 1187 | + | tuf | WP_424526613.1 |
| GCF_021228795.1 | NZ_CP089309.1 | 895866 | 897050 | 1184 | - | tuf | WP_338969653.1 |
| GCF_000227745.2 | NC_018012.1 | 1257295 | 1258548 | 1253 | - | tuf | WP_338980156.1 |
| GCF_000227745.2 | NC_018012.1 | 61748 | 63001 | 1253 | + | tuf | WP_338961656.1 |
| GCF_000327045.1 | NC_019940.1 | 1274172 | 1275356 | 1184 | + | tuf | WP_338969016.1 |
| GCF_000327045.1 | NC_019940.1 | 1536965 | 1538218 | 1253 | + | tuf | WP_425382428.1 |
| GCF_004001325.1 | NZ_AP018722.1 | 1264278 | 1265531 | 1253 | - | tuf | WP_338961656.1 |
| GCF_004001325.1 | NZ_AP018722.1 | 1323541 | 1324782 | 1241 | + | tuf | WP_342276281.1 |
| GCF_021654855.1 | NZ_AP024202.1 | 828481 | 829665 | 1184 | - | tuf | WP_215825561.1 |
| GCF_021654855.1 | NZ_AP024202.1 | 264579 | 265763 | 1184 | + | tuf | WP_338969016.1 |
| GCF_004293625.1 | NZ_CP033040.1 | 50472 | 51725 | 1253 | + | tuf | WP_342257783.1 |
| GCF_004293625.1 | NZ_CP033040.1 | 217448 | 218701 | 1253 | - | tuf | WP_395473388.1 |
| GCF_029201445.1 | NZ_CP102381.1 | 852409 | 853662 | 1253 | + | tuf | WP_286641788.1 |
| GCF_029201445.1 | NZ_CP102381.1 | 739629 | 740813 | 1184 | - | tuf | WP_338983384.1 |
| GCF_005885815.1 | NZ_CP040602.1 | 253250 | 254434 | 1184 | + | tuf | WP_115557861.1 |
| GCF_005885815.1 | NZ_CP040602.1 | 1311807 | 1313060 | 1253 | - | tuf | WP_338978343.1 |
| GCF_013282625.1 | NZ_CP054020.1 | 622405 | 623592 | 1187 | + | tuf | WP_253301309.1 |
| GCF_013282625.1 | NZ_CP054020.1 | 690847 | 692034 | 1187 | + | tuf | WP_342256781.1 |
| GCF_000227665.2 | NZ_CP007030.1 | 46768 | 47955 | 1187 | + | tuf | WP_338955468.1 |
| GCF_000227665.2 | NZ_CP007030.1 | 238763 | 239950 | 1187 | + | tuf | WP_374696016.1 |
| GCF_000214825.1 | NC_015581.1 | 81252 | 82505 | 1253 | + | tuf | WP_425381329.1 |
| GCF_000214825.1 | NC_015581.1 | 926629 | 927816 | 1187 | + | tuf | WP_338964859.1 |
| GCF_001267175.1 | NZ_CP012358.1 | 863198 | 864385 | 1187 | - | tuf | WP_338990251.1 |
| GCF_001267175.1 | NZ_CP012358.1 | 100963 | 102150 | 1187 | + | tuf | WP_425380101.1 |
| GCF_033954835.1 | NZ_CP121471.1 | 60755 | 62008 | 1253 | + | tuf | WP_338961656.1 |
| GCF_033954835.1 | NZ_CP121471.1 | 110441 | 111694 | 1253 | + | tuf | WP_286641788.1 |
| GCF_033954455.1 | NZ_CP121473.1 | 1240811 | 1241995 | 1184 | - | tuf | WP_375315363.1 |
| GCF_033954455.1 | NZ_CP121473.1 | 78643 | 79896 | 1253 | + | tuf | WP_338961656.1 |
| GCF_036208045.1 | NZ_CP121472.1 | 92902 | 94155 | 1253 | - | tuf | WP_342266616.1 |

| | | | | | | | |
|---|---|---|---|---|---|---|---|
| GCF_036208045.1 | NZ_CP121472.1 | 1446269 | 1447510 | 1241 | - | tuf | WP_375317533.1 |
| GCF_002009425.1 | NZ_CP019936.1 | 190707 | 191960 | 1253 | + | tuf | WP_342254104.1 |
| GCF_002009425.1 | NZ_CP019936.1 | 93206 | 94387 | 1181 | + | tuf | WP_047790972.1 |
| GCF_011398355.1 | NZ_AP021889.1 | 1216756 | 1217940 | 1184 | - | tuf | WP_100917066.1 |
| GCF_011398355.1 | NZ_AP021889.1 | 1100231 | 1101415 | 1184 | - | tuf | WP_134298274.1 |
| GCF_011398155.1 | NZ_AP021888.1 | 1242647 | 1243831 | 1184 | - | tuf | WP_069117434.1 |
| GCF_011398155.1 | NZ_AP021888.1 | 96600 | 97853 | 1253 | + | tuf | WP_286641788.1 |
| GCF_017901135.1 | NZ_CP072801.1 | 1367090 | 1368277 | 1187 | - | tuf | WP_053391487.1 |
| GCF_017901135.1 | NZ_CP072801.1 | 1118797 | 1119981 | 1184 | - | tuf | WP_075058720.1 |
| GCF_016772315.1 | NZ_CP053482.1 | 71313 | 72494 | 1181 | + | tuf | WP_025316900.1 |
| GCF_021650935.1 | NZ_CP091244.1 | 831130 | 832314 | 1184 | - | tuf | WP_101781045.1 |
| GCF_021650935.1 | NZ_CP091244.1 | 874257 | 875444 | 1187 | - | tuf | WP_114564848.1 |
| GCF_000023065.1 | NC_012691.1 | 661995 | 663236 | 1241 | - | tuf | WP_168104950.1 |
| GCF_000023065.1 | NC_012691.1 | 1783040 | 1784227 | 1187 | - | tuf | WP_040094444.1 |
| GCF_030053895.1 | NZ_CP125781.1 | 207957 | 209141 | 1184 | + | tuf | WP_025250726.1 |
| GCF_030053895.1 | NZ_CP125781.1 | 94199 | 95383 | 1184 | + | tuf | WP_016340335.1 |
| GCF_036251705.1 | NZ_CP133959.1 | 1207216 | 1208400 | 1184 | - | tuf | WP_156007341.1 |
| GCF_036251705.1 | NZ_CP133959.1 | 1018381 | 1019565 | 1184 | - | tuf | WP_020834770.1 |
| GCF_016132445.1 | NZ_CP046056.1 | 1198953 | 1200137 | 1184 | - | tuf | WP_075048838.1 |
| GCF_016132445.1 | NZ_CP046056.1 | 397378 | 398559 | 1181 | - | tuf | WP_015637948.1 |
| GCF_016132445.1 | NZ_CP046056.1 | 317780 | 318964 | 1184 | + | tuf | WP_025734492.1 |
| GCF_013377275.1 | NZ_CP016615.1 | 131939 | 133123 | 1184 | + | tuf | WP_024863815.1 |
| GCF_013377275.1 | NZ_CP016615.1 | 2271319 | 2272506 | 1187 | + | tuf | WP_012525973.1 |
| GCF_024346755.1 | NZ_AP024861.1 | 2624210 | 2625397 | 1187 | - | tuf | WP_012525973.1 |
| GCF_024346755.1 | NZ_AP024861.1 | 6223086 | 6224273 | 1187 | + | tuf | WP_248356329.1 |
| GCF_019670485.1 | NZ_AP024165.1 | 6579145 | 6580332 | 1187 | - | tuf | WP_248357248.1 |
| GCF_019670485.1 | NZ_AP024165.1 | 706997 | 708184 | 1187 | + | tuf | WP_248344263.1 |
| GCF_009601765.2 | NZ_CP045699.1 | 975137 | 976324 | 1187 | - | tuf | WP_248344263.1 |
| GCF_009601765.2 | NZ_CP045699.1 | 4382330 | 4383502 | 1172 | - | tuf | WP_047856422.1 |
| GCF_023650915.1 | NZ_CP098034.1 | 5897829 | 5899016 | 1187 | + | tuf | WP_047857257.1 |
| GCF_023650915.1 | NZ_CP098034.1 | 8244434 | 8245621 | 1187 | - | tuf | WP_047857257.1 |

| | | | | | | | |
|---|---|---|---|---|---|---|---|
| GCF_000024825.1 | NC_013456.1 | 4626438 | 4627625 | 1187 | + | tuf | WP_376762789.1 |
| GCF_000024825.1 | NC_013456.1 | 7322349 | 7323536 | 1187 | - | tuf | WP_376754909.1 |
| GCF_002157735.2 | NZ_AP018689.1 | 8932554 | 8933726 | 1172 | + | tuf | WP_376755997.1 |
| GCF_002157735.2 | NZ_AP018689.1 | 3746748 | 3747935 | 1187 | + | tuf | WP_014395860.1 |
| GCF_030297155.1 | NZ_AP028067.1 | 6265624 | 6266811 | 1187 | - | tuf | WP_014395860.1 |
| GCF_030297155.1 | NZ_AP028067.1 | 3606375 | 3607562 | 1187 | + | tuf | WP_013939874.1 |
| GCF_009363415.1 | NZ_CP045350.1 | 3893321 | 3894508 | 1187 | + | tuf | WP_013939874.1 |
| GCF_009363415.1 | NZ_CP045350.1 | 113329 | 114516 | 1187 | - | tuf | WP_095975577.1 |
| GCF_024347295.1 | NZ_AP025458.1 | 7984275 | 7985462 | 1187 | + | tuf | WP_095975577.1 |
| GCF_024347295.1 | NZ_AP025458.1 | 912743 | 913930 | 1187 | + | tuf | WP_206716907.1 |
| GCF_010587385.1 | NZ_CP047475.1 | 2492490 | 2493677 | 1187 | - | tuf | WP_206716907.1 |
| GCF_010587385.1 | NZ_CP047475.1 | 4436527 | 4437714 | 1187 | + | tuf | WP_015349098.1 |
| GCF_024347315.1 | NZ_AP025460.1 | 5968872 | 5970059 | 1187 | - | tuf | WP_015350187.1 |
| GCF_024347315.1 | NZ_AP025460.1 | 3593508 | 3594695 | 1187 | + | tuf | WP_011553119.1 |
| GCF_002849855.1 | NZ_CP018616.1 | 3853234 | 3854421 | 1187 | + | tuf | WP_011553342.1 |
| GCF_002849855.1 | NZ_CP018616.1 | 1509272 | 1510459 | 1187 | + | tuf | WP_394826485.1 |
| GCF_014879975.1 | NZ_CP062500.1 | 1563107 | 1564294 | 1187 | + | tuf | WP_394826527.1 |
| GCF_014879975.1 | NZ_CP062500.1 | 230007 | 231176 | 1169 | - | tuf | WP_394846018.1 |
| GCF_024346795.1 | NZ_AP024864.1 | 9874593 | 9875780 | 1187 | - | tuf | WP_394842977.1 |
| GCF_024346795.1 | NZ_AP024864.1 | 9919514 | 9920701 | 1187 | - | tuf | WP_394843008.1 |
| GCF_002906475.1 | NZ_CP026321.1 | 9987624 | 9988811 | 1187 | - | tuf | WP_394832415.1 |
| GCF_002906475.1 | NZ_CP026321.1 | 10036834 | 10038021 | 1187 | - | tuf | WP_394832450.1 |
| GCF_040412085.2 | NZ_CP115920.1 | 4601917 | 4603104 | 1187 | - | tuf | WP_049872326.1 |
| GCF_040412085.2 | NZ_CP115920.1 | 4940561 | 4941748 | 1187 | - | tuf | WP_049872326.1 |
| GCF_024347355.1 | NZ_AP025465.1 | 5208073 | 5209260 | 1187 | - | tuf | WP_206728474.1 |
| GCF_024347355.1 | NZ_AP025465.1 | 5676372 | 5677559 | 1187 | - | tuf | WP_206728474.1 |
| GCF_008369605.1 | NZ_CP043554.1 | 2346565 | 2347752 | 1187 | - | tuf | WP_050725882.1 |
| GCF_008369605.1 | NZ_CP043554.1 | 2460794 | 2461981 | 1187 | - | tuf | WP_050725882.1 |
| GCF_009665415.1 | NZ_CP035921.1 | 311623 | 312813 | 1190 | + | tuf | WP_012937652.1 |
| GCF_009665415.1 | NZ_CP035921.1 | 2134933 | 2136123 | 1190 | - | tuf | WP_012937652.1 |
| GCF_024346835.1 | NZ_AP024866.1 | 2055211 | 2056395 | 1184 | - | tuf | WP_014129083.1 |

| | | | | | | | |
|---|---|---|---|---|---|---|---|
| GCF_024346835.1 | NZ_AP024866.1 | 2174214 | 2175398 | 1184 | - | tuf | WP_014129083.1 |
| GCF_029541605.1 | NZ_CP120719.1 | 1467741 | 1468925 | 1184 | - | tuf | WP_108850696.1 |
| GCF_029541605.1 | NZ_CP120719.1 | 1636038 | 1637222 | 1184 | - | tuf | WP_143332667.1 |
| GCF_024347395.1 | NZ_AP025472.1 | 2000420 | 2001604 | 1184 | + | tuf | WP_108850696.1 |
| GCF_024347395.1 | NZ_AP025472.1 | 2169250 | 2170434 | 1184 | + | tuf | WP_108850696.1 |
| GCF_024347415.1 | NZ_AP025476.1 | 251285 | 252469 | 1184 | - | tuf | WP_273187623.1 |
| GCF_024347415.1 | NZ_AP025476.1 | 1259628 | 1260812 | 1184 | + | tuf | WP_273185634.1 |
| GCF_005144905.1 | NZ_CP039700.1 | 412526 | 413710 | 1184 | + | tuf | WP_008538884.1 |
| GCF_005144905.1 | NZ_CP039700.1 | 1525109 | 1526293 | 1184 | - | tuf | WP_027889348.1 |
| GCF_011801455.1 | NZ_CP042451.1 | 1123788 | 1124972 | 1184 | + | tuf | WP_014015702.1 |
| GCF_011801455.1 | NZ_CP042451.1 | 1168689 | 1169879 | 1190 | + | tuf | WP_014015733.1 |
| GCF_038452265.1 | NZ_CP151842.1 | 2594617 | 2595807 | 1190 | - | tuf | WP_113855078.1 |
| GCF_038452265.1 | NZ_CP151842.1 | 2647990 | 2649180 | 1190 | - | tuf | WP_113855078.1 |
| GCF_024347455.1 | NZ_AP025483.1 | 811504 | 812688 | 1184 | - | tuf | WP_115759885.1 |
| GCF_024347455.1 | NZ_AP025483.1 | 902891 | 904075 | 1184 | - | tuf | WP_115759885.1 |
| GCF_001558415.2 | NZ_CP014035.2 | 1626668 | 1627855 | 1187 | - | tuf | WP_427111902.1 |
| GCF_001558415.2 | NZ_CP014035.2 | 1721607 | 1722794 | 1187 | - | tuf | WP_427111902.1 |
| GCF_024347475.1 | NZ_AP025487.1 | 2222756 | 2223955 | 1199 | + | tuf | WP_126308372.1 |
| GCF_024347475.1 | NZ_AP025487.1 | 2240820 | 2242019 | 1199 | + | tuf | WP_126308372.1 |
| GCF_006364355.1 | NZ_CP040990.1 | 323598 | 324782 | 1184 | + | tuf | WP_182187666.1 |
| GCF_006364355.1 | NZ_CP040990.1 | 1056339 | 1057523 | 1184 | + | tuf | WP_113077255.1 |
| GCF_024347495.1 | NZ_AP025490.1 | 1094990 | 1096174 | 1184 | + | tuf | WP_113077255.1 |
| GCF_024347495.1 | NZ_AP025490.1 | 1770959 | 1772146 | 1187 | - | tuf | WP_009145072.1 |
| GCF_024346875.1 | NZ_AP024871.1 | 1838793 | 1839980 | 1187 | - | tuf | WP_009145072.1 |
| GCF_024346875.1 | NZ_AP024871.1 | 1933984 | 1935168 | 1184 | - | tuf | WP_147670314.1 |
| GCF_026001925.1 | NZ_AP021869.1 | 1853863 | 1855047 | 1184 | - | tuf | WP_304986868.1 |
| GCF_026001925.1 | NZ_AP021869.1 | 1642739 | 1643923 | 1184 | - | tuf | WP_006191715.1 |
| GCF_023920225.1 | NZ_CP092587.1 | 1081609 | 1082793 | 1184 | + | tuf | WP_009441263.1 |
| GCF_023920225.1 | NZ_CP092587.1 | 485998 | 487197 | 1199 | + | tuf | WP_094607976.1 |
| GCF_022371215.1 | NZ_CP092384.1 | 502891 | 504090 | 1199 | + | tuf | WP_094607977.1 |
| GCF_022371215.1 | NZ_CP092384.1 | 446354 | 447553 | 1199 | + | tuf | WP_371376757.1 |

| | | | | | | | |
|---|---|---|---|---|---|---|---|
| GCF_030060435.1 | NZ_CP125875.1 | 463962 | 465161 | 1199 | + | tuf | WP_371376796.1 |
| GCF_030060435.1 | NZ_CP125875.1 | 452732 | 453931 | 1199 | + | tuf | WP_425059795.1 |
| GCF_002906655.1 | NZ_CP025794.1 | 469579 | 470778 | 1199 | + | tuf | WP_425059795.1 |
| GCF_002906655.1 | NZ_CP025794.1 | 4782820 | 4784019 | 1199 | - | tuf | WP_371360906.1 |
| GCF_024346935.1 | NZ_AP024878.1 | 4799580 | 4800779 | 1199 | - | tuf | WP_371360906.1 |
| GCF_024346935.1 | NZ_AP024878.1 | 399558 | 400757 | 1199 | + | tuf | WP_021168675.1 |
| GCF_024347675.1 | NZ_AP024881.1 | 417133 | 418332 | 1199 | + | tuf | WP_021168675.1 |
| GCF_024347675.1 | NZ_AP024881.1 | 475753 | 476952 | 1199 | + | tuf | WP_188399590.1 |
| GCF_024582835.1 | NZ_CP102096.1 | 492782 | 493981 | 1199 | + | tuf | WP_188399586.1 |
| GCF_024582835.1 | NZ_CP102096.1 | 480088 | 481287 | 1199 | + | tuf | WP_371365795.1 |
| GCF_002887615.1 | NZ_CP025792.1 | 498220 | 499419 | 1199 | + | tuf | WP_371365795.1 |
| GCF_002887615.1 | NZ_CP025792.1 | 439963 | 441162 | 1199 | + | tuf | WP_094607438.1 |
| GCF_024347535.1 | NZ_AP025497.1 | 456780 | 457979 | 1199 | + | tuf | WP_094607438.1 |
| GCF_024347535.1 | NZ_AP025497.1 | 499604 | 500803 | 1199 | + | tuf | WP_075752792.1 |
| GCF_024346955.1 | NZ_AP024883.1 | 516510 | 517709 | 1199 | + | tuf | WP_075752792.1 |
| GCF_024346955.1 | NZ_AP024883.1 | 37184 | 38383 | 1199 | + | tuf | WP_144348517.1 |
| GCF_030580635.1 | NZ_CP129421.1 | 54370 | 55569 | 1199 | + | tuf | WP_144348517.1 |
| GCF_030580635.1 | NZ_CP129421.1 | 1674969 | 1676153 | 1184 | - | tuf | WP_126413463.1 |
| GCF_029635775.1 | NZ_CP113241.1 | 1698155 | 1699339 | 1184 | - | tuf | WP_039968974.1 |
| GCF_029635775.1 | NZ_CP113241.1 | 1618373 | 1619557 | 1184 | - | tuf | WP_105084932.1 |
| GCF_001456255.1 | NZ_CP009977.1 | 1642800 | 1643984 | 1184 | - | tuf | WP_105084932.1 |
| GCF_001456255.1 | NZ_CP009977.1 | 1634337 | 1635521 | 1184 | - | tuf | WP_105084932.1 |
| GCF_024346975.1 | NZ_AP024885.1 | 1658246 | 1659430 | 1184 | - | tuf | WP_105084932.1 |
| GCF_024346975.1 | NZ_AP024885.1 | 1698100 | 1699284 | 1184 | - | tuf | WP_012864763.1 |
| GCF_027924785.1 | NZ_AP027049.1 | 1723615 | 1724799 | 1184 | - | tuf | WP_012864763.1 |
| GCF_027924785.1 | NZ_AP027049.1 | 1676123 | 1677307 | 1184 | - | tuf | WP_095066419.1 |
| GCF_019226825.1 | NZ_CP076643.1 | 1701700 | 1702884 | 1184 | - | tuf | WP_095066429.1 |
| GCF_019226825.1 | NZ_CP076643.1 | 2836559 | 2837743 | 1184 | + | tuf | WP_131155432.1 |
| GCF_002021755.1 | NZ_CP019959.1 | 2856777 | 2857961 | 1184 | + | tuf | WP_131155432.1 |
| GCF_002021755.1 | NZ_CP019959.1 | 650372 | 651556 | 1184 | + | tuf | WP_130648539.1 |
| GCF_024346995.1 | NZ_AP024887.1 | 672261 | 673445 | 1184 | + | tuf | WP_130648539.1 |

| | | | | | | | |
|---|---|---|---|---|---|---|---|
| GCF_024346995.1 | NZ_AP024887.1 | 5102309 | 5103493 | 1184 | - | tuf | WP_114593377.1 |
| GCF_037672085.1 | NZ_JBBHLM0100000C | 5147778 | 5148962 | 1184 | - | tuf | WP_114593377.1 |
| GCF_037672085.1 | NZ_JBBHLM0100000C | 2686470 | 2687657 | 1187 | - | tuf | WP_282012274.1 |
| GCF_000196095.1 | NC_004603.1 | 1579383 | 1580585 | 1202 | + | tuf | WP_312646243.1 |
| GCF_000196095.1 | NC_004603.1 | 1595460 | 1596662 | 1202 | + | tuf | WP_312646243.1 |
| GCF_024347015.1 | NZ_AP024889.1 | 1656833 | 1658035 | 1202 | + | tuf | WP_062484667.1 |
| GCF_024347015.1 | NZ_AP024889.1 | 1673271 | 1674473 | 1202 | + | tuf | WP_062484667.1 |
| GCF_024347575.1 | NZ_AP025503.1 | 4186138 | 4187340 | 1202 | + | tuf | WP_312744067.1 |
| GCF_024347575.1 | NZ_AP025503.1 | 4202211 | 4203413 | 1202 | + | tuf | WP_312744067.1 |
| GCF_019977755.1 | NZ_AP025144.1 | 852136 | 853332 | 1196 | + | tuf | WP_014449063.1 |
| GCF_019977755.1 | NZ_AP025144.1 | 869933 | 871129 | 1196 | + | tuf | WP_014449075.1 |
| GCF_024347595.1 | NZ_AP025506.1 | 2196805 | 2198007 | 1202 | + | tuf | WP_080886867.1 |
| GCF_024347595.1 | NZ_AP025506.1 | 2213112 | 2214314 | 1202 | + | tuf | WP_080886867.1 |
| GCF_009938225.1 | NZ_AP019657.1 | 1128737 | 1129939 | 1202 | + | tuf | WP_053378932.1 |
| GCF_009938225.1 | NZ_AP019657.1 | 1145239 | 1146441 | 1202 | + | tuf | WP_053378932.1 |
| GCF_024347055.1 | NZ_AP024895.1 | 784383 | 785567 | 1184 | + | tuf | WP_130606560.1 |
| GCF_024347055.1 | NZ_AP024895.1 | 792924 | 794108 | 1184 | + | tuf | WP_148696735.1 |
| GCF_002257545.1 | NZ_CP022741.1 | 699525 | 700709 | 1184 | + | tuf | WP_338636938.1 |
| GCF_002257545.1 | NZ_CP022741.1 | 2461599 | 2462786 | 1187 | - | tuf | WP_221032147.1 |
| GCF_024529975.1 | NZ_AP024897.1 | 3811642 | 3812829 | 1187 | - | tuf | WP_283395005.1 |
| GCF_024529975.1 | NZ_AP024897.1 | 3269245 | 3270432 | 1187 | + | tuf | WP_013044440.1 |
| GCF_024347075.1 | NZ_AP024900.1 | 1194518 | 1195705 | 1187 | - | tuf | WP_308949775.1 |
| GCF_024347075.1 | NZ_AP024900.1 | 1830751 | 1831938 | 1187 | + | tuf | WP_108824809.1 |
| GCF_024347095.1 | NZ_AP024903.1 | 937419 | 938606 | 1187 | + | tuf | WP_220163947.1 |
| GCF_024347095.1 | NZ_AP024903.1 | 546720 | 547907 | 1187 | - | tuf | WP_069960798.1 |
| GCF_002214395.1 | NZ_CP018312.1 | 1251153 | 1252340 | 1187 | - | tuf | WP_096055034.1 |
| GCF_002214395.1 | NZ_CP018312.1 | 267887 | 269074 | 1187 | - | tuf | WP_012373068.1 |
| GCF_024347115.1 | NZ_AP024905.1 | 5092490 | 5093677 | 1187 | - | tuf | WP_317833148.1 |
| GCF_024347115.1 | NZ_AP024905.1 | 504485 | 505672 | 1187 | + | tuf | WP_330930135.1 |
| GCF_002218045.2 | NZ_AP018685.1 | 5789462 | 5790649 | 1187 | + | tuf | WP_068769889.1 |
| GCF_002218045.2 | NZ_AP018685.1 | 3673920 | 3675128 | 1208 | - | tuf | WP_146663511.1 |

| | | | | | | | |
|---|---|---|---|---|---|---|---|
| GCF_040869025.1 | NZ_CP134271.1 | 2813955 | 2815163 | 1208 | - | tuf | WP_206295066.1 |
| GCF_040869025.1 | NZ_CP134271.1 | 6332611 | 6333804 | 1193 | + | tuf | WP_206292228.1 |
| GCF_024347135.1 | NZ_AP024907.1 | 2218635 | 2219840 | 1205 | - | tuf | WP_146683527.1 |
| GCF_024347135.1 | NZ_AP024907.1 | 175147 | 176349 | 1202 | + | tuf | WP_145444366.1 |
| GCF_022812045.1 | NZ_CP089203.1 | 965412 | 966614 | 1202 | + | tuf | WP_014436210.1 |
| GCF_022812045.1 | NZ_CP089203.1 | 730207 | 731409 | 1202 | - | tuf | WP_145074377.1 |
| GCF_002119525.1 | NZ_CP017916.1 | 2693834 | 2695039 | 1205 | + | tuf | WP_077541384.1 |
| GCF_002119525.1 | NZ_CP017916.1 | 1321109 | 1322314 | 1205 | - | tuf | WP_085755291.1 |
| GCF_024347635.1 | NZ_AP025510.1 | 5871498 | 5872694 | 1196 | - | tuf | WP_145063750.1 |
| GCF_024347635.1 | NZ_AP025510.1 | 5360257 | 5361453 | 1196 | + | tuf | WP_145249774.1 |
| GCF_024347655.1 | NZ_AP025514.1 | 2207767 | 2208960 | 1193 | + | tuf | WP_145358565.1 |
| GCF_024347655.1 | NZ_AP025514.1 | 7149430 | 7150623 | 1193 | + | tuf | WP_145096067.1 |
| GCF_001547935.1 | NZ_AP014635.1 | 1326595 | 1327803 | 1208 | - | tuf | WP_148591679.1 |
| GCF_001547935.1 | NZ_AP014635.1 | 1366303 | 1367493 | 1190 | + | tuf | WP_145074797.1 |
| GCF_000772105.1 | NZ_CP009354.1 | 5540250 | 5541428 | 1178 | - | tuf | WP_145081626.1 |
| GCF_000772105.1 | NZ_CP009354.1 | 5659874 | 5661070 | 1196 | - | tuf | WP_145116715.1 |
| GCF_002224265.1 | NZ_CP012881.1 | 5881004 | 5882197 | 1193 | - | tuf | WP_144976845.1 |
| GCF_002224265.1 | NZ_CP012881.1 | 1217581 | 1218777 | 1196 | - | tuf | WP_148072474.1 |
| GCF_003716875.1 | NZ_CP033078.1 | 2155746 | 2156939 | 1193 | - | tuf | WP_145261948.1 |
| GCF_003716875.1 | NZ_CP033078.1 | 1859378 | 1860571 | 1193 | + | tuf | WP_145028733.1 |
| GCF_011064285.1 | NZ_CP049331.1 | 1635905 | 1637098 | 1193 | + | tuf | WP_145294820.1 |
| GCF_011064285.1 | NZ_CP049331.1 | 2080440 | 2081639 | 1199 | + | tuf | WP_145295681.1 |
| GCF_034479615.1 | NZ_CP140255.1 | 463644 | 464837 | 1193 | - | tuf | WP_168565055.1 |
| GCF_012427705.1 | NZ_CP048602.1 | 6994246 | 6995445 | 1199 | - | tuf | WP_146414702.1 |
| GCF_019722725.1 | NZ_CP077941.1 | 7850208 | 7851506 | 1298 | + | tuf | WP_171474553.1 |
| GCF_043585715.1 | NZ_CP162600.1 | 7645431 | 7646624 | 1193 | - | tuf | WP_077027142.1 |
| GCF_900142895.1 | NZ_LT670847.1 | 7040868 | 7042166 | 1298 | - | tuf | WP_162670852.1 |
| GCF_014697095.1 | NZ_CP059082.1 | 7911280 | 7912452 | 1172 | + | tuf | WP_162671514.1 |
| GCF_015689495.1 | NZ_CP065135.1 | 705795 | 706973 | 1178 | + | tuf | WP_145210578.1 |
| GCF_001187785.1 | NZ_CP012154.1 | 7171499 | 7172692 | 1193 | - | tuf | WP_145222138.1 |
| GCF_001187785.1 | NZ_CP012154.1 | 7324372 | 7325565 | 1193 | - | tuf | WP_145231972.1 |

| | | | | | | |
|---|---|---|---|---|---|---|
| GCF_000247565.1 | NC_016893.1 | 616904 | 618097 | 1193 - | tuf | WP_144980955.1 |
| GCF_032164335.1 | NZ_CP134152.1 | 6633649 | 6634827 | 1178 - | tuf | WP_144989493.1 |
| GCF_032164335.1 | NZ_CP134152.1 | 3102169 | 3103347 | 1178 - | tuf | WP_149340871.1 |
| GCF_001677435.1 | NZ_CP016268.1 | 4296472 | 4297665 | 1193 + | tuf | WP_155365048.1 |
| GCF_001677435.1 | NZ_CP016268.1 | 806940 | 808118 | 1178 + | tuf | WP_145305783.1 |
| GCF_009931595.1 | NZ_CP046570.1 | 7131051 | 7132244 | 1193 - | tuf | WP_145313163.1 |
| GCF_009931595.1 | NZ_CP046570.1 | 7260101 | 7261294 | 1193 - | tuf | WP_002647429.1 |
| GCF_905367715.1 | NZ_HG999362.1 | 729813 | 730991 | 1178 + | tuf | WP_145448086.1 |
| GCF_905367715.1 | NZ_HG999362.1 | 7231729 | 7232922 | 1193 - | tuf | WP_145114916.1 |
| GCF_041519315.1 | NZ_CP167998.1 | 164044 | 165261 | 1217 - | tuf | WP_013562999.1 |
| GCF_041519315.1 | NZ_CP167998.1 | 6664573 | 6665769 | 1196 - | tuf | WP_145436040.1 |
| GCF_041240605.1 | NZ_CP162487.1 | 6723605 | 6724801 | 1196 - | tuf | WP_152101595.1 |
| GCF_041240605.1 | NZ_CP162487.1 | 8262074 | 8263267 | 1193 - | tuf | WP_145057465.1 |
| GCF_013388375.1 | NZ_CP058243.1 | 2470763 | 2472001 | 1238 + | tuf | WP_149110010.1 |
| GCF_013388375.1 | NZ_CP058243.1 | 2484046 | 2485230 | 1184 - | tuf | WP_246173683.1 |
| GCF_040616535.1 | NZ_CP136580.1 | 5531996 | 5533189 | 1193 + | tuf | WP_145370969.1 |
| GCF_040616535.1 | NZ_CP136580.1 | 4991340 | 4992533 | 1193 - | tuf | WP_075082429.1 |
| GCF_000454545.1 | NZ_CM002139.1 | 3483816 | 3485024 | 1208 + | tuf | WP_076346386.1 |
| GCF_021906995.1 | NZ_CP083804.1 | 6149243 | 6150439 | 1196 + | tuf | WP_145291523.1 |
| GCF_021906995.1 | NZ_CP083804.1 | 1448367 | 1449560 | 1193 - | tuf | WP_336522550.1 |
| GCF_000961215.1 | NZ_CP008998.1 | 1476709 | 1477902 | 1193 + | tuf | WP_145297035.1 |
| GCF_000961215.1 | NZ_CP008998.1 | 545039 | 546232 | 1193 + | tuf | WP_013108716.1 |
| GCF_009883735.1 | NZ_CP033326.1 | 4571376 | 4572569 | 1193 - | tuf | WP_144997378.1 |
| GCF_009883735.1 | NZ_CP033326.1 | 4217484 | 4218677 | 1193 + | tuf | WP_011121621.1 |
| GCF_032698475.1 | NZ_CP103837.1 | 1257801 | 1258994 | 1193 + | tuf | WP_145350547.1 |
| GCF_032698475.1 | NZ_CP103837.1 | 1974542 | 1975735 | 1193 + | tuf | WP_068132479.1 |
| GCF_903989455.1 | NZ_LR861803.1 | 5969103 | 5970296 | 1193 - | tuf | WP_145100551.1 |
| GCF_903989455.1 | NZ_LR861803.1 | 5971366 | 5972559 | 1193 + | tuf | WP_145123152.1 |
| GCF_017724035.1 | NZ_CP072268.1 | 6361519 | 6362712 | 1193 - | tuf | WP_145348323.1 |
| GCF_017724035.1 | NZ_CP072268.1 | 2381560 | 2382753 | 1193 - | tuf | WP_013628343.1 |
| GCF_900183975.1 | NZ_LT853882.1 | 6897175 | 6898368 | 1193 - | tuf | WP_145173956.1 |

| | | | | | | | |
|---|---|---|---|---|---|---|---|
| GCF_900183975.1 | NZ_LT853882.1 | 654614 | 655822 | 1208 | - | tuf | WP_015244129.1 |
| GCF_030168915.1 | NZ_CP076255.1 | 1565643 | 1566836 | 1193 | - | tuf | WP_147866775.1 |
| GCF_030168915.1 | NZ_CP076255.1 | 9076617 | 9077816 | 1199 | - | tuf | WP_147871259.1 |
| GCF_031834635.1 | NZ_CP115873.1 | 601769 | 602968 | 1199 | + | tuf | WP_145384488.1 |
| GCF_031834635.1 | NZ_CP115873.1 | 8278682 | 8279875 | 1193 | + | tuf | WP_145390320.1 |
| GCF_002285515.1 | NZ_CP016878.1 | 4046542 | 4047741 | 1199 | - | tuf | WP_146521149.1 |
| GCF_002285515.1 | NZ_CP016878.1 | 6079624 | 6080817 | 1193 | + | tuf | WP_146520884.1 |
| GCF_009769165.1 | NZ_CP043476.1 | 2186304 | 2187497 | 1193 | + | tuf | WP_145363503.1 |
| GCF_009769165.1 | NZ_CP043476.1 | 101115 | 102323 | 1208 | + | tuf | WP_145266250.1 |
| GCF_032697525.1 | NZ_CP103836.1 | 3627839 | 3629047 | 1208 | - | tuf | WP_213493926.1 |
| GCF_032697525.1 | NZ_CP103836.1 | 5009717 | 5010910 | 1193 | - | tuf | WP_145202864.1 |
| GCF_040529045.1 | NZ_CP131914.1 | 3714976 | 3716172 | 1196 | + | tuf | WP_095415973.1 |
| GCF_040529045.1 | NZ_CP131914.1 | 2435268 | 2436485 | 1217 | + | tuf | WP_162657664.1 |
| GCF_041242975.1 | NZ_CP162488.1 | 2743462 | 2744784 | 1322 | + | tuf | WP_145237654.1 |
| GCF_041242975.1 | NZ_CP162488.1 | 3228598 | 3229770 | 1172 | - | tuf | WP_145238797.1 |
| GCF_023337365.3 | NZ_CP142084.2 | 1764506 | 1765699 | 1193 | - | tuf | WP_228350410.1 |
| GCF_023337365.3 | NZ_CP142084.2 | 4220235 | 4221428 | 1193 | + | tuf | WP_228350410.1 |
| GCF_008370835.2 | NZ_CP043403.1 | 963703 | 964896 | 1193 | + | tuf | WP_012843349.1 |
| GCF_008370835.2 | NZ_CP043403.1 | 1272860 | 1274053 | 1193 | + | tuf | WP_012843604.1 |
| GCF_008639345.1 | NZ_CP041380.1 | 1434623 | 1435810 | 1187 | + | tuf | WP_011403793.1 |
| GCF_008639345.1 | NZ_CP041380.1 | 2284982 | 2286169 | 1187 | - | tuf | WP_118828945.1 |
| GCF_025266575.1 | NZ_CP096142.1 | 807202 | 808401 | 1199 | + | tuf | WP_119066825.1 |
| GCF_025266575.1 | NZ_CP096142.1 | 822739 | 823938 | 1199 | + | tuf | WP_119066825.1 |
| GCF_033170385.1 | NZ_CP126170.1 | 2168956 | 2170155 | 1199 | - | tuf | WP_011565116.1 |
| GCF_033170385.1 | NZ_CP126170.1 | 2184314 | 2185513 | 1199 | - | tuf | WP_011565116.1 |
| GCF_040529065.1 | NZ_CP132343.1 | 5736930 | 5738114 | 1184 | - | tuf | WP_264789346.1 |
| GCF_040529065.1 | NZ_CP132343.1 | 5775055 | 5776248 | 1193 | + | tuf | WP_013766944.1 |
| GCF_040529055.1 | NZ_CP132342.1 | 2339568 | 2340752 | 1184 | + | tuf | WP_015692647.1 |
| GCF_040529055.1 | NZ_CP132342.1 | 2673761 | 2674945 | 1184 | - | tuf | WP_317896814.1 |
| GCF_041245805.1 | NZ_CP162490.1 | 4512146 | 4513330 | 1184 | - | tuf | WP_022833148.1 |
| GCF_041245805.1 | NZ_CP162490.1 | 3995057 | 3996241 | 1184 | - | tuf | WP_157539718.1 |

| | | | | | | | |
|---|---|---|---|---|---|---|---|
| GCF_014236795.1 | NZ_CP049017.1 | 1894684 | 1895868 | 1184 | - | tuf | WP_157524301.1 |
| GCF_014236795.1 | NZ_CP049017.1 | 4812595 | 4813779 | 1184 | - | tuf | WP_147033947.1 |
| GCF_017301775.1 | NZ_CP063993.1 | 4718036 | 4719220 | 1184 | + | tuf | WP_090642950.1 |
| GCF_017301775.1 | NZ_CP063993.1 | 58303 | 59487 | 1184 | + | tuf | WP_091166595.1 |
| GCF_000277895.2 | NZ_CP034655.1 | 3991183 | 3992367 | 1184 | - | tuf | WP_096353764.1 |
| GCF_000277895.2 | NZ_CP034655.1 | 1684182 | 1685366 | 1184 | + | tuf | WP_166584404.1 |
| GCF_001908725.1 | NZ_CP018725.1 | 4211497 | 4212681 | 1184 | - | tuf | WP_173416168.1 |
| GCF_001908725.1 | NZ_CP018725.1 | 4419267 | 4420451 | 1184 | - | tuf | WP_091376052.1 |
| GCF_030272115.1 | NZ_CP119194.1 | 5055333 | 5056517 | 1184 | + | tuf | WP_008509243.1 |
| GCF_030272115.1 | NZ_CP119194.1 | 3183414 | 3184598 | 1184 | + | tuf | WP_158824927.1 |
| GCF_017743015.1 | NZ_CP072455.1 | 2045393 | 2046577 | 1184 | + | tuf | WP_321564707.1 |
| GCF_017743015.1 | NZ_CP072455.1 | 1866987 | 1868171 | 1184 | + | tuf | WP_094570055.1 |
| GCF_000968195.1 | NZ_FO704550.1 | 317058 | 318242 | 1184 | + | tuf | WP_039450248.1 |
| GCF_000968195.1 | NZ_FO704550.1 | 4727108 | 4728292 | 1184 | + | tuf | WP_010601091.1 |
| GCF_030016155.2 | NZ_CP133479.1 | 3531622 | 3532806 | 1184 | - | tuf | WP_173090975.1 |
| GCF_030016155.2 | NZ_CP133479.1 | 993442 | 994626 | 1184 | + | tuf | WP_012781010.1 |
| GCF_001721185.1 | NZ_CP016176.1 | 2715877 | 2717061 | 1184 | - | tuf | WP_041878245.1 |
| GCF_001721185.1 | NZ_CP016176.1 | 2740722 | 2741906 | 1184 | - | tuf | WP_238414037.1 |
| GCF_000252955.1 | NC_014228.1 | 765088 | 766272 | 1184 | + | tuf | WP_086545150.1 |
| GCF_000968175.1 | NZ_FO704551.1 | 844535 | 845719 | 1184 | + | tuf | WP_187593732.1 |
| GCF_000968175.1 | NZ_FO704551.1 | 6187940 | 6189124 | 1184 | - | tuf | WP_121273950.1 |
| GCF_036512875.1 | NZ_AP028978.1 | 1569964 | 1571148 | 1184 | - | tuf | WP_010601091.1 |
| GCF_036512875.1 | NZ_AP028978.1 | 2642371 | 2643555 | 1184 | - | tuf | WP_013633306.1 |
| GCF_028891345.1 | NZ_CP118539.1 | 514727 | 515911 | 1184 | + | tuf | WP_014678779.1 |
| GCF_028891345.1 | NZ_CP118539.1 | 1442737 | 1443921 | 1184 | + | tuf | WP_103787696.1 |
| GCF_013177435.1 | NZ_CP053627.1 | 1710154 | 1711338 | 1184 | - | tuf | WP_320185785.1 |
| GCF_013177435.1 | NZ_CP053627.1 | 689553 | 690737 | 1184 | + | tuf | WP_094255410.1 |
| GCF_000834395.1 | NZ_CP009781.1 | 3681057 | 3682241 | 1184 | - | tuf | WP_149526999.1 |
| GCF_000834395.1 | NZ_CP009781.1 | 129796 | 130980 | 1184 | - | tuf | WP_103907701.1 |
| GCF_025133195.1 | NZ_CP104006.1 | 832784 | 833968 | 1184 | + | tuf | WP_093098550.1 |
| GCF_025133195.1 | NZ_CP104006.1 | 558090 | 559271 | 1181 | + | tuf | WP_075991174.1 |

| | | | | | | | |
|---|---|---|---|---|---|---|---|
| GCF_037060685.1 | NZ_CP124240.1 | 4183506 | 4184690 | 1184 | - | tuf | WP_274266751.1 |
| GCF_037060685.1 | NZ_CP124240.1 | 3851513 | 3852694 | 1181 | - | tuf | WP_153847177.1 |
| GCF_009831415.1 | NZ_CP043727.1 | 635531 | 636715 | 1184 | + | tuf | WP_085472253.1 |
| GCF_009831415.1 | NZ_CP043727.1 | 1168448 | 1169629 | 1181 | + | tuf | WP_075991174.1 |
| GCF_000253175.1 | NC_017564.1 | 1059095 | 1060276 | 1181 | - | tuf | WP_002992774.1 |
| GCF_000253175.1 | NC_017564.1 | 720028 | 721212 | 1184 | - | tuf | WP_028072656.1 |
| GCF_001656035.1 | NZ_CP010029.1 | 810313 | 811497 | 1184 | + | tuf | WP_153509776.1 |
| GCF_001656035.1 | NZ_CP010029.1 | 491360 | 492541 | 1181 | + | tuf | WP_025419655.1 |
| GCF_004124235.1 | NZ_CP032487.1 | 498323 | 499504 | 1181 | + | tuf | WP_025408099.1 |
| GCF_004124235.1 | NZ_CP032487.1 | 508098 | 509279 | 1181 | + | tuf | WP_012538231.1 |
| GCF_009730055.1 | NZ_CP046293.1 | 500652 | 501833 | 1181 | + | tuf | WP_012422241.1 |
| GCF_009730055.1 | NZ_CP046293.1 | 493252 | 494433 | 1181 | + | tuf | WP_151552237.1 |
| GCF_013282725.1 | NZ_CP054043.1 | 495486 | 496667 | 1181 | + | tuf | WP_020954851.1 |
| GCF_013282725.1 | NZ_CP054043.1 | 499877 | 501058 | 1181 | + | tuf | WP_025399802.1 |
| GCF_000222975.1 | NC_017168.1 | 497576 | 498757 | 1181 | + | tuf | WP_025375490.1 |
| GCF_000222975.1 | NC_017168.1 | 499157 | 500338 | 1181 | + | tuf | WP_247065628.1 |
| GCF_900637475.1 | NZ_LR134373.1 | 514819 | 516000 | 1181 | + | tuf | WP_012538231.1 |
| GCF_900637475.1 | NZ_LR134373.1 | 514743 | 515924 | 1181 | + | tuf | WP_120104267.1 |
| GCF_003600645.1 | NZ_CP032482.1 | 497704 | 498885 | 1181 | + | tuf | WP_011772423.1 |
| GCF_003600645.1 | NZ_CP032482.1 | 497356 | 498537 | 1181 | + | tuf | WP_004789670.1 |
| GCF_000834455.1 | NZ_CP009787.1 | 494670 | 495851 | 1181 | + | tuf | WKD00810.1 |
| GCF_000834455.1 | NZ_CP009787.1 | 492416 | 493597 | 1181 | + | tuf | WP_316255282.1 |
| GCF_017498685.1 | NZ_CP071802.1 | 497347 | 498528 | 1181 | + | tuf | WP_011193796.1 |
| GCF_017498685.1 | NZ_CP071802.1 | 493710 | 494891 | 1181 | + | tuf | WP_014023706.1 |
| GCF_000582515.1 | NZ_CP007230.1 | 495156 | 496337 | 1181 | + | tuf | WP_184123904.1 |
| GCF_000582515.1 | NZ_CP007230.1 | 494096 | 495277 | 1181 | + | tuf | WP_014023706.1 |
| GCF_044905075.1 | NZ_CP173260.1 | 494171 | 495352 | 1181 | + | tuf | WP_091972162.1 |
| GCF_044905075.1 | NZ_CP173260.1 | 497107 | 498288 | 1181 | + | tuf | WP_004791386.1 |
| GCF_001586255.1 | NZ_CP014544.1 | 499504 | 500685 | 1181 | + | tuf | WP_301393080.1 |
| GCF_001586255.1 | NZ_CP014544.1 | 493847 | 495028 | 1181 | + | tuf | WP_014023706.1 |
| GCF_002407485.1 | NZ_CP012621.1 | 494483 | 495664 | 1181 | + | tuf | WP_184107272.1 |

| | | | | | | | |
|---|---|---|---|---|---|---|---|
| GCF_002407485.1 | NZ_CP012621.1 | 496711 | 497892 | 1181 | + | tuf | WP_301395093.1 |
| GCF_011745665.1 | NZ_CP050253.1 | 496143 | 497324 | 1181 | + | tuf | WP_002657015.1 |
| GCF_011745665.1 | NZ_CP050253.1 | 407267 | 408448 | 1181 | - | tuf | WP_320292361.1 |
| GCF_003610015.1 | NZ_AP018933.1 | 496023 | 497204 | 1181 | + | tuf | WP_006433994.1 |
| GCF_003574215.1 | NZ_AP018558.1 | 495069 | 496250 | 1181 | + | tuf | WP_301353770.1 |
| GCF_003574215.1 | NZ_AP018558.1 | 495372 | 496553 | 1181 | + | tuf | WP_301394185.1 |
| GCF_002795805.1 | NZ_CP018799.1 | 496263 | 497444 | 1181 | + | tuf | WP_006068879.1 |
| GCF_002795805.1 | NZ_CP018799.1 | 494906 | 496087 | 1181 | + | tuf | WP_183219270.1 |
| GCF_002795825.1 | NZ_CP018800.1 | 2242775 | 2243998 | 1223 | + | tuf | WP_012671618.1 |
| GCF_002795825.1 | NZ_CP018800.1 | 2877858 | 2879081 | 1223 | - | tuf | WP_014488947.1 |
| GCF_011682235.2 | NZ_CP076718.1 | 2098935 | 2100158 | 1223 | + | tuf | WP_013114301.1 |
| GCF_011682235.2 | NZ_CP076718.1 | 1132364 | 1133587 | 1223 | - | tuf | WP_013243972.1 |
| GCF_000024845.1 | NC_013501.1 | 3861743 | 3862930 | 1187 | + | tuf | WP_215627937.1 |
| GCF_000024845.1 | NC_013501.1 | 2637381 | 2638568 | 1187 | + | tuf | WP_255946254.1 |
| GCF_003491405.1 | NZ_CP030361.1 | 452733 | 453914 | 1181 | - | tuf | WP_025375490.1 |
| GCF_003491405.1 | NZ_CP030361.1 | 511918 | 513099 | 1181 | + | tuf | WP_108729249.1 |
| GCF_001936255.1 | NZ_CP013704.1 | 2819299 | 2820480 | 1181 | - | tuf | WP_326927121.1 |
| GCF_023035295.1 | NZ_CP075076.1 | 1449610 | 1450791 | 1181 | - | tuf | WP_167696003.1 |
| GCF_000019685.1 | NC_011229.1 | 377606 | 378790 | 1184 | - | tuf | WP_167699859.1 |
| GCF_020422925.2 | NZ_CP084623.1 | 371723 | 372904 | 1181 | - | tuf | WP_167703659.1 |
| GCF_008931845.1 | NZ_CP044535.1 | 437973 | 439160 | 1187 | + | tuf | WP_013967859.1 |
| GCF_019668505.1 | NZ_AP024371.1 | 3545192 | 3546388 | 1196 | - | tuf | WP_015712882.1 |
| GCF_023035795.1 | NZ_CP073148.1 | 2042036 | 2043238 | 1202 | - | tuf | WP_012388941.1 |
| GCF_000512145.1 | NZ_CP007022.1 | 876546 | 877748 | 1202 | + | tuf | WP_002748403.1 |
| GCF_023035875.1 | NZ_CP075379.1 | 3005953 | 3007155 | 1202 | - | tuf | WP_020770357.1 |
| GCF_000019705.1 | NC_011244.1 | 1479049 | 1480251 | 1202 | - | tuf | WP_001040571.1 |
| GCF_003606285.1 | NZ_CP028884.1 | 1620840 | 1622042 | 1202 | - | tuf | WP_010573748.1 |
| GCF_000012085.2 | NC_008710.1 | 2316128 | 2317330 | 1202 | - | tuf | WP_109019452.1 |
| GCF_000741005.1 | NZ_CP009212.1 | 3306024 | 3307226 | 1202 | - | tuf | WP_002748403.1 |
| GCA_030445085.2 | CP124113.1 | 3160296 | 3161498 | 1202 | - | tuf | WP_004457701.1 |
| GCF_032595975.2 | NZ_CP132472.1 | 3363344 | 3364546 | 1202 | - | tuf | WP_069608386.1 |

| | | | | | | | |
|---|---|---|---|---|---|---|---|
| GCF_000196215.1 | NC_006156.1 | 3381043 | 3382245 | 1202 | - | tuf | WP_004496941.1 |
| GCF_000222305.1 | NC_015921.1 | 2510760 | 2511944 | 1184 | - | tuf | WP_149486647.1 |
| GCF_030438475.2 | NZ_CP124076.1 | 2020853 | 2022037 | 1184 | - | tuf | WP_013740309.1 |
| GCF_032595855.2 | NZ_CP132453.1 | 936191 | 937378 | 1187 | + | tuf | WP_041401147.1 |
| GCF_030438365.2 | NZ_CP124070.1 | 1010057 | 1011244 | 1187 | + | tuf | WP_013253551.1 |
| GCF_026572315.1 | NZ_CP075232.1 | 3290341 | 3291528 | 1187 | - | tuf | WP_244772298.1 |
| GCF_030438115.2 | NZ_CP124062.1 | 3078313 | 3079500 | 1187 | - | tuf | WP_013608504.1 |
| GCF_030437945.2 | NZ_CP124058.1 | 2342616 | 2343803 | 1187 | + | tuf | WP_014270749.1 |
| GCF_030437705.2 | NZ_CP124054.1 | 658156 | 659346 | 1190 | + | tuf | WP_014454650.1 |
| GCF_030440365.2 | NZ_CP124050.1 | 616417 | 617604 | 1187 | + | tuf | WP_014624187.1 |
| GCF_001945665.1 | NZ_CP015780.1 | 1250121 | 1251323 | 1202 | + | tuf | WP_271436454.1 |
| GCF_033969665.2 | NZ_CP135180.1 | 2350948 | 2352135 | 1187 | + | tuf | WP_149568395.1 |
| GCF_030436445.2 | NZ_CP124042.1 | 2700975 | 2702159 | 1184 | - | tuf | WP_013759407.1 |
| GCF_030439725.2 | NZ_CP124034.1 | 479547 | 480728 | 1181 | + | tuf | WP_074641252.1 |
| GCF_030439285.2 | NZ_CP124018.1 | 804872 | 806056 | 1184 | + | tuf | WP_002669993.1 |
| GCF_000170955.2 | NZ_ABCY02000001.1 | 694509 | 695693 | 1184 | - | tuf | WP_016522599.1 |
| GCF_030436165.1 | NZ_CP123998.1 | 204532 | 205716 | 1184 | + | tuf | WP_010881634.1 |
| GCF_017751225.1 | NZ_CP046932.1 | 201946 | 203130 | 1184 | + | tuf | WP_010881634.1 |
| GCF_000223215.1 | NC_017243.1 | 341953 | 343137 | 1184 | + | tuf | WP_210117240.1 |
| GCF_000092845.1 | NC_014150.1 | 2720575 | 2721759 | 1184 | - | tuf | WP_020966413.1 |
| GCF_000143725.1 | NC_014330.1 | 2554454 | 2555638 | 1184 | - | tuf | WP_177528781.1 |
| GCF_016616095.2 | NZ_CP067089.2 | 3301743 | 3302927 | 1184 | + | tuf | WP_024752674.1 |
| GCF_022376835.1 | NZ_CP084606.1 | 2814294 | 2815481 | 1187 | - | tuf | WP_015708630.1 |
| GCF_027925085.1 | NZ_AP027070.1 | 1964459 | 1965643 | 1184 | - | tuf | WP_044977712.1 |
| GCF_003076595.1 | NZ_CP025785.1 | 373723 | 374907 | 1184 | + | tuf | WP_002706022.1 |
| GCF_029867005.1 | NZ_CP123443.1 | 510080 | 511273 | 1193 | + | tuf | WP_180486862.1 |
| GCF_028748145.1 | NZ_CP118181.1 | 2614292 | 2615476 | 1184 | - | tuf | WP_013702570.1 |
| GCF_028748125.1 | NZ_CP118174.1 | 2630092 | 2631294 | 1202 | + | tuf | WP_014803692.1 |
| GCF_028748105.1 | NZ_CP118168.1 | 658858 | 660054 | 1196 | + | tuf | WP_274372696.1 |
| GCF_000219725.1 | NC_015732.1 | 1966262 | 1967458 | 1196 | - | tuf | WP_274372696.1 |
| GCF_000214355.1 | NC_015577.1 | 330583 | 331782 | 1199 | + | tuf | WP_013047721.1 |

| | | | | | | | |
|---|---|---|---|---|---|---|---|
| GCF_000017685.1 | NC_010602.1 | 663744 | 664943 | 1199 | + | tuf | WP_013048026.1 |
| GCF_003516145.1 | NZ_CP026671.1 | 1742683 | 1743873 | 1190 | - | tuf | WP_006301283.1 |
| GCF_040833995.1 | NZ_CP162137.1 | 2045121 | 2046311 | 1190 | - | tuf | WP_006301283.1 |
| GCF_002073495.2 | NZ_CP020414.2 | 1131664 | 1132851 | 1187 | + | tuf | WP_066743727.1 |
| GCF_003722295.1 | NZ_CP033614.1 | 3564266 | 3565453 | 1187 | - | tuf | WP_066743727.1 |
| GCF_003114835.2 | NZ_AP025028.1 | 89113 | 90312 | 1199 | - | tuf | WP_409152299.1 |
| GCF_000306675.2 | NZ_CP024871.1 | 1761315 | 1762514 | 1199 | + | tuf | WP_298779972.1 |
| GCF_000313175.2 | NZ_CP006694.1 | 652225 | 653421 | 1196 | + | tuf | WP_009165642.1 |
| GCF_001729245.1 | NZ_CP015217.1 | 2134537 | 2135733 | 1196 | - | tuf | WP_009165642.1 |
| GCF_006874765.1 | NZ_CP040840.1 | 1225381 | 1226571 | 1190 | - | tuf | WP_012869930.1 |
| GCF_008329965.1 | NZ_CP036150.1 | 1584025 | 1585215 | 1190 | - | tuf | WP_012869930.1 |
| GCF_000208385.1 | NC_015436.1 | 285942 | 287132 | 1190 | + | tuf | WP_006582921.1 |
| GCF_000507245.1 | NC_023035.1 | 631627 | 632817 | 1190 | + | tuf | WP_006582921.1 |
| GCF_000143985.1 | NC_014364.1 | 312244 | 313434 | 1190 | + | tuf | WP_011416212.1 |
| GCF_022869165.1 | NZ_CP094929.1 | 525796 | 526986 | 1190 | + | tuf | WP_408998886.1 |
| GCF_000190435.1 | NC_015152.1 | 370034 | 371224 | 1190 | - | tuf | WP_137423029.1 |
| GCF_000236685.1 | NC_016633.1 | 384608 | 385798 | 1190 | - | tuf | WP_137423029.1 |
| GCF_000242595.2 | NC_017098.1 | 1901006 | 1902196 | 1190 | + | tuf | WP_011698399.1 |
| GCF_000184345.1 | NC_017583.1 | 1915762 | 1916952 | 1190 | + | tuf | WP_011698399.1 |
| GCF_023525255.1 | NZ_CP073355.1 | 270419 | 271615 | 1196 | + | tuf | WP_158065949.1 |
| GCF_008329945.1 | NZ_CP035807.1 | 1864459 | 1865655 | 1196 | + | tuf | WP_158065949.1 |
| GCF_000212415.1 | NC_015500.1 | 1526292 | 1527476 | 1184 | + | tuf | WP_114206378.1 |
| GCF_036492245.1 | NZ_AP025286.1 | 5262826 | 5264010 | 1184 | + | tuf | WP_114206378.1 |
| GCF_000008185.1 | NC_002967.9 | 1717513 | 1718697 | 1184 | - | tuf | WP_015896586.1 |
| GCF_017161265.1 | NZ_CP031393.1 | 3384187 | 3385371 | 1184 | + | tuf | WP_015896586.1 |
| GCF_000410535.2 | NC_021490.2 | 1510559 | 1511743 | 1184 | + | tuf | WP_011522028.1 |
| GCF_000217655.1 | NC_015714.1 | 5543905 | 5545089 | 1184 | - | tuf | WP_011522028.1 |
| GCF_017893965.1 | NZ_CP054142.1 | 1221248 | 1222432 | 1184 | - | tuf | WP_260703923.1 |
| GCF_000447675.1 | NC_022097.1 | 3618277 | 3619461 | 1184 | + | tuf | WP_260703923.1 |
| GCF_016117655.1 | NZ_CP064936.1 | 750844 | 752028 | 1184 | + | tuf | WP_348268173.1 |
| GCF_008153345.1 | NZ_CP042818.1 | 955467 | 956651 | 1184 | + | tuf | WP_348268173.1 |

| | | | | | | | |
|---|---|---|---|---|---|---|---|
| GCF_000214375.1 | NC_015578.1 | 665096 | 666280 | 1184 | + | tuf | WP_014263805.1 |
| GCF_024401155.1 | NZ_CP038804.1 | 875175 | 876359 | 1184 | + | tuf | WP_014263805.1 |
| GCF_036250655.1 | NZ_AP025288.1 | 1095706 | 1096890 | 1184 | + | tuf | WP_013579308.1 |
| GCF_024181585.1 | NZ_CP054258.1 | 3427501 | 3428685 | 1184 | - | tuf | WP_013579308.1 |
| GCF_000195275.1 | NC_015385.1 | 2514366 | 2515550 | 1184 | + | tuf | WP_260795908.1 |
| GCF_000266885.1 | NC_018020.1 | 2604746 | 2605930 | 1184 | + | tuf | WP_260795984.1 |
| GCF_018069705.1 | NZ_CP072943.1 | 2822789 | 2823976 | 1187 | - | tuf | WP_194447947.1 |
| GCF_018069705.1 | NZ_CP072943.1 | 6657067 | 6658254 | 1187 | + | tuf | WP_194447947.1 |
| GCF_000025885.1 | NC_014011.1 | 1698123 | 1699307 | 1184 | + | tuf | WP_420237592.1 |
| GCF_000025885.1 | NC_014011.1 | 4491844 | 4493028 | 1184 | - | tuf | WP_420237592.1 |
| GCF_000165795.1 | NZ_CM001022.1 | 1023453 | 1024637 | 1184 | - | tuf | WP_014784803.1 |
| GCF_000165795.1 | NZ_CM001022.1 | 4441746 | 4442930 | 1184 | - | tuf | WP_014784803.1 |
| GCF_001701045.1 | NZ_CP016757.1 | 3941097 | 3942281 | 1184 | - | tuf | WP_013569725.1 |
| GCF_001701045.1 | NZ_CP016757.1 | 4167029 | 4168213 | 1184 | - | tuf | WP_013569725.1 |
| GCF_964642095.1 | NZ_OZ209244.1 | 2995909 | 2997093 | 1184 | + | tuf | WP_179636944.1 |
| GCF_964642095.1 | NZ_OZ209244.1 | 3386599 | 3387783 | 1184 | + | tuf | WP_353068428.1 |
| GCF_022846135.1 | NZ_AP025585.1 | 4774625 | 4775809 | 1184 | - | tuf | WP_179636944.1 |
| GCF_022846135.1 | NZ_AP025585.1 | 5414528 | 5415712 | 1184 | - | tuf | WP_179636944.1 |
| GCF_000024905.1 | NC_013522.1 | 2484609 | 2485793 | 1184 | - | tuf | WP_260736975.1 |
| GCF_000024905.1 | NC_013522.1 | 5509306 | 5510490 | 1184 | + | tuf | WP_260736975.1 |
| GCF_000237825.1 | NZ_CM001377.1 | 1796723 | 1797907 | 1184 | + | tuf | WP_353063398.1 |
| GCF_000237825.1 | NZ_CM001377.1 | 4644754 | 4645938 | 1184 | - | tuf | WP_353063398.1 |
| GCF_001577525.1 | NZ_CP013015.1 | 124241 | 125437 | 1196 | + | tuf | WP_068511825.1 |
| GCF_001577525.1 | NZ_CP013015.1 | 1676894 | 1678090 | 1196 | + | tuf | WP_068511825.1 |
| GCF_000143965.1 | NC_014365.1 | 299023 | 300219 | 1196 | - | tuf | WP_013906911.1 |
| GCF_000143965.1 | NC_014365.1 | 1538368 | 1539564 | 1196 | + | tuf | WP_013908065.1 |
| GCF_037076455.1 | NZ_AP028679.1 | 1041173 | 1042369 | 1196 | - | tuf | WP_038549660.1 |
| GCF_037076455.1 | NZ_AP028679.1 | 1700597 | 1701793 | 1196 | + | tuf | WP_038549660.1 |
| GCF_000195295.1 | NC_015388.1 | 162437 | 163633 | 1196 | - | tuf | WP_013908987.1 |
| GCF_000195295.1 | NC_015388.1 | 744915 | 746111 | 1196 | + | tuf | WP_013908987.1 |
| GCF_000021905.1 | NC_011768.1 | 480803 | 481999 | 1196 | - | tuf | WP_168719102.1 |

| | | | | | | | |
|---|---|---|---|---|---|---|---|
| GCF_000021905.1 | NC_011768.1 | 575578 | 576774 | 1196 | - | tuf | WP_168719102.1 |
| GCF_004319545.1 | NZ_CP036313.1 | 652643 | 653839 | 1196 | - | tuf | WP_166031559.1 |
| GCF_000233695.2 | NZ_CM001488.1 | 668752 | 669948 | 1196 | - | tuf | WP_166031559.1 |
| GCF_000307105.1 | NC_018645.1 | 467942 | 469156 | 1214 | + | tuf | WP_108308694.1 |
| GCF_000307105.1 | NC_018645.1 | 1029113 | 1030327 | 1214 | - | tuf | WP_108308694.1 |
| GCF_002009335.2 | NZ_CP019913.2 | 503663 | 504877 | 1214 | + | tuf | WP_013755870.1 |
| GCF_002009335.2 | NZ_CP019913.2 | 780424 | 781638 | 1214 | + | tuf | WP_013755870.1 |
| GCF_021654575.1 | NZ_AP024488.1 | 439003 | 440199 | 1196 | + | tuf | WP_203473001.1 |
| GCF_021654575.1 | NZ_AP024488.1 | 639376 | 640572 | 1196 | + | tuf | WP_203473001.1 |
| GCF_017377355.1 | NZ_CP061799.1 | 36 | 1232 | 1196 | + | tuf | WP_353684206.1 |
| GCF_017377355.1 | NZ_CP061799.1 | 1739600 | 1740796 | 1196 | + | tuf | WP_353683996.1 |
| GCF_017377395.1 | NZ_CP061800.1 | 1440238 | 1441434 | 1196 | - | tuf | WP_353685803.1 |
| GCF_000020365.1 | NC_012108.1 | 1601853 | 1603049 | 1196 | - | tuf | WP_353685803.1 |
| GCF_000020365.1 | NC_012108.1 | 1319439 | 1320635 | 1196 | - | tuf | WP_012545401.1 |
| GCF_009688945.1 | NZ_AP021874.1 | 1383641 | 1384837 | 1196 | - | tuf | WP_012545401.1 |
| GCF_009688945.1 | NZ_AP021874.1 | 4863523 | 4864704 | 1181 | - | tuf | WP_272474261.1 |
| GCF_009688985.1 | NZ_AP021876.1 | 4880632 | 4881813 | 1181 | - | tuf | WP_146922788.1 |
| GCF_009688985.1 | NZ_AP021876.1 | 4806323 | 4807504 | 1181 | - | tuf | WP_259312295.1 |
| GCF_048565275.1 | NZ_CP159846.1 | 1416598 | 1417779 | 1181 | + | tuf | WP_012932818.1 |
| GCF_048565275.1 | NZ_CP159846.1 | 2759138 | 2760328 | 1190 | - | tuf | WP_217913729.1 |
| GCF_009688965.1 | NZ_AP021875.1 | 752902 | 754086 | 1184 | + | tuf | WP_217923220.1 |
| GCF_009688965.1 | NZ_AP021875.1 | 988697 | 989878 | 1181 | + | tuf | WP_254572221.1 |
| GCF_000018405.1 | NC_009943.1 | 4779022 | 4780203 | 1181 | - | tuf | WP_353864194.1 |
| GCF_000018405.1 | NC_009943.1 | 1148907 | 1150103 | 1196 | + | tuf | WP_012871517.1 |
| GCF_016446285.1 | NZ_CP054140.1 | 1770324 | 1771520 | 1196 | - | tuf | WP_012872092.1 |
| GCF_016446285.1 | NZ_CP054140.1 | 670981 | 672180 | 1199 | + | tuf | WP_012642069.1 |
| GCF_002952055.1 | NZ_CP021255.1 | 946167 | 947366 | 1199 | - | tuf | WP_012642069.1 |
| GCF_002952055.1 | NZ_CP021255.1 | 1458551 | 1459750 | 1199 | - | tuf | WP_068550216.1 |
| GCF_000186885.1 | NC_014972.1 | 1474704 | 1475903 | 1199 | - | tuf | WP_068550216.1 |
| GCF_000186885.1 | NC_014972.1 | 94693 | 95886 | 1193 | - | tuf | WP_161847242.1 |
| GCF_000341395.1 | NC_020304.1 | 170125 | 171321 | 1196 | + | tuf | WP_045087099.1 |

| | | | | | | | |
|---|---|---|---|---|---|---|---|
| GCF_000341395.1 | NC_020304.1 | 948657 | 949856 | 1199 | - | tuf | WP_090221999.1 |
| GCF_023169885.1 | NZ_AP025516.1 | 1493188 | 1494387 | 1199 | - | tuf | WP_033192026.1 |
| GCF_023169885.1 | NZ_AP025516.1 | 1224404 | 1225603 | 1199 | - | tuf | WP_011994300.1 |
| GCF_025998535.1 | NZ_AP024233.1 | 1451013 | 1452209 | 1196 | - | tuf | WP_014451930.1 |
| GCF_025998535.1 | NZ_AP024233.1 | 2024532 | 2025728 | 1196 | - | tuf | WP_015869228.1 |
| GCF_019703855.1 | NZ_AP024086.1 | 1866904 | 1868100 | 1196 | + | tuf | WP_047755020.1 |
| GCF_019703855.1 | NZ_AP024086.1 | 439309 | 440505 | 1196 | - | tuf | WP_129409932.1 |
| GCF_005116655.2 | NZ_CP050698.1 | 876642 | 877838 | 1196 | - | tuf | WP_014296370.1 |
| GCF_005116655.2 | NZ_CP050698.1 | 361481 | 362674 | 1193 | - | tuf | WP_169698241.1 |
| GCF_005116675.2 | NZ_CP050699.1 | 728564 | 729757 | 1193 | + | tuf | WP_006492341.1 |
| GCF_005116675.2 | NZ_CP050699.1 | 830502 | 831698 | 1196 | - | tuf | WP_012208619.1 |
| GCF_000025945.1 | NC_006138.1 | 600797 | 601993 | 1196 | + | tuf | WP_012002624.1 |
| GCF_000025945.1 | NC_006138.1 | 1690332 | 1691528 | 1196 | + | tuf | WP_031503255.1 |
| GCF_000092205.1 | NC_014216.1 | 607634 | 608830 | 1196 | + | tuf | WP_012002624.1 |
| GCF_000092205.1 | NC_014216.1 | 730729 | 731925 | 1196 | - | tuf | WP_013932053.1 |
| GCF_000266945.1 | NC_018025.1 | 1153614 | 1154810 | 1196 | + | tuf | WP_190615961.1 |
| GCF_000266945.1 | NC_018025.1 | 1234215 | 1235414 | 1199 | + | tuf | WP_004101434.1 |
| GCF_002355955.1 | NZ_AP017368.1 | 824940 | 826139 | 1199 | + | tuf | WP_207567398.1 |
| GCF_000212675.2 | NC_016629.1 | 987037 | 988236 | 1199 | + | tuf | WP_012057173.1 |
| GCF_000212675.2 | NC_016629.1 | 806798 | 807994 | 1196 | + | tuf | WP_041076658.1 |
| GCF_000024325.1 | NC_013223.1 | 1315207 | 1316406 | 1199 | + | tuf | WP_004081839.1 |
| GCF_000024325.1 | NC_013223.1 | 981235 | 982434 | 1199 | + | tuf | WP_015919484.1 |
| GCF_013376455.1 | NZ_CP045504.1 | 1301020 | 1302219 | 1199 | + | tuf | WP_011943784.1 |
| GCF_013376455.1 | NZ_CP045504.1 | 1670137 | 1671333 | 1196 | - | tuf | WP_041083330.1 |
| GCF_013376475.1 | NZ_CP045508.1 | 678449 | 679648 | 1199 | - | tuf | WP_160626763.1 |
| GCF_013376475.1 | NZ_CP045508.1 | 2983637 | 2984827 | 1190 | - | tuf | WP_114218409.1 |
| GCF_000023225.1 | NC_013173.1 | 2999671 | 3000861 | 1190 | - | tuf | WP_128753056.1 |
| GCF_000023225.1 | NC_013173.1 | 49272 | 50465 | 1193 | - | tuf | WP_072536332.1 |
| GCF_001553625.1 | NZ_CP014230.1 | 1479391 | 1480584 | 1193 | - | tuf | WP_072536332.1 |
| GCF_001553625.1 | NZ_CP014230.1 | 910790 | 911983 | 1193 | + | tuf | WP_117522332.1 |
| GCF_047716005.1 | NZ_CP168306.1 | 1392385 | 1393578 | 1193 | - | tuf | WP_117522332.1 |

| | | | | | | | |
|---|---|---|---|---|---|---|---|
| GCF_017815575.1 | NZ_CP072608.1 | 999965 | 1001158 | 1193 | + | tuf | WP_198482170.1 |
| GCF_001553605.1 | NZ_CP014229.1 | 1745022 | 1746215 | 1193 | - | tuf | WP_200225826.1 |
| GCF_003966735.1 | NZ_AP017378.1 | 524134 | 525327 | 1193 | + | tuf | WP_015777418.1 |
| GCF_003966735.1 | NZ_AP017378.1 | 1226109 | 1227302 | 1193 | - | tuf | WP_015777418.1 |
| GCF_026230175.1 | NZ_CP104208.1 | 384712 | 385905 | 1193 | + | tuf | WP_198482170.1 |
| GCF_027558435.1 | NZ_CP087381.1 | 1130140 | 1131333 | 1193 | - | tuf | WP_198482135.1 |
| GCF_000331715.1 | NC_020127.1 | 633013 | 634203 | 1190 | - | tuf | WP_075573968.1 |
| GCF_000331025.1 | NC_020055.1 | 1198752 | 1199942 | 1190 | + | tuf | WP_075573968.1 |
| GCF_000023445.1 | NC_012881.1 | 2375656 | 2376846 | 1190 | - | tuf | WP_014968404.1 |
| GCF_000468495.1 | NC_022444.1 | 2391789 | 2392979 | 1190 | - | tuf | WP_014968404.1 |
| GCF_000015485.1 | NC_008751.1 | 1828150 | 1829340 | 1190 | - | tuf | WP_128426460.1 |
| GCF_013085545.1 | NZ_CP039543.1 | 1844044 | 1845234 | 1190 | - | tuf | WP_128426471.1 |
| GCF_013085545.1 | NZ_CP039543.1 | 451923 | 453113 | 1190 | + | tuf | WP_134710457.1 |
| GCF_000012665.1 | NC_007519.1 | 1942697 | 1943887 | 1190 | - | tuf | WP_202690501.1 |
| GCF_000176915.2 | NC_014844.1 | 1958591 | 1959781 | 1190 | - | tuf | WP_202690503.1 |
| GCF_009762795.1 | NZ_CP046400.1 | 703600 | 704790 | 1190 | + | tuf | WP_425539730.1 |
| GCF_000189295.2 | NC_016803.1 | 726648 | 727838 | 1190 | + | tuf | WP_425539730.1 |
| GCF_037094465.1 | NZ_CP146609.1 | 118937 | 120127 | 1190 | - | tuf | WP_243661114.1 |
| GCF_027923765.1 | NZ_AP026709.1 | 464474 | 465664 | 1190 | - | tuf | WP_243661426.1 |
| GCF_000341895.1 | NC_020409.1 | 1426042 | 1427229 | 1187 | - | tuf | WP_071705823.1 |
| GCF_026000375.1 | NZ_AP026708.1 | 482444 | 483634 | 1190 | - | tuf | WP_004831804.1 |
| GCF_900217235.1 | NZ_LT907975.1 | 1216618 | 1217808 | 1190 | - | tuf | WP_068474487.1 |
| GCF_020886695.1 | NZ_AP024485.1 | 337625 | 338815 | 1190 | + | tuf | WP_271191327.1 |
| GCF_021353295.2 | NZ_CP120635.1 | 1486363 | 1487553 | 1190 | - | tuf | WP_271191327.1 |
| GCF_022809775.1 | NZ_CP094380.1 | 264043 | 265233 | 1190 | - | tuf | WP_250342211.1 |
| GCF_008001235.1 | NZ_CP040751.1 | 759193 | 760383 | 1190 | - | tuf | WP_250342484.1 |
| GCF_004135975.1 | NZ_CP026538.1 | 821571 | 822761 | 1190 | - | tuf | WP_112889499.1 |
| GCF_004135975.1 | NZ_CP026538.1 | 1624395 | 1625585 | 1190 | + | tuf | WP_112890057.1 |
| GCF_000177215.2 | NZ_CM001368.1 | 1501937 | 1503118 | 1181 | - | tuf | WP_067570716.1 |
| GCF_000177215.2 | NZ_CM001368.1 | 571451 | 572629 | 1178 | - | tuf | WP_067776340.1 |
| GCF_000010665.1 | NC_012796.1 | 1304661 | 1305842 | 1181 | - | tuf | WP_012420089.1 |

| | | | | | | | |
|---|---|---|---|---|---|---|---|
| GCF_000010665.1 | NC_012796.1 | 17474 | 18661 | 1187 | + | tuf | WP_283846412.1 |
| GCF_025232395.1 | NZ_CP065938.1 | 824232 | 825413 | 1181 | - | tuf | WP_338688457.1 |
| GCF_000020725.1 | NC_011146.1 | 3324776 | 3325957 | 1181 | - | tuf | WP_211629804.1 |
| GCF_000020725.1 | NC_011146.1 | 4267586 | 4268764 | 1178 | + | tuf | WP_169455947.1 |
| GCF_014218275.1 | NZ_AP023213.1 | 1033918 | 1035099 | 1181 | + | tuf | WP_164365479.1 |
| GCF_014218275.1 | NZ_AP023213.1 | 6667098 | 6668279 | 1181 | - | tuf | WP_009964691.1 |
| GCF_001278055.1 | NZ_CP010802.1 | 5782914 | 5784098 | 1184 | - | tuf | WP_110173164.1 |
| GCF_001278055.1 | NZ_CP010802.1 | 5806160 | 5807344 | 1184 | - | tuf | WP_110173164.1 |
| GCF_963662975.1 | NZ_OY759968.1 | 2019461 | 2020660 | 1199 | - | tuf | WP_317994440.1 |
| GCF_963662975.1 | NZ_OY759968.1 | 320988 | 322175 | 1187 | + | tuf | WP_100276750.1 |
| GCF_019704135.1 | NZ_AP024355.1 | 2521274 | 2522461 | 1187 | - | tuf | WP_100278580.1 |
| GCF_019704135.1 | NZ_AP024355.1 | 1948956 | 1950143 | 1187 | - | tuf | WP_100266088.1 |
| GCF_025263685.1 | NZ_CP092109.1 | 2285443 | 2286630 | 1187 | - | tuf | WP_100266088.1 |
| GCF_025263685.1 | NZ_CP092109.1 | 22509 | 23708 | 1199 | - | tuf | WP_026137265.1 |
| GCF_000827125.1 | NZ_CP010311.1 | 1716960 | 1718159 | 1199 | + | tuf | WP_026137265.1 |
| GCF_000827125.1 | NZ_CP010311.1 | 1958754 | 1959953 | 1199 | - | tuf | WP_013433111.1 |
| GCF_017338855.1 | NZ_CP071382.1 | 906467 | 907666 | 1199 | + | tuf | WP_013403741.1 |
| GCF_017338855.1 | NZ_CP071382.1 | 2067115 | 2068314 | 1199 | - | tuf | WP_011916518.1 |
| GCF_000012925.1 | NC_007517.1 | 1706830 | 1708029 | 1199 | - | tuf | WP_013403741.1 |
| GCF_000012925.1 | NC_007517.1 | 1904879 | 1906078 | 1199 | - | tuf | WP_013403741.1 |
| GCF_000817955.1 | NZ_CP009788.1 | 2005094 | 2006293 | 1199 | - | tuf | WP_013403741.1 |
| GCF_000817955.1 | NZ_CP009788.1 | 662954 | 664153 | 1199 | + | tuf | WP_013290055.1 |
| GCF_000007985.2 | NC_002939.5 | 1881908 | 1883107 | 1199 | - | tuf | WP_045165673.1 |
| GCF_000007985.2 | NC_002939.5 | 863683 | 864882 | 1199 | + | tuf | WP_013290055.1 |
| GCF_019063885.1 | NZ_CP077684.1 | 758230 | 759429 | 1199 | + | tuf | WP_013411633.1 |
| GCF_019063885.1 | NZ_CP077684.1 | 1082330 | 1083529 | 1199 | + | tuf | WP_011916518.1 |
| GCF_018986915.1 | NZ_CP076723.1 | 510104 | 511306 | 1202 | + | tuf | WP_088369561.1 |
| GCF_018986915.1 | NZ_CP076723.1 | 526951 | 528153 | 1202 | + | tuf | WP_088369561.1 |
| GCF_023221575.1 | NZ_CP096574.1 | 383604 | 384791 | 1187 | + | tuf | WP_072513986.1 |
| GCF_023221575.1 | NZ_CP096574.1 | 994647 | 995834 | 1187 | - | tuf | WP_072514507.1 |
| GCF_019203205.1 | NZ_CP078096.1 | 1327565 | 1328752 | 1187 | - | tuf | WP_240526869.1 |

| | | | | | | | |
|---|---|---|---|---|---|---|---|
| GCF_019203205.1 | NZ_CP078096.1 | 298333 | 299508 | 1175 | - | tuf | WP_231179770.1 |
| GCF_000022265.1 | NC_011979.1 | 2533099 | 2534319 | 1220 | + | tuf | WP_259095499.1 |
| GCF_000022265.1 | NC_011979.1 | 730921 | 732120 | 1199 | + | tuf | WP_128699425.1 |
| GCF_000016745.1 | NC_009483.1 | 566453 | 567631 | 1178 | - | tuf | WP_260762394.1 |
| GCF_000016745.1 | NC_009483.1 | 779677 | 780876 | 1199 | + | tuf | WP_012874665.1 |
| GCF_000015045.1 | NC_008609.1 | 1964125 | 1965324 | 1199 | - | tuf | WP_012874665.1 |
| GCF_000015045.1 | NC_008609.1 | 2895574 | 2896800 | 1226 | + | tuf | WP_281152645.1 |
| GCF_001887715.1 | NZ_CP015518.1 | 2723526 | 2724752 | 1226 | + | tuf | WP_044107442.1 |
| GCF_001887715.1 | NZ_CP015518.1 | 1742735 | 1743961 | 1226 | - | tuf | WP_119260197.1 |
| GCF_001887775.1 | NZ_CP015519.1 | 471151 | 472341 | 1190 | + | tuf | WP_405279861.1 |
| GCF_001887775.1 | NZ_CP015519.1 | 487932 | 489122 | 1190 | + | tuf | WP_405279861.1 |
| GCF_000012885.1 | NC_007498.2 | 190058 | 191248 | 1190 | + | tuf | WP_033178719.1 |
| GCF_000012885.1 | NC_007498.2 | 455934 | 457115 | 1181 | + | tuf | WP_391592197.1 |
| GCF_933509905.1 | NZ_OW150024.1 | 102816 | 104000 | 1184 | + | tuf | WP_205499581.1 |
| GCF_933509905.1 | NZ_OW150024.1 | 145124 | 146314 | 1190 | + | tuf | WP_114190709.1 |
| GCF_000020385.1 | NC_010814.1 | 1188982 | 1190163 | 1181 | - | tuf | WP_004025345.1 |
| GCF_000020385.1 | NC_010814.1 | 90061 | 91242 | 1181 | + | tuf | WP_412032368.1 |
| GCF_000013405.1 | NC_007759.1 | 42293 | 43474 | 1181 | + | tuf | WP_011076858.1 |
| GCF_046097065.1 | NZ_CP175563.1 | 403293 | 404474 | 1181 | - | tuf | WP_022934762.1 |
| GCF_005377625.1 | NZ_CP040098.1 | 640228 | 641415 | 1187 | - | tuf | WP_041594541.1 |
| GCF_005377625.1 | NZ_CP040098.1 | 587115 | 588320 | 1205 | - | tuf | WP_044635440.1 |
| GCF_000014965.1 | NC_008554.1 | 180302 | 181507 | 1205 | + | tuf | WP_002557626.1 |
| GCF_000014965.1 | NC_008554.1 | 676058 | 677263 | 1205 | - | tuf | WP_011284269.1 |
| GCF_001548275.1 | NZ_AP014945.1 | 687637 | 688842 | 1205 | - | tuf | WP_020104673.1 |
| GCF_001548275.1 | NZ_AP014945.1 | 897394 | 898581 | 1187 | - | tuf | WP_280101896.1 |
| GCF_000217795.1 | NC_015681.1 | 513065 | 514258 | 1193 | + | tuf | WP_022934608.1 |
| GCF_000217795.1 | NC_015681.1 | 758534 | 759721 | 1187 | - | tuf | WP_027123418.1 |
| GCF_000734015.1 | NZ_CP008796.1 | 728376 | 729563 | 1187 | - | tuf | WP_129719989.1 |
| GCF_000734015.1 | NZ_CP008796.1 | 331930 | 333135 | 1205 | - | tuf | WP_010321121.1 |
| GCF_000215975.1 | NC_015682.1 | 293394 | 294584 | 1190 | - | tuf | WP_002880918.1 |
| GCF_000215975.1 | NC_015682.1 | 472140 | 473330 | 1190 | - | tuf | WP_012498300.1 |

| | | | | | | | |
|---|---|---|---|---|---|---|---|
| GCF_012317585.1 | NZ_CP042909.1 | 651407 | 652597 | 1190 | - | tuf | WP_412031472.1 |
| GCF_012317585.1 | NZ_CP042909.1 | 537091 | 538281 | 1190 | + | tuf | WP_045434054.1 |
| GCF_011207455.1 | NZ_CP048877.1 | 585119 | 586309 | 1190 | - | tuf | WP_029330109.1 |
| GCF_011207455.1 | NZ_CP048877.1 | 192079 | 193269 | 1190 | + | tuf | WP_140031565.1 |
| GCF_003057965.1 | NZ_CP020921.1 | 476386 | 477576 | 1190 | + | tuf | WP_405311214.1 |
| GCF_003057965.1 | NZ_CP020921.1 | 320332 | 321522 | 1190 | - | tuf | WP_330463683.1 |
| GCF_000212395.1 | NC_015499.1 | 62811 | 64001 | 1190 | + | tuf | WP_012855329.1 |
| GCF_000212395.1 | NC_015499.1 | 655953 | 657143 | 1190 | + | tuf | WP_036441217.1 |
| GCF_000024985.1 | NC_013523.1 | 616304 | 617494 | 1190 | + | tuf | WP_022935742.1 |
| GCF_000024985.1 | NC_013523.1 | 181995 | 183185 | 1190 | + | tuf | WP_116171923.1 |
| GCF_000021685.1 | NC_011959.1 | 224886 | 226076 | 1190 | - | tuf | WP_024544157.1 |
| GCF_000021685.1 | NC_011959.1 | 83263 | 84453 | 1190 | + | tuf | WP_027122526.1 |
| GCF_001547735.1 | NZ_AP013035.1 | 657221 | 658414 | 1193 | - | tuf | WP_406602450.1 |
| GCF_001547735.1 | NZ_AP013035.1 | 496389 | 497579 | 1190 | + | tuf | WP_121941015.1 |
| GCF_009936215.1 | NZ_AP019551.1 | 291542 | 292741 | 1199 | - | tuf | WP_011264744.1 |
| GCF_000953715.1 | NZ_LN824141.1 | 519788 | 520969 | 1181 | + | tuf | WP_256547790.1 |
| GCF_004117075.1 | NZ_CP026721.1 | 306682 | 307863 | 1181 | + | tuf | WP_011113468.1 |
| GCF_000767275.3 | NZ_CP014334.2 | 554433 | 555614 | 1181 | - | tuf | WP_009885583.1 |
| GCF_000017545.1 | NC_009718.1 | 25477 | 26658 | 1181 | + | tuf | WP_010875022.1 |
| GCF_000235405.2 | NC_017095.1 | 824354 | 825541 | 1187 | - | tuf | WP_406616881.1 |
| GCF_000023325.1 | NC_012785.1 | 382492 | 383679 | 1187 | + | tuf | WP_004024476.1 |
| GCF_001027025.1 | NZ_CP011232.1 | 633037 | 634221 | 1184 | - | tuf | WP_006886429.1 |
| GCF_029910535.1 | NZ_CP069362.1 | 630446 | 631636 | 1190 | + | tuf | WP_004415386.1 |
| GCF_000255135.1 | NC_016751.1 | 497006 | 498193 | 1187 | + | tuf | WP_004418907.1 |
| GCF_900157305.1 | NZ_LS974202.1 | 523168 | 524352 | 1184 | + | tuf | WP_099309096.1 |
| GCF_000147715.2 | NC_017934.1 | 474522 | 475709 | 1187 | + | tuf | WP_004024476.1 |
| GCF_000018605.1 | NC_010003.1 | 517235 | 518422 | 1187 | + | tuf | WP_038561836.1 |
| GCF_000504085.1 | NC_022792.1 | 809338 | 810522 | 1184 | - | tuf | WP_004795211.1 |
| GCF_000504105.1 | NC_022795.1 | 901917 | 903101 | 1184 | - | tuf | WP_126118367.1 |
| GCF_000017865.1 | NC_009828.1 | 523539 | 524723 | 1184 | - | tuf | WP_129725323.1 |
| GCF_000217815.1 | NC_015707.1 | 664815 | 666005 | 1190 | + | tuf | WP_006608829.1 |

| | | | | | | | |
|---|---|---|---|---|---|---|---|
| GCF_014701195.1 | NZ_AP018712.1 | 141378 | 142559 | 1181 | - | tuf | WP_129622736.1 |
| GCF_000021285.1 | NC_011653.1 | 792125 | 793309 | 1184 | + | tuf | WP_036434301.1 |
| GCF_017358165.1 | NZ_CP071446.1 | 929364 | 930548 | 1184 | - | tuf | WP_015287681.1 |
| GCF_000016905.1 | NC_009616.1 | 721901 | 723085 | 1184 | - | tuf | WP_117275722.1 |
| GCF_000828655.1 | NZ_AP014509.1 | 791508 | 792689 | 1181 | - | tuf | WP_129722737.1 |
| GCF_000230655.2 | NC_023151.1 | 848729 | 849913 | 1184 | + | tuf | WP_036430197.1 |
| GCF_000018945.1 | NC_011978.1 | 254968 | 256152 | 1184 | + | tuf | WP_013526754.1 |
| GCF_000016785.1 | NC_009486.1 | 603639 | 604823 | 1184 | - | tuf | WP_129620326.1 |
| GCF_000828675.1 | NZ_AP014510.1 | 19271 | 20455 | 1184 | - | tuf | WP_119572352.1 |
| GCF_949774925.1 | NZ_OX458932.1 | 573519 | 574703 | 1184 | + | tuf | WP_119572352.1 |
| GCF_017310505.1 | NZ_CP065957.1 | 921567 | 922751 | 1184 | + | tuf | WP_027333868.1 |
| GCF_000019665.1 | NC_010794.1 | 696677 | 697858 | 1181 | - | tuf | WP_406615180.1 |
| GCF_007475525.1 | NZ_CP037899.1 | 695989 | 697170 | 1181 | - | tuf | WP_406617845.1 |
| GCF_019679435.2 | NZ_CP139781.1 | 809262 | 810449 | 1187 | - | tuf | WP_129647105.1 |
| GCF_014529625.1 | NZ_CP076039.1 | 111719 | 112900 | 1181 | + | tuf | WP_046097128.1 |
| GCF_000025905.1 | NC_014008.1 | 553142 | 554329 | 1187 | - | tuf | WP_130429491.1 |
| GCF_033878955.1 | NZ_CP138858.1 | 452473 | 453660 | 1187 | - | tuf | WP_129690203.1 |
| GCF_003096195.1 | NZ_CP023004.1 | 806629 | 807810 | 1181 | + | tuf | WP_020003176.1 |
| GCF_019464535.1 | NZ_CP080507.1 | 96860 | 98044 | 1184 | + | tuf | WP_078747171.1 |
| GCF_001746835.1 | NZ_CP016094.1 | 53141 | 54319 | 1178 | - | tuf | WP_208895042.1 |
| GCF_002310495.1 | NZ_CP023344.1 | 647230 | 648411 | 1181 | - | tuf | WP_006688507.1 |
| GCF_000019965.1 | NC_010571.1 | 2022524 | 2023711 | 1187 | + | tuf | WP_050429774.1 |
| GCF_033100135.1 | NZ_CP136920.1 | 2483212 | 2484399 | 1187 | + | tuf | WP_050430052.1 |
| GCF_029023725.1 | NZ_CP119075.1 | 5498178 | 5499368 | 1190 | - | tuf | WP_012829105.1 |
| GCF_038725785.1 | NZ_CP109796.1 | 5522510 | 5523700 | 1190 | - | tuf | WP_012829105.1 |
| GCF_026072915.1 | NZ_AP025943.1 | 338748 | 339935 | 1187 | - | tuf | WP_146645413.1 |
| GCF_900097105.1 | NZ_LT629973.1 | 630005 | 631192 | 1187 | - | tuf | WP_146645413.1 |
| GCF_009731575.1 | NZ_AP021898.1 | 11313684 | 11314880 | 1196 | - | tuf | WP_267775467.1 |
| GCF_024832785.1 | NZ_CP116615.1 | 11331397 | 11332593 | 1196 | - | tuf | WP_267775467.1 |
| GCF_037076345.1 | NZ_AP024702.1 | 1411873 | 1413060 | 1187 | - | tuf | WP_075277110.1 |
| GCF_018137965.1 | NZ_CP073100.1 | 78931 | 80118 | 1187 | - | tuf | WP_248913849.1 |

| | | | | | | | |
|---|---|---|---|---|---|---|---|
| GCF_012913485.1 | NZ_CP051774.1 | 4828287 | 4829474 | 1187 | + | tuf | WP_248913849.1 |
| GCF_010681825.2 | NZ_CP066776.1 | 3144607 | 3145794 | 1187 | - | tuf | WP_053232793.1 |
| GCF_000172155.1 | NZ_ABIZ01000001.1 | 9062129 | 9063316 | 1187 | - | tuf | WP_053232793.1 |
| GCF_027923555.1 | NZ_AP025523.1 | 581602 | 582789 | 1187 | + | tuf | WP_012233037.1 |
| GCF_000513475.1 | NC_023003.1 | 1193098 | 1194285 | 1187 | + | tuf | WP_012233037.1 |
| GCF_003366055.1 | NZ_CP025544.1 | 5205800 | 5207005 | 1205 | - | tuf | WP_145068219.1 |
| GCF_003258315.1 | NZ_CP030032.1 | 6361852 | 6363048 | 1196 | + | tuf | WP_145261940.1 |
| GCF_003258315.1 | NZ_CP030032.1 | 3841214 | 3842419 | 1205 | - | tuf | WP_145189870.1 |
| GCF_022026155.1 | NZ_CP085041.1 | 5445993 | 5447195 | 1202 | - | tuf | WP_145201143.1 |
| GCF_022026155.1 | NZ_CP085041.1 | 1713552 | 1714745 | 1193 | - | tuf | WP_145270019.1 |
| GCF_007993755.1 | NZ_CP042467.1 | 5738464 | 5739663 | 1199 | + | tuf | WP_145214566.1 |
| GCF_007993775.1 | NZ_CP042468.1 | 7796098 | 7797291 | 1193 | - | tuf | WP_145218177.1 |
| GCF_007993775.1 | NZ_CP042468.1 | 5325082 | 5326275 | 1193 | + | tuf | WP_145419747.1 |
| GCF_030866785.1 | NZ_CP133018.1 | 3132965 | 3134152 | 1187 | - | tuf | WP_308797144.1 |
| GCF_044772865.1 | NZ_CP173029.1 | 2959359 | 2960531 | 1172 | - | tuf | WP_420861269.1 |
| GCF_044772865.1 | NZ_CP173029.1 | 3113786 | 3114958 | 1172 | - | tuf | WP_420861269.1 |
| GCF_018398915.1 | NZ_CP074691.1 | 640048 | 641244 | 1196 | + | tuf | WP_274372696.1 |
| GCF_018398915.1 | NZ_CP074691.1 | 1955132 | 1956328 | 1196 | - | tuf | WP_274372696.1 |
| GCF_047275885.1 | NZ_CP178454.1 | 2144007 | 2145233 | 1226 | - | tuf | WP_413175196.1 |
| GCF_015243575.1 | NZ_CP042469.1 | 4551468 | 4552658 | 1190 | + | tuf | WP_419822540.1 |
| GCF_015243575.1 | NZ_CP042469.1 | 4568771 | 4569961 | 1190 | + | tuf | WP_419822540.1 |
| GCF_048270625.1 | NZ_CP158164.1 | 3848031 | 3849218 | 1187 | - | tuf | WP_386645268.1 |
| GCF_049676565.1 | NZ_CP152277.1 | 394723 | 395907 | 1184 | - | tuf | WP_424245440.1 |
| GCF_049676565.1 | NZ_CP152277.1 | 435583 | 436767 | 1184 | - | tuf | WP_424245440.1 |
| GCF_047159375.1 | NZ_AP031381.1 | 62088 | 63272 | 1184 | + | tuf | WP_412728816.1 |
| GCF_001644665.1 | NZ_CP011393.1 | 24734 | 25930 | 1196 | - | tuf | WP_064011014.1 |
| GCF_041154365.1 | NZ_AP035449.1 | 1886442 | 1887632 | 1190 | - | tuf | WP_400166285.1 |
| GCF_041154365.1 | NZ_AP035449.1 | 1902029 | 1903219 | 1190 | - | tuf | WP_400166285.1 |
| GCF_044054575.1 | NZ_CP168716.1 | 1563216 | 1564400 | 1184 | - | tuf | WP_395318978.1 |
| GCF_019720755.1 | NZ_CP081303.1 | 2804280 | 2805458 | 1178 | - | tuf | WP_394654743.1 |
| GCF_019856515.1 | NZ_CP082230.1 | 4805469 | 4806647 | 1178 | + | tuf | WP_394668687.1 |

| | | | | | | | |
|---|---|---|---|---|---|---|---|
| GCF_964332355.1 | NZ_OZ201293.1 | 2971620 | 2972807 | 1187 | - | tuf | WP_009523912.1 |
| GCF_964332355.1 | NZ_OZ201293.1 | 2990945 | 2992132 | 1187 | - | tuf | WP_009523912.1 |
| GCF_043950205.1 | NZ_CP171642.1 | 4904760 | 4905950 | 1190 | + | tuf | WP_395295779.1 |
| GCF_043633585.1 | NZ_CP141832.1 | 548607 | 549797 | 1190 | + | tuf | WP_394524930.1 |
| GCF_044053885.1 | NZ_CP168707.1 | 1277581 | 1278768 | 1187 | + | tuf | WP_125762958.1 |
| GCF_050500245.1 | NZ_CP191408.1 | 1858780 | 1860006 | 1226 | - | tuf | WP_430014430.1 |
| GCF_050500235.1 | NZ_CP191407.1 | 1854904 | 1856130 | 1226 | - | tuf | WP_430014430.1 |
| GCF_044704065.1 | NZ_CP162613.1 | 146731 | 147915 | 1184 | + | tuf | WP_400244848.1 |
| GCF_048213115.1 | NZ_CP172404.1 | 560789 | 561976 | 1187 | + | tuf | WP_417069438.1 |
| GCF_048213115.1 | NZ_CP172404.1 | 577406 | 578593 | 1187 | + | tuf | WP_417069450.1 |
| GCF_031656915.1 | NZ_CP134227.1 | 2164483 | 2165673 | 1190 | - | tuf | WP_310961104.1 |
| GCF_050389785.1 | NZ_CP015019.1 | 1020631 | 1021818 | 1187 | - | tuf | WP_428897108.1 |
| GCF_050389785.1 | NZ_CP015019.1 | 1208565 | 1209752 | 1187 | + | tuf | WP_428897108.1 |
| GCF_042466665.1 | NZ_CP162102.1 | 5009672 | 5010862 | 1190 | + | tuf | WP_377270828.1 |
| GCF_046254905.1 | NZ_CP158016.1 | 3326858 | 3328039 | 1181 | - | tuf | WP_409421178.1 |
| GCF_046254905.1 | NZ_CP158016.1 | 3347581 | 3348762 | 1181 | - | tuf | WP_409421178.1 |
| GCF_046352735.2 | NZ_CP110853.2 | 4802682 | 4803872 | 1190 | - | tuf | WP_003176426.1 |
| GCF_964291045.1 | NZ_OZ197277.1 | 1067805 | 1068986 | 1181 | - | tuf | WP_404665074.1 |
| GCF_047160125.1 | NZ_AP038788.1 | 642884 | 644065 | 1181 | - | tuf | WP_226692005.1 |
| GCF_047160125.1 | NZ_AP038788.1 | 654995 | 656176 | 1181 | - | tuf | WP_226692005.1 |
| GCF_042140045.1 | NZ_CP152419.1 | 1630025 | 1631218 | 1193 | - | tuf | WP_001040718.1 |
| GCF_045519255.1 | NZ_CP169548.1 | 1159278 | 1160456 | 1178 | - | tuf | WP_365899139.1 |
| GCF_045519255.1 | NZ_CP169548.1 | 4614115 | 4615305 | 1190 | + | tuf | WP_103546614.1 |
| GCF_045822005.1 | NZ_CP169550.1 | 3768899 | 3770089 | 1190 | - | tuf | WP_407837694.1 |
| GCF_048537595.1 | NZ_CP175923.1 | 190530 | 191699 | 1169 | + | tuf | WP_419247305.1 |
| GCF_048537595.1 | NZ_CP175923.1 | 431158 | 432327 | 1169 | - | tuf | WP_419247305.1 |

| locus_tag | product | Organism Taxonomic ID |
|---|---|---|
| AFER_RS02050 | elongation factor Tu | 525909 |
| GH723_RS01210 | elongation factor Tu | 2789776 |
| GH723_RS01290 | elongation factor Tu | 2789776 |
| ACE2AJ_RS08250 | elongation factor Tu | 1052257 |
| ACE2AJ_RS08430 | elongation factor Tu | 1052257 |
| LH044_RS11055 | elongation factor Tu | 2884263 |
| LH044_RS11135 | elongation factor Tu | 2884263 |
| PO878_RS19475 | elongation factor Tu | 467976 |
| PO878_RS19565 | elongation factor Tu | 467976 |
| YM304_RS03955 | elongation factor Tu | 1313172 |
| YM304_RS04110 | elongation factor Tu | 1313172 |
| ATC_RS02925 | elongation factor Tu | 990288 |
| ATC_RS02990 | elongation factor Tu | 990288 |
| GL267_RS13050 | elongation factor Tu | 2678518 |
| GL267_RS13115 | elongation factor Tu | 2678518 |
| AFERRID_RS05410 | elongation factor Tu | 1232575 |
| AFERRID_RS05475 | elongation factor Tu | 1232575 |
| PJU76_RS11095 | elongation factor Tu | 1689834 |
| PJU76_RS11160 | elongation factor Tu | 1689834 |
| H2515_RS02505 | elongation factor Tu | 160808 |
| H2515_RS02570 | elongation factor Tu | 160808 |
| AFE_RS01470 | elongation factor Tu | 243159 |
| AFE_RS01535 | elongation factor Tu | 243159 |
| HF292_RS02335 | elongation factor Tu | 3063951 |
| HF292_RS02400 | elongation factor Tu | 3063951 |
| HHS34_RS12720 | elongation factor Tu | 2729135 |
| HHS34_RS12785 | elongation factor Tu | 2729135 |
| EC580_RS02755 | elongation factor Tu | 1867749 |
| EC580_RS02820 | elongation factor Tu | 1867749 |
| GCD22_RS01930 | elongation factor Tu | 637390 |

| | | |
|---|---|---|
| GCD22_RS01995 | elongation factor Tu | 637390 |
| RG903_RS11290 | elongation factor Tu | 1123368 |
| RG903_RS11355 | elongation factor Tu | 1123368 |
| BKM31_RS24970 | elongation factor Tu | 1909395 |
| BKM31_RS40965 | elongation factor Tu | 1909395 |
| OG362_RS12565 | elongation factor Tu | 1464011 |
| OG362_RS27815 | elongation factor Tu | 1464011 |
| RCP80_RS19220 | elongation factor Tu | 3064286 |
| RCP37_RS03605 | elongation factor Tu | 3064288 |
| RCP38_RS03250 | elongation factor Tu | 3064287 |
| MSTE_RS19710 | elongation factor Tu | 1520670 |
| R2K23_RS18290 | elongation factor Tu | 3064284 |
| ASQ49_RS09125 | elongation factor Tu | 1748 |
| FEZ32_RS07905 | elongation factor Tu | 1749 |
| JS278_RS04680 | elongation factor Tu | 2057246 |
| ACEL_RS01590 | elongation factor Tu | 351607 |
| OIE69_RS17735 | elongation factor Tu | 235986 |
| OIE69_RS34115 | elongation factor Tu | 235986 |
| RVR_RS22145 | elongation factor Tu | 659352 |
| UA74_RS02920 | elongation factor Tu | 1612552 |
| AHOG_RS02825 | elongation factor Tu | 1470176 |
| TL08_RS02915 | elongation factor Tu | 340345 |
| Asera_RS10490 | elongation factor Tu | 390989 |
| Athai_RS26555 | elongation factor Tu | 227318 |
| V5P93_RS34350 | elongation factor Tu | 155976 |
| AGRA3207_RS20230 | elongation factor Tu | 2750812 |
| AGRA3207_RS24460 | elongation factor Tu | 2750812 |
| V6738_RS32795 | elongation factor Tu | 46163 |
| V6738_RS37760 | elongation factor Tu | 46163 |
| K8Z49_RS29480 | elongation factor Tu | 1993 |
| K8Z49_RS34575 | elongation factor Tu | 1993 |

| | | |
|---|---|---|
| MANAM107_RS08240 | elongation factor Tu | 2755559 |
| HRL51_RS09145 | elongation factor Tu | 2722820 |
| B6G06_RS07265 | elongation factor Tu | 1960083 |
| EL245_RS05545 | elongation factor Tu | 52771 |
| AIF0345_RS12580 | elongation factor Tu | 1659 |
| D5R93_RS03160 | elongation factor Tu | 2321394 |
| HPC72_RS02345 | elongation factor Tu | 2737173 |
| I6H94_RS13170 | elongation factor Tu | 1655 |
| I6I08_RS08700 | elongation factor Tu | 544580 |
| BQ7111_RS07450 | elongation factor Tu | 1852377 |
| E4J16_RS11410 | elongation factor Tu | 2560010 |
| CWT10_RS13185 | elongation factor Tu | 2057800 |
| AXF14_RS07170 | elongation factor Tu | 111015 |
| ID810_RS09600 | elongation factor Tu | 2744574 |
| EL266_RS05155 | elongation factor Tu | 52774 |
| I2V18_RS02140 | elongation factor Tu | 2763540 |
| EL340_RS14390 | elongation factor Tu | 1656 |
| JG540_RS02325 | elongation factor Tu | 675090 |
| CWS50_RS09255 | elongation factor Tu | 2057743 |
| BLU81_RS20220 | elongation factor Tu | 113562 |
| BLU81_RS30265 | elongation factor Tu | 113562 |
| AFR_RS04550 | elongation factor Tu | 1246995 |
| AFR_RS32580 | elongation factor Tu | 1246995 |
| Aiant_RS07405 | elongation factor Tu | 122358 |
| Aiant_RS24490 | elongation factor Tu | 122358 |
| AMIS_RS03065 | elongation factor Tu | 512565 |
| AMIS_RS28490 | elongation factor Tu | 512565 |
| Actob_RS03085 | elongation factor Tu | 3040509 |
| Actob_RS30950 | elongation factor Tu | 3040509 |
| Q0Z83_RS31460 | elongation factor Tu | 512349 |
| Q0Z83_RS40630 | elongation factor Tu | 512349 |

| | | |
|---|---|---|
| ACTEI_RS02965 | elongation factor Tu | 457423 |
| BLU27_RS13460 | elongation factor Tu | 117157 |
| CDG81_RS19995 | elongation factor Tu | 414996 |
| AMIR_RS32770 | elongation factor Tu | 446462 |
| CNX65_RS32460 | elongation factor Tu | 42197 |
| FB03_RS03885 | elongation factor Tu | 59505 |
| CYJ17_RS08760 | elongation factor Tu | 1870995 |
| C3E78_RS14925 | elongation factor Tu | 2079793 |
| B5D60_RS05005 | elongation factor Tu | 1736691 |
| NP095_RS02300 | elongation factor Tu | 2968086 |
| Aeryth_RS13695 | elongation factor Tu | 2041 |
| HMPREF0063_RS10465 | elongation factor Tu | 585531 |
| H9L21_RS02330 | elongation factor Tu | 2663859 |
| GEV26_RS15030 | elongation factor Tu | 2662028 |
| Q9250_RS02520 | elongation factor Tu | 3068634 |
| BLT67_RS09295 | elongation factor Tu | 684552 |
| BLQ67_RS05095 | elongation factor Tu | 399736 |
| G127AT_RS04565 | elongation factor Tu | 2781962 |
| ATC03_RS15340 | elongation factor Tu | 453304 |
| DCE93_RS11570 | elongation factor Tu | 2080742 |
| BLT99_RS00940 | elongation factor Tu | 589382 |
| FLP10_RS12505 | elongation factor Tu | 2592652 |
| FHG54_RS05385 | elongation factor Tu | 2585717 |
| MTO99_RS10530 | elongation factor Tu | 2929802 |
| QUE38_RS04140 | elongation factor Tu | 1858653 |
| DSM26151_RS11785 | elongation factor Tu | 1389020 |
| ET445_RS07130 | elongation factor Tu | 2509455 |
| MTP13_RS09280 | elongation factor Tu | 659012 |
| BLT28_RS14675 | elongation factor Tu | 211114 |
| BLT28_RS25340 | elongation factor Tu | 211114 |
| GIY23_RS20240 | elongation factor Tu | 2665642 |

| | | |
|---|---|---|
| AADG42_RS03670 | elongation factor Tu | 3138240 |
| LWP59_RS03085 | elongation factor Tu | 715473 |
| KOI47_RS31200 | elongation factor Tu | 2842453 |
| A4R43_RS34280 | elongation factor Tu | 1804986 |
| AMYBAR_RS03120 | elongation factor Tu | 941986 |
| QRX50_RS04240 | elongation factor Tu | 715471 |
| ORV05_RS30255 | elongation factor Tu | 2995223 |
| AJAP_RS35865 | elongation factor Tu | 208439 |
| AORI_RS44275 | elongation factor Tu | 129921 |
| RAM_RS03445 | elongation factor Tu | 713604 |
| RAM_RS21185 | elongation factor Tu | 713604 |
| AMETH_RS03245 | elongation factor Tu | 1068978 |
| QRX60_RS07955 | elongation factor Tu | 715475 |
| QRX60_RS35125 | elongation factor Tu | 715475 |
| QP939_RS09220 | elongation factor Tu | 715472 |
| QP939_RS43185 | elongation factor Tu | 715472 |
| SD37_RS37890 | elongation factor Tu | 31958 |
| VSH64_RS40915 | elongation factor Tu | 1448059 |
| NQK81_RS19530 | elongation factor Tu | 700274 |
| LH407_RS05585 | elongation factor Tu | 2880260 |
| HRU87_RS00730 | elongation factor Tu | 2499157 |
| EL272_RS10900 | elongation factor Tu | 1750 |
| V7R85_RS04455 | elongation factor Tu | 1547448 |
| HC352_RS07175 | elongation factor Tu | 2722751 |
| P7079_RS02060 | elongation factor Tu | 999183 |
| DQN53_RS03160 | elongation factor Tu | 28264 |
| BLT51_RS04200 | elongation factor Tu | 131112 |
| JTE88_RS07035 | elongation factor Tu | 1302235 |
| NG665_RS06805 | elongation factor Tu | 1503041 |
| P8A24_RS02360 | elongation factor Tu | 1803177 |
| ACF3NS_RS00840 | elongation factor Tu | 2663847 |

| | | |
|---|---|---|
| QK900_RS12990 | elongation factor Tu | 1076331 |
| PTW37_RS12500 | elongation factor Tu | 37921 |
| MB46_RS13455 | elongation factor Tu | 656366 |
| MWM45_RS13695 | elongation factor Tu | 2895818 |
| NF551_RS12805 | elongation factor Tu | 2886935 |
| AC20117_RS01580 | elongation factor Tu | 37928 |
| DMB86_RS19110 | elongation factor Tu | 2211210 |
| MUG94_RS13775 | elongation factor Tu | 2886940 |
| KKR91_RS13485 | elongation factor Tu | 2817475 |
| N2K98_RS13400 | elongation factor Tu | 2964609 |
| ACJJV6_RS01980 | elongation factor Tu | 683150 |
| J0916_RS12360 | elongation factor Tu | 2813727 |
| MNQ99_RS05210 | elongation factor Tu | 2486855 |
| KG104_RS14385 | elongation factor Tu | 2816859 |
| MUK71_RS12750 | elongation factor Tu | 2886936 |
| N2K95_RS12695 | elongation factor Tu | 2964616 |
| AUMI_RS06265 | elongation factor Tu | 708131 |
| AURMO_RS00880 | elongation factor Tu | 1987356 |
| BLT52_RS07925 | elongation factor Tu | 675864 |
| QDX22_RS11465 | elongation factor Tu | 678932 |
| QUE40_RS02935 | elongation factor Tu | 1535208 |
| BCAV_RS15835 | elongation factor Tu | 471853 |
| AB656_RS03575 | elongation factor Tu | 1437605 |
| BAD_RS02840 | elongation factor Tu | 367928 |
| BBAG_RS02525 | elongation factor Tu | 518635 |
| BALAT_RS03070 | elongation factor Tu | 555970 |
| QN215_RS03975 | elongation factor Tu | 3059038 |
| BA20089_RS04905 | elongation factor Tu | 1437594 |
| CNCMI4319_RS06365 | elongation factor Tu | 1681 |
| BBBR_RS02720 | elongation factor Tu | 518634 |
| BBCT_RS02745 | elongation factor Tu | 566552 |

| | | |
|---|---|---|
| BcFMB_RS05590 | elongation factor Tu | 35760 |
| bcor_RS04720 | elongation factor Tu | 1687 |
| BBDE_RS03585 | elongation factor Tu | 1150423 |
| BE0216_RS03615 | elongation factor Tu | 1765219 |
| QN062_RS00105 | elongation factor Tu | 3059035 |
| BLI708_RS00960 | elongation factor Tu | 2020965 |
| BL8807_RS07445 | elongation factor Tu | 1603886 |
| BLLJ_RS02635 | elongation factor Tu | 565042 |
| BBPC_RS02985 | elongation factor Tu | 547043 |
| BP20092_RS06170 | elongation factor Tu | 1410605 |
| BSD967_RS08635 | elongation factor Tu | 762210 |
| BBSC_RS03995 | elongation factor Tu | 1150461 |
| BS3272_RS03970 | elongation factor Tu | 77635 |
| U6N30_RS25190 | elongation factor Tu | 1564165 |
| ABDB74_RS18110 | elongation factor Tu | 3144973 |
| R8377_RS05270 | elongation factor Tu | 2932182 |
| BK816_RS02040 | elongation factor Tu | 1912795 |
| CFK39_RS04330 | elongation factor Tu | 2017485 |
| CFK41_RS04545 | elongation factor Tu | 1331682 |
| Bra3105_RS15860 | elongation factor Tu | 2887346 |
| BRM3_RS00755 | elongation factor Tu | 1652028 |
| M4486_RS00030 | elongation factor Tu | 2942909 |
| DWV08_RS09425 | elongation factor Tu | 556288 |
| CFK38_RS06835 | elongation factor Tu | 2017484 |
| GUY23_RS06120 | elongation factor Tu | 2697563 |
| CXR25_RS05845 | elongation factor Tu | 273384 |
| I6G93_RS00290 | elongation factor Tu | 33889 |
| AAFP32_RS04835 | elongation factor Tu | 3140787 |
| GUY37_RS12815 | elongation factor Tu | 2697565 |
| CXR29_RS05265 | elongation factor Tu | 1703 |
| J9314_RS06155 | elongation factor Tu | 358099 |

| | | |
|---|---|---|
| GUY30_RS05175 | elongation factor Tu | 1496080 |
| BLT92_RS02950 | elongation factor Tu | 629680 |
| BLU88_RS15030 | elongation factor Tu | 1136497 |
| L1F31_RS05975 | elongation factor Tu | 2909672 |
| LQ788_RS06690 | elongation factor Tu | 2898795 |
| QUE25_RS10895 | elongation factor Tu | 3041175 |
| ABYF32_RS00265 | elongation factor Tu | 3231492 |
| M5I08_RS03620 | elongation factor Tu | 2943498 |
| PT015_RS22270 | elongation factor Tu | 3027808 |
| B1sIIB91_RS00555 | elongation factor Tu | 1884916 |
| B1s21160_RS00630 | elongation factor Tu | 1884915 |
| B1s21122_RS05655 | elongation factor Tu | 1884634 |
| A1s21155_RS05710 | elongation factor Tu | 1884914 |
| A1sIIB106_RS05780 | elongation factor Tu | 1884913 |
| PHILAsVB114_RS01205 | elongation factor Tu | 573600 |
| A1sIA56_RS05625 | elongation factor Tu | 1884904 |
| A7sIIA15_RS01015 | elongation factor Tu | 1884907 |
| A1sIA105_RS05650 | elongation factor Tu | 1884905 |
| A4Z71_RS06045 | elongation factor Tu | 535712 |
| CACI_RS04575 | elongation factor Tu | 479433 |
| CACI_RS43835 | elongation factor Tu | 479433 |
| NP064_RS13695 | elongation factor Tu | 2968084 |
| KKR89_RS13180 | elongation factor Tu | 2819979 |
| KG102_RS13455 | elongation factor Tu | 2819978 |
| CELF_RS05030 | elongation factor Tu | 590998 |
| CFLA_RS13435 | elongation factor Tu | 446466 |
| CELGI_RS04090 | elongation factor Tu | 593907 |
| F1D97_RS16415 | elongation factor Tu | 2608402 |
| E5225_RS04370 | elongation factor Tu | 2566013 |
| HGK68_RS13050 | elongation factor Tu | 2729175 |
| KG103_RS13430 | elongation factor Tu | 2816956 |

| | | |
|---|---|---|
| NP075_RS13845 | elongation factor Tu | 2968085 |
| NP048_RS13445 | elongation factor Tu | 2968083 |
| J5A69_RS10940 | elongation factor Tu | 1710 |
| FIC82_RS16735 | elongation factor Tu | 2587808 |
| H2O17_RS02885 | elongation factor Tu | 2758440 |
| IMT31_RS04970 | elongation factor Tu | 863575 |
| FGD68_RS00335 | elongation factor Tu | 1401995 |
| GW571_RS12730 | elongation factor Tu | 1874630 |
| K0V08_RS09010 | elongation factor Tu | 28447 |
| CMN_RS12750 | elongation factor Tu | 1097677 |
| FGI33_RS00180 | elongation factor Tu | 1734031 |
| KYT88_RS13615 | elongation factor Tu | 2860285 |
| CMS_RS01400 | elongation factor Tu | 31964 |
| H9X71_RS13060 | elongation factor Tu | 2768071 |
| B5808_RS04630 | elongation factor Tu | 1619308 |
| CACC_RS02075 | elongation factor Tu | 38284 |
| CAFEL_RS01625 | elongation factor Tu | 144184 |
| CAMM_RS02335 | elongation factor Tu | 649754 |
| I6J19_RS01200 | elongation factor Tu | 43765 |
| GP473_RS07960 | elongation factor Tu | 2684406 |
| CAPP_RS01450 | elongation factor Tu | 1161099 |
| CAQUA_RS09845 | elongation factor Tu | 1190508 |
| CAQU_RS01635 | elongation factor Tu | 1431546 |
| CARG_RS01210 | elongation factor Tu | 1348662 |
| CATRI_RS01930 | elongation factor Tu | 1336740 |
| CATYP_RS01875 | elongation factor Tu | 191610 |
| CAURI_RS02070 | elongation factor Tu | 548476 |
| CAURIS_RS01895 | elongation factor Tu | 44750 |
| CAURIC_RS09455 | elongation factor Tu | 99807 |
| ABMV07_RS02345 | elongation factor Tu | 2014537 |
| QP027_RS01355 | elongation factor Tu | 3049799 |

| | | |
|---|---|---|
| H924_RS02450 | elongation factor Tu | 1121353 |
| UL81_RS01720 | elongation factor Tu | 161896 |
| CCANI_RS01675 | elongation factor Tu | 679663 |
| CAPI_RS01320 | elongation factor Tu | 1121354 |
| CCASEI_RS02730 | elongation factor Tu | 1285583 |
| CCASP_RS07340 | elongation factor Tu | 1121355 |
| CCHOA_RS09070 | elongation factor Tu | 1862358 |
| CCICO_RS01325 | elongation factor Tu | 1121356 |
| CETAM_RS02175 | elongation factor Tu | 2675218 |
| CCONF_RS01895 | elongation factor Tu | 71254 |
| I6L55_RS06660 | elongation factor Tu | 53374 |
| ccrud_RS02590 | elongation factor Tu | 1652495 |
| CKV99_RS09950 | elongation factor Tu | 35757 |
| CDES_RS02565 | elongation factor Tu | 931089 |
| AT687_RS01910 | elongation factor Tu | 1717 |
| CDOO_RS02655 | elongation factor Tu | 558173 |
| CDUR_RS01980 | elongation factor Tu | 61592 |
| CE_RS02745 | elongation factor Tu | 196164 |
| CENDO_RS01820 | elongation factor Tu | 2488819 |
| CEPID_RS01820 | elongation factor Tu | 1050174 |
| CFAEC_RS02090 | elongation factor Tu | 1758466 |
| LA343_RS00395 | elongation factor Tu | 108486 |
| CFELI_RS01640 | elongation factor Tu | 131318 |
| CFLV_RS02075 | elongation factor Tu | 28028 |
| CFOUR_RS01650 | elongation factor Tu | 1852390 |
| CFRA_RS01930 | elongation factor Tu | 1437875 |
| CFREI_RS01575 | elongation factor Tu | 556548 |
| I6L56_RS10045 | elongation factor Tu | 134034 |
| KBP53_RS01610 | elongation factor Tu | 585529 |
| CGERO_RS09225 | elongation factor Tu | 2079234 |
| CGLAUT_RS01840 | elongation factor Tu | 187491 |

| | | |
|---|---|---|
| CGLUCO_RS02235 | elongation factor Tu | 1121361 |
| CGL_RS02540 | elongation factor Tu | 196627 |
| CGLY_RS03460 | elongation factor Tu | 1404245 |
| CGOTT_RS01700 | elongation factor Tu | 2041036 |
| CGUA_RS02400 | elongation factor Tu | 1783348 |
| A605_RS02460 | elongation factor Tu | 1121362 |
| CHAN_RS01785 | elongation factor Tu | 394964 |
| CHEID_RS08945 | elongation factor Tu | 2055947 |
| R3O64_RS01890 | elongation factor Tu | 2913502 |
| HW450_RS05500 | elongation factor Tu | 699041 |
| CIHUM_RS01795 | elongation factor Tu | 1232427 |
| CKV97_RS02075 | elongation factor Tu | 156978 |
| H0194_RS07375 | elongation factor Tu | 2754725 |
| CJEDD_RS01945 | elongation factor Tu | 1414719 |
| CJEIK_RS09940 | elongation factor Tu | 38289 |
| CKALI_RS10140 | elongation factor Tu | 2675216 |
| I6I12_RS11000 | elongation factor Tu | 1979527 |
| CKROP_RS09110 | elongation factor Tu | 645127 |
| UL82_RS01490 | elongation factor Tu | 35755 |
| CLAC_RS01900 | elongation factor Tu | 1408189 |
| C3E79_RS01700 | elongation factor Tu | 2079535 |
| G7Y31_RS02155 | elongation factor Tu | 2709394 |
| IAU68_RS01600 | elongation factor Tu | 2763010 |
| I6J25_RS02245 | elongation factor Tu | 38290 |
| B840_RS01800 | elongation factor Tu | 1224162 |
| B841_RS02520 | elongation factor Tu | 1224163 |
| CMASS_RS01650 | elongation factor Tu | 1121364 |
| VLL26_RS10390 | elongation factor Tu | 3062461 |
| DQN65_RS10670 | elongation factor Tu | 38301 |
| CMUST_RS02075 | elongation factor Tu | 571915 |
| BLS40_RS09330 | elongation factor Tu | 38302 |

| | | |
|---|---|---|
| FSW06_RS06975 | elongation factor Tu | 863239 |
| COCCU_RS02245 | elongation factor Tu | 2675219 |
| CPELA_RS09490 | elongation factor Tu | 1471400 |
| CPHO_RS10700 | elongation factor Tu | 161895 |
| GP475_RS02235 | elongation factor Tu | 2684405 |
| I6I69_RS03635 | elongation factor Tu | 43769 |
| KBP54_RS01875 | elongation factor Tu | 38303 |
| Q0N40_RS09495 | elongation factor Tu | 2804917 |
| CPPEL_RS09615 | elongation factor Tu | 2080757 |
| CpATCC19410_RS01745 | elongation factor Tu | 1719 |
| G7Y29_RS01530 | elongation factor Tu | 2709392 |
| CULFRC11_RS01885 | elongation factor Tu | 3026968 |
| DQN72_RS05955 | elongation factor Tu | 1724 |
| CRES_RS09865 | elongation factor Tu | 662755 |
| AK829_RS08140 | elongation factor Tu | 156976 |
| CIP100161_RS02455 | elongation factor Tu | 2719119 |
| E3227_RS02480 | elongation factor Tu | 2594913 |
| CBE74_RS14465 | elongation factor Tu | 2320431 |
| AWU68_RS05310 | elongation factor Tu | 146827 |
| CSING_RS02100 | elongation factor Tu | 161899 |
| CSPHI_RS01430 | elongation factor Tu | 1437874 |
| CSTAT_RS02355 | elongation factor Tu | 1705 |
| I6I72_RS04895 | elongation factor Tu | 43770 |
| QP029_RS05835 | elongation factor Tu | 3049801 |
| N24_RS03295 | elongation factor Tu | 2506452 |
| A606_RS10105 | elongation factor Tu | 1200352 |
| CTEST_RS01955 | elongation factor Tu | 136857 |
| I6I74_RS07005 | elongation factor Tu | 38304 |
| LH390_RS01735 | elongation factor Tu | 2883169 |
| CULC809_RS02000 | elongation factor Tu | 945711 |
| CKV82_RS09000 | elongation factor Tu | 43771 |

| | | |
|---|---|---|
| CUREI_RS01480 | elongation factor Tu | 401472 |
| CUROG_RS08760 | elongation factor Tu | 2487892 |
| CUTER_RS01520 | elongation factor Tu | 1072256 |
| B843_RS02185 | elongation factor Tu | 1224164 |
| C3B44_RS01685 | elongation factor Tu | 2080740 |
| IAU67_RS08215 | elongation factor Tu | 2768834 |
| DOE79_RS15935 | elongation factor Tu | 2220095 |
| KM842_RS14440 | elongation factor Tu | 2841594 |
| NY057_RS16785 | elongation factor Tu | 2035 |
| ACL00U_RS15845 | elongation factor Tu | 159612 |
| FNX46_RS09930 | elongation factor Tu | 1734925 |
| PALO_RS01620 | elongation factor Tu | 1170318 |
| O6R08_RS08545 | elongation factor Tu | 3016342 |
| TPCG7_RS07190 | elongation factor Tu | 33011 |
| Daura_RS43215 | elongation factor Tu | 35754 |
| Daura_RS48350 | elongation factor Tu | 35754 |
| Dfulv_RS39330 | elongation factor Tu | 53359 |
| Dfulv_RS44400 | elongation factor Tu | 53359 |
| Drose_RS29945 | elongation factor Tu | 47989 |
| Drose_RS34405 | elongation factor Tu | 47989 |
| Dvina_RS44565 | elongation factor Tu | 53362 |
| Dvina_RS49875 | elongation factor Tu | 53362 |
| RN607_RS11865 | elongation factor Tu | 3075620 |
| FXY94_RS05435 | elongation factor Tu | 1667168 |
| DAD186_RS02520 | elongation factor Tu | 1630135 |
| HX89_RS03575 | elongation factor Tu | 1274 |
| CKV89_RS09980 | elongation factor Tu | 1863 |
| BN1724_RS10450 | elongation factor Tu | 472569 |
| VYL96_RS11770 | elongation factor Tu | 321318 |
| VZF99_RS11150 | elongation factor Tu | 322509 |
| A6035_RS11615 | elongation factor Tu | 546160 |

| | | |
|---|---|---|
| A6048_RS12585 | elongation factor Tu | 139021 |
| BJL86_RS12075 | elongation factor Tu | 499555 |
| EK0264_RS14175 | elongation factor Tu | 1891644 |
| H2O75_RS02985 | elongation factor Tu | 2758573 |
| EJO69_RS11910 | elongation factor Tu | 1282737 |
| FRAAL_RS04680 | elongation factor Tu | 326424 |
| FRANCCI3_RS02915 | elongation factor Tu | 106370 |
| BLT72_RS17635 | elongation factor Tu | 546871 |
| QUE79_RS06375 | elongation factor Tu | 1268041 |
| DOD25_RS04075 | elongation factor Tu | 2792978 |
| GAVG_RS01675 | elongation factor Tu | 585528 |
| GOBS_RS22070 | elongation factor Tu | 526225 |
| EBO36_RS12565 | elongation factor Tu | 2483799 |
| FE251_RS13345 | elongation factor Tu | 2585135 |
| FE374_RS15855 | elongation factor Tu | 2589797 |
| G9V96_RS05350 | elongation factor Tu | 2716227 |
| CJ185_RS07585 | elongation factor Tu | 66228 |
| AARI_RS12745 | elongation factor Tu | 861360 |
| GcLGCM259_RS10755 | elongation factor Tu | 162496 |
| QMQ05_RS11650 | elongation factor Tu | 3046593 |
| ACJD00_RS11525 | elongation factor Tu | 1522174 |
| AOZ07_RS12135 | elongation factor Tu | 1933880 |
| D3791_RS03065 | elongation factor Tu | 1775880 |
| JQN66_RS12070 | elongation factor Tu | 37930 |
| C5O27_RS08855 | elongation factor Tu | 84096 |
| GII30_RS18435 | elongation factor Tu | 36821 |
| IHQ52_RS22185 | elongation factor Tu | 89053 |
| GBRO_RS18900 | elongation factor Tu | 526226 |
| P9A14_RS18305 | elongation factor Tu | 1701090 |
| RVF87_RS12080 | elongation factor Tu | 40516 |
| D7316_RS24155 | elongation factor Tu | 2420509 |

| | | |
|---|---|---|
| C6V83_RS16320 | elongation factor Tu | 1004901 |
| H1R19_RS18030 | elongation factor Tu | 2758710 |
| NWF22_RS03660 | elongation factor Tu | 2665643 |
| LK459_RS15700 | elongation factor Tu | 249058 |
| ACH46_RS16130 | elongation factor Tu | 1136941 |
| GPOL_RS18415 | elongation factor Tu | 1112204 |
| GII31_RS17500 | elongation factor Tu | 2831662 |
| H3V45_RS00395 | elongation factor Tu | 36822 |
| P5P27_RS02320 | elongation factor Tu | 173559 |
| BCM27_RS19675 | elongation factor Tu | 2055 |
| FO044_RS12080 | elongation factor Tu | 2597659 |
| D7I44_RS05265 | elongation factor Tu | 2419771 |
| GMOLON4_RS14750 | elongation factor Tu | 256821 |
| R3I40_RS07590 | elongation factor Tu | 521392 |
| AS9A_RS02210 | elongation factor Tu | 443218 |
| FPZ11_RS09715 | elongation factor Tu | 2599293 |
| INTCA_RS13940 | elongation factor Tu | 710696 |
| I598_RS08160 | elongation factor Tu | 1300344 |
| ISOVA_RS03295 | elongation factor Tu | 743718 |
| V1351_RS02760 | elongation factor Tu | 3115209 |
| EEW87_RS08320 | elongation factor Tu | 262209 |
| V3C34_RS05720 | elongation factor Tu | 103817 |
| M6B22_RS08915 | elongation factor Tu | 2944128 |
| M6D93_RS17140 | elongation factor Tu | 2038343 |
| BLV05_RS35300 | elongation factor Tu | 419479 |
| DQN74_RS02935 | elongation factor Tu | 43674 |
| AOZ06_RS12360 | elongation factor Tu | 860235 |
| AOZ06_RS49785 | elongation factor Tu | 860235 |
| KRAD_RS07680 | elongation factor Tu | 266940 |
| ABWK59_RS14800 | elongation factor Tu | 3156397 |
| O1G21_RS22760 | elongation factor Tu | 3004092 |

| | | |
|---|---|---|
| QMQ26_RS14770 | elongation factor Tu | 2991545 |
| ACI1MP_RS22660 | elongation factor Tu | 2064 |
| OG267_RS16500 | elongation factor Tu | 68217 |
| OG715_RS18725 | elongation factor Tu | 67352 |
| OG715_RS36230 | elongation factor Tu | 67352 |
| KSE_RS15705 | elongation factor Tu | 452652 |
| AS188_RS14050 | elongation factor Tu | 446860 |
| OG312_RS08995 | elongation factor Tu | 72000 |
| AYX06_RS05750 | elongation factor Tu | 388357 |
| KFLA_RS29530 | elongation factor Tu | 479435 |
| KFLA_RS30440 | elongation factor Tu | 479435 |
| KALB_RS40435 | elongation factor Tu | 1449976 |
| Q4V65_RS29180 | elongation factor Tu | 1045776 |
| M3Q35_RS30730 | elongation factor Tu | 1483604 |
| I6I18_RS08170 | elongation factor Tu | 1276 |
| IY73_RS02890 | elongation factor Tu | 1528099 |
| O159_RS01835 | elongation factor Tu | 1389489 |
| BBK82_RS17550 | elongation factor Tu | 1586287 |
| BBK82_RS20290 | elongation factor Tu | 1586287 |
| MUN78_RS03660 | elongation factor Tu | 2932247 |
| G7066_RS04665 | elongation factor Tu | 2714933 |
| H9L06_RS10985 | elongation factor Tu | 683042 |
| G7067_RS05990 | elongation factor Tu | 2714934 |
| Leucomu_RS04025 | elongation factor Tu | 1935379 |
| MUN76_RS05305 | elongation factor Tu | 2932245 |
| KVY00_RS04960 | elongation factor Tu | 2873270 |
| EVS81_RS04140 | elongation factor Tu | 1784719 |
| G7068_RS14140 | elongation factor Tu | 2714935 |
| KDY119_RS04030 | elongation factor Tu | 1133546 |
| VV02_RS21270 | elongation factor Tu | 571913 |
| BHD05_RS04610 | elongation factor Tu | 674079 |

| | | |
|---|---|---|
| IM776_RS03560 | elongation factor Tu | 2782166 |
| QSU92_RS11410 | elongation factor Tu | 3050384 |
| G6N81_RS09705 | elongation factor Tu | 936337 |
| QUC20_RS14065 | elongation factor Tu | 33883 |
| IR212_RS02585 | elongation factor Tu | 2782168 |
| KV397_RS03490 | elongation factor Tu | 2851642 |
| BOH66_RS14305 | elongation factor Tu | 36805 |
| NFX31_RS13905 | elongation factor Tu | 582680 |
| N8K70_RS13560 | elongation factor Tu | 2981139 |
| F6J84_RS02075 | elongation factor Tu | 2614638 |
| A8L33_RS05805 | elongation factor Tu | 84292 |
| IM778_RS13820 | elongation factor Tu | 2782169 |
| G6N83_RS04635 | elongation factor Tu | 1526412 |
| DXT68_RS13310 | elongation factor Tu | 104336 |
| OB895_RS15840 | elongation factor Tu | 2982533 |
| KV396_RS02870 | elongation factor Tu | 2851646 |
| IC745_RS03340 | elongation factor Tu | 162426 |
| PU630_RS02385 | elongation factor Tu | 3028316 |
| CVS47_RS02140 | elongation factor Tu | 1072463 |
| RYJ27_RS11535 | elongation factor Tu | 3079935 |
| F6J85_RS02485 | elongation factor Tu | 2614639 |
| IM777_RS14150 | elongation factor Tu | 2782167 |
| NMQ05_RS14620 | elongation factor Tu | 33918 |
| JSY13_RS10495 | elongation factor Tu | 2810535 |
| BWL13_RS12370 | elongation factor Tu | 273677 |
| D7D94_RS07905 | elongation factor Tu | 743009 |
| BLU02_RS06705 | elongation factor Tu | 199592 |
| VC184_RS14005 | elongation factor Tu | 1816425 |
| ET475_RS09745 | elongation factor Tu | 2509458 |
| BLT19_RS00295 | elongation factor Tu | 370764 |
| K8F61_RS17435 | elongation factor Tu | 156977 |

| | | |
|---|---|---|
| SM116_RS04395 | elongation factor Tu | 1678237 |
| E3O41_RS11685 | elongation factor Tu | 904291 |
| KV394_RS01230 | elongation factor Tu | 2851649 |
| OAU46_RS12715 | elongation factor Tu | 344163 |
| QU603_RS02455 | elongation factor Tu | 3242192 |
| E4K62_RS02660 | elongation factor Tu | 2541726 |
| AAH991_RS10940 | elongation factor Tu | 3144104 |
| OVN18_RS06560 | elongation factor Tu | 2994971 |
| HGB54_RS01585 | elongation factor Tu | 1804990 |
| NNL39_RS06550 | elongation factor Tu | 2963406 |
| HUJ41_RS01025 | elongation factor Tu | 2750620 |
| LG274_RS05240 | elongation factor Tu | 86171 |
| DQN81_RS01945 | elongation factor Tu | 465515 |
| KW076_RS05445 | elongation factor Tu | 2856555 |
| FOE78_RS05930 | elongation factor Tu | 2596828 |
| MLP_RS05305 | elongation factor Tu | 1032480 |
| BLU42_RS14335 | elongation factor Tu | 546874 |
| BLU38_RS18365 | elongation factor Tu | 630515 |
| MICAU_RS27570 | elongation factor Tu | 644283 |
| GA0070611_RS19125 | elongation factor Tu | 261654 |
| HXZ27_RS27645 | elongation factor Tu | 47853 |
| PVK37_RS02550 | elongation factor Tu | 3028804 |
| PVK74_RS17795 | elongation factor Tu | 1874 |
| GA0070612_RS09450 | elongation factor Tu | 356851 |
| GA0070607_RS16065 | elongation factor Tu | 285665 |
| GA0070614_RS12550 | elongation factor Tu | 356852 |
| ID554_RS19015 | elongation factor Tu | 2294034 |
| GA0070609_RS32820 | elongation factor Tu | 47857 |
| GA0070610_RS01455 | elongation factor Tu | 47858 |
| OG994_RS28770 | elongation factor Tu | 1894969 |
| RMN56_RS15240 | elongation factor Tu | 709879 |

| | | |
|---|---|---|
| ABUL04_RS02325 | elongation factor Tu | 1490225 |
| GA0070613_RS12725 | elongation factor Tu | 47865 |
| GA0070620_RS25460 | elongation factor Tu | 307121 |
| VAB18032_RS26235 | elongation factor Tu | 263358 |
| GA0070621_RS20270 | elongation factor Tu | 299146 |
| GA0074696_RS06455 | elongation factor Tu | 47872 |
| GA0070623_RS22035 | elongation factor Tu | 291594 |
| H1D33_RS28000 | elongation factor Tu | 2749844 |
| IGA62_RS08835 | elongation factor Tu | 47875 |
| GA0074704_RS18250 | elongation factor Tu | 299152 |
| GA0074704_RS28020 | elongation factor Tu | 299152 |
| GCE86_RS22170 | elongation factor Tu | 1914461 |
| GA0074695_RS05125 | elongation factor Tu | 1881 |
| GA0070619_RS24635 | elongation factor Tu | 709883 |
| MPLG2_RS01900 | elongation factor Tu | 75385 |
| BLT62_RS04830 | elongation factor Tu | 412690 |
| HMPREF0573_RS01725 | elongation factor Tu | 548479 |
| MODMU_RS23310 | elongation factor Tu | 2732864 |
| AB3M34_RS18690 | elongation factor Tu | 3234852 |
| KXZ72_RS11450 | elongation factor Tu | 2859226 |
| C3E77_RS11605 | elongation factor Tu | 2079792 |
| A7U43_RS25400 | elongation factor Tu | 1682113 |
| BJP76_RS44960 | elongation factor Tu | 439334 |
| MB901379_RS04230 | elongation factor Tu | 2094119 |
| G6N47_RS03260 | elongation factor Tu | 43348 |
| MCAN_RS03565 | elongation factor Tu | 1048245 |
| G6N66_RS02870 | elongation factor Tu | 44010 |
| G6N27_RS20900 | elongation factor Tu | 1775 |
| BTO20_RS28775 | elongation factor Tu | 482462 |
| U8D42_RS17820 | elongation factor Tu | 761804 |
| G6N55_RS12790 | elongation factor Tu | 292462 |

| | | |
|---|---|---|
| G6N42_RS27190 | elongation factor Tu | 39689 |
| B586_RS03530 | elongation factor Tu | 1202450 |
| JUH53_RS19310 | elongation factor Tu | 110505 |
| G6N25_RS05890 | elongation factor Tu | 53376 |
| LK403_RS05525 | elongation factor Tu | 1767 |
| MKAN_RS18660 | elongation factor Tu | 557599 |
| GAN17_RS04790 | elongation factor Tu | 120959 |
| G6N24_RS17135 | elongation factor Tu | 169765 |
| MJO58_RS04640 | elongation factor Tu | 141349 |
| DIJ64_RS10305 | elongation factor Tu | 1769 |
| MLPF_RS11035 | elongation factor Tu | 480418 |
| K3U93_RS04225 | elongation factor Tu | 1780 |
| G6N50_RS15745 | elongation factor Tu | 560555 |
| MMARE11_RS04800 | elongation factor Tu | 1131442 |
| CKJ54_RS20850 | elongation factor Tu | 701042 |
| G6N15_RS12490 | elongation factor Tu | 459858 |
| G6N19_RS05660 | elongation factor Tu | 2492438 |
| LTS72_RS20615 | elongation factor Tu | 2738409 |
| C0J29_RS05535 | elongation factor Tu | 1389713 |
| OCQ_RS21905 | elongation factor Tu | 1138383 |
| G6N51_RS23120 | elongation factor Tu | 590652 |
| MKK62_RS14465 | elongation factor Tu | 577492 |
| G6N48_RS15460 | elongation factor Tu | 185642 |
| MPSD_RS05700 | elongation factor Tu | 1136880 |
| AADZ78_RS04815 | elongation factor Tu | 486698 |
| G6N56_RS22070 | elongation factor Tu | 220927 |
| MTY59_RS03590 | elongation factor Tu | 2775496 |
| G6N37_RS24000 | elongation factor Tu | 386911 |
| MSG_RS04295 | elongation factor Tu | 722731 |
| G6N20_RS13645 | elongation factor Tu | 398694 |
| G6N41_RS12600 | elongation factor Tu | 133549 |

| | | |
|---|---|---|
| G6N33_RS07980 | elongation factor Tu | 1784 |
| F6B93_RS04230 | elongation factor Tu | 886343 |
| G6N54_RS14065 | elongation factor Tu | 470076 |
| Rv0685 | elongation factor Tu | 83332 |
| SHTP_RS21000 | elongation factor Tu | 1124626 |
| H0P51_RS05025 | elongation factor Tu | 1719132 |
| MAB_RS19495 | elongation factor Tu | 561007 |
| BB28_RS19310 | elongation factor Tu | 1460372 |
| ABG82_RS21815 | elongation factor Tu | 83262 |
| DSM43276_RS18225 | elongation factor Tu | 404941 |
| MYCSP_RS18035 | elongation factor Tu | 1578165 |
| G6N11_RS11530 | elongation factor Tu | 1069220 |
| MIU77_RS15125 | elongation factor Tu | 1069221 |
| K3U94_RS03885 | elongation factor Tu | 512402 |
| G6N14_RS01010 | elongation factor Tu | 29314 |
| G6N09_RS11675 | elongation factor Tu | 1118379 |
| JDM601_RS03460 | elongation factor Tu | 875328 |
| G6N23_RS16735 | elongation factor Tu | 1788 |
| MJO54_RS03905 | elongation factor Tu | 1795032 |
| G6N32_RS22840 | elongation factor Tu | 1799 |
| G6N44_RS12080 | elongation factor Tu | 67081 |
| G6N35_RS24520 | elongation factor Tu | 1431246 |
| G6N61_RS13515 | elongation factor Tu | 1286181 |
| G6N59_RS11065 | elongation factor Tu | 319707 |
| EL337_RS05260 | elongation factor Tu | 1791 |
| JN090_RS05155 | elongation factor Tu | 39687 |
| I7X18_RS22770 | elongation factor Tu | 2761578 |
| G6N57_RS31070 | elongation factor Tu | 146017 |
| L2Z93_RS16330 | elongation factor Tu | 85968 |
| G6N18_RS11600 | elongation factor Tu | 1249101 |
| EL338_RS19210 | elongation factor Tu | 1792 |

| | | |
|---|---|---|
| G6N34_RS19085 | elongation factor Tu | 28047 |
| MI149_RS05755 | elongation factor Tu | 388459 |
| MI149_RS08930 | elongation factor Tu | 388459 |
| K0O62_RS05875 | elongation factor Tu | 1801 |
| G6N07_RS03090 | elongation factor Tu | 126673 |
| G6N31_RS16905 | elongation factor Tu | 39688 |
| G6N10_RS10640 | elongation factor Tu | 1793 |
| K6L26_RS25590 | elongation factor Tu | 1802 |
| HZU40_RS09955 | elongation factor Tu | 258505 |
| MFTT_RS07190 | elongation factor Tu | 144549 |
| EXE63_RS09515 | elongation factor Tu | 117567 |
| G6N36_RS14040 | elongation factor Tu | 1794 |
| MSPYR1_RS22080 | elongation factor Tu | 278137 |
| MI170_RS21435 | elongation factor Tu | 134601 |
| FZ046_RS11285 | elongation factor Tu | 1552759 |
| MHAS_RS01020 | elongation factor Tu | 1122247 |
| G6N38_RS05965 | elongation factor Tu | 1534349 |
| K3U96_RS21315 | elongation factor Tu | 1226752 |
| G6N16_RS06455 | elongation factor Tu | 444597 |
| G6N30_RS15740 | elongation factor Tu | 758802 |
| G6N60_RS05455 | elongation factor Tu | 212765 |
| G6N67_RS20940 | elongation factor Tu | 53462 |
| I5054_RS20265 | elongation factor Tu | 2736649 |
| G6N49_RS18180 | elongation factor Tu | 85693 |
| G6N43_RS26920 | elongation factor Tu | 39691 |
| C1S78_RS23060 | elongation factor Tu | 1226753 |
| D174_RS06400 | elongation factor Tu | 700508 |
| EH231_RS25130 | elongation factor Tu | 2487344 |
| K0O64_RS05005 | elongation factor Tu | 370524 |
| NTM_RS01460 | elongation factor Tu | 39692 |
| G6N46_RS23980 | elongation factor Tu | 319706 |

| | | |
|---|---|---|
| G6N39_RS07770 | elongation factor Tu | 39694 |
| G6N45_RS26825 | elongation factor Tu | 216929 |
| G6N28_RS05870 | elongation factor Tu | 36813 |
| MJO55_RS08990 | elongation factor Tu | 318424 |
| BLW81_RS00135 | elongation factor Tu | 370526 |
| G6N13_RS12390 | elongation factor Tu | 1534348 |
| G6N64_RS11650 | elongation factor Tu | 1286180 |
| K3U95_RS06140 | elongation factor Tu | 1796 |
| JVX95_RS11725 | elongation factor Tu | 98668 |
| KZ780_RS06605 | elongation factor Tu | 1772 |
| CKW28_RS04430 | elongation factor Tu | 1797 |
| G6N58_RS15875 | elongation factor Tu | 39695 |
| MYVA_RS05295 | elongation factor Tu | 1354275 |
| MVAN_RS06480 | elongation factor Tu | 350058 |
| QUE51_RS04720 | elongation factor Tu | 1162966 |
| EH165_RS11675 | elongation factor Tu | 1902245 |
| NAMU_RS05580 | elongation factor Tu | 479431 |
| BLS97_RS12585 | elongation factor Tu | 1090615 |
| JQS30_RS12280 | elongation factor Tu | 2811108 |
| JQS43_RS03965 | elongation factor Tu | 2811111 |
| EL493_RS31995 | elongation factor Tu | 1824 |
| CEQ30_RS33380 | elongation factor Tu | 37326 |
| CEQ30_RS36350 | elongation factor Tu | 37326 |
| C5B73_RS12070 | elongation factor Tu | 135487 |
| NFA_RS25155 | elongation factor Tu | 247156 |
| LTT66_RS08175 | elongation factor Tu | 2749991 |
| H0264_RS09040 | elongation factor Tu | 2755382 |
| H0264_RS15715 | elongation factor Tu | 2755382 |
| H0264_RS18990 | elongation factor Tu | 2755382 |
| KV110_RS34390 | elongation factor Tu | 204891 |
| KV110_RS37370 | elongation factor Tu | 204891 |

| | | |
|---|---|---|
| BOX37_RS29590 | elongation factor Tu | 2213200 |
| FOH10_RS00625 | elongation factor Tu | 1823 |
| FOH10_RS20740 | elongation factor Tu | 1823 |
| OHB21_RS07205 | elongation factor Tu | 208602 |
| OHB21_RS11900 | elongation factor Tu | 208602 |
| OHS03_RS24905 | elongation factor Tu | 53431 |
| QMG86_RS27860 | elongation factor Tu | 2984338 |
| CRH09_RS10325 | elongation factor Tu | 455432 |
| CRH09_RS36390 | elongation factor Tu | 455432 |
| NWFMUON74_RS03590 | elongation factor Tu | 480035 |
| NWFMUON74_RS31755 | elongation factor Tu | 480035 |
| LTV02_RS15535 | elongation factor Tu | 209247 |
| LTV02_RS17355 | elongation factor Tu | 209247 |
| LTV02_RS23975 | elongation factor Tu | 209247 |
| D7D52_RS06985 | elongation factor Tu | 2382165 |
| D7D52_RS21290 | elongation factor Tu | 2382165 |
| D7D52_RS33075 | elongation factor Tu | 2382165 |
| G5V58_RS01645 | elongation factor Tu | 2712223 |
| ENKNEFLB_RS05090 | elongation factor Tu | 160826 |
| KCTC_RS10795 | elongation factor Tu | 1804624 |
| SHK19_RS17885 | elongation factor Tu | 3095076 |
| SHK19_RS20360 | elongation factor Tu | 3095076 |
| H8838_RS15335 | elongation factor Tu | 2736757 |
| K6T13_RS14445 | elongation factor Tu | 2872154 |
| E3N83_RS17620 | elongation factor Tu | 2558918 |
| I601_RS02355 | elongation factor Tu | 1300347 |
| H4O22_RS17270 | elongation factor Tu | 2760089 |
| EXE57_RS15520 | elongation factor Tu | 2518370 |
| KG111_RS15025 | elongation factor Tu | 2803858 |
| FIV44_RS08295 | elongation factor Tu | 449461 |
| FIV44_RS22040 | elongation factor Tu | 449461 |

| | | |
|---|---|---|
| HPC71_RS17540 | elongation factor Tu | 2663857 |
| H9L09_RS12300 | elongation factor Tu | 433659 |
| HBO46_RS03125 | elongation factor Tu | 2685869 |
| LN652_RS17215 | elongation factor Tu | 2894081 |
| KRR39_RS15900 | elongation factor Tu | 2849501 |
| EBO35_RS16315 | elongation factor Tu | 2483798 |
| SHK17_RS03840 | elongation factor Tu | 3095075 |
| K8W59_RS05135 | elongation factor Tu | 1774216 |
| FIV43_RS00215 | elongation factor Tu | 2589074 |
| BLU55_RS10890 | elongation factor Tu | 642780 |
| EXE58_RS04235 | elongation factor Tu | 2518371 |
| EOV43_RS12295 | elongation factor Tu | 2500546 |
| KGD83_RS26130 | elongation factor Tu | 2831968 |
| KGD84_RS28400 | elongation factor Tu | 2831969 |
| EL646_RS20930 | elongation factor Tu | 2014 |
| NE857_RS05235 | elongation factor Tu | 163604 |
| CDO52_RS25245 | elongation factor Tu | 1235441 |
| Nocox_RS04280 | elongation factor Tu | 1122611 |
| Nocox_RS18350 | elongation factor Tu | 1122611 |
| ACFMQL_RS17645 | elongation factor Tu | 46173 |
| ACFMQL_RS30495 | elongation factor Tu | 46173 |
| OIE67_RS24560 | elongation factor Tu | 1291556 |
| OIE67_RS42260 | elongation factor Tu | 1291556 |
| LCN96_RS04685 | elongation factor Tu | 93944 |
| LCN96_RS21995 | elongation factor Tu | 93944 |
| OHA68_RS18890 | elongation factor Tu | 2047744 |
| OHA68_RS33715 | elongation factor Tu | 2047744 |
| GBF35_RS04745 | elongation factor Tu | 2219224 |
| GKS42_RS05380 | elongation factor Tu | 2675754 |
| DV701_RS16875 | elongation factor Tu | 2283195 |
| FNH13_RS05060 | elongation factor Tu | 2594265 |

| | | |
|---|---|---|
| NF557_RS03575 | elongation factor Tu | 2934161 |
| NF556_RS04430 | elongation factor Tu | 2934158 |
| NF556_RS08240 | elongation factor Tu | 2934158 |
| E3Z34_RS13250 | elongation factor Tu | 1288636 |
| FY030_RS11970 | elongation factor Tu | 2593973 |
| ESZ52_RS14550 | elongation factor Tu | 2508882 |
| ACFLFB_RS15390 | elongation factor Tu | 3149179 |
| ACFWN3_RS14210 | elongation factor Tu | 1620385 |
| ABI796_RS13930 | elongation factor Tu | 43663 |
| JMY29_RS14225 | elongation factor Tu | 29320 |
| AYX19_RS09275 | elongation factor Tu | 37931 |
| HCR76_RS03380 | elongation factor Tu | 2769067 |
| HCR84_RS03865 | elongation factor Tu | 2708079 |
| BLU20_RS12175 | elongation factor Tu | 545619 |
| QUE47_RS02440 | elongation factor Tu | 1138587 |
| PSDT_RS06870 | elongation factor Tu | 864564 |
| BH719_RS04570 | elongation factor Tu | 178339 |
| BLQ34_RS18475 | elongation factor Tu | 443156 |
| VTA80_RS12360 | elongation factor Tu | 3114394 |
| J4E96_RS17650 | elongation factor Tu | 2819238 |
| C7M71_RS11915 | elongation factor Tu | 2126346 |
| LG324_RS17580 | elongation factor Tu | 367299 |
| Psuf_RS05860 | elongation factor Tu | 624315 |
| Psuf_RS19020 | elongation factor Tu | 624315 |
| KR76_RS02880 | elongation factor Tu | 2045 |
| KR76_RS06275 | elongation factor Tu | 2045 |
| Prubr_RS02110 | elongation factor Tu | 338584 |
| Prubr_RS07840 | elongation factor Tu | 338584 |
| C3B54_RS01530 | elongation factor Tu | 1159327 |
| BAY61_RS02760 | elongation factor Tu | 530584 |
| C7H75_RS18975 | elongation factor Tu | 43767 |

| | | |
|---|---|---|
| ABI214_RS19720 | elongation factor Tu | 1543852 |
| EGX94_RS09710 | elongation factor Tu | 556499 |
| EL282_RS09640 | elongation factor Tu | 119981 |
| CR251_RS08330 | elongation factor Tu | 1744 |
| G7070_RS03025 | elongation factor Tu | 2714937 |
| PCC79_RS04560 | elongation factor Tu | 2775081 |
| D7I47_RS04375 | elongation factor Tu | 2419774 |
| FLP23_RS03935 | elongation factor Tu | 2592654 |
| ACHL_RS13595 | elongation factor Tu | 452863 |
| BLT71_RS15500 | elongation factor Tu | 728066 |
| WHH00_RS04995 | elongation factor Tu | 3128980 |
| FRAEUI1C_RS01395 | elongation factor Tu | 298654 |
| FRAEUI1C_RS30955 | elongation factor Tu | 298654 |
| I4I81_RS28185 | elongation factor Tu | 2792008 |
| Pdca_RS28935 | elongation factor Tu | 2074 |
| HOP40_RS12135 | elongation factor Tu | 2736640 |
| PSED_RS27280 | elongation factor Tu | 675635 |
| H6H00_RS12140 | elongation factor Tu | 37331 |
| UM93_RS09760 | elongation factor Tu | 1618207 |
| C1I64_RS00740 | elongation factor Tu | 1328866 |
| C7V51_RS09485 | elongation factor Tu | 59737 |
| C1O28_RS01560 | elongation factor Tu | 33887 |
| QU604_RS03895 | elongation factor Tu | 3144168 |
| GSU10_RS12075 | elongation factor Tu | 1671680 |
| AYW78_RS09135 | elongation factor Tu | 145458 |
| A6122_RS02955 | elongation factor Tu | 33888 |
| RSAL33209_RS09595 | elongation factor Tu | 288705 |
| OCS65_RS09995 | elongation factor Tu | 191292 |
| RHODO2019_RS02245 | elongation factor Tu | 2987751 |
| DQO56_RS18190 | elongation factor Tu | 38310 |
| H351_RS23180 | elongation factor Tu | 1289591 |

| | | |
|---|---|---|
| KYT97_RS09705 | elongation factor Tu | 33008 |
| ACM26Y_RS17780 | elongation factor Tu | 3055869 |
| JWS14_RS09840 | elongation factor Tu | 99653 |
| K2Z90_RS28790 | elongation factor Tu | 543736 |
| CBI38_RS08500 | elongation factor Tu | 1990687 |
| JWS13_RS35990 | elongation factor Tu | 2811421 |
| Y013_RS08650 | elongation factor Tu | 1435356 |
| RQCS_RS08575 | elongation factor Tu | 334542 |
| PVE95_RS17270 | elongation factor Tu | 1829 |
| F1734_RS19540 | elongation factor Tu | 1830 |
| OED52_RS15540 | elongation factor Tu | 2962047 |
| WDS16_RS16445 | elongation factor Tu | 1805840 |
| G4H72_RS14170 | elongation factor Tu | 300028 |
| WKC72_RS13365 | elongation factor Tu | 44752 |
| RHOLA_RS06190 | elongation factor Tu | 529884 |
| FFA38_RS00810 | elongation factor Tu | 232537 |
| I6I94_RS09075 | elongation factor Tu | 172042 |
| IDM48_RS03110 | elongation factor Tu | 169480 |
| HMPREF0733_RS01445 | elongation factor Tu | 762948 |
| I6G21_RS01395 | elongation factor Tu | 37923 |
| IDM49_RS03215 | elongation factor Tu | 396015 |
| IM660_RS15265 | elongation factor Tu | 2779775 |
| LQF10_RS15165 | elongation factor Tu | 2897773 |
| LQF12_RS13180 | elongation factor Tu | 2897774 |
| FU260_RS19535 | elongation factor Tu | 2603206 |
| SACAZDRAFT_RS15530 | elongation factor Tu | 882081 |
| SACCYDRAFT_RS02410 | elongation factor Tu | 882082 |
| SACGLDRAFT_RS02080 | elongation factor Tu | 928724 |
| SACMADRAFT_RS03385 | elongation factor Tu | 882083 |
| SVIR_RS01955 | elongation factor Tu | 471857 |
| SACE_RS32925 | elongation factor Tu | 405948 |

| | | |
|---|---|---|
| H1226_RS25875 | elongation factor Tu | 33914 |
| DL519_RS33355 | elongation factor Tu | 333966 |
| NV263_RS28295 | elongation factor Tu | 524884 |
| BN6_RS38415 | elongation factor Tu | 1179773 |
| EKG83_RS44090 | elongation factor Tu | 103733 |
| STROP_RS19745 | elongation factor Tu | 369723 |
| SANBI_RS14505 | elongation factor Tu | 3030830 |
| SKED_RS14615 | elongation factor Tu | 446469 |
| LWF01_RS12385 | elongation factor Tu | 2909229 |
| SCIP_RS00890 | elongation factor Tu | 1150468 |
| FFA61_RS01430 | elongation factor Tu | 888050 |
| I6H42_RS06105 | elongation factor Tu | 52773 |
| RDV55_RS06860 | elongation factor Tu | 1660 |
| BLT69_RS07995 | elongation factor Tu | 131110 |
| SAC06_RS08520 | elongation factor Tu | 3079930 |
| SROT_RS02760 | elongation factor Tu | 640132 |
| FHD63_RS11365 | elongation factor Tu | 767452 |
| SGUI_RS14790 | elongation factor Tu | 1758689 |
| SA2016_RS15000 | elongation factor Tu | 37927 |
| SCMU_RS14895 | elongation factor Tu | 322009 |
| AB5L97_RS13885 | elongation factor Tu | 3238584 |
| KV203_RS03570 | elongation factor Tu | 39122 |
| L0C25_RS14505 | elongation factor Tu | 2908642 |
| ACOZ38_RS00795 | elongation factor Tu | 46189 |
| ACOZ38_RS05390 | elongation factor Tu | 46189 |
| J4H86_RS19705 | elongation factor Tu | 2820884 |
| SNAS_RS05850 | elongation factor Tu | 446470 |
| SCAT_RS23130 | elongation factor Tu | 1003195 |
| SCAT_RS25035 | elongation factor Tu | 1003195 |
| EKD16_RS20545 | elongation factor Tu | 2498135 |
| OG350_RS22655 | elongation factor Tu | 67255 |

| | | |
|---|---|---|
| K1J60_RS17360 | elongation factor Tu | 2865673 |
| K1J60_RS37875 | elongation factor Tu | 2865673 |
| N8I86_RS07300 | elongation factor Tu | 2981135 |
| N8I86_RS17020 | elongation factor Tu | 2981135 |
| OG330_RS19590 | elongation factor Tu | 1886 |
| OG330_RS28680 | elongation factor Tu | 1886 |
| CP975_RS04660 | elongation factor Tu | 132473 |
| CP975_RS20940 | elongation factor Tu | 132473 |
| NR995_RS12955 | elongation factor Tu | 1888 |
| H9W91_RS06485 | elongation factor Tu | 1642299 |
| H9W91_RS22700 | elongation factor Tu | 1642299 |
| OG872_RS13545 | elongation factor Tu | 83380 |
| OG872_RS28665 | elongation factor Tu | 83380 |
| SAM23877_RS06380 | elongation factor Tu | 278992 |
| SAM23877_RS21240 | elongation factor Tu | 278992 |
| K7396_RS03390 | elongation factor Tu | 285578 |
| K7396_RS20425 | elongation factor Tu | 285578 |
| OIE72_RS14940 | elongation factor Tu | 68174 |
| OIE72_RS31655 | elongation factor Tu | 68174 |
| OG751_RS27435 | elongation factor Tu | 68175 |
| OG751_RS39370 | elongation factor Tu | 68175 |
| JI76_RS04340 | elongation factor Tu | 1892 |
| JI76_RS22800 | elongation factor Tu | 1892 |
| EJC51_RS09080 | elongation factor Tu | 2548456 |
| EJC51_RS29030 | elongation factor Tu | 2548456 |
| EJC51_RS41250 | elongation factor Tu | 2548456 |
| HKX69_RS13810 | elongation factor Tu | 2726118 |
| HKX69_RS29150 | elongation factor Tu | 2726118 |
| DVA86_RS05060 | elongation factor Tu | 83291 |
| DVA86_RS14840 | elongation factor Tu | 83291 |
| C5746_RS07025 | elongation factor Tu | 1893 |

| | | |
|---|---|---|
| C5746_RS24765 | elongation factor Tu | 1893 |
| IGS71_RS24030 | elongation factor Tu | 47760 |
| IGS71_RS34575 | elongation factor Tu | 47760 |
| SU9_RS14830 | elongation factor Tu | 1160718 |
| SU9_RS31130 | elongation factor Tu | 1160718 |
| SAVERM_RS25400 | elongation factor Tu | 227882 |
| SAVERM_RS36110 | elongation factor Tu | 227882 |
| OG373_RS16995 | elongation factor Tu | 1895 |
| OG373_RS32205 | elongation factor Tu | 1895 |
| G4Z16_RS12170 | elongation factor Tu | 2710756 |
| G4Z16_RS30895 | elongation factor Tu | 2710756 |
| OHT53_RS17085 | elongation factor Tu | 67280 |
| OHT53_RS34600 | elongation factor Tu | 67280 |
| ACNNMR_RS03500 | elongation factor Tu | 2039455 |
| ACNNMR_RS26640 | elongation factor Tu | 2039455 |
| GQF42_RS01640 | elongation factor Tu | 2686304 |
| GQF42_RS09140 | elongation factor Tu | 2686304 |
| GQF42_RS26020 | elongation factor Tu | 2686304 |
| GQF42_RS39190 | elongation factor Tu | 2686304 |
| H7H31_RS13215 | elongation factor Tu | 2763006 |
| DN051_RS16905 | elongation factor Tu | 2184053 |
| I6J39_RS03500 | elongation factor Tu | 67351 |
| I6J39_RS21445 | elongation factor Tu | 67351 |
| CD934_RS18650 | elongation factor Tu | 67282 |
| CD934_RS29365 | elongation factor Tu | 67282 |
| O1G22_RS17135 | elongation factor Tu | 3004093 |
| O1G22_RS35210 | elongation factor Tu | 3004093 |
| OG436_RS17315 | elongation factor Tu | 285557 |
| OG436_RS36240 | elongation factor Tu | 285557 |
| IHE65_RS17550 | elongation factor Tu | 2746961 |
| IHE65_RS37920 | elongation factor Tu | 2746961 |

| | | |
|---|---|---|
| OHT68_RS19295 | elongation factor Tu | 58343 |
| OHT68_RS39775 | elongation factor Tu | 58343 |
| CES90_RS13845 | elongation factor Tu | 2014920 |
| CES90_RS33520 | elongation factor Tu | 2014920 |
| N0X72_RS02445 | elongation factor Tu | 285558 |
| N0X72_RS11585 | elongation factor Tu | 285558 |
| P8A18_RS03100 | elongation factor Tu | 3033811 |
| P8A18_RS20700 | elongation factor Tu | 3033811 |
| PYS65_RS14935 | elongation factor Tu | 3031124 |
| NRO40_RS03560 | elongation factor Tu | 2964669 |
| NRO40_RS17880 | elongation factor Tu | 2964669 |
| CP983_RS17455 | elongation factor Tu | 1969 |
| CP983_RS36780 | elongation factor Tu | 1969 |
| QRN89_RS04740 | elongation factor Tu | 3053466 |
| QRN89_RS20655 | elongation factor Tu | 3053466 |
| IPT68_RS05680 | elongation factor Tu | 42881 |
| IPT68_RS21440 | elongation factor Tu | 42881 |
| CP977_RS20580 | elongation factor Tu | 67260 |
| STRCI_RS06160 | elongation factor Tu | 67287 |
| STRCI_RS24180 | elongation factor Tu | 67287 |
| STRCI_RS36725 | elongation factor Tu | 67287 |
| V2W30_RS05710 | elongation factor Tu | 3118173 |
| V2W30_RS16385 | elongation factor Tu | 3118173 |
| OG927_RS04205 | elongation factor Tu | 68188 |
| OG927_RS19675 | elongation factor Tu | 68188 |
| CRV15_RS10300 | elongation factor Tu | 1901 |
| OHA99_RS14985 | elongation factor Tu | 285562 |
| OHA99_RS31560 | elongation factor Tu | 285562 |
| RFN52_RS05860 | elongation factor Tu | 42684 |
| RFN52_RS24285 | elongation factor Tu | 42684 |
| EJ357_RS03420 | elongation factor Tu | 2496836 |

| | | |
|---|---|---|
| EJ357_RS07245 | elongation factor Tu | 2496836 |
| EJ357_RS27615 | elongation factor Tu | 2496836 |
| R2B67_RS13500 | elongation factor Tu | 66883 |
| R2B67_RS32055 | elongation factor Tu | 66883 |
| S1361_RS07305 | elongation factor Tu | 80860 |
| S1361_RS23415 | elongation factor Tu | 80860 |
| N8I84_RS07225 | elongation factor Tu | 2981134 |
| N8I84_RS23850 | elongation factor Tu | 2981134 |
| D9753_RS14405 | elongation factor Tu | 1442032 |
| BN159_RS05240 | elongation factor Tu | 1214101 |
| BN159_RS18250 | elongation factor Tu | 1214101 |
| BN159_RS36085 | elongation factor Tu | 1214101 |
| L3078_RS07660 | elongation factor Tu | 424188 |
| L3078_RS28035 | elongation factor Tu | 424188 |
| OH780_RS04380 | elongation factor Tu | 249567 |
| OH780_RS22665 | elongation factor Tu | 249567 |
| C4B68_RS03400 | elongation factor Tu | 2049881 |
| C4B68_RS16280 | elongation factor Tu | 2049881 |
| C4B68_RS34945 | elongation factor Tu | 2049881 |
| M4V62_RS18355 | elongation factor Tu | 319947 |
| M4V62_RS35590 | elongation factor Tu | 319947 |
| OHA91_RS15870 | elongation factor Tu | 285574 |
| OHA91_RS31375 | elongation factor Tu | 285574 |
| OG858_RS07280 | elongation factor Tu | 146819 |
| OG858_RS27960 | elongation factor Tu | 146819 |
| GFH48_RS17265 | elongation factor Tu | 2662397 |
| GFH48_RS33465 | elongation factor Tu | 2662397 |
| IM697_RS16190 | elongation factor Tu | 1413221 |
| IM697_RS40780 | elongation factor Tu | 1413221 |
| EIZ62_RS12945 | elongation factor Tu | 1977088 |
| EIZ62_RS28335 | elongation factor Tu | 1977088 |

| | | |
|---|---|---|
| BFF78_RS17535 | elongation factor Tu | 1904616 |
| BFF78_RS35050 | elongation factor Tu | 1904616 |
| J4032_RS03825 | elongation factor Tu | 1616117 |
| J4032_RS22090 | elongation factor Tu | 1616117 |
| CP974_RS03140 | elongation factor Tu | 1319510 |
| CP974_RS18095 | elongation factor Tu | 1319510 |
| LL044_RS05250 | elongation factor Tu | 68203 |
| LL044_RS20895 | elongation factor Tu | 68203 |
| H4W23_RS24190 | elongation factor Tu | 66892 |
| IAG43_RS04035 | elongation factor Tu | 2768068 |
| IAG43_RS18705 | elongation factor Tu | 2768068 |
| B1H19_RS18590 | elongation factor Tu | 553510 |
| B1H19_RS36465 | elongation factor Tu | 553510 |
| WQO_RS04395 | elongation factor Tu | 1172567 |
| WQO_RS21340 | elongation factor Tu | 1172567 |
| P8A20_RS14440 | elongation factor Tu | 3033808 |
| P8A20_RS32160 | elongation factor Tu | 3033808 |
| test1122_RS14780 | elongation factor Tu | 2875706 |
| test1122_RS16205 | elongation factor Tu | 2875706 |
| test1122_RS25920 | elongation factor Tu | 2875706 |
| PET44_RS19420 | elongation factor Tu | 1942 |
| PET44_RS26900 | elongation factor Tu | 1942 |
| OG378_RS17395 | elongation factor Tu | 53448 |
| OG378_RS25510 | elongation factor Tu | 53448 |
| SGFS_RS35210 | elongation factor Tu | 68212 |
| SGFS_RS46415 | elongation factor Tu | 68212 |
| SGFS_RS49610 | elongation factor Tu | 68212 |
| V9K83_RS13245 | elongation factor Tu | 68213 |
| V9K83_RS27985 | elongation factor Tu | 68213 |
| AVL59_RS04140 | elongation factor Tu | 68214 |
| AVL59_RS32975 | elongation factor Tu | 68214 |

| | | |
|---|---|---|
| HEP81_RS16235 | elongation factor Tu | 146922 |
| HEP81_RS32190 | elongation factor Tu | 146922 |
| DWG14_RS17605 | elongation factor Tu | 67304 |
| DWG14_RS36370 | elongation factor Tu | 67304 |
| K9S39_RS03230 | elongation factor Tu | 2879846 |
| K9S39_RS24955 | elongation factor Tu | 2879846 |
| OG923_RS14100 | elongation factor Tu | 1944 |
| OG923_RS30235 | elongation factor Tu | 1944 |
| HUT13_RS01960 | elongation factor Tu | 1176198 |
| HUT13_RS09275 | elongation factor Tu | 1176198 |
| CEB94_RS06795 | elongation factor Tu | 67305 |
| CEB94_RS24290 | elongation factor Tu | 67305 |
| OIE73_RS05630 | elongation factor Tu | 35620 |
| OIE73_RS23095 | elongation factor Tu | 35620 |
| J2N69_RS04605 | elongation factor Tu | 1854574 |
| J2N69_RS22000 | elongation factor Tu | 1854574 |
| V8J11_RS22280 | elongation factor Tu | 68042 |
| V8J11_RS33390 | elongation factor Tu | 68042 |
| RFN58_RS06700 | elongation factor Tu | 68219 |
| RFN58_RS25635 | elongation factor Tu | 68219 |
| ABB07_RS15900 | elongation factor Tu | 665007 |
| ABB07_RS32715 | elongation factor Tu | 665007 |
| KI385_RS20450 | elongation factor Tu | 2293571 |
| KI385_RS39340 | elongation factor Tu | 2293571 |
| RY074_RS08485 | elongation factor Tu | 576784 |
| RY074_RS21815 | elongation factor Tu | 576784 |
| RI060_RS16375 | elongation factor Tu | 67308 |
| RI060_RS35950 | elongation factor Tu | 67308 |
| OHB09_RS04815 | elongation factor Tu | 235987 |
| OHB09_RS19465 | elongation factor Tu | 235987 |
| CP970_RS17255 | elongation factor Tu | 1967 |

| | | |
|---|---|---|
| CP970_RS37445 | elongation factor Tu | 1967 |
| KJK29_RS14270 | elongation factor Tu | 2838015 |
| KJK29_RS32805 | elongation factor Tu | 2838015 |
| D0C37_RS20240 | elongation factor Tu | 188770 |
| P8A22_RS15640 | elongation factor Tu | 887464 |
| P8A22_RS33865 | elongation factor Tu | 887464 |
| SLAV_RS15800 | elongation factor Tu | 58340 |
| SLAV_RS30955 | elongation factor Tu | 58340 |
| BN2145_RS16295 | elongation factor Tu | 1437453 |
| BN2145_RS30655 | elongation factor Tu | 1437453 |
| HA039_RS13495 | elongation factor Tu | 2717324 |
| HA039_RS29635 | elongation factor Tu | 2717324 |
| STRLI_RS04390 | elongation factor Tu | 68044 |
| STRLI_RS21835 | elongation factor Tu | 68044 |
| BJ961_RS03385 | elongation factor Tu | 284035 |
| BJ961_RS23475 | elongation factor Tu | 284035 |
| JEQ17_RS27840 | elongation factor Tu | 2797636 |
| JEQ17_RS39245 | elongation factor Tu | 2797636 |
| SLINC_RS08025 | elongation factor Tu | 1915 |
| SLINC_RS26875 | elongation factor Tu | 1915 |
| SLINC_RS40645 | elongation factor Tu | 1915 |
| LGI35_RS09765 | elongation factor Tu | 2880933 |
| LGI35_RS26390 | elongation factor Tu | 2880933 |
| SLUN_RS05895 | elongation factor Tu | 1535768 |
| SLUN_RS23865 | elongation factor Tu | 1535768 |
| PS467_RS05865 | elongation factor Tu | 3026192 |
| PS467_RS23960 | elongation factor Tu | 3026192 |
| AB8735_RS17515 | elongation factor Tu | 173858 |
| BXT88_RS17390 | elongation factor Tu | 47763 |
| BXT88_RS34215 | elongation factor Tu | 47763 |
| SMALA_RS22515 | elongation factor Tu | 92644 |

| | | |
|---|---|---|
| SMALA_RS34380 | elongation factor Tu | 92644 |
| ACPU2J_RS05755 | elongation factor Tu | 1077945 |
| ACPU2J_RS20345 | elongation factor Tu | 1077945 |
| LC193_RS18945 | elongation factor Tu | 2878388 |
| HUT09_RS03965 | elongation factor Tu | 1919 |
| HUT09_RS21335 | elongation factor Tu | 1919 |
| NLG24_RS07975 | elongation factor Tu | 476552 |
| NLG24_RS22490 | elongation factor Tu | 476552 |
| OG981_RS07015 | elongation factor Tu | 68239 |
| OG981_RS12505 | elongation factor Tu | 68239 |
| OG981_RS31775 | elongation factor Tu | 68239 |
| K7I03_RS18530 | elongation factor Tu | 35621 |
| GO605_RS03155 | elongation factor Tu | 33900 |
| GO605_RS24245 | elongation factor Tu | 33900 |
| ACLIN4_RS05370 | elongation factor Tu | 67332 |
| ACLIN4_RS20955 | elongation factor Tu | 67332 |
| WEB32_RS15195 | elongation factor Tu | 55404 |
| DC008_RS04685 | elongation factor Tu | 1827580 |
| DC008_RS20765 | elongation factor Tu | 1827580 |
| STRNI_RS04345 | elongation factor Tu | 1920 |
| STRNI_RS23860 | elongation factor Tu | 1920 |
| CP967_RS13400 | elongation factor Tu | 28894 |
| CP967_RS30010 | elongation factor Tu | 28894 |
| OG275_RS03750 | elongation factor Tu | 193462 |
| OG275_RS22855 | elongation factor Tu | 193462 |
| CP978_RS06260 | elongation factor Tu | 40318 |
| CP978_RS20005 | elongation factor Tu | 40318 |
| JYK04_RS09695 | elongation factor Tu | 66374 |
| JYK04_RS25290 | elongation factor Tu | 66374 |
| JNO44_RS20310 | elongation factor Tu | 1971 |
| JNO44_RS38180 | elongation factor Tu | 1971 |

| | | |
|---|---|---|
| FYC74_RS15375 | elongation factor Tu | 47716 |
| FYC74_RS31375 | elongation factor Tu | 47716 |
| DVK44_RS02335 | elongation factor Tu | 2282738 |
| DVK44_RS12095 | elongation factor Tu | 2282738 |
| Spa2297_RS04855 | elongation factor Tu | 146923 |
| Spa2297_RS19680 | elongation factor Tu | 146923 |
| OG437_RS09285 | elongation factor Tu | 1923 |
| OG437_RS20805 | elongation factor Tu | 1923 |
| F9278_RS06080 | elongation factor Tu | 2653200 |
| F9278_RS30535 | elongation factor Tu | 2653200 |
| NFX46_RS03725 | elongation factor Tu | 114686 |
| NFX46_RS27010 | elongation factor Tu | 114686 |
| ABXR15_RS09935 | elongation factor Tu | 1150371 |
| CP981_RS17410 | elongation factor Tu | 58346 |
| CP981_RS34920 | elongation factor Tu | 58346 |
| LK06_RS03080 | elongation factor Tu | 1355015 |
| LK06_RS12400 | elongation factor Tu | 1355015 |
| P8A21_RS05110 | elongation factor Tu | 2798799 |
| P8A21_RS23890 | elongation factor Tu | 2798799 |
| CP972_RS05695 | elongation factor Tu | 67345 |
| CP972_RS21280 | elongation factor Tu | 67345 |
| HED23_RS03495 | elongation factor Tu | 1169025 |
| HED23_RS20695 | elongation factor Tu | 1169025 |
| H3L99_RS05545 | elongation factor Tu | 38300 |
| H3L99_RS21045 | elongation factor Tu | 38300 |
| K4G22_RS00525 | elongation factor Tu | 2867410 |
| K4G22_RS10950 | elongation factor Tu | 2867410 |
| OG588_RS07480 | elongation factor Tu | 67348 |
| OG588_RS29865 | elongation factor Tu | 67348 |
| OG588_RS45455 | elongation factor Tu | 67348 |
| OHO81_RS18395 | elongation factor Tu | 67350 |

| | | |
|---|---|---|
| OHO81_RS36275 | elongation factor Tu | 67350 |
| OHU35_RS06820 | elongation factor Tu | 1924 |
| OHU35_RS25820 | elongation factor Tu | 1924 |
| A4E84_RS06435 | elongation factor Tu | 1783515 |
| A4E84_RS24195 | elongation factor Tu | 1783515 |
| FQU76_RS03440 | elongation factor Tu | 2599401 |
| FQU76_RS20675 | elongation factor Tu | 2599401 |
| LIV37_RS08300 | elongation factor Tu | 1343740 |
| LIV37_RS21775 | elongation factor Tu | 1343740 |
| FGW37_RS14100 | elongation factor Tu | 173860 |
| SRIM_RS04555 | elongation factor Tu | 1265868 |
| SRIM_RS18760 | elongation factor Tu | 1265868 |
| P7W03_RS06475 | elongation factor Tu | 1928 |
| P7W03_RS21575 | elongation factor Tu | 1928 |
| IAG44_RS16070 | elongation factor Tu | 2768066 |
| IAG44_RS33980 | elongation factor Tu | 2768066 |
| IAG44_RS36755 | elongation factor Tu | 2768066 |
| M878_RS65940 | elongation factor Tu | 1352936 |
| M878_RS85090 | elongation factor Tu | 1352936 |
| Sru02f_RS13165 | elongation factor Tu | 194673 |
| Sru02f_RS34405 | elongation factor Tu | 194673 |
| F0345_RS02755 | elongation factor Tu | 53451 |
| F0345_RS10800 | elongation factor Tu | 53451 |
| SCAB_RS17585 | elongation factor Tu | 680198 |
| SCAB_RS37175 | elongation factor Tu | 680198 |
| AAC944_RS02725 | elongation factor Tu | 1957 |
| AAC944_RS17065 | elongation factor Tu | 1957 |
| OG949_RS15110 | elongation factor Tu | 452529 |
| OG949_RS32080 | elongation factor Tu | 452529 |
| D0Z67_RS03520 | elongation factor Tu | 73044 |
| D0Z67_RS17195 | elongation factor Tu | 73044 |

| | | |
|---|---|---|
| WAB15_RS18655 | elongation factor Tu | 3127701 |
| WAB15_RS36465 | elongation factor Tu | 3127701 |
| ABZO31_RS04060 | elongation factor Tu | 3231716 |
| ABZO31_RS19660 | elongation factor Tu | 3231716 |
| R2D22_RS15285 | elongation factor Tu | 3081768 |
| R2D22_RS31465 | elongation factor Tu | 3081768 |
| I1A49_RS26265 | elongation factor Tu | 1929477 |
| I1A49_RS37750 | elongation factor Tu | 1929477 |
| CP982_RS07050 | elongation factor Tu | 68270 |
| CP982_RS25170 | elongation factor Tu | 68270 |
| LXH13_RS06010 | elongation factor Tu | 2906474 |
| LXH13_RS17415 | elongation factor Tu | 2906474 |
| LXH13_RS35780 | elongation factor Tu | 2906474 |
| DDQ41_RS10735 | elongation factor Tu | 1690221 |
| DDQ41_RS18875 | elongation factor Tu | 1690221 |
| CP968_RS04965 | elongation factor Tu | 36818 |
| CP968_RS13955 | elongation factor Tu | 36818 |
| MW084_RS05135 | elongation factor Tu | 436397 |
| SSEG_RS07015 | elongation factor Tu | 463191 |
| SSEG_RS25175 | elongation factor Tu | 463191 |
| ABII15_RS05780 | elongation factor Tu | 3156398 |
| ABII15_RS22880 | elongation factor Tu | 3156398 |
| LDH80_RS16705 | elongation factor Tu | 67367 |
| OG288_RS05920 | elongation factor Tu | 68274 |
| OG288_RS25210 | elongation factor Tu | 68274 |
| OG288_RS35580 | elongation factor Tu | 68274 |
| AY578_RS04240 | elongation factor Tu | 59299 |
| AY578_RS19580 | elongation factor Tu | 59299 |
| ACI1V9_RS14835 | elongation factor Tu | 44033 |
| OG899_RS10585 | elongation factor Tu | 1952 |
| DDW44_RS10705 | elongation factor Tu | 2174846 |

| | | |
|---|---|---|
| DDW44_RS17975 | elongation factor Tu | 2174846 |
| ACNJ93_RS03485 | elongation factor Tu | 2054410 |
| ACNJ93_RS18770 | elongation factor Tu | 2054410 |
| B7R87_RS03340 | elongation factor Tu | 83656 |
| B7R87_RS20055 | elongation factor Tu | 83656 |
| MMF93_RS19670 | elongation factor Tu | 2923272 |
| STRTU_RS02230 | elongation factor Tu | 47759 |
| STRTU_RS19945 | elongation factor Tu | 47759 |
| IGS69_RS04340 | elongation factor Tu | 68278 |
| IGS69_RS20970 | elongation factor Tu | 68278 |
| OG711_RS23595 | elongation factor Tu | 1048205 |
| OG711_RS33275 | elongation factor Tu | 1048205 |
| DEJ43_RS21910 | elongation factor Tu | 953739 |
| SVTN_RS22805 | elongation factor Tu | 362257 |
| FDM97_RS13625 | elongation factor Tu | 3051092 |
| FDM97_RS31755 | elongation factor Tu | 3051092 |
| CP980_RS13895 | elongation factor Tu | 1960 |
| CP980_RS28200 | elongation factor Tu | 1960 |
| R2E43_RS15185 | elongation factor Tu | 1935 |
| R2E43_RS31850 | elongation factor Tu | 1935 |
| OG542_RS15975 | elongation factor Tu | 1936 |
| OG542_RS34410 | elongation factor Tu | 1936 |
| CP969_RS07570 | elongation factor Tu | 665577 |
| CP969_RS23025 | elongation factor Tu | 665577 |
| IAG42_RS14335 | elongation factor Tu | 2768069 |
| IAG42_RS30480 | elongation factor Tu | 2768069 |
| KPP03845_RS07585 | elongation factor Tu | 67385 |
| KPP03845_RS22490 | elongation factor Tu | 67385 |
| SXIM_RS10340 | elongation factor Tu | 408015 |
| SXIM_RS26165 | elongation factor Tu | 408015 |
| SXIN_RS01065 | elongation factor Tu | 1038929 |

| | | |
|---|---|---|
| SXIN_RS12550 | elongation factor Tu | 1038929 |
| NRK68_RS05600 | elongation factor Tu | 1648182 |
| NRK68_RS20115 | elongation factor Tu | 1648182 |
| J8403_RS06585 | elongation factor Tu | 155177 |
| J8403_RS18975 | elongation factor Tu | 155177 |
| MOV08_RS06135 | elongation factor Tu | 156453 |
| MOV08_RS23640 | elongation factor Tu | 156453 |
| OG567_RS24425 | elongation factor Tu | 68286 |
| SROS_RS05180 | elongation factor Tu | 479432 |
| ACMATS_RS21505 | elongation factor Tu | 1433415 |
| LQ955_RS00430 | elongation factor Tu | 2895559 |
| BW730_RS09085 | elongation factor Tu | 1332264 |
| BW733_RS03225 | elongation factor Tu | 399497 |
| RPIT_RS02930 | elongation factor Tu | 1610493 |
| QH948_RS03695 | elongation factor Tu | 3041766 |
| KDB89_RS10425 | elongation factor Tu | 2829499 |
| DHT94_RS01445 | elongation factor Tu | 2161816 |
| FOF52_RS13685 | elongation factor Tu | 53522 |
| TFU_RS13170 | elongation factor Tu | 269800 |
| NI17_RS16305 | elongation factor Tu | 483545 |
| TBIS_RS02800 | elongation factor Tu | 469371 |
| D3U04_RS12830 | elongation factor Tu | 1411117 |
| TCUR_RS21230 | elongation factor Tu | 471852 |
| FO059_RS03595 | elongation factor Tu | 2597660 |
| H4F70_RS04325 | elongation factor Tu | 2760083 |
| TW_RS00685 | elongation factor Tu | 218496 |
| QPC17_RS01940 | elongation factor Tu | 59561 |
| EL234_RS02760 | elongation factor Tu | 312285 |
| HLG82_RS02140 | elongation factor Tu | 2733571 |
| AN946_RS05520 | elongation factor Tu | 1661 |
| TPAU_RS17315 | elongation factor Tu | 521096 |

| | | |
|---|---|---|
| CZ356_RS05320 | elongation factor Tu | 1871034 |
| KO216_RS08020 | elongation factor Tu | 2811781 |
| OG579_RS07635 | elongation factor Tu | 1603258 |
| ET495_RS15785 | elongation factor Tu | 2509459 |
| XCEL_RS03140 | elongation factor Tu | 446471 |
| ET471_RS08555 | elongation factor Tu | 2509457 |
| AFIC_RS06640 | elongation factor Tu | 290 |
| LGT41_RS06795 | elongation factor Tu | 2881234 |
| LGT41_RS06955 | elongation factor Tu | 2881234 |
| S101447_RS01525 | elongation factor Tu | 481146 |
| CBI36_RS05010 | elongation factor Tu | 1633874 |
| EOV40_RS04200 | elongation factor Tu | 2500548 |
| FCN51_RS14935 | elongation factor Tu | 438 |
| ASN_RS00735 | elongation factor Tu | 446692 |
| FLP30_RS00620 | elongation factor Tu | 2592655 |
| DEF76_RS05500 | elongation factor Tu | 1969806 |
| ACMV_RS10910 | elongation factor Tu | 926570 |
| QP803_RS20220 | elongation factor Tu | 3040935 |
| HHL28_RS11565 | elongation factor Tu | 1709002 |
| HHL28_RS11640 | elongation factor Tu | 1709002 |
| IF205_RS16230 | elongation factor Tu | 2775080 |
| IF205_RS16300 | elongation factor Tu | 2775080 |
| OCA5_RS11715 | elongation factor Tu | 504832 |
| KZ699_RS08355 | elongation factor Tu | 2862866 |
| KZ699_RS08440 | elongation factor Tu | 2862866 |
| ATU_RS09525 | elongation factor Tu | 176299 |
| ATU_RS09610 | elongation factor Tu | 176299 |
| CFBP5477_RS06385 | elongation factor Tu | 160699 |
| CFBP5477_RS06505 | elongation factor Tu | 160699 |
| CFBP4996_RS08065 | elongation factor Tu | 2792015 |
| CFBP4996_RS08150 | elongation factor Tu | 2792015 |

| | | |
|---|---|---|
| HQN82_RS07640 | elongation factor Tu | 648995 |
| HQN82_RS07725 | elongation factor Tu | 648995 |
| G6L97_RS07815 | elongation factor Tu | 358 |
| G6L97_RS07900 | elongation factor Tu | 358 |
| HRR99_RS05290 | elongation factor Tu | 2735528 |
| HRR99_RS05380 | elongation factor Tu | 2735528 |
| GO499_RS00680 | elongation factor Tu | 2683284 |
| GO499_RS11600 | elongation factor Tu | 2683284 |
| K3J57_RS01170 | elongation factor Tu | 154981 |
| K3J57_RS01240 | elongation factor Tu | 154981 |
| QEZ52_RS03575 | elongation factor Tu | 1524259 |
| QEZ52_RS20320 | elongation factor Tu | 1524259 |
| AVI_RS07085 | elongation factor Tu | 311402 |
| AVI_RS07175 | elongation factor Tu | 311402 |
| G6N78_RS15910 | elongation factor Tu | 379684 |
| G6N78_RS15985 | elongation factor Tu | 379684 |
| B9N75_RS07295 | elongation factor Tu | 941907 |
| AAHM83_RS01090 | elongation factor Tu | 3077944 |
| P7228_RS11655 | elongation factor Tu | 3032592 |
| AMC99_RS07555 | elongation factor Tu | 361183 |
| A6F69_RS06465 | elongation factor Tu | 476157 |
| QQX03_RS05520 | elongation factor Tu | 2173831 |
| QQW98_RS09815 | elongation factor Tu | 3053610 |
| RB602_RS10980 | elongation factor Tu | 3071408 |
| O1Y97_RS16565 | elongation factor Tu | 83265 |
| O1Y97_RS16650 | elongation factor Tu | 83265 |
| ACIS_RS01805 | elongation factor Tu | 574556 |
| ACIS_RS04325 | elongation factor Tu | 574556 |
| AMF_RS05985 | elongation factor Tu | 320483 |
| AMF_RS08485 | elongation factor Tu | 320483 |
| AOV_RS00755 | elongation factor Tu | 1248439 |

| | | |
|---|---|---|
| AOV_RS03355 | elongation factor Tu | 1248439 |
| ANPL_RS03715 | elongation factor Tu | 949 |
| SNOV_RS07135 | elongation factor Tu | 639283 |
| SNOV_RS07225 | elongation factor Tu | 639283 |
| G3A50_RS00635 | elongation factor Tu | 1745854 |
| G3A50_RS00730 | elongation factor Tu | 1745854 |
| IAI54_RS04550 | elongation factor Tu | 675281 |
| IAI54_RS09110 | elongation factor Tu | 675281 |
| BSQ44_RS11375 | elongation factor Tu | 1670800 |
| BSQ44_RS16250 | elongation factor Tu | 1670800 |
| K0O24_RS05010 | elongation factor Tu | 1550733 |
| DM15PD_RS03035 | elongation factor Tu | 2697033 |
| Asbog_RS03600 | elongation factor Tu | 1231624 |
| ASTEX_RS04645 | elongation factor Tu | 573065 |
| CP97_RS07050 | elongation factor Tu | 1648404 |
| BMF35_RS07455 | elongation factor Tu | 502682 |
| H1343_RS07710 | elongation factor Tu | 2758041 |
| H1343_RS10905 | elongation factor Tu | 2758041 |
| ACLNGM_RS16560 | elongation factor Tu | 1166078 |
| ACLNGM_RS16635 | elongation factor Tu | 1166078 |
| J7654_RS12235 | elongation factor Tu | 1701758 |
| J7654_RS12310 | elongation factor Tu | 1701758 |
| AZC_RS04515 | elongation factor Tu | 438753 |
| AZC_RS13215 | elongation factor Tu | 438753 |
| ABAZ39_RS03490 | elongation factor Tu | 2970906 |
| ABAZ39_RS03560 | elongation factor Tu | 2970906 |
| Sp245p_RS05660 | elongation factor Tu | 1064539 |
| Sp245p_RS05730 | elongation factor Tu | 1064539 |
| A6A40_RS01775 | elongation factor Tu | 1226968 |
| A6A40_RS01845 | elongation factor Tu | 1226968 |
| HUE56_RS20865 | elongation factor Tu | 286727 |

| | | |
|---|---|---|
| HUE56_RS20930 | elongation factor Tu | 286727 |
| DM194_RS01515 | elongation factor Tu | 682998 |
| DM194_RS01580 | elongation factor Tu | 682998 |
| DEW08_RS10320 | elongation factor Tu | 2202148 |
| DEW08_RS10385 | elongation factor Tu | 2202148 |
| AL072_RS01485 | elongation factor Tu | 528244 |
| AL072_RS01550 | elongation factor Tu | 528244 |
| HWV54_RS01040 | elongation factor Tu | 52764 |
| HWV54_RS06450 | elongation factor Tu | 52764 |
| PU02_RS04050 | elongation factor Tu | 1318743 |
| PU02_RS05290 | elongation factor Tu | 1318743 |
| BANH1_RS02760 | elongation factor Tu | 1094489 |
| BANH1_RS04185 | elongation factor Tu | 1094489 |
| BARBAKC583_RS02710 | elongation factor Tu | 360095 |
| BARBAKC583_RS03215 | elongation factor Tu | 360095 |
| BARBAKC583_RS03375 | elongation factor Tu | 360095 |
| BARCL_RS04160 | elongation factor Tu | 696125 |
| BARCL_RS04350 | elongation factor Tu | 696125 |
| EL285_RS04180 | elongation factor Tu | 807 |
| EL285_RS04845 | elongation factor Tu | 807 |
| QHG54_RS00590 | elongation factor Tu | 3004109 |
| QHG54_RS02600 | elongation factor Tu | 3004109 |
| BGR_RS03290 | elongation factor Tu | 634504 |
| BGR_RS06370 | elongation factor Tu | 634504 |
| NMK50_RS02020 | elongation factor Tu | 2961895 |
| NMK50_RS02855 | elongation factor Tu | 2961895 |
| ACKC4Y_RS02435 | elongation factor Tu | 1461000 |
| ACKC4Y_RS04730 | elongation factor Tu | 1461000 |
| K8O99_RS02235 | elongation factor Tu | 38323 |
| K8O99_RS04360 | elongation factor Tu | 38323 |
| D1093_RS03600 | elongation factor Tu | 2133959 |

| | | |
|---|---|---|
| D1093_RS05935 | elongation factor Tu | 2133959 |
| D1092_RS02460 | elongation factor Tu | 2267275 |
| D1092_RS05260 | elongation factor Tu | 2267275 |
| LNM86_RS02255 | elongation factor Tu | 2893471 |
| LNM86_RS04815 | elongation factor Tu | 2893471 |
| MF1_RS02180 | elongation factor Tu | 803 |
| MF1_RS02700 | elongation factor Tu | 803 |
| AAJP84_RS01425 | elongation factor Tu | 165694 |
| AAJP84_RS02850 | elongation factor Tu | 165694 |
| LAJ60_RS00630 | elongation factor Tu | 33046 |
| LAJ60_RS02275 | elongation factor Tu | 33046 |
| BTR_RS04220 | elongation factor Tu | 382640 |
| BTR_RS07260 | elongation factor Tu | 382640 |
| EL291_RS05810 | elongation factor Tu | 33047 |
| EL291_RS08535 | elongation factor Tu | 33047 |
| BIND_RS06865 | elongation factor Tu | 395963 |
| BIND_RS09860 | elongation factor Tu | 395963 |
| BLTE_RS08865 | elongation factor Tu | 2233851 |
| BVIR_RS06670 | elongation factor Tu | 1079 |
| B5J99_RS00460 | elongation factor Tu | 1550728 |
| T8S45_RS01760 | elongation factor Tu | 1867408 |
| BHK69_RS23150 | elongation factor Tu | 1526658 |
| BHK69_RS23235 | elongation factor Tu | 1526658 |
| CIT40_RS20735 | elongation factor Tu | 1404768 |
| WN72_RS29405 | elongation factor Tu | 858423 |
| G6321_RS19715 | elongation factor Tu | 2823807 |
| F8237_RS29345 | elongation factor Tu | 244734 |
| QA635_RS18785 | elongation factor Tu | 1419277 |
| P6U31_RS15355 | elongation factor Tu | 1300039 |
| IC761_RS14520 | elongation factor Tu | 83627 |
| IC761_RS28050 | elongation factor Tu | 83627 |

| | | |
|---|---|---|
| S23_RS14075 | elongation factor Tu | 1404864 |
| LPJ38_RS28595 | elongation factor Tu | 993502 |
| AAV28_RS23820 | elongation factor Tu | 224911 |
| JEY66_RS20125 | elongation factor Tu | 398525 |
| ACLFLU_RS26540 | elongation factor Tu | 1179767 |
| X265_RS22775 | elongation factor Tu | 1325090 |
| X268_RS19015 | elongation factor Tu | 1325115 |
| XH91_RS14495 | elongation factor Tu | 1325095 |
| N2603_RS18245 | elongation factor Tu | 990078 |
| LMTR13_RS16735 | elongation factor Tu | 1274631 |
| HXT67_RS25795 | elongation factor Tu | 375 |
| ACFKHW_RS13160 | elongation factor Tu | 136996 |
| S58_RS22215 | elongation factor Tu | 1245469 |
| LQG66_RS09225 | elongation factor Tu | 2898149 |
| CIT37_RS16865 | elongation factor Tu | 931866 |
| QA639_RS26210 | elongation factor Tu | 280333 |
| LMTR21_RS14065 | elongation factor Tu | 190148 |
| RS897_RS28060 | elongation factor Tu | 2854389 |
| RS897_RS28170 | elongation factor Tu | 2854389 |
| J4P68_RS03180 | elongation factor Tu | 2748629 |
| QUH67_RS13965 | elongation factor Tu | 3056648 |
| KMZ29_RS11900 | elongation factor Tu | 2840469 |
| HAP48_RS07570 | elongation factor Tu | 1404411 |
| HAP48_RS43945 | elongation factor Tu | 1404411 |
| HAV00_RS20500 | elongation factor Tu | 1404367 |
| BRAD3257_RS16515 | elongation factor Tu | 1549949 |
| I3J27_RS23225 | elongation factor Tu | 2736875 |
| N2605_RS20940 | elongation factor Tu | 108015 |
| XH85_RS19810 | elongation factor Tu | 1325107 |
| M8231_RS08225 | elongation factor Tu | 1312364 |
| M8231_RS09430 | elongation factor Tu | 1312364 |

| | | |
|---|---|---|
| U0030_RS04515 | elongation factor Tu | 13160 |
| U0030_RS04685 | elongation factor Tu | 13160 |
| JX001_RS08635 | elongation factor Tu | 2813778 |
| JX001_RS11970 | elongation factor Tu | 2813778 |
| IFJ75_RS01920 | elongation factor Tu | 2774190 |
| IFJ75_RS05950 | elongation factor Tu | 2774190 |
| ACETKC_RS05800 | elongation factor Tu | 74315 |
| ACETKC_RS10685 | elongation factor Tu | 74315 |
| KWG56_RS02575 | elongation factor Tu | 172043 |
| KWG56_RS02745 | elongation factor Tu | 172043 |
| V8J38_RS06730 | elongation factor Tu | 657642 |
| V8J38_RS12545 | elongation factor Tu | 657642 |
| PIB44_RS05910 | elongation factor Tu | 3021811 |
| PIB44_RS06075 | elongation factor Tu | 3021811 |
| ACETK8_RS12455 | elongation factor Tu | 74326 |
| ACETK8_RS16315 | elongation factor Tu | 74326 |
| BRESU_RS07855 | elongation factor Tu | 633149 |
| BRESU_RS08885 | elongation factor Tu | 633149 |
| CEP68_RS18270 | elongation factor Tu | 41276 |
| CEP68_RS31955 | elongation factor Tu | 41276 |
| JIP62_RS00455 | elongation factor Tu | 2800818 |
| JIP62_RS14230 | elongation factor Tu | 2800818 |
| IR196_RS16470 | elongation factor Tu | 529 |
| IR196_RS16540 | elongation factor Tu | 529 |
| BCAN_RS05750 | elongation factor Tu | 483179 |
| BCAN_RS05820 | elongation factor Tu | 483179 |
| ACL594_RS00785 | elongation factor Tu | 120577 |
| ACL594_RS00855 | elongation factor Tu | 120577 |
| BR141012304_RS07130 | elongation factor Tu | 1218315 |
| BR141012304_RS07200 | elongation factor Tu | 1218315 |
| BME_RS03700 | elongation factor Tu | 224914 |

| | | |
|---|---|---|
| BME_RS03770 | elongation factor Tu | 224914 |
| BMI_RS05780 | elongation factor Tu | 568815 |
| BMI_RS05850 | elongation factor Tu | 568815 |
| BOV_RS05940 | elongation factor Tu | 444178 |
| BOV_RS06010 | elongation factor Tu | 444178 |
| NIK97_RS03930 | elongation factor Tu | 370111 |
| NIK97_RS04005 | elongation factor Tu | 370111 |
| L7D45_RS04240 | elongation factor Tu | 419475 |
| L7D45_RS04310 | elongation factor Tu | 419475 |
| BR_RS05730 | elongation factor Tu | 204722 |
| BR_RS05800 | elongation factor Tu | 204722 |
| EGO55_RS01150 | elongation factor Tu | 1219035 |
| N3Z17_RS01400 | elongation factor Tu | 2570586 |
| N3Z17_RS02050 | elongation factor Tu | 2570586 |
| Bandiella_RS04095 | elongation factor Tu | 1664265 |
| Bandiella_RS05280 | elongation factor Tu | 1664265 |
| Bealeia1_RS08025 | elongation factor Tu | 1921001 |
| Bealeia1_RS09150 | elongation factor Tu | 1921001 |
| CPBP_RS00375 | elongation factor Tu | 2741701 |
| FZC35_RS00780 | elongation factor Tu | 2601575 |
| QQM06_RS02710 | elongation factor Tu | 2066024 |
| deia_RS03780 | elongation factor Tu | 2163644 |
| A1OE_RS00800 | elongation factor Tu | 1193729 |
| AAGD37_RS00985 | elongation factor Tu | 3077946 |
| Fokcrypt_RS01255 | elongation factor Tu | 1920990 |
| Fsol_RS00810 | elongation factor Tu | 1802984 |
| CR143_RS02225 | elongation factor Tu | 1925548 |
| Gromo_RS02285 | elongation factor Tu | 2806609 |
| MAGTDC_RS02570 | elongation factor Tu | 573658 |
| MAGTDC_RS02860 | elongation factor Tu | 573658 |
| HYD_RS03740 | elongation factor Tu | 2558174 |

| | | |
|---|---|---|
| N5W20_RS00855 | elongation factor Tu | 2984322 |
| AACL20_RS01710 | elongation factor Tu | 3066224 |
| AACK98_RS02405 | elongation factor Tu | 3077949 |
| G293_RS00925 | elongation factor Tu | 1277257 |
| G293_RS04145 | elongation factor Tu | 1277257 |
| LAM_RS02215 | elongation factor Tu | 1261131 |
| LAM_RS04605 | elongation factor Tu | 1261131 |
| CD16_RS00105 | elongation factor Tu | 34021 |
| CD16_RS00675 | elongation factor Tu | 34021 |
| CKC_RS04865 | elongation factor Tu | 658172 |
| CKC_RS05205 | elongation factor Tu | 658172 |
| N3Z16_RS03110 | elongation factor Tu | 988779 |
| AACL19_RS06080 | elongation factor Tu | 3077948 |
| AACL19_RS06590 | elongation factor Tu | 3077948 |
| AACL09_RS02290 | elongation factor Tu | 3066221 |
| AACL09_RS04935 | elongation factor Tu | 3066221 |
| MIDI_RS02800 | elongation factor Tu | 696127 |
| Q5948_RS00525 | elongation factor Tu | 2601668 |
| GQ61_RS08220 | elongation factor Tu | 1414854 |
| GQ61_RS08285 | elongation factor Tu | 1414854 |
| ID47_RS08080 | elongation factor Tu | 91604 |
| ID47_RS08145 | elongation factor Tu | 91604 |
| E5R92_RS00380 | elongation factor Tu | 2563896 |
| SAR11_RS05650 | elongation factor Tu | 335992 |
| phytr_RS03940 | elongation factor Tu | 2115978 |
| SAR116_RS12155 | elongation factor Tu | 488538 |
| SAR116_RS12235 | elongation factor Tu | 488538 |
| FZC37_RS02410 | elongation factor Tu | 2601646 |
| AAGD44_RS06835 | elongation factor Tu | 3066262 |
| ACQH1V_RS02590 | elongation factor Tu | 3077928 |
| AAGD19_RS02345 | elongation factor Tu | 3066253 |

| | | |
|---|---|---|
| AAGD42_RS02945 | elongation factor Tu | 3066254 |
| AAGD31_RS02300 | elongation factor Tu | 3066255 |
| AAGD29_RS07935 | elongation factor Tu | 3066272 |
| AAHM86_RS03585 | elongation factor Tu | 3066259 |
| AAGD53_RS06180 | elongation factor Tu | 3066261 |
| AAHM80_RS01135 | elongation factor Tu | 3066274 |
| ACN1AG_RS02840 | elongation factor Tu | 3066271 |
| AAHN03_RS04270 | elongation factor Tu | 3066257 |
| ACQH6S_RS04185 | elongation factor Tu | 3139330 |
| AB3211_RS01395 | elongation factor Tu | 3139332 |
| AAGD39_RS04410 | elongation factor Tu | 3066263 |
| ACFELW_RS06665 | elongation factor Tu | 3077926 |
| ACFE3K_RS01635 | elongation factor Tu | 3139336 |
| ACFEL1_RS03725 | elongation factor Tu | 3077925 |
| AAHN00_RS11090 | elongation factor Tu | 3066275 |
| AAGD26_RS05140 | elongation factor Tu | 3066273 |
| ACFEL0_RS07490 | elongation factor Tu | 3139333 |
| AAGD36_RS05945 | elongation factor Tu | 3066260 |
| AAGD20_RS02000 | elongation factor Tu | 2029164 |
| ACQH1W_RS05585 | elongation factor Tu | 3139337 |
| ACFEL2_RS00930 | elongation factor Tu | 3139335 |
| AAGD45_RS00470 | elongation factor Tu | 3066258 |
| ACFELY_RS03970 | elongation factor Tu | 3139334 |
| R2I74_RS07340 | elongation factor Tu | 1346319 |
| ATE48_RS18865 | elongation factor Tu | 1759059 |
| C1707_RS13045 | elongation factor Tu | 1679497 |
| C1707_RS24230 | elongation factor Tu | 1679497 |
| AQ619_RS06185 | elongation factor Tu | 69395 |
| AQ619_RS14755 | elongation factor Tu | 69395 |
| CSW64_RS03150 | elongation factor Tu | 69666 |
| CSW64_RS09075 | elongation factor Tu | 69666 |

| | | |
|---|---|---|
| G3M57_RS16325 | elongation factor Tu | 2010972 |
| G3M57_RS21965 | elongation factor Tu | 2010972 |
| B7G68_RS03865 | elongation factor Tu | 88688 |
| B7G68_RS07615 | elongation factor Tu | 88688 |
| G3M62_RS04270 | elongation factor Tu | 2708539 |
| G3M62_RS08325 | elongation factor Tu | 2708539 |
| CEW89_RS14965 | elongation factor Tu | 1758178 |
| CEW89_RS15055 | elongation factor Tu | 1758178 |
| P73_RS03230 | elongation factor Tu | 1208324 |
| P73_RS03295 | elongation factor Tu | 1208324 |
| IMCC12053_RS07190 | elongation factor Tu | 1397108 |
| IMCC12053_RS07280 | elongation factor Tu | 1397108 |
| LV780_RS11625 | elongation factor Tu | 43057 |
| LV780_RS11695 | elongation factor Tu | 43057 |
| RSP_RS01435 | elongation factor Tu | 272943 |
| RSP_RS01510 | elongation factor Tu | 272943 |
| H4I97_RS09305 | elongation factor Tu | 1969821 |
| H4I97_RS09385 | elongation factor Tu | 1969821 |
| SLU04_RS03375 | elongation factor Tu | 655353 |
| SLU04_RS03450 | elongation factor Tu | 655353 |
| D9V35_RS07025 | elongation factor Tu | 2070537 |
| RA157_RS01440 | elongation factor Tu | 3066369 |
| RA157_RS01510 | elongation factor Tu | 3066369 |
| Q7I88_RS09190 | elongation factor Tu | 3064139 |
| WYH_RS00370 | elongation factor Tu | 1267766 |
| H4O24_RS11605 | elongation factor Tu | 450378 |
| AB433_RS05680 | elongation factor Tu | 1348774 |
| HQ394_RS04935 | elongation factor Tu | 111831 |
| HQ394_RS04990 | elongation factor Tu | 111831 |
| GDR53_RS17070 | elongation factor Tu | 2657486 |
| FPZ08_RS06740 | elongation factor Tu | 400770 |

| | | |
|---|---|---|
| MF606_RS11300 | elongation factor Tu | 2917991 |
| N8A98_RS18315 | elongation factor Tu | 191302 |
| JI749_RS10750 | elongation factor Tu | 2801335 |
| JI748_RS05430 | elongation factor Tu | 2774137 |
| QOV41_RS08595 | elongation factor Tu | 3049774 |
| PSQ90_RS00455 | elongation factor Tu | 3026423 |
| K1X15_RS09580 | elongation factor Tu | 2860336 |
| DSHI_RS01110 | elongation factor Tu | 398580 |
| DSHI_RS01365 | elongation factor Tu | 398580 |
| ECAJ_RS00850 | elongation factor Tu | 269484 |
| ECAJ_RS03165 | elongation factor Tu | 269484 |
| ECH_RS01660 | elongation factor Tu | 205920 |
| ECH_RS03930 | elongation factor Tu | 205920 |
| EHF_RS01550 | elongation factor Tu | 391036 |
| EHF_RS03765 | elongation factor Tu | 391036 |
| MR76_RS00760 | elongation factor Tu | 1423892 |
| MR76_RS03060 | elongation factor Tu | 1423892 |
| ERUM_RS00900 | elongation factor Tu | 254945 |
| ERUM_RS03305 | elongation factor Tu | 254945 |
| KO353_RS12070 | elongation factor Tu | 2843330 |
| KO353_RS12135 | elongation factor Tu | 2843330 |
| I862_RS03605 | elongation factor Tu | 86106 |
| I862_RS05945 | elongation factor Tu | 86106 |
| FA04_RS05420 | elongation factor Tu | 106592 |
| FA04_RS05495 | elongation factor Tu | 106592 |
| J3R84_RS05365 | elongation factor Tu | 555315 |
| J3R84_RS05440 | elongation factor Tu | 555315 |
| JGUZn3_RS00305 | elongation factor Tu | 2762277 |
| L1K66_RS12260 | elongation factor Tu | 2909249 |
| DVR09_RS11900 | elongation factor Tu | 2182384 |
| Ga0102493_RS05925 | elongation factor Tu | 39960 |

| | | |
|---|---|---|
| HQR01_RS11545 | elongation factor Tu | 2739433 |
| A9D12_RS13900 | elongation factor Tu | 1112 |
| RC74_RS04300 | elongation factor Tu | 1579316 |
| RC74_RS04390 | elongation factor Tu | 1579316 |
| GR316_RS09915 | elongation factor Tu | 2692330 |
| GR316_RS09990 | elongation factor Tu | 2692330 |
| FNB15_RS15450 | elongation factor Tu | 2594003 |
| FNB15_RS17465 | elongation factor Tu | 2594003 |
| QTA57_RS07220 | elongation factor Tu | 2978973 |
| QTA57_RS07325 | elongation factor Tu | 2978973 |
| E3E12_RS01925 | elongation factor Tu | 2558361 |
| AB6B37_RS03110 | elongation factor Tu | 570162 |
| AB6B37_RS04245 | elongation factor Tu | 570162 |
| AKL17_RS01855 | elongation factor Tu | 1335048 |
| AKL17_RS01935 | elongation factor Tu | 1335048 |
| QF092_RS15535 | elongation factor Tu | 3042299 |
| QF092_RS15620 | elongation factor Tu | 3042299 |
| KM031_RS11990 | elongation factor Tu | 2840474 |
| KM031_RS12080 | elongation factor Tu | 2840474 |
| D5400_RS12005 | elongation factor Tu | 2341112 |
| D5400_RS12075 | elongation factor Tu | 2341112 |
| PH603_RS07585 | elongation factor Tu | 3024400 |
| PH603_RS07650 | elongation factor Tu | 3024400 |
| GDIA_RS14865 | elongation factor Tu | 272568 |
| A0U94_RS09035 | elongation factor Tu | 318683 |
| GLS_RS02030 | elongation factor Tu | 1288313 |
| IGS75_RS02405 | elongation factor Tu | 574987 |
| AB6B38_RS01100 | elongation factor Tu | 1433403 |
| AB6B38_RS01180 | elongation factor Tu | 1433403 |
| X907_RS04190 | elongation factor Tu | 1434191 |
| X907_RS04260 | elongation factor Tu | 1434191 |

| | | |
|---|---|---|
| GbCGDNIH7_RS02860 | elongation factor Tu | 364410 |
| HmaOT1_RS10810 | elongation factor Tu | 195105 |
| HmaOT1_RS10900 | elongation factor Tu | 195105 |
| AY555_RS00975 | elongation factor Tu | 1549855 |
| AY555_RS01040 | elongation factor Tu | 1549855 |
| E6W36_RS13120 | elongation factor Tu | 1763828 |
| HDIA_RS07540 | elongation factor Tu | 1482074 |
| HDIA_RS12690 | elongation factor Tu | 1482074 |
| HBAL_RS10465 | elongation factor Tu | 582402 |
| HPDFL43_RS09440 | elongation factor Tu | 411684 |
| HPDFL43_RS09525 | elongation factor Tu | 411684 |
| FRZ61_RS08945 | elongation factor Tu | 2602016 |
| FRZ61_RS09010 | elongation factor Tu | 2602016 |
| FRZ44_RS14610 | elongation factor Tu | 2602015 |
| FRZ44_RS14675 | elongation factor Tu | 2602015 |
| PUV54_RS13580 | elongation factor Tu | 1866326 |
| ABFZ83_RS03535 | elongation factor Tu | 3143534 |
| HDEN_RS05010 | elongation factor Tu | 582899 |
| W911_RS13675 | elongation factor Tu | 1029756 |
| HF931_RS13195 | elongation factor Tu | 1280948 |
| HF931_RS16515 | elongation factor Tu | 1280948 |
| HNE_RS08560 | elongation factor Tu | 228405 |
| HNE_RS14120 | elongation factor Tu | 228405 |
| DOL89_RS01810 | elongation factor Tu | 2220096 |
| DOL89_RS01875 | elongation factor Tu | 2220096 |
| P8627_RS02565 | elongation factor Tu | 3038773 |
| P8627_RS02660 | elongation factor Tu | 3038773 |
| HW532_RS00365 | elongation factor Tu | 2593653 |
| HW532_RS00430 | elongation factor Tu | 2593653 |
| BVG79_RS10990 | elongation factor Tu | 92947 |
| BVG79_RS11070 | elongation factor Tu | 92947 |

| | | |
|---|---|---|
| KVH_RS13270 | elongation factor Tu | 92945 |
| KVH_RS13360 | elongation factor Tu | 92945 |
| ACE2AA_RS08150 | elongation factor Tu | 1932662 |
| S101446_RS07610 | elongation factor Tu | 33995 |
| GLX_RS05585 | elongation factor Tu | 634177 |
| B0W47_RS05015 | elongation factor Tu | 265960 |
| GWK63_RS11215 | elongation factor Tu | 215221 |
| KSAC_RS06285 | elongation factor Tu | 265959 |
| R5N89_RS03265 | elongation factor Tu | 1307942 |
| CXP35_RS06535 | elongation factor Tu | 28448 |
| ICL80_RS08520 | elongation factor Tu | 2161677 |
| ICL80_RS08585 | elongation factor Tu | 2161677 |
| A0U90_RS08125 | elongation factor Tu | 153496 |
| ACMDCR_RS14005 | elongation factor Tu | 346911 |
| ACMDCR_RS14080 | elongation factor Tu | 346911 |
| R2C4_RS16060 | elongation factor Tu | 1396826 |
| R2C4_RS16155 | elongation factor Tu | 1396826 |
| K3721_RS01075 | elongation factor Tu | 506591 |
| K3721_RS16040 | elongation factor Tu | 506591 |
| METH_RS00905 | elongation factor Tu | 999552 |
| METH_RS00995 | elongation factor Tu | 999552 |
| OKQ63_RS01120 | elongation factor Tu | 2930199 |
| OKQ63_RS01215 | elongation factor Tu | 2930199 |
| RL73_RS02180 | elongation factor Tu | 1273132 |
| RL73_RS04325 | elongation factor Tu | 1273132 |
| HN018_RS05190 | elongation factor Tu | 1484109 |
| T8T21_RS12290 | elongation factor Tu | 1492771 |
| T8T21_RS12380 | elongation factor Tu | 1492771 |
| KBW81_RS00790 | elongation factor Tu | 195913 |
| KBW81_RS00870 | elongation factor Tu | 195913 |
| FE788_RS04320 | elongation factor Tu | 2582913 |

| | | |
|---|---|---|
| FE788_RS04385 | elongation factor Tu | 2582913 |
| MMC1_RS04260 | elongation factor Tu | 156889 |
| MMC1_RS04330 | elongation factor Tu | 156889 |
| MGMSRV2_RS00105 | elongation factor Tu | 1430440 |
| MGMSRV2_RS00185 | elongation factor Tu | 1430440 |
| L1P08_RS04715 | elongation factor Tu | 2909283 |
| L1P08_RS04800 | elongation factor Tu | 2909283 |
| JW792_RS06600 | elongation factor Tu | 2029849 |
| JW792_RS06680 | elongation factor Tu | 2029849 |
| MXMO3_RS08630 | elongation factor Tu | 454601 |
| MXMO3_RS09185 | elongation factor Tu | 454601 |
| TM49_RS18825 | elongation factor Tu | 1486262 |
| TM49_RS19975 | elongation factor Tu | 1486262 |
| Mame_RS11895 | elongation factor Tu | 1122214 |
| Mame_RS19950 | elongation factor Tu | 1122214 |
| A6B35_RS19875 | elongation factor Tu | 1082933 |
| A6B35_RS19975 | elongation factor Tu | 1082933 |
| MESAU_RS20730 | elongation factor Tu | 754035 |
| MESAU_RS20820 | elongation factor Tu | 754035 |
| EB233_RS20160 | elongation factor Tu | 1777866 |
| EB233_RS20255 | elongation factor Tu | 1777866 |
| U0R22_RS15460 | elongation factor Tu | 28104 |
| U0R22_RS15575 | elongation factor Tu | 28104 |
| R7A2020_RS19610 | elongation factor Tu | 935547 |
| R7A2020_RS19705 | elongation factor Tu | 935547 |
| EB229_RS23395 | elongation factor Tu | 1777867 |
| EB229_RS23485 | elongation factor Tu | 1777867 |
| RBH77_RS18165 | elongation factor Tu | 3074855 |
| RBH77_RS18235 | elongation factor Tu | 3074855 |
| LRP31_RS22425 | elongation factor Tu | 43617 |
| LRP31_RS22510 | elongation factor Tu | 43617 |

| | | |
|---|---|---|
| IHQ72_RS22705 | elongation factor Tu | 2775404 |
| IHQ72_RS22800 | elongation factor Tu | 2775404 |
| MESOP_RS22865 | elongation factor Tu | 536019 |
| MESOP_RS22970 | elongation factor Tu | 536019 |
| AB3461_RS06685 | elongation factor Tu | 570505 |
| AB3461_RS06775 | elongation factor Tu | 570505 |
| DA075_RS13845 | elongation factor Tu | 2051553 |
| DA075_RS13915 | elongation factor Tu | 2051553 |
| DK389_RS30995 | elongation factor Tu | 2202825 |
| DK389_RS31075 | elongation factor Tu | 2202825 |
| OF380_RS26785 | elongation factor Tu | 2984843 |
| OF380_RS26865 | elongation factor Tu | 2984843 |
| miv_RS06290 | elongation factor Tu | 1775910 |
| miv_RS15860 | elongation factor Tu | 1775910 |
| MNOD_RS09135 | elongation factor Tu | 460265 |
| MNOD_RS09205 | elongation factor Tu | 460265 |
| MMB17_RS16140 | elongation factor Tu | 410 |
| MMB17_RS23380 | elongation factor Tu | 410 |
| MOC_RS11805 | elongation factor Tu | 693986 |
| MOC_RS19770 | elongation factor Tu | 693986 |
| MCBMB27_RS08665 | elongation factor Tu | 418223 |
| MCBMB27_RS16075 | elongation factor Tu | 418223 |
| DK427_RS18835 | elongation factor Tu | 2202828 |
| DK427_RS22350 | elongation factor Tu | 2202828 |
| MRAD2831_RS42945 | elongation factor Tu | 426355 |
| MRAD2831_RS51060 | elongation factor Tu | 426355 |
| DK419_RS19060 | elongation factor Tu | 2202827 |
| DK419_RS19130 | elongation factor Tu | 2202827 |
| RZS28_RS16430 | elongation factor Tu | 3080828 |
| GL4_RS11185 | elongation factor Tu | 1384459 |
| MSIL_RS02960 | elongation factor Tu | 395965 |

| | | |
|---|---|---|
| SIN04_RS08670 | elongation factor Tu | 227605 |
| QMG80_RS06785 | elongation factor Tu | 655015 |
| QMG80_RS08245 | elongation factor Tu | 655015 |
| RVU70_RS08840 | elongation factor Tu | 29468 |
| RVU70_RS16590 | elongation factor Tu | 29468 |
| H2LOC_RS06525 | elongation factor Tu | 391905 |
| H2LOC_RS17650 | elongation factor Tu | 391905 |
| QMG84_RS10375 | elongation factor Tu | 2885079 |
| QMG84_RS16805 | elongation factor Tu | 2885079 |
| MMG94_RS04210 | elongation factor Tu | 1134912 |
| MMG94_RS10205 | elongation factor Tu | 1134912 |
| ABS361_RS04415 | elongation factor Tu | 3162530 |
| ABS361_RS04485 | elongation factor Tu | 3162530 |
| METD_RS13010 | elongation factor Tu | 661410 |
| METD_RS24825 | elongation factor Tu | 661410 |
| MPOP_RS10595 | elongation factor Tu | 441620 |
| MPOP_RS24250 | elongation factor Tu | 441620 |
| CWB41_RS15665 | elongation factor Tu | 569860 |
| U0023_RS00655 | elongation factor Tu | 864069 |
| U0023_RS00735 | elongation factor Tu | 864069 |
| BB934_RS21050 | elongation factor Tu | 1882682 |
| BB934_RS21130 | elongation factor Tu | 1882682 |
| HPT29_RS09145 | elongation factor Tu | 2740529 |
| HPT29_RS09220 | elongation factor Tu | 2740529 |
| ACP6CK_RS07700 | elongation factor Tu | 2231721 |
| ACP6CK_RS07785 | elongation factor Tu | 2231721 |
| A0U93_RS06360 | elongation factor Tu | 320497 |
| LUA81_RS00510 | elongation factor Tu | 89586 |
| LUA81_RS02940 | elongation factor Tu | 89586 |
| D5366_RS01785 | elongation factor Tu | 661191 |
| RG540_RS06360 | elongation factor Tu | 1028800 |

| | | |
|---|---|---|
| RG540_RS06430 | elongation factor Tu | 1028800 |
| QEO92_RS07145 | elongation factor Tu | 515361 |
| QEO92_RS07230 | elongation factor Tu | 515361 |
| GP480_RS02780 | elongation factor Tu | 2686014 |
| NHE_RS02895 | elongation factor Tu | 1286528 |
| NRI_RS02865 | elongation factor Tu | 434131 |
| NSE_RS02775 | elongation factor Tu | 222891 |
| JO391_RS00595 | elongation factor Tu | 2807096 |
| JO391_RS13465 | elongation factor Tu | 2807096 |
| OLX08_RS09095 | elongation factor Tu | 282199 |
| OLX08_RS09180 | elongation factor Tu | 282199 |
| AAC691_RS20265 | elongation factor Tu | 1216886 |
| NUH88_RS14650 | elongation factor Tu | 1862145 |
| NUH88_RS14715 | elongation factor Tu | 1862145 |
| ACLIR7_RS02910 | elongation factor Tu | 889300 |
| ACLIR7_RS02985 | elongation factor Tu | 889300 |
| KW403_RS17965 | elongation factor Tu | 430679 |
| KW403_RS18035 | elongation factor Tu | 430679 |
| FQ775_RS04535 | elongation factor Tu | 2599600 |
| FQ775_RS04610 | elongation factor Tu | 2599600 |
| NTH_RS19230 | elongation factor Tu | 2267699 |
| NTH_RS19325 | elongation factor Tu | 2267699 |
| NHAM_RS07900 | elongation factor Tu | 323097 |
| NWI_RS07065 | elongation factor Tu | 323098 |
| Y958_RS04045 | elongation factor Tu | 1441467 |
| Y958_RS09240 | elongation factor Tu | 1441467 |
| C0V82_RS15045 | elongation factor Tu | 1612173 |
| C0V82_RS15140 | elongation factor Tu | 1612173 |
| IFJ82_RS05095 | elongation factor Tu | 436 |
| SARO_RS06240 | elongation factor Tu | 279238 |
| HT578_RS19325 | elongation factor Tu | 2698673 |

| | | |
|---|---|---|
| FRF71_RS15235 | elongation factor Tu | 1176536 |
| PQ457_RS11595 | elongation factor Tu | 2282397 |
| LO787_RS09410 | elongation factor Tu | 2896849 |
| AACH40_RS04240 | elongation factor Tu | 2728851 |
| JI59_RS14850 | elongation factor Tu | 1088721 |
| BES08_RS09290 | elongation factor Tu | 158500 |
| H7X45_RS00550 | elongation factor Tu | 2762729 |
| ACOQOU_RS17485 | elongation factor Tu | 2184161 |
| ACOQOU_RS17575 | elongation factor Tu | 2184161 |
| CES85_RS15820 | elongation factor Tu | 271865 |
| CES85_RS15895 | elongation factor Tu | 271865 |
| OAN307_RS22280 | elongation factor Tu | 391626 |
| OA238_RS01310 | elongation factor Tu | 391616 |
| OA238_RS01405 | elongation factor Tu | 391616 |
| OSB_RS01195 | elongation factor Tu | 1458307 |
| OSB_RS01305 | elongation factor Tu | 1458307 |
| E3E11_RS01835 | elongation factor Tu | 2558360 |
| HTY61_RS07105 | elongation factor Tu | 2742145 |
| HTY61_RS07175 | elongation factor Tu | 2742145 |
| DK406_RS04130 | elongation factor Tu | 784 |
| DK406_RS10240 | elongation factor Tu | 784 |
| CBW24_RS13675 | elongation factor Tu | 1411902 |
| CBW24_RS13765 | elongation factor Tu | 1411902 |
| Q5701_RS22760 | elongation factor Tu | 3064531 |
| PAE61_RS06810 | elongation factor Tu | 3017781 |
| PAE61_RS12290 | elongation factor Tu | 3017781 |
| PAF20_RS02215 | elongation factor Tu | 3017784 |
| PAF20_RS05670 | elongation factor Tu | 3017784 |
| JHW40_RS00060 | elongation factor Tu | 34002 |
| JHW40_RS16360 | elongation factor Tu | 34002 |
| JCM7686_RS16255 | elongation factor Tu | 1367847 |

| | | |
|---|---|---|
| JCM7686_RS16350 | elongation factor Tu | 1367847 |
| JCM7685_RS02940 | elongation factor Tu | 34004 |
| JCM7685_RS03050 | elongation factor Tu | 34004 |
| B0A89_RS01895 | elongation factor Tu | 1945662 |
| B0A89_RS07485 | elongation factor Tu | 1945662 |
| EO213_RS11610 | elongation factor Tu | 266 |
| EO213_RS11725 | elongation factor Tu | 266 |
| LZ585_RS12370 | elongation factor Tu | 2903900 |
| LZ585_RS12490 | elongation factor Tu | 2903900 |
| JHX87_RS15300 | elongation factor Tu | 658446 |
| JHX87_RS15420 | elongation factor Tu | 658446 |
| CYR75_RS02075 | elongation factor Tu | 2065379 |
| CYR75_RS08200 | elongation factor Tu | 2065379 |
| F8A10_RS09065 | elongation factor Tu | 135740 |
| F8A10_RS09170 | elongation factor Tu | 135740 |
| E4191_RS02890 | elongation factor Tu | 2560053 |
| E4191_RS13150 | elongation factor Tu | 2560053 |
| CP157_RS02650 | elongation factor Tu | 59779 |
| CP157_RS12155 | elongation factor Tu | 59779 |
| JWJ88_RS02695 | elongation factor Tu | 2812658 |
| JWJ88_RS02800 | elongation factor Tu | 2812658 |
| ESD82_RS18085 | elongation factor Tu | 82367 |
| ESD82_RS18195 | elongation factor Tu | 82367 |
| JHX88_RS15635 | elongation factor Tu | 405559 |
| JHX88_RS15735 | elongation factor Tu | 405559 |
| HGN31_RS09290 | elongation factor Tu | 1545044 |
| HGN31_RS09420 | elongation factor Tu | 1545044 |
| PAF18_RS01145 | elongation factor Tu | 3017783 |
| PAF18_RS03930 | elongation factor Tu | 3017783 |
| JHW44_RS03090 | elongation factor Tu | 184748 |
| JHW44_RS03180 | elongation factor Tu | 184748 |

| | | |
|---|---|---|
| JHW45_RS15265 | elongation factor Tu | 659350 |
| JHW45_RS15370 | elongation factor Tu | 659350 |
| DRW48_RS06220 | elongation factor Tu | 2259340 |
| DRW48_RS06330 | elongation factor Tu | 2259340 |
| CUV01_RS05045 | elongation factor Tu | 1529068 |
| CUV01_RS10160 | elongation factor Tu | 1529068 |
| E3U25_RS16710 | elongation factor Tu | 34007 |
| E3U25_RS16825 | elongation factor Tu | 34007 |
| PYTT13_RS01770 | elongation factor Tu | 147645 |
| PYTT13_RS03210 | elongation factor Tu | 147645 |
| CX676_RS08985 | elongation factor Tu | 1077935 |
| CX676_RS10065 | elongation factor Tu | 1077935 |
| JNE37_RS17140 | elongation factor Tu | 1335043 |
| FNA67_RS13350 | elongation factor Tu | 1447062 |
| HYN69_RS01815 | elongation factor Tu | 2169400 |
| HYN69_RS01935 | elongation factor Tu | 2169400 |
| AMB_RS15780 | elongation factor Tu | 342108 |
| AMB_RS15855 | elongation factor Tu | 342108 |
| PY308_RS09915 | elongation factor Tu | 1395951 |
| PY308_RS10000 | elongation factor Tu | 1395951 |
| PYR65_RS14815 | elongation factor Tu | 3031126 |
| PYR65_RS14890 | elongation factor Tu | 3031126 |
| EBB79_RS01215 | elongation factor Tu | 2483033 |
| EBB79_RS17625 | elongation factor Tu | 2483033 |
| QPJ95_RS05895 | elongation factor Tu | 2997337 |
| QPJ95_RS10205 | elongation factor Tu | 2997337 |
| HFP57_RS15075 | elongation factor Tu | 2026624 |
| J4G78_RS15810 | elongation factor Tu | 2806553 |
| HF685_RS08005 | elongation factor Tu | 2725558 |
| BS29_RS12085 | elongation factor Tu | 1123271 |
| A6F65_RS06400 | elongation factor Tu | 645517 |

| | | |
|---|---|---|
| FIV45_RS14525 | elongation factor Tu | 2043170 |
| FIV45_RS14590 | elongation factor Tu | 2043170 |
| FDP22_RS01720 | elongation factor Tu | 2579971 |
| FDP22_RS01785 | elongation factor Tu | 2579971 |
| PLAV_RS13810 | elongation factor Tu | 402881 |
| PLAV_RS13875 | elongation factor Tu | 402881 |
| PB2503_RS08890 | elongation factor Tu | 314260 |
| AM2010_RS02500 | elongation factor Tu | 543877 |
| T8T98_RS04850 | elongation factor Tu | 538382 |
| OF122_RS09245 | elongation factor Tu | 2984530 |
| OF122_RS09315 | elongation factor Tu | 2984530 |
| HKM20_RS08540 | elongation factor Tu | 531813 |
| HKM20_RS08610 | elongation factor Tu | 531813 |
| V6617_RS08355 | elongation factor Tu | 1046114 |
| V6617_RS08425 | elongation factor Tu | 1046114 |
| I8N54_RS15960 | elongation factor Tu | 2588711 |
| I8N54_RS16065 | elongation factor Tu | 2588711 |
| QTL56_RS02950 | elongation factor Tu | 2919917 |
| QTL56_RS03025 | elongation factor Tu | 2919917 |
| FE840_RS11955 | elongation factor Tu | 1813451 |
| FE840_RS12030 | elongation factor Tu | 1813451 |
| GAL_RS01300 | elongation factor Tu | 1423144 |
| GAL_RS01415 | elongation factor Tu | 1423144 |
| PhaeoP54_RS14920 | elongation factor Tu | 221822 |
| PhaeoP54_RS15045 | elongation factor Tu | 221822 |
| OL67_RS15060 | elongation factor Tu | 1580596 |
| OL67_RS15160 | elongation factor Tu | 1580596 |
| PhaeoP97_RS01760 | elongation factor Tu | 1844006 |
| PhaeoP97_RS01860 | elongation factor Tu | 1844006 |
| KCG34_RS23450 | elongation factor Tu | 2823693 |
| KCG34_RS23540 | elongation factor Tu | 2823693 |

| | | |
|---|---|---|
| HYN04_RS05335 | elongation factor Tu | 2201350 |
| HYN04_RS05420 | elongation factor Tu | 2201350 |
| PHZ_RS06700 | elongation factor Tu | 450851 |
| PHZ_RS06795 | elongation factor Tu | 450851 |
| E8L99_RS15745 | elongation factor Tu | 2570229 |
| E8L99_RS15820 | elongation factor Tu | 2570229 |
| C6569_RS06050 | elongation factor Tu | 1868589 |
| C6569_RS06120 | elongation factor Tu | 1868589 |
| E8M01_RS33330 | elongation factor Tu | 1940610 |
| E8M01_RS33405 | elongation factor Tu | 1940610 |
| BLM14_RS09685 | elongation factor Tu | 1867719 |
| BLM14_RS09760 | elongation factor Tu | 1867719 |
| G5B40_RS16725 | elongation factor Tu | 2748098 |
| G5B40_RS20235 | elongation factor Tu | 2748098 |
| KTC28_RS08660 | elongation factor Tu | 2835787 |
| SL003B_RS08720 | elongation factor Tu | 991905 |
| SL003B_RS08790 | elongation factor Tu | 991905 |
| GQA70_RS02340 | elongation factor Tu | 1943633 |
| GQA70_RS17155 | elongation factor Tu | 1943633 |
| G8E03_RS04615 | elongation factor Tu | 2758038 |
| G8E03_RS04700 | elongation factor Tu | 2758038 |
| I0K15_RS11550 | elongation factor Tu | 2789856 |
| I0K15_RS11635 | elongation factor Tu | 2789856 |
| BAR1_RS00550 | elongation factor Tu | 2171755 |
| BAR1_RS16770 | elongation factor Tu | 2171755 |
| KW060_RS03035 | elongation factor Tu | 2857064 |
| KW060_RS03100 | elongation factor Tu | 2857064 |
| KHQ09_RS11245 | elongation factor Tu | 2834768 |
| KHQ09_RS11315 | elongation factor Tu | 2834768 |
| B6K69_RS12860 | elongation factor Tu | 1075 |
| B6K69_RS12940 | elongation factor Tu | 1075 |

| | | |
|---|---|---|
| DW352_RS02945 | elongation factor Tu | 331696 |
| PSAL_RS10220 | elongation factor Tu | 1537215 |
| PSAL_RS10295 | elongation factor Tu | 1537215 |
| HF937_RS13590 | elongation factor Tu | 1461694 |
| HF937_RS13735 | elongation factor Tu | 1461694 |
| HF934_RS04800 | elongation factor Tu | 517719 |
| F3W81_RS19540 | elongation factor Tu | 2613965 |
| F3W81_RS19625 | elongation factor Tu | 2613965 |
| GFD27_RS07470 | elongation factor Tu | 1335061 |
| GFD27_RS07545 | elongation factor Tu | 1335061 |
| EOK75_RS05545 | elongation factor Tu | 2500533 |
| EOK75_RS15115 | elongation factor Tu | 2500533 |
| CAK95_RS21255 | elongation factor Tu | 1235591 |
| HT745_RS03800 | elongation factor Tu | 1402135 |
| HT745_RS03910 | elongation factor Tu | 1402135 |
| KGB56_RS13535 | elongation factor Tu | 1898042 |
| KGB56_RS13605 | elongation factor Tu | 1898042 |
| C6Y53_RS10420 | elongation factor Tu | 2605946 |
| C6Y53_RS12620 | elongation factor Tu | 2605946 |
| HG718_RS04985 | elongation factor Tu | 1712261 |
| HG718_RS05060 | elongation factor Tu | 1712261 |
| FPZ52_RS00710 | elongation factor Tu | 2599296 |
| FPZ52_RS00785 | elongation factor Tu | 2599296 |
| K3148_RS11155 | elongation factor Tu | 2867233 |
| D0Y83_RS12970 | elongation factor Tu | 192812 |
| K3136_RS02690 | elongation factor Tu | 2867231 |
| SD421_RS08770 | elongation factor Tu | 3113984 |
| K3166_RS07455 | elongation factor Tu | 2867238 |
| E2O00_RS06135 | elongation factor Tu | 1532023 |
| CVE41_RS01185 | elongation factor Tu | 266951 |
| IRL76_RS07830 | elongation factor Tu | 2782568 |

| | | |
|---|---|---|
| L1F33_RS07835 | elongation factor Tu | 2909673 |
| CO657_RS06620 | elongation factor Tu | 1538158 |
| CO657_RS06695 | elongation factor Tu | 1538158 |
| NE851_RS18590 | elongation factor Tu | 2018689 |
| NE851_RS18670 | elongation factor Tu | 2018689 |
| J2J98_RS08260 | elongation factor Tu | 1138189 |
| J2J98_RS08335 | elongation factor Tu | 1138189 |
| J2J99_RS08495 | elongation factor Tu | 1138190 |
| J2J99_RS08570 | elongation factor Tu | 1138190 |
| RLCC275e_RS07205 | elongation factor Tu | 3019932 |
| RLCC275e_RS07280 | elongation factor Tu | 3019932 |
| ACLBKR_RS06480 | elongation factor Tu | 2813827 |
| ACLBKR_RS06560 | elongation factor Tu | 2813827 |
| GT349_RS21425 | elongation factor Tu | 240521 |
| GT349_RS21505 | elongation factor Tu | 240521 |
| AMJ99_RS08680 | elongation factor Tu | 1967781 |
| AMJ99_RS08755 | elongation factor Tu | 1967781 |
| RHE_RS08490 | elongation factor Tu | 347834 |
| RHE_RS08565 | elongation factor Tu | 347834 |
| LPU83_RS46015 | elongation factor Tu | 348824 |
| LPU83_RS46095 | elongation factor Tu | 348824 |
| RGR602_RS07695 | elongation factor Tu | 1041138 |
| RGR602_RS07775 | elongation factor Tu | 1041138 |
| FFM81_RS12730 | elongation factor Tu | 1538159 |
| FFM81_RS12805 | elongation factor Tu | 1538159 |
| FFM53_RS03920 | elongation factor Tu | 2583231 |
| FFM53_RS03995 | elongation factor Tu | 2583231 |
| U5G49_RS07270 | elongation factor Tu | 158891 |
| U5G49_RS07345 | elongation factor Tu | 158891 |
| CCGE525_RS08575 | elongation factor Tu | 1312183 |
| CCGE525_RS08670 | elongation factor Tu | 1312183 |

| | | |
|---|---|---|
| RL_RS09160 | elongation factor Tu | 216596 |
| RL_RS09235 | elongation factor Tu | 216596 |
| J0663_RS17225 | elongation factor Tu | 1138194 |
| J0663_RS17300 | elongation factor Tu | 1138194 |
| G3A56_RS00290 | elongation factor Tu | 2267833 |
| G3A56_RS00375 | elongation factor Tu | 2267833 |
| AMC88_RS08510 | elongation factor Tu | 396 |
| AMC88_RS08585 | elongation factor Tu | 396 |
| F3X89_RS05410 | elongation factor Tu | 359 |
| F3X89_RS05500 | elongation factor Tu | 359 |
| G6N80_RS19770 | elongation factor Tu | 451876 |
| G6N80_RS19845 | elongation factor Tu | 451876 |
| PR018_RS05275 | elongation factor Tu | 2506430 |
| PR018_RS05500 | elongation factor Tu | 2506430 |
| J3P73_RS09860 | elongation factor Tu | 2081791 |
| J3P73_RS09935 | elongation factor Tu | 2081791 |
| N2597_RS06685 | elongation factor Tu | 1535245 |
| N2597_RS06765 | elongation factor Tu | 1535245 |
| N2599_RS10060 | elongation factor Tu | 50338 |
| N2599_RS10145 | elongation factor Tu | 50338 |
| RTCIAT899_RS07245 | elongation factor Tu | 698761 |
| RTCIAT899_RS07340 | elongation factor Tu | 698761 |
| PR017_RS05495 | elongation factor Tu | 2041385 |
| PR017_RS05615 | elongation factor Tu | 2041385 |
| CMV14_RS07965 | elongation factor Tu | 1850238 |
| G6P88_RS18245 | elongation factor Tu | 2711156 |
| HRJ34_RS26445 | elongation factor Tu | 160791 |
| RCAP_RS00740 | elongation factor Tu | 272942 |
| RCAP_RS01530 | elongation factor Tu | 272942 |
| LPB142_RS13880 | elongation factor Tu | 1850250 |
| LPB142_RS13965 | elongation factor Tu | 1850250 |

| | | |
|---|---|---|
| RVAN_RS14960 | elongation factor Tu | 648757 |
| E0H22_RS16935 | elongation factor Tu | 475937 |
| E0H22_RS17065 | elongation factor Tu | 475937 |
| I8G32_RS16710 | elongation factor Tu | 1076 |
| I8G32_RS16870 | elongation factor Tu | 1076 |
| RC1_RS03265 | elongation factor Tu | 414684 |
| RC1_RS03345 | elongation factor Tu | 414684 |
| RRU_RS13890 | elongation factor Tu | 269796 |
| RRU_RS13955 | elongation factor Tu | 269796 |
| NBY65_RS14355 | elongation factor Tu | 504468 |
| A6W98_RS01255 | elongation factor Tu | 1188256 |
| A6W98_RS01330 | elongation factor Tu | 1188256 |
| A1C_RS04585 | elongation factor Tu | 293614 |
| RAS_RS05990 | elongation factor Tu | 238800 |
| MC5_RS02560 | elongation factor Tu | 1105110 |
| RBE_RS05525 | elongation factor Tu | 336407 |
| RCA_RS03800 | elongation factor Tu | 1105107 |
| RC_RS05040 | elongation factor Tu | 272944 |
| AAHM75_RS09110 | elongation factor Tu | 3066247 |
| AAGD46_RS02130 | elongation factor Tu | 3066248 |
| AAGD64_RS01935 | elongation factor Tu | 3066249 |
| AAGD55_RS08255 | elongation factor Tu | 3066266 |
| AAGD49_RS05700 | elongation factor Tu | 3066269 |
| ACOAL3_RS01570 | elongation factor Tu | 3066268 |
| NBW66_RS10515 | elongation factor Tu | 931099 |
| ACICQC_RS07440 | elongation factor Tu | 3066267 |
| AAGD48_RS06895 | elongation factor Tu | 3066251 |
| AAGD54_RS06930 | elongation factor Tu | 3066252 |
| AAHF54_RS05585 | elongation factor Tu | 3066270 |
| AB1146_RS00530 | elongation factor Tu | 35789 |
| RJP_RS05240 | elongation factor Tu | 652620 |

| | | |
|---|---|---|
| RMONA_RS06875 | elongation factor Tu | 109232 |
| M9W_RS03340 | elongation factor Tu | 1105094 |
| MCC_RS05715 | elongation factor Tu | 1105113 |
| A1G_RS04990 | elongation factor Tu | 392021 |
| RSIB_RS01220 | elongation factor Tu | 272951 |
| RSL_RS05210 | elongation factor Tu | 941638 |
| H6P87_RS05360 | elongation factor Tu | 2761623 |
| RT_RS03230 | elongation factor Tu | 257363 |
| EF513_RS04765 | elongation factor Tu | 2486578 |
| EF513_RS05165 | elongation factor Tu | 2486578 |
| SADFL11_RS02370 | elongation factor Tu | 244592 |
| SADFL11_RS02445 | elongation factor Tu | 244592 |
| B0E33_RS24450 | elongation factor Tu | 2857014 |
| B0E33_RS24520 | elongation factor Tu | 2857014 |
| K1718_RS13455 | elongation factor Tu | 2866279 |
| K1718_RS13525 | elongation factor Tu | 2866279 |
| ROSELON_RS07190 | elongation factor Tu | 1294273 |
| ROSELON_RS07310 | elongation factor Tu | 1294273 |
| E0E05_RS08185 | elongation factor Tu | 1852022 |
| E0E05_RS08295 | elongation factor Tu | 1852022 |
| CEP88_RS09295 | elongation factor Tu | 2434 |
| CEP88_RS16910 | elongation factor Tu | 2434 |
| ROLI_RS16035 | elongation factor Tu | 1537242 |
| ROLI_RS17250 | elongation factor Tu | 1537242 |
| RLO149_RS02105 | elongation factor Tu | 391595 |
| RLO149_RS16475 | elongation factor Tu | 391595 |
| G3256_RS02785 | elongation factor Tu | 1891787 |
| G3256_RS02875 | elongation factor Tu | 1891787 |
| ICW72_RS17835 | elongation factor Tu | 2771361 |
| ICW72_RS17900 | elongation factor Tu | 2771361 |
| LPC08_RS06285 | elongation factor Tu | 2897332 |

| | | |
|---|---|---|
| LPC08_RS14810 | elongation factor Tu | 2897332 |
| MWM08_RS20550 | elongation factor Tu | 1750527 |
| MWM08_RS20635 | elongation factor Tu | 1750527 |
| RGI145_RS13900 | elongation factor Tu | 257708 |
| RGI145_RS13980 | elongation factor Tu | 257708 |
| IAI59_RS01040 | elongation factor Tu | 2768162 |
| IAI59_RS01105 | elongation factor Tu | 2768162 |
| IAI58_RS04695 | elongation factor Tu | 2768161 |
| IAI58_RS04765 | elongation factor Tu | 2768161 |
| HVIM_RS20535 | elongation factor Tu | 207340 |
| HVIM_RS20620 | elongation factor Tu | 207340 |
| EI983_RS04825 | elongation factor Tu | 2494550 |
| RIdsm_RS01030 | elongation factor Tu | 540747 |
| RIdsm_RS19305 | elongation factor Tu | 540747 |
| N7U68_RS07255 | elongation factor Tu | 2980108 |
| N7U68_RS07350 | elongation factor Tu | 2980108 |
| RZ517_RS15660 | elongation factor Tu | 3080976 |
| RZ517_RS15915 | elongation factor Tu | 3080976 |
| RZS32_RS10910 | elongation factor Tu | 3080827 |
| RZS32_RS11130 | elongation factor Tu | 3080827 |
| PVU19_RS04570 | elongation factor Tu | 74031 |
| PVU19_RS13185 | elongation factor Tu | 74031 |
| SPO_RS03675 | elongation factor Tu | 246200 |
| SPO_RS17715 | elongation factor Tu | 246200 |
| ABFK29_RS21030 | elongation factor Tu | 388399 |
| ABFK29_RS21150 | elongation factor Tu | 388399 |
| BVL55_RS06020 | elongation factor Tu | 2712698 |
| BVL55_RS06325 | elongation factor Tu | 2712698 |
| Ga0080574_RS07840 | elongation factor Tu | 1250539 |
| Ga0080574_RS07945 | elongation factor Tu | 1250539 |
| Ga0080559_RS10040 | elongation factor Tu | 1229727 |

| | | |
|---|---|---|
| Ga0080559_RS10135 | elongation factor Tu | 1229727 |
| H3309_RS01290 | elongation factor Tu | 2759707 |
| R9Z33_RS07825 | elongation factor Tu | 1225128 |
| R9Z33_RS07910 | elongation factor Tu | 1225128 |
| K6301_RS05855 | elongation factor Tu | 2871820 |
| K6301_RS05930 | elongation factor Tu | 2871820 |
| Q9313_RS07760 | elongation factor Tu | 1967501 |
| Q9313_RS07835 | elongation factor Tu | 1967501 |
| K8M09_RS06635 | elongation factor Tu | 352475 |
| K8M09_RS06705 | elongation factor Tu | 352475 |
| EF888_RS00500 | elongation factor Tu | 1826607 |
| EF888_RS20300 | elongation factor Tu | 1826607 |
| EKH55_RS05325 | elongation factor Tu | 1752398 |
| EKH55_RS05405 | elongation factor Tu | 1752398 |
| SAMCFNEI73_RS06990 | elongation factor Tu | 194963 |
| SAMCFNEI73_RS07065 | elongation factor Tu | 194963 |
| RB548_RS05870 | elongation factor Tu | 501572 |
| RB548_RS05950 | elongation factor Tu | 501572 |
| AOX55_RS06160 | elongation factor Tu | 1128330 |
| AOX55_RS06235 | elongation factor Tu | 1128330 |
| PZN02_RS18820 | elongation factor Tu | 680247 |
| PZN02_RS18900 | elongation factor Tu | 680247 |
| PZL22_RS07465 | elongation factor Tu | 158892 |
| PZL22_RS07545 | elongation factor Tu | 158892 |
| V7S97_RS16390 | elongation factor Tu | 382 |
| V7S97_RS16470 | elongation factor Tu | 382 |
| FKV68_RS07100 | elongation factor Tu | 375549 |
| FKV68_RS07180 | elongation factor Tu | 375549 |
| PYH37_RS17925 | elongation factor Tu | 680248 |
| PYH37_RS18005 | elongation factor Tu | 680248 |
| SJ05684_RS05205 | elongation factor Tu | 716928 |

| | | |
|---|---|---|
| SJ05684_RS05610 | elongation factor Tu | 716928 |
| QA637_RS04860 | elongation factor Tu | 110322 |
| QA637_RS04940 | elongation factor Tu | 110322 |
| IGS68_RS07595 | elongation factor Tu | 2775420 |
| IGS68_RS20115 | elongation factor Tu | 2775420 |
| JL100_RS06025 | elongation factor Tu | 1789672 |
| JL100_RS19965 | elongation factor Tu | 1789672 |
| DPR14_RS05590 | elongation factor Tu | 2233999 |
| DPR14_RS18490 | elongation factor Tu | 2233999 |
| JL101_RS05895 | elongation factor Tu | 1817965 |
| JL101_RS18725 | elongation factor Tu | 1817965 |
| OIR97_RS13250 | elongation factor Tu | 2599305 |
| OIR97_RS13315 | elongation factor Tu | 2599305 |
| NBZ79_RS13915 | elongation factor Tu | 2950108 |
| NBZ79_RS13980 | elongation factor Tu | 2950108 |
| SAMIE_RS14215 | elongation factor Tu | 135719 |
| SCLO_RS10695 | elongation factor Tu | 120107 |
| H5V43_RS12290 | elongation factor Tu | 336203 |
| SJA_RS02950 | elongation factor Tu | 452662 |
| ACGGKE_RS09725 | elongation factor Tu | 1886786 |
| KFK14_RS01550 | elongation factor Tu | 1592790 |
| CJD35_RS08540 | elongation factor Tu | 121428 |
| A6768_RS18830 | elongation factor Tu | 13690 |
| NUW81_RS04685 | elongation factor Tu | 1582356 |
| NVV54_RS07545 | elongation factor Tu | 1229164 |
| PBT88_RS04125 | elongation factor Tu | 3012344 |
| NEF64_RS06595 | elongation factor Tu | 185951 |
| AB3474_RS12850 | elongation factor Tu | 2762591 |
| H5J25_RS05990 | elongation factor Tu | 2759526 |
| SBA_RS05300 | elongation factor Tu | 296544 |
| LZK98_RS18190 | elongation factor Tu | 2899123 |

| | | |
|---|---|---|
| ACLN6N_RS14940 | elongation factor Tu | 1166323 |
| GVO57_RS00290 | elongation factor Tu | 2698679 |
| M9980_RS02535 | elongation factor Tu | 2949093 |
| NV382_RS17525 | elongation factor Tu | 2972485 |
| QU596_RS05725 | elongation factor Tu | 3132797 |
| GCU42_RS05775 | elongation factor Tu | 363835 |
| M1K48_RS13440 | elongation factor Tu | 2938948 |
| TS85_RS15400 | elongation factor Tu | 1609977 |
| GTH33_RS17390 | elongation factor Tu | 424800 |
| BRX40_RS18015 | elongation factor Tu | 93064 |
| GV829_RS05255 | elongation factor Tu | 2698828 |
| J3P39_RS10940 | elongation factor Tu | 1045317 |
| LHA26_RS10965 | elongation factor Tu | 1541170 |
| PQ455_RS15165 | elongation factor Tu | 1813468 |
| K8P63_RS19510 | elongation factor Tu | 2872652 |
| D3Y57_RS09075 | elongation factor Tu | 2319844 |
| J0A91_RS22795 | elongation factor Tu | 1560345 |
| FPZ24_RS05700 | elongation factor Tu | 1813879 |
| I6G65_RS06110 | elongation factor Tu | 13689 |
| G7077_RS10630 | elongation factor Tu | 2714943 |
| GNT64_RS01405 | elongation factor Tu | 2681549 |
| CVN68_RS20655 | elongation factor Tu | 1327635 |
| NMP03_RS01300 | elongation factor Tu | 2918765 |
| LLW23_RS03175 | elongation factor Tu | 2890321 |
| H9L12_RS08580 | elongation factor Tu | 2071607 |
| H8M03_RS05200 | elongation factor Tu | 2764186 |
| NX02_RS19635 | elongation factor Tu | 1123269 |
| NX02_RS27400 | elongation factor Tu | 1123269 |
| H9L14_RS09925 | elongation factor Tu | 386874 |
| G7078_RS04115 | elongation factor Tu | 2714944 |
| FPZ54_RS01120 | elongation factor Tu | 2599297 |

| | | |
|---|---|---|
| MC45_RS11865 | elongation factor Tu | 1549858 |
| FMM02_RS03440 | elongation factor Tu | 2594473 |
| SALA_RS14355 | elongation factor Tu | 317655 |
| VSX79_RS17075 | elongation factor Tu | 180400 |
| SKP52_RS20675 | elongation factor Tu | 1515612 |
| SGRAN_RS01495 | elongation factor Tu | 267128 |
| KNJ79_RS21880 | elongation factor Tu | 436663 |
| VSX77_RS07515 | elongation factor Tu | 3114367 |
| CVO77_RS11055 | elongation factor Tu | 2054227 |
| LH19_RS23515 | elongation factor Tu | 33050 |
| EUU25_RS14485 | elongation factor Tu | 392610 |
| LPB140_RS11720 | elongation factor Tu | 1913578 |
| IC614_RS06995 | elongation factor Tu | 2771430 |
| G5C33_RS16830 | elongation factor Tu | 2711215 |
| RPR59_RS06305 | elongation factor Tu | 3075605 |
| GH266_RS04825 | elongation factor Tu | 538381 |
| GH266_RS04890 | elongation factor Tu | 538381 |
| STVA_RS04145 | elongation factor Tu | 94 |
| STVA_RS04210 | elongation factor Tu | 94 |
| KDD17_RS12155 | elongation factor Tu | 2829501 |
| KDD17_RS12280 | elongation factor Tu | 2829501 |
| BOO69_RS01860 | elongation factor Tu | 1917485 |
| BOO69_RS01960 | elongation factor Tu | 1917485 |
| T7987_RS02695 | elongation factor Tu | 1775881 |
| T7987_RS14775 | elongation factor Tu | 1775881 |
| DSM14862_RS02500 | elongation factor Tu | 225422 |
| DSM14862_RS14490 | elongation factor Tu | 225422 |
| JNX02_RS10720 | elongation factor Tu | 83219 |
| JNX02_RS10845 | elongation factor Tu | 83219 |
| AB1495_RS03455 | elongation factor Tu | 60137 |
| AB1495_RS03530 | elongation factor Tu | 60137 |

| | | |
|---|---|---|
| E3D00_RS02015 | elongation factor Tu | 1293412 |
| EI545_RS08100 | elongation factor Tu | 2494374 |
| EI545_RS08185 | elongation factor Tu | 2494374 |
| BSL82_RS16925 | elongation factor Tu | 1921510 |
| RPMA_RS17805 | elongation factor Tu | 340268 |
| RPMA_RS18010 | elongation factor Tu | 340268 |
| QGN29_RS00245 | elongation factor Tu | 1456591 |
| QGN29_RS00315 | elongation factor Tu | 1456591 |
| FZF13_RS25360 | elongation factor Tu | 2725666 |
| FZF13_RS25465 | elongation factor Tu | 2725666 |
| DSM104635_RS09090 | elongation factor Tu | 2686094 |
| IZ6_RS09770 | elongation factor Tu | 708113 |
| IZ6_RS09850 | elongation factor Tu | 708113 |
| L8F45_RS05700 | elongation factor Tu | 2895980 |
| DY252_RS08810 | elongation factor Tu | 1891279 |
| DY252_RS08875 | elongation factor Tu | 1891279 |
| CSC3H3_RS05755 | elongation factor Tu | 2048283 |
| CSC3H3_RS05820 | elongation factor Tu | 2048283 |
| TH3_RS05975 | elongation factor Tu | 1123366 |
| TH3_RS06040 | elongation factor Tu | 1123366 |
| HFZ77_RS07920 | elongation factor Tu | 53501 |
| HFZ77_RS13340 | elongation factor Tu | 53501 |
| AKL02_RS18315 | elongation factor Tu | 1549850 |
| AKL02_RS18480 | elongation factor Tu | 1549850 |
| RPE78_RS03390 | elongation factor Tu | 3076557 |
| RPE78_RS03475 | elongation factor Tu | 3076557 |
| BMG03_RS17835 | elongation factor Tu | 1915078 |
| BMG03_RS17990 | elongation factor Tu | 1915078 |
| P7L64_RS07015 | elongation factor Tu | 657419 |
| P7L64_RS07080 | elongation factor Tu | 657419 |
| PJ900_RS23335 | elongation factor Tu | 171437 |

| | | |
|---|---|---|
| PJ900_RS23405 | elongation factor Tu | 171437 |
| K529_RS00830 | elongation factor Tu | 1265309 |
| K529_RS00925 | elongation factor Tu | 1265309 |
| R1T40_RS14300 | elongation factor Tu | 483013 |
| R1T40_RS19130 | elongation factor Tu | 483013 |
| QF118_RS06520 | elongation factor Tu | 3058420 |
| QF118_RS06625 | elongation factor Tu | 3058420 |
| D4766_RS02205 | elongation factor Tu | 2338327 |
| A6F68_RS06290 | elongation factor Tu | 692370 |
| CJO11_RS05600 | elongation factor Tu | 1982042 |
| GJW30_RS17980 | elongation factor Tu | 1333996 |
| GJW30_RS18115 | elongation factor Tu | 1333996 |
| P3M64_RS12630 | elongation factor Tu | 1773489 |
| P3M64_RS12695 | elongation factor Tu | 1773489 |
| OOL03_RS04490 | elongation factor Tu | 2953971 |
| OOL03_RS05440 | elongation factor Tu | 2953971 |
| AAGD56_RS03085 | elongation factor Tu | 3066130 |
| AAGD56_RS04340 | elongation factor Tu | 3066130 |
| ABWU67_RS02310 | elongation factor Tu | 3077919 |
| ABWU67_RS06045 | elongation factor Tu | 3077919 |
| ABWU31_RS01565 | elongation factor Tu | 3066188 |
| ABWU31_RS05050 | elongation factor Tu | 3066188 |
| OOK90_RS01615 | elongation factor Tu | 2953974 |
| OOK90_RS04825 | elongation factor Tu | 2953974 |
| AAGD71_RS01955 | elongation factor Tu | 3066189 |
| AAGD71_RS05590 | elongation factor Tu | 3066189 |
| AAGD80_RS00295 | elongation factor Tu | 3066190 |
| AAGD80_RS04240 | elongation factor Tu | 3066190 |
| OPR53_RS01825 | elongation factor Tu | 2953975 |
| OPR53_RS05435 | elongation factor Tu | 2953975 |
| AAGD84_RS02460 | elongation factor Tu | 3066191 |

| | | |
|---|---|---|
| AAGD84_RS05735 | elongation factor Tu | 3066191 |
| OOK94_RS02565 | elongation factor Tu | 2953976 |
| OOK94_RS05410 | elongation factor Tu | 2953976 |
| ABWU73_RS02135 | elongation factor Tu | 3066192 |
| ABWU73_RS06905 | elongation factor Tu | 3066192 |
| ABWU57_RS01535 | elongation factor Tu | 3066193 |
| ABWU57_RS05980 | elongation factor Tu | 3066193 |
| OPR57_RS02555 | elongation factor Tu | 2953978 |
| OPR57_RS03095 | elongation factor Tu | 2953978 |
| OPR26_RS01515 | elongation factor Tu | 2953979 |
| OPR26_RS02440 | elongation factor Tu | 2953979 |
| ABWU75_RS01520 | elongation factor Tu | 3066194 |
| ABWU75_RS04190 | elongation factor Tu | 3066194 |
| OPR43_RS03130 | elongation factor Tu | 2953980 |
| OPR43_RS04380 | elongation factor Tu | 2953980 |
| ACICP7_RS01390 | elongation factor Tu | 3066195 |
| ACICP7_RS03995 | elongation factor Tu | 3066195 |
| AAGD50_RS01560 | elongation factor Tu | 3066165 |
| AAGD50_RS05160 | elongation factor Tu | 3066165 |
| ACQH1X_RS02590 | elongation factor Tu | 3139322 |
| ACQH1X_RS04130 | elongation factor Tu | 3139322 |
| ABWU69_RS02905 | elongation factor Tu | 3066139 |
| ABWU69_RS04190 | elongation factor Tu | 3066139 |
| OPR48_RS03740 | elongation factor Tu | 2953987 |
| OPR48_RS05000 | elongation factor Tu | 2953987 |
| OPR49_RS02045 | elongation factor Tu | 2953988 |
| OPR49_RS02510 | elongation factor Tu | 2953988 |
| AAGD93_RS00810 | elongation factor Tu | 3066140 |
| AAGD93_RS04970 | elongation factor Tu | 3066140 |
| OPR55_RS01430 | elongation factor Tu | 2953989 |
| OPR55_RS05575 | elongation factor Tu | 2953989 |

| | | |
|---|---|---|
| AAGD74_RS02020 | elongation factor Tu | 3066196 |
| AAGD74_RS05415 | elongation factor Tu | 3066196 |
| AAGD59_RS01435 | elongation factor Tu | 3077920 |
| AAGD59_RS04985 | elongation factor Tu | 3077920 |
| ACQH2I_RS01980 | elongation factor Tu | 3139309 |
| ACQH2I_RS02750 | elongation factor Tu | 3139309 |
| AAGD72_RS01585 | elongation factor Tu | 3066197 |
| AAGD72_RS02830 | elongation factor Tu | 3066197 |
| OOK91_RS01460 | elongation factor Tu | 2953995 |
| OOK91_RS04955 | elongation factor Tu | 2953995 |
| AAGD99_RS01530 | elongation factor Tu | 3066133 |
| AAGD99_RS05295 | elongation factor Tu | 3066133 |
| ACFELZ_RS01715 | elongation factor Tu | 3139321 |
| ACFELZ_RS05900 | elongation factor Tu | 3139321 |
| AAGD69_RS02600 | elongation factor Tu | 3066143 |
| AAGD69_RS04605 | elongation factor Tu | 3066143 |
| OPR62_RS01500 | elongation factor Tu | 2953999 |
| OPR62_RS02790 | elongation factor Tu | 2953999 |
| ABWU55_RS03565 | elongation factor Tu | 3066144 |
| ABWU55_RS04440 | elongation factor Tu | 3066144 |
| ABWU40_RS01460 | elongation factor Tu | 3066169 |
| ABWU40_RS04650 | elongation factor Tu | 3066169 |
| AAGD76_RS02475 | elongation factor Tu | 3066170 |
| AAGD76_RS02930 | elongation factor Tu | 3066170 |
| AAGD66_RS02750 | elongation factor Tu | 3066171 |
| AAGD66_RS05040 | elongation factor Tu | 3066171 |
| OPR31_RS03565 | elongation factor Tu | 2954002 |
| OPR31_RS04505 | elongation factor Tu | 2954002 |
| OPQ79_RS05070 | elongation factor Tu | 2954004 |
| OPQ79_RS05600 | elongation factor Tu | 2954004 |
| ABWU41_RS03175 | elongation factor Tu | 3066174 |

| | | |
|---|---|---|
| ABWU41_RS06135 | elongation factor Tu | 3066174 |
| OPR50_RS02460 | elongation factor Tu | 2954006 |
| OPR50_RS06555 | elongation factor Tu | 2954006 |
| OPR51_RS01490 | elongation factor Tu | 2954007 |
| OPR51_RS04325 | elongation factor Tu | 2954007 |
| OPR28_RS03070 | elongation factor Tu | 2954008 |
| OPR28_RS06405 | elongation factor Tu | 2954008 |
| AAGD62_RS02705 | elongation factor Tu | 3066145 |
| AAGD62_RS04805 | elongation factor Tu | 3066145 |
| OPR40_RS01455 | elongation factor Tu | 2954015 |
| OPR40_RS05720 | elongation factor Tu | 2954015 |
| ABWU47_RS02490 | elongation factor Tu | 3066200 |
| ABWU47_RS05440 | elongation factor Tu | 3066200 |
| OOK97_RS02045 | elongation factor Tu | 2954016 |
| OOK97_RS04245 | elongation factor Tu | 2954016 |
| ABWU23_RS02410 | elongation factor Tu | 3077921 |
| ABWU23_RS06550 | elongation factor Tu | 3077921 |
| OPR52_RS01775 | elongation factor Tu | 2954018 |
| OPR52_RS03140 | elongation factor Tu | 2954018 |
| OPR17_RS02465 | elongation factor Tu | 2954019 |
| OPR17_RS03780 | elongation factor Tu | 2954019 |
| AAGD90_RS01780 | elongation factor Tu | 3077922 |
| AAGD90_RS05300 | elongation factor Tu | 3077922 |
| ABWU71_RS02125 | elongation factor Tu | 3066201 |
| ABWU71_RS04880 | elongation factor Tu | 3066201 |
| OPR16_RS03035 | elongation factor Tu | 2954023 |
| OPR16_RS04675 | elongation factor Tu | 2954023 |
| OOK89_RS02170 | elongation factor Tu | 2954025 |
| OOK89_RS04720 | elongation factor Tu | 2954025 |
| AAGD58_RS01290 | elongation factor Tu | 3066202 |
| AAGD58_RS02115 | elongation factor Tu | 3066202 |

| | | |
|---|---|---|
| ACE38P_RS03565 | elongation factor Tu | 3139318 |
| ACE38P_RS07025 | elongation factor Tu | 3139318 |
| AABM58_RS01110 | elongation factor Tu | 3066134 |
| AABM58_RS02215 | elongation factor Tu | 3066134 |
| ABWU72_RS03205 | elongation factor Tu | 3066146 |
| ABWU72_RS07600 | elongation factor Tu | 3066146 |
| AAGD68_RS02315 | elongation factor Tu | 3066147 |
| AAGD68_RS03485 | elongation factor Tu | 3066147 |
| OPR61_RS02460 | elongation factor Tu | 2954028 |
| OPR61_RS03185 | elongation factor Tu | 2954028 |
| ACQH2Q_RS00940 | elongation factor Tu | 3139311 |
| ACQH2Q_RS02235 | elongation factor Tu | 3139311 |
| OPR23_RS01620 | elongation factor Tu | 2954031 |
| OPR23_RS04880 | elongation factor Tu | 2954031 |
| ACQH2M_RS03735 | elongation factor Tu | 3139310 |
| ACQH2M_RS05650 | elongation factor Tu | 3139310 |
| AAGD91_RS00815 | elongation factor Tu | 3066148 |
| AAGD91_RS03290 | elongation factor Tu | 3066148 |
| OPQ78_RS01585 | elongation factor Tu | 2954033 |
| OPQ78_RS05250 | elongation factor Tu | 2954033 |
| AAGD57_RS00690 | elongation factor Tu | 3066204 |
| AAGD57_RS04100 | elongation factor Tu | 3066204 |
| AAGD70_RS04415 | elongation factor Tu | 3066205 |
| AAGD70_RS05730 | elongation factor Tu | 3066205 |
| ABWU43_RS01505 | elongation factor Tu | 3066206 |
| ABWU43_RS04430 | elongation factor Tu | 3066206 |
| AAGD98_RS03720 | elongation factor Tu | 3066207 |
| AAGD98_RS04480 | elongation factor Tu | 3066207 |
| AACH14_RS03475 | elongation factor Tu | 3066208 |
| AACH14_RS06730 | elongation factor Tu | 3066208 |
| ACICP3_RS01590 | elongation factor Tu | 3077923 |

| | | |
|---|---|---|
| ACICP3_RS05065 | elongation factor Tu | 3077923 |
| ACFZP3_RS00530 | elongation factor Tu | 3066150 |
| ACFZP3_RS03125 | elongation factor Tu | 3066150 |
| AACH21_RS02125 | elongation factor Tu | 3066209 |
| AACH21_RS05385 | elongation factor Tu | 3066209 |
| AACK80_RS01935 | elongation factor Tu | 3066210 |
| AACK80_RS05435 | elongation factor Tu | 3066210 |
| ACFEL3_RS00575 | elongation factor Tu | 3077917 |
| ACFEL3_RS04160 | elongation factor Tu | 3077917 |
| AAGD79_RS04440 | elongation factor Tu | 3066211 |
| AAGD79_RS05245 | elongation factor Tu | 3066211 |
| ABWU74_RS03945 | elongation factor Tu | 3066212 |
| ABWU74_RS05905 | elongation factor Tu | 3066212 |
| OOK78_RS01305 | elongation factor Tu | 2954042 |
| OOK78_RS04590 | elongation factor Tu | 2954042 |
| ABWU29_RS02770 | elongation factor Tu | 3066152 |
| ABWU29_RS05635 | elongation factor Tu | 3066152 |
| OPR38_RS01505 | elongation factor Tu | 2954044 |
| OPR38_RS04960 | elongation factor Tu | 2954044 |
| AAGD87_RS05485 | elongation factor Tu | 3066153 |
| AAGD87_RS05860 | elongation factor Tu | 3066153 |
| ABWU68_RS01195 | elongation factor Tu | 3077924 |
| ABWU68_RS04610 | elongation factor Tu | 3077924 |
| OPR21_RS01755 | elongation factor Tu | 2954046 |
| OPR21_RS06835 | elongation factor Tu | 2954046 |
| ABWU50_RS01750 | elongation factor Tu | 3066154 |
| ABWU50_RS03110 | elongation factor Tu | 3066154 |
| ABWU54_RS00130 | elongation factor Tu | 3066136 |
| ABWU54_RS04020 | elongation factor Tu | 3066136 |
| ABWU36_RS01395 | elongation factor Tu | 3066155 |
| ABWU36_RS04665 | elongation factor Tu | 3066155 |

| | | |
|---|---|---|
| OPR63_RS01500 | elongation factor Tu | 2954050 |
| OPR63_RS04985 | elongation factor Tu | 2954050 |
| OOK92_RS00380 | elongation factor Tu | 2954053 |
| OOK92_RS04295 | elongation factor Tu | 2954053 |
| ABWU27_RS01805 | elongation factor Tu | 3066213 |
| ABWU27_RS04865 | elongation factor Tu | 3066213 |
| OPR46_RS01850 | elongation factor Tu | 2954055 |
| OPR46_RS04945 | elongation factor Tu | 2954055 |
| ACQH2N_RS01245 | elongation factor Tu | 3139314 |
| ACQH2N_RS02410 | elongation factor Tu | 3139314 |
| ACQH1Y_RS02295 | elongation factor Tu | 3139319 |
| ACQH1Y_RS05360 | elongation factor Tu | 3139319 |
| OPR24_RS02595 | elongation factor Tu | 2954056 |
| OPR24_RS05000 | elongation factor Tu | 2954056 |
| OOK84_RS02005 | elongation factor Tu | 2954057 |
| OOK84_RS05320 | elongation factor Tu | 2954057 |
| OPR25_RS01575 | elongation factor Tu | 2954060 |
| OPR25_RS04290 | elongation factor Tu | 2954060 |
| ACQH3L_RS01525 | elongation factor Tu | 3139315 |
| ACQH3L_RS04460 | elongation factor Tu | 3139315 |
| OOK87_RS00390 | elongation factor Tu | 2954061 |
| OOK87_RS05175 | elongation factor Tu | 2954061 |
| ACICQD_RS01280 | elongation factor Tu | 3066214 |
| ACICQD_RS03990 | elongation factor Tu | 3066214 |
| OPR34_RS01640 | elongation factor Tu | 2954063 |
| OPR34_RS04785 | elongation factor Tu | 2954063 |
| ABWU48_RS04255 | elongation factor Tu | 3066215 |
| ABWU48_RS05205 | elongation factor Tu | 3066215 |
| OPR60_RS02880 | elongation factor Tu | 2954064 |
| OPR60_RS05865 | elongation factor Tu | 2954064 |
| ABWU32_RS02080 | elongation factor Tu | 3066183 |

| | | |
|---|---|---|
| ABWU32_RS05980 | elongation factor Tu | 3066183 |
| ABWU38_RS05305 | elongation factor Tu | 3066156 |
| ABWU38_RS05800 | elongation factor Tu | 3066156 |
| AAGD51_RS02200 | elongation factor Tu | 3066158 |
| AAGD51_RS02495 | elongation factor Tu | 3066158 |
| OPR39_RS03810 | elongation factor Tu | 2954068 |
| OPR39_RS05730 | elongation factor Tu | 2954068 |
| ACICP1_RS01195 | elongation factor Tu | 3077915 |
| ACICP1_RS04375 | elongation factor Tu | 3077915 |
| ABWU66_RS00695 | elongation factor Tu | 3077918 |
| ABWU66_RS03710 | elongation factor Tu | 3077918 |
| OOL10_RS01615 | elongation factor Tu | 2953972 |
| OOL10_RS07005 | elongation factor Tu | 2953972 |
| AAGD77_RS01625 | elongation factor Tu | 3066161 |
| AAGD77_RS05470 | elongation factor Tu | 3066161 |
| ABWU46_RS01430 | elongation factor Tu | 3066162 |
| ABWU46_RS04390 | elongation factor Tu | 3066162 |
| ABWU53_RS01245 | elongation factor Tu | 3066163 |
| ABWU53_RS04545 | elongation factor Tu | 3066163 |
| ACQH1A_RS01435 | elongation factor Tu | 3139312 |
| ACQH1A_RS04745 | elongation factor Tu | 3139312 |
| ABWU39_RS02135 | elongation factor Tu | 3066164 |
| ABWU39_RS05785 | elongation factor Tu | 3066164 |
| OOK79_RS00690 | elongation factor Tu | 2953981 |
| OOK79_RS03585 | elongation factor Tu | 2953981 |
| OOK82_RS00180 | elongation factor Tu | 2953982 |
| OOK82_RS03675 | elongation factor Tu | 2953982 |
| OPR20_RS01370 | elongation factor Tu | 2953983 |
| OPR20_RS05690 | elongation factor Tu | 2953983 |
| OOL02_RS01795 | elongation factor Tu | 2953984 |
| OOL02_RS07020 | elongation factor Tu | 2953984 |

| | | |
|---|---|---|
| AAGE00_RS01760 | elongation factor Tu | 3066166 |
| AAGE00_RS06190 | elongation factor Tu | 3066166 |
| OPR30_RS01620 | elongation factor Tu | 2953986 |
| OPR30_RS04940 | elongation factor Tu | 2953986 |
| OOL08_RS01580 | elongation factor Tu | 2953990 |
| OOL08_RS05535 | elongation factor Tu | 2953990 |
| OPR44_RS06065 | elongation factor Tu | 2953991 |
| OPR44_RS06960 | elongation factor Tu | 2953991 |
| OOK86_RS02880 | elongation factor Tu | 2953993 |
| OOK86_RS06185 | elongation factor Tu | 2953993 |
| AAGD65_RS01420 | elongation factor Tu | 3066167 |
| AAGD65_RS05410 | elongation factor Tu | 3066167 |
| OPS21_RS01865 | elongation factor Tu | 2953997 |
| OPS21_RS02605 | elongation factor Tu | 2953997 |
| OOK80_RS02210 | elongation factor Tu | 2953998 |
| OOK80_RS03865 | elongation factor Tu | 2953998 |
| ABWU33_RS01855 | elongation factor Tu | 3066168 |
| ABWU33_RS06505 | elongation factor Tu | 3066168 |
| AAGD67_RS01600 | elongation factor Tu | 3066172 |
| AAGD67_RS05975 | elongation factor Tu | 3066172 |
| AAGD75_RS03130 | elongation factor Tu | 3066173 |
| AAGD75_RS06375 | elongation factor Tu | 3066173 |
| OPR56_RS01265 | elongation factor Tu | 2954003 |
| OPR56_RS04975 | elongation factor Tu | 2954003 |
| OPR54_RS01360 | elongation factor Tu | 2954005 |
| OPR54_RS05965 | elongation factor Tu | 2954005 |
| AAGD95_RS03460 | elongation factor Tu | 3066175 |
| AAGD95_RS04830 | elongation factor Tu | 3066175 |
| OOL05_RS01165 | elongation factor Tu | 2954009 |
| OOL05_RS04320 | elongation factor Tu | 2954009 |
| OOL04_RS01470 | elongation factor Tu | 2954011 |

| | | |
|---|---|---|
| OOL04_RS04295 | elongation factor Tu | 2954011 |
| OOK99_RS01510 | elongation factor Tu | 2954012 |
| OOK99_RS04940 | elongation factor Tu | 2954012 |
| OOK93_RS01645 | elongation factor Tu | 2954013 |
| OOK93_RS04440 | elongation factor Tu | 2954013 |
| OOK83_RS01805 | elongation factor Tu | 2954014 |
| OOK83_RS04825 | elongation factor Tu | 2954014 |
| ACQH21_RS06805 | elongation factor Tu | 3139316 |
| ACQH21_RS07295 | elongation factor Tu | 3139316 |
| ABWU61_RS01245 | elongation factor Tu | 3066176 |
| ABWU61_RS04780 | elongation factor Tu | 3066176 |
| AAGD63_RS01335 | elongation factor Tu | 3077916 |
| AAGD63_RS01715 | elongation factor Tu | 3077916 |
| ABWU62_RS03625 | elongation factor Tu | 3066159 |
| ABWU62_RS07425 | elongation factor Tu | 3066159 |
| ABWU24_RS02615 | elongation factor Tu | 3066177 |
| ABWU24_RS05605 | elongation factor Tu | 3066177 |
| ABWU56_RS01755 | elongation factor Tu | 3066178 |
| ABWU56_RS05600 | elongation factor Tu | 3066178 |
| OPR32_RS01420 | elongation factor Tu | 2954017 |
| OPR32_RS04680 | elongation factor Tu | 2954017 |
| AAGD97_RS01910 | elongation factor Tu | 3066179 |
| AAGD97_RS06605 | elongation factor Tu | 3066179 |
| OPR58_RS03875 | elongation factor Tu | 2954021 |
| OPR58_RS06185 | elongation factor Tu | 2954021 |
| ACE38O_RS02295 | elongation factor Tu | 3139317 |
| ACE38O_RS04065 | elongation factor Tu | 3139317 |
| ABWU45_RS05940 | elongation factor Tu | 3066135 |
| ABWU45_RS06670 | elongation factor Tu | 3066135 |
| OPR37_RS01310 | elongation factor Tu | 2954027 |
| OPR37_RS04615 | elongation factor Tu | 2954027 |

| | | |
|---|---|---|
| AAHF52_RS03025 | elongation factor Tu | 2954029 |
| AAHF52_RS06120 | elongation factor Tu | 2954029 |
| OPR42_RS02450 | elongation factor Tu | 2954030 |
| OPR42_RS05855 | elongation factor Tu | 2954030 |
| ACICQB_RS00490 | elongation factor Tu | 3066180 |
| ACICQB_RS04520 | elongation factor Tu | 3066180 |
| OOK76_RS01295 | elongation factor Tu | 2954034 |
| OOK76_RS04240 | elongation factor Tu | 2954034 |
| ACN1AB_RS01095 | elongation factor Tu | 2954036 |
| ACN1AB_RS03965 | elongation factor Tu | 2954036 |
| OPR29_RS01330 | elongation factor Tu | 2954037 |
| OPR29_RS03980 | elongation factor Tu | 2954037 |
| OOK98_RS01465 | elongation factor Tu | 2954039 |
| OOK98_RS05550 | elongation factor Tu | 2954039 |
| OOT12_RS01665 | elongation factor Tu | 2954040 |
| OOT12_RS05680 | elongation factor Tu | 2954040 |
| OOK88_RS01730 | elongation factor Tu | 2954041 |
| OOK88_RS04675 | elongation factor Tu | 2954041 |
| OPR70_RS00885 | elongation factor Tu | 2954043 |
| OPR70_RS03710 | elongation factor Tu | 2954043 |
| AAGD82_RS02255 | elongation factor Tu | 3066151 |
| AAGD82_RS05075 | elongation factor Tu | 3066151 |
| OPR22_RS01390 | elongation factor Tu | 2954045 |
| OPR22_RS04355 | elongation factor Tu | 2954045 |
| ABWU28_RS00940 | elongation factor Tu | 3066160 |
| ABWU28_RS05565 | elongation factor Tu | 3066160 |
| OOL06_RS02450 | elongation factor Tu | 2954049 |
| OOL06_RS03935 | elongation factor Tu | 2954049 |
| OPR35_RS01695 | elongation factor Tu | 2954051 |
| OPR35_RS02335 | elongation factor Tu | 2954051 |
| OOK81_RS01450 | elongation factor Tu | 2954052 |

| | | |
|---|---|---|
| OOK81_RS04545 | elongation factor Tu | 2954052 |
| OOL01_RS06550 | elongation factor Tu | 2954054 |
| OOL01_RS06925 | elongation factor Tu | 2954054 |
| AB3210_RS01595 | elongation factor Tu | 3139320 |
| AB3210_RS05190 | elongation factor Tu | 3139320 |
| ABWU65_RS03015 | elongation factor Tu | 3066137 |
| ABWU65_RS05895 | elongation factor Tu | 3066137 |
| ABWU60_RS01375 | elongation factor Tu | 3066181 |
| ABWU60_RS04375 | elongation factor Tu | 3066181 |
| OPR36_RS02115 | elongation factor Tu | 2954062 |
| OPR36_RS04945 | elongation factor Tu | 2954062 |
| ABWU63_RS01760 | elongation factor Tu | 3066182 |
| ABWU63_RS04680 | elongation factor Tu | 3066182 |
| ABWU52_RS01400 | elongation factor Tu | 3066157 |
| ABWU52_RS05730 | elongation factor Tu | 3066157 |
| OOL00_RS02045 | elongation factor Tu | 2954066 |
| OOL00_RS06130 | elongation factor Tu | 2954066 |
| ABWU64_RS01300 | elongation factor Tu | 3066184 |
| ABWU64_RS04815 | elongation factor Tu | 3066184 |
| OOK95_RS01580 | elongation factor Tu | 2954067 |
| OOK95_RS04930 | elongation factor Tu | 2954067 |
| ABWU35_RS01435 | elongation factor Tu | 3066185 |
| ABWU35_RS03370 | elongation factor Tu | 3066185 |
| AAGD73_RS01175 | elongation factor Tu | 3066186 |
| AAGD73_RS04055 | elongation factor Tu | 3066186 |
| AAGD89_RS02130 | elongation factor Tu | 3066149 |
| AAGD89_RS02460 | elongation factor Tu | 3066149 |
| J8Z11_RS00080 | elongation factor Tu | 546165 |
| J8Z11_RS03350 | elongation factor Tu | 546165 |
| NHG98_RS00200 | elongation factor Tu | 167957 |
| NHG98_RS05060 | elongation factor Tu | 167957 |

| | | |
|---|---|---|
| NOX90_RS01595 | elongation factor Tu | 2850562 |
| NOX90_RS06130 | elongation factor Tu | 2850562 |
| ABLO99_RS01380 | elongation factor Tu | 3158571 |
| ABLO99_RS02865 | elongation factor Tu | 3158571 |
| BBB02_RS01385 | elongation factor Tu | 215173 |
| BBB02_RS04430 | elongation factor Tu | 215173 |
| WBP_RS02075 | elongation factor Tu | 96495 |
| WBP_RS03750 | elongation factor Tu | 96495 |
| ACICP6_RS01345 | elongation factor Tu | 3066132 |
| ACICP6_RS05120 | elongation factor Tu | 3066132 |
| wCauA_RS02865 | elongation factor Tu | 2591635 |
| wCauA_RS03670 | elongation factor Tu | 2591635 |
| HUB95_RS02275 | elongation factor Tu | 497299 |
| HUB95_RS04945 | elongation factor Tu | 497299 |
| CAI20_RS00180 | elongation factor Tu | 1335053 |
| CAI20_RS03390 | elongation factor Tu | 1335053 |
| WCLE_RS01075 | elongation factor Tu | 246273 |
| WCLE_RS04815 | elongation factor Tu | 246273 |
| GOY13_RS01305 | elongation factor Tu | 1812111 |
| GOY13_RS03055 | elongation factor Tu | 1812111 |
| HF196_RS02655 | elongation factor Tu | 2732594 |
| HF196_RS05475 | elongation factor Tu | 2732594 |
| HF197_RS00650 | elongation factor Tu | 2732593 |
| HF197_RS03545 | elongation factor Tu | 2732593 |
| WP_RS02770 | elongation factor Tu | 570417 |
| WP_RS05830 | elongation factor Tu | 570417 |
| L2227_RS01490 | elongation factor Tu | 502352 |
| L2227_RS06875 | elongation factor Tu | 502352 |
| FK497_RS01590 | elongation factor Tu | 116598 |
| FK497_RS04535 | elongation factor Tu | 116598 |
| GOY14_RS00865 | elongation factor Tu | 1812112 |

| | | |
|---|---|---|
| GOY14_RS02445 | elongation factor Tu | 1812112 |
| GOY12_RS01015 | elongation factor Tu | 1812115 |
| GOY12_RS02755 | elongation factor Tu | 1812115 |
| R0F62_RS00735 | elongation factor Tu | 3080329 |
| R0F62_RS04970 | elongation factor Tu | 3080329 |
| EA652_RS00075 | elongation factor Tu | 307502 |
| EA652_RS02735 | elongation factor Tu | 307502 |
| WG67_RS00075 | elongation factor Tu | 1633785 |
| WG67_RS02900 | elongation factor Tu | 1633785 |
| EJB00_RS01550 | elongation factor Tu | 109663 |
| EJB00_RS04930 | elongation factor Tu | 109663 |
| GQX67_RS00085 | elongation factor Tu | 163164 |
| GQX67_RS03325 | elongation factor Tu | 163164 |
| ONI09_RS00080 | elongation factor Tu | 375919 |
| ONI09_RS02850 | elongation factor Tu | 375919 |
| wSan_RS00080 | elongation factor Tu | 260917 |
| wSan_RS02490 | elongation factor Tu | 260917 |
| WHA_RS00075 | elongation factor Tu | 1236909 |
| WHA_RS03120 | elongation factor Tu | 1236909 |
| ABL518_RS02075 | elongation factor Tu | 295084 |
| ABL518_RS05315 | elongation factor Tu | 295084 |
| wYak_RS00080 | elongation factor Tu | 260915 |
| wYak_RS02530 | elongation factor Tu | 260915 |
| AAE962_RS01075 | elongation factor Tu | 77125 |
| AAE962_RS04305 | elongation factor Tu | 77125 |
| ASM33_RS00255 | elongation factor Tu | 169402 |
| ASM33_RS00710 | elongation factor Tu | 169402 |
| M3L71_RS03200 | elongation factor Tu | 2902422 |
| M3L71_RS06560 | elongation factor Tu | 2902422 |
| MC061_RS01525 | elongation factor Tu | 2916816 |
| ACFDSB_RS00090 | elongation factor Tu | 2867522 |

| | | |
|---|---|---|
| ACFDSB_RS05155 | elongation factor Tu | 2867522 |
| GOY07_RS02735 | elongation factor Tu | 80850 |
| NBW37_RS01305 | elongation factor Tu | 931100 |
| NBW37_RS05660 | elongation factor Tu | 931100 |
| WOO_RS00535 | elongation factor Tu | 100901 |
| WOO_RS03565 | elongation factor Tu | 100901 |
| WOVOC_RS02415 | elongation factor Tu | 1410384 |
| WOVOC_RS03090 | elongation factor Tu | 1410384 |
| MJ631_RS04230 | elongation factor Tu | 573241 |
| MJ631_RS07945 | elongation factor Tu | 573241 |
| M1L25_RS01705 | elongation factor Tu | 154048 |
| M1L25_RS05195 | elongation factor Tu | 154048 |
| M1L26_RS01740 | elongation factor Tu | 183754 |
| M1L26_RS05230 | elongation factor Tu | 183754 |
| NL281_RS00245 | elongation factor Tu | 2886377 |
| NL281_RS06035 | elongation factor Tu | 2886377 |
| NL277_RS00820 | elongation factor Tu | 2883236 |
| NL277_RS02170 | elongation factor Tu | 2883236 |
| ACJZL3_RS00075 | elongation factor Tu | 1220542 |
| ACJZL3_RS01430 | elongation factor Tu | 1220542 |
| JKF54_RS02240 | elongation factor Tu | 2769078 |
| JKF54_RS05110 | elongation factor Tu | 2769078 |
| O1Q06_RS00165 | elongation factor Tu | 169184 |
| O1Q06_RS04030 | elongation factor Tu | 169184 |
| wTpre_RS02330 | elongation factor Tu | 125593 |
| wTpre_RS05230 | elongation factor Tu | 125593 |
| HUB92_RS01855 | elongation factor Tu | 2742717 |
| HUB92_RS02170 | elongation factor Tu | 2742717 |
| R0F63_RS03445 | elongation factor Tu | 2603209 |
| R0F63_RS05915 | elongation factor Tu | 2603209 |
| WBM_RS02015 | elongation factor Tu | 292805 |

| | | |
|---|---|---|
| WBM_RS03935 | elongation factor Tu | 292805 |
| FRT62_RS00075 | elongation factor Tu | 955 |
| FRT62_RS03340 | elongation factor Tu | 955 |
| EZH22_RS16210 | elongation factor Tu | 2528964 |
| EZH22_RS17870 | elongation factor Tu | 2528964 |
| AABB28_RS15735 | elongation factor Tu | 3137368 |
| AABB28_RS17520 | elongation factor Tu | 3137368 |
| AABB29_RS03130 | elongation factor Tu | 3137369 |
| AABB29_RS03265 | elongation factor Tu | 3137369 |
| AABB31_RS16765 | elongation factor Tu | 3137370 |
| AABB31_RS16870 | elongation factor Tu | 3137370 |
| ZCP4_RS03830 | elongation factor Tu | 627343 |
| ANT_RS05005 | elongation factor Tu | 926569 |
| ANT_RS09540 | elongation factor Tu | 926569 |
| CFX1CAM_RS09785 | elongation factor Tu | 1986204 |
| Pelsub_RS00070 | elongation factor Tu | 913107 |
| Pelsub_RS12845 | elongation factor Tu | 913107 |
| AQ_RS00010 | elongation factor Tu | 224324 |
| AQ_RS07585 | elongation factor Tu | 224324 |
| DESTER_RS00820 | elongation factor Tu | 868864 |
| DESTER_RS00985 | elongation factor Tu | 868864 |
| HYDTH_RS04285 | elongation factor Tu | 608538 |
| HYDTH_RS04355 | elongation factor Tu | 608538 |
| PERMA_RS05855 | elongation factor Tu | 123214 |
| PERMA_RS05925 | elongation factor Tu | 123214 |
| SULAZ_RS08585 | elongation factor Tu | 204536 |
| THAL_RS03100 | elongation factor Tu | 638303 |
| THAL_RS03170 | elongation factor Tu | 638303 |
| B5444_RS03260 | elongation factor Tu | 381751 |
| B5444_RS03330 | elongation factor Tu | 381751 |
| THERU_RS07045 | elongation factor Tu | 75906 |

| | | |
|---|---|---|
| THERU_RS07115 | elongation factor Tu | 75906 |
| THEAM_RS01305 | elongation factor Tu | 648996 |
| THEAM_RS08020 | elongation factor Tu | 648996 |
| CFX0092_RS16180 | elongation factor Tu | 1806508 |
| CFX0092_RS20675 | elongation factor Tu | 1806508 |
| D5261_RS17020 | elongation factor Tu | 2219043 |
| D5261_RS19320 | elongation factor Tu | 2219043 |
| RT761_RS01590 | elongation factor Tu | 2847778 |
| RT761_RS13115 | elongation factor Tu | 2847778 |
| ACKTT1_RS01695 | elongation factor Tu | 3064161 |
| ACKTT1_RS08185 | elongation factor Tu | 3064161 |
| QBE54_RS02885 | elongation factor Tu | 3039681 |
| QBE54_RS08800 | elongation factor Tu | 3039681 |
| BSEL_RS00665 | elongation factor Tu | 439292 |
| FOC79_RS04505 | elongation factor Tu | 46125 |
| BVH56_RS07925 | elongation factor Tu | 1817405 |
| ELX58_RS04380 | elongation factor Tu | 1720083 |
| QR721_RS00695 | elongation factor Tu | 3054939 |
| RI196_RS00750 | elongation factor Tu | 1868734 |
| AWM71_RS04480 | elongation factor Tu | 87541 |
| AWM72_RS03235 | elongation factor Tu | 119206 |
| F6I27_RS16255 | elongation factor Tu | 3078812 |
| AWM73_RS06990 | elongation factor Tu | 1376 |
| AWM74_RS05790 | elongation factor Tu | 51665 |
| AWM75_RS03665 | elongation factor Tu | 128944 |
| AWM76_RS01890 | elongation factor Tu | 1377 |
| ACNAN0_RS07365 | elongation factor Tu | 2586909 |
| AACI_RS13455 | elongation factor Tu | 521098 |
| K1I37_RS01670 | elongation factor Tu | 1450 |
| JI721_RS07730 | elongation factor Tu | 1457 |
| NZD86_RS02285 | elongation factor Tu | 1475485 |

| | | |
|---|---|---|
| NZD89_RS01785 | elongation factor Tu | 392011 |
| JZ786_RS02870 | elongation factor Tu | 2931921 |
| AB4027_RS01015 | elongation factor Tu | 426703 |
| FTX54_RS00655 | elongation factor Tu | 1667239 |
| PQ478_RS00695 | elongation factor Tu | 79885 |
| PT602_RS13805 | elongation factor Tu | 570278 |
| AXY_RS00680 | elongation factor Tu | 698758 |
| LA20533_RS07725 | elongation factor Tu | 1423721 |
| AWH56_RS00860 | elongation factor Tu | 1532552 |
| CB4_RS01100 | elongation factor Tu | 1500254 |
| K3F53_RS00935 | elongation factor Tu | 143495 |
| PO771_RS18635 | elongation factor Tu | 2966586 |
| AFLV_RS00600 | elongation factor Tu | 491915 |
| AFK25_RS00595 | elongation factor Tu | 198467 |
| GFC30_RS00385 | elongation factor Tu | 294699 |
| OH548_RS04385 | elongation factor Tu | 1218495 |
| MOO46_RS02670 | elongation factor Tu | 2923364 |
| D7I45_RS02795 | elongation factor Tu | 2419772 |
| K2W83_RS04350 | elongation factor Tu | 148814 |
| QRY64_RS03015 | elongation factor Tu | 2026189 |
| ID12_RS05545 | elongation factor Tu | 293387 |
| J5X95_RS09950 | elongation factor Tu | 1390 |
| GBAA_RS00610 | elongation factor Tu | 261594 |
| K6959_RS17410 | elongation factor Tu | 2672567 |
| QRY57_RS02300 | elongation factor Tu | 2819290 |
| AAHT65_RS07175 | elongation factor Tu | 1452 |
| EFK13_RS00715 | elongation factor Tu | 2487276 |
| BCER98_RS00650 | elongation factor Tu | 315749 |
| HHK01_RS00350 | elongation factor Tu | 2728853 |
| EQZ20_RS00735 | elongation factor Tu | 1664069 |
| AM592_RS18605 | elongation factor Tu | 1441095 |

| | | |
|---|---|---|
| DIC78_RS09510 | elongation factor Tu | 260554 |
| H2R00_RS14360 | elongation factor Tu | 1925021 |
| M1M80_RS00710 | elongation factor Tu | 483913 |
| ABXS71_RS24075 | elongation factor Tu | 324767 |
| WDJ61_RS00670 | elongation factor Tu | 3129297 |
| SR852_RS02385 | elongation factor Tu | 1402 |
| EUC40_RS25255 | elongation factor Tu | 2026191 |
| ABVJ71_RS10675 | elongation factor Tu | 3229854 |
| HC660_RS00710 | elongation factor Tu | 72360 |
| EXW63_RS13400 | elongation factor Tu | 1405 |
| LMD38_RS25615 | elongation factor Tu | 2026187 |
| CP943_RS00885 | elongation factor Tu | 1648923 |
| NLJ82_RS00660 | elongation factor Tu | 2026186 |
| BPMYX0001_RS00605 | elongation factor Tu | 527000 |
| BPUM_RS00660 | elongation factor Tu | 315750 |
| M0696_RS00710 | elongation factor Tu | 2715209 |
| FX981_RS06990 | elongation factor Tu | 561879 |
| LGQ02_RS00735 | elongation factor Tu | 1983719 |
| BSM4216_RS00640 | elongation factor Tu | 1479 |
| SK061_RS00730 | elongation factor Tu | 119858 |
| GYO_RS22280 | elongation factor Tu | 1052585 |
| QA442_RS21880 | elongation factor Tu | 2054641 |
| BSU_01130 | elongation factor Tu | 224308 |
| RBH91_RS00730 | elongation factor Tu | 1925020 |
| G4P54_RS00710 | elongation factor Tu | 227866 |
| BTHUR0008_RS00565 | elongation factor Tu | 527031 |
| I0K03_RS00660 | elongation factor Tu | 155322 |
| P3F89_RS26570 | elongation factor Tu | 2026188 |
| D9779_RS01440 | elongation factor Tu | 72361 |
| RBAM_RS00695 | elongation factor Tu | 326423 |
| A9C19_RS19620 | elongation factor Tu | 1547283 |

| | | |
|---|---|---|
| D4A37_RS00660 | elongation factor Tu | 1890302 |
| BK049_RS07515 | elongation factor Tu | 1178537 |
| ACK2UD_RS03980 | elongation factor Tu | 1303590 |
| MOO45_RS03650 | elongation factor Tu | 2923362 |
| MOO47_RS00670 | elongation factor Tu | 2923363 |
| BA6348_RS01550 | elongation factor Tu | 51101 |
| Q7F15_RS19520 | elongation factor Tu | 927786 |
| V5G20_RS26770 | elongation factor Tu | 45462 |
| EL268_RS01490 | elongation factor Tu | 1393 |
| ACLIN2_RS01320 | elongation factor Tu | 54910 |
| JD108_RS01340 | elongation factor Tu | 2796470 |
| LOK74_RS22000 | elongation factor Tu | 2895522 |
| BrL25_RS11015 | elongation factor Tu | 1121121 |
| EJ378_RS01075 | elongation factor Tu | 2496837 |
| IRJ17_RS03095 | elongation factor Tu | 54914 |
| NDK47_RS01470 | elongation factor Tu | 2950604 |
| ACJYYY_RS00520 | elongation factor Tu | 2757 |
| BFC19_RS06545 | elongation factor Tu | 2756 |
| HUR95_RS03215 | elongation factor Tu | 986075 |
| NSQ14_RS00670 | elongation factor Tu | 996558 |
| A5888_RS02495 | elongation factor Tu | 1834193 |
| A5888_RS11075 | elongation factor Tu | 1834193 |
| A5889_RS01615 | elongation factor Tu | 1834192 |
| A5889_RS11370 | elongation factor Tu | 1834192 |
| A5866_RS03030 | elongation factor Tu | 1834167 |
| A5866_RS09145 | elongation factor Tu | 1834167 |
| DOK78_RS03460 | elongation factor Tu | 2230877 |
| DOK78_RS13425 | elongation factor Tu | 2230877 |
| DOK79_RS06405 | elongation factor Tu | 2230878 |
| DOK79_RS11390 | elongation factor Tu | 2230878 |
| A5821_RS02960 | elongation factor Tu | 1834189 |

| | | |
|---|---|---|
| A5821_RS13870 | elongation factor Tu | 1834189 |
| Q783_RS10810 | elongation factor Tu | 1266845 |
| QOS94_RS01055 | elongation factor Tu | 2751 |
| QOS94_RS04185 | elongation factor Tu | 2751 |
| QOS94_RS07580 | elongation factor Tu | 2751 |
| ACF3OH_RS01445 | elongation factor Tu | 1441723 |
| KCTCHS21_RS26975 | elongation factor Tu | 2507935 |
| EAV92_RS16655 | elongation factor Tu | 2674991 |
| FPL14_RS10605 | elongation factor Tu | 2598458 |
| HH215_RS19210 | elongation factor Tu | 2728023 |
| LA20249_RS01875 | elongation factor Tu | 1423720 |
| BTM29_RS10815 | elongation factor Tu | 1847728 |
| BI355_RS06260 | elongation factor Tu | 392416 |
| LF20184_RS07760 | elongation factor Tu | 936140 |
| FG051_RS07320 | elongation factor Tu | 938155 |
| ABM34_RS02055 | elongation factor Tu | 1007676 |
| JP39_RS04910 | elongation factor Tu | 1074467 |
| ABWI26_RS07785 | elongation factor Tu | 1903258 |
| G6534_RS04135 | elongation factor Tu | 2714036 |
| ACI6PB_RS07570 | elongation factor Tu | 83526 |
| D1B17_RS07380 | elongation factor Tu | 2304606 |
| FSZ17_RS00685 | elongation factor Tu | 1742359 |
| J1899_RS00710 | elongation factor Tu | 859144 |
| LG276_RS00625 | elongation factor Tu | 859143 |
| AADC60_RS00705 | elongation factor Tu | 3051614 |
| FI644_RS09075 | elongation factor Tu | 1637975 |
| LS684_RS00710 | elongation factor Tu | 2901381 |
| VUQ06_RS04870 | elongation factor Tu | 3110288 |
| skT53_RS11960 | elongation factor Tu | 2743772 |
| ECBG_RS01155 | elongation factor Tu | 565655 |
| ECBG_RS10875 | elongation factor Tu | 565655 |

| | | |
|---|---|---|
| DQL78_RS01225 | elongation factor Tu | 44008 |
| CJZ72_RS05315 | elongation factor Tu | 53345 |
| CJZ72_RS09895 | elongation factor Tu | 53345 |
| E6A31_RS00325 | elongation factor Tu | 1352 |
| E6A31_RS05005 | elongation factor Tu | 1352 |
| HED61_RS04320 | elongation factor Tu | 1353 |
| HED61_RS07770 | elongation factor Tu | 1353 |
| EGCR1_RS06345 | elongation factor Tu | 160453 |
| EGCR1_RS13455 | elongation factor Tu | 160453 |
| A6J73_RS02785 | elongation factor Tu | 1354 |
| A6J73_RS11210 | elongation factor Tu | 1354 |
| ENLAB_RS10515 | elongation factor Tu | 2839759 |
| ENLAB_RS14105 | elongation factor Tu | 2839759 |
| KU781_RS00275 | elongation factor Tu | 357441 |
| KU781_RS08000 | elongation factor Tu | 357441 |
| P3T75_RS01305 | elongation factor Tu | 3031993 |
| P3T75_RS05245 | elongation factor Tu | 3031993 |
| EM4838_RS04845 | elongation factor Tu | 53346 |
| EM4838_RS14650 | elongation factor Tu | 53346 |
| J9537_RS01325 | elongation factor Tu | 71452 |
| J9537_RS08600 | elongation factor Tu | 71452 |
| ATZ35_RS03295 | elongation factor Tu | 118060 |
| ATZ35_RS13605 | elongation factor Tu | 118060 |
| PYW32_RS01435 | elongation factor Tu | 1814218 |
| EsVE80_RS08005 | elongation factor Tu | 1805431 |
| EsVE80_RS11785 | elongation factor Tu | 1805431 |
| CK496_RS00320 | elongation factor Tu | 417368 |
| CK496_RS04210 | elongation factor Tu | 417368 |
| CC204_RS03235 | elongation factor Tu | 2005703 |
| CC204_RS10250 | elongation factor Tu | 2005703 |
| BCELL_RS00680 | elongation factor Tu | 649639 |

| | | |
|---|---|---|
| KKI46_RS00665 | elongation factor Tu | 41170 |
| KB235_RS02310 | elongation factor Tu | 1428684 |
| EXIG_RS00660 | elongation factor Tu | 262543 |
| OE104_RS12890 | elongation factor Tu | 2980026 |
| OE105_RS12730 | elongation factor Tu | 2980027 |
| LSG31_RS06545 | elongation factor Tu | 2901836 |
| M3M37_RS01640 | elongation factor Tu | 2940297 |
| M3M40_RS01545 | elongation factor Tu | 2940299 |
| M3M39_RS04035 | elongation factor Tu | 2940300 |
| M8332_RS04315 | elongation factor Tu | 1750535 |
| AYR60_RS04515 | elongation factor Tu | 53444 |
| M3M35_RS02130 | elongation factor Tu | 2940301 |
| PY770_RS05030 | elongation factor Tu | 1625 |
| R4B61_RS04625 | elongation factor Tu | 1249630 |
| M3M36_RS06425 | elongation factor Tu | 2940302 |
| NRE15_RS08040 | elongation factor Tu | 2969821 |
| LROSL1_RS07515 | elongation factor Tu | 231049 |
| DQN46_RS05665 | elongation factor Tu | 29391 |
| FOC50_RS01650 | elongation factor Tu | 84135 |
| GK_RS00875 | elongation factor Tu | 235909 |
| GS3922_RS15110 | elongation factor Tu | 129338 |
| GD3902_RS08610 | elongation factor Tu | 33940 |
| GT3570_RS00580 | elongation factor Tu | 33941 |
| QWV57_RS00640 | elongation factor Tu | 213419 |
| CYJ72_RS07125 | elongation factor Tu | 13076 |
| MUN88_RS04965 | elongation factor Tu | 2932256 |
| MUN87_RS14345 | elongation factor Tu | 2932255 |
| GI584_RS00910 | elongation factor Tu | 2663022 |
| K8O88_RS03175 | elongation factor Tu | 46124 |
| FED51_RS00820 | elongation factor Tu | 86665 |
| MUO15_RS13190 | elongation factor Tu | 2932259 |

| | | |
|---|---|---|
| HBHAL_RS00725 | elongation factor Tu | 866895 |
| HM131_RS00440 | elongation factor Tu | 402384 |
| P9989_RS00855 | elongation factor Tu | 554880 |
| MUN89_RS01155 | elongation factor Tu | 2932257 |
| MUO14_RS12510 | elongation factor Tu | 2932258 |
| BF29_RS11745 | elongation factor Tu | 1121088 |
| ABR335_RS16615 | elongation factor Tu | 2824910 |
| I5776_RS00665 | elongation factor Tu | 1476025 |
| M0R79_RS00925 | elongation factor Tu | 89093 |
| G7057_RS02980 | elongation factor Tu | 1868794 |
| EJN90_RS03605 | elongation factor Tu | 2496265 |
| BW727_RS07320 | elongation factor Tu | 708126 |
| G7058_RS08455 | elongation factor Tu | 1868793 |
| RZ44_RS08870 | elongation factor Tu | 1461582 |
| GXN75_RS01265 | elongation factor Tu | 714067 |
| GXN76_RS14895 | elongation factor Tu | 1380685 |
| CVV65_RS00775 | elongation factor Tu | 2055160 |
| BTUS_RS00790 | elongation factor Tu | 562970 |
| NYR52_RS01195 | elongation factor Tu | 37482 |
| LBCZ_RS06200 | elongation factor Tu | 1423732 |
| LHUE1_RS11990 | elongation factor Tu | 3035291 |
| LM010_RS06875 | elongation factor Tu | 88233 |
| PQ472_RS06720 | elongation factor Tu | 3025672 |
| LBPG_RS06480 | elongation factor Tu | 537973 |
| LACPH_RS06190 | elongation factor Tu | 3068305 |
| FG342_RS09270 | elongation factor Tu | 47715 |
| LACSTY_RS06595 | elongation factor Tu | 3068306 |
| LACZS1_RS08670 | elongation factor Tu | 3068307 |
| V7R88_RS08175 | elongation factor Tu | 271881 |
| ACKTTY_RS08000 | elongation factor Tu | 3069269 |
| P6S16_RS14725 | elongation factor Tu | 2993456 |

| | | |
|---|---|---|
| PWO93_RS08735 | elongation factor Tu | 60520 |
| LP314_RS10010 | elongation factor Tu | 1589 |
| C7M33_RS00305 | elongation factor Tu | 1590 |
| ACI6O9_RS08960 | elongation factor Tu | 942150 |
| LA749_RS05010 | elongation factor Tu | 1600 |
| SO785_RS04440 | elongation factor Tu | 1579 |
| D0Z64_RS03920 | elongation factor Tu | 83683 |
| LAB52_RS04320 | elongation factor Tu | 695562 |
| DKL56_RS03580 | elongation factor Tu | 303541 |
| E6A57_RS04450 | elongation factor Tu | 47770 |
| LJ046_RS05345 | elongation factor Tu | 1217420 |
| LGAS_RS05740 | elongation factor Tu | 324831 |
| DLD54_RS04110 | elongation factor Tu | 1218494 |
| AADV52_RS04350 | elongation factor Tu | 1587 |
| G6Z82_RS02960 | elongation factor Tu | 147802 |
| KBW87_RS04635 | elongation factor Tu | 151781 |
| QM512_RS04595 | elongation factor Tu | 2993455 |
| BUE77_RS05655 | elongation factor Tu | 109790 |
| H0I41_RS05640 | elongation factor Tu | 33959 |
| ICI50_RS06385 | elongation factor Tu | 267818 |
| DKL58_RS04400 | elongation factor Tu | 1218493 |
| ABZU12_RS04580 | elongation factor Tu | 2508708 |
| GYM71_RS04720 | elongation factor Tu | 2012495 |
| LpgJCM5343_RS05845 | elongation factor Tu | 2107999 |
| H1A07_RS05605 | elongation factor Tu | 508451 |
| CS873_RS04525 | elongation factor Tu | 2269374 |
| H4B44_RS06885 | elongation factor Tu | 227945 |
| R8389_RS03710 | elongation factor Tu | 2976676 |
| D7I46_RS07260 | elongation factor Tu | 2419773 |
| BHS00_RS07580 | elongation factor Tu | 2749961 |
| KW2_RS09535 | elongation factor Tu | 1295826 |

| | | |
|---|---|---|
| VNN39_RS05295 | elongation factor Tu | 1281486 |
| I6G86_RS04090 | elongation factor Tu | 1363 |
| LL14B4_RS09850 | elongation factor Tu | 1360 |
| BHS01_RS08350 | elongation factor Tu | 2749962 |
| lgb_RS08465 | elongation factor Tu | 1940789 |
| FLP15_RS12410 | elongation factor Tu | 2592653 |
| G3T13_RS05720 | elongation factor Tu | 1366 |
| JW886_RS08755 | elongation factor Tu | 1151742 |
| LH506_RS07400 | elongation factor Tu | 51664 |
| LCU_RS01905 | elongation factor Tu | 1293592 |
| LFUMFP_RS06070 | elongation factor Tu | 164393 |
| LG542_RS05455 | elongation factor Tu | 60519 |
| GJ664_RS09280 | elongation factor Tu | 1599 |
| DQN84_RS00665 | elongation factor Tu | 1467 |
| AOX59_RS12870 | elongation factor Tu | 1472767 |
| O2S85_RS00640 | elongation factor Tu | 2911514 |
| KE627_RS09775 | elongation factor Tu | 1581 |
| PL11_RS00515 | elongation factor Tu | 1138822 |
| GQR93_RS05945 | elongation factor Tu | 1588 |
| DNL43_RS01425 | elongation factor Tu | 33962 |
| LHL71_RS02030 | elongation factor Tu | 2883998 |
| ILQ00_RS00800 | elongation factor Tu | 152331 |
| O0236_RS07750 | elongation factor Tu | 3003483 |
| FGL89_RS07690 | elongation factor Tu | 1252 |
| FGL72_RS05520 | elongation factor Tu | 33964 |
| ACI3ES_RS03295 | elongation factor Tu | 2766470 |
| LEGAS_RS06125 | elongation factor Tu | 762550 |
| C269_RS06085 | elongation factor Tu | 1229756 |
| LKI_RS02420 | elongation factor Tu | 762051 |
| FGL83_RS00670 | elongation factor Tu | 1246 |
| LEUM_RS03065 | elongation factor Tu | 203120 |

| | | |
|---|---|---|
| I6H60_RS03410 | elongation factor Tu | 33968 |
| A6B45_RS03095 | elongation factor Tu | 1511761 |
| DQN51_RS06910 | elongation factor Tu | 1580 |
| AB3234_RS05260 | elongation factor Tu | 1704076 |
| ABN16_RS11620 | elongation factor Tu | 637971 |
| ACEPSU_RS05980 | elongation factor Tu | 357278 |
| H3M12_RS08355 | elongation factor Tu | 2292255 |
| ACI6PE_RS05430 | elongation factor Tu | 267363 |
| BQ5903_RS06715 | elongation factor Tu | 89059 |
| N4597_RS06800 | elongation factor Tu | 1601 |
| GSR62_RS03755 | elongation factor Tu | 1605 |
| QPK35_RS03920 | elongation factor Tu | 3050131 |
| QFX10_RS10235 | elongation factor Tu | 762833 |
| CPS94_RS04650 | elongation factor Tu | 1622 |
| LRC_RS06505 | elongation factor Tu | 1069534 |
| V3C97_RS04295 | elongation factor Tu | 228229 |
| PTA89_RS03590 | elongation factor Tu | 1624 |
| P8770_RS04335 | elongation factor Tu | 1613 |
| LF145_RS03405 | elongation factor Tu | 104955 |
| H0G69_RS03070 | elongation factor Tu | 97478 |
| N4599_RS00540 | elongation factor Tu | 1632 |
| LP475_RS03405 | elongation factor Tu | 35787 |
| PRK60_RS05435 | elongation factor Tu | 2742601 |
| N4627_RS03620 | elongation factor Tu | 1633 |
| G6O70_RS09335 | elongation factor Tu | 1423759 |
| LM596_RS07155 | elongation factor Tu | 1618 |
| G6O73_RS02585 | elongation factor Tu | 1122150 |
| EL241_RS13065 | elongation factor Tu | 1641 |
| PQQ29_RS13490 | elongation factor Tu | 1642 |
| JL52_RS13555 | elongation factor Tu | 202751 |
| lmo2653 | elongation factor Tu | 169963 |

| | | |
|---|---|---|
| OTR81_RS08310 | elongation factor Tu | 1497681 |
| LSE_RS13015 | elongation factor Tu | 683837 |
| QJV29_RS13205 | elongation factor Tu | 2713501 |
| UE46_RS01685 | elongation factor Tu | 1006155 |
| CKV90_RS13480 | elongation factor Tu | 1643 |
| AYR53_RS10190 | elongation factor Tu | 375175 |
| LB003_RS01475 | elongation factor Tu | 1607 |
| LC20001_RS06545 | elongation factor Tu | 913848 |
| FJQ98_RS25635 | elongation factor Tu | 2590012 |
| LCP48_RS22700 | elongation factor Tu | 2115968 |
| HR49_RS00855 | elongation factor Tu | 28031 |
| OU989_RS21650 | elongation factor Tu | 2998077 |
| R6U77_RS05790 | elongation factor Tu | 1470088 |
| I6G82_RS19830 | elongation factor Tu | 33935 |
| QNH24_RS25340 | elongation factor Tu | 759811 |
| C9J31_RS22780 | elongation factor Tu | 436516 |
| LS41612_RS01100 | elongation factor Tu | 1421 |
| MKY04_RS21380 | elongation factor Tu | 1714840 |
| C9963_RS10755 | elongation factor Tu | 2086577 |
| T479_RS21930 | elongation factor Tu | 1145276 |
| HT586_RS11680 | elongation factor Tu | 1903056 |
| MCCS_RS11770 | elongation factor Tu | 1855823 |
| I6G25_RS08585 | elongation factor Tu | 69966 |
| LAU42_RS11095 | elongation factor Tu | 2875764 |
| QSV55_RS10660 | elongation factor Tu | 69968 |
| MGG13_RS10525 | elongation factor Tu | 198483 |
| KFV12_RS12505 | elongation factor Tu | 1902580 |
| KFV11_RS10735 | elongation factor Tu | 69967 |
| QA539_RS10025 | elongation factor Tu | 3039389 |
| ACNMOQ_RS13185 | elongation factor Tu | 42858 |
| DQL71_RS12955 | elongation factor Tu | 1296 |

| | | |
|---|---|---|
| CKV64_RS11180 | elongation factor Tu | 643214 |
| I6J10_RS11965 | elongation factor Tu | 71237 |
| G8O30_RS14370 | elongation factor Tu | 2593652 |
| DAT869_RS01065 | elongation factor Tu | 33970 |
| CD004_RS22960 | elongation factor Tu | 129985 |
| LG336_RS00670 | elongation factor Tu | 1643336 |
| K8L98_RS00735 | elongation factor Tu | 2874282 |
| HUW50_RS09900 | elongation factor Tu | 2745198 |
| QLQ22_RS00735 | elongation factor Tu | 3048891 |
| ACK1LH_RS00740 | elongation factor Tu | 246786 |
| WCV66_RS02795 | elongation factor Tu | 3117747 |
| GMB29_RS00715 | elongation factor Tu | 2567941 |
| WCV65_RS00745 | elongation factor Tu | 3117746 |
| ACN6MT_RS03420 | elongation factor Tu | 86668 |
| FOC77_RS01275 | elongation factor Tu | 1397 |
| R4Z09_RS00640 | elongation factor Tu | 1737571 |
| MOO44_RS06195 | elongation factor Tu | 2925830 |
| KPF49_RS07260 | elongation factor Tu | 489910 |
| B0W44_RS00670 | elongation factor Tu | 1471761 |
| OB_RS00735 | elongation factor Tu | 221109 |
| LG273_RS00740 | elongation factor Tu | 481353 |
| C794_RS00135 | elongation factor Tu | 1238184 |
| CUC15_RS00650 | elongation factor Tu | 2052660 |
| OKIT_RS08110 | elongation factor Tu | 1045004 |
| J3U91_RS06420 | elongation factor Tu | 1247 |
| DLJ48_RS06395 | elongation factor Tu | 2203724 |
| FLK61_RS01100 | elongation factor Tu | 2592382 |
| KJW29_RS30300 | elongation factor Tu | 171404 |
| HGI30_RS03040 | elongation factor Tu | 2726185 |
| EJC50_RS01160 | elongation factor Tu | 2495582 |
| E6C60_RS18490 | elongation factor Tu | 2565926 |

| | | |
|---|---|---|
| V6668_RS26835 | elongation factor Tu | 1451 |
| NYR53_RS32290 | elongation factor Tu | 2975482 |
| F0T56_RS07855 | elongation factor Tu | 253703 |
| MJA45_RS01320 | elongation factor Tu | 2918900 |
| L6442_RS29980 | elongation factor Tu | 116718 |
| EKN16_RS26495 | elongation factor Tu | 1712516 |
| HUB98_RS13685 | elongation factor Tu | 198119 |
| VN24_RS12630 | elongation factor Tu | 1126833 |
| PBOR_RS31560 | elongation factor Tu | 160799 |
| AR543_RS22135 | elongation factor Tu | 1616788 |
| GCU48_RS07385 | elongation factor Tu | 128574 |
| HUB94_RS26320 | elongation factor Tu | 375489 |
| GCU45_RS19520 | elongation factor Tu | 562959 |
| PC41400_RS29015 | elongation factor Tu | 79263 |
| LPB68_RS14800 | elongation factor Tu | 1763538 |
| L6439_RS24930 | elongation factor Tu | 130049 |
| B9T62_RS28565 | elongation factor Tu | 414771 |
| PDUR_RS23480 | elongation factor Tu | 44251 |
| OF512_RS01390 | elongation factor Tu | 189691 |
| WDY75_RS03395 | elongation factor Tu | 59843 |
| PGRAT_RS27990 | elongation factor Tu | 189425 |
| GCU39_RS21855 | elongation factor Tu | 1473112 |
| L0M14_RS30160 | elongation factor Tu | 2908839 |
| SK066_RS20345 | elongation factor Tu | 539262 |
| BBD41_RS17500 | elongation factor Tu | 1870820 |
| GT351_RS05095 | elongation factor Tu | 682956 |
| B4V02_RS02855 | elongation factor Tu | 172713 |
| PQ456_RS19240 | elongation factor Tu | 1452732 |
| ERICIV_RS20665 | elongation factor Tu | 147375 |
| EIM92_RS07240 | elongation factor Tu | 1338368 |
| EI981_RS25515 | elongation factor Tu | 2707005 |

| | | |
|---|---|---|
| GXP70_RS27740 | elongation factor Tu | 2704462 |
| PTQ21_RS01710 | elongation factor Tu | 3026433 |
| B2K_RS34485 | elongation factor Tu | 997761 |
| IJ22_RS22445 | elongation factor Tu | 162209 |
| RCP16_RS22295 | elongation factor Tu | 1886670 |
| PODO_RS27035 | elongation factor Tu | 189426 |
| QU597_RS25865 | elongation factor Tu | 3242193 |
| CBE73_RS11655 | elongation factor Tu | 1619311 |
| QPK24_RS21175 | elongation factor Tu | 3050112 |
| CUU60_RS13355 | elongation factor Tu | 1036171 |
| ET464_RS17115 | elongation factor Tu | 2509456 |
| EHS13_RS01125 | elongation factor Tu | 1778678 |
| GZH47_RS18855 | elongation factor Tu | 2704463 |
| PRIO_RS30220 | elongation factor Tu | 1073571 |
| MJB10_RS24985 | elongation factor Tu | 2918901 |
| PSAB_RS21790 | elongation factor Tu | 1268072 |
| QU599_RS28545 | elongation factor Tu | 3110539 |
| KP014_RS00195 | elongation factor Tu | 1333845 |
| L1F29_RS31360 | elongation factor Tu | 2909671 |
| PSTEL_RS23220 | elongation factor Tu | 169760 |
| SY83_RS11380 | elongation factor Tu | 1178515 |
| HPL003_RS03890 | elongation factor Tu | 985665 |
| FLT43_RS13940 | elongation factor Tu | 49283 |
| JRJ22_RS26245 | elongation factor Tu | 2810347 |
| KDC22_RS29785 | elongation factor Tu | 1873425 |
| V6W81_RS25650 | elongation factor Tu | 528187 |
| B9N86_RS02340 | elongation factor Tu | 1313296 |
| PUW25_RS22235 | elongation factor Tu | 521520 |
| QNH46_RS21565 | elongation factor Tu | 307580 |
| BS614_RS30295 | elongation factor Tu | 528191 |
| AWM70_RS16590 | elongation factor Tu | 1462996 |

| | | |
|---|---|---|
| E2636_RS00760 | elongation factor Tu | 417367 |
| D3873_RS12575 | elongation factor Tu | 2320858 |
| BCV53_RS02425 | elongation factor Tu | 1426 |
| JTI59_RS01810 | elongation factor Tu | 153151 |
| DM447_RS00840 | elongation factor Tu | 2213194 |
| LOOC260_RS06415 | elongation factor Tu | 1291742 |
| F0161_RS07420 | elongation factor Tu | 1296540 |
| LACOL_RS05735 | elongation factor Tu | 1423778 |
| ABTQ33_RS07940 | elongation factor Tu | 152335 |
| HN015_RS06035 | elongation factor Tu | 1254 |
| AYR57_RS08270 | elongation factor Tu | 187452 |
| ADU73_RS00415 | elongation factor Tu | 51663 |
| SIN07_RS04115 | elongation factor Tu | 114090 |
| AB3217_RS04870 | elongation factor Tu | 54062 |
| PEPE_RS05520 | elongation factor Tu | 278197 |
| ACLJJ6_RS03975 | elongation factor Tu | 381829 |
| L8956_RS00690 | elongation factor Tu | 450367 |
| GW626_RS00560 | elongation factor Tu | 264697 |
| I6J18_RS13810 | elongation factor Tu | 1407 |
| BS1321_RS11630 | elongation factor Tu | 1349754 |
| EQG49_RS11290 | elongation factor Tu | 2506420 |
| BBH88_RS00645 | elongation factor Tu | 1185653 |
| BCM40_RS00705 | elongation factor Tu | 414778 |
| AJGP001_RS00620 | elongation factor Tu | 1598147 |
| HF394_RS07540 | elongation factor Tu | 459472 |
| BBI08_RS00610 | elongation factor Tu | 1215089 |
| AUO94_RS08145 | elongation factor Tu | 1374 |
| B0X71_RS00560 | elongation factor Tu | 2213202 |
| QWY22_RS00625 | elongation factor Tu | 3058394 |
| BBI11_RS00610 | elongation factor Tu | 192421 |
| BBI15_RS00610 | elongation factor Tu | 1038856 |

| | | |
|---|---|---|
| AUC31_RS16425 | elongation factor Tu | 200991 |
| I858_RS00575 | elongation factor Tu | 1302659 |
| KI215_RS00905 | elongation factor Tu | 1125966 |
| CR091_RS00665 | elongation factor Tu | 412384 |
| LZP85_RS19920 | elongation factor Tu | 86664 |
| EQG57_RS00660 | elongation factor Tu | 1348623 |
| PB01_RS19675 | elongation factor Tu | 2283160 |
| FN924_RS00750 | elongation factor Tu | 2594883 |
| KO561_RS00430 | elongation factor Tu | 2844358 |
| K6T23_RS00760 | elongation factor Tu | 189381 |
| FFV09_RS07710 | elongation factor Tu | 2583377 |
| BV455_RS08150 | elongation factor Tu | 81408 |
| HUG15_RS01060 | elongation factor Tu | 2743000 |
| HUG20_RS00785 | elongation factor Tu | 2743001 |
| EPH95_RS10185 | elongation factor Tu | 2502791 |
| DT065_RS11915 | elongation factor Tu | 2099786 |
| AAV35_RS13195 | elongation factor Tu | 1230341 |
| RQP18_RS12535 | elongation factor Tu | 3136731 |
| BBEV_RS16325 | elongation factor Tu | 632773 |
| FMM01_RS03855 | elongation factor Tu | 304207 |
| FGL90_RS00685 | elongation factor Tu | 176292 |
| AYR61_RS10955 | elongation factor Tu | 240427 |
| ERJ70_RS00105 | elongation factor Tu | 2729421 |
| SR921_RS00630 | elongation factor Tu | 79880 |
| PQ477_RS08430 | elongation factor Tu | 766894 |
| FPV17_RS00830 | elongation factor Tu | 2598861 |
| V5G21_RS10060 | elongation factor Tu | 866786 |
| O7776_RS19760 | elongation factor Tu | 2912187 |
| K7399_RS13915 | elongation factor Tu | 269673 |
| PGH26_RS14775 | elongation factor Tu | 3020056 |
| QWT69_RS15800 | elongation factor Tu | 3056444 |

| | | |
|---|---|---|
| AB1H92_RS15220 | elongation factor Tu | 1474 |
| AZE41_RS01425 | elongation factor Tu | 1476 |
| QWT68_RS13285 | elongation factor Tu | 3056445 |
| SporoS204_RS10395 | elongation factor Tu | 1571 |
| BI350_RS02090 | elongation factor Tu | 298596 |
| GJE18_RS01295 | elongation factor Tu | 985762 |
| SAMSHR1132_RS02625 | elongation factor Tu | 985002 |
| MUA37_RS11630 | elongation factor Tu | 29378 |
| I6G39_RS01805 | elongation factor Tu | 29379 |
| V6C80_RS10930 | elongation factor Tu | 74703 |
| JMUB898_RS11540 | elongation factor Tu | 29380 |
| I6G40_RS09480 | elongation factor Tu | 1281 |
| QMO72_RS11680 | elongation factor Tu | 201828 |
| DWB97_RS10540 | elongation factor Tu | 46126 |
| KM149_RS11075 | elongation factor Tu | 74706 |
| EL167_RS11560 | elongation factor Tu | 70255 |
| CNQ82_RS12940 | elongation factor Tu | 2044912 |
| MUA44_RS11360 | elongation factor Tu | 53344 |
| EQW00_RS11075 | elongation factor Tu | 1282 |
| I6I25_RS10055 | elongation factor Tu | 246432 |
| C7J90_RS01895 | elongation factor Tu | 46127 |
| JN168_RS11930 | elongation factor Tu | 1293 |
| EQ029_RS10970 | elongation factor Tu | 1283 |
| EGX58_RS03525 | elongation factor Tu | 1290 |
| QQM35_RS04330 | elongation factor Tu | 3051183 |
| SHYC_RS10630 | elongation factor Tu | 1284 |
| C7J89_RS12920 | elongation factor Tu | 29384 |
| ISP08_RS11115 | elongation factor Tu | 2781774 |
| AL499_RS05690 | elongation factor Tu | 28035 |
| B5P37_RS07170 | elongation factor Tu | 155085 |
| ACF3N8_RS01120 | elongation factor Tu | 555791 |

| | | |
|---|---|---|
| CKV93_RS01005 | elongation factor Tu | 1294 |
| BJG89_RS12485 | elongation factor Tu | 214473 |
| I6I26_RS10995 | elongation factor Tu | 45972 |
| CEP67_RS09565 | elongation factor Tu | 170573 |
| CKV71_RS11305 | elongation factor Tu | 70258 |
| I6G87_RS07935 | elongation factor Tu | 283734 |
| LG295_RS12080 | elongation factor Tu | 2282419 |
| LN051_RS10220 | elongation factor Tu | 2892440 |
| OVY33_RS07705 | elongation factor Tu | 522262 |
| I6I31_RS05495 | elongation factor Tu | 33028 |
| SSP_RS11030 | elongation factor Tu | 342451 |
| MUA40_RS10885 | elongation factor Tu | 1295 |
| EL116_RS02670 | elongation factor Tu | 1654388 |
| CKV88_RS02435 | elongation factor Tu | 308354 |
| MUA87_RS11440 | elongation factor Tu | 1286 |
| PYW31_RS11330 | elongation factor Tu | 61015 |
| MUA21_RS01235 | elongation factor Tu | 94138 |
| D3P10_RS11230 | elongation factor Tu | 1292 |
| SXYLSMQ121_RS11175 | elongation factor Tu | 1288 |
| EG886_RS00870 | elongation factor Tu | 2490858 |
| AMM49_RS04165 | elongation factor Tu | 1311 |
| EL079_RS01340 | elongation factor Tu | 1328 |
| DQL20_RS07545 | elongation factor Tu | 113107 |
| EL097_RS10015 | elongation factor Tu | 1329 |
| DDV21_RS08080 | elongation factor Tu | 2173853 |
| I6J13_RS03810 | elongation factor Tu | 76860 |
| DQM67_RS03470 | elongation factor Tu | 889201 |
| SM121_RS05220 | elongation factor Tu | 3098075 |
| I6H74_RS09240 | elongation factor Tu | 1334 |
| I2432_RS06930 | elongation factor Tu | 40041 |
| DQL21_RS06650 | elongation factor Tu | 1345 |

| | | |
|---|---|---|
| FOC63_RS00565 | elongation factor Tu | 315405 |
| I6L84_RS01850 | elongation factor Tu | 1302 |
| D7D53_RS04820 | elongation factor Tu | 1433513 |
| K5J94_RS05510 | elongation factor Tu | 254785 |
| A2G56_RS00140 | elongation factor Tu | 1814128 |
| AB6M99_RS08290 | elongation factor Tu | 2282420 |
| BFM96_RS05685 | elongation factor Tu | 1888195 |
| N596_RS04145 | elongation factor Tu | 1156431 |
| PW252_RS08515 | elongation factor Tu | 3028084 |
| I6H76_RS08820 | elongation factor Tu | 102684 |
| SI82_RS06360 | elongation factor Tu | 1346 |
| DQN42_RS03475 | elongation factor Tu | 1338 |
| D7D50_RS06390 | elongation factor Tu | 2382163 |
| J4854_RS07450 | elongation factor Tu | 684066 |
| SM123_RS03990 | elongation factor Tu | 3098076 |
| DQN23_RS06240 | elongation factor Tu | 150055 |
| PY824_RS08165 | elongation factor Tu | 59310 |
| A4H00_RS08050 | elongation factor Tu | 1825069 |
| CKV83_RS01945 | elongation factor Tu | 400065 |
| RN80_RS07290 | elongation factor Tu | 28037 |
| I6L88_RS05590 | elongation factor Tu | 1309 |
| EL140_RS03970 | elongation factor Tu | 655813 |
| INT76_RS10775 | elongation factor Tu | 2781599 |
| A0O21_RS02720 | elongation factor Tu | 1811193 |
| SP4011_RS04090 | elongation factor Tu | 2993430 |
| HMPREF0833_RS04260 | elongation factor Tu | 760570 |
| L6410_RS03105 | elongation factor Tu | 1501662 |
| SPB_RS00745 | elongation factor Tu | 873447 |
| E8M05_RS07600 | elongation factor Tu | 197614 |
| EHW89_RS05925 | elongation factor Tu | 2490633 |
| C0J00_RS06930 | elongation factor Tu | 82348 |

| | | |
|---|---|---|
| AT689_RS06645 | elongation factor Tu | 1313 |
| DQM45_RS06605 | elongation factor Tu | 1340 |
| SPPN_RS07180 | elongation factor Tu | 1054460 |
| EL147_RS09525 | elongation factor Tu | 361101 |
| DQM35_RS06470 | elongation factor Tu | 1314 |
| FY406_RS06905 | elongation factor Tu | 1341 |
| CHF41_RS08405 | elongation factor Tu | 2021971 |
| SR187_RS07200 | elongation factor Tu | 1917441 |
| GPZ88_RS01550 | elongation factor Tu | 2686210 |
| SALIVA_RS02330 | elongation factor Tu | 347253 |
| DQM63_RS07365 | elongation factor Tu | 1305 |
| DK181_RS07710 | elongation factor Tu | 1310 |
| SSUBM407_RS02950 | elongation factor Tu | 568814 |
| PXH68_RS02800 | elongation factor Tu | 3028082 |
| H0506_RS02485 | elongation factor Tu | 1308 |
| STYK_RS04505 | elongation factor Tu | 2819619 |
| SRT_RS06880 | elongation factor Tu | 1111760 |
| DQM95_RS03530 | elongation factor Tu | 1349 |
| EL133_RS06970 | elongation factor Tu | 149016 |
| EL076_RS02655 | elongation factor Tu | 1343 |
| J5M87_RS06500 | elongation factor Tu | 2664091 |
| CL176_RS11330 | elongation factor Tu | 2036206 |
| BC6307_RS23675 | elongation factor Tu | 33932 |
| B4U37_RS00645 | elongation factor Tu | 79883 |
| WDJ59_RS13150 | elongation factor Tu | 3131995 |
| GZ22_RS17135 | elongation factor Tu | 361277 |
| AC806_RS10370 | elongation factor Tu | 51669 |
| C7K43_RS07965 | elongation factor Tu | 290335 |
| C7K38_RS05570 | elongation factor Tu | 526944 |
| ISX45_RS16310 | elongation factor Tu | 247480 |
| FA954_RS00415 | elongation factor Tu | 2026 |

| | | |
|---|---|---|
| THECO_RS01335 | elongation factor Tu | 717605 |
| U2959_RS12750 | elongation factor Tu | 82803 |
| SNQ41_RS04420 | elongation factor Tu | 2094020 |
| CIG75_RS19745 | elongation factor Tu | 1214604 |
| CBW65_RS08320 | elongation factor Tu | 1903704 |
| LG296_RS18750 | elongation factor Tu | 1202712 |
| DKZ56_RS03005 | elongation factor Tu | 367743 |
| UtA1_RS03495 | elongation factor Tu | 51173 |
| H9L18_RS01055 | elongation factor Tu | 218144 |
| H9L18_RS13720 | elongation factor Tu | 218144 |
| G7081_RS02025 | elongation factor Tu | 2714946 |
| G7082_RS05600 | elongation factor Tu | 2714947 |
| OL234_RS00205 | elongation factor Tu | 2991418 |
| VSF34_RS00295 | elongation factor Tu | 3109030 |
| G314FT_RS02885 | elongation factor Tu | 2920380 |
| M2919_RS06720 | elongation factor Tu | 81947 |
| BHY08_RS10245 | elongation factor Tu | 519472 |
| E4Z98_RS09370 | elongation factor Tu | 2562451 |
| FA707_RS00295 | elongation factor Tu | 2571750 |
| A21D_RS06385 | elongation factor Tu | 302167 |
| BME96_RS00720 | elongation factor Tu | 1482 |
| OLD84_RS01375 | elongation factor Tu | 1216297 |
| CFK40_RS00975 | elongation factor Tu | 163877 |
| KBP50_RS00880 | elongation factor Tu | 1473 |
| CFK37_RS05475 | elongation factor Tu | 2017483 |
| KHQ31_RS02380 | elongation factor Tu | 759620 |
| AUC62_RS04515 | elongation factor Tu | 137591 |
| G7084_RS06210 | elongation factor Tu | 2714949 |
| C6P13_RS04170 | elongation factor Tu | 1583 |
| H9L19_RS01060 | elongation factor Tu | 1162633 |
| FOL01_RS06435 | elongation factor Tu | 1631871 |

| | | |
|---|---|---|
| WKK_RS05850 | elongation factor Tu | 1045854 |
| QRX23_RS05670 | elongation factor Tu | 1249 |
| WSWS_RS03915 | elongation factor Tu | 155866 |
| KHQ32_RS02350 | elongation factor Tu | 887702 |
| IE337_RS03465 | elongation factor Tu | 1629 |
| R8495_RS08565 | elongation factor Tu | 2932184 |
| R8749_RS08730 | elongation factor Tu | 2932183 |
| C0V70_RS18135 | elongation factor Tu | 960 |
| C0V70_RS18200 | elongation factor Tu | 960 |
| BMS_RS15760 | elongation factor Tu | 862908 |
| BMS_RS15825 | elongation factor Tu | 862908 |
| SOO65_RS00660 | elongation factor Tu | 28202 |
| SOO65_RS00725 | elongation factor Tu | 28202 |
| FMF02_RS03560 | elongation factor Tu | 2585118 |
| FME97_RS04150 | elongation factor Tu | 2585119 |
| NQ491_RS06090 | elongation factor Tu | 1211813 |
| NQ495_RS06580 | elongation factor Tu | 742725 |
| ED734_RS07105 | elongation factor Tu | 2364787 |
| Alistipes_RS02745 | elongation factor Tu | 2585117 |
| NQ492_RS04360 | elongation factor Tu | 717959 |
| CDL62_RS11905 | elongation factor Tu | 889453 |
| AQPE_RS16210 | elongation factor Tu | 2010828 |
| INE72_RS02405 | elongation factor Tu | 997873 |
| A4V03_RS16790 | elongation factor Tu | 1796613 |
| INE78_RS01710 | elongation factor Tu | 246787 |
| ACNOGP_RS02530 | elongation factor Tu | 2763021 |
| NQ546_RS01265 | elongation factor Tu | 28111 |
| K6V30_RS02625 | elongation factor Tu | 674529 |
| BacF7301_RS02335 | elongation factor Tu | 2715212 |
| BF9343_RS19510 | elongation factor Tu | 272559 |
| SLT85_RS05370 | elongation factor Tu | 477666 |

| | | |
|---|---|---|
| BACHE_RS05225 | elongation factor Tu | 693979 |
| C3V43_RS06450 | elongation factor Tu | 28113 |
| R3O62_RS03290 | elongation factor Tu | 2763023 |
| JXR92_RS05910 | elongation factor Tu | 2792859 |
| GD631_RS02685 | elongation factor Tu | 2650158 |
| VYM24_RS08180 | elongation factor Tu | 3117552 |
| K8P02_RS16055 | elongation factor Tu | 291645 |
| Bovatus_RS13610 | elongation factor Tu | 28116 |
| M1B74_RS01995 | elongation factor Tu | 310300 |
| RB604_RS02150 | elongation factor Tu | 291644 |
| ABWU87_RS10640 | elongation factor Tu | 2136147 |
| FE838_RS03195 | elongation factor Tu | 818 |
| QWU49_RS06500 | elongation factor Tu | 820 |
| INE92_RS16415 | elongation factor Tu | 371601 |
| GD630_RS17630 | elongation factor Tu | 2650157 |
| C4H11_RS06065 | elongation factor Tu | 28119 |
| BARVI_RS10765 | elongation factor Tu | 880074 |
| D8S85_RS09725 | elongation factor Tu | 2093856 |
| F1611_RS19970 | elongation factor Tu | 1472416 |
| F1644_RS16575 | elongation factor Tu | 1472417 |
| NQ494_RS02810 | elongation factor Tu | 544645 |
| CFPG_RS00070 | elongation factor Tu | 511995 |
| QUE35_RS07540 | elongation factor Tu | 1099853 |
| Cop2CBH44_RS12985 | elongation factor Tu | 2751153 |
| G0Q07_RS17505 | elongation factor Tu | 2706887 |
| E7747_RS06610 | elongation factor Tu | 2518971 |
| AABJ51_RS15915 | elongation factor Tu | 45254 |
| NQ518_RS11875 | elongation factor Tu | 679190 |
| AS203_RS00705 | elongation factor Tu | 76123 |
| ALGA_RS13250 | elongation factor Tu | 1717717 |
| ABNK31_RS04355 | elongation factor Tu | 3229863 |

| | | |
|---|---|---|
| GM418_RS05295 | elongation factor Tu | 2681766 |
| U2921_RS11400 | elongation factor Tu | 570161 |
| E7746_RS01480 | elongation factor Tu | 2530390 |
| A4V02_RS10585 | elongation factor Tu | 1796646 |
| CKW01_RS10020 | elongation factor Tu | 28118 |
| PALPR_RS08585 | elongation factor Tu | 694427 |
| NEE14_RS08305 | elongation factor Tu | 2951805 |
| P3L47_RS01285 | elongation factor Tu | 2685834 |
| FQN59_RS03940 | elongation factor Tu | 823 |
| K6V26_RS28395 | elongation factor Tu | 328812 |
| NQ564_RS19120 | elongation factor Tu | 537006 |
| NF347_RS10940 | elongation factor Tu | 2094151 |
| E0E47_RS09540 | elongation factor Tu | 762982 |
| ING2E5A_RS11710 | elongation factor Tu | 1642646 |
| I6J50_RS14150 | elongation factor Tu | 387090 |
| GKD17_RS20235 | elongation factor Tu | 357276 |
| BACSA_RS12625 | elongation factor Tu | 667015 |
| BVU_RS04165 | elongation factor Tu | 435590 |
| PORAS_RS03710 | elongation factor Tu | 879243 |
| EL262_RS08620 | elongation factor Tu | 36874 |
| DQN70_RS01225 | elongation factor Tu | 393921 |
| LA319_RS03390 | elongation factor Tu | 28124 |
| PGN_RS07520 | elongation factor Tu | 431947 |
| VYJ22_RS10890 | elongation factor Tu | 867595 |
| MWL85_RS10930 | elongation factor Tu | 322095 |
| L6468_RS01805 | elongation factor Tu | 2913614 |
| PREDE_RS01045 | elongation factor Tu | 908937 |
| J4866_RS05525 | elongation factor Tu | 28129 |
| ADJ77_RS04905 | elongation factor Tu | 1236517 |
| prwr041_RS04390 | elongation factor Tu | 2801997 |
| J5A62_RS06770 | elongation factor Tu | 470565 |

| | | |
|---|---|---|
| CRM71_RS01470 | elongation factor Tu | 1177574 |
| HMPREF0659_RS01495 | elongation factor Tu | 553174 |
| J4864_RS03895 | elongation factor Tu | 282402 |
| J5A59_RS12065 | elongation factor Tu | 28133 |
| J4856_RS06755 | elongation factor Tu | 1236518 |
| J5A55_RS06950 | elongation factor Tu | 28137 |
| KDN43_RS01790 | elongation factor Tu | 2829812 |
| PSM36_RS14365 | elongation factor Tu | 1642647 |
| C7Y71_RS09325 | elongation factor Tu | 2133944 |
| L21SP5_RS05535 | elongation factor Tu | 1307839 |
| L6471_RS13065 | elongation factor Tu | 77095 |
| LK433_RS00580 | elongation factor Tu | 537011 |
| EL210_RS12135 | elongation factor Tu | 28135 |
| GF423_RS04425 | elongation factor Tu | 2606626 |
| BFO_RS08440 | elongation factor Tu | 203275 |
| BCB71_RS00725 | elongation factor Tu | 712710 |
| FHG85_RS04410 | elongation factor Tu | 2590900 |
| M9189_RS08885 | elongation factor Tu | 2949635 |
| PRU_RS11045 | elongation factor Tu | 264731 |
| CYPRO_RS04000 | elongation factor Tu | 1457365 |
| CYPRO_RS05010 | elongation factor Tu | 1457365 |
| BD_RS13700 | elongation factor Tu | 264462 |
| MNR06_RS15990 | elongation factor Tu | 2835041 |
| A11Q_RS09680 | elongation factor Tu | 1184267 |
| DTPSY_RS01335 | elongation factor Tu | 535289 |
| DTPSY_RS16295 | elongation factor Tu | 535289 |
| FOC80_RS04245 | elongation factor Tu | 32002 |
| FOC80_RS04340 | elongation factor Tu | 32002 |
| EL339_RS29535 | elongation factor Tu | 217204 |
| EL339_RS29630 | elongation factor Tu | 217204 |
| QEP15_RS25605 | elongation factor Tu | 1389922 |

| | | |
|---|---|---|
| QEP15_RS25700 | elongation factor Tu | 1389922 |
| AruCF_RS28595 | elongation factor Tu | 72557 |
| AruCF_RS28690 | elongation factor Tu | 72557 |
| RAS12_RS21975 | elongation factor Tu | 3047478 |
| RAS12_RS22070 | elongation factor Tu | 3047478 |
| CVS48_RS06235 | elongation factor Tu | 217203 |
| CVS48_RS06335 | elongation factor Tu | 217203 |
| WHX56_RS04410 | elongation factor Tu | 2069367 |
| WHX56_RS04505 | elongation factor Tu | 2069367 |
| I6I48_RS03875 | elongation factor Tu | 85698 |
| I6I48_RS03970 | elongation factor Tu | 85698 |
| CBP33_RS01340 | elongation factor Tu | 553814 |
| CBP33_RS01605 | elongation factor Tu | 553814 |
| ACGHC0_RS02125 | elongation factor Tu | 3230736 |
| ACGHC0_RS23285 | elongation factor Tu | 3230736 |
| PQV96_RS18925 | elongation factor Tu | 80878 |
| PQV96_RS19175 | elongation factor Tu | 80878 |
| MIM_RS21165 | elongation factor Tu | 1247726 |
| MIM_RS21505 | elongation factor Tu | 1247726 |
| FE795_RS16905 | elongation factor Tu | 2582914 |
| FE795_RS17000 | elongation factor Tu | 2582914 |
| MTR80_RS17215 | elongation factor Tu | 323284 |
| MTR80_RS17310 | elongation factor Tu | 323284 |
| ALIDE2_RS01750 | elongation factor Tu | 596154 |
| ALIDE2_RS02010 | elongation factor Tu | 596154 |
| LVJ80_RS07695 | elongation factor Tu | 153491 |
| LVJ80_RS07775 | elongation factor Tu | 153491 |
| JF568_RS05900 | elongation factor Tu | 1120981 |
| JF568_RS05985 | elongation factor Tu | 1120981 |
| ACMYR3_RS15455 | elongation factor Tu | 1341549 |
| ACMYR3_RS15520 | elongation factor Tu | 1341549 |

| | | |
|---|---|---|
| DEH84_RS00810 | elongation factor Tu | 1296669 |
| DEH84_RS00905 | elongation factor Tu | 1296669 |
| CUN60_RS02020 | elongation factor Tu | 2067065 |
| CUN60_RS11080 | elongation factor Tu | 2067065 |
| MW290_RS30555 | elongation factor Tu | 391953 |
| MW290_RS30625 | elongation factor Tu | 391953 |
| FAZ30_RS07165 | elongation factor Tu | 1537400 |
| FAZ30_RS07235 | elongation factor Tu | 1537400 |
| GSR16_RS18370 | elongation factor Tu | 519081 |
| GSR16_RS18440 | elongation factor Tu | 519081 |
| EBN1_RS10640 | elongation factor Tu | 76114 |
| EBN1_RS10705 | elongation factor Tu | 76114 |
| pbN1_RS11420 | elongation factor Tu | 76115 |
| pbN1_RS11485 | elongation factor Tu | 76115 |
| U5817_RS22375 | elongation factor Tu | 59406 |
| U5817_RS22440 | elongation factor Tu | 59406 |
| ToN1_RS09595 | elongation factor Tu | 76116 |
| ToN1_RS09660 | elongation factor Tu | 76116 |
| dqs_RS17540 | elongation factor Tu | 418699 |
| dqs_RS17605 | elongation factor Tu | 418699 |
| Azoinq_RS05315 | elongation factor Tu | 2785627 |
| Azoinq_RS05380 | elongation factor Tu | 2785627 |
| DSUI_RS01720 | elongation factor Tu | 640081 |
| DSUI_RS01785 | elongation factor Tu | 640081 |
| IWH25_RS05965 | elongation factor Tu | 404405 |
| IWH25_RS06030 | elongation factor Tu | 404405 |
| IX83_RS00370 | elongation factor Tu | 1072685 |
| IX83_RS08425 | elongation factor Tu | 1072685 |
| ACM67C_RS00235 | elongation factor Tu | 494 |
| ACM67C_RS00405 | elongation factor Tu | 494 |
| U0029_RS17030 | elongation factor Tu | 521 |

| | | |
|---|---|---|
| U0029_RS17120 | elongation factor Tu | 521 |
| BAU06_RS25580 | elongation factor Tu | 463025 |
| BAU06_RS25690 | elongation factor Tu | 463025 |
| EL288_RS24050 | elongation factor Tu | 518 |
| EL288_RS24155 | elongation factor Tu | 518 |
| BAU07_RS25705 | elongation factor Tu | 463014 |
| BAU07_RS25835 | elongation factor Tu | 463014 |
| EL326_RS16400 | elongation factor Tu | 103855 |
| EL326_RS16495 | elongation factor Tu | 103855 |
| LK440_RS16515 | elongation factor Tu | 35814 |
| LK440_RS16615 | elongation factor Tu | 35814 |
| LK409_RS20415 | elongation factor Tu | 519 |
| LK409_RS20525 | elongation factor Tu | 519 |
| CXD86_RS00050 | elongation factor Tu | 520 |
| CXD86_RS01320 | elongation factor Tu | 520 |
| BPET_RS25055 | elongation factor Tu | 94624 |
| BPET_RS25150 | elongation factor Tu | 94624 |
| BBN53_RS20580 | elongation factor Tu | 1331258 |
| BBN53_RS20675 | elongation factor Tu | 1331258 |
| ACN8ZM_RS01455 | elongation factor Tu | 2015348 |
| ACN8ZM_RS01525 | elongation factor Tu | 2015348 |
| CH72_RS06630 | elongation factor Tu | 339670 |
| CH72_RS06700 | elongation factor Tu | 339670 |
| OHZ10_RS01495 | elongation factor Tu | 488730 |
| OHZ10_RS01565 | elongation factor Tu | 488730 |
| WL90_RS01395 | elongation factor Tu | 95486 |
| WL90_RS01465 | elongation factor Tu | 95486 |
| BURCE16_RS01390 | elongation factor Tu | 292 |
| BURCE16_RS01460 | elongation factor Tu | 292 |
| ABNN95_RS26820 | elongation factor Tu | 488447 |
| ABNN95_RS26890 | elongation factor Tu | 488447 |

| | | |
|---|---|---|
| K8O94_RS04500 | elongation factor Tu | 152500 |
| K8O94_RS04580 | elongation factor Tu | 152500 |
| JKG63_RS01460 | elongation factor Tu | 28095 |
| JKG63_RS01580 | elongation factor Tu | 28095 |
| KS03_RS18435 | elongation factor Tu | 1176492 |
| KS03_RS18510 | elongation factor Tu | 1176492 |
| AQ610_RS01470 | elongation factor Tu | 430531 |
| AQ610_RS01540 | elongation factor Tu | 430531 |
| BCEP18194_RS07255 | elongation factor Tu | 482957 |
| BCEP18194_RS07325 | elongation factor Tu | 482957 |
| atcc23344_RS22740 | elongation factor Tu | 13373 |
| atcc23344_RS22815 | elongation factor Tu | 13373 |
| WS70_RS01680 | elongation factor Tu | 1385591 |
| WS70_RS01750 | elongation factor Tu | 1385591 |
| WJ16_RS01290 | elongation factor Tu | 488729 |
| WJ16_RS01360 | elongation factor Tu | 488729 |
| V1241_RS11835 | elongation factor Tu | 87883 |
| V1241_RS11905 | elongation factor Tu | 87883 |
| BG90_RS08880 | elongation factor Tu | 441162 |
| BG90_RS08950 | elongation factor Tu | 441162 |
| BpN12_RS15770 | elongation factor Tu | 2615214 |
| BpN12_RS15845 | elongation factor Tu | 2615214 |
| bpln_RS01220 | elongation factor Tu | 41899 |
| bpln_RS01295 | elongation factor Tu | 41899 |
| QUF21_RS16705 | elongation factor Tu | 28450 |
| QUF21_RS16780 | elongation factor Tu | 28450 |
| ABD05_RS07320 | elongation factor Tu | 60550 |
| ABD05_RS07390 | elongation factor Tu | 60550 |
| WS78_RS00505 | elongation factor Tu | 1637837 |
| WS78_RS00580 | elongation factor Tu | 1637837 |
| WJ12_RS01345 | elongation factor Tu | 488731 |

| | | |
|---|---|---|
| WJ12_RS01415 | elongation factor Tu | 488731 |
| AQ611_RS02565 | elongation factor Tu | 1503053 |
| AQ611_RS02640 | elongation factor Tu | 1503053 |
| BSTAB16_RS11270 | elongation factor Tu | 95485 |
| BSTAB16_RS11345 | elongation factor Tu | 95485 |
| ABFO07_RS00910 | elongation factor Tu | 1503054 |
| ABFO07_RS00985 | elongation factor Tu | 1503054 |
| WS51_RS12105 | elongation factor Tu | 1503055 |
| WS51_RS12175 | elongation factor Tu | 1503055 |
| BTH_RS27835 | elongation factor Tu | 271848 |
| BTH_RS27905 | elongation factor Tu | 271848 |
| ACN22W_RS01635 | elongation factor Tu | 3143496 |
| ACN22W_RS01715 | elongation factor Tu | 3143496 |
| B7P44_RS01515 | elongation factor Tu | 265293 |
| B7P44_RS01585 | elongation factor Tu | 265293 |
| AK36_RS12380 | elongation factor Tu | 1449978 |
| AK36_RS12450 | elongation factor Tu | 1449978 |
| LDZ21_RS01260 | elongation factor Tu | 1071679 |
| LDZ21_RS01330 | elongation factor Tu | 1071679 |
| BRPE64_RS12460 | elongation factor Tu | 758793 |
| BRPE64_RS12530 | elongation factor Tu | 758793 |
| LDZ23_RS12900 | elongation factor Tu | 871203 |
| LDZ23_RS12970 | elongation factor Tu | 871203 |
| OMP39_RS13540 | elongation factor Tu | 376175 |
| OMP39_RS13670 | elongation factor Tu | 376175 |
| AAW51_RS24465 | elongation factor Tu | 413882 |
| AAW51_RS24540 | elongation factor Tu | 413882 |
| IS481_RS15910 | elongation factor Tu | 215580 |
| IS481_RS15985 | elongation factor Tu | 215580 |
| CAP2UW1_RS17330 | elongation factor Tu | 327160 |
| CAP2UW1_RS17395 | elongation factor Tu | 327160 |

| | | |
|---|---|---|
| Arg41_RS02310 | elongation factor Tu | 1561003 |
| Arg41_RS02375 | elongation factor Tu | 1561003 |
| CKBE_RS00770 | elongation factor Tu | 1208923 |
| CKBE_RS00835 | elongation factor Tu | 1208923 |
| CDEE_RS03790 | elongation factor Tu | 1208918 |
| CDEE_RS03855 | elongation factor Tu | 1208918 |
| CDSE_RS03835 | elongation factor Tu | 1208919 |
| CDSE_RS03900 | elongation factor Tu | 1208919 |
| CKSOR_RS03490 | elongation factor Tu | 1576550 |
| CONE_RS03640 | elongation factor Tu | 1208920 |
| CONE_RS03705 | elongation factor Tu | 1208920 |
| FIT68_RS00960 | elongation factor Tu | 1581557 |
| FIT68_RS01025 | elongation factor Tu | 1581557 |
| FIT61_RS05815 | elongation factor Tu | 2588535 |
| FIT61_RS05880 | elongation factor Tu | 2588535 |
| FIT94_RS00910 | elongation factor Tu | 2588536 |
| FIT94_RS00975 | elongation factor Tu | 2588536 |
| MP021_RS00655 | elongation factor Tu | 1160784 |
| MKZ32_RS13950 | elongation factor Tu | 453162 |
| MKZ32_RS14015 | elongation factor Tu | 453162 |
| SSDC_RS00105 | elongation factor Tu | 669502 |
| CENROD_RS01670 | elongation factor Tu | 946483 |
| CENROD_RS04685 | elongation factor Tu | 946483 |
| GKR41_RS00350 | elongation factor Tu | 2018052 |
| GKR41_RS00415 | elongation factor Tu | 2018052 |
| GJV44_RS00215 | elongation factor Tu | 1177211 |
| GJV44_RS00285 | elongation factor Tu | 1177211 |
| KW136_RS03750 | elongation factor Tu | 1177213 |
| KW136_RS03825 | elongation factor Tu | 1177213 |
| AACL00_RS00575 | elongation factor Tu | 3077951 |
| FKL89_RS04595 | elongation factor Tu | 2591109 |

| | | |
|---|---|---|
| FKL89_RS04660 | elongation factor Tu | 2591109 |
| BN940_RS00605 | elongation factor Tu | 1437824 |
| BN940_RS00770 | elongation factor Tu | 1437824 |
| ABRZ00_RS05350 | elongation factor Tu | 546114 |
| ABRZ00_RS05465 | elongation factor Tu | 546114 |
| HQ393_RS10950 | elongation factor Tu | 2739434 |
| HQ393_RS11015 | elongation factor Tu | 2739434 |
| HZU75_RS10330 | elongation factor Tu | 1737446 |
| HZU75_RS10395 | elongation factor Tu | 1737446 |
| FNU76_RS08410 | elongation factor Tu | 2594795 |
| FNU76_RS08475 | elongation factor Tu | 2594795 |
| ABFG87_RS20190 | elongation factor Tu | 2571148 |
| ABFG87_RS20255 | elongation factor Tu | 2571148 |
| N8I74_RS05855 | elongation factor Tu | 2981137 |
| N8I74_RS05920 | elongation factor Tu | 2981137 |
| FLM21_RS05085 | elongation factor Tu | 682798 |
| FLM21_RS05150 | elongation factor Tu | 682798 |
| GYH29_RS01880 | elongation factor Tu | 394935 |
| GYH29_RS01950 | elongation factor Tu | 394935 |
| FYK34_RS19260 | elongation factor Tu | 2605945 |
| FYK34_RS19330 | elongation factor Tu | 2605945 |
| DK842_RS10365 | elongation factor Tu | 2202141 |
| DK842_RS10435 | elongation factor Tu | 2202141 |
| JC616_RS02530 | elongation factor Tu | 1778675 |
| JC616_RS02600 | elongation factor Tu | 1778675 |
| U6151_RS21635 | elongation factor Tu | 251747 |
| U6151_RS21705 | elongation factor Tu | 251747 |
| BKX93_RS16460 | elongation factor Tu | 1108595 |
| BKX93_RS16530 | elongation factor Tu | 1108595 |
| I6K03_RS09570 | elongation factor Tu | 536 |
| I6K03_RS09640 | elongation factor Tu | 536 |

| | | |
|---|---|---|
| CAter282_RS01405 | elongation factor Tu | 279058 |
| CAter282_RS01475 | elongation factor Tu | 279058 |
| CFter6_RS02835 | elongation factor Tu | 158899 |
| CFter6_RS02915 | elongation factor Tu | 158899 |
| CPter291_RS01815 | elongation factor Tu | 279113 |
| CPter291_RS01890 | elongation factor Tu | 279113 |
| HUK68_RS17805 | elongation factor Tu | 2743470 |
| HUK68_RS18025 | elongation factor Tu | 2743470 |
| KAQ61_RS17615 | elongation factor Tu | 225991 |
| KAQ61_RS17855 | elongation factor Tu | 225991 |
| M9799_RS07120 | elongation factor Tu | 2949090 |
| M9799_RS07340 | elongation factor Tu | 2949090 |
| FOZ74_RS05435 | elongation factor Tu | 2597701 |
| FOZ74_RS06355 | elongation factor Tu | 2597701 |
| JDW18_RS01490 | elongation factor Tu | 2796366 |
| JDW18_RS01790 | elongation factor Tu | 2796366 |
| B5M06_RS04670 | elongation factor Tu | 225992 |
| B5M06_RS04910 | elongation factor Tu | 225992 |
| LAD35_RS02005 | elongation factor Tu | 379895 |
| LAD35_RS02335 | elongation factor Tu | 379895 |
| HS961_RS00455 | elongation factor Tu | 1562974 |
| HS961_RS00730 | elongation factor Tu | 1562974 |
| QMY55_RS01410 | elongation factor Tu | 3046670 |
| QMY55_RS01665 | elongation factor Tu | 3046670 |
| CCO03_RS01010 | elongation factor Tu | 1082851 |
| CCO03_RS19220 | elongation factor Tu | 1082851 |
| N4T19_RS14580 | elongation factor Tu | 2977320 |
| N4T19_RS14855 | elongation factor Tu | 2977320 |
| CT3_RS01190 | elongation factor Tu | 1219032 |
| CT3_RS01630 | elongation factor Tu | 1219032 |
| Cthiooxydans_RS01540 | elongation factor Tu | 363952 |

| | | |
|---|---|---|
| Cthiooxydans_RS01795 | elongation factor Tu | 363952 |
| H3L98_RS01870 | elongation factor Tu | 153493 |
| H3L98_RS02010 | elongation factor Tu | 153493 |
| DWG20_RS03525 | elongation factor Tu | 2290923 |
| DWG20_RS03595 | elongation factor Tu | 2290923 |
| F7R26_RS19095 | elongation factor Tu | 68895 |
| FOB47_RS00630 | elongation factor Tu | 82541 |
| FOB47_RS00705 | elongation factor Tu | 82541 |
| BKK80_RS19750 | elongation factor Tu | 367825 |
| BKK80_RS19820 | elongation factor Tu | 367825 |
| CNE_RS16795 | elongation factor Tu | 1042878 |
| CNE_RS16865 | elongation factor Tu | 1042878 |
| CBM2589_RS15490 | elongation factor Tu | 1040979 |
| CBM2589_RS15565 | elongation factor Tu | 1040979 |
| JTE92_RS28990 | elongation factor Tu | 96344 |
| JTE92_RS29060 | elongation factor Tu | 96344 |
| K8O84_RS06925 | elongation factor Tu | 82633 |
| K8O84_RS06995 | elongation factor Tu | 82633 |
| K2O51_RS29315 | elongation factor Tu | 248026 |
| K2O51_RS29390 | elongation factor Tu | 248026 |
| RALTA_RS14390 | elongation factor Tu | 977880 |
| RALTA_RS14460 | elongation factor Tu | 977880 |
| VX159_RS14400 | elongation factor Tu | 2664436 |
| VX159_RS14465 | elongation factor Tu | 2664436 |
| HQN60_RS03990 | elongation factor Tu | 2739061 |
| HQN60_RS04055 | elongation factor Tu | 2739061 |
| K4H28_RS12435 | elongation factor Tu | 2865840 |
| K4H28_RS12500 | elongation factor Tu | 2865840 |
| DACI_RS01945 | elongation factor Tu | 398578 |
| DACI_RS02545 | elongation factor Tu | 398578 |
| U6958_RS03990 | elongation factor Tu | 558537 |

| | | |
|---|---|---|
| U6958_RS04240 | elongation factor Tu | 558537 |
| PW274_RS29375 | elongation factor Tu | 180282 |
| PW274_RS29625 | elongation factor Tu | 180282 |
| DENOEST_RS14885 | elongation factor Tu | 311182 |
| DENOEST_RS14950 | elongation factor Tu | 311182 |
| H9K75_RS11715 | elongation factor Tu | 1288495 |
| H9K75_RS15395 | elongation factor Tu | 1288495 |
| P4826_RS11125 | elongation factor Tu | 3036128 |
| P4826_RS11350 | elongation factor Tu | 3036128 |
| BA022_RS13880 | elongation factor Tu | 164759 |
| BA022_RS16705 | elongation factor Tu | 164759 |
| H9K76_RS00820 | elongation factor Tu | 1715720 |
| H9K76_RS22265 | elongation factor Tu | 1715720 |
| HH213_RS12755 | elongation factor Tu | 2728021 |
| HH213_RS13135 | elongation factor Tu | 2728021 |
| SR858_RS24395 | elongation factor Tu | 75659 |
| SR858_RS24695 | elongation factor Tu | 75659 |
| NXF97_RS06170 | elongation factor Tu | 1852381 |
| NXF97_RS06245 | elongation factor Tu | 1852381 |
| CKV94_RS05675 | elongation factor Tu | 539 |
| CKV94_RS05710 | elongation factor Tu | 539 |
| EZJ17_RS00090 | elongation factor Tu | 2528037 |
| EZJ17_RS00125 | elongation factor Tu | 2528037 |
| N7L95_RS17395 | elongation factor Tu | 1597781 |
| N7L95_RS17475 | elongation factor Tu | 1597781 |
| DTO96_RS00655 | elongation factor Tu | 2268024 |
| DTO96_RS01380 | elongation factor Tu | 2268024 |
| KI610_RS01665 | elongation factor Tu | 76259 |
| KI610_RS01730 | elongation factor Tu | 76259 |
| FGKAn22_RS09565 | elongation factor Tu | 1682490 |
| FGKAn22_RS09630 | elongation factor Tu | 1682490 |

| | | |
|---|---|---|
| OYT1_RS02280 | elongation factor Tu | 1188319 |
| OYT1_RS02345 | elongation factor Tu | 1188319 |
| HO274_RS08935 | elongation factor Tu | 416213 |
| HO274_RS09000 | elongation factor Tu | 416213 |
| SHINM1_RS09645 | elongation factor Tu | 1751046 |
| SHINM1_RS09710 | elongation factor Tu | 1751046 |
| GALF_RS02765 | elongation factor Tu | 395494 |
| GALF_RS02830 | elongation factor Tu | 395494 |
| ACEOAC_RS00345 | elongation factor Tu | 92645 |
| ACEOAC_RS00420 | elongation factor Tu | 92645 |
| F506_RS21655 | elongation factor Tu | 1262470 |
| F506_RS21730 | elongation factor Tu | 1262470 |
| E2K99_RS00695 | elongation factor Tu | 863372 |
| E2K99_RS00770 | elongation factor Tu | 863372 |
| BUQ72_RS15155 | elongation factor Tu | 2014887 |
| BUQ72_RS15230 | elongation factor Tu | 2014887 |
| RC54_RS00550 | elongation factor Tu | 80842 |
| RC54_RS00630 | elongation factor Tu | 80842 |
| ACP92_RS00480 | elongation factor Tu | 964 |
| ACP92_RS00555 | elongation factor Tu | 964 |
| BQ6873_RS13005 | elongation factor Tu | 1809410 |
| BQ6873_RS13080 | elongation factor Tu | 1809410 |
| LPB072_RS01910 | elongation factor Tu | 1763535 |
| LPB072_RS21480 | elongation factor Tu | 1763535 |
| G9Q37_RS16130 | elongation factor Tu | 2716225 |
| G9Q37_RS18795 | elongation factor Tu | 2716225 |
| HPF_RS02040 | elongation factor Tu | 47421 |
| HPF_RS21695 | elongation factor Tu | 47421 |
| LRM40_RS15490 | elongation factor Tu | 36863 |
| LRM40_RS15560 | elongation factor Tu | 36863 |
| FF090_RS03165 | elongation factor Tu | 392593 |

| | | |
|---|---|---|
| FF090_RS03235 | elongation factor Tu | 392593 |
| EJO50_RS11425 | elongation factor Tu | 2496266 |
| EJO50_RS11490 | elongation factor Tu | 2496266 |
| OPV09_RS03160 | elongation factor Tu | 2985511 |
| OPV09_RS03400 | elongation factor Tu | 2985511 |
| GJA_RS00005 | elongation factor Tu | 1349767 |
| G3257_RS02830 | elongation factor Tu | 29581 |
| G3257_RS03055 | elongation factor Tu | 29581 |
| P9875_RS02925 | elongation factor Tu | 2751478 |
| P9875_RS03165 | elongation factor Tu | 2751478 |
| CNX70_RS02995 | elongation factor Tu | 368607 |
| CNX70_RS03205 | elongation factor Tu | 368607 |
| FJQ89_RS19960 | elongation factor Tu | 2590869 |
| FJQ89_RS20165 | elongation factor Tu | 2590869 |
| ACEW7X_RS16145 | elongation factor Tu | 206506 |
| ACEW7X_RS16850 | elongation factor Tu | 206506 |
| I6I17_RS00910 | elongation factor Tu | 502 |
| I6I17_RS01285 | elongation factor Tu | 502 |
| DQN52_RS02040 | elongation factor Tu | 504 |
| DQN52_RS02790 | elongation factor Tu | 504 |
| QEO93_RS07860 | elongation factor Tu | 1522312 |
| QEO93_RS08525 | elongation factor Tu | 1522312 |
| EL249_RS03595 | elongation factor Tu | 47671 |
| EL249_RS03665 | elongation factor Tu | 47671 |
| LCHO_RS19225 | elongation factor Tu | 395495 |
| LCHO_RS19285 | elongation factor Tu | 395495 |
| HKT17_RS01720 | elongation factor Tu | 2732163 |
| HKT17_RS01785 | elongation factor Tu | 2732163 |
| RGQ30_RS14315 | elongation factor Tu | 131080 |
| RGQ30_RS14380 | elongation factor Tu | 131080 |
| IV454_RS09835 | elongation factor Tu | 2765360 |

| | | |
|---|---|---|
| IV454_RS10215 | elongation factor Tu | 2765360 |
| HF313_RS03610 | elongation factor Tu | 2495579 |
| HF313_RS04020 | elongation factor Tu | 2495579 |
| ACEN9H_RS28440 | elongation factor Tu | 2683234 |
| ACEN9H_RS28865 | elongation factor Tu | 2683234 |
| LSQ66_RS20245 | elongation factor Tu | 2899220 |
| LSQ66_RS23570 | elongation factor Tu | 2899220 |
| HH212_RS06400 | elongation factor Tu | 2728020 |
| HH212_RS06705 | elongation factor Tu | 2728020 |
| LPB04_RS03995 | elongation factor Tu | 2769491 |
| LPB04_RS07320 | elongation factor Tu | 2769491 |
| DIR46_RS11990 | elongation factor Tu | 945844 |
| DIR46_RS12355 | elongation factor Tu | 945844 |
| BVG12_RS21460 | elongation factor Tu | 1141883 |
| BVG12_RS21755 | elongation factor Tu | 1141883 |
| CR152_RS03725 | elongation factor Tu | 2045208 |
| CR152_RS04200 | elongation factor Tu | 2045208 |
| LVC68_RS01255 | elongation factor Tu | 1267217 |
| LVC68_RS15400 | elongation factor Tu | 1267217 |
| C6568_RS07630 | elongation factor Tu | 2109913 |
| C6568_RS10550 | elongation factor Tu | 2109913 |
| MPE_RS17405 | elongation factor Tu | 420662 |
| MPE_RS17475 | elongation factor Tu | 420662 |
| MFLA_RS01380 | elongation factor Tu | 265072 |
| MFLA_RS01445 | elongation factor Tu | 265072 |
| FIU01_RS01620 | elongation factor Tu | 2588534 |
| FIU01_RS01685 | elongation factor Tu | 2588534 |
| ZMTM_RS01400 | elongation factor Tu | 2778876 |
| ZMTM_RS01470 | elongation factor Tu | 2778876 |
| MMOL_RS01700 | elongation factor Tu | 583345 |
| MMOL_RS01765 | elongation factor Tu | 583345 |

| | | |
|---|---|---|
| M301_RS01685 | elongation factor Tu | 666681 |
| M301_RS01750 | elongation factor Tu | 666681 |
| MSIP34_RS01505 | elongation factor Tu | 582744 |
| MSIP34_RS01570 | elongation factor Tu | 582744 |
| DAI18_RS04985 | elongation factor Tu | 57480 |
| DAI18_RS05050 | elongation factor Tu | 57480 |
| MON37_RS00355 | elongation factor Tu | 1056807 |
| MON37_RS00420 | elongation factor Tu | 1056807 |
| IHE33_RS01275 | elongation factor Tu | 417203 |
| IHE33_RS01360 | elongation factor Tu | 417203 |
| IHE31_RS04615 | elongation factor Tu | 412963 |
| IHE31_RS04700 | elongation factor Tu | 412963 |
| MCB1EB_RS01140 | elongation factor Tu | 1553431 |
| D0T90_RS04820 | elongation factor Tu | 492 |
| D0T90_RS05030 | elongation factor Tu | 492 |
| EL216_RS07830 | elongation factor Tu | 326522 |
| EL216_RS08120 | elongation factor Tu | 326522 |
| LVJ86_RS00770 | elongation factor Tu | 1470200 |
| LVJ86_RS10655 | elongation factor Tu | 1470200 |
| H3L91_RS02305 | elongation factor Tu | 267212 |
| H3L91_RS02640 | elongation factor Tu | 267212 |
| GJV52_RS06350 | elongation factor Tu | 2666100 |
| GJV52_RS06430 | elongation factor Tu | 2666100 |
| EL143_RS00830 | elongation factor Tu | 493 |
| EL143_RS05280 | elongation factor Tu | 493 |
| BG910_RS04535 | elongation factor Tu | 1853278 |
| BG910_RS05160 | elongation factor Tu | 1853278 |
| DQM57_RS08640 | elongation factor Tu | 483 |
| DQM57_RS08715 | elongation factor Tu | 483 |
| H3L92_RS10765 | elongation factor Tu | 194197 |
| H3L92_RS12525 | elongation factor Tu | 194197 |

| | | |
|---|---|---|
| LVJ88_RS01105 | elongation factor Tu | 1931275 |
| LVJ88_RS10350 | elongation factor Tu | 1931275 |
| FAH67_RS09905 | elongation factor Tu | 484 |
| FAH67_RS09985 | elongation factor Tu | 484 |
| HT085_RS10225 | elongation factor Tu | 485 |
| HT085_RS10320 | elongation factor Tu | 485 |
| DV311_RS10745 | elongation factor Tu | 486 |
| DV311_RS10820 | elongation factor Tu | 486 |
| ORY85_RS06680 | elongation factor Tu | 2995413 |
| ORY85_RS06985 | elongation factor Tu | 2995413 |
| PJU73_RS08645 | elongation factor Tu | 2912188 |
| PJU73_RS08710 | elongation factor Tu | 2912188 |
| MON40_RS01700 | elongation factor Tu | 997348 |
| MON40_RS01765 | elongation factor Tu | 997348 |
| IUJ45_RS06845 | elongation factor Tu | 487 |
| IUJ45_RS06915 | elongation factor Tu | 487 |
| H7A79_RS10255 | elongation factor Tu | 1815583 |
| H7A79_RS11005 | elongation factor Tu | 1815583 |
| H3L94_RS10405 | elongation factor Tu | 607712 |
| H3L94_RS11170 | elongation factor Tu | 607712 |
| J7445_RS10965 | elongation factor Tu | 490 |
| J7445_RS11030 | elongation factor Tu | 490 |
| FAH66_RS08795 | elongation factor Tu | 28449 |
| FAH66_RS08865 | elongation factor Tu | 28449 |
| H3L96_RS01485 | elongation factor Tu | 607711 |
| H3L96_RS05535 | elongation factor Tu | 607711 |
| EL309_RS06490 | elongation factor Tu | 28091 |
| EL309_RS06555 | elongation factor Tu | 28091 |
| H4O27_RS00515 | elongation factor Tu | 2709396 |
| H4O27_RS00600 | elongation factor Tu | 2709396 |
| D0T92_RS02620 | elongation factor Tu | 640030 |

| | | |
|---|---|---|
| D0T92_RS03425 | elongation factor Tu | 640030 |
| CKV66_RS00160 | elongation factor Tu | 326523 |
| CKV66_RS03650 | elongation factor Tu | 326523 |
| N7E01_RS08265 | elongation factor Tu | 2979868 |
| N7E01_RS08395 | elongation factor Tu | 2979868 |
| J0W34_RS18290 | elongation factor Tu | 2815343 |
| J0W34_RS18355 | elongation factor Tu | 2815343 |
| G3580_RS16420 | elongation factor Tu | 2016596 |
| G3580_RS16485 | elongation factor Tu | 2016596 |
| AAW31_RS01335 | elongation factor Tu | 44574 |
| AAW31_RS16710 | elongation factor Tu | 44574 |
| ALW85_RS02125 | elongation factor Tu | 228410 |
| ALW85_RS10695 | elongation factor Tu | 228410 |
| NEUT_RS02805 | elongation factor Tu | 335283 |
| NEUT_RS09130 | elongation factor Tu | 335283 |
| ATY38_RS08540 | elongation factor Tu | 44577 |
| ATY38_RS10040 | elongation factor Tu | 44577 |
| F822_RS04355 | elongation factor Tu | 1266925 |
| F822_RS04425 | elongation factor Tu | 1266925 |
| EBAPG3_RS05990 | elongation factor Tu | 1288494 |
| EBAPG3_RS06070 | elongation factor Tu | 1288494 |
| NMUL_RS03985 | elongation factor Tu | 323848 |
| NMUL_RS04055 | elongation factor Tu | 323848 |
| JY500_RS03280 | elongation factor Tu | 2811415 |
| JY500_RS03345 | elongation factor Tu | 2811415 |
| I6G29_RS06900 | elongation factor Tu | 90244 |
| I6G29_RS07125 | elongation factor Tu | 90244 |
| DHf2319_RS00790 | elongation factor Tu | 2782176 |
| DHf2319_RS00855 | elongation factor Tu | 2782176 |
| ODI_RS00475 | elongation factor Tu | 1851544 |
| ODI_RS00695 | elongation factor Tu | 1851544 |

| | | |
|---|---|---|
| DBV39_RS14180 | elongation factor Tu | 2163011 |
| DBV39_RS14245 | elongation factor Tu | 2163011 |
| OTERR_RS14060 | elongation factor Tu | 1735038 |
| OTERR_RS14125 | elongation factor Tu | 1735038 |
| C6570_RS01610 | elongation factor Tu | 2109914 |
| C6570_RS02500 | elongation factor Tu | 2109914 |
| BRW83_RS00385 | elongation factor Tu | 847 |
| BRW83_RS00450 | elongation factor Tu | 847 |
| NB640_RS07770 | elongation factor Tu | 933080 |
| NB640_RS07835 | elongation factor Tu | 933080 |
| N7U67_RS12625 | elongation factor Tu | 2980099 |
| N7U67_RS12705 | elongation factor Tu | 2980099 |
| JNO50_RS17780 | elongation factor Tu | 1225769 |
| JNO50_RS17855 | elongation factor Tu | 1225769 |
| JNO49_RS19465 | elongation factor Tu | 400061 |
| JNO49_RS19535 | elongation factor Tu | 400061 |
| AT395_RS01390 | elongation factor Tu | 93218 |
| AT395_RS01555 | elongation factor Tu | 93218 |
| NTU39_RS25120 | elongation factor Tu | 2508289 |
| NTU39_RS25300 | elongation factor Tu | 2508289 |
| AB870_RS21415 | elongation factor Tu | 656179 |
| AB870_RS21585 | elongation factor Tu | 656179 |
| PI93_RS04870 | elongation factor Tu | 1891094 |
| PI93_RS05045 | elongation factor Tu | 1891094 |
| AT302_RS25480 | elongation factor Tu | 93219 |
| AT302_RS25660 | elongation factor Tu | 93219 |
| MB84_RS23130 | elongation factor Tu | 573737 |
| MB84_RS23315 | elongation factor Tu | 573737 |
| X636_RS22740 | elongation factor Tu | 93220 |
| X636_RS22900 | elongation factor Tu | 93220 |
| RO07_RS23775 | elongation factor Tu | 93221 |

| | | |
|---|---|---|
| RO07_RS23935 | elongation factor Tu | 93221 |
| CKW10_RS23775 | elongation factor Tu | 93222 |
| CKW10_RS23950 | elongation factor Tu | 93222 |
| ABW99_RS01070 | elongation factor Tu | 445709 |
| ABW99_RS01135 | elongation factor Tu | 445709 |
| UC34_RS22990 | elongation factor Tu | 656178 |
| UC34_RS23170 | elongation factor Tu | 656178 |
| CJU94_RS06640 | elongation factor Tu | 2026199 |
| CJU94_RS06710 | elongation factor Tu | 2026199 |
| BC1002_RS14510 | elongation factor Tu | 2654982 |
| BC1002_RS14580 | elongation factor Tu | 2654982 |
| NDK50_RS19495 | elongation factor Tu | 420952 |
| NDK50_RS19570 | elongation factor Tu | 420952 |
| DSC91_RS24355 | elongation factor Tu | 1761016 |
| DSC91_RS24425 | elongation factor Tu | 1761016 |
| K788_RS04795 | elongation factor Tu | 1323664 |
| K788_RS04865 | elongation factor Tu | 1323664 |
| PDMSB3_RS18865 | elongation factor Tu | 2604047 |
| PDMSB3_RS18935 | elongation factor Tu | 2604047 |
| DMB87_RS12745 | elongation factor Tu | 2211211 |
| DMB87_RS12905 | elongation factor Tu | 2211211 |
| KZJ38_RS02870 | elongation factor Tu | 2861782 |
| KZJ38_RS02950 | elongation factor Tu | 2861782 |
| OI25_RS14045 | elongation factor Tu | 134537 |
| OI25_RS14115 | elongation factor Tu | 134537 |
| I6I06_RS13110 | elongation factor Tu | 311231 |
| I6I06_RS13185 | elongation factor Tu | 311231 |
| CUJ91_RS16635 | elongation factor Tu | 60548 |
| CUJ91_RS16705 | elongation factor Tu | 60548 |
| C2L64_RS16865 | elongation factor Tu | 169430 |
| C2L64_RS16935 | elongation factor Tu | 169430 |

| | | |
|---|---|---|
| RW095_RS33855 | elongation factor Tu | 1245436 |
| RW095_RS33925 | elongation factor Tu | 1245436 |
| ACMFWW_RS16660 | elongation factor Tu | 984307 |
| ACMFWW_RS16730 | elongation factor Tu | 984307 |
| PPGU16_RS15200 | elongation factor Tu | 3014751 |
| PPGU16_RS15270 | elongation factor Tu | 3014751 |
| E1956_RS04140 | elongation factor Tu | 2547399 |
| E1956_RS04300 | elongation factor Tu | 2547399 |
| BPHY_RS14440 | elongation factor Tu | 391038 |
| BPHY_RS14510 | elongation factor Tu | 391038 |
| BPHYT_RS18125 | elongation factor Tu | 398527 |
| BPHYT_RS18195 | elongation factor Tu | 398527 |
| QEN71_RS28275 | elongation factor Tu | 273251 |
| QEN71_RS28345 | elongation factor Tu | 273251 |
| BJG93_RS15295 | elongation factor Tu | 754502 |
| BJG93_RS15365 | elongation factor Tu | 754502 |
| C2L65_RS14700 | elongation factor Tu | 311230 |
| C2L65_RS14770 | elongation factor Tu | 311230 |
| CUJ90_RS16760 | elongation factor Tu | 169427 |
| CUJ90_RS16830 | elongation factor Tu | 169427 |
| G5S35_RS13435 | elongation factor Tu | 92647 |
| G5S35_RS13510 | elongation factor Tu | 92647 |
| BXE_RS20335 | elongation factor Tu | 266265 |
| BXE_RS20405 | elongation factor Tu | 266265 |
| ACAV_RS01960 | elongation factor Tu | 643561 |
| ACAV_RS22110 | elongation factor Tu | 643561 |
| FRC97_RS03740 | elongation factor Tu | 80869 |
| FRC97_RS06860 | elongation factor Tu | 80869 |
| JQU52_RS03930 | elongation factor Tu | 2809020 |
| JQU52_RS04040 | elongation factor Tu | 2809020 |
| PFX98_RS10150 | elongation factor Tu | 3019553 |

| | | |
|---|---|---|
| PFX98_RS10225 | elongation factor Tu | 3019553 |
| F9B76_RS06910 | elongation factor Tu | 2652177 |
| F9B76_RS07060 | elongation factor Tu | 2652177 |
| FXN63_RS25235 | elongation factor Tu | 1940612 |
| FXN63_RS25270 | elongation factor Tu | 1940612 |
| CPZ87_RS02585 | elongation factor Tu | 946333 |
| CPZ87_RS02655 | elongation factor Tu | 946333 |
| ACNH6H_RS10165 | elongation factor Tu | 1547922 |
| ACNH6H_RS10235 | elongation factor Tu | 1547922 |
| ACMDW7_RS01760 | elongation factor Tu | 1793872 |
| ACMDW7_RS16195 | elongation factor Tu | 1793872 |
| ABLV49_RS01030 | elongation factor Tu | 335476 |
| ABLV49_RS02910 | elongation factor Tu | 335476 |
| PNAP_RS00985 | elongation factor Tu | 365044 |
| PNAP_RS18215 | elongation factor Tu | 365044 |
| HC248_RS00790 | elongation factor Tu | 37448 |
| HC248_RS16475 | elongation factor Tu | 37448 |
| CKA81_RS14445 | elongation factor Tu | 2028345 |
| CKA81_RS14525 | elongation factor Tu | 2028345 |
| AOC32_RS00190 | elongation factor Tu | 556053 |
| AOC32_RS00255 | elongation factor Tu | 556053 |
| DCO16_RS00195 | elongation factor Tu | 1743162 |
| DCO16_RS00260 | elongation factor Tu | 1743162 |
| DN92_RS00195 | elongation factor Tu | 1743165 |
| DN92_RS00260 | elongation factor Tu | 1743165 |
| PNUC_RS00240 | elongation factor Tu | 312153 |
| PNUC_RS00305 | elongation factor Tu | 312153 |
| C2747_RS00220 | elongation factor Tu | 2081042 |
| C2747_RS00285 | elongation factor Tu | 2081042 |
| AOC34_RS00190 | elongation factor Tu | 556054 |
| AOC34_RS00255 | elongation factor Tu | 556054 |

| | | |
|---|---|---|
| CL55_RS00225 | elongation factor Tu | 1835254 |
| CL55_RS00290 | elongation factor Tu | 1835254 |
| AOC20_RS00230 | elongation factor Tu | 1819725 |
| AOC20_RS00295 | elongation factor Tu | 1819725 |
| DXE33_RS08100 | elongation factor Tu | 576610 |
| DXE33_RS08165 | elongation factor Tu | 576610 |
| AOC06_RS00225 | elongation factor Tu | 1855895 |
| AOC06_RS00290 | elongation factor Tu | 1855895 |
| C2750_RS00225 | elongation factor Tu | 2527775 |
| C2750_RS00290 | elongation factor Tu | 2527775 |
| DCO17_RS00235 | elongation factor Tu | 1743174 |
| DCO17_RS00300 | elongation factor Tu | 1743174 |
| A8O14_RS00230 | elongation factor Tu | 1743168 |
| A8O14_RS00295 | elongation factor Tu | 1743168 |
| QMF28_RS00230 | elongation factor Tu | 1743159 |
| QMF28_RS00295 | elongation factor Tu | 1743159 |
| C0099_RS02915 | elongation factor Tu | 2067960 |
| C0099_RS02980 | elongation factor Tu | 2067960 |
| EYF70_RS26860 | elongation factor Tu | 321983 |
| EYF70_RS27075 | elongation factor Tu | 321983 |
| KT882_RS03675 | elongation factor Tu | 2660641 |
| KT882_RS03980 | elongation factor Tu | 2660641 |
| C9I28_RS26440 | elongation factor Tu | 2072590 |
| C9I28_RS26775 | elongation factor Tu | 2072590 |
| PX653_RS23855 | elongation factor Tu | 34070 |
| PX653_RS24210 | elongation factor Tu | 34070 |
| GO485_RS17895 | elongation factor Tu | 871742 |
| GO485_RS18205 | elongation factor Tu | 871742 |
| EWM63_RS11075 | elongation factor Tu | 321985 |
| EWM63_RS11285 | elongation factor Tu | 321985 |
| E1742_RS18150 | elongation factor Tu | 321984 |

| | | |
|---|---|---|
| E1742_RS18415 | elongation factor Tu | 321984 |
| FCL38_RS22355 | elongation factor Tu | 864828 |
| FCL38_RS22575 | elongation factor Tu | 864828 |
| DIE29_RS02165 | elongation factor Tu | 2184083 |
| DIE29_RS02230 | elongation factor Tu | 2184083 |
| C7H73_RS02665 | elongation factor Tu | 2116657 |
| C7H73_RS03225 | elongation factor Tu | 2116657 |
| KI612_RS01540 | elongation factor Tu | 138118 |
| KI612_RS01605 | elongation factor Tu | 138118 |
| A9Y76_RS16050 | elongation factor Tu | 190721 |
| A9Y76_RS16165 | elongation factor Tu | 190721 |
| TK49_RS00745 | elongation factor Tu | 105219 |
| TK49_RS00860 | elongation factor Tu | 105219 |
| GO999_RS01705 | elongation factor Tu | 3037696 |
| GO999_RS01810 | elongation factor Tu | 3037696 |
| RALBFv3_RS07200 | elongation factor Tu | 305 |
| RALBFv3_RS07305 | elongation factor Tu | 305 |
| AACJ33_RS01780 | elongation factor Tu | 1310167 |
| AACJ33_RS01885 | elongation factor Tu | 1310167 |
| KOL96_RS22010 | elongation factor Tu | 2842456 |
| KOL96_RS22125 | elongation factor Tu | 2842456 |
| RTA_RS01620 | elongation factor Tu | 365046 |
| RTA_RS18295 | elongation factor Tu | 365046 |
| RA876_RS03110 | elongation factor Tu | 81479 |
| RA876_RS18815 | elongation factor Tu | 81479 |
| EXZ61_RS19950 | elongation factor Tu | 2527691 |
| EXZ61_RS20560 | elongation factor Tu | 2527691 |
| RFER_RS18235 | elongation factor Tu | 338969 |
| RFER_RS19295 | elongation factor Tu | 338969 |
| RD110_RS01905 | elongation factor Tu | 1842727 |
| RD110_RS03745 | elongation factor Tu | 1842727 |

| | | |
|---|---|---|
| RAN89_RS00645 | elongation factor Tu | 3068341 |
| RAN89_RS01590 | elongation factor Tu | 3068341 |
| RS694_RS01495 | elongation factor Tu | 1484693 |
| RS694_RS02400 | elongation factor Tu | 1484693 |
| EUB48_RS01455 | elongation factor Tu | 2509614 |
| EUB48_RS20250 | elongation factor Tu | 2509614 |
| SJQ66_RS28510 | elongation factor Tu | 28092 |
| SJQ66_RS28595 | elongation factor Tu | 28092 |
| N4261_RS02935 | elongation factor Tu | 2978473 |
| N4261_RS03005 | elongation factor Tu | 2978473 |
| RD2015_RS02655 | elongation factor Tu | 76731 |
| RD2015_RS02725 | elongation factor Tu | 76731 |
| ACNFWO_RS23555 | elongation factor Tu | 28068 |
| ACNFWO_RS23625 | elongation factor Tu | 28068 |
| PG1C_RS12790 | elongation factor Tu | 1565605 |
| PG1C_RS12855 | elongation factor Tu | 1565605 |
| SMCB_RS10195 | elongation factor Tu | 1458426 |
| SMCB_RS10650 | elongation factor Tu | 1458426 |
| SRAA_RS10180 | elongation factor Tu | 1458425 |
| SRAA_RS10630 | elongation factor Tu | 1458425 |
| L6418_RS03395 | elongation factor Tu | 2764705 |
| L6418_RS03460 | elongation factor Tu | 2764705 |
| SLIT_RS03765 | elongation factor Tu | 580332 |
| SLIT_RS03830 | elongation factor Tu | 580332 |
| N7220_RS14255 | elongation factor Tu | 2980100 |
| N7220_RS14320 | elongation factor Tu | 2980100 |
| BWP33_RS11380 | elongation factor Tu | 641147 |
| BWP33_RS11535 | elongation factor Tu | 641147 |
| C6571_RS05885 | elongation factor Tu | 2109915 |
| C6571_RS06105 | elongation factor Tu | 2109915 |
| SALWKB2_RS01290 | elongation factor Tu | 1196094 |

| | | |
|---|---|---|
| SALWKB2_RS01355 | elongation factor Tu | 1196094 |
| NGK70_RS05225 | elongation factor Tu | 2914710 |
| NGK70_RS05290 | elongation factor Tu | 2914710 |
| EWH46_RS00860 | elongation factor Tu | 639200 |
| EWH46_RS15400 | elongation factor Tu | 639200 |
| SCD_RS03030 | elongation factor Tu | 1163617 |
| SCD_RS03095 | elongation factor Tu | 1163617 |
| EL335_RS02515 | elongation factor Tu | 2211108 |
| EL335_RS02580 | elongation factor Tu | 2211108 |
| M52SOB_RS02760 | elongation factor Tu | 2496847 |
| M52SOB_RS02825 | elongation factor Tu | 2496847 |
| SFSGTM_RS02915 | elongation factor Tu | 2675298 |
| SFSGTM_RS02980 | elongation factor Tu | 2675298 |
| GZH91_RS03535 | elongation factor Tu | 171865 |
| GZH91_RS03600 | elongation factor Tu | 171865 |
| SKTS_RS02725 | elongation factor Tu | 2715678 |
| SKTS_RS02790 | elongation factor Tu | 2715678 |
| SUTH_RS03200 | elongation factor Tu | 1223802 |
| SUTH_RS03265 | elongation factor Tu | 1223802 |
| EL388_RS02215 | elongation factor Tu | 1914471 |
| EL388_RS02285 | elongation factor Tu | 1914471 |
| FG381_RS08330 | elongation factor Tu | 2584944 |
| FG381_RS08395 | elongation factor Tu | 2584944 |
| S6FBBBH3_RS01425 | elongation factor Tu | 2494234 |
| S6FBBBH3_RS01490 | elongation factor Tu | 2494234 |
| I6J16_RS10055 | elongation factor Tu | 40545 |
| I6J16_RS10120 | elongation factor Tu | 40545 |
| TASI_RS07520 | elongation factor Tu | 1008459 |
| TASI_RS07665 | elongation factor Tu | 1008459 |
| KPH58_RS07705 | elongation factor Tu | 29575 |
| KPH58_RS07865 | elongation factor Tu | 29575 |

| | | |
|---|---|---|
| Q9246_RS26465 | translation elongation fa | 3068633 |
| Q9246_RS26480 | elongation factor Tu | 3068633 |
| P0M04_RS07180 | elongation factor Tu | 34071 |
| P0M04_RS07565 | elongation factor Tu | 34071 |
| LCC91_RS13275 | elongation factor Tu | 307486 |
| LCC91_RS13545 | elongation factor Tu | 307486 |
| TMZ1T_RS16100 | elongation factor Tu | 164330 |
| TMZ1T_RS16165 | elongation factor Tu | 164330 |
| Tharo_RS13815 | elongation factor Tu | 44139 |
| Tharo_RS13880 | elongation factor Tu | 44139 |
| ACNQFN_RS18525 | elongation factor Tu | 86174 |
| ACNQFN_RS18590 | elongation factor Tu | 86174 |
| Tchl_RS08660 | elongation factor Tu | 96773 |
| Tchl_RS08725 | elongation factor Tu | 96773 |
| AC731_RS11605 | elongation factor Tu | 1134435 |
| AC731_RS11670 | elongation factor Tu | 1134435 |
| VA613_RS02255 | elongation factor Tu | 3110231 |
| VA613_RS02320 | elongation factor Tu | 3110231 |
| THI_RS15865 | elongation factor Tu | 426114 |
| THI_RS17145 | elongation factor Tu | 426114 |
| BVH73_RS06980 | elongation factor Tu | 926 |
| BVH73_RS07775 | elongation factor Tu | 926 |
| U0034_RS08095 | elongation factor Tu | 28094 |
| U0034_RS08200 | elongation factor Tu | 28094 |
| FAZ95_RS20200 | elongation factor Tu | 2571746 |
| FAZ95_RS20270 | elongation factor Tu | 2571746 |
| I5Q81_RS07085 | elongation factor Tu | 1796652 |
| I5Q81_RS07150 | elongation factor Tu | 1796652 |
| RF679_RS02665 | elongation factor Tu | 3073561 |
| RF679_RS03575 | elongation factor Tu | 3073561 |
| EJN92_RS05975 | elongation factor Tu | 401471 |

| | | |
|---|---|---|
| EJN92_RS11440 | elongation factor Tu | 401471 |
| LVJ78_RS11375 | elongation factor Tu | 252130 |
| LVJ78_RS11440 | elongation factor Tu | 252130 |
| LVJ83_RS12865 | elongation factor Tu | 1282863 |
| LVJ83_RS12930 | elongation factor Tu | 1282863 |
| DSM104440_RS16000 | elongation factor Tu | 2732487 |
| DSM104440_RS16065 | elongation factor Tu | 2732487 |
| DSM104443_RS18390 | elongation factor Tu | 2732067 |
| DSM104443_RS18455 | elongation factor Tu | 2732067 |
| ABL837_RS02935 | elongation factor Tu | 1321610 |
| ABL837_RS24280 | elongation factor Tu | 1321610 |
| VEIS_RS06160 | elongation factor Tu | 391735 |
| VEIS_RS11035 | elongation factor Tu | 391735 |
| VITFI_RS03950 | elongation factor Tu | 63 |
| VITFI_RS06170 | elongation factor Tu | 63 |
| LVJ81_RS11640 | elongation factor Tu | 61 |
| LVJ81_RS11705 | elongation factor Tu | 61 |
| MIS45_RS00605 | elongation factor Tu | 2917790 |
| MIS45_RS01435 | elongation factor Tu | 2917790 |
| GT347_RS11880 | elongation factor Tu | 2697032 |
| GT347_RS14425 | elongation factor Tu | 2697032 |
| J8C02_RS01130 | elongation factor Tu | 2821542 |
| J8C02_RS07355 | elongation factor Tu | 2821542 |
| CABTHER_RS01150 | elongation factor Tu | 981222 |
| CABTHER_RS07340 | elongation factor Tu | 981222 |
| J8C06_RS01805 | elongation factor Tu | 2821543 |
| J8C06_RS03245 | elongation factor Tu | 2821543 |
| CLDAP_RS02075 | elongation factor Tu | 926550 |
| CLDAP_RS07550 | elongation factor Tu | 926550 |
| CSE_RS05335 | elongation factor Tu | 511051 |
| CSE_RS05575 | elongation factor Tu | 511051 |

| | | |
|---|---|---|
| Cabys_RS04150 | elongation factor Tu | 880073 |
| Cabys_RS04220 | elongation factor Tu | 880073 |
| BABL1_RS02055 | elongation factor Tu | 673862 |
| C0J27_RS01755 | elongation factor Tu | 2066483 |
| KsCSTR_RS05335 | elongation factor Tu | 174633 |
| KsCSTR_RS05405 | elongation factor Tu | 174633 |
| CLOAM_RS02220 | elongation factor Tu | 459349 |
| HS1_RS02465 | elongation factor Tu | 1621989 |
| HS1_RS02530 | elongation factor Tu | 1621989 |
| G4Z02_RS06240 | elongation factor Tu | 2782163 |
| HF295_RS05180 | elongation factor Tu | 2725268 |
| ACMCNE_RS01945 | elongation factor Tu | 3238481 |
| FBF37_RS02025 | elongation factor Tu | 2572088 |
| TM7x_RS01975 | elongation factor Tu | 2093824 |
| L336_RS05325 | elongation factor Tu | 1332188 |
| SEML1_RS02365 | elongation factor Tu | 3043284 |
| G4Y79_RS10610 | elongation factor Tu | 2710758 |
| UABAM_RS19445 | elongation factor Tu | 2596890 |
| UABHE_RS02370 | elongation factor Tu | 3095367 |
| D6B99_RS03455 | elongation factor Tu | 2341117 |
| K9M52_RS04995 | elongation factor Tu | 2875539 |
| GWR21_RS23955 | elongation factor Tu | 2703787 |
| COR50_RS10160 | elongation factor Tu | 2029983 |
| WJU22_RS25980 | elongation factor Tu | 2267521 |
| MKQ68_RS01430 | elongation factor Tu | 2920382 |
| HF329_RS20330 | elongation factor Tu | 2725414 |
| KTO58_RS18180 | elongation factor Tu | 2849666 |
| CPIN_RS08560 | elongation factor Tu | 485918 |
| WJU16_RS18870 | elongation factor Tu | 3133966 |
| U0033_RS26075 | elongation factor Tu | 1004 |
| K9M53_RS07550 | elongation factor Tu | 2875540 |

| | | |
|---|---|---|
| LK994_RS10270 | elongation factor Tu | 563172 |
| FLA_RS03360 | elongation factor Tu | 477680 |
| KJS93_RS10685 | elongation factor Tu | 2838157 |
| KJS94_RS08070 | elongation factor Tu | 2838156 |
| FSB75_RS21265 | elongation factor Tu | 661481 |
| SY85_RS22310 | elongation factor Tu | 1492898 |
| H4075_RS06425 | elongation factor Tu | 2760713 |
| PIECOFPK_RS06205 | elongation factor Tu | 2962693 |
| A8C56_RS16045 | elongation factor Tu | 1176587 |
| ABDK00_RS12300 | elongation factor Tu | 3144874 |
| NIASO_RS10285 | elongation factor Tu | 929713 |
| U0035_RS07385 | elongation factor Tu | 577386 |
| NIAKO_RS24230 | elongation factor Tu | 700598 |
| FRZ67_RS03605 | elongation factor Tu | 1813871 |
| D3H65_RS16190 | elongation factor Tu | 2315862 |
| RS881_RS45310 | elongation factor Tu | 3076558 |
| EPD17_RS16855 | elongation factor Tu | 2502779 |
| E0W69_RS15970 | elongation factor Tu | 2545455 |
| Cs308_RS03330 | elongation factor Tu | 1806891 |
| PC_RS02855 | elongation factor Tu | 264201 |
| PNK_RS08525 | elongation factor Tu | 389348 |
| RHABOEDO_RS04600 | elongation factor Tu | 2720720 |
| RHAB15C_RS02365 | elongation factor Tu | 225148 |
| ABCH23_RS01120 | elongation factor Tu | 83555 |
| HBN95_RS02075 | elongation factor Tu | 2494525 |
| CCA_RS03510 | elongation factor Tu | 227941 |
| H9Q19_RS00695 | elongation factor Tu | 2766982 |
| CF_RS01580 | elongation factor Tu | 264202 |
| M787_RS00360 | elongation factor Tu | 1143323 |
| TC_RS03020 | elongation factor Tu | 243161 |
| MarsBar_RS04745 | elongation factor Tu | 85991 |

| | | |
|---|---|---|
| CPB_RS00370 | elongation factor Tu | 182082 |
| C10C_RS03525 | elongation factor Tu | 1967783 |
| G5O_RS08630 | elongation factor Tu | 331636 |
| C834KP_RS00260 | elongation factor Tu | 1967782 |
| B6E82_RS01725 | elongation factor Tu | 83559 |
| CT_322 | elongation factor Tu | 272561 |
| KJA58_RS00665 | elongation factor Tu | 2681469 |
| KJA62_RS01075 | elongation factor Tu | 2681470 |
| PUV_RS09295 | elongation factor Tu | 765952 |
| SNE_RS07855 | elongation factor Tu | 331113 |
| WCW_RS02790 | elongation factor Tu | 716544 |
| BIU88_RS12210 | elongation factor Tu | 274537 |
| CPAR_RS00920 | elongation factor Tu | 517417 |
| AYT24_RS09945 | elongation factor Tu | 194439 |
| CLIM_RS11210 | elongation factor Tu | 290315 |
| CPHA266_RS12555 | elongation factor Tu | 290317 |
| CTHA_RS05665 | elongation factor Tu | 517418 |
| PLUT_RS00940 | elongation factor Tu | 319225 |
| PPHA_RS01455 | elongation factor Tu | 324925 |
| PAES_RS10305 | elongation factor Tu | 290512 |
| OZ401_RS06970 | elongation factor Tu | 3003348 |
| OZ401_RS07020 | elongation factor Tu | 3003348 |
| CAGG_RS07325 | elongation factor Tu | 326427 |
| CAGG_RS15080 | elongation factor Tu | 326427 |
| CAUR_RS11410 | elongation factor Tu | 324602 |
| CAUR_RS12415 | elongation factor Tu | 324602 |
| RCAS_RS20910 | elongation factor Tu | 383372 |
| RCAS_RS22385 | elongation factor Tu | 383372 |
| SELIN_RS10050 | elongation factor Tu | 653733 |
| CCALI_RS02955 | elongation factor Tu | 1303518 |
| CCALI_RS14150 | elongation factor Tu | 1303518 |

| | | |
|---|---|---|
| NQ535_RS04620 | elongation factor Tu | 518636 |
| NBW53_RS02560 | elongation factor Tu | 36835 |
| MWL92_RS16260 | elongation factor Tu | 89153 |
| CLOSA_RS19020 | elongation factor Tu | 610130 |
| HDCHBGLK_RS03860 | elongation factor Tu | 411468 |
| NQ541_RS10720 | elongation factor Tu | 471875 |
| ACH9UM_RS12015 | elongation factor Tu | 33039 |
| ACETAC_RS10075 | elongation factor Tu | 2755338 |
| ACETAC_RS10145 | elongation factor Tu | 2755338 |
| CLOCL_RS03220 | elongation factor Tu | 720554 |
| HVS_RS12925 | elongation factor Tu | 1677857 |
| CTHE_RS14255 | elongation factor Tu | 203119 |
| CLOST_RS11095 | elongation factor Tu | 1511 |
| CLOST_RS11160 | elongation factor Tu | 1511 |
| ACIZ62_RS03340 | elongation factor Tu | 52690 |
| ACIZ62_RS15220 | elongation factor Tu | 52690 |
| ACIUZZ_RS03550 | elongation factor Tu | 1068975 |
| ACIUZZ_RS15995 | elongation factor Tu | 1068975 |
| CHL1_RS03200 | elongation factor Tu | 52694 |
| CHL1_RS14310 | elongation factor Tu | 52694 |
| AWO_RS04640 | elongation factor Tu | 931626 |
| AWO_RS14735 | elongation factor Tu | 931626 |
| acear_RS00865 | elongation factor Tu | 574087 |
| acear_RS00930 | elongation factor Tu | 574087 |
| MNU25_RS03850 | elongation factor Tu | 1561005 |
| ADH66_RS01785 | elongation factor Tu | 1796620 |
| MIO91_RS01755 | elongation factor Tu | 39491 |
| J0B03_RS06495 | elongation factor Tu | 2815577 |
| J0B03_RS06565 | elongation factor Tu | 2815577 |
| HYG86_RS08310 | elongation factor Tu | 2731377 |
| HYG86_RS08380 | elongation factor Tu | 2731377 |

| | | |
|---|---|---|
| AMET_RS22385 | elongation factor Tu | 293826 |
| AMET_RS22450 | elongation factor Tu | 293826 |
| CLOS_RS02475 | elongation factor Tu | 350688 |
| CLOS_RS02545 | elongation factor Tu | 350688 |
| Ami103574_RS13405 | elongation factor Tu | 433296 |
| Ami103574_RS13495 | elongation factor Tu | 433296 |
| EQM06_RS09955 | elongation factor Tu | 2507160 |
| EQM06_RS10045 | elongation factor Tu | 2507160 |
| Ami3637_RS10760 | elongation factor Tu | 2697030 |
| Ami3637_RS10860 | elongation factor Tu | 2697030 |
| ADEG_RS02075 | elongation factor Tu | 429009 |
| ADEG_RS07700 | elongation factor Tu | 429009 |
| PYS60_RS02220 | elongation factor Tu | 3029272 |
| KDJ91_RS02390 | elongation factor Tu | 39488 |
| acsn021_RS21180 | elongation factor Tu | 433286 |
| bsdcttw_RS01815 | elongation factor Tu | 1727145 |
| Ana3638_RS03850 | elongation factor Tu | 2696063 |
| bsdtb5_RS20410 | elongation factor Tu | 2785025 |
| NBX03_RS13435 | elongation factor Tu | 2934315 |
| LCQ53_RS06720 | elongation factor Tu | 105841 |
| RBI15_RS01725 | elongation factor Tu | 649756 |
| AR1Y2_RS17105 | elongation factor Tu | 1229621 |
| CPRO_RS12620 | elongation factor Tu | 991789 |
| K5I23_RS11595 | elongation factor Tu | 169435 |
| BBF96_RS16025 | elongation factor Tu | 1323375 |
| BBF96_RS16100 | elongation factor Tu | 1323375 |
| JYE49_RS06235 | elongation factor Tu | 3046382 |
| JYE49_RS13240 | elongation factor Tu | 3046382 |
| JYE50_RS05840 | elongation factor Tu | 3046383 |
| JYE50_RS12860 | elongation factor Tu | 3046383 |
| RFF05_RS16770 | elongation factor Tu | 3073143 |

| | | |
|---|---|---|
| D2962_RS01280 | elongation factor Tu | 2316383 |
| D2962_RS15980 | elongation factor Tu | 2316383 |
| DQQ01_RS01620 | elongation factor Tu | 1912897 |
| BLCOC_RS01655 | elongation factor Tu | 1532 |
| CGC63_RS14390 | elongation factor Tu | 537007 |
| BLHYD_RS00935 | elongation factor Tu | 476272 |
| INP51_RS01265 | elongation factor Tu | 2779518 |
| R8695_RS15460 | elongation factor Tu | 89014 |
| NQ503_RS15445 | elongation factor Tu | 411459 |
| LC508_RS16830 | elongation factor Tu | 2877527 |
| GXM18_RS16395 | elongation factor Tu | 1121114 |
| A4V09_RS18070 | elongation factor Tu | 1796616 |
| NQ550_RS18925 | elongation factor Tu | 1121115 |
| WAA20_RS17010 | elongation factor Tu | 831 |
| bhn_RS03595 | elongation factor Tu | 185008 |
| BPR_RS11950 | elongation factor Tu | 515622 |
| TTE_RS10955 | elongation factor Tu | 273068 |
| TTE_RS11025 | elongation factor Tu | 273068 |
| caldi_RS16375 | elongation factor Tu | 671077 |
| Daudx_RS01150 | elongation factor Tu | 471827 |
| Daudx_RS01230 | elongation factor Tu | 471827 |
| PCY70_RS07925 | elongation factor Tu | 420336 |
| DCMF_RS22215 | elongation factor Tu | 1761012 |
| DCMF_RS22285 | elongation factor Tu | 1761012 |
| HH195_RS09765 | elongation factor Tu | 2726954 |
| HH195_RS09830 | elongation factor Tu | 2726954 |
| SYNTR_RS10785 | elongation factor Tu | 2293317 |
| SYNTR_RS10860 | elongation factor Tu | 2293317 |
| HCR03_RS02980 | elongation factor Tu | 2576756 |
| H6X83_RS08080 | elongation factor Tu | 2766537 |
| PXC00_RS08095 | elongation factor Tu | 3030016 |

| | | |
|---|---|---|
| GJQ69_RS08065 | elongation factor Tu | 2666138 |
| CHY_RS10720 | elongation factor Tu | 246194 |
| CHY_RS10790 | elongation factor Tu | 246194 |
| CLOLE_RS17820 | elongation factor Tu | 642492 |
| MJY40_RS10085 | elongation factor Tu | 2918914 |
| G4C92_RS10195 | elongation factor Tu | 2709410 |
| PUP29_RS04590 | elongation factor Tu | 1805714 |
| PUP29_RS09790 | elongation factor Tu | 1805714 |
| B1H56_RS06745 | elongation factor Tu | 626937 |
| B1H56_RS10290 | elongation factor Tu | 626937 |
| R2J37_RS02605 | elongation factor Tu | 3028070 |
| KNZ77_RS00585 | elongation factor Tu | 1496 |
| KNZ77_RS00655 | elongation factor Tu | 1496 |
| CACET_RS17645 | elongation factor Tu | 84022 |
| CACET_RS17715 | elongation factor Tu | 84022 |
| SMB_RS16040 | elongation factor Tu | 991791 |
| SMB_RS16110 | elongation factor Tu | 991791 |
| RSJ17_RS03530 | elongation factor Tu | 29341 |
| RSJ17_RS03600 | elongation factor Tu | 29341 |
| CLAU_RS09400 | elongation factor Tu | 1341692 |
| CLAU_RS09470 | elongation factor Tu | 1341692 |
| NPD11_RS01790 | elongation factor Tu | 1561 |
| NPD11_RS01855 | elongation factor Tu | 1561 |
| CM240_RS12130 | elongation factor Tu | 1216932 |
| CM240_RS12200 | elongation factor Tu | 1216932 |
| CBO_RS17235 | elongation factor Tu | 413999 |
| CBO_RS17305 | elongation factor Tu | 413999 |
| FF104_RS17150 | elongation factor Tu | 1492 |
| FF104_RS17215 | elongation factor Tu | 1492 |
| KQH81_RS15585 | elongation factor Tu | 1529 |
| KQH81_RS15655 | elongation factor Tu | 1529 |

| | | |
|---|---|---|
| Ccar_RS19830 | elongation factor Tu | 536227 |
| Ccar_RS19900 | elongation factor Tu | 536227 |
| CLOCEL_RS18615 | elongation factor Tu | 573061 |
| CLOCEL_RS18685 | elongation factor Tu | 573061 |
| BTM20_RS01085 | elongation factor Tu | 46867 |
| BTM20_RS01150 | elongation factor Tu | 46867 |
| CKV72_RS11165 | elongation factor Tu | 1494 |
| CKV72_RS11235 | elongation factor Tu | 1494 |
| F3K33_RS00845 | elongation factor Tu | 223919 |
| F3K33_RS00910 | elongation factor Tu | 223919 |
| B9W14_RS23745 | elongation factor Tu | 332101 |
| B9W14_RS23815 | elongation factor Tu | 332101 |
| KTC99_RS01650 | elongation factor Tu | 238834 |
| KTC99_RS01720 | elongation factor Tu | 238834 |
| CLFE_RS03845 | elongation factor Tu | 1121310 |
| CLFE_RS03915 | elongation factor Tu | 1121310 |
| D4Z93_RS01230 | elongation factor Tu | 2068654 |
| D4Z93_RS01300 | elongation factor Tu | 2068654 |
| CLFO_RS19715 | elongation factor Tu | 1497 |
| CLFO_RS19785 | elongation factor Tu | 1497 |
| J1C67_RS04835 | elongation factor Tu | 94869 |
| J1C67_RS04905 | elongation factor Tu | 94869 |
| psyc5s11_RS00910 | elongation factor Tu | 704125 |
| psyc5s11_RS00975 | elongation factor Tu | 704125 |
| P8F83_RS15915 | elongation factor Tu | 36845 |
| P8F83_RS15985 | elongation factor Tu | 36845 |
| BEN51_RS01130 | elongation factor Tu | 182773 |
| BEN51_RS01195 | elongation factor Tu | 182773 |
| CKR_RS01025 | elongation factor Tu | 583346 |
| CKR_RS01095 | elongation factor Tu | 583346 |
| CLJU_RS20225 | elongation factor Tu | 748727 |

| | | |
|---|---|---|
| CLJU_RS20295 | elongation factor Tu | 748727 |
| C1I91_RS03540 | elongation factor Tu | 2320868 |
| C1I91_RS03620 | elongation factor Tu | 2320868 |
| DFH04_RS02800 | elongation factor Tu | 1542 |
| DFH04_RS02870 | elongation factor Tu | 1542 |
| AQ983_RS17930 | elongation factor Tu | 1262449 |
| AQ983_RS18000 | elongation factor Tu | 1262449 |
| I6G60_RS02980 | elongation factor Tu | 1502 |
| I6G60_RS03050 | elongation factor Tu | 1502 |
| CLSA_RS00930 | elongation factor Tu | 1345695 |
| CLSA_RS00995 | elongation factor Tu | 1345695 |
| CSPA_RS00850 | elongation factor Tu | 931276 |
| CSPA_RS00915 | elongation factor Tu | 931276 |
| Csca_RS20725 | elongation factor Tu | 1548 |
| Csca_RS20795 | elongation factor Tu | 1548 |
| CP523_RS08910 | elongation factor Tu | 1504 |
| CP523_RS08975 | elongation factor Tu | 1504 |
| CLSPO_RS18235 | elongation factor Tu | 1509 |
| CLSPO_RS18305 | elongation factor Tu | 1509 |
| BGI42_RS01000 | elongation factor Tu | 394958 |
| BGI42_RS01065 | elongation factor Tu | 394958 |
| LL095_RS20930 | elongation factor Tu | 360422 |
| LL095_RS21000 | elongation factor Tu | 360422 |
| CTC_RS13380 | elongation factor Tu | 212717 |
| CTC_RS13450 | elongation factor Tu | 212717 |
| FHY60_RS15430 | elongation factor Tu | 1716543 |
| FHY60_RS15500 | elongation factor Tu | 1716543 |
| CDO33_RS15610 | elongation factor Tu | 84032 |
| CTK_RS14030 | elongation factor Tu | 1519 |
| CTK_RS14100 | elongation factor Tu | 1519 |
| NQ556_RS13990 | elongation factor Tu | 470146 |

| | | |
|---|---|---|
| NQ536_RS12375 | elongation factor Tu | 33043 |
| KVH43_RS06315 | elongation factor Tu | 2855394 |
| KVH43_RS06385 | elongation factor Tu | 2855394 |
| K7H06_RS16540 | elongation factor Tu | 1286698 |
| K7H06_RS16610 | elongation factor Tu | 1286698 |
| FQB35_RS01555 | elongation factor Tu | 2599308 |
| FQB35_RS01625 | elongation factor Tu | 2599308 |
| QBE51_RS07715 | elongation factor Tu | 879970 |
| GQ588_RS02750 | elongation factor Tu | 55583 |
| CEQ75_RS05635 | elongation factor Tu | 51515 |
| CEQ75_RS05705 | elongation factor Tu | 51515 |
| DESDE_RS02055 | elongation factor Tu | 756499 |
| DESDI_RS02095 | elongation factor Tu | 871963 |
| DHAF_RS02190 | elongation factor Tu | 272564 |
| DESME_RS01375 | elongation factor Tu | 871968 |
| DTOX_RS01395 | elongation factor Tu | 485916 |
| DTOX_RS01460 | elongation factor Tu | 485916 |
| DESKU_RS15040 | elongation factor Tu | 760568 |
| DESKU_RS15110 | elongation factor Tu | 760568 |
| B0537_RS01865 | elongation factor Tu | 1833852 |
| B0537_RS01935 | elongation factor Tu | 1833852 |
| DRED_RS01105 | elongation factor Tu | 349161 |
| DRED_RS01170 | elongation factor Tu | 349161 |
| DESRU_RS01490 | elongation factor Tu | 696281 |
| DESRU_RS01560 | elongation factor Tu | 696281 |
| DESGI_RS01680 | elongation factor Tu | 767817 |
| DESGI_RS01745 | elongation factor Tu | 767817 |
| DESACI_RS02385 | elongation factor Tu | 646529 |
| DESMER_RS01170 | elongation factor Tu | 768704 |
| DESOR_RS01200 | elongation factor Tu | 768706 |
| VZF35_RS12090 | elongation factor Tu | 3115649 |

| | | |
|---|---|---|
| DESYODRAFT_RS01305 | elongation factor Tu | 768710 |
| DESCA_RS01120 | elongation factor Tu | 868595 |
| DESCA_RS01185 | elongation factor Tu | 868595 |
| NQ560_RS00740 | elongation factor Tu | 39486 |
| NQ508_RS12285 | elongation factor Tu | 88431 |
| FOC47_RS19505 | elongation factor Tu | 1531 |
| NQ527_RS11025 | elongation factor Tu | 45851 |
| ETHHA_RS05455 | elongation factor Tu | 663278 |
| ELI_RS18070 | elongation factor Tu | 53442 |
| ELI_RS18645 | elongation factor Tu | 53442 |
| B2M23_RS05735 | elongation factor Tu | 1736 |
| B2M23_RS06310 | elongation factor Tu | 1736 |
| CPZ25_RS10440 | elongation factor Tu | 2041044 |
| CPZ25_RS11010 | elongation factor Tu | 2041044 |
| ACIRUQ_RS10855 | elongation factor Tu | 39490 |
| GXM22_RS10035 | elongation factor Tu | 411483 |
| PXT33_RS07105 | elongation factor Tu | 3030638 |
| ACGG81_RS02730 | elongation factor Tu | 1912855 |
| C5Q98_RS04510 | elongation factor Tu | 236753 |
| M1R53_RS06510 | elongation factor Tu | 938288 |
| KO172_RS01240 | elongation factor Tu | 2811778 |
| HMPREF0389_RS03090 | elongation factor Tu | 546269 |
| HMPREF0389_RS06140 | elongation factor Tu | 546269 |
| GXM20_RS09150 | elongation factor Tu | 292800 |
| Gferi_RS17305 | elongation factor Tu | 1424294 |
| Gferi_RS17385 | elongation factor Tu | 1424294 |
| HALSA_RS09780 | elongation factor Tu | 656519 |
| HALSA_RS09850 | elongation factor Tu | 656519 |
| HALHA_RS00885 | elongation factor Tu | 748449 |
| HALHA_RS00955 | elongation factor Tu | 748449 |
| HORE_RS00610 | elongation factor Tu | 373903 |

| | | |
|---|---|---|
| HORE_RS00685 | elongation factor Tu | 373903 |
| FGL08_RS11720 | elongation factor Tu | 1498 |
| FGL08_RS11790 | elongation factor Tu | 1498 |
| AACH12_RS12595 | elongation factor Tu | 3065157 |
| AACH12_RS12665 | elongation factor Tu | 3065157 |
| HM1_RS06255 | elongation factor Tu | 498761 |
| FTV88_RS14200 | elongation factor Tu | 356322 |
| SD1D_RS10970 | elongation factor Tu | 1679721 |
| NQ487_RS27055 | elongation factor Tu | 154046 |
| NQ514_RS01515 | elongation factor Tu | 261299 |
| NQ514_RS01580 | elongation factor Tu | 261299 |
| SRB521_RS05120 | elongation factor Tu | 1297617 |
| GM661_RS15725 | elongation factor Tu | 2682810 |
| GM661_RS15795 | elongation factor Tu | 2682810 |
| V6984_RS04075 | elongation factor Tu | 3123317 |
| H0A61_RS06550 | elongation factor Tu | 2751313 |
| H0A61_RS06620 | elongation factor Tu | 2751313 |
| D4A81_RS00400 | elongation factor Tu | 617123 |
| CPHY_RS01305 | elongation factor Tu | 357809 |
| EUBELI_RS01385 | elongation factor Tu | 515620 |
| OW255_RS18250 | elongation factor Tu | 29375 |
| HMPREF0868_RS02280 | elongation factor Tu | 699246 |
| MAHAU_RS10755 | elongation factor Tu | 697281 |
| MAHAU_RS10825 | elongation factor Tu | 697281 |
| NQ534_RS13300 | elongation factor Tu | 478749 |
| ACGHPO_RS13020 | elongation factor Tu | 342942 |
| RGna_RS08815 | elongation factor Tu | 411470 |
| MFMK1_RS00625 | elongation factor Tu | 3039180 |
| MFMK1_RS00695 | elongation factor Tu | 3039180 |
| C5Q96_RS00400 | elongation factor Tu | 114527 |
| C5Q96_RS06305 | elongation factor Tu | 114527 |

| | | |
|---|---|---|
| QU661_RS01690 | elongation factor Tu | 114528 |
| QU661_RS07375 | elongation factor Tu | 114528 |
| B9O19_RS06435 | elongation factor Tu | 1981510 |
| NTHER_RS01010 | elongation factor Tu | 457570 |
| NTHER_RS01085 | elongation factor Tu | 457570 |
| ACONDI_RS15055 | elongation factor Tu | 2742683 |
| ACONDI_RS15125 | elongation factor Tu | 2742683 |
| MCACP_RS13840 | elongation factor Tu | 3062783 |
| MCACP_RS13910 | elongation factor Tu | 3062783 |
| MGLY_RS05905 | elongation factor Tu | 55779 |
| MGLY_RS05975 | elongation factor Tu | 55779 |
| MHOCP_RS12245 | elongation factor Tu | 676965 |
| MHOCP_RS12315 | elongation factor Tu | 676965 |
| MOTHA_RS12715 | elongation factor Tu | 1525 |
| KNL20_RS01755 | elongation factor Tu | 2068655 |
| H8790_RS01630 | elongation factor Tu | 2763056 |
| ACKTS9_RS16125 | elongation factor Tu | 3136601 |
| KXZ80_RS00505 | elongation factor Tu | 1490 |
| KXZ80_RS00570 | elongation factor Tu | 1490 |
| ACN6I1_RS15050 | elongation factor Tu | 2662455 |
| ACN6I1_RS15115 | elongation factor Tu | 2662455 |
| LZ906_RS00505 | elongation factor Tu | 29358 |
| LZ906_RS00570 | elongation factor Tu | 29358 |
| ATCC9714_RS00140 | elongation factor Tu | 1505 |
| ATCC9714_RS00205 | elongation factor Tu | 1505 |
| KGNDJEFE_RS00265 | elongation factor Tu | 89152 |
| KGNDJEFE_RS00330 | elongation factor Tu | 89152 |
| EAL2_RS00665 | elongation factor Tu | 1286171 |
| EAL2_RS00735 | elongation factor Tu | 1286171 |
| O0R46_RS09645 | elongation factor Tu | 3003601 |
| O0R46_RS09815 | elongation factor Tu | 3003601 |

| | | |
|---|---|---|
| PATL70BA_RS12540 | elongation factor Tu | 2173034 |
| PRVXH_RS00855 | elongation factor Tu | 1826727 |
| PRVXT_RS00830 | elongation factor Tu | 1260629 |
| KJS55_RS03125 | elongation factor Tu | 2714358 |
| H9Q78_RS06290 | elongation factor Tu | 2763669 |
| CRIB_RS12210 | elongation factor Tu | 1115758 |
| CRIB_RS12275 | elongation factor Tu | 1115758 |
| RHOM_RS01035 | elongation factor Tu | 585394 |
| RIL182_RS18450 | elongation factor Tu | 536231 |
| H8S51_RS02630 | elongation factor Tu | 2763062 |
| CCEL_RS01590 | elongation factor Tu | 394503 |
| EHE19_RS17265 | elongation factor Tu | 2488810 |
| P0092_RS01870 | elongation factor Tu | 588581 |
| RUMAL_RS14775 | elongation factor Tu | 697329 |
| E5Z56_RS06515 | elongation factor Tu | 2564099 |
| NQ502_RS08505 | elongation factor Tu | 438033 |
| ACI3DM_RS07685 | elongation factor Tu | 2592333 |
| ACI3DL_RS13105 | elongation factor Tu | 1653434 |
| HZR23_RS06375 | elongation factor Tu | 1734049 |
| HZR23_RS06445 | elongation factor Tu | 1734049 |
| H9Q77_RS12845 | elongation factor Tu | 2763672 |
| C12CBH8_RS08845 | elongation factor Tu | 2780922 |
| NQ490_RS14045 | elongation factor Tu | 214851 |
| STH_RS15295 | elongation factor Tu | 292459 |
| SGLY_RS02045 | elongation factor Tu | 645991 |
| SWOL_RS12095 | elongation factor Tu | 335541 |
| SWOL_RS12170 | elongation factor Tu | 335541 |
| SLIP_RS11090 | elongation factor Tu | 643648 |
| M2214_RS16590 | elongation factor Tu | 2925782 |
| M2214_RS16660 | elongation factor Tu | 2925782 |
| P4S50_RS19100 | elongation factor Tu | 3036126 |

| | | |
|---|---|---|
| P4S50_RS19170 | elongation factor Tu | 3036126 |
| TEGL_RS00520 | elongation factor Tu | 1121315 |
| TEGL_RS00585 | elongation factor Tu | 1121315 |
| JW646_RS08805 | elongation factor Tu | 2813371 |
| JW646_RS08870 | elongation factor Tu | 2813371 |
| TEMA_RS14725 | elongation factor Tu | 1541 |
| TEMA_RS14790 | elongation factor Tu | 1541 |
| TEPE_RS00520 | elongation factor Tu | 1460447 |
| TEPE_RS00585 | elongation factor Tu | 1460447 |
| TPH_RS12830 | elongation factor Tu | 1089553 |
| B9A14_RS00285 | elongation factor Tu | 698762 |
| BR63_RS14800 | elongation factor Tu | 1712410 |
| BR63_RS14870 | elongation factor Tu | 1712410 |
| THEBR_RS01825 | elongation factor Tu | 509193 |
| THEBR_RS01890 | elongation factor Tu | 509193 |
| EB239_RS13845 | elongation factor Tu | 509192 |
| EB239_RS13915 | elongation factor Tu | 509192 |
| THIT_RS09935 | elongation factor Tu | 580331 |
| THIT_RS10000 | elongation factor Tu | 580331 |
| TKV_RS10045 | elongation factor Tu | 2325 |
| TKV_RS10115 | elongation factor Tu | 2325 |
| TMATH_RS09730 | elongation factor Tu | 583358 |
| TETH39_RS01825 | elongation factor Tu | 340099 |
| TETH39_RS01890 | elongation factor Tu | 340099 |
| THEWI_RS11285 | elongation factor Tu | 697303 |
| THEWI_RS11350 | elongation factor Tu | 697303 |
| TSAC_RS05020 | elongation factor Tu | 1094508 |
| TSAC_RS05090 | elongation factor Tu | 1094508 |
| TTHE_RS02070 | elongation factor Tu | 580327 |
| TTHE_RS02135 | elongation factor Tu | 580327 |
| THEXY_RS01660 | elongation factor Tu | 858215 |

| | | |
|---|---|---|
| THEXY_RS01725 | elongation factor Tu | 858215 |
| NOG13_RS05285 | elongation factor Tu | 2966552 |
| CST_RS12465 | elongation factor Tu | 1121335 |
| TOCE_RS00605 | elongation factor Tu | 555079 |
| TOCE_RS10595 | elongation factor Tu | 555079 |
| HYG85_RS08545 | elongation factor Tu | 1185412 |
| HZI73_RS03860 | elongation factor Tu | 1348613 |
| KJS28_RS08250 | elongation factor Tu | 2714355 |
| KI236_RS10680 | elongation factor Tu | 2714353 |
| H9Q79_RS16175 | elongation factor Tu | 2763667 |
| H9Q76_RS10410 | elongation factor Tu | 2763670 |
| EYS13_RS14435 | elongation factor Tu | 2528593 |
| EYS13_RS14505 | elongation factor Tu | 2528593 |
| COPRO5265_RS04535 | elongation factor Tu | 309798 |
| COPRO5265_RS04805 | elongation factor Tu | 309798 |
| AEQU_RS07390 | elongation factor Tu | 1384484 |
| HRR28_RS04650 | elongation factor Tu | 2707299 |
| FX134_RS14120 | elongation factor Tu | 1870985 |
| JI75_RS06280 | elongation factor Tu | 1531429 |
| HLV38_RS07170 | elongation factor Tu | 2734634 |
| GXM19_RS05635 | elongation factor Tu | 411903 |
| NQ498_RS01025 | elongation factor Tu | 147206 |
| CORGL_RS07035 | elongation factor Tu | 700015 |
| CCUR_RS04795 | elongation factor Tu | 469378 |
| AAY81_RS07130 | elongation factor Tu | 79604 |
| GS424_RS12095 | elongation factor Tu | 2690837 |
| LUA64_RS10325 | elongation factor Tu | 84112 |
| I6G91_RS05310 | elongation factor Tu | 82135 |
| APAR_RS06030 | elongation factor Tu | 521095 |
| OR601_RS02300 | elongation factor Tu | 2894156 |
| BQ5347_RS00355 | elongation factor Tu | 1805478 |

| | | |
|---|---|---|
| OLSU_RS08135 | elongation factor Tu | 633147 |
| BLT96_RS08720 | elongation factor Tu | 604330 |
| Pcatena_RS00925 | elongation factor Tu | 2003188 |
| BQ7373_RS07735 | elongation factor Tu | 1871022 |
| B5449_RS02235 | elongation factor Tu | 1944646 |
| MWL89_RS10490 | elongation factor Tu | 1907662 |
| EL225_RS03845 | elongation factor Tu | 84110 |
| INP52_RS08905 | elongation factor Tu | 2779519 |
| J7S26_RS05635 | elongation factor Tu | 2682776 |
| AM1_RS08015 | elongation factor Tu | 329726 |
| MIC7113_RS29940 | elongation factor Tu | 1173027 |
| ANACY_RS12320 | elongation factor Tu | 272123 |
| IM676_RS08480 | elongation factor Tu | 2779889 |
| IL331_RS05700 | elongation factor Tu | 2778476 |
| DP114_RS06735 | elongation factor Tu | 415709 |
| UCYN_RS03890 | elongation factor Tu | 1453429 |
| ACL6C3_RS30040 | elongation factor Tu | 1972598 |
| CHA6605_RS20650 | elongation factor Tu | 1173020 |
| CHRO_RS23415 | elongation factor Tu | 251229 |
| CRI9333_RS14790 | elongation factor Tu | 1173022 |
| CCE_RS19300 | elongation factor Tu | 43989 |
| CYAN10605_RS02100 | elongation factor Tu | 755178 |
| CYAGR_RS08800 | elongation factor Tu | 292564 |
| U9970_RS11025 | elongation factor Tu | 2116684 |
| IAR63_RS15170 | elongation factor Tu | 2666332 |
| C6N34_RS08000 | elongation factor Tu | 2108055 |
| CYLST_RS23740 | elongation factor Tu | 56107 |
| DACSA_RS03025 | elongation factor Tu | 13035 |
| CA730_RS22495 | elongation factor Tu | 1973481 |
| HGD76_RS03790 | elongation factor Tu | 315271 |
| NG743_RS20440 | elongation factor Tu | 747523 |

| | | |
|---|---|---|
| FRE64_RS11650 | elongation factor Tu | 522448 |
| SYN6308_RS05160 | elongation factor Tu | 113355 |
| GKIL_RS12925 | elongation factor Tu | 1183438 |
| ISF26_RS17385 | elongation factor Tu | 2781736 |
| GLL_RS20235 | elongation factor Tu | 251221 |
| P0S91_RS08410 | elongation factor Tu | 2859516 |
| GlitD10_RS06075 | elongation factor Tu | 1188229 |
| PCC7424_RS00250 | elongation factor Tu | 65393 |
| CYAN7822_RS10425 | elongation factor Tu | 497965 |
| XM38_RS04350 | elongation factor Tu | 1641165 |
| QI031_RS02635 | elongation factor Tu | 1539056 |
| KIK02_RS11065 | elongation factor Tu | 2835870 |
| LBWT_RS16470 | elongation factor Tu | 411966 |
| J5X98_RS15705 | elongation factor Tu | 2016068 |
| XZ46_RS20715 | elongation factor Tu | 376219 |
| P2A85_RS25235 | elongation factor Tu | 2705319 |
| BDGGKGIB_RS10690 | elongation factor Tu | 2761619 |
| BMF81_RS10265 | elongation factor Tu | 1914872 |
| AAZO_RS03245 | elongation factor Tu | 551115 |
| HUN01_RS14800 | elongation factor Tu | 1472755 |
| COO91_RS19430 | elongation factor Tu | 2038116 |
| ACX27_RS27160 | elongation factor Tu | 224013 |
| NPUN_RS19630 | elongation factor Tu | 63737 |
| D1367_RS24655 | elongation factor Tu | 446679 |
| WJM97_RS01735 | elongation factor Tu | 3133117 |
| OSCIL6304_RS20615 | elongation factor Tu | 56110 |
| OSC7112_RS00315 | elongation factor Tu | 179408 |
| OSC7112_RS02070 | elongation factor Tu | 179408 |
| HCG48_RS19935 | elongation factor Tu | 2064643 |
| TX72_RS10750 | elongation factor Tu | 84588 |
| BRW62_RS04045 | elongation factor Tu | 1917166 |

| | | |
|---|---|---|
| NEA10_RS02380 | elongation factor Tu | 2940671 |
| ABWT76_RS25450 | elongation factor Tu | 2792601 |
| NMG88_RS05345 | elongation factor Tu | 1160 |
| NMK37_RS20735 | elongation factor Tu | 1678309 |
| P9515_RS08055 | elongation factor Tu | 167542 |
| OA858_RS13665 | elongation factor Tu | 2602720 |
| M4D78_RS11570 | elongation factor Tu | 2942274 |
| B6N60_RS23200 | elongation factor Tu | 1357546 |
| PCC8801_RS07465 | elongation factor Tu | 41431 |
| K2F26_RS08270 | elongation factor Tu | 984208 |
| STA7437_RS11200 | elongation factor Tu | 111780 |
| PCC6301pg_RS04625 | elongation factor Tu | 269084 |
| OXH18_RS06105 | elongation factor Tu | 2016057 |
| HNI00_RS12490 | elongation factor Tu | 2547457 |
| HPC62_RS19850 | elongation factor Tu | 2737172 |
| D3A95_RS10095 | elongation factor Tu | 2016101 |
| TLL_RS08885 | elongation factor Tu | 197221 |
| PH595_RS03635 | elongation factor Tu | 3023083 |
| TERY_RS02245 | elongation factor Tu | 203124 |
| GSQ19_RS11695 | elongation factor Tu | 375899 |
| OOK60_RS11350 | elongation factor Tu | 2016064 |
| HUW48_RS20630 | elongation factor Tu | 2745197 |
| HUW51_RS03410 | elongation factor Tu | 2086471 |
| OM944_RS15710 | elongation factor Tu | 2991499 |
| ALPR1_RS18765 | elongation factor Tu | 388413 |
| AO498_RS05450 | elongation factor Tu | 1727163 |
| B9A52_RS00980 | elongation factor Tu | 758820 |
| G9X63_RS01060 | elongation factor Tu | 2516559 |
| G9X62_RS01065 | elongation factor Tu | 2696560 |
| HME7025_RS01625 | elongation factor Tu | 2516557 |
| DJ013_RS14545 | elongation factor Tu | 1784714 |

| | | |
|---|---|---|
| AABK36_RS02360 | elongation factor Tu | 866807 |
| BELBA_RS08305 | elongation factor Tu | 866536 |
| FLELI_RS13040 | elongation factor Tu | 880071 |
| AASI_RS07180 | elongation factor Tu | 452471 |
| DK880_RS00110 | elongation factor Tu | 247481 |
| AL022_RS01520 | elongation factor Tu | 1231626 |
| ACOALF_RS02865 | elongation factor Tu | 3066217 |
| NBW36_RS03650 | elongation factor Tu | 931101 |
| AAHM81_RS01125 | elongation factor Tu | 3077941 |
| CE557_RS04655 | elongation factor Tu | 650378 |
| AACK88_RS04480 | elongation factor Tu | 3066216 |
| PZB72_RS09055 | elongation factor Tu | 1397264 |
| OQ292_RS06225 | elongation factor Tu | 2994555 |
| D4L85_RS12025 | elongation factor Tu | 2321403 |
| CA2015_RS12450 | elongation factor Tu | 320787 |
| CYCMA_RS12745 | elongation factor Tu | 880070 |
| CHU_RS15450 | elongation factor Tu | 269798 |
| NFI80_RS20050 | elongation factor Tu | 2909339 |
| MUK70_RS25175 | elongation factor Tu | 2906456 |
| NFI81_RS21210 | elongation factor Tu | 2906443 |
| DFER_RS17120 | elongation factor Tu | 471854 |
| ON006_RS23170 | elongation factor Tu | 2910158 |
| HWI92_RS20695 | elongation factor Tu | 2747268 |
| KZP23_RS01280 | elongation factor Tu | 2859768 |
| FDP09_RS01295 | elongation factor Tu | 1807691 |
| FKX85_RS01325 | elongation factor Tu | 2591634 |
| DN752_RS07890 | elongation factor Tu | 1795355 |
| ECHVI_RS01325 | elongation factor Tu | 926556 |
| EMTOL_RS05955 | elongation factor Tu | 929562 |
| FAES_RS05450 | elongation factor Tu | 1166018 |
| KM029_RS15045 | elongation factor Tu | 373891 |

| | | |
|---|---|---|
| EI427_RS14360 | elongation factor Tu | 2494373 |
| KMW28_RS18125 | elongation factor Tu | 367791 |
| AABK39_RS16765 | elongation factor Tu | 1182444 |
| LVD16_RS00960 | elongation factor Tu | 2904246 |
| JR347_RS00045 | elongation factor Tu | 2810512 |
| LVD15_RS04910 | elongation factor Tu | 2904247 |
| LVD17_RS05895 | elongation factor Tu | 2904245 |
| MUN82_RS20740 | elongation factor Tu | 2932251 |
| F6X24_RS01745 | elongation factor Tu | 2615203 |
| O3303_RS12210 | elongation factor Tu | 2999067 |
| MUN80_RS01080 | elongation factor Tu | 2932249 |
| FHG12_RS11640 | elongation factor Tu | 2502781 |
| MTP16_RS02365 | elongation factor Tu | 1705399 |
| DDQ68_RS12915 | elongation factor Tu | 1850093 |
| D3Y59_RS16060 | elongation factor Tu | 2319843 |
| EPD59_RS03895 | elongation factor Tu | 2496028 |
| AUC43_RS07885 | elongation factor Tu | 1411621 |
| H4317_RS03390 | elongation factor Tu | 2761579 |
| MWH26_RS15000 | elongation factor Tu | 2933777 |
| HSW_RS18885 | elongation factor Tu | 1227739 |
| HMJ29_RS04090 | elongation factor Tu | 2735321 |
| MTX78_RS19555 | elongation factor Tu | 497967 |
| MUN86_RS19445 | elongation factor Tu | 2932254 |
| O9Z63_RS14190 | elongation factor Tu | 748197 |
| RT717_RS19185 | elongation factor Tu | 1324217 |
| OQ371_RS09935 | elongation factor Tu | 332158 |
| LBYS_RS11825 | elongation factor Tu | 649349 |
| DCC35_RS04205 | elongation factor Tu | 2715131 |
| LAG90_RS09950 | elongation factor Tu | 2975157 |
| QYS47_RS05055 | elongation factor Tu | 3059076 |
| Q3Y49_RS09070 | elongation factor Tu | 2010992 |

| | | |
|---|---|---|
| QYS49_RS18685 | elongation factor Tu | 3059078 |
| FTRAC_RS15095 | elongation factor Tu | 643867 |
| IPZ59_RS09055 | elongation factor Tu | 2782006 |
| GU926_RS15250 | elongation factor Tu | 2698458 |
| AABK40_RS12040 | elongation factor Tu | 387638 |
| CA264_RS19320 | elongation factor Tu | 323450 |
| A0W33_RS10630 | elongation factor Tu | 388950 |
| OH144_RS19805 | elongation factor Tu | 2592049 |
| PKOR_RS08400 | elongation factor Tu | 400092 |
| GSQ66_RS12175 | elongation factor Tu | 2694930 |
| GSQ62_RS13440 | elongation factor Tu | 2694929 |
| PZB74_RS01490 | elongation factor Tu | 930972 |
| N6H18_RS02510 | elongation factor Tu | 2979464 |
| N7E81_RS08525 | elongation factor Tu | 2979869 |
| GXP67_RS21680 | elongation factor Tu | 2704465 |
| T7867_RS10095 | elongation factor Tu | 333140 |
| TH63_RS01025 | elongation factor Tu | 1379910 |
| DC20_RS11540 | elongation factor Tu | 512763 |
| DR864_RS17975 | elongation factor Tu | 2259595 |
| RUNSL_RS08930 | elongation factor Tu | 761193 |
| B5M13_RS27580 | elongation factor Tu | 1211326 |
| G8759_RS00845 | elongation factor Tu | 2692134 |
| GJR95_RS25255 | elongation factor Tu | 2666025 |
| H3H32_RS25250 | elongation factor Tu | 2710596 |
| SLIN_RS22010 | elongation factor Tu | 504472 |
| AWR27_RS21540 | elongation factor Tu | 1178516 |
| LQ777_RS23350 | elongation factor Tu | 2898794 |
| CWM47_RS04840 | elongation factor Tu | 2057025 |
| SD10_RS24220 | elongation factor Tu | 1379870 |
| HH216_RS07640 | elongation factor Tu | 2728024 |
| B5M14_RS02850 | elongation factor Tu | 564064 |

| | | |
|---|---|---|
| HNV11_RS00255 | elongation factor Tu | 2735870 |
| L0Y31_RS06975 | elongation factor Tu | 2907205 |
| ORG26_RS12750 | elongation factor Tu | 2674997 |
| P0M28_RS24705 | elongation factor Tu | 931531 |
| CALNI_RS07600 | elongation factor Tu | 768670 |
| CALNI_RS07745 | elongation factor Tu | 768670 |
| DEFDS_RS08920 | elongation factor Tu | 639282 |
| DEFDS_RS08990 | elongation factor Tu | 639282 |
| DACET_RS01795 | elongation factor Tu | 522772 |
| DACET_RS01865 | elongation factor Tu | 522772 |
| FLEXSI_RS02230 | elongation factor Tu | 717231 |
| FLEXSI_RS02300 | elongation factor Tu | 717231 |
| EP073_RS04920 | elongation factor Tu | 139438 |
| EP073_RS04990 | elongation factor Tu | 139438 |
| N508_RS04770 | elongation factor Tu | 1379858 |
| V8247_RS06285 | elongation factor Tu | 3127115 |
| GKN94_RS02870 | elongation factor Tu | 3038979 |
| DET_RS05180 | elongation factor Tu | 243164 |
| HX448_RS05835 | elongation factor Tu | 1536648 |
| Dform_RS06085 | elongation factor Tu | 1839801 |
| MESIL_RS13495 | elongation factor Tu | 526227 |
| MESIL_RS14630 | elongation factor Tu | 526227 |
| DAETH_RS01920 | elongation factor Tu | 200252 |
| DAETH_RS10810 | elongation factor Tu | 200252 |
| QYS50_RS00355 | elongation factor Tu | 468914 |
| QYS50_RS10635 | elongation factor Tu | 468914 |
| M8445_RS01880 | elongation factor Tu | 328692 |
| M8445_RS08810 | elongation factor Tu | 328692 |
| DEIDE_RS03525 | elongation factor Tu | 546414 |
| DEIDE_RS10865 | elongation factor Tu | 546414 |
| DFI_RS02160 | elongation factor Tu | 317577 |

| | | |
|---|---|---|
| DFI_RS02985 | elongation factor Tu | 317577 |
| DGEO_RS05915 | elongation factor Tu | 319795 |
| DGEO_RS12105 | elongation factor Tu | 319795 |
| DGO_RS05810 | elongation factor Tu | 745776 |
| DGO_RS13110 | elongation factor Tu | 745776 |
| DKM44_RS01505 | elongation factor Tu | 2202254 |
| DKM44_RS05395 | elongation factor Tu | 2202254 |
| DEIMA_RS04200 | elongation factor Tu | 709986 |
| DEIMA_RS14755 | elongation factor Tu | 709986 |
| E5F05_RS15765 | elongation factor Tu | 1211322 |
| E5F05_RS17865 | elongation factor Tu | 1211322 |
| DEIPE_RS04785 | elongation factor Tu | 937777 |
| DEIPE_RS14825 | elongation factor Tu | 937777 |
| DEIPR_RS02045 | elongation factor Tu | 693977 |
| DEIPR_RS07275 | elongation factor Tu | 693977 |
| EHF33_RS02130 | elongation factor Tu | 2489213 |
| EHF33_RS08690 | elongation factor Tu | 2489213 |
| SU48_RS10060 | elongation factor Tu | 1182568 |
| SU48_RS11425 | elongation factor Tu | 1182568 |
| Q0M94_RS03245 | elongation factor Tu | 468916 |
| Q0M94_RS08625 | elongation factor Tu | 468916 |
| QR90_RS01070 | elongation factor Tu | 57497 |
| QR90_RS05440 | elongation factor Tu | 57497 |
| N0D28_RS03095 | elongation factor Tu | 1889240 |
| N0D28_RS04645 | elongation factor Tu | 1889240 |
| ABOD76_RS09240 | elongation factor Tu | 694439 |
| ABOD76_RS19070 | elongation factor Tu | 694439 |
| LAJ19_RS02385 | elongation factor Tu | 2737050 |
| LAJ19_RS09670 | elongation factor Tu | 2737050 |
| G6R31_RS01015 | elongation factor Tu | 1288484 |
| G6R31_RS10105 | elongation factor Tu | 1288484 |

| | | |
|---|---|---|
| ACLF6M_RS08850 | elongation factor Tu | 367282 |
| ACLF6M_RS10895 | elongation factor Tu | 367282 |
| MARKY_RS02260 | elongation factor Tu | 869210 |
| MARKY_RS08690 | elongation factor Tu | 869210 |
| MRUB_RS05995 | elongation factor Tu | 504728 |
| MRUB_RS13180 | elongation factor Tu | 504728 |
| MTAI_RS02635 | elongation factor Tu | 1339250 |
| MTAI_RS08845 | elongation factor Tu | 1339250 |
| KNN15_RS02840 | elongation factor Tu | 88190 |
| KNN15_RS05085 | elongation factor Tu | 88190 |
| TO73_RS00345 | elongation factor Tu | 498848 |
| TO73_RS01795 | elongation factor Tu | 498848 |
| MWM05_RS09560 | elongation factor Tu | 56956 |
| MWM05_RS09930 | elongation factor Tu | 56956 |
| EBI04_RS08200 | elongation factor Tu | 2483360 |
| EBI04_RS10405 | elongation factor Tu | 2483360 |
| THEOS_RS01380 | elongation factor Tu | 751945 |
| THEOS_RS09665 | elongation factor Tu | 751945 |
| AV541_RS00175 | elongation factor Tu | 456163 |
| AV541_RS02290 | elongation factor Tu | 456163 |
| TTH_RS01315 | elongation factor Tu | 300852 |
| TTH_RS08570 | elongation factor Tu | 300852 |
| TRAD_RS05170 | elongation factor Tu | 649638 |
| TRAD_RS12365 | elongation factor Tu | 649638 |
| DN745_RS01000 | elongation factor Tu | 1548548 |
| DN745_RS17985 | elongation factor Tu | 1548548 |
| LGS26_RS03525 | elongation factor Tu | 1750598 |
| LGS26_RS04275 | elongation factor Tu | 1750598 |
| FRD01_RS18700 | elongation factor Tu | 2600177 |
| FRD00_RS17695 | elongation factor Tu | 2292766 |
| FRD00_RS18390 | elongation factor Tu | 2292766 |

| | | |
|---|---|---|
| DEBA_RS14670 | elongation factor Tu | 644282 |
| DEBA_RS14740 | elongation factor Tu | 644282 |
| AACH32_RS17830 | elongation factor Tu | 3058419 |
| AACH32_RS17900 | elongation factor Tu | 3058419 |
| DESAC_RS07260 | elongation factor Tu | 880072 |
| DESAC_RS07325 | elongation factor Tu | 880072 |
| DALK_RS09895 | elongation factor Tu | 218208 |
| DALK_RS09960 | elongation factor Tu | 218208 |
| EYB58_RS09090 | elongation factor Tu | 2291 |
| DESPODRAFT_RS12100 | elongation factor Tu | 879212 |
| TOL2_RS18565 | elongation factor Tu | 651182 |
| TOL2_RS18630 | elongation factor Tu | 651182 |
| B2D07_RS14795 | elongation factor Tu | 897 |
| B2D07_RS14860 | elongation factor Tu | 897 |
| L6434_RS19785 | elongation factor Tu | 2810562 |
| L6434_RS19850 | elongation factor Tu | 2810562 |
| dnl_RS04900 | elongation factor Tu | 45656 |
| dnl_RS04960 | elongation factor Tu | 45656 |
| dnm_RS04120 | elongation factor Tu | 45655 |
| HRM2_RS17910 | elongation factor Tu | 177437 |
| HRM2_RS17970 | elongation factor Tu | 177437 |
| DSCA_RS10550 | elongation factor Tu | 571177 |
| DSCA_RS10615 | elongation factor Tu | 571177 |
| GN113_RS30215 | elongation factor Tu | 885957 |
| GN113_RS30280 | elongation factor Tu | 885957 |
| Dvar_RS09830 | elongation factor Tu | 859321 |
| Dvar_RS09895 | elongation factor Tu | 859321 |
| GN111_RS11385 | elongation factor Tu | 947919 |
| GN111_RS11450 | elongation factor Tu | 947919 |
| DOLE_RS03585 | elongation factor Tu | 96561 |
| DOLE_RS03645 | elongation factor Tu | 96561 |

| | | |
|---|---|---|
| HP555_RS07480 | elongation factor Tu | 1909699 |
| HP555_RS07550 | elongation factor Tu | 1909699 |
| CAY53_RS10835 | elongation factor Tu | 1986146 |
| CAY53_RS10905 | elongation factor Tu | 1986146 |
| DESPR_RS03835 | elongation factor Tu | 577650 |
| DESPR_RS03900 | elongation factor Tu | 577650 |
| UWK_RS10105 | elongation factor Tu | 1167006 |
| UWK_RS10170 | elongation factor Tu | 1167006 |
| DPPLL_RS02405 | elongation factor Tu | 2740163 |
| DPPLL_RS02475 | elongation factor Tu | 2740163 |
| GF1_RS11575 | elongation factor Tu | 2795293 |
| GF1_RS11645 | elongation factor Tu | 2795293 |
| LO777_RS02255 | elongation factor Tu | 2772557 |
| LO777_RS02325 | elongation factor Tu | 2772557 |
| FCL46_RS02085 | elongation factor Tu | 2569541 |
| FCL46_RS02155 | elongation factor Tu | 2569541 |
| FCL45_RS21910 | elongation factor Tu | 2569540 |
| FCL45_RS21980 | elongation factor Tu | 2569540 |
| DP_RS05585 | elongation factor Tu | 177439 |
| DP_RS05655 | elongation factor Tu | 177439 |
| DAAHT2_RS07415 | elongation factor Tu | 589865 |
| DAAHT2_RS07485 | elongation factor Tu | 589865 |
| DESTI_RS26025 | elongation factor Tu | 706587 |
| DESTI_RS26095 | elongation factor Tu | 706587 |
| RSDT_RS01870 | elongation factor Tu | 1725232 |
| DESAF_RS01135 | elongation factor Tu | 690850 |
| DESAF_RS11005 | elongation factor Tu | 690850 |
| DRET_RS10310 | elongation factor Tu | 485915 |
| DRET_RS10380 | elongation factor Tu | 485915 |
| GD606_RS00875 | elongation factor Tu | 1121464 |
| GD606_RS07150 | elongation factor Tu | 1121464 |

| | | |
|---|---|---|
| GD604_RS05400 | elongation factor Tu | 2773299 |
| GD604_RS07610 | elongation factor Tu | 2773299 |
| DBAC_RS13870 | elongation factor Tu | 525897 |
| DBAC_RS13935 | elongation factor Tu | 525897 |
| AXF15_RS01035 | elongation factor Tu | 888061 |
| AXF15_RS01105 | elongation factor Tu | 888061 |
| ACEB74_RS16385 | elongation factor Tu | 81425 |
| J8J02_RS10045 | elongation factor Tu | 876 |
| AXF13_RS03755 | elongation factor Tu | 44742 |
| EL361_RS01860 | elongation factor Tu | 241368 |
| EL361_RS01960 | elongation factor Tu | 241368 |
| N1030_RS16170 | elongation factor Tu | 2976983 |
| LN040_RS14915 | elongation factor Tu | 2718620 |
| LAW_RS05100 | elongation factor Tu | 1234378 |
| DESAM_RS15350 | elongation factor Tu | 1121451 |
| DESAL_RS05895 | elongation factor Tu | 526222 |
| DGI_RS08240 | elongation factor Tu | 1121448 |
| DVUL_RS02320 | elongation factor Tu | 391774 |
| E8L03_RS01345 | elongation factor Tu | 370038 |
| E8L03_RS17030 | elongation factor Tu | 370038 |
| DDE_RS13700 | elongation factor Tu | 207559 |
| DAES_RS11435 | elongation factor Tu | 643562 |
| GM415_RS11615 | elongation factor Tu | 2678688 |
| DND132_RS17040 | elongation factor Tu | 641491 |
| V8V93_RS08975 | elongation factor Tu | 3126690 |
| SYK_RS17260 | elongation factor Tu | 2973072 |
| BN4_RS16515 | elongation factor Tu | 1322246 |
| OO730_RS14375 | elongation factor Tu | 231439 |
| DPRO_RS19220 | elongation factor Tu | 57320 |
| SRBAKS_RS17430 | elongation factor Tu | 2810563 |
| LF599_RS05715 | elongation factor Tu | 2880651 |

| | | |
|---|---|---|
| MPN23_RS00690 | elongation factor Tu | 463192 |
| FGL65_RS14980 | elongation factor Tu | 2583823 |
| C3Y92_RS06645 | elongation factor Tu | 296842 |
| C3Y92_RS15205 | elongation factor Tu | 296842 |
| DFW101_RS01440 | elongation factor Tu | 694327 |
| DFW101_RS15900 | elongation factor Tu | 694327 |
| DMR_RS05575 | elongation factor Tu | 573370 |
| DMR_RS14925 | elongation factor Tu | 573370 |
| JBF11_RS09845 | elongation factor Tu | 2787751 |
| HIPMA_RS07115 | elongation factor Tu | 760142 |
| HIPMA_RS07185 | elongation factor Tu | 760142 |
| GBEM_RS04585 | elongation factor Tu | 404380 |
| GBEM_RS04655 | elongation factor Tu | 404380 |
| GEOBRER4_RS16315 | elongation factor Tu | 60035 |
| GEOBRER4_RS16385 | elongation factor Tu | 60035 |
| DSOUD_RS13955 | elongation factor Tu | 1603606 |
| DSOUD_RS14025 | elongation factor Tu | 1603606 |
| SNR20_RS06885 | elongation factor Tu | 57664 |
| SNR20_RS06955 | elongation factor Tu | 57664 |
| DESUT3_RS05565 | elongation factor Tu | 2802975 |
| DESUT3_RS05635 | elongation factor Tu | 2802975 |
| L9S41_RS14885 | elongation factor Tu | 2847045 |
| L9S41_RS14955 | elongation factor Tu | 2847045 |
| GSUB_RS12660 | elongation factor Tu | 483547 |
| GSUB_RS12725 | elongation factor Tu | 483547 |
| JZM60_RS08680 | elongation factor Tu | 2815309 |
| JZM60_RS08750 | elongation factor Tu | 2815309 |
| GMET_RS03075 | elongation factor Tu | 269799 |
| GMET_RS03145 | elongation factor Tu | 269799 |
| GPICK_RS03250 | elongation factor Tu | 345632 |
| GPICK_RS03320 | elongation factor Tu | 345632 |

| | | |
|---|---|---|
| GS_RS14355 | elongation factor Tu | 243231 |
| GS_RS14425 | elongation factor Tu | 243231 |
| KP003_RS15485 | elongation factor Tu | 2847991 |
| KP003_RS15555 | elongation factor Tu | 2847991 |
| KP004_RS15605 | elongation factor Tu | 2847992 |
| KP004_RS15675 | elongation factor Tu | 2847992 |
| M1B72_RS04325 | elongation factor Tu | 2740185 |
| M1B72_RS04395 | elongation factor Tu | 2740185 |
| KP002_RS08635 | elongation factor Tu | 2847989 |
| KP002_RS08705 | elongation factor Tu | 2847989 |
| GEOB_RS18050 | elongation factor Tu | 316067 |
| GEOB_RS18120 | elongation factor Tu | 316067 |
| GURA_RS05350 | elongation factor Tu | 351605 |
| GURA_RS05420 | elongation factor Tu | 351605 |
| PPRO_RS03265 | elongation factor Tu | 338966 |
| PPRO_RS03335 | elongation factor Tu | 338966 |
| A7E75_RS03485 | elongation factor Tu | 29542 |
| A7E75_RS03555 | elongation factor Tu | 29542 |
| A7E78_RS07615 | elongation factor Tu | 1842532 |
| A7E78_RS07685 | elongation factor Tu | 1842532 |
| PCAR_RS04205 | elongation factor Tu | 338963 |
| PCAR_RS04275 | elongation factor Tu | 338963 |
| RAK07_RS03665 | elongation factor Tu | 2916410 |
| RAK07_RS03735 | elongation factor Tu | 2916410 |
| GLOV_RS06560 | elongation factor Tu | 398767 |
| GLOV_RS06625 | elongation factor Tu | 398767 |
| HUE98_RS01870 | elongation factor Tu | 338645 |
| DICTH_RS04010 | elongation factor Tu | 309799 |
| DICTH_RS06095 | elongation factor Tu | 309799 |
| DTUR_RS04995 | elongation factor Tu | 515635 |
| DTUR_RS07135 | elongation factor Tu | 515635 |

| | | |
|---|---|---|
| EMIN_RS07230 | elongation factor Tu | 445932 |
| EMIN_RS07640 | elongation factor Tu | 445932 |
| RSTT_RS00675 | elongation factor Tu | 1408204 |
| RSTT_RS01650 | elongation factor Tu | 1408204 |
| Epro_RS04110 | elongation factor Tu | 1408281 |
| Epro_RS06670 | elongation factor Tu | 1408281 |
| ABU_RS09455 | elongation factor Tu | 367737 |
| ACBT_RS02785 | elongation factor Tu | 255507 |
| ACRYA_RS02175 | elongation factor Tu | 1032070 |
| AFAEC_RS01595 | elongation factor Tu | 1564138 |
| ALANTH_RS09110 | elongation factor Tu | 1355374 |
| ASKIR_RS02415 | elongation factor Tu | 1032239 |
| ATH_RS07740 | elongation factor Tu | 1032240 |
| ATR_RS07765 | elongation factor Tu | 1032241 |
| ACNSOS_RS04625 | elongation factor Tu | 2011099 |
| AACT_RS12920 | elongation factor Tu | 1849015 |
| AAQM_RS10795 | elongation factor Tu | 1315211 |
| ACLO_RS11495 | elongation factor Tu | 1054034 |
| ADFLV_RS13010 | elongation factor Tu | 873191 |
| AELL_RS12280 | elongation factor Tu | 913109 |
| ARNIT_RS13125 | elongation factor Tu | 572480 |
| D3Z07_RS12430 | elongation factor Tu | 2320140 |
| APORC_RS08180 | elongation factor Tu | 1935204 |
| NJU99_RS12430 | elongation factor Tu | 2961520 |
| ASUIS_RS11530 | elongation factor Tu | 663365 |
| AVENP_RS13300 | elongation factor Tu | 1054033 |
| FE773_RS00105 | elongation factor Tu | 391592 |
| C6V80_RS08365 | elongation factor Tu | 1424653 |
| CARM_RS02135 | elongation factor Tu | 2505970 |
| CAV_RS05675 | elongation factor Tu | 522484 |
| ACEN41_RS01870 | elongation factor Tu | 2691918 |

| | | |
|---|---|---|
| CBLAS_RS02195 | elongation factor Tu | 2042961 |
| CCAL_RS06580 | elongation factor Tu | 1032243 |
| CCANL266_RS00230 | elongation factor Tu | 449520 |
| FOC43_RS05085 | elongation factor Tu | 195 |
| CCON33237_RS03450 | elongation factor Tu | 199 |
| CCORG_RS01975 | elongation factor Tu | 1448857 |
| CCUN_RS03165 | elongation factor Tu | 1121267 |
| CCVT_RS06870 | elongation factor Tu | 200 |
| GZ985_RS06500 | elongation factor Tu | 196 |
| CGEO_RS02470 | elongation factor Tu | 1780362 |
| CGRAC_RS08290 | elongation factor Tu | 824 |
| CHELV3228_RS01755 | elongation factor Tu | 28898 |
| A2J15_RS00300 | elongation factor Tu | 1813019 |
| CHAB381_RS07820 | elongation factor Tu | 360107 |
| CHLWT_RS02330 | elongation factor Tu | 91353 |
| CIG1485E_RS06695 | elongation factor Tu | 1244531 |
| CINS_RS02105 | elongation factor Tu | 1031564 |
| Cj0470 | elongation factor Tu | 192222 |
| CLAN_RS02535 | elongation factor Tu | 1031753 |
| CLCT_RS02200 | elongation factor Tu | 488545 |
| CMCT_RS02295 | elongation factor Tu | 202 |
| CNZW441b_RS02675 | elongation factor Tu | 2267891 |
| CORN_RS02245 | elongation factor Tu | 1848766 |
| CPEL_RS02255 | elongation factor Tu | 1388753 |
| CPIN18020_RS06850 | elongation factor Tu | 1874362 |
| CSUIS_RS02935 | elongation factor Tu | 1660073 |
| CRECT_RS09260 | elongation factor Tu | 203 |
| CSHOW_RS02550 | elongation factor Tu | 204 |
| CSPT_RS02260 | elongation factor Tu | 32024 |
| CSUB8523_RS02290 | elongation factor Tu | 1388752 |
| CUP3940_RS06665 | elongation factor Tu | 1031733 |

| | | |
|---|---|---|
| CURT_RS07235 | elongation factor Tu | 827 |
| CVIC9261_RS02260 | elongation factor Tu | 1660076 |
| CVOLT_RS02180 | elongation factor Tu | 1031542 |
| CVULP_RS01705 | elongation factor Tu | 1655500 |
| CINF_RS03835 | elongation factor Tu | 2561898 |
| HUE88_RS11365 | elongation factor Tu | 2740404 |
| HUE87_RS01475 | elongation factor Tu | 2740405 |
| AANAER_RS12440 | elongation factor Tu | 877500 |
| ABIV_RS01850 | elongation factor Tu | 663364 |
| AEBR_RS12625 | elongation factor Tu | 1462615 |
| HAC_RS06690 | elongation factor Tu | 382638 |
| CDV25_RS08210 | elongation factor Tu | 135569 |
| HCAN_RS01105 | elongation factor Tu | 537970 |
| HCD_RS02955 | elongation factor Tu | 1163745 |
| EL253_RS03655 | elongation factor Tu | 45498 |
| HCN_RS07625 | elongation factor Tu | 1172562 |
| BBW65_RS04320 | elongation factor Tu | 222136 |
| HFELIS_RS02590 | elongation factor Tu | 936155 |
| QUE23_RS03590 | elongation factor Tu | 3040201 |
| K6J72_RS05945 | elongation factor Tu | 2849641 |
| HH_RS01830 | elongation factor Tu | 235279 |
| A3217_RS03735 | elongation factor Tu | 1591088 |
| V3I05_RS01220 | elongation factor Tu | 287948 |
| DQN48_RS01750 | elongation factor Tu | 217 |
| EL247_RS07440 | elongation factor Tu | 35818 |
| OW489_RS06980 | elongation factor Tu | 210 |
| HXS61_RS07370 | elongation factor Tu | 104628 |
| BN2458_RS05935 | elongation factor Tu | 76936 |
| IP358_RS00950 | elongation factor Tu | 556267 |
| QUD54_RS08015 | elongation factor Tu | 2993563 |
| ABXS81_RS07225 | elongation factor Tu | 2954236 |

| | | |
|---|---|---|
| RZR97_RS00895 | elongation factor Tu | 223786 |
| JMG82_RS06655 | elongation factor Tu | 2740515 |
| ACAN_RS11635 | elongation factor Tu | 1912871 |
| AHALO_RS11120 | elongation factor Tu | 197482 |
| AMRN_RS11600 | elongation factor Tu | 505249 |
| AMOL_RS11595 | elongation factor Tu | 870501 |
| AMYT_RS11460 | elongation factor Tu | 1032238 |
| APAC_RS10790 | elongation factor Tu | 1080223 |
| NAMH_RS00840 | elongation factor Tu | 598659 |
| NITSA_RS09915 | elongation factor Tu | 749222 |
| NITER_RS07160 | elongation factor Tu | 1069081 |
| EPR_RS01080 | elongation factor Tu | 2714855 |
| QML81_RS04345 | elongation factor Tu | 2930200 |
| NIL_RS00915 | elongation factor Tu | 2706014 |
| ALEK_RS14635 | elongation factor Tu | 1904463 |
| LPB137_RS11650 | elongation factor Tu | 1850254 |
| SULKU_RS11455 | elongation factor Tu | 709032 |
| GJV85_RS01645 | elongation factor Tu | 2672570 |
| SAUT_RS01570 | elongation factor Tu | 563040 |
| SUDEN_RS01805 | elongation factor Tu | 326298 |
| WCY31_RS01950 | elongation factor Tu | 3131939 |
| SMGD1_RS04495 | elongation factor Tu | 929558 |
| ETP70_RS10620 | elongation factor Tu | 2509341 |
| FJR48_RS10090 | elongation factor Tu | 2590022 |
| FJR03_RS01350 | elongation factor Tu | 2590551 |
| FM071_RS01270 | elongation factor Tu | 317658 |
| FJR45_RS01880 | elongation factor Tu | 2590020 |
| FJR47_RS01380 | elongation factor Tu | 2590021 |
| SULBA_RS01890 | elongation factor Tu | 760154 |
| SDEL_RS01780 | elongation factor Tu | 525898 |
| SJPD1_RS02145 | elongation factor Tu | 1854492 |

| | | |
|---|---|---|
| SHALO_RS02395 | elongation factor Tu | 1193502 |
| SMUL_RS02390 | elongation factor Tu | 1150621 |
| IMZ28_RS10195 | elongation factor Tu | 2779528 |
| YH65_RS01170 | elongation factor Tu | 206403 |
| LN246_RS00750 | elongation factor Tu | 2893889 |
| EL168_RS07470 | elongation factor Tu | 844 |
| G4D54_RS03735 | elongation factor Tu | 1522 |
| KWG62_RS07980 | elongation factor Tu | 2834459 |
| NMU03_RS09515 | elongation factor Tu | 2944914 |
| A9CBEGH2_RS11510 | elongation factor Tu | 2583452 |
| NQ499_RS03200 | elongation factor Tu | 100886 |
| NMG63_RS06390 | elongation factor Tu | 2929157 |
| LLS04_RS00720 | elongation factor Tu | 2890314 |
| H9L01_RS05910 | elongation factor Tu | 225084 |
| AOC36_RS03375 | elongation factor Tu | 1514105 |
| EEI45_RS04205 | elongation factor Tu | 2485784 |
| ERH_RS06050 | elongation factor Tu | 650150 |
| Fi14EGH31_RS02505 | elongation factor Tu | 1982626 |
| aalo17_RS01915 | elongation factor Tu | 1702221 |
| SG0102_RS03660 | elongation factor Tu | 2487118 |
| KJS44_RS01520 | elongation factor Tu | 1796635 |
| ACGHJ8_RS11710 | elongation factor Tu | 1776392 |
| RGT18_RS12005 | elongation factor Tu | 102148 |
| LRR82_RS10850 | elongation factor Tu | 2899121 |
| I6I62_RS15355 | elongation factor Tu | 1547 |
| NQ543_RS11515 | elongation factor Tu | 428126 |
| J0J80_RS04790 | elongation factor Tu | 2735723 |
| AACH31_RS00465 | elongation factor Tu | 2963365 |
| HLK68_RS05535 | elongation factor Tu | 154288 |
| FSU_RS08265 | elongation factor Tu | 59374 |
| FSU_RS09005 | elongation factor Tu | 59374 |

| | | |
|---|---|---|
| OP10G_RS21615 | elongation factor Tu | 661478 |
| MQE35_RS14160 | elongation factor Tu | 2926907 |
| ISU00_RS01325 | elongation factor Tu | 2893202 |
| EI546_RS00045 | elongation factor Tu | 2494375 |
| JK629_RS09390 | elongation factor Tu | 2803865 |
| QCQ61_RS08825 | elongation factor Tu | 3040348 |
| Q2U82_RS01110 | elongation factor Tu | 1287690 |
| AEQSU_RS09915 | elongation factor Tu | 746697 |
| R1X58_RS01820 | elongation factor Tu | 2920935 |
| FGM00_RS03535 | elongation factor Tu | 2583587 |
| O5O44_RS12615 | elongation factor Tu | 1178825 |
| MURRU_RS10665 | elongation factor Tu | 886377 |
| KFZ70_RS00230 | elongation factor Tu | 2583814 |
| FK178_RS07965 | elongation factor Tu | 2585771 |
| FHG64_RS15390 | elongation factor Tu | 2058175 |
| EOV51_RS00365 | elongation factor Tu | 2500547 |
| AREALGSMS7_RS00075 | elongation factor Tu | 616991 |
| KCTC52924_RS11260 | elongation factor Tu | 2040469 |
| FF125_RS15005 | elongation factor Tu | 2584122 |
| DBX24_RS03835 | elongation factor Tu | 1585976 |
| DM817_RS00825 | elongation factor Tu | 164516 |
| H0H70_RS02370 | elongation factor Tu | 1653831 |
| DM780_RS00820 | elongation factor Tu | 164514 |
| NG655_RS07525 | elongation factor Tu | 2820661 |
| J4771_RS06930 | elongation factor Tu | 2820270 |
| ASU30_RS01075 | elongation factor Tu | 336810 |
| QOX03_RS01875 | elongation factor Tu | 2497989 |
| N2590_RS00030 | elongation factor Tu | 2943807 |
| AAHJ01_RS00270 | elongation factor Tu | 3066219 |
| FNIIJ_RS00830 | elongation factor Tu | 1415657 |
| CGC47_RS04250 | elongation factor Tu | 28188 |

| | | |
|---|---|---|
| ACF3OB_RS09505 | elongation factor Tu | 1848903 |
| CGC48_RS08930 | elongation factor Tu | 28189 |
| CBG49_RS00220 | elongation factor Tu | 2708117 |
| CGC50_RS06925 | elongation factor Tu | 1017 |
| CKV80_RS06120 | elongation factor Tu | 45243 |
| COCH_RS06090 | elongation factor Tu | 521097 |
| CGC57_RS08490 | elongation factor Tu | 1019 |
| CGC58_RS03210 | elongation factor Tu | 1848904 |
| CELAL_RS11780 | elongation factor Tu | 688270 |
| M666_RS09175 | elongation factor Tu | 1348584 |
| CELLY_RS02670 | elongation factor Tu | 867900 |
| PBN93_RS02650 | elongation factor Tu | 1888890 |
| FAF07_RS02950 | elongation factor Tu | 2049305 |
| GRFL_RS12035 | elongation factor Tu | 1229726 |
| GFO_RS11215 | elongation factor Tu | 411154 |
| C7S20_RS11050 | elongation factor Tu | 2126553 |
| LPB144_RS06575 | elongation factor Tu | 1913577 |
| J8N07_RS12915 | elongation factor Tu | 651561 |
| EKK86_RS11340 | elongation factor Tu | 2497456 |
| EB354_RS00050 | elongation factor Tu | 246 |
| EG339_RS02415 | elongation factor Tu | 1241978 |
| CGB83_RS09870 | elongation factor Tu | 1265445 |
| EG346_RS09485 | elongation factor Tu | 1124835 |
| ODZ84_RS11425 | elongation factor Tu | 2983606 |
| OK18_RS04960 | elongation factor Tu | 1324352 |
| PFY09_RS09895 | elongation factor Tu | 373672 |
| A0O34_RS12115 | elongation factor Tu | 1685010 |
| I6I60_RS07090 | elongation factor Tu | 250 |
| QGN23_RS13165 | elongation factor Tu | 3042315 |
| EG358_RS18495 | elongation factor Tu | 254 |
| EG359_RS11845 | elongation factor Tu | 112234 |

| | | |
|---|---|---|
| EG342_RS02280 | elongation factor Tu | 1241981 |
| H0S70_RS12975 | elongation factor Tu | 2754694 |
| EL260_RS22950 | elongation factor Tu | 1241982 |
| M0D58_RS08750 | elongation factor Tu | 1854498 |
| MTP08_RS12070 | elongation factor Tu | 2929799 |
| NG806_RS12925 | elongation factor Tu | 2956784 |
| CJF12_RS17015 | elongation factor Tu | 558152 |
| NBC122_RS07240 | elongation factor Tu | 2547600 |
| EG350_RS02620 | elongation factor Tu | 1493872 |
| MTP09_RS02885 | elongation factor Tu | 2929800 |
| E0E43_RS14135 | elongation factor Tu | 1141221 |
| CKV81_RS06075 | elongation factor Tu | 536441 |
| KKQ76_RS08270 | elongation factor Tu | 2004710 |
| EB819_RS01005 | elongation factor Tu | 237258 |
| ALE3EI_RS08610 | elongation factor Tu | 2562285 |
| HYG79_RS02400 | elongation factor Tu | 343403 |
| CA2559_RS00065 | elongation factor Tu | 216432 |
| H4K34_RS03035 | elongation factor Tu | 2761580 |
| IMZ16_RS04675 | elongation factor Tu | 1118202 |
| I597_RS13335 | elongation factor Tu | 1300343 |
| BAZ09_RS13325 | elongation factor Tu | 1246994 |
| AYC65_RS15445 | elongation factor Tu | 1756149 |
| DQN19_RS10510 | elongation factor Tu | 238 |
| BBD34_RS11195 | elongation factor Tu | 1756150 |
| FH779_RS01555 | elongation factor Tu | 343874 |
| HNV03_RS00080 | elongation factor Tu | 1628248 |
| EIB74_RS05755 | elongation factor Tu | 2487072 |
| D1013_RS06450 | elongation factor Tu | 1761453 |
| J9309_RS06770 | elongation factor Tu | 2898187 |
| VC82_RS11175 | elongation factor Tu | 516051 |
| HME9304_RS12150 | elongation factor Tu | 1383885 |

| | | |
|---|---|---|
| GVT53_RS01660 | elongation factor Tu | 2698672 |
| Q4Q34_RS12680 | elongation factor Tu | 3063533 |
| C1H87_RS05840 | elongation factor Tu | 1803846 |
| LNP19_RS07435 | elongation factor Tu | 2893883 |
| OZP08_RS07890 | elongation factor Tu | 3003257 |
| K5I29_RS08310 | elongation factor Tu | 2870839 |
| HYN59_RS08785 | elongation factor Tu | 2175091 |
| FUA48_RS00470 | elongation factor Tu | 2602769 |
| LPC20_RS00165 | elongation factor Tu | 1751056 |
| LPC21_RS00165 | elongation factor Tu | 1751095 |
| BB050_RS03240 | elongation factor Tu | 459526 |
| OZP12_RS18360 | elongation factor Tu | 3003261 |
| DVK85_RS00765 | elongation factor Tu | 1784713 |
| SBO79_RS00265 | elongation factor Tu | 2038737 |
| M0M57_RS02370 | elongation factor Tu | 1871076 |
| L0669_RS10200 | elongation factor Tu | 2704140 |
| FBFL15_RS03965 | elongation factor Tu | 1034807 |
| RN605_RS07025 | elongation factor Tu | 3075618 |
| LOS89_RS09910 | elongation factor Tu | 2897181 |
| OGZ34_RS09925 | elongation factor Tu | 1114861 |
| HUE53_RS02450 | elongation factor Tu | 996 |
| BIW12_RS10960 | elongation factor Tu | 1306519 |
| AWN65_RS09210 | elongation factor Tu | 2906076 |
| LB076_RS00170 | elongation factor Tu | 1763534 |
| HYN56_RS10470 | elongation factor Tu | 2183896 |
| LNP23_RS09875 | elongation factor Tu | 2893885 |
| LOS86_RS03580 | elongation factor Tu | 2895947 |
| NU10_RS12220 | elongation factor Tu | 1563479 |
| NAB20_RS00205 | elongation factor Tu | 2906077 |
| OZP15_RS13740 | elongation factor Tu | 3003263 |
| J0383_RS05860 | elongation factor Tu | 2816357 |

| | | |
|---|---|---|
| FFWV33_RS00190 | elongation factor Tu | 1355330 |
| OZP09_RS17270 | elongation factor Tu | 3003258 |
| LNP22_RS09580 | elongation factor Tu | 2893884 |
| HYN86_RS09270 | elongation factor Tu | 2249356 |
| LNP27_RS00630 | elongation factor Tu | 2893886 |
| OZP11_RS09730 | elongation factor Tu | 3003260 |
| EM308_RS08020 | elongation factor Tu | 1492737 |
| V6624_RS04835 | elongation factor Tu | 687844 |
| P5P89_RS13905 | elongation factor Tu | 1310418 |
| LJY17_RS08755 | elongation factor Tu | 1176532 |
| M0M44_RS10080 | elongation factor Tu | 2937442 |
| KQS_RS10670 | elongation factor Tu | 1094466 |
| HW120_RS00215 | elongation factor Tu | 2748320 |
| FJOH_RS10070 | elongation factor Tu | 376686 |
| L2Z92_RS11285 | elongation factor Tu | 998845 |
| MG292_RS00180 | elongation factor Tu | 1306518 |
| FK004_RS00145 | elongation factor Tu | 1678728 |
| O6P34_RS01150 | elongation factor Tu | 3016339 |
| OZP13_RS00190 | elongation factor Tu | 3003262 |
| K1I41_RS00750 | elongation factor Tu | 2856519 |
| OZP10_RS13790 | elongation factor Tu | 3003259 |
| HYN48_RS06385 | elongation factor Tu | 2162713 |
| OZP07_RS17175 | elongation factor Tu | 3003256 |
| E1750_RS14765 | elongation factor Tu | 2547394 |
| RF683_RS02470 | elongation factor Tu | 3073563 |
| P5P90_RS12945 | elongation factor Tu | 992289 |
| F0T65_RS02400 | elongation factor Tu | 1617283 |
| KK2020170_RS06695 | elongation factor Tu | 2830782 |
| ABZP37_RS00180 | elongation factor Tu | 1928857 |
| HYN49_RS06710 | elongation factor Tu | 2172098 |
| P2W65_RS10545 | elongation factor Tu | 3034146 |

| | | |
|---|---|---|
| NLJ00_RS10550 | elongation factor Tu | 2654843 |
| NLG42_RS10030 | elongation factor Tu | 1114867 |
| OYT91_RS09305 | elongation factor Tu | 3002900 |
| E5165_RS08200 | elongation factor Tu | 96345 |
| DYH63_RS13255 | elongation factor Tu | 2294119 |
| P7V56_RS13470 | elongation factor Tu | 2732161 |
| GS03_RS00390 | elongation factor Tu | 2518177 |
| LXD69_RS09250 | elongation factor Tu | 2024526 |
| DI487_RS10815 | elongation factor Tu | 2201181 |
| AABD74_RS09935 | elongation factor Tu | 2903098 |
| FLUTA_RS10790 | elongation factor Tu | 755732 |
| BN863_RS12460 | elongation factor Tu | 1347342 |
| FNB79_RS00485 | elongation factor Tu | 2594004 |
| RNZ46_RS11155 | elongation factor Tu | 3075823 |
| JBKA6_RS00600 | elongation factor Tu | 242600 |
| PT603_RS00780 | elongation factor Tu | 1165090 |
| EL218_RS08175 | elongation factor Tu | 266748 |
| EIB73_RS13065 | elongation factor Tu | 1241979 |
| LL667_RS08625 | elongation factor Tu | 2852098 |
| EL270_RS12440 | elongation factor Tu | 266749 |
| LC814_RS10620 | elongation factor Tu | 2878534 |
| IMCC3317_RS20195 | elongation factor Tu | 1218801 |
| N7U66_RS07165 | elongation factor Tu | 2980107 |
| BWR22_RS11450 | elongation factor Tu | 1486034 |
| LDL79_RS03185 | elongation factor Tu | 573501 |
| Lupro_RS04445 | elongation factor Tu | 1622118 |
| QZH61_RS11980 | elongation factor Tu | 3060215 |
| R3L16_RS12985 | elongation factor Tu | 3080055 |
| N8A89_RS02455 | elongation factor Tu | 1130723 |
| CJ263_RS14350 | elongation factor Tu | 1178778 |
| BLT83_RS02470 | elongation factor Tu | 320912 |

| | | |
|---|---|---|
| N8A86_RS02940 | elongation factor Tu | 2980554 |
| BTR34_RS03340 | elongation factor Tu | 1836467 |
| NYZ99_RS13605 | elongation factor Tu | 2976127 |
| QLS71_RS12670 | elongation factor Tu | 3045158 |
| IFB02_RS01370 | elongation factor Tu | 2708110 |
| EQY75_RS02705 | elongation factor Tu | 2507538 |
| NWE55_RS05245 | elongation factor Tu | 2562892 |
| FBR08_RS09065 | elongation factor Tu | 2572594 |
| E0Z07_RS02915 | elongation factor Tu | 76832 |
| I6I88_RS15015 | elongation factor Tu | 256 |
| MPR_RS15865 | elongation factor Tu | 480520 |
| DDD_RS09875 | elongation factor Tu | 592029 |
| NMS_RS07115 | elongation factor Tu | 1454201 |
| EJ995_RS05185 | elongation factor Tu | 2496866 |
| BST97_RS15005 | elongation factor Tu | 331648 |
| BST91_RS09400 | elongation factor Tu | 323273 |
| R1T42_RS14295 | elongation factor Tu | 906888 |
| ORNRH_RS07445 | elongation factor Tu | 867902 |
| OWEHO_RS07285 | elongation factor Tu | 926562 |
| NPX36_RS07235 | elongation factor Tu | 2968490 |
| K6119_RS14945 | elongation factor Tu | 2862509 |
| NYQ84_RS13275 | elongation factor Tu | 2829795 |
| WG945_RS05545 | elongation factor Tu | 1333662 |
| JL193_RS14150 | elongation factor Tu | 2808900 |
| WG951_RS06665 | elongation factor Tu | 218490 |
| J3359_RS16425 | elongation factor Tu | 2818493 |
| LPB302_RS05465 | elongation factor Tu | 326329 |
| WHA43_RS04755 | elongation factor Tu | 574710 |
| H9I45_RS14880 | elongation factor Tu | 1888915 |
| KCTC32516_RS08055 | elongation factor Tu | 1354726 |
| K8354_RS03115 | elongation factor Tu | 1908341 |

| | | |
|---|---|---|
| WG950_RS04600 | elongation factor Tu | 1642819 |
| WHC90_RS04685 | elongation factor Tu | 1775173 |
| H9W90_RS02185 | elongation factor Tu | 2738844 |
| BW723_RS10160 | elongation factor Tu | 996801 |
| NQP51_RS07090 | elongation factor Tu | 985043 |
| WHD07_RS05905 | elongation factor Tu | 2022337 |
| WHD54_RS02890 | elongation factor Tu | 1920034 |
| LPB03_RS05175 | elongation factor Tu | 1774273 |
| APS56_RS05570 | elongation factor Tu | 1736674 |
| C1A40_RS10680 | elongation factor Tu | 2069432 |
| P700755_RS11800 | elongation factor Tu | 313595 |
| MUN68_RS12050 | elongation factor Tu | 2932268 |
| C5O00_RS01445 | elongation factor Tu | 2094025 |
| G5B37_RS05580 | elongation factor Tu | 2744027 |
| RIEAN_RS07645 | elongation factor Tu | 693978 |
| RB2501_RS14500 | elongation factor Tu | 313596 |
| LDL76_RS01540 | elongation factor Tu | 270918 |
| B5488_RS15395 | elongation factor Tu | 143223 |
| JRG66_RS03825 | elongation factor Tu | 2809707 |
| RA156_RS02445 | elongation factor Tu | 3069118 |
| AW14_RS00605 | elongation factor Tu | 1454006 |
| LS482_RS04170 | elongation factor Tu | 2900738 |
| ACN5J4_RS04220 | elongation factor Tu | 2006131 |
| ABNT14_RS09055 | elongation factor Tu | 669041 |
| ACN5J5_RS02365 | elongation factor Tu | 2029677 |
| AQ1685_RS06170 | elongation factor Tu | 584609 |
| MARIT_RS04720 | elongation factor Tu | 1349785 |
| F9Y86_RS03930 | elongation factor Tu | 104268 |
| PG913_RS03055 | elongation factor Tu | 3018314 |
| ABNT61_RS06435 | elongation factor Tu | 3137852 |
| ABNT22_RS14410 | elongation factor Tu | 3137857 |

| | | |
|---|---|---|
| PG912_RS10330 | elongation factor Tu | 3018315 |
| D6T69_RS06390 | elongation factor Tu | 2358479 |
| P8625_RS06725 | elongation factor Tu | 3038772 |
| LPB136_RS06195 | elongation factor Tu | 1850252 |
| ABNT65_RS10730 | elongation factor Tu | 3137860 |
| RHP49_RS11675 | elongation factor Tu | 3074124 |
| LPB138_RS05660 | elongation factor Tu | 1850246 |
| EL204_RS00390 | elongation factor Tu | 1014 |
| AXE80_RS02260 | elongation factor Tu | 1790137 |
| HM987_RS13245 | elongation factor Tu | 2686077 |
| HM990_RS12900 | elongation factor Tu | 2686078 |
| BLT74_RS11935 | elongation factor Tu | 1382466 |
| MQE36_RS00985 | elongation factor Tu | 2202721 |
| P0077_RS13805 | elongation factor Tu | 3032586 |
| ZOBGAL_RS03030 | elongation factor Tu | 63186 |
| SJY45_RS08300 | elongation factor Tu | 248907 |
| ZPR_RS05565 | elongation factor Tu | 655815 |
| MKD34_RS07750 | elongation factor Tu | 188913 |
| MKD34_RS08380 | elongation factor Tu | 188913 |
| RO08_RS00715 | elongation factor Tu | 76859 |
| I6I83_RS10745 | elongation factor Tu | 285729 |
| C4N16_RS00270 | elongation factor Tu | 469615 |
| C4N16_RS01020 | elongation factor Tu | 469615 |
| H9Q81_RS08375 | elongation factor Tu | 2764326 |
| H9Q81_RS09070 | elongation factor Tu | 2764326 |
| RN92_RS04190 | elongation factor Tu | 1307443 |
| C4N19_RS02125 | elongation factor Tu | 469616 |
| C4N19_RS03740 | elongation factor Tu | 469616 |
| BWX37_RS00515 | elongation factor Tu | 143387 |
| BWX37_RS01350 | elongation factor Tu | 143387 |
| C7Y58_RS00300 | elongation factor Tu | 190304 |

| | | |
|---|---|---|
| AT688_RS10600 | elongation factor Tu | 76857 |
| C4N20_RS05060 | elongation factor Tu | 861 |
| C4N20_RS06635 | elongation factor Tu | 861 |
| C4N18_RS03830 | elongation factor Tu | 469618 |
| C4N18_RS05250 | elongation factor Tu | 469618 |
| JY399_RS02965 | elongation factor Tu | 155615 |
| ACM8CJ_RS03465 | elongation factor Tu | 2686067 |
| RDY08_RS00430 | elongation factor Tu | 2996794 |
| RDY08_RS03170 | elongation factor Tu | 2996794 |
| ILYOP_RS07475 | elongation factor Tu | 572544 |
| ILYOP_RS09130 | elongation factor Tu | 572544 |
| LEBU_RS01200 | elongation factor Tu | 523794 |
| LEBU_RS09505 | elongation factor Tu | 523794 |
| FVE77_RS01510 | elongation factor Tu | 157688 |
| FVE77_RS11360 | elongation factor Tu | 157688 |
| FVE75_RS00575 | elongation factor Tu | 554406 |
| FVE75_RS08835 | elongation factor Tu | 554406 |
| AB8B23_RS00610 | elongation factor Tu | 3239303 |
| AB8B23_RS01450 | elongation factor Tu | 3239303 |
| JMUB4039_RS11220 | elongation factor Tu | 109328 |
| JMUB4039_RS12140 | elongation factor Tu | 109328 |
| FVE73_RS08965 | elongation factor Tu | 157687 |
| FVE73_RS09830 | elongation factor Tu | 157687 |
| VC03_RS00510 | elongation factor Tu | 187101 |
| VC03_RS04190 | elongation factor Tu | 187101 |
| SMON_RS00940 | elongation factor Tu | 519441 |
| SMON_RS05345 | elongation factor Tu | 519441 |
| ENTCL_RS01945 | elongation factor Tu | 701347 |
| ENTCL_RS20920 | elongation factor Tu | 701347 |
| A6037_RS02710 | elongation factor Tu | 730 |
| A6037_RS03590 | elongation factor Tu | 730 |

| | | |
|---|---|---|
| MS_RS00775 | elongation factor Tu | 221988 |
| MS_RS09935 | elongation factor Tu | 221988 |
| J1C60_RS01755 | elongation factor Tu | 1324864 |
| J1C60_RS17235 | elongation factor Tu | 1324864 |
| CVPH_RS06310 | elongation factor Tu | 1235283 |
| CVPH_RS07820 | elongation factor Tu | 1235283 |
| A9R16_RS02640 | elongation factor Tu | 163359 |
| A9R16_RS02705 | elongation factor Tu | 163359 |
| BJI67_RS11475 | elongation factor Tu | 2792603 |
| BJI67_RS11540 | elongation factor Tu | 2792603 |
| BW247_RS14095 | elongation factor Tu | 1765967 |
| BW247_RS14160 | elongation factor Tu | 1765967 |
| BI364_RS12680 | elongation factor Tu | 2819280 |
| BI364_RS12745 | elongation factor Tu | 2819280 |
| FQU82_RS01885 | elongation factor Tu | 470 |
| FQU82_RS04500 | elongation factor Tu | 470 |
| ACIAD_RS01420 | elongation factor Tu | 62977 |
| ACIAD_RS04085 | elongation factor Tu | 62977 |
| I9192_RS17760 | elongation factor Tu | 106648 |
| I9192_RS20595 | elongation factor Tu | 106648 |
| KI216_RS03765 | elongation factor Tu | 202951 |
| KI216_RS14775 | elongation factor Tu | 202951 |
| CDG60_RS02220 | elongation factor Tu | 2004650 |
| CDG60_RS04550 | elongation factor Tu | 2004650 |
| QLH32_RS09440 | elongation factor Tu | 3045147 |
| QLH32_RS15410 | elongation factor Tu | 3045147 |
| C9E88_RS12835 | elongation factor Tu | 2136182 |
| C9E88_RS14900 | elongation factor Tu | 2136182 |
| DJ533_RS04375 | elongation factor Tu | 1871111 |
| DJ533_RS15285 | elongation factor Tu | 1871111 |
| FPL17_RS09780 | elongation factor Tu | 70348 |

| | | |
|---|---|---|
| FPL17_RS12680 | elongation factor Tu | 70348 |
| AOY20_RS02800 | elongation factor Tu | 1324350 |
| AOY20_RS06940 | elongation factor Tu | 1324350 |
| AS4_RS01240 | elongation factor Tu | 106649 |
| AS4_RS17375 | elongation factor Tu | 106649 |
| DS737_RS11795 | elongation factor Tu | 29430 |
| DS737_RS14620 | elongation factor Tu | 29430 |
| GA693_RS08945 | elongation factor Tu | 756892 |
| GA693_RS13660 | elongation factor Tu | 756892 |
| H2677_RS04100 | elongation factor Tu | 40215 |
| H2677_RS14245 | elongation factor Tu | 40215 |
| HLV59_RS14350 | elongation factor Tu | 1785128 |
| HLV59_RS17095 | elongation factor Tu | 1785128 |
| G8D99_RS11925 | elongation factor Tu | 2715163 |
| G8D99_RS14200 | elongation factor Tu | 2715163 |
| BFG52_RS02675 | elongation factor Tu | 1789224 |
| BFG52_RS15065 | elongation factor Tu | 1789224 |
| J7649_RS07220 | elongation factor Tu | 28090 |
| J7649_RS11610 | elongation factor Tu | 28090 |
| ABLC00_RS01580 | elongation factor Tu | 106654 |
| ABLC00_RS04390 | elongation factor Tu | 106654 |
| AOLE_RS15340 | elongation factor Tu | 436717 |
| AOLE_RS18070 | elongation factor Tu | 436717 |
| BDGL_000132 | elongation factor Tu | 871585 |
| BDGL_003185 | -- | 871585 |
| DOM24_RS00985 | elongation factor Tu | 40216 |
| DOM24_RS11225 | elongation factor Tu | 40216 |
| FSC11_RS11410 | elongation factor Tu | 108981 |
| FSC11_RS13525 | elongation factor Tu | 108981 |
| I8T81_RS14690 | elongation factor Tu | 1530123 |
| I8T81_RS17345 | elongation factor Tu | 1530123 |

| | | |
|---|---|---|
| I6L31_RS08790 | elongation factor Tu | 465797 |
| I6L31_RS13750 | elongation factor Tu | 465797 |
| KCV09_RS00515 | elongation factor Tu | 487316 |
| KCV09_RS10415 | elongation factor Tu | 487316 |
| F2A31_RS11430 | elongation factor Tu | 2609668 |
| F2A31_RS14260 | elongation factor Tu | 2609668 |
| ABEF82_RS03665 | elongation factor Tu | 3151487 |
| ABEF82_RS13555 | elongation factor Tu | 3151487 |
| M5E07_RS03310 | elongation factor Tu | 2943497 |
| M5E07_RS14880 | elongation factor Tu | 2943497 |
| J4G44_RS03745 | elongation factor Tu | 202956 |
| J4G44_RS12085 | elongation factor Tu | 202956 |
| Q3F87_RS12615 | elongation factor Tu | 70346 |
| Q3F87_RS14845 | elongation factor Tu | 70346 |
| BEN71_RS01750 | elongation factor Tu | 1879050 |
| BEN71_RS04920 | elongation factor Tu | 1879050 |
| EL259_RS03945 | elongation factor Tu | 51161 |
| EL259_RS06640 | elongation factor Tu | 51161 |
| EL121_RS06390 | elongation factor Tu | 718 |
| EL121_RS06810 | elongation factor Tu | 718 |
| EXH44_RS06255 | elongation factor Tu | 51049 |
| EXH44_RS06475 | elongation factor Tu | 51049 |
| DRF63_RS08100 | elongation factor Tu | 715 |
| DRF63_RS08250 | elongation factor Tu | 715 |
| DDU33_RS01860 | elongation factor Tu | 189834 |
| DDU33_RS03850 | elongation factor Tu | 189834 |
| ASUC_RS00505 | elongation factor Tu | 339671 |
| ASUC_RS01450 | elongation factor Tu | 339671 |
| ASU1_RS08915 | elongation factor Tu | 743972 |
| ASU1_RS10685 | elongation factor Tu | 743972 |
| EYZ66_RS01805 | elongation factor Tu | 2518989 |

| | | |
|---|---|---|
| EYZ66_RS01870 | elongation factor Tu | 2518989 |
| I6G90_RS17375 | elongation factor Tu | 656 |
| I6G90_RS17465 | elongation factor Tu | 656 |
| IU367_RS20425 | elongation factor Tu | 105751 |
| IU367_RS20515 | elongation factor Tu | 105751 |
| DQN91_RS01325 | elongation factor Tu | 648 |
| DQN91_RS01415 | elongation factor Tu | 648 |
| CK627_RS02335 | elongation factor Tu | 196024 |
| CK627_RS02425 | elongation factor Tu | 196024 |
| EL255_RS01170 | elongation factor Tu | 73010 |
| EL255_RS01260 | elongation factor Tu | 73010 |
| LA321_RS07835 | elongation factor Tu | 29489 |
| LA321_RS07925 | elongation factor Tu | 29489 |
| AHA_RS20300 | elongation factor Tu | 380703 |
| AHA_RS20390 | elongation factor Tu | 380703 |
| I6H43_RS09235 | elongation factor Tu | 650 |
| I6H43_RS09325 | elongation factor Tu | 650 |
| KI797_RS19335 | elongation factor Tu | 651 |
| KI797_RS19425 | elongation factor Tu | 651 |
| WJM78_RS02200 | elongation factor Tu | 70856 |
| WJM78_RS02290 | elongation factor Tu | 70856 |
| KYK33_RS01035 | elongation factor Tu | 648794 |
| KYK33_RS01125 | elongation factor Tu | 648794 |
| ONZ54_RS20865 | elongation factor Tu | 645 |
| ONZ54_RS20955 | elongation factor Tu | 645 |
| FE240_RS18555 | elongation factor Tu | 218936 |
| FE240_RS18645 | elongation factor Tu | 218936 |
| FOB40_RS16300 | elongation factor Tu | 654 |
| FOB40_RS16390 | elongation factor Tu | 654 |
| ALSL_RS10140 | elongation factor Tu | 2010829 |
| ALSL_RS10205 | elongation factor Tu | 2010829 |

| | | |
|---|---|---|
| BVC89_RS21660 | elongation factor Tu | 1737490 |
| BVC89_RS21725 | elongation factor Tu | 1737490 |
| BVC89_RS21790 | elongation factor Tu | 1737490 |
| K5609_RS18690 | elongation factor Tu | 1563703 |
| K5609_RS18775 | elongation factor Tu | 1563703 |
| K5620_RS18710 | elongation factor Tu | 182262 |
| K5620_RS18795 | elongation factor Tu | 182262 |
| K5620_RS18880 | elongation factor Tu | 182262 |
| LIN98_RS01875 | elongation factor Tu | 714 |
| LIN98_RS04240 | elongation factor Tu | 714 |
| EL144_RS00545 | elongation factor Tu | 985008 |
| EL144_RS06460 | elongation factor Tu | 985008 |
| DQN58_RS00690 | elongation factor Tu | 888057 |
| DQN58_RS01955 | elongation factor Tu | 888057 |
| H8B22_RS10350 | elongation factor Tu | 2865112 |
| H8B22_RS10415 | elongation factor Tu | 2865112 |
| C7S18_RS16160 | elongation factor Tu | 2021234 |
| C7S18_RS16225 | elongation factor Tu | 2021234 |
| ABO_RS01935 | elongation factor Tu | 393595 |
| ABO_RS02000 | elongation factor Tu | 393595 |
| OCU49_RS21675 | elongation factor Tu | 1524258 |
| OCU49_RS21740 | elongation factor Tu | 1524258 |
| PCI15_RS21610 | elongation factor Tu | 2895352 |
| PCI15_RS21675 | elongation factor Tu | 2895352 |
| VF_RS01140 | elongation factor Tu | 312309 |
| VF_RS12545 | elongation factor Tu | 312309 |
| VSAL_RS01835 | elongation factor Tu | 316275 |
| VSAL_RS15080 | elongation factor Tu | 316275 |
| MLG_RS02275 | elongation factor Tu | 187272 |
| MLG_RS02340 | elongation factor Tu | 187272 |
| NKI27_RS17550 | elongation factor Tu | 2961619 |

| | | |
|---|---|---|
| MY523_RS20530 | elongation factor Tu | 2935863 |
| NNL22_RS02015 | elongation factor Tu | 1632866 |
| B5T_RS18780 | elongation factor Tu | 930169 |
| B5T_RS18845 | elongation factor Tu | 930169 |
| ACMG4O_RS16060 | elongation factor Tu | 172371 |
| ACMG4O_RS16125 | elongation factor Tu | 172371 |
| RYH70_RS17560 | elongation factor Tu | 1094342 |
| RYH70_RS17625 | elongation factor Tu | 1094342 |
| Atep_RS06635 | elongation factor Tu | 553982 |
| Atep_RS06700 | elongation factor Tu | 553982 |
| ALVIN_RS11785 | elongation factor Tu | 572477 |
| ALVIN_RS11850 | elongation factor Tu | 572477 |
| E3E15_RS06525 | elongation factor Tu | 1085644 |
| SD28_RS02565 | elongation factor Tu | 594679 |
| E4K63_RS01265 | elongation factor Tu | 1085647 |
| TE101_RS17125 | elongation factor Tu | 28108 |
| TE101_RS18020 | elongation factor Tu | 28108 |
| AMBT_RS00255 | elongation factor Tu | 715451 |
| AMBT_RS19235 | elongation factor Tu | 715451 |
| CA267_RS11845 | elongation factor Tu | 1858656 |
| CA267_RS12545 | elongation factor Tu | 1858656 |
| AVL56_RS17950 | elongation factor Tu | 233316 |
| OCU43_RS20825 | elongation factor Tu | 355243 |
| OCU43_RS20890 | elongation factor Tu | 355243 |
| PT3_RS16450 | elongation factor Tu | 452627 |
| PT3_RS16515 | elongation factor Tu | 452627 |
| AQUSIP_RS07980 | elongation factor Tu | 254246 |
| AQUSIP_RS08045 | elongation factor Tu | 254246 |
| AQULUS_RS02165 | elongation factor Tu | 254247 |
| AQULUS_RS02230 | elongation factor Tu | 254247 |
| A0T30_RS19795 | elongation factor Tu | 43263 |

| | | |
|---|---|---|
| A0T30_RS19860 | elongation factor Tu | 43263 |
| HNE05_RS02445 | elongation factor Tu | 2731681 |
| HNE05_RS02510 | elongation factor Tu | 2731681 |
| HYN46_RS08225 | elongation factor Tu | 2283318 |
| HYN46_RS11275 | elongation factor Tu | 2283318 |
| LDL57_RS14585 | elongation factor Tu | 2879119 |
| LDL57_RS14765 | elongation factor Tu | 2879119 |
| WBAF_RS02160 | elongation factor Tu | 235559 |
| E3U36_RS02045 | elongation factor Tu | 1231049 |
| E3U36_RS02560 | elongation factor Tu | 1231049 |
| AACL31_RS05465 | elongation factor Tu | 3066235 |
| AACL31_RS20385 | elongation factor Tu | 3066235 |
| ACICP2_RS03075 | elongation factor Tu | 363258 |
| ArsFIN_RS19160 | elongation factor Tu | 638 |
| ArsFIN_RS19355 | elongation factor Tu | 638 |
| ABNS14_RS00805 | elongation factor Tu | 634121 |
| I6G45_RS00605 | elongation factor Tu | 565 |
| I6G45_RS18540 | elongation factor Tu | 565 |
| NLZ15_RS01820 | elongation factor Tu | 255519 |
| NLZ15_RS20965 | elongation factor Tu | 255519 |
| HBL79_RS09850 | elongation factor Tu | 728 |
| HBL79_RS11555 | elongation factor Tu | 728 |
| ELZ61_RS01200 | elongation factor Tu | 762 |
| ELZ61_RS02190 | elongation factor Tu | 762 |
| ACGI6K_RS18740 | elongation factor Tu | 69963 |
| ACGI6K_RS18805 | elongation factor Tu | 69963 |
| ACG10_RS18605 | elongation factor Tu | 353 |
| ACG10_RS18670 | elongation factor Tu | 353 |
| GCU53_RS15790 | elongation factor Tu | 69964 |
| GCU53_RS15855 | elongation factor Tu | 69964 |
| AVIN_RS02850 | elongation factor Tu | 322710 |

| | | |
|---|---|---|
| AVIN_RS02915 | elongation factor Tu | 322710 |
| A4G13_RS07955 | elongation factor Tu | 653940 |
| A4G13_RS09415 | elongation factor Tu | 653940 |
| MS2017_RS02265 | elongation factor Tu | 2360 |
| MS2017_RS09215 | elongation factor Tu | 2360 |
| AL038_RS09695 | elongation factor Tu | 288004 |
| AL038_RS13990 | elongation factor Tu | 288004 |
| HF888_RS14495 | elongation factor Tu | 207949 |
| HF888_RS14565 | elongation factor Tu | 207949 |
| F542_RS09070 | elongation factor Tu | 1263829 |
| F542_RS09575 | elongation factor Tu | 1263829 |
| HNO51_RS18355 | elongation factor Tu | 2733484 |
| A6A11_RS01140 | elongation factor Tu | 109472 |
| A6A11_RS08865 | elongation factor Tu | 109472 |
| BOBLI757_RS02845 | elongation factor Tu | 1505597 |
| M9396_RS02850 | elongation factor Tu | 2945587 |
| GN161_RS02100 | elongation factor Tu | 2681986 |
| M9401_RS01145 | elongation factor Tu | 700220 |
| M9409_RS03170 | elongation factor Tu | 2945589 |
| GN160_RS02675 | elongation factor Tu | 2681987 |
| BTURN675_RS02795 | elongation factor Tu | 1505596 |
| AWC36_RS17590 | elongation factor Tu | 1109412 |
| AWC36_RS21395 | elongation factor Tu | 1109412 |
| HC231_RS02195 | elongation factor Tu | 2722756 |
| HC231_RS22400 | elongation factor Tu | 2722756 |
| EH206_RS02200 | elongation factor Tu | 1121120 |
| EH206_RS21210 | elongation factor Tu | 1121120 |
| RS889_RS01940 | elongation factor Tu | 3057057 |
| RS889_RS21945 | elongation factor Tu | 3057057 |
| GNIT_RS03240 | elongation factor Tu | 1085623 |
| GNIT_RS14420 | elongation factor Tu | 1085623 |

| | | |
|---|---|---|
| BUSG_RS02665 | elongation factor Tu | 198804 |
| MNO13_RS01805 | elongation factor Tu | 82989 |
| MNO13_RS23175 | elongation factor Tu | 82989 |
| RHD99_RS01805 | elongation factor Tu | 3073902 |
| RHD99_RS22595 | elongation factor Tu | 3073902 |
| GWK36_RS03425 | elongation factor Tu | 2699430 |
| GWK36_RS03490 | elongation factor Tu | 2699430 |
| C3B56_RS00920 | elongation factor Tu | 1302411 |
| GFK87_RS00150 | elongation factor Tu | 1345117 |
| AUT07_RS01435 | elongation factor Tu | 634113 |
| WH390_RS12810 | elongation factor Tu | 1247023 |
| WH390_RS13080 | elongation factor Tu | 1247023 |
| ESZ_RS01610 | elongation factor Tu | 2652803 |
| M9404_RS02565 | elongation factor Tu | 108080 |
| BCHRO640_RS02895 | elongation factor Tu | 1240471 |
| BPEN_RS02880 | elongation factor Tu | 291272 |
| BVAF_RS02805 | elongation factor Tu | 859654 |
| A353_RS00915 | elongation factor Tu | 1202538 |
| MMH89_RS03580 | elongation factor Tu | 2919598 |
| MMH89_RS03665 | elongation factor Tu | 2919598 |
| CleRt_RS06915 | elongation factor Tu | 2054173 |
| CleRt_RS06980 | elongation factor Tu | 2054173 |
| ACICP0_RS02035 | elongation factor Tu | 1191162 |
| A4A68_RS01635 | elongation factor Tu | 1778262 |
| L0Y14_RS01695 | elongation factor Tu | 393765 |
| L0Y14_RS01760 | elongation factor Tu | 393765 |
| BTN50_RS05765 | elongation factor Tu | 1927128 |
| HACICURV3402_RS01245 | elongation factor Tu | 1922217 |
| QI37_RS05170 | elongation factor Tu | 653937 |
| AAHH42_RS01710 | elongation factor Tu | 3077957 |
| FVIR_RS00180 | elongation factor Tu | 1070130 |

| | | |
|---|---|---|
| FVIR_RS01625 | elongation factor Tu | 1070130 |
| HDEF_RS03430 | elongation factor Tu | 572265 |
| A4A70_RS01595 | elongation factor Tu | 1778263 |
| ICMP_RS01245 | elongation factor Tu | 476281 |
| ICMP_RS02620 | elongation factor Tu | 476281 |
| CCU22_RS00815 | elongation factor Tu | 2005262 |
| ABNT83_RS10575 | elongation factor Tu | 3108543 |
| ABNT83_RS11635 | elongation factor Tu | 3108543 |
| F6R98_RS14430 | elongation factor Tu | 1808979 |
| F6R98_RS14945 | elongation factor Tu | 1808979 |
| A4A67_RS00515 | elongation factor Tu | 1778264 |
| MPC_RS01370 | elongation factor Tu | 1234603 |
| NSCAC_RS01675 | elongation factor Tu | 553981 |
| NSCAC_RS01740 | elongation factor Tu | 553981 |
| TAO_RS03295 | elongation factor Tu | 1630141 |
| TAO_RS03360 | elongation factor Tu | 1630141 |
| IM45_RS02735 | elongation factor Tu | 186490 |
| BMSBPS_RS03475 | elongation factor Tu | 1235990 |
| D8B20_RS00935 | elongation factor Tu | 3098669 |
| D8B20_RS15800 | elongation factor Tu | 3098669 |
| E0F26_RS04675 | elongation factor Tu | 2518994 |
| B186_RS00380 | elongation factor Tu | 1206109 |
| GII40_RS01935 | elongation factor Tu | 1987921 |
| KTT71_RS02055 | elongation factor Tu | 1177216 |
| AAGD61_RS01075 | elongation factor Tu | 1715285 |
| C3B55_RS01225 | elongation factor Tu | 1302376 |
| W908_RS01355 | elongation factor Tu | 1125411 |
| W908_RS02220 | elongation factor Tu | 1125411 |
| C9I82_RS01095 | elongation factor Tu | 472834 |
| AACL30_RS08950 | elongation factor Tu | 3077956 |
| HUE57_RS07065 | elongation factor Tu | 2738883 |

| | | |
|---|---|---|
| HUE57_RS07130 | elongation factor Tu | 2738883 |
| DMP02_RS05340 | elongation factor Tu | 676208 |
| RIEPE_RS01765 | elongation factor Tu | 515618 |
| AOQ87_RS00845 | elongation factor Tu | 428411 |
| HUE58_RS00010 | elongation factor Tu | 2738852 |
| HUE58_RS03990 | elongation factor Tu | 2738852 |
| RMAG_RS00870 | elongation factor Tu | 413404 |
| RMAG_RS04035 | elongation factor Tu | 413404 |
| KEC37_RS00075 | elongation factor Tu | 1081631 |
| SOPEG_RS01220 | elongation factor Tu | 2342 |
| SOPEG_RS19345 | elongation factor Tu | 2342 |
| GFK82_RS01840 | elongation factor Tu | 1076626 |
| TGUWTKB_RS01695 | elongation factor Tu | 1410383 |
| TGUWTKB_RS01895 | elongation factor Tu | 1410383 |
| SOJ49_RS16605 | elongation factor Tu | 3100113 |
| SOJ49_RS16725 | elongation factor Tu | 3100113 |
| THSYN_RS05680 | elongation factor Tu | 1166950 |
| THSYN_RS23925 | elongation factor Tu | 1166950 |
| AB8525_RS07655 | elongation factor Tu | 2738886 |
| AB8525_RS08250 | elongation factor Tu | 2738886 |
| SP60_RS05245 | elongation factor Tu | 1705394 |
| SP60_RS06005 | elongation factor Tu | 1705394 |
| J8380_RS07380 | elongation factor Tu | 2823902 |
| J8380_RS07445 | elongation factor Tu | 2823902 |
| COSY_RS00820 | elongation factor Tu | 412965 |
| COSY_RS03735 | elongation factor Tu | 412965 |
| WEOB_RS00565 | elongation factor Tu | 1594731 |
| AACL23_RS08420 | elongation factor Tu | 3077954 |
| EL199_RS03830 | elongation factor Tu | 2718 |
| EL199_RS03895 | elongation factor Tu | 2718 |
| OLW01_RS02190 | elongation factor Tu | 2984846 |

| | | |
|---|---|---|
| OLW01_RS02255 | elongation factor Tu | 2984846 |
| FNT16_RS08790 | elongation factor Tu | 1740262 |
| FNT16_RS08855 | elongation factor Tu | 1740262 |
| ACG1VR_RS01535 | elongation factor Tu | 158484 |
| ACG1VR_RS21810 | elongation factor Tu | 158484 |
| EL098_RS01105 | elongation factor Tu | 158823 |
| EL098_RS21250 | elongation factor Tu | 158823 |
| CO704_RS01070 | elongation factor Tu | 158822 |
| CO704_RS22975 | elongation factor Tu | 158822 |
| LDO33_RS17205 | elongation factor Tu | 412034 |
| LDO33_RS17295 | elongation factor Tu | 412034 |
| CJA_RS03340 | elongation factor Tu | 498211 |
| CJA_RS03405 | elongation factor Tu | 498211 |
| Z042_RS02550 | elongation factor Tu | 1441930 |
| Z042_RS05975 | elongation factor Tu | 1441930 |
| SR908_RS09310 | elongation factor Tu | 141389 |
| CSAL_RS02145 | elongation factor Tu | 290398 |
| LA348_RS17115 | elongation factor Tu | 35703 |
| LA348_RS20755 | elongation factor Tu | 35703 |
| E1B03_RS02320 | elongation factor Tu | 2546350 |
| E1B03_RS24170 | elongation factor Tu | 2546350 |
| GFC06_RS03210 | elongation factor Tu | 57706 |
| GFC06_RS06365 | elongation factor Tu | 57706 |
| JSN25_RS03920 | elongation factor Tu | 1748967 |
| JSN25_RS06595 | elongation factor Tu | 1748967 |
| WHR59_RS01915 | elongation factor Tu | 1914243 |
| WHR59_RS22635 | elongation factor Tu | 1914243 |
| I6L53_RS00400 | elongation factor Tu | 67824 |
| I6L53_RS20760 | elongation factor Tu | 67824 |
| JMV71_RS01875 | elongation factor Tu | 546 |
| JMV71_RS22315 | elongation factor Tu | 546 |

| | | |
|---|---|---|
| CKO_RS12850 | elongation factor Tu | 290338 |
| CKO_RS20240 | elongation factor Tu | 290338 |
| NMD08_RS02085 | elongation factor Tu | 2894201 |
| NMD08_RS24315 | elongation factor Tu | 2894201 |
| I6L54_RS00180 | elongation factor Tu | 1563222 |
| I6L54_RS03255 | elongation factor Tu | 1563222 |
| LY264_RS01970 | elongation factor Tu | 1639133 |
| LY264_RS23285 | elongation factor Tu | 1639133 |
| K7R23_RS04105 | elongation factor Tu | 1218085 |
| K7R23_RS07680 | elongation factor Tu | 1218085 |
| JY391_RS02180 | elongation factor Tu | 67826 |
| JY391_RS21175 | elongation factor Tu | 67826 |
| E4Z61_RS15790 | elongation factor Tu | 2562449 |
| E4Z61_RS18915 | elongation factor Tu | 2562449 |
| FOB24_RS02855 | elongation factor Tu | 67827 |
| FOB24_RS23120 | elongation factor Tu | 67827 |
| R1T44_RS00760 | elongation factor Tu | 1055104 |
| BFX80_RS00590 | elongation factor Tu | 28258 |
| B5D82_RS09195 | elongation factor Tu | 1967665 |
| B5D82_RS09265 | elongation factor Tu | 1967665 |
| CPS_RS21430 | elongation factor Tu | 167879 |
| CPS_RS21510 | elongation factor Tu | 167879 |
| R0137_RS16030 | elongation factor Tu | 3081201 |
| R0137_RS16100 | elongation factor Tu | 3081201 |
| KT71_RS04530 | elongation factor Tu | 314285 |
| KT71_RS04600 | elongation factor Tu | 314285 |
| R0135_RS15705 | elongation factor Tu | 3081200 |
| R0135_RS15775 | elongation factor Tu | 3081200 |
| CBU_0236 | protein translation elong | 227377 |
| Z664_RS01505 | elongation factor Tu | 325775 |
| Z664_RS01575 | elongation factor Tu | 325775 |

| | | |
|---|---|---|
| CEAn_RS01145 | elongation factor Tu | 2749996 |
| CEAn_RS01210 | elongation factor Tu | 2749996 |
| EGQ50_RS01490 | elongation factor Tu | 2487929 |
| B1F76_RS01530 | elongation factor Tu | 1987500 |
| B1F76_RS01600 | elongation factor Tu | 1987500 |
| AFK62_RS01650 | elongation factor Tu | 1073999 |
| AFK62_RS18270 | elongation factor Tu | 1073999 |
| AFK67_RS01620 | elongation factor Tu | 1159554 |
| AFK67_RS18855 | elongation factor Tu | 1159554 |
| AFK66_RS00975 | elongation factor Tu | 1159491 |
| AFK66_RS17885 | elongation factor Tu | 1159491 |
| AFK63_RS00370 | elongation factor Tu | 1159613 |
| AFK63_RS16970 | elongation factor Tu | 1159613 |
| CsakCS931_RS14650 | elongation factor Tu | 28141 |
| CsakCS931_RS17880 | elongation factor Tu | 28141 |
| AB8W27_RS01715 | elongation factor Tu | 413502 |
| AB8W27_RS19305 | elongation factor Tu | 413502 |
| AFK65_RS01560 | elongation factor Tu | 1074000 |
| AFK65_RS17995 | elongation factor Tu | 1074000 |
| DAQ1742_RS01325 | elongation factor Tu | 1401087 |
| DAQ1742_RS18385 | elongation factor Tu | 1401087 |
| DD1591_RS01615 | elongation factor Tu | 561229 |
| DD1591_RS19315 | elongation factor Tu | 561229 |
| CO076_RS01150 | elongation factor Tu | 204038 |
| CO076_RS20095 | elongation factor Tu | 204038 |
| DZA65_RS01120 | elongation factor Tu | 204039 |
| DZA65_RS20525 | elongation factor Tu | 204039 |
| CVE23_RS01145 | elongation factor Tu | 1778540 |
| CVE23_RS20280 | elongation factor Tu | 1778540 |
| LF929_RS01915 | elongation factor Tu | 1240404 |
| LF929_RS19645 | elongation factor Tu | 1240404 |

| | | |
|---|---|---|
| DD586_RS01080 | elongation factor Tu | 590409 |
| DD586_RS18615 | elongation factor Tu | 590409 |
| Dpoa569_RS01100 | elongation factor Tu | 568768 |
| Dpoa569_RS02065 | elongation factor Tu | 568768 |
| A4U42_RS06855 | elongation factor Tu | 1225786 |
| A4U42_RS10230 | elongation factor Tu | 1225786 |
| C1O30_RS01135 | elongation factor Tu | 204042 |
| C1O30_RS18905 | elongation factor Tu | 204042 |
| I596_RS12320 | elongation factor Tu | 1300342 |
| I596_RS12385 | elongation factor Tu | 1300342 |
| DB847_RS20990 | elongation factor Tu | 2047966 |
| J9874_RS00960 | elongation factor Tu | 1619313 |
| J9874_RS15335 | elongation factor Tu | 1619313 |
| RA164_RS04300 | elongation factor Tu | 3069105 |
| RA164_RS04365 | elongation factor Tu | 3069105 |
| ISN74_RS14455 | elongation factor Tu | 1849581 |
| ISN74_RS14520 | elongation factor Tu | 1849581 |
| H8F01_RS04885 | elongation factor Tu | 2763498 |
| H8F01_RS04950 | elongation factor Tu | 2763498 |
| ATSB10_RS09390 | elongation factor Tu | 445710 |
| ATSB10_RS09455 | elongation factor Tu | 445710 |
| L1F06_RS21545 | elongation factor Tu | 2493633 |
| L1F06_RS21610 | elongation factor Tu | 2493633 |
| JLK41_RS21645 | elongation factor Tu | 2502979 |
| JLK41_RS21710 | elongation factor Tu | 2502979 |
| DW68_RS03315 | elongation factor Tu | 1225174 |
| DW68_RS03380 | elongation factor Tu | 1225174 |
| JWV26_RS06835 | elongation factor Tu | 554344 |
| JWV26_RS06900 | elongation factor Tu | 554344 |
| ETEE_RS06420 | elongation factor Tu | 667120 |
| ETEE_RS08710 | elongation factor Tu | 667120 |

| | | |
|---|---|---|
| I6G97_RS02040 | elongation factor Tu | 93378 |
| I6G97_RS04385 | elongation factor Tu | 93378 |
| DB741_RS00755 | elongation factor Tu | 67780 |
| DB741_RS16580 | elongation factor Tu | 67780 |
| L1P05_RS00875 | elongation factor Tu | 1263550 |
| L1P05_RS15535 | elongation factor Tu | 1263550 |
| K8O98_RS00265 | elongation factor Tu | 636 |
| K8O98_RS02860 | elongation factor Tu | 636 |
| BSEPE_RS01395 | elongation factor Tu | 1303921 |
| BSEPE_RS02190 | elongation factor Tu | 1303921 |
| NAREPO1_RS00940 | elongation factor Tu | 650377 |
| NARPIN1_RS00435 | elongation factor Tu | 1971488 |
| NARRFE1_RS00725 | elongation factor Tu | 1972133 |
| NARSGI1_RS00635 | elongation factor Tu | 1972134 |
| EBS_RS00895 | elongation factor Tu | 1248727 |
| EBS_RS00960 | elongation factor Tu | 1248727 |
| NX720_RS13290 | elongation factor Tu | 1234143 |
| EZMO1_RS23265 | elongation factor Tu | 570277 |
| AAHF05_RS01060 | elongation factor Tu | 3142537 |
| AAHF05_RS20660 | elongation factor Tu | 3142537 |
| FOY94_RS00960 | elongation factor Tu | 61645 |
| FOY94_RS20435 | elongation factor Tu | 61645 |
| DG357_RS01075 | elongation factor Tu | 881260 |
| DG357_RS20850 | elongation factor Tu | 881260 |
| K3T75_RS00135 | elongation factor Tu | 69218 |
| K3T75_RS21310 | elongation factor Tu | 69218 |
| FY206_RS01110 | elongation factor Tu | 2494701 |
| FY206_RS22515 | elongation factor Tu | 2494701 |
| MX657_RS01070 | elongation factor Tu | 2497875 |
| MX657_RS20410 | elongation factor Tu | 2497875 |
| LQ108_RS01150 | elongation factor Tu | 550 |

| | | |
|---|---|---|
| LQ108_RS21060 | elongation factor Tu | 550 |
| F0320_RS00910 | elongation factor Tu | 2797506 |
| F0320_RS19780 | elongation factor Tu | 2797506 |
| I6L63_RS00730 | elongation factor Tu | 158836 |
| I6L63_RS20100 | elongation factor Tu | 158836 |
| D5067_RS01970 | elongation factor Tu | 2494702 |
| D5067_RS21970 | elongation factor Tu | 2494702 |
| ACN6BD_RS01185 | elongation factor Tu | 3133180 |
| ACN6BD_RS20545 | elongation factor Tu | 3133180 |
| LCD42_RS00930 | elongation factor Tu | 208224 |
| LCD42_RS20140 | elongation factor Tu | 208224 |
| BH714_RS13450 | elongation factor Tu | 299767 |
| BH714_RS16185 | elongation factor Tu | 299767 |
| L6Y89_RS01050 | elongation factor Tu | 539813 |
| L6Y89_RS20265 | elongation factor Tu | 539813 |
| EoCCA6_RS09860 | elongation factor Tu | 2478464 |
| EoCCA6_RS13000 | elongation factor Tu | 2478464 |
| QVH37_RS00955 | elongation factor Tu | 2996112 |
| QVH37_RS21070 | elongation factor Tu | 2996112 |
| V1572_RS00910 | elongation factor Tu | 2497436 |
| V1572_RS03870 | elongation factor Tu | 2497436 |
| BFV67_RS00970 | elongation factor Tu | 1812935 |
| BFV67_RS20205 | elongation factor Tu | 1812935 |
| C1N69_RS01025 | elongation factor Tu | 2071710 |
| C1N69_RS20880 | elongation factor Tu | 2071710 |
| ENTAS_RS01080 | elongation factor Tu | 885040 |
| ENTAS_RS20430 | elongation factor Tu | 885040 |
| AB3N39_RS00955 | elongation factor Tu | 2797505 |
| AB3N39_RS19085 | elongation factor Tu | 2797505 |
| VPX56_RS01375 | elongation factor Tu | 2529381 |
| VPX56_RS20900 | elongation factor Tu | 2529381 |

| | | |
|---|---|---|
| GJU03_RS01620 | elongation factor Tu | 2675772 |
| GJT99_RS01610 | elongation factor Tu | 2675773 |
| GJT94_RS01625 | elongation factor Tu | 2675774 |
| GJT92_RS01605 | elongation factor Tu | 2675775 |
| GJT95_RS01615 | elongation factor Tu | 2675776 |
| GJT90_RS01595 | elongation factor Tu | 2675777 |
| GJU05_RS01635 | elongation factor Tu | 2675778 |
| GJU04_RS01630 | elongation factor Tu | 2675779 |
| GJT96_RS01650 | elongation factor Tu | 2675780 |
| GJT93_RS01620 | elongation factor Tu | 2675781 |
| GJT97_RS01590 | elongation factor Tu | 2675782 |
| GJT91_RS01615 | elongation factor Tu | 2675783 |
| GJU00_RS01620 | elongation factor Tu | 2675784 |
| GJT98_RS01640 | elongation factor Tu | 2675785 |
| GJU02_RS01615 | elongation factor Tu | 2675786 |
| GJT88_RS01685 | elongation factor Tu | 2675787 |
| GJT89_RS01605 | elongation factor Tu | 2675788 |
| GJU01_RS01615 | elongation factor Tu | 2675789 |
| GJT86_RS01650 | elongation factor Tu | 2675790 |
| GJT87_RS01655 | elongation factor Tu | 2675791 |
| GJT85_RS01670 | elongation factor Tu | 2675792 |
| GJT80_RS01750 | elongation factor Tu | 2675793 |
| GJT81_RS01740 | elongation factor Tu | 2675794 |
| GJT83_RS01745 | elongation factor Tu | 2675795 |
| GJT82_RS01715 | elongation factor Tu | 2675796 |
| GJT84_RS01765 | elongation factor Tu | 2675797 |
| JHT90_RS13330 | elongation factor Tu | 2785331 |
| DM558_RS14190 | elongation factor Tu | 2213226 |
| ACFXQC_RS01425 | elongation factor Tu | 552 |
| ACFXQC_RS16150 | elongation factor Tu | 552 |
| EBC_RS02750 | elongation factor Tu | 634500 |

| | | |
|---|---|---|
| EBC_RS22380 | elongation factor Tu | 634500 |
| K6L24_RS01685 | elongation factor Tu | 55211 |
| K6L24_RS20125 | elongation factor Tu | 55211 |
| Q3V30_RS01985 | elongation factor Tu | 3062598 |
| Q3V30_RS19980 | elongation factor Tu | 3062598 |
| CPI84_RS01460 | elongation factor Tu | 79967 |
| CPI84_RS17775 | elongation factor Tu | 79967 |
| LJN55_RS02020 | elongation factor Tu | 55212 |
| LJN55_RS21765 | elongation factor Tu | 55212 |
| GN242_RS01765 | elongation factor Tu | 2681984 |
| GN242_RS19800 | elongation factor Tu | 2681984 |
| ETA_RS01765 | elongation factor Tu | 465817 |
| ETA_RS17035 | elongation factor Tu | 465817 |
| LU633_RS01375 | elongation factor Tu | 65700 |
| LU633_RS02830 | elongation factor Tu | 65700 |
| PS053_RS12090 | elongation factor Tu | 208962 |
| PS053_RS15855 | elongation factor Tu | 208962 |
| ECs_4190 | translation elongation fa | 386585 |
| ECs_4903 | translation elongation fa | 386585 |
| b3339 | translation elongation fa | 511145 |
| b3980 | translation elongation fa | 511145 |
| LA347_RS16160 | elongation factor Tu | 564 |
| LA347_RS19675 | elongation factor Tu | 564 |
| J8I93_RS01025 | elongation factor Tu | 1499973 |
| J8I93_RS19375 | elongation factor Tu | 1499973 |
| ACGTJS_RS00730 | elongation factor Tu | 1309795 |
| ACGTJS_RS03265 | elongation factor Tu | 1309795 |
| FBAL_RS01290 | elongation factor Tu | 550540 |
| FBAL_RS01350 | elongation factor Tu | 550540 |
| HER31_RS16930 | elongation factor Tu | 2724191 |
| HER31_RS16990 | elongation factor Tu | 2724191 |

| | | |
|---|---|---|
| JN531_RS03420 | elongation factor Tu | 2806210 |
| JN531_RS03485 | elongation factor Tu | 2806210 |
| HUU81_RS02285 | elongation factor Tu | 2744479 |
| HUU81_RS15435 | elongation factor Tu | 2744479 |
| CDH04_RS01945 | elongation factor Tu | 2007306 |
| KX01_RS03255 | elongation factor Tu | 1542390 |
| CDV26_RS01280 | elongation factor Tu | 549298 |
| FN3523_RS08240 | elongation factor Tu | 622488 |
| F0R75_RS10055 | elongation factor Tu | 2249302 |
| CGC45_RS07870 | elongation factor Tu | 2016517 |
| M973_RS08590 | elongation factor Tu | 1356861 |
| ACH24_RS00170 | elongation factor Tu | 1086726 |
| IMS64_RS08040 | elongation factor Tu | 28110 |
| KQR59_RS08320 | elongation factor Tu | 2599927 |
| F7308_RS03175 | elongation factor Tu | 573569 |
| AQ14_RS05415 | elongation factor Tu | 1450527 |
| F7310_RS09080 | elongation factor Tu | 573570 |
| FRAAU_RS03150 | elongation factor Tu | 767434 |
| FRAAU_RS03215 | elongation factor Tu | 767434 |
| LQ772_RS03840 | elongation factor Tu | 2898793 |
| LQ772_RS03905 | elongation factor Tu | 2898793 |
| LQ771_RS03155 | elongation factor Tu | 1542730 |
| LQ771_RS03220 | elongation factor Tu | 1542730 |
| A4G17_RS03805 | elongation factor Tu | 123824 |
| A4G17_RS03815 | elongation factor Tu | 123824 |
| FPB0191_RS09810 | elongation factor Tu | 1267021 |
| FPB0191_RS09850 | elongation factor Tu | 1267021 |
| PVT67_RS01520 | elongation factor Tu | 3029055 |
| PVT67_RS01585 | elongation factor Tu | 3029055 |
| DW350_RS01310 | elongation factor Tu | 2291597 |
| DW350_RS01370 | elongation factor Tu | 2291597 |

| | | |
|---|---|---|
| NYR30_RS10775 | elongation factor Tu | 505341 |
| NYR30_RS13135 | elongation factor Tu | 505341 |
| BLU22_RS06475 | elongation factor Tu | 1245526 |
| AWC35_RS10450 | elongation factor Tu | 929813 |
| AWC35_RS13505 | elongation factor Tu | 929813 |
| GAPWK_RS01955 | elongation factor Tu | 1196095 |
| GAPWK_RS12910 | elongation factor Tu | 1196095 |
| RAM03_RS01965 | elongation factor Tu | 1970738 |
| RAM03_RS03020 | elongation factor Tu | 1970738 |
| ACGHAD_RS02745 | elongation factor Tu | 1796469 |
| ACGHAD_RS02810 | elongation factor Tu | 1796469 |
| J1G47_RS01300 | elongation factor Tu | 738 |
| J1G47_RS02855 | elongation factor Tu | 738 |
| IMCC3135_RS08165 | elongation factor Tu | 1192854 |
| IMCC3135_RS08455 | elongation factor Tu | 1192854 |
| AL542_RS08275 | elongation factor Tu | 673 |
| AL542_RS10630 | elongation factor Tu | 673 |
| K6Q96_RS01230 | elongation factor Tu | 2872157 |
| K6Q96_RS16020 | elongation factor Tu | 2872157 |
| SR882_RS10685 | elongation factor Tu | 47960 |
| SR882_RS10750 | elongation factor Tu | 47960 |
| DQN49_RS02305 | elongation factor Tu | 197575 |
| DQN49_RS02755 | elongation factor Tu | 197575 |
| DQN30_RS02050 | elongation factor Tu | 726 |
| DQN30_RS02935 | elongation factor Tu | 726 |
| LK401_RS03300 | elongation factor Tu | 727 |
| LK401_RS03500 | elongation factor Tu | 727 |
| I6J29_RS06725 | elongation factor Tu | 735 |
| I6J29_RS08450 | elongation factor Tu | 735 |
| I6H57_RS08465 | elongation factor Tu | 729 |
| I6H57_RS09495 | elongation factor Tu | 729 |

| | | |
|---|---|---|
| CKV74_RS01595 | elongation factor Tu | 249188 |
| CKV74_RS03230 | elongation factor Tu | 249188 |
| HBA19_RS01315 | elongation factor Tu | 569 |
| HBA19_RS19540 | elongation factor Tu | 569 |
| K9N75_RS01285 | elongation factor Tu | 546367 |
| K9N75_RS20080 | elongation factor Tu | 546367 |
| HCH_RS27970 | elongation factor Tu | 349521 |
| HCH_RS28035 | elongation factor Tu | 349521 |
| EY643_RS03100 | elongation factor Tu | 2601894 |
| EY643_RS03185 | elongation factor Tu | 2601894 |
| ACGHM3_RS03930 | elongation factor Tu | 1306992 |
| ACGHM3_RS04005 | elongation factor Tu | 1306992 |
| BOX17_RS09045 | elongation factor Tu | 1897729 |
| QEN58_RS00765 | elongation factor Tu | 232346 |
| QYS58_RS16785 | elongation factor Tu | 1930622 |
| BEI_RS17025 | elongation factor Tu | 475662 |
| E4T21_RS01880 | elongation factor Tu | 2562282 |
| LOKO_RS00575 | elongation factor Tu | 507626 |
| HELO_RS01235 | elongation factor Tu | 768066 |
| QWG60_RS15445 | elongation factor Tu | 29573 |
| AR456_RS01685 | elongation factor Tu | 1178482 |
| HHSLTHF2_RS00895 | elongation factor Tu | 115561 |
| R5M92_RS14435 | elongation factor Tu | 3084920 |
| P1P91_RS02240 | elongation factor Tu | 3031727 |
| K1Y77_RS00900 | elongation factor Tu | 2866211 |
| M0220_RS06790 | elongation factor Tu | 2937790 |
| HNO53_RS18660 | elongation factor Tu | 2733488 |
| QUD69_RS06810 | elongation factor Tu | 857252 |
| QUD69_RS06875 | elongation factor Tu | 857252 |
| BLU11_RS18205 | elongation factor Tu | 797277 |
| BLU11_RS18270 | elongation factor Tu | 797277 |

| | | |
|---|---|---|
| HV822_RS07060 | elongation factor Tu | 2918528 |
| HV822_RS07125 | elongation factor Tu | 2918528 |
| KEM63_RS15475 | elongation factor Tu | 2830842 |
| KEM63_RS15540 | elongation factor Tu | 2830842 |
| U2971_RS04245 | elongation factor Tu | 1708783 |
| U2971_RS04310 | elongation factor Tu | 1708783 |
| BVH74_RS03435 | elongation factor Tu | 1931241 |
| BVH74_RS03500 | elongation factor Tu | 1931241 |
| BLU26_RS13725 | elongation factor Tu | 472181 |
| BLU26_RS13790 | elongation factor Tu | 472181 |
| BLU07_RS13845 | elongation factor Tu | 1434072 |
| BLU07_RS13910 | elongation factor Tu | 1434072 |
| BLT85_RS13200 | elongation factor Tu | 487184 |
| BLT85_RS13265 | elongation factor Tu | 487184 |
| HH1059_RS11130 | elongation factor Tu | 1052 |
| HH1059_RS11195 | elongation factor Tu | 1052 |
| HHAL_RS04355 | elongation factor Tu | 349124 |
| HHAL_RS04420 | elongation factor Tu | 349124 |
| A9404_RS10485 | elongation factor Tu | 1860122 |
| A9404_RS10550 | elongation factor Tu | 1860122 |
| HNEAP_RS01530 | elongation factor Tu | 555778 |
| HNEAP_RS01595 | elongation factor Tu | 555778 |
| HSM_RS00310 | elongation factor Tu | 228400 |
| HSM_RS09310 | elongation factor Tu | 228400 |
| soil367_RS02720 | elongation factor Tu | 2259620 |
| soil367_RS02785 | elongation factor Tu | 2259620 |
| GHNINEIG_RS01445 | elongation factor Tu | 39765 |
| GHNINEIG_RS01510 | elongation factor Tu | 39765 |
| HVMH_RS01615 | elongation factor Tu | 28885 |
| HVMH_RS01680 | elongation factor Tu | 28885 |
| EPV75_RS01925 | elongation factor Tu | 265883 |

| | | |
|---|---|---|
| EPV75_RS01990 | elongation factor Tu | 265883 |
| K5X84_RS01545 | elongation factor Tu | 86102 |
| K5X84_RS01610 | elongation factor Tu | 86102 |
| IL_RS01725 | elongation factor Tu | 283942 |
| IL_RS01790 | elongation factor Tu | 283942 |
| WMO13_RS01035 | elongation factor Tu | 1111732 |
| WMO13_RS08180 | elongation factor Tu | 1111732 |
| MMG00_RS08695 | elongation factor Tu | 2923279 |
| MMG00_RS13755 | elongation factor Tu | 2923279 |
| PG2T_RS11600 | elongation factor Tu | 1810504 |
| PG2T_RS11665 | elongation factor Tu | 1810504 |
| S7S_RS02795 | elongation factor Tu | 391936 |
| S7S_RS02860 | elongation factor Tu | 391936 |
| CVFO_RS00690 | elongation factor Tu | 1235284 |
| CVFO_RS08000 | elongation factor Tu | 1235284 |
| FEM41_RS03395 | elongation factor Tu | 2579935 |
| FEM41_RS06945 | elongation factor Tu | 2579935 |
| TQ33_RS09785 | elongation factor Tu | 914150 |
| TQ33_RS09850 | elongation factor Tu | 914150 |
| KKOR_RS02125 | elongation factor Tu | 523791 |
| KKOR_RS02190 | elongation factor Tu | 523791 |
| KS2013_RS10030 | elongation factor Tu | 1144748 |
| KS2013_RS10095 | elongation factor Tu | 1144748 |
| Kalk_RS06355 | elongation factor Tu | 1917421 |
| Kalk_RS06420 | elongation factor Tu | 1917421 |
| I6L72_RS00260 | elongation factor Tu | 548 |
| I6L72_RS22075 | elongation factor Tu | 548 |
| LGL98_RS01995 | elongation factor Tu | 2489010 |
| LGL98_RS23880 | elongation factor Tu | 2489010 |
| Electrica_RS01900 | elongation factor Tu | 1259973 |
| Electrica_RS24020 | elongation factor Tu | 1259973 |

| | | |
|---|---|---|
| ACE3MN_RS02045 | elongation factor Tu | 2058152 |
| ACE3MN_RS25370 | elongation factor Tu | 2058152 |
| DA718_RS02490 | elongation factor Tu | 2153354 |
| DA718_RS27910 | elongation factor Tu | 2153354 |
| JLG71_RS02155 | elongation factor Tu | 1134687 |
| JLG71_RS26280 | elongation factor Tu | 1134687 |
| EL226_RS00175 | elongation factor Tu | 571 |
| EL226_RS03615 | elongation factor Tu | 571 |
| KCG39_RS16260 | elongation factor Tu | 2587529 |
| KCG39_RS19845 | elongation factor Tu | 2587529 |
| LA349_RS10820 | elongation factor Tu | 1463165 |
| LA349_RS14090 | elongation factor Tu | 1463165 |
| LGM30_RS02015 | elongation factor Tu | 2026240 |
| LGM30_RS25120 | elongation factor Tu | 2026240 |
| LGN93_RS01980 | elongation factor Tu | 2590157 |
| LGN93_RS25420 | elongation factor Tu | 2590157 |
| MY052_RS01970 | elongation factor Tu | 51288 |
| MY052_RS22825 | elongation factor Tu | 51288 |
| GHC21_RS02080 | elongation factor Tu | 61648 |
| GHC21_RS20585 | elongation factor Tu | 61648 |
| AB2B48_RS15845 | elongation factor Tu | 2725494 |
| AB2B48_RS19210 | elongation factor Tu | 2725494 |
| GCU50_RS14390 | elongation factor Tu | 551989 |
| GCU50_RS17515 | elongation factor Tu | 551989 |
| AAEY27_RS02065 | elongation factor Tu | 3139408 |
| AAEY27_RS20990 | elongation factor Tu | 3139408 |
| ODS25_RS01940 | elongation factor Tu | 208223 |
| ODS25_RS20730 | elongation factor Tu | 208223 |
| AWR26_RS02045 | elongation factor Tu | 497725 |
| AWR26_RS24365 | elongation factor Tu | 497725 |
| O9K67_RS02290 | elongation factor Tu | 1005665 |

| | | |
|---|---|---|
| O9K67_RS23355 | elongation factor Tu | 1005665 |
| IP581_RS01100 | elongation factor Tu | 1646340 |
| IP581_RS21210 | elongation factor Tu | 1646340 |
| FEI17_RS02010 | elongation factor Tu | 283686 |
| FEI17_RS25500 | elongation factor Tu | 283686 |
| C813_RS24925 | elongation factor Tu | 1235834 |
| C813_RS44860 | elongation factor Tu | 1235834 |
| B9G99_RS03615 | elongation factor Tu | 698828 |
| B9H00_RS11520 | elongation factor Tu | 157779 |
| FY550_RS14015 | elongation factor Tu | 657387 |
| APT61_RS02035 | elongation factor Tu | 83655 |
| APT61_RS21230 | elongation factor Tu | 83655 |
| JZ655_RS00915 | elongation factor Tu | 2815358 |
| JZ655_RS19040 | elongation factor Tu | 2815358 |
| DLD14_RS17550 | elongation factor Tu | 28082 |
| DLD14_RS17615 | elongation factor Tu | 28082 |
| HRS36_RS01795 | elongation factor Tu | 2708020 |
| HRS36_RS01860 | elongation factor Tu | 2708020 |
| PXX05_RS12060 | elongation factor Tu | 1071983 |
| PXX05_RS12125 | elongation factor Tu | 1071983 |
| clem_RS13255 | elongation factor Tu | 1867846 |
| clem_RS13320 | elongation factor Tu | 1867846 |
| LFA_RS12815 | elongation factor Tu | 1212491 |
| LFA_RS12880 | elongation factor Tu | 1212491 |
| E4T54_RS07435 | elongation factor Tu | 45065 |
| E4T54_RS07500 | elongation factor Tu | 45065 |
| LHA_RS01475 | elongation factor Tu | 449 |
| LHA_RS01540 | elongation factor Tu | 449 |
| FOG18_RS01180 | elongation factor Tu | 454 |
| FOG18_RS01245 | elongation factor Tu | 454 |
| EL203_RS01185 | elongation factor Tu | 456 |

| | | |
|---|---|---|
| EL203_RS01250 | elongation factor Tu | 456 |
| CKV79_RS01100 | elongation factor Tu | 45067 |
| CKV79_RS01165 | elongation factor Tu | 45067 |
| LLO_RS02455 | elongation factor Tu | 661367 |
| LLO_RS02520 | elongation factor Tu | 661367 |
| J2N86_RS01340 | elongation factor Tu | 96232 |
| J2N86_RS01405 | elongation factor Tu | 96232 |
| LMI_RS01950 | elongation factor Tu | 451 |
| LMI_RS02015 | elongation factor Tu | 451 |
| AVR58_RS01600 | elongation factor Tu | 446 |
| AVR58_RS01665 | elongation factor Tu | 446 |
| CAB17_RS18620 | elongation factor Tu | 28087 |
| CAB17_RS18685 | elongation factor Tu | 28087 |
| CKW05_RS01885 | elongation factor Tu | 452 |
| CKW05_RS01950 | elongation factor Tu | 452 |
| CKW04_RS03075 | elongation factor Tu | 66969 |
| CKW04_RS03140 | elongation factor Tu | 66969 |
| I6L74_RS06465 | elongation factor Tu | 61646 |
| I6L74_RS09695 | elongation factor Tu | 61646 |
| ACB496_RS00975 | elongation factor Tu | 69220 |
| ACB496_RS20350 | elongation factor Tu | 69220 |
| V8L73_RS02075 | elongation factor Tu | 82981 |
| V8L73_RS17500 | elongation factor Tu | 82981 |
| DQM29_RS01740 | elongation factor Tu | 158841 |
| DQM29_RS15865 | elongation factor Tu | 158841 |
| HYN51_RS01065 | elongation factor Tu | 2172103 |
| HYN51_RS14055 | elongation factor Tu | 2172103 |
| EKN56_RS17960 | elongation factor Tu | 2498113 |
| EKN56_RS18585 | elongation factor Tu | 2498113 |
| EMK97_RS09805 | elongation factor Tu | 718192 |
| EMK97_RS09870 | elongation factor Tu | 718192 |

| | | |
|---|---|---|
| GH975_RS09885 | elongation factor Tu | 418701 |
| GH975_RS09950 | elongation factor Tu | 418701 |
| O1Q83_RS00695 | elongation factor Tu | 53417 |
| O1Q83_RS11355 | elongation factor Tu | 53417 |
| CKQ53_RS03590 | elongation factor Tu | 1082704 |
| CKQ53_RS06230 | elongation factor Tu | 1082704 |
| I6N93_RS01440 | elongation factor Tu | 1172565 |
| I6N93_RS16030 | elongation factor Tu | 1172565 |
| L2Y96_RS05560 | elongation factor Tu | 2911538 |
| L2Y96_RS05625 | elongation factor Tu | 2911538 |
| L2Y94_RS16820 | elongation factor Tu | 2911537 |
| L2Y94_RS16885 | elongation factor Tu | 2911537 |
| L2Y97_RS05025 | elongation factor Tu | 2911539 |
| L2Y97_RS05090 | elongation factor Tu | 2911539 |
| KR767_RS02120 | elongation factor Tu | 564369 |
| KR767_RS02185 | elongation factor Tu | 564369 |
| IM816_RS03775 | elongation factor Tu | 2780091 |
| IM816_RS03840 | elongation factor Tu | 2780091 |
| FIV34_RS04985 | elongation factor Tu | 2589080 |
| FIV34_RS05050 | elongation factor Tu | 2589080 |
| BJI69_RS12060 | elongation factor Tu | 1440763 |
| BJI69_RS12125 | elongation factor Tu | 1440763 |
| CNR27_RS06120 | elongation factor Tu | 2006110 |
| CNR27_RS06185 | elongation factor Tu | 2006110 |
| LU699_RS03775 | elongation factor Tu | 2901869 |
| LU699_RS03840 | elongation factor Tu | 2901869 |
| FPZ22_RS10655 | elongation factor Tu | 1176533 |
| FPZ22_RS10720 | elongation factor Tu | 1176533 |
| E5843_RS10885 | elongation factor Tu | 2565782 |
| E5843_RS10950 | elongation factor Tu | 2565782 |
| LA76x_RS17925 | elongation factor Tu | 84531 |

| | | |
|---|---|---|
| LA76x_RS17990 | elongation factor Tu | 84531 |
| HIV01_RS08800 | elongation factor Tu | 2795387 |
| HIV01_RS08865 | elongation factor Tu | 2795387 |
| LA521A_RS04720 | elongation factor Tu | 2992573 |
| LA521A_RS04785 | elongation factor Tu | 2992573 |
| IEQ11_RS18180 | elongation factor Tu | 435897 |
| IEQ11_RS18245 | elongation factor Tu | 435897 |
| LEN_RS06550 | elongation factor Tu | 69 |
| LEN_RS06615 | elongation factor Tu | 69 |
| ABU614_RS12185 | elongation factor Tu | 1792846 |
| ABU614_RS12250 | elongation factor Tu | 1792846 |
| LG3211_RS17085 | elongation factor Tu | 262324 |
| LG3211_RS17150 | elongation factor Tu | 262324 |
| LYSHEL_RS02580 | elongation factor Tu | 2675059 |
| LYSHEL_RS02645 | elongation factor Tu | 2675059 |
| GOY17_RS04900 | elongation factor Tu | 453783 |
| GOY17_RS04965 | elongation factor Tu | 453783 |
| H8L67_RS03305 | elongation factor Tu | 2764185 |
| H8L67_RS03370 | elongation factor Tu | 2764185 |
| RDV84_RS13910 | elongation factor Tu | 1003114 |
| RDV84_RS13975 | elongation factor Tu | 1003114 |
| MPHASIOC01_RS19095 | elongation factor Tu | 1691903 |
| MPHASIOC01_RS26030 | elongation factor Tu | 1691903 |
| ICJ55_RS00975 | elongation factor Tu | 2770636 |
| ICJ55_RS03130 | elongation factor Tu | 2770636 |
| ACJ7Z2_RS00735 | elongation factor Tu | 85401 |
| ACJ7Z2_RS03290 | elongation factor Tu | 85401 |
| CKG22_RS02180 | elongation factor Tu | 75985 |
| CKG22_RS13515 | elongation factor Tu | 75985 |
| HV560_RS01430 | elongation factor Tu | 111844 |
| HV560_RS03780 | elongation factor Tu | 111844 |

| | | |
|---|---|---|
| A6B40_RS04870 | elongation factor Tu | 85404 |
| A6B40_RS05780 | elongation factor Tu | 85404 |
| MARPU_RS03610 | elongation factor Tu | 765910 |
| MARPU_RS03675 | elongation factor Tu | 765910 |
| ACGHZ6_RS02155 | elongation factor Tu | 1008307 |
| ACGHZ6_RS02220 | elongation factor Tu | 1008307 |
| FKV23_RS11830 | elongation factor Tu | 2591633 |
| FKV23_RS11895 | elongation factor Tu | 2591633 |
| C9I47_RS12750 | elongation factor Tu | 1605891 |
| C9I47_RS12815 | elongation factor Tu | 1605891 |
| MARGE09_RS01720 | elongation factor Tu | 2721545 |
| TYM08_RS04230 | elongation factor Tu | 2724896 |
| TYM08_RS17820 | elongation factor Tu | 2724896 |
| KQ249_RS05940 | elongation factor Tu | 1033846 |
| KQ249_RS06005 | elongation factor Tu | 1033846 |
| AAGT77_RS02750 | elongation factor Tu | 254719 |
| AAGT77_RS02815 | elongation factor Tu | 254719 |
| QUE89_RS02880 | elongation factor Tu | 3035454 |
| QUE89_RS02945 | elongation factor Tu | 3035454 |
| ACGK9W_RS20490 | elongation factor Tu | 262076 |
| ACGK9W_RS20555 | elongation factor Tu | 262076 |
| FXO11_RS03440 | elongation factor Tu | 2603215 |
| FXO11_RS03505 | elongation factor Tu | 2603215 |
| NLK58_RS16215 | elongation factor Tu | 2961995 |
| NLK58_RS16280 | elongation factor Tu | 2961995 |
| MARHY_RS02725 | elongation factor Tu | 1163748 |
| MARHY_RS02790 | elongation factor Tu | 1163748 |
| ABA45_RS03135 | elongation factor Tu | 330734 |
| ABA45_RS03200 | elongation factor Tu | 330734 |
| MARSALSMR5_RS17390 | elongation factor Tu | 1420917 |
| MARSALSMR5_RS17455 | elongation factor Tu | 1420917 |

| | | |
|---|---|---|
| LPB19_RS06220 | elongation factor Tu | 2769486 |
| LPB19_RS06285 | elongation factor Tu | 2769486 |
| BKP64_RS12455 | elongation factor Tu | 1874317 |
| BKP64_RS12520 | elongation factor Tu | 1874317 |
| QUE75_RS03500 | elongation factor Tu | 1389223 |
| QUE75_RS03565 | elongation factor Tu | 1389223 |
| AU14_RS18430 | elongation factor Tu | 1420916 |
| A8C75_RS21455 | elongation factor Tu | 1821621 |
| A8C75_RS21520 | elongation factor Tu | 1821621 |
| CFI10_RS01160 | elongation factor Tu | 48076 |
| CFI10_RS01225 | elongation factor Tu | 48076 |
| IEZ33_RS01025 | elongation factor Tu | 2773454 |
| IBG28_RS20025 | elongation factor Tu | 383750 |
| JSY38_RS08305 | elongation factor Tu | 491950 |
| MARME_RS01195 | elongation factor Tu | 717774 |
| AB8E32_RS15560 | elongation factor Tu | 293552 |
| AABA11_RS15635 | elongation factor Tu | 264739 |
| MAR181_RS17045 | elongation factor Tu | 491952 |
| MP3633_RS17965 | elongation factor Tu | 178399 |
| J8N69_RS07070 | elongation factor Tu | 2726122 |
| CEP47_RS00030 | elongation factor Tu | 221402 |
| CEP47_RS01215 | elongation factor Tu | 221402 |
| OXH74_RS10840 | elongation factor Tu | 2822138 |
| OXH74_RS10905 | elongation factor Tu | 2822138 |
| KF707C_RS02760 | elongation factor Tu | 1149133 |
| KF707C_RS02825 | elongation factor Tu | 1149133 |
| FXN65_RS02845 | elongation factor Tu | 2604832 |
| FXN65_RS02910 | elongation factor Tu | 2604832 |
| PotMrB4_RS03290 | elongation factor Tu | 319939 |
| PotMrB4_RS03355 | elongation factor Tu | 319939 |
| sS8_RS13175 | elongation factor Tu | 1432792 |

| | | |
|---|---|---|
| sS8_RS15995 | elongation factor Tu | 1432792 |
| QEN43_RS11420 | elongation factor Tu | 73780 |
| QEN43_RS14905 | elongation factor Tu | 73780 |
| MCA_RS05205 | elongation factor Tu | 243233 |
| MCA_RS11625 | elongation factor Tu | 243233 |
| GNH96_RS12735 | elongation factor Tu | 2681310 |
| GNH96_RS14725 | elongation factor Tu | 2681310 |
| OOT43_RS16795 | elongation factor Tu | 2993564 |
| OOT43_RS18225 | elongation factor Tu | 2993564 |
| K5607_RS01905 | elongation factor Tu | 1295382 |
| K5607_RS01970 | elongation factor Tu | 1295382 |
| K5658_RS04675 | elongation factor Tu | 1760988 |
| K5658_RS18555 | elongation factor Tu | 1760988 |
| MCIT9_RS00560 | elongation factor Tu | 438856 |
| MCIT9_RS00625 | elongation factor Tu | 438856 |
| MIN45_RS06055 | elongation factor Tu | 2840183 |
| MIN45_RS06120 | elongation factor Tu | 2840183 |
| Q9L42_RS01620 | elongation factor Tu | 3067653 |
| Q9L42_RS01685 | elongation factor Tu | 3067653 |
| METAL_RS16065 | elongation factor Tu | 686340 |
| METAL_RS16130 | elongation factor Tu | 686340 |
| JT25_RS14655 | elongation factor Tu | 1538553 |
| JT25_RS14720 | elongation factor Tu | 1538553 |
| MKFW12EY_RS18265 | elongation factor Tu | 702114 |
| MKFW12EY_RS18330 | elongation factor Tu | 702114 |
| QZJ86_RS17080 | elongation factor Tu | 3058963 |
| QZJ86_RS17145 | elongation factor Tu | 3058963 |
| KEF85_RS13165 | elongation factor Tu | 1173101 |
| KEF85_RS13230 | elongation factor Tu | 1173101 |
| NM686_RS18425 | elongation factor Tu | 2963939 |
| NM686_RS18490 | elongation factor Tu | 2963939 |

| | | |
|---|---|---|
| F1E05_RS02425 | elongation factor Tu | 2608981 |
| F1E05_RS02490 | elongation factor Tu | 2608981 |
| Q7C_RS11400 | elongation factor Tu | 754477 |
| Q7C_RS11465 | elongation factor Tu | 754477 |
| QUE24_RS05135 | elongation factor Tu | 45495 |
| QUE24_RS05200 | elongation factor Tu | 45495 |
| Q7A_RS13995 | elongation factor Tu | 754476 |
| Q7A_RS14060 | elongation factor Tu | 754476 |
| LGT42_RS13730 | elongation factor Tu | 2881052 |
| LGT42_RS13795 | elongation factor Tu | 2881052 |
| MEALZ_RS16380 | elongation factor Tu | 1091494 |
| MEALZ_RS16445 | elongation factor Tu | 1091494 |
| EQU24_RS03800 | elongation factor Tu | 95641 |
| EQU24_RS03865 | elongation factor Tu | 95641 |
| CEK71_RS05235 | elongation factor Tu | 1704499 |
| CEK71_RS05300 | elongation factor Tu | 1704499 |
| Mag101_RS14825 | elongation factor Tu | 260552 |
| Mag101_RS14890 | elongation factor Tu | 260552 |
| AUP74_RS06745 | elongation factor Tu | 1769779 |
| AUP74_RS06810 | elongation factor Tu | 1769779 |
| PVT68_RS08260 | elongation factor Tu | 3029061 |
| PVT68_RS08325 | elongation factor Tu | 3029061 |
| LPW13_RS12015 | elongation factor Tu | 435905 |
| LPW13_RS12080 | elongation factor Tu | 435905 |
| LRR79_RS06015 | elongation factor Tu | 86173 |
| LRR79_RS06080 | elongation factor Tu | 86173 |
| GTQ55_RS14765 | elongation factor Tu | 48074 |
| GTQ55_RS14830 | elongation factor Tu | 48074 |
| R5R33_RS06425 | elongation factor Tu | 407164 |
| R5R33_RS06490 | elongation factor Tu | 407164 |
| M8T91_RS16015 | elongation factor Tu | 2944933 |

| | | |
|---|---|---|
| M8T91_RS16080 | elongation factor Tu | 2944933 |
| A3224_RS02370 | elongation factor Tu | 252514 |
| A3224_RS02435 | elongation factor Tu | 252514 |
| C2E15_RS02050 | elongation factor Tu | 665914 |
| C2E15_RS19990 | elongation factor Tu | 665914 |
| K6958_RS01960 | elongation factor Tu | 2872648 |
| K6958_RS18850 | elongation factor Tu | 2872648 |
| C7M51_RS15725 | elongation factor Tu | 1615494 |
| C7M51_RS19530 | elongation factor Tu | 1615494 |
| MNY66_RS01225 | elongation factor Tu | 158849 |
| MNY66_RS15160 | elongation factor Tu | 158849 |
| LP107_RS01640 | elongation factor Tu | 476 |
| LP107_RS12665 | elongation factor Tu | 476 |
| AAX10_RS08140 | elongation factor Tu | 386891 |
| AAX10_RS08295 | elongation factor Tu | 386891 |
| U0021_RS06140 | elongation factor Tu | 90239 |
| U0021_RS06255 | elongation factor Tu | 90239 |
| MC195_RS07105 | elongation factor Tu | 480 |
| MC195_RS07220 | elongation factor Tu | 480 |
| EL162_RS03845 | elongation factor Tu | 34061 |
| EL162_RS03960 | elongation factor Tu | 34061 |
| LU276_RS01280 | elongation factor Tu | 2904119 |
| LU276_RS08655 | elongation factor Tu | 2904119 |
| WDM69_RS02370 | elongation factor Tu | 90241 |
| WDM69_RS07270 | elongation factor Tu | 90241 |
| LU290_RS01165 | elongation factor Tu | 2904120 |
| LU290_RS09375 | elongation factor Tu | 2904120 |
| LU297_RS03700 | elongation factor Tu | 2904122 |
| LU297_RS06275 | elongation factor Tu | 2904122 |
| LU293_RS03755 | elongation factor Tu | 2904121 |
| LU293_RS06355 | elongation factor Tu | 2904121 |

| | | |
|---|---|---|
| I6G26_RS02720 | elongation factor Tu | 478 |
| I6G26_RS10660 | elongation factor Tu | 478 |
| AXE82_RS03690 | elongation factor Tu | 34062 |
| AXE82_RS09080 | elongation factor Tu | 34062 |
| MOVS_RS08790 | elongation factor Tu | 29433 |
| MOVS_RS08930 | elongation factor Tu | 29433 |
| ACE1C5_RS00735 | elongation factor Tu | 3344666 |
| ACE1C5_RS00845 | elongation factor Tu | 3344666 |
| EO986_RS01000 | elongation factor Tu | 180434 |
| EO986_RS17660 | elongation factor Tu | 180434 |
| FR932_RS16320 | elongation factor Tu | 1202962 |
| MORIYA_RS12635 | elongation factor Tu | 69539 |
| DD703_RS02010 | elongation factor Tu | 579405 |
| DD703_RS18635 | elongation factor Tu | 579405 |
| QUD59_RS06690 | elongation factor Tu | 651666 |
| QUD59_RS06755 | elongation factor Tu | 651666 |
| ACOQOE_RS00020 | elongation factor Tu | 1806670 |
| ACOQOE_RS00085 | elongation factor Tu | 1806670 |
| F0U83_RS15775 | elongation factor Tu | 1031538 |
| F0U83_RS15840 | elongation factor Tu | 1031538 |
| FLM49_RS00330 | elongation factor Tu | 1572645 |
| FLM49_RS00395 | elongation factor Tu | 1572645 |
| F5I99_RS01130 | elongation factor Tu | 2614693 |
| F5I99_RS01195 | elongation factor Tu | 2614693 |
| NHAL_RS11550 | elongation factor Tu | 472759 |
| NHAL_RS11615 | elongation factor Tu | 472759 |
| NOC_RS12420 | elongation factor Tu | 323261 |
| NOC_RS12485 | elongation factor Tu | 323261 |
| E3U44_RS09090 | elongation factor Tu | 1814290 |
| E3U44_RS09155 | elongation factor Tu | 1814290 |
| NWAT_RS10585 | elongation factor Tu | 105559 |

| | | |
|---|---|---|
| NWAT_RS10650 | elongation factor Tu | 105559 |
| NWAT_RS10715 | elongation factor Tu | 105559 |
| LYSCAS_RS02580 | elongation factor Tu | 2675101 |
| LYSCAS_RS02645 | elongation factor Tu | 2675101 |
| INQ42_RS02485 | elongation factor Tu | 2781023 |
| INQ42_RS02550 | elongation factor Tu | 2781023 |
| INQ41_RS02415 | elongation factor Tu | 2781022 |
| INQ41_RS02480 | elongation factor Tu | 2781022 |
| KOD61_RS10355 | elongation factor Tu | 2822368 |
| KOD61_RS10420 | elongation factor Tu | 2822368 |
| BST96_RS04740 | elongation factor Tu | 716816 |
| BST96_RS10155 | elongation factor Tu | 716816 |
| PU634_RS00935 | elongation factor Tu | 3028314 |
| PU634_RS14335 | elongation factor Tu | 3028314 |
| CBP12_RS08620 | elongation factor Tu | 1903694 |
| CBP31_RS05475 | elongation factor Tu | 1416627 |
| OLMES_RS03330 | elongation factor Tu | 141451 |
| RHO11_RS11125 | elongation factor Tu | 3074107 |
| RHO11_RS11290 | elongation factor Tu | 3074107 |
| RHO12_RS09735 | elongation factor Tu | 3074109 |
| RHO12_RS09785 | elongation factor Tu | 3074109 |
| RHO14_RS12425 | elongation factor Tu | 3074111 |
| RHO14_RS12465 | elongation factor Tu | 3074111 |
| A6A10_RS01215 | elongation factor Tu | 1032623 |
| A6A10_RS08515 | elongation factor Tu | 1032623 |
| I6L77_RS01655 | elongation factor Tu | 549 |
| I6L77_RS04485 | elongation factor Tu | 549 |
| K6R05_RS02115 | elongation factor Tu | 3074822 |
| K6R05_RS17645 | elongation factor Tu | 3074822 |
| B1H58_RS05655 | elongation factor Tu | 1891675 |
| B1H58_RS08860 | elongation factor Tu | 1891675 |

| | | |
|---|---|---|
| QNH99_RS02145 | elongation factor Tu | 574096 |
| QNH99_RS20670 | elongation factor Tu | 574096 |
| PAGR_RS02095 | elongation factor Tu | 1095774 |
| PAGR_RS20885 | elongation factor Tu | 1095774 |
| OJ965_RS04635 | elongation factor Tu | 470931 |
| OJ965_RS19825 | elongation factor Tu | 470931 |
| J1C59_RS02035 | elongation factor Tu | 470932 |
| J1C59_RS17895 | elongation factor Tu | 470932 |
| K7H94_RS01435 | elongation factor Tu | 59814 |
| K7H94_RS17335 | elongation factor Tu | 59814 |
| EE896_RS02095 | elongation factor Tu | 470933 |
| EE896_RS18025 | elongation factor Tu | 470933 |
| LAC65_RS00735 | elongation factor Tu | 472693 |
| LAC65_RS14625 | elongation factor Tu | 472693 |
| ACF2G4_RS12600 | elongation factor Tu | 2769024 |
| ACF2G4_RS15580 | elongation factor Tu | 2769024 |
| G5574_RS11725 | elongation factor Tu | 66269 |
| G5574_RS14500 | elongation factor Tu | 66269 |
| EGO56_RS02195 | elongation factor Tu | 470934 |
| EGO56_RS18125 | elongation factor Tu | 470934 |
| PMPD1_RS01885 | elongation factor Tu | 2740817 |
| PMPD1_RS20675 | elongation factor Tu | 2740817 |
| CF386_RS02545 | elongation factor Tu | 1755811 |
| CF386_RS03895 | elongation factor Tu | 1755811 |
| E2I05_RS00615 | elongation factor Tu | 342950 |
| E2I05_RS21435 | elongation factor Tu | 342950 |
| E2H97_RS00445 | elongation factor Tu | 2547970 |
| E2H97_RS00545 | elongation factor Tu | 2547970 |
| U9966_RS01550 | elongation factor Tu | 2827233 |
| U9966_RS03485 | elongation factor Tu | 2827233 |
| K7G93_RS07685 | elongation factor Tu | 753 |

| | | |
|---|---|---|
| K7G93_RS09690 | elongation factor Tu | 753 |
| CKV78_RS07365 | elongation factor Tu | 754 |
| CKV78_RS08080 | elongation factor Tu | 754 |
| A6J57_RS03165 | elongation factor Tu | 747 |
| A6J57_RS05255 | elongation factor Tu | 747 |
| DMB82_RS00955 | elongation factor Tu | 2204145 |
| DMB82_RS18355 | elongation factor Tu | 2204145 |
| AACH44_RS01760 | elongation factor Tu | 3073862 |
| AACH44_RS19235 | elongation factor Tu | 3073862 |
| I2D83_RS02225 | elongation factor Tu | 1201031 |
| I2D83_RS20870 | elongation factor Tu | 1201031 |
| GZ59_RS01150 | elongation factor Tu | 29471 |
| GZ59_RS19630 | elongation factor Tu | 29471 |
| GT391_RS08290 | elongation factor Tu | 180957 |
| GT391_RS09485 | elongation factor Tu | 180957 |
| RFN81_RS01470 | elongation factor Tu | 69221 |
| RFN81_RS16770 | elongation factor Tu | 69221 |
| HER17_RS02045 | elongation factor Tu | 554 |
| HER17_RS20465 | elongation factor Tu | 554 |
| LCF41_RS01000 | elongation factor Tu | 2878098 |
| LCF41_RS19755 | elongation factor Tu | 2878098 |
| EO763_RS01045 | elongation factor Tu | 78398 |
| EO763_RS19880 | elongation factor Tu | 78398 |
| A8F97_RS17375 | elongation factor Tu | 1905730 |
| A8F97_RS21475 | elongation factor Tu | 1905730 |
| LOZ86_RS01950 | elongation factor Tu | 2778550 |
| LOZ86_RS03225 | elongation factor Tu | 2778550 |
| DUT67_RS02265 | elongation factor Tu | 2066479 |
| DUT67_RS20895 | elongation factor Tu | 2066479 |
| BJK05_RS14605 | elongation factor Tu | 2042057 |
| BJK05_RS18225 | elongation factor Tu | 2042057 |

| | | |
|---|---|---|
| E2566_RS01210 | elongation factor Tu | 2108399 |
| E2566_RS19270 | elongation factor Tu | 2108399 |
| IG609_RS01020 | elongation factor Tu | 2774015 |
| IG609_RS18095 | elongation factor Tu | 2774015 |
| EIP93_RS01220 | elongation factor Tu | 2488639 |
| EIP93_RS20105 | elongation factor Tu | 2488639 |
| A7983_RS05150 | elongation factor Tu | 1175631 |
| A7983_RS06450 | elongation factor Tu | 1175631 |
| A6B44_RS00440 | elongation factor Tu | 97481 |
| A6B44_RS09470 | elongation factor Tu | 97481 |
| NNL38_RS01285 | elongation factor Tu | 865757 |
| NNL38_RS15145 | elongation factor Tu | 865757 |
| WAX86_RS01145 | elongation factor Tu | 85581 |
| WAX86_RS13830 | elongation factor Tu | 85581 |
| FH974_RS01260 | elongation factor Tu | 320778 |
| FH974_RS14990 | elongation factor Tu | 320778 |
| ACN08M_RS05725 | elongation factor Tu | 48408 |
| ACN08M_RS07965 | elongation factor Tu | 48408 |
| ACJZU6_RS00685 | elongation factor Tu | 1378299 |
| ACJZU6_RS02950 | elongation factor Tu | 1378299 |
| OCU87_RS01270 | elongation factor Tu | 875932 |
| OCU87_RS16135 | elongation factor Tu | 875932 |
| OCU77_RS01350 | elongation factor Tu | 680026 |
| OCU77_RS16330 | elongation factor Tu | 680026 |
| OC457_RS01195 | elongation factor Tu | 1935446 |
| OC457_RS13115 | elongation factor Tu | 1935446 |
| B0X70_RS01995 | elongation factor Tu | 171438 |
| B0X70_RS23785 | elongation factor Tu | 171438 |
| PAU_RS01695 | elongation factor Tu | 291112 |
| PAU_RS20980 | elongation factor Tu | 291112 |
| PluTT01m_RS02200 | elongation factor Tu | 141679 |

| | | |
|---|---|---|
| PluTT01m_RS24310 | elongation factor Tu | 141679 |
| VY86_RS05950 | elongation factor Tu | 230089 |
| VY86_RS06965 | elongation factor Tu | 230089 |
| AB182_RS23845 | elongation factor Tu | 1972431 |
| AB182_RS27710 | elongation factor Tu | 1972431 |
| Psal070_RS14760 | elongation factor Tu | 1238 |
| FGL86_RS02490 | elongation factor Tu | 1883414 |
| Q9312_RS09295 | elongation factor Tu | 3070815 |
| Q9312_RS09360 | elongation factor Tu | 3070815 |
| LPH35_RS04285 | elongation factor Tu | 703 |
| LPH35_RS06990 | elongation factor Tu | 703 |
| A8H26_RS02535 | elongation factor Tu | 61647 |
| A8H26_RS05590 | elongation factor Tu | 61647 |
| ACLIJP_RS24030 | elongation factor Tu | 1635171 |
| EL341_RS01190 | elongation factor Tu | 82985 |
| EL341_RS15290 | elongation factor Tu | 82985 |
| SER39006_RS02385 | elongation factor Tu | 104623 |
| SER39006_RS04695 | elongation factor Tu | 104623 |
| QQS39_RS00820 | elongation factor Tu | 3034648 |
| QQS39_RS17125 | elongation factor Tu | 3034648 |
| PH4a_RS05525 | elongation factor Tu | 183417 |
| PH4a_RS07690 | elongation factor Tu | 183417 |
| PMI_RS13790 | elongation factor Tu | 529507 |
| PMI_RS16130 | elongation factor Tu | 529507 |
| H1225_RS14245 | elongation factor Tu | 102862 |
| H1225_RS16545 | elongation factor Tu | 102862 |
| GTH25_RS00620 | elongation factor Tu | 626774 |
| GTH25_RS16485 | elongation factor Tu | 626774 |
| N1711_RS01470 | elongation factor Tu | 585 |
| N1711_RS17370 | elongation factor Tu | 585 |
| PZ638_RS01605 | elongation factor Tu | 3031799 |

| | | |
|---|---|---|
| PZ638_RS19935 | elongation factor Tu | 3031799 |
| DQN80_RS01900 | elongation factor Tu | 333962 |
| DQN80_RS18845 | elongation factor Tu | 333962 |
| V6066_RS01885 | elongation factor Tu | 2027290 |
| V6066_RS19340 | elongation factor Tu | 2027290 |
| JI723_RS02060 | elongation factor Tu | 2923283 |
| JI723_RS19090 | elongation factor Tu | 2923283 |
| I6L80_RS01315 | elongation factor Tu | 587 |
| I6L80_RS03825 | elongation factor Tu | 587 |
| EL244_RS01945 | elongation factor Tu | 158850 |
| EL244_RS17190 | elongation factor Tu | 158850 |
| OO7_RS03100 | elongation factor Tu | 1141660 |
| VQ573_RS01810 | elongation factor Tu | 588 |
| VQ573_RS19520 | elongation factor Tu | 588 |
| ACE2AL_RS01695 | elongation factor Tu | 3342830 |
| ACE2AL_RS19580 | elongation factor Tu | 3342830 |
| ACE2AM_RS01720 | elongation factor Tu | 3342829 |
| ACE2AM_RS18490 | elongation factor Tu | 3342829 |
| QS795_RS01515 | elongation factor Tu | 3053982 |
| QS795_RS16705 | elongation factor Tu | 3053982 |
| FN633_RS00875 | elongation factor Tu | 566 |
| B0W48_RS07480 | elongation factor Tu | 247523 |
| B0W48_RS09620 | elongation factor Tu | 247523 |
| QUE86_RS06730 | elongation factor Tu | 3035452 |
| QUE86_RS09300 | elongation factor Tu | 3035452 |
| PARC_RS00625 | elongation factor Tu | 1117313 |
| PARC_RS01325 | elongation factor Tu | 1117313 |
| ALFOR1_RS00625 | elongation factor Tu | 1314868 |
| ALFOR1_RS15020 | elongation factor Tu | 1314868 |
| FFU37_RS00035 | elongation factor Tu | 77608 |
| FFU37_RS16105 | elongation factor Tu | 77608 |

| | | |
|---|---|---|
| PESP_RS01375 | elongation factor Tu | 1314869 |
| PESP_RS16170 | elongation factor Tu | 1314869 |
| PISS_RS01240 | elongation factor Tu | 152297 |
| PISS_RS15130 | elongation factor Tu | 152297 |
| D0907_RS01980 | elongation factor Tu | 570156 |
| D0907_RS15770 | elongation factor Tu | 570156 |
| FLM44_RS20045 | elongation factor Tu | 43657 |
| FLM44_RS20615 | elongation factor Tu | 43657 |
| CTT31_RS01105 | elongation factor Tu | 184924 |
| CTT31_RS01655 | elongation factor Tu | 184924 |
| PNIG_RS01210 | elongation factor Tu | 28109 |
| PNIG_RS15985 | elongation factor Tu | 28109 |
| PWO49_RS01410 | elongation factor Tu | 2774154 |
| PWO49_RS16510 | elongation factor Tu | 2774154 |
| PP2015_RS00770 | elongation factor Tu | 161398 |
| PP2015_RS01055 | elongation factor Tu | 161398 |
| OM33_RS03830 | elongation factor Tu | 1348114 |
| OM33_RS03900 | elongation factor Tu | 1348114 |
| K5642_RS01070 | elongation factor Tu | 43662 |
| K5642_RS01610 | elongation factor Tu | 43662 |
| PPRY_RS00025 | elongation factor Tu | 1315282 |
| PPRY_RS04515 | elongation factor Tu | 1315282 |
| PPRY_RS07190 | elongation factor Tu | 1315282 |
| PVK79_RS00820 | elongation factor Tu | 2518973 |
| PVK79_RS02850 | elongation factor Tu | 2518973 |
| PSPO_RS00625 | elongation factor Tu | 1117319 |
| PSPO_RS00695 | elongation factor Tu | 1117319 |
| PSHA_RS01120 | elongation factor Tu | 166935 |
| PSHA_RS14280 | elongation factor Tu | 166935 |
| D1819_RS13270 | elongation factor Tu | 314281 |
| D1819_RS13340 | elongation factor Tu | 314281 |

| | | |
|---|---|---|
| PUND_RS01215 | elongation factor Tu | 43660 |
| PUND_RS14505 | elongation factor Tu | 43660 |
| J5X90_RS04650 | elongation factor Tu | 339617 |
| J5X90_RS07060 | elongation factor Tu | 339617 |
| G163CM_RS16760 | elongation factor Tu | 2891570 |
| G163CM_RS19925 | elongation factor Tu | 2891570 |
| PS2015_RS12310 | elongation factor Tu | 1249552 |
| PS2015_RS12370 | elongation factor Tu | 1249552 |
| ELE36_RS15065 | elongation factor Tu | 2511995 |
| ELE36_RS15130 | elongation factor Tu | 2511995 |
| L9B60_RS14710 | elongation factor Tu | 2931382 |
| PA4265 | elongation factor Tu | 208964 |
| PA4277 | elongation factor Tu | 208964 |
| AABM56_RS02040 | elongation factor Tu | 3136739 |
| AABM56_RS02105 | elongation factor Tu | 3136739 |
| GPJ81_RS03135 | elongation factor Tu | 237609 |
| GPJ81_RS03200 | elongation factor Tu | 237609 |
| IF199_RS27005 | elongation factor Tu | 2774460 |
| IF199_RS27070 | elongation factor Tu | 2774460 |
| LU682_RS02240 | elongation factor Tu | 1940621 |
| LU682_RS02305 | elongation factor Tu | 1940621 |
| KSS97_RS26125 | elongation factor Tu | 2842348 |
| KSS97_RS26190 | elongation factor Tu | 2842348 |
| PSYTB_RS25230 | elongation factor Tu | 573066 |
| BLQ27_RS06805 | elongation factor Tu | 219572 |
| HU763_RS02175 | elongation factor Tu | 485870 |
| HU763_RS02240 | elongation factor Tu | 485870 |
| BLQ41_RS07995 | elongation factor Tu | 702115 |
| KSS96_RS25795 | elongation factor Tu | 2842349 |
| PsasTeo6_RS02470 | elongation factor Tu | 2219225 |
| PsasTeo6_RS02535 | elongation factor Tu | 2219225 |

| | | |
|---|---|---|
| BLU37_RS10560 | elongation factor Tu | 53407 |
| JWU58_RS03575 | elongation factor Tu | 2565368 |
| JWU58_RS03640 | elongation factor Tu | 2565368 |
| KSS91_RS25950 | elongation factor Tu | 2842350 |
| BLR69_RS18340 | elongation factor Tu | 47878 |
| NUH87_RS20420 | elongation factor Tu | 226910 |
| NUH87_RS20485 | elongation factor Tu | 226910 |
| PSH92_RS25575 | elongation factor Tu | 2954101 |
| PSH92_RS25640 | elongation factor Tu | 2954101 |
| SBP02_RS17960 | elongation factor Tu | 556533 |
| SBP02_RS18025 | elongation factor Tu | 556533 |
| V6W80_RS21780 | elongation factor Tu | 2993566 |
| V6W80_RS21845 | elongation factor Tu | 2993566 |
| HS968_RS22175 | elongation factor Tu | 2726956 |
| HS968_RS22240 | elongation factor Tu | 2726956 |
| M3M50_RS27530 | elongation factor Tu | 2681983 |
| M3M50_RS27595 | elongation factor Tu | 2681983 |
| ELZ14_RS28195 | elongation factor Tu | 930166 |
| ELZ14_RS28260 | elongation factor Tu | 930166 |
| V6L81_RS05690 | elongation factor Tu | 2316085 |
| RB216_RS26280 | elongation factor Tu | 915099 |
| JGS08_RS02340 | elongation factor Tu | 757414 |
| D3880_RS02730 | elongation factor Tu | 2320867 |
| D3880_RS02795 | elongation factor Tu | 2320867 |
| BLU91_RS23105 | elongation factor Tu | 651740 |
| V4Y05_RS28880 | elongation factor Tu | 333 |
| V4Y05_RS28945 | elongation factor Tu | 333 |
| KGD89_RS22840 | elongation factor Tu | 36746 |
| PcP3B5_RS03645 | elongation factor Tu | 53408 |
| PcP3B5_RS03710 | elongation factor Tu | 53408 |
| POR16_RS22055 | elongation factor Tu | 563797 |

| | | |
|---|---|---|
| AXG94_RS12415 | elongation factor Tu | 47879 |
| AXG94_RS12480 | elongation factor Tu | 47879 |
| PSH97_RS25140 | elongation factor Tu | 2954082 |
| PSH97_RS25205 | elongation factor Tu | 2954082 |
| P7C01_RS10385 | elongation factor Tu | 2497146 |
| P7C01_RS10450 | elongation factor Tu | 2497146 |
| BV82_RS24205 | elongation factor Tu | 1163398 |
| BV82_RS24270 | elongation factor Tu | 1163398 |
| PSEEN_RS02200 | elongation factor Tu | 384676 |
| PSEEN_RS02265 | elongation factor Tu | 384676 |
| HWQ56_RS25900 | elongation factor Tu | 2599595 |
| BLR63_RS25665 | elongation factor Tu | 359110 |
| LUU92_RS25955 | elongation factor Tu | 169669 |
| KSS94_RS02530 | elongation factor Tu | 2842355 |
| KSS94_RS02595 | elongation factor Tu | 2842355 |
| LJU29_RS03195 | elongation factor Tu | 53410 |
| KJY40_RS27270 | elongation factor Tu | 2837969 |
| KJY40_RS27335 | elongation factor Tu | 2837969 |
| CPH89_RS06960 | elongation factor Tu | 294 |
| OZ911_RS02415 | elongation factor Tu | 3233375 |
| OZ911_RS02480 | elongation factor Tu | 3233375 |
| IS178_RS21585 | elongation factor Tu | 296 |
| BLL42_RS17025 | elongation factor Tu | 104087 |
| BLL42_RS17090 | elongation factor Tu | 104087 |
| Q9323_RS19955 | elongation factor Tu | 47880 |
| Q9323_RS20020 | elongation factor Tu | 47880 |
| J0G10_RS27720 | elongation factor Tu | 2815720 |
| J0G10_RS27785 | elongation factor Tu | 2815720 |
| AWU82_RS27515 | elongation factor Tu | 1785145 |
| U771_RS27925 | elongation factor Tu | 3240790 |
| IHQ43_RS26165 | elongation factor Tu | 2774461 |

| | | |
|---|---|---|
| IHQ43_RS26230 | elongation factor Tu | 2774461 |
| BLU52_RS21905 | elongation factor Tu | 1421430 |
| BLU52_RS21970 | elongation factor Tu | 1421430 |
| AABM52_RS27580 | elongation factor Tu | 3136736 |
| AB1N67_RS22235 | elongation factor Tu | 1288410 |
| AB1N67_RS22300 | elongation factor Tu | 1288410 |
| HU739_RS26110 | elongation factor Tu | 2745504 |
| HU739_RS26175 | elongation factor Tu | 2745504 |
| ACJJU9_RS02245 | elongation factor Tu | 1608996 |
| AABM55_RS26315 | elongation factor Tu | 3136738 |
| AABM55_RS26380 | elongation factor Tu | 3136738 |
| KJF94_RS23565 | elongation factor Tu | 3093767 |
| OYW20_RS23375 | elongation factor Tu | 3002903 |
| WIN67_RS27690 | elongation factor Tu | 2942628 |
| WIN67_RS27755 | elongation factor Tu | 2942628 |
| RIN61_RS14065 | elongation factor Tu | 2078786 |
| RIN61_RS14130 | elongation factor Tu | 2078786 |
| HU724_RS25355 | elongation factor Tu | 2745503 |
| HU724_RS25420 | elongation factor Tu | 2745503 |
| LAB08_RS27380 | elongation factor Tu | 2684212 |
| L1P09_RS02110 | elongation factor Tu | 2666183 |
| L1P09_RS02180 | elongation factor Tu | 2666183 |
| PZ739_RS02290 | elongation factor Tu | 2745482 |
| PZ739_RS02360 | elongation factor Tu | 2745482 |
| QYQ93_RS26285 | elongation factor Tu | 2842351 |
| Q1W70_RS13075 | elongation factor Tu | 2762577 |
| Q1W70_RS13140 | elongation factor Tu | 2762577 |
| GVI47_RS03115 | elongation factor Tu | 65741 |
| GVI47_RS03180 | elongation factor Tu | 65741 |
| A8L59_RS23740 | elongation factor Tu | 198620 |
| A8L59_RS23805 | elongation factor Tu | 198620 |

| | | |
|---|---|---|
| DLD99_RS25900 | elongation factor Tu | 1628086 |
| DLD99_RS25965 | elongation factor Tu | 1628086 |
| U5A05_RS02775 | elongation factor Tu | 3043408 |
| QQ992_RS02505 | elongation factor Tu | 485880 |
| QQ992_RS02570 | elongation factor Tu | 485880 |
| SLU14_RS14240 | elongation factor Tu | 2364649 |
| SLU14_RS14305 | elongation factor Tu | 2364649 |
| ACMYQ0_RS21595 | elongation factor Tu | 2448482 |
| ACMYQ0_RS21660 | elongation factor Tu | 2448482 |
| KQP88_RS21870 | elongation factor Tu | 2995658 |
| BLU65_RS08275 | elongation factor Tu | 163011 |
| BLU65_RS08340 | elongation factor Tu | 163011 |
| IF654_RS03195 | elongation factor Tu | 86185 |
| BLU63_RS13720 | elongation factor Tu | 75612 |
| BLU63_RS13785 | elongation factor Tu | 75612 |
| QMK57_RS28030 | elongation factor Tu | 298 |
| PMYSY11_RS03305 | elongation factor Tu | 437900 |
| PMYSY11_RS03370 | elongation factor Tu | 437900 |
| KSS90_RS02410 | elongation factor Tu | 2842354 |
| KSS90_RS02475 | elongation factor Tu | 2842354 |
| BLU23_RS05825 | elongation factor Tu | 183795 |
| BLU23_RS05890 | elongation factor Tu | 183795 |
| ACMDB8_RS24970 | elongation factor Tu | 3060212 |
| HV782_RS26220 | elongation factor Tu | 2745509 |
| HV782_RS26285 | elongation factor Tu | 2745509 |
| CHR29_RS02620 | elongation factor Tu | 76759 |
| CHR29_RS02685 | elongation factor Tu | 76759 |
| ACGJAX_RS03230 | elongation factor Tu | 395599 |
| CSH50_RS23940 | elongation factor Tu | 78327 |
| CSH50_RS24005 | elongation factor Tu | 78327 |
| DQO04_RS24140 | elongation factor Tu | 46679 |

| | | |
|---|---|---|
| G4G71_RS29400 | elongation factor Tu | 95301 |
| G4G71_RS29465 | elongation factor Tu | 95301 |
| KSS95_RS00600 | elongation factor Tu | 2842357 |
| KSS95_RS00665 | elongation factor Tu | 2842357 |
| NK667_RS30665 | elongation factor Tu | 2961896 |
| NK667_RS30730 | elongation factor Tu | 2961896 |
| HU749_RS26900 | elongation factor Tu | 1114970 |
| HU749_RS26965 | elongation factor Tu | 1114970 |
| C4J98_RS23920 | elongation factor Tu | 76758 |
| BLT78_RS15830 | elongation factor Tu | 1392877 |
| BLT78_RS15895 | elongation factor Tu | 1392877 |
| EJA05_RS02470 | elongation factor Tu | 2894079 |
| EJA05_RS02535 | elongation factor Tu | 2894079 |
| VCJ09_RS02415 | elongation factor Tu | 3110772 |
| VCJ09_RS02480 | elongation factor Tu | 3110772 |
| MJP36_RS05295 | elongation factor Tu | 191390 |
| ACK2V3_RS02830 | elongation factor Tu | 2750625 |
| B7D75_RS03345 | elongation factor Tu | 2994495 |
| B7D75_RS03410 | elongation factor Tu | 2994495 |
| U5A08_RS26555 | elongation factor Tu | 3040598 |
| KUA23_RS27070 | elongation factor Tu | 2853159 |
| ACI512_RS32010 | elongation factor Tu | 2899822 |
| HU825_RS07750 | elongation factor Tu | 2746321 |
| HU825_RS07815 | elongation factor Tu | 2746321 |
| DVB73_RS13445 | elongation factor Tu | 70775 |
| DVB73_RS13510 | elongation factor Tu | 70775 |
| H045_RS18010 | elongation factor Tu | 1282356 |
| BLT89_RS00865 | elongation factor Tu | 364197 |
| BLT89_RS00930 | elongation factor Tu | 364197 |
| HU725_RS02220 | elongation factor Tu | 485898 |
| HU725_RS02285 | elongation factor Tu | 485898 |

| | | |
|---|---|---|
| BLU01_RS16010 | elongation factor Tu | 1148509 |
| BLU01_RS16075 | elongation factor Tu | 1148509 |
| PPRCHA0_RS28300 | elongation factor Tu | 1124983 |
| PPRCHA0_RS28365 | elongation factor Tu | 1124983 |
| G5J76_RS05480 | elongation factor Tu | 122355 |
| AABM54_RS23335 | elongation factor Tu | 3136737 |
| AABM54_RS23400 | elongation factor Tu | 3136737 |
| PP4_RS02430 | elongation factor Tu | 1211579 |
| PP4_RS02495 | elongation factor Tu | 1211579 |
| KH389_RS24305 | elongation factor Tu | 2056231 |
| KH389_RS24370 | elongation factor Tu | 2056231 |
| OSC50_RS01740 | elongation factor Tu | 2995174 |
| BLS73_RS15905 | elongation factor Tu | 395598 |
| CRX69_RS24140 | elongation factor Tu | 2045200 |
| LT40_RS18345 | elongation factor Tu | 216142 |
| HRH33_RS24165 | elongation factor Tu | 76760 |
| HU731_RS12745 | elongation factor Tu | 2745514 |
| KP808_RS22625 | elongation factor Tu | 29438 |
| HNQ25_RS04835 | elongation factor Tu | 1691904 |
| HNQ25_RS04900 | elongation factor Tu | 1691904 |
| NN484_RS13390 | elongation factor Tu | 2964670 |
| NN484_RS13455 | elongation factor Tu | 2964670 |
| C4K39_RS20095 | elongation factor Tu | 658629 |
| C4K39_RS20160 | elongation factor Tu | 658629 |
| HU773_RS25155 | elongation factor Tu | 2745512 |
| ACLIMG_RS02325 | elongation factor Tu | 2745494 |
| ACLIMG_RS02390 | elongation factor Tu | 2745494 |
| IMX18_RS22055 | elongation factor Tu | 1940636 |
| IMX18_RS22120 | elongation factor Tu | 1940636 |
| LOY51_RS02495 | elongation factor Tu | 2213015 |
| LOY51_RS02560 | elongation factor Tu | 2213015 |

| | | |
|---|---|---|
| BLT86_RS15645 | elongation factor Tu | 1274359 |
| BLT86_RS15710 | elongation factor Tu | 1274359 |
| PMA3_RS27275 | elongation factor Tu | 1853130 |
| NF676_RS24300 | elongation factor Tu | 2842346 |
| NF676_RS24365 | elongation factor Tu | 2842346 |
| PF1751_RS24985 | elongation factor Tu | 321846 |
| LRO94_RS25650 | elongation factor Tu | 1880678 |
| ABS648_RS03675 | elongation factor Tu | 2731552 |
| ABS648_RS03740 | elongation factor Tu | 2731552 |
| LU688_RS02170 | elongation factor Tu | 1306993 |
| LU688_RS02235 | elongation factor Tu | 1306993 |
| QFX16_RS26145 | elongation factor Tu | 3042029 |
| QFX16_RS26210 | elongation factor Tu | 3042029 |
| C4K01_RS27200 | elongation factor Tu | 47883 |
| KIJ28_RS22065 | elongation factor Tu | 317 |
| MME58_RS23225 | elongation factor Tu | 129140 |
| PSPTO_RS03305 | elongation factor Tu | 223283 |
| DQN55_RS02555 | elongation factor Tu | 47884 |
| HU718_RS27400 | elongation factor Tu | 2745511 |
| HU718_RS27465 | elongation factor Tu | 2745511 |
| HSX14_RS03975 | elongation factor Tu | 2725477 |
| HSX14_RS04040 | elongation factor Tu | 2725477 |
| I9H09_RS03265 | elongation factor Tu | 200454 |
| OU419_RS03255 | elongation factor Tu | 2961893 |
| OU419_RS03320 | elongation factor Tu | 2961893 |
| HU722_RS26615 | elongation factor Tu | 2745518 |
| VRB54_RS21385 | elongation factor Tu | 200450 |
| EXN22_RS24860 | elongation factor Tu | 2518644 |
| EXN22_RS24925 | elongation factor Tu | 2518644 |
| HGP31_RS28215 | elongation factor Tu | 198618 |
| BLV08_RS28155 | elongation factor Tu | 95300 |

| | | |
|---|---|---|
| HU752_RS29400 | elongation factor Tu | 2745495 |
| PverR02_RS28475 | elongation factor Tu | 76761 |
| AOC04_RS22670 | elongation factor Tu | 1788301 |
| EPZ47_RS26790 | elongation factor Tu | 2505979 |
| EPZ47_RS26855 | elongation factor Tu | 2505979 |
| CFBP1590_RS03355 | elongation factor Tu | 33069 |
| ABVN19_RS25365 | elongation factor Tu | 321661 |
| ABVN19_RS25430 | elongation factor Tu | 321661 |
| J7655_RS18430 | elongation factor Tu | 2906062 |
| J7655_RS18495 | elongation factor Tu | 2906062 |
| PSH88_RS03195 | elongation factor Tu | 2954098 |
| PSH88_RS03260 | elongation factor Tu | 2954098 |
| HU772_RS02395 | elongation factor Tu | 2745490 |
| HU772_RS02460 | elongation factor Tu | 2745490 |
| KSS93_RS05655 | elongation factor Tu | 2842356 |
| KSS93_RS05720 | elongation factor Tu | 2842356 |
| BLU46_RS19430 | elongation factor Tu | 515393 |
| HU754_RS27130 | elongation factor Tu | 2745510 |
| HU754_RS27195 | elongation factor Tu | 2745510 |
| MNQ95_RS02595 | elongation factor Tu | 266062 |
| MNQ95_RS02660 | elongation factor Tu | 266062 |
| ACGHX3_RS14345 | elongation factor Tu | 69284 |
| ACGHX3_RS14410 | elongation factor Tu | 69284 |
| IAE60_RS08420 | elongation factor Tu | 128785 |
| IAE60_RS08485 | elongation factor Tu | 128785 |
| DSC_RS11730 | elongation factor Tu | 1045855 |
| DSC_RS11795 | elongation factor Tu | 1045855 |
| MWN52_RS05230 | elongation factor Tu | 2480810 |
| MWN52_RS05295 | elongation factor Tu | 2480810 |
| A3K91_RS02490 | elongation factor Tu | 261164 |
| A3K91_RS12195 | elongation factor Tu | 261164 |

| | | |
|---|---|---|
| PSYC_RS02005 | elongation factor Tu | 259536 |
| PSYC_RS09805 | elongation factor Tu | 259536 |
| JMV70_RS05295 | elongation factor Tu | 256326 |
| JMV70_RS10165 | elongation factor Tu | 256326 |
| PCRYO_RS02215 | elongation factor Tu | 335284 |
| PCRYO_RS11160 | elongation factor Tu | 335284 |
| GYK50_RS01820 | elongation factor Tu | 112002 |
| GYK50_RS10420 | elongation factor Tu | 112002 |
| MN210_RS10125 | elongation factor Tu | 2565531 |
| MN210_RS11165 | elongation factor Tu | 2565531 |
| PING_RS17770 | elongation factor Tu | 357804 |
| J9318_RS00330 | elongation factor Tu | 1266052 |
| J9318_RS00410 | elongation factor Tu | 1266052 |
| J1N51_RS08440 | elongation factor Tu | 2820710 |
| J1N51_RS08520 | elongation factor Tu | 2820710 |
| MNO10_RS15100 | elongation factor Tu | 2703885 |
| MNO10_RS18225 | elongation factor Tu | 2703885 |
| RAHAQ2_RS01645 | elongation factor Tu | 745277 |
| RAHAQ2_RS20450 | elongation factor Tu | 745277 |
| I2123_RS00360 | elongation factor Tu | 58169 |
| I2123_RS03475 | elongation factor Tu | 58169 |
| BV494_RS15125 | elongation factor Tu | 1805933 |
| BV494_RS18350 | elongation factor Tu | 1805933 |
| J9880_RS06185 | elongation factor Tu | 1510570 |
| J9880_RS09325 | elongation factor Tu | 1510570 |
| Q1L34_RS02065 | elongation factor Tu | 54291 |
| Q1L34_RS25010 | elongation factor Tu | 54291 |
| MC50_RS06185 | elongation factor Tu | 575 |
| MC50_RS09625 | elongation factor Tu | 575 |
| HCK03_RS01905 | elongation factor Tu | 577 |
| HCK03_RS25040 | elongation factor Tu | 577 |

| | | |
|---|---|---|
| REIFOR_RS15600 | elongation factor Tu | 1336806 |
| EK374_RS01415 | elongation factor Tu | 2498451 |
| EK374_RS01480 | elongation factor Tu | 2498451 |
| LRK52_RS04845 | elongation factor Tu | 666685 |
| LRK52_RS04910 | elongation factor Tu | 666685 |
| AAHF87_RS06980 | elongation factor Tu | 3077936 |
| KX723_RS02095 | elongation factor Tu | 2856608 |
| ACICP9_RS04800 | elongation factor Tu | 3066237 |
| AAHH40_RS05530 | elongation factor Tu | 3066239 |
| AAHI99_RS05450 | elongation factor Tu | 3077937 |
| AACL18_RS05790 | elongation factor Tu | 3066238 |
| FEE42_RS07545 | elongation factor Tu | 1796644 |
| FEE42_RS08565 | elongation factor Tu | 1796644 |
| IHV77_RS05925 | elongation factor Tu | 2778911 |
| IHV77_RS06025 | elongation factor Tu | 2778911 |
| O1V65_RS17375 | elongation factor Tu | 1646377 |
| O1V65_RS20185 | elongation factor Tu | 1646377 |
| C2869_RS09865 | elongation factor Tu | 2172099 |
| C2869_RS09930 | elongation factor Tu | 2172099 |
| SDE_RS04905 | elongation factor Tu | 203122 |
| SDE_RS04920 | elongation factor Tu | 203122 |
| DW349_RS02515 | elongation factor Tu | 2161747 |
| DW349_RS02580 | elongation factor Tu | 2161747 |
| FBQ74_RS01925 | elongation factor Tu | 2572577 |
| FBQ74_RS02575 | elongation factor Tu | 2572577 |
| EZV72_RS02315 | elongation factor Tu | 914153 |
| EZV72_RS15640 | elongation factor Tu | 914153 |
| IT774_RS01350 | elongation factor Tu | 2785918 |
| IT774_RS01980 | elongation factor Tu | 2785918 |
| D0Y50_RS16525 | elongation factor Tu | 2303538 |
| D0Y50_RS17390 | elongation factor Tu | 2303538 |

| | | |
|---|---|---|
| HMF8227_RS02105 | elongation factor Tu | 2183582 |
| HMF8227_RS02180 | elongation factor Tu | 2183582 |
| O4546_RS01380 | elongation factor Tu | 1908198 |
| O4546_RS12210 | elongation factor Tu | 1908198 |
| N7E60_RS01415 | elongation factor Tu | 334715 |
| N7E60_RS13215 | elongation factor Tu | 334715 |
| SBG_RS15865 | elongation factor Tu | 218493 |
| SBG_RS18960 | elongation factor Tu | 218493 |
| STM3445 | protein chain elongation | 99287 |
| STM4146 | protein chain elongation | 99287 |
| A8O29_RS02440 | elongation factor Tu | 1851514 |
| A8O29_RS21410 | elongation factor Tu | 1851514 |
| A359_RS01650 | elongation factor Tu | 1199245 |
| A35E_RS02110 | elongation factor Tu | 134287 |
| TREMTM_RS01435 | elongation factor Tu | 1835721 |
| AAY24_RS14415 | elongation factor Tu | 1543721 |
| AAY24_RS14480 | elongation factor Tu | 1543721 |
| SAQUA_RS01670 | elongation factor Tu | 1737515 |
| SAQUA_RS22700 | elongation factor Tu | 1737515 |
| KHA73_RS00985 | elongation factor Tu | 42906 |
| KHA73_RS21950 | elongation factor Tu | 42906 |
| CKW09_RS01270 | elongation factor Tu | 61651 |
| CKW09_RS22575 | elongation factor Tu | 61651 |
| WN53_RS05865 | elongation factor Tu | 47917 |
| WN53_RS09865 | elongation factor Tu | 47917 |
| M495_RS22530 | elongation factor Tu | 1346614 |
| M495_RS24565 | elongation factor Tu | 1346614 |
| QRD25_RS01220 | elongation factor Tu | 615 |
| QRD25_RS22060 | elongation factor Tu | 615 |
| E5D60_RS01125 | elongation factor Tu | 458197 |
| E5D60_RS23155 | elongation factor Tu | 458197 |

| | | |
|---|---|---|
| V8N38_RS01145 | elongation factor Tu | 2703794 |
| V8N38_RS23635 | elongation factor Tu | 2703794 |
| SERAS9_RS01185 | elongation factor Tu | 768492 |
| SERAS9_RS23230 | elongation factor Tu | 768492 |
| QOT07_RS01295 | elongation factor Tu | 28151 |
| QOT07_RS23225 | elongation factor Tu | 28151 |
| EL336_RS01300 | elongation factor Tu | 137545 |
| EL336_RS23430 | elongation factor Tu | 137545 |
| FO014_RS08555 | elongation factor Tu | 2597702 |
| FO014_RS11530 | elongation factor Tu | 2597702 |
| I6G83_RS05225 | elongation factor Tu | 61652 |
| I6G83_RS08345 | elongation factor Tu | 61652 |
| VOT19_RS01180 | elongation factor Tu | 3020826 |
| VOT19_RS22080 | elongation factor Tu | 3020826 |
| ATE40_RS19125 | elongation factor Tu | 2741499 |
| ATE40_RS22110 | elongation factor Tu | 2741499 |
| SYMBAF_RS14630 | elongation factor Tu | 138074 |
| SYMBAF_RS16210 | elongation factor Tu | 138074 |
| J0X03_RS01530 | elongation factor Tu | 300181 |
| J0X03_RS22650 | elongation factor Tu | 300181 |
| K0H61_RS00695 | elongation factor Tu | 2864212 |
| K0H61_RS00755 | elongation factor Tu | 2864212 |
| K0H80_RS00755 | elongation factor Tu | 2864203 |
| K0H80_RS00815 | elongation factor Tu | 2864203 |
| E1N14_RS01145 | elongation factor Tu | 38313 |
| E1N14_RS01205 | elongation factor Tu | 38313 |
| K0J45_RS00760 | elongation factor Tu | 2864209 |
| K0J45_RS00820 | elongation factor Tu | 2864209 |
| SAMA_RS01055 | elongation factor Tu | 326297 |
| SAMA_RS01115 | elongation factor Tu | 326297 |
| JYB87_RS17435 | elongation factor Tu | 2814294 |

| | | |
|---|---|---|
| JYB87_RS17495 | elongation factor Tu | 2814294 |
| SBAL678_RS24320 | elongation factor Tu | 693973 |
| SBAL678_RS24380 | elongation factor Tu | 693973 |
| K5621_RS00960 | elongation factor Tu | 1987584 |
| K5621_RS01020 | elongation factor Tu | 1987584 |
| WKR14_RS00975 | elongation factor Tu | 558541 |
| WKR14_RS19675 | elongation factor Tu | 558541 |
| JYB84_RS00955 | elongation factor Tu | 2814292 |
| JYB84_RS01015 | elongation factor Tu | 2814292 |
| EYR25_RS19845 | elongation factor Tu | 256839 |
| SDEN_RS00910 | elongation factor Tu | 318161 |
| FPK91_RS14530 | elongation factor Tu | 238836 |
| FPK91_RS14590 | elongation factor Tu | 238836 |
| FM038_RS24065 | elongation factor Tu | 2593656 |
| FM038_RS24125 | elongation factor Tu | 2593656 |
| SFRI_RS00725 | elongation factor Tu | 318167 |
| SFRI_RS00785 | elongation factor Tu | 318167 |
| FJ709_RS18795 | elongation factor Tu | 2590884 |
| FJ709_RS18855 | elongation factor Tu | 2590884 |
| QPX86_RS00940 | elongation factor Tu | 3050637 |
| QPX86_RS01000 | elongation factor Tu | 3050637 |
| SHAL_RS21385 | elongation factor Tu | 458817 |
| SHAL_RS21445 | elongation factor Tu | 458817 |
| K0H81_RS00755 | elongation factor Tu | 2864204 |
| K0H81_RS00815 | elongation factor Tu | 2864204 |
| K8B83_RS02905 | elongation factor Tu | 1738770 |
| K8B83_RS02965 | elongation factor Tu | 1738770 |
| SJ2017_RS20280 | elongation factor Tu | 93973 |
| SJ2017_RS20340 | elongation factor Tu | 93973 |
| STH12_RS19370 | elongation factor Tu | 1965282 |
| STH12_RS19430 | elongation factor Tu | 1965282 |

| | | |
|---|---|---|
| JQC75_RS00945 | elongation factor Tu | 1173586 |
| JQC75_RS01005 | elongation factor Tu | 1173586 |
| EGC82_RS01860 | elongation factor Tu | 150120 |
| EGC82_RS01920 | elongation factor Tu | 150120 |
| SHEW_RS00760 | elongation factor Tu | 323850 |
| SHEW_RS00820 | elongation factor Tu | 323850 |
| K0H60_RS01225 | elongation factor Tu | 2864211 |
| K0H60_RS01285 | elongation factor Tu | 2864211 |
| CFF01_RS00715 | elongation factor Tu | 260364 |
| CFF01_RS00775 | elongation factor Tu | 260364 |
| EXU30_RS11085 | elongation factor Tu | 2520507 |
| EXU30_RS11145 | elongation factor Tu | 2520507 |
| K0I73_RS00675 | elongation factor Tu | 2864208 |
| K0I73_RS00735 | elongation factor Tu | 2864208 |
| RA178_RS01025 | elongation factor Tu | 2726434 |
| RA178_RS01085 | elongation factor Tu | 2726434 |
| SO_RS01030 | elongation factor Tu | 211586 |
| SO_RS01090 | elongation factor Tu | 211586 |
| SPEA_RS00935 | elongation factor Tu | 398579 |
| SPEA_RS00995 | elongation factor Tu | 398579 |
| SWP_RS08880 | elongation factor Tu | 225849 |
| SWP_RS08940 | elongation factor Tu | 225849 |
| FH971_RS19415 | elongation factor Tu | 2588449 |
| FH971_RS19475 | elongation factor Tu | 2588449 |
| EGC80_RS05730 | elongation factor Tu | 2487742 |
| EGC80_RS05790 | elongation factor Tu | 2487742 |
| FM037_RS26850 | elongation factor Tu | 2593655 |
| FM037_RS26910 | elongation factor Tu | 2593655 |
| K0I62_RS00690 | elongation factor Tu | 2864206 |
| K0I62_RS00750 | elongation factor Tu | 2864206 |
| JEZ96_RS00770 | elongation factor Tu | 24 |

| | | |
|---|---|---|
| JEZ96_RS00830 | elongation factor Tu | 24 |
| K0I63_RS00765 | elongation factor Tu | 2864207 |
| K0I63_RS00825 | elongation factor Tu | 2864207 |
| JYB85_RS01005 | elongation factor Tu | 2814293 |
| JYB85_RS01065 | elongation factor Tu | 2814293 |
| K0I31_RS00760 | elongation factor Tu | 2864205 |
| K0I31_RS00820 | elongation factor Tu | 2864205 |
| SVI_RS19695 | elongation factor Tu | 637905 |
| SVI_RS19755 | elongation factor Tu | 637905 |
| SWOO_RS24065 | elongation factor Tu | 392500 |
| SWOO_RS24125 | elongation factor Tu | 392500 |
| KXJ80_RS22265 | elongation factor Tu | 332186 |
| KXJ80_RS22325 | elongation factor Tu | 332186 |
| RGE70_RS17965 | elongation factor Tu | 2999068 |
| RGE70_RS18025 | elongation factor Tu | 2999068 |
| KDN34_RS00885 | elongation factor Tu | 2829809 |
| KDN34_RS00945 | elongation factor Tu | 2829809 |
| K0H63_RS01020 | elongation factor Tu | 2864213 |
| K0H63_RS01080 | elongation factor Tu | 2864213 |
| HUZ68_RS00525 | elongation factor Tu | 622 |
| HUZ68_RS20880 | elongation factor Tu | 622 |
| SF3357 | elongation factor Tu | 198214 |
| SF4053 | elongation factor Tu | 198214 |
| C1P89_RS18075 | elongation factor Tu | 624 |
| C1P89_RS21965 | elongation factor Tu | 624 |
| EBL_RS01330 | elongation factor Tu | 630626 |
| EBL_RS18060 | elongation factor Tu | 630626 |
| KFZ77_RS01640 | elongation factor Tu | 1505757 |
| KFZ77_RS18075 | elongation factor Tu | 1505757 |
| ACGHTT_RS02355 | elongation factor Tu | 1435348 |
| ACGHTT_RS02420 | elongation factor Tu | 1435348 |

| | | |
|---|---|---|
| HBA_RS00345 | elongation factor Tu | 1929246 |
| SGP1_RS01090 | elongation factor Tu | 343509 |
| SGP1_RS20870 | elongation factor Tu | 343509 |
| AB4W47_RS00460 | elongation factor Tu | 2994493 |
| DCD74_RS01460 | elongation factor Tu | 2698682 |
| DCD74_RS01525 | elongation factor Tu | 2698682 |
| G4Y78_RS25480 | elongation factor Tu | 2683272 |
| G4Y78_RS25545 | elongation factor Tu | 2683272 |
| LMH63_RS14860 | elongation factor Tu | 2182432 |
| LMH63_RS14925 | elongation factor Tu | 2182432 |
| BBH56_RS07535 | elongation factor Tu | 1855875 |
| BBH56_RS07600 | elongation factor Tu | 1855875 |
| SPISAL_RS07600 | elongation factor Tu | 1260251 |
| I6N98_RS14380 | elongation factor Tu | 2794344 |
| I6N98_RS14445 | elongation factor Tu | 2794344 |
| NCG89_RS07595 | elongation factor Tu | 1748242 |
| NCG89_RS07660 | elongation factor Tu | 1748242 |
| F0P95_RS07440 | elongation factor Tu | 128780 |
| F0P95_RS07505 | elongation factor Tu | 128780 |
| PDM28_RS03910 | elongation factor Tu | 3003272 |
| PDM28_RS03975 | elongation factor Tu | 3003272 |
| K5L94_RS03845 | elongation factor Tu | 2871169 |
| K5L94_RS03910 | elongation factor Tu | 2871169 |
| RI032_RS05110 | elongation factor Tu | 86188 |
| RI032_RS05175 | elongation factor Tu | 86188 |
| DAIF1_RS04010 | elongation factor Tu | 2045451 |
| DAIF1_RS04080 | elongation factor Tu | 2045451 |
| AB9L18_RS16470 | elongation factor Tu | 2045214 |
| AB9L18_RS16535 | elongation factor Tu | 2045214 |
| CKW06_RS04150 | elongation factor Tu | 40324 |
| CKW06_RS04220 | elongation factor Tu | 40324 |

| | | |
|---|---|---|
| PDM29_RS11415 | elongation factor Tu | 3003271 |
| PDM29_RS11480 | elongation factor Tu | 3003271 |
| Q5Z11_RS05160 | elongation factor Tu | 3055080 |
| Q5Z11_RS05225 | elongation factor Tu | 3055080 |
| DX03_RS15825 | elongation factor Tu | 216778 |
| DX03_RS15890 | elongation factor Tu | 216778 |
| SriyCCGPB3_RS03720 | elongation factor Tu | 2859893 |
| SriyCCGPB3_RS03785 | elongation factor Tu | 2859893 |
| AB9H29_RS03665 | elongation factor Tu | 2860290 |
| AB9H29_RS03730 | elongation factor Tu | 2860290 |
| ACG33_RS05170 | elongation factor Tu | 465721 |
| ACG33_RS05235 | elongation factor Tu | 465721 |
| CL52_RS17410 | elongation factor Tu | 1123016 |
| CL52_RS17475 | elongation factor Tu | 1123016 |
| LLJ08_RS17290 | elongation factor Tu | 203192 |
| LLJ08_RS17355 | elongation factor Tu | 203192 |
| UIB01_RS17840 | elongation factor Tu | 3022791 |
| UIB01_RS17905 | elongation factor Tu | 3022791 |
| I6G33_RS12090 | elongation factor Tu | 2968968 |
| I6G33_RS12155 | elongation factor Tu | 2968968 |
| I6G34_RS08670 | elongation factor Tu | 2968969 |
| I6G34_RS08735 | elongation factor Tu | 2968969 |
| PszF2a_RS03420 | elongation factor Tu | 316 |
| PszF2a_RS03485 | elongation factor Tu | 316 |
| BLW61_RS04410 | elongation factor Tu | 271420 |
| BLW61_RS04475 | elongation factor Tu | 271420 |
| JKJ12_RS00470 | elongation factor Tu | 83771 |
| JKJ12_RS00545 | elongation factor Tu | 83771 |
| SCL_RS11320 | elongation factor Tu | 1620215 |
| SCL_RS11385 | elongation factor Tu | 1620215 |
| EL386_RS11845 | elongation factor Tu | 1811807 |

| | | |
|---|---|---|
| EL386_RS11910 | elongation factor Tu | 1811807 |
| SVA_RS16495 | elongation factor Tu | 1675686 |
| SVA_RS16560 | elongation factor Tu | 1675686 |
| EP379_RS03290 | elongation factor Tu | 1972068 |
| EP379_RS03355 | elongation factor Tu | 1972068 |
| K6K13_RS21340 | elongation factor Tu | 2871826 |
| K6K13_RS22565 | elongation factor Tu | 2871826 |
| N4264_RS22020 | elongation factor Tu | 2976241 |
| N4264_RS22085 | elongation factor Tu | 2976241 |
| A7K98_RS00535 | elongation factor Tu | 53336 |
| A7K98_RS18590 | elongation factor Tu | 53336 |
| H5715_RS00820 | elongation factor Tu | 2731755 |
| H5647_RS19825 | elongation factor Tu | 2731756 |
| WKI13_RS17165 | elongation factor Tu | 2426 |
| HUF19_RS16115 | elongation factor Tu | 2742796 |
| HUF19_RS16125 | elongation factor Tu | 2742796 |
| HUF19_RS16190 | elongation factor Tu | 2742796 |
| TOL_RS16670 | elongation factor Tu | 1298593 |
| TOL_RS16735 | elongation factor Tu | 1298593 |
| TOL_RS16745 | elongation factor Tu | 1298593 |
| SG35_RS26635 | elongation factor Tu | 485447 |
| SG35_RS26705 | elongation factor Tu | 485447 |
| H3N35_RS25650 | elongation factor Tu | 485448 |
| H3N35_RS25720 | elongation factor Tu | 485448 |
| SG34_RS01790 | elongation factor Tu | 137584 |
| SG34_RS01855 | elongation factor Tu | 137584 |
| QUD85_RS01595 | elongation factor Tu | 349064 |
| QUD85_RS02135 | elongation factor Tu | 349064 |
| ACAY00_RS13015 | elongation factor Tu | 3242415 |
| ACAY00_RS13035 | elongation factor Tu | 3242415 |
| ACAY00_RS13105 | elongation factor Tu | 3242415 |

| | | |
|---|---|---|
| QUE03_RS02395 | elongation factor Tu | 2054316 |
| QUE03_RS02465 | elongation factor Tu | 2054316 |
| LT090_RS01275 | elongation factor Tu | 1763536 |
| LT090_RS02025 | elongation factor Tu | 1763536 |
| RI844_RS16280 | elongation factor Tu | 3065649 |
| RI844_RS16350 | elongation factor Tu | 3065649 |
| RI844_RS16430 | elongation factor Tu | 3065649 |
| QUE72_RS01780 | elongation factor Tu | 3030492 |
| QUE72_RS01850 | elongation factor Tu | 3030492 |
| ACAX20_RS13570 | elongation factor Tu | 3242416 |
| ACAX20_RS13605 | elongation factor Tu | 3242416 |
| ACAX20_RS13690 | elongation factor Tu | 3242416 |
| RI845_RS02295 | elongation factor Tu | 3065648 |
| RI845_RS02365 | elongation factor Tu | 3065648 |
| RI845_RS02435 | elongation factor Tu | 3065648 |
| QUD79_RS15835 | elongation factor Tu | 1230533 |
| QUD79_RS15900 | elongation factor Tu | 1230533 |
| ACAY30_RS13165 | elongation factor Tu | 1523392 |
| ACAY30_RS13210 | elongation factor Tu | 1523392 |
| ACAY30_RS13285 | elongation factor Tu | 1523392 |
| RGQ13_RS17375 | elongation factor Tu | 3065647 |
| RGQ13_RS17445 | elongation factor Tu | 3065647 |
| RGQ13_RS17515 | elongation factor Tu | 3065647 |
| QUE09_RS15550 | elongation factor Tu | 1759089 |
| QUE09_RS15625 | elongation factor Tu | 1759089 |
| TSUB_RS01245 | elongation factor Tu | 1891207 |
| TSUB_RS15315 | elongation factor Tu | 1891207 |
| E6P07_RS07290 | elongation factor Tu | 316276 |
| E6P07_RS07355 | elongation factor Tu | 316276 |
| FHQ07_RS04865 | elongation factor Tu | 2202149 |
| FHQ07_RS04930 | elongation factor Tu | 2202149 |

| | | |
|---|---|---|
| H9L17_RS12025 | elongation factor Tu | 215691 |
| H9L17_RS12090 | elongation factor Tu | 215691 |
| LIW09_RS02505 | elongation factor Tu | 2884874 |
| LIW09_RS02570 | elongation factor Tu | 2884874 |
| TVNIR_RS12140 | elongation factor Tu | 1255043 |
| TVNIR_RS12205 | elongation factor Tu | 1255043 |
| THITH_RS13850 | elongation factor Tu | 713585 |
| THITH_RS13915 | elongation factor Tu | 713585 |
| TGR7_RS11645 | elongation factor Tu | 396588 |
| TGR7_RS11710 | elongation factor Tu | 396588 |
| TVD_RS02285 | elongation factor Tu | 106634 |
| TVD_RS02350 | elongation factor Tu | 106634 |
| LT988_RS20375 | elongation factor Tu | 521689 |
| LT988_RS20440 | elongation factor Tu | 521689 |
| THIVI_RS20225 | elongation factor Tu | 765911 |
| THIVI_RS20290 | elongation factor Tu | 765911 |
| THIMO_RS13640 | elongation factor Tu | 765912 |
| THIMO_RS13705 | elongation factor Tu | 765912 |
| EP181_RS01425 | elongation factor Tu | 2211106 |
| EP181_RS01490 | elongation factor Tu | 2211106 |
| L6421_RS01695 | elongation factor Tu | 2791037 |
| L6421_RS01760 | elongation factor Tu | 2791037 |
| D9T12_RS01765 | elongation factor Tu | 2267253 |
| D9T12_RS01830 | elongation factor Tu | 2267253 |
| NR989_RS01725 | elongation factor Tu | 2949997 |
| NR989_RS01790 | elongation factor Tu | 2949997 |
| FE785_RS01490 | elongation factor Tu | 2580412 |
| FE785_RS01555 | elongation factor Tu | 2580412 |
| HQN79_RS01910 | elongation factor Tu | 2739063 |
| HQN79_RS01975 | elongation factor Tu | 2739063 |
| THIAE_RS00665 | elongation factor Tu | 717772 |

| | | |
|---|---|---|
| THIAE_RS00730 | elongation factor Tu | 717772 |
| THICY_RS00570 | elongation factor Tu | 717773 |
| THICY_RS00635 | elongation factor Tu | 717773 |
| AKN87_RS09925 | elongation factor Tu | 1697053 |
| AKN87_RS09990 | elongation factor Tu | 1697053 |
| Thiofri_RS08810 | elongation factor Tu | 631362 |
| Thiofri_RS08875 | elongation factor Tu | 631362 |
| Thiosp_RS16885 | elongation factor Tu | 2952932 |
| Thiosp_RS16950 | elongation factor Tu | 2952932 |
| Thiowin_RS08225 | elongation factor Tu | 77007 |
| Thiowin_RS08300 | elongation factor Tu | 77007 |
| B1781_RS20495 | elongation factor Tu | 1973002 |
| B1781_RS20560 | elongation factor Tu | 1973002 |
| HRR27_RS11310 | elongation factor Tu | 2675054 |
| HRR27_RS11375 | elongation factor Tu | 2675054 |
| THMIRH_RS02255 | elongation factor Tu | 2675053 |
| THMIRH_RS02320 | elongation factor Tu | 2675053 |
| J9253_RS19710 | elongation factor Tu | 2891210 |
| J9253_RS19775 | elongation factor Tu | 2891210 |
| HMY34_RS06765 | elongation factor Tu | 2735563 |
| L2Y54_RS07240 | elongation factor Tu | 96472 |
| L2Y54_RS07305 | elongation factor Tu | 96472 |
| TOLA_RS14205 | elongation factor Tu | 595494 |
| TOLA_RS14290 | elongation factor Tu | 595494 |
| QMG90_RS01605 | elongation factor Tu | 379893 |
| QMG90_RS20805 | elongation factor Tu | 379893 |
| RHO15_RS02750 | elongation factor Tu | 3074108 |
| RHO15_RS02800 | elongation factor Tu | 3074108 |
| GJQ55_RS01010 | elongation factor Tu | 2661630 |
| GJQ55_RS01075 | elongation factor Tu | 2661630 |
| GJQ55_RS01085 | elongation factor Tu | 2661630 |

| | | |
|---|---|---|
| A6B43_RS02770 | elongation factor Tu | 1443036 |
| A6B43_RS05505 | elongation factor Tu | 1443036 |
| OCV29_RS15810 | elongation factor Tu | 92172 |
| OCV29_RS16055 | elongation factor Tu | 92172 |
| VA249_RS13065 | elongation factor Tu | 1074311 |
| VA249_RS13840 | elongation factor Tu | 1074311 |
| GFB47_RS00495 | elongation factor Tu | 2662262 |
| GFB47_RS10545 | elongation factor Tu | 2662262 |
| NAL94_RS08655 | elongation factor Tu | 663 |
| NAL94_RS09400 | elongation factor Tu | 663 |
| VEA_RS02855 | elongation factor Tu | 150340 |
| VEA_RS03625 | elongation factor Tu | 150340 |
| VCA1004_RS01305 | elongation factor Tu | 1891186 |
| VCA1004_RS11000 | elongation factor Tu | 1891186 |
| QUF19_RS15455 | elongation factor Tu | 3035453 |
| QUF19_RS16150 | elongation factor Tu | 3035453 |
| FIV01_RS01280 | elongation factor Tu | 2587862 |
| FIV01_RS01525 | elongation factor Tu | 2587862 |
| OCU36_RS12360 | elongation factor Tu | 446374 |
| OCU36_RS12980 | elongation factor Tu | 446374 |
| GT360_RS12910 | elongation factor Tu | 1481923 |
| GT360_RS13170 | elongation factor Tu | 1481923 |
| OCV30_RS14015 | elongation factor Tu | 693153 |
| OCV30_RS14670 | elongation factor Tu | 693153 |
| BS333_RS12660 | elongation factor Tu | 512649 |
| BS333_RS13320 | elongation factor Tu | 512649 |
| IHV80_RS14735 | elongation factor Tu | 2777577 |
| IHV80_RS15385 | elongation factor Tu | 2777577 |
| OCU35_RS00580 | elongation factor Tu | 553239 |
| OCU35_RS12090 | elongation factor Tu | 553239 |
| C1N50_RS00810 | elongation factor Tu | 680 |

| | | |
|---|---|---|
| C1N50_RS01610 | elongation factor Tu | 680 |
| PG915_RS01540 | elongation factor Tu | 3016528 |
| PG915_RS15410 | elongation factor Tu | 3016528 |
| OCV52_RS14225 | elongation factor Tu | 170679 |
| OCV52_RS14870 | elongation factor Tu | 170679 |
| F0316_RS12170 | elongation factor Tu | 666 |
| F0316_RS12425 | elongation factor Tu | 666 |
| EWA64_RS09585 | elongation factor Tu | 1763883 |
| EWA64_RS10290 | elongation factor Tu | 1763883 |
| OCU46_RS00585 | elongation factor Tu | 413401 |
| OCU46_RS12690 | elongation factor Tu | 413401 |
| P6988_RS04170 | elongation factor Tu | 190893 |
| P6988_RS04435 | elongation factor Tu | 190893 |
| OCV39_RS01390 | elongation factor Tu | 1328770 |
| OCV39_RS13615 | elongation factor Tu | 1328770 |
| OC193_RS14165 | elongation factor Tu | 246167 |
| OC193_RS14810 | elongation factor Tu | 246167 |
| FAZ90_RS00790 | elongation factor Tu | 47951 |
| FAZ90_RS01445 | elongation factor Tu | 47951 |
| FR741_RS00895 | elongation factor Tu | 50719 |
| FR741_RS01650 | elongation factor Tu | 50719 |
| AAGA51_RS13735 | elongation factor Tu | 685 |
| AAGA51_RS13995 | elongation factor Tu | 685 |
| OCV36_RS00765 | elongation factor Tu | 2100116 |
| OCV36_RS01410 | elongation factor Tu | 2100116 |
| AL536_RS41930 | elongation factor Tu | 676 |
| AL536_RS42190 | elongation factor Tu | 676 |
| OCV50_RS13035 | elongation factor Tu | 212667 |
| OCV50_RS13650 | elongation factor Tu | 212667 |
| FIU11_RS03980 | elongation factor Tu | 29494 |
| FIU11_RS04235 | elongation factor Tu | 29494 |

| | | |
|---|---|---|
| OCU78_RS12960 | elongation factor Tu | 552386 |
| OCU78_RS13585 | elongation factor Tu | 552386 |
| OCU28_RS00980 | elongation factor Tu | 190897 |
| OCU28_RS11115 | elongation factor Tu | 190897 |
| Vgang_RS00390 | elongation factor Tu | 2077090 |
| Vgang_RS10865 | elongation factor Tu | 2077090 |
| MKS89_RS13920 | elongation factor Tu | 687 |
| MKS89_RS14145 | elongation factor Tu | 687 |
| MID13_RS14055 | elongation factor Tu | 296199 |
| MID13_RS14710 | elongation factor Tu | 296199 |
| QMY49_RS15425 | elongation factor Tu | 669 |
| QMY49_RS16190 | elongation factor Tu | 669 |
| C1S74_RS11660 | elongation factor Tu | 1534743 |
| C1S74_RS12515 | elongation factor Tu | 1534743 |
| OCU48_RS00580 | elongation factor Tu | 413402 |
| OCU48_RS12045 | elongation factor Tu | 413402 |
| Pcarn_RS01090 | elongation factor Tu | 1481914 |
| Pcarn_RS13015 | elongation factor Tu | 1481914 |
| NP165_RS01230 | elongation factor Tu | 1824638 |
| NP165_RS01515 | elongation factor Tu | 1824638 |
| C1S73_RS11495 | elongation factor Tu | 1280002 |
| C1S73_RS12325 | elongation factor Tu | 1280002 |
| OCV24_RS00725 | elongation factor Tu | 170673 |
| OCV24_RS01460 | elongation factor Tu | 170673 |
| OCU74_RS01625 | elongation factor Tu | 474394 |
| OCU74_RS01850 | elongation factor Tu | 474394 |
| QYQ96_RS12020 | elongation factor Tu | 1481663 |
| QYQ96_RS12275 | elongation factor Tu | 1481663 |
| OOK74_RS01040 | elongation factor Tu | 2993714 |
| OOK74_RS01290 | elongation factor Tu | 2993714 |
| PN96_RS00085 | elongation factor Tu | 1219067 |

| | | |
|---|---|---|
| PN96_RS14945 | elongation factor Tu | 1219067 |
| OCU38_RS01065 | elongation factor Tu | 278860 |
| OCU38_RS12065 | elongation factor Tu | 278860 |
| QMG91_RS00900 | elongation factor Tu | 28173 |
| QMG91_RS01655 | elongation factor Tu | 28173 |
| KNV97_RS17240 | elongation factor Tu | 2841925 |
| KNV97_RS17495 | elongation factor Tu | 2841925 |
| A9237_RS09085 | elongation factor Tu | 696485 |
| A9237_RS09860 | elongation factor Tu | 696485 |
| OCU30_RS01100 | elongation factor Tu | 1918946 |
| OCU30_RS01350 | elongation factor Tu | 1918946 |
| R4537_RS18325 | elongation factor Tu | 650003 |
| R4537_RS18570 | elongation factor Tu | 650003 |
| VP_RS13600 | elongation factor Tu | 223926 |
| VP_RS14450 | elongation factor Tu | 223926 |
| OCV27_RS01400 | elongation factor Tu | 62763 |
| OCV27_RS01640 | elongation factor Tu | 62763 |
| vsple_RS12705 | elongation factor Tu | 28169 |
| vsple_RS13310 | elongation factor Tu | 28169 |
| LDO37_RS01085 | elongation factor Tu | 104609 |
| LDO37_RS01990 | elongation factor Tu | 104609 |
| OCV12_RS14620 | elongation factor Tu | 198832 |
| OCV12_RS15265 | elongation factor Tu | 198832 |
| GZN30_RS11495 | elongation factor Tu | 265668 |
| GZN30_RS11770 | elongation factor Tu | 265668 |
| OCV11_RS01355 | elongation factor Tu | 1123496 |
| OCV11_RS01620 | elongation factor Tu | 1123496 |
| CCZ37_RS01015 | elongation factor Tu | 2025808 |
| CCZ37_RS01260 | elongation factor Tu | 2025808 |
| OC443_RS01690 | elongation factor Tu | 1117707 |
| OC443_RS01950 | elongation factor Tu | 1117707 |

| | | |
|---|---|---|
| OCU56_RS01030 | elongation factor Tu | 413403 |
| OCU56_RS12225 | elongation factor Tu | 413403 |
| OCV37_RS13650 | elongation factor Tu | 398736 |
| OCV37_RS13875 | elongation factor Tu | 398736 |
| BSZ04_RS09610 | elongation factor Tu | 190895 |
| BSZ04_RS24555 | elongation factor Tu | 190895 |
| OCU42_RS13625 | elongation factor Tu | 184755 |
| OCU42_RS13850 | elongation factor Tu | 184755 |
| VRUMOI_RS01410 | elongation factor Tu | 76258 |
| VRUMOI_RS12305 | elongation factor Tu | 76258 |
| RJD38_RS04890 | elongation factor Tu | 45658 |
| RJD38_RS05145 | elongation factor Tu | 45658 |
| OCU60_RS15865 | elongation factor Tu | 1918945 |
| OCU60_RS16100 | elongation factor Tu | 1918945 |
| LTQ54_RS09590 | elongation factor Tu | 29497 |
| LTQ54_RS10250 | elongation factor Tu | 29497 |
| K08M4_RS00795 | elongation factor Tu | 3034029 |
| K08M4_RS01455 | elongation factor Tu | 3034029 |
| OCV44_RS00795 | elongation factor Tu | 212663 |
| OCV44_RS01465 | elongation factor Tu | 212663 |
| OCU50_RS12900 | elongation factor Tu | 1194427 |
| OCU50_RS13540 | elongation factor Tu | 1194427 |
| JCM16456_RS01280 | elongation factor Tu | 1435069 |
| JCM16456_RS01540 | elongation factor Tu | 1435069 |
| IX91_RS13460 | elongation factor Tu | 1051646 |
| IX91_RS13725 | elongation factor Tu | 1051646 |
| AOT11_RS07765 | elongation factor Tu | 1219061 |
| AOT11_RS08455 | elongation factor Tu | 1219061 |
| EAE30_RS06215 | elongation factor Tu | 2479546 |
| EAE30_RS06460 | elongation factor Tu | 2479546 |
| G5S32_RS13260 | elongation factor Tu | 2711221 |

| | | |
|---|---|---|
| G5S32_RS13545 | elongation factor Tu | 2711221 |
| SR894_RS00575 | elongation factor Tu | 115551 |
| GYM47_RS00930 | elongation factor Tu | 2609667 |
| KUO20_RS00450 | elongation factor Tu | 2852117 |
| AB4Y29_RS04280 | elongation factor Tu | 502821 |
| B5495_RS07925 | elongation factor Tu | 29571 |
| HZS52_RS03445 | elongation factor Tu | 664683 |
| IR195_RS00895 | elongation factor Tu | 44935 |
| WM2015_RS11595 | elongation factor Tu | 1579979 |
| WM2015_RS11695 | elongation factor Tu | 1579979 |
| WIGMOR_RS01185 | elongation factor Tu | 1142511 |
| RIN69_RS01905 | elongation factor Tu | 92490 |
| RIN69_RS21010 | elongation factor Tu | 92490 |
| BA177_RS04820 | elongation factor Tu | 1548547 |
| BA177_RS04890 | elongation factor Tu | 1548547 |
| XaFJ1_RS02170 | elongation factor Tu | 29447 |
| XaFJ1_RS02240 | elongation factor Tu | 29447 |
| KHN93_RS05030 | elongation factor Tu | 195709 |
| KHN93_RS05095 | elongation factor Tu | 195709 |
| ACDH77_RS04600 | elongation factor Tu | 487832 |
| ACDH77_RS04665 | elongation factor Tu | 487832 |
| AB3X10_RS17715 | elongation factor Tu | 3160842 |
| AB3X10_RS17780 | elongation factor Tu | 3160842 |
| AD14011_RS16165 | elongation factor Tu | 359385 |
| AD14011_RS16230 | elongation factor Tu | 359385 |
| RZV17_RS04945 | elongation factor Tu | 1885674 |
| RZV17_RS05010 | elongation factor Tu | 1885674 |
| XCSCFBP4642_RS0100480 | elongation factor Tu | 1219375 |
| K8O61_RS15235 | elongation factor Tu | 152263 |
| K8O61_RS15300 | elongation factor Tu | 152263 |
| AMD14_RS05375 | elongation factor Tu | 611301 |

| | | |
|---|---|---|
| AMD14_RS05440 | elongation factor Tu | 611301 |
| EBN15_RS03735 | elongation factor Tu | 56453 |
| EBN15_RS03800 | elongation factor Tu | 56453 |
| NYR95_RS05475 | elongation factor Tu | 743699 |
| NYR95_RS05540 | elongation factor Tu | 743699 |
| H7A86_RS04665 | elongation factor Tu | 2259622 |
| H7A86_RS04730 | elongation factor Tu | 2259622 |
| XeaCFBP3836p_RS05000 | elongation factor Tu | 359387 |
| XeaCFBP3836p_RS05065 | elongation factor Tu | 359387 |
| PD885_RS14650 | elongation factor Tu | 48664 |
| PD885_RS14720 | elongation factor Tu | 48664 |
| KHF85_RS04030 | elongation factor Tu | 134874 |
| KHF85_RS04095 | elongation factor Tu | 134874 |
| PG878_RS04175 | elongation factor Tu | 3003247 |
| PG878_RS04240 | elongation factor Tu | 3003247 |
| XJ27_RS11055 | elongation factor Tu | 56454 |
| XJ27_RS11120 | elongation factor Tu | 56454 |
| FZ025_RS03850 | elongation factor Tu | 56455 |
| FZ025_RS03920 | elongation factor Tu | 56455 |
| NYR97_RS13975 | elongation factor Tu | 2775159 |
| NYR97_RS14040 | elongation factor Tu | 2775159 |
| Q7W82_RS07340 | elongation factor Tu | 2912242 |
| Q7W82_RS07405 | elongation factor Tu | 2912242 |
| AB3X07_RS17890 | elongation factor Tu | 3160841 |
| AB3X07_RS17955 | elongation factor Tu | 3160841 |
| M3O59_RS04720 | elongation factor Tu | 1843581 |
| M3O59_RS04785 | elongation factor Tu | 1843581 |
| XOCgx_RS18000 | elongation factor Tu | 129394 |
| XOCgx_RS18065 | elongation factor Tu | 129394 |
| FLG15_RS04425 | elongation factor Tu | 92828 |
| FLG15_RS04490 | elongation factor Tu | 92828 |

| | | |
|---|---|---|
| M0D43_RS06245 | elongation factor Tu | 2053930 |
| M0D43_RS06310 | elongation factor Tu | 2053930 |
| QN245_RS16785 | elongation factor Tu | 3046274 |
| QN245_RS16850 | elongation factor Tu | 3046274 |
| RAB71_RS17650 | elongation factor Tu | 56458 |
| RAB71_RS17715 | elongation factor Tu | 56458 |
| RAB70_RS20100 | elongation factor Tu | 2650745 |
| RAB70_RS20165 | elongation factor Tu | 2650745 |
| AB3X08_RS17210 | elongation factor Tu | 3160839 |
| AB3X08_RS17275 | elongation factor Tu | 3160839 |
| G4Q83_RS14875 | elongation factor Tu | 56464 |
| G4Q83_RS14940 | elongation factor Tu | 56464 |
| ISN38_RS06385 | elongation factor Tu | 487909 |
| ISN38_RS06450 | elongation factor Tu | 487909 |
| KWO_RS05075 | elongation factor Tu | 1094184 |
| KWO_RS05140 | elongation factor Tu | 1094184 |
| BJD12_RS17735 | elongation factor Tu | 925775 |
| BJD12_RS17800 | elongation factor Tu | 925775 |
| PXH59_RS14205 | elongation factor Tu | 3026967 |
| PXH59_RS16310 | elongation factor Tu | 3026967 |
| HGO23_RS01470 | elongation factor Tu | 290110 |
| HGO23_RS18625 | elongation factor Tu | 290110 |
| XDD1_RS01320 | elongation factor Tu | 351671 |
| XDD1_RS16875 | elongation factor Tu | 351671 |
| QL128_RS01665 | elongation factor Tu | 351672 |
| QL128_RS20045 | elongation factor Tu | 351672 |
| A9255_RS14565 | elongation factor Tu | 351679 |
| A9255_RS16650 | elongation factor Tu | 351679 |
| XNC1_RS01285 | elongation factor Tu | 406817 |
| XPG1_RS14305 | elongation factor Tu | 1354304 |
| XPG1_RS16105 | elongation factor Tu | 1354304 |

| | | |
|---|---|---|
| ACE1SE_RS01320 | elongation factor Tu | 3085177 |
| ACE1SE_RS17975 | elongation factor Tu | 3085177 |
| PXH81_RS10875 | elongation factor Tu | 2371 |
| PXH81_RS10940 | elongation factor Tu | 2371 |
| PLS229_RS10320 | elongation factor Tu | 1444770 |
| PLS229_RS10385 | elongation factor Tu | 1444770 |
| AT01_RS10745 | elongation factor Tu | 1453495 |
| AT01_RS13515 | elongation factor Tu | 1453495 |
| N0H69_RS01230 | elongation factor Tu | 2890317 |
| N0H69_RS20485 | elongation factor Tu | 2890317 |
| QJR59_RS18385 | elongation factor Tu | 634 |
| QJR59_RS18685 | elongation factor Tu | 634 |
| F0T03_RS19800 | elongation factor Tu | 2607663 |
| F0T03_RS20090 | elongation factor Tu | 2607663 |
| Y11_RS15485 | elongation factor Tu | 930944 |
| Y11_RS16525 | elongation factor Tu | 930944 |
| PL78_RS06945 | elongation factor Tu | 935293 |
| PL78_RS11375 | elongation factor Tu | 935293 |
| D5F51_RS01515 | elongation factor Tu | 2339259 |
| D5F51_RS20645 | elongation factor Tu | 2339259 |
| FOC38_RS00560 | elongation factor Tu | 631 |
| FOC38_RS20185 | elongation factor Tu | 631 |
| HRD69_RS03740 | elongation factor Tu | 349967 |
| HRD69_RS06835 | elongation factor Tu | 349967 |
| A1122_RS04095 | elongation factor Tu | 1035377 |
| A1122_RS07140 | elongation factor Tu | 1035377 |
| EL213_RS01310 | elongation factor Tu | 633 |
| EL213_RS19390 | elongation factor Tu | 633 |
| DXZ79_RS18750 | elongation factor Tu | 1604335 |
| DXZ79_RS19050 | elongation factor Tu | 1604335 |
| CH64_RS11580 | elongation factor Tu | 29485 |

| | | |
|---|---|---|
| CH64_RS14195 | elongation factor Tu | 29485 |
| YR821_RS01110 | elongation factor Tu | 29486 |
| YR821_RS15525 | elongation factor Tu | 29486 |
| BF17_RS05695 | elongation factor Tu | 367190 |
| BF17_RS08785 | elongation factor Tu | 367190 |
| ACIOG0_RS01130 | elongation factor Tu | 2890318 |
| ACIOG0_RS20320 | elongation factor Tu | 2890318 |
| AZF00_RS02070 | elongation factor Tu | 1470434 |
| AZF00_RS02135 | elongation factor Tu | 1470434 |
| AN401_RS01265 | elongation factor Tu | 347534 |
| AN401_RS01710 | elongation factor Tu | 347534 |
| IPMB12_RS03465 | elongation factor Tu | 1635327 |
| IPMB12_RS03510 | elongation factor Tu | 1635327 |
| ZBT109_RS01455 | elongation factor Tu | 33074 |
| GAU_RS04455 | elongation factor Tu | 379066 |
| HKW67_RS10525 | elongation factor Tu | 2732249 |
| GEMMAAP_RS05145 | elongation factor Tu | 1379270 |
| J421_RS13890 | elongation factor Tu | 861299 |
| Strain318_RS03575 | elongation factor Tu | 3062599 |
| QUD34_RS13340 | elongation factor Tu | 2927975 |
| R2J75_RS01190 | elongation factor Tu | 2927980 |
| R2J76_RS01200 | elongation factor Tu | 2927979 |
| J3U87_RS10665 | elongation factor Tu | 2818388 |
| TTHT_RS06840 | elongation factor Tu | 981385 |
| TTHT_RS06910 | elongation factor Tu | 981385 |
| HPTL_RS01935 | elongation factor Tu | 297 |
| HPTL_RS02000 | elongation factor Tu | 297 |
| IALB_RS04545 | elongation factor Tu | 945713 |
| MROS_RS01140 | elongation factor Tu | 1191523 |
| L21SP4_RS08480 | elongation factor Tu | 1307763 |
| EPA93_RS30540 | elongation factor Tu | 2509675 |

| | | |
|---|---|---|
| EPA93_RS31820 | elongation factor Tu | 2509675 |
| PQO03_RS12860 | elongation factor Tu | 1658616 |
| U7230_RS01505 | elongation factor Tu | 3109565 |
| VLY81_RS12130 | elongation factor Tu | 3109564 |
| LIP_RS15460 | elongation factor Tu | 1555112 |
| LIP_RS15530 | elongation factor Tu | 1555112 |
| QOL44_RS04610 | elongation factor Tu | 591154 |
| IT6_RS02400 | elongation factor Tu | 2795386 |
| MINF_RS03555 | elongation factor Tu | 481448 |
| kam1_RS04040 | elongation factor Tu | 1202785 |
| EXC59_RS04605 | elongation factor Tu | 264636 |
| ACL_RS00920 | elongation factor Tu | 441768 |
| BN854_RS06360 | elongation factor Tu | 1318466 |
| X271_RS02530 | elongation factor Tu | 1427984 |
| O3I44_RS01120 | elongation factor Tu | 3004105 |
| QN326_RS02090 | elongation factor Tu | 85620 |
| AADV48_RS02105 | elongation factor Tu | 35780 |
| LFWB_RS01375 | elongation factor Tu | 35773 |
| ATP_RS02165 | elongation factor Tu | 37692 |
| RS022_RS03240 | elongation factor Tu | 399025 |
| PSOL_RS01355 | elongation factor Tu | 69896 |
| CWO85_RS02445 | elongation factor Tu | 135727 |
| EFREU_RS00525 | elongation factor Tu | 74700 |
| EXC62_RS02755 | elongation factor Tu | 29552 |
| MPAN_RS01640 | elongation factor Tu | 2735436 |
| CK556_RS03510 | elongation factor Tu | 216427 |
| MCOLE_RS03355 | elongation factor Tu | 324078 |
| CS528_RS03500 | elongation factor Tu | 2149 |
| MFL_RS03310 | elongation factor Tu | 265311 |
| CP520_RS03510 | elongation factor Tu | 81460 |
| EMELA_RS04075 | elongation factor Tu | 81459 |

| | | |
|---|---|---|
| CXP39_RS00525 | elongation factor Tu | 225999 |
| MTABA_RS03500 | elongation factor Tu | 219745 |
| ABDB96_RS00205 | elongation factor Tu | 572419 |
| FRW55_RS03695 | elongation factor Tu | 519450 |
| ACM0IS_RS00425 | elongation factor Tu | 547986 |
| NMG68_RS01105 | elongation factor Tu | 2963128 |
| MCAP_RS00780 | elongation factor Tu | 340047 |
| ACJA23_RS03595 | elongation factor Tu | 2106 |
| NX779_RS00460 | elongation factor Tu | 51364 |
| MCRO_RS00210 | elongation factor Tu | 512564 |
| MFERI15220_RS00710 | elongation factor Tu | 1179777 |
| ACPYPZ_RS03480 | elongation factor Tu | 525886 |
| NV226_RS02600 | elongation factor Tu | 292461 |
| MSB_RS00990 | elongation factor Tu | 880447 |
| HLA87_RS03265 | elongation factor Tu | 754515 |
| HLA92_RS02575 | elongation factor Tu | 754516 |
| I7639_RS03670 | elongation factor Tu | 40477 |
| FG904_RS02865 | elongation factor Tu | 353852 |
| MR07_RS00895 | elongation factor Tu | 1415773 |
| PRV_RS02425 | elongation factor Tu | 1403316 |
| HGG69_RS00640 | elongation factor Tu | 754517 |
| HGG64_RS00510 | elongation factor Tu | 2726117 |
| D2843_RS03340 | elongation factor Tu | 2123 |
| Q8852_RS04135 | elongation factor Tu | 512749 |
| EGN60_RS01910 | elongation factor Tu | 538220 |
| MSU_RS01420 | elongation factor Tu | 768700 |
| JS510_RS01550 | elongation factor Tu | 547987 |
| H3143_RS01100 | elongation factor Tu | 1612150 |
| WEN_RS01505 | elongation factor Tu | 1197325 |
| WEN_RS02680 | elongation factor Tu | 1197325 |
| MYE_RS00505 | elongation factor Tu | 743967 |

| | | |
|---|---|---|
| BN853_RS01605 | elongation factor Tu | 61635 |
| SALLE_RS01420 | elongation factor Tu | 216931 |
| SAPIS_RS04860 | elongation factor Tu | 1276258 |
| SCANT_RS04925 | elongation factor Tu | 362837 |
| SCHIN_RS05970 | elongation factor Tu | 216932 |
| SCHRY_RS00290 | elongation factor Tu | 1276227 |
| SCITRI_RS00445 | elongation factor Tu | 2133 |
| SCLAR_RS00855 | elongation factor Tu | 2139 |
| SCORR_RS04890 | elongation factor Tu | 216934 |
| SCULI_RS05130 | elongation factor Tu | 1276246 |
| SDIMI_RS04125 | elongation factor Tu | 1276221 |
| AAHM96_RS00310 | elongation factor Tu | 3066306 |
| AACK93_RS07065 | elongation factor Tu | 3077930 |
| ACQH20_RS01075 | elongation factor Tu | 3139329 |
| ABNT63_RS01030 | elongation factor Tu | 3066315 |
| AAHH39_RS02590 | elongation factor Tu | 3066288 |
| AACL03_RS03675 | elongation factor Tu | 3066316 |
| NPX79_RS02195 | elongation factor Tu | 2967972 |
| AACL02_RS00380 | elongation factor Tu | 3066307 |
| AAHJ00_RS07315 | elongation factor Tu | 3066281 |
| AAHM97_RS04785 | elongation factor Tu | 3066276 |
| AACK92_RS05425 | elongation factor Tu | 3066291 |
| AACL04_RS04950 | elongation factor Tu | 3066278 |
| AACL15_RS05165 | elongation factor Tu | 3066279 |
| AAHM82_RS11825 | elongation factor Tu | 3066282 |
| AACL24_RS02275 | elongation factor Tu | 3066310 |
| AACL01_RS03095 | elongation factor Tu | 3066286 |
| AACL11_RS02405 | elongation factor Tu | 3066299 |
| AAHF53_RS04600 | elongation factor Tu | 3066292 |
| AAHM84_RS04910 | elongation factor Tu | 3066290 |
| AACL10_RS04460 | elongation factor Tu | 3066322 |

| | | |
|---|---|---|
| AAHM88_RS07055 | elongation factor Tu | 3066283 |
| AACK86_RS05825 | elongation factor Tu | 3066300 |
| ACOBWH_RS00535 | elongation factor Tu | 2004463 |
| AACK85_RS04565 | elongation factor Tu | 3066305 |
| AACK95_RS06840 | elongation factor Tu | 3066319 |
| AACK81_RS00325 | elongation factor Tu | 3066320 |
| AACK97_RS07245 | elongation factor Tu | 3066297 |
| ACQH26_RS10090 | elongation factor Tu | 3139324 |
| AACK94_RS06745 | elongation factor Tu | 3066312 |
| AAHM98_RS07855 | elongation factor Tu | 3066284 |
| I5Q57_RS05330 | elongation factor Tu | 2792084 |
| AAHN02_RS01530 | elongation factor Tu | 3066302 |
| AAHM79_RS00240 | elongation factor Tu | 3066325 |
| ACICQF_RS01230 | elongation factor Tu | 3077933 |
| AAHM85_RS05145 | elongation factor Tu | 3066285 |
| AACL07_RS03525 | elongation factor Tu | 3066289 |
| AACK87_RS01185 | elongation factor Tu | 3066314 |
| AACK89_RS07305 | elongation factor Tu | 3066301 |
| NL276_RS04115 | elongation factor Tu | 2886375 |
| AAHM76_RS03985 | elongation factor Tu | 1209220 |
| AACK78_RS00205 | elongation factor Tu | 3066287 |
| ACE38S_RS01235 | elongation factor Tu | 3139326 |
| ACQH25_RS00390 | elongation factor Tu | 3139327 |
| AACK91_RS05310 | elongation factor Tu | 3066303 |
| AACL22_RS05540 | elongation factor Tu | 3066321 |
| ACQH24_RS00465 | elongation factor Tu | 3139328 |
| AACL05_RS00315 | elongation factor Tu | 3066294 |
| AACK83_RS00700 | elongation factor Tu | 3066326 |
| ACFEL5_RS07475 | elongation factor Tu | 3066295 |
| AACK96_RS00415 | elongation factor Tu | 3066296 |
| AAHM89_RS00530 | elongation factor Tu | 3066293 |

| | | |
|---|---|---|
| ACFEL4_RS09400 | elongation factor Tu | 3139323 |
| AAHM78_RS01265 | elongation factor Tu | 3066313 |
| SERIO_RS00410 | elongation factor Tu | 315358 |
| SFLOR_RS05510 | elongation factor Tu | 1336749 |
| SGLAD_RS04975 | elongation factor Tu | 2143 |
| SHELI_RS05515 | elongation factor Tu | 216938 |
| SIXOD_RS00555 | elongation factor Tu | 1276219 |
| SKUN_RS07430 | elongation factor Tu | 273035 |
| SLITO_RS05300 | elongation factor Tu | 216942 |
| SATRI_RS00310 | elongation factor Tu | 2144 |
| SMONO_RS03960 | elongation factor Tu | 1336748 |
| SDAV_RS05490 | elongation factor Tu | 1276259 |
| SPLAT_RS03150 | elongation factor Tu | 1276218 |
| MSROBK_RS10135 | elongation factor Tu | 2138 |
| SSABA_RS00955 | elongation factor Tu | 1276257 |
| SSYRP_RS00370 | elongation factor Tu | 1276229 |
| STABA_RS05455 | elongation factor Tu | 324079 |
| STAIW_RS05160 | elongation factor Tu | 1276220 |
| STURON_RS05340 | elongation factor Tu | 216946 |
| SLY_RS01950 | elongation factor Tu | 980422 |
| ELUMI_RS01345 | elongation factor Tu | 214888 |
| ESOMN_RS00565 | elongation factor Tu | 215578 |
| A2CP1_RS10130 | elongation factor Tu | 455488 |
| A2CP1_RS11900 | elongation factor Tu | 455488 |
| AMOR_RS27335 | elongation factor Tu | 2918170 |
| AMOR_RS29040 | elongation factor Tu | 2918170 |
| AMPC_RS03120 | elongation factor Tu | 2918171 |
| AMPC_RS04510 | elongation factor Tu | 2918171 |
| AA314_RS17610 | elongation factor Tu | 48 |
| AA314_RS23170 | elongation factor Tu | 48 |
| AA314_RS32055 | elongation factor Tu | 48 |

| | | |
|---|---|---|
| KYC5002_RS19040 | elongation factor Tu | 83451 |
| KYC5002_RS29100 | elongation factor Tu | 83451 |
| KYC5002_RS35035 | elongation factor Tu | 83451 |
| COCOR_RS15175 | elongation factor Tu | 1144275 |
| COCOR_RS24830 | elongation factor Tu | 1144275 |
| MYMAC_RS15175 | elongation factor Tu | 1189310 |
| MYMAC_RS16410 | elongation factor Tu | 1189310 |
| MEBOL_RS00490 | elongation factor Tu | 1294270 |
| MEBOL_RS33095 | elongation factor Tu | 1294270 |
| JY572_RS03535 | elongation factor Tu | 2813577 |
| JY572_RS09000 | elongation factor Tu | 2813577 |
| MYSTI_RS17465 | elongation factor Tu | 1278073 |
| MYSTI_RS23000 | elongation factor Tu | 1278073 |
| MXAN_RS14870 | elongation factor Tu | 246197 |
| MXAN_RS15985 | elongation factor Tu | 246197 |
| LZC94_RS06195 | elongation factor Tu | 2741071 |
| LZC94_RS06420 | elongation factor Tu | 2741071 |
| LZC95_RS00965 | elongation factor Tu | 2905690 |
| LZC95_RS38960 | elongation factor Tu | 2905690 |
| LZC95_RS39145 | elongation factor Tu | 2905690 |
| LVJ94_RS38855 | elongation factor Tu | 2741070 |
| LVJ94_RS39050 | elongation factor Tu | 2741070 |
| A176_RS17555 | elongation factor Tu | 1297742 |
| A176_RS18980 | elongation factor Tu | 1297742 |
| JY651_RS19380 | elongation factor Tu | 2813578 |
| JY651_RS21320 | elongation factor Tu | 2813578 |
| AKJ08_RS09670 | elongation factor Tu | 1391653 |
| AKJ08_RS10170 | elongation factor Tu | 1391653 |
| ACFER_RS01365 | elongation factor Tu | 591001 |
| ACFER_RS09890 | elongation factor Tu | 591001 |
| NQ562_RS09655 | elongation factor Tu | 187327 |

| | | |
|---|---|---|
| NQ562_RS10245 | elongation factor Tu | 187327 |
| Dia5BBH33_RS06775 | elongation factor Tu | 2582419 |
| Dia5BBH33_RS07625 | elongation factor Tu | 2582419 |
| DHT67_RS09220 | elongation factor Tu | 2161821 |
| DHT67_RS10085 | elongation factor Tu | 2161821 |
| NW747_RS01080 | elongation factor Tu | 487173 |
| NW747_RS05180 | elongation factor Tu | 487173 |
| GXM21_RS01975 | elongation factor Tu | 437897 |
| CKV65_RS07340 | elongation factor Tu | 158847 |
| C6362_RS05270 | elongation factor Tu | 1064535 |
| C6362_RS05430 | elongation factor Tu | 1064535 |
| ACGTIC_RS12600 | elongation factor Tu | 1675036 |
| ACGTIC_RS12945 | elongation factor Tu | 1675036 |
| DKB62_RS03930 | elongation factor Tu | 2144175 |
| DKB62_RS04285 | elongation factor Tu | 2144175 |
| NXG62_RS07895 | elongation factor Tu | 349094 |
| NXG62_RS08275 | elongation factor Tu | 349094 |
| MAMMFC1_RS09965 | elongation factor Tu | 1930071 |
| MAMMFC1_RS10040 | elongation factor Tu | 1930071 |
| F8M42_RS01540 | elongation factor Tu | 865 |
| PFJ30894_RS05100 | elongation factor Tu | 33025 |
| PFJ30894_RS05265 | elongation factor Tu | 33025 |
| M5E81_RS08370 | elongation factor Tu | 626940 |
| M5E81_RS08815 | elongation factor Tu | 626940 |
| P3F81_RS09320 | elongation factor Tu | 3054208 |
| QU667_RS09135 | elongation factor Tu | 135079 |
| SELSP_RS07535 | elongation factor Tu | 546271 |
| H1B31_RS04990 | elongation factor Tu | 2754044 |
| SPACI_RS02180 | elongation factor Tu | 1123286 |
| SPACI_RS02255 | elongation factor Tu | 1123286 |
| SATMO3_RS01995 | elongation factor Tu | 204936 |

| | | |
|---|---|---|
| SATMO3_RS02075 | elongation factor Tu | 204936 |
| SCACP_RS02105 | elongation factor Tu | 3076075 |
| SCACP_RS02180 | elongation factor Tu | 3076075 |
| SMWOGL2_RS22870 | elongation factor Tu | 112901 |
| SMWOGL2_RS22940 | elongation factor Tu | 112901 |
| SOV_RS01910 | elongation factor Tu | 1123288 |
| SOV_RS01995 | elongation factor Tu | 1123288 |
| SPX_RS02110 | elongation factor Tu | 2376 |
| SPX_RS02185 | elongation factor Tu | 2376 |
| SRRS_RS02120 | elongation factor Tu | 357999 |
| SRRS_RS02205 | elongation factor Tu | 357999 |
| SPSIL_RS02035 | elongation factor Tu | 1123289 |
| SPSIL_RS02110 | elongation factor Tu | 1123289 |
| SPSPH_RS02230 | elongation factor Tu | 1337886 |
| SPSPH_RS02305 | elongation factor Tu | 1337886 |
| SPTER_RS00210 | elongation factor Tu | 2377 |
| SPTER_RS00280 | elongation factor Tu | 2377 |
| EL171_RS07910 | elongation factor Tu | 39778 |
| EL171_RS07995 | elongation factor Tu | 39778 |
| VEIT17_RS07480 | elongation factor Tu | 2682456 |
| VEIT17_RS07570 | elongation factor Tu | 2682456 |
| KJS32_RS07540 | elongation factor Tu | 2682455 |
| KJS32_RS07630 | elongation factor Tu | 2682455 |
| CKV63_RS07690 | elongation factor Tu | 29466 |
| CKV63_RS07780 | elongation factor Tu | 29466 |
| CKV62_RS07815 | elongation factor Tu | 248315 |
| CKV62_RS07905 | elongation factor Tu | 248315 |
| ER308_RS13255 | elongation factor Tu | 1670831 |
| ER308_RS13345 | elongation factor Tu | 1670831 |
| ELR47_RS03025 | elongation factor Tu | 1670830 |
| ELR47_RS03115 | elongation factor Tu | 1670830 |

| | | |
|---|---|---|
| DVS28_RS22025 | elongation factor Tu | 1608957 |
| DVS28_RS22205 | elongation factor Tu | 1608957 |
| QML71_RS12565 | elongation factor Tu | 1323948 |
| PP769_RS06920 | elongation factor Tu | 3020900 |
| PP769_RS06990 | elongation factor Tu | 3020900 |
| NITINOP_RS07900 | elongation factor Tu | 1715989 |
| NITINOP_RS07965 | elongation factor Tu | 1715989 |
| PQG83_RS18040 | elongation factor Tu | 3020899 |
| PQG83_RS18110 | elongation factor Tu | 3020899 |
| LFE_RS04450 | elongation factor Tu | 1162668 |
| LFE_RS04515 | elongation factor Tu | 1162668 |
| NSJP_RS10570 | elongation factor Tu | 1325564 |
| NSJP_RS10635 | elongation factor Tu | 1325564 |
| NITMOv2_RS05525 | elongation factor Tu | 42253 |
| NITMOv2_RS05590 | elongation factor Tu | 42253 |
| EZS29_RS03435 | elongation factor Tu | 2493639 |
| AXG55_RS03425 | elongation factor Tu | 1915309 |
| Spiro2_RS03080 | elongation factor Tu | 495820 |
| K1X11_RS09725 | elongation factor Tu | 2866162 |
| GA004_RS16645 | elongation factor Tu | 2684113 |
| CAKA_RS13435 | elongation factor Tu | 583355 |
| SH580_RS04615 | elongation factor Tu | 3092156 |
| CKA38_RS06820 | elongation factor Tu | 1796921 |
| K0B96_RS03620 | elongation factor Tu | 2862869 |
| Verru16B_RS02430 | elongation factor Tu | 1838286 |
| CMV30_RS05230 | elongation factor Tu | 2576891 |
| OTER_RS01195 | elongation factor Tu | 452637 |
| RZN69_RS19940 | elongation factor Tu | 3080537 |
| PXH66_RS01905 | elongation factor Tu | 2909285 |
| OH491_RS20265 | elongation factor Tu | 1184151 |
| STSP2_RS14830 | elongation factor Tu | 1936003 |

| | | |
|---|---|---|
| IPV69_RS10510 | elongation factor Tu | 2807512 |
| IPV69_RS23820 | elongation factor Tu | 2807512 |
| SMSP2_RS08445 | elongation factor Tu | 1851148 |
| Pan265_RS00750 | elongation factor Tu | 2527982 |
| PSMK_RS03955 | elongation factor Tu | 1142394 |
| KS4_RS02895 | elongation factor Tu | 2528020 |
| L21SP3_RS10745 | elongation factor Tu | 1940790 |
| STSP1_RS04965 | elongation factor Tu | 1941349 |
| HG15A2_RS23230 | elongation factor Tu | 1930276 |
| Pan181_RS21010 | elongation factor Tu | 2527972 |
| CA12_RS08630 | elongation factor Tu | 2527989 |
| ETAA8_RS27005 | elongation factor Tu | 2528021 |
| OJF2_RS04580 | elongation factor Tu | 406548 |
| Q31a_RS04815 | elongation factor Tu | 2527968 |
| Q31a_RS19335 | elongation factor Tu | 2527968 |
| Spa11_RS21840 | elongation factor Tu | 2528022 |
| Pan97_RS23435 | elongation factor Tu | 2527984 |
| Pr1d_RS04855 | elongation factor Tu | 1400387 |
| V22_RS08965 | elongation factor Tu | 2528027 |
| Pan44_RS07355 | elongation factor Tu | 2527983 |
| Mal65_RS06015 | elongation factor Tu | 2527996 |
| Mal65_RS07900 | elongation factor Tu | 2527996 |
| ACLV6H_RS02320 | elongation factor Tu | 2724528 |
| ACLV6H_RS26210 | elongation factor Tu | 2724528 |
| FTUN_RS32485 | elongation factor Tu | 2774151 |
| Fuma_RS28710 | elongation factor Tu | 1891926 |
| SOIL9_RS29060 | elongation factor Tu | 1210884 |
| SOIL9_RS32810 | elongation factor Tu | 1210884 |
| Pan241w_RS02660 | elongation factor Tu | 2527973 |
| Pan241w_RS26810 | elongation factor Tu | 2527973 |
| Pan161_RS27685 | elongation factor Tu | 2527971 |

| | | |
|---|---|---|
| V144x_RS02565 | elongation factor Tu | 2527964 |
| V144x_RS25310 | elongation factor Tu | 2527964 |
| F1728_RS11730 | elongation factor Tu | 2608982 |
| F1728_RS16560 | elongation factor Tu | 2608982 |
| Enr17x_RS03140 | elongation factor Tu | 2527976 |
| Enr17x_RS26955 | elongation factor Tu | 2527976 |
| GmarT_RS27095 | elongation factor Tu | 122 |
| Enr10x_RS02800 | elongation factor Tu | 2527978 |
| Enr10x_RS27270 | elongation factor Tu | 2527978 |
| ISOP_RS00605 | elongation factor Tu | 575540 |
| I41_RS26220 | elongation factor Tu | 2528024 |
| PLANPX_RS26295 | elongation factor Tu | 2650471 |
| Pla8534_RS30630 | elongation factor Tu | 2528010 |
| PX52LOC_RS10265 | elongation factor Tu | 2598579 |
| PX52LOC_RS10320 | elongation factor Tu | 2598579 |
| Mal4_RS20265 | elongation factor Tu | 2528026 |
| MFFC18_RS18550 | elongation factor Tu | 980251 |
| BSF38_RS13495 | elongation factor Tu | 1387353 |
| Pla175_RS23890 | elongation factor Tu | 2528009 |
| SH412_RS05700 | elongation factor Tu | 3064265 |
| Spb1_RS05600 | elongation factor Tu | 2528019 |
| PLIM_RS02285 | elongation factor Tu | 521674 |
| Pla110_RS16940 | elongation factor Tu | 2528007 |
| RB_RS16140 | elongation factor Tu | 243090 |
| FF011L_RS04730 | elongation factor Tu | 1930274 |
| UC8_RS06900 | elongation factor Tu | 980254 |
| Poly24_RS21635 | elongation factor Tu | 2528017 |
| CA51_RS21155 | elongation factor Tu | 2527990 |
| EC9_RS22605 | elongation factor Tu | 1930277 |
| PLABR_RS10125 | elongation factor Tu | 756272 |
| K227x_RS24140 | elongation factor Tu | 1930273 |

| | | |
|---|---|---|
| SINAC_RS02490 | elongation factor Tu | 886293 |
| Mal15_RS05095 | elongation factor Tu | 2795974 |
| Mal15_RS30750 | elongation factor Tu | 2795974 |
| Enr13x_RS02235 | elongation factor Tu | 2527979 |
| Enr13x_RS28865 | elongation factor Tu | 2527979 |
| Pla52nx_RS13395 | elongation factor Tu | 2528005 |
| Pla52nx_RS20585 | elongation factor Tu | 2528005 |
| Pan189_RS08525 | elongation factor Tu | 2527980 |
| ElP_RS00490 | elongation factor Tu | 2527974 |
| KIH39_RS14335 | elongation factor Tu | 1123043 |
| Mal48_RS18115 | elongation factor Tu | 2527994 |
| THTE_RS13795 | elongation factor Tu | 1331910 |
| GMBLW1_RS09465 | elongation factor Tu | 692036 |
| ETAA1_RS11095 | elongation factor Tu | 2528023 |
| ETAA1_RS13210 | elongation factor Tu | 2528023 |
| GQ464_RS07300 | elongation factor Tu | 2558931 |
| GQ464_RS17495 | elongation factor Tu | 2558931 |
| RMAR_RS04190 | elongation factor Tu | 518766 |
| RMAR_RS05485 | elongation factor Tu | 518766 |
| DRG95_RS06180 | elongation factor Tu | 146919 |
| DRG95_RS09920 | elongation factor Tu | 146919 |
| DU509_RS03980 | elongation factor Tu | 2051957 |
| DU509_RS04050 | elongation factor Tu | 2051957 |
| RXYL_RS10830 | elongation factor Tu | 266117 |
| RXYL_RS10895 | elongation factor Tu | 266117 |
| AsAng_RS22455 | elongation factor Tu | 2864201 |
| HALHY_RS22900 | elongation factor Tu | 760192 |
| SGRA_RS10445 | elongation factor Tu | 984262 |
| P2086_RS11095 | elongation factor Tu | 3036359 |
| HYN43_RS18305 | elongation factor Tu | 2305508 |
| LLH06_RS17060 | elongation factor Tu | 398049 |

| | | |
|---|---|---|
| GO620_RS08720 | elongation factor Tu | 2682091 |
| FRZ54_RS21910 | elongation factor Tu | 398053 |
| FSB76_RS19760 | elongation factor Tu | 862126 |
| J3L18_RS00285 | elongation factor Tu | 551996 |
| MgSA37_RS17610 | elongation factor Tu | 1550579 |
| PQO05_RS07705 | elongation factor Tu | 1176721 |
| HQ865_RS17620 | elongation factor Tu | 2740462 |
| BLU33_RS17730 | elongation factor Tu | 652787 |
| MUCPA_RS21420 | elongation factor Tu | 714943 |
| HH214_RS14190 | elongation factor Tu | 2728022 |
| SNE25_RS08720 | elongation factor Tu | 1173583 |
| MuYL_RS08120 | elongation factor Tu | 1234841 |
| FYC62_RS01450 | elongation factor Tu | 2605747 |
| IZT61_RS19350 | elongation factor Tu | 2789740 |
| AB3465_RS14480 | elongation factor Tu | 2739058 |
| PHEP_RS04235 | elongation factor Tu | 485917 |
| PL_RS11505 | elongation factor Tu | 1503925 |
| LOK61_RS11420 | elongation factor Tu | 2895286 |
| H9N25_RS03120 | elongation factor Tu | 2766984 |
| H9L23_RS03780 | elongation factor Tu | 336820 |
| EAO65_RS25075 | elongation factor Tu | 2201271 |
| DV942_RS06390 | elongation factor Tu | 332999 |
| PEDSA_RS11345 | elongation factor Tu | 762903 |
| SOLCA_RS02015 | elongation factor Tu | 929556 |
| L2B55_RS06110 | elongation factor Tu | 2911172 |
| SCB77_RS07360 | elongation factor Tu | 662858 |
| ACF3OC_RS02980 | elongation factor Tu | 1768011 |
| DSM08_RS15505 | elongation factor Tu | 649196 |
| G6N79_RS00590 | elongation factor Tu | 797291 |
| CKW03_RS03600 | elongation factor Tu | 1010 |
| I6J33_RS02375 | elongation factor Tu | 28454 |

| | | |
|---|---|---|
| PQ465_RS17145 | elongation factor Tu | 3025669 |
| HS960_RS16215 | elongation factor Tu | 2886510 |
| E2P86_RS02720 | elongation factor Tu | 561061 |
| K2F45_RS04760 | elongation factor Tu | 459529 |
| I6J01_RS04430 | elongation factor Tu | 258 |
| FGL37_RS03160 | elongation factor Tu | 259 |
| GFH32_RS03585 | elongation factor Tu | 2662364 |
| N187_RS02360 | elongation factor Tu | 1365188 |
| bcCo53_RS02390 | elongation factor Tu | 144 |
| BDU_RS02380 | elongation factor Tu | 412419 |
| K9R62_RS07600 | elongation factor Tu | 140 |
| DB723_RS02380 | elongation factor Tu | 2761123 |
| K5Q05_RS02350 | elongation factor Tu | 47466 |
| bhYOR_RS02395 | elongation factor Tu | 1293576 |
| X966_RS02370 | elongation factor Tu | 1435045 |
| bpuSUM_RS02395 | elongation factor Tu | 2756107 |
| BRE_RS02430 | elongation factor Tu | 412418 |
| DB313_RS02500 | elongation factor Tu | 1104446 |
| BT0_RS02395 | elongation factor Tu | 314724 |
| P612_RS02395 | elongation factor Tu | 1398490 |
| QIA01_02370 | elongation factor Tu | 478807 |
| QIA43_RS02375 | elongation factor Tu | 42109 |
| BG_RS02375 | elongation factor Tu | 290434 |
| BBIDN127_RS02365 | elongation factor Tu | 521010 |
| QIA17_RS02920 | elongation factor Tu | 373543 |
| QIA46_RS02380 | elongation factor Tu | 478174 |
| QIA19_RS02935 | elongation factor Tu | 498741 |
| KCI44_RS02385 | elongation factor Tu | 29519 |
| QIA21_RS02995 | elongation factor Tu | 34095 |
| QIA22_RS02945 | elongation factor Tu | 1196056 |
| QIA23_RS02920 | elongation factor Tu | 373540 |

| | | |
|---|---|---|
| QIA24_RS03105 | elongation factor Tu | 100177 |
| Bmayo_RS02400 | elongation factor Tu | 1674146 |
| QIA41_RS01885 | elongation factor Tu | 87162 |
| QIA25_RS02970 | elongation factor Tu | 88916 |
| QIA27_RS02905 | elongation factor Tu | 56146 |
| QIA31_RS02920 | elongation factor Tu | 57863 |
| BVAVS116_RS02380 | elongation factor Tu | 445987 |
| QIA35_RS02940 | elongation factor Tu | 683292 |
| GQX59_RS09665 | elongation factor Tu | 159 |
| BINT_RS12560 | elongation factor Tu | 1045858 |
| BMUR_RS09210 | elongation factor Tu | 526224 |
| BP951000_RS05050 | elongation factor Tu | 759914 |
| JFL75_RS17365 | elongation factor Tu | 2798577 |
| K7I13_RS12675 | elongation factor Tu | 2880008 |
| QMG63_RS02040 | elongation factor Tu | 2518322 |
| CR532_RS02510 | elongation factor Tu | 1964448 |
| P0082_RS10670 | elongation factor Tu | 3034016 |
| PVA46_RS06900 | elongation factor Tu | 2719989 |
| PVA45_RS01850 | elongation factor Tu | 2719988 |
| PVA44_RS01780 | elongation factor Tu | 2719987 |
| SPICA_RS01945 | elongation factor Tu | 744872 |
| TREAZ_RS15705 | elongation factor Tu | 545695 |
| LEPBI_RS09715 | elongation factor Tu | 456481 |
| C4Q31_RS04130 | elongation factor Tu | 508536 |
| AB3N61_RS13810 | elongation factor Tu | 3235173 |
| A6J42_RS08275 | elongation factor Tu | 44275 |
| EFP84_RS07595 | elongation factor Tu | 408139 |
| DI077_RS10400 | elongation factor Tu | 1917830 |
| LEP1GSC190_RS15200 | elongation factor Tu | 1192864 |
| LSS_RS14995 | elongation factor Tu | 758847 |
| A0128_RS15790 | elongation factor Tu | 2564040 |

| | | |
|---|---|---|
| FHG67_RS16460 | elongation factor Tu | 28184 |
| EXM22_RS11425 | elongation factor Tu | 2518645 |
| SPICO_RS08750 | elongation factor Tu | 760011 |
| L21SP2_RS04070 | elongation factor Tu | 1307761 |
| SPIRS_RS04825 | elongation factor Tu | 573413 |
| MUG09_RS15265 | elongation factor Tu | 1129264 |
| SPIBUDDY_RS14395 | elongation factor Tu | 158189 |
| SPIGRAPES_RS10625 | elongation factor Tu | 158190 |
| SPIAF_RS02785 | elongation factor Tu | 889378 |
| SPITH_RS02620 | elongation factor Tu | 869211 |
| KDW03_RS05875 | elongation factor Tu | 2828656 |
| EW093_RS10680 | elongation factor Tu | 252967 |
| TREBR_RS11840 | elongation factor Tu | 906968 |
| AABJ44_RS02500 | elongation factor Tu | 163 |
| TDE_RS03700 | elongation factor Tu | 243275 |
| DWB79_RS03170 | elongation factor Tu | 58231 |
| TPANIC_RS00930 | elongation factor Tu | 243276 |
| TPCCA_RS00930 | elongation factor Tu | 545776 |
| HRQ91_RS01560 | elongation factor Tu | 138851 |
| TPE_RS12435 | elongation factor Tu | 1291379 |
| IWA51_RS11825 | elongation factor Tu | 2787628 |
| FUT79_RS14560 | elongation factor Tu | 162 |
| TREPR_RS12205 | elongation factor Tu | 545694 |
| E4N74_RS09170 | elongation factor Tu | 221027 |
| AABB03_RS01550 | elongation factor Tu | 165 |
| HRI97_RS02205 | elongation factor Tu | 69713 |
| TRESU_RS12545 | elongation factor Tu | 869209 |
| TURPA_RS12615 | elongation factor Tu | 869212 |
| KAR29_RS02815 | elongation factor Tu | 3029084 |
| KAR29_RS09145 | elongation factor Tu | 3029084 |
| AMICO_RS01550 | elongation factor Tu | 572547 |

| | | |
|---|---|---|
| AMICO_RS03150 | elongation factor Tu | 572547 |
| APAU_RS08315 | elongation factor Tu | 584708 |
| APAU_RS09725 | elongation factor Tu | 584708 |
| BED41_RS05070 | elongation factor Tu | 1197717 |
| BED41_RS15935 | elongation factor Tu | 1197717 |
| ACK28C_RS00400 | elongation factor Tu | 651822 |
| ACK28C_RS07805 | elongation factor Tu | 651822 |
| MWL86_RS02945 | elongation factor Tu | 638849 |
| MWL86_RS10285 | elongation factor Tu | 638849 |
| TACI_RS06115 | elongation factor Tu | 525903 |
| TACI_RS07835 | elongation factor Tu | 525903 |
| THEVEDRAFT_RS01275 | elongation factor Tu | 926567 |
| THEVEDRAFT_RS02905 | elongation factor Tu | 926567 |
| SYN_RS01460 | elongation factor Tu | 56780 |
| ACJ77P_RS02355 | elongation factor Tu | 43774 |
| FDQ92_RS01880 | elongation factor Tu | 980445 |
| FDQ92_RS01940 | elongation factor Tu | 980445 |
| SFUM_RS07990 | elongation factor Tu | 335543 |
| SFUM_RS08045 | elongation factor Tu | 335543 |
| Tbon_RS01390 | elongation factor Tu | 2601677 |
| Tbon_RS09375 | elongation factor Tu | 2601677 |
| ACPOL_RS06735 | elongation factor Tu | 2211140 |
| ACPOL_RS22310 | elongation factor Tu | 2211140 |
| ACP_RS06955 | elongation factor Tu | 240015 |
| ACP_RS13885 | elongation factor Tu | 240015 |
| ACID345_RS06305 | elongation factor Tu | 204669 |
| ACID345_RS24370 | elongation factor Tu | 204669 |
| KFE13_RS05055 | elongation factor Tu | 1933044 |
| KFE13_RS15115 | elongation factor Tu | 1933044 |
| P4G45_RS02785 | elongation factor Tu | 3035702 |
| P4G45_RS03780 | elongation factor Tu | 3035702 |

| | | |
|---|---|---|
| ACIX8_RS02825 | elongation factor Tu | 682795 |
| ACIX8_RS03790 | elongation factor Tu | 682795 |
| ACIX9_RS04610 | elongation factor Tu | 1198114 |
| ACIX9_RS14755 | elongation factor Tu | 1198114 |
| MOP44_RS10065 | elongation factor Tu | 1002689 |
| MOP44_RS10495 | elongation factor Tu | 1002689 |
| IRI77_RS11025 | elongation factor Tu | 1473598 |
| IRI77_RS26185 | elongation factor Tu | 1473598 |
| ACOBR2_RS07345 | elongation factor Tu | 474953 |
| ACOBR2_RS18640 | elongation factor Tu | 474953 |
| TERRO_RS04440 | elongation factor Tu | 926566 |
| TERRO_RS18365 | elongation factor Tu | 926566 |
| ACIPR4_RS16110 | elongation factor Tu | 401053 |
| ACIPR4_RS17130 | elongation factor Tu | 401053 |
| RBB75_RS12305 | elongation factor Tu | 3069691 |
| RBB75_RS14015 | elongation factor Tu | 3069691 |
| RBB81_RS20610 | elongation factor Tu | 3069689 |
| RBB81_RS23290 | elongation factor Tu | 3069689 |
| KFE12_RS10530 | elongation factor Tu | 2051959 |
| KFE12_RS23040 | elongation factor Tu | 2051959 |
| RBB77_RS07290 | elongation factor Tu | 3069686 |
| RBB77_RS19525 | elongation factor Tu | 3069686 |
| THC_RS00580 | elongation factor Tu | 1653476 |
| THC_RS08370 | elongation factor Tu | 1653476 |
| THEIN_RS01425 | elongation factor Tu | 667014 |
| THEIN_RS07440 | elongation factor Tu | 667014 |
| HL41_RS05275 | elongation factor Tu | 289377 |
| HL41_RS08565 | elongation factor Tu | 289377 |
| TOPB45_RS00870 | elongation factor Tu | 795359 |
| TOPB45_RS03890 | elongation factor Tu | 795359 |
| FVE67_RS02555 | elongation factor Tu | 2047767 |

| | | |
|---|---|---|
| FVE67_RS03040 | elongation factor Tu | 2047767 |
| G4V39_RS03160 | elongation factor Tu | 1936021 |
| G4V39_RS03230 | elongation factor Tu | 1936021 |
| TDSAC_RS02365 | elongation factor Tu | 1794699 |
| TDSAC_RS05250 | elongation factor Tu | 1794699 |
| THENA_RS02535 | elongation factor Tu | 747365 |
| THENA_RS04070 | elongation factor Tu | 747365 |
| JTV28_RS02230 | elongation factor Tu | 2715679 |
| JTV28_RS03300 | elongation factor Tu | 2715679 |
| V4D30_RS00005 | elongation factor Tu | 3118333 |
| V4D30_RS09025 | elongation factor Tu | 3118333 |
| V4D31_RS07380 | elongation factor Tu | 3118332 |
| V4D31_RS08365 | elongation factor Tu | 3118332 |
| THEYE_RS06700 | elongation factor Tu | 289376 |
| THEYE_RS07155 | elongation factor Tu | 289376 |
| DSM104299_RS24360 | elongation factor Tu | 2997333 |
| FSW04_RS23760 | elongation factor Tu | 496014 |
| DSM104329_RS23540 | elongation factor Tu | 2884022 |
| CWOE_RS06705 | elongation factor Tu | 469383 |
| ITJ85_RS14040 | elongation factor Tu | 2843215 |
| IU369_RS03690 | elongation factor Tu | 2843216 |
| NBH00_RS04830 | elongation factor Tu | 2949664 |
| LRS13_RS24300 | elongation factor Tu | 2898438 |
| STHE_RS05180 | elongation factor Tu | 479434 |
| STHE_RS08160 | elongation factor Tu | 479434 |
| TRD_RS03095 | elongation factor Tu | 309801 |
| TRD_RS04535 | elongation factor Tu | 309801 |
| AXH83_RS07540 | elongation factor Tu | 1298851 |
| AXH83_RS07610 | elongation factor Tu | 1298851 |
| ATHSA_RS00500 | elongation factor Tu | 1441386 |
| DTL3_RS00775 | elongation factor Tu | 1006576 |

| | | |
|---|---|---|
| CBS1_RS04550 | elongation factor Tu | 310769 |
| NA23_RS07230 | elongation factor Tu | 2423 |
| FNOD_RS05965 | elongation factor Tu | 381764 |
| FERPE_RS06955 | elongation factor Tu | 771875 |
| KOLE_RS09620 | elongation factor Tu | 521045 |
| IX53_RS08730 | elongation factor Tu | 1330330 |
| JRV97_RS02105 | elongation factor Tu | 2809031 |
| MARPI_RS04340 | elongation factor Tu | 443254 |
| MESINF_RS01730 | elongation factor Tu | 1236046 |
| THEBA_RS03490 | elongation factor Tu | 660470 |
| PMOB_RS04085 | elongation factor Tu | 403833 |
| TEL01S_RS02865 | elongation factor Tu | 1123383 |
| THY01S_RS08110 | elongation factor Tu | 1123384 |
| TLET_RS02955 | elongation factor Tu | 416591 |
| THETH_RS03765 | elongation factor Tu | 688269 |
| IGS63_RS05300 | elongation factor Tu | 2108365 |
| THA_RS06110 | elongation factor Tu | 484019 |
| JYK00_RS04005 | elongation factor Tu | 2571116 |
| TMEL_RS04965 | elongation factor Tu | 391009 |
| TSP01S_RS03960 | elongation factor Tu | 1408159 |
| THEMA_RS06710 | elongation factor Tu | 243274 |
| CTN_RS04900 | elongation factor Tu | 309803 |
| TPET_RS06585 | elongation factor Tu | 390874 |
| TSP02S_RS08170 | elongation factor Tu | 1408160 |
| GT409_RS02885 | elongation factor Tu | 2697043 |
| EQM13_RS14415 | elongation factor Tu | 2507161 |
| EQM13_RS14485 | elongation factor Tu | 2507161 |
| BQ4451_RS00225 | elongation factor Tu | 1870984 |
| BQ4451_RS07250 | elongation factor Tu | 1870984 |
| ABXG81_RS04425 | elongation factor Tu | 1755241 |
| ABXG81_RS06820 | elongation factor Tu | 1755241 |

| | | |
|---|---|---|
| I6H46_RS05025 | elongation factor Tu | 1287640 |
| I6H46_RS08805 | elongation factor Tu | 1287640 |
| APRE_RS02440 | elongation factor Tu | 525919 |
| APRE_RS05805 | elongation factor Tu | 525919 |
| I6H45_RS01995 | elongation factor Tu | 33037 |
| I6H45_RS05500 | elongation factor Tu | 33037 |
| BQ4440_RS03040 | elongation factor Tu | 1852374 |
| BQ4440_RS05770 | elongation factor Tu | 1852374 |
| CURI_RS11335 | elongation factor Tu | 1128398 |
| CURI_RS11405 | elongation factor Tu | 1128398 |
| EC328_RS08875 | elongation factor Tu | 1582259 |
| EC328_RS08945 | elongation factor Tu | 1582259 |
| EQF90_RS02095 | elongation factor Tu | 72026 |
| JL105_RS09520 | elongation factor Tu | 1278232 |
| JL105_RS09590 | elongation factor Tu | 1278232 |
| VZL98_RS03245 | elongation factor Tu | 3118105 |
| VZL98_RS03360 | elongation factor Tu | 3118105 |
| JFY71_RS00610 | elongation factor Tu | 2931922 |
| JFY71_RS02235 | elongation factor Tu | 2931922 |
| BN8034_RS06455 | elongation factor Tu | 1852373 |
| EL196_RS02320 | elongation factor Tu | 33033 |
| ACKTSI_RS06010 | elongation factor Tu | 2769485 |
| O6R05_RS01725 | elongation factor Tu | 3016343 |
| O6R05_RS07275 | elongation factor Tu | 3016343 |
| M9426_RS01275 | elongation factor Tu | 3036303 |
| M9426_RS03650 | elongation factor Tu | 3036303 |
| I6I93_RS03780 | elongation factor Tu | 54005 |
| I6I93_RS07685 | elongation factor Tu | 54005 |
| OQH67_RS06160 | elongation factor Tu | 2946555 |
| PYTT_RS02330 | elongation factor Tu | 1679444 |
| GOZ73_RS05570 | elongation factor Tu | 239935 |

| | | |
|---|---|---|
| NXS98_RS00080 | elongation factor Tu | 2974023 |
| HAHE_RS02875 | elongation factor Tu | 490095 |
| KBB96_RS12625 | elongation factor Tu | 2824561 |
| HHL09_RS17605 | elongation factor Tu | 2728835 |
| G3M56_RS04005 | elongation factor Tu | 2707525 |
| VSP_RS27100 | elongation factor Tu | 240016 |
| LuPra_RS24310 | elongation factor Tu | 1855912 |
| LuPra_RS24400 | elongation factor Tu | 1855912 |
| WPS_RS10435 | elongation factor Tu | 3016050 |
| Ga0123461_RS01630 | elongation factor Tu | 1921086 |
| Ga0123461_RS12170 | elongation factor Tu | 1921086 |
| Ga0123462_RS09575 | elongation factor Tu | 1921087 |
| Ga0123462_RS11200 | elongation factor Tu | 1921087 |
| ACPT9Q_RS00150 | elongation factor Tu | 1454753 |
| ACPT9Q_RS08675 | elongation factor Tu | 1454753 |
| CALKR_RS09520 | elongation factor Tu | 632335 |
| ATHE_RS04055 | elongation factor Tu | 521460 |
| ELD05_RS10110 | elongation factor Tu | 1222016 |
| CaldiYA01_RS08300 | elongation factor Tu | 2806205 |
| CALHY_RS09485 | elongation factor Tu | 632292 |
| CALKRO_RS09290 | elongation factor Tu | 632348 |
| OTK00_RS03130 | elongation factor Tu | 1387555 |
| OTJ99_RS09460 | elongation factor Tu | 29324 |
| COB47_RS03670 | elongation factor Tu | 608506 |
| CALOW_RS03295 | elongation factor Tu | 632518 |
| CSAC_RS04860 | elongation factor Tu | 351627 |
| AZF37_RS03255 | elongation factor Tu | 243899 |
| AZF37_RS03335 | elongation factor Tu | 243899 |
| BQ7385_RS01880 | elongation factor Tu | 1871025 |
| BQ7385_RS04845 | elongation factor Tu | 1871025 |
| LA303_RS05200 | elongation factor Tu | 2875962 |

| | | |
|---|---|---|
| HLG78_RS01420 | elongation factor Tu | 2735562 |
| Q2T83_RS11690 | elongation factor Tu | 1448929 |
| BU251_RS03025 | elongation factor Tu | 1930593 |
| GII36_RS03215 | elongation factor Tu | 2665156 |
| TTER_RS03575 | elongation factor Tu | 525904 |
| TTER_RS09280 | elongation factor Tu | 525904 |
| P3F56_RS09685 | elongation factor Tu | 3034674 |
| ETSB_RS08505 | elongation factor Tu | 1228987 |
| RGRSB_RS05600 | elongation factor Tu | 1763363 |
| V3331_RS01945 | elongation factor Tu | 3119007 |
| V3331_RS02010 | elongation factor Tu | 3119007 |
| DP065_RS01040 | elongation factor Tu | 92400 |
| MCAV_RS01895 | elongation factor Tu | 92757 |
| WG616_RS00440 | elongation factor Tu | 92404 |
| DA803_RS00585 | elongation factor Tu | 142651 |
| EXC57_RS04765 | elongation factor Tu | 2116 |
| ACLFO7_RS00370 | elongation factor Tu | 2119 |
| MYPE_RS00160 | elongation factor Tu | 272633 |
| MYB_RS01585 | elongation factor Tu | 743966 |
| MCJ_RS02620 | elongation factor Tu | 45361 |
| MDIS_RS02140 | elongation factor Tu | 86660 |
| MYF_RS00755 | elongation factor Tu | 743971 |
| MHJ_RS02785 | elongation factor Tu | 262719 |
| D2830_RS03050 | elongation factor Tu | 2100 |
| QEG99_RS03985 | elongation factor Tu | 171286 |
| ACJA28_RS02525 | elongation factor Tu | 171287 |
| NX772_RS03540 | elongation factor Tu | 171288 |
| EXC65_RS02870 | elongation factor Tu | 2120 |
| PWA39_RS01405 | elongation factor Tu | 743968 |
| D2845_RS01240 | elongation factor Tu | 45363 |
| EXC42_RS01960 | elongation factor Tu | 2111 |

| | | |
|---|---|---|
| ACLFP3_RS02980 | elongation factor Tu | 55602 |
| MCAN360_RS02170 | elongation factor Tu | 29554 |
| DK849_RS02605 | elongation factor Tu | 92401 |
| R9B83_RS00805 | elongation factor Tu | 92402 |
| LQ356_RS02245 | elongation factor Tu | 56142 |
| V2E26_RS01525 | elongation factor Tu | 35769 |
| MLBD4_RS00255 | elongation factor Tu | 1267000 |
| NMG93_RS02730 | elongation factor Tu | 29559 |
| EXC29_RS02785 | elongation factor Tu | 2121 |
| DMC14_RS00725 | elongation factor Tu | 142649 |
| EXC60_RS01120 | elongation factor Tu | 2124 |
| AB5V95_RS00340 | elongation factor Tu | 92406 |
| ACJA29_RS03200 | elongation factor Tu | 2125 |
| R9C05_RS02275 | elongation factor Tu | 92407 |
| MMOB_RS01215 | elongation factor Tu | 267748 |
| NPA10_RS01575 | elongation factor Tu | 92758 |
| D2833_RS01325 | elongation factor Tu | 2096 |
| ABS333_RS02725 | elongation factor Tu | 2097 |
| EXC43_RS00115 | elongation factor Tu | 2104 |
| ACJA22_RS03460 | elongation factor Tu | 51362 |
| MAG_RS01670 | elongation factor Tu | 347257 |
| DP067_RS02605 | elongation factor Tu | 171279 |
| MARG145_RS02885 | elongation factor Tu | 2094 |
| EXC34_RS02110 | elongation factor Tu | 2112 |
| EXC44_RS02130 | elongation factor Tu | 29553 |
| MBOVPG45_RS02055 | elongation factor Tu | 289397 |
| MCFN_RS02075 | elongation factor Tu | 2113 |
| AXW82_RS03190 | elongation factor Tu | 1117644 |
| NPA07_RS04415 | elongation factor Tu | 55603 |
| EXC58_RS01695 | elongation factor Tu | 171281 |
| EXC37_RS02725 | elongation factor Tu | 114881 |

| | | |
|---|---|---|
| EXC55_RS00455 | elongation factor Tu | 114880 |
| EXC45_RS03275 | elongation factor Tu | 171282 |
| MCYN_RS03800 | elongation factor Tu | 1246955 |
| D2846_RS03075 | elongation factor Tu | 53558 |
| NPA09_RS03515 | elongation factor Tu | 114883 |
| RRG57_RS03565 | elongation factor Tu | 33923 |
| MFEM64YM_RS01205 | elongation factor Tu | 943945 |
| EXC51_RS02260 | elongation factor Tu | 29556 |
| EXC53_RS00080 | elongation factor Tu | 76629 |
| EXC53_RS02365 | elongation factor Tu | 76629 |
| EXC46_RS03610 | elongation factor Tu | 171285 |
| ACJA25_RS02875 | elongation factor Tu | 29558 |
| ACJA27_RS03150 | elongation factor Tu | 2117 |
| EXC47_RS03305 | elongation factor Tu | 114885 |
| EXC33_RS00545 | elongation factor Tu | 29561 |
| EG856_RS02175 | elongation factor Tu | 142650 |
| EXC36_RS01925 | elongation factor Tu | 2107 |
| VY93_RS03630 | elongation factor Tu | 1267001 |
| SAM46_RS00395 | elongation factor Tu | 171291 |
| JM47_RS00245 | elongation factor Tu | 42094 |
| UPA3_RS02795 | elongation factor Tu | 505682 |
| CMC5_RS07600 | elongation factor Tu | 52 |
| CMC5_RS09235 | elongation factor Tu | 52 |
| HOCH_RS19720 | elongation factor Tu | 502025 |
| HOCH_RS19800 | elongation factor Tu | 502025 |
| AKJ09_RS01645 | elongation factor Tu | 1391654 |
| AKJ09_RS02930 | elongation factor Tu | 1391654 |
| O0S08_RS43405 | elongation factor Tu | 2995304 |
| O0S08_RS43475 | elongation factor Tu | 2995304 |
| BCY86_RS07125 | elongation factor Tu | 1882918 |
| E8A73_RS00280 | elongation factor Tu | 2567896 |

| | | |
|---|---|---|
| E8A73_RS19460 | elongation factor Tu | 2567896 |
| DB32_RS13185 | elongation factor Tu | 927083 |
| DB32_RS38220 | elongation factor Tu | 927083 |
| SCE_RS02115 | elongation factor Tu | 448385 |
| SCE_RS04410 | elongation factor Tu | 448385 |
| Pla133_RS19175 | elongation factor Tu | 2528011 |
| Pan216_RS23440 | elongation factor Tu | 2527975 |
| Pla163_RS14365 | elongation factor Tu | 2528008 |
| Poly30_RS19550 | elongation factor Tu | 2528018 |
| SV7mr_RS05710 | elongation factor Tu | 2528025 |
| TBK1r_RS20040 | elongation factor Tu | 2527963 |
| TBK1r_RS27710 | elongation factor Tu | 2527963 |
| K239x_RS18885 | elongation factor Tu | 1930275 |
| QU602_RS14385 | elongation factor Tu | 3388266 |
| ACIRQ6_RS14245 | elongation factor Tu | 3377108 |
| ACIRQ6_RS15050 | elongation factor Tu | 3377108 |
| KIH16_RS02755 | elongation factor Tu | 3415223 |
| KIH16_RS09040 | elongation factor Tu | 3415223 |
| ACLLO5_RS09430 | elongation factor Tu | 3392114 |
| FRZ06_RS20275 | elongation factor Tu | 3408580 |
| FRZ06_RS20345 | elongation factor Tu | 3408580 |
| ABN702_RS19640 | elongation factor Tu | 3160967 |
| Dip510_RS01905 | elongation factor Tu | 3116653 |
| Dip510_RS02115 | elongation factor Tu | 3116653 |
| ACMCL3_RS00315 | elongation factor Tu | 3132096 |
| JM64_RS00205 | elongation factor Tu | 3389972 |
| strainIA91_RS08125 | elongation factor Tu | 3377529 |
| strainIA91_RS08195 | elongation factor Tu | 3377529 |
| ACEPSS_RS08000 | elongation factor Tu | 3242688 |
| K4L44_RS10970 | elongation factor Tu | 3374600 |
| K5X82_RS17675 | elongation factor Tu | 3374599 |

| | | |
|---|---|---|
| ACQ3C8_RS13740 | elongation factor Tu | 3058603 |
| ACQ3C8_RS13810 | elongation factor Tu | 3058603 |
| ACF9IK_RS21445 | elongation factor Tu | 3369522 |
| VIG27_RS02410 | elongation factor Tu | 3111456 |
| ACEPSZ_RS05880 | elongation factor Tu | 3242689 |
| ACQY1Y_RS08980 | elongation factor Tu | 3422495 |
| ACQY1X_RS09025 | elongation factor Tu | 3422496 |
| AB3U99_RS00735 | elongation factor Tu | 3237105 |
| ACHFB9_RS02685 | elongation factor Tu | 3373598 |
| ACHFB9_RS02750 | elongation factor Tu | 3373598 |
| Q9R13_RS10410 | elongation factor Tu | 3388267 |
| Dip518_RS04555 | elongation factor Tu | 3228953 |
| Dip518_RS05480 | elongation factor Tu | 3228953 |
| AB0761_RS22040 | elongation factor Tu | 3232308 |
| ABHF91_RS15680 | elongation factor Tu | 3153928 |
| ABHF91_RS15775 | elongation factor Tu | 3153928 |
| OQ460_RS21935 | elongation factor Tu | 2992837 |
| ACI7T6_RS03710 | elongation factor Tu | 3342349 |
| ACMCNJ_RS02970 | elongation factor Tu | 3381777 |
| ACMCNJ_RS03055 | elongation factor Tu | 3381777 |
| AAFQ93_RS07790 | elongation factor Tu | 3140788 |
| ACE1N8_RS05165 | elongation factor Tu | 3344655 |
| ACE1N8_RS20885 | elongation factor Tu | 3344655 |
| ACE1OC_RS16850 | elongation factor Tu | 3344656 |
| ACJZL1_RS00975 | elongation factor Tu | 3383250 |
| ACJZL1_RS02205 | elongation factor Tu | 3383250 |

| Organism Name | Phylum | Class |
|---|---|---|
| Acidimicrobium ferrooxidans DSM 10331 | Actinomycetota | Acidimicrobiia |
| Actinomarinicola tropica | Actinomycetota | Acidimicrobiia |
| Actinomarinicola tropica | Actinomycetota | Acidimicrobiia |
| Aquihabitans daechungensis | Actinomycetota | Acidimicrobiia |
| Aquihabitans daechungensis | Actinomycetota | Acidimicrobiia |
| Dermatobacter hominis | Actinomycetota | Acidimicrobiia |
| Dermatobacter hominis | Actinomycetota | Acidimicrobiia |
| Iamia majanohamensis | Actinomycetota | Acidimicrobiia |
| Iamia majanohamensis | Actinomycetota | Acidimicrobiia |
| Ilumatobacter coccineus YM16-304 | Actinomycetota | Acidimicrobiia |
| Ilumatobacter coccineus YM16-304 | Actinomycetota | Acidimicrobiia |
| Acidithiobacillus caldus SM-1 | Pseudomonadota | Acidithiobacillia |
| Acidithiobacillus caldus SM-1 | Pseudomonadota | Acidithiobacillia |
| Acidithiobacillus ferrianus | Pseudomonadota | Acidithiobacillia |
| Acidithiobacillus ferrianus | Pseudomonadota | Acidithiobacillia |
| Acidithiobacillus ferridurans | Pseudomonadota | Acidithiobacillia |
| Acidithiobacillus ferridurans | Pseudomonadota | Acidithiobacillia |
| Acidithiobacillus ferriphilus | Pseudomonadota | Acidithiobacillia |
| Acidithiobacillus ferriphilus | Pseudomonadota | Acidithiobacillia |
| Acidithiobacillus ferrivorans | Pseudomonadota | Acidithiobacillia |
| Acidithiobacillus ferrivorans | Pseudomonadota | Acidithiobacillia |
| Acidithiobacillus ferrooxidans ATCC 23270 | Pseudomonadota | Acidithiobacillia |
| Acidithiobacillus ferrooxidans ATCC 23270 | Pseudomonadota | Acidithiobacillia |
| Acidithiobacillus ferruginosus | Pseudomonadota | Acidithiobacillia |
| Acidithiobacillus ferruginosus | Pseudomonadota | Acidithiobacillia |
| Acidithiobacillus montserratensis | Pseudomonadota | Acidithiobacillia |
| Acidithiobacillus montserratensis | Pseudomonadota | Acidithiobacillia |
| Acidithiobacillus sulfuriphilus | Pseudomonadota | Acidithiobacillia |
| Acidithiobacillus sulfuriphilus | Pseudomonadota | Acidithiobacillia |
| Acidithiobacillus thiooxidans ATCC 19377 | Pseudomonadota | Acidithiobacillia |

| | | |
|---|---|---|
| Acidithiobacillus thiooxidans ATCC 19377 | Pseudomonadota | Acidithiobacillia |
| Thermithiobacillus tepidarius DSM 3134 | Pseudomonadota | Acidithiobacillia |
| Thermithiobacillus tepidarius DSM 3134 | Pseudomonadota | Acidithiobacillia |
| [Actinomadura] parvosata subsp. kistnae | Actinomycetota | Actinomycetes |
| [Actinomadura] parvosata subsp. kistnae | Actinomycetota | Actinomycetes |
| [Kitasatospora] papulosa | Actinomycetota | Actinomycetes |
| [Kitasatospora] papulosa | Actinomycetota | Actinomycetes |
| [Mycobacterium] burgundiense | Actinomycetota | Actinomycetes |
| [Mycobacterium] holstebronense | Actinomycetota | Actinomycetes |
| [Mycobacterium] kokjensenii | Actinomycetota | Actinomycetes |
| [Mycobacterium] stephanolepidis | Actinomycetota | Actinomycetes |
| [Mycobacterium] wendilense | Actinomycetota | Actinomycetes |
| Acidipropionibacterium acidipropionici | Actinomycetota | Actinomycetes |
| Acidipropionibacterium jensenii | Actinomycetota | Actinomycetes |
| Acidipropionibacterium virtanenii | Actinomycetota | Actinomycetes |
| Acidothermus cellulolyticus 11B | Actinomycetota | Actinomycetes |
| Actinacidiphila glaucinigra | Actinomycetota | Actinomycetes |
| Actinacidiphila glaucinigra | Actinomycetota | Actinomycetes |
| Actinacidiphila reveromycinica | Actinomycetota | Actinomycetes |
| Actinoalloteichus fjordicus | Actinomycetota | Actinomycetes |
| Actinoalloteichus hoggarensis | Actinomycetota | Actinomycetes |
| Actinoalloteichus hymeniacidonis | Actinomycetota | Actinomycetes |
| Actinocatenispora sera | Actinomycetota | Actinomycetes |
| Actinocatenispora thailandica | Actinomycetota | Actinomycetes |
| Actinokineospora auranticolor | Actinomycetota | Actinomycetes |
| Actinomadura graeca | Actinomycetota | Actinomycetes |
| Actinomadura graeca | Actinomycetota | Actinomycetes |
| Actinomadura luteofluorescens | Actinomycetota | Actinomycetes |
| Actinomadura luteofluorescens | Actinomycetota | Actinomycetes |
| Actinomadura madurae | Actinomycetota | Actinomycetes |
| Actinomadura madurae | Actinomycetota | Actinomycetes |

| | | |
|---|---|---|
| Actinomyces capricornis | Actinomycetota | Actinomycetes |
| Actinomyces faecalis | Actinomycetota | Actinomycetes |
| Actinomyces gaoshouyii | Actinomycetota | Actinomycetes |
| Actinomyces howellii | Actinomycetota | Actinomycetes |
| Actinomyces israelii | Actinomycetota | Actinomycetes |
| Actinomyces lilanjuaniae | Actinomycetota | Actinomycetes |
| Actinomyces marmotae | Actinomycetota | Actinomycetes |
| Actinomyces naeslundii | Actinomycetota | Actinomycetes |
| Actinomyces oris | Actinomycetota | Actinomycetes |
| Actinomyces pacaensis | Actinomycetota | Actinomycetes |
| Actinomyces procaprae | Actinomycetota | Actinomycetes |
| Actinomyces qiguomingii | Actinomycetota | Actinomycetes |
| Actinomyces radicidentis | Actinomycetota | Actinomycetes |
| Actinomyces respiraculi | Actinomycetota | Actinomycetes |
| Actinomyces slackii | Actinomycetota | Actinomycetes |
| Actinomyces trachealis | Actinomycetota | Actinomycetes |
| Actinomyces viscosus | Actinomycetota | Actinomycetes |
| Actinomyces weissii | Actinomycetota | Actinomycetes |
| Actinomyces wuliandei | Actinomycetota | Actinomycetes |
| Actinoplanes derwentensis | Actinomycetota | Actinomycetes |
| Actinoplanes derwentensis | Actinomycetota | Actinomycetes |
| Actinoplanes friuliensis DSM 7358 | Actinomycetota | Actinomycetes |
| Actinoplanes friuliensis DSM 7358 | Actinomycetota | Actinomycetes |
| Actinoplanes ianthinogenes | Actinomycetota | Actinomycetes |
| Actinoplanes ianthinogenes | Actinomycetota | Actinomycetes |
| Actinoplanes missouriensis 431 | Actinomycetota | Actinomycetes |
| Actinoplanes missouriensis 431 | Actinomycetota | Actinomycetes |
| Actinoplanes oblitus | Actinomycetota | Actinomycetes |
| Actinoplanes oblitus | Actinomycetota | Actinomycetes |
| Actinoplanes sichuanensis | Actinomycetota | Actinomycetes |
| Actinoplanes sichuanensis | Actinomycetota | Actinomycetes |

| | | |
|---|---|---|
| Actinoplanes teichomyceticus ATCC 31121 | Actinomycetota | Actinomycetes |
| Actinopolymorpha singaporensis | Actinomycetota | Actinomycetes |
| Actinopolyspora erythraea | Actinomycetota | Actinomycetes |
| Actinosynnema mirum DSM 43827 | Actinomycetota | Actinomycetes |
| Actinosynnema pretiosum | Actinomycetota | Actinomycetes |
| Actinotignum schaalii | Actinomycetota | Actinomycetes |
| Actinotignum timonense | Actinomycetota | Actinomycetes |
| Aeromicrobium chenweiae | Actinomycetota | Actinomycetes |
| Aeromicrobium choanae | Actinomycetota | Actinomycetes |
| Aeromicrobium duanguangcaii | Actinomycetota | Actinomycetes |
| Aeromicrobium erythreum | Actinomycetota | Actinomycetes |
| Aeromicrobium marinum DSM 15272 | Actinomycetota | Actinomycetes |
| Aeromicrobium senzhongii | Actinomycetota | Actinomycetes |
| Aeromicrobium yanjiei | Actinomycetota | Actinomycetes |
| Agrococcus beijingensis | Actinomycetota | Actinomycetes |
| Agrococcus carbonis | Actinomycetota | Actinomycetes |
| Agrococcus jejuensis | Actinomycetota | Actinomycetes |
| Agromyces archimandritae | Actinomycetota | Actinomycetes |
| Agromyces aureus | Actinomycetota | Actinomycetes |
| Agromyces badenianii | Actinomycetota | Actinomycetes |
| Agromyces flavus | Actinomycetota | Actinomycetes |
| Agromyces intestinalis | Actinomycetota | Actinomycetes |
| Agromyces laixinhei | Actinomycetota | Actinomycetes |
| Agromyces larvae | Actinomycetota | Actinomycetes |
| Agromyces mangrovi | Actinomycetota | Actinomycetes |
| Agromyces marinus | Actinomycetota | Actinomycetes |
| Agromyces protaetiae | Actinomycetota | Actinomycetes |
| Agromyces soli | Actinomycetota | Actinomycetes |
| Allokutzneria albata | Actinomycetota | Actinomycetes |
| Allokutzneria albata | Actinomycetota | Actinomycetes |
| Allosaccharopolyspora coralli | Actinomycetota | Actinomycetes |

| | | |
|---|---|---|
| Ammonicoccus fulvus | Actinomycetota | Actinomycetes |
| Amycolatopsis acidiphila | Actinomycetota | Actinomycetes |
| Amycolatopsis aidingensis | Actinomycetota | Actinomycetes |
| Amycolatopsis albispora | Actinomycetota | Actinomycetes |
| Amycolatopsis bartoniae | Actinomycetota | Actinomycetes |
| Amycolatopsis carbonis | Actinomycetota | Actinomycetes |
| Amycolatopsis cynarae | Actinomycetota | Actinomycetes |
| Amycolatopsis japonica | Actinomycetota | Actinomycetes |
| Amycolatopsis keratiniphila | Actinomycetota | Actinomycetes |
| Amycolatopsis mediterranei S699 | Actinomycetota | Actinomycetes |
| Amycolatopsis mediterranei S699 | Actinomycetota | Actinomycetes |
| Amycolatopsis methanolica 239 | Actinomycetota | Actinomycetes |
| Amycolatopsis mongoliensis | Actinomycetota | Actinomycetes |
| Amycolatopsis mongoliensis | Actinomycetota | Actinomycetes |
| Amycolatopsis nalaikhensis | Actinomycetota | Actinomycetes |
| Amycolatopsis nalaikhensis | Actinomycetota | Actinomycetes |
| Amycolatopsis orientalis | Actinomycetota | Actinomycetes |
| Amycolatopsis rhabdoformis | Actinomycetota | Actinomycetes |
| Amycolatopsis roodepoortensis | Actinomycetota | Actinomycetes |
| Antiquaquibacter oligotrophicus | Actinomycetota | Actinomycetes |
| Aquiluna borgnonia | Actinomycetota | Actinomycetes |
| Arachnia propionica | Actinomycetota | Actinomycetes |
| Arachnia rubra | Actinomycetota | Actinomycetes |
| Arcanobacterium buesumense | Actinomycetota | Actinomycetes |
| Arcanobacterium canis | Actinomycetota | Actinomycetes |
| Arcanobacterium haemolyticum | Actinomycetota | Actinomycetes |
| Arcanobacterium phocae | Actinomycetota | Actinomycetes |
| Arcanobacterium phocisimile | Actinomycetota | Actinomycetes |
| Arcanobacterium pinnipediorum | Actinomycetota | Actinomycetes |
| Arcanobacterium wilhelmae | Actinomycetota | Actinomycetes |
| Arsenicicoccus cauae | Actinomycetota | Actinomycetes |

| | | |
|---|---|---|
| Arsenicicoccus dermatophilus | Actinomycetota | Actinomycetes |
| Arthrobacter agilis | Actinomycetota | Actinomycetes |
| Arthrobacter alpinus | Actinomycetota | Actinomycetes |
| Arthrobacter antioxidans | Actinomycetota | Actinomycetes |
| Arthrobacter caoxuetaonis | Actinomycetota | Actinomycetes |
| Arthrobacter crystallopoietes | Actinomycetota | Actinomycetes |
| Arthrobacter dokdonellae | Actinomycetota | Actinomycetes |
| Arthrobacter gengyunqii | Actinomycetota | Actinomycetes |
| Arthrobacter jiangjiafuii | Actinomycetota | Actinomycetes |
| Arthrobacter jinronghuae | Actinomycetota | Actinomycetes |
| Arthrobacter nitrophenolicus | Actinomycetota | Actinomycetes |
| Arthrobacter polaris | Actinomycetota | Actinomycetes |
| Arthrobacter sulfonylureivorans | Actinomycetota | Actinomycetes |
| Arthrobacter sunyaminii | Actinomycetota | Actinomycetes |
| Arthrobacter zhangbolii | Actinomycetota | Actinomycetes |
| Arthrobacter zhaoxinii | Actinomycetota | Actinomycetes |
| Aurantimicrobium minutum | Actinomycetota | Actinomycetes |
| Aurantimicrobium photophilum | Actinomycetota | Actinomycetes |
| Auraticoccus monumenti | Actinomycetota | Actinomycetes |
| Auritidibacter ignavus | Actinomycetota | Actinomycetes |
| Barrientosiimonas endolithica | Actinomycetota | Actinomycetes |
| Beutenbergia cavernae DSM 12333 | Actinomycetota | Actinomycetes |
| Bifidobacterium actinocoloniiforme DSM 22766 | Actinomycetota | Actinomycetes |
| Bifidobacterium adolescentis ATCC 15703 | Actinomycetota | Actinomycetes |
| Bifidobacterium angulatum DSM 20098 = JCM 7096 | Actinomycetota | Actinomycetes |
| Bifidobacterium animalis subsp. lactis DSM 10140 | Actinomycetota | Actinomycetes |
| Bifidobacterium aquikefiricola | Actinomycetota | Actinomycetes |
| Bifidobacterium asteroides DSM 20089 | Actinomycetota | Actinomycetes |
| Bifidobacterium bifidum | Actinomycetota | Actinomycetes |
| Bifidobacterium breve DSM 20213 = JCM 1192 | Actinomycetota | Actinomycetes |
| Bifidobacterium catenulatum DSM 16992 = JCM 1194 = LMG 11043 | Actinomycetota | Actinomycetes |

| | | |
|---|---|---|
| Bifidobacterium choerinum | Actinomycetota | Actinomycetes |
| Bifidobacterium coryneforme | Actinomycetota | Actinomycetes |
| Bifidobacterium dentium JCM 1195 = DSM 20436 | Actinomycetota | Actinomycetes |
| Bifidobacterium eulemuris | Actinomycetota | Actinomycetes |
| Bifidobacterium fermentum | Actinomycetota | Actinomycetes |
| Bifidobacterium imperatoris | Actinomycetota | Actinomycetes |
| Bifidobacterium lemurum | Actinomycetota | Actinomycetes |
| Bifidobacterium longum subsp. longum JCM 1217 | Actinomycetota | Actinomycetes |
| Bifidobacterium pseudocatenulatum DSM 20438 = JCM 1200 = LMG 10505 | Actinomycetota | Actinomycetes |
| Bifidobacterium pseudolongum subsp. globosum DSM 20092 | Actinomycetota | Actinomycetes |
| Bifidobacterium saguini | Actinomycetota | Actinomycetes |
| Bifidobacterium scardovii JCM 12489 = DSM 13734 | Actinomycetota | Actinomycetes |
| Bifidobacterium subtile | Actinomycetota | Actinomycetes |
| Blastococcus brunescens | Actinomycetota | Actinomycetes |
| Blastococcus montanus | Actinomycetota | Actinomycetes |
| Bombiscardovia apis | Actinomycetota | Actinomycetes |
| Boudabousia tangfeifanii | Actinomycetota | Actinomycetes |
| Brachybacterium avium | Actinomycetota | Actinomycetes |
| Brachybacterium ginsengisoli | Actinomycetota | Actinomycetes |
| Brachybacterium halotolerans subsp. kimchii | Actinomycetota | Actinomycetes |
| Brachybacterium huguangmaarense | Actinomycetota | Actinomycetes |
| Brachybacterium kimchii | Actinomycetota | Actinomycetes |
| Brachybacterium saurashtrense | Actinomycetota | Actinomycetes |
| Brachybacterium vulturis | Actinomycetota | Actinomycetes |
| Brevibacterium atlanticum | Actinomycetota | Actinomycetes |
| Brevibacterium aurantiacum | Actinomycetota | Actinomycetes |
| Brevibacterium casei | Actinomycetota | Actinomycetes |
| Brevibacterium koreense | Actinomycetota | Actinomycetes |
| Brevibacterium limosum | Actinomycetota | Actinomycetes |
| Brevibacterium linens | Actinomycetota | Actinomycetes |
| Brevibacterium oceani | Actinomycetota | Actinomycetes |

| | | |
|---|---|---|
| Brevibacterium pigmentatum | Actinomycetota | Actinomycetes |
| Brevibacterium sandarakinum | Actinomycetota | Actinomycetes |
| Brevibacterium siliguriense | Actinomycetota | Actinomycetes |
| Brevibacterium spongiae | Actinomycetota | Actinomycetes |
| Brevibacterium zhoupengii | Actinomycetota | Actinomycetes |
| Brooklawnia propionicigenes | Actinomycetota | Actinomycetes |
| Buchananella felis | Actinomycetota | Actinomycetes |
| Candidatus Mycobacterium methanotrophicum | Actinomycetota | Actinomycetes |
| Candidatus Mycobacterium wuenschmannii | Actinomycetota | Actinomycetes |
| Candidatus Nanopelagicus abundans | Actinomycetota | Actinomycetes |
| Candidatus Nanopelagicus hibericus | Actinomycetota | Actinomycetes |
| Candidatus Nanopelagicus limnes | Actinomycetota | Actinomycetes |
| Candidatus Planktophila dulcis | Actinomycetota | Actinomycetes |
| Candidatus Planktophila lacus | Actinomycetota | Actinomycetes |
| Candidatus Planktophila limnetica | Actinomycetota | Actinomycetes |
| Candidatus Planktophila sulfonica | Actinomycetota | Actinomycetes |
| Candidatus Planktophila vernalis | Actinomycetota | Actinomycetes |
| Candidatus Planktophila versatilis | Actinomycetota | Actinomycetes |
| Candidatus Rhodoluna planktonica | Actinomycetota | Actinomycetes |
| Catenulispora acidiphila DSM 44928 | Actinomycetota | Actinomycetes |
| Catenulispora acidiphila DSM 44928 | Actinomycetota | Actinomycetes |
| Cellulomonas chengniuliangii | Actinomycetota | Actinomycetes |
| Cellulomonas dongxiuzhuiae | Actinomycetota | Actinomycetes |
| Cellulomonas fengjieae | Actinomycetota | Actinomycetes |
| Cellulomonas fimi ATCC 484 | Actinomycetota | Actinomycetes |
| Cellulomonas flavigena DSM 20109 | Actinomycetota | Actinomycetes |
| Cellulomonas gilvus ATCC 13127 | Actinomycetota | Actinomycetes |
| Cellulomonas palmilytica | Actinomycetota | Actinomycetes |
| Cellulomonas shaoxiangyii | Actinomycetota | Actinomycetes |
| Cellulomonas taurus | Actinomycetota | Actinomycetes |
| Cellulomonas wangleii | Actinomycetota | Actinomycetes |

| | | |
|---|---|---|
| Cellulomonas wangsupingiae | Actinomycetota | Actinomycetes |
| Cellulomonas xiejunii | Actinomycetota | Actinomycetes |
| Cellulosimicrobium cellulans | Actinomycetota | Actinomycetes |
| Cellulosimicrobium protaetiae | Actinomycetota | Actinomycetes |
| Changpingibacter yushuensis | Actinomycetota | Actinomycetes |
| Citricoccus nitrophenolicus | Actinomycetota | Actinomycetes |
| Clavibacter californiensis | Actinomycetota | Actinomycetes |
| Clavibacter capsici | Actinomycetota | Actinomycetes |
| Clavibacter michiganensis | Actinomycetota | Actinomycetes |
| Clavibacter nebraskensis NCPPB 2581 | Actinomycetota | Actinomycetes |
| Clavibacter phaseoli | Actinomycetota | Actinomycetes |
| Clavibacter seminis | Actinomycetota | Actinomycetes |
| Clavibacter sepedonicus | Actinomycetota | Actinomycetes |
| Clavibacter zhangzhiyongii | Actinomycetota | Actinomycetes |
| Cnuibacter physcomitrellae | Actinomycetota | Actinomycetes |
| Corynebacterium accolens | Actinomycetota | Actinomycetes |
| Corynebacterium afermentans subsp. lipophilum | Actinomycetota | Actinomycetes |
| Corynebacterium ammoniagenes DSM 20306 | Actinomycetota | Actinomycetes |
| Corynebacterium amycolatum | Actinomycetota | Actinomycetes |
| Corynebacterium anserum | Actinomycetota | Actinomycetes |
| Corynebacterium appendicis CIP 107643 | Actinomycetota | Actinomycetes |
| Corynebacterium aquatimens | Actinomycetota | Actinomycetes |
| Corynebacterium aquilae DSM 44791 | Actinomycetota | Actinomycetes |
| Corynebacterium argentoratense DSM 44202 | Actinomycetota | Actinomycetes |
| Corynebacterium atrinae | Actinomycetota | Actinomycetes |
| Corynebacterium atypicum | Actinomycetota | Actinomycetes |
| Corynebacterium aurimucosum ATCC 700975 | Actinomycetota | Actinomycetes |
| Corynebacterium auris | Actinomycetota | Actinomycetes |
| Corynebacterium auriscanis | Actinomycetota | Actinomycetes |
| Corynebacterium belfantii | Actinomycetota | Actinomycetes |
| Corynebacterium breve | Actinomycetota | Actinomycetes |

| | | |
|---|---|---|
| Corynebacterium callunae DSM 20147 | Actinomycetota | Actinomycetes |
| Corynebacterium camporealensis | Actinomycetota | Actinomycetes |
| Corynebacterium canis | Actinomycetota | Actinomycetes |
| Corynebacterium capitovis DSM 44611 | Actinomycetota | Actinomycetes |
| Corynebacterium casei LMG S-19264 | Actinomycetota | Actinomycetes |
| Corynebacterium caspium DSM 44850 | Actinomycetota | Actinomycetes |
| Corynebacterium choanae | Actinomycetota | Actinomycetes |
| Corynebacterium ciconiae DSM 44920 | Actinomycetota | Actinomycetes |
| Corynebacterium comes | Actinomycetota | Actinomycetes |
| Corynebacterium confusum | Actinomycetota | Actinomycetes |
| Corynebacterium coyleae | Actinomycetota | Actinomycetes |
| Corynebacterium crudilactis | Actinomycetota | Actinomycetes |
| Corynebacterium cystitidis | Actinomycetota | Actinomycetes |
| Corynebacterium deserti GIMN1.010 | Actinomycetota | Actinomycetes |
| Corynebacterium diphtheriae | Actinomycetota | Actinomycetes |
| Corynebacterium doosanense CAU 212 = DSM 45436 | Actinomycetota | Actinomycetes |
| Corynebacterium durum | Actinomycetota | Actinomycetes |
| Corynebacterium efficiens YS-314 | Actinomycetota | Actinomycetes |
| Corynebacterium endometrii | Actinomycetota | Actinomycetes |
| Corynebacterium epidermidicanis | Actinomycetota | Actinomycetes |
| Corynebacterium faecale | Actinomycetota | Actinomycetes |
| Corynebacterium falsenii | Actinomycetota | Actinomycetes |
| Corynebacterium felinum | Actinomycetota | Actinomycetes |
| Corynebacterium flavescens | Actinomycetota | Actinomycetes |
| Corynebacterium fournieri | Actinomycetota | Actinomycetes |
| Corynebacterium frankenforstense DSM 45800 | Actinomycetota | Actinomycetes |
| Corynebacterium freiburgense | Actinomycetota | Actinomycetes |
| Corynebacterium freneyi | Actinomycetota | Actinomycetes |
| Corynebacterium genitalium ATCC 33030 | Actinomycetota | Actinomycetes |
| Corynebacterium gerontici | Actinomycetota | Actinomycetes |
| Corynebacterium glaucum | Actinomycetota | Actinomycetes |

| | | |
|---|---|---|
| Corynebacterium glucuronolyticum DSM 44120 | Actinomycetota | Actinomycetes |
| Corynebacterium glutamicum ATCC 13032 | Actinomycetota | Actinomycetes |
| Corynebacterium glyciniphilum AJ 3170 | Actinomycetota | Actinomycetes |
| Corynebacterium gottingense | Actinomycetota | Actinomycetes |
| Corynebacterium guangdongense | Actinomycetota | Actinomycetes |
| Corynebacterium halotolerans YIM 70093 = DSM 44683 | Actinomycetota | Actinomycetes |
| Corynebacterium hansenii | Actinomycetota | Actinomycetes |
| Corynebacterium heidelbergense | Actinomycetota | Actinomycetes |
| Corynebacterium hesseae | Actinomycetota | Actinomycetes |
| Corynebacterium hindlerae | Actinomycetota | Actinomycetes |
| Corynebacterium ihumii | Actinomycetota | Actinomycetes |
| Corynebacterium imitans | Actinomycetota | Actinomycetes |
| Corynebacterium incognita | Actinomycetota | Actinomycetes |
| Corynebacterium jeddahense | Actinomycetota | Actinomycetes |
| Corynebacterium jeikeium | Actinomycetota | Actinomycetes |
| Corynebacterium kalinowskii | Actinomycetota | Actinomycetes |
| Corynebacterium kefirresidentii | Actinomycetota | Actinomycetes |
| Corynebacterium kroppenstedtii DSM 44385 | Actinomycetota | Actinomycetes |
| Corynebacterium kutscheri | Actinomycetota | Actinomycetes |
| Corynebacterium lactis RW2-5 | Actinomycetota | Actinomycetes |
| Corynebacterium liangguodongii | Actinomycetota | Actinomycetes |
| Corynebacterium lizhenjunii | Actinomycetota | Actinomycetes |
| Corynebacterium lujinxingii | Actinomycetota | Actinomycetes |
| Corynebacterium macginleyi | Actinomycetota | Actinomycetes |
| Corynebacterium marinum DSM 44953 | Actinomycetota | Actinomycetes |
| Corynebacterium maris DSM 45190 | Actinomycetota | Actinomycetes |
| Corynebacterium massiliense DSM 45435 | Actinomycetota | Actinomycetes |
| Corynebacterium mayonis | Actinomycetota | Actinomycetes |
| Corynebacterium minutissimum | Actinomycetota | Actinomycetes |
| Corynebacterium mustelae | Actinomycetota | Actinomycetes |
| Corynebacterium mycetoides | Actinomycetota | Actinomycetes |

| | | |
|---|---|---|
| Corynebacterium nuruki S6-4 | Actinomycetota | Actinomycetes |
| Corynebacterium occultum | Actinomycetota | Actinomycetes |
| Corynebacterium pelargi | Actinomycetota | Actinomycetes |
| Corynebacterium phocae | Actinomycetota | Actinomycetes |
| Corynebacterium poyangense | Actinomycetota | Actinomycetes |
| Corynebacterium propinquum | Actinomycetota | Actinomycetes |
| Corynebacterium pseudogenitalium | Actinomycetota | Actinomycetes |
| Corynebacterium pseudokroppenstedtii | Actinomycetota | Actinomycetes |
| Corynebacterium pseudopelargi | Actinomycetota | Actinomycetes |
| Corynebacterium pseudotuberculosis | Actinomycetota | Actinomycetes |
| Corynebacterium qintianiae | Actinomycetota | Actinomycetes |
| Corynebacterium ramonii | Actinomycetota | Actinomycetes |
| Corynebacterium renale | Actinomycetota | Actinomycetes |
| Corynebacterium resistens DSM 45100 | Actinomycetota | Actinomycetes |
| Corynebacterium riegelii | Actinomycetota | Actinomycetes |
| Corynebacterium rouxii | Actinomycetota | Actinomycetes |
| Corynebacterium sanguinis | Actinomycetota | Actinomycetes |
| Corynebacterium silvaticum | Actinomycetota | Actinomycetes |
| Corynebacterium simulans | Actinomycetota | Actinomycetes |
| Corynebacterium singulare | Actinomycetota | Actinomycetes |
| Corynebacterium sphenisci DSM 44792 | Actinomycetota | Actinomycetes |
| Corynebacterium stationis | Actinomycetota | Actinomycetes |
| Corynebacterium striatum | Actinomycetota | Actinomycetes |
| Corynebacterium suedekumii | Actinomycetota | Actinomycetes |
| Corynebacterium suranareeae | Actinomycetota | Actinomycetes |
| Corynebacterium terpenotabidum Y-11 | Actinomycetota | Actinomycetes |
| Corynebacterium testudinoris | Actinomycetota | Actinomycetes |
| Corynebacterium tuberculostearicum | Actinomycetota | Actinomycetes |
| Corynebacterium uberis | Actinomycetota | Actinomycetes |
| Corynebacterium ulcerans 809 | Actinomycetota | Actinomycetes |
| Corynebacterium urealyticum | Actinomycetota | Actinomycetes |

| | | |
|---|---|---|
| Corynebacterium ureicelerivorans | Actinomycetota | Actinomycetes |
| Corynebacterium urogenitale | Actinomycetota | Actinomycetes |
| Corynebacterium uterequi | Actinomycetota | Actinomycetes |
| Corynebacterium vitaeruminis DSM 20294 | Actinomycetota | Actinomycetes |
| Corynebacterium yudongzhengii | Actinomycetota | Actinomycetes |
| Corynebacterium zhongnanshanii | Actinomycetota | Actinomycetes |
| Cryobacterium soli | Actinomycetota | Actinomycetes |
| Curtobacterium aetherium | Actinomycetota | Actinomycetes |
| Curtobacterium flaccumfaciens | Actinomycetota | Actinomycetes |
| Curtobacterium poinsettiae | Actinomycetota | Actinomycetes |
| Cutibacterium acnes subsp. acnes | Actinomycetota | Actinomycetes |
| Cutibacterium avidum 44067 | Actinomycetota | Actinomycetes |
| Cutibacterium equinum | Actinomycetota | Actinomycetes |
| Cutibacterium granulosum | Actinomycetota | Actinomycetes |
| Dactylosporangium aurantiacum | Actinomycetota | Actinomycetes |
| Dactylosporangium aurantiacum | Actinomycetota | Actinomycetes |
| Dactylosporangium fulvum | Actinomycetota | Actinomycetes |
| Dactylosporangium fulvum | Actinomycetota | Actinomycetes |
| Dactylosporangium roseum | Actinomycetota | Actinomycetes |
| Dactylosporangium roseum | Actinomycetota | Actinomycetes |
| Dactylosporangium vinaceum | Actinomycetota | Actinomycetes |
| Dactylosporangium vinaceum | Actinomycetota | Actinomycetes |
| Demequina capsici | Actinomycetota | Actinomycetes |
| Dermabacter jinjuensis | Actinomycetota | Actinomycetes |
| Dermabacter vaginalis | Actinomycetota | Actinomycetes |
| Dermacoccus nishinomiyaensis | Actinomycetota | Actinomycetes |
| Dermatophilus congolensis | Actinomycetota | Actinomycetes |
| Devriesea agamarum | Actinomycetota | Actinomycetes |
| Dietzia cinnamea | Actinomycetota | Actinomycetes |
| Dietzia kunjamensis | Actinomycetota | Actinomycetes |
| Dietzia lutea | Actinomycetota | Actinomycetes |

| | | |
|---|---|---|
| Dietzia psychralcaliphila | Actinomycetota | Actinomycetes |
| Dietzia timorensis | Actinomycetota | Actinomycetes |
| Epidermidibacterium keratini | Actinomycetota | Actinomycetes |
| Flaviflexus equikiangi | Actinomycetota | Actinomycetes |
| Flaviflexus salsibiostraticola | Actinomycetota | Actinomycetes |
| Frankia alni ACN14a | Actinomycetota | Actinomycetes |
| Frankia casuarinae | Actinomycetota | Actinomycetes |
| Friedmanniella luteola | Actinomycetota | Actinomycetes |
| Frondihabitans sucicola | Actinomycetota | Actinomycetes |
| Gardnerella leopoldii | Actinomycetota | Actinomycetes |
| Gardnerella vaginalis ATCC 14018 = JCM 11026 | Actinomycetota | Actinomycetes |
| Geodermatophilus obscurus DSM 43160 | Actinomycetota | Actinomycetes |
| Georgenia faecalis | Actinomycetota | Actinomycetes |
| Georgenia wutianyii | Actinomycetota | Actinomycetes |
| Georgenia yuyongxinii | Actinomycetota | Actinomycetes |
| Gephyromycinifex aptenodytis | Actinomycetota | Actinomycetes |
| Gleimia europaea | Actinomycetota | Actinomycetes |
| Glutamicibacter arilaitensis Re117 | Actinomycetota | Actinomycetes |
| Glutamicibacter creatinolyticus | Actinomycetota | Actinomycetes |
| Glutamicibacter ectropisis | Actinomycetota | Actinomycetes |
| Glutamicibacter endophyticus | Actinomycetota | Actinomycetes |
| Glutamicibacter halophytocola | Actinomycetota | Actinomycetes |
| Glutamicibacter mishrai | Actinomycetota | Actinomycetes |
| Glutamicibacter protophormiae | Actinomycetota | Actinomycetes |
| Gordonia alkanivorans | Actinomycetota | Actinomycetes |
| Gordonia amarae | Actinomycetota | Actinomycetes |
| Gordonia amicalis | Actinomycetota | Actinomycetes |
| Gordonia bronchialis DSM 43247 | Actinomycetota | Actinomycetes |
| Gordonia hongkongensis | Actinomycetota | Actinomycetes |
| Gordonia hydrophobica | Actinomycetota | Actinomycetes |
| Gordonia insulae | Actinomycetota | Actinomycetes |

| | | |
|---|---|---|
| Gordonia iterans | Actinomycetota | Actinomycetes |
| Gordonia jinghuaiqii | Actinomycetota | Actinomycetes |
| Gordonia mangrovi | Actinomycetota | Actinomycetes |
| Gordonia otitidis | Actinomycetota | Actinomycetes |
| Gordonia phthalatica | Actinomycetota | Actinomycetes |
| Gordonia polyisoprenivorans VH2 | Actinomycetota | Actinomycetes |
| Gordonia pseudamarae | Actinomycetota | Actinomycetes |
| Gordonia rubripertincta | Actinomycetota | Actinomycetes |
| Gordonia sihwensis | Actinomycetota | Actinomycetes |
| Gordonia terrae | Actinomycetota | Actinomycetes |
| Gordonia zhaorongruii | Actinomycetota | Actinomycetes |
| Gryllotalpicola protaetiae | Actinomycetota | Actinomycetes |
| Gulosibacter molinativorax | Actinomycetota | Actinomycetes |
| Helcobacillus massiliensis | Actinomycetota | Actinomycetes |
| Hoyosella subflava DQS3-9A1 | Actinomycetota | Actinomycetes |
| Humibacter ginsenosidimutans | Actinomycetota | Actinomycetes |
| Intrasporangium calvum DSM 43043 | Actinomycetota | Actinomycetes |
| Isoptericola dokdonensis DS-3 | Actinomycetota | Actinomycetes |
| Isoptericola variabilis 225 | Actinomycetota | Actinomycetes |
| Janibacter alittae | Actinomycetota | Actinomycetes |
| Janibacter melonis | Actinomycetota | Actinomycetes |
| Janibacter terrae | Actinomycetota | Actinomycetes |
| Jatrophihabitans cynanchi | Actinomycetota | Actinomycetes |
| Jatrophihabitans telluris | Actinomycetota | Actinomycetes |
| Jiangella alkaliphila | Actinomycetota | Actinomycetes |
| Jonesia denitrificans | Actinomycetota | Actinomycetes |
| Kibdelosporangium phytohabitans | Actinomycetota | Actinomycetes |
| Kibdelosporangium phytohabitans | Actinomycetota | Actinomycetes |
| Kineococcus radiotolerans SRS30216 = ATCC BAA-149 | Actinomycetota | Actinomycetes |
| Kitasatospora camelliae | Actinomycetota | Actinomycetes |
| Kitasatospora cathayae | Actinomycetota | Actinomycetes |

| | | |
|---|---|---|
| Kitasatospora fiedleri | Actinomycetota | Actinomycetes |
| Kitasatospora griseola | Actinomycetota | Actinomycetes |
| Kitasatospora herbaricolor | Actinomycetota | Actinomycetes |
| Kitasatospora purpeofusca | Actinomycetota | Actinomycetes |
| Kitasatospora purpeofusca | Actinomycetota | Actinomycetes |
| Kitasatospora setae KM-6054 | Actinomycetota | Actinomycetes |
| Kocuria flava | Actinomycetota | Actinomycetes |
| Kocuria rhizophila | Actinomycetota | Actinomycetes |
| Kocuria turfanensis | Actinomycetota | Actinomycetes |
| Kribbella flavida DSM 17836 | Actinomycetota | Actinomycetes |
| Kribbella flavida DSM 17836 | Actinomycetota | Actinomycetes |
| Kutzneria albida DSM 43870 | Actinomycetota | Actinomycetes |
| Kutzneria buriramensis | Actinomycetota | Actinomycetes |
| Kutzneria chonburiensis | Actinomycetota | Actinomycetes |
| Kytococcus sedentarius | Actinomycetota | Actinomycetes |
| Lawsonella clevelandensis | Actinomycetota | Actinomycetes |
| Leifsonia xyli subsp. cynodontis DSM 46306 | Actinomycetota | Actinomycetes |
| Lentzea guizhouensis | Actinomycetota | Actinomycetes |
| Lentzea guizhouensis | Actinomycetota | Actinomycetes |
| Leucobacter allii | Actinomycetota | Actinomycetes |
| Leucobacter coleopterorum | Actinomycetota | Actinomycetes |
| Leucobacter denitrificans | Actinomycetota | Actinomycetes |
| Leucobacter insecticola | Actinomycetota | Actinomycetes |
| Leucobacter muris | Actinomycetota | Actinomycetes |
| Leucobacter rhizosphaerae | Actinomycetota | Actinomycetes |
| Leucobacter tenebrionis | Actinomycetota | Actinomycetes |
| Leucobacter triazinivorans | Actinomycetota | Actinomycetes |
| Leucobacter viscericola | Actinomycetota | Actinomycetes |
| Luteimicrobium xylanilyticum | Actinomycetota | Actinomycetes |
| Luteipulveratus mongoliensis | Actinomycetota | Actinomycetes |
| Marisediminicola antarctica | Actinomycetota | Actinomycetes |

| | | |
|---|---|---|
| Microbacterium abyssi | Actinomycetota | Actinomycetes |
| Microbacterium albipurpureum | Actinomycetota | Actinomycetes |
| Microbacterium amylolyticum | Actinomycetota | Actinomycetes |
| Microbacterium arborescens | Actinomycetota | Actinomycetes |
| Microbacterium atlanticum | Actinomycetota | Actinomycetes |
| Microbacterium aurugineum | Actinomycetota | Actinomycetes |
| Microbacterium aurum | Actinomycetota | Actinomycetes |
| Microbacterium azadirachtae | Actinomycetota | Actinomycetes |
| Microbacterium betulae | Actinomycetota | Actinomycetes |
| Microbacterium caowuchunii | Actinomycetota | Actinomycetes |
| Microbacterium chocolatum | Actinomycetota | Actinomycetes |
| Microbacterium cremeum | Actinomycetota | Actinomycetes |
| Microbacterium endophyticum | Actinomycetota | Actinomycetes |
| Microbacterium foliorum | Actinomycetota | Actinomycetes |
| Microbacterium forte | Actinomycetota | Actinomycetes |
| Microbacterium galbinum | Actinomycetota | Actinomycetes |
| Microbacterium hominis | Actinomycetota | Actinomycetes |
| Microbacterium horticulturae | Actinomycetota | Actinomycetes |
| Microbacterium lemovicicum | Actinomycetota | Actinomycetes |
| Microbacterium limosum | Actinomycetota | Actinomycetes |
| Microbacterium lushaniae | Actinomycetota | Actinomycetes |
| Microbacterium luteum | Actinomycetota | Actinomycetes |
| Microbacterium maritypicum | Actinomycetota | Actinomycetes |
| Microbacterium neungamense | Actinomycetota | Actinomycetes |
| Microbacterium oleivorans | Actinomycetota | Actinomycetes |
| Microbacterium oryzae | Actinomycetota | Actinomycetes |
| Microbacterium paraoxydans | Actinomycetota | Actinomycetes |
| Microbacterium plantarum | Actinomycetota | Actinomycetes |
| Microbacterium protaetiae | Actinomycetota | Actinomycetes |
| Microbacterium pygmaeum | Actinomycetota | Actinomycetes |
| Microbacterium resistens | Actinomycetota | Actinomycetes |

| | | |
|---|---|---|
| Microbacterium rhizosphaerae | Actinomycetota | Actinomycetes |
| Microbacterium sediminis | Actinomycetota | Actinomycetes |
| Microbacterium sufflavum | Actinomycetota | Actinomycetes |
| Microbacterium terricola | Actinomycetota | Actinomycetes |
| Microbacterium terrisoli | Actinomycetota | Actinomycetes |
| Microbacterium wangchenii | Actinomycetota | Actinomycetes |
| Microbispora maris | Actinomycetota | Actinomycetes |
| Microcella daejeonensis | Actinomycetota | Actinomycetes |
| Microcella flavibacter | Actinomycetota | Actinomycetes |
| Microcella humidisoli | Actinomycetota | Actinomycetes |
| Microcella indica | Actinomycetota | Actinomycetes |
| Micrococcus antarcticus | Actinomycetota | Actinomycetes |
| Micrococcus luteus NCTC 2665 | Actinomycetota | Actinomycetes |
| Micrococcus porci | Actinomycetota | Actinomycetes |
| Microlunatus elymi | Actinomycetota | Actinomycetes |
| Microlunatus phosphovorus NM-1 | Actinomycetota | Actinomycetes |
| Microlunatus sagamiharensis | Actinomycetota | Actinomycetes |
| Microlunatus soli | Actinomycetota | Actinomycetes |
| Micromonospora aurantiaca ATCC 27029 | Actinomycetota | Actinomycetes |
| Micromonospora auratinigra | Actinomycetota | Actinomycetes |
| Micromonospora carbonacea | Actinomycetota | Actinomycetes |
| Micromonospora cathayae | Actinomycetota | Actinomycetes |
| Micromonospora chalcea | Actinomycetota | Actinomycetes |
| Micromonospora chokoriensis | Actinomycetota | Actinomycetes |
| Micromonospora coriariae | Actinomycetota | Actinomycetes |
| Micromonospora coxensis | Actinomycetota | Actinomycetes |
| Micromonospora craniellae | Actinomycetota | Actinomycetes |
| Micromonospora echinaurantiaca | Actinomycetota | Actinomycetes |
| Micromonospora echinofusca | Actinomycetota | Actinomycetes |
| Micromonospora globbae | Actinomycetota | Actinomycetes |
| Micromonospora halotolerans | Actinomycetota | Actinomycetes |

| | | |
|---|---|---|
| Micromonospora harpali | Actinomycetota | Actinomycetes |
| Micromonospora inositola | Actinomycetota | Actinomycetes |
| Micromonospora krabiensis | Actinomycetota | Actinomycetes |
| Micromonospora maris AB-18-032 | Actinomycetota | Actinomycetes |
| Micromonospora narathiwatensis | Actinomycetota | Actinomycetes |
| Micromonospora purpureochromogenes | Actinomycetota | Actinomycetes |
| Micromonospora rifamycinica | Actinomycetota | Actinomycetes |
| Micromonospora robiginosa | Actinomycetota | Actinomycetes |
| Micromonospora sagamiensis | Actinomycetota | Actinomycetes |
| Micromonospora siamensis | Actinomycetota | Actinomycetes |
| Micromonospora siamensis | Actinomycetota | Actinomycetes |
| Micromonospora terminaliae | Actinomycetota | Actinomycetes |
| Micromonospora viridifaciens | Actinomycetota | Actinomycetes |
| Micromonospora zamorensis | Actinomycetota | Actinomycetes |
| Micropruina glycogenica | Actinomycetota | Actinomycetes |
| Microterricola viridarii | Actinomycetota | Actinomycetes |
| Mobiluncus curtisii ATCC 43063 | Actinomycetota | Actinomycetes |
| Modestobacter italicus | Actinomycetota | Actinomycetes |
| Mumia qirimensis | Actinomycetota | Actinomycetes |
| Mycetocola spongiae | Actinomycetota | Actinomycetes |
| Mycetocola zhujimingii | Actinomycetota | Actinomycetes |
| Mycobacterium adipatum | Actinomycetota | Actinomycetes |
| Mycobacterium avium subsp. hominissuis | Actinomycetota | Actinomycetes |
| Mycobacterium basiliense | Actinomycetota | Actinomycetes |
| Mycobacterium branderi | Actinomycetota | Actinomycetes |
| Mycobacterium canettii CIPT 140010059 | Actinomycetota | Actinomycetes |
| Mycobacterium conspicuum | Actinomycetota | Actinomycetes |
| Mycobacterium cookii | Actinomycetota | Actinomycetes |
| Mycobacterium dioxanotrophicus | Actinomycetota | Actinomycetes |
| Mycobacterium europaeum | Actinomycetota | Actinomycetes |
| Mycobacterium florentinum | Actinomycetota | Actinomycetes |

| | | |
|---|---|---|
| Mycobacterium gallinarum | Actinomycetota | Actinomycetes |
| Mycobacterium haemophilum DSM 44634 | Actinomycetota | Actinomycetes |
| Mycobacterium heckeshornense | Actinomycetota | Actinomycetes |
| Mycobacterium heidelbergense | Actinomycetota | Actinomycetes |
| Mycobacterium intracellulare | Actinomycetota | Actinomycetes |
| Mycobacterium kansasii ATCC 12478 | Actinomycetota | Actinomycetes |
| Mycobacterium kubicae | Actinomycetota | Actinomycetes |
| Mycobacterium lacus | Actinomycetota | Actinomycetes |
| Mycobacterium lentiflavum | Actinomycetota | Actinomycetes |
| Mycobacterium leprae | Actinomycetota | Actinomycetes |
| Mycobacterium lepromatosis | Actinomycetota | Actinomycetes |
| Mycobacterium malmoense | Actinomycetota | Actinomycetes |
| Mycobacterium mantenii | Actinomycetota | Actinomycetes |
| Mycobacterium marinum E11 | Actinomycetota | Actinomycetes |
| Mycobacterium marseillense | Actinomycetota | Actinomycetes |
| Mycobacterium noviomagense | Actinomycetota | Actinomycetes |
| Mycobacterium novum | Actinomycetota | Actinomycetes |
| Mycobacterium ostraviense | Actinomycetota | Actinomycetes |
| Mycobacterium paragordonae | Actinomycetota | Actinomycetes |
| Mycobacterium paraintracellulare | Actinomycetota | Actinomycetes |
| Mycobacterium paraseoulense | Actinomycetota | Actinomycetes |
| Mycobacterium paraterrae | Actinomycetota | Actinomycetes |
| Mycobacterium parmense | Actinomycetota | Actinomycetes |
| Mycobacterium pseudoshottsii JCM 15466 | Actinomycetota | Actinomycetes |
| Mycobacterium riyadhense | Actinomycetota | Actinomycetes |
| Mycobacterium saskatchewanense | Actinomycetota | Actinomycetes |
| Mycobacterium senriense | Actinomycetota | Actinomycetes |
| Mycobacterium seoulense | Actinomycetota | Actinomycetes |
| Mycobacterium shigaense | Actinomycetota | Actinomycetes |
| Mycobacterium shinjukuense | Actinomycetota | Actinomycetes |
| Mycobacterium shottsii | Actinomycetota | Actinomycetes |

| | | |
|---|---|---|
| Mycobacterium simiae | Actinomycetota | Actinomycetes |
| Mycobacterium spongiae | Actinomycetota | Actinomycetes |
| Mycobacterium stomatepiae | Actinomycetota | Actinomycetes |
| Mycobacterium tuberculosis H37Rv | Actinomycetota | Actinomycetes |
| Mycobacterium ulcerans subsp. shinshuense | Actinomycetota | Actinomycetes |
| Mycobacterium vicinigordonae | Actinomycetota | Actinomycetes |
| Mycobacteroides abscessus ATCC 19977 | Actinomycetota | Actinomycetes |
| Mycobacteroides chelonae CCUG 47445 | Actinomycetota | Actinomycetes |
| Mycobacteroides immunogenum | Actinomycetota | Actinomycetes |
| Mycobacteroides salmoniphilum | Actinomycetota | Actinomycetes |
| Mycobacteroides saopaulense | Actinomycetota | Actinomycetes |
| Mycolicibacillus koreensis | Actinomycetota | Actinomycetes |
| Mycolicibacillus parakoreensis | Actinomycetota | Actinomycetes |
| Mycolicibacter heraklionensis | Actinomycetota | Actinomycetes |
| Mycolicibacter hiberniae | Actinomycetota | Actinomycetes |
| Mycolicibacter minnesotensis | Actinomycetota | Actinomycetes |
| Mycolicibacter sinensis | Actinomycetota | Actinomycetes |
| Mycolicibacter terrae | Actinomycetota | Actinomycetes |
| Mycolicibacter virginiensis | Actinomycetota | Actinomycetes |
| Mycolicibacterium aichiense | Actinomycetota | Actinomycetes |
| Mycolicibacterium alvei | Actinomycetota | Actinomycetes |
| Mycolicibacterium anyangense | Actinomycetota | Actinomycetes |
| Mycolicibacterium arabiense | Actinomycetota | Actinomycetes |
| Mycolicibacterium aubagnense | Actinomycetota | Actinomycetes |
| Mycolicibacterium aurum | Actinomycetota | Actinomycetes |
| Mycolicibacterium austroafricanum | Actinomycetota | Actinomycetes |
| Mycolicibacterium baixiangningiae | Actinomycetota | Actinomycetes |
| Mycolicibacterium boenickei | Actinomycetota | Actinomycetes |
| Mycolicibacterium brumae | Actinomycetota | Actinomycetes |
| Mycolicibacterium celeriflavum | Actinomycetota | Actinomycetes |
| Mycolicibacterium chitae | Actinomycetota | Actinomycetes |

| | | |
|---|---|---|
| Mycolicibacterium confluentis | Actinomycetota | Actinomycetes |
| Mycolicibacterium crocinum | Actinomycetota | Actinomycetes |
| Mycolicibacterium crocinum | Actinomycetota | Actinomycetes |
| Mycolicibacterium diernhoferi | Actinomycetota | Actinomycetes |
| Mycolicibacterium doricum | Actinomycetota | Actinomycetes |
| Mycolicibacterium duvalii | Actinomycetota | Actinomycetes |
| Mycolicibacterium fallax | Actinomycetota | Actinomycetes |
| Mycolicibacterium farcinogenes | Actinomycetota | Actinomycetes |
| Mycolicibacterium fluoranthenivorans | Actinomycetota | Actinomycetes |
| Mycolicibacterium fortuitum subsp. fortuitum | Actinomycetota | Actinomycetes |
| Mycolicibacterium frederiksbergense | Actinomycetota | Actinomycetes |
| Mycolicibacterium gadium | Actinomycetota | Actinomycetes |
| Mycolicibacterium gilvum Spyr1 | Actinomycetota | Actinomycetes |
| Mycolicibacterium goodii | Actinomycetota | Actinomycetes |
| Mycolicibacterium grossiae | Actinomycetota | Actinomycetes |
| Mycolicibacterium hassiacum DSM 44199 | Actinomycetota | Actinomycetes |
| Mycolicibacterium helvum | Actinomycetota | Actinomycetes |
| Mycolicibacterium holsaticum DSM 44478 = JCM 12374 | Actinomycetota | Actinomycetes |
| Mycolicibacterium insubricum | Actinomycetota | Actinomycetes |
| Mycolicibacterium litorale | Actinomycetota | Actinomycetes |
| Mycolicibacterium madagascariense | Actinomycetota | Actinomycetes |
| Mycolicibacterium mageritense | Actinomycetota | Actinomycetes |
| Mycolicibacterium mengxianglii | Actinomycetota | Actinomycetes |
| Mycolicibacterium monacense | Actinomycetota | Actinomycetes |
| Mycolicibacterium moriokaense | Actinomycetota | Actinomycetes |
| Mycolicibacterium mucogenicum DSM 44124 | Actinomycetota | Actinomycetes |
| Mycolicibacterium neoaurum VKM Ac-1815D | Actinomycetota | Actinomycetes |
| Mycolicibacterium nivoides | Actinomycetota | Actinomycetes |
| Mycolicibacterium pallens | Actinomycetota | Actinomycetes |
| Mycolicibacterium parafortuitum | Actinomycetota | Actinomycetes |
| Mycolicibacterium phocaicum | Actinomycetota | Actinomycetes |

| | | |
|---|---|---|
| Mycolicibacterium poriferae | Actinomycetota | Actinomycetes |
| Mycolicibacterium psychrotolerans | Actinomycetota | Actinomycetes |
| Mycolicibacterium pulveris | Actinomycetota | Actinomycetes |
| Mycolicibacterium rufum | Actinomycetota | Actinomycetes |
| Mycolicibacterium rutilum | Actinomycetota | Actinomycetes |
| Mycolicibacterium sarraceniae | Actinomycetota | Actinomycetes |
| Mycolicibacterium sediminis | Actinomycetota | Actinomycetes |
| Mycolicibacterium senegalense | Actinomycetota | Actinomycetes |
| Mycolicibacterium septicum | Actinomycetota | Actinomycetes |
| Mycolicibacterium smegmatis | Actinomycetota | Actinomycetes |
| Mycolicibacterium thermoresistibile | Actinomycetota | Actinomycetes |
| Mycolicibacterium tokaiense | Actinomycetota | Actinomycetes |
| Mycolicibacterium vaccae 95051 | Actinomycetota | Actinomycetes |
| Mycolicibacterium vanbaalenii PYR-1 | Actinomycetota | Actinomycetes |
| Naasia aerilata | Actinomycetota | Actinomycetes |
| Nakamurella antarctica | Actinomycetota | Actinomycetes |
| Nakamurella multipartita DSM 44233 | Actinomycetota | Actinomycetes |
| Nakamurella panacisegetis | Actinomycetota | Actinomycetes |
| Natronoglycomyces albus | Actinomycetota | Actinomycetes |
| Natronosporangium hydrolyticum | Actinomycetota | Actinomycetes |
| Nocardia asteroides | Actinomycetota | Actinomycetes |
| Nocardia brasiliensis | Actinomycetota | Actinomycetes |
| Nocardia brasiliensis | Actinomycetota | Actinomycetes |
| Nocardia cyriacigeorgica | Actinomycetota | Actinomycetes |
| Nocardia farcinica IFM 10152 | Actinomycetota | Actinomycetes |
| Nocardia gipuzkoensis | Actinomycetota | Actinomycetes |
| Nocardia huaxiensis | Actinomycetota | Actinomycetes |
| Nocardia huaxiensis | Actinomycetota | Actinomycetes |
| Nocardia huaxiensis | Actinomycetota | Actinomycetes |
| Nocardia iowensis | Actinomycetota | Actinomycetes |
| Nocardia iowensis | Actinomycetota | Actinomycetes |

| | | |
|---|---|---|
| Nocardia mangyaensis | Actinomycetota | Actinomycetes |
| Nocardia otitidiscaviarum | Actinomycetota | Actinomycetes |
| Nocardia otitidiscaviarum | Actinomycetota | Actinomycetes |
| Nocardia puris | Actinomycetota | Actinomycetes |
| Nocardia puris | Actinomycetota | Actinomycetes |
| Nocardia salmonicida | Actinomycetota | Actinomycetes |
| Nocardia sputorum | Actinomycetota | Actinomycetes |
| Nocardia terpenica | Actinomycetota | Actinomycetes |
| Nocardia terpenica | Actinomycetota | Actinomycetes |
| Nocardia wallacei | Actinomycetota | Actinomycetes |
| Nocardia wallacei | Actinomycetota | Actinomycetes |
| Nocardia yamanashiensis | Actinomycetota | Actinomycetes |
| Nocardia yamanashiensis | Actinomycetota | Actinomycetes |
| Nocardia yamanashiensis | Actinomycetota | Actinomycetes |
| Nocardia yunnanensis | Actinomycetota | Actinomycetes |
| Nocardia yunnanensis | Actinomycetota | Actinomycetes |
| Nocardia yunnanensis | Actinomycetota | Actinomycetes |
| Nocardioides anomalus | Actinomycetota | Actinomycetes |
| Nocardioides aquaticus | Actinomycetota | Actinomycetes |
| Nocardioides baekrokdamisoli | Actinomycetota | Actinomycetes |
| Nocardioides bizhenqiangii | Actinomycetota | Actinomycetes |
| Nocardioides bizhenqiangii | Actinomycetota | Actinomycetes |
| Nocardioides campestrisoli | Actinomycetota | Actinomycetes |
| Nocardioides coralli | Actinomycetota | Actinomycetes |
| Nocardioides cynanchi | Actinomycetota | Actinomycetes |
| Nocardioides dokdonensis FR1436 | Actinomycetota | Actinomycetes |
| Nocardioides dongkuii | Actinomycetota | Actinomycetes |
| Nocardioides euryhalodurans | Actinomycetota | Actinomycetes |
| Nocardioides faecalis | Actinomycetota | Actinomycetes |
| Nocardioides humi | Actinomycetota | Actinomycetes |
| Nocardioides humi | Actinomycetota | Actinomycetes |

| | | |
|---|---|---|
| Nocardioides marmotae | Actinomycetota | Actinomycetes |
| Nocardioides mesophilus | Actinomycetota | Actinomycetes |
| Nocardioides ochotonae | Actinomycetota | Actinomycetes |
| Nocardioides okcheonensis | Actinomycetota | Actinomycetes |
| Nocardioides panacis | Actinomycetota | Actinomycetes |
| Nocardioides pantholopis | Actinomycetota | Actinomycetes |
| Nocardioides renjunii | Actinomycetota | Actinomycetes |
| Nocardioides rotundus | Actinomycetota | Actinomycetes |
| Nocardioides sambongensis | Actinomycetota | Actinomycetes |
| Nocardioides scoriae | Actinomycetota | Actinomycetes |
| Nocardioides seonyuensis | Actinomycetota | Actinomycetes |
| Nocardioides yefusunii | Actinomycetota | Actinomycetes |
| Nocardiopsis akebiae | Actinomycetota | Actinomycetes |
| Nocardiopsis changdeensis | Actinomycetota | Actinomycetes |
| Nocardiopsis dassonvillei | Actinomycetota | Actinomycetes |
| Nocardiopsis exhalans | Actinomycetota | Actinomycetes |
| Nocardiopsis gilva YIM 90087 | Actinomycetota | Actinomycetes |
| Nonomuraea coxensis DSM 45129 | Actinomycetota | Actinomycetes |
| Nonomuraea coxensis DSM 45129 | Actinomycetota | Actinomycetes |
| Nonomuraea fastidiosa | Actinomycetota | Actinomycetes |
| Nonomuraea fastidiosa | Actinomycetota | Actinomycetes |
| Nonomuraea fuscirosea | Actinomycetota | Actinomycetes |
| Nonomuraea fuscirosea | Actinomycetota | Actinomycetes |
| Nonomuraea gerenzanensis | Actinomycetota | Actinomycetes |
| Nonomuraea gerenzanensis | Actinomycetota | Actinomycetes |
| Nonomuraea glycinis | Actinomycetota | Actinomycetes |
| Nonomuraea glycinis | Actinomycetota | Actinomycetes |
| Nonomuraea phyllanthi | Actinomycetota | Actinomycetes |
| Occultella kanbiaonis | Actinomycetota | Actinomycetes |
| Ornithinimicrobium avium | Actinomycetota | Actinomycetes |
| Ornithinimicrobium ciconiae | Actinomycetota | Actinomycetes |

| | | |
|---|---|---|
| Ornithinimicrobium cryptoxanthini | Actinomycetota | Actinomycetes |
| Ornithinimicrobium faecis | Actinomycetota | Actinomycetes |
| Ornithinimicrobium faecis | Actinomycetota | Actinomycetes |
| Ornithinimicrobium flavum | Actinomycetota | Actinomycetes |
| Ornithinimicrobium pratense | Actinomycetota | Actinomycetes |
| Ornithinimicrobium sufpigmenti | Actinomycetota | Actinomycetes |
| Oryzobacter telluris | Actinomycetota | Actinomycetes |
| Oryzobacter terrae | Actinomycetota | Actinomycetes |
| Paenarthrobacter aurescens | Actinomycetota | Actinomycetes |
| Paenarthrobacter nicotinovorans | Actinomycetota | Actinomycetes |
| Paenarthrobacter ureafaciens | Actinomycetota | Actinomycetes |
| Paramicrobacterium chengjingii | Actinomycetota | Actinomycetes |
| Paramicrobacterium fandaimingii | Actinomycetota | Actinomycetes |
| Paraoerskovia marina | Actinomycetota | Actinomycetes |
| Paraoerskovia sediminicola | Actinomycetota | Actinomycetes |
| Parascardovia denticolens DSM 10105 = JCM 12538 | Actinomycetota | Actinomycetes |
| Pauljensenia hongkongensis | Actinomycetota | Actinomycetes |
| Pedococcus dokdonensis | Actinomycetota | Actinomycetes |
| Pengzhenrongella phosphoraccumulans | Actinomycetota | Actinomycetes |
| Pengzhenrongella sicca | Actinomycetota | Actinomycetes |
| Peterkaempfera bronchialis | Actinomycetota | Actinomycetes |
| Phycicoccus jejuensis | Actinomycetota | Actinomycetes |
| Phytohabitans suffuscus | Actinomycetota | Actinomycetes |
| Phytohabitans suffuscus | Actinomycetota | Actinomycetes |
| Pimelobacter simplex | Actinomycetota | Actinomycetes |
| Pimelobacter simplex | Actinomycetota | Actinomycetes |
| Polymorphospora rubra | Actinomycetota | Actinomycetes |
| Polymorphospora rubra | Actinomycetota | Actinomycetes |
| Pontimonas salivibrio | Actinomycetota | Actinomycetes |
| Prauserella marina | Actinomycetota | Actinomycetes |
| Prescottella equi | Actinomycetota | Actinomycetes |

| | | |
|---|---|---|
| Prescottella soli | Actinomycetota | Actinomycetes |
| Propionibacterium acidifaciens | Actinomycetota | Actinomycetes |
| Propionibacterium australiense | Actinomycetota | Actinomycetes |
| Propionibacterium freudenreichii | Actinomycetota | Actinomycetes |
| Propioniciclava coleopterorum | Actinomycetota | Actinomycetes |
| Propioniciclava soli | Actinomycetota | Actinomycetes |
| Protaetiibacter intestinalis | Actinomycetota | Actinomycetes |
| Protaetiibacter larvae | Actinomycetota | Actinomycetes |
| Pseudarthrobacter chlorophenolicus A6 | Actinomycetota | Actinomycetes |
| Pseudarthrobacter equi | Actinomycetota | Actinomycetes |
| Pseudarthrobacter quantipunctorum | Actinomycetota | Actinomycetes |
| Pseudofrankia inefficax | Actinomycetota | Actinomycetes |
| Pseudofrankia inefficax | Actinomycetota | Actinomycetes |
| Pseudonocardia abyssalis | Actinomycetota | Actinomycetes |
| Pseudonocardia autotrophica | Actinomycetota | Actinomycetes |
| Pseudonocardia broussonetiae | Actinomycetota | Actinomycetes |
| Pseudonocardia dioxanivorans CB1190 | Actinomycetota | Actinomycetes |
| Pseudonocardia petroleophila | Actinomycetota | Actinomycetes |
| Psychromicrobium lacuslunae | Actinomycetota | Actinomycetes |
| Rathayibacter festucae DSM 15932 | Actinomycetota | Actinomycetes |
| Rathayibacter iranicus | Actinomycetota | Actinomycetes |
| Rathayibacter rathayi | Actinomycetota | Actinomycetes |
| Rathayibacter soli | Actinomycetota | Actinomycetes |
| Rathayibacter tanaceti | Actinomycetota | Actinomycetes |
| Rathayibacter toxicus | Actinomycetota | Actinomycetes |
| Rathayibacter tritici | Actinomycetota | Actinomycetes |
| Renibacterium salmoninarum ATCC 33209 | Actinomycetota | Actinomycetes |
| Rhodococcus aetherivorans | Actinomycetota | Actinomycetes |
| Rhodococcus antarcticus | Actinomycetota | Actinomycetes |
| Rhodococcus coprophilus | Actinomycetota | Actinomycetes |
| Rhodococcus erythropolis R138 | Actinomycetota | Actinomycetes |

| | | |
|---|---|---|
| Rhodococcus globerulus | Actinomycetota | Actinomycetes |
| Rhodococcus indonesiensis | Actinomycetota | Actinomycetes |
| Rhodococcus koreensis | Actinomycetota | Actinomycetes |
| Rhodococcus opacus PD630 | Actinomycetota | Actinomycetes |
| Rhodococcus oxybenzonivorans | Actinomycetota | Actinomycetes |
| Rhodococcus pseudokoreensis | Actinomycetota | Actinomycetes |
| Rhodococcus pyridinivorans SB3094 | Actinomycetota | Actinomycetes |
| Rhodococcus qingshengii | Actinomycetota | Actinomycetes |
| Rhodococcus rhodochrous | Actinomycetota | Actinomycetes |
| Rhodococcus ruber | Actinomycetota | Actinomycetes |
| Rhodococcus sacchari | Actinomycetota | Actinomycetes |
| Rhodococcus sovatensis | Actinomycetota | Actinomycetes |
| Rhodococcus triatomae | Actinomycetota | Actinomycetes |
| Rhodococcus wratislaviensis | Actinomycetota | Actinomycetes |
| Rhodoluna lacicola | Actinomycetota | Actinomycetes |
| Rhodoluna limnophila | Actinomycetota | Actinomycetes |
| Rothia aeria | Actinomycetota | Actinomycetes |
| Rothia amarae | Actinomycetota | Actinomycetes |
| Rothia dentocariosa ATCC 17931 | Actinomycetota | Actinomycetes |
| Rothia kristinae | Actinomycetota | Actinomycetes |
| Rothia terrae | Actinomycetota | Actinomycetes |
| Ruania alkalisoli | Actinomycetota | Actinomycetes |
| Ruania halotolerans | Actinomycetota | Actinomycetes |
| Ruania suaedae | Actinomycetota | Actinomycetes |
| Ruania zhangjianzhongii | Actinomycetota | Actinomycetes |
| Saccharomonospora azurea NA-128 | Actinomycetota | Actinomycetes |
| Saccharomonospora cyanea NA-134 | Actinomycetota | Actinomycetes |
| Saccharomonospora glauca K62 | Actinomycetota | Actinomycetes |
| Saccharomonospora marina XMU15 | Actinomycetota | Actinomycetes |
| Saccharomonospora viridis DSM 43017 | Actinomycetota | Actinomycetes |
| Saccharopolyspora erythraea NRRL 2338 | Actinomycetota | Actinomycetes |

| | | |
|---|---|---|
| Saccharopolyspora gregorii | Actinomycetota | Actinomycetes |
| Saccharopolyspora pogona | Actinomycetota | Actinomycetes |
| Saccharopolyspora rosea | Actinomycetota | Actinomycetes |
| Saccharothrix espanaensis DSM 44229 | Actinomycetota | Actinomycetes |
| Saccharothrix syringae | Actinomycetota | Actinomycetes |
| Salinispora tropica CNB-440 | Actinomycetota | Actinomycetes |
| Sanguibacter biliveldensis | Actinomycetota | Actinomycetes |
| Sanguibacter keddieii DSM 10542 | Actinomycetota | Actinomycetes |
| Saxibacter everestensis | Actinomycetota | Actinomycetes |
| Scardovia inopinata JCM 12537 | Actinomycetota | Actinomycetes |
| Schaalia cardiffensis F0333 | Actinomycetota | Actinomycetes |
| Schaalia meyeri | Actinomycetota | Actinomycetes |
| Schaalia odontolytica | Actinomycetota | Actinomycetes |
| Schaalia radingae | Actinomycetota | Actinomycetes |
| Scrofimicrobium appendicitidis | Actinomycetota | Actinomycetes |
| Segniliparus rotundus DSM 44985 | Actinomycetota | Actinomycetes |
| Serinicoccus chungangensis | Actinomycetota | Actinomycetes |
| Serinicoccus hydrothermalis | Actinomycetota | Actinomycetes |
| Sinomonas atrocyanea | Actinomycetota | Actinomycetes |
| Sinomonas cyclohexanicum | Actinomycetota | Actinomycetes |
| Sinomonas puerhi | Actinomycetota | Actinomycetes |
| Skermania piniformis | Actinomycetota | Actinomycetes |
| Solicola gregarius | Actinomycetota | Actinomycetes |
| Sphaerisporangium viridialbum | Actinomycetota | Actinomycetes |
| Sphaerisporangium viridialbum | Actinomycetota | Actinomycetes |
| Spiractinospora alimapuensis | Actinomycetota | Actinomycetes |
| Stackebrandtia nassauensis DSM 44728 | Actinomycetota | Actinomycetes |
| Streptantibioticus cattleyicolor NRRL 8057 = DSM 46488 | Actinomycetota | Actinomycetes |
| Streptantibioticus cattleyicolor NRRL 8057 = DSM 46488 | Actinomycetota | Actinomycetes |
| Streptomonospora litoralis | Actinomycetota | Actinomycetes |
| Streptomyces achromogenes | Actinomycetota | Actinomycetes |

| | | |
|---|---|---|
| Streptomyces akebiae | Actinomycetota | Actinomycetes |
| Streptomyces akebiae | Actinomycetota | Actinomycetes |
| Streptomyces albidocamelliae | Actinomycetota | Actinomycetes |
| Streptomyces albidocamelliae | Actinomycetota | Actinomycetes |
| Streptomyces albidoflavus | Actinomycetota | Actinomycetes |
| Streptomyces albidoflavus | Actinomycetota | Actinomycetes |
| Streptomyces alboniger | Actinomycetota | Actinomycetes |
| Streptomyces alboniger | Actinomycetota | Actinomycetes |
| Streptomyces albus | Actinomycetota | Actinomycetes |
| Streptomyces alfalfae | Actinomycetota | Actinomycetes |
| Streptomyces alfalfae | Actinomycetota | Actinomycetes |
| Streptomyces althioticus | Actinomycetota | Actinomycetes |
| Streptomyces althioticus | Actinomycetota | Actinomycetes |
| Streptomyces ambofaciens ATCC 23877 | Actinomycetota | Actinomycetes |
| Streptomyces ambofaciens ATCC 23877 | Actinomycetota | Actinomycetes |
| Streptomyces angustmyceticus | Actinomycetota | Actinomycetes |
| Streptomyces angustmyceticus | Actinomycetota | Actinomycetes |
| Streptomyces anthocyanicus | Actinomycetota | Actinomycetes |
| Streptomyces anthocyanicus | Actinomycetota | Actinomycetes |
| Streptomyces antimycoticus | Actinomycetota | Actinomycetes |
| Streptomyces antimycoticus | Actinomycetota | Actinomycetes |
| Streptomyces anulatus | Actinomycetota | Actinomycetes |
| Streptomyces anulatus | Actinomycetota | Actinomycetes |
| Streptomyces aquilus | Actinomycetota | Actinomycetes |
| Streptomyces aquilus | Actinomycetota | Actinomycetes |
| Streptomyces aquilus | Actinomycetota | Actinomycetes |
| Streptomyces argyrophylli | Actinomycetota | Actinomycetes |
| Streptomyces argyrophylli | Actinomycetota | Actinomycetes |
| Streptomyces armeniacus | Actinomycetota | Actinomycetes |
| Streptomyces armeniacus | Actinomycetota | Actinomycetes |
| Streptomyces atratus | Actinomycetota | Actinomycetes |

| | | |
|---|---|---|
| Streptomyces atratus | Actinomycetota | Actinomycetes |
| Streptomyces aurantiacus | Actinomycetota | Actinomycetes |
| Streptomyces aurantiacus | Actinomycetota | Actinomycetes |
| Streptomyces auratus AGR0001 | Actinomycetota | Actinomycetes |
| Streptomyces auratus AGR0001 | Actinomycetota | Actinomycetes |
| Streptomyces avermitilis MA-4680 = NBRC 14893 | Actinomycetota | Actinomycetes |
| Streptomyces avermitilis MA-4680 = NBRC 14893 | Actinomycetota | Actinomycetes |
| Streptomyces avidinii | Actinomycetota | Actinomycetes |
| Streptomyces avidinii | Actinomycetota | Actinomycetes |
| Streptomyces bathyalis | Actinomycetota | Actinomycetes |
| Streptomyces bathyalis | Actinomycetota | Actinomycetes |
| Streptomyces bobili | Actinomycetota | Actinomycetes |
| Streptomyces bobili | Actinomycetota | Actinomycetes |
| Streptomyces boninensis | Actinomycetota | Actinomycetes |
| Streptomyces boninensis | Actinomycetota | Actinomycetes |
| Streptomyces broussonetiae | Actinomycetota | Actinomycetes |
| Streptomyces broussonetiae | Actinomycetota | Actinomycetes |
| Streptomyces broussonetiae | Actinomycetota | Actinomycetes |
| Streptomyces broussonetiae | Actinomycetota | Actinomycetes |
| Streptomyces buecherae | Actinomycetota | Actinomycetes |
| Streptomyces cadmiisoli | Actinomycetota | Actinomycetes |
| Streptomyces californicus | Actinomycetota | Actinomycetes |
| Streptomyces californicus | Actinomycetota | Actinomycetes |
| Streptomyces calvus | Actinomycetota | Actinomycetes |
| Streptomyces calvus | Actinomycetota | Actinomycetes |
| Streptomyces camelliae | Actinomycetota | Actinomycetes |
| Streptomyces camelliae | Actinomycetota | Actinomycetes |
| Streptomyces caniferus | Actinomycetota | Actinomycetes |
| Streptomyces caniferus | Actinomycetota | Actinomycetes |
| Streptomyces caniscabiei | Actinomycetota | Actinomycetes |
| Streptomyces caniscabiei | Actinomycetota | Actinomycetes |

| | | |
|---|---|---|
| Streptomyces canus | Actinomycetota | Actinomycetes |
| Streptomyces canus | Actinomycetota | Actinomycetes |
| Streptomyces capitiformicae | Actinomycetota | Actinomycetes |
| Streptomyces capitiformicae | Actinomycetota | Actinomycetes |
| Streptomyces carpaticus | Actinomycetota | Actinomycetes |
| Streptomyces carpaticus | Actinomycetota | Actinomycetes |
| Streptomyces castrisilvae | Actinomycetota | Actinomycetes |
| Streptomyces castrisilvae | Actinomycetota | Actinomycetes |
| Streptomyces cathayae | Actinomycetota | Actinomycetes |
| Streptomyces changanensis | Actinomycetota | Actinomycetes |
| Streptomyces changanensis | Actinomycetota | Actinomycetes |
| Streptomyces chartreusis | Actinomycetota | Actinomycetes |
| Streptomyces chartreusis | Actinomycetota | Actinomycetes |
| Streptomyces chengbuensis | Actinomycetota | Actinomycetes |
| Streptomyces chengbuensis | Actinomycetota | Actinomycetes |
| Streptomyces chromofuscus | Actinomycetota | Actinomycetes |
| Streptomyces chromofuscus | Actinomycetota | Actinomycetes |
| Streptomyces cinereoruber | Actinomycetota | Actinomycetes |
| Streptomyces cinnabarinus | Actinomycetota | Actinomycetes |
| Streptomyces cinnabarinus | Actinomycetota | Actinomycetes |
| Streptomyces cinnabarinus | Actinomycetota | Actinomycetes |
| Streptomyces citrinus | Actinomycetota | Actinomycetes |
| Streptomyces citrinus | Actinomycetota | Actinomycetes |
| Streptomyces clavifer | Actinomycetota | Actinomycetes |
| Streptomyces clavifer | Actinomycetota | Actinomycetes |
| Streptomyces clavuligerus | Actinomycetota | Actinomycetes |
| Streptomyces coelicoflavus | Actinomycetota | Actinomycetes |
| Streptomyces coelicoflavus | Actinomycetota | Actinomycetes |
| Streptomyces collinus | Actinomycetota | Actinomycetes |
| Streptomyces collinus | Actinomycetota | Actinomycetes |
| Streptomyces cyaneochromogenes | Actinomycetota | Actinomycetes |

| | | |
|---|---|---|
| Streptomyces cyaneochromogenes | Actinomycetota | Actinomycetes |
| Streptomyces cyaneochromogenes | Actinomycetota | Actinomycetes |
| Streptomyces cyaneofuscatus | Actinomycetota | Actinomycetes |
| Streptomyces cyaneofuscatus | Actinomycetota | Actinomycetes |
| Streptomyces cyanogenus | Actinomycetota | Actinomycetes |
| Streptomyces cyanogenus | Actinomycetota | Actinomycetes |
| Streptomyces cynarae | Actinomycetota | Actinomycetes |
| Streptomyces cynarae | Actinomycetota | Actinomycetes |
| Streptomyces dangxiongensis | Actinomycetota | Actinomycetes |
| Streptomyces davaonensis JCM 4913 | Actinomycetota | Actinomycetes |
| Streptomyces davaonensis JCM 4913 | Actinomycetota | Actinomycetes |
| Streptomyces davaonensis JCM 4913 | Actinomycetota | Actinomycetes |
| Streptomyces deccanensis | Actinomycetota | Actinomycetes |
| Streptomyces deccanensis | Actinomycetota | Actinomycetes |
| Streptomyces decoyicus | Actinomycetota | Actinomycetes |
| Streptomyces decoyicus | Actinomycetota | Actinomycetes |
| Streptomyces dengpaensis | Actinomycetota | Actinomycetes |
| Streptomyces dengpaensis | Actinomycetota | Actinomycetes |
| Streptomyces dengpaensis | Actinomycetota | Actinomycetes |
| Streptomyces durmitorensis | Actinomycetota | Actinomycetes |
| Streptomyces durmitorensis | Actinomycetota | Actinomycetes |
| Streptomyces erythrochromogenes | Actinomycetota | Actinomycetes |
| Streptomyces erythrochromogenes | Actinomycetota | Actinomycetes |
| Streptomyces europaeiscabiei | Actinomycetota | Actinomycetes |
| Streptomyces europaeiscabiei | Actinomycetota | Actinomycetes |
| Streptomyces fagopyri | Actinomycetota | Actinomycetes |
| Streptomyces fagopyri | Actinomycetota | Actinomycetes |
| Streptomyces ferrugineus | Actinomycetota | Actinomycetes |
| Streptomyces ferrugineus | Actinomycetota | Actinomycetes |
| Streptomyces ficellus | Actinomycetota | Actinomycetes |
| Streptomyces ficellus | Actinomycetota | Actinomycetes |

| | | |
|---|---|---|
| Streptomyces fodineus | Actinomycetota | Actinomycetes |
| Streptomyces fodineus | Actinomycetota | Actinomycetes |
| Streptomyces formicae | Actinomycetota | Actinomycetes |
| Streptomyces formicae | Actinomycetota | Actinomycetes |
| Streptomyces fradiae ATCC 10745 = DSM 40063 | Actinomycetota | Actinomycetes |
| Streptomyces fradiae ATCC 10745 = DSM 40063 | Actinomycetota | Actinomycetes |
| Streptomyces fungicidicus | Actinomycetota | Actinomycetes |
| Streptomyces fungicidicus | Actinomycetota | Actinomycetes |
| Streptomyces gardneri | Actinomycetota | Actinomycetes |
| Streptomyces genisteinicus | Actinomycetota | Actinomycetes |
| Streptomyces genisteinicus | Actinomycetota | Actinomycetes |
| Streptomyces gilvosporeus | Actinomycetota | Actinomycetes |
| Streptomyces gilvosporeus | Actinomycetota | Actinomycetes |
| Streptomyces globisporus C-1027 | Actinomycetota | Actinomycetes |
| Streptomyces globisporus C-1027 | Actinomycetota | Actinomycetes |
| Streptomyces glycanivorans | Actinomycetota | Actinomycetes |
| Streptomyces glycanivorans | Actinomycetota | Actinomycetes |
| Streptomyces gobiensis | Actinomycetota | Actinomycetes |
| Streptomyces gobiensis | Actinomycetota | Actinomycetes |
| Streptomyces gobiensis | Actinomycetota | Actinomycetes |
| Streptomyces goshikiensis | Actinomycetota | Actinomycetes |
| Streptomyces goshikiensis | Actinomycetota | Actinomycetes |
| Streptomyces gougerotii | Actinomycetota | Actinomycetes |
| Streptomyces gougerotii | Actinomycetota | Actinomycetes |
| Streptomyces graminofaciens | Actinomycetota | Actinomycetes |
| Streptomyces graminofaciens | Actinomycetota | Actinomycetes |
| Streptomyces graminofaciens | Actinomycetota | Actinomycetes |
| Streptomyces griseoaurantiacus | Actinomycetota | Actinomycetes |
| Streptomyces griseoaurantiacus | Actinomycetota | Actinomycetes |
| Streptomyces griseochromogenes | Actinomycetota | Actinomycetes |
| Streptomyces griseochromogenes | Actinomycetota | Actinomycetes |

| | | |
|---|---|---|
| Streptomyces griseofuscus | Actinomycetota | Actinomycetes |
| Streptomyces griseofuscus | Actinomycetota | Actinomycetes |
| Streptomyces griseorubiginosus | Actinomycetota | Actinomycetes |
| Streptomyces griseorubiginosus | Actinomycetota | Actinomycetes |
| Streptomyces halobius | Actinomycetota | Actinomycetes |
| Streptomyces halobius | Actinomycetota | Actinomycetes |
| Streptomyces halstedii | Actinomycetota | Actinomycetes |
| Streptomyces halstedii | Actinomycetota | Actinomycetes |
| Streptomyces harbinensis | Actinomycetota | Actinomycetes |
| Streptomyces harbinensis | Actinomycetota | Actinomycetes |
| Streptomyces hawaiiensis | Actinomycetota | Actinomycetes |
| Streptomyces hawaiiensis | Actinomycetota | Actinomycetes |
| Streptomyces hirsutus | Actinomycetota | Actinomycetes |
| Streptomyces hirsutus | Actinomycetota | Actinomycetes |
| Streptomyces huasconensis | Actinomycetota | Actinomycetes |
| Streptomyces huasconensis | Actinomycetota | Actinomycetes |
| Streptomyces hygroscopicus subsp. hygroscopicus | Actinomycetota | Actinomycetes |
| Streptomyces hygroscopicus subsp. hygroscopicus | Actinomycetota | Actinomycetes |
| Streptomyces iakyrus | Actinomycetota | Actinomycetes |
| Streptomyces iakyrus | Actinomycetota | Actinomycetes |
| Streptomyces incarnatus | Actinomycetota | Actinomycetes |
| Streptomyces incarnatus | Actinomycetota | Actinomycetes |
| Streptomyces inhibens | Actinomycetota | Actinomycetes |
| Streptomyces inhibens | Actinomycetota | Actinomycetes |
| Streptomyces iranensis | Actinomycetota | Actinomycetes |
| Streptomyces iranensis | Actinomycetota | Actinomycetes |
| Streptomyces janthinus | Actinomycetota | Actinomycetes |
| Streptomyces janthinus | Actinomycetota | Actinomycetes |
| Streptomyces jietaisiensis | Actinomycetota | Actinomycetes |
| Streptomyces jietaisiensis | Actinomycetota | Actinomycetes |
| Streptomyces kanamyceticus | Actinomycetota | Actinomycetes |

| | | |
|---|---|---|
| Streptomyces kanamyceticus | Actinomycetota | Actinomycetes |
| Streptomyces koelreuteriae | Actinomycetota | Actinomycetes |
| Streptomyces koelreuteriae | Actinomycetota | Actinomycetes |
| Streptomyces koyangensis | Actinomycetota | Actinomycetes |
| Streptomyces laculatispora | Actinomycetota | Actinomycetes |
| Streptomyces laculatispora | Actinomycetota | Actinomycetes |
| Streptomyces lavendulae subsp. lavendulae | Actinomycetota | Actinomycetes |
| Streptomyces lavendulae subsp. lavendulae | Actinomycetota | Actinomycetes |
| Streptomyces leeuwenhoekii | Actinomycetota | Actinomycetes |
| Streptomyces leeuwenhoekii | Actinomycetota | Actinomycetes |
| Streptomyces liangshanensis | Actinomycetota | Actinomycetes |
| Streptomyces liangshanensis | Actinomycetota | Actinomycetes |
| Streptomyces libani subsp. libani | Actinomycetota | Actinomycetes |
| Streptomyces libani subsp. libani | Actinomycetota | Actinomycetes |
| Streptomyces lienomycini | Actinomycetota | Actinomycetes |
| Streptomyces lienomycini | Actinomycetota | Actinomycetes |
| Streptomyces liliifuscus | Actinomycetota | Actinomycetes |
| Streptomyces liliifuscus | Actinomycetota | Actinomycetes |
| Streptomyces lincolnensis | Actinomycetota | Actinomycetes |
| Streptomyces lincolnensis | Actinomycetota | Actinomycetes |
| Streptomyces lincolnensis | Actinomycetota | Actinomycetes |
| Streptomyces longhuiensis | Actinomycetota | Actinomycetes |
| Streptomyces longhuiensis | Actinomycetota | Actinomycetes |
| Streptomyces lunaelactis | Actinomycetota | Actinomycetes |
| Streptomyces lunaelactis | Actinomycetota | Actinomycetes |
| Streptomyces luomodiensis | Actinomycetota | Actinomycetes |
| Streptomyces luomodiensis | Actinomycetota | Actinomycetes |
| Streptomyces luteireticuli | Actinomycetota | Actinomycetes |
| Streptomyces lydicus | Actinomycetota | Actinomycetes |
| Streptomyces lydicus | Actinomycetota | Actinomycetes |
| Streptomyces malaysiensis | Actinomycetota | Actinomycetes |

| | | |
|---|---|---|
| Streptomyces malaysiensis | Actinomycetota | Actinomycetes |
| Streptomyces manipurensis | Actinomycetota | Actinomycetes |
| Streptomyces manipurensis | Actinomycetota | Actinomycetes |
| Streptomyces marincola | Actinomycetota | Actinomycetes |
| Streptomyces microflavus | Actinomycetota | Actinomycetes |
| Streptomyces microflavus | Actinomycetota | Actinomycetes |
| Streptomyces milbemycinicus | Actinomycetota | Actinomycetes |
| Streptomyces milbemycinicus | Actinomycetota | Actinomycetes |
| Streptomyces mirabilis | Actinomycetota | Actinomycetes |
| Streptomyces mirabilis | Actinomycetota | Actinomycetes |
| Streptomyces mirabilis | Actinomycetota | Actinomycetes |
| Streptomyces mobaraensis | Actinomycetota | Actinomycetes |
| Streptomyces murinus | Actinomycetota | Actinomycetes |
| Streptomyces murinus | Actinomycetota | Actinomycetes |
| Streptomyces mutabilis | Actinomycetota | Actinomycetes |
| Streptomyces mutabilis | Actinomycetota | Actinomycetes |
| Streptomyces netropsis | Actinomycetota | Actinomycetes |
| Streptomyces nigra | Actinomycetota | Actinomycetes |
| Streptomyces nigra | Actinomycetota | Actinomycetes |
| Streptomyces nigrescens | Actinomycetota | Actinomycetes |
| Streptomyces nigrescens | Actinomycetota | Actinomycetes |
| Streptomyces nitrosporeus | Actinomycetota | Actinomycetes |
| Streptomyces nitrosporeus | Actinomycetota | Actinomycetes |
| Streptomyces niveus | Actinomycetota | Actinomycetes |
| Streptomyces niveus | Actinomycetota | Actinomycetes |
| Streptomyces nodosus | Actinomycetota | Actinomycetes |
| Streptomyces nodosus | Actinomycetota | Actinomycetes |
| Streptomyces nojiriensis | Actinomycetota | Actinomycetes |
| Streptomyces nojiriensis | Actinomycetota | Actinomycetes |
| Streptomyces noursei | Actinomycetota | Actinomycetes |
| Streptomyces noursei | Actinomycetota | Actinomycetes |

| | | |
|---|---|---|
| Streptomyces olivaceus | Actinomycetota | Actinomycetes |
| Streptomyces olivaceus | Actinomycetota | Actinomycetes |
| Streptomyces paludis | Actinomycetota | Actinomycetes |
| Streptomyces paludis | Actinomycetota | Actinomycetes |
| Streptomyces parvulus | Actinomycetota | Actinomycetes |
| Streptomyces parvulus | Actinomycetota | Actinomycetes |
| Streptomyces phaeochromogenes | Actinomycetota | Actinomycetes |
| Streptomyces phaeochromogenes | Actinomycetota | Actinomycetes |
| Streptomyces phaeolivaceus | Actinomycetota | Actinomycetes |
| Streptomyces phaeolivaceus | Actinomycetota | Actinomycetes |
| Streptomyces phaeoluteigriseus | Actinomycetota | Actinomycetes |
| Streptomyces phaeoluteigriseus | Actinomycetota | Actinomycetes |
| Streptomyces phytohabitans | Actinomycetota | Actinomycetes |
| Streptomyces platensis | Actinomycetota | Actinomycetes |
| Streptomyces platensis | Actinomycetota | Actinomycetes |
| Streptomyces pluripotens | Actinomycetota | Actinomycetes |
| Streptomyces pluripotens | Actinomycetota | Actinomycetes |
| Streptomyces poriferorum | Actinomycetota | Actinomycetes |
| Streptomyces poriferorum | Actinomycetota | Actinomycetes |
| Streptomyces prasinus | Actinomycetota | Actinomycetes |
| Streptomyces prasinus | Actinomycetota | Actinomycetes |
| Streptomyces pratensis | Actinomycetota | Actinomycetes |
| Streptomyces pratensis | Actinomycetota | Actinomycetes |
| Streptomyces pristinaespiralis | Actinomycetota | Actinomycetes |
| Streptomyces pristinaespiralis | Actinomycetota | Actinomycetes |
| Streptomyces profundus | Actinomycetota | Actinomycetes |
| Streptomyces profundus | Actinomycetota | Actinomycetes |
| Streptomyces prunicolor | Actinomycetota | Actinomycetes |
| Streptomyces prunicolor | Actinomycetota | Actinomycetes |
| Streptomyces prunicolor | Actinomycetota | Actinomycetes |
| Streptomyces pseudovenezuelae | Actinomycetota | Actinomycetes |

| | | |
|---|---|---|
| Streptomyces pseudovenezuelae | Actinomycetota | Actinomycetes |
| Streptomyces purpurascens | Actinomycetota | Actinomycetes |
| Streptomyces purpurascens | Actinomycetota | Actinomycetes |
| Streptomyces qaidamensis | Actinomycetota | Actinomycetes |
| Streptomyces qaidamensis | Actinomycetota | Actinomycetes |
| Streptomyces qinzhouensis | Actinomycetota | Actinomycetes |
| Streptomyces qinzhouensis | Actinomycetota | Actinomycetes |
| Streptomyces rapamycinicus NRRL 5491 | Actinomycetota | Actinomycetes |
| Streptomyces rapamycinicus NRRL 5491 | Actinomycetota | Actinomycetes |
| Streptomyces rectiverticillatus | Actinomycetota | Actinomycetes |
| Streptomyces rimosus subsp. rimosus ATCC 10970 | Actinomycetota | Actinomycetes |
| Streptomyces rimosus subsp. rimosus ATCC 10970 | Actinomycetota | Actinomycetes |
| Streptomyces rochei | Actinomycetota | Actinomycetes |
| Streptomyces rochei | Actinomycetota | Actinomycetes |
| Streptomyces roseirectus | Actinomycetota | Actinomycetes |
| Streptomyces roseirectus | Actinomycetota | Actinomycetes |
| Streptomyces roseirectus | Actinomycetota | Actinomycetes |
| Streptomyces roseochromogenus subsp. oscitans DS 12.976 | Actinomycetota | Actinomycetes |
| Streptomyces roseochromogenus subsp. oscitans DS 12.976 | Actinomycetota | Actinomycetes |
| Streptomyces rubrogriseus | Actinomycetota | Actinomycetes |
| Streptomyces rubrogriseus | Actinomycetota | Actinomycetes |
| Streptomyces rutgersensis | Actinomycetota | Actinomycetes |
| Streptomyces rutgersensis | Actinomycetota | Actinomycetes |
| Streptomyces scabiei 87.22 | Actinomycetota | Actinomycetes |
| Streptomyces scabiei 87.22 | Actinomycetota | Actinomycetes |
| Streptomyces sclerotialus | Actinomycetota | Actinomycetes |
| Streptomyces sclerotialus | Actinomycetota | Actinomycetes |
| Streptomyces scopuliridis | Actinomycetota | Actinomycetes |
| Streptomyces scopuliridis | Actinomycetota | Actinomycetes |
| Streptomyces seoulensis | Actinomycetota | Actinomycetes |
| Streptomyces seoulensis | Actinomycetota | Actinomycetes |

| | | |
|---|---|---|
| Streptomyces sirii | Actinomycetota | Actinomycetes |
| Streptomyces sirii | Actinomycetota | Actinomycetes |
| Streptomyces solicamelliae | Actinomycetota | Actinomycetes |
| Streptomyces solicamelliae | Actinomycetota | Actinomycetes |
| Streptomyces solicathayae | Actinomycetota | Actinomycetes |
| Streptomyces solicathayae | Actinomycetota | Actinomycetes |
| Streptomyces solisilvae | Actinomycetota | Actinomycetes |
| Streptomyces solisilvae | Actinomycetota | Actinomycetes |
| Streptomyces spectabilis | Actinomycetota | Actinomycetes |
| Streptomyces spectabilis | Actinomycetota | Actinomycetes |
| Streptomyces spinosirectus | Actinomycetota | Actinomycetes |
| Streptomyces spinosirectus | Actinomycetota | Actinomycetes |
| Streptomyces spinosirectus | Actinomycetota | Actinomycetes |
| Streptomyces spongiicola | Actinomycetota | Actinomycetes |
| Streptomyces spongiicola | Actinomycetota | Actinomycetes |
| Streptomyces subrutilus | Actinomycetota | Actinomycetes |
| Streptomyces subrutilus | Actinomycetota | Actinomycetes |
| Streptomyces sudanensis | Actinomycetota | Actinomycetes |
| Streptomyces sviceus ATCC 29083 | Actinomycetota | Actinomycetes |
| Streptomyces sviceus ATCC 29083 | Actinomycetota | Actinomycetes |
| Streptomyces tabacisoli | Actinomycetota | Actinomycetes |
| Streptomyces tabacisoli | Actinomycetota | Actinomycetes |
| Streptomyces tanashiensis | Actinomycetota | Actinomycetes |
| Streptomyces tauricus | Actinomycetota | Actinomycetes |
| Streptomyces tauricus | Actinomycetota | Actinomycetes |
| Streptomyces tauricus | Actinomycetota | Actinomycetes |
| Streptomyces thermocarboxydus | Actinomycetota | Actinomycetes |
| Streptomyces thermocarboxydus | Actinomycetota | Actinomycetes |
| Streptomyces thermolineatus | Actinomycetota | Actinomycetes |
| Streptomyces thermoviolaceus | Actinomycetota | Actinomycetes |
| Streptomyces tirandamycinicus | Actinomycetota | Actinomycetes |

| | | |
|---|---|---|
| Streptomyces tirandamycinicus | Actinomycetota | Actinomycetes |
| Streptomyces tritici | Actinomycetota | Actinomycetes |
| Streptomyces tritici | Actinomycetota | Actinomycetes |
| Streptomyces tsukubensis | Actinomycetota | Actinomycetes |
| Streptomyces tsukubensis | Actinomycetota | Actinomycetes |
| Streptomyces tubbatahanensis | Actinomycetota | Actinomycetes |
| Streptomyces tubercidicus | Actinomycetota | Actinomycetes |
| Streptomyces tubercidicus | Actinomycetota | Actinomycetes |
| Streptomyces tuirus | Actinomycetota | Actinomycetes |
| Streptomyces tuirus | Actinomycetota | Actinomycetes |
| Streptomyces uncialis | Actinomycetota | Actinomycetes |
| Streptomyces uncialis | Actinomycetota | Actinomycetes |
| Streptomyces venezuelae ATCC 10712 | Actinomycetota | Actinomycetes |
| Streptomyces vietnamensis | Actinomycetota | Actinomycetes |
| Streptomyces vilmorinianum | Actinomycetota | Actinomycetes |
| Streptomyces vilmorinianum | Actinomycetota | Actinomycetes |
| Streptomyces vinaceus | Actinomycetota | Actinomycetes |
| Streptomyces vinaceus | Actinomycetota | Actinomycetes |
| Streptomyces violaceoruber | Actinomycetota | Actinomycetes |
| Streptomyces violaceoruber | Actinomycetota | Actinomycetes |
| Streptomyces violaceus | Actinomycetota | Actinomycetes |
| Streptomyces violaceus | Actinomycetota | Actinomycetes |
| Streptomyces viridosporus T7A | Actinomycetota | Actinomycetes |
| Streptomyces viridosporus T7A | Actinomycetota | Actinomycetes |
| Streptomyces xanthii | Actinomycetota | Actinomycetes |
| Streptomyces xanthii | Actinomycetota | Actinomycetes |
| Streptomyces xanthophaeus | Actinomycetota | Actinomycetes |
| Streptomyces xanthophaeus | Actinomycetota | Actinomycetes |
| Streptomyces xiamenensis | Actinomycetota | Actinomycetes |
| Streptomyces xiamenensis | Actinomycetota | Actinomycetes |
| Streptomyces xinghaiensis S187 | Actinomycetota | Actinomycetes |

| | | |
|---|---|---|
| Streptomyces xinghaiensis S187 | Actinomycetota | Actinomycetes |
| Streptomyces yangpuensis | Actinomycetota | Actinomycetes |
| Streptomyces yangpuensis | Actinomycetota | Actinomycetes |
| Streptomyces yatensis | Actinomycetota | Actinomycetes |
| Streptomyces yatensis | Actinomycetota | Actinomycetes |
| Streptomyces yunnanensis | Actinomycetota | Actinomycetes |
| Streptomyces yunnanensis | Actinomycetota | Actinomycetes |
| Streptomyces zaomyceticus | Actinomycetota | Actinomycetes |
| Streptosporangium roseum DSM 43021 | Actinomycetota | Actinomycetes |
| Streptoverticillium reticulum | Actinomycetota | Actinomycetes |
| Subtercola endophyticus | Actinomycetota | Actinomycetes |
| Tessaracoccus aquimaris | Actinomycetota | Actinomycetes |
| Tessaracoccus flavescens | Actinomycetota | Actinomycetes |
| Tessaracoccus flavus | Actinomycetota | Actinomycetes |
| Tessaracoccus lacteus | Actinomycetota | Actinomycetes |
| Tessaracoccus palaemonis | Actinomycetota | Actinomycetes |
| Tessaracoccus timonensis | Actinomycetota | Actinomycetes |
| Thermobifida alba | Actinomycetota | Actinomycetes |
| Thermobifida fusca YX | Actinomycetota | Actinomycetes |
| Thermobifida halotolerans | Actinomycetota | Actinomycetes |
| Thermobispora bispora DSM 43833 | Actinomycetota | Actinomycetes |
| Thermomonospora amylolytica | Actinomycetota | Actinomycetes |
| Thermomonospora curvata DSM 43183 | Actinomycetota | Actinomycetes |
| Tomitella fengzijianii | Actinomycetota | Actinomycetes |
| Tomitella gaofuii | Actinomycetota | Actinomycetes |
| Tropheryma whipplei TW08/27 | Actinomycetota | Actinomycetes |
| Trueperella bernardiae | Actinomycetota | Actinomycetes |
| Trueperella bialowiezensis | Actinomycetota | Actinomycetes |
| Trueperella pecoris | Actinomycetota | Actinomycetes |
| Trueperella pyogenes | Actinomycetota | Actinomycetes |
| Tsukamurella paurometabola DSM 20162 | Actinomycetota | Actinomycetes |

| | | |
|---|---|---|
| Vaginimicrobium propionicum | Actinomycetota | Actinomycetes |
| Varibaculum prostatecancerukia | Actinomycetota | Actinomycetes |
| Williamsia herbipolensis | Actinomycetota | Actinomycetes |
| Xylanimonas allomyrinae | Actinomycetota | Actinomycetes |
| Xylanimonas cellulosilytica DSM 15894 | Actinomycetota | Actinomycetes |
| Xylanimonas protaetiae | Actinomycetota | Actinomycetes |
| [Pseudomonas] carboxydohydrogena | Pseudomonadota | Alphaproteobacteria |
| Abyssibius alkaniclasticus | Pseudomonadota | Alphaproteobacteria |
| Abyssibius alkaniclasticus | Pseudomonadota | Alphaproteobacteria |
| Acetobacter ascendens | Pseudomonadota | Alphaproteobacteria |
| Acetobacter oryzifermentans | Pseudomonadota | Alphaproteobacteria |
| Acetobacter oryzoeni | Pseudomonadota | Alphaproteobacteria |
| Acetobacter pasteurianus | Pseudomonadota | Alphaproteobacteria |
| Acetobacter senegalensis | Pseudomonadota | Alphaproteobacteria |
| Acetobacter vaccinii | Pseudomonadota | Alphaproteobacteria |
| Acidibrevibacterium fodinaquatile | Pseudomonadota | Alphaproteobacteria |
| Acidiphilium multivorum AIU301 | Pseudomonadota | Alphaproteobacteria |
| Acidisoma cladoniae | Pseudomonadota | Alphaproteobacteria |
| Aerophototrophica crusticola | Pseudomonadota | Alphaproteobacteria |
| Aerophototrophica crusticola | Pseudomonadota | Alphaproteobacteria |
| Aestuariispira ectoiniformans | Pseudomonadota | Alphaproteobacteria |
| Aestuariispira ectoiniformans | Pseudomonadota | Alphaproteobacteria |
| Afipia carboxidovorans OM5 | Pseudomonadota | Alphaproteobacteria |
| Agrobacterium cucumeris | Pseudomonadota | Alphaproteobacteria |
| Agrobacterium cucumeris | Pseudomonadota | Alphaproteobacteria |
| Agrobacterium fabrum str. C58 | Pseudomonadota | Alphaproteobacteria |
| Agrobacterium fabrum str. C58 | Pseudomonadota | Alphaproteobacteria |
| Agrobacterium larrymoorei | Pseudomonadota | Alphaproteobacteria |
| Agrobacterium larrymoorei | Pseudomonadota | Alphaproteobacteria |
| Agrobacterium leguminum | Pseudomonadota | Alphaproteobacteria |
| Agrobacterium leguminum | Pseudomonadota | Alphaproteobacteria |

| | | |
|---|---|---|
| Agrobacterium pusense | Pseudomonadota | Alphaproteobacteria |
| Agrobacterium pusense | Pseudomonadota | Alphaproteobacteria |
| Agrobacterium tumefaciens | Pseudomonadota | Alphaproteobacteria |
| Agrobacterium tumefaciens | Pseudomonadota | Alphaproteobacteria |
| Agrobacterium vaccinii | Pseudomonadota | Alphaproteobacteria |
| Agrobacterium vaccinii | Pseudomonadota | Alphaproteobacteria |
| Algicella marina | Pseudomonadota | Alphaproteobacteria |
| Algicella marina | Pseudomonadota | Alphaproteobacteria |
| Aliiroseovarius crassostreae | Pseudomonadota | Alphaproteobacteria |
| Aliiroseovarius crassostreae | Pseudomonadota | Alphaproteobacteria |
| Aliisedimentitalea scapharcae | Pseudomonadota | Alphaproteobacteria |
| Aliisedimentitalea scapharcae | Pseudomonadota | Alphaproteobacteria |
| Allorhizobium ampelinum S4 | Pseudomonadota | Alphaproteobacteria |
| Allorhizobium ampelinum S4 | Pseudomonadota | Alphaproteobacteria |
| Allorhizobium pseudoryzae | Pseudomonadota | Alphaproteobacteria |
| Allorhizobium pseudoryzae | Pseudomonadota | Alphaproteobacteria |
| Allosphingosinicella indica | Pseudomonadota | Alphaproteobacteria |
| Alphaproteobacteria bacterium endosymbiont of Tiliacea citrago | Pseudomonadota | Alphaproteobacteria |
| Altererythrobacter arenosus | Pseudomonadota | Alphaproteobacteria |
| Altererythrobacter epoxidivorans | Pseudomonadota | Alphaproteobacteria |
| Altererythrobacter ishigakiensis | Pseudomonadota | Alphaproteobacteria |
| Altererythrobacter rubellus | Pseudomonadota | Alphaproteobacteria |
| Alteriqipengyuania flavescens | Pseudomonadota | Alphaproteobacteria |
| Alterisphingorhabdus coralli | Pseudomonadota | Alphaproteobacteria |
| Aminobacter niigataensis | Pseudomonadota | Alphaproteobacteria |
| Aminobacter niigataensis | Pseudomonadota | Alphaproteobacteria |
| Anaplasma centrale str. Israel | Pseudomonadota | Alphaproteobacteria |
| Anaplasma centrale str. Israel | Pseudomonadota | Alphaproteobacteria |
| Anaplasma marginale str. Florida | Pseudomonadota | Alphaproteobacteria |
| Anaplasma marginale str. Florida | Pseudomonadota | Alphaproteobacteria |
| Anaplasma ovis str. Haibei | Pseudomonadota | Alphaproteobacteria |

| | | |
|---|---|---|
| Anaplasma ovis str. Haibei | Pseudomonadota | Alphaproteobacteria |
| Anaplasma platys | Pseudomonadota | Alphaproteobacteria |
| Ancylobacter novellus DSM 506 | Pseudomonadota | Alphaproteobacteria |
| Ancylobacter novellus DSM 506 | Pseudomonadota | Alphaproteobacteria |
| Ancylobacter pratisalsi | Pseudomonadota | Alphaproteobacteria |
| Ancylobacter pratisalsi | Pseudomonadota | Alphaproteobacteria |
| Aquibium microcysteis | Pseudomonadota | Alphaproteobacteria |
| Aquibium microcysteis | Pseudomonadota | Alphaproteobacteria |
| Aquibium oceanicum | Pseudomonadota | Alphaproteobacteria |
| Aquibium oceanicum | Pseudomonadota | Alphaproteobacteria |
| Aquisediminimonas profunda | Pseudomonadota | Alphaproteobacteria |
| Aristophania vespae | Pseudomonadota | Alphaproteobacteria |
| Asaia bogorensis NBRC 16594 | Pseudomonadota | Alphaproteobacteria |
| Asticcacaulis excentricus CB 48 | Pseudomonadota | Alphaproteobacteria |
| Aurantiacibacter atlanticus | Pseudomonadota | Alphaproteobacteria |
| Aurantiacibacter gangjinensis | Pseudomonadota | Alphaproteobacteria |
| Aureimonas mangrovi | Pseudomonadota | Alphaproteobacteria |
| Aureimonas mangrovi | Pseudomonadota | Alphaproteobacteria |
| Aureimonas phyllosphaerae | Pseudomonadota | Alphaproteobacteria |
| Aureimonas phyllosphaerae | Pseudomonadota | Alphaproteobacteria |
| Aureimonas populi | Pseudomonadota | Alphaproteobacteria |
| Aureimonas populi | Pseudomonadota | Alphaproteobacteria |
| Azorhizobium caulinodans ORS 571 | Pseudomonadota | Alphaproteobacteria |
| Azorhizobium caulinodans ORS 571 | Pseudomonadota | Alphaproteobacteria |
| Azospirillum argentinense | Pseudomonadota | Alphaproteobacteria |
| Azospirillum argentinense | Pseudomonadota | Alphaproteobacteria |
| Azospirillum baldaniorum | Pseudomonadota | Alphaproteobacteria |
| Azospirillum baldaniorum | Pseudomonadota | Alphaproteobacteria |
| Azospirillum humicireducens | Pseudomonadota | Alphaproteobacteria |
| Azospirillum humicireducens | Pseudomonadota | Alphaproteobacteria |
| Azospirillum oryzae | Pseudomonadota | Alphaproteobacteria |

| | | |
|---|---|---|
| Azospirillum oryzae | Pseudomonadota | Alphaproteobacteria |
| Azospirillum ramasamyi | Pseudomonadota | Alphaproteobacteria |
| Azospirillum ramasamyi | Pseudomonadota | Alphaproteobacteria |
| Azospirillum thermophilum | Pseudomonadota | Alphaproteobacteria |
| Azospirillum thermophilum | Pseudomonadota | Alphaproteobacteria |
| Azospirillum thiophilum | Pseudomonadota | Alphaproteobacteria |
| Azospirillum thiophilum | Pseudomonadota | Alphaproteobacteria |
| Bartonella alsatica | Pseudomonadota | Alphaproteobacteria |
| Bartonella alsatica | Pseudomonadota | Alphaproteobacteria |
| Bartonella ancashensis | Pseudomonadota | Alphaproteobacteria |
| Bartonella ancashensis | Pseudomonadota | Alphaproteobacteria |
| Bartonella australis AUST/NH1 | Pseudomonadota | Alphaproteobacteria |
| Bartonella australis AUST/NH1 | Pseudomonadota | Alphaproteobacteria |
| Bartonella bacilliformis KC583 | Pseudomonadota | Alphaproteobacteria |
| Bartonella bacilliformis KC583 | Pseudomonadota | Alphaproteobacteria |
| Bartonella bacilliformis KC583 | Pseudomonadota | Alphaproteobacteria |
| Bartonella clarridgeiae 73 | Pseudomonadota | Alphaproteobacteria |
| Bartonella clarridgeiae 73 | Pseudomonadota | Alphaproteobacteria |
| Bartonella elizabethae | Pseudomonadota | Alphaproteobacteria |
| Bartonella elizabethae | Pseudomonadota | Alphaproteobacteria |
| Bartonella gliris | Pseudomonadota | Alphaproteobacteria |
| Bartonella gliris | Pseudomonadota | Alphaproteobacteria |
| Bartonella grahamii as4aup | Pseudomonadota | Alphaproteobacteria |
| Bartonella grahamii as4aup | Pseudomonadota | Alphaproteobacteria |
| Bartonella harrusi | Pseudomonadota | Alphaproteobacteria |
| Bartonella harrusi | Pseudomonadota | Alphaproteobacteria |
| Bartonella heixiaziensis | Pseudomonadota | Alphaproteobacteria |
| Bartonella heixiaziensis | Pseudomonadota | Alphaproteobacteria |
| Bartonella henselae | Pseudomonadota | Alphaproteobacteria |
| Bartonella henselae | Pseudomonadota | Alphaproteobacteria |
| Bartonella kosoyi | Pseudomonadota | Alphaproteobacteria |

| | | |
|---|---|---|
| Bartonella kosoyi | Pseudomonadota | Alphaproteobacteria |
| Bartonella krasnovii | Pseudomonadota | Alphaproteobacteria |
| Bartonella krasnovii | Pseudomonadota | Alphaproteobacteria |
| Bartonella machadoae | Pseudomonadota | Alphaproteobacteria |
| Bartonella machadoae | Pseudomonadota | Alphaproteobacteria |
| Bartonella quintana | Pseudomonadota | Alphaproteobacteria |
| Bartonella quintana | Pseudomonadota | Alphaproteobacteria |
| Bartonella schoenbuchensis | Pseudomonadota | Alphaproteobacteria |
| Bartonella schoenbuchensis | Pseudomonadota | Alphaproteobacteria |
| Bartonella taylorii | Pseudomonadota | Alphaproteobacteria |
| Bartonella taylorii | Pseudomonadota | Alphaproteobacteria |
| Bartonella tribocorum CIP 105476 | Pseudomonadota | Alphaproteobacteria |
| Bartonella tribocorum CIP 105476 | Pseudomonadota | Alphaproteobacteria |
| Bartonella vinsonii | Pseudomonadota | Alphaproteobacteria |
| Bartonella vinsonii | Pseudomonadota | Alphaproteobacteria |
| Beijerinckia indica subsp. indica ATCC 9039 | Pseudomonadota | Alphaproteobacteria |
| Beijerinckia indica subsp. indica ATCC 9039 | Pseudomonadota | Alphaproteobacteria |
| Blastochloris tepida | Pseudomonadota | Alphaproteobacteria |
| Blastochloris viridis | Pseudomonadota | Alphaproteobacteria |
| Blastomonas fulva | Pseudomonadota | Alphaproteobacteria |
| Blastomonas marina | Pseudomonadota | Alphaproteobacteria |
| Bosea vaviloviae | Pseudomonadota | Alphaproteobacteria |
| Bosea vaviloviae | Pseudomonadota | Alphaproteobacteria |
| Bradyrhizobium amphicarpaeae | Pseudomonadota | Alphaproteobacteria |
| Bradyrhizobium arachidis | Pseudomonadota | Alphaproteobacteria |
| Bradyrhizobium barranii subsp. barranii | Pseudomonadota | Alphaproteobacteria |
| Bradyrhizobium betae | Pseudomonadota | Alphaproteobacteria |
| Bradyrhizobium brasilense | Pseudomonadota | Alphaproteobacteria |
| Bradyrhizobium centrosematis | Pseudomonadota | Alphaproteobacteria |
| Bradyrhizobium commune | Pseudomonadota | Alphaproteobacteria |
| Bradyrhizobium commune | Pseudomonadota | Alphaproteobacteria |

| | | |
|---|---|---|
| Bradyrhizobium cosmicum | Pseudomonadota | Alphaproteobacteria |
| Bradyrhizobium daqingense | Pseudomonadota | Alphaproteobacteria |
| Bradyrhizobium diazoefficiens USDA 110 | Pseudomonadota | Alphaproteobacteria |
| Bradyrhizobium elkanii USDA 76 | Pseudomonadota | Alphaproteobacteria |
| Bradyrhizobium ganzhouense | Pseudomonadota | Alphaproteobacteria |
| Bradyrhizobium guangdongense | Pseudomonadota | Alphaproteobacteria |
| Bradyrhizobium guangxiense | Pseudomonadota | Alphaproteobacteria |
| Bradyrhizobium guangzhouense | Pseudomonadota | Alphaproteobacteria |
| Bradyrhizobium huanghuaihaiense | Pseudomonadota | Alphaproteobacteria |
| Bradyrhizobium icense | Pseudomonadota | Alphaproteobacteria |
| Bradyrhizobium japonicum | Pseudomonadota | Alphaproteobacteria |
| Bradyrhizobium lupini | Pseudomonadota | Alphaproteobacteria |
| Bradyrhizobium oligotrophicum S58 | Pseudomonadota | Alphaproteobacteria |
| Bradyrhizobium ontarionense | Pseudomonadota | Alphaproteobacteria |
| Bradyrhizobium ottawaense | Pseudomonadota | Alphaproteobacteria |
| Bradyrhizobium pachyrhizi | Pseudomonadota | Alphaproteobacteria |
| Bradyrhizobium paxllaeri | Pseudomonadota | Alphaproteobacteria |
| Bradyrhizobium prioriisuperbiae | Pseudomonadota | Alphaproteobacteria |
| Bradyrhizobium prioriisuperbiae | Pseudomonadota | Alphaproteobacteria |
| Bradyrhizobium quebecense | Pseudomonadota | Alphaproteobacteria |
| Bradyrhizobium roseum | Pseudomonadota | Alphaproteobacteria |
| Bradyrhizobium sediminis | Pseudomonadota | Alphaproteobacteria |
| Bradyrhizobium septentrionale | Pseudomonadota | Alphaproteobacteria |
| Bradyrhizobium septentrionale | Pseudomonadota | Alphaproteobacteria |
| Bradyrhizobium symbiodeficiens | Pseudomonadota | Alphaproteobacteria |
| Bradyrhizobium vignae | Pseudomonadota | Alphaproteobacteria |
| Bradyrhizobium xenonodulans | Pseudomonadota | Alphaproteobacteria |
| Bradyrhizobium yuanmingense | Pseudomonadota | Alphaproteobacteria |
| Bradyrhizobium zhanjiangense | Pseudomonadota | Alphaproteobacteria |
| Brevundimonas albigilva | Pseudomonadota | Alphaproteobacteria |
| Brevundimonas albigilva | Pseudomonadota | Alphaproteobacteria |

| | | |
|---|---|---|
| Brevundimonas bullata | Pseudomonadota | Alphaproteobacteria |
| Brevundimonas bullata | Pseudomonadota | Alphaproteobacteria |
| Brevundimonas fontaquae | Pseudomonadota | Alphaproteobacteria |
| Brevundimonas fontaquae | Pseudomonadota | Alphaproteobacteria |
| Brevundimonas goettingensis | Pseudomonadota | Alphaproteobacteria |
| Brevundimonas goettingensis | Pseudomonadota | Alphaproteobacteria |
| Brevundimonas intermedia | Pseudomonadota | Alphaproteobacteria |
| Brevundimonas intermedia | Pseudomonadota | Alphaproteobacteria |
| Brevundimonas nasdae | Pseudomonadota | Alphaproteobacteria |
| Brevundimonas nasdae | Pseudomonadota | Alphaproteobacteria |
| Brevundimonas olei | Pseudomonadota | Alphaproteobacteria |
| Brevundimonas olei | Pseudomonadota | Alphaproteobacteria |
| Brevundimonas sanguinis | Pseudomonadota | Alphaproteobacteria |
| Brevundimonas sanguinis | Pseudomonadota | Alphaproteobacteria |
| Brevundimonas staleyi | Pseudomonadota | Alphaproteobacteria |
| Brevundimonas staleyi | Pseudomonadota | Alphaproteobacteria |
| Brevundimonas subvibrioides ATCC 15264 | Pseudomonadota | Alphaproteobacteria |
| Brevundimonas subvibrioides ATCC 15264 | Pseudomonadota | Alphaproteobacteria |
| Brevundimonas vesicularis | Pseudomonadota | Alphaproteobacteria |
| Brevundimonas vesicularis | Pseudomonadota | Alphaproteobacteria |
| Brevundimonas vitisensis | Pseudomonadota | Alphaproteobacteria |
| Brevundimonas vitisensis | Pseudomonadota | Alphaproteobacteria |
| Brucella anthropi | Pseudomonadota | Alphaproteobacteria |
| Brucella anthropi | Pseudomonadota | Alphaproteobacteria |
| Brucella canis ATCC 23365 | Pseudomonadota | Alphaproteobacteria |
| Brucella canis ATCC 23365 | Pseudomonadota | Alphaproteobacteria |
| Brucella ceti | Pseudomonadota | Alphaproteobacteria |
| Brucella ceti | Pseudomonadota | Alphaproteobacteria |
| Brucella inopinata | Pseudomonadota | Alphaproteobacteria |
| Brucella inopinata | Pseudomonadota | Alphaproteobacteria |
| Brucella melitensis bv. 1 str. 16M | Pseudomonadota | Alphaproteobacteria |

| | | |
|---|---|---|
| Brucella melitensis bv. 1 str. 16M | Pseudomonadota | Alphaproteobacteria |
| Brucella microti CCM 4915 | Pseudomonadota | Alphaproteobacteria |
| Brucella microti CCM 4915 | Pseudomonadota | Alphaproteobacteria |
| Brucella ovis ATCC 25840 | Pseudomonadota | Alphaproteobacteria |
| Brucella ovis ATCC 25840 | Pseudomonadota | Alphaproteobacteria |
| Brucella pseudintermedia | Pseudomonadota | Alphaproteobacteria |
| Brucella pseudintermedia | Pseudomonadota | Alphaproteobacteria |
| Brucella pseudogrignonensis | Pseudomonadota | Alphaproteobacteria |
| Brucella pseudogrignonensis | Pseudomonadota | Alphaproteobacteria |
| Brucella suis 1330 | Pseudomonadota | Alphaproteobacteria |
| Brucella suis 1330 | Pseudomonadota | Alphaproteobacteria |
| Caenibius tardaugens NBRC 16725 | Pseudomonadota | Alphaproteobacteria |
| Candidatus Bandiella numerosa | Pseudomonadota | Alphaproteobacteria |
| Candidatus Bandiella numerosa | Pseudomonadota | Alphaproteobacteria |
| Candidatus Bandiella woodruffii | Pseudomonadota | Alphaproteobacteria |
| Candidatus Bandiella woodruffii | Pseudomonadota | Alphaproteobacteria |
| Candidatus Bealeia paramacronuclearis | Pseudomonadota | Alphaproteobacteria |
| Candidatus Bealeia paramacronuclearis | Pseudomonadota | Alphaproteobacteria |
| Candidatus Bodocaedibacter vickermanii | Pseudomonadota | Alphaproteobacteria |
| Candidatus Cytomitobacter indipagum | Pseudomonadota | Alphaproteobacteria |
| Candidatus Cytomitobacter primus | Pseudomonadota | Alphaproteobacteria |
| Candidatus Deianiraea vastatrix | Pseudomonadota | Alphaproteobacteria |
| Candidatus Endolissoclinum faulkneri L2 | Pseudomonadota | Alphaproteobacteria |
| Candidatus Endowatersipora endosymbiont of Watersipora subatra | Pseudomonadota | Alphaproteobacteria |
| Candidatus Fokinia cryptica | Pseudomonadota | Alphaproteobacteria |
| Candidatus Fokinia solitaria | Pseudomonadota | Alphaproteobacteria |
| Candidatus Fonsibacter ubiquis | Pseudomonadota | Alphaproteobacteria |
| Candidatus Gromoviella agglomerans | Pseudomonadota | Alphaproteobacteria |
| Candidatus Hodgkinia cicadicola | Pseudomonadota | Alphaproteobacteria |
| Candidatus Hodgkinia cicadicola | Pseudomonadota | Alphaproteobacteria |
| Candidatus Hydrogenosomobacter endosymbioticus | Pseudomonadota | Alphaproteobacteria |

| | | |
|---|---|---|
| Candidatus Kirkpatrickella diaphorinae | Pseudomonadota | Alphaproteobacteria |
| Candidatus Lariskella endosymbiont of Epinotia ramella | Pseudomonadota | Alphaproteobacteria |
| Candidatus Lariskella endosymbiont of Hedychridium roseum | Pseudomonadota | Alphaproteobacteria |
| Candidatus Liberibacter africanus PTSAPSY | Pseudomonadota | Alphaproteobacteria |
| Candidatus Liberibacter africanus PTSAPSY | Pseudomonadota | Alphaproteobacteria |
| Candidatus Liberibacter americanus str. Sao Paulo | Pseudomonadota | Alphaproteobacteria |
| Candidatus Liberibacter americanus str. Sao Paulo | Pseudomonadota | Alphaproteobacteria |
| Candidatus Liberibacter asiaticus | Pseudomonadota | Alphaproteobacteria |
| Candidatus Liberibacter asiaticus | Pseudomonadota | Alphaproteobacteria |
| Candidatus Liberibacter solanacearum CLso-ZC1 | Pseudomonadota | Alphaproteobacteria |
| Candidatus Liberibacter solanacearum CLso-ZC1 | Pseudomonadota | Alphaproteobacteria |
| Candidatus Megaera polyxenophila | Pseudomonadota | Alphaproteobacteria |
| Candidatus Mesenet endosymbiont of Agriotes lineatus | Pseudomonadota | Alphaproteobacteria |
| Candidatus Mesenet endosymbiont of Agriotes lineatus | Pseudomonadota | Alphaproteobacteria |
| Candidatus Mesenet endosymbiont of Phosphuga atrata | Pseudomonadota | Alphaproteobacteria |
| Candidatus Mesenet endosymbiont of Phosphuga atrata | Pseudomonadota | Alphaproteobacteria |
| Candidatus Midichloria mitochondrii IricVA | Pseudomonadota | Alphaproteobacteria |
| Candidatus Nesciobacter abundans | Pseudomonadota | Alphaproteobacteria |
| Candidatus Nucleicultrix amoebiphila FS5 | Pseudomonadota | Alphaproteobacteria |
| Candidatus Nucleicultrix amoebiphila FS5 | Pseudomonadota | Alphaproteobacteria |
| Candidatus Odyssella acanthamoebae | Pseudomonadota | Alphaproteobacteria |
| Candidatus Odyssella acanthamoebae | Pseudomonadota | Alphaproteobacteria |
| Candidatus Pelagibacter giovannonii | Pseudomonadota | Alphaproteobacteria |
| Candidatus Pelagibacter ubique HTCC1062 | Pseudomonadota | Alphaproteobacteria |
| Candidatus Phycorickettsia trachydisci | Pseudomonadota | Alphaproteobacteria |
| Candidatus Puniceispirillum marinum IMCC1322 | Pseudomonadota | Alphaproteobacteria |
| Candidatus Puniceispirillum marinum IMCC1322 | Pseudomonadota | Alphaproteobacteria |
| Candidatus Sneabacter namystus | Pseudomonadota | Alphaproteobacteria |
| Candidatus Tisiphia endosymbiont of Beris chalybata | Pseudomonadota | Alphaproteobacteria |
| Candidatus Tisiphia endosymbiont of Ceraclea dissimilis | Pseudomonadota | Alphaproteobacteria |
| Candidatus Tisiphia endosymbiont of Dascillus cervinus | Pseudomonadota | Alphaproteobacteria |

| | | |
|---|---|---|
| Candidatus Tisiphia endosymbiont of Dioctria linearis | Pseudomonadota | Alphaproteobacteria |
| Candidatus Tisiphia endosymbiont of Dioctria rufipes | Pseudomonadota | Alphaproteobacteria |
| Candidatus Tisiphia endosymbiont of Ditula angustiorana | Pseudomonadota | Alphaproteobacteria |
| Candidatus Tisiphia endosymbiont of Empis tessellata | Pseudomonadota | Alphaproteobacteria |
| Candidatus Tisiphia endosymbiont of Melanophora roralis | Pseudomonadota | Alphaproteobacteria |
| Candidatus Tisiphia endosymbiont of Metellina segmentata | Pseudomonadota | Alphaproteobacteria |
| Candidatus Tisiphia endosymbiont of Micropterix aruncella | Pseudomonadota | Alphaproteobacteria |
| Candidatus Tisiphia endosymbiont of Myopa tessellatipennis | Pseudomonadota | Alphaproteobacteria |
| Candidatus Tisiphia endosymbiont of Mystacides longicornis | Pseudomonadota | Alphaproteobacteria |
| Candidatus Tisiphia endosymbiont of Nedyus quadrimaculatus | Pseudomonadota | Alphaproteobacteria |
| Candidatus Tisiphia endosymbiont of Nemotelus nigrinus | Pseudomonadota | Alphaproteobacteria |
| Candidatus Tisiphia endosymbiont of Nemotelus uliginosus | Pseudomonadota | Alphaproteobacteria |
| Candidatus Tisiphia endosymbiont of Neophilaenus lineatus | Pseudomonadota | Alphaproteobacteria |
| Candidatus Tisiphia endosymbiont of Oplodontha viridula | Pseudomonadota | Alphaproteobacteria |
| Candidatus Tisiphia endosymbiont of Parasteatoda lunata | Pseudomonadota | Alphaproteobacteria |
| Candidatus Tisiphia endosymbiont of Piscicola geometra | Pseudomonadota | Alphaproteobacteria |
| Candidatus Tisiphia endosymbiont of Psammoecus bipunctatus | Pseudomonadota | Alphaproteobacteria |
| Candidatus Tisiphia endosymbiont of Ptychoptera albimana | Pseudomonadota | Alphaproteobacteria |
| Candidatus Tisiphia endosymbiont of Sialis lutaria | Pseudomonadota | Alphaproteobacteria |
| Candidatus Tisiphia endosymbiont of Stenodema calcarata | Pseudomonadota | Alphaproteobacteria |
| Candidatus Tisiphia endosymbiont of Temnostethus pusillus | Pseudomonadota | Alphaproteobacteria |
| Candidatus Tisiphia endosymbiont of Thecophora atra | Pseudomonadota | Alphaproteobacteria |
| Candidatus Tisiphia endosymbiont of Xenochironomus xenolabis | Pseudomonadota | Alphaproteobacteria |
| Candidatus Trichorickettsia mobilis | Pseudomonadota | Alphaproteobacteria |
| Candidatus Viadribacter manganicus | Pseudomonadota | Alphaproteobacteria |
| Caulobacter flavus | Pseudomonadota | Alphaproteobacteria |
| Caulobacter flavus | Pseudomonadota | Alphaproteobacteria |
| Caulobacter henricii | Pseudomonadota | Alphaproteobacteria |
| Caulobacter henricii | Pseudomonadota | Alphaproteobacteria |
| Caulobacter mirabilis | Pseudomonadota | Alphaproteobacteria |
| Caulobacter mirabilis | Pseudomonadota | Alphaproteobacteria |

| | | |
|---|---|---|
| Caulobacter rhizosphaerae | Pseudomonadota | Alphaproteobacteria |
| Caulobacter rhizosphaerae | Pseudomonadota | Alphaproteobacteria |
| Caulobacter segnis | Pseudomonadota | Alphaproteobacteria |
| Caulobacter segnis | Pseudomonadota | Alphaproteobacteria |
| Caulobacter soli | Pseudomonadota | Alphaproteobacteria |
| Caulobacter soli | Pseudomonadota | Alphaproteobacteria |
| Celeribacter ethanolicus | Pseudomonadota | Alphaproteobacteria |
| Celeribacter ethanolicus | Pseudomonadota | Alphaproteobacteria |
| Celeribacter indicus | Pseudomonadota | Alphaproteobacteria |
| Celeribacter indicus | Pseudomonadota | Alphaproteobacteria |
| Celeribacter marinus | Pseudomonadota | Alphaproteobacteria |
| Celeribacter marinus | Pseudomonadota | Alphaproteobacteria |
| Cereibacter azotoformans | Pseudomonadota | Alphaproteobacteria |
| Cereibacter azotoformans | Pseudomonadota | Alphaproteobacteria |
| Cereibacter sphaeroides 2.4.1 | Pseudomonadota | Alphaproteobacteria |
| Cereibacter sphaeroides 2.4.1 | Pseudomonadota | Alphaproteobacteria |
| Ciceribacter thiooxidans | Pseudomonadota | Alphaproteobacteria |
| Ciceribacter thiooxidans | Pseudomonadota | Alphaproteobacteria |
| Cohaesibacter marisflavi | Pseudomonadota | Alphaproteobacteria |
| Cohaesibacter marisflavi | Pseudomonadota | Alphaproteobacteria |
| Commensalibacter melissae | Pseudomonadota | Alphaproteobacteria |
| Coralliovum pocilloporae | Pseudomonadota | Alphaproteobacteria |
| Coralliovum pocilloporae | Pseudomonadota | Alphaproteobacteria |
| Croceibacterium aestuarii | Pseudomonadota | Alphaproteobacteria |
| Croceibacterium atlanticum | Pseudomonadota | Alphaproteobacteria |
| Croceicoccus marinus | Pseudomonadota | Alphaproteobacteria |
| Croceicoccus naphthovorans | Pseudomonadota | Alphaproteobacteria |
| Defluviicoccus vanus | Pseudomonadota | Alphaproteobacteria |
| Defluviicoccus vanus | Pseudomonadota | Alphaproteobacteria |
| Devosia beringensis | Pseudomonadota | Alphaproteobacteria |
| Devosia ginsengisoli | Pseudomonadota | Alphaproteobacteria |

| | | |
|---|---|---|
| Devosia lacusdianchii | Pseudomonadota | Alphaproteobacteria |
| Devosia neptuniae | Pseudomonadota | Alphaproteobacteria |
| Devosia oryziradicis | Pseudomonadota | Alphaproteobacteria |
| Devosia rhizoryzae | Pseudomonadota | Alphaproteobacteria |
| Devosia rhizosphaerae | Pseudomonadota | Alphaproteobacteria |
| Devosia rhodophyticola | Pseudomonadota | Alphaproteobacteria |
| Devosia salina | Pseudomonadota | Alphaproteobacteria |
| Dinoroseobacter shibae DFL 12 = DSM 16493 | Pseudomonadota | Alphaproteobacteria |
| Dinoroseobacter shibae DFL 12 = DSM 16493 | Pseudomonadota | Alphaproteobacteria |
| Ehrlichia canis str. Jake | Pseudomonadota | Alphaproteobacteria |
| Ehrlichia canis str. Jake | Pseudomonadota | Alphaproteobacteria |
| Ehrlichia chaffeensis str. Arkansas | Pseudomonadota | Alphaproteobacteria |
| Ehrlichia chaffeensis str. Arkansas | Pseudomonadota | Alphaproteobacteria |
| Ehrlichia japonica | Pseudomonadota | Alphaproteobacteria |
| Ehrlichia japonica | Pseudomonadota | Alphaproteobacteria |
| Ehrlichia muris AS145 | Pseudomonadota | Alphaproteobacteria |
| Ehrlichia muris AS145 | Pseudomonadota | Alphaproteobacteria |
| Ehrlichia ruminantium str. Welgevonden | Pseudomonadota | Alphaproteobacteria |
| Ehrlichia ruminantium str. Welgevonden | Pseudomonadota | Alphaproteobacteria |
| Elioraea tepida | Pseudomonadota | Alphaproteobacteria |
| Elioraea tepida | Pseudomonadota | Alphaproteobacteria |
| endosymbiont of Acanthamoeba sp. UWC8 | Pseudomonadota | Alphaproteobacteria |
| endosymbiont of Acanthamoeba sp. UWC8 | Pseudomonadota | Alphaproteobacteria |
| Ensifer adhaerens | Pseudomonadota | Alphaproteobacteria |
| Ensifer adhaerens | Pseudomonadota | Alphaproteobacteria |
| Ensifer canadensis | Pseudomonadota | Alphaproteobacteria |
| Ensifer canadensis | Pseudomonadota | Alphaproteobacteria |
| Entomobacter blattae | Pseudomonadota | Alphaproteobacteria |
| Erythrobacter aurantius | Pseudomonadota | Alphaproteobacteria |
| Erythrobacter aureus | Pseudomonadota | Alphaproteobacteria |
| Erythrobacter litoralis | Pseudomonadota | Alphaproteobacteria |

| | | |
|---|---|---|
| Erythrobacter mangrovi | Pseudomonadota | Alphaproteobacteria |
| Erythrobacter neustonensis | Pseudomonadota | Alphaproteobacteria |
| Falsihalocynthiibacter arcticus | Pseudomonadota | Alphaproteobacteria |
| Falsihalocynthiibacter arcticus | Pseudomonadota | Alphaproteobacteria |
| Falsirhodobacter algicola | Pseudomonadota | Alphaproteobacteria |
| Falsirhodobacter algicola | Pseudomonadota | Alphaproteobacteria |
| Ferrovibrio terrae | Pseudomonadota | Alphaproteobacteria |
| Ferrovibrio terrae | Pseudomonadota | Alphaproteobacteria |
| Fontisubflavum oceani | Pseudomonadota | Alphaproteobacteria |
| Fontisubflavum oceani | Pseudomonadota | Alphaproteobacteria |
| Formicincola oecophyllae | Pseudomonadota | Alphaproteobacteria |
| Fretibacter rubidus | Pseudomonadota | Alphaproteobacteria |
| Fretibacter rubidus | Pseudomonadota | Alphaproteobacteria |
| Frigidibacter mobilis | Pseudomonadota | Alphaproteobacteria |
| Frigidibacter mobilis | Pseudomonadota | Alphaproteobacteria |
| Fuscovulum ytuae | Pseudomonadota | Alphaproteobacteria |
| Fuscovulum ytuae | Pseudomonadota | Alphaproteobacteria |
| Gemmobacter fulvus | Pseudomonadota | Alphaproteobacteria |
| Gemmobacter fulvus | Pseudomonadota | Alphaproteobacteria |
| Georhizobium profundi | Pseudomonadota | Alphaproteobacteria |
| Georhizobium profundi | Pseudomonadota | Alphaproteobacteria |
| Gimibacter soli | Pseudomonadota | Alphaproteobacteria |
| Gimibacter soli | Pseudomonadota | Alphaproteobacteria |
| Gluconacetobacter diazotrophicus PA1 5 | Pseudomonadota | Alphaproteobacteria |
| Gluconobacter albidus | Pseudomonadota | Alphaproteobacteria |
| Gluconobacter oxydans DSM 3504 | Pseudomonadota | Alphaproteobacteria |
| Gluconobacter sphaericus | Pseudomonadota | Alphaproteobacteria |
| Glycocaulis abyssi | Pseudomonadota | Alphaproteobacteria |
| Glycocaulis abyssi | Pseudomonadota | Alphaproteobacteria |
| Glycocaulis alkaliphilus | Pseudomonadota | Alphaproteobacteria |
| Glycocaulis alkaliphilus | Pseudomonadota | Alphaproteobacteria |

| | | |
|---|---|---|
| Granulibacter bethesdensis | Pseudomonadota | Alphaproteobacteria |
| Haematobacter massiliensis | Pseudomonadota | Alphaproteobacteria |
| Haematobacter massiliensis | Pseudomonadota | Alphaproteobacteria |
| Haematospirillum jordaniae | Pseudomonadota | Alphaproteobacteria |
| Haematospirillum jordaniae | Pseudomonadota | Alphaproteobacteria |
| Hankyongella ginsenosidimutans | Pseudomonadota | Alphaproteobacteria |
| Hartmannibacter diazotrophicus | Pseudomonadota | Alphaproteobacteria |
| Hartmannibacter diazotrophicus | Pseudomonadota | Alphaproteobacteria |
| Hirschia baltica ATCC 49814 | Pseudomonadota | Alphaproteobacteria |
| Hoeflea phototrophica DFL-43 | Pseudomonadota | Alphaproteobacteria |
| Hoeflea phototrophica DFL-43 | Pseudomonadota | Alphaproteobacteria |
| Hypericibacter adhaerens | Pseudomonadota | Alphaproteobacteria |
| Hypericibacter adhaerens | Pseudomonadota | Alphaproteobacteria |
| Hypericibacter terrae | Pseudomonadota | Alphaproteobacteria |
| Hypericibacter terrae | Pseudomonadota | Alphaproteobacteria |
| Hyphococcus flavus | Pseudomonadota | Alphaproteobacteria |
| Hyphococcus formosus | Pseudomonadota | Alphaproteobacteria |
| Hyphomicrobium denitrificans ATCC 51888 | Pseudomonadota | Alphaproteobacteria |
| Hyphomicrobium nitrativorans NL23 | Pseudomonadota | Alphaproteobacteria |
| Hyphomonas atlantica | Pseudomonadota | Alphaproteobacteria |
| Hyphomonas atlantica | Pseudomonadota | Alphaproteobacteria |
| Hyphomonas neptunium ATCC 15444 | Pseudomonadota | Alphaproteobacteria |
| Hyphomonas neptunium ATCC 15444 | Pseudomonadota | Alphaproteobacteria |
| Indioceanicola profundi | Pseudomonadota | Alphaproteobacteria |
| Indioceanicola profundi | Pseudomonadota | Alphaproteobacteria |
| Jannaschia ovalis | Pseudomonadota | Alphaproteobacteria |
| Jannaschia ovalis | Pseudomonadota | Alphaproteobacteria |
| Kaustia mangrovi | Pseudomonadota | Alphaproteobacteria |
| Kaustia mangrovi | Pseudomonadota | Alphaproteobacteria |
| Ketogulonicigenium robustum | Pseudomonadota | Alphaproteobacteria |
| Ketogulonicigenium robustum | Pseudomonadota | Alphaproteobacteria |

| | | |
|---|---|---|
| Ketogulonicigenium vulgare | Pseudomonadota | Alphaproteobacteria |
| Ketogulonicigenium vulgare | Pseudomonadota | Alphaproteobacteria |
| Komagataeibacter diospyri | Pseudomonadota | Alphaproteobacteria |
| Komagataeibacter europaeus | Pseudomonadota | Alphaproteobacteria |
| Komagataeibacter medellinensis NBRC 3288 | Pseudomonadota | Alphaproteobacteria |
| Komagataeibacter nataicola | Pseudomonadota | Alphaproteobacteria |
| Komagataeibacter rhaeticus | Pseudomonadota | Alphaproteobacteria |
| Komagataeibacter saccharivorans | Pseudomonadota | Alphaproteobacteria |
| Komagataeibacter sucrofermentans DSM 15973 | Pseudomonadota | Alphaproteobacteria |
| Komagataeibacter xylinus | Pseudomonadota | Alphaproteobacteria |
| Kordiimonas pumila | Pseudomonadota | Alphaproteobacteria |
| Kordiimonas pumila | Pseudomonadota | Alphaproteobacteria |
| Kozakia baliensis | Pseudomonadota | Alphaproteobacteria |
| Labrys okinawensis | Pseudomonadota | Alphaproteobacteria |
| Labrys okinawensis | Pseudomonadota | Alphaproteobacteria |
| Leisingera aquaemixtae | Pseudomonadota | Alphaproteobacteria |
| Leisingera aquaemixtae | Pseudomonadota | Alphaproteobacteria |
| Leisingera caerulea | Pseudomonadota | Alphaproteobacteria |
| Leisingera caerulea | Pseudomonadota | Alphaproteobacteria |
| Leisingera methylohalidivorans DSM 14336 | Pseudomonadota | Alphaproteobacteria |
| Leisingera methylohalidivorans DSM 14336 | Pseudomonadota | Alphaproteobacteria |
| Leisingera thetidis | Pseudomonadota | Alphaproteobacteria |
| Leisingera thetidis | Pseudomonadota | Alphaproteobacteria |
| Liberibacter crescens | Pseudomonadota | Alphaproteobacteria |
| Liberibacter crescens | Pseudomonadota | Alphaproteobacteria |
| Lichenicola cladoniae | Pseudomonadota | Alphaproteobacteria |
| Limimaricola variabilis | Pseudomonadota | Alphaproteobacteria |
| Limimaricola variabilis | Pseudomonadota | Alphaproteobacteria |
| Loktanella salsilacus | Pseudomonadota | Alphaproteobacteria |
| Loktanella salsilacus | Pseudomonadota | Alphaproteobacteria |
| Luteithermobacter gelatinilyticus | Pseudomonadota | Alphaproteobacteria |

| | | |
|---|---|---|
| Luteithermobacter gelatinilyticus | Pseudomonadota | Alphaproteobacteria |
| Magnetococcus marinus MC-1 | Pseudomonadota | Alphaproteobacteria |
| Magnetococcus marinus MC-1 | Pseudomonadota | Alphaproteobacteria |
| Magnetospirillum gryphiswaldense MSR-1 v2 | Pseudomonadota | Alphaproteobacteria |
| Magnetospirillum gryphiswaldense MSR-1 v2 | Pseudomonadota | Alphaproteobacteria |
| Mariluticola halotolerans | Pseudomonadota | Alphaproteobacteria |
| Mariluticola halotolerans | Pseudomonadota | Alphaproteobacteria |
| Marinicauda algicola | Pseudomonadota | Alphaproteobacteria |
| Marinicauda algicola | Pseudomonadota | Alphaproteobacteria |
| Maritalea myrionectae | Pseudomonadota | Alphaproteobacteria |
| Maritalea myrionectae | Pseudomonadota | Alphaproteobacteria |
| Martelella endophytica | Pseudomonadota | Alphaproteobacteria |
| Martelella endophytica | Pseudomonadota | Alphaproteobacteria |
| Martelella mediterranea DSM 17316 | Pseudomonadota | Alphaproteobacteria |
| Martelella mediterranea DSM 17316 | Pseudomonadota | Alphaproteobacteria |
| Mesorhizobium amorphae CCNWGS0123 | Pseudomonadota | Alphaproteobacteria |
| Mesorhizobium amorphae CCNWGS0123 | Pseudomonadota | Alphaproteobacteria |
| Mesorhizobium australicum WSM2073 | Pseudomonadota | Alphaproteobacteria |
| Mesorhizobium australicum WSM2073 | Pseudomonadota | Alphaproteobacteria |
| Mesorhizobium erdmanii | Pseudomonadota | Alphaproteobacteria |
| Mesorhizobium erdmanii | Pseudomonadota | Alphaproteobacteria |
| Mesorhizobium huakuii | Pseudomonadota | Alphaproteobacteria |
| Mesorhizobium huakuii | Pseudomonadota | Alphaproteobacteria |
| Mesorhizobium japonicum R7A | Pseudomonadota | Alphaproteobacteria |
| Mesorhizobium japonicum R7A | Pseudomonadota | Alphaproteobacteria |
| Mesorhizobium jarvisii | Pseudomonadota | Alphaproteobacteria |
| Mesorhizobium jarvisii | Pseudomonadota | Alphaproteobacteria |
| Mesorhizobium koreense | Pseudomonadota | Alphaproteobacteria |
| Mesorhizobium koreense | Pseudomonadota | Alphaproteobacteria |
| Mesorhizobium mediterraneum | Pseudomonadota | Alphaproteobacteria |
| Mesorhizobium mediterraneum | Pseudomonadota | Alphaproteobacteria |

| | | |
|---|---|---|
| Mesorhizobium onobrychidis | Pseudomonadota | Alphaproteobacteria |
| Mesorhizobium onobrychidis | Pseudomonadota | Alphaproteobacteria |
| Mesorhizobium opportunistum WSM2075 | Pseudomonadota | Alphaproteobacteria |
| Mesorhizobium opportunistum WSM2075 | Pseudomonadota | Alphaproteobacteria |
| Methylobacterium bullatum | Pseudomonadota | Alphaproteobacteria |
| Methylobacterium bullatum | Pseudomonadota | Alphaproteobacteria |
| Methylobacterium currus | Pseudomonadota | Alphaproteobacteria |
| Methylobacterium currus | Pseudomonadota | Alphaproteobacteria |
| Methylobacterium durans | Pseudomonadota | Alphaproteobacteria |
| Methylobacterium durans | Pseudomonadota | Alphaproteobacteria |
| Methylobacterium flocculans | Pseudomonadota | Alphaproteobacteria |
| Methylobacterium flocculans | Pseudomonadota | Alphaproteobacteria |
| Methylobacterium indicum | Pseudomonadota | Alphaproteobacteria |
| Methylobacterium indicum | Pseudomonadota | Alphaproteobacteria |
| Methylobacterium nodulans ORS 2060 | Pseudomonadota | Alphaproteobacteria |
| Methylobacterium nodulans ORS 2060 | Pseudomonadota | Alphaproteobacteria |
| Methylobacterium organophilum | Pseudomonadota | Alphaproteobacteria |
| Methylobacterium organophilum | Pseudomonadota | Alphaproteobacteria |
| Methylobacterium oryzae CBMB20 | Pseudomonadota | Alphaproteobacteria |
| Methylobacterium oryzae CBMB20 | Pseudomonadota | Alphaproteobacteria |
| Methylobacterium phyllosphaerae | Pseudomonadota | Alphaproteobacteria |
| Methylobacterium phyllosphaerae | Pseudomonadota | Alphaproteobacteria |
| Methylobacterium radiodurans | Pseudomonadota | Alphaproteobacteria |
| Methylobacterium radiodurans | Pseudomonadota | Alphaproteobacteria |
| Methylobacterium radiotolerans JCM 2831 | Pseudomonadota | Alphaproteobacteria |
| Methylobacterium radiotolerans JCM 2831 | Pseudomonadota | Alphaproteobacteria |
| Methylobacterium terrae | Pseudomonadota | Alphaproteobacteria |
| Methylobacterium terrae | Pseudomonadota | Alphaproteobacteria |
| Methylocapsa polymorpha | Pseudomonadota | Alphaproteobacteria |
| Methyloceanibacter caenitepidi | Pseudomonadota | Alphaproteobacteria |
| Methylocella silvestris BL2 | Pseudomonadota | Alphaproteobacteria |

| | | |
|---|---|---|
| Methylocella tundrae | Pseudomonadota | Alphaproteobacteria |
| Methylocystis bryophila | Pseudomonadota | Alphaproteobacteria |
| Methylocystis bryophila | Pseudomonadota | Alphaproteobacteria |
| Methylocystis echinoides | Pseudomonadota | Alphaproteobacteria |
| Methylocystis echinoides | Pseudomonadota | Alphaproteobacteria |
| Methylocystis heyeri | Pseudomonadota | Alphaproteobacteria |
| Methylocystis heyeri | Pseudomonadota | Alphaproteobacteria |
| Methylocystis iwaonis | Pseudomonadota | Alphaproteobacteria |
| Methylocystis iwaonis | Pseudomonadota | Alphaproteobacteria |
| Methylocystis parvus OBBP | Pseudomonadota | Alphaproteobacteria |
| Methylocystis parvus OBBP | Pseudomonadota | Alphaproteobacteria |
| Methyloraptor flagellatus | Pseudomonadota | Alphaproteobacteria |
| Methyloraptor flagellatus | Pseudomonadota | Alphaproteobacteria |
| Methylorubrum extorquens DM4 | Pseudomonadota | Alphaproteobacteria |
| Methylorubrum extorquens DM4 | Pseudomonadota | Alphaproteobacteria |
| Methylorubrum populi BJ001 | Pseudomonadota | Alphaproteobacteria |
| Methylorubrum populi BJ001 | Pseudomonadota | Alphaproteobacteria |
| Methylovirgula ligni | Pseudomonadota | Alphaproteobacteria |
| Microvirga lotononidis | Pseudomonadota | Alphaproteobacteria |
| Microvirga lotononidis | Pseudomonadota | Alphaproteobacteria |
| Microvirga ossetica | Pseudomonadota | Alphaproteobacteria |
| Microvirga ossetica | Pseudomonadota | Alphaproteobacteria |
| Microvirga terrae | Pseudomonadota | Alphaproteobacteria |
| Microvirga terrae | Pseudomonadota | Alphaproteobacteria |
| Muricoccus radiodurans | Pseudomonadota | Alphaproteobacteria |
| Muricoccus radiodurans | Pseudomonadota | Alphaproteobacteria |
| Neoasaia chiangmaiensis | Pseudomonadota | Alphaproteobacteria |
| Neoehrlichia mikurensis | Pseudomonadota | Alphaproteobacteria |
| Neoehrlichia mikurensis | Pseudomonadota | Alphaproteobacteria |
| Neokomagataea tanensis | Pseudomonadota | Alphaproteobacteria |
| Neorhizobium galegae bv. orientalis str. HAMBI 540 | Pseudomonadota | Alphaproteobacteria |

| | | |
|---|---|---|
| Neorhizobium galegae bv. orientalis str. HAMBI 540 | Pseudomonadota | Alphaproteobacteria |
| Neorhizobium petrolearium | Pseudomonadota | Alphaproteobacteria |
| Neorhizobium petrolearium | Pseudomonadota | Alphaproteobacteria |
| Neorickettsia findlayensis | Pseudomonadota | Alphaproteobacteria |
| Neorickettsia helminthoeca str. Oregon | Pseudomonadota | Alphaproteobacteria |
| Neorickettsia risticii str. Illinois | Pseudomonadota | Alphaproteobacteria |
| Neorickettsia sennetsu str. Miyayama | Pseudomonadota | Alphaproteobacteria |
| Neotabrizicola shimadae | Pseudomonadota | Alphaproteobacteria |
| Neotabrizicola shimadae | Pseudomonadota | Alphaproteobacteria |
| Nereida ignava | Pseudomonadota | Alphaproteobacteria |
| Nereida ignava | Pseudomonadota | Alphaproteobacteria |
| Nguyenibacter vanlangensis | Pseudomonadota | Alphaproteobacteria |
| Nisaea acidiphila | Pseudomonadota | Alphaproteobacteria |
| Nisaea acidiphila | Pseudomonadota | Alphaproteobacteria |
| Nitratireductor aquimarinus | Pseudomonadota | Alphaproteobacteria |
| Nitratireductor aquimarinus | Pseudomonadota | Alphaproteobacteria |
| Nitratireductor kimnyeongensis | Pseudomonadota | Alphaproteobacteria |
| Nitratireductor kimnyeongensis | Pseudomonadota | Alphaproteobacteria |
| Nitratireductor mangrovi | Pseudomonadota | Alphaproteobacteria |
| Nitratireductor mangrovi | Pseudomonadota | Alphaproteobacteria |
| Nitratireductor thuwali | Pseudomonadota | Alphaproteobacteria |
| Nitratireductor thuwali | Pseudomonadota | Alphaproteobacteria |
| Nitrobacter hamburgensis X14 | Pseudomonadota | Alphaproteobacteria |
| Nitrobacter winogradskyi Nb-255 | Pseudomonadota | Alphaproteobacteria |
| Nitrospirillum viridazoti CBAmc | Pseudomonadota | Alphaproteobacteria |
| Nitrospirillum viridazoti CBAmc | Pseudomonadota | Alphaproteobacteria |
| Niveispirillum cyanobacteriorum | Pseudomonadota | Alphaproteobacteria |
| Niveispirillum cyanobacteriorum | Pseudomonadota | Alphaproteobacteria |
| Novacetimonas hansenii | Pseudomonadota | Alphaproteobacteria |
| Novosphingobium aromaticivorans DSM 12444 | Pseudomonadota | Alphaproteobacteria |
| Novosphingobium decolorationis | Pseudomonadota | Alphaproteobacteria |

| | | |
|---|---|---|
| Novosphingobium ginsenosidimutans | Pseudomonadota | Alphaproteobacteria |
| Novosphingobium humi | Pseudomonadota | Alphaproteobacteria |
| Novosphingobium kaempferiae | Pseudomonadota | Alphaproteobacteria |
| Novosphingobium olei | Pseudomonadota | Alphaproteobacteria |
| Novosphingobium pentaromativorans US6-1 | Pseudomonadota | Alphaproteobacteria |
| Novosphingobium resinovorum | Pseudomonadota | Alphaproteobacteria |
| Novosphingopyxis iocasae | Pseudomonadota | Alphaproteobacteria |
| Oceaniradius stylonematis | Pseudomonadota | Alphaproteobacteria |
| Oceaniradius stylonematis | Pseudomonadota | Alphaproteobacteria |
| Ochrobactrum quorumnocens | Pseudomonadota | Alphaproteobacteria |
| Ochrobactrum quorumnocens | Pseudomonadota | Alphaproteobacteria |
| Octadecabacter antarcticus 307 | Pseudomonadota | Alphaproteobacteria |
| Octadecabacter arcticus 238 | Pseudomonadota | Alphaproteobacteria |
| Octadecabacter arcticus 238 | Pseudomonadota | Alphaproteobacteria |
| Octadecabacter temperatus | Pseudomonadota | Alphaproteobacteria |
| Octadecabacter temperatus | Pseudomonadota | Alphaproteobacteria |
| Oecophyllibacter saccharovorans | Pseudomonadota | Alphaproteobacteria |
| Oricola thermophila | Pseudomonadota | Alphaproteobacteria |
| Oricola thermophila | Pseudomonadota | Alphaproteobacteria |
| Orientia tsutsugamushi | Pseudomonadota | Alphaproteobacteria |
| Orientia tsutsugamushi | Pseudomonadota | Alphaproteobacteria |
| Pacificitalea manganoxidans | Pseudomonadota | Alphaproteobacteria |
| Pacificitalea manganoxidans | Pseudomonadota | Alphaproteobacteria |
| Paracoccus actinidiae | Pseudomonadota | Alphaproteobacteria |
| Paracoccus aerodenitrificans | Pseudomonadota | Alphaproteobacteria |
| Paracoccus aerodenitrificans | Pseudomonadota | Alphaproteobacteria |
| Paracoccus albus | Pseudomonadota | Alphaproteobacteria |
| Paracoccus albus | Pseudomonadota | Alphaproteobacteria |
| Paracoccus alcaliphilus | Pseudomonadota | Alphaproteobacteria |
| Paracoccus alcaliphilus | Pseudomonadota | Alphaproteobacteria |
| Paracoccus aminophilus JCM 7686 | Pseudomonadota | Alphaproteobacteria |

| | | |
|---|---|---|
| Paracoccus aminophilus JCM 7686 | Pseudomonadota | Alphaproteobacteria |
| Paracoccus aminovorans | Pseudomonadota | Alphaproteobacteria |
| Paracoccus aminovorans | Pseudomonadota | Alphaproteobacteria |
| Paracoccus contaminans | Pseudomonadota | Alphaproteobacteria |
| Paracoccus contaminans | Pseudomonadota | Alphaproteobacteria |
| Paracoccus denitrificans | Pseudomonadota | Alphaproteobacteria |
| Paracoccus denitrificans | Pseudomonadota | Alphaproteobacteria |
| Paracoccus everestensis | Pseudomonadota | Alphaproteobacteria |
| Paracoccus everestensis | Pseudomonadota | Alphaproteobacteria |
| Paracoccus fistulariae | Pseudomonadota | Alphaproteobacteria |
| Paracoccus fistulariae | Pseudomonadota | Alphaproteobacteria |
| Paracoccus jeotgali | Pseudomonadota | Alphaproteobacteria |
| Paracoccus jeotgali | Pseudomonadota | Alphaproteobacteria |
| Paracoccus kondratievae | Pseudomonadota | Alphaproteobacteria |
| Paracoccus kondratievae | Pseudomonadota | Alphaproteobacteria |
| Paracoccus liaowanqingii | Pseudomonadota | Alphaproteobacteria |
| Paracoccus liaowanqingii | Pseudomonadota | Alphaproteobacteria |
| Paracoccus marcusii | Pseudomonadota | Alphaproteobacteria |
| Paracoccus marcusii | Pseudomonadota | Alphaproteobacteria |
| Paracoccus methylovorus | Pseudomonadota | Alphaproteobacteria |
| Paracoccus methylovorus | Pseudomonadota | Alphaproteobacteria |
| Paracoccus pantotrophus | Pseudomonadota | Alphaproteobacteria |
| Paracoccus pantotrophus | Pseudomonadota | Alphaproteobacteria |
| Paracoccus saliphilus | Pseudomonadota | Alphaproteobacteria |
| Paracoccus saliphilus | Pseudomonadota | Alphaproteobacteria |
| Paracoccus sanguinis | Pseudomonadota | Alphaproteobacteria |
| Paracoccus sanguinis | Pseudomonadota | Alphaproteobacteria |
| Paracoccus sediminicola | Pseudomonadota | Alphaproteobacteria |
| Paracoccus sediminicola | Pseudomonadota | Alphaproteobacteria |
| Paracoccus seriniphilus | Pseudomonadota | Alphaproteobacteria |
| Paracoccus seriniphilus | Pseudomonadota | Alphaproteobacteria |

| | | |
|---|---|---|
| Paracoccus stylophorae | Pseudomonadota | Alphaproteobacteria |
| Paracoccus stylophorae | Pseudomonadota | Alphaproteobacteria |
| Paracoccus suum | Pseudomonadota | Alphaproteobacteria |
| Paracoccus suum | Pseudomonadota | Alphaproteobacteria |
| Paracoccus tegillarcae | Pseudomonadota | Alphaproteobacteria |
| Paracoccus tegillarcae | Pseudomonadota | Alphaproteobacteria |
| Paracoccus versutus | Pseudomonadota | Alphaproteobacteria |
| Paracoccus versutus | Pseudomonadota | Alphaproteobacteria |
| Paracoccus yeei | Pseudomonadota | Alphaproteobacteria |
| Paracoccus yeei | Pseudomonadota | Alphaproteobacteria |
| Paracoccus zhejiangensis | Pseudomonadota | Alphaproteobacteria |
| Paracoccus zhejiangensis | Pseudomonadota | Alphaproteobacteria |
| Paradevosia shaoguanensis | Pseudomonadota | Alphaproteobacteria |
| Paradevosia tibetensis | Pseudomonadota | Alphaproteobacteria |
| Paragemmobacter aquarius | Pseudomonadota | Alphaproteobacteria |
| Paragemmobacter aquarius | Pseudomonadota | Alphaproteobacteria |
| Paramagnetospirillum magneticum AMB-1 | Pseudomonadota | Alphaproteobacteria |
| Paramagnetospirillum magneticum AMB-1 | Pseudomonadota | Alphaproteobacteria |
| Pararhizobium gei | Pseudomonadota | Alphaproteobacteria |
| Pararhizobium gei | Pseudomonadota | Alphaproteobacteria |
| Pararhizobium qamdonense | Pseudomonadota | Alphaproteobacteria |
| Pararhizobium qamdonense | Pseudomonadota | Alphaproteobacteria |
| Parasedimentitalea marina | Pseudomonadota | Alphaproteobacteria |
| Parasedimentitalea marina | Pseudomonadota | Alphaproteobacteria |
| Parasedimentitalea psychrophila | Pseudomonadota | Alphaproteobacteria |
| Parasedimentitalea psychrophila | Pseudomonadota | Alphaproteobacteria |
| Parasphingopyxis algicola | Pseudomonadota | Alphaproteobacteria |
| Parasphingorhabdus cellanae | Pseudomonadota | Alphaproteobacteria |
| Parasphingorhabdus halotolerans | Pseudomonadota | Alphaproteobacteria |
| Parasphingorhabdus litoris DSM 22379 | Pseudomonadota | Alphaproteobacteria |
| Paraurantiacibacter namhicola | Pseudomonadota | Alphaproteobacteria |

| | | |
|---|---|---|
| Paremcibacter congregatus | Pseudomonadota | Alphaproteobacteria |
| Paremcibacter congregatus | Pseudomonadota | Alphaproteobacteria |
| Paroceanicella profunda | Pseudomonadota | Alphaproteobacteria |
| Paroceanicella profunda | Pseudomonadota | Alphaproteobacteria |
| Parvibaculum lavamentivorans DS-1 | Pseudomonadota | Alphaproteobacteria |
| Parvibaculum lavamentivorans DS-1 | Pseudomonadota | Alphaproteobacteria |
| Parvularcula bermudensis HTCC2503 | Pseudomonadota | Alphaproteobacteria |
| Pelagerythrobacter marensis | Pseudomonadota | Alphaproteobacteria |
| Pelagerythrobacter marinus | Pseudomonadota | Alphaproteobacteria |
| Pelagibacterium flavum | Pseudomonadota | Alphaproteobacteria |
| Pelagibacterium flavum | Pseudomonadota | Alphaproteobacteria |
| Pelagibacterium halotolerans | Pseudomonadota | Alphaproteobacteria |
| Pelagibacterium halotolerans | Pseudomonadota | Alphaproteobacteria |
| Pelagibacterium nitratireducens | Pseudomonadota | Alphaproteobacteria |
| Pelagibacterium nitratireducens | Pseudomonadota | Alphaproteobacteria |
| Pelagovum pacificum | Pseudomonadota | Alphaproteobacteria |
| Pelagovum pacificum | Pseudomonadota | Alphaproteobacteria |
| Peteryoungia algae | Pseudomonadota | Alphaproteobacteria |
| Peteryoungia algae | Pseudomonadota | Alphaproteobacteria |
| Peteryoungia desertarenae | Pseudomonadota | Alphaproteobacteria |
| Peteryoungia desertarenae | Pseudomonadota | Alphaproteobacteria |
| Phaeobacter gallaeciensis DSM 26640 | Pseudomonadota | Alphaproteobacteria |
| Phaeobacter gallaeciensis DSM 26640 | Pseudomonadota | Alphaproteobacteria |
| Phaeobacter inhibens | Pseudomonadota | Alphaproteobacteria |
| Phaeobacter inhibens | Pseudomonadota | Alphaproteobacteria |
| Phaeobacter piscinae | Pseudomonadota | Alphaproteobacteria |
| Phaeobacter piscinae | Pseudomonadota | Alphaproteobacteria |
| Phaeobacter porticola | Pseudomonadota | Alphaproteobacteria |
| Phaeobacter porticola | Pseudomonadota | Alphaproteobacteria |
| Phenylobacterium montanum | Pseudomonadota | Alphaproteobacteria |
| Phenylobacterium montanum | Pseudomonadota | Alphaproteobacteria |

| | | |
|---|---|---|
| Phenylobacterium parvum | Pseudomonadota | Alphaproteobacteria |
| Phenylobacterium parvum | Pseudomonadota | Alphaproteobacteria |
| Phenylobacterium zucineum HLK1 | Pseudomonadota | Alphaproteobacteria |
| Phenylobacterium zucineum HLK1 | Pseudomonadota | Alphaproteobacteria |
| Phreatobacter aquaticus | Pseudomonadota | Alphaproteobacteria |
| Phreatobacter aquaticus | Pseudomonadota | Alphaproteobacteria |
| Phreatobacter cathodiphilus | Pseudomonadota | Alphaproteobacteria |
| Phreatobacter cathodiphilus | Pseudomonadota | Alphaproteobacteria |
| Phreatobacter stygius | Pseudomonadota | Alphaproteobacteria |
| Phreatobacter stygius | Pseudomonadota | Alphaproteobacteria |
| Phyllobacterium zundukense | Pseudomonadota | Alphaproteobacteria |
| Phyllobacterium zundukense | Pseudomonadota | Alphaproteobacteria |
| Pikeienuella piscinae | Pseudomonadota | Alphaproteobacteria |
| Pikeienuella piscinae | Pseudomonadota | Alphaproteobacteria |
| Polymorphobacter megasporae | Pseudomonadota | Alphaproteobacteria |
| Polymorphum gilvum SL003B-26A1 | Pseudomonadota | Alphaproteobacteria |
| Polymorphum gilvum SL003B-26A1 | Pseudomonadota | Alphaproteobacteria |
| Ponticoccus alexandrii | Pseudomonadota | Alphaproteobacteria |
| Ponticoccus alexandrii | Pseudomonadota | Alphaproteobacteria |
| Pontivivens nitratireducens | Pseudomonadota | Alphaproteobacteria |
| Pontivivens nitratireducens | Pseudomonadota | Alphaproteobacteria |
| Pontivivens ytuae | Pseudomonadota | Alphaproteobacteria |
| Pontivivens ytuae | Pseudomonadota | Alphaproteobacteria |
| Profundibacter amoris | Pseudomonadota | Alphaproteobacteria |
| Profundibacter amoris | Pseudomonadota | Alphaproteobacteria |
| Pseudemcibacter aquimaris | Pseudomonadota | Alphaproteobacteria |
| Pseudemcibacter aquimaris | Pseudomonadota | Alphaproteobacteria |
| Pseudochrobactrum algeriensis | Pseudomonadota | Alphaproteobacteria |
| Pseudochrobactrum algeriensis | Pseudomonadota | Alphaproteobacteria |
| Pseudogemmobacter blasticus | Pseudomonadota | Alphaproteobacteria |
| Pseudogemmobacter blasticus | Pseudomonadota | Alphaproteobacteria |

| | | |
|---|---|---|
| Pseudolabrys taiwanensis | Pseudomonadota | Alphaproteobacteria |
| Pseudooceanicola algae | Pseudomonadota | Alphaproteobacteria |
| Pseudooceanicola algae | Pseudomonadota | Alphaproteobacteria |
| Pseudooceanicola atlanticus | Pseudomonadota | Alphaproteobacteria |
| Pseudooceanicola atlanticus | Pseudomonadota | Alphaproteobacteria |
| Pseudooceanicola nitratireducens | Pseudomonadota | Alphaproteobacteria |
| Pseudooceanicola spongiae | Pseudomonadota | Alphaproteobacteria |
| Pseudooceanicola spongiae | Pseudomonadota | Alphaproteobacteria |
| Pseudorhizobium flavum | Pseudomonadota | Alphaproteobacteria |
| Pseudorhizobium flavum | Pseudomonadota | Alphaproteobacteria |
| Pseudorhodobacter turbinis | Pseudomonadota | Alphaproteobacteria |
| Pseudorhodobacter turbinis | Pseudomonadota | Alphaproteobacteria |
| Pseudorhodoplanes sinuspersici | Pseudomonadota | Alphaproteobacteria |
| Pseudosulfitobacter pseudonitzschiae | Pseudomonadota | Alphaproteobacteria |
| Pseudosulfitobacter pseudonitzschiae | Pseudomonadota | Alphaproteobacteria |
| Pseudovibrio brasiliensis | Pseudomonadota | Alphaproteobacteria |
| Pseudovibrio brasiliensis | Pseudomonadota | Alphaproteobacteria |
| Pukyongiella litopenaei | Pseudomonadota | Alphaproteobacteria |
| Pukyongiella litopenaei | Pseudomonadota | Alphaproteobacteria |
| Pyruvatibacter mobilis | Pseudomonadota | Alphaproteobacteria |
| Pyruvatibacter mobilis | Pseudomonadota | Alphaproteobacteria |
| Qingshengfaniella alkalisoli | Pseudomonadota | Alphaproteobacteria |
| Qingshengfaniella alkalisoli | Pseudomonadota | Alphaproteobacteria |
| Qipengyuania aurantiaca | Pseudomonadota | Alphaproteobacteria |
| Qipengyuania flava | Pseudomonadota | Alphaproteobacteria |
| Qipengyuania gelatinilytica | Pseudomonadota | Alphaproteobacteria |
| Qipengyuania profunda | Pseudomonadota | Alphaproteobacteria |
| Qipengyuania psychrotolerans | Pseudomonadota | Alphaproteobacteria |
| Qipengyuania sediminis | Pseudomonadota | Alphaproteobacteria |
| Qipengyuania seohaensis | Pseudomonadota | Alphaproteobacteria |
| Qipengyuania soli | Pseudomonadota | Alphaproteobacteria |

| | | |
|---|---|---|
| Qipengyuania spongiae | Pseudomonadota | Alphaproteobacteria |
| Rhizobium acidisoli | Pseudomonadota | Alphaproteobacteria |
| Rhizobium acidisoli | Pseudomonadota | Alphaproteobacteria |
| Rhizobium anhuiense bv. trifolii | Pseudomonadota | Alphaproteobacteria |
| Rhizobium anhuiense bv. trifolii | Pseudomonadota | Alphaproteobacteria |
| Rhizobium bangladeshense | Pseudomonadota | Alphaproteobacteria |
| Rhizobium bangladeshense | Pseudomonadota | Alphaproteobacteria |
| Rhizobium binae | Pseudomonadota | Alphaproteobacteria |
| Rhizobium binae | Pseudomonadota | Alphaproteobacteria |
| Rhizobium brockwellii | Pseudomonadota | Alphaproteobacteria |
| Rhizobium brockwellii | Pseudomonadota | Alphaproteobacteria |
| Rhizobium cremeum | Pseudomonadota | Alphaproteobacteria |
| Rhizobium cremeum | Pseudomonadota | Alphaproteobacteria |
| Rhizobium daejeonense | Pseudomonadota | Alphaproteobacteria |
| Rhizobium daejeonense | Pseudomonadota | Alphaproteobacteria |
| Rhizobium esperanzae | Pseudomonadota | Alphaproteobacteria |
| Rhizobium esperanzae | Pseudomonadota | Alphaproteobacteria |
| Rhizobium etli CFN 42 | Pseudomonadota | Alphaproteobacteria |
| Rhizobium etli CFN 42 | Pseudomonadota | Alphaproteobacteria |
| Rhizobium favelukesii | Pseudomonadota | Alphaproteobacteria |
| Rhizobium favelukesii | Pseudomonadota | Alphaproteobacteria |
| Rhizobium gallicum bv. gallicum R602sp | Pseudomonadota | Alphaproteobacteria |
| Rhizobium gallicum bv. gallicum R602sp | Pseudomonadota | Alphaproteobacteria |
| Rhizobium hidalgonense | Pseudomonadota | Alphaproteobacteria |
| Rhizobium hidalgonense | Pseudomonadota | Alphaproteobacteria |
| Rhizobium indicum | Pseudomonadota | Alphaproteobacteria |
| Rhizobium indicum | Pseudomonadota | Alphaproteobacteria |
| Rhizobium indigoferae | Pseudomonadota | Alphaproteobacteria |
| Rhizobium indigoferae | Pseudomonadota | Alphaproteobacteria |
| Rhizobium jaguaris | Pseudomonadota | Alphaproteobacteria |
| Rhizobium jaguaris | Pseudomonadota | Alphaproteobacteria |

| | | |
|---|---|---|
| Rhizobium johnstonii 3841 | Pseudomonadota | Alphaproteobacteria |
| Rhizobium johnstonii 3841 | Pseudomonadota | Alphaproteobacteria |
| Rhizobium lentis | Pseudomonadota | Alphaproteobacteria |
| Rhizobium lentis | Pseudomonadota | Alphaproteobacteria |
| Rhizobium oryzihabitans | Pseudomonadota | Alphaproteobacteria |
| Rhizobium oryzihabitans | Pseudomonadota | Alphaproteobacteria |
| Rhizobium phaseoli | Pseudomonadota | Alphaproteobacteria |
| Rhizobium phaseoli | Pseudomonadota | Alphaproteobacteria |
| Rhizobium rhizogenes | Pseudomonadota | Alphaproteobacteria |
| Rhizobium rhizogenes | Pseudomonadota | Alphaproteobacteria |
| Rhizobium rhizoryzae | Pseudomonadota | Alphaproteobacteria |
| Rhizobium rhizoryzae | Pseudomonadota | Alphaproteobacteria |
| Rhizobium rhododendri | Pseudomonadota | Alphaproteobacteria |
| Rhizobium rhododendri | Pseudomonadota | Alphaproteobacteria |
| Rhizobium ruizarguesonis | Pseudomonadota | Alphaproteobacteria |
| Rhizobium ruizarguesonis | Pseudomonadota | Alphaproteobacteria |
| Rhizobium sophoriradicis | Pseudomonadota | Alphaproteobacteria |
| Rhizobium sophoriradicis | Pseudomonadota | Alphaproteobacteria |
| Rhizobium sullae | Pseudomonadota | Alphaproteobacteria |
| Rhizobium sullae | Pseudomonadota | Alphaproteobacteria |
| Rhizobium tropici CIAT 899 | Pseudomonadota | Alphaproteobacteria |
| Rhizobium tropici CIAT 899 | Pseudomonadota | Alphaproteobacteria |
| Rhizobium tumorigenes | Pseudomonadota | Alphaproteobacteria |
| Rhizobium tumorigenes | Pseudomonadota | Alphaproteobacteria |
| Rhizorhabdus dicambivorans | Pseudomonadota | Alphaproteobacteria |
| Rhizorhabdus phycosphaerae | Pseudomonadota | Alphaproteobacteria |
| Rhizorhabdus wittichii | Pseudomonadota | Alphaproteobacteria |
| Rhodobacter capsulatus SB 1003 | Pseudomonadota | Alphaproteobacteria |
| Rhodobacter capsulatus SB 1003 | Pseudomonadota | Alphaproteobacteria |
| Rhodobacter xanthinilyticus | Pseudomonadota | Alphaproteobacteria |
| Rhodobacter xanthinilyticus | Pseudomonadota | Alphaproteobacteria |

| | | |
|---|---|---|
| Rhodomicrobium vannielii ATCC 17100 | Pseudomonadota | Alphaproteobacteria |
| Rhodopseudomonas boonkerdii | Pseudomonadota | Alphaproteobacteria |
| Rhodopseudomonas boonkerdii | Pseudomonadota | Alphaproteobacteria |
| Rhodopseudomonas palustris | Pseudomonadota | Alphaproteobacteria |
| Rhodopseudomonas palustris | Pseudomonadota | Alphaproteobacteria |
| Rhodospirillum centenum SW | Pseudomonadota | Alphaproteobacteria |
| Rhodospirillum centenum SW | Pseudomonadota | Alphaproteobacteria |
| Rhodospirillum rubrum ATCC 11170 | Pseudomonadota | Alphaproteobacteria |
| Rhodospirillum rubrum ATCC 11170 | Pseudomonadota | Alphaproteobacteria |
| Rhodovastum atsumiense | Pseudomonadota | Alphaproteobacteria |
| Rhodovulum sulfidophilum DSM 1374 | Pseudomonadota | Alphaproteobacteria |
| Rhodovulum sulfidophilum DSM 1374 | Pseudomonadota | Alphaproteobacteria |
| Rickettsia akari str. Hartford | Pseudomonadota | Alphaproteobacteria |
| Rickettsia asiatica | Pseudomonadota | Alphaproteobacteria |
| Rickettsia australis str. Cutlack | Pseudomonadota | Alphaproteobacteria |
| Rickettsia bellii RML369-C | Pseudomonadota | Alphaproteobacteria |
| Rickettsia canadensis str. CA410 | Pseudomonadota | Alphaproteobacteria |
| Rickettsia conorii str. Malish 7 | Pseudomonadota | Alphaproteobacteria |
| Rickettsia endosymbiont of Aspidapion aeneum | Pseudomonadota | Alphaproteobacteria |
| Rickettsia endosymbiont of Cantharis rufa | Pseudomonadota | Alphaproteobacteria |
| Rickettsia endosymbiont of Ceutorhynchus obstrictus | Pseudomonadota | Alphaproteobacteria |
| Rickettsia endosymbiont of Gonocerus acuteangulatus | Pseudomonadota | Alphaproteobacteria |
| Rickettsia endosymbiont of Lasioglossum villosulum | Pseudomonadota | Alphaproteobacteria |
| Rickettsia endosymbiont of Nabis limbatus | Pseudomonadota | Alphaproteobacteria |
| Rickettsia endosymbiont of Oedothorax gibbosus | Pseudomonadota | Alphaproteobacteria |
| Rickettsia endosymbiont of Pantilius tunicatus | Pseudomonadota | Alphaproteobacteria |
| Rickettsia endosymbiont of Polydrusus tereticollis | Pseudomonadota | Alphaproteobacteria |
| Rickettsia endosymbiont of Rhinocyllus conicus | Pseudomonadota | Alphaproteobacteria |
| Rickettsia endosymbiont of Seladonia tumulorum | Pseudomonadota | Alphaproteobacteria |
| Rickettsia helvetica | Pseudomonadota | Alphaproteobacteria |
| Rickettsia japonica YH | Pseudomonadota | Alphaproteobacteria |

| | | |
|---|---|---|
| Rickettsia monacensis | Pseudomonadota | Alphaproteobacteria |
| Rickettsia prowazekii str. Chernikova | Pseudomonadota | Alphaproteobacteria |
| Rickettsia rhipicephali str. 3-7-female6-CWPP | Pseudomonadota | Alphaproteobacteria |
| Rickettsia rickettsii str. 'Sheila Smith' | Pseudomonadota | Alphaproteobacteria |
| Rickettsia sibirica 246 | Pseudomonadota | Alphaproteobacteria |
| Rickettsia slovaca 13-B | Pseudomonadota | Alphaproteobacteria |
| Rickettsia tillamookensis | Pseudomonadota | Alphaproteobacteria |
| Rickettsia typhi str. Wilmington | Pseudomonadota | Alphaproteobacteria |
| Rickettsiales endosymbiont of Stachyamoeba lipophora | Pseudomonadota | Alphaproteobacteria |
| Rickettsiales endosymbiont of Stachyamoeba lipophora | Pseudomonadota | Alphaproteobacteria |
| Roseibium alexandrii DFL-11 | Pseudomonadota | Alphaproteobacteria |
| Roseibium alexandrii DFL-11 | Pseudomonadota | Alphaproteobacteria |
| Roseibium algicola | Pseudomonadota | Alphaproteobacteria |
| Roseibium algicola | Pseudomonadota | Alphaproteobacteria |
| Roseibium porphyridii | Pseudomonadota | Alphaproteobacteria |
| Roseibium porphyridii | Pseudomonadota | Alphaproteobacteria |
| Roseicyclus elongatus DSM 19469 | Pseudomonadota | Alphaproteobacteria |
| Roseicyclus elongatus DSM 19469 | Pseudomonadota | Alphaproteobacteria |
| Roseitalea porphyridii | Pseudomonadota | Alphaproteobacteria |
| Roseitalea porphyridii | Pseudomonadota | Alphaproteobacteria |
| Roseobacter denitrificans | Pseudomonadota | Alphaproteobacteria |
| Roseobacter denitrificans | Pseudomonadota | Alphaproteobacteria |
| Roseobacter fucihabitans | Pseudomonadota | Alphaproteobacteria |
| Roseobacter fucihabitans | Pseudomonadota | Alphaproteobacteria |
| Roseobacter litoralis Och 149 | Pseudomonadota | Alphaproteobacteria |
| Roseobacter litoralis Och 149 | Pseudomonadota | Alphaproteobacteria |
| Roseobacter ponti | Pseudomonadota | Alphaproteobacteria |
| Roseobacter ponti | Pseudomonadota | Alphaproteobacteria |
| Roseococcus microcysteis | Pseudomonadota | Alphaproteobacteria |
| Roseococcus microcysteis | Pseudomonadota | Alphaproteobacteria |
| Roseomonas cutis | Pseudomonadota | Alphaproteobacteria |

| | | |
|---|---|---|
| Roseomonas cutis | Pseudomonadota | Alphaproteobacteria |
| Roseomonas fluvialis | Pseudomonadota | Alphaproteobacteria |
| Roseomonas fluvialis | Pseudomonadota | Alphaproteobacteria |
| Roseomonas gilardii | Pseudomonadota | Alphaproteobacteria |
| Roseomonas gilardii | Pseudomonadota | Alphaproteobacteria |
| Roseomonas haemaphysalidis | Pseudomonadota | Alphaproteobacteria |
| Roseomonas haemaphysalidis | Pseudomonadota | Alphaproteobacteria |
| Roseomonas marmotae | Pseudomonadota | Alphaproteobacteria |
| Roseomonas marmotae | Pseudomonadota | Alphaproteobacteria |
| Roseomonas mucosa | Pseudomonadota | Alphaproteobacteria |
| Roseomonas mucosa | Pseudomonadota | Alphaproteobacteria |
| Roseovarius faecimaris | Pseudomonadota | Alphaproteobacteria |
| Roseovarius indicus | Pseudomonadota | Alphaproteobacteria |
| Roseovarius indicus | Pseudomonadota | Alphaproteobacteria |
| Roseovarius pelagicus | Pseudomonadota | Alphaproteobacteria |
| Roseovarius pelagicus | Pseudomonadota | Alphaproteobacteria |
| Roseovarius phycicola | Pseudomonadota | Alphaproteobacteria |
| Roseovarius phycicola | Pseudomonadota | Alphaproteobacteria |
| Roseovarius rhodophyticola | Pseudomonadota | Alphaproteobacteria |
| Roseovarius rhodophyticola | Pseudomonadota | Alphaproteobacteria |
| Roseovarius tolerans | Pseudomonadota | Alphaproteobacteria |
| Roseovarius tolerans | Pseudomonadota | Alphaproteobacteria |
| Ruegeria pomeroyi DSS-3 | Pseudomonadota | Alphaproteobacteria |
| Ruegeria pomeroyi DSS-3 | Pseudomonadota | Alphaproteobacteria |
| Sagittula stellata E-37 | Pseudomonadota | Alphaproteobacteria |
| Sagittula stellata E-37 | Pseudomonadota | Alphaproteobacteria |
| Salaquimonas pukyongi | Pseudomonadota | Alphaproteobacteria |
| Salaquimonas pukyongi | Pseudomonadota | Alphaproteobacteria |
| Salipiger abyssi | Pseudomonadota | Alphaproteobacteria |
| Salipiger abyssi | Pseudomonadota | Alphaproteobacteria |
| Salipiger profundus | Pseudomonadota | Alphaproteobacteria |

| | | |
|---|---|---|
| Salipiger profundus | Pseudomonadota | Alphaproteobacteria |
| Sandaracinobacteroides saxicola | Pseudomonadota | Alphaproteobacteria |
| Sediminicoccus rosea | Pseudomonadota | Alphaproteobacteria |
| Sediminicoccus rosea | Pseudomonadota | Alphaproteobacteria |
| Shinella oryzae | Pseudomonadota | Alphaproteobacteria |
| Shinella oryzae | Pseudomonadota | Alphaproteobacteria |
| Shinella sumterensis | Pseudomonadota | Alphaproteobacteria |
| Shinella sumterensis | Pseudomonadota | Alphaproteobacteria |
| Shinella zoogloeoides | Pseudomonadota | Alphaproteobacteria |
| Shinella zoogloeoides | Pseudomonadota | Alphaproteobacteria |
| Silicimonas algicola | Pseudomonadota | Alphaproteobacteria |
| Silicimonas algicola | Pseudomonadota | Alphaproteobacteria |
| Sinorhizobium alkalisoli | Pseudomonadota | Alphaproteobacteria |
| Sinorhizobium alkalisoli | Pseudomonadota | Alphaproteobacteria |
| Sinorhizobium americanum | Pseudomonadota | Alphaproteobacteria |
| Sinorhizobium americanum | Pseudomonadota | Alphaproteobacteria |
| Sinorhizobium chiapasense | Pseudomonadota | Alphaproteobacteria |
| Sinorhizobium chiapasense | Pseudomonadota | Alphaproteobacteria |
| Sinorhizobium fredii CCBAU 25509 | Pseudomonadota | Alphaproteobacteria |
| Sinorhizobium fredii CCBAU 25509 | Pseudomonadota | Alphaproteobacteria |
| Sinorhizobium garamanticum | Pseudomonadota | Alphaproteobacteria |
| Sinorhizobium garamanticum | Pseudomonadota | Alphaproteobacteria |
| Sinorhizobium kummerowiae | Pseudomonadota | Alphaproteobacteria |
| Sinorhizobium kummerowiae | Pseudomonadota | Alphaproteobacteria |
| Sinorhizobium meliloti | Pseudomonadota | Alphaproteobacteria |
| Sinorhizobium meliloti | Pseudomonadota | Alphaproteobacteria |
| Sinorhizobium mexicanum | Pseudomonadota | Alphaproteobacteria |
| Sinorhizobium mexicanum | Pseudomonadota | Alphaproteobacteria |
| Sinorhizobium numidicum | Pseudomonadota | Alphaproteobacteria |
| Sinorhizobium numidicum | Pseudomonadota | Alphaproteobacteria |
| Sinorhizobium sojae CCBAU 05684 | Pseudomonadota | Alphaproteobacteria |

| | | |
|---|---|---|
| Sinorhizobium sojae CCBAU 05684 | Pseudomonadota | Alphaproteobacteria |
| Sinorhizobium terangae | Pseudomonadota | Alphaproteobacteria |
| Sinorhizobium terangae | Pseudomonadota | Alphaproteobacteria |
| Skermanella cutis | Pseudomonadota | Alphaproteobacteria |
| Skermanella cutis | Pseudomonadota | Alphaproteobacteria |
| Skermanella mucosa | Pseudomonadota | Alphaproteobacteria |
| Skermanella mucosa | Pseudomonadota | Alphaproteobacteria |
| Skermanella pratensis | Pseudomonadota | Alphaproteobacteria |
| Skermanella pratensis | Pseudomonadota | Alphaproteobacteria |
| Skermanella rosea | Pseudomonadota | Alphaproteobacteria |
| Skermanella rosea | Pseudomonadota | Alphaproteobacteria |
| Sneathiella aquimaris | Pseudomonadota | Alphaproteobacteria |
| Sneathiella aquimaris | Pseudomonadota | Alphaproteobacteria |
| Sneathiella marina | Pseudomonadota | Alphaproteobacteria |
| Sneathiella marina | Pseudomonadota | Alphaproteobacteria |
| Sphingobium amiense | Pseudomonadota | Alphaproteobacteria |
| Sphingobium cloacae | Pseudomonadota | Alphaproteobacteria |
| Sphingobium fuliginis | Pseudomonadota | Alphaproteobacteria |
| Sphingobium indicum UT26S | Pseudomonadota | Alphaproteobacteria |
| Sphingobium naphthae | Pseudomonadota | Alphaproteobacteria |
| Sphingobium phenoxybenzoativorans | Pseudomonadota | Alphaproteobacteria |
| Sphingobium xenophagum | Pseudomonadota | Alphaproteobacteria |
| Sphingobium yanoikuyae | Pseudomonadota | Alphaproteobacteria |
| Sphingomicrobium aestuariivivum | Pseudomonadota | Alphaproteobacteria |
| Sphingomicrobium flavum | Pseudomonadota | Alphaproteobacteria |
| Sphingomonas abietis | Pseudomonadota | Alphaproteobacteria |
| Sphingomonas aerolata | Pseudomonadota | Alphaproteobacteria |
| Sphingomonas albertensis | Pseudomonadota | Alphaproteobacteria |
| Sphingomonas aliaeris | Pseudomonadota | Alphaproteobacteria |
| Sphingomonas bisphenolicum | Pseudomonadota | Alphaproteobacteria |
| Sphingomonas cannabina | Pseudomonadota | Alphaproteobacteria |

| | | |
|---|---|---|
| Sphingomonas carotinifaciens | Pseudomonadota | Alphaproteobacteria |
| Sphingomonas changnyeongensis | Pseudomonadota | Alphaproteobacteria |
| Sphingomonas donggukensis | Pseudomonadota | Alphaproteobacteria |
| Sphingomonas endolithica | Pseudomonadota | Alphaproteobacteria |
| Sphingomonas flavescens | Pseudomonadota | Alphaproteobacteria |
| Sphingomonas ginsengisoli (ex An et al. 2013) | Pseudomonadota | Alphaproteobacteria |
| Sphingomonas glaciei | Pseudomonadota | Alphaproteobacteria |
| Sphingomonas hengshuiensis | Pseudomonadota | Alphaproteobacteria |
| Sphingomonas insulae | Pseudomonadota | Alphaproteobacteria |
| Sphingomonas koreensis | Pseudomonadota | Alphaproteobacteria |
| Sphingomonas lacunae | Pseudomonadota | Alphaproteobacteria |
| Sphingomonas lutea | Pseudomonadota | Alphaproteobacteria |
| Sphingomonas morindae | Pseudomonadota | Alphaproteobacteria |
| Sphingomonas naphthae | Pseudomonadota | Alphaproteobacteria |
| Sphingomonas nostoxanthinifaciens | Pseudomonadota | Alphaproteobacteria |
| Sphingomonas paeninsulae | Pseudomonadota | Alphaproteobacteria |
| Sphingomonas panacis | Pseudomonadota | Alphaproteobacteria |
| Sphingomonas panacisoli | Pseudomonadota | Alphaproteobacteria |
| Sphingomonas paucimobilis | Pseudomonadota | Alphaproteobacteria |
| Sphingomonas piscis | Pseudomonadota | Alphaproteobacteria |
| Sphingomonas profundi | Pseudomonadota | Alphaproteobacteria |
| Sphingomonas psychrotolerans | Pseudomonadota | Alphaproteobacteria |
| Sphingomonas qomolangmaensis | Pseudomonadota | Alphaproteobacteria |
| Sphingomonas radiodurans | Pseudomonadota | Alphaproteobacteria |
| Sphingomonas rhizophila | Pseudomonadota | Alphaproteobacteria |
| Sphingomonas sabuli | Pseudomonadota | Alphaproteobacteria |
| Sphingomonas sanxanigenens DSM 19645 = NX02 | Pseudomonadota | Alphaproteobacteria |
| Sphingomonas sanxanigenens DSM 19645 = NX02 | Pseudomonadota | Alphaproteobacteria |
| Sphingomonas sediminicola | Pseudomonadota | Alphaproteobacteria |
| Sphingomonas sinipercae | Pseudomonadota | Alphaproteobacteria |
| Sphingomonas suaedae | Pseudomonadota | Alphaproteobacteria |

| | | |
|---|---|---|
| Sphingomonas taxi | Pseudomonadota | Alphaproteobacteria |
| Sphingomonas xanthus | Pseudomonadota | Alphaproteobacteria |
| Sphingopyxis alaskensis RB2256 | Pseudomonadota | Alphaproteobacteria |
| Sphingopyxis chilensis | Pseudomonadota | Alphaproteobacteria |
| Sphingopyxis fribergensis | Pseudomonadota | Alphaproteobacteria |
| Sphingopyxis granuli | Pseudomonadota | Alphaproteobacteria |
| Sphingopyxis indica | Pseudomonadota | Alphaproteobacteria |
| Sphingopyxis kveilinensis | Pseudomonadota | Alphaproteobacteria |
| Sphingopyxis lindanitolerans | Pseudomonadota | Alphaproteobacteria |
| Sphingopyxis macrogoltabida | Pseudomonadota | Alphaproteobacteria |
| Sphingorhabdus lacus | Pseudomonadota | Alphaproteobacteria |
| Sphingorhabdus lutea | Pseudomonadota | Alphaproteobacteria |
| Sphingosinicella flava | Pseudomonadota | Alphaproteobacteria |
| Sphingosinithalassobacter tenebrarum | Pseudomonadota | Alphaproteobacteria |
| Stakelama saccharophila | Pseudomonadota | Alphaproteobacteria |
| Stappia indica | Pseudomonadota | Alphaproteobacteria |
| Stappia indica | Pseudomonadota | Alphaproteobacteria |
| Stella humosa | Pseudomonadota | Alphaproteobacteria |
| Stella humosa | Pseudomonadota | Alphaproteobacteria |
| Sulfitobacter albidus | Pseudomonadota | Alphaproteobacteria |
| Sulfitobacter albidus | Pseudomonadota | Alphaproteobacteria |
| Sulfitobacter alexandrii | Pseudomonadota | Alphaproteobacteria |
| Sulfitobacter alexandrii | Pseudomonadota | Alphaproteobacteria |
| Sulfitobacter faviae | Pseudomonadota | Alphaproteobacteria |
| Sulfitobacter faviae | Pseudomonadota | Alphaproteobacteria |
| Sulfitobacter indolifex | Pseudomonadota | Alphaproteobacteria |
| Sulfitobacter indolifex | Pseudomonadota | Alphaproteobacteria |
| Sulfitobacter mediterraneus | Pseudomonadota | Alphaproteobacteria |
| Sulfitobacter mediterraneus | Pseudomonadota | Alphaproteobacteria |
| Sulfitobacter pontiacus | Pseudomonadota | Alphaproteobacteria |
| Sulfitobacter pontiacus | Pseudomonadota | Alphaproteobacteria |

| | | |
|---|---|---|
| Swingsia samuiensis | Pseudomonadota | Alphaproteobacteria |
| Tabrizicola piscis | Pseudomonadota | Alphaproteobacteria |
| Tabrizicola piscis | Pseudomonadota | Alphaproteobacteria |
| Tardibacter chloracetimidivorans | Pseudomonadota | Alphaproteobacteria |
| Tardiphaga alba | Pseudomonadota | Alphaproteobacteria |
| Tardiphaga alba | Pseudomonadota | Alphaproteobacteria |
| Temperatibacter marinus | Pseudomonadota | Alphaproteobacteria |
| Temperatibacter marinus | Pseudomonadota | Alphaproteobacteria |
| Terribium terrae | Pseudomonadota | Alphaproteobacteria |
| Terribium terrae | Pseudomonadota | Alphaproteobacteria |
| Terricaulis silvestris | Pseudomonadota | Alphaproteobacteria |
| Terrihabitans soli | Pseudomonadota | Alphaproteobacteria |
| Terrihabitans soli | Pseudomonadota | Alphaproteobacteria |
| Terrirubrum flagellatum | Pseudomonadota | Alphaproteobacteria |
| Thalassospira indica | Pseudomonadota | Alphaproteobacteria |
| Thalassospira indica | Pseudomonadota | Alphaproteobacteria |
| Thalassospira marina | Pseudomonadota | Alphaproteobacteria |
| Thalassospira marina | Pseudomonadota | Alphaproteobacteria |
| Thalassospira xiamenensis M-5 = DSM 17429 | Pseudomonadota | Alphaproteobacteria |
| Thalassospira xiamenensis M-5 = DSM 17429 | Pseudomonadota | Alphaproteobacteria |
| Thalassovita gelatinovora | Pseudomonadota | Alphaproteobacteria |
| Thalassovita gelatinovora | Pseudomonadota | Alphaproteobacteria |
| Thioclava electrotropha | Pseudomonadota | Alphaproteobacteria |
| Thioclava electrotropha | Pseudomonadota | Alphaproteobacteria |
| Thioclava litoralis | Pseudomonadota | Alphaproteobacteria |
| Thioclava litoralis | Pseudomonadota | Alphaproteobacteria |
| Thioclava nitratireducens | Pseudomonadota | Alphaproteobacteria |
| Thioclava nitratireducens | Pseudomonadota | Alphaproteobacteria |
| Tistrella bauzanensis | Pseudomonadota | Alphaproteobacteria |
| Tistrella bauzanensis | Pseudomonadota | Alphaproteobacteria |
| Tistrella mobilis | Pseudomonadota | Alphaproteobacteria |

| | | |
|---|---|---|
| Tistrella mobilis | Pseudomonadota | Alphaproteobacteria |
| Tritonibacter mobilis F1926 | Pseudomonadota | Alphaproteobacteria |
| Tritonibacter mobilis F1926 | Pseudomonadota | Alphaproteobacteria |
| Tritonibacter scottomollicae | Pseudomonadota | Alphaproteobacteria |
| Tritonibacter scottomollicae | Pseudomonadota | Alphaproteobacteria |
| Tropicibacter oceani | Pseudomonadota | Alphaproteobacteria |
| Tropicibacter oceani | Pseudomonadota | Alphaproteobacteria |
| Tsuneonella amylolytica | Pseudomonadota | Alphaproteobacteria |
| Tsuneonella dongtanensis | Pseudomonadota | Alphaproteobacteria |
| Tsuneonella mangrovi | Pseudomonadota | Alphaproteobacteria |
| Variibacter gotjawalensis | Pseudomonadota | Alphaproteobacteria |
| Variibacter gotjawalensis | Pseudomonadota | Alphaproteobacteria |
| Varunaivibrio sulfuroxidans | Pseudomonadota | Alphaproteobacteria |
| Varunaivibrio sulfuroxidans | Pseudomonadota | Alphaproteobacteria |
| Wolbachia endosymbiont (group A) of Acrocera orbiculus | Pseudomonadota | Alphaproteobacteria |
| Wolbachia endosymbiont (group A) of Acrocera orbiculus | Pseudomonadota | Alphaproteobacteria |
| Wolbachia endosymbiont (group A) of Agelastica alni | Pseudomonadota | Alphaproteobacteria |
| Wolbachia endosymbiont (group A) of Agelastica alni | Pseudomonadota | Alphaproteobacteria |
| Wolbachia endosymbiont (group A) of Aleiodes leptofemur | Pseudomonadota | Alphaproteobacteria |
| Wolbachia endosymbiont (group A) of Aleiodes leptofemur | Pseudomonadota | Alphaproteobacteria |
| Wolbachia endosymbiont (group A) of Alloplasta piceator | Pseudomonadota | Alphaproteobacteria |
| Wolbachia endosymbiont (group A) of Alloplasta piceator | Pseudomonadota | Alphaproteobacteria |
| Wolbachia endosymbiont (group A) of Ancistrocerus nigricornis | Pseudomonadota | Alphaproteobacteria |
| Wolbachia endosymbiont (group A) of Ancistrocerus nigricornis | Pseudomonadota | Alphaproteobacteria |
| Wolbachia endosymbiont (group A) of Andrena bucephala | Pseudomonadota | Alphaproteobacteria |
| Wolbachia endosymbiont (group A) of Andrena bucephala | Pseudomonadota | Alphaproteobacteria |
| Wolbachia endosymbiont (group A) of Andrena carantonica | Pseudomonadota | Alphaproteobacteria |
| Wolbachia endosymbiont (group A) of Andrena carantonica | Pseudomonadota | Alphaproteobacteria |
| Wolbachia endosymbiont (group A) of Andrena dorsata | Pseudomonadota | Alphaproteobacteria |
| Wolbachia endosymbiont (group A) of Andrena dorsata | Pseudomonadota | Alphaproteobacteria |
| Wolbachia endosymbiont (group A) of Andrena fulva | Pseudomonadota | Alphaproteobacteria |

| | | |
|---|---|---|
| Wolbachia endosymbiont (group A) of Andrena fulva | Pseudomonadota | Alphaproteobacteria |
| Wolbachia endosymbiont (group A) of Andrena haemorrhoa | Pseudomonadota | Alphaproteobacteria |
| Wolbachia endosymbiont (group A) of Andrena haemorrhoa | Pseudomonadota | Alphaproteobacteria |
| Wolbachia endosymbiont (group A) of Andrena helvola | Pseudomonadota | Alphaproteobacteria |
| Wolbachia endosymbiont (group A) of Andrena helvola | Pseudomonadota | Alphaproteobacteria |
| Wolbachia endosymbiont (group A) of Andrena trimmerana | Pseudomonadota | Alphaproteobacteria |
| Wolbachia endosymbiont (group A) of Andrena trimmerana | Pseudomonadota | Alphaproteobacteria |
| Wolbachia endosymbiont (group A) of Anomoia purmunda | Pseudomonadota | Alphaproteobacteria |
| Wolbachia endosymbiont (group A) of Anomoia purmunda | Pseudomonadota | Alphaproteobacteria |
| Wolbachia endosymbiont (group A) of Anoplius nigerrimus | Pseudomonadota | Alphaproteobacteria |
| Wolbachia endosymbiont (group A) of Anoplius nigerrimus | Pseudomonadota | Alphaproteobacteria |
| Wolbachia endosymbiont (group A) of Anthophora plumipes | Pseudomonadota | Alphaproteobacteria |
| Wolbachia endosymbiont (group A) of Anthophora plumipes | Pseudomonadota | Alphaproteobacteria |
| Wolbachia endosymbiont (group A) of Apoderus coryli | Pseudomonadota | Alphaproteobacteria |
| Wolbachia endosymbiont (group A) of Apoderus coryli | Pseudomonadota | Alphaproteobacteria |
| Wolbachia endosymbiont (group A) of Aporus unicolor | Pseudomonadota | Alphaproteobacteria |
| Wolbachia endosymbiont (group A) of Aporus unicolor | Pseudomonadota | Alphaproteobacteria |
| Wolbachia endosymbiont (group A) of Apotomis capreana | Pseudomonadota | Alphaproteobacteria |
| Wolbachia endosymbiont (group A) of Apotomis capreana | Pseudomonadota | Alphaproteobacteria |
| Wolbachia endosymbiont (group A) of Barypeithes pellucidus | Pseudomonadota | Alphaproteobacteria |
| Wolbachia endosymbiont (group A) of Barypeithes pellucidus | Pseudomonadota | Alphaproteobacteria |
| Wolbachia endosymbiont (group A) of Beris morrisii | Pseudomonadota | Alphaproteobacteria |
| Wolbachia endosymbiont (group A) of Beris morrisii | Pseudomonadota | Alphaproteobacteria |
| Wolbachia endosymbiont (group A) of Bibio marci | Pseudomonadota | Alphaproteobacteria |
| Wolbachia endosymbiont (group A) of Bibio marci | Pseudomonadota | Alphaproteobacteria |
| Wolbachia endosymbiont (group A) of Bombylius major | Pseudomonadota | Alphaproteobacteria |
| Wolbachia endosymbiont (group A) of Bombylius major | Pseudomonadota | Alphaproteobacteria |
| Wolbachia endosymbiont (group A) of Brachyopa scutellaris | Pseudomonadota | Alphaproteobacteria |
| Wolbachia endosymbiont (group A) of Brachyopa scutellaris | Pseudomonadota | Alphaproteobacteria |
| Wolbachia endosymbiont (group A) of Calamotropha paludella | Pseudomonadota | Alphaproteobacteria |
| Wolbachia endosymbiont (group A) of Calamotropha paludella | Pseudomonadota | Alphaproteobacteria |

| | | |
|---|---|---|
| Wolbachia endosymbiont (group A) of Campoletis raptor | Pseudomonadota | Alphaproteobacteria |
| Wolbachia endosymbiont (group A) of Campoletis raptor | Pseudomonadota | Alphaproteobacteria |
| Wolbachia endosymbiont (group A) of Cephus spinipes | Pseudomonadota | Alphaproteobacteria |
| Wolbachia endosymbiont (group A) of Cephus spinipes | Pseudomonadota | Alphaproteobacteria |
| Wolbachia endosymbiont (group A) of Cerceris ruficornis | Pseudomonadota | Alphaproteobacteria |
| Wolbachia endosymbiont (group A) of Cerceris ruficornis | Pseudomonadota | Alphaproteobacteria |
| Wolbachia endosymbiont (group A) of Chalcis sispes | Pseudomonadota | Alphaproteobacteria |
| Wolbachia endosymbiont (group A) of Chalcis sispes | Pseudomonadota | Alphaproteobacteria |
| Wolbachia endosymbiont (group A) of Cheilosia soror | Pseudomonadota | Alphaproteobacteria |
| Wolbachia endosymbiont (group A) of Cheilosia soror | Pseudomonadota | Alphaproteobacteria |
| Wolbachia endosymbiont (group A) of Clivina fossor | Pseudomonadota | Alphaproteobacteria |
| Wolbachia endosymbiont (group A) of Clivina fossor | Pseudomonadota | Alphaproteobacteria |
| Wolbachia endosymbiont (group A) of Colletes cunicularius | Pseudomonadota | Alphaproteobacteria |
| Wolbachia endosymbiont (group A) of Colletes cunicularius | Pseudomonadota | Alphaproteobacteria |
| Wolbachia endosymbiont (group A) of Conops quadrifasciatus | Pseudomonadota | Alphaproteobacteria |
| Wolbachia endosymbiont (group A) of Conops quadrifasciatus | Pseudomonadota | Alphaproteobacteria |
| Wolbachia endosymbiont (group A) of Coremacera marginata | Pseudomonadota | Alphaproteobacteria |
| Wolbachia endosymbiont (group A) of Coremacera marginata | Pseudomonadota | Alphaproteobacteria |
| Wolbachia endosymbiont (group A) of Crataerina pallida | Pseudomonadota | Alphaproteobacteria |
| Wolbachia endosymbiont (group A) of Crataerina pallida | Pseudomonadota | Alphaproteobacteria |
| Wolbachia endosymbiont (group A) of Cydia amplana | Pseudomonadota | Alphaproteobacteria |
| Wolbachia endosymbiont (group A) of Cydia amplana | Pseudomonadota | Alphaproteobacteria |
| Wolbachia endosymbiont (group A) of Cydia strobilella | Pseudomonadota | Alphaproteobacteria |
| Wolbachia endosymbiont (group A) of Cydia strobilella | Pseudomonadota | Alphaproteobacteria |
| Wolbachia endosymbiont (group A) of Dendrolimus pini | Pseudomonadota | Alphaproteobacteria |
| Wolbachia endosymbiont (group A) of Dendrolimus pini | Pseudomonadota | Alphaproteobacteria |
| Wolbachia endosymbiont (group A) of Ectemnius continuus | Pseudomonadota | Alphaproteobacteria |
| Wolbachia endosymbiont (group A) of Ectemnius continuus | Pseudomonadota | Alphaproteobacteria |
| Wolbachia endosymbiont (group A) of Endotricha flammealis | Pseudomonadota | Alphaproteobacteria |
| Wolbachia endosymbiont (group A) of Endotricha flammealis | Pseudomonadota | Alphaproteobacteria |
| Wolbachia endosymbiont (group A) of Ennomos erosarius | Pseudomonadota | Alphaproteobacteria |

| | | |
|---|---|---|
| Wolbachia endosymbiont (group A) of Ennomos erosarius | Pseudomonadota | Alphaproteobacteria |
| Wolbachia endosymbiont (group A) of Epagoge grotiana | Pseudomonadota | Alphaproteobacteria |
| Wolbachia endosymbiont (group A) of Epagoge grotiana | Pseudomonadota | Alphaproteobacteria |
| Wolbachia endosymbiont (group A) of Epirrhoe alternata | Pseudomonadota | Alphaproteobacteria |
| Wolbachia endosymbiont (group A) of Epirrhoe alternata | Pseudomonadota | Alphaproteobacteria |
| Wolbachia endosymbiont (group A) of Epistrophe grossularia | Pseudomonadota | Alphaproteobacteria |
| Wolbachia endosymbiont (group A) of Epistrophe grossularia | Pseudomonadota | Alphaproteobacteria |
| Wolbachia endosymbiont (group A) of Eristalinus aeneus | Pseudomonadota | Alphaproteobacteria |
| Wolbachia endosymbiont (group A) of Eristalinus aeneus | Pseudomonadota | Alphaproteobacteria |
| Wolbachia endosymbiont (group A) of Eupithecia tripunctaria | Pseudomonadota | Alphaproteobacteria |
| Wolbachia endosymbiont (group A) of Eupithecia tripunctaria | Pseudomonadota | Alphaproteobacteria |
| Wolbachia endosymbiont (group A) of Gastracanthus pulcherrimus | Pseudomonadota | Alphaproteobacteria |
| Wolbachia endosymbiont (group A) of Gastracanthus pulcherrimus | Pseudomonadota | Alphaproteobacteria |
| Wolbachia endosymbiont (group A) of Gymnosoma rotundatum | Pseudomonadota | Alphaproteobacteria |
| Wolbachia endosymbiont (group A) of Gymnosoma rotundatum | Pseudomonadota | Alphaproteobacteria |
| Wolbachia endosymbiont (group A) of Hedychridium roseum | Pseudomonadota | Alphaproteobacteria |
| Wolbachia endosymbiont (group A) of Hedychridium roseum | Pseudomonadota | Alphaproteobacteria |
| Wolbachia endosymbiont (group A) of Hylaeus communis | Pseudomonadota | Alphaproteobacteria |
| Wolbachia endosymbiont (group A) of Hylaeus communis | Pseudomonadota | Alphaproteobacteria |
| Wolbachia endosymbiont (group A) of Icerya purchasi | Pseudomonadota | Alphaproteobacteria |
| Wolbachia endosymbiont (group A) of Icerya purchasi | Pseudomonadota | Alphaproteobacteria |
| Wolbachia endosymbiont (group A) of Ischnus inquisitorius | Pseudomonadota | Alphaproteobacteria |
| Wolbachia endosymbiont (group A) of Ischnus inquisitorius | Pseudomonadota | Alphaproteobacteria |
| Wolbachia endosymbiont (group A) of Lasioglossum fulvicorne | Pseudomonadota | Alphaproteobacteria |
| Wolbachia endosymbiont (group A) of Lasioglossum fulvicorne | Pseudomonadota | Alphaproteobacteria |
| Wolbachia endosymbiont (group A) of Lasioglossum lativentre | Pseudomonadota | Alphaproteobacteria |
| Wolbachia endosymbiont (group A) of Lasioglossum lativentre | Pseudomonadota | Alphaproteobacteria |
| Wolbachia endosymbiont (group A) of Lasioglossum morio | Pseudomonadota | Alphaproteobacteria |
| Wolbachia endosymbiont (group A) of Lasioglossum morio | Pseudomonadota | Alphaproteobacteria |
| Wolbachia endosymbiont (group A) of Lasioglossum villosulum | Pseudomonadota | Alphaproteobacteria |
| Wolbachia endosymbiont (group A) of Lasioglossum villosulum | Pseudomonadota | Alphaproteobacteria |

| | | |
|---|---|---|
| Wolbachia endosymbiont (group A) of Limnophora tigrina | Pseudomonadota | Alphaproteobacteria |
| Wolbachia endosymbiont (group A) of Limnophora tigrina | Pseudomonadota | Alphaproteobacteria |
| Wolbachia endosymbiont (group A) of Longitarsus flavicornis | Pseudomonadota | Alphaproteobacteria |
| Wolbachia endosymbiont (group A) of Longitarsus flavicornis | Pseudomonadota | Alphaproteobacteria |
| Wolbachia endosymbiont (group A) of Lypha dubia | Pseudomonadota | Alphaproteobacteria |
| Wolbachia endosymbiont (group A) of Lypha dubia | Pseudomonadota | Alphaproteobacteria |
| Wolbachia endosymbiont (group A) of Machimus atricapillus | Pseudomonadota | Alphaproteobacteria |
| Wolbachia endosymbiont (group A) of Machimus atricapillus | Pseudomonadota | Alphaproteobacteria |
| Wolbachia endosymbiont (group A) of Macropis europaea | Pseudomonadota | Alphaproteobacteria |
| Wolbachia endosymbiont (group A) of Macropis europaea | Pseudomonadota | Alphaproteobacteria |
| Wolbachia endosymbiont (group A) of Melieria omissa | Pseudomonadota | Alphaproteobacteria |
| Wolbachia endosymbiont (group A) of Melieria omissa | Pseudomonadota | Alphaproteobacteria |
| Wolbachia endosymbiont (group A) of Merzomyia westermanni | Pseudomonadota | Alphaproteobacteria |
| Wolbachia endosymbiont (group A) of Merzomyia westermanni | Pseudomonadota | Alphaproteobacteria |
| Wolbachia endosymbiont (group A) of Microdon myrmicae | Pseudomonadota | Alphaproteobacteria |
| Wolbachia endosymbiont (group A) of Microdon myrmicae | Pseudomonadota | Alphaproteobacteria |
| Wolbachia endosymbiont (group A) of Myopa testacea | Pseudomonadota | Alphaproteobacteria |
| Wolbachia endosymbiont (group A) of Myopa testacea | Pseudomonadota | Alphaproteobacteria |
| Wolbachia endosymbiont (group A) of Nomada fabriciana | Pseudomonadota | Alphaproteobacteria |
| Wolbachia endosymbiont (group A) of Nomada fabriciana | Pseudomonadota | Alphaproteobacteria |
| Wolbachia endosymbiont (group A) of Nomada ferruginata | Pseudomonadota | Alphaproteobacteria |
| Wolbachia endosymbiont (group A) of Nomada ferruginata | Pseudomonadota | Alphaproteobacteria |
| Wolbachia endosymbiont (group A) of Nomada flava | Pseudomonadota | Alphaproteobacteria |
| Wolbachia endosymbiont (group A) of Nomada flava | Pseudomonadota | Alphaproteobacteria |
| Wolbachia endosymbiont (group A) of Nomada fucata | Pseudomonadota | Alphaproteobacteria |
| Wolbachia endosymbiont (group A) of Nomada fucata | Pseudomonadota | Alphaproteobacteria |
| Wolbachia endosymbiont (group A) of Nomada goodeniana | Pseudomonadota | Alphaproteobacteria |
| Wolbachia endosymbiont (group A) of Nomada goodeniana | Pseudomonadota | Alphaproteobacteria |
| Wolbachia endosymbiont (group A) of Nomada hirtipes | Pseudomonadota | Alphaproteobacteria |
| Wolbachia endosymbiont (group A) of Nomada hirtipes | Pseudomonadota | Alphaproteobacteria |
| Wolbachia endosymbiont (group A) of Nomada rufipes | Pseudomonadota | Alphaproteobacteria |

| | | |
|---|---|---|
| Wolbachia endosymbiont (group A) of Nomada rufipes | Pseudomonadota | Alphaproteobacteria |
| Wolbachia endosymbiont (group A) of Norellia spinipes | Pseudomonadota | Alphaproteobacteria |
| Wolbachia endosymbiont (group A) of Norellia spinipes | Pseudomonadota | Alphaproteobacteria |
| Wolbachia endosymbiont (group A) of Ophion costatus | Pseudomonadota | Alphaproteobacteria |
| Wolbachia endosymbiont (group A) of Ophion costatus | Pseudomonadota | Alphaproteobacteria |
| Wolbachia endosymbiont (group A) of Ophion ellenae | Pseudomonadota | Alphaproteobacteria |
| Wolbachia endosymbiont (group A) of Ophion ellenae | Pseudomonadota | Alphaproteobacteria |
| Wolbachia endosymbiont (group A) of Opomyza germinationis | Pseudomonadota | Alphaproteobacteria |
| Wolbachia endosymbiont (group A) of Opomyza germinationis | Pseudomonadota | Alphaproteobacteria |
| Wolbachia endosymbiont (group A) of Oxytorus armatus | Pseudomonadota | Alphaproteobacteria |
| Wolbachia endosymbiont (group A) of Oxytorus armatus | Pseudomonadota | Alphaproteobacteria |
| Wolbachia endosymbiont (group A) of Paraperithous gnathaulax | Pseudomonadota | Alphaproteobacteria |
| Wolbachia endosymbiont (group A) of Paraperithous gnathaulax | Pseudomonadota | Alphaproteobacteria |
| Wolbachia endosymbiont (group A) of Phalera bucephala | Pseudomonadota | Alphaproteobacteria |
| Wolbachia endosymbiont (group A) of Phalera bucephala | Pseudomonadota | Alphaproteobacteria |
| Wolbachia endosymbiont (group A) of Phasia obesa | Pseudomonadota | Alphaproteobacteria |
| Wolbachia endosymbiont (group A) of Phasia obesa | Pseudomonadota | Alphaproteobacteria |
| Wolbachia endosymbiont (group A) of Pheosia gnoma | Pseudomonadota | Alphaproteobacteria |
| Wolbachia endosymbiont (group A) of Pheosia gnoma | Pseudomonadota | Alphaproteobacteria |
| Wolbachia endosymbiont (group A) of Pherbina coryleti | Pseudomonadota | Alphaproteobacteria |
| Wolbachia endosymbiont (group A) of Pherbina coryleti | Pseudomonadota | Alphaproteobacteria |
| Wolbachia endosymbiont (group A) of Philanthus triangulum | Pseudomonadota | Alphaproteobacteria |
| Wolbachia endosymbiont (group A) of Philanthus triangulum | Pseudomonadota | Alphaproteobacteria |
| Wolbachia endosymbiont (group A) of Philonthus cognatus | Pseudomonadota | Alphaproteobacteria |
| Wolbachia endosymbiont (group A) of Philonthus cognatus | Pseudomonadota | Alphaproteobacteria |
| Wolbachia endosymbiont (group A) of Pipizella viduata | Pseudomonadota | Alphaproteobacteria |
| Wolbachia endosymbiont (group A) of Pipizella viduata | Pseudomonadota | Alphaproteobacteria |
| Wolbachia endosymbiont (group A) of Pogonocherus hispidulus | Pseudomonadota | Alphaproteobacteria |
| Wolbachia endosymbiont (group A) of Pogonocherus hispidulus | Pseudomonadota | Alphaproteobacteria |
| Wolbachia endosymbiont (group A) of Portevinia maculata | Pseudomonadota | Alphaproteobacteria |
| Wolbachia endosymbiont (group A) of Portevinia maculata | Pseudomonadota | Alphaproteobacteria |

| | | |
|---|---|---|
| Wolbachia endosymbiont (group A) of Protocalliphora azurea | Pseudomonadota | Alphaproteobacteria |
| Wolbachia endosymbiont (group A) of Protocalliphora azurea | Pseudomonadota | Alphaproteobacteria |
| Wolbachia endosymbiont (group A) of Rhinocyllus conicus | Pseudomonadota | Alphaproteobacteria |
| Wolbachia endosymbiont (group A) of Rhinocyllus conicus | Pseudomonadota | Alphaproteobacteria |
| Wolbachia endosymbiont (group A) of Rhorus exstirpatorius | Pseudomonadota | Alphaproteobacteria |
| Wolbachia endosymbiont (group A) of Rhorus exstirpatorius | Pseudomonadota | Alphaproteobacteria |
| Wolbachia endosymbiont (group A) of Scambus nigricans | Pseudomonadota | Alphaproteobacteria |
| Wolbachia endosymbiont (group A) of Scambus nigricans | Pseudomonadota | Alphaproteobacteria |
| Wolbachia endosymbiont (group A) of Schoenobius gigantella | Pseudomonadota | Alphaproteobacteria |
| Wolbachia endosymbiont (group A) of Schoenobius gigantella | Pseudomonadota | Alphaproteobacteria |
| Wolbachia endosymbiont (group A) of Scoparia ambigualis | Pseudomonadota | Alphaproteobacteria |
| Wolbachia endosymbiont (group A) of Scoparia ambigualis | Pseudomonadota | Alphaproteobacteria |
| Wolbachia endosymbiont (group A) of Sicus ferrugineus | Pseudomonadota | Alphaproteobacteria |
| Wolbachia endosymbiont (group A) of Sicus ferrugineus | Pseudomonadota | Alphaproteobacteria |
| Wolbachia endosymbiont (group A) of Sphaerophoria taeniata | Pseudomonadota | Alphaproteobacteria |
| Wolbachia endosymbiont (group A) of Sphaerophoria taeniata | Pseudomonadota | Alphaproteobacteria |
| Wolbachia endosymbiont (group A) of Sphecodes monilicornis | Pseudomonadota | Alphaproteobacteria |
| Wolbachia endosymbiont (group A) of Sphecodes monilicornis | Pseudomonadota | Alphaproteobacteria |
| Wolbachia endosymbiont (group A) of Sturmia bella | Pseudomonadota | Alphaproteobacteria |
| Wolbachia endosymbiont (group A) of Sturmia bella | Pseudomonadota | Alphaproteobacteria |
| Wolbachia endosymbiont (group A) of Sympetrum striolatum | Pseudomonadota | Alphaproteobacteria |
| Wolbachia endosymbiont (group A) of Sympetrum striolatum | Pseudomonadota | Alphaproteobacteria |
| Wolbachia endosymbiont (group A) of Therophilus tumidulus | Pseudomonadota | Alphaproteobacteria |
| Wolbachia endosymbiont (group A) of Therophilus tumidulus | Pseudomonadota | Alphaproteobacteria |
| Wolbachia endosymbiont (group A) of Tiphia femorata | Pseudomonadota | Alphaproteobacteria |
| Wolbachia endosymbiont (group A) of Tiphia femorata | Pseudomonadota | Alphaproteobacteria |
| Wolbachia endosymbiont (group A) of Tromatobia lineatoria | Pseudomonadota | Alphaproteobacteria |
| Wolbachia endosymbiont (group A) of Tromatobia lineatoria | Pseudomonadota | Alphaproteobacteria |
| Wolbachia endosymbiont (group A) of Trypoxylon clavicerum | Pseudomonadota | Alphaproteobacteria |
| Wolbachia endosymbiont (group A) of Trypoxylon clavicerum | Pseudomonadota | Alphaproteobacteria |
| Wolbachia endosymbiont (group A) of Udea olivalis | Pseudomonadota | Alphaproteobacteria |

| | | |
|---|---|---|
| Wolbachia endosymbiont (group A) of Udea olivalis | Pseudomonadota | Alphaproteobacteria |
| Wolbachia endosymbiont (group A) of Urophora cardui | Pseudomonadota | Alphaproteobacteria |
| Wolbachia endosymbiont (group A) of Urophora cardui | Pseudomonadota | Alphaproteobacteria |
| Wolbachia endosymbiont (group A) of Volucella bombylans | Pseudomonadota | Alphaproteobacteria |
| Wolbachia endosymbiont (group A) of Volucella bombylans | Pseudomonadota | Alphaproteobacteria |
| Wolbachia endosymbiont (group A) of Yponomeuta plumbellus | Pseudomonadota | Alphaproteobacteria |
| Wolbachia endosymbiont (group A) of Yponomeuta plumbellus | Pseudomonadota | Alphaproteobacteria |
| Wolbachia endosymbiont (group B) of Ablattaria laevigata | Pseudomonadota | Alphaproteobacteria |
| Wolbachia endosymbiont (group B) of Ablattaria laevigata | Pseudomonadota | Alphaproteobacteria |
| Wolbachia endosymbiont (group B) of Agriphila inquinatella | Pseudomonadota | Alphaproteobacteria |
| Wolbachia endosymbiont (group B) of Agriphila inquinatella | Pseudomonadota | Alphaproteobacteria |
| Wolbachia endosymbiont (group B) of Agriphila straminella | Pseudomonadota | Alphaproteobacteria |
| Wolbachia endosymbiont (group B) of Agriphila straminella | Pseudomonadota | Alphaproteobacteria |
| Wolbachia endosymbiont (group B) of Agrotis exclamationis | Pseudomonadota | Alphaproteobacteria |
| Wolbachia endosymbiont (group B) of Agrotis exclamationis | Pseudomonadota | Alphaproteobacteria |
| Wolbachia endosymbiont (group B) of Agrotis puta | Pseudomonadota | Alphaproteobacteria |
| Wolbachia endosymbiont (group B) of Agrotis puta | Pseudomonadota | Alphaproteobacteria |
| Wolbachia endosymbiont (group B) of Alcis repandata | Pseudomonadota | Alphaproteobacteria |
| Wolbachia endosymbiont (group B) of Alcis repandata | Pseudomonadota | Alphaproteobacteria |
| Wolbachia endosymbiont (group B) of Anania hortulata | Pseudomonadota | Alphaproteobacteria |
| Wolbachia endosymbiont (group B) of Anania hortulata | Pseudomonadota | Alphaproteobacteria |
| Wolbachia endosymbiont (group B) of Apomyelois bistriatella | Pseudomonadota | Alphaproteobacteria |
| Wolbachia endosymbiont (group B) of Apomyelois bistriatella | Pseudomonadota | Alphaproteobacteria |
| Wolbachia endosymbiont (group B) of Aporia crataegi | Pseudomonadota | Alphaproteobacteria |
| Wolbachia endosymbiont (group B) of Aporia crataegi | Pseudomonadota | Alphaproteobacteria |
| Wolbachia endosymbiont (group B) of Apotomis betuletana | Pseudomonadota | Alphaproteobacteria |
| Wolbachia endosymbiont (group B) of Apotomis betuletana | Pseudomonadota | Alphaproteobacteria |
| Wolbachia endosymbiont (group B) of Apotomis turbidana | Pseudomonadota | Alphaproteobacteria |
| Wolbachia endosymbiont (group B) of Apotomis turbidana | Pseudomonadota | Alphaproteobacteria |
| Wolbachia endosymbiont (group B) of Archips podanus | Pseudomonadota | Alphaproteobacteria |
| Wolbachia endosymbiont (group B) of Archips podanus | Pseudomonadota | Alphaproteobacteria |

| | | |
|---|---|---|
| Wolbachia endosymbiont (group B) of Aricia artaxerxes | Pseudomonadota | Alphaproteobacteria |
| Wolbachia endosymbiont (group B) of Aricia artaxerxes | Pseudomonadota | Alphaproteobacteria |
| Wolbachia endosymbiont (group B) of Athalia cordata | Pseudomonadota | Alphaproteobacteria |
| Wolbachia endosymbiont (group B) of Athalia cordata | Pseudomonadota | Alphaproteobacteria |
| Wolbachia endosymbiont (group B) of Campaea margaritaria | Pseudomonadota | Alphaproteobacteria |
| Wolbachia endosymbiont (group B) of Campaea margaritaria | Pseudomonadota | Alphaproteobacteria |
| Wolbachia endosymbiont (group B) of Camptogramma bilineatum | Pseudomonadota | Alphaproteobacteria |
| Wolbachia endosymbiont (group B) of Camptogramma bilineatum | Pseudomonadota | Alphaproteobacteria |
| Wolbachia endosymbiont (group B) of Catoptria pinella | Pseudomonadota | Alphaproteobacteria |
| Wolbachia endosymbiont (group B) of Catoptria pinella | Pseudomonadota | Alphaproteobacteria |
| Wolbachia endosymbiont (group B) of Chesias legatella | Pseudomonadota | Alphaproteobacteria |
| Wolbachia endosymbiont (group B) of Chesias legatella | Pseudomonadota | Alphaproteobacteria |
| Wolbachia endosymbiont (group B) of Chorthippus parallelus | Pseudomonadota | Alphaproteobacteria |
| Wolbachia endosymbiont (group B) of Chorthippus parallelus | Pseudomonadota | Alphaproteobacteria |
| Wolbachia endosymbiont (group B) of Colias croceus | Pseudomonadota | Alphaproteobacteria |
| Wolbachia endosymbiont (group B) of Colias croceus | Pseudomonadota | Alphaproteobacteria |
| Wolbachia endosymbiont (group B) of Cyclophora punctaria | Pseudomonadota | Alphaproteobacteria |
| Wolbachia endosymbiont (group B) of Cyclophora punctaria | Pseudomonadota | Alphaproteobacteria |
| Wolbachia endosymbiont (group B) of Dendrolimus pini | Pseudomonadota | Alphaproteobacteria |
| Wolbachia endosymbiont (group B) of Dendrolimus pini | Pseudomonadota | Alphaproteobacteria |
| Wolbachia endosymbiont (group B) of Elophila nymphaeata | Pseudomonadota | Alphaproteobacteria |
| Wolbachia endosymbiont (group B) of Elophila nymphaeata | Pseudomonadota | Alphaproteobacteria |
| Wolbachia endosymbiont (group B) of Emmelina monodactyla | Pseudomonadota | Alphaproteobacteria |
| Wolbachia endosymbiont (group B) of Emmelina monodactyla | Pseudomonadota | Alphaproteobacteria |
| Wolbachia endosymbiont (group B) of Endotricha flammealis | Pseudomonadota | Alphaproteobacteria |
| Wolbachia endosymbiont (group B) of Endotricha flammealis | Pseudomonadota | Alphaproteobacteria |
| Wolbachia endosymbiont (group B) of Ennomos erosarius | Pseudomonadota | Alphaproteobacteria |
| Wolbachia endosymbiont (group B) of Ennomos erosarius | Pseudomonadota | Alphaproteobacteria |
| Wolbachia endosymbiont (group B) of Episyrphus balteatus | Pseudomonadota | Alphaproteobacteria |
| Wolbachia endosymbiont (group B) of Episyrphus balteatus | Pseudomonadota | Alphaproteobacteria |
| Wolbachia endosymbiont (group B) of Erynnis tages | Pseudomonadota | Alphaproteobacteria |

| | | |
|---|---|---|
| Wolbachia endosymbiont (group B) of Erynnis tages | Pseudomonadota | Alphaproteobacteria |
| Wolbachia endosymbiont (group B) of Eucosma cana | Pseudomonadota | Alphaproteobacteria |
| Wolbachia endosymbiont (group B) of Eucosma cana | Pseudomonadota | Alphaproteobacteria |
| Wolbachia endosymbiont (group B) of Eupeodes latifasciatus | Pseudomonadota | Alphaproteobacteria |
| Wolbachia endosymbiont (group B) of Eupeodes latifasciatus | Pseudomonadota | Alphaproteobacteria |
| Wolbachia endosymbiont (group B) of Euphydryas aurinia | Pseudomonadota | Alphaproteobacteria |
| Wolbachia endosymbiont (group B) of Euphydryas aurinia | Pseudomonadota | Alphaproteobacteria |
| Wolbachia endosymbiont (group B) of Eupithecia inturbata | Pseudomonadota | Alphaproteobacteria |
| Wolbachia endosymbiont (group B) of Eupithecia inturbata | Pseudomonadota | Alphaproteobacteria |
| Wolbachia endosymbiont (group B) of Falcaria lacertinaria | Pseudomonadota | Alphaproteobacteria |
| Wolbachia endosymbiont (group B) of Falcaria lacertinaria | Pseudomonadota | Alphaproteobacteria |
| Wolbachia endosymbiont (group B) of Germaria angustata | Pseudomonadota | Alphaproteobacteria |
| Wolbachia endosymbiont (group B) of Germaria angustata | Pseudomonadota | Alphaproteobacteria |
| Wolbachia endosymbiont (group B) of Gerris lacustris | Pseudomonadota | Alphaproteobacteria |
| Wolbachia endosymbiont (group B) of Gerris lacustris | Pseudomonadota | Alphaproteobacteria |
| Wolbachia endosymbiont (group B) of Hofmannophila pseudospretella | Pseudomonadota | Alphaproteobacteria |
| Wolbachia endosymbiont (group B) of Hofmannophila pseudospretella | Pseudomonadota | Alphaproteobacteria |
| Wolbachia endosymbiont (group B) of Horisme vitalbata | Pseudomonadota | Alphaproteobacteria |
| Wolbachia endosymbiont (group B) of Horisme vitalbata | Pseudomonadota | Alphaproteobacteria |
| Wolbachia endosymbiont (group B) of Hylaea fasciaria | Pseudomonadota | Alphaproteobacteria |
| Wolbachia endosymbiont (group B) of Hylaea fasciaria | Pseudomonadota | Alphaproteobacteria |
| Wolbachia endosymbiont (group B) of Idaea biselata | Pseudomonadota | Alphaproteobacteria |
| Wolbachia endosymbiont (group B) of Idaea biselata | Pseudomonadota | Alphaproteobacteria |
| Wolbachia endosymbiont (group B) of Ischnura elegans | Pseudomonadota | Alphaproteobacteria |
| Wolbachia endosymbiont (group B) of Ischnura elegans | Pseudomonadota | Alphaproteobacteria |
| Wolbachia endosymbiont (group B) of Limnophora tigrina | Pseudomonadota | Alphaproteobacteria |
| Wolbachia endosymbiont (group B) of Limnophora tigrina | Pseudomonadota | Alphaproteobacteria |
| Wolbachia endosymbiont (group B) of Longitarsus flavicornis | Pseudomonadota | Alphaproteobacteria |
| Wolbachia endosymbiont (group B) of Longitarsus flavicornis | Pseudomonadota | Alphaproteobacteria |
| Wolbachia endosymbiont (group B) of Lycaena phlaeas | Pseudomonadota | Alphaproteobacteria |
| Wolbachia endosymbiont (group B) of Lycaena phlaeas | Pseudomonadota | Alphaproteobacteria |

| | | |
|---|---|---|
| Wolbachia endosymbiont (group B) of Melanargia galathea | Pseudomonadota | Alphaproteobacteria |
| Wolbachia endosymbiont (group B) of Melanargia galathea | Pseudomonadota | Alphaproteobacteria |
| Wolbachia endosymbiont (group B) of Melanostoma mellinum | Pseudomonadota | Alphaproteobacteria |
| Wolbachia endosymbiont (group B) of Melanostoma mellinum | Pseudomonadota | Alphaproteobacteria |
| Wolbachia endosymbiont (group B) of Myelois circumvoluta | Pseudomonadota | Alphaproteobacteria |
| Wolbachia endosymbiont (group B) of Myelois circumvoluta | Pseudomonadota | Alphaproteobacteria |
| Wolbachia endosymbiont (group B) of Nymphalis c-album | Pseudomonadota | Alphaproteobacteria |
| Wolbachia endosymbiont (group B) of Nymphalis c-album | Pseudomonadota | Alphaproteobacteria |
| Wolbachia endosymbiont (group B) of Orgyia antiqua | Pseudomonadota | Alphaproteobacteria |
| Wolbachia endosymbiont (group B) of Orgyia antiqua | Pseudomonadota | Alphaproteobacteria |
| Wolbachia endosymbiont (group B) of Pammene fasciana | Pseudomonadota | Alphaproteobacteria |
| Wolbachia endosymbiont (group B) of Pammene fasciana | Pseudomonadota | Alphaproteobacteria |
| Wolbachia endosymbiont (group B) of Pandemis corylana | Pseudomonadota | Alphaproteobacteria |
| Wolbachia endosymbiont (group B) of Pandemis corylana | Pseudomonadota | Alphaproteobacteria |
| Wolbachia endosymbiont (group B) of Parapoynx stratiotata | Pseudomonadota | Alphaproteobacteria |
| Wolbachia endosymbiont (group B) of Parapoynx stratiotata | Pseudomonadota | Alphaproteobacteria |
| Wolbachia endosymbiont (group B) of Pararge aegeria | Pseudomonadota | Alphaproteobacteria |
| Wolbachia endosymbiont (group B) of Pararge aegeria | Pseudomonadota | Alphaproteobacteria |
| Wolbachia endosymbiont (group B) of Phalera bucephala | Pseudomonadota | Alphaproteobacteria |
| Wolbachia endosymbiont (group B) of Phalera bucephala | Pseudomonadota | Alphaproteobacteria |
| Wolbachia endosymbiont (group B) of Phasia obesa | Pseudomonadota | Alphaproteobacteria |
| Wolbachia endosymbiont (group B) of Phasia obesa | Pseudomonadota | Alphaproteobacteria |
| Wolbachia endosymbiont (group B) of Pheosia tremula | Pseudomonadota | Alphaproteobacteria |
| Wolbachia endosymbiont (group B) of Pheosia tremula | Pseudomonadota | Alphaproteobacteria |
| Wolbachia endosymbiont (group B) of Pilophorus perplexus | Pseudomonadota | Alphaproteobacteria |
| Wolbachia endosymbiont (group B) of Pilophorus perplexus | Pseudomonadota | Alphaproteobacteria |
| Wolbachia endosymbiont (group B) of Polyommatus icarus | Pseudomonadota | Alphaproteobacteria |
| Wolbachia endosymbiont (group B) of Polyommatus icarus | Pseudomonadota | Alphaproteobacteria |
| Wolbachia endosymbiont (group B) of Protocalliphora azurea | Pseudomonadota | Alphaproteobacteria |
| Wolbachia endosymbiont (group B) of Protocalliphora azurea | Pseudomonadota | Alphaproteobacteria |
| Wolbachia endosymbiont (group B) of Pyrgus malvae | Pseudomonadota | Alphaproteobacteria |

| | | |
|---|---|---|
| Wolbachia endosymbiont (group B) of Pyrgus malvae | Pseudomonadota | Alphaproteobacteria |
| Wolbachia endosymbiont (group B) of Rhopobota naevana | Pseudomonadota | Alphaproteobacteria |
| Wolbachia endosymbiont (group B) of Rhopobota naevana | Pseudomonadota | Alphaproteobacteria |
| Wolbachia endosymbiont (group B) of Scrobipalpa ocellatella | Pseudomonadota | Alphaproteobacteria |
| Wolbachia endosymbiont (group B) of Scrobipalpa ocellatella | Pseudomonadota | Alphaproteobacteria |
| Wolbachia endosymbiont (group B) of Silvanus unidentatus | Pseudomonadota | Alphaproteobacteria |
| Wolbachia endosymbiont (group B) of Silvanus unidentatus | Pseudomonadota | Alphaproteobacteria |
| Wolbachia endosymbiont (group B) of Tholera decimalis | Pseudomonadota | Alphaproteobacteria |
| Wolbachia endosymbiont (group B) of Tholera decimalis | Pseudomonadota | Alphaproteobacteria |
| Wolbachia endosymbiont (group B) of Thymelicus sylvestris | Pseudomonadota | Alphaproteobacteria |
| Wolbachia endosymbiont (group B) of Thymelicus sylvestris | Pseudomonadota | Alphaproteobacteria |
| Wolbachia endosymbiont (group B) of Udea ferrugalis | Pseudomonadota | Alphaproteobacteria |
| Wolbachia endosymbiont (group B) of Udea ferrugalis | Pseudomonadota | Alphaproteobacteria |
| Wolbachia endosymbiont (group B) of Villa cingulata | Pseudomonadota | Alphaproteobacteria |
| Wolbachia endosymbiont (group B) of Villa cingulata | Pseudomonadota | Alphaproteobacteria |
| Wolbachia endosymbiont (group B) of Watsonalla binaria | Pseudomonadota | Alphaproteobacteria |
| Wolbachia endosymbiont (group B) of Watsonalla binaria | Pseudomonadota | Alphaproteobacteria |
| Wolbachia endosymbiont (group B) of Xanthorhoe designata | Pseudomonadota | Alphaproteobacteria |
| Wolbachia endosymbiont (group B) of Xanthorhoe designata | Pseudomonadota | Alphaproteobacteria |
| Wolbachia endosymbiont (group B) of Xestia c-nigrum | Pseudomonadota | Alphaproteobacteria |
| Wolbachia endosymbiont (group B) of Xestia c-nigrum | Pseudomonadota | Alphaproteobacteria |
| Wolbachia endosymbiont (group B) of Xylocampa areola | Pseudomonadota | Alphaproteobacteria |
| Wolbachia endosymbiont (group B) of Xylocampa areola | Pseudomonadota | Alphaproteobacteria |
| Wolbachia endosymbiont (group B) of Yponomeuta malinellus | Pseudomonadota | Alphaproteobacteria |
| Wolbachia endosymbiont (group B) of Yponomeuta malinellus | Pseudomonadota | Alphaproteobacteria |
| Wolbachia endosymbiont (group E) of Neria commutata | Pseudomonadota | Alphaproteobacteria |
| Wolbachia endosymbiont (group E) of Neria commutata | Pseudomonadota | Alphaproteobacteria |
| Wolbachia endosymbiont of Aedes aegypti | Pseudomonadota | Alphaproteobacteria |
| Wolbachia endosymbiont of Aedes aegypti | Pseudomonadota | Alphaproteobacteria |
| Wolbachia endosymbiont of Aedes albopictus | Pseudomonadota | Alphaproteobacteria |
| Wolbachia endosymbiont of Aedes albopictus | Pseudomonadota | Alphaproteobacteria |

| | | |
|---|---|---|
| Wolbachia endosymbiont of Anurida maritima | Pseudomonadota | Alphaproteobacteria |
| Wolbachia endosymbiont of Anurida maritima | Pseudomonadota | Alphaproteobacteria |
| Wolbachia endosymbiont of Armadillidium arcangelii | Pseudomonadota | Alphaproteobacteria |
| Wolbachia endosymbiont of Armadillidium arcangelii | Pseudomonadota | Alphaproteobacteria |
| Wolbachia endosymbiont of Bemisia tabaci | Pseudomonadota | Alphaproteobacteria |
| Wolbachia endosymbiont of Bemisia tabaci | Pseudomonadota | Alphaproteobacteria |
| Wolbachia endosymbiont of Brugia pahangi | Pseudomonadota | Alphaproteobacteria |
| Wolbachia endosymbiont of Brugia pahangi | Pseudomonadota | Alphaproteobacteria |
| Wolbachia endosymbiont of Cantharis cryptica | Pseudomonadota | Alphaproteobacteria |
| Wolbachia endosymbiont of Cantharis cryptica | Pseudomonadota | Alphaproteobacteria |
| Wolbachia endosymbiont of Carposina sasakii | Pseudomonadota | Alphaproteobacteria |
| Wolbachia endosymbiont of Carposina sasakii | Pseudomonadota | Alphaproteobacteria |
| Wolbachia endosymbiont of Ceratosolen solmsi | Pseudomonadota | Alphaproteobacteria |
| Wolbachia endosymbiont of Ceratosolen solmsi | Pseudomonadota | Alphaproteobacteria |
| Wolbachia endosymbiont of Chrysomya megacephala | Pseudomonadota | Alphaproteobacteria |
| Wolbachia endosymbiont of Chrysomya megacephala | Pseudomonadota | Alphaproteobacteria |
| Wolbachia endosymbiont of Cimex lectularius | Pseudomonadota | Alphaproteobacteria |
| Wolbachia endosymbiont of Cimex lectularius | Pseudomonadota | Alphaproteobacteria |
| Wolbachia endosymbiont of Cruorifilaria tuberocauda | Pseudomonadota | Alphaproteobacteria |
| Wolbachia endosymbiont of Cruorifilaria tuberocauda | Pseudomonadota | Alphaproteobacteria |
| Wolbachia endosymbiont of Ctenocephalides felis wCfeJ | Pseudomonadota | Alphaproteobacteria |
| Wolbachia endosymbiont of Ctenocephalides felis wCfeJ | Pseudomonadota | Alphaproteobacteria |
| Wolbachia endosymbiont of Ctenocephalides felis wCfeT | Pseudomonadota | Alphaproteobacteria |
| Wolbachia endosymbiont of Ctenocephalides felis wCfeT | Pseudomonadota | Alphaproteobacteria |
| Wolbachia endosymbiont of Culex quinquefasciatus Pel | Pseudomonadota | Alphaproteobacteria |
| Wolbachia endosymbiont of Culex quinquefasciatus Pel | Pseudomonadota | Alphaproteobacteria |
| Wolbachia endosymbiont of Delia radicum | Pseudomonadota | Alphaproteobacteria |
| Wolbachia endosymbiont of Delia radicum | Pseudomonadota | Alphaproteobacteria |
| Wolbachia endosymbiont of Diaphorina citri | Pseudomonadota | Alphaproteobacteria |
| Wolbachia endosymbiont of Diaphorina citri | Pseudomonadota | Alphaproteobacteria |
| Wolbachia endosymbiont of Dipetalonema caudispina | Pseudomonadota | Alphaproteobacteria |

| | | |
|---|---|---|
| Wolbachia endosymbiont of Dipetalonema caudispina | Pseudomonadota | Alphaproteobacteria |
| Wolbachia endosymbiont of Dirofilaria (Dirofilaria) immitis | Pseudomonadota | Alphaproteobacteria |
| Wolbachia endosymbiont of Dirofilaria (Dirofilaria) immitis | Pseudomonadota | Alphaproteobacteria |
| Wolbachia endosymbiont of Drosophila aff. chauvacae BK-2020 | Pseudomonadota | Alphaproteobacteria |
| Wolbachia endosymbiont of Drosophila aff. chauvacae BK-2020 | Pseudomonadota | Alphaproteobacteria |
| Wolbachia endosymbiont of Drosophila ananassae | Pseudomonadota | Alphaproteobacteria |
| Wolbachia endosymbiont of Drosophila ananassae | Pseudomonadota | Alphaproteobacteria |
| Wolbachia endosymbiont of Drosophila incompta | Pseudomonadota | Alphaproteobacteria |
| Wolbachia endosymbiont of Drosophila incompta | Pseudomonadota | Alphaproteobacteria |
| Wolbachia endosymbiont of Drosophila mauritiana | Pseudomonadota | Alphaproteobacteria |
| Wolbachia endosymbiont of Drosophila mauritiana | Pseudomonadota | Alphaproteobacteria |
| Wolbachia endosymbiont of Drosophila melanogaster | Pseudomonadota | Alphaproteobacteria |
| Wolbachia endosymbiont of Drosophila melanogaster | Pseudomonadota | Alphaproteobacteria |
| Wolbachia endosymbiont of Drosophila pseudotakahashii | Pseudomonadota | Alphaproteobacteria |
| Wolbachia endosymbiont of Drosophila pseudotakahashii | Pseudomonadota | Alphaproteobacteria |
| Wolbachia endosymbiont of Drosophila santomea | Pseudomonadota | Alphaproteobacteria |
| Wolbachia endosymbiont of Drosophila santomea | Pseudomonadota | Alphaproteobacteria |
| Wolbachia endosymbiont of Drosophila simulans wHa | Pseudomonadota | Alphaproteobacteria |
| Wolbachia endosymbiont of Drosophila simulans wHa | Pseudomonadota | Alphaproteobacteria |
| Wolbachia endosymbiont of Drosophila willistoni | Pseudomonadota | Alphaproteobacteria |
| Wolbachia endosymbiont of Drosophila willistoni | Pseudomonadota | Alphaproteobacteria |
| Wolbachia endosymbiont of Drosophila yakuba | Pseudomonadota | Alphaproteobacteria |
| Wolbachia endosymbiont of Drosophila yakuba | Pseudomonadota | Alphaproteobacteria |
| Wolbachia endosymbiont of Encarsia formosa | Pseudomonadota | Alphaproteobacteria |
| Wolbachia endosymbiont of Encarsia formosa | Pseudomonadota | Alphaproteobacteria |
| Wolbachia endosymbiont of Folsomia candida | Pseudomonadota | Alphaproteobacteria |
| Wolbachia endosymbiont of Folsomia candida | Pseudomonadota | Alphaproteobacteria |
| Wolbachia endosymbiont of Frankliniella intonsa | Pseudomonadota | Alphaproteobacteria |
| Wolbachia endosymbiont of Frankliniella intonsa | Pseudomonadota | Alphaproteobacteria |
| Wolbachia endosymbiont of Howardula sp. | Pseudomonadota | Alphaproteobacteria |
| Wolbachia endosymbiont of Liriomyza huidobrensis | Pseudomonadota | Alphaproteobacteria |

| | | |
|---|---|---|
| Wolbachia endosymbiont of Liriomyza huidobrensis | Pseudomonadota | Alphaproteobacteria |
| Wolbachia endosymbiont of Litomosoides sigmodontis | Pseudomonadota | Alphaproteobacteria |
| Wolbachia endosymbiont of Oedothorax gibbosus | Pseudomonadota | Alphaproteobacteria |
| Wolbachia endosymbiont of Oedothorax gibbosus | Pseudomonadota | Alphaproteobacteria |
| Wolbachia endosymbiont of Onchocerca ochengi | Pseudomonadota | Alphaproteobacteria |
| Wolbachia endosymbiont of Onchocerca ochengi | Pseudomonadota | Alphaproteobacteria |
| Wolbachia endosymbiont of Onchocerca volvulus str. Cameroon | Pseudomonadota | Alphaproteobacteria |
| Wolbachia endosymbiont of Onchocerca volvulus str. Cameroon | Pseudomonadota | Alphaproteobacteria |
| Wolbachia endosymbiont of Oryzaephilus surinamensis | Pseudomonadota | Alphaproteobacteria |
| Wolbachia endosymbiont of Oryzaephilus surinamensis | Pseudomonadota | Alphaproteobacteria |
| Wolbachia endosymbiont of Ostrinia furnacalis | Pseudomonadota | Alphaproteobacteria |
| Wolbachia endosymbiont of Ostrinia furnacalis | Pseudomonadota | Alphaproteobacteria |
| Wolbachia endosymbiont of Ostrinia scapulalis | Pseudomonadota | Alphaproteobacteria |
| Wolbachia endosymbiont of Ostrinia scapulalis | Pseudomonadota | Alphaproteobacteria |
| Wolbachia endosymbiont of Phyllotreta cruciferae | Pseudomonadota | Alphaproteobacteria |
| Wolbachia endosymbiont of Phyllotreta cruciferae | Pseudomonadota | Alphaproteobacteria |
| Wolbachia endosymbiont of Psylliodes chrysocephala | Pseudomonadota | Alphaproteobacteria |
| Wolbachia endosymbiont of Psylliodes chrysocephala | Pseudomonadota | Alphaproteobacteria |
| Wolbachia endosymbiont of Rhagoletis cingulata | Pseudomonadota | Alphaproteobacteria |
| Wolbachia endosymbiont of Rhagoletis cingulata | Pseudomonadota | Alphaproteobacteria |
| Wolbachia endosymbiont of Spodoptera picta | Pseudomonadota | Alphaproteobacteria |
| Wolbachia endosymbiont of Spodoptera picta | Pseudomonadota | Alphaproteobacteria |
| Wolbachia endosymbiont of Tetranychus urticae | Pseudomonadota | Alphaproteobacteria |
| Wolbachia endosymbiont of Tetranychus urticae | Pseudomonadota | Alphaproteobacteria |
| Wolbachia endosymbiont of Trichogramma pretiosum | Pseudomonadota | Alphaproteobacteria |
| Wolbachia endosymbiont of Trichogramma pretiosum | Pseudomonadota | Alphaproteobacteria |
| Wolbachia endosymbiont of Wiebesia pumilae | Pseudomonadota | Alphaproteobacteria |
| Wolbachia endosymbiont of Wiebesia pumilae | Pseudomonadota | Alphaproteobacteria |
| Wolbachia endosymbiont of Zaprionus tsacasi | Pseudomonadota | Alphaproteobacteria |
| Wolbachia endosymbiont of Zaprionus tsacasi | Pseudomonadota | Alphaproteobacteria |
| Wolbachia endosymbiont strain TRS of Brugia malayi | Pseudomonadota | Alphaproteobacteria |

| | | |
|---|---|---|
| Wolbachia endosymbiont strain TRS of Brugia malayi | Pseudomonadota | Alphaproteobacteria |
| Wolbachia pipientis | Pseudomonadota | Alphaproteobacteria |
| Wolbachia pipientis | Pseudomonadota | Alphaproteobacteria |
| Xanthobacter dioxanivorans | Pseudomonadota | Alphaproteobacteria |
| Xanthobacter dioxanivorans | Pseudomonadota | Alphaproteobacteria |
| Yoonia algicola | Pseudomonadota | Alphaproteobacteria |
| Yoonia algicola | Pseudomonadota | Alphaproteobacteria |
| Yoonia phaeophyticola | Pseudomonadota | Alphaproteobacteria |
| Yoonia phaeophyticola | Pseudomonadota | Alphaproteobacteria |
| Yoonia rhodophyticola | Pseudomonadota | Alphaproteobacteria |
| Yoonia rhodophyticola | Pseudomonadota | Alphaproteobacteria |
| Zymomonas mobilis subsp. mobilis str. CP4 = NRRL B-14023 | Pseudomonadota | Alphaproteobacteria |
| Anaerolinea thermophila UNI-1 | Chloroflexota | Anaerolineae |
| Anaerolinea thermophila UNI-1 | Chloroflexota | Anaerolineae |
| Brevefilum fermentans | Chloroflexota | Anaerolineae |
| Pelolinea submarina | Chloroflexota | Anaerolineae |
| Pelolinea submarina | Chloroflexota | Anaerolineae |
| Aquifex aeolicus VF5 | Aquificota | Aquificia |
| Aquifex aeolicus VF5 | Aquificota | Aquificia |
| Desulfurobacterium thermolithotrophum DSM 11699 | Aquificota | Aquificia |
| Desulfurobacterium thermolithotrophum DSM 11699 | Aquificota | Aquificia |
| Hydrogenobacter thermophilus TK-6 | Aquificota | Aquificia |
| Hydrogenobacter thermophilus TK-6 | Aquificota | Aquificia |
| Persephonella marina EX-H1 | Aquificota | Aquificia |
| Persephonella marina EX-H1 | Aquificota | Aquificia |
| Sulfurihydrogenibium azorense Az-Fu1 | Aquificota | Aquificia |
| Thermocrinis albus DSM 14484 | Aquificota | Aquificia |
| Thermocrinis albus DSM 14484 | Aquificota | Aquificia |
| Thermocrinis minervae | Aquificota | Aquificia |
| Thermocrinis minervae | Aquificota | Aquificia |
| Thermocrinis ruber | Aquificota | Aquificia |

| | | |
|---|---|---|
| Thermocrinis ruber | Aquificota | Aquificia |
| Thermovibrio ammonificans HB-1 | Aquificota | Aquificia |
| Thermovibrio ammonificans HB-1 | Aquificota | Aquificia |
| Candidatus Promineifilum breve | Chloroflexota | Ardenticatenia |
| Candidatus Promineifilum breve | Chloroflexota | Ardenticatenia |
| Capsulimonas corticalis | Armatimonadota | Armatimonadia |
| Capsulimonas corticalis | Armatimonadota | Armatimonadia |
| Atribacter laminatus | Atribacterota | Atribacteria |
| Atribacter laminatus | Atribacterota | Atribacteria |
| Atrimonas thermophila | Atribacterota | Atribacteria |
| Atrimonas thermophila | Atribacterota | Atribacteria |
| Thermatribacter velox | Atribacterota | Atribacteria |
| Thermatribacter velox | Atribacterota | Atribacteria |
| [Bacillus] selenitireducens MLS10 | Bacillota | Bacilli |
| Abiotrophia defectiva | Bacillota | Bacilli |
| Abyssicoccus albus | Bacillota | Bacilli |
| Acetilactobacillus jinshanensis | Bacillota | Bacilli |
| Aciduricibacillus chroicocephali | Bacillota | Bacilli |
| Aeribacillus composti | Bacillota | Bacilli |
| Aerococcus christensenii | Bacillota | Bacilli |
| Aerococcus sanguinicola | Bacillota | Bacilli |
| Aerococcus tenax | Bacillota | Bacilli |
| Aerococcus urinae | Bacillota | Bacilli |
| Aerococcus urinaeequi | Bacillota | Bacilli |
| Aerococcus urinaehominis | Bacillota | Bacilli |
| Aerococcus viridans | Bacillota | Bacilli |
| Agrilactobacillus fermenti | Bacillota | Bacilli |
| Alicyclobacillus acidocaldarius subsp. acidocaldarius DSM 446 | Bacillota | Bacilli |
| Alicyclobacillus acidoterrestris | Bacillota | Bacilli |
| Alicyclobacillus cycloheptanicus | Bacillota | Bacilli |
| Alicyclobacillus dauci | Bacillota | Bacilli |

| | | |
|---|---|---|
| Alicyclobacillus fastidiosus | Bacillota | Bacilli |
| Alicyclobacillus mengziensis | Bacillota | Bacilli |
| Alkalibacterium putridalgicola | Bacillota | Bacilli |
| Alkalicoccus halolimnae | Bacillota | Bacilli |
| Alkalihalophilus pseudofirmus | Bacillota | Bacilli |
| Allobacillus halotolerans | Bacillota | Bacilli |
| Amphibacillus xylanus NBRC 15112 | Bacillota | Bacilli |
| Amylolactobacillus amylophilus DSM 20533 = JCM 1125 | Bacillota | Bacilli |
| Anaerobacillus isosaccharinicus | Bacillota | Bacilli |
| Aneurinibacillus soli | Bacillota | Bacilli |
| Aneurinibacillus thermoaerophilus | Bacillota | Bacilli |
| Aneurinibacillus uraniidurans | Bacillota | Bacilli |
| Anoxybacillus flavithermus WK1 | Bacillota | Bacilli |
| Anoxybacillus gonensis | Bacillota | Bacilli |
| Anoxybacteroides amylolyticum | Bacillota | Bacilli |
| Apilactobacillus apinorum | Bacillota | Bacilli |
| Apilactobacillus apisilvae | Bacillota | Bacilli |
| Apilactobacillus bombintestini | Bacillota | Bacilli |
| Apilactobacillus kunkeei | Bacillota | Bacilli |
| Bacillus albus | Bacillota | Bacilli |
| Bacillus altitudinis | Bacillota | Bacilli |
| Bacillus amyloliquefaciens | Bacillota | Bacilli |
| Bacillus anthracis str. 'Ames Ancestor' | Bacillota | Bacilli |
| Bacillus aquiflavi | Bacillota | Bacilli |
| Bacillus arachidis | Bacillota | Bacilli |
| Bacillus atrophaeus | Bacillota | Bacilli |
| Bacillus cabrialesii | Bacillota | Bacilli |
| Bacillus cytotoxicus NVH 391-98 | Bacillota | Bacilli |
| Bacillus fonticola | Bacillota | Bacilli |
| Bacillus glycinifermentans | Bacillota | Bacilli |
| Bacillus gobiensis | Bacillota | Bacilli |

| | | |
|---|---|---|
| Bacillus halotolerans | Bacillota | Bacilli |
| Bacillus haynesii | Bacillota | Bacilli |
| Bacillus inaquosorum | Bacillota | Bacilli |
| Bacillus infantis | Bacillota | Bacilli |
| Bacillus kandeliae | Bacillota | Bacilli |
| Bacillus licheniformis | Bacillota | Bacilli |
| Bacillus luti | Bacillota | Bacilli |
| Bacillus maqinnsis | Bacillota | Bacilli |
| Bacillus mojavensis | Bacillota | Bacilli |
| Bacillus mycoides | Bacillota | Bacilli |
| Bacillus pacificus | Bacillota | Bacilli |
| Bacillus paralicheniformis | Bacillota | Bacilli |
| Bacillus paranthracis | Bacillota | Bacilli |
| Bacillus pseudomycoides DSM 12442 | Bacillota | Bacilli |
| Bacillus pumilus SAFR-032 | Bacillota | Bacilli |
| Bacillus rugosus | Bacillota | Bacilli |
| Bacillus safensis | Bacillota | Bacilli |
| Bacillus shivajii | Bacillota | Bacilli |
| Bacillus smithii | Bacillota | Bacilli |
| Bacillus sonorensis | Bacillota | Bacilli |
| Bacillus spizizenii TU-B-10 | Bacillota | Bacilli |
| Bacillus stercoris | Bacillota | Bacilli |
| Bacillus subtilis subsp. subtilis str. 168 | Bacillota | Bacilli |
| Bacillus swezeyi | Bacillota | Bacilli |
| Bacillus tequilensis | Bacillota | Bacilli |
| Bacillus thuringiensis serovar berliner ATCC 10792 | Bacillota | Bacilli |
| Bacillus toyonensis | Bacillota | Bacilli |
| Bacillus tropicus | Bacillota | Bacilli |
| Bacillus vallismortis | Bacillota | Bacilli |
| Bacillus velezensis FZB42 | Bacillota | Bacilli |
| Bacillus weihaiensis | Bacillota | Bacilli |

| | | |
|---|---|---|
| Bacillus wiedmannii | Bacillota | Bacilli |
| Bacillus xiamenensis | Bacillota | Bacilli |
| Bombilactobacillus bombi | Bacillota | Bacilli |
| Bombilactobacillus folatiphilus | Bacillota | Bacilli |
| Bombilactobacillus thymidiniphilus | Bacillota | Bacilli |
| Brevibacillus agri | Bacillota | Bacilli |
| Brevibacillus aydinogluensis | Bacillota | Bacilli |
| Brevibacillus borstelensis | Bacillota | Bacilli |
| Brevibacillus brevis | Bacillota | Bacilli |
| Brevibacillus centrosporus | Bacillota | Bacilli |
| Brevibacillus composti | Bacillota | Bacilli |
| Brevibacillus humidisoli | Bacillota | Bacilli |
| Brevibacillus laterosporus DSM 25 | Bacillota | Bacilli |
| Brevibacillus marinus | Bacillota | Bacilli |
| Brevibacillus parabrevis | Bacillota | Bacilli |
| Brevibacillus ruminantium | Bacillota | Bacilli |
| Brochothrix campestris | Bacillota | Bacilli |
| Brochothrix thermosphacta | Bacillota | Bacilli |
| Caldalkalibacillus thermarum TA2.A1 | Bacillota | Bacilli |
| Caldifermentibacillus hisashii | Bacillota | Bacilli |
| Candidatus Enterococcus clewellii | Bacillota | Bacilli |
| Candidatus Enterococcus clewellii | Bacillota | Bacilli |
| Candidatus Enterococcus dunnyi | Bacillota | Bacilli |
| Candidatus Enterococcus dunnyi | Bacillota | Bacilli |
| Candidatus Enterococcus lemimoniae | Bacillota | Bacilli |
| Candidatus Enterococcus lemimoniae | Bacillota | Bacilli |
| Candidatus Enterococcus lowellii | Bacillota | Bacilli |
| Candidatus Enterococcus lowellii | Bacillota | Bacilli |
| Candidatus Enterococcus mangumiae | Bacillota | Bacilli |
| Candidatus Enterococcus mangumiae | Bacillota | Bacilli |
| Candidatus Enterococcus palustris | Bacillota | Bacilli |

| | | |
|---|---|---|
| Candidatus Enterococcus palustris | Bacillota | Bacilli |
| Carnobacterium inhibens subsp. gilichinskyi | Bacillota | Bacilli |
| Carnobacterium maltaromaticum | Bacillota | Bacilli |
| Carnobacterium maltaromaticum | Bacillota | Bacilli |
| Carnobacterium maltaromaticum | Bacillota | Bacilli |
| Chryseomicrobium aureum | Bacillota | Bacilli |
| Cohnella abietis | Bacillota | Bacilli |
| Cohnella candidum | Bacillota | Bacilli |
| Cohnella cholangitidis | Bacillota | Bacilli |
| Cohnella herbarum | Bacillota | Bacilli |
| Companilactobacillus alimentarius DSM 20249 | Bacillota | Bacilli |
| Companilactobacillus allii | Bacillota | Bacilli |
| Companilactobacillus crustorum | Bacillota | Bacilli |
| Companilactobacillus farciminis KCTC 3681 = DSM 20184 | Bacillota | Bacilli |
| Companilactobacillus futsaii | Bacillota | Bacilli |
| Companilactobacillus ginsenosidimutans | Bacillota | Bacilli |
| Companilactobacillus heilongjiangensis | Bacillota | Bacilli |
| Companilactobacillus musae | Bacillota | Bacilli |
| Companilactobacillus pabuli | Bacillota | Bacilli |
| Companilactobacillus paralimentarius | Bacillota | Bacilli |
| Companilactobacillus zhachilii | Bacillota | Bacilli |
| Cytobacillus dafuensis | Bacillota | Bacilli |
| Cytobacillus gottheilii | Bacillota | Bacilli |
| Cytobacillus kochii | Bacillota | Bacilli |
| Cytobacillus pseudoceanisediminis | Bacillota | Bacilli |
| Cytobacillus solani | Bacillota | Bacilli |
| Cytobacillus spongiae | Bacillota | Bacilli |
| Dolosigranulum savutiense | Bacillota | Bacilli |
| Effusibacillus dendaii | Bacillota | Bacilli |
| Enterococcus casseliflavus EC20 | Bacillota | Bacilli |
| Enterococcus casseliflavus EC20 | Bacillota | Bacilli |

| | | |
|---|---|---|
| Enterococcus cecorum | Bacillota | Bacilli |
| Enterococcus durans | Bacillota | Bacilli |
| Enterococcus durans | Bacillota | Bacilli |
| Enterococcus faecium | Bacillota | Bacilli |
| Enterococcus faecium | Bacillota | Bacilli |
| Enterococcus gallinarum | Bacillota | Bacilli |
| Enterococcus gallinarum | Bacillota | Bacilli |
| Enterococcus gilvus | Bacillota | Bacilli |
| Enterococcus gilvus | Bacillota | Bacilli |
| Enterococcus hirae | Bacillota | Bacilli |
| Enterococcus hirae | Bacillota | Bacilli |
| Enterococcus innesii | Bacillota | Bacilli |
| Enterococcus innesii | Bacillota | Bacilli |
| Enterococcus lactis | Bacillota | Bacilli |
| Enterococcus lactis | Bacillota | Bacilli |
| Enterococcus montenegrensis | Bacillota | Bacilli |
| Enterococcus montenegrensis | Bacillota | Bacilli |
| Enterococcus mundtii | Bacillota | Bacilli |
| Enterococcus mundtii | Bacillota | Bacilli |
| Enterococcus raffinosus | Bacillota | Bacilli |
| Enterococcus raffinosus | Bacillota | Bacilli |
| Enterococcus rotai | Bacillota | Bacilli |
| Enterococcus rotai | Bacillota | Bacilli |
| Enterococcus saccharolyticus subsp. saccharolyticus | Bacillota | Bacilli |
| Enterococcus saigonensis | Bacillota | Bacilli |
| Enterococcus saigonensis | Bacillota | Bacilli |
| Enterococcus thailandicus | Bacillota | Bacilli |
| Enterococcus thailandicus | Bacillota | Bacilli |
| Enterococcus wangshanyuanii | Bacillota | Bacilli |
| Enterococcus wangshanyuanii | Bacillota | Bacilli |
| Evansella cellulosilytica DSM 2522 | Bacillota | Bacilli |

| | | |
|---|---|---|
| Exiguobacterium acetylicum | Bacillota | Bacilli |
| Exiguobacterium alkaliphilum | Bacillota | Bacilli |
| Exiguobacterium sibiricum 255-15 | Bacillota | Bacilli |
| Fervidibacillus albus | Bacillota | Bacilli |
| Fervidibacillus halotolerans | Bacillota | Bacilli |
| Fodinisporobacter ferrooxydans | Bacillota | Bacilli |
| Fructilactobacillus carniphilus | Bacillota | Bacilli |
| Fructilactobacillus cliffordii | Bacillota | Bacilli |
| Fructilactobacillus hinvesii | Bacillota | Bacilli |
| Fructilactobacillus ixorae | Bacillota | Bacilli |
| Fructilactobacillus lindneri | Bacillota | Bacilli |
| Fructilactobacillus myrtifloralis | Bacillota | Bacilli |
| Fructilactobacillus sanfranciscensis | Bacillota | Bacilli |
| Fructilactobacillus vespulae | Bacillota | Bacilli |
| Fructobacillus americanaquae | Bacillota | Bacilli |
| Fundicoccus culcitae | Bacillota | Bacilli |
| Furfurilactobacillus rossiae | Bacillota | Bacilli |
| Gemella morbillorum | Bacillota | Bacilli |
| Gemella sanguinis | Bacillota | Bacilli |
| Geobacillus kaustophilus HTA426 | Bacillota | Bacilli |
| Geobacillus subterraneus | Bacillota | Bacilli |
| Geobacillus thermodenitrificans | Bacillota | Bacilli |
| Geobacillus thermoleovorans | Bacillota | Bacilli |
| Geobacillus zalihae | Bacillota | Bacilli |
| Globicatella sanguinis | Bacillota | Bacilli |
| Gracilibacillus caseinilyticus | Bacillota | Bacilli |
| Gracilibacillus salinarum | Bacillota | Bacilli |
| Gracilibacillus salitolerans | Bacillota | Bacilli |
| Granulicatella adiacens | Bacillota | Bacilli |
| Halalkalibacterium halodurans | Bacillota | Bacilli |
| Halobacillus amylolyticus | Bacillota | Bacilli |

| | | |
|---|---|---|
| Halobacillus halophilus DSM 2266 | Bacillota | Bacilli |
| Halobacillus mangrovi | Bacillota | Bacilli |
| Halobacillus naozhouensis | Bacillota | Bacilli |
| Halobacillus salinarum | Bacillota | Bacilli |
| Halobacillus shinanisalinarum | Bacillota | Bacilli |
| Heyndrickxia coagulans DSM 1 = ATCC 7050 | Bacillota | Bacilli |
| Heyndrickxia faecalis | Bacillota | Bacilli |
| Heyndrickxia vini | Bacillota | Bacilli |
| Ignavigranum ruoffiae | Bacillota | Bacilli |
| Jeotgalibaca arthritidis | Bacillota | Bacilli |
| Jeotgalibaca ciconiae | Bacillota | Bacilli |
| Jeotgalibaca dankookensis | Bacillota | Bacilli |
| Jeotgalibaca porci | Bacillota | Bacilli |
| Jeotgalicoccus saudimassiliensis | Bacillota | Bacilli |
| Kroppenstedtia eburnea | Bacillota | Bacilli |
| Kroppenstedtia pulmonis | Bacillota | Bacilli |
| Kyrpidia spormannii | Bacillota | Bacilli |
| Kyrpidia tusciae DSM 2912 | Bacillota | Bacilli |
| Laceyella sacchari | Bacillota | Bacilli |
| Lacticaseibacillus casei DSM 20011 = JCM 1134 = ATCC 393 | Bacillota | Bacilli |
| Lacticaseibacillus huelsenbergensis | Bacillota | Bacilli |
| Lacticaseibacillus manihotivorans | Bacillota | Bacilli |
| Lacticaseibacillus pabuli | Bacillota | Bacilli |
| Lacticaseibacillus paracasei subsp. paracasei 8700:2 | Bacillota | Bacilli |
| Lacticaseibacillus parahuelsenbergensis | Bacillota | Bacilli |
| Lacticaseibacillus rhamnosus | Bacillota | Bacilli |
| Lacticaseibacillus styriensis | Bacillota | Bacilli |
| Lacticaseibacillus zeae subsp. silagei | Bacillota | Bacilli |
| Lactiplantibacillus argentoratensis | Bacillota | Bacilli |
| Lactiplantibacillus brownii | Bacillota | Bacilli |
| Lactiplantibacillus carotarum | Bacillota | Bacilli |

| | | |
|---|---|---|
| Lactiplantibacillus paraplantarum | Bacillota | Bacilli |
| Lactiplantibacillus pentosus | Bacillota | Bacilli |
| Lactiplantibacillus plantarum | Bacillota | Bacilli |
| Lactiplantibacillus xiangfangensis | Bacillota | Bacilli |
| Lactobacillus acetotolerans | Bacillota | Bacilli |
| Lactobacillus acidophilus | Bacillota | Bacilli |
| Lactobacillus amylolyticus | Bacillota | Bacilli |
| Lactobacillus amylovorus GRL1118 | Bacillota | Bacilli |
| Lactobacillus apis | Bacillota | Bacilli |
| Lactobacillus crispatus | Bacillota | Bacilli |
| Lactobacillus delbrueckii subsp. jakobsenii ZN7a-9 = DSM 26046 | Bacillota | Bacilli |
| Lactobacillus gasseri ATCC 33323 = JCM 1131 | Bacillota | Bacilli |
| Lactobacillus helsingborgensis | Bacillota | Bacilli |
| Lactobacillus helveticus | Bacillota | Bacilli |
| Lactobacillus iners | Bacillota | Bacilli |
| Lactobacillus intestinalis | Bacillota | Bacilli |
| Lactobacillus isalae | Bacillota | Bacilli |
| Lactobacillus jensenii | Bacillota | Bacilli |
| Lactobacillus johnsonii | Bacillota | Bacilli |
| Lactobacillus kefiranofaciens | Bacillota | Bacilli |
| Lactobacillus kullabergensis | Bacillota | Bacilli |
| Lactobacillus mulieris | Bacillota | Bacilli |
| Lactobacillus panisapium | Bacillota | Bacilli |
| Lactobacillus paragasseri | Bacillota | Bacilli |
| Lactobacillus taiwanensis | Bacillota | Bacilli |
| Lactobacillus terrae | Bacillota | Bacilli |
| Lactobacillus ultunensis | Bacillota | Bacilli |
| Lactobacillus xylocopicola | Bacillota | Bacilli |
| Lactococcus allomyrinae | Bacillota | Bacilli |
| Lactococcus carnosus | Bacillota | Bacilli |
| Lactococcus cremoris subsp. cremoris KW2 | Bacillota | Bacilli |

| | | |
|---|---|---|
| Lactococcus formosensis | Bacillota | Bacilli |
| Lactococcus garvieae | Bacillota | Bacilli |
| Lactococcus lactis subsp. lactis | Bacillota | Bacilli |
| Lactococcus paracarnosus | Bacillota | Bacilli |
| Lactococcus petauri | Bacillota | Bacilli |
| Lactococcus protaetiae | Bacillota | Bacilli |
| Lactococcus raffinolactis | Bacillota | Bacilli |
| Lactococcus taiwanensis | Bacillota | Bacilli |
| Lapidilactobacillus dextrinicus | Bacillota | Bacilli |
| Latilactobacillus curvatus JCM 1096 = DSM 20019 | Bacillota | Bacilli |
| Latilactobacillus fuchuensis | Bacillota | Bacilli |
| Latilactobacillus graminis | Bacillota | Bacilli |
| Latilactobacillus sakei | Bacillota | Bacilli |
| Lederbergia lenta | Bacillota | Bacilli |
| Lentibacillus amyloliquefaciens | Bacillota | Bacilli |
| Lentibacillus daqui | Bacillota | Bacilli |
| Lentilactobacillus buchneri | Bacillota | Bacilli |
| Lentilactobacillus curieae | Bacillota | Bacilli |
| Lentilactobacillus hilgardii | Bacillota | Bacilli |
| Lentilactobacillus kefiri | Bacillota | Bacilli |
| Lentilactobacillus laojiaonis | Bacillota | Bacilli |
| Lentilactobacillus parabuchneri | Bacillota | Bacilli |
| Lentilactobacillus terminaliae | Bacillota | Bacilli |
| Leuconostoc carnosum | Bacillota | Bacilli |
| Leuconostoc citreum | Bacillota | Bacilli |
| Leuconostoc falkenbergense | Bacillota | Bacilli |
| Leuconostoc gasicomitatum LMG 18811 | Bacillota | Bacilli |
| Leuconostoc gelidum JB7 | Bacillota | Bacilli |
| Leuconostoc kimchii IMSNU 11154 | Bacillota | Bacilli |
| Leuconostoc lactis | Bacillota | Bacilli |
| Leuconostoc mesenteroides subsp. mesenteroides ATCC 8293 | Bacillota | Bacilli |

| | | |
|---|---|---|
| Leuconostoc pseudomesenteroides | Bacillota | Bacilli |
| Leuconostoc suionicum | Bacillota | Bacilli |
| Levilactobacillus brevis | Bacillota | Bacilli |
| Levilactobacillus cerevisiae | Bacillota | Bacilli |
| Levilactobacillus koreensis | Bacillota | Bacilli |
| Levilactobacillus parabrevis | Bacillota | Bacilli |
| Levilactobacillus suantsaii | Bacillota | Bacilli |
| Levilactobacillus zymae | Bacillota | Bacilli |
| Ligilactobacillus acidipiscis | Bacillota | Bacilli |
| Ligilactobacillus agilis | Bacillota | Bacilli |
| Ligilactobacillus animalis | Bacillota | Bacilli |
| Ligilactobacillus cholophilus | Bacillota | Bacilli |
| Ligilactobacillus faecis | Bacillota | Bacilli |
| Ligilactobacillus murinus | Bacillota | Bacilli |
| Ligilactobacillus ruminis ATCC 27782 | Bacillota | Bacilli |
| Ligilactobacillus saerimneri | Bacillota | Bacilli |
| Ligilactobacillus salivarius | Bacillota | Bacilli |
| Limosilactobacillus fermentum | Bacillota | Bacilli |
| Limosilactobacillus frumenti | Bacillota | Bacilli |
| Limosilactobacillus mucosae | Bacillota | Bacilli |
| Limosilactobacillus oris | Bacillota | Bacilli |
| Limosilactobacillus pontis | Bacillota | Bacilli |
| Limosilactobacillus portuensis | Bacillota | Bacilli |
| Limosilactobacillus vaginalis | Bacillota | Bacilli |
| Liquorilactobacillus hordei DSM 19519 | Bacillota | Bacilli |
| Liquorilactobacillus mali | Bacillota | Bacilli |
| Liquorilactobacillus nagelii DSM 13675 | Bacillota | Bacilli |
| Listeria grayi | Bacillota | Bacilli |
| Listeria innocua | Bacillota | Bacilli |
| Listeria ivanovii subsp. ivanovii | Bacillota | Bacilli |
| Listeria monocytogenes EGD-e | Bacillota | Bacilli |

| | | |
|---|---|---|
| Listeria newyorkensis | Bacillota | Bacilli |
| Listeria seeligeri serovar 1/2b str. SLCC3954 | Bacillota | Bacilli |
| Listeria swaminathanii | Bacillota | Bacilli |
| Listeria weihenstephanensis | Bacillota | Bacilli |
| Listeria welshimeri | Bacillota | Bacilli |
| Loigolactobacillus backii | Bacillota | Bacilli |
| Loigolactobacillus bifermentans | Bacillota | Bacilli |
| Loigolactobacillus coryniformis subsp. coryniformis KCTC 3167 = DSM 20001 | Bacillota | Bacilli |
| Lysinibacillus agricola | Bacillota | Bacilli |
| Lysinibacillus capsici | Bacillota | Bacilli |
| Lysinibacillus fusiformis | Bacillota | Bacilli |
| Lysinibacillus irui | Bacillota | Bacilli |
| Lysinibacillus louembei | Bacillota | Bacilli |
| Lysinibacillus macroides | Bacillota | Bacilli |
| Lysinibacillus pakistanensis | Bacillota | Bacilli |
| Lysinibacillus parviboronicapiens | Bacillota | Bacilli |
| Lysinibacillus sphaericus | Bacillota | Bacilli |
| Lysinibacillus telephonicus | Bacillota | Bacilli |
| Lysinibacillus timonensis | Bacillota | Bacilli |
| Lysinibacillus varians | Bacillota | Bacilli |
| Macrococcoides bohemicum | Bacillota | Bacilli |
| Macrococcoides canis | Bacillota | Bacilli |
| Macrococcoides caseolyticum | Bacillota | Bacilli |
| Macrococcus armenti | Bacillota | Bacilli |
| Macrococcus bovicus | Bacillota | Bacilli |
| Macrococcus brunensis | Bacillota | Bacilli |
| Macrococcus epidermidis | Bacillota | Bacilli |
| Macrococcus equipercicus | Bacillota | Bacilli |
| Macrococcus psychrotolerans | Bacillota | Bacilli |
| Mammaliicoccus lentus | Bacillota | Bacilli |
| Mammaliicoccus sciuri | Bacillota | Bacilli |

| | | |
|---|---|---|
| Mammaliicoccus stepanovicii | Bacillota | Bacilli |
| Mammaliicoccus vitulinus | Bacillota | Bacilli |
| Mangrovibacillus cuniculi | Bacillota | Bacilli |
| Melissococcus plutonius | Bacillota | Bacilli |
| Mesobacillus jeotgali | Bacillota | Bacilli |
| Mesobacillus maritimus | Bacillota | Bacilli |
| Metabacillus dongyingensis | Bacillota | Bacilli |
| Metabacillus elymi | Bacillota | Bacilli |
| Metabacillus hrfriensis | Bacillota | Bacilli |
| Metabacillus indicus | Bacillota | Bacilli |
| Metabacillus rhizosphaerae | Bacillota | Bacilli |
| Metabacillus sediminilitoris | Bacillota | Bacilli |
| Metabacillus sediminis | Bacillota | Bacilli |
| Neobacillus niacini | Bacillota | Bacilli |
| Niallia circulans | Bacillota | Bacilli |
| Niallia oryzisoli | Bacillota | Bacilli |
| Nicoliella spurrieriana | Bacillota | Bacilli |
| Nosocomiicoccus ampullae | Bacillota | Bacilli |
| Novibacillus thermophilus | Bacillota | Bacilli |
| Oceanobacillus iheyensis HTE831 | Bacillota | Bacilli |
| Oceanobacillus kapialis | Bacillota | Bacilli |
| Oceanobacillus kimchii X50 | Bacillota | Bacilli |
| Oceanobacillus zhaokaii | Bacillota | Bacilli |
| Oenococcus kitaharae DSM 17330 | Bacillota | Bacilli |
| Oenococcus oeni | Bacillota | Bacilli |
| Oenococcus sicerae | Bacillota | Bacilli |
| Paenalkalicoccus suaedae | Bacillota | Bacilli |
| Paenibacillus agaridevorans | Bacillota | Bacilli |
| Paenibacillus albicereus | Bacillota | Bacilli |
| Paenibacillus albus | Bacillota | Bacilli |
| Paenibacillus algicola | Bacillota | Bacilli |

| | | |
|---|---|---|
| Paenibacillus amylolyticus | Bacillota | Bacilli |
| Paenibacillus andongensis | Bacillota | Bacilli |
| Paenibacillus antarcticus | Bacillota | Bacilli |
| Paenibacillus aurantius | Bacillota | Bacilli |
| Paenibacillus azoreducens | Bacillota | Bacilli |
| Paenibacillus baekrokdamisoli | Bacillota | Bacilli |
| Paenibacillus barcinonensis | Bacillota | Bacilli |
| Paenibacillus beijingensis | Bacillota | Bacilli |
| Paenibacillus borealis | Bacillota | Bacilli |
| Paenibacillus bovis | Bacillota | Bacilli |
| Paenibacillus brasilensis | Bacillota | Bacilli |
| Paenibacillus cellulosilyticus | Bacillota | Bacilli |
| Paenibacillus cellulositrophicus | Bacillota | Bacilli |
| Paenibacillus chitinolyticus | Bacillota | Bacilli |
| Paenibacillus crassostreae | Bacillota | Bacilli |
| Paenibacillus dendritiformis | Bacillota | Bacilli |
| Paenibacillus donghaensis | Bacillota | Bacilli |
| Paenibacillus durus | Bacillota | Bacilli |
| Paenibacillus elgii | Bacillota | Bacilli |
| Paenibacillus glucanolyticus | Bacillota | Bacilli |
| Paenibacillus graminis | Bacillota | Bacilli |
| Paenibacillus guangzhouensis | Bacillota | Bacilli |
| Paenibacillus hexagrammi | Bacillota | Bacilli |
| Paenibacillus hunanensis | Bacillota | Bacilli |
| Paenibacillus ihbetae | Bacillota | Bacilli |
| Paenibacillus jilunlii | Bacillota | Bacilli |
| Paenibacillus kribbensis | Bacillota | Bacilli |
| Paenibacillus kyungheensis | Bacillota | Bacilli |
| Paenibacillus larvae subsp. larvae | Bacillota | Bacilli |
| Paenibacillus lentus | Bacillota | Bacilli |
| Paenibacillus lutimineralis | Bacillota | Bacilli |

| | | |
|---|---|---|
| Paenibacillus lycopersici | Bacillota | Bacilli |
| Paenibacillus marchantiae | Bacillota | Bacilli |
| Paenibacillus mucilaginosus K02 | Bacillota | Bacilli |
| Paenibacillus naphthalenovorans | Bacillota | Bacilli |
| Paenibacillus nuruki | Bacillota | Bacilli |
| Paenibacillus odorifer | Bacillota | Bacilli |
| Paenibacillus pedocola | Bacillota | Bacilli |
| Paenibacillus physcomitrellae | Bacillota | Bacilli |
| Paenibacillus polygoni | Bacillota | Bacilli |
| Paenibacillus polymyxa ATCC 842 | Bacillota | Bacilli |
| Paenibacillus protaetiae | Bacillota | Bacilli |
| Paenibacillus psychroresistens | Bacillota | Bacilli |
| Paenibacillus rhizovicinus | Bacillota | Bacilli |
| Paenibacillus riograndensis SBR5 | Bacillota | Bacilli |
| Paenibacillus roseopurpureus | Bacillota | Bacilli |
| Paenibacillus sabinae T27 | Bacillota | Bacilli |
| Paenibacillus silvisoli | Bacillota | Bacilli |
| Paenibacillus sophorae | Bacillota | Bacilli |
| Paenibacillus spongiae | Bacillota | Bacilli |
| Paenibacillus stellifer | Bacillota | Bacilli |
| Paenibacillus swuensis | Bacillota | Bacilli |
| Paenibacillus terrae HPL-003 | Bacillota | Bacilli |
| Paenibacillus thiaminolyticus | Bacillota | Bacilli |
| Paenibacillus tianjinensis | Bacillota | Bacilli |
| Paenibacillus tritici | Bacillota | Bacilli |
| Paenibacillus tundrae | Bacillota | Bacilli |
| Paenibacillus uliginis N3/975 | Bacillota | Bacilli |
| Paenibacillus urinalis | Bacillota | Bacilli |
| Paenibacillus woosongensis | Bacillota | Bacilli |
| Paenibacillus xylanexedens | Bacillota | Bacilli |
| Paenibacillus yonginensis | Bacillota | Bacilli |

| | | |
|---|---|---|
| Paenisporosarcina antarctica | Bacillota | Bacilli |
| Paenisporosarcina cavernae | Bacillota | Bacilli |
| Parageobacillus thermoglucosidasius | Bacillota | Bacilli |
| Parageobacillus toebii | Bacillota | Bacilli |
| Paraliobacillus zengyii | Bacillota | Bacilli |
| Paucilactobacillus hokkaidonensis JCM 18461 | Bacillota | Bacilli |
| Paucilactobacillus nenjiangensis | Bacillota | Bacilli |
| Paucilactobacillus oligofermentans DSM 15707 = LMG 22743 | Bacillota | Bacilli |
| Paucilactobacillus suebicus | Bacillota | Bacilli |
| Pediococcus acidilactici | Bacillota | Bacilli |
| Pediococcus claussenii | Bacillota | Bacilli |
| Pediococcus damnosus | Bacillota | Bacilli |
| Pediococcus inopinatus | Bacillota | Bacilli |
| Pediococcus parvulus | Bacillota | Bacilli |
| Pediococcus pentosaceus ATCC 25745 | Bacillota | Bacilli |
| Pediococcus siamensis | Bacillota | Bacilli |
| Peribacillus frigoritolerans | Bacillota | Bacilli |
| Peribacillus muralis | Bacillota | Bacilli |
| Peribacillus psychrosaccharolyticus | Bacillota | Bacilli |
| Peribacillus simplex NBRC 15720 = DSM 1321 | Bacillota | Bacilli |
| Periweissella cryptocerci | Bacillota | Bacilli |
| Planococcus antarcticus DSM 14505 | Bacillota | Bacilli |
| Planococcus donghaensis | Bacillota | Bacilli |
| Planococcus faecalis | Bacillota | Bacilli |
| Planococcus glaciei | Bacillota | Bacilli |
| Planococcus halocryophilus | Bacillota | Bacilli |
| Planococcus kocurii | Bacillota | Bacilli |
| Planococcus lenghuensis | Bacillota | Bacilli |
| Planococcus liqunii | Bacillota | Bacilli |
| Planococcus maritimus | Bacillota | Bacilli |
| Planococcus plakortidis | Bacillota | Bacilli |

| | | |
|---|---|---|
| Planococcus rifietoensis | Bacillota | Bacilli |
| Planococcus versutus | Bacillota | Bacilli |
| Polycladomyces abyssicola | Bacillota | Bacilli |
| Priestia aryabhattai | Bacillota | Bacilli |
| Priestia flexa | Bacillota | Bacilli |
| Priestia megaterium NBRC 15308 = ATCC 14581 | Bacillota | Bacilli |
| Psychrobacillus glaciei | Bacillota | Bacilli |
| Radiobacillus deserti | Bacillota | Bacilli |
| Radiobacillus kanasensis | Bacillota | Bacilli |
| Rossellomorea marisflavi | Bacillota | Bacilli |
| Saccharibacillus brassicae | Bacillota | Bacilli |
| Saccharococcus caldoxylosilyticus | Bacillota | Bacilli |
| Salicibibacter cibarius | Bacillota | Bacilli |
| Salicibibacter cibi | Bacillota | Bacilli |
| Salicibibacter halophilus | Bacillota | Bacilli |
| Salicibibacter kimchii | Bacillota | Bacilli |
| Salimicrobium jeotgali | Bacillota | Bacilli |
| Salinicoccus bachuensis | Bacillota | Bacilli |
| Salisediminibacterium beveridgei | Bacillota | Bacilli |
| Schleiferilactobacillus harbinensis | Bacillota | Bacilli |
| Secundilactobacillus malefermentans | Bacillota | Bacilli |
| Secundilactobacillus paracollinoides | Bacillota | Bacilli |
| Sediminibacillus dalangtanensis | Bacillota | Bacilli |
| Shouchella clausii | Bacillota | Bacilli |
| Shouchella hunanensis | Bacillota | Bacilli |
| Shouchella miscanthi | Bacillota | Bacilli |
| Shouchella rhizosphaerae | Bacillota | Bacilli |
| Solibacillus daqui | Bacillota | Bacilli |
| Sporolactobacillus terrae | Bacillota | Bacilli |
| Sporosarcina jeotgali | Bacillota | Bacilli |
| Sporosarcina oncorhynchi | Bacillota | Bacilli |

| | | |
|---|---|---|
| Sporosarcina pasteurii | Bacillota | Bacilli |
| Sporosarcina psychrophila | Bacillota | Bacilli |
| Sporosarcina trichiuri | Bacillota | Bacilli |
| Sporosarcina ureae | Bacillota | Bacilli |
| Sporosarcina ureilytica | Bacillota | Bacilli |
| Staphylococcus agnetis | Bacillota | Bacilli |
| Staphylococcus argenteus | Bacillota | Bacilli |
| Staphylococcus arlettae | Bacillota | Bacilli |
| Staphylococcus auricularis | Bacillota | Bacilli |
| Staphylococcus capitis subsp. urealyticus | Bacillota | Bacilli |
| Staphylococcus caprae | Bacillota | Bacilli |
| Staphylococcus carnosus | Bacillota | Bacilli |
| Staphylococcus casei | Bacillota | Bacilli |
| Staphylococcus chromogenes | Bacillota | Bacilli |
| Staphylococcus coagulans | Bacillota | Bacilli |
| Staphylococcus condimenti | Bacillota | Bacilli |
| Staphylococcus debuckii | Bacillota | Bacilli |
| Staphylococcus delphini | Bacillota | Bacilli |
| Staphylococcus epidermidis | Bacillota | Bacilli |
| Staphylococcus equorum | Bacillota | Bacilli |
| Staphylococcus felis | Bacillota | Bacilli |
| Staphylococcus gallinarum | Bacillota | Bacilli |
| Staphylococcus haemolyticus | Bacillota | Bacilli |
| Staphylococcus hominis | Bacillota | Bacilli |
| Staphylococcus hsinchuensis | Bacillota | Bacilli |
| Staphylococcus hyicus | Bacillota | Bacilli |
| Staphylococcus kloosii | Bacillota | Bacilli |
| Staphylococcus lloydii | Bacillota | Bacilli |
| Staphylococcus lugdunensis | Bacillota | Bacilli |
| Staphylococcus lutrae | Bacillota | Bacilli |
| Staphylococcus massiliensis | Bacillota | Bacilli |

| | | |
|---|---|---|
| Staphylococcus muscae | Bacillota | Bacilli |
| Staphylococcus nepalensis | Bacillota | Bacilli |
| Staphylococcus pasteuri | Bacillota | Bacilli |
| Staphylococcus pettenkoferi | Bacillota | Bacilli |
| Staphylococcus piscifermentans | Bacillota | Bacilli |
| Staphylococcus pseudintermedius | Bacillota | Bacilli |
| Staphylococcus pseudoxylosus | Bacillota | Bacilli |
| Staphylococcus ratti | Bacillota | Bacilli |
| Staphylococcus rostri | Bacillota | Bacilli |
| Staphylococcus saccharolyticus | Bacillota | Bacilli |
| Staphylococcus saprophyticus subsp. saprophyticus ATCC 15305 = NCTC 729[illegible] | Bacillota | Bacilli |
| Staphylococcus schleiferi | Bacillota | Bacilli |
| Staphylococcus schweitzeri | Bacillota | Bacilli |
| Staphylococcus simiae | Bacillota | Bacilli |
| Staphylococcus simulans | Bacillota | Bacilli |
| Staphylococcus succinus | Bacillota | Bacilli |
| Staphylococcus ureilyticus | Bacillota | Bacilli |
| Staphylococcus warneri | Bacillota | Bacilli |
| Staphylococcus xylosus | Bacillota | Bacilli |
| Staphylospora marina | Bacillota | Bacilli |
| Streptococcus agalactiae | Bacillota | Bacilli |
| Streptococcus anginosus | Bacillota | Bacilli |
| Streptococcus australis | Bacillota | Bacilli |
| Streptococcus canis | Bacillota | Bacilli |
| Streptococcus chenjunshii | Bacillota | Bacilli |
| Streptococcus constellatus | Bacillota | Bacilli |
| Streptococcus cristatus ATCC 51100 | Bacillota | Bacilli |
| Streptococcus dentalis | Bacillota | Bacilli |
| Streptococcus dysgalactiae | Bacillota | Bacilli |
| Streptococcus equi subsp. zooepidemicus | Bacillota | Bacilli |
| Streptococcus ferus | Bacillota | Bacilli |

| | | |
|---|---|---|
| Streptococcus gallolyticus | Bacillota | Bacilli |
| Streptococcus gordonii | Bacillota | Bacilli |
| Streptococcus gwangjuensis | Bacillota | Bacilli |
| Streptococcus halichoeri | Bacillota | Bacilli |
| Streptococcus halotolerans | Bacillota | Bacilli |
| Streptococcus hillyeri | Bacillota | Bacilli |
| Streptococcus himalayensis | Bacillota | Bacilli |
| Streptococcus ilei | Bacillota | Bacilli |
| Streptococcus iners | Bacillota | Bacilli |
| Streptococcus infantarius | Bacillota | Bacilli |
| Streptococcus iniae | Bacillota | Bacilli |
| Streptococcus intermedius | Bacillota | Bacilli |
| Streptococcus koreensis | Bacillota | Bacilli |
| Streptococcus lactarius | Bacillota | Bacilli |
| Streptococcus lingualis | Bacillota | Bacilli |
| Streptococcus lutetiensis | Bacillota | Bacilli |
| Streptococcus macedonicus | Bacillota | Bacilli |
| Streptococcus marmotae | Bacillota | Bacilli |
| Streptococcus merionis | Bacillota | Bacilli |
| Streptococcus mitis | Bacillota | Bacilli |
| Streptococcus mutans | Bacillota | Bacilli |
| Streptococcus oralis ATCC 35037 | Bacillota | Bacilli |
| Streptococcus oriscaviae | Bacillota | Bacilli |
| Streptococcus pantholopis | Bacillota | Bacilli |
| Streptococcus parapneumoniae | Bacillota | Bacilli |
| Streptococcus parasanguinis ATCC 15912 | Bacillota | Bacilli |
| Streptococcus parasuis | Bacillota | Bacilli |
| Streptococcus parauberis NCFD 2020 | Bacillota | Bacilli |
| Streptococcus pasteurianus | Bacillota | Bacilli |
| Streptococcus periodonticum | Bacillota | Bacilli |
| Streptococcus pluranimalium | Bacillota | Bacilli |

| | | |
|---|---|---|
| Streptococcus pneumoniae | Bacillota | Bacilli |
| Streptococcus porcinus | Bacillota | Bacilli |
| Streptococcus pseudopneumoniae IS7493 | Bacillota | Bacilli |
| Streptococcus pseudoporcinus | Bacillota | Bacilli |
| Streptococcus pyogenes | Bacillota | Bacilli |
| Streptococcus ratti | Bacillota | Bacilli |
| Streptococcus respiraculi | Bacillota | Bacilli |
| Streptococcus ruminantium | Bacillota | Bacilli |
| Streptococcus ruminicola | Bacillota | Bacilli |
| Streptococcus salivarius JIM8777 | Bacillota | Bacilli |
| Streptococcus sanguinis | Bacillota | Bacilli |
| Streptococcus sobrinus | Bacillota | Bacilli |
| Streptococcus suis BM407 | Bacillota | Bacilli |
| Streptococcus suivaginalis | Bacillota | Bacilli |
| Streptococcus thermophilus | Bacillota | Bacilli |
| Streptococcus toyakuensis | Bacillota | Bacilli |
| Streptococcus troglodytae | Bacillota | Bacilli |
| Streptococcus uberis | Bacillota | Bacilli |
| Streptococcus urinalis | Bacillota | Bacilli |
| Streptococcus vestibularis | Bacillota | Bacilli |
| Streptococcus zhangguiae | Bacillota | Bacilli |
| Suicoccus acidiformans | Bacillota | Bacilli |
| Sutcliffiella cohnii | Bacillota | Bacilli |
| Sutcliffiella horikoshii | Bacillota | Bacilli |
| Tepidibacillus marianensis | Bacillota | Bacilli |
| Terribacillus saccharophilus | Bacillota | Bacilli |
| Tetragenococcus halophilus | Bacillota | Bacilli |
| Tetragenococcus koreensis | Bacillota | Bacilli |
| Tetragenococcus osmophilus | Bacillota | Bacilli |
| Thermaerobacillus caldiproteolyticus | Bacillota | Bacilli |
| Thermoactinomyces vulgaris | Bacillota | Bacilli |

| | | |
|---|---|---|
| Thermobacillus composti KWC4 | Bacillota | Bacilli |
| Trichococcus flocculiformis | Bacillota | Bacilli |
| Trichococcus shcherbakoviae | Bacillota | Bacilli |
| Tumebacillus algifaecis | Bacillota | Bacilli |
| Tumebacillus avium | Bacillota | Bacilli |
| Ureibacillus chungkukjangi | Bacillota | Bacilli |
| Ureibacillus thermophilus | Bacillota | Bacilli |
| Ureibacillus thermosphaericus | Bacillota | Bacilli |
| Vagococcus carniphilus | Bacillota | Bacilli |
| Vagococcus carniphilus | Bacillota | Bacilli |
| Vagococcus coleopterorum | Bacillota | Bacilli |
| Vagococcus hydrophili | Bacillota | Bacilli |
| Vagococcus intermedius | Bacillota | Bacilli |
| Vagococcus jeotgali | Bacillota | Bacilli |
| Vagococcus luciliae | Bacillota | Bacilli |
| Vagococcus lutrae | Bacillota | Bacilli |
| Vagococcus teuberi | Bacillota | Bacilli |
| Vagococcus xieshaowenii | Bacillota | Bacilli |
| Vagococcus zengguangii | Bacillota | Bacilli |
| Virgibacillus dokdonensis | Bacillota | Bacilli |
| Virgibacillus halodenitrificans | Bacillota | Bacilli |
| Virgibacillus natechei | Bacillota | Bacilli |
| Virgibacillus necropolis | Bacillota | Bacilli |
| Virgibacillus pantothenticus | Bacillota | Bacilli |
| Virgibacillus phasianinus | Bacillota | Bacilli |
| Weissella ceti | Bacillota | Bacilli |
| Weissella cibaria | Bacillota | Bacilli |
| Weissella coleopterorum | Bacillota | Bacilli |
| Weissella confusa | Bacillota | Bacilli |
| Weissella diestrammenae | Bacillota | Bacilli |
| Weissella jogaejeotgali | Bacillota | Bacilli |

| | | |
|---|---|---|
| Weissella koreensis KACC 15510 | Bacillota | Bacilli |
| Weissella paramesenteroides | Bacillota | Bacilli |
| Weissella soli | Bacillota | Bacilli |
| Weissella tructae | Bacillota | Bacilli |
| Weissella viridescens | Bacillota | Bacilli |
| Xylocopilactobacillus apicola | Bacillota | Bacilli |
| Xylocopilactobacillus apis | Bacillota | Bacilli |
| Bacteriovorax stolpii | Bdellovibrionota | Bacteriovoracia |
| Bacteriovorax stolpii | Bdellovibrionota | Bacteriovoracia |
| Halobacteriovorax marinus SJ | Bdellovibrionota | Bacteriovoracia |
| Halobacteriovorax marinus SJ | Bdellovibrionota | Bacteriovoracia |
| Peredibacter starrii | Bdellovibrionota | Bacteriovoracia |
| Peredibacter starrii | Bdellovibrionota | Bacteriovoracia |
| Alistipes communis | Bacteroidota | Bacteroidia |
| Alistipes dispar | Bacteroidota | Bacteroidia |
| Alistipes ihumii AP11 | Bacteroidota | Bacteroidia |
| Alistipes indistinctus YIT 12060 | Bacteroidota | Bacteroidia |
| Alistipes megaguti | Bacteroidota | Bacteroidia |
| Alistipes onderdonkii subsp. vulgaris | Bacteroidota | Bacteroidia |
| Alistipes shahii WAL 8301 | Bacteroidota | Bacteroidia |
| Alkalitalea saponilacus | Bacteroidota | Bacteroidia |
| Aquipluma nitroreducens | Bacteroidota | Bacteroidia |
| Bacteroides caccae CL03T12C61 | Bacteroidota | Bacteroidia |
| Bacteroides caecimuris | Bacteroidota | Bacteroidia |
| Bacteroides cellulosilyticus | Bacteroidota | Bacteroidia |
| Bacteroides difficilis | Bacteroidota | Bacteroidia |
| Bacteroides eggerthii | Bacteroidota | Bacteroidia |
| Bacteroides faecis | Bacteroidota | Bacteroidia |
| Bacteroides faecium | Bacteroidota | Bacteroidia |
| Bacteroides fragilis NCTC 9343 | Bacteroidota | Bacteroidia |
| Bacteroides graminisolvens | Bacteroidota | Bacteroidia |

| | | |
|---|---|---|
| Bacteroides helcogenes P 36-108 | Bacteroidota | Bacteroidia |
| Bacteroides heparinolyticus | Bacteroidota | Bacteroidia |
| Bacteroides hominis | Bacteroidota | Bacteroidia |
| Bacteroides humanifaecis | Bacteroidota | Bacteroidia |
| Bacteroides luhongzhouii | Bacteroidota | Bacteroidia |
| Bacteroides maternus | Bacteroidota | Bacteroidia |
| Bacteroides nordii | Bacteroidota | Bacteroidia |
| Bacteroides ovatus | Bacteroidota | Bacteroidia |
| Bacteroides pyogenes | Bacteroidota | Bacteroidia |
| Bacteroides salyersiae | Bacteroidota | Bacteroidia |
| Bacteroides sedimenti | Bacteroidota | Bacteroidia |
| Bacteroides thetaiotaomicron | Bacteroidota | Bacteroidia |
| Bacteroides uniformis | Bacteroidota | Bacteroidia |
| Bacteroides xylanisolvens | Bacteroidota | Bacteroidia |
| Bacteroides zhangwenhongii | Bacteroidota | Bacteroidia |
| Bacteroides zoogleoformans | Bacteroidota | Bacteroidia |
| Barnesiella viscericola DSM 18177 | Bacteroidota | Bacteroidia |
| Butyricimonas faecalis | Bacteroidota | Bacteroidia |
| Butyricimonas faecihominis | Bacteroidota | Bacteroidia |
| Butyricimonas paravirosa | Bacteroidota | Bacteroidia |
| Butyricimonas virosa | Bacteroidota | Bacteroidia |
| Candidatus Azobacteroides pseudotrichonymphae genomovar. CFP2 | Bacteroidota | Bacteroidia |
| Coprobacter fastidiosus | Bacteroidota | Bacteroidia |
| Coprobacter secundus subsp. similis | Bacteroidota | Bacteroidia |
| Draconibacterium halophilum | Bacteroidota | Bacteroidia |
| Duncaniella dubosii | Bacteroidota | Bacteroidia |
| Dysgonomonas capnocytophagoides | Bacteroidota | Bacteroidia |
| Hoylesella buccalis ATCC 35310 | Bacteroidota | Bacteroidia |
| Hoylesella enoeca | Bacteroidota | Bacteroidia |
| Labilibaculum antarcticum | Bacteroidota | Bacteroidia |
| Limibacterium fermenti | Bacteroidota | Bacteroidia |

| | | |
|---|---|---|
| Maribellus comscasis | Bacteroidota | Bacteroidia |
| Marinifilum fragile | Bacteroidota | Bacteroidia |
| Muribaculum gordoncarteri | Bacteroidota | Bacteroidia |
| Muribaculum intestinale | Bacteroidota | Bacteroidia |
| Odoribacter splanchnicus | Bacteroidota | Bacteroidia |
| Paludibacter propionicigenes WB4 | Bacteroidota | Bacteroidia |
| Parabacteroides absconsus | Bacteroidota | Bacteroidia |
| Parabacteroides chongii | Bacteroidota | Bacteroidia |
| Parabacteroides distasonis | Bacteroidota | Bacteroidia |
| Parabacteroides goldsteinii | Bacteroidota | Bacteroidia |
| Parabacteroides johnsonii DSM 18315 | Bacteroidota | Bacteroidia |
| Paramuribaculum intestinale | Bacteroidota | Bacteroidia |
| Paraprevotella xylaniphila YIT 11841 | Bacteroidota | Bacteroidia |
| Petrimonas mucosa | Bacteroidota | Bacteroidia |
| Phocaeicola coprophilus | Bacteroidota | Bacteroidia |
| Phocaeicola dorei | Bacteroidota | Bacteroidia |
| Phocaeicola salanitronis DSM 18170 | Bacteroidota | Bacteroidia |
| Phocaeicola vulgatus ATCC 8482 | Bacteroidota | Bacteroidia |
| Porphyromonas asaccharolytica DSM 20707 | Bacteroidota | Bacteroidia |
| Porphyromonas cangingivalis | Bacteroidota | Bacteroidia |
| Porphyromonas crevioricanis | Bacteroidota | Bacteroidia |
| Porphyromonas endodontalis | Bacteroidota | Bacteroidia |
| Porphyromonas gingivalis ATCC 33277 | Bacteroidota | Bacteroidia |
| Porphyromonas pogonae | Bacteroidota | Bacteroidia |
| Porphyromonas somerae | Bacteroidota | Bacteroidia |
| Prevotella communis | Bacteroidota | Bacteroidia |
| Prevotella dentalis DSM 3688 | Bacteroidota | Bacteroidia |
| Prevotella denticola | Bacteroidota | Bacteroidia |
| Prevotella fusca JCM 17724 | Bacteroidota | Bacteroidia |
| Prevotella herbatica | Bacteroidota | Bacteroidia |
| Prevotella histicola | Bacteroidota | Bacteroidia |

| | | |
|---|---|---|
| Prevotella jejuni | Bacteroidota | Bacteroidia |
| Prevotella melaninogenica ATCC 25845 | Bacteroidota | Bacteroidia |
| Prevotella multiformis | Bacteroidota | Bacteroidia |
| Prevotella nigrescens | Bacteroidota | Bacteroidia |
| Prevotella scopos JCM 17725 | Bacteroidota | Bacteroidia |
| Prevotella veroralis | Bacteroidota | Bacteroidia |
| Proteiniphilum propionicum | Bacteroidota | Bacteroidia |
| Proteiniphilum saccharofermentans | Bacteroidota | Bacteroidia |
| Pseudoprevotella muciniphila | Bacteroidota | Bacteroidia |
| Salinivirga cyanobacteriivorans | Bacteroidota | Bacteroidia |
| Segatella bryantii | Bacteroidota | Bacteroidia |
| Segatella copri DSM 18205 | Bacteroidota | Bacteroidia |
| Segatella oris | Bacteroidota | Bacteroidia |
| Sodaliphilus pleomorphus | Bacteroidota | Bacteroidia |
| Tannerella forsythia 92A2 | Bacteroidota | Bacteroidia |
| Tannerella serpentiformis | Bacteroidota | Bacteroidia |
| Tenuifilum thalassicum | Bacteroidota | Bacteroidia |
| Xiashengella succiniciproducens | Bacteroidota | Bacteroidia |
| Xylanibacter ruminicola 23 | Bacteroidota | Bacteroidia |
| Cyclonatronum proteinivorum | Balneolota | Balneolia |
| Cyclonatronum proteinivorum | Balneolota | Balneolia |
| Bdellovibrio bacteriovorus HD100 | Bdellovibrionota | Bdellovibrionia |
| Bdellovibrio reynosensis | Bdellovibrionota | Bdellovibrionia |
| Pseudobdellovibrio exovorus JSS | Bdellovibrionota | Bdellovibrionia |
| [Acidovorax] ebreus TPSY | Pseudomonadota | Betaproteobacteria |
| [Acidovorax] ebreus TPSY | Pseudomonadota | Betaproteobacteria |
| Achromobacter denitrificans | Pseudomonadota | Betaproteobacteria |
| Achromobacter denitrificans | Pseudomonadota | Betaproteobacteria |
| Achromobacter insolitus | Pseudomonadota | Betaproteobacteria |
| Achromobacter insolitus | Pseudomonadota | Betaproteobacteria |
| Achromobacter mucicolens | Pseudomonadota | Betaproteobacteria |

| | | |
|---|---|---|
| Achromobacter mucicolens | Pseudomonadota | Betaproteobacteria |
| Achromobacter ruhlandii | Pseudomonadota | Betaproteobacteria |
| Achromobacter ruhlandii | Pseudomonadota | Betaproteobacteria |
| Achromobacter seleniivolatilans | Pseudomonadota | Betaproteobacteria |
| Achromobacter seleniivolatilans | Pseudomonadota | Betaproteobacteria |
| Achromobacter spanius | Pseudomonadota | Betaproteobacteria |
| Achromobacter spanius | Pseudomonadota | Betaproteobacteria |
| Achromobacter veterisilvae | Pseudomonadota | Betaproteobacteria |
| Achromobacter veterisilvae | Pseudomonadota | Betaproteobacteria |
| Achromobacter xylosoxidans | Pseudomonadota | Betaproteobacteria |
| Achromobacter xylosoxidans | Pseudomonadota | Betaproteobacteria |
| Acidovorax carolinensis | Pseudomonadota | Betaproteobacteria |
| Acidovorax carolinensis | Pseudomonadota | Betaproteobacteria |
| Acidovorax sacchari | Pseudomonadota | Betaproteobacteria |
| Acidovorax sacchari | Pseudomonadota | Betaproteobacteria |
| Acidovorax temperans | Pseudomonadota | Betaproteobacteria |
| Acidovorax temperans | Pseudomonadota | Betaproteobacteria |
| Advenella mimigardefordensis DPN7 | Pseudomonadota | Betaproteobacteria |
| Advenella mimigardefordensis DPN7 | Pseudomonadota | Betaproteobacteria |
| Alcaligenes ammonioxydans | Pseudomonadota | Betaproteobacteria |
| Alcaligenes ammonioxydans | Pseudomonadota | Betaproteobacteria |
| Alcaligenes aquatilis | Pseudomonadota | Betaproteobacteria |
| Alcaligenes aquatilis | Pseudomonadota | Betaproteobacteria |
| Alicycliphilus denitrificans K601 | Pseudomonadota | Betaproteobacteria |
| Alicycliphilus denitrificans K601 | Pseudomonadota | Betaproteobacteria |
| Alysiella crassa | Pseudomonadota | Betaproteobacteria |
| Alysiella crassa | Pseudomonadota | Betaproteobacteria |
| Alysiella filiformis DSM 16848 | Pseudomonadota | Betaproteobacteria |
| Alysiella filiformis DSM 16848 | Pseudomonadota | Betaproteobacteria |
| Ampullimonas aquatilis | Pseudomonadota | Betaproteobacteria |
| Ampullimonas aquatilis | Pseudomonadota | Betaproteobacteria |

| | | |
|---|---|---|
| Aquabacterium olei | Pseudomonadota | Betaproteobacteria |
| Aquabacterium olei | Pseudomonadota | Betaproteobacteria |
| Aquella oligotrophica | Pseudomonadota | Betaproteobacteria |
| Aquella oligotrophica | Pseudomonadota | Betaproteobacteria |
| Aquincola tertiaricarbonis | Pseudomonadota | Betaproteobacteria |
| Aquincola tertiaricarbonis | Pseudomonadota | Betaproteobacteria |
| Aquitalea aquatilis | Pseudomonadota | Betaproteobacteria |
| Aquitalea aquatilis | Pseudomonadota | Betaproteobacteria |
| Aquitalea denitrificans | Pseudomonadota | Betaproteobacteria |
| Aquitalea denitrificans | Pseudomonadota | Betaproteobacteria |
| Aromatoleum aromaticum EbN1 | Pseudomonadota | Betaproteobacteria |
| Aromatoleum aromaticum EbN1 | Pseudomonadota | Betaproteobacteria |
| Aromatoleum bremense | Pseudomonadota | Betaproteobacteria |
| Aromatoleum bremense | Pseudomonadota | Betaproteobacteria |
| Aromatoleum evansii | Pseudomonadota | Betaproteobacteria |
| Aromatoleum evansii | Pseudomonadota | Betaproteobacteria |
| Aromatoleum petrolei | Pseudomonadota | Betaproteobacteria |
| Aromatoleum petrolei | Pseudomonadota | Betaproteobacteria |
| Azoarcus olearius | Pseudomonadota | Betaproteobacteria |
| Azoarcus olearius | Pseudomonadota | Betaproteobacteria |
| Azospira inquinata | Pseudomonadota | Betaproteobacteria |
| Azospira inquinata | Pseudomonadota | Betaproteobacteria |
| Azospira oryzae PS | Pseudomonadota | Betaproteobacteria |
| Azospira oryzae PS | Pseudomonadota | Betaproteobacteria |
| Azospira restricta | Pseudomonadota | Betaproteobacteria |
| Azospira restricta | Pseudomonadota | Betaproteobacteria |
| Basilea psittacipulmonis DSM 24701 | Pseudomonadota | Betaproteobacteria |
| Basilea psittacipulmonis DSM 24701 | Pseudomonadota | Betaproteobacteria |
| Bergeriella denitrificans | Pseudomonadota | Betaproteobacteria |
| Bergeriella denitrificans | Pseudomonadota | Betaproteobacteria |
| Bordetella avium | Pseudomonadota | Betaproteobacteria |

| | | |
|---|---|---|
| Bordetella avium | Pseudomonadota | Betaproteobacteria |
| Bordetella bronchialis | Pseudomonadota | Betaproteobacteria |
| Bordetella bronchialis | Pseudomonadota | Betaproteobacteria |
| Bordetella bronchiseptica | Pseudomonadota | Betaproteobacteria |
| Bordetella bronchiseptica | Pseudomonadota | Betaproteobacteria |
| Bordetella flabilis | Pseudomonadota | Betaproteobacteria |
| Bordetella flabilis | Pseudomonadota | Betaproteobacteria |
| Bordetella hinzii | Pseudomonadota | Betaproteobacteria |
| Bordetella hinzii | Pseudomonadota | Betaproteobacteria |
| Bordetella holmesii | Pseudomonadota | Betaproteobacteria |
| Bordetella holmesii | Pseudomonadota | Betaproteobacteria |
| Bordetella parapertussis | Pseudomonadota | Betaproteobacteria |
| Bordetella parapertussis | Pseudomonadota | Betaproteobacteria |
| Bordetella pertussis | Pseudomonadota | Betaproteobacteria |
| Bordetella pertussis | Pseudomonadota | Betaproteobacteria |
| Bordetella petrii | Pseudomonadota | Betaproteobacteria |
| Bordetella petrii | Pseudomonadota | Betaproteobacteria |
| Bordetella pseudohinzii | Pseudomonadota | Betaproteobacteria |
| Bordetella pseudohinzii | Pseudomonadota | Betaproteobacteria |
| Burkholderia aenigmatica | Pseudomonadota | Betaproteobacteria |
| Burkholderia aenigmatica | Pseudomonadota | Betaproteobacteria |
| Burkholderia ambifaria AMMD | Pseudomonadota | Betaproteobacteria |
| Burkholderia ambifaria AMMD | Pseudomonadota | Betaproteobacteria |
| Burkholderia arboris | Pseudomonadota | Betaproteobacteria |
| Burkholderia arboris | Pseudomonadota | Betaproteobacteria |
| Burkholderia cenocepacia | Pseudomonadota | Betaproteobacteria |
| Burkholderia cenocepacia | Pseudomonadota | Betaproteobacteria |
| Burkholderia cepacia | Pseudomonadota | Betaproteobacteria |
| Burkholderia cepacia | Pseudomonadota | Betaproteobacteria |
| Burkholderia contaminans | Pseudomonadota | Betaproteobacteria |
| Burkholderia contaminans | Pseudomonadota | Betaproteobacteria |

| | | |
|---|---|---|
| Burkholderia dolosa | Pseudomonadota | Betaproteobacteria |
| Burkholderia dolosa | Pseudomonadota | Betaproteobacteria |
| Burkholderia gladioli | Pseudomonadota | Betaproteobacteria |
| Burkholderia gladioli | Pseudomonadota | Betaproteobacteria |
| Burkholderia glumae LMG 2196 = ATCC 33617 | Pseudomonadota | Betaproteobacteria |
| Burkholderia glumae LMG 2196 = ATCC 33617 | Pseudomonadota | Betaproteobacteria |
| Burkholderia humptydooensis | Pseudomonadota | Betaproteobacteria |
| Burkholderia humptydooensis | Pseudomonadota | Betaproteobacteria |
| Burkholderia lata | Pseudomonadota | Betaproteobacteria |
| Burkholderia lata | Pseudomonadota | Betaproteobacteria |
| Burkholderia mallei | Pseudomonadota | Betaproteobacteria |
| Burkholderia mallei | Pseudomonadota | Betaproteobacteria |
| Burkholderia mayonis | Pseudomonadota | Betaproteobacteria |
| Burkholderia mayonis | Pseudomonadota | Betaproteobacteria |
| Burkholderia metallica | Pseudomonadota | Betaproteobacteria |
| Burkholderia metallica | Pseudomonadota | Betaproteobacteria |
| Burkholderia multivorans | Pseudomonadota | Betaproteobacteria |
| Burkholderia multivorans | Pseudomonadota | Betaproteobacteria |
| Burkholderia oklahomensis C6786 | Pseudomonadota | Betaproteobacteria |
| Burkholderia oklahomensis C6786 | Pseudomonadota | Betaproteobacteria |
| Burkholderia perseverans | Pseudomonadota | Betaproteobacteria |
| Burkholderia perseverans | Pseudomonadota | Betaproteobacteria |
| Burkholderia plantarii | Pseudomonadota | Betaproteobacteria |
| Burkholderia plantarii | Pseudomonadota | Betaproteobacteria |
| Burkholderia pseudomallei | Pseudomonadota | Betaproteobacteria |
| Burkholderia pseudomallei | Pseudomonadota | Betaproteobacteria |
| Burkholderia pyrrocinia | Pseudomonadota | Betaproteobacteria |
| Burkholderia pyrrocinia | Pseudomonadota | Betaproteobacteria |
| Burkholderia savannae | Pseudomonadota | Betaproteobacteria |
| Burkholderia savannae | Pseudomonadota | Betaproteobacteria |
| Burkholderia seminalis | Pseudomonadota | Betaproteobacteria |

| | | |
|---|---|---|
| Burkholderia seminalis | Pseudomonadota | Betaproteobacteria |
| Burkholderia singularis | Pseudomonadota | Betaproteobacteria |
| Burkholderia singularis | Pseudomonadota | Betaproteobacteria |
| Burkholderia stabilis | Pseudomonadota | Betaproteobacteria |
| Burkholderia stabilis | Pseudomonadota | Betaproteobacteria |
| Burkholderia stagnalis | Pseudomonadota | Betaproteobacteria |
| Burkholderia stagnalis | Pseudomonadota | Betaproteobacteria |
| Burkholderia territorii | Pseudomonadota | Betaproteobacteria |
| Burkholderia territorii | Pseudomonadota | Betaproteobacteria |
| Burkholderia thailandensis E264 | Pseudomonadota | Betaproteobacteria |
| Burkholderia thailandensis E264 | Pseudomonadota | Betaproteobacteria |
| Burkholderia theae | Pseudomonadota | Betaproteobacteria |
| Burkholderia theae | Pseudomonadota | Betaproteobacteria |
| Burkholderia ubonensis subsp. mesacidophila | Pseudomonadota | Betaproteobacteria |
| Burkholderia ubonensis subsp. mesacidophila | Pseudomonadota | Betaproteobacteria |
| Burkholderia vietnamiensis LMG 10929 | Pseudomonadota | Betaproteobacteria |
| Burkholderia vietnamiensis LMG 10929 | Pseudomonadota | Betaproteobacteria |
| Caballeronia grimmiae | Pseudomonadota | Betaproteobacteria |
| Caballeronia grimmiae | Pseudomonadota | Betaproteobacteria |
| Caballeronia insecticola | Pseudomonadota | Betaproteobacteria |
| Caballeronia insecticola | Pseudomonadota | Betaproteobacteria |
| Caballeronia zhejiangensis | Pseudomonadota | Betaproteobacteria |
| Caballeronia zhejiangensis | Pseudomonadota | Betaproteobacteria |
| Caldimonas aquatica | Pseudomonadota | Betaproteobacteria |
| Caldimonas aquatica | Pseudomonadota | Betaproteobacteria |
| Caldimonas brevitalea | Pseudomonadota | Betaproteobacteria |
| Caldimonas brevitalea | Pseudomonadota | Betaproteobacteria |
| Caldimonas thermodepolymerans | Pseudomonadota | Betaproteobacteria |
| Caldimonas thermodepolymerans | Pseudomonadota | Betaproteobacteria |
| Candidatus Accumulibacter phosphatis | Pseudomonadota | Betaproteobacteria |
| Candidatus Accumulibacter phosphatis | Pseudomonadota | Betaproteobacteria |

| | | |
|---|---|---|
| Candidatus Ichthyocystis hellenicum | Pseudomonadota | Betaproteobacteria |
| Candidatus Ichthyocystis hellenicum | Pseudomonadota | Betaproteobacteria |
| Candidatus Kinetoplastidibacterium blastocrithidiae (ex Strigomonas culicis) | Pseudomonadota | Betaproteobacteria |
| Candidatus Kinetoplastidibacterium blastocrithidiae (ex Strigomonas culicis) | Pseudomonadota | Betaproteobacteria |
| Candidatus Kinetoplastidibacterium crithidiae TCC036E | Pseudomonadota | Betaproteobacteria |
| Candidatus Kinetoplastidibacterium crithidiae TCC036E | Pseudomonadota | Betaproteobacteria |
| Candidatus Kinetoplastidibacterium desouzai TCC079E | Pseudomonadota | Betaproteobacteria |
| Candidatus Kinetoplastidibacterium desouzai TCC079E | Pseudomonadota | Betaproteobacteria |
| Candidatus Kinetoplastidibacterium kentomonadis | Pseudomonadota | Betaproteobacteria |
| Candidatus Kinetoplastidibacterium stringomonadis TCC290E | Pseudomonadota | Betaproteobacteria |
| Candidatus Kinetoplastidibacterium stringomonadis TCC290E | Pseudomonadota | Betaproteobacteria |
| Candidatus Methylopumilus planktonicus | Pseudomonadota | Betaproteobacteria |
| Candidatus Methylopumilus planktonicus | Pseudomonadota | Betaproteobacteria |
| Candidatus Methylopumilus rimovensis | Pseudomonadota | Betaproteobacteria |
| Candidatus Methylopumilus rimovensis | Pseudomonadota | Betaproteobacteria |
| Candidatus Methylopumilus universalis | Pseudomonadota | Betaproteobacteria |
| Candidatus Methylopumilus universalis | Pseudomonadota | Betaproteobacteria |
| Candidatus Nasuia deltocephalinicola | Pseudomonadota | Betaproteobacteria |
| Candidatus Nitrotoga arctica | Pseudomonadota | Betaproteobacteria |
| Candidatus Nitrotoga arctica | Pseudomonadota | Betaproteobacteria |
| Candidatus Profftella armatura | Pseudomonadota | Betaproteobacteria |
| Candidatus Symbiobacter mobilis CR | Pseudomonadota | Betaproteobacteria |
| Candidatus Symbiobacter mobilis CR | Pseudomonadota | Betaproteobacteria |
| Candidatus Vallotia laricis | Pseudomonadota | Betaproteobacteria |
| Candidatus Vallotia laricis | Pseudomonadota | Betaproteobacteria |
| Candidatus Vallotiella adelgis | Pseudomonadota | Betaproteobacteria |
| Candidatus Vallotiella adelgis | Pseudomonadota | Betaproteobacteria |
| Candidatus Vallotiella hemipterorum | Pseudomonadota | Betaproteobacteria |
| Candidatus Vallotiella hemipterorum | Pseudomonadota | Betaproteobacteria |
| Candidatus Zinderia endosymbiont of Aphrophora alni | Pseudomonadota | Betaproteobacteria |
| Casimicrobium huifangae | Pseudomonadota | Betaproteobacteria |

| | | |
|---|---|---|
| Casimicrobium huifangae | Pseudomonadota | Betaproteobacteria |
| Castellaniella defragrans 65Phen | Pseudomonadota | Betaproteobacteria |
| Castellaniella defragrans 65Phen | Pseudomonadota | Betaproteobacteria |
| Castellaniella ginsengisoli | Pseudomonadota | Betaproteobacteria |
| Castellaniella ginsengisoli | Pseudomonadota | Betaproteobacteria |
| Chitinibacter bivalviorum | Pseudomonadota | Betaproteobacteria |
| Chitinibacter bivalviorum | Pseudomonadota | Betaproteobacteria |
| Chitinibacter fontanus | Pseudomonadota | Betaproteobacteria |
| Chitinibacter fontanus | Pseudomonadota | Betaproteobacteria |
| Chitinimonas arctica | Pseudomonadota | Betaproteobacteria |
| Chitinimonas arctica | Pseudomonadota | Betaproteobacteria |
| Chitiniphilus eburneus | Pseudomonadota | Betaproteobacteria |
| Chitiniphilus eburneus | Pseudomonadota | Betaproteobacteria |
| Chitiniphilus purpureus | Pseudomonadota | Betaproteobacteria |
| Chitiniphilus purpureus | Pseudomonadota | Betaproteobacteria |
| Chitinolyticbacter meiyuanensis | Pseudomonadota | Betaproteobacteria |
| Chitinolyticbacter meiyuanensis | Pseudomonadota | Betaproteobacteria |
| Chromobacterium haemolyticum | Pseudomonadota | Betaproteobacteria |
| Chromobacterium haemolyticum | Pseudomonadota | Betaproteobacteria |
| Chromobacterium paludis | Pseudomonadota | Betaproteobacteria |
| Chromobacterium paludis | Pseudomonadota | Betaproteobacteria |
| Chromobacterium phragmitis | Pseudomonadota | Betaproteobacteria |
| Chromobacterium phragmitis | Pseudomonadota | Betaproteobacteria |
| Chromobacterium rhizoryzae | Pseudomonadota | Betaproteobacteria |
| Chromobacterium rhizoryzae | Pseudomonadota | Betaproteobacteria |
| Chromobacterium subtsugae | Pseudomonadota | Betaproteobacteria |
| Chromobacterium subtsugae | Pseudomonadota | Betaproteobacteria |
| Chromobacterium vaccinii | Pseudomonadota | Betaproteobacteria |
| Chromobacterium vaccinii | Pseudomonadota | Betaproteobacteria |
| Chromobacterium violaceum | Pseudomonadota | Betaproteobacteria |
| Chromobacterium violaceum | Pseudomonadota | Betaproteobacteria |

| | | |
|---|---|---|
| Collimonas arenae | Pseudomonadota | Betaproteobacteria |
| Collimonas arenae | Pseudomonadota | Betaproteobacteria |
| Collimonas fungivorans | Pseudomonadota | Betaproteobacteria |
| Collimonas fungivorans | Pseudomonadota | Betaproteobacteria |
| Collimonas pratensis | Pseudomonadota | Betaproteobacteria |
| Collimonas pratensis | Pseudomonadota | Betaproteobacteria |
| Comamonas antarctica | Pseudomonadota | Betaproteobacteria |
| Comamonas antarctica | Pseudomonadota | Betaproteobacteria |
| Comamonas aquatica | Pseudomonadota | Betaproteobacteria |
| Comamonas aquatica | Pseudomonadota | Betaproteobacteria |
| Comamonas endophytica | Pseudomonadota | Betaproteobacteria |
| Comamonas endophytica | Pseudomonadota | Betaproteobacteria |
| Comamonas flocculans | Pseudomonadota | Betaproteobacteria |
| Comamonas flocculans | Pseudomonadota | Betaproteobacteria |
| Comamonas fluminis | Pseudomonadota | Betaproteobacteria |
| Comamonas fluminis | Pseudomonadota | Betaproteobacteria |
| Comamonas kerstersii | Pseudomonadota | Betaproteobacteria |
| Comamonas kerstersii | Pseudomonadota | Betaproteobacteria |
| Comamonas odontotermitis | Pseudomonadota | Betaproteobacteria |
| Comamonas odontotermitis | Pseudomonadota | Betaproteobacteria |
| Comamonas piscis | Pseudomonadota | Betaproteobacteria |
| Comamonas piscis | Pseudomonadota | Betaproteobacteria |
| Comamonas resistens | Pseudomonadota | Betaproteobacteria |
| Comamonas resistens | Pseudomonadota | Betaproteobacteria |
| Comamonas serinivorans | Pseudomonadota | Betaproteobacteria |
| Comamonas serinivorans | Pseudomonadota | Betaproteobacteria |
| Comamonas squillarum | Pseudomonadota | Betaproteobacteria |
| Comamonas squillarum | Pseudomonadota | Betaproteobacteria |
| Comamonas terrigena NBRC 13299 | Pseudomonadota | Betaproteobacteria |
| Comamonas terrigena NBRC 13299 | Pseudomonadota | Betaproteobacteria |
| Comamonas thiooxydans | Pseudomonadota | Betaproteobacteria |

| | | |
|---|---|---|
| Comamonas thiooxydans | Pseudomonadota | Betaproteobacteria |
| Conchiformibius steedae | Pseudomonadota | Betaproteobacteria |
| Conchiformibius steedae | Pseudomonadota | Betaproteobacteria |
| Crenobacter cavernae | Pseudomonadota | Betaproteobacteria |
| Crenobacter cavernae | Pseudomonadota | Betaproteobacteria |
| Cupriavidus basilensis | Pseudomonadota | Betaproteobacteria |
| Cupriavidus gilardii | Pseudomonadota | Betaproteobacteria |
| Cupriavidus gilardii | Pseudomonadota | Betaproteobacteria |
| Cupriavidus malaysiensis | Pseudomonadota | Betaproteobacteria |
| Cupriavidus malaysiensis | Pseudomonadota | Betaproteobacteria |
| Cupriavidus necator N-1 | Pseudomonadota | Betaproteobacteria |
| Cupriavidus necator N-1 | Pseudomonadota | Betaproteobacteria |
| Cupriavidus neocaledonicus | Pseudomonadota | Betaproteobacteria |
| Cupriavidus neocaledonicus | Pseudomonadota | Betaproteobacteria |
| Cupriavidus oxalaticus | Pseudomonadota | Betaproteobacteria |
| Cupriavidus oxalaticus | Pseudomonadota | Betaproteobacteria |
| Cupriavidus pauculus | Pseudomonadota | Betaproteobacteria |
| Cupriavidus pauculus | Pseudomonadota | Betaproteobacteria |
| Cupriavidus pinatubonensis | Pseudomonadota | Betaproteobacteria |
| Cupriavidus pinatubonensis | Pseudomonadota | Betaproteobacteria |
| Cupriavidus taiwanensis LMG 19424 | Pseudomonadota | Betaproteobacteria |
| Cupriavidus taiwanensis LMG 19424 | Pseudomonadota | Betaproteobacteria |
| Dechloromonas aquae | Pseudomonadota | Betaproteobacteria |
| Dechloromonas aquae | Pseudomonadota | Betaproteobacteria |
| Deefgea piscis | Pseudomonadota | Betaproteobacteria |
| Deefgea piscis | Pseudomonadota | Betaproteobacteria |
| Deefgea tanakiae | Pseudomonadota | Betaproteobacteria |
| Deefgea tanakiae | Pseudomonadota | Betaproteobacteria |
| Delftia acidovorans SPH-1 | Pseudomonadota | Betaproteobacteria |
| Delftia acidovorans SPH-1 | Pseudomonadota | Betaproteobacteria |
| Delftia lacustris | Pseudomonadota | Betaproteobacteria |

| | | |
|---|---|---|
| Delftia lacustris | Pseudomonadota | Betaproteobacteria |
| Delftia tsuruhatensis | Pseudomonadota | Betaproteobacteria |
| Delftia tsuruhatensis | Pseudomonadota | Betaproteobacteria |
| Denitratisoma oestradiolicum | Pseudomonadota | Betaproteobacteria |
| Denitratisoma oestradiolicum | Pseudomonadota | Betaproteobacteria |
| Diaphorobacter aerolatus | Pseudomonadota | Betaproteobacteria |
| Diaphorobacter aerolatus | Pseudomonadota | Betaproteobacteria |
| Diaphorobacter limosus | Pseudomonadota | Betaproteobacteria |
| Diaphorobacter limosus | Pseudomonadota | Betaproteobacteria |
| Diaphorobacter nitroreducens | Pseudomonadota | Betaproteobacteria |
| Diaphorobacter nitroreducens | Pseudomonadota | Betaproteobacteria |
| Diaphorobacter ruginosibacter | Pseudomonadota | Betaproteobacteria |
| Diaphorobacter ruginosibacter | Pseudomonadota | Betaproteobacteria |
| Duganella dendranthematis | Pseudomonadota | Betaproteobacteria |
| Duganella dendranthematis | Pseudomonadota | Betaproteobacteria |
| Duganella zoogloeoides | Pseudomonadota | Betaproteobacteria |
| Duganella zoogloeoides | Pseudomonadota | Betaproteobacteria |
| Duodenibacillus massiliensis | Pseudomonadota | Betaproteobacteria |
| Duodenibacillus massiliensis | Pseudomonadota | Betaproteobacteria |
| Eikenella corrodens | Pseudomonadota | Betaproteobacteria |
| Eikenella corrodens | Pseudomonadota | Betaproteobacteria |
| Eikenella exigua | Pseudomonadota | Betaproteobacteria |
| Eikenella exigua | Pseudomonadota | Betaproteobacteria |
| Eleftheria terrae | Pseudomonadota | Betaproteobacteria |
| Eleftheria terrae | Pseudomonadota | Betaproteobacteria |
| Ephemeroptericola cinctiostellae | Pseudomonadota | Betaproteobacteria |
| Ephemeroptericola cinctiostellae | Pseudomonadota | Betaproteobacteria |
| Ferribacterium limneticum | Pseudomonadota | Betaproteobacteria |
| Ferribacterium limneticum | Pseudomonadota | Betaproteobacteria |
| Ferrigenium kumadai | Pseudomonadota | Betaproteobacteria |
| Ferrigenium kumadai | Pseudomonadota | Betaproteobacteria |

| | | |
|---|---|---|
| Ferriphaselus amnicola | Pseudomonadota | Betaproteobacteria |
| Ferriphaselus amnicola | Pseudomonadota | Betaproteobacteria |
| Ferrovum myxofaciens | Pseudomonadota | Betaproteobacteria |
| Ferrovum myxofaciens | Pseudomonadota | Betaproteobacteria |
| Fluviibacter phosphoraccumulans | Pseudomonadota | Betaproteobacteria |
| Fluviibacter phosphoraccumulans | Pseudomonadota | Betaproteobacteria |
| Gallionella capsiferriformans ES-2 | Pseudomonadota | Betaproteobacteria |
| Gallionella capsiferriformans ES-2 | Pseudomonadota | Betaproteobacteria |
| Herbaspirillum frisingense | Pseudomonadota | Betaproteobacteria |
| Herbaspirillum frisingense | Pseudomonadota | Betaproteobacteria |
| Herbaspirillum hiltneri N3 | Pseudomonadota | Betaproteobacteria |
| Herbaspirillum hiltneri N3 | Pseudomonadota | Betaproteobacteria |
| Herbaspirillum huttiense | Pseudomonadota | Betaproteobacteria |
| Herbaspirillum huttiense | Pseudomonadota | Betaproteobacteria |
| Herbaspirillum robiniae | Pseudomonadota | Betaproteobacteria |
| Herbaspirillum robiniae | Pseudomonadota | Betaproteobacteria |
| Herbaspirillum rubrisubalbicans | Pseudomonadota | Betaproteobacteria |
| Herbaspirillum rubrisubalbicans | Pseudomonadota | Betaproteobacteria |
| Herbaspirillum seropedicae | Pseudomonadota | Betaproteobacteria |
| Herbaspirillum seropedicae | Pseudomonadota | Betaproteobacteria |
| Herminiimonas arsenitoxidans | Pseudomonadota | Betaproteobacteria |
| Herminiimonas arsenitoxidans | Pseudomonadota | Betaproteobacteria |
| Hydrogenophaga crassostreae | Pseudomonadota | Betaproteobacteria |
| Hydrogenophaga crassostreae | Pseudomonadota | Betaproteobacteria |
| Hydrogenophaga crocea | Pseudomonadota | Betaproteobacteria |
| Hydrogenophaga crocea | Pseudomonadota | Betaproteobacteria |
| Hydrogenophaga pseudoflava | Pseudomonadota | Betaproteobacteria |
| Hydrogenophaga pseudoflava | Pseudomonadota | Betaproteobacteria |
| Ideonella dechloratans | Pseudomonadota | Betaproteobacteria |
| Ideonella dechloratans | Pseudomonadota | Betaproteobacteria |
| Inhella inkyongensis | Pseudomonadota | Betaproteobacteria |

| | | |
|---|---|---|
| Inhella inkyongensis | Pseudomonadota | Betaproteobacteria |
| Iodobacter ciconiae | Pseudomonadota | Betaproteobacteria |
| Iodobacter ciconiae | Pseudomonadota | Betaproteobacteria |
| Janthinobacterium aestuarii | Pseudomonadota | Betaproteobacteria |
| Janthinobacterium aestuarii | Pseudomonadota | Betaproteobacteria |
| Janthinobacterium agaricidamnosum NBRC 102515 = DSM 9628 | Pseudomonadota | Betaproteobacteria |
| Janthinobacterium lividum | Pseudomonadota | Betaproteobacteria |
| Janthinobacterium lividum | Pseudomonadota | Betaproteobacteria |
| Janthinobacterium rivuli | Pseudomonadota | Betaproteobacteria |
| Janthinobacterium rivuli | Pseudomonadota | Betaproteobacteria |
| Janthinobacterium svalbardensis | Pseudomonadota | Betaproteobacteria |
| Janthinobacterium svalbardensis | Pseudomonadota | Betaproteobacteria |
| Janthinobacterium tructae | Pseudomonadota | Betaproteobacteria |
| Janthinobacterium tructae | Pseudomonadota | Betaproteobacteria |
| Kerstersia gyiorum | Pseudomonadota | Betaproteobacteria |
| Kerstersia gyiorum | Pseudomonadota | Betaproteobacteria |
| Kingella denitrificans | Pseudomonadota | Betaproteobacteria |
| Kingella denitrificans | Pseudomonadota | Betaproteobacteria |
| Kingella kingae | Pseudomonadota | Betaproteobacteria |
| Kingella kingae | Pseudomonadota | Betaproteobacteria |
| Kingella negevensis | Pseudomonadota | Betaproteobacteria |
| Kingella negevensis | Pseudomonadota | Betaproteobacteria |
| Lautropia mirabilis | Pseudomonadota | Betaproteobacteria |
| Lautropia mirabilis | Pseudomonadota | Betaproteobacteria |
| Leptothrix cholodnii SP-6 | Pseudomonadota | Betaproteobacteria |
| Leptothrix cholodnii SP-6 | Pseudomonadota | Betaproteobacteria |
| Limnobacter profundi | Pseudomonadota | Betaproteobacteria |
| Limnobacter profundi | Pseudomonadota | Betaproteobacteria |
| Limnobacter thiooxidans | Pseudomonadota | Betaproteobacteria |
| Limnobacter thiooxidans | Pseudomonadota | Betaproteobacteria |
| Massilia antarctica | Pseudomonadota | Betaproteobacteria |

| | | |
|---|---|---|
| Massilia antarctica | Pseudomonadota | Betaproteobacteria |
| Massilia atriviolacea | Pseudomonadota | Betaproteobacteria |
| Massilia atriviolacea | Pseudomonadota | Betaproteobacteria |
| Massilia cellulosiltytica | Pseudomonadota | Betaproteobacteria |
| Massilia cellulosiltytica | Pseudomonadota | Betaproteobacteria |
| Massilia endophytica | Pseudomonadota | Betaproteobacteria |
| Massilia endophytica | Pseudomonadota | Betaproteobacteria |
| Massilia forsythiae | Pseudomonadota | Betaproteobacteria |
| Massilia forsythiae | Pseudomonadota | Betaproteobacteria |
| Massilia litorea | Pseudomonadota | Betaproteobacteria |
| Massilia litorea | Pseudomonadota | Betaproteobacteria |
| Massilia oculi | Pseudomonadota | Betaproteobacteria |
| Massilia oculi | Pseudomonadota | Betaproteobacteria |
| Massilia putida | Pseudomonadota | Betaproteobacteria |
| Massilia putida | Pseudomonadota | Betaproteobacteria |
| Massilia violaceinigra | Pseudomonadota | Betaproteobacteria |
| Massilia violaceinigra | Pseudomonadota | Betaproteobacteria |
| Melaminivora jejuensis | Pseudomonadota | Betaproteobacteria |
| Melaminivora jejuensis | Pseudomonadota | Betaproteobacteria |
| Melaminivora suipulveris | Pseudomonadota | Betaproteobacteria |
| Melaminivora suipulveris | Pseudomonadota | Betaproteobacteria |
| Methylibium petroleiphilum PM1 | Pseudomonadota | Betaproteobacteria |
| Methylibium petroleiphilum PM1 | Pseudomonadota | Betaproteobacteria |
| Methylobacillus flagellatus KT | Pseudomonadota | Betaproteobacteria |
| Methylobacillus flagellatus KT | Pseudomonadota | Betaproteobacteria |
| Methylophilus medardicus | Pseudomonadota | Betaproteobacteria |
| Methylophilus medardicus | Pseudomonadota | Betaproteobacteria |
| Methyloradius palustris | Pseudomonadota | Betaproteobacteria |
| Methyloradius palustris | Pseudomonadota | Betaproteobacteria |
| Methylotenera mobilis JLW8 | Pseudomonadota | Betaproteobacteria |
| Methylotenera mobilis JLW8 | Pseudomonadota | Betaproteobacteria |

| | | |
|---|---|---|
| Methylotenera versatilis 301 | Pseudomonadota | Betaproteobacteria |
| Methylotenera versatilis 301 | Pseudomonadota | Betaproteobacteria |
| Methylovorus glucosotrophus SIP3-4 | Pseudomonadota | Betaproteobacteria |
| Methylovorus glucosotrophus SIP3-4 | Pseudomonadota | Betaproteobacteria |
| Microvirgula aerodenitrificans | Pseudomonadota | Betaproteobacteria |
| Microvirgula aerodenitrificans | Pseudomonadota | Betaproteobacteria |
| Morococcus cerebrosus | Pseudomonadota | Betaproteobacteria |
| Morococcus cerebrosus | Pseudomonadota | Betaproteobacteria |
| Mycetohabitans endofungorum | Pseudomonadota | Betaproteobacteria |
| Mycetohabitans endofungorum | Pseudomonadota | Betaproteobacteria |
| Mycetohabitans rhizoxinica | Pseudomonadota | Betaproteobacteria |
| Mycetohabitans rhizoxinica | Pseudomonadota | Betaproteobacteria |
| Mycoavidus cysteinexigens | Pseudomonadota | Betaproteobacteria |
| Neisseria animalis | Pseudomonadota | Betaproteobacteria |
| Neisseria animalis | Pseudomonadota | Betaproteobacteria |
| Neisseria animaloris | Pseudomonadota | Betaproteobacteria |
| Neisseria animaloris | Pseudomonadota | Betaproteobacteria |
| Neisseria arctica | Pseudomonadota | Betaproteobacteria |
| Neisseria arctica | Pseudomonadota | Betaproteobacteria |
| Neisseria bacilliformis | Pseudomonadota | Betaproteobacteria |
| Neisseria bacilliformis | Pseudomonadota | Betaproteobacteria |
| Neisseria brasiliensis | Pseudomonadota | Betaproteobacteria |
| Neisseria brasiliensis | Pseudomonadota | Betaproteobacteria |
| Neisseria canis | Pseudomonadota | Betaproteobacteria |
| Neisseria canis | Pseudomonadota | Betaproteobacteria |
| Neisseria chenwenguii | Pseudomonadota | Betaproteobacteria |
| Neisseria chenwenguii | Pseudomonadota | Betaproteobacteria |
| Neisseria cinerea | Pseudomonadota | Betaproteobacteria |
| Neisseria cinerea | Pseudomonadota | Betaproteobacteria |
| Neisseria dentiae | Pseudomonadota | Betaproteobacteria |
| Neisseria dentiae | Pseudomonadota | Betaproteobacteria |

| | | |
|---|---|---|
| Neisseria dumasiana | Pseudomonadota | Betaproteobacteria |
| Neisseria dumasiana | Pseudomonadota | Betaproteobacteria |
| Neisseria flavescens | Pseudomonadota | Betaproteobacteria |
| Neisseria flavescens | Pseudomonadota | Betaproteobacteria |
| Neisseria gonorrhoeae | Pseudomonadota | Betaproteobacteria |
| Neisseria gonorrhoeae | Pseudomonadota | Betaproteobacteria |
| Neisseria lactamica | Pseudomonadota | Betaproteobacteria |
| Neisseria lactamica | Pseudomonadota | Betaproteobacteria |
| Neisseria leonii | Pseudomonadota | Betaproteobacteria |
| Neisseria leonii | Pseudomonadota | Betaproteobacteria |
| Neisseria lisongii | Pseudomonadota | Betaproteobacteria |
| Neisseria lisongii | Pseudomonadota | Betaproteobacteria |
| Neisseria macacae ATCC 33926 | Pseudomonadota | Betaproteobacteria |
| Neisseria macacae ATCC 33926 | Pseudomonadota | Betaproteobacteria |
| Neisseria meningitidis | Pseudomonadota | Betaproteobacteria |
| Neisseria meningitidis | Pseudomonadota | Betaproteobacteria |
| Neisseria musculi | Pseudomonadota | Betaproteobacteria |
| Neisseria musculi | Pseudomonadota | Betaproteobacteria |
| Neisseria shayeganii | Pseudomonadota | Betaproteobacteria |
| Neisseria shayeganii | Pseudomonadota | Betaproteobacteria |
| Neisseria sicca | Pseudomonadota | Betaproteobacteria |
| Neisseria sicca | Pseudomonadota | Betaproteobacteria |
| Neisseria subflava | Pseudomonadota | Betaproteobacteria |
| Neisseria subflava | Pseudomonadota | Betaproteobacteria |
| Neisseria wadsworthii | Pseudomonadota | Betaproteobacteria |
| Neisseria wadsworthii | Pseudomonadota | Betaproteobacteria |
| Neisseria weaveri | Pseudomonadota | Betaproteobacteria |
| Neisseria weaveri | Pseudomonadota | Betaproteobacteria |
| Neisseria yangbaofengii | Pseudomonadota | Betaproteobacteria |
| Neisseria yangbaofengii | Pseudomonadota | Betaproteobacteria |
| Neisseria zalophi | Pseudomonadota | Betaproteobacteria |

| | | |
|---|---|---|
| Neisseria zalophi | Pseudomonadota | Betaproteobacteria |
| Neisseria zoodegmatis | Pseudomonadota | Betaproteobacteria |
| Neisseria zoodegmatis | Pseudomonadota | Betaproteobacteria |
| Neopusillimonas aromaticivorans | Pseudomonadota | Betaproteobacteria |
| Neopusillimonas aromaticivorans | Pseudomonadota | Betaproteobacteria |
| Nitrogeniibacter aestuarii | Pseudomonadota | Betaproteobacteria |
| Nitrogeniibacter aestuarii | Pseudomonadota | Betaproteobacteria |
| Nitrogeniibacter mangrovi | Pseudomonadota | Betaproteobacteria |
| Nitrogeniibacter mangrovi | Pseudomonadota | Betaproteobacteria |
| Nitrosomonas communis | Pseudomonadota | Betaproteobacteria |
| Nitrosomonas communis | Pseudomonadota | Betaproteobacteria |
| Nitrosomonas europaea ATCC 19718 | Pseudomonadota | Betaproteobacteria |
| Nitrosomonas europaea ATCC 19718 | Pseudomonadota | Betaproteobacteria |
| Nitrosomonas eutropha C91 | Pseudomonadota | Betaproteobacteria |
| Nitrosomonas eutropha C91 | Pseudomonadota | Betaproteobacteria |
| Nitrosomonas ureae | Pseudomonadota | Betaproteobacteria |
| Nitrosomonas ureae | Pseudomonadota | Betaproteobacteria |
| Nitrosospira briensis C-128 | Pseudomonadota | Betaproteobacteria |
| Nitrosospira briensis C-128 | Pseudomonadota | Betaproteobacteria |
| Nitrosospira lacus | Pseudomonadota | Betaproteobacteria |
| Nitrosospira lacus | Pseudomonadota | Betaproteobacteria |
| Nitrosospira multiformis ATCC 25196 | Pseudomonadota | Betaproteobacteria |
| Nitrosospira multiformis ATCC 25196 | Pseudomonadota | Betaproteobacteria |
| Niveibacterium microcysteis | Pseudomonadota | Betaproteobacteria |
| Niveibacterium microcysteis | Pseudomonadota | Betaproteobacteria |
| Oligella ureolytica | Pseudomonadota | Betaproteobacteria |
| Oligella ureolytica | Pseudomonadota | Betaproteobacteria |
| Orrella daihaiensis | Pseudomonadota | Betaproteobacteria |
| Orrella daihaiensis | Pseudomonadota | Betaproteobacteria |
| Orrella dioscoreae | Pseudomonadota | Betaproteobacteria |
| Orrella dioscoreae | Pseudomonadota | Betaproteobacteria |

| | | |
|---|---|---|
| Orrella marina | Pseudomonadota | Betaproteobacteria |
| Orrella marina | Pseudomonadota | Betaproteobacteria |
| Oryzomicrobium terrae | Pseudomonadota | Betaproteobacteria |
| Oryzomicrobium terrae | Pseudomonadota | Betaproteobacteria |
| Ottowia oryzae | Pseudomonadota | Betaproteobacteria |
| Ottowia oryzae | Pseudomonadota | Betaproteobacteria |
| Oxalobacter formigenes | Pseudomonadota | Betaproteobacteria |
| Oxalobacter formigenes | Pseudomonadota | Betaproteobacteria |
| Oxalobacter vibrioformis | Pseudomonadota | Betaproteobacteria |
| Oxalobacter vibrioformis | Pseudomonadota | Betaproteobacteria |
| Paenalcaligenes faecalis | Pseudomonadota | Betaproteobacteria |
| Paenalcaligenes faecalis | Pseudomonadota | Betaproteobacteria |
| Paludibacterium paludis | Pseudomonadota | Betaproteobacteria |
| Paludibacterium paludis | Pseudomonadota | Betaproteobacteria |
| Paludibacterium yongneupense | Pseudomonadota | Betaproteobacteria |
| Paludibacterium yongneupense | Pseudomonadota | Betaproteobacteria |
| Pandoraea apista | Pseudomonadota | Betaproteobacteria |
| Pandoraea apista | Pseudomonadota | Betaproteobacteria |
| Pandoraea commovens | Pseudomonadota | Betaproteobacteria |
| Pandoraea commovens | Pseudomonadota | Betaproteobacteria |
| Pandoraea faecigallinarum | Pseudomonadota | Betaproteobacteria |
| Pandoraea faecigallinarum | Pseudomonadota | Betaproteobacteria |
| Pandoraea fibrosis | Pseudomonadota | Betaproteobacteria |
| Pandoraea fibrosis | Pseudomonadota | Betaproteobacteria |
| Pandoraea norimbergensis | Pseudomonadota | Betaproteobacteria |
| Pandoraea norimbergensis | Pseudomonadota | Betaproteobacteria |
| Pandoraea oxalativorans | Pseudomonadota | Betaproteobacteria |
| Pandoraea oxalativorans | Pseudomonadota | Betaproteobacteria |
| Pandoraea pnomenusa | Pseudomonadota | Betaproteobacteria |
| Pandoraea pnomenusa | Pseudomonadota | Betaproteobacteria |
| Pandoraea pulmonicola | Pseudomonadota | Betaproteobacteria |

| | | |
|---|---|---|
| Pandoraea pulmonicola | Pseudomonadota | Betaproteobacteria |
| Pandoraea sputorum | Pseudomonadota | Betaproteobacteria |
| Pandoraea sputorum | Pseudomonadota | Betaproteobacteria |
| Pandoraea thiooxydans | Pseudomonadota | Betaproteobacteria |
| Pandoraea thiooxydans | Pseudomonadota | Betaproteobacteria |
| Pandoraea vervacti | Pseudomonadota | Betaproteobacteria |
| Pandoraea vervacti | Pseudomonadota | Betaproteobacteria |
| Paraburkholderia aromaticivorans | Pseudomonadota | Betaproteobacteria |
| Paraburkholderia aromaticivorans | Pseudomonadota | Betaproteobacteria |
| Paraburkholderia atlantica | Pseudomonadota | Betaproteobacteria |
| Paraburkholderia atlantica | Pseudomonadota | Betaproteobacteria |
| Paraburkholderia bryophila | Pseudomonadota | Betaproteobacteria |
| Paraburkholderia bryophila | Pseudomonadota | Betaproteobacteria |
| Paraburkholderia caffeinilytica | Pseudomonadota | Betaproteobacteria |
| Paraburkholderia caffeinilytica | Pseudomonadota | Betaproteobacteria |
| Paraburkholderia caribensis MBA4 | Pseudomonadota | Betaproteobacteria |
| Paraburkholderia caribensis MBA4 | Pseudomonadota | Betaproteobacteria |
| Paraburkholderia dioscoreae | Pseudomonadota | Betaproteobacteria |
| Paraburkholderia dioscoreae | Pseudomonadota | Betaproteobacteria |
| Paraburkholderia dokdonensis | Pseudomonadota | Betaproteobacteria |
| Paraburkholderia dokdonensis | Pseudomonadota | Betaproteobacteria |
| Paraburkholderia edwinii | Pseudomonadota | Betaproteobacteria |
| Paraburkholderia edwinii | Pseudomonadota | Betaproteobacteria |
| Paraburkholderia fungorum | Pseudomonadota | Betaproteobacteria |
| Paraburkholderia fungorum | Pseudomonadota | Betaproteobacteria |
| Paraburkholderia ginsengisoli | Pseudomonadota | Betaproteobacteria |
| Paraburkholderia ginsengisoli | Pseudomonadota | Betaproteobacteria |
| Paraburkholderia graminis | Pseudomonadota | Betaproteobacteria |
| Paraburkholderia graminis | Pseudomonadota | Betaproteobacteria |
| Paraburkholderia hospita | Pseudomonadota | Betaproteobacteria |
| Paraburkholderia hospita | Pseudomonadota | Betaproteobacteria |

| | | |
|---|---|---|
| Paraburkholderia kirstenboschensis | Pseudomonadota | Betaproteobacteria |
| Paraburkholderia kirstenboschensis | Pseudomonadota | Betaproteobacteria |
| Paraburkholderia kururiensis | Pseudomonadota | Betaproteobacteria |
| Paraburkholderia kururiensis | Pseudomonadota | Betaproteobacteria |
| Paraburkholderia largidicola | Pseudomonadota | Betaproteobacteria |
| Paraburkholderia largidicola | Pseudomonadota | Betaproteobacteria |
| Paraburkholderia pallida | Pseudomonadota | Betaproteobacteria |
| Paraburkholderia pallida | Pseudomonadota | Betaproteobacteria |
| Paraburkholderia phymatum STM815 | Pseudomonadota | Betaproteobacteria |
| Paraburkholderia phymatum STM815 | Pseudomonadota | Betaproteobacteria |
| Paraburkholderia phytofirmans PsJN | Pseudomonadota | Betaproteobacteria |
| Paraburkholderia phytofirmans PsJN | Pseudomonadota | Betaproteobacteria |
| Paraburkholderia sabiae | Pseudomonadota | Betaproteobacteria |
| Paraburkholderia sabiae | Pseudomonadota | Betaproteobacteria |
| Paraburkholderia sprentiae WSM5005 | Pseudomonadota | Betaproteobacteria |
| Paraburkholderia sprentiae WSM5005 | Pseudomonadota | Betaproteobacteria |
| Paraburkholderia terrae | Pseudomonadota | Betaproteobacteria |
| Paraburkholderia terrae | Pseudomonadota | Betaproteobacteria |
| Paraburkholderia terricola | Pseudomonadota | Betaproteobacteria |
| Paraburkholderia terricola | Pseudomonadota | Betaproteobacteria |
| Paraburkholderia tropica | Pseudomonadota | Betaproteobacteria |
| Paraburkholderia tropica | Pseudomonadota | Betaproteobacteria |
| Paraburkholderia xenovorans LB400 | Pseudomonadota | Betaproteobacteria |
| Paraburkholderia xenovorans LB400 | Pseudomonadota | Betaproteobacteria |
| Paracidovorax avenae ATCC 19860 | Pseudomonadota | Betaproteobacteria |
| Paracidovorax avenae ATCC 19860 | Pseudomonadota | Betaproteobacteria |
| Paracidovorax citrulli | Pseudomonadota | Betaproteobacteria |
| Paracidovorax citrulli | Pseudomonadota | Betaproteobacteria |
| Paralysiella testudinis | Pseudomonadota | Betaproteobacteria |
| Paralysiella testudinis | Pseudomonadota | Betaproteobacteria |
| Paucibacter sediminis | Pseudomonadota | Betaproteobacteria |

| | | |
|---|---|---|
| Paucibacter sediminis | Pseudomonadota | Betaproteobacteria |
| Pelistega ratti | Pseudomonadota | Betaproteobacteria |
| Pelistega ratti | Pseudomonadota | Betaproteobacteria |
| Pigmentiphaga aceris | Pseudomonadota | Betaproteobacteria |
| Pigmentiphaga aceris | Pseudomonadota | Betaproteobacteria |
| Piscinibacter gummiphilus | Pseudomonadota | Betaproteobacteria |
| Piscinibacter gummiphilus | Pseudomonadota | Betaproteobacteria |
| Piscinibacter sakaiensis | Pseudomonadota | Betaproteobacteria |
| Piscinibacter sakaiensis | Pseudomonadota | Betaproteobacteria |
| Piscinibacterium candidicorallinum | Pseudomonadota | Betaproteobacteria |
| Piscinibacterium candidicorallinum | Pseudomonadota | Betaproteobacteria |
| Polaromonas hydrogenivorans | Pseudomonadota | Betaproteobacteria |
| Polaromonas hydrogenivorans | Pseudomonadota | Betaproteobacteria |
| Polaromonas naphthalenivorans CJ2 | Pseudomonadota | Betaproteobacteria |
| Polaromonas naphthalenivorans CJ2 | Pseudomonadota | Betaproteobacteria |
| Polaromonas vacuolata | Pseudomonadota | Betaproteobacteria |
| Polaromonas vacuolata | Pseudomonadota | Betaproteobacteria |
| Pollutimonas thiosulfatoxidans | Pseudomonadota | Betaproteobacteria |
| Pollutimonas thiosulfatoxidans | Pseudomonadota | Betaproteobacteria |
| Polynucleobacter acidiphobus | Pseudomonadota | Betaproteobacteria |
| Polynucleobacter acidiphobus | Pseudomonadota | Betaproteobacteria |
| Polynucleobacter antarcticus | Pseudomonadota | Betaproteobacteria |
| Polynucleobacter antarcticus | Pseudomonadota | Betaproteobacteria |
| Polynucleobacter arcticus | Pseudomonadota | Betaproteobacteria |
| Polynucleobacter arcticus | Pseudomonadota | Betaproteobacteria |
| Polynucleobacter asymbioticus QLW-P1DMWA-1 | Pseudomonadota | Betaproteobacteria |
| Polynucleobacter asymbioticus QLW-P1DMWA-1 | Pseudomonadota | Betaproteobacteria |
| Polynucleobacter corsicus | Pseudomonadota | Betaproteobacteria |
| Polynucleobacter corsicus | Pseudomonadota | Betaproteobacteria |
| Polynucleobacter difficilis | Pseudomonadota | Betaproteobacteria |
| Polynucleobacter difficilis | Pseudomonadota | Betaproteobacteria |

| | | |
|---|---|---|
| Polynucleobacter duraquae | Pseudomonadota | Betaproteobacteria |
| Polynucleobacter duraquae | Pseudomonadota | Betaproteobacteria |
| Polynucleobacter ibericus | Pseudomonadota | Betaproteobacteria |
| Polynucleobacter ibericus | Pseudomonadota | Betaproteobacteria |
| Polynucleobacter necessarius | Pseudomonadota | Betaproteobacteria |
| Polynucleobacter necessarius | Pseudomonadota | Betaproteobacteria |
| Polynucleobacter paludilacus | Pseudomonadota | Betaproteobacteria |
| Polynucleobacter paludilacus | Pseudomonadota | Betaproteobacteria |
| Polynucleobacter paneuropaeus | Pseudomonadota | Betaproteobacteria |
| Polynucleobacter paneuropaeus | Pseudomonadota | Betaproteobacteria |
| Polynucleobacter tropicus | Pseudomonadota | Betaproteobacteria |
| Polynucleobacter tropicus | Pseudomonadota | Betaproteobacteria |
| Polynucleobacter wuianus | Pseudomonadota | Betaproteobacteria |
| Polynucleobacter wuianus | Pseudomonadota | Betaproteobacteria |
| Polynucleobacter yangtzensis | Pseudomonadota | Betaproteobacteria |
| Polynucleobacter yangtzensis | Pseudomonadota | Betaproteobacteria |
| Pseudazoarcus pumilus | Pseudomonadota | Betaproteobacteria |
| Pseudazoarcus pumilus | Pseudomonadota | Betaproteobacteria |
| Pseudoduganella albidiflava | Pseudomonadota | Betaproteobacteria |
| Pseudoduganella albidiflava | Pseudomonadota | Betaproteobacteria |
| Pseudoduganella aquatica | Pseudomonadota | Betaproteobacteria |
| Pseudoduganella aquatica | Pseudomonadota | Betaproteobacteria |
| Pseudoduganella armeniaca | Pseudomonadota | Betaproteobacteria |
| Pseudoduganella armeniaca | Pseudomonadota | Betaproteobacteria |
| Pseudoduganella chitinolytica | Pseudomonadota | Betaproteobacteria |
| Pseudoduganella chitinolytica | Pseudomonadota | Betaproteobacteria |
| Pseudoduganella flava | Pseudomonadota | Betaproteobacteria |
| Pseudoduganella flava | Pseudomonadota | Betaproteobacteria |
| Pseudoduganella lutea | Pseudomonadota | Betaproteobacteria |
| Pseudoduganella lutea | Pseudomonadota | Betaproteobacteria |
| Pseudoduganella plicata | Pseudomonadota | Betaproteobacteria |

| | | |
|---|---|---|
| Pseudoduganella plicata | Pseudomonadota | Betaproteobacteria |
| Pseudoduganella umbonata | Pseudomonadota | Betaproteobacteria |
| Pseudoduganella umbonata | Pseudomonadota | Betaproteobacteria |
| Pseudothauera hydrothermalis | Pseudomonadota | Betaproteobacteria |
| Pseudothauera hydrothermalis | Pseudomonadota | Betaproteobacteria |
| Pulveribacter suum | Pseudomonadota | Betaproteobacteria |
| Pulveribacter suum | Pseudomonadota | Betaproteobacteria |
| Quatrionicoccus australiensis | Pseudomonadota | Betaproteobacteria |
| Quatrionicoccus australiensis | Pseudomonadota | Betaproteobacteria |
| Ralstonia insidiosa | Pseudomonadota | Betaproteobacteria |
| Ralstonia insidiosa | Pseudomonadota | Betaproteobacteria |
| Ralstonia mannitolilytica | Pseudomonadota | Betaproteobacteria |
| Ralstonia mannitolilytica | Pseudomonadota | Betaproteobacteria |
| Ralstonia nicotianae | Pseudomonadota | Betaproteobacteria |
| Ralstonia nicotianae | Pseudomonadota | Betaproteobacteria |
| Ralstonia solanacearum | Pseudomonadota | Betaproteobacteria |
| Ralstonia solanacearum | Pseudomonadota | Betaproteobacteria |
| Ralstonia syzygii subsp. indonesiensis | Pseudomonadota | Betaproteobacteria |
| Ralstonia syzygii subsp. indonesiensis | Pseudomonadota | Betaproteobacteria |
| Ralstonia wenshanensis | Pseudomonadota | Betaproteobacteria |
| Ralstonia wenshanensis | Pseudomonadota | Betaproteobacteria |
| Ramlibacter tataouinensis TTB310 | Pseudomonadota | Betaproteobacteria |
| Ramlibacter tataouinensis TTB310 | Pseudomonadota | Betaproteobacteria |
| Rhodoferax antarcticus | Pseudomonadota | Betaproteobacteria |
| Rhodoferax antarcticus | Pseudomonadota | Betaproteobacteria |
| Rhodoferax aquaticus | Pseudomonadota | Betaproteobacteria |
| Rhodoferax aquaticus | Pseudomonadota | Betaproteobacteria |
| Rhodoferax ferrireducens T118 | Pseudomonadota | Betaproteobacteria |
| Rhodoferax ferrireducens T118 | Pseudomonadota | Betaproteobacteria |
| Rhodoferax koreensis | Pseudomonadota | Betaproteobacteria |
| Rhodoferax koreensis | Pseudomonadota | Betaproteobacteria |

| | | |
|---|---|---|
| Rhodoferax mekongensis | Pseudomonadota | Betaproteobacteria |
| Rhodoferax mekongensis | Pseudomonadota | Betaproteobacteria |
| Rhodoferax saidenbachensis | Pseudomonadota | Betaproteobacteria |
| Rhodoferax saidenbachensis | Pseudomonadota | Betaproteobacteria |
| Rhodoferax sediminis | Pseudomonadota | Betaproteobacteria |
| Rhodoferax sediminis | Pseudomonadota | Betaproteobacteria |
| Robbsia andropogonis | Pseudomonadota | Betaproteobacteria |
| Robbsia andropogonis | Pseudomonadota | Betaproteobacteria |
| Roseateles amylovorans | Pseudomonadota | Betaproteobacteria |
| Roseateles amylovorans | Pseudomonadota | Betaproteobacteria |
| Roseateles depolymerans | Pseudomonadota | Betaproteobacteria |
| Roseateles depolymerans | Pseudomonadota | Betaproteobacteria |
| Rubrivivax gelatinosus | Pseudomonadota | Betaproteobacteria |
| Rubrivivax gelatinosus | Pseudomonadota | Betaproteobacteria |
| Rugosibacter aromaticivorans | Pseudomonadota | Betaproteobacteria |
| Rugosibacter aromaticivorans | Pseudomonadota | Betaproteobacteria |
| Serpentinimonas maccroryi | Pseudomonadota | Betaproteobacteria |
| Serpentinimonas maccroryi | Pseudomonadota | Betaproteobacteria |
| Serpentinimonas raichei | Pseudomonadota | Betaproteobacteria |
| Serpentinimonas raichei | Pseudomonadota | Betaproteobacteria |
| Sideroxyarcus emersonii | Pseudomonadota | Betaproteobacteria |
| Sideroxyarcus emersonii | Pseudomonadota | Betaproteobacteria |
| Sideroxydans lithotrophicus ES-1 | Pseudomonadota | Betaproteobacteria |
| Sideroxydans lithotrophicus ES-1 | Pseudomonadota | Betaproteobacteria |
| Silvimonas soli | Pseudomonadota | Betaproteobacteria |
| Silvimonas soli | Pseudomonadota | Betaproteobacteria |
| Simonsiella muelleri ATCC 29453 | Pseudomonadota | Betaproteobacteria |
| Simonsiella muelleri ATCC 29453 | Pseudomonadota | Betaproteobacteria |
| Simplicispira suum | Pseudomonadota | Betaproteobacteria |
| Simplicispira suum | Pseudomonadota | Betaproteobacteria |
| Snodgrassella alvi wkB2 | Pseudomonadota | Betaproteobacteria |

| | | |
|---|---|---|
| Snodgrassella alvi wkB2 | Pseudomonadota | Betaproteobacteria |
| Sphaerotilus microaerophilus | Pseudomonadota | Betaproteobacteria |
| Sphaerotilus microaerophilus | Pseudomonadota | Betaproteobacteria |
| Sphaerotilus sulfidivorans | Pseudomonadota | Betaproteobacteria |
| Sphaerotilus sulfidivorans | Pseudomonadota | Betaproteobacteria |
| Sulfuricella denitrificans skB26 | Pseudomonadota | Betaproteobacteria |
| Sulfuricella denitrificans skB26 | Pseudomonadota | Betaproteobacteria |
| Sulfuricystis multivorans | Pseudomonadota | Betaproteobacteria |
| Sulfuricystis multivorans | Pseudomonadota | Betaproteobacteria |
| Sulfuricystis thermophila | Pseudomonadota | Betaproteobacteria |
| Sulfuricystis thermophila | Pseudomonadota | Betaproteobacteria |
| Sulfuriferula nivalis | Pseudomonadota | Betaproteobacteria |
| Sulfuriferula nivalis | Pseudomonadota | Betaproteobacteria |
| Sulfuriferula plumbiphila | Pseudomonadota | Betaproteobacteria |
| Sulfuriferula plumbiphila | Pseudomonadota | Betaproteobacteria |
| Sulfurimicrobium lacus | Pseudomonadota | Betaproteobacteria |
| Sulfurimicrobium lacus | Pseudomonadota | Betaproteobacteria |
| Sulfuritalea hydrogenivorans sk43H | Pseudomonadota | Betaproteobacteria |
| Sulfuritalea hydrogenivorans sk43H | Pseudomonadota | Betaproteobacteria |
| Sulfuritortus calidifontis | Pseudomonadota | Betaproteobacteria |
| Sulfuritortus calidifontis | Pseudomonadota | Betaproteobacteria |
| Sutterella faecalis | Pseudomonadota | Betaproteobacteria |
| Sutterella faecalis | Pseudomonadota | Betaproteobacteria |
| Sutterella megalosphaeroides | Pseudomonadota | Betaproteobacteria |
| Sutterella megalosphaeroides | Pseudomonadota | Betaproteobacteria |
| Sutterella wadsworthensis | Pseudomonadota | Betaproteobacteria |
| Sutterella wadsworthensis | Pseudomonadota | Betaproteobacteria |
| Taylorella asinigenitalis MCE3 | Pseudomonadota | Betaproteobacteria |
| Taylorella asinigenitalis MCE3 | Pseudomonadota | Betaproteobacteria |
| Taylorella equigenitalis | Pseudomonadota | Betaproteobacteria |
| Taylorella equigenitalis | Pseudomonadota | Betaproteobacteria |

| | | |
|---|---|---|
| Telluria beijingensis | Pseudomonadota | Betaproteobacteria |
| Telluria beijingensis | Pseudomonadota | Betaproteobacteria |
| Telluria mixta | Pseudomonadota | Betaproteobacteria |
| Telluria mixta | Pseudomonadota | Betaproteobacteria |
| Tepidimonas taiwanensis | Pseudomonadota | Betaproteobacteria |
| Tepidimonas taiwanensis | Pseudomonadota | Betaproteobacteria |
| Thauera aminoaromatica | Pseudomonadota | Betaproteobacteria |
| Thauera aminoaromatica | Pseudomonadota | Betaproteobacteria |
| Thauera aromatica K172 | Pseudomonadota | Betaproteobacteria |
| Thauera aromatica K172 | Pseudomonadota | Betaproteobacteria |
| Thauera butanivorans | Pseudomonadota | Betaproteobacteria |
| Thauera butanivorans | Pseudomonadota | Betaproteobacteria |
| Thauera chlorobenzoica | Pseudomonadota | Betaproteobacteria |
| Thauera chlorobenzoica | Pseudomonadota | Betaproteobacteria |
| Thauera humireducens | Pseudomonadota | Betaproteobacteria |
| Thauera humireducens | Pseudomonadota | Betaproteobacteria |
| Thiobacillus sedimenti | Pseudomonadota | Betaproteobacteria |
| Thiobacillus sedimenti | Pseudomonadota | Betaproteobacteria |
| Thiomonas arsenitoxydans | Pseudomonadota | Betaproteobacteria |
| Thiomonas arsenitoxydans | Pseudomonadota | Betaproteobacteria |
| Thiomonas intermedia | Pseudomonadota | Betaproteobacteria |
| Thiomonas intermedia | Pseudomonadota | Betaproteobacteria |
| Trinickia caryophylli | Pseudomonadota | Betaproteobacteria |
| Trinickia caryophylli | Pseudomonadota | Betaproteobacteria |
| Trinickia violacea | Pseudomonadota | Betaproteobacteria |
| Trinickia violacea | Pseudomonadota | Betaproteobacteria |
| Turicimonas muris | Pseudomonadota | Betaproteobacteria |
| Turicimonas muris | Pseudomonadota | Betaproteobacteria |
| Undibacterium cyanobacteriorum | Pseudomonadota | Betaproteobacteria |
| Undibacterium cyanobacteriorum | Pseudomonadota | Betaproteobacteria |
| Undibacterium parvum | Pseudomonadota | Betaproteobacteria |

| | | |
|---|---|---|
| Undibacterium parvum | Pseudomonadota | Betaproteobacteria |
| Uruburuella suis | Pseudomonadota | Betaproteobacteria |
| Uruburuella suis | Pseudomonadota | Betaproteobacteria |
| Uruburuella testudinis | Pseudomonadota | Betaproteobacteria |
| Uruburuella testudinis | Pseudomonadota | Betaproteobacteria |
| Usitatibacter palustris | Pseudomonadota | Betaproteobacteria |
| Usitatibacter palustris | Pseudomonadota | Betaproteobacteria |
| Usitatibacter rugosus | Pseudomonadota | Betaproteobacteria |
| Usitatibacter rugosus | Pseudomonadota | Betaproteobacteria |
| Variovorax paradoxus NBRC 15149 | Pseudomonadota | Betaproteobacteria |
| Variovorax paradoxus NBRC 15149 | Pseudomonadota | Betaproteobacteria |
| Verminephrobacter eiseniae EF01-2 | Pseudomonadota | Betaproteobacteria |
| Verminephrobacter eiseniae EF01-2 | Pseudomonadota | Betaproteobacteria |
| Vitreoscilla filiformis | Pseudomonadota | Betaproteobacteria |
| Vitreoscilla filiformis | Pseudomonadota | Betaproteobacteria |
| Vitreoscilla stercoraria | Pseudomonadota | Betaproteobacteria |
| Vitreoscilla stercoraria | Pseudomonadota | Betaproteobacteria |
| Wielerella bovis | Pseudomonadota | Betaproteobacteria |
| Wielerella bovis | Pseudomonadota | Betaproteobacteria |
| Xylophilus rhododendri | Pseudomonadota | Betaproteobacteria |
| Xylophilus rhododendri | Pseudomonadota | Betaproteobacteria |
| Chloracidobacterium sp. MS 40/45 | Acidobacteriota | Blastocatellia |
| Chloracidobacterium sp. MS 40/45 | Acidobacteriota | Blastocatellia |
| Chloracidobacterium thermophilum B | Acidobacteriota | Blastocatellia |
| Chloracidobacterium thermophilum B | Acidobacteriota | Blastocatellia |
| Chloracidobacterium validum | Acidobacteriota | Blastocatellia |
| Chloracidobacterium validum | Acidobacteriota | Blastocatellia |
| Caldilinea aerophila DSM 14535 = NBRC 104270 | Chloroflexota | Caldilineae |
| Caldilinea aerophila DSM 14535 = NBRC 104270 | Chloroflexota | Caldilineae |
| Caldisericum exile AZM16c01 | Caldisericota | Caldisericia |
| Caldisericum exile AZM16c01 | Caldisericota | Caldisericia |

| | | |
|---|---|---|
| Caldithrix abyssi DSM 13497 | Calditrichota | Calditrichia |
| Caldithrix abyssi DSM 13497 | Calditrichota | Calditrichia |
| Candidatus Babela massiliensis | | Candidatus Babeliae |
| Candidatus Chromulinivorax destructor | | Candidatus Babeliae |
| Candidatus Kuenenia stuttgartensis | Planctomycetota | Candidatus Brocadiia |
| Candidatus Kuenenia stuttgartensis | Planctomycetota | Candidatus Brocadiia |
| Candidatus Cloacimonas acidaminovorans str. Evry | Candidatus Cloacimonadota | Candidatus Cloacimonadia |
| Candidatus Desulfofervidus auxilii | Thermodesulfobacteriota | Candidatus Desulfofervidia |
| Candidatus Desulfofervidus auxilii | Thermodesulfobacteriota | Candidatus Desulfofervidia |
| Candidatus Xianfuyuplasma coldseepsis | Mycoplasmatota | Candidatus Izimaplasma |
| Hujiaoplasma nucleasis | Mycoplasmatota | Candidatus Izimaplasma |
| Minisyncoccus archaeiphilus | Candidatus Paceibacterota | Candidatus Paceibacteria |
| Candidatus Nanosynbacter featherlites | Candidatus Saccharibacteria | Candidatus Saccharimonadia |
| Candidatus Nanosynbacter lyticus | Candidatus Saccharibacteria | Candidatus Saccharimonadia |
| Candidatus Saccharimonas aalborgensis | Candidatus Saccharibacteria | Candidatus Saccharimonadia |
| Candidatus Southlakia epibionticum | Candidatus Saccharibacteria | Candidatus Saccharimonadia |
| Phototrophicus methaneseepsis | Chloroflexota | Candidatus Thermofonsia |
| Candidatus Uabimicrobium amorphum | Planctomycetota | Candidatus Uabimicrobiia |
| Candidatus Uabimicrobium helgolandensis | Planctomycetota | Candidatus Uabimicrobiia |
| Arachidicoccus soli | Bacteroidota | Chitinophagia |
| Arachidicoccus terrestris | Bacteroidota | Chitinophagia |
| Chitinophaga agri | Bacteroidota | Chitinophagia |
| Chitinophaga caeni | Bacteroidota | Chitinophagia |
| Chitinophaga caseinilytica | Bacteroidota | Chitinophagia |
| Chitinophaga horti | Bacteroidota | Chitinophagia |
| Chitinophaga oryzae | Bacteroidota | Chitinophagia |
| Chitinophaga pendula | Bacteroidota | Chitinophagia |
| Chitinophaga pinensis DSM 2588 | Bacteroidota | Chitinophagia |
| Chitinophaga pollutisoli | Bacteroidota | Chitinophagia |
| Chitinophaga sancti | Bacteroidota | Chitinophagia |
| Ferruginibacter albus | Bacteroidota | Chitinophagia |

| | | |
|---|---|---|
| Ferruginibacter lapsinanis | Bacteroidota | Chitinophagia |
| Filimonas lacunae | Bacteroidota | Chitinophagia |
| Flavihumibacter fluvii | Bacteroidota | Chitinophagia |
| Flavihumibacter rivuli | Bacteroidota | Chitinophagia |
| Flavisolibacter ginsenosidimutans | Bacteroidota | Chitinophagia |
| Flavisolibacter tropicus | Bacteroidota | Chitinophagia |
| Lacibacter sediminis | Bacteroidota | Chitinophagia |
| Mycovorax composti | Bacteroidota | Chitinophagia |
| Niabella ginsenosidivorans | Bacteroidota | Chitinophagia |
| Niabella insulamsoli | Bacteroidota | Chitinophagia |
| Niabella soli DSM 19437 | Bacteroidota | Chitinophagia |
| Niabella yanshanensis | Bacteroidota | Chitinophagia |
| Niastella koreensis GR20-10 | Bacteroidota | Chitinophagia |
| Panacibacter ginsenosidivorans | Bacteroidota | Chitinophagia |
| Paraflavitalea soli | Bacteroidota | Chitinophagia |
| Paraflavitalea speifideaquila | Bacteroidota | Chitinophagia |
| Pseudocnuella soli | Bacteroidota | Chitinophagia |
| Rhizosphaericola mali | Bacteroidota | Chitinophagia |
| Candidatus Chlamydia sanziniae | Chlamydiota | Chlamydiia |
| Candidatus Protochlamydia amoebophila UWE25 | Chlamydiota | Chlamydiia |
| Candidatus Protochlamydia naegleriophila | Chlamydiota | Chlamydiia |
| Candidatus Rhabdochlamydia oedothoracis | Chlamydiota | Chlamydiia |
| Candidatus Rhabdochlamydia porcellionis | Chlamydiota | Chlamydiia |
| Chlamydia abortus | Chlamydiota | Chlamydiia |
| Chlamydia buteonis | Chlamydiota | Chlamydiia |
| Chlamydia caviae GPIC | Chlamydiota | Chlamydiia |
| Chlamydia crocodili | Chlamydiota | Chlamydiia |
| Chlamydia felis Fe/C-56 | Chlamydiota | Chlamydiia |
| Chlamydia gallinacea 08-1274/3 | Chlamydiota | Chlamydiia |
| Chlamydia muridarum str. Nigg | Chlamydiota | Chlamydiia |
| Chlamydia pecorum | Chlamydiota | Chlamydiia |

| | | |
|---|---|---|
| Chlamydia pneumoniae TW-183 | Chlamydiota | Chlamydiia |
| Chlamydia poikilotherma | Chlamydiota | Chlamydiia |
| Chlamydia psittaci 6BC | Chlamydiota | Chlamydiia |
| Chlamydia serpentis | Chlamydiota | Chlamydiia |
| Chlamydia suis | Chlamydiota | Chlamydiia |
| Chlamydia trachomatis D/UW-3/CX | Chlamydiota | Chlamydiia |
| Chlamydiifrater phoenicopteri | Chlamydiota | Chlamydiia |
| Chlamydiifrater volucris | Chlamydiota | Chlamydiia |
| Parachlamydia acanthamoebae UV-7 | Chlamydiota | Chlamydiia |
| Simkania negevensis Z | Chlamydiota | Chlamydiia |
| Waddlia chondrophila WSU 86-1044 | Chlamydiota | Chlamydiia |
| Chlorobaculum limnaeum | Chlorobiota | Chlorobiia |
| Chlorobaculum parvum NCIB 8327 | Chlorobiota | Chlorobiia |
| Chlorobaculum tepidum TLS | Chlorobiota | Chlorobiia |
| Chlorobium limicola DSM 245 | Chlorobiota | Chlorobiia |
| Chlorobium phaeobacteroides DSM 266 | Chlorobiota | Chlorobiia |
| Chloroherpeton thalassium ATCC 35110 | Chlorobiota | Chlorobiia |
| Pelodictyon luteolum DSM 273 | Chlorobiota | Chlorobiia |
| Pelodictyon phaeoclathratiforme BU-1 | Chlorobiota | Chlorobiia |
| Prosthecochloris aestuarii DSM 271 | Chlorobiota | Chlorobiia |
| Candidatus Chlorohelix allophototropha | Chloroflexota | Chloroflexia |
| Candidatus Chlorohelix allophototropha | Chloroflexota | Chloroflexia |
| Chloroflexus aggregans DSM 9485 | Chloroflexota | Chloroflexia |
| Chloroflexus aggregans DSM 9485 | Chloroflexota | Chloroflexia |
| Chloroflexus aurantiacus J-10-fl | Chloroflexota | Chloroflexia |
| Chloroflexus aurantiacus J-10-fl | Chloroflexota | Chloroflexia |
| Roseiflexus castenholzii DSM 13941 | Chloroflexota | Chloroflexia |
| Roseiflexus castenholzii DSM 13941 | Chloroflexota | Chloroflexia |
| Desulfurispirillum indicum S5 | Chrysiogenota | Chrysiogenia |
| Chthonomonas calidirosea T49 | Armatimonadota | Chthonomonadia |
| Chthonomonas calidirosea T49 | Armatimonadota | Chthonomonadia |

| | | |
|---|---|---|
| [Clostridium] asparagiforme DSM 15981 | Bacillota | Clostridia |
| [Clostridium] colinum | Bacillota | Clostridia |
| [Clostridium] hylemonae | Bacillota | Clostridia |
| [Clostridium] saccharolyticum WM1 | Bacillota | Clostridia |
| [Clostridium] scindens ATCC 35704 | Bacillota | Clostridia |
| [Ruminococcus] lactaris ATCC 29176 | Bacillota | Clostridia |
| [Ruminococcus] torques | Bacillota | Clostridia |
| Aceticella autotrophica | Bacillota | Clostridia |
| Aceticella autotrophica | Bacillota | Clostridia |
| Acetivibrio clariflavus DSM 19732 | Bacillota | Clostridia |
| Acetivibrio saccincola | Bacillota | Clostridia |
| Acetivibrio thermocellus ATCC 27405 | Bacillota | Clostridia |
| Acetoanaerobium sticklandii | Bacillota | Clostridia |
| Acetoanaerobium sticklandii | Bacillota | Clostridia |
| Acetobacterium carbinolicum | Bacillota | Clostridia |
| Acetobacterium carbinolicum | Bacillota | Clostridia |
| Acetobacterium malicum subsp. dehalogenans DSM 11527 | Bacillota | Clostridia |
| Acetobacterium malicum subsp. dehalogenans DSM 11527 | Bacillota | Clostridia |
| Acetobacterium wieringae | Bacillota | Clostridia |
| Acetobacterium wieringae | Bacillota | Clostridia |
| Acetobacterium woodii DSM 1030 | Bacillota | Clostridia |
| Acetobacterium woodii DSM 1030 | Bacillota | Clostridia |
| Acetohalobium arabaticum DSM 5501 | Bacillota | Clostridia |
| Acetohalobium arabaticum DSM 5501 | Bacillota | Clostridia |
| Acididesulfobacillus acetoxydans | Bacillota | Clostridia |
| Acutalibacter muris | Bacillota | Clostridia |
| Agathobacter rectalis | Bacillota | Clostridia |
| Alkalibacter rhizosphaerae | Bacillota | Clostridia |
| Alkalibacter rhizosphaerae | Bacillota | Clostridia |
| Alkalicella caledoniensis | Bacillota | Clostridia |
| Alkalicella caledoniensis | Bacillota | Clostridia |

| | | |
|---|---|---|
| Alkaliphilus metalliredigens QYMF | Bacillota | Clostridia |
| Alkaliphilus metalliredigens QYMF | Bacillota | Clostridia |
| Alkaliphilus oremlandii OhILAs | Bacillota | Clostridia |
| Alkaliphilus oremlandii OhILAs | Bacillota | Clostridia |
| Aminipila butyrica | Bacillota | Clostridia |
| Aminipila butyrica | Bacillota | Clostridia |
| Aminipila luticellarii | Bacillota | Clostridia |
| Aminipila luticellarii | Bacillota | Clostridia |
| Aminipila terrae | Bacillota | Clostridia |
| Aminipila terrae | Bacillota | Clostridia |
| Ammonifex degensii KC4 | Bacillota | Clostridia |
| Ammonifex degensii KC4 | Bacillota | Clostridia |
| Amygdalobacter indicium | Bacillota | Clostridia |
| Anaerobutyricum hallii | Bacillota | Clostridia |
| Anaerocolumna cellulosilytica | Bacillota | Clostridia |
| Anaerocolumna chitinilytica | Bacillota | Clostridia |
| Anaerocolumna sedimenticola | Bacillota | Clostridia |
| Anaeromicropila herbilytica | Bacillota | Clostridia |
| Anaeropeptidivorans aminofermentans | Bacillota | Clostridia |
| Anaerostipes caccae | Bacillota | Clostridia |
| Anaerostipes hadrus | Bacillota | Clostridia |
| Anaerostipes rhamnosivorans | Bacillota | Clostridia |
| Anaerotignum propionicum DSM 1682 | Bacillota | Clostridia |
| Anaerotruncus colihominis | Bacillota | Clostridia |
| Anoxybacter fermentans | Bacillota | Clostridia |
| Anoxybacter fermentans | Bacillota | Clostridia |
| Aristaeella hokkaidonensis | Bacillota | Clostridia |
| Aristaeella hokkaidonensis | Bacillota | Clostridia |
| Aristaeella lactis | Bacillota | Clostridia |
| Aristaeella lactis | Bacillota | Clostridia |
| Bengtsoniella intestinalis | Bacillota | Clostridia |

| | | |
|---|---|---|
| Biomaibacter acetigenes | Bacillota | Clostridia |
| Biomaibacter acetigenes | Bacillota | Clostridia |
| Blautia argi | Bacillota | Clostridia |
| Blautia coccoides | Bacillota | Clostridia |
| Blautia hansenii DSM 20583 | Bacillota | Clostridia |
| Blautia hydrogenotrophica DSM 10507 | Bacillota | Clostridia |
| Blautia liquoris | Bacillota | Clostridia |
| Blautia luti | Bacillota | Clostridia |
| Blautia obeum ATCC 29174 | Bacillota | Clostridia |
| Blautia parvula | Bacillota | Clostridia |
| Blautia producta ATCC 27340 = DSM 2950 | Bacillota | Clostridia |
| Blautia pseudococcoides | Bacillota | Clostridia |
| Blautia wexlerae DSM 19850 | Bacillota | Clostridia |
| Butyrivibrio fibrisolvens | Bacillota | Clostridia |
| Butyrivibrio hungatei | Bacillota | Clostridia |
| Butyrivibrio proteoclasticus B316 | Bacillota | Clostridia |
| Caldanaerobacter subterraneus subsp. tengcongensis MB4 | Bacillota | Clostridia |
| Caldanaerobacter subterraneus subsp. tengcongensis MB4 | Bacillota | Clostridia |
| Caldinitratiruptor microaerophilus | Bacillota | Clostridia |
| Candidatus Desulforudis audaxviator | Bacillota | Clostridia |
| Candidatus Desulforudis audaxviator | Bacillota | Clostridia |
| Candidatus Epulonipiscium viviparus | Bacillota | Clostridia |
| Candidatus Formimonas warabiya | Bacillota | Clostridia |
| Candidatus Formimonas warabiya | Bacillota | Clostridia |
| Candidatus Sarcina troglodytae | Bacillota | Clostridia |
| Candidatus Sarcina troglodytae | Bacillota | Clostridia |
| Candidatus Syntrophocurvum alkaliphilum | Bacillota | Clostridia |
| Candidatus Syntrophocurvum alkaliphilum | Bacillota | Clostridia |
| Caproicibacter fermentans | Bacillota | Clostridia |
| Caproicibacterium amylolyticum | Bacillota | Clostridia |
| Caproicibacterium argilliputei | Bacillota | Clostridia |

| | | |
|---|---|---|
| Caproicibacterium lactatifermentans | Bacillota | Clostridia |
| Carboxydothermus hydrogenoformans Z-2901 | Bacillota | Clostridia |
| Carboxydothermus hydrogenoformans Z-2901 | Bacillota | Clostridia |
| Cellulosilyticum lentocellum DSM 5427 | Bacillota | Clostridia |
| Chakrabartyella piscis | Bacillota | Clostridia |
| Chordicoccus furentiruminis | Bacillota | Clostridia |
| Christensenella massiliensis | Bacillota | Clostridia |
| Christensenella massiliensis | Bacillota | Clostridia |
| Christensenella minuta | Bacillota | Clostridia |
| Christensenella minuta | Bacillota | Clostridia |
| Claveliimonas bilis | Bacillota | Clostridia |
| Clostridioides difficile | Bacillota | Clostridia |
| Clostridioides difficile | Bacillota | Clostridia |
| Clostridium aceticum | Bacillota | Clostridia |
| Clostridium aceticum | Bacillota | Clostridia |
| Clostridium acetobutylicum DSM 1731 | Bacillota | Clostridia |
| Clostridium acetobutylicum DSM 1731 | Bacillota | Clostridia |
| Clostridium argentinense | Bacillota | Clostridia |
| Clostridium argentinense | Bacillota | Clostridia |
| Clostridium autoethanogenum DSM 10061 | Bacillota | Clostridia |
| Clostridium autoethanogenum DSM 10061 | Bacillota | Clostridia |
| Clostridium baratii | Bacillota | Clostridia |
| Clostridium baratii | Bacillota | Clostridia |
| Clostridium bornimense | Bacillota | Clostridia |
| Clostridium bornimense | Bacillota | Clostridia |
| Clostridium botulinum A str. ATCC 3502 | Bacillota | Clostridia |
| Clostridium botulinum A str. ATCC 3502 | Bacillota | Clostridia |
| Clostridium butyricum | Bacillota | Clostridia |
| Clostridium butyricum | Bacillota | Clostridia |
| Clostridium cadaveris | Bacillota | Clostridia |
| Clostridium cadaveris | Bacillota | Clostridia |

| | | |
|---|---|---|
| Clostridium carboxidivorans P7 | Bacillota | Clostridia |
| Clostridium carboxidivorans P7 | Bacillota | Clostridia |
| Clostridium cellulovorans 743B | Bacillota | Clostridia |
| Clostridium cellulovorans 743B | Bacillota | Clostridia |
| Clostridium chauvoei | Bacillota | Clostridia |
| Clostridium chauvoei | Bacillota | Clostridia |
| Clostridium cochlearium | Bacillota | Clostridia |
| Clostridium cochlearium | Bacillota | Clostridia |
| Clostridium diolis | Bacillota | Clostridia |
| Clostridium diolis | Bacillota | Clostridia |
| Clostridium drakei | Bacillota | Clostridia |
| Clostridium drakei | Bacillota | Clostridia |
| Clostridium estertheticum | Bacillota | Clostridia |
| Clostridium estertheticum | Bacillota | Clostridia |
| Clostridium felsineum DSM 794 | Bacillota | Clostridia |
| Clostridium felsineum DSM 794 | Bacillota | Clostridia |
| Clostridium fermenticellae | Bacillota | Clostridia |
| Clostridium fermenticellae | Bacillota | Clostridia |
| Clostridium formicaceticum | Bacillota | Clostridia |
| Clostridium formicaceticum | Bacillota | Clostridia |
| Clostridium gasigenes | Bacillota | Clostridia |
| Clostridium gasigenes | Bacillota | Clostridia |
| Clostridium gelidum | Bacillota | Clostridia |
| Clostridium gelidum | Bacillota | Clostridia |
| Clostridium intestinale | Bacillota | Clostridia |
| Clostridium intestinale | Bacillota | Clostridia |
| Clostridium isatidis | Bacillota | Clostridia |
| Clostridium isatidis | Bacillota | Clostridia |
| Clostridium kluyveri NBRC 12016 | Bacillota | Clostridia |
| Clostridium kluyveri NBRC 12016 | Bacillota | Clostridia |
| Clostridium ljungdahlii DSM 13528 | Bacillota | Clostridia |

| | | |
|---|---|---|
| Clostridium ljungdahlii DSM 13528 | Bacillota | Clostridia |
| Clostridium manihotivorum | Bacillota | Clostridia |
| Clostridium manihotivorum | Bacillota | Clostridia |
| Clostridium novyi | Bacillota | Clostridia |
| Clostridium novyi | Bacillota | Clostridia |
| Clostridium pasteurianum DSM 525 = ATCC 6013 | Bacillota | Clostridia |
| Clostridium pasteurianum DSM 525 = ATCC 6013 | Bacillota | Clostridia |
| Clostridium perfringens | Bacillota | Clostridia |
| Clostridium perfringens | Bacillota | Clostridia |
| Clostridium saccharobutylicum DSM 13864 | Bacillota | Clostridia |
| Clostridium saccharobutylicum DSM 13864 | Bacillota | Clostridia |
| Clostridium saccharoperbutylacetonicum N1-4(HMT) | Bacillota | Clostridia |
| Clostridium saccharoperbutylacetonicum N1-4(HMT) | Bacillota | Clostridia |
| Clostridium scatologenes | Bacillota | Clostridia |
| Clostridium scatologenes | Bacillota | Clostridia |
| Clostridium septicum | Bacillota | Clostridia |
| Clostridium septicum | Bacillota | Clostridia |
| Clostridium sporogenes | Bacillota | Clostridia |
| Clostridium sporogenes | Bacillota | Clostridia |
| Clostridium taeniosporum | Bacillota | Clostridia |
| Clostridium taeniosporum | Bacillota | Clostridia |
| Clostridium tagluense | Bacillota | Clostridia |
| Clostridium tagluense | Bacillota | Clostridia |
| Clostridium tetani E88 | Bacillota | Clostridia |
| Clostridium tetani E88 | Bacillota | Clostridia |
| Clostridium thermarum | Bacillota | Clostridia |
| Clostridium thermarum | Bacillota | Clostridia |
| Clostridium thermosuccinogenes | Bacillota | Clostridia |
| Clostridium tyrobutyricum | Bacillota | Clostridia |
| Clostridium tyrobutyricum | Bacillota | Clostridia |
| Coprococcus comes ATCC 27758 | Bacillota | Clostridia |

| | | |
|---|---|---|
| Coprococcus eutactus | Bacillota | Clostridia |
| Crassaminicella indica | Bacillota | Clostridia |
| Crassaminicella indica | Bacillota | Clostridia |
| Crassaminicella profunda | Bacillota | Clostridia |
| Crassaminicella profunda | Bacillota | Clostridia |
| Crassaminicella thermophila | Bacillota | Clostridia |
| Crassaminicella thermophila | Bacillota | Clostridia |
| Defluviitalea saccharophila | Bacillota | Clostridia |
| Dehalobacter restrictus | Bacillota | Clostridia |
| Dehalobacterium formicoaceticum | Bacillota | Clostridia |
| Dehalobacterium formicoaceticum | Bacillota | Clostridia |
| Desulfitobacterium dehalogenans ATCC 51507 | Bacillota | Clostridia |
| Desulfitobacterium dichloroeliminans LMG P-21439 | Bacillota | Clostridia |
| Desulfitobacterium hafniense DCB-2 | Bacillota | Clostridia |
| Desulfitobacterium metallireducens DSM 15288 | Bacillota | Clostridia |
| Desulfofarcimen acetoxidans DSM 771 | Bacillota | Clostridia |
| Desulfofarcimen acetoxidans DSM 771 | Bacillota | Clostridia |
| Desulfofundulus kuznetsovii DSM 6115 | Bacillota | Clostridia |
| Desulfofundulus kuznetsovii DSM 6115 | Bacillota | Clostridia |
| Desulforamulus ferrireducens | Bacillota | Clostridia |
| Desulforamulus ferrireducens | Bacillota | Clostridia |
| Desulforamulus reducens MI-1 | Bacillota | Clostridia |
| Desulforamulus reducens MI-1 | Bacillota | Clostridia |
| Desulforamulus ruminis DSM 2154 | Bacillota | Clostridia |
| Desulforamulus ruminis DSM 2154 | Bacillota | Clostridia |
| Desulfoscipio gibsoniae DSM 7213 | Bacillota | Clostridia |
| Desulfoscipio gibsoniae DSM 7213 | Bacillota | Clostridia |
| Desulfosporosinus acidiphilus SJ4 | Bacillota | Clostridia |
| Desulfosporosinus meridiei DSM 13257 | Bacillota | Clostridia |
| Desulfosporosinus orientis DSM 765 | Bacillota | Clostridia |
| Desulfosporosinus paludis | Bacillota | Clostridia |

| | | |
|---|---|---|
| Desulfosporosinus youngiae DSM 17734 | Bacillota | Clostridia |
| Desulfotomaculum nigrificans CO-1-SRB | Bacillota | Clostridia |
| Desulfotomaculum nigrificans CO-1-SRB | Bacillota | Clostridia |
| Dorea formicigenerans | Bacillota | Clostridia |
| Dorea longicatena | Bacillota | Clostridia |
| Enterocloster clostridioformis | Bacillota | Clostridia |
| Eshraghiella crossota | Bacillota | Clostridia |
| Ethanoligenens harbinense YUAN-3 | Bacillota | Clostridia |
| Eubacterium callanderi | Bacillota | Clostridia |
| Eubacterium callanderi | Bacillota | Clostridia |
| Eubacterium limosum | Bacillota | Clostridia |
| Eubacterium limosum | Bacillota | Clostridia |
| Eubacterium maltosivorans | Bacillota | Clostridia |
| Eubacterium maltosivorans | Bacillota | Clostridia |
| Eubacterium ramulus | Bacillota | Clostridia |
| Faecalibacterium duncaniae | Bacillota | Clostridia |
| Faecalibacterium taiwanense | Bacillota | Clostridia |
| Faecalimonas umbilicata | Bacillota | Clostridia |
| Fastidiosipila sanguinis | Bacillota | Clostridia |
| Fenollaria massiliensis | Bacillota | Clostridia |
| Fenollaria sporofastidiosus | Bacillota | Clostridia |
| Filifactor alocis ATCC 35896 | Bacillota | Clostridia |
| Filifactor alocis ATCC 35896 | Bacillota | Clostridia |
| Flavonifractor plautii | Bacillota | Clostridia |
| Geosporobacter ferrireducens | Bacillota | Clostridia |
| Geosporobacter ferrireducens | Bacillota | Clostridia |
| Halanaerobium hydrogeniformans | Bacillota | Clostridia |
| Halanaerobium hydrogeniformans | Bacillota | Clostridia |
| Halobacteroides halobius DSM 5150 | Bacillota | Clostridia |
| Halobacteroides halobius DSM 5150 | Bacillota | Clostridia |
| Halothermothrix orenii H 168 | Bacillota | Clostridia |

| | | |
|---|---|---|
| Halothermothrix orenii H 168 | Bacillota | Clostridia |
| Hathewaya histolytica | Bacillota | Clostridia |
| Hathewaya histolytica | Bacillota | Clostridia |
| Helicovermis profundi | Bacillota | Clostridia |
| Helicovermis profundi | Bacillota | Clostridia |
| Heliomicrobium modesticaldum Ice1 | Bacillota | Clostridia |
| Heliorestis convoluta | Bacillota | Clostridia |
| Herbinix luporum | Bacillota | Clostridia |
| Hungatella hathewayi | Bacillota | Clostridia |
| Intestinibacter bartlettii | Bacillota | Clostridia |
| Intestinibacter bartlettii | Bacillota | Clostridia |
| Intestinimonas butyriciproducens | Bacillota | Clostridia |
| Iocasia fonsfrigidae | Bacillota | Clostridia |
| Iocasia fonsfrigidae | Bacillota | Clostridia |
| Kineothrix sedimenti | Bacillota | Clostridia |
| Koleobacter methoxysyntrophicus | Bacillota | Clostridia |
| Koleobacter methoxysyntrophicus | Bacillota | Clostridia |
| Lachnoanaerobaculum umeaense | Bacillota | Clostridia |
| Lachnoclostridium phytofermentans ISDg | Bacillota | Clostridia |
| Lachnospira eligens ATCC 27750 | Bacillota | Clostridia |
| Lacrimispora xylanolytica | Bacillota | Clostridia |
| Mageeibacillus indolicus UPII9-5 | Bacillota | Clostridia |
| Mahella australiensis 50-1 BON | Bacillota | Clostridia |
| Mahella australiensis 50-1 BON | Bacillota | Clostridia |
| Marvinbryantia formatexigens DSM 14469 | Bacillota | Clostridia |
| Mediterraneibacter glycyrrhizinilyticus | Bacillota | Clostridia |
| Mediterraneibacter gnavus ATCC 29149 | Bacillota | Clostridia |
| Metallumcola ferriviriculae | Bacillota | Clostridia |
| Metallumcola ferriviriculae | Bacillota | Clostridia |
| Mogibacterium diversum | Bacillota | Clostridia |
| Mogibacterium diversum | Bacillota | Clostridia |

| | | |
|---|---|---|
| Mogibacterium neglectum | Bacillota | Clostridia |
| Mogibacterium neglectum | Bacillota | Clostridia |
| Monoglobus pectinilyticus | Bacillota | Clostridia |
| Natranaerobius thermophilus JW/NM-WN-LF | Bacillota | Clostridia |
| Natranaerobius thermophilus JW/NM-WN-LF | Bacillota | Clostridia |
| Natranaerofaba carboxydovora | Bacillota | Clostridia |
| Natranaerofaba carboxydovora | Bacillota | Clostridia |
| Neomoorella carbonis | Bacillota | Clostridia |
| Neomoorella carbonis | Bacillota | Clostridia |
| Neomoorella glycerini | Bacillota | Clostridia |
| Neomoorella glycerini | Bacillota | Clostridia |
| Neomoorella humiferrea | Bacillota | Clostridia |
| Neomoorella humiferrea | Bacillota | Clostridia |
| Neomoorella thermoacetica | Bacillota | Clostridia |
| Novisyntrophococcus fermenticellae | Bacillota | Clostridia |
| Oscillibacter hominis | Bacillota | Clostridia |
| Owariibacterium komagatae | Bacillota | Clostridia |
| Paraclostridium bifermentans | Bacillota | Clostridia |
| Paraclostridium bifermentans | Bacillota | Clostridia |
| Paraclostridium dentum | Bacillota | Clostridia |
| Paraclostridium dentum | Bacillota | Clostridia |
| Paraclostridium ghonii | Bacillota | Clostridia |
| Paraclostridium ghonii | Bacillota | Clostridia |
| Paraclostridium sordellii | Bacillota | Clostridia |
| Paraclostridium sordellii | Bacillota | Clostridia |
| Peptacetobacter hiranonis | Bacillota | Clostridia |
| Peptacetobacter hiranonis | Bacillota | Clostridia |
| Peptoclostridium acidaminophilum DSM 3953 | Bacillota | Clostridia |
| Peptoclostridium acidaminophilum DSM 3953 | Bacillota | Clostridia |
| Peptostreptococcus equinus | Bacillota | Clostridia |
| Peptostreptococcus equinus | Bacillota | Clostridia |

| | | |
|---|---|---|
| Petrocella atlantisensis | Bacillota | Clostridia |
| Proteinivorax hydrogeniformans | Bacillota | Clostridia |
| Proteinivorax tanatarense | Bacillota | Clostridia |
| Pusillibacter faecalis | Bacillota | Clostridia |
| Qiania dongpingensis | Bacillota | Clostridia |
| Romboutsia ilealis | Bacillota | Clostridia |
| Romboutsia ilealis | Bacillota | Clostridia |
| Roseburia hominis A2-183 | Bacillota | Clostridia |
| Roseburia intestinalis L1-82 | Bacillota | Clostridia |
| Roseburia rectibacter | Bacillota | Clostridia |
| Ruminiclostridium cellulolyticum H10 | Bacillota | Clostridia |
| Ruminiclostridium herbifermentans | Bacillota | Clostridia |
| Ruminiclostridium papyrosolvens DSM 2782 | Bacillota | Clostridia |
| Ruminococcus albus 7 = DSM 20455 | Bacillota | Clostridia |
| Ruminococcus bovis | Bacillota | Clostridia |
| Ruminococcus gauvreauii | Bacillota | Clostridia |
| Sellimonas caecigallum | Bacillota | Clostridia |
| Sellimonas intestinalis | Bacillota | Clostridia |
| Serpentinicella alkaliphila | Bacillota | Clostridia |
| Serpentinicella alkaliphila | Bacillota | Clostridia |
| Simiaoa sunii | Bacillota | Clostridia |
| Solibaculum mannosilyticum | Bacillota | Clostridia |
| Subdoligranulum variabile | Bacillota | Clostridia |
| Symbiobacterium thermophilum IAM 14863 | Bacillota | Clostridia |
| Syntrophobotulus glycolicus DSM 8271 | Bacillota | Clostridia |
| Syntrophomonas wolfei subsp. wolfei str. Goettingen G311 | Bacillota | Clostridia |
| Syntrophomonas wolfei subsp. wolfei str. Goettingen G311 | Bacillota | Clostridia |
| Syntrophothermus lipocalidus DSM 12680 | Bacillota | Clostridia |
| Tepidibacter aestuarii | Bacillota | Clostridia |
| Tepidibacter aestuarii | Bacillota | Clostridia |
| Tepidibacter hydrothermalis | Bacillota | Clostridia |

| | | |
|---|---|---|
| Tepidibacter hydrothermalis | Bacillota | Clostridia |
| Terrisporobacter glycolicus ATCC 14880 = DSM 1288 | Bacillota | Clostridia |
| Terrisporobacter glycolicus ATCC 14880 = DSM 1288 | Bacillota | Clostridia |
| Terrisporobacter hibernicus | Bacillota | Clostridia |
| Terrisporobacter hibernicus | Bacillota | Clostridia |
| Terrisporobacter mayombei | Bacillota | Clostridia |
| Terrisporobacter mayombei | Bacillota | Clostridia |
| Terrisporobacter petrolearius | Bacillota | Clostridia |
| Terrisporobacter petrolearius | Bacillota | Clostridia |
| Thermacetogenium phaeum DSM 12270 | Bacillota | Clostridia |
| Thermanaeromonas toyohensis ToBE | Bacillota | Clostridia |
| Thermanaerosceptrum fracticalcis | Bacillota | Clostridia |
| Thermanaerosceptrum fracticalcis | Bacillota | Clostridia |
| Thermoanaerobacter brockii subsp. finnii Ako-1 | Bacillota | Clostridia |
| Thermoanaerobacter brockii subsp. finnii Ako-1 | Bacillota | Clostridia |
| Thermoanaerobacter ethanolicus JW 200 | Bacillota | Clostridia |
| Thermoanaerobacter ethanolicus JW 200 | Bacillota | Clostridia |
| Thermoanaerobacter italicus Ab9 | Bacillota | Clostridia |
| Thermoanaerobacter italicus Ab9 | Bacillota | Clostridia |
| Thermoanaerobacter kivui | Bacillota | Clostridia |
| Thermoanaerobacter kivui | Bacillota | Clostridia |
| Thermoanaerobacter mathranii subsp. mathranii str. A3 | Bacillota | Clostridia |
| Thermoanaerobacter pseudethanolicus ATCC 33223 | Bacillota | Clostridia |
| Thermoanaerobacter pseudethanolicus ATCC 33223 | Bacillota | Clostridia |
| Thermoanaerobacter wiegelii Rt8.B1 | Bacillota | Clostridia |
| Thermoanaerobacter wiegelii Rt8.B1 | Bacillota | Clostridia |
| Thermoanaerobacterium saccharolyticum JW/SL-YS485 | Bacillota | Clostridia |
| Thermoanaerobacterium saccharolyticum JW/SL-YS485 | Bacillota | Clostridia |
| Thermoanaerobacterium thermosaccharolyticum DSM 571 | Bacillota | Clostridia |
| Thermoanaerobacterium thermosaccharolyticum DSM 571 | Bacillota | Clostridia |
| Thermoanaerobacterium xylanolyticum LX-11 | Bacillota | Clostridia |

| | | |
|---|---|---|
| Thermoanaerobacterium xylanolyticum LX-11 | Bacillota | Clostridia |
| Thermocaproicibacter melissae | Bacillota | Clostridia |
| Thermoclostridium stercorarium subsp. stercorarium DSM 8532 | Bacillota | Clostridia |
| Thermosediminibacter oceani DSM 16646 | Bacillota | Clostridia |
| Thermosediminibacter oceani DSM 16646 | Bacillota | Clostridia |
| Vallitalea guaymasensis | Bacillota | Clostridia |
| Vallitalea pronyensis | Bacillota | Clostridia |
| Vescimonas coprocola | Bacillota | Clostridia |
| Vescimonas fastidiosa | Bacillota | Clostridia |
| Wansuia hejianensis | Bacillota | Clostridia |
| Wujia chipingensis | Bacillota | Clostridia |
| Zhaonella formicivorans | Bacillota | Clostridia |
| Zhaonella formicivorans | Bacillota | Clostridia |
| Coprothermobacter proteolyticus DSM 5265 | Coprothermobacterota | Coprothermobacteria |
| Coprothermobacter proteolyticus DSM 5265 | Coprothermobacterota | Coprothermobacteria |
| Adlercreutzia equolifaciens DSM 19450 | Actinomycetota | Coriobacteriia |
| Adlercreutzia hattorii | Actinomycetota | Coriobacteriia |
| Arabiibacter massiliensis | Actinomycetota | Coriobacteriia |
| Berryella intestinalis | Actinomycetota | Coriobacteriia |
| Berryella wangjianweii | Actinomycetota | Coriobacteriia |
| Collinsella aerofaciens ATCC 25986 | Actinomycetota | Coriobacteriia |
| Collinsella stercoris | Actinomycetota | Coriobacteriia |
| Coriobacterium glomerans PW2 | Actinomycetota | Coriobacteriia |
| Cryptobacterium curtum DSM 15641 | Actinomycetota | Coriobacteriia |
| Denitrobacterium detoxificans | Actinomycetota | Coriobacteriia |
| Eggerthella guodeyinii | Actinomycetota | Coriobacteriia |
| Eggerthella lenta | Actinomycetota | Coriobacteriia |
| Fannyhessea vaginae | Actinomycetota | Coriobacteriia |
| Lancefieldella parvula DSM 20469 | Actinomycetota | Coriobacteriia |
| Leptogranulimonas caecicola | Actinomycetota | Coriobacteriia |
| Olsenella timonensis | Actinomycetota | Coriobacteriia |

| | | |
|---|---|---|
| Olsenella uli DSM 7084 | Actinomycetota | Coriobacteriia |
| Parafannyhessea umbonata | Actinomycetota | Coriobacteriia |
| Parolsenella catena | Actinomycetota | Coriobacteriia |
| Parolsenella massiliensis | Actinomycetota | Coriobacteriia |
| Phoenicibacter congonensis | Actinomycetota | Coriobacteriia |
| Raoultibacter timonensis | Actinomycetota | Coriobacteriia |
| Slackia heliotrinireducens | Actinomycetota | Coriobacteriia |
| Thermophilibacter immobilis | Actinomycetota | Coriobacteriia |
| Xiamenia xianingshaonis | Actinomycetota | Coriobacteriia |
| Acaryochloris marina MBIC11017 | Cyanobacteriota | Cyanophyceae |
| Allocoleopsis franciscana PCC 7113 | Cyanobacteriota | Cyanophyceae |
| Anabaena cylindrica PCC 7122 | Cyanobacteriota | Cyanophyceae |
| Anabaenopsis elenkinii CCIBt3563 | Cyanobacteriota | Cyanophyceae |
| Anthocerotibacter panamensis C109 | Cyanobacteriota | Cyanophyceae |
| Brasilonema sennae CENA114 | Cyanobacteriota | Cyanophyceae |
| Candidatus Atelocyanobacterium thalassa isolate ALOHA | Cyanobacteriota | Cyanophyceae |
| Capilliphycus salinus ALCB114379 | Cyanobacteriota | Cyanophyceae |
| Chamaesiphon minutus PCC 6605 | Cyanobacteriota | Cyanophyceae |
| Chroococcidiopsis thermalis PCC 7203 | Cyanobacteriota | Cyanophyceae |
| Crinalium epipsammum PCC 9333 | Cyanobacteriota | Cyanophyceae |
| Crocosphaera subtropica ATCC 51142 | Cyanobacteriota | Cyanophyceae |
| Cyanobacterium aponinum PCC 10605 | Cyanobacteriota | Cyanophyceae |
| Cyanobium gracile PCC 6307 | Cyanobacteriota | Cyanophyceae |
| Cyanobium usitatum str. Tous | Cyanobacteriota | Cyanophyceae |
| Cylindrospermopsis curvispora GIHE-G1 | Cyanobacteriota | Cyanophyceae |
| Cylindrospermopsis raciborskii Cr2010 | Cyanobacteriota | Cyanophyceae |
| Cylindrospermum stagnale PCC 7417 | Cyanobacteriota | Cyanophyceae |
| Dactylococcopsis salina PCC 8305 | Cyanobacteriota | Cyanophyceae |
| Dolichospermum compactum NIES-806 | Cyanobacteriota | Cyanophyceae |
| Dolichospermum flos-aquae CCAP 1403/13F | Cyanobacteriota | Cyanophyceae |
| Dolichospermum heterosporum TAC447 | Cyanobacteriota | Cyanophyceae |

| | | |
|---|---|---|
| Euhalothece natronophila Z-M001 | Cyanobacteriota | Cyanophyceae |
| Geminocystis herdmanii PCC 6308 | Cyanobacteriota | Cyanophyceae |
| Gloeobacter kilaueensis JS1 | Cyanobacteriota | Cyanophyceae |
| Gloeobacter morelensis MG652769 | Cyanobacteriota | Cyanophyceae |
| Gloeobacter violaceus PCC 7421 | Cyanobacteriota | Cyanophyceae |
| Gloeocapsopsis dulcis | Cyanobacteriota | Cyanophyceae |
| Gloeomargarita lithophora Alchichica-D10 | Cyanobacteriota | Cyanophyceae |
| Gloeothece citriformis PCC 7424 | Cyanobacteriota | Cyanophyceae |
| Gloeothece verrucosa PCC 7822 | Cyanobacteriota | Cyanophyceae |
| Halomicronema hongdechloris C2206 | Cyanobacteriota | Cyanophyceae |
| Halotia branconii CENA392 | Cyanobacteriota | Cyanophyceae |
| Leptodesmis sichuanensis A121 | Cyanobacteriota | Cyanophyceae |
| Leptolyngbya boryana IAM M-101 | Cyanobacteriota | Cyanophyceae |
| Leptothermofonsia sichuanensis E412 | Cyanobacteriota | Cyanophyceae |
| Limnospira indica PCC 8005 | Cyanobacteriota | Cyanophyceae |
| Microcoleus anatoxicus | Cyanobacteriota | Cyanophyceae |
| Nodularia sphaerocarpa UHCC 0038 | Cyanobacteriota | Cyanophyceae |
| Nodularia spumigena UHCC 0039 | Cyanobacteriota | Cyanophyceae |
| 'Nostoc azollae' 0708 | Cyanobacteriota | Cyanophyceae |
| Nostoc edaphicum CCNP1411 | Cyanobacteriota | Cyanophyceae |
| Nostoc flagelliforme CCNUN1 | Cyanobacteriota | Cyanophyceae |
| Nostoc piscinale CENA21 | Cyanobacteriota | Cyanophyceae |
| Nostoc punctiforme PCC 73102 | Cyanobacteriota | Cyanophyceae |
| Nostoc sphaeroides | Cyanobacteriota | Cyanophyceae |
| Okeanomitos corallinicola TIOX110 | Cyanobacteriota | Cyanophyceae |
| Oscillatoria acuminata PCC 6304 | Cyanobacteriota | Cyanophyceae |
| Oscillatoria nigro-viridis PCC 7112 | Cyanobacteriota | Cyanophyceae |
| Oscillatoria nigro-viridis PCC 7112 | Cyanobacteriota | Cyanophyceae |
| Oxynema aestuarii AP17 | Cyanobacteriota | Cyanophyceae |
| Parasynechococcus marenigrum WH 8102 | Cyanobacteriota | Cyanophyceae |
| Parathermosynechococcus lividus PCC 6715 | Cyanobacteriota | Cyanophyceae |

| | | |
|---|---|---|
| Phormidium yuhuli AB48 | Cyanobacteriota | Cyanophyceae |
| Planktothricoides raciborskii GIHE-MW2 | Cyanobacteriota | Cyanophyceae |
| Planktothrix agardhii | Cyanobacteriota | Cyanophyceae |
| Planktothrix tepida | Cyanobacteriota | Cyanophyceae |
| Prochlorococcus marinus str. MIT 9515 | Cyanobacteriota | Cyanophyceae |
| Pseudanabaena galeata CCNP1313 | Cyanobacteriota | Cyanophyceae |
| Pseudanabaena mucicola str. Chao 1806 | Cyanobacteriota | Cyanophyceae |
| Richelia sinica FACHB-800 | Cyanobacteriota | Cyanophyceae |
| Rippkaea orientalis PCC 8801 | Cyanobacteriota | Cyanophyceae |
| Sphaerospermopsis torques-reginae ITEP-024 | Cyanobacteriota | Cyanophyceae |
| Stanieria cyanosphaera PCC 7437 | Cyanobacteriota | Cyanophyceae |
| Synechococcus elongatus PCC 6301 | Cyanobacteriota | Cyanophyceae |
| Thermocoleostomius sinensis A174 | Cyanobacteriota | Cyanophyceae |
| Thermoleptolyngbya oregonensis NK1-22 | Cyanobacteriota | Cyanophyceae |
| Thermoleptolyngbya sichuanensis A183 | Cyanobacteriota | Cyanophyceae |
| Thermosynechococcus sichuanensis E542 | Cyanobacteriota | Cyanophyceae |
| Thermosynechococcus vestitus BP-1 | Cyanobacteriota | Cyanophyceae |
| Trichocoleus desertorum NBK24 | Cyanobacteriota | Cyanophyceae |
| Trichodesmium erythraeum IMS101 | Cyanobacteriota | Cyanophyceae |
| Trichormus variabilis 0441 | Cyanobacteriota | Cyanophyceae |
| Trichothermofontia sichuanensis B231 | Cyanobacteriota | Cyanophyceae |
| Adhaeribacter radiodurans | Bacteroidota | Cytophagia |
| Adhaeribacter swui | Bacteroidota | Cytophagia |
| Algoriphagus halophytocola | Bacteroidota | Cytophagia |
| Algoriphagus machipongonensis | Bacteroidota | Cytophagia |
| Algoriphagus sanaruensis | Bacteroidota | Cytophagia |
| Aquiflexum balticum DSM 16537 | Bacteroidota | Cytophagia |
| Aquirufa antheringensis | Bacteroidota | Cytophagia |
| Aquirufa lenticrescens | Bacteroidota | Cytophagia |
| Aquirufa nivalisilvae | Bacteroidota | Cytophagia |
| Arcticibacterium luteifluviistationis | Bacteroidota | Cytophagia |

| | | |
|---|---|---|
| Aureibacter tunicatorum | Bacteroidota | Cytophagia |
| Belliella baltica DSM 15883 | Bacteroidota | Cytophagia |
| Bernardetia litoralis DSM 6794 | Bacteroidota | Cytophagia |
| Candidatus Amoebophilus asiaticus 5a2 | Bacteroidota | Cytophagia |
| Candidatus Cardinium hertigii | Bacteroidota | Cytophagia |
| Cardinium endosymbiont cEper1 of Encarsia pergandiella | Bacteroidota | Cytophagia |
| Cardinium endosymbiont of Nabis limbatus | Bacteroidota | Cytophagia |
| Cardinium endosymbiont of Oedothorax gibbosus | Bacteroidota | Cytophagia |
| Cardinium endosymbiont of Philonthus spinipes | Bacteroidota | Cytophagia |
| Cardinium endosymbiont of Sogatella furcifera | Bacteroidota | Cytophagia |
| Cardinium endosymbiont of Tipula unca | Bacteroidota | Cytophagia |
| Catalinimonas niigatensis | Bacteroidota | Cytophagia |
| Chondrinema litorale | Bacteroidota | Cytophagia |
| Chryseolinea soli | Bacteroidota | Cytophagia |
| Cyclobacterium amurskyense | Bacteroidota | Cytophagia |
| Cyclobacterium marinum DSM 745 | Bacteroidota | Cytophagia |
| Cytophaga hutchinsonii ATCC 33406 | Bacteroidota | Cytophagia |
| Dyadobacter chenhuakuii | Bacteroidota | Cytophagia |
| Dyadobacter chenwenxiniae | Bacteroidota | Cytophagia |
| Dyadobacter fanqingshengii | Bacteroidota | Cytophagia |
| Dyadobacter fermentans DSM 18053 | Bacteroidota | Cytophagia |
| Dyadobacter pollutisoli | Bacteroidota | Cytophagia |
| Dyadobacter sandarakinus | Bacteroidota | Cytophagia |
| Echinicola marina | Bacteroidota | Cytophagia |
| Echinicola rosea | Bacteroidota | Cytophagia |
| Echinicola soli | Bacteroidota | Cytophagia |
| Echinicola strongylocentroti | Bacteroidota | Cytophagia |
| Echinicola vietnamensis DSM 17526 | Bacteroidota | Cytophagia |
| Emticicia oligotrophica DSM 17448 | Bacteroidota | Cytophagia |
| Fibrella aestuarina BUZ 2 | Bacteroidota | Cytophagia |
| Flammeovirga kamogawensis | Bacteroidota | Cytophagia |

| | | |
|---|---|---|
| Flammeovirga pectinis | Bacteroidota | Cytophagia |
| Flammeovirga yaeyamensis | Bacteroidota | Cytophagia |
| Fulvitalea axinellae | Bacteroidota | Cytophagia |
| Fulvivirga ligni | Bacteroidota | Cytophagia |
| Fulvivirga lutea | Bacteroidota | Cytophagia |
| Fulvivirga maritima | Bacteroidota | Cytophagia |
| Fulvivirga ulvae | Bacteroidota | Cytophagia |
| Hymenobacter aerilatus | Bacteroidota | Cytophagia |
| Hymenobacter baengnokdamensis | Bacteroidota | Cytophagia |
| Hymenobacter canadensis | Bacteroidota | Cytophagia |
| Hymenobacter cellulosivorans | Bacteroidota | Cytophagia |
| Hymenobacter jejuensis | Bacteroidota | Cytophagia |
| Hymenobacter monticola | Bacteroidota | Cytophagia |
| Hymenobacter nivis | Bacteroidota | Cytophagia |
| Hymenobacter oligotrophus | Bacteroidota | Cytophagia |
| Hymenobacter radiodurans | Bacteroidota | Cytophagia |
| Hymenobacter sedentarius | Bacteroidota | Cytophagia |
| Hymenobacter sediminicola | Bacteroidota | Cytophagia |
| Hymenobacter sublimis | Bacteroidota | Cytophagia |
| Hymenobacter swuensis DY53 | Bacteroidota | Cytophagia |
| Hymenobacter taeanensis | Bacteroidota | Cytophagia |
| Hymenobacter tibetensis | Bacteroidota | Cytophagia |
| Hymenobacter volaticus | Bacteroidota | Cytophagia |
| Hymenobacter yonginensis | Bacteroidota | Cytophagia |
| Imperialibacter roseus | Bacteroidota | Cytophagia |
| Larkinella insperata | Bacteroidota | Cytophagia |
| Leadbetterella byssophila DSM 17132 | Bacteroidota | Cytophagia |
| Mangrovivirga cuniculi | Bacteroidota | Cytophagia |
| Marinilongibacter aquaticus | Bacteroidota | Cytophagia |
| Marivirga arenosa | Bacteroidota | Cytophagia |
| Marivirga harenae | Bacteroidota | Cytophagia |

| | | |
|---|---|---|
| Marivirga salinarum | Bacteroidota | Cytophagia |
| Marivirga tractuosa DSM 4126 | Bacteroidota | Cytophagia |
| Mongoliitalea daihaiensis | Bacteroidota | Cytophagia |
| Nibribacter ruber | Bacteroidota | Cytophagia |
| Persicobacter psychrovividus | Bacteroidota | Cytophagia |
| Pontibacter actiniarum | Bacteroidota | Cytophagia |
| Pontibacter akesuensis | Bacteroidota | Cytophagia |
| Pontibacter kalidii | Bacteroidota | Cytophagia |
| Pontibacter korlensis | Bacteroidota | Cytophagia |
| Pontibacter pudoricolor | Bacteroidota | Cytophagia |
| Pontibacter russatus | Bacteroidota | Cytophagia |
| Porifericola rhodea | Bacteroidota | Cytophagia |
| Reichenbachiella agarivorans | Bacteroidota | Cytophagia |
| Reichenbachiella carrageenanivorans | Bacteroidota | Cytophagia |
| Rhodocytophaga rosea | Bacteroidota | Cytophagia |
| Roseivirga spongicola | Bacteroidota | Cytophagia |
| Rufibacter radiotolerans | Bacteroidota | Cytophagia |
| Rufibacter tibetensis | Bacteroidota | Cytophagia |
| Runella rosea | Bacteroidota | Cytophagia |
| Runella slithyformis DSM 19594 | Bacteroidota | Cytophagia |
| Spirosoma aerolatum | Bacteroidota | Cytophagia |
| Spirosoma aureum | Bacteroidota | Cytophagia |
| Spirosoma endbachense | Bacteroidota | Cytophagia |
| Spirosoma foliorum | Bacteroidota | Cytophagia |
| Spirosoma linguale DSM 74 | Bacteroidota | Cytophagia |
| Spirosoma montaniterrae | Bacteroidota | Cytophagia |
| Spirosoma oryzicola | Bacteroidota | Cytophagia |
| Spirosoma pollinicola | Bacteroidota | Cytophagia |
| Spirosoma radiotolerans | Bacteroidota | Cytophagia |
| Spirosoma rhododendri | Bacteroidota | Cytophagia |
| Spirosoma rigui | Bacteroidota | Cytophagia |

| | | |
|---|---|---|
| Spirosoma taeanense | Bacteroidota | Cytophagia |
| Tellurirhabdus bombi | Bacteroidota | Cytophagia |
| Tellurirhabdus rosea | Bacteroidota | Cytophagia |
| Tunicatimonas pelagia | Bacteroidota | Cytophagia |
| Calditerrivibrio nitroreducens DSM 19672 | Deferribacterota | Deferribacteres |
| Calditerrivibrio nitroreducens DSM 19672 | Deferribacterota | Deferribacteres |
| Deferribacter desulfuricans SSM1 | Deferribacterota | Deferribacteres |
| Deferribacter desulfuricans SSM1 | Deferribacterota | Deferribacteres |
| Denitrovibrio acetiphilus DSM 12809 | Deferribacterota | Deferribacteres |
| Denitrovibrio acetiphilus DSM 12809 | Deferribacterota | Deferribacteres |
| Flexistipes sinusarabici DSM 4947 | Deferribacterota | Deferribacteres |
| Flexistipes sinusarabici DSM 4947 | Deferribacterota | Deferribacteres |
| Geovibrio thiophilus | Deferribacterota | Deferribacteres |
| Geovibrio thiophilus | Deferribacterota | Deferribacteres |
| Mucispirillum schaedleri ASF457 | Deferribacterota | Deferribacteres |
| Candidatus Dehalogenimonas loeffleri | Chloroflexota | Dehalococcoidia |
| Candidatus Lucifugimonas marina | Chloroflexota | Dehalococcoidia |
| Dehalococcoides mccartyi 195 | Chloroflexota | Dehalococcoidia |
| Dehalogenimonas etheniformans | Chloroflexota | Dehalococcoidia |
| Dehalogenimonas formicexedens | Chloroflexota | Dehalococcoidia |
| Allomeiothermus silvanus DSM 9946 | Deinococcota | Deinococci |
| Allomeiothermus silvanus DSM 9946 | Deinococcota | Deinococci |
| Deinococcus aetherius | Deinococcota | Deinococci |
| Deinococcus aetherius | Deinococcota | Deinococci |
| Deinococcus altitudinis | Deinococcota | Deinococci |
| Deinococcus altitudinis | Deinococcota | Deinococci |
| Deinococcus aquaticus | Deinococcota | Deinococci |
| Deinococcus aquaticus | Deinococcota | Deinococci |
| Deinococcus deserti VCD115 | Deinococcota | Deinococci |
| Deinococcus deserti VCD115 | Deinococcota | Deinococci |
| Deinococcus ficus | Deinococcota | Deinococci |

| | | |
|---|---|---|
| Deinococcus ficus | Deinococcota | Deinococci |
| Deinococcus geothermalis DSM 11300 | Deinococcota | Deinococci |
| Deinococcus geothermalis DSM 11300 | Deinococcota | Deinococci |
| Deinococcus gobiensis I-0 | Deinococcota | Deinococci |
| Deinococcus gobiensis I-0 | Deinococcota | Deinococci |
| Deinococcus irradiatisoli | Deinococcota | Deinococci |
| Deinococcus irradiatisoli | Deinococcota | Deinococci |
| Deinococcus maricopensis DSM 21211 | Deinococcota | Deinococci |
| Deinococcus maricopensis DSM 21211 | Deinococcota | Deinococci |
| Deinococcus metallilatus | Deinococcota | Deinococci |
| Deinococcus metallilatus | Deinococcota | Deinococci |
| Deinococcus peraridilitoris DSM 19664 | Deinococcota | Deinococci |
| Deinococcus peraridilitoris DSM 19664 | Deinococcota | Deinococci |
| Deinococcus proteolyticus MRP | Deinococcota | Deinococci |
| Deinococcus proteolyticus MRP | Deinococcota | Deinococci |
| Deinococcus psychrotolerans | Deinococcota | Deinococci |
| Deinococcus psychrotolerans | Deinococcota | Deinococci |
| Deinococcus puniceus | Deinococcota | Deinococci |
| Deinococcus puniceus | Deinococcota | Deinococci |
| Deinococcus radiomollis | Deinococcota | Deinococci |
| Deinococcus radiomollis | Deinococcota | Deinococci |
| Deinococcus radiopugnans | Deinococcota | Deinococci |
| Deinococcus radiopugnans | Deinococcota | Deinococci |
| Deinococcus rubellus | Deinococcota | Deinococci |
| Deinococcus rubellus | Deinococcota | Deinococci |
| Deinococcus sonorensis KR-87 | Deinococcota | Deinococci |
| Deinococcus sonorensis KR-87 | Deinococcota | Deinococci |
| Deinococcus taeanensis | Deinococcota | Deinococci |
| Deinococcus taeanensis | Deinococcota | Deinococci |
| Deinococcus wulumuqiensis R12 | Deinococcota | Deinococci |
| Deinococcus wulumuqiensis R12 | Deinococcota | Deinococci |

| | | |
|---|---|---|
| Deinococcus yunweiensis | Deinococcota | Deinococci |
| Deinococcus yunweiensis | Deinococcota | Deinococci |
| Marinithermus hydrothermalis DSM 14884 | Deinococcota | Deinococci |
| Marinithermus hydrothermalis DSM 14884 | Deinococcota | Deinococci |
| Meiothermus ruber DSM 1279 | Deinococcota | Deinococci |
| Meiothermus ruber DSM 1279 | Deinococcota | Deinococci |
| Meiothermus taiwanensis WR-220 | Deinococcota | Deinococci |
| Meiothermus taiwanensis WR-220 | Deinococcota | Deinococci |
| Thermus antranikianii | Deinococcota | Deinococci |
| Thermus antranikianii | Deinococcota | Deinococci |
| Thermus aquaticus Y51MC23 | Deinococcota | Deinococci |
| Thermus aquaticus Y51MC23 | Deinococcota | Deinococci |
| Thermus brockianus | Deinococcota | Deinococci |
| Thermus brockianus | Deinococcota | Deinococci |
| Thermus caldilimi | Deinococcota | Deinococci |
| Thermus caldilimi | Deinococcota | Deinococci |
| Thermus oshimai JL-2 | Deinococcota | Deinococci |
| Thermus oshimai JL-2 | Deinococcota | Deinococci |
| Thermus parvatiensis | Deinococcota | Deinococci |
| Thermus parvatiensis | Deinococcota | Deinococci |
| Thermus thermophilus HB8 | Deinococcota | Deinococci |
| Thermus thermophilus HB8 | Deinococcota | Deinococci |
| Truepera radiovictrix DSM 17093 | Deinococcota | Deinococci |
| Truepera radiovictrix DSM 17093 | Deinococcota | Deinococci |
| Bradymonas sediminis | | Deltaproteobacteria |
| Bradymonas sediminis | | Deltaproteobacteria |
| Dissulfurimicrobium hydrothermale | | Deltaproteobacteria |
| Dissulfurimicrobium hydrothermale | | Deltaproteobacteria |
| Microvenator marinus | | Deltaproteobacteria |
| Persicimonas caeni | | Deltaproteobacteria |
| Persicimonas caeni | | Deltaproteobacteria |

| | | |
|---|---|---|
| Desulfarculus baarsii DSM 2075 | Thermodesulfobacteriota | Desulfarculia |
| Desulfarculus baarsii DSM 2075 | Thermodesulfobacteriota | Desulfarculia |
| Desulfoferula mesophila | Thermodesulfobacteriota | Desulfarculia |
| Desulfoferula mesophila | Thermodesulfobacteriota | Desulfarculia |
| Desulfobacca acetoxidans DSM 11109 | Thermodesulfobacteriota | Desulfobaccia |
| Desulfobacca acetoxidans DSM 11109 | Thermodesulfobacteriota | Desulfobaccia |
| Desulfatibacillum aliphaticivorans | Thermodesulfobacteriota | Desulfobacteria |
| Desulfatibacillum aliphaticivorans | Thermodesulfobacteriota | Desulfobacteria |
| Desulfobacter hydrogenophilus | Thermodesulfobacteriota | Desulfobacteria |
| Desulfobacter postgatei 2ac9 | Thermodesulfobacteriota | Desulfobacteria |
| Desulfobacula toluolica Tol2 | Thermodesulfobacteriota | Desulfobacteria |
| Desulfobacula toluolica Tol2 | Thermodesulfobacteriota | Desulfobacteria |
| Desulfococcus multivorans | Thermodesulfobacteriota | Desulfobacteria |
| Desulfococcus multivorans | Thermodesulfobacteriota | Desulfobacteria |
| Desulfoluna limicola | Thermodesulfobacteriota | Desulfobacteria |
| Desulfoluna limicola | Thermodesulfobacteriota | Desulfobacteria |
| Desulfonema limicola | Thermodesulfobacteriota | Desulfobacteria |
| Desulfonema limicola | Thermodesulfobacteriota | Desulfobacteria |
| Desulfonema magnum | Thermodesulfobacteriota | Desulfobacteria |
| Desulforapulum autotrophicum HRM2 | Thermodesulfobacteriota | Desulfobacteria |
| Desulforapulum autotrophicum HRM2 | Thermodesulfobacteriota | Desulfobacteria |
| Desulfosarcina alkanivorans | Thermodesulfobacteriota | Desulfobacteria |
| Desulfosarcina alkanivorans | Thermodesulfobacteriota | Desulfobacteria |
| Desulfosarcina ovata subsp. sediminis | Thermodesulfobacteriota | Desulfobacteria |
| Desulfosarcina ovata subsp. sediminis | Thermodesulfobacteriota | Desulfobacteria |
| Desulfosarcina variabilis str. Montpellier | Thermodesulfobacteriota | Desulfobacteria |
| Desulfosarcina variabilis str. Montpellier | Thermodesulfobacteriota | Desulfobacteria |
| Desulfosarcina widdelii | Thermodesulfobacteriota | Desulfobacteria |
| Desulfosarcina widdelii | Thermodesulfobacteriota | Desulfobacteria |
| Desulfosudis oleivorans Hxd3 | Thermodesulfobacteriota | Desulfobacteria |
| Desulfosudis oleivorans Hxd3 | Thermodesulfobacteriota | Desulfobacteria |

| | | |
|---|---|---|
| Desulfobulbus oligotrophicus | Thermodesulfobacteriota | Desulfobulbia |
| Desulfobulbus oligotrophicus | Thermodesulfobacteriota | Desulfobulbia |
| Desulfobulbus oralis | Thermodesulfobacteriota | Desulfobulbia |
| Desulfobulbus oralis | Thermodesulfobacteriota | Desulfobulbia |
| Desulfobulbus propionicus DSM 2032 | Thermodesulfobacteriota | Desulfobulbia |
| Desulfobulbus propionicus DSM 2032 | Thermodesulfobacteriota | Desulfobulbia |
| Desulfocapsa sulfexigens DSM 10523 | Thermodesulfobacteriota | Desulfobulbia |
| Desulfocapsa sulfexigens DSM 10523 | Thermodesulfobacteriota | Desulfobulbia |
| Desulfofustis limnaeus | Thermodesulfobacteriota | Desulfobulbia |
| Desulfofustis limnaeus | Thermodesulfobacteriota | Desulfobulbia |
| Desulfolithobacter dissulfuricans | Thermodesulfobacteriota | Desulfobulbia |
| Desulfolithobacter dissulfuricans | Thermodesulfobacteriota | Desulfobulbia |
| Desulfomarina profundi | Thermodesulfobacteriota | Desulfobulbia |
| Desulfomarina profundi | Thermodesulfobacteriota | Desulfobulbia |
| Desulfosediminicola flagellatus | Thermodesulfobacteriota | Desulfobulbia |
| Desulfosediminicola flagellatus | Thermodesulfobacteriota | Desulfobulbia |
| Desulfosediminicola ganghwensis | Thermodesulfobacteriota | Desulfobulbia |
| Desulfosediminicola ganghwensis | Thermodesulfobacteriota | Desulfobulbia |
| Desulfotalea psychrophila LSv54 | Thermodesulfobacteriota | Desulfobulbia |
| Desulfotalea psychrophila LSv54 | Thermodesulfobacteriota | Desulfobulbia |
| Desulfurivibrio alkaliphilus AHT 2 | Thermodesulfobacteriota | Desulfobulbia |
| Desulfurivibrio alkaliphilus AHT 2 | Thermodesulfobacteriota | Desulfobulbia |
| Desulfomonile tiedjei DSM 6799 | Thermodesulfobacteriota | Desulfomonilia |
| Desulfomonile tiedjei DSM 6799 | Thermodesulfobacteriota | Desulfomonilia |
| Candidatus Desulfovibrio trichonymphae | Thermodesulfobacteriota | Desulfovibrionia |
| Desulfocurvibacter africanus subsp. africanus str. Walvis Bay | Thermodesulfobacteriota | Desulfovibrionia |
| Desulfocurvibacter africanus subsp. africanus str. Walvis Bay | Thermodesulfobacteriota | Desulfovibrionia |
| Desulfohalobium retbaense DSM 5692 | Thermodesulfobacteriota | Desulfovibrionia |
| Desulfohalobium retbaense DSM 5692 | Thermodesulfobacteriota | Desulfovibrionia |
| Desulfolutivibrio sulfodismutans DSM 3696 | Thermodesulfobacteriota | Desulfovibrionia |
| Desulfolutivibrio sulfodismutans DSM 3696 | Thermodesulfobacteriota | Desulfovibrionia |

| | | |
|---|---|---|
| Desulfolutivibrio sulfoxidireducens | Thermodesulfobacteriota | Desulfovibrionia |
| Desulfolutivibrio sulfoxidireducens | Thermodesulfobacteriota | Desulfovibrionia |
| Desulfomicrobium baculatum DSM 4028 | Thermodesulfobacteriota | Desulfovibrionia |
| Desulfomicrobium baculatum DSM 4028 | Thermodesulfobacteriota | Desulfovibrionia |
| Desulfomicrobium orale DSM 12838 | Thermodesulfobacteriota | Desulfovibrionia |
| Desulfomicrobium orale DSM 12838 | Thermodesulfobacteriota | Desulfovibrionia |
| Desulfovibrio aminophilus | Thermodesulfobacteriota | Desulfovibrionia |
| Desulfovibrio desulfuricans | Thermodesulfobacteriota | Desulfovibrionia |
| Desulfovibrio fairfieldensis | Thermodesulfobacteriota | Desulfovibrionia |
| Desulfovibrio ferrophilus | Thermodesulfobacteriota | Desulfovibrionia |
| Desulfovibrio ferrophilus | Thermodesulfobacteriota | Desulfovibrionia |
| Desulfovibrio mangrovi | Thermodesulfobacteriota | Desulfovibrionia |
| Desulfovibrio subterraneus | Thermodesulfobacteriota | Desulfovibrionia |
| Lawsonia intracellularis N343 | Thermodesulfobacteriota | Desulfovibrionia |
| Maridesulfovibrio hydrothermalis AM13 = DSM 14728 | Thermodesulfobacteriota | Desulfovibrionia |
| Maridesulfovibrio salexigens DSM 2638 | Thermodesulfobacteriota | Desulfovibrionia |
| Megalodesulfovibrio gigas DSM 1382 = ATCC 19364 | Thermodesulfobacteriota | Desulfovibrionia |
| Nitratidesulfovibrio vulgaris DP4 | Thermodesulfobacteriota | Desulfovibrionia |
| Oceanidesulfovibrio marinus | Thermodesulfobacteriota | Desulfovibrionia |
| Oceanidesulfovibrio marinus | Thermodesulfobacteriota | Desulfovibrionia |
| Oleidesulfovibrio alaskensis G20 | Thermodesulfobacteriota | Desulfovibrionia |
| Pseudodesulfovibrio aespoeensis Aspo-2 | Thermodesulfobacteriota | Desulfovibrionia |
| Pseudodesulfovibrio cashew | Thermodesulfobacteriota | Desulfovibrionia |
| Pseudodesulfovibrio mercurii | Thermodesulfobacteriota | Desulfovibrionia |
| Pseudodesulfovibrio methanolicus | Thermodesulfobacteriota | Desulfovibrionia |
| Pseudodesulfovibrio nedwellii | Thermodesulfobacteriota | Desulfovibrionia |
| Pseudodesulfovibrio piezophilus C1TLV30 | Thermodesulfobacteriota | Desulfovibrionia |
| Pseudodesulfovibrio portus | Thermodesulfobacteriota | Desulfovibrionia |
| Pseudodesulfovibrio profundus | Thermodesulfobacteriota | Desulfovibrionia |
| Pseudodesulfovibrio sediminis | Thermodesulfobacteriota | Desulfovibrionia |
| Pseudodesulfovibrio thermohalotolerans | Thermodesulfobacteriota | Desulfovibrionia |

| | | |
|---|---|---|
| Pseudodesulfovibrio tunisiensis | Thermodesulfobacteriota | Desulfovibrionia |
| Salidesulfovibrio onnuriiensis | Thermodesulfobacteriota | Desulfovibrionia |
| Solidesulfovibrio carbinolicus | Thermodesulfobacteriota | Desulfovibrionia |
| Solidesulfovibrio carbinolicus | Thermodesulfobacteriota | Desulfovibrionia |
| Solidesulfovibrio carbinoliphilus subsp. oakridgensis | Thermodesulfobacteriota | Desulfovibrionia |
| Solidesulfovibrio carbinoliphilus subsp. oakridgensis | Thermodesulfobacteriota | Desulfovibrionia |
| Solidesulfovibrio magneticus RS-1 | Thermodesulfobacteriota | Desulfovibrionia |
| Solidesulfovibrio magneticus RS-1 | Thermodesulfobacteriota | Desulfovibrionia |
| Taurinivorans muris | Thermodesulfobacteriota | Desulfovibrionia |
| Hippea maritima DSM 10411 | Campylobacterota | Desulfurellia |
| Hippea maritima DSM 10411 | Campylobacterota | Desulfurellia |
| Citrifermentans bemidjiense Bem | Thermodesulfobacteriota | Desulfuromonadia |
| Citrifermentans bemidjiense Bem | Thermodesulfobacteriota | Desulfuromonadia |
| Citrifermentans bremense | Thermodesulfobacteriota | Desulfuromonadia |
| Citrifermentans bremense | Thermodesulfobacteriota | Desulfuromonadia |
| Desulfuromonas soudanensis | Thermodesulfobacteriota | Desulfuromonadia |
| Desulfuromonas soudanensis | Thermodesulfobacteriota | Desulfuromonadia |
| Desulfuromonas thiophila | Thermodesulfobacteriota | Desulfuromonadia |
| Desulfuromonas thiophila | Thermodesulfobacteriota | Desulfuromonadia |
| Desulfuromonas versatilis | Thermodesulfobacteriota | Desulfuromonadia |
| Desulfuromonas versatilis | Thermodesulfobacteriota | Desulfuromonadia |
| Geoalkalibacter halelectricus | Thermodesulfobacteriota | Desulfuromonadia |
| Geoalkalibacter halelectricus | Thermodesulfobacteriota | Desulfuromonadia |
| Geoalkalibacter subterraneus | Thermodesulfobacteriota | Desulfuromonadia |
| Geoalkalibacter subterraneus | Thermodesulfobacteriota | Desulfuromonadia |
| Geobacter benzoatilyticus | Thermodesulfobacteriota | Desulfuromonadia |
| Geobacter benzoatilyticus | Thermodesulfobacteriota | Desulfuromonadia |
| Geobacter metallireducens GS-15 | Thermodesulfobacteriota | Desulfuromonadia |
| Geobacter metallireducens GS-15 | Thermodesulfobacteriota | Desulfuromonadia |
| Geobacter pickeringii | Thermodesulfobacteriota | Desulfuromonadia |
| Geobacter pickeringii | Thermodesulfobacteriota | Desulfuromonadia |

| | | |
|---|---|---|
| Geobacter sulfurreducens PCA | Thermodesulfobacteriota | Desulfuromonadia |
| Geobacter sulfurreducens PCA | Thermodesulfobacteriota | Desulfuromonadia |
| Geomonas nitrogeniifigens | Thermodesulfobacteriota | Desulfuromonadia |
| Geomonas nitrogeniifigens | Thermodesulfobacteriota | Desulfuromonadia |
| Geomonas oryzisoli | Thermodesulfobacteriota | Desulfuromonadia |
| Geomonas oryzisoli | Thermodesulfobacteriota | Desulfuromonadia |
| Geomonas paludis | Thermodesulfobacteriota | Desulfuromonadia |
| Geomonas paludis | Thermodesulfobacteriota | Desulfuromonadia |
| Geomonas subterranea | Thermodesulfobacteriota | Desulfuromonadia |
| Geomonas subterranea | Thermodesulfobacteriota | Desulfuromonadia |
| Geotalea daltonii FRC-32 | Thermodesulfobacteriota | Desulfuromonadia |
| Geotalea daltonii FRC-32 | Thermodesulfobacteriota | Desulfuromonadia |
| Geotalea uraniireducens Rf4 | Thermodesulfobacteriota | Desulfuromonadia |
| Geotalea uraniireducens Rf4 | Thermodesulfobacteriota | Desulfuromonadia |
| Pelobacter propionicus DSM 2379 | Thermodesulfobacteriota | Desulfuromonadia |
| Pelobacter propionicus DSM 2379 | Thermodesulfobacteriota | Desulfuromonadia |
| Syntrophotalea acetylenica | Thermodesulfobacteriota | Desulfuromonadia |
| Syntrophotalea acetylenica | Thermodesulfobacteriota | Desulfuromonadia |
| Syntrophotalea acetylenivorans | Thermodesulfobacteriota | Desulfuromonadia |
| Syntrophotalea acetylenivorans | Thermodesulfobacteriota | Desulfuromonadia |
| Syntrophotalea carbinolica DSM 2380 | Thermodesulfobacteriota | Desulfuromonadia |
| Syntrophotalea carbinolica DSM 2380 | Thermodesulfobacteriota | Desulfuromonadia |
| Trichlorobacter ammonificans | Thermodesulfobacteriota | Desulfuromonadia |
| Trichlorobacter ammonificans | Thermodesulfobacteriota | Desulfuromonadia |
| Trichlorobacter lovleyi SZ | Thermodesulfobacteriota | Desulfuromonadia |
| Trichlorobacter lovleyi SZ | Thermodesulfobacteriota | Desulfuromonadia |
| Candidatus Contubernalis alkaliaceticus | Bacillota | Dethiobacteria |
| Dictyoglomus thermophilum H-6-12 | Dictyoglomota | Dictyoglomia |
| Dictyoglomus thermophilum H-6-12 | Dictyoglomota | Dictyoglomia |
| Dictyoglomus turgidum DSM 6724 | Dictyoglomota | Dictyoglomia |
| Dictyoglomus turgidum DSM 6724 | Dictyoglomota | Dictyoglomia |

| | | |
|---|---|---|
| Elusimicrobium minutum Pei191 | Elusimicrobiota | Elusimicrobia |
| Elusimicrobium minutum Pei191 | Elusimicrobiota | Elusimicrobia |
| Candidatus Endomicrobiellum trichonymphae | Elusimicrobiota | Endomicrobiia |
| Candidatus Endomicrobiellum trichonymphae | Elusimicrobiota | Endomicrobiia |
| Endomicrobium proavitum | Elusimicrobiota | Endomicrobiia |
| Endomicrobium proavitum | Elusimicrobiota | Endomicrobiia |
| Aliarcobacter butzleri RM4018 | Campylobacterota | Epsilonproteobacteria |
| Aliarcobacter cibarius | Campylobacterota | Epsilonproteobacteria |
| Aliarcobacter cryaerophilus ATCC 43158 | Campylobacterota | Epsilonproteobacteria |
| Aliarcobacter faecis | Campylobacterota | Epsilonproteobacteria |
| Aliarcobacter lanthieri | Campylobacterota | Epsilonproteobacteria |
| Aliarcobacter skirrowii CCUG 10374 | Campylobacterota | Epsilonproteobacteria |
| Aliarcobacter thereius LMG 24486 | Campylobacterota | Epsilonproteobacteria |
| Aliarcobacter trophiarum LMG 25534 | Campylobacterota | Epsilonproteobacteria |
| Aliarcobacter vitoriensis | Campylobacterota | Epsilonproteobacteria |
| Arcobacter acticola | Campylobacterota | Epsilonproteobacteria |
| Arcobacter aquimarinus | Campylobacterota | Epsilonproteobacteria |
| Arcobacter cloacae | Campylobacterota | Epsilonproteobacteria |
| Arcobacter defluvii | Campylobacterota | Epsilonproteobacteria |
| Arcobacter ellisii | Campylobacterota | Epsilonproteobacteria |
| Arcobacter nitrofigilis DSM 7299 | Campylobacterota | Epsilonproteobacteria |
| Arcobacter peruensis | Campylobacterota | Epsilonproteobacteria |
| Arcobacter porcinus | Campylobacterota | Epsilonproteobacteria |
| Arcobacter roscoffensis | Campylobacterota | Epsilonproteobacteria |
| Arcobacter suis CECT 7833 | Campylobacterota | Epsilonproteobacteria |
| Arcobacter venerupis | Campylobacterota | Epsilonproteobacteria |
| Caminibacter mediatlanticus TB-2 | Campylobacterota | Epsilonproteobacteria |
| Caminibacter pacificus | Campylobacterota | Epsilonproteobacteria |
| Campylobacter armoricus | Campylobacterota | Epsilonproteobacteria |
| Campylobacter avium LMG 24591 | Campylobacterota | Epsilonproteobacteria |
| Campylobacter bilis | Campylobacterota | Epsilonproteobacteria |

| | | |
|---|---|---|
| Campylobacter blaseri | Campylobacterota | Epsilonproteobacteria |
| Campylobacter californiensis | Campylobacterota | Epsilonproteobacteria |
| Campylobacter canadensis | Campylobacterota | Epsilonproteobacteria |
| Campylobacter coli | Campylobacterota | Epsilonproteobacteria |
| Campylobacter concisus | Campylobacterota | Epsilonproteobacteria |
| Campylobacter corcagiensis | Campylobacterota | Epsilonproteobacteria |
| Campylobacter cuniculorum DSM 23162 = LMG 24588 | Campylobacterota | Epsilonproteobacteria |
| Campylobacter curvus | Campylobacterota | Epsilonproteobacteria |
| Campylobacter fetus | Campylobacterota | Epsilonproteobacteria |
| Campylobacter geochelonis | Campylobacterota | Epsilonproteobacteria |
| Campylobacter gracilis | Campylobacterota | Epsilonproteobacteria |
| Campylobacter helveticus | Campylobacterota | Epsilonproteobacteria |
| Campylobacter hepaticus | Campylobacterota | Epsilonproteobacteria |
| Campylobacter hominis ATCC BAA-381 | Campylobacterota | Epsilonproteobacteria |
| Campylobacter hyointestinalis subsp. lawsonii | Campylobacterota | Epsilonproteobacteria |
| Campylobacter iguaniorum | Campylobacterota | Epsilonproteobacteria |
| Campylobacter insulaenigrae NCTC 12927 | Campylobacterota | Epsilonproteobacteria |
| Campylobacter jejuni subsp. jejuni NCTC 11168 = ATCC 700819 | Campylobacterota | Epsilonproteobacteria |
| Campylobacter lanienae NCTC 13004 | Campylobacterota | Epsilonproteobacteria |
| Campylobacter lari subsp. concheus | Campylobacterota | Epsilonproteobacteria |
| Campylobacter mucosalis | Campylobacterota | Epsilonproteobacteria |
| Campylobacter novaezeelandiae | Campylobacterota | Epsilonproteobacteria |
| Campylobacter ornithocola | Campylobacterota | Epsilonproteobacteria |
| Campylobacter peloridis LMG 23910 | Campylobacterota | Epsilonproteobacteria |
| Campylobacter pinnipediorum subsp. caledonicus | Campylobacterota | Epsilonproteobacteria |
| Campylobacter porcelli | Campylobacterota | Epsilonproteobacteria |
| Campylobacter rectus | Campylobacterota | Epsilonproteobacteria |
| Campylobacter showae | Campylobacterota | Epsilonproteobacteria |
| Campylobacter sputorum subsp. sputorum | Campylobacterota | Epsilonproteobacteria |
| Campylobacter subantarcticus LMG 24377 | Campylobacterota | Epsilonproteobacteria |
| Campylobacter upsaliensis RM3940 | Campylobacterota | Epsilonproteobacteria |

| | | |
|---|---|---|
| Campylobacter ureolyticus | Campylobacterota | Epsilonproteobacteria |
| Campylobacter vicugnae | Campylobacterota | Epsilonproteobacteria |
| Campylobacter volucris | Campylobacterota | Epsilonproteobacteria |
| Campylobacter vulpis | Campylobacterota | Epsilonproteobacteria |
| Candidatus Campylobacter infans | Campylobacterota | Epsilonproteobacteria |
| Candidatus Sulfurimonas baltica | Campylobacterota | Epsilonproteobacteria |
| Candidatus Sulfurimonas marisnigri | Campylobacterota | Epsilonproteobacteria |
| Halarcobacter anaerophilus | Campylobacterota | Epsilonproteobacteria |
| Halarcobacter bivalviorum | Campylobacterota | Epsilonproteobacteria |
| Halarcobacter ebronensis | Campylobacterota | Epsilonproteobacteria |
| Helicobacter acinonychis str. Sheeba | Campylobacterota | Epsilonproteobacteria |
| Helicobacter apodemus | Campylobacterota | Epsilonproteobacteria |
| Helicobacter canadensis MIT 98-5491 | Campylobacterota | Epsilonproteobacteria |
| Helicobacter cetorum MIT 99-5656 | Campylobacterota | Epsilonproteobacteria |
| Helicobacter cholecystus | Campylobacterota | Epsilonproteobacteria |
| Helicobacter cinaedi PAGU611 | Campylobacterota | Epsilonproteobacteria |
| Helicobacter enhydrae | Campylobacterota | Epsilonproteobacteria |
| Helicobacter felis ATCC 49179 | Campylobacterota | Epsilonproteobacteria |
| Helicobacter felistomachi | Campylobacterota | Epsilonproteobacteria |
| Helicobacter gastrocanis | Campylobacterota | Epsilonproteobacteria |
| Helicobacter hepaticus ATCC 51449 | Campylobacterota | Epsilonproteobacteria |
| Helicobacter himalayensis | Campylobacterota | Epsilonproteobacteria |
| Helicobacter mastomyrinus | Campylobacterota | Epsilonproteobacteria |
| Helicobacter mustelae | Campylobacterota | Epsilonproteobacteria |
| Helicobacter pullorum | Campylobacterota | Epsilonproteobacteria |
| Helicobacter pylori | Campylobacterota | Epsilonproteobacteria |
| Helicobacter suis | Campylobacterota | Epsilonproteobacteria |
| Helicobacter typhlonius | Campylobacterota | Epsilonproteobacteria |
| Helicobacter winghamensis ATCC BAA-430 | Campylobacterota | Epsilonproteobacteria |
| Hydrogenimonas cancrithermarum | Campylobacterota | Epsilonproteobacteria |
| Hydrogenimonas leucolamina | Campylobacterota | Epsilonproteobacteria |

| | | |
|---|---|---|
| Hydrogenimonas thermophila | Campylobacterota | Epsilonproteobacteria |
| Hydrogenimonas urashimensis | Campylobacterota | Epsilonproteobacteria |
| Malaciobacter canalis | Campylobacterota | Epsilonproteobacteria |
| Malaciobacter halophilus | Campylobacterota | Epsilonproteobacteria |
| Malaciobacter marinus | Campylobacterota | Epsilonproteobacteria |
| Malaciobacter molluscorum LMG 25693 | Campylobacterota | Epsilonproteobacteria |
| Malaciobacter mytili LMG 24559 | Campylobacterota | Epsilonproteobacteria |
| Malaciobacter pacificus | Campylobacterota | Epsilonproteobacteria |
| Nautilia profundicola AmH | Campylobacterota | Epsilonproteobacteria |
| Nitratifractor salsuginis DSM 16511 | Campylobacterota | Epsilonproteobacteria |
| Nitratiruptor tergarcus DSM 16512 | Campylobacterota | Epsilonproteobacteria |
| Nitrosophilus alvini | Campylobacterota | Epsilonproteobacteria |
| Nitrosophilus kaiyonis | Campylobacterota | Epsilonproteobacteria |
| Nitrosophilus labii | Campylobacterota | Epsilonproteobacteria |
| Poseidonibacter lekithochrous | Campylobacterota | Epsilonproteobacteria |
| Poseidonibacter parvus | Campylobacterota | Epsilonproteobacteria |
| Sulfuricurvum kujiense DSM 16994 | Campylobacterota | Epsilonproteobacteria |
| Sulfurimonas aquatica | Campylobacterota | Epsilonproteobacteria |
| Sulfurimonas autotrophica DSM 16294 | Campylobacterota | Epsilonproteobacteria |
| Sulfurimonas denitrificans DSM 1251 | Campylobacterota | Epsilonproteobacteria |
| Sulfurimonas diazotrophicus | Campylobacterota | Epsilonproteobacteria |
| Sulfurimonas gotlandica GD1 | Campylobacterota | Epsilonproteobacteria |
| Sulfurimonas hydrogeniphila | Campylobacterota | Epsilonproteobacteria |
| Sulfurimonas lithotrophica | Campylobacterota | Epsilonproteobacteria |
| Sulfurimonas marina | Campylobacterota | Epsilonproteobacteria |
| Sulfurimonas paralvinellae | Campylobacterota | Epsilonproteobacteria |
| Sulfurimonas sediminis | Campylobacterota | Epsilonproteobacteria |
| Sulfurimonas xiamenensis | Campylobacterota | Epsilonproteobacteria |
| Sulfurospirillum barnesii SES-3 | Campylobacterota | Epsilonproteobacteria |
| Sulfurospirillum deleyianum DSM 6946 | Campylobacterota | Epsilonproteobacteria |
| Sulfurospirillum diekertiae | Campylobacterota | Epsilonproteobacteria |

| | | |
|---|---|---|
| Sulfurospirillum halorespirans DSM 13726 | Campylobacterota | Epsilonproteobacteria |
| Sulfurospirillum multivorans DSM 12446 | Campylobacterota | Epsilonproteobacteria |
| Sulfurovum indicum | Campylobacterota | Epsilonproteobacteria |
| Sulfurovum lithotrophicum | Campylobacterota | Epsilonproteobacteria |
| Sulfurovum mangrovi | Campylobacterota | Epsilonproteobacteria |
| Wolinella succinogenes | Campylobacterota | Epsilonproteobacteria |
| [Clostridium] innocuum | Bacillota | Erysipelotrichia |
| Allobaculum mucilyticum | Bacillota | Erysipelotrichia |
| Allocoprobacillus halotolerans | Bacillota | Erysipelotrichia |
| Amedibacterium intestinale | Bacillota | Erysipelotrichia |
| Catenibacterium mitsuokai | Bacillota | Erysipelotrichia |
| Erysipelothrix amsterdamensis | Bacillota | Erysipelotrichia |
| Erysipelothrix enhydrae | Bacillota | Erysipelotrichia |
| Erysipelothrix inopinata | Bacillota | Erysipelotrichia |
| Erysipelothrix larvae | Bacillota | Erysipelotrichia |
| Erysipelothrix piscisicarius | Bacillota | Erysipelotrichia |
| Erysipelothrix rhusiopathiae str. Fujisawa | Bacillota | Erysipelotrichia |
| Faecalibacillus intestinalis | Bacillota | Erysipelotrichia |
| Faecalibaculum rodentium | Bacillota | Erysipelotrichia |
| Intestinibaculum porci | Bacillota | Erysipelotrichia |
| Longicatena caecimuris | Bacillota | Erysipelotrichia |
| Massilimicrobiota timonensis | Bacillota | Erysipelotrichia |
| Solobacterium moorei | Bacillota | Erysipelotrichia |
| Tannockella kyphosi | Bacillota | Erysipelotrichia |
| Thomasclavelia ramosa | Bacillota | Erysipelotrichia |
| Thomasclavelia spiroformis DSM 1552 | Bacillota | Erysipelotrichia |
| Turicibacter bilis | Bacillota | Erysipelotrichia |
| Turicibacter faecis | Bacillota | Erysipelotrichia |
| Turicibacter sanguinis | Bacillota | Erysipelotrichia |
| Fibrobacter succinogenes subsp. succinogenes S85 | Fibrobacterota | Fibrobacteria |
| Fibrobacter succinogenes subsp. succinogenes S85 | Fibrobacterota | Fibrobacteria |

| | | |
|---|---|---|
| Fimbriimonas ginsengisoli Gsoil 348 | Armatimonadota | Fimbriimonadia |
| Abyssalbus ytuae | Bacteroidota | Flavobacteriia |
| Aegicerativicinus sediminis | Bacteroidota | Flavobacteriia |
| Aequorivita ciconiae | Bacteroidota | Flavobacteriia |
| Aequorivita iocasae | Bacteroidota | Flavobacteriia |
| Aequorivita marisscotiae | Bacteroidota | Flavobacteriia |
| Aequorivita nionensis | Bacteroidota | Flavobacteriia |
| Aequorivita sublithincola DSM 14238 | Bacteroidota | Flavobacteriia |
| Aestuariibaculum lutulentum | Bacteroidota | Flavobacteriia |
| Aggregatimonas sangjinii | Bacteroidota | Flavobacteriia |
| Algibacter luteus | Bacteroidota | Flavobacteriia |
| Allomuricauda ruestringensis DSM 13258 | Bacteroidota | Flavobacteriia |
| Allotamlana fucoidanivorans | Bacteroidota | Flavobacteriia |
| Antarcticibacterium arcticum | Bacteroidota | Flavobacteriia |
| Antarcticibacterium flavum | Bacteroidota | Flavobacteriia |
| Apibacter raozihei | Bacteroidota | Flavobacteriia |
| Arenibacter algicola | Bacteroidota | Flavobacteriia |
| Arenibacter antarcticus | Bacteroidota | Flavobacteriia |
| Aureibaculum algae | Bacteroidota | Flavobacteriia |
| Bergeyella cardium | Bacteroidota | Flavobacteriia |
| Blattabacterium clevelandi | Bacteroidota | Flavobacteriia |
| Blattabacterium cuenoti | Bacteroidota | Flavobacteriia |
| Blattabacterium punctulatus | Bacteroidota | Flavobacteriia |
| Candidatus Arcticimaribacter forsetii | Bacteroidota | Flavobacteriia |
| Candidatus Kaistella beijingensis | Bacteroidota | Flavobacteriia |
| Candidatus Karelsulcia muelleri | Bacteroidota | Flavobacteriia |
| Candidatus Ornithobacterium hominis | Bacteroidota | Flavobacteriia |
| Candidatus Shikimatogenerans bostrichidophilus | Bacteroidota | Flavobacteriia |
| Candidatus Walczuchella endosymbiont of Icerya purchasi | Bacteroidota | Flavobacteriia |
| Candidatus Walczuchella monophlebidarum | Bacteroidota | Flavobacteriia |
| Capnocytophaga canimorsus | Bacteroidota | Flavobacteriia |

| | | |
|---|---|---|
| Capnocytophaga canis | Bacteroidota | Flavobacteriia |
| Capnocytophaga cynodegmi | Bacteroidota | Flavobacteriia |
| Capnocytophaga endodontalis | Bacteroidota | Flavobacteriia |
| Capnocytophaga gingivalis | Bacteroidota | Flavobacteriia |
| Capnocytophaga haemolytica | Bacteroidota | Flavobacteriia |
| Capnocytophaga ochracea DSM 7271 | Bacteroidota | Flavobacteriia |
| Capnocytophaga sputigena | Bacteroidota | Flavobacteriia |
| Capnocytophaga stomatis | Bacteroidota | Flavobacteriia |
| Cellulophaga algicola DSM 14237 | Bacteroidota | Flavobacteriia |
| Cellulophaga baltica 18 | Bacteroidota | Flavobacteriia |
| Cellulophaga lytica DSM 7489 | Bacteroidota | Flavobacteriia |
| Cellulophaga omnivescoria | Bacteroidota | Flavobacteriia |
| Changchengzhania lutea | Bacteroidota | Flavobacteriia |
| Christiangramia flava JLT2011 | Bacteroidota | Flavobacteriia |
| Christiangramia forsetii KT0803 | Bacteroidota | Flavobacteriia |
| Christiangramia fulva | Bacteroidota | Flavobacteriia |
| Christiangramia salexigens | Bacteroidota | Flavobacteriia |
| Chryseobacterium arthrosphaerae | Bacteroidota | Flavobacteriia |
| Chryseobacterium aureum | Bacteroidota | Flavobacteriia |
| Chryseobacterium balustinum | Bacteroidota | Flavobacteriia |
| Chryseobacterium bernardetii | Bacteroidota | Flavobacteriia |
| Chryseobacterium camelliae | Bacteroidota | Flavobacteriia |
| Chryseobacterium carnipullorum | Bacteroidota | Flavobacteriia |
| Chryseobacterium fluminis | Bacteroidota | Flavobacteriia |
| Chryseobacterium gallinarum | Bacteroidota | Flavobacteriia |
| Chryseobacterium gambrini | Bacteroidota | Flavobacteriia |
| Chryseobacterium glaciei | Bacteroidota | Flavobacteriia |
| Chryseobacterium gleum | Bacteroidota | Flavobacteriia |
| Chryseobacterium gotjawalense | Bacteroidota | Flavobacteriia |
| Chryseobacterium indoltheticum | Bacteroidota | Flavobacteriia |
| Chryseobacterium joostei | Bacteroidota | Flavobacteriia |

| | | |
|---|---|---|
| Chryseobacterium lactis | Bacteroidota | Flavobacteriia |
| Chryseobacterium manosquense | Bacteroidota | Flavobacteriia |
| Chryseobacterium nakagawai | Bacteroidota | Flavobacteriia |
| Chryseobacterium nepalense | Bacteroidota | Flavobacteriia |
| Chryseobacterium oryzae | Bacteroidota | Flavobacteriia |
| Chryseobacterium paludis | Bacteroidota | Flavobacteriia |
| Chryseobacterium piperi | Bacteroidota | Flavobacteriia |
| Chryseobacterium salivictor | Bacteroidota | Flavobacteriia |
| Chryseobacterium shandongense | Bacteroidota | Flavobacteriia |
| Chryseobacterium suipulveris | Bacteroidota | Flavobacteriia |
| Chryseobacterium taihuense | Bacteroidota | Flavobacteriia |
| Chryseobacterium taklimakanense | Bacteroidota | Flavobacteriia |
| Cloacibacterium caeni | Bacteroidota | Flavobacteriia |
| Cloacibacterium normanense | Bacteroidota | Flavobacteriia |
| Constantimarinum furrinae | Bacteroidota | Flavobacteriia |
| Costertonia aggregata | Bacteroidota | Flavobacteriia |
| Croceibacter atlanticus HTCC2559 | Bacteroidota | Flavobacteriia |
| Croceimicrobium hydrocarbonivorans | Bacteroidota | Flavobacteriia |
| Cruoricaptor ignavus | Bacteroidota | Flavobacteriia |
| Dokdonia donghaensis DSW-1 | Bacteroidota | Flavobacteriia |
| Elizabethkingia anophelis R26 | Bacteroidota | Flavobacteriia |
| Elizabethkingia bruuniana | Bacteroidota | Flavobacteriia |
| Elizabethkingia meningoseptica | Bacteroidota | Flavobacteriia |
| Elizabethkingia ursingii | Bacteroidota | Flavobacteriia |
| Empedobacter falsenii | Bacteroidota | Flavobacteriia |
| Empedobacter stercoris | Bacteroidota | Flavobacteriia |
| Epilithonimonas vandammei | Bacteroidota | Flavobacteriia |
| Euzebyella marina | Bacteroidota | Flavobacteriia |
| Faecalibacter bovis | Bacteroidota | Flavobacteriia |
| Flagellimonas lutaonensis | Bacteroidota | Flavobacteriia |
| Flagellimonas maritima | Bacteroidota | Flavobacteriia |

| | | |
|---|---|---|
| Flagellimonas oceani | Bacteroidota | Flavobacteriia |
| Flavivirga abyssicola | Bacteroidota | Flavobacteriia |
| Flavivirga eckloniae | Bacteroidota | Flavobacteriia |
| Flavobacterium acetivorans | Bacteroidota | Flavobacteriia |
| Flavobacterium aestivum | Bacteroidota | Flavobacteriia |
| Flavobacterium agricola | Bacteroidota | Flavobacteriia |
| Flavobacterium album | Bacteroidota | Flavobacteriia |
| Flavobacterium alkalisoli | Bacteroidota | Flavobacteriia |
| Flavobacterium ammonificans | Bacteroidota | Flavobacteriia |
| Flavobacterium ammoniigenes | Bacteroidota | Flavobacteriia |
| Flavobacterium anhuiense | Bacteroidota | Flavobacteriia |
| Flavobacterium aquiphilum | Bacteroidota | Flavobacteriia |
| Flavobacterium arcticum | Bacteroidota | Flavobacteriia |
| Flavobacterium ardleyense | Bacteroidota | Flavobacteriia |
| Flavobacterium azooxidireducens | Bacteroidota | Flavobacteriia |
| Flavobacterium bizetiae | Bacteroidota | Flavobacteriia |
| Flavobacterium branchiophilum FL-15 | Bacteroidota | Flavobacteriia |
| Flavobacterium capsici | Bacteroidota | Flavobacteriia |
| Flavobacterium channae | Bacteroidota | Flavobacteriia |
| Flavobacterium collinsii | Bacteroidota | Flavobacteriia |
| Flavobacterium columnare | Bacteroidota | Flavobacteriia |
| Flavobacterium commune | Bacteroidota | Flavobacteriia |
| Flavobacterium covae | Bacteroidota | Flavobacteriia |
| Flavobacterium crassostreae | Bacteroidota | Flavobacteriia |
| Flavobacterium crocinum | Bacteroidota | Flavobacteriia |
| Flavobacterium cupriresistens | Bacteroidota | Flavobacteriia |
| Flavobacterium cyclinae | Bacteroidota | Flavobacteriia |
| Flavobacterium dauae | Bacteroidota | Flavobacteriia |
| Flavobacterium davisii | Bacteroidota | Flavobacteriia |
| Flavobacterium eburneipallidum | Bacteroidota | Flavobacteriia |
| Flavobacterium endoglycinae | Bacteroidota | Flavobacteriia |

| | | |
|---|---|---|
| Flavobacterium faecale | Bacteroidota | Flavobacteriia |
| Flavobacterium flavigenum | Bacteroidota | Flavobacteriia |
| Flavobacterium flavipigmentatum | Bacteroidota | Flavobacteriia |
| Flavobacterium fluviale | Bacteroidota | Flavobacteriia |
| Flavobacterium galactosidilyticum | Bacteroidota | Flavobacteriia |
| Flavobacterium gelatinilyticum | Bacteroidota | Flavobacteriia |
| Flavobacterium gilvum | Bacteroidota | Flavobacteriia |
| Flavobacterium ginsenosidimutans | Bacteroidota | Flavobacteriia |
| Flavobacterium gyeonganense | Bacteroidota | Flavobacteriia |
| Flavobacterium hankyongi | Bacteroidota | Flavobacteriia |
| Flavobacterium humidisoli | Bacteroidota | Flavobacteriia |
| Flavobacterium indicum GPTSA100-9 = DSM 17447 | Bacteroidota | Flavobacteriia |
| Flavobacterium inviolabile | Bacteroidota | Flavobacteriia |
| Flavobacterium johnsoniae UW101 | Bacteroidota | Flavobacteriia |
| Flavobacterium jumunjinense | Bacteroidota | Flavobacteriia |
| Flavobacterium keumense | Bacteroidota | Flavobacteriia |
| Flavobacterium kingsejongi | Bacteroidota | Flavobacteriia |
| Flavobacterium lacustre | Bacteroidota | Flavobacteriia |
| Flavobacterium limnophilum | Bacteroidota | Flavobacteriia |
| Flavobacterium litorale | Bacteroidota | Flavobacteriia |
| Flavobacterium luteolum | Bacteroidota | Flavobacteriia |
| Flavobacterium magnum | Bacteroidota | Flavobacteriia |
| Flavobacterium marginilacus | Bacteroidota | Flavobacteriia |
| Flavobacterium nackdongense | Bacteroidota | Flavobacteriia |
| Flavobacterium nakdongensis | Bacteroidota | Flavobacteriia |
| Flavobacterium nitratireducens | Bacteroidota | Flavobacteriia |
| Flavobacterium nitrogenifigens | Bacteroidota | Flavobacteriia |
| Flavobacterium okayamense | Bacteroidota | Flavobacteriia |
| Flavobacterium ovatum | Bacteroidota | Flavobacteriia |
| Flavobacterium pallidum | Bacteroidota | Flavobacteriia |
| Flavobacterium panacagri | Bacteroidota | Flavobacteriia |

| | | |
|---|---|---|
| Flavobacterium panici | Bacteroidota | Flavobacteriia |
| Flavobacterium plurextorum | Bacteroidota | Flavobacteriia |
| Flavobacterium praedii | Bacteroidota | Flavobacteriia |
| Flavobacterium psychrophilum | Bacteroidota | Flavobacteriia |
| Flavobacterium psychrotrophum | Bacteroidota | Flavobacteriia |
| Flavobacterium rivulicola | Bacteroidota | Flavobacteriia |
| Flavobacterium sangjuense | Bacteroidota | Flavobacteriia |
| Flavobacterium sediminilitoris | Bacteroidota | Flavobacteriia |
| Flavobacterium sediminis | Bacteroidota | Flavobacteriia |
| Flavobacterium soyae | Bacteroidota | Flavobacteriia |
| Fluviicola taffensis DSM 16823 | Bacteroidota | Flavobacteriia |
| Formosa agariphila KMM 3901 | Bacteroidota | Flavobacteriia |
| Formosa sediminum | Bacteroidota | Flavobacteriia |
| Hwangdonia lutea | Bacteroidota | Flavobacteriia |
| Ichthyobacterium seriolicida | Bacteroidota | Flavobacteriia |
| Imtechella halotolerans | Bacteroidota | Flavobacteriia |
| Kaistella antarctica | Bacteroidota | Flavobacteriia |
| Kaistella carnis | Bacteroidota | Flavobacteriia |
| Kaistella faecalis | Bacteroidota | Flavobacteriia |
| Kaistella jeonii | Bacteroidota | Flavobacteriia |
| Kaistella polysaccharea | Bacteroidota | Flavobacteriia |
| Kordia antarctica | Bacteroidota | Flavobacteriia |
| Lacinutrix neustonica | Bacteroidota | Flavobacteriia |
| Lacinutrix venerupis | Bacteroidota | Flavobacteriia |
| Leeuwenhoekiella palythoae | Bacteroidota | Flavobacteriia |
| Lutibacter profundi | Bacteroidota | Flavobacteriia |
| Lutimonas zeaxanthinifaciens | Bacteroidota | Flavobacteriia |
| Mangrovimonas cancribranchiae | Bacteroidota | Flavobacteriia |
| Maribacter aestuarii | Bacteroidota | Flavobacteriia |
| Maribacter cobaltidurans | Bacteroidota | Flavobacteriia |
| Maribacter dokdonensis | Bacteroidota | Flavobacteriia |

| | | |
|---|---|---|
| Maribacter halichondriae | Bacteroidota | Flavobacteriia |
| Maribacter hydrothermalis | Bacteroidota | Flavobacteriia |
| Maribacter litopenaei | Bacteroidota | Flavobacteriia |
| Mariniflexile litorale | Bacteroidota | Flavobacteriia |
| Mesoflavibacter profundi | Bacteroidota | Flavobacteriia |
| Muriicola soli | Bacteroidota | Flavobacteriia |
| Myroides albus | Bacteroidota | Flavobacteriia |
| Myroides fluvii | Bacteroidota | Flavobacteriia |
| Myroides odoratimimus | Bacteroidota | Flavobacteriia |
| Myroides odoratus | Bacteroidota | Flavobacteriia |
| Myroides profundi | Bacteroidota | Flavobacteriia |
| Nonlabens dokdonensis DSW-6 | Bacteroidota | Flavobacteriia |
| Nonlabens marinus S1-08 | Bacteroidota | Flavobacteriia |
| Nonlabens ponticola | Bacteroidota | Flavobacteriia |
| Nonlabens spongiae | Bacteroidota | Flavobacteriia |
| Nonlabens tegetincola | Bacteroidota | Flavobacteriia |
| Nonlabens ulvanivorans | Bacteroidota | Flavobacteriia |
| Ornithobacterium rhinotracheale DSM 15997 | Bacteroidota | Flavobacteriia |
| Owenweeksia hongkongensis DSM 17368 | Bacteroidota | Flavobacteriia |
| Paenimyroides aestuarii | Bacteroidota | Flavobacteriia |
| Paracrocinitomix mangrovi | Bacteroidota | Flavobacteriia |
| Parvicella tangerina | Bacteroidota | Flavobacteriia |
| Polaribacter atrinae | Bacteroidota | Flavobacteriia |
| Polaribacter batillariae | Bacteroidota | Flavobacteriia |
| Polaribacter butkevichii | Bacteroidota | Flavobacteriia |
| Polaribacter cellanae | Bacteroidota | Flavobacteriia |
| Polaribacter dokdonensis | Bacteroidota | Flavobacteriia |
| Polaribacter gangjinensis | Bacteroidota | Flavobacteriia |
| Polaribacter haliotis | Bacteroidota | Flavobacteriia |
| Polaribacter huanghezhanensis | Bacteroidota | Flavobacteriia |
| Polaribacter litorisediminis | Bacteroidota | Flavobacteriia |

| | | |
|---|---|---|
| Polaribacter marinaquae | Bacteroidota | Flavobacteriia |
| Polaribacter pacificus | Bacteroidota | Flavobacteriia |
| Polaribacter pectinis | Bacteroidota | Flavobacteriia |
| Polaribacter reichenbachii | Bacteroidota | Flavobacteriia |
| Polaribacter sejongensis | Bacteroidota | Flavobacteriia |
| Polaribacter staleyi | Bacteroidota | Flavobacteriia |
| Polaribacter tangerinus | Bacteroidota | Flavobacteriia |
| Polaribacter vadi | Bacteroidota | Flavobacteriia |
| Pseudalgibacter alginicilyticus | Bacteroidota | Flavobacteriia |
| Pseudotamlana carrageenivorans | Bacteroidota | Flavobacteriia |
| Psychroflexus torquis ATCC 700755 | Bacteroidota | Flavobacteriia |
| Psychroserpens ponticola | Bacteroidota | Flavobacteriia |
| Pukyongia salina | Bacteroidota | Flavobacteriia |
| Rasiella rasia | Bacteroidota | Flavobacteriia |
| Riemerella anatipestifer ATCC 11845 = DSM 15868 | Bacteroidota | Flavobacteriia |
| Robiginitalea biformata HTCC2501 | Bacteroidota | Flavobacteriia |
| Salegentibacter mishustinae | Bacteroidota | Flavobacteriia |
| Salegentibacter salegens | Bacteroidota | Flavobacteriia |
| Salinimicrobium tongyeongense | Bacteroidota | Flavobacteriia |
| Sanyastnella coralliicola | Bacteroidota | Flavobacteriia |
| Siansivirga zeaxanthinifaciens CC-SAMT-1 | Bacteroidota | Flavobacteriia |
| Sinomicrobium kalidii | Bacteroidota | Flavobacteriia |
| Tenacibaculum aestuariivivum | Bacteroidota | Flavobacteriia |
| Tenacibaculum dicentrarchi | Bacteroidota | Flavobacteriia |
| Tenacibaculum insulae | Bacteroidota | Flavobacteriia |
| Tenacibaculum jejuense | Bacteroidota | Flavobacteriia |
| Tenacibaculum maritimum NCIMB 2154 | Bacteroidota | Flavobacteriia |
| Tenacibaculum mesophilum | Bacteroidota | Flavobacteriia |
| Tenacibaculum pacificus | Bacteroidota | Flavobacteriia |
| Tenacibaculum platacis | Bacteroidota | Flavobacteriia |
| Tenacibaculum polynesiense | Bacteroidota | Flavobacteriia |

| | | |
|---|---|---|
| Tenacibaculum retecalamus | Bacteroidota | Flavobacteriia |
| Tenacibaculum singaporense | Bacteroidota | Flavobacteriia |
| Tenacibaculum tangerinum | Bacteroidota | Flavobacteriia |
| Tenacibaculum todarodis | Bacteroidota | Flavobacteriia |
| Tenacibaculum vairaonense | Bacteroidota | Flavobacteriia |
| Thalassobellus suaedae | Bacteroidota | Flavobacteriia |
| Urechidicola croceus | Bacteroidota | Flavobacteriia |
| Weeksella virosa | Bacteroidota | Flavobacteriia |
| Wenyingzhuangia fucanilytica | Bacteroidota | Flavobacteriia |
| Winogradskyella forsetii | Bacteroidota | Flavobacteriia |
| Winogradskyella schleiferi | Bacteroidota | Flavobacteriia |
| Winogradskyella sediminis | Bacteroidota | Flavobacteriia |
| Zhouia spongiae | Bacteroidota | Flavobacteriia |
| Zobellia alginiliquefaciens | Bacteroidota | Flavobacteriia |
| Zobellia galactanivorans | Bacteroidota | Flavobacteriia |
| Zobellia russellii | Bacteroidota | Flavobacteriia |
| Zunongwangia profunda SM-A87 | Bacteroidota | Flavobacteriia |
| Cetobacterium somerae | Fusobacteriota | Fusobacteriia |
| Cetobacterium somerae | Fusobacteriota | Fusobacteriia |
| Fusobacterium animalis | Fusobacteriota | Fusobacteriia |
| Fusobacterium canifelinum | Fusobacteriota | Fusobacteriia |
| Fusobacterium gonidiaformans ATCC 25563 | Fusobacteriota | Fusobacteriia |
| Fusobacterium gonidiaformans ATCC 25563 | Fusobacteriota | Fusobacteriia |
| Fusobacterium hominis | Fusobacteriota | Fusobacteriia |
| Fusobacterium hominis | Fusobacteriota | Fusobacteriia |
| Fusobacterium hwasookii ChDC F206 | Fusobacteriota | Fusobacteriia |
| Fusobacterium mortiferum ATCC 9817 | Fusobacteriota | Fusobacteriia |
| Fusobacterium mortiferum ATCC 9817 | Fusobacteriota | Fusobacteriia |
| Fusobacterium necrophorum subsp. funduliforme | Fusobacteriota | Fusobacteriia |
| Fusobacterium necrophorum subsp. funduliforme | Fusobacteriota | Fusobacteriia |
| Fusobacterium nucleatum subsp. nucleatum ATCC 25586 | Fusobacteriota | Fusobacteriia |

| | | |
|---|---|---|
| Fusobacterium polymorphum | Fusobacteriota | Fusobacteriia |
| Fusobacterium ulcerans | Fusobacteriota | Fusobacteriia |
| Fusobacterium ulcerans | Fusobacteriota | Fusobacteriia |
| Fusobacterium varium ATCC 27725 | Fusobacteriota | Fusobacteriia |
| Fusobacterium varium ATCC 27725 | Fusobacteriota | Fusobacteriia |
| Fusobacterium vincentii | Fusobacteriota | Fusobacteriia |
| Fusobacterium watanabei | Fusobacteriota | Fusobacteriia |
| Haliovirga abyssi | Fusobacteriota | Fusobacteriia |
| Haliovirga abyssi | Fusobacteriota | Fusobacteriia |
| Ilyobacter polytropus DSM 2926 | Fusobacteriota | Fusobacteriia |
| Ilyobacter polytropus DSM 2926 | Fusobacteriota | Fusobacteriia |
| Leptotrichia buccalis C-1013-b | Fusobacteriota | Fusobacteriia |
| Leptotrichia buccalis C-1013-b | Fusobacteriota | Fusobacteriia |
| Leptotrichia hofstadii | Fusobacteriota | Fusobacteriia |
| Leptotrichia hofstadii | Fusobacteriota | Fusobacteriia |
| Leptotrichia hongkongensis | Fusobacteriota | Fusobacteriia |
| Leptotrichia hongkongensis | Fusobacteriota | Fusobacteriia |
| Leptotrichia mesophila | Fusobacteriota | Fusobacteriia |
| Leptotrichia mesophila | Fusobacteriota | Fusobacteriia |
| Leptotrichia trevisanii | Fusobacteriota | Fusobacteriia |
| Leptotrichia trevisanii | Fusobacteriota | Fusobacteriia |
| Leptotrichia wadei | Fusobacteriota | Fusobacteriia |
| Leptotrichia wadei | Fusobacteriota | Fusobacteriia |
| Sneathia vaginalis | Fusobacteriota | Fusobacteriia |
| Sneathia vaginalis | Fusobacteriota | Fusobacteriia |
| Streptobacillus moniliformis DSM 12112 | Fusobacteriota | Fusobacteriia |
| Streptobacillus moniliformis DSM 12112 | Fusobacteriota | Fusobacteriia |
| [Enterobacter] lignolyticus SCF1 | Pseudomonadota | Gammaproteobacteria |
| [Enterobacter] lignolyticus SCF1 | Pseudomonadota | Gammaproteobacteria |
| [Haemophilus] ducreyi | Pseudomonadota | Gammaproteobacteria |
| [Haemophilus] ducreyi | Pseudomonadota | Gammaproteobacteria |

| | | |
|---|---|---|
| [Mannheimia] succiniciproducens MBEL55E | Pseudomonadota | Gammaproteobacteria |
| [Mannheimia] succiniciproducens MBEL55E | Pseudomonadota | Gammaproteobacteria |
| [Pantoea] beijingensis | Pseudomonadota | Gammaproteobacteria |
| [Pantoea] beijingensis | Pseudomonadota | Gammaproteobacteria |
| Abyssogena phaseoliformis symbiont OG214 | Pseudomonadota | Gammaproteobacteria |
| Abyssogena phaseoliformis symbiont OG214 | Pseudomonadota | Gammaproteobacteria |
| Acidiferrobacter thiooxydans | Pseudomonadota | Gammaproteobacteria |
| Acidiferrobacter thiooxydans | Pseudomonadota | Gammaproteobacteria |
| Acidihalobacter aeolianus | Pseudomonadota | Gammaproteobacteria |
| Acidihalobacter aeolianus | Pseudomonadota | Gammaproteobacteria |
| Acidihalobacter ferrooxydans | Pseudomonadota | Gammaproteobacteria |
| Acidihalobacter ferrooxydans | Pseudomonadota | Gammaproteobacteria |
| Acidihalobacter yilgarnensis | Pseudomonadota | Gammaproteobacteria |
| Acidihalobacter yilgarnensis | Pseudomonadota | Gammaproteobacteria |
| Acinetobacter baumannii | Pseudomonadota | Gammaproteobacteria |
| Acinetobacter baumannii | Pseudomonadota | Gammaproteobacteria |
| Acinetobacter baylyi ADP1 | Pseudomonadota | Gammaproteobacteria |
| Acinetobacter baylyi ADP1 | Pseudomonadota | Gammaproteobacteria |
| Acinetobacter bereziniae | Pseudomonadota | Gammaproteobacteria |
| Acinetobacter bereziniae | Pseudomonadota | Gammaproteobacteria |
| Acinetobacter bouvetii | Pseudomonadota | Gammaproteobacteria |
| Acinetobacter bouvetii | Pseudomonadota | Gammaproteobacteria |
| Acinetobacter chinensis | Pseudomonadota | Gammaproteobacteria |
| Acinetobacter chinensis | Pseudomonadota | Gammaproteobacteria |
| Acinetobacter corruptisaponis | Pseudomonadota | Gammaproteobacteria |
| Acinetobacter corruptisaponis | Pseudomonadota | Gammaproteobacteria |
| Acinetobacter cumulans | Pseudomonadota | Gammaproteobacteria |
| Acinetobacter cumulans | Pseudomonadota | Gammaproteobacteria |
| Acinetobacter defluvii | Pseudomonadota | Gammaproteobacteria |
| Acinetobacter defluvii | Pseudomonadota | Gammaproteobacteria |
| Acinetobacter dispersus | Pseudomonadota | Gammaproteobacteria |

| | | |
|---|---|---|
| Acinetobacter dispersus | Pseudomonadota | Gammaproteobacteria |
| Acinetobacter equi | Pseudomonadota | Gammaproteobacteria |
| Acinetobacter equi | Pseudomonadota | Gammaproteobacteria |
| Acinetobacter guillouiae | Pseudomonadota | Gammaproteobacteria |
| Acinetobacter guillouiae | Pseudomonadota | Gammaproteobacteria |
| Acinetobacter haemolyticus | Pseudomonadota | Gammaproteobacteria |
| Acinetobacter haemolyticus | Pseudomonadota | Gammaproteobacteria |
| Acinetobacter indicus | Pseudomonadota | Gammaproteobacteria |
| Acinetobacter indicus | Pseudomonadota | Gammaproteobacteria |
| Acinetobacter junii | Pseudomonadota | Gammaproteobacteria |
| Acinetobacter junii | Pseudomonadota | Gammaproteobacteria |
| Acinetobacter lactucae | Pseudomonadota | Gammaproteobacteria |
| Acinetobacter lactucae | Pseudomonadota | Gammaproteobacteria |
| Acinetobacter lanii | Pseudomonadota | Gammaproteobacteria |
| Acinetobacter lanii | Pseudomonadota | Gammaproteobacteria |
| Acinetobacter larvae | Pseudomonadota | Gammaproteobacteria |
| Acinetobacter larvae | Pseudomonadota | Gammaproteobacteria |
| Acinetobacter lwoffii | Pseudomonadota | Gammaproteobacteria |
| Acinetobacter lwoffii | Pseudomonadota | Gammaproteobacteria |
| Acinetobacter nosocomialis | Pseudomonadota | Gammaproteobacteria |
| Acinetobacter nosocomialis | Pseudomonadota | Gammaproteobacteria |
| Acinetobacter oleivorans DR1 | Pseudomonadota | Gammaproteobacteria |
| Acinetobacter oleivorans DR1 | Pseudomonadota | Gammaproteobacteria |
| Acinetobacter pittii PHEA-2 | Pseudomonadota | Gammaproteobacteria |
| Acinetobacter pittii PHEA-2 | Pseudomonadota | Gammaproteobacteria |
| Acinetobacter radioresistens | Pseudomonadota | Gammaproteobacteria |
| Acinetobacter radioresistens | Pseudomonadota | Gammaproteobacteria |
| Acinetobacter schindleri | Pseudomonadota | Gammaproteobacteria |
| Acinetobacter schindleri | Pseudomonadota | Gammaproteobacteria |
| Acinetobacter seifertii | Pseudomonadota | Gammaproteobacteria |
| Acinetobacter seifertii | Pseudomonadota | Gammaproteobacteria |

| | | |
|---|---|---|
| Acinetobacter septicus | Pseudomonadota | Gammaproteobacteria |
| Acinetobacter septicus | Pseudomonadota | Gammaproteobacteria |
| Acinetobacter soli | Pseudomonadota | Gammaproteobacteria |
| Acinetobacter soli | Pseudomonadota | Gammaproteobacteria |
| Acinetobacter suaedae | Pseudomonadota | Gammaproteobacteria |
| Acinetobacter suaedae | Pseudomonadota | Gammaproteobacteria |
| Acinetobacter thermotolerans | Pseudomonadota | Gammaproteobacteria |
| Acinetobacter thermotolerans | Pseudomonadota | Gammaproteobacteria |
| Acinetobacter tibetensis | Pseudomonadota | Gammaproteobacteria |
| Acinetobacter tibetensis | Pseudomonadota | Gammaproteobacteria |
| Acinetobacter towneri | Pseudomonadota | Gammaproteobacteria |
| Acinetobacter towneri | Pseudomonadota | Gammaproteobacteria |
| Acinetobacter variabilis | Pseudomonadota | Gammaproteobacteria |
| Acinetobacter variabilis | Pseudomonadota | Gammaproteobacteria |
| Acinetobacter wuhouensis | Pseudomonadota | Gammaproteobacteria |
| Acinetobacter wuhouensis | Pseudomonadota | Gammaproteobacteria |
| Actinobacillus delphinicola | Pseudomonadota | Gammaproteobacteria |
| Actinobacillus delphinicola | Pseudomonadota | Gammaproteobacteria |
| Actinobacillus equuli | Pseudomonadota | Gammaproteobacteria |
| Actinobacillus equuli | Pseudomonadota | Gammaproteobacteria |
| Actinobacillus indolicus | Pseudomonadota | Gammaproteobacteria |
| Actinobacillus indolicus | Pseudomonadota | Gammaproteobacteria |
| Actinobacillus pleuropneumoniae | Pseudomonadota | Gammaproteobacteria |
| Actinobacillus pleuropneumoniae | Pseudomonadota | Gammaproteobacteria |
| Actinobacillus porcitonsillarum | Pseudomonadota | Gammaproteobacteria |
| Actinobacillus porcitonsillarum | Pseudomonadota | Gammaproteobacteria |
| Actinobacillus succinogenes 130Z | Pseudomonadota | Gammaproteobacteria |
| Actinobacillus succinogenes 130Z | Pseudomonadota | Gammaproteobacteria |
| Actinobacillus suis ATCC 33415 | Pseudomonadota | Gammaproteobacteria |
| Actinobacillus suis ATCC 33415 | Pseudomonadota | Gammaproteobacteria |
| Aequoribacter fuscus | Pseudomonadota | Gammaproteobacteria |

| | | |
|---|---|---|
| Aequoribacter fuscus | Pseudomonadota | Gammaproteobacteria |
| Aeromonas allosaccharophila | Pseudomonadota | Gammaproteobacteria |
| Aeromonas allosaccharophila | Pseudomonadota | Gammaproteobacteria |
| Aeromonas bestiarum | Pseudomonadota | Gammaproteobacteria |
| Aeromonas bestiarum | Pseudomonadota | Gammaproteobacteria |
| Aeromonas caviae | Pseudomonadota | Gammaproteobacteria |
| Aeromonas caviae | Pseudomonadota | Gammaproteobacteria |
| Aeromonas dhakensis | Pseudomonadota | Gammaproteobacteria |
| Aeromonas dhakensis | Pseudomonadota | Gammaproteobacteria |
| Aeromonas encheleia | Pseudomonadota | Gammaproteobacteria |
| Aeromonas encheleia | Pseudomonadota | Gammaproteobacteria |
| Aeromonas enteropelogenes | Pseudomonadota | Gammaproteobacteria |
| Aeromonas enteropelogenes | Pseudomonadota | Gammaproteobacteria |
| Aeromonas hydrophila subsp. hydrophila ATCC 7966 | Pseudomonadota | Gammaproteobacteria |
| Aeromonas hydrophila subsp. hydrophila ATCC 7966 | Pseudomonadota | Gammaproteobacteria |
| Aeromonas jandaei | Pseudomonadota | Gammaproteobacteria |
| Aeromonas jandaei | Pseudomonadota | Gammaproteobacteria |
| Aeromonas media | Pseudomonadota | Gammaproteobacteria |
| Aeromonas media | Pseudomonadota | Gammaproteobacteria |
| Aeromonas popoffii | Pseudomonadota | Gammaproteobacteria |
| Aeromonas popoffii | Pseudomonadota | Gammaproteobacteria |
| Aeromonas rivuli | Pseudomonadota | Gammaproteobacteria |
| Aeromonas rivuli | Pseudomonadota | Gammaproteobacteria |
| Aeromonas salmonicida | Pseudomonadota | Gammaproteobacteria |
| Aeromonas salmonicida | Pseudomonadota | Gammaproteobacteria |
| Aeromonas simiae | Pseudomonadota | Gammaproteobacteria |
| Aeromonas simiae | Pseudomonadota | Gammaproteobacteria |
| Aeromonas veronii | Pseudomonadota | Gammaproteobacteria |
| Aeromonas veronii | Pseudomonadota | Gammaproteobacteria |
| Aerosticca soli | Pseudomonadota | Gammaproteobacteria |
| Aerosticca soli | Pseudomonadota | Gammaproteobacteria |

| | | |
|---|---|---|
| Agarilytica rhodophyticola | Pseudomonadota | Gammaproteobacteria |
| Agarilytica rhodophyticola | Pseudomonadota | Gammaproteobacteria |
| Agarilytica rhodophyticola | Pseudomonadota | Gammaproteobacteria |
| Agarivorans aestuarii | Pseudomonadota | Gammaproteobacteria |
| Agarivorans aestuarii | Pseudomonadota | Gammaproteobacteria |
| Agarivorans albus | Pseudomonadota | Gammaproteobacteria |
| Agarivorans albus | Pseudomonadota | Gammaproteobacteria |
| Agarivorans albus | Pseudomonadota | Gammaproteobacteria |
| Aggregatibacter actinomycetemcomitans | Pseudomonadota | Gammaproteobacteria |
| Aggregatibacter actinomycetemcomitans | Pseudomonadota | Gammaproteobacteria |
| Aggregatibacter aphrophilus ATCC 33389 | Pseudomonadota | Gammaproteobacteria |
| Aggregatibacter aphrophilus ATCC 33389 | Pseudomonadota | Gammaproteobacteria |
| Aggregatibacter segnis ATCC 33393 | Pseudomonadota | Gammaproteobacteria |
| Aggregatibacter segnis ATCC 33393 | Pseudomonadota | Gammaproteobacteria |
| Agrilutibacter terrestris | Pseudomonadota | Gammaproteobacteria |
| Agrilutibacter terrestris | Pseudomonadota | Gammaproteobacteria |
| Ahniella affigens | Pseudomonadota | Gammaproteobacteria |
| Ahniella affigens | Pseudomonadota | Gammaproteobacteria |
| Alcanivorax borkumensis SK2 | Pseudomonadota | Gammaproteobacteria |
| Alcanivorax borkumensis SK2 | Pseudomonadota | Gammaproteobacteria |
| Aliamphritea ceti | Pseudomonadota | Gammaproteobacteria |
| Aliamphritea ceti | Pseudomonadota | Gammaproteobacteria |
| Aliamphritea hakodatensis | Pseudomonadota | Gammaproteobacteria |
| Aliamphritea hakodatensis | Pseudomonadota | Gammaproteobacteria |
| Aliivibrio fischeri ES114 | Pseudomonadota | Gammaproteobacteria |
| Aliivibrio fischeri ES114 | Pseudomonadota | Gammaproteobacteria |
| Aliivibrio salmonicida LFI1238 | Pseudomonadota | Gammaproteobacteria |
| Aliivibrio salmonicida LFI1238 | Pseudomonadota | Gammaproteobacteria |
| Alkalilimnicola ehrlichii MLHE-1 | Pseudomonadota | Gammaproteobacteria |
| Alkalilimnicola ehrlichii MLHE-1 | Pseudomonadota | Gammaproteobacteria |
| Alkalimarinus alittae | Pseudomonadota | Gammaproteobacteria |

| | | |
|---|---|---|
| Alkalimarinus coralli | Pseudomonadota | Gammaproteobacteria |
| Alkalimarinus sediminis | Pseudomonadota | Gammaproteobacteria |
| Alloalcanivorax dieselolei B5 | Pseudomonadota | Gammaproteobacteria |
| Alloalcanivorax dieselolei B5 | Pseudomonadota | Gammaproteobacteria |
| Alloalcanivorax venustensis | Pseudomonadota | Gammaproteobacteria |
| Alloalcanivorax venustensis | Pseudomonadota | Gammaproteobacteria |
| Alloalcanivorax xenomutans | Pseudomonadota | Gammaproteobacteria |
| Alloalcanivorax xenomutans | Pseudomonadota | Gammaproteobacteria |
| Allochromatium tepidum | Pseudomonadota | Gammaproteobacteria |
| Allochromatium tepidum | Pseudomonadota | Gammaproteobacteria |
| Allochromatium vinosum DSM 180 | Pseudomonadota | Gammaproteobacteria |
| Allochromatium vinosum DSM 180 | Pseudomonadota | Gammaproteobacteria |
| Allofrancisella frigidaquae | Pseudomonadota | Gammaproteobacteria |
| Allofrancisella guangzhouensis | Pseudomonadota | Gammaproteobacteria |
| Allofrancisella inopinata | Pseudomonadota | Gammaproteobacteria |
| Alteromonas macleodii | Pseudomonadota | Gammaproteobacteria |
| Alteromonas macleodii | Pseudomonadota | Gammaproteobacteria |
| Alteromonas naphthalenivorans | Pseudomonadota | Gammaproteobacteria |
| Alteromonas naphthalenivorans | Pseudomonadota | Gammaproteobacteria |
| Alteromonas pelagimontana | Pseudomonadota | Gammaproteobacteria |
| Alteromonas pelagimontana | Pseudomonadota | Gammaproteobacteria |
| Alteromonas stellipolaris | Pseudomonadota | Gammaproteobacteria |
| Amphritea atlantica | Pseudomonadota | Gammaproteobacteria |
| Amphritea atlantica | Pseudomonadota | Gammaproteobacteria |
| Amphritea japonica | Pseudomonadota | Gammaproteobacteria |
| Amphritea japonica | Pseudomonadota | Gammaproteobacteria |
| Aquicella lusitana | Pseudomonadota | Gammaproteobacteria |
| Aquicella lusitana | Pseudomonadota | Gammaproteobacteria |
| Aquicella siphonis | Pseudomonadota | Gammaproteobacteria |
| Aquicella siphonis | Pseudomonadota | Gammaproteobacteria |
| Aquipseudomonas alcaligenes | Pseudomonadota | Gammaproteobacteria |

| | | |
|---|---|---|
| Aquipseudomonas alcaligenes | Pseudomonadota | Gammaproteobacteria |
| Aquipseudomonas campi | Pseudomonadota | Gammaproteobacteria |
| Aquipseudomonas campi | Pseudomonadota | Gammaproteobacteria |
| Aquirhabdus parva | Pseudomonadota | Gammaproteobacteria |
| Aquirhabdus parva | Pseudomonadota | Gammaproteobacteria |
| Arsenophonus apicola | Pseudomonadota | Gammaproteobacteria |
| Arsenophonus apicola | Pseudomonadota | Gammaproteobacteria |
| Arsenophonus endosymbiont of Aleurodicus dispersus | Pseudomonadota | Gammaproteobacteria |
| Arsenophonus endosymbiont of Aphis craccivora | Pseudomonadota | Gammaproteobacteria |
| Arsenophonus endosymbiont of Aphis craccivora | Pseudomonadota | Gammaproteobacteria |
| Arsenophonus endosymbiont of Crataerina pallida | Pseudomonadota | Gammaproteobacteria |
| Arsenophonus endosymbiont of Crataerina pallida | Pseudomonadota | Gammaproteobacteria |
| Arsenophonus endosymbiont of Lipoptena cervi | Pseudomonadota | Gammaproteobacteria |
| Arsenophonus nasoniae | Pseudomonadota | Gammaproteobacteria |
| Arsenophonus nasoniae | Pseudomonadota | Gammaproteobacteria |
| Arsenophonus symbiont of Ornithomya chloropus | Pseudomonadota | Gammaproteobacteria |
| Atlantibacter hermannii | Pseudomonadota | Gammaproteobacteria |
| Atlantibacter hermannii | Pseudomonadota | Gammaproteobacteria |
| Atlantibacter subterraneus | Pseudomonadota | Gammaproteobacteria |
| Atlantibacter subterraneus | Pseudomonadota | Gammaproteobacteria |
| Avibacterium paragallinarum | Pseudomonadota | Gammaproteobacteria |
| Avibacterium paragallinarum | Pseudomonadota | Gammaproteobacteria |
| Avibacterium volantium | Pseudomonadota | Gammaproteobacteria |
| Avibacterium volantium | Pseudomonadota | Gammaproteobacteria |
| Azorhizophilus paspali | Pseudomonadota | Gammaproteobacteria |
| Azorhizophilus paspali | Pseudomonadota | Gammaproteobacteria |
| Azotobacter chroococcum | Pseudomonadota | Gammaproteobacteria |
| Azotobacter chroococcum | Pseudomonadota | Gammaproteobacteria |
| Azotobacter salinestris | Pseudomonadota | Gammaproteobacteria |
| Azotobacter salinestris | Pseudomonadota | Gammaproteobacteria |
| Azotobacter vinelandii DJ | Pseudomonadota | Gammaproteobacteria |

| | | |
|---|---|---|
| Azotobacter vinelandii DJ | Pseudomonadota | Gammaproteobacteria |
| Basfia succiniciproducens | Pseudomonadota | Gammaproteobacteria |
| Basfia succiniciproducens | Pseudomonadota | Gammaproteobacteria |
| Bathymodiolus thermophilus thioautotrophic gill symbiont | Pseudomonadota | Gammaproteobacteria |
| Bathymodiolus thermophilus thioautotrophic gill symbiont | Pseudomonadota | Gammaproteobacteria |
| Beggiatoa leptomitoformis | Pseudomonadota | Gammaproteobacteria |
| Beggiatoa leptomitoformis | Pseudomonadota | Gammaproteobacteria |
| Bermanella marisrubri | Pseudomonadota | Gammaproteobacteria |
| Bermanella marisrubri | Pseudomonadota | Gammaproteobacteria |
| Bibersteinia trehalosi USDA-ARS-USMARC-188 | Pseudomonadota | Gammaproteobacteria |
| Bibersteinia trehalosi USDA-ARS-USMARC-188 | Pseudomonadota | Gammaproteobacteria |
| Billgrantia sulfidoxydans | Pseudomonadota | Gammaproteobacteria |
| Bisgaardia hudsonensis | Pseudomonadota | Gammaproteobacteria |
| Bisgaardia hudsonensis | Pseudomonadota | Gammaproteobacteria |
| Blochmannia endosymbiont of Camponotus (Colobopsis) obliquus | Pseudomonadota | Gammaproteobacteria |
| Blochmannia endosymbiont of Camponotus modoc | Pseudomonadota | Gammaproteobacteria |
| Blochmannia endosymbiont of Camponotus nipponensis | Pseudomonadota | Gammaproteobacteria |
| Blochmannia endosymbiont of Camponotus sp. | Pseudomonadota | Gammaproteobacteria |
| Blochmannia endosymbiont of Camponotus sp. C-046 | Pseudomonadota | Gammaproteobacteria |
| Blochmannia endosymbiont of Colobopsis nipponica | Pseudomonadota | Gammaproteobacteria |
| Blochmannia endosymbiont of Polyrhachis (Hedomyrma) turneri | Pseudomonadota | Gammaproteobacteria |
| Brenneria goodwinii | Pseudomonadota | Gammaproteobacteria |
| Brenneria goodwinii | Pseudomonadota | Gammaproteobacteria |
| Brenneria izadpanahii | Pseudomonadota | Gammaproteobacteria |
| Brenneria izadpanahii | Pseudomonadota | Gammaproteobacteria |
| Brenneria nigrifluens DSM 30175 = ATCC 13028 | Pseudomonadota | Gammaproteobacteria |
| Brenneria nigrifluens DSM 30175 = ATCC 13028 | Pseudomonadota | Gammaproteobacteria |
| Brenneria uluponensis | Pseudomonadota | Gammaproteobacteria |
| Brenneria uluponensis | Pseudomonadota | Gammaproteobacteria |
| Brumicola nitratireducens FR1064 | Pseudomonadota | Gammaproteobacteria |
| Brumicola nitratireducens FR1064 | Pseudomonadota | Gammaproteobacteria |

| | | |
|---|---|---|
| Buchnera aphidicola str. Sg (Schizaphis graminum) | Pseudomonadota | Gammaproteobacteria |
| Buttiauxella ferragutiae | Pseudomonadota | Gammaproteobacteria |
| Buttiauxella ferragutiae | Pseudomonadota | Gammaproteobacteria |
| Buttiauxella selenatireducens | Pseudomonadota | Gammaproteobacteria |
| Buttiauxella selenatireducens | Pseudomonadota | Gammaproteobacteria |
| Caldichromatium japonicum | Pseudomonadota | Gammaproteobacteria |
| Caldichromatium japonicum | Pseudomonadota | Gammaproteobacteria |
| Candidatus Annandia adelgestsuga | Pseudomonadota | Gammaproteobacteria |
| Candidatus Annandia pinicola | Pseudomonadota | Gammaproteobacteria |
| Candidatus Arsenophonus lipoptenae | Pseudomonadota | Gammaproteobacteria |
| Candidatus Arsenophonus nilaparvatae | Pseudomonadota | Gammaproteobacteria |
| Candidatus Arsenophonus nilaparvatae | Pseudomonadota | Gammaproteobacteria |
| Candidatus Azoamicus ciliaticola | Pseudomonadota | Gammaproteobacteria |
| Candidatus Blochmanniella camponoti | Pseudomonadota | Gammaproteobacteria |
| Candidatus Blochmanniella chromaiodes str. 640 | Pseudomonadota | Gammaproteobacteria |
| Candidatus Blochmanniella pennsylvanica str. BPEN | Pseudomonadota | Gammaproteobacteria |
| Candidatus Blochmanniella vafra str. BVAF | Pseudomonadota | Gammaproteobacteria |
| Candidatus Carsonella ruddii HC isolate Thao2000 | Pseudomonadota | Gammaproteobacteria |
| Candidatus Comchoanobacter bicostacola | Pseudomonadota | Gammaproteobacteria |
| Candidatus Comchoanobacter bicostacola | Pseudomonadota | Gammaproteobacteria |
| Candidatus Coxiella mudrowiae | Pseudomonadota | Gammaproteobacteria |
| Candidatus Coxiella mudrowiae | Pseudomonadota | Gammaproteobacteria |
| Candidatus Curculioniphilus buchneri endosymbiont of Curculio glandium | Pseudomonadota | Gammaproteobacteria |
| Candidatus Doolittlea endobia | Pseudomonadota | Gammaproteobacteria |
| Candidatus Endoriftia persephonae | Pseudomonadota | Gammaproteobacteria |
| Candidatus Endoriftia persephonae | Pseudomonadota | Gammaproteobacteria |
| Candidatus Enterovibrio altilux | Pseudomonadota | Gammaproteobacteria |
| Candidatus Erwinia haradaeae | Pseudomonadota | Gammaproteobacteria |
| Candidatus Francisella endociliophora | Pseudomonadota | Gammaproteobacteria |
| Candidatus Fukatsuia endosymbiont of Tuberolachnus salignus | Pseudomonadota | Gammaproteobacteria |
| Candidatus Gullanella endobia | Pseudomonadota | Gammaproteobacteria |

| | | |
|---|---|---|
| Candidatus Gullanella endobia | Pseudomonadota | Gammaproteobacteria |
| Candidatus Hamiltonella defensa 5AT (Acyrthosiphon pisum) | Pseudomonadota | Gammaproteobacteria |
| Candidatus Hoaglandella endobia | Pseudomonadota | Gammaproteobacteria |
| Candidatus Ishikawaella capsulata Mpkobe | Pseudomonadota | Gammaproteobacteria |
| Candidatus Ishikawaella capsulata Mpkobe | Pseudomonadota | Gammaproteobacteria |
| Candidatus Legionella polyplacis | Pseudomonadota | Gammaproteobacteria |
| Candidatus Methylocalor cossyra | Pseudomonadota | Gammaproteobacteria |
| Candidatus Methylocalor cossyra | Pseudomonadota | Gammaproteobacteria |
| Candidatus Methylospira mobilis | Pseudomonadota | Gammaproteobacteria |
| Candidatus Methylospira mobilis | Pseudomonadota | Gammaproteobacteria |
| Candidatus Mikella endobia | Pseudomonadota | Gammaproteobacteria |
| Candidatus Moranella endobia PCVAL | Pseudomonadota | Gammaproteobacteria |
| Candidatus Nitrosacidococcus tergens | Pseudomonadota | Gammaproteobacteria |
| Candidatus Nitrosacidococcus tergens | Pseudomonadota | Gammaproteobacteria |
| Candidatus Nitrosoglobus terrae | Pseudomonadota | Gammaproteobacteria |
| Candidatus Nitrosoglobus terrae | Pseudomonadota | Gammaproteobacteria |
| Candidatus Palibaumannia cicadellinicola | Pseudomonadota | Gammaproteobacteria |
| Candidatus Pantoea carbekii | Pseudomonadota | Gammaproteobacteria |
| Candidatus Pantoea soli | Pseudomonadota | Gammaproteobacteria |
| Candidatus Pantoea soli | Pseudomonadota | Gammaproteobacteria |
| Candidatus Paraluminiphilus aquimaris | Pseudomonadota | Gammaproteobacteria |
| Candidatus Portiera aleyrodidarum BT-B-HRs | Pseudomonadota | Gammaproteobacteria |
| Candidatus Profftia lariciata | Pseudomonadota | Gammaproteobacteria |
| Candidatus Profftia tarda | Pseudomonadota | Gammaproteobacteria |
| Candidatus Providencia siddallii | Pseudomonadota | Gammaproteobacteria |
| Candidatus Pseudomonas adelgestsugas | Pseudomonadota | Gammaproteobacteria |
| Candidatus Pseudothioglobus singularis PS1 | Pseudomonadota | Gammaproteobacteria |
| Candidatus Pseudothioglobus singularis PS1 | Pseudomonadota | Gammaproteobacteria |
| Candidatus Purcelliella pentastirinorum | Pseudomonadota | Gammaproteobacteria |
| Candidatus Regiella endosymbiont of Tuberolachnus salignus | Pseudomonadota | Gammaproteobacteria |
| Candidatus Reidiella endopervernicosa | Pseudomonadota | Gammaproteobacteria |

| | | |
|---|---|---|
| Candidatus Reidiella endopervernicosa | Pseudomonadota | Gammaproteobacteria |
| Candidatus Rickettsiella viridis | Pseudomonadota | Gammaproteobacteria |
| Candidatus Riesia pediculicola USDA | Pseudomonadota | Gammaproteobacteria |
| Candidatus Riesia pediculischaeffi | Pseudomonadota | Gammaproteobacteria |
| Candidatus Ruthia endofausta | Pseudomonadota | Gammaproteobacteria |
| Candidatus Ruthia endofausta | Pseudomonadota | Gammaproteobacteria |
| Candidatus Ruthia magnifica str. Cm (Calyptogena magnifica) | Pseudomonadota | Gammaproteobacteria |
| Candidatus Ruthia magnifica str. Cm (Calyptogena magnifica) | Pseudomonadota | Gammaproteobacteria |
| Candidatus Schneideria nysicola | Pseudomonadota | Gammaproteobacteria |
| Candidatus Sodalis pierantonii str. SOPE | Pseudomonadota | Gammaproteobacteria |
| Candidatus Sodalis pierantonii str. SOPE | Pseudomonadota | Gammaproteobacteria |
| Candidatus Steffania adelgidicola | Pseudomonadota | Gammaproteobacteria |
| Candidatus Tachikawaea gelatinosa | Pseudomonadota | Gammaproteobacteria |
| Candidatus Tachikawaea gelatinosa | Pseudomonadota | Gammaproteobacteria |
| Candidatus Thalassolituus haligoni | Pseudomonadota | Gammaproteobacteria |
| Candidatus Thalassolituus haligoni | Pseudomonadota | Gammaproteobacteria |
| Candidatus Thiodictyon syntrophicum | Pseudomonadota | Gammaproteobacteria |
| Candidatus Thiodictyon syntrophicum | Pseudomonadota | Gammaproteobacteria |
| Candidatus Thiodubiliella endoseptemdiera | Pseudomonadota | Gammaproteobacteria |
| Candidatus Thiodubiliella endoseptemdiera | Pseudomonadota | Gammaproteobacteria |
| Candidatus Thioglobus autotrophicus | Pseudomonadota | Gammaproteobacteria |
| Candidatus Thioglobus autotrophicus | Pseudomonadota | Gammaproteobacteria |
| Candidatus Thiothrix anitrata | Pseudomonadota | Gammaproteobacteria |
| Candidatus Thiothrix anitrata | Pseudomonadota | Gammaproteobacteria |
| Candidatus Vesicomyidisocius calyptogenae | Pseudomonadota | Gammaproteobacteria |
| Candidatus Vesicomyidisocius calyptogenae | Pseudomonadota | Gammaproteobacteria |
| Candidatus Westeberhardia cardiocondylae | Pseudomonadota | Gammaproteobacteria |
| Candidatus Williamhamiltonella endosymbiont of Tuberolachnus salignus | Pseudomonadota | Gammaproteobacteria |
| Cardiobacterium hominis | Pseudomonadota | Gammaproteobacteria |
| Cardiobacterium hominis | Pseudomonadota | Gammaproteobacteria |
| Catenovulum adriaticum | Pseudomonadota | Gammaproteobacteria |

| | | |
|---|---|---|
| Catenovulum adriaticum | Pseudomonadota | Gammaproteobacteria |
| Catenovulum sediminis | Pseudomonadota | Gammaproteobacteria |
| Catenovulum sediminis | Pseudomonadota | Gammaproteobacteria |
| Cedecea davisae | Pseudomonadota | Gammaproteobacteria |
| Cedecea davisae | Pseudomonadota | Gammaproteobacteria |
| Cedecea lapagei | Pseudomonadota | Gammaproteobacteria |
| Cedecea lapagei | Pseudomonadota | Gammaproteobacteria |
| Cedecea neteri | Pseudomonadota | Gammaproteobacteria |
| Cedecea neteri | Pseudomonadota | Gammaproteobacteria |
| Celerinatantimonas diazotrophica | Pseudomonadota | Gammaproteobacteria |
| Celerinatantimonas diazotrophica | Pseudomonadota | Gammaproteobacteria |
| Cellvibrio japonicus Ueda107 | Pseudomonadota | Gammaproteobacteria |
| Cellvibrio japonicus Ueda107 | Pseudomonadota | Gammaproteobacteria |
| Chania multitudinisentens RB-25 | Pseudomonadota | Gammaproteobacteria |
| Chania multitudinisentens RB-25 | Pseudomonadota | Gammaproteobacteria |
| Chromohalobacter canadensis | Pseudomonadota | Gammaproteobacteria |
| Chromohalobacter israelensis DSM 3043 | Pseudomonadota | Gammaproteobacteria |
| Citrobacter amalonaticus | Pseudomonadota | Gammaproteobacteria |
| Citrobacter amalonaticus | Pseudomonadota | Gammaproteobacteria |
| Citrobacter arsenatis | Pseudomonadota | Gammaproteobacteria |
| Citrobacter arsenatis | Pseudomonadota | Gammaproteobacteria |
| Citrobacter braakii | Pseudomonadota | Gammaproteobacteria |
| Citrobacter braakii | Pseudomonadota | Gammaproteobacteria |
| Citrobacter cronae | Pseudomonadota | Gammaproteobacteria |
| Citrobacter cronae | Pseudomonadota | Gammaproteobacteria |
| Citrobacter europaeus | Pseudomonadota | Gammaproteobacteria |
| Citrobacter europaeus | Pseudomonadota | Gammaproteobacteria |
| Citrobacter farmeri | Pseudomonadota | Gammaproteobacteria |
| Citrobacter farmeri | Pseudomonadota | Gammaproteobacteria |
| Citrobacter freundii | Pseudomonadota | Gammaproteobacteria |
| Citrobacter freundii | Pseudomonadota | Gammaproteobacteria |

| | | |
|---|---|---|
| Citrobacter koseri ATCC BAA-895 | Pseudomonadota | Gammaproteobacteria |
| Citrobacter koseri ATCC BAA-895 | Pseudomonadota | Gammaproteobacteria |
| Citrobacter meridianamericanus | Pseudomonadota | Gammaproteobacteria |
| Citrobacter meridianamericanus | Pseudomonadota | Gammaproteobacteria |
| Citrobacter pasteurii | Pseudomonadota | Gammaproteobacteria |
| Citrobacter pasteurii | Pseudomonadota | Gammaproteobacteria |
| Citrobacter portucalensis | Pseudomonadota | Gammaproteobacteria |
| Citrobacter portucalensis | Pseudomonadota | Gammaproteobacteria |
| Citrobacter rodentium NBRC 105723 = DSM 16636 | Pseudomonadota | Gammaproteobacteria |
| Citrobacter rodentium NBRC 105723 = DSM 16636 | Pseudomonadota | Gammaproteobacteria |
| Citrobacter sedlakii | Pseudomonadota | Gammaproteobacteria |
| Citrobacter sedlakii | Pseudomonadota | Gammaproteobacteria |
| Citrobacter tructae | Pseudomonadota | Gammaproteobacteria |
| Citrobacter tructae | Pseudomonadota | Gammaproteobacteria |
| Citrobacter werkmanii | Pseudomonadota | Gammaproteobacteria |
| Citrobacter werkmanii | Pseudomonadota | Gammaproteobacteria |
| Cobetia amphilecti | Pseudomonadota | Gammaproteobacteria |
| Cobetia marina | Pseudomonadota | Gammaproteobacteria |
| Cognaticolwellia beringensis | Pseudomonadota | Gammaproteobacteria |
| Cognaticolwellia beringensis | Pseudomonadota | Gammaproteobacteria |
| Colwellia psychrerythraea 34H | Pseudomonadota | Gammaproteobacteria |
| Colwellia psychrerythraea 34H | Pseudomonadota | Gammaproteobacteria |
| Congregibacter brevis | Pseudomonadota | Gammaproteobacteria |
| Congregibacter brevis | Pseudomonadota | Gammaproteobacteria |
| Congregibacter litoralis KT71 | Pseudomonadota | Gammaproteobacteria |
| Congregibacter litoralis KT71 | Pseudomonadota | Gammaproteobacteria |
| Congregibacter variabilis | Pseudomonadota | Gammaproteobacteria |
| Congregibacter variabilis | Pseudomonadota | Gammaproteobacteria |
| Coxiella burnetii RSA 493 | Pseudomonadota | Gammaproteobacteria |
| Coxiella endosymbiont of Amblyomma americanum | Pseudomonadota | Gammaproteobacteria |
| Coxiella endosymbiont of Amblyomma americanum | Pseudomonadota | Gammaproteobacteria |

| | | |
|---|---|---|
| Coxiella endosymbiont of Amblyomma nuttalli | Pseudomonadota | Gammaproteobacteria |
| Coxiella endosymbiont of Amblyomma nuttalli | Pseudomonadota | Gammaproteobacteria |
| Coxiella endosymbiont of Amblyomma sculptum | Pseudomonadota | Gammaproteobacteria |
| Coxiella-like endosymbiont of Amblyomma americanum | Pseudomonadota | Gammaproteobacteria |
| Coxiella-like endosymbiont of Amblyomma americanum | Pseudomonadota | Gammaproteobacteria |
| Cronobacter condimenti 1330 | Pseudomonadota | Gammaproteobacteria |
| Cronobacter condimenti 1330 | Pseudomonadota | Gammaproteobacteria |
| Cronobacter dublinensis subsp. dublinensis LMG 23823 | Pseudomonadota | Gammaproteobacteria |
| Cronobacter dublinensis subsp. dublinensis LMG 23823 | Pseudomonadota | Gammaproteobacteria |
| Cronobacter malonaticus LMG 23826 | Pseudomonadota | Gammaproteobacteria |
| Cronobacter malonaticus LMG 23826 | Pseudomonadota | Gammaproteobacteria |
| Cronobacter muytjensii ATCC 51329 | Pseudomonadota | Gammaproteobacteria |
| Cronobacter muytjensii ATCC 51329 | Pseudomonadota | Gammaproteobacteria |
| Cronobacter sakazakii | Pseudomonadota | Gammaproteobacteria |
| Cronobacter sakazakii | Pseudomonadota | Gammaproteobacteria |
| Cronobacter turicensis | Pseudomonadota | Gammaproteobacteria |
| Cronobacter turicensis | Pseudomonadota | Gammaproteobacteria |
| Cronobacter universalis NCTC 9529 | Pseudomonadota | Gammaproteobacteria |
| Cronobacter universalis NCTC 9529 | Pseudomonadota | Gammaproteobacteria |
| Dickeya aquatica | Pseudomonadota | Gammaproteobacteria |
| Dickeya aquatica | Pseudomonadota | Gammaproteobacteria |
| Dickeya chrysanthemi Ech1591 | Pseudomonadota | Gammaproteobacteria |
| Dickeya chrysanthemi Ech1591 | Pseudomonadota | Gammaproteobacteria |
| Dickeya dadantii | Pseudomonadota | Gammaproteobacteria |
| Dickeya dadantii | Pseudomonadota | Gammaproteobacteria |
| Dickeya dianthicola | Pseudomonadota | Gammaproteobacteria |
| Dickeya dianthicola | Pseudomonadota | Gammaproteobacteria |
| Dickeya fangzhongdai | Pseudomonadota | Gammaproteobacteria |
| Dickeya fangzhongdai | Pseudomonadota | Gammaproteobacteria |
| Dickeya oryzae | Pseudomonadota | Gammaproteobacteria |
| Dickeya oryzae | Pseudomonadota | Gammaproteobacteria |

| | | |
|---|---|---|
| Dickeya parazeae Ech586 | Pseudomonadota | Gammaproteobacteria |
| Dickeya parazeae Ech586 | Pseudomonadota | Gammaproteobacteria |
| Dickeya poaceiphila | Pseudomonadota | Gammaproteobacteria |
| Dickeya poaceiphila | Pseudomonadota | Gammaproteobacteria |
| Dickeya solani IPO 2222 | Pseudomonadota | Gammaproteobacteria |
| Dickeya solani IPO 2222 | Pseudomonadota | Gammaproteobacteria |
| Dickeya zeae | Pseudomonadota | Gammaproteobacteria |
| Dickeya zeae | Pseudomonadota | Gammaproteobacteria |
| Dokdonella koreensis DS-123 | Pseudomonadota | Gammaproteobacteria |
| Dokdonella koreensis DS-123 | Pseudomonadota | Gammaproteobacteria |
| Dongshaea marina | Pseudomonadota | Gammaproteobacteria |
| Duffyella gerundensis | Pseudomonadota | Gammaproteobacteria |
| Duffyella gerundensis | Pseudomonadota | Gammaproteobacteria |
| Dyella aluminiiresistens | Pseudomonadota | Gammaproteobacteria |
| Dyella aluminiiresistens | Pseudomonadota | Gammaproteobacteria |
| Dyella caseinilytica | Pseudomonadota | Gammaproteobacteria |
| Dyella caseinilytica | Pseudomonadota | Gammaproteobacteria |
| Dyella telluris | Pseudomonadota | Gammaproteobacteria |
| Dyella telluris | Pseudomonadota | Gammaproteobacteria |
| Dyella thiooxydans | Pseudomonadota | Gammaproteobacteria |
| Dyella thiooxydans | Pseudomonadota | Gammaproteobacteria |
| Ectopseudomonas hydrolytica | Pseudomonadota | Gammaproteobacteria |
| Ectopseudomonas hydrolytica | Pseudomonadota | Gammaproteobacteria |
| Ectopseudomonas khazarica | Pseudomonadota | Gammaproteobacteria |
| Ectopseudomonas khazarica | Pseudomonadota | Gammaproteobacteria |
| Ectopseudomonas mendocina S5.2 | Pseudomonadota | Gammaproteobacteria |
| Ectopseudomonas mendocina S5.2 | Pseudomonadota | Gammaproteobacteria |
| Ectopseudomonas toyotomiensis | Pseudomonadota | Gammaproteobacteria |
| Ectopseudomonas toyotomiensis | Pseudomonadota | Gammaproteobacteria |
| Edwardsiella anguillarum ET080813 | Pseudomonadota | Gammaproteobacteria |
| Edwardsiella anguillarum ET080813 | Pseudomonadota | Gammaproteobacteria |

| | | |
|---|---|---|
| Edwardsiella hoshinae | Pseudomonadota | Gammaproteobacteria |
| Edwardsiella hoshinae | Pseudomonadota | Gammaproteobacteria |
| Edwardsiella ictaluri | Pseudomonadota | Gammaproteobacteria |
| Edwardsiella ictaluri | Pseudomonadota | Gammaproteobacteria |
| Edwardsiella piscicida | Pseudomonadota | Gammaproteobacteria |
| Edwardsiella piscicida | Pseudomonadota | Gammaproteobacteria |
| Edwardsiella tarda | Pseudomonadota | Gammaproteobacteria |
| Edwardsiella tarda | Pseudomonadota | Gammaproteobacteria |
| endosymbiont of Bathymodiolus septemdierum str. Myojin knoll | Pseudomonadota | Gammaproteobacteria |
| endosymbiont of Bathymodiolus septemdierum str. Myojin knoll | Pseudomonadota | Gammaproteobacteria |
| endosymbiont of Euscepes postfasciatus | Pseudomonadota | Gammaproteobacteria |
| endosymbiont of Pachyrhynchus infernalis | Pseudomonadota | Gammaproteobacteria |
| endosymbiont of Rhynchophorus ferrugineus | Pseudomonadota | Gammaproteobacteria |
| endosymbiont of Sipalinus gigas | Pseudomonadota | Gammaproteobacteria |
| endosymbiont of unidentified scaly snail isolate Monju | Pseudomonadota | Gammaproteobacteria |
| endosymbiont of unidentified scaly snail isolate Monju | Pseudomonadota | Gammaproteobacteria |
| Endozoicomonas euniceicola | Pseudomonadota | Gammaproteobacteria |
| Endozoicomonas montiporae CL-33 | Pseudomonadota | Gammaproteobacteria |
| Enterobacter adelaidei | Pseudomonadota | Gammaproteobacteria |
| Enterobacter adelaidei | Pseudomonadota | Gammaproteobacteria |
| Enterobacter asburiae | Pseudomonadota | Gammaproteobacteria |
| Enterobacter asburiae | Pseudomonadota | Gammaproteobacteria |
| Enterobacter bugandensis | Pseudomonadota | Gammaproteobacteria |
| Enterobacter bugandensis | Pseudomonadota | Gammaproteobacteria |
| Enterobacter cancerogenus | Pseudomonadota | Gammaproteobacteria |
| Enterobacter cancerogenus | Pseudomonadota | Gammaproteobacteria |
| Enterobacter chengduensis | Pseudomonadota | Gammaproteobacteria |
| Enterobacter chengduensis | Pseudomonadota | Gammaproteobacteria |
| Enterobacter chuandaensis | Pseudomonadota | Gammaproteobacteria |
| Enterobacter chuandaensis | Pseudomonadota | Gammaproteobacteria |
| Enterobacter cloacae | Pseudomonadota | Gammaproteobacteria |

| | | |
|---|---|---|
| Enterobacter cloacae | Pseudomonadota | Gammaproteobacteria |
| Enterobacter dykesii | Pseudomonadota | Gammaproteobacteria |
| Enterobacter dykesii | Pseudomonadota | Gammaproteobacteria |
| Enterobacter hormaechei | Pseudomonadota | Gammaproteobacteria |
| Enterobacter hormaechei | Pseudomonadota | Gammaproteobacteria |
| Enterobacter huaxiensis | Pseudomonadota | Gammaproteobacteria |
| Enterobacter huaxiensis | Pseudomonadota | Gammaproteobacteria |
| Enterobacter intestinihominis | Pseudomonadota | Gammaproteobacteria |
| Enterobacter intestinihominis | Pseudomonadota | Gammaproteobacteria |
| Enterobacter kobei | Pseudomonadota | Gammaproteobacteria |
| Enterobacter kobei | Pseudomonadota | Gammaproteobacteria |
| Enterobacter ludwigii | Pseudomonadota | Gammaproteobacteria |
| Enterobacter ludwigii | Pseudomonadota | Gammaproteobacteria |
| Enterobacter mori | Pseudomonadota | Gammaproteobacteria |
| Enterobacter mori | Pseudomonadota | Gammaproteobacteria |
| Enterobacter oligotrophicus | Pseudomonadota | Gammaproteobacteria |
| Enterobacter oligotrophicus | Pseudomonadota | Gammaproteobacteria |
| Enterobacter pseudoroggenkampii | Pseudomonadota | Gammaproteobacteria |
| Enterobacter pseudoroggenkampii | Pseudomonadota | Gammaproteobacteria |
| Enterobacter quasiroggenkampii | Pseudomonadota | Gammaproteobacteria |
| Enterobacter quasiroggenkampii | Pseudomonadota | Gammaproteobacteria |
| Enterobacter roggenkampii | Pseudomonadota | Gammaproteobacteria |
| Enterobacter roggenkampii | Pseudomonadota | Gammaproteobacteria |
| Enterobacter sichuanensis | Pseudomonadota | Gammaproteobacteria |
| Enterobacter sichuanensis | Pseudomonadota | Gammaproteobacteria |
| Enterobacter soli | Pseudomonadota | Gammaproteobacteria |
| Enterobacter soli | Pseudomonadota | Gammaproteobacteria |
| Enterobacter vonholyi | Pseudomonadota | Gammaproteobacteria |
| Enterobacter vonholyi | Pseudomonadota | Gammaproteobacteria |
| Enterobacter wuhouensis | Pseudomonadota | Gammaproteobacteria |
| Enterobacter wuhouensis | Pseudomonadota | Gammaproteobacteria |

| | | |
|---|---|---|
| Enterobacteriaceae endosymbiont of Donacia bicoloricornis | Pseudomonadota | Gammaproteobacteria |
| Enterobacteriaceae endosymbiont of Donacia cincticornis | Pseudomonadota | Gammaproteobacteria |
| Enterobacteriaceae endosymbiont of Donacia cinerea | Pseudomonadota | Gammaproteobacteria |
| Enterobacteriaceae endosymbiont of Donacia clavipes | Pseudomonadota | Gammaproteobacteria |
| Enterobacteriaceae endosymbiont of Donacia crassipes | Pseudomonadota | Gammaproteobacteria |
| Enterobacteriaceae endosymbiont of Donacia dentata | Pseudomonadota | Gammaproteobacteria |
| Enterobacteriaceae endosymbiont of Donacia fulgens | Pseudomonadota | Gammaproteobacteria |
| Enterobacteriaceae endosymbiont of Donacia marginata | Pseudomonadota | Gammaproteobacteria |
| Enterobacteriaceae endosymbiont of Donacia piscatrix | Pseudomonadota | Gammaproteobacteria |
| Enterobacteriaceae endosymbiont of Donacia provostii | Pseudomonadota | Gammaproteobacteria |
| Enterobacteriaceae endosymbiont of Donacia proxima | Pseudomonadota | Gammaproteobacteria |
| Enterobacteriaceae endosymbiont of Donacia semicuprea | Pseudomonadota | Gammaproteobacteria |
| Enterobacteriaceae endosymbiont of Donacia simplex | Pseudomonadota | Gammaproteobacteria |
| Enterobacteriaceae endosymbiont of Donacia sparganii | Pseudomonadota | Gammaproteobacteria |
| Enterobacteriaceae endosymbiont of Donacia thalassina | Pseudomonadota | Gammaproteobacteria |
| Enterobacteriaceae endosymbiont of Donacia tomentosa | Pseudomonadota | Gammaproteobacteria |
| Enterobacteriaceae endosymbiont of Donacia versicolorea | Pseudomonadota | Gammaproteobacteria |
| Enterobacteriaceae endosymbiont of Donacia vulgaris | Pseudomonadota | Gammaproteobacteria |
| Enterobacteriaceae endosymbiont of Macroplea appendiculata | Pseudomonadota | Gammaproteobacteria |
| Enterobacteriaceae endosymbiont of Macroplea mutica | Pseudomonadota | Gammaproteobacteria |
| Enterobacteriaceae endosymbiont of Neohaemonia nigricornis | Pseudomonadota | Gammaproteobacteria |
| Enterobacteriaceae endosymbiont of Plateumaris braccata | Pseudomonadota | Gammaproteobacteria |
| Enterobacteriaceae endosymbiont of Plateumaris consimilis | Pseudomonadota | Gammaproteobacteria |
| Enterobacteriaceae endosymbiont of Plateumaris pusilla | Pseudomonadota | Gammaproteobacteria |
| Enterobacteriaceae endosymbiont of Plateumaris rustica | Pseudomonadota | Gammaproteobacteria |
| Enterobacteriaceae endosymbiont of Plateumaris sericea | Pseudomonadota | Gammaproteobacteria |
| Entomomonas asaccharolytica | Pseudomonadota | Gammaproteobacteria |
| Entomomonas moraniae | Pseudomonadota | Gammaproteobacteria |
| Erwinia amylovora | Pseudomonadota | Gammaproteobacteria |
| Erwinia amylovora | Pseudomonadota | Gammaproteobacteria |
| Erwinia billingiae Eb661 | Pseudomonadota | Gammaproteobacteria |

| | | |
|---|---|---|
| Erwinia billingiae Eb661 | Pseudomonadota | Gammaproteobacteria |
| Erwinia persicina | Pseudomonadota | Gammaproteobacteria |
| Erwinia persicina | Pseudomonadota | Gammaproteobacteria |
| Erwinia pyri | Pseudomonadota | Gammaproteobacteria |
| Erwinia pyri | Pseudomonadota | Gammaproteobacteria |
| Erwinia pyrifoliae | Pseudomonadota | Gammaproteobacteria |
| Erwinia pyrifoliae | Pseudomonadota | Gammaproteobacteria |
| Erwinia rhapontici | Pseudomonadota | Gammaproteobacteria |
| Erwinia rhapontici | Pseudomonadota | Gammaproteobacteria |
| Erwinia sorbitola | Pseudomonadota | Gammaproteobacteria |
| Erwinia sorbitola | Pseudomonadota | Gammaproteobacteria |
| Erwinia tasmaniensis Et1/99 | Pseudomonadota | Gammaproteobacteria |
| Erwinia tasmaniensis Et1/99 | Pseudomonadota | Gammaproteobacteria |
| Erwinia tracheiphila | Pseudomonadota | Gammaproteobacteria |
| Erwinia tracheiphila | Pseudomonadota | Gammaproteobacteria |
| Escherichia albertii | Pseudomonadota | Gammaproteobacteria |
| Escherichia albertii | Pseudomonadota | Gammaproteobacteria |
| Escherichia coli O157:H7 str. Sakai | Pseudomonadota | Gammaproteobacteria |
| Escherichia coli O157:H7 str. Sakai | Pseudomonadota | Gammaproteobacteria |
| Escherichia coli str. K-12 substr. MG1655 | Pseudomonadota | Gammaproteobacteria |
| Escherichia coli str. K-12 substr. MG1655 | Pseudomonadota | Gammaproteobacteria |
| Escherichia fergusonii | Pseudomonadota | Gammaproteobacteria |
| Escherichia fergusonii | Pseudomonadota | Gammaproteobacteria |
| Escherichia marmotae | Pseudomonadota | Gammaproteobacteria |
| Escherichia marmotae | Pseudomonadota | Gammaproteobacteria |
| Faucicola mancuniensis | Pseudomonadota | Gammaproteobacteria |
| Faucicola mancuniensis | Pseudomonadota | Gammaproteobacteria |
| Ferrimonas balearica DSM 9799 | Pseudomonadota | Gammaproteobacteria |
| Ferrimonas balearica DSM 9799 | Pseudomonadota | Gammaproteobacteria |
| Ferrimonas lipolytica | Pseudomonadota | Gammaproteobacteria |
| Ferrimonas lipolytica | Pseudomonadota | Gammaproteobacteria |

| | | |
|---|---|---|
| Flagellatimonas centrodinii | Pseudomonadota | Gammaproteobacteria |
| Flagellatimonas centrodinii | Pseudomonadota | Gammaproteobacteria |
| Flocculibacter collagenilyticus | Pseudomonadota | Gammaproteobacteria |
| Flocculibacter collagenilyticus | Pseudomonadota | Gammaproteobacteria |
| Francisella adeliensis | Pseudomonadota | Gammaproteobacteria |
| Francisella frigiditurris | Pseudomonadota | Gammaproteobacteria |
| Francisella halioticida | Pseudomonadota | Gammaproteobacteria |
| Francisella hispaniensis | Pseudomonadota | Gammaproteobacteria |
| Francisella marina | Pseudomonadota | Gammaproteobacteria |
| Francisella opportunistica | Pseudomonadota | Gammaproteobacteria |
| Francisella orientalis LADL 07-285A | Pseudomonadota | Gammaproteobacteria |
| Francisella persica ATCC VR-331 | Pseudomonadota | Gammaproteobacteria |
| Francisella philomiragia | Pseudomonadota | Gammaproteobacteria |
| Francisella salimarina | Pseudomonadota | Gammaproteobacteria |
| Francisella salina | Pseudomonadota | Gammaproteobacteria |
| Francisella tularensis subsp. novicida D9876 | Pseudomonadota | Gammaproteobacteria |
| Francisella uliginis | Pseudomonadota | Gammaproteobacteria |
| Frateuria aurantia DSM 6220 | Pseudomonadota | Gammaproteobacteria |
| Frateuria aurantia DSM 6220 | Pseudomonadota | Gammaproteobacteria |
| Frateuria edaphi | Pseudomonadota | Gammaproteobacteria |
| Frateuria edaphi | Pseudomonadota | Gammaproteobacteria |
| Frateuria soli | Pseudomonadota | Gammaproteobacteria |
| Frateuria soli | Pseudomonadota | Gammaproteobacteria |
| Frederiksenia canicola | Pseudomonadota | Gammaproteobacteria |
| Frederiksenia canicola | Pseudomonadota | Gammaproteobacteria |
| Frischella perrara | Pseudomonadota | Gammaproteobacteria |
| Frischella perrara | Pseudomonadota | Gammaproteobacteria |
| Gallaecimonas kandeliae | Pseudomonadota | Gammaproteobacteria |
| Gallaecimonas kandeliae | Pseudomonadota | Gammaproteobacteria |
| Gallaecimonas mangrovi | Pseudomonadota | Gammaproteobacteria |
| Gallaecimonas mangrovi | Pseudomonadota | Gammaproteobacteria |

| | | |
|---|---|---|
| Gallibacterium salpingitidis | Pseudomonadota | Gammaproteobacteria |
| Gallibacterium salpingitidis | Pseudomonadota | Gammaproteobacteria |
| Geopseudomonas guangdongensis | Pseudomonadota | Gammaproteobacteria |
| Gibbsiella quercinecans | Pseudomonadota | Gammaproteobacteria |
| Gibbsiella quercinecans | Pseudomonadota | Gammaproteobacteria |
| Gilliamella apicola | Pseudomonadota | Gammaproteobacteria |
| Gilliamella apicola | Pseudomonadota | Gammaproteobacteria |
| Gilliamella apis | Pseudomonadota | Gammaproteobacteria |
| Gilliamella apis | Pseudomonadota | Gammaproteobacteria |
| Gilvimarinus japonicus | Pseudomonadota | Gammaproteobacteria |
| Gilvimarinus japonicus | Pseudomonadota | Gammaproteobacteria |
| Glaesserella parasuis | Pseudomonadota | Gammaproteobacteria |
| Glaesserella parasuis | Pseudomonadota | Gammaproteobacteria |
| Granulosicoccus antarcticus IMCC3135 | Pseudomonadota | Gammaproteobacteria |
| Granulosicoccus antarcticus IMCC3135 | Pseudomonadota | Gammaproteobacteria |
| Grimontia hollisae | Pseudomonadota | Gammaproteobacteria |
| Grimontia hollisae | Pseudomonadota | Gammaproteobacteria |
| Grimontia kaedaensis | Pseudomonadota | Gammaproteobacteria |
| Grimontia kaedaensis | Pseudomonadota | Gammaproteobacteria |
| Guyparkeria halophila | Pseudomonadota | Gammaproteobacteria |
| Guyparkeria halophila | Pseudomonadota | Gammaproteobacteria |
| Haemophilus aegyptius | Pseudomonadota | Gammaproteobacteria |
| Haemophilus aegyptius | Pseudomonadota | Gammaproteobacteria |
| Haemophilus haemolyticus | Pseudomonadota | Gammaproteobacteria |
| Haemophilus haemolyticus | Pseudomonadota | Gammaproteobacteria |
| Haemophilus influenzae | Pseudomonadota | Gammaproteobacteria |
| Haemophilus influenzae | Pseudomonadota | Gammaproteobacteria |
| Haemophilus parahaemolyticus | Pseudomonadota | Gammaproteobacteria |
| Haemophilus parahaemolyticus | Pseudomonadota | Gammaproteobacteria |
| Haemophilus parainfluenzae | Pseudomonadota | Gammaproteobacteria |
| Haemophilus parainfluenzae | Pseudomonadota | Gammaproteobacteria |

| | | |
|---|---|---|
| Haemophilus pittmaniae | Pseudomonadota | Gammaproteobacteria |
| Haemophilus pittmaniae | Pseudomonadota | Gammaproteobacteria |
| Hafnia alvei | Pseudomonadota | Gammaproteobacteria |
| Hafnia alvei | Pseudomonadota | Gammaproteobacteria |
| Hafnia paralvei | Pseudomonadota | Gammaproteobacteria |
| Hafnia paralvei | Pseudomonadota | Gammaproteobacteria |
| Hahella chejuensis KCTC 2396 | Pseudomonadota | Gammaproteobacteria |
| Hahella chejuensis KCTC 2396 | Pseudomonadota | Gammaproteobacteria |
| Halioglobus maricola | Pseudomonadota | Gammaproteobacteria |
| Halioglobus maricola | Pseudomonadota | Gammaproteobacteria |
| Halioxenophilus aromaticivorans | Pseudomonadota | Gammaproteobacteria |
| Halioxenophilus aromaticivorans | Pseudomonadota | Gammaproteobacteria |
| Halomonas aestuarii | Pseudomonadota | Gammaproteobacteria |
| Halomonas alkaliantarctica | Pseudomonadota | Gammaproteobacteria |
| Halomonas alkalicola | Pseudomonadota | Gammaproteobacteria |
| Halomonas beimenensis | Pseudomonadota | Gammaproteobacteria |
| Halomonas binhaiensis | Pseudomonadota | Gammaproteobacteria |
| Halomonas chromatireducens | Pseudomonadota | Gammaproteobacteria |
| Halomonas elongata DSM 2581 | Pseudomonadota | Gammaproteobacteria |
| Halomonas halophila | Pseudomonadota | Gammaproteobacteria |
| Halomonas huangheensis | Pseudomonadota | Gammaproteobacteria |
| Halomonas hydrothermalis | Pseudomonadota | Gammaproteobacteria |
| Halomonas kashgarensis | Pseudomonadota | Gammaproteobacteria |
| Halomonas piscis | Pseudomonadota | Gammaproteobacteria |
| Halomonas qaidamensis | Pseudomonadota | Gammaproteobacteria |
| Halomonas qinghailakensis | Pseudomonadota | Gammaproteobacteria |
| Halomonas sulfidivorans | Pseudomonadota | Gammaproteobacteria |
| Halopseudomonas aestusnigri | Pseudomonadota | Gammaproteobacteria |
| Halopseudomonas aestusnigri | Pseudomonadota | Gammaproteobacteria |
| Halopseudomonas litoralis | Pseudomonadota | Gammaproteobacteria |
| Halopseudomonas litoralis | Pseudomonadota | Gammaproteobacteria |

| | | |
|---|---|---|
| Halopseudomonas maritima | Pseudomonadota | Gammaproteobacteria |
| Halopseudomonas maritima | Pseudomonadota | Gammaproteobacteria |
| Halopseudomonas nanhaiensis | Pseudomonadota | Gammaproteobacteria |
| Halopseudomonas nanhaiensis | Pseudomonadota | Gammaproteobacteria |
| Halopseudomonas oceani | Pseudomonadota | Gammaproteobacteria |
| Halopseudomonas oceani | Pseudomonadota | Gammaproteobacteria |
| Halopseudomonas phragmitis | Pseudomonadota | Gammaproteobacteria |
| Halopseudomonas phragmitis | Pseudomonadota | Gammaproteobacteria |
| Halopseudomonas sabulinigri | Pseudomonadota | Gammaproteobacteria |
| Halopseudomonas sabulinigri | Pseudomonadota | Gammaproteobacteria |
| Halopseudomonas salegens | Pseudomonadota | Gammaproteobacteria |
| Halopseudomonas salegens | Pseudomonadota | Gammaproteobacteria |
| Halopseudomonas xinjiangensis | Pseudomonadota | Gammaproteobacteria |
| Halopseudomonas xinjiangensis | Pseudomonadota | Gammaproteobacteria |
| Halorhodospira halochloris | Pseudomonadota | Gammaproteobacteria |
| Halorhodospira halochloris | Pseudomonadota | Gammaproteobacteria |
| Halorhodospira halophila SL1 | Pseudomonadota | Gammaproteobacteria |
| Halorhodospira halophila SL1 | Pseudomonadota | Gammaproteobacteria |
| Halothiobacillus diazotrophicus | Pseudomonadota | Gammaproteobacteria |
| Halothiobacillus diazotrophicus | Pseudomonadota | Gammaproteobacteria |
| Halothiobacillus neapolitanus c2 | Pseudomonadota | Gammaproteobacteria |
| Halothiobacillus neapolitanus c2 | Pseudomonadota | Gammaproteobacteria |
| Histophilus somni 2336 | Pseudomonadota | Gammaproteobacteria |
| Histophilus somni 2336 | Pseudomonadota | Gammaproteobacteria |
| Hydrocarboniclastica marina | Pseudomonadota | Gammaproteobacteria |
| Hydrocarboniclastica marina | Pseudomonadota | Gammaproteobacteria |
| Hydrogenovibrio crunogenus | Pseudomonadota | Gammaproteobacteria |
| Hydrogenovibrio crunogenus | Pseudomonadota | Gammaproteobacteria |
| Hydrogenovibrio marinus | Pseudomonadota | Gammaproteobacteria |
| Hydrogenovibrio marinus | Pseudomonadota | Gammaproteobacteria |
| Hydrogenovibrio thermophilus | Pseudomonadota | Gammaproteobacteria |

| | | |
|---|---|---|
| Hydrogenovibrio thermophilus | Pseudomonadota | Gammaproteobacteria |
| Idiomarina abyssalis | Pseudomonadota | Gammaproteobacteria |
| Idiomarina abyssalis | Pseudomonadota | Gammaproteobacteria |
| Idiomarina loihiensis L2TR | Pseudomonadota | Gammaproteobacteria |
| Idiomarina loihiensis L2TR | Pseudomonadota | Gammaproteobacteria |
| Ignatzschineria larvae DSM 13226 | Pseudomonadota | Gammaproteobacteria |
| Ignatzschineria larvae DSM 13226 | Pseudomonadota | Gammaproteobacteria |
| Ignatzschineria rhizosphaerae | Pseudomonadota | Gammaproteobacteria |
| Ignatzschineria rhizosphaerae | Pseudomonadota | Gammaproteobacteria |
| Immundisolibacter cernigliae | Pseudomonadota | Gammaproteobacteria |
| Immundisolibacter cernigliae | Pseudomonadota | Gammaproteobacteria |
| Isoalcanivorax pacificus W11-5 | Pseudomonadota | Gammaproteobacteria |
| Isoalcanivorax pacificus W11-5 | Pseudomonadota | Gammaproteobacteria |
| Isorropodon fossajaponicum endosymbiont JTNG4 | Pseudomonadota | Gammaproteobacteria |
| Isorropodon fossajaponicum endosymbiont JTNG4 | Pseudomonadota | Gammaproteobacteria |
| Jejubacter calystegiae | Pseudomonadota | Gammaproteobacteria |
| Jejubacter calystegiae | Pseudomonadota | Gammaproteobacteria |
| Kangiella geojedonensis | Pseudomonadota | Gammaproteobacteria |
| Kangiella geojedonensis | Pseudomonadota | Gammaproteobacteria |
| Kangiella koreensis DSM 16069 | Pseudomonadota | Gammaproteobacteria |
| Kangiella koreensis DSM 16069 | Pseudomonadota | Gammaproteobacteria |
| Kangiella sediminilitoris | Pseudomonadota | Gammaproteobacteria |
| Kangiella sediminilitoris | Pseudomonadota | Gammaproteobacteria |
| Ketobacter alkanivorans | Pseudomonadota | Gammaproteobacteria |
| Ketobacter alkanivorans | Pseudomonadota | Gammaproteobacteria |
| Klebsiella aerogenes | Pseudomonadota | Gammaproteobacteria |
| Klebsiella aerogenes | Pseudomonadota | Gammaproteobacteria |
| Klebsiella africana | Pseudomonadota | Gammaproteobacteria |
| Klebsiella africana | Pseudomonadota | Gammaproteobacteria |
| Klebsiella electrica | Pseudomonadota | Gammaproteobacteria |
| Klebsiella electrica | Pseudomonadota | Gammaproteobacteria |

| | | |
|---|---|---|
| Klebsiella grimontii | Pseudomonadota | Gammaproteobacteria |
| Klebsiella grimontii | Pseudomonadota | Gammaproteobacteria |
| Klebsiella huaxiensis | Pseudomonadota | Gammaproteobacteria |
| Klebsiella huaxiensis | Pseudomonadota | Gammaproteobacteria |
| Klebsiella michiganensis | Pseudomonadota | Gammaproteobacteria |
| Klebsiella michiganensis | Pseudomonadota | Gammaproteobacteria |
| Klebsiella oxytoca | Pseudomonadota | Gammaproteobacteria |
| Klebsiella oxytoca | Pseudomonadota | Gammaproteobacteria |
| Klebsiella pasteurii | Pseudomonadota | Gammaproteobacteria |
| Klebsiella pasteurii | Pseudomonadota | Gammaproteobacteria |
| Klebsiella quasipneumoniae | Pseudomonadota | Gammaproteobacteria |
| Klebsiella quasipneumoniae | Pseudomonadota | Gammaproteobacteria |
| Klebsiella quasivariicola | Pseudomonadota | Gammaproteobacteria |
| Klebsiella quasivariicola | Pseudomonadota | Gammaproteobacteria |
| Klebsiella variicola subsp. variicola | Pseudomonadota | Gammaproteobacteria |
| Klebsiella variicola subsp. variicola | Pseudomonadota | Gammaproteobacteria |
| Kluyvera ascorbata | Pseudomonadota | Gammaproteobacteria |
| Kluyvera ascorbata | Pseudomonadota | Gammaproteobacteria |
| Kluyvera intermedia | Pseudomonadota | Gammaproteobacteria |
| Kluyvera intermedia | Pseudomonadota | Gammaproteobacteria |
| Kluyvera sichuanensis | Pseudomonadota | Gammaproteobacteria |
| Kluyvera sichuanensis | Pseudomonadota | Gammaproteobacteria |
| Kosakonia arachidis | Pseudomonadota | Gammaproteobacteria |
| Kosakonia arachidis | Pseudomonadota | Gammaproteobacteria |
| Kosakonia calanthes | Pseudomonadota | Gammaproteobacteria |
| Kosakonia calanthes | Pseudomonadota | Gammaproteobacteria |
| Kosakonia cowanii | Pseudomonadota | Gammaproteobacteria |
| Kosakonia cowanii | Pseudomonadota | Gammaproteobacteria |
| Kosakonia oryzae | Pseudomonadota | Gammaproteobacteria |
| Kosakonia oryzae | Pseudomonadota | Gammaproteobacteria |
| Kosakonia oryzendophytica | Pseudomonadota | Gammaproteobacteria |

| | | |
|---|---|---|
| Kosakonia oryzendophytica | Pseudomonadota | Gammaproteobacteria |
| Kosakonia pseudosacchari | Pseudomonadota | Gammaproteobacteria |
| Kosakonia pseudosacchari | Pseudomonadota | Gammaproteobacteria |
| Kosakonia radicincitans | Pseudomonadota | Gammaproteobacteria |
| Kosakonia radicincitans | Pseudomonadota | Gammaproteobacteria |
| Kosakonia sacchari SP1 | Pseudomonadota | Gammaproteobacteria |
| Kosakonia sacchari SP1 | Pseudomonadota | Gammaproteobacteria |
| Kushneria konosiri | Pseudomonadota | Gammaproteobacteria |
| Kushneria marisflavi | Pseudomonadota | Gammaproteobacteria |
| Kushneria phosphatilytica | Pseudomonadota | Gammaproteobacteria |
| Leclercia adecarboxylata | Pseudomonadota | Gammaproteobacteria |
| Leclercia adecarboxylata | Pseudomonadota | Gammaproteobacteria |
| Leclercia pneumoniae | Pseudomonadota | Gammaproteobacteria |
| Leclercia pneumoniae | Pseudomonadota | Gammaproteobacteria |
| Legionella anisa | Pseudomonadota | Gammaproteobacteria |
| Legionella anisa | Pseudomonadota | Gammaproteobacteria |
| Legionella antarctica | Pseudomonadota | Gammaproteobacteria |
| Legionella antarctica | Pseudomonadota | Gammaproteobacteria |
| Legionella cardiaca | Pseudomonadota | Gammaproteobacteria |
| Legionella cardiaca | Pseudomonadota | Gammaproteobacteria |
| Legionella clemsonensis | Pseudomonadota | Gammaproteobacteria |
| Legionella clemsonensis | Pseudomonadota | Gammaproteobacteria |
| Legionella fallonii LLAP-10 | Pseudomonadota | Gammaproteobacteria |
| Legionella fallonii LLAP-10 | Pseudomonadota | Gammaproteobacteria |
| Legionella geestiana | Pseudomonadota | Gammaproteobacteria |
| Legionella geestiana | Pseudomonadota | Gammaproteobacteria |
| Legionella hackeliae | Pseudomonadota | Gammaproteobacteria |
| Legionella hackeliae | Pseudomonadota | Gammaproteobacteria |
| Legionella israelensis | Pseudomonadota | Gammaproteobacteria |
| Legionella israelensis | Pseudomonadota | Gammaproteobacteria |
| Legionella jordanis | Pseudomonadota | Gammaproteobacteria |

| | | |
|---|---|---|
| Legionella jordanis | Pseudomonadota | Gammaproteobacteria |
| Legionella lansingensis | Pseudomonadota | Gammaproteobacteria |
| Legionella lansingensis | Pseudomonadota | Gammaproteobacteria |
| Legionella longbeachae NSW150 | Pseudomonadota | Gammaproteobacteria |
| Legionella longbeachae NSW150 | Pseudomonadota | Gammaproteobacteria |
| Legionella lytica | Pseudomonadota | Gammaproteobacteria |
| Legionella lytica | Pseudomonadota | Gammaproteobacteria |
| Legionella micdadei | Pseudomonadota | Gammaproteobacteria |
| Legionella micdadei | Pseudomonadota | Gammaproteobacteria |
| Legionella pneumophila | Pseudomonadota | Gammaproteobacteria |
| Legionella pneumophila | Pseudomonadota | Gammaproteobacteria |
| Legionella sainthelensi | Pseudomonadota | Gammaproteobacteria |
| Legionella sainthelensi | Pseudomonadota | Gammaproteobacteria |
| Legionella spiritensis | Pseudomonadota | Gammaproteobacteria |
| Legionella spiritensis | Pseudomonadota | Gammaproteobacteria |
| Legionella waltersii | Pseudomonadota | Gammaproteobacteria |
| Legionella waltersii | Pseudomonadota | Gammaproteobacteria |
| Lelliottia amnigena | Pseudomonadota | Gammaproteobacteria |
| Lelliottia amnigena | Pseudomonadota | Gammaproteobacteria |
| Lelliottia nimipressuralis | Pseudomonadota | Gammaproteobacteria |
| Lelliottia nimipressuralis | Pseudomonadota | Gammaproteobacteria |
| Leminorella grimontii | Pseudomonadota | Gammaproteobacteria |
| Leminorella grimontii | Pseudomonadota | Gammaproteobacteria |
| Leminorella richardii | Pseudomonadota | Gammaproteobacteria |
| Leminorella richardii | Pseudomonadota | Gammaproteobacteria |
| Limnobaculum parvum | Pseudomonadota | Gammaproteobacteria |
| Limnobaculum parvum | Pseudomonadota | Gammaproteobacteria |
| Limnobaculum zhutongyuii | Pseudomonadota | Gammaproteobacteria |
| Limnobaculum zhutongyuii | Pseudomonadota | Gammaproteobacteria |
| Litorilituus sediminis | Pseudomonadota | Gammaproteobacteria |
| Litorilituus sediminis | Pseudomonadota | Gammaproteobacteria |

| | | |
|---|---|---|
| Litorivicinus lipolyticus | Pseudomonadota | Gammaproteobacteria |
| Litorivicinus lipolyticus | Pseudomonadota | Gammaproteobacteria |
| Lonepinella koalarum | Pseudomonadota | Gammaproteobacteria |
| Lonepinella koalarum | Pseudomonadota | Gammaproteobacteria |
| Lonsdalea britannica | Pseudomonadota | Gammaproteobacteria |
| Lonsdalea britannica | Pseudomonadota | Gammaproteobacteria |
| Lonsdalea populi | Pseudomonadota | Gammaproteobacteria |
| Lonsdalea populi | Pseudomonadota | Gammaproteobacteria |
| Luteibacter aegosomaticola | Pseudomonadota | Gammaproteobacteria |
| Luteibacter aegosomaticola | Pseudomonadota | Gammaproteobacteria |
| Luteibacter aegosomatis | Pseudomonadota | Gammaproteobacteria |
| Luteibacter aegosomatis | Pseudomonadota | Gammaproteobacteria |
| Luteibacter aegosomatissinici | Pseudomonadota | Gammaproteobacteria |
| Luteibacter aegosomatissinici | Pseudomonadota | Gammaproteobacteria |
| Luteibacter anthropi | Pseudomonadota | Gammaproteobacteria |
| Luteibacter anthropi | Pseudomonadota | Gammaproteobacteria |
| Luteibacter flocculans | Pseudomonadota | Gammaproteobacteria |
| Luteibacter flocculans | Pseudomonadota | Gammaproteobacteria |
| Luteibacter pinisoli | Pseudomonadota | Gammaproteobacteria |
| Luteibacter pinisoli | Pseudomonadota | Gammaproteobacteria |
| Luteibacter rhizovicinus DSM 16549 | Pseudomonadota | Gammaproteobacteria |
| Luteibacter rhizovicinus DSM 16549 | Pseudomonadota | Gammaproteobacteria |
| Luteimonas chenhongjianii | Pseudomonadota | Gammaproteobacteria |
| Luteimonas chenhongjianii | Pseudomonadota | Gammaproteobacteria |
| Luteimonas fraxinea | Pseudomonadota | Gammaproteobacteria |
| Luteimonas fraxinea | Pseudomonadota | Gammaproteobacteria |
| Luteimonas granuli | Pseudomonadota | Gammaproteobacteria |
| Luteimonas granuli | Pseudomonadota | Gammaproteobacteria |
| Luteimonas yindakuii | Pseudomonadota | Gammaproteobacteria |
| Luteimonas yindakuii | Pseudomonadota | Gammaproteobacteria |
| Lysobacter antibioticus | Pseudomonadota | Gammaproteobacteria |

| | | |
|---|---|---|
| Lysobacter antibioticus | Pseudomonadota | Gammaproteobacteria |
| Lysobacter arenosi | Pseudomonadota | Gammaproteobacteria |
| Lysobacter arenosi | Pseudomonadota | Gammaproteobacteria |
| Lysobacter auxotrophicus | Pseudomonadota | Gammaproteobacteria |
| Lysobacter auxotrophicus | Pseudomonadota | Gammaproteobacteria |
| Lysobacter capsici | Pseudomonadota | Gammaproteobacteria |
| Lysobacter capsici | Pseudomonadota | Gammaproteobacteria |
| Lysobacter enzymogenes | Pseudomonadota | Gammaproteobacteria |
| Lysobacter enzymogenes | Pseudomonadota | Gammaproteobacteria |
| Lysobacter firmicutimachus | Pseudomonadota | Gammaproteobacteria |
| Lysobacter firmicutimachus | Pseudomonadota | Gammaproteobacteria |
| Lysobacter gummosus | Pseudomonadota | Gammaproteobacteria |
| Lysobacter gummosus | Pseudomonadota | Gammaproteobacteria |
| Lysobacter helvus | Pseudomonadota | Gammaproteobacteria |
| Lysobacter helvus | Pseudomonadota | Gammaproteobacteria |
| Lysobacter soli | Pseudomonadota | Gammaproteobacteria |
| Lysobacter soli | Pseudomonadota | Gammaproteobacteria |
| Lysobacter soyae | Pseudomonadota | Gammaproteobacteria |
| Lysobacter soyae | Pseudomonadota | Gammaproteobacteria |
| Lysobacter yananisis | Pseudomonadota | Gammaproteobacteria |
| Lysobacter yananisis | Pseudomonadota | Gammaproteobacteria |
| Mangrovibacter phragmitis | Pseudomonadota | Gammaproteobacteria |
| Mangrovibacter phragmitis | Pseudomonadota | Gammaproteobacteria |
| Mannheimia bovis | Pseudomonadota | Gammaproteobacteria |
| Mannheimia bovis | Pseudomonadota | Gammaproteobacteria |
| Mannheimia glucosida | Pseudomonadota | Gammaproteobacteria |
| Mannheimia glucosida | Pseudomonadota | Gammaproteobacteria |
| Mannheimia haemolytica | Pseudomonadota | Gammaproteobacteria |
| Mannheimia haemolytica | Pseudomonadota | Gammaproteobacteria |
| Mannheimia pernigra | Pseudomonadota | Gammaproteobacteria |
| Mannheimia pernigra | Pseudomonadota | Gammaproteobacteria |

| | | |
|---|---|---|
| Mannheimia varigena | Pseudomonadota | Gammaproteobacteria |
| Mannheimia varigena | Pseudomonadota | Gammaproteobacteria |
| Marichromatium purpuratum 984 | Pseudomonadota | Gammaproteobacteria |
| Marichromatium purpuratum 984 | Pseudomonadota | Gammaproteobacteria |
| Maricurvus nonylphenolicus | Pseudomonadota | Gammaproteobacteria |
| Maricurvus nonylphenolicus | Pseudomonadota | Gammaproteobacteria |
| Marilutibacter alkalisoli | Pseudomonadota | Gammaproteobacteria |
| Marilutibacter alkalisoli | Pseudomonadota | Gammaproteobacteria |
| Marilutibacter maris | Pseudomonadota | Gammaproteobacteria |
| Marilutibacter maris | Pseudomonadota | Gammaproteobacteria |
| Marinagarivorans cellulosilyticus | Pseudomonadota | Gammaproteobacteria |
| Marinicellulosiphila megalodicopiae | Pseudomonadota | Gammaproteobacteria |
| Marinicellulosiphila megalodicopiae | Pseudomonadota | Gammaproteobacteria |
| Marinobacter adhaerens | Pseudomonadota | Gammaproteobacteria |
| Marinobacter adhaerens | Pseudomonadota | Gammaproteobacteria |
| Marinobacter alkaliphilus | Pseudomonadota | Gammaproteobacteria |
| Marinobacter alkaliphilus | Pseudomonadota | Gammaproteobacteria |
| Marinobacter apostichopi | Pseudomonadota | Gammaproteobacteria |
| Marinobacter apostichopi | Pseudomonadota | Gammaproteobacteria |
| Marinobacter flavimaris | Pseudomonadota | Gammaproteobacteria |
| Marinobacter flavimaris | Pseudomonadota | Gammaproteobacteria |
| Marinobacter fonticola | Pseudomonadota | Gammaproteobacteria |
| Marinobacter fonticola | Pseudomonadota | Gammaproteobacteria |
| Marinobacter metalliresistant | Pseudomonadota | Gammaproteobacteria |
| Marinobacter metalliresistant | Pseudomonadota | Gammaproteobacteria |
| Marinobacter nauticus ATCC 49840 | Pseudomonadota | Gammaproteobacteria |
| Marinobacter nauticus ATCC 49840 | Pseudomonadota | Gammaproteobacteria |
| Marinobacter psychrophilus | Pseudomonadota | Gammaproteobacteria |
| Marinobacter psychrophilus | Pseudomonadota | Gammaproteobacteria |
| Marinobacter salarius | Pseudomonadota | Gammaproteobacteria |
| Marinobacter salarius | Pseudomonadota | Gammaproteobacteria |

| | | |
|---|---|---|
| Marinobacter salinisoli | Pseudomonadota | Gammaproteobacteria |
| Marinobacter salinisoli | Pseudomonadota | Gammaproteobacteria |
| Marinobacter salinus | Pseudomonadota | Gammaproteobacteria |
| Marinobacter salinus | Pseudomonadota | Gammaproteobacteria |
| Marinobacter shengliensis | Pseudomonadota | Gammaproteobacteria |
| Marinobacter shengliensis | Pseudomonadota | Gammaproteobacteria |
| Marinobacter similis | Pseudomonadota | Gammaproteobacteria |
| Marinobacterium aestuarii | Pseudomonadota | Gammaproteobacteria |
| Marinobacterium aestuarii | Pseudomonadota | Gammaproteobacteria |
| Marinobacterium iners | Pseudomonadota | Gammaproteobacteria |
| Marinobacterium iners | Pseudomonadota | Gammaproteobacteria |
| Marinomonas algicola | Pseudomonadota | Gammaproteobacteria |
| Marinomonas arctica | Pseudomonadota | Gammaproteobacteria |
| Marinomonas foliarum | Pseudomonadota | Gammaproteobacteria |
| Marinomonas mediterranea MMB-1 | Pseudomonadota | Gammaproteobacteria |
| Marinomonas polaris | Pseudomonadota | Gammaproteobacteria |
| Marinomonas pontica | Pseudomonadota | Gammaproteobacteria |
| Marinomonas posidonica IVIA-Po-181 | Pseudomonadota | Gammaproteobacteria |
| Marinomonas primoryensis | Pseudomonadota | Gammaproteobacteria |
| Marinomonas profundi | Pseudomonadota | Gammaproteobacteria |
| Mergibacter septicus | Pseudomonadota | Gammaproteobacteria |
| Mergibacter septicus | Pseudomonadota | Gammaproteobacteria |
| Metapseudomonas boanensis | Pseudomonadota | Gammaproteobacteria |
| Metapseudomonas boanensis | Pseudomonadota | Gammaproteobacteria |
| Metapseudomonas furukawaii | Pseudomonadota | Gammaproteobacteria |
| Metapseudomonas furukawaii | Pseudomonadota | Gammaproteobacteria |
| Metapseudomonas lalkuanensis | Pseudomonadota | Gammaproteobacteria |
| Metapseudomonas lalkuanensis | Pseudomonadota | Gammaproteobacteria |
| Metapseudomonas otitidis | Pseudomonadota | Gammaproteobacteria |
| Metapseudomonas otitidis | Pseudomonadota | Gammaproteobacteria |
| Methylocaldum marinum | Pseudomonadota | Gammaproteobacteria |

| | | |
|---|---|---|
| Methylocaldum marinum | Pseudomonadota | Gammaproteobacteria |
| Methylocaldum szegediense | Pseudomonadota | Gammaproteobacteria |
| Methylocaldum szegediense | Pseudomonadota | Gammaproteobacteria |
| Methylococcus capsulatus str. Bath | Pseudomonadota | Gammaproteobacteria |
| Methylococcus capsulatus str. Bath | Pseudomonadota | Gammaproteobacteria |
| Methylococcus geothermalis | Pseudomonadota | Gammaproteobacteria |
| Methylococcus geothermalis | Pseudomonadota | Gammaproteobacteria |
| Methylococcus mesophilus | Pseudomonadota | Gammaproteobacteria |
| Methylococcus mesophilus | Pseudomonadota | Gammaproteobacteria |
| Methylogaea oryzae | Pseudomonadota | Gammaproteobacteria |
| Methylogaea oryzae | Pseudomonadota | Gammaproteobacteria |
| Methylomagnum ishizawai | Pseudomonadota | Gammaproteobacteria |
| Methylomagnum ishizawai | Pseudomonadota | Gammaproteobacteria |
| Methylomarinovum caldicuralii | Pseudomonadota | Gammaproteobacteria |
| Methylomarinovum caldicuralii | Pseudomonadota | Gammaproteobacteria |
| Methylomarinovum tepidoasis | Pseudomonadota | Gammaproteobacteria |
| Methylomarinovum tepidoasis | Pseudomonadota | Gammaproteobacteria |
| Methylomarinum roseum | Pseudomonadota | Gammaproteobacteria |
| Methylomarinum roseum | Pseudomonadota | Gammaproteobacteria |
| Methylomicrobium album BG8 | Pseudomonadota | Gammaproteobacteria |
| Methylomicrobium album BG8 | Pseudomonadota | Gammaproteobacteria |
| Methylomonas denitrificans | Pseudomonadota | Gammaproteobacteria |
| Methylomonas denitrificans | Pseudomonadota | Gammaproteobacteria |
| Methylomonas koyamae | Pseudomonadota | Gammaproteobacteria |
| Methylomonas koyamae | Pseudomonadota | Gammaproteobacteria |
| Methylomonas montana | Pseudomonadota | Gammaproteobacteria |
| Methylomonas montana | Pseudomonadota | Gammaproteobacteria |
| Methylomonas paludis | Pseudomonadota | Gammaproteobacteria |
| Methylomonas paludis | Pseudomonadota | Gammaproteobacteria |
| Methylomonas rapida | Pseudomonadota | Gammaproteobacteria |
| Methylomonas rapida | Pseudomonadota | Gammaproteobacteria |

| | | |
|---|---|---|
| Methylomonas rhizoryzae | Pseudomonadota | Gammaproteobacteria |
| Methylomonas rhizoryzae | Pseudomonadota | Gammaproteobacteria |
| Methylophaga frappieri | Pseudomonadota | Gammaproteobacteria |
| Methylophaga frappieri | Pseudomonadota | Gammaproteobacteria |
| Methylophaga marina | Pseudomonadota | Gammaproteobacteria |
| Methylophaga marina | Pseudomonadota | Gammaproteobacteria |
| Methylophaga nitratireducenticrescens | Pseudomonadota | Gammaproteobacteria |
| Methylophaga nitratireducenticrescens | Pseudomonadota | Gammaproteobacteria |
| Methylophaga pinxianii | Pseudomonadota | Gammaproteobacteria |
| Methylophaga pinxianii | Pseudomonadota | Gammaproteobacteria |
| Methylotuvimicrobium alcaliphilum 20Z | Pseudomonadota | Gammaproteobacteria |
| Methylotuvimicrobium alcaliphilum 20Z | Pseudomonadota | Gammaproteobacteria |
| Methylotuvimicrobium buryatense | Pseudomonadota | Gammaproteobacteria |
| Methylotuvimicrobium buryatense | Pseudomonadota | Gammaproteobacteria |
| Methylovulum psychrotolerans | Pseudomonadota | Gammaproteobacteria |
| Methylovulum psychrotolerans | Pseudomonadota | Gammaproteobacteria |
| Microbulbifer agarilyticus | Pseudomonadota | Gammaproteobacteria |
| Microbulbifer agarilyticus | Pseudomonadota | Gammaproteobacteria |
| Microbulbifer aggregans | Pseudomonadota | Gammaproteobacteria |
| Microbulbifer aggregans | Pseudomonadota | Gammaproteobacteria |
| Microbulbifer bruguierae | Pseudomonadota | Gammaproteobacteria |
| Microbulbifer bruguierae | Pseudomonadota | Gammaproteobacteria |
| Microbulbifer celer | Pseudomonadota | Gammaproteobacteria |
| Microbulbifer celer | Pseudomonadota | Gammaproteobacteria |
| Microbulbifer elongatus | Pseudomonadota | Gammaproteobacteria |
| Microbulbifer elongatus | Pseudomonadota | Gammaproteobacteria |
| Microbulbifer hydrolyticus | Pseudomonadota | Gammaproteobacteria |
| Microbulbifer hydrolyticus | Pseudomonadota | Gammaproteobacteria |
| Microbulbifer pacificus | Pseudomonadota | Gammaproteobacteria |
| Microbulbifer pacificus | Pseudomonadota | Gammaproteobacteria |
| Microbulbifer spongiae | Pseudomonadota | Gammaproteobacteria |

| | | |
|---|---|---|
| Microbulbifer spongiae | Pseudomonadota | Gammaproteobacteria |
| Microbulbifer thermotolerans | Pseudomonadota | Gammaproteobacteria |
| Microbulbifer thermotolerans | Pseudomonadota | Gammaproteobacteria |
| Mixta gaviniae | Pseudomonadota | Gammaproteobacteria |
| Mixta gaviniae | Pseudomonadota | Gammaproteobacteria |
| Mixta hanseatica | Pseudomonadota | Gammaproteobacteria |
| Mixta hanseatica | Pseudomonadota | Gammaproteobacteria |
| Mixta intestinalis | Pseudomonadota | Gammaproteobacteria |
| Mixta intestinalis | Pseudomonadota | Gammaproteobacteria |
| Moellerella wisconsensis | Pseudomonadota | Gammaproteobacteria |
| Moellerella wisconsensis | Pseudomonadota | Gammaproteobacteria |
| Moraxella bovis | Pseudomonadota | Gammaproteobacteria |
| Moraxella bovis | Pseudomonadota | Gammaproteobacteria |
| Moraxella bovoculi | Pseudomonadota | Gammaproteobacteria |
| Moraxella bovoculi | Pseudomonadota | Gammaproteobacteria |
| Moraxella canis | Pseudomonadota | Gammaproteobacteria |
| Moraxella canis | Pseudomonadota | Gammaproteobacteria |
| Moraxella catarrhalis | Pseudomonadota | Gammaproteobacteria |
| Moraxella catarrhalis | Pseudomonadota | Gammaproteobacteria |
| Moraxella cuniculi | Pseudomonadota | Gammaproteobacteria |
| Moraxella cuniculi | Pseudomonadota | Gammaproteobacteria |
| Moraxella haemolytica | Pseudomonadota | Gammaproteobacteria |
| Moraxella haemolytica | Pseudomonadota | Gammaproteobacteria |
| Moraxella lincolnii | Pseudomonadota | Gammaproteobacteria |
| Moraxella lincolnii | Pseudomonadota | Gammaproteobacteria |
| Moraxella nasibovis | Pseudomonadota | Gammaproteobacteria |
| Moraxella nasibovis | Pseudomonadota | Gammaproteobacteria |
| Moraxella nasicaprae | Pseudomonadota | Gammaproteobacteria |
| Moraxella nasicaprae | Pseudomonadota | Gammaproteobacteria |
| Moraxella nasovis | Pseudomonadota | Gammaproteobacteria |
| Moraxella nasovis | Pseudomonadota | Gammaproteobacteria |

| | | |
|---|---|---|
| Moraxella nonliquefaciens | Pseudomonadota | Gammaproteobacteria |
| Moraxella nonliquefaciens | Pseudomonadota | Gammaproteobacteria |
| Moraxella osloensis | Pseudomonadota | Gammaproteobacteria |
| Moraxella osloensis | Pseudomonadota | Gammaproteobacteria |
| Moraxella ovis | Pseudomonadota | Gammaproteobacteria |
| Moraxella ovis | Pseudomonadota | Gammaproteobacteria |
| Moraxella veridica | Pseudomonadota | Gammaproteobacteria |
| Moraxella veridica | Pseudomonadota | Gammaproteobacteria |
| Morganella morganii subsp. morganii | Pseudomonadota | Gammaproteobacteria |
| Morganella morganii subsp. morganii | Pseudomonadota | Gammaproteobacteria |
| Moritella marina ATCC 15381 | Pseudomonadota | Gammaproteobacteria |
| Moritella yayanosii | Pseudomonadota | Gammaproteobacteria |
| Musicola paradisiaca Ech703 | Pseudomonadota | Gammaproteobacteria |
| Musicola paradisiaca Ech703 | Pseudomonadota | Gammaproteobacteria |
| Neptuniibacter halophilus | Pseudomonadota | Gammaproteobacteria |
| Neptuniibacter halophilus | Pseudomonadota | Gammaproteobacteria |
| Neptuniibacter marinus | Pseudomonadota | Gammaproteobacteria |
| Neptuniibacter marinus | Pseudomonadota | Gammaproteobacteria |
| Neptunomonas concharum | Pseudomonadota | Gammaproteobacteria |
| Neptunomonas concharum | Pseudomonadota | Gammaproteobacteria |
| Neptunomonas phycophila | Pseudomonadota | Gammaproteobacteria |
| Neptunomonas phycophila | Pseudomonadota | Gammaproteobacteria |
| Nitrincola iocasae | Pseudomonadota | Gammaproteobacteria |
| Nitrincola iocasae | Pseudomonadota | Gammaproteobacteria |
| Nitrosococcus halophilus Nc 4 | Pseudomonadota | Gammaproteobacteria |
| Nitrosococcus halophilus Nc 4 | Pseudomonadota | Gammaproteobacteria |
| Nitrosococcus oceani ATCC 19707 | Pseudomonadota | Gammaproteobacteria |
| Nitrosococcus oceani ATCC 19707 | Pseudomonadota | Gammaproteobacteria |
| Nitrosococcus wardiae | Pseudomonadota | Gammaproteobacteria |
| Nitrosococcus wardiae | Pseudomonadota | Gammaproteobacteria |
| Nitrosococcus watsonii C-113 | Pseudomonadota | Gammaproteobacteria |

| | | |
|---|---|---|
| Nitrosococcus watsonii C-113 | Pseudomonadota | Gammaproteobacteria |
| Nitrosococcus watsonii C-113 | Pseudomonadota | Gammaproteobacteria |
| Noviluteimonas caseinilytica | Pseudomonadota | Gammaproteobacteria |
| Noviluteimonas caseinilytica | Pseudomonadota | Gammaproteobacteria |
| Novilysobacter avium | Pseudomonadota | Gammaproteobacteria |
| Novilysobacter avium | Pseudomonadota | Gammaproteobacteria |
| Novilysobacter ciconiae | Pseudomonadota | Gammaproteobacteria |
| Novilysobacter ciconiae | Pseudomonadota | Gammaproteobacteria |
| Novilysobacter luteus | Pseudomonadota | Gammaproteobacteria |
| Novilysobacter luteus | Pseudomonadota | Gammaproteobacteria |
| Oceanicoccus sagamiensis | Pseudomonadota | Gammaproteobacteria |
| Oceanicoccus sagamiensis | Pseudomonadota | Gammaproteobacteria |
| Oceanimonas pelagia | Pseudomonadota | Gammaproteobacteria |
| Oceanimonas pelagia | Pseudomonadota | Gammaproteobacteria |
| Oceanisphaera avium | Pseudomonadota | Gammaproteobacteria |
| Oceanisphaera profunda | Pseudomonadota | Gammaproteobacteria |
| Oleiphilus messinensis | Pseudomonadota | Gammaproteobacteria |
| Orbus mooreae | Pseudomonadota | Gammaproteobacteria |
| Orbus mooreae | Pseudomonadota | Gammaproteobacteria |
| Orbus sturtevantii | Pseudomonadota | Gammaproteobacteria |
| Orbus sturtevantii | Pseudomonadota | Gammaproteobacteria |
| Orbus wheelerorum | Pseudomonadota | Gammaproteobacteria |
| Orbus wheelerorum | Pseudomonadota | Gammaproteobacteria |
| Otariodibacter oris | Pseudomonadota | Gammaproteobacteria |
| Otariodibacter oris | Pseudomonadota | Gammaproteobacteria |
| Pantoea agglomerans | Pseudomonadota | Gammaproteobacteria |
| Pantoea agglomerans | Pseudomonadota | Gammaproteobacteria |
| Pantoea alfalfae | Pseudomonadota | Gammaproteobacteria |
| Pantoea alfalfae | Pseudomonadota | Gammaproteobacteria |
| Pantoea alhagi | Pseudomonadota | Gammaproteobacteria |
| Pantoea alhagi | Pseudomonadota | Gammaproteobacteria |

| | | |
|---|---|---|
| Pantoea allii | Pseudomonadota | Gammaproteobacteria |
| Pantoea allii | Pseudomonadota | Gammaproteobacteria |
| Pantoea ananatis PA13 | Pseudomonadota | Gammaproteobacteria |
| Pantoea ananatis PA13 | Pseudomonadota | Gammaproteobacteria |
| Pantoea anthophila | Pseudomonadota | Gammaproteobacteria |
| Pantoea anthophila | Pseudomonadota | Gammaproteobacteria |
| Pantoea deleyi | Pseudomonadota | Gammaproteobacteria |
| Pantoea deleyi | Pseudomonadota | Gammaproteobacteria |
| Pantoea dispersa | Pseudomonadota | Gammaproteobacteria |
| Pantoea dispersa | Pseudomonadota | Gammaproteobacteria |
| Pantoea eucalypti | Pseudomonadota | Gammaproteobacteria |
| Pantoea eucalypti | Pseudomonadota | Gammaproteobacteria |
| Pantoea eucrina | Pseudomonadota | Gammaproteobacteria |
| Pantoea eucrina | Pseudomonadota | Gammaproteobacteria |
| Pantoea phytostimulans | Pseudomonadota | Gammaproteobacteria |
| Pantoea phytostimulans | Pseudomonadota | Gammaproteobacteria |
| Pantoea stewartii | Pseudomonadota | Gammaproteobacteria |
| Pantoea stewartii | Pseudomonadota | Gammaproteobacteria |
| Pantoea vagans | Pseudomonadota | Gammaproteobacteria |
| Pantoea vagans | Pseudomonadota | Gammaproteobacteria |
| Paramixta manurensis | Pseudomonadota | Gammaproteobacteria |
| Paramixta manurensis | Pseudomonadota | Gammaproteobacteria |
| Paraphotobacterium marinum | Pseudomonadota | Gammaproteobacteria |
| Paraphotobacterium marinum | Pseudomonadota | Gammaproteobacteria |
| Parashewanella spongiae | Pseudomonadota | Gammaproteobacteria |
| Parashewanella spongiae | Pseudomonadota | Gammaproteobacteria |
| Parashewanella tropica | Pseudomonadota | Gammaproteobacteria |
| Parashewanella tropica | Pseudomonadota | Gammaproteobacteria |
| Pasteurella atlantica | Pseudomonadota | Gammaproteobacteria |
| Pasteurella atlantica | Pseudomonadota | Gammaproteobacteria |
| Pasteurella canis | Pseudomonadota | Gammaproteobacteria |

| | | |
|---|---|---|
| Pasteurella canis | Pseudomonadota | Gammaproteobacteria |
| Pasteurella dagmatis | Pseudomonadota | Gammaproteobacteria |
| Pasteurella dagmatis | Pseudomonadota | Gammaproteobacteria |
| Pasteurella multocida | Pseudomonadota | Gammaproteobacteria |
| Pasteurella multocida | Pseudomonadota | Gammaproteobacteria |
| Pectobacterium aquaticum | Pseudomonadota | Gammaproteobacteria |
| Pectobacterium aquaticum | Pseudomonadota | Gammaproteobacteria |
| Pectobacterium araliae | Pseudomonadota | Gammaproteobacteria |
| Pectobacterium araliae | Pseudomonadota | Gammaproteobacteria |
| Pectobacterium aroidearum | Pseudomonadota | Gammaproteobacteria |
| Pectobacterium aroidearum | Pseudomonadota | Gammaproteobacteria |
| Pectobacterium atrosepticum | Pseudomonadota | Gammaproteobacteria |
| Pectobacterium atrosepticum | Pseudomonadota | Gammaproteobacteria |
| Pectobacterium brasiliense | Pseudomonadota | Gammaproteobacteria |
| Pectobacterium brasiliense | Pseudomonadota | Gammaproteobacteria |
| Pectobacterium cacticida | Pseudomonadota | Gammaproteobacteria |
| Pectobacterium cacticida | Pseudomonadota | Gammaproteobacteria |
| Pectobacterium carotovorum | Pseudomonadota | Gammaproteobacteria |
| Pectobacterium carotovorum | Pseudomonadota | Gammaproteobacteria |
| Pectobacterium colocasium | Pseudomonadota | Gammaproteobacteria |
| Pectobacterium colocasium | Pseudomonadota | Gammaproteobacteria |
| Pectobacterium odoriferum | Pseudomonadota | Gammaproteobacteria |
| Pectobacterium odoriferum | Pseudomonadota | Gammaproteobacteria |
| Pectobacterium parmentieri | Pseudomonadota | Gammaproteobacteria |
| Pectobacterium parmentieri | Pseudomonadota | Gammaproteobacteria |
| Pectobacterium parvum | Pseudomonadota | Gammaproteobacteria |
| Pectobacterium parvum | Pseudomonadota | Gammaproteobacteria |
| Pectobacterium peruviense | Pseudomonadota | Gammaproteobacteria |
| Pectobacterium peruviense | Pseudomonadota | Gammaproteobacteria |
| Pectobacterium polaris | Pseudomonadota | Gammaproteobacteria |
| Pectobacterium polaris | Pseudomonadota | Gammaproteobacteria |

| | | |
|---|---|---|
| Pectobacterium punjabense | Pseudomonadota | Gammaproteobacteria |
| Pectobacterium punjabense | Pseudomonadota | Gammaproteobacteria |
| Pectobacterium quasiaquaticum | Pseudomonadota | Gammaproteobacteria |
| Pectobacterium quasiaquaticum | Pseudomonadota | Gammaproteobacteria |
| Pectobacterium versatile | Pseudomonadota | Gammaproteobacteria |
| Pectobacterium versatile | Pseudomonadota | Gammaproteobacteria |
| Pectobacterium wasabiae CFBP 3304 | Pseudomonadota | Gammaproteobacteria |
| Pectobacterium wasabiae CFBP 3304 | Pseudomonadota | Gammaproteobacteria |
| Phocoenobacter skyensis | Pseudomonadota | Gammaproteobacteria |
| Phocoenobacter skyensis | Pseudomonadota | Gammaproteobacteria |
| Photobacterium atrarenae | Pseudomonadota | Gammaproteobacteria |
| Photobacterium atrarenae | Pseudomonadota | Gammaproteobacteria |
| Photobacterium damselae subsp. damselae | Pseudomonadota | Gammaproteobacteria |
| Photobacterium damselae subsp. damselae | Pseudomonadota | Gammaproteobacteria |
| Photobacterium ganghwense | Pseudomonadota | Gammaproteobacteria |
| Photobacterium ganghwense | Pseudomonadota | Gammaproteobacteria |
| Photobacterium leiognathi subsp. mandapamensis | Pseudomonadota | Gammaproteobacteria |
| Photobacterium leiognathi subsp. mandapamensis | Pseudomonadota | Gammaproteobacteria |
| Photobacterium piscicola | Pseudomonadota | Gammaproteobacteria |
| Photobacterium piscicola | Pseudomonadota | Gammaproteobacteria |
| Photobacterium sanguinicancri | Pseudomonadota | Gammaproteobacteria |
| Photobacterium sanguinicancri | Pseudomonadota | Gammaproteobacteria |
| Photobacterium swingsii | Pseudomonadota | Gammaproteobacteria |
| Photobacterium swingsii | Pseudomonadota | Gammaproteobacteria |
| Photobacterium toruni | Pseudomonadota | Gammaproteobacteria |
| Photobacterium toruni | Pseudomonadota | Gammaproteobacteria |
| Photorhabdus akhurstii | Pseudomonadota | Gammaproteobacteria |
| Photorhabdus akhurstii | Pseudomonadota | Gammaproteobacteria |
| Photorhabdus asymbiotica | Pseudomonadota | Gammaproteobacteria |
| Photorhabdus asymbiotica | Pseudomonadota | Gammaproteobacteria |
| Photorhabdus laumondii subsp. laumondii | Pseudomonadota | Gammaproteobacteria |

| | | |
|---|---|---|
| Photorhabdus laumondii subsp. laumondii | Pseudomonadota | Gammaproteobacteria |
| Photorhabdus thracensis | Pseudomonadota | Gammaproteobacteria |
| Photorhabdus thracensis | Pseudomonadota | Gammaproteobacteria |
| Phytobacter ursingii | Pseudomonadota | Gammaproteobacteria |
| Phytobacter ursingii | Pseudomonadota | Gammaproteobacteria |
| Piscirickettsia salmonis | Pseudomonadota | Gammaproteobacteria |
| Pistricoccus aurantiacus | Pseudomonadota | Gammaproteobacteria |
| Pleionea litopenaei | Pseudomonadota | Gammaproteobacteria |
| Pleionea litopenaei | Pseudomonadota | Gammaproteobacteria |
| Plesiomonas shigelloides | Pseudomonadota | Gammaproteobacteria |
| Plesiomonas shigelloides | Pseudomonadota | Gammaproteobacteria |
| Pluralibacter gergoviae | Pseudomonadota | Gammaproteobacteria |
| Pluralibacter gergoviae | Pseudomonadota | Gammaproteobacteria |
| Pokkaliibacter plantistimulans | Pseudomonadota | Gammaproteobacteria |
| Pragia fontium | Pseudomonadota | Gammaproteobacteria |
| Pragia fontium | Pseudomonadota | Gammaproteobacteria |
| Prodigiosinella confusarubida | Pseudomonadota | Gammaproteobacteria |
| Prodigiosinella confusarubida | Pseudomonadota | Gammaproteobacteria |
| Proteus appendicitidis | Pseudomonadota | Gammaproteobacteria |
| Proteus appendicitidis | Pseudomonadota | Gammaproteobacteria |
| Proteus hauseri | Pseudomonadota | Gammaproteobacteria |
| Proteus hauseri | Pseudomonadota | Gammaproteobacteria |
| Proteus mirabilis HI4320 | Pseudomonadota | Gammaproteobacteria |
| Proteus mirabilis HI4320 | Pseudomonadota | Gammaproteobacteria |
| Proteus penneri | Pseudomonadota | Gammaproteobacteria |
| Proteus penneri | Pseudomonadota | Gammaproteobacteria |
| Proteus terrae subsp. cibarius | Pseudomonadota | Gammaproteobacteria |
| Proteus terrae subsp. cibarius | Pseudomonadota | Gammaproteobacteria |
| Proteus vulgaris | Pseudomonadota | Gammaproteobacteria |
| Proteus vulgaris | Pseudomonadota | Gammaproteobacteria |
| Providencia hangzhouensis | Pseudomonadota | Gammaproteobacteria |

| | | |
|---|---|---|
| Providencia hangzhouensis | Pseudomonadota | Gammaproteobacteria |
| Providencia heimbachae | Pseudomonadota | Gammaproteobacteria |
| Providencia heimbachae | Pseudomonadota | Gammaproteobacteria |
| Providencia huaxiensis | Pseudomonadota | Gammaproteobacteria |
| Providencia huaxiensis | Pseudomonadota | Gammaproteobacteria |
| Providencia manganoxydans | Pseudomonadota | Gammaproteobacteria |
| Providencia manganoxydans | Pseudomonadota | Gammaproteobacteria |
| Providencia rettgeri | Pseudomonadota | Gammaproteobacteria |
| Providencia rettgeri | Pseudomonadota | Gammaproteobacteria |
| Providencia rustigianii | Pseudomonadota | Gammaproteobacteria |
| Providencia rustigianii | Pseudomonadota | Gammaproteobacteria |
| Providencia sneebia DSM 19967 | Pseudomonadota | Gammaproteobacteria |
| Providencia stuartii | Pseudomonadota | Gammaproteobacteria |
| Providencia stuartii | Pseudomonadota | Gammaproteobacteria |
| Providencia xihuensis | Pseudomonadota | Gammaproteobacteria |
| Providencia xihuensis | Pseudomonadota | Gammaproteobacteria |
| Providencia zhejiangensis | Pseudomonadota | Gammaproteobacteria |
| Providencia zhejiangensis | Pseudomonadota | Gammaproteobacteria |
| Providencia zhijiangensis | Pseudomonadota | Gammaproteobacteria |
| Providencia zhijiangensis | Pseudomonadota | Gammaproteobacteria |
| Pseudescherichia vulneris | Pseudomonadota | Gammaproteobacteria |
| Pseudoalteromonas aliena | Pseudomonadota | Gammaproteobacteria |
| Pseudoalteromonas aliena | Pseudomonadota | Gammaproteobacteria |
| Pseudoalteromonas apostichopi | Pseudomonadota | Gammaproteobacteria |
| Pseudoalteromonas apostichopi | Pseudomonadota | Gammaproteobacteria |
| Pseudoalteromonas arctica A 37-1-2 | Pseudomonadota | Gammaproteobacteria |
| Pseudoalteromonas arctica A 37-1-2 | Pseudomonadota | Gammaproteobacteria |
| Pseudoalteromonas carrageenovora IAM 12662 | Pseudomonadota | Gammaproteobacteria |
| Pseudoalteromonas carrageenovora IAM 12662 | Pseudomonadota | Gammaproteobacteria |
| Pseudoalteromonas distincta | Pseudomonadota | Gammaproteobacteria |
| Pseudoalteromonas distincta | Pseudomonadota | Gammaproteobacteria |

| | | |
|---|---|---|
| Pseudoalteromonas espejiana DSM 9414 | Pseudomonadota | Gammaproteobacteria |
| Pseudoalteromonas espejiana DSM 9414 | Pseudomonadota | Gammaproteobacteria |
| Pseudoalteromonas issachenkonii | Pseudomonadota | Gammaproteobacteria |
| Pseudoalteromonas issachenkonii | Pseudomonadota | Gammaproteobacteria |
| Pseudoalteromonas lipolytica | Pseudomonadota | Gammaproteobacteria |
| Pseudoalteromonas lipolytica | Pseudomonadota | Gammaproteobacteria |
| Pseudoalteromonas luteoviolacea | Pseudomonadota | Gammaproteobacteria |
| Pseudoalteromonas luteoviolacea | Pseudomonadota | Gammaproteobacteria |
| Pseudoalteromonas maricaloris | Pseudomonadota | Gammaproteobacteria |
| Pseudoalteromonas maricaloris | Pseudomonadota | Gammaproteobacteria |
| Pseudoalteromonas nigrifaciens | Pseudomonadota | Gammaproteobacteria |
| Pseudoalteromonas nigrifaciens | Pseudomonadota | Gammaproteobacteria |
| Pseudoalteromonas ostreae | Pseudomonadota | Gammaproteobacteria |
| Pseudoalteromonas ostreae | Pseudomonadota | Gammaproteobacteria |
| Pseudoalteromonas phenolica | Pseudomonadota | Gammaproteobacteria |
| Pseudoalteromonas phenolica | Pseudomonadota | Gammaproteobacteria |
| Pseudoalteromonas piratica | Pseudomonadota | Gammaproteobacteria |
| Pseudoalteromonas piratica | Pseudomonadota | Gammaproteobacteria |
| Pseudoalteromonas piscicida | Pseudomonadota | Gammaproteobacteria |
| Pseudoalteromonas piscicida | Pseudomonadota | Gammaproteobacteria |
| Pseudoalteromonas prydzensis ACAM 620 | Pseudomonadota | Gammaproteobacteria |
| Pseudoalteromonas prydzensis ACAM 620 | Pseudomonadota | Gammaproteobacteria |
| Pseudoalteromonas prydzensis ACAM 620 | Pseudomonadota | Gammaproteobacteria |
| Pseudoalteromonas rhizosphaerae | Pseudomonadota | Gammaproteobacteria |
| Pseudoalteromonas rhizosphaerae | Pseudomonadota | Gammaproteobacteria |
| Pseudoalteromonas spongiae UST010723-006 | Pseudomonadota | Gammaproteobacteria |
| Pseudoalteromonas spongiae UST010723-006 | Pseudomonadota | Gammaproteobacteria |
| Pseudoalteromonas translucida | Pseudomonadota | Gammaproteobacteria |
| Pseudoalteromonas translucida | Pseudomonadota | Gammaproteobacteria |
| Pseudoalteromonas tunicata | Pseudomonadota | Gammaproteobacteria |
| Pseudoalteromonas tunicata | Pseudomonadota | Gammaproteobacteria |

| | | |
|---|---|---|
| Pseudoalteromonas undina | Pseudomonadota | Gammaproteobacteria |
| Pseudoalteromonas undina | Pseudomonadota | Gammaproteobacteria |
| Pseudoalteromonas viridis | Pseudomonadota | Gammaproteobacteria |
| Pseudoalteromonas viridis | Pseudomonadota | Gammaproteobacteria |
| Pseudocitrobacter corydidari | Pseudomonadota | Gammaproteobacteria |
| Pseudocitrobacter corydidari | Pseudomonadota | Gammaproteobacteria |
| Pseudohongiella spirulinae | Pseudomonadota | Gammaproteobacteria |
| Pseudohongiella spirulinae | Pseudomonadota | Gammaproteobacteria |
| Pseudolysobacter antarcticus | Pseudomonadota | Gammaproteobacteria |
| Pseudolysobacter antarcticus | Pseudomonadota | Gammaproteobacteria |
| Pseudomonas abieticivorans | Pseudomonadota | Gammaproteobacteria |
| Pseudomonas aeruginosa PAO1 | Pseudomonadota | Gammaproteobacteria |
| Pseudomonas aeruginosa PAO1 | Pseudomonadota | Gammaproteobacteria |
| Pseudomonas aestiva | Pseudomonadota | Gammaproteobacteria |
| Pseudomonas aestiva | Pseudomonadota | Gammaproteobacteria |
| Pseudomonas alkylphenolica | Pseudomonadota | Gammaproteobacteria |
| Pseudomonas alkylphenolica | Pseudomonadota | Gammaproteobacteria |
| Pseudomonas allokribbensis | Pseudomonadota | Gammaproteobacteria |
| Pseudomonas allokribbensis | Pseudomonadota | Gammaproteobacteria |
| Pseudomonas alloputida | Pseudomonadota | Gammaproteobacteria |
| Pseudomonas alloputida | Pseudomonadota | Gammaproteobacteria |
| Pseudomonas alvandae | Pseudomonadota | Gammaproteobacteria |
| Pseudomonas alvandae | Pseudomonadota | Gammaproteobacteria |
| Pseudomonas amygdali pv. tabaci str. ATCC 11528 | Pseudomonadota | Gammaproteobacteria |
| Pseudomonas antarctica | Pseudomonadota | Gammaproteobacteria |
| Pseudomonas anuradhapurensis | Pseudomonadota | Gammaproteobacteria |
| Pseudomonas anuradhapurensis | Pseudomonadota | Gammaproteobacteria |
| Pseudomonas arsenicoxydans | Pseudomonadota | Gammaproteobacteria |
| Pseudomonas asgharzadehiana | Pseudomonadota | Gammaproteobacteria |
| Pseudomonas asiatica | Pseudomonadota | Gammaproteobacteria |
| Pseudomonas asiatica | Pseudomonadota | Gammaproteobacteria |

| | | |
|---|---|---|
| Pseudomonas asplenii | Pseudomonadota | Gammaproteobacteria |
| Pseudomonas atacamensis | Pseudomonadota | Gammaproteobacteria |
| Pseudomonas atacamensis | Pseudomonadota | Gammaproteobacteria |
| Pseudomonas azerbaijanorientalis | Pseudomonadota | Gammaproteobacteria |
| Pseudomonas azotoformans | Pseudomonadota | Gammaproteobacteria |
| Pseudomonas batumici | Pseudomonadota | Gammaproteobacteria |
| Pseudomonas batumici | Pseudomonadota | Gammaproteobacteria |
| Pseudomonas beijingensis | Pseudomonadota | Gammaproteobacteria |
| Pseudomonas beijingensis | Pseudomonadota | Gammaproteobacteria |
| Pseudomonas benzenivorans | Pseudomonadota | Gammaproteobacteria |
| Pseudomonas benzenivorans | Pseudomonadota | Gammaproteobacteria |
| Pseudomonas benzopyrenica | Pseudomonadota | Gammaproteobacteria |
| Pseudomonas benzopyrenica | Pseudomonadota | Gammaproteobacteria |
| Pseudomonas berkeleyensis | Pseudomonadota | Gammaproteobacteria |
| Pseudomonas berkeleyensis | Pseudomonadota | Gammaproteobacteria |
| Pseudomonas bijieensis | Pseudomonadota | Gammaproteobacteria |
| Pseudomonas bijieensis | Pseudomonadota | Gammaproteobacteria |
| Pseudomonas brassicacearum | Pseudomonadota | Gammaproteobacteria |
| Pseudomonas brassicacearum | Pseudomonadota | Gammaproteobacteria |
| Pseudomonas bubulae | Pseudomonadota | Gammaproteobacteria |
| Pseudomonas canadensis | Pseudomonadota | Gammaproteobacteria |
| Pseudomonas cannabina pv. alisalensis | Pseudomonadota | Gammaproteobacteria |
| Pseudomonas cavernae | Pseudomonadota | Gammaproteobacteria |
| Pseudomonas cavernae | Pseudomonadota | Gammaproteobacteria |
| Pseudomonas cedrina | Pseudomonadota | Gammaproteobacteria |
| Pseudomonas chlororaphis subsp. chlororaphis | Pseudomonadota | Gammaproteobacteria |
| Pseudomonas chlororaphis subsp. chlororaphis | Pseudomonadota | Gammaproteobacteria |
| Pseudomonas cichorii | Pseudomonadota | Gammaproteobacteria |
| Pseudomonas citronellolis | Pseudomonadota | Gammaproteobacteria |
| Pseudomonas citronellolis | Pseudomonadota | Gammaproteobacteria |
| Pseudomonas coronafaciens pv. oryzae str. 1_6 | Pseudomonadota | Gammaproteobacteria |

| | | |
|---|---|---|
| Pseudomonas corrugata | Pseudomonadota | Gammaproteobacteria |
| Pseudomonas corrugata | Pseudomonadota | Gammaproteobacteria |
| Pseudomonas cucumis | Pseudomonadota | Gammaproteobacteria |
| Pseudomonas cucumis | Pseudomonadota | Gammaproteobacteria |
| Pseudomonas danubii | Pseudomonadota | Gammaproteobacteria |
| Pseudomonas danubii | Pseudomonadota | Gammaproteobacteria |
| Pseudomonas donghuensis | Pseudomonadota | Gammaproteobacteria |
| Pseudomonas donghuensis | Pseudomonadota | Gammaproteobacteria |
| Pseudomonas entomophila L48 | Pseudomonadota | Gammaproteobacteria |
| Pseudomonas entomophila L48 | Pseudomonadota | Gammaproteobacteria |
| Pseudomonas eucalypticola | Pseudomonadota | Gammaproteobacteria |
| Pseudomonas extremaustralis | Pseudomonadota | Gammaproteobacteria |
| Pseudomonas extremorientalis | Pseudomonadota | Gammaproteobacteria |
| Pseudomonas fakonensis | Pseudomonadota | Gammaproteobacteria |
| Pseudomonas fakonensis | Pseudomonadota | Gammaproteobacteria |
| Pseudomonas ficuserectae | Pseudomonadota | Gammaproteobacteria |
| Pseudomonas fitomaticsae | Pseudomonadota | Gammaproteobacteria |
| Pseudomonas fitomaticsae | Pseudomonadota | Gammaproteobacteria |
| Pseudomonas fluorescens | Pseudomonadota | Gammaproteobacteria |
| Pseudomonas fortuita | Pseudomonadota | Gammaproteobacteria |
| Pseudomonas fortuita | Pseudomonadota | Gammaproteobacteria |
| Pseudomonas fragi | Pseudomonadota | Gammaproteobacteria |
| Pseudomonas frederiksbergensis | Pseudomonadota | Gammaproteobacteria |
| Pseudomonas frederiksbergensis | Pseudomonadota | Gammaproteobacteria |
| Pseudomonas fulva | Pseudomonadota | Gammaproteobacteria |
| Pseudomonas fulva | Pseudomonadota | Gammaproteobacteria |
| Pseudomonas germanica | Pseudomonadota | Gammaproteobacteria |
| Pseudomonas germanica | Pseudomonadota | Gammaproteobacteria |
| Pseudomonas glycinae | Pseudomonadota | Gammaproteobacteria |
| Pseudomonas gorinensis | Pseudomonadota | Gammaproteobacteria |
| Pseudomonas gozinkensis | Pseudomonadota | Gammaproteobacteria |

| | | |
|---|---|---|
| Pseudomonas gozinkensis | Pseudomonadota | Gammaproteobacteria |
| Pseudomonas granadensis | Pseudomonadota | Gammaproteobacteria |
| Pseudomonas granadensis | Pseudomonadota | Gammaproteobacteria |
| Pseudomonas grandcourensis | Pseudomonadota | Gammaproteobacteria |
| Pseudomonas guariconensis | Pseudomonadota | Gammaproteobacteria |
| Pseudomonas guariconensis | Pseudomonadota | Gammaproteobacteria |
| Pseudomonas hamedanensis | Pseudomonadota | Gammaproteobacteria |
| Pseudomonas hamedanensis | Pseudomonadota | Gammaproteobacteria |
| Pseudomonas helleri | Pseudomonadota | Gammaproteobacteria |
| Pseudomonas helvetica | Pseudomonadota | Gammaproteobacteria |
| Pseudomonas helvetica | Pseudomonadota | Gammaproteobacteria |
| Pseudomonas hormoni | Pseudomonadota | Gammaproteobacteria |
| Pseudomonas huanghezhanensis | Pseudomonadota | Gammaproteobacteria |
| Pseudomonas idahonensis | Pseudomonadota | Gammaproteobacteria |
| Pseudomonas idahonensis | Pseudomonadota | Gammaproteobacteria |
| Pseudomonas inefficax | Pseudomonadota | Gammaproteobacteria |
| Pseudomonas inefficax | Pseudomonadota | Gammaproteobacteria |
| Pseudomonas iranensis | Pseudomonadota | Gammaproteobacteria |
| Pseudomonas iranensis | Pseudomonadota | Gammaproteobacteria |
| Pseudomonas izuensis | Pseudomonadota | Gammaproteobacteria |
| Pseudomonas juntendi | Pseudomonadota | Gammaproteobacteria |
| Pseudomonas juntendi | Pseudomonadota | Gammaproteobacteria |
| Pseudomonas kermanshahensis | Pseudomonadota | Gammaproteobacteria |
| Pseudomonas kermanshahensis | Pseudomonadota | Gammaproteobacteria |
| Pseudomonas khavaziana | Pseudomonadota | Gammaproteobacteria |
| Pseudomonas kielensis | Pseudomonadota | Gammaproteobacteria |
| Pseudomonas kielensis | Pseudomonadota | Gammaproteobacteria |
| Pseudomonas knackmussii | Pseudomonadota | Gammaproteobacteria |
| Pseudomonas knackmussii | Pseudomonadota | Gammaproteobacteria |
| Pseudomonas koreensis | Pseudomonadota | Gammaproteobacteria |
| Pseudomonas koreensis | Pseudomonadota | Gammaproteobacteria |

| | | |
|---|---|---|
| Pseudomonas kribbensis | Pseudomonadota | Gammaproteobacteria |
| Pseudomonas kribbensis | Pseudomonadota | Gammaproteobacteria |
| Pseudomonas kulmbachensis | Pseudomonadota | Gammaproteobacteria |
| Pseudomonas kurunegalensis | Pseudomonadota | Gammaproteobacteria |
| Pseudomonas kurunegalensis | Pseudomonadota | Gammaproteobacteria |
| Pseudomonas laurentiana | Pseudomonadota | Gammaproteobacteria |
| Pseudomonas laurentiana | Pseudomonadota | Gammaproteobacteria |
| Pseudomonas leptonychotis | Pseudomonadota | Gammaproteobacteria |
| Pseudomonas leptonychotis | Pseudomonadota | Gammaproteobacteria |
| Pseudomonas lijiangensis | Pseudomonadota | Gammaproteobacteria |
| Pseudomonas lini | Pseudomonadota | Gammaproteobacteria |
| Pseudomonas lini | Pseudomonadota | Gammaproteobacteria |
| Pseudomonas lundensis | Pseudomonadota | Gammaproteobacteria |
| Pseudomonas mandelii | Pseudomonadota | Gammaproteobacteria |
| Pseudomonas mandelii | Pseudomonadota | Gammaproteobacteria |
| Pseudomonas marginalis | Pseudomonadota | Gammaproteobacteria |
| Pseudomonas marincola | Pseudomonadota | Gammaproteobacteria |
| Pseudomonas marincola | Pseudomonadota | Gammaproteobacteria |
| Pseudomonas maumuensis | Pseudomonadota | Gammaproteobacteria |
| Pseudomonas maumuensis | Pseudomonadota | Gammaproteobacteria |
| Pseudomonas mediterranea | Pseudomonadota | Gammaproteobacteria |
| Pseudomonas mediterranea | Pseudomonadota | Gammaproteobacteria |
| Pseudomonas monachiensis | Pseudomonadota | Gammaproteobacteria |
| Pseudomonas monsensis | Pseudomonadota | Gammaproteobacteria |
| Pseudomonas monsensis | Pseudomonadota | Gammaproteobacteria |
| Pseudomonas monteilii | Pseudomonadota | Gammaproteobacteria |
| Pseudomonas monteilii | Pseudomonadota | Gammaproteobacteria |
| Pseudomonas moorei | Pseudomonadota | Gammaproteobacteria |
| Pseudomonas mosselii | Pseudomonadota | Gammaproteobacteria |
| Pseudomonas mosselii | Pseudomonadota | Gammaproteobacteria |
| Pseudomonas mucidolens | Pseudomonadota | Gammaproteobacteria |

| | | |
|---|---|---|
| Pseudomonas multiresinivorans | Pseudomonadota | Gammaproteobacteria |
| Pseudomonas multiresinivorans | Pseudomonadota | Gammaproteobacteria |
| Pseudomonas muyukensis | Pseudomonadota | Gammaproteobacteria |
| Pseudomonas muyukensis | Pseudomonadota | Gammaproteobacteria |
| Pseudomonas nunensis | Pseudomonadota | Gammaproteobacteria |
| Pseudomonas nunensis | Pseudomonadota | Gammaproteobacteria |
| Pseudomonas ogarae | Pseudomonadota | Gammaproteobacteria |
| Pseudomonas ogarae | Pseudomonadota | Gammaproteobacteria |
| Pseudomonas orientalis | Pseudomonadota | Gammaproteobacteria |
| Pseudomonas oryzae | Pseudomonadota | Gammaproteobacteria |
| Pseudomonas oryzae | Pseudomonadota | Gammaproteobacteria |
| Pseudomonas oryziphila | Pseudomonadota | Gammaproteobacteria |
| Pseudomonas oryziphila | Pseudomonadota | Gammaproteobacteria |
| Pseudomonas paeninsulae | Pseudomonadota | Gammaproteobacteria |
| Pseudomonas paeninsulae | Pseudomonadota | Gammaproteobacteria |
| Pseudomonas palleroniana | Pseudomonadota | Gammaproteobacteria |
| Pseudomonas paracarnis | Pseudomonadota | Gammaproteobacteria |
| Pseudomonas paraeruginosa | Pseudomonadota | Gammaproteobacteria |
| Pseudomonas paraeruginosa | Pseudomonadota | Gammaproteobacteria |
| Pseudomonas paraveronii | Pseudomonadota | Gammaproteobacteria |
| Pseudomonas pergaminensis | Pseudomonadota | Gammaproteobacteria |
| Pseudomonas petroselini | Pseudomonadota | Gammaproteobacteria |
| Pseudomonas phenolilytica | Pseudomonadota | Gammaproteobacteria |
| Pseudomonas phenolilytica | Pseudomonadota | Gammaproteobacteria |
| Pseudomonas plecoglossicida | Pseudomonadota | Gammaproteobacteria |
| Pseudomonas plecoglossicida | Pseudomonadota | Gammaproteobacteria |
| Pseudomonas poae RE*1-1-14 | Pseudomonadota | Gammaproteobacteria |
| Pseudomonas pohangensis | Pseudomonadota | Gammaproteobacteria |
| Pseudomonas pohangensis | Pseudomonadota | Gammaproteobacteria |
| Pseudomonas promysalinigenes | Pseudomonadota | Gammaproteobacteria |
| Pseudomonas promysalinigenes | Pseudomonadota | Gammaproteobacteria |

| | | |
|---|---|---|
| Pseudomonas prosekii | Pseudomonadota | Gammaproteobacteria |
| Pseudomonas prosekii | Pseudomonadota | Gammaproteobacteria |
| Pseudomonas protegens CHA0 | Pseudomonadota | Gammaproteobacteria |
| Pseudomonas protegens CHA0 | Pseudomonadota | Gammaproteobacteria |
| Pseudomonas psychrophila | Pseudomonadota | Gammaproteobacteria |
| Pseudomonas purpurea | Pseudomonadota | Gammaproteobacteria |
| Pseudomonas purpurea | Pseudomonadota | Gammaproteobacteria |
| Pseudomonas putida NBRC 14164 | Pseudomonadota | Gammaproteobacteria |
| Pseudomonas putida NBRC 14164 | Pseudomonadota | Gammaproteobacteria |
| Pseudomonas qingdaonensis | Pseudomonadota | Gammaproteobacteria |
| Pseudomonas qingdaonensis | Pseudomonadota | Gammaproteobacteria |
| Pseudomonas quebecensis | Pseudomonadota | Gammaproteobacteria |
| Pseudomonas reinekei | Pseudomonadota | Gammaproteobacteria |
| Pseudomonas rhizophila | Pseudomonadota | Gammaproteobacteria |
| Pseudomonas rhizosphaerae | Pseudomonadota | Gammaproteobacteria |
| Pseudomonas rhodesiae | Pseudomonadota | Gammaproteobacteria |
| Pseudomonas salmasensis | Pseudomonadota | Gammaproteobacteria |
| Pseudomonas savastanoi | Pseudomonadota | Gammaproteobacteria |
| Pseudomonas sediminis | Pseudomonadota | Gammaproteobacteria |
| Pseudomonas sediminis | Pseudomonadota | Gammaproteobacteria |
| Pseudomonas serboccidentalis | Pseudomonadota | Gammaproteobacteria |
| Pseudomonas serboccidentalis | Pseudomonadota | Gammaproteobacteria |
| Pseudomonas sessilinigenes | Pseudomonadota | Gammaproteobacteria |
| Pseudomonas sessilinigenes | Pseudomonadota | Gammaproteobacteria |
| Pseudomonas shahriarae | Pseudomonadota | Gammaproteobacteria |
| Pseudomonas shirazensis | Pseudomonadota | Gammaproteobacteria |
| Pseudomonas shirazensis | Pseudomonadota | Gammaproteobacteria |
| Pseudomonas shirazica | Pseudomonadota | Gammaproteobacteria |
| Pseudomonas shirazica | Pseudomonadota | Gammaproteobacteria |
| Pseudomonas sichuanensis | Pseudomonadota | Gammaproteobacteria |
| Pseudomonas sichuanensis | Pseudomonadota | Gammaproteobacteria |

| | | |
|---|---|---|
| Pseudomonas sihuiensis | Pseudomonadota | Gammaproteobacteria |
| Pseudomonas sihuiensis | Pseudomonadota | Gammaproteobacteria |
| Pseudomonas silesiensis | Pseudomonadota | Gammaproteobacteria |
| Pseudomonas siliginis | Pseudomonadota | Gammaproteobacteria |
| Pseudomonas siliginis | Pseudomonadota | Gammaproteobacteria |
| Pseudomonas simiae | Pseudomonadota | Gammaproteobacteria |
| Pseudomonas sivasensis | Pseudomonadota | Gammaproteobacteria |
| Pseudomonas solani | Pseudomonadota | Gammaproteobacteria |
| Pseudomonas solani | Pseudomonadota | Gammaproteobacteria |
| Pseudomonas soli | Pseudomonadota | Gammaproteobacteria |
| Pseudomonas soli | Pseudomonadota | Gammaproteobacteria |
| Pseudomonas svalbardensis | Pseudomonadota | Gammaproteobacteria |
| Pseudomonas svalbardensis | Pseudomonadota | Gammaproteobacteria |
| Pseudomonas synxantha | Pseudomonadota | Gammaproteobacteria |
| Pseudomonas syringae | Pseudomonadota | Gammaproteobacteria |
| Pseudomonas syringae pv. tagetis | Pseudomonadota | Gammaproteobacteria |
| Pseudomonas syringae pv. tomato str. DC3000 | Pseudomonadota | Gammaproteobacteria |
| Pseudomonas taetrolens | Pseudomonadota | Gammaproteobacteria |
| Pseudomonas tensinigenes | Pseudomonadota | Gammaproteobacteria |
| Pseudomonas tensinigenes | Pseudomonadota | Gammaproteobacteria |
| Pseudomonas tohonis | Pseudomonadota | Gammaproteobacteria |
| Pseudomonas tohonis | Pseudomonadota | Gammaproteobacteria |
| Pseudomonas tremae | Pseudomonadota | Gammaproteobacteria |
| Pseudomonas triclosanedens | Pseudomonadota | Gammaproteobacteria |
| Pseudomonas triclosanedens | Pseudomonadota | Gammaproteobacteria |
| Pseudomonas tritici | Pseudomonadota | Gammaproteobacteria |
| Pseudomonas trivialis | Pseudomonadota | Gammaproteobacteria |
| Pseudomonas tructae | Pseudomonadota | Gammaproteobacteria |
| Pseudomonas tructae | Pseudomonadota | Gammaproteobacteria |
| Pseudomonas umsongensis | Pseudomonadota | Gammaproteobacteria |
| Pseudomonas vancouverensis | Pseudomonadota | Gammaproteobacteria |

| | | |
|---|---|---|
| Pseudomonas vanderleydeniana | Pseudomonadota | Gammaproteobacteria |
| Pseudomonas veronii | Pseudomonadota | Gammaproteobacteria |
| Pseudomonas versuta | Pseudomonadota | Gammaproteobacteria |
| Pseudomonas viciae | Pseudomonadota | Gammaproteobacteria |
| Pseudomonas viciae | Pseudomonadota | Gammaproteobacteria |
| Pseudomonas viridiflava | Pseudomonadota | Gammaproteobacteria |
| Pseudomonas vranovensis | Pseudomonadota | Gammaproteobacteria |
| Pseudomonas vranovensis | Pseudomonadota | Gammaproteobacteria |
| Pseudomonas wenzhouensis | Pseudomonadota | Gammaproteobacteria |
| Pseudomonas wenzhouensis | Pseudomonadota | Gammaproteobacteria |
| Pseudomonas wuhanensis | Pseudomonadota | Gammaproteobacteria |
| Pseudomonas wuhanensis | Pseudomonadota | Gammaproteobacteria |
| Pseudomonas xantholysinigenes | Pseudomonadota | Gammaproteobacteria |
| Pseudomonas xantholysinigenes | Pseudomonadota | Gammaproteobacteria |
| Pseudomonas xanthosomatis | Pseudomonadota | Gammaproteobacteria |
| Pseudomonas xanthosomatis | Pseudomonadota | Gammaproteobacteria |
| Pseudomonas yamanorum | Pseudomonadota | Gammaproteobacteria |
| Pseudomonas zeae | Pseudomonadota | Gammaproteobacteria |
| Pseudomonas zeae | Pseudomonadota | Gammaproteobacteria |
| Pseudoxanthomonas daejeonensis | Pseudomonadota | Gammaproteobacteria |
| Pseudoxanthomonas daejeonensis | Pseudomonadota | Gammaproteobacteria |
| Pseudoxanthomonas japonensis | Pseudomonadota | Gammaproteobacteria |
| Pseudoxanthomonas japonensis | Pseudomonadota | Gammaproteobacteria |
| Pseudoxanthomonas mexicana | Pseudomonadota | Gammaproteobacteria |
| Pseudoxanthomonas mexicana | Pseudomonadota | Gammaproteobacteria |
| Pseudoxanthomonas spadix BD-a59 | Pseudomonadota | Gammaproteobacteria |
| Pseudoxanthomonas spadix BD-a59 | Pseudomonadota | Gammaproteobacteria |
| Pseudoxanthomonas winnipegensis | Pseudomonadota | Gammaproteobacteria |
| Pseudoxanthomonas winnipegensis | Pseudomonadota | Gammaproteobacteria |
| Psychrobacter alimentarius | Pseudomonadota | Gammaproteobacteria |
| Psychrobacter alimentarius | Pseudomonadota | Gammaproteobacteria |

| | | |
|---|---|---|
| Psychrobacter arcticus 273-4 | Pseudomonadota | Gammaproteobacteria |
| Psychrobacter arcticus 273-4 | Pseudomonadota | Gammaproteobacteria |
| Psychrobacter arenosus | Pseudomonadota | Gammaproteobacteria |
| Psychrobacter arenosus | Pseudomonadota | Gammaproteobacteria |
| Psychrobacter cryohalolentis K5 | Pseudomonadota | Gammaproteobacteria |
| Psychrobacter cryohalolentis K5 | Pseudomonadota | Gammaproteobacteria |
| Psychrobacter pacificensis | Pseudomonadota | Gammaproteobacteria |
| Psychrobacter pacificensis | Pseudomonadota | Gammaproteobacteria |
| Psychrobacter raelei | Pseudomonadota | Gammaproteobacteria |
| Psychrobacter raelei | Pseudomonadota | Gammaproteobacteria |
| Psychromonas ingrahamii 37 | Pseudomonadota | Gammaproteobacteria |
| Psychrosphaera aestuarii | Pseudomonadota | Gammaproteobacteria |
| Psychrosphaera aestuarii | Pseudomonadota | Gammaproteobacteria |
| Psychrosphaera ytuae | Pseudomonadota | Gammaproteobacteria |
| Psychrosphaera ytuae | Pseudomonadota | Gammaproteobacteria |
| Rahnella aceris | Pseudomonadota | Gammaproteobacteria |
| Rahnella aceris | Pseudomonadota | Gammaproteobacteria |
| Rahnella aquatilis CIP 78.65 = ATCC 33071 | Pseudomonadota | Gammaproteobacteria |
| Rahnella aquatilis CIP 78.65 = ATCC 33071 | Pseudomonadota | Gammaproteobacteria |
| Rahnella inusitata | Pseudomonadota | Gammaproteobacteria |
| Rahnella inusitata | Pseudomonadota | Gammaproteobacteria |
| Rahnella sikkimica | Pseudomonadota | Gammaproteobacteria |
| Rahnella sikkimica | Pseudomonadota | Gammaproteobacteria |
| Rahnella victoriana | Pseudomonadota | Gammaproteobacteria |
| Rahnella victoriana | Pseudomonadota | Gammaproteobacteria |
| Raoultella ornithinolytica | Pseudomonadota | Gammaproteobacteria |
| Raoultella ornithinolytica | Pseudomonadota | Gammaproteobacteria |
| Raoultella planticola | Pseudomonadota | Gammaproteobacteria |
| Raoultella planticola | Pseudomonadota | Gammaproteobacteria |
| Raoultella terrigena | Pseudomonadota | Gammaproteobacteria |
| Raoultella terrigena | Pseudomonadota | Gammaproteobacteria |

| | | |
|---|---|---|
| Reinekea forsetii | Pseudomonadota | Gammaproteobacteria |
| Rheinheimera mangrovi | Pseudomonadota | Gammaproteobacteria |
| Rheinheimera mangrovi | Pseudomonadota | Gammaproteobacteria |
| Rhodanobacter denitrificans | Pseudomonadota | Gammaproteobacteria |
| Rhodanobacter denitrificans | Pseudomonadota | Gammaproteobacteria |
| Rickettsiella endosymbiont of Aleochara curtula | Pseudomonadota | Gammaproteobacteria |
| Rickettsiella endosymbiont of Dermanyssus gallinae | Pseudomonadota | Gammaproteobacteria |
| Rickettsiella endosymbiont of Litargus connexus | Pseudomonadota | Gammaproteobacteria |
| Rickettsiella endosymbiont of Miltochrista miniata | Pseudomonadota | Gammaproteobacteria |
| Rickettsiella endosymbiont of Rhagonycha lignosa | Pseudomonadota | Gammaproteobacteria |
| Rickettsiella endosymbiont of Xylota segnis | Pseudomonadota | Gammaproteobacteria |
| Rodentibacter caecimuris | Pseudomonadota | Gammaproteobacteria |
| Rodentibacter caecimuris | Pseudomonadota | Gammaproteobacteria |
| Rodentibacter haemolyticus | Pseudomonadota | Gammaproteobacteria |
| Rodentibacter haemolyticus | Pseudomonadota | Gammaproteobacteria |
| Rouxiella badensis | Pseudomonadota | Gammaproteobacteria |
| Rouxiella badensis | Pseudomonadota | Gammaproteobacteria |
| Saccharobesus litoralis | Pseudomonadota | Gammaproteobacteria |
| Saccharobesus litoralis | Pseudomonadota | Gammaproteobacteria |
| Saccharophagus degradans 2-40 | Pseudomonadota | Gammaproteobacteria |
| Saccharophagus degradans 2-40 | Pseudomonadota | Gammaproteobacteria |
| Saccharospirillum mangrovi | Pseudomonadota | Gammaproteobacteria |
| Saccharospirillum mangrovi | Pseudomonadota | Gammaproteobacteria |
| Salinimonas iocasae | Pseudomonadota | Gammaproteobacteria |
| Salinimonas iocasae | Pseudomonadota | Gammaproteobacteria |
| Salinimonas lutimaris | Pseudomonadota | Gammaproteobacteria |
| Salinimonas lutimaris | Pseudomonadota | Gammaproteobacteria |
| Salinimonas marina | Pseudomonadota | Gammaproteobacteria |
| Salinimonas marina | Pseudomonadota | Gammaproteobacteria |
| Salinimonas sediminis | Pseudomonadota | Gammaproteobacteria |
| Salinimonas sediminis | Pseudomonadota | Gammaproteobacteria |

| | | |
|---|---|---|
| Saliniradius amylolyticus | Pseudomonadota | Gammaproteobacteria |
| Saliniradius amylolyticus | Pseudomonadota | Gammaproteobacteria |
| Salinivibrio kushneri | Pseudomonadota | Gammaproteobacteria |
| Salinivibrio kushneri | Pseudomonadota | Gammaproteobacteria |
| Salinivibrio proteolyticus | Pseudomonadota | Gammaproteobacteria |
| Salinivibrio proteolyticus | Pseudomonadota | Gammaproteobacteria |
| Salmonella bongori NCTC 12419 | Pseudomonadota | Gammaproteobacteria |
| Salmonella bongori NCTC 12419 | Pseudomonadota | Gammaproteobacteria |
| Salmonella enterica subsp. enterica serovar Typhimurium str. LT2 | Pseudomonadota | Gammaproteobacteria |
| Salmonella enterica subsp. enterica serovar Typhimurium str. LT2 | Pseudomonadota | Gammaproteobacteria |
| Scandinavium goeteborgense | Pseudomonadota | Gammaproteobacteria |
| Scandinavium goeteborgense | Pseudomonadota | Gammaproteobacteria |
| secondary endosymbiont of Ctenarytaina eucalypti | Pseudomonadota | Gammaproteobacteria |
| secondary endosymbiont of Heteropsylla cubana | Pseudomonadota | Gammaproteobacteria |
| secondary endosymbiont of Trabutina mannipara | Pseudomonadota | Gammaproteobacteria |
| Sedimenticola thiotaurini | Pseudomonadota | Gammaproteobacteria |
| Sedimenticola thiotaurini | Pseudomonadota | Gammaproteobacteria |
| Serratia aquatilis | Pseudomonadota | Gammaproteobacteria |
| Serratia aquatilis | Pseudomonadota | Gammaproteobacteria |
| Serratia entomophila | Pseudomonadota | Gammaproteobacteria |
| Serratia entomophila | Pseudomonadota | Gammaproteobacteria |
| Serratia ficaria | Pseudomonadota | Gammaproteobacteria |
| Serratia ficaria | Pseudomonadota | Gammaproteobacteria |
| Serratia fonticola | Pseudomonadota | Gammaproteobacteria |
| Serratia fonticola | Pseudomonadota | Gammaproteobacteria |
| Serratia liquefaciens ATCC 27592 | Pseudomonadota | Gammaproteobacteria |
| Serratia liquefaciens ATCC 27592 | Pseudomonadota | Gammaproteobacteria |
| Serratia marcescens | Pseudomonadota | Gammaproteobacteria |
| Serratia marcescens | Pseudomonadota | Gammaproteobacteria |
| Serratia nematodiphila | Pseudomonadota | Gammaproteobacteria |
| Serratia nematodiphila | Pseudomonadota | Gammaproteobacteria |

| | | |
|---|---|---|
| Serratia nevei | Pseudomonadota | Gammaproteobacteria |
| Serratia nevei | Pseudomonadota | Gammaproteobacteria |
| Serratia plymuthica AS9 | Pseudomonadota | Gammaproteobacteria |
| Serratia plymuthica AS9 | Pseudomonadota | Gammaproteobacteria |
| Serratia proteamaculans | Pseudomonadota | Gammaproteobacteria |
| Serratia proteamaculans | Pseudomonadota | Gammaproteobacteria |
| Serratia quinivorans | Pseudomonadota | Gammaproteobacteria |
| Serratia quinivorans | Pseudomonadota | Gammaproteobacteria |
| Serratia rhizosphaerae | Pseudomonadota | Gammaproteobacteria |
| Serratia rhizosphaerae | Pseudomonadota | Gammaproteobacteria |
| Serratia rubidaea | Pseudomonadota | Gammaproteobacteria |
| Serratia rubidaea | Pseudomonadota | Gammaproteobacteria |
| Serratia sarumanii | Pseudomonadota | Gammaproteobacteria |
| Serratia sarumanii | Pseudomonadota | Gammaproteobacteria |
| Serratia surfactantfaciens | Pseudomonadota | Gammaproteobacteria |
| Serratia surfactantfaciens | Pseudomonadota | Gammaproteobacteria |
| Serratia symbiotica | Pseudomonadota | Gammaproteobacteria |
| Serratia symbiotica | Pseudomonadota | Gammaproteobacteria |
| Serratia ureilytica | Pseudomonadota | Gammaproteobacteria |
| Serratia ureilytica | Pseudomonadota | Gammaproteobacteria |
| Shewanella acanthi | Pseudomonadota | Gammaproteobacteria |
| Shewanella acanthi | Pseudomonadota | Gammaproteobacteria |
| Shewanella aegiceratis | Pseudomonadota | Gammaproteobacteria |
| Shewanella aegiceratis | Pseudomonadota | Gammaproteobacteria |
| Shewanella algae | Pseudomonadota | Gammaproteobacteria |
| Shewanella algae | Pseudomonadota | Gammaproteobacteria |
| Shewanella alkalitolerans | Pseudomonadota | Gammaproteobacteria |
| Shewanella alkalitolerans | Pseudomonadota | Gammaproteobacteria |
| Shewanella amazonensis SB2B | Pseudomonadota | Gammaproteobacteria |
| Shewanella amazonensis SB2B | Pseudomonadota | Gammaproteobacteria |
| Shewanella avicenniae | Pseudomonadota | Gammaproteobacteria |

| | | |
|---|---|---|
| Shewanella avicenniae | Pseudomonadota | Gammaproteobacteria |
| Shewanella baltica OS678 | Pseudomonadota | Gammaproteobacteria |
| Shewanella baltica OS678 | Pseudomonadota | Gammaproteobacteria |
| Shewanella carassii | Pseudomonadota | Gammaproteobacteria |
| Shewanella carassii | Pseudomonadota | Gammaproteobacteria |
| Shewanella chilikensis | Pseudomonadota | Gammaproteobacteria |
| Shewanella chilikensis | Pseudomonadota | Gammaproteobacteria |
| Shewanella cyperi | Pseudomonadota | Gammaproteobacteria |
| Shewanella cyperi | Pseudomonadota | Gammaproteobacteria |
| Shewanella decolorationis | Pseudomonadota | Gammaproteobacteria |
| Shewanella denitrificans OS217 | Pseudomonadota | Gammaproteobacteria |
| Shewanella donghaensis | Pseudomonadota | Gammaproteobacteria |
| Shewanella donghaensis | Pseudomonadota | Gammaproteobacteria |
| Shewanella eurypsychrophilus | Pseudomonadota | Gammaproteobacteria |
| Shewanella eurypsychrophilus | Pseudomonadota | Gammaproteobacteria |
| Shewanella frigidimarina NCIMB 400 | Pseudomonadota | Gammaproteobacteria |
| Shewanella frigidimarina NCIMB 400 | Pseudomonadota | Gammaproteobacteria |
| Shewanella glacialimarina | Pseudomonadota | Gammaproteobacteria |
| Shewanella glacialimarina | Pseudomonadota | Gammaproteobacteria |
| Shewanella goraebulensis | Pseudomonadota | Gammaproteobacteria |
| Shewanella goraebulensis | Pseudomonadota | Gammaproteobacteria |
| Shewanella halifaxensis HAW-EB4 | Pseudomonadota | Gammaproteobacteria |
| Shewanella halifaxensis HAW-EB4 | Pseudomonadota | Gammaproteobacteria |
| Shewanella halotolerans | Pseudomonadota | Gammaproteobacteria |
| Shewanella halotolerans | Pseudomonadota | Gammaproteobacteria |
| Shewanella inventionis | Pseudomonadota | Gammaproteobacteria |
| Shewanella inventionis | Pseudomonadota | Gammaproteobacteria |
| Shewanella japonica | Pseudomonadota | Gammaproteobacteria |
| Shewanella japonica | Pseudomonadota | Gammaproteobacteria |
| Shewanella khirikhana | Pseudomonadota | Gammaproteobacteria |
| Shewanella khirikhana | Pseudomonadota | Gammaproteobacteria |

| | | |
|---|---|---|
| Shewanella litorisediminis | Pseudomonadota | Gammaproteobacteria |
| Shewanella litorisediminis | Pseudomonadota | Gammaproteobacteria |
| Shewanella livingstonensis | Pseudomonadota | Gammaproteobacteria |
| Shewanella livingstonensis | Pseudomonadota | Gammaproteobacteria |
| Shewanella loihica PV-4 | Pseudomonadota | Gammaproteobacteria |
| Shewanella loihica PV-4 | Pseudomonadota | Gammaproteobacteria |
| Shewanella mangrovisoli | Pseudomonadota | Gammaproteobacteria |
| Shewanella mangrovisoli | Pseudomonadota | Gammaproteobacteria |
| Shewanella marisflavi | Pseudomonadota | Gammaproteobacteria |
| Shewanella marisflavi | Pseudomonadota | Gammaproteobacteria |
| Shewanella maritima | Pseudomonadota | Gammaproteobacteria |
| Shewanella maritima | Pseudomonadota | Gammaproteobacteria |
| Shewanella mesophila | Pseudomonadota | Gammaproteobacteria |
| Shewanella mesophila | Pseudomonadota | Gammaproteobacteria |
| Shewanella oncorhynchi | Pseudomonadota | Gammaproteobacteria |
| Shewanella oncorhynchi | Pseudomonadota | Gammaproteobacteria |
| Shewanella oneidensis MR-1 | Pseudomonadota | Gammaproteobacteria |
| Shewanella oneidensis MR-1 | Pseudomonadota | Gammaproteobacteria |
| Shewanella pealeana ATCC 700345 | Pseudomonadota | Gammaproteobacteria |
| Shewanella pealeana ATCC 700345 | Pseudomonadota | Gammaproteobacteria |
| Shewanella piezotolerans WP3 | Pseudomonadota | Gammaproteobacteria |
| Shewanella piezotolerans WP3 | Pseudomonadota | Gammaproteobacteria |
| Shewanella polaris | Pseudomonadota | Gammaproteobacteria |
| Shewanella polaris | Pseudomonadota | Gammaproteobacteria |
| Shewanella psychromarinicola | Pseudomonadota | Gammaproteobacteria |
| Shewanella psychromarinicola | Pseudomonadota | Gammaproteobacteria |
| Shewanella psychropiezotolerans | Pseudomonadota | Gammaproteobacteria |
| Shewanella psychropiezotolerans | Pseudomonadota | Gammaproteobacteria |
| Shewanella psychrotolerans | Pseudomonadota | Gammaproteobacteria |
| Shewanella psychrotolerans | Pseudomonadota | Gammaproteobacteria |
| Shewanella putrefaciens | Pseudomonadota | Gammaproteobacteria |

| | | |
|---|---|---|
| Shewanella putrefaciens | Pseudomonadota | Gammaproteobacteria |
| Shewanella rhizosphaerae | Pseudomonadota | Gammaproteobacteria |
| Shewanella rhizosphaerae | Pseudomonadota | Gammaproteobacteria |
| Shewanella sedimentimangrovi | Pseudomonadota | Gammaproteobacteria |
| Shewanella sedimentimangrovi | Pseudomonadota | Gammaproteobacteria |
| Shewanella spartinae | Pseudomonadota | Gammaproteobacteria |
| Shewanella spartinae | Pseudomonadota | Gammaproteobacteria |
| Shewanella violacea DSS12 | Pseudomonadota | Gammaproteobacteria |
| Shewanella violacea DSS12 | Pseudomonadota | Gammaproteobacteria |
| Shewanella woodyi ATCC 51908 | Pseudomonadota | Gammaproteobacteria |
| Shewanella woodyi ATCC 51908 | Pseudomonadota | Gammaproteobacteria |
| Shewanella xiamenensis | Pseudomonadota | Gammaproteobacteria |
| Shewanella xiamenensis | Pseudomonadota | Gammaproteobacteria |
| Shewanella youngdeokensis | Pseudomonadota | Gammaproteobacteria |
| Shewanella youngdeokensis | Pseudomonadota | Gammaproteobacteria |
| Shewanella yunxiaonensis | Pseudomonadota | Gammaproteobacteria |
| Shewanella yunxiaonensis | Pseudomonadota | Gammaproteobacteria |
| Shewanella zhangzhouensis | Pseudomonadota | Gammaproteobacteria |
| Shewanella zhangzhouensis | Pseudomonadota | Gammaproteobacteria |
| Shigella dysenteriae | Pseudomonadota | Gammaproteobacteria |
| Shigella dysenteriae | Pseudomonadota | Gammaproteobacteria |
| Shigella flexneri 2a str. 301 | Pseudomonadota | Gammaproteobacteria |
| Shigella flexneri 2a str. 301 | Pseudomonadota | Gammaproteobacteria |
| Shigella sonnei | Pseudomonadota | Gammaproteobacteria |
| Shigella sonnei | Pseudomonadota | Gammaproteobacteria |
| Shimwellia blattae DSM 4481 = NBRC 105725 | Pseudomonadota | Gammaproteobacteria |
| Shimwellia blattae DSM 4481 = NBRC 105725 | Pseudomonadota | Gammaproteobacteria |
| Siccibacter colletis | Pseudomonadota | Gammaproteobacteria |
| Siccibacter colletis | Pseudomonadota | Gammaproteobacteria |
| Simiduia litorea | Pseudomonadota | Gammaproteobacteria |
| Simiduia litorea | Pseudomonadota | Gammaproteobacteria |

| | | |
|---|---|---|
| Sodalis endosymbiont of Henestaris halophilus | Pseudomonadota | Gammaproteobacteria |
| Sodalis glossinidius str. 'morsitans' | Pseudomonadota | Gammaproteobacteria |
| Sodalis glossinidius str. 'morsitans' | Pseudomonadota | Gammaproteobacteria |
| Sodalis-like secondary symbiont of Drepanosiphum platanoidis | Pseudomonadota | Gammaproteobacteria |
| Solilutibacter oculi | Pseudomonadota | Gammaproteobacteria |
| Solilutibacter oculi | Pseudomonadota | Gammaproteobacteria |
| Spartinivicinus ruber | Pseudomonadota | Gammaproteobacteria |
| Spartinivicinus ruber | Pseudomonadota | Gammaproteobacteria |
| Spiribacter halobius | Pseudomonadota | Gammaproteobacteria |
| Spiribacter halobius | Pseudomonadota | Gammaproteobacteria |
| Spiribacter roseus | Pseudomonadota | Gammaproteobacteria |
| Spiribacter roseus | Pseudomonadota | Gammaproteobacteria |
| Spiribacter salinus M19-40 | Pseudomonadota | Gammaproteobacteria |
| Spongiibacter nanhainus | Pseudomonadota | Gammaproteobacteria |
| Spongiibacter nanhainus | Pseudomonadota | Gammaproteobacteria |
| Spongiibacter taiwanensis | Pseudomonadota | Gammaproteobacteria |
| Spongiibacter taiwanensis | Pseudomonadota | Gammaproteobacteria |
| Stenotrophomonas acidaminiphila | Pseudomonadota | Gammaproteobacteria |
| Stenotrophomonas acidaminiphila | Pseudomonadota | Gammaproteobacteria |
| Stenotrophomonas aracearum | Pseudomonadota | Gammaproteobacteria |
| Stenotrophomonas aracearum | Pseudomonadota | Gammaproteobacteria |
| Stenotrophomonas forensis | Pseudomonadota | Gammaproteobacteria |
| Stenotrophomonas forensis | Pseudomonadota | Gammaproteobacteria |
| Stenotrophomonas geniculata | Pseudomonadota | Gammaproteobacteria |
| Stenotrophomonas geniculata | Pseudomonadota | Gammaproteobacteria |
| Stenotrophomonas indicatrix | Pseudomonadota | Gammaproteobacteria |
| Stenotrophomonas indicatrix | Pseudomonadota | Gammaproteobacteria |
| Stenotrophomonas lactitubi | Pseudomonadota | Gammaproteobacteria |
| Stenotrophomonas lactitubi | Pseudomonadota | Gammaproteobacteria |
| Stenotrophomonas maltophilia | Pseudomonadota | Gammaproteobacteria |
| Stenotrophomonas maltophilia | Pseudomonadota | Gammaproteobacteria |

| | | |
|---|---|---|
| Stenotrophomonas oahuensis | Pseudomonadota | Gammaproteobacteria |
| Stenotrophomonas oahuensis | Pseudomonadota | Gammaproteobacteria |
| Stenotrophomonas pigmentata | Pseudomonadota | Gammaproteobacteria |
| Stenotrophomonas pigmentata | Pseudomonadota | Gammaproteobacteria |
| Stenotrophomonas rhizophila | Pseudomonadota | Gammaproteobacteria |
| Stenotrophomonas rhizophila | Pseudomonadota | Gammaproteobacteria |
| Stenotrophomonas riyadhensis | Pseudomonadota | Gammaproteobacteria |
| Stenotrophomonas riyadhensis | Pseudomonadota | Gammaproteobacteria |
| Stenotrophomonas sepilia | Pseudomonadota | Gammaproteobacteria |
| Stenotrophomonas sepilia | Pseudomonadota | Gammaproteobacteria |
| Steroidobacter denitrificans | Pseudomonadota | Gammaproteobacteria |
| Steroidobacter denitrificans | Pseudomonadota | Gammaproteobacteria |
| Stutzerimonas balearica DSM 6083 | Pseudomonadota | Gammaproteobacteria |
| Stutzerimonas balearica DSM 6083 | Pseudomonadota | Gammaproteobacteria |
| Stutzerimonas chloritidismutans | Pseudomonadota | Gammaproteobacteria |
| Stutzerimonas chloritidismutans | Pseudomonadota | Gammaproteobacteria |
| Stutzerimonas decontaminans | Pseudomonadota | Gammaproteobacteria |
| Stutzerimonas decontaminans | Pseudomonadota | Gammaproteobacteria |
| Stutzerimonas degradans | Pseudomonadota | Gammaproteobacteria |
| Stutzerimonas degradans | Pseudomonadota | Gammaproteobacteria |
| Stutzerimonas frequens | Pseudomonadota | Gammaproteobacteria |
| Stutzerimonas frequens | Pseudomonadota | Gammaproteobacteria |
| Stutzerimonas stutzeri | Pseudomonadota | Gammaproteobacteria |
| Stutzerimonas stutzeri | Pseudomonadota | Gammaproteobacteria |
| Stutzerimonas xanthomarina | Pseudomonadota | Gammaproteobacteria |
| Stutzerimonas xanthomarina | Pseudomonadota | Gammaproteobacteria |
| Succinivibrio dextrinosolvens | Pseudomonadota | Gammaproteobacteria |
| Succinivibrio dextrinosolvens | Pseudomonadota | Gammaproteobacteria |
| Sulfuricaulis limicola | Pseudomonadota | Gammaproteobacteria |
| Sulfuricaulis limicola | Pseudomonadota | Gammaproteobacteria |
| Sulfuriflexus mobilis | Pseudomonadota | Gammaproteobacteria |

| | | |
|---|---|---|
| Sulfuriflexus mobilis | Pseudomonadota | Gammaproteobacteria |
| Sulfurifustis variabilis | Pseudomonadota | Gammaproteobacteria |
| Sulfurifustis variabilis | Pseudomonadota | Gammaproteobacteria |
| Sulfurivermis fontis | Pseudomonadota | Gammaproteobacteria |
| Sulfurivermis fontis | Pseudomonadota | Gammaproteobacteria |
| Symbiopectobacterium purcellii | Pseudomonadota | Gammaproteobacteria |
| Symbiopectobacterium purcellii | Pseudomonadota | Gammaproteobacteria |
| Tahibacter amnicola | Pseudomonadota | Gammaproteobacteria |
| Tahibacter amnicola | Pseudomonadota | Gammaproteobacteria |
| Tatumella citrea | Pseudomonadota | Gammaproteobacteria |
| Tatumella citrea | Pseudomonadota | Gammaproteobacteria |
| Teredinibacter haidensis | Pseudomonadota | Gammaproteobacteria |
| Teredinibacter purpureus | Pseudomonadota | Gammaproteobacteria |
| Teredinibacter turnerae | Pseudomonadota | Gammaproteobacteria |
| Thalassolituus hydrocarboniclasticus | Pseudomonadota | Gammaproteobacteria |
| Thalassolituus hydrocarboniclasticus | Pseudomonadota | Gammaproteobacteria |
| Thalassolituus hydrocarboniclasticus | Pseudomonadota | Gammaproteobacteria |
| Thalassolituus oleivorans MIL-1 | Pseudomonadota | Gammaproteobacteria |
| Thalassolituus oleivorans MIL-1 | Pseudomonadota | Gammaproteobacteria |
| Thalassolituus oleivorans MIL-1 | Pseudomonadota | Gammaproteobacteria |
| Thalassomonas actiniarum | Pseudomonadota | Gammaproteobacteria |
| Thalassomonas actiniarum | Pseudomonadota | Gammaproteobacteria |
| Thalassomonas haliotis | Pseudomonadota | Gammaproteobacteria |
| Thalassomonas haliotis | Pseudomonadota | Gammaproteobacteria |
| Thalassomonas viridans | Pseudomonadota | Gammaproteobacteria |
| Thalassomonas viridans | Pseudomonadota | Gammaproteobacteria |
| Thalassotalea agarivorans | Pseudomonadota | Gammaproteobacteria |
| Thalassotalea agarivorans | Pseudomonadota | Gammaproteobacteria |
| Thalassotalea aquiviva | Pseudomonadota | Gammaproteobacteria |
| Thalassotalea aquiviva | Pseudomonadota | Gammaproteobacteria |
| Thalassotalea aquiviva | Pseudomonadota | Gammaproteobacteria |

| | | |
|---|---|---|
| Thalassotalea atypica | Pseudomonadota | Gammaproteobacteria |
| Thalassotalea atypica | Pseudomonadota | Gammaproteobacteria |
| Thalassotalea crassostreae | Pseudomonadota | Gammaproteobacteria |
| Thalassotalea crassostreae | Pseudomonadota | Gammaproteobacteria |
| Thalassotalea fonticola | Pseudomonadota | Gammaproteobacteria |
| Thalassotalea fonticola | Pseudomonadota | Gammaproteobacteria |
| Thalassotalea fonticola | Pseudomonadota | Gammaproteobacteria |
| Thalassotalea hakodatensis | Pseudomonadota | Gammaproteobacteria |
| Thalassotalea hakodatensis | Pseudomonadota | Gammaproteobacteria |
| Thalassotalea maritima | Pseudomonadota | Gammaproteobacteria |
| Thalassotalea maritima | Pseudomonadota | Gammaproteobacteria |
| Thalassotalea maritima | Pseudomonadota | Gammaproteobacteria |
| Thalassotalea nanhaiensis | Pseudomonadota | Gammaproteobacteria |
| Thalassotalea nanhaiensis | Pseudomonadota | Gammaproteobacteria |
| Thalassotalea nanhaiensis | Pseudomonadota | Gammaproteobacteria |
| Thalassotalea piscium | Pseudomonadota | Gammaproteobacteria |
| Thalassotalea piscium | Pseudomonadota | Gammaproteobacteria |
| Thalassotalea ponticola | Pseudomonadota | Gammaproteobacteria |
| Thalassotalea ponticola | Pseudomonadota | Gammaproteobacteria |
| Thalassotalea ponticola | Pseudomonadota | Gammaproteobacteria |
| Thalassotalea psychrophila | Pseudomonadota | Gammaproteobacteria |
| Thalassotalea psychrophila | Pseudomonadota | Gammaproteobacteria |
| Thalassotalea psychrophila | Pseudomonadota | Gammaproteobacteria |
| Thalassotalea sediminis | Pseudomonadota | Gammaproteobacteria |
| Thalassotalea sediminis | Pseudomonadota | Gammaproteobacteria |
| Thaumasiovibrio subtropicus | Pseudomonadota | Gammaproteobacteria |
| Thaumasiovibrio subtropicus | Pseudomonadota | Gammaproteobacteria |
| Thermochromatium tepidum ATCC 43061 | Pseudomonadota | Gammaproteobacteria |
| Thermochromatium tepidum ATCC 43061 | Pseudomonadota | Gammaproteobacteria |
| Thermomonas aquatica | Pseudomonadota | Gammaproteobacteria |
| Thermomonas aquatica | Pseudomonadota | Gammaproteobacteria |

| | | |
|---|---|---|
| Thermomonas brevis | Pseudomonadota | Gammaproteobacteria |
| Thermomonas brevis | Pseudomonadota | Gammaproteobacteria |
| Thermomonas paludicola | Pseudomonadota | Gammaproteobacteria |
| Thermomonas paludicola | Pseudomonadota | Gammaproteobacteria |
| Thioalkalivibrio nitratireducens DSM 14787 | Pseudomonadota | Gammaproteobacteria |
| Thioalkalivibrio nitratireducens DSM 14787 | Pseudomonadota | Gammaproteobacteria |
| Thioalkalivibrio paradoxus ARh 1 | Pseudomonadota | Gammaproteobacteria |
| Thioalkalivibrio paradoxus ARh 1 | Pseudomonadota | Gammaproteobacteria |
| Thioalkalivibrio sulfidiphilus HL-EbGr7 | Pseudomonadota | Gammaproteobacteria |
| Thioalkalivibrio sulfidiphilus HL-EbGr7 | Pseudomonadota | Gammaproteobacteria |
| Thioalkalivibrio versutus | Pseudomonadota | Gammaproteobacteria |
| Thioalkalivibrio versutus | Pseudomonadota | Gammaproteobacteria |
| Thiocapsa bogorovii | Pseudomonadota | Gammaproteobacteria |
| Thiocapsa bogorovii | Pseudomonadota | Gammaproteobacteria |
| Thiocystis violascens DSM 198 | Pseudomonadota | Gammaproteobacteria |
| Thiocystis violascens DSM 198 | Pseudomonadota | Gammaproteobacteria |
| Thioflavicoccus mobilis 8321 | Pseudomonadota | Gammaproteobacteria |
| Thioflavicoccus mobilis 8321 | Pseudomonadota | Gammaproteobacteria |
| Thiomicrorhabdus aquaedulcis | Pseudomonadota | Gammaproteobacteria |
| Thiomicrorhabdus aquaedulcis | Pseudomonadota | Gammaproteobacteria |
| Thiomicrorhabdus immobilis | Pseudomonadota | Gammaproteobacteria |
| Thiomicrorhabdus immobilis | Pseudomonadota | Gammaproteobacteria |
| Thiomicrorhabdus indica | Pseudomonadota | Gammaproteobacteria |
| Thiomicrorhabdus indica | Pseudomonadota | Gammaproteobacteria |
| Thiomicrorhabdus lithotrophica | Pseudomonadota | Gammaproteobacteria |
| Thiomicrorhabdus lithotrophica | Pseudomonadota | Gammaproteobacteria |
| Thiomicrorhabdus sediminis | Pseudomonadota | Gammaproteobacteria |
| Thiomicrorhabdus sediminis | Pseudomonadota | Gammaproteobacteria |
| Thiomicrorhabdus xiamenensis | Pseudomonadota | Gammaproteobacteria |
| Thiomicrorhabdus xiamenensis | Pseudomonadota | Gammaproteobacteria |
| Thiomicrospira aerophila AL3 | Pseudomonadota | Gammaproteobacteria |

| | | |
|---|---|---|
| Thiomicrospira aerophila AL3 | Pseudomonadota | Gammaproteobacteria |
| Thiomicrospira cyclica ALM1 | Pseudomonadota | Gammaproteobacteria |
| Thiomicrospira cyclica ALM1 | Pseudomonadota | Gammaproteobacteria |
| Thiopseudomonas alkaliphila | Pseudomonadota | Gammaproteobacteria |
| Thiopseudomonas alkaliphila | Pseudomonadota | Gammaproteobacteria |
| Thiorhodovibrio frisius | Pseudomonadota | Gammaproteobacteria |
| Thiorhodovibrio frisius | Pseudomonadota | Gammaproteobacteria |
| Thiorhodovibrio litoralis | Pseudomonadota | Gammaproteobacteria |
| Thiorhodovibrio litoralis | Pseudomonadota | Gammaproteobacteria |
| Thiorhodovibrio winogradskyi | Pseudomonadota | Gammaproteobacteria |
| Thiorhodovibrio winogradskyi | Pseudomonadota | Gammaproteobacteria |
| Thiosocius teredinicola | Pseudomonadota | Gammaproteobacteria |
| Thiosocius teredinicola | Pseudomonadota | Gammaproteobacteria |
| Thiosulfatimonas sediminis | Pseudomonadota | Gammaproteobacteria |
| Thiosulfatimonas sediminis | Pseudomonadota | Gammaproteobacteria |
| Thiosulfativibrio zosterae | Pseudomonadota | Gammaproteobacteria |
| Thiosulfativibrio zosterae | Pseudomonadota | Gammaproteobacteria |
| Thiothrix litoralis | Pseudomonadota | Gammaproteobacteria |
| Thiothrix litoralis | Pseudomonadota | Gammaproteobacteria |
| Thiothrix subterranea | Pseudomonadota | Gammaproteobacteria |
| Thiothrix winogradskyi | Pseudomonadota | Gammaproteobacteria |
| Thiothrix winogradskyi | Pseudomonadota | Gammaproteobacteria |
| Tolumonas auensis DSM 9187 | Pseudomonadota | Gammaproteobacteria |
| Tolumonas auensis DSM 9187 | Pseudomonadota | Gammaproteobacteria |
| Trabulsiella odontotermitis | Pseudomonadota | Gammaproteobacteria |
| Trabulsiella odontotermitis | Pseudomonadota | Gammaproteobacteria |
| Utexia brackfieldae | Pseudomonadota | Gammaproteobacteria |
| Utexia brackfieldae | Pseudomonadota | Gammaproteobacteria |
| Venatoribacter cucullus | Pseudomonadota | Gammaproteobacteria |
| Venatoribacter cucullus | Pseudomonadota | Gammaproteobacteria |
| Venatoribacter cucullus | Pseudomonadota | Gammaproteobacteria |

| | | |
|---|---|---|
| Vespertiliibacter pulmonis | Pseudomonadota | Gammaproteobacteria |
| Vespertiliibacter pulmonis | Pseudomonadota | Gammaproteobacteria |
| Vibrio aerogenes | Pseudomonadota | Gammaproteobacteria |
| Vibrio aerogenes | Pseudomonadota | Gammaproteobacteria |
| Vibrio alfacsensis | Pseudomonadota | Gammaproteobacteria |
| Vibrio alfacsensis | Pseudomonadota | Gammaproteobacteria |
| Vibrio algicola | Pseudomonadota | Gammaproteobacteria |
| Vibrio algicola | Pseudomonadota | Gammaproteobacteria |
| Vibrio alginolyticus | Pseudomonadota | Gammaproteobacteria |
| Vibrio alginolyticus | Pseudomonadota | Gammaproteobacteria |
| Vibrio antiquarius | Pseudomonadota | Gammaproteobacteria |
| Vibrio antiquarius | Pseudomonadota | Gammaproteobacteria |
| Vibrio aphrogenes | Pseudomonadota | Gammaproteobacteria |
| Vibrio aphrogenes | Pseudomonadota | Gammaproteobacteria |
| Vibrio apostichopi | Pseudomonadota | Gammaproteobacteria |
| Vibrio apostichopi | Pseudomonadota | Gammaproteobacteria |
| Vibrio aquimaris | Pseudomonadota | Gammaproteobacteria |
| Vibrio aquimaris | Pseudomonadota | Gammaproteobacteria |
| Vibrio artabrorum | Pseudomonadota | Gammaproteobacteria |
| Vibrio artabrorum | Pseudomonadota | Gammaproteobacteria |
| Vibrio astriarenae | Pseudomonadota | Gammaproteobacteria |
| Vibrio astriarenae | Pseudomonadota | Gammaproteobacteria |
| Vibrio atlanticus | Pseudomonadota | Gammaproteobacteria |
| Vibrio atlanticus | Pseudomonadota | Gammaproteobacteria |
| Vibrio azureus | Pseudomonadota | Gammaproteobacteria |
| Vibrio azureus | Pseudomonadota | Gammaproteobacteria |
| Vibrio bathopelagicus | Pseudomonadota | Gammaproteobacteria |
| Vibrio bathopelagicus | Pseudomonadota | Gammaproteobacteria |
| Vibrio breoganii | Pseudomonadota | Gammaproteobacteria |
| Vibrio breoganii | Pseudomonadota | Gammaproteobacteria |
| Vibrio campbellii | Pseudomonadota | Gammaproteobacteria |

| | | |
|---|---|---|
| Vibrio campbellii | Pseudomonadota | Gammaproteobacteria |
| Vibrio chaetopteri | Pseudomonadota | Gammaproteobacteria |
| Vibrio chaetopteri | Pseudomonadota | Gammaproteobacteria |
| Vibrio chagasii | Pseudomonadota | Gammaproteobacteria |
| Vibrio chagasii | Pseudomonadota | Gammaproteobacteria |
| Vibrio cholerae | Pseudomonadota | Gammaproteobacteria |
| Vibrio cholerae | Pseudomonadota | Gammaproteobacteria |
| Vibrio cidicii | Pseudomonadota | Gammaproteobacteria |
| Vibrio cidicii | Pseudomonadota | Gammaproteobacteria |
| Vibrio comitans | Pseudomonadota | Gammaproteobacteria |
| Vibrio comitans | Pseudomonadota | Gammaproteobacteria |
| Vibrio coralliilyticus | Pseudomonadota | Gammaproteobacteria |
| Vibrio coralliilyticus | Pseudomonadota | Gammaproteobacteria |
| Vibrio cortegadensis | Pseudomonadota | Gammaproteobacteria |
| Vibrio cortegadensis | Pseudomonadota | Gammaproteobacteria |
| Vibrio crassostreae | Pseudomonadota | Gammaproteobacteria |
| Vibrio crassostreae | Pseudomonadota | Gammaproteobacteria |
| Vibrio cyclitrophicus | Pseudomonadota | Gammaproteobacteria |
| Vibrio cyclitrophicus | Pseudomonadota | Gammaproteobacteria |
| Vibrio diabolicus | Pseudomonadota | Gammaproteobacteria |
| Vibrio diabolicus | Pseudomonadota | Gammaproteobacteria |
| Vibrio diazotrophicus | Pseudomonadota | Gammaproteobacteria |
| Vibrio diazotrophicus | Pseudomonadota | Gammaproteobacteria |
| Vibrio echinoideorum | Pseudomonadota | Gammaproteobacteria |
| Vibrio echinoideorum | Pseudomonadota | Gammaproteobacteria |
| Vibrio fluvialis | Pseudomonadota | Gammaproteobacteria |
| Vibrio fluvialis | Pseudomonadota | Gammaproteobacteria |
| Vibrio fortis | Pseudomonadota | Gammaproteobacteria |
| Vibrio fortis | Pseudomonadota | Gammaproteobacteria |
| Vibrio furnissii | Pseudomonadota | Gammaproteobacteria |
| Vibrio furnissii | Pseudomonadota | Gammaproteobacteria |

| | | |
|---|---|---|
| Vibrio gallaecicus | Pseudomonadota | Gammaproteobacteria |
| Vibrio gallaecicus | Pseudomonadota | Gammaproteobacteria |
| Vibrio gallicus | Pseudomonadota | Gammaproteobacteria |
| Vibrio gallicus | Pseudomonadota | Gammaproteobacteria |
| Vibrio gangliei | Pseudomonadota | Gammaproteobacteria |
| Vibrio gangliei | Pseudomonadota | Gammaproteobacteria |
| Vibrio gazogenes | Pseudomonadota | Gammaproteobacteria |
| Vibrio gazogenes | Pseudomonadota | Gammaproteobacteria |
| Vibrio gigantis | Pseudomonadota | Gammaproteobacteria |
| Vibrio gigantis | Pseudomonadota | Gammaproteobacteria |
| Vibrio harveyi | Pseudomonadota | Gammaproteobacteria |
| Vibrio harveyi | Pseudomonadota | Gammaproteobacteria |
| Vibrio hyugaensis | Pseudomonadota | Gammaproteobacteria |
| Vibrio hyugaensis | Pseudomonadota | Gammaproteobacteria |
| Vibrio inusitatus | Pseudomonadota | Gammaproteobacteria |
| Vibrio inusitatus | Pseudomonadota | Gammaproteobacteria |
| Vibrio ishigakensis | Pseudomonadota | Gammaproteobacteria |
| Vibrio ishigakensis | Pseudomonadota | Gammaproteobacteria |
| Vibrio japonicus | Pseudomonadota | Gammaproteobacteria |
| Vibrio japonicus | Pseudomonadota | Gammaproteobacteria |
| Vibrio jasicida 090810c | Pseudomonadota | Gammaproteobacteria |
| Vibrio jasicida 090810c | Pseudomonadota | Gammaproteobacteria |
| Vibrio kanaloae | Pseudomonadota | Gammaproteobacteria |
| Vibrio kanaloae | Pseudomonadota | Gammaproteobacteria |
| Vibrio mangrovi | Pseudomonadota | Gammaproteobacteria |
| Vibrio mangrovi | Pseudomonadota | Gammaproteobacteria |
| Vibrio metoecus | Pseudomonadota | Gammaproteobacteria |
| Vibrio metoecus | Pseudomonadota | Gammaproteobacteria |
| Vibrio misgurnus | Pseudomonadota | Gammaproteobacteria |
| Vibrio misgurnus | Pseudomonadota | Gammaproteobacteria |
| Vibrio natriegens NBRC 15636 = ATCC 14048 = DSM 759 | Pseudomonadota | Gammaproteobacteria |

| | | |
|---|---|---|
| Vibrio natriegens NBRC 15636 = ATCC 14048 = DSM 759 | Pseudomonadota | Gammaproteobacteria |
| Vibrio neonatus | Pseudomonadota | Gammaproteobacteria |
| Vibrio neonatus | Pseudomonadota | Gammaproteobacteria |
| Vibrio nigripulchritudo | Pseudomonadota | Gammaproteobacteria |
| Vibrio nigripulchritudo | Pseudomonadota | Gammaproteobacteria |
| Vibrio ostreae | Pseudomonadota | Gammaproteobacteria |
| Vibrio ostreae | Pseudomonadota | Gammaproteobacteria |
| Vibrio owensii | Pseudomonadota | Gammaproteobacteria |
| Vibrio owensii | Pseudomonadota | Gammaproteobacteria |
| Vibrio palustris | Pseudomonadota | Gammaproteobacteria |
| Vibrio palustris | Pseudomonadota | Gammaproteobacteria |
| Vibrio paracholerae | Pseudomonadota | Gammaproteobacteria |
| Vibrio paracholerae | Pseudomonadota | Gammaproteobacteria |
| Vibrio parahaemolyticus RIMD 2210633 | Pseudomonadota | Gammaproteobacteria |
| Vibrio parahaemolyticus RIMD 2210633 | Pseudomonadota | Gammaproteobacteria |
| Vibrio pectenicida | Pseudomonadota | Gammaproteobacteria |
| Vibrio pectenicida | Pseudomonadota | Gammaproteobacteria |
| Vibrio pelagius | Pseudomonadota | Gammaproteobacteria |
| Vibrio pelagius | Pseudomonadota | Gammaproteobacteria |
| Vibrio penaeicida | Pseudomonadota | Gammaproteobacteria |
| Vibrio penaeicida | Pseudomonadota | Gammaproteobacteria |
| Vibrio pomeroyi | Pseudomonadota | Gammaproteobacteria |
| Vibrio pomeroyi | Pseudomonadota | Gammaproteobacteria |
| Vibrio ponticus | Pseudomonadota | Gammaproteobacteria |
| Vibrio ponticus | Pseudomonadota | Gammaproteobacteria |
| Vibrio porteresiae DSM 19223 | Pseudomonadota | Gammaproteobacteria |
| Vibrio porteresiae DSM 19223 | Pseudomonadota | Gammaproteobacteria |
| Vibrio qinghaiensis | Pseudomonadota | Gammaproteobacteria |
| Vibrio qinghaiensis | Pseudomonadota | Gammaproteobacteria |
| Vibrio quintilis | Pseudomonadota | Gammaproteobacteria |
| Vibrio quintilis | Pseudomonadota | Gammaproteobacteria |

| | | |
|---|---|---|
| Vibrio rarus | Pseudomonadota | Gammaproteobacteria |
| Vibrio rarus | Pseudomonadota | Gammaproteobacteria |
| Vibrio rhizosphaerae | Pseudomonadota | Gammaproteobacteria |
| Vibrio rhizosphaerae | Pseudomonadota | Gammaproteobacteria |
| Vibrio rotiferianus | Pseudomonadota | Gammaproteobacteria |
| Vibrio rotiferianus | Pseudomonadota | Gammaproteobacteria |
| Vibrio ruber | Pseudomonadota | Gammaproteobacteria |
| Vibrio ruber | Pseudomonadota | Gammaproteobacteria |
| Vibrio rumoiensis | Pseudomonadota | Gammaproteobacteria |
| Vibrio rumoiensis | Pseudomonadota | Gammaproteobacteria |
| Vibrio scophthalmi | Pseudomonadota | Gammaproteobacteria |
| Vibrio scophthalmi | Pseudomonadota | Gammaproteobacteria |
| Vibrio spartinae | Pseudomonadota | Gammaproteobacteria |
| Vibrio spartinae | Pseudomonadota | Gammaproteobacteria |
| Vibrio splendidus | Pseudomonadota | Gammaproteobacteria |
| Vibrio splendidus | Pseudomonadota | Gammaproteobacteria |
| Vibrio syngnathi | Pseudomonadota | Gammaproteobacteria |
| Vibrio syngnathi | Pseudomonadota | Gammaproteobacteria |
| Vibrio tasmaniensis | Pseudomonadota | Gammaproteobacteria |
| Vibrio tasmaniensis | Pseudomonadota | Gammaproteobacteria |
| Vibrio toranzoniae | Pseudomonadota | Gammaproteobacteria |
| Vibrio toranzoniae | Pseudomonadota | Gammaproteobacteria |
| Vibrio tritonius | Pseudomonadota | Gammaproteobacteria |
| Vibrio tritonius | Pseudomonadota | Gammaproteobacteria |
| Vibrio tubiashii ATCC 19109 | Pseudomonadota | Gammaproteobacteria |
| Vibrio tubiashii ATCC 19109 | Pseudomonadota | Gammaproteobacteria |
| Vibrio vulnificus NBRC 15645 = ATCC 27562 | Pseudomonadota | Gammaproteobacteria |
| Vibrio vulnificus NBRC 15645 = ATCC 27562 | Pseudomonadota | Gammaproteobacteria |
| Vibrio zhugei | Pseudomonadota | Gammaproteobacteria |
| Vibrio zhugei | Pseudomonadota | Gammaproteobacteria |
| Vibrio ziniensis | Pseudomonadota | Gammaproteobacteria |

| | | |
|---|---|---|
| Vibrio ziniensis | Pseudomonadota | Gammaproteobacteria |
| Vreelandella neptunia | Pseudomonadota | Gammaproteobacteria |
| Vreelandella piezotolerans | Pseudomonadota | Gammaproteobacteria |
| Vreelandella profundi | Pseudomonadota | Gammaproteobacteria |
| Vreelandella stevensii | Pseudomonadota | Gammaproteobacteria |
| Vreelandella subglaciescola | Pseudomonadota | Gammaproteobacteria |
| Vreelandella titanicae | Pseudomonadota | Gammaproteobacteria |
| Vreelandella venusta | Pseudomonadota | Gammaproteobacteria |
| Wenzhouxiangella marina | Pseudomonadota | Gammaproteobacteria |
| Wenzhouxiangella marina | Pseudomonadota | Gammaproteobacteria |
| Wigglesworthia glossinidia endosymbiont of Glossina morsitans morsitans (Yale | Pseudomonadota | Gammaproteobacteria |
| Winslowiella toletana | Pseudomonadota | Gammaproteobacteria |
| Winslowiella toletana | Pseudomonadota | Gammaproteobacteria |
| Woeseia oceani | Pseudomonadota | Gammaproteobacteria |
| Woeseia oceani | Pseudomonadota | Gammaproteobacteria |
| Xanthomonas albilineans | Pseudomonadota | Gammaproteobacteria |
| Xanthomonas albilineans | Pseudomonadota | Gammaproteobacteria |
| Xanthomonas arboricola pv. juglandis | Pseudomonadota | Gammaproteobacteria |
| Xanthomonas arboricola pv. juglandis | Pseudomonadota | Gammaproteobacteria |
| Xanthomonas axonopodis pv. maculifoliigardeniae | Pseudomonadota | Gammaproteobacteria |
| Xanthomonas axonopodis pv. maculifoliigardeniae | Pseudomonadota | Gammaproteobacteria |
| Xanthomonas bundabergensis | Pseudomonadota | Gammaproteobacteria |
| Xanthomonas bundabergensis | Pseudomonadota | Gammaproteobacteria |
| Xanthomonas campestris pv. raphani | Pseudomonadota | Gammaproteobacteria |
| Xanthomonas campestris pv. raphani | Pseudomonadota | Gammaproteobacteria |
| Xanthomonas cannabis | Pseudomonadota | Gammaproteobacteria |
| Xanthomonas cannabis | Pseudomonadota | Gammaproteobacteria |
| Xanthomonas cassavae CFBP 4642 | Pseudomonadota | Gammaproteobacteria |
| Xanthomonas cerealis pv. cerealis | Pseudomonadota | Gammaproteobacteria |
| Xanthomonas cerealis pv. cerealis | Pseudomonadota | Gammaproteobacteria |
| Xanthomonas citri pv. citri | Pseudomonadota | Gammaproteobacteria |

| | | |
|---|---|---|
| Xanthomonas citri pv. citri | Pseudomonadota | Gammaproteobacteria |
| Xanthomonas cucurbitae | Pseudomonadota | Gammaproteobacteria |
| Xanthomonas cucurbitae | Pseudomonadota | Gammaproteobacteria |
| Xanthomonas dyei | Pseudomonadota | Gammaproteobacteria |
| Xanthomonas dyei | Pseudomonadota | Gammaproteobacteria |
| Xanthomonas euroxanthea | Pseudomonadota | Gammaproteobacteria |
| Xanthomonas euroxanthea | Pseudomonadota | Gammaproteobacteria |
| Xanthomonas euvesicatoria pv. alfalfae | Pseudomonadota | Gammaproteobacteria |
| Xanthomonas euvesicatoria pv. alfalfae | Pseudomonadota | Gammaproteobacteria |
| Xanthomonas fragariae | Pseudomonadota | Gammaproteobacteria |
| Xanthomonas fragariae | Pseudomonadota | Gammaproteobacteria |
| Xanthomonas graminis pv. graminis | Pseudomonadota | Gammaproteobacteria |
| Xanthomonas graminis pv. graminis | Pseudomonadota | Gammaproteobacteria |
| Xanthomonas hawaiiensis | Pseudomonadota | Gammaproteobacteria |
| Xanthomonas hawaiiensis | Pseudomonadota | Gammaproteobacteria |
| Xanthomonas hortorum | Pseudomonadota | Gammaproteobacteria |
| Xanthomonas hortorum | Pseudomonadota | Gammaproteobacteria |
| Xanthomonas hyacinthi | Pseudomonadota | Gammaproteobacteria |
| Xanthomonas hyacinthi | Pseudomonadota | Gammaproteobacteria |
| Xanthomonas hydrangeae | Pseudomonadota | Gammaproteobacteria |
| Xanthomonas hydrangeae | Pseudomonadota | Gammaproteobacteria |
| Xanthomonas indica | Pseudomonadota | Gammaproteobacteria |
| Xanthomonas indica | Pseudomonadota | Gammaproteobacteria |
| Xanthomonas medicagonis | Pseudomonadota | Gammaproteobacteria |
| Xanthomonas medicagonis | Pseudomonadota | Gammaproteobacteria |
| Xanthomonas nasturtii | Pseudomonadota | Gammaproteobacteria |
| Xanthomonas nasturtii | Pseudomonadota | Gammaproteobacteria |
| Xanthomonas oryzae pv. oryzicola | Pseudomonadota | Gammaproteobacteria |
| Xanthomonas oryzae pv. oryzicola | Pseudomonadota | Gammaproteobacteria |
| Xanthomonas phaseoli pv. dieffenbachiae | Pseudomonadota | Gammaproteobacteria |
| Xanthomonas phaseoli pv. dieffenbachiae | Pseudomonadota | Gammaproteobacteria |

| | | |
|---|---|---|
| Xanthomonas prunicola | Pseudomonadota | Gammaproteobacteria |
| Xanthomonas prunicola | Pseudomonadota | Gammaproteobacteria |
| Xanthomonas rydalmerensis | Pseudomonadota | Gammaproteobacteria |
| Xanthomonas rydalmerensis | Pseudomonadota | Gammaproteobacteria |
| Xanthomonas sacchari | Pseudomonadota | Gammaproteobacteria |
| Xanthomonas sacchari | Pseudomonadota | Gammaproteobacteria |
| Xanthomonas sontii | Pseudomonadota | Gammaproteobacteria |
| Xanthomonas sontii | Pseudomonadota | Gammaproteobacteria |
| Xanthomonas tesorieronis | Pseudomonadota | Gammaproteobacteria |
| Xanthomonas tesorieronis | Pseudomonadota | Gammaproteobacteria |
| Xanthomonas theicola | Pseudomonadota | Gammaproteobacteria |
| Xanthomonas theicola | Pseudomonadota | Gammaproteobacteria |
| Xanthomonas translucens pv. undulosa | Pseudomonadota | Gammaproteobacteria |
| Xanthomonas translucens pv. undulosa | Pseudomonadota | Gammaproteobacteria |
| Xanthomonas vasicola pv. musacearum NCPPB 4379 | Pseudomonadota | Gammaproteobacteria |
| Xanthomonas vasicola pv. musacearum NCPPB 4379 | Pseudomonadota | Gammaproteobacteria |
| Xanthomonas vesicatoria ATCC 35937 | Pseudomonadota | Gammaproteobacteria |
| Xanthomonas vesicatoria ATCC 35937 | Pseudomonadota | Gammaproteobacteria |
| Xenorhabdus bakwenae | Pseudomonadota | Gammaproteobacteria |
| Xenorhabdus bakwenae | Pseudomonadota | Gammaproteobacteria |
| Xenorhabdus budapestensis | Pseudomonadota | Gammaproteobacteria |
| Xenorhabdus budapestensis | Pseudomonadota | Gammaproteobacteria |
| Xenorhabdus doucetiae | Pseudomonadota | Gammaproteobacteria |
| Xenorhabdus doucetiae | Pseudomonadota | Gammaproteobacteria |
| Xenorhabdus griffiniae | Pseudomonadota | Gammaproteobacteria |
| Xenorhabdus griffiniae | Pseudomonadota | Gammaproteobacteria |
| Xenorhabdus hominickii | Pseudomonadota | Gammaproteobacteria |
| Xenorhabdus hominickii | Pseudomonadota | Gammaproteobacteria |
| Xenorhabdus nematophila ATCC 19061 | Pseudomonadota | Gammaproteobacteria |
| Xenorhabdus poinarii G6 | Pseudomonadota | Gammaproteobacteria |
| Xenorhabdus poinarii G6 | Pseudomonadota | Gammaproteobacteria |

| | | |
|---|---|---|
| Xenorhabdus taiwanensis | Pseudomonadota | Gammaproteobacteria |
| Xenorhabdus taiwanensis | Pseudomonadota | Gammaproteobacteria |
| Xylella fastidiosa | Pseudomonadota | Gammaproteobacteria |
| Xylella fastidiosa | Pseudomonadota | Gammaproteobacteria |
| Xylella taiwanensis | Pseudomonadota | Gammaproteobacteria |
| Xylella taiwanensis | Pseudomonadota | Gammaproteobacteria |
| Yersinia aldovae 670-83 | Pseudomonadota | Gammaproteobacteria |
| Yersinia aldovae 670-83 | Pseudomonadota | Gammaproteobacteria |
| Yersinia alsatica | Pseudomonadota | Gammaproteobacteria |
| Yersinia alsatica | Pseudomonadota | Gammaproteobacteria |
| Yersinia bercovieri | Pseudomonadota | Gammaproteobacteria |
| Yersinia bercovieri | Pseudomonadota | Gammaproteobacteria |
| Yersinia canariae | Pseudomonadota | Gammaproteobacteria |
| Yersinia canariae | Pseudomonadota | Gammaproteobacteria |
| Yersinia enterocolitica subsp. palearctica Y11 | Pseudomonadota | Gammaproteobacteria |
| Yersinia enterocolitica subsp. palearctica Y11 | Pseudomonadota | Gammaproteobacteria |
| Yersinia entomophaga | Pseudomonadota | Gammaproteobacteria |
| Yersinia entomophaga | Pseudomonadota | Gammaproteobacteria |
| Yersinia hibernica | Pseudomonadota | Gammaproteobacteria |
| Yersinia hibernica | Pseudomonadota | Gammaproteobacteria |
| Yersinia intermedia | Pseudomonadota | Gammaproteobacteria |
| Yersinia intermedia | Pseudomonadota | Gammaproteobacteria |
| Yersinia mollaretii ATCC 43969 | Pseudomonadota | Gammaproteobacteria |
| Yersinia mollaretii ATCC 43969 | Pseudomonadota | Gammaproteobacteria |
| Yersinia pestis A1122 | Pseudomonadota | Gammaproteobacteria |
| Yersinia pestis A1122 | Pseudomonadota | Gammaproteobacteria |
| Yersinia pseudotuberculosis | Pseudomonadota | Gammaproteobacteria |
| Yersinia pseudotuberculosis | Pseudomonadota | Gammaproteobacteria |
| Yersinia rochesterensis | Pseudomonadota | Gammaproteobacteria |
| Yersinia rochesterensis | Pseudomonadota | Gammaproteobacteria |
| Yersinia rohdei | Pseudomonadota | Gammaproteobacteria |

| | | |
|---|---|---|
| Yersinia rohdei | Pseudomonadota | Gammaproteobacteria |
| Yersinia ruckeri | Pseudomonadota | Gammaproteobacteria |
| Yersinia ruckeri | Pseudomonadota | Gammaproteobacteria |
| Yersinia similis | Pseudomonadota | Gammaproteobacteria |
| Yersinia similis | Pseudomonadota | Gammaproteobacteria |
| Yersinia vastinensis | Pseudomonadota | Gammaproteobacteria |
| Yersinia vastinensis | Pseudomonadota | Gammaproteobacteria |
| Zhongshania aliphaticivorans | Pseudomonadota | Gammaproteobacteria |
| Zhongshania aliphaticivorans | Pseudomonadota | Gammaproteobacteria |
| Zobellella denitrificans | Pseudomonadota | Gammaproteobacteria |
| Zobellella denitrificans | Pseudomonadota | Gammaproteobacteria |
| Zophobihabitans entericus | Pseudomonadota | Gammaproteobacteria |
| Zophobihabitans entericus | Pseudomonadota | Gammaproteobacteria |
| Zymobacter palmae | Pseudomonadota | Gammaproteobacteria |
| Gemmatimonas aurantiaca T-27 | Gemmatimonadota | Gemmatimonadia |
| Gemmatimonas groenlandica | Gemmatimonadota | Gemmatimonadia |
| Gemmatimonas phototrophica | Gemmatimonadota | Gemmatimonadia |
| Gemmatirosa kalamazoonensis | Gemmatimonadota | Gemmatimonadia |
| Pseudogemmatithrix spongiicola | Gemmatimonadota | Gemmatimonadia |
| Geothrix oryzae | Acidobacteriota | Holophagae |
| Mesoterricola sediminis | Acidobacteriota | Holophagae |
| Mesoterricola silvestris | Acidobacteriota | Holophagae |
| Sulfidibacter corallicola | Acidobacteriota | Holophagae |
| Thermotomaculum hydrothermale | Acidobacteriota | Holophagae |
| Thermotomaculum hydrothermale | Acidobacteriota | Holophagae |
| Hydrogenophilus thermoluteolus | Pseudomonadota | Hydrogenophilia |
| Hydrogenophilus thermoluteolus | Pseudomonadota | Hydrogenophilia |
| Ignavibacterium album JCM 16511 | Ignavibacteriota | Ignavibacteria |
| Melioribacter roseus P3M-2 | Ignavibacteriota | Ignavibacteria |
| Kiritimatiella glycovorans | Kiritimatiellota | Kiritimatiellia |
| Ktedonosporobacter rubrisoli | Chloroflexota | Ktedonobacteria |

| | | |
|---|---|---|
| Ktedonosporobacter rubrisoli | Chloroflexota | Ktedonobacteria |
| Lentisphaera profundi | Lentisphaerota | Lentisphaeria |
| Carboxydichorda subterranea | Bacillota | Limnochordia |
| Geochorda subterranea | Bacillota | Limnochordia |
| Limnochorda pilosa | Bacillota | Limnochordia |
| Limnochorda pilosa | Bacillota | Limnochordia |
| Candidatus Methylacidiphilum fumarolicum | Verrucomicrobiota | Methylacidiphilae |
| Methylacidiphilum caldifontis | Verrucomicrobiota | Methylacidiphilae |
| Methylacidiphilum infernorum V4 | Verrucomicrobiota | Methylacidiphilae |
| Methylacidiphilum kamchatkense Kam1 | Verrucomicrobiota | Methylacidiphilae |
| Acholeplasma hippikon | Mycoplasmatota | Mollicutes |
| Acholeplasma laidlawii PG-8A | Mycoplasmatota | Mollicutes |
| Alteracholeplasma palmae J233 | Mycoplasmatota | Mollicutes |
| Candidatus Hepatoplasma crinochetorum Av | Mycoplasmatota | Mollicutes |
| Candidatus Mycoplasma mahonii | Mycoplasmatota | Mollicutes |
| Candidatus Phytoplasma asteris | Mycoplasmatota | Mollicutes |
| Candidatus Phytoplasma fraxini | Mycoplasmatota | Mollicutes |
| Candidatus Phytoplasma luffae | Mycoplasmatota | Mollicutes |
| Candidatus Phytoplasma mali | Mycoplasmatota | Mollicutes |
| Candidatus Phytoplasma rubi | Mycoplasmatota | Mollicutes |
| Candidatus Phytoplasma solani | Mycoplasmatota | Mollicutes |
| Candidatus Phytoplasma ziziphi | Mycoplasmatota | Mollicutes |
| Entomoplasma freundtii | Mycoplasmatota | Mollicutes |
| Haploplasma axanthum | Mycoplasmatota | Mollicutes |
| Mariniplasma anaerobium | Mycoplasmatota | Mollicutes |
| Mesoplasma chauliocola | Mycoplasmatota | Mollicutes |
| Mesoplasma coleopterae | Mycoplasmatota | Mollicutes |
| Mesoplasma entomophilum | Mycoplasmatota | Mollicutes |
| Mesoplasma florum L1 | Mycoplasmatota | Mollicutes |
| Mesoplasma lactucae ATCC 49193 | Mycoplasmatota | Mollicutes |
| Mesoplasma melaleucae | Mycoplasmatota | Mollicutes |

| | | |
|---|---|---|
| Mesoplasma syrphidae | Mycoplasmatota | Mollicutes |
| Mesoplasma tabanidae | Mycoplasmatota | Mollicutes |
| Mycoplasma amphoriforme A39 | Mycoplasmatota | Mollicutes |
| Mycoplasma anserisalpingitidis | Mycoplasmatota | Mollicutes |
| Mycoplasma aquilae ATCC BAA-1896 | Mycoplasmatota | Mollicutes |
| Mycoplasma bradburyae | Mycoplasmatota | Mollicutes |
| Mycoplasma capricolum subsp. capricolum ATCC 27343 | Mycoplasmatota | Mollicutes |
| Mycoplasma corogypsi | Mycoplasmatota | Mollicutes |
| Mycoplasma cottewii | Mycoplasmatota | Mollicutes |
| Mycoplasma crocodyli MP145 | Mycoplasmatota | Mollicutes |
| Mycoplasma feriruminatoris | Mycoplasmatota | Mollicutes |
| Mycoplasma hafezii | Mycoplasmatota | Mollicutes |
| Mycoplasma iguanae | Mycoplasmatota | Mollicutes |
| Mycoplasma leachii PG50 | Mycoplasmatota | Mollicutes |
| Mycoplasma miroungigenitalium | Mycoplasmatota | Mollicutes |
| Mycoplasma miroungirhinis | Mycoplasmatota | Mollicutes |
| Mycoplasma mycoides subsp. capri | Mycoplasmatota | Mollicutes |
| Mycoplasma nasistruthionis | Mycoplasmatota | Mollicutes |
| Mycoplasma ovis str. Michigan | Mycoplasmatota | Mollicutes |
| Mycoplasma parvum str. Indiana | Mycoplasmatota | Mollicutes |
| Mycoplasma phocoenae | Mycoplasmatota | Mollicutes |
| Mycoplasma phocoeninasale | Mycoplasmatota | Mollicutes |
| Mycoplasma putrefaciens | Mycoplasmatota | Mollicutes |
| Mycoplasma seminis | Mycoplasmatota | Mollicutes |
| Mycoplasma struthionis | Mycoplasmatota | Mollicutes |
| Mycoplasma suis str. Illinois | Mycoplasmatota | Mollicutes |
| Mycoplasma tauri | Mycoplasmatota | Mollicutes |
| Mycoplasma tullyi | Mycoplasmatota | Mollicutes |
| Mycoplasma wenyonii str. Massachusetts | Mycoplasmatota | Mollicutes |
| Mycoplasma wenyonii str. Massachusetts | Mycoplasmatota | Mollicutes |
| Mycoplasma yeatsii GM274B | Mycoplasmatota | Mollicutes |

| | | |
|---|---|---|
| Paracholeplasma brassicae | Mycoplasmatota | Mollicutes |
| Spiroplasma alleghenense | Mycoplasmatota | Mollicutes |
| Spiroplasma apis B31 | Mycoplasmatota | Mollicutes |
| Spiroplasma cantharicola | Mycoplasmatota | Mollicutes |
| Spiroplasma chinense | Mycoplasmatota | Mollicutes |
| Spiroplasma chrysopicola DF-1 | Mycoplasmatota | Mollicutes |
| Spiroplasma citri | Mycoplasmatota | Mollicutes |
| Spiroplasma clarkii | Mycoplasmatota | Mollicutes |
| Spiroplasma corruscae | Mycoplasmatota | Mollicutes |
| Spiroplasma culicicola AES-1 | Mycoplasmatota | Mollicutes |
| Spiroplasma diminutum CUAS-1 | Mycoplasmatota | Mollicutes |
| Spiroplasma endosymbiont of Acasis viretata | Mycoplasmatota | Mollicutes |
| Spiroplasma endosymbiont of Agriotes lineatus | Mycoplasmatota | Mollicutes |
| Spiroplasma endosymbiont of Aleiodes alternator | Mycoplasmatota | Mollicutes |
| Spiroplasma endosymbiont of Ammophila pubescens | Mycoplasmatota | Mollicutes |
| Spiroplasma endosymbiont of Amphimallon solstitiale | Mycoplasmatota | Mollicutes |
| Spiroplasma endosymbiont of Andrena trimmerana | Mycoplasmatota | Mollicutes |
| Spiroplasma endosymbiont of Anurida maritima | Mycoplasmatota | Mollicutes |
| Spiroplasma endosymbiont of Apeira syringaria | Mycoplasmatota | Mollicutes |
| Spiroplasma endosymbiont of Asaphidion curtum | Mycoplasmatota | Mollicutes |
| Spiroplasma endosymbiont of Aspidapion aeneum | Mycoplasmatota | Mollicutes |
| Spiroplasma endosymbiont of Atherix ibis | Mycoplasmatota | Mollicutes |
| Spiroplasma endosymbiont of Cantharis nigra | Mycoplasmatota | Mollicutes |
| Spiroplasma endosymbiont of Cantharis rufa | Mycoplasmatota | Mollicutes |
| Spiroplasma endosymbiont of Clivina fossor | Mycoplasmatota | Mollicutes |
| Spiroplasma endosymbiont of Colias croceus | Mycoplasmatota | Mollicutes |
| Spiroplasma endosymbiont of Crioceris asparagi | Mycoplasmatota | Mollicutes |
| Spiroplasma endosymbiont of Dasysyrphus albostriatus | Mycoplasmatota | Mollicutes |
| Spiroplasma endosymbiont of Dilophus febrilis | Mycoplasmatota | Mollicutes |
| Spiroplasma endosymbiont of Dioctria linearis | Mycoplasmatota | Mollicutes |
| Spiroplasma endosymbiont of Diplazon laetatorius | Mycoplasmatota | Mollicutes |

| | | |
|---|---|---|
| Spiroplasma endosymbiont of Dromius quadrimaculatus | Mycoplasmatota | Mollicutes |
| Spiroplasma endosymbiont of Eupeodes luniger | Mycoplasmatota | Mollicutes |
| Spiroplasma endosymbiont of Glossina fuscipes fuscipes | Mycoplasmatota | Mollicutes |
| Spiroplasma endosymbiont of Labia minor | Mycoplasmatota | Mollicutes |
| Spiroplasma endosymbiont of Lasioglossum malachurum | Mycoplasmatota | Mollicutes |
| Spiroplasma endosymbiont of Lasioglossum villosulum | Mycoplasmatota | Mollicutes |
| Spiroplasma endosymbiont of Lonchoptera lutea | Mycoplasmatota | Mollicutes |
| Spiroplasma endosymbiont of Melieria omissa | Mycoplasmatota | Mollicutes |
| Spiroplasma endosymbiont of Monopis laevigella | Mycoplasmatota | Mollicutes |
| Spiroplasma endosymbiont of Nebria brevicollis | Mycoplasmatota | Mollicutes |
| Spiroplasma endosymbiont of 'Nebria riversi' | Mycoplasmatota | Mollicutes |
| Spiroplasma endosymbiont of Nephrotoma flavescens | Mycoplasmatota | Mollicutes |
| Spiroplasma endosymbiont of Nomada ruficornis | Mycoplasmatota | Mollicutes |
| Spiroplasma endosymbiont of Nomada rufipes | Mycoplasmatota | Mollicutes |
| Spiroplasma endosymbiont of Notiophilus biguttatus | Mycoplasmatota | Mollicutes |
| Spiroplasma endosymbiont of Othius punctulatus | Mycoplasmatota | Mollicutes |
| Spiroplasma endosymbiont of Panorpa germanica | Mycoplasmatota | Mollicutes |
| Spiroplasma endosymbiont of Panzeria rudis | Mycoplasmatota | Mollicutes |
| Spiroplasma endosymbiont of Phyllotreta cruciferae | Mycoplasmatota | Mollicutes |
| Spiroplasma endosymbiont of Poecilobothrus nobilitatus | Mycoplasmatota | Mollicutes |
| Spiroplasma endosymbiont of Polydrusus cervinus | Mycoplasmatota | Mollicutes |
| Spiroplasma endosymbiont of Polydrusus formosus | Mycoplasmatota | Mollicutes |
| Spiroplasma endosymbiont of Polydrusus pterygomalis | Mycoplasmatota | Mollicutes |
| Spiroplasma endosymbiont of Sarcophaga carnaria | Mycoplasmatota | Mollicutes |
| Spiroplasma endosymbiont of Seladonia tumulorum | Mycoplasmatota | Mollicutes |
| Spiroplasma endosymbiont of Stenodema calcarata | Mycoplasmatota | Mollicutes |
| Spiroplasma endosymbiont of Thecophora atra | Mycoplasmatota | Mollicutes |
| Spiroplasma endosymbiont of Tiphia femorata | Mycoplasmatota | Mollicutes |
| Spiroplasma endosymbiont of Tipula paludosa | Mycoplasmatota | Mollicutes |
| Spiroplasma endosymbiont of Tricholauxania praeusta | Mycoplasmatota | Mollicutes |
| Spiroplasma endosymbiont of Villa modesta | Mycoplasmatota | Mollicutes |

| | | |
|---|---|---|
| Spiroplasma endosymbiont of Virgichneumon dumeticola | Mycoplasmatota | Mollicutes |
| Spiroplasma endosymbiont of Zeiraphera isertana | Mycoplasmatota | Mollicutes |
| Spiroplasma eriocheiris | Mycoplasmatota | Mollicutes |
| Spiroplasma floricola 23-6 | Mycoplasmatota | Mollicutes |
| Spiroplasma gladiatoris | Mycoplasmatota | Mollicutes |
| Spiroplasma helicoides | Mycoplasmatota | Mollicutes |
| Spiroplasma ixodetis Y32 | Mycoplasmatota | Mollicutes |
| Spiroplasma kunkelii CR2-3x | Mycoplasmatota | Mollicutes |
| Spiroplasma litorale | Mycoplasmatota | Mollicutes |
| Spiroplasma mirum | Mycoplasmatota | Mollicutes |
| Spiroplasma monobiae MQ-1 | Mycoplasmatota | Mollicutes |
| Spiroplasma phoeniceum P40 | Mycoplasmatota | Mollicutes |
| Spiroplasma platyhelix PALS-1 | Mycoplasmatota | Mollicutes |
| Spiroplasma poulsonii | Mycoplasmatota | Mollicutes |
| Spiroplasma sabaudiense Ar-1343 | Mycoplasmatota | Mollicutes |
| Spiroplasma syrphidicola EA-1 | Mycoplasmatota | Mollicutes |
| Spiroplasma tabanidicola | Mycoplasmatota | Mollicutes |
| Spiroplasma taiwanense CT-1 | Mycoplasmatota | Mollicutes |
| Spiroplasma turonicum | Mycoplasmatota | Mollicutes |
| Strawberry lethal yellows phytoplasma (CPA) str. NZSb11 | Mycoplasmatota | Mollicutes |
| Williamsoniiplasma luminosum | Mycoplasmatota | Mollicutes |
| Williamsoniiplasma somnilux | Mycoplasmatota | Mollicutes |
| Anaeromyxobacter dehalogenans 2CP-1 | Myxococcota | Myxococcia |
| Anaeromyxobacter dehalogenans 2CP-1 | Myxococcota | Myxococcia |
| Anaeromyxobacter oryzae | Myxococcota | Myxococcia |
| Anaeromyxobacter oryzae | Myxococcota | Myxococcia |
| Anaeromyxobacter paludicola | Myxococcota | Myxococcia |
| Anaeromyxobacter paludicola | Myxococcota | Myxococcia |
| Archangium gephyra | Myxococcota | Myxococcia |
| Archangium gephyra | Myxococcota | Myxococcia |
| Archangium gephyra | Myxococcota | Myxococcia |

| | | |
|---|---|---|
| Archangium violaceum | Myxococcota | Myxococcia |
| Archangium violaceum | Myxococcota | Myxococcia |
| Archangium violaceum | Myxococcota | Myxococcia |
| Corallococcus coralloides DSM 2259 | Myxococcota | Myxococcia |
| Corallococcus coralloides DSM 2259 | Myxococcota | Myxococcia |
| Corallococcus macrosporus DSM 14697 | Myxococcota | Myxococcia |
| Corallococcus macrosporus DSM 14697 | Myxococcota | Myxococcia |
| Melittangium boletus DSM 14713 | Myxococcota | Myxococcia |
| Melittangium boletus DSM 14713 | Myxococcota | Myxococcia |
| Myxococcus landrumensis | Myxococcota | Myxococcia |
| Myxococcus landrumensis | Myxococcota | Myxococcia |
| Myxococcus stipitatus DSM 14675 | Myxococcota | Myxococcia |
| Myxococcus stipitatus DSM 14675 | Myxococcota | Myxococcia |
| Myxococcus xanthus DK 1622 | Myxococcota | Myxococcia |
| Myxococcus xanthus DK 1622 | Myxococcota | Myxococcia |
| Pendulispora albinea | Myxococcota | Myxococcia |
| Pendulispora albinea | Myxococcota | Myxococcia |
| Pendulispora brunnea | Myxococcota | Myxococcia |
| Pendulispora brunnea | Myxococcota | Myxococcia |
| Pendulispora brunnea | Myxococcota | Myxococcia |
| Pendulispora rubella | Myxococcota | Myxococcia |
| Pendulispora rubella | Myxococcota | Myxococcia |
| Pseudomyxococcus hansupus | Myxococcota | Myxococcia |
| Pseudomyxococcus hansupus | Myxococcota | Myxococcia |
| Pyxidicoccus parkwayensis | Myxococcota | Myxococcia |
| Pyxidicoccus parkwayensis | Myxococcota | Myxococcia |
| Vulgatibacter incomptus | Myxococcota | Myxococcia |
| Vulgatibacter incomptus | Myxococcota | Myxococcia |
| Acidaminococcus fermentans DSM 20731 | Bacillota | Negativicutes |
| Acidaminococcus fermentans DSM 20731 | Bacillota | Negativicutes |
| Acidaminococcus intestini | Bacillota | Negativicutes |

| | | |
|---|---|---|
| Acidaminococcus intestini | Bacillota | Negativicutes |
| Dialister hominis | Bacillota | Negativicutes |
| Dialister hominis | Bacillota | Negativicutes |
| Dialister massiliensis | Bacillota | Negativicutes |
| Dialister massiliensis | Bacillota | Negativicutes |
| Dialister succinatiphilus | Bacillota | Negativicutes |
| Dialister succinatiphilus | Bacillota | Negativicutes |
| Megamonas funiformis | Bacillota | Negativicutes |
| Megamonas hypermegale | Bacillota | Negativicutes |
| Megasphaera elsdenii DSM 20460 | Bacillota | Negativicutes |
| Megasphaera elsdenii DSM 20460 | Bacillota | Negativicutes |
| Megasphaera hexanoica | Bacillota | Negativicutes |
| Megasphaera hexanoica | Bacillota | Negativicutes |
| Megasphaera stantonii | Bacillota | Negativicutes |
| Megasphaera stantonii | Bacillota | Negativicutes |
| Megasphaera sueciensis | Bacillota | Negativicutes |
| Megasphaera sueciensis | Bacillota | Negativicutes |
| Methylomusa anaerophila | Bacillota | Negativicutes |
| Methylomusa anaerophila | Bacillota | Negativicutes |
| Pectinatus frisingensis | Bacillota | Negativicutes |
| Phascolarctobacterium faecium | Bacillota | Negativicutes |
| Phascolarctobacterium faecium | Bacillota | Negativicutes |
| Phascolarctobacterium succinatutens | Bacillota | Negativicutes |
| Phascolarctobacterium succinatutens | Bacillota | Negativicutes |
| Selenobaculum gibii | Bacillota | Negativicutes |
| Selenomonas dianae | Bacillota | Negativicutes |
| Selenomonas sputigena ATCC 35185 | Bacillota | Negativicutes |
| Selenomonas timonae | Bacillota | Negativicutes |
| Sporomusa acidovorans DSM 3132 | Bacillota | Negativicutes |
| Sporomusa acidovorans DSM 3132 | Bacillota | Negativicutes |
| Sporomusa aerivorans | Bacillota | Negativicutes |

| | | |
|---|---|---|
| Sporomusa aerivorans | Bacillota | Negativicutes |
| Sporomusa carbonis | Bacillota | Negativicutes |
| Sporomusa carbonis | Bacillota | Negativicutes |
| Sporomusa malonica | Bacillota | Negativicutes |
| Sporomusa malonica | Bacillota | Negativicutes |
| Sporomusa ovata DSM 2662 | Bacillota | Negativicutes |
| Sporomusa ovata DSM 2662 | Bacillota | Negativicutes |
| Sporomusa paucivorans | Bacillota | Negativicutes |
| Sporomusa paucivorans | Bacillota | Negativicutes |
| Sporomusa rhizae | Bacillota | Negativicutes |
| Sporomusa rhizae | Bacillota | Negativicutes |
| Sporomusa silvacetica DSM 10669 | Bacillota | Negativicutes |
| Sporomusa silvacetica DSM 10669 | Bacillota | Negativicutes |
| Sporomusa sphaeroides DSM 2875 | Bacillota | Negativicutes |
| Sporomusa sphaeroides DSM 2875 | Bacillota | Negativicutes |
| Sporomusa termitida | Bacillota | Negativicutes |
| Sporomusa termitida | Bacillota | Negativicutes |
| Veillonella dispar | Bacillota | Negativicutes |
| Veillonella dispar | Bacillota | Negativicutes |
| Veillonella nakazawae | Bacillota | Negativicutes |
| Veillonella nakazawae | Bacillota | Negativicutes |
| Veillonella orientalis | Bacillota | Negativicutes |
| Veillonella orientalis | Bacillota | Negativicutes |
| Veillonella parvula | Bacillota | Negativicutes |
| Veillonella parvula | Bacillota | Negativicutes |
| Veillonella rodentium | Bacillota | Negativicutes |
| Veillonella rodentium | Bacillota | Negativicutes |
| Egibacter rhizosphaerae | Actinomycetota | Nitriliruptoria |
| Egibacter rhizosphaerae | Actinomycetota | Nitriliruptoria |
| Egicoccus halophilus | Actinomycetota | Nitriliruptoria |
| Egicoccus halophilus | Actinomycetota | Nitriliruptoria |

| | | |
|---|---|---|
| Euzebya pacifica | Actinomycetota | Nitriliruptoria |
| Euzebya pacifica | Actinomycetota | Nitriliruptoria |
| Nitrospina watsonii | Nitrospinota | Nitrospinia |
| Candidatus Nitrospira allomarina | Nitrospirota | Nitrospiria |
| Candidatus Nitrospira allomarina | Nitrospirota | Nitrospiria |
| Candidatus Nitrospira inopinata | Nitrospirota | Nitrospiria |
| Candidatus Nitrospira inopinata | Nitrospirota | Nitrospiria |
| Candidatus Nitrospira neomarina | Nitrospirota | Nitrospiria |
| Candidatus Nitrospira neomarina | Nitrospirota | Nitrospiria |
| Leptospirillum ferrooxidans C2-3 | Nitrospirota | Nitrospiria |
| Leptospirillum ferrooxidans C2-3 | Nitrospirota | Nitrospiria |
| Nitrospira japonica | Nitrospirota | Nitrospiria |
| Nitrospira japonica | Nitrospirota | Nitrospiria |
| Nitrospira moscoviensis | Nitrospirota | Nitrospiria |
| Nitrospira moscoviensis | Nitrospirota | Nitrospiria |
| Fluviispira sanaruensis | Bdellovibrionota | Oligoflexia |
| Silvanigrella aquatica | Bdellovibrionota | Oligoflexia |
| Spirobacillus cienkowskii | Bdellovibrionota | Oligoflexia |
| Actomonas aquatica | Verrucomicrobiota | Opitutia |
| Candidatus Pelagisphaera phototrophica | Verrucomicrobiota | Opitutia |
| Coraliomargarita akajimensis DSM 45221 | Verrucomicrobiota | Opitutia |
| Coraliomargarita algicola | Verrucomicrobiota | Opitutia |
| Ereboglobus luteus | Verrucomicrobiota | Opitutia |
| Horticoccus luteus | Verrucomicrobiota | Opitutia |
| Lacunisphaera limnophila | Verrucomicrobiota | Opitutia |
| Nibricoccus aquaticus | Verrucomicrobiota | Opitutia |
| Opitutus terrae PB90-1 | Verrucomicrobiota | Opitutia |
| Rubellicoccus peritrichatus | Verrucomicrobiota | Opitutia |
| Synoicihabitans lomoniglobus | Verrucomicrobiota | Opitutia |
| Termitidicoccus mucosus | Verrucomicrobiota | Opitutia |
| Anaerohalosphaera lusitana | Planctomycetota | Phycisphaerae |

| | | |
|---|---|---|
| Humisphaera borealis | Planctomycetota | Phycisphaerae |
| Humisphaera borealis | Planctomycetota | Phycisphaerae |
| Limihaloglobus sulfuriphilus | Planctomycetota | Phycisphaerae |
| Mucisphaera calidilacus | Planctomycetota | Phycisphaerae |
| Phycisphaera mikurensis NBRC 102666 | Planctomycetota | Phycisphaerae |
| Poriferisphaera corsica | Planctomycetota | Phycisphaerae |
| Sedimentisphaera cyanobacteriorum | Planctomycetota | Phycisphaerae |
| Sedimentisphaera salicampi | Planctomycetota | Phycisphaerae |
| Adhaeretor mobilis | Planctomycetota | Planctomycetia |
| Aeoliella mucimassa | Planctomycetota | Planctomycetia |
| Alienimonas californiensis | Planctomycetota | Planctomycetia |
| Anatilimnocola aggregata | Planctomycetota | Planctomycetia |
| Aquisphaera giovannonii | Planctomycetota | Planctomycetia |
| Aureliella helgolandensis | Planctomycetota | Planctomycetia |
| Aureliella helgolandensis | Planctomycetota | Planctomycetia |
| Botrimarina mediterranea | Planctomycetota | Planctomycetia |
| Bremerella volcania | Planctomycetota | Planctomycetia |
| Bythopirellula goksoeyrii | Planctomycetota | Planctomycetia |
| Calycomorphotria hydatis | Planctomycetota | Planctomycetia |
| Caulifigura coniformis | Planctomycetota | Planctomycetia |
| Crateriforma conspicua | Planctomycetota | Planctomycetia |
| Crateriforma conspicua | Planctomycetota | Planctomycetia |
| Crateriforma spongiae | Planctomycetota | Planctomycetia |
| Crateriforma spongiae | Planctomycetota | Planctomycetia |
| Frigoriglobus tundricola | Planctomycetota | Planctomycetia |
| Fuerstiella marisgermanici | Planctomycetota | Planctomycetia |
| Gemmata massiliana | Planctomycetota | Planctomycetia |
| Gemmata massiliana | Planctomycetota | Planctomycetia |
| Gimesia alba | Planctomycetota | Planctomycetia |
| Gimesia alba | Planctomycetota | Planctomycetia |
| Gimesia algae | Planctomycetota | Planctomycetia |

| | | |
|---|---|---|
| Gimesia aquarii | Planctomycetota | Planctomycetia |
| Gimesia aquarii | Planctomycetota | Planctomycetia |
| Gimesia benthica | Planctomycetota | Planctomycetia |
| Gimesia benthica | Planctomycetota | Planctomycetia |
| Gimesia fumaroli | Planctomycetota | Planctomycetia |
| Gimesia fumaroli | Planctomycetota | Planctomycetia |
| Gimesia maris | Planctomycetota | Planctomycetia |
| Gimesia panareensis | Planctomycetota | Planctomycetia |
| Gimesia panareensis | Planctomycetota | Planctomycetia |
| Isosphaera pallida ATCC 43644 | Planctomycetota | Planctomycetia |
| Lacipirellula limnantheis | Planctomycetota | Planctomycetia |
| Lacipirellula parvula | Planctomycetota | Planctomycetia |
| Lignipirellula cremea | Planctomycetota | Planctomycetia |
| Limnoglobus roseus | Planctomycetota | Planctomycetia |
| Limnoglobus roseus | Planctomycetota | Planctomycetia |
| Maioricimonas rarisocia | Planctomycetota | Planctomycetia |
| Mariniblastus fucicola | Planctomycetota | Planctomycetia |
| Paludisphaera borealis | Planctomycetota | Planctomycetia |
| Pirellulimonas nuda | Planctomycetota | Planctomycetia |
| Planctellipticum variicoloris | Planctomycetota | Planctomycetia |
| Planctopirus ephydatiae | Planctomycetota | Planctomycetia |
| Planctopirus limnophila DSM 3776 | Planctomycetota | Planctomycetia |
| Polystyrenella longa | Planctomycetota | Planctomycetia |
| Rhodopirellula baltica SH 1 | Planctomycetota | Planctomycetia |
| Roseimaritima multifibrata | Planctomycetota | Planctomycetia |
| Roseimaritima ulvae | Planctomycetota | Planctomycetia |
| Rosistilla carotiformis | Planctomycetota | Planctomycetia |
| Rosistilla oblonga | Planctomycetota | Planctomycetia |
| Rosistilla ulvae | Planctomycetota | Planctomycetia |
| Rubinisphaera brasiliensis DSM 5305 | Planctomycetota | Planctomycetia |
| Rubripirellula lacrimiformis | Planctomycetota | Planctomycetia |

| | | |
|---|---|---|
| Singulisphaera acidiphila DSM 18658 | Planctomycetota | Planctomycetia |
| Stieleria maiorica | Planctomycetota | Planctomycetia |
| Stieleria maiorica | Planctomycetota | Planctomycetia |
| Stieleria neptunia | Planctomycetota | Planctomycetia |
| Stieleria neptunia | Planctomycetota | Planctomycetia |
| Stieleria varia | Planctomycetota | Planctomycetia |
| Stieleria varia | Planctomycetota | Planctomycetia |
| Stratiformator vulcanicus | Planctomycetota | Planctomycetia |
| Tautonia plasticadhaerens | Planctomycetota | Planctomycetia |
| Telmatocola sphagniphila | Planctomycetota | Planctomycetia |
| Thalassoglobus polymorphus | Planctomycetota | Planctomycetia |
| Thermogutta terrifontis | Planctomycetota | Planctomycetia |
| Tuwongella immobilis | Planctomycetota | Planctomycetia |
| Urbifossiella limnaea | Planctomycetota | Planctomycetia |
| Urbifossiella limnaea | Planctomycetota | Planctomycetia |
| Rhodocaloribacter litoris | Rhodothermota | Rhodothermia |
| Rhodocaloribacter litoris | Rhodothermota | Rhodothermia |
| Rhodothermus marinus DSM 4252 | Rhodothermota | Rhodothermia |
| Rhodothermus marinus DSM 4252 | Rhodothermota | Rhodothermia |
| Salinibacter ruber | Rhodothermota | Rhodothermia |
| Salinibacter ruber | Rhodothermota | Rhodothermia |
| Rubrobacter indicoceani | Actinomycetota | Rubrobacteria |
| Rubrobacter indicoceani | Actinomycetota | Rubrobacteria |
| Rubrobacter xylanophilus DSM 9941 | Actinomycetota | Rubrobacteria |
| Rubrobacter xylanophilus DSM 9941 | Actinomycetota | Rubrobacteria |
| Aureispira anguillae | Bacteroidota | Saprospiria |
| Haliscomenobacter hydrossis DSM 1100 | Bacteroidota | Saprospiria |
| Saprospira grandis str. Lewin | Bacteroidota | Saprospiria |
| Aurantibacillus circumpalustris | Bacteroidota | Sphingobacteriia |
| Mucilaginibacter celer | Bacteroidota | Sphingobacteriia |
| Mucilaginibacter daejeonensis | Bacteroidota | Sphingobacteriia |

| | | |
|---|---|---|
| Mucilaginibacter ginkgonis | Bacteroidota | Sphingobacteriia |
| Mucilaginibacter ginsenosidivorans | Bacteroidota | Sphingobacteriia |
| Mucilaginibacter ginsenosidivorax | Bacteroidota | Sphingobacteriia |
| Mucilaginibacter gossypii | Bacteroidota | Sphingobacteriia |
| Mucilaginibacter gotjawali | Bacteroidota | Sphingobacteriia |
| Mucilaginibacter jinjuensis | Bacteroidota | Sphingobacteriia |
| Mucilaginibacter mali | Bacteroidota | Sphingobacteriia |
| Mucilaginibacter mallensis | Bacteroidota | Sphingobacteriia |
| Mucilaginibacter paludis DSM 18603 | Bacteroidota | Sphingobacteriia |
| Mucilaginibacter robiniae | Bacteroidota | Sphingobacteriia |
| Mucilaginibacter sabulilitoris | Bacteroidota | Sphingobacteriia |
| Mucilaginibacter xinganensis | Bacteroidota | Sphingobacteriia |
| Pedobacter aquae | Bacteroidota | Sphingobacteriia |
| Pedobacter endophyticus | Bacteroidota | Sphingobacteriia |
| Pedobacter foliorum | Bacteroidota | Sphingobacteriia |
| Pedobacter heparinus DSM 2366 | Bacteroidota | Sphingobacteriia |
| Pedobacter lusitanus | Bacteroidota | Sphingobacteriia |
| Pedobacter mucosus | Bacteroidota | Sphingobacteriia |
| Pedobacter riviphilus | Bacteroidota | Sphingobacteriia |
| Pedobacter roseus | Bacteroidota | Sphingobacteriia |
| Pedobacter schmidteae | Bacteroidota | Sphingobacteriia |
| Pedobacter suwonensis | Bacteroidota | Sphingobacteriia |
| Pseudopedobacter saltans DSM 12145 | Bacteroidota | Sphingobacteriia |
| Solitalea canadensis DSM 3403 | Bacteroidota | Sphingobacteriia |
| Solitalea lacus | Bacteroidota | Sphingobacteriia |
| Sphingobacterium bambusae | Bacteroidota | Sphingobacteriia |
| Sphingobacterium cellulitidis | Bacteroidota | Sphingobacteriia |
| Sphingobacterium hotanense | Bacteroidota | Sphingobacteriia |
| Sphingobacterium lactis | Bacteroidota | Sphingobacteriia |
| Sphingobacterium mizutaii | Bacteroidota | Sphingobacteriia |
| Sphingobacterium multivorum | Bacteroidota | Sphingobacteriia |

| | | |
|---|---|---|
| Sphingobacterium oryzagri | Bacteroidota | Sphingobacteriia |
| Sphingobacterium paramultivorum | Bacteroidota | Sphingobacteriia |
| Sphingobacterium psychroaquaticum | Bacteroidota | Sphingobacteriia |
| Sphingobacterium siyangense | Bacteroidota | Sphingobacteriia |
| Sphingobacterium spiritivorum | Bacteroidota | Sphingobacteriia |
| Sphingobacterium thalpophilum | Bacteroidota | Sphingobacteriia |
| Sphingobacterium zhuxiongii | Bacteroidota | Sphingobacteriia |
| Borrelia anserina Es | Spirochaetota | Spirochaetia |
| Borrelia coriaceae | Spirochaetota | Spirochaetia |
| Borrelia duttonii Ly | Spirochaetota | Spirochaetia |
| Borrelia hermsii | Spirochaetota | Spirochaetia |
| Borrelia maritima | Spirochaetota | Spirochaetia |
| Borrelia miyamotoi | Spirochaetota | Spirochaetia |
| Borrelia nietonii YOR | Spirochaetota | Spirochaetia |
| Borrelia parkeri HR1 | Spirochaetota | Spirochaetia |
| Borrelia puertoricensis | Spirochaetota | Spirochaetia |
| Borrelia recurrentis A1 | Spirochaetota | Spirochaetia |
| Borrelia turcica IST7 | Spirochaetota | Spirochaetia |
| Borrelia turicatae 91E135 | Spirochaetota | Spirochaetia |
| Borreliella afzelii Tom3107 | Spirochaetota | Spirochaetia |
| Borreliella americana | Spirochaetota | Spirochaetia |
| Borreliella andersonii | Spirochaetota | Spirochaetia |
| Borreliella bavariensis PBi | Spirochaetota | Spirochaetia |
| Borreliella bissettiae DN127 | Spirochaetota | Spirochaetia |
| Borreliella californiensis | Spirochaetota | Spirochaetia |
| Borreliella carolinensis | Spirochaetota | Spirochaetia |
| Borreliella finlandensis | Spirochaetota | Spirochaetia |
| Borreliella garinii | Spirochaetota | Spirochaetia |
| Borreliella japonica | Spirochaetota | Spirochaetia |
| Borreliella kurtenbachii | Spirochaetota | Spirochaetia |
| Borreliella lanei | Spirochaetota | Spirochaetia |

| | | |
|---|---|---|
| Borreliella lusitaniae | Spirochaetota | Spirochaetia |
| Borreliella mayonii | Spirochaetota | Spirochaetia |
| Borreliella sinica | Spirochaetota | Spirochaetia |
| Borreliella spielmanii | Spirochaetota | Spirochaetia |
| Borreliella tanukii | Spirochaetota | Spirochaetia |
| Borreliella turdi | Spirochaetota | Spirochaetia |
| Borreliella valaisiana VS116 | Spirochaetota | Spirochaetia |
| Borreliella yangtzensis | Spirochaetota | Spirochaetia |
| Brachyspira hyodysenteriae | Spirochaetota | Spirochaetia |
| Brachyspira intermedia PWS/A | Spirochaetota | Spirochaetia |
| Brachyspira murdochii DSM 12563 | Spirochaetota | Spirochaetia |
| Brachyspira pilosicoli 95/1000 | Spirochaetota | Spirochaetia |
| Breznakiella homolactica | Spirochaetota | Spirochaetia |
| Brucepastera parasyntrophica | Spirochaetota | Spirochaetia |
| Candidatus Borrelia fainii | Spirochaetota | Spirochaetia |
| Candidatus Borreliella tachyglossi | Spirochaetota | Spirochaetia |
| Candidatus Haliotispira prima | Spirochaetota | Spirochaetia |
| Entomospira culicis | Spirochaetota | Spirochaetia |
| Entomospira entomophila | Spirochaetota | Spirochaetia |
| Entomospira nematocerorum | Spirochaetota | Spirochaetia |
| Gracilinema caldarium DSM 7334 | Spirochaetota | Spirochaetia |
| Leadbettera azotonutricia ZAS-9 | Spirochaetota | Spirochaetia |
| Leptospira biflexa serovar Patoc strain 'Patoc 1 (Paris)' | Spirochaetota | Spirochaetia |
| Leptospira borgpetersenii serovar Ceylonica | Spirochaetota | Spirochaetia |
| Leptospira cinconiae | Spirochaetota | Spirochaetia |
| Leptospira interrogans serovar Copenhageni | Spirochaetota | Spirochaetia |
| Leptospira kmetyi | Spirochaetota | Spirochaetia |
| Leptospira kobayashii | Spirochaetota | Spirochaetia |
| Leptospira mayottensis 200901116 | Spirochaetota | Spirochaetia |
| Leptospira santarosai serovar Shermani str. LT 821 | Spirochaetota | Spirochaetia |
| Leptospira tipperaryensis | Spirochaetota | Spirochaetia |

| | | |
|---|---|---|
| Leptospira weilii | Spirochaetota | Spirochaetia |
| Oceanispirochaeta crateris | Spirochaetota | Spirochaetia |
| Parasphaerochaeta coccoides DSM 17374 | Spirochaetota | Spirochaetia |
| Salinispira pacifica | Spirochaetota | Spirochaetia |
| Sediminispirochaeta smaragdinae DSM 11293 | Spirochaetota | Spirochaetia |
| Sphaerochaeta associata | Spirochaetota | Spirochaetia |
| Sphaerochaeta globosa str. Buddy | Spirochaetota | Spirochaetia |
| Sphaerochaeta pleomorpha str. Grapes | Spirochaetota | Spirochaetia |
| Spirochaeta africana DSM 8902 | Spirochaetota | Spirochaetia |
| Spirochaeta thermophila DSM 6578 | Spirochaetota | Spirochaetia |
| Thermospira aquatica | Spirochaetota | Spirochaetia |
| Thiospirochaeta perfilievii | Spirochaetota | Spirochaetia |
| Treponema brennaborense DSM 12168 | Spirochaetota | Spirochaetia |
| Treponema bryantii | Spirochaetota | Spirochaetia |
| Treponema denticola ATCC 35405 | Spirochaetota | Spirochaetia |
| Treponema medium | Spirochaetota | Spirochaetia |
| Treponema pallidum subsp. pallidum str. Nichols | Spirochaetota | Spirochaetia |
| Treponema paraluiscuniculi Cuniculi A | Spirochaetota | Spirochaetia |
| Treponema parvum | Spirochaetota | Spirochaetia |
| Treponema pedis str. T A4 | Spirochaetota | Spirochaetia |
| Treponema peruense | Spirochaetota | Spirochaetia |
| Treponema phagedenis | Spirochaetota | Spirochaetia |
| Treponema primitia ZAS-2 | Spirochaetota | Spirochaetia |
| Treponema putidum | Spirochaetota | Spirochaetia |
| Treponema saccharophilum | Spirochaetota | Spirochaetia |
| Treponema socranskii subsp. buccale | Spirochaetota | Spirochaetia |
| Treponema succinifaciens DSM 2489 | Spirochaetota | Spirochaetia |
| Turneriella parva DSM 21527 | Spirochaetota | Spirochaetia |
| Aminithiophilus ramosus | Synergistota | Synergistia |
| Aminithiophilus ramosus | Synergistota | Synergistia |
| Aminobacterium colombiense DSM 12261 | Synergistota | Synergistia |

| | | |
|---|---|---|
| Aminobacterium colombiense DSM 12261 | Synergistota | Synergistia |
| Aminomonas paucivorans DSM 12260 | Synergistota | Synergistia |
| Aminomonas paucivorans DSM 12260 | Synergistota | Synergistia |
| Cloacibacillus porcorum | Synergistota | Synergistia |
| Cloacibacillus porcorum | Synergistota | Synergistia |
| Fretibacterium fastidiosum | Synergistota | Synergistia |
| Fretibacterium fastidiosum | Synergistota | Synergistia |
| Pyramidobacter piscolens | Synergistota | Synergistia |
| Pyramidobacter piscolens | Synergistota | Synergistia |
| Thermanaerovibrio acidaminovorans DSM 6589 | Synergistota | Synergistia |
| Thermanaerovibrio acidaminovorans DSM 6589 | Synergistota | Synergistia |
| Thermanaerovibrio velox DSM 12556 | Synergistota | Synergistia |
| Thermanaerovibrio velox DSM 12556 | Synergistota | Synergistia |
| Syntrophus aciditrophicus SB | Thermodesulfobacteriota | Syntrophia |
| Syntrophus buswellii | Thermodesulfobacteriota | Syntrophia |
| Desulfoglaeba alkanexedens ALDC | Thermodesulfobacteriota | Syntrophobacteria |
| Desulfoglaeba alkanexedens ALDC | Thermodesulfobacteriota | Syntrophobacteria |
| Syntrophobacter fumaroxidans MPOB | Thermodesulfobacteriota | Syntrophobacteria |
| Syntrophobacter fumaroxidans MPOB | Thermodesulfobacteriota | Syntrophobacteria |
| Tepidiforma bonchosmolovskayae | Chloroflexota | Tepidiformia |
| Tepidiforma bonchosmolovskayae | Chloroflexota | Tepidiformia |
| Acidisarcina polymorpha | Acidobacteriota | Terriglobia |
| Acidisarcina polymorpha | Acidobacteriota | Terriglobia |
| Acidobacterium capsulatum ATCC 51196 | Acidobacteriota | Terriglobia |
| Acidobacterium capsulatum ATCC 51196 | Acidobacteriota | Terriglobia |
| Candidatus Koribacter versatilis Ellin345 | Acidobacteriota | Terriglobia |
| Candidatus Koribacter versatilis Ellin345 | Acidobacteriota | Terriglobia |
| Edaphobacter flagellatus | Acidobacteriota | Terriglobia |
| Edaphobacter flagellatus | Acidobacteriota | Terriglobia |
| Edaphobacter paludis | Acidobacteriota | Terriglobia |
| Edaphobacter paludis | Acidobacteriota | Terriglobia |

| | | |
|---|---|---|
| Granulicella mallensis MP5ACTX8 | Acidobacteriota | Terriglobia |
| Granulicella mallensis MP5ACTX8 | Acidobacteriota | Terriglobia |
| Granulicella tundricola MP5ACTX9 | Acidobacteriota | Terriglobia |
| Granulicella tundricola MP5ACTX9 | Acidobacteriota | Terriglobia |
| Occallatibacter riparius | Acidobacteriota | Terriglobia |
| Occallatibacter riparius | Acidobacteriota | Terriglobia |
| Paludibaculum fermentans | Acidobacteriota | Terriglobia |
| Paludibaculum fermentans | Acidobacteriota | Terriglobia |
| Telmatobacter bradus | Acidobacteriota | Terriglobia |
| Telmatobacter bradus | Acidobacteriota | Terriglobia |
| Terriglobus roseus DSM 18391 | Acidobacteriota | Terriglobia |
| Terriglobus roseus DSM 18391 | Acidobacteriota | Terriglobia |
| Terriglobus saanensis SP1PR4 | Acidobacteriota | Terriglobia |
| Terriglobus saanensis SP1PR4 | Acidobacteriota | Terriglobia |
| Tunturiibacter empetritectus | Acidobacteriota | Terriglobia |
| Tunturiibacter empetritectus | Acidobacteriota | Terriglobia |
| Tunturiibacter gelidiferens | Acidobacteriota | Terriglobia |
| Tunturiibacter gelidiferens | Acidobacteriota | Terriglobia |
| Tunturiibacter lichenicola | Acidobacteriota | Terriglobia |
| Tunturiibacter lichenicola | Acidobacteriota | Terriglobia |
| Tunturiibacter psychrotolerans | Acidobacteriota | Terriglobia |
| Tunturiibacter psychrotolerans | Acidobacteriota | Terriglobia |
| Caldimicrobium thiodismutans | Thermodesulfobacteriota | Thermodesulfobacteria |
| Caldimicrobium thiodismutans | Thermodesulfobacteriota | Thermodesulfobacteria |
| Thermodesulfatator indicus DSM 15286 | Thermodesulfobacteriota | Thermodesulfobacteria |
| Thermodesulfatator indicus DSM 15286 | Thermodesulfobacteriota | Thermodesulfobacteria |
| Thermodesulfobacterium commune DSM 2178 | Thermodesulfobacteriota | Thermodesulfobacteria |
| Thermodesulfobacterium commune DSM 2178 | Thermodesulfobacteriota | Thermodesulfobacteria |
| Thermodesulfobacterium geofontis OPF15 | Thermodesulfobacteriota | Thermodesulfobacteria |
| Thermodesulfobacterium geofontis OPF15 | Thermodesulfobacteriota | Thermodesulfobacteria |
| Thermosulfurimonas marina | Thermodesulfobacteriota | Thermodesulfobacteria |

| | | |
|---|---|---|
| Thermosulfurimonas marina | Thermodesulfobacteriota | Thermodesulfobacteria |
| Thermosulfuriphilus ammonigenes | Thermodesulfobacteriota | Thermodesulfobacteria |
| Thermosulfuriphilus ammonigenes | Thermodesulfobacteriota | Thermodesulfobacteria |
| Thermodesulfobium acidiphilum | Thermodesulfobiota | Thermodesulfobiia |
| Thermodesulfobium acidiphilum | Thermodesulfobiota | Thermodesulfobiia |
| Thermodesulfobium narugense DSM 14796 | Thermodesulfobiota | Thermodesulfobiia |
| Thermodesulfobium narugense DSM 14796 | Thermodesulfobiota | Thermodesulfobiia |
| Dissulfurispira thermophila | Nitrospirota | Thermodesulfovibrionia |
| Dissulfurispira thermophila | Nitrospirota | Thermodesulfovibrionia |
| Thermodesulfovibrio autotrophicus | Nitrospirota | Thermodesulfovibrionia |
| Thermodesulfovibrio autotrophicus | Nitrospirota | Thermodesulfovibrionia |
| Thermodesulfovibrio obliviosus | Nitrospirota | Thermodesulfovibrionia |
| Thermodesulfovibrio obliviosus | Nitrospirota | Thermodesulfovibrionia |
| Thermodesulfovibrio yellowstonii DSM 11347 | Nitrospirota | Thermodesulfovibrionia |
| Thermodesulfovibrio yellowstonii DSM 11347 | Nitrospirota | Thermodesulfovibrionia |
| Baekduia alba | Actinomycetota | Thermoleophilia |
| Baekduia soli | Actinomycetota | Thermoleophilia |
| Capillimicrobium parvum | Actinomycetota | Thermoleophilia |
| Conexibacter woesei DSM 14684 | Actinomycetota | Thermoleophilia |
| Miltoncostaea marina | Actinomycetota | Thermoleophilia |
| Miltoncostaea oceani | Actinomycetota | Thermoleophilia |
| Paraconexibacter antarcticus | Actinomycetota | Thermoleophilia |
| Svornostia abyssi | Actinomycetota | Thermoleophilia |
| Sphaerobacter thermophilus DSM 20745 | Thermomicrobiota | Thermomicrobia |
| Sphaerobacter thermophilus DSM 20745 | Thermomicrobiota | Thermomicrobia |
| Thermomicrobium roseum DSM 5159 | Thermomicrobiota | Thermomicrobia |
| Thermomicrobium roseum DSM 5159 | Thermomicrobiota | Thermomicrobia |
| Thermosulfidibacter takaii ABI70S6 | Thermosulfidibacterota | Thermosulfidibacteria |
| Thermosulfidibacter takaii ABI70S6 | Thermosulfidibacterota | Thermosulfidibacteria |
| Athalassotoga saccharophila | Thermotogota | Thermotogae |
| Defluviitoga tunisiensis | Thermotogota | Thermotogae |

| | | |
|---|---|---|
| Fervidobacterium changbaicum | Thermotogota | Thermotogae |
| Fervidobacterium islandicum | Thermotogota | Thermotogae |
| Fervidobacterium nodosum Rt17-B1 | Thermotogota | Thermotogae |
| Fervidobacterium pennivorans DSM 9078 | Thermotogota | Thermotogae |
| Kosmotoga olearia TBF 19.5.1 | Thermotogota | Thermotogae |
| Kosmotoga pacifica | Thermotogota | Thermotogae |
| Marinitoga aeolica | Thermotogota | Thermotogae |
| Marinitoga piezophila KA3 | Thermotogota | Thermotogae |
| Mesotoga infera | Thermotogota | Thermotogae |
| Mesotoga prima MesG1.Ag.4.2 | Thermotogota | Thermotogae |
| Petrotoga mobilis SJ95 | Thermotogota | Thermotogae |
| Pseudothermotoga elfii DSM 9442 = NBRC 107921 | Thermotogota | Thermotogae |
| Pseudothermotoga hypogea DSM 11164 = NBRC 106472 | Thermotogota | Thermotogae |
| Pseudothermotoga lettingae TMO | Thermotogota | Thermotogae |
| Pseudothermotoga thermarum DSM 5069 | Thermotogota | Thermotogae |
| Tepiditoga spiralis | Thermotogota | Thermotogae |
| Thermosipho africanus TCF52B | Thermotogota | Thermotogae |
| Thermosipho ferrireducens | Thermotogota | Thermotogae |
| Thermosipho melanesiensis BI429 | Thermotogota | Thermotogae |
| Thermotoga caldifontis AZM44c09 | Thermotogota | Thermotogae |
| Thermotoga maritima MSB8 | Thermotogota | Thermotogae |
| Thermotoga neapolitana DSM 4359 | Thermotogota | Thermotogae |
| Thermotoga petrophila RKU-1 | Thermotogota | Thermotogae |
| Thermotoga profunda AZM34c06 | Thermotogota | Thermotogae |
| Tichowtungia aerotolerans | Kiritimatiellota | Tichowtungiia |
| Acidilutibacter cellobiosedens | Bacillota | Tissierellia |
| Acidilutibacter cellobiosedens | Bacillota | Tissierellia |
| Anaerococcus mediterraneensis | Bacillota | Tissierellia |
| Anaerococcus mediterraneensis | Bacillota | Tissierellia |
| Anaerococcus nagyae | Bacillota | Tissierellia |
| Anaerococcus nagyae | Bacillota | Tissierellia |

| | | |
|---|---|---|
| Anaerococcus obesiensis | Bacillota | Tissierellia |
| Anaerococcus obesiensis | Bacillota | Tissierellia |
| Anaerococcus prevotii DSM 20548 | Bacillota | Tissierellia |
| Anaerococcus prevotii DSM 20548 | Bacillota | Tissierellia |
| Anaerococcus vaginalis | Bacillota | Tissierellia |
| Anaerococcus vaginalis | Bacillota | Tissierellia |
| Ezakiella massiliensis | Bacillota | Tissierellia |
| Ezakiella massiliensis | Bacillota | Tissierellia |
| Gottschalkia acidurici 9a | Bacillota | Tissierellia |
| Gottschalkia acidurici 9a | Bacillota | Tissierellia |
| Gudongella oleilytica | Bacillota | Tissierellia |
| Gudongella oleilytica | Bacillota | Tissierellia |
| Helcococcus ovis | Bacillota | Tissierellia |
| Keratinibaculum paraultunense | Bacillota | Tissierellia |
| Keratinibaculum paraultunense | Bacillota | Tissierellia |
| Microaceticoccus formicicus | Bacillota | Tissierellia |
| Microaceticoccus formicicus | Bacillota | Tissierellia |
| Miniphocaeibacter halophilus | Bacillota | Tissierellia |
| Miniphocaeibacter halophilus | Bacillota | Tissierellia |
| Murdochiella vaginalis | Bacillota | Tissierellia |
| Parvimonas micra | Bacillota | Tissierellia |
| Parvimonas parva | Bacillota | Tissierellia |
| Peptoniphilus equinus | Bacillota | Tissierellia |
| Peptoniphilus equinus | Bacillota | Tissierellia |
| Peptoniphilus genitalis | Bacillota | Tissierellia |
| Peptoniphilus genitalis | Bacillota | Tissierellia |
| Peptoniphilus harei | Bacillota | Tissierellia |
| Peptoniphilus harei | Bacillota | Tissierellia |
| Akkermansia biwaensis | Verrucomicrobiota | Verrucomicrobiia |
| Akkermansia glycaniphila | Verrucomicrobiota | Verrucomicrobiia |
| Akkermansia muciniphila | Verrucomicrobiota | Verrucomicrobiia |

| | | |
|---|---|---|
| Fontisphaera persica | Verrucomicrobiota | Verrucomicrobiia |
| Haloferula helveola | Verrucomicrobiota | Verrucomicrobiia |
| Luteolibacter ambystomatis | Verrucomicrobiota | Verrucomicrobiia |
| Luteolibacter luteus | Verrucomicrobiota | Verrucomicrobiia |
| Sulfuriroseicoccus oceanibius | Verrucomicrobiota | Verrucomicrobiia |
| Verrucomicrobium spinosum DSM 4136 = JCM 18804 | Verrucomicrobiota | Verrucomicrobiia |
| Luteitalea pratensis | Acidobacteriota | Vicinamibacteria |
| Luteitalea pratensis | Acidobacteriota | Vicinamibacteria |
| Vulcanimicrobium alpinum | Vulcanimicrobiota | Vulcanimicrobiia |
| Mariprofundus aestuarium | Pseudomonadota | Zetaproteobacteria |
| Mariprofundus aestuarium | Pseudomonadota | Zetaproteobacteria |
| Mariprofundus ferrinatatus | Pseudomonadota | Zetaproteobacteria |
| Mariprofundus ferrinatatus | Pseudomonadota | Zetaproteobacteria |
| Candidatus Caldatribacterium saccharofermentans | Atribacterota | |
| Candidatus Caldatribacterium saccharofermentans | Atribacterota | |
| Caldicellulosiruptor acetigenus I77R1B | Bacillota | |
| Caldicellulosiruptor bescii DSM 6725 | Bacillota | |
| Caldicellulosiruptor changbaiensis | Bacillota | |
| Caldicellulosiruptor diazotrophicus | Bacillota | |
| Caldicellulosiruptor hydrothermalis 108 | Bacillota | |
| Caldicellulosiruptor kronotskyensis 2002 | Bacillota | |
| Caldicellulosiruptor morganii | Bacillota | |
| Caldicellulosiruptor naganoensis | Bacillota | |
| Caldicellulosiruptor obsidiansis OB47 | Bacillota | |
| Caldicellulosiruptor owensensis OL | Bacillota | |
| Caldicellulosiruptor saccharolyticus DSM 8903 | Bacillota | |
| endosymbiont 'TC1' of Trimyema compressum | Bacillota | |
| endosymbiont 'TC1' of Trimyema compressum | Bacillota | |
| Ndongobacter massiliensis | Bacillota | |
| Ndongobacter massiliensis | Bacillota | |
| Candidatus Sulfidibacterium hydrothermale | Bacteroidota | |

| | |
|---|---|
| Candidatus Absconditicoccus praedator | Candidatus Absconditabacteria |
| Candidatus Fervidibacter sacchari | Candidatus Fervidibacterota |
| Candidatus Velamenicoccus archaeovorus | Candidatus Omnitrophota |
| Candidatus Mycosynbacter amalyticus | Candidatus Saccharibacteria |
| Thermobaculum terrenum ATCC BAA-798 | Chloroflexota |
| Thermobaculum terrenum ATCC BAA-798 | Chloroflexota |
| cyanobacterium endosymbiont of Epithemia clementina EcSB | Cyanobacteriota |
| cyanobacterium endosymbiont of Epithemia turgida isolate EtSB Lake Yunoko | Cyanobacteriota |
| cyanobacterium endosymbiont of Rhopalodia gibberula | Cyanobacteriota |
| Gaopeijia maritima | Gemmatimonadota |
| Gaopeijia maritima | Gemmatimonadota |
| [Mycoplasma] anseris | Mycoplasmatota |
| [Mycoplasma] cavipharyngis | Mycoplasmatota |
| [Mycoplasma] gypis | Mycoplasmatota |
| [Mycoplasma] phocae | Mycoplasmatota |
| Malacoplasma iowae | Mycoplasmatota |
| Malacoplasma muris | Mycoplasmatota |
| Malacoplasma penetrans HF-2 | Mycoplasmatota |
| Mesomycoplasma bovoculi M165/69 | Mycoplasmatota |
| Mesomycoplasma conjunctivae | Mycoplasmatota |
| Mesomycoplasma dispar | Mycoplasmatota |
| Mesomycoplasma flocculare ATCC 27399 | Mycoplasmatota |
| Mesomycoplasma hyopneumoniae J | Mycoplasmatota |
| Mesomycoplasma hyorhinis | Mycoplasmatota |
| Mesomycoplasma lagogenitalium | Mycoplasmatota |
| Mesomycoplasma moatsii | Mycoplasmatota |
| Mesomycoplasma molare | Mycoplasmatota |
| Mesomycoplasma neurolyticum | Mycoplasmatota |
| Mesomycoplasma ovipneumoniae ATCC 29419 | Mycoplasmatota |
| Metamycoplasma alkalescens | Mycoplasmatota |
| Metamycoplasma arthritidis | Mycoplasmatota |

| | |
|---|---|
| Metamycoplasma buccale | Mycoplasmatota |
| Metamycoplasma canadense | Mycoplasmatota |
| Metamycoplasma cloacale | Mycoplasmatota |
| Metamycoplasma equirhinis | Mycoplasmatota |
| Metamycoplasma faucium | Mycoplasmatota |
| Metamycoplasma gateae | Mycoplasmatota |
| Metamycoplasma hominis ATCC 27545 | Mycoplasmatota |
| Metamycoplasma hyosynoviae | Mycoplasmatota |
| Metamycoplasma orale | Mycoplasmatota |
| Metamycoplasma phocicerebrale | Mycoplasmatota |
| Metamycoplasma salivarium | Mycoplasmatota |
| Metamycoplasma spumans | Mycoplasmatota |
| Metamycoplasma sualvi | Mycoplasmatota |
| Metamycoplasma subdolum | Mycoplasmatota |
| Mycoplasma mobile 163K | Mycoplasmatota |
| Mycoplasmoides fastidiosum | Mycoplasmatota |
| Mycoplasmoides gallisepticum | Mycoplasmatota |
| Mycoplasmoides genitalium | Mycoplasmatota |
| Mycoplasmoides pneumoniae | Mycoplasmatota |
| Mycoplasmopsis adleri | Mycoplasmatota |
| Mycoplasmopsis agalactiae PG2 | Mycoplasmatota |
| Mycoplasmopsis anatis | Mycoplasmatota |
| Mycoplasmopsis arginini | Mycoplasmatota |
| Mycoplasmopsis bovigenitalium | Mycoplasmatota |
| Mycoplasmopsis bovirhinis | Mycoplasmatota |
| Mycoplasmopsis bovis PG45 | Mycoplasmatota |
| Mycoplasmopsis californica | Mycoplasmatota |
| Mycoplasmopsis canis PG 14 | Mycoplasmatota |
| Mycoplasmopsis caviae | Mycoplasmatota |
| Mycoplasmopsis citelli | Mycoplasmatota |
| Mycoplasmopsis columbina | Mycoplasmatota |

| | |
|---|---|
| Mycoplasmopsis columbinasalis | Mycoplasmatota |
| Mycoplasmopsis columboralis | Mycoplasmatota |
| Mycoplasmopsis cynos C142 | Mycoplasmatota |
| Mycoplasmopsis edwardii | Mycoplasmatota |
| Mycoplasmopsis equigenitalium | Mycoplasmatota |
| Mycoplasmopsis felis | Mycoplasmatota |
| Mycoplasmopsis fermentans M64 | Mycoplasmatota |
| Mycoplasmopsis gallinacea | Mycoplasmatota |
| Mycoplasmopsis gallopavonis | Mycoplasmatota |
| Mycoplasmopsis gallopavonis | Mycoplasmatota |
| Mycoplasmopsis glycophila | Mycoplasmatota |
| Mycoplasmopsis hyopharyngis | Mycoplasmatota |
| Mycoplasmopsis lipophila | Mycoplasmatota |
| Mycoplasmopsis maculosa | Mycoplasmatota |
| Mycoplasmopsis meleagridis | Mycoplasmatota |
| Mycoplasmopsis phocirhinis | Mycoplasmatota |
| Mycoplasmopsis pulmonis | Mycoplasmatota |
| Mycoplasmopsis synoviae ATCC 25204 | Mycoplasmatota |
| Mycoplasmopsis verecunda | Mycoplasmatota |
| Ureaplasma diversum | Mycoplasmatota |
| Ureaplasma parvum serovar 3 str. ATCC 27815 | Mycoplasmatota |
| Chondromyces crocatus | Myxococcota |
| Chondromyces crocatus | Myxococcota |
| Haliangium ochraceum DSM 14365 | Myxococcota |
| Haliangium ochraceum DSM 14365 | Myxococcota |
| Labilithrix luteola | Myxococcota |
| Labilithrix luteola | Myxococcota |
| Nannocystis punicea | Myxococcota |
| Nannocystis punicea | Myxococcota |
| Pajaroellobacter abortibovis | Myxococcota |
| Polyangium aurulentum | Myxococcota |

| | |
|---|---|
| Polyangium aurulentum | Myxococcota |
| Sandaracinus amylolyticus | Myxococcota |
| Sandaracinus amylolyticus | Myxococcota |
| Sorangium cellulosum So ce56 | Myxococcota |
| Sorangium cellulosum So ce56 | Myxococcota |
| Engelhardtia mirabilis | Planctomycetota |
| Kolteria novifilia | Planctomycetota |
| Rohdeia mirabilis | Planctomycetota |
| Saltatorellus ferox | Planctomycetota |
| Stieleria bergensis | Planctomycetota |
| Stieleria magnilauensis | Planctomycetota |
| Stieleria magnilauensis | Planctomycetota |
| Stieleria marina | Planctomycetota |
| Agromyces silvae | |
| Algirhabdus cladophorae | |
| Algirhabdus cladophorae | |
| Aminirod propionatiphilus | |
| Aminirod propionatiphilus | |
| Anabaena azotica SJ-1 | |
| Anoxybacterium hadale | |
| Anoxybacterium hadale | |
| Bacillus haimaensis | |
| Elusimicrobium posterum | |
| Elusimicrobium posterum | |
| Enterobacterales bacterium endosymbiont of Anomoneura mori | |
| Fervidobacterium ngatamarikiense | |
| Fidelibacter multiformis | |
| Fidelibacter multiformis | |
| Fructilactobacillus frigidiflavus | |
| Halosquirtibacter laminarini | |
| Halosquirtibacter xylanolyticus | |

Imbroritus primus
Imbroritus primus
Kitasatospora hibisci
Lacrimispora sinapis
Levilactobacillus lettrarii
Microcystis ichthyoblabe FBCC-A1114
Microcystis protocystis FBCC-A270
Niallia tiangongensis
Niveibacterium terrae
Niveibacterium terrae
Nocardioides terrisoli
Parelusimicrobium proximum
Parelusimicrobium proximum
Peterkaempfera podocarpi
Pseudaeromonas aegiceratis
Pseudaeromonas aegiceratis
Pseudomonas imrae
Rhabdochlamydiaceae symbiont of Dictyostelium giganteum
Rodentibacter abscessus
Rodentibacter abscessus
Streptococcus nidrosiense
Streptomyces okerensis
Streptomyces okerensis
Streptomyces stoeckheimensis
Wolbachia endosymbiont of Rhagoletis indifferens
Wolbachia endosymbiont of Rhagoletis indifferens

| Organism Name | Phylum | Class | Refseq_ID | tuf_count |
|---|---|---|---|---|
| Chloracidobacterium sp. MS 40/45 | Acidobacteriota | Blastocatellia | NZ_CP072646.1 | 2 |
| Chloracidobacterium thermophilum B | Acidobacteriota | Blastocatellia | NC_016024.1 | 2 |
| Chloracidobacterium validum | Acidobacteriota | Blastocatellia | NZ_CP072648.1 | 2 |
| Geothrix oryzae | Acidobacteriota | Holophagae | NZ_AP027079.1 | 1 |
| Mesoterricola sediminis | Acidobacteriota | Holophagae | NZ_AP027081.1 | 1 |
| Mesoterricola silvestris | Acidobacteriota | Holophagae | NZ_AP027080.1 | 1 |
| Sulfidibacter corallicola | Acidobacteriota | Holophagae | NZ_CP071793.1 | 1 |
| Thermotomaculum hydrothermale | Acidobacteriota | Holophagae | NZ_AP017470.1 | 2 |
| Acidisarcina polymorpha | Acidobacteriota | Terriglobia | NZ_CP030840.1 | 2 |
| Acidobacterium capsulatum ATCC 51196 | Acidobacteriota | Terriglobia | NC_012483.1 | 2 |
| Candidatus Koribacter versatilis Ellin345 | Acidobacteriota | Terriglobia | NC_008009.1 | 2 |
| Edaphobacter flagellatus | Acidobacteriota | Terriglobia | NZ_CP073697.1 | 2 |
| Edaphobacter paludis | Acidobacteriota | Terriglobia | NZ_CP121194.1 | 2 |
| Granulicella mallensis MP5ACTX8 | Acidobacteriota | Terriglobia | NC_016631.1 | 2 |
| Granulicella tundricola MP5ACTX9 | Acidobacteriota | Terriglobia | NC_015064.1 | 2 |
| Occallatibacter riparius | Acidobacteriota | Terriglobia | NZ_CP093313.1 | 2 |
| Paludibaculum fermentans | Acidobacteriota | Terriglobia | NZ_CP063849.1 | 2 |
| Telmatobacter bradus | Acidobacteriota | Terriglobia | NZ_CP184532.1 | 2 |
| Terriglobus roseus DSM 18391 | Acidobacteriota | Terriglobia | NC_018014.1 | 2 |
| Terriglobus saanensis SP1PR4 | Acidobacteriota | Terriglobia | NC_014963.1 | 2 |
| Tunturiibacter empetritectus | Acidobacteriota | Terriglobia | NZ_CP132932.1 | 2 |
| Tunturiibacter gelidiferens | Acidobacteriota | Terriglobia | NZ_CP132938.1 | 2 |
| Tunturiibacter lichenicola | Acidobacteriota | Terriglobia | NZ_CP073696.1 | 2 |
| Tunturiibacter psychrotolerans | Acidobacteriota | Terriglobia | NZ_CP132942.1 | 2 |
| Luteitalea pratensis | Acidobacteriota | Vicinamibacteria | NZ_CP015136.1 | 2 |
| Acidimicrobium ferrooxidans DSM 10331 | Actinomycetota | Acidimicrobiia | NC_013124.1 | 1 |
| Actinomarinicola tropica | Actinomycetota | Acidimicrobiia | NZ_CP045851.1 | 2 |
| Aquihabitans daechungensis | Actinomycetota | Acidimicrobiia | NZ_CP172265.1 | 2 |
| Dermatobacter hominis | Actinomycetota | Acidimicrobiia | NZ_CP085840.1 | 2 |
| Iamia majanohamensis | Actinomycetota | Acidimicrobiia | NZ_CP116942.1 | 2 |
| Ilumatobacter coccineus YM16-304 | Actinomycetota | Acidimicrobiia | NC_020520.1 | 2 |

| | | | | |
|---|---|---|---|---|
| [Actinomadura] parvosata subsp. kistnae | Actinomycetota | Actinomycetes | NZ_CP017717.1 | 2 |
| [Kitasatospora] papulosa | Actinomycetota | Actinomycetes | NZ_CP108452.1 | 2 |
| [Mycobacterium] burgundiense | Actinomycetota | Actinomycetes | NZ_OY726397.1 | 1 |
| [Mycobacterium] holstebronense | Actinomycetota | Actinomycetes | NZ_OY726398.1 | 1 |
| [Mycobacterium] kokjensenii | Actinomycetota | Actinomycetes | NZ_OY726394.1 | 1 |
| [Mycobacterium] stephanolepidis | Actinomycetota | Actinomycetes | NZ_AP018165.1 | 1 |
| [Mycobacterium] wendilense | Actinomycetota | Actinomycetes | NZ_OY726395.1 | 1 |
| Acidipropionibacterium acidipropionici | Actinomycetota | Actinomycetes | NZ_CP013126.1 | 1 |
| Acidipropionibacterium jensenii | Actinomycetota | Actinomycetes | NZ_CP040635.1 | 1 |
| Acidipropionibacterium virtanenii | Actinomycetota | Actinomycetes | NZ_CP025198.1 | 1 |
| Acidothermus cellulolyticus 11B | Actinomycetota | Actinomycetes | NC_008578.1 | 1 |
| Actinacidiphila glaucinigra | Actinomycetota | Actinomycetes | NZ_CP109144.1 | 2 |
| Actinacidiphila reveromycinica | Actinomycetota | Actinomycetes | NZ_AP018365.1 | 1 |
| Actinoalloteichus fjordicus | Actinomycetota | Actinomycetes | NZ_CP016076.1 | 1 |
| Actinoalloteichus hoggarensis | Actinomycetota | Actinomycetes | NZ_CP022521.1 | 1 |
| Actinoalloteichus hymeniacidonis | Actinomycetota | Actinomycetes | NZ_CP014859.1 | 1 |
| Actinocatenispora sera | Actinomycetota | Actinomycetes | NZ_AP023354.1 | 1 |
| Actinocatenispora thailandica | Actinomycetota | Actinomycetes | NZ_AP023355.1 | 1 |
| Actinokineospora auranticolor | Actinomycetota | Actinomycetes | NZ_CP154825.1 | 1 |
| Actinomadura graeca | Actinomycetota | Actinomycetes | NZ_CP059572.1 | 2 |
| Actinomadura luteofluorescens | Actinomycetota | Actinomycetes | NZ_CP146086.1 | 2 |
| Actinomadura madurae | Actinomycetota | Actinomycetes | NZ_CP083237.1 | 2 |
| Actinomyces capricornis | Actinomycetota | Actinomycetes | NZ_AP025017.1 | 1 |
| Actinomyces faecalis | Actinomycetota | Actinomycetes | NZ_CP063418.1 | 1 |
| Actinomyces gaoshouyii | Actinomycetota | Actinomycetes | NZ_CP020468.1 | 1 |
| Actinomyces howellii | Actinomycetota | Actinomycetes | NZ_LR134350.1 | 1 |
| Actinomyces israelii | Actinomycetota | Actinomycetes | NZ_CP124548.1 | 1 |
| Actinomyces lilanjuaniae | Actinomycetota | Actinomycetes | NZ_CP032514.1 | 1 |
| Actinomyces marmotae | Actinomycetota | Actinomycetes | NZ_CP053642.1 | 1 |
| Actinomyces naeslundii | Actinomycetota | Actinomycetes | NZ_CP066049.1 | 1 |
| Actinomyces oris | Actinomycetota | Actinomycetes | NZ_CP066060.1 | 1 |
| Actinomyces pacaensis | Actinomycetota | Actinomycetes | NZ_LT635457.1 | 1 |

| | | | | |
|---|---|---|---|---|
| Actinomyces procaprae | Actinomycetota | Actinomycetes | NZ_CP039292.1 | 1 |
| Actinomyces qiguomingii | Actinomycetota | Actinomycetes | NZ_CP025228.1 | 1 |
| Actinomyces radicidentis | Actinomycetota | Actinomycetes | NZ_CP014228.1 | 1 |
| Actinomyces respiraculi | Actinomycetota | Actinomycetes | NZ_CP063989.1 | 1 |
| Actinomyces slackii | Actinomycetota | Actinomycetes | NZ_LR134363.1 | 1 |
| Actinomyces trachealis | Actinomycetota | Actinomycetes | NZ_CP065027.1 | 1 |
| Actinomyces viscosus | Actinomycetota | Actinomycetes | NZ_LR134477.1 | 1 |
| Actinomyces weissii | Actinomycetota | Actinomycetes | NZ_CP066802.1 | 1 |
| Actinomyces wuliandei | Actinomycetota | Actinomycetes | NZ_CP025227.1 | 1 |
| Actinoplanes derwentensis | Actinomycetota | Actinomycetes | NZ_LT629758.1 | 2 |
| Actinoplanes friuliensis DSM 7358 | Actinomycetota | Actinomycetes | NC_022657.1 | 2 |
| Actinoplanes ianthinogenes | Actinomycetota | Actinomycetes | NZ_AP023356.1 | 2 |
| Actinoplanes missouriensis 431 | Actinomycetota | Actinomycetes | NC_017093.1 | 2 |
| Actinoplanes oblitus | Actinomycetota | Actinomycetes | NZ_CP126980.1 | 2 |
| Actinoplanes sichuanensis | Actinomycetota | Actinomycetes | NZ_AP028461.1 | 2 |
| Actinoplanes teichomyceticus ATCC 31121 | Actinomycetota | Actinomycetes | NZ_CP023865.1 | 1 |
| Actinopolymorpha singaporensis | Actinomycetota | Actinomycetes | NZ_LT629732.1 | 1 |
| Actinopolyspora erythraea | Actinomycetota | Actinomycetes | NZ_CP022752.1 | 1 |
| Actinosynnema mirum DSM 43827 | Actinomycetota | Actinomycetes | NC_013093.1 | 1 |
| Actinosynnema pretiosum | Actinomycetota | Actinomycetes | NZ_CP023445.1 | 1 |
| Actinotignum schaalii | Actinomycetota | Actinomycetes | NZ_CP008802.1 | 1 |
| Actinotignum timonense | Actinomycetota | Actinomycetes | NZ_CP136960.1 | 1 |
| Aeromicrobium chenweiae | Actinomycetota | Actinomycetes | NZ_CP026952.1 | 1 |
| Aeromicrobium choanae | Actinomycetota | Actinomycetes | NZ_LT796768.1 | 1 |
| Aeromicrobium duanguangcaii | Actinomycetota | Actinomycetes | NZ_CP101990.1 | 1 |
| Aeromicrobium erythreum | Actinomycetota | Actinomycetes | NZ_CP011502.1 | 1 |
| Aeromicrobium marinum DSM 15272 | Actinomycetota | Actinomycetes | NZ_CM001024.1 | 1 |
| Aeromicrobium senzhongii | Actinomycetota | Actinomycetes | NZ_CP060587.1 | 1 |
| Aeromicrobium yanjiei | Actinomycetota | Actinomycetes | NZ_CP045737.1 | 1 |
| Agrococcus beijingensis | Actinomycetota | Actinomycetes | NZ_CP132360.1 | 1 |
| Agrococcus carbonis | Actinomycetota | Actinomycetes | NZ_LT629734.1 | 1 |
| Agrococcus jejuensis | Actinomycetota | Actinomycetes | NZ_LT629695.1 | 1 |

| | | | | |
|---|---|---|---|---|
| Agromyces archimandritae | Actinomycetota | Actinomycetes | NZ_CP071696.1 | 1 |
| Agromyces aureus | Actinomycetota | Actinomycetes | NZ_CP013979.1 | 1 |
| Agromyces badenianii | Actinomycetota | Actinomycetes | NZ_CP028913.1 | 1 |
| Agromyces flavus | Actinomycetota | Actinomycetes | NZ_LT629755.1 | 1 |
| Agromyces intestinalis | Actinomycetota | Actinomycetes | NZ_CP043505.1 | 1 |
| Agromyces laixinhei | Actinomycetota | Actinomycetes | NZ_CP040872.1 | 1 |
| Agromyces larvae | Actinomycetota | Actinomycetes | NZ_CP094528.1 | 1 |
| Agromyces mangrovi | Actinomycetota | Actinomycetes | NZ_AP027737.1 | 1 |
| Agromyces marinus | Actinomycetota | Actinomycetes | NZ_CP087879.1 | 1 |
| Agromyces protaetiae | Actinomycetota | Actinomycetes | NZ_CP035491.1 | 1 |
| Agromyces soli | Actinomycetota | Actinomycetes | NZ_CP094533.1 | 1 |
| Allokutzneria albata | Actinomycetota | Actinomycetes | NZ_LT629701.1 | 2 |
| Allosaccharopolyspora coralli | Actinomycetota | Actinomycetes | NZ_CP045929.1 | 1 |
| Ammonicoccus fulvus | Actinomycetota | Actinomycetes | NZ_CP154795.1 | 1 |
| Amycolatopsis acidiphila | Actinomycetota | Actinomycetes | NZ_CP090063.1 | 1 |
| Amycolatopsis aidingensis | Actinomycetota | Actinomycetes | NZ_CP076538.1 | 1 |
| Amycolatopsis albispora | Actinomycetota | Actinomycetes | NZ_CP015163.1 | 1 |
| Amycolatopsis bartoniae | Actinomycetota | Actinomycetes | NZ_CP174150.1 | 1 |
| Amycolatopsis carbonis | Actinomycetota | Actinomycetes | NZ_CP127294.1 | 1 |
| Amycolatopsis cynarae | Actinomycetota | Actinomycetes | NZ_CP113836.1 | 1 |
| Amycolatopsis japonica | Actinomycetota | Actinomycetes | NZ_CP008953.1 | 1 |
| Amycolatopsis keratiniphila | Actinomycetota | Actinomycetes | NC_021252.1 | 1 |
| Amycolatopsis mediterranei S699 | Actinomycetota | Actinomycetes | NC_017186.1 | 2 |
| Amycolatopsis methanolica 239 | Actinomycetota | Actinomycetes | NZ_CP009110.1 | 1 |
| Amycolatopsis mongoliensis | Actinomycetota | Actinomycetes | NZ_CP127295.1 | 2 |
| Amycolatopsis nalaikhensis | Actinomycetota | Actinomycetes | NZ_CP127173.1 | 2 |
| Amycolatopsis orientalis | Actinomycetota | Actinomycetes | NZ_CP016174.1 | 1 |
| Amycolatopsis rhabdoformis | Actinomycetota | Actinomycetes | NZ_CP142149.1 | 1 |
| Amycolatopsis roodepoortensis | Actinomycetota | Actinomycetes | NZ_CP102415.1 | 1 |
| Antiquaquibacter oligotrophicus | Actinomycetota | Actinomycetes | NZ_CP085036.1 | 1 |
| Aquiluna borgnonia | Actinomycetota | Actinomycetes | NZ_CP054056.1 | 1 |
| Arachnia propionica | Actinomycetota | Actinomycetes | NZ_LR134406.1 | 1 |

| | | | | |
|---|---|---|---|---|
| Arachnia rubra | Actinomycetota | Actinomycetes | NZ_CP146372.1 | 1 |
| Arcanobacterium buesumense | Actinomycetota | Actinomycetes | NZ_CP050804.1 | 1 |
| Arcanobacterium canis | Actinomycetota | Actinomycetes | NZ_CP121208.1 | 1 |
| Arcanobacterium haemolyticum | Actinomycetota | Actinomycetes | NZ_LS483427.1 | 1 |
| Arcanobacterium phocae | Actinomycetota | Actinomycetes | NZ_LT629804.1 | 1 |
| Arcanobacterium phocisimile | Actinomycetota | Actinomycetes | NZ_CP070228.1 | 1 |
| Arcanobacterium pinnipediorum | Actinomycetota | Actinomycetes | NZ_CP099547.1 | 1 |
| Arcanobacterium wilhelmae | Actinomycetota | Actinomycetes | NZ_CP121247.1 | 1 |
| Arsenicicoccus cauae | Actinomycetota | Actinomycetes | NZ_CP171001.1 | 1 |
| Arsenicicoccus dermatophilus | Actinomycetota | Actinomycetes | NZ_CP124757.1 | 1 |
| Arthrobacter agilis | Actinomycetota | Actinomycetes | NZ_CP117869.1 | 1 |
| Arthrobacter alpinus | Actinomycetota | Actinomycetes | NZ_CP013745.1 | 1 |
| Arthrobacter antioxidans | Actinomycetota | Actinomycetes | NZ_CP095501.1 | 1 |
| Arthrobacter caoxuetaonis | Actinomycetota | Actinomycetes | NZ_CP099466.1 | 1 |
| Arthrobacter crystallopoietes | Actinomycetota | Actinomycetes | NZ_CP018863.1 | 1 |
| Arthrobacter dokdonellae | Actinomycetota | Actinomycetes | NZ_CP029642.1 | 1 |
| Arthrobacter gengyunqii | Actinomycetota | Actinomycetes | NZ_CP095461.1 | 1 |
| Arthrobacter jiangjiafuii | Actinomycetota | Actinomycetes | NZ_CP076022.1 | 1 |
| Arthrobacter jinronghuae | Actinomycetota | Actinomycetes | NZ_CP104263.1 | 1 |
| Arthrobacter nitrophenolicus | Actinomycetota | Actinomycetes | NZ_CP174504.1 | 1 |
| Arthrobacter polaris | Actinomycetota | Actinomycetes | NZ_CP071516.1 | 1 |
| Arthrobacter sulfonylureivorans | Actinomycetota | Actinomycetes | NZ_CP093326.1 | 1 |
| Arthrobacter sunyaminii | Actinomycetota | Actinomycetes | NZ_CP076456.1 | 1 |
| Arthrobacter zhangbolii | Actinomycetota | Actinomycetes | NZ_CP094984.1 | 1 |
| Arthrobacter zhaoxinii | Actinomycetota | Actinomycetes | NZ_CP104275.1 | 1 |
| Aurantimicrobium minutum | Actinomycetota | Actinomycetes | NZ_AP017457.1 | 1 |
| Aurantimicrobium photophilum | Actinomycetota | Actinomycetes | NZ_CP023994.1 | 1 |
| Auraticoccus monumenti | Actinomycetota | Actinomycetes | NZ_LT629688.1 | 1 |
| Auritidibacter ignavus | Actinomycetota | Actinomycetes | NZ_CP122564.1 | 1 |
| Barrientosiimonas endolithica | Actinomycetota | Actinomycetes | NZ_AP027735.1 | 1 |
| Beutenbergia cavernae DSM 12333 | Actinomycetota | Actinomycetes | NC_012669.1 | 1 |
| Bifidobacterium actinocoloniiforme DSM 22766 | Actinomycetota | Actinomycetes | NZ_CP011786.1 | 1 |

| | | | | |
|---|---|---|---|---|
| Bifidobacterium adolescentis ATCC 15703 | Actinomycetota | Actinomycetes | NC_008618.1 | 1 |
| Bifidobacterium angulatum DSM 20098 = JCM 7096 | Actinomycetota | Actinomycetes | NZ_AP012322.1 | 1 |
| Bifidobacterium animalis subsp. lactis DSM 10140 | Actinomycetota | Actinomycetes | NC_012815.1 | 1 |
| Bifidobacterium aquikefiricola | Actinomycetota | Actinomycetes | NZ_CP129674.1 | 1 |
| Bifidobacterium asteroides DSM 20089 | Actinomycetota | Actinomycetes | NZ_CP017696.1 | 1 |
| Bifidobacterium bifidum | Actinomycetota | Actinomycetes | NZ_CP058603.1 | 1 |
| Bifidobacterium breve DSM 20213 = JCM 1192 | Actinomycetota | Actinomycetes | NZ_AP012324.1 | 1 |
| Bifidobacterium catenulatum DSM 16992 = JCM 1194 = LM | Actinomycetota | Actinomycetes | NZ_AP012325.1 | 1 |
| Bifidobacterium choerinum | Actinomycetota | Actinomycetes | NZ_CP018044.1 | 1 |
| Bifidobacterium coryneforme | Actinomycetota | Actinomycetes | NZ_CP007287.1 | 1 |
| Bifidobacterium dentium JCM 1195 = DSM 20436 | Actinomycetota | Actinomycetes | NZ_AP012326.1 | 1 |
| Bifidobacterium eulemuris | Actinomycetota | Actinomycetes | NZ_CP062938.1 | 1 |
| Bifidobacterium fermentum | Actinomycetota | Actinomycetes | NZ_CP129683.1 | 1 |
| Bifidobacterium imperatoris | Actinomycetota | Actinomycetes | NZ_CP071591.1 | 1 |
| Bifidobacterium lemurum | Actinomycetota | Actinomycetes | NZ_CP062948.1 | 1 |
| Bifidobacterium longum subsp. longum JCM 1217 | Actinomycetota | Actinomycetes | NC_015067.1 | 1 |
| Bifidobacterium pseudocatenulatum DSM 20438 = JCM 12 | Actinomycetota | Actinomycetes | NZ_AP012330.1 | 1 |
| Bifidobacterium pseudolongum subsp. globosum DSM 200 | Actinomycetota | Actinomycetes | NZ_CP017695.1 | 1 |
| Bifidobacterium saguini | Actinomycetota | Actinomycetes | NZ_CP071732.1 | 1 |
| Bifidobacterium scardovii JCM 12489 = DSM 13734 | Actinomycetota | Actinomycetes | NZ_AP012331.1 | 1 |
| Bifidobacterium subtile | Actinomycetota | Actinomycetes | NZ_CP062939.1 | 1 |
| Blastococcus brunescens | Actinomycetota | Actinomycetes | NZ_CP141261.1 | 1 |
| Blastococcus montanus | Actinomycetota | Actinomycetes | NZ_CP155551.1 | 1 |
| Bombiscardovia apis | Actinomycetota | Actinomycetes | NZ_AP026800.1 | 1 |
| Boudabousia tangfeifanii | Actinomycetota | Actinomycetes | NZ_CP017812.1 | 1 |
| Brachybacterium avium | Actinomycetota | Actinomycetes | NZ_CP022316.1 | 1 |
| Brachybacterium ginsengisoli | Actinomycetota | Actinomycetes | NZ_CP023564.1 | 1 |
| Brachybacterium halotolerans subsp. kimchii | Actinomycetota | Actinomycetes | NZ_CP086254.1 | 1 |
| Brachybacterium huguangmaarense | Actinomycetota | Actinomycetes | NZ_CP107020.1 | 1 |
| Brachybacterium kimchii | Actinomycetota | Actinomycetes | NZ_CP097218.1 | 1 |
| Brachybacterium saurashtrense | Actinomycetota | Actinomycetes | NZ_CP031356.1 | 1 |
| Brachybacterium vulturis | Actinomycetota | Actinomycetes | NZ_CP023563.1 | 1 |

| | | | | |
|---|---|---|---|---|
| Brevibacterium atlanticum | Actinomycetota | Actinomycetes | NZ_CP050152.1 | 1 |
| Brevibacterium aurantiacum | Actinomycetota | Actinomycetes | NZ_CP025333.1 | 1 |
| Brevibacterium casei | Actinomycetota | Actinomycetes | NZ_CP065630.1 | 1 |
| Brevibacterium koreense | Actinomycetota | Actinomycetes | NZ_CP158281.1 | 1 |
| Brevibacterium limosum | Actinomycetota | Actinomycetes | NZ_CP050154.1 | 1 |
| Brevibacterium linens | Actinomycetota | Actinomycetes | NZ_CP026734.1 | 1 |
| Brevibacterium oceani | Actinomycetota | Actinomycetes | NZ_CP072846.1 | 1 |
| Brevibacterium pigmentatum | Actinomycetota | Actinomycetes | NZ_CP050153.1 | 1 |
| Brevibacterium sandarakinum | Actinomycetota | Actinomycetes | NZ_LT629739.1 | 1 |
| Brevibacterium siliguriense | Actinomycetota | Actinomycetes | NZ_LT629766.1 | 1 |
| Brevibacterium spongiae | Actinomycetota | Actinomycetes | NZ_CP093443.1 | 1 |
| Brevibacterium zhoupengii | Actinomycetota | Actinomycetes | NZ_CP088298.1 | 1 |
| Brooklawnia propionicigenes | Actinomycetota | Actinomycetes | NZ_AP028056.1 | 1 |
| Buchananella felis | Actinomycetota | Actinomycetes | NZ_CP159980.1 | 1 |
| Candidatus Mycobacterium methanotrophicum | Actinomycetota | Actinomycetes | NZ_CP097320.1 | 1 |
| Candidatus Mycobacterium wuenschmannii | Actinomycetota | Actinomycetes | NZ_CP126981.1 | 1 |
| Candidatus Nanopelagicus abundans | Actinomycetota | Actinomycetes | NZ_CP016779.1 | 1 |
| Candidatus Nanopelagicus hibericus | Actinomycetota | Actinomycetes | NZ_CP016771.1 | 1 |
| Candidatus Nanopelagicus limnes | Actinomycetota | Actinomycetes | NZ_CP016768.2 | 1 |
| Candidatus Planktophila dulcis | Actinomycetota | Actinomycetes | NZ_CP016770.1 | 1 |
| Candidatus Planktophila lacus | Actinomycetota | Actinomycetes | NZ_CP016780.1 | 1 |
| Candidatus Planktophila limnetica | Actinomycetota | Actinomycetes | NZ_CP016782.1 | 1 |
| Candidatus Planktophila sulfonica | Actinomycetota | Actinomycetes | NZ_CP016773.1 | 1 |
| Candidatus Planktophila vernalis | Actinomycetota | Actinomycetes | NZ_CP016776.1 | 1 |
| Candidatus Planktophila versatilis | Actinomycetota | Actinomycetes | NZ_CP016775.1 | 1 |
| Candidatus Rhodoluna planktonica | Actinomycetota | Actinomycetes | NZ_CP015208.1 | 1 |
| Catenulispora acidiphila DSM 44928 | Actinomycetota | Actinomycetes | NC_013131.1 | 2 |
| Cellulomonas chengniuliangii | Actinomycetota | Actinomycetes | NZ_CP101988.1 | 1 |
| Cellulomonas dongxiuzhuiae | Actinomycetota | Actinomycetes | NZ_CP076023.1 | 1 |
| Cellulomonas fengjieae | Actinomycetota | Actinomycetes | NZ_CP074404.1 | 1 |
| Cellulomonas fimi ATCC 484 | Actinomycetota | Actinomycetes | NC_015514.1 | 1 |
| Cellulomonas flavigena DSM 20109 | Actinomycetota | Actinomycetes | NC_014151.1 | 1 |

| | | | | |
|---|---|---|---|---|
| Cellulomonas gilvus ATCC 13127 | Actinomycetota | Actinomycetes | NC_015671.1 | 1 |
| Cellulomonas palmilytica | Actinomycetota | Actinomycetes | NZ_CP062221.1 | 1 |
| Cellulomonas shaoxiangyii | Actinomycetota | Actinomycetes | NZ_CP039291.1 | 1 |
| Cellulomonas taurus | Actinomycetota | Actinomycetes | NZ_CP051884.1 | 1 |
| Cellulomonas wangleii | Actinomycetota | Actinomycetes | NZ_CP074405.1 | 1 |
| Cellulomonas wangsupingiae | Actinomycetota | Actinomycetes | NZ_CP101989.1 | 1 |
| Cellulomonas xiejunii | Actinomycetota | Actinomycetes | NZ_CP101987.1 | 1 |
| Cellulosimicrobium cellulans | Actinomycetota | Actinomycetes | NZ_CP072387.1 | 1 |
| Cellulosimicrobium protaetiae | Actinomycetota | Actinomycetes | NZ_CP052757.1 | 1 |
| Changpingibacter yushuensis | Actinomycetota | Actinomycetes | NZ_CP059492.1 | 1 |
| Citricoccus nitrophenolicus | Actinomycetota | Actinomycetes | NZ_CP063070.1 | 1 |
| Clavibacter californiensis | Actinomycetota | Actinomycetes | NZ_CP040792.1 | 1 |
| Clavibacter capsici | Actinomycetota | Actinomycetes | NZ_CP048047.1 | 1 |
| Clavibacter michiganensis | Actinomycetota | Actinomycetes | NZ_CP080437.1 | 1 |
| Clavibacter nebraskensis NCPPB 2581 | Actinomycetota | Actinomycetes | NC_020891.1 | 1 |
| Clavibacter phaseoli | Actinomycetota | Actinomycetes | NZ_CP040786.1 | 1 |
| Clavibacter seminis | Actinomycetota | Actinomycetes | NZ_CP083439.1 | 1 |
| Clavibacter sepedonicus | Actinomycetota | Actinomycetes | NC_010407.1 | 1 |
| Clavibacter zhangzhiyongii | Actinomycetota | Actinomycetes | NZ_CP061274.1 | 1 |
| Cnuibacter physcomitrellae | Actinomycetota | Actinomycetes | NZ_CP020715.1 | 1 |
| Corynebacterium accolens | Actinomycetota | Actinomycetes | NZ_CP046605.1 | 1 |
| Corynebacterium afermentans subsp. lipophilum | Actinomycetota | Actinomycetes | NZ_CP046530.1 | 1 |
| Corynebacterium ammoniagenes DSM 20306 | Actinomycetota | Actinomycetes | NZ_CP009244.1 | 1 |
| Corynebacterium amycolatum | Actinomycetota | Actinomycetes | NZ_CP069513.1 | 1 |
| Corynebacterium anserum | Actinomycetota | Actinomycetes | NZ_CP046883.1 | 1 |
| Corynebacterium appendicis CIP 107643 | Actinomycetota | Actinomycetes | NZ_CP046976.1 | 1 |
| Corynebacterium aquatimens | Actinomycetota | Actinomycetes | NZ_CP046980.1 | 1 |
| Corynebacterium aquilae DSM 44791 | Actinomycetota | Actinomycetes | NZ_CP009245.1 | 1 |
| Corynebacterium argentoratense DSM 44202 | Actinomycetota | Actinomycetes | NC_022198.1 | 1 |
| Corynebacterium atrinae | Actinomycetota | Actinomycetes | NZ_CP046977.1 | 1 |
| Corynebacterium atypicum | Actinomycetota | Actinomycetes | NZ_CP008944.1 | 1 |
| Corynebacterium aurimucosum ATCC 700975 | Actinomycetota | Actinomycetes | NC_012590.1 | 1 |

| | | | | |
|---|---|---|---|---|
| Corynebacterium auris | Actinomycetota | Actinomycetes | NZ_CP047047.1 | 1 |
| Corynebacterium auriscanis | Actinomycetota | Actinomycetes | NZ_CP047046.1 | 1 |
| Corynebacterium belfantii | Actinomycetota | Actinomycetes | NZ_CP157823.1 | 1 |
| Corynebacterium breve | Actinomycetota | Actinomycetes | NZ_CP126969.1 | 1 |
| Corynebacterium callunae DSM 20147 | Actinomycetota | Actinomycetes | NC_020506.1 | 1 |
| Corynebacterium camporealensis | Actinomycetota | Actinomycetes | NZ_CP011311.1 | 1 |
| Corynebacterium canis | Actinomycetota | Actinomycetes | NZ_CP047080.1 | 1 |
| Corynebacterium capitovis DSM 44611 | Actinomycetota | Actinomycetes | NZ_CP047117.1 | 1 |
| Corynebacterium casei LMG S-19264 | Actinomycetota | Actinomycetes | NZ_CP004350.1 | 1 |
| Corynebacterium caspium DSM 44850 | Actinomycetota | Actinomycetes | NZ_CP047118.1 | 1 |
| Corynebacterium choanae | Actinomycetota | Actinomycetes | NZ_CP033896.1 | 1 |
| Corynebacterium ciconiae DSM 44920 | Actinomycetota | Actinomycetes | NZ_CP047189.1 | 1 |
| Corynebacterium comes | Actinomycetota | Actinomycetes | NZ_CP046453.1 | 1 |
| Corynebacterium confusum | Actinomycetota | Actinomycetes | NZ_CP047202.1 | 1 |
| Corynebacterium coyleae | Actinomycetota | Actinomycetes | NZ_CP077302.1 | 1 |
| Corynebacterium crudilactis | Actinomycetota | Actinomycetes | NZ_CP015622.1 | 1 |
| Corynebacterium cystitidis | Actinomycetota | Actinomycetes | NZ_LT906473.1 | 1 |
| Corynebacterium deserti GIMN1.010 | Actinomycetota | Actinomycetes | NZ_CP009220.1 | 1 |
| Corynebacterium diphtheriae | Actinomycetota | Actinomycetes | NZ_LN831026.1 | 1 |
| Corynebacterium doosanense CAU 212 = DSM 45436 | Actinomycetota | Actinomycetes | NZ_CP006764.1 | 1 |
| Corynebacterium durum | Actinomycetota | Actinomycetes | NZ_CP047200.1 | 1 |
| Corynebacterium efficiens YS-314 | Actinomycetota | Actinomycetes | NC_004369.1 | 1 |
| Corynebacterium endometrii | Actinomycetota | Actinomycetes | NZ_CP039247.1 | 1 |
| Corynebacterium epidermidicanis | Actinomycetota | Actinomycetes | NZ_CP011541.1 | 1 |
| Corynebacterium faecale | Actinomycetota | Actinomycetes | NZ_CP047204.1 | 1 |
| Corynebacterium falsenii | Actinomycetota | Actinomycetes | NZ_CP083646.1 | 1 |
| Corynebacterium felinum | Actinomycetota | Actinomycetes | NZ_CP047209.1 | 1 |
| Corynebacterium flavescens | Actinomycetota | Actinomycetes | NZ_CP009246.1 | 1 |
| Corynebacterium fournieri | Actinomycetota | Actinomycetes | NZ_CP047210.1 | 1 |
| Corynebacterium frankenforstense DSM 45800 | Actinomycetota | Actinomycetes | NZ_CP009247.1 | 1 |
| Corynebacterium freiburgense | Actinomycetota | Actinomycetes | NZ_CP047355.1 | 1 |
| Corynebacterium freneyi | Actinomycetota | Actinomycetes | NZ_CP077258.1 | 1 |

| | | | | |
|---|---|---|---|---|
| Corynebacterium genitalium ATCC 33030 | Actinomycetota | Actinomycetes | NZ_CP072935.1 | 1 |
| Corynebacterium gerontici | Actinomycetota | Actinomycetes | NZ_CP033897.1 | 1 |
| Corynebacterium glaucum | Actinomycetota | Actinomycetes | NZ_CP047358.1 | 1 |
| Corynebacterium glucuronolyticum DSM 44120 | Actinomycetota | Actinomycetes | NZ_CP047452.1 | 1 |
| Corynebacterium glutamicum ATCC 13032 | Actinomycetota | Actinomycetes | NC_003450.3 | 1 |
| Corynebacterium glyciniphilum AJ 3170 | Actinomycetota | Actinomycetes | NZ_CP006842.1 | 1 |
| Corynebacterium gottingense | Actinomycetota | Actinomycetes | NZ_CP047454.1 | 1 |
| Corynebacterium guangdongense | Actinomycetota | Actinomycetes | NZ_CP047654.1 | 1 |
| Corynebacterium halotolerans YIM 70093 = DSM 44683 | Actinomycetota | Actinomycetes | NC_020302.1 | 1 |
| Corynebacterium hansenii | Actinomycetota | Actinomycetes | NZ_CP047211.1 | 1 |
| Corynebacterium heidelbergense | Actinomycetota | Actinomycetes | NZ_CP063191.1 | 1 |
| Corynebacterium hesseae | Actinomycetota | Actinomycetes | NZ_CP137211.1 | 1 |
| Corynebacterium hindlerae | Actinomycetota | Actinomycetes | NZ_CP059833.1 | 1 |
| Corynebacterium ihumii | Actinomycetota | Actinomycetes | NZ_CP063190.1 | 1 |
| Corynebacterium imitans | Actinomycetota | Actinomycetes | NZ_LT906467.1 | 1 |
| Corynebacterium incognita | Actinomycetota | Actinomycetes | NZ_CP059404.1 | 1 |
| Corynebacterium jeddahense | Actinomycetota | Actinomycetes | NZ_CP063194.1 | 1 |
| Corynebacterium jeikeium | Actinomycetota | Actinomycetes | NZ_CP063195.1 | 1 |
| Corynebacterium kalinowskii | Actinomycetota | Actinomycetes | NZ_CP046452.1 | 1 |
| Corynebacterium kefirresidentii | Actinomycetota | Actinomycetes | NZ_CP067012.1 | 1 |
| Corynebacterium kroppenstedtii DSM 44385 | Actinomycetota | Actinomycetes | NC_012704.1 | 1 |
| Corynebacterium kutscheri | Actinomycetota | Actinomycetes | NZ_CP011312.1 | 1 |
| Corynebacterium lactis RW2-5 | Actinomycetota | Actinomycetes | NZ_CP006841.1 | 1 |
| Corynebacterium liangguodongii | Actinomycetota | Actinomycetes | NZ_CP026948.1 | 1 |
| Corynebacterium lizhenjunii | Actinomycetota | Actinomycetes | NZ_CP064954.1 | 1 |
| Corynebacterium lujinxingii | Actinomycetota | Actinomycetes | NZ_CP061032.1 | 1 |
| Corynebacterium macginleyi | Actinomycetota | Actinomycetes | NZ_CP069516.1 | 1 |
| Corynebacterium marinum DSM 44953 | Actinomycetota | Actinomycetes | NZ_CP007790.1 | 1 |
| Corynebacterium maris DSM 45190 | Actinomycetota | Actinomycetes | NC_021915.1 | 1 |
| Corynebacterium massiliense DSM 45435 | Actinomycetota | Actinomycetes | NZ_CP063189.1 | 1 |
| Corynebacterium mayonis | Actinomycetota | Actinomycetes | NZ_CP141643.1 | 1 |
| Corynebacterium minutissimum | Actinomycetota | Actinomycetes | NZ_LS483460.1 | 1 |

| | | | | |
|---|---|---|---|---|
| Corynebacterium mustelae | Actinomycetota | Actinomycetes | NZ_CP011542.1 | 1 |
| Corynebacterium mycetoides | Actinomycetota | Actinomycetes | NZ_LT629700.1 | 1 |
| Corynebacterium nuruki S6-4 | Actinomycetota | Actinomycetes | NZ_CP042429.1 | 1 |
| Corynebacterium occultum | Actinomycetota | Actinomycetes | NZ_CP046455.1 | 1 |
| Corynebacterium pelargi | Actinomycetota | Actinomycetes | NZ_CP035299.1 | 1 |
| Corynebacterium phocae | Actinomycetota | Actinomycetes | NZ_CP009249.1 | 1 |
| Corynebacterium poyangense | Actinomycetota | Actinomycetes | NZ_CP046884.1 | 1 |
| Corynebacterium propinquum | Actinomycetota | Actinomycetes | NZ_CP068161.1 | 1 |
| Corynebacterium pseudogenitalium | Actinomycetota | Actinomycetes | NZ_CP072934.1 | 1 |
| Corynebacterium pseudokroppenstedtii | Actinomycetota | Actinomycetes | NZ_CP137757.1 | 1 |
| Corynebacterium pseudopelargi | Actinomycetota | Actinomycetes | NZ_CP033898.1 | 1 |
| Corynebacterium pseudotuberculosis | Actinomycetota | Actinomycetes | NZ_CP021251.1 | 1 |
| Corynebacterium qintianiae | Actinomycetota | Actinomycetes | NZ_CP064955.1 | 1 |
| Corynebacterium ramonii | Actinomycetota | Actinomycetes | NZ_CP009622.1 | 1 |
| Corynebacterium renale | Actinomycetota | Actinomycetes | NZ_LS483464.1 | 1 |
| Corynebacterium resistens DSM 45100 | Actinomycetota | Actinomycetes | NC_015673.1 | 1 |
| Corynebacterium riegelii | Actinomycetota | Actinomycetes | NZ_CP012342.1 | 1 |
| Corynebacterium rouxii | Actinomycetota | Actinomycetes | NZ_LR738855.1 | 1 |
| Corynebacterium sanguinis | Actinomycetota | Actinomycetes | NZ_CP038157.1 | 1 |
| Corynebacterium silvaticum | Actinomycetota | Actinomycetes | NZ_CP021417.2 | 1 |
| Corynebacterium simulans | Actinomycetota | Actinomycetes | NZ_CP014635.1 | 1 |
| Corynebacterium singulare | Actinomycetota | Actinomycetes | NZ_CP010827.1 | 1 |
| Corynebacterium sphenisci DSM 44792 | Actinomycetota | Actinomycetes | NZ_CP009248.1 | 1 |
| Corynebacterium stationis | Actinomycetota | Actinomycetes | NZ_CP009251.1 | 1 |
| Corynebacterium striatum | Actinomycetota | Actinomycetes | NZ_CP068158.1 | 1 |
| Corynebacterium suedekumii | Actinomycetota | Actinomycetes | NZ_CP126970.1 | 1 |
| Corynebacterium suranareeae | Actinomycetota | Actinomycetes | NZ_AP017369.1 | 1 |
| Corynebacterium terpenotabidum Y-11 | Actinomycetota | Actinomycetes | NC_021663.1 | 1 |
| Corynebacterium testudinoris | Actinomycetota | Actinomycetes | NZ_CP011545.1 | 1 |
| Corynebacterium tuberculostearicum | Actinomycetota | Actinomycetes | NZ_CP068156.1 | 1 |
| Corynebacterium uberis | Actinomycetota | Actinomycetes | NZ_CP085051.1 | 1 |
| Corynebacterium ulcerans 809 | Actinomycetota | Actinomycetes | NC_017317.1 | 1 |

| | | | | |
|---|---|---|---|---|
| Corynebacterium urealyticum | Actinomycetota | Actinomycetes | NZ_LT906481.1 | 1 |
| Corynebacterium ureicelerivorans | Actinomycetota | Actinomycetes | NZ_CP009215.1 | 1 |
| Corynebacterium urogenitale | Actinomycetota | Actinomycetes | NZ_CP045032.1 | 1 |
| Corynebacterium uterequi | Actinomycetota | Actinomycetes | NZ_CP011546.1 | 1 |
| Corynebacterium vitaeruminis DSM 20294 | Actinomycetota | Actinomycetes | NZ_CP004353.1 | 1 |
| Corynebacterium yudongzhengii | Actinomycetota | Actinomycetes | NZ_CP026947.1 | 1 |
| Corynebacterium zhongnanshanii | Actinomycetota | Actinomycetes | NZ_CP061033.1 | 1 |
| Cryobacterium soli | Actinomycetota | Actinomycetes | NZ_CP030033.1 | 1 |
| Curtobacterium aetherium | Actinomycetota | Actinomycetes | NZ_CP076544.1 | 1 |
| Curtobacterium flaccumfaciens | Actinomycetota | Actinomycetes | NZ_CP103853.1 | 1 |
| Curtobacterium poinsettiae | Actinomycetota | Actinomycetes | NZ_CP178434.1 | 1 |
| Cutibacterium acnes subsp. acnes | Actinomycetota | Actinomycetes | NZ_AP019723.1 | 1 |
| Cutibacterium avidum 44067 | Actinomycetota | Actinomycetes | NC_021064.1 | 1 |
| Cutibacterium equinum | Actinomycetota | Actinomycetes | NZ_CP115668.1 | 1 |
| Cutibacterium granulosum | Actinomycetota | Actinomycetes | NZ_AP026710.1 | 1 |
| Dactylosporangium aurantiacum | Actinomycetota | Actinomycetes | NZ_CP073767.1 | 2 |
| Dactylosporangium fulvum | Actinomycetota | Actinomycetes | NZ_CP073720.1 | 2 |
| Dactylosporangium roseum | Actinomycetota | Actinomycetes | NZ_CP073721.1 | 2 |
| Dactylosporangium vinaceum | Actinomycetota | Actinomycetes | NZ_CP061913.1 | 2 |
| Demequina capsici | Actinomycetota | Actinomycetes | NZ_CP134880.1 | 1 |
| Dermabacter jinjuensis | Actinomycetota | Actinomycetes | NZ_LR698963.1 | 1 |
| Dermabacter vaginalis | Actinomycetota | Actinomycetes | NZ_CP012117.1 | 1 |
| Dermacoccus nishinomiyaensis | Actinomycetota | Actinomycetes | NZ_CP008889.1 | 1 |
| Dermatophilus congolensis | Actinomycetota | Actinomycetes | NZ_LT906453.1 | 1 |
| Devriesea agamarum | Actinomycetota | Actinomycetes | NZ_LN849456.1 | 1 |
| Dietzia cinnamea | Actinomycetota | Actinomycetes | NZ_CP143053.1 | 1 |
| Dietzia kunjamensis | Actinomycetota | Actinomycetes | NZ_CP143297.1 | 1 |
| Dietzia lutea | Actinomycetota | Actinomycetes | NZ_CP015449.1 | 1 |
| Dietzia psychralcaliphila | Actinomycetota | Actinomycetes | NZ_CP015453.1 | 1 |
| Dietzia timorensis | Actinomycetota | Actinomycetes | NZ_CP015961.1 | 1 |
| Epidermidibacterium keratini | Actinomycetota | Actinomycetes | NZ_CP047156.1 | 1 |
| Flaviflexus equikiangi | Actinomycetota | Actinomycetes | NZ_CP059676.1 | 1 |

| | | | | |
|---|---|---|---|---|
| Flaviflexus salsibiostraticola | Actinomycetota | Actinomycetes | NZ_CP034438.1 | 1 |
| Frankia alni ACN14a | Actinomycetota | Actinomycetes | NC_008278.1 | 1 |
| Frankia casuarinae | Actinomycetota | Actinomycetes | NC_007777.1 | 1 |
| Friedmanniella luteola | Actinomycetota | Actinomycetes | NZ_LT629749.1 | 1 |
| Frondihabitans sucicola | Actinomycetota | Actinomycetes | NZ_AP027732.1 | 1 |
| Gardnerella leopoldii | Actinomycetota | Actinomycetes | NZ_CP029984.1 | 1 |
| Gardnerella vaginalis ATCC 14018 = JCM 11026 | Actinomycetota | Actinomycetes | NZ_AP012332.1 | 1 |
| Geodermatophilus obscurus DSM 43160 | Actinomycetota | Actinomycetes | NC_013757.1 | 1 |
| Georgenia faecalis | Actinomycetota | Actinomycetes | NZ_CP033325.1 | 1 |
| Georgenia wutianyii | Actinomycetota | Actinomycetes | NZ_CP040899.1 | 1 |
| Georgenia yuyongxinii | Actinomycetota | Actinomycetes | NZ_CP040915.1 | 1 |
| Gephyromycinifex aptenodytis | Actinomycetota | Actinomycetes | NZ_CP051155.1 | 1 |
| Gleimia europaea | Actinomycetota | Actinomycetes | NZ_CP126966.1 | 1 |
| Glutamicibacter arilaitensis Re117 | Actinomycetota | Actinomycetes | NC_014550.1 | 1 |
| Glutamicibacter creatinolyticus | Actinomycetota | Actinomycetes | NZ_CP034412.1 | 1 |
| Glutamicibacter ectropisis | Actinomycetota | Actinomycetes | NZ_CP125942.1 | 1 |
| Glutamicibacter endophyticus | Actinomycetota | Actinomycetes | NZ_CP174200.1 | 1 |
| Glutamicibacter halophytocola | Actinomycetota | Actinomycetes | NZ_CP012750.1 | 1 |
| Glutamicibacter mishrai | Actinomycetota | Actinomycetes | NZ_CP032549.1 | 1 |
| Glutamicibacter protophormiae | Actinomycetota | Actinomycetes | NZ_CP069806.1 | 1 |
| Gordonia alkanivorans | Actinomycetota | Actinomycetes | NZ_CP027114.1 | 1 |
| Gordonia amarae | Actinomycetota | Actinomycetes | NZ_CP045810.1 | 1 |
| Gordonia amicalis | Actinomycetota | Actinomycetes | NZ_CP091855.1 | 1 |
| Gordonia bronchialis DSM 43247 | Actinomycetota | Actinomycetes | NC_013441.1 | 1 |
| Gordonia hongkongensis | Actinomycetota | Actinomycetes | NZ_CP121270.1 | 1 |
| Gordonia hydrophobica | Actinomycetota | Actinomycetes | NZ_CP136137.1 | 1 |
| Gordonia insulae | Actinomycetota | Actinomycetes | NZ_CP033972.1 | 1 |
| Gordonia iterans | Actinomycetota | Actinomycetes | NZ_CP027433.1 | 1 |
| Gordonia jinghuaiqii | Actinomycetota | Actinomycetes | NZ_CP059491.1 | 1 |
| Gordonia mangrovi | Actinomycetota | Actinomycetes | NZ_CP102850.1 | 1 |
| Gordonia otitidis | Actinomycetota | Actinomycetes | NZ_CP085974.1 | 1 |
| Gordonia phthalatica | Actinomycetota | Actinomycetes | NZ_CP011853.1 | 1 |

| | | | | |
|---|---|---|---|---|
| Gordonia polyisoprenivorans VH2 | Actinomycetota | Actinomycetes | NC_016906.1 | 1 |
| Gordonia pseudamarae | Actinomycetota | Actinomycetes | NZ_CP045809.1 | 1 |
| Gordonia rubripertincta | Actinomycetota | Actinomycetes | NZ_CP059694.1 | 1 |
| Gordonia sihwensis | Actinomycetota | Actinomycetes | NZ_CP121248.1 | 1 |
| Gordonia terrae | Actinomycetota | Actinomycetes | NZ_CP016594.1 | 1 |
| Gordonia zhaorongruii | Actinomycetota | Actinomycetes | NZ_CP041763.1 | 1 |
| Gryllotalpicola protaetiae | Actinomycetota | Actinomycetes | NZ_CP032624.1 | 1 |
| Gulosibacter molinativorax | Actinomycetota | Actinomycetes | NZ_CP028426.1 | 1 |
| Helcobacillus massiliensis | Actinomycetota | Actinomycetes | NZ_CP136959.1 | 1 |
| Hoyosella subflava DQS3-9A1 | Actinomycetota | Actinomycetes | NC_015564.1 | 1 |
| Humibacter ginsenosidimutans | Actinomycetota | Actinomycetes | NZ_CP042305.1 | 1 |
| Intrasporangium calvum DSM 43043 | Actinomycetota | Actinomycetes | NC_014830.1 | 1 |
| Isoptericola dokdonensis DS-3 | Actinomycetota | Actinomycetes | NZ_CP014209.1 | 1 |
| Isoptericola variabilis 225 | Actinomycetota | Actinomycetes | NC_015588.1 | 1 |
| Janibacter alittae | Actinomycetota | Actinomycetes | NZ_CP144913.1 | 1 |
| Janibacter melonis | Actinomycetota | Actinomycetes | NZ_CP044548.2 | 1 |
| Janibacter terrae | Actinomycetota | Actinomycetes | NZ_CP144167.1 | 1 |
| Jatrophihabitans cynanchi | Actinomycetota | Actinomycetes | NZ_CP097463.1 | 1 |
| Jatrophihabitans telluris | Actinomycetota | Actinomycetes | NZ_CP097332.1 | 1 |
| Jiangella alkaliphila | Actinomycetota | Actinomycetes | NZ_LT629791.1 | 1 |
| Jonesia denitrificans | Actinomycetota | Actinomycetes | NZ_LS483423.1 | 1 |
| Kibdelosporangium phytohabitans | Actinomycetota | Actinomycetes | NZ_CP012752.1 | 2 |
| Kineococcus radiotolerans SRS30216 = ATCC BAA-149 | Actinomycetota | Actinomycetes | NC_009664.2 | 1 |
| Kitasatospora camelliae | Actinomycetota | Actinomycetes | NZ_CP159872.1 | 1 |
| Kitasatospora cathayae | Actinomycetota | Actinomycetes | NZ_CP115450.1 | 1 |
| Kitasatospora fiedleri | Actinomycetota | Actinomycetes | NZ_OX419519.1 | 1 |
| Kitasatospora griseola | Actinomycetota | Actinomycetes | NZ_CP173360.1 | 1 |
| Kitasatospora herbaricolor | Actinomycetota | Actinomycetes | NZ_CP108460.1 | 1 |
| Kitasatospora purpeofusca | Actinomycetota | Actinomycetes | NZ_CP108561.1 | 2 |
| Kitasatospora setae KM-6054 | Actinomycetota | Actinomycetes | NC_016109.1 | 1 |
| Kocuria flava | Actinomycetota | Actinomycetes | NZ_CP013254.1 | 1 |
| Kocuria rhizophila | Actinomycetota | Actinomycetes | NZ_CP108528.1 | 1 |

| | | | | |
|---|---|---|---|---|
| Kocuria turfanensis | Actinomycetota | Actinomycetes | NZ_CP014480.1 | 1 |
| Kribbella flavida DSM 17836 | Actinomycetota | Actinomycetes | NC_013729.1 | 2 |
| Kutzneria albida DSM 43870 | Actinomycetota | Actinomycetes | NZ_CP007155.1 | 1 |
| Kutzneria buriramensis | Actinomycetota | Actinomycetes | NZ_CP144375.1 | 1 |
| Kutzneria chonburiensis | Actinomycetota | Actinomycetes | NZ_CP097263.1 | 1 |
| Kytococcus sedentarius | Actinomycetota | Actinomycetes | NZ_CP066015.1 | 1 |
| Lawsonella clevelandensis | Actinomycetota | Actinomycetes | NZ_CP009312.1 | 1 |
| Leifsonia xyli subsp. cynodontis DSM 46306 | Actinomycetota | Actinomycetes | NC_022438.1 | 1 |
| Lentzea guizhouensis | Actinomycetota | Actinomycetes | NZ_CP016793.1 | 2 |
| Leucobacter allii | Actinomycetota | Actinomycetes | NZ_CP095045.1 | 1 |
| Leucobacter coleopterorum | Actinomycetota | Actinomycetes | NZ_CP049933.1 | 1 |
| Leucobacter denitrificans | Actinomycetota | Actinomycetes | NZ_CP060716.1 | 1 |
| Leucobacter insecticola | Actinomycetota | Actinomycetes | NZ_CP049934.1 | 1 |
| Leucobacter muris | Actinomycetota | Actinomycetes | NZ_CP035037.1 | 1 |
| Leucobacter rhizosphaerae | Actinomycetota | Actinomycetes | NZ_CP095043.1 | 1 |
| Leucobacter tenebrionis | Actinomycetota | Actinomycetes | NZ_CP082322.1 | 1 |
| Leucobacter triazinivorans | Actinomycetota | Actinomycetes | NZ_CP035806.1 | 1 |
| Leucobacter viscericola | Actinomycetota | Actinomycetes | NZ_CP049863.1 | 1 |
| Luteimicrobium xylanilyticum | Actinomycetota | Actinomycetes | NZ_CP045529.1 | 1 |
| Luteipulveratus mongoliensis | Actinomycetota | Actinomycetes | NZ_CP011112.1 | 1 |
| Marisediminicola antarctica | Actinomycetota | Actinomycetes | NZ_CP017146.1 | 1 |
| Microbacterium abyssi | Actinomycetota | Actinomycetes | NZ_CP063815.1 | 1 |
| Microbacterium albipurpureum | Actinomycetota | Actinomycetes | NZ_CP128170.1 | 1 |
| Microbacterium amylolyticum | Actinomycetota | Actinomycetes | NZ_CP049253.1 | 1 |
| Microbacterium arborescens | Actinomycetota | Actinomycetes | NZ_CP128474.1 | 1 |
| Microbacterium atlanticum | Actinomycetota | Actinomycetes | NZ_CP063813.1 | 1 |
| Microbacterium aurugineum | Actinomycetota | Actinomycetes | NZ_CP078078.1 | 1 |
| Microbacterium aurum | Actinomycetota | Actinomycetes | NZ_CP018762.1 | 1 |
| Microbacterium azadirachtae | Actinomycetota | Actinomycetes | NZ_CP099706.1 | 1 |
| Microbacterium betulae | Actinomycetota | Actinomycetes | NZ_CP118157.1 | 1 |
| Microbacterium caowuchunii | Actinomycetota | Actinomycetes | NZ_CP044231.1 | 1 |
| Microbacterium chocolatum | Actinomycetota | Actinomycetes | NZ_CP015810.1 | 1 |

| | | | | |
|---|---|---|---|---|
| Microbacterium cremeum | Actinomycetota | Actinomycetes | NZ_CP063812.1 | 1 |
| Microbacterium endophyticum | Actinomycetota | Actinomycetes | NZ_CP049255.1 | 1 |
| Microbacterium foliorum | Actinomycetota | Actinomycetes | NZ_CP031425.1 | 1 |
| Microbacterium forte | Actinomycetota | Actinomycetes | NZ_CP116871.1 | 1 |
| Microbacterium galbinum | Actinomycetota | Actinomycetes | NZ_CP078077.1 | 1 |
| Microbacterium hominis | Actinomycetota | Actinomycetes | NZ_CP061344.1 | 1 |
| Microbacterium horticulturae | Actinomycetota | Actinomycetes | NZ_CP119108.1 | 1 |
| Microbacterium lemovicicum | Actinomycetota | Actinomycetes | NZ_CP031423.1 | 1 |
| Microbacterium limosum | Actinomycetota | Actinomycetes | NZ_CP137080.1 | 1 |
| Microbacterium lushaniae | Actinomycetota | Actinomycetes | NZ_CP044232.1 | 1 |
| Microbacterium luteum | Actinomycetota | Actinomycetes | NZ_CP063814.1 | 1 |
| Microbacterium maritypicum | Actinomycetota | Actinomycetes | NZ_CP101471.1 | 1 |
| Microbacterium neungamense | Actinomycetota | Actinomycetes | NZ_CP069717.1 | 1 |
| Microbacterium oleivorans | Actinomycetota | Actinomycetes | NZ_CP031421.1 | 1 |
| Microbacterium oryzae | Actinomycetota | Actinomycetes | NZ_CP032550.1 | 1 |
| Microbacterium paraoxydans | Actinomycetota | Actinomycetes | NZ_LT629770.1 | 1 |
| Microbacterium plantarum | Actinomycetota | Actinomycetes | NZ_CP141117.1 | 1 |
| Microbacterium protaetiae | Actinomycetota | Actinomycetes | NZ_CP035494.1 | 1 |
| Microbacterium pygmaeum | Actinomycetota | Actinomycetes | NZ_LT629692.1 | 1 |
| Microbacterium resistens | Actinomycetota | Actinomycetes | NZ_CP082781.1 | 1 |
| Microbacterium rhizosphaerae | Actinomycetota | Actinomycetes | NZ_CP139368.1 | 1 |
| Microbacterium sediminis | Actinomycetota | Actinomycetes | NZ_CP038256.1 | 1 |
| Microbacterium sufflavum | Actinomycetota | Actinomycetes | NZ_CP078076.1 | 1 |
| Microbacterium terricola | Actinomycetota | Actinomycetes | NZ_CP107222.1 | 1 |
| Microbacterium terrisoli | Actinomycetota | Actinomycetes | NZ_CP133019.1 | 1 |
| Microbacterium wangchenii | Actinomycetota | Actinomycetes | NZ_CP038266.1 | 1 |
| Microbispora maris | Actinomycetota | Actinomycetes | NZ_JBDJAW01000 | 1 |
| Microcella daejeonensis | Actinomycetota | Actinomycetes | NZ_CP113089.1 | 1 |
| Microcella flavibacter | Actinomycetota | Actinomycetes | NZ_CP051299.1 | 1 |
| Microcella humidisoli | Actinomycetota | Actinomycetes | NZ_CP101497.1 | 1 |
| Microcella indica | Actinomycetota | Actinomycetes | NZ_CP058670.1 | 1 |
| Micrococcus antarcticus | Actinomycetota | Actinomycetes | NZ_CP085253.1 | 1 |

| | | | | |
|---|---|---|---|---|
| Micrococcus luteus NCTC 2665 | Actinomycetota | Actinomycetes | NZ_LS483396.1 | 1 |
| Micrococcus porci | Actinomycetota | Actinomycetes | NZ_CP083691.1 | 1 |
| Microlunatus elymi | Actinomycetota | Actinomycetes | NZ_CP041692.1 | 1 |
| Microlunatus phosphovorus NM-1 | Actinomycetota | Actinomycetes | NC_015635.1 | 1 |
| Microlunatus sagamiharensis | Actinomycetota | Actinomycetes | NZ_LT629799.1 | 1 |
| Microlunatus soli | Actinomycetota | Actinomycetes | NZ_LT629772.1 | 1 |
| Micromonospora aurantiaca ATCC 27029 | Actinomycetota | Actinomycetes | NC_014391.1 | 1 |
| Micromonospora auratinigra | Actinomycetota | Actinomycetes | NZ_LT594323.1 | 1 |
| Micromonospora carbonacea | Actinomycetota | Actinomycetes | NZ_CP058322.1 | 1 |
| Micromonospora cathayae | Actinomycetota | Actinomycetes | NZ_CP118615.1 | 1 |
| Micromonospora chalcea | Actinomycetota | Actinomycetes | NZ_CP118233.1 | 1 |
| Micromonospora chokoriensis | Actinomycetota | Actinomycetes | NZ_LT607409.1 | 1 |
| Micromonospora coriariae | Actinomycetota | Actinomycetes | NZ_LT607412.1 | 1 |
| Micromonospora coxensis | Actinomycetota | Actinomycetes | NZ_LT607753.1 | 1 |
| Micromonospora craniellae | Actinomycetota | Actinomycetes | NZ_CP061725.1 | 1 |
| Micromonospora echinaurantiaca | Actinomycetota | Actinomycetes | NZ_LT607750.1 | 1 |
| Micromonospora echinofusca | Actinomycetota | Actinomycetes | NZ_LT607733.1 | 1 |
| Micromonospora globbae | Actinomycetota | Actinomycetes | NZ_CP108084.1 | 1 |
| Micromonospora halotolerans | Actinomycetota | Actinomycetes | NZ_CP134876.1 | 1 |
| Micromonospora harpali | Actinomycetota | Actinomycetes | NZ_CP160047.1 | 1 |
| Micromonospora inositola | Actinomycetota | Actinomycetes | NZ_LT607754.1 | 1 |
| Micromonospora krabiensis | Actinomycetota | Actinomycetes | NZ_LT598496.1 | 1 |
| Micromonospora maris AB-18-032 | Actinomycetota | Actinomycetes | NC_015434.1 | 1 |
| Micromonospora narathiwatensis | Actinomycetota | Actinomycetes | NZ_LT594324.1 | 1 |
| Micromonospora purpureochromogenes | Actinomycetota | Actinomycetes | NZ_LT607410.1 | 1 |
| Micromonospora rifamycinica | Actinomycetota | Actinomycetes | NZ_LT607752.1 | 1 |
| Micromonospora robiginosa | Actinomycetota | Actinomycetes | NZ_CP059322.2 | 1 |
| Micromonospora sagamiensis | Actinomycetota | Actinomycetes | NZ_AP023438.1 | 1 |
| Micromonospora siamensis | Actinomycetota | Actinomycetes | NZ_LT607751.1 | 2 |
| Micromonospora terminaliae | Actinomycetota | Actinomycetes | NZ_CP045309.1 | 1 |
| Micromonospora viridifaciens | Actinomycetota | Actinomycetes | NZ_LT607411.1 | 1 |
| Micromonospora zamorensis | Actinomycetota | Actinomycetes | NZ_LT607755.1 | 1 |

| | | | | |
|---|---|---|---|---|
| Micropruina glycogenica | Actinomycetota | Actinomycetes | NZ_LT985188.1 | 1 |
| Microterricola viridarii | Actinomycetota | Actinomycetes | NZ_LT629742.1 | 1 |
| Mobiluncus curtisii ATCC 43063 | Actinomycetota | Actinomycetes | NC_014246.1 | 1 |
| Modestobacter italicus | Actinomycetota | Actinomycetes | NC_017955.1 | 1 |
| Mumia qirimensis | Actinomycetota | Actinomycetes | NZ_CP162023.1 | 1 |
| Mycetocola spongiae | Actinomycetota | Actinomycetes | NZ_CP080203.1 | 1 |
| Mycetocola zhujimingii | Actinomycetota | Actinomycetes | NZ_CP026949.1 | 1 |
| Mycobacterium adipatum | Actinomycetota | Actinomycetes | NZ_CP015596.1 | 1 |
| Mycobacterium avium subsp. hominissuis | Actinomycetota | Actinomycetes | NZ_CP018019.2 | 1 |
| Mycobacterium basiliense | Actinomycetota | Actinomycetes | NZ_LR130759.1 | 1 |
| Mycobacterium branderi | Actinomycetota | Actinomycetes | NZ_AP022606.1 | 1 |
| Mycobacterium canettii CIPT 140010059 | Actinomycetota | Actinomycetes | NC_015848.1 | 1 |
| Mycobacterium conspicuum | Actinomycetota | Actinomycetes | NZ_AP022613.1 | 1 |
| Mycobacterium cookii | Actinomycetota | Actinomycetes | NZ_AP022569.1 | 1 |
| Mycobacterium dioxanotrophicus | Actinomycetota | Actinomycetes | NZ_CP020809.1 | 1 |
| Mycobacterium europaeum | Actinomycetota | Actinomycetes | NZ_CP157315.1 | 1 |
| Mycobacterium florentinum | Actinomycetota | Actinomycetes | NZ_AP022576.1 | 1 |
| Mycobacterium gallinarum | Actinomycetota | Actinomycetes | NZ_AP022601.1 | 1 |
| Mycobacterium haemophilum DSM 44634 | Actinomycetota | Actinomycetes | NZ_CP011883.2 | 1 |
| Mycobacterium heckeshornense | Actinomycetota | Actinomycetes | NZ_AP024310.1 | 1 |
| Mycobacterium heidelbergense | Actinomycetota | Actinomycetes | NZ_AP022615.1 | 1 |
| Mycobacterium intracellulare | Actinomycetota | Actinomycetes | NZ_CP085945.1 | 1 |
| Mycobacterium kansasii ATCC 12478 | Actinomycetota | Actinomycetes | NC_022663.1 | 1 |
| Mycobacterium kubicae | Actinomycetota | Actinomycetes | NZ_CP045081.1 | 1 |
| Mycobacterium lacus | Actinomycetota | Actinomycetes | NZ_AP022581.1 | 1 |
| Mycobacterium lentiflavum | Actinomycetota | Actinomycetes | NZ_CP092423.2 | 1 |
| Mycobacterium leprae | Actinomycetota | Actinomycetes | NZ_CP029543.1 | 1 |
| Mycobacterium lepromatosis | Actinomycetota | Actinomycetes | NZ_CP083405.1 | 1 |
| Mycobacterium malmoense | Actinomycetota | Actinomycetes | NZ_CP080999.1 | 1 |
| Mycobacterium mantenii | Actinomycetota | Actinomycetes | NZ_AP022590.1 | 1 |
| Mycobacterium marinum E11 | Actinomycetota | Actinomycetes | NZ_HG917972.2 | 1 |
| Mycobacterium marseillense | Actinomycetota | Actinomycetes | NZ_CP023147.1 | 1 |

| | | | | |
|---|---|---|---|---|
| Mycobacterium noviomagense | Actinomycetota | Actinomycetes | NZ_AP022583.1 | 1 |
| Mycobacterium novum | Actinomycetota | Actinomycetes | NZ_AP022562.1 | 1 |
| Mycobacterium ostraviense | Actinomycetota | Actinomycetes | NZ_CP089224.1 | 1 |
| Mycobacterium paragordonae | Actinomycetota | Actinomycetes | NZ_CP025546.1 | 1 |
| Mycobacterium paraintracellulare | Actinomycetota | Actinomycetes | NC_016948.1 | 1 |
| Mycobacterium paraseoulense | Actinomycetota | Actinomycetes | NZ_AP022619.1 | 1 |
| Mycobacterium paraterrae | Actinomycetota | Actinomycetes | NZ_CP092488.2 | 1 |
| Mycobacterium parmense | Actinomycetota | Actinomycetes | NZ_AP022614.1 | 1 |
| Mycobacterium pseudoshottsii JCM 15466 | Actinomycetota | Actinomycetes | NZ_AP018410.1 | 1 |
| Mycobacterium riyadhense | Actinomycetota | Actinomycetes | NZ_OY970456.1 | 1 |
| Mycobacterium saskatchewanense | Actinomycetota | Actinomycetes | NZ_AP022573.1 | 1 |
| Mycobacterium senriense | Actinomycetota | Actinomycetes | NZ_AP024828.1 | 1 |
| Mycobacterium seoulense | Actinomycetota | Actinomycetes | NZ_AP022582.1 | 1 |
| Mycobacterium shigaense | Actinomycetota | Actinomycetes | NZ_AP018164.1 | 1 |
| Mycobacterium shinjukuense | Actinomycetota | Actinomycetes | NZ_AP022575.1 | 1 |
| Mycobacterium shottsii | Actinomycetota | Actinomycetes | NZ_AP022572.1 | 1 |
| Mycobacterium simiae | Actinomycetota | Actinomycetes | NZ_AP022568.1 | 1 |
| Mycobacterium spongiae | Actinomycetota | Actinomycetes | NZ_CP046600.1 | 1 |
| Mycobacterium stomatepiae | Actinomycetota | Actinomycetes | NZ_AP022587.1 | 1 |
| Mycobacterium tuberculosis H37Rv | Actinomycetota | Actinomycetes | NC_000962.3 | 1 |
| Mycobacterium ulcerans subsp. shinshuense | Actinomycetota | Actinomycetes | NZ_AP017624.1 | 1 |
| Mycobacterium vicinigordonae | Actinomycetota | Actinomycetes | NZ_CP059165.1 | 1 |
| Mycobacteroides abscessus ATCC 19977 | Actinomycetota | Actinomycetes | NC_010397.1 | 1 |
| Mycobacteroides chelonae CCUG 47445 | Actinomycetota | Actinomycetes | NZ_CP007220.1 | 1 |
| Mycobacteroides immunogenum | Actinomycetota | Actinomycetes | NZ_CP011530.1 | 1 |
| Mycobacteroides salmoniphilum | Actinomycetota | Actinomycetes | NZ_CP024633.1 | 1 |
| Mycobacteroides saopaulense | Actinomycetota | Actinomycetes | NZ_CP010271.1 | 1 |
| Mycolicibacillus koreensis | Actinomycetota | Actinomycetes | NZ_AP022594.1 | 1 |
| Mycolicibacillus parakoreensis | Actinomycetota | Actinomycetes | NZ_CP092365.1 | 1 |
| Mycolicibacter heraklionensis | Actinomycetota | Actinomycetes | NZ_CP080997.1 | 1 |
| Mycolicibacter hiberniae | Actinomycetota | Actinomycetes | NZ_AP022609.1 | 1 |
| Mycolicibacter minnesotensis | Actinomycetota | Actinomycetes | NZ_AP022589.1 | 1 |

| | | | | |
|---|---|---|---|---|
| Mycolicibacter sinensis | Actinomycetota | Actinomycetes | NC_015576.1 | 1 |
| Mycolicibacter terrae | Actinomycetota | Actinomycetes | NZ_AP022564.1 | 1 |
| Mycolicibacter virginiensis | Actinomycetota | Actinomycetes | NZ_CP092430.2 | 1 |
| Mycolicibacterium aichiense | Actinomycetota | Actinomycetes | NZ_AP022561.1 | 1 |
| Mycolicibacterium alvei | Actinomycetota | Actinomycetes | NZ_AP022565.1 | 1 |
| Mycolicibacterium anyangense | Actinomycetota | Actinomycetes | NZ_AP022620.1 | 1 |
| Mycolicibacterium arabiense | Actinomycetota | Actinomycetes | NZ_AP022593.1 | 1 |
| Mycolicibacterium aubagnense | Actinomycetota | Actinomycetes | NZ_AP022577.1 | 1 |
| Mycolicibacterium aurum | Actinomycetota | Actinomycetes | NZ_LR134356.1 | 1 |
| Mycolicibacterium austroafricanum | Actinomycetota | Actinomycetes | NZ_CP070380.1 | 1 |
| Mycolicibacterium baixiangningiae | Actinomycetota | Actinomycetes | NZ_CP066218.1 | 1 |
| Mycolicibacterium boenickei | Actinomycetota | Actinomycetes | NZ_AP022579.1 | 1 |
| Mycolicibacterium brumae | Actinomycetota | Actinomycetes | NZ_CP104302.1 | 1 |
| Mycolicibacterium celeriflavum | Actinomycetota | Actinomycetes | NZ_AP022591.1 | 1 |
| Mycolicibacterium chitae | Actinomycetota | Actinomycetes | NZ_LR134355.1 | 1 |
| Mycolicibacterium confluentis | Actinomycetota | Actinomycetes | NZ_AP022612.1 | 1 |
| Mycolicibacterium crocinum | Actinomycetota | Actinomycetes | NZ_CP092362.2 | 2 |
| Mycolicibacterium diernhoferi | Actinomycetota | Actinomycetes | NZ_CP080332.1 | 1 |
| Mycolicibacterium doricum | Actinomycetota | Actinomycetes | NZ_AP022605.1 | 1 |
| Mycolicibacterium duvalii | Actinomycetota | Actinomycetes | NZ_AP022563.1 | 1 |
| Mycolicibacterium fallax | Actinomycetota | Actinomycetes | NZ_AP022603.1 | 1 |
| Mycolicibacterium farcinogenes | Actinomycetota | Actinomycetes | NZ_CP081673.1 | 1 |
| Mycolicibacterium fluoranthenivorans | Actinomycetota | Actinomycetes | NZ_CP059894.1 | 1 |
| Mycolicibacterium fortuitum subsp. fortuitum | Actinomycetota | Actinomycetes | NZ_AP025518.1 | 1 |
| Mycolicibacterium frederiksbergense | Actinomycetota | Actinomycetes | NZ_CP038799.1 | 1 |
| Mycolicibacterium gadium | Actinomycetota | Actinomycetes | NZ_AP022608.1 | 1 |
| Mycolicibacterium gilvum Spyr1 | Actinomycetota | Actinomycetes | NC_014814.1 | 1 |
| Mycolicibacterium goodii | Actinomycetota | Actinomycetes | NZ_CP092364.2 | 1 |
| Mycolicibacterium grossiae | Actinomycetota | Actinomycetes | NZ_CP043474.1 | 1 |
| Mycolicibacterium hassiacum DSM 44199 | Actinomycetota | Actinomycetes | NZ_LR026975.1 | 1 |
| Mycolicibacterium helvum | Actinomycetota | Actinomycetes | NZ_AP022596.1 | 1 |
| Mycolicibacterium holsaticum DSM 44478 = JCM 12374 | Actinomycetota | Actinomycetes | NZ_CP080998.1 | 1 |

| | | | | |
|---|---|---|---|---|
| Mycolicibacterium insubricum | Actinomycetota | Actinomycetes | NZ_AP022618.1 | 1 |
| Mycolicibacterium litorale | Actinomycetota | Actinomycetes | NZ_AP022586.1 | 1 |
| Mycolicibacterium madagascariense | Actinomycetota | Actinomycetes | NZ_AP022610.1 | 1 |
| Mycolicibacterium mageritense | Actinomycetota | Actinomycetes | NZ_AP022567.1 | 1 |
| Mycolicibacterium mengxianglii | Actinomycetota | Actinomycetes | NZ_CP065373.1 | 1 |
| Mycolicibacterium monacense | Actinomycetota | Actinomycetes | NZ_AP022617.1 | 1 |
| Mycolicibacterium moriokaense | Actinomycetota | Actinomycetes | NZ_AP022560.1 | 1 |
| Mycolicibacterium mucogenicum DSM 44124 | Actinomycetota | Actinomycetes | NZ_CP062008.1 | 1 |
| Mycolicibacterium neoaurum VKM Ac-1815D | Actinomycetota | Actinomycetes | NC_023036.2 | 1 |
| Mycolicibacterium nivoides | Actinomycetota | Actinomycetes | NZ_CP034072.1 | 1 |
| Mycolicibacterium pallens | Actinomycetota | Actinomycetes | NZ_CP080333.1 | 1 |
| Mycolicibacterium parafortuitum | Actinomycetota | Actinomycetes | NZ_AP022598.1 | 1 |
| Mycolicibacterium phocaicum | Actinomycetota | Actinomycetes | NZ_AP022616.1 | 1 |
| Mycolicibacterium poriferae | Actinomycetota | Actinomycetes | NZ_AP022570.1 | 1 |
| Mycolicibacterium psychrotolerans | Actinomycetota | Actinomycetes | NZ_AP022574.1 | 1 |
| Mycolicibacterium pulveris | Actinomycetota | Actinomycetes | NZ_AP022599.1 | 1 |
| Mycolicibacterium rufum | Actinomycetota | Actinomycetes | NZ_CP092427.2 | 1 |
| Mycolicibacterium rutilum | Actinomycetota | Actinomycetes | NZ_LT629971.1 | 1 |
| Mycolicibacterium sarraceniae | Actinomycetota | Actinomycetes | NZ_AP022595.1 | 1 |
| Mycolicibacterium sediminis | Actinomycetota | Actinomycetes | NZ_AP022588.1 | 1 |
| Mycolicibacterium senegalense | Actinomycetota | Actinomycetes | NZ_CP081000.1 | 1 |
| Mycolicibacterium septicum | Actinomycetota | Actinomycetes | NZ_CP070349.1 | 1 |
| Mycolicibacterium smegmatis | Actinomycetota | Actinomycetes | NZ_CP080274.1 | 1 |
| Mycolicibacterium thermoresistibile | Actinomycetota | Actinomycetes | NZ_LT906483.1 | 1 |
| Mycolicibacterium tokaiense | Actinomycetota | Actinomycetes | NZ_AP022600.1 | 1 |
| Mycolicibacterium vaccae 95051 | Actinomycetota | Actinomycetes | NZ_CP011491.1 | 1 |
| Mycolicibacterium vanbaalenii PYR-1 | Actinomycetota | Actinomycetes | NC_008726.1 | 1 |
| Naasia aerilata | Actinomycetota | Actinomycetes | NZ_AP027731.1 | 1 |
| Nakamurella antarctica | Actinomycetota | Actinomycetes | NZ_CP034170.1 | 1 |
| Nakamurella multipartita DSM 44233 | Actinomycetota | Actinomycetes | NC_013235.1 | 1 |
| Nakamurella panacisegetis | Actinomycetota | Actinomycetes | NZ_LT629710.1 | 1 |
| Natronoglycomyces albus | Actinomycetota | Actinomycetes | NZ_CP070496.1 | 1 |

| | | | | |
|---|---|---|---|---|
| Natronosporangium hydrolyticum | Actinomycetota | Actinomycetes | NZ_CP070499.1 | 1 |
| Nocardia asteroides | Actinomycetota | Actinomycetes | NZ_LR134352.1 | 1 |
| Nocardia brasiliensis | Actinomycetota | Actinomycetes | NZ_CP022088.2 | 2 |
| Nocardia cyriacigeorgica | Actinomycetota | Actinomycetes | NZ_CP026746.1 | 1 |
| Nocardia farcinica IFM 10152 | Actinomycetota | Actinomycetes | NC_006361.1 | 1 |
| Nocardia gipuzkoensis | Actinomycetota | Actinomycetes | NZ_CP089225.1 | 1 |
| Nocardia huaxiensis | Actinomycetota | Actinomycetes | NZ_CP059399.1 | 3 |
| Nocardia iowensis | Actinomycetota | Actinomycetes | NZ_CP078145.1 | 2 |
| Nocardia mangyaensis | Actinomycetota | Actinomycetes | NZ_CP018082.1 | 1 |
| Nocardia otitidiscaviarum | Actinomycetota | Actinomycetes | NZ_CP041695.1 | 2 |
| Nocardia puris | Actinomycetota | Actinomycetes | NZ_CP107943.1 | 2 |
| Nocardia salmonicida | Actinomycetota | Actinomycetes | NZ_CP108014.1 | 1 |
| Nocardia sputorum | Actinomycetota | Actinomycetes | NZ_AP026978.1 | 1 |
| Nocardia terpenica | Actinomycetota | Actinomycetes | NZ_CP023778.1 | 2 |
| Nocardia wallacei | Actinomycetota | Actinomycetes | NZ_AP023396.1 | 2 |
| Nocardia yamanashiensis | Actinomycetota | Actinomycetes | NZ_CP089226.1 | 3 |
| Nocardia yunnanensis | Actinomycetota | Actinomycetes | NZ_CP032568.1 | 3 |
| Nocardioides anomalus | Actinomycetota | Actinomycetes | NZ_CP049257.1 | 1 |
| Nocardioides aquaticus | Actinomycetota | Actinomycetes | NZ_CP075371.1 | 1 |
| Nocardioides baekrokdamisoli | Actinomycetota | Actinomycetes | NZ_AP019307.1 | 1 |
| Nocardioides bizhenqiangii | Actinomycetota | Actinomycetes | NZ_CP141059.1 | 2 |
| Nocardioides campestrisoli | Actinomycetota | Actinomycetes | NZ_CP061768.1 | 1 |
| Nocardioides coralli | Actinomycetota | Actinomycetes | NZ_CP082273.1 | 1 |
| Nocardioides cynanchi | Actinomycetota | Actinomycetes | NZ_CP044344.1 | 1 |
| Nocardioides dokdonensis FR1436 | Actinomycetota | Actinomycetes | NZ_CP015079.1 | 1 |
| Nocardioides dongkuii | Actinomycetota | Actinomycetes | NZ_CP059903.1 | 1 |
| Nocardioides euryhalodurans | Actinomycetota | Actinomycetes | NZ_CP038267.1 | 1 |
| Nocardioides faecalis | Actinomycetota | Actinomycetes | NZ_CP074406.1 | 1 |
| Nocardioides humi | Actinomycetota | Actinomycetes | NZ_CP041146.1 | 2 |
| Nocardioides marmotae | Actinomycetota | Actinomycetes | NZ_CP053660.1 | 1 |
| Nocardioides mesophilus | Actinomycetota | Actinomycetes | NZ_CP060713.1 | 1 |
| Nocardioides ochotonae | Actinomycetota | Actinomycetes | NZ_CP061769.1 | 1 |

| | | | | |
|---|---|---|---|---|
| Nocardioides okcheonensis | Actinomycetota | Actinomycetes | NZ_CP087710.1 | 1 |
| Nocardioides panacis | Actinomycetota | Actinomycetes | NZ_CP077062.1 | 1 |
| Nocardioides pantholopis | Actinomycetota | Actinomycetes | NZ_CP033324.1 | 1 |
| Nocardioides renjunii | Actinomycetota | Actinomycetes | NZ_CP141058.1 | 1 |
| Nocardioides rotundus | Actinomycetota | Actinomycetes | NZ_CP082922.1 | 1 |
| Nocardioides sambongensis | Actinomycetota | Actinomycetes | NZ_CP041091.1 | 1 |
| Nocardioides scoriae | Actinomycetota | Actinomycetes | NZ_LT629757.1 | 1 |
| Nocardioides seonyuensis | Actinomycetota | Actinomycetes | NZ_CP038436.1 | 1 |
| Nocardioides yefusunii | Actinomycetota | Actinomycetes | NZ_CP034929.1 | 1 |
| Nocardiopsis akebiae | Actinomycetota | Actinomycetes | NZ_CP074132.1 | 1 |
| Nocardiopsis changdeensis | Actinomycetota | Actinomycetes | NZ_CP074133.1 | 1 |
| Nocardiopsis dassonvillei | Actinomycetota | Actinomycetes | NZ_LR134501.1 | 1 |
| Nocardiopsis exhalans | Actinomycetota | Actinomycetes | NZ_CP099837.1 | 1 |
| Nocardiopsis gilva YIM 90087 | Actinomycetota | Actinomycetes | NZ_CP022753.1 | 1 |
| Nonomuraea coxensis DSM 45129 | Actinomycetota | Actinomycetes | NZ_CP068985.1 | 2 |
| Nonomuraea fastidiosa | Actinomycetota | Actinomycetes | NZ_CP170570.1 | 2 |
| Nonomuraea fuscirosea | Actinomycetota | Actinomycetes | NZ_CP109074.1 | 2 |
| Nonomuraea gerenzanensis | Actinomycetota | Actinomycetes | NZ_CP084058.1 | 2 |
| Nonomuraea glycinis | Actinomycetota | Actinomycetes | NZ_CP109166.1 | 2 |
| Nonomuraea phyllanthi | Actinomycetota | Actinomycetes | NZ_CP045572.1 | 1 |
| Occultella kanbiaonis | Actinomycetota | Actinomycetes | NZ_CP046175.1 | 1 |
| Ornithinimicrobium avium | Actinomycetota | Actinomycetes | NZ_CP031229.1 | 1 |
| Ornithinimicrobium ciconiae | Actinomycetota | Actinomycetes | NZ_CP041616.1 | 1 |
| Ornithinimicrobium cryptoxanthini | Actinomycetota | Actinomycetes | NZ_CP099490.1 | 1 |
| Ornithinimicrobium faecis | Actinomycetota | Actinomycetes | NZ_CP099489.1 | 2 |
| Ornithinimicrobium flavum | Actinomycetota | Actinomycetes | NZ_CP038213.1 | 1 |
| Ornithinimicrobium pratense | Actinomycetota | Actinomycetes | NZ_CP044427.1 | 1 |
| Ornithinimicrobium sufpigmenti | Actinomycetota | Actinomycetes | NZ_CP036403.1 | 1 |
| Oryzobacter telluris | Actinomycetota | Actinomycetes | NZ_CP170723.1 | 1 |
| Oryzobacter terrae | Actinomycetota | Actinomycetes | NZ_CP170578.1 | 1 |
| Paenarthrobacter aurescens | Actinomycetota | Actinomycetes | NZ_CP157456.1 | 1 |
| Paenarthrobacter nicotinovorans | Actinomycetota | Actinomycetes | NZ_CP089293.1 | 1 |

| | | | | |
|---|---|---|---|---|
| Paenarthrobacter ureafaciens | Actinomycetota | Actinomycetes | NZ_CP014574.1 | 1 |
| Paramicrobacterium chengjingii | Actinomycetota | Actinomycetes | NZ_CP061169.1 | 1 |
| Paramicrobacterium fandaimingii | Actinomycetota | Actinomycetes | NZ_CP061170.1 | 1 |
| Paraoerskovia marina | Actinomycetota | Actinomycetes | NZ_LT629776.1 | 1 |
| Paraoerskovia sediminicola | Actinomycetota | Actinomycetes | NZ_AP027729.1 | 1 |
| Parascardovia denticolens DSM 10105 = JCM 12538 | Actinomycetota | Actinomycetes | NZ_AP012333.1 | 1 |
| Pauljensenia hongkongensis | Actinomycetota | Actinomycetes | NZ_CP017298.1 | 1 |
| Pedococcus dokdonensis | Actinomycetota | Actinomycetes | NZ_LT629711.1 | 1 |
| Pengzhenrongella phosphoraccumulans | Actinomycetota | Actinomycetes | NZ_CP144210.1 | 1 |
| Pengzhenrongella sicca | Actinomycetota | Actinomycetes | NZ_CP071868.1 | 1 |
| Peterkaempfera bronchialis | Actinomycetota | Actinomycetes | NZ_CP031264.1 | 1 |
| Phycicoccus jejuensis | Actinomycetota | Actinomycetes | NZ_CP085209.1 | 1 |
| Phytohabitans suffuscus | Actinomycetota | Actinomycetes | NZ_AP022871.1 | 2 |
| Pimelobacter simplex | Actinomycetota | Actinomycetes | NZ_CP009896.1 | 2 |
| Polymorphospora rubra | Actinomycetota | Actinomycetes | NZ_AP023359.1 | 2 |
| Pontimonas salivibrio | Actinomycetota | Actinomycetes | NZ_CP026923.1 | 1 |
| Prauserella marina | Actinomycetota | Actinomycetes | NZ_CP016353.1 | 1 |
| Prescottella equi | Actinomycetota | Actinomycetes | NZ_CP027793.1 | 1 |
| Prescottella soli | Actinomycetota | Actinomycetes | NZ_CP157276.1 | 1 |
| Propionibacterium acidifaciens | Actinomycetota | Actinomycetes | NZ_CP033719.1 | 1 |
| Propionibacterium australiense | Actinomycetota | Actinomycetes | NZ_LR134442.1 | 1 |
| Propionibacterium freudenreichii | Actinomycetota | Actinomycetes | NZ_LT593929.1 | 1 |
| Propioniciclava coleopterorum | Actinomycetota | Actinomycetes | NZ_CP049865.1 | 1 |
| Propioniciclava soli | Actinomycetota | Actinomycetes | NZ_CP115965.1 | 1 |
| Protaetiibacter intestinalis | Actinomycetota | Actinomycetes | NZ_CP032630.1 | 1 |
| Protaetiibacter larvae | Actinomycetota | Actinomycetes | NZ_CP043504.1 | 1 |
| Pseudarthrobacter chlorophenolicus A6 | Actinomycetota | Actinomycetes | NC_011886.1 | 1 |
| Pseudarthrobacter equi | Actinomycetota | Actinomycetes | NZ_LT629779.1 | 1 |
| Pseudarthrobacter quantipunctorum | Actinomycetota | Actinomycetes | NZ_CP148033.1 | 1 |
| Pseudofrankia inefficax | Actinomycetota | Actinomycetes | NC_014666.1 | 2 |
| Pseudonocardia abyssalis | Actinomycetota | Actinomycetes | NZ_JADQDK01000 | 1 |
| Pseudonocardia autotrophica | Actinomycetota | Actinomycetes | NZ_AP018920.1 | 1 |

| | | | | |
|---|---|---|---|---|
| Pseudonocardia broussonetiae | Actinomycetota | Actinomycetes | NZ_CP053564.1 | 1 |
| Pseudonocardia dioxanivorans CB1190 | Actinomycetota | Actinomycetes | NC_015312.1 | 1 |
| Pseudonocardia petroleophila | Actinomycetota | Actinomycetes | NZ_CP060131.1 | 1 |
| Psychromicrobium lacuslunae | Actinomycetota | Actinomycetes | NZ_CP011005.1 | 1 |
| Rathayibacter festucae DSM 15932 | Actinomycetota | Actinomycetes | NZ_CP028137.1 | 1 |
| Rathayibacter iranicus | Actinomycetota | Actinomycetes | NZ_CP028130.1 | 1 |
| Rathayibacter rathayi | Actinomycetota | Actinomycetes | NZ_CP028129.1 | 1 |
| Rathayibacter soli | Actinomycetota | Actinomycetes | NZ_CP133020.1 | 1 |
| Rathayibacter tanaceti | Actinomycetota | Actinomycetes | NZ_CP047186.1 | 1 |
| Rathayibacter toxicus | Actinomycetota | Actinomycetes | NZ_CP061742.1 | 1 |
| Rathayibacter tritici | Actinomycetota | Actinomycetes | NZ_CP015515.1 | 1 |
| Renibacterium salmoninarum ATCC 33209 | Actinomycetota | Actinomycetes | NC_010168.1 | 1 |
| Rhodococcus aetherivorans | Actinomycetota | Actinomycetes | NZ_CP106982.1 | 1 |
| Rhodococcus antarcticus | Actinomycetota | Actinomycetes | NZ_CP110615.1 | 1 |
| Rhodococcus coprophilus | Actinomycetota | Actinomycetes | NZ_LS483468.1 | 1 |
| Rhodococcus erythropolis R138 | Actinomycetota | Actinomycetes | NZ_CP007255.1 | 1 |
| Rhodococcus globerulus | Actinomycetota | Actinomycetes | NZ_CP079698.1 | 1 |
| Rhodococcus indonesiensis | Actinomycetota | Actinomycetes | NZ_CP180630.1 | 1 |
| Rhodococcus koreensis | Actinomycetota | Actinomycetes | NZ_CP070609.1 | 1 |
| Rhodococcus opacus PD630 | Actinomycetota | Actinomycetes | NZ_CP080954.1 | 1 |
| Rhodococcus oxybenzonivorans | Actinomycetota | Actinomycetes | NZ_CP021354.1 | 1 |
| Rhodococcus pseudokoreensis | Actinomycetota | Actinomycetes | NZ_CP070619.1 | 1 |
| Rhodococcus pyridinivorans SB3094 | Actinomycetota | Actinomycetes | NC_023150.1 | 1 |
| Rhodococcus qingshengii | Actinomycetota | Actinomycetes | NZ_AP023172.1 | 1 |
| Rhodococcus rhodochrous | Actinomycetota | Actinomycetes | NZ_CP120356.1 | 1 |
| Rhodococcus ruber | Actinomycetota | Actinomycetes | NZ_CP044211.1 | 1 |
| Rhodococcus sacchari | Actinomycetota | Actinomycetes | NZ_CP107551.1 | 1 |
| Rhodococcus sovatensis | Actinomycetota | Actinomycetes | NZ_CP147846.1 | 1 |
| Rhodococcus triatomae | Actinomycetota | Actinomycetes | NZ_CP048813.1 | 1 |
| Rhodococcus wratislaviensis | Actinomycetota | Actinomycetes | NZ_CP150488.1 | 1 |
| Rhodoluna lacicola | Actinomycetota | Actinomycetes | NZ_CP007490.1 | 1 |
| Rhodoluna limnophila | Actinomycetota | Actinomycetes | NZ_CP040509.1 | 1 |

| | | | | |
|---|---|---|---|---|
| Rothia aeria | Actinomycetota | Actinomycetes | NZ_CP068102.1 | 1 |
| Rothia amarae | Actinomycetota | Actinomycetes | NZ_CP061538.1 | 1 |
| Rothia dentocariosa ATCC 17931 | Actinomycetota | Actinomycetes | NC_014643.1 | 1 |
| Rothia kristinae | Actinomycetota | Actinomycetes | NZ_CP065738.1 | 1 |
| Rothia terrae | Actinomycetota | Actinomycetes | NZ_CP061539.1 | 1 |
| Ruania alkalisoli | Actinomycetota | Actinomycetes | NZ_CP063169.1 | 1 |
| Ruania halotolerans | Actinomycetota | Actinomycetes | NZ_CP088017.1 | 1 |
| Ruania suaedae | Actinomycetota | Actinomycetes | NZ_CP088018.1 | 1 |
| Ruania zhangjianzhongii | Actinomycetota | Actinomycetes | NZ_CP042828.1 | 1 |
| Saccharomonospora azurea NA-128 | Actinomycetota | Actinomycetes | NZ_CM001466.1 | 1 |
| Saccharomonospora cyanea NA-134 | Actinomycetota | Actinomycetes | NZ_CM001440.1 | 1 |
| Saccharomonospora glauca K62 | Actinomycetota | Actinomycetes | NZ_CM001484.1 | 1 |
| Saccharomonospora marina XMU15 | Actinomycetota | Actinomycetes | NZ_CM001439.1 | 1 |
| Saccharomonospora viridis DSM 43017 | Actinomycetota | Actinomycetes | NC_013159.1 | 1 |
| Saccharopolyspora erythraea NRRL 2338 | Actinomycetota | Actinomycetes | NC_009142.1 | 1 |
| Saccharopolyspora gregorii | Actinomycetota | Actinomycetes | NZ_CP059556.1 | 1 |
| Saccharopolyspora pogona | Actinomycetota | Actinomycetes | NZ_CP031142.1 | 1 |
| Saccharopolyspora rosea | Actinomycetota | Actinomycetes | NZ_CP102826.1 | 1 |
| Saccharothrix espanaensis DSM 44229 | Actinomycetota | Actinomycetes | NC_019673.1 | 1 |
| Saccharothrix syringae | Actinomycetota | Actinomycetes | NZ_CP034550.1 | 1 |
| Salinispora tropica CNB-440 | Actinomycetota | Actinomycetes | NC_009380.1 | 1 |
| Sanguibacter biliveldensis | Actinomycetota | Actinomycetes | NZ_CP138359.1 | 1 |
| Sanguibacter keddieii DSM 10542 | Actinomycetota | Actinomycetes | NC_013521.1 | 1 |
| Saxibacter everestensis | Actinomycetota | Actinomycetes | NZ_CP090958.1 | 1 |
| Scardovia inopinata JCM 12537 | Actinomycetota | Actinomycetes | NZ_AP012334.1 | 1 |
| Schaalia cardiffensis F0333 | Actinomycetota | Actinomycetes | NZ_CP040505.1 | 1 |
| Schaalia meyeri | Actinomycetota | Actinomycetes | NZ_CP066065.1 | 1 |
| Schaalia odontolytica | Actinomycetota | Actinomycetes | NZ_CP133472.1 | 1 |
| Schaalia radingae | Actinomycetota | Actinomycetes | NZ_LT629792.1 | 1 |
| Scrofimicrobium appendicitidis | Actinomycetota | Actinomycetes | NZ_CP138335.1 | 1 |
| Segniliparus rotundus DSM 44985 | Actinomycetota | Actinomycetes | NC_014168.1 | 1 |
| Serinicoccus chungangensis | Actinomycetota | Actinomycetes | NZ_CP040887.1 | 1 |

| | | | | |
|---|---|---|---|---|
| Serinicoccus hydrothermalis | Actinomycetota | Actinomycetes | NZ_CP014989.1 | 1 |
| Sinomonas atrocyanea | Actinomycetota | Actinomycetes | NZ_CP014518.1 | 1 |
| Sinomonas cyclohexanicum | Actinomycetota | Actinomycetes | NZ_AP024525.1 | 1 |
| Sinomonas puerhi | Actinomycetota | Actinomycetes | NZ_CP163302.1 | 1 |
| Skermania piniformis | Actinomycetota | Actinomycetes | NZ_CP079105.1 | 1 |
| Solicola gregarius | Actinomycetota | Actinomycetes | NZ_CP094970.1 | 1 |
| Sphaerisporangium viridialbum | Actinomycetota | Actinomycetes | NZ_CP187232.1 | 2 |
| Spiractinospora alimapuensis | Actinomycetota | Actinomycetes | NZ_CP072467.1 | 1 |
| Stackebrandtia nassauensis DSM 44728 | Actinomycetota | Actinomycetes | NC_013947.1 | 1 |
| Streptantibioticus cattleyicolor NRRL 8057 = DSM 46488 | Actinomycetota | Actinomycetes | NC_016111.1 | 2 |
| Streptomonospora litoralis | Actinomycetota | Actinomycetes | NZ_CP036455.1 | 1 |
| Streptomyces achromogenes | Actinomycetota | Actinomycetes | NZ_CP108164.1 | 1 |
| Streptomyces akebiae | Actinomycetota | Actinomycetes | NZ_CP080647.1 | 2 |
| Streptomyces albidocamelliae | Actinomycetota | Actinomycetes | NZ_CP106795.1 | 2 |
| Streptomyces albidoflavus | Actinomycetota | Actinomycetes | NZ_CP108647.1 | 2 |
| Streptomyces alboniger | Actinomycetota | Actinomycetes | NZ_CP023695.1 | 2 |
| Streptomyces albus | Actinomycetota | Actinomycetes | NZ_CP103060.1 | 1 |
| Streptomyces alfalfae | Actinomycetota | Actinomycetes | NZ_CP060742.1 | 2 |
| Streptomyces althioticus | Actinomycetota | Actinomycetes | NZ_CP109234.1 | 2 |
| Streptomyces ambofaciens ATCC 23877 | Actinomycetota | Actinomycetes | NZ_CP012382.1 | 2 |
| Streptomyces angustmyceticus | Actinomycetota | Actinomycetes | NZ_CP082945.1 | 2 |
| Streptomyces anthocyanicus | Actinomycetota | Actinomycetes | NZ_CP109102.1 | 2 |
| Streptomyces antimycoticus | Actinomycetota | Actinomycetes | NZ_CP108856.1 | 2 |
| Streptomyces anulatus | Actinomycetota | Actinomycetes | NZ_CM003601.1 | 2 |
| Streptomyces aquilus | Actinomycetota | Actinomycetes | NZ_CP034463.1 | 3 |
| Streptomyces argyrophylli | Actinomycetota | Actinomycetes | NZ_CP053189.1 | 2 |
| Streptomyces armeniacus | Actinomycetota | Actinomycetes | NZ_CP031320.1 | 2 |
| Streptomyces atratus | Actinomycetota | Actinomycetes | NZ_CP027306.1 | 2 |
| Streptomyces aurantiacus | Actinomycetota | Actinomycetes | NZ_AP023440.1 | 2 |
| Streptomyces auratus AGR0001 | Actinomycetota | Actinomycetes | NZ_CP072931.1 | 2 |
| Streptomyces avermitilis MA-4680 = NBRC 14893 | Actinomycetota | Actinomycetes | NC_003155.5 | 2 |
| Streptomyces avidinii | Actinomycetota | Actinomycetes | NZ_CP108863.1 | 2 |

| | | | | |
|---|---|---|---|---|
| Streptomyces bathyalis | Actinomycetota | Actinomycetes | NZ_CP048882.1 | 2 |
| Streptomyces bobili | Actinomycetota | Actinomycetes | NZ_CP108038.1 | 2 |
| Streptomyces boninensis | Actinomycetota | Actinomycetes | NZ_CP183837.1 | 2 |
| Streptomyces broussonetiae | Actinomycetota | Actinomycetes | NZ_CP047020.1 | 4 |
| Streptomyces buecherae | Actinomycetota | Actinomycetes | NZ_CP060404.1 | 1 |
| Streptomyces cadmiisoli | Actinomycetota | Actinomycetes | NZ_CP030073.1 | 1 |
| Streptomyces californicus | Actinomycetota | Actinomycetes | NZ_CP070242.1 | 2 |
| Streptomyces calvus | Actinomycetota | Actinomycetes | NZ_CP022310.1 | 2 |
| Streptomyces camelliae | Actinomycetota | Actinomycetes | NZ_CP115300.1 | 2 |
| Streptomyces caniferus | Actinomycetota | Actinomycetes | NZ_CP108073.1 | 2 |
| Streptomyces caniscabiei | Actinomycetota | Actinomycetes | NZ_CP119182.1 | 2 |
| Streptomyces canus | Actinomycetota | Actinomycetes | NZ_CP108065.1 | 2 |
| Streptomyces capitiformicae | Actinomycetota | Actinomycetes | NZ_CP022161.1 | 2 |
| Streptomyces carpaticus | Actinomycetota | Actinomycetes | NZ_CP104005.1 | 2 |
| Streptomyces castrisilvae | Actinomycetota | Actinomycetes | NZ_CP120997.1 | 2 |
| Streptomyces cathayae | Actinomycetota | Actinomycetes | NZ_CP121682.1 | 1 |
| Streptomyces changanensis | Actinomycetota | Actinomycetes | NZ_CP102332.1 | 2 |
| Streptomyces chartreusis | Actinomycetota | Actinomycetes | NZ_CP023689.1 | 2 |
| Streptomyces chengbuensis | Actinomycetota | Actinomycetes | NZ_CP129137.1 | 2 |
| Streptomyces chromofuscus | Actinomycetota | Actinomycetes | NZ_CP063374.1 | 2 |
| Streptomyces cinereoruber | Actinomycetota | Actinomycetes | NZ_CP023693.1 | 1 |
| Streptomyces cinnabarinus | Actinomycetota | Actinomycetes | NZ_CP114413.1 | 3 |
| Streptomyces citrinus | Actinomycetota | Actinomycetes | NZ_CP146022.1 | 2 |
| Streptomyces clavifer | Actinomycetota | Actinomycetes | NZ_CP107793.1 | 2 |
| Streptomyces clavuligerus | Actinomycetota | Actinomycetes | NZ_CP027858.1 | 1 |
| Streptomyces coelicoflavus | Actinomycetota | Actinomycetes | NZ_CP108218.1 | 2 |
| Streptomyces collinus | Actinomycetota | Actinomycetes | NZ_CP133771.1 | 2 |
| Streptomyces cyaneochromogenes | Actinomycetota | Actinomycetes | NZ_CP034539.1 | 3 |
| Streptomyces cyaneofuscatus | Actinomycetota | Actinomycetes | NZ_CP137002.1 | 2 |
| Streptomyces cyanogenus | Actinomycetota | Actinomycetes | NZ_CP071839.1 | 2 |
| Streptomyces cynarae | Actinomycetota | Actinomycetes | NZ_CP106793.1 | 2 |
| Streptomyces dangxiongensis | Actinomycetota | Actinomycetes | NZ_CP033073.1 | 1 |

| | | | | |
|---|---|---|---|---|
| Streptomyces davaonensis JCM 4913 | Actinomycetota | Actinomycetes | NC_020504.1 | 3 |
| Streptomyces deccanensis | Actinomycetota | Actinomycetes | NZ_CP092431.1 | 2 |
| Streptomyces decoyicus | Actinomycetota | Actinomycetes | NZ_CP109339.1 | 2 |
| Streptomyces dengpaensis | Actinomycetota | Actinomycetes | NZ_CP026652.1 | 3 |
| Streptomyces durmitorensis | Actinomycetota | Actinomycetes | NZ_CP097289.1 | 2 |
| Streptomyces erythrochromogenes | Actinomycetota | Actinomycetes | NZ_CP108036.1 | 2 |
| Streptomyces europaeiscabiei | Actinomycetota | Actinomycetes | NZ_CP107841.1 | 2 |
| Streptomyces fagopyri | Actinomycetota | Actinomycetes | NZ_CP045643.1 | 2 |
| Streptomyces ferrugineus | Actinomycetota | Actinomycetes | NZ_CP063373.1 | 2 |
| Streptomyces ficellus | Actinomycetota | Actinomycetes | NZ_CP034279.1 | 2 |
| Streptomyces fodineus | Actinomycetota | Actinomycetes | NZ_CP017248.1 | 2 |
| Streptomyces formicae | Actinomycetota | Actinomycetes | NZ_CP071872.1 | 2 |
| Streptomyces fradiae ATCC 10745 = DSM 40063 | Actinomycetota | Actinomycetes | NZ_CP023696.1 | 2 |
| Streptomyces fungicidicus | Actinomycetota | Actinomycetes | NZ_CP086173.1 | 2 |
| Streptomyces gardneri | Actinomycetota | Actinomycetes | NZ_CP059991.1 | 1 |
| Streptomyces genisteinicus | Actinomycetota | Actinomycetes | NZ_CP060825.1 | 2 |
| Streptomyces gilvosporeus | Actinomycetota | Actinomycetes | NZ_CP020569.1 | 2 |
| Streptomyces globisporus C-1027 | Actinomycetota | Actinomycetes | NZ_CP013738.1 | 2 |
| Streptomyces glycanivorans | Actinomycetota | Actinomycetes | NZ_CP120983.1 | 2 |
| Streptomyces gobiensis | Actinomycetota | Actinomycetes | NZ_CP086120.1 | 3 |
| Streptomyces goshikiensis | Actinomycetota | Actinomycetes | NZ_CP115936.1 | 2 |
| Streptomyces gougerotii | Actinomycetota | Actinomycetes | NZ_CP108653.1 | 2 |
| Streptomyces graminofaciens | Actinomycetota | Actinomycetes | NZ_AP018448.1 | 3 |
| Streptomyces griseoaurantiacus | Actinomycetota | Actinomycetes | NZ_CP146837.1 | 2 |
| Streptomyces griseochromogenes | Actinomycetota | Actinomycetes | NZ_CP016279.1 | 2 |
| Streptomyces griseofuscus | Actinomycetota | Actinomycetes | NZ_CP051006.1 | 2 |
| Streptomyces griseorubiginosus | Actinomycetota | Actinomycetes | NZ_CP032427.1 | 2 |
| Streptomyces halobius | Actinomycetota | Actinomycetes | NZ_CP086322.1 | 2 |
| Streptomyces halstedii | Actinomycetota | Actinomycetes | NZ_CP109044.1 | 2 |
| Streptomyces harbinensis | Actinomycetota | Actinomycetes | NZ_CP054938.1 | 2 |
| Streptomyces hawaiiensis | Actinomycetota | Actinomycetes | NZ_CP021978.1 | 2 |
| Streptomyces hirsutus | Actinomycetota | Actinomycetes | NZ_CP109134.1 | 2 |

| | | | | |
|---|---|---|---|---|
| Streptomyces huasconensis | Actinomycetota | Actinomycetes | NZ_CP086119.1 | 2 |
| Streptomyces hygroscopicus subsp. hygroscopicus | Actinomycetota | Actinomycetes | NZ_CP146988.1 | 2 |
| Streptomyces iakyrus | Actinomycetota | Actinomycetes | NZ_CP143088.1 | 2 |
| Streptomyces incarnatus | Actinomycetota | Actinomycetes | NZ_CP011497.1 | 2 |
| Streptomyces inhibens | Actinomycetota | Actinomycetes | NZ_CP084681.1 | 2 |
| Streptomyces iranensis | Actinomycetota | Actinomycetes | NZ_CP136563.1 | 2 |
| Streptomyces janthinus | Actinomycetota | Actinomycetes | NZ_CP134213.1 | 2 |
| Streptomyces jietaisiensis | Actinomycetota | Actinomycetes | NZ_CP108326.1 | 2 |
| Streptomyces kanamyceticus | Actinomycetota | Actinomycetes | NZ_CP023699.1 | 2 |
| Streptomyces koelreuteriae | Actinomycetota | Actinomycetes | NZ_CP075896.1 | 2 |
| Streptomyces koyangensis | Actinomycetota | Actinomycetes | NZ_CP031742.1 | 1 |
| Streptomyces laculatispora | Actinomycetota | Actinomycetes | NZ_CP120992.1 | 2 |
| Streptomyces lavendulae subsp. lavendulae | Actinomycetota | Actinomycetes | NZ_CP024985.1 | 2 |
| Streptomyces leeuwenhoekii | Actinomycetota | Actinomycetes | NZ_LN831790.1 | 2 |
| Streptomyces liangshanensis | Actinomycetota | Actinomycetes | NZ_CP050177.1 | 2 |
| Streptomyces libani subsp. libani | Actinomycetota | Actinomycetes | NZ_CP114202.1 | 2 |
| Streptomyces lienomycini | Actinomycetota | Actinomycetes | NZ_CP116257.1 | 2 |
| Streptomyces liliifuscus | Actinomycetota | Actinomycetes | NZ_CP066831.1 | 2 |
| Streptomyces lincolnensis | Actinomycetota | Actinomycetes | NZ_CP016438.1 | 3 |
| Streptomyces longhuiensis | Actinomycetota | Actinomycetes | NZ_CP085173.1 | 2 |
| Streptomyces lunaelactis | Actinomycetota | Actinomycetes | NZ_CP026304.1 | 2 |
| Streptomyces luomodiensis | Actinomycetota | Actinomycetes | NZ_CP117522.1 | 2 |
| Streptomyces luteireticuli | Actinomycetota | Actinomycetes | NZ_CP167927.1 | 1 |
| Streptomyces lydicus | Actinomycetota | Actinomycetes | NZ_CP019457.1 | 2 |
| Streptomyces malaysiensis | Actinomycetota | Actinomycetes | NZ_CP023992.1 | 2 |
| Streptomyces manipurensis | Actinomycetota | Actinomycetes | NZ_CP186950.1 | 2 |
| Streptomyces marincola | Actinomycetota | Actinomycetes | NZ_CP084541.1 | 1 |
| Streptomyces microflavus | Actinomycetota | Actinomycetes | NZ_CP054926.1 | 2 |
| Streptomyces milbemycinicus | Actinomycetota | Actinomycetes | NZ_CP109672.1 | 2 |
| Streptomyces mirabilis | Actinomycetota | Actinomycetes | NZ_CP107957.1 | 3 |
| Streptomyces mobaraensis | Actinomycetota | Actinomycetes | NZ_CP083590.1 | 1 |
| Streptomyces murinus | Actinomycetota | Actinomycetes | NZ_CP046623.1 | 2 |

| | | | | |
|---|---|---|---|---|
| Streptomyces mutabilis | Actinomycetota | Actinomycetes | NZ_CP176874.1 | 2 |
| Streptomyces netropsis | Actinomycetota | Actinomycetes | NZ_CP147867.1 | 1 |
| Streptomyces nigra | Actinomycetota | Actinomycetes | NZ_CP029043.1 | 2 |
| Streptomyces nigrescens | Actinomycetota | Actinomycetes | NZ_CP114203.1 | 2 |
| Streptomyces nitrosporeus | Actinomycetota | Actinomycetes | NZ_CP023702.1 | 2 |
| Streptomyces niveus | Actinomycetota | Actinomycetes | NZ_CP109483.1 | 2 |
| Streptomyces nodosus | Actinomycetota | Actinomycetes | NZ_CP023747.1 | 2 |
| Streptomyces nojiriensis | Actinomycetota | Actinomycetes | NZ_CP071139.1 | 2 |
| Streptomyces noursei | Actinomycetota | Actinomycetes | NZ_CP070326.1 | 2 |
| Streptomyces olivaceus | Actinomycetota | Actinomycetes | NZ_CP043317.1 | 2 |
| Streptomyces paludis | Actinomycetota | Actinomycetes | NZ_CP031194.1 | 2 |
| Streptomyces parvulus | Actinomycetota | Actinomycetes | NZ_CP015866.1 | 2 |
| Streptomyces phaeochromogenes | Actinomycetota | Actinomycetes | NZ_CP108382.1 | 2 |
| Streptomyces phaeolivaceus | Actinomycetota | Actinomycetes | NZ_CP045096.1 | 2 |
| Streptomyces phaeoluteigriseus | Actinomycetota | Actinomycetes | NZ_CP099468.1 | 2 |
| Streptomyces phytohabitans | Actinomycetota | Actinomycetes | NZ_CP160458.1 | 1 |
| Streptomyces platensis | Actinomycetota | Actinomycetes | NZ_CP023691.1 | 2 |
| Streptomyces pluripotens | Actinomycetota | Actinomycetes | NZ_CP021080.1 | 2 |
| Streptomyces poriferorum | Actinomycetota | Actinomycetes | NZ_CP120990.1 | 2 |
| Streptomyces prasinus | Actinomycetota | Actinomycetes | NZ_CP023697.1 | 2 |
| Streptomyces pratensis | Actinomycetota | Actinomycetes | NZ_CP051486.1 | 2 |
| Streptomyces pristinaespiralis | Actinomycetota | Actinomycetes | NZ_CP059696.1 | 2 |
| Streptomyces profundus | Actinomycetota | Actinomycetes | NZ_CP082362.1 | 2 |
| Streptomyces prunicolor | Actinomycetota | Actinomycetes | NZ_CP108676.1 | 3 |
| Streptomyces pseudovenezuelae | Actinomycetota | Actinomycetes | NZ_CP107755.1 | 2 |
| Streptomyces purpurascens | Actinomycetota | Actinomycetes | NZ_CP108341.1 | 2 |
| Streptomyces qaidamensis | Actinomycetota | Actinomycetes | NZ_CP015098.1 | 2 |
| Streptomyces qinzhouensis | Actinomycetota | Actinomycetes | NZ_CP042266.1 | 2 |
| Streptomyces rapamycinicus NRRL 5491 | Actinomycetota | Actinomycetes | NZ_CP085193.1 | 2 |
| Streptomyces rectiverticillatus | Actinomycetota | Actinomycetes | NZ_CP040752.1 | 1 |
| Streptomyces rimosus subsp. rimosus ATCC 10970 | Actinomycetota | Actinomycetes | NZ_CP048261.1 | 2 |
| Streptomyces rochei | Actinomycetota | Actinomycetes | NZ_CP121271.1 | 2 |

| | | | | |
|---|---|---|---|---|
| Streptomyces roseirectus | Actinomycetota | Actinomycetes | NZ_CP060828.1 | 3 |
| Streptomyces roseochromogenus subsp. oscitans DS 12.9 | Actinomycetota | Actinomycetes | NZ_CM002285.1 | 2 |
| Streptomyces rubrogriseus | Actinomycetota | Actinomycetes | NZ_CP116256.1 | 2 |
| Streptomyces rutgersensis | Actinomycetota | Actinomycetes | NZ_CP045704.1 | 2 |
| Streptomyces scabiei 87.22 | Actinomycetota | Actinomycetes | NC_013929.1 | 2 |
| Streptomyces sclerotialus | Actinomycetota | Actinomycetes | NZ_JBFOHP01000 | 2 |
| Streptomyces scopuliridis | Actinomycetota | Actinomycetes | NZ_CP109091.1 | 2 |
| Streptomyces seoulensis | Actinomycetota | Actinomycetes | NZ_CP032229.1 | 2 |
| Streptomyces sirii | Actinomycetota | Actinomycetes | NZ_CP147982.1 | 2 |
| Streptomyces solicamelliae | Actinomycetota | Actinomycetes | NZ_CP160404.1 | 2 |
| Streptomyces solicathayae | Actinomycetota | Actinomycetes | NZ_CP137573.1 | 2 |
| Streptomyces solisilvae | Actinomycetota | Actinomycetes | NZ_CP065050.1 | 2 |
| Streptomyces spectabilis | Actinomycetota | Actinomycetes | NZ_CP023690.1 | 2 |
| Streptomyces spinosirectus | Actinomycetota | Actinomycetes | NZ_CP090447.1 | 3 |
| Streptomyces spongiicola | Actinomycetota | Actinomycetes | NZ_CP029254.1 | 2 |
| Streptomyces subrutilus | Actinomycetota | Actinomycetes | NZ_CP023701.1 | 2 |
| Streptomyces sudanensis | Actinomycetota | Actinomycetes | NZ_CP095474.1 | 1 |
| Streptomyces sviceus ATCC 29083 | Actinomycetota | Actinomycetes | NZ_CM000951.1 | 2 |
| Streptomyces tabacisoli | Actinomycetota | Actinomycetes | NZ_CP159534.1 | 2 |
| Streptomyces tanashiensis | Actinomycetota | Actinomycetes | NZ_CP084204.1 | 1 |
| Streptomyces tauricus | Actinomycetota | Actinomycetes | NZ_CP108133.1 | 3 |
| Streptomyces thermocarboxydus | Actinomycetota | Actinomycetes | NZ_CP058273.1 | 2 |
| Streptomyces thermolineatus | Actinomycetota | Actinomycetes | NZ_CP173276.1 | 1 |
| Streptomyces thermoviolaceus | Actinomycetota | Actinomycetes | NZ_CP109280.1 | 1 |
| Streptomyces tirandamycinicus | Actinomycetota | Actinomycetes | NZ_CP029188.1 | 2 |
| Streptomyces tritici | Actinomycetota | Actinomycetes | NZ_CP183183.1 | 2 |
| Streptomyces tsukubensis | Actinomycetota | Actinomycetes | NZ_CP020700.1 | 2 |
| Streptomyces tubbatahanensis | Actinomycetota | Actinomycetes | NZ_CP093846.1 | 1 |
| Streptomyces tubercidicus | Actinomycetota | Actinomycetes | NZ_CP114205.1 | 2 |
| Streptomyces tuirus | Actinomycetota | Actinomycetes | NZ_AP023439.1 | 2 |
| Streptomyces uncialis | Actinomycetota | Actinomycetes | NZ_CP109583.1 | 2 |
| Streptomyces venezuelae ATCC 10712 | Actinomycetota | Actinomycetes | NZ_CP029197.1 | 1 |

| | | | | |
|---|---|---|---|---|
| Streptomyces vietnamensis | Actinomycetota | Actinomycetes | NZ_CP010407.1 | 1 |
| Streptomyces vilmorinianum | Actinomycetota | Actinomycetes | NZ_CP040244.1 | 2 |
| Streptomyces vinaceus | Actinomycetota | Actinomycetes | NZ_CP023692.1 | 2 |
| Streptomyces violaceoruber | Actinomycetota | Actinomycetes | NZ_CP137734.1 | 2 |
| Streptomyces violaceus | Actinomycetota | Actinomycetes | NZ_CP107911.1 | 2 |
| Streptomyces viridosporus T7A | Actinomycetota | Actinomycetes | NZ_CP023700.1 | 2 |
| Streptomyces xanthii | Actinomycetota | Actinomycetes | NZ_CP061281.1 | 2 |
| Streptomyces xanthophaeus | Actinomycetota | Actinomycetes | NZ_CP116355.1 | 2 |
| Streptomyces xiamenensis | Actinomycetota | Actinomycetes | NZ_CP009922.3 | 2 |
| Streptomyces xinghaiensis S187 | Actinomycetota | Actinomycetes | NZ_CP023202.1 | 2 |
| Streptomyces yangpuensis | Actinomycetota | Actinomycetes | NZ_CP102514.1 | 2 |
| Streptomyces yatensis | Actinomycetota | Actinomycetes | NZ_CP072941.1 | 2 |
| Streptomyces yunnanensis | Actinomycetota | Actinomycetes | NZ_CP095749.1 | 2 |
| Streptomyces zaomyceticus | Actinomycetota | Actinomycetes | NZ_CP107933.1 | 1 |
| Streptosporangium roseum DSM 43021 | Actinomycetota | Actinomycetes | NC_013595.1 | 1 |
| Streptoverticillium reticulum | Actinomycetota | Actinomycetes | NZ_CP179755.1 | 1 |
| Subtercola endophyticus | Actinomycetota | Actinomycetes | NZ_CP087997.1 | 1 |
| Tessaracoccus aquimaris | Actinomycetota | Actinomycetes | NZ_CP019606.1 | 1 |
| Tessaracoccus flavescens | Actinomycetota | Actinomycetes | NZ_CP019607.1 | 1 |
| Tessaracoccus flavus | Actinomycetota | Actinomycetes | NZ_CP019605.1 | 1 |
| Tessaracoccus lacteus | Actinomycetota | Actinomycetes | NZ_CP123967.1 | 1 |
| Tessaracoccus palaemonis | Actinomycetota | Actinomycetes | NZ_CP079216.1 | 1 |
| Tessaracoccus timonensis | Actinomycetota | Actinomycetes | NZ_LT996886.1 | 1 |
| Thermobifida alba | Actinomycetota | Actinomycetes | NZ_CP051627.1 | 1 |
| Thermobifida fusca YX | Actinomycetota | Actinomycetes | NC_007333.1 | 1 |
| Thermobifida halotolerans | Actinomycetota | Actinomycetes | NZ_CP063196.1 | 1 |
| Thermobispora bispora DSM 43833 | Actinomycetota | Actinomycetes | NC_014165.1 | 1 |
| Thermomonospora amylolytica | Actinomycetota | Actinomycetes | NZ_CP032402.1 | 1 |
| Thermomonospora curvata DSM 43183 | Actinomycetota | Actinomycetes | NC_013510.1 | 1 |
| Tomitella fengzijianii | Actinomycetota | Actinomycetes | NZ_CP041765.1 | 1 |
| Tomitella gaofuii | Actinomycetota | Actinomycetes | NZ_CP059900.1 | 1 |
| Tropheryma whipplei TW08/27 | Actinomycetota | Actinomycetes | NC_004551.1 | 1 |

| | | | | |
|---|---|---|---|---|
| Trueperella bernardiae | Actinomycetota | Actinomycetes | NZ_CP127099.1 | 1 |
| Trueperella bialowiezensis | Actinomycetota | Actinomycetes | NZ_LR134476.1 | 1 |
| Trueperella pecoris | Actinomycetota | Actinomycetes | NZ_CP053291.1 | 1 |
| Trueperella pyogenes | Actinomycetota | Actinomycetes | NZ_CP012649.1 | 1 |
| Tsukamurella paurometabola DSM 20162 | Actinomycetota | Actinomycetes | NC_014158.1 | 1 |
| Vaginimicrobium propionicum | Actinomycetota | Actinomycetes | NZ_LT706985.1 | 1 |
| Varibaculum prostatecancerukia | Actinomycetota | Actinomycetes | NZ_OW968402.1 | 1 |
| Williamsia herbipolensis | Actinomycetota | Actinomycetes | NZ_CP108021.1 | 1 |
| Xylanimonas allomyrinae | Actinomycetota | Actinomycetes | NZ_CP035495.1 | 1 |
| Xylanimonas cellulosilytica DSM 15894 | Actinomycetota | Actinomycetes | NC_013530.1 | 1 |
| Xylanimonas protaetiae | Actinomycetota | Actinomycetes | NZ_CP035493.1 | 1 |
| Adlercreutzia equolifaciens DSM 19450 | Actinomycetota | Coriobacteriia | NC_022567.1 | 1 |
| Adlercreutzia hattorii | Actinomycetota | Coriobacteriia | NZ_AP022829.1 | 1 |
| Arabiibacter massiliensis | Actinomycetota | Coriobacteriia | NZ_LR698996.1 | 1 |
| Berryella intestinalis | Actinomycetota | Coriobacteriia | NZ_CP009302.1 | 1 |
| Berryella wangjianweii | Actinomycetota | Coriobacteriia | NZ_CP053716.1 | 1 |
| Collinsella aerofaciens ATCC 25986 | Actinomycetota | Coriobacteriia | NZ_CP048433.1 | 1 |
| Collinsella stercoris | Actinomycetota | Coriobacteriia | NZ_CP102276.1 | 1 |
| Coriobacterium glomerans PW2 | Actinomycetota | Coriobacteriia | NC_015389.1 | 1 |
| Cryptobacterium curtum DSM 15641 | Actinomycetota | Coriobacteriia | NC_013170.1 | 1 |
| Denitrobacterium detoxificans | Actinomycetota | Coriobacteriia | NZ_CP011402.1 | 1 |
| Eggerthella guodeyinii | Actinomycetota | Coriobacteriia | NZ_CP063310.1 | 1 |
| Eggerthella lenta | Actinomycetota | Coriobacteriia | NZ_CP089331.1 | 1 |
| Fannyhessea vaginae | Actinomycetota | Coriobacteriia | NZ_CP065631.1 | 1 |
| Lancefieldella parvula DSM 20469 | Actinomycetota | Coriobacteriia | NC_013203.1 | 1 |
| Leptogranulimonas caecicola | Actinomycetota | Coriobacteriia | NZ_AP025285.1 | 1 |
| Olsenella timonensis | Actinomycetota | Coriobacteriia | NZ_LT635455.1 | 1 |
| Olsenella uli DSM 7084 | Actinomycetota | Coriobacteriia | NC_014363.1 | 1 |
| Parafannyhessea umbonata | Actinomycetota | Coriobacteriia | NZ_LT629759.1 | 1 |
| Parolsenella catena | Actinomycetota | Coriobacteriia | NZ_AP019367.1 | 1 |
| Parolsenella massiliensis | Actinomycetota | Coriobacteriia | NZ_LT671675.1 | 1 |
| Phoenicibacter congonensis | Actinomycetota | Coriobacteriia | NZ_LT821227.1 | 1 |

| | | | | |
|---|---|---|---|---|
| Raoultibacter timonensis | Actinomycetota | Coriobacteriia | NZ_AP025564.1 | 1 |
| Slackia heliotrinireducens | Actinomycetota | Coriobacteriia | NZ_LR134379.1 | 1 |
| Thermophilibacter immobilis | Actinomycetota | Coriobacteriia | NZ_CP063767.1 | 1 |
| Xiamenia xianingshaonis | Actinomycetota | Coriobacteriia | NZ_CP072829.1 | 1 |
| Egibacter rhizosphaerae | Actinomycetota | Nitriliruptoria | NZ_CP036402.1 | 2 |
| Egicoccus halophilus | Actinomycetota | Nitriliruptoria | NZ_CP036250.1 | 2 |
| Euzebya pacifica | Actinomycetota | Nitriliruptoria | NZ_CP031165.1 | 2 |
| Rubrobacter indicoceani | Actinomycetota | Rubrobacteria | NZ_CP031115.1 | 2 |
| Rubrobacter xylanophilus DSM 9941 | Actinomycetota | Rubrobacteria | NC_008148.1 | 2 |
| Baekduia alba | Actinomycetota | Thermoleophilia | NZ_CP114013.1 | 1 |
| Baekduia soli | Actinomycetota | Thermoleophilia | NZ_CP042430.1 | 1 |
| Capillimicrobium parvum | Actinomycetota | Thermoleophilia | NZ_CP087164.1 | 1 |
| Conexibacter woesei DSM 14684 | Actinomycetota | Thermoleophilia | NC_013739.1 | 1 |
| Miltoncostaea marina | Actinomycetota | Thermoleophilia | NZ_CP064655.1 | 1 |
| Miltoncostaea oceani | Actinomycetota | Thermoleophilia | NZ_CP064356.1 | 1 |
| Paraconexibacter antarcticus | Actinomycetota | Thermoleophilia | NZ_CP098502.1 | 1 |
| Svornostia abyssi | Actinomycetota | Thermoleophilia | NZ_CP088295.1 | 1 |
| Aquifex aeolicus VF5 | Aquificota | Aquificia | NC_000918.1 | 2 |
| Desulfurobacterium thermolithotrophum DSM 11699 | Aquificota | Aquificia | NC_015185.1 | 2 |
| Hydrogenobacter thermophilus TK-6 | Aquificota | Aquificia | NC_017161.1 | 2 |
| Persephonella marina EX-H1 | Aquificota | Aquificia | NC_012440.1 | 2 |
| Sulfurihydrogenibium azorense Az-Fu1 | Aquificota | Aquificia | NC_012438.1 | 1 |
| Thermocrinis albus DSM 14484 | Aquificota | Aquificia | NC_013894.1 | 2 |
| Thermocrinis minervae | Aquificota | Aquificia | NZ_LT670846.1 | 2 |
| Thermocrinis ruber | Aquificota | Aquificia | NZ_CP007028.1 | 2 |
| Thermovibrio ammonificans HB-1 | Aquificota | Aquificia | NC_014926.1 | 2 |
| Capsulimonas corticalis | Armatimonadota | Armatimonadia | NZ_AP025739.1 | 2 |
| Chthonomonas calidirosea T49 | Armatimonadota | Chthonomonadia | NC_021487.1 | 2 |
| Fimbriimonas ginsengisoli Gsoil 348 | Armatimonadota | Fimbriimonadia | NZ_CP007139.1 | 1 |
| Atribacter laminatus | Atribacterota | Atribacteria | NZ_CP065383.1 | 2 |
| Atrimonas thermophila | Atribacterota | Atribacteria | NZ_AP028803.1 | 2 |
| Thermatribacter velox | Atribacterota | Atribacteria | NZ_CP121689.1 | 2 |

| | | | | |
|---|---|---|---|---|
| [Bacillus] selenitireducens MLS10 | Bacillota | Bacilli | NC_014219.1 | 1 |
| Abiotrophia defectiva | Bacillota | Bacilli | NZ_CP053988.1 | 1 |
| Abyssicoccus albus | Bacillota | Bacilli | NZ_CP019573.1 | 1 |
| Acetilactobacillus jinshanensis | Bacillota | Bacilli | NZ_CP034726.1 | 1 |
| Aciduricibacillus chroicocephali | Bacillota | Bacilli | NZ_CP129113.1 | 1 |
| Aeribacillus composti | Bacillota | Bacilli | NZ_CP134501.1 | 1 |
| Aerococcus christensenii | Bacillota | Bacilli | NZ_CP014159.1 | 1 |
| Aerococcus sanguinicola | Bacillota | Bacilli | NZ_CP014160.1 | 1 |
| Aerococcus tenax | Bacillota | Bacilli | NZ_CP145134.1 | 1 |
| Aerococcus urinae | Bacillota | Bacilli | NZ_CP014161.1 | 1 |
| Aerococcus urinaeequi | Bacillota | Bacilli | NZ_CP014162.1 | 1 |
| Aerococcus urinaehominis | Bacillota | Bacilli | NZ_CP014163.1 | 1 |
| Aerococcus viridans | Bacillota | Bacilli | NZ_CP014164.1 | 1 |
| Agrilactobacillus fermenti | Bacillota | Bacilli | NZ_CP181347.1 | 1 |
| Alicyclobacillus acidocaldarius subsp. acidocaldarius DSM | Bacillota | Bacilli | NC_013205.1 | 1 |
| Alicyclobacillus acidoterrestris | Bacillota | Bacilli | NZ_CP080467.1 | 1 |
| Alicyclobacillus cycloheptanicus | Bacillota | Bacilli | NZ_CP067097.1 | 1 |
| Alicyclobacillus dauci | Bacillota | Bacilli | NZ_CP104064.1 | 1 |
| Alicyclobacillus fastidiosus | Bacillota | Bacilli | NZ_CP104067.1 | 1 |
| Alicyclobacillus mengziensis | Bacillota | Bacilli | NZ_CP071182.1 | 1 |
| Alkalibacterium putridalgicola | Bacillota | Bacilli | NZ_CP162615.1 | 1 |
| Alkalicoccus halolimnae | Bacillota | Bacilli | NZ_CP144914.1 | 1 |
| Alkalihalophilus pseudofirmus | Bacillota | Bacilli | NZ_CP117835.1 | 1 |
| Allobacillus halotolerans | Bacillota | Bacilli | NZ_CP117968.1 | 1 |
| Amphibacillus xylanus NBRC 15112 | Bacillota | Bacilli | NC_018704.1 | 1 |
| Amylolactobacillus amylophilus DSM 20533 = JCM 1125 | Bacillota | Bacilli | NZ_CP018888.1 | 1 |
| Anaerobacillus isosaccharinicus | Bacillota | Bacilli | NZ_CP063356.2 | 1 |
| Aneurinibacillus soli | Bacillota | Bacilli | NZ_AP017312.1 | 1 |
| Aneurinibacillus thermoaerophilus | Bacillota | Bacilli | NZ_CP080764.1 | 1 |
| Aneurinibacillus uraniidurans | Bacillota | Bacilli | NZ_CP116902.1 | 1 |
| Anoxybacillus flavithermus WK1 | Bacillota | Bacilli | NC_011567.1 | 1 |
| Anoxybacillus gonensis | Bacillota | Bacilli | NZ_CP012152.1 | 1 |

| | | | | |
|---|---|---|---|---|
| Anoxybacteroides amylolyticum | Bacillota | Bacilli | NZ_CP015438.1 | 1 |
| Apilactobacillus apinorum | Bacillota | Bacilli | NZ_OX335302.1 | 1 |
| Apilactobacillus apisilvae | Bacillota | Bacilli | NZ_CP093362.1 | 1 |
| Apilactobacillus bombintestini | Bacillota | Bacilli | NZ_CP032626.1 | 1 |
| Apilactobacillus kunkeei | Bacillota | Bacilli | NZ_CP080568.1 | 1 |
| Bacillus albus | Bacillota | Bacilli | NZ_CP128152.1 | 1 |
| Bacillus altitudinis | Bacillota | Bacilli | NZ_CP009108.1 | 1 |
| Bacillus amyloliquefaciens | Bacillota | Bacilli | NZ_CP072120.1 | 1 |
| Bacillus anthracis str. 'Ames Ancestor' | Bacillota | Bacilli | NC_007530.2 | 1 |
| Bacillus aquiflavi | Bacillota | Bacilli | NZ_CP082780.1 | 1 |
| Bacillus arachidis | Bacillota | Bacilli | NZ_CP127376.1 | 1 |
| Bacillus atrophaeus | Bacillota | Bacilli | NZ_CP154443.1 | 1 |
| Bacillus cabrialesii | Bacillota | Bacilli | NZ_CP096889.1 | 1 |
| Bacillus cytotoxicus NVH 391-98 | Bacillota | Bacilli | NC_009674.1 | 1 |
| Bacillus fonticola | Bacillota | Bacilli | NZ_CP052135.1 | 1 |
| Bacillus glycinifermentans | Bacillota | Bacilli | NZ_CP035232.1 | 1 |
| Bacillus gobiensis | Bacillota | Bacilli | NZ_CP012600.1 | 1 |
| Bacillus halotolerans | Bacillota | Bacilli | NZ_CP029364.1 | 1 |
| Bacillus haynesii | Bacillota | Bacilli | NZ_CP059494.1 | 1 |
| Bacillus inaquosorum | Bacillota | Bacilli | NZ_CP096592.1 | 1 |
| Bacillus infantis | Bacillota | Bacilli | NZ_CP160000.1 | 1 |
| Bacillus kandeliae | Bacillota | Bacilli | NZ_CP147404.1 | 1 |
| Bacillus licheniformis | Bacillota | Bacilli | NZ_CP140161.1 | 1 |
| Bacillus luti | Bacillota | Bacilli | NZ_CP040336.1 | 1 |
| Bacillus maqinnsis | Bacillota | Bacilli | NZ_CP159358.1 | 1 |
| Bacillus mojavensis | Bacillota | Bacilli | NZ_CP051464.1 | 1 |
| Bacillus mycoides | Bacillota | Bacilli | NZ_CP035997.1 | 1 |
| Bacillus pacificus | Bacillota | Bacilli | NZ_CP086328.1 | 1 |
| Bacillus paralicheniformis | Bacillota | Bacilli | NZ_CP023665.1 | 1 |
| Bacillus paranthracis | Bacillota | Bacilli | NZ_CP101135.1 | 1 |
| Bacillus pseudomycoides DSM 12442 | Bacillota | Bacilli | NZ_CM000745.1 | 1 |
| Bacillus pumilus SAFR-032 | Bacillota | Bacilli | NC_009848.4 | 1 |

| | | | | |
|---|---|---|---|---|
| Bacillus rugosus | Bacillota | Bacilli | NZ_CP096590.1 | 1 |
| Bacillus safensis | Bacillota | Bacilli | NZ_CP043404.1 | 1 |
| Bacillus shivajii | Bacillota | Bacilli | NZ_CP084703.1 | 1 |
| Bacillus smithii | Bacillota | Bacilli | NZ_CP012024.1 | 1 |
| Bacillus sonorensis | Bacillota | Bacilli | NZ_CP139190.1 | 1 |
| Bacillus spizizenii TU-B-10 | Bacillota | Bacilli | NC_016047.1 | 1 |
| Bacillus stercoris | Bacillota | Bacilli | NZ_CP121861.1 | 1 |
| Bacillus subtilis subsp. subtilis str. 168 | Bacillota | Bacilli | NC_000964.3 | 1 |
| Bacillus swezeyi | Bacillota | Bacilli | NZ_CP133085.1 | 1 |
| Bacillus tequilensis | Bacillota | Bacilli | NZ_CP048852.1 | 1 |
| Bacillus thuringiensis serovar berliner ATCC 10792 | Bacillota | Bacilli | NZ_CM000753.1 | 1 |
| Bacillus toyonensis | Bacillota | Bacilli | NZ_CP064875.1 | 1 |
| Bacillus tropicus | Bacillota | Bacilli | NZ_CP119875.1 | 1 |
| Bacillus vallismortis | Bacillota | Bacilli | NZ_CP033052.1 | 1 |
| Bacillus velezensis FZB42 | Bacillota | Bacilli | NC_009725.2 | 1 |
| Bacillus weihaiensis | Bacillota | Bacilli | NZ_CP016020.1 | 1 |
| Bacillus wiedmannii | Bacillota | Bacilli | NZ_CP032365.1 | 1 |
| Bacillus xiamenensis | Bacillota | Bacilli | NZ_CP017786.1 | 1 |
| Bombilactobacillus bombi | Bacillota | Bacilli | NZ_OZ204251.1 | 1 |
| Bombilactobacillus folatiphilus | Bacillota | Bacilli | NZ_CP093366.1 | 1 |
| Bombilactobacillus thymidiniphilus | Bacillota | Bacilli | NZ_CP093365.1 | 1 |
| Brevibacillus agri | Bacillota | Bacilli | NZ_CP026363.1 | 1 |
| Brevibacillus aydinogluensis | Bacillota | Bacilli | NZ_OY569118.1 | 1 |
| Brevibacillus borstelensis | Bacillota | Bacilli | NZ_CP144922.1 | 1 |
| Brevibacillus brevis | Bacillota | Bacilli | NZ_LR134338.1 | 1 |
| Brevibacillus centrosporus | Bacillota | Bacilli | NZ_CP176856.1 | 1 |
| Brevibacillus composti | Bacillota | Bacilli | NZ_CP066308.1 | 1 |
| Brevibacillus humidisoli | Bacillota | Bacilli | NZ_CP087263.1 | 1 |
| Brevibacillus laterosporus DSM 25 | Bacillota | Bacilli | NZ_CP017705.1 | 1 |
| Brevibacillus marinus | Bacillota | Bacilli | NZ_CP034541.1 | 1 |
| Brevibacillus parabrevis | Bacillota | Bacilli | NZ_CP064090.1 | 1 |
| Brevibacillus ruminantium | Bacillota | Bacilli | NZ_CP098755.1 | 1 |

| | | | | |
|---|---|---|---|---|
| Brochothrix campestris | Bacillota | Bacilli | NZ_CP175511.1 | 1 |
| Brochothrix thermosphacta | Bacillota | Bacilli | NZ_CP016839.1 | 1 |
| Caldalkalibacillus thermarum TA2.A1 | Bacillota | Bacilli | NZ_CP082237.1 | 1 |
| Caldifermentibacillus hisashii | Bacillota | Bacilli | NZ_CP155467.1 | 1 |
| Candidatus Enterococcus clewellii | Bacillota | Bacilli | NZ_CP147247.1 | 2 |
| Candidatus Enterococcus dunnyi | Bacillota | Bacilli | NZ_CP147246.1 | 2 |
| Candidatus Enterococcus lemimoniae | Bacillota | Bacilli | NZ_CP147248.1 | 2 |
| Candidatus Enterococcus lowellii | Bacillota | Bacilli | NZ_CP147251.1 | 2 |
| Candidatus Enterococcus mangumiae | Bacillota | Bacilli | NZ_CP147250.1 | 2 |
| Candidatus Enterococcus palustris | Bacillota | Bacilli | NZ_CP147244.1 | 2 |
| Carnobacterium inhibens subsp. gilichinskyi | Bacillota | Bacilli | NC_022606.1 | 1 |
| Carnobacterium maltaromaticum | Bacillota | Bacilli | NZ_OX460976.1 | 3 |
| Chryseomicrobium aureum | Bacillota | Bacilli | NZ_CP171117.1 | 1 |
| Cohnella abietis | Bacillota | Bacilli | NZ_AP019400.1 | 1 |
| Cohnella candidum | Bacillota | Bacilli | NZ_CP033433.1 | 1 |
| Cohnella cholangitidis | Bacillota | Bacilli | NZ_CP041969.1 | 1 |
| Cohnella herbarum | Bacillota | Bacilli | NZ_CP051680.1 | 1 |
| Companilactobacillus alimentarius DSM 20249 | Bacillota | Bacilli | NZ_CP018867.1 | 1 |
| Companilactobacillus allii | Bacillota | Bacilli | NZ_CP019323.1 | 1 |
| Companilactobacillus crustorum | Bacillota | Bacilli | NZ_CP017996.1 | 1 |
| Companilactobacillus farciminis KCTC 3681 = DSM 20184 | Bacillota | Bacilli | NZ_CP017702.1 | 1 |
| Companilactobacillus futsaii | Bacillota | Bacilli | NZ_CP040736.1 | 1 |
| Companilactobacillus ginsenosidimutans | Bacillota | Bacilli | NZ_CP012034.1 | 1 |
| Companilactobacillus heilongjiangensis | Bacillota | Bacilli | NZ_CP012559.1 | 1 |
| Companilactobacillus musae | Bacillota | Bacilli | NZ_CP159983.1 | 1 |
| Companilactobacillus pabuli | Bacillota | Bacilli | NZ_CP049366.1 | 1 |
| Companilactobacillus paralimentarius | Bacillota | Bacilli | NZ_OY724131.1 | 1 |
| Companilactobacillus zhachilii | Bacillota | Bacilli | NZ_CP031933.2 | 1 |
| Cytobacillus dafuensis | Bacillota | Bacilli | NZ_CP042593.1 | 1 |
| Cytobacillus gottheilii | Bacillota | Bacilli | NZ_CP071709.1 | 1 |
| Cytobacillus kochii | Bacillota | Bacilli | NZ_CP085254.1 | 1 |
| Cytobacillus pseudoceanisediminis | Bacillota | Bacilli | NZ_CP151651.1 | 1 |

| | | | | |
|---|---|---|---|---|
| Cytobacillus solani | Bacillota | Bacilli | NZ_CP041305.1 | 1 |
| Cytobacillus spongiae | Bacillota | Bacilli | NZ_CP089997.1 | 1 |
| Dolosigranulum savutiense | Bacillota | Bacilli | NZ_CP142435.1 | 1 |
| Effusibacillus dendaii | Bacillota | Bacilli | NZ_AP023366.1 | 1 |
| Enterococcus casseliflavus EC20 | Bacillota | Bacilli | NC_020995.1 | 2 |
| Enterococcus cecorum | Bacillota | Bacilli | NZ_LS483306.1 | 1 |
| Enterococcus durans | Bacillota | Bacilli | NZ_CP022930.1 | 2 |
| Enterococcus faecium | Bacillota | Bacilli | NZ_CP038996.1 | 2 |
| Enterococcus gallinarum | Bacillota | Bacilli | NZ_CP050816.1 | 2 |
| Enterococcus gilvus | Bacillota | Bacilli | NZ_CP030932.1 | 2 |
| Enterococcus hirae | Bacillota | Bacilli | NZ_CP023011.2 | 2 |
| Enterococcus innesii | Bacillota | Bacilli | NZ_AP025635.1 | 2 |
| Enterococcus lactis | Bacillota | Bacilli | NZ_CP079880.1 | 2 |
| Enterococcus montenegrensis | Bacillota | Bacilli | NZ_CP120467.1 | 2 |
| Enterococcus mundtii | Bacillota | Bacilli | NZ_CP018061.1 | 2 |
| Enterococcus raffinosus | Bacillota | Bacilli | NZ_CP072888.1 | 2 |
| Enterococcus rotai | Bacillota | Bacilli | NZ_CP013655.1 | 2 |
| Enterococcus saccharolyticus subsp. saccharolyticus | Bacillota | Bacilli | NZ_CP118957.1 | 1 |
| Enterococcus saigonensis | Bacillota | Bacilli | NZ_AP022822.1 | 2 |
| Enterococcus thailandicus | Bacillota | Bacilli | NZ_CP023074.1 | 2 |
| Enterococcus wangshanyuanii | Bacillota | Bacilli | NZ_CP021874.1 | 2 |
| Evansella cellulosilytica DSM 2522 | Bacillota | Bacilli | NC_014829.1 | 1 |
| Exiguobacterium acetylicum | Bacillota | Bacilli | NZ_CP075897.1 | 1 |
| Exiguobacterium alkaliphilum | Bacillota | Bacilli | NZ_CP073101.1 | 1 |
| Exiguobacterium sibiricum 255-15 | Bacillota | Bacilli | NC_010556.1 | 1 |
| Fervidibacillus albus | Bacillota | Bacilli | NZ_CP106878.1 | 1 |
| Fervidibacillus halotolerans | Bacillota | Bacilli | NZ_CP106877.1 | 1 |
| Fodinisporobacter ferrooxydans | Bacillota | Bacilli | NZ_CP089291.1 | 1 |
| Fructilactobacillus carniphilus | Bacillota | Bacilli | NZ_CP097121.1 | 1 |
| Fructilactobacillus cliffordii | Bacillota | Bacilli | NZ_CP097119.1 | 1 |
| Fructilactobacillus hinvesii | Bacillota | Bacilli | NZ_CP097118.1 | 1 |
| Fructilactobacillus ixorae | Bacillota | Bacilli | NZ_CP097478.1 | 1 |

| | | | | |
|---|---|---|---|---|
| Fructilactobacillus lindneri | Bacillota | Bacilli | NZ_CP014872.1 | 1 |
| Fructilactobacillus myrtifloralis | Bacillota | Bacilli | NZ_CP097116.1 | 1 |
| Fructilactobacillus sanfranciscensis | Bacillota | Bacilli | NZ_CP118925.1 | 1 |
| Fructilactobacillus vespulae | Bacillota | Bacilli | NZ_CP137138.1 | 1 |
| Fructobacillus americanaquae | Bacillota | Bacilli | NZ_CP097122.1 | 1 |
| Fundicoccus culcitae | Bacillota | Bacilli | NZ_CP102453.1 | 1 |
| Furfurilactobacillus rossiae | Bacillota | Bacilli | NZ_CP040586.1 | 1 |
| Gemella morbillorum | Bacillota | Bacilli | NZ_LS483440.1 | 1 |
| Gemella sanguinis | Bacillota | Bacilli | NZ_CP046313.1 | 1 |
| Geobacillus kaustophilus HTA426 | Bacillota | Bacilli | NC_006510.1 | 1 |
| Geobacillus subterraneus | Bacillota | Bacilli | NZ_CP014342.1 | 1 |
| Geobacillus thermodenitrificans | Bacillota | Bacilli | NZ_CP017694.1 | 1 |
| Geobacillus thermoleovorans | Bacillota | Bacilli | NZ_CP014335.1 | 1 |
| Geobacillus zalihae | Bacillota | Bacilli | NZ_CP129217.1 | 1 |
| Globicatella sanguinis | Bacillota | Bacilli | NZ_CP126962.1 | 1 |
| Gracilibacillus caseinilyticus | Bacillota | Bacilli | NZ_CP095072.1 | 1 |
| Gracilibacillus salinarum | Bacillota | Bacilli | NZ_CP095071.1 | 1 |
| Gracilibacillus salitolerans | Bacillota | Bacilli | NZ_CP045915.1 | 1 |
| Granulicatella adiacens | Bacillota | Bacilli | NZ_CP082858.1 | 1 |
| Halalkalibacterium halodurans | Bacillota | Bacilli | NZ_CP040441.1 | 1 |
| Halobacillus amylolyticus | Bacillota | Bacilli | NZ_CP095075.1 | 1 |
| Halobacillus halophilus DSM 2266 | Bacillota | Bacilli | NC_017668.1 | 1 |
| Halobacillus mangrovi | Bacillota | Bacilli | NZ_CP020772.1 | 1 |
| Halobacillus naozhouensis | Bacillota | Bacilli | NZ_CP121671.1 | 1 |
| Halobacillus salinarum | Bacillota | Bacilli | NZ_CP095073.1 | 1 |
| Halobacillus shinanisalinarum | Bacillota | Bacilli | NZ_CP095074.1 | 1 |
| Heyndrickxia coagulans DSM 1 = ATCC 7050 | Bacillota | Bacilli | NZ_CP009709.1 | 1 |
| Heyndrickxia faecalis | Bacillota | Bacilli | NZ_CP158453.1 | 1 |
| Heyndrickxia vini | Bacillota | Bacilli | NZ_CP065425.1 | 1 |
| Ignavigranum ruoffiae | Bacillota | Bacilli | NZ_CP096206.2 | 1 |
| Jeotgalibaca arthritidis | Bacillota | Bacilli | NZ_CP049740.1 | 1 |
| Jeotgalibaca ciconiae | Bacillota | Bacilli | NZ_CP034465.1 | 1 |

| | | | | |
|---|---|---|---|---|
| Jeotgalibaca dankookensis | Bacillota | Bacilli | NZ_CP019728.1 | 1 |
| Jeotgalibaca porci | Bacillota | Bacilli | NZ_CP049889.1 | 1 |
| Jeotgalicoccus saudimassiliensis | Bacillota | Bacilli | NZ_CCSE0100000 | 1 |
| Kroppenstedtia eburnea | Bacillota | Bacilli | NZ_CP048103.1 | 1 |
| Kroppenstedtia pulmonis | Bacillota | Bacilli | NZ_CP048104.1 | 1 |
| Kyrpidia spormannii | Bacillota | Bacilli | NZ_CP024955.1 | 1 |
| Kyrpidia tusciae DSM 2912 | Bacillota | Bacilli | NC_014098.1 | 1 |
| Laceyella sacchari | Bacillota | Bacilli | NZ_CP103866.1 | 1 |
| Lacticaseibacillus casei DSM 20011 = JCM 1134 = ATCC : | Bacillota | Bacilli | NZ_AP012544.1 | 1 |
| Lacticaseibacillus huelsenbergensis | Bacillota | Bacilli | NZ_CP120687.1 | 1 |
| Lacticaseibacillus manihotivorans | Bacillota | Bacilli | NZ_CP045068.1 | 1 |
| Lacticaseibacillus pabuli | Bacillota | Bacilli | NZ_CP117884.1 | 1 |
| Lacticaseibacillus paracasei subsp. paracasei 8700:2 | Bacillota | Bacilli | NC_022112.1 | 1 |
| Lacticaseibacillus parahuelsenbergensis | Bacillota | Bacilli | NZ_CP132482.1 | 1 |
| Lacticaseibacillus rhamnosus | Bacillota | Bacilli | NZ_CP040780.1 | 1 |
| Lacticaseibacillus styriensis | Bacillota | Bacilli | NZ_CP132483.1 | 1 |
| Lacticaseibacillus zeae subsp. silagei | Bacillota | Bacilli | NZ_CP132484.1 | 1 |
| Lactiplantibacillus argentoratensis | Bacillota | Bacilli | NZ_CP147647.1 | 1 |
| Lactiplantibacillus brownii | Bacillota | Bacilli | NZ_AP027463.1 | 1 |
| Lactiplantibacillus carotarum | Bacillota | Bacilli | NZ_OX443570.1 | 1 |
| Lactiplantibacillus paraplantarum | Bacillota | Bacilli | NZ_CP118745.1 | 1 |
| Lactiplantibacillus pentosus | Bacillota | Bacilli | NZ_CP032757.1 | 1 |
| Lactiplantibacillus plantarum | Bacillota | Bacilli | NZ_CP028221.1 | 1 |
| Lactiplantibacillus xiangfangensis | Bacillota | Bacilli | NZ_OY725316.1 | 1 |
| Lactobacillus acetotolerans | Bacillota | Bacilli | NZ_CP044496.1 | 1 |
| Lactobacillus acidophilus | Bacillota | Bacilli | NZ_CP139575.1 | 1 |
| Lactobacillus amylolyticus | Bacillota | Bacilli | NZ_CP031835.1 | 1 |
| Lactobacillus amylovorus GRL1118 | Bacillota | Bacilli | NC_017470.1 | 1 |
| Lactobacillus apis | Bacillota | Bacilli | NZ_CP029476.1 | 1 |
| Lactobacillus crispatus | Bacillota | Bacilli | NZ_CP039266.1 | 1 |
| Lactobacillus delbrueckii subsp. jakobsenii ZN7a-9 = DSM | Bacillota | Bacilli | NZ_CP018218.1 | 1 |
| Lactobacillus gasseri ATCC 33323 = JCM 1131 | Bacillota | Bacilli | NC_008530.1 | 1 |

| | | | | |
|---|---|---|---|---|
| Lactobacillus helsingborgensis | Bacillota | Bacilli | NZ_CP029544.1 | 1 |
| Lactobacillus helveticus | Bacillota | Bacilli | NZ_CP151471.1 | 1 |
| Lactobacillus iners | Bacillota | Bacilli | NZ_CP049230.1 | 1 |
| Lactobacillus intestinalis | Bacillota | Bacilli | NZ_CP072983.1 | 1 |
| Lactobacillus isalae | Bacillota | Bacilli | NZ_OX443569.1 | 1 |
| Lactobacillus jensenii | Bacillota | Bacilli | NZ_CP018809.1 | 1 |
| Lactobacillus johnsonii | Bacillota | Bacilli | NZ_CP059055.1 | 1 |
| Lactobacillus kefiranofaciens | Bacillota | Bacilli | NZ_CP061341.1 | 1 |
| Lactobacillus kullabergensis | Bacillota | Bacilli | NZ_CP029477.1 | 1 |
| Lactobacillus mulieris | Bacillota | Bacilli | NZ_CP160088.1 | 1 |
| Lactobacillus panisapium | Bacillota | Bacilli | NZ_CP048268.1 | 1 |
| Lactobacillus paragasseri | Bacillota | Bacilli | NZ_AP018549.1 | 1 |
| Lactobacillus taiwanensis | Bacillota | Bacilli | NZ_CP059276.1 | 1 |
| Lactobacillus terrae | Bacillota | Bacilli | NZ_CP024610.1 | 1 |
| Lactobacillus ultunensis | Bacillota | Bacilli | NZ_CP059829.1 | 1 |
| Lactobacillus xylocopicola | Bacillota | Bacilli | NZ_AP026803.1 | 1 |
| Lactococcus allomyrinae | Bacillota | Bacilli | NZ_CP032627.1 | 1 |
| Lactococcus carnosus | Bacillota | Bacilli | NZ_CP017194.1 | 1 |
| Lactococcus cremoris subsp. cremoris KW2 | Bacillota | Bacilli | NC_022369.1 | 1 |
| Lactococcus formosensis | Bacillota | Bacilli | NZ_CP141721.1 | 1 |
| Lactococcus garvieae | Bacillota | Bacilli | NZ_CP065637.1 | 1 |
| Lactococcus lactis subsp. lactis | Bacillota | Bacilli | NZ_CP028160.1 | 1 |
| Lactococcus paracarnosus | Bacillota | Bacilli | NZ_CP017195.1 | 1 |
| Lactococcus petauri | Bacillota | Bacilli | NZ_CP094882.1 | 1 |
| Lactococcus protaetiae | Bacillota | Bacilli | NZ_CP041356.1 | 1 |
| Lactococcus raffinolactis | Bacillota | Bacilli | NZ_CP048686.1 | 1 |
| Lactococcus taiwanensis | Bacillota | Bacilli | NZ_CP070872.1 | 1 |
| Lapidilactobacillus dextrinicus | Bacillota | Bacilli | NZ_CP044499.1 | 1 |
| Latilactobacillus curvatus JCM 1096 = DSM 20019 | Bacillota | Bacilli | NZ_CP026116.1 | 1 |
| Latilactobacillus fuchuensis | Bacillota | Bacilli | NZ_LT984417.1 | 1 |
| Latilactobacillus graminis | Bacillota | Bacilli | NZ_CP045007.1 | 1 |
| Latilactobacillus sakei | Bacillota | Bacilli | NZ_CP046037.1 | 1 |

| | | | | |
|---|---|---|---|---|
| Lederbergia lenta | Bacillota | Bacilli | NZ_LS483476.1 | 1 |
| Lentibacillus amyloliquefaciens | Bacillota | Bacilli | NZ_CP013862.1 | 1 |
| Lentibacillus daqui | Bacillota | Bacilli | NZ_CP114176.1 | 1 |
| Lentilactobacillus buchneri | Bacillota | Bacilli | NZ_CP073066.1 | 1 |
| Lentilactobacillus curieae | Bacillota | Bacilli | NZ_CP018906.1 | 1 |
| Lentilactobacillus hilgardii | Bacillota | Bacilli | NZ_CP047121.1 | 1 |
| Lentilactobacillus kefiri | Bacillota | Bacilli | NZ_CP029971.1 | 1 |
| Lentilactobacillus laojiaonis | Bacillota | Bacilli | NZ_CP085278.1 | 1 |
| Lentilactobacillus parabuchneri | Bacillota | Bacilli | NZ_CP062473.1 | 1 |
| Lentilactobacillus terminaliae | Bacillota | Bacilli | NZ_CP168151.1 | 1 |
| Leuconostoc carnosum | Bacillota | Bacilli | NZ_CP042374.1 | 1 |
| Leuconostoc citreum | Bacillota | Bacilli | NZ_CP042410.1 | 1 |
| Leuconostoc falkenbergense | Bacillota | Bacilli | NZ_CP174386.1 | 1 |
| Leuconostoc gasicomitatum LMG 18811 | Bacillota | Bacilli | NC_014319.1 | 1 |
| Leuconostoc gelidum JB7 | Bacillota | Bacilli | NC_018631.1 | 1 |
| Leuconostoc kimchii IMSNU 11154 | Bacillota | Bacilli | NC_014136.1 | 1 |
| Leuconostoc lactis | Bacillota | Bacilli | NZ_CP042387.1 | 1 |
| Leuconostoc mesenteroides subsp. mesenteroides ATCC | Bacillota | Bacilli | NC_008531.1 | 1 |
| Leuconostoc pseudomesenteroides | Bacillota | Bacilli | NZ_CP065993.1 | 1 |
| Leuconostoc suionicum | Bacillota | Bacilli | NZ_CP015247.1 | 1 |
| Levilactobacillus brevis | Bacillota | Bacilli | NZ_LS483405.1 | 1 |
| Levilactobacillus cerevisiae | Bacillota | Bacilli | NZ_OZ061257.1 | 1 |
| Levilactobacillus koreensis | Bacillota | Bacilli | NZ_CP012033.1 | 1 |
| Levilactobacillus parabrevis | Bacillota | Bacilli | NZ_CP168688.1 | 1 |
| Levilactobacillus suantsaii | Bacillota | Bacilli | NZ_CP059603.1 | 1 |
| Levilactobacillus zymae | Bacillota | Bacilli | NZ_OY725319.1 | 1 |
| Ligilactobacillus acidipiscis | Bacillota | Bacilli | NZ_LT630287.1 | 1 |
| Ligilactobacillus agilis | Bacillota | Bacilli | NZ_CP104397.1 | 1 |
| Ligilactobacillus animalis | Bacillota | Bacilli | NZ_CP047141.1 | 1 |
| Ligilactobacillus cholophilus | Bacillota | Bacilli | NZ_CP127832.1 | 1 |
| Ligilactobacillus faecis | Bacillota | Bacilli | NZ_CP123639.1 | 1 |
| Ligilactobacillus murinus | Bacillota | Bacilli | NZ_CP023565.1 | 1 |

| | | | | |
|---|---|---|---|---|
| Ligilactobacillus ruminis ATCC 27782 | Bacillota | Bacilli | NC_015975.1 | 1 |
| Ligilactobacillus saerimneri | Bacillota | Bacilli | NZ_CP144759.1 | 1 |
| Ligilactobacillus salivarius | Bacillota | Bacilli | NZ_CP117983.1 | 1 |
| Limosilactobacillus fermentum | Bacillota | Bacilli | NZ_CP124737.1 | 1 |
| Limosilactobacillus frumenti | Bacillota | Bacilli | NZ_CP044534.1 | 1 |
| Limosilactobacillus mucosae | Bacillota | Bacilli | NZ_CP058954.1 | 1 |
| Limosilactobacillus oris | Bacillota | Bacilli | NZ_CP104398.1 | 1 |
| Limosilactobacillus pontis | Bacillota | Bacilli | NZ_CP045530.1 | 1 |
| Limosilactobacillus portuensis | Bacillota | Bacilli | NZ_CP117296.1 | 1 |
| Limosilactobacillus vaginalis | Bacillota | Bacilli | NZ_CP104399.1 | 1 |
| Liquorilactobacillus hordei DSM 19519 | Bacillota | Bacilli | NZ_CP049303.1 | 1 |
| Liquorilactobacillus mali | Bacillota | Bacilli | NZ_CP045035.1 | 1 |
| Liquorilactobacillus nagelii DSM 13675 | Bacillota | Bacilli | NZ_CP049304.1 | 1 |
| Listeria grayi | Bacillota | Bacilli | NZ_LR134483.1 | 1 |
| Listeria innocua | Bacillota | Bacilli | NZ_CP117229.1 | 1 |
| Listeria ivanovii subsp. ivanovii | Bacillota | Bacilli | NZ_CP009577.1 | 1 |
| Listeria monocytogenes EGD-e | Bacillota | Bacilli | NC_003210.1 | 1 |
| Listeria newyorkensis | Bacillota | Bacilli | NZ_CP113980.1 | 1 |
| Listeria seeligeri serovar 1/2b str. SLCC3954 | Bacillota | Bacilli | NC_013891.1 | 1 |
| Listeria swaminathanii | Bacillota | Bacilli | NZ_CP156021.1 | 1 |
| Listeria weihenstephanensis | Bacillota | Bacilli | NZ_CP011102.1 | 1 |
| Listeria welshimeri | Bacillota | Bacilli | NZ_LT906444.1 | 1 |
| Loigolactobacillus backii | Bacillota | Bacilli | NZ_CP014873.1 | 1 |
| Loigolactobacillus bifermentans | Bacillota | Bacilli | NZ_CP045872.1 | 1 |
| Loigolactobacillus coryniformis subsp. coryniformis KCTC 3 | Bacillota | Bacilli | NZ_CP017713.1 | 1 |
| Lysinibacillus agricola | Bacillota | Bacilli | NZ_CP067341.1 | 1 |
| Lysinibacillus capsici | Bacillota | Bacilli | NZ_CP084108.1 | 1 |
| Lysinibacillus fusiformis | Bacillota | Bacilli | NZ_CP010820.1 | 1 |
| Lysinibacillus irui | Bacillota | Bacilli | NZ_CP113527.1 | 1 |
| Lysinibacillus louembei | Bacillota | Bacilli | NZ_CP137624.1 | 1 |
| Lysinibacillus macroides | Bacillota | Bacilli | NZ_CP065643.1 | 1 |
| Lysinibacillus pakistanensis | Bacillota | Bacilli | NZ_CP126101.1 | 1 |

| | | | | |
|---|---|---|---|---|
| Lysinibacillus parviboronicapiens | Bacillota | Bacilli | NZ_CP073713.1 | 1 |
| Lysinibacillus sphaericus | Bacillota | Bacilli | NZ_CP019980.1 | 1 |
| Lysinibacillus telephonicus | Bacillota | Bacilli | NZ_CP155468.1 | 1 |
| Lysinibacillus timonensis | Bacillota | Bacilli | NZ_LT985980.1 | 1 |
| Lysinibacillus varians | Bacillota | Bacilli | NZ_CP006837.1 | 1 |
| Macrococcoides bohemicum | Bacillota | Bacilli | NZ_CP054482.1 | 1 |
| Macrococcoides canis | Bacillota | Bacilli | NZ_CP021059.1 | 1 |
| Macrococcoides caseolyticum | Bacillota | Bacilli | NZ_CP065729.1 | 1 |
| Macrococcus armenti | Bacillota | Bacilli | NZ_CP083608.1 | 1 |
| Macrococcus bovicus | Bacillota | Bacilli | NZ_CP128470.1 | 1 |
| Macrococcus brunensis | Bacillota | Bacilli | NZ_CP092179.1 | 1 |
| Macrococcus epidermidis | Bacillota | Bacilli | NZ_CP073819.1 | 1 |
| Macrococcus equipercicus | Bacillota | Bacilli | NZ_CP073809.1 | 1 |
| Macrococcus psychrotolerans | Bacillota | Bacilli | NZ_CP124591.1 | 1 |
| Mammaliicoccus lentus | Bacillota | Bacilli | NZ_OX344834.1 | 1 |
| Mammaliicoccus sciuri | Bacillota | Bacilli | NZ_LS483305.1 | 1 |
| Mammaliicoccus stepanovicii | Bacillota | Bacilli | NZ_LT906462.1 | 1 |
| Mammaliicoccus vitulinus | Bacillota | Bacilli | NZ_CP068061.1 | 1 |
| Mangrovibacillus cuniculi | Bacillota | Bacilli | NZ_CP049742.1 | 1 |
| Melissococcus plutonius | Bacillota | Bacilli | NZ_AP018492.1 | 1 |
| Mesobacillus jeotgali | Bacillota | Bacilli | NZ_CP025025.1 | 1 |
| Mesobacillus maritimus | Bacillota | Bacilli | NZ_CP085225.1 | 1 |
| Metabacillus dongyingensis | Bacillota | Bacilli | NZ_CP082944.1 | 1 |
| Metabacillus elymi | Bacillota | Bacilli | NZ_CP055263.1 | 1 |
| Metabacillus hrfriensis | Bacillota | Bacilli | NZ_CP126116.1 | 1 |
| Metabacillus indicus | Bacillota | Bacilli | NZ_CP176757.1 | 1 |
| Metabacillus rhizosphaerae | Bacillota | Bacilli | NZ_CP147403.1 | 1 |
| Metabacillus sediminilitoris | Bacillota | Bacilli | NZ_CP046266.1 | 1 |
| Metabacillus sediminis | Bacillota | Bacilli | NZ_CP147407.1 | 1 |
| Neobacillus niacini | Bacillota | Bacilli | NZ_CP183883.1 | 1 |
| Niallia circulans | Bacillota | Bacilli | NZ_CP053989.1 | 1 |
| Niallia oryzisoli | Bacillota | Bacilli | NZ_CP137640.1 | 1 |

| | | | | |
|---|---|---|---|---|
| Nicoliella spurrieriana | Bacillota | Bacilli | NZ_CP093361.1 | 1 |
| Nosocomiicoccus ampullae | Bacillota | Bacilli | NZ_CP079110.1 | 1 |
| Novibacillus thermophilus | Bacillota | Bacilli | NZ_CP019699.1 | 1 |
| Oceanobacillus iheyensis HTE831 | Bacillota | Bacilli | NC_004193.1 | 1 |
| Oceanobacillus kapialis | Bacillota | Bacilli | NZ_CP085256.1 | 1 |
| Oceanobacillus kimchii X50 | Bacillota | Bacilli | NZ_CM001792.1 | 1 |
| Oceanobacillus zhaokaii | Bacillota | Bacilli | NZ_CP024848.1 | 1 |
| Oenococcus kitaharae DSM 17330 | Bacillota | Bacilli | NZ_CM001398.1 | 1 |
| Oenococcus oeni | Bacillota | Bacilli | NZ_CP084701.1 | 1 |
| Oenococcus sicerae | Bacillota | Bacilli | NZ_CP029684.2 | 1 |
| Paenalkalicoccus suaedae | Bacillota | Bacilli | NZ_CP041372.2 | 1 |
| Paenibacillus agaridevorans | Bacillota | Bacilli | NZ_CP075889.1 | 1 |
| Paenibacillus albicereus | Bacillota | Bacilli | NZ_CP051428.1 | 1 |
| Paenibacillus albus | Bacillota | Bacilli | NZ_CP034437.1 | 1 |
| Paenibacillus algicola | Bacillota | Bacilli | NZ_CP040396.1 | 1 |
| Paenibacillus amylolyticus | Bacillota | Bacilli | NZ_CP145892.1 | 1 |
| Paenibacillus andongensis | Bacillota | Bacilli | NZ_CP104467.1 | 1 |
| Paenibacillus antarcticus | Bacillota | Bacilli | NZ_CP043611.1 | 1 |
| Paenibacillus aurantius | Bacillota | Bacilli | NZ_CP130318.1 | 1 |
| Paenibacillus azoreducens | Bacillota | Bacilli | NZ_AP025343.1 | 1 |
| Paenibacillus baekrokdamisoli | Bacillota | Bacilli | NZ_AP019308.1 | 1 |
| Paenibacillus barcinonensis | Bacillota | Bacilli | NZ_CP054614.1 | 1 |
| Paenibacillus beijingensis | Bacillota | Bacilli | NZ_CP011058.1 | 1 |
| Paenibacillus borealis | Bacillota | Bacilli | NZ_CP009285.1 | 1 |
| Paenibacillus bovis | Bacillota | Bacilli | NZ_CP013023.1 | 1 |
| Paenibacillus brasilensis | Bacillota | Bacilli | NZ_CP045298.1 | 1 |
| Paenibacillus cellulosilyticus | Bacillota | Bacilli | NZ_CP054613.1 | 1 |
| Paenibacillus cellulositrophicus | Bacillota | Bacilli | NZ_CP045295.1 | 1 |
| Paenibacillus chitinolyticus | Bacillota | Bacilli | NZ_CP026520.1 | 1 |
| Paenibacillus crassostreae | Bacillota | Bacilli | NZ_CP017770.1 | 1 |
| Paenibacillus dendritiformis | Bacillota | Bacilli | NZ_AP025344.1 | 1 |
| Paenibacillus donghaensis | Bacillota | Bacilli | NZ_CP021780.1 | 1 |

| | | | | |
|---|---|---|---|---|
| Paenibacillus durus | Bacillota | Bacilli | NZ_CP009288.1 | 1 |
| Paenibacillus elgii | Bacillota | Bacilli | NZ_CP121215.1 | 1 |
| Paenibacillus glucanolyticus | Bacillota | Bacilli | NZ_CP147845.1 | 1 |
| Paenibacillus graminis | Bacillota | Bacilli | NZ_CP009287.1 | 1 |
| Paenibacillus guangzhouensis | Bacillota | Bacilli | NZ_CP045293.1 | 1 |
| Paenibacillus hexagrammi | Bacillota | Bacilli | NZ_CP090978.1 | 1 |
| Paenibacillus hunanensis | Bacillota | Bacilli | NZ_CP139194.1 | 1 |
| Paenibacillus ihbetae | Bacillota | Bacilli | NZ_CP016809.1 | 1 |
| Paenibacillus jilunlii | Bacillota | Bacilli | NZ_CP048429.1 | 1 |
| Paenibacillus kribbensis | Bacillota | Bacilli | NZ_CP020028.1 | 1 |
| Paenibacillus kyungheensis | Bacillota | Bacilli | NZ_CP117416.1 | 1 |
| Paenibacillus larvae subsp. larvae | Bacillota | Bacilli | NZ_CP019659.1 | 1 |
| Paenibacillus lentus | Bacillota | Bacilli | NZ_CP034248.1 | 1 |
| Paenibacillus lutimineralis | Bacillota | Bacilli | NZ_CP034346.1 | 1 |
| Paenibacillus lycopersici | Bacillota | Bacilli | NZ_CP048209.1 | 1 |
| Paenibacillus marchantiae | Bacillota | Bacilli | NZ_CP118270.1 | 1 |
| Paenibacillus mucilaginosus K02 | Bacillota | Bacilli | NC_017672.3 | 1 |
| Paenibacillus naphthalenovorans | Bacillota | Bacilli | NZ_CP013652.1 | 1 |
| Paenibacillus nuruki | Bacillota | Bacilli | NZ_OY725677.1 | 1 |
| Paenibacillus odorifer | Bacillota | Bacilli | NZ_CP009428.1 | 1 |
| Paenibacillus pedocola | Bacillota | Bacilli | NZ_CP134223.1 | 1 |
| Paenibacillus physcomitrellae | Bacillota | Bacilli | NZ_CP022584.1 | 1 |
| Paenibacillus polygoni | Bacillota | Bacilli | NZ_CP127162.1 | 1 |
| Paenibacillus polymyxa ATCC 842 | Bacillota | Bacilli | NZ_CP024795.1 | 1 |
| Paenibacillus protaetiae | Bacillota | Bacilli | NZ_CP035492.1 | 1 |
| Paenibacillus psychroresistens | Bacillota | Bacilli | NZ_CP034235.1 | 1 |
| Paenibacillus rhizovicinus | Bacillota | Bacilli | NZ_CP048286.1 | 1 |
| Paenibacillus riograndensis SBR5 | Bacillota | Bacilli | NZ_LN831776.1 | 1 |
| Paenibacillus roseopurpureus | Bacillota | Bacilli | NZ_CP130319.1 | 1 |
| Paenibacillus sabinae T27 | Bacillota | Bacilli | NZ_CP004078.1 | 1 |
| Paenibacillus silvisoli | Bacillota | Bacilli | NZ_CP133017.1 | 1 |
| Paenibacillus sophorae | Bacillota | Bacilli | NZ_CP076607.1 | 1 |

| | | | | |
|---|---|---|---|---|
| Paenibacillus spongiae | Bacillota | Bacilli | NZ_CP091430.1 | 1 |
| Paenibacillus stellifer | Bacillota | Bacilli | NZ_CP009286.1 | 1 |
| Paenibacillus swuensis | Bacillota | Bacilli | NZ_CP011388.1 | 1 |
| Paenibacillus terrae HPL-003 | Bacillota | Bacilli | NC_016641.1 | 1 |
| Paenibacillus thiaminolyticus | Bacillota | Bacilli | NZ_CP041405.1 | 1 |
| Paenibacillus tianjinensis | Bacillota | Bacilli | NZ_CP070969.1 | 1 |
| Paenibacillus tritici | Bacillota | Bacilli | NZ_CP073365.1 | 1 |
| Paenibacillus tundrae | Bacillota | Bacilli | NZ_CP145605.1 | 1 |
| Paenibacillus uliginis N3/975 | Bacillota | Bacilli | NZ_LT840184.1 | 1 |
| Paenibacillus urinalis | Bacillota | Bacilli | NZ_CP118108.1 | 1 |
| Paenibacillus woosongensis | Bacillota | Bacilli | NZ_CP126084.1 | 1 |
| Paenibacillus xylanexedens | Bacillota | Bacilli | NZ_CP018620.1 | 1 |
| Paenibacillus yonginensis | Bacillota | Bacilli | NZ_CP014167.1 | 1 |
| Paenisporosarcina antarctica | Bacillota | Bacilli | NZ_CP038015.1 | 1 |
| Paenisporosarcina cavernae | Bacillota | Bacilli | NZ_CP032418.1 | 1 |
| Parageobacillus thermoglucosidasius | Bacillota | Bacilli | NZ_CP016622.1 | 1 |
| Parageobacillus toebii | Bacillota | Bacilli | NZ_CP070511.1 | 1 |
| Paraliobacillus zengyii | Bacillota | Bacilli | NZ_CP029797.1 | 1 |
| Paucilactobacillus hokkaidonensis JCM 18461 | Bacillota | Bacilli | NZ_AP014680.1 | 1 |
| Paucilactobacillus nenjiangensis | Bacillota | Bacilli | NZ_CP043939.1 | 1 |
| Paucilactobacillus oligofermentans DSM 15707 = LMG 227 | Bacillota | Bacilli | NZ_LN898144.1 | 1 |
| Paucilactobacillus suebicus | Bacillota | Bacilli | NZ_CP158670.1 | 1 |
| Pediococcus acidilactici | Bacillota | Bacilli | NZ_CP053421.1 | 1 |
| Pediococcus claussenii | Bacillota | Bacilli | NZ_CP014933.1 | 1 |
| Pediococcus damnosus | Bacillota | Bacilli | NZ_CP012294.1 | 1 |
| Pediococcus inopinatus | Bacillota | Bacilli | NZ_CP139080.1 | 1 |
| Pediococcus parvulus | Bacillota | Bacilli | NZ_OZ061292.1 | 1 |
| Pediococcus pentosaceus ATCC 25745 | Bacillota | Bacilli | NC_008525.1 | 1 |
| Pediococcus siamensis | Bacillota | Bacilli | NZ_CP178375.1 | 1 |
| Peribacillus frigoritolerans | Bacillota | Bacilli | NZ_CP091882.1 | 1 |
| Peribacillus muralis | Bacillota | Bacilli | NZ_CP048037.1 | 1 |
| Peribacillus psychrosaccharolyticus | Bacillota | Bacilli | NZ_CP068053.1 | 1 |

| | | | | |
|---|---|---|---|---|
| Peribacillus simplex NBRC 15720 = DSM 1321 | Bacillota | Bacilli | NZ_CP017704.1 | 1 |
| Periweissella cryptocerci | Bacillota | Bacilli | NZ_CP037940.1 | 1 |
| Planococcus antarcticus DSM 14505 | Bacillota | Bacilli | NZ_CP016534.2 | 1 |
| Planococcus donghaensis | Bacillota | Bacilli | NZ_CP016543.2 | 1 |
| Planococcus faecalis | Bacillota | Bacilli | NZ_CP019401.1 | 1 |
| Planococcus glaciei | Bacillota | Bacilli | NZ_CP051177.1 | 1 |
| Planococcus halocryophilus | Bacillota | Bacilli | NZ_CP016537.2 | 1 |
| Planococcus kocurii | Bacillota | Bacilli | NZ_CP013661.2 | 1 |
| Planococcus lenghuensis | Bacillota | Bacilli | NZ_CP019640.1 | 1 |
| Planococcus liqunii | Bacillota | Bacilli | NZ_CP129238.1 | 1 |
| Planococcus maritimus | Bacillota | Bacilli | NZ_CP016538.2 | 1 |
| Planococcus plakortidis | Bacillota | Bacilli | NZ_CP016539.2 | 1 |
| Planococcus rifietoensis | Bacillota | Bacilli | NZ_CP013659.2 | 1 |
| Planococcus versutus | Bacillota | Bacilli | NZ_CP016540.2 | 1 |
| Polycladomyces abyssicola | Bacillota | Bacilli | NZ_AP024601.1 | 1 |
| Priestia aryabhattai | Bacillota | Bacilli | NZ_CP024035.1 | 1 |
| Priestia flexa | Bacillota | Bacilli | NZ_CP090431.1 | 1 |
| Priestia megaterium NBRC 15308 = ATCC 14581 | Bacillota | Bacilli | NZ_CP035094.1 | 1 |
| Psychrobacillus glaciei | Bacillota | Bacilli | NZ_CP031223.1 | 1 |
| Radiobacillus deserti | Bacillota | Bacilli | NZ_CP041666.1 | 1 |
| Radiobacillus kanasensis | Bacillota | Bacilli | NZ_CP088020.1 | 1 |
| Rossellomorea marisflavi | Bacillota | Bacilli | NZ_CP081870.1 | 1 |
| Saccharibacillus brassicae | Bacillota | Bacilli | NZ_CP041217.1 | 1 |
| Saccharococcus caldoxylosilyticus | Bacillota | Bacilli | NZ_CP040553.1 | 1 |
| Salicibibacter cibarius | Bacillota | Bacilli | NZ_CP054705.1 | 1 |
| Salicibibacter cibi | Bacillota | Bacilli | NZ_CP054706.1 | 1 |
| Salicibibacter halophilus | Bacillota | Bacilli | NZ_CP035485.1 | 1 |
| Salicibibacter kimchii | Bacillota | Bacilli | NZ_CP031092.1 | 1 |
| Salimicrobium jeotgali | Bacillota | Bacilli | NZ_CP011361.2 | 1 |
| Salinicoccus bachuensis | Bacillota | Bacilli | NZ_CP138333.2 | 1 |
| Salisediminibacterium beveridgei | Bacillota | Bacilli | NZ_CP012502.1 | 1 |
| Schleiferilactobacillus harbinensis | Bacillota | Bacilli | NZ_CP041364.1 | 1 |

| | | | | |
|---|---|---|---|---|
| Secundilactobacillus malefermentans | Bacillota | Bacilli | NZ_CP042371.1 | 1 |
| Secundilactobacillus paracollinoides | Bacillota | Bacilli | NZ_CP014912.1 | 1 |
| Sediminibacillus dalangtanensis | Bacillota | Bacilli | NZ_CP046956.1 | 1 |
| Shouchella clausii | Bacillota | Bacilli | NZ_CP140150.1 | 1 |
| Shouchella hunanensis | Bacillota | Bacilli | NZ_CP117834.1 | 1 |
| Shouchella miscanthi | Bacillota | Bacilli | NZ_CP042163.1 | 1 |
| Shouchella rhizosphaerae | Bacillota | Bacilli | NZ_CP144921.1 | 1 |
| Solibacillus daqui | Bacillota | Bacilli | NZ_CP114887.1 | 1 |
| Sporolactobacillus terrae | Bacillota | Bacilli | NZ_CP082837.1 | 1 |
| Sporosarcina jeotgali | Bacillota | Bacilli | NZ_CP116341.1 | 1 |
| Sporosarcina oncorhynchi | Bacillota | Bacilli | NZ_CP129118.1 | 1 |
| Sporosarcina pasteurii | Bacillota | Bacilli | NZ_CP160452.1 | 1 |
| Sporosarcina psychrophila | Bacillota | Bacilli | NZ_CP014616.1 | 1 |
| Sporosarcina trichiuri | Bacillota | Bacilli | NZ_CP129119.1 | 1 |
| Sporosarcina ureae | Bacillota | Bacilli | NZ_CP015108.1 | 1 |
| Sporosarcina ureilytica | Bacillota | Bacilli | NZ_CP017560.1 | 1 |
| Staphylococcus agnetis | Bacillota | Bacilli | NZ_CP045927.1 | 1 |
| Staphylococcus argenteus | Bacillota | Bacilli | NC_016941.1 | 1 |
| Staphylococcus arlettae | Bacillota | Bacilli | NZ_CP094803.1 | 1 |
| Staphylococcus auricularis | Bacillota | Bacilli | NZ_CP065712.1 | 1 |
| Staphylococcus capitis subsp. urealyticus | Bacillota | Bacilli | NZ_CP145258.1 | 1 |
| Staphylococcus caprae | Bacillota | Bacilli | NZ_AP018587.1 | 1 |
| Staphylococcus carnosus | Bacillota | Bacilli | NZ_CP065711.1 | 1 |
| Staphylococcus casei | Bacillota | Bacilli | NZ_CP125718.1 | 1 |
| Staphylococcus chromogenes | Bacillota | Bacilli | NZ_CP031274.1 | 1 |
| Staphylococcus coagulans | Bacillota | Bacilli | NZ_CP092966.1 | 1 |
| Staphylococcus condimenti | Bacillota | Bacilli | NZ_LR134360.1 | 1 |
| Staphylococcus debuckii | Bacillota | Bacilli | NZ_CP033460.1 | 1 |
| Staphylococcus delphini | Bacillota | Bacilli | NZ_CP094740.1 | 1 |
| Staphylococcus epidermidis | Bacillota | Bacilli | NZ_CP035288.1 | 1 |
| Staphylococcus equorum | Bacillota | Bacilli | NZ_CP066013.1 | 1 |
| Staphylococcus felis | Bacillota | Bacilli | NZ_CP027770.1 | 1 |

| | | | | |
|---|---|---|---|---|
| Staphylococcus gallinarum | Bacillota | Bacilli | NZ_CP069082.1 | 1 |
| Staphylococcus haemolyticus | Bacillota | Bacilli | NZ_CP035291.1 | 1 |
| Staphylococcus hominis | Bacillota | Bacilli | NZ_CP033732.1 | 1 |
| Staphylococcus hsinchuensis | Bacillota | Bacilli | NZ_CP128355.1 | 1 |
| Staphylococcus hyicus | Bacillota | Bacilli | NZ_CP008747.1 | 1 |
| Staphylococcus kloosii | Bacillota | Bacilli | NZ_CP027846.1 | 1 |
| Staphylococcus lloydii | Bacillota | Bacilli | NZ_CP064056.1 | 1 |
| Staphylococcus lugdunensis | Bacillota | Bacilli | NZ_CP014022.1 | 1 |
| Staphylococcus lutrae | Bacillota | Bacilli | NZ_CP020773.1 | 1 |
| Staphylococcus massiliensis | Bacillota | Bacilli | NZ_CP171119.1 | 1 |
| Staphylococcus muscae | Bacillota | Bacilli | NZ_LT906464.1 | 1 |
| Staphylococcus nepalensis | Bacillota | Bacilli | NZ_CP017466.1 | 1 |
| Staphylococcus pasteuri | Bacillota | Bacilli | NZ_CP067014.1 | 1 |
| Staphylococcus pettenkoferi | Bacillota | Bacilli | NZ_CP022096.2 | 1 |
| Staphylococcus piscifermentans | Bacillota | Bacilli | NZ_LT906447.1 | 1 |
| Staphylococcus pseudintermedius | Bacillota | Bacilli | NZ_CP065635.1 | 1 |
| Staphylococcus pseudoxylosus | Bacillota | Bacilli | NZ_CP085024.1 | 1 |
| Staphylococcus ratti | Bacillota | Bacilli | NZ_CP086654.1 | 1 |
| Staphylococcus rostri | Bacillota | Bacilli | NZ_CP113107.1 | 1 |
| Staphylococcus saccharolyticus | Bacillota | Bacilli | NZ_CP066042.1 | 1 |
| Staphylococcus saprophyticus subsp. saprophyticus ATCC | Bacillota | Bacilli | NC_007350.1 | 1 |
| Staphylococcus schleiferi | Bacillota | Bacilli | NZ_CP094712.1 | 1 |
| Staphylococcus schweitzeri | Bacillota | Bacilli | NZ_LR134304.1 | 1 |
| Staphylococcus simiae | Bacillota | Bacilli | NZ_LT906460.1 | 1 |
| Staphylococcus simulans | Bacillota | Bacilli | NZ_CP095100.1 | 1 |
| Staphylococcus succinus | Bacillota | Bacilli | NZ_CP118976.1 | 1 |
| Staphylococcus ureilyticus | Bacillota | Bacilli | NZ_CP094825.1 | 1 |
| Staphylococcus warneri | Bacillota | Bacilli | NZ_CP032159.1 | 1 |
| Staphylococcus xylosus | Bacillota | Bacilli | NZ_CP008724.1 | 1 |
| Staphylospora marina | Bacillota | Bacilli | NZ_CP034118.1 | 1 |
| Streptococcus agalactiae | Bacillota | Bacilli | NZ_CP012480.1 | 1 |
| Streptococcus anginosus | Bacillota | Bacilli | NZ_LR134283.1 | 1 |

| | | | | |
|---|---|---|---|---|
| Streptococcus australis | Bacillota | Bacilli | NZ_LS483444.1 | 1 |
| Streptococcus canis | Bacillota | Bacilli | NZ_LR134293.1 | 1 |
| Streptococcus chenjunshii | Bacillota | Bacilli | NZ_CP031733.1 | 1 |
| Streptococcus constellatus | Bacillota | Bacilli | NZ_CP068213.1 | 1 |
| Streptococcus cristatus ATCC 51100 | Bacillota | Bacilli | NZ_LS483383.1 | 1 |
| Streptococcus dentalis | Bacillota | Bacilli | NZ_CP139418.1 | 1 |
| Streptococcus dysgalactiae | Bacillota | Bacilli | NZ_CP066069.1 | 1 |
| Streptococcus equi subsp. zooepidemicus | Bacillota | Bacilli | NZ_CP065054.1 | 1 |
| Streptococcus ferus | Bacillota | Bacilli | NZ_LS483343.1 | 1 |
| Streptococcus gallolyticus | Bacillota | Bacilli | NZ_CP054015.1 | 1 |
| Streptococcus gordonii | Bacillota | Bacilli | NZ_CP077224.1 | 1 |
| Streptococcus gwangjuensis | Bacillota | Bacilli | NZ_CP032621.1 | 1 |
| Streptococcus halichoeri | Bacillota | Bacilli | NZ_CP081502.1 | 1 |
| Streptococcus halotolerans | Bacillota | Bacilli | NZ_CP014835.1 | 1 |
| Streptococcus hillyeri | Bacillota | Bacilli | NZ_CP163513.1 | 1 |
| Streptococcus himalayensis | Bacillota | Bacilli | NZ_CP016953.1 | 1 |
| Streptococcus ilei | Bacillota | Bacilli | NC_022584.1 | 1 |
| Streptococcus iners | Bacillota | Bacilli | NZ_CP118735.1 | 1 |
| Streptococcus infantarius | Bacillota | Bacilli | NZ_CP065994.1 | 1 |
| Streptococcus iniae | Bacillota | Bacilli | NZ_CP010783.1 | 1 |
| Streptococcus intermedius | Bacillota | Bacilli | NZ_LS483436.1 | 1 |
| Streptococcus koreensis | Bacillota | Bacilli | NZ_CP032620.1 | 1 |
| Streptococcus lactarius | Bacillota | Bacilli | NZ_CP072329.1 | 1 |
| Streptococcus lingualis | Bacillota | Bacilli | NZ_CP139419.1 | 1 |
| Streptococcus lutetiensis | Bacillota | Bacilli | NZ_LS483403.1 | 1 |
| Streptococcus macedonicus | Bacillota | Bacilli | NZ_CP119172.1 | 1 |
| Streptococcus marmotae | Bacillota | Bacilli | NZ_CP015196.1 | 1 |
| Streptococcus merionis | Bacillota | Bacilli | NZ_LT906439.1 | 1 |
| Streptococcus mitis | Bacillota | Bacilli | NZ_CP012646.1 | 1 |
| Streptococcus mutans | Bacillota | Bacilli | NZ_CP077404.1 | 1 |
| Streptococcus oralis ATCC 35037 | Bacillota | Bacilli | NZ_LR134336.1 | 1 |
| Streptococcus oriscaviae | Bacillota | Bacilli | NZ_CP073084.1 | 1 |

| | | | | |
|---|---|---|---|---|
| Streptococcus pantholopis | Bacillota | Bacilli | NZ_CP014699.1 | 1 |
| Streptococcus parapneumoniae | Bacillota | Bacilli | NZ_AP026968.1 | 1 |
| Streptococcus parasanguinis ATCC 15912 | Bacillota | Bacilli | NC_015678.1 | 1 |
| Streptococcus parasuis | Bacillota | Bacilli | NZ_AP024276.1 | 1 |
| Streptococcus parauberis NCFD 2020 | Bacillota | Bacilli | NZ_AEUT0200000[illegible] | 1 |
| Streptococcus pasteurianus | Bacillota | Bacilli | NZ_CP039457.1 | 1 |
| Streptococcus periodonticum | Bacillota | Bacilli | NZ_CP034543.1 | 1 |
| Streptococcus pluranimalium | Bacillota | Bacilli | NZ_CP025536.1 | 1 |
| Streptococcus pneumoniae | Bacillota | Bacilli | NZ_LN831051.1 | 1 |
| Streptococcus porcinus | Bacillota | Bacilli | NZ_LS483388.1 | 1 |
| Streptococcus pseudopneumoniae IS7493 | Bacillota | Bacilli | NC_015875.1 | 1 |
| Streptococcus pseudoporcinus | Bacillota | Bacilli | NZ_LR134341.1 | 1 |
| Streptococcus pyogenes | Bacillota | Bacilli | NZ_LS483338.1 | 1 |
| Streptococcus ratti | Bacillota | Bacilli | NZ_CP043405.1 | 1 |
| Streptococcus respiraculi | Bacillota | Bacilli | NZ_CP022680.1 | 1 |
| Streptococcus ruminantium | Bacillota | Bacilli | NZ_AP018400.1 | 1 |
| Streptococcus ruminicola | Bacillota | Bacilli | NZ_CP046919.1 | 1 |
| Streptococcus salivarius JIM8777 | Bacillota | Bacilli | NC_017595.1 | 1 |
| Streptococcus sanguinis | Bacillota | Bacilli | NZ_LS483385.1 | 1 |
| Streptococcus sobrinus | Bacillota | Bacilli | NZ_CP029490.1 | 1 |
| Streptococcus suis BM407 | Bacillota | Bacilli | NC_012926.1 | 1 |
| Streptococcus suivaginalis | Bacillota | Bacilli | NZ_CP118733.1 | 1 |
| Streptococcus thermophilus | Bacillota | Bacilli | NZ_LR822015.1 | 1 |
| Streptococcus toyakuensis | Bacillota | Bacilli | NZ_AP024523.1 | 1 |
| Streptococcus troglodytae | Bacillota | Bacilli | NZ_AP014612.1 | 1 |
| Streptococcus uberis | Bacillota | Bacilli | NZ_LS483397.1 | 1 |
| Streptococcus urinalis | Bacillota | Bacilli | NZ_LR134323.1 | 1 |
| Streptococcus vestibularis | Bacillota | Bacilli | NZ_LR134275.1 | 1 |
| Streptococcus zhangguiae | Bacillota | Bacilli | NZ_CP072115.1 | 1 |
| Suicoccus acidiformans | Bacillota | Bacilli | NZ_CP023434.1 | 1 |
| Sutcliffiella cohnii | Bacillota | Bacilli | NZ_CP018866.1 | 1 |
| Sutcliffiella horikoshii | Bacillota | Bacilli | NZ_CP020880.1 | 1 |

| | | | | |
|---|---|---|---|---|
| Tepidibacillus marianensis | Bacillota | Bacilli | NZ_CP149939.1 | 1 |
| Terribacillus saccharophilus | Bacillota | Bacilli | NZ_CP008876.1 | 1 |
| Tetragenococcus halophilus | Bacillota | Bacilli | NZ_CP012047.1 | 1 |
| Tetragenococcus koreensis | Bacillota | Bacilli | NZ_CP027786.1 | 1 |
| Tetragenococcus osmophilus | Bacillota | Bacilli | NZ_CP027783.1 | 1 |
| Thermaerobacillus caldiproteolyticus | Bacillota | Bacilli | NZ_CP064060.1 | 1 |
| Thermoactinomyces vulgaris | Bacillota | Bacilli | NZ_CP039710.1 | 1 |
| Thermobacillus composti KWC4 | Bacillota | Bacilli | NC_019897.1 | 1 |
| Trichococcus flocculiformis | Bacillota | Bacilli | NZ_OY759718.1 | 1 |
| Trichococcus shcherbakoviae | Bacillota | Bacilli | NZ_OY762653.1 | 1 |
| Tumebacillus algifaecis | Bacillota | Bacilli | NZ_CP022657.1 | 1 |
| Tumebacillus avium | Bacillota | Bacilli | NZ_CP021434.1 | 1 |
| Ureibacillus chungkukjangi | Bacillota | Bacilli | NZ_CP085009.1 | 1 |
| Ureibacillus thermophilus | Bacillota | Bacilli | NZ_CP036528.1 | 1 |
| Ureibacillus thermosphaericus | Bacillota | Bacilli | NZ_AP018335.1 | 1 |
| Vagococcus carniphilus | Bacillota | Bacilli | NZ_CP060720.1 | 2 |
| Vagococcus coleopterorum | Bacillota | Bacilli | NZ_CP049886.1 | 1 |
| Vagococcus hydrophili | Bacillota | Bacilli | NZ_CP049887.1 | 1 |
| Vagococcus intermedius | Bacillota | Bacilli | NZ_CP110232.1 | 1 |
| Vagococcus jeotgali | Bacillota | Bacilli | NZ_CP142146.1 | 1 |
| Vagococcus luciliae | Bacillota | Bacilli | NZ_CP102451.1 | 1 |
| Vagococcus lutrae | Bacillota | Bacilli | NZ_CP097017.1 | 1 |
| Vagococcus teuberi | Bacillota | Bacilli | NZ_CP017267.1 | 1 |
| Vagococcus xieshaowenii | Bacillota | Bacilli | NZ_CP038865.1 | 1 |
| Vagococcus zengguangii | Bacillota | Bacilli | NZ_CP039712.1 | 1 |
| Virgibacillus dokdonensis | Bacillota | Bacilli | NZ_CP018622.1 | 1 |
| Virgibacillus halodenitrificans | Bacillota | Bacilli | NZ_CP017962.1 | 1 |
| Virgibacillus natechei | Bacillota | Bacilli | NZ_CP110224.1 | 1 |
| Virgibacillus necropolis | Bacillota | Bacilli | NZ_CP022437.1 | 1 |
| Virgibacillus pantothenticus | Bacillota | Bacilli | NZ_CP073011.1 | 1 |
| Virgibacillus phasianinus | Bacillota | Bacilli | NZ_CP022315.1 | 1 |
| Weissella ceti | Bacillota | Bacilli | NZ_CP074441.1 | 1 |

| | | | | |
|---|---|---|---|---|
| Weissella cibaria | Bacillota | Bacilli | NZ_CP013934.1 | 1 |
| Weissella coleopterorum | Bacillota | Bacilli | NZ_CP049888.1 | 1 |
| Weissella confusa | Bacillota | Bacilli | NZ_CP027563.1 | 1 |
| Weissella diestrammenae | Bacillota | Bacilli | NZ_CP060724.1 | 1 |
| Weissella jogaejeotgali | Bacillota | Bacilli | NZ_CP014332.1 | 1 |
| Weissella koreensis KACC 15510 | Bacillota | Bacilli | NC_015759.1 | 1 |
| Weissella paramesenteroides | Bacillota | Bacilli | NZ_CP130559.1 | 1 |
| Weissella soli | Bacillota | Bacilli | NZ_CP017326.1 | 1 |
| Weissella tructae | Bacillota | Bacilli | NZ_CP075544.1 | 1 |
| Weissella viridescens | Bacillota | Bacilli | NZ_CP061835.1 | 1 |
| Xylocopilactobacillus apicola | Bacillota | Bacilli | NZ_AP026802.1 | 1 |
| Xylocopilactobacillus apis | Bacillota | Bacilli | NZ_AP026801.1 | 1 |
| [Clostridium] asparagiforme DSM 15981 | Bacillota | Clostridia | NZ_CP102272.1 | 1 |
| [Clostridium] colinum | Bacillota | Clostridia | NZ_OW712331.1 | 1 |
| [Clostridium] hylemonae | Bacillota | Clostridia | NZ_AP025566.1 | 1 |
| [Clostridium] saccharolyticum WM1 | Bacillota | Clostridia | NC_014376.1 | 1 |
| [Clostridium] scindens ATCC 35704 | Bacillota | Clostridia | NZ_CP036170.1 | 1 |
| [Ruminococcus] lactaris ATCC 29176 | Bacillota | Clostridia | NZ_CP102292.1 | 1 |
| [Ruminococcus] torques | Bacillota | Clostridia | NZ_CP172594.1 | 1 |
| Aceticella autotrophica | Bacillota | Clostridia | NZ_CP060096.1 | 2 |
| Acetivibrio clariflavus DSM 19732 | Bacillota | Clostridia | NC_016627.1 | 1 |
| Acetivibrio saccincola | Bacillota | Clostridia | NZ_CP025197.1 | 1 |
| Acetivibrio thermocellus ATCC 27405 | Bacillota | Clostridia | NC_009012.1 | 1 |
| Acetoanaerobium sticklandii | Bacillota | Clostridia | NC_014614.1 | 2 |
| Acetobacterium carbinolicum | Bacillota | Clostridia | NZ_CP173357.1 | 2 |
| Acetobacterium malicum subsp. dehalogenans DSM 1152 | Bacillota | Clostridia | NZ_CP174122.1 | 2 |
| Acetobacterium wieringae | Bacillota | Clostridia | NZ_CP097897.1 | 2 |
| Acetobacterium woodii DSM 1030 | Bacillota | Clostridia | NC_016894.1 | 2 |
| Acetohalobium arabaticum DSM 5501 | Bacillota | Clostridia | NC_014378.1 | 2 |
| Acididesulfobacillus acetoxydans | Bacillota | Clostridia | NZ_LR746496.1 | 1 |
| Acutalibacter muris | Bacillota | Clostridia | NZ_CP021422.1 | 1 |
| Agathobacter rectalis | Bacillota | Clostridia | NZ_CP092643.1 | 1 |

| | | | | |
|---|---|---|---|---|
| Alkalibacter rhizosphaerae | Bacillota | Clostridia | NZ_CP071444.1 | 2 |
| Alkalicella caledoniensis | Bacillota | Clostridia | NZ_CP058559.1 | 2 |
| Alkaliphilus metalliredigens QYMF | Bacillota | Clostridia | NC_009633.1 | 2 |
| Alkaliphilus oremlandii OhILAs | Bacillota | Clostridia | NC_009922.1 | 2 |
| Aminipila butyrica | Bacillota | Clostridia | NZ_CP048649.1 | 2 |
| Aminipila luticellarii | Bacillota | Clostridia | NZ_CP035281.1 | 2 |
| Aminipila terrae | Bacillota | Clostridia | NZ_CP047591.1 | 2 |
| Ammonifex degensii KC4 | Bacillota | Clostridia | NC_013385.1 | 2 |
| Amygdalobacter indicium | Bacillota | Clostridia | NZ_CP118866.1 | 1 |
| Anaerobutyricum hallii | Bacillota | Clostridia | NZ_CP073280.1 | 1 |
| Anaerocolumna cellulosilytica | Bacillota | Clostridia | NZ_AP023367.1 | 1 |
| Anaerocolumna chitinilytica | Bacillota | Clostridia | NZ_AP023368.1 | 1 |
| Anaerocolumna sedimenticola | Bacillota | Clostridia | NZ_CP048000.1 | 1 |
| Anaeromicropila herbilytica | Bacillota | Clostridia | NZ_AP024169.1 | 1 |
| Anaeropeptidivorans aminofermentans | Bacillota | Clostridia | NZ_OW711693.1 | 1 |
| Anaerostipes caccae | Bacillota | Clostridia | NZ_CP084016.1 | 1 |
| Anaerostipes hadrus | Bacillota | Clostridia | NZ_CP132968.1 | 1 |
| Anaerostipes rhamnosivorans | Bacillota | Clostridia | NZ_CP040058.1 | 1 |
| Anaerotignum propionicum DSM 1682 | Bacillota | Clostridia | NZ_CP014223.1 | 1 |
| Anaerotruncus colihominis | Bacillota | Clostridia | NZ_CP094682.1 | 1 |
| Anoxybacter fermentans | Bacillota | Clostridia | NZ_CP016379.1 | 2 |
| Aristaeella hokkaidonensis | Bacillota | Clostridia | NZ_CP068393.1 | 2 |
| Aristaeella lactis | Bacillota | Clostridia | NZ_CP069421.1 | 2 |
| Bengtsoniella intestinalis | Bacillota | Clostridia | NZ_CP133584.1 | 1 |
| Biomaibacter acetigenes | Bacillota | Clostridia | NZ_CP033169.1 | 2 |
| Blautia argi | Bacillota | Clostridia | NZ_CP030280.1 | 1 |
| Blautia coccoides | Bacillota | Clostridia | NZ_CP136422.1 | 1 |
| Blautia hansenii DSM 20583 | Bacillota | Clostridia | NZ_CP022413.2 | 1 |
| Blautia hydrogenotrophica DSM 10507 | Bacillota | Clostridia | NZ_CP136423.1 | 1 |
| Blautia liquoris | Bacillota | Clostridia | NZ_CP063304.1 | 1 |
| Blautia luti | Bacillota | Clostridia | NZ_AP028156.1 | 1 |
| Blautia obeum ATCC 29174 | Bacillota | Clostridia | NZ_CP102265.1 | 1 |

| | | | | |
|---|---|---|---|---|
| Blautia parvula | Bacillota | Clostridia | NZ_CP084061.1 | 1 |
| Blautia producta ATCC 27340 = DSM 2950 | Bacillota | Clostridia | NZ_CP048626.1 | 1 |
| Blautia pseudococcoides | Bacillota | Clostridia | NZ_CP015405.2 | 1 |
| Blautia wexlerae DSM 19850 | Bacillota | Clostridia | NZ_CP102267.1 | 1 |
| Butyrivibrio fibrisolvens | Bacillota | Clostridia | NZ_CP146963.1 | 1 |
| Butyrivibrio hungatei | Bacillota | Clostridia | NZ_CP017831.1 | 1 |
| Butyrivibrio proteoclasticus B316 | Bacillota | Clostridia | NC_014387.1 | 1 |
| Caldanaerobacter subterraneus subsp. tengcongensis MB4 | Bacillota | Clostridia | NC_003869.1 | 2 |
| Caldinitratiruptor microaerophilus | Bacillota | Clostridia | NZ_AP025628.1 | 1 |
| Candidatus Desulforudis audaxviator | Bacillota | Clostridia | NZ_CP034260.1 | 2 |
| Candidatus Epulonipiscium viviparus | Bacillota | Clostridia | NZ_CP117982.1 | 1 |
| Candidatus Formimonas warabiya | Bacillota | Clostridia | NZ_CP017634.1 | 2 |
| Candidatus Sarcina troglodytae | Bacillota | Clostridia | NZ_CP051754.1 | 2 |
| Candidatus Syntrophocurvum alkaliphilum | Bacillota | Clostridia | NZ_CP046457.1 | 2 |
| Caproicibacter fermentans | Bacillota | Clostridia | NZ_CP060286.1 | 1 |
| Caproicibacterium amylolyticum | Bacillota | Clostridia | NZ_CP060696.1 | 1 |
| Caproicibacterium argilliputei | Bacillota | Clostridia | NZ_CP135996.1 | 1 |
| Caproicibacterium lactatifermentans | Bacillota | Clostridia | NZ_CP046051.1 | 1 |
| Carboxydothermus hydrogenoformans Z-2901 | Bacillota | Clostridia | NC_007503.1 | 2 |
| Cellulosilyticum lentocellum DSM 5427 | Bacillota | Clostridia | NC_015275.1 | 1 |
| Chakrabartyella piscis | Bacillota | Clostridia | NZ_CP092441.1 | 1 |
| Chordicoccus furentiruminis | Bacillota | Clostridia | NZ_CP048829.1 | 1 |
| Christensenella massiliensis | Bacillota | Clostridia | NZ_CP117826.1 | 2 |
| Christensenella minuta | Bacillota | Clostridia | NZ_CP029256.1 | 2 |
| Claveliimonas bilis | Bacillota | Clostridia | NZ_AP027742.1 | 1 |
| Clostridioides difficile | Bacillota | Clostridia | NZ_CP076401.1 | 2 |
| Clostridium aceticum | Bacillota | Clostridia | NZ_CP009687.1 | 2 |
| Clostridium acetobutylicum DSM 1731 | Bacillota | Clostridia | NC_015687.1 | 2 |
| Clostridium argentinense | Bacillota | Clostridia | NZ_CP014176.1 | 2 |
| Clostridium autoethanogenum DSM 10061 | Bacillota | Clostridia | NZ_CP012395.1 | 2 |
| Clostridium baratii | Bacillota | Clostridia | NZ_CP014204.2 | 2 |
| Clostridium bornimense | Bacillota | Clostridia | NZ_HG917868.1 | 2 |

| | | | | |
|---|---|---|---|---|
| Clostridium botulinum A str. ATCC 3502 | Bacillota | Clostridia | NC_009495.1 | 2 |
| Clostridium butyricum | Bacillota | Clostridia | NZ_CP040626.1 | 2 |
| Clostridium cadaveris | Bacillota | Clostridia | NZ_CP076620.1 | 2 |
| Clostridium carboxidivorans P7 | Bacillota | Clostridia | NZ_CP011803.1 | 2 |
| Clostridium cellulovorans 743B | Bacillota | Clostridia | NC_014393.1 | 2 |
| Clostridium chauvoei | Bacillota | Clostridia | NZ_CP018630.1 | 2 |
| Clostridium cochlearium | Bacillota | Clostridia | NZ_LT906477.1 | 2 |
| Clostridium diolis | Bacillota | Clostridia | NZ_CP043998.1 | 2 |
| Clostridium drakei | Bacillota | Clostridia | NZ_CP020953.1 | 2 |
| Clostridium estertheticum | Bacillota | Clostridia | NZ_CP077615.1 | 2 |
| Clostridium felsineum DSM 794 | Bacillota | Clostridia | NZ_CP096980.1 | 2 |
| Clostridium fermenticellae | Bacillota | Clostridia | NZ_CP032416.1 | 2 |
| Clostridium formicaceticum | Bacillota | Clostridia | NZ_CP020559.1 | 2 |
| Clostridium gasigenes | Bacillota | Clostridia | NZ_CP071376.1 | 2 |
| Clostridium gelidum | Bacillota | Clostridia | NZ_AP024849.1 | 2 |
| Clostridium intestinale | Bacillota | Clostridia | NZ_CP121166.1 | 2 |
| Clostridium isatidis | Bacillota | Clostridia | NZ_CP016786.1 | 2 |
| Clostridium kluyveri NBRC 12016 | Bacillota | Clostridia | NC_011837.1 | 2 |
| Clostridium ljungdahlii DSM 13528 | Bacillota | Clostridia | NC_014328.1 | 2 |
| Clostridium manihotivorum | Bacillota | Clostridia | NZ_CP025746.1 | 2 |
| Clostridium novyi | Bacillota | Clostridia | NZ_CP029458.1 | 2 |
| Clostridium pasteurianum DSM 525 = ATCC 6013 | Bacillota | Clostridia | NZ_CP013018.1 | 2 |
| Clostridium perfringens | Bacillota | Clostridia | NZ_CP065681.1 | 2 |
| Clostridium saccharobutylicum DSM 13864 | Bacillota | Clostridia | NC_022571.1 | 2 |
| Clostridium saccharoperbutylacetonicum N1-4(HMT) | Bacillota | Clostridia | NC_020291.1 | 2 |
| Clostridium scatologenes | Bacillota | Clostridia | NZ_CP009933.1 | 2 |
| Clostridium septicum | Bacillota | Clostridia | NZ_CP023671.1 | 2 |
| Clostridium sporogenes | Bacillota | Clostridia | NZ_CP009225.1 | 2 |
| Clostridium taeniosporum | Bacillota | Clostridia | NZ_CP017253.2 | 2 |
| Clostridium tagluense | Bacillota | Clostridia | NZ_CP086155.1 | 2 |
| Clostridium tetani E88 | Bacillota | Clostridia | NC_004557.1 | 2 |
| Clostridium thermarum | Bacillota | Clostridia | NZ_CP040924.1 | 2 |

| | | | | |
|---|---|---|---|---|
| Clostridium thermosuccinogenes | Bacillota | Clostridia | NZ_CP021850.1 | 1 |
| Clostridium tyrobutyricum | Bacillota | Clostridia | NZ_CP014170.1 | 2 |
| Coprococcus comes ATCC 27758 | Bacillota | Clostridia | NZ_CP102277.1 | 1 |
| Coprococcus eutactus | Bacillota | Clostridia | NZ_CP102278.1 | 1 |
| Crassaminicella indica | Bacillota | Clostridia | NZ_CP078093.1 | 2 |
| Crassaminicella profunda | Bacillota | Clostridia | NZ_CP082326.1 | 2 |
| Crassaminicella thermophila | Bacillota | Clostridia | NZ_CP042243.1 | 2 |
| Defluviitalea saccharophila | Bacillota | Clostridia | NZ_CP121687.1 | 1 |
| Dehalobacter restrictus | Bacillota | Clostridia | NZ_CP046996.1 | 1 |
| Dehalobacterium formicoaceticum | Bacillota | Clostridia | NZ_CP022121.1 | 2 |
| Desulfitobacterium dehalogenans ATCC 51507 | Bacillota | Clostridia | NC_018017.1 | 1 |
| Desulfitobacterium dichloroeliminans LMG P-21439 | Bacillota | Clostridia | NC_019903.1 | 1 |
| Desulfitobacterium hafniense DCB-2 | Bacillota | Clostridia | NC_011830.1 | 1 |
| Desulfitobacterium metallireducens DSM 15288 | Bacillota | Clostridia | NZ_CP007032.1 | 1 |
| Desulfofarcimen acetoxidans DSM 771 | Bacillota | Clostridia | NC_013216.1 | 2 |
| Desulfofundulus kuznetsovii DSM 6115 | Bacillota | Clostridia | NC_015573.1 | 2 |
| Desulforamulus ferrireducens | Bacillota | Clostridia | NZ_CP019698.1 | 2 |
| Desulforamulus reducens MI-1 | Bacillota | Clostridia | NC_009253.1 | 2 |
| Desulforamulus ruminis DSM 2154 | Bacillota | Clostridia | NC_015589.1 | 2 |
| Desulfoscipio gibsoniae DSM 7213 | Bacillota | Clostridia | NC_021184.1 | 2 |
| Desulfosporosinus acidiphilus SJ4 | Bacillota | Clostridia | NC_018068.1 | 1 |
| Desulfosporosinus meridiei DSM 13257 | Bacillota | Clostridia | NC_018515.1 | 1 |
| Desulfosporosinus orientis DSM 765 | Bacillota | Clostridia | NC_016584.1 | 1 |
| Desulfosporosinus paludis | Bacillota | Clostridia | NZ_CP144211.1 | 1 |
| Desulfosporosinus youngiae DSM 17734 | Bacillota | Clostridia | NZ_CM001441.1 | 1 |
| Desulfotomaculum nigrificans CO-1-SRB | Bacillota | Clostridia | NC_015565.1 | 2 |
| Dorea formicigenerans | Bacillota | Clostridia | NZ_CP102279.1 | 1 |
| Dorea longicatena | Bacillota | Clostridia | NZ_CP102280.1 | 1 |
| Enterocloster clostridioformis | Bacillota | Clostridia | NZ_CP050964.1 | 1 |
| Eshraghiella crossota | Bacillota | Clostridia | NZ_CP102270.1 | 1 |
| Ethanoligenens harbinense YUAN-3 | Bacillota | Clostridia | NC_014828.1 | 1 |
| Eubacterium callanderi | Bacillota | Clostridia | NC_014624.2 | 2 |

| | | | | |
|---|---|---|---|---|
| Eubacterium limosum | Bacillota | Clostridia | NZ_CP019962.1 | 2 |
| Eubacterium maltosivorans | Bacillota | Clostridia | NZ_CP029487.1 | 2 |
| Eubacterium ramulus | Bacillota | Clostridia | NZ_CP173382.1 | 1 |
| Faecalibacterium duncaniae | Bacillota | Clostridia | NZ_CP048437.1 | 1 |
| Faecalibacterium taiwanense | Bacillota | Clostridia | NZ_CP155552.1 | 1 |
| Faecalimonas umbilicata | Bacillota | Clostridia | NZ_AP031411.1 | 1 |
| Fastidiosipila sanguinis | Bacillota | Clostridia | NZ_CP027226.1 | 1 |
| Fenollaria massiliensis | Bacillota | Clostridia | NZ_CP096649.1 | 1 |
| Fenollaria sporofastidiosus | Bacillota | Clostridia | NZ_OW968186.1 | 1 |
| Filifactor alocis ATCC 35896 | Bacillota | Clostridia | NC_016630.1 | 2 |
| Flavonifractor plautii | Bacillota | Clostridia | NZ_CP048436.1 | 1 |
| Geosporobacter ferrireducens | Bacillota | Clostridia | NZ_CP017269.1 | 2 |
| Halanaerobium hydrogeniformans | Bacillota | Clostridia | NC_014654.1 | 2 |
| Halobacteroides halobius DSM 5150 | Bacillota | Clostridia | NC_019978.1 | 2 |
| Halothermothrix orenii H 168 | Bacillota | Clostridia | NC_011899.1 | 2 |
| Hathewaya histolytica | Bacillota | Clostridia | NZ_LR590481.1 | 2 |
| Helicovermis profundi | Bacillota | Clostridia | NZ_AP028654.1 | 2 |
| Heliomicrobium modesticaldum Ice1 | Bacillota | Clostridia | NC_010337.2 | 1 |
| Heliorestis convoluta | Bacillota | Clostridia | NZ_CP045875.1 | 1 |
| Herbinix luporum | Bacillota | Clostridia | NZ_LN879430.1 | 1 |
| Hungatella hathewayi | Bacillota | Clostridia | NZ_CP102274.1 | 1 |
| Intestinibacter bartlettii | Bacillota | Clostridia | NZ_CP102273.1 | 2 |
| Intestinimonas butyriciproducens | Bacillota | Clostridia | NZ_CP011524.1 | 1 |
| Iocasia fonsfrigidae | Bacillota | Clostridia | NZ_CP046640.1 | 2 |
| Kineothrix sedimenti | Bacillota | Clostridia | NZ_CP146256.1 | 1 |
| Koleobacter methoxysyntrophicus | Bacillota | Clostridia | NZ_CP059066.1 | 2 |
| Lachnoanaerobaculum umeaense | Bacillota | Clostridia | NZ_CP032364.1 | 1 |
| Lachnoclostridium phytofermentans ISDg | Bacillota | Clostridia | NC_010001.1 | 1 |
| Lachnospira eligens ATCC 27750 | Bacillota | Clostridia | NC_012778.1 | 1 |
| Lacrimispora xylanolytica | Bacillota | Clostridia | NZ_CP113524.1 | 1 |
| Mageeibacillus indolicus UPII9-5 | Bacillota | Clostridia | NC_013895.2 | 1 |
| Mahella australiensis 50-1 BON | Bacillota | Clostridia | NC_015520.1 | 2 |

| | | | | |
|---|---|---|---|---|
| Marvinbryantia formatexigens DSM 14469 | Bacillota | Clostridia | NZ_CP102268.1 | 1 |
| Mediterraneibacter glycyrrhizinilyticus | Bacillota | Clostridia | NZ_AP031414.1 | 1 |
| Mediterraneibacter gnavus ATCC 29149 | Bacillota | Clostridia | NZ_CP027002.1 | 1 |
| Metallumcola ferriviriculae | Bacillota | Clostridia | NZ_CP121694.1 | 2 |
| Mogibacterium diversum | Bacillota | Clostridia | NZ_CP027228.1 | 2 |
| Mogibacterium neglectum | Bacillota | Clostridia | NZ_CP128647.1 | 2 |
| Monoglobus pectinilyticus | Bacillota | Clostridia | NZ_CP020991.1 | 1 |
| Natranaerobius thermophilus JW/NM-WN-LF | Bacillota | Clostridia | NC_010718.1 | 2 |
| Natranaerofaba carboxydovora | Bacillota | Clostridia | NZ_CP054394.1 | 2 |
| Neomoorella carbonis | Bacillota | Clostridia | NZ_CP136420.1 | 2 |
| Neomoorella glycerini | Bacillota | Clostridia | NZ_CP046244.1 | 2 |
| Neomoorella humiferrea | Bacillota | Clostridia | NZ_CP136421.1 | 2 |
| Neomoorella thermoacetica | Bacillota | Clostridia | NZ_CP012370.1 | 1 |
| Novisyntrophococcus fermenticellae | Bacillota | Clostridia | NZ_CP076458.1 | 1 |
| Oscillibacter hominis | Bacillota | Clostridia | NZ_CP060490.1 | 1 |
| Owariibacterium komagatae | Bacillota | Clostridia | NZ_AP031406.1 | 1 |
| Paraclostridium bifermentans | Bacillota | Clostridia | NZ_CP079737.1 | 2 |
| Paraclostridium dentum | Bacillota | Clostridia | NZ_CP183813.1 | 2 |
| Paraclostridium ghonii | Bacillota | Clostridia | NZ_CP161000.1 | 2 |
| Paraclostridium sordellii | Bacillota | Clostridia | NZ_LN679998.1 | 2 |
| Peptacetobacter hiranonis | Bacillota | Clostridia | NZ_CP036523.1 | 2 |
| Peptoclostridium acidaminophilum DSM 3953 | Bacillota | Clostridia | NZ_CP007452.1 | 2 |
| Peptostreptococcus equinus | Bacillota | Clostridia | NZ_CP114052.1 | 2 |
| Petrocella atlantisensis | Bacillota | Clostridia | NZ_LR130778.1 | 1 |
| Proteinivorax hydrogeniformans | Bacillota | Clostridia | NZ_CP159485.1 | 1 |
| Proteinivorax tanatarense | Bacillota | Clostridia | NZ_CP158367.1 | 1 |
| Pusillibacter faecalis | Bacillota | Clostridia | NZ_AP023420.1 | 1 |
| Qiania dongpingensis | Bacillota | Clostridia | NZ_CP060634.1 | 1 |
| Romboutsia ilealis | Bacillota | Clostridia | NZ_LN555523.1 | 2 |
| Roseburia hominis A2-183 | Bacillota | Clostridia | NC_015977.1 | 1 |
| Roseburia intestinalis L1-82 | Bacillota | Clostridia | NZ_LR027880.1 | 1 |
| Roseburia rectibacter | Bacillota | Clostridia | NZ_CP092473.1 | 1 |

| | | | | |
|---|---|---|---|---|
| Ruminiclostridium cellulolyticum H10 | Bacillota | Clostridia | NC_011898.1 | 1 |
| Ruminiclostridium herbifermentans | Bacillota | Clostridia | NZ_CP061336.1 | 1 |
| Ruminiclostridium papyrosolvens DSM 2782 | Bacillota | Clostridia | NZ_CP119677.1 | 1 |
| Ruminococcus albus 7 = DSM 20455 | Bacillota | Clostridia | NC_014833.1 | 1 |
| Ruminococcus bovis | Bacillota | Clostridia | NZ_CP039381.1 | 1 |
| Ruminococcus gauvreauii | Bacillota | Clostridia | NZ_CP102290.1 | 1 |
| Sellimonas caecigallum | Bacillota | Clostridia | NZ_CP173660.1 | 1 |
| Sellimonas intestinalis | Bacillota | Clostridia | NZ_CP173694.1 | 1 |
| Serpentinicella alkaliphila | Bacillota | Clostridia | NZ_CP058648.1 | 2 |
| Simiaoa sunii | Bacillota | Clostridia | NZ_CP060633.1 | 1 |
| Solibaculum mannosilyticum | Bacillota | Clostridia | NZ_AP023321.1 | 1 |
| Subdoligranulum variabile | Bacillota | Clostridia | NZ_CP102293.1 | 1 |
| Symbiobacterium thermophilum IAM 14863 | Bacillota | Clostridia | NC_006177.1 | 1 |
| Syntrophobotulus glycolicus DSM 8271 | Bacillota | Clostridia | NC_015172.1 | 1 |
| Syntrophomonas wolfei subsp. wolfei str. Goettingen G311 | Bacillota | Clostridia | NC_008346.1 | 2 |
| Syntrophothermus lipocalidus DSM 12680 | Bacillota | Clostridia | NC_014220.1 | 1 |
| Tepidibacter aestuarii | Bacillota | Clostridia | NZ_OW235315.1 | 2 |
| Tepidibacter hydrothermalis | Bacillota | Clostridia | NZ_CP120733.1 | 2 |
| Terrisporobacter glycolicus ATCC 14880 = DSM 1288 | Bacillota | Clostridia | NZ_CP117523.1 | 2 |
| Terrisporobacter hibernicus | Bacillota | Clostridia | NZ_CP081135.1 | 2 |
| Terrisporobacter mayombei | Bacillota | Clostridia | NZ_CP101637.1 | 2 |
| Terrisporobacter petrolearius | Bacillota | Clostridia | NZ_CP102984.1 | 2 |
| Thermacetogenium phaeum DSM 12270 | Bacillota | Clostridia | NC_018870.1 | 1 |
| Thermanaeromonas toyohensis ToBE | Bacillota | Clostridia | NZ_LT838272.1 | 1 |
| Thermanaerosceptrum fracticalcis | Bacillota | Clostridia | NZ_CP045798.1 | 2 |
| Thermoanaerobacter brockii subsp. finnii Ako-1 | Bacillota | Clostridia | NC_014964.1 | 2 |
| Thermoanaerobacter ethanolicus JW 200 | Bacillota | Clostridia | NZ_CP033580.1 | 2 |
| Thermoanaerobacter italicus Ab9 | Bacillota | Clostridia | NC_013921.1 | 2 |
| Thermoanaerobacter kivui | Bacillota | Clostridia | NZ_CP009170.1 | 2 |
| Thermoanaerobacter mathranii subsp. mathranii str. A3 | Bacillota | Clostridia | NC_014209.1 | 1 |
| Thermoanaerobacter pseudethanolicus ATCC 33223 | Bacillota | Clostridia | NC_010321.1 | 2 |
| Thermoanaerobacter wiegelii Rt8.B1 | Bacillota | Clostridia | NC_015958.1 | 2 |

| | | | | |
|---|---|---|---|---|
| Thermoanaerobacterium saccharolyticum JW/SL-YS485 | Bacillota | Clostridia | NC_017992.1 | 2 |
| Thermoanaerobacterium thermosaccharolyticum DSM 571 | Bacillota | Clostridia | NC_014410.1 | 2 |
| Thermoanaerobacterium xylanolyticum LX-11 | Bacillota | Clostridia | NC_015555.1 | 2 |
| Thermocaproicibacter melissae | Bacillota | Clostridia | NZ_CP101827.1 | 1 |
| Thermoclostridium stercorarium subsp. stercorarium DSM | Bacillota | Clostridia | NC_020134.1 | 1 |
| Thermosediminibacter oceani DSM 16646 | Bacillota | Clostridia | NC_014377.1 | 2 |
| Vallitalea guaymasensis | Bacillota | Clostridia | NZ_CP058561.1 | 1 |
| Vallitalea pronyensis | Bacillota | Clostridia | NZ_CP058649.1 | 1 |
| Vescimonas coprocola | Bacillota | Clostridia | NZ_AP023418.1 | 1 |
| Vescimonas fastidiosa | Bacillota | Clostridia | NZ_AP023416.1 | 1 |
| Wansuia hejianensis | Bacillota | Clostridia | NZ_CP060635.1 | 1 |
| Wujia chipingensis | Bacillota | Clostridia | NZ_CP060632.1 | 1 |
| Zhaonella formicivorans | Bacillota | Clostridia | NZ_CP085524.1 | 2 |
| Candidatus Contubernalis alkaliaceticus | Bacillota | Dethiobacteria | NZ_CP054699.1 | 1 |
| [Clostridium] innocuum | Bacillota | Erysipelotrichia | NZ_CP048838.1 | 1 |
| Allobaculum mucilyticum | Bacillota | Erysipelotrichia | NZ_CP078089.1 | 1 |
| Allocoprobacillus halotolerans | Bacillota | Erysipelotrichia | NZ_CP101620.1 | 1 |
| Amedibacterium intestinale | Bacillota | Erysipelotrichia | NZ_AP019711.1 | 1 |
| Catenibacterium mitsuokai | Bacillota | Erysipelotrichia | NZ_CP102271.1 | 1 |
| Erysipelothrix amsterdamensis | Bacillota | Erysipelotrichia | NZ_OW659496.1 | 1 |
| Erysipelothrix enhydrae | Bacillota | Erysipelotrichia | NZ_CP125807.1 | 1 |
| Erysipelothrix inopinata | Bacillota | Erysipelotrichia | NZ_CP060715.1 | 1 |
| Erysipelothrix larvae | Bacillota | Erysipelotrichia | NZ_CP013213.1 | 1 |
| Erysipelothrix piscisicarius | Bacillota | Erysipelotrichia | NZ_CP034234.1 | 1 |
| Erysipelothrix rhusiopathiae str. Fujisawa | Bacillota | Erysipelotrichia | NC_015601.1 | 1 |
| Faecalibacillus intestinalis | Bacillota | Erysipelotrichia | NZ_AP024085.1 | 1 |
| Faecalibaculum rodentium | Bacillota | Erysipelotrichia | NZ_CP011391.1 | 1 |
| Intestinibaculum porci | Bacillota | Erysipelotrichia | NZ_AP019309.1 | 1 |
| Longicatena caecimuris | Bacillota | Erysipelotrichia | NZ_AP024510.1 | 1 |
| Massilimicrobiota timonensis | Bacillota | Erysipelotrichia | NZ_AP031415.1 | 1 |
| Solobacterium moorei | Bacillota | Erysipelotrichia | NZ_AP028934.1 | 1 |
| Tannockella kyphosi | Bacillota | Erysipelotrichia | NZ_CP088239.1 | 1 |

| | | | | |
|---|---|---|---|---|
| Thomasclavelia ramosa | Bacillota | Erysipelotrichia | NZ_CP068170.1 | 1 |
| Thomasclavelia spiroformis DSM 1552 | Bacillota | Erysipelotrichia | NZ_CP102275.1 | 1 |
| Turicibacter bilis | Bacillota | Erysipelotrichia | NZ_CP071251.1 | 1 |
| Turicibacter faecis | Bacillota | Erysipelotrichia | NZ_AP028127.1 | 1 |
| Turicibacter sanguinis | Bacillota | Erysipelotrichia | NZ_CP053187.1 | 1 |
| Carboxydichorda subterranea | Bacillota | Limnochordia | NZ_CP141615.1 | 1 |
| Geochorda subterranea | Bacillota | Limnochordia | NZ_CP141614.1 | 1 |
| Limnochorda pilosa | Bacillota | Limnochordia | NZ_AP014924.1 | 2 |
| Acidaminococcus fermentans DSM 20731 | Bacillota | Negativicutes | NC_013740.1 | 2 |
| Acidaminococcus intestini | Bacillota | Negativicutes | NZ_CP102249.1 | 2 |
| Dialister hominis | Bacillota | Negativicutes | NZ_AP019697.1 | 2 |
| Dialister massiliensis | Bacillota | Negativicutes | NZ_LT996885.1 | 2 |
| Dialister succinatiphilus | Bacillota | Negativicutes | NZ_OV788641.1 | 2 |
| Megamonas funiformis | Bacillota | Negativicutes | NZ_CP048627.1 | 1 |
| Megamonas hypermegale | Bacillota | Negativicutes | NZ_LT906446.1 | 1 |
| Megasphaera elsdenii DSM 20460 | Bacillota | Negativicutes | NZ_CP027570.1 | 2 |
| Megasphaera hexanoica | Bacillota | Negativicutes | NZ_CP171361.1 | 2 |
| Megasphaera stantonii | Bacillota | Negativicutes | NZ_CP029462.1 | 2 |
| Megasphaera sueciensis | Bacillota | Negativicutes | NZ_CP116939.1 | 2 |
| Methylomusa anaerophila | Bacillota | Negativicutes | NZ_AP018449.1 | 2 |
| Pectinatus frisingensis | Bacillota | Negativicutes | NZ_CP045424.1 | 1 |
| Phascolarctobacterium faecium | Bacillota | Negativicutes | NZ_AP019004.1 | 2 |
| Phascolarctobacterium succinatutens | Bacillota | Negativicutes | NZ_CP097284.1 | 2 |
| Selenobaculum gibii | Bacillota | Negativicutes | NZ_CP120678.1 | 1 |
| Selenomonas dianae | Bacillota | Negativicutes | NZ_CP128650.1 | 1 |
| Selenomonas sputigena ATCC 35185 | Bacillota | Negativicutes | NC_015437.1 | 1 |
| Selenomonas timonae | Bacillota | Negativicutes | NZ_CP060204.1 | 1 |
| Sporomusa acidovorans DSM 3132 | Bacillota | Negativicutes | NZ_CP155571.1 | 2 |
| Sporomusa aerivorans | Bacillota | Negativicutes | NZ_CP156926.1 | 2 |
| Sporomusa carbonis | Bacillota | Negativicutes | NZ_CP155570.1 | 2 |
| Sporomusa malonica | Bacillota | Negativicutes | NZ_CP155572.1 | 2 |
| Sporomusa ovata DSM 2662 | Bacillota | Negativicutes | NZ_CP146301.1 | 2 |

| | | | | |
|---|---|---|---|---|
| Sporomusa paucivorans | Bacillota | Negativicutes | NZ_CP155574.1 | 2 |
| Sporomusa rhizae | Bacillota | Negativicutes | NZ_CP156925.1 | 2 |
| Sporomusa silvacetica DSM 10669 | Bacillota | Negativicutes | NZ_CP155573.1 | 2 |
| Sporomusa sphaeroides DSM 2875 | Bacillota | Negativicutes | NZ_CP146991.1 | 2 |
| Sporomusa termitida | Bacillota | Negativicutes | NZ_CP036259.1 | 2 |
| Veillonella dispar | Bacillota | Negativicutes | NZ_LR134375.1 | 2 |
| Veillonella nakazawae | Bacillota | Negativicutes | NZ_AP022321.1 | 2 |
| Veillonella orientalis | Bacillota | Negativicutes | NZ_AP022322.1 | 2 |
| Veillonella parvula | Bacillota | Negativicutes | NZ_LT906445.1 | 2 |
| Veillonella rodentium | Bacillota | Negativicutes | NZ_LT906470.1 | 2 |
| Acidilutibacter cellobiosedens | Bacillota | Tissierellia | NZ_CP035282.1 | 2 |
| Anaerococcus mediterraneensis | Bacillota | Tissierellia | NZ_LT635772.1 | 2 |
| Anaerococcus nagyae | Bacillota | Tissierellia | NZ_AP031484.1 | 2 |
| Anaerococcus obesiensis | Bacillota | Tissierellia | NZ_CP067016.1 | 2 |
| Anaerococcus prevotii DSM 20548 | Bacillota | Tissierellia | NC_013171.1 | 2 |
| Anaerococcus vaginalis | Bacillota | Tissierellia | NZ_CP066014.1 | 2 |
| Ezakiella massiliensis | Bacillota | Tissierellia | NZ_LT635475.1 | 2 |
| Gottschalkia acidurici 9a | Bacillota | Tissierellia | NC_018664.1 | 2 |
| Gudongella oleilytica | Bacillota | Tissierellia | NZ_CP035130.1 | 2 |
| Helcococcus ovis | Bacillota | Tissierellia | NZ_CP119081.1 | 1 |
| Keratinibaculum paraultunense | Bacillota | Tissierellia | NZ_CP068564.1 | 2 |
| Microaceticoccus formicicus | Bacillota | Tissierellia | NZ_CP143259.1 | 2 |
| Miniphocaeibacter halophilus | Bacillota | Tissierellia | NZ_CP066744.1 | 2 |
| Murdochiella vaginalis | Bacillota | Tissierellia | NZ_LT632322.1 | 1 |
| Parvimonas micra | Bacillota | Tissierellia | NZ_LR134472.1 | 1 |
| Parvimonas parva | Bacillota | Tissierellia | NZ_AP038371.1 | 1 |
| Peptoniphilus equinus | Bacillota | Tissierellia | NZ_CP115667.1 | 2 |
| Peptoniphilus genitalis | Bacillota | Tissierellia | NZ_CP097885.1 | 2 |
| Peptoniphilus harei | Bacillota | Tissierellia | NZ_CP068103.1 | 2 |
| Alistipes communis | Bacteroidota | Bacteroidia | NZ_AP019735.1 | 1 |
| Alistipes dispar | Bacteroidota | Bacteroidia | NZ_AP019736.1 | 1 |
| Alistipes ihumii AP11 | Bacteroidota | Bacteroidia | NZ_CP102294.1 | 1 |

| | | | | |
|---|---|---|---|---|
| Alistipes indistinctus YIT 12060 | Bacteroidota | Bacteroidia | NZ_CP102250.1 | 1 |
| Alistipes megaguti | Bacteroidota | Bacteroidia | NZ_LR027382.1 | 1 |
| Alistipes onderdonkii subsp. vulgaris | Bacteroidota | Bacteroidia | NZ_AP019734.1 | 1 |
| Alistipes shahii WAL 8301 | Bacteroidota | Bacteroidia | NZ_CP102253.1 | 1 |
| Alkalitalea saponilacus | Bacteroidota | Bacteroidia | NZ_CP021904.1 | 1 |
| Aquipluma nitroreducens | Bacteroidota | Bacteroidia | NZ_AP018694.1 | 1 |
| Bacteroides caccae CL03T12C61 | Bacteroidota | Bacteroidia | NZ_CP072258.1 | 1 |
| Bacteroides caecimuris | Bacteroidota | Bacteroidia | NZ_CP015401.2 | 1 |
| Bacteroides cellulosilyticus | Bacteroidota | Bacteroidia | NZ_CP072251.1 | 1 |
| Bacteroides difficilis | Bacteroidota | Bacteroidia | NZ_CP182814.1 | 1 |
| Bacteroides eggerthii | Bacteroidota | Bacteroidia | NZ_CP102258.1 | 1 |
| Bacteroides faecis | Bacteroidota | Bacteroidia | NZ_CP081916.1 | 1 |
| Bacteroides faecium | Bacteroidota | Bacteroidia | NZ_CP050831.1 | 1 |
| Bacteroides fragilis NCTC 9343 | Bacteroidota | Bacteroidia | NC_003228.3 | 1 |
| Bacteroides graminisolvens | Bacteroidota | Bacteroidia | NZ_OY762556.1 | 1 |
| Bacteroides helcogenes P 36-108 | Bacteroidota | Bacteroidia | NC_014933.1 | 1 |
| Bacteroides heparinolyticus | Bacteroidota | Bacteroidia | NZ_CP027234.1 | 1 |
| Bacteroides hominis | Bacteroidota | Bacteroidia | NZ_CP178829.1 | 1 |
| Bacteroides humanifaecis | Bacteroidota | Bacteroidia | NZ_CP084680.1 | 1 |
| Bacteroides luhongzhouii | Bacteroidota | Bacteroidia | NZ_CP059973.1 | 1 |
| Bacteroides maternus | Bacteroidota | Bacteroidia | NZ_CP143867.1 | 1 |
| Bacteroides nordii | Bacteroidota | Bacteroidia | NZ_CP082886.1 | 1 |
| Bacteroides ovatus | Bacteroidota | Bacteroidia | NZ_CP012938.1 | 1 |
| Bacteroides pyogenes | Bacteroidota | Bacteroidia | NZ_CP184347.1 | 1 |
| Bacteroides salyersiae | Bacteroidota | Bacteroidia | NZ_CP133452.1 | 1 |
| Bacteroides sedimenti | Bacteroidota | Bacteroidia | NZ_AP028055.1 | 1 |
| Bacteroides thetaiotaomicron | Bacteroidota | Bacteroidia | NZ_CP040530.1 | 1 |
| Bacteroides uniformis | Bacteroidota | Bacteroidia | NZ_CP129143.1 | 1 |
| Bacteroides xylanisolvens | Bacteroidota | Bacteroidia | NZ_CP072216.1 | 1 |
| Bacteroides zhangwenhongii | Bacteroidota | Bacteroidia | NZ_CP059856.1 | 1 |
| Bacteroides zoogleoformans | Bacteroidota | Bacteroidia | NZ_CP027231.1 | 1 |
| Barnesiella viscericola DSM 18177 | Bacteroidota | Bacteroidia | NZ_CP007034.1 | 1 |

| | | | | |
|---|---|---|---|---|
| Butyricimonas faecalis | Bacteroidota | Bacteroidia | NZ_CP032819.1 | 1 |
| Butyricimonas faecihominis | Bacteroidota | Bacteroidia | NZ_CP043836.1 | 1 |
| Butyricimonas paravirosa | Bacteroidota | Bacteroidia | NZ_CP043839.1 | 1 |
| Butyricimonas virosa | Bacteroidota | Bacteroidia | NZ_CP102269.1 | 1 |
| Candidatus Azobacteroides pseudotrichonymphae genom | Bacteroidota | Bacteroidia | NC_011565.1 | 1 |
| Coprobacter fastidiosus | Bacteroidota | Bacteroidia | NZ_AP028032.1 | 1 |
| Coprobacter secundus subsp. similis | Bacteroidota | Bacteroidia | NZ_AP023322.1 | 1 |
| Draconibacterium halophilum | Bacteroidota | Bacteroidia | NZ_CP048409.1 | 1 |
| Duncaniella dubosii | Bacteroidota | Bacteroidia | NZ_CP039396.1 | 1 |
| Dysgonomonas capnocytophagoides | Bacteroidota | Bacteroidia | NZ_AP028867.1 | 1 |
| Hoylesella buccalis ATCC 35310 | Bacteroidota | Bacteroidia | NZ_CP102287.1 | 1 |
| Hoylesella enoeca | Bacteroidota | Bacteroidia | NZ_CP013195.1 | 1 |
| Labilibaculum antarcticum | Bacteroidota | Bacteroidia | NZ_AP018042.1 | 1 |
| Limibacterium fermenti | Bacteroidota | Bacteroidia | NZ_CP159486.1 | 1 |
| Maribellus comscasis | Bacteroidota | Bacteroidia | NZ_CP046401.1 | 1 |
| Marinifilum fragile | Bacteroidota | Bacteroidia | NZ_OY781962.1 | 1 |
| Muribaculum gordoncarteri | Bacteroidota | Bacteroidia | NZ_CP039393.1 | 1 |
| Muribaculum intestinale | Bacteroidota | Bacteroidia | NZ_CP015402.2 | 1 |
| Odoribacter splanchnicus | Bacteroidota | Bacteroidia | NZ_LT906459.1 | 1 |
| Paludibacter propionicigenes WB4 | Bacteroidota | Bacteroidia | NC_014734.1 | 1 |
| Parabacteroides absconsus | Bacteroidota | Bacteroidia | NZ_CP146284.1 | 1 |
| Parabacteroides chongii | Bacteroidota | Bacteroidia | NZ_CP120849.1 | 1 |
| Parabacteroides distasonis | Bacteroidota | Bacteroidia | NZ_CP042285.1 | 1 |
| Parabacteroides goldsteinii | Bacteroidota | Bacteroidia | NZ_CP081906.1 | 1 |
| Parabacteroides johnsonii DSM 18315 | Bacteroidota | Bacteroidia | NZ_CP102285.1 | 1 |
| Paramuribaculum intestinale | Bacteroidota | Bacteroidia | NZ_CP098825.1 | 1 |
| Paraprevotella xylaniphila YIT 11841 | Bacteroidota | Bacteroidia | NZ_LR215980.1 | 1 |
| Petrimonas mucosa | Bacteroidota | Bacteroidia | NZ_LT608328.1 | 1 |
| Phocaeicola coprophilus | Bacteroidota | Bacteroidia | NZ_CP069440.1 | 1 |
| Phocaeicola dorei | Bacteroidota | Bacteroidia | NZ_CP046176.1 | 1 |
| Phocaeicola salanitronis DSM 18170 | Bacteroidota | Bacteroidia | NC_015164.1 | 1 |
| Phocaeicola vulgatus ATCC 8482 | Bacteroidota | Bacteroidia | NC_009614.1 | 1 |

| | | | | |
|---|---|---|---|---|
| Porphyromonas asaccharolytica DSM 20707 | Bacteroidota | Bacteroidia | NC_015501.1 | 1 |
| Porphyromonas cangingivalis | Bacteroidota | Bacteroidia | NZ_LR134506.1 | 1 |
| Porphyromonas crevioricanis | Bacteroidota | Bacteroidia | NZ_LS483447.1 | 1 |
| Porphyromonas endodontalis | Bacteroidota | Bacteroidia | NZ_CP083629.1 | 1 |
| Porphyromonas gingivalis ATCC 33277 | Bacteroidota | Bacteroidia | NC_010729.1 | 1 |
| Porphyromonas pogonae | Bacteroidota | Bacteroidia | NZ_CP143258.1 | 1 |
| Porphyromonas somerae | Bacteroidota | Bacteroidia | NZ_AP025559.1 | 1 |
| Prevotella communis | Bacteroidota | Bacteroidia | NZ_CP091792.1 | 1 |
| Prevotella dentalis DSM 3688 | Bacteroidota | Bacteroidia | NC_019960.1 | 1 |
| Prevotella denticola | Bacteroidota | Bacteroidia | NZ_CP072374.1 | 1 |
| Prevotella fusca JCM 17724 | Bacteroidota | Bacteroidia | NZ_CP012074.1 | 1 |
| Prevotella herbatica | Bacteroidota | Bacteroidia | NZ_AP024484.1 | 1 |
| Prevotella histicola | Bacteroidota | Bacteroidia | NZ_CP072367.1 | 1 |
| Prevotella jejuni | Bacteroidota | Bacteroidia | NZ_CP023863.1 | 1 |
| Prevotella melaninogenica ATCC 25845 | Bacteroidota | Bacteroidia | NC_014370.1 | 1 |
| Prevotella multiformis | Bacteroidota | Bacteroidia | NZ_CP072357.1 | 1 |
| Prevotella nigrescens | Bacteroidota | Bacteroidia | NZ_CP072339.1 | 1 |
| Prevotella scopos JCM 17725 | Bacteroidota | Bacteroidia | NZ_CP072390.1 | 1 |
| Prevotella veroralis | Bacteroidota | Bacteroidia | NZ_CP072330.1 | 1 |
| Proteiniphilum propionicum | Bacteroidota | Bacteroidia | NZ_CP073586.1 | 1 |
| Proteiniphilum saccharofermentans | Bacteroidota | Bacteroidia | NZ_LT605205.1 | 1 |
| Pseudoprevotella muciniphila | Bacteroidota | Bacteroidia | NZ_CP033459.1 | 1 |
| Salinivirga cyanobacteriivorans | Bacteroidota | Bacteroidia | NZ_CP013118.1 | 1 |
| Segatella bryantii | Bacteroidota | Bacteroidia | NZ_CP091798.1 | 1 |
| Segatella copri DSM 18205 | Bacteroidota | Bacteroidia | NZ_CP085932.1 | 1 |
| Segatella oris | Bacteroidota | Bacteroidia | NZ_LR134384.1 | 1 |
| Sodaliphilus pleomorphus | Bacteroidota | Bacteroidia | NZ_CP045696.1 | 1 |
| Tannerella forsythia 92A2 | Bacteroidota | Bacteroidia | NC_016610.1 | 1 |
| Tannerella serpentiformis | Bacteroidota | Bacteroidia | NZ_CP017038.2 | 1 |
| Tenuifilum thalassicum | Bacteroidota | Bacteroidia | NZ_CP041345.1 | 1 |
| Xiashengella succiniciproducens | Bacteroidota | Bacteroidia | NZ_CP098400.1 | 1 |
| Xylanibacter ruminicola 23 | Bacteroidota | Bacteroidia | NC_014033.1 | 1 |

| | | | | |
|---|---|---|---|---|
| Arachidicoccus soli | Bacteroidota | Chitinophagia | NZ_CP032489.1 | 1 |
| Arachidicoccus terrestris | Bacteroidota | Chitinophagia | NZ_CP083387.1 | 1 |
| Chitinophaga agri | Bacteroidota | Chitinophagia | NZ_CP048113.1 | 1 |
| Chitinophaga caeni | Bacteroidota | Chitinophagia | NZ_CP023777.1 | 1 |
| Chitinophaga caseinilytica | Bacteroidota | Chitinophagia | NZ_CP150096.1 | 1 |
| Chitinophaga horti | Bacteroidota | Chitinophagia | NZ_CP107006.1 | 1 |
| Chitinophaga oryzae | Bacteroidota | Chitinophagia | NZ_CP051205.1 | 1 |
| Chitinophaga pendula | Bacteroidota | Chitinophagia | NZ_CP077769.1 | 1 |
| Chitinophaga pinensis DSM 2588 | Bacteroidota | Chitinophagia | NC_013132.1 | 1 |
| Chitinophaga pollutisoli | Bacteroidota | Chitinophagia | NZ_CP149822.1 | 1 |
| Chitinophaga sancti | Bacteroidota | Chitinophagia | NZ_CP139972.1 | 1 |
| Ferruginibacter albus | Bacteroidota | Chitinophagia | NZ_CP083388.1 | 1 |
| Ferruginibacter lapsinanis | Bacteroidota | Chitinophagia | NZ_CP086063.1 | 1 |
| Filimonas lacunae | Bacteroidota | Chitinophagia | NZ_AP017422.1 | 1 |
| Flavihumibacter fluvii | Bacteroidota | Chitinophagia | NZ_CP092333.1 | 1 |
| Flavihumibacter rivuli | Bacteroidota | Chitinophagia | NZ_CP092334.1 | 1 |
| Flavisolibacter ginsenosidimutans | Bacteroidota | Chitinophagia | NZ_CP042433.1 | 1 |
| Flavisolibacter tropicus | Bacteroidota | Chitinophagia | NZ_CP011390.1 | 1 |
| Lacibacter sediminis | Bacteroidota | Chitinophagia | NZ_CP060007.1 | 1 |
| Mycovorax composti | Bacteroidota | Chitinophagia | NZ_CP144143.1 | 1 |
| Niabella ginsenosidivorans | Bacteroidota | Chitinophagia | NZ_CP015772.1 | 1 |
| Niabella insulamsoli | Bacteroidota | Chitinophagia | NZ_CP190246.1 | 1 |
| Niabella soli DSM 19437 | Bacteroidota | Chitinophagia | NZ_CP007035.1 | 1 |
| Niabella yanshanensis | Bacteroidota | Chitinophagia | NZ_CP139960.1 | 1 |
| Niastella koreensis GR20-10 | Bacteroidota | Chitinophagia | NC_016609.1 | 1 |
| Panacibacter ginsenosidivorans | Bacteroidota | Chitinophagia | NZ_CP042435.1 | 1 |
| Paraflavitalea soli | Bacteroidota | Chitinophagia | NZ_CP032157.1 | 1 |
| Paraflavitalea speifideaquila | Bacteroidota | Chitinophagia | NZ_CP135920.1 | 1 |
| Pseudocnuella soli | Bacteroidota | Chitinophagia | NZ_CP037755.1 | 1 |
| Rhizosphaericola mali | Bacteroidota | Chitinophagia | NZ_CP044016.1 | 1 |
| Adhaeribacter radiodurans | Bacteroidota | Cytophagia | NZ_CP055153.1 | 1 |
| Adhaeribacter swui | Bacteroidota | Cytophagia | NZ_CP055156.1 | 1 |

| | | | | |
|---|---|---|---|---|
| Algoriphagus halophytocola | Bacteroidota | Cytophagia | NZ_CP110226.1 | 1 |
| Algoriphagus machipongonensis | Bacteroidota | Cytophagia | NZ_CM001023.1 | 1 |
| Algoriphagus sanaruensis | Bacteroidota | Cytophagia | NZ_CP012836.1 | 1 |
| Aquiflexum balticum DSM 16537 | Bacteroidota | Cytophagia | NZ_LT838813.1 | 1 |
| Aquirufa antheringensis | Bacteroidota | Cytophagia | NZ_CP049835.1 | 1 |
| Aquirufa lenticrescens | Bacteroidota | Cytophagia | NZ_CP049834.1 | 1 |
| Aquirufa nivalisilvae | Bacteroidota | Cytophagia | NZ_CP029346.1 | 1 |
| Arcticibacterium luteifluviistationis | Bacteroidota | Cytophagia | NZ_CP029480.1 | 1 |
| Aureibacter tunicatorum | Bacteroidota | Cytophagia | NZ_AP025305.1 | 1 |
| Belliella baltica DSM 15883 | Bacteroidota | Cytophagia | NC_018010.1 | 1 |
| Bernardetia litoralis DSM 6794 | Bacteroidota | Cytophagia | NC_018018.1 | 1 |
| Candidatus Amoebophilus asiaticus 5a2 | Bacteroidota | Cytophagia | NC_010830.1 | 1 |
| Candidatus Cardinium hertigii | Bacteroidota | Cytophagia | NZ_CP029619.1 | 1 |
| Cardinium endosymbiont cEper1 of Encarsia pergandiella | Bacteroidota | Cytophagia | NC_018605.1 | 1 |
| Cardinium endosymbiont of Nabis limbatus | Bacteroidota | Cytophagia | NZ_OZ239769.1 | 1 |
| Cardinium endosymbiont of Oedothorax gibbosus | Bacteroidota | Cytophagia | NZ_OW441264.1 | 1 |
| Cardinium endosymbiont of Philonthus spinipes | Bacteroidota | Cytophagia | NZ_OZ034918.1 | 1 |
| Cardinium endosymbiont of Sogatella furcifera | Bacteroidota | Cytophagia | NZ_CP022339.1 | 1 |
| Cardinium endosymbiont of Tipula unca | Bacteroidota | Cytophagia | NZ_OZ026476.1 | 1 |
| Catalinimonas niigatensis | Bacteroidota | Cytophagia | NZ_CP119422.1 | 1 |
| Chondrinema litorale | Bacteroidota | Cytophagia | NZ_CP111043.1 | 1 |
| Chryseolinea soli | Bacteroidota | Cytophagia | NZ_CP032382.1 | 1 |
| Cyclobacterium amurskyense | Bacteroidota | Cytophagia | NZ_CP012040.1 | 1 |
| Cyclobacterium marinum DSM 745 | Bacteroidota | Cytophagia | NC_015914.1 | 1 |
| Cytophaga hutchinsonii ATCC 33406 | Bacteroidota | Cytophagia | NC_008255.1 | 1 |
| Dyadobacter chenhuakuii | Bacteroidota | Cytophagia | NZ_CP098805.1 | 1 |
| Dyadobacter chenwenxiniae | Bacteroidota | Cytophagia | NZ_CP094997.1 | 1 |
| Dyadobacter fanqingshengii | Bacteroidota | Cytophagia | NZ_CP098806.1 | 1 |
| Dyadobacter fermentans DSM 18053 | Bacteroidota | Cytophagia | NC_013037.1 | 1 |
| Dyadobacter pollutisoli | Bacteroidota | Cytophagia | NZ_CP112998.1 | 1 |
| Dyadobacter sandarakinus | Bacteroidota | Cytophagia | NZ_CP056775.1 | 1 |
| Echinicola marina | Bacteroidota | Cytophagia | NZ_CP080025.1 | 1 |

| | | | | |
|---|---|---|---|---|
| Echinicola rosea | Bacteroidota | Cytophagia | NZ_CP040106.1 | 1 |
| Echinicola soli | Bacteroidota | Cytophagia | NZ_CP041253.1 | 1 |
| Echinicola strongylocentroti | Bacteroidota | Cytophagia | NZ_CP030041.1 | 1 |
| Echinicola vietnamensis DSM 17526 | Bacteroidota | Cytophagia | NC_019904.1 | 1 |
| Emticicia oligotrophica DSM 17448 | Bacteroidota | Cytophagia | NC_018748.1 | 1 |
| Fibrella aestuarina BUZ 2 | Bacteroidota | Cytophagia | NC_020054.1 | 1 |
| Flammeovirga kamogawensis | Bacteroidota | Cytophagia | NZ_CP076128.1 | 1 |
| Flammeovirga pectinis | Bacteroidota | Cytophagia | NZ_CP034562.1 | 1 |
| Flammeovirga yaeyamensis | Bacteroidota | Cytophagia | NZ_CP076132.1 | 1 |
| Fulvitalea axinellae | Bacteroidota | Cytophagia | NZ_AP025314.1 | 1 |
| Fulvivirga ligni | Bacteroidota | Cytophagia | NZ_CP089979.1 | 1 |
| Fulvivirga lutea | Bacteroidota | Cytophagia | NZ_CP070608.1 | 1 |
| Fulvivirga maritima | Bacteroidota | Cytophagia | NZ_CP089980.1 | 1 |
| Fulvivirga ulvae | Bacteroidota | Cytophagia | NZ_CP089981.1 | 1 |
| Hymenobacter aerilatus | Bacteroidota | Cytophagia | NZ_CP095053.1 | 1 |
| Hymenobacter baengnokdamensis | Bacteroidota | Cytophagia | NZ_CP044285.1 | 1 |
| Hymenobacter canadensis | Bacteroidota | Cytophagia | NZ_CP114767.1 | 1 |
| Hymenobacter cellulosivorans | Bacteroidota | Cytophagia | NZ_CP095049.1 | 1 |
| Hymenobacter jejuensis | Bacteroidota | Cytophagia | NZ_CP040896.1 | 1 |
| Hymenobacter monticola | Bacteroidota | Cytophagia | NZ_CP094534.1 | 1 |
| Hymenobacter nivis | Bacteroidota | Cytophagia | NZ_CP029145.1 | 1 |
| Hymenobacter oligotrophus | Bacteroidota | Cytophagia | NZ_CP032317.1 | 1 |
| Hymenobacter radiodurans | Bacteroidota | Cytophagia | NZ_CP037922.1 | 1 |
| Hymenobacter sedentarius | Bacteroidota | Cytophagia | NZ_CP013909.1 | 1 |
| Hymenobacter sediminicola | Bacteroidota | Cytophagia | NZ_CP060202.1 | 1 |
| Hymenobacter sublimis | Bacteroidota | Cytophagia | NZ_CP095848.1 | 1 |
| Hymenobacter swuensis DY53 | Bacteroidota | Cytophagia | NZ_CP007145.1 | 1 |
| Hymenobacter taeanensis | Bacteroidota | Cytophagia | NZ_CP053538.1 | 1 |
| Hymenobacter tibetensis | Bacteroidota | Cytophagia | NZ_CP094669.1 | 1 |
| Hymenobacter volaticus | Bacteroidota | Cytophagia | NZ_CP095061.1 | 1 |
| Hymenobacter yonginensis | Bacteroidota | Cytophagia | NZ_CP115396.1 | 1 |
| Imperialibacter roseus | Bacteroidota | Cytophagia | NZ_CP136051.1 | 1 |

| | | | | |
|---|---|---|---|---|
| Larkinella insperata | Bacteroidota | Cytophagia | NZ_CP110973.1 | 1 |
| Leadbetterella byssophila DSM 17132 | Bacteroidota | Cytophagia | NC_014655.1 | 1 |
| Mangrovivirga cuniculi | Bacteroidota | Cytophagia | NZ_CP028923.1 | 1 |
| Marinilongibacter aquaticus | Bacteroidota | Cytophagia | NZ_CP083757.1 | 1 |
| Marivirga arenosa | Bacteroidota | Cytophagia | NZ_CP129968.2 | 1 |
| Marivirga harenae | Bacteroidota | Cytophagia | NZ_CP130565.1 | 1 |
| Marivirga salinarum | Bacteroidota | Cytophagia | NZ_CP129971.1 | 1 |
| Marivirga tractuosa DSM 4126 | Bacteroidota | Cytophagia | NC_014759.1 | 1 |
| Mongoliitalea daihaiensis | Bacteroidota | Cytophagia | NZ_CP063779.1 | 1 |
| Nibribacter ruber | Bacteroidota | Cytophagia | NZ_CP047897.1 | 1 |
| Persicobacter psychrovividus | Bacteroidota | Cytophagia | NZ_AP025292.1 | 1 |
| Pontibacter actiniarum | Bacteroidota | Cytophagia | NZ_CP021235.1 | 1 |
| Pontibacter akesuensis | Bacteroidota | Cytophagia | NZ_CP014766.1 | 1 |
| Pontibacter kalidii | Bacteroidota | Cytophagia | NZ_CP111079.1 | 1 |
| Pontibacter korlensis | Bacteroidota | Cytophagia | NZ_CP009621.1 | 1 |
| Pontibacter pudoricolor | Bacteroidota | Cytophagia | NZ_CP048106.1 | 1 |
| Pontibacter russatus | Bacteroidota | Cytophagia | NZ_CP047984.1 | 1 |
| Porifericola rhodea | Bacteroidota | Cytophagia | NZ_CP119421.1 | 1 |
| Reichenbachiella agarivorans | Bacteroidota | Cytophagia | NZ_CP106679.1 | 1 |
| Reichenbachiella carrageenanivorans | Bacteroidota | Cytophagia | NZ_CP106735.1 | 1 |
| Rhodocytophaga rosea | Bacteroidota | Cytophagia | NZ_CP048222.1 | 1 |
| Roseivirga spongicola | Bacteroidota | Cytophagia | NZ_CP139724.1 | 1 |
| Rufibacter radiotolerans | Bacteroidota | Cytophagia | NZ_CP010777.1 | 1 |
| Rufibacter tibetensis | Bacteroidota | Cytophagia | NZ_CP012643.1 | 1 |
| Runella rosea | Bacteroidota | Cytophagia | NZ_CP030850.1 | 1 |
| Runella slithyformis DSM 19594 | Bacteroidota | Cytophagia | NC_015703.1 | 1 |
| Spirosoma aerolatum | Bacteroidota | Cytophagia | NZ_CP020104.1 | 1 |
| Spirosoma aureum | Bacteroidota | Cytophagia | NZ_CP050063.1 | 1 |
| Spirosoma endbachense | Bacteroidota | Cytophagia | NZ_CP045997.1 | 1 |
| Spirosoma foliorum | Bacteroidota | Cytophagia | NZ_CP059732.1 | 1 |
| Spirosoma linguale DSM 74 | Bacteroidota | Cytophagia | NC_013730.1 | 1 |
| Spirosoma montaniterrae | Bacteroidota | Cytophagia | NZ_CP014263.1 | 1 |

| | | | | |
|---|---|---|---|---|
| Spirosoma oryzicola | Bacteroidota | Cytophagia | NZ_CP089538.1 | 1 |
| Spirosoma pollinicola | Bacteroidota | Cytophagia | NZ_CP025096.1 | 1 |
| Spirosoma radiotolerans | Bacteroidota | Cytophagia | NZ_CP010429.1 | 1 |
| Spirosoma rhododendri | Bacteroidota | Cytophagia | NZ_CP051677.1 | 1 |
| Spirosoma rigui | Bacteroidota | Cytophagia | NZ_CP020105.1 | 1 |
| Spirosoma taeanense | Bacteroidota | Cytophagia | NZ_CP053435.1 | 1 |
| Tellurirhabdus bombi | Bacteroidota | Cytophagia | NZ_CP090557.1 | 1 |
| Tellurirhabdus rosea | Bacteroidota | Cytophagia | NZ_CP111085.1 | 1 |
| Tunicatimonas pelagia | Bacteroidota | Cytophagia | NZ_CP120683.1 | 1 |
| Abyssalbus ytuae | Bacteroidota | Flavobacteriia | NZ_CP094358.1 | 1 |
| Aegicerativicinus sediminis | Bacteroidota | Flavobacteriia | NZ_CP064295.1 | 1 |
| Aequorivita ciconiae | Bacteroidota | Flavobacteriia | NZ_CP034951.1 | 1 |
| Aequorivita iocasae | Bacteroidota | Flavobacteriia | NZ_CP068439.1 | 1 |
| Aequorivita marisscotiae | Bacteroidota | Flavobacteriia | NZ_CP122379.1 | 1 |
| Aequorivita nionensis | Bacteroidota | Flavobacteriia | NZ_CP130281.1 | 1 |
| Aequorivita sublithincola DSM 14238 | Bacteroidota | Flavobacteriia | NC_018013.1 | 1 |
| Aestuariibaculum lutulentum | Bacteroidota | Flavobacteriia | NZ_CP136709.1 | 1 |
| Aggregatimonas sangjinii | Bacteroidota | Flavobacteriia | NZ_CP040710.1 | 1 |
| Algibacter luteus | Bacteroidota | Flavobacteriia | NZ_CP128313.1 | 1 |
| Allomuricauda ruestringensis DSM 13258 | Bacteroidota | Flavobacteriia | NC_015945.1 | 1 |
| Allotamlana fucoidanivorans | Bacteroidota | Flavobacteriia | NZ_CP074074.1 | 1 |
| Antarcticibacterium arcticum | Bacteroidota | Flavobacteriia | NZ_CP042476.1 | 1 |
| Antarcticibacterium flavum | Bacteroidota | Flavobacteriia | NZ_CP040812.1 | 1 |
| Apibacter raozihei | Bacteroidota | Flavobacteriia | NZ_CP034930.1 | 1 |
| Arenibacter algicola | Bacteroidota | Flavobacteriia | NZ_CP022515.1 | 1 |
| Arenibacter antarcticus | Bacteroidota | Flavobacteriia | NZ_CP166679.1 | 1 |
| Aureibaculum algae | Bacteroidota | Flavobacteriia | NZ_CP040749.1 | 1 |
| Bergeyella cardium | Bacteroidota | Flavobacteriia | NZ_CP029149.1 | 1 |
| Blattabacterium clevelandi | Bacteroidota | Flavobacteriia | NZ_CP029844.1 | 1 |
| Blattabacterium cuenoti | Bacteroidota | Flavobacteriia | NZ_CP059188.1 | 1 |
| Blattabacterium punctulatus | Bacteroidota | Flavobacteriia | NZ_CP029808.1 | 1 |
| Candidatus Arcticimaribacter forsetii | Bacteroidota | Flavobacteriia | NZ_OU342399.1 | 1 |

| | | | | |
|---|---|---|---|---|
| Candidatus Kaistella beijingensis | Bacteroidota | Flavobacteriia | NZ_CP071953.1 | 1 |
| Candidatus Karelsulcia muelleri | Bacteroidota | Flavobacteriia | NZ_CP013212.1 | 1 |
| Candidatus Ornithobacterium hominis | Bacteroidota | Flavobacteriia | NZ_OX579588.1 | 1 |
| Candidatus Shikimatogenerans bostrichidophilus | Bacteroidota | Flavobacteriia | NZ_CP104094.1 | 1 |
| Candidatus Walczuchella endosymbiont of Icerya purchasi | Bacteroidota | Flavobacteriia | NZ_OZ034981.1 | 1 |
| Candidatus Walczuchella monophlebidarum | Bacteroidota | Flavobacteriia | NZ_CP006873.1 | 1 |
| Capnocytophaga canimorsus | Bacteroidota | Flavobacteriia | NZ_CP022382.1 | 1 |
| Capnocytophaga canis | Bacteroidota | Flavobacteriia | NZ_CP171121.1 | 1 |
| Capnocytophaga cynodegmi | Bacteroidota | Flavobacteriia | NZ_CP022378.1 | 1 |
| Capnocytophaga endodontalis | Bacteroidota | Flavobacteriia | NZ_CP022022.1 | 1 |
| Capnocytophaga gingivalis | Bacteroidota | Flavobacteriia | NZ_CP022386.1 | 1 |
| Capnocytophaga haemolytica | Bacteroidota | Flavobacteriia | NZ_LT906449.1 | 1 |
| Capnocytophaga ochracea DSM 7271 | Bacteroidota | Flavobacteriia | NC_013162.1 | 1 |
| Capnocytophaga sputigena | Bacteroidota | Flavobacteriia | NZ_CP022379.1 | 1 |
| Capnocytophaga stomatis | Bacteroidota | Flavobacteriia | NZ_CP022387.1 | 1 |
| Cellulophaga algicola DSM 14237 | Bacteroidota | Flavobacteriia | NC_014934.1 | 1 |
| Cellulophaga baltica 18 | Bacteroidota | Flavobacteriia | NZ_CP009976.1 | 1 |
| Cellulophaga lytica DSM 7489 | Bacteroidota | Flavobacteriia | NC_015167.1 | 1 |
| Cellulophaga omnivescoria | Bacteroidota | Flavobacteriia | NZ_CP115819.1 | 1 |
| Changchengzhania lutea | Bacteroidota | Flavobacteriia | NZ_CP039456.1 | 1 |
| Christiangramia flava JLT2011 | Bacteroidota | Flavobacteriia | NZ_CP016359.1 | 1 |
| Christiangramia forsetii KT0803 | Bacteroidota | Flavobacteriia | NC_008571.1 | 1 |
| Christiangramia fulva | Bacteroidota | Flavobacteriia | NZ_CP028136.1 | 1 |
| Christiangramia salexigens | Bacteroidota | Flavobacteriia | NZ_CP018153.1 | 1 |
| Chryseobacterium arthrosphaerae | Bacteroidota | Flavobacteriia | NZ_CP073074.1 | 1 |
| Chryseobacterium aureum | Bacteroidota | Flavobacteriia | NZ_CP034661.1 | 1 |
| Chryseobacterium balustinum | Bacteroidota | Flavobacteriia | NZ_CP033934.1 | 1 |
| Chryseobacterium bernardetii | Bacteroidota | Flavobacteriia | NZ_CP033932.1 | 1 |
| Chryseobacterium camelliae | Bacteroidota | Flavobacteriia | NZ_CP022986.1 | 1 |
| Chryseobacterium carnipullorum | Bacteroidota | Flavobacteriia | NZ_CP033920.1 | 1 |
| Chryseobacterium fluminis | Bacteroidota | Flavobacteriia | NZ_CP111121.1 | 1 |
| Chryseobacterium gallinarum | Bacteroidota | Flavobacteriia | NZ_CP009928.1 | 1 |

| | | | | |
|---|---|---|---|---|
| Chryseobacterium gambrini | Bacteroidota | Flavobacteriia | NZ_CP115857.1 | 1 |
| Chryseobacterium glaciei | Bacteroidota | Flavobacteriia | NZ_CP015199.1 | 1 |
| Chryseobacterium gleum | Bacteroidota | Flavobacteriia | NZ_CP068486.1 | 1 |
| Chryseobacterium gotjawalense | Bacteroidota | Flavobacteriia | NZ_CP124855.1 | 1 |
| Chryseobacterium indoltheticum | Bacteroidota | Flavobacteriia | NZ_CP033929.1 | 1 |
| Chryseobacterium joostei | Bacteroidota | Flavobacteriia | NZ_CP033926.1 | 1 |
| Chryseobacterium lactis | Bacteroidota | Flavobacteriia | NZ_CP033924.1 | 1 |
| Chryseobacterium manosquense | Bacteroidota | Flavobacteriia | NZ_CP060203.1 | 1 |
| Chryseobacterium nakagawai | Bacteroidota | Flavobacteriia | NZ_LR134386.1 | 1 |
| Chryseobacterium nepalense | Bacteroidota | Flavobacteriia | NZ_CP096203.1 | 1 |
| Chryseobacterium oryzae | Bacteroidota | Flavobacteriia | NZ_CP094529.1 | 1 |
| Chryseobacterium paludis | Bacteroidota | Flavobacteriia | NZ_CP099966.1 | 1 |
| Chryseobacterium piperi | Bacteroidota | Flavobacteriia | NZ_CP023049.2 | 1 |
| Chryseobacterium salivictor | Bacteroidota | Flavobacteriia | NZ_CP037954.1 | 1 |
| Chryseobacterium shandongense | Bacteroidota | Flavobacteriia | NZ_CP033914.1 | 1 |
| Chryseobacterium suipulveris | Bacteroidota | Flavobacteriia | NZ_CP094532.1 | 1 |
| Chryseobacterium taihuense | Bacteroidota | Flavobacteriia | NZ_LR215974.1 | 1 |
| Chryseobacterium taklimakanense | Bacteroidota | Flavobacteriia | NZ_LT906465.1 | 1 |
| Cloacibacterium caeni | Bacteroidota | Flavobacteriia | NZ_OU015321.1 | 1 |
| Cloacibacterium normanense | Bacteroidota | Flavobacteriia | NZ_CP034157.1 | 1 |
| Constantimarinum furrinae | Bacteroidota | Flavobacteriia | NZ_CP052909.1 | 1 |
| Costertonia aggregata | Bacteroidota | Flavobacteriia | NZ_CP058595.1 | 1 |
| Croceibacter atlanticus HTCC2559 | Bacteroidota | Flavobacteriia | NC_014230.1 | 1 |
| Croceimicrobium hydrocarbonivorans | Bacteroidota | Flavobacteriia | NZ_CP060139.1 | 1 |
| Cruoricaptor ignavus | Bacteroidota | Flavobacteriia | NZ_CP063145.1 | 1 |
| Dokdonia donghaensis DSW-1 | Bacteroidota | Flavobacteriia | NZ_CP015125.1 | 1 |
| Elizabethkingia anophelis R26 | Bacteroidota | Flavobacteriia | NZ_CP023401.1 | 1 |
| Elizabethkingia bruuniana | Bacteroidota | Flavobacteriia | NZ_CP014337.1 | 1 |
| Elizabethkingia meningoseptica | Bacteroidota | Flavobacteriia | NZ_LS483376.1 | 1 |
| Elizabethkingia ursingii | Bacteroidota | Flavobacteriia | NZ_CP016377.1 | 1 |
| Empedobacter falsenii | Bacteroidota | Flavobacteriia | NZ_CP040908.1 | 1 |
| Empedobacter stercoris | Bacteroidota | Flavobacteriia | NZ_CP053698.1 | 1 |

| | | | | |
|---|---|---|---|---|
| Epilithonimonas vandammei | Bacteroidota | Flavobacteriia | NZ_CP034161.1 | 1 |
| Euzebyella marina | Bacteroidota | Flavobacteriia | NZ_CP032050.1 | 1 |
| Faecalibacter bovis | Bacteroidota | Flavobacteriia | NZ_CP072842.1 | 1 |
| Flagellimonas lutaonensis | Bacteroidota | Flavobacteriia | NZ_CP011071.1 | 1 |
| Flagellimonas maritima | Bacteroidota | Flavobacteriia | NZ_CP030104.1 | 1 |
| Flagellimonas oceani | Bacteroidota | Flavobacteriia | NZ_CP049616.1 | 1 |
| Flavivirga abyssicola | Bacteroidota | Flavobacteriia | NZ_CP141266.1 | 1 |
| Flavivirga eckloniae | Bacteroidota | Flavobacteriia | NZ_CP025791.1 | 1 |
| Flavobacterium acetivorans | Bacteroidota | Flavobacteriia | NZ_CP087132.1 | 1 |
| Flavobacterium aestivum | Bacteroidota | Flavobacteriia | NZ_CP113977.2 | 1 |
| Flavobacterium agricola | Bacteroidota | Flavobacteriia | NZ_CP081495.1 | 1 |
| Flavobacterium album | Bacteroidota | Flavobacteriia | NZ_CP029186.1 | 1 |
| Flavobacterium alkalisoli | Bacteroidota | Flavobacteriia | NZ_CP042831.1 | 1 |
| Flavobacterium ammonificans | Bacteroidota | Flavobacteriia | NZ_AP025185.1 | 1 |
| Flavobacterium ammoniigenes | Bacteroidota | Flavobacteriia | NZ_AP025184.1 | 1 |
| Flavobacterium anhuiense | Bacteroidota | Flavobacteriia | NZ_CP016907.1 | 1 |
| Flavobacterium aquiphilum | Bacteroidota | Flavobacteriia | NZ_CP114288.1 | 1 |
| Flavobacterium arcticum | Bacteroidota | Flavobacteriia | NZ_CP031188.1 | 1 |
| Flavobacterium ardleyense | Bacteroidota | Flavobacteriia | NZ_CP137891.1 | 1 |
| Flavobacterium azooxidireducens | Bacteroidota | Flavobacteriia | NZ_CP096205.1 | 1 |
| Flavobacterium bizetiae | Bacteroidota | Flavobacteriia | NZ_CP091861.1 | 1 |
| Flavobacterium branchiophilum FL-15 | Bacteroidota | Flavobacteriia | NC_016001.1 | 1 |
| Flavobacterium capsici | Bacteroidota | Flavobacteriia | NZ_CP134890.1 | 1 |
| Flavobacterium channae | Bacteroidota | Flavobacteriia | NZ_CP089096.1 | 1 |
| Flavobacterium collinsii | Bacteroidota | Flavobacteriia | NZ_OX336425.1 | 1 |
| Flavobacterium columnare | Bacteroidota | Flavobacteriia | NZ_CP054494.1 | 1 |
| Flavobacterium commune | Bacteroidota | Flavobacteriia | NZ_CP017774.1 | 1 |
| Flavobacterium covae | Bacteroidota | Flavobacteriia | NZ_CP013992.1 | 1 |
| Flavobacterium crassostreae | Bacteroidota | Flavobacteriia | NZ_CP017688.1 | 1 |
| Flavobacterium crocinum | Bacteroidota | Flavobacteriia | NZ_CP029255.1 | 1 |
| Flavobacterium cupriresistens | Bacteroidota | Flavobacteriia | NZ_CP087134.1 | 1 |
| Flavobacterium cyclinae | Bacteroidota | Flavobacteriia | NZ_CP089095.1 | 1 |

| | | | | |
|---|---|---|---|---|
| Flavobacterium dauae | Bacteroidota | Flavobacteriia | NZ_CP130821.1 | 1 |
| Flavobacterium davisii | Bacteroidota | Flavobacteriia | NZ_CP097869.1 | 1 |
| Flavobacterium eburneipallidum | Bacteroidota | Flavobacteriia | NZ_CP114291.2 | 1 |
| Flavobacterium endoglycinae | Bacteroidota | Flavobacteriia | NZ_CP071448.1 | 1 |
| Flavobacterium faecale | Bacteroidota | Flavobacteriia | NZ_CP020918.1 | 1 |
| Flavobacterium flavigenum | Bacteroidota | Flavobacteriia | NZ_CP114285.2 | 1 |
| Flavobacterium flavipigmentatum | Bacteroidota | Flavobacteriia | NZ_CP087133.1 | 1 |
| Flavobacterium fluviale | Bacteroidota | Flavobacteriia | NZ_CP030261.1 | 1 |
| Flavobacterium galactosidilyticum | Bacteroidota | Flavobacteriia | NZ_CP087135.1 | 1 |
| Flavobacterium gelatinilyticum | Bacteroidota | Flavobacteriia | NZ_CP114287.1 | 1 |
| Flavobacterium gilvum | Bacteroidota | Flavobacteriia | NZ_CP017479.1 | 1 |
| Flavobacterium ginsenosidimutans | Bacteroidota | Flavobacteriia | NZ_CP147988.1 | 1 |
| Flavobacterium gyeonganense | Bacteroidota | Flavobacteriia | NZ_CP121112.1 | 1 |
| Flavobacterium hankyongi | Bacteroidota | Flavobacteriia | NZ_CP085725.1 | 1 |
| Flavobacterium humidisoli | Bacteroidota | Flavobacteriia | NZ_CP096829.1 | 1 |
| Flavobacterium indicum GPTSA100-9 = DSM 17447 | Bacteroidota | Flavobacteriia | NC_017025.1 | 1 |
| Flavobacterium inviolabile | Bacteroidota | Flavobacteriia | NZ_CP058278.1 | 1 |
| Flavobacterium johnsoniae UW101 | Bacteroidota | Flavobacteriia | NC_009441.1 | 1 |
| Flavobacterium jumunjinense | Bacteroidota | Flavobacteriia | NZ_CP091285.1 | 1 |
| Flavobacterium keumense | Bacteroidota | Flavobacteriia | NZ_CP092332.1 | 1 |
| Flavobacterium kingsejongi | Bacteroidota | Flavobacteriia | NZ_CP020919.1 | 1 |
| Flavobacterium lacustre | Bacteroidota | Flavobacteriia | NZ_CP114882.2 | 1 |
| Flavobacterium limnophilum | Bacteroidota | Flavobacteriia | NZ_CP114289.2 | 1 |
| Flavobacterium litorale | Bacteroidota | Flavobacteriia | NZ_CP080429.1 | 1 |
| Flavobacterium luteolum | Bacteroidota | Flavobacteriia | NZ_CP114286.1 | 1 |
| Flavobacterium magnum | Bacteroidota | Flavobacteriia | NZ_CP028811.1 | 1 |
| Flavobacterium marginilacus | Bacteroidota | Flavobacteriia | NZ_CP113975.1 | 1 |
| Flavobacterium nackdongense | Bacteroidota | Flavobacteriia | NZ_CP037933.1 | 1 |
| Flavobacterium nakdongensis | Bacteroidota | Flavobacteriia | NZ_CP133721.1 | 1 |
| Flavobacterium nitratireducens | Bacteroidota | Flavobacteriia | NZ_CP121111.1 | 1 |
| Flavobacterium nitrogenifigens | Bacteroidota | Flavobacteriia | NZ_CP043612.1 | 1 |
| Flavobacterium okayamense | Bacteroidota | Flavobacteriia | NZ_AP024749.1 | 1 |

| | | | | |
|---|---|---|---|---|
| Flavobacterium ovatum | Bacteroidota | Flavobacteriia | NZ_CP160035.1 | 1 |
| Flavobacterium pallidum | Bacteroidota | Flavobacteriia | NZ_CP029187.1 | 1 |
| Flavobacterium panacagri | Bacteroidota | Flavobacteriia | NZ_CP119766.1 | 1 |
| Flavobacterium panici | Bacteroidota | Flavobacteriia | NZ_CP100437.1 | 1 |
| Flavobacterium plurextorum | Bacteroidota | Flavobacteriia | NZ_CP100442.1 | 1 |
| Flavobacterium praedii | Bacteroidota | Flavobacteriia | NZ_CP113948.1 | 1 |
| Flavobacterium psychrophilum | Bacteroidota | Flavobacteriia | NZ_CP039120.2 | 1 |
| Flavobacterium psychrotrophum | Bacteroidota | Flavobacteriia | NZ_CP031557.1 | 1 |
| Flavobacterium rivulicola | Bacteroidota | Flavobacteriia | NZ_CP121446.1 | 1 |
| Flavobacterium sangjuense | Bacteroidota | Flavobacteriia | NZ_CP038810.1 | 1 |
| Flavobacterium sediminilitoris | Bacteroidota | Flavobacteriia | NZ_CP090145.1 | 1 |
| Flavobacterium sediminis | Bacteroidota | Flavobacteriia | NZ_CP029463.1 | 1 |
| Flavobacterium soyae | Bacteroidota | Flavobacteriia | NZ_CP150845.1 | 1 |
| Fluviicola taffensis DSM 16823 | Bacteroidota | Flavobacteriia | NC_015321.1 | 1 |
| Formosa agariphila KMM 3901 | Bacteroidota | Flavobacteriia | NZ_HG315671.1 | 1 |
| Formosa sediminum | Bacteroidota | Flavobacteriia | NZ_CP041637.1 | 1 |
| Hwangdonia lutea | Bacteroidota | Flavobacteriia | NZ_CP136521.1 | 1 |
| Ichthyobacterium seriolicida | Bacteroidota | Flavobacteriia | NZ_AP014564.1 | 1 |
| Imtechella halotolerans | Bacteroidota | Flavobacteriia | NZ_CP117969.2 | 1 |
| Kaistella antarctica | Bacteroidota | Flavobacteriia | NZ_LR134441.1 | 1 |
| Kaistella carnis | Bacteroidota | Flavobacteriia | NZ_CP034159.1 | 1 |
| Kaistella faecalis | Bacteroidota | Flavobacteriia | NZ_CP087583.1 | 1 |
| Kaistella jeonii | Bacteroidota | Flavobacteriia | NZ_LR134503.1 | 1 |
| Kaistella polysaccharea | Bacteroidota | Flavobacteriia | NZ_CP084528.1 | 1 |
| Kordia antarctica | Bacteroidota | Flavobacteriia | NZ_CP019288.1 | 1 |
| Lacinutrix neustonica | Bacteroidota | Flavobacteriia | NZ_CP113088.1 | 1 |
| Lacinutrix venerupis | Bacteroidota | Flavobacteriia | NZ_CP019352.1 | 1 |
| Leeuwenhoekiella palythoae | Bacteroidota | Flavobacteriia | NZ_CP084318.1 | 1 |
| Lutibacter profundi | Bacteroidota | Flavobacteriia | NZ_CP013355.1 | 1 |
| Lutimonas zeaxanthinifaciens | Bacteroidota | Flavobacteriia | NZ_CP129964.1 | 1 |
| Mangrovimonas cancribranchiae | Bacteroidota | Flavobacteriia | NZ_CP136924.1 | 1 |
| Maribacter aestuarii | Bacteroidota | Flavobacteriia | NZ_CP107031.2 | 1 |

| | | | | |
|---|---|---|---|---|
| Maribacter cobaltidurans | Bacteroidota | Flavobacteriia | NZ_CP022957.1 | 1 |
| Maribacter dokdonensis | Bacteroidota | Flavobacteriia | NZ_LT629754.1 | 1 |
| Maribacter halichondriae | Bacteroidota | Flavobacteriia | NZ_CP107030.1 | 1 |
| Maribacter hydrothermalis | Bacteroidota | Flavobacteriia | NZ_CP018760.1 | 1 |
| Maribacter litopenaei | Bacteroidota | Flavobacteriia | NZ_CP104205.1 | 1 |
| Mariniflexile litorale | Bacteroidota | Flavobacteriia | NZ_CP155618.1 | 1 |
| Mesoflavibacter profundi | Bacteroidota | Flavobacteriia | NZ_CP061703.1 | 1 |
| Muriicola soli | Bacteroidota | Flavobacteriia | NZ_CP035544.1 | 1 |
| Myroides albus | Bacteroidota | Flavobacteriia | NZ_CP102754.1 | 1 |
| Myroides fluvii | Bacteroidota | Flavobacteriia | NZ_CP039934.1 | 1 |
| Myroides odoratimimus | Bacteroidota | Flavobacteriia | NZ_CP037427.1 | 1 |
| Myroides odoratus | Bacteroidota | Flavobacteriia | NZ_CP068108.1 | 1 |
| Myroides profundi | Bacteroidota | Flavobacteriia | NZ_CP010817.1 | 1 |
| Nonlabens dokdonensis DSW-6 | Bacteroidota | Flavobacteriia | NC_020156.1 | 1 |
| Nonlabens marinus S1-08 | Bacteroidota | Flavobacteriia | NZ_AP014548.1 | 1 |
| Nonlabens ponticola | Bacteroidota | Flavobacteriia | NZ_CP034549.1 | 1 |
| Nonlabens spongiae | Bacteroidota | Flavobacteriia | NZ_CP019344.1 | 1 |
| Nonlabens tegetincola | Bacteroidota | Flavobacteriia | NZ_CP019342.1 | 1 |
| Nonlabens ulvanivorans | Bacteroidota | Flavobacteriia | NZ_CP136694.1 | 1 |
| Ornithobacterium rhinotracheale DSM 15997 | Bacteroidota | Flavobacteriia | NC_018016.1 | 1 |
| Owenweeksia hongkongensis DSM 17368 | Bacteroidota | Flavobacteriia | NC_016599.1 | 1 |
| Paenimyroides aestuarii | Bacteroidota | Flavobacteriia | NZ_CP102382.1 | 1 |
| Paracrocinitomix mangrovi | Bacteroidota | Flavobacteriia | NZ_CP091819.1 | 1 |
| Parvicella tangerina | Bacteroidota | Flavobacteriia | NZ_OU015584.1 | 1 |
| Polaribacter atrinae | Bacteroidota | Flavobacteriia | NZ_CP150660.1 | 1 |
| Polaribacter batillariae | Bacteroidota | Flavobacteriia | NZ_CP071795.1 | 1 |
| Polaribacter butkevichii | Bacteroidota | Flavobacteriia | NZ_CP150661.1 | 1 |
| Polaribacter cellanae | Bacteroidota | Flavobacteriia | NZ_CP071869.1 | 1 |
| Polaribacter dokdonensis | Bacteroidota | Flavobacteriia | NZ_CP101505.1 | 1 |
| Polaribacter gangjinensis | Bacteroidota | Flavobacteriia | NZ_CP150662.1 | 1 |
| Polaribacter haliotis | Bacteroidota | Flavobacteriia | NZ_CP061813.1 | 1 |
| Polaribacter huanghezhanensis | Bacteroidota | Flavobacteriia | NZ_CP128595.1 | 1 |

| | | | | |
|---|---|---|---|---|
| Polaribacter litorisediminis | Bacteroidota | Flavobacteriia | NZ_CP082966.1 | 1 |
| Polaribacter marinaquae | Bacteroidota | Flavobacteriia | NZ_CP150496.1 | 1 |
| Polaribacter pacificus | Bacteroidota | Flavobacteriia | NZ_CP150664.1 | 1 |
| Polaribacter pectinis | Bacteroidota | Flavobacteriia | NZ_CP060695.1 | 1 |
| Polaribacter reichenbachii | Bacteroidota | Flavobacteriia | NZ_CP019419.1 | 1 |
| Polaribacter sejongensis | Bacteroidota | Flavobacteriia | NZ_CP103460.1 | 1 |
| Polaribacter staleyi | Bacteroidota | Flavobacteriia | NZ_CP150665.1 | 1 |
| Polaribacter tangerinus | Bacteroidota | Flavobacteriia | NZ_CP150668.1 | 1 |
| Polaribacter vadi | Bacteroidota | Flavobacteriia | NZ_CP017477.1 | 1 |
| Pseudalgibacter alginicilyticus | Bacteroidota | Flavobacteriia | NZ_CP012898.1 | 1 |
| Pseudotamlana carrageenivorans | Bacteroidota | Flavobacteriia | NZ_CP025938.1 | 1 |
| Psychroflexus torquis ATCC 700755 | Bacteroidota | Flavobacteriia | NC_018721.1 | 1 |
| Psychroserpens ponticola | Bacteroidota | Flavobacteriia | NZ_CP116221.1 | 1 |
| Pukyongia salina | Bacteroidota | Flavobacteriia | NZ_CP027062.1 | 1 |
| Rasiella rasia | Bacteroidota | Flavobacteriia | NZ_CP049057.1 | 1 |
| Riemerella anatipestifer ATCC 11845 = DSM 15868 | Bacteroidota | Flavobacteriia | NC_014738.1 | 1 |
| Robiginitalea biformata HTCC2501 | Bacteroidota | Flavobacteriia | NC_013222.1 | 1 |
| Salegentibacter mishustinae | Bacteroidota | Flavobacteriia | NZ_CP084316.1 | 1 |
| Salegentibacter salegens | Bacteroidota | Flavobacteriia | NZ_LT670848.1 | 1 |
| Salinimicrobium tongyeongense | Bacteroidota | Flavobacteriia | NZ_CP069620.1 | 1 |
| Sanyastnella coralliicola | Bacteroidota | Flavobacteriia | NZ_CP132543.1 | 1 |
| Siansivirga zeaxanthinifaciens CC-SAMT-1 | Bacteroidota | Flavobacteriia | NZ_CP007202.1 | 1 |
| Sinomicrobium kalidii | Bacteroidota | Flavobacteriia | NZ_CP089211.1 | 1 |
| Tenacibaculum aestuariivivum | Bacteroidota | Flavobacteriia | NZ_OZ208414.1 | 1 |
| Tenacibaculum dicentrarchi | Bacteroidota | Flavobacteriia | NZ_OZ038524.1 | 1 |
| Tenacibaculum insulae | Bacteroidota | Flavobacteriia | NZ_OZ208512.1 | 1 |
| Tenacibaculum jejuense | Bacteroidota | Flavobacteriia | NZ_LT899436.1 | 1 |
| Tenacibaculum maritimum NCIMB 2154 | Bacteroidota | Flavobacteriia | NZ_LT634361.1 | 1 |
| Tenacibaculum mesophilum | Bacteroidota | Flavobacteriia | NZ_CP045192.1 | 1 |
| Tenacibaculum pacificus | Bacteroidota | Flavobacteriia | NZ_CP115917.1 | 1 |
| Tenacibaculum platacis | Bacteroidota | Flavobacteriia | NZ_OZ038523.1 | 1 |
| Tenacibaculum polynesiense | Bacteroidota | Flavobacteriia | NZ_OZ040189.1 | 1 |

| | | | | |
|---|---|---|---|---|
| Tenacibaculum retecalamus | Bacteroidota | Flavobacteriia | NZ_CP115916.1 | 1 |
| Tenacibaculum singaporense | Bacteroidota | Flavobacteriia | NZ_CP032548.1 | 1 |
| Tenacibaculum tangerinum | Bacteroidota | Flavobacteriia | NZ_CP122539.1 | 1 |
| Tenacibaculum todarodis | Bacteroidota | Flavobacteriia | NZ_CP018155.1 | 1 |
| Tenacibaculum vairaonense | Bacteroidota | Flavobacteriia | NZ_OZ038525.1 | 1 |
| Thalassobellus suaedae | Bacteroidota | Flavobacteriia | NZ_CP134536.1 | 1 |
| Urechidicola croceus | Bacteroidota | Flavobacteriia | NZ_CP017478.1 | 1 |
| Weeksella virosa | Bacteroidota | Flavobacteriia | NZ_LR134404.1 | 1 |
| Wenyingzhuangia fucanilytica | Bacteroidota | Flavobacteriia | NZ_CP014224.1 | 1 |
| Winogradskyella forsetii | Bacteroidota | Flavobacteriia | NZ_CP053348.1 | 1 |
| Winogradskyella schleiferi | Bacteroidota | Flavobacteriia | NZ_CP053351.1 | 1 |
| Winogradskyella sediminis | Bacteroidota | Flavobacteriia | NZ_LT629774.1 | 1 |
| Zhouia spongiae | Bacteroidota | Flavobacteriia | NZ_CP094326.1 | 1 |
| Zobellia alginiliquefaciens | Bacteroidota | Flavobacteriia | NZ_CP119758.1 | 1 |
| Zobellia galactanivorans | Bacteroidota | Flavobacteriia | NC_015844.1 | 1 |
| Zobellia russellii | Bacteroidota | Flavobacteriia | NZ_CP139000.1 | 1 |
| Zunongwangia profunda SM-A87 | Bacteroidota | Flavobacteriia | NC_014041.1 | 1 |
| Aureispira anguillae | Bacteroidota | Saprospiria | NZ_AP026867.1 | 1 |
| Haliscomenobacter hydrossis DSM 1100 | Bacteroidota | Saprospiria | NC_015510.1 | 1 |
| Saprospira grandis str. Lewin | Bacteroidota | Saprospiria | NC_016940.1 | 1 |
| Aurantibacillus circumpalustris | Bacteroidota | Sphingobacteriia | NZ_CP121197.1 | 1 |
| Mucilaginibacter celer | Bacteroidota | Sphingobacteriia | NZ_CP032869.1 | 1 |
| Mucilaginibacter daejeonensis | Bacteroidota | Sphingobacteriia | NZ_CP086068.1 | 1 |
| Mucilaginibacter ginkgonis | Bacteroidota | Sphingobacteriia | NZ_CP066775.1 | 1 |
| Mucilaginibacter ginsenosidivorans | Bacteroidota | Sphingobacteriia | NZ_CP042436.1 | 1 |
| Mucilaginibacter ginsenosidivorax | Bacteroidota | Sphingobacteriia | NZ_CP042437.1 | 1 |
| Mucilaginibacter gossypii | Bacteroidota | Sphingobacteriia | NZ_CP071878.2 | 1 |
| Mucilaginibacter gotjawali | Bacteroidota | Sphingobacteriia | NZ_AP017313.1 | 1 |
| Mucilaginibacter jinjuensis | Bacteroidota | Sphingobacteriia | NZ_CP117167.1 | 1 |
| Mucilaginibacter mali | Bacteroidota | Sphingobacteriia | NZ_CP054139.1 | 1 |
| Mucilaginibacter mallensis | Bacteroidota | Sphingobacteriia | NZ_LT629740.1 | 1 |
| Mucilaginibacter paludis DSM 18603 | Bacteroidota | Sphingobacteriia | NZ_CM001403.1 | 1 |

| | | | | |
|---|---|---|---|---|
| Mucilaginibacter robiniae | Bacteroidota | Sphingobacteriia | NZ_CP051682.1 | 1 |
| Mucilaginibacter sabulilitoris | Bacteroidota | Sphingobacteriia | NZ_CP139558.1 | 1 |
| Mucilaginibacter xinganensis | Bacteroidota | Sphingobacteriia | NZ_CP022743.1 | 1 |
| Pedobacter aquae | Bacteroidota | Sphingobacteriia | NZ_CP043329.1 | 1 |
| Pedobacter endophyticus | Bacteroidota | Sphingobacteriia | NZ_CP064939.1 | 1 |
| Pedobacter foliorum | Bacteroidota | Sphingobacteriia | NZ_CP162529.1 | 1 |
| Pedobacter heparinus DSM 2366 | Bacteroidota | Sphingobacteriia | NC_013061.1 | 1 |
| Pedobacter lusitanus | Bacteroidota | Sphingobacteriia | NZ_CP157278.1 | 1 |
| Pedobacter mucosus | Bacteroidota | Sphingobacteriia | NZ_CP087585.1 | 1 |
| Pedobacter riviphilus | Bacteroidota | Sphingobacteriia | NZ_CP061171.1 | 1 |
| Pedobacter roseus | Bacteroidota | Sphingobacteriia | NZ_CP060723.1 | 1 |
| Pedobacter schmidteae | Bacteroidota | Sphingobacteriia | NZ_LS999839.1 | 1 |
| Pedobacter suwonensis | Bacteroidota | Sphingobacteriia | NZ_CP031708.1 | 1 |
| Pseudopedobacter saltans DSM 12145 | Bacteroidota | Sphingobacteriia | NC_015177.1 | 1 |
| Solitalea canadensis DSM 3403 | Bacteroidota | Sphingobacteriia | NC_017770.1 | 1 |
| Solitalea lacus | Bacteroidota | Sphingobacteriia | NZ_CP091740.1 | 1 |
| Sphingobacterium bambusae | Bacteroidota | Sphingobacteriia | NZ_CP138332.1 | 1 |
| Sphingobacterium cellulitidis | Bacteroidota | Sphingobacteriia | NZ_CP171136.1 | 1 |
| Sphingobacterium hotanense | Bacteroidota | Sphingobacteriia | NZ_CP030848.1 | 1 |
| Sphingobacterium lactis | Bacteroidota | Sphingobacteriia | NZ_CP049246.1 | 1 |
| Sphingobacterium mizutaii | Bacteroidota | Sphingobacteriia | NZ_LT906468.1 | 1 |
| Sphingobacterium multivorum | Bacteroidota | Sphingobacteriia | NZ_CP069793.1 | 1 |
| Sphingobacterium oryzagri | Bacteroidota | Sphingobacteriia | NZ_CP117880.1 | 1 |
| Sphingobacterium paramultivorum | Bacteroidota | Sphingobacteriia | NZ_CP058555.1 | 1 |
| Sphingobacterium psychroaquaticum | Bacteroidota | Sphingobacteriia | NZ_CP038029.1 | 1 |
| Sphingobacterium siyangense | Bacteroidota | Sphingobacteriia | NZ_CP080574.1 | 1 |
| Sphingobacterium spiritivorum | Bacteroidota | Sphingobacteriia | NZ_CP068084.1 | 1 |
| Sphingobacterium thalpophilum | Bacteroidota | Sphingobacteriia | NZ_LR590484.1 | 1 |
| Sphingobacterium zhuxiongii | Bacteroidota | Sphingobacteriia | NZ_CP045652.1 | 1 |
| Cyclonatronum proteinivorum | Balneolota | Balneolia | NZ_CP027806.1 | 2 |
| Bacteriovorax stolpii | Bdellovibrionota | Bacteriovoracia | NZ_CP025704.1 | 2 |
| Halobacteriovorax marinus SJ | Bdellovibrionota | Bacteriovoracia | NC_016620.1 | 2 |

| | | | | |
|---|---|---|---|---|
| Peredibacter starrii | Bdellovibrionota | Bacteriovoracia | NZ_CP139487.1 | 2 |
| Bdellovibrio bacteriovorus HD100 | Bdellovibrionota | Bdellovibrionia | NC_005363.1 | 1 |
| Bdellovibrio reynosensis | Bdellovibrionota | Bdellovibrionia | NZ_CP093442.1 | 1 |
| Pseudobdellovibrio exovorus JSS | Bdellovibrionota | Bdellovibrionia | NC_020813.1 | 1 |
| Fluviispira sanaruensis | Bdellovibrionota | Oligoflexia | NZ_AP019368.1 | 1 |
| Silvanigrella aquatica | Bdellovibrionota | Oligoflexia | NZ_CP017834.1 | 1 |
| Spirobacillus cienkowskii | Bdellovibrionota | Oligoflexia | NZ_CP146516.1 | 1 |
| Caldisericum exile AZM16c01 | Caldisericota | Caldisericia | NC_017096.1 | 2 |
| Caldithrix abyssi DSM 13497 | Calditrichota | Calditrichia | NZ_CP018099.1 | 2 |
| Hippea maritima DSM 10411 | Campylobacterota | Desulfurellia | NC_015318.1 | 2 |
| Aliarcobacter butzleri RM4018 | Campylobacterota | Epsilonproteobacteria | NC_009850.1 | 1 |
| Aliarcobacter cibarius | Campylobacterota | Epsilonproteobacteria | NZ_CP054051.1 | 1 |
| Aliarcobacter cryaerophilus ATCC 43158 | Campylobacterota | Epsilonproteobacteria | NZ_CP032823.1 | 1 |
| Aliarcobacter faecis | Campylobacterota | Epsilonproteobacteria | NZ_CP053837.1 | 1 |
| Aliarcobacter lanthieri | Campylobacterota | Epsilonproteobacteria | NZ_CP053839.1 | 1 |
| Aliarcobacter skirrowii CCUG 10374 | Campylobacterota | Epsilonproteobacteria | NZ_CP032099.1 | 1 |
| Aliarcobacter thereius LMG 24486 | Campylobacterota | Epsilonproteobacteria | NZ_CP035926.1 | 1 |
| Aliarcobacter trophiarum LMG 25534 | Campylobacterota | Epsilonproteobacteria | NZ_CP031367.1 | 1 |
| Aliarcobacter vitoriensis | Campylobacterota | Epsilonproteobacteria | NZ_CP182882.1 | 1 |
| Arcobacter acticola | Campylobacterota | Epsilonproteobacteria | NZ_CP042652.1 | 1 |
| Arcobacter aquimarinus | Campylobacterota | Epsilonproteobacteria | NZ_CP030944.1 | 1 |
| Arcobacter cloacae | Campylobacterota | Epsilonproteobacteria | NZ_CP053833.1 | 1 |
| Arcobacter defluvii | Campylobacterota | Epsilonproteobacteria | NZ_CP053835.1 | 1 |
| Arcobacter ellisii | Campylobacterota | Epsilonproteobacteria | NZ_CP032097.1 | 1 |
| Arcobacter nitrofigilis DSM 7299 | Campylobacterota | Epsilonproteobacteria | NC_014166.1 | 1 |
| Arcobacter peruensis | Campylobacterota | Epsilonproteobacteria | NZ_CP032363.1 | 1 |
| Arcobacter porcinus | Campylobacterota | Epsilonproteobacteria | NZ_CP036246.2 | 1 |
| Arcobacter roscoffensis | Campylobacterota | Epsilonproteobacteria | NZ_CP100595.1 | 1 |
| Arcobacter suis CECT 7833 | Campylobacterota | Epsilonproteobacteria | NZ_CP032100.1 | 1 |
| Arcobacter venerupis | Campylobacterota | Epsilonproteobacteria | NZ_CP053840.1 | 1 |
| Caminibacter mediatlanticus TB-2 | Campylobacterota | Epsilonproteobacteria | NZ_CP040463.1 | 1 |
| Caminibacter pacificus | Campylobacterota | Epsilonproteobacteria | NZ_CP027432.2 | 1 |

| | | | | |
|---|---|---|---|---|
| Campylobacter armoricus | Campylobacterota | Epsilonproteobacteria | NZ_CP053825.1 | 1 |
| Campylobacter avium LMG 24591 | Campylobacterota | Epsilonproteobacteria | NZ_CP022347.1 | 1 |
| Campylobacter bilis | Campylobacterota | Epsilonproteobacteria | NZ_CP168723.1 | 1 |
| Campylobacter blaseri | Campylobacterota | Epsilonproteobacteria | NZ_CP053841.1 | 1 |
| Campylobacter californiensis | Campylobacterota | Epsilonproteobacteria | NZ_CP012545.1 | 1 |
| Campylobacter canadensis | Campylobacterota | Epsilonproteobacteria | NZ_CP035946.1 | 1 |
| Campylobacter coli | Campylobacterota | Epsilonproteobacteria | NZ_CP046317.1 | 1 |
| Campylobacter concisus | Campylobacterota | Epsilonproteobacteria | NZ_CP012541.1 | 1 |
| Campylobacter corcagiensis | Campylobacterota | Epsilonproteobacteria | NZ_CP053842.1 | 1 |
| Campylobacter cuniculorum DSM 23162 = LMG 24588 | Campylobacterota | Epsilonproteobacteria | NZ_CP020867.1 | 1 |
| Campylobacter curvus | Campylobacterota | Epsilonproteobacteria | NZ_CP053826.1 | 1 |
| Campylobacter fetus | Campylobacterota | Epsilonproteobacteria | NZ_CP059443.1 | 1 |
| Campylobacter geochelonis | Campylobacterota | Epsilonproteobacteria | NZ_CP053844.1 | 1 |
| Campylobacter gracilis | Campylobacterota | Epsilonproteobacteria | NZ_CP012196.1 | 1 |
| Campylobacter helveticus | Campylobacterota | Epsilonproteobacteria | NZ_CP020478.1 | 1 |
| Campylobacter hepaticus | Campylobacterota | Epsilonproteobacteria | NZ_CP031611.1 | 1 |
| Campylobacter hominis ATCC BAA-381 | Campylobacterota | Epsilonproteobacteria | NC_009714.1 | 1 |
| Campylobacter hyointestinalis subsp. lawsonii | Campylobacterota | Epsilonproteobacteria | NZ_CP053828.1 | 1 |
| Campylobacter iguaniorum | Campylobacterota | Epsilonproteobacteria | NZ_CP009043.1 | 1 |
| Campylobacter insulaenigrae NCTC 12927 | Campylobacterota | Epsilonproteobacteria | NZ_CP007770.1 | 1 |
| Campylobacter jejuni subsp. jejuni NCTC 11168 = ATCC 7 | Campylobacterota | Epsilonproteobacteria | NC_002163.1 | 1 |
| Campylobacter lanienae NCTC 13004 | Campylobacterota | Epsilonproteobacteria | NZ_CP015578.1 | 1 |
| Campylobacter lari subsp. concheus | Campylobacterota | Epsilonproteobacteria | NZ_CP043426.1 | 1 |
| Campylobacter mucosalis | Campylobacterota | Epsilonproteobacteria | NZ_CP053831.1 | 1 |
| Campylobacter novaezeelandiae | Campylobacterota | Epsilonproteobacteria | NZ_CP076657.1 | 1 |
| Campylobacter ornithocola | Campylobacterota | Epsilonproteobacteria | NZ_CP053848.1 | 1 |
| Campylobacter peloridis LMG 23910 | Campylobacterota | Epsilonproteobacteria | NZ_CP007766.1 | 1 |
| Campylobacter pinnipediorum subsp. caledonicus | Campylobacterota | Epsilonproteobacteria | NZ_CP017018.1 | 1 |
| Campylobacter porcelli | Campylobacterota | Epsilonproteobacteria | NZ_CP018789.1 | 1 |
| Campylobacter rectus | Campylobacterota | Epsilonproteobacteria | NZ_CP012543.1 | 1 |
| Campylobacter showae | Campylobacterota | Epsilonproteobacteria | NZ_CP012544.1 | 1 |
| Campylobacter sputorum subsp. sputorum | Campylobacterota | Epsilonproteobacteria | NZ_CP043427.1 | 1 |

| | | | | |
|---|---|---|---|---|
| Campylobacter subantarcticus LMG 24377 | Campylobacterota | Epsilonproteobacteria | NZ_CP007773.1 | 1 |
| Campylobacter upsaliensis RM3940 | Campylobacterota | Epsilonproteobacteria | NZ_CP053849.1 | 1 |
| Campylobacter ureolyticus | Campylobacterota | Epsilonproteobacteria | NZ_CP053832.1 | 1 |
| Campylobacter vicugnae | Campylobacterota | Epsilonproteobacteria | NZ_CP144916.1 | 1 |
| Campylobacter volucris | Campylobacterota | Epsilonproteobacteria | NZ_CP043428.1 | 1 |
| Campylobacter vulpis | Campylobacterota | Epsilonproteobacteria | NZ_CP041617.1 | 1 |
| Candidatus Campylobacter infans | Campylobacterota | Epsilonproteobacteria | NZ_CP049075.1 | 1 |
| Candidatus Sulfurimonas baltica | Campylobacterota | Epsilonproteobacteria | NZ_CP054492.1 | 1 |
| Candidatus Sulfurimonas marisnigri | Campylobacterota | Epsilonproteobacteria | NZ_CP054493.1 | 1 |
| Halarcobacter anaerophilus | Campylobacterota | Epsilonproteobacteria | NZ_CP041070.1 | 1 |
| Halarcobacter bivalviorum | Campylobacterota | Epsilonproteobacteria | NZ_CP031217.1 | 1 |
| Halarcobacter ebronensis | Campylobacterota | Epsilonproteobacteria | NZ_CP053836.1 | 1 |
| Helicobacter acinonychis str. Sheeba | Campylobacterota | Epsilonproteobacteria | NC_008229.1 | 1 |
| Helicobacter apodemus | Campylobacterota | Epsilonproteobacteria | NZ_CP021886.1 | 1 |
| Helicobacter canadensis MIT 98-5491 | Campylobacterota | Epsilonproteobacteria | NZ_CM000776.2 | 1 |
| Helicobacter cetorum MIT 99-5656 | Campylobacterota | Epsilonproteobacteria | NC_017735.1 | 1 |
| Helicobacter cholecystus | Campylobacterota | Epsilonproteobacteria | NZ_LR134518.1 | 1 |
| Helicobacter cinaedi PAGU611 | Campylobacterota | Epsilonproteobacteria | NC_017761.1 | 1 |
| Helicobacter enhydrae | Campylobacterota | Epsilonproteobacteria | NZ_CP016503.1 | 1 |
| Helicobacter felis ATCC 49179 | Campylobacterota | Epsilonproteobacteria | NC_014810.2 | 1 |
| Helicobacter felistomachi | Campylobacterota | Epsilonproteobacteria | NZ_AP028022.1 | 1 |
| Helicobacter gastrocanis | Campylobacterota | Epsilonproteobacteria | NZ_AP024814.1 | 1 |
| Helicobacter hepaticus ATCC 51449 | Campylobacterota | Epsilonproteobacteria | NC_004917.1 | 1 |
| Helicobacter himalayensis | Campylobacterota | Epsilonproteobacteria | NZ_CP014991.1 | 1 |
| Helicobacter mastomyrinus | Campylobacterota | Epsilonproteobacteria | NZ_CP145316.1 | 1 |
| Helicobacter mustelae | Campylobacterota | Epsilonproteobacteria | NZ_LS483446.1 | 1 |
| Helicobacter pullorum | Campylobacterota | Epsilonproteobacteria | NZ_LR134509.1 | 1 |
| Helicobacter pylori | Campylobacterota | Epsilonproteobacteria | NZ_AP026446.1 | 1 |
| Helicobacter suis | Campylobacterota | Epsilonproteobacteria | NZ_AP023046.1 | 1 |
| Helicobacter typhlonius | Campylobacterota | Epsilonproteobacteria | NZ_LN907858.1 | 1 |
| Helicobacter winghamensis ATCC BAA-430 | Campylobacterota | Epsilonproteobacteria | NZ_CP063533.1 | 1 |
| Hydrogenimonas cancrithermarum | Campylobacterota | Epsilonproteobacteria | NZ_AP027370.1 | 1 |

| | | | | |
|---|---|---|---|---|
| Hydrogenimonas leucolamina | Campylobacterota | Epsilonproteobacteria | NZ_AP026369.1 | 1 |
| Hydrogenimonas thermophila | Campylobacterota | Epsilonproteobacteria | NZ_CP136593.1 | 1 |
| Hydrogenimonas urashimensis | Campylobacterota | Epsilonproteobacteria | NZ_AP023212.1 | 1 |
| Malaciobacter canalis | Campylobacterota | Epsilonproteobacteria | NZ_CP042812.1 | 1 |
| Malaciobacter halophilus | Campylobacterota | Epsilonproteobacteria | NZ_CP031218.1 | 1 |
| Malaciobacter marinus | Campylobacterota | Epsilonproteobacteria | NZ_CP032101.1 | 1 |
| Malaciobacter molluscorum LMG 25693 | Campylobacterota | Epsilonproteobacteria | NZ_CP032098.1 | 1 |
| Malaciobacter mytili LMG 24559 | Campylobacterota | Epsilonproteobacteria | NZ_CP031219.1 | 1 |
| Malaciobacter pacificus | Campylobacterota | Epsilonproteobacteria | NZ_CP035928.1 | 1 |
| Nautilia profundicola AmH | Campylobacterota | Epsilonproteobacteria | NC_012115.1 | 1 |
| Nitratifractor salsuginis DSM 16511 | Campylobacterota | Epsilonproteobacteria | NC_014935.1 | 1 |
| Nitratiruptor tergarcus DSM 16512 | Campylobacterota | Epsilonproteobacteria | NZ_AP026671.1 | 1 |
| Nitrosophilus alvini | Campylobacterota | Epsilonproteobacteria | NZ_AP022847.1 | 1 |
| Nitrosophilus kaiyonis | Campylobacterota | Epsilonproteobacteria | NZ_AP025696.1 | 1 |
| Nitrosophilus labii | Campylobacterota | Epsilonproteobacteria | NZ_AP022826.1 | 1 |
| Poseidonibacter lekithochrous | Campylobacterota | Epsilonproteobacteria | NZ_CP054052.1 | 1 |
| Poseidonibacter parvus | Campylobacterota | Epsilonproteobacteria | NZ_CP019070.1 | 1 |
| Sulfuricurvum kujiense DSM 16994 | Campylobacterota | Epsilonproteobacteria | NC_014762.1 | 1 |
| Sulfurimonas aquatica | Campylobacterota | Epsilonproteobacteria | NZ_CP046072.1 | 1 |
| Sulfurimonas autotrophica DSM 16294 | Campylobacterota | Epsilonproteobacteria | NC_014506.1 | 1 |
| Sulfurimonas denitrificans DSM 1251 | Campylobacterota | Epsilonproteobacteria | NC_007575.1 | 1 |
| Sulfurimonas diazotrophicus | Campylobacterota | Epsilonproteobacteria | NZ_CP147920.1 | 1 |
| Sulfurimonas gotlandica GD1 | Campylobacterota | Epsilonproteobacteria | NZ_AFRZ0100000[illegible] | 1 |
| Sulfurimonas hydrogeniphila | Campylobacterota | Epsilonproteobacteria | NZ_CP035534.1 | 1 |
| Sulfurimonas lithotrophica | Campylobacterota | Epsilonproteobacteria | NZ_CP043617.1 | 1 |
| Sulfurimonas marina | Campylobacterota | Epsilonproteobacteria | NZ_CP041165.1 | 1 |
| Sulfurimonas paralvinellae | Campylobacterota | Epsilonproteobacteria | NZ_CP041406.1 | 1 |
| Sulfurimonas sediminis | Campylobacterota | Epsilonproteobacteria | NZ_CP041235.1 | 1 |
| Sulfurimonas xiamenensis | Campylobacterota | Epsilonproteobacteria | NZ_CP041166.1 | 1 |
| Sulfurospirillum barnesii SES-3 | Campylobacterota | Epsilonproteobacteria | NC_018002.1 | 1 |
| Sulfurospirillum deleyianum DSM 6946 | Campylobacterota | Epsilonproteobacteria | NC_013512.1 | 1 |
| Sulfurospirillum diekertiae | Campylobacterota | Epsilonproteobacteria | NZ_CP023275.1 | 1 |

| | | | | |
|---|---|---|---|---|
| Sulfurospirillum halorespirans DSM 13726 | Campylobacterota | Epsilonproteobacteria | NZ_CP017111.1 | 1 |
| Sulfurospirillum multivorans DSM 12446 | Campylobacterota | Epsilonproteobacteria | NZ_CP007201.1 | 1 |
| Sulfurovum indicum | Campylobacterota | Epsilonproteobacteria | NZ_CP063164.1 | 1 |
| Sulfurovum lithotrophicum | Campylobacterota | Epsilonproteobacteria | NZ_CP011308.1 | 1 |
| Sulfurovum mangrovi | Campylobacterota | Epsilonproteobacteria | NZ_CP087124.1 | 1 |
| Wolinella succinogenes | Campylobacterota | Epsilonproteobacteria | NZ_LR134366.1 | 1 |
| Candidatus Cloacimonas acidaminovorans str. Evry | Candidatus Cloacimonad | Candidatus Cloacimona | NC_020449.1 | 1 |
| Minisyncoccus archaeiphilus | Candidatus Paceibactero | Candidatus Paceibacte | NZ_AP038758.1 | 1 |
| Candidatus Nanosynbacter featherlites | Candidatus Saccharibact | Candidatus Saccharimc | NZ_CP040004.1 | 1 |
| Candidatus Nanosynbacter lyticus | Candidatus Saccharibact | Candidatus Saccharimc | NZ_CP007496.1 | 1 |
| Candidatus Saccharimonas aalborgensis | Candidatus Saccharibact | Candidatus Saccharimc | NC_021219.1 | 1 |
| Candidatus Southlakia epibionticum | Candidatus Saccharibact | Candidatus Saccharimc | NZ_CP124550.1 | 1 |
| Candidatus Chlamydia sanziniae | Chlamydiota | Chlamydiia | NZ_CP014639.1 | 1 |
| Candidatus Protochlamydia amoebophila UWE25 | Chlamydiota | Chlamydiia | NC_005861.2 | 1 |
| Candidatus Protochlamydia naegleriophila | Chlamydiota | Chlamydiia | NZ_LN879502.1 | 1 |
| Candidatus Rhabdochlamydia oedothoracis | Chlamydiota | Chlamydiia | NZ_CP075587.1 | 1 |
| Candidatus Rhabdochlamydia porcellionis | Chlamydiota | Chlamydiia | NZ_CP075585.1 | 1 |
| Chlamydia abortus | Chlamydiota | Chlamydiia | NZ_CP158101.1 | 1 |
| Chlamydia buteonis | Chlamydiota | Chlamydiia | NZ_CP050318.1 | 1 |
| Chlamydia caviae GPIC | Chlamydiota | Chlamydiia | NC_003361.3 | 1 |
| Chlamydia crocodili | Chlamydiota | Chlamydiia | NZ_CP060791.1 | 1 |
| Chlamydia felis Fe/C-56 | Chlamydiota | Chlamydiia | NC_007899.1 | 1 |
| Chlamydia gallinacea 08-1274/3 | Chlamydiota | Chlamydiia | NZ_CP015840.1 | 1 |
| Chlamydia muridarum str. Nigg | Chlamydiota | Chlamydiia | NC_002620.2 | 1 |
| Chlamydia pecorum | Chlamydiota | Chlamydiia | NZ_CP080403.1 | 1 |
| Chlamydia pneumoniae TW-183 | Chlamydiota | Chlamydiia | NC_005043.1 | 1 |
| Chlamydia poikilotherma | Chlamydiota | Chlamydiia | NZ_LS992154.1 | 1 |
| Chlamydia psittaci 6BC | Chlamydiota | Chlamydiia | NC_017287.1 | 1 |
| Chlamydia serpentis | Chlamydiota | Chlamydiia | NZ_LT993738.1 | 1 |
| Chlamydia suis | Chlamydiota | Chlamydiia | NZ_LT999999.1 | 1 |
| Chlamydia trachomatis D/UW-3/CX | Chlamydiota | Chlamydiia | NC_000117.1 | 1 |
| Chlamydiifrater phoenicopteri | Chlamydiota | Chlamydiia | NZ_LR777658.1 | 1 |

| | | | | |
|---|---|---|---|---|
| Chlamydiifrater volucris | Chlamydiota | Chlamydiia | NZ_LR777654.1 | 1 |
| Parachlamydia acanthamoebae UV-7 | Chlamydiota | Chlamydiia | NC_015702.1 | 1 |
| Simkania negevensis Z | Chlamydiota | Chlamydiia | NC_015713.1 | 1 |
| Waddlia chondrophila WSU 86-1044 | Chlamydiota | Chlamydiia | NC_014225.1 | 1 |
| Chlorobaculum limnaeum | Chlorobiota | Chlorobiia | NZ_CP017305.1 | 1 |
| Chlorobaculum parvum NCIB 8327 | Chlorobiota | Chlorobiia | NC_011027.1 | 1 |
| Chlorobaculum tepidum TLS | Chlorobiota | Chlorobiia | NC_002932.3 | 1 |
| Chlorobium limicola DSM 245 | Chlorobiota | Chlorobiia | NC_010803.1 | 1 |
| Chlorobium phaeobacteroides DSM 266 | Chlorobiota | Chlorobiia | NC_008639.1 | 1 |
| Chloroherpeton thalassium ATCC 35110 | Chlorobiota | Chlorobiia | NC_011026.1 | 1 |
| Pelodictyon luteolum DSM 273 | Chlorobiota | Chlorobiia | NC_007512.1 | 1 |
| Pelodictyon phaeoclathratiforme BU-1 | Chlorobiota | Chlorobiia | NC_011060.1 | 1 |
| Prosthecochloris aestuarii DSM 271 | Chlorobiota | Chlorobiia | NC_011059.1 | 1 |
| Anaerolinea thermophila UNI-1 | Chloroflexota | Anaerolineae | NC_014960.1 | 2 |
| Brevefilum fermentans | Chloroflexota | Anaerolineae | NZ_LT859958.1 | 1 |
| Pelolinea submarina | Chloroflexota | Anaerolineae | NZ_AP018437.1 | 2 |
| Candidatus Promineifilum breve | Chloroflexota | Ardenticatenia | NZ_LN890656.1 | 1 |
| Candidatus Promineifilum breve | Chloroflexota | Ardenticatenia | NZ_LN890655.1 | 1 |
| Caldilinea aerophila DSM 14535 = NBRC 104270 | Chloroflexota | Caldilineae | NC_017079.1 | 2 |
| Phototrophicus methaneseepsis | Chloroflexota | Candidatus Thermofons | NZ_CP062983.1 | 1 |
| Candidatus Chlorohelix allophototropha | Chloroflexota | Chloroflexia | NZ_CP128399.1 | 2 |
| Chloroflexus aggregans DSM 9485 | Chloroflexota | Chloroflexia | NC_011831.1 | 2 |
| Chloroflexus aurantiacus J-10-fl | Chloroflexota | Chloroflexia | NC_010175.1 | 2 |
| Roseiflexus castenholzii DSM 13941 | Chloroflexota | Chloroflexia | NC_009767.1 | 2 |
| Candidatus Dehalogenimonas loeffleri | Chloroflexota | Dehalococcoidia | NZ_CP146612.1 | 1 |
| Candidatus Lucifugimonas marina | Chloroflexota | Dehalococcoidia | NZ_CP046146.1 | 1 |
| Dehalococcoides mccartyi 195 | Chloroflexota | Dehalococcoidia | NC_002936.3 | 1 |
| Dehalogenimonas etheniformans | Chloroflexota | Dehalococcoidia | NZ_CP058566.2 | 1 |
| Dehalogenimonas formicexedens | Chloroflexota | Dehalococcoidia | NZ_CP018258.1 | 1 |
| Ktedonosporobacter rubrisoli | Chloroflexota | Ktedonobacteria | NZ_CP035758.1 | 2 |
| Tepidiforma bonchosmolovskayae | Chloroflexota | Tepidiformia | NZ_CP042829.1 | 2 |
| Desulfurispirillum indicum S5 | Chrysiogenota | Chrysiogenia | NC_014836.1 | 1 |

| | | | | |
|---|---|---|---|---|
| Coprothermobacter proteolyticus DSM 5265 | Coprothermobacterota | Coprothermobacteria | NC_011295.1 | 2 |
| Acaryochloris marina MBIC11017 | Cyanobacteriota | Cyanophyceae | NC_009925.1 | 1 |
| Allocoleopsis franciscana PCC 7113 | Cyanobacteriota | Cyanophyceae | NC_019738.1 | 1 |
| Anabaena cylindrica PCC 7122 | Cyanobacteriota | Cyanophyceae | NC_019771.1 | 1 |
| Anabaenopsis elenkinii CCIBt3563 | Cyanobacteriota | Cyanophyceae | NZ_CP063311.1 | 1 |
| Anthocerotibacter panamensis C109 | Cyanobacteriota | Cyanophyceae | NZ_CP062698.1 | 1 |
| Brasilonema sennae CENA114 | Cyanobacteriota | Cyanophyceae | NZ_CP030118.1 | 1 |
| Candidatus Atelocyanobacterium thalassa isolate ALOHA | Cyanobacteriota | Cyanophyceae | NC_013771.1 | 1 |
| Capilliphycus salinus ALCB114379 | Cyanobacteriota | Cyanophyceae | NZ_CP178543.1 | 1 |
| Chamaesiphon minutus PCC 6605 | Cyanobacteriota | Cyanophyceae | NC_019697.1 | 1 |
| Chroococcidiopsis thermalis PCC 7203 | Cyanobacteriota | Cyanophyceae | NC_019695.1 | 1 |
| Crinalium epipsammum PCC 9333 | Cyanobacteriota | Cyanophyceae | NC_019753.1 | 1 |
| Crocosphaera subtropica ATCC 51142 | Cyanobacteriota | Cyanophyceae | NC_010546.1 | 1 |
| Cyanobacterium aponinum PCC 10605 | Cyanobacteriota | Cyanophyceae | NC_019776.1 | 1 |
| Cyanobium gracile PCC 6307 | Cyanobacteriota | Cyanophyceae | NC_019675.1 | 1 |
| Cyanobium usitatum str. Tous | Cyanobacteriota | Cyanophyceae | NZ_OY986431.1 | 1 |
| Cylindrospermopsis curvispora GIHE-G1 | Cyanobacteriota | Cyanophyceae | NZ_CP060822.1 | 1 |
| Cylindrospermopsis raciborskii Cr2010 | Cyanobacteriota | Cyanophyceae | NZ_CP065936.1 | 1 |
| Cylindrospermum stagnale PCC 7417 | Cyanobacteriota | Cyanophyceae | NC_019757.1 | 1 |
| Dactylococcopsis salina PCC 8305 | Cyanobacteriota | Cyanophyceae | NC_019780.1 | 1 |
| Dolichospermum compactum NIES-806 | Cyanobacteriota | Cyanophyceae | NZ_AP018316.1 | 1 |
| Dolichospermum flos-aquae CCAP 1403/13F | Cyanobacteriota | Cyanophyceae | NZ_CP051206.1 | 1 |
| Dolichospermum heterosporum TAC447 | Cyanobacteriota | Cyanophyceae | NZ_CP099464.1 | 1 |
| Euhalothece natronophila Z-M001 | Cyanobacteriota | Cyanophyceae | NZ_CP042326.1 | 1 |
| Geminocystis herdmanii PCC 6308 | Cyanobacteriota | Cyanophyceae | NZ_CM001775.1 | 1 |
| Gloeobacter kilaueensis JS1 | Cyanobacteriota | Cyanophyceae | NC_022600.1 | 1 |
| Gloeobacter morelensis MG652769 | Cyanobacteriota | Cyanophyceae | NZ_CP063845.1 | 1 |
| Gloeobacter violaceus PCC 7421 | Cyanobacteriota | Cyanophyceae | NC_005125.1 | 1 |
| Gloeocapsopsis dulcis | Cyanobacteriota | Cyanophyceae | NZ_CP119968.1 | 1 |
| Gloeomargarita lithophora Alchichica-D10 | Cyanobacteriota | Cyanophyceae | NZ_CP017675.1 | 1 |
| Gloeothece citriformis PCC 7424 | Cyanobacteriota | Cyanophyceae | NC_011729.1 | 1 |
| Gloeothece verrucosa PCC 7822 | Cyanobacteriota | Cyanophyceae | NC_014501.1 | 1 |

| | | | | |
|---|---|---|---|---|
| Halomicronema hongdechloris C2206 | Cyanobacteriota | Cyanophyceae | NZ_CP021983.2 | 1 |
| Halotia branconii CENA392 | Cyanobacteriota | Cyanophyceae | NZ_CP124543.1 | 1 |
| Leptodesmis sichuanensis A121 | Cyanobacteriota | Cyanophyceae | NZ_CP075171.1 | 1 |
| Leptolyngbya boryana IAM M-101 | Cyanobacteriota | Cyanophyceae | NZ_AP014638.1 | 1 |
| Leptothermofonsia sichuanensis E412 | Cyanobacteriota | Cyanophyceae | NZ_CP072600.1 | 1 |
| Limnospira indica PCC 8005 | Cyanobacteriota | Cyanophyceae | NZ_FO818640.1 | 1 |
| Microcoleus anatoxicus | Cyanobacteriota | Cyanophyceae | NZ_CP120853.1 | 1 |
| Nodularia sphaerocarpa UHCC 0038 | Cyanobacteriota | Cyanophyceae | NZ_CP060140.1 | 1 |
| Nodularia spumigena UHCC 0039 | Cyanobacteriota | Cyanophyceae | NZ_CP020114.1 | 1 |
| 'Nostoc azollae' 0708 | Cyanobacteriota | Cyanophyceae | NC_014248.1 | 1 |
| Nostoc edaphicum CCNP1411 | Cyanobacteriota | Cyanophyceae | NZ_CP054698.1 | 1 |
| Nostoc flagelliforme CCNUN1 | Cyanobacteriota | Cyanophyceae | NZ_CP024785.1 | 1 |
| Nostoc piscinale CENA21 | Cyanobacteriota | Cyanophyceae | NZ_CP012036.1 | 1 |
| Nostoc punctiforme PCC 73102 | Cyanobacteriota | Cyanophyceae | NC_010628.1 | 1 |
| Nostoc sphaeroides | Cyanobacteriota | Cyanophyceae | NZ_CP031941.1 | 1 |
| Okeanomitos corallinicola TIOX110 | Cyanobacteriota | Cyanophyceae | NZ_CP150886.1 | 1 |
| Oscillatoria acuminata PCC 6304 | Cyanobacteriota | Cyanophyceae | NC_019693.1 | 1 |
| Oscillatoria nigro-viridis PCC 7112 | Cyanobacteriota | Cyanophyceae | NC_019729.1 | 2 |
| Oxynema aestuarii AP17 | Cyanobacteriota | Cyanophyceae | NZ_CP051167.1 | 1 |
| Parasynechococcus marenigrum WH 8102 | Cyanobacteriota | Cyanophyceae | NC_005070.1 | 1 |
| Parathermosynechococcus lividus PCC 6715 | Cyanobacteriota | Cyanophyceae | NZ_CP018092.1 | 1 |
| Phormidium yuhuli AB48 | Cyanobacteriota | Cyanophyceae | NZ_CP098611.1 | 1 |
| Planktothricoides raciborskii GIHE-MW2 | Cyanobacteriota | Cyanophyceae | NZ_CP159837.1 | 1 |
| Planktothrix agardhii | Cyanobacteriota | Cyanophyceae | NZ_LR882952.1 | 1 |
| Planktothrix tepida | Cyanobacteriota | Cyanophyceae | NZ_LR882970.1 | 1 |
| Prochlorococcus marinus str. MIT 9515 | Cyanobacteriota | Cyanophyceae | NC_008817.1 | 1 |
| Pseudanabaena galeata CCNP1313 | Cyanobacteriota | Cyanophyceae | NZ_CP112874.1 | 1 |
| Pseudanabaena mucicola str. Chao 1806 | Cyanobacteriota | Cyanophyceae | NZ_CP097329.1 | 1 |
| Richelia sinica FACHB-800 | Cyanobacteriota | Cyanophyceae | NZ_CP021056.1 | 1 |
| Rippkaea orientalis PCC 8801 | Cyanobacteriota | Cyanophyceae | NC_011726.1 | 1 |
| Sphaerospermopsis torques-reginae ITEP-024 | Cyanobacteriota | Cyanophyceae | NZ_CP080598.1 | 1 |
| Stanieria cyanosphaera PCC 7437 | Cyanobacteriota | Cyanophyceae | NC_019748.1 | 1 |

| | | | | |
|---|---|---|---|---|
| Synechococcus elongatus PCC 6301 | Cyanobacteriota | Cyanophyceae | NZ_CP085785.1 | 1 |
| Thermocoleostomius sinensis A174 | Cyanobacteriota | Cyanophyceae | NZ_CP113797.1 | 1 |
| Thermoleptolyngbya oregonensis NK1-22 | Cyanobacteriota | Cyanophyceae | NZ_CP053540.1 | 1 |
| Thermoleptolyngbya sichuanensis A183 | Cyanobacteriota | Cyanophyceae | NZ_CP053661.1 | 1 |
| Thermosynechococcus sichuanensis E542 | Cyanobacteriota | Cyanophyceae | NZ_CP032152.1 | 1 |
| Thermosynechococcus vestitus BP-1 | Cyanobacteriota | Cyanophyceae | NC_004113.1 | 1 |
| Trichocoleus desertorum NBK24 | Cyanobacteriota | Cyanophyceae | NZ_CP116619.1 | 1 |
| Trichodesmium erythraeum IMS101 | Cyanobacteriota | Cyanophyceae | NC_008312.1 | 1 |
| Trichormus variabilis 0441 | Cyanobacteriota | Cyanophyceae | NZ_CP047242.1 | 1 |
| Trichothermofontia sichuanensis B231 | Cyanobacteriota | Cyanophyceae | NZ_CP110848.1 | 1 |
| Calditerrivibrio nitroreducens DSM 19672 | Deferribacterota | Deferribacteres | NC_014758.1 | 2 |
| Deferribacter desulfuricans SSM1 | Deferribacterota | Deferribacteres | NC_013939.1 | 2 |
| Denitrovibrio acetiphilus DSM 12809 | Deferribacterota | Deferribacteres | NC_013943.1 | 2 |
| Flexistipes sinusarabici DSM 4947 | Deferribacterota | Deferribacteres | NC_015672.1 | 2 |
| Geovibrio thiophilus | Deferribacterota | Deferribacteres | NZ_CP035108.1 | 2 |
| Mucispirillum schaedleri ASF457 | Deferribacterota | Deferribacteres | NZ_CP097562.1 | 1 |
| Allomeiothermus silvanus DSM 9946 | Deinococcota | Deinococci | NC_014212.1 | 2 |
| Deinococcus aetherius | Deinococcota | Deinococci | NZ_AP026560.1 | 2 |
| Deinococcus altitudinis | Deinococcota | Deinococci | NZ_CP129415.1 | 2 |
| Deinococcus aquaticus | Deinococcota | Deinococci | NZ_CP115165.1 | 2 |
| Deinococcus deserti VCD115 | Deinococcota | Deinococci | NC_012526.1 | 2 |
| Deinococcus ficus | Deinococcota | Deinococci | NZ_CP021081.1 | 2 |
| Deinococcus geothermalis DSM 11300 | Deinococcota | Deinococci | NC_008025.1 | 2 |
| Deinococcus gobiensis I-0 | Deinococcota | Deinococci | NC_017790.1 | 2 |
| Deinococcus irradiatisoli | Deinococcota | Deinococci | NZ_CP029494.1 | 2 |
| Deinococcus maricopensis DSM 21211 | Deinococcota | Deinococci | NC_014958.1 | 2 |
| Deinococcus metallilatus | Deinococcota | Deinococci | NZ_CP038512.1 | 2 |
| Deinococcus pераridilitoris DSM 19664 | Deinococcota | Deinococci | NC_019793.1 | 2 |
| Deinococcus proteolyticus MRP | Deinococcota | Deinococci | NC_015161.1 | 2 |
| Deinococcus psychrotolerans | Deinococcota | Deinococci | NZ_CP034183.1 | 2 |
| Deinococcus puniceus | Deinococcota | Deinococci | NZ_CP011387.1 | 2 |
| Deinococcus radiomollis | Deinococcota | Deinococci | NZ_CP129906.1 | 2 |

| | | | | |
|---|---|---|---|---|
| Deinococcus radiopugnans | Deinococcota | Deinococci | NZ_CP010028.1 | 2 |
| Deinococcus rubellus | Deinococcota | Deinococci | NZ_CP104213.1 | 2 |
| Deinococcus sonorensis KR-87 | Deinococcota | Deinococci | NZ_CP158299.1 | 2 |
| Deinococcus taeanensis | Deinococcota | Deinococci | NZ_CP083455.1 | 2 |
| Deinococcus wulumuqiensis R12 | Deinococcota | Deinococci | NZ_CP049357.1 | 2 |
| Deinococcus yunweiensis | Deinococcota | Deinococci | NZ_CP177252.1 | 2 |
| Marinithermus hydrothermalis DSM 14884 | Deinococcota | Deinococci | NC_015387.1 | 2 |
| Meiothermus ruber DSM 1279 | Deinococcota | Deinococci | NC_013946.1 | 2 |
| Meiothermus taiwanensis WR-220 | Deinococcota | Deinococci | NZ_CP021130.1 | 2 |
| Thermus antranikianii | Deinococcota | Deinococci | NZ_CP076325.1 | 2 |
| Thermus aquaticus Y51MC23 | Deinococcota | Deinococci | NZ_CP010822.1 | 2 |
| Thermus brockianus | Deinococcota | Deinococci | NZ_AP025593.1 | 2 |
| Thermus caldilimi | Deinococcota | Deinococci | NZ_CP038452.1 | 2 |
| Thermus oshimai JL-2 | Deinococcota | Deinococci | NC_019386.1 | 2 |
| Thermus parvatiensis | Deinococcota | Deinococci | NZ_CP014141.1 | 2 |
| Thermus thermophilus HB8 | Deinococcota | Deinococci | NC_006461.1 | 2 |
| Truepera radiovictrix DSM 17093 | Deinococcota | Deinococci | NC_014221.1 | 2 |
| Dictyoglomus thermophilum H-6-12 | Dictyoglomota | Dictyoglomia | NC_011297.1 | 2 |
| Dictyoglomus turgidum DSM 6724 | Dictyoglomota | Dictyoglomia | NC_011661.1 | 2 |
| Elusimicrobium minutum Pei191 | Elusimicrobiota | Elusimicrobia | NC_010644.1 | 2 |
| Candidatus Endomicrobiellum trichonymphae | Elusimicrobiota | Endomicrobiia | NZ_AP017459.1 | 2 |
| Endomicrobium proavitum | Elusimicrobiota | Endomicrobiia | NZ_CP009498.1 | 2 |
| Fibrobacter succinogenes subsp. succinogenes S85 | Fibrobacterota | Fibrobacteria | NC_017448.1 | 2 |
| Cetobacterium somerae | Fusobacteriota | Fusobacteriia | NZ_CP092519.1 | 2 |
| Fusobacterium animalis | Fusobacteriota | Fusobacteriia | NZ_CP012715.1 | 1 |
| Fusobacterium canifelinum | Fusobacteriota | Fusobacteriia | NZ_CP068114.1 | 1 |
| Fusobacterium gonidiaformans ATCC 25563 | Fusobacteriota | Fusobacteriia | NZ_CP028106.1 | 2 |
| Fusobacterium hominis | Fusobacteriota | Fusobacteriia | NZ_CP060637.1 | 2 |
| Fusobacterium hwasookii ChDC F206 | Fusobacteriota | Fusobacteriia | NZ_CP013336.1 | 1 |
| Fusobacterium mortiferum ATCC 9817 | Fusobacteriota | Fusobacteriia | NZ_CP028102.1 | 2 |
| Fusobacterium necrophorum subsp. funduliforme | Fusobacteriota | Fusobacteriia | NZ_CP019306.1 | 2 |
| Fusobacterium nucleatum subsp. nucleatum ATCC 25586 | Fusobacteriota | Fusobacteriia | NZ_CP028101.1 | 1 |

| | | | | |
|---|---|---|---|---|
| Fusobacterium polymorphum | Fusobacteriota | Fusobacteriia | NZ_LN831027.1 | 1 |
| Fusobacterium ulcerans | Fusobacteriota | Fusobacteriia | NZ_CP028105.1 | 2 |
| Fusobacterium varium ATCC 27725 | Fusobacteriota | Fusobacteriia | NZ_CP028103.1 | 2 |
| Fusobacterium vincentii | Fusobacteriota | Fusobacteriia | NZ_CP071093.1 | 1 |
| Fusobacterium watanabei | Fusobacteriota | Fusobacteriia | NZ_OZ221964.1 | 1 |
| Haliovirga abyssi | Fusobacteriota | Fusobacteriia | NZ_AP027059.1 | 2 |
| Ilyobacter polytropus DSM 2926 | Fusobacteriota | Fusobacteriia | NC_014632.1 | 2 |
| Leptotrichia buccalis C-1013-b | Fusobacteriota | Fusobacteriia | NC_013192.1 | 2 |
| Leptotrichia hofstadii | Fusobacteriota | Fusobacteriia | NZ_AP019823.1 | 2 |
| Leptotrichia hongkongensis | Fusobacteriota | Fusobacteriia | NZ_AP019846.1 | 2 |
| Leptotrichia mesophila | Fusobacteriota | Fusobacteriia | NZ_CP165646.1 | 2 |
| Leptotrichia trevisanii | Fusobacteriota | Fusobacteriia | NZ_AP019831.1 | 2 |
| Leptotrichia wadei | Fusobacteriota | Fusobacteriia | NZ_AP019829.2 | 2 |
| Sneathia vaginalis | Fusobacteriota | Fusobacteriia | NZ_CP011280.1 | 2 |
| Streptobacillus moniliformis DSM 12112 | Fusobacteriota | Fusobacteriia | NC_013515.1 | 2 |
| Gemmatimonas aurantiaca T-27 | Gemmatimonadota | Gemmatimonadia | NC_012489.1 | 1 |
| Gemmatimonas groenlandica | Gemmatimonadota | Gemmatimonadia | NZ_CP053085.1 | 1 |
| Gemmatimonas phototrophica | Gemmatimonadota | Gemmatimonadia | NZ_CP011454.1 | 1 |
| Gemmatirosa kalamazoonensis | Gemmatimonadota | Gemmatimonadia | NZ_CP007128.1 | 1 |
| Pseudogemmatithrix spongiicola | Gemmatimonadota | Gemmatimonadia | NZ_CP130613.1 | 1 |
| Ignavibacterium album JCM 16511 | Ignavibacteriota | Ignavibacteria | NC_017464.1 | 1 |
| Melioribacter roseus P3M-2 | Ignavibacteriota | Ignavibacteria | NC_018178.1 | 1 |
| Kiritimatiella glycovorans | Kiritimatiellota | Kiritimatiellia | NZ_CP010904.1 | 1 |
| Tichowtungia aerotolerans | Kiritimatiellota | Tichowtungiia | NZ_CP047593.1 | 1 |
| Lentisphaera profundi | Lentisphaerota | Lentisphaeria | NZ_CP117812.1 | 1 |
| Candidatus Xianfuyuplasma coldseepsis | Mycoplasmatota | Candidatus Izimaplasma | NZ_CP048914.1 | 1 |
| Hujiaoplasma nucleasis | Mycoplasmatota | Candidatus Izimaplasma | NZ_CP051151.1 | 1 |
| Acholeplasma hippikon | Mycoplasmatota | Mollicutes | NZ_LR215050.1 | 1 |
| Acholeplasma laidlawii PG-8A | Mycoplasmatota | Mollicutes | NC_010163.1 | 1 |
| Alteracholeplasma palmae J233 | Mycoplasmatota | Mollicutes | NC_022538.1 | 1 |
| Candidatus Hepatoplasma crinochetorum Av | Mycoplasmatota | Mollicutes | NZ_CP006932.1 | 1 |
| Candidatus Mycoplasma mahonii | Mycoplasmatota | Mollicutes | NZ_CP114583.1 | 1 |

| | | | | |
|---|---|---|---|---|
| Candidatus Phytoplasma asteris | Mycoplasmatota | Mollicutes | NZ_CP128414.1 | 1 |
| Candidatus Phytoplasma fraxini | Mycoplasmatota | Mollicutes | NZ_CP146843.1 | 1 |
| Candidatus Phytoplasma luffae | Mycoplasmatota | Mollicutes | NZ_CP054393.1 | 1 |
| Candidatus Phytoplasma mali | Mycoplasmatota | Mollicutes | NC_011047.1 | 1 |
| Candidatus Phytoplasma rubi | Mycoplasmatota | Mollicutes | NZ_CP114006.1 | 1 |
| Candidatus Phytoplasma solani | Mycoplasmatota | Mollicutes | NZ_CP155828.1 | 1 |
| Candidatus Phytoplasma ziziphi | Mycoplasmatota | Mollicutes | NZ_CP025121.1 | 1 |
| Entomoplasma freundtii | Mycoplasmatota | Mollicutes | NZ_CP024962.1 | 1 |
| Haploplasma axanthum | Mycoplasmatota | Mollicutes | NZ_LR215048.1 | 1 |
| Mariniplasma anaerobium | Mycoplasmatota | Mollicutes | NZ_AP024412.1 | 1 |
| Mesoplasma chauliocola | Mycoplasmatota | Mollicutes | NZ_CP023173.1 | 1 |
| Mesoplasma coleopterae | Mycoplasmatota | Mollicutes | NZ_CP024968.1 | 1 |
| Mesoplasma entomophilum | Mycoplasmatota | Mollicutes | NZ_CP024411.1 | 1 |
| Mesoplasma florum L1 | Mycoplasmatota | Mollicutes | NC_006055.1 | 1 |
| Mesoplasma lactucae ATCC 49193 | Mycoplasmatota | Mollicutes | NZ_CP023668.1 | 1 |
| Mesoplasma melaleucae | Mycoplasmatota | Mollicutes | NZ_CP024964.1 | 1 |
| Mesoplasma syrphidae | Mycoplasmatota | Mollicutes | NZ_CP025257.1 | 1 |
| Mesoplasma tabanidae | Mycoplasmatota | Mollicutes | NZ_CP024969.1 | 1 |
| Mycoplasma amphoriforme A39 | Mycoplasmatota | Mollicutes | NZ_HG937516.1 | 1 |
| Mycoplasma anserisalpingitidis | Mycoplasmatota | Mollicutes | NZ_CP042295.1 | 1 |
| Mycoplasma aquilae ATCC BAA-1896 | Mycoplasmatota | Mollicutes | NZ_CP182208.1 | 1 |
| Mycoplasma bradburyae | Mycoplasmatota | Mollicutes | NZ_CP101414.1 | 1 |
| Mycoplasma capricolum subsp. capricolum ATCC 27343 | Mycoplasmatota | Mollicutes | NC_007633.1 | 1 |
| Mycoplasma corogypsi | Mycoplasmatota | Mollicutes | NZ_CP174163.1 | 1 |
| Mycoplasma cottewii | Mycoplasmatota | Mollicutes | NZ_CP103424.1 | 1 |
| Mycoplasma crocodyli MP145 | Mycoplasmatota | Mollicutes | NC_014014.1 | 1 |
| Mycoplasma feriruminatoris | Mycoplasmatota | Mollicutes | NZ_CP104009.1 | 1 |
| Mycoplasma hafezii | Mycoplasmatota | Mollicutes | NZ_CP187940.1 | 1 |
| Mycoplasma iguanae | Mycoplasmatota | Mollicutes | NZ_CP102734.1 | 1 |
| Mycoplasma leachii PG50 | Mycoplasmatota | Mollicutes | NC_014751.1 | 1 |
| Mycoplasma miroungigenitalium | Mycoplasmatota | Mollicutes | NZ_CP053096.1 | 1 |
| Mycoplasma miroungirhinis | Mycoplasmatota | Mollicutes | NZ_CP053097.1 | 1 |

| | | | | |
|---|---|---|---|---|
| Mycoplasma mycoides subsp. capri | Mycoplasmatota | Mollicutes | NZ_CP065581.1 | 1 |
| Mycoplasma nasistruthionis | Mycoplasmatota | Mollicutes | NZ_CP040825.1 | 1 |
| Mycoplasma ovis str. Michigan | Mycoplasmatota | Mollicutes | NC_023062.1 | 1 |
| Mycoplasma parvum str. Indiana | Mycoplasmatota | Mollicutes | NC_022575.1 | 1 |
| Mycoplasma phocoenae | Mycoplasmatota | Mollicutes | NZ_CP051481.1 | 1 |
| Mycoplasma phocoeninasale | Mycoplasmatota | Mollicutes | NZ_CP051480.1 | 1 |
| Mycoplasma putrefaciens | Mycoplasmatota | Mollicutes | NZ_LS991954.1 | 1 |
| Mycoplasma seminis | Mycoplasmatota | Mollicutes | NZ_CP132191.1 | 1 |
| Mycoplasma struthionis | Mycoplasmatota | Mollicutes | NZ_CP034044.1 | 1 |
| Mycoplasma suis str. Illinois | Mycoplasmatota | Mollicutes | NC_015155.1 | 1 |
| Mycoplasma tauri | Mycoplasmatota | Mollicutes | NZ_CP070479.1 | 1 |
| Mycoplasma tullyi | Mycoplasmatota | Mollicutes | NZ_CP059674.1 | 1 |
| Mycoplasma wenyonii str. Massachusetts | Mycoplasmatota | Mollicutes | NC_018149.1 | 2 |
| Mycoplasma yeatsii GM274B | Mycoplasmatota | Mollicutes | NZ_CP007520.1 | 1 |
| Paracholeplasma brassicae | Mycoplasmatota | Mollicutes | NC_022549.1 | 1 |
| Spiroplasma alleghenense | Mycoplasmatota | Mollicutes | NZ_CP031376.1 | 1 |
| Spiroplasma apis B31 | Mycoplasmatota | Mollicutes | NC_022998.1 | 1 |
| Spiroplasma cantharicola | Mycoplasmatota | Mollicutes | NZ_CP012622.1 | 1 |
| Spiroplasma chinense | Mycoplasmatota | Mollicutes | NZ_CP043026.1 | 1 |
| Spiroplasma chrysopicola DF-1 | Mycoplasmatota | Mollicutes | NC_021280.1 | 1 |
| Spiroplasma citri | Mycoplasmatota | Mollicutes | NZ_CP013197.1 | 1 |
| Spiroplasma clarkii | Mycoplasmatota | Mollicutes | NZ_CP024870.1 | 1 |
| Spiroplasma corruscae | Mycoplasmatota | Mollicutes | NZ_CP022535.1 | 1 |
| Spiroplasma culicicola AES-1 | Mycoplasmatota | Mollicutes | NZ_CP006681.1 | 1 |
| Spiroplasma diminutum CUAS-1 | Mycoplasmatota | Mollicutes | NC_021833.1 | 1 |
| Spiroplasma endosymbiont of Acasis viretata | Mycoplasmatota | Mollicutes | NZ_OZ035019.1 | 1 |
| Spiroplasma endosymbiont of Agriotes lineatus | Mycoplasmatota | Mollicutes | NZ_OZ026448.1 | 1 |
| Spiroplasma endosymbiont of Aleiodes alternator | Mycoplasmatota | Mollicutes | NZ_OZ035008.1 | 1 |
| Spiroplasma endosymbiont of Ammophila pubescens | Mycoplasmatota | Mollicutes | NZ_OZ026611.1 | 1 |
| Spiroplasma endosymbiont of Amphimallon solstitiale | Mycoplasmatota | Mollicutes | NZ_OZ034999.1 | 1 |
| Spiroplasma endosymbiont of Andrena trimmerana | Mycoplasmatota | Mollicutes | NZ_OZ026455.1 | 1 |
| Spiroplasma endosymbiont of Anurida maritima | Mycoplasmatota | Mollicutes | NZ_CP102059.1 | 1 |

| | | | | |
|---|---|---|---|---|
| Spiroplasma endosymbiont of Apeira syringaria | Mycoplasmatota | Mollicutes | NZ_OZ026543.1 | 1 |
| Spiroplasma endosymbiont of Asaphidion curtum | Mycoplasmatota | Mollicutes | NZ_OZ035001.1 | 1 |
| Spiroplasma endosymbiont of Aspidapion aeneum | Mycoplasmatota | Mollicutes | NZ_OZ034994.1 | 1 |
| Spiroplasma endosymbiont of Atherix ibis | Mycoplasmatota | Mollicutes | NZ_OZ026474.1 | 1 |
| Spiroplasma endosymbiont of Cantharis nigra | Mycoplasmatota | Mollicutes | NZ_OZ026470.1 | 1 |
| Spiroplasma endosymbiont of Cantharis rufa | Mycoplasmatota | Mollicutes | NZ_OZ026459.1 | 1 |
| Spiroplasma endosymbiont of Clivina fossor | Mycoplasmatota | Mollicutes | NZ_OZ035006.1 | 1 |
| Spiroplasma endosymbiont of Colias croceus | Mycoplasmatota | Mollicutes | NZ_OZ026536.1 | 1 |
| Spiroplasma endosymbiont of Crioceris asparagi | Mycoplasmatota | Mollicutes | NZ_OZ026475.1 | 1 |
| Spiroplasma endosymbiont of Dasysyrphus albostriatus | Mycoplasmatota | Mollicutes | NZ_OZ026447.1 | 1 |
| Spiroplasma endosymbiont of Dilophus febrilis | Mycoplasmatota | Mollicutes | NZ_OZ034912.1 | 1 |
| Spiroplasma endosymbiont of Dioctria linearis | Mycoplasmatota | Mollicutes | NZ_OZ034984.1 | 1 |
| Spiroplasma endosymbiont of Diplazon laetatorius | Mycoplasmatota | Mollicutes | NZ_OZ026458.1 | 1 |
| Spiroplasma endosymbiont of Dromius quadrimaculatus | Mycoplasmatota | Mollicutes | NZ_OZ035010.1 | 1 |
| Spiroplasma endosymbiont of Eupeodes luniger | Mycoplasmatota | Mollicutes | NZ_OZ026453.1 | 1 |
| Spiroplasma endosymbiont of Glossina fuscipes fuscipes | Mycoplasmatota | Mollicutes | NZ_CP187162.1 | 1 |
| Spiroplasma endosymbiont of Labia minor | Mycoplasmatota | Mollicutes | NZ_OZ026465.1 | 1 |
| Spiroplasma endosymbiont of Lasioglossum malachurum | Mycoplasmatota | Mollicutes | NZ_OZ026534.1 | 1 |
| Spiroplasma endosymbiont of Lasioglossum villosulum | Mycoplasmatota | Mollicutes | NZ_OZ026539.1 | 1 |
| Spiroplasma endosymbiont of Lonchoptera lutea | Mycoplasmatota | Mollicutes | NZ_OZ026463.1 | 1 |
| Spiroplasma endosymbiont of Melieria omissa | Mycoplasmatota | Mollicutes | NZ_OZ035012.1 | 1 |
| Spiroplasma endosymbiont of Monopis laevigella | Mycoplasmatota | Mollicutes | NZ_OZ026473.1 | 1 |
| Spiroplasma endosymbiont of Nebria brevicollis | Mycoplasmatota | Mollicutes | NZ_OZ034986.1 | 1 |
| Spiroplasma endosymbiont of 'Nebria riversi' | Mycoplasmatota | Mollicutes | NZ_CP065193.1 | 1 |
| Spiroplasma endosymbiont of Nephrotoma flavescens | Mycoplasmatota | Mollicutes | NZ_OZ034982.1 | 1 |
| Spiroplasma endosymbiont of Nomada ruficornis | Mycoplasmatota | Mollicutes | NZ_OZ034962.1 | 1 |
| Spiroplasma endosymbiont of Nomada rufipes | Mycoplasmatota | Mollicutes | NZ_OZ195517.1 | 1 |
| Spiroplasma endosymbiont of Notiophilus biguttatus | Mycoplasmatota | Mollicutes | NZ_OZ034993.1 | 1 |
| Spiroplasma endosymbiont of Othius punctulatus | Mycoplasmatota | Mollicutes | NZ_OZ026449.1 | 1 |
| Spiroplasma endosymbiont of Panorpa germanica | Mycoplasmatota | Mollicutes | NZ_OZ026461.1 | 1 |
| Spiroplasma endosymbiont of Panzeria rudis | Mycoplasmatota | Mollicutes | NZ_OZ026460.1 | 1 |
| Spiroplasma endosymbiont of Phyllotreta cruciferae | Mycoplasmatota | Mollicutes | NZ_OU953854.1 | 1 |

| | | | | |
|---|---|---|---|---|
| Spiroplasma endosymbiont of Poecilobothrus nobilitatus | Mycoplasmatota | Mollicutes | NZ_OZ034915.1 | 1 |
| Spiroplasma endosymbiont of Polydrusus cervinus | Mycoplasmatota | Mollicutes | NZ_OZ026469.1 | 1 |
| Spiroplasma endosymbiont of Polydrusus formosus | Mycoplasmatota | Mollicutes | NZ_OZ181656.1 | 1 |
| Spiroplasma endosymbiont of Polydrusus pterygomalis | Mycoplasmatota | Mollicutes | NZ_OZ034964.1 | 1 |
| Spiroplasma endosymbiont of Sarcophaga carnaria | Mycoplasmatota | Mollicutes | NZ_OZ026456.1 | 1 |
| Spiroplasma endosymbiont of Seladonia tumulorum | Mycoplasmatota | Mollicutes | NZ_OZ026545.1 | 1 |
| Spiroplasma endosymbiont of Stenodema calcarata | Mycoplasmatota | Mollicutes | NZ_OZ035018.1 | 1 |
| Spiroplasma endosymbiont of Thecophora atra | Mycoplasmatota | Mollicutes | NZ_OZ026464.1 | 1 |
| Spiroplasma endosymbiont of Tiphia femorata | Mycoplasmatota | Mollicutes | NZ_OZ026544.1 | 1 |
| Spiroplasma endosymbiont of Tipula paludosa | Mycoplasmatota | Mollicutes | NZ_OZ183484.1 | 1 |
| Spiroplasma endosymbiont of Tricholauxania praeusta | Mycoplasmatota | Mollicutes | NZ_OZ026450.1 | 1 |
| Spiroplasma endosymbiont of Villa modesta | Mycoplasmatota | Mollicutes | NZ_OZ034992.1 | 1 |
| Spiroplasma endosymbiont of Virgichneumon dumeticola | Mycoplasmatota | Mollicutes | NZ_OZ183478.1 | 1 |
| Spiroplasma endosymbiont of Zeiraphera isertana | Mycoplasmatota | Mollicutes | NZ_OZ034965.1 | 1 |
| Spiroplasma eriocheiris | Mycoplasmatota | Mollicutes | NZ_CP011856.1 | 1 |
| Spiroplasma floricola 23-6 | Mycoplasmatota | Mollicutes | NZ_CP025057.1 | 1 |
| Spiroplasma gladiatoris | Mycoplasmatota | Mollicutes | NZ_CP038013.1 | 1 |
| Spiroplasma helicoides | Mycoplasmatota | Mollicutes | NZ_CP017015.1 | 1 |
| Spiroplasma ixodetis Y32 | Mycoplasmatota | Mollicutes | NZ_CP127039.1 | 1 |
| Spiroplasma kunkelii CR2-3x | Mycoplasmatota | Mollicutes | NZ_CP010899.1 | 1 |
| Spiroplasma litorale | Mycoplasmatota | Mollicutes | NZ_CP012357.1 | 1 |
| Spiroplasma mirum | Mycoplasmatota | Mollicutes | NZ_CP011855.1 | 1 |
| Spiroplasma monobiae MQ-1 | Mycoplasmatota | Mollicutes | NZ_CP025543.1 | 1 |
| Spiroplasma phoeniceum P40 | Mycoplasmatota | Mollicutes | NZ_CP031088.1 | 1 |
| Spiroplasma platyhelix PALS-1 | Mycoplasmatota | Mollicutes | NZ_CP051215.1 | 1 |
| Spiroplasma poulsonii | Mycoplasmatota | Mollicutes | NZ_CM020866.1 | 1 |
| Spiroplasma sabaudiense Ar-1343 | Mycoplasmatota | Mollicutes | NZ_CP006934.1 | 1 |
| Spiroplasma syrphidicola EA-1 | Mycoplasmatota | Mollicutes | NC_021284.1 | 1 |
| Spiroplasma tabanidicola | Mycoplasmatota | Mollicutes | NZ_CP046276.1 | 1 |
| Spiroplasma taiwanense CT-1 | Mycoplasmatota | Mollicutes | NC_021846.1 | 1 |
| Spiroplasma turonicum | Mycoplasmatota | Mollicutes | NZ_CP012328.1 | 1 |
| Strawberry lethal yellows phytoplasma (CPA) str. NZSb11 | Mycoplasmatota | Mollicutes | NC_021236.1 | 1 |

| | | | | |
|---|---|---|---|---|
| Williamsoniiplasma luminosum | Mycoplasmatota | Mollicutes | NZ_CP024963.1 | 1 |
| Williamsoniiplasma somnilux | Mycoplasmatota | Mollicutes | NZ_CP024965.1 | 1 |
| Anaeromyxobacter dehalogenans 2CP-1 | Myxococcota | Myxococcia | NC_011891.1 | 2 |
| Anaeromyxobacter oryzae | Myxococcota | Myxococcia | NZ_AP025591.1 | 2 |
| Anaeromyxobacter paludicola | Myxococcota | Myxococcia | NZ_AP025592.1 | 2 |
| Archangium gephyra | Myxococcota | Myxococcia | NZ_CP011509.1 | 3 |
| Archangium violaceum | Myxococcota | Myxococcia | NZ_CP137851.1 | 3 |
| Corallococcus coralloides DSM 2259 | Myxococcota | Myxococcia | NC_017030.1 | 2 |
| Corallococcus macrosporus DSM 14697 | Myxococcota | Myxococcia | NZ_CP022203.1 | 2 |
| Melittangium boletus DSM 14713 | Myxococcota | Myxococcia | NZ_CP022163.1 | 2 |
| Myxococcus landrumensis | Myxococcota | Myxococcia | NZ_CP071091.1 | 2 |
| Myxococcus stipitatus DSM 14675 | Myxococcota | Myxococcia | NC_020126.1 | 2 |
| Myxococcus xanthus DK 1622 | Myxococcota | Myxococcia | NC_008095.1 | 2 |
| Pendulispora albinea | Myxococcota | Myxococcia | NZ_CP089984.1 | 2 |
| Pendulispora brunnea | Myxococcota | Myxococcia | NZ_CP089982.1 | 3 |
| Pendulispora rubella | Myxococcota | Myxococcia | NZ_CP089983.1 | 2 |
| Pseudomyxococcus hansupus | Myxococcota | Myxococcia | NZ_CP012109.1 | 2 |
| Pyxidicoccus parkwayensis | Myxococcota | Myxococcia | NZ_CP071090.1 | 2 |
| Vulgatibacter incomptus | Myxococcota | Myxococcia | NZ_CP012332.1 | 2 |
| Nitrospina watsonii | Nitrospinota | Nitrospinia | NZ_OX336137.1 | 1 |
| Candidatus Nitrospira allomarina | Nitrospirota | Nitrospiria | NZ_CP116967.1 | 2 |
| Candidatus Nitrospira inopinata | Nitrospirota | Nitrospiria | NZ_LN885086.1 | 2 |
| Candidatus Nitrospira neomarina | Nitrospirota | Nitrospiria | NZ_CP116968.1 | 2 |
| Leptospirillum ferrooxidans C2-3 | Nitrospirota | Nitrospiria | NC_017094.1 | 2 |
| Nitrospira japonica | Nitrospirota | Nitrospiria | NZ_LT828648.1 | 2 |
| Nitrospira moscoviensis | Nitrospirota | Nitrospiria | NZ_CP011801.1 | 2 |
| Dissulfurispira thermophila | Nitrospirota | Thermodesulfovibrionia | NZ_AP022873.1 | 2 |
| Thermodesulfovibrio autotrophicus | Nitrospirota | Thermodesulfovibrionia | NZ_CP144373.1 | 2 |
| Thermodesulfovibrio obliviosus | Nitrospirota | Thermodesulfovibrionia | NZ_CP144374.1 | 2 |
| Thermodesulfovibrio yellowstonii DSM 11347 | Nitrospirota | Thermodesulfovibrionia | NC_011296.1 | 2 |
| Candidatus Kuenenia stuttgartensis | Planctomycetota | Candidatus Brocadiia | NZ_CP049055.1 | 2 |
| Candidatus Uabimicrobium amorphum | Planctomycetota | Candidatus Uabimicrobi | NZ_AP019860.1 | 1 |

| | | | | |
|---|---|---|---|---|
| Candidatus Uabimicrobium helgolandensis | Planctomycetota | Candidatus Uabimicrobi | NZ_CP165719.1 | 1 |
| Anaerohalosphaera lusitana | Planctomycetota | Phycisphaerae | NZ_CP019791.1 | 1 |
| Humisphaera borealis | Planctomycetota | Phycisphaerae | NZ_CP063458.1 | 2 |
| Limihaloglobus sulfuriphilus | Planctomycetota | Phycisphaerae | NZ_CP019646.1 | 1 |
| Mucisphaera calidilacus | Planctomycetota | Phycisphaerae | NZ_CP036280.1 | 1 |
| Phycisphaera mikurensis NBRC 102666 | Planctomycetota | Phycisphaerae | NC_017080.1 | 1 |
| Poriferisphaera corsica | Planctomycetota | Phycisphaerae | NZ_CP036425.1 | 1 |
| Sedimentisphaera cyanobacteriorum | Planctomycetota | Phycisphaerae | NZ_CP019633.1 | 1 |
| Sedimentisphaera salicampi | Planctomycetota | Phycisphaerae | NZ_CP021023.1 | 1 |
| Adhaeretor mobilis | Planctomycetota | Planctomycetia | NZ_CP036263.1 | 1 |
| Aeoliella mucimassa | Planctomycetota | Planctomycetia | NZ_CP036278.1 | 1 |
| Alienimonas californiensis | Planctomycetota | Planctomycetia | NZ_CP036265.1 | 1 |
| Anatilimnocola aggregata | Planctomycetota | Planctomycetia | NZ_CP036274.1 | 1 |
| Aquisphaera giovannonii | Planctomycetota | Planctomycetia | NZ_CP042997.1 | 1 |
| Aureliella helgolandensis | Planctomycetota | Planctomycetia | NZ_CP036298.1 | 2 |
| Botrimarina mediterranea | Planctomycetota | Planctomycetia | NZ_CP036349.1 | 1 |
| Bremerella volcania | Planctomycetota | Planctomycetia | NZ_CP036289.1 | 1 |
| Bythopirellula goksoeyrii | Planctomycetota | Planctomycetia | NZ_CP042913.1 | 1 |
| Calycomorphotria hydatis | Planctomycetota | Planctomycetia | NZ_CP036316.1 | 1 |
| Caulifigura coniformis | Planctomycetota | Planctomycetia | NZ_CP036271.1 | 1 |
| Crateriforma conspicua | Planctomycetota | Planctomycetia | NZ_CP036319.1 | 2 |
| Crateriforma spongiae | Planctomycetota | Planctomycetia | NZ_CP178615.1 | 2 |
| Frigoriglobus tundricola | Planctomycetota | Planctomycetia | NZ_CP053452.2 | 1 |
| Fuerstiella marisgermanici | Planctomycetota | Planctomycetia | NZ_CP017641.1 | 1 |
| Gemmata massiliana | Planctomycetota | Planctomycetia | NZ_LR593886.1 | 2 |
| Gimesia alba | Planctomycetota | Planctomycetia | NZ_CP036269.1 | 2 |
| Gimesia algae | Planctomycetota | Planctomycetia | NZ_CP036343.1 | 1 |
| Gimesia aquarii | Planctomycetota | Planctomycetia | NZ_CP037920.1 | 2 |
| Gimesia benthica | Planctomycetota | Planctomycetia | NZ_CP043930.1 | 2 |
| Gimesia fumaroli | Planctomycetota | Planctomycetia | NZ_CP037452.1 | 2 |
| Gimesia maris | Planctomycetota | Planctomycetia | NZ_CP042910.1 | 1 |
| Gimesia panareensis | Planctomycetota | Planctomycetia | NZ_CP037421.1 | 2 |

| | | | | |
|---|---|---|---|---|
| Isosphaera pallida ATCC 43644 | Planctomycetota | Planctomycetia | NC_014962.1 | 1 |
| Lacipirellula limnantheis | Planctomycetota | Planctomycetia | NZ_CP036339.1 | 1 |
| Lacipirellula parvula | Planctomycetota | Planctomycetia | NZ_AP021861.1 | 1 |
| Lignipirellula cremea | Planctomycetota | Planctomycetia | NZ_CP036433.1 | 1 |
| Limnoglobus roseus | Planctomycetota | Planctomycetia | NZ_CP042425.1 | 2 |
| Maioricimonas rarisocia | Planctomycetota | Planctomycetia | NZ_CP036275.1 | 1 |
| Mariniblastus fucicola | Planctomycetota | Planctomycetia | NZ_CP042912.1 | 1 |
| Paludisphaera borealis | Planctomycetota | Planctomycetia | NZ_CP019082.1 | 1 |
| Pirellulimonas nuda | Planctomycetota | Planctomycetia | NZ_CP036291.1 | 1 |
| Planctellipticum variicoloris | Planctomycetota | Planctomycetia | NZ_CP130886.1 | 1 |
| Planctopirus ephydatiae | Planctomycetota | Planctomycetia | NZ_CP036299.1 | 1 |
| Planctopirus limnophila DSM 3776 | Planctomycetota | Planctomycetia | NC_014148.1 | 1 |
| Polystyrenella longa | Planctomycetota | Planctomycetia | NZ_CP036281.1 | 1 |
| Rhodopirellula baltica SH 1 | Planctomycetota | Planctomycetia | NC_005027.1 | 1 |
| Roseimaritima multifibrata | Planctomycetota | Planctomycetia | NZ_CP036262.1 | 1 |
| Roseimaritima ulvae | Planctomycetota | Planctomycetia | NZ_CP042914.1 | 1 |
| Rosistilla carotiformis | Planctomycetota | Planctomycetia | NZ_CP036348.1 | 1 |
| Rosistilla oblonga | Planctomycetota | Planctomycetia | NZ_CP036292.1 | 1 |
| Rosistilla ulvae | Planctomycetota | Planctomycetia | NZ_CP036261.1 | 1 |
| Rubinisphaera brasiliensis DSM 5305 | Planctomycetota | Planctomycetia | NC_015174.1 | 1 |
| Rubripirellula lacrimiformis | Planctomycetota | Planctomycetia | NZ_CP036525.1 | 1 |
| Singulisphaera acidiphila DSM 18658 | Planctomycetota | Planctomycetia | NC_019892.1 | 1 |
| Stieleria maiorica | Planctomycetota | Planctomycetia | NZ_CP036264.1 | 2 |
| Stieleria neptunia | Planctomycetota | Planctomycetia | NZ_CP037423.1 | 2 |
| Stieleria varia | Planctomycetota | Planctomycetia | NZ_CP151726.1 | 2 |
| Stratiformator vulcanicus | Planctomycetota | Planctomycetia | NZ_CP036268.1 | 1 |
| Tautonia plasticadhaerens | Planctomycetota | Planctomycetia | NZ_CP036426.1 | 1 |
| Telmatocola sphagniphila | Planctomycetota | Planctomycetia | NZ_CP074694.1 | 1 |
| Thalassoglobus polymorphus | Planctomycetota | Planctomycetia | NZ_CP036267.1 | 1 |
| Thermogutta terrifontis | Planctomycetota | Planctomycetia | NZ_CP018477.1 | 1 |
| Tuwongella immobilis | Planctomycetota | Planctomycetia | NZ_LR593887.1 | 1 |
| Urbifossiella limnaea | Planctomycetota | Planctomycetia | NZ_CP036273.1 | 2 |

| | | | | |
|---|---|---|---|---|
| Acidithiobacillus caldus SM-1 | Pseudomonadota | Acidithiobacillia | NC_015850.1 | 2 |
| Acidithiobacillus ferrianus | Pseudomonadota | Acidithiobacillia | NZ_CP127523.1 | 2 |
| Acidithiobacillus ferridurans | Pseudomonadota | Acidithiobacillia | NZ_AP018795.1 | 2 |
| Acidithiobacillus ferriphilus | Pseudomonadota | Acidithiobacillia | NZ_CP116458.1 | 2 |
| Acidithiobacillus ferrivorans | Pseudomonadota | Acidithiobacillia | NZ_CP059488.1 | 2 |
| Acidithiobacillus ferrooxidans ATCC 23270 | Pseudomonadota | Acidithiobacillia | NC_011761.1 | 2 |
| Acidithiobacillus ferruginosus | Pseudomonadota | Acidithiobacillia | NZ_CP130946.1 | 2 |
| Acidithiobacillus montserratensis | Pseudomonadota | Acidithiobacillia | NZ_CP127526.1 | 2 |
| Acidithiobacillus sulfuriphilus | Pseudomonadota | Acidithiobacillia | NZ_CP127527.1 | 2 |
| Acidithiobacillus thiooxidans ATCC 19377 | Pseudomonadota | Acidithiobacillia | NZ_CP045571.1 | 2 |
| Thermithiobacillus tepidarius DSM 3134 | Pseudomonadota | Acidithiobacillia | NZ_CP133759.1 | 2 |
| [Pseudomonas] carboxydohydrogena | Pseudomonadota | Alphaproteobacteria | NZ_CP113162.1 | 1 |
| Abyssibius alkaniclasticus | Pseudomonadota | Alphaproteobacteria | NZ_CP095732.1 | 2 |
| Acetobacter ascendens | Pseudomonadota | Alphaproteobacteria | NZ_CP021524.1 | 1 |
| Acetobacter oryzifermentans | Pseudomonadota | Alphaproteobacteria | NZ_CP022374.1 | 1 |
| Acetobacter oryzoeni | Pseudomonadota | Alphaproteobacteria | NZ_CP042808.1 | 1 |
| Acetobacter pasteurianus | Pseudomonadota | Alphaproteobacteria | NZ_CP039846.2 | 1 |
| Acetobacter senegalensis | Pseudomonadota | Alphaproteobacteria | NZ_LN606600.1 | 1 |
| Acetobacter vaccinii | Pseudomonadota | Alphaproteobacteria | NZ_CP043506.1 | 1 |
| Acidibrevibacterium fodinaquatile | Pseudomonadota | Alphaproteobacteria | NZ_CP029176.1 | 1 |
| Acidiphilium multivorum AIU301 | Pseudomonadota | Alphaproteobacteria | NC_015186.1 | 1 |
| Acidisoma cladoniae | Pseudomonadota | Alphaproteobacteria | NZ_CP126994.1 | 1 |
| Aerophototrophica crusticola | Pseudomonadota | Alphaproteobacteria | NZ_CP051775.1 | 2 |
| Aestuariispira ectoiniformans | Pseudomonadota | Alphaproteobacteria | NZ_CP062788.1 | 2 |
| Afipia carboxidovorans OM5 | Pseudomonadota | Alphaproteobacteria | NC_015684.1 | 1 |
| Agrobacterium cucumeris | Pseudomonadota | Alphaproteobacteria | NZ_CP080387.1 | 2 |
| Agrobacterium fabrum str. C58 | Pseudomonadota | Alphaproteobacteria | NC_003062.2 | 2 |
| Agrobacterium larrymoorei | Pseudomonadota | Alphaproteobacteria | NZ_CP124733.1 | 2 |
| Agrobacterium leguminum | Pseudomonadota | Alphaproteobacteria | NZ_CP120211.1 | 2 |
| Agrobacterium pusense | Pseudomonadota | Alphaproteobacteria | NZ_CP053856.1 | 2 |
| Agrobacterium tumefaciens | Pseudomonadota | Alphaproteobacteria | NZ_CP115841.1 | 2 |
| Agrobacterium vaccinii | Pseudomonadota | Alphaproteobacteria | NZ_CP054150.1 | 2 |

| | | | | |
|---|---|---|---|---|
| Algicella marina | Pseudomonadota | Alphaproteobacteria | NZ_CP046620.1 | 2 |
| Aliiroseovarius crassostreae | Pseudomonadota | Alphaproteobacteria | NZ_CP080772.1 | 2 |
| Aliisedimentitalea scapharcae | Pseudomonadota | Alphaproteobacteria | NZ_CP123584.1 | 2 |
| Allorhizobium ampelinum S4 | Pseudomonadota | Alphaproteobacteria | NC_011989.1 | 2 |
| Allorhizobium pseudoryzae | Pseudomonadota | Alphaproteobacteria | NZ_CP049241.1 | 2 |
| Allosphingosinicella indica | Pseudomonadota | Alphaproteobacteria | NZ_LT840185.1 | 1 |
| Alphaproteobacteria bacterium endosymbiont of Tiliacea ci | Pseudomonadota | Alphaproteobacteria | NZ_OZ034995.1 | 1 |
| Altererythrobacter arenosus | Pseudomonadota | Alphaproteobacteria | NZ_CP121106.1 | 1 |
| Altererythrobacter epoxidivorans | Pseudomonadota | Alphaproteobacteria | NZ_CP012669.1 | 1 |
| Altererythrobacter ishigakiensis | Pseudomonadota | Alphaproteobacteria | NZ_CP015963.1 | 1 |
| Altererythrobacter rubellus | Pseudomonadota | Alphaproteobacteria | NZ_CP127221.1 | 1 |
| Alteriqipengyuania flavescens | Pseudomonadota | Alphaproteobacteria | NZ_CP129107.1 | 1 |
| Alterisphingorhabdus coralli | Pseudomonadota | Alphaproteobacteria | NZ_CP136594.1 | 1 |
| Aminobacter niigataensis | Pseudomonadota | Alphaproteobacteria | NZ_OX341517.1 | 2 |
| Anaplasma centrale str. Israel | Pseudomonadota | Alphaproteobacteria | NC_013532.1 | 2 |
| Anaplasma marginale str. Florida | Pseudomonadota | Alphaproteobacteria | NC_012026.1 | 2 |
| Anaplasma ovis str. Haibei | Pseudomonadota | Alphaproteobacteria | NZ_CP015994.1 | 2 |
| Anaplasma platys | Pseudomonadota | Alphaproteobacteria | NZ_CP046391.1 | 1 |
| Ancylobacter novellus DSM 506 | Pseudomonadota | Alphaproteobacteria | NC_014217.1 | 2 |
| Ancylobacter pratisalsi | Pseudomonadota | Alphaproteobacteria | NZ_CP048630.1 | 2 |
| Aquibium microcysteis | Pseudomonadota | Alphaproteobacteria | NZ_CP061080.1 | 2 |
| Aquibium oceanicum | Pseudomonadota | Alphaproteobacteria | NZ_CP018171.1 | 2 |
| Aquisediminimonas profunda | Pseudomonadota | Alphaproteobacteria | NZ_CP080327.1 | 1 |
| Aristophania vespae | Pseudomonadota | Alphaproteobacteria | NZ_CP092734.1 | 1 |
| Asaia bogorensis NBRC 16594 | Pseudomonadota | Alphaproteobacteria | NZ_AP014690.1 | 1 |
| Asticcacaulis excentricus CB 48 | Pseudomonadota | Alphaproteobacteria | NC_014816.1 | 1 |
| Aurantiacibacter atlanticus | Pseudomonadota | Alphaproteobacteria | NZ_CP011310.1 | 1 |
| Aurantiacibacter gangjinensis | Pseudomonadota | Alphaproteobacteria | NZ_CP018097.1 | 1 |
| Aureimonas mangrovi | Pseudomonadota | Alphaproteobacteria | NZ_CP059692.1 | 2 |
| Aureimonas phyllosphaerae | Pseudomonadota | Alphaproteobacteria | NZ_CP181348.1 | 2 |
| Aureimonas populi | Pseudomonadota | Alphaproteobacteria | NZ_CP072611.1 | 2 |
| Azorhizobium caulinodans ORS 571 | Pseudomonadota | Alphaproteobacteria | NC_009937.1 | 2 |

| | | | | |
|---|---|---|---|---|
| Azospirillum argentinense | Pseudomonadota | Alphaproteobacteria | NZ_CP007793.1 | 2 |
| Azospirillum baldaniorum | Pseudomonadota | Alphaproteobacteria | NZ_CP022253.1 | 2 |
| Azospirillum humicireducens | Pseudomonadota | Alphaproteobacteria | NZ_CP015285.1 | 2 |
| Azospirillum oryzae | Pseudomonadota | Alphaproteobacteria | NZ_CP054619.1 | 2 |
| Azospirillum ramasamyi | Pseudomonadota | Alphaproteobacteria | NZ_CP029829.1 | 2 |
| Azospirillum thermophilum | Pseudomonadota | Alphaproteobacteria | NZ_CP029353.1 | 2 |
| Azospirillum thiophilum | Pseudomonadota | Alphaproteobacteria | NZ_CP012401.1 | 2 |
| Bartonella alsatica | Pseudomonadota | Alphaproteobacteria | NZ_CP058235.1 | 2 |
| Bartonella ancashensis | Pseudomonadota | Alphaproteobacteria | NZ_CP010401.1 | 2 |
| Bartonella australis AUST/NH1 | Pseudomonadota | Alphaproteobacteria | NC_020300.1 | 2 |
| Bartonella bacilliformis KC583 | Pseudomonadota | Alphaproteobacteria | NC_008783.1 | 3 |
| Bartonella clarridgeiae 73 | Pseudomonadota | Alphaproteobacteria | NC_014932.1 | 2 |
| Bartonella elizabethae | Pseudomonadota | Alphaproteobacteria | NZ_LR134527.1 | 2 |
| Bartonella gliris | Pseudomonadota | Alphaproteobacteria | NZ_CP123962.1 | 2 |
| Bartonella grahamii as4aup | Pseudomonadota | Alphaproteobacteria | NC_012846.1 | 2 |
| Bartonella harrusi | Pseudomonadota | Alphaproteobacteria | NZ_CP101114.1 | 2 |
| Bartonella heixiaziensis | Pseudomonadota | Alphaproteobacteria | NZ_CP176404.1 | 2 |
| Bartonella henselae | Pseudomonadota | Alphaproteobacteria | NZ_CP082885.1 | 2 |
| Bartonella kosoyi | Pseudomonadota | Alphaproteobacteria | NZ_CP031843.2 | 2 |
| Bartonella krasnovii | Pseudomonadota | Alphaproteobacteria | NZ_CP031844.2 | 2 |
| Bartonella machadoae | Pseudomonadota | Alphaproteobacteria | NZ_CP087114.1 | 2 |
| Bartonella quintana | Pseudomonadota | Alphaproteobacteria | NZ_AP019773.1 | 2 |
| Bartonella schoenbuchensis | Pseudomonadota | Alphaproteobacteria | NZ_CP154603.1 | 2 |
| Bartonella taylorii | Pseudomonadota | Alphaproteobacteria | NZ_CP083444.1 | 2 |
| Bartonella tribocorum CIP 105476 | Pseudomonadota | Alphaproteobacteria | NC_010161.1 | 2 |
| Bartonella vinsonii | Pseudomonadota | Alphaproteobacteria | NZ_LR134529.1 | 2 |
| Beijerinckia indica subsp. indica ATCC 9039 | Pseudomonadota | Alphaproteobacteria | NC_010581.1 | 2 |
| Blastochloris tepida | Pseudomonadota | Alphaproteobacteria | NZ_AP018907.1 | 1 |
| Blastochloris viridis | Pseudomonadota | Alphaproteobacteria | NZ_CP012946.1 | 1 |
| Blastomonas fulva | Pseudomonadota | Alphaproteobacteria | NZ_CP020083.1 | 1 |
| Blastomonas marina | Pseudomonadota | Alphaproteobacteria | NZ_CP139718.1 | 1 |
| Bosea vaviloviae | Pseudomonadota | Alphaproteobacteria | NZ_CP017147.1 | 2 |

| | | | | |
|---|---|---|---|---|
| Bradyrhizobium amphicarpaeae | Pseudomonadota | Alphaproteobacteria | NZ_CP029426.2 | 1 |
| Bradyrhizobium arachidis | Pseudomonadota | Alphaproteobacteria | NZ_CP030050.1 | 1 |
| Bradyrhizobium barranii subsp. barranii | Pseudomonadota | Alphaproteobacteria | NZ_CP088280.1 | 1 |
| Bradyrhizobium betae | Pseudomonadota | Alphaproteobacteria | NZ_CP044543.1 | 1 |
| Bradyrhizobium brasilense | Pseudomonadota | Alphaproteobacteria | NZ_CP121669.1 | 1 |
| Bradyrhizobium centrosematis | Pseudomonadota | Alphaproteobacteria | NZ_CP120703.1 | 1 |
| Bradyrhizobium commune | Pseudomonadota | Alphaproteobacteria | NZ_CP061379.1 | 2 |
| Bradyrhizobium cosmicum | Pseudomonadota | Alphaproteobacteria | NC_017082.1 | 1 |
| Bradyrhizobium daqingense | Pseudomonadota | Alphaproteobacteria | NZ_CP088014.1 | 1 |
| Bradyrhizobium diazoefficiens USDA 110 | Pseudomonadota | Alphaproteobacteria | NZ_CP011360.1 | 1 |
| Bradyrhizobium elkanii USDA 76 | Pseudomonadota | Alphaproteobacteria | NZ_CP066356.1 | 1 |
| Bradyrhizobium ganzhouense | Pseudomonadota | Alphaproteobacteria | NZ_CP181318.1 | 1 |
| Bradyrhizobium guangdongense | Pseudomonadota | Alphaproteobacteria | NZ_CP030051.1 | 1 |
| Bradyrhizobium guangxiense | Pseudomonadota | Alphaproteobacteria | NZ_CP022219.1 | 1 |
| Bradyrhizobium guangzhouense | Pseudomonadota | Alphaproteobacteria | NZ_CP030053.1 | 1 |
| Bradyrhizobium huanghuaihaiense | Pseudomonadota | Alphaproteobacteria | NZ_CP104172.1 | 1 |
| Bradyrhizobium icense | Pseudomonadota | Alphaproteobacteria | NZ_CP016428.1 | 1 |
| Bradyrhizobium japonicum | Pseudomonadota | Alphaproteobacteria | NZ_CP058354.1 | 1 |
| Bradyrhizobium lupini | Pseudomonadota | Alphaproteobacteria | NZ_CP170504.1 | 1 |
| Bradyrhizobium oligotrophicum S58 | Pseudomonadota | Alphaproteobacteria | NC_020453.1 | 1 |
| Bradyrhizobium ontarionense | Pseudomonadota | Alphaproteobacteria | NZ_CP088156.1 | 1 |
| Bradyrhizobium ottawaense | Pseudomonadota | Alphaproteobacteria | NZ_CP029425.2 | 1 |
| Bradyrhizobium pachyrhizi | Pseudomonadota | Alphaproteobacteria | NZ_CP121667.1 | 1 |
| Bradyrhizobium paxllaeri | Pseudomonadota | Alphaproteobacteria | NZ_CP042968.1 | 1 |
| Bradyrhizobium prioriisuperbiae | Pseudomonadota | Alphaproteobacteria | NZ_CP135921.1 | 2 |
| Bradyrhizobium quebecense | Pseudomonadota | Alphaproteobacteria | NZ_CP088282.1 | 1 |
| Bradyrhizobium roseum | Pseudomonadota | Alphaproteobacteria | NZ_CP129212.1 | 1 |
| Bradyrhizobium sediminis | Pseudomonadota | Alphaproteobacteria | NZ_CP076134.1 | 1 |
| Bradyrhizobium septentrionale | Pseudomonadota | Alphaproteobacteria | NZ_CP088285.1 | 2 |
| Bradyrhizobium symbiodeficiens | Pseudomonadota | Alphaproteobacteria | NZ_CP050066.2 | 1 |
| Bradyrhizobium vignae | Pseudomonadota | Alphaproteobacteria | NZ_LS398110.1 | 1 |
| Bradyrhizobium xenonodulans | Pseudomonadota | Alphaproteobacteria | NZ_CP089391.1 | 1 |

| | | | | |
|---|---|---|---|---|
| Bradyrhizobium yuanmingense | Pseudomonadota | Alphaproteobacteria | NZ_CP104173.1 | 1 |
| Bradyrhizobium zhanjiangense | Pseudomonadota | Alphaproteobacteria | NZ_CP022221.1 | 1 |
| Brevundimonas albigilva | Pseudomonadota | Alphaproteobacteria | NZ_CP097649.1 | 2 |
| Brevundimonas bullata | Pseudomonadota | Alphaproteobacteria | NZ_CP139996.1 | 2 |
| Brevundimonas fontaquae | Pseudomonadota | Alphaproteobacteria | NZ_CP070968.1 | 2 |
| Brevundimonas goettingensis | Pseudomonadota | Alphaproteobacteria | NZ_CP062222.1 | 2 |
| Brevundimonas intermedia | Pseudomonadota | Alphaproteobacteria | NZ_CP169067.1 | 2 |
| Brevundimonas nasdae | Pseudomonadota | Alphaproteobacteria | NZ_CP080034.1 | 2 |
| Brevundimonas olei | Pseudomonadota | Alphaproteobacteria | NZ_CP146369.1 | 2 |
| Brevundimonas sanguinis | Pseudomonadota | Alphaproteobacteria | NZ_CP116770.1 | 2 |
| Brevundimonas staleyi | Pseudomonadota | Alphaproteobacteria | NZ_CP169082.1 | 2 |
| Brevundimonas subvibrioides ATCC 15264 | Pseudomonadota | Alphaproteobacteria | NC_014375.1 | 2 |
| Brevundimonas vesicularis | Pseudomonadota | Alphaproteobacteria | NZ_CP022048.2 | 2 |
| Brevundimonas vitisensis | Pseudomonadota | Alphaproteobacteria | NZ_CP067977.1 | 2 |
| Brucella anthropi | Pseudomonadota | Alphaproteobacteria | NZ_CP064063.1 | 2 |
| Brucella canis ATCC 23365 | Pseudomonadota | Alphaproteobacteria | NC_010103.1 | 2 |
| Brucella ceti | Pseudomonadota | Alphaproteobacteria | NZ_CP178745.1 | 2 |
| Brucella inopinata | Pseudomonadota | Alphaproteobacteria | NZ_LT605585.1 | 2 |
| Brucella melitensis bv. 1 str. 16M | Pseudomonadota | Alphaproteobacteria | NC_003317.1 | 2 |
| Brucella microti CCM 4915 | Pseudomonadota | Alphaproteobacteria | NC_013119.1 | 2 |
| Brucella ovis ATCC 25840 | Pseudomonadota | Alphaproteobacteria | NC_009505.1 | 2 |
| Brucella pseudintermedia | Pseudomonadota | Alphaproteobacteria | NZ_CP099967.1 | 2 |
| Brucella pseudogrignonensis | Pseudomonadota | Alphaproteobacteria | NZ_CP091783.1 | 2 |
| Brucella suis 1330 | Pseudomonadota | Alphaproteobacteria | NC_004310.3 | 2 |
| Caenibius tardaugens NBRC 16725 | Pseudomonadota | Alphaproteobacteria | NZ_CP034179.1 | 1 |
| Candidatus Bandiella numerosa | Pseudomonadota | Alphaproteobacteria | NZ_CP104164.1 | 2 |
| Candidatus Bandiella woodruffii | Pseudomonadota | Alphaproteobacteria | NZ_CP110820.1 | 2 |
| Candidatus Bealeia paramacronuclearis | Pseudomonadota | Alphaproteobacteria | NZ_CP133270.1 | 2 |
| Candidatus Bodocaedibacter vickermanii | Pseudomonadota | Alphaproteobacteria | NZ_CP054719.1 | 1 |
| Candidatus Cytomitobacter indipagum | Pseudomonadota | Alphaproteobacteria | NZ_CP043315.1 | 1 |
| Candidatus Cytomitobacter primus | Pseudomonadota | Alphaproteobacteria | NZ_OX638200.1 | 1 |
| Candidatus Deianiraea vastatrix | Pseudomonadota | Alphaproteobacteria | NZ_CP029077.1 | 1 |

| | | | | |
|---|---|---|---|---|
| Candidatus Endolissoclinum faulkneri L2 | Pseudomonadota | Alphaproteobacteria | NC_019566.1 | 1 |
| Candidatus Endowatersipora endosymbiont of Watersipora | Pseudomonadota | Alphaproteobacteria | NZ_OZ032160.1 | 1 |
| Candidatus Fokinia cryptica | Pseudomonadota | Alphaproteobacteria | NZ_CP110343.1 | 1 |
| Candidatus Fokinia solitaria | Pseudomonadota | Alphaproteobacteria | NZ_CP025989.1 | 1 |
| Candidatus Fonsibacter ubiquis | Pseudomonadota | Alphaproteobacteria | NZ_CP024034.1 | 1 |
| Candidatus Gromoviella agglomerans | Pseudomonadota | Alphaproteobacteria | NZ_CP069247.1 | 1 |
| Candidatus Hodgkinia cicadicola | Pseudomonadota | Alphaproteobacteria | NZ_NXGO0100007 | 1 |
| Candidatus Hodgkinia cicadicola | Pseudomonadota | Alphaproteobacteria | NZ_NXGO0100005 | 1 |
| Candidatus Hydrogenosomobacter endosymbioticus | Pseudomonadota | Alphaproteobacteria | NZ_AP025225.1 | 1 |
| Candidatus Kirkpatrickella diaphorinae | Pseudomonadota | Alphaproteobacteria | NZ_CP107052.1 | 1 |
| Candidatus Lariskella endosymbiont of Epinotia ramella | Pseudomonadota | Alphaproteobacteria | NZ_OZ026472.1 | 1 |
| Candidatus Lariskella endosymbiont of Hedychridium roseu | Pseudomonadota | Alphaproteobacteria | NZ_OZ026538.1 | 1 |
| Candidatus Liberibacter africanus PTSAPSY | Pseudomonadota | Alphaproteobacteria | NZ_CP004021.1 | 2 |
| Candidatus Liberibacter americanus str. Sao Paulo | Pseudomonadota | Alphaproteobacteria | NC_022793.1 | 2 |
| Candidatus Liberibacter asiaticus | Pseudomonadota | Alphaproteobacteria | NZ_CP010804.2 | 2 |
| Candidatus Liberibacter solanacearum CLso-ZC1 | Pseudomonadota | Alphaproteobacteria | NC_014774.1 | 2 |
| Candidatus Megaera polyxenophila | Pseudomonadota | Alphaproteobacteria | NZ_CP104166.1 | 1 |
| Candidatus Mesenet endosymbiont of Agriotes lineatus | Pseudomonadota | Alphaproteobacteria | NZ_OZ026454.1 | 2 |
| Candidatus Mesenet endosymbiont of Phosphuga atrata | Pseudomonadota | Alphaproteobacteria | NZ_OZ026541.1 | 2 |
| Candidatus Midichloria mitochondrii IricVA | Pseudomonadota | Alphaproteobacteria | NC_015722.1 | 1 |
| Candidatus Nesciobacter abundans | Pseudomonadota | Alphaproteobacteria | NZ_OY490516.1 | 1 |
| Candidatus Nucleicultrix amoebiphila FS5 | Pseudomonadota | Alphaproteobacteria | NZ_CP008743.1 | 2 |
| Candidatus Odyssella acanthamoebae | Pseudomonadota | Alphaproteobacteria | NZ_CP008941.1 | 2 |
| Candidatus Pelagibacter giovannonii | Pseudomonadota | Alphaproteobacteria | NZ_CP038852.1 | 1 |
| Candidatus Pelagibacter ubique HTCC1062 | Pseudomonadota | Alphaproteobacteria | NC_007205.1 | 1 |
| Candidatus Phycorickettsia trachydisci | Pseudomonadota | Alphaproteobacteria | NZ_CP027845.1 | 1 |
| Candidatus Puniceispirillum marinum IMCC1322 | Pseudomonadota | Alphaproteobacteria | NC_014010.1 | 2 |
| Candidatus Sneabacter namystus | Pseudomonadota | Alphaproteobacteria | NZ_CP043312.1 | 1 |
| Candidatus Tisiphia endosymbiont of Beris chalybata | Pseudomonadota | Alphaproteobacteria | NZ_OZ032159.1 | 1 |
| Candidatus Tisiphia endosymbiont of Ceraclea dissimilis | Pseudomonadota | Alphaproteobacteria | NZ_OZ034917.1 | 1 |
| Candidatus Tisiphia endosymbiont of Dascillus cervinus | Pseudomonadota | Alphaproteobacteria | NZ_OZ032146.1 | 1 |
| Candidatus Tisiphia endosymbiont of Dioctria linearis | Pseudomonadota | Alphaproteobacteria | NZ_OZ032156.1 | 1 |

| | | | | |
|---|---|---|---|---|
| Candidatus Tisiphia endosymbiont of Dioctria rufipes | Pseudomonadota | Alphaproteobacteria | NZ_OZ032163.1 | 1 |
| Candidatus Tisiphia endosymbiont of Ditula angustiorana | Pseudomonadota | Alphaproteobacteria | NZ_OZ032143.1 | 1 |
| Candidatus Tisiphia endosymbiont of Empis tessellata | Pseudomonadota | Alphaproteobacteria | NZ_OZ034997.1 | 1 |
| Candidatus Tisiphia endosymbiont of Melanophora roralis | Pseudomonadota | Alphaproteobacteria | NZ_OZ032161.1 | 1 |
| Candidatus Tisiphia endosymbiont of Metellina segmentata | Pseudomonadota | Alphaproteobacteria | NZ_OZ034919.1 | 1 |
| Candidatus Tisiphia endosymbiont of Micropterix aruncella | Pseudomonadota | Alphaproteobacteria | NZ_OZ238268.1 | 1 |
| Candidatus Tisiphia endosymbiont of Myopa tessellatipenn | Pseudomonadota | Alphaproteobacteria | NZ_OZ035005.1 | 1 |
| Candidatus Tisiphia endosymbiont of Mystacides longicorn | Pseudomonadota | Alphaproteobacteria | NZ_OZ035004.1 | 1 |
| Candidatus Tisiphia endosymbiont of Nedyus quadrimacula | Pseudomonadota | Alphaproteobacteria | NZ_OZ060452.1 | 1 |
| Candidatus Tisiphia endosymbiont of Nemotelus nigrinus | Pseudomonadota | Alphaproteobacteria | NZ_OZ032151.1 | 1 |
| Candidatus Tisiphia endosymbiont of Nemotelus uliginosus | Pseudomonadota | Alphaproteobacteria | NZ_OZ183481.1 | 1 |
| Candidatus Tisiphia endosymbiont of Neophilaenus lineatu | Pseudomonadota | Alphaproteobacteria | NZ_OZ183649.1 | 1 |
| Candidatus Tisiphia endosymbiont of Oplodontha viridula | Pseudomonadota | Alphaproteobacteria | NZ_OZ183486.1 | 1 |
| Candidatus Tisiphia endosymbiont of Parasteatoda lunata | Pseudomonadota | Alphaproteobacteria | NZ_OZ035003.1 | 1 |
| Candidatus Tisiphia endosymbiont of Piscicola geometra | Pseudomonadota | Alphaproteobacteria | NZ_OZ032144.1 | 1 |
| Candidatus Tisiphia endosymbiont of Psammoecus bipunc | Pseudomonadota | Alphaproteobacteria | NZ_OZ183487.1 | 1 |
| Candidatus Tisiphia endosymbiont of Ptychoptera albimana | Pseudomonadota | Alphaproteobacteria | NZ_OZ032164.1 | 1 |
| Candidatus Tisiphia endosymbiont of Sialis lutaria | Pseudomonadota | Alphaproteobacteria | NZ_OZ032153.1 | 1 |
| Candidatus Tisiphia endosymbiont of Stenodema calcarata | Pseudomonadota | Alphaproteobacteria | NZ_OZ034911.1 | 1 |
| Candidatus Tisiphia endosymbiont of Temnostethus pusillu | Pseudomonadota | Alphaproteobacteria | NZ_OZ183472.1 | 1 |
| Candidatus Tisiphia endosymbiont of Thecophora atra | Pseudomonadota | Alphaproteobacteria | NZ_OZ032145.1 | 1 |
| Candidatus Tisiphia endosymbiont of Xenochironomus xen | Pseudomonadota | Alphaproteobacteria | NZ_OZ183473.1 | 1 |
| Candidatus Trichorickettsia mobilis | Pseudomonadota | Alphaproteobacteria | NZ_OY728607.1 | 1 |
| Candidatus Viadribacter manganicus | Pseudomonadota | Alphaproteobacteria | NZ_CP013244.1 | 1 |
| Caulobacter flavus | Pseudomonadota | Alphaproteobacteria | NZ_CP026100.1 | 2 |
| Caulobacter henricii | Pseudomonadota | Alphaproteobacteria | NZ_CP013002.1 | 2 |
| Caulobacter mirabilis | Pseudomonadota | Alphaproteobacteria | NZ_CP024201.1 | 2 |
| Caulobacter rhizosphaerae | Pseudomonadota | Alphaproteobacteria | NZ_CP048815.1 | 2 |
| Caulobacter segnis | Pseudomonadota | Alphaproteobacteria | NZ_CP027850.1 | 2 |
| Caulobacter soli | Pseudomonadota | Alphaproteobacteria | NZ_CP049199.1 | 2 |
| Celeribacter ethanolicus | Pseudomonadota | Alphaproteobacteria | NZ_CP022196.1 | 2 |
| Celeribacter indicus | Pseudomonadota | Alphaproteobacteria | NZ_CP004393.1 | 2 |

| | | | | |
|---|---|---|---|---|
| Celeribacter marinus | Pseudomonadota | Alphaproteobacteria | NZ_CP012023.1 | 2 |
| Cereibacter azotoformans | Pseudomonadota | Alphaproteobacteria | NZ_CP090021.1 | 2 |
| Cereibacter sphaeroides 2.4.1 | Pseudomonadota | Alphaproteobacteria | NC_007493.2 | 2 |
| Ciceribacter thiooxidans | Pseudomonadota | Alphaproteobacteria | NZ_CP059896.1 | 2 |
| Cohaesibacter marisflavi | Pseudomonadota | Alphaproteobacteria | NZ_OY762534.1 | 2 |
| Commensalibacter melissae | Pseudomonadota | Alphaproteobacteria | NZ_CP033087.1 | 1 |
| Coralliovum pocilloporae | Pseudomonadota | Alphaproteobacteria | NZ_CP132542.1 | 2 |
| Croceibacterium aestuarii | Pseudomonadota | Alphaproteobacteria | NZ_CP131039.1 | 1 |
| Croceibacterium atlanticum | Pseudomonadota | Alphaproteobacteria | NZ_CP011452.2 | 1 |
| Croceicoccus marinus | Pseudomonadota | Alphaproteobacteria | NZ_CP060052.1 | 1 |
| Croceicoccus naphthovorans | Pseudomonadota | Alphaproteobacteria | NZ_CP011770.1 | 1 |
| Defluviicoccus vanus | Pseudomonadota | Alphaproteobacteria | NZ_CP053923.1 | 2 |
| Devosia beringensis | Pseudomonadota | Alphaproteobacteria | NZ_CP045422.1 | 1 |
| Devosia ginsengisoli | Pseudomonadota | Alphaproteobacteria | NZ_CP042304.1 | 1 |
| Devosia lacusdianchii | Pseudomonadota | Alphaproteobacteria | NZ_CP092483.1 | 1 |
| Devosia neptuniae | Pseudomonadota | Alphaproteobacteria | NZ_CP104965.1 | 1 |
| Devosia oryziradicis | Pseudomonadota | Alphaproteobacteria | NZ_CP068047.1 | 1 |
| Devosia rhizoryzae | Pseudomonadota | Alphaproteobacteria | NZ_CP068046.1 | 1 |
| Devosia rhizosphaerae | Pseudomonadota | Alphaproteobacteria | NZ_CP126539.1 | 1 |
| Devosia rhodophyticola | Pseudomonadota | Alphaproteobacteria | NZ_CP118247.1 | 1 |
| Devosia salina | Pseudomonadota | Alphaproteobacteria | NZ_CP080590.1 | 1 |
| Dinoroseobacter shibae DFL 12 = DSM 16493 | Pseudomonadota | Alphaproteobacteria | NC_009952.1 | 2 |
| Ehrlichia canis str. Jake | Pseudomonadota | Alphaproteobacteria | NC_007354.1 | 2 |
| Ehrlichia chaffeensis str. Arkansas | Pseudomonadota | Alphaproteobacteria | NC_007799.1 | 2 |
| Ehrlichia japonica | Pseudomonadota | Alphaproteobacteria | NZ_CP007474.1 | 2 |
| Ehrlichia muris AS145 | Pseudomonadota | Alphaproteobacteria | NC_023063.1 | 2 |
| Ehrlichia ruminantium str. Welgevonden | Pseudomonadota | Alphaproteobacteria | NC_005295.2 | 2 |
| Elioraea tepida | Pseudomonadota | Alphaproteobacteria | NZ_CP076448.1 | 2 |
| endosymbiont of Acanthamoeba sp. UWC8 | Pseudomonadota | Alphaproteobacteria | NZ_CP004403.1 | 2 |
| Ensifer adhaerens | Pseudomonadota | Alphaproteobacteria | NZ_CP015880.1 | 2 |
| Ensifer canadensis | Pseudomonadota | Alphaproteobacteria | NZ_CP083370.1 | 2 |
| Entomobacter blattae | Pseudomonadota | Alphaproteobacteria | NZ_CP060244.1 | 1 |

| | | | | |
|---|---|---|---|---|
| Erythrobacter aurantius | Pseudomonadota | Alphaproteobacteria | NZ_CP090949.1 | 1 |
| Erythrobacter aureus | Pseudomonadota | Alphaproteobacteria | NZ_CP031357.1 | 1 |
| Erythrobacter litoralis | Pseudomonadota | Alphaproteobacteria | NZ_CP017057.1 | 1 |
| Erythrobacter mangrovi | Pseudomonadota | Alphaproteobacteria | NZ_CP053921.1 | 1 |
| Erythrobacter neustonensis | Pseudomonadota | Alphaproteobacteria | NZ_CP016033.1 | 1 |
| Falsihalocynthiibacter arcticus | Pseudomonadota | Alphaproteobacteria | NZ_CP014327.1 | 2 |
| Falsirhodobacter algicola | Pseudomonadota | Alphaproteobacteria | NZ_CP047289.1 | 2 |
| Ferrovibrio terrae | Pseudomonadota | Alphaproteobacteria | NZ_CP041636.1 | 2 |
| Fontisubflavum oceani | Pseudomonadota | Alphaproteobacteria | NZ_CP129111.1 | 2 |
| Formicincola oecophyllae | Pseudomonadota | Alphaproteobacteria | NZ_CP038231.1 | 1 |
| Fretibacter rubidus | Pseudomonadota | Alphaproteobacteria | NZ_CP163423.1 | 2 |
| Frigidibacter mobilis | Pseudomonadota | Alphaproteobacteria | NZ_CP012661.1 | 2 |
| Fuscovulum ytuae | Pseudomonadota | Alphaproteobacteria | NZ_CP124535.1 | 2 |
| Gemmobacter fulvus | Pseudomonadota | Alphaproteobacteria | NZ_CP076361.1 | 2 |
| Georhizobium profundi | Pseudomonadota | Alphaproteobacteria | NZ_CP032509.1 | 2 |
| Gimibacter soli | Pseudomonadota | Alphaproteobacteria | NZ_CP116805.1 | 2 |
| Gluconacetobacter diazotrophicus PA1 5 | Pseudomonadota | Alphaproteobacteria | NC_011365.1 | 1 |
| Gluconobacter albidus | Pseudomonadota | Alphaproteobacteria | NZ_CP014689.1 | 1 |
| Gluconobacter oxydans DSM 3504 | Pseudomonadota | Alphaproteobacteria | NZ_CP004373.1 | 1 |
| Gluconobacter sphaericus | Pseudomonadota | Alphaproteobacteria | NZ_CP068419.1 | 1 |
| Glycocaulis abyssi | Pseudomonadota | Alphaproteobacteria | NZ_CP163421.1 | 2 |
| Glycocaulis alkaliphilus | Pseudomonadota | Alphaproteobacteria | NZ_CP018911.1 | 2 |
| Granulibacter bethesdensis | Pseudomonadota | Alphaproteobacteria | NZ_CP018193.1 | 1 |
| Haematobacter massiliensis | Pseudomonadota | Alphaproteobacteria | NZ_CP035510.1 | 2 |
| Haematospirillum jordaniae | Pseudomonadota | Alphaproteobacteria | NZ_CP014525.1 | 2 |
| Hankyongella ginsenosidimutans | Pseudomonadota | Alphaproteobacteria | NZ_CP039704.1 | 1 |
| Hartmannibacter diazotrophicus | Pseudomonadota | Alphaproteobacteria | NZ_LT960614.1 | 2 |
| Hirschia baltica ATCC 49814 | Pseudomonadota | Alphaproteobacteria | NC_012982.1 | 1 |
| Hoeflea phototrophica DFL-43 | Pseudomonadota | Alphaproteobacteria | NZ_CM002917.1 | 2 |
| Hypericibacter adhaerens | Pseudomonadota | Alphaproteobacteria | NZ_CP042582.1 | 2 |
| Hypericibacter terrae | Pseudomonadota | Alphaproteobacteria | NZ_CP042906.1 | 2 |
| Hyphococcus flavus | Pseudomonadota | Alphaproteobacteria | NZ_CP118166.1 | 1 |

| | | | | |
|---|---|---|---|---|
| Hyphococcus formosus | Pseudomonadota | Alphaproteobacteria | NZ_CP157210.1 | 1 |
| Hyphomicrobium denitrificans ATCC 51888 | Pseudomonadota | Alphaproteobacteria | NC_014313.1 | 1 |
| Hyphomicrobium nitrativorans NL23 | Pseudomonadota | Alphaproteobacteria | NC_022997.1 | 1 |
| Hyphomonas atlantica | Pseudomonadota | Alphaproteobacteria | NZ_CP051254.1 | 2 |
| Hyphomonas neptunium ATCC 15444 | Pseudomonadota | Alphaproteobacteria | NC_008358.1 | 2 |
| Indioceanicola profundi | Pseudomonadota | Alphaproteobacteria | NZ_CP030126.1 | 2 |
| Jannaschia ovalis | Pseudomonadota | Alphaproteobacteria | NZ_CP122537.1 | 2 |
| Kaustia mangrovi | Pseudomonadota | Alphaproteobacteria | NZ_CP058214.1 | 2 |
| Ketogulonicigenium robustum | Pseudomonadota | Alphaproteobacteria | NZ_CP019937.1 | 2 |
| Ketogulonicigenium vulgare | Pseudomonadota | Alphaproteobacteria | NZ_CP012908.1 | 2 |
| Komagataeibacter diospyri | Pseudomonadota | Alphaproteobacteria | NZ_CP171836.1 | 1 |
| Komagataeibacter europaeus | Pseudomonadota | Alphaproteobacteria | NZ_CP021467.1 | 1 |
| Komagataeibacter medellinensis NBRC 3288 | Pseudomonadota | Alphaproteobacteria | NC_016027.1 | 1 |
| Komagataeibacter nataicola | Pseudomonadota | Alphaproteobacteria | NZ_CP019875.1 | 1 |
| Komagataeibacter rhaeticus | Pseudomonadota | Alphaproteobacteria | NZ_CP050139.1 | 1 |
| Komagataeibacter saccharivorans | Pseudomonadota | Alphaproteobacteria | NZ_CP036404.1 | 1 |
| Komagataeibacter sucrofermentans DSM 15973 | Pseudomonadota | Alphaproteobacteria | NZ_CP137157.1 | 1 |
| Komagataeibacter xylinus | Pseudomonadota | Alphaproteobacteria | NZ_CP025269.1 | 1 |
| Kordiimonas pumila | Pseudomonadota | Alphaproteobacteria | NZ_CP061205.1 | 2 |
| Kozakia baliensis | Pseudomonadota | Alphaproteobacteria | NZ_CP014681.1 | 1 |
| Labrys okinawensis | Pseudomonadota | Alphaproteobacteria | NZ_CP178864.1 | 2 |
| Leisingera aquaemixtae | Pseudomonadota | Alphaproteobacteria | NZ_CP041159.1 | 2 |
| Leisingera caerulea | Pseudomonadota | Alphaproteobacteria | NZ_CP081070.1 | 2 |
| Leisingera methylohalidivorans DSM 14336 | Pseudomonadota | Alphaproteobacteria | NC_023135.1 | 2 |
| Leisingera thetidis | Pseudomonadota | Alphaproteobacteria | NZ_CP109787.1 | 2 |
| Liberibacter crescens | Pseudomonadota | Alphaproteobacteria | NZ_CP010522.1 | 2 |
| Lichenicola cladoniae | Pseudomonadota | Alphaproteobacteria | NZ_CP053708.1 | 1 |
| Limimaricola variabilis | Pseudomonadota | Alphaproteobacteria | NZ_CP139691.1 | 2 |
| Loktanella salsilacus | Pseudomonadota | Alphaproteobacteria | NZ_CP072994.1 | 2 |
| Luteithermobacter gelatinilyticus | Pseudomonadota | Alphaproteobacteria | NZ_CP040517.1 | 2 |
| Magnetococcus marinus MC-1 | Pseudomonadota | Alphaproteobacteria | NC_008576.1 | 2 |
| Magnetospirillum gryphiswaldense MSR-1 v2 | Pseudomonadota | Alphaproteobacteria | NC_023065.1 | 2 |

| | | | | |
|---|---|---|---|---|
| Mariluticola halotolerans | Pseudomonadota | Alphaproteobacteria | NZ_CP090960.1 | 2 |
| Marinicauda algicola | Pseudomonadota | Alphaproteobacteria | NZ_CP071057.1 | 2 |
| Maritalea myrionectae | Pseudomonadota | Alphaproteobacteria | NZ_CP021330.1 | 2 |
| Martelella endophytica | Pseudomonadota | Alphaproteobacteria | NZ_CP010803.1 | 2 |
| Martelella mediterranea DSM 17316 | Pseudomonadota | Alphaproteobacteria | NZ_CP020330.1 | 2 |
| Mesorhizobium amorphae CCNWGS0123 | Pseudomonadota | Alphaproteobacteria | NZ_CP015318.1 | 2 |
| Mesorhizobium australicum WSM2073 | Pseudomonadota | Alphaproteobacteria | NC_019973.1 | 2 |
| Mesorhizobium erdmanii | Pseudomonadota | Alphaproteobacteria | NZ_CP033361.1 | 2 |
| Mesorhizobium huakuii | Pseudomonadota | Alphaproteobacteria | NZ_CP139858.1 | 2 |
| Mesorhizobium japonicum R7A | Pseudomonadota | Alphaproteobacteria | NZ_CP051772.1 | 2 |
| Mesorhizobium jarvisii | Pseudomonadota | Alphaproteobacteria | NZ_CP033507.1 | 2 |
| Mesorhizobium koreense | Pseudomonadota | Alphaproteobacteria | NZ_CP134228.1 | 2 |
| Mesorhizobium mediterraneum | Pseudomonadota | Alphaproteobacteria | NZ_CP088151.1 | 2 |
| Mesorhizobium onobrychidis | Pseudomonadota | Alphaproteobacteria | NZ_CP062229.1 | 2 |
| Mesorhizobium opportunistum WSM2075 | Pseudomonadota | Alphaproteobacteria | NC_015675.1 | 2 |
| Methylobacterium bullatum | Pseudomonadota | Alphaproteobacteria | NZ_CP162530.1 | 2 |
| Methylobacterium currus | Pseudomonadota | Alphaproteobacteria | NZ_CP028843.1 | 2 |
| Methylobacterium durans | Pseudomonadota | Alphaproteobacteria | NZ_CP029550.1 | 2 |
| Methylobacterium flocculans | Pseudomonadota | Alphaproteobacteria | NZ_CP107532.1 | 2 |
| Methylobacterium indicum | Pseudomonadota | Alphaproteobacteria | NZ_AP024145.1 | 2 |
| Methylobacterium nodulans ORS 2060 | Pseudomonadota | Alphaproteobacteria | NC_011894.1 | 2 |
| Methylobacterium organophilum | Pseudomonadota | Alphaproteobacteria | NZ_CP092848.1 | 2 |
| Methylobacterium oryzae CBMB20 | Pseudomonadota | Alphaproteobacteria | NZ_CP003811.1 | 2 |
| Methylobacterium phyllosphaerae | Pseudomonadota | Alphaproteobacteria | NZ_CP015367.1 | 2 |
| Methylobacterium radiodurans | Pseudomonadota | Alphaproteobacteria | NZ_CP029551.1 | 2 |
| Methylobacterium radiotolerans JCM 2831 | Pseudomonadota | Alphaproteobacteria | NC_010505.1 | 2 |
| Methylobacterium terrae | Pseudomonadota | Alphaproteobacteria | NZ_CP029553.1 | 2 |
| Methylocapsa polymorpha | Pseudomonadota | Alphaproteobacteria | NZ_CP136862.1 | 1 |
| Methyloceanibacter caenitepidi | Pseudomonadota | Alphaproteobacteria | NZ_AP014648.1 | 1 |
| Methylocella silvestris BL2 | Pseudomonadota | Alphaproteobacteria | NC_011666.1 | 1 |
| Methylocella tundrae | Pseudomonadota | Alphaproteobacteria | NZ_CP139089.1 | 1 |
| Methylocystis bryophila | Pseudomonadota | Alphaproteobacteria | NZ_AP027149.1 | 2 |

| | | | | |
|---|---|---|---|---|
| Methylocystis echinoides | Pseudomonadota | Alphaproteobacteria | NZ_CP156084.1 | 2 |
| Methylocystis heyeri | Pseudomonadota | Alphaproteobacteria | NZ_CP046052.1 | 2 |
| Methylocystis iwaonis | Pseudomonadota | Alphaproteobacteria | NZ_AP027142.1 | 2 |
| Methylocystis parvus OBBP | Pseudomonadota | Alphaproteobacteria | NZ_CP092968.1 | 2 |
| Methyloraptor flagellatus | Pseudomonadota | Alphaproteobacteria | NZ_CP158568.1 | 2 |
| Methylorubrum extorquens DM4 | Pseudomonadota | Alphaproteobacteria | NC_012988.1 | 2 |
| Methylorubrum populi BJ001 | Pseudomonadota | Alphaproteobacteria | NC_010725.1 | 2 |
| Methylovirgula ligni | Pseudomonadota | Alphaproteobacteria | NZ_CP025086.1 | 1 |
| Microvirga lotononidis | Pseudomonadota | Alphaproteobacteria | NZ_CP141048.1 | 2 |
| Microvirga ossetica | Pseudomonadota | Alphaproteobacteria | NZ_CP016616.1 | 2 |
| Microvirga terrae | Pseudomonadota | Alphaproteobacteria | NZ_CP102845.1 | 2 |
| Muricoccus radiodurans | Pseudomonadota | Alphaproteobacteria | NZ_CP189561.1 | 2 |
| Neoasaia chiangmaiensis | Pseudomonadota | Alphaproteobacteria | NZ_CP014691.1 | 1 |
| Neoehrlichia mikurensis | Pseudomonadota | Alphaproteobacteria | NZ_CP089285.1 | 2 |
| Neokomagataea tanensis | Pseudomonadota | Alphaproteobacteria | NZ_CP032485.1 | 1 |
| Neorhizobium galegae bv. orientalis str. HAMBI 540 | Pseudomonadota | Alphaproteobacteria | NZ_HG938353.1 | 2 |
| Neorhizobium petrolearium | Pseudomonadota | Alphaproteobacteria | NZ_CP123000.1 | 2 |
| Neorickettsia findlayensis | Pseudomonadota | Alphaproteobacteria | NZ_CP047224.1 | 1 |
| Neorickettsia helminthoeca str. Oregon | Pseudomonadota | Alphaproteobacteria | NZ_CP007481.1 | 1 |
| Neorickettsia risticii str. Illinois | Pseudomonadota | Alphaproteobacteria | NC_013009.1 | 1 |
| Neorickettsia sennetsu str. Miyayama | Pseudomonadota | Alphaproteobacteria | NC_007798.1 | 1 |
| Neotabrizicola shimadae | Pseudomonadota | Alphaproteobacteria | NZ_CP069370.1 | 2 |
| Nereida ignava | Pseudomonadota | Alphaproteobacteria | NZ_CP112885.1 | 2 |
| Nguyenibacter vanlangensis | Pseudomonadota | Alphaproteobacteria | NZ_CP152276.1 | 1 |
| Nisaea acidiphila | Pseudomonadota | Alphaproteobacteria | NZ_CP102480.1 | 2 |
| Nitratireductor aquimarinus | Pseudomonadota | Alphaproteobacteria | NZ_CP177239.1 | 2 |
| Nitratireductor kimnyeongensis | Pseudomonadota | Alphaproteobacteria | NZ_CP078143.1 | 2 |
| Nitratireductor mangrovi | Pseudomonadota | Alphaproteobacteria | NZ_CP042301.2 | 2 |
| Nitratireductor thuwali | Pseudomonadota | Alphaproteobacteria | NZ_CP030941.1 | 2 |
| Nitrobacter hamburgensis X14 | Pseudomonadota | Alphaproteobacteria | NC_007964.1 | 1 |
| Nitrobacter winogradskyi Nb-255 | Pseudomonadota | Alphaproteobacteria | NC_007406.1 | 1 |
| Nitrospirillum viridazoti CBAmc | Pseudomonadota | Alphaproteobacteria | NZ_CP022110.1 | 2 |

| | | | | |
|---|---|---|---|---|
| Niveispirillum cyanobacteriorum | Pseudomonadota | Alphaproteobacteria | NZ_CP025611.1 | 2 |
| Novacetimonas hansenii | Pseudomonadota | Alphaproteobacteria | NZ_CP062147.1 | 1 |
| Novosphingobium aromaticivorans DSM 12444 | Pseudomonadota | Alphaproteobacteria | NC_007794.1 | 1 |
| Novosphingobium decolorationis | Pseudomonadota | Alphaproteobacteria | NZ_CP054856.1 | 1 |
| Novosphingobium ginsenosidimutans | Pseudomonadota | Alphaproteobacteria | NZ_CP042345.1 | 1 |
| Novosphingobium humi | Pseudomonadota | Alphaproteobacteria | NZ_CP117417.1 | 1 |
| Novosphingobium kaempferiae | Pseudomonadota | Alphaproteobacteria | NZ_CP089301.1 | 1 |
| Novosphingobium olei | Pseudomonadota | Alphaproteobacteria | NZ_AP029021.1 | 1 |
| Novosphingobium pentaromativorans US6-1 | Pseudomonadota | Alphaproteobacteria | NZ_CP009291.1 | 1 |
| Novosphingobium resinovorum | Pseudomonadota | Alphaproteobacteria | NZ_CP017075.1 | 1 |
| Novosphingopyxis iocasae | Pseudomonadota | Alphaproteobacteria | NZ_CP060495.1 | 1 |
| Oceaniradius stylonematis | Pseudomonadota | Alphaproteobacteria | NZ_OZ244994.1 | 2 |
| Ochrobactrum quorumnocens | Pseudomonadota | Alphaproteobacteria | NZ_CP022604.1 | 2 |
| Octadecabacter antarcticus 307 | Pseudomonadota | Alphaproteobacteria | NC_020911.1 | 1 |
| Octadecabacter arcticus 238 | Pseudomonadota | Alphaproteobacteria | NC_020908.1 | 2 |
| Octadecabacter temperatus | Pseudomonadota | Alphaproteobacteria | NZ_CP012160.1 | 2 |
| Oecophyllibacter saccharovorans | Pseudomonadota | Alphaproteobacteria | NZ_CP038143.1 | 1 |
| Oricola thermophila | Pseudomonadota | Alphaproteobacteria | NZ_CP054836.1 | 2 |
| Orientia tsutsugamushi | Pseudomonadota | Alphaproteobacteria | NZ_LS398552.1 | 2 |
| Pacificitalea manganoxidans | Pseudomonadota | Alphaproteobacteria | NZ_CP021404.1 | 2 |
| Paracoccus actinidiae | Pseudomonadota | Alphaproteobacteria | NZ_CP132956.1 | 1 |
| Paracoccus aerodenitrificans | Pseudomonadota | Alphaproteobacteria | NZ_CP115784.1 | 2 |
| Paracoccus albus | Pseudomonadota | Alphaproteobacteria | NZ_CP115775.1 | 2 |
| Paracoccus alcaliphilus | Pseudomonadota | Alphaproteobacteria | NZ_CP067124.1 | 2 |
| Paracoccus aminophilus JCM 7686 | Pseudomonadota | Alphaproteobacteria | NC_022041.1 | 2 |
| Paracoccus aminovorans | Pseudomonadota | Alphaproteobacteria | NZ_LN832559.1 | 2 |
| Paracoccus contaminans | Pseudomonadota | Alphaproteobacteria | NZ_CP020612.1 | 2 |
| Paracoccus denitrificans | Pseudomonadota | Alphaproteobacteria | NZ_CP035090.1 | 2 |
| Paracoccus everestensis | Pseudomonadota | Alphaproteobacteria | NZ_CP090836.1 | 2 |
| Paracoccus fistulariae | Pseudomonadota | Alphaproteobacteria | NZ_CP067136.1 | 2 |
| Paracoccus jeotgali | Pseudomonadota | Alphaproteobacteria | NZ_CP025583.1 | 2 |
| Paracoccus kondratievae | Pseudomonadota | Alphaproteobacteria | NZ_CP045072.1 | 2 |

| | | | | |
|---|---|---|---|---|
| Paracoccus liaowanqingii | Pseudomonadota | Alphaproteobacteria | NZ_CP038439.1 | 2 |
| Paracoccus marcusii | Pseudomonadota | Alphaproteobacteria | NZ_CP065892.1 | 2 |
| Paracoccus methylovorus | Pseudomonadota | Alphaproteobacteria | NZ_CP070368.1 | 2 |
| Paracoccus pantotrophus | Pseudomonadota | Alphaproteobacteria | NZ_CP044426.1 | 2 |
| Paracoccus saliphilus | Pseudomonadota | Alphaproteobacteria | NZ_CP067140.1 | 2 |
| Paracoccus sanguinis | Pseudomonadota | Alphaproteobacteria | NZ_CP051542.1 | 2 |
| Paracoccus sediminicola | Pseudomonadota | Alphaproteobacteria | NZ_CP115768.1 | 2 |
| Paracoccus seriniphilus | Pseudomonadota | Alphaproteobacteria | NZ_CP067129.1 | 2 |
| Paracoccus stylophorae | Pseudomonadota | Alphaproteobacteria | NZ_CP067134.1 | 2 |
| Paracoccus suum | Pseudomonadota | Alphaproteobacteria | NZ_CP030918.1 | 2 |
| Paracoccus tegillarcae | Pseudomonadota | Alphaproteobacteria | NZ_CP025408.1 | 2 |
| Paracoccus versutus | Pseudomonadota | Alphaproteobacteria | NZ_CP038197.1 | 2 |
| Paracoccus yeei | Pseudomonadota | Alphaproteobacteria | NZ_CP024422.1 | 2 |
| Paracoccus zhejiangensis | Pseudomonadota | Alphaproteobacteria | NZ_CP025430.1 | 2 |
| Paradevosia shaoguanensis | Pseudomonadota | Alphaproteobacteria | NZ_CP068983.1 | 1 |
| Paradevosia tibetensis | Pseudomonadota | Alphaproteobacteria | NZ_CP041690.1 | 1 |
| Paragemmobacter aquarius | Pseudomonadota | Alphaproteobacteria | NZ_CP028918.1 | 2 |
| Paramagnetospirillum magneticum AMB-1 | Pseudomonadota | Alphaproteobacteria | NC_007626.1 | 2 |
| Pararhizobium gei | Pseudomonadota | Alphaproteobacteria | NZ_CP119409.1 | 2 |
| Pararhizobium qamdonense | Pseudomonadota | Alphaproteobacteria | NZ_CP119566.1 | 2 |
| Parasedimentitalea marina | Pseudomonadota | Alphaproteobacteria | NZ_CP033219.1 | 2 |
| Parasedimentitalea psychrophila | Pseudomonadota | Alphaproteobacteria | NZ_CP127247.1 | 2 |
| Parasphingopyxis algicola | Pseudomonadota | Alphaproteobacteria | NZ_CP051131.1 | 1 |
| Parasphingorhabdus cellanae | Pseudomonadota | Alphaproteobacteria | NZ_CP071794.1 | 1 |
| Parasphingorhabdus halotolerans | Pseudomonadota | Alphaproteobacteria | NZ_CP051217.1 | 1 |
| Parasphingorhabdus litoris DSM 22379 | Pseudomonadota | Alphaproteobacteria | NZ_CP086727.1 | 1 |
| Paraurantiacibacter namhicola | Pseudomonadota | Alphaproteobacteria | NZ_CP016545.1 | 1 |
| Paremcibacter congregatus | Pseudomonadota | Alphaproteobacteria | NZ_CP041025.1 | 2 |
| Paroceanicella profunda | Pseudomonadota | Alphaproteobacteria | NZ_CP040818.1 | 2 |
| Parvibaculum lavamentivorans DS-1 | Pseudomonadota | Alphaproteobacteria | NC_009719.1 | 2 |
| Parvularcula bermudensis HTCC2503 | Pseudomonadota | Alphaproteobacteria | NC_014414.1 | 1 |
| Pelagerythrobacter marensis | Pseudomonadota | Alphaproteobacteria | NZ_CP011805.1 | 1 |

| | | | | |
|---|---|---|---|---|
| Pelagerythrobacter marinus | Pseudomonadota | Alphaproteobacteria | NZ_CP139719.1 | 1 |
| Pelagibacterium flavum | Pseudomonadota | Alphaproteobacteria | NZ_CP107716.1 | 2 |
| Pelagibacterium halotolerans | Pseudomonadota | Alphaproteobacteria | NZ_CP053086.1 | 2 |
| Pelagibacterium nitratireducens | Pseudomonadota | Alphaproteobacteria | NZ_CP146275.1 | 2 |
| Pelagovum pacificum | Pseudomonadota | Alphaproteobacteria | NZ_CP065915.1 | 2 |
| Peteryoungia algae | Pseudomonadota | Alphaproteobacteria | NZ_CP128477.1 | 2 |
| Peteryoungia desertarenae | Pseudomonadota | Alphaproteobacteria | NZ_CP058350.1 | 2 |
| Phaeobacter gallaeciensis DSM 26640 | Pseudomonadota | Alphaproteobacteria | NC_023137.1 | 2 |
| Phaeobacter inhibens | Pseudomonadota | Alphaproteobacteria | NZ_CP010650.1 | 2 |
| Phaeobacter piscinae | Pseudomonadota | Alphaproteobacteria | NZ_CP080275.1 | 2 |
| Phaeobacter porticola | Pseudomonadota | Alphaproteobacteria | NZ_CP016364.1 | 2 |
| Phenylobacterium montanum | Pseudomonadota | Alphaproteobacteria | NZ_CP073078.1 | 2 |
| Phenylobacterium parvum | Pseudomonadota | Alphaproteobacteria | NZ_CP029479.1 | 2 |
| Phenylobacterium zucineum HLK1 | Pseudomonadota | Alphaproteobacteria | NC_011144.1 | 2 |
| Phreatobacter aquaticus | Pseudomonadota | Alphaproteobacteria | NZ_CP039865.1 | 2 |
| Phreatobacter cathodiphilus | Pseudomonadota | Alphaproteobacteria | NZ_CP027668.1 | 2 |
| Phreatobacter stygius | Pseudomonadota | Alphaproteobacteria | NZ_CP039690.1 | 2 |
| Phyllobacterium zundukense | Pseudomonadota | Alphaproteobacteria | NZ_CP017940.1 | 2 |
| Pikeienuella piscinae | Pseudomonadota | Alphaproteobacteria | NZ_CP049056.1 | 2 |
| Polymorphobacter megasporae | Pseudomonadota | Alphaproteobacteria | NZ_CP081848.1 | 1 |
| Polymorphum gilvum SL003B-26A1 | Pseudomonadota | Alphaproteobacteria | NC_015259.1 | 2 |
| Ponticoccus alexandrii | Pseudomonadota | Alphaproteobacteria | NZ_CP047166.1 | 2 |
| Pontivivens nitratireducens | Pseudomonadota | Alphaproteobacteria | NZ_CP049811.1 | 2 |
| Pontivivens ytuae | Pseudomonadota | Alphaproteobacteria | NZ_CP064942.1 | 2 |
| Profundibacter amoris | Pseudomonadota | Alphaproteobacteria | NZ_CP032125.1 | 2 |
| Pseudemcibacter aquimaris | Pseudomonadota | Alphaproteobacteria | NZ_CP079800.1 | 2 |
| Pseudochrobactrum algeriensis | Pseudomonadota | Alphaproteobacteria | NZ_CP075361.1 | 2 |
| Pseudogemmobacter blasticus | Pseudomonadota | Alphaproteobacteria | NZ_CP020470.1 | 2 |
| Pseudolabrys taiwanensis | Pseudomonadota | Alphaproteobacteria | NZ_CP031417.1 | 1 |
| Pseudooceanicola algae | Pseudomonadota | Alphaproteobacteria | NZ_CP060436.1 | 2 |
| Pseudooceanicola atlanticus | Pseudomonadota | Alphaproteobacteria | NZ_CP051248.1 | 2 |
| Pseudooceanicola nitratireducens | Pseudomonadota | Alphaproteobacteria | NZ_CP051251.1 | 1 |

| | | | | |
|---|---|---|---|---|
| Pseudooceanicola spongiae | Pseudomonadota | Alphaproteobacteria | NZ_CP045201.1 | 2 |
| Pseudorhizobium flavum | Pseudomonadota | Alphaproteobacteria | NZ_LR723670.1 | 2 |
| Pseudorhodobacter turbinis | Pseudomonadota | Alphaproteobacteria | NZ_CP039965.1 | 1 |
| Pseudorhodobacter turbinis | Pseudomonadota | Alphaproteobacteria | NZ_CP039964.1 | 1 |
| Pseudorhodoplanes sinuspersici | Pseudomonadota | Alphaproteobacteria | NZ_CP021112.1 | 1 |
| Pseudosulfitobacter pseudonitzschiae | Pseudomonadota | Alphaproteobacteria | NZ_CP054599.1 | 2 |
| Pseudovibrio brasiliensis | Pseudomonadota | Alphaproteobacteria | NZ_CP074126.1 | 2 |
| Pukyongiella litopenaei | Pseudomonadota | Alphaproteobacteria | NZ_CP027665.1 | 2 |
| Pyruvatibacter mobilis | Pseudomonadota | Alphaproteobacteria | NZ_CP051630.1 | 2 |
| Qingshengfaniella alkalisoli | Pseudomonadota | Alphaproteobacteria | NZ_CP042261.1 | 2 |
| Qipengyuania aurantiaca | Pseudomonadota | Alphaproteobacteria | NZ_CP081295.1 | 1 |
| Qipengyuania flava | Pseudomonadota | Alphaproteobacteria | NZ_CP032228.1 | 1 |
| Qipengyuania gelatinilytica | Pseudomonadota | Alphaproteobacteria | NZ_CP081294.1 | 1 |
| Qipengyuania profunda | Pseudomonadota | Alphaproteobacteria | NZ_CP138853.1 | 1 |
| Qipengyuania psychrotolerans | Pseudomonadota | Alphaproteobacteria | NZ_CP081297.1 | 1 |
| Qipengyuania sediminis | Pseudomonadota | Alphaproteobacteria | NZ_CP037948.1 | 1 |
| Qipengyuania seohaensis | Pseudomonadota | Alphaproteobacteria | NZ_CP024920.1 | 1 |
| Qipengyuania soli | Pseudomonadota | Alphaproteobacteria | NZ_CP064654.1 | 1 |
| Qipengyuania spongiae | Pseudomonadota | Alphaproteobacteria | NZ_CP092471.1 | 1 |
| Rhizobium acidisoli | Pseudomonadota | Alphaproteobacteria | NZ_CP034998.1 | 2 |
| Rhizobium anhuiense bv. trifolii | Pseudomonadota | Alphaproteobacteria | NZ_CP098803.1 | 2 |
| Rhizobium bangladeshense | Pseudomonadota | Alphaproteobacteria | NZ_CP071612.1 | 2 |
| Rhizobium binae | Pseudomonadota | Alphaproteobacteria | NZ_CP071604.1 | 2 |
| Rhizobium brockwellii | Pseudomonadota | Alphaproteobacteria | NZ_CP053439.1 | 2 |
| Rhizobium cremeum | Pseudomonadota | Alphaproteobacteria | NZ_CP180217.1 | 2 |
| Rhizobium daejeonense | Pseudomonadota | Alphaproteobacteria | NZ_CP048427.1 | 2 |
| Rhizobium esperanzae | Pseudomonadota | Alphaproteobacteria | NZ_CP013500.1 | 2 |
| Rhizobium etli CFN 42 | Pseudomonadota | Alphaproteobacteria | NC_007761.1 | 2 |
| Rhizobium favelukesii | Pseudomonadota | Alphaproteobacteria | NZ_HG916852.1 | 2 |
| Rhizobium gallicum bv. gallicum R602sp | Pseudomonadota | Alphaproteobacteria | NZ_CP006877.1 | 2 |
| Rhizobium hidalgonense | Pseudomonadota | Alphaproteobacteria | NZ_CP054027.1 | 2 |
| Rhizobium indicum | Pseudomonadota | Alphaproteobacteria | NZ_CP054021.1 | 2 |

| | | | | |
|---|---|---|---|---|
| Rhizobium indigoferae | Pseudomonadota | Alphaproteobacteria | NZ_CP140635.1 | 2 |
| Rhizobium jaguaris | Pseudomonadota | Alphaproteobacteria | NZ_CP032694.1 | 2 |
| Rhizobium johnstonii 3841 | Pseudomonadota | Alphaproteobacteria | NC_008380.1 | 2 |
| Rhizobium lentis | Pseudomonadota | Alphaproteobacteria | NZ_CP071454.1 | 2 |
| Rhizobium oryzihabitans | Pseudomonadota | Alphaproteobacteria | NZ_CP048632.1 | 2 |
| Rhizobium phaseoli | Pseudomonadota | Alphaproteobacteria | NZ_CP013532.1 | 2 |
| Rhizobium rhizogenes | Pseudomonadota | Alphaproteobacteria | NZ_CP044122.1 | 2 |
| Rhizobium rhizoryzae | Pseudomonadota | Alphaproteobacteria | NZ_CP049250.1 | 2 |
| Rhizobium rhododendri | Pseudomonadota | Alphaproteobacteria | NZ_CP117267.1 | 2 |
| Rhizobium ruizarguesonis | Pseudomonadota | Alphaproteobacteria | NZ_CP071678.1 | 2 |
| Rhizobium sophoriradicis | Pseudomonadota | Alphaproteobacteria | NZ_CP104152.1 | 2 |
| Rhizobium sullae | Pseudomonadota | Alphaproteobacteria | NZ_CP104143.1 | 2 |
| Rhizobium tropici CIAT 899 | Pseudomonadota | Alphaproteobacteria | NC_020059.1 | 2 |
| Rhizobium tumorigenes | Pseudomonadota | Alphaproteobacteria | NZ_CP117255.1 | 2 |
| Rhizorhabdus dicambivorans | Pseudomonadota | Alphaproteobacteria | NZ_CP023449.1 | 1 |
| Rhizorhabdus phycosphaerae | Pseudomonadota | Alphaproteobacteria | NZ_CP049107.1 | 1 |
| Rhizorhabdus wittichii | Pseudomonadota | Alphaproteobacteria | NZ_CP059319.1 | 1 |
| Rhodobacter capsulatus SB 1003 | Pseudomonadota | Alphaproteobacteria | NC_014034.1 | 2 |
| Rhodobacter xanthinilyticus | Pseudomonadota | Alphaproteobacteria | NZ_CP017781.1 | 2 |
| Rhodomicrobium vannielii ATCC 17100 | Pseudomonadota | Alphaproteobacteria | NC_014664.1 | 1 |
| Rhodopseudomonas boonkerdii | Pseudomonadota | Alphaproteobacteria | NZ_CP036537.1 | 2 |
| Rhodopseudomonas palustris | Pseudomonadota | Alphaproteobacteria | NZ_CP066699.1 | 2 |
| Rhodospirillum centenum SW | Pseudomonadota | Alphaproteobacteria | NC_011420.2 | 2 |
| Rhodospirillum rubrum ATCC 11170 | Pseudomonadota | Alphaproteobacteria | NC_007643.1 | 2 |
| Rhodovastum atsumiense | Pseudomonadota | Alphaproteobacteria | NZ_OW485601.1 | 1 |
| Rhodovulum sulfidophilum DSM 1374 | Pseudomonadota | Alphaproteobacteria | NZ_CP015418.1 | 2 |
| Rickettsia akari str. Hartford | Pseudomonadota | Alphaproteobacteria | NC_009881.1 | 1 |
| Rickettsia asiatica | Pseudomonadota | Alphaproteobacteria | NZ_AP019563.1 | 1 |
| Rickettsia australis str. Cutlack | Pseudomonadota | Alphaproteobacteria | NC_017058.1 | 1 |
| Rickettsia bellii RML369-C | Pseudomonadota | Alphaproteobacteria | NC_007940.1 | 1 |
| Rickettsia canadensis str. CA410 | Pseudomonadota | Alphaproteobacteria | NC_016929.1 | 1 |
| Rickettsia conorii str. Malish 7 | Pseudomonadota | Alphaproteobacteria | NC_003103.1 | 1 |

| | | | | |
|---|---|---|---|---|
| Rickettsia endosymbiont of Aspidapion aeneum | Pseudomonadota | Alphaproteobacteria | NZ_OZ034920.1 | 1 |
| Rickettsia endosymbiont of Cantharis rufa | Pseudomonadota | Alphaproteobacteria | NZ_OZ032150.1 | 1 |
| Rickettsia endosymbiont of Ceutorhynchus obstrictus | Pseudomonadota | Alphaproteobacteria | NZ_OZ032162.1 | 1 |
| Rickettsia endosymbiont of Gonocerus acuteangulatus | Pseudomonadota | Alphaproteobacteria | NZ_OZ032147.1 | 1 |
| Rickettsia endosymbiont of Lasioglossum villosulum | Pseudomonadota | Alphaproteobacteria | NZ_OZ032152.1 | 1 |
| Rickettsia endosymbiont of Nabis limbatus | Pseudomonadota | Alphaproteobacteria | NZ_OZ239771.1 | 1 |
| Rickettsia endosymbiont of Oedothorax gibbosus | Pseudomonadota | Alphaproteobacteria | NZ_OW370493.1 | 1 |
| Rickettsia endosymbiont of Pantilius tunicatus | Pseudomonadota | Alphaproteobacteria | NZ_OZ198175.1 | 1 |
| Rickettsia endosymbiont of Polydrusus tereticollis | Pseudomonadota | Alphaproteobacteria | NZ_OZ032142.1 | 1 |
| Rickettsia endosymbiont of Rhinocyllus conicus | Pseudomonadota | Alphaproteobacteria | NZ_OZ032155.1 | 1 |
| Rickettsia endosymbiont of Seladonia tumulorum | Pseudomonadota | Alphaproteobacteria | NZ_OZ034963.1 | 1 |
| Rickettsia helvetica | Pseudomonadota | Alphaproteobacteria | NZ_OZ018776.1 | 1 |
| Rickettsia japonica YH | Pseudomonadota | Alphaproteobacteria | NC_016050.1 | 1 |
| Rickettsia monacensis | Pseudomonadota | Alphaproteobacteria | NZ_LN794217.1 | 1 |
| Rickettsia prowazekii str. Chernikova | Pseudomonadota | Alphaproteobacteria | NC_017049.1 | 1 |
| Rickettsia rhipicephali str. 3-7-female6-CWPP | Pseudomonadota | Alphaproteobacteria | NC_017042.1 | 1 |
| Rickettsia rickettsii str. 'Sheila Smith' | Pseudomonadota | Alphaproteobacteria | NC_009882.1 | 1 |
| Rickettsia sibirica 246 | Pseudomonadota | Alphaproteobacteria | NZ_AABW0100000 | 1 |
| Rickettsia slovaca 13-B | Pseudomonadota | Alphaproteobacteria | NC_016639.1 | 1 |
| Rickettsia tillamookensis | Pseudomonadota | Alphaproteobacteria | NZ_CP060138.2 | 1 |
| Rickettsia typhi str. Wilmington | Pseudomonadota | Alphaproteobacteria | NC_006142.1 | 1 |
| Rickettsiales endosymbiont of Stachyamoeba lipophora | Pseudomonadota | Alphaproteobacteria | NZ_CP033611.1 | 2 |
| Roseibium alexandrii DFL-11 | Pseudomonadota | Alphaproteobacteria | NZ_CM011002.1 | 2 |
| Roseibium algicola | Pseudomonadota | Alphaproteobacteria | NZ_CP019630.1 | 2 |
| Roseibium porphyridii | Pseudomonadota | Alphaproteobacteria | NZ_CP120863.1 | 2 |
| Roseicyclus elongatus DSM 19469 | Pseudomonadota | Alphaproteobacteria | NZ_CP004372.1 | 2 |
| Roseitalea porphyridii | Pseudomonadota | Alphaproteobacteria | NZ_CP036532.1 | 2 |
| Roseobacter denitrificans | Pseudomonadota | Alphaproteobacteria | NZ_CP027407.1 | 2 |
| Roseobacter fucihabitans | Pseudomonadota | Alphaproteobacteria | NZ_CP143423.1 | 2 |
| Roseobacter litoralis Och 149 | Pseudomonadota | Alphaproteobacteria | NC_015730.1 | 2 |
| Roseobacter ponti | Pseudomonadota | Alphaproteobacteria | NZ_CP048788.1 | 2 |
| Roseococcus microcysteis | Pseudomonadota | Alphaproteobacteria | NZ_CP061718.1 | 2 |

| | | | | |
|---|---|---|---|---|
| Roseomonas cutis | Pseudomonadota | Alphaproteobacteria | NZ_CP087719.1 | 2 |
| Roseomonas fluvialis | Pseudomonadota | Alphaproteobacteria | NZ_AP025637.1 | 2 |
| Roseomonas gilardii | Pseudomonadota | Alphaproteobacteria | NZ_CP015583.1 | 2 |
| Roseomonas haemaphysalidis | Pseudomonadota | Alphaproteobacteria | NZ_CP061177.1 | 2 |
| Roseomonas marmotae | Pseudomonadota | Alphaproteobacteria | NZ_CP061091.1 | 2 |
| Roseomonas mucosa | Pseudomonadota | Alphaproteobacteria | NZ_CP025060.1 | 2 |
| Roseovarius faecimaris | Pseudomonadota | Alphaproteobacteria | NZ_CP034348.1 | 1 |
| Roseovarius indicus | Pseudomonadota | Alphaproteobacteria | NZ_CP031598.1 | 2 |
| Roseovarius pelagicus | Pseudomonadota | Alphaproteobacteria | NZ_CP106738.1 | 2 |
| Roseovarius phycicola | Pseudomonadota | Alphaproteobacteria | NZ_CP146069.1 | 2 |
| Roseovarius rhodophyticola | Pseudomonadota | Alphaproteobacteria | NZ_CP146606.1 | 2 |
| Roseovarius tolerans | Pseudomonadota | Alphaproteobacteria | NZ_CP118494.1 | 2 |
| Ruegeria pomeroyi DSS-3 | Pseudomonadota | Alphaproteobacteria | NC_003911.12 | 2 |
| Sagittula stellata E-37 | Pseudomonadota | Alphaproteobacteria | NZ_CP155729.1 | 2 |
| Salaquimonas pukyongi | Pseudomonadota | Alphaproteobacteria | NZ_CP019044.1 | 2 |
| Salipiger abyssi | Pseudomonadota | Alphaproteobacteria | NZ_CP015093.1 | 2 |
| Salipiger profundus | Pseudomonadota | Alphaproteobacteria | NZ_CP014796.1 | 2 |
| Sandaracinobacteroides saxicola | Pseudomonadota | Alphaproteobacteria | NZ_CP059851.1 | 1 |
| Sediminicoccus rosea | Pseudomonadota | Alphaproteobacteria | NZ_CP137852.1 | 2 |
| Shinella oryzae | Pseudomonadota | Alphaproteobacteria | NZ_CP081987.1 | 2 |
| Shinella sumterensis | Pseudomonadota | Alphaproteobacteria | NZ_CP132302.1 | 2 |
| Shinella zoogloeoides | Pseudomonadota | Alphaproteobacteria | NZ_CP086610.1 | 2 |
| Silicimonas algicola | Pseudomonadota | Alphaproteobacteria | NZ_CP034588.1 | 2 |
| Sinorhizobium alkalisoli | Pseudomonadota | Alphaproteobacteria | NZ_CP034909.1 | 2 |
| Sinorhizobium americanum | Pseudomonadota | Alphaproteobacteria | NZ_CP013107.1 | 2 |
| Sinorhizobium chiapasense | Pseudomonadota | Alphaproteobacteria | NZ_CP133148.1 | 2 |
| Sinorhizobium fredii CCBAU 25509 | Pseudomonadota | Alphaproteobacteria | NZ_CP029451.1 | 2 |
| Sinorhizobium garamanticum | Pseudomonadota | Alphaproteobacteria | NZ_CP120373.1 | 2 |
| Sinorhizobium kummerowiae | Pseudomonadota | Alphaproteobacteria | NZ_CP120365.1 | 2 |
| Sinorhizobium meliloti | Pseudomonadota | Alphaproteobacteria | NZ_CP146208.1 | 2 |
| Sinorhizobium mexicanum | Pseudomonadota | Alphaproteobacteria | NZ_CP041238.1 | 2 |
| Sinorhizobium numidicum | Pseudomonadota | Alphaproteobacteria | NZ_CP120368.1 | 2 |

| | | | | |
|---|---|---|---|---|
| Sinorhizobium sojae CCBAU 05684 | Pseudomonadota | Alphaproteobacteria | NZ_CP023067.1 | 2 |
| Sinorhizobium terangae | Pseudomonadota | Alphaproteobacteria | NZ_CP121659.1 | 2 |
| Skermanella cutis | Pseudomonadota | Alphaproteobacteria | NZ_CP067420.1 | 2 |
| Skermanella mucosa | Pseudomonadota | Alphaproteobacteria | NZ_CP086106.1 | 2 |
| Skermanella pratensis | Pseudomonadota | Alphaproteobacteria | NZ_CP030265.1 | 2 |
| Skermanella rosea | Pseudomonadota | Alphaproteobacteria | NZ_CP086111.1 | 2 |
| Sneathiella aquimaris | Pseudomonadota | Alphaproteobacteria | NZ_CP112881.1 | 2 |
| Sneathiella marina | Pseudomonadota | Alphaproteobacteria | NZ_CP098747.1 | 2 |
| Sphingobium amiense | Pseudomonadota | Alphaproteobacteria | NZ_AP018664.1 | 1 |
| Sphingobium cloacae | Pseudomonadota | Alphaproteobacteria | NZ_AP017655.1 | 1 |
| Sphingobium fuliginis | Pseudomonadota | Alphaproteobacteria | NZ_CP060035.1 | 1 |
| Sphingobium indicum UT26S | Pseudomonadota | Alphaproteobacteria | NC_014006.1 | 1 |
| Sphingobium naphthae | Pseudomonadota | Alphaproteobacteria | NZ_CP171454.1 | 1 |
| Sphingobium phenoxybenzoativorans | Pseudomonadota | Alphaproteobacteria | NZ_CP073910.1 | 1 |
| Sphingobium xenophagum | Pseudomonadota | Alphaproteobacteria | NZ_CP022745.1 | 1 |
| Sphingobium yanoikuyae | Pseudomonadota | Alphaproteobacteria | NZ_CP023741.1 | 1 |
| Sphingomicrobium aestuariivivum | Pseudomonadota | Alphaproteobacteria | NZ_CP102629.1 | 1 |
| Sphingomicrobium flavum | Pseudomonadota | Alphaproteobacteria | NZ_CP102630.1 | 1 |
| Sphingomonas abietis | Pseudomonadota | Alphaproteobacteria | NZ_CP115174.1 | 1 |
| Sphingomonas aerolata | Pseudomonadota | Alphaproteobacteria | NZ_CP098762.1 | 1 |
| Sphingomonas albertensis | Pseudomonadota | Alphaproteobacteria | NZ_CP162536.1 | 1 |
| Sphingomonas aliaeris | Pseudomonadota | Alphaproteobacteria | NZ_CP061035.1 | 1 |
| Sphingomonas bisphenolicum | Pseudomonadota | Alphaproteobacteria | NZ_AP018817.1 | 1 |
| Sphingomonas cannabina | Pseudomonadota | Alphaproteobacteria | NZ_CP090059.1 | 1 |
| Sphingomonas carotinifaciens | Pseudomonadota | Alphaproteobacteria | NZ_CP178397.1 | 1 |
| Sphingomonas changnyeongensis | Pseudomonadota | Alphaproteobacteria | NZ_CP047895.1 | 1 |
| Sphingomonas donggukensis | Pseudomonadota | Alphaproteobacteria | NZ_CP098401.1 | 1 |
| Sphingomonas endolithica | Pseudomonadota | Alphaproteobacteria | NZ_CP103057.1 | 1 |
| Sphingomonas flavescens | Pseudomonadota | Alphaproteobacteria | NZ_CP133016.1 | 1 |
| Sphingomonas ginsengisoli (ex An et al. 2013) | Pseudomonadota | Alphaproteobacteria | NZ_CP045434.1 | 1 |
| Sphingomonas glaciei | Pseudomonadota | Alphaproteobacteria | NZ_CP097253.1 | 1 |
| Sphingomonas hengshuiensis | Pseudomonadota | Alphaproteobacteria | NZ_CP010836.1 | 1 |

| | | | | |
|---|---|---|---|---|
| Sphingomonas insulae | Pseudomonadota | Alphaproteobacteria | NZ_CP048422.1 | 1 |
| Sphingomonas koreensis | Pseudomonadota | Alphaproteobacteria | NZ_CP018820.1 | 1 |
| Sphingomonas lacunae | Pseudomonadota | Alphaproteobacteria | NZ_CP053015.1 | 1 |
| Sphingomonas lutea | Pseudomonadota | Alphaproteobacteria | NZ_CP071573.1 | 1 |
| Sphingomonas morindae | Pseudomonadota | Alphaproteobacteria | NZ_CP084930.1 | 1 |
| Sphingomonas naphthae | Pseudomonadota | Alphaproteobacteria | NZ_CP117411.1 | 1 |
| Sphingomonas nostoxanthinifaciens | Pseudomonadota | Alphaproteobacteria | NZ_CP082839.1 | 1 |
| Sphingomonas paeninsulae | Pseudomonadota | Alphaproteobacteria | NZ_CP032829.1 | 1 |
| Sphingomonas panacis | Pseudomonadota | Alphaproteobacteria | NZ_CP014168.1 | 1 |
| Sphingomonas panacisoli | Pseudomonadota | Alphaproteobacteria | NZ_CP042306.1 | 1 |
| Sphingomonas paucimobilis | Pseudomonadota | Alphaproteobacteria | NZ_CP065670.1 | 1 |
| Sphingomonas piscis | Pseudomonadota | Alphaproteobacteria | NZ_CP049869.1 | 1 |
| Sphingomonas profundi | Pseudomonadota | Alphaproteobacteria | NZ_CP046535.1 | 1 |
| Sphingomonas psychrotolerans | Pseudomonadota | Alphaproteobacteria | NZ_CP024923.1 | 1 |
| Sphingomonas qomolangmaensis | Pseudomonadota | Alphaproteobacteria | NZ_CP101740.1 | 1 |
| Sphingomonas radiodurans | Pseudomonadota | Alphaproteobacteria | NZ_CP086594.1 | 1 |
| Sphingomonas rhizophila | Pseudomonadota | Alphaproteobacteria | NZ_CP060717.1 | 1 |
| Sphingomonas sabuli | Pseudomonadota | Alphaproteobacteria | NZ_CP060697.1 | 1 |
| Sphingomonas sanxanigenens DSM 19645 = NX02 | Pseudomonadota | Alphaproteobacteria | NZ_CP006644.1 | 2 |
| Sphingomonas sediminicola | Pseudomonadota | Alphaproteobacteria | NZ_CP060782.1 | 1 |
| Sphingomonas sinipercae | Pseudomonadota | Alphaproteobacteria | NZ_CP049871.1 | 1 |
| Sphingomonas suaedae | Pseudomonadota | Alphaproteobacteria | NZ_CP042239.1 | 1 |
| Sphingomonas taxi | Pseudomonadota | Alphaproteobacteria | NZ_CP009571.1 | 1 |
| Sphingomonas xanthus | Pseudomonadota | Alphaproteobacteria | NZ_CP041659.1 | 1 |
| Sphingopyxis alaskensis RB2256 | Pseudomonadota | Alphaproteobacteria | NC_008048.1 | 1 |
| Sphingopyxis chilensis | Pseudomonadota | Alphaproteobacteria | NZ_CP142394.1 | 1 |
| Sphingopyxis fribergensis | Pseudomonadota | Alphaproteobacteria | NZ_CP009122.1 | 1 |
| Sphingopyxis granuli | Pseudomonadota | Alphaproteobacteria | NZ_CP012199.1 | 1 |
| Sphingopyxis indica | Pseudomonadota | Alphaproteobacteria | NZ_CP076395.1 | 1 |
| Sphingopyxis kveilinensis | Pseudomonadota | Alphaproteobacteria | NZ_CP144683.1 | 1 |
| Sphingopyxis lindanitolerans | Pseudomonadota | Alphaproteobacteria | NZ_CM009578.1 | 1 |
| Sphingopyxis macrogoltabida | Pseudomonadota | Alphaproteobacteria | NZ_CP009429.1 | 1 |

| | | | | |
|---|---|---|---|---|
| Sphingorhabdus lacus | Pseudomonadota | Alphaproteobacteria | NZ_CP035733.1 | 1 |
| Sphingorhabdus lutea | Pseudomonadota | Alphaproteobacteria | NZ_CP018154.1 | 1 |
| Sphingosinicella flava | Pseudomonadota | Alphaproteobacteria | NZ_CP065592.1 | 1 |
| Sphingosinithalassobacter tenebrarum | Pseudomonadota | Alphaproteobacteria | NZ_CP049109.1 | 1 |
| Stakelama saccharophila | Pseudomonadota | Alphaproteobacteria | NZ_CP135076.1 | 1 |
| Stappia indica | Pseudomonadota | Alphaproteobacteria | NZ_CP046908.1 | 2 |
| Stella humosa | Pseudomonadota | Alphaproteobacteria | NZ_AP019700.1 | 2 |
| Sulfitobacter albidus | Pseudomonadota | Alphaproteobacteria | NZ_CP073581.1 | 2 |
| Sulfitobacter alexandrii | Pseudomonadota | Alphaproteobacteria | NZ_CP018076.1 | 2 |
| Sulfitobacter faviae | Pseudomonadota | Alphaproteobacteria | NZ_CP139725.1 | 2 |
| Sulfitobacter indolifex | Pseudomonadota | Alphaproteobacteria | NZ_CP084951.1 | 2 |
| Sulfitobacter mediterraneus | Pseudomonadota | Alphaproteobacteria | NZ_CP068998.1 | 2 |
| Sulfitobacter pontiacus | Pseudomonadota | Alphaproteobacteria | NZ_CP160849.1 | 2 |
| Swingsia samuiensis | Pseudomonadota | Alphaproteobacteria | NZ_CP038141.1 | 1 |
| Tabrizicola piscis | Pseudomonadota | Alphaproteobacteria | NZ_CP034328.1 | 2 |
| Tardibacter chloracetimidivorans | Pseudomonadota | Alphaproteobacteria | NZ_CP018221.1 | 1 |
| Tardiphaga alba | Pseudomonadota | Alphaproteobacteria | NZ_CP036498.1 | 2 |
| Temperatibacter marinus | Pseudomonadota | Alphaproteobacteria | NZ_CP123872.1 | 2 |
| Terribium terrae | Pseudomonadota | Alphaproteobacteria | NZ_CP044218.1 | 2 |
| Terricaulis silvestris | Pseudomonadota | Alphaproteobacteria | NZ_CP047045.1 | 1 |
| Terrihabitans soli | Pseudomonadota | Alphaproteobacteria | NZ_AP023361.1 | 2 |
| Terrirubrum flagellatum | Pseudomonadota | Alphaproteobacteria | NZ_CP091851.1 | 1 |
| Thalassospira indica | Pseudomonadota | Alphaproteobacteria | NZ_CP031555.1 | 2 |
| Thalassospira marina | Pseudomonadota | Alphaproteobacteria | NZ_CP024199.1 | 2 |
| Thalassospira xiamenensis M-5 = DSM 17429 | Pseudomonadota | Alphaproteobacteria | NZ_CP004388.1 | 2 |
| Thalassovita gelatinovora | Pseudomonadota | Alphaproteobacteria | NZ_CP051181.1 | 2 |
| Thioclava electrotropha | Pseudomonadota | Alphaproteobacteria | NZ_CP053562.1 | 2 |
| Thioclava litoralis | Pseudomonadota | Alphaproteobacteria | NZ_CP135443.1 | 2 |
| Thioclava nitratireducens | Pseudomonadota | Alphaproteobacteria | NZ_CP019437.1 | 2 |
| Tistrella bauzanensis | Pseudomonadota | Alphaproteobacteria | NZ_CP121009.1 | 2 |
| Tistrella mobilis | Pseudomonadota | Alphaproteobacteria | NZ_CP121045.1 | 2 |
| Tritonibacter mobilis F1926 | Pseudomonadota | Alphaproteobacteria | NZ_CP015230.1 | 2 |

| | | | | |
|---|---|---|---|---|
| Tritonibacter scottomollicae | Pseudomonadota | Alphaproteobacteria | NZ_CP136704.1 | 2 |
| Tropicibacter oceani | Pseudomonadota | Alphaproteobacteria | NZ_CP124616.1 | 2 |
| Tsuneonella amylolytica | Pseudomonadota | Alphaproteobacteria | NZ_CP032570.1 | 1 |
| Tsuneonella dongtanensis | Pseudomonadota | Alphaproteobacteria | NZ_CP016591.1 | 1 |
| Tsuneonella mangrovi | Pseudomonadota | Alphaproteobacteria | NZ_CP022889.1 | 1 |
| Variibacter gotjawalensis | Pseudomonadota | Alphaproteobacteria | NZ_AP014946.1 | 2 |
| Varunaivibrio sulfuroxidans | Pseudomonadota | Alphaproteobacteria | NZ_CP119676.1 | 2 |
| Wolbachia endosymbiont (group A) of Acrocera orbiculus | Pseudomonadota | Alphaproteobacteria | NZ_OX366272.1 | 2 |
| Wolbachia endosymbiont (group A) of Agelastica alni | Pseudomonadota | Alphaproteobacteria | NZ_OZ034699.1 | 2 |
| Wolbachia endosymbiont (group A) of Aleiodes leptofemur | Pseudomonadota | Alphaproteobacteria | NZ_OZ034782.1 | 2 |
| Wolbachia endosymbiont (group A) of Alloplasta piceator | Pseudomonadota | Alphaproteobacteria | NZ_OZ034793.1 | 2 |
| Wolbachia endosymbiont (group A) of Ancistrocerus nigrico | Pseudomonadota | Alphaproteobacteria | NZ_OX366348.1 | 2 |
| Wolbachia endosymbiont (group A) of Andrena bucephala | Pseudomonadota | Alphaproteobacteria | NZ_OZ034690.1 | 2 |
| Wolbachia endosymbiont (group A) of Andrena carantonica | Pseudomonadota | Alphaproteobacteria | NZ_OZ034726.1 | 2 |
| Wolbachia endosymbiont (group A) of Andrena dorsata | Pseudomonadota | Alphaproteobacteria | NZ_OX366389.1 | 2 |
| Wolbachia endosymbiont (group A) of Andrena fulva | Pseudomonadota | Alphaproteobacteria | NZ_OZ034701.1 | 2 |
| Wolbachia endosymbiont (group A) of Andrena haemorrho | Pseudomonadota | Alphaproteobacteria | NZ_OX366326.1 | 2 |
| Wolbachia endosymbiont (group A) of Andrena helvola | Pseudomonadota | Alphaproteobacteria | NZ_OZ034779.1 | 2 |
| Wolbachia endosymbiont (group A) of Andrena trimmerana | Pseudomonadota | Alphaproteobacteria | NZ_OZ034776.1 | 2 |
| Wolbachia endosymbiont (group A) of Anomoia purmunda | Pseudomonadota | Alphaproteobacteria | NZ_OX366362.1 | 2 |
| Wolbachia endosymbiont (group A) of Anoplius nigerrimus | Pseudomonadota | Alphaproteobacteria | NZ_OX366385.1 | 2 |
| Wolbachia endosymbiont (group A) of Anthophora plumipe | Pseudomonadota | Alphaproteobacteria | NZ_OZ034756.1 | 2 |
| Wolbachia endosymbiont (group A) of Apoderus coryli | Pseudomonadota | Alphaproteobacteria | NZ_OX366367.1 | 2 |
| Wolbachia endosymbiont (group A) of Aporus unicolor | Pseudomonadota | Alphaproteobacteria | NZ_OZ195518.1 | 2 |
| Wolbachia endosymbiont (group A) of Apotomis capreana | Pseudomonadota | Alphaproteobacteria | NZ_OZ034721.1 | 2 |
| Wolbachia endosymbiont (group A) of Barypeithes pellucid | Pseudomonadota | Alphaproteobacteria | NZ_OZ034985.1 | 2 |
| Wolbachia endosymbiont (group A) of Beris morrisii | Pseudomonadota | Alphaproteobacteria | NZ_OZ034760.1 | 2 |
| Wolbachia endosymbiont (group A) of Bibio marci | Pseudomonadota | Alphaproteobacteria | NZ_OX366364.1 | 2 |
| Wolbachia endosymbiont (group A) of Bombylius major | Pseudomonadota | Alphaproteobacteria | NZ_OX366395.1 | 2 |
| Wolbachia endosymbiont (group A) of Brachyopa scutellari | Pseudomonadota | Alphaproteobacteria | NZ_OZ034720.1 | 2 |
| Wolbachia endosymbiont (group A) of Calamotropha paluc | Pseudomonadota | Alphaproteobacteria | NZ_OX366397.1 | 2 |
| Wolbachia endosymbiont (group A) of Campoletis raptor | Pseudomonadota | Alphaproteobacteria | NZ_OZ034695.1 | 2 |

| | | | | |
|---|---|---|---|---|
| Wolbachia endosymbiont (group A) of Cephus spinipes | Pseudomonadota | Alphaproteobacteria | NZ_OZ034713.1 | 2 |
| Wolbachia endosymbiont (group A) of Cerceris ruficornis | Pseudomonadota | Alphaproteobacteria | NZ_OZ035014.1 | 2 |
| Wolbachia endosymbiont (group A) of Chalcis sispes | Pseudomonadota | Alphaproteobacteria | NZ_OZ034697.1 | 2 |
| Wolbachia endosymbiont (group A) of Cheilosia soror | Pseudomonadota | Alphaproteobacteria | NZ_OX366332.1 | 2 |
| Wolbachia endosymbiont (group A) of Clivina fossor | Pseudomonadota | Alphaproteobacteria | NZ_OZ034707.1 | 2 |
| Wolbachia endosymbiont (group A) of Colletes cunicularius | Pseudomonadota | Alphaproteobacteria | NZ_OZ183471.1 | 2 |
| Wolbachia endosymbiont (group A) of Conops quadrifascia | Pseudomonadota | Alphaproteobacteria | NZ_OZ034722.1 | 2 |
| Wolbachia endosymbiont (group A) of Coremacera margin | Pseudomonadota | Alphaproteobacteria | NZ_OX366399.1 | 2 |
| Wolbachia endosymbiont (group A) of Crataerina pallida | Pseudomonadota | Alphaproteobacteria | NZ_OZ034785.1 | 2 |
| Wolbachia endosymbiont (group A) of Cydia amplana | Pseudomonadota | Alphaproteobacteria | NZ_OZ034792.1 | 2 |
| Wolbachia endosymbiont (group A) of Cydia strobilella | Pseudomonadota | Alphaproteobacteria | NZ_OZ034694.1 | 2 |
| Wolbachia endosymbiont (group A) of Dendrolimus pini | Pseudomonadota | Alphaproteobacteria | NZ_OZ034702.1 | 2 |
| Wolbachia endosymbiont (group A) of Ectemnius continuus | Pseudomonadota | Alphaproteobacteria | NZ_OX366378.1 | 2 |
| Wolbachia endosymbiont (group A) of Endotricha flammea | Pseudomonadota | Alphaproteobacteria | NZ_OX366352.1 | 2 |
| Wolbachia endosymbiont (group A) of Ennomos erosarius | Pseudomonadota | Alphaproteobacteria | NZ_OZ034790.1 | 2 |
| Wolbachia endosymbiont (group A) of Epagoge grotiana | Pseudomonadota | Alphaproteobacteria | NZ_OX366387.1 | 2 |
| Wolbachia endosymbiont (group A) of Epirrhoe alternata | Pseudomonadota | Alphaproteobacteria | NZ_OX366359.1 | 2 |
| Wolbachia endosymbiont (group A) of Epistrophe grossula | Pseudomonadota | Alphaproteobacteria | NZ_OX366363.1 | 2 |
| Wolbachia endosymbiont (group A) of Eristalinus aeneus | Pseudomonadota | Alphaproteobacteria | NZ_OZ034712.1 | 2 |
| Wolbachia endosymbiont (group A) of Eupithecia tripuncta | Pseudomonadota | Alphaproteobacteria | NZ_OX366369.1 | 2 |
| Wolbachia endosymbiont (group A) of Gastracanthus pulcl | Pseudomonadota | Alphaproteobacteria | NZ_OZ034763.1 | 2 |
| Wolbachia endosymbiont (group A) of Gymnosoma rotund | Pseudomonadota | Alphaproteobacteria | NZ_OX366347.1 | 2 |
| Wolbachia endosymbiont (group A) of Hedychridium roseu | Pseudomonadota | Alphaproteobacteria | NZ_OZ034755.1 | 2 |
| Wolbachia endosymbiont (group A) of Hylaeus communis | Pseudomonadota | Alphaproteobacteria | NZ_OX366396.1 | 2 |
| Wolbachia endosymbiont (group A) of Icerya purchasi | Pseudomonadota | Alphaproteobacteria | NZ_OX366357.1 | 2 |
| Wolbachia endosymbiont (group A) of Ischnus inquisitorius | Pseudomonadota | Alphaproteobacteria | NZ_OZ034715.1 | 2 |
| Wolbachia endosymbiont (group A) of Lasioglossum fulvic | Pseudomonadota | Alphaproteobacteria | NZ_OZ034798.1 | 2 |
| Wolbachia endosymbiont (group A) of Lasioglossum lative | Pseudomonadota | Alphaproteobacteria | NZ_OX366398.1 | 2 |
| Wolbachia endosymbiont (group A) of Lasioglossum morio | Pseudomonadota | Alphaproteobacteria | NZ_OX366327.1 | 2 |
| Wolbachia endosymbiont (group A) of Lasioglossum villosu | Pseudomonadota | Alphaproteobacteria | NZ_OZ034716.1 | 2 |
| Wolbachia endosymbiont (group A) of Limnophora tigrina | Pseudomonadota | Alphaproteobacteria | NZ_OZ181654.1 | 2 |
| Wolbachia endosymbiont (group A) of Longitarsus flavicorn | Pseudomonadota | Alphaproteobacteria | NZ_OZ008337.1 | 2 |

| | | | | |
|---|---|---|---|---|
| Wolbachia endosymbiont (group A) of Lypha dubia | Pseudomonadota | Alphaproteobacteria | NZ_OZ034796.1 | 2 |
| Wolbachia endosymbiont (group A) of Machimus atricapillu | Pseudomonadota | Alphaproteobacteria | NZ_OZ034727.1 | 2 |
| Wolbachia endosymbiont (group A) of Macropis europaea | Pseudomonadota | Alphaproteobacteria | NZ_OX366372.1 | 2 |
| Wolbachia endosymbiont (group A) of Melieria omissa | Pseudomonadota | Alphaproteobacteria | NZ_OZ035007.1 | 2 |
| Wolbachia endosymbiont (group A) of Merzomyia westerm | Pseudomonadota | Alphaproteobacteria | NZ_OX366375.1 | 2 |
| Wolbachia endosymbiont (group A) of Microdon myrmicae | Pseudomonadota | Alphaproteobacteria | NZ_OZ034998.1 | 2 |
| Wolbachia endosymbiont (group A) of Myopa testacea | Pseudomonadota | Alphaproteobacteria | NZ_OZ034729.1 | 2 |
| Wolbachia endosymbiont (group A) of Nomada fabriciana | Pseudomonadota | Alphaproteobacteria | NZ_OX366351.1 | 2 |
| Wolbachia endosymbiont (group A) of Nomada ferruginata | Pseudomonadota | Alphaproteobacteria | NZ_OZ034728.1 | 2 |
| Wolbachia endosymbiont (group A) of Nomada flava | Pseudomonadota | Alphaproteobacteria | NZ_OZ034725.1 | 2 |
| Wolbachia endosymbiont (group A) of Nomada fucata | Pseudomonadota | Alphaproteobacteria | NZ_OZ034762.1 | 2 |
| Wolbachia endosymbiont (group A) of Nomada goodeniana | Pseudomonadota | Alphaproteobacteria | NZ_OZ034693.1 | 2 |
| Wolbachia endosymbiont (group A) of Nomada hirtipes | Pseudomonadota | Alphaproteobacteria | NZ_OZ014602.1 | 2 |
| Wolbachia endosymbiont (group A) of Nomada rufipes | Pseudomonadota | Alphaproteobacteria | NZ_OZ195525.1 | 2 |
| Wolbachia endosymbiont (group A) of Norellia spinipes | Pseudomonadota | Alphaproteobacteria | NZ_OZ187006.1 | 2 |
| Wolbachia endosymbiont (group A) of Ophion costatus | Pseudomonadota | Alphaproteobacteria | NZ_OZ017714.1 | 2 |
| Wolbachia endosymbiont (group A) of Ophion ellenae | Pseudomonadota | Alphaproteobacteria | NZ_OZ025779.1 | 2 |
| Wolbachia endosymbiont (group A) of Opomyza germinatio | Pseudomonadota | Alphaproteobacteria | NZ_OZ183482.1 | 2 |
| Wolbachia endosymbiont (group A) of Oxytorus armatus | Pseudomonadota | Alphaproteobacteria | NZ_OZ034719.1 | 2 |
| Wolbachia endosymbiont (group A) of Paraperithous gnath | Pseudomonadota | Alphaproteobacteria | NZ_OZ034784.1 | 2 |
| Wolbachia endosymbiont (group A) of Phalera bucephala | Pseudomonadota | Alphaproteobacteria | NZ_OX366319.1 | 2 |
| Wolbachia endosymbiont (group A) of Phasia obesa | Pseudomonadota | Alphaproteobacteria | NZ_OZ034764.1 | 2 |
| Wolbachia endosymbiont (group A) of Pheosia gnoma | Pseudomonadota | Alphaproteobacteria | NZ_OX366379.1 | 2 |
| Wolbachia endosymbiont (group A) of Pherbina coryleti | Pseudomonadota | Alphaproteobacteria | NZ_OZ034706.1 | 2 |
| Wolbachia endosymbiont (group A) of Philanthus triangulu | Pseudomonadota | Alphaproteobacteria | NZ_OZ034757.1 | 2 |
| Wolbachia endosymbiont (group A) of Philonthus cognatus | Pseudomonadota | Alphaproteobacteria | NZ_OX366388.1 | 2 |
| Wolbachia endosymbiont (group A) of Pipizella viduata | Pseudomonadota | Alphaproteobacteria | NZ_OZ034787.1 | 2 |
| Wolbachia endosymbiont (group A) of Pogonocherus hispi | Pseudomonadota | Alphaproteobacteria | NZ_OZ034770.1 | 2 |
| Wolbachia endosymbiont (group A) of Portevinia maculata | Pseudomonadota | Alphaproteobacteria | NZ_OZ034786.1 | 2 |
| Wolbachia endosymbiont (group A) of Protocalliphora azur | Pseudomonadota | Alphaproteobacteria | NZ_OX366368.1 | 2 |
| Wolbachia endosymbiont (group A) of Rhinocyllus conicus | Pseudomonadota | Alphaproteobacteria | NZ_OX366349.1 | 2 |
| Wolbachia endosymbiont (group A) of Rhorus exstirpatoriu | Pseudomonadota | Alphaproteobacteria | NZ_OZ034769.1 | 2 |

| | | | | |
|---|---|---|---|---|
| Wolbachia endosymbiont (group A) of Scambus nigricans | Pseudomonadota | Alphaproteobacteria | NZ_OX366356.1 | 2 |
| Wolbachia endosymbiont (group A) of Schoenobius gigant | Pseudomonadota | Alphaproteobacteria | NZ_OZ034991.1 | 2 |
| Wolbachia endosymbiont (group A) of Scoparia ambigualis | Pseudomonadota | Alphaproteobacteria | NZ_OZ034913.1 | 2 |
| Wolbachia endosymbiont (group A) of Sicus ferrugineus | Pseudomonadota | Alphaproteobacteria | NZ_OX366370.1 | 2 |
| Wolbachia endosymbiont (group A) of Sphaerophoria taen | Pseudomonadota | Alphaproteobacteria | NZ_OX366344.1 | 2 |
| Wolbachia endosymbiont (group A) of Sphecodes monilico | Pseudomonadota | Alphaproteobacteria | NZ_OX366400.1 | 2 |
| Wolbachia endosymbiont (group A) of Sturmia bella | Pseudomonadota | Alphaproteobacteria | NZ_OZ034988.1 | 2 |
| Wolbachia endosymbiont (group A) of Sympetrum striolatu | Pseudomonadota | Alphaproteobacteria | NZ_OX366342.1 | 2 |
| Wolbachia endosymbiont (group A) of Therophilus tumidul | Pseudomonadota | Alphaproteobacteria | NZ_OZ195515.1 | 2 |
| Wolbachia endosymbiont (group A) of Tiphia femorata | Pseudomonadota | Alphaproteobacteria | NZ_OX366380.1 | 2 |
| Wolbachia endosymbiont (group A) of Tromatobia lineatori | Pseudomonadota | Alphaproteobacteria | NZ_OZ034791.1 | 2 |
| Wolbachia endosymbiont (group A) of Trypoxylon claviceru | Pseudomonadota | Alphaproteobacteria | NZ_OX366374.1 | 2 |
| Wolbachia endosymbiont (group A) of Udea olivalis | Pseudomonadota | Alphaproteobacteria | NZ_OZ034773.1 | 2 |
| Wolbachia endosymbiont (group A) of Urophora cardui | Pseudomonadota | Alphaproteobacteria | NZ_OZ034774.1 | 2 |
| Wolbachia endosymbiont (group A) of Volucella bombylans | Pseudomonadota | Alphaproteobacteria | NZ_OZ034717.1 | 2 |
| Wolbachia endosymbiont (group A) of Yponomeuta plumbe | Pseudomonadota | Alphaproteobacteria | NZ_OX366353.1 | 2 |
| Wolbachia endosymbiont (group B) of Ablattaria laevigata | Pseudomonadota | Alphaproteobacteria | NZ_OZ195519.1 | 2 |
| Wolbachia endosymbiont (group B) of Agriphila inquinatella | Pseudomonadota | Alphaproteobacteria | NZ_OZ034789.1 | 2 |
| Wolbachia endosymbiont (group B) of Agriphila straminella | Pseudomonadota | Alphaproteobacteria | NZ_OX366345.1 | 2 |
| Wolbachia endosymbiont (group B) of Agrotis exclamationi | Pseudomonadota | Alphaproteobacteria | NZ_OZ034705.1 | 2 |
| Wolbachia endosymbiont (group B) of Agrotis puta | Pseudomonadota | Alphaproteobacteria | NZ_OZ034765.1 | 2 |
| Wolbachia endosymbiont (group B) of Alcis repandata | Pseudomonadota | Alphaproteobacteria | NZ_OZ034751.1 | 2 |
| Wolbachia endosymbiont (group B) of Anania hortulata | Pseudomonadota | Alphaproteobacteria | NZ_OZ034914.1 | 2 |
| Wolbachia endosymbiont (group B) of Apomyelois bistriate | Pseudomonadota | Alphaproteobacteria | NZ_OZ034783.1 | 2 |
| Wolbachia endosymbiont (group B) of Aporia crataegi | Pseudomonadota | Alphaproteobacteria | NZ_OX366322.1 | 2 |
| Wolbachia endosymbiont (group B) of Apotomis betuletana | Pseudomonadota | Alphaproteobacteria | NZ_OX366320.1 | 2 |
| Wolbachia endosymbiont (group B) of Apotomis turbidana | Pseudomonadota | Alphaproteobacteria | NZ_OX366384.1 | 2 |
| Wolbachia endosymbiont (group B) of Archips podanus | Pseudomonadota | Alphaproteobacteria | NZ_OX366341.1 | 2 |
| Wolbachia endosymbiont (group B) of Aricia artaxerxes | Pseudomonadota | Alphaproteobacteria | NZ_OZ034718.1 | 2 |
| Wolbachia endosymbiont (group B) of Athalia cordata | Pseudomonadota | Alphaproteobacteria | NZ_OX366383.1 | 2 |
| Wolbachia endosymbiont (group B) of Campaea margarita | Pseudomonadota | Alphaproteobacteria | NZ_OX366330.1 | 2 |
| Wolbachia endosymbiont (group B) of Camptogramma bilii | Pseudomonadota | Alphaproteobacteria | NZ_OX366391.1 | 2 |

| | | | | |
|---|---|---|---|---|
| Wolbachia endosymbiont (group B) of Catoptria pinella | Pseudomonadota | Alphaproteobacteria | NZ_OX366321.1 | 2 |
| Wolbachia endosymbiont (group B) of Chesias legatella | Pseudomonadota | Alphaproteobacteria | NZ_OZ034711.1 | 2 |
| Wolbachia endosymbiont (group B) of Chorthippus parallel | Pseudomonadota | Alphaproteobacteria | NZ_OX366390.1 | 2 |
| Wolbachia endosymbiont (group B) of Colias croceus | Pseudomonadota | Alphaproteobacteria | NZ_OX366323.1 | 2 |
| Wolbachia endosymbiont (group B) of Cyclophora punctari | Pseudomonadota | Alphaproteobacteria | NZ_OZ034749.1 | 2 |
| Wolbachia endosymbiont (group B) of Dendrolimus pini | Pseudomonadota | Alphaproteobacteria | NZ_OZ034689.1 | 2 |
| Wolbachia endosymbiont (group B) of Elophila nymphaeat | Pseudomonadota | Alphaproteobacteria | NZ_OZ034703.1 | 2 |
| Wolbachia endosymbiont (group B) of Emmelina monodac | Pseudomonadota | Alphaproteobacteria | NZ_OX366361.1 | 2 |
| Wolbachia endosymbiont (group B) of Endotricha flammea | Pseudomonadota | Alphaproteobacteria | NZ_OX366382.1 | 2 |
| Wolbachia endosymbiont (group B) of Ennomos erosarius | Pseudomonadota | Alphaproteobacteria | NZ_OZ034708.1 | 2 |
| Wolbachia endosymbiont (group B) of Episyrphus balteatu | Pseudomonadota | Alphaproteobacteria | NZ_OX366325.1 | 2 |
| Wolbachia endosymbiont (group B) of Erynnis tages | Pseudomonadota | Alphaproteobacteria | NZ_OX366331.1 | 2 |
| Wolbachia endosymbiont (group B) of Eucosma cana | Pseudomonadota | Alphaproteobacteria | NZ_OX366334.1 | 2 |
| Wolbachia endosymbiont (group B) of Eupeodes latifasciat | Pseudomonadota | Alphaproteobacteria | NZ_OX366337.1 | 2 |
| Wolbachia endosymbiont (group B) of Euphydryas aurinia | Pseudomonadota | Alphaproteobacteria | NZ_OX366324.1 | 2 |
| Wolbachia endosymbiont (group B) of Eupithecia inturbata | Pseudomonadota | Alphaproteobacteria | NZ_OZ034987.1 | 2 |
| Wolbachia endosymbiont (group B) of Falcaria lacertinaria | Pseudomonadota | Alphaproteobacteria | NZ_OZ034799.1 | 2 |
| Wolbachia endosymbiont (group B) of Germaria angustata | Pseudomonadota | Alphaproteobacteria | NZ_OZ034691.1 | 2 |
| Wolbachia endosymbiont (group B) of Gerris lacustris | Pseudomonadota | Alphaproteobacteria | NZ_OZ034761.1 | 2 |
| Wolbachia endosymbiont (group B) of Hofmannophila pseu | Pseudomonadota | Alphaproteobacteria | NZ_OZ034788.1 | 2 |
| Wolbachia endosymbiont (group B) of Horisme vitalbata | Pseudomonadota | Alphaproteobacteria | NZ_OZ034797.1 | 2 |
| Wolbachia endosymbiont (group B) of Hylaea fasciaria | Pseudomonadota | Alphaproteobacteria | NZ_OX366401.1 | 2 |
| Wolbachia endosymbiont (group B) of Idaea biselata | Pseudomonadota | Alphaproteobacteria | NZ_OZ034700.1 | 2 |
| Wolbachia endosymbiont (group B) of Ischnura elegans | Pseudomonadota | Alphaproteobacteria | NZ_OX366371.1 | 2 |
| Wolbachia endosymbiont (group B) of Limnophora tigrina | Pseudomonadota | Alphaproteobacteria | NZ_OZ181655.1 | 2 |
| Wolbachia endosymbiont (group B) of Longitarsus flavicorn | Pseudomonadota | Alphaproteobacteria | NZ_OZ034758.1 | 2 |
| Wolbachia endosymbiont (group B) of Lycaena phlaeas | Pseudomonadota | Alphaproteobacteria | NZ_OX366392.1 | 2 |
| Wolbachia endosymbiont (group B) of Melanargia galathea | Pseudomonadota | Alphaproteobacteria | NZ_OZ034910.1 | 2 |
| Wolbachia endosymbiont (group B) of Melanostoma mellin | Pseudomonadota | Alphaproteobacteria | NZ_OX366366.1 | 2 |
| Wolbachia endosymbiont (group B) of Myelois circumvoluta | Pseudomonadota | Alphaproteobacteria | NZ_OZ195524.1 | 2 |
| Wolbachia endosymbiont (group B) of Nymphalis c-album | Pseudomonadota | Alphaproteobacteria | NZ_OX366317.1 | 2 |
| Wolbachia endosymbiont (group B) of Orgyia antiqua | Pseudomonadota | Alphaproteobacteria | NZ_OZ238265.1 | 2 |

| | | | | |
|---|---|---|---|---|
| Wolbachia endosymbiont (group B) of Pammene fasciana | Pseudomonadota | Alphaproteobacteria | NZ_OX366360.1 | 2 |
| Wolbachia endosymbiont (group B) of Pandemis corylana | Pseudomonadota | Alphaproteobacteria | NZ_OX366328.1 | 2 |
| Wolbachia endosymbiont (group B) of Parapoynx stratiotat | Pseudomonadota | Alphaproteobacteria | NZ_OX366335.1 | 2 |
| Wolbachia endosymbiont (group B) of Pararge aegeria | Pseudomonadota | Alphaproteobacteria | NZ_OX366343.1 | 2 |
| Wolbachia endosymbiont (group B) of Phalera bucephala | Pseudomonadota | Alphaproteobacteria | NZ_OX366358.1 | 2 |
| Wolbachia endosymbiont (group B) of Phasia obesa | Pseudomonadota | Alphaproteobacteria | NZ_OZ034723.1 | 2 |
| Wolbachia endosymbiont (group B) of Pheosia tremula | Pseudomonadota | Alphaproteobacteria | NZ_OX366373.1 | 2 |
| Wolbachia endosymbiont (group B) of Pilophorus perplexu | Pseudomonadota | Alphaproteobacteria | NZ_OZ034754.1 | 2 |
| Wolbachia endosymbiont (group B) of Polyommatus icarus | Pseudomonadota | Alphaproteobacteria | NZ_OX366346.1 | 2 |
| Wolbachia endosymbiont (group B) of Protocalliphora azur | Pseudomonadota | Alphaproteobacteria | NZ_OX366394.1 | 2 |
| Wolbachia endosymbiont (group B) of Pyrgus malvae | Pseudomonadota | Alphaproteobacteria | NZ_OX366329.1 | 2 |
| Wolbachia endosymbiont (group B) of Rhopobota naevana | Pseudomonadota | Alphaproteobacteria | NZ_OX366333.1 | 2 |
| Wolbachia endosymbiont (group B) of Scrobipalpa ocellate | Pseudomonadota | Alphaproteobacteria | NZ_OZ077190.1 | 2 |
| Wolbachia endosymbiont (group B) of Silvanus unidentatus | Pseudomonadota | Alphaproteobacteria | NZ_OZ034795.1 | 2 |
| Wolbachia endosymbiont (group B) of Tholera decimalis | Pseudomonadota | Alphaproteobacteria | NZ_OZ034777.1 | 2 |
| Wolbachia endosymbiont (group B) of Thymelicus sylvestri | Pseudomonadota | Alphaproteobacteria | NZ_OX366365.1 | 2 |
| Wolbachia endosymbiont (group B) of Udea ferrugalis | Pseudomonadota | Alphaproteobacteria | NZ_OZ034752.1 | 2 |
| Wolbachia endosymbiont (group B) of Villa cingulata | Pseudomonadota | Alphaproteobacteria | NZ_OZ034766.1 | 2 |
| Wolbachia endosymbiont (group B) of Watsonalla binaria | Pseudomonadota | Alphaproteobacteria | NZ_OX366336.1 | 2 |
| Wolbachia endosymbiont (group B) of Xanthorhoe designa | Pseudomonadota | Alphaproteobacteria | NZ_OZ034794.1 | 2 |
| Wolbachia endosymbiont (group B) of Xestia c-nigrum | Pseudomonadota | Alphaproteobacteria | NZ_OX366340.1 | 2 |
| Wolbachia endosymbiont (group B) of Xylocampa areola | Pseudomonadota | Alphaproteobacteria | NZ_OZ034771.1 | 2 |
| Wolbachia endosymbiont (group B) of Yponomeuta maline | Pseudomonadota | Alphaproteobacteria | NZ_OZ034698.1 | 2 |
| Wolbachia endosymbiont (group E) of Neria commutata | Pseudomonadota | Alphaproteobacteria | NZ_OZ034692.1 | 2 |
| Wolbachia endosymbiont of Aedes aegypti | Pseudomonadota | Alphaproteobacteria | NZ_CP072669.1 | 2 |
| Wolbachia endosymbiont of Aedes albopictus | Pseudomonadota | Alphaproteobacteria | NZ_CP101657.1 | 2 |
| Wolbachia endosymbiont of Anurida maritima | Pseudomonadota | Alphaproteobacteria | NZ_CP102058.1 | 2 |
| Wolbachia endosymbiont of Armadillidium arcangelii | Pseudomonadota | Alphaproteobacteria | NZ_CP157942.1 | 2 |
| Wolbachia endosymbiont of Bemisia tabaci | Pseudomonadota | Alphaproteobacteria | NZ_CP016430.1 | 2 |
| Wolbachia endosymbiont of Brugia pahangi | Pseudomonadota | Alphaproteobacteria | NZ_CP050521.1 | 2 |
| Wolbachia endosymbiont of Cantharis cryptica | Pseudomonadota | Alphaproteobacteria | NZ_OZ195521.1 | 2 |
| Wolbachia endosymbiont of Carposina sasakii | Pseudomonadota | Alphaproteobacteria | NZ_CP041215.1 | 2 |

| | | | | |
|---|---|---|---|---|
| Wolbachia endosymbiont of Ceratosolen solmsi | Pseudomonadota | Alphaproteobacteria | NZ_CP054598.1 | 2 |
| Wolbachia endosymbiont of Chrysomya megacephala | Pseudomonadota | Alphaproteobacteria | NZ_CP021120.1 | 2 |
| Wolbachia endosymbiont of Cimex lectularius | Pseudomonadota | Alphaproteobacteria | NZ_AP013028.1 | 2 |
| Wolbachia endosymbiont of Cruorifilaria tuberocauda | Pseudomonadota | Alphaproteobacteria | NZ_CP046579.1 | 2 |
| Wolbachia endosymbiont of Ctenocephalides felis wCfeJ | Pseudomonadota | Alphaproteobacteria | NZ_CP051157.1 | 2 |
| Wolbachia endosymbiont of Ctenocephalides felis wCfeT | Pseudomonadota | Alphaproteobacteria | NZ_CP051156.1 | 2 |
| Wolbachia endosymbiont of Culex quinquefasciatus Pel | Pseudomonadota | Alphaproteobacteria | NC_010981.1 | 2 |
| Wolbachia endosymbiont of Delia radicum | Pseudomonadota | Alphaproteobacteria | NZ_CP091195.1 | 2 |
| Wolbachia endosymbiont of Diaphorina citri | Pseudomonadota | Alphaproteobacteria | NZ_CP051608.1 | 2 |
| Wolbachia endosymbiont of Dipetalonema caudispina | Pseudomonadota | Alphaproteobacteria | NZ_CP046580.1 | 2 |
| Wolbachia endosymbiont of Dirofilaria (Dirofilaria) immitis | Pseudomonadota | Alphaproteobacteria | NZ_CP046578.1 | 2 |
| Wolbachia endosymbiont of Drosophila aff. chauvacae BK | Pseudomonadota | Alphaproteobacteria | NZ_CP136586.1 | 2 |
| Wolbachia endosymbiont of Drosophila ananassae | Pseudomonadota | Alphaproteobacteria | NZ_CP042904.1 | 2 |
| Wolbachia endosymbiont of Drosophila incompta | Pseudomonadota | Alphaproteobacteria | NZ_CP011148.1 | 2 |
| Wolbachia endosymbiont of Drosophila mauritiana | Pseudomonadota | Alphaproteobacteria | NZ_CP034335.1 | 2 |
| Wolbachia endosymbiont of Drosophila melanogaster | Pseudomonadota | Alphaproteobacteria | NZ_CP046925.1 | 2 |
| Wolbachia endosymbiont of Drosophila pseudotakahashii | Pseudomonadota | Alphaproteobacteria | NZ_CP110359.1 | 2 |
| Wolbachia endosymbiont of Drosophila santomea | Pseudomonadota | Alphaproteobacteria | NZ_CP069060.1 | 2 |
| Wolbachia endosymbiont of Drosophila simulans wHa | Pseudomonadota | Alphaproteobacteria | NC_021089.1 | 2 |
| Wolbachia endosymbiont of Drosophila willistoni | Pseudomonadota | Alphaproteobacteria | NZ_CP157591.1 | 2 |
| Wolbachia endosymbiont of Drosophila yakuba | Pseudomonadota | Alphaproteobacteria | NZ_CP069053.1 | 2 |
| Wolbachia endosymbiont of Encarsia formosa | Pseudomonadota | Alphaproteobacteria | NZ_CP154278.1 | 2 |
| Wolbachia endosymbiont of Folsomia candida | Pseudomonadota | Alphaproteobacteria | NZ_CP015510.2 | 2 |
| Wolbachia endosymbiont of Frankliniella intonsa | Pseudomonadota | Alphaproteobacteria | NZ_CP097148.1 | 2 |
| Wolbachia endosymbiont of Howardula sp. | Pseudomonadota | Alphaproteobacteria | NZ_CP092368.1 | 1 |
| Wolbachia endosymbiont of Liriomyza huidobrensis | Pseudomonadota | Alphaproteobacteria | NZ_CP170632.1 | 2 |
| Wolbachia endosymbiont of Litomosoides sigmodontis | Pseudomonadota | Alphaproteobacteria | NZ_CP046577.1 | 1 |
| Wolbachia endosymbiont of Oedothorax gibbosus | Pseudomonadota | Alphaproteobacteria | NZ_OW370537.1 | 2 |
| Wolbachia endosymbiont of Onchocerca ochengi | Pseudomonadota | Alphaproteobacteria | NC_018267.1 | 2 |
| Wolbachia endosymbiont of Onchocerca volvulus str. Cam | Pseudomonadota | Alphaproteobacteria | NZ_HG810405.1 | 2 |
| Wolbachia endosymbiont of Oryzaephilus surinamensis | Pseudomonadota | Alphaproteobacteria | NZ_CP092526.1 | 2 |
| Wolbachia endosymbiont of Ostrinia furnacalis | Pseudomonadota | Alphaproteobacteria | NZ_CP096925.1 | 2 |

| | | | | |
|---|---|---|---|---|
| Wolbachia endosymbiont of Ostrinia scapulalis | Pseudomonadota | Alphaproteobacteria | NZ_CP096926.1 | 2 |
| Wolbachia endosymbiont of Phyllotreta cruciferae | Pseudomonadota | Alphaproteobacteria | NZ_OU953855.1 | 2 |
| Wolbachia endosymbiont of Psylliodes chrysocephala | Pseudomonadota | Alphaproteobacteria | NZ_OU909470.1 | 2 |
| Wolbachia endosymbiont of Rhagoletis cingulata | Pseudomonadota | Alphaproteobacteria | NZ_CP175927.1 | 2 |
| Wolbachia endosymbiont of Spodoptera picta | Pseudomonadota | Alphaproteobacteria | NZ_CP067976.1 | 2 |
| Wolbachia endosymbiont of Tetranychus urticae | Pseudomonadota | Alphaproteobacteria | NZ_CP114181.1 | 2 |
| Wolbachia endosymbiont of Trichogramma pretiosum | Pseudomonadota | Alphaproteobacteria | NZ_CM003641.1 | 2 |
| Wolbachia endosymbiont of Wiebesia pumilae | Pseudomonadota | Alphaproteobacteria | NZ_CP054557.1 | 2 |
| Wolbachia endosymbiont of Zaprionus tsacasi | Pseudomonadota | Alphaproteobacteria | NZ_CP136587.1 | 2 |
| Wolbachia endosymbiont strain TRS of Brugia malayi | Pseudomonadota | Alphaproteobacteria | NC_006833.1 | 2 |
| Wolbachia pipientis | Pseudomonadota | Alphaproteobacteria | NZ_CP042446.1 | 2 |
| Xanthobacter dioxanivorans | Pseudomonadota | Alphaproteobacteria | NZ_CP063362.1 | 2 |
| Yoonia algicola | Pseudomonadota | Alphaproteobacteria | NZ_CP151762.1 | 2 |
| Yoonia phaeophyticola | Pseudomonadota | Alphaproteobacteria | NZ_CP150951.2 | 2 |
| Yoonia rhodophyticola | Pseudomonadota | Alphaproteobacteria | NZ_CP151767.2 | 2 |
| Zymomonas mobilis subsp. mobilis str. CP4 = NRRL B-14( | Pseudomonadota | Alphaproteobacteria | NC_022900.1 | 1 |
| [Acidovorax] ebreus TPSY | Pseudomonadota | Betaproteobacteria | NC_011992.1 | 2 |
| Achromobacter denitrificans | Pseudomonadota | Betaproteobacteria | NZ_CP053987.1 | 2 |
| Achromobacter insolitus | Pseudomonadota | Betaproteobacteria | NZ_LR134361.1 | 2 |
| Achromobacter mucicolens | Pseudomonadota | Betaproteobacteria | NZ_CP123364.1 | 2 |
| Achromobacter ruhlandii | Pseudomonadota | Betaproteobacteria | NZ_CP017433.1 | 2 |
| Achromobacter seleniivolatilans | Pseudomonadota | Betaproteobacteria | NZ_CP132976.1 | 2 |
| Achromobacter spanius | Pseudomonadota | Betaproteobacteria | NZ_CP025030.1 | 2 |
| Achromobacter veterisilvae | Pseudomonadota | Betaproteobacteria | NZ_CP148753.1 | 2 |
| Achromobacter xylosoxidans | Pseudomonadota | Betaproteobacteria | NZ_CP068211.1 | 2 |
| Acidovorax carolinensis | Pseudomonadota | Betaproteobacteria | NZ_CP021359.1 | 2 |
| Acidovorax sacchari | Pseudomonadota | Betaproteobacteria | NZ_AP035783.1 | 2 |
| Acidovorax temperans | Pseudomonadota | Betaproteobacteria | NZ_CP117193.1 | 2 |
| Advenella mimigardefordensis DPN7 | Pseudomonadota | Betaproteobacteria | NZ_CP003915.1 | 2 |
| Alcaligenes ammonioxydans | Pseudomonadota | Betaproteobacteria | NZ_CP049362.1 | 2 |
| Alcaligenes aquatilis | Pseudomonadota | Betaproteobacteria | NZ_CP094619.1 | 2 |
| Alicycliphilus denitrificans K601 | Pseudomonadota | Betaproteobacteria | NC_015422.1 | 2 |

| | | | | |
|---|---|---|---|---|
| Alysiella crassa | Pseudomonadota | Betaproteobacteria | NZ_CP091519.2 | 2 |
| Alysiella filiformis DSM 16848 | Pseudomonadota | Betaproteobacteria | NZ_CP083931.1 | 2 |
| Ampullimonas aquatilis | Pseudomonadota | Betaproteobacteria | NZ_CP183834.1 | 2 |
| Aquabacterium olei | Pseudomonadota | Betaproteobacteria | NZ_CP029210.1 | 2 |
| Aquella oligotrophica | Pseudomonadota | Betaproteobacteria | NZ_CP024847.1 | 2 |
| Aquincola tertiaricarbonis | Pseudomonadota | Betaproteobacteria | NZ_CP097636.1 | 2 |
| Aquitalea aquatilis | Pseudomonadota | Betaproteobacteria | NZ_CP039731.1 | 2 |
| Aquitalea denitrificans | Pseudomonadota | Betaproteobacteria | NZ_CP047241.1 | 2 |
| Aromatoleum aromaticum EbN1 | Pseudomonadota | Betaproteobacteria | NC_006513.1 | 2 |
| Aromatoleum bremense | Pseudomonadota | Betaproteobacteria | NZ_CP059467.1 | 2 |
| Aromatoleum evansii | Pseudomonadota | Betaproteobacteria | NZ_CP141259.1 | 2 |
| Aromatoleum petrolei | Pseudomonadota | Betaproteobacteria | NZ_CP059560.1 | 2 |
| Azoarcus olearius | Pseudomonadota | Betaproteobacteria | NZ_CP016210.1 | 2 |
| Azospira inquinata | Pseudomonadota | Betaproteobacteria | NZ_CP064782.1 | 2 |
| Azospira oryzae PS | Pseudomonadota | Betaproteobacteria | NC_016616.1 | 2 |
| Azospira restricta | Pseudomonadota | Betaproteobacteria | NZ_CP064781.1 | 2 |
| Basilea psittacipulmonis DSM 24701 | Pseudomonadota | Betaproteobacteria | NZ_CP009238.1 | 2 |
| Bergeriella denitrificans | Pseudomonadota | Betaproteobacteria | NZ_CP181246.1 | 2 |
| Bordetella avium | Pseudomonadota | Betaproteobacteria | NZ_CP139969.1 | 2 |
| Bordetella bronchialis | Pseudomonadota | Betaproteobacteria | NZ_CP016170.1 | 2 |
| Bordetella bronchiseptica | Pseudomonadota | Betaproteobacteria | NZ_LR134326.1 | 2 |
| Bordetella flabilis | Pseudomonadota | Betaproteobacteria | NZ_CP016172.1 | 2 |
| Bordetella hinzii | Pseudomonadota | Betaproteobacteria | NZ_LR134382.1 | 2 |
| Bordetella holmesii | Pseudomonadota | Betaproteobacteria | NZ_CP085973.1 | 2 |
| Bordetella parapertussis | Pseudomonadota | Betaproteobacteria | NZ_CP085971.1 | 2 |
| Bordetella pertussis | Pseudomonadota | Betaproteobacteria | NZ_CP025371.1 | 2 |
| Bordetella petrii | Pseudomonadota | Betaproteobacteria | NC_010170.1 | 2 |
| Bordetella pseudohinzii | Pseudomonadota | Betaproteobacteria | NZ_CP016440.1 | 2 |
| Burkholderia aenigmatica | Pseudomonadota | Betaproteobacteria | NZ_CP184468.1 | 2 |
| Burkholderia ambifaria AMMD | Pseudomonadota | Betaproteobacteria | NZ_CP009798.1 | 2 |
| Burkholderia arboris | Pseudomonadota | Betaproteobacteria | NZ_CP109821.1 | 2 |
| Burkholderia cenocepacia | Pseudomonadota | Betaproteobacteria | NZ_CP013450.1 | 2 |

| | | | | |
|---|---|---|---|---|
| Burkholderia cepacia | Pseudomonadota | Betaproteobacteria | NZ_CP045235.1 | 2 |
| Burkholderia contaminans | Pseudomonadota | Betaproteobacteria | NZ_CP157953.1 | 2 |
| Burkholderia dolosa | Pseudomonadota | Betaproteobacteria | NZ_CP082882.1 | 2 |
| Burkholderia gladioli | Pseudomonadota | Betaproteobacteria | NZ_CP068049.1 | 2 |
| Burkholderia glumae LMG 2196 = ATCC 33617 | Pseudomonadota | Betaproteobacteria | NZ_CP009435.1 | 2 |
| Burkholderia humptydooensis | Pseudomonadota | Betaproteobacteria | NZ_CP013380.1 | 2 |
| Burkholderia lata | Pseudomonadota | Betaproteobacteria | NC_007510.1 | 2 |
| Burkholderia mallei | Pseudomonadota | Betaproteobacteria | NZ_CP124608.1 | 2 |
| Burkholderia mayonis | Pseudomonadota | Betaproteobacteria | NZ_CP013386.1 | 2 |
| Burkholderia metallica | Pseudomonadota | Betaproteobacteria | NZ_CP013401.1 | 2 |
| Burkholderia multivorans | Pseudomonadota | Betaproteobacteria | NZ_CP143552.1 | 2 |
| Burkholderia oklahomensis C6786 | Pseudomonadota | Betaproteobacteria | NZ_CP009555.1 | 2 |
| Burkholderia perseverans | Pseudomonadota | Betaproteobacteria | NZ_CP045094.1 | 2 |
| Burkholderia plantarii | Pseudomonadota | Betaproteobacteria | NZ_CP007212.1 | 2 |
| Burkholderia pseudomallei | Pseudomonadota | Betaproteobacteria | NZ_AP028077.1 | 2 |
| Burkholderia pyrrocinia | Pseudomonadota | Betaproteobacteria | NZ_CP011503.1 | 2 |
| Burkholderia savannae | Pseudomonadota | Betaproteobacteria | NZ_CP013417.1 | 2 |
| Burkholderia seminalis | Pseudomonadota | Betaproteobacteria | NZ_CP013398.1 | 2 |
| Burkholderia singularis | Pseudomonadota | Betaproteobacteria | NZ_CP013448.1 | 2 |
| Burkholderia stabilis | Pseudomonadota | Betaproteobacteria | NZ_LR025742.1 | 2 |
| Burkholderia stagnalis | Pseudomonadota | Betaproteobacteria | NZ_CP156685.1 | 2 |
| Burkholderia territorii | Pseudomonadota | Betaproteobacteria | NZ_CP013366.1 | 2 |
| Burkholderia thailandensis E264 | Pseudomonadota | Betaproteobacteria | NC_007651.1 | 2 |
| Burkholderia theae | Pseudomonadota | Betaproteobacteria | NZ_CP183461.1 | 2 |
| Burkholderia ubonensis subsp. mesacidophila | Pseudomonadota | Betaproteobacteria | NZ_CP020737.1 | 2 |
| Burkholderia vietnamiensis LMG 10929 | Pseudomonadota | Betaproteobacteria | NZ_CP009631.1 | 2 |
| Caballeronia grimmiae | Pseudomonadota | Betaproteobacteria | NZ_CP084271.1 | 2 |
| Caballeronia insecticola | Pseudomonadota | Betaproteobacteria | NC_021287.1 | 2 |
| Caballeronia zhejiangensis | Pseudomonadota | Betaproteobacteria | NZ_CP084283.1 | 2 |
| Caldimonas aquatica | Pseudomonadota | Betaproteobacteria | NZ_CP110257.1 | 2 |
| Caldimonas brevitalea | Pseudomonadota | Betaproteobacteria | NZ_CP011371.1 | 2 |
| Caldimonas thermodepolymerans | Pseudomonadota | Betaproteobacteria | NZ_CP064338.1 | 2 |

| | | | | |
|---|---|---|---|---|
| Candidatus Accumulibacter phosphatis | Pseudomonadota | Betaproteobacteria | NC_013194.1 | 2 |
| Candidatus Ichthyocystis hellenicum | Pseudomonadota | Betaproteobacteria | NZ_LN906597.1 | 2 |
| Candidatus Kinetoplastidibacterium blastocrithidiae (ex Stri | Pseudomonadota | Betaproteobacteria | NC_019814.1 | 2 |
| Candidatus Kinetoplastidibacterium crithidiae TCC036E | Pseudomonadota | Betaproteobacteria | NC_020283.1 | 2 |
| Candidatus Kinetoplastidibacterium desouzai TCC079E | Pseudomonadota | Betaproteobacteria | NC_020294.1 | 2 |
| Candidatus Kinetoplastidibacterium kentomonadis | Pseudomonadota | Betaproteobacteria | NZ_CP025628.1 | 1 |
| Candidatus Kinetoplastidibacterium stringomonadis TCC2( | Pseudomonadota | Betaproteobacteria | NC_020299.1 | 2 |
| Candidatus Methylopumilus planktonicus | Pseudomonadota | Betaproteobacteria | NZ_CP040979.1 | 2 |
| Candidatus Methylopumilus rimovensis | Pseudomonadota | Betaproteobacteria | NZ_CP040986.1 | 2 |
| Candidatus Methylopumilus universalis | Pseudomonadota | Betaproteobacteria | NZ_CP040953.1 | 2 |
| Candidatus Nasuia deltocephalinicola | Pseudomonadota | Betaproteobacteria | NZ_CP094197.1 | 1 |
| Candidatus Nitrotoga arctica | Pseudomonadota | Betaproteobacteria | NZ_OU912926.1 | 2 |
| Candidatus Profftella armatura | Pseudomonadota | Betaproteobacteria | NC_021885.1 | 1 |
| Candidatus Symbiobacter mobilis CR | Pseudomonadota | Betaproteobacteria | NC_022576.1 | 2 |
| Candidatus Vallotia laricis | Pseudomonadota | Betaproteobacteria | NZ_CP047399.1 | 2 |
| Candidatus Vallotiella adelgis | Pseudomonadota | Betaproteobacteria | NZ_CP047396.1 | 2 |
| Candidatus Vallotiella hemipterorum | Pseudomonadota | Betaproteobacteria | NZ_OU343031.1 | 2 |
| Candidatus Zinderia endosymbiont of Aphrophora alni | Pseudomonadota | Betaproteobacteria | NZ_OZ026462.1 | 1 |
| Casimicrobium huifangae | Pseudomonadota | Betaproteobacteria | NZ_CP041352.1 | 2 |
| Castellaniella defragrans 65Phen | Pseudomonadota | Betaproteobacteria | NZ_HG916765.1 | 2 |
| Castellaniella ginsengisoli | Pseudomonadota | Betaproteobacteria | NZ_CP158257.1 | 2 |
| Chitinibacter bivalviorum | Pseudomonadota | Betaproteobacteria | NZ_CP058627.1 | 2 |
| Chitinibacter fontanus | Pseudomonadota | Betaproteobacteria | NZ_CP058952.1 | 2 |
| Chitinimonas arctica | Pseudomonadota | Betaproteobacteria | NZ_CP041730.1 | 2 |
| Chitiniphilus eburneus | Pseudomonadota | Betaproteobacteria | NZ_CP156074.1 | 2 |
| Chitiniphilus purpureus | Pseudomonadota | Betaproteobacteria | NZ_CP106753.1 | 2 |
| Chitinolyticbacter meiyuanensis | Pseudomonadota | Betaproteobacteria | NZ_CP041335.1 | 2 |
| Chromobacterium haemolyticum | Pseudomonadota | Betaproteobacteria | NZ_AP019312.1 | 2 |
| Chromobacterium paludis | Pseudomonadota | Betaproteobacteria | NZ_CP043473.1 | 2 |
| Chromobacterium phragmitis | Pseudomonadota | Betaproteobacteria | NZ_CP029495.1 | 2 |
| Chromobacterium rhizoryzae | Pseudomonadota | Betaproteobacteria | NZ_CP066126.1 | 2 |
| Chromobacterium subtsugae | Pseudomonadota | Betaproteobacteria | NZ_CP143257.1 | 2 |

| | | | | |
|---|---|---|---|---|
| Chromobacterium vaccinii | Pseudomonadota | Betaproteobacteria | NZ_CP017707.1 | 2 |
| Chromobacterium violaceum | Pseudomonadota | Betaproteobacteria | NZ_CP069587.1 | 2 |
| Collimonas arenae | Pseudomonadota | Betaproteobacteria | NZ_CP013235.1 | 2 |
| Collimonas fungivorans | Pseudomonadota | Betaproteobacteria | NZ_CP013232.1 | 2 |
| Collimonas pratensis | Pseudomonadota | Betaproteobacteria | NZ_CP013236.1 | 2 |
| Comamonas antarctica | Pseudomonadota | Betaproteobacteria | NZ_CP054840.1 | 2 |
| Comamonas aquatica | Pseudomonadota | Betaproteobacteria | NZ_CP072916.1 | 2 |
| Comamonas endophytica | Pseudomonadota | Betaproteobacteria | NZ_CP106881.1 | 2 |
| Comamonas flocculans | Pseudomonadota | Betaproteobacteria | NZ_CP042344.1 | 2 |
| Comamonas fluminis | Pseudomonadota | Betaproteobacteria | NZ_CP066783.1 | 2 |
| Comamonas kerstersii | Pseudomonadota | Betaproteobacteria | NZ_CP020121.1 | 2 |
| Comamonas odontotermitis | Pseudomonadota | Betaproteobacteria | NZ_CP083451.1 | 2 |
| Comamonas piscis | Pseudomonadota | Betaproteobacteria | NZ_CP058554.1 | 2 |
| Comamonas resistens | Pseudomonadota | Betaproteobacteria | NZ_CP125947.1 | 2 |
| Comamonas serinivorans | Pseudomonadota | Betaproteobacteria | NZ_CP021455.1 | 2 |
| Comamonas squillarum | Pseudomonadota | Betaproteobacteria | NZ_CP104377.1 | 2 |
| Comamonas terrigena NBRC 13299 | Pseudomonadota | Betaproteobacteria | NZ_AP019749.1 | 2 |
| Comamonas thiooxydans | Pseudomonadota | Betaproteobacteria | NZ_AP025193.1 | 2 |
| Conchiformibius steedae | Pseudomonadota | Betaproteobacteria | NZ_CP059563.1 | 2 |
| Crenobacter cavernae | Pseudomonadota | Betaproteobacteria | NZ_CP031337.1 | 2 |
| Cupriavidus basilensis | Pseudomonadota | Betaproteobacteria | NZ_CP062803.1 | 1 |
| Cupriavidus gilardii | Pseudomonadota | Betaproteobacteria | NZ_CP054624.1 | 2 |
| Cupriavidus malaysiensis | Pseudomonadota | Betaproteobacteria | NZ_CP017754.1 | 2 |
| Cupriavidus necator N-1 | Pseudomonadota | Betaproteobacteria | NC_015726.1 | 2 |
| Cupriavidus neocaledonicus | Pseudomonadota | Betaproteobacteria | NZ_LT984806.1 | 2 |
| Cupriavidus oxalaticus | Pseudomonadota | Betaproteobacteria | NZ_CP069812.1 | 2 |
| Cupriavidus pauculus | Pseudomonadota | Betaproteobacteria | NZ_CP082862.1 | 2 |
| Cupriavidus pinatubonensis | Pseudomonadota | Betaproteobacteria | NZ_CP080767.1 | 2 |
| Cupriavidus taiwanensis LMG 19424 | Pseudomonadota | Betaproteobacteria | NC_010528.1 | 2 |
| Dechloromonas aquae | Pseudomonadota | Betaproteobacteria | NZ_CP144212.1 | 2 |
| Deefgea piscis | Pseudomonadota | Betaproteobacteria | NZ_CP054143.1 | 2 |
| Deefgea tanakiae | Pseudomonadota | Betaproteobacteria | NZ_CP081150.1 | 2 |

| | | | | |
|---|---|---|---|---|
| Delftia acidovorans SPH-1 | Pseudomonadota | Betaproteobacteria | NC_010002.1 | 2 |
| Delftia lacustris | Pseudomonadota | Betaproteobacteria | NZ_CP141274.1 | 2 |
| Delftia tsuruhatensis | Pseudomonadota | Betaproteobacteria | NZ_CP118775.1 | 2 |
| Denitratisoma oestradiolicum | Pseudomonadota | Betaproteobacteria | NZ_LR778301.1 | 2 |
| Diaphorobacter aerolatus | Pseudomonadota | Betaproteobacteria | NZ_CP060783.1 | 2 |
| Diaphorobacter limosus | Pseudomonadota | Betaproteobacteria | NZ_CP136921.1 | 2 |
| Diaphorobacter nitroreducens | Pseudomonadota | Betaproteobacteria | NZ_CP016278.1 | 2 |
| Diaphorobacter ruginosibacter | Pseudomonadota | Betaproteobacteria | NZ_CP060714.1 | 2 |
| Duganella dendranthematis | Pseudomonadota | Betaproteobacteria | NZ_CP051684.1 | 2 |
| Duganella zoogloeoides | Pseudomonadota | Betaproteobacteria | NZ_CP140152.1 | 2 |
| Duodenibacillus massiliensis | Pseudomonadota | Betaproteobacteria | NZ_OV788642.1 | 2 |
| Eikenella corrodens | Pseudomonadota | Betaproteobacteria | NZ_LT906482.1 | 2 |
| Eikenella exigua | Pseudomonadota | Betaproteobacteria | NZ_CP038018.1 | 2 |
| Eleftheria terrae | Pseudomonadota | Betaproteobacteria | NZ_CP106951.1 | 2 |
| Ephemeroptericola cinctiostellae | Pseudomonadota | Betaproteobacteria | NZ_CP031124.1 | 2 |
| Ferribacterium limneticum | Pseudomonadota | Betaproteobacteria | NZ_CP075189.1 | 2 |
| Ferrigenium kumadai | Pseudomonadota | Betaproteobacteria | NZ_AP019536.1 | 2 |
| Ferriphaselus amnicola | Pseudomonadota | Betaproteobacteria | NZ_AP018738.1 | 2 |
| Ferrovum myxofaciens | Pseudomonadota | Betaproteobacteria | NZ_CP053676.1 | 2 |
| Fluviibacter phosphoraccumulans | Pseudomonadota | Betaproteobacteria | NZ_AP019011.1 | 2 |
| Gallionella capsiferriformans ES-2 | Pseudomonadota | Betaproteobacteria | NC_014394.1 | 2 |
| Herbaspirillum frisingense | Pseudomonadota | Betaproteobacteria | NZ_CP168558.1 | 2 |
| Herbaspirillum hiltneri N3 | Pseudomonadota | Betaproteobacteria | NZ_CP011409.1 | 2 |
| Herbaspirillum huttiense | Pseudomonadota | Betaproteobacteria | NZ_CP037993.1 | 2 |
| Herbaspirillum robiniae | Pseudomonadota | Betaproteobacteria | NZ_CP018845.1 | 2 |
| Herbaspirillum rubrisubalbicans | Pseudomonadota | Betaproteobacteria | NZ_CP024996.1 | 2 |
| Herbaspirillum seropedicae | Pseudomonadota | Betaproteobacteria | NZ_CP011930.1 | 2 |
| Herminiimonas arsenitoxidans | Pseudomonadota | Betaproteobacteria | NZ_LT671418.1 | 2 |
| Hydrogenophaga crassostreae | Pseudomonadota | Betaproteobacteria | NZ_CP017476.1 | 2 |
| Hydrogenophaga crocea | Pseudomonadota | Betaproteobacteria | NZ_CP049989.1 | 2 |
| Hydrogenophaga pseudoflava | Pseudomonadota | Betaproteobacteria | NZ_CP037867.1 | 2 |
| Ideonella dechloratans | Pseudomonadota | Betaproteobacteria | NZ_CP088081.1 | 2 |

| | | | | |
|---|---|---|---|---|
| Inhella inkyongensis | Pseudomonadota | Betaproteobacteria | NZ_CP040709.1 | 2 |
| Iodobacter ciconiae | Pseudomonadota | Betaproteobacteria | NZ_CP034433.1 | 2 |
| Janthinobacterium aestuarii | Pseudomonadota | Betaproteobacteria | NZ_CP142523.1 | 2 |
| Janthinobacterium agaricidamnosum NBRC 102515 = DSI | Pseudomonadota | Betaproteobacteria | NZ_HG322949.1 | 1 |
| Janthinobacterium lividum | Pseudomonadota | Betaproteobacteria | NZ_CP048832.1 | 2 |
| Janthinobacterium rivuli | Pseudomonadota | Betaproteobacteria | NZ_CP121464.1 | 2 |
| Janthinobacterium svalbardensis | Pseudomonadota | Betaproteobacteria | NZ_CP023422.1 | 2 |
| Janthinobacterium tructae | Pseudomonadota | Betaproteobacteria | NZ_CP041185.1 | 2 |
| Kerstersia gyiorum | Pseudomonadota | Betaproteobacteria | NZ_CP169556.1 | 2 |
| Kingella denitrificans | Pseudomonadota | Betaproteobacteria | NZ_CP065999.1 | 2 |
| Kingella kingae | Pseudomonadota | Betaproteobacteria | NZ_LS483426.1 | 2 |
| Kingella negevensis | Pseudomonadota | Betaproteobacteria | NZ_CP123448.1 | 2 |
| Lautropia mirabilis | Pseudomonadota | Betaproteobacteria | NZ_LR134378.1 | 2 |
| Leptothrix cholodnii SP-6 | Pseudomonadota | Betaproteobacteria | NC_010524.1 | 2 |
| Limnobacter profundi | Pseudomonadota | Betaproteobacteria | NZ_CP053084.1 | 2 |
| Limnobacter thiooxidans | Pseudomonadota | Betaproteobacteria | NZ_AP028947.1 | 2 |
| Massilia antarctica | Pseudomonadota | Betaproteobacteria | NZ_CP065053.1 | 2 |
| Massilia atriviolacea | Pseudomonadota | Betaproteobacteria | NZ_CP051166.1 | 2 |
| Massilia cellulosiltytica | Pseudomonadota | Betaproteobacteria | NZ_CP168562.1 | 2 |
| Massilia endophytica | Pseudomonadota | Betaproteobacteria | NZ_CP088952.1 | 2 |
| Massilia forsythiae | Pseudomonadota | Betaproteobacteria | NZ_CP051685.1 | 2 |
| Massilia litorea | Pseudomonadota | Betaproteobacteria | NZ_CP062941.1 | 2 |
| Massilia oculi | Pseudomonadota | Betaproteobacteria | NZ_CP029343.1 | 2 |
| Massilia putida | Pseudomonadota | Betaproteobacteria | NZ_CP019038.1 | 2 |
| Massilia violaceinigra | Pseudomonadota | Betaproteobacteria | NZ_CP024608.1 | 2 |
| Melaminivora jejuensis | Pseudomonadota | Betaproteobacteria | NZ_CP089742.1 | 2 |
| Melaminivora suipulveris | Pseudomonadota | Betaproteobacteria | NZ_CP027667.1 | 2 |
| Methylibium petroleiphilum PM1 | Pseudomonadota | Betaproteobacteria | NC_008825.1 | 2 |
| Methylobacillus flagellatus KT | Pseudomonadota | Betaproteobacteria | NC_007947.1 | 2 |
| Methylophilus medardicus | Pseudomonadota | Betaproteobacteria | NZ_CP040946.1 | 2 |
| Methyloradius palustris | Pseudomonadota | Betaproteobacteria | NZ_AP024110.1 | 2 |
| Methylotenera mobilis JLW8 | Pseudomonadota | Betaproteobacteria | NC_012968.1 | 2 |

| | | | | |
|---|---|---|---|---|
| Methylotenera versatilis 301 | Pseudomonadota | Betaproteobacteria | NC_014207.1 | 2 |
| Methylovorus glucosotrophus SIP3-4 | Pseudomonadota | Betaproteobacteria | NC_012969.1 | 2 |
| Microvirgula aerodenitrificans | Pseudomonadota | Betaproteobacteria | NZ_CP028519.1 | 2 |
| Morococcus cerebrosus | Pseudomonadota | Betaproteobacteria | NZ_CP094242.1 | 2 |
| Mycetohabitans endofungorum | Pseudomonadota | Betaproteobacteria | NZ_CP062168.1 | 2 |
| Mycetohabitans rhizoxinica | Pseudomonadota | Betaproteobacteria | NZ_CP062174.1 | 2 |
| Mycoavidus cysteinexigens | Pseudomonadota | Betaproteobacteria | NZ_AP018150.1 | 1 |
| Neisseria animalis | Pseudomonadota | Betaproteobacteria | NZ_CP031699.1 | 2 |
| Neisseria animaloris | Pseudomonadota | Betaproteobacteria | NZ_LR134440.1 | 2 |
| Neisseria arctica | Pseudomonadota | Betaproteobacteria | NZ_CP091510.1 | 2 |
| Neisseria bacilliformis | Pseudomonadota | Betaproteobacteria | NZ_CP059571.1 | 2 |
| Neisseria brasiliensis | Pseudomonadota | Betaproteobacteria | NZ_CP046027.1 | 2 |
| Neisseria canis | Pseudomonadota | Betaproteobacteria | NZ_LR134313.1 | 2 |
| Neisseria chenwenguii | Pseudomonadota | Betaproteobacteria | NZ_CP022278.1 | 2 |
| Neisseria cinerea | Pseudomonadota | Betaproteobacteria | NZ_LS483369.1 | 2 |
| Neisseria dentiae | Pseudomonadota | Betaproteobacteria | NZ_CP059570.1 | 2 |
| Neisseria dumasiana | Pseudomonadota | Betaproteobacteria | NZ_CP091509.1 | 2 |
| Neisseria flavescens | Pseudomonadota | Betaproteobacteria | NZ_CP039886.1 | 2 |
| Neisseria gonorrhoeae | Pseudomonadota | Betaproteobacteria | NZ_AP023069.1 | 2 |
| Neisseria lactamica | Pseudomonadota | Betaproteobacteria | NZ_CP031253.1 | 2 |
| Neisseria leonii | Pseudomonadota | Betaproteobacteria | NZ_CP145606.1 | 2 |
| Neisseria lisongii | Pseudomonadota | Betaproteobacteria | NZ_CP116766.1 | 2 |
| Neisseria macacae ATCC 33926 | Pseudomonadota | Betaproteobacteria | NZ_CP094241.1 | 2 |
| Neisseria meningitidis | Pseudomonadota | Betaproteobacteria | NZ_CP064367.1 | 2 |
| Neisseria musculi | Pseudomonadota | Betaproteobacteria | NZ_CP060414.2 | 2 |
| Neisseria shayeganii | Pseudomonadota | Betaproteobacteria | NZ_CP059567.1 | 2 |
| Neisseria sicca | Pseudomonadota | Betaproteobacteria | NZ_CP072524.1 | 2 |
| Neisseria subflava | Pseudomonadota | Betaproteobacteria | NZ_CP039887.1 | 2 |
| Neisseria wadsworthii | Pseudomonadota | Betaproteobacteria | NZ_CP059565.1 | 2 |
| Neisseria weaveri | Pseudomonadota | Betaproteobacteria | NZ_LR134533.1 | 2 |
| Neisseria yangbaofengii | Pseudomonadota | Betaproteobacteria | NZ_CP062976.1 | 2 |
| Neisseria zalophi | Pseudomonadota | Betaproteobacteria | NZ_CP031700.1 | 2 |

| | | | | |
|---|---|---|---|---|
| Neisseria zoodegmatis | Pseudomonadota | Betaproteobacteria | NZ_LT906434.1 | 2 |
| Neopusillimonas aromaticivorans | Pseudomonadota | Betaproteobacteria | NZ_CP128233.1 | 2 |
| Nitrogeniibacter aestuarii | Pseudomonadota | Betaproteobacteria | NZ_CP071321.1 | 2 |
| Nitrogeniibacter mangrovi | Pseudomonadota | Betaproteobacteria | NZ_CP048836.1 | 2 |
| Nitrosomonas communis | Pseudomonadota | Betaproteobacteria | NZ_CP011451.1 | 2 |
| Nitrosomonas europaea ATCC 19718 | Pseudomonadota | Betaproteobacteria | NC_004757.1 | 2 |
| Nitrosomonas eutropha C91 | Pseudomonadota | Betaproteobacteria | NC_008344.1 | 2 |
| Nitrosomonas ureae | Pseudomonadota | Betaproteobacteria | NZ_CP013341.1 | 2 |
| Nitrosospira briensis C-128 | Pseudomonadota | Betaproteobacteria | NZ_CP012371.1 | 2 |
| Nitrosospira lacus | Pseudomonadota | Betaproteobacteria | NZ_CP021106.3 | 2 |
| Nitrosospira multiformis ATCC 25196 | Pseudomonadota | Betaproteobacteria | NC_007614.1 | 2 |
| Niveibacterium microcysteis | Pseudomonadota | Betaproteobacteria | NZ_CP071060.1 | 2 |
| Oligella ureolytica | Pseudomonadota | Betaproteobacteria | NZ_CP065725.1 | 2 |
| Orrella daihaiensis | Pseudomonadota | Betaproteobacteria | NZ_CP063982.1 | 2 |
| Orrella dioscoreae | Pseudomonadota | Betaproteobacteria | NZ_LT907988.1 | 2 |
| Orrella marina | Pseudomonadota | Betaproteobacteria | NZ_CP028901.1 | 2 |
| Oryzomicrobium terrae | Pseudomonadota | Betaproteobacteria | NZ_CP022579.1 | 2 |
| Ottowia oryzae | Pseudomonadota | Betaproteobacteria | NZ_CP027666.1 | 2 |
| Oxalobacter formigenes | Pseudomonadota | Betaproteobacteria | NZ_CP018787.1 | 2 |
| Oxalobacter vibrioformis | Pseudomonadota | Betaproteobacteria | NZ_CP098242.1 | 2 |
| Paenalcaligenes faecalis | Pseudomonadota | Betaproteobacteria | NZ_CP106841.1 | 2 |
| Paludibacterium paludis | Pseudomonadota | Betaproteobacteria | NZ_CP069161.1 | 2 |
| Paludibacterium yongneupense | Pseudomonadota | Betaproteobacteria | NZ_CP069162.1 | 2 |
| Pandoraea apista | Pseudomonadota | Betaproteobacteria | NZ_CP013481.2 | 2 |
| Pandoraea commovens | Pseudomonadota | Betaproteobacteria | NZ_CP102780.1 | 2 |
| Pandoraea faecigallinarum | Pseudomonadota | Betaproteobacteria | NZ_CP011807.3 | 2 |
| Pandoraea fibrosis | Pseudomonadota | Betaproteobacteria | NZ_CP047385.1 | 2 |
| Pandoraea norimbergensis | Pseudomonadota | Betaproteobacteria | NZ_CP013480.3 | 2 |
| Pandoraea oxalativorans | Pseudomonadota | Betaproteobacteria | NZ_CP011253.3 | 2 |
| Pandoraea pnomenusa | Pseudomonadota | Betaproteobacteria | NC_023018.2 | 2 |
| Pandoraea pulmonicola | Pseudomonadota | Betaproteobacteria | NZ_CP010310.2 | 2 |
| Pandoraea sputorum | Pseudomonadota | Betaproteobacteria | NZ_LT906435.1 | 2 |

| | | | | |
|---|---|---|---|---|
| Pandoraea thiooxydans | Pseudomonadota | Betaproteobacteria | NZ_CP011568.3 | 2 |
| Pandoraea vervacti | Pseudomonadota | Betaproteobacteria | NZ_CP010897.2 | 2 |
| Paraburkholderia aromaticivorans | Pseudomonadota | Betaproteobacteria | NZ_CP022989.1 | 2 |
| Paraburkholderia atlantica | Pseudomonadota | Betaproteobacteria | NC_014117.1 | 2 |
| Paraburkholderia bryophila | Pseudomonadota | Betaproteobacteria | NZ_CP099394.1 | 2 |
| Paraburkholderia caffeinilytica | Pseudomonadota | Betaproteobacteria | NZ_CP031467.1 | 2 |
| Paraburkholderia caribensis MBA4 | Pseudomonadota | Betaproteobacteria | NZ_CP012746.1 | 2 |
| Paraburkholderia dioscoreae | Pseudomonadota | Betaproteobacteria | NZ_LR699553.1 | 2 |
| Paraburkholderia dokdonensis | Pseudomonadota | Betaproteobacteria | NZ_CP029640.1 | 2 |
| Paraburkholderia edwinii | Pseudomonadota | Betaproteobacteria | NZ_CP080095.1 | 2 |
| Paraburkholderia fungorum | Pseudomonadota | Betaproteobacteria | NZ_CP010026.1 | 2 |
| Paraburkholderia ginsengisoli | Pseudomonadota | Betaproteobacteria | NZ_CP066075.1 | 2 |
| Paraburkholderia graminis | Pseudomonadota | Betaproteobacteria | NZ_CP024934.1 | 2 |
| Paraburkholderia hospita | Pseudomonadota | Betaproteobacteria | NZ_CP026105.1 | 2 |
| Paraburkholderia kirstenboschensis | Pseudomonadota | Betaproteobacteria | NZ_CP136513.1 | 2 |
| Paraburkholderia kururiensis | Pseudomonadota | Betaproteobacteria | NZ_OZ221290.1 | 2 |
| Paraburkholderia largidicola | Pseudomonadota | Betaproteobacteria | NZ_AP023174.1 | 2 |
| Paraburkholderia pallida | Pseudomonadota | Betaproteobacteria | NZ_CP038148.1 | 2 |
| Paraburkholderia phymatum STM815 | Pseudomonadota | Betaproteobacteria | NC_010622.1 | 2 |
| Paraburkholderia phytofirmans PsJN | Pseudomonadota | Betaproteobacteria | NC_010681.1 | 2 |
| Paraburkholderia sabiae | Pseudomonadota | Betaproteobacteria | NZ_CP125295.1 | 2 |
| Paraburkholderia sprentiae WSM5005 | Pseudomonadota | Betaproteobacteria | NZ_CP017561.2 | 2 |
| Paraburkholderia terrae | Pseudomonadota | Betaproteobacteria | NZ_CP026111.1 | 2 |
| Paraburkholderia terricola | Pseudomonadota | Betaproteobacteria | NZ_CP024941.1 | 2 |
| Paraburkholderia tropica | Pseudomonadota | Betaproteobacteria | NZ_CP049134.1 | 2 |
| Paraburkholderia xenovorans LB400 | Pseudomonadota | Betaproteobacteria | NC_007951.1 | 2 |
| Paracidovorax avenae ATCC 19860 | Pseudomonadota | Betaproteobacteria | NC_015138.1 | 2 |
| Paracidovorax citrulli | Pseudomonadota | Betaproteobacteria | NZ_CP042323.1 | 2 |
| Paralysiella testudinis | Pseudomonadota | Betaproteobacteria | NZ_CP069798.1 | 2 |
| Paucibacter sediminis | Pseudomonadota | Betaproteobacteria | NZ_CP116346.1 | 2 |
| Pelistega ratti | Pseudomonadota | Betaproteobacteria | NZ_CP047165.1 | 2 |
| Pigmentiphaga aceris | Pseudomonadota | Betaproteobacteria | NZ_CP043046.1 | 2 |

| | | | | |
|---|---|---|---|---|
| Piscinibacter gummiphilus | Pseudomonadota | Betaproteobacteria | NZ_CP024645.1 | 2 |
| Piscinibacter sakaiensis | Pseudomonadota | Betaproteobacteria | NZ_CP182947.1 | 2 |
| Piscinibacterium candidicorallinum | Pseudomonadota | Betaproteobacteria | NZ_CP180191.1 | 2 |
| Polaromonas hydrogenivorans | Pseudomonadota | Betaproteobacteria | NZ_CP157675.1 | 2 |
| Polaromonas naphthalenivorans CJ2 | Pseudomonadota | Betaproteobacteria | NC_008781.1 | 2 |
| Polaromonas vacuolata | Pseudomonadota | Betaproteobacteria | NZ_CP051461.1 | 2 |
| Pollutimonas thiosulfatoxidans | Pseudomonadota | Betaproteobacteria | NZ_CP022987.1 | 2 |
| Polynucleobacter acidiphobus | Pseudomonadota | Betaproteobacteria | NZ_CP023277.1 | 2 |
| Polynucleobacter antarcticus | Pseudomonadota | Betaproteobacteria | NZ_CP028941.1 | 2 |
| Polynucleobacter arcticus | Pseudomonadota | Betaproteobacteria | NZ_CP028940.1 | 2 |
| Polynucleobacter asymbioticus QLW-P1DMWA-1 | Pseudomonadota | Betaproteobacteria | NC_009379.1 | 2 |
| Polynucleobacter corsicus | Pseudomonadota | Betaproteobacteria | NZ_CP061314.1 | 2 |
| Polynucleobacter difficilis | Pseudomonadota | Betaproteobacteria | NZ_CP023276.1 | 2 |
| Polynucleobacter duraquae | Pseudomonadota | Betaproteobacteria | NZ_CP007501.1 | 2 |
| Polynucleobacter ibericus | Pseudomonadota | Betaproteobacteria | NZ_CP061309.1 | 2 |
| Polynucleobacter necessarius | Pseudomonadota | Betaproteobacteria | NZ_LT615228.1 | 2 |
| Polynucleobacter paludilacus | Pseudomonadota | Betaproteobacteria | NZ_CP061298.1 | 2 |
| Polynucleobacter paneuropaeus | Pseudomonadota | Betaproteobacteria | NZ_CP049682.1 | 2 |
| Polynucleobacter tropicus | Pseudomonadota | Betaproteobacteria | NZ_CP028942.1 | 2 |
| Polynucleobacter wuianus | Pseudomonadota | Betaproteobacteria | NZ_CP015922.1 | 2 |
| Polynucleobacter yangtzensis | Pseudomonadota | Betaproteobacteria | NZ_AP026973.1 | 2 |
| Pseudazoarcus pumilus | Pseudomonadota | Betaproteobacteria | NZ_CP025682.1 | 2 |
| Pseudoduganella albidiflava | Pseudomonadota | Betaproteobacteria | NZ_CP036401.1 | 2 |
| Pseudoduganella aquatica | Pseudomonadota | Betaproteobacteria | NZ_CP086370.1 | 2 |
| Pseudoduganella armeniaca | Pseudomonadota | Betaproteobacteria | NZ_CP028324.1 | 2 |
| Pseudoduganella chitinolytica | Pseudomonadota | Betaproteobacteria | NZ_CP119083.1 | 2 |
| Pseudoduganella flava | Pseudomonadota | Betaproteobacteria | NZ_CP046904.1 | 2 |
| Pseudoduganella lutea | Pseudomonadota | Betaproteobacteria | NZ_CP035913.1 | 2 |
| Pseudoduganella plicata | Pseudomonadota | Betaproteobacteria | NZ_CP038026.1 | 2 |
| Pseudoduganella umbonata | Pseudomonadota | Betaproteobacteria | NZ_CP040017.1 | 2 |
| Pseudothauera hydrothermalis | Pseudomonadota | Betaproteobacteria | NZ_CP029331.1 | 2 |
| Pulveribacter suum | Pseudomonadota | Betaproteobacteria | NZ_CP027792.1 | 2 |

| | | | | |
|---|---|---|---|---|
| Quatrionicoccus australiensis | Pseudomonadota | Betaproteobacteria | NZ_CP075188.1 | 2 |
| Ralstonia insidiosa | Pseudomonadota | Betaproteobacteria | NZ_CP016022.1 | 2 |
| Ralstonia mannitolilytica | Pseudomonadota | Betaproteobacteria | NZ_CP010799.1 | 2 |
| Ralstonia nicotianae | Pseudomonadota | Betaproteobacteria | NZ_CP046674.1 | 2 |
| Ralstonia solanacearum | Pseudomonadota | Betaproteobacteria | NZ_CP012943.1 | 2 |
| Ralstonia syzygii subsp. indonesiensis | Pseudomonadota | Betaproteobacteria | NZ_AP029014.1 | 2 |
| Ralstonia wenshanensis | Pseudomonadota | Betaproteobacteria | NZ_CP076413.1 | 2 |
| Ramlibacter tataouinensis TTB310 | Pseudomonadota | Betaproteobacteria | NC_015677.1 | 2 |
| Rhodoferax antarcticus | Pseudomonadota | Betaproteobacteria | NZ_CP019240.1 | 2 |
| Rhodoferax aquaticus | Pseudomonadota | Betaproteobacteria | NZ_CP036282.1 | 2 |
| Rhodoferax ferrireducens T118 | Pseudomonadota | Betaproteobacteria | NC_007908.1 | 2 |
| Rhodoferax koreensis | Pseudomonadota | Betaproteobacteria | NZ_CP019236.1 | 2 |
| Rhodoferax mekongensis | Pseudomonadota | Betaproteobacteria | NZ_CP132507.1 | 2 |
| Rhodoferax saidenbachensis | Pseudomonadota | Betaproteobacteria | NZ_CP019239.1 | 2 |
| Rhodoferax sediminis | Pseudomonadota | Betaproteobacteria | NZ_CP035503.1 | 2 |
| Robbsia andropogonis | Pseudomonadota | Betaproteobacteria | NZ_CP139170.1 | 2 |
| Roseateles amylovorans | Pseudomonadota | Betaproteobacteria | NZ_CP104562.2 | 2 |
| Roseateles depolymerans | Pseudomonadota | Betaproteobacteria | NZ_CP013729.1 | 2 |
| Rubrivivax gelatinosus | Pseudomonadota | Betaproteobacteria | NZ_CP181386.1 | 2 |
| Rugosibacter aromaticivorans | Pseudomonadota | Betaproteobacteria | NZ_CP010554.1 | 2 |
| Serpentinimonas maccroryi | Pseudomonadota | Betaproteobacteria | NZ_AP014569.1 | 2 |
| Serpentinimonas raichei | Pseudomonadota | Betaproteobacteria | NZ_AP014568.1 | 2 |
| Sideroxyarcus emersonii | Pseudomonadota | Betaproteobacteria | NZ_AP023423.1 | 2 |
| Sideroxydans lithotrophicus ES-1 | Pseudomonadota | Betaproteobacteria | NC_013959.1 | 2 |
| Silvimonas soli | Pseudomonadota | Betaproteobacteria | NZ_CP106736.1 | 2 |
| Simonsiella muelleri ATCC 29453 | Pseudomonadota | Betaproteobacteria | NZ_CP019448.1 | 2 |
| Simplicispira suum | Pseudomonadota | Betaproteobacteria | NZ_CP027669.1 | 2 |
| Snodgrassella alvi wkB2 | Pseudomonadota | Betaproteobacteria | NZ_CP007446.1 | 2 |
| Sphaerotilus microaerophilus | Pseudomonadota | Betaproteobacteria | NZ_AP025730.1 | 2 |
| Sphaerotilus sulfidivorans | Pseudomonadota | Betaproteobacteria | NZ_CP035708.1 | 2 |
| Sulfuricella denitrificans skB26 | Pseudomonadota | Betaproteobacteria | NC_022357.1 | 2 |
| Sulfuricystis multivorans | Pseudomonadota | Betaproteobacteria | NZ_AP018718.1 | 2 |

| | | | | |
|---|---|---|---|---|
| Sulfuricystis thermophila | Pseudomonadota | Betaproteobacteria | NZ_AP019373.1 | 2 |
| Sulfuriferula nivalis | Pseudomonadota | Betaproteobacteria | NZ_AP021881.1 | 2 |
| Sulfuriferula plumbiphila | Pseudomonadota | Betaproteobacteria | NZ_AP021884.1 | 2 |
| Sulfurimicrobium lacus | Pseudomonadota | Betaproteobacteria | NZ_AP022853.1 | 2 |
| Sulfuritalea hydrogenivorans sk43H | Pseudomonadota | Betaproteobacteria | NZ_AP012547.1 | 2 |
| Sulfuritortus calidifontis | Pseudomonadota | Betaproteobacteria | NZ_AP018721.1 | 2 |
| Sutterella faecalis | Pseudomonadota | Betaproteobacteria | NZ_CP040882.1 | 2 |
| Sutterella megalosphaeroides | Pseudomonadota | Betaproteobacteria | NZ_AP018786.1 | 2 |
| Sutterella wadsworthensis | Pseudomonadota | Betaproteobacteria | NZ_CP068055.1 | 2 |
| Taylorella asinigenitalis MCE3 | Pseudomonadota | Betaproteobacteria | NC_016043.1 | 2 |
| Taylorella equigenitalis | Pseudomonadota | Betaproteobacteria | NZ_CP076516.1 | 2 |
| Telluria beijingensis | Pseudomonadota | Betaproteobacteria | NZ_CP132480.1 | 2 |
| Telluria mixta | Pseudomonadota | Betaproteobacteria | NZ_CP119520.1 | 2 |
| Tepidimonas taiwanensis | Pseudomonadota | Betaproteobacteria | NZ_CP083911.1 | 2 |
| Thauera aminoaromatica | Pseudomonadota | Betaproteobacteria | NC_011662.2 | 2 |
| Thauera aromatica K172 | Pseudomonadota | Betaproteobacteria | NZ_CP028339.1 | 2 |
| Thauera butanivorans | Pseudomonadota | Betaproteobacteria | NZ_CP183038.1 | 2 |
| Thauera chlorobenzoica | Pseudomonadota | Betaproteobacteria | NZ_CP018839.1 | 2 |
| Thauera humireducens | Pseudomonadota | Betaproteobacteria | NZ_CP014646.1 | 2 |
| Thiobacillus sedimenti | Pseudomonadota | Betaproteobacteria | NZ_CP141769.1 | 2 |
| Thiomonas arsenitoxydans | Pseudomonadota | Betaproteobacteria | NC_014145.1 | 2 |
| Thiomonas intermedia | Pseudomonadota | Betaproteobacteria | NZ_CP020046.1 | 2 |
| Trinickia caryophylli | Pseudomonadota | Betaproteobacteria | NZ_CP139970.1 | 2 |
| Trinickia violacea | Pseudomonadota | Betaproteobacteria | NZ_CP040077.1 | 2 |
| Turicimonas muris | Pseudomonadota | Betaproteobacteria | NZ_CP065313.1 | 2 |
| Undibacterium cyanobacteriorum | Pseudomonadota | Betaproteobacteria | NZ_CP133720.1 | 2 |
| Undibacterium parvum | Pseudomonadota | Betaproteobacteria | NZ_CP034464.1 | 2 |
| Uruburuella suis | Pseudomonadota | Betaproteobacteria | NZ_CP091507.1 | 2 |
| Uruburuella testudinis | Pseudomonadota | Betaproteobacteria | NZ_CP091508.1 | 2 |
| Usitatibacter palustris | Pseudomonadota | Betaproteobacteria | NZ_CP053073.1 | 2 |
| Usitatibacter rugosus | Pseudomonadota | Betaproteobacteria | NZ_CP053069.1 | 2 |
| Variovorax paradoxus NBRC 15149 | Pseudomonadota | Betaproteobacteria | NZ_CP157626.1 | 2 |

| | | | | |
|---|---|---|---|---|
| Verminephrobacter eiseniae EF01-2 | Pseudomonadota | Betaproteobacteria | NC_008786.1 | 2 |
| Vitreoscilla filiformis | Pseudomonadota | Betaproteobacteria | NZ_CP022423.1 | 2 |
| Vitreoscilla stercoraria | Pseudomonadota | Betaproteobacteria | NZ_CP091512.1 | 2 |
| Wielerella bovis | Pseudomonadota | Betaproteobacteria | NZ_CP092361.1 | 2 |
| Xylophilus rhododendri | Pseudomonadota | Betaproteobacteria | NZ_CP047650.1 | 2 |
| [Enterobacter] lignolyticus SCF1 | Pseudomonadota | Gammaproteobacteria | NC_014618.1 | 2 |
| [Haemophilus] ducreyi | Pseudomonadota | Gammaproteobacteria | NZ_CP015425.1 | 2 |
| [Mannheimia] succiniciproducens MBEL55E | Pseudomonadota | Gammaproteobacteria | NC_006300.1 | 2 |
| [Pantoea] beijingensis | Pseudomonadota | Gammaproteobacteria | NZ_CP071409.1 | 2 |
| Abyssogena phaseoliformis symbiont OG214 | Pseudomonadota | Gammaproteobacteria | NZ_AP012977.1 | 2 |
| Acidiferrobacter thiooxydans | Pseudomonadota | Gammaproteobacteria | NZ_CP080624.1 | 2 |
| Acidihalobacter aeolianus | Pseudomonadota | Gammaproteobacteria | NZ_CP017448.1 | 2 |
| Acidihalobacter ferrooxydans | Pseudomonadota | Gammaproteobacteria | NZ_CP019434.1 | 2 |
| Acidihalobacter yilgarnensis | Pseudomonadota | Gammaproteobacteria | NZ_CP017415.1 | 2 |
| Acinetobacter baumannii | Pseudomonadota | Gammaproteobacteria | NZ_CP045110.1 | 2 |
| Acinetobacter baylyi ADP1 | Pseudomonadota | Gammaproteobacteria | NC_005966.1 | 2 |
| Acinetobacter bereziniae | Pseudomonadota | Gammaproteobacteria | NZ_CP066119.1 | 2 |
| Acinetobacter bouvetii | Pseudomonadota | Gammaproteobacteria | NZ_AP024595.1 | 2 |
| Acinetobacter chinensis | Pseudomonadota | Gammaproteobacteria | NZ_CP032134.1 | 2 |
| Acinetobacter corruptisaponis | Pseudomonadota | Gammaproteobacteria | NZ_CP125669.1 | 2 |
| Acinetobacter cumulans | Pseudomonadota | Gammaproteobacteria | NZ_CP035934.2 | 2 |
| Acinetobacter defluvii | Pseudomonadota | Gammaproteobacteria | NZ_CP029397.2 | 2 |
| Acinetobacter dispersus | Pseudomonadota | Gammaproteobacteria | NZ_CP041970.1 | 2 |
| Acinetobacter equi | Pseudomonadota | Gammaproteobacteria | NZ_CP012808.1 | 2 |
| Acinetobacter guillouiae | Pseudomonadota | Gammaproteobacteria | NZ_AP014630.1 | 2 |
| Acinetobacter haemolyticus | Pseudomonadota | Gammaproteobacteria | NZ_CP030880.1 | 2 |
| Acinetobacter indicus | Pseudomonadota | Gammaproteobacteria | NZ_CP045198.1 | 2 |
| Acinetobacter junii | Pseudomonadota | Gammaproteobacteria | NZ_CP059558.1 | 2 |
| Acinetobacter lactucae | Pseudomonadota | Gammaproteobacteria | NZ_CP053391.1 | 2 |
| Acinetobacter lanii | Pseudomonadota | Gammaproteobacteria | NZ_CP049916.1 | 2 |
| Acinetobacter larvae | Pseudomonadota | Gammaproteobacteria | NZ_CP016895.1 | 2 |
| Acinetobacter lwoffii | Pseudomonadota | Gammaproteobacteria | NZ_CP072549.1 | 2 |

| | | | | |
|---|---|---|---|---|
| Acinetobacter nosocomialis | Pseudomonadota | Gammaproteobacteria | NZ_CP157432.1 | 2 |
| Acinetobacter oleivorans DR1 | Pseudomonadota | Gammaproteobacteria | NC_014259.1 | 2 |
| Acinetobacter pittii PHEA-2 | Pseudomonadota | Gammaproteobacteria | NC_016603.1 | 2 |
| Acinetobacter radioresistens | Pseudomonadota | Gammaproteobacteria | NZ_CP030031.1 | 2 |
| Acinetobacter schindleri | Pseudomonadota | Gammaproteobacteria | NZ_CP044483.1 | 2 |
| Acinetobacter seifertii | Pseudomonadota | Gammaproteobacteria | NZ_CP065820.1 | 2 |
| Acinetobacter septicus | Pseudomonadota | Gammaproteobacteria | NZ_CP079898.1 | 2 |
| Acinetobacter soli | Pseudomonadota | Gammaproteobacteria | NZ_CP073241.1 | 2 |
| Acinetobacter suaedae | Pseudomonadota | Gammaproteobacteria | NZ_CP043909.1 | 2 |
| Acinetobacter thermotolerans | Pseudomonadota | Gammaproteobacteria | NZ_CP156787.1 | 2 |
| Acinetobacter tibetensis | Pseudomonadota | Gammaproteobacteria | NZ_CP098732.1 | 2 |
| Acinetobacter towneri | Pseudomonadota | Gammaproteobacteria | NZ_CP071766.1 | 2 |
| Acinetobacter variabilis | Pseudomonadota | Gammaproteobacteria | NZ_CP130328.1 | 2 |
| Acinetobacter wuhouensis | Pseudomonadota | Gammaproteobacteria | NZ_CP031716.1 | 2 |
| Actinobacillus delphinicola | Pseudomonadota | Gammaproteobacteria | NZ_LR134510.1 | 2 |
| Actinobacillus equuli | Pseudomonadota | Gammaproteobacteria | NZ_LR134310.1 | 2 |
| Actinobacillus indolicus | Pseudomonadota | Gammaproteobacteria | NZ_CP038145.1 | 2 |
| Actinobacillus pleuropneumoniae | Pseudomonadota | Gammaproteobacteria | NZ_CP030753.1 | 2 |
| Actinobacillus porcitonsillarum | Pseudomonadota | Gammaproteobacteria | NZ_CP029206.1 | 2 |
| Actinobacillus succinogenes 130Z | Pseudomonadota | Gammaproteobacteria | NC_009655.1 | 2 |
| Actinobacillus suis ATCC 33415 | Pseudomonadota | Gammaproteobacteria | NZ_CP009159.1 | 2 |
| Aequoribacter fuscus | Pseudomonadota | Gammaproteobacteria | NZ_CP036423.1 | 2 |
| Aeromonas allosaccharophila | Pseudomonadota | Gammaproteobacteria | NZ_CP065745.1 | 2 |
| Aeromonas bestiarum | Pseudomonadota | Gammaproteobacteria | NZ_CP064746.1 | 2 |
| Aeromonas caviae | Pseudomonadota | Gammaproteobacteria | NZ_LS483441.1 | 2 |
| Aeromonas dhakensis | Pseudomonadota | Gammaproteobacteria | NZ_CP023141.1 | 2 |
| Aeromonas encheleia | Pseudomonadota | Gammaproteobacteria | NZ_LR134376.1 | 2 |
| Aeromonas enteropelogenes | Pseudomonadota | Gammaproteobacteria | NZ_CP083625.1 | 2 |
| Aeromonas hydrophila subsp. hydrophila ATCC 7966 | Pseudomonadota | Gammaproteobacteria | NC_008570.1 | 2 |
| Aeromonas jandaei | Pseudomonadota | Gammaproteobacteria | NZ_CP066092.1 | 2 |
| Aeromonas media | Pseudomonadota | Gammaproteobacteria | NZ_CP075564.1 | 2 |
| Aeromonas popoffii | Pseudomonadota | Gammaproteobacteria | NZ_CP149129.1 | 2 |

| | | | | |
|---|---|---|---|---|
| Aeromonas rivuli | Pseudomonadota | Gammaproteobacteria | NZ_CP079742.1 | 2 |
| Aeromonas salmonicida | Pseudomonadota | Gammaproteobacteria | NZ_CP110645.1 | 2 |
| Aeromonas simiae | Pseudomonadota | Gammaproteobacteria | NZ_CP040449.1 | 2 |
| Aeromonas veronii | Pseudomonadota | Gammaproteobacteria | NZ_CP044060.1 | 2 |
| Aerosticca soli | Pseudomonadota | Gammaproteobacteria | NZ_AP018560.1 | 2 |
| Agarilytica rhodophyticola | Pseudomonadota | Gammaproteobacteria | NZ_CP020038.1 | 3 |
| Agarivorans aestuarii | Pseudomonadota | Gammaproteobacteria | NZ_AP023033.1 | 2 |
| Agarivorans albus | Pseudomonadota | Gammaproteobacteria | NZ_AP023032.1 | 3 |
| Aggregatibacter actinomycetemcomitans | Pseudomonadota | Gammaproteobacteria | NZ_CP085096.1 | 2 |
| Aggregatibacter aphrophilus ATCC 33389 | Pseudomonadota | Gammaproteobacteria | NZ_LR134327.1 | 2 |
| Aggregatibacter segnis ATCC 33393 | Pseudomonadota | Gammaproteobacteria | NZ_LS483443.1 | 2 |
| Agrilutibacter terrestris | Pseudomonadota | Gammaproteobacteria | NZ_CP060820.1 | 2 |
| Ahniella affigens | Pseudomonadota | Gammaproteobacteria | NZ_CP027860.1 | 2 |
| Alcanivorax borkumensis SK2 | Pseudomonadota | Gammaproteobacteria | NC_008260.1 | 2 |
| Aliamphritea ceti | Pseudomonadota | Gammaproteobacteria | NZ_AP025282.1 | 2 |
| Aliamphritea hakodatensis | Pseudomonadota | Gammaproteobacteria | NZ_AP025281.1 | 2 |
| Aliivibrio fischeri ES114 | Pseudomonadota | Gammaproteobacteria | NC_006840.2 | 2 |
| Aliivibrio salmonicida LFI1238 | Pseudomonadota | Gammaproteobacteria | NC_011312.1 | 2 |
| Alkalilimnicola ehrlichii MLHE-1 | Pseudomonadota | Gammaproteobacteria | NC_008340.1 | 2 |
| Alkalimarinus alittae | Pseudomonadota | Gammaproteobacteria | NZ_CP100390.1 | 1 |
| Alkalimarinus coralli | Pseudomonadota | Gammaproteobacteria | NZ_CP096016.1 | 1 |
| Alkalimarinus sediminis | Pseudomonadota | Gammaproteobacteria | NZ_CP101527.1 | 1 |
| Alloalcanivorax dieselolei B5 | Pseudomonadota | Gammaproteobacteria | NC_018691.1 | 2 |
| Alloalcanivorax venustensis | Pseudomonadota | Gammaproteobacteria | NZ_CP179934.1 | 2 |
| Alloalcanivorax xenomutans | Pseudomonadota | Gammaproteobacteria | NZ_CP136538.1 | 2 |
| Allochromatium tepidum | Pseudomonadota | Gammaproteobacteria | NZ_AP024563.1 | 2 |
| Allochromatium vinosum DSM 180 | Pseudomonadota | Gammaproteobacteria | NC_013851.1 | 2 |
| Allofrancisella frigidaquae | Pseudomonadota | Gammaproteobacteria | NZ_CP038017.1 | 1 |
| Allofrancisella guangzhouensis | Pseudomonadota | Gammaproteobacteria | NZ_CP010427.1 | 1 |
| Allofrancisella inopinata | Pseudomonadota | Gammaproteobacteria | NZ_CP038241.1 | 1 |
| Alteromonas macleodii | Pseudomonadota | Gammaproteobacteria | NZ_CP018321.1 | 2 |
| Alteromonas naphthalenivorans | Pseudomonadota | Gammaproteobacteria | NC_015554.1 | 2 |

| | | | | |
|---|---|---|---|---|
| Alteromonas pelagimontana | Pseudomonadota | Gammaproteobacteria | NZ_CP052766.1 | 2 |
| Alteromonas stellipolaris | Pseudomonadota | Gammaproteobacteria | NZ_CP014322.1 | 1 |
| Amphritea atlantica | Pseudomonadota | Gammaproteobacteria | NZ_AP025284.1 | 2 |
| Amphritea japonica | Pseudomonadota | Gammaproteobacteria | NZ_AP025761.1 | 2 |
| Aquicella lusitana | Pseudomonadota | Gammaproteobacteria | NZ_LR699114.1 | 2 |
| Aquicella siphonis | Pseudomonadota | Gammaproteobacteria | NZ_LR699119.1 | 2 |
| Aquipseudomonas alcaligenes | Pseudomonadota | Gammaproteobacteria | NZ_CP014784.1 | 2 |
| Aquipseudomonas campi | Pseudomonadota | Gammaproteobacteria | NZ_CP053697.2 | 2 |
| Aquirhabdus parva | Pseudomonadota | Gammaproteobacteria | NZ_CP031222.1 | 2 |
| Arsenophonus apicola | Pseudomonadota | Gammaproteobacteria | NZ_CP084222.1 | 2 |
| Arsenophonus endosymbiont of Aleurodicus dispersus | Pseudomonadota | Gammaproteobacteria | NZ_LR025108.1 | 1 |
| Arsenophonus endosymbiont of Aphis craccivora | Pseudomonadota | Gammaproteobacteria | NZ_CP038155.1 | 2 |
| Arsenophonus endosymbiont of Crataerina pallida | Pseudomonadota | Gammaproteobacteria | NZ_OZ026540.1 | 2 |
| Arsenophonus endosymbiont of Lipoptena cervi | Pseudomonadota | Gammaproteobacteria | NZ_OZ195522.1 | 1 |
| Arsenophonus nasoniae | Pseudomonadota | Gammaproteobacteria | NZ_CP038613.1 | 2 |
| Arsenophonus symbiont of Ornithomya chloropus | Pseudomonadota | Gammaproteobacteria | NZ_OZ032148.1 | 1 |
| Atlantibacter hermannii | Pseudomonadota | Gammaproteobacteria | NZ_CP065700.1 | 2 |
| Atlantibacter subterraneus | Pseudomonadota | Gammaproteobacteria | NZ_CP100494.1 | 2 |
| Avibacterium paragallinarum | Pseudomonadota | Gammaproteobacteria | NZ_CP050316.1 | 2 |
| Avibacterium volantium | Pseudomonadota | Gammaproteobacteria | NZ_LR134167.1 | 2 |
| Azorhizophilus paspali | Pseudomonadota | Gammaproteobacteria | NZ_CP171449.1 | 2 |
| Azotobacter chroococcum | Pseudomonadota | Gammaproteobacteria | NZ_CP011835.1 | 2 |
| Azotobacter salinestris | Pseudomonadota | Gammaproteobacteria | NZ_CP045302.1 | 2 |
| Azotobacter vinelandii DJ | Pseudomonadota | Gammaproteobacteria | NC_012560.1 | 2 |
| Basfia succiniciproducens | Pseudomonadota | Gammaproteobacteria | NZ_CP015031.1 | 2 |
| Bathymodiolus thermophilus thioautotrophic gill symbiont | Pseudomonadota | Gammaproteobacteria | NZ_CP024634.1 | 2 |
| Beggiatoa leptomitoformis | Pseudomonadota | Gammaproteobacteria | NZ_CP012373.2 | 2 |
| Bermanella marisrubri | Pseudomonadota | Gammaproteobacteria | NZ_CP051183.1 | 2 |
| Bibersteinia trehalosi USDA-ARS-USMARC-188 | Pseudomonadota | Gammaproteobacteria | NZ_CP006954.1 | 2 |
| Billgrantia sulfidoxydans | Pseudomonadota | Gammaproteobacteria | NZ_CP053381.1 | 1 |
| Bisgaardia hudsonensis | Pseudomonadota | Gammaproteobacteria | NZ_CP016605.1 | 2 |
| Blochmannia endosymbiont of Camponotus (Colobopsis) c | Pseudomonadota | Gammaproteobacteria | NZ_CP010049.1 | 1 |

| | | | | |
|---|---|---|---|---|
| Blochmannia endosymbiont of Camponotus modoc | Pseudomonadota | Gammaproteobacteria | NZ_CP097765.1 | 1 |
| Blochmannia endosymbiont of Camponotus nipponensis | Pseudomonadota | Gammaproteobacteria | NZ_CP046534.1 | 1 |
| Blochmannia endosymbiont of Camponotus sp. | Pseudomonadota | Gammaproteobacteria | NZ_CP097760.1 | 1 |
| Blochmannia endosymbiont of Camponotus sp. C-046 | Pseudomonadota | Gammaproteobacteria | NZ_CP097752.1 | 1 |
| Blochmannia endosymbiont of Colobopsis nipponica | Pseudomonadota | Gammaproteobacteria | NZ_CP046533.1 | 1 |
| Blochmannia endosymbiont of Polyrhachis (Hedomyrma) t | Pseudomonadota | Gammaproteobacteria | NZ_CP010048.1 | 1 |
| Brenneria goodwinii | Pseudomonadota | Gammaproteobacteria | NZ_CP014137.1 | 2 |
| Brenneria izadpanahii | Pseudomonadota | Gammaproteobacteria | NZ_CP050854.1 | 2 |
| Brenneria nigrifluens DSM 30175 = ATCC 13028 | Pseudomonadota | Gammaproteobacteria | NZ_CP034036.1 | 2 |
| Brenneria uluponensis | Pseudomonadota | Gammaproteobacteria | NZ_CP135917.1 | 2 |
| Brumicola nitratireducens FR1064 | Pseudomonadota | Gammaproteobacteria | NC_016041.1 | 2 |
| Buchnera aphidicola str. Sg (Schizaphis graminum) | Pseudomonadota | Gammaproteobacteria | NC_004061.1 | 1 |
| Buttiauxella ferragutiae | Pseudomonadota | Gammaproteobacteria | NZ_CP093332.1 | 2 |
| Buttiauxella selenatireducens | Pseudomonadota | Gammaproteobacteria | NZ_CP133838.1 | 2 |
| Caldichromatium japonicum | Pseudomonadota | Gammaproteobacteria | NZ_CP048029.1 | 2 |
| Candidatus Annandia adelgestsuga | Pseudomonadota | Gammaproteobacteria | NZ_CP026513.1 | 1 |
| Candidatus Annandia pinicola | Pseudomonadota | Gammaproteobacteria | NZ_CP045876.1 | 1 |
| Candidatus Arsenophonus lipoptenae | Pseudomonadota | Gammaproteobacteria | NZ_CP013920.1 | 1 |
| Candidatus Arsenophonus nilaparvatae | Pseudomonadota | Gammaproteobacteria | NZ_CP158507.1 | 2 |
| Candidatus Azoamicus ciliaticola | Pseudomonadota | Gammaproteobacteria | NZ_LR794158.1 | 1 |
| Candidatus Blochmanniella camponoti | Pseudomonadota | Gammaproteobacteria | NZ_CP097750.1 | 1 |
| Candidatus Blochmanniella chromaiodes str. 640 | Pseudomonadota | Gammaproteobacteria | NC_020075.1 | 1 |
| Candidatus Blochmanniella pennsylvanica str. BPEN | Pseudomonadota | Gammaproteobacteria | NC_007292.1 | 1 |
| Candidatus Blochmanniella vafra str. BVAF | Pseudomonadota | Gammaproteobacteria | NC_014909.2 | 1 |
| Candidatus Carsonella ruddii HC isolate Thao2000 | Pseudomonadota | Gammaproteobacteria | NC_018416.1 | 1 |
| Candidatus Comchoanobacter bicostacola | Pseudomonadota | Gammaproteobacteria | NZ_CP092900.1 | 2 |
| Candidatus Coxiella mudrowiae | Pseudomonadota | Gammaproteobacteria | NZ_CP011126.1 | 2 |
| Candidatus Curculioniphilus buchneri endosymbiont of Cur | Pseudomonadota | Gammaproteobacteria | NZ_OZ193413.1 | 1 |
| Candidatus Doolittlea endobia | Pseudomonadota | Gammaproteobacteria | NZ_LN999833.1 | 1 |
| Candidatus Endoriftia persephonae | Pseudomonadota | Gammaproteobacteria | NZ_CP090569.1 | 2 |
| Candidatus Enterovibrio altilux | Pseudomonadota | Gammaproteobacteria | NZ_CP020660.1 | 1 |
| Candidatus Erwinia haradaeae | Pseudomonadota | Gammaproteobacteria | NZ_LR217713.1 | 1 |

| | | | | |
|---|---|---|---|---|
| Candidatus Francisella endociliophora | Pseudomonadota | Gammaproteobacteria | NZ_CP009574.1 | 1 |
| Candidatus Fukatsuia endosymbiont of Tuberolachnus sali | Pseudomonadota | Gammaproteobacteria | NZ_OZ034983.1 | 1 |
| Candidatus Gullanella endobia | Pseudomonadota | Gammaproteobacteria | NZ_LN999832.1 | 2 |
| Candidatus Hamiltonella defensa 5AT (Acyrthosiphon pisu | Pseudomonadota | Gammaproteobacteria | NC_012751.1 | 1 |
| Candidatus Hoaglandella endobia | Pseudomonadota | Gammaproteobacteria | NZ_LN999835.1 | 1 |
| Candidatus Ishikawaella capsulata Mpkobe | Pseudomonadota | Gammaproteobacteria | NZ_AP010872.1 | 2 |
| Candidatus Legionella polyplacis | Pseudomonadota | Gammaproteobacteria | NZ_CP021497.1 | 1 |
| Candidatus Methylocalor cossyra | Pseudomonadota | Gammaproteobacteria | NZ_OZ026884.1 | 2 |
| Candidatus Methylospira mobilis | Pseudomonadota | Gammaproteobacteria | NZ_CP044205.1 | 2 |
| Candidatus Mikella endobia | Pseudomonadota | Gammaproteobacteria | NZ_LN999831.1 | 1 |
| Candidatus Moranella endobia PCVAL | Pseudomonadota | Gammaproteobacteria | NC_021057.1 | 1 |
| Candidatus Nitrosacidococcus tergens | Pseudomonadota | Gammaproteobacteria | NZ_LR778175.1 | 2 |
| Candidatus Nitrosoglobus terrae | Pseudomonadota | Gammaproteobacteria | NZ_AP014836.1 | 2 |
| Candidatus Palibaumannia cicadellinicola | Pseudomonadota | Gammaproteobacteria | NZ_CP008985.1 | 1 |
| Candidatus Pantoea carbekii | Pseudomonadota | Gammaproteobacteria | NZ_CP010907.1 | 1 |
| Candidatus Pantoea soli | Pseudomonadota | Gammaproteobacteria | NZ_CP032702.1 | 2 |
| Candidatus Paraluminiphilus aquimaris | Pseudomonadota | Gammaproteobacteria | NZ_CP036501.1 | 1 |
| Candidatus Portiera aleyrodidarum BT-B-HRs | Pseudomonadota | Gammaproteobacteria | NC_018507.1 | 1 |
| Candidatus Profftia lariciata | Pseudomonadota | Gammaproteobacteria | NZ_CP045925.1 | 1 |
| Candidatus Profftia tarda | Pseudomonadota | Gammaproteobacteria | NZ_LR890047.1 | 1 |
| Candidatus Providencia siddallii | Pseudomonadota | Gammaproteobacteria | NZ_OZ034688.1 | 1 |
| Candidatus Pseudomonas adelgestsugas | Pseudomonadota | Gammaproteobacteria | NZ_CP026512.1 | 1 |
| Candidatus Pseudothioglobus singularis PS1 | Pseudomonadota | Gammaproteobacteria | NZ_CP006911.1 | 2 |
| Candidatus Purcelliella pentastirinorum | Pseudomonadota | Gammaproteobacteria | NZ_CP028374.1 | 1 |
| Candidatus Regiella endosymbiont of Tuberolachnus saligı | Pseudomonadota | Gammaproteobacteria | NZ_OZ026542.1 | 1 |
| Candidatus Reidiella endopervernicosa | Pseudomonadota | Gammaproteobacteria | NZ_CP054491.1 | 2 |
| Candidatus Rickettsiella viridis | Pseudomonadota | Gammaproteobacteria | NZ_AP018005.1 | 1 |
| Candidatus Riesia pediculicola USDA | Pseudomonadota | Gammaproteobacteria | NC_014109.1 | 1 |
| Candidatus Riesia pediculischaeffi | Pseudomonadota | Gammaproteobacteria | NZ_CP012839.1 | 1 |
| Candidatus Ruthia endofausta | Pseudomonadota | Gammaproteobacteria | NZ_CP054490.1 | 2 |
| Candidatus Ruthia magnifica str. Cm (Calyptogena magnifi | Pseudomonadota | Gammaproteobacteria | NC_008610.1 | 2 |
| Candidatus Schneideria nysicola | Pseudomonadota | Gammaproteobacteria | NZ_CP074435.1 | 1 |

| | | | | |
|---|---|---|---|---|
| Candidatus Sodalis pierantonii str. SOPE | Pseudomonadota | Gammaproteobacteria | NZ_CP006568.1 | 2 |
| Candidatus Steffania adelgidicola | Pseudomonadota | Gammaproteobacteria | NZ_CP045889.1 | 1 |
| Candidatus Tachikawaea gelatinosa | Pseudomonadota | Gammaproteobacteria | NZ_AP014521.1 | 2 |
| Candidatus Thalassolituus haligoni | Pseudomonadota | Gammaproteobacteria | NZ_CP139482.1 | 2 |
| Candidatus Thiodictyon syntrophicum | Pseudomonadota | Gammaproteobacteria | NZ_CP020370.1 | 2 |
| Candidatus Thiodubiliella endoseptemdiera | Pseudomonadota | Gammaproteobacteria | NZ_OZ156463.1 | 2 |
| Candidatus Thioglobus autotrophicus | Pseudomonadota | Gammaproteobacteria | NZ_CP010552.1 | 2 |
| Candidatus Thiothrix anitrata | Pseudomonadota | Gammaproteobacteria | NZ_CP072800.1 | 2 |
| Candidatus Vesicomyidisocius calyptogenae | Pseudomonadota | Gammaproteobacteria | NC_009465.1 | 2 |
| Candidatus Westeberhardia cardiocondylae | Pseudomonadota | Gammaproteobacteria | NZ_LN774881.1 | 1 |
| Candidatus Williamhamiltonella endosymbiont of Tuberola | Pseudomonadota | Gammaproteobacteria | NZ_OZ026471.1 | 1 |
| Cardiobacterium hominis | Pseudomonadota | Gammaproteobacteria | NZ_LR134365.1 | 2 |
| Catenovulum adriaticum | Pseudomonadota | Gammaproteobacteria | NZ_CP109965.1 | 2 |
| Catenovulum sediminis | Pseudomonadota | Gammaproteobacteria | NZ_CP041660.1 | 2 |
| Cedecea davisae | Pseudomonadota | Gammaproteobacteria | NZ_CP172054.1 | 2 |
| Cedecea lapagei | Pseudomonadota | Gammaproteobacteria | NZ_LR134201.1 | 2 |
| Cedecea neteri | Pseudomonadota | Gammaproteobacteria | NZ_CP023525.1 | 2 |
| Celerinatantimonas diazotrophica | Pseudomonadota | Gammaproteobacteria | NZ_OU594967.1 | 2 |
| Cellvibrio japonicus Ueda107 | Pseudomonadota | Gammaproteobacteria | NC_010995.1 | 2 |
| Chania multitudinisentens RB-25 | Pseudomonadota | Gammaproteobacteria | NZ_CP007044.2 | 2 |
| Chromohalobacter canadensis | Pseudomonadota | Gammaproteobacteria | NZ_CP140151.1 | 1 |
| Chromohalobacter israelensis DSM 3043 | Pseudomonadota | Gammaproteobacteria | NC_007963.1 | 1 |
| Citrobacter amalonaticus | Pseudomonadota | Gammaproteobacteria | NZ_CP083651.1 | 2 |
| Citrobacter arsenatis | Pseudomonadota | Gammaproteobacteria | NZ_CP037864.1 | 2 |
| Citrobacter braakii | Pseudomonadota | Gammaproteobacteria | NZ_CP045771.1 | 2 |
| Citrobacter cronae | Pseudomonadota | Gammaproteobacteria | NZ_CP069763.1 | 2 |
| Citrobacter europaeus | Pseudomonadota | Gammaproteobacteria | NZ_CP148084.1 | 2 |
| Citrobacter farmeri | Pseudomonadota | Gammaproteobacteria | NZ_CP077291.1 | 2 |
| Citrobacter freundii | Pseudomonadota | Gammaproteobacteria | NZ_LR890181.1 | 2 |
| Citrobacter koseri ATCC BAA-895 | Pseudomonadota | Gammaproteobacteria | NC_009792.1 | 2 |
| Citrobacter meridianamericanus | Pseudomonadota | Gammaproteobacteria | NZ_CP101036.1 | 2 |
| Citrobacter pasteurii | Pseudomonadota | Gammaproteobacteria | NZ_CP077262.1 | 2 |

| | | | | |
|---|---|---|---|---|
| Citrobacter portucalensis | Pseudomonadota | Gammaproteobacteria | NZ_CP092466.1 | 2 |
| Citrobacter rodentium NBRC 105723 = DSM 16636 | Pseudomonadota | Gammaproteobacteria | NZ_CP082833.1 | 2 |
| Citrobacter sedlakii | Pseudomonadota | Gammaproteobacteria | NZ_CP071070.1 | 2 |
| Citrobacter tructae | Pseudomonadota | Gammaproteobacteria | NZ_CP038469.1 | 2 |
| Citrobacter werkmanii | Pseudomonadota | Gammaproteobacteria | NZ_CP044101.1 | 2 |
| Cobetia amphilecti | Pseudomonadota | Gammaproteobacteria | NZ_CP136695.1 | 1 |
| Cobetia marina | Pseudomonadota | Gammaproteobacteria | NZ_CP017114.1 | 1 |
| Cognaticolwellia beringensis | Pseudomonadota | Gammaproteobacteria | NZ_CP020465.1 | 2 |
| Colwellia psychrerythraea 34H | Pseudomonadota | Gammaproteobacteria | NC_003910.7 | 2 |
| Congregibacter brevis | Pseudomonadota | Gammaproteobacteria | NZ_CP136865.1 | 2 |
| Congregibacter litoralis KT71 | Pseudomonadota | Gammaproteobacteria | NZ_CM002299.1 | 2 |
| Congregibacter variabilis | Pseudomonadota | Gammaproteobacteria | NZ_CP136864.1 | 2 |
| Coxiella burnetii RSA 493 | Pseudomonadota | Gammaproteobacteria | NC_002971.4 | 1 |
| Coxiella endosymbiont of Amblyomma americanum | Pseudomonadota | Gammaproteobacteria | NZ_CP007541.1 | 2 |
| Coxiella endosymbiont of Amblyomma nuttalli | Pseudomonadota | Gammaproteobacteria | NZ_CP064834.1 | 2 |
| Coxiella endosymbiont of Amblyomma sculptum | Pseudomonadota | Gammaproteobacteria | NZ_CP033868.1 | 1 |
| Coxiella-like endosymbiont of Amblyomma americanum | Pseudomonadota | Gammaproteobacteria | NZ_CP021379.1 | 2 |
| Cronobacter condimenti 1330 | Pseudomonadota | Gammaproteobacteria | NZ_CP012264.1 | 2 |
| Cronobacter dublinensis subsp. dublinensis LMG 23823 | Pseudomonadota | Gammaproteobacteria | NZ_CP012266.1 | 2 |
| Cronobacter malonaticus LMG 23826 | Pseudomonadota | Gammaproteobacteria | NZ_CP013940.1 | 2 |
| Cronobacter muytjensii ATCC 51329 | Pseudomonadota | Gammaproteobacteria | NZ_CP012268.1 | 2 |
| Cronobacter sakazakii | Pseudomonadota | Gammaproteobacteria | NZ_CP027107.1 | 2 |
| Cronobacter turicensis | Pseudomonadota | Gammaproteobacteria | NZ_CP165997.1 | 2 |
| Cronobacter universalis NCTC 9529 | Pseudomonadota | Gammaproteobacteria | NZ_CP012257.1 | 2 |
| Dickeya aquatica | Pseudomonadota | Gammaproteobacteria | NZ_LT615367.1 | 2 |
| Dickeya chrysanthemi Ech1591 | Pseudomonadota | Gammaproteobacteria | NC_012912.1 | 2 |
| Dickeya dadantii | Pseudomonadota | Gammaproteobacteria | NZ_CP023467.1 | 2 |
| Dickeya dianthicola | Pseudomonadota | Gammaproteobacteria | NZ_CP031560.1 | 2 |
| Dickeya fangzhongdai | Pseudomonadota | Gammaproteobacteria | NZ_CP025003.1 | 2 |
| Dickeya oryzae | Pseudomonadota | Gammaproteobacteria | NZ_CP162670.1 | 2 |
| Dickeya parazeae Ech586 | Pseudomonadota | Gammaproteobacteria | NC_013592.1 | 2 |
| Dickeya poaceiphila | Pseudomonadota | Gammaproteobacteria | NZ_CP042220.2 | 2 |

| | | | | |
|---|---|---|---|---|
| Dickeya solani IPO 2222 | Pseudomonadota | Gammaproteobacteria | NZ_CP015137.1 | 2 |
| Dickeya zeae | Pseudomonadota | Gammaproteobacteria | NZ_CP025799.1 | 2 |
| Dokdonella koreensis DS-123 | Pseudomonadota | Gammaproteobacteria | NZ_CP015249.1 | 2 |
| Dongshaea marina | Pseudomonadota | Gammaproteobacteria | NZ_CP028897.1 | 1 |
| Duffyella gerundensis | Pseudomonadota | Gammaproteobacteria | NZ_CP073262.1 | 2 |
| Dyella aluminiiresistens | Pseudomonadota | Gammaproteobacteria | NZ_CP132911.1 | 2 |
| Dyella caseinilytica | Pseudomonadota | Gammaproteobacteria | NZ_CP064030.1 | 2 |
| Dyella telluris | Pseudomonadota | Gammaproteobacteria | NZ_CP060412.1 | 2 |
| Dyella thiooxydans | Pseudomonadota | Gammaproteobacteria | NZ_CP014841.1 | 2 |
| Ectopseudomonas hydrolytica | Pseudomonadota | Gammaproteobacteria | NZ_CP099397.1 | 2 |
| Ectopseudomonas khazarica | Pseudomonadota | Gammaproteobacteria | NZ_CP068551.1 | 2 |
| Ectopseudomonas mendocina S5.2 | Pseudomonadota | Gammaproteobacteria | NZ_CP013124.1 | 2 |
| Ectopseudomonas toyotomiensis | Pseudomonadota | Gammaproteobacteria | NZ_CP070505.1 | 2 |
| Edwardsiella anguillarum ET080813 | Pseudomonadota | Gammaproteobacteria | NZ_CP006664.1 | 2 |
| Edwardsiella hoshinae | Pseudomonadota | Gammaproteobacteria | NZ_CP065626.1 | 2 |
| Edwardsiella ictaluri | Pseudomonadota | Gammaproteobacteria | NZ_CP084521.1 | 2 |
| Edwardsiella piscicida | Pseudomonadota | Gammaproteobacteria | NZ_CP090968.1 | 2 |
| Edwardsiella tarda | Pseudomonadota | Gammaproteobacteria | NZ_CP082939.1 | 2 |
| endosymbiont of Bathymodiolus septemdierum str. Myojin | Pseudomonadota | Gammaproteobacteria | NZ_AP013042.1 | 2 |
| endosymbiont of Euscepes postfasciatus | Pseudomonadota | Gammaproteobacteria | NZ_AP018159.1 | 1 |
| endosymbiont of Pachyrhynchus infernalis | Pseudomonadota | Gammaproteobacteria | NZ_AP018160.1 | 1 |
| endosymbiont of Rhynchophorus ferrugineus | Pseudomonadota | Gammaproteobacteria | NZ_AP018161.1 | 1 |
| endosymbiont of Sipalinus gigas | Pseudomonadota | Gammaproteobacteria | NZ_AP018162.1 | 1 |
| endosymbiont of unidentified scaly snail isolate Monju | Pseudomonadota | Gammaproteobacteria | NZ_AP012978.1 | 2 |
| Endozoicomonas euniceicola | Pseudomonadota | Gammaproteobacteria | NZ_CP103300.1 | 1 |
| Endozoicomonas montiporae CL-33 | Pseudomonadota | Gammaproteobacteria | NZ_CP013251.1 | 1 |
| Enterobacter adelaidei | Pseudomonadota | Gammaproteobacteria | NZ_CP154353.1 | 2 |
| Enterobacter asburiae | Pseudomonadota | Gammaproteobacteria | NZ_AP019630.1 | 2 |
| Enterobacter bugandensis | Pseudomonadota | Gammaproteobacteria | NZ_LT992502.1 | 2 |
| Enterobacter cancerogenus | Pseudomonadota | Gammaproteobacteria | NZ_CP081105.1 | 2 |
| Enterobacter chengduensis | Pseudomonadota | Gammaproteobacteria | NZ_CP043318.1 | 2 |
| Enterobacter chuandaensis | Pseudomonadota | Gammaproteobacteria | NZ_CP097173.1 | 2 |

| | | | | |
|---|---|---|---|---|
| Enterobacter cloacae | Pseudomonadota | Gammaproteobacteria | NZ_OW968328.1 | 2 |
| Enterobacter dykesii | Pseudomonadota | Gammaproteobacteria | NZ_CP126604.1 | 2 |
| Enterobacter hormaechei | Pseudomonadota | Gammaproteobacteria | NZ_CP077308.1 | 2 |
| Enterobacter huaxiensis | Pseudomonadota | Gammaproteobacteria | NZ_CP043342.1 | 2 |
| Enterobacter intestinihominis | Pseudomonadota | Gammaproteobacteria | NZ_CP183772.1 | 2 |
| Enterobacter kobei | Pseudomonadota | Gammaproteobacteria | NZ_CP083828.1 | 2 |
| Enterobacter ludwigii | Pseudomonadota | Gammaproteobacteria | NZ_CP017279.1 | 2 |
| Enterobacter mori | Pseudomonadota | Gammaproteobacteria | NZ_CP091779.1 | 2 |
| Enterobacter oligotrophicus | Pseudomonadota | Gammaproteobacteria | NZ_AP019007.1 | 2 |
| Enterobacter pseudoroggenkampii | Pseudomonadota | Gammaproteobacteria | NZ_CP129026.1 | 2 |
| Enterobacter quasiroggenkampii | Pseudomonadota | Gammaproteobacteria | NZ_CP143745.1 | 2 |
| Enterobacter roggenkampii | Pseudomonadota | Gammaproteobacteria | NZ_CP017184.1 | 2 |
| Enterobacter sichuanensis | Pseudomonadota | Gammaproteobacteria | NZ_CP027986.1 | 2 |
| Enterobacter soli | Pseudomonadota | Gammaproteobacteria | NC_015968.1 | 2 |
| Enterobacter vonholyi | Pseudomonadota | Gammaproteobacteria | NZ_CP162152.1 | 2 |
| Enterobacter wuhouensis | Pseudomonadota | Gammaproteobacteria | NZ_CP142124.1 | 2 |
| Enterobacteriaceae endosymbiont of Donacia bicoloricorni | Pseudomonadota | Gammaproteobacteria | NZ_CP046186.1 | 1 |
| Enterobacteriaceae endosymbiont of Donacia cincticornis | Pseudomonadota | Gammaproteobacteria | NZ_CP046194.1 | 1 |
| Enterobacteriaceae endosymbiont of Donacia cinerea | Pseudomonadota | Gammaproteobacteria | NZ_CP046204.1 | 1 |
| Enterobacteriaceae endosymbiont of Donacia clavipes | Pseudomonadota | Gammaproteobacteria | NZ_CP046208.1 | 1 |
| Enterobacteriaceae endosymbiont of Donacia crassipes | Pseudomonadota | Gammaproteobacteria | NZ_CP046202.1 | 1 |
| Enterobacteriaceae endosymbiont of Donacia dentata | Pseudomonadota | Gammaproteobacteria | NZ_CP046212.1 | 1 |
| Enterobacteriaceae endosymbiont of Donacia fulgens | Pseudomonadota | Gammaproteobacteria | NZ_CP046182.1 | 1 |
| Enterobacteriaceae endosymbiont of Donacia marginata | Pseudomonadota | Gammaproteobacteria | NZ_CP046184.1 | 1 |
| Enterobacteriaceae endosymbiont of Donacia piscatrix | Pseudomonadota | Gammaproteobacteria | NZ_CP046200.1 | 1 |
| Enterobacteriaceae endosymbiont of Donacia provostii | Pseudomonadota | Gammaproteobacteria | NZ_CP046206.1 | 1 |
| Enterobacteriaceae endosymbiont of Donacia proxima | Pseudomonadota | Gammaproteobacteria | NZ_CP046198.1 | 1 |
| Enterobacteriaceae endosymbiont of Donacia semicuprea | Pseudomonadota | Gammaproteobacteria | NZ_CP046210.1 | 1 |
| Enterobacteriaceae endosymbiont of Donacia simplex | Pseudomonadota | Gammaproteobacteria | NZ_CP046192.1 | 1 |
| Enterobacteriaceae endosymbiont of Donacia sparganii | Pseudomonadota | Gammaproteobacteria | NZ_CP046196.1 | 1 |
| Enterobacteriaceae endosymbiont of Donacia thalassina | Pseudomonadota | Gammaproteobacteria | NZ_CP046188.1 | 1 |
| Enterobacteriaceae endosymbiont of Donacia tomentosa | Pseudomonadota | Gammaproteobacteria | NZ_CP046216.1 | 1 |

| | | | | |
|---|---|---|---|---|
| Enterobacteriaceae endosymbiont of Donacia versicolorea | Pseudomonadota | Gammaproteobacteria | NZ_CP046214.1 | 1 |
| Enterobacteriaceae endosymbiont of Donacia vulgaris | Pseudomonadota | Gammaproteobacteria | NZ_CP046190.1 | 1 |
| Enterobacteriaceae endosymbiont of Macroplea appendicu | Pseudomonadota | Gammaproteobacteria | NZ_CP046220.1 | 1 |
| Enterobacteriaceae endosymbiont of Macroplea mutica | Pseudomonadota | Gammaproteobacteria | NZ_CP046218.1 | 1 |
| Enterobacteriaceae endosymbiont of Neohaemonia nigrico | Pseudomonadota | Gammaproteobacteria | NZ_CP046222.1 | 1 |
| Enterobacteriaceae endosymbiont of Plateumaris braccata | Pseudomonadota | Gammaproteobacteria | NZ_CP046232.1 | 1 |
| Enterobacteriaceae endosymbiont of Plateumaris consimili | Pseudomonadota | Gammaproteobacteria | NZ_CP046230.1 | 1 |
| Enterobacteriaceae endosymbiont of Plateumaris pusilla | Pseudomonadota | Gammaproteobacteria | NZ_CP046226.1 | 1 |
| Enterobacteriaceae endosymbiont of Plateumaris rustica | Pseudomonadota | Gammaproteobacteria | NZ_CP046228.1 | 1 |
| Enterobacteriaceae endosymbiont of Plateumaris sericea | Pseudomonadota | Gammaproteobacteria | NZ_CP046224.1 | 1 |
| Entomomonas asaccharolytica | Pseudomonadota | Gammaproteobacteria | NZ_CP067393.1 | 1 |
| Entomomonas moraniae | Pseudomonadota | Gammaproteobacteria | NZ_CP029822.1 | 1 |
| Erwinia amylovora | Pseudomonadota | Gammaproteobacteria | NZ_CP171267.1 | 2 |
| Erwinia billingiae Eb661 | Pseudomonadota | Gammaproteobacteria | NC_014306.1 | 2 |
| Erwinia persicina | Pseudomonadota | Gammaproteobacteria | NZ_CP082141.1 | 2 |
| Erwinia pyri | Pseudomonadota | Gammaproteobacteria | NZ_CP132353.1 | 2 |
| Erwinia pyrifoliae | Pseudomonadota | Gammaproteobacteria | NZ_CP023567.1 | 2 |
| Erwinia rhapontici | Pseudomonadota | Gammaproteobacteria | NZ_CP085627.1 | 2 |
| Erwinia sorbitola | Pseudomonadota | Gammaproteobacteria | NZ_CP046509.1 | 2 |
| Erwinia tasmaniensis Et1/99 | Pseudomonadota | Gammaproteobacteria | NC_010694.1 | 2 |
| Erwinia tracheiphila | Pseudomonadota | Gammaproteobacteria | NZ_CP089932.1 | 2 |
| Escherichia albertii | Pseudomonadota | Gammaproteobacteria | NZ_CP117568.1 | 2 |
| Escherichia coli O157:H7 str. Sakai | Pseudomonadota | Gammaproteobacteria | NC_002695.2 | 2 |
| Escherichia coli str. K-12 substr. MG1655 | Pseudomonadota | Gammaproteobacteria | NC_000913.3 | 2 |
| Escherichia fergusonii | Pseudomonadota | Gammaproteobacteria | NZ_CP083638.1 | 2 |
| Escherichia marmotae | Pseudomonadota | Gammaproteobacteria | NZ_CP072689.1 | 2 |
| Faucicola mancuniensis | Pseudomonadota | Gammaproteobacteria | NZ_CP171363.1 | 2 |
| Ferrimonas balearica DSM 9799 | Pseudomonadota | Gammaproteobacteria | NC_014541.1 | 2 |
| Ferrimonas lipolytica | Pseudomonadota | Gammaproteobacteria | NZ_CP051180.1 | 2 |
| Flagellatimonas centrodinii | Pseudomonadota | Gammaproteobacteria | NZ_CP092104.1 | 2 |
| Flocculibacter collagenilyticus | Pseudomonadota | Gammaproteobacteria | NZ_CP059888.1 | 2 |
| Francisella adeliensis | Pseudomonadota | Gammaproteobacteria | NZ_CP021781.1 | 1 |

| | | | | |
|---|---|---|---|---|
| Francisella frigiditurris | Pseudomonadota | Gammaproteobacteria | NZ_CP009654.1 | 1 |
| Francisella halioticida | Pseudomonadota | Gammaproteobacteria | NZ_CP022132.1 | 1 |
| Francisella hispaniensis | Pseudomonadota | Gammaproteobacteria | NC_017449.1 | 1 |
| Francisella marina | Pseudomonadota | Gammaproteobacteria | NZ_CP043552.1 | 1 |
| Francisella opportunistica | Pseudomonadota | Gammaproteobacteria | NZ_CP022377.1 | 1 |
| Francisella orientalis LADL 07-285A | Pseudomonadota | Gammaproteobacteria | NC_023029.1 | 1 |
| Francisella persica ATCC VR-331 | Pseudomonadota | Gammaproteobacteria | NZ_CP012505.1 | 1 |
| Francisella philomiragia | Pseudomonadota | Gammaproteobacteria | NZ_CP063138.1 | 1 |
| Francisella salimarina | Pseudomonadota | Gammaproteobacteria | NZ_CP076680.1 | 1 |
| Francisella salina | Pseudomonadota | Gammaproteobacteria | NC_015696.1 | 1 |
| Francisella tularensis subsp. novicida D9876 | Pseudomonadota | Gammaproteobacteria | NZ_CP009607.1 | 1 |
| Francisella uliginis | Pseudomonadota | Gammaproteobacteria | NZ_CP016796.1 | 1 |
| Frateuria aurantia DSM 6220 | Pseudomonadota | Gammaproteobacteria | NC_017033.1 | 2 |
| Frateuria edaphi | Pseudomonadota | Gammaproteobacteria | NZ_CP088251.1 | 2 |
| Frateuria soli | Pseudomonadota | Gammaproteobacteria | NZ_CP088252.1 | 2 |
| Frederiksenia canicola | Pseudomonadota | Gammaproteobacteria | NZ_CP015029.1 | 2 |
| Frischella perrara | Pseudomonadota | Gammaproteobacteria | NZ_CP009056.1 | 2 |
| Gallaecimonas kandeliae | Pseudomonadota | Gammaproteobacteria | NZ_CP118480.1 | 2 |
| Gallaecimonas mangrovi | Pseudomonadota | Gammaproteobacteria | NZ_CP031416.1 | 2 |
| Gallibacterium salpingitidis | Pseudomonadota | Gammaproteobacteria | NZ_CP103875.1 | 2 |
| Geopseudomonas guangdongensis | Pseudomonadota | Gammaproteobacteria | NZ_LT629780.1 | 1 |
| Gibbsiella quercinecans | Pseudomonadota | Gammaproteobacteria | NZ_CP014136.1 | 2 |
| Gilliamella apicola | Pseudomonadota | Gammaproteobacteria | NZ_CP007445.1 | 2 |
| Gilliamella apis | Pseudomonadota | Gammaproteobacteria | NZ_CP132382.1 | 2 |
| Gilvimarinus japonicus | Pseudomonadota | Gammaproteobacteria | NZ_AP031500.1 | 2 |
| Glaesserella parasuis | Pseudomonadota | Gammaproteobacteria | NZ_CP071487.1 | 2 |
| Granulosicoccus antarcticus IMCC3135 | Pseudomonadota | Gammaproteobacteria | NZ_CP018632.1 | 2 |
| Grimontia hollisae | Pseudomonadota | Gammaproteobacteria | NZ_CP014056.2 | 2 |
| Grimontia kaedaensis | Pseudomonadota | Gammaproteobacteria | NZ_CP082275.1 | 2 |
| Guyparkeria halophila | Pseudomonadota | Gammaproteobacteria | NZ_CP140153.1 | 2 |
| Haemophilus aegyptius | Pseudomonadota | Gammaproteobacteria | NZ_LS483429.1 | 2 |
| Haemophilus haemolyticus | Pseudomonadota | Gammaproteobacteria | NZ_LS483458.1 | 2 |

| | | | | |
|---|---|---|---|---|
| Haemophilus influenzae | Pseudomonadota | Gammaproteobacteria | NZ_CP085952.1 | 2 |
| Haemophilus parahaemolyticus | Pseudomonadota | Gammaproteobacteria | NZ_CP069510.1 | 2 |
| Haemophilus parainfluenzae | Pseudomonadota | Gammaproteobacteria | NZ_CP065991.1 | 2 |
| Haemophilus pittmaniae | Pseudomonadota | Gammaproteobacteria | NZ_LT906463.1 | 2 |
| Hafnia alvei | Pseudomonadota | Gammaproteobacteria | NZ_CP050150.1 | 2 |
| Hafnia paralvei | Pseudomonadota | Gammaproteobacteria | NZ_CP083737.1 | 2 |
| Hahella chejuensis KCTC 2396 | Pseudomonadota | Gammaproteobacteria | NC_007645.1 | 2 |
| Halioglobus maricola | Pseudomonadota | Gammaproteobacteria | NZ_CP036422.1 | 2 |
| Halioxenophilus aromaticivorans | Pseudomonadota | Gammaproteobacteria | NZ_AP031496.1 | 2 |
| Halomonas aestuarii | Pseudomonadota | Gammaproteobacteria | NZ_CP018139.1 | 1 |
| Halomonas alkaliantarctica | Pseudomonadota | Gammaproteobacteria | NZ_CP122961.1 | 1 |
| Halomonas alkalicola | Pseudomonadota | Gammaproteobacteria | NZ_CP130143.1 | 1 |
| Halomonas beimenensis | Pseudomonadota | Gammaproteobacteria | NZ_CP021435.1 | 1 |
| Halomonas binhaiensis | Pseudomonadota | Gammaproteobacteria | NZ_CP038437.2 | 1 |
| Halomonas chromatireducens | Pseudomonadota | Gammaproteobacteria | NZ_CP014226.1 | 1 |
| Halomonas elongata DSM 2581 | Pseudomonadota | Gammaproteobacteria | NC_014532.2 | 1 |
| Halomonas halophila | Pseudomonadota | Gammaproteobacteria | NZ_CP129121.1 | 1 |
| Halomonas huangheensis | Pseudomonadota | Gammaproteobacteria | NZ_CP013106.1 | 1 |
| Halomonas hydrothermalis | Pseudomonadota | Gammaproteobacteria | NZ_AP022843.1 | 1 |
| Halomonas kashgarensis | Pseudomonadota | Gammaproteobacteria | NZ_CP137552.1 | 1 |
| Halomonas piscis | Pseudomonadota | Gammaproteobacteria | NZ_CP119391.1 | 1 |
| Halomonas qaidamensis | Pseudomonadota | Gammaproteobacteria | NZ_CP080627.1 | 1 |
| Halomonas qinghailakensis | Pseudomonadota | Gammaproteobacteria | NZ_CP096973.1 | 1 |
| Halomonas sulfidivorans | Pseudomonadota | Gammaproteobacteria | NZ_CP053383.1 | 1 |
| Halopseudomonas aestusnigri | Pseudomonadota | Gammaproteobacteria | NZ_AP027273.1 | 2 |
| Halopseudomonas litoralis | Pseudomonadota | Gammaproteobacteria | NZ_LT629748.1 | 2 |
| Halopseudomonas maritima | Pseudomonadota | Gammaproteobacteria | NZ_CP079801.1 | 2 |
| Halopseudomonas nanhaiensis | Pseudomonadota | Gammaproteobacteria | NZ_CP073751.1 | 2 |
| Halopseudomonas oceani | Pseudomonadota | Gammaproteobacteria | NZ_OY781932.1 | 2 |
| Halopseudomonas phragmitis | Pseudomonadota | Gammaproteobacteria | NZ_CP020100.1 | 2 |
| Halopseudomonas sabulinigri | Pseudomonadota | Gammaproteobacteria | NZ_LT629763.1 | 2 |
| Halopseudomonas salegens | Pseudomonadota | Gammaproteobacteria | NZ_LT629787.1 | 2 |

| | | | | |
|---|---|---|---|---|
| Halopseudomonas xinjiangensis | Pseudomonadota | Gammaproteobacteria | NZ_LT629736.1 | 2 |
| Halorhodospira halochloris | Pseudomonadota | Gammaproteobacteria | NZ_AP017372.2 | 2 |
| Halorhodospira halophila SL1 | Pseudomonadota | Gammaproteobacteria | NC_008789.1 | 2 |
| Halothiobacillus diazotrophicus | Pseudomonadota | Gammaproteobacteria | NZ_CP016027.1 | 2 |
| Halothiobacillus neapolitanus c2 | Pseudomonadota | Gammaproteobacteria | NC_013422.1 | 2 |
| Histophilus somni 2336 | Pseudomonadota | Gammaproteobacteria | NC_010519.1 | 2 |
| Hydrocarboniclastica marina | Pseudomonadota | Gammaproteobacteria | NZ_CP031093.1 | 2 |
| Hydrogenovibrio crunogenus | Pseudomonadota | Gammaproteobacteria | NZ_CP032096.1 | 2 |
| Hydrogenovibrio marinus | Pseudomonadota | Gammaproteobacteria | NZ_AP020335.1 | 2 |
| Hydrogenovibrio thermophilus | Pseudomonadota | Gammaproteobacteria | NZ_CP035033.1 | 2 |
| Idiomarina abyssalis | Pseudomonadota | Gammaproteobacteria | NZ_CP081832.1 | 2 |
| Idiomarina loihiensis L2TR | Pseudomonadota | Gammaproteobacteria | NC_006512.1 | 2 |
| Ignatzschineria larvae DSM 13226 | Pseudomonadota | Gammaproteobacteria | NZ_CP150637.1 | 2 |
| Ignatzschineria rhizosphaerae | Pseudomonadota | Gammaproteobacteria | NZ_CP093379.1 | 2 |
| Immundisolibacter cernigliae | Pseudomonadota | Gammaproteobacteria | NZ_CP014671.1 | 2 |
| Isoalcanivorax pacificus W11-5 | Pseudomonadota | Gammaproteobacteria | NZ_CP004387.1 | 2 |
| Isorropodon fossajaponicum endosymbiont JTNG4 | Pseudomonadota | Gammaproteobacteria | NZ_AP013043.1 | 2 |
| Jejubacter calystegiae | Pseudomonadota | Gammaproteobacteria | NZ_CP040428.1 | 2 |
| Kangiella geojedonensis | Pseudomonadota | Gammaproteobacteria | NZ_CP010975.1 | 2 |
| Kangiella koreensis DSM 16069 | Pseudomonadota | Gammaproteobacteria | NC_013166.1 | 2 |
| Kangiella sediminilitoris | Pseudomonadota | Gammaproteobacteria | NZ_CP012418.1 | 2 |
| Ketobacter alkanivorans | Pseudomonadota | Gammaproteobacteria | NZ_CP022684.1 | 2 |
| Klebsiella aerogenes | Pseudomonadota | Gammaproteobacteria | NZ_CP077293.1 | 2 |
| Klebsiella africana | Pseudomonadota | Gammaproteobacteria | NZ_CP084874.1 | 2 |
| Klebsiella electrica | Pseudomonadota | Gammaproteobacteria | NZ_CP041247.1 | 2 |
| Klebsiella grimontii | Pseudomonadota | Gammaproteobacteria | NZ_CP169721.1 | 2 |
| Klebsiella huaxiensis | Pseudomonadota | Gammaproteobacteria | NZ_CP036175.1 | 2 |
| Klebsiella michiganensis | Pseudomonadota | Gammaproteobacteria | NZ_AP022547.1 | 2 |
| Klebsiella oxytoca | Pseudomonadota | Gammaproteobacteria | NZ_LR134333.1 | 2 |
| Klebsiella pasteurii | Pseudomonadota | Gammaproteobacteria | NZ_CP073236.1 | 2 |
| Klebsiella quasipneumoniae | Pseudomonadota | Gammaproteobacteria | NZ_CP083633.1 | 2 |
| Klebsiella quasivariicola | Pseudomonadota | Gammaproteobacteria | NZ_CP084768.1 | 2 |

| | | | | |
|---|---|---|---|---|
| Klebsiella variicola subsp. variicola | Pseudomonadota | Gammaproteobacteria | NZ_CP084765.1 | 2 |
| Kluyvera ascorbata | Pseudomonadota | Gammaproteobacteria | NZ_CP096201.1 | 2 |
| Kluyvera intermedia | Pseudomonadota | Gammaproteobacteria | NZ_CP045845.1 | 2 |
| Kluyvera sichuanensis | Pseudomonadota | Gammaproteobacteria | NZ_CP162271.1 | 2 |
| Kosakonia arachidis | Pseudomonadota | Gammaproteobacteria | NZ_CP045300.1 | 2 |
| Kosakonia calanthes | Pseudomonadota | Gammaproteobacteria | NZ_CP151800.1 | 2 |
| Kosakonia cowanii | Pseudomonadota | Gammaproteobacteria | NZ_CP107077.1 | 2 |
| Kosakonia oryzae | Pseudomonadota | Gammaproteobacteria | NZ_CP014007.2 | 2 |
| Kosakonia oryzendophytica | Pseudomonadota | Gammaproteobacteria | NZ_CP115659.1 | 2 |
| Kosakonia pseudosacchari | Pseudomonadota | Gammaproteobacteria | NZ_CP063425.1 | 2 |
| Kosakonia radicincitans | Pseudomonadota | Gammaproteobacteria | NZ_CP040392.1 | 2 |
| Kosakonia sacchari SP1 | Pseudomonadota | Gammaproteobacteria | NZ_CP007215.2 | 2 |
| Kushneria konosiri | Pseudomonadota | Gammaproteobacteria | NZ_CP021323.1 | 1 |
| Kushneria marisflavi | Pseudomonadota | Gammaproteobacteria | NZ_CP021358.1 | 1 |
| Kushneria phosphatilytica | Pseudomonadota | Gammaproteobacteria | NZ_CP043420.1 | 1 |
| Leclercia adecarboxylata | Pseudomonadota | Gammaproteobacteria | NZ_CP013990.1 | 2 |
| Leclercia pneumoniae | Pseudomonadota | Gammaproteobacteria | NZ_CP071383.1 | 2 |
| Legionella anisa | Pseudomonadota | Gammaproteobacteria | NZ_CP029563.1 | 2 |
| Legionella antarctica | Pseudomonadota | Gammaproteobacteria | NZ_AP022839.1 | 2 |
| Legionella cardiaca | Pseudomonadota | Gammaproteobacteria | NZ_CP119078.1 | 2 |
| Legionella clemsonensis | Pseudomonadota | Gammaproteobacteria | NZ_CP016397.1 | 2 |
| Legionella fallonii LLAP-10 | Pseudomonadota | Gammaproteobacteria | NZ_LN614827.1 | 2 |
| Legionella geestiana | Pseudomonadota | Gammaproteobacteria | NZ_CP038271.1 | 2 |
| Legionella hackeliae | Pseudomonadota | Gammaproteobacteria | NZ_LN681225.1 | 2 |
| Legionella israelensis | Pseudomonadota | Gammaproteobacteria | NZ_CP041668.1 | 2 |
| Legionella jordanis | Pseudomonadota | Gammaproteobacteria | NZ_LR134383.1 | 2 |
| Legionella lansingensis | Pseudomonadota | Gammaproteobacteria | NZ_LT906451.1 | 2 |
| Legionella longbeachae NSW150 | Pseudomonadota | Gammaproteobacteria | NC_013861.1 | 2 |
| Legionella lytica | Pseudomonadota | Gammaproteobacteria | NZ_CP071527.1 | 2 |
| Legionella micdadei | Pseudomonadota | Gammaproteobacteria | NZ_LN614830.1 | 2 |
| Legionella pneumophila | Pseudomonadota | Gammaproteobacteria | NZ_CP013742.1 | 2 |
| Legionella sainthelensi | Pseudomonadota | Gammaproteobacteria | NZ_CP025491.2 | 2 |

| | | | | |
|---|---|---|---|---|
| Legionella spiritensis | Pseudomonadota | Gammaproteobacteria | NZ_LT906457.1 | 2 |
| Legionella waltersii | Pseudomonadota | Gammaproteobacteria | NZ_LT906442.1 | 2 |
| Lelliottia amnigena | Pseudomonadota | Gammaproteobacteria | NZ_CP077236.1 | 2 |
| Lelliottia nimipressuralis | Pseudomonadota | Gammaproteobacteria | NZ_CP167172.1 | 2 |
| Leminorella grimontii | Pseudomonadota | Gammaproteobacteria | NZ_CP146357.1 | 2 |
| Leminorella richardii | Pseudomonadota | Gammaproteobacteria | NZ_LS483470.1 | 2 |
| Limnobaculum parvum | Pseudomonadota | Gammaproteobacteria | NZ_CP029185.2 | 2 |
| Limnobaculum zhutongyuii | Pseudomonadota | Gammaproteobacteria | NZ_CP034752.1 | 2 |
| Litorilituus sediminis | Pseudomonadota | Gammaproteobacteria | NZ_CP034759.1 | 2 |
| Litorivicinus lipolyticus | Pseudomonadota | Gammaproteobacteria | NZ_CP045871.1 | 2 |
| Lonepinella koalarum | Pseudomonadota | Gammaproteobacteria | NZ_CP170642.1 | 2 |
| Lonsdalea britannica | Pseudomonadota | Gammaproteobacteria | NZ_CP023009.1 | 2 |
| Lonsdalea populi | Pseudomonadota | Gammaproteobacteria | NZ_CP065534.1 | 2 |
| Luteibacter aegosomaticola | Pseudomonadota | Gammaproteobacteria | NZ_CP095741.1 | 2 |
| Luteibacter aegosomatis | Pseudomonadota | Gammaproteobacteria | NZ_CP095740.1 | 2 |
| Luteibacter aegosomatissinici | Pseudomonadota | Gammaproteobacteria | NZ_CP095742.1 | 2 |
| Luteibacter anthropi | Pseudomonadota | Gammaproteobacteria | NZ_CP077072.1 | 2 |
| Luteibacter flocculans | Pseudomonadota | Gammaproteobacteria | NZ_CP063231.1 | 2 |
| Luteibacter pinisoli | Pseudomonadota | Gammaproteobacteria | NZ_CP041046.1 | 2 |
| Luteibacter rhizovicinus DSM 16549 | Pseudomonadota | Gammaproteobacteria | NZ_CP017480.1 | 2 |
| Luteimonas chenhongjianii | Pseudomonadota | Gammaproteobacteria | NZ_CP023406.1 | 2 |
| Luteimonas fraxinea | Pseudomonadota | Gammaproteobacteria | NZ_CP089507.1 | 2 |
| Luteimonas granuli | Pseudomonadota | Gammaproteobacteria | NZ_CP042218.1 | 2 |
| Luteimonas yindakuii | Pseudomonadota | Gammaproteobacteria | NZ_CP039383.2 | 2 |
| Lysobacter antibioticus | Pseudomonadota | Gammaproteobacteria | NZ_CP011129.1 | 2 |
| Lysobacter arenosi | Pseudomonadota | Gammaproteobacteria | NZ_CP071517.1 | 2 |
| Lysobacter auxotrophicus | Pseudomonadota | Gammaproteobacteria | NZ_AP027041.1 | 2 |
| Lysobacter capsici | Pseudomonadota | Gammaproteobacteria | NZ_CP094357.1 | 2 |
| Lysobacter enzymogenes | Pseudomonadota | Gammaproteobacteria | NZ_AP014940.1 | 2 |
| Lysobacter firmicutimachus | Pseudomonadota | Gammaproteobacteria | NZ_CP159925.1 | 2 |
| Lysobacter gummosus | Pseudomonadota | Gammaproteobacteria | NZ_CP011131.1 | 2 |
| Lysobacter helvus | Pseudomonadota | Gammaproteobacteria | NZ_AP024546.1 | 2 |

| | | | | |
|---|---|---|---|---|
| Lysobacter soli | Pseudomonadota | Gammaproteobacteria | NZ_CP046603.1 | 2 |
| Lysobacter soyae | Pseudomonadota | Gammaproteobacteria | NZ_CP080544.1 | 2 |
| Lysobacter yananisis | Pseudomonadota | Gammaproteobacteria | NZ_CP133568.1 | 2 |
| Mangrovibacter phragmitis | Pseudomonadota | Gammaproteobacteria | NZ_CP134782.1 | 2 |
| Mannheimia bovis | Pseudomonadota | Gammaproteobacteria | NZ_CP061280.1 | 2 |
| Mannheimia glucosida | Pseudomonadota | Gammaproteobacteria | NZ_CP176502.1 | 2 |
| Mannheimia haemolytica | Pseudomonadota | Gammaproteobacteria | NZ_CP023044.1 | 2 |
| Mannheimia pernigra | Pseudomonadota | Gammaproteobacteria | NZ_CP055305.1 | 2 |
| Mannheimia varigena | Pseudomonadota | Gammaproteobacteria | NZ_CP016226.1 | 2 |
| Marichromatium purpuratum 984 | Pseudomonadota | Gammaproteobacteria | NZ_CP007031.1 | 2 |
| Maricurvus nonylphenolicus | Pseudomonadota | Gammaproteobacteria | NZ_AP031498.1 | 2 |
| Marilutibacter alkalisoli | Pseudomonadota | Gammaproteobacteria | NZ_CP041242.1 | 2 |
| Marilutibacter maris | Pseudomonadota | Gammaproteobacteria | NZ_CP029843.1 | 2 |
| Marinagarivorans cellulosilyticus | Pseudomonadota | Gammaproteobacteria | NZ_AP023086.1 | 1 |
| Marinicellulosiphila megalodicopiae | Pseudomonadota | Gammaproteobacteria | NZ_AP023087.1 | 2 |
| Marinobacter adhaerens | Pseudomonadota | Gammaproteobacteria | NZ_CP076686.1 | 2 |
| Marinobacter alkaliphilus | Pseudomonadota | Gammaproteobacteria | NZ_CP152380.1 | 2 |
| Marinobacter apostichopi | Pseudomonadota | Gammaproteobacteria | NZ_AP028070.1 | 2 |
| Marinobacter flavimaris | Pseudomonadota | Gammaproteobacteria | NZ_CP171359.1 | 2 |
| Marinobacter fonticola | Pseudomonadota | Gammaproteobacteria | NZ_CP043042.1 | 2 |
| Marinobacter metalliresistant | Pseudomonadota | Gammaproteobacteria | NZ_CP101118.1 | 2 |
| Marinobacter nauticus ATCC 49840 | Pseudomonadota | Gammaproteobacteria | NC_017067.1 | 2 |
| Marinobacter psychrophilus | Pseudomonadota | Gammaproteobacteria | NZ_CP011494.1 | 2 |
| Marinobacter salarius | Pseudomonadota | Gammaproteobacteria | NZ_CP020931.1 | 2 |
| Marinobacter salinisoli | Pseudomonadota | Gammaproteobacteria | NZ_CP071247.1 | 2 |
| Marinobacter salinus | Pseudomonadota | Gammaproteobacteria | NZ_CP017715.1 | 2 |
| Marinobacter shengliensis | Pseudomonadota | Gammaproteobacteria | NZ_AP028062.1 | 2 |
| Marinobacter similis | Pseudomonadota | Gammaproteobacteria | NZ_CP007151.1 | 1 |
| Marinobacterium aestuarii | Pseudomonadota | Gammaproteobacteria | NZ_CP015839.1 | 2 |
| Marinobacterium iners | Pseudomonadota | Gammaproteobacteria | NZ_CP022297.1 | 2 |
| Marinomonas algicola | Pseudomonadota | Gammaproteobacteria | NZ_CP061941.1 | 1 |
| Marinomonas arctica | Pseudomonadota | Gammaproteobacteria | NZ_CP061081.1 | 1 |

| | | | | |
|---|---|---|---|---|
| Marinomonas foliarum | Pseudomonadota | Gammaproteobacteria | NZ_CP070273.1 | 1 |
| Marinomonas mediterranea MMB-1 | Pseudomonadota | Gammaproteobacteria | NC_015276.1 | 1 |
| Marinomonas polaris | Pseudomonadota | Gammaproteobacteria | NZ_CP165966.1 | 1 |
| Marinomonas pontica | Pseudomonadota | Gammaproteobacteria | NZ_AP027271.1 | 1 |
| Marinomonas posidonica IVIA-Po-181 | Pseudomonadota | Gammaproteobacteria | NC_015559.1 | 1 |
| Marinomonas primoryensis | Pseudomonadota | Gammaproteobacteria | NZ_CP054301.1 | 1 |
| Marinomonas profundi | Pseudomonadota | Gammaproteobacteria | NZ_CP073013.1 | 1 |
| Mergibacter septicus | Pseudomonadota | Gammaproteobacteria | NZ_CP022013.1 | 2 |
| Metapseudomonas boanensis | Pseudomonadota | Gammaproteobacteria | NZ_CP113889.1 | 2 |
| Metapseudomonas furukawaii | Pseudomonadota | Gammaproteobacteria | NZ_AP014862.1 | 2 |
| Metapseudomonas lalkuanensis | Pseudomonadota | Gammaproteobacteria | NZ_CP043311.1 | 2 |
| Metapseudomonas otitidis | Pseudomonadota | Gammaproteobacteria | NZ_AP022642.1 | 2 |
| Methylocaldum marinum | Pseudomonadota | Gammaproteobacteria | NZ_AP017928.1 | 2 |
| Methylocaldum szegediense | Pseudomonadota | Gammaproteobacteria | NZ_OX458333.1 | 2 |
| Methylococcus capsulatus str. Bath | Pseudomonadota | Gammaproteobacteria | NC_002977.6 | 2 |
| Methylococcus geothermalis | Pseudomonadota | Gammaproteobacteria | NZ_CP046565.1 | 2 |
| Methylococcus mesophilus | Pseudomonadota | Gammaproteobacteria | NZ_CP110921.1 | 2 |
| Methylogaea oryzae | Pseudomonadota | Gammaproteobacteria | NZ_AP019782.1 | 2 |
| Methylomagnum ishizawai | Pseudomonadota | Gammaproteobacteria | NZ_AP019783.1 | 2 |
| Methylomarinovum caldicuralii | Pseudomonadota | Gammaproteobacteria | NZ_AP024714.1 | 2 |
| Methylomarinovum tepidoasis | Pseudomonadota | Gammaproteobacteria | NZ_AP024718.1 | 2 |
| Methylomarinum roseum | Pseudomonadota | Gammaproteobacteria | NZ_CP157743.1 | 2 |
| Methylomicrobium album BG8 | Pseudomonadota | Gammaproteobacteria | NZ_CM001475.1 | 2 |
| Methylomonas denitrificans | Pseudomonadota | Gammaproteobacteria | NZ_CP014476.1 | 2 |
| Methylomonas koyamae | Pseudomonadota | Gammaproteobacteria | NZ_AP019777.1 | 2 |
| Methylomonas montana | Pseudomonadota | Gammaproteobacteria | NZ_CP129884.1 | 2 |
| Methylomonas paludis | Pseudomonadota | Gammaproteobacteria | NZ_CP073754.1 | 2 |
| Methylomonas rapida | Pseudomonadota | Gammaproteobacteria | NZ_CP113517.1 | 2 |
| Methylomonas rhizoryzae | Pseudomonadota | Gammaproteobacteria | NZ_CP043929.1 | 2 |
| Methylophaga frappieri | Pseudomonadota | Gammaproteobacteria | NC_017856.1 | 2 |
| Methylophaga marina | Pseudomonadota | Gammaproteobacteria | NZ_AP027741.1 | 2 |
| Methylophaga nitratireducenticrescens | Pseudomonadota | Gammaproteobacteria | NC_017857.3 | 2 |

| | | | | |
|---|---|---|---|---|
| Methylophaga pinxianii | Pseudomonadota | Gammaproteobacteria | NZ_CP095733.1 | 2 |
| Methylotuvimicrobium alcaliphilum 20Z | Pseudomonadota | Gammaproteobacteria | NC_016112.1 | 2 |
| Methylotuvimicrobium buryatense | Pseudomonadota | Gammaproteobacteria | NZ_CP035467.1 | 2 |
| Methylovulum psychrotolerans | Pseudomonadota | Gammaproteobacteria | NZ_CP022129.1 | 2 |
| Microbulbifer agarilyticus | Pseudomonadota | Gammaproteobacteria | NZ_CP019650.1 | 2 |
| Microbulbifer aggregans | Pseudomonadota | Gammaproteobacteria | NZ_CP014143.1 | 2 |
| Microbulbifer bruguierae | Pseudomonadota | Gammaproteobacteria | NZ_CP118605.1 | 2 |
| Microbulbifer celer | Pseudomonadota | Gammaproteobacteria | NZ_CP087715.1 | 2 |
| Microbulbifer elongatus | Pseudomonadota | Gammaproteobacteria | NZ_CP088953.1 | 2 |
| Microbulbifer hydrolyticus | Pseudomonadota | Gammaproteobacteria | NZ_CP047491.1 | 2 |
| Microbulbifer pacificus | Pseudomonadota | Gammaproteobacteria | NZ_CP137555.1 | 2 |
| Microbulbifer spongiae | Pseudomonadota | Gammaproteobacteria | NZ_CP098023.1 | 2 |
| Microbulbifer thermotolerans | Pseudomonadota | Gammaproteobacteria | NZ_CP014864.1 | 2 |
| Mixta gaviniae | Pseudomonadota | Gammaproteobacteria | NZ_CP026377.1 | 2 |
| Mixta hanseatica | Pseudomonadota | Gammaproteobacteria | NZ_CP082904.1 | 2 |
| Mixta intestinalis | Pseudomonadota | Gammaproteobacteria | NZ_CP028271.1 | 2 |
| Moellerella wisconsensis | Pseudomonadota | Gammaproteobacteria | NZ_CP093242.1 | 2 |
| Moraxella bovis | Pseudomonadota | Gammaproteobacteria | NZ_CP087840.1 | 2 |
| Moraxella bovoculi | Pseudomonadota | Gammaproteobacteria | NZ_CP011381.2 | 2 |
| Moraxella canis | Pseudomonadota | Gammaproteobacteria | NZ_CP139961.1 | 2 |
| Moraxella catarrhalis | Pseudomonadota | Gammaproteobacteria | NZ_CP018059.1 | 2 |
| Moraxella cuniculi | Pseudomonadota | Gammaproteobacteria | NZ_LR134343.1 | 2 |
| Moraxella haemolytica | Pseudomonadota | Gammaproteobacteria | NZ_CP089974.1 | 2 |
| Moraxella lincolnii | Pseudomonadota | Gammaproteobacteria | NZ_CP147511.1 | 2 |
| Moraxella nasibovis | Pseudomonadota | Gammaproteobacteria | NZ_CP089975.1 | 2 |
| Moraxella nasicaprae | Pseudomonadota | Gammaproteobacteria | NZ_CP089977.1 | 2 |
| Moraxella nasovis | Pseudomonadota | Gammaproteobacteria | NZ_CP089976.1 | 2 |
| Moraxella nonliquefaciens | Pseudomonadota | Gammaproteobacteria | NZ_CP065728.1 | 2 |
| Moraxella osloensis | Pseudomonadota | Gammaproteobacteria | NZ_CP014234.1 | 2 |
| Moraxella ovis | Pseudomonadota | Gammaproteobacteria | NZ_CP011158.1 | 2 |
| Moraxella veridica | Pseudomonadota | Gammaproteobacteria | NZ_CP170124.1 | 2 |
| Morganella morganii subsp. morganii | Pseudomonadota | Gammaproteobacteria | NZ_CP034944.1 | 2 |

| | | | | |
|---|---|---|---|---|
| Moritella marina ATCC 15381 | Pseudomonadota | Gammaproteobacteria | NZ_CP044399.1 | 1 |
| Moritella yayanosii | Pseudomonadota | Gammaproteobacteria | NZ_LS483250.1 | 1 |
| Musicola paradisiaca Ech703 | Pseudomonadota | Gammaproteobacteria | NC_012880.1 | 2 |
| Neptuniibacter halophilus | Pseudomonadota | Gammaproteobacteria | NZ_AP027292.1 | 2 |
| Neptuniibacter marinus | Pseudomonadota | Gammaproteobacteria | NZ_OZ245004.1 | 2 |
| Neptunomonas concharum | Pseudomonadota | Gammaproteobacteria | NZ_CP043869.1 | 2 |
| Neptunomonas phycophila | Pseudomonadota | Gammaproteobacteria | NZ_CP041336.1 | 2 |
| Nitrincola iocasae | Pseudomonadota | Gammaproteobacteria | NZ_CP044222.1 | 2 |
| Nitrosococcus halophilus Nc 4 | Pseudomonadota | Gammaproteobacteria | NC_013960.1 | 2 |
| Nitrosococcus oceani ATCC 19707 | Pseudomonadota | Gammaproteobacteria | NC_007484.1 | 2 |
| Nitrosococcus wardiae | Pseudomonadota | Gammaproteobacteria | NZ_CP038033.1 | 2 |
| Nitrosococcus watsonii C-113 | Pseudomonadota | Gammaproteobacteria | NC_014315.1 | 3 |
| Noviluteimonas caseinilytica | Pseudomonadota | Gammaproteobacteria | NZ_AP024545.1 | 2 |
| Novilysobacter avium | Pseudomonadota | Gammaproteobacteria | NZ_CP063657.1 | 2 |
| Novilysobacter ciconiae | Pseudomonadota | Gammaproteobacteria | NZ_CP063656.1 | 2 |
| Novilysobacter luteus | Pseudomonadota | Gammaproteobacteria | NZ_OU015430.1 | 2 |
| Oceanicoccus sagamiensis | Pseudomonadota | Gammaproteobacteria | NZ_CP019343.1 | 2 |
| Oceanimonas pelagia | Pseudomonadota | Gammaproteobacteria | NZ_CP118224.1 | 2 |
| Oceanisphaera avium | Pseudomonadota | Gammaproteobacteria | NZ_CP021376.1 | 1 |
| Oceanisphaera profunda | Pseudomonadota | Gammaproteobacteria | NZ_CP021377.1 | 1 |
| Oleiphilus messinensis | Pseudomonadota | Gammaproteobacteria | NZ_CP021425.1 | 1 |
| Orbus mooreae | Pseudomonadota | Gammaproteobacteria | NZ_CP133958.1 | 2 |
| Orbus sturtevantii | Pseudomonadota | Gammaproteobacteria | NZ_CP133960.1 | 2 |
| Orbus wheelerorum | Pseudomonadota | Gammaproteobacteria | NZ_CP133967.1 | 2 |
| Otariodibacter oris | Pseudomonadota | Gammaproteobacteria | NZ_CP016604.1 | 2 |
| Pantoea agglomerans | Pseudomonadota | Gammaproteobacteria | NZ_CP077366.1 | 2 |
| Pantoea alfalfae | Pseudomonadota | Gammaproteobacteria | NZ_CP082292.1 | 2 |
| Pantoea alhagi | Pseudomonadota | Gammaproteobacteria | NZ_CP019706.1 | 2 |
| Pantoea allii | Pseudomonadota | Gammaproteobacteria | NZ_CP125958.1 | 2 |
| Pantoea ananatis PA13 | Pseudomonadota | Gammaproteobacteria | NC_017554.1 | 2 |
| Pantoea anthophila | Pseudomonadota | Gammaproteobacteria | NZ_CP110473.1 | 2 |
| Pantoea deleyi | Pseudomonadota | Gammaproteobacteria | NZ_CP071405.1 | 2 |

| | | | | |
|---|---|---|---|---|
| Pantoea dispersa | Pseudomonadota | Gammaproteobacteria | NZ_CP082346.1 | 2 |
| Pantoea eucalypti | Pseudomonadota | Gammaproteobacteria | NZ_CP045720.1 | 2 |
| Pantoea eucrina | Pseudomonadota | Gammaproteobacteria | NZ_CP083448.1 | 2 |
| Pantoea phytostimulans | Pseudomonadota | Gammaproteobacteria | NZ_CP173698.1 | 2 |
| Pantoea stewartii | Pseudomonadota | Gammaproteobacteria | NZ_CP049115.1 | 2 |
| Pantoea vagans | Pseudomonadota | Gammaproteobacteria | NZ_CP038853.1 | 2 |
| Paramixta manurensis | Pseudomonadota | Gammaproteobacteria | NZ_CP054212.1 | 2 |
| Paraphotobacterium marinum | Pseudomonadota | Gammaproteobacteria | NZ_CP022355.1 | 2 |
| Parashewanella spongiae | Pseudomonadota | Gammaproteobacteria | NZ_CP037952.1 | 2 |
| Parashewanella tropica | Pseudomonadota | Gammaproteobacteria | NZ_CP037951.1 | 2 |
| Pasteurella atlantica | Pseudomonadota | Gammaproteobacteria | NZ_OY856306.1 | 2 |
| Pasteurella canis | Pseudomonadota | Gammaproteobacteria | NZ_CP085871.1 | 2 |
| Pasteurella dagmatis | Pseudomonadota | Gammaproteobacteria | NZ_LT906448.1 | 2 |
| Pasteurella multocida | Pseudomonadota | Gammaproteobacteria | NZ_CP020405.2 | 2 |
| Pectobacterium aquaticum | Pseudomonadota | Gammaproteobacteria | NZ_CP086253.1 | 2 |
| Pectobacterium araliae | Pseudomonadota | Gammaproteobacteria | NZ_AP028908.1 | 2 |
| Pectobacterium aroidearum | Pseudomonadota | Gammaproteobacteria | NZ_CP065044.1 | 2 |
| Pectobacterium atrosepticum | Pseudomonadota | Gammaproteobacteria | NZ_CP009125.1 | 2 |
| Pectobacterium brasiliense | Pseudomonadota | Gammaproteobacteria | NZ_CP047495.1 | 2 |
| Pectobacterium cacticida | Pseudomonadota | Gammaproteobacteria | NZ_CP133656.1 | 2 |
| Pectobacterium carotovorum | Pseudomonadota | Gammaproteobacteria | NZ_CP051652.1 | 2 |
| Pectobacterium colocasium | Pseudomonadota | Gammaproteobacteria | NZ_CP084032.1 | 2 |
| Pectobacterium odoriferum | Pseudomonadota | Gammaproteobacteria | NZ_CP034938.1 | 2 |
| Pectobacterium parmentieri | Pseudomonadota | Gammaproteobacteria | NZ_CP015749.1 | 2 |
| Pectobacterium parvum | Pseudomonadota | Gammaproteobacteria | NZ_CP087392.1 | 2 |
| Pectobacterium peruviense | Pseudomonadota | Gammaproteobacteria | NZ_CP161826.1 | 2 |
| Pectobacterium polaris | Pseudomonadota | Gammaproteobacteria | NZ_CP017482.1 | 2 |
| Pectobacterium punjabense | Pseudomonadota | Gammaproteobacteria | NZ_CP038498.1 | 2 |
| Pectobacterium quasiaquaticum | Pseudomonadota | Gammaproteobacteria | NZ_CP065177.1 | 2 |
| Pectobacterium versatile | Pseudomonadota | Gammaproteobacteria | NZ_CP034276.1 | 2 |
| Pectobacterium wasabiae CFBP 3304 | Pseudomonadota | Gammaproteobacteria | NZ_CP015750.1 | 2 |
| Phocoenobacter skyensis | Pseudomonadota | Gammaproteobacteria | NZ_CP016180.1 | 2 |

| | | | | |
|---|---|---|---|---|
| Photobacterium atrarenae | Pseudomonadota | Gammaproteobacteria | NZ_CP101508.1 | 2 |
| Photobacterium damselae subsp. damselae | Pseudomonadota | Gammaproteobacteria | NZ_CP146970.1 | 2 |
| Photobacterium ganghwense | Pseudomonadota | Gammaproteobacteria | NZ_CP071325.1 | 2 |
| Photobacterium leiognathi subsp. mandapamensis | Pseudomonadota | Gammaproteobacteria | NZ_CP183250.1 | 2 |
| Photobacterium piscicola | Pseudomonadota | Gammaproteobacteria | NZ_CP175534.1 | 2 |
| Photobacterium sanguinicancri | Pseudomonadota | Gammaproteobacteria | NZ_AP024850.1 | 2 |
| Photobacterium swingsii | Pseudomonadota | Gammaproteobacteria | NZ_AP024852.1 | 2 |
| Photobacterium toruni | Pseudomonadota | Gammaproteobacteria | NZ_AP024854.1 | 2 |
| Photorhabdus akhurstii | Pseudomonadota | Gammaproteobacteria | NZ_CP020335.1 | 2 |
| Photorhabdus asymbiotica | Pseudomonadota | Gammaproteobacteria | NC_012962.1 | 2 |
| Photorhabdus laumondii subsp. laumondii | Pseudomonadota | Gammaproteobacteria | NZ_CP024901.1 | 2 |
| Photorhabdus thracensis | Pseudomonadota | Gammaproteobacteria | NZ_CP011104.1 | 2 |
| Phytobacter ursingii | Pseudomonadota | Gammaproteobacteria | NZ_CP011602.1 | 2 |
| Piscirickettsia salmonis | Pseudomonadota | Gammaproteobacteria | NZ_CP039032.1 | 1 |
| Pistricoccus aurantiacus | Pseudomonadota | Gammaproteobacteria | NZ_CP042382.1 | 1 |
| Pleionea litopenaei | Pseudomonadota | Gammaproteobacteria | NZ_CP133548.1 | 2 |
| Plesiomonas shigelloides | Pseudomonadota | Gammaproteobacteria | NZ_CP087711.1 | 2 |
| Pluralibacter gergoviae | Pseudomonadota | Gammaproteobacteria | NZ_CP020388.1 | 2 |
| Pokkaliibacter plantistimulans | Pseudomonadota | Gammaproteobacteria | NZ_CP177354.1 | 1 |
| Pragia fontium | Pseudomonadota | Gammaproteobacteria | NZ_LR134531.1 | 2 |
| Prodigiosinella confusarubida | Pseudomonadota | Gammaproteobacteria | NZ_CP025084.1 | 2 |
| Proteus appendicitidis | Pseudomonadota | Gammaproteobacteria | NZ_CP127389.1 | 2 |
| Proteus hauseri | Pseudomonadota | Gammaproteobacteria | NZ_CP026364.1 | 2 |
| Proteus mirabilis HI4320 | Pseudomonadota | Gammaproteobacteria | NC_010554.1 | 2 |
| Proteus penneri | Pseudomonadota | Gammaproteobacteria | NZ_CP059690.1 | 2 |
| Proteus terrae subsp. cibarius | Pseudomonadota | Gammaproteobacteria | NZ_CP047349.1 | 2 |
| Proteus vulgaris | Pseudomonadota | Gammaproteobacteria | NZ_CP104121.1 | 2 |
| Providencia hangzhouensis | Pseudomonadota | Gammaproteobacteria | NZ_CP135052.1 | 2 |
| Providencia heimbachae | Pseudomonadota | Gammaproteobacteria | NZ_LS483422.1 | 2 |
| Providencia huaxiensis | Pseudomonadota | Gammaproteobacteria | NZ_CP145931.1 | 2 |
| Providencia manganoxydans | Pseudomonadota | Gammaproteobacteria | NZ_CP067099.1 | 2 |
| Providencia rettgeri | Pseudomonadota | Gammaproteobacteria | NZ_CP077317.1 | 2 |

| | | | | |
|---|---|---|---|---|
| Providencia rustigianii | Pseudomonadota | Gammaproteobacteria | NZ_LR134396.1 | 2 |
| Providencia sneebia DSM 19967 | Pseudomonadota | Gammaproteobacteria | NZ_CM001773.1 | 1 |
| Providencia stuartii | Pseudomonadota | Gammaproteobacteria | NZ_CP142095.1 | 2 |
| Providencia xihuensis | Pseudomonadota | Gammaproteobacteria | NZ_CP169565.1 | 2 |
| Providencia zhejiangensis | Pseudomonadota | Gammaproteobacteria | NZ_CP169566.1 | 2 |
| Providencia zhijiangensis | Pseudomonadota | Gammaproteobacteria | NZ_CP135990.1 | 2 |
| Pseudescherichia vulneris | Pseudomonadota | Gammaproteobacteria | NZ_LR607338.1 | 1 |
| Pseudoalteromonas aliena | Pseudomonadota | Gammaproteobacteria | NZ_CP019628.1 | 2 |
| Pseudoalteromonas apostichopi | Pseudomonadota | Gammaproteobacteria | NZ_AP028065.1 | 2 |
| Pseudoalteromonas arctica A 37-1-2 | Pseudomonadota | Gammaproteobacteria | NZ_CP011025.1 | 2 |
| Pseudoalteromonas carrageenovora IAM 12662 | Pseudomonadota | Gammaproteobacteria | NZ_LT965928.1 | 2 |
| Pseudoalteromonas distincta | Pseudomonadota | Gammaproteobacteria | NZ_CP040558.1 | 2 |
| Pseudoalteromonas espejiana DSM 9414 | Pseudomonadota | Gammaproteobacteria | NZ_CP011028.1 | 2 |
| Pseudoalteromonas issachenkonii | Pseudomonadota | Gammaproteobacteria | NZ_CP011030.1 | 2 |
| Pseudoalteromonas lipolytica | Pseudomonadota | Gammaproteobacteria | NZ_CP032090.1 | 2 |
| Pseudoalteromonas luteoviolacea | Pseudomonadota | Gammaproteobacteria | NZ_VIGJ01000001 | 2 |
| Pseudoalteromonas maricaloris | Pseudomonadota | Gammaproteobacteria | NZ_CP024625.1 | 2 |
| Pseudoalteromonas nigrifaciens | Pseudomonadota | Gammaproteobacteria | NZ_CP011036.1 | 2 |
| Pseudoalteromonas ostreae | Pseudomonadota | Gammaproteobacteria | NZ_CP118916.1 | 2 |
| Pseudoalteromonas phenolica | Pseudomonadota | Gammaproteobacteria | NZ_CP013187.1 | 2 |
| Pseudoalteromonas piratica | Pseudomonadota | Gammaproteobacteria | NZ_CP009888.1 | 2 |
| Pseudoalteromonas piscicida | Pseudomonadota | Gammaproteobacteria | NZ_CP081860.1 | 2 |
| Pseudoalteromonas prydzensis ACAM 620 | Pseudomonadota | Gammaproteobacteria | NZ_AQHH0100001 | 1 |
| Pseudoalteromonas prydzensis ACAM 620 | Pseudomonadota | Gammaproteobacteria | NZ_AQHH0100000 | 1 |
| Pseudoalteromonas prydzensis ACAM 620 | Pseudomonadota | Gammaproteobacteria | NZ_AQHH0100000 | 1 |
| Pseudoalteromonas rhizosphaerae | Pseudomonadota | Gammaproteobacteria | NZ_CP118496.1 | 2 |
| Pseudoalteromonas spongiae UST010723-006 | Pseudomonadota | Gammaproteobacteria | NZ_CP011039.1 | 2 |
| Pseudoalteromonas translucida | Pseudomonadota | Gammaproteobacteria | NC_007481.1 | 2 |
| Pseudoalteromonas tunicata | Pseudomonadota | Gammaproteobacteria | NZ_CP031961.1 | 2 |
| Pseudoalteromonas undina | Pseudomonadota | Gammaproteobacteria | NZ_AHCF0300000[illegible] | 1 |
| Pseudoalteromonas undina | Pseudomonadota | Gammaproteobacteria | NZ_AHCF0300000[illegible] | 1 |
| Pseudoalteromonas viridis | Pseudomonadota | Gammaproteobacteria | NZ_CP072425.1 | 2 |

| | | | | |
|---|---|---|---|---|
| Pseudocitrobacter corydidari | Pseudomonadota | Gammaproteobacteria | NZ_CP087880.1 | 2 |
| Pseudohongiella spirulinae | Pseudomonadota | Gammaproteobacteria | NZ_CP013189.1 | 2 |
| Pseudolysobacter antarcticus | Pseudomonadota | Gammaproteobacteria | NZ_CP035704.1 | 2 |
| Pseudomonas abieticivorans | Pseudomonadota | Gammaproteobacteria | NZ_CP094975.1 | 1 |
| Pseudomonas aeruginosa PAO1 | Pseudomonadota | Gammaproteobacteria | NC_002516.2 | 2 |
| Pseudomonas aestiva | Pseudomonadota | Gammaproteobacteria | NZ_CP150916.1 | 2 |
| Pseudomonas alkylphenolica | Pseudomonadota | Gammaproteobacteria | NZ_CP046621.1 | 2 |
| Pseudomonas allokribbensis | Pseudomonadota | Gammaproteobacteria | NZ_CP062252.1 | 2 |
| Pseudomonas alloputida | Pseudomonadota | Gammaproteobacteria | NZ_CP128540.1 | 2 |
| Pseudomonas alvandae | Pseudomonadota | Gammaproteobacteria | NZ_CP077080.1 | 2 |
| Pseudomonas amygdali pv. tabaci str. ATCC 11528 | Pseudomonadota | Gammaproteobacteria | NZ_CP042804.1 | 1 |
| Pseudomonas antarctica | Pseudomonadota | Gammaproteobacteria | NZ_LT629704.1 | 1 |
| Pseudomonas anuradhapurensis | Pseudomonadota | Gammaproteobacteria | NZ_CP077097.1 | 2 |
| Pseudomonas arsenicoxydans | Pseudomonadota | Gammaproteobacteria | NZ_LT629705.1 | 1 |
| Pseudomonas asgharzadehiana | Pseudomonadota | Gammaproteobacteria | NZ_CP077079.1 | 1 |
| Pseudomonas asiatica | Pseudomonadota | Gammaproteobacteria | NZ_CP139776.1 | 2 |
| Pseudomonas asplenii | Pseudomonadota | Gammaproteobacteria | NZ_LT629777.1 | 1 |
| Pseudomonas atacamensis | Pseudomonadota | Gammaproteobacteria | NZ_CP070503.1 | 2 |
| Pseudomonas azerbaijanorientalis | Pseudomonadota | Gammaproteobacteria | NZ_CP077078.1 | 1 |
| Pseudomonas azotoformans | Pseudomonadota | Gammaproteobacteria | NZ_LT629702.1 | 1 |
| Pseudomonas batumici | Pseudomonadota | Gammaproteobacteria | NZ_CP144470.1 | 2 |
| Pseudomonas beijingensis | Pseudomonadota | Gammaproteobacteria | NZ_CP117451.1 | 2 |
| Pseudomonas benzenivorans | Pseudomonadota | Gammaproteobacteria | NZ_CP137892.1 | 2 |
| Pseudomonas benzopyrenica | Pseudomonadota | Gammaproteobacteria | NZ_CP145723.1 | 2 |
| Pseudomonas berkeleyensis | Pseudomonadota | Gammaproteobacteria | NZ_CP059139.1 | 2 |
| Pseudomonas bijieensis | Pseudomonadota | Gammaproteobacteria | NZ_CP097108.1 | 2 |
| Pseudomonas brassicacearum | Pseudomonadota | Gammaproteobacteria | NZ_CP034725.1 | 2 |
| Pseudomonas bubulae | Pseudomonadota | Gammaproteobacteria | NZ_CP146077.1 | 1 |
| Pseudomonas canadensis | Pseudomonadota | Gammaproteobacteria | NZ_CP133298.1 | 1 |
| Pseudomonas cannabina pv. alisalensis | Pseudomonadota | Gammaproteobacteria | NZ_CP067022.1 | 1 |
| Pseudomonas cavernae | Pseudomonadota | Gammaproteobacteria | NZ_CP032419.1 | 2 |
| Pseudomonas cedrina | Pseudomonadota | Gammaproteobacteria | NZ_LT629753.1 | 1 |

| | | | | |
|---|---|---|---|---|
| Pseudomonas chlororaphis subsp. chlororaphis | Pseudomonadota | Gammaproteobacteria | NZ_CP144767.1 | 2 |
| Pseudomonas cichorii | Pseudomonadota | Gammaproteobacteria | NZ_CP074349.1 | 1 |
| Pseudomonas citronellolis | Pseudomonadota | Gammaproteobacteria | NZ_CP014158.1 | 2 |
| Pseudomonas coronafaciens pv. oryzae str. 1_6 | Pseudomonadota | Gammaproteobacteria | NZ_CP046035.1 | 1 |
| Pseudomonas corrugata | Pseudomonadota | Gammaproteobacteria | NZ_CP014262.1 | 2 |
| Pseudomonas cucumis | Pseudomonadota | Gammaproteobacteria | NZ_CP117454.1 | 2 |
| Pseudomonas danubii | Pseudomonadota | Gammaproteobacteria | NZ_CP120725.1 | 2 |
| Pseudomonas donghuensis | Pseudomonadota | Gammaproteobacteria | NZ_CP071706.1 | 2 |
| Pseudomonas entomophila L48 | Pseudomonadota | Gammaproteobacteria | NC_008027.1 | 2 |
| Pseudomonas eucalypticola | Pseudomonadota | Gammaproteobacteria | NZ_CP056030.1 | 1 |
| Pseudomonas extremaustralis | Pseudomonadota | Gammaproteobacteria | NZ_LT629689.1 | 1 |
| Pseudomonas extremorientalis | Pseudomonadota | Gammaproteobacteria | NZ_CP089519.1 | 1 |
| Pseudomonas fakonensis | Pseudomonadota | Gammaproteobacteria | NZ_CP077076.1 | 2 |
| Pseudomonas ficuserectae | Pseudomonadota | Gammaproteobacteria | NZ_CP085684.1 | 1 |
| Pseudomonas fitomaticsae | Pseudomonadota | Gammaproteobacteria | NZ_CP075567.1 | 2 |
| Pseudomonas fluorescens | Pseudomonadota | Gammaproteobacteria | NZ_LT907842.1 | 1 |
| Pseudomonas fortuita | Pseudomonadota | Gammaproteobacteria | NZ_CP114035.2 | 2 |
| Pseudomonas fragi | Pseudomonadota | Gammaproteobacteria | NZ_CP064354.1 | 1 |
| Pseudomonas frederiksbergensis | Pseudomonadota | Gammaproteobacteria | NZ_CP017886.1 | 2 |
| Pseudomonas fulva | Pseudomonadota | Gammaproteobacteria | NZ_CP132347.1 | 2 |
| Pseudomonas germanica | Pseudomonadota | Gammaproteobacteria | NZ_CP071586.1 | 2 |
| Pseudomonas glycinae | Pseudomonadota | Gammaproteobacteria | NZ_CP014205.2 | 1 |
| Pseudomonas gorinensis | Pseudomonadota | Gammaproteobacteria | NC_023064.1 | 1 |
| Pseudomonas gozinkensis | Pseudomonadota | Gammaproteobacteria | NZ_CP062253.1 | 2 |
| Pseudomonas granadensis | Pseudomonadota | Gammaproteobacteria | NZ_LT629778.1 | 2 |
| Pseudomonas grandcourensis | Pseudomonadota | Gammaproteobacteria | NZ_CP150919.1 | 1 |
| Pseudomonas guariconensis | Pseudomonadota | Gammaproteobacteria | NZ_CP162012.1 | 2 |
| Pseudomonas hamedanensis | Pseudomonadota | Gammaproteobacteria | NZ_CP077091.1 | 2 |
| Pseudomonas helleri | Pseudomonadota | Gammaproteobacteria | NZ_CP174501.1 | 1 |
| Pseudomonas helvetica | Pseudomonadota | Gammaproteobacteria | NZ_CP150917.1 | 2 |
| Pseudomonas hormoni | Pseudomonadota | Gammaproteobacteria | NZ_CP075566.1 | 1 |
| Pseudomonas huanghezhanensis | Pseudomonadota | Gammaproteobacteria | NZ_CP113949.1 | 1 |

| | | | | |
|---|---|---|---|---|
| Pseudomonas idahonensis | Pseudomonadota | Gammaproteobacteria | NZ_CP149586.1 | 2 |
| Pseudomonas inefficax | Pseudomonadota | Gammaproteobacteria | NZ_CP134401.1 | 2 |
| Pseudomonas iranensis | Pseudomonadota | Gammaproteobacteria | NZ_CP077092.1 | 2 |
| Pseudomonas izuensis | Pseudomonadota | Gammaproteobacteria | NZ_AP017423.2 | 1 |
| Pseudomonas juntendi | Pseudomonadota | Gammaproteobacteria | NZ_CP091088.1 | 2 |
| Pseudomonas kermanshahensis | Pseudomonadota | Gammaproteobacteria | NZ_CP119382.1 | 2 |
| Pseudomonas khavaziana | Pseudomonadota | Gammaproteobacteria | NZ_CP129946.1 | 1 |
| Pseudomonas kielensis | Pseudomonadota | Gammaproteobacteria | NZ_CP130043.1 | 2 |
| Pseudomonas knackmussii | Pseudomonadota | Gammaproteobacteria | NZ_CP047698.1 | 2 |
| Pseudomonas koreensis | Pseudomonadota | Gammaproteobacteria | NZ_CP015852.1 | 2 |
| Pseudomonas kribbensis | Pseudomonadota | Gammaproteobacteria | NZ_CP029608.1 | 2 |
| Pseudomonas kulmbachensis | Pseudomonadota | Gammaproteobacteria | NZ_OY817088.1 | 1 |
| Pseudomonas kurunegalensis | Pseudomonadota | Gammaproteobacteria | NZ_CP127871.1 | 2 |
| Pseudomonas laurentiana | Pseudomonadota | Gammaproteobacteria | NZ_OY782661.1 | 2 |
| Pseudomonas leptonychotis | Pseudomonadota | Gammaproteobacteria | NZ_CP180477.1 | 2 |
| Pseudomonas lijiangensis | Pseudomonadota | Gammaproteobacteria | NZ_CP076668.1 | 1 |
| Pseudomonas lini | Pseudomonadota | Gammaproteobacteria | NZ_LT629746.1 | 2 |
| Pseudomonas lundensis | Pseudomonadota | Gammaproteobacteria | NZ_CP062158.2 | 1 |
| Pseudomonas mandelii | Pseudomonadota | Gammaproteobacteria | NZ_LT629796.1 | 2 |
| Pseudomonas marginalis | Pseudomonadota | Gammaproteobacteria | NZ_CP125381.1 | 1 |
| Pseudomonas marincola | Pseudomonadota | Gammaproteobacteria | NZ_LR215729.2 | 2 |
| Pseudomonas maumuensis | Pseudomonadota | Gammaproteobacteria | NZ_CP077077.1 | 2 |
| Pseudomonas mediterranea | Pseudomonadota | Gammaproteobacteria | NZ_LT629790.1 | 2 |
| Pseudomonas monachiensis | Pseudomonadota | Gammaproteobacteria | NZ_CP178857.1 | 1 |
| Pseudomonas monsensis | Pseudomonadota | Gammaproteobacteria | NZ_CP077087.1 | 2 |
| Pseudomonas monteilii | Pseudomonadota | Gammaproteobacteria | NZ_CP022562.1 | 2 |
| Pseudomonas moorei | Pseudomonadota | Gammaproteobacteria | NZ_AP035776.1 | 1 |
| Pseudomonas mosselii | Pseudomonadota | Gammaproteobacteria | NZ_CP024159.1 | 2 |
| Pseudomonas mucidolens | Pseudomonadota | Gammaproteobacteria | NZ_LS483433.1 | 1 |
| Pseudomonas multiresinivorans | Pseudomonadota | Gammaproteobacteria | NZ_CP048833.1 | 2 |
| Pseudomonas muyukensis | Pseudomonadota | Gammaproteobacteria | NZ_CP077073.1 | 2 |
| Pseudomonas nunensis | Pseudomonadota | Gammaproteobacteria | NZ_CP101125.1 | 2 |

| | | | | |
|---|---|---|---|---|
| Pseudomonas ogarae | Pseudomonadota | Gammaproteobacteria | NZ_CP077086.1 | 2 |
| Pseudomonas orientalis | Pseudomonadota | Gammaproteobacteria | NZ_CP027723.1 | 1 |
| Pseudomonas oryzae | Pseudomonadota | Gammaproteobacteria | NZ_LT629751.1 | 2 |
| Pseudomonas oryziphila | Pseudomonadota | Gammaproteobacteria | NZ_CP034338.1 | 2 |
| Pseudomonas paeninsulae | Pseudomonadota | Gammaproteobacteria | NZ_CP141799.1 | 2 |
| Pseudomonas palleroniana | Pseudomonadota | Gammaproteobacteria | NZ_CP092411.1 | 1 |
| Pseudomonas paracarnis | Pseudomonadota | Gammaproteobacteria | NZ_CP176709.1 | 1 |
| Pseudomonas paraeruginosa | Pseudomonadota | Gammaproteobacteria | NZ_CP020560.1 | 2 |
| Pseudomonas paraveronii | Pseudomonadota | Gammaproteobacteria | NZ_OY817130.1 | 1 |
| Pseudomonas pergaminensis | Pseudomonadota | Gammaproteobacteria | NZ_CP078013.2 | 1 |
| Pseudomonas petroselini | Pseudomonadota | Gammaproteobacteria | NZ_CP173614.1 | 1 |
| Pseudomonas phenolilytica | Pseudomonadota | Gammaproteobacteria | NZ_CP058908.1 | 2 |
| Pseudomonas plecoglossicida | Pseudomonadota | Gammaproteobacteria | NZ_CP031146.1 | 2 |
| Pseudomonas poae RE*1-1-14 | Pseudomonadota | Gammaproteobacteria | NC_020209.1 | 1 |
| Pseudomonas pohangensis | Pseudomonadota | Gammaproteobacteria | NZ_LT629785.1 | 2 |
| Pseudomonas promysalinigenes | Pseudomonadota | Gammaproteobacteria | NZ_CP077094.1 | 2 |
| Pseudomonas prosekii | Pseudomonadota | Gammaproteobacteria | NZ_LT629762.1 | 2 |
| Pseudomonas protegens CHA0 | Pseudomonadota | Gammaproteobacteria | NZ_LS999205.1 | 2 |
| Pseudomonas psychrophila | Pseudomonadota | Gammaproteobacteria | NZ_CP049044.1 | 1 |
| Pseudomonas purpurea | Pseudomonadota | Gammaproteobacteria | NZ_CP150918.1 | 2 |
| Pseudomonas putida NBRC 14164 | Pseudomonadota | Gammaproteobacteria | NC_021505.1 | 2 |
| Pseudomonas qingdaonensis | Pseudomonadota | Gammaproteobacteria | NZ_CP074676.1 | 2 |
| Pseudomonas quebecensis | Pseudomonadota | Gammaproteobacteria | NZ_CP112866.1 | 1 |
| Pseudomonas reinekei | Pseudomonadota | Gammaproteobacteria | NZ_LT629709.1 | 1 |
| Pseudomonas rhizophila | Pseudomonadota | Gammaproteobacteria | NZ_CP024081.1 | 1 |
| Pseudomonas rhizosphaerae | Pseudomonadota | Gammaproteobacteria | NZ_CP009533.1 | 1 |
| Pseudomonas rhodesiae | Pseudomonadota | Gammaproteobacteria | NZ_CP054205.1 | 1 |
| Pseudomonas salmasensis | Pseudomonadota | Gammaproteobacteria | NZ_CP077083.1 | 1 |
| Pseudomonas savastanoi | Pseudomonadota | Gammaproteobacteria | NZ_CP076652.1 | 1 |
| Pseudomonas sediminis | Pseudomonadota | Gammaproteobacteria | NZ_CP060009.1 | 2 |
| Pseudomonas serboccidentalis | Pseudomonadota | Gammaproteobacteria | NZ_CP101655.1 | 2 |
| Pseudomonas sessilinigenes | Pseudomonadota | Gammaproteobacteria | NZ_CP027706.1 | 2 |

| | | | | |
|---|---|---|---|---|
| Pseudomonas shahriarae | Pseudomonadota | Gammaproteobacteria | NZ_CP077085.1 | 1 |
| Pseudomonas shirazensis | Pseudomonadota | Gammaproteobacteria | NZ_CP177041.1 | 2 |
| Pseudomonas shirazica | Pseudomonadota | Gammaproteobacteria | NZ_CP063456.1 | 2 |
| Pseudomonas sichuanensis | Pseudomonadota | Gammaproteobacteria | NZ_CP087185.1 | 2 |
| Pseudomonas sihuiensis | Pseudomonadota | Gammaproteobacteria | NZ_LT629797.1 | 2 |
| Pseudomonas silesiensis | Pseudomonadota | Gammaproteobacteria | NZ_CP014870.1 | 1 |
| Pseudomonas siliginis | Pseudomonadota | Gammaproteobacteria | NZ_CP099598.1 | 2 |
| Pseudomonas simiae | Pseudomonadota | Gammaproteobacteria | NZ_CP010896.1 | 1 |
| Pseudomonas sivasensis | Pseudomonadota | Gammaproteobacteria | NZ_CP090029.1 | 1 |
| Pseudomonas solani | Pseudomonadota | Gammaproteobacteria | NZ_CP158373.1 | 2 |
| Pseudomonas soli | Pseudomonadota | Gammaproteobacteria | NZ_CP128543.1 | 2 |
| Pseudomonas svalbardensis | Pseudomonadota | Gammaproteobacteria | NZ_CP125619.1 | 2 |
| Pseudomonas synxantha | Pseudomonadota | Gammaproteobacteria | NZ_CP027756.1 | 1 |
| Pseudomonas syringae | Pseudomonadota | Gammaproteobacteria | NZ_CP074578.1 | 1 |
| Pseudomonas syringae pv. tagetis | Pseudomonadota | Gammaproteobacteria | NZ_CP092923.1 | 1 |
| Pseudomonas syringae pv. tomato str. DC3000 | Pseudomonadota | Gammaproteobacteria | NC_004578.1 | 1 |
| Pseudomonas taetrolens | Pseudomonadota | Gammaproteobacteria | NZ_LS483370.1 | 1 |
| Pseudomonas tensinigenes | Pseudomonadota | Gammaproteobacteria | NZ_CP077089.1 | 2 |
| Pseudomonas tohonis | Pseudomonadota | Gammaproteobacteria | NZ_AP023189.1 | 2 |
| Pseudomonas tremae | Pseudomonadota | Gammaproteobacteria | NZ_CP066270.1 | 1 |
| Pseudomonas triclosanedens | Pseudomonadota | Gammaproteobacteria | NZ_CP113432.1 | 2 |
| Pseudomonas tritici | Pseudomonadota | Gammaproteobacteria | NZ_CP077084.1 | 1 |
| Pseudomonas trivialis | Pseudomonadota | Gammaproteobacteria | NZ_CP142151.1 | 1 |
| Pseudomonas tructae | Pseudomonadota | Gammaproteobacteria | NZ_CP035952.1 | 2 |
| Pseudomonas umsongensis | Pseudomonadota | Gammaproteobacteria | NZ_CP051487.1 | 1 |
| Pseudomonas vancouverensis | Pseudomonadota | Gammaproteobacteria | NZ_LT629803.1 | 1 |
| Pseudomonas vanderleydeniana | Pseudomonadota | Gammaproteobacteria | NZ_CP077093.1 | 1 |
| Pseudomonas veronii | Pseudomonadota | Gammaproteobacteria | NZ_CP018420.1 | 1 |
| Pseudomonas versuta | Pseudomonadota | Gammaproteobacteria | NZ_CP012676.1 | 1 |
| Pseudomonas viciae | Pseudomonadota | Gammaproteobacteria | NZ_CP035088.1 | 2 |
| Pseudomonas viridiflava | Pseudomonadota | Gammaproteobacteria | NZ_LT855380.1 | 1 |
| Pseudomonas vranovensis | Pseudomonadota | Gammaproteobacteria | NZ_CP159259.1 | 2 |

| | | | | |
|---|---|---|---|---|
| Pseudomonas wenzhouensis | Pseudomonadota | Gammaproteobacteria | NZ_CP072610.1 | 2 |
| Pseudomonas wuhanensis | Pseudomonadota | Gammaproteobacteria | NZ_CP117430.1 | 2 |
| Pseudomonas xantholysinigenes | Pseudomonadota | Gammaproteobacteria | NZ_CP077095.1 | 2 |
| Pseudomonas xanthosomatis | Pseudomonadota | Gammaproteobacteria | NZ_CP077075.1 | 2 |
| Pseudomonas yamanorum | Pseudomonadota | Gammaproteobacteria | NZ_LT629793.1 | 1 |
| Pseudomonas zeae | Pseudomonadota | Gammaproteobacteria | NZ_CP077090.1 | 2 |
| Pseudoxanthomonas daejeonensis | Pseudomonadota | Gammaproteobacteria | NZ_CP093331.1 | 2 |
| Pseudoxanthomonas japonensis | Pseudomonadota | Gammaproteobacteria | NZ_CP171632.1 | 2 |
| Pseudoxanthomonas mexicana | Pseudomonadota | Gammaproteobacteria | NZ_CP060731.1 | 2 |
| Pseudoxanthomonas spadix BD-a59 | Pseudomonadota | Gammaproteobacteria | NC_016147.2 | 2 |
| Pseudoxanthomonas winnipegensis | Pseudomonadota | Gammaproteobacteria | NZ_CP095475.1 | 2 |
| Psychrobacter alimentarius | Pseudomonadota | Gammaproteobacteria | NZ_CP014945.1 | 2 |
| Psychrobacter arcticus 273-4 | Pseudomonadota | Gammaproteobacteria | NC_007204.1 | 2 |
| Psychrobacter arenosus | Pseudomonadota | Gammaproteobacteria | NZ_LR884459.1 | 2 |
| Psychrobacter cryohalolentis K5 | Pseudomonadota | Gammaproteobacteria | NC_007969.1 | 2 |
| Psychrobacter pacificensis | Pseudomonadota | Gammaproteobacteria | NZ_CP048752.1 | 2 |
| Psychrobacter raelei | Pseudomonadota | Gammaproteobacteria | NZ_CP093310.2 | 2 |
| Psychromonas ingrahamii 37 | Pseudomonadota | Gammaproteobacteria | NC_008709.1 | 1 |
| Psychrosphaera aestuarii | Pseudomonadota | Gammaproteobacteria | NZ_CP072844.1 | 2 |
| Psychrosphaera ytuae | Pseudomonadota | Gammaproteobacteria | NZ_CP072110.1 | 2 |
| Rahnella aceris | Pseudomonadota | Gammaproteobacteria | NZ_CP093328.1 | 2 |
| Rahnella aquatilis CIP 78.65 = ATCC 33071 | Pseudomonadota | Gammaproteobacteria | NC_016818.1 | 2 |
| Rahnella inusitata | Pseudomonadota | Gammaproteobacteria | NZ_CP065024.1 | 2 |
| Rahnella sikkimica | Pseudomonadota | Gammaproteobacteria | NZ_CP019062.1 | 2 |
| Rahnella victoriana | Pseudomonadota | Gammaproteobacteria | NZ_CP089919.1 | 2 |
| Raoultella ornithinolytica | Pseudomonadota | Gammaproteobacteria | NZ_CP130153.1 | 2 |
| Raoultella planticola | Pseudomonadota | Gammaproteobacteria | NZ_CP026047.1 | 2 |
| Raoultella terrigena | Pseudomonadota | Gammaproteobacteria | NZ_CP050508.1 | 2 |
| Reinekea forsetii | Pseudomonadota | Gammaproteobacteria | NZ_CP011797.1 | 1 |
| Rheinheimera mangrovi | Pseudomonadota | Gammaproteobacteria | NZ_CP034683.1 | 2 |
| Rhodanobacter denitrificans | Pseudomonadota | Gammaproteobacteria | NZ_CP088980.1 | 2 |
| Rickettsiella endosymbiont of Aleochara curtula | Pseudomonadota | Gammaproteobacteria | NZ_OZ034990.1 | 1 |

| | | | | |
|---|---|---|---|---|
| Rickettsiella endosymbiont of Dermanyssus gallinae | Pseudomonadota | Gammaproteobacteria | NZ_CP079094.1 | 1 |
| Rickettsiella endosymbiont of Litargus connexus | Pseudomonadota | Gammaproteobacteria | NZ_OZ195516.1 | 1 |
| Rickettsiella endosymbiont of Miltochrista miniata | Pseudomonadota | Gammaproteobacteria | NZ_OZ035017.1 | 1 |
| Rickettsiella endosymbiont of Rhagonycha lignosa | Pseudomonadota | Gammaproteobacteria | NZ_OZ035011.1 | 1 |
| Rickettsiella endosymbiont of Xylota segnis | Pseudomonadota | Gammaproteobacteria | NZ_OZ026451.1 | 1 |
| Rodentibacter caecimuris | Pseudomonadota | Gammaproteobacteria | NZ_CP040863.1 | 2 |
| Rodentibacter haemolyticus | Pseudomonadota | Gammaproteobacteria | NZ_CP063056.1 | 2 |
| Rouxiella badensis | Pseudomonadota | Gammaproteobacteria | NZ_CP114062.1 | 2 |
| Saccharobesus litoralis | Pseudomonadota | Gammaproteobacteria | NZ_CP026604.1 | 2 |
| Saccharophagus degradans 2-40 | Pseudomonadota | Gammaproteobacteria | NC_007912.1 | 2 |
| Saccharospirillum mangrovi | Pseudomonadota | Gammaproteobacteria | NZ_CP031415.1 | 2 |
| Salinimonas iocasae | Pseudomonadota | Gammaproteobacteria | NZ_CP039852.1 | 2 |
| Salinimonas lutimaris | Pseudomonadota | Gammaproteobacteria | NZ_CP036536.1 | 2 |
| Salinimonas marina | Pseudomonadota | Gammaproteobacteria | NZ_CP064795.1 | 2 |
| Salinimonas sediminis | Pseudomonadota | Gammaproteobacteria | NZ_CP031769.1 | 2 |
| Saliniradius amylolyticus | Pseudomonadota | Gammaproteobacteria | NZ_CP029347.1 | 2 |
| Salinivibrio kushneri | Pseudomonadota | Gammaproteobacteria | NZ_CP114590.1 | 2 |
| Salinivibrio proteolyticus | Pseudomonadota | Gammaproteobacteria | NZ_CP114584.1 | 2 |
| Salmonella bongori NCTC 12419 | Pseudomonadota | Gammaproteobacteria | NC_015761.1 | 2 |
| Salmonella enterica subsp. enterica serovar Typhimurium s | Pseudomonadota | Gammaproteobacteria | NC_003197.2 | 2 |
| Scandinavium goeteborgense | Pseudomonadota | Gammaproteobacteria | NZ_CP054058.1 | 2 |
| secondary endosymbiont of Ctenarytaina eucalypti | Pseudomonadota | Gammaproteobacteria | NC_018419.1 | 1 |
| secondary endosymbiont of Heteropsylla cubana | Pseudomonadota | Gammaproteobacteria | NC_018420.1 | 1 |
| secondary endosymbiont of Trabutina mannipara | Pseudomonadota | Gammaproteobacteria | NZ_LT594522.1 | 1 |
| Sedimenticola thiotaurini | Pseudomonadota | Gammaproteobacteria | NZ_CP011412.1 | 2 |
| Serratia aquatilis | Pseudomonadota | Gammaproteobacteria | NZ_CP173186.1 | 2 |
| Serratia entomophila | Pseudomonadota | Gammaproteobacteria | NZ_CP082787.1 | 2 |
| Serratia ficaria | Pseudomonadota | Gammaproteobacteria | NZ_LT906479.1 | 2 |
| Serratia fonticola | Pseudomonadota | Gammaproteobacteria | NZ_CP011254.1 | 2 |
| Serratia liquefaciens ATCC 27592 | Pseudomonadota | Gammaproteobacteria | NC_021742.1 | 1 |
| Serratia liquefaciens ATCC 27592 | Pseudomonadota | Gammaproteobacteria | NC_021741.1 | 1 |
| Serratia marcescens | Pseudomonadota | Gammaproteobacteria | NZ_CP127881.1 | 2 |

| | | | | |
|---|---|---|---|---|
| Serratia nematodiphila | Pseudomonadota | Gammaproteobacteria | NZ_CP038662.1 | 2 |
| Serratia nevei | Pseudomonadota | Gammaproteobacteria | NZ_CP149940.1 | 2 |
| Serratia plymuthica AS9 | Pseudomonadota | Gammaproteobacteria | NC_015567.1 | 2 |
| Serratia proteamaculans | Pseudomonadota | Gammaproteobacteria | NZ_OX461105.1 | 2 |
| Serratia quinivorans | Pseudomonadota | Gammaproteobacteria | NZ_LR134494.1 | 2 |
| Serratia rhizosphaerae | Pseudomonadota | Gammaproteobacteria | NZ_CP041764.1 | 2 |
| Serratia rubidaea | Pseudomonadota | Gammaproteobacteria | NZ_CP065640.1 | 2 |
| Serratia sarumanii | Pseudomonadota | Gammaproteobacteria | NZ_CP142089.1 | 2 |
| Serratia surfactantfaciens | Pseudomonadota | Gammaproteobacteria | NZ_CP016948.1 | 2 |
| Serratia symbiotica | Pseudomonadota | Gammaproteobacteria | NZ_CP050855.1 | 2 |
| Serratia ureilytica | Pseudomonadota | Gammaproteobacteria | NZ_CP071320.1 | 2 |
| Shewanella acanthi | Pseudomonadota | Gammaproteobacteria | NZ_CP080413.1 | 2 |
| Shewanella aegiceratis | Pseudomonadota | Gammaproteobacteria | NZ_CP080416.1 | 2 |
| Shewanella algae | Pseudomonadota | Gammaproteobacteria | NZ_CP068230.1 | 2 |
| Shewanella alkalitolerans | Pseudomonadota | Gammaproteobacteria | NZ_CP080422.1 | 2 |
| Shewanella amazonensis SB2B | Pseudomonadota | Gammaproteobacteria | NC_008700.1 | 2 |
| Shewanella avicenniae | Pseudomonadota | Gammaproteobacteria | NZ_CP071503.1 | 2 |
| Shewanella baltica OS678 | Pseudomonadota | Gammaproteobacteria | NC_016901.1 | 2 |
| Shewanella carassii | Pseudomonadota | Gammaproteobacteria | NZ_AP024618.1 | 2 |
| Shewanella chilikensis | Pseudomonadota | Gammaproteobacteria | NZ_CP149966.1 | 2 |
| Shewanella cyperi | Pseudomonadota | Gammaproteobacteria | NZ_CP071501.1 | 2 |
| Shewanella decolorationis | Pseudomonadota | Gammaproteobacteria | NZ_CP037898.1 | 1 |
| Shewanella denitrificans OS217 | Pseudomonadota | Gammaproteobacteria | NC_007954.1 | 1 |
| Shewanella donghaensis | Pseudomonadota | Gammaproteobacteria | NZ_CP041783.1 | 2 |
| Shewanella eurypsychrophilus | Pseudomonadota | Gammaproteobacteria | NZ_CP045503.2 | 2 |
| Shewanella frigidimarina NCIMB 400 | Pseudomonadota | Gammaproteobacteria | NC_008345.1 | 2 |
| Shewanella glacialimarina | Pseudomonadota | Gammaproteobacteria | NZ_CP041216.1 | 2 |
| Shewanella goraebulensis | Pseudomonadota | Gammaproteobacteria | NZ_CP126972.1 | 2 |
| Shewanella halifaxensis HAW-EB4 | Pseudomonadota | Gammaproteobacteria | NC_010334.1 | 2 |
| Shewanella halotolerans | Pseudomonadota | Gammaproteobacteria | NZ_CP080417.1 | 2 |
| Shewanella inventionis | Pseudomonadota | Gammaproteobacteria | NZ_CP082926.1 | 2 |
| Shewanella japonica | Pseudomonadota | Gammaproteobacteria | NZ_CP020472.1 | 2 |

| | | | | |
|---|---|---|---|---|
| Shewanella khirikhana | Pseudomonadota | Gammaproteobacteria | NZ_CP020373.1 | 2 |
| Shewanella litorisediminis | Pseudomonadota | Gammaproteobacteria | NZ_CP069213.1 | 2 |
| Shewanella livingstonensis | Pseudomonadota | Gammaproteobacteria | NZ_CP034015.1 | 2 |
| Shewanella loihica PV-4 | Pseudomonadota | Gammaproteobacteria | NC_009092.1 | 2 |
| Shewanella mangrovisoli | Pseudomonadota | Gammaproteobacteria | NZ_CP080412.1 | 2 |
| Shewanella marisflavi | Pseudomonadota | Gammaproteobacteria | NZ_CP022272.1 | 2 |
| Shewanella maritima | Pseudomonadota | Gammaproteobacteria | NZ_CP036200.1 | 2 |
| Shewanella mesophila | Pseudomonadota | Gammaproteobacteria | NZ_CP080421.1 | 2 |
| Shewanella oncorhynchi | Pseudomonadota | Gammaproteobacteria | NZ_CP132914.1 | 2 |
| Shewanella oneidensis MR-1 | Pseudomonadota | Gammaproteobacteria | NC_004347.2 | 2 |
| Shewanella pealeana ATCC 700345 | Pseudomonadota | Gammaproteobacteria | NC_009901.1 | 2 |
| Shewanella piezotolerans WP3 | Pseudomonadota | Gammaproteobacteria | NC_011566.1 | 2 |
| Shewanella polaris | Pseudomonadota | Gammaproteobacteria | NZ_CP041036.1 | 2 |
| Shewanella psychromarinicola | Pseudomonadota | Gammaproteobacteria | NZ_CP034073.1 | 2 |
| Shewanella psychropiezotolerans | Pseudomonadota | Gammaproteobacteria | NZ_CP041614.1 | 2 |
| Shewanella psychrotolerans | Pseudomonadota | Gammaproteobacteria | NZ_CP080419.1 | 2 |
| Shewanella putrefaciens | Pseudomonadota | Gammaproteobacteria | NZ_CP066370.1 | 2 |
| Shewanella rhizosphaerae | Pseudomonadota | Gammaproteobacteria | NZ_CP080420.1 | 2 |
| Shewanella sedimentimangrovi | Pseudomonadota | Gammaproteobacteria | NZ_CP071502.1 | 2 |
| Shewanella spartinae | Pseudomonadota | Gammaproteobacteria | NZ_CP080418.1 | 2 |
| Shewanella violacea DSS12 | Pseudomonadota | Gammaproteobacteria | NC_014012.1 | 2 |
| Shewanella woodyi ATCC 51908 | Pseudomonadota | Gammaproteobacteria | NC_010506.1 | 2 |
| Shewanella xiamenensis | Pseudomonadota | Gammaproteobacteria | NZ_CP079718.1 | 2 |
| Shewanella youngdeokensis | Pseudomonadota | Gammaproteobacteria | NZ_CP136522.1 | 2 |
| Shewanella yunxiaonensis | Pseudomonadota | Gammaproteobacteria | NZ_CP073587.1 | 2 |
| Shewanella zhangzhouensis | Pseudomonadota | Gammaproteobacteria | NZ_CP080414.1 | 2 |
| Shigella dysenteriae | Pseudomonadota | Gammaproteobacteria | NZ_CP055055.1 | 2 |
| Shigella flexneri 2a str. 301 | Pseudomonadota | Gammaproteobacteria | NC_004337.2 | 2 |
| Shigella sonnei | Pseudomonadota | Gammaproteobacteria | NZ_CP026802.1 | 2 |
| Shimwellia blattae DSM 4481 = NBRC 105725 | Pseudomonadota | Gammaproteobacteria | NC_017910.1 | 2 |
| Siccibacter colletis | Pseudomonadota | Gammaproteobacteria | NZ_CP074352.1 | 2 |
| Simiduia litorea | Pseudomonadota | Gammaproteobacteria | NZ_AP031497.1 | 2 |

| | | | | |
|---|---|---|---|---|
| Sodalis endosymbiont of Henestaris halophilus | Pseudomonadota | Gammaproteobacteria | NZ_LT897836.1 | 1 |
| Sodalis glossinidius str. 'morsitans' | Pseudomonadota | Gammaproteobacteria | NC_007712.1 | 2 |
| Sodalis-like secondary symbiont of Drepanosiphum platan | Pseudomonadota | Gammaproteobacteria | NZ_OZ060924.1 | 1 |
| Solilutibacter oculi | Pseudomonadota | Gammaproteobacteria | NZ_CP029556.1 | 2 |
| Spartinivicinus ruber | Pseudomonadota | Gammaproteobacteria | NZ_CP048878.1 | 2 |
| Spiribacter halobius | Pseudomonadota | Gammaproteobacteria | NZ_CP086615.1 | 2 |
| Spiribacter roseus | Pseudomonadota | Gammaproteobacteria | NZ_CP016382.1 | 2 |
| Spiribacter salinus M19-40 | Pseudomonadota | Gammaproteobacteria | NC_021291.1 | 1 |
| Spongiibacter nanhainus | Pseudomonadota | Gammaproteobacteria | NZ_CP066167.1 | 2 |
| Spongiibacter taiwanensis | Pseudomonadota | Gammaproteobacteria | NZ_CP098455.1 | 2 |
| Stenotrophomonas acidaminiphila | Pseudomonadota | Gammaproteobacteria | NZ_CP043567.1 | 2 |
| Stenotrophomonas aracearum | Pseudomonadota | Gammaproteobacteria | NZ_CP115543.1 | 2 |
| Stenotrophomonas forensis | Pseudomonadota | Gammaproteobacteria | NZ_CP082270.1 | 2 |
| Stenotrophomonas geniculata | Pseudomonadota | Gammaproteobacteria | NZ_CP140571.1 | 2 |
| Stenotrophomonas indicatrix | Pseudomonadota | Gammaproteobacteria | NZ_CP037883.1 | 2 |
| Stenotrophomonas lactitubi | Pseudomonadota | Gammaproteobacteria | NZ_CP167120.1 | 2 |
| Stenotrophomonas maltophilia | Pseudomonadota | Gammaproteobacteria | NZ_LT906480.1 | 2 |
| Stenotrophomonas oahuensis | Pseudomonadota | Gammaproteobacteria | NZ_CP115541.1 | 2 |
| Stenotrophomonas pigmentata | Pseudomonadota | Gammaproteobacteria | NZ_CP130832.1 | 2 |
| Stenotrophomonas rhizophila | Pseudomonadota | Gammaproteobacteria | NZ_CP007597.1 | 2 |
| Stenotrophomonas riyadhensis | Pseudomonadota | Gammaproteobacteria | NZ_CP155568.1 | 2 |
| Stenotrophomonas sepilia | Pseudomonadota | Gammaproteobacteria | NZ_CP168059.1 | 2 |
| Steroidobacter denitrificans | Pseudomonadota | Gammaproteobacteria | NZ_CP011971.1 | 2 |
| Stutzerimonas balearica DSM 6083 | Pseudomonadota | Gammaproteobacteria | NZ_CP007511.1 | 2 |
| Stutzerimonas chloritidismutans | Pseudomonadota | Gammaproteobacteria | NZ_CP086067.1 | 2 |
| Stutzerimonas decontaminans | Pseudomonadota | Gammaproteobacteria | NZ_CP007509.1 | 2 |
| Stutzerimonas degradans | Pseudomonadota | Gammaproteobacteria | NZ_CP065721.1 | 2 |
| Stutzerimonas frequens | Pseudomonadota | Gammaproteobacteria | NZ_CP065720.1 | 2 |
| Stutzerimonas stutzeri | Pseudomonadota | Gammaproteobacteria | NZ_AP024722.1 | 2 |
| Stutzerimonas xanthomarina | Pseudomonadota | Gammaproteobacteria | NZ_LT629970.1 | 2 |
| Succinivibrio dextrinosolvens | Pseudomonadota | Gammaproteobacteria | NZ_CP068345.1 | 2 |
| Sulfuricaulis limicola | Pseudomonadota | Gammaproteobacteria | NZ_AP014879.1 | 2 |

| | | | | |
|---|---|---|---|---|
| Sulfuriflexus mobilis | Pseudomonadota | Gammaproteobacteria | NZ_AP018725.1 | 2 |
| Sulfurifustis variabilis | Pseudomonadota | Gammaproteobacteria | NZ_AP014936.1 | 2 |
| Sulfurivermis fontis | Pseudomonadota | Gammaproteobacteria | NZ_AP018724.1 | 2 |
| Symbiopectobacterium purcellii | Pseudomonadota | Gammaproteobacteria | NZ_CP081864.1 | 2 |
| Tahibacter amnicola | Pseudomonadota | Gammaproteobacteria | NZ_CP104694.1 | 2 |
| Tatumella citrea | Pseudomonadota | Gammaproteobacteria | NZ_CP015579.1 | 2 |
| Teredinibacter haidensis | Pseudomonadota | Gammaproteobacteria | NZ_CP060084.1 | 1 |
| Teredinibacter purpureus | Pseudomonadota | Gammaproteobacteria | NZ_CP060092.1 | 1 |
| Teredinibacter turnerae | Pseudomonadota | Gammaproteobacteria | NZ_CP149817.1 | 1 |
| Thalassolituus hydrocarboniclasticus | Pseudomonadota | Gammaproteobacteria | NZ_CP054475.1 | 3 |
| Thalassolituus oleivorans MIL-1 | Pseudomonadota | Gammaproteobacteria | NC_020888.1 | 3 |
| Thalassomonas actiniarum | Pseudomonadota | Gammaproteobacteria | NZ_CP059735.1 | 2 |
| Thalassomonas haliotis | Pseudomonadota | Gammaproteobacteria | NZ_CP059693.1 | 2 |
| Thalassomonas viridans | Pseudomonadota | Gammaproteobacteria | NZ_CP059733.1 | 2 |
| Thalassotalea agarivorans | Pseudomonadota | Gammaproteobacteria | NZ_AP027363.1 | 2 |
| Thalassotalea aquiviva | Pseudomonadota | Gammaproteobacteria | NZ_CP166961.1 | 3 |
| Thalassotalea atypica | Pseudomonadota | Gammaproteobacteria | NZ_AP027364.1 | 2 |
| Thalassotalea crassostreae | Pseudomonadota | Gammaproteobacteria | NZ_CP017689.1 | 2 |
| Thalassotalea fonticola | Pseudomonadota | Gammaproteobacteria | NZ_CP136600.1 | 3 |
| Thalassotalea hakodatensis | Pseudomonadota | Gammaproteobacteria | NZ_AP027365.1 | 2 |
| Thalassotalea maritima | Pseudomonadota | Gammaproteobacteria | NZ_CP166919.1 | 3 |
| Thalassotalea nanhaiensis | Pseudomonadota | Gammaproteobacteria | NZ_CP134146.1 | 3 |
| Thalassotalea piscium | Pseudomonadota | Gammaproteobacteria | NZ_AP027362.1 | 2 |
| Thalassotalea ponticola | Pseudomonadota | Gammaproteobacteria | NZ_CP166871.1 | 3 |
| Thalassotalea psychrophila | Pseudomonadota | Gammaproteobacteria | NZ_CP134145.1 | 3 |
| Thalassotalea sediminis | Pseudomonadota | Gammaproteobacteria | NZ_AP027361.1 | 2 |
| Thaumasiovibrio subtropicus | Pseudomonadota | Gammaproteobacteria | NZ_AP023054.1 | 2 |
| Thermochromatium tepidum ATCC 43061 | Pseudomonadota | Gammaproteobacteria | NZ_CP039268.1 | 2 |
| Thermomonas aquatica | Pseudomonadota | Gammaproteobacteria | NZ_CP040871.1 | 2 |
| Thermomonas brevis | Pseudomonadota | Gammaproteobacteria | NZ_CP060711.1 | 2 |
| Thermomonas paludicola | Pseudomonadota | Gammaproteobacteria | NZ_CP093311.1 | 2 |
| Thioalkalivibrio nitratireducens DSM 14787 | Pseudomonadota | Gammaproteobacteria | NC_019902.2 | 2 |

| | | | | |
|---|---|---|---|---|
| Thioalkalivibrio paradoxus ARh 1 | Pseudomonadota | Gammaproteobacteria | NZ_CP007029.1 | 2 |
| Thioalkalivibrio sulfidiphilus HL-EbGr7 | Pseudomonadota | Gammaproteobacteria | NC_011901.1 | 2 |
| Thioalkalivibrio versutus | Pseudomonadota | Gammaproteobacteria | NZ_CP011367.1 | 2 |
| Thiocapsa bogorovii | Pseudomonadota | Gammaproteobacteria | NZ_CP089309.1 | 2 |
| Thiocystis violascens DSM 198 | Pseudomonadota | Gammaproteobacteria | NC_018012.1 | 2 |
| Thioflavicoccus mobilis 8321 | Pseudomonadota | Gammaproteobacteria | NC_019940.1 | 2 |
| Thiomicrorhabdus aquaedulcis | Pseudomonadota | Gammaproteobacteria | NZ_AP018722.1 | 2 |
| Thiomicrorhabdus immobilis | Pseudomonadota | Gammaproteobacteria | NZ_AP024202.1 | 2 |
| Thiomicrorhabdus indica | Pseudomonadota | Gammaproteobacteria | NZ_CP033040.1 | 2 |
| Thiomicrorhabdus lithotrophica | Pseudomonadota | Gammaproteobacteria | NZ_CP102381.1 | 2 |
| Thiomicrorhabdus sediminis | Pseudomonadota | Gammaproteobacteria | NZ_CP040602.1 | 2 |
| Thiomicrorhabdus xiamenensis | Pseudomonadota | Gammaproteobacteria | NZ_CP054020.1 | 2 |
| Thiomicrospira aerophila AL3 | Pseudomonadota | Gammaproteobacteria | NZ_CP007030.1 | 2 |
| Thiomicrospira cyclica ALM1 | Pseudomonadota | Gammaproteobacteria | NC_015581.1 | 2 |
| Thiopseudomonas alkaliphila | Pseudomonadota | Gammaproteobacteria | NZ_CP012358.1 | 2 |
| Thiorhodovibrio frisius | Pseudomonadota | Gammaproteobacteria | NZ_CP121471.1 | 2 |
| Thiorhodovibrio litoralis | Pseudomonadota | Gammaproteobacteria | NZ_CP121473.1 | 2 |
| Thiorhodovibrio winogradskyi | Pseudomonadota | Gammaproteobacteria | NZ_CP121472.1 | 2 |
| Thiosocius teredinicola | Pseudomonadota | Gammaproteobacteria | NZ_CP019936.1 | 2 |
| Thiosulfatimonas sediminis | Pseudomonadota | Gammaproteobacteria | NZ_AP021889.1 | 2 |
| Thiosulfativibrio zosterae | Pseudomonadota | Gammaproteobacteria | NZ_AP021888.1 | 2 |
| Thiothrix litoralis | Pseudomonadota | Gammaproteobacteria | NZ_CP072801.1 | 2 |
| Thiothrix subterranea | Pseudomonadota | Gammaproteobacteria | NZ_CP053482.1 | 1 |
| Thiothrix winogradskyi | Pseudomonadota | Gammaproteobacteria | NZ_CP091244.1 | 2 |
| Tolumonas auensis DSM 9187 | Pseudomonadota | Gammaproteobacteria | NC_012691.1 | 2 |
| Trabulsiella odontotermitis | Pseudomonadota | Gammaproteobacteria | NZ_CP125781.1 | 2 |
| Utexia brackfieldae | Pseudomonadota | Gammaproteobacteria | NZ_CP133959.1 | 2 |
| Venatoribacter cucullus | Pseudomonadota | Gammaproteobacteria | NZ_CP046056.1 | 3 |
| Vespertiliibacter pulmonis | Pseudomonadota | Gammaproteobacteria | NZ_CP016615.1 | 2 |
| Vibrio aerogenes | Pseudomonadota | Gammaproteobacteria | NZ_AP024861.1 | 2 |
| Vibrio alfacsensis | Pseudomonadota | Gammaproteobacteria | NZ_AP024165.1 | 2 |
| Vibrio algicola | Pseudomonadota | Gammaproteobacteria | NZ_CP045699.1 | 2 |

| | | | | |
|---|---|---|---|---|
| Vibrio alginolyticus | Pseudomonadota | Gammaproteobacteria | NZ_CP098034.1 | 2 |
| Vibrio antiquarius | Pseudomonadota | Gammaproteobacteria | NC_013456.1 | 2 |
| Vibrio aphrogenes | Pseudomonadota | Gammaproteobacteria | NZ_AP018689.1 | 2 |
| Vibrio apostichopi | Pseudomonadota | Gammaproteobacteria | NZ_AP028067.1 | 2 |
| Vibrio aquimaris | Pseudomonadota | Gammaproteobacteria | NZ_CP045350.1 | 2 |
| Vibrio artabrorum | Pseudomonadota | Gammaproteobacteria | NZ_AP025458.1 | 2 |
| Vibrio astriarenae | Pseudomonadota | Gammaproteobacteria | NZ_CP047475.1 | 2 |
| Vibrio atlanticus | Pseudomonadota | Gammaproteobacteria | NZ_AP025460.1 | 2 |
| Vibrio azureus | Pseudomonadota | Gammaproteobacteria | NZ_CP018616.1 | 2 |
| Vibrio bathopelagicus | Pseudomonadota | Gammaproteobacteria | NZ_CP062500.1 | 2 |
| Vibrio breoganii | Pseudomonadota | Gammaproteobacteria | NZ_AP024864.1 | 2 |
| Vibrio campbellii | Pseudomonadota | Gammaproteobacteria | NZ_CP026321.1 | 2 |
| Vibrio chaetopteri | Pseudomonadota | Gammaproteobacteria | NZ_CP115920.1 | 2 |
| Vibrio chagasii | Pseudomonadota | Gammaproteobacteria | NZ_AP025465.1 | 2 |
| Vibrio cholerae | Pseudomonadota | Gammaproteobacteria | NZ_CP043554.1 | 2 |
| Vibrio cidicii | Pseudomonadota | Gammaproteobacteria | NZ_CP035921.1 | 2 |
| Vibrio comitans | Pseudomonadota | Gammaproteobacteria | NZ_AP024866.1 | 2 |
| Vibrio coralliilyticus | Pseudomonadota | Gammaproteobacteria | NZ_CP120719.1 | 2 |
| Vibrio cortegadensis | Pseudomonadota | Gammaproteobacteria | NZ_AP025472.1 | 2 |
| Vibrio crassostreae | Pseudomonadota | Gammaproteobacteria | NZ_AP025476.1 | 2 |
| Vibrio cyclitrophicus | Pseudomonadota | Gammaproteobacteria | NZ_CP039700.1 | 2 |
| Vibrio diabolicus | Pseudomonadota | Gammaproteobacteria | NZ_CP042451.1 | 2 |
| Vibrio diazotrophicus | Pseudomonadota | Gammaproteobacteria | NZ_CP151842.1 | 2 |
| Vibrio echinoideorum | Pseudomonadota | Gammaproteobacteria | NZ_AP025483.1 | 2 |
| Vibrio fluvialis | Pseudomonadota | Gammaproteobacteria | NZ_CP014035.2 | 2 |
| Vibrio fortis | Pseudomonadota | Gammaproteobacteria | NZ_AP025487.1 | 2 |
| Vibrio furnissii | Pseudomonadota | Gammaproteobacteria | NZ_CP040990.1 | 2 |
| Vibrio gallaecicus | Pseudomonadota | Gammaproteobacteria | NZ_AP025490.1 | 2 |
| Vibrio gallicus | Pseudomonadota | Gammaproteobacteria | NZ_AP024871.1 | 2 |
| Vibrio gangliei | Pseudomonadota | Gammaproteobacteria | NZ_AP021869.1 | 2 |
| Vibrio gazogenes | Pseudomonadota | Gammaproteobacteria | NZ_CP092587.1 | 2 |
| Vibrio gigantis | Pseudomonadota | Gammaproteobacteria | NZ_CP092384.1 | 2 |

| | | | | |
|---|---|---|---|---|
| Vibrio harveyi | Pseudomonadota | Gammaproteobacteria | NZ_CP125875.1 | 2 |
| Vibrio hyugaensis | Pseudomonadota | Gammaproteobacteria | NZ_CP025794.1 | 2 |
| Vibrio inusitatus | Pseudomonadota | Gammaproteobacteria | NZ_AP024878.1 | 2 |
| Vibrio ishigakensis | Pseudomonadota | Gammaproteobacteria | NZ_AP024881.1 | 2 |
| Vibrio japonicus | Pseudomonadota | Gammaproteobacteria | NZ_CP102096.1 | 2 |
| Vibrio jasicida 090810c | Pseudomonadota | Gammaproteobacteria | NZ_CP025792.1 | 2 |
| Vibrio kanaloae | Pseudomonadota | Gammaproteobacteria | NZ_AP025497.1 | 2 |
| Vibrio mangrovi | Pseudomonadota | Gammaproteobacteria | NZ_AP024883.1 | 2 |
| Vibrio metoecus | Pseudomonadota | Gammaproteobacteria | NZ_CP129421.1 | 2 |
| Vibrio misgurnus | Pseudomonadota | Gammaproteobacteria | NZ_CP113241.1 | 2 |
| Vibrio natriegens NBRC 15636 = ATCC 14048 = DSM 759 | Pseudomonadota | Gammaproteobacteria | NZ_CP009977.1 | 2 |
| Vibrio neonatus | Pseudomonadota | Gammaproteobacteria | NZ_AP024885.1 | 2 |
| Vibrio nigripulchritudo | Pseudomonadota | Gammaproteobacteria | NZ_AP027049.1 | 2 |
| Vibrio ostreae | Pseudomonadota | Gammaproteobacteria | NZ_CP076643.1 | 2 |
| Vibrio owensii | Pseudomonadota | Gammaproteobacteria | NZ_CP019959.1 | 2 |
| Vibrio palustris | Pseudomonadota | Gammaproteobacteria | NZ_AP024887.1 | 2 |
| Vibrio paracholerae | Pseudomonadota | Gammaproteobacteria | NZ_JBBHLM01000 | 2 |
| Vibrio parahaemolyticus RIMD 2210633 | Pseudomonadota | Gammaproteobacteria | NC_004603.1 | 2 |
| Vibrio pectenicida | Pseudomonadota | Gammaproteobacteria | NZ_AP024889.1 | 2 |
| Vibrio pelagius | Pseudomonadota | Gammaproteobacteria | NZ_AP025503.1 | 2 |
| Vibrio penaeicida | Pseudomonadota | Gammaproteobacteria | NZ_AP025144.1 | 2 |
| Vibrio pomeroyi | Pseudomonadota | Gammaproteobacteria | NZ_AP025506.1 | 2 |
| Vibrio ponticus | Pseudomonadota | Gammaproteobacteria | NZ_AP019657.1 | 2 |
| Vibrio porteresiae DSM 19223 | Pseudomonadota | Gammaproteobacteria | NZ_AP024895.1 | 2 |
| Vibrio qinghaiensis | Pseudomonadota | Gammaproteobacteria | NZ_CP022741.1 | 2 |
| Vibrio quintilis | Pseudomonadota | Gammaproteobacteria | NZ_AP024897.1 | 2 |
| Vibrio rarus | Pseudomonadota | Gammaproteobacteria | NZ_AP024900.1 | 2 |
| Vibrio rhizosphaerae | Pseudomonadota | Gammaproteobacteria | NZ_AP024903.1 | 2 |
| Vibrio rotiferianus | Pseudomonadota | Gammaproteobacteria | NZ_CP018312.1 | 2 |
| Vibrio ruber | Pseudomonadota | Gammaproteobacteria | NZ_AP024905.1 | 2 |
| Vibrio rumoiensis | Pseudomonadota | Gammaproteobacteria | NZ_AP018685.1 | 2 |
| Vibrio scophthalmi | Pseudomonadota | Gammaproteobacteria | NZ_CP134271.1 | 2 |

| | | | | |
|---|---|---|---|---|
| Vibrio spartinae | Pseudomonadota | Gammaproteobacteria | NZ_AP024907.1 | 2 |
| Vibrio splendidus | Pseudomonadota | Gammaproteobacteria | NZ_CP089203.1 | 2 |
| Vibrio syngnathi | Pseudomonadota | Gammaproteobacteria | NZ_CP017916.1 | 2 |
| Vibrio tasmaniensis | Pseudomonadota | Gammaproteobacteria | NZ_AP025510.1 | 2 |
| Vibrio toranzoniae | Pseudomonadota | Gammaproteobacteria | NZ_AP025514.1 | 2 |
| Vibrio tritonius | Pseudomonadota | Gammaproteobacteria | NZ_AP014635.1 | 2 |
| Vibrio tubiashii ATCC 19109 | Pseudomonadota | Gammaproteobacteria | NZ_CP009354.1 | 2 |
| Vibrio vulnificus NBRC 15645 = ATCC 27562 | Pseudomonadota | Gammaproteobacteria | NZ_CP012881.1 | 2 |
| Vibrio zhugei | Pseudomonadota | Gammaproteobacteria | NZ_CP033078.1 | 2 |
| Vibrio ziniensis | Pseudomonadota | Gammaproteobacteria | NZ_CP049331.1 | 2 |
| Vreelandella neptunia | Pseudomonadota | Gammaproteobacteria | NZ_CP140255.1 | 1 |
| Vreelandella piezotolerans | Pseudomonadota | Gammaproteobacteria | NZ_CP048602.1 | 1 |
| Vreelandella profundi | Pseudomonadota | Gammaproteobacteria | NZ_CP077941.1 | 1 |
| Vreelandella stevensii | Pseudomonadota | Gammaproteobacteria | NZ_CP162600.1 | 1 |
| Vreelandella subglaciescola | Pseudomonadota | Gammaproteobacteria | NZ_LT670847.1 | 1 |
| Vreelandella titanicae | Pseudomonadota | Gammaproteobacteria | NZ_CP059082.1 | 1 |
| Vreelandella venusta | Pseudomonadota | Gammaproteobacteria | NZ_CP065135.1 | 1 |
| Wenzhouxiangella marina | Pseudomonadota | Gammaproteobacteria | NZ_CP012154.1 | 2 |
| Wigglesworthia glossinidia endosymbiont of Glossina mors | Pseudomonadota | Gammaproteobacteria | NC_016893.1 | 1 |
| Winslowiella toletana | Pseudomonadota | Gammaproteobacteria | NZ_CP134152.1 | 2 |
| Woeseia oceani | Pseudomonadota | Gammaproteobacteria | NZ_CP016268.1 | 2 |
| Xanthomonas albilineans | Pseudomonadota | Gammaproteobacteria | NZ_CP046570.1 | 2 |
| Xanthomonas arboricola pv. juglandis | Pseudomonadota | Gammaproteobacteria | NZ_HG999362.1 | 2 |
| Xanthomonas axonopodis pv. maculifoliigardeniae | Pseudomonadota | Gammaproteobacteria | NZ_CP167998.1 | 2 |
| Xanthomonas bundabergensis | Pseudomonadota | Gammaproteobacteria | NZ_CP162487.1 | 2 |
| Xanthomonas campestris pv. raphani | Pseudomonadota | Gammaproteobacteria | NZ_CP058243.1 | 2 |
| Xanthomonas cannabis | Pseudomonadota | Gammaproteobacteria | NZ_CP136580.1 | 2 |
| Xanthomonas cassavae CFBP 4642 | Pseudomonadota | Gammaproteobacteria | NZ_CM002139.1 | 1 |
| Xanthomonas cerealis pv. cerealis | Pseudomonadota | Gammaproteobacteria | NZ_CP083804.1 | 2 |
| Xanthomonas citri pv. citri | Pseudomonadota | Gammaproteobacteria | NZ_CP008998.1 | 2 |
| Xanthomonas cucurbitae | Pseudomonadota | Gammaproteobacteria | NZ_CP033326.1 | 2 |
| Xanthomonas dyei | Pseudomonadota | Gammaproteobacteria | NZ_CP103837.1 | 2 |

| | | | | |
|---|---|---|---|---|
| Xanthomonas euroxanthea | Pseudomonadota | Gammaproteobacteria | NZ_LR861803.1 | 2 |
| Xanthomonas euvesicatoria pv. alfalfae | Pseudomonadota | Gammaproteobacteria | NZ_CP072268.1 | 2 |
| Xanthomonas fragariae | Pseudomonadota | Gammaproteobacteria | NZ_LT853882.1 | 2 |
| Xanthomonas graminis pv. graminis | Pseudomonadota | Gammaproteobacteria | NZ_CP076255.1 | 2 |
| Xanthomonas hawaiiensis | Pseudomonadota | Gammaproteobacteria | NZ_CP115873.1 | 2 |
| Xanthomonas hortorum | Pseudomonadota | Gammaproteobacteria | NZ_CP016878.1 | 2 |
| Xanthomonas hyacinthi | Pseudomonadota | Gammaproteobacteria | NZ_CP043476.1 | 2 |
| Xanthomonas hydrangeae | Pseudomonadota | Gammaproteobacteria | NZ_CP103836.1 | 2 |
| Xanthomonas indica | Pseudomonadota | Gammaproteobacteria | NZ_CP131914.1 | 2 |
| Xanthomonas medicagonis | Pseudomonadota | Gammaproteobacteria | NZ_CP162488.1 | 2 |
| Xanthomonas nasturtii | Pseudomonadota | Gammaproteobacteria | NZ_CP142084.2 | 2 |
| Xanthomonas oryzae pv. oryzicola | Pseudomonadota | Gammaproteobacteria | NZ_CP043403.1 | 2 |
| Xanthomonas phaseoli pv. dieffenbachiae | Pseudomonadota | Gammaproteobacteria | NZ_CP041380.1 | 2 |
| Xanthomonas prunicola | Pseudomonadota | Gammaproteobacteria | NZ_CP096142.1 | 2 |
| Xanthomonas rydalmerensis | Pseudomonadota | Gammaproteobacteria | NZ_CP126170.1 | 2 |
| Xanthomonas sacchari | Pseudomonadota | Gammaproteobacteria | NZ_CP132343.1 | 2 |
| Xanthomonas sontii | Pseudomonadota | Gammaproteobacteria | NZ_CP132342.1 | 2 |
| Xanthomonas tesorieronis | Pseudomonadota | Gammaproteobacteria | NZ_CP162490.1 | 2 |
| Xanthomonas theicola | Pseudomonadota | Gammaproteobacteria | NZ_CP049017.1 | 2 |
| Xanthomonas translucens pv. undulosa | Pseudomonadota | Gammaproteobacteria | NZ_CP063993.1 | 2 |
| Xanthomonas vasicola pv. musacearum NCPPB 4379 | Pseudomonadota | Gammaproteobacteria | NZ_CP034655.1 | 2 |
| Xanthomonas vesicatoria ATCC 35937 | Pseudomonadota | Gammaproteobacteria | NZ_CP018725.1 | 2 |
| Xenorhabdus bakwenae | Pseudomonadota | Gammaproteobacteria | NZ_CP119194.1 | 2 |
| Xenorhabdus budapestensis | Pseudomonadota | Gammaproteobacteria | NZ_CP072455.1 | 2 |
| Xenorhabdus doucetiae | Pseudomonadota | Gammaproteobacteria | NZ_FO704550.1 | 2 |
| Xenorhabdus griffiniae | Pseudomonadota | Gammaproteobacteria | NZ_CP133479.1 | 2 |
| Xenorhabdus hominickii | Pseudomonadota | Gammaproteobacteria | NZ_CP016176.1 | 2 |
| Xenorhabdus nematophila ATCC 19061 | Pseudomonadota | Gammaproteobacteria | NC_014228.1 | 1 |
| Xenorhabdus poinarii G6 | Pseudomonadota | Gammaproteobacteria | NZ_FO704551.1 | 2 |
| Xenorhabdus taiwanensis | Pseudomonadota | Gammaproteobacteria | NZ_AP028978.1 | 2 |
| Xylella fastidiosa | Pseudomonadota | Gammaproteobacteria | NZ_CP118539.1 | 2 |
| Xylella taiwanensis | Pseudomonadota | Gammaproteobacteria | NZ_CP053627.1 | 2 |

| | | | | |
|---|---|---|---|---|
| Yersinia aldovae 670-83 | Pseudomonadota | Gammaproteobacteria | NZ_CP009781.1 | 2 |
| Yersinia alsatica | Pseudomonadota | Gammaproteobacteria | NZ_CP104006.1 | 2 |
| Yersinia bercovieri | Pseudomonadota | Gammaproteobacteria | NZ_CP124240.1 | 2 |
| Yersinia canariae | Pseudomonadota | Gammaproteobacteria | NZ_CP043727.1 | 2 |
| Yersinia enterocolitica subsp. palearctica Y11 | Pseudomonadota | Gammaproteobacteria | NC_017564.1 | 2 |
| Yersinia entomophaga | Pseudomonadota | Gammaproteobacteria | NZ_CP010029.1 | 2 |
| Yersinia hibernica | Pseudomonadota | Gammaproteobacteria | NZ_CP032487.1 | 2 |
| Yersinia intermedia | Pseudomonadota | Gammaproteobacteria | NZ_CP046293.1 | 2 |
| Yersinia mollaretii ATCC 43969 | Pseudomonadota | Gammaproteobacteria | NZ_CP054043.1 | 2 |
| Yersinia pestis A1122 | Pseudomonadota | Gammaproteobacteria | NC_017168.1 | 2 |
| Yersinia pseudotuberculosis | Pseudomonadota | Gammaproteobacteria | NZ_LR134373.1 | 2 |
| Yersinia rochesterensis | Pseudomonadota | Gammaproteobacteria | NZ_CP032482.1 | 2 |
| Yersinia rohdei | Pseudomonadota | Gammaproteobacteria | NZ_CP009787.1 | 2 |
| Yersinia ruckeri | Pseudomonadota | Gammaproteobacteria | NZ_CP071802.1 | 2 |
| Yersinia similis | Pseudomonadota | Gammaproteobacteria | NZ_CP007230.1 | 2 |
| Yersinia vastinensis | Pseudomonadota | Gammaproteobacteria | NZ_CP173260.1 | 2 |
| Zhongshania aliphaticivorans | Pseudomonadota | Gammaproteobacteria | NZ_CP014544.1 | 2 |
| Zobellella denitrificans | Pseudomonadota | Gammaproteobacteria | NZ_CP012621.1 | 2 |
| Zophobihabitans entericus | Pseudomonadota | Gammaproteobacteria | NZ_CP050253.1 | 2 |
| Zymobacter palmae | Pseudomonadota | Gammaproteobacteria | NZ_AP018933.1 | 1 |
| Hydrogenophilus thermoluteolus | Pseudomonadota | Hydrogenophilia | NZ_AP018558.1 | 2 |
| Mariprofundus aestuarium | Pseudomonadota | Zetaproteobacteria | NZ_CP018799.1 | 2 |
| Mariprofundus ferrinatatus | Pseudomonadota | Zetaproteobacteria | NZ_CP018800.1 | 2 |
| Rhodocaloribacter litoris | Rhodothermota | Rhodothermia | NZ_CP076718.1 | 2 |
| Rhodothermus marinus DSM 4252 | Rhodothermota | Rhodothermia | NC_013501.1 | 2 |
| Salinibacter ruber | Rhodothermota | Rhodothermia | NZ_CP030361.1 | 2 |
| Borrelia anserina Es | Spirochaetota | Spirochaetia | NZ_CP013704.1 | 1 |
| Borrelia coriaceae | Spirochaetota | Spirochaetia | NZ_CP075076.1 | 1 |
| Borrelia duttonii Ly | Spirochaetota | Spirochaetia | NC_011229.1 | 1 |
| Borrelia hermsii | Spirochaetota | Spirochaetia | NZ_CP084623.1 | 1 |
| Borrelia maritima | Spirochaetota | Spirochaetia | NZ_CP044535.1 | 1 |
| Borrelia miyamotoi | Spirochaetota | Spirochaetia | NZ_AP024371.1 | 1 |

| | | | | |
|---|---|---|---|---|
| Borrelia nietonii YOR | Spirochaetota | Spirochaetia | NZ_CP073148.1 | 1 |
| Borrelia parkeri HR1 | Spirochaetota | Spirochaetia | NZ_CP007022.1 | 1 |
| Borrelia puertoricensis | Spirochaetota | Spirochaetia | NZ_CP075379.1 | 1 |
| Borrelia recurrentis A1 | Spirochaetota | Spirochaetia | NC_011244.1 | 1 |
| Borrelia turcica IST7 | Spirochaetota | Spirochaetia | NZ_CP028884.1 | 1 |
| Borrelia turicatae 91E135 | Spirochaetota | Spirochaetia | NC_008710.1 | 1 |
| Borreliella afzelii Tom3107 | Spirochaetota | Spirochaetia | NZ_CP009212.1 | 1 |
| Borreliella americana | Spirochaetota | Spirochaetia | CP124113.1 | 1 |
| Borreliella andersonii | Spirochaetota | Spirochaetia | NZ_CP132472.1 | 1 |
| Borreliella bavariensis PBi | Spirochaetota | Spirochaetia | NC_006156.1 | 1 |
| Borreliella bissettiae DN127 | Spirochaetota | Spirochaetia | NC_015921.1 | 1 |
| Borreliella californiensis | Spirochaetota | Spirochaetia | NZ_CP124076.1 | 1 |
| Borreliella carolinensis | Spirochaetota | Spirochaetia | NZ_CP132453.1 | 1 |
| Borreliella finlandensis | Spirochaetota | Spirochaetia | NZ_CP124070.1 | 1 |
| Borreliella garinii | Spirochaetota | Spirochaetia | NZ_CP075232.1 | 1 |
| Borreliella japonica | Spirochaetota | Spirochaetia | NZ_CP124062.1 | 1 |
| Borreliella kurtenbachii | Spirochaetota | Spirochaetia | NZ_CP124058.1 | 1 |
| Borreliella lanei | Spirochaetota | Spirochaetia | NZ_CP124054.1 | 1 |
| Borreliella lusitaniae | Spirochaetota | Spirochaetia | NZ_CP124050.1 | 1 |
| Borreliella mayonii | Spirochaetota | Spirochaetia | NZ_CP015780.1 | 1 |
| Borreliella sinica | Spirochaetota | Spirochaetia | NZ_CP135180.1 | 1 |
| Borreliella spielmanii | Spirochaetota | Spirochaetia | NZ_CP124042.1 | 1 |
| Borreliella tanukii | Spirochaetota | Spirochaetia | NZ_CP124034.1 | 1 |
| Borreliella turdi | Spirochaetota | Spirochaetia | NZ_CP124018.1 | 1 |
| Borreliella valaisiana VS116 | Spirochaetota | Spirochaetia | NZ_ABCY0200000 | 1 |
| Borreliella yangtzensis | Spirochaetota | Spirochaetia | NZ_CP123998.1 | 1 |
| Brachyspira hyodysenteriae | Spirochaetota | Spirochaetia | NZ_CP046932.1 | 1 |
| Brachyspira intermedia PWS/A | Spirochaetota | Spirochaetia | NC_017243.1 | 1 |
| Brachyspira murdochii DSM 12563 | Spirochaetota | Spirochaetia | NC_014150.1 | 1 |
| Brachyspira pilosicoli 95/1000 | Spirochaetota | Spirochaetia | NC_014330.1 | 1 |
| Breznakiella homolactica | Spirochaetota | Spirochaetia | NZ_CP067089.2 | 1 |
| Brucepastera parasyntrophica | Spirochaetota | Spirochaetia | NZ_CP084606.1 | 1 |

| | | | | |
|---|---|---|---|---|
| Candidatus Borrelia fainii | Spirochaetota | Spirochaetia | NZ_AP027070.1 | 1 |
| Candidatus Borreliella tachyglossi | Spirochaetota | Spirochaetia | NZ_CP025785.1 | 1 |
| Candidatus Haliotispira prima | Spirochaetota | Spirochaetia | NZ_CP123443.1 | 1 |
| Entomospira culicis | Spirochaetota | Spirochaetia | NZ_CP118181.1 | 1 |
| Entomospira entomophila | Spirochaetota | Spirochaetia | NZ_CP118174.1 | 1 |
| Entomospira nematocerorum | Spirochaetota | Spirochaetia | NZ_CP118168.1 | 1 |
| Gracilinema caldarium DSM 7334 | Spirochaetota | Spirochaetia | NC_015732.1 | 1 |
| Leadbettera azotonutricia ZAS-9 | Spirochaetota | Spirochaetia | NC_015577.1 | 1 |
| Leptospira biflexa serovar Patoc strain 'Patoc 1 (Paris)' | Spirochaetota | Spirochaetia | NC_010602.1 | 1 |
| Leptospira borgpetersenii serovar Ceylonica | Spirochaetota | Spirochaetia | NZ_CP026671.1 | 1 |
| Leptospira cinconiae | Spirochaetota | Spirochaetia | NZ_CP162137.1 | 1 |
| Leptospira interrogans serovar Copenhageni | Spirochaetota | Spirochaetia | NZ_CP020414.2 | 1 |
| Leptospira kmetyi | Spirochaetota | Spirochaetia | NZ_CP033614.1 | 1 |
| Leptospira kobayashii | Spirochaetota | Spirochaetia | NZ_AP025028.1 | 1 |
| Leptospira mayottensis 200901116 | Spirochaetota | Spirochaetia | NZ_CP024871.1 | 1 |
| Leptospira santarosai serovar Shermani str. LT 821 | Spirochaetota | Spirochaetia | NZ_CP006694.1 | 1 |
| Leptospira tipperaryensis | Spirochaetota | Spirochaetia | NZ_CP015217.1 | 1 |
| Leptospira weilii | Spirochaetota | Spirochaetia | NZ_CP040840.1 | 1 |
| Oceanispirochaeta crateris | Spirochaetota | Spirochaetia | NZ_CP036150.1 | 1 |
| Parasphaerochaeta coccoides DSM 17374 | Spirochaetota | Spirochaetia | NC_015436.1 | 1 |
| Salinispira pacifica | Spirochaetota | Spirochaetia | NC_023035.1 | 1 |
| Sediminispirochaeta smaragdinae DSM 11293 | Spirochaetota | Spirochaetia | NC_014364.1 | 1 |
| Sphaerochaeta associata | Spirochaetota | Spirochaetia | NZ_CP094929.1 | 1 |
| Sphaerochaeta globosa str. Buddy | Spirochaetota | Spirochaetia | NC_015152.1 | 1 |
| Sphaerochaeta pleomorpha str. Grapes | Spirochaetota | Spirochaetia | NC_016633.1 | 1 |
| Spirochaeta africana DSM 8902 | Spirochaetota | Spirochaetia | NC_017098.1 | 1 |
| Spirochaeta thermophila DSM 6578 | Spirochaetota | Spirochaetia | NC_017583.1 | 1 |
| Thermospira aquatica | Spirochaetota | Spirochaetia | NZ_CP073355.1 | 1 |
| Thiospirochaeta perfilievii | Spirochaetota | Spirochaetia | NZ_CP035807.1 | 1 |
| Treponema brennaborense DSM 12168 | Spirochaetota | Spirochaetia | NC_015500.1 | 1 |
| Treponema bryantii | Spirochaetota | Spirochaetia | NZ_AP025286.1 | 1 |
| Treponema denticola ATCC 35405 | Spirochaetota | Spirochaetia | NC_002967.9 | 1 |

| | | | | |
|---|---|---|---|---|
| Treponema medium | Spirochaetota | Spirochaetia | NZ_CP031393.1 | 1 |
| Treponema pallidum subsp. pallidum str. Nichols | Spirochaetota | Spirochaetia | NC_021490.2 | 1 |
| Treponema paraluiscuniculi Cuniculi A | Spirochaetota | Spirochaetia | NC_015714.1 | 1 |
| Treponema parvum | Spirochaetota | Spirochaetia | NZ_CP054142.1 | 1 |
| Treponema pedis str. T A4 | Spirochaetota | Spirochaetia | NC_022097.1 | 1 |
| Treponema peruense | Spirochaetota | Spirochaetia | NZ_CP064936.1 | 1 |
| Treponema phagedenis | Spirochaetota | Spirochaetia | NZ_CP042818.1 | 1 |
| Treponema primitia ZAS-2 | Spirochaetota | Spirochaetia | NC_015578.1 | 1 |
| Treponema putidum | Spirochaetota | Spirochaetia | NZ_CP038804.1 | 1 |
| Treponema saccharophilum | Spirochaetota | Spirochaetia | NZ_AP025288.1 | 1 |
| Treponema socranskii subsp. buccale | Spirochaetota | Spirochaetia | NZ_CP054258.1 | 1 |
| Treponema succinifaciens DSM 2489 | Spirochaetota | Spirochaetia | NC_015385.1 | 1 |
| Turneriella parva DSM 21527 | Spirochaetota | Spirochaetia | NC_018020.1 | 1 |
| Aminithiophilus ramosus | Synergistota | Synergistia | NZ_CP072943.1 | 2 |
| Aminobacterium colombiense DSM 12261 | Synergistota | Synergistia | NC_014011.1 | 2 |
| Aminomonas paucivorans DSM 12260 | Synergistota | Synergistia | NZ_CM001022.1 | 2 |
| Cloacibacillus porcorum | Synergistota | Synergistia | NZ_CP016757.1 | 2 |
| Fretibacterium fastidiosum | Synergistota | Synergistia | NZ_OZ209244.1 | 2 |
| Pyramidobacter piscolens | Synergistota | Synergistia | NZ_AP025585.1 | 2 |
| Thermanaerovibrio acidaminovorans DSM 6589 | Synergistota | Synergistia | NC_013522.1 | 2 |
| Thermanaerovibrio velox DSM 12556 | Synergistota | Synergistia | NZ_CM001377.1 | 2 |
| Candidatus Desulfofervidus auxilii | Thermodesulfobacteriota | Candidatus Desulfoferv | NZ_CP013015.1 | 2 |
| Desulfarculus baarsii DSM 2075 | Thermodesulfobacteriota | Desulfarculia | NC_014365.1 | 2 |
| Desulfoferula mesophila | Thermodesulfobacteriota | Desulfarculia | NZ_AP028679.1 | 2 |
| Desulfobacca acetoxidans DSM 11109 | Thermodesulfobacteriota | Desulfobaccia | NC_015388.1 | 2 |
| Desulfatibacillum aliphaticivorans | Thermodesulfobacteriota | Desulfobacteria | NC_011768.1 | 2 |
| Desulfobacter hydrogenophilus | Thermodesulfobacteriota | Desulfobacteria | NZ_CP036313.1 | 1 |
| Desulfobacter postgatei 2ac9 | Thermodesulfobacteriota | Desulfobacteria | NZ_CM001488.1 | 1 |
| Desulfobacula toluolica Tol2 | Thermodesulfobacteriota | Desulfobacteria | NC_018645.1 | 2 |
| Desulfococcus multivorans | Thermodesulfobacteriota | Desulfobacteria | NZ_CP019913.2 | 2 |
| Desulfoluna limicola | Thermodesulfobacteriota | Desulfobacteria | NZ_AP024488.1 | 2 |
| Desulfonema limicola | Thermodesulfobacteriota | Desulfobacteria | NZ_CP061799.1 | 2 |

| | | | | |
|---|---|---|---|---|
| Desulfonema magnum | Thermodesulfobacteriota | Desulfobacteria | NZ_CP061800.1 | 1 |
| Desulforapulum autotrophicum HRM2 | Thermodesulfobacteriota | Desulfobacteria | NC_012108.1 | 2 |
| Desulfosarcina alkanivorans | Thermodesulfobacteriota | Desulfobacteria | NZ_AP021874.1 | 2 |
| Desulfosarcina ovata subsp. sediminis | Thermodesulfobacteriota | Desulfobacteria | NZ_AP021876.1 | 2 |
| Desulfosarcina variabilis str. Montpellier | Thermodesulfobacteriota | Desulfobacteria | NZ_CP159846.1 | 2 |
| Desulfosarcina widdelii | Thermodesulfobacteriota | Desulfobacteria | NZ_AP021875.1 | 2 |
| Desulfosudis oleivorans Hxd3 | Thermodesulfobacteriota | Desulfobacteria | NC_009943.1 | 2 |
| Desulfobulbus oligotrophicus | Thermodesulfobacteriota | Desulfobulbia | NZ_CP054140.1 | 2 |
| Desulfobulbus oralis | Thermodesulfobacteriota | Desulfobulbia | NZ_CP021255.1 | 2 |
| Desulfobulbus propionicus DSM 2032 | Thermodesulfobacteriota | Desulfobulbia | NC_014972.1 | 2 |
| Desulfocapsa sulfexigens DSM 10523 | Thermodesulfobacteriota | Desulfobulbia | NC_020304.1 | 2 |
| Desulfofustis limnaeus | Thermodesulfobacteriota | Desulfobulbia | NZ_AP025516.1 | 2 |
| Desulfolithobacter dissulfuricans | Thermodesulfobacteriota | Desulfobulbia | NZ_AP024233.1 | 2 |
| Desulfomarina profundi | Thermodesulfobacteriota | Desulfobulbia | NZ_AP024086.1 | 2 |
| Desulfosediminicola flagellatus | Thermodesulfobacteriota | Desulfobulbia | NZ_CP050698.1 | 2 |
| Desulfosediminicola ganghwensis | Thermodesulfobacteriota | Desulfobulbia | NZ_CP050699.1 | 2 |
| Desulfotalea psychrophila LSv54 | Thermodesulfobacteriota | Desulfobulbia | NC_006138.1 | 2 |
| Desulfurivibrio alkaliphilus AHT 2 | Thermodesulfobacteriota | Desulfobulbia | NC_014216.1 | 2 |
| Desulfomonile tiedjei DSM 6799 | Thermodesulfobacteriota | Desulfomonilia | NC_018025.1 | 2 |
| Candidatus Desulfovibrio trichonymphae | Thermodesulfobacteriota | Desulfovibrionia | NZ_AP017368.1 | 1 |
| Desulfocurvibacter africanus subsp. africanus str. Walvis E | Thermodesulfobacteriota | Desulfovibrionia | NC_016629.1 | 2 |
| Desulfohalobium retbaense DSM 5692 | Thermodesulfobacteriota | Desulfovibrionia | NC_013223.1 | 2 |
| Desulfolutivibrio sulfodismutans DSM 3696 | Thermodesulfobacteriota | Desulfovibrionia | NZ_CP045504.1 | 2 |
| Desulfolutivibrio sulfoxidireducens | Thermodesulfobacteriota | Desulfovibrionia | NZ_CP045508.1 | 2 |
| Desulfomicrobium baculatum DSM 4028 | Thermodesulfobacteriota | Desulfovibrionia | NC_013173.1 | 2 |
| Desulfomicrobium orale DSM 12838 | Thermodesulfobacteriota | Desulfovibrionia | NZ_CP014230.1 | 2 |
| Desulfovibrio aminophilus | Thermodesulfobacteriota | Desulfovibrionia | NZ_CP168306.1 | 1 |
| Desulfovibrio desulfuricans | Thermodesulfobacteriota | Desulfovibrionia | NZ_CP072608.1 | 1 |
| Desulfovibrio fairfieldensis | Thermodesulfobacteriota | Desulfovibrionia | NZ_CP014229.1 | 1 |
| Desulfovibrio ferrophilus | Thermodesulfobacteriota | Desulfovibrionia | NZ_AP017378.1 | 2 |
| Desulfovibrio mangrovi | Thermodesulfobacteriota | Desulfovibrionia | NZ_CP104208.1 | 1 |
| Desulfovibrio subterraneus | Thermodesulfobacteriota | Desulfovibrionia | NZ_CP087381.1 | 1 |

| | | | |
|---|---|---|---|
| Lawsonia intracellularis N343 | Thermodesulfobacteriota Desulfovibrionia | NC_020127.1 | 1 |
| Maridesulfovibrio hydrothermalis AM13 = DSM 14728 | Thermodesulfobacteriota Desulfovibrionia | NC_020055.1 | 1 |
| Maridesulfovibrio salexigens DSM 2638 | Thermodesulfobacteriota Desulfovibrionia | NC_012881.1 | 1 |
| Megalodesulfovibrio gigas DSM 1382 = ATCC 19364 | Thermodesulfobacteriota Desulfovibrionia | NC_022444.1 | 1 |
| Nitratidesulfovibrio vulgaris DP4 | Thermodesulfobacteriota Desulfovibrionia | NC_008751.1 | 1 |
| Oceanidesulfovibrio marinus | Thermodesulfobacteriota Desulfovibrionia | NZ_CP039543.1 | 2 |
| Oleidesulfovibrio alaskensis G20 | Thermodesulfobacteriota Desulfovibrionia | NC_007519.1 | 1 |
| Pseudodesulfovibrio aespoeensis Aspo-2 | Thermodesulfobacteriota Desulfovibrionia | NC_014844.1 | 1 |
| Pseudodesulfovibrio cashew | Thermodesulfobacteriota Desulfovibrionia | NZ_CP046400.1 | 1 |
| Pseudodesulfovibrio mercurii | Thermodesulfobacteriota Desulfovibrionia | NC_016803.1 | 1 |
| Pseudodesulfovibrio methanolicus | Thermodesulfobacteriota Desulfovibrionia | NZ_CP146609.1 | 1 |
| Pseudodesulfovibrio nedwellii | Thermodesulfobacteriota Desulfovibrionia | NZ_AP026709.1 | 1 |
| Pseudodesulfovibrio piezophilus C1TLV30 | Thermodesulfobacteriota Desulfovibrionia | NC_020409.1 | 1 |
| Pseudodesulfovibrio portus | Thermodesulfobacteriota Desulfovibrionia | NZ_AP026708.1 | 1 |
| Pseudodesulfovibrio profundus | Thermodesulfobacteriota Desulfovibrionia | NZ_LT907975.1 | 1 |
| Pseudodesulfovibrio sediminis | Thermodesulfobacteriota Desulfovibrionia | NZ_AP024485.1 | 1 |
| Pseudodesulfovibrio thermohalotolerans | Thermodesulfobacteriota Desulfovibrionia | NZ_CP120635.1 | 1 |
| Pseudodesulfovibrio tunisiensis | Thermodesulfobacteriota Desulfovibrionia | NZ_CP094380.1 | 1 |
| Salidesulfovibrio onnuriiensis | Thermodesulfobacteriota Desulfovibrionia | NZ_CP040751.1 | 1 |
| Solidesulfovibrio carbinolicus | Thermodesulfobacteriota Desulfovibrionia | NZ_CP026538.1 | 2 |
| Solidesulfovibrio carbinoliphilus subsp. oakridgensis | Thermodesulfobacteriota Desulfovibrionia | NZ_CM001368.1 | 2 |
| Solidesulfovibrio magneticus RS-1 | Thermodesulfobacteriota Desulfovibrionia | NC_012796.1 | 2 |
| Taurinivorans muris | Thermodesulfobacteriota Desulfovibrionia | NZ_CP065938.1 | 1 |
| Citrifermentans bemidjiense Bem | Thermodesulfobacteriota Desulfuromonadia | NC_011146.1 | 2 |
| Citrifermentans bremense | Thermodesulfobacteriota Desulfuromonadia | NZ_AP023213.1 | 2 |
| Desulfuromonas soudanensis | Thermodesulfobacteriota Desulfuromonadia | NZ_CP010802.1 | 2 |
| Desulfuromonas thiophila | Thermodesulfobacteriota Desulfuromonadia | NZ_OY759968.1 | 2 |
| Desulfuromonas versatilis | Thermodesulfobacteriota Desulfuromonadia | NZ_AP024355.1 | 2 |
| Geoalkalibacter halelectricus | Thermodesulfobacteriota Desulfuromonadia | NZ_CP092109.1 | 2 |
| Geoalkalibacter subterraneus | Thermodesulfobacteriota Desulfuromonadia | NZ_CP010311.1 | 2 |
| Geobacter benzoatilyticus | Thermodesulfobacteriota Desulfuromonadia | NZ_CP071382.1 | 2 |
| Geobacter metallireducens GS-15 | Thermodesulfobacteriota Desulfuromonadia | NC_007517.1 | 2 |

| | | | | |
|---|---|---|---|---|
| Geobacter pickeringii | Thermodesulfobacteriota | Desulfuromonadia | NZ_CP009788.1 | 2 |
| Geobacter sulfurreducens PCA | Thermodesulfobacteriota | Desulfuromonadia | NC_002939.5 | 2 |
| Geomonas nitrogeniifigens | Thermodesulfobacteriota | Desulfuromonadia | NZ_CP077684.1 | 2 |
| Geomonas oryzisoli | Thermodesulfobacteriota | Desulfuromonadia | NZ_CP076723.1 | 2 |
| Geomonas paludis | Thermodesulfobacteriota | Desulfuromonadia | NZ_CP096574.1 | 2 |
| Geomonas subterranea | Thermodesulfobacteriota | Desulfuromonadia | NZ_CP078096.1 | 2 |
| Geotalea daltonii FRC-32 | Thermodesulfobacteriota | Desulfuromonadia | NC_011979.1 | 2 |
| Geotalea uraniireducens Rf4 | Thermodesulfobacteriota | Desulfuromonadia | NC_009483.1 | 2 |
| Pelobacter propionicus DSM 2379 | Thermodesulfobacteriota | Desulfuromonadia | NC_008609.1 | 2 |
| Syntrophotalea acetylenica | Thermodesulfobacteriota | Desulfuromonadia | NZ_CP015518.1 | 2 |
| Syntrophotalea acetylenivorans | Thermodesulfobacteriota | Desulfuromonadia | NZ_CP015519.1 | 2 |
| Syntrophotalea carbinolica DSM 2380 | Thermodesulfobacteriota | Desulfuromonadia | NC_007498.2 | 2 |
| Trichlorobacter ammonificans | Thermodesulfobacteriota | Desulfuromonadia | NZ_OW150024.1 | 2 |
| Trichlorobacter lovleyi SZ | Thermodesulfobacteriota | Desulfuromonadia | NC_010814.1 | 2 |
| Syntrophus aciditrophicus SB | Thermodesulfobacteriota | Syntrophia | NC_007759.1 | 1 |
| Syntrophus buswellii | Thermodesulfobacteriota | Syntrophia | NZ_CP175563.1 | 1 |
| Desulfoglaeba alkanexedens ALDC | Thermodesulfobacteriota | Syntrophobacteria | NZ_CP040098.1 | 2 |
| Syntrophobacter fumaroxidans MPOB | Thermodesulfobacteriota | Syntrophobacteria | NC_008554.1 | 2 |
| Caldimicrobium thiodismutans | Thermodesulfobacteriota | Thermodesulfobacteria | NZ_AP014945.1 | 2 |
| Thermodesulfatator indicus DSM 15286 | Thermodesulfobacteriota | Thermodesulfobacteria | NC_015681.1 | 2 |
| Thermodesulfobacterium commune DSM 2178 | Thermodesulfobacteriota | Thermodesulfobacteria | NZ_CP008796.1 | 2 |
| Thermodesulfobacterium geofontis OPF15 | Thermodesulfobacteriota | Thermodesulfobacteria | NC_015682.1 | 2 |
| Thermosulfurimonas marina | Thermodesulfobacteriota | Thermodesulfobacteria | NZ_CP042909.1 | 2 |
| Thermosulfuriphilus ammonigenes | Thermodesulfobacteriota | Thermodesulfobacteria | NZ_CP048877.1 | 2 |
| Thermodesulfobium acidiphilum | Thermodesulfobiota | Thermodesulfobiia | NZ_CP020921.1 | 2 |
| Thermodesulfobium narugense DSM 14796 | Thermodesulfobiota | Thermodesulfobiia | NC_015499.1 | 2 |
| Sphaerobacter thermophilus DSM 20745 | Thermomicrobiota | Thermomicrobia | NC_013523.1 | 2 |
| Thermomicrobium roseum DSM 5159 | Thermomicrobiota | Thermomicrobia | NC_011959.1 | 2 |
| Thermosulfidibacter takaii ABI70S6 | Thermosulfidibacterota | Thermosulfidibacteria | NZ_AP013035.1 | 2 |
| Athalassotoga saccharophila | Thermotogota | Thermotogae | NZ_AP019551.1 | 1 |
| Defluviitoga tunisiensis | Thermotogota | Thermotogae | NZ_LN824141.1 | 1 |
| Fervidobacterium changbaicum | Thermotogota | Thermotogae | NZ_CP026721.1 | 1 |

| | | | | |
|---|---|---|---|---|
| Fervidobacterium islandicum | Thermotogota | Thermotogae | NZ_CP014334.2 | 1 |
| Fervidobacterium nodosum Rt17-B1 | Thermotogota | Thermotogae | NC_009718.1 | 1 |
| Fervidobacterium pennivorans DSM 9078 | Thermotogota | Thermotogae | NC_017095.1 | 1 |
| Kosmotoga olearia TBF 19.5.1 | Thermotogota | Thermotogae | NC_012785.1 | 1 |
| Kosmotoga pacifica | Thermotogota | Thermotogae | NZ_CP011232.1 | 1 |
| Marinitoga aeolica | Thermotogota | Thermotogae | NZ_CP069362.1 | 1 |
| Marinitoga piezophila KA3 | Thermotogota | Thermotogae | NC_016751.1 | 1 |
| Mesotoga infera | Thermotogota | Thermotogae | NZ_LS974202.1 | 1 |
| Mesotoga prima MesG1.Ag.4.2 | Thermotogota | Thermotogae | NC_017934.1 | 1 |
| Petrotoga mobilis SJ95 | Thermotogota | Thermotogae | NC_010003.1 | 1 |
| Pseudothermotoga elfii DSM 9442 = NBRC 107921 | Thermotogota | Thermotogae | NC_022792.1 | 1 |
| Pseudothermotoga hypogea DSM 11164 = NBRC 106472 | Thermotogota | Thermotogae | NC_022795.1 | 1 |
| Pseudothermotoga lettingae TMO | Thermotogota | Thermotogae | NC_009828.1 | 1 |
| Pseudothermotoga thermarum DSM 5069 | Thermotogota | Thermotogae | NC_015707.1 | 1 |
| Tepiditoga spiralis | Thermotogota | Thermotogae | NZ_AP018712.1 | 1 |
| Thermosipho africanus TCF52B | Thermotogota | Thermotogae | NC_011653.1 | 1 |
| Thermosipho ferrireducens | Thermotogota | Thermotogae | NZ_CP071446.1 | 1 |
| Thermosipho melanesiensis BI429 | Thermotogota | Thermotogae | NC_009616.1 | 1 |
| Thermotoga caldifontis AZM44c09 | Thermotogota | Thermotogae | NZ_AP014509.1 | 1 |
| Thermotoga maritima MSB8 | Thermotogota | Thermotogae | NC_023151.1 | 1 |
| Thermotoga neapolitana DSM 4359 | Thermotogota | Thermotogae | NC_011978.1 | 1 |
| Thermotoga petrophila RKU-1 | Thermotogota | Thermotogae | NC_009486.1 | 1 |
| Thermotoga profunda AZM34c06 | Thermotogota | Thermotogae | NZ_AP014510.1 | 1 |
| Candidatus Methylacidiphilum fumarolicum | Verrucomicrobiota | Methylacidiphilae | NZ_OX458932.1 | 1 |
| Methylacidiphilum caldifontis | Verrucomicrobiota | Methylacidiphilae | NZ_CP065957.1 | 1 |
| Methylacidiphilum infernorum V4 | Verrucomicrobiota | Methylacidiphilae | NC_010794.1 | 1 |
| Methylacidiphilum kamchatkense Kam1 | Verrucomicrobiota | Methylacidiphilae | NZ_CP037899.1 | 1 |
| Actomonas aquatica | Verrucomicrobiota | Opitutia | NZ_CP139781.1 | 1 |
| Candidatus Pelagisphaera phototrophica | Verrucomicrobiota | Opitutia | NZ_CP076039.1 | 1 |
| Coraliomargarita akajimensis DSM 45221 | Verrucomicrobiota | Opitutia | NC_014008.1 | 1 |
| Coraliomargarita algicola | Verrucomicrobiota | Opitutia | NZ_CP138858.1 | 1 |
| Ereboglobus luteus | Verrucomicrobiota | Opitutia | NZ_CP023004.1 | 1 |

| | | | | |
|---|---|---|---|---|
| Horticoccus luteus | Verrucomicrobiota | Opitutia | NZ_CP080507.1 | 1 |
| Lacunisphaera limnophila | Verrucomicrobiota | Opitutia | NZ_CP016094.1 | 1 |
| Nibricoccus aquaticus | Verrucomicrobiota | Opitutia | NZ_CP023344.1 | 1 |
| Opitutus terrae PB90-1 | Verrucomicrobiota | Opitutia | NC_010571.1 | 1 |
| Rubellicoccus peritrichatus | Verrucomicrobiota | Opitutia | NZ_CP136920.1 | 1 |
| Synoicihabitans lomoniglobus | Verrucomicrobiota | Opitutia | NZ_CP119075.1 | 1 |
| Termitidicoccus mucosus | Verrucomicrobiota | Opitutia | NZ_CP109796.1 | 1 |
| Akkermansia biwaensis | Verrucomicrobiota | Verrucomicrobiia | NZ_AP025943.1 | 1 |
| Akkermansia glycaniphila | Verrucomicrobiota | Verrucomicrobiia | NZ_LT629973.1 | 1 |
| Akkermansia muciniphila | Verrucomicrobiota | Verrucomicrobiia | NZ_AP021898.1 | 1 |
| Fontisphaera persica | Verrucomicrobiota | Verrucomicrobiia | NZ_CP116615.1 | 1 |
| Haloferula helveola | Verrucomicrobiota | Verrucomicrobiia | NZ_AP024702.1 | 1 |
| Luteolibacter ambystomatis | Verrucomicrobiota | Verrucomicrobiia | NZ_CP073100.1 | 1 |
| Luteolibacter luteus | Verrucomicrobiota | Verrucomicrobiia | NZ_CP051774.1 | 1 |
| Sulfuriroseicoccus oceanibius | Verrucomicrobiota | Verrucomicrobiia | NZ_CP066776.1 | 1 |
| Verrucomicrobium spinosum DSM 4136 = JCM 18804 | Verrucomicrobiota | Verrucomicrobiia | NZ_ABIZ01000001 | 1 |
| Vulcanimicrobium alpinum | Vulcanimicrobiota | Vulcanimicrobiia | NZ_AP025523.1 | 1 |

| | # of tuf genes in the genome | | | | |
|---|---|---|---|---|---|
| **Phylum\Class** | **1** | **2** | **3** | **4** | **Total species count:** |
| **Acidobacteriota** | 4 | 21 | | | 25 |
| Blastocatellia | | 3 | | | 3 |
| Holophagae | 4 | 1 | | | 5 |
| Terriglobia | | 16 | | | 16 |
| Vicinamibacteria | | 1 | | | 1 |
| **Actinomycetota** | 848 | 214 | 16 | 1 | 1079 |
| Acidimicrobiia | 1 | 5 | | | 6 |
| Actinomycetes | 814 | 204 | 16 | 1 | 1035 |
| Coriobacteriia | 25 | | | | 25 |
| Nitriliruptoria | | 3 | | | 3 |
| Rubrobacteria | | 2 | | | 2 |
| Thermoleophilia | 8 | | | | 8 |
| **Aquificota** | 1 | 8 | | | 9 |
| Aquificia | 1 | 8 | | | 9 |
| **Armatimonadota** | 1 | 2 | | | 3 |
| Armatimonadia | | 1 | | | 1 |
| Chthonomonadia | | 1 | | | 1 |
| Fimbriimonadia | 1 | | | | 1 |
| **Atribacterota** | | 3 | | | 3 |
| Atribacteria | | 3 | | | 3 |
| **Bacillota** | 795 | 193 | 1 | | 989 |
| Bacilli | 629 | 22 | 1 | | 652 |
| Clostridia | 129 | 128 | | | 257 |
| Dethiobacteria | 1 | | | | 1 |
| Erysipelotrichia | 23 | | | | 23 |
| Limnochordia | 2 | 1 | | | 3 |
| Negativicutes | 7 | 27 | | | 34 |
| Tissierellia | 4 | 15 | | | 19 |
| **Bacteroidota** | 544 | | | | 544 |
| Bacteroidia | 99 | | | | 99 |
| Chitinophagia | 30 | | | | 30 |

| | | | | | |
|---|---|---|---|---|---|
| Cytophagia | 107 | | | | 107 |
| Flavobacteriia | 264 | | | | 264 |
| Saprospiria | 3 | | | | 3 |
| Sphingobacteriia | 41 | | | | 41 |
| **Balneolota** | | 1 | | | 1 |
| Balneolia | | 1 | | | 1 |
| **Bdellovibrionota** | 6 | 3 | | | 9 |
| Bacteriovoracia | | 3 | | | 3 |
| Bdellovibrionia | 3 | | | | 3 |
| Oligoflexia | 3 | | | | 3 |
| **Caldisericota** | | 1 | | | 1 |
| Caldisericia | | 1 | | | 1 |
| **Calditrichota** | | 1 | | | 1 |
| Calditrichia | | 1 | | | 1 |
| **Campylobacterota** | 124 | 1 | | | 125 |
| Desulfurellia | | 1 | | | 1 |
| Epsilonproteobacteria | 124 | | | | 124 |
| **Candidatus Cloacimonadota** | 1 | | | | 1 |
| Candidatus Cloacimonadia | 1 | | | | 1 |
| **Candidatus Paceibacterota** | 1 | | | | 1 |
| Candidatus Paceibacteria | 1 | | | | 1 |
| **Candidatus Saccharibacteria** | 4 | | | | 4 |
| Candidatus Saccharimonadia | 4 | | | | 4 |
| **Chlamydiota** | 24 | | | | 24 |
| Chlamydiia | 24 | | | | 24 |
| **Chlorobiota** | 9 | | | | 9 |
| Chlorobiia | 9 | | | | 9 |
| **Chloroflexota** | 9 | 9 | | | 18 |
| Anaerolineae | 1 | 2 | | | 3 |
| Ardenticatenia | 2 | | | | 2 |
| Caldilineae | | 1 | | | 1 |
| Candidatus Thermofonsia | 1 | | | | 1 |
| Chloroflexia | | 4 | | | 4 |
| Dehalococcoidia | 5 | | | | 5 |

| | | | | | |
|---|---|---|---|---|---|
| Ktedonobacteria | | 1 | | | 1 |
| Tepidiformia | | 1 | | | 1 |
| **Chrysiogenota** | 1 | | | | 1 |
| Chrysiogenia | 1 | | | | 1 |
| **Coprothermobacterota** | | 1 | | | 1 |
| Coprothermobacteria | | 1 | | | 1 |
| **Cyanobacteriota** | 72 | 1 | | | 73 |
| Cyanophyceae | 72 | 1 | | | 73 |
| **Deferribacterota** | 1 | 5 | | | 6 |
| Deferribacteres | 1 | 5 | | | 6 |
| **Deinococcota** | | 33 | | | 33 |
| Deinococci | | 33 | | | 33 |
| **Dictyoglomota** | | 2 | | | 2 |
| Dictyoglomia | | 2 | | | 2 |
| **Elusimicrobiota** | | 3 | | | 3 |
| Elusimicrobia | | 1 | | | 1 |
| Endomicrobiia | | 2 | | | 2 |
| **Fibrobacterota** | | 1 | | | 1 |
| Fibrobacteria | | 1 | | | 1 |
| **Fusobacteriota** | 7 | 17 | | | 24 |
| Fusobacteriia | 7 | 17 | | | 24 |
| **Gemmatimonadota** | 5 | | | | 5 |
| Gemmatimonadia | 5 | | | | 5 |
| **Ignavibacteriota** | 2 | | | | 2 |
| Ignavibacteria | 2 | | | | 2 |
| **Kiritimatiellota** | 2 | | | | 2 |
| Kiritimatiellia | 1 | | | | 1 |
| Tichowtungiia | 1 | | | | 1 |
| **Lentisphaerota** | 1 | | | | 1 |
| Lentisphaeria | 1 | | | | 1 |
| **Mycoplasmatota** | 136 | 1 | | | 137 |
| Candidatus Izimaplasma | 2 | | | | 2 |
| Mollicutes | 134 | 1 | | | 135 |
| **Myxococcota** | | 14 | 3 | | 17 |

| | | | | | |
|---|---|---|---|---|---|
| Myxococcia | | 14 | 3 | | 17 |
| **Nitrospinota** | 1 | | | | 1 |
| Nitrospinia | 1 | | | | 1 |
| **Nitrospirota** | | 10 | | | 10 |
| Nitrospiria | | 6 | | | 6 |
| Thermodesulfovibrionia | | 4 | | | 4 |
| **Planctomycetota** | 50 | 16 | | | 66 |
| Candidatus Brocadiia | | 1 | | | 1 |
| Candidatus Uabimicrobiia | 2 | | | | 2 |
| Phycisphaerae | 7 | 1 | | | 8 |
| Planctomycetia | 41 | 14 | | | 55 |
| **Pseudomonadota** | 575 | 2027 | 13 | | 2615 |
| Acidithiobacillia | | 11 | | | 11 |
| Alphaproteobacteria | 311 | 621 | 1 | | 933 |
| Betaproteobacteria | 7 | 398 | | | 405 |
| Gammaproteobacteria | 257 | 994 | 12 | | 1263 |
| Hydrogenophilia | | 1 | | | 1 |
| Zetaproteobacteria | | 2 | | | 2 |
| **Rhodothermota** | | 3 | | | 3 |
| Rhodothermia | | 3 | | | 3 |
| **Spirochaetota** | 83 | | | | 83 |
| Spirochaetia | 83 | | | | 83 |
| **Synergistota** | | 8 | | | 8 |
| Synergistia | | 8 | | | 8 |
| **Thermodesulfobacteriota** | 30 | 69 | | | 99 |
| Candidatus Desulfofervidia | | 1 | | | 1 |
| Desulfarculia | | 2 | | | 2 |
| Desulfobaccia | | 1 | | | 1 |
| Desulfobacteria | 3 | 11 | | | 14 |
| Desulfobulbia | | 11 | | | 11 |
| Desulfomonilia | | 1 | | | 1 |
| Desulfovibrionia | 25 | 11 | | | 36 |
| Desulfuromonadia | | 23 | | | 23 |
| Syntrophia | 2 | | | | 2 |

| | | | | | |
|---|---|---|---|---|---|
| Syntrophobacteria | | 2 | | | 2 |
| Thermodesulfobacteria | | 6 | | | 6 |
| **Thermodesulfobiota** | | 2 | | | 2 |
| Thermodesulfobiia | | 2 | | | 2 |
| **Thermomicrobiota** | | 2 | | | 2 |
| Thermomicrobia | | 2 | | | 2 |
| **Thermosulfidibacterota** | | 1 | | | 1 |
| Thermosulfidibacteria | | 1 | | | 1 |
| **Thermotogota** | 26 | | | | 26 |
| Thermotogae | 26 | | | | 26 |
| **Verrucomicrobiota** | 25 | | | | 25 |
| Methylacidiphilae | 4 | | | | 4 |
| Opitutia | 12 | | | | 12 |
| Verrucomicrobiia | 9 | | | | 9 |
| **Vulcanimicrobiota** | 1 | | | | 1 |
| Vulcanimicrobiia | 1 | | | | 1 |
| **Total species count:** | **3389** | **2673** | **33** | **1** | **6096** |
